# Leaf movements as a quantitative metric for early stress detection

**DOI:** 10.64898/2026.06.16.732190

**Authors:** Eva Herrero, Samalka Wijeweera, Alison R. Gill, Charlotte Bampton, Wendy Sullivan, John D. Stamford, Jennifer Bromley, Andreas Z. Antoniades, Jenny C. Mortimer, Alex A.R. Webb, Matthew Gilliham, A. Harvey Millar

## Abstract

Early, precise, and non-destructive stress detection is essential for maintaining crop productivity, particularly in high-density plant growth systems like controlled environment agriculture (CEA), where manual monitoring is often impractical. Using plant motion as a proxy for growth and plant health, we demonstrate a method for early, non-invasive stress detection through quantitative leaf-movement analysis in lettuce and five other CEA relevant crops. Leaf-movement dynamics under stress were imaged with a low-cost, scalable Raspberry Pi imaging setup and quantified using a repurposed open-source motion estimation algorithm; Tracking Rhythms in Plants (TRiP). Our system detected stress-induced changes in leaf-movement within 1 hour of stress, with the timing dependent on the nature of the stress. Sustained reductions in leaf-movement coincide with decreased biomass accumulation. This approach offers a non-invasive, rapid, scalable, and cost-effective solution for continuous crop monitoring, with potential for application in both terrestrial and space farming CEA systems.

Graphical abstract:
Quantification of leaf-movement dynamics as a high-throughput proxy for plant physiological status, enabling early stress detection and timely intervention to mitigate yield penalties in CEA settings (image made with biorender.org).

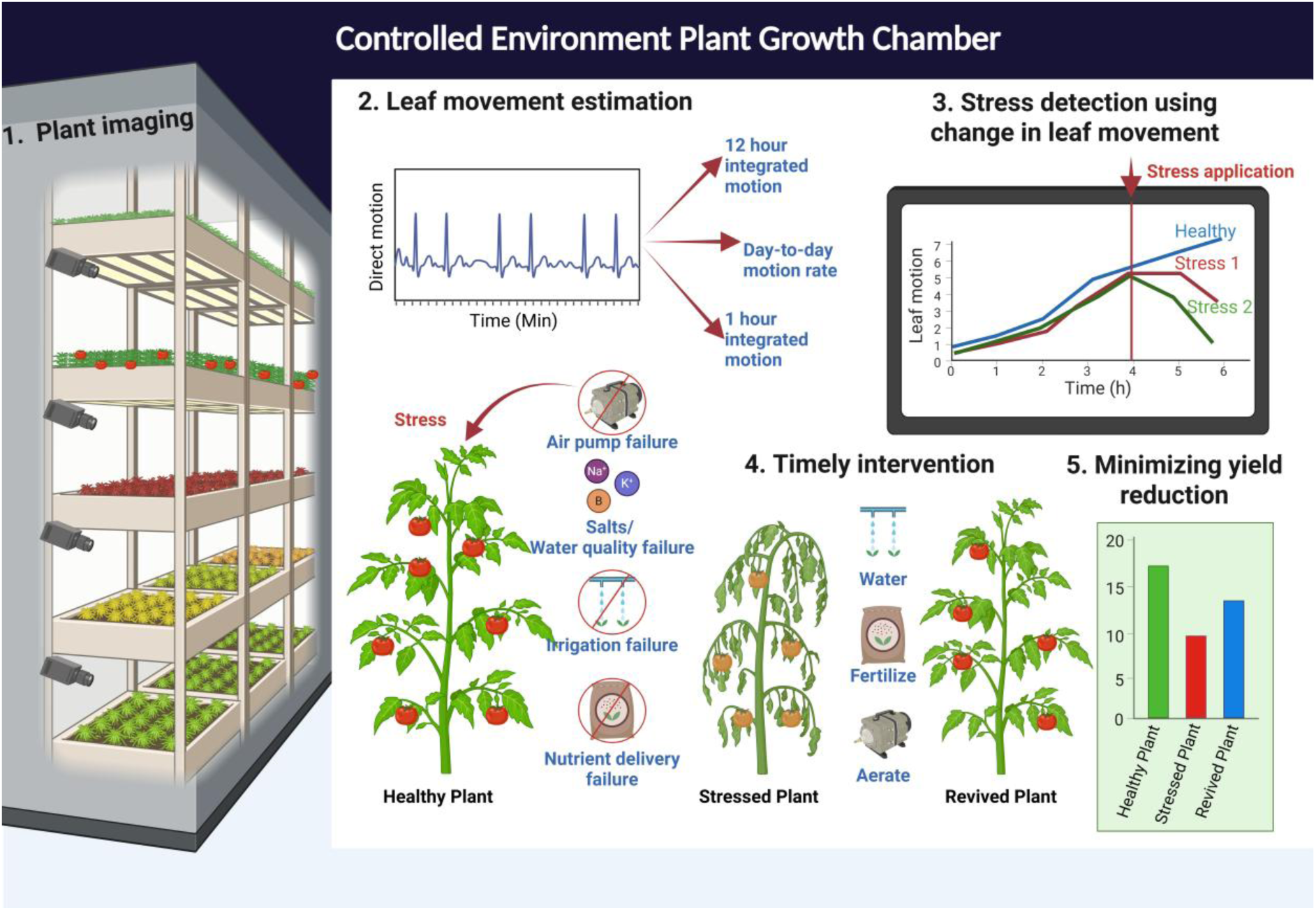

## Introduction

Environmental stresses are responsible for up to 80% of annual crop yield losses (Junaid & Gökçe, 2024), reducing productivity and posing a major challenge to global food security (Humplík et al., 2015). Although modern large-scale controlled-environment agriculture (CEA) systems provide precise regulation of growth conditions, crops remain vulnerable to a range of stresses, including irrigation failures, nutrient imbalances, salinity, suboptimal lighting, equipment or system faults, which can each rapidly impair plant performance and yield (Ragaveena, et al 2021). Addressing these challenges requires high-throughput non-invasive methods capable of detecting plant stress at early stages. In this context, automated phenotyping tools based on imaging technologies offer significant potential for continuous monitoring and early stress detection in large-scale agricultural production systems (Humplík et al., 2015).

A wide range of advanced imaging-based tools has been developed to detect plant stress and monitor plant health in controlled environments (CE), including infrared, near-infrared, ultraviolet, fluorescence, and multi or hyperspectral imaging approaches (Zubler & Yoon, 2020). These techniques can provide rich physiological information and are increasingly integrated into multi-sensor platforms to improve precision and automation (Walsh et al., 2024). However, their cost, technical complexity, and substantial data-processing requirements limit their routine deployment in commercial settings.

The majority of plant imaging for health monitoring in CE is still performed using the visible range of light (red-green-blue/RGB) spectrum, owing to simplicity, accessibility, and compatibility with low-cost imaging hardware. But despite major advances in plant phenotyping (Pieruschka & Schurr, 2019; Sarić et al., 2022), many RGB-based platforms remain primarily focused on growth assessment rather than early stress detection and is only carried out during the daylight period. Many existing RGB-imaging systems require complex infrastructure, lack robustness across applications, do not support continuous monitoring, or rely on narrowly defined detection parameters that vary with plant stress condition (Sarić et al., 2022).

Many dicotyledonous plants have diel leaf-movements (nyctinasty) regulated by the circadian clock and light signalling, while higher-frequency movements, known as circumnutations, are also observed and result from differential growth at the base of the petiole (Bours et al., 2012; Geldhof et al., 2021). Beyond endogenous rhythms, changes in environmental conditions including light, temperature, salinity, and water availability, can induce pronounced nastic responses, such as pronounced epinastic (downward) and hyponastic (upward) leaf-movements (Park et al., 2019; Polko et al., 2011). As many abiotic stresses inhibit plant growth, associated reductions in the amplitude or dynamics of leaf-movement may provide early indicators of stress (Geldhof et al., 2021).

Leaf-movement monitoring has been explored as a proxy for plant stress detection (Geldhof et al., 2021; Humplík et al., 2015; Rehman et al., 2020; Vantoai & Roberts, 2003) and for estimating plant growth (Nagano et al., 2019). While contact-based digital sensors mounted on leaves or stems have been used to track leaf-angle dynamics across developmental stages and under stress (Geldhof et al., 2021), non-invasive imaging approaches offer higher-throughput, commercially deployable alternatives. These methods combine time-lapse imaging with algorithms to extract movement-related image features, including top projected canopy area (Vantoai & Roberts, 2003), mean pixel displacement or composite motion metrices (Nagano et al., 2019). For leaf-movement dynamics to be widely adopted as indicators of plant stress, imaging and data-analysis solutions must be both low-cost and operationally streamlined (Walsh et al., 2024), while also providing quantitative metrics capable of early and reliable stress detection.

We recently established an inexpensive and straightforward imaging system that enables quantitative analysis of diel leaf-movement dynamics in lettuce and Arabidopsis grown in a laboratory facsimile of a commercial vertical farm (Herrero et al., 2026). The system uses a Raspberry Pi Camera Module 3 NoIR to acquire time-lapse images under very low-intensity green light (dimG < 0.3 μmol m⁻² s⁻¹), allowing continuous day and night imaging under stable illumination conditions (Herrero et al., 2026). Using a motion quantification approach based on the Tracking Rhythms in Plants (TRiP) algorithm (Greenham et al., 2015), modified to derive a direct movement profile (Herrero et al., 2026), we captured diel movements driven by light–dark cycles, higher-frequency fluctuations predominantly occurring at night, and a progressive increase in overall movement consistent with growth.

In this study, we extend our leaf-movement quantification framework into a versatile and broadly applicable platform for plant stress detection. The system enables simultaneous monitoring of stress responses at both the individual-plant level and across plant arrays. Using this approach, we detect both early stress-induced alterations in leaf-movement dynamics and sustained inhibition of movement align with reduced growth. We validate the platform across multiple stress conditions in lettuce, across independent experimental locations, and in several other horticultural crop species, demonstrating its suitability for deployment in CEA.

## Materials and Methods

### Plant material and growth conditions

All experiments were conducted in CE facilities across three sites with consistent environmental conditions (temperature daytime 22°C, nighttime 20°C; humidity 60%) (Supplementary Information S1). At the University of Western Australia (UWA); a closed Plant Growth Chamber Flex (Conviron, Manitoba, Canada) with precise temperature and humidity control, at Adelaide University (AU); a walk-in plant growth room with temperature and humidity control and at the University of Cambridge (UoC); a laboratory room with broad temperature and humidity control were used. Commercially developed horticultural LED fixtures (Vertical Future, UK) were used in all three locations to provide light conditions representative of typical production settings. LED arrays were placed 30 cm above the plant canopy provided 160 µmolm^-2^s^-1^ light in the form of 105 µmolm^-2^s^-1^ red (wavelength 600-700 nm), 20 μmolm^-2^s^-1^ green (wavelength 500-600 nm), and 35 μmolm^-2^s^-1^ blue (wavelength 400-500 nm) at plant height. The photoperiod was 12 h of light and 12 h of dark.

Lettuce (*Lactuca sativa* var. Outredgeous), radish (*Raphanus sativus* var. cherry belle), mizuna (*Brassica juncea* var. japonica), tomato (*Solanum lycopersicum* var. tommy toe), rocket *(Eruca sativa* var. runway), and amaranth (*Amaranthus hypochondiacus* AGG acc. 132876) were the horticultural crops used. Seeds were germinated in coco coir for 17 days before transfer to hydroponics using ‘Ionic Grow’ nutrient solution (Growth Technology, Australia) added at 5 mL/L (half strength) and adjusted to pH 6 using pH UP (Growth Technology, Australia). Hydroponic tanks consisted of 30 L plastic tubs with twelve round holes of diameter 5 cm in the lid, for a total of twelve plants per tank. 24 L of full strength ‘Ionic Grow’ nutrient solution (10 mL/L) with pH adjusted to 6 using pH UP was added to each tank, with basal nutrients as follows: nitrate 2.20% (w/v), phosphorus 0.35%, potassium 3.19%, calcium 1.16%, magnesium 0.44%, sulphur 0.12%, iron 0.04%, manganese 0.007%, boron 0.003%, zinc 0.003%, copper 0.002%, and molybdenum 0.001%. An aquarium pump continuously aerated the nutrient solution in each tank.

On day 17, seedlings were removed from the coco coir, washed with DI water, put into a foam collar, and placed into the holes in the lid. Salinity, nutrient withdrawal and hypoxia stress treatments were imposed 7 days after the plants were moved to hydroponics (day 24 post sowing), and the plants were grown for an additional 14 days after stress imposition (Figure 1a). As an exception, water withdrawal stress was imposed after 12 days, but plants were grown for only another 8 days after water re-addition. Stresses imposed were as follows: Salinity (100 mM NaCl added in one dose [140 g NaCl per tank], or 150 mM NaCl added in 3 doses over 3 days [210 g NaCl per tank] or 100 mM KCl [179 g per tank]) added in one dose, Nutrient withdrawal (NW) (nutrient solution replaced with water), Hypoxia (N_2(g)_ bubbled through solution at 2-3 L/min), 100 mM Boron (as boric acid, 370.71 mg per tank), and water withdrawal (all solution was removed from the tub for 24 h). For timeline of the experiments refer to experiment log (Supplementary Information S2).

**Figure 1.**
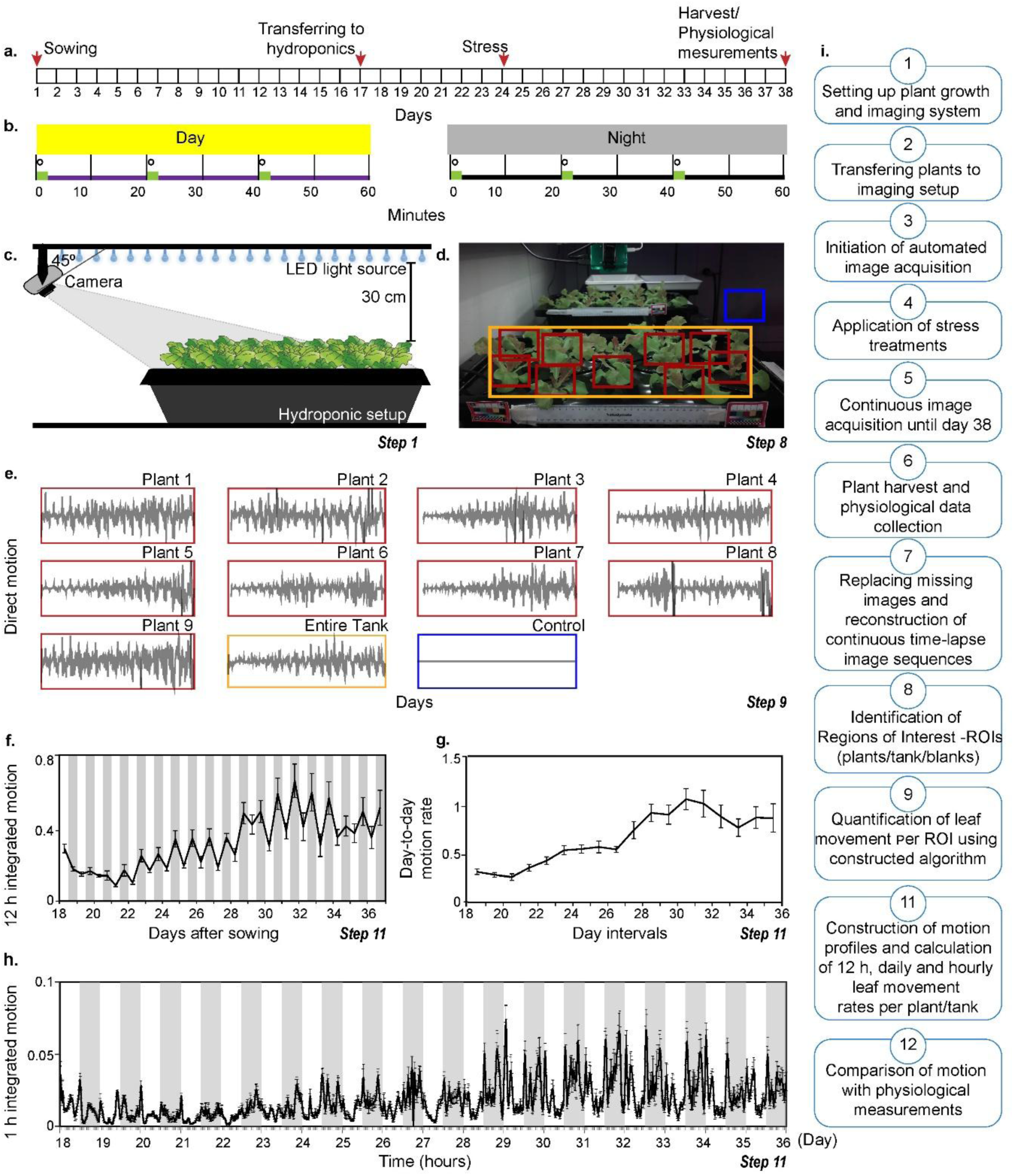
| Timeline, imaging setup, and motion-based outputs for quantifying leaf-movement dynamics during stress exposure. **(a)** representative timeline of imaging, stress application, and harvest physiological measurements. **(b)** imaging and illumination schedule. During the daytime, full-spectrum RGB illumination (purple line) was used to support plant growth, while darkness (black line) was maintained at night. The day–night illumination regime was interrupted every 20 minutes by a 2-minute pulse of dimG illumination (green lines), synchronised with image acquisition. **(c)** overview of the plant growth and imaging setup (step 1). **(d)** representative image showing regions of interest (ROIs) defined for motion estimation: individual plants (red), the entire tank (yellow), and a control region without plants (blue) (step 8). **(e)** representative direct motion time series for each ROI (step 9). **(f)** 12 h integrated motion per tank (step 11). **(g)** day-to-day motion rate per tank (step 11). **(h)** 1 h integrated motion shown as a moving average of three frames per tank (step 11). Error bars represent SEM (n = 9) (see Supplementary Table S1a-e for numerical data). **(i)** schematic overview of key steps from plant growth to stress detection.

#### Leaf-movement imaging system

The imaging was performed as described previously (Herrero et al. 2026). Briefly, images were captured from day 17 to day 38 under a 2 min pulse of dim green light (dimG, 0.3 µmol m^-2^ s^-1^, wavelength 500-600 nm) (Figure 1b), using a Raspberry Pi Camera Module 3 NoIR (Herrero et al., 2026) positioned 20 cm above the plants at a 45° angle centred on the longest side (Figure 1c). Each image consisted of the average of 90 consecutive frames to minimise high frequency leaf-movements caused by airflow and reduce illumination variability during dimG pulses. Image data can be accessed online (Mortimer & Gilliham, 2025).

### Harvest measurements

Plants were harvested 38 days after sowing (Figure 1a). Chlorophyll content was measured using a chlorophyll meter (CCM-300, Opti-Sciences, United States, or SPAD-502, Konica Minolta, United States) at four positions per leaf on the third youngest leaf. Six to twelve plants were measured per tank. Plants were cut at the intersection between the shoot and root, with both parts patted dried and weighed for fresh weight (FW). Shoot and root dry weight (DW) was measured after drying at 60°C for at least 48 h.

### Quantification of leaf-movement dynamics based on TRiP

The analysis of leaf-movement dynamics was performed as previously described (Herrero et al., 2026). The analysis initiated on day 18 at ZT0 (Zeitgeber time, 0 hours after dawn). Regions of interest (ROI) covering 9-12 individual plants per tank and one ROI covering the entire tank were selected, with plant growth over time considered to ensure plants remained within the ROI. In addition, one ROI covering a control region with no plants (and therefore no movement) were also selected (Figure 1d). ROI movement was estimated with the TRiP algorithm (Greenham et al., 2015), with a modification to reduce the averaging window of movement visualization (line 83: taps = 13 replaced by taps = 1) to obtain the ‘*direct motion’* (Figure 1e). We calculated the trapezoidal area under the curve (trapz AUC) to estimate the ‘*12 h integrated motion’* for the day or night interval (Figure 1f) and to estimate ‘*day-to-day motion rate*’ (Figure 1g), which was calculated as the slope of cumulative area of direct motion between consecutive days. To increase sensitivity of the movement estimate we calculated ‘*1 h integrated motion’* using hourly trapezoidal area under the curve (Figure 1h; Supplementary Table S1a-e). The TRiP output contains seven fewer measurements than the image sequence, leading to a forward shift in the motion curve when plotting 1 h integrated motion. This offset was corrected prior to the integration step. For each tank, average values and standard error of the mean (SEM) were calculated (Figure 1f-h). The schematic of the sequential process from plant growth to motion estimation is also visualised (Figure 1i).

### Statistical analysis

For the harvest data, two-sample *t*-tests (Non-stressed (Control) vs. Stressed (Treatment)) and 1-way ANOVA Tukey’s HSD tests were performed, and graphs were generated, in GraphPad Prism (version 10.5.0). For motion data, 12 h integrated motion and day-to-day motion rate following stress imposition were compared to the corresponding values at day 23 using two-sample t-tests to identify significant differences in leaf movement in response to stress.

## Results

### Quantification of leaf-movement dynamics for stress detection

To test the applicably of our system for early stress detection, we applied a range of stresses relevant to CEA on lettuce plants during their mid-growth cycle and leaf movement was quantified (Supplementary Figure S1a-b; Supplementary Table S2a-b). Salinity, a common constraint in hydroponic and substrate based systems that impairs growth and plant physiology (Liu et al., 2024), was tested using three regimes: (i) a single step application of 100 mM NaCl (Figure 2a-c), and (ii) a single step application of 100 mM KCl (Figure 2d-f) and (iii) a gradual increase of 50 mM NaCl per day to a final concentration of 150 mM (Supplementary Figure S2a-c). These regimes were designed to distinguish between progressively accumulating stress and strongly imposed acute stress (Munns, 1993). KCl was used at the same concentration as NaCl to check general osmotic effects from Na⁺ specific ion toxicity (Munns, 2002).

**Figure 2.**
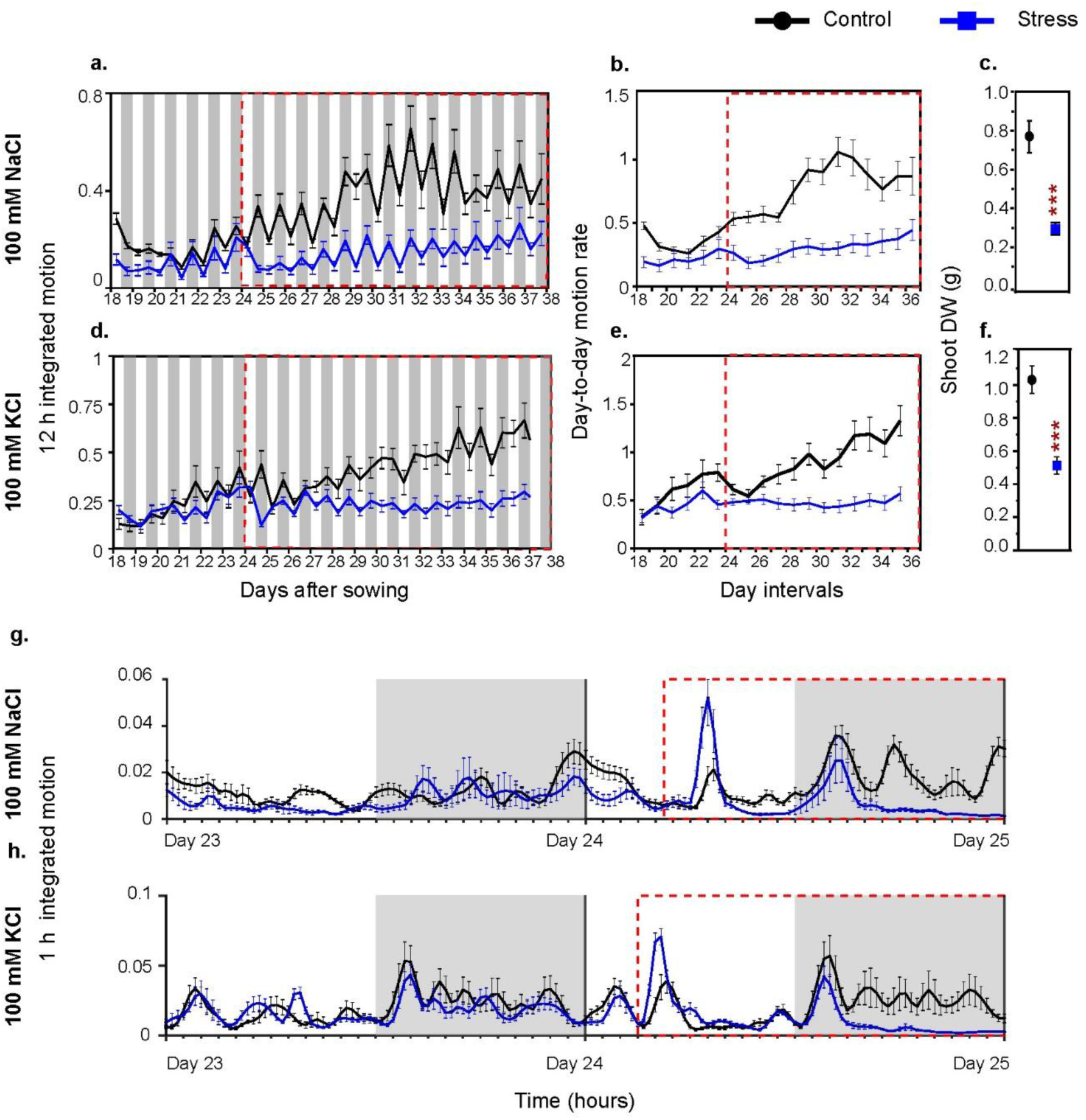
| Salinity stress reduces lettuce leaf-movement dynamics and biomass accumulation. Responses to salinity treatments: **(a–c)** 100 mM NaCl and **(d–f)** 100 mM KCl. Panels show **(a,d)** 12 h integrated motion, **(b,e)** day-to-day motion rate, and **(c,f)** shoot dry weight (DW). Panels **(g-h)** show 1 h integrated motion following the application of **(g)** 100 mM NaCl and **(h)** 100 mM KCl, displayed from day 23 to day 25 to highlight the immediate changes in leaf-movement dynamics after stress application on day 24. Data represent means of 9-12 lettuce plants for non-stressed (black) and stress (blue) treatments; error bars indicate SEM (see Supplementary Table S3b, S4b, S5b and S6a for numerical data). The red dashed line marks the timing of stress application. White and grey backgrounds indicate day and night, respectively. Asterisks denote significant differences between stress and non-stressed treatments (\**p* < 0.05, \*\**p* < 0.01, \*\*\**p* < 0.001; two sample *t*-tests).

Hypoxic conditions (Supplementary Figure S2d-f) were induced to mimic air-pump failures frequently observed in CE settings. Excess boron (B) (Supplementary Figure S2g-i) supply was applied to assess ion toxicity arising from nutrient mis-delivery; a known risk in hydroponic systems given the variability in irrigation water quality (Brdar-jokanovi, 2020; Song et al., 2022). Plants were subjected to nutrient withdrawal (NW) (Figure 3a-c), to model nutrient delivery failures encountered in CEA systems. A transient 24 h water-withdrawal treatment was used to mimic irrigation failures associated with emitter clogging (Shi et al., 2022) and to assess post-stress recovery (Figure 3d-f).

**Figure 3.**
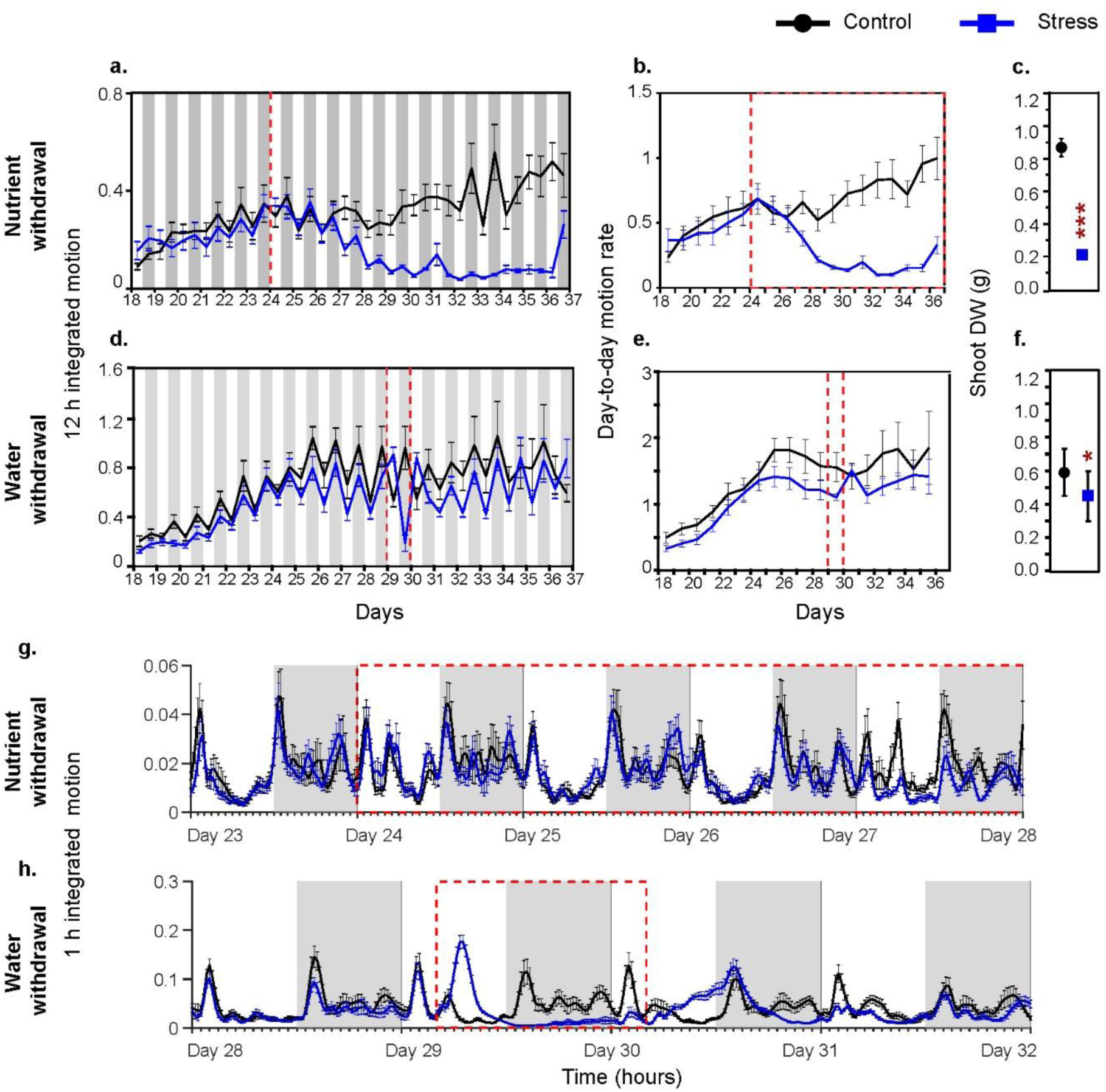
| Nutrient and water delivery failures alter lettuce leaf-movement dynamics and biomass accumulation. Responses to **(a–c)** nutrient withdrawal and **(d–f)** water withdrawal. Panels show **(a,d)** 12 h integrated motion, **(b,e)** day-to-day motion rate, and **(c,f)** shoot dry weight (DW) measured at day 38. Panels **(g-h)** show 1 h integrated motion following **(g)** nutrient withdrawal and **(h)** water withdrawal, displayed from the day prior to stress application to the onset of changes in leaf-movement dynamics after stress application. Data represent means of 9–12 lettuce plants for non-stressed (black) and stress (blue) treatments; error bars indicate SEM (see Supplementary Table S3b, S4b, S5b and S6a for numerical data). Red dashed lines mark the timing of stress application. White and grey backgrounds indicate day and night periods, respectively. Asterisks denote significant differences between stressed and non-stressed plants (*p < 0.05, **p < 0.01, ***p < 0.001; two sample t-tests).

### Acute and long-term changes in leaf-movement dynamics upon stress application

Under non-stressed conditions (control), lettuce plants had greater leaf movement during night than during the day, as visualised with 12 h integrated motion. The day-to-day motion rate progressively increased over time coinciding with visible increases in plant size observed in the image sequences (Figures 2-3; Supplementary Figure S2-S3; Supplementary Table S3a-b and S4a-b). To assess reproducibility across CE settings, we compared non-stressed experiments conducted at three locations with equivalent CEs (UWA, AU and UoC; Supplementary Information S1). Across sites, lettuce plants had consistent leaf-movement dynamics with only minor differences, accompanied by comparable shoot dry weight (DW) at the end of the experiment (Supplementary Figure S3a-c; Supplementary Table S2b, S3b, S4b, S5b). These results indicate robust and comparable performance of the imaging approach across all three locations despite their minor variations in room set up, temperature and humidity control.

In stressed plants, the baseline leaf-movement dynamics prior to stress application were comparable to non-stressed plants, showing an overall increase in motion over time (Figure 2; Supplementary Figure S2a-c, S3). During stepwise NaCl application at UWA (50 mM added per day to a final concentration of 150 mM to limit osmotic shock), nighttime leaf movement was already reduced after the first 50 mM increment on day 24 (Supplementary Figure S2a, S3d). After completion of salt application, from day 27 (1 day after stress completion) onwards, the increase in day-to-day motion plateaued and remained below non-stressed levels for the remainder of the experiment (Supplementary Figure S2b, S3e), consistent with a strong reduction in shoot DW (Supplementary Figure S2c, S3f). Single-step application of 100 mM NaCl on day 24 (day of stress) at AU caused a sharp increase in daytime leaf movement (113%), consistent with acute turgor loss, followed by a 62% reduction during the subsequent night (Figure 2a; Supplementary Figure S2d; Supplementary Table S3a-b). From day 25 (1 day after stress) onwards, the day-to-day motion rate remained below non-stressed levels until the end of the experiment (Figure 2b; Supplementary Figure S2e; Supplementary Table S4a-b). A similar response was observed at UoC on day 24 (day of stress), although the sustained reduction in motion was less pronounced than at AU (Supplementary Figure S2d-f). Although leaf-movement dynamics varied slightly between sites, all experiments demonstrated a consistent association between stress-induced changes in leaf-motion dynamics and reduced biomass accumulation (Supplementary Figure S3). Single-step application of 100 mM KCl elicited leaf-movement responses like those observed under NaCl treatment, although a transient recovery was evident on days 25–26 (Figure 2d). Nevertheless, from day 24 (day of stress) onwards, the day-to-day motion rate did not increase (Figure 2e). This response is consistent with the deceleration of leaf-movement dynamics observed across all salinity treatments and with the significantly reduced shoot DW accumulated by stressed plants compared to non-stressed (Figure 2f).

The rapid increase in lettuce leaf movement following one-step salt application prompted us to assess how rapidly changes in leaf-movement dynamics could be detected. Hourly integrated direct motion was therefore quantified (Figure 2g-h; Supplementary Figure S3j; Supplementary Table S6a). An epinastic response detected as increased motion was visible within 1 h of both one-step 100 mM NaCl and KCl application (Figure 2g-h), demonstrating the sensitivity of our imaging system for early stress detection. In both treatments, high-frequency nighttime leaf movements were reduced starting from day 24 (the day of stress). Taken together, these results show that both one-step and ramped salt treatments disrupted leaf-movement trajectories by preventing the progressive increase in leaf movement observed in non-stressed plants (Figure 2; Supplementary Figure S2a-c, S3).

Nutrient withdrawal caused a gradual reduction in both day and nighttime 12 h integrated motion from day 27 (3 days after stress) onwards (Figure 3a). From this point, the day-to-day motion rate declined and remained low (Figure 3b), consistent with strong, significant reduction in biomass accumulation relative to non-stressed plants (Figure 3c). To evaluate whether rapid stress detection combined with timely intervention could mitigate yield loss, we performed a transient 24 h water withdrawal to simulate irrigation interruption starting ZT4 (4 h after dawn) on day 29 (Figure 3d-f). This treatment triggered a sharp increase in motion within 1 h (Figure 3h), coincident with rapid dehydration and wilting. This response was also evident in the 12 h integrated motion profiles, which showed a pronounced increase (110%) in daytime motion, followed by a marked reduction in nighttime motion (Figure 3d; Supplementary Table S3b). Restoring irrigation on day 30 resulted in a rapid increase of motion as plants recovered turgor, although nighttime motion remained reduced (Figure 3d). By day 31 (1 day after stress recovery), normal rhythmic motion resumed (Figure 3d,e,h). End-point shoot DW showed a significant but relatively modest reduction from this transient water withdrawal (Figure 3f) compared to treatments with continuous stress exposure such as 100 mM NaCl and 100 mM KCl (Figure 2c,f), or nutrient withdrawal (Figure 3c). Overall, acute irrigation stress was detected within an hour, which could allow timely intervention to minimize yield loss (Figure 3h).

In contrast, hypoxia did not induce detectable changes in lettuce leaf-movement dynamics and had no effect on shoot DW (Supplementary Figure S3d-f), suggesting that this treatment did not induce measurable changes under the conditions tested. Excess boron treatment caused a transient reduction of leaf-movement between days 24 (day of stress) and 29 (5 days after stress), associated with a significant reduction in shoot DW (Supplementary Figure S3g-i). For several stress conditions, we performed replicate experiments within the same site, including hypoxia and nutrient withdrawal at UWA and 100 mM NaCl at AU (Supplementary Figure S4). These replicates confirmed the reproducibility of the observed leaf-movement dynamics and associated biomass responses.

Taken together, our system captured both rapid, acute increases in lettuce leaf movement occurring within hours of stress application and sustained reductions in motion across multiple stresses, with prolonged motion suppression associated with reduced biomass accumulation.

### Altered leaf movement in response to stress can be detected in different lighting regimes

In our experiments, image acquisition was synchonized with brief dimG illumination pulses to maintain consistent pixel intensity values and minimize image analysis artifacts associated with light-dark transitions. However, such pulsed dimG illumination might not be practical in many CEA settings, particularly in commercial applications. We demonstrated that image sequences acquired using full-spectrum RGB light during the day combined with dimG illumination at night (RGBday and dimG^night^) are sufficient to accurately quantify diel and high frequency leaf-movement dynamics (Herrero *et al*., 2026). To determine whether this simplified illumination regime affects stress detection, we compared leaf-movement dynamics in the one-step 100 mM KCl treatment under continuous dimG (dimG^day/night^) and RGB^day^/dimG^night^ illumination (Supplementary Figure S1d; Supplementary Table S2d). Motion profiles were similar between the two lighting regimes. For example, the reduction in nighttime motion on day 24 (day of stress) was comparable under both illumination conditions (Figure 4a,c), and reduced motion persisted during days 30–32 (Figure 4b,d). Although RGB^day^/dimG^night^ imaging consistently produced higher motion peaks at light–dark transitions and higher absolute 12 h integrated motion values, the stress-induced inhibition of leaf movement was equally visible in stressed plants under both regimes (Figure 4e). Together, these results demonstrate that stress-induced changes in leaf-movement dynamics can be robustly detected using simplified and more practical illumination conditions, such as RGB^day^/dimG^night^ rather than consistent dimG^day/night^ illumination.

**Figure 4.**
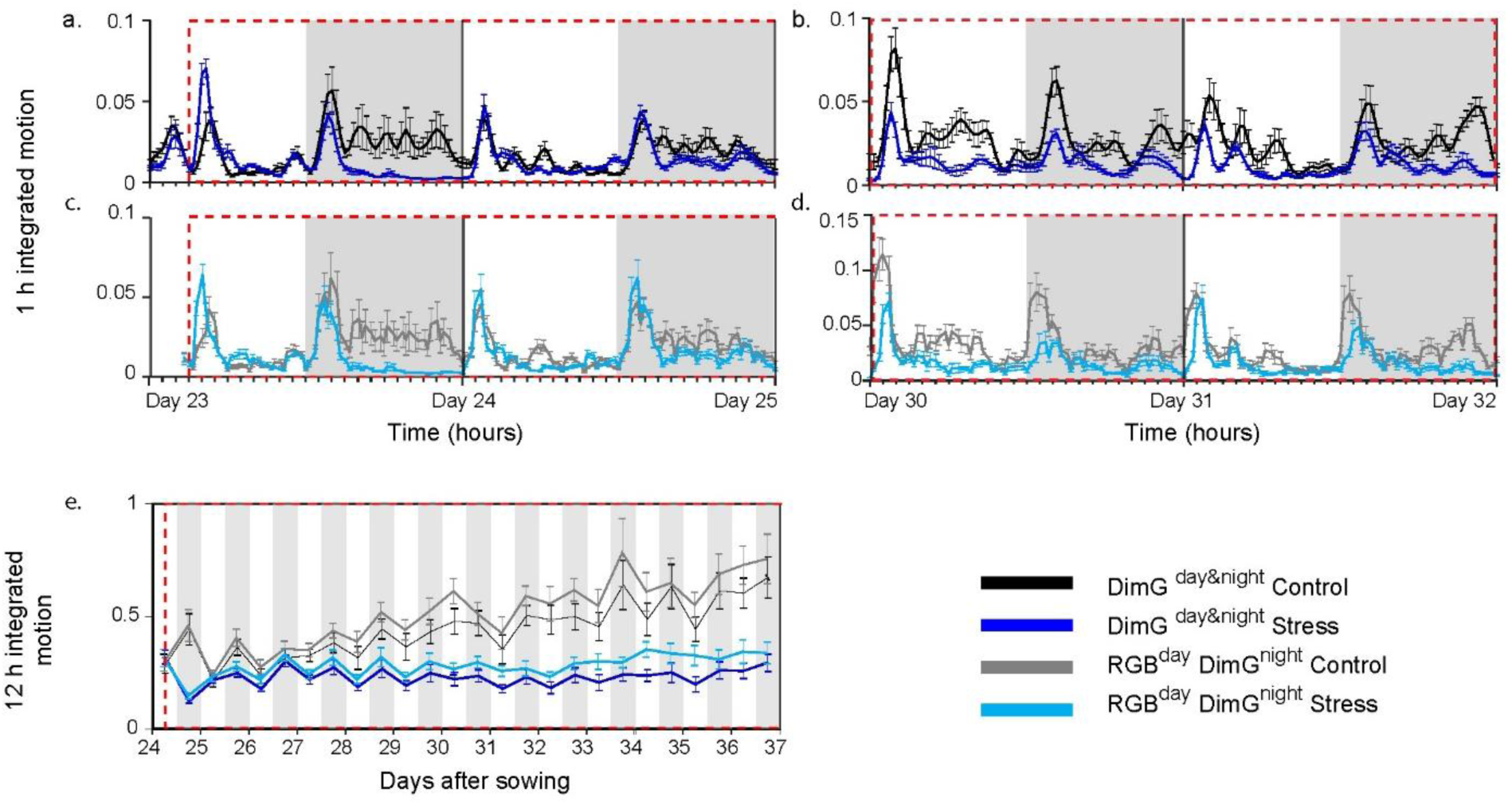
| Leaf-movement dynamics under salinity stress are robust to changes in illumination regime. Lettuce plants were exposed to 100 mM KCl from day 24 onwards (same experiment as Figure 2d-f). Leaf-movement dynamics were quantified in non-stressed and stressed plants under two imaging regimes: continuous dim green pulse illumination during both day and night (dimG^day & night^; non-stressed (black) and stress (dark blue)), and RGB illumination during the day with dim green illumination at night (RGB^day^ dimG^night^; non-stressed (grey) and stress (light blue)). Panels **(a-d)** show 1 h integrated motion during **(a,c)** days 23–25 and **(b,d)** days 30–32. Curves represent values calculated using a centred 1 h moving integration window. Panel **(e)** shows 12 h integrated motion. Data represent means of 9–10 plants; error bars indicate SEM (see Supplementary Table S3e and S6c for numerical data)

### Leaf-movement-based stress detection in plant arrays

Imaging plant arrays is particularly advantageous in dense production systems, where individual plants are closely spaced and leaf overlap can limit reliable segmentation. By enabling the derivation of stress metrics at the canopy or tray level, this approach might provide a more representative basis for assessing crop status in CEs. We therefore tested whether our stress detection system can capture stress-induced leaf-movement dynamics when applied to entire hydroponic tanks, rather than individual plants under both dimG^day/night^ and RGB^day^/dimG^night^ imaging regimes (Supplementary Figure S5a-d). Under dimG^day/night^, direct motion analysis revealed clear diel rhythms of leaf movement in both non-stressed and stressed canopies. Day-to-day motion rates (Supplementary Figure S5c) derived at the canopy level closely matched the mean of individual plants (Supplementary Figure S5d), with stress-treated plants showing a sustained reduction in leaf movement following stress application. Under RGB^day^/dimG^night^, higher motion peaks were detected at multiple light-dark transitions in both non-stressed and stressed canopies (Supplementary Figure S5b). Although the difference in day-to-day motion rate remained evident, these peaks increased the overall motion estimates (Supplementary Figure S5c). Excluding light-dark transition periods from the daily integrated motion yielded comparable motion rates across illumination regimes and consistent with those derived from the mean of individual plants (Supplementary Figure S5d). Together, these results indicate that the system can be reliably applied to quantify stress responses at the level of whole plant arrays and across illumination regimes.

### Leaf movement changes under stress can be observed across various plant species

Our approach can be potentially expanded to other plants species with similar leaf architecture to lettuce that exhibit pronounced, reversible petiole-mediated leaf movements amenable to robust imaging-based quantification. To assess this, we applied salinity stress to five additional horticultural crops commonly grown in CEA: amaranth, mizuna, radish, rocket, and tomato; and successfully quantified leaf-movement dynamics (Supplementary Figure S1c, Supplementary Table S2c) as 12 h integrated motion (Supplementary Table S3d), day-to-day motion rate (Supplementary Table S4d) and 1 h integrated motion (Supplementary Figure S6c).

In non-stressed conditions, each species had a gradual increase in leaf movement over time, consistent with visible growth. Day–night differences in 12 h integrated motion was most pronounced in amaranth, rocket, and radish, but were less evident in mizuna (Figure 5a,d,g,j,m). Notably, amaranth had an inverted diel pattern relative to lettuce, with higher daytime than nighttime motion, consistent with increased high frequency movements during the day (Supplementary Figure S6a).

**Figure 5.**
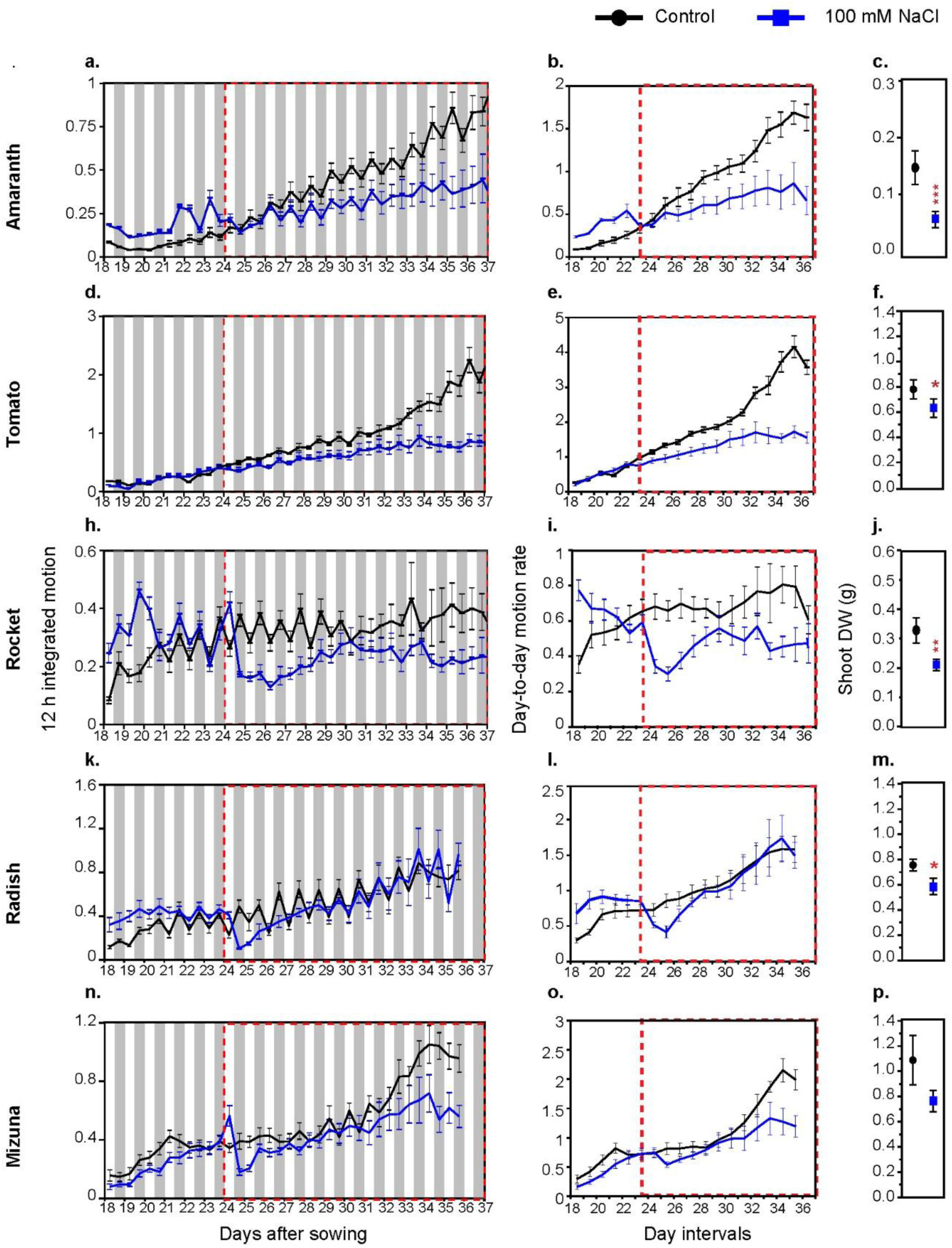
| Salinity-induced reductions in leaf-movement dynamics and biomass accumulation vary between crop species. Responses of **(a–c)** amaranth, **(d–f)** tomato, **(g–i)** rocket, **(j–l)** radish, and **(m–o)** mizuna to salinity stress. For each species, panels show **(a,d,g,j,m)** 12 h integrated motion, **(b,e,h,k,n)** day-to-day motion rate, and **(c,f,i,l,o)** shoot dry weight (DW) measured at day 38 for non-stressed control plants (black) and plants exposed to 100 mM NaCl (blue) from day 24 onwards. Data represent means of 9–12 plants; error bars indicate SEM (see Supplementary Table S3d, S4d and S5d for numerical data). Red dashed lines mark the timing of stress application. White and grey backgrounds indicate day and night periods, respectively. Asterisks denote significant differences between stressed and non-stressed plants (*p < 0.05, **p < 0.01, ***p < 0.001; two sample t-tests). 1 h integrated motion is shown in Supplementary Figure S5.

In both amaranth and tomato, stress application led to a rapid reduction in leaf movement from day 24 (day of stress), followed by sustained inhibition of the day-to-day motion rate below non-stressed levels (Figure 5b,e), consistent with a significant reduction in shoot DW (Figure 5c,f). We confirmed that the RGB^day^ dimG^night^ light regime is also applicable to amaranth and tomato, enabling detection of stress-induced leaf-movement dynamics in both individual plants and whole tank arrays (Supplementary Figure S5e-l).

Similar to lettuce (Figure 2), salt application triggered a transient increase in leaf movement in rocket, radish and mizuna (Figure 5g-o; Supplementary Figure S6c-e). Comparison of sustained salinity stress effects across species revealed a graded tolerance to the stress, with lettuce showing the strongest and most sustained reductions in motion and shoot DW (Figure 2), rocket displaying intermediate sensitivity, and radish and mizuna largely maintaining motion dynamics and biomass similar to non-stressed levels (Figure 5g-o).

Taken together, these results show that salinity stress induces species-specific changes in leaf movement that were consistently detected between plants by our system. Rocket, radish and mizuna had rapid changes in motion that were detectable shortly after stress imposition, while tomato and amaranth had delayed motion changes (Supplementary Figure S6a-e). Radish and mizuna showed eventual partial stress recovery with relatively lower reduction in shoot DW (Figure 5c,f,i,l,o) while other crops showed sustained reductions in motion coincide with significant reduction in shoot DW. This indicates the leaf-movement system can detect differences in stress susceptibility but further characterisation of each species individually, is required for precise outcomes. Overall, these findings support the broad applicability of leaf movement monitoring for stress detection in broad leaved CEA crops.

## Discussion

Despite the high level of control in CEA (Gill et al., 2025), stress continues to limit yield and product consistency, underscoring the need for rapid, low-cost, autonomous, and scalable stress detection approaches that can trigger repair or maintenance protocols.

## Addressing key limitations in plant stress detection through scalable quantitative leaf-movement dynamics

Our plant stress detection system directly addresses several key limitations of existing image-based stress detection approaches (Humplík et al., 2015). Whereas leaf angle sensors rely on costly, plant-level instrumentation that limits scalability and throughput (Geldhof et al., 2021), our platform enables simultaneous, non-contact monitoring of leaf movement across an array of plants using a low-cost imaging framework that could be expanded by replication to cover large plant populations.

While most leaf movement tracking algorithms assume stable visible-light conditions (Nagano et al., 2019), we show that stress-induced changes in leaf-movement dynamics can be detected consistently under both RGB and dimG imaging, relaxing a key practical constraint for diel, population-scale stress detection (Figure 4).

Canopy-level growth and area-based stress metrics are sensitive to plant overlap and leaf elevation, leading to increased noise (Vantoai & Roberts, 2003; Rehman et al., 2020), whereas we show that dynamic leaf-movement patterns remain detectable under these conditions and reveal stress earlier than growth-based measures.

Other image-based systems are stress specific and lack demonstrated generalisability across stresses or crops (Vantoai & Roberts, 2003), whereas by quantifying leaf-movement dynamics as a general physiological response, the system we have devised detects conserved early and sustained stress signatures across multiple stresses (Figure 2-3, Supplementary Figure S2, S4), species (Figure 5), and locations (Supplementary Figure S3). While we find species difference exist in the timing and duration of leaf movement changes, tests could be tuned for each species to enable stress detection. Other stress-detection approaches rely on visible phenotypic symptoms that may take days or weeks to emerge (Ro et al., 2023; Sakamoto et al., 2021), whereas our system detects stress-induced changes as early as one hour after stress application (Figure 2g-h, Figure 3h), or within 1–3 days thereafter, depending on the stress (Figure 2-4, Supplementary Figure S2–S4), preceding visible symptoms like yellowing, reddening, and stunted growth. The timing of stress induced changes was dependent on the nature of the stress, varying from 1 hour to few days. This early detection provides a meaningful window for informed management intervention.

Diel dynamics of leaf movement provided a temporal baseline for early stress detection. Under non-stress conditions, leaf motion followed a reproducible day–night pattern, with higher night-time activity in most species but elevated daytime motion in amaranth (Figure 2-5). Deviations from these patterns were evident on the day of stress application, including increased daytime motion consistent with a stress-associated hyponastic response and reduced night-time motion. These findings suggest that sensitivity to stress detection may depend on the timing of stress application, warranting further time-of-day analyses. Overall, diel signatures support time-resolved leaf-motion analysis as a sensitive, non-invasive approach for early stress detection.

Our results further support dimG as a practical solution for continuous diel leaf movement monitoring. DimG can be readily implemented using dimmable LED systems widely used in CEA, providing a simple alternative to IR for night-time imaging. Although transitions from RGB to dimG still introduce some artefacts in motion detection, these do not affect stress detection sensitivity (Figure 4, S5) and are smaller than those introduced by IR night-time imaging (Herrero et al., 2026).

Collectively, these results position our platform as a robust and scalable alternative to existing image-based plant stress detection technologies, capturing early and long-term, dynamic leaf-movement responses that are broadly informative across stress types and crop species. This positions leaf-movement based phenotyping as a practical and generalisable framework for non-invasive stress monitoring in CEA.

## Stress specific leaf-movement signatures

We identified multiple, distinct stress signatures in leaf-movement dynamics. The first signature was a transient increase in leaf movement, observed in lettuce following abrupt osmotic stress and water withdrawal. Under one-step NaCl and KCl addition (Figure 2a-f and Figure 5), this response is consistent with rapid turgor loss followed by recovery as ionic homeostasis is re-established (Graus et al., 2023), whereas during water withdrawal (Figure 3d-f) it reflected acute turgor loss leading to wilting and motion arrest (Zhang et al., 2010). These increases were not prominently evident in gradual NaCl addition (Supplementary Figure S2D) or nutrient withdrawal from the irrigation line (Figure 3a-c), consistent with turgor change being the driver. This transient response, decoupled from baseline diel dynamics, provides a robust early indicator of acute stress.

A second, broadly conserved signature was a reduction in nighttime motion on the day of stress application (Figure 2g–h, 3h and Supplementary Figure S6c–e), likely reflecting an early attenuation of growth-associated leaf movement that represents a consistent early response to stress across conditions and crops. This reduction is in line with previous observations that stress alters leaf-movement dynamics, including reductions in nutation amplitude (Geldhof et al., 2021).

A third signature was a sustained suppression of day-to-day motion rate emerging within 1–3 days of stress onset and coinciding with reduced biomass accumulation. Under nutrient withdrawal, this effect appeared later (Figure 3a-c), consistent with transient buffering via nutrient remobilisation before growth limitation became evident (Maillard et al., 2015). In 150 mM NaCl (step wise) this appeared starting from day 3 when the stress reached its maximum also result in reduced biomass (Supplementary Figure S2a-c). Across species, variation in the severity of motion inhibition revealed differential stress sensitivity, highlighting the potential of leaf-movement dynamics as a phenotyping readout of stress tolerance.

Overall, our findings highlight the potential for developing early stress-detection systems based on specific leaf-movement signatures, with training datasets spanning stress types, severities, durations, and developmental stages enabling improved calibration. Further gains in precision and sensitivity could be achieved by integrating leaf-movement analysis with existing environmental sensors (Paul et al., 2022), enabling multimodal analytics that jointly capture plant physiological status and environmental context, thereby enhancing system versatility and decision relevance in commercial settings (Zandi et al., 2025).

## Limitations and future opportunities

Our leaf-movement–based stress-detection system supports early detection and may, with further development, be extended to broader applications, including commercial settings. In this study, quantitative leaf-movement analysis was performed retrospectively at the end of the experiments; extending the approach to real-time processing through an automated image acquisition and analysis pipeline represents an important next step to enable interventions. The current pipeline, implemented in Python scripts, should be compatible with automated execution, supporting future development towards real-time operation.

Leaf-movement dynamics have been shown to predict plant yield in lettuce (Nagano, 2019), indicating that motion signals capture biologically relevant information associated with growth. Increases in plant size and leaf area lead to a larger visible plant structure and greater detectable motion (Mylo & Poppinga, 2024) that can be captured through analysis of motion trends. Imaging platforms such as Phytotyping4D capture plant geometry over time, enabling direct separation of growth and movement components (Apelt et al., 2015). In our stress detection system, TRiP motion outputs integrate multiple components of leaf movement, including circadian oscillations (Greenham et al., 2015), transient stress responses, and growth-related changes. Then, growth-related contributions can be estimated indirectly in the day-to-day motion rate through the integration of cumulative motion. This suggests that simplified, low-cost imaging approaches provide a scalable alternative for capturing growth-associated signals. The development of motion analysis approaches that explicitly separate growth-related components, or calibration of motion increases against independent measures of plant size, such as pixel-based area estimates, could further enable the use of motion data as a proxy for growth in high-throughput phenotyping applications.

Taken together, our results support leaf-movement analysis as a practical, low-cost and extensible framework for early, non-invasive stress monitoring, with further development and calibration expected to enable real-time implementation to trigger repair or maintenance protocols, and extension to growth and yield estimation.

## Acknowledgements

The authors acknowledge that this work was conceived, conducted, and prepared as part of a collaborative project funded by the United Kingdom Space Agency International Bilateral Fund and the Australian Space Agency. All authors gratefully acknowledge support from this funding. The authors thank project partners from Vertical Future, Axiom Space, Saber Astronautics, and the University of Southern Queensland for their contributions to the collaboration. We are grateful to Rosemary Farthing at The University of Western Australia for her assistance with plant growth experiments and to Dr. Laura Wilkinson at Adelaide University for generously providing amaranth seeds used in this study.

## Data availability

Supporting datasets, and analysis scripts will be shared via the University of Cambridge Apollo repository and will be made publicly available at the time of publication via the DOIs 10.17863/CAM.130855 and 10.17863/CAM.130856.

## Author contributions

**EHS:** Writing – original draft, review & editing, Methodology, Investigation, Formal analysis, Conceptualization; **SW:** Writing – original draft, review & editing, Methodology, Investigation, Formal analysis, Conceptualization; **ARG:** Writing – original draft, review & editing, Methodology, Investigation, Formal analysis, Conceptualization**; JDS:** Writing – review & editing, Methodology, Investigation, Formal analysis, Conceptualization; **CB:** Writing – review & editing, Investigation; **WS:** Writing – review & editing, Investigation; **AZA:** Writing –review & editing, Methodology, Conceptualization; **JRB:** Writing –review & editing, Methodology, Conceptualization; **JCM:** Writing – review & editing, Methodology, Conceptualization; Supervision; **AARW:** Writing – review & editing, Methodology, Conceptualization; Supervision**; MG:** Writing – review & editing, Methodology, Conceptualization; Supervision**; AHM:** Writing – original draft, review & editing, Methodology, Conceptualization; Supervision.

## Supplementary documents

**Supplementary Table S1.** Data corresponding to the control experiment presented in Figure 1, including: **(a)** raw motion data, **(b)** area under the motion curve (AUC), **(c)** 12 h integrated motion, **(d)** day-to-day motion rate, and **(e)** 1 h integrated motion.

**Supplementary Table S2.** Raw motion data from: **(a)** different stress treatments in lettuce, **(b)** salt stress applied at different locations on lettuce, **(c)** 100 mM NaCl treatment on different CEA crops and, **(d)** 100 mM KCl treatment under dimG^day/night^ and RGB^day/^dimG^night^ with their respective non-stressed plants.

**Supplementary Table S3.** Mean, SEM, fold changes, *p*-values (t-test), and percentage differences in motion obtained from the statistical comparison of 12 h integrated motion following stress imposition with the corresponding time point on day before stress imposition in **(a)** different stress treatments in lettuce, **(b)** salt stress applied at different locations on lettuce, **(c)** within location replication of different stresses, **(d)** 100 mM NaCl treatment on different CEA crops and **(e)** 100 mM KCl treatment under dimG^day/night^ and RGB^day/^dimG^night^ with their respective non-stressed plants.

**Supplementary Table S4.** Mean, SEM, fold changes, *p*-values (t-test), and percentage differences in motion obtained from the statistical comparison of day-today motion following stress imposition in **(a)** different stress treatments in lettuce, **(b)** salt stress applied at different locations on lettuce, **(c)** within location replication of different stresses and **(d)** 100 mM NaCl treatment on different CEA crops, with their respective non-stressed plants.

**Supplementary Table S5.** The end point root and shoot fresh weights, dry weights and chlorophyll content data of **(a)** different stress treatments in lettuce, **(b)** salt stress applied at different locations on lettuce, **(c)** within location replication of different stresses and **(d)** 100 mM NaCl treatment on different CEA crops, with their respective non-stressed plants.

**Supplementary Table S6.** 1 h integrated motion data in **(a)** lettuce subjected to different stresses, **(b)** 100 mM NaCl treatment on different CEA crops and, **(c)** 100 mM KCl treatment under dimG^day/night^ and RGB^day/^dimG^night^ with their respective non-stressed plants.

**Supplementary Table S7.** Quantification of 100 mM NaCl stress-induced leaf-movement dynamics in **(a-d)** lettuce, **(e-h)** Amaranth, **(i-l)** Tomato plant arrays and the respective controls. **(a,e,i)** Direct motion under RGB^day/^dimG^night^ lighting conditions **(b,f,j)** Direct motion under dimG^day/night^ lighting conditions **(c,g,k)** Mean day-to-day motion rate in measured under dimG^day/night^ and RGB^day/^dimG^night^, calculated using either 24 h motion integration or 22 h motion integration excluding day/night transitions. **(d,h,l)** Mean day-to-day motion rate of individual plants under dimG^day/night^ and RGB^day/^dimG^night^. Data represent means of 9–10 plants.

**Supplementary Figure S1.** Raw motion curves corresponding to: **(a)** different stress treatments in lettuce, **(b)** salt stress applied at different locations on lettuce, and **(c)** 100 mM NaCl stress in various crop species and their respective controls (see Supplementary Table S2a-d for numerical data).

**Supplementary Figure S2.** Leaf-movement responses to **(a-c)** 150 mM NaCl, **(d-f)** Hypoxia and **(g-i)** 100 mM Boron. For each treatment, panels show **(a,d,g)** 12 h integrated motion, **(b,e,h)** day-to-day motion rate, **(c,f,i)** Shoot DW. **(j-l)** 1 h integrated motion following application of **(j)** 150 mM NaCl, **(k)** Hypoxia and **(l)** 100 mM Boron, shown from day 23 to the onset of changes in leaf-movement dynamics after stress application on day 24. Data represent means of 9-12 plants for control (black) and stress (blue) treatments; error bars indicate SEM (see Supplementary Table S3a, S4a, S5a and S6a for numerical data). The red dashed line marks the timing of stress application. White and grey backgrounds indicate day and night, respectively. Asterisks denote significant differences between stress and control treatments (\**p* < 0.05, \*\**p* < 0.01, \*\*\**p* < 0.001; two-sample *t*-tests).

**Supplementary Figure S3.** Leaf movement in a closed Conviron growth cabinet (UWA), in a controlled environment plant growth walk-in room (AU), and in a laboratory room (UoC) under **(a-c)** control and **(d-f)** NaCl (150 mM in UWA; 100 mM in AU and UoC) conditions. For each treatment, panels show **(a,d)** 12 h integrated motion, **(b,e)** day-to-day motion rate, **(c,f)** Shoot DW. Data represent means of 9-12 plants for control (black) and stress (blue) treatments; error bars indicate SEM (see Supplementary Table S3b, S4b, S5b for numerical data). The red dashed line marks the timing of stress application. White and grey backgrounds indicate day and night, respectively. Asterisks denote significant differences between stress and control treatments (\**p* < 0.05, \*\**p* < 0.01, \*\*\**p* < 0.001; two-sample *t*-tests).

**Supplementary Figure S4.** Leaf movement in replicate experiments conducted in the same location under **(a-c)** Hypoxia, **(d-f)** Control (UWA) **(g-i)** Nutrient withdrawal, **(j-l)** 100 mM NaCl, and **(m-o)** Control (AU) conditions. For each treatment, panels show **(a,d,g,j,m)** 12 h integrated motion, **(b,e,h,k,n)** day-to-day motion rate, **(c,f,i,l,o)** Shoot DW. Data represent means of 9-12 plants for control (black) and stress (blue) treatments; error bars indicate SEM (see Supplementary Table S3c, S4c, S5c for numerical data). The red dashed line marks the timing of stress application. White and grey backgrounds indicate day and night, respectively. Different letters in shoot dry weight denote significant differences between locations within a treatment (\**p* < 0.05) as determined by 1-way ANOVA analysis, Tukey’s HSD test.

**Supplementary Figure S5.** Quantification of stress-induced leaf-movement dynamics in **(a-d)** lettuce, **(e-h)** Amaranth, **(i-l)** Tomato plant arrays. **(a,e,i)** Region of interest used for leaf-movement quantification in control and stress treatment tank, showing lettuce plants at the start (day 18) and end (day 38) of the experiment. **(b,f,j)** Mean direct motion across days for Control and Stress tanks under dimG^day/night^ and RGB^day^ dimG^night^ imaging regimes. **(c,g,k)** Mean day-to-day motion rate in Control and Stress tanks measured under dimG^day/night^ and RGB^day^ dimG^night^, calculated using either 24 h motion integration or 22 h motion integration excluding day/night transitions. **(d,h,l)** Mean day-to-day motion rate of individual plants under dimG^day/night^ and RGB^day^ dimG^night.^ Data represent means of 9–10 plants, shaded areas depict SEM. Red dashed lines indicate stress application. See Supplementary Table S7 for numerical data.

**Supplementary Figure S6.** 1 h integrated motion in **(a)** amaranth, **(b)** mizuna, **(c)** radish, **(d)** rocket, and **(e)** tomato subjected to 100 mM NaCl stress, with their respective non-stressed plants. (see Supplementary Table S6c for numerical data) The red dashed line marks the timing of stress application. White and grey backgrounds indicate day and night, respectively.

**Supplementary Information S1**. Plant growth and imaging setups across locations.

**Supplementary Information S2.** Experimental logs indicating key stages of experimental process.

## References

1. Apelt, F., Breuer, D., Nikoloski, Z., Stitt, M. and Kragler, F. (2015), Phytotyping4D: a light-field imaging system for non-invasive and accurate monitoring of spatio-temporal plant growth. Plant J, 82: 693–706. 10.1111/tpj.12833

2. Bours, R., Muthuraman, M., Bouwmeester, H., & van der Krol, A. (2012). Oscillator: A system for analysis of diurnal leaf growth using infrared photography combined with wavelet transformation. Plant Methods, 8(1), 1–12. 10.1186/1746-4811-8-29

3. Brdar-jokanovi, M. (2020). Boron Toxicity and Deficiency in Agricultural Plants. International Journal of Molecular Sciences, 21(1424), 1–20.

4. Geldhof, B., Pattyn, J., Eyland, D., Carpentier, S., & Van de Poel, B. (2021). A digital sensor to measure real-time leaf-movements and detect abiotic stress in plants. Plant Physiology, 187(3), 1131–1148. 10.1093/plphys/kiab407

5. Gill, A. R., Miller, T. K., Wijeweera, S., Herrero, E., Massa, G. D., Mortimer, J. C., Webb, A. A. R., Millar, A. H., & Gilliham, M. (2025). Turbocharging fundamental science translation through controlled environment agriculture. Trends in Plant Science, xx(xx), 1–14. 10.1016/j.tplants.2025.08.014

6. Graus, D., Li, K., Rathje, J. M., Ding, M., Krischke, M., Cuin, T. A., Al-rasheid, K. A. S., Marten, I., Hedrich, R., Konrad, K. R., & Konrad, K. R. (2023). Tobacco leaf tissue rapidly detoxifies direct salt loads without activation of calcium and SOS signaling. New Phytologist, 237, 217–231. 10.1111/nph.18501

7. Greenham, K., Lou, P., Remsen, S. E., Farid, H., & McClung, C. R. (2015). TRiP: Tracking Rhythms in Plants, an automated leaf-movement analysis program for circadian period estimation. Plant Methods, 11(1), 1–11. 10.1186/s13007-015-0075-5

8. Herrero, E., Gill, A. R., Wijeweera, S., Ginzburg, D. N., Stamford, J. D., Antoniades, A., Bromley, J. R., Mortimer, J. C., Gilliham, M., Millar, A. H., & Webb, A. A. R. (2026). Dim Green Light Enables Day-and-Night Monitoring of Leaf Movements Author list and affiliations. BioRxiv, May. 10.64898/2026.05.08.723725

9. Humplík, J. F., Lazár, D., Husičková, A., & Spíchal, L. (2015). Automated phenotyping of plant shoots using imaging methods for analysis of plant stress responses – A review. Plant Methods, 11(1), 1–10. 10.1186/s13007-015-0072-8

10. Junaid, M. D., & Gökçe, A. F. (2024). Global Agricultural Losses And Their Causes. Bulletin of Biological and Allied Sciences Research, 9(66), 1–11.

11. Liu, C., Jiang, X., & Yuan, Z. (2024). Plant Responses and Adaptations to Salt Stress: A Review. Horticulturae, 10(11), 1–19. 10.3390/horticulturae10111221

12. Maillard, A., Diquélou, S., Billard, V., Laîné, P., Garnica, M., Prudent, M., Garcia-Mina, J. M., Yvin, J. C., & Ourry, A. (2015). Leaf mineral nutrient remobilization during leaf senescence and modulation by nutrient deficiency. Frontiers in Plant Science, 6(MAY), 1–15. 10.3389/fpls.2015.00317

13. Mortimer, J., & Gilliham, M. (2025). Autonomous Agriculture to Support Space Exploration. The University of Adelaide. 10.25909/29595234.v1

14. Munns, R. (1993). Physiological processes limiting plant growth in saline soils: some dogmas and hypotheses. Plant Cell and Environment, 16(1), 15–24.

15. Munns, R. (2002). Comparative physiology of salt and water stress. Plant, Cell and Environment, 25, 239–250. 10.1046/j.0016-8025.2001.00808.x

16. Mylo MD and Poppinga S (2024) Digital image correlation techniques for motion analysis and biomechanical characterization of plants. Frontiers in Plant Science, 14(1335445), 10.3389/fpls.2023.1335445

17. Nagano, S., Moriyuki, S., Wakamori, K., Mineno, H., & Fukuda, H. (2019). Leaf-movement-based growth prediction model using optical flow analysis and machine learning in plant factory. Frontiers in Plant Science, 10(March), 1–10. 10.3389/fpls.2019.00227

18. Park, Y. J., Lee, H. J., Gil, K. E., Kim, J. Y., Lee, J. H., Lee, H., Cho, H. T., Vu, L. D., Smet, I. De, & Park, C. M. (2019). Developmental programming of thermonastic leaf-movement. Plant Physiology, 180(2), 1185–1197. 10.1104/pp.19.00139

19. Paul, K., Chatterjee, S. S., Pai, P., Varshney, A., Juikar, S., Prasad, V., Bhadra, B., & Dasgupta, S. (2022). Viable smart sensors and their application in data driven agriculture. Computers and Electronics in Agriculture, 198(April), 107096. 10.1016/j.compag.2022.107096

20. Pieruschka, R., & Schurr, U. (2019). Plant Phenotyping: Past, Present, and Future. Plant Phenomics, 2019(7507131), 1–6. 10.34133/2019/7507131

21. Polko, J. K., Voesenek, L. A. C. J., Peeters, A. J. M., & Pierik, R. (2011). Petiole hyponasty: An ethylene-driven, adaptive response to changes in the environment. AoB PLANTS, 11(1), 1–11. 10.1093/aobpla/plr031

22. Ragaveena, S., Shirly Edward, A., & Surendran, U. (2021). Smart controlled environment agriculture methods: a holistic review. Reviews in Environmental Science and Biotechnology, 20(4), 887–913. 10.1007/s11157-021-09591-z

23. Rehman, T. U., Zhang, L., Wang, L., Ma, D., Maki, H., Sánchez-Gallego, J. A., Mickelbart, M. V., & Jin, J. (2020). Automated leaf-movement tracking in time-lapse imaging for plant phenotyping. Computers and Electronics in Agriculture, 175(765), 1–33. 10.1016/j.compag.2020.105623

24. Ro, M., Mihalache, G., & Stoleru, V. (2023). Tomato responses to salinity stress: From morphological traits to genetic changes. Frontiers in Plant Science, 14, 1–26. 10.3389/fpls.2023.1118383

25. Sakamoto, M., Komatsu, Y., & Suzuki, T. (2021). Nutrient Deficiency Affects the Growth and Nitrate Concentration of Hydroponic Radish. Horticulturae, 7(525), 1–12.

26. Sarić, R., Nguyen, V. D., Burge, T., Berkowitz, O., Trtílek, M., Whelan, J., Lewsey, M. G., & Čustović, E. (2022). Applications of hyperspectral imaging in plant phenotyping. Trends in Plant Science, 27(3), 301–315. 10.1016/j.tplants.2021.12.003

27. Shi, K., Lu, T., Zheng, W., Zhang, X., & Zhangzhong, L. (2022). A Review of the Category, Mechanism, and Controlling Methods of Chemical Clogging in Drip Irrigation System. Agriculture, 12(202), 1–20. doi10.3390/agriculture12020202

28. Song, X., Hao, X., Song, B., Zhao, X., Wu, Z., Wang, X., & Du, J. (2022). The Oxidative Damage and Morphological Changes of Sugar Beet ( Beta vulgaris L.) Leaves at Seedlings Stage Exposed to Boron Deficiency in Hydroponics. Sugar Tech, 24(2), 532–541. 10.1007/s12355-021-01064-5

29. Vantoai, T. T., & Roberts, G. (2003). Monitoring Soybean’s Tolerance to Flood Stress using an Image Processing Technique. In Digital Imaging and Spectral Techniques: Applications to Precision Agriculture and Crop Physiology. (Issue 66, pp. 43–51). ASA Special Publication.

30. Walsh, J. J., Mangina, E., & Negrão, S. (2024). Advancements in Imaging Sensors and AI for Plant Stress Detection: A Systematic Literature Review. Plant Phenomics, 6. 10.34133/plantphenomics.0153

31. Zandi, A., Hosseinirad, S., Kashani Zadeh, H., Tavakolian, K., Cho, B. K., Vasefi, F., Kim, M. S., & Tavakolian, P. (2025). A systematic review of multi-mode analytics for enhanced plant stress evaluation. Frontiers in Plant Science, 16(April), 1–17. 10.3389/fpls.2025.1545025

32. Zhang, Y. L., Zhang, H. Z., Du, M. W., Li, W., Luo, H. H., Chow, W. S., & Zhang, W. F. (2010). Leaf wilting movement can protect water-stressed cotton (Gossypium hirsutum L.) plants against photoinhibition of photosynthesis and maintain carbon assimilation in the field. Journal of Plant Biology, 53(1), 52–60. 10.1007/s12374-009-9085-z

33. Zubler, A. V., & Yoon, J. Y. (2020). Proximal Methods for Plant Stress Detection Using Optical Sensors and Machine Learning. Biosensors, 10(12), 1–27. 10.3390/BIOS10120193

