## Supplemental Figure S1 for "Leaf movements as a quantitative metric for early stress detection"

**Supporting Figure S1.** Raw motion curves corresponding to: (**a**) different stress treatments in lettuce and their respective controls.

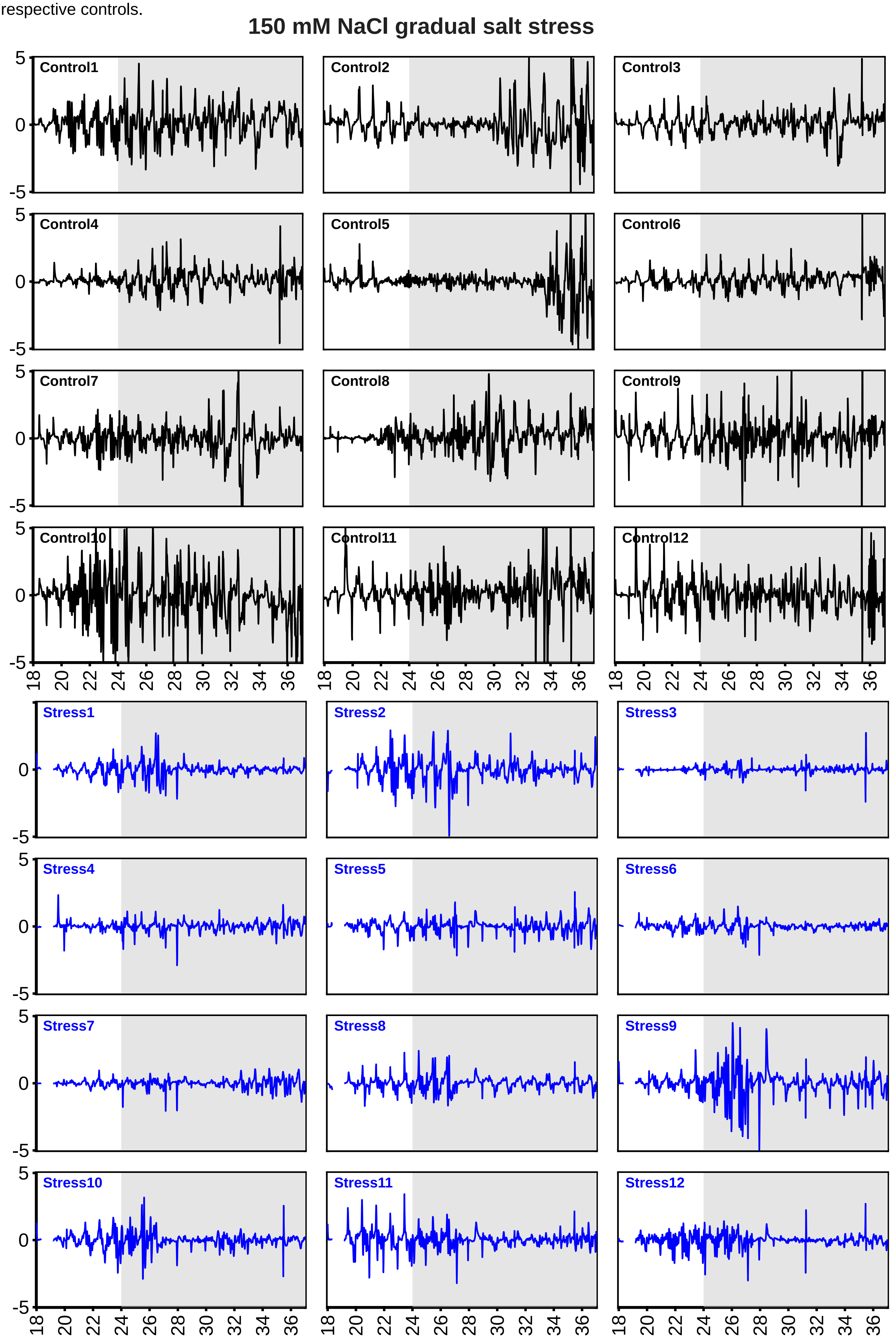

### 100 mM NaCl

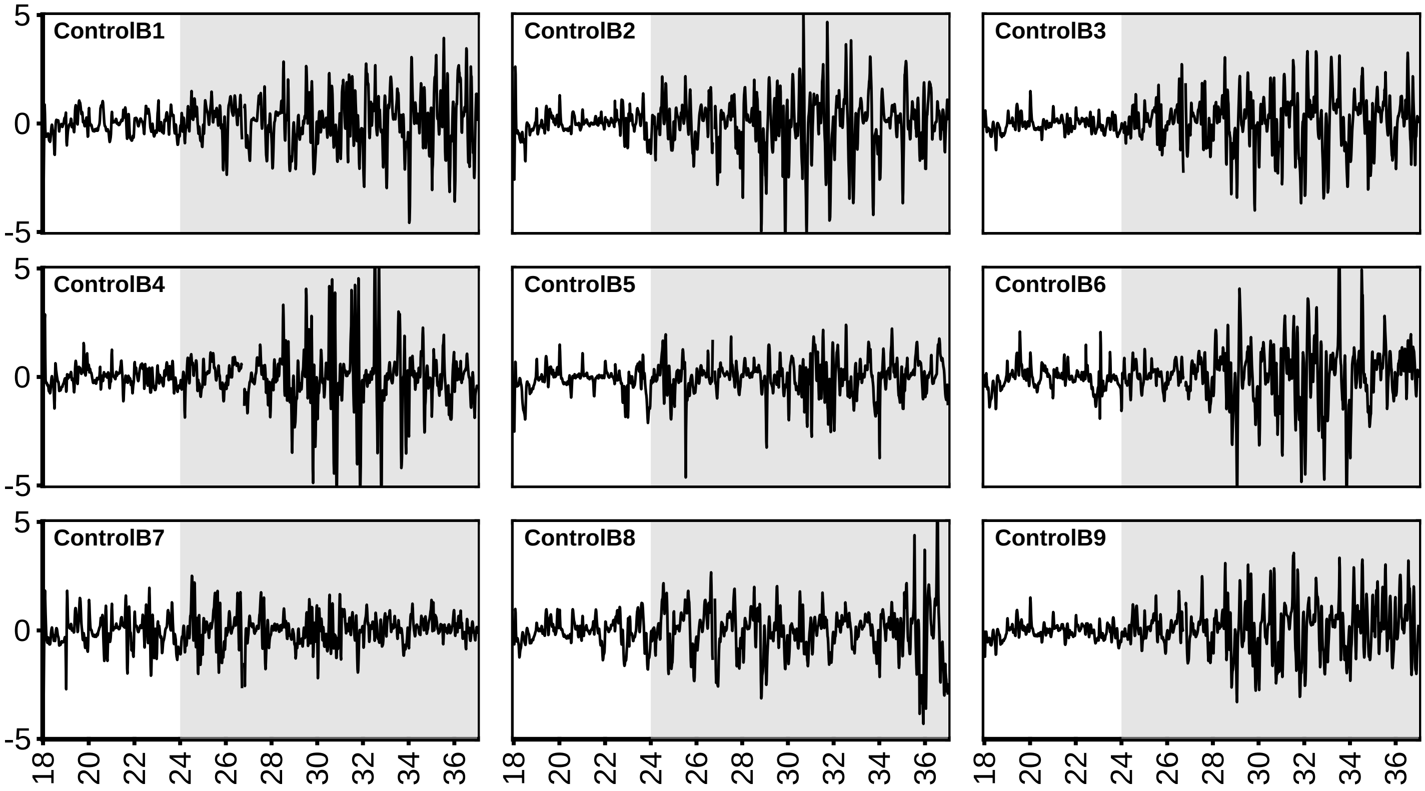
ControlB1
ControlB2
ControlB3
ControlB4
ControlB5
ControlB6
ControlB7
ControlB8
ControlB9
5
0
-5
18 20 22 24 26 28 30 32 34 36

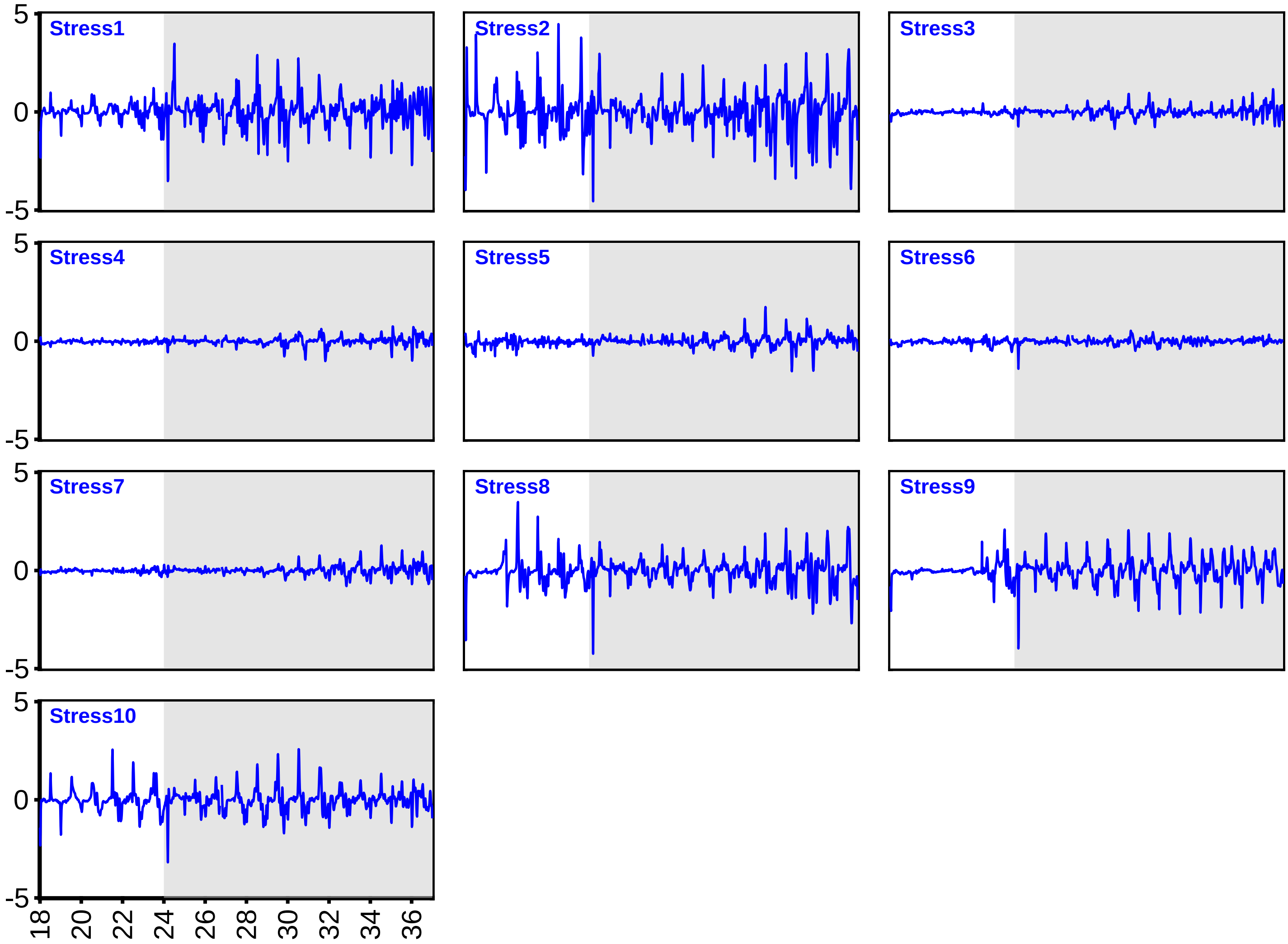
Stress1
Stress2
Stress3
Stress4
Stress5
Stress6
Stress7
Stress8
Stress9
Stress10
5
0
-5
18 20 22 24 26 28 30 32 34 36

### 100 mM KCl

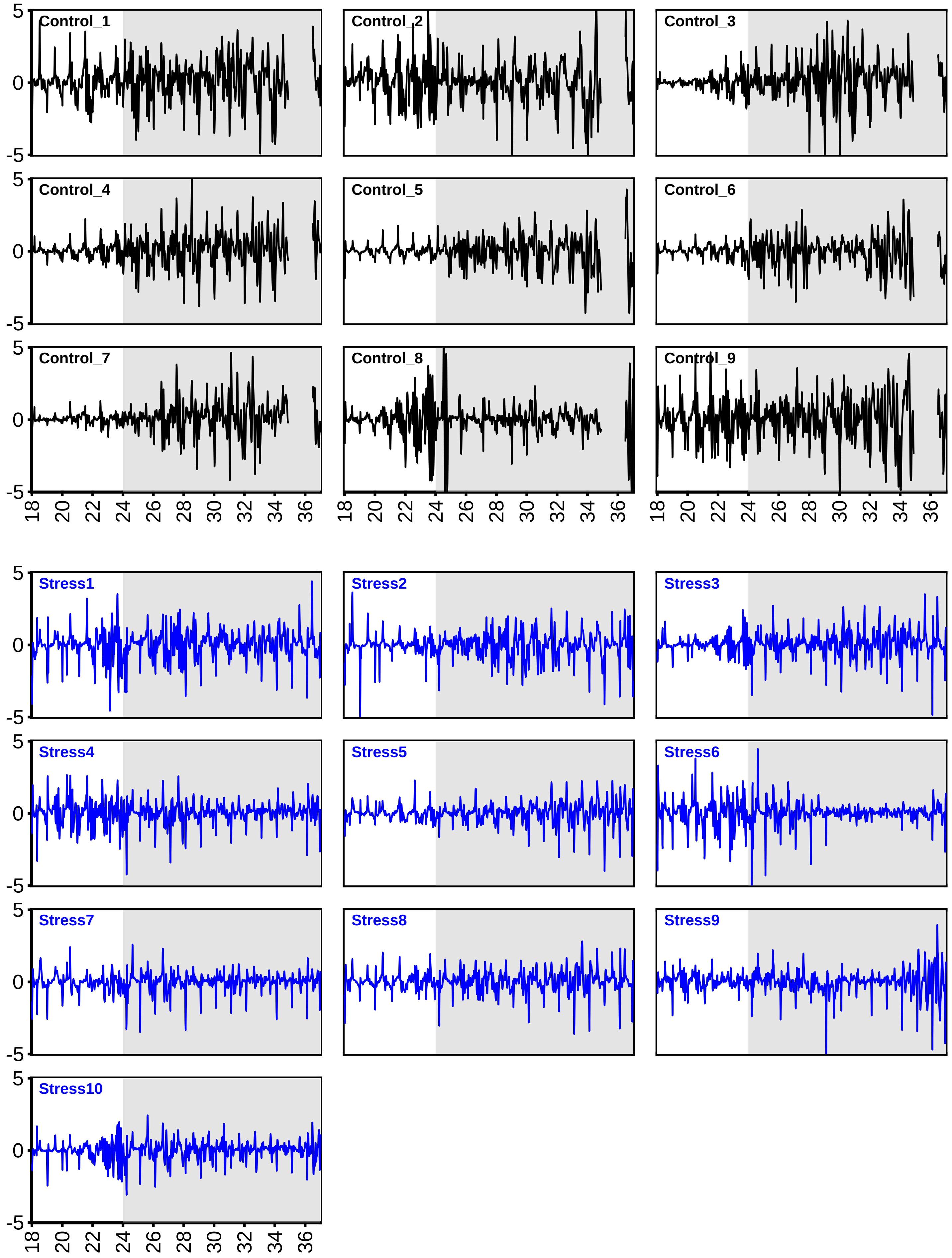

Control_1
Control_2
Control_3
Control_4
Control_5
Control_6
Control_7
Control_8
Control_9
Stress1
Stress2
Stress3
Stress4
Stress5
Stress6
Stress7
Stress8
Stress9
Stress10
5
0
-5
18
20
22
24
26
28
30
32
34
36

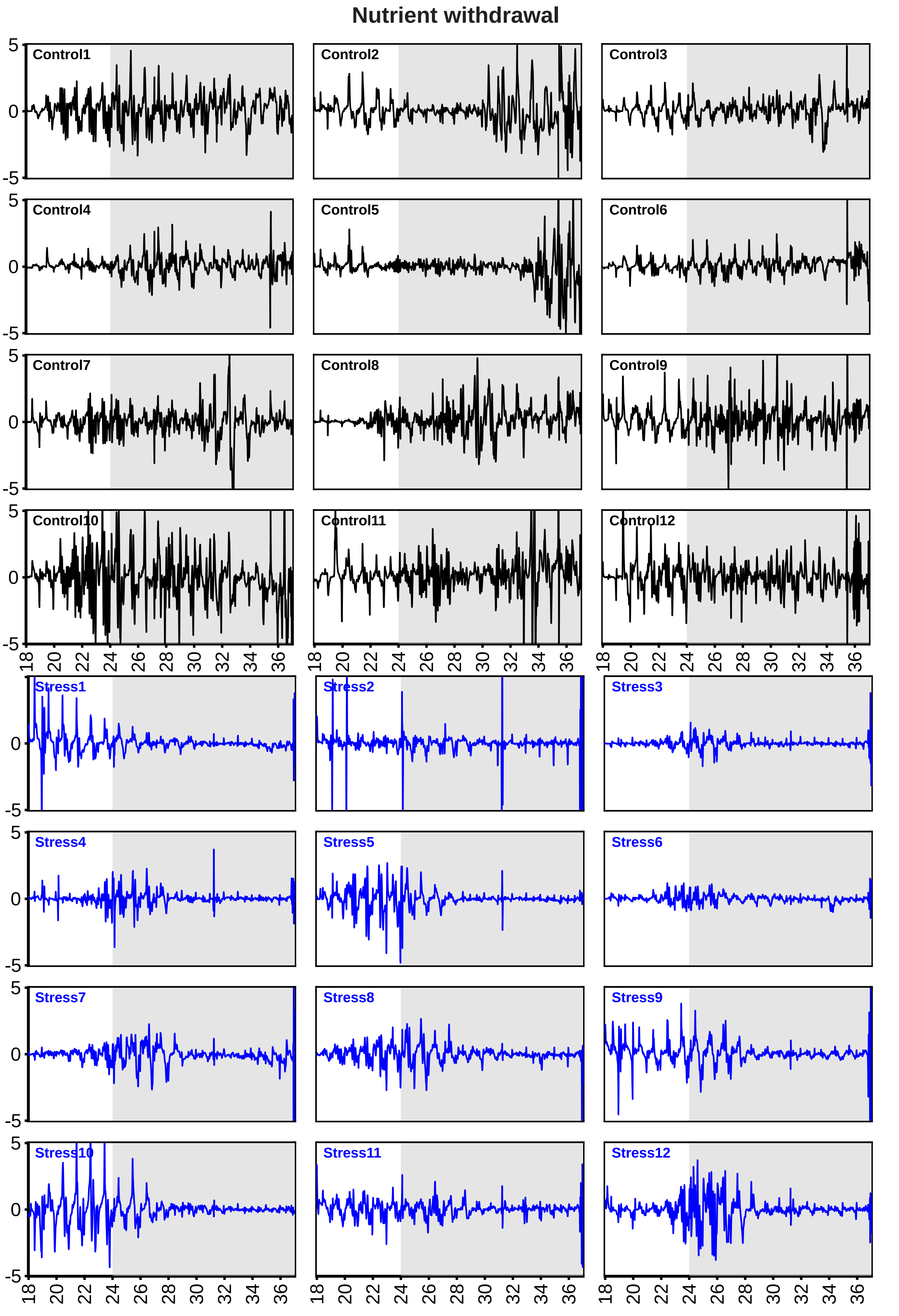
Nutrient withdrawal
Control1
Control2
Control3
Control4
Control5
Control6
Control7
Control8
Control9
Control10
Control11
Control12
Stress1
Stress2
Stress3
Stress4
Stress5
Stress6
Stress7
Stress8
Stress9
Stress10
Stress11
Stress12
5
0
-5
18 20 22 24 26 28 30 32 34 36

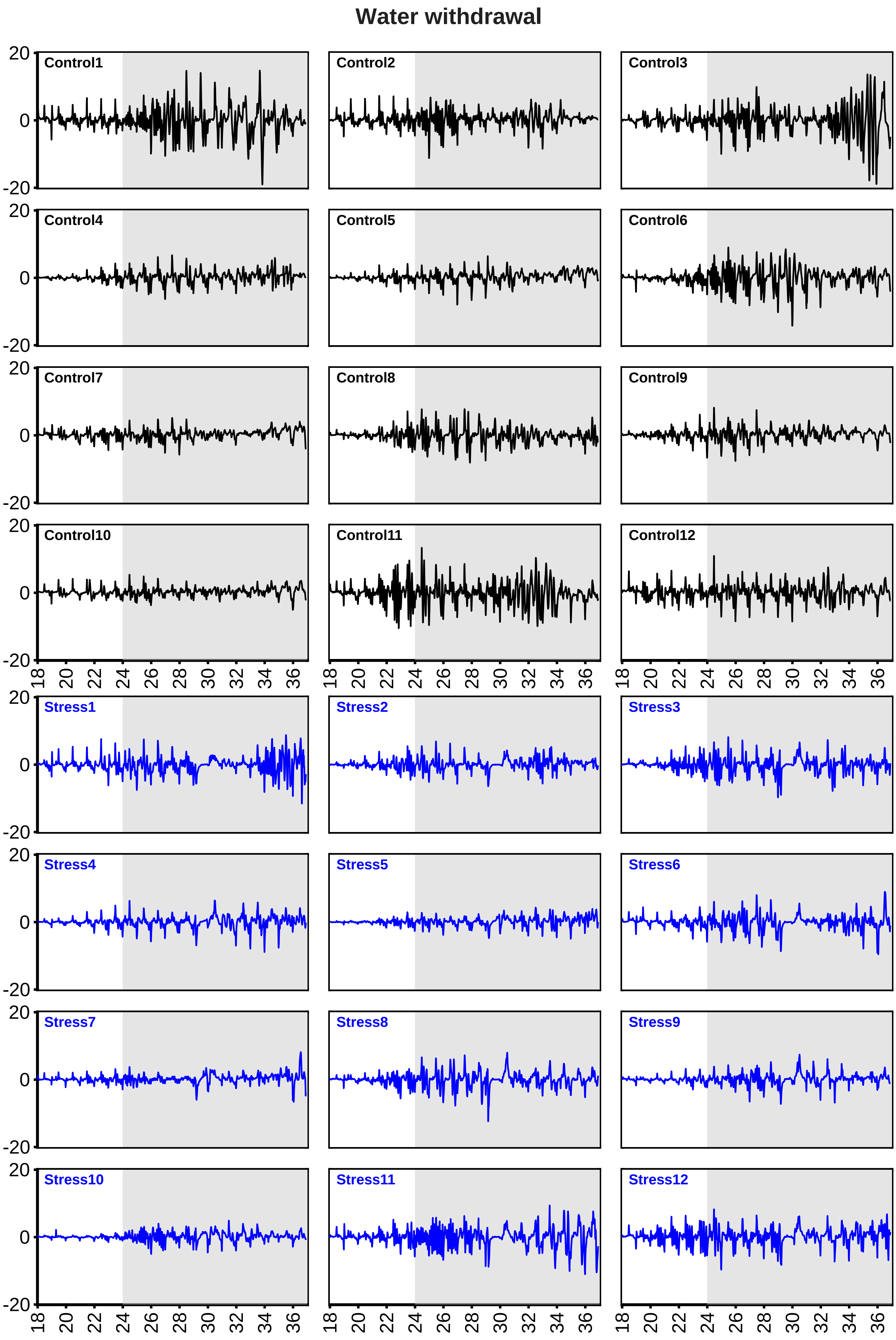

Water withdrawal
Control1
Control2
Control3
Control4
Control5
Control6
Control7
Control8
Control9
Control10
Control11
Control12
Stress1
Stress2
Stress3
Stress4
Stress5
Stress6
Stress7
Stress8
Stress9
Stress10
Stress11
Stress12
20
0
-20
18
20
22
24
26
28
30
32
34
36

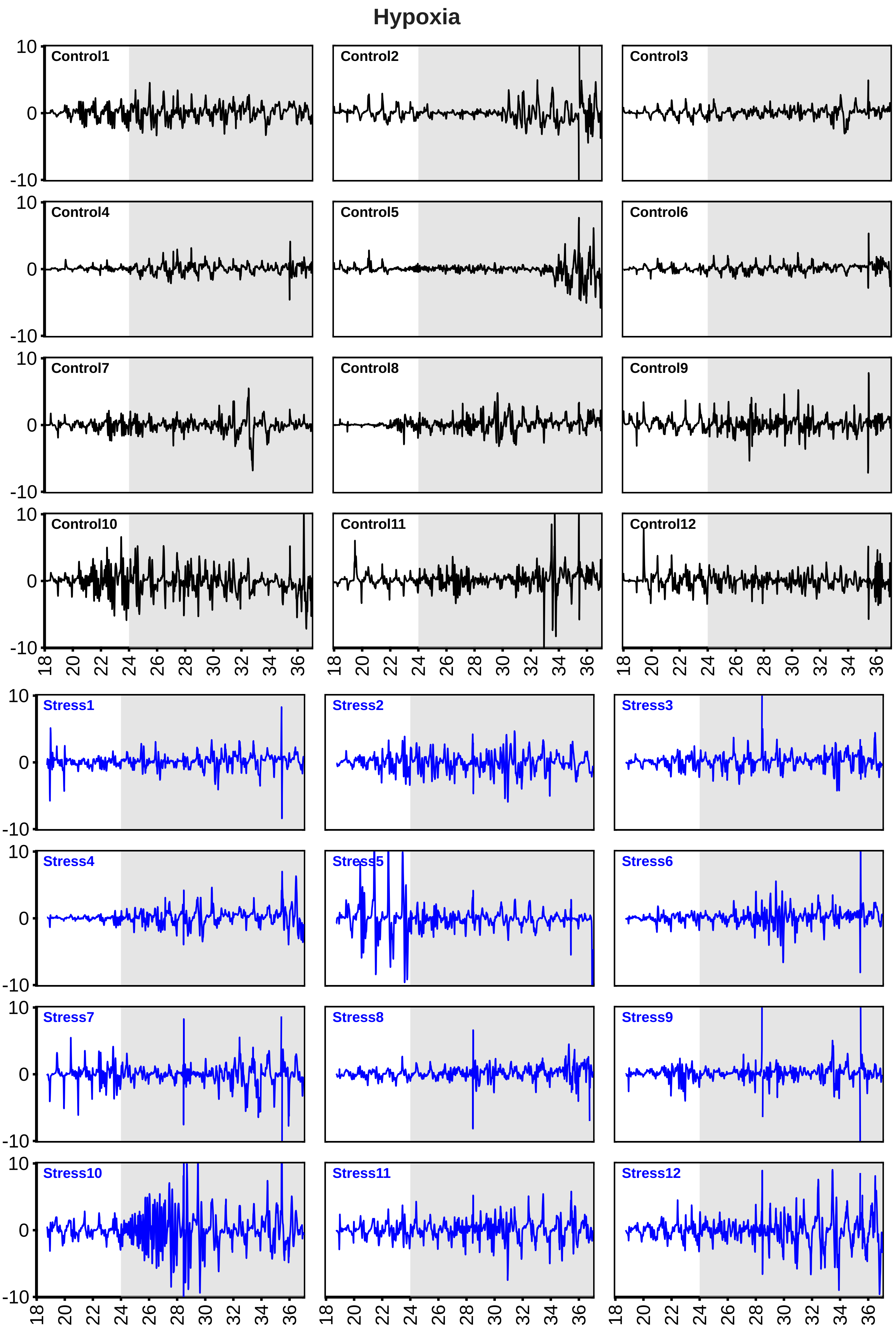
Hypoxia
Control1
Control2
Control3
Control4
Control5
Control6
Control7
Control8
Control9
Control10
Control11
Control12
Stress1
Stress2
Stress3
Stress4
Stress5
Stress6
Stress7
Stress8
Stress9
Stress10
Stress11
Stress12
10
0
-10
18 20 22 24 26 28 30 32 34 36

### 100 mM Boron

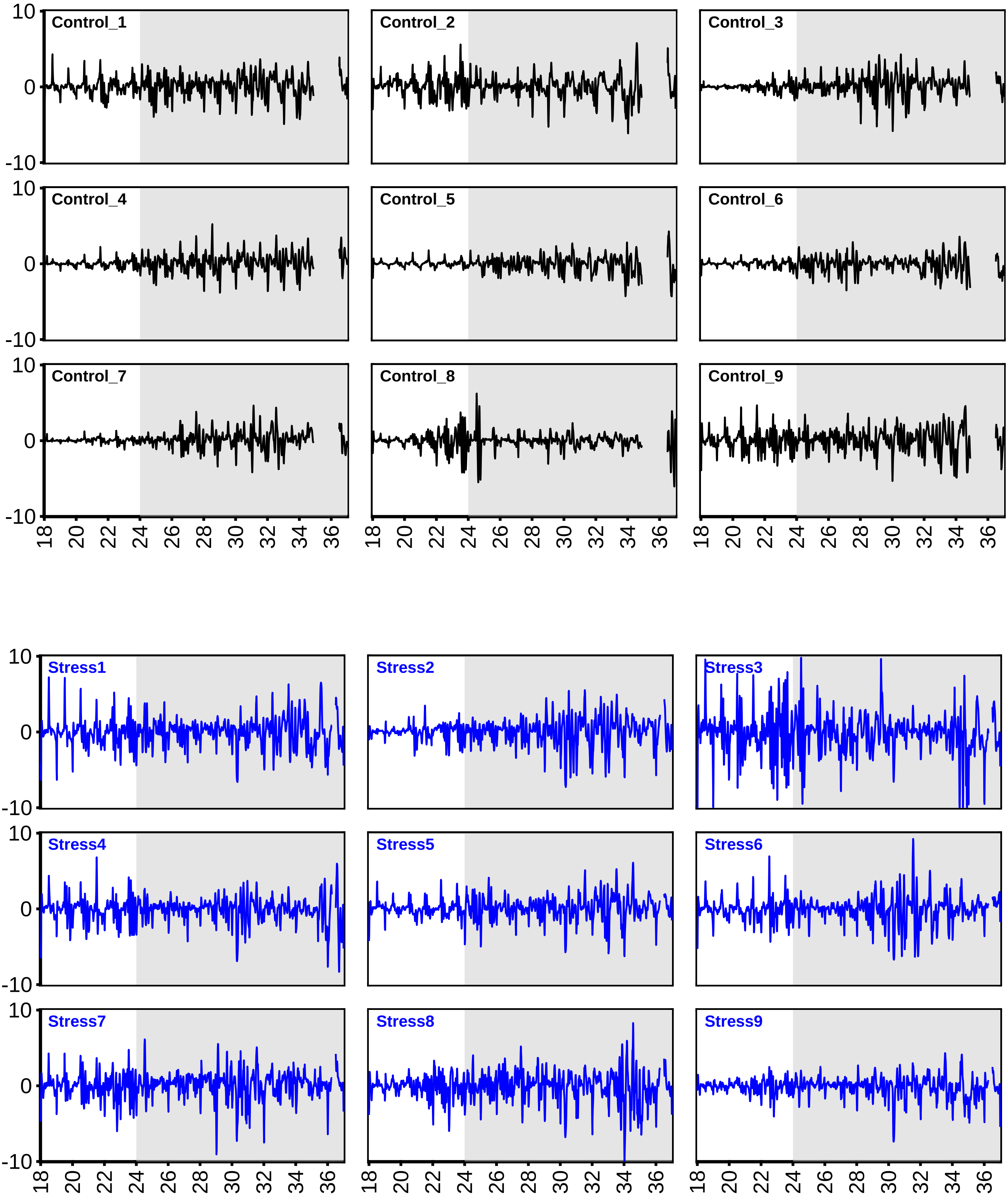
Control_1
Control_2
Control_3
Control_4
Control_5
Control_6
Control_7
Control_8
Control_9
Stress1
Stress2
Stress3
Stress4
Stress5
Stress6
Stress7
Stress8
Stress9
10
0
-10
18
20
22
24
26
28
30
32
34
36

**Supporting Figure S1**. Raw motion curves corresponding to: **(b)** salt stress applied at different locations on lettuce and their respective controls.

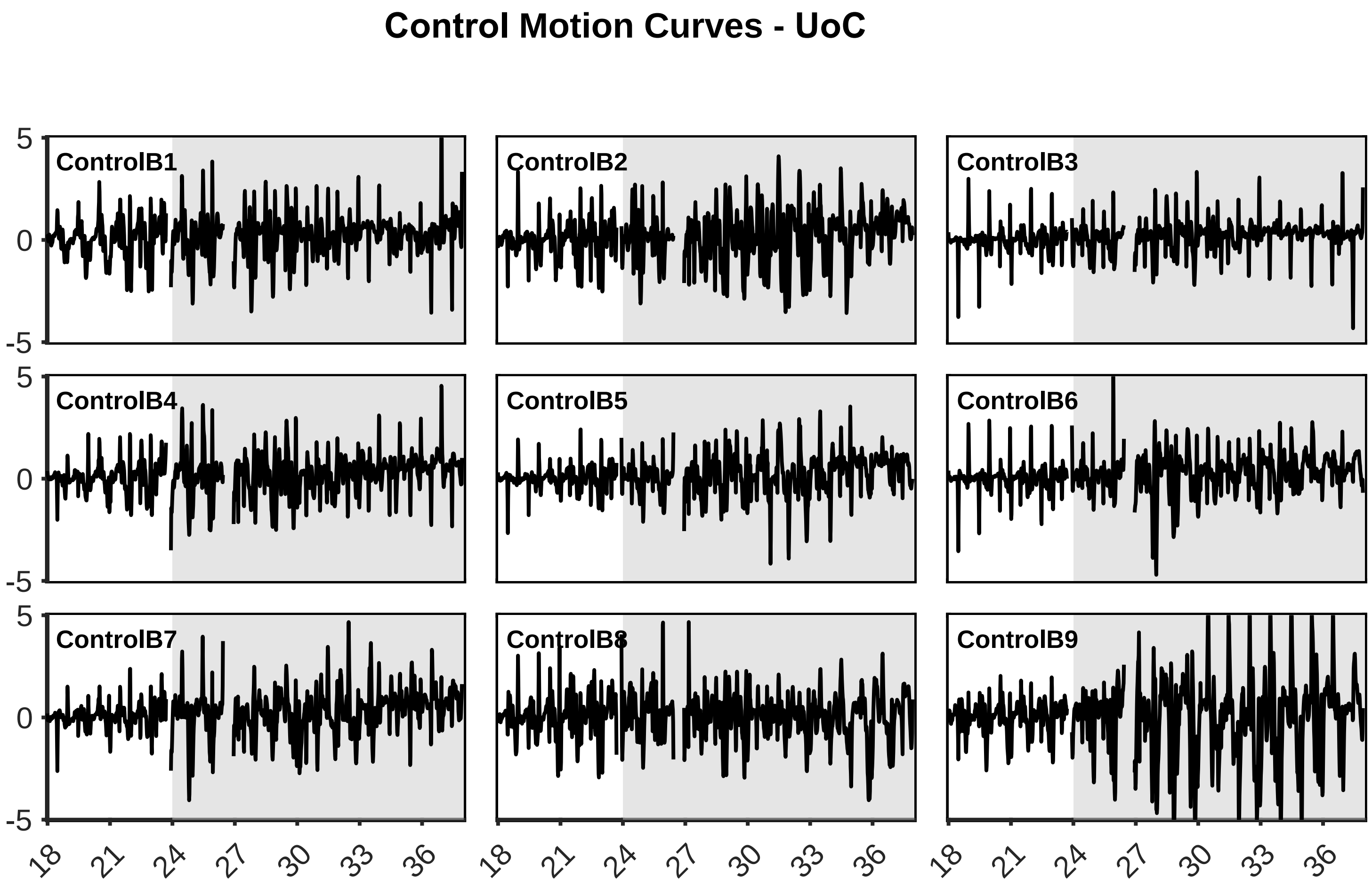

### 100 mM NaCl Motion Curves - UoC

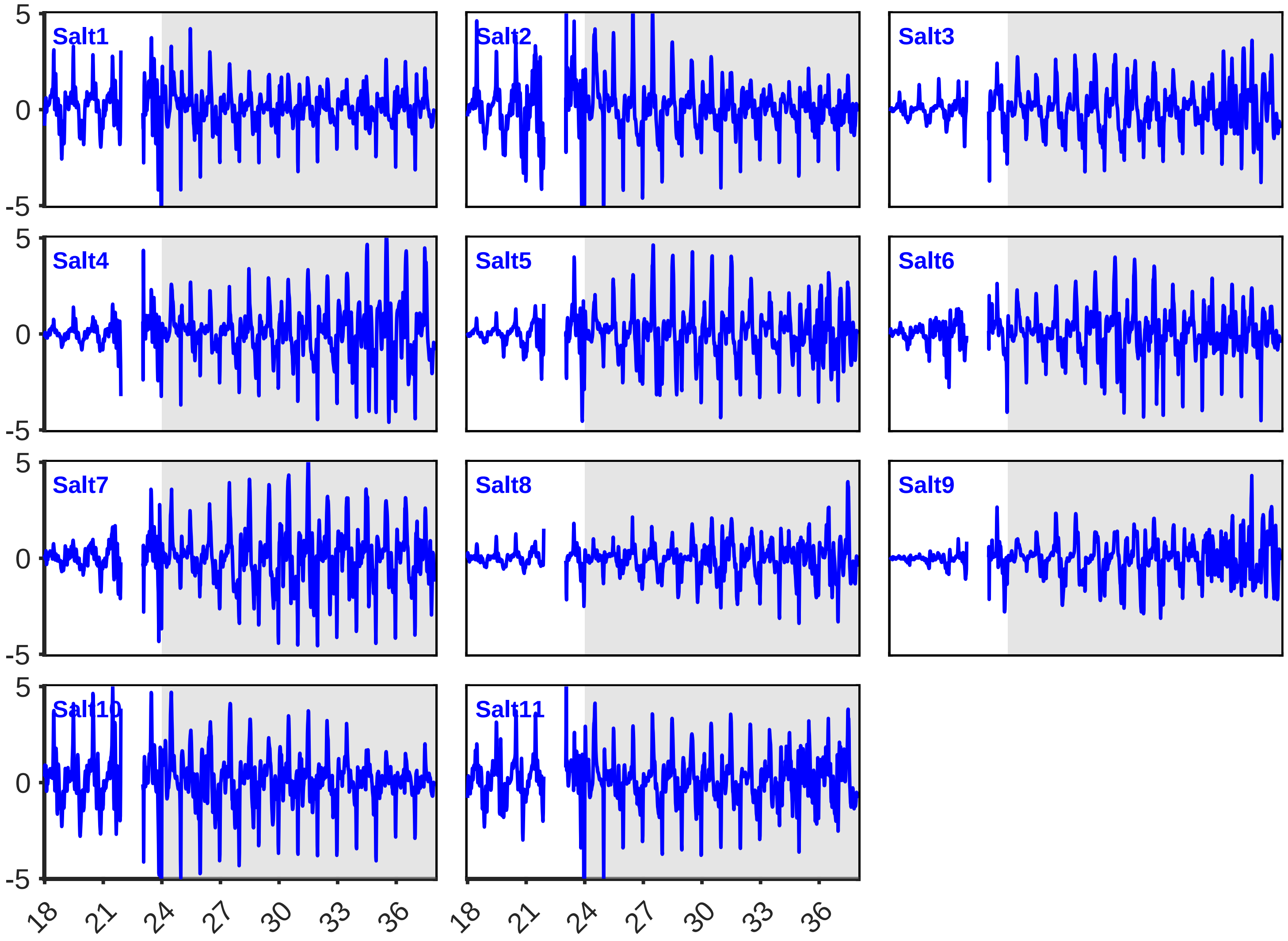

### Control Motion Curves: UWA

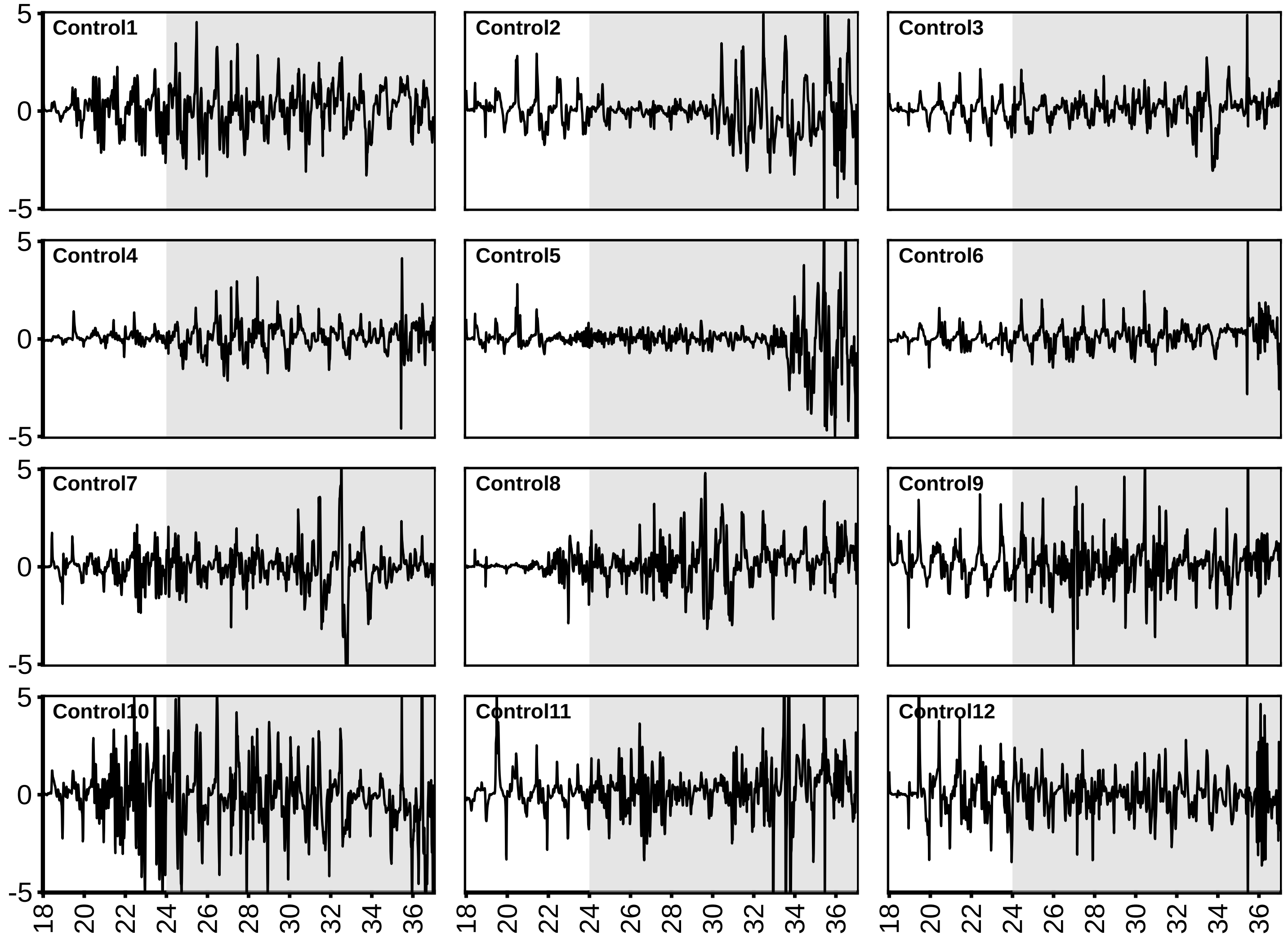

### 150 mM Gradual NaCl Motion Curves: UWA

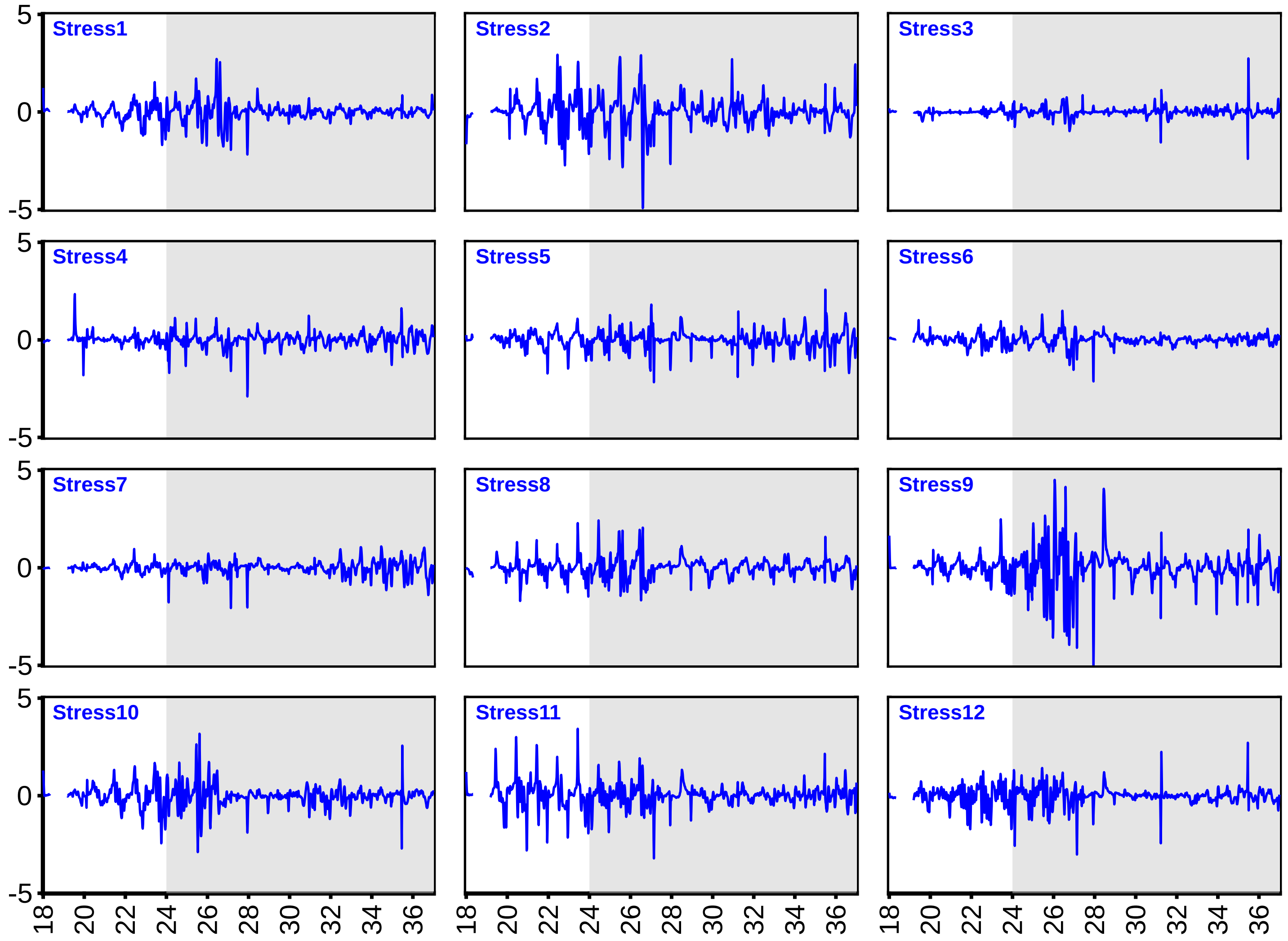

#### Control Motion Curves: UoA

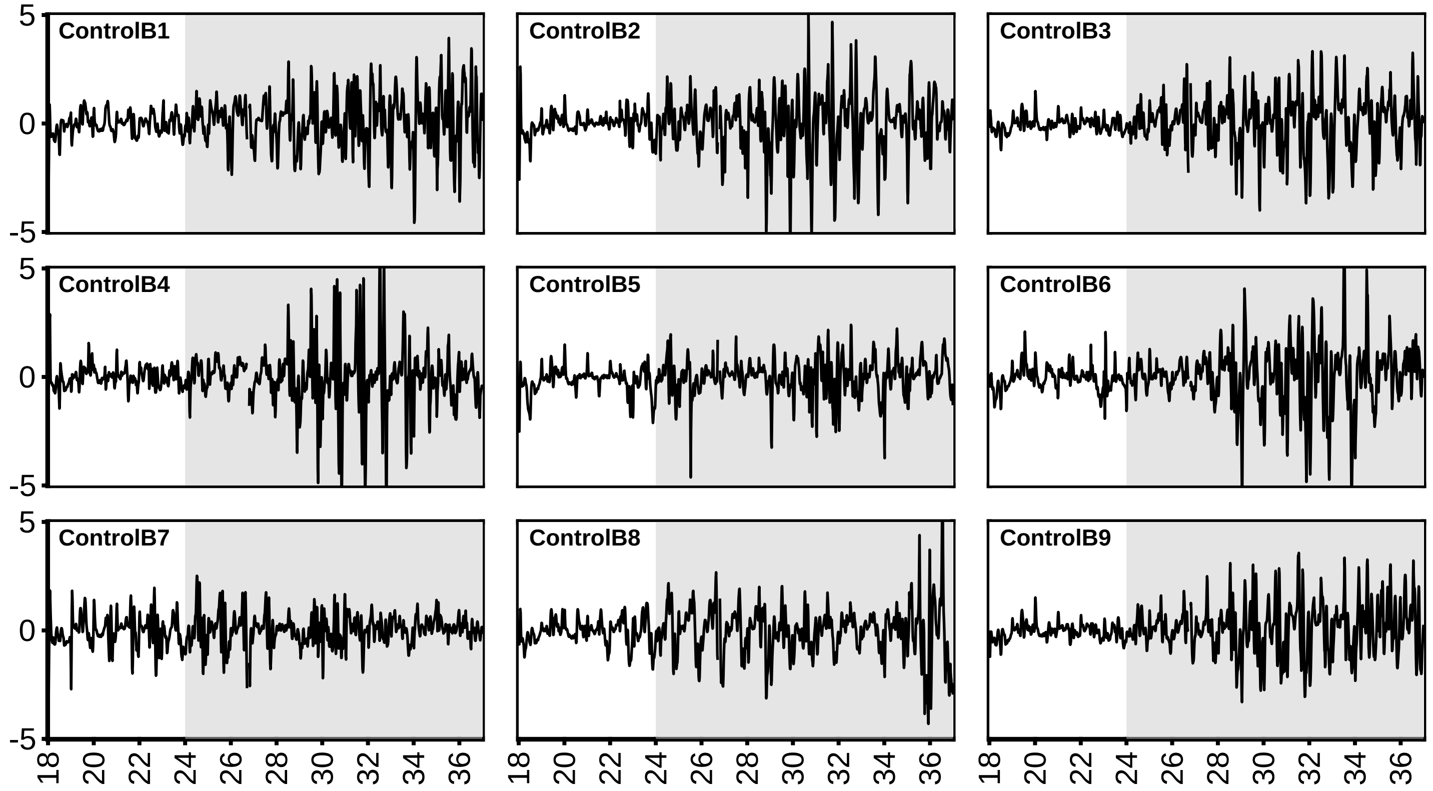

### 100 mM NaCl Motion Curves: UoA

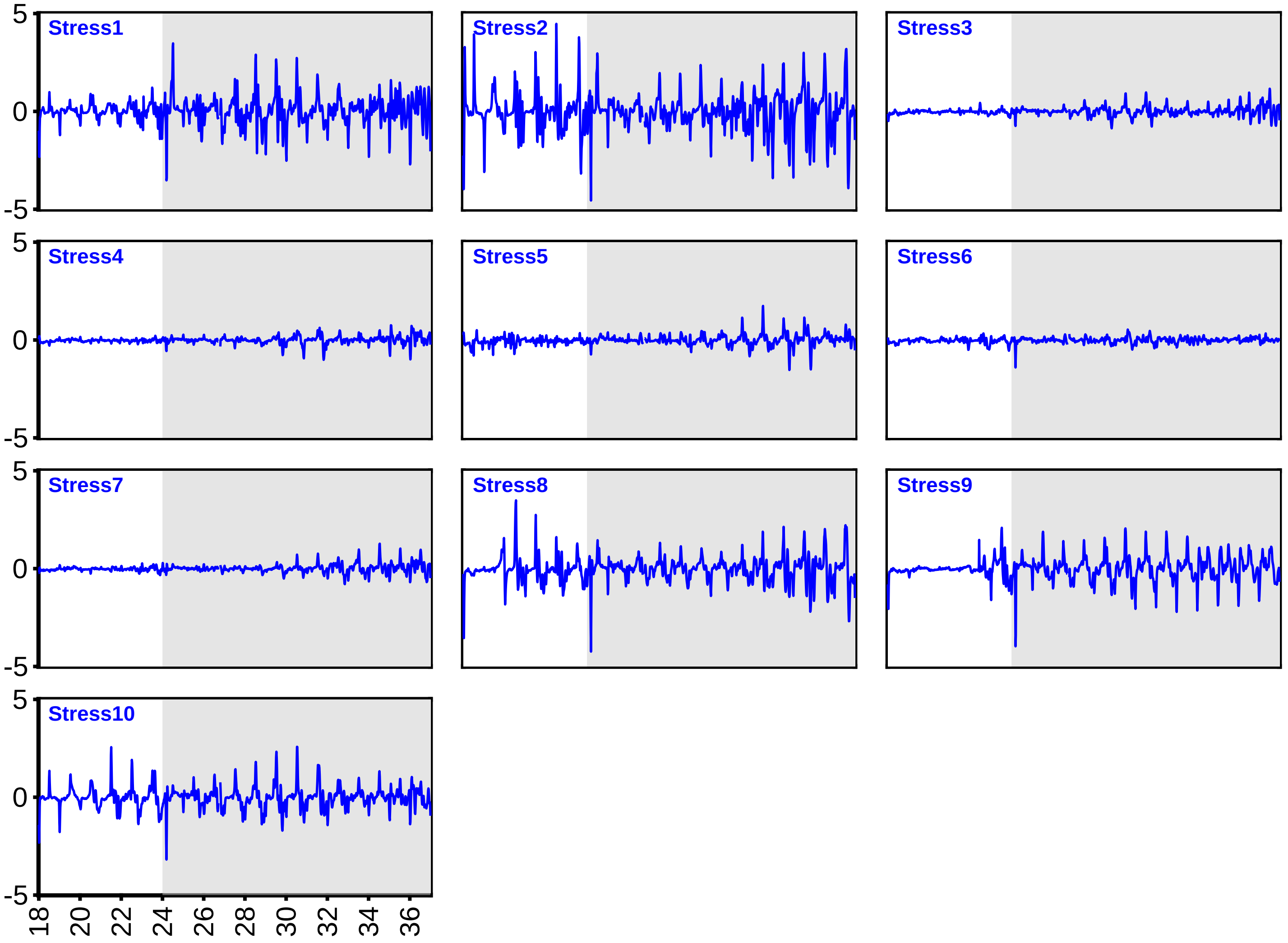

**Supporting Figure S1.** Raw motion curves corresponding to: **(c)**100 mM NaCl stress in various crop species and their respective controls.

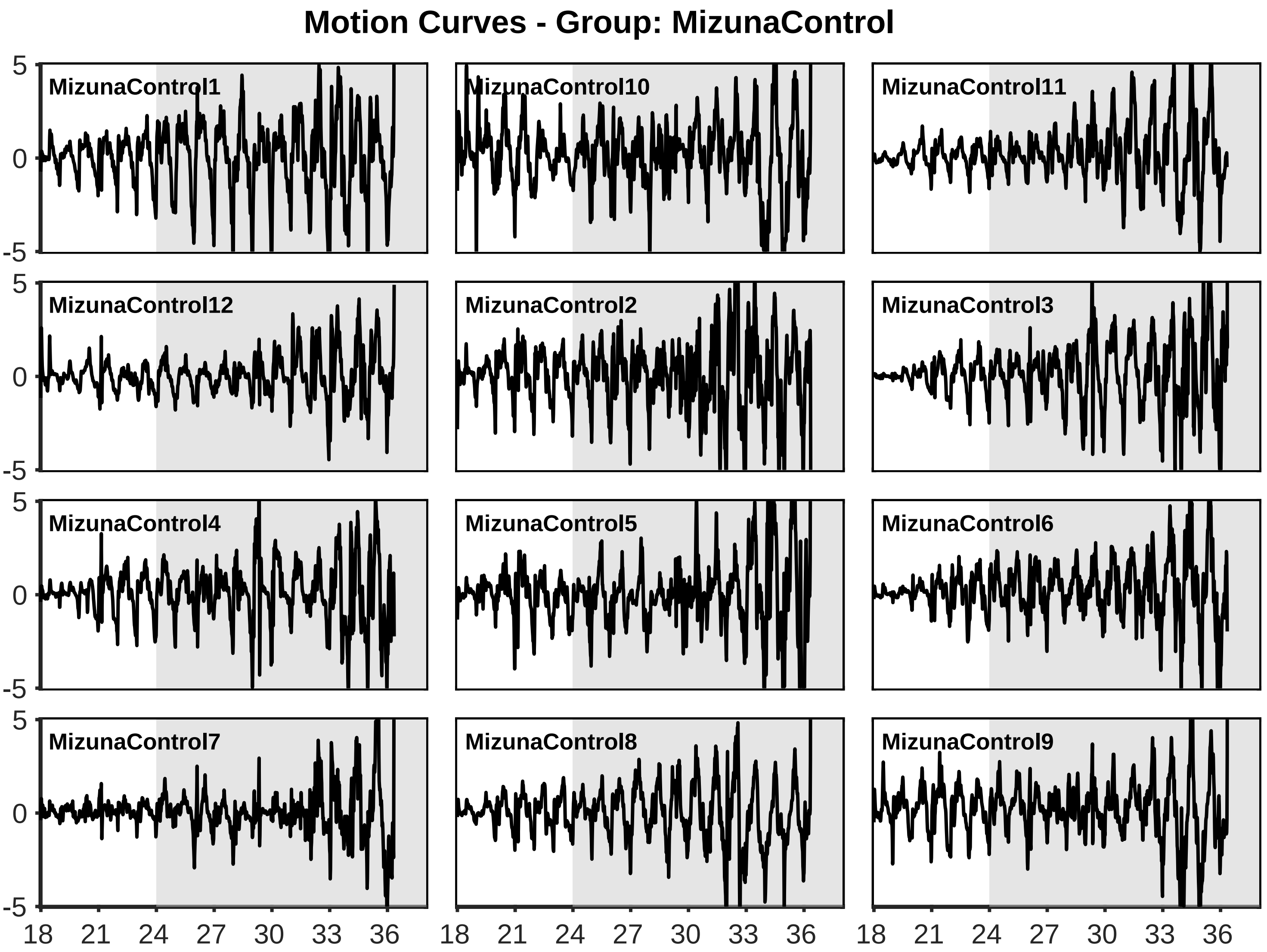

### Motion Curves - Group: Mizuna 100 mM NaCl

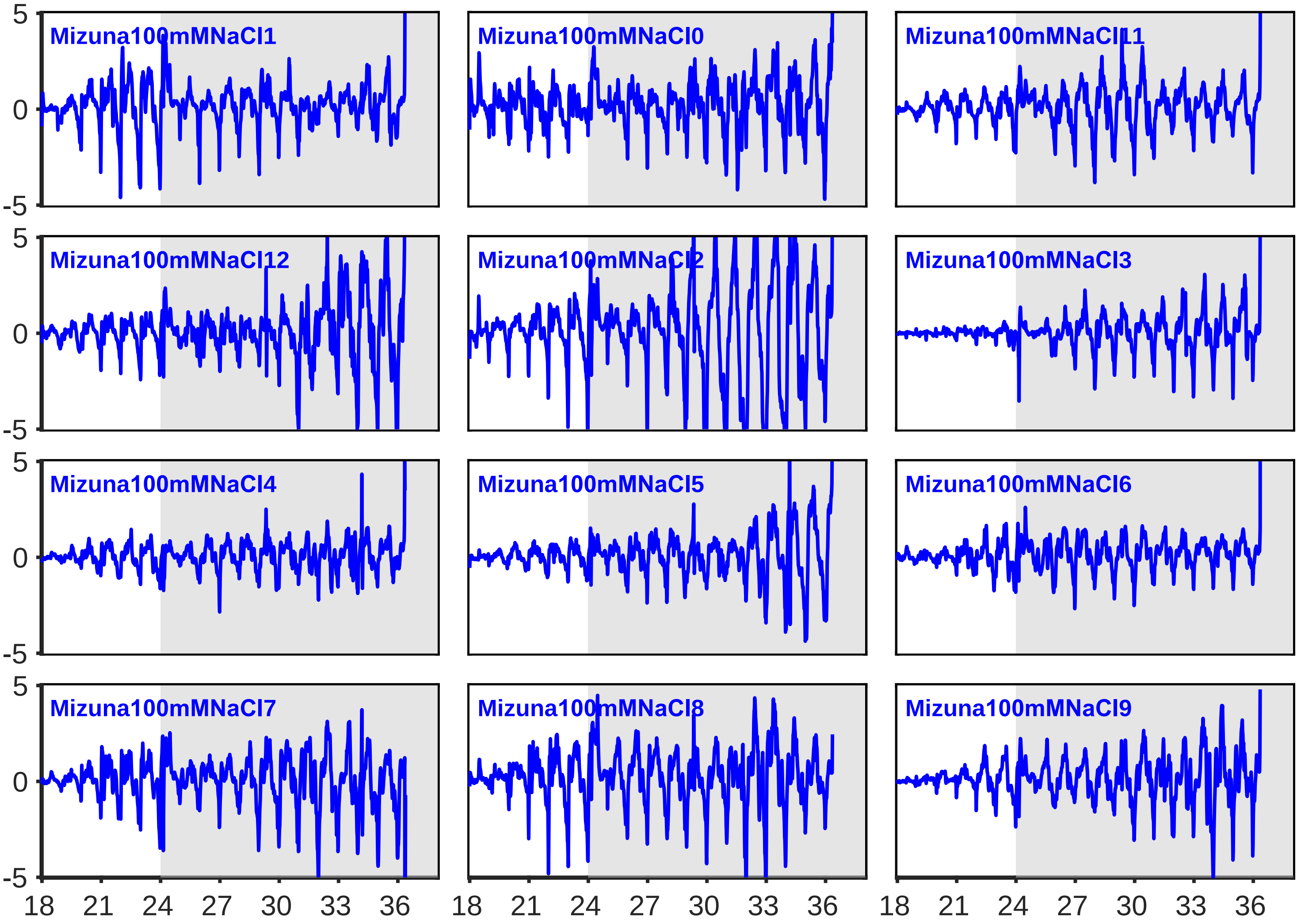

### Motion Curves - Group: RadishControl

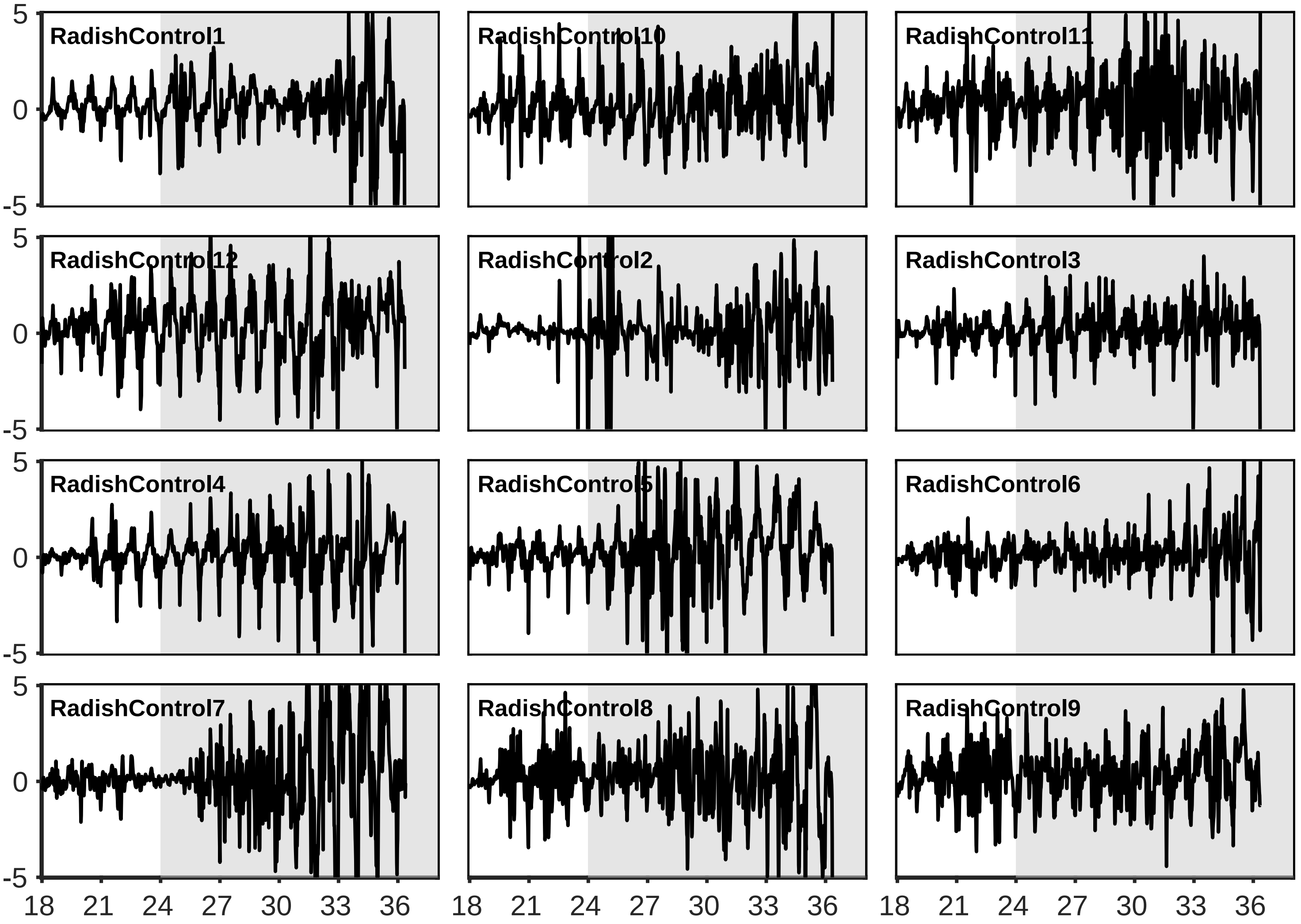

#### Motion Curves - Group: Radish 100 mM NaCl

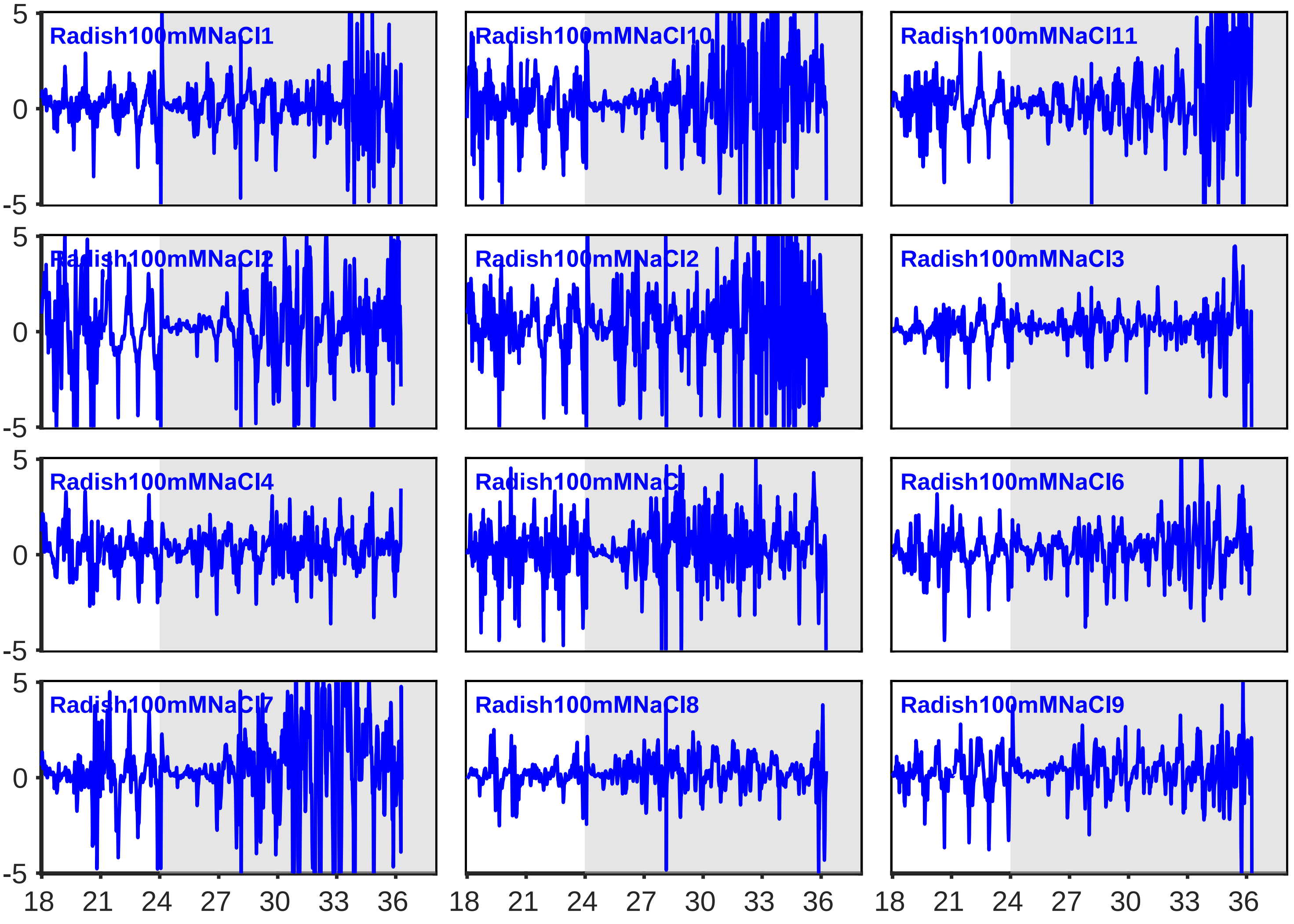

### Motion Curves - Group: AmaranthControl

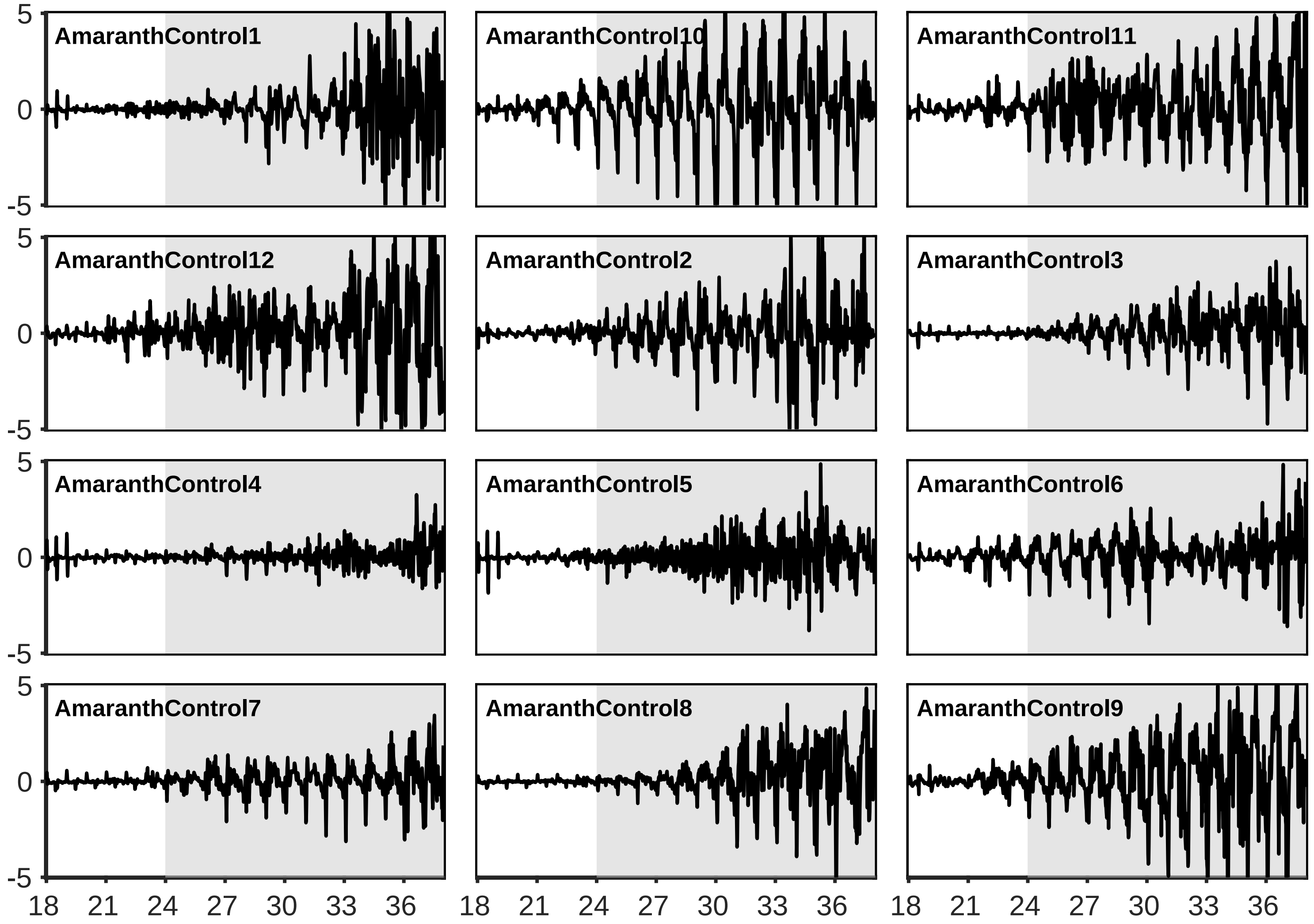

### Motion Curves - Group: Amaranth100mMNaCl

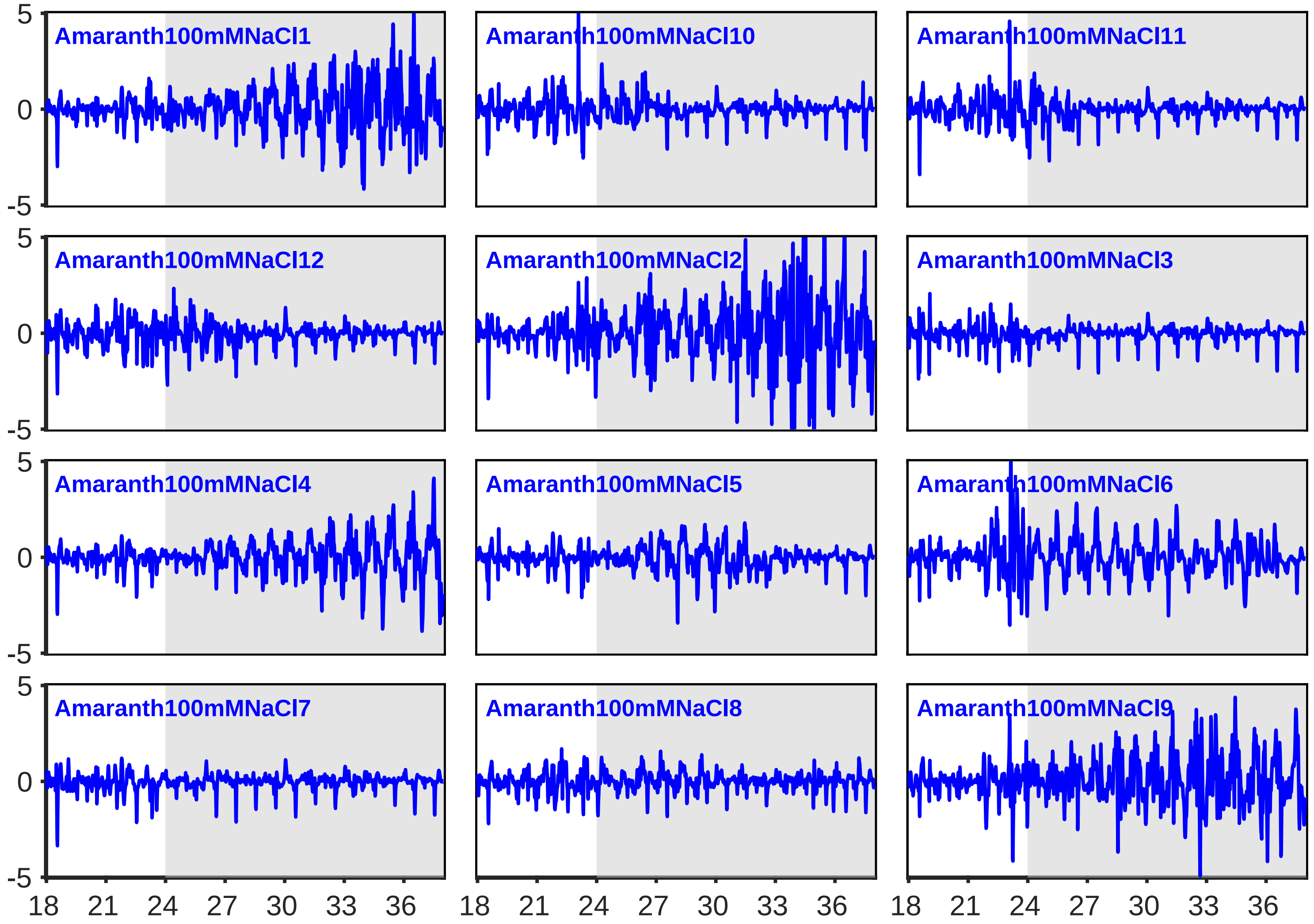

### Motion Curves - Group: TomatoControl

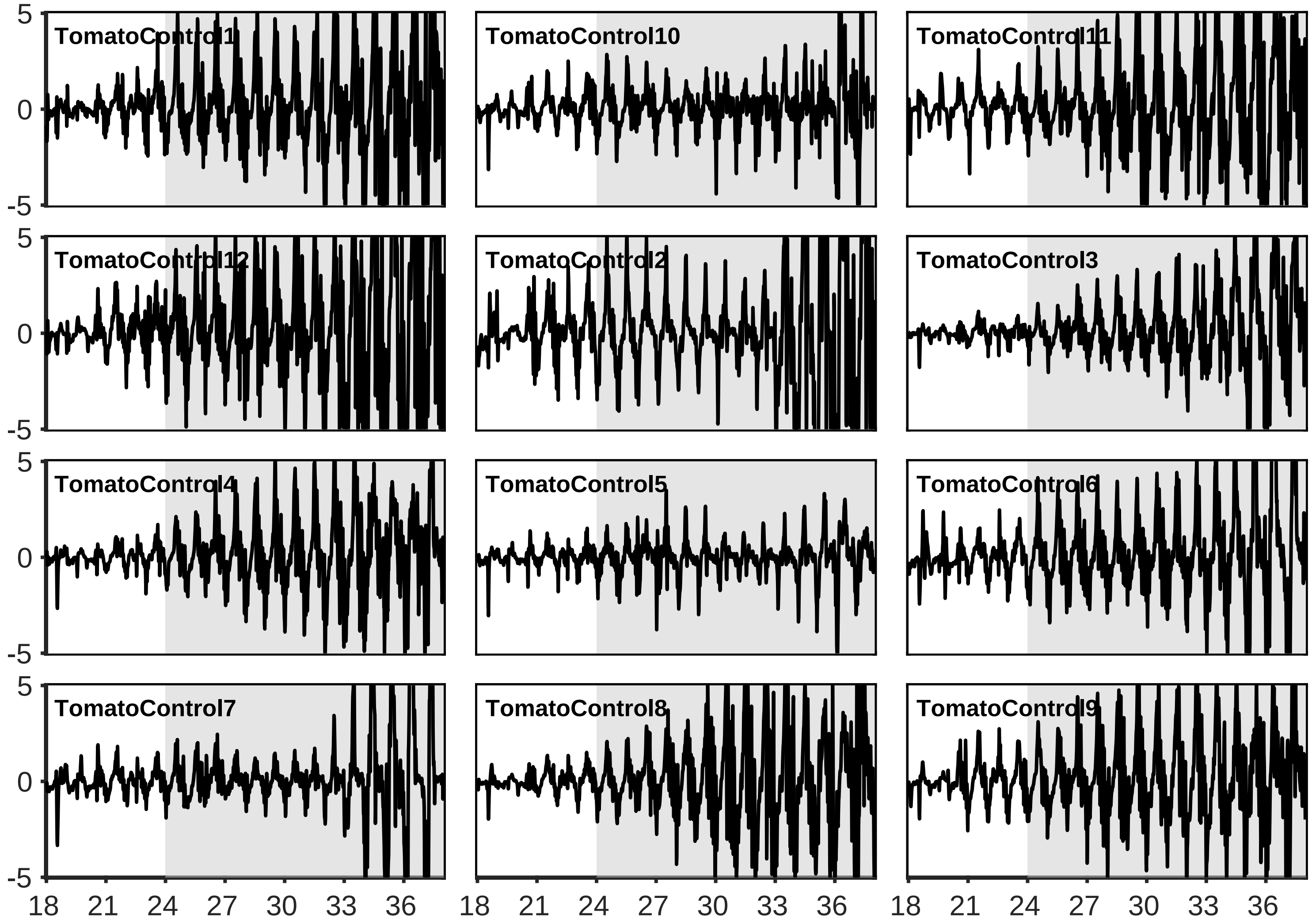

### Motion Curves - Group: Tomato100mMNaCl

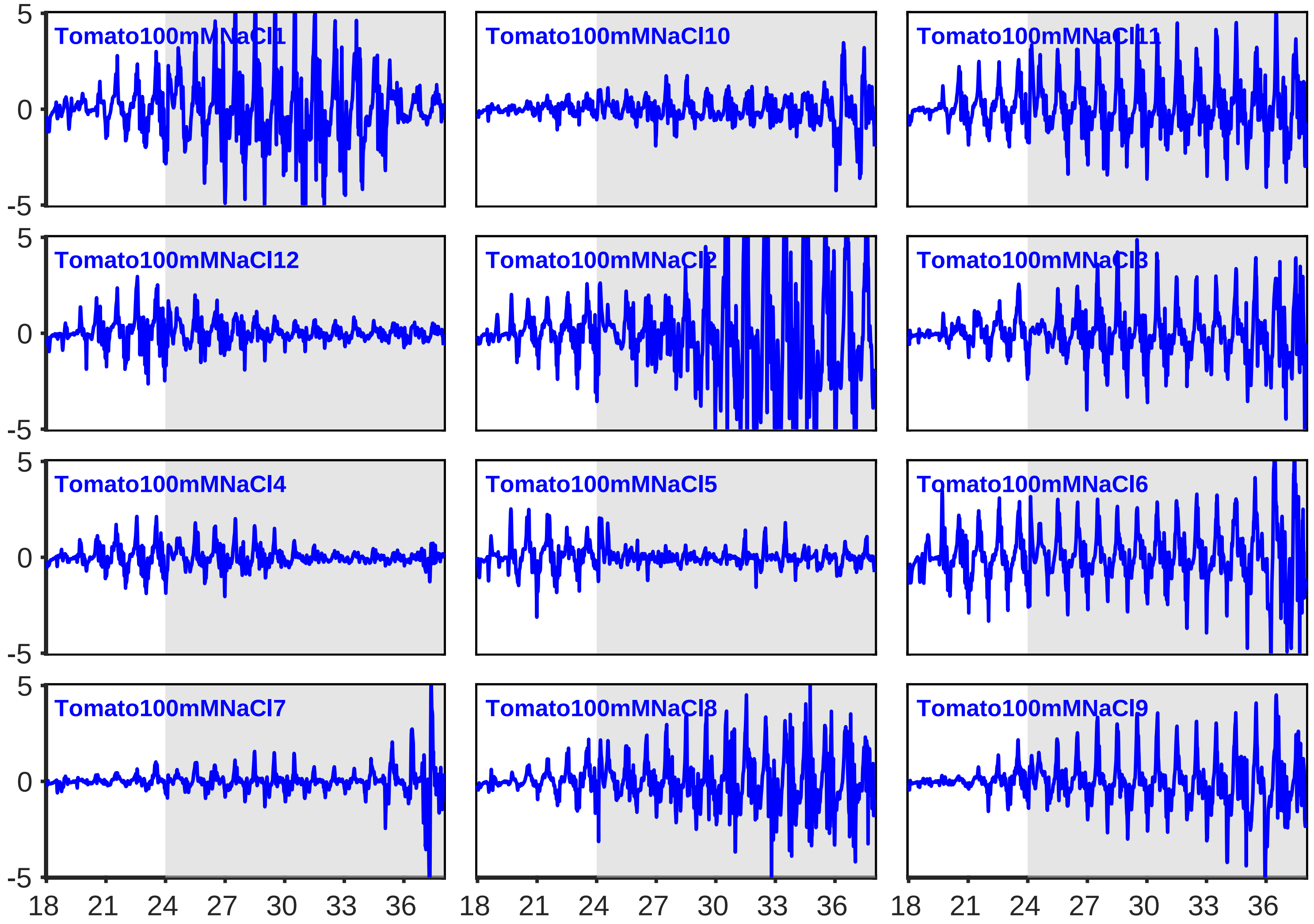

#### Motion Curves - Group: RocketControl

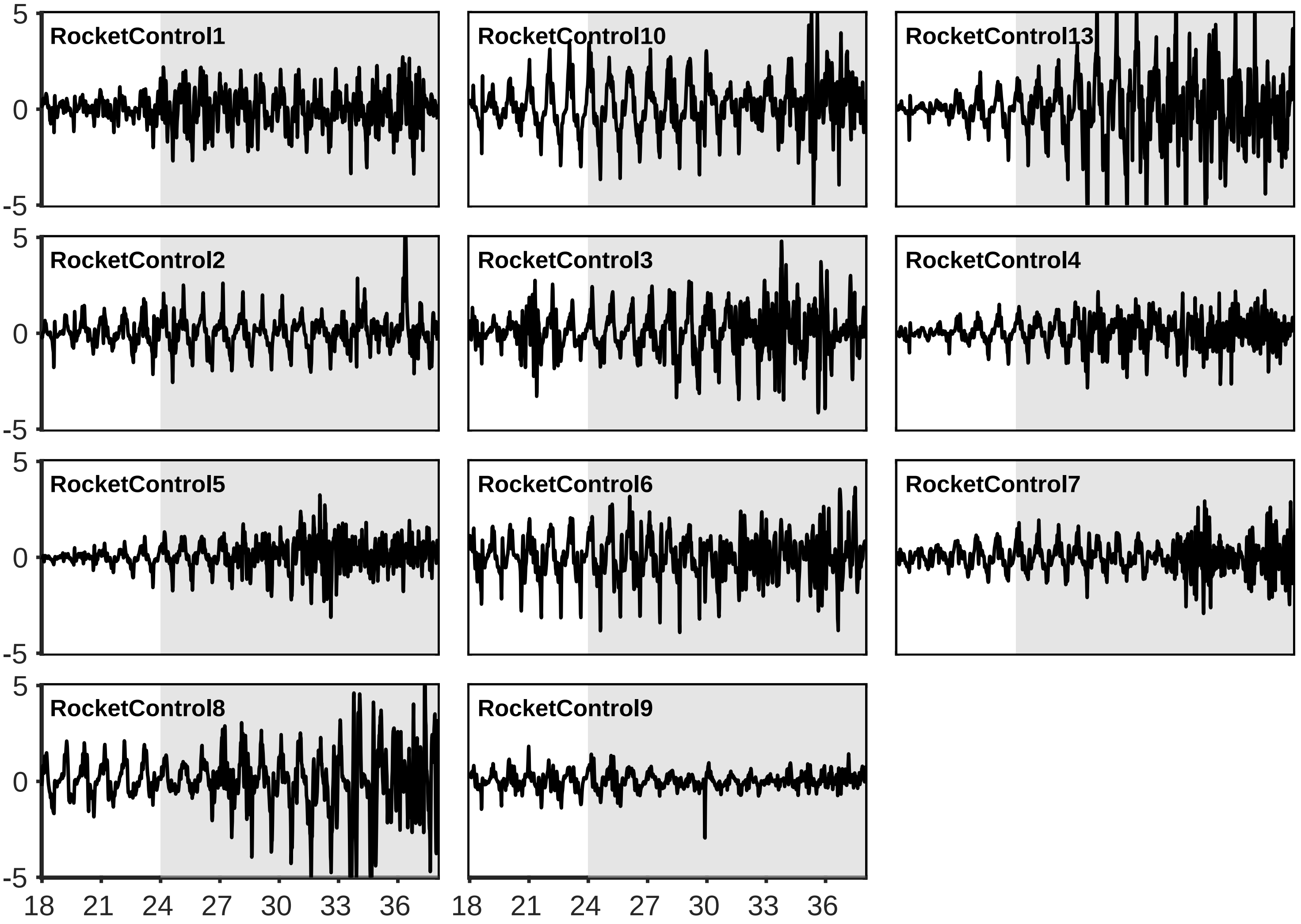

#### Motion Curves - Group: Rocket100mMNaCl

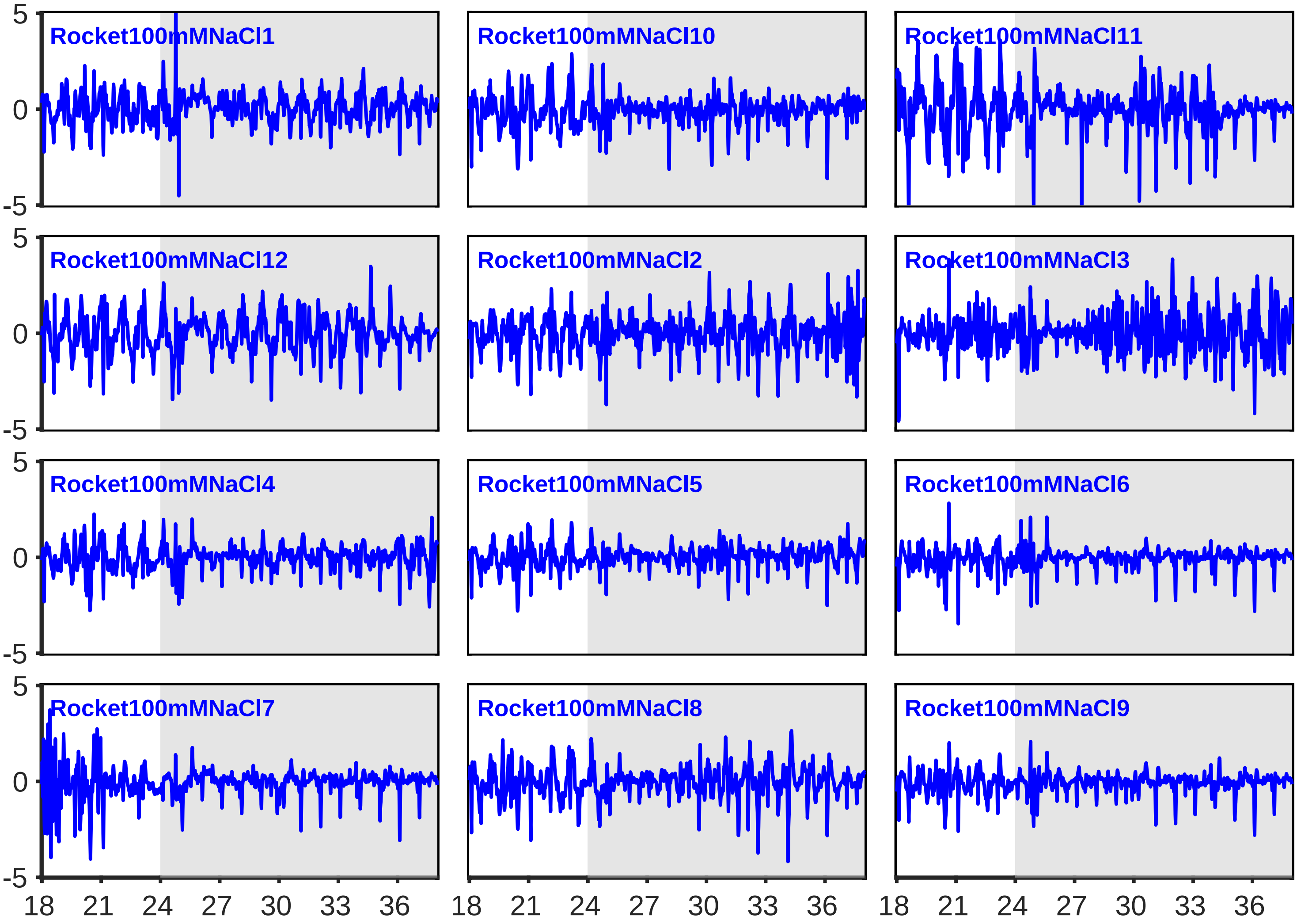

**Supporting Figure S1**. Raw motion curves corresponding to: **(d)** lettuce plants exposed to 100 mM KCl under two lighting regimes; dimG$^{day\ \&\ night}$ and RGB$^{day}$ dimG$^{night}$

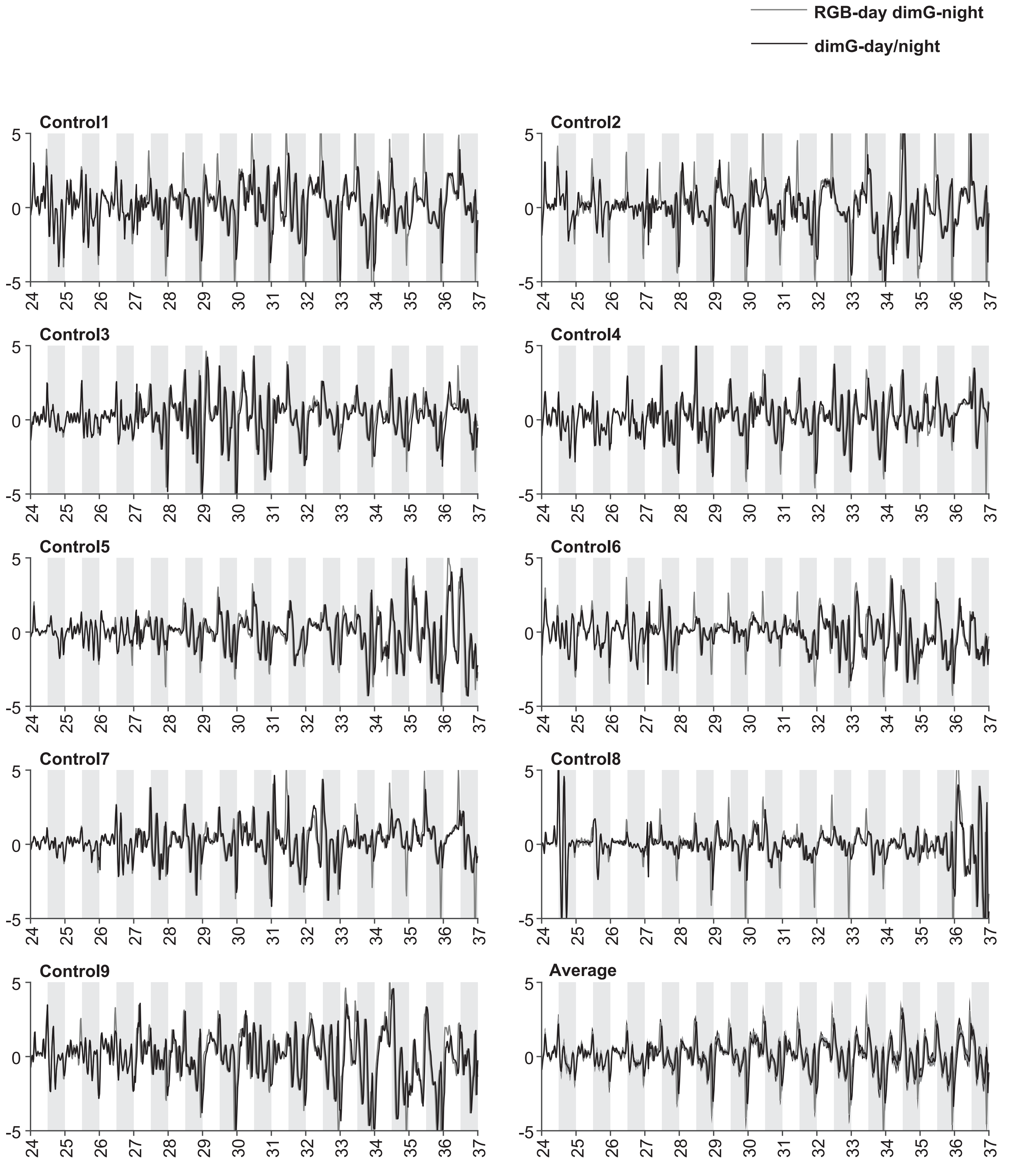

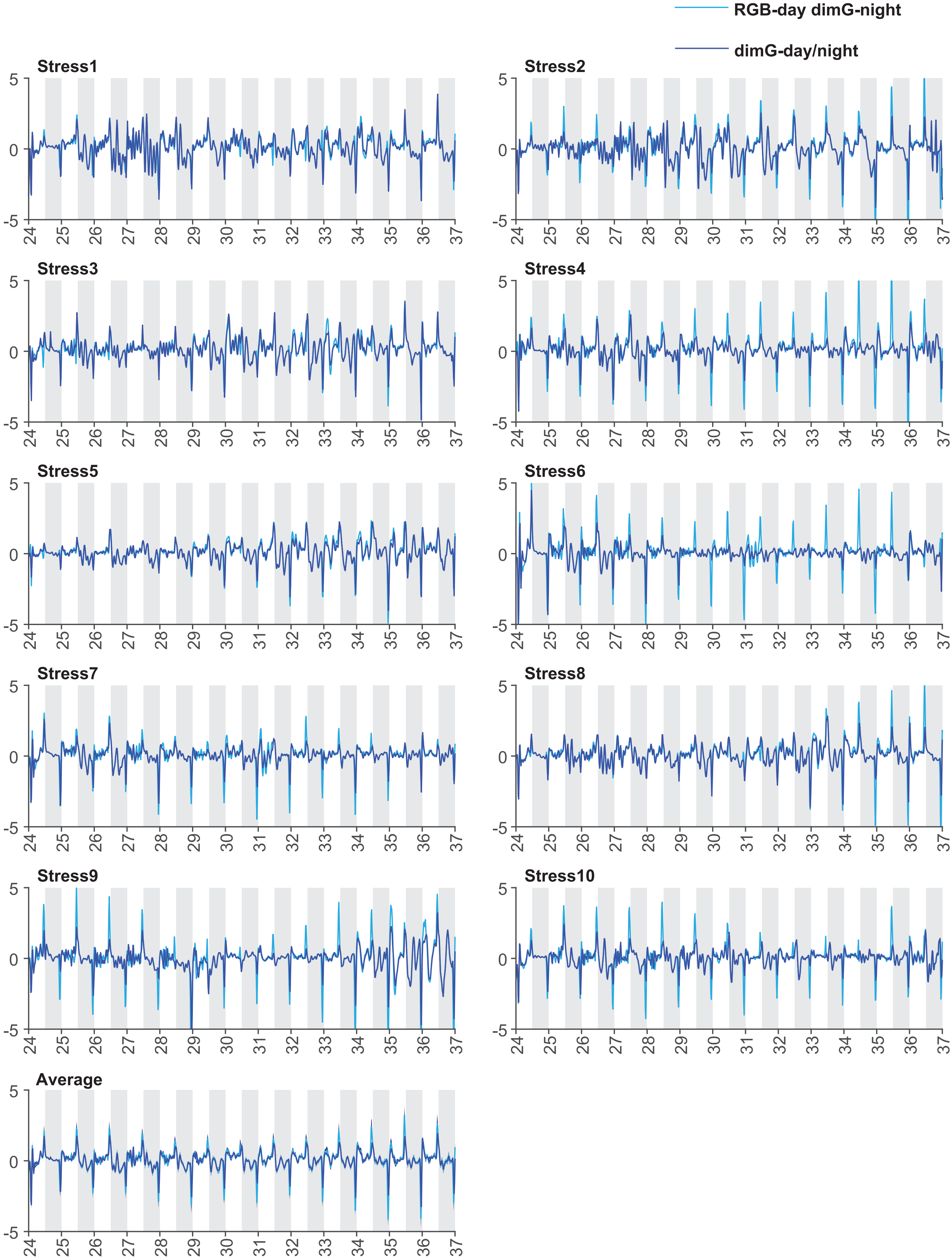
RGB-day dimG-night
dimG-day/night
Stress1
Stress2
Stress3
Stress4
Stress5
Stress6
Stress7
Stress8
Stress9
Stress10
Average
5
0
-5
