## Supplemental Figure S2 for "Leaf movements as a quantitative metric for early stress detection"

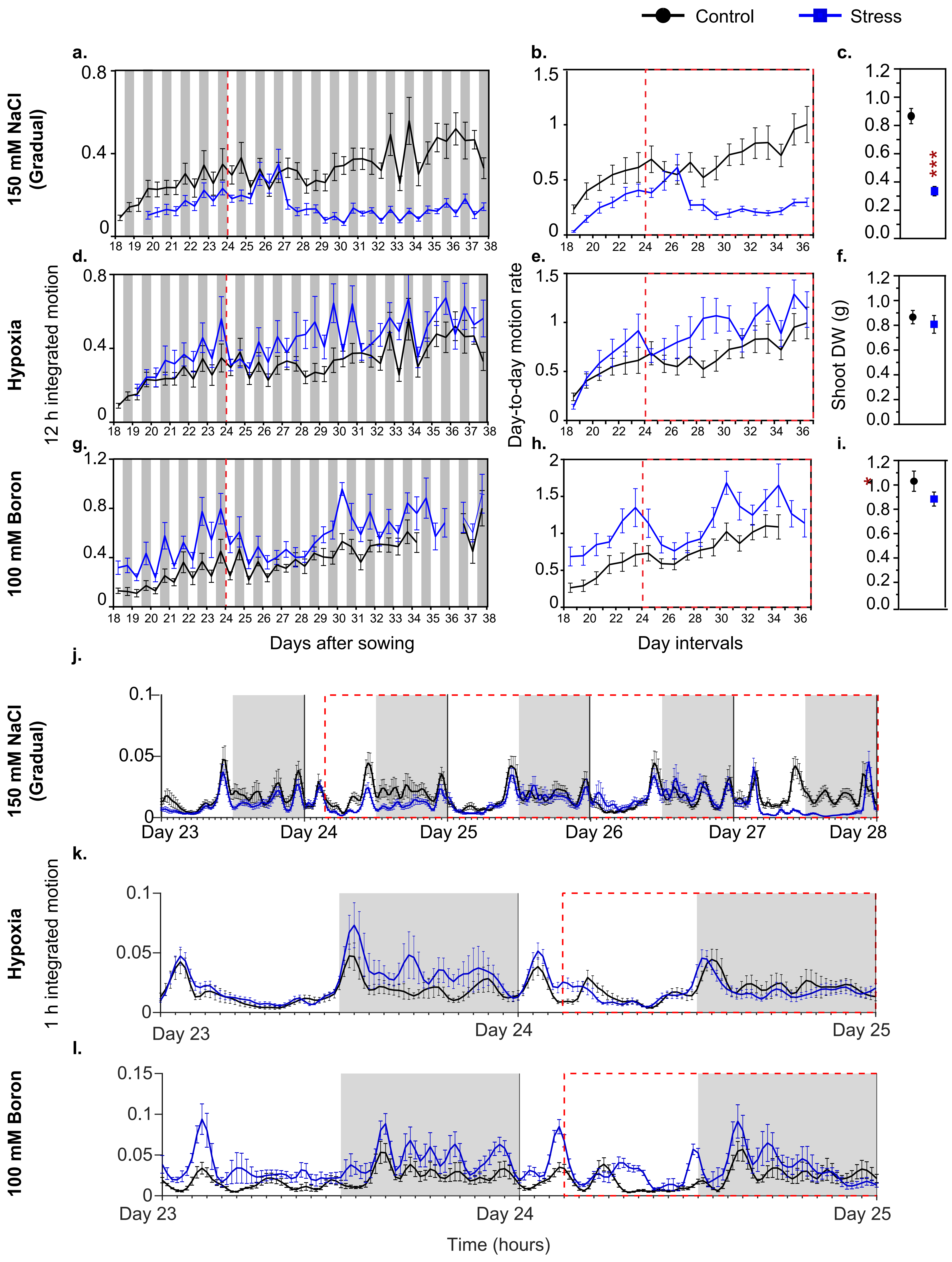


Supplementary Figure S2. Leaf movement responses to (a-c) 150 mM NaCl, (d-f) Hypoxia and (g-i) 100 mM Boron. For each treatment, panels show (a,d,g) 12 h integrated motion, (b,e,h) day-to-day motion rate, (c,f,i) Shoot DW. (j-l) 1 h integrated motion following application of (j) 150 mM NaCl, (k) Hypoxia and (l) 100 mM Boron, shown from day 23 to the onset of changes in leaf-movement dynamics after stress application on day 24. Data represent means of 9-12 plants for control (black) and stress (blue) treatments; error bars indicate SEM (see Supplementary Table S3a, S4a, S5a and S6a for numerical data). The red dashed line marks the timing of stress application. White and grey backgrounds indicate day and night, respectively. Asterisks denote significant differences between stress and control treatments (*$p < 0.05$, **$p < 0.01$, ***$p < 0.001$; two-sample t-tests).
