## Supplemental Figure S3 for "Leaf movements as a quantitative metric for early stress detection"

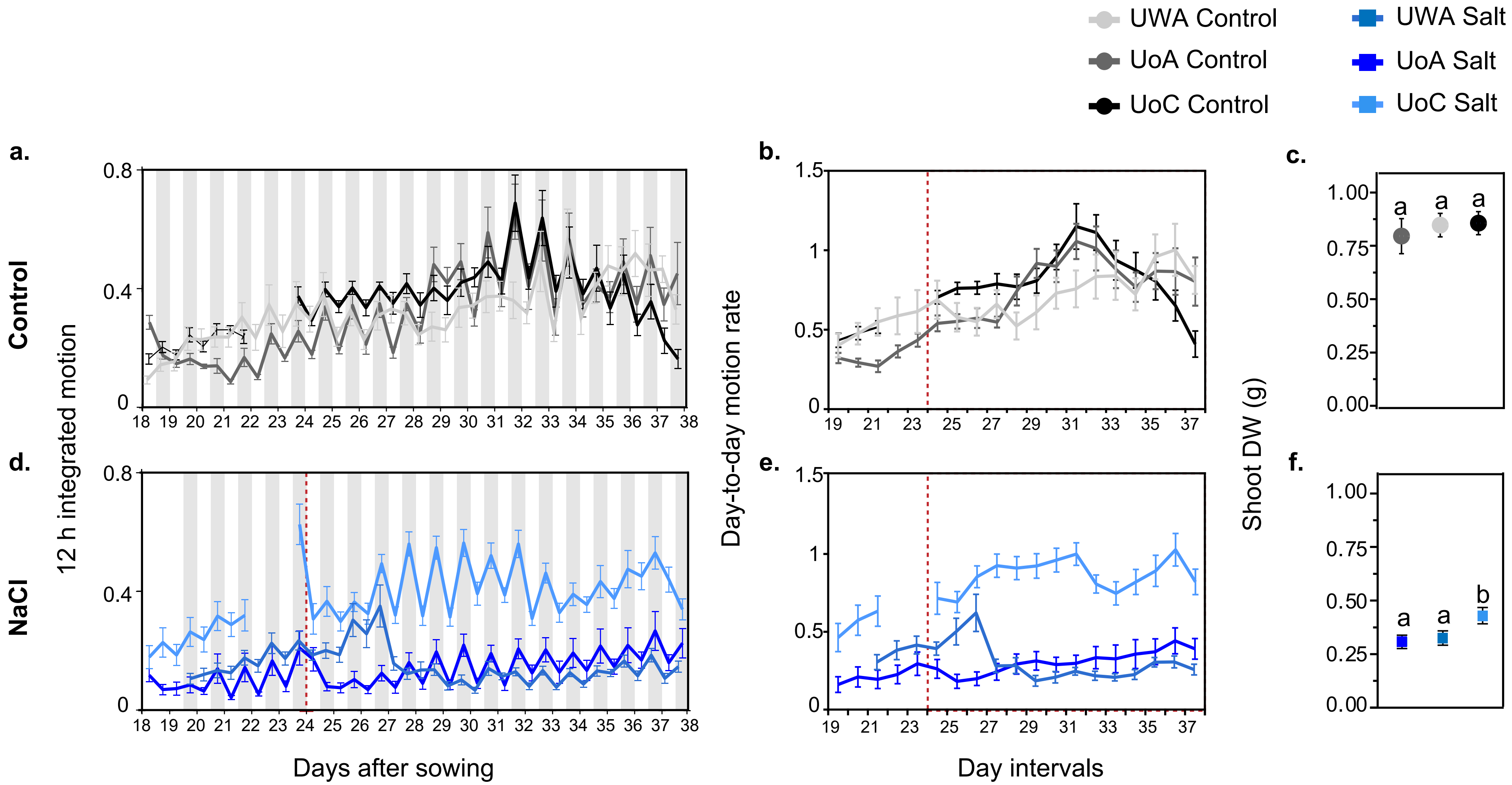


Supplementary Figure S3. Leaf-movement in a closed Conviron growth cabinet (UWA), in a controlled environment plant growth walk-in room (AU), and in a laboratory room (UoC) under (a-c) control and (d-f) NaCl (150 mM in UWA; 100 mM in AU and UoC) conditions. For each treatment, panels show (a,d) 12 h integrated motion, (b,e) day-to-day motion rate, (c,f) Shoot DW. Data represent means of 9-12 plants for control (black) and stress (blue) treatments; error bars indicate SEM (see Supplementary Table S3b, S4b, S5b for numerical data). The red dashed line marks the timing of stress application. White and grey backgrounds indicate day and night, respectively. Asterisks denote significant differences between stress and control treatments (*p < 0.05, **p < 0.01, ***p < 0.001; two-sample t-tests).
