## Supplemental Figure S4 for "Leaf movements as a quantitative metric for early stress detection"

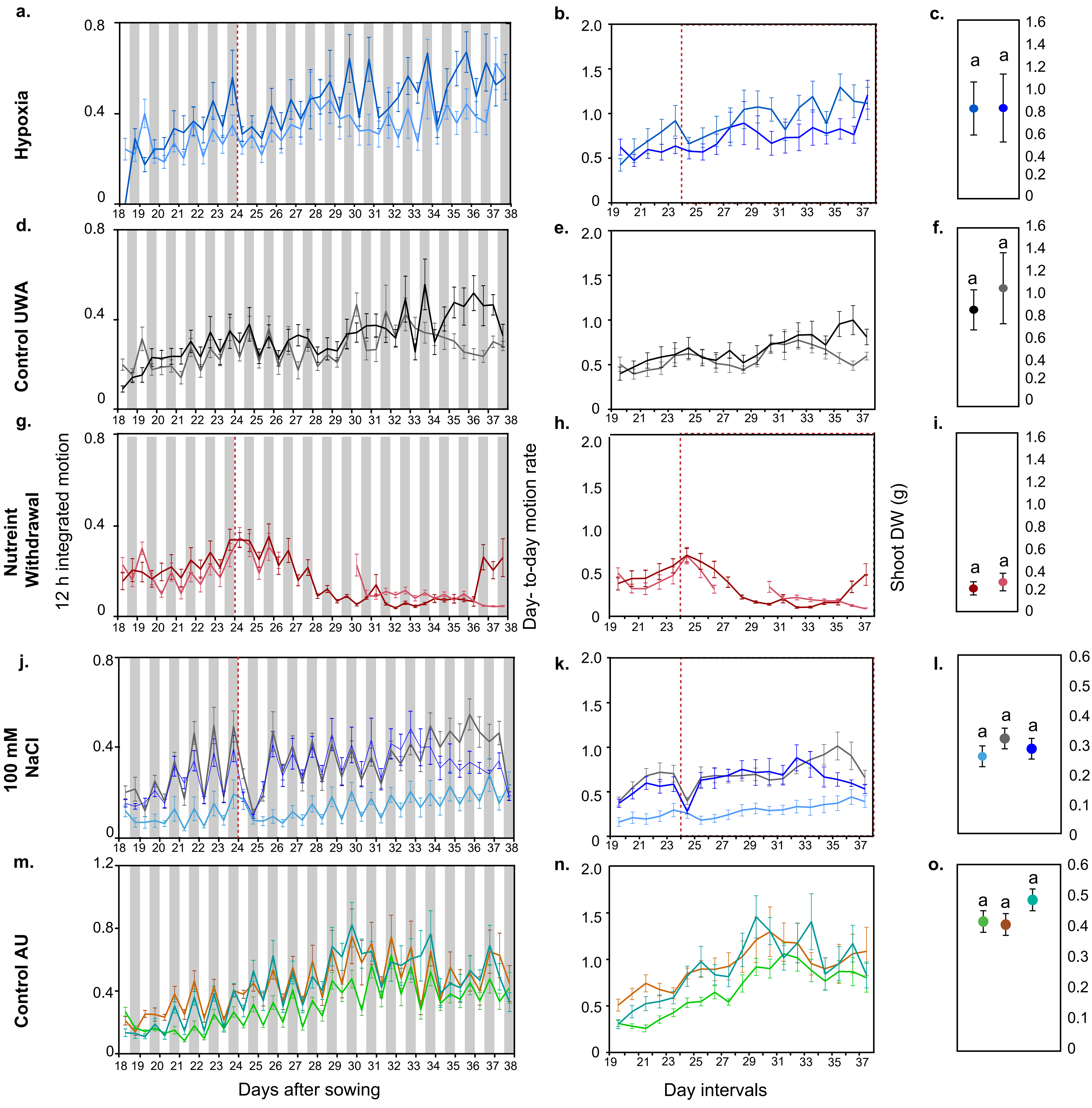


Supplementary Figure S4. Leaf-movement in replicate experiments conducted in the same location under (a-c) Hypoxia, (d-f) Control (UWA) (g-i) Nutrient withdrawal, (j-l) 100 mM NaCl, and (m-o) Control (AU) conditions. For each treatment, panels show (a,d,g,j,m) 12 h integrated motion, (b,e,h,k,n) day-to-day motion rate, (c,f,i,l,o) Shoot DW. Data represent means of 9-12 plants for control (black) and stress (blue) treatments; error bars indicate SEM (see Supplementary Table S3c, S4c, S5c for numerical data). The red dashed line marks the timing of stress application. White and grey backgrounds indicate day and night, respectively. Different letters in shoot dry weight denote significant differences between locations within a treatment (*p < 0.05) as determined by 1-way ANOVA analysis, Tukey's HSD test.
