## Supplemental Figure S5 for "Leaf movements as a quantitative metric for early stress detection"

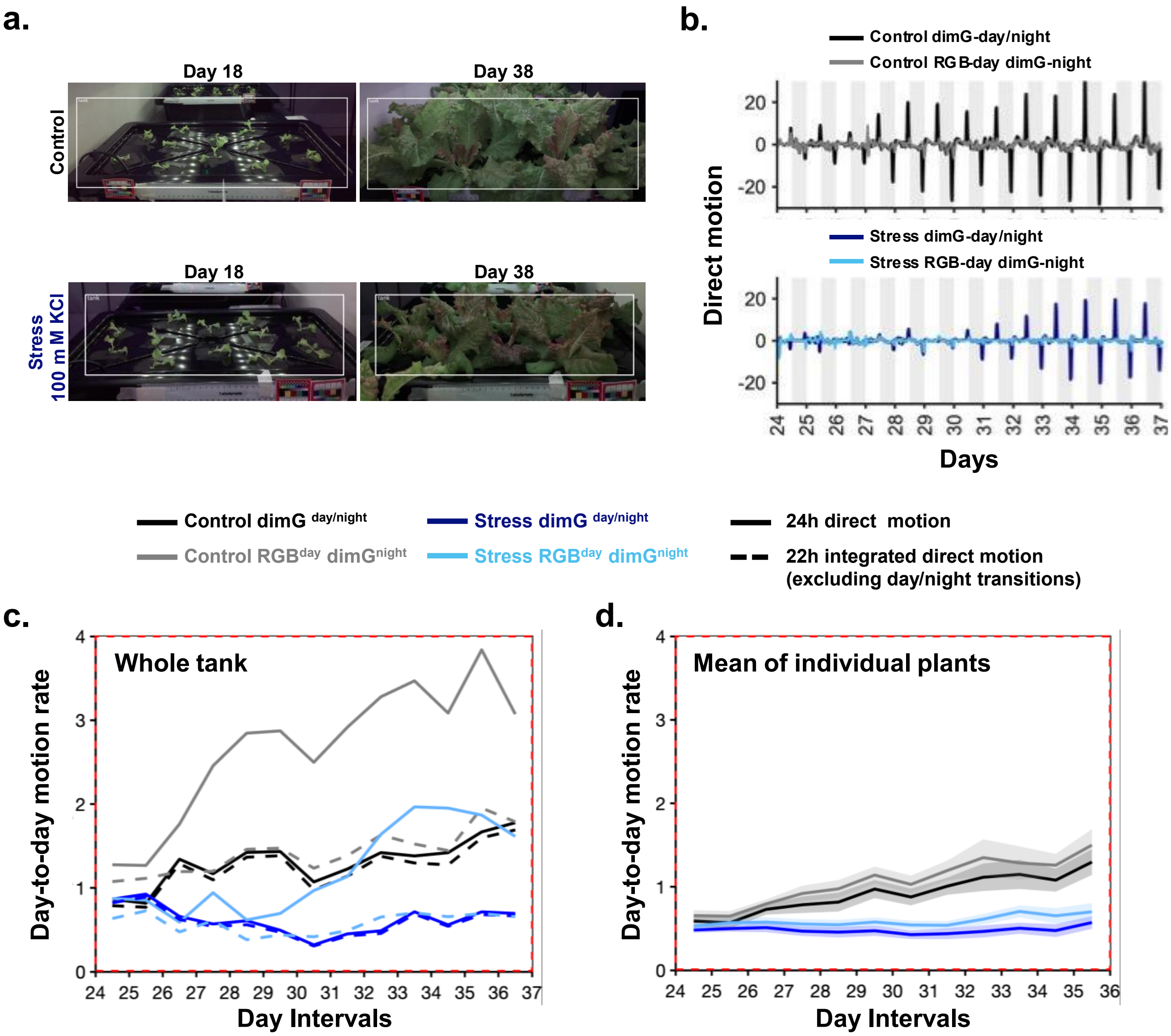


**Fig. S5 Quantification of stress-induced leaf movement dynamics in lettuce plant arrays.**

**a.** Region of interest used for leaf-movement quantification in control and stress treatment tank, showing lettuce plants at the start (day 18) and end (day 38) of the experiment.

**b.** Direct motion across days for Control and Stress tanks under dimG $^{day/night}$ and RGB$^{day}$ dimG$^{night}$ imaging regimes.

**c.** Day-to-day motion rate in Control and Stress tanks measured under dimG $^{day/night}$ and RGB$^{day}$ dimG$^{night}$ , calculated using either 24 h motion integration or 22 h motion integration excluding day/night transitions.

**d.** Mean day-to-day motion rate of individual plants under dimG $^{day/night}$ and RGB$^{day}$ dimG$^{night}$ Data represent means of 9–10 plants, shaded areas depicts SEM. Red dashed lines indicate stress application.

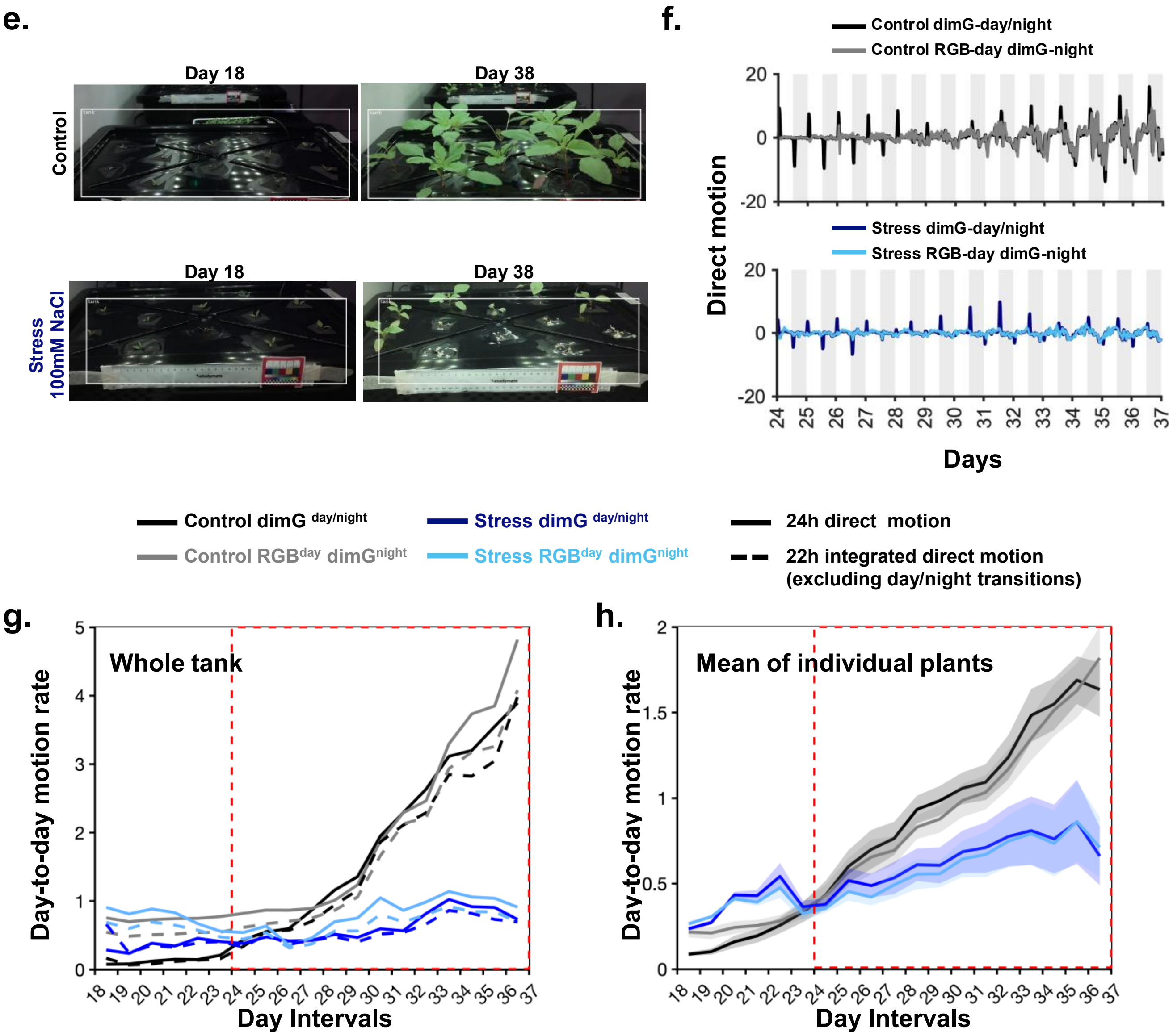


**Fig. S5 Quantification of stress-induced leaf movement dynamics in amaranth plant arrays.**

**e.** Region of interest used for leaf-movement quantification in control and stress treatment tank, showing lettuce plants at the start (day 18) and end (day 38) of the experiment.
**f.** Direct motion across days for Control and Stress tanks under dimG $^{day/night}$ and RGB$^{day}$ dimG$^{night}$ imaging regimes.
**g.** Day-to-day motion rate in Control and Stress tanks measured under dimG $^{day/night}$ and RGB$^{day}$ dimG$^{night}$ , calculated using either 24 h motion integration or 22 h motion integration excluding day/night transitions.
**h.** Mean day-to-day motion rate of individual plants under dimG $^{day/night}$ and RGB$^{day}$ dimG$^{night}$ Data represent means of 9–10 plants, shaded areas depicts SEM. Red dashed lines indicate stress application.

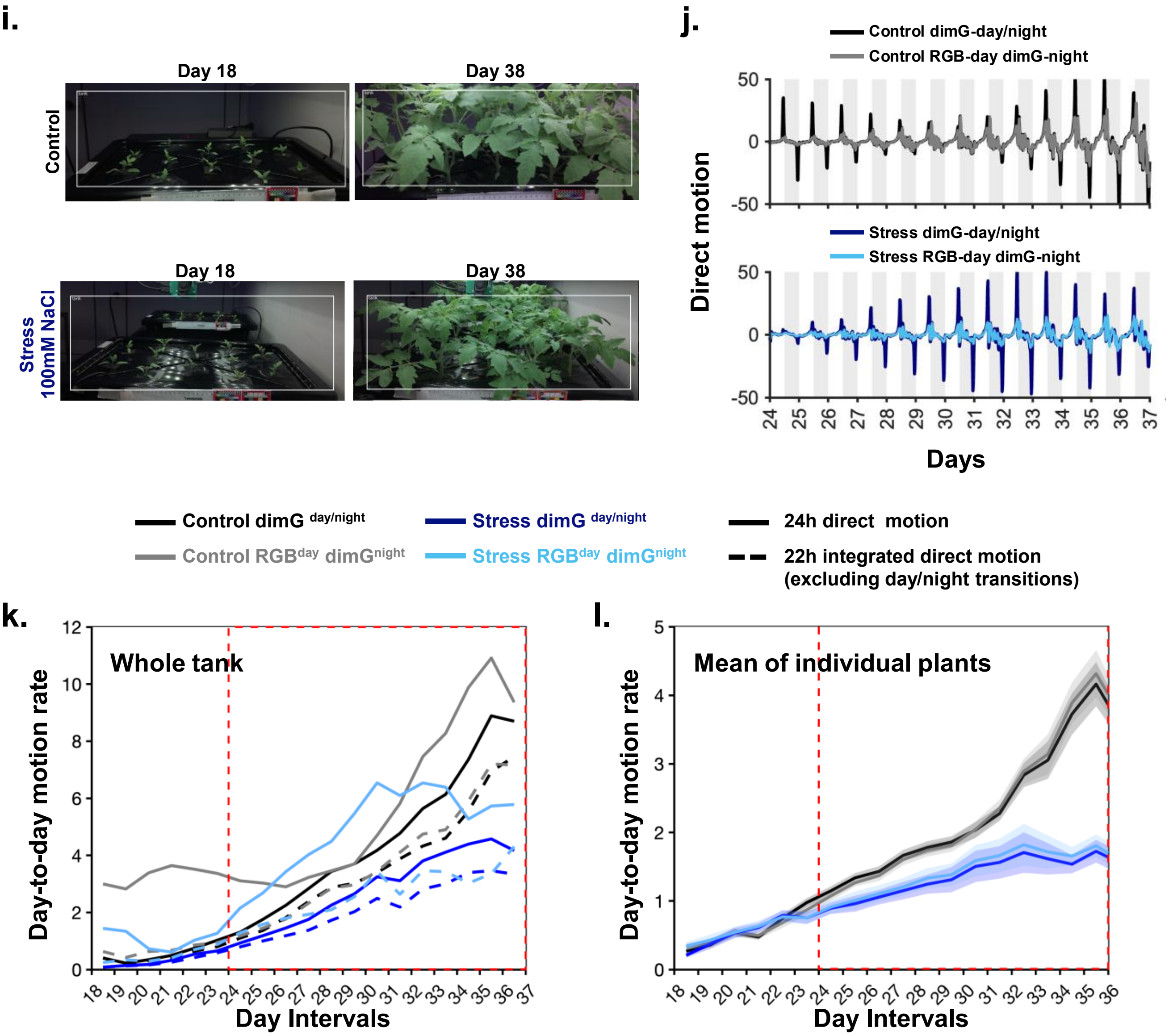


**Fig. S5 Quantification of stress-induced leaf movement dynamics in tomato plant arrays.**

**i.** Region of interest used for leaf-movement quantification in control and stress treatment tank, showing lettuce plants at the start (day 18) and end (day 38) of the experiment.
**j.** Direct motion across days for Control and Stress tanks under dimG $^{day/night}$ and RGB$^{day}$ dimG$^{night}$ imaging regimes.
**k.** Day-to-day motion rate in Control and Stress tanks measured under dimG $^{day/night}$ and RGB$^{day}$ dimG$^{night}$, calculated using either 24 h motion integration or 22 h motion integration excluding day/night transitions.
**l.** Mean day-to-day motion rate of individual plants under dimG $^{day/night}$ and RGB$^{day}$ dimG$^{night}$ Data represent means of 9–10 plants, shaded areas depicts SEM. Red dashed lines indicate stress application.
