## Supplemental Figure S6 for "Leaf movements as a quantitative metric for early stress detection"

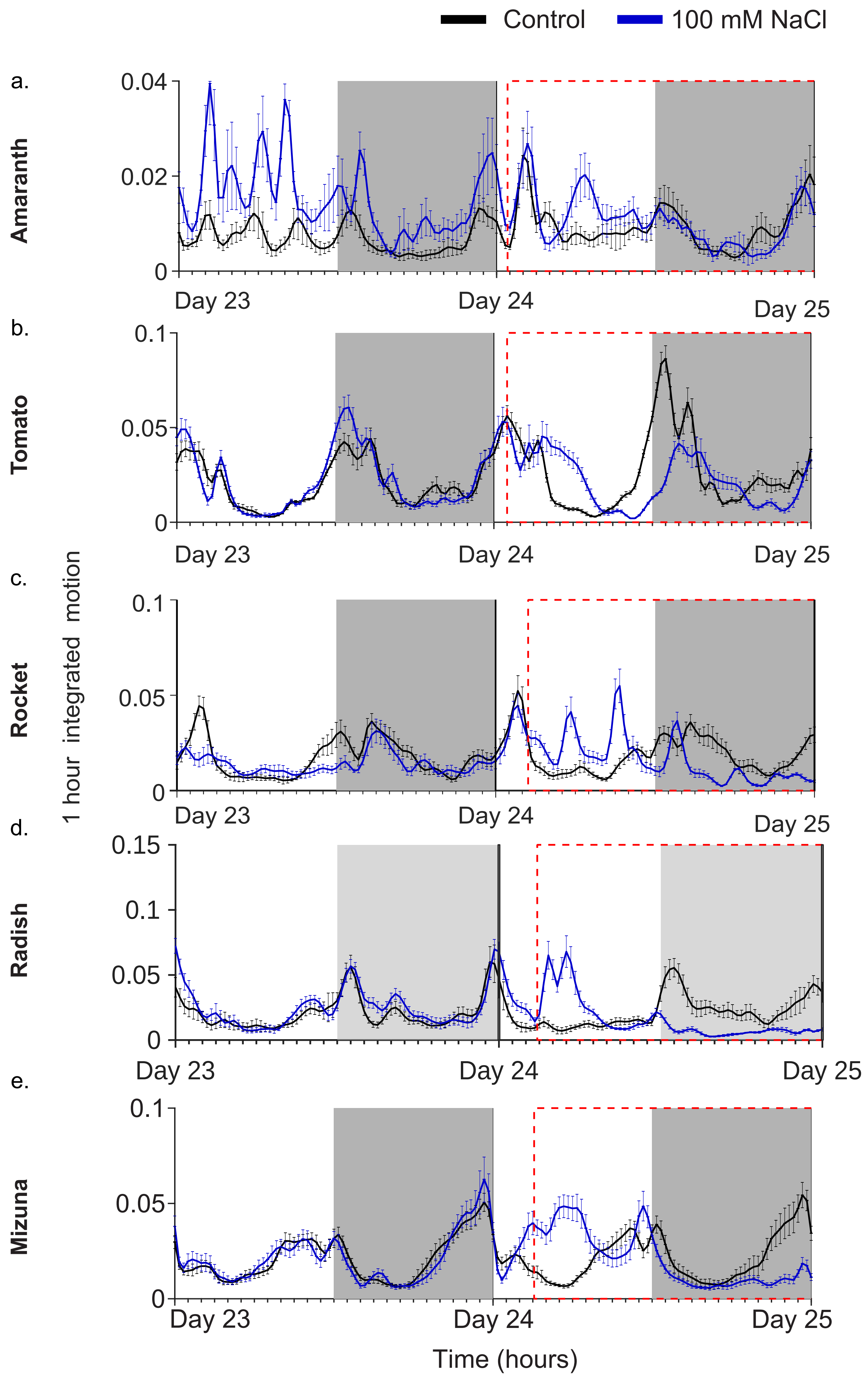


Supplementary Figure S6. 1 h integrated motion in (a) amaranth, (b) mizuna, (c) radish, (d) rocket, and (e) tomato subjected to 100 mM NaCl stress, with their respective non-stressed plants. (see Supplementary Table S6c for numerical data) The red dashed line marks the timing of stress application. White and grey backgrounds indicate day and nighrespectively.
