## Supplemental Information S1 for "Leaf movements as a quantitative metric for early stress detection"

### Supporting information S1. Plant growth and imaging setups across locations

University of Western Australia, Australia - Plant Growth Chamber (PGC) Flex closed growth cabinet with tight temperature and humidity control (Conviron, Manitoba, Canada)

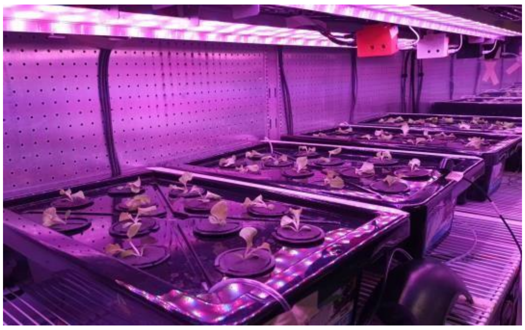

Adeleide University, Australia- a controlled environment plant growth walk-in room with temperature and humidity control

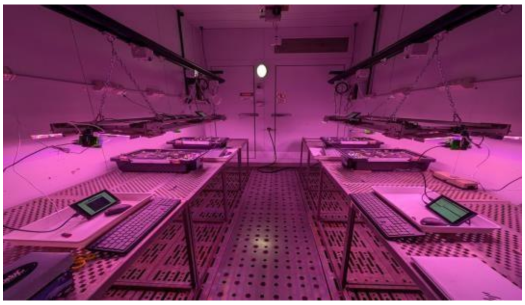

University of Cambridge, UK - A laboratory room with broad temperature and humidity control.

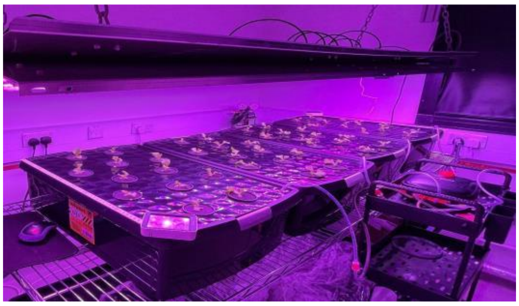

LED lights 30 cm above the plant canopy provided 160 µmol $m^{-2}$ $s^{-1}$ in the form of 105 µmol $m^{-2}$ $s^{-1}$ red (wavelength 600-700 nm), 20 µmol $m^{-2}$ $s^{-1}$ green (wavelength 500-600 nm), and 35 µmol $m^{-2}$ $s^{-1}$ blue (wavelength 400-500 nm) at plant height. Photoperiod was 12 h of light at 22°C and 12 h of night at 20°C and humidity set at 60%.

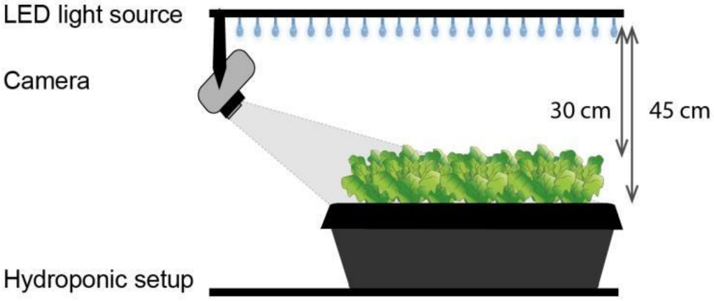
