## Supplemental Information S2 for "Leaf movements as a quantitative metric for early stress detection"

**Supporting Information S2.** Experimental logs indicating key stages of experimental process.

| Experiment | Time lights on | Time lights off | Sowing day | Transfer to tanks (day 17) | Analysis start (day 18) | Day of stress application | Time of stress application | Day of harvest (day38) | Tank 1 | Tank2 | Tank3 | Tank4 |
|---|---|---|---|---|---|---|---|---|---|---|---|---|
| UA-Exp4 | 7:00 | 19:00 | 9/9/2024 | 25/9/2024 | 27/9/2024 | 3/10/2024 | 100 mM NaCl at 12:40:00 | 17/10/204 | 100 mM NaCl lettuce | Control lettuce | 100 mM NaCl lettuce | Control lettuce |
| UA-Exp5 | 8:00 | 20:00 | 1/10/2024 | 18/10/2024 | 19/10/2024 | 25/10/2024 | 100 mM NaCl at 11:00 | 8/11/2024 | 100 mM NaCl lettuce | Control lettuce | Control lettuce | 100 mM NaCl lettuce |
| UA-Exp6 | 8:00 | 20:00 | 2/11/2024 | 19/11/2024 | 20/11/2024 | 26/11/2025 | Boron 11:10, KCl 10:50, nutrients replenished 10:50 | 10/12/2024 | Control lettuce | Boron lettuce | 100 mM KCl lettuce | Nutrient replenish lettuce |
| UA_Exp7 | 8:00 | 20:00 | 11/11/2024 | 28/11/2024 | 29/11/2024 | 4/12/2024 | Two nozzles closed off at 15:55 | 19/12/2024 | Canal experiment | | | |
| UA_Exp8 | 21:30 | 9:30 | 24/1/2025 | 9/2/2025 | 10/2/2025 | 17/2/2025 | Fixed floating collars (particularly tub 2, rocket) between 12.30 and 12.50pm local time (02:00 and 02:20 on images) Some disruption to imaging expected at this time. Added salt (140 g salt per tub) at 3.50 pm (05:20 on images) | 03/03/2025 | Rocket control | Rocket 100 mM NaCl | Lettuce control | Lettuce 100 mM NaCl |
| UA-Exp9 | 8:00 | 20:00 | 22/02/2025 | 11/3/2025 | 12/03/2025 | 18/3/2025 | Added salt at 9.10am | 01/04/2025 | Amaranth control | Amaranth 100 mM NaCl | Tomato 100 mM NaCl | Tomato control |
| UoC_Exp1 | 10:00 | 22:00 | 10/9/2024 | 26/9/2024 | 27/9/2024 | 4/10/2024 | 100mM between image 10:20 and image 10:40 | 18/10/2024 | 100 mM NaCL lettuce | Control lettuce | Control Lettuce | Transient water withdrawal |
| | | | | | | 10/10/2024 | Removed nutrient solution at 14:00 | | | | | |
| | | | | | | 11/10/2024 | Restored nutrient solution at 14:00 | | | | | |
| UoC_Exp8 | 10:00 | 22:00 | 20/7/2025 | 5/8/2025 | 6/8/2025 | 17/8/2025 | Removed nutrient solution at 14:00 | 25/8/2025 | Control lettuce | Transient water withdrawal | | |
| | | | | | | 18/8/2025 | Restored nutrient solution at 14:00 | | | | | |
| UWA_Exp4 | 06:00 | 18:00 | 08/09/2024 | 25/09/2024 | 26/09/2024 | 02/10/2024 | Stresses applied 9-9.30 am. | 16/10/2024 | Nutrient Withdrawal Lettuce | 150 mM NaCl lettuce | Control lettuce | Hypoxia lettuce |
| UWA_Epx6 | 06:00 | 18:00 | 25/10/2024 | 11/11/2024 | 12/11/2024 | 18/11/2024 | Stresses applied 9-9.30 am. | 2/12/2024 | Nutrient Withdrawal Lettuce | 150 mM NaCl lettuce | Control lettuce | Hypoxia lettuce |
| UWA_Exp9 | 06:00 | 18:00 | 11/02/2026 | 02/03/2025 | 03/03/2025 | 09/03/2025 | Stresses applied 9-9.30 am. | 23/03/2025 | 100 mM NaCl Mizuna | Control Mizuna | Control Radish | 100 mM NaCl radish |
