## Supplemental Table S1 for "Leaf movements as a quantitative metric for early stress detection"

**Supplementary Table S1.** Data corresponding to the control experiment presented in Figure 1, including: **(a)** raw motion data, **(b)** area under the motion curve (AUC), **(c)** 12 h integrated motion, **(d)** day-to-day motion rate, and **(e)** 1 h integrated motion.

**S1a Raw motion data for control experiment**

| Day | Plant1 | Plant2 | Plant3 | Plant4 | Plant5 | Plant6 | Plant7 | Plant8 | Plant9 | Whole Tank ROI | Empty ROI |
|---|---|---|---|---|---|---|---|---|---|---|---|
| 18 | -0.45008 | -0.29144 | -0.20011 | 0.351355 | 0.030443 | 0.129136 | -2.74893 | -0.01682 | -0.0011 | -0.405243 | 0 |
| 18.01389 | -1.85246 | -1.74027 | 0.078379 | 0.089679 | -1.19837 | 0.092218 | -4.75128 | -0.41286 | -0.7913 | -1.599851 | 0 |
| 18.02778 | -2.64248 | -2.58533 | 0.552324 | -0.4473 | -2.51619 | -0.24431 | -2.99979 | -0.64264 | -1.21287 | -3.015904 | 0 |
| 18.04167 | -1.30252 | -1.28108 | 0.60062 | -0.18439 | -1.39165 | -0.44058 | -1.40841 | -0.33426 | -0.55164 | -2.143187 | 0 |
| 18.05556 | -0.10311 | 0.62603 | 0.370327 | 0.92628 | -0.07692 | -0.36827 | -0.04257 | 0.292843 | -0.2009 | 0.199701 | 0 |
| 18.06944 | 0.872961 | 2.621784 | 0.185485 | 2.646077 | 0.697789 | 0.04493 | 1.330566 | 0.988953 | -0.06051 | 1.649717 | 0 |
| 18.08333 | 0.451911 | 2.263823 | -0.05021 | 2.884997 | 0.669681 | 0.17018 | 1.835847 | 0.805798 | -0.0591 | 1.045999 | 0 |
| 18.09722 | -0.25444 | 0.607823 | -0.30504 | 0.951926 | 0.053135 | -0.08597 | 1.348679 | 0.012386 | -0.19891 | -0.353299 | 0 |
| 18.11111 | -0.51426 | -0.28351 | -0.39472 | -0.13283 | -0.31932 | -0.33474 | 0.769237 | -0.42833 | -0.30889 | -0.536401 | 0 |
| 18.125 | -0.58612 | -0.42189 | -0.35019 | -0.22559 | -0.39524 | -0.46877 | 0.304266 | -0.50404 | -0.42028 | -0.156197 | 0 |
| 18.13889 | -0.6147 | -0.44913 | -0.33448 | -0.26541 | -0.44022 | -0.57275 | -0.10134 | -0.50744 | -0.5459 | -0.149032 | 0 |
| 18.15278 | -0.55274 | -0.3946 | -0.26964 | -0.29116 | -0.40215 | -0.54497 | -0.33458 | -0.43025 | -0.51551 | -0.352378 | 0 |
| 18.16667 | -0.48449 | -0.3152 | -0.19548 | -0.27743 | -0.31437 | -0.48229 | -0.45054 | -0.35946 | -0.39854 | -0.35719 | 0 |
| 18.18056 | -0.51959 | -0.36309 | -0.24327 | -0.32578 | -0.33386 | -0.56714 | -0.53449 | -0.41783 | -0.36018 | -0.298919 | 0 |
| 18.19444 | -0.56988 | -0.46488 | -0.32859 | -0.39756 | -0.41124 | -0.78915 | -0.5718 | -0.52551 | -0.41599 | -0.344947 | 0 |
| 18.20833 | -0.52957 | -0.49138 | -0.31743 | -0.39051 | -0.38174 | -1.08219 | -0.46373 | -0.59059 | -0.39531 | -0.32645 | 0 |
| 18.22222 | -0.45958 | -0.43557 | -0.2569 | -0.36917 | -0.27941 | -1.39929 | -0.32178 | -0.71782 | -0.34285 | -0.379536 | 0 |
| 18.23611 | -0.51077 | -0.46538 | -0.28444 | -0.48056 | -0.22295 | -1.35949 | -0.34307 | -1.008 | -0.39314 | -0.592202 | 0 |
| 18.25 | -0.66102 | -0.6068 | -0.41813 | -0.63503 | -0.14038 | -1.21176 | -0.50614 | -1.24317 | -0.53635 | -0.758685 | 0 |
| 18.26389 | -0.77658 | -0.71395 | -0.5409 | -0.67011 | -0.0523 | -1.10134 | -0.61938 | -1.17162 | -0.65093 | -0.649808 | 0 |
| 18.27778 | -0.80507 | -0.75367 | -0.59676 | -0.64498 | -0.07077 | -1.05507 | -0.64116 | -1.00211 | -0.67638 | -0.555865 | 0 |
| 18.29167 | -0.81841 | -0.8055 | -0.60851 | -0.67985 | -0.11475 | -1.01986 | -0.64831 | -0.8856 | -0.68015 | -0.588805 | 0 |
| 18.30556 | -0.8463 | -0.87045 | -0.61227 | -0.75338 | -0.25136 | -1.00473 | -0.64176 | -0.84398 | -0.68753 | -0.662296 | 0 |
| 18.31944 | -0.80041 | -0.85327 | -0.57317 | -0.7216 | -0.31862 | -0.91292 | -0.58284 | -0.76386 | -0.63334 | -0.668323 | 0 |
| 18.33333 | -0.63256 | -0.7103 | -0.45206 | -0.54258 | -0.37259 | -0.70958 | -0.37272 | -0.61217 | -0.48976 | -0.551497 | 0 |
| 18.34722 | -0.43473 | -0.54234 | -0.28046 | -0.32995 | -0.47723 | -0.50768 | -0.15622 | -0.45142 | -0.31212 | -0.370946 | 0 |
| 18.36111 | -0.2513 | -0.4257 | -0.1118 | -0.15358 | -0.6625 | -0.33669 | -0.01031 | -0.30963 | -0.14751 | -0.267325 | 0 |
| 18.375 | -0.10479 | -0.39266 | 0.025774 | -0.01021 | -0.99953 | -0.1949 | 0.085753 | -0.17972 | -0.01598 | -0.291913 | 0 |
| 18.38889 | -0.05418 | -0.48273 | 0.077156 | 0.037297 | -1.20807 | -0.17671 | 0.135281 | -0.0842 | 0.027759 | -0.484505 | 0 |
| 18.40278 | -0.12839 | -0.70091 | -0.00059 | -0.05949 | -1.34856 | -0.28091 | 0.040963 | -0.09742 | -0.05219 | -0.651095 | 0 |
| 18.41667 | -0.2588 | -0.86634 | -0.1288 | -0.21561 | -1.48935 | -0.4233 | -0.10736 | -0.18786 | -0.18888 | -0.834594 | 0 |
| 18.43056 | -0.34373 | -0.87551 | -0.21281 | -0.32555 | -1.55477 | -0.51273 | -0.20493 | -0.26552 | -0.28926 | -0.966138 | 0 |
| 18.44444 | -0.38315 | -0.7751 | -0.25791 | -0.39446 | -1.56566 | -0.57627 | -0.2403 | -0.30883 | -0.34804 | -1.01644 | 0 |
| 18.45833 | -0.46966 | -0.72436 | -0.35355 | -0.49811 | -1.6544 | -0.66866 | -0.3476 | -0.38324 | -0.4387 | -1.026604 | 0 |
| 18.47222 | -0.65628 | -0.84471 | -0.53838 | -0.68885 | -1.69978 | -0.83309 | -0.48085 | -0.51521 | -0.60719 | -1.068763 | 0 |
| 18.48611 | -1.00586 | -1.18984 | -0.82736 | -1.08127 | -1.80751 | -1.12026 | -0.4122 | -0.58667 | -0.85666 | -1.198163 | 0 |
| 18.5 | -1.4255 | -1.67522 | -1.15231 | -1.45328 | -1.95841 | -1.47838 | -0.23466 | -0.58142 | -1.11662 | -1.265365 | 0 |
| 18.51389 | -1.44272 | -1.73987 | -1.22594 | -1.07011 | -1.83204 | -1.45007 | -0.3304 | -0.66888 | -1.09991 | -1.351471 | 0 |
| 18.52778 | -0.93271 | -1.20817 | -0.90706 | 0.028367 | -1.29005 | -0.9461 | -0.48645 | -0.57961 | -0.74806 | -1.096237 | 0 |
| 18.54167 | -0.345 | -0.54478 | -0.39385 | 0.631093 | -0.75269 | -0.38113 | -0.33325 | -0.35036 | -0.29352 | -0.697195 | 0 |
| 18.55556 | -0.00336 | -0.15649 | -0.06099 | 0.545835 | -0.4246 | -0.05776 | -0.11005 | -0.09565 | -0.02739 | -0.33619 | 0 |
| 18.56944 | 0.11948 | -0.01544 | 0.01425 | 0.323366 | -0.30057 | 0.042018 | 0.024472 | 0.045501 | 0.050723 | -0.189651 | 0 |
| 18.58333 | 0.169435 | 0.044972 | 0.021301 | 0.221573 | -0.21645 | 0.074608 | 0.103459 | 0.111091 | 0.05976 | -0.147609 | 0 |
| 18.59722 | 0.16233 | 0.027467 | -0.00366 | 0.100926 | -0.21115 | 0.038957 | 0.095838 | 0.082456 | 0.008346 | -0.236757 | 0 |
| 18.61111 | 0.158704 | -0.00367 | -0.03175 | 0.005068 | -0.26204 | -0.04965 | 0.040897 | 0.010025 | -0.0571 | -0.330407 | 0 |
| 18.625 | 0.148763 | -0.02354 | -0.06449 | -0.0758 | -0.30771 | -0.14349 | -0.05289 | -0.0565 | -0.11532 | -0.297954 | 0 |
| 18.63889 | 0.097338 | -0.08843 | -0.14037 | -0.18716 | -0.36262 | -0.20486 | -0.18395 | -0.12347 | -0.19086 | -0.276088 | 0 |
| 18.65278 | 0.075596 | -0.10376 | -0.16588 | -0.22557 | -0.36196 | -0.18987 | -0.26669 | -0.15044 | -0.20329 | -0.288482 | 0 |
| 18.66667 | 0.140047 | -0.07851 | -0.17722 | -0.21681 | -0.34161 | -0.17947 | -0.29119 | -0.17662 | -0.19628 | -0.289692 | 0 |
| 18.68056 | 0.032109 | -0.17011 | -0.25584 | -0.27521 | -0.41016 | -0.25887 | -0.35402 | -0.26068 | -0.27794 | -0.369119 | 0 |
| 18.69444 | -0.20174 | -0.37798 | -0.40406 | -0.4503 | -0.59301 | -0.45111 | -0.51948 | -0.42463 | -0.46454 | -0.595079 | 0 |
| 18.70833 | -0.36584 | -0.53968 | -0.55078 | -0.64619 | -0.78311 | -0.64215 | -0.69754 | -0.59276 | -0.65025 | -0.805553 | 0 |
| 18.72222 | -0.24265 | -0.4485 | -0.51926 | -0.62393 | -0.7422 | -0.59929 | -0.66772 | -0.55475 | -0.60551 | -0.75171 | 0 |
| 18.73611 | -0.06213 | -0.29124 | -0.45717 | -0.52329 | -0.62493 | -0.50703 | -0.57319 | -0.46789 | -0.49074 | -0.575352 | 0 |
| 18.75 | -0.09372 | -0.32121 | -0.55449 | -0.57029 | -0.67457 | -0.56283 | -0.62725 | -0.53249 | -0.5456 | -0.544007 | 0 |

| | | | | | | | | | | | |
|---|---|---|---|---|---|---|---|---|---|---|---|
| 18.76389 | -0.1382 | -0.36928 | -0.62456 | -0.61917 | -0.71476 | -0.57976 | -0.66479 | -0.58434 | -0.60277 | -0.571637 | 0 |
| 18.77778 | -0.12157 | -0.33901 | -0.59645 | -0.59274 | -0.66766 | -0.52144 | -0.62321 | -0.55454 | -0.58398 | -0.53219 | 0 |
| 18.79167 | -0.05797 | -0.29078 | -0.51834 | -0.53697 | -0.60577 | -0.45818 | -0.57349 | -0.48869 | -0.51954 | -0.458944 | 0 |
| 18.80556 | -0.02163 | -0.24982 | -0.45187 | -0.48103 | -0.53847 | -0.42061 | -0.53114 | -0.43339 | -0.45172 | -0.392232 | 0 |
| 18.81944 | -0.04971 | -0.26269 | -0.45946 | -0.44744 | -0.51738 | -0.41922 | -0.49835 | -0.41272 | -0.43714 | -0.42336 | 0 |
| 18.83333 | -0.11409 | -0.29348 | -0.51484 | -0.45322 | -0.54607 | -0.43081 | -0.50707 | -0.43368 | -0.46542 | -0.427211 | 0 |
| 18.84722 | -0.15814 | -0.31865 | -0.56034 | -0.45728 | -0.55571 | -0.42089 | -0.55103 | -0.44983 | -0.48539 | -0.402395 | 0 |
| 18.86111 | -0.10887 | -0.27373 | -0.52738 | -0.39211 | -0.46803 | -0.3591 | -0.54964 | -0.37897 | -0.43541 | -0.375104 | 0 |
| 18.875 | -0.057 | -0.22106 | -0.48765 | -0.33811 | -0.39679 | -0.30818 | -0.5313 | -0.3154 | -0.3926 | -0.375438 | 0 |
| 18.88889 | -0.03717 | -0.1903 | -0.46311 | -0.31604 | -0.37739 | -0.28552 | -0.53118 | -0.28913 | -0.39176 | -0.374857 | 0 |
| 18.90278 | 0.022014 | -0.13199 | -0.41775 | -0.27707 | -0.33276 | -0.23838 | -0.52957 | -0.24983 | -0.36292 | -0.279438 | 0 |
| 18.91667 | 0.113485 | -0.0795 | -0.36622 | -0.24048 | -0.28112 | -0.20034 | -0.50716 | -0.2127 | -0.32592 | -0.155751 | 0 |
| 18.93056 | 0.211626 | -0.00594 | -0.27953 | -0.17539 | -0.20594 | -0.1455 | -0.44451 | -0.15687 | -0.2604 | -0.042532 | 0 |
| 18.94444 | 0.265533 | 0.050821 | -0.19539 | -0.10711 | -0.12136 | -0.06373 | -0.35216 | -0.09805 | -0.19446 | 0.010228 | 0 |
| 18.95833 | 0.259156 | 0.058638 | -0.16226 | -0.06874 | -0.07211 | 0.011902 | -0.30896 | -0.05482 | -0.16542 | 0.054756 | 0 |
| 18.97222 | 0.250736 | 0.08331 | -0.1424 | -0.06036 | -0.00204 | 0.087334 | -0.32892 | -0.01206 | -0.1367 | 0.088428 | 0 |
| 18.98611 | 0.392572 | 0.3124 | -0.07884 | -0.10744 | 0.274263 | 0.33505 | -0.56982 | 0.126899 | 0.033004 | 0.375584 | 0 |
| 19 | 0.502171 | 0.695654 | 0.012951 | -0.29765 | 0.760329 | 0.743491 | -1.67203 | 0.349166 | 0.387644 | 0.924012 | 0 |
| 19.01389 | -0.00463 | 0.565596 | 0.190845 | -0.6764 | 0.829387 | 0.71198 | -2.7098 | 0.339384 | 0.557476 | 0.506913 | 0 |
| 19.02778 | -0.84735 | -0.10119 | 0.425951 | -0.64263 | 0.36932 | 0.194903 | -0.98533 | 0.166441 | 0.354492 | 0.010083 | 0 |
| 19.04167 | -1.01108 | -0.45656 | 0.589008 | -0.12009 | 0.015229 | -0.18277 | 0.979658 | 0.129612 | 0.159455 | 0.075845 | 0 |
| 19.05556 | -0.77048 | -0.49633 | 0.488965 | 0.123146 | -0.08924 | -0.2149 | 1.827609 | 0.109066 | 0.090416 | 0.059132 | 0 |
| 19.06944 | -0.57712 | -0.43418 | 0.280815 | 0.073311 | -0.11445 | -0.18518 | 1.425666 | 0.047118 | 0.04739 | 0.032027 | 0 |
| 19.08333 | -0.38689 | -0.33485 | 0.112278 | -0.03524 | -0.1417 | -0.14359 | 0.67937 | -0.00735 | 0.01685 | -0.015533 | 0 |
| 19.09722 | -0.26881 | -0.27199 | 0.026835 | -0.0716 | -0.15883 | -0.10741 | 0.250602 | -0.03291 | 0.005148 | -0.027855 | 0 |
| 19.11111 | -0.20879 | -0.23561 | 0.012815 | -0.04073 | -0.14855 | -0.05114 | 0.140064 | -0.0273 | 0.014066 | -0.046595 | 0 |
| 19.125 | -0.15147 | -0.17773 | 0.049308 | 0.021446 | -0.10113 | 0.042398 | 0.177839 | 0.01979 | 0.065088 | -0.011432 | 0 |
| 19.13889 | -0.07047 | -0.0943 | 0.09931 | 0.091117 | -0.03134 | 0.130353 | 0.216914 | 0.072588 | 0.134994 | 0.08063 | 0 |
| 19.15278 | -0.0376 | -0.08571 | 0.092844 | 0.099114 | -0.01632 | 0.146002 | 0.18134 | 0.052387 | 0.134449 | 0.093322 | 0 |
| 19.16667 | -0.06403 | -0.1226 | 0.041976 | 0.036958 | -0.06246 | 0.111393 | 0.114181 | -0.01245 | 0.081299 | 0.042472 | 0 |
| 19.18056 | -0.06602 | -0.10503 | 0.028679 | -0.0003 | -0.0873 | 0.113895 | 0.081575 | -0.03174 | 0.063057 | 0.032615 | 0 |
| 19.19444 | -0.00406 | -0.03167 | 0.07655 | 0.053647 | -0.04462 | 0.157265 | 0.118411 | 0.012992 | 0.113128 | 0.044109 | 0 |
| 19.20833 | 0.113348 | 0.070516 | 0.174266 | 0.15971 | 0.04913 | 0.235112 | 0.216807 | 0.112916 | 0.212172 | 0.097384 | 0 |
| 19.22222 | 0.220317 | 0.160937 | 0.271716 | 0.229017 | 0.121022 | 0.317618 | 0.299509 | 0.20356 | 0.292928 | 0.139985 | 0 |
| 19.23611 | 0.19155 | 0.126538 | 0.244798 | 0.171101 | 0.082531 | 0.298341 | 0.265722 | 0.163711 | 0.244841 | 0.12762 | 0 |
| 19.25 | -0.00568 | -0.06083 | 0.044118 | -0.01802 | -0.08283 | 0.142562 | 0.108664 | -0.02089 | 0.048755 | -0.022415 | 0 |
| 19.26389 | -0.16841 | -0.20078 | -0.12604 | -0.14114 | -0.21015 | 0.003932 | -0.00438 | -0.15067 | -0.10464 | -0.170467 | 0 |
| 19.27778 | -0.1387 | -0.16532 | -0.10118 | -0.11261 | -0.15621 | 0.012793 | 0.00516 | -0.10618 | -0.08384 | -0.154382 | 0 |
| 19.29167 | -0.04749 | -0.08654 | -0.00321 | -0.07766 | -0.03325 | 0.076709 | 0.049459 | -0.01102 | 0.002699 | -0.053554 | 0 |
| 19.30556 | -0.02066 | -0.06229 | 0.03697 | -0.10445 | -0.00711 | 0.096007 | 0.077686 | 0.01992 | 0.036218 | -0.031293 | 0 |
| 19.31944 | -0.07211 | -0.11077 | -0.00528 | -0.17789 | -0.07996 | 0.045731 | 0.058724 | -0.02802 | -0.01377 | -0.048043 | 0 |
| 19.33333 | -0.20203 | -0.24397 | -0.12488 | -0.32016 | -0.18725 | -0.05494 | -0.02003 | -0.13136 | -0.12717 | -0.062074 | 0 |
| 19.34722 | -0.35194 | -0.39528 | -0.26104 | -0.55321 | -0.23843 | -0.09089 | -0.09685 | -0.23162 | -0.23966 | -0.083652 | 0 |
| 19.36111 | -0.39942 | -0.4533 | -0.29645 | -0.70616 | -0.16279 | 0.015105 | -0.06656 | -0.24504 | -0.25159 | -0.074819 | 0 |
| 19.375 | -0.18289 | -0.24855 | -0.06454 | -0.52444 | 0.127321 | 0.297728 | 0.184299 | -0.01019 | -0.02283 | 0.184969 | 0 |
| 19.38889 | 0.260224 | 0.197852 | 0.386382 | -0.03325 | 0.560763 | 0.703866 | 0.60489 | 0.428224 | 0.427215 | 0.670932 | 0 |
| 19.40278 | 0.605611 | 0.552504 | 0.717714 | 0.38703 | 0.866528 | 0.983404 | 0.915986 | 0.767606 | 0.751357 | 0.998921 | 0 |
| 19.41667 | 0.76934 | 0.724289 | 0.853156 | 0.636877 | 0.958876 | 1.091256 | 1.027731 | 0.92258 | 0.897029 | 1.098043 | 0 |
| 19.43056 | 0.843225 | 0.808757 | 0.902014 | 0.742909 | 0.912216 | 1.113134 | 1.025071 | 0.972281 | 0.940476 | 1.033501 | 0 |
| 19.44444 | 0.77835 | 0.74848 | 0.841716 | 0.668836 | 0.75471 | 1.003389 | 0.896636 | 0.868987 | 0.837233 | 0.854813 | 0 |
| 19.45833 | 0.612276 | 0.581755 | 0.699404 | 0.510811 | 0.557215 | 0.817097 | 0.724248 | 0.677272 | 0.653549 | 0.617294 | 0 |
| 19.47222 | 0.43929 | 0.406565 | 0.539315 | 0.347386 | 0.365063 | 0.607616 | 0.569611 | 0.501896 | 0.480467 | 0.392509 | 0 |
| 19.48611 | 0.239129 | 0.184114 | 0.323746 | 0.113787 | 0.109985 | 0.392541 | 0.425387 | 0.328024 | 0.28235 | 0.197264 | 0 |
| 19.5 | -0.0009 | -0.14345 | -0.02005 | -0.20016 | -0.24154 | 0.154819 | 0.190247 | 0.060501 | -0.03391 | -0.053812 | 0 |
| 19.51389 | -0.04271 | -0.30684 | -0.33905 | -0.32091 | -0.46678 | 0.068258 | -0.07198 | -0.21139 | -0.3269 | -0.279445 | 0 |
| 19.52778 | 0.25761 | -0.08171 | -0.31213 | -0.07069 | -0.32168 | 0.531118 | -0.03158 | -0.18985 | -0.28537 | -0.211784 | 0 |
| 19.54167 | 0.639425 | 0.297608 | -0.05875 | 0.251427 | -0.00872 | 1.539607 | 0.230739 | 0.088394 | -0.04468 | 0.062512 | 0 |

| | | | | | | | | | | | |
|---|---|---|---|---|---|---|---|---|---|---|---|
| 19.55556 | 0.849341 | 0.523941 | 0.140159 | 0.420379 | 0.21006 | 2.089206 | 0.389583 | 0.330723 | 0.136261 | 0.204931 | 0 |
| 19.56944 | 0.994014 | 0.658643 | 0.347074 | 0.46817 | 0.373842 | 1.705497 | 0.602935 | 0.496313 | 0.280149 | 0.331483 | 0 |
| 19.58333 | 1.061109 | 0.669198 | 0.5365 | 0.43842 | 0.454682 | 1.182557 | 0.960503 | 0.588153 | 0.368031 | 0.571507 | 0 |
| 19.59722 | 0.997398 | 0.550153 | 0.585866 | 0.375082 | 0.400537 | 0.741838 | 1.27931 | 0.599281 | 0.368504 | 0.522139 | 0 |
| 19.61111 | 0.97771 | 0.515193 | 0.590077 | 0.385949 | 0.410385 | 0.559961 | 1.495238 | 0.685778 | 0.435694 | 0.720429 | 0 |
| 19.625 | 0.943753 | 0.51065 | 0.575157 | 0.420019 | 0.474994 | 0.602486 | 1.46793 | 0.737005 | 0.505659 | 0.735538 | 0 |
| 19.63889 | 0.832832 | 0.429099 | 0.424336 | 0.358068 | 0.423173 | 0.560413 | 1.322349 | 0.638282 | 0.425979 | 0.768307 | 0 |
| 19.65278 | 0.703271 | 0.321955 | 0.162066 | 0.247216 | 0.294494 | 0.393887 | 0.998653 | 0.436386 | 0.255665 | 0.657544 | 0 |
| 19.66667 | 0.544897 | 0.239279 | -0.05499 | 0.15505 | 0.168994 | 0.186504 | 0.594101 | 0.242962 | 0.099115 | 0.465328 | 0 |
| 19.68056 | 0.430457 | 0.227323 | -0.15082 | 0.141752 | 0.113456 | 0.038667 | 0.295244 | 0.170085 | 0.019263 | 0.334843 | 0 |
| 19.69444 | 0.442147 | 0.292432 | -0.1178 | 0.220222 | 0.191528 | 0.024208 | 0.122846 | 0.220261 | 0.064933 | 0.324403 | 0 |
| 19.70833 | 0.526579 | 0.360072 | -0.04138 | 0.317224 | 0.323506 | 0.083582 | 0.006605 | 0.290846 | 0.175978 | 0.383095 | 0 |
| 19.72222 | 0.481143 | 0.326917 | 0.033306 | 0.334224 | 0.322009 | 0.040524 | -0.1964 | 0.288238 | 0.207966 | 0.374548 | 0 |
| 19.73611 | 0.265583 | 0.176964 | -0.00582 | 0.216598 | 0.140554 | -0.08448 | -0.3987 | 0.141298 | 0.091601 | 0.168714 | 0 |
| 19.75 | 0.062165 | 0.048673 | -0.04757 | 0.228369 | -0.01587 | -0.08004 | -0.44565 | -0.03825 | -0.02038 | 0.103855 | 0 |
| 19.76389 | 0.062199 | 0.111547 | 0.03518 | 0.788335 | 0.055292 | 0.159513 | -0.26981 | 0.004493 | 0.058518 | 0.178702 | 0 |
| 19.77778 | 0.121544 | 0.216643 | 0.090209 | 1.562426 | 0.186808 | 0.331264 | -0.12395 | 0.130346 | 0.148239 | 0.173791 | 0 |
| 19.79167 | 0.064714 | 0.132549 | -0.05201 | 1.451657 | 0.150156 | 0.205197 | -0.19836 | 0.074029 | 0.060897 | 0.050895 | 0 |
| 19.80556 | -0.04249 | -0.01782 | -0.21486 | 0.691346 | 0.045215 | 0.0014 | -0.34185 | -0.09062 | -0.08915 | -0.040537 | 0 |
| 19.81944 | -0.12094 | -0.07033 | -0.24157 | 0.238604 | -0.00236 | -0.12057 | -0.55221 | -0.20289 | -0.19064 | -0.087573 | 0 |
| 19.83333 | -0.12514 | 0.008236 | -0.19005 | 0.155149 | 0.021008 | -0.16605 | -0.71357 | -0.183 | -0.17814 | -0.103802 | 0 |
| 19.84722 | -0.09936 | 0.087152 | -0.16435 | 0.120231 | 0.040388 | -0.15942 | -0.80708 | -0.12401 | -0.11736 | -0.116419 | 0 |
| 19.86111 | -0.06278 | 0.132653 | -0.16424 | 0.067852 | 0.066388 | -0.05609 | -0.71301 | -0.06492 | -0.06616 | -0.135564 | 0 |
| 19.875 | 0.073546 | 0.25174 | -0.06023 | 0.169262 | 0.203488 | 0.187029 | -0.49867 | 0.074331 | 0.074322 | 0.000551 | 0 |
| 19.88889 | 0.228383 | 0.360896 | 0.048208 | 0.37179 | 0.352894 | 0.372209 | -0.35271 | 0.324029 | 0.210591 | 0.187238 | 0 |
| 19.90278 | 0.27788 | 0.396761 | 0.086184 | 0.608244 | 0.423504 | 0.319722 | -0.37357 | 0.70421 | 0.264607 | 0.251713 | 0 |
| 19.91667 | 0.18275 | 0.374955 | 0.102159 | 0.927769 | 0.453044 | 0.129694 | -0.59283 | 0.981338 | 0.266195 | 0.216343 | 0 |
| 19.93056 | -0.04243 | 0.257474 | 0.096214 | 1.087491 | 0.410535 | -0.07431 | -0.78089 | 0.817294 | 0.217912 | 0.122652 | 0 |
| 19.94444 | -0.34141 | 0.045726 | 0.009304 | 0.754034 | 0.277536 | -0.26224 | -0.93862 | 0.48366 | 0.093888 | -0.0842 | 0 |
| 19.95833 | -0.60645 | -0.15214 | -0.1294 | 0.404934 | 0.108018 | -0.4157 | -0.97992 | 0.16372 | -0.05935 | -0.27419 | 0 |
| 19.97222 | -0.57765 | -0.13399 | -0.09883 | 0.331974 | 0.117315 | -0.325 | -0.70307 | 0.108527 | -0.02253 | -0.20829 | 0 |
| 19.98611 | -0.08443 | 0.274522 | 0.286059 | 0.429176 | 0.492199 | 0.106868 | -0.03514 | 0.36563 | 0.372789 | 0.143339 | 0 |
| 20 | 0.535604 | 0.943924 | 0.997806 | 0.56228 | 1.147367 | 0.74406 | 0.847783 | 0.826011 | 1.068178 | 0.714364 | 0 |
| 20.01389 | 0.712157 | 1.300856 | 1.492397 | 0.454274 | 1.490426 | 1.124365 | 1.404994 | 0.947112 | 1.515951 | 1.039005 | 0 |
| 20.02778 | 0.359975 | 1.030767 | 1.28889 | 0.209716 | 1.183985 | 0.965811 | 1.224369 | 0.528722 | 1.276119 | 0.862711 | 0 |
| 20.04167 | 0.090383 | 0.600515 | 0.83312 | 0.282383 | 0.680885 | 0.632793 | 0.769629 | 0.179299 | 0.80926 | 0.493728 | 0 |
| 20.05556 | -0.00125 | 0.292647 | 0.525521 | 0.376193 | 0.311168 | 0.396767 | 0.444112 | 0.128237 | 0.475281 | 0.185156 | 0 |
| 20.06944 | -0.06588 | 0.078606 | 0.289784 | 0.308843 | 0.041718 | 0.186917 | 0.192769 | 0.090516 | 0.234474 | -0.063101 | 0 |
| 20.08333 | -0.0871 | -0.03613 | 0.131453 | 0.214716 | -0.07898 | 0.041764 | 0.038685 | 0.027876 | 0.105413 | -0.108002 | 0 |
| 20.09722 | -0.06625 | -0.05147 | 0.085523 | 0.20117 | -0.07435 | 0.006352 | 0.021447 | 0.035446 | 0.097261 | -0.014543 | 0 |
| 20.11111 | -0.0549 | -0.04927 | 0.078155 | 0.193631 | -0.05793 | -0.02212 | 0.031923 | 0.055537 | 0.093993 | 0.046879 | 0 |
| 20.125 | -0.09503 | -0.08855 | 0.036177 | 0.129448 | -0.10144 | -0.11942 | 0.000809 | 0.026338 | 0.036995 | 0.018394 | 0 |
| 20.13889 | -0.17144 | -0.15411 | -0.0444 | 0.035398 | -0.17879 | -0.24474 | -0.07263 | -0.0527 | -0.04922 | -0.07458 | 0 |
| 20.15278 | -0.23294 | -0.21395 | -0.10964 | -0.05445 | -0.24177 | -0.32023 | -0.14608 | -0.12624 | -0.12894 | -0.189013 | 0 |
| 20.16667 | -0.27678 | -0.2598 | -0.15583 | -0.13041 | -0.28009 | -0.37793 | -0.19501 | -0.17687 | -0.17736 | -0.259968 | 0 |
| 20.18056 | -0.33149 | -0.31951 | -0.23325 | -0.22991 | -0.32325 | -0.46692 | -0.25276 | -0.25184 | -0.23006 | -0.305447 | 0 |
| 20.19444 | -0.35041 | -0.33799 | -0.28198 | -0.27574 | -0.33176 | -0.48829 | -0.26841 | -0.28914 | -0.25357 | -0.332322 | 0 |
| 20.20833 | -0.29911 | -0.28093 | -0.24346 | -0.23521 | -0.27395 | -0.38964 | -0.20668 | -0.22496 | -0.20653 | -0.291241 | 0 |
| 20.22222 | -0.29349 | -0.2478 | -0.22258 | -0.23289 | -0.25279 | -0.34356 | -0.19575 | -0.19341 | -0.1926 | -0.257473 | 0 |
| 20.23611 | -0.34902 | -0.27169 | -0.27149 | -0.27117 | -0.29216 | -0.40962 | -0.26681 | -0.25808 | -0.24489 | -0.288144 | 0 |
| 20.25 | -0.36277 | -0.28231 | -0.29787 | -0.24391 | -0.30484 | -0.44596 | -0.29347 | -0.30476 | -0.26472 | -0.320842 | 0 |
| 20.26389 | -0.30492 | -0.2552 | -0.25394 | -0.16159 | -0.26776 | -0.40401 | -0.24112 | -0.28077 | -0.219 | -0.323488 | 0 |
| 20.27778 | -0.26169 | -0.24649 | -0.22291 | -0.14273 | -0.26814 | -0.43878 | -0.22011 | -0.28779 | -0.20197 | -0.333442 | 0 |
| 20.29167 | -0.25822 | -0.256 | -0.23195 | -0.20386 | -0.32107 | -0.60999 | -0.24953 | -0.33531 | -0.23141 | -0.321503 | 0 |
| 20.30556 | -0.27184 | -0.26562 | -0.24235 | -0.26179 | -0.35309 | -0.70417 | -0.27221 | -0.35917 | -0.25334 | -0.311794 | 0 |
| 20.31944 | -0.31726 | -0.30484 | -0.26784 | -0.30909 | -0.36526 | -0.6844 | -0.29617 | -0.37253 | -0.2724 | -0.328286 | 0 |
| 20.33333 | -0.33129 | -0.31561 | -0.26877 | -0.27669 | -0.33468 | -0.59276 | -0.28111 | -0.35289 | -0.25805 | -0.31755 | 0 |

| | | | | | | | | | | | |
|---|---|---|---|---|---|---|---|---|---|---|---|
| 20.34722 | -0.2729 | -0.2635 | -0.20976 | -0.14457 | -0.2555 | -0.44202 | -0.20639 | -0.28194 | -0.18908 | -0.221782 | 0 |
| 20.36111 | -0.23093 | -0.2344 | -0.16577 | -0.04459 | -0.2063 | -0.29972 | -0.1544 | -0.24253 | -0.14017 | -0.140126 | 0 |
| 20.375 | -0.27049 | -0.28294 | -0.2038 | -0.05872 | -0.24785 | -0.26939 | -0.18339 | -0.29013 | -0.17673 | -0.187233 | 0 |
| 20.38889 | -0.3309 | -0.34388 | -0.2861 | -0.1225 | -0.32452 | -0.31644 | -0.22502 | -0.36077 | -0.25418 | -0.295562 | 0 |
| 20.40278 | -0.30376 | -0.32976 | -0.29715 | -0.11233 | -0.31962 | -0.30433 | -0.16956 | -0.35472 | -0.25514 | -0.30317 | 0 |
| 20.41667 | -0.20951 | -0.26459 | -0.23334 | -0.06127 | -0.21353 | -0.23403 | -0.06306 | -0.29075 | -0.17336 | -0.205087 | 0 |
| 20.43056 | -0.1405 | -0.23283 | -0.1972 | -0.05747 | -0.03541 | -0.18583 | -0.01657 | -0.25287 | -0.11855 | -0.156748 | 0 |
| 20.44444 | -0.1012 | -0.21286 | -0.19809 | -0.04189 | 0.072826 | -0.1597 | 0.033044 | -0.23539 | -0.10474 | -0.184077 | 0 |
| 20.45833 | -0.066 | -0.18028 | -0.18975 | -0.00433 | -0.08941 | -0.12803 | 0.092561 | -0.20545 | -0.0877 | -0.126292 | 0 |
| 20.47222 | -0.00267 | -0.14742 | -0.16485 | 0.067619 | -0.37739 | -0.05206 | 0.140893 | -0.16861 | -0.07781 | -0.074393 | 0 |
| 20.48611 | 0.153977 | -0.17595 | -0.22713 | 0.27456 | -0.53248 | 0.167949 | 0.158009 | -0.1841 | -0.13835 | -0.001336 | 0 |
| 20.5 | 0.41575 | -0.33506 | -0.50918 | 0.465608 | -0.73996 | 0.539758 | 0.214475 | -0.34741 | -0.34262 | 0.184055 | 0 |
| 20.51389 | 0.703618 | -0.48356 | -0.75664 | -0.14977 | -0.94862 | 0.670797 | 0.406034 | -0.57878 | -0.55211 | 0.169049 | 0 |
| 20.52778 | 0.75231 | -0.41185 | -0.65914 | -0.57776 | -0.83589 | 0.547326 | 0.580407 | -0.54041 | -0.5361 | 0.002922 | 0 |
| 20.54167 | 0.82657 | -0.02222 | -0.32467 | -0.486 | -0.50557 | 0.5586 | 0.512817 | -0.10141 | -0.37898 | 0.126323 | 0 |
| 20.55556 | 0.831223 | 0.299048 | -0.04544 | -0.24125 | -0.201 | 0.609362 | 0.32796 | 0.442177 | -0.22189 | 0.224482 | 0 |
| 20.56944 | 0.852664 | 0.477717 | 0.101988 | -0.05784 | -0.02029 | 0.530923 | 0.265131 | 0.843749 | -0.06361 | 0.379604 | 0 |
| 20.58333 | 0.837783 | 0.534732 | 0.188923 | 0.058708 | 0.043438 | 0.385257 | 0.478093 | 0.974777 | 0.120458 | 0.572002 | 0 |
| 20.59722 | 0.966928 | 0.549091 | 0.192074 | 0.029103 | 0.022507 | 0.30017 | 0.711811 | 0.92121 | 0.280836 | 0.707825 | 0 |
| 20.61111 | 1.033653 | 0.54194 | 0.137438 | -0.1279 | -0.02653 | 0.276473 | 0.589521 | 0.747646 | 0.369492 | 0.651543 | 0 |
| 20.625 | 0.966464 | 0.501138 | 0.102271 | -0.18474 | -0.04488 | 0.362676 | 0.177342 | 0.578375 | 0.344492 | 0.412015 | 0 |
| 20.63889 | 0.738054 | 0.406199 | 0.093397 | -0.05292 | -0.0176 | 0.461087 | -0.31916 | 0.447706 | 0.210778 | 0.323772 | 0 |
| 20.65278 | 0.507387 | 0.387848 | 0.043156 | 0.136217 | 0.075171 | 0.525667 | -0.70483 | 0.393801 | 0.136368 | 0.231378 | 0 |
| 20.66667 | 0.273708 | 0.328485 | -0.07574 | 0.218382 | 0.125507 | 0.491636 | -1.08322 | 0.333101 | 0.067119 | 0.048458 | 0 |
| 20.68056 | 0.048995 | 0.168181 | -0.22026 | 0.129764 | 0.050497 | 0.31385 | -1.40422 | 0.206522 | -0.10289 | -0.202486 | 0 |
| 20.69444 | -0.06723 | 0.016998 | -0.26308 | 0.047916 | -0.03648 | 0.204805 | -1.28361 | 0.099848 | -0.22879 | -0.304453 | 0 |
| 20.70833 | -0.12572 | -0.06029 | -0.1646 | 0.027485 | -0.05642 | 0.218592 | -0.69217 | 0.051604 | -0.17735 | -0.188952 | 0 |
| 20.72222 | -0.13737 | -0.02643 | 0.02353 | 0.059182 | 0.003515 | 0.342167 | 0.175603 | 0.066465 | -0.01022 | 0.013191 | 0 |
| 20.73611 | -0.15826 | 0.007346 | 0.123091 | 0.093093 | 0.038016 | 0.360688 | 0.882078 | 0.065049 | 0.084844 | 0.090307 | 0 |
| 20.75 | -0.24261 | -0.02721 | 0.044618 | 0.110065 | 0.007257 | 0.174251 | 1.136871 | 0.047188 | 0.033921 | -0.00821 | 0 |
| 20.76389 | -0.32299 | -0.05406 | -0.0753 | 0.122224 | 0.002829 | 0.012731 | 0.91507 | 0.10375 | -0.04291 | -0.082162 | 0 |
| 20.77778 | -0.34609 | -0.03446 | -0.11189 | 0.12851 | 0.049664 | 0.04314 | 0.336506 | 0.204986 | -0.08146 | -0.111211 | 0 |
| 20.79167 | -0.31891 | -0.02315 | -0.11169 | 0.070018 | 0.095667 | 0.158365 | -0.52227 | 0.2073 | -0.07088 | -0.219027 | 0 |
| 20.80556 | -0.30178 | -0.0731 | -0.1793 | -0.0294 | 0.053719 | 0.129618 | -1.28585 | -0.00013 | -0.10371 | -0.297724 | 0 |
| 20.81944 | -0.27422 | -0.06174 | -0.1938 | -0.00884 | 0.019569 | 0.033702 | -1.40227 | -0.2266 | -0.09766 | -0.284846 | 0 |
| 20.83333 | -0.26791 | -0.00356 | -0.11788 | 0.106101 | 0.055746 | 0.00618 | -0.80619 | -0.30989 | -0.00847 | -0.196761 | 0 |
| 20.84722 | -0.31695 | 0.026792 | -0.03791 | 0.209457 | 0.114945 | 0.04197 | -0.32094 | -0.28189 | 0.073446 | -0.184971 | 0 |
| 20.86111 | -0.36597 | 0.004552 | 0.022868 | 0.19411 | 0.154947 | 0.057432 | -0.17425 | -0.20639 | 0.076047 | -0.196992 | 0 |
| 20.875 | -0.45132 | -0.09773 | 0.014136 | 0.062962 | 0.093899 | -0.07011 | -0.14014 | -0.09609 | -0.04524 | -0.24656 | 0 |
| 20.88889 | -0.59573 | -0.19786 | -0.04328 | -0.16 | -0.00396 | -0.2773 | -0.1832 | 0.078178 | -0.18315 | -0.344207 | 0 |
| 20.90278 | -0.75914 | -0.27554 | -0.11791 | -0.44178 | -0.05952 | -0.46712 | -0.34173 | 0.212468 | -0.28143 | -0.434023 | 0 |
| 20.91667 | -0.85198 | -0.31447 | -0.15834 | -0.52286 | -0.0662 | -0.52601 | -0.48498 | 0.15453 | -0.30128 | -0.420144 | 0 |
| 20.93056 | -0.81329 | -0.27444 | -0.11873 | -0.35956 | -0.02885 | -0.43868 | -0.52793 | -0.02601 | -0.21781 | -0.360617 | 0 |
| 20.94444 | -0.73262 | -0.22532 | -0.03472 | -0.19807 | 0.03748 | -0.3812 | -0.46614 | -0.19828 | -0.10336 | -0.297882 | 0 |
| 20.95833 | -0.64477 | -0.18316 | 0.031247 | -0.23433 | 0.087917 | -0.40768 | -0.34728 | -0.4095 | -0.04544 | -0.320778 | 0 |
| 20.97222 | -0.47861 | -0.1268 | 0.086514 | -0.20269 | 0.134423 | -0.46046 | -0.11836 | -0.47186 | 0.036717 | -0.125908 | 0 |
| 20.98611 | -0.09056 | 0.110064 | 0.270793 | 0.281096 | 0.33761 | -0.60782 | 0.343586 | -0.11912 | 0.283069 | 0.648508 | 0 |
| 21 | 0.365355 | 0.485755 | 0.582041 | 1.040232 | 0.705367 | -0.97702 | 0.958662 | 0.435066 | 0.646476 | 1.764691 | 0 |
| 21.01389 | 0.534744 | 0.642985 | 0.797044 | 1.255609 | 1.093487 | -0.86941 | 1.225858 | 0.666154 | 0.844969 | 1.984743 | 0 |
| 21.02778 | 0.347314 | 0.496643 | 0.721861 | 0.741767 | 0.805745 | -0.06401 | 0.83082 | 0.515134 | 0.774985 | 1.364087 | 0 |
| 21.04167 | 0.166478 | 0.275079 | 0.415007 | 0.243097 | 0.401573 | 0.348948 | 0.354496 | 0.123854 | 0.44982 | 0.34195 | 0 |
| 21.05556 | 0.081047 | 0.058569 | 0.155669 | 0.107282 | 0.128675 | 0.242743 | 0.087318 | -0.1956 | 0.18275 | 0.028081 | 0 |
| 21.06944 | 0.045292 | -0.03975 | 0.048907 | 0.087196 | 0.034358 | 0.086132 | 0.002597 | -0.25394 | 0.078753 | -0.002087 | 0 |
| 21.08333 | 0.069347 | 0.009198 | 0.081241 | 0.055206 | 0.063111 | 0.127186 | 0.044428 | -0.04391 | 0.106948 | -0.071114 | 0 |
| 21.09722 | 0.104844 | 0.095677 | 0.155999 | 0.00235 | 0.107102 | 0.304408 | 0.110123 | 0.110513 | 0.167644 | -0.308192 | 0 |
| 21.11111 | 0.097972 | 0.101866 | 0.179526 | -0.01799 | 0.114678 | 0.287969 | 0.135068 | 0.113249 | 0.176009 | -0.349529 | 0 |
| 21.125 | 0.088952 | 0.072236 | 0.158388 | 0.01906 | 0.101114 | 0.107248 | 0.145049 | 0.055654 | 0.154585 | -0.084459 | 0 |

| | | | | | | | | | | | |
|---|---|---|---|---|---|---|---|---|---|---|---|
| 21.13889 | 0.070099 | 0.04233 | 0.118629 | -0.0016 | 0.083692 | -0.00484 | 0.133334 | 0.02567 | 0.122456 | 0.05993 | 0 |
| 21.15278 | 0.059269 | 0.051604 | 0.111864 | -0.08909 | 0.054809 | 0.115708 | 0.099834 | 0.071746 | 0.121222 | -0.163514 | 0 |
| 21.16667 | 0.114295 | 0.12731 | 0.180992 | -0.10199 | 0.075737 | 0.402149 | 0.113889 | 0.191667 | 0.183899 | -0.421724 | 0 |
| 21.18056 | 0.181456 | 0.183909 | 0.23072 | -0.01137 | 0.115642 | 0.410213 | 0.174904 | 0.239476 | 0.233984 | -0.226424 | 0 |
| 21.19444 | 0.213356 | 0.183265 | 0.215831 | 0.142294 | 0.188937 | 0.141434 | 0.267606 | 0.15593 | 0.236585 | 0.231311 | 0 |
| 21.20833 | 0.193677 | 0.14313 | 0.175044 | 0.238482 | 0.188582 | -0.0414 | 0.317779 | 0.074476 | 0.205982 | 0.479485 | 0 |
| 21.22222 | 0.093383 | 0.087007 | 0.124152 | 0.157191 | 0.084929 | 0.064975 | 0.233847 | 0.047124 | 0.114386 | -0.019297 | 0 |
| 21.23611 | -0.00705 | 0.014217 | 0.051904 | 0.057552 | -0.03485 | 0.197733 | 0.121356 | 0.005444 | -0.0069 | -0.483597 | 0 |
| 21.25 | -0.01066 | -0.03345 | 0.018185 | 0.115067 | -0.05318 | 0.093708 | 0.125369 | -0.04363 | -0.0174 | -0.18604 | 0 |
| 21.26389 | 0.026654 | -0.01995 | 0.034699 | 0.185271 | -0.01581 | 0.030438 | 0.165049 | -0.03878 | 0.054816 | 0.084791 | 0 |
| 21.27778 | 0.023028 | -0.00466 | 0.047567 | 0.160394 | 0.001528 | 0.060383 | 0.162561 | -0.02554 | 0.090081 | -0.029807 | 0 |
| 21.29167 | -0.01907 | -0.0297 | 0.038537 | 0.145824 | -0.02415 | 0.050051 | 0.133404 | -0.05878 | 0.054033 | -0.108127 | 0 |
| 21.30556 | -0.06691 | -0.09465 | 0.028801 | 0.143704 | -0.08697 | -0.02228 | 0.076258 | -0.13392 | 0.01337 | -0.110055 | 0 |
| 21.31944 | -0.07171 | -0.12366 | 0.071041 | 0.17461 | -0.1049 | -0.04901 | 0.056338 | -0.1699 | 0.024279 | -0.038957 | 0 |
| 21.33333 | -0.05776 | -0.09081 | 0.146654 | 0.199692 | -0.07402 | -0.01403 | 0.073544 | -0.14429 | 0.068504 | 0.029651 | 0 |
| 21.34722 | -0.05212 | -0.05152 | 0.208608 | 0.171162 | -0.0591 | 0.012406 | 0.073012 | -0.10539 | 0.100132 | -0.014978 | 0 |
| 21.36111 | -0.00925 | -0.01696 | 0.299241 | 0.208134 | -0.0319 | 0.064605 | 0.111638 | -0.05097 | 0.141712 | -0.029499 | 0 |
| 21.375 | 0.049074 | 0.019615 | 0.382033 | 0.350034 | 0.005038 | 0.141543 | 0.207088 | 0.006747 | 0.22255 | 0.114015 | 0 |
| 21.38889 | 0.064922 | 0.038589 | 0.393554 | 0.472384 | 0.015758 | 0.181956 | 0.325335 | 0.03557 | 0.282299 | 0.231202 | 0 |
| 21.40278 | 0.066352 | 0.023414 | 0.323284 | 0.552509 | 0.005774 | 0.097398 | 0.467378 | 0.024519 | 0.265836 | 0.385526 | 0 |
| 21.41667 | 0.11456 | 0.009107 | 0.236745 | 0.733313 | 0.004393 | -0.07702 | 0.564142 | 0.027401 | 0.278006 | 0.646136 | 0 |
| 21.43056 | 0.17321 | 0.012065 | 0.175066 | 0.751955 | 0.021015 | -0.10379 | 0.608741 | 0.077895 | 0.285177 | 0.520558 | 0 |
| 21.44444 | 0.176698 | 0.01265 | 0.107501 | 0.537588 | -0.03741 | -0.01572 | 0.607361 | 0.133578 | 0.193024 | 0.239797 | 0 |
| 21.45833 | 0.146325 | -0.04203 | -0.00531 | 0.277353 | -0.12812 | 0.102086 | 0.41456 | 0.173974 | 0.069899 | 0.001699 | 0 |
| 21.47222 | 0.123098 | -0.04966 | -0.02501 | 0.074148 | -0.15923 | 0.349701 | 0.231226 | 0.294325 | 0.036613 | -0.111739 | 0 |
| 21.48611 | 0.190656 | 0.024374 | -0.06064 | -0.24094 | -0.25064 | 0.901862 | 0.266913 | 0.419591 | 0.095053 | -0.086027 | 0 |
| 21.5 | 0.39268 | -0.00815 | -0.29419 | -0.78787 | -0.61325 | 1.030215 | 0.339916 | 0.309955 | -0.04813 | -0.20787 | 0 |
| 21.51389 | 0.631683 | -0.02562 | -0.61871 | -1.11996 | -0.87603 | 0.476506 | 0.19367 | -0.02341 | -0.38191 | 0.208112 | 0 |
| 21.52778 | 0.478323 | -0.03621 | -0.63009 | -1.0174 | -0.88716 | -0.23469 | 0.177986 | -0.18732 | -0.63562 | -0.007562 | 0 |
| 21.54167 | 0.331625 | 0.046866 | -0.50445 | -0.60901 | -0.63018 | -0.40048 | 0.339012 | 0.097408 | -0.66085 | -0.146154 | 0 |
| 21.55556 | 0.275703 | 0.1947 | -0.39332 | -0.32156 | -0.31469 | -0.27228 | 0.296899 | 0.420017 | -0.61778 | -0.1873 | 0 |
| 21.56944 | 0.326305 | 0.282516 | -0.36011 | -0.28813 | -0.13729 | -0.14097 | 0.13129 | 0.642184 | -0.48377 | -0.063496 | 0 |
| 21.58333 | 0.525663 | 0.338796 | -0.2459 | -0.27204 | -0.06798 | 0.012301 | 0.336939 | 0.820711 | -0.2306 | 0.218763 | 0 |
| 21.59722 | 0.739402 | 0.332046 | -0.0296 | -0.1051 | -0.03527 | 0.171395 | 0.916107 | 0.963343 | 0.018059 | 0.514879 | 0 |
| 21.61111 | 0.674507 | 0.216401 | 0.246925 | 0.058685 | -0.04932 | 0.357698 | 1.417768 | 0.888905 | 0.181786 | 0.605551 | 0 |
| 21.625 | 0.393694 | 0.084988 | 0.443212 | 0.057184 | -0.04484 | 0.612319 | 1.606454 | 0.651869 | 0.29638 | 0.493139 | 0 |
| 21.63889 | 0.112125 | -0.01923 | 0.346304 | -0.06613 | -0.02012 | 0.855857 | 1.273113 | 0.416311 | 0.31243 | 0.253947 | 0 |
| 21.65278 | -0.21358 | -0.05537 | 0.015078 | -0.06751 | 0.029131 | 0.922745 | 0.416062 | 0.237187 | 0.179385 | -0.088617 | 0 |
| 21.66667 | -0.53165 | -0.04665 | -0.32292 | -0.02 | 0.054933 | 0.727462 | -0.74435 | 0.089045 | -0.06209 | -0.525038 | 0 |
| 21.68056 | -0.70385 | -0.06674 | -0.48205 | -0.10598 | 0.012481 | 0.396397 | -1.76079 | -0.04186 | -0.29465 | -0.770765 | 0 |
| 21.69444 | -0.65343 | -0.09271 | -0.45789 | -0.25006 | -0.04948 | 0.107842 | -1.98477 | -0.13127 | -0.35027 | -0.714444 | 0 |
| 21.70833 | -0.38831 | -0.05962 | -0.31076 | -0.26818 | -0.06745 | -0.08004 | -1.46101 | -0.11666 | -0.21894 | -0.445184 | 0 |
| 21.72222 | 0.073046 | 0.098758 | -0.03374 | -0.09122 | 0.02187 | -0.11211 | -0.63719 | 0.028242 | -0.04319 | -0.077149 | 0 |
| 21.73611 | 0.432909 | 0.251122 | 0.185564 | 0.054251 | 0.095195 | -0.09766 | 0.156812 | 0.079983 | 0.013585 | 0.122198 | 0 |
| 21.75 | 0.428684 | 0.250606 | 0.242385 | 0.010353 | 0.068089 | -0.12026 | 0.787352 | -0.04136 | -0.03626 | 0.089822 | 0 |
| 21.76389 | 0.173146 | 0.171541 | 0.236007 | -0.06519 | 0.050183 | -0.08806 | 1.099608 | -0.2054 | -0.0659 | 0.000806 | 0 |
| 21.77778 | -0.16903 | 0.086008 | 0.132501 | -0.07441 | 0.021944 | -0.02992 | 0.90573 | -0.40604 | -0.10943 | -0.064728 | 0 |
| 21.79167 | -0.46492 | 0.023328 | -0.10019 | -0.05113 | -0.01639 | 0.018184 | 0.424949 | -0.5845 | -0.1778 | -0.231313 | 0 |
| 21.80556 | -0.64138 | 0.12093 | -0.33539 | -0.06123 | -0.00512 | 0.044264 | -0.01887 | -0.68954 | -0.2012 | -0.419313 | 0 |
| 21.81944 | -0.69324 | 0.415244 | -0.38682 | -0.10234 | 0.057435 | 0.052904 | -0.11701 | -0.66605 | -0.14652 | -0.516158 | 0 |
| 21.83333 | -0.67132 | 0.540937 | -0.29805 | -0.24791 | 0.071226 | 0.043191 | -0.10121 | -0.5582 | -0.08944 | -0.487919 | 0 |
| 21.84722 | -0.71301 | 0.329041 | -0.18537 | -0.36854 | 0.008996 | -0.01665 | -0.29551 | -0.48653 | -0.10847 | -0.451899 | 0 |
| 21.86111 | -0.75584 | 0.133714 | -0.12474 | -0.29324 | -0.04292 | -0.13197 | -0.68151 | -0.57225 | -0.18895 | -0.514813 | 0 |
| 21.875 | -0.78523 | 0.042139 | -0.12582 | -0.20864 | -0.05117 | -0.23085 | -0.92489 | -0.84362 | -0.27421 | -0.613015 | 0 |
| 21.88889 | -0.78858 | 0.003663 | -0.15961 | -0.32594 | -0.00665 | -0.25064 | -0.8191 | -1.16614 | -0.29787 | -0.669141 | 0 |
| 21.90278 | -0.75209 | -0.04169 | -0.20987 | -0.57935 | 0.042242 | -0.23896 | -0.65613 | -1.37286 | -0.27567 | -0.676569 | 0 |
| 21.91667 | -0.65899 | -0.0483 | -0.2092 | -0.75254 | 0.07169 | -0.18851 | -0.60184 | -1.30834 | -0.2306 | -0.725985 | 0 |

| | | | | | | | | | | | |
|---|---|---|---|---|---|---|---|---|---|---|---|
| 21.93056 | -0.56826 | -0.06262 | -0.196 | -0.5943 | 0.055126 | -0.15532 | -0.77845 | -1.1532 | -0.20502 | -0.729219 | 0 |
| 21.94444 | -0.59823 | -0.17989 | -0.23985 | -0.40811 | -0.02338 | -0.19564 | -1.14544 | -1.08259 | -0.24022 | -0.795773 | 0 |
| 21.95833 | -0.65712 | -0.2716 | -0.23978 | -0.30993 | -0.05148 | -0.22275 | -1.21461 | -1.06842 | -0.25013 | -0.745052 | 0 |
| 21.97222 | -0.54675 | -0.25535 | -0.14728 | -0.20521 | 0.041699 | -0.21056 | -0.81056 | -0.98292 | -0.16046 | -0.450438 | 0 |
| 21.98611 | -0.26472 | -0.07381 | 0.138807 | 0.155021 | 0.305833 | -0.12778 | -0.08677 | -0.61401 | 0.126198 | 0.013979 | 0 |
| 22 | 0.089369 | 0.222922 | 0.566339 | 0.602217 | 0.691444 | -0.00185 | 0.654311 | -0.01693 | 0.538496 | 0.512719 | 0 |
| 22.01389 | 0.243359 | 0.288064 | 0.737689 | 0.684566 | 0.735636 | 0.070437 | 0.864784 | 0.407427 | 0.703457 | 0.755762 | 0 |
| 22.02778 | 0.185923 | 0.151244 | 0.512232 | 0.378668 | 0.569184 | 0.031017 | 0.539475 | 0.396468 | 0.471961 | 0.551748 | 0 |
| 22.04167 | 0.184895 | 0.072775 | 0.278079 | 0.218084 | 0.377317 | 0.179202 | 0.254293 | 0.25763 | 0.249967 | 0.352008 | 0 |
| 22.05556 | 0.252881 | 0.112461 | 0.205297 | 0.381842 | 0.281091 | 0.302857 | 0.213326 | 0.152013 | 0.207978 | 0.320937 | 0 |
| 22.06944 | 0.261862 | 0.105928 | 0.179847 | 0.429125 | 0.213954 | 0.23998 | 0.212214 | 0.054374 | 0.209302 | 0.265478 | 0 |
| 22.08333 | 0.243456 | 0.12496 | 0.196307 | 0.274442 | 0.173673 | 0.22834 | 0.205355 | 0.060437 | 0.228693 | 0.175525 | 0 |
| 22.09722 | 0.230214 | 0.160907 | 0.232463 | 0.158209 | 0.192916 | 0.242155 | 0.206943 | 0.124897 | 0.25103 | 0.140619 | 0 |
| 22.11111 | 0.173628 | 0.122335 | 0.194754 | 0.10233 | 0.16398 | 0.146659 | 0.157176 | 0.12949 | 0.197365 | 0.152609 | 0 |
| 22.125 | 0.089195 | 0.046513 | 0.11614 | 0.022053 | 0.087453 | 0.034917 | 0.072044 | 0.065331 | 0.099316 | 0.112664 | 0 |
| 22.13889 | 0.00253 | -0.02493 | 0.048637 | -0.06998 | 0.032747 | -0.0046 | 0.00045 | -0.018 | 0.020746 | 0.059115 | 0 |
| 22.15278 | -0.07344 | -0.08406 | -0.00083 | -0.15027 | 0.003719 | -0.00851 | -0.04753 | -0.07018 | -0.02252 | 0.018955 | 0 |
| 22.16667 | -0.12358 | -0.11671 | -0.02496 | -0.19155 | -0.02112 | -0.00064 | -0.07606 | -0.06893 | -0.03566 | -0.046276 | 0 |
| 22.18056 | -0.12645 | -0.10792 | -0.01724 | -0.15094 | -0.02986 | 0.01218 | -0.07311 | -0.02901 | -0.01991 | -0.083708 | 0 |
| 22.19444 | -0.06477 | -0.04993 | 0.021839 | -0.0117 | -0.00558 | 0.052721 | -0.01303 | 0.032064 | 0.028081 | -0.026231 | 0 |
| 22.20833 | 0.002654 | 0.008662 | 0.051178 | 0.127529 | 0.015301 | 0.06932 | 0.051514 | 0.063364 | 0.058185 | 0.018645 | 0 |
| 22.22222 | 0.045964 | 0.034456 | 0.060365 | 0.251934 | 0.027018 | -0.03258 | 0.097664 | 0.04666 | 0.063432 | 0.052307 | 0 |
| 22.23611 | 0.102878 | 0.064727 | 0.091295 | 0.379796 | 0.053466 | -0.1173 | 0.151721 | 0.056841 | 0.102234 | 0.115014 | 0 |
| 22.25 | 0.175724 | 0.162382 | 0.175041 | 0.432668 | 0.106755 | 0.078747 | 0.203709 | 0.163118 | 0.208089 | 0.164 | 0 |
| 22.26389 | 0.228359 | 0.250066 | 0.254075 | 0.487875 | 0.170943 | 0.225427 | 0.234542 | 0.257966 | 0.304086 | 0.178437 | 0 |
| 22.27778 | 0.225755 | 0.228924 | 0.251032 | 0.564395 | 0.163239 | 0.071206 | 0.221495 | 0.252979 | 0.308726 | 0.207023 | 0 |
| 22.29167 | 0.186662 | 0.191504 | 0.20718 | 0.557832 | 0.085855 | 0.011508 | 0.163583 | 0.253538 | 0.275889 | 0.180472 | 0 |
| 22.30556 | 0.146111 | 0.153274 | 0.152066 | 0.555144 | 0.01268 | 0.06091 | 0.109645 | 0.262355 | 0.240566 | 0.140551 | 0 |
| 22.31944 | 0.121163 | 0.072661 | 0.101579 | 0.602076 | 0.004914 | -0.11156 | 0.118508 | 0.232935 | 0.206461 | 0.19926 | 0 |
| 22.33333 | 0.105229 | 0.034191 | 0.117514 | 0.617891 | 0.022814 | -0.1836 | 0.245331 | 0.262642 | 0.205819 | 0.236504 | 0 |
| 22.34722 | 0.068472 | 0.07264 | 0.182858 | 0.445563 | 0.001244 | 0.019429 | 0.503753 | 0.39318 | 0.216474 | 0.216281 | 0 |
| 22.36111 | 0.014192 | 0.076896 | 0.206507 | 0.371205 | -0.04117 | 0.230348 | 0.722619 | 0.473837 | 0.212832 | 0.151852 | 0 |
| 22.375 | -0.044 | -0.00839 | 0.169295 | 0.4394 | -0.08012 | 0.101489 | 0.684553 | 0.467088 | 0.177808 | 0.240058 | 0 |
| 22.38889 | -0.12256 | -0.11462 | 0.109316 | 0.54881 | -0.13316 | -0.04742 | 0.446497 | 0.468497 | 0.134402 | 0.248796 | 0 |
| 22.40278 | -0.2353 | -0.15589 | -0.0024 | 0.52045 | -0.18731 | -0.08771 | 0.16634 | 0.49278 | 0.055641 | 0.09863 | 0 |
| 22.41667 | -0.44124 | 0.070632 | -0.13472 | 0.041776 | -0.32647 | -0.10177 | -0.07272 | 0.266577 | -0.16125 | -0.58737 | 0 |
| 22.43056 | -0.48423 | 1.042086 | -0.16367 | -0.54367 | -0.19386 | 0.227224 | -0.47571 | -0.03658 | -0.19081 | -0.392831 | 0 |
| 22.44444 | -0.12997 | 0.252017 | 0.027122 | -0.34145 | 0.581928 | 1.481288 | -0.78651 | 0.123558 | -0.06909 | -0.370165 | 0 |
| 22.45833 | 0.126013 | 0.07924 | 0.235778 | 0.407756 | 0.444159 | 0.648864 | -0.31752 | 0.600241 | 0.31941 | 0.119679 | 0 |
| 22.47222 | 0.275227 | 0.278349 | 0.25681 | 0.593156 | 0.189552 | 0.28591 | 0.000717 | 0.643553 | 0.277955 | 0.270301 | 0 |
| 22.48611 | 0.500505 | 0.449605 | 0.215403 | 0.246205 | 0.070407 | 0.27806 | 0.416567 | 0.462098 | 0.159979 | 0.402603 | 0 |
| 22.5 | 0.809124 | 0.629289 | 0.171957 | -0.16793 | 0.005341 | 0.54428 | 1.008841 | 0.536137 | 0.238647 | 0.678045 | 0 |
| 22.51389 | 0.804517 | 0.48097 | 0.074804 | -0.27746 | -0.10309 | 0.536098 | 1.191736 | 0.759019 | 0.322797 | 0.914917 | 0 |
| 22.52778 | 0.641588 | 0.225282 | -0.02037 | 0.05442 | -0.16172 | 0.246905 | 1.026643 | 0.908265 | 0.296685 | 0.58087 | 0 |
| 22.54167 | 0.553522 | 0.17363 | -0.02263 | 0.381187 | -0.12598 | 0.061164 | 0.887834 | 0.932327 | 0.189098 | 0.300851 | 0 |
| 22.55556 | 0.52099 | 0.203441 | -0.03227 | 0.146341 | -0.11066 | -0.04534 | 0.488039 | 0.657998 | 0.033344 | -0.002783 | 0 |
| 22.56944 | 0.462143 | 0.171523 | -0.11195 | -0.24306 | -0.14873 | -0.18373 | -0.08305 | 0.388065 | -0.19925 | -0.252033 | 0 |
| 22.58333 | 0.537702 | 0.157778 | -0.11634 | -0.40017 | -0.11163 | -0.21038 | -0.47397 | 0.329202 | -0.30899 | -0.180761 | 0 |
| 22.59722 | 0.656648 | 0.15874 | 0.011258 | -0.32574 | 0.065585 | -0.0872 | -0.58341 | 0.409636 | -0.22285 | 0.023035 | 0 |
| 22.61111 | 0.672723 | 0.074032 | 0.1898 | -0.20084 | 0.267513 | 0.025485 | -0.27464 | 0.587918 | -0.05694 | 0.117213 | 0 |
| 22.625 | 0.59876 | -0.0338 | 0.419663 | 0.016321 | 0.289172 | 0.085869 | 0.505921 | 0.836894 | 0.107665 | 0.179282 | 0 |
| 22.63889 | 0.499476 | -0.10326 | 0.535527 | 0.32862 | 0.091715 | 0.128057 | 1.52756 | 1.079682 | 0.22583 | 0.30234 | 0 |
| 22.65278 | 0.412969 | -0.26291 | 0.383791 | 0.424711 | -0.00081 | 0.067192 | 1.963006 | 1.039163 | 0.272166 | 0.291237 | 0 |
| 22.66667 | 0.264133 | -0.55804 | -0.02165 | 0.058876 | -0.06709 | -0.10537 | 1.507299 | 0.718058 | 0.227268 | 0.087016 | 0 |
| 22.68056 | 0.027314 | -0.4605 | -0.36913 | -0.4749 | -0.16531 | -0.35945 | 0.525299 | 0.292942 | 0.10135 | -0.013168 | 0 |
| 22.69444 | -0.16719 | 0.223123 | -0.4926 | -0.71577 | -0.29299 | -0.53333 | -0.64066 | -0.15286 | -0.05471 | 0.002193 | 0 |
| 22.70833 | -0.2432 | 0.856379 | -0.46939 | -0.71812 | -0.31617 | -0.56733 | -1.63479 | -0.50774 | -0.1271 | 0.043312 | 0 |

| | | | | | | | | | | | |
|---|---|---|---|---|---|---|---|---|---|---|---|
| 22.72222 | -0.28528 | 1.098018 | -0.30451 | -0.71161 | -0.27122 | -0.49376 | -2.08524 | -0.64868 | -0.11148 | 0.076637 | 0 |
| 22.73611 | -0.37729 | 1.081132 | -0.11781 | -0.56733 | -0.31884 | -0.46816 | -2.01953 | -0.51634 | -0.10398 | -0.050997 | 0 |
| 22.75 | -0.38092 | 1.076711 | 0.116972 | -0.09685 | -0.61869 | -0.60354 | -1.34963 | -0.20903 | -0.06545 | -0.034443 | 0 |
| 22.76389 | -0.21854 | 1.07535 | 0.300135 | 0.346469 | -1.07172 | -0.80726 | -0.34227 | 0.028873 | -0.01069 | 0.157337 | 0 |
| 22.77778 | -0.03137 | 0.987633 | 0.288986 | 0.429843 | -1.18734 | -0.88999 | 0.450711 | -0.06136 | 0.02333 | 0.335402 | 0 |
| 22.79167 | -0.00826 | 0.710588 | 0.103809 | 0.198043 | -1.03228 | -0.91631 | 0.721774 | -0.4734 | 0.030206 | 0.33522 | 0 |
| 22.80556 | -0.15534 | 0.318278 | -0.09855 | -0.08382 | -0.83283 | -0.89584 | 0.557682 | -0.97182 | -0.06998 | 0.074028 | 0 |
| 22.81944 | -0.32858 | -0.12895 | -0.231 | -0.30274 | -0.5392 | -0.82058 | 0.07027 | -1.35663 | -0.2559 | -0.359398 | 0 |
| 22.83333 | -0.46446 | -0.5746 | -0.3324 | -0.48331 | -0.42384 | -0.81471 | -0.52576 | -1.54143 | -0.41892 | -0.897504 | 0 |
| 22.84722 | -0.56116 | -0.91597 | -0.42482 | -0.50059 | -0.66241 | -1.05517 | -1.0326 | -1.63156 | -0.51259 | -1.244551 | 0 |
| 22.86111 | -0.59756 | -1.07119 | -0.46918 | -0.42531 | -1.13516 | -1.28402 | -1.27333 | -1.58899 | -0.59381 | -1.362499 | 0 |
| 22.875 | -0.56902 | -0.95012 | -0.43815 | -0.44376 | -1.57187 | -1.37068 | -1.0888 | -1.45341 | -0.5792 | -1.154226 | 0 |
| 22.88889 | -0.52108 | -0.53768 | -0.3854 | -0.57574 | -1.84075 | -1.35774 | -0.56494 | -1.36195 | -0.51822 | -0.860854 | 0 |
| 22.90278 | -0.45513 | -0.05265 | -0.31029 | -0.60357 | -1.79879 | -1.15999 | -0.02895 | -1.39429 | -0.43411 | -0.686762 | 0 |
| 22.91667 | -0.37784 | 0.250535 | -0.25942 | -0.59722 | -1.50567 | -0.98145 | 0.0777 | -1.38084 | -0.38084 | -0.569372 | 0 |
| 22.93056 | -0.32345 | 0.172142 | -0.25371 | -0.54283 | -1.10208 | -0.86105 | -0.25966 | -1.21679 | -0.41499 | -0.546004 | 0 |
| 22.94444 | -0.34067 | -0.21942 | -0.20934 | -0.43443 | -0.74767 | -0.726 | -0.73225 | -0.95259 | -0.46478 | -0.509237 | 0 |
| 22.95833 | -0.40642 | -0.68767 | -0.10307 | -0.28685 | -0.50077 | -0.60713 | -1.11177 | -0.71482 | -0.41564 | -0.579321 | 0 |
| 22.97222 | -0.4193 | -0.98941 | -0.00137 | -0.11079 | -0.39208 | -0.49926 | -1.31371 | -0.49457 | -0.32854 | -0.645585 | 0 |
| 22.98611 | -0.24973 | -1.13325 | 0.134791 | 0.11571 | -0.8371 | -0.46224 | -1.04387 | -0.12144 | -0.33332 | -0.585422 | 0 |
| 23 | 0.118059 | -1.06544 | 0.431939 | 0.427856 | -1.86755 | -0.41319 | -0.27018 | 0.360014 | -0.46399 | -0.381269 | 0 |
| 23.01389 | 0.472208 | -0.34082 | 0.630198 | 0.529576 | -1.51001 | 0.250737 | 0.313624 | 0.566709 | -0.23751 | -0.057337 | 0 |
| 23.02778 | 0.607771 | 0.456622 | 0.446346 | 0.2809 | -0.12959 | 0.257213 | 0.415389 | 0.267795 | 0.261717 | 0.39085 | 0 |
| 23.04167 | 0.896375 | 0.42757 | 0.071703 | 0.170182 | 0.232959 | -1.05417 | 0.234001 | -0.383 | 0.540766 | 1.181424 | 0 |
| 23.05556 | 1.06331 | 0.096645 | -0.16646 | 0.131942 | -0.07894 | -1.92123 | -0.02435 | -0.73774 | 0.439714 | 1.179763 | 0 |
| 23.06944 | 0.37927 | 0.398327 | -0.16497 | -0.17327 | -0.15058 | -0.02696 | 0.027964 | -0.10058 | 0.172217 | 0.089059 | 0 |
| 23.08333 | -0.26356 | 0.911498 | -0.04185 | -0.30127 | 0.054913 | 2.069573 | 0.208575 | 0.636226 | -0.02188 | -0.928268 | 0 |
| 23.09722 | -0.33718 | 0.625108 | -0.08112 | -0.04474 | 0.040449 | 1.488533 | 0.09134 | 0.442255 | -0.12441 | -0.780255 | 0 |
| 23.11111 | -0.24399 | 0.230545 | -0.21781 | 0.086436 | -0.11358 | 0.16575 | -0.10343 | -0.0323 | -0.22991 | -0.167936 | 0 |
| 23.125 | -0.24016 | 0.140214 | -0.31775 | -0.01724 | -0.23587 | -0.25629 | -0.26928 | -0.17924 | -0.33723 | -0.105386 | 0 |
| 23.13889 | -0.13246 | 0.185928 | -0.32274 | -0.10676 | -0.294 | -0.60795 | -0.3335 | -0.1702 | -0.34217 | 0.063474 | 0 |
| 23.15278 | 0.011133 | 0.303888 | -0.25291 | -0.10782 | -0.26092 | -0.88486 | -0.31926 | -0.1233 | -0.24705 | 0.325497 | 0 |
| 23.16667 | 0.064049 | 0.421601 | -0.12917 | -0.02987 | -0.06307 | -0.80378 | -0.27495 | -0.13701 | -0.12781 | 0.515599 | 0 |
| 23.18056 | -0.07415 | 0.38399 | -0.08195 | -0.02912 | 0.026899 | -0.41934 | -0.35466 | -0.18392 | -0.13581 | 0.286412 | 0 |
| 23.19444 | -0.33246 | 0.174656 | -0.20822 | -0.16299 | -0.14758 | -0.17957 | -0.6948 | -0.25357 | -0.28771 | -0.108598 | 0 |
| 23.20833 | -0.39738 | -0.03992 | -0.39005 | -0.2597 | -0.39077 | -0.42779 | -1.04308 | -0.37005 | -0.41702 | -0.189001 | 0 |
| 23.22222 | -0.28452 | -0.09643 | -0.49904 | -0.27937 | -0.48737 | -0.65806 | -1.01805 | -0.48507 | -0.47026 | -0.105778 | 0 |
| 23.23611 | -0.24709 | -0.072 | -0.53083 | -0.31948 | -0.48791 | -0.55269 | -0.81311 | -0.4895 | -0.45922 | -0.219935 | 0 |
| 23.25 | -0.3215 | -0.11549 | -0.53879 | -0.3873 | -0.46799 | -0.45383 | -0.60443 | -0.38561 | -0.43339 | -0.367097 | 0 |
| 23.26389 | -0.43341 | -0.2645 | -0.55669 | -0.42686 | -0.41973 | -0.47195 | -0.50693 | -0.31503 | -0.43174 | -0.428515 | 0 |
| 23.27778 | -0.45907 | -0.36792 | -0.5373 | -0.35766 | -0.4035 | -0.48431 | -0.46563 | -0.3159 | -0.41548 | -0.393156 | 0 |
| 23.29167 | -0.27062 | -0.27885 | -0.42036 | -0.10541 | -0.38041 | -0.49872 | -0.4098 | -0.31922 | -0.31647 | -0.180329 | 0 |
| 23.30556 | 0.01299 | -0.12246 | -0.29058 | 0.206215 | -0.28925 | -0.617 | -0.33737 | -0.28918 | -0.22613 | 0.122278 | 0 |
| 23.31944 | 0.171551 | -0.05448 | -0.21078 | 0.303406 | -0.14771 | -0.62415 | -0.2504 | -0.19434 | -0.19705 | 0.166488 | 0 |
| 23.33333 | 0.155378 | -0.03744 | -0.1588 | 0.177001 | 0.027655 | -0.43926 | -0.13525 | -0.09145 | -0.17502 | 0.008962 | 0 |
| 23.34722 | 0.116236 | -0.00587 | -0.10894 | 0.044109 | 0.213703 | -0.29366 | -0.01065 | -0.05521 | -0.09317 | -0.018399 | 0 |
| 23.36111 | 0.120537 | 0.061801 | -0.04477 | 0.07536 | 0.370112 | -0.20273 | 0.059355 | 0.086952 | 0.070118 | 0.138703 | 0 |
| 23.375 | 0.222737 | 0.236609 | 0.097771 | 0.226252 | 0.45976 | -0.05729 | 0.093077 | 0.374749 | 0.264579 | 0.332111 | 0 |
| 23.38889 | 0.383703 | 0.483584 | 0.234893 | 0.435948 | 0.509314 | 0.065908 | 0.116069 | 0.659357 | 0.356337 | 0.486769 | 0 |
| 23.40278 | 0.489563 | 0.660831 | 0.252481 | 0.610072 | 0.503324 | 0.053191 | 0.131408 | 0.865002 | 0.292576 | 0.564061 | 0 |
| 23.41667 | 0.516265 | 0.70295 | 0.220368 | 0.673662 | 0.540728 | -0.05919 | 0.181686 | 0.928265 | 0.173242 | 0.67776 | 0 |
| 23.43056 | 0.47075 | 0.636208 | 0.193443 | 0.617327 | 0.614923 | -0.18408 | 0.336891 | 1.040358 | 0.144006 | 0.833193 | 0 |
| 23.44444 | 0.348143 | 0.592356 | 0.215746 | 0.457893 | 0.592473 | -0.10664 | 0.621235 | 1.061908 | 0.211998 | 0.875656 | 0 |
| 23.45833 | 0.248899 | 0.556179 | 0.252331 | 0.339532 | 0.476698 | -0.01201 | 0.884163 | 0.904884 | 0.262251 | 0.775496 | 0 |
| 23.47222 | 0.179312 | 0.474397 | 0.226787 | 0.200458 | 0.356052 | 0.084349 | 0.934602 | 0.55105 | 0.230049 | 0.529341 | 0 |
| 23.48611 | -0.00884 | 0.422229 | 0.048386 | 0.028437 | 0.254622 | 0.472531 | 0.821058 | 0.22159 | 0.070827 | -0.024354 | 0 |
| 23.5 | -0.35636 | 0.394986 | -0.24665 | -0.1462 | 0.324331 | 1.146637 | 0.825702 | -0.0301 | -0.07463 | -0.35604 | 0 |

| | | | | | | | | | | | |
|---|---|---|---|---|---|---|---|---|---|---|---|
| 23.51389 | -0.51194 | 0.343308 | -0.29055 | 0.072432 | 0.701265 | 0.806721 | 0.646857 | -0.07772 | -0.01588 | -0.056751 | 0 |
| 23.52778 | -0.25209 | 0.095982 | -0.05979 | 0.398134 | 0.526293 | 0.11139 | 0.506921 | 0.103625 | 0.122923 | 0.316632 | 0 |
| 23.54167 | 0.058118 | -0.07739 | 0.099829 | 0.49803 | 0.275716 | -0.21884 | 0.345985 | 0.336336 | -0.05424 | 0.286945 | 0 |
| 23.55556 | 0.228244 | -0.11025 | 0.042266 | 0.253144 | -0.14611 | -0.17271 | 0.232973 | 0.579843 | -0.3917 | -0.00937 | 0 |
| 23.56944 | 0.381085 | -0.00297 | 0.047663 | 0.043221 | -0.49731 | -0.08416 | 0.11361 | 0.892098 | -0.54354 | -0.189573 | 0 |
| 23.58333 | 0.646887 | 0.238604 | 0.244825 | -0.0354 | -0.56271 | 0.015287 | 0.020247 | 1.171574 | -0.38946 | -0.021425 | 0 |
| 23.59722 | 0.86818 | 0.438051 | 0.465995 | -0.10394 | -0.28941 | 0.04402 | -0.00188 | 1.267003 | -0.11094 | 0.288877 | 0 |
| 23.61111 | 0.80433 | 0.402306 | 0.535724 | -0.09606 | 0.05487 | 0.008173 | 0.195835 | 1.237505 | 0.219285 | 0.473336 | 0 |
| 23.625 | 0.579465 | 0.397318 | 0.428423 | 0.017637 | 0.425202 | -0.04724 | 0.737333 | 1.252868 | 0.521073 | 0.61535 | 0 |
| 23.63889 | 0.441341 | 0.682414 | 0.191844 | 0.243547 | 0.657622 | -0.01697 | 1.293049 | 1.210805 | 0.599689 | 0.891692 | 0 |
| 23.65278 | 0.400389 | 1.157168 | -0.11513 | 0.581909 | 1.02233 | 0.09817 | 1.235231 | 1.016517 | 0.580429 | 1.074165 | 0 |
| 23.66667 | 0.354399 | 1.38432 | -0.32039 | 0.738271 | 0.961697 | 0.111257 | 0.828884 | 0.704041 | 0.426389 | 0.921822 | 0 |
| 23.68056 | 0.351233 | 1.162101 | -0.27963 | 0.636614 | 0.386509 | -0.03095 | 0.381843 | 0.440687 | 0.20479 | 0.509084 | 0 |
| 23.69444 | 0.393775 | 0.634957 | -0.04769 | 0.285008 | -0.07188 | -0.09782 | 0.091648 | 0.181382 | 0.036329 | 0.124684 | 0 |
| 23.70833 | 0.384489 | 0.066376 | 0.241589 | -0.11781 | 0.026473 | -0.11183 | -0.19598 | -0.25125 | -0.1054 | -0.219088 | 0 |
| 23.72222 | 0.114067 | -0.50249 | 0.391774 | -0.53252 | 0.177694 | -0.14723 | -0.58234 | -0.71927 | -0.25258 | -0.513226 | 0 |
| 23.73611 | -0.17168 | -0.73938 | 0.337785 | -0.75683 | 0.043391 | -0.12783 | -0.63028 | -1.07103 | -0.35476 | -0.727803 | 0 |
| 23.75 | -0.33543 | -0.56301 | 0.133735 | -0.63622 | -0.27718 | -0.0175 | -0.39976 | -1.02384 | -0.35781 | -0.620286 | 0 |
| 23.76389 | -0.45374 | -0.33133 | -0.13965 | -0.31219 | -0.46915 | 0.040794 | -0.16469 | -0.7966 | -0.31233 | -0.379218 | 0 |
| 23.77778 | -0.4586 | -0.18497 | -0.35068 | -0.07762 | -0.60362 | -0.01289 | -0.21593 | -0.44691 | -0.22813 | -0.1418 | 0 |
| 23.79167 | -0.24173 | -0.00219 | -0.44694 | -0.02897 | -0.75444 | -0.05682 | -0.31908 | -0.12489 | -0.04151 | 0.005468 | 0 |
| 23.80556 | -0.04483 | 0.035723 | -0.51557 | -0.11706 | -0.84439 | -0.08899 | -0.41873 | -0.13984 | 0.098189 | -0.019131 | 0 |
| 23.81944 | -0.09495 | -0.30764 | -0.60773 | -0.24398 | -1.01099 | -0.18999 | -0.75569 | -0.60055 | -0.03274 | -0.340063 | 0 |
| 23.83333 | -0.31363 | -0.82829 | -0.61839 | -0.36886 | -1.29818 | -0.28386 | -1.06813 | -1.20307 | -0.35469 | -0.85472 | 0 |
| 23.84722 | -0.52187 | -1.17081 | -0.54423 | -0.42518 | -1.58909 | -0.31281 | -1.21187 | -1.58133 | -0.5824 | -1.223974 | 0 |
| 23.86111 | -0.6749 | -1.32703 | -0.4277 | -0.48639 | -1.91656 | -0.30615 | -1.27102 | -1.7838 | -0.70832 | -1.449794 | 0 |
| 23.875 | -0.83935 | -1.32982 | -0.43733 | -0.62232 | -2.11881 | -0.35662 | -1.35924 | -1.80085 | -0.81038 | -1.575696 | 0 |
| 23.88889 | -0.97861 | -1.3054 | -0.47976 | -0.68576 | -1.95646 | -0.38497 | -1.1869 | -1.55999 | -0.81884 | -1.450974 | 0 |
| 23.90278 | -0.97659 | -1.2137 | -0.45094 | -0.54541 | -1.70921 | -0.3817 | -0.7825 | -1.30831 | -0.73638 | -1.147652 | 0 |
| 23.91667 | -0.87782 | -1.01991 | -0.39927 | -0.43039 | -1.52355 | -0.39417 | -0.75172 | -1.27761 | -0.67819 | -0.93218 | 0 |
| 23.93056 | -0.71384 | -0.82733 | -0.36178 | -0.43866 | -1.41828 | -0.45725 | -1.01242 | -1.2575 | -0.61405 | -0.925075 | 0 |
| 23.94444 | -0.53888 | -0.75361 | -0.31286 | -0.41909 | -1.40322 | -0.48688 | -1.05418 | -1.13631 | -0.50776 | -0.922852 | 0 |
| 23.95833 | -0.51043 | -0.86848 | -0.27634 | -0.38387 | -1.46193 | -0.51246 | -0.90095 | -0.98626 | -0.45941 | -0.867329 | 0 |
| 23.97222 | -0.59948 | -1.07153 | -0.25986 | -0.43808 | -1.3477 | -0.56769 | -0.85438 | -0.83196 | -0.51214 | -0.781171 | 0 |
| 23.98611 | -0.49832 | -1.28585 | -0.19654 | -0.51031 | -0.90521 | -0.90434 | -0.86655 | -0.54756 | -0.52969 | -0.138848 | 0 |
| 24 | -0.19741 | -1.39282 | -0.05205 | -0.44656 | -0.40041 | -1.56528 | -0.98575 | -0.26736 | -0.3577 | 1.049073 | 0 |
| 24.01389 | -0.08944 | -1.09244 | 0.008749 | -0.29912 | -0.14999 | -1.44573 | -1.1025 | -0.0696 | -0.10639 | 1.149911 | 0 |
| 24.02778 | -0.29305 | -0.67978 | -0.10247 | -0.14538 | -0.316 | -0.29653 | -1.01428 | -0.01615 | -0.06122 | -0.038622 | 0 |
| 24.04167 | -0.2171 | -0.38549 | -0.17034 | -0.22304 | -0.29431 | 0.181849 | -0.73044 | -0.27023 | -0.16671 | -0.689533 | 0 |
| 24.05556 | 0.187441 | 0.01895 | -0.11875 | -0.48429 | -0.06959 | 0.017359 | -0.5073 | -0.49595 | -0.21605 | -0.396216 | 0 |
| 24.06944 | 0.482919 | 0.386938 | -0.07896 | -0.58059 | 0.003247 | -0.07816 | -0.36171 | -0.31073 | -0.17982 | -0.09119 | 0 |
| 24.08333 | 0.349913 | 0.3854 | -0.10098 | -0.45394 | -0.07663 | -0.0331 | -0.28801 | -0.05464 | -0.13114 | -0.119552 | 0 |
| 24.09722 | -0.03538 | 0.139608 | -0.25415 | -0.26364 | -0.20031 | 0.248997 | -0.29514 | 0.076835 | -0.17077 | -0.69329 | 0 |
| 24.11111 | -0.37311 | -0.00521 | -0.4282 | -0.11939 | -0.23585 | 0.723481 | -0.30334 | 0.000831 | -0.27593 | -1.617811 | 0 |
| 24.125 | -0.26624 | 0.073489 | -0.27589 | 0.071282 | 0.030371 | 0.875489 | -0.12159 | 0.045995 | -0.19902 | -1.518037 | 0 |
| 24.13889 | 0.09323 | 0.249927 | 0.006783 | 0.236784 | 0.358441 | 0.486247 | 0.122949 | 0.132114 | 0.013103 | -0.4469 | 0 |
| 24.15278 | 0.267468 | 0.320862 | 0.05602 | 0.215304 | 0.320002 | 0.133439 | 0.216302 | 0.063378 | 0.084949 | 0.106677 | 0 |
| 24.16667 | 0.200468 | 0.290821 | -0.25823 | 0.089904 | 0.137361 | 0.010679 | 0.196068 | -0.07509 | -0.01031 | 0.036723 | 0 |
| 24.18056 | -0.17263 | -0.23741 | -0.78154 | -0.43161 | 0.114495 | -0.1152 | -0.06185 | -0.39026 | -0.15226 | -0.354305 | 0 |
| 24.19444 | -0.74714 | -1.40732 | -0.91993 | -1.52896 | 0.040458 | -0.28511 | -0.69262 | -0.98666 | -0.22109 | -0.960373 | 0 |
| 24.20833 | -0.90564 | -1.69505 | -0.55947 | -1.86961 | -0.08284 | -0.27054 | -1.00806 | -1.15869 | -0.26298 | -1.103398 | 0 |
| 24.22222 | -0.60677 | -0.88223 | -0.22908 | -1.08817 | -0.17788 | -0.19833 | -0.7887 | -0.69781 | -0.23696 | -0.710082 | 0 |
| 24.23611 | -0.3753 | -0.36459 | -0.1537 | -0.50295 | -0.20755 | -0.27115 | -0.61743 | -0.38902 | -0.1887 | -0.353735 | 0 |
| 24.25 | -0.20886 | -0.22215 | -0.22547 | -0.31981 | -0.17249 | -0.39786 | -0.61326 | -0.3688 | -0.17543 | -0.184009 | 0 |
| 24.26389 | -0.13184 | -0.17208 | -0.39867 | -0.22496 | -0.1531 | -0.36205 | -0.67912 | -0.33395 | -0.29248 | -0.356447 | 0 |
| 24.27778 | -0.18755 | -0.21863 | -0.50088 | -0.10182 | -0.12817 | -0.29534 | -0.69945 | -0.1766 | -0.42356 | -0.55168 | 0 |
| 24.29167 | -0.145 | -0.19947 | -0.37259 | 0.183072 | 0.029999 | -0.27829 | -0.54251 | 0.039917 | -0.33942 | -0.313653 | 0 |

| | | | | | | | | | | | |
|---|---|---|---|---|---|---|---|---|---|---|---|
| 24.30556 | 0.050237 | -0.04961 | -0.13591 | 0.518264 | 0.243166 | -0.27587 | -0.29639 | 0.147927 | -0.14474 | 0.094783 | 0 |
| 24.31944 | 0.361907 | 0.19445 | 0.091615 | 0.767736 | 0.435876 | -0.15367 | -0.03827 | 0.208898 | 0.046932 | 0.326891 | 0 |
| 24.33333 | 0.691259 | 0.442766 | 0.291628 | 0.884179 | 0.550832 | 0.098496 | 0.180217 | 0.318124 | 0.221251 | 0.287702 | 0 |
| 24.34722 | 0.797539 | 0.434107 | 0.315156 | 0.773159 | 0.456463 | 0.125438 | 0.194964 | 0.327084 | 0.24597 | 0.156768 | 0 |
| 24.36111 | 0.510162 | 0.083222 | 0.131871 | 0.533139 | 0.247806 | -0.17402 | 0.037525 | 0.230571 | 0.062765 | 0.021588 | 0 |
| 24.375 | 0.07544 | -0.29274 | -0.08719 | 0.363412 | 0.152569 | -0.51458 | -0.10983 | 0.163941 | -0.19024 | -0.08351 | 0 |
| 24.38889 | -0.25594 | -0.51973 | -0.35463 | 0.074271 | 0.142433 | -0.86131 | -0.22931 | -0.02911 | -0.43728 | -0.165681 | 0 |
| 24.40278 | -0.36102 | -0.52392 | -0.68437 | -0.29196 | 0.079686 | -1.06589 | -0.35693 | -0.42009 | -0.63276 | -0.313244 | 0 |
| 24.41667 | 0.003181 | -0.04734 | -0.63433 | -0.16763 | 0.152571 | -0.86069 | -0.16955 | -0.36952 | -0.46288 | -0.060032 | 0 |
| 24.43056 | 0.598901 | 0.654828 | -0.17049 | 0.452984 | 0.397692 | -0.33428 | 0.353826 | 0.113348 | 0.022884 | 0.548086 | 0 |
| 24.44444 | 0.513833 | 0.742605 | 0.173705 | 0.875672 | 0.610188 | 0.027371 | 0.659965 | 0.471749 | 0.3243 | 0.824993 | 0 |
| 24.45833 | 0.216434 | 0.609183 | 0.341978 | 0.905258 | 0.76223 | 0.244026 | 0.665712 | 0.667824 | 0.453836 | 0.881589 | 0 |
| 24.47222 | 0.46076 | 0.80862 | 0.60929 | 0.971545 | 0.940587 | 0.581587 | 0.85169 | 1.008417 | 0.669619 | 1.192181 | 0 |
| 24.48611 | 1.179072 | 1.524972 | 0.833962 | 1.048084 | 1.108262 | 0.90669 | 1.363496 | 1.33816 | 0.953064 | 1.843633 | 0 |
| 24.5 | 1.495018 | 2.164418 | 0.741755 | 0.8766 | 1.208825 | 1.117874 | 2.082601 | 1.437605 | 1.198608 | 2.439048 | 0 |
| 24.51389 | 1.03674 | 1.848269 | 0.695725 | 0.621901 | 1.416465 | 0.898886 | 2.515457 | 1.391499 | 1.085975 | 2.566885 | 0 |
| 24.52778 | 0.732955 | 1.161753 | 0.863493 | 0.568085 | 1.67351 | 0.483421 | 2.482348 | 1.620287 | 0.727807 | 2.119294 | 0 |
| 24.54167 | 0.590713 | 0.631908 | 0.728374 | 0.639394 | 0.963922 | 0.312279 | 1.797918 | 2.069714 | 0.510116 | 1.357435 | 0 |
| 24.55556 | 0.40267 | 0.230254 | 0.208943 | 0.650381 | -0.044 | 0.349219 | 0.759351 | 2.175103 | 0.195103 | 0.484332 | 0 |
| 24.56944 | 0.167237 | 0.003591 | -0.24407 | 0.365779 | -0.81844 | 0.474028 | -0.12062 | 1.73609 | -0.10701 | -0.18136 | 0 |
| 24.58333 | 0.250836 | -0.02329 | -0.23933 | 0.089659 | -1.22802 | 0.577086 | -0.46524 | 1.03918 | -0.25193 | -0.402213 | 0 |
| 24.59722 | 0.530143 | 0.06378 | 0.176936 | 0.057738 | -0.96861 | 0.559092 | -0.04969 | 0.764415 | -0.22693 | -0.029086 | 0 |
| 24.61111 | 0.928888 | 0.330398 | 0.822488 | 0.255133 | -0.09894 | 0.654939 | 1.048061 | 0.968313 | 0.061504 | 0.828688 | 0 |
| 24.625 | 1.15316 | 0.789946 | 1.347562 | 0.424172 | 1.048785 | 0.816671 | 2.20907 | 1.24052 | 0.602891 | 1.59854 | 0 |
| 24.63889 | 1.104006 | 1.390924 | 1.181623 | 0.436956 | 1.871392 | 0.824614 | 2.072327 | 1.179744 | 0.983491 | 1.723753 | 0 |
| 24.65278 | 0.817657 | 1.850985 | 0.576893 | 0.3836 | 1.961315 | 0.723803 | 1.064555 | 1.135483 | 1.124677 | 1.340438 | 0 |
| 24.66667 | 0.487591 | 1.778282 | 0.017093 | 0.375906 | 1.219871 | 0.519177 | 0.045853 | 1.392076 | 1.135787 | 0.846891 | 0 |
| 24.68056 | 0.32781 | 1.159754 | -0.21021 | 0.586969 | 0.319012 | 0.254442 | -0.53069 | 1.686777 | 0.956226 | 0.483617 | 0 |
| 24.69444 | 0.387817 | 0.645456 | -0.15568 | 0.760407 | -0.44855 | 0.056988 | -0.77225 | 1.727168 | 0.673734 | 0.392424 | 0 |
| 24.70833 | 0.563812 | 0.34163 | 0.037764 | 0.750911 | -0.80408 | -0.04345 | -0.42451 | 1.376279 | 0.370174 | 0.428822 | 0 |
| 24.72222 | 0.771512 | 0.101622 | 0.312673 | 0.604035 | -0.85318 | -0.05455 | 0.182872 | 0.790192 | 0.122527 | 0.300853 | 0 |
| 24.73611 | 0.7205 | -0.18855 | 0.504671 | 0.201466 | -0.72753 | -0.04793 | 0.576916 | -0.04833 | -0.01003 | -0.029681 | 0 |
| 24.75 | 0.385542 | -0.55209 | 0.389745 | -0.36722 | -0.57034 | -0.09563 | 0.111739 | -0.97574 | -0.09865 | -0.53889 | 0 |
| 24.76389 | 0.062329 | -0.71993 | 0.059946 | -0.64598 | -0.24298 | -0.0832 | -0.89895 | -1.82552 | -0.16906 | -1.001842 | 0 |
| 24.77778 | -0.11017 | -0.4804 | -0.23609 | -0.38211 | 0.185576 | 0.046576 | -1.78267 | -2.00643 | -0.15154 | -1.057298 | 0 |
| 24.79167 | -0.30564 | 0.015502 | -0.47309 | -0.13307 | 0.515113 | 0.143872 | -2.00243 | -1.5359 | -0.11721 | -0.702661 | 0 |
| 24.80556 | -0.47259 | 0.517387 | -0.58765 | -0.27083 | 0.506967 | 0.120512 | -1.03667 | -0.95829 | -0.09857 | -0.167956 | 0 |
| 24.81944 | -0.3462 | 0.67795 | -0.59754 | -0.38233 | 0.221935 | 0.056714 | 0.294002 | -0.39604 | 0.019553 | 0.234377 | 0 |
| 24.83333 | -0.14363 | 0.340381 | -0.53252 | -0.29497 | -0.23372 | -0.05601 | 0.704828 | -0.15188 | 0.115981 | 0.11186 | 0 |
| 24.84722 | -0.22913 | -0.18748 | -0.58117 | -0.32638 | -0.84872 | -0.31747 | -0.01201 | -0.48894 | 0.010295 | -0.54429 | 0 |
| 24.86111 | -0.47561 | -0.61883 | -0.61924 | -0.49039 | -1.50877 | -0.62749 | -1.02921 | -1.01922 | -0.25571 | -1.237625 | 0 |
| 24.875 | -0.68243 | -0.83539 | -0.64206 | -0.66502 | -1.91247 | -0.78525 | -1.49997 | -1.42723 | -0.60112 | -1.646922 | 0 |
| 24.88889 | -0.78394 | -1.05873 | -0.73595 | -0.88377 | -1.95502 | -0.85475 | -1.58348 | -1.66122 | -0.94168 | -1.732569 | 0 |
| 24.90278 | -0.81257 | -1.23824 | -0.7352 | -0.91257 | -1.73667 | -0.80313 | -1.38977 | -1.773 | -0.93368 | -1.487708 | 0 |
| 24.91667 | -0.68159 | -1.21715 | -0.47799 | -0.76561 | -1.18568 | -0.54981 | -0.85653 | -1.63601 | -0.64513 | -1.012076 | 0 |
| 24.93056 | -0.65733 | -1.19804 | -0.26482 | -0.61598 | -0.67407 | -0.4204 | -0.30744 | -1.51804 | -0.44431 | -0.758968 | 0 |
| 24.94444 | -0.8549 | -1.24282 | -0.33108 | -0.51967 | -0.38187 | -0.38776 | -0.21519 | -1.47994 | -0.42139 | -0.817887 | 0 |
| 24.95833 | -1.05988 | -1.12434 | -0.39494 | -0.45262 | -0.33882 | -0.30837 | -0.53274 | -1.13469 | -0.41063 | -0.939476 | 0 |
| 24.97222 | -1.0801 | -0.82292 | -0.31479 | -0.4828 | -0.47911 | -0.22133 | -1.0296 | -0.56775 | -0.38588 | -0.958325 | 0 |
| 24.98611 | -0.78882 | -0.39446 | 0.007911 | -0.28112 | -0.38609 | 0.040998 | -1.11659 | 0.115405 | -0.14579 | -0.66032 | 0 |
| 25 | -0.22655 | 0.083957 | 0.645382 | 0.345547 | -0.10919 | 0.726844 | -0.47264 | 0.809673 | 0.488545 | -0.095187 | 0 |
| 25.01389 | -0.05124 | 0.093039 | 1.061176 | 0.819911 | -0.42035 | 1.285939 | 0.26095 | 0.87632 | 0.91588 | 0.162071 | 0 |
| 25.02778 | -0.42791 | -0.39382 | 0.907942 | 0.831164 | -1.19791 | 1.183852 | 0.564026 | 0.304104 | 0.678133 | -0.028753 | 0 |
| 25.04167 | -0.47387 | -0.76139 | 0.619675 | 0.571897 | -1.37292 | 0.749994 | 0.550501 | -0.1068 | 0.177835 | -0.228326 | 0 |
| 25.05556 | -0.18194 | -0.68883 | 0.362719 | 0.153785 | -0.84301 | 0.302256 | 0.337391 | -0.15975 | -0.21148 | -0.321831 | 0 |
| 25.06944 | 0.336525 | -0.23548 | 0.147743 | -0.26614 | -0.43617 | -0.02 | 0.046606 | -0.09096 | -0.36291 | -0.33652 | 0 |
| 25.08333 | 0.825495 | 0.331654 | 0.075068 | -0.35226 | -0.26926 | -0.0968 | -0.09527 | 0.086881 | -0.3025 | -0.190841 | 0 |

| | | | | | | | | | | | |
|---|---|---|---|---|---|---|---|---|---|---|---|
| 25.09722 | 1.069078 | 0.685317 | 0.149572 | -0.15259 | -0.18029 | -0.02517 | -0.06634 | 0.200018 | -0.18033 | 0.06306 | 0 |
| 25.11111 | 0.997489 | 0.729738 | 0.296217 | 0.047654 | -0.04385 | -0.00404 | 0.003467 | 0.166118 | -0.07674 | 0.254793 | 0 |
| 25.125 | 0.871048 | 0.620105 | 0.436948 | 0.157207 | 0.172692 | -0.05701 | 0.121171 | 0.108414 | 0.037046 | 0.382252 | 0 |
| 25.13889 | 0.776432 | 0.490492 | 0.468877 | 0.186112 | 0.268235 | -0.10433 | 0.291966 | 0.081372 | 0.133846 | 0.385255 | 0 |
| 25.15278 | 0.688813 | 0.409388 | 0.357227 | 0.174419 | 0.245126 | -0.0953 | 0.406165 | 0.090003 | 0.161498 | 0.288114 | 0 |
| 25.16667 | 0.621143 | 0.357562 | 0.19966 | 0.137418 | 0.260267 | -0.0259 | 0.387822 | 0.135624 | 0.161232 | 0.202395 | 0 |
| 25.18056 | 0.574274 | 0.325189 | 0.077437 | 0.130945 | 0.322085 | 0.047202 | 0.297292 | 0.19364 | 0.123346 | 0.179349 | 0 |
| 25.19444 | 0.514266 | 0.298506 | 0.052044 | 0.194628 | 0.278866 | 0.108162 | 0.220576 | 0.221008 | 0.073375 | 0.193758 | 0 |
| 25.20833 | 0.451001 | 0.249188 | 0.044236 | 0.255082 | -0.01807 | 0.148652 | 0.149884 | 0.168231 | 0.073012 | 0.155013 | 0 |
| 25.22222 | 0.368564 | 0.154045 | -0.01494 | 0.255976 | -0.26886 | 0.17265 | 0.087051 | 0.068393 | 0.154021 | 0.047927 | 0 |
| 25.23611 | 0.222248 | 0.016492 | -0.10047 | 0.215745 | -0.23505 | 0.180573 | -0.03195 | -0.01776 | 0.190454 | -0.080154 | 0 |
| 25.25 | 0.032411 | -0.15487 | -0.14247 | 0.188067 | 0.02476 | 0.15847 | -0.24346 | -0.039 | 0.082461 | -0.195619 | 0 |
| 25.26389 | -0.05915 | -0.28832 | -0.09319 | 0.187812 | 0.244787 | 0.087315 | -0.34363 | 0.06393 | -0.02717 | -0.123006 | 0 |
| 25.27778 | 0.056791 | -0.30178 | 0.043942 | 0.216284 | 0.29397 | -0.06847 | -0.1964 | 0.223546 | 0.035532 | 0.26286 | 0 |
| 25.29167 | 0.321986 | -0.11459 | 0.170207 | 0.347811 | 0.169164 | -0.02526 | 0.054495 | 0.390643 | 0.198294 | 0.53818 | 0 |
| 25.30556 | 0.51167 | 0.234789 | 0.220918 | 0.680549 | -0.10087 | 0.461392 | 0.219858 | 0.619035 | 0.296445 | 0.292787 | 0 |
| 25.31944 | 0.64744 | 0.495527 | 0.281069 | 1.008654 | -0.19496 | 0.852391 | 0.300578 | 0.79578 | 0.384718 | 0.076873 | 0 |
| 25.33333 | 0.802662 | 0.47611 | 0.364898 | 1.128582 | -0.02842 | 0.661716 | 0.325392 | 0.822279 | 0.468476 | 0.296986 | 0 |
| 25.34722 | 0.952654 | 0.446968 | 0.445099 | 1.035732 | 0.145179 | 0.378087 | 0.215323 | 0.70583 | 0.414388 | 0.565414 | 0 |
| 25.36111 | 1.177279 | 0.579764 | 0.548729 | 0.900816 | 0.394442 | 0.266332 | 0.085808 | 0.567748 | 0.339838 | 0.652747 | 0 |
| 25.375 | 1.466545 | 0.858426 | 0.610344 | 0.849719 | 0.824342 | 0.210163 | 0.108891 | 0.513141 | 0.407503 | 0.680581 | 0 |
| 25.38889 | 1.262452 | 0.794426 | 0.596997 | 0.775722 | 1.127716 | 0.238964 | 0.232145 | 0.573249 | 0.608502 | 0.711834 | 0 |
| 25.40278 | 0.918052 | 0.538638 | 0.595474 | 0.704319 | 1.270296 | 0.3114 | 0.327084 | 0.580705 | 0.752974 | 0.755304 | 0 |
| 25.41667 | 0.558276 | 0.305587 | 0.60555 | 0.741548 | 1.151709 | 0.339804 | 0.391561 | 0.534322 | 0.779317 | 0.723964 | 0 |
| 25.43056 | 0.202035 | 0.080283 | 0.644699 | 0.907114 | 0.806119 | 0.325661 | 0.503614 | 0.451829 | 0.68817 | 0.61377 | 0 |
| 25.44444 | 0.051388 | -0.00683 | 0.737914 | 1.094674 | 0.441509 | 0.320853 | 0.720457 | 0.487508 | 0.564583 | 0.607262 | 0 |
| 25.45833 | 0.185796 | 0.094564 | 0.747318 | 1.012771 | 0.377509 | 0.337977 | 0.934832 | 0.663647 | 0.472472 | 0.695424 | 0 |
| 25.47222 | 0.350919 | 0.226528 | 0.601731 | 0.701367 | 0.534751 | 0.349328 | 1.004855 | 0.858212 | 0.466486 | 0.752979 | 0 |
| 25.48611 | 0.465757 | 0.599283 | 0.510929 | 0.435331 | 0.68228 | 0.340276 | 1.060639 | 1.094685 | 0.688548 | 1.026274 | 0 |
| 25.5 | 0.574863 | 1.52586 | 0.76852 | 0.391611 | 0.322023 | 0.166144 | 1.365817 | 1.419396 | 1.395158 | 1.266678 | 0 |
| 25.51389 | 0.55173 | 2.177132 | 1.140066 | 0.438769 | -2.28485 | -0.30204 | 1.605193 | 1.360126 | 1.605578 | 0.546994 | 0 |
| 25.52778 | 0.361291 | 1.66872 | 1.204751 | 0.408761 | -4.63003 | -0.40574 | 1.728947 | 1.050375 | 0.971178 | -0.094298 | 0 |
| 25.54167 | 0.304469 | 1.233528 | 0.517408 | 0.30852 | -2.45493 | -0.06689 | 1.432247 | 0.887673 | 0.324875 | 0.161087 | 0 |
| 25.55556 | 0.359822 | 0.857425 | -0.45498 | 0.038871 | -1.47499 | 0.162327 | 0.445909 | 0.876592 | -0.15364 | 0.024546 | 0 |
| 25.56944 | 0.478195 | 0.562989 | -0.97818 | -0.12977 | -0.99133 | 0.315674 | -0.5442 | 0.813797 | -0.48978 | -0.346201 | 0 |
| 25.58333 | 0.634077 | 0.409782 | -0.59209 | -0.16069 | -0.99089 | 0.381355 | -1.00949 | 0.787841 | -0.54168 | -0.455165 | 0 |
| 25.59722 | 0.715143 | 0.278656 | 0.37253 | -0.08908 | -1.07668 | 0.319634 | -0.69097 | 0.8804 | -0.23532 | -0.149976 | 0 |
| 25.61111 | 0.630601 | 0.043081 | 1.345592 | 0.058244 | -0.92762 | 0.235809 | 0.330055 | 1.28448 | 0.206889 | 0.425247 | 0 |
| 25.625 | 0.436925 | -0.18571 | 1.779455 | 0.119231 | -0.60336 | 0.236531 | 1.451139 | 1.755431 | 0.396004 | 0.792907 | 0 |
| 25.63889 | 0.253223 | -0.40508 | 1.049745 | 0.065087 | -0.38632 | 0.191758 | 1.818618 | 1.823529 | 0.3029 | 0.702626 | 0 |
| 25.65278 | 0.018212 | -0.51188 | -0.08173 | 0.057865 | -0.29375 | 0.059005 | 1.194889 | 1.555669 | 0.041945 | 0.27822 | 0 |
| 25.66667 | -0.09621 | -0.25624 | -0.97176 | 0.32875 | -0.34774 | 0.015577 | -0.19716 | 1.394238 | -0.16789 | -0.176384 | 0 |
| 25.68056 | -0.08292 | 0.293202 | -1.21308 | 0.524783 | -0.61879 | 0.113288 | -1.45663 | 1.294175 | 0.020208 | -0.425304 | 0 |
| 25.69444 | -0.03502 | 0.877658 | -0.765 | 0.364887 | -0.8718 | 0.177739 | -1.94646 | 0.877587 | 0.490197 | -0.282333 | 0 |
| 25.70833 | 0.054945 | 1.360043 | -0.06843 | 0.076432 | -0.75082 | 0.178181 | -1.40823 | 0.323237 | 0.741774 | 0.201393 | 0 |
| 25.72222 | 0.456805 | 1.564737 | 0.671173 | 0.094989 | -0.37575 | 0.239009 | -0.01848 | -0.16315 | 0.57394 | 0.792865 | 0 |
| 25.73611 | 0.833571 | 1.290497 | 0.875053 | 0.022888 | -0.12873 | 0.248398 | 1.246195 | -0.72469 | 0.183294 | 0.911533 | 0 |
| 25.75 | 0.790052 | 0.732157 | 0.23513 | -0.36105 | -0.05452 | 0.105788 | 1.26891 | -1.3546 | -0.23166 | 0.364819 | 0 |
| 25.76389 | 0.518575 | 0.261013 | -0.68503 | -0.53511 | 0.008063 | -0.02302 | 0.470017 | -1.67802 | -0.36503 | -0.411451 | 0 |
| 25.77778 | 0.100971 | -0.16586 | -1.45778 | -0.48842 | 0.078742 | -0.12826 | -0.48775 | -1.68553 | -0.34109 | -0.944029 | 0 |
| 25.79167 | -0.22231 | -0.58312 | -1.38316 | -0.4459 | 0.089719 | -0.32156 | -1.20037 | -1.40343 | -0.33818 | -1.134055 | 0 |
| 25.80556 | -0.16454 | -0.8712 | -0.7816 | -0.48357 | 0.024315 | -0.55541 | -1.45747 | -1.00239 | -0.30446 | -0.990991 | 0 |
| 25.81944 | -0.04178 | -1.12176 | -0.28573 | -0.55949 | -0.13744 | -0.68551 | -1.49368 | -0.6978 | -0.35167 | -0.933957 | 0 |
| 25.83333 | -0.17293 | -1.38258 | -0.37737 | -0.47221 | -0.34633 | -0.75471 | -1.4996 | -0.60959 | -0.44365 | -1.135925 | 0 |
| 25.84722 | -0.4854 | -1.33883 | -0.70023 | -0.26241 | -0.46234 | -0.83585 | -1.28098 | -0.83724 | -0.43975 | -1.359501 | 0 |
| 25.86111 | -1.05883 | -1.48013 | -0.91705 | -0.30921 | -0.53055 | -1.01299 | -1.07598 | -1.47688 | -0.49484 | -1.650178 | 0 |
| 25.875 | -2.01902 | -1.98131 | -1.02652 | -0.63661 | -0.69336 | -1.1078 | -1.1436 | -2.1094 | -0.55573 | -1.928642 | 0 |

| | | | | | | | | | | | |
|---|---|---|---|---|---|---|---|---|---|---|---|
| 25.88889 | -2.15381 | -1.86531 | -0.86324 | -0.98705 | -0.76521 | -0.94914 | -1.24678 | -2.31073 | -0.59638 | -2.047206 | 0 |
| 25.90278 | -1.59714 | -1.32675 | -0.68192 | -1.12096 | -0.75467 | -0.59364 | -0.93613 | -2.33251 | -0.7898 | -1.89384 | 0 |
| 25.91667 | -0.66368 | -0.8272 | -0.41422 | -1.06193 | -0.7403 | -0.31524 | -0.62069 | -2.16407 | -1.00138 | -1.442159 | 0 |
| 25.93056 | 0.119893 | -0.46425 | -0.17651 | -0.70787 | -0.73163 | -0.26734 | -0.50291 | -1.6056 | -1.06764 | -0.969278 | 0 |
| 25.94444 | 0.318737 | -0.37598 | -0.09202 | -0.52104 | -0.72429 | -0.431 | -0.72447 | -1.10825 | -0.98373 | -0.772171 | 0 |
| 25.95833 | 0.045375 | -0.42831 | -0.13183 | -0.41421 | -0.5751 | -0.56886 | -0.88883 | -0.66435 | -0.61445 | -0.745757 | 0 |
| 25.97222 | -0.40827 | -0.47122 | -0.24637 | -0.36104 | -0.38621 | -0.52752 | -0.70791 | -0.449 | -0.26528 | -0.78468 | 0 |
| 25.98611 | -0.81803 | -0.61832 | -0.31268 | -0.3342 | -0.20209 | -0.31883 | -0.28797 | -0.35531 | -0.14766 | -0.896626 | 0 |
| 26 | -1.08871 | -0.8621 | -0.17243 | -0.2711 | 0.120416 | 0.072316 | 0.095025 | -0.2851 | -0.06678 | -1.005784 | 0 |
| 26.01389 | -1.52391 | -1.08643 | 0.063716 | -0.39981 | 0.387097 | 0.228255 | 0.10521 | -0.46603 | 0.009764 | -1.145488 | 0 |
| 26.02778 | -2.06445 | -1.19594 | 0.09712 | -0.55214 | 0.356491 | -0.07745 | -0.16193 | -0.83719 | -0.12612 | -1.161832 | 0 |
| 26.04167 | -2.36306 | -1.18942 | 0.079629 | -0.23672 | 0.237128 | -0.3524 | -0.2358 | -0.91923 | -0.31672 | -0.964501 | 0 |
| 26.05556 | -1.93602 | -0.95956 | 0.078521 | 0.056606 | 0.150411 | -0.28249 | -0.13324 | -0.72352 | -0.39581 | -0.744729 | 0 |
| 26.06944 | -1.14699 | -0.65174 | 0.011085 | -0.11324 | 0.020747 | -0.13709 | -0.09012 | -0.55098 | -0.40249 | -0.525777 | 0 |
| 26.08333 | -0.37925 | -0.38477 | -0.02887 | -0.38008 | -0.18865 | -0.09913 | -0.08925 | -0.36235 | -0.37905 | -0.384577 | 0 |
| 26.09722 | 0.227925 | -0.12541 | 0.047658 | -0.36868 | -0.29473 | -0.12407 | -0.01084 | -0.09036 | -0.29497 | -0.253625 | 0 |
| 26.11111 | 0.654666 | 0.146218 | 0.221159 | -0.1601 | -0.30524 | -0.13475 | 0.223156 | 0.109433 | -0.20315 | -0.084667 | 0 |
| 26.125 | 0.839648 | 0.372308 | 0.381745 | 0.115307 | -0.23158 | -0.12674 | 0.532786 | 0.249519 | -0.09078 | 0.193328 | 0 |
| 26.13889 | 0.877325 | 0.501232 | 0.483324 | 0.404148 | -0.04578 | -0.10425 | 0.793049 | 0.386306 | 0.055749 | 0.500196 | 0 |
| 26.15278 | 0.940313 | 0.589677 | 0.602447 | 0.533354 | 0.191283 | -0.00364 | 0.882432 | 0.481956 | 0.236182 | 0.690208 | 0 |
| 26.16667 | 1.015259 | 0.684616 | 0.677935 | 0.381154 | 0.395552 | 0.154957 | 0.805359 | 0.480095 | 0.404773 | 0.760852 | 0 |
| 26.18056 | 1.132921 | 0.786666 | 0.682317 | 0.14214 | 0.499695 | 0.228871 | 0.596194 | 0.396448 | 0.470247 | 0.72857 | 0 |
| 26.19444 | 1.237608 | 0.840371 | 0.557409 | -0.00836 | 0.53624 | 0.17176 | 0.374469 | 0.312126 | 0.41189 | 0.609099 | 0 |
| 26.20833 | 1.253485 | 0.870486 | 0.375233 | 0.053381 | 0.523182 | 0.097063 | 0.134435 | 0.281388 | 0.288612 | 0.390548 | 0 |
| 26.22222 | 1.175749 | 0.789781 | 0.224913 | 0.263652 | 0.430571 | 0.114839 | -0.07502 | 0.227399 | 0.151715 | 0.175858 | 0 |
| 26.23611 | 1.004209 | 0.506086 | 0.114481 | 0.34047 | 0.247637 | 0.188314 | -0.14242 | 0.067918 | 0.00562 | -0.023722 | 0 |
| 26.25 | 0.745308 | 0.216533 | 0.012402 | 0.162266 | 0.076589 | 0.236046 | -0.11581 | -0.07111 | -0.09808 | -0.174707 | 0 |
| 26.26389 | 0.570833 | 0.142211 | -0.02367 | -0.01954 | 0.07603 | 0.270685 | -0.02743 | -0.04982 | -0.0825 | -0.161894 | 0 |
| 26.27778 | 0.510536 | 0.166789 | 0.006043 | 0.117861 | 0.197925 | 0.281492 | 0.045281 | 0.105506 | -0.00058 | -0.071116 | 0 |
| 26.29167 | 0.456277 | 0.136171 | 0.026795 | 0.533 | 0.264732 | 0.275905 | 0.079904 | 0.291264 | 0.068806 | 0.016304 | 0 |
| 26.30556 | 0.393963 | 0.121976 | 0.075712 | 0.894368 | 0.30081 | 0.37444 | 0.105851 | 0.423638 | 0.135373 | 0.133964 | 0 |
| 26.31944 | 0.36252 | 0.187954 | 0.207864 | 0.853304 | 0.283278 | 0.587637 | 0.151618 | 0.580555 | 0.211711 | 0.263024 | 0 |
| 26.33333 | 0.37921 | 0.273237 | 0.371412 | 0.620975 | 0.218635 | 0.781841 | 0.184025 | 0.857279 | 0.274349 | 0.34201 | 0 |
| 26.34722 | 0.546247 | 0.362357 | 0.536216 | 0.557964 | 0.162855 | 0.806034 | 0.236613 | 1.176387 | 0.336082 | 0.459564 | 0 |
| 26.36111 | 0.81858 | 0.495836 | 0.626812 | 0.739242 | 0.14874 | 0.861853 | 0.358756 | 1.350032 | 0.393117 | 0.66098 | 0 |
| 26.375 | 1.105066 | 0.645893 | 0.618567 | 0.917312 | 0.172691 | 0.914858 | 0.520293 | 1.397364 | 0.430314 | 0.8733 | 0 |
| 26.38889 | 1.28928 | 0.724682 | 0.559516 | 0.917571 | 0.144775 | 0.910574 | 0.601081 | 1.405316 | 0.407759 | 1.021072 | 0 |
| 26.40278 | 1.231564 | 0.706873 | 0.553039 | 0.816846 | 0.039635 | 0.831878 | 0.547185 | 1.296912 | 0.356037 | 1.027423 | 0 |
| 26.41667 | 1.029685 | 0.649057 | 0.642657 | 0.680007 | -0.0164 | 0.646299 | 0.339808 | 1.112154 | 0.30602 | 0.899713 | 0 |
| 26.43056 | 0.793391 | 0.566045 | 0.819101 | 0.637151 | -0.00058 | 0.495646 | 0.224962 | 0.874103 | 0.400258 | 0.833623 | 0 |
| 26.44444 | 0.674357 | 0.432802 | 0.879243 | 0.661072 | 0.039671 | 0.382534 | 0.307005 | 0.730312 | 0.651003 | 0.85618 | 0 |
| 26.45833 | 0.539506 | 0.256415 | 0.637123 | 0.634121 | 0.103827 | 0.278052 | 0.429065 | 0.488705 | 0.757099 | 0.734607 | 0 |
| 26.47222 | 0.334148 | 0.022124 | 0.198231 | 0.516584 | 0.283124 | 0.134244 | 0.498525 | 0.208063 | 0.662417 | 0.550468 | 0 |
| 26.48611 | 0.105473 | -0.28439 | 0.044318 | 0.535306 | 0.469774 | -0.14079 | 0.815321 | 0.084856 | 0.68775 | 0.749983 | 0 |
| 26.5 | -0.06463 | -0.61601 | 0.600446 | 0.638898 | 1.207062 | -0.39546 | 1.319524 | 0.240477 | 1.244076 | 1.431129 | 0 |
| 26.51389 | -0.0704 | -0.44373 | 1.529096 | 0.553423 | 1.047573 | -0.29828 | 1.730918 | 0.719488 | 1.813804 | 1.816415 | 0 |
| 26.52778 | 0.276553 | 0.626069 | 2.067539 | 0.620981 | 0.218673 | 0.093044 | 1.548873 | 1.1626 | 1.611272 | 1.704973 | 0 |
| 26.54167 | 0.639572 | 1.525046 | 1.49739 | 0.592076 | -0.03661 | 0.226852 | 0.917028 | 1.271489 | 1.179514 | 1.462607 | 0 |
| 26.55556 | 0.643397 | 1.339206 | 0.277686 | 0.448836 | -0.11839 | 0.150793 | 0.064271 | 1.064726 | 0.633677 | 0.860053 | 0 |
| 26.56944 | 0.487697 | 0.697654 | -0.75744 | 0.317367 | -0.17197 | -0.07162 | -0.62249 | 0.814561 | 0.030401 | 0.271627 | 0 |
| 26.58333 | 0.373714 | 0.640604 | -1.22654 | 0.27036 | -0.26184 | -0.24305 | -0.91282 | 0.851884 | -0.47599 | 0.096018 | 0 |
| 26.59722 | 0.595239 | 1.017214 | -0.95908 | 0.327605 | -0.30466 | -0.11643 | -0.58196 | 1.320871 | -0.57664 | 0.380651 | 0 |
| 26.61111 | 0.851933 | 1.518005 | 0.085403 | 0.364111 | -0.31944 | 0.328571 | 0.501805 | 1.851434 | -0.17454 | 0.986086 | 0 |
| 26.625 | 1.164446 | 1.668559 | 1.575245 | 0.355782 | -0.24398 | 0.772149 | 1.56066 | 2.423247 | 0.372329 | 1.613779 | 0 |
| 26.63889 | 1.31211 | 1.399559 | 2.731779 | 0.372396 | -0.04741 | 0.924054 | 1.758814 | 2.675568 | 0.86769 | 1.83355 | 0 |
| 26.65278 | 1.006684 | 0.505381 | 2.394605 | 0.390252 | -0.14069 | 0.783736 | 1.252739 | 2.051977 | 0.744428 | 1.317111 | 0 |
| 26.66667 | 0.432076 | -0.59036 | 0.937587 | 0.421946 | -0.22601 | 0.462989 | 0.320898 | 0.994542 | 0.163478 | 0.332038 | 0 |

| | | | | | | | | | | | |
|---|---|---|---|---|---|---|---|---|---|---|---|
| 26.68056 | -0.01525 | -1.34167 | -0.66438 | 0.602832 | -0.15452 | 0.174847 | -0.56577 | -0.10989 | -0.01085 | -0.624116 | 0 |
| 26.69444 | -0.45295 | -1.35659 | -1.74694 | 0.527982 | 0.496996 | -0.06125 | -1.82783 | -1.14017 | 0.047512 | -1.311532 | 0 |
| 26.70833 | -0.72981 | -0.87184 | -2.27158 | 0.528954 | 1.711509 | 0.025661 | -2.66269 | -1.35887 | 0.057034 | -1.617102 | 0 |
| 26.72222 | -0.86273 | 0.465437 | -3.12645 | 1.740315 | 1.242413 | 0.703595 | -0.91544 | -1.20685 | -0.89358 | -2.227811 | 0 |
| 26.73611 | -1.39967 | 2.123333 | -5.40473 | 3.813266 | 0.655664 | -0.01416 | 1.304045 | -2.07308 | -3.06363 | -3.458413 | 0 |
| 26.75 | NaN | NaN | NaN | NaN | NaN | NaN | NaN | NaN | NaN | NaN | 0 |
| 26.76389 | NaN | NaN | NaN | NaN | NaN | NaN | NaN | NaN | NaN | NaN | 0 |
| 26.77778 | NaN | NaN | NaN | NaN | NaN | NaN | NaN | NaN | NaN | NaN | 0 |
| 26.79167 | 1.550495 | -1.51844 | 4.342729 | -3.3047 | 0.785358 | 0.224181 | -2.64306 | 2.606838 | 2.678288 | 1.599619 | 0 |
| 26.80556 | 0.703733 | -0.25882 | 1.858359 | -1.33187 | 0.296094 | -0.45255 | -1.49587 | 1.469213 | 1.154355 | 1.911017 | 0 |
| 26.81944 | 0.798988 | 0.80386 | 0.786572 | -0.55366 | -0.1807 | -0.23268 | -1.80624 | 0.400797 | 1.261113 | 0.441451 | 0 |
| 26.83333 | 0.859118 | 0.641029 | 0.099939 | -0.80291 | -0.30933 | -0.28551 | -2.5611 | -0.77719 | 0.987896 | -0.067298 | 0 |
| 26.84722 | 0.570138 | 0.235545 | -0.65852 | -1.01729 | -0.26701 | -0.36904 | -1.51585 | -1.85646 | 0.07028 | -0.740845 | 0 |
| 26.86111 | 0.043015 | -0.15338 | -1.24812 | -1.00009 | -0.44404 | -0.61094 | -0.3714 | -2.4162 | -0.55564 | -0.906894 | 0 |
| 26.875 | -0.59345 | -0.69874 | -1.50519 | -0.80132 | -0.66508 | -0.62149 | 0.0412 | -2.33113 | -1.06621 | -1.005675 | 0 |
| 26.88889 | -1.07822 | -1.5659 | -1.52281 | -0.70557 | -0.87811 | -0.44962 | 0.098922 | -2.0605 | -1.3646 | -1.096384 | 0 |
| 26.90278 | -1.11324 | -2.41918 | -1.33171 | -0.8335 | -0.9888 | -0.50649 | -0.0383 | -2.29262 | -1.51613 | -1.24891 | 0 |
| 26.91667 | -0.80477 | -2.82354 | -0.92678 | -1.09569 | -0.79703 | -0.79225 | -0.22788 | -2.45599 | -1.45934 | -1.525489 | 0 |
| 26.93056 | -0.53809 | -2.66079 | -0.51482 | -1.34285 | -0.50329 | -1.01968 | -0.46304 | -2.55259 | -1.42138 | -1.872543 | 0 |
| 26.94444 | -0.61625 | -2.15971 | -0.20868 | -1.67164 | -0.2083 | -1.02696 | -0.59482 | -2.58234 | -1.32117 | -2.028179 | 0 |
| 26.95833 | -0.80797 | -1.44593 | -0.0479 | -1.55888 | 0.151546 | -0.77976 | -0.46415 | -2.11193 | -0.97593 | -1.677433 | 0 |
| 26.97222 | -1.00186 | -0.78856 | -0.08864 | -1.14082 | 0.358098 | -0.57311 | -0.30018 | -1.38437 | -0.60147 | -1.092878 | 0 |
| 26.98611 | -1.14265 | -0.61919 | -0.25199 | -0.81372 | 0.303868 | -0.44247 | -0.17199 | -0.87076 | -0.3748 | -0.742051 | 0 |
| 27 | -1.15302 | -1.07634 | -0.41309 | -0.58616 | 0.098364 | -0.41418 | 0.081546 | -0.53093 | -0.2661 | -0.610147 | 0 |
| 27.01389 | -1.25707 | -2.02377 | -0.54418 | -0.49446 | 0.003199 | -0.64538 | 0.348426 | -0.37356 | -0.26671 | -0.654134 | 0 |
| 27.02778 | -1.46336 | -2.24472 | -0.16032 | -0.37972 | 0.296963 | -0.43878 | 0.555785 | -0.4752 | -0.09858 | -0.470819 | 0 |
| 27.04167 | -1.68221 | -0.76642 | 0.488148 | -0.24129 | 0.646287 | 0.089385 | 0.674166 | -0.53073 | 0.373054 | -0.039827 | 0 |
| 27.05556 | -1.71771 | 0.402182 | 0.952504 | -0.21908 | 0.661419 | 0.488193 | 0.593062 | -0.37862 | 0.635848 | 0.194016 | 0 |
| 27.06944 | -1.2617 | 0.514125 | 0.820212 | -0.1798 | 0.505148 | 0.633489 | 0.374478 | -0.06434 | 0.445801 | 0.097827 | 0 |
| 27.08333 | -0.62159 | 0.615185 | 0.475376 | -0.11318 | 0.433021 | 0.722679 | 0.22225 | 0.265201 | 0.169035 | -0.086474 | 0 |
| 27.09722 | -0.15516 | 0.346895 | 0.335009 | 0.047255 | 0.405354 | 0.754953 | 0.238798 | 0.384743 | 0.093854 | -0.144677 | 0 |
| 27.11111 | 0.082229 | 0.181724 | 0.415318 | 0.185521 | 0.43384 | 0.68773 | 0.320804 | 0.336691 | 0.235259 | 0.103249 | 0 |
| 27.125 | 0.174042 | 0.206692 | 0.500634 | 0.195972 | 0.409726 | 0.47987 | 0.321597 | 0.314134 | 0.323273 | 0.320216 | 0 |
| 27.13889 | 0.188539 | 0.322817 | 0.461728 | 0.114054 | 0.299226 | 0.27307 | 0.240617 | 0.318992 | 0.136894 | 0.289956 | 0 |
| 27.15278 | 0.178308 | 0.435517 | 0.377808 | -0.03409 | 0.221942 | 0.149345 | 0.15279 | 0.260623 | -0.09484 | 0.168429 | 0 |
| 27.16667 | 0.166898 | 0.492105 | 0.350449 | -0.19885 | 0.182896 | 0.190777 | 0.158391 | 0.137785 | -0.18769 | 0.012583 | 0 |
| 27.18056 | 0.168629 | 0.460701 | 0.346463 | -0.24725 | 0.161053 | 0.421055 | 0.250461 | -0.02425 | -0.13378 | -0.121729 | 0 |
| 27.19444 | 0.216318 | 0.350924 | 0.291914 | -0.16073 | 0.14276 | 0.568173 | 0.312314 | -0.18606 | -0.02422 | -0.165413 | 0 |
| 27.20833 | 0.256425 | 0.274502 | 0.204917 | -0.10936 | 0.10399 | 0.480529 | 0.247816 | -0.26449 | -0.00952 | -0.173934 | 0 |
| 27.22222 | 0.249802 | 0.203605 | 0.128864 | -0.18513 | 0.070918 | 0.239031 | 0.097027 | -0.14373 | -0.12131 | -0.223145 | 0 |
| 27.23611 | 0.202129 | 0.064232 | 0.086631 | -0.29658 | 0.033953 | 0.020442 | -0.0491 | 0.081784 | -0.23836 | -0.252653 | 0 |
| 27.25 | 0.172932 | -0.11664 | 0.066287 | -0.3291 | 0.073792 | -0.1036 | -0.13064 | 0.139681 | -0.30429 | -0.220751 | 0 |
| 27.26389 | 0.158728 | -0.22971 | 0.064819 | -0.26571 | 0.148464 | -0.03142 | -0.14797 | 0.035241 | -0.29606 | -0.136862 | 0 |
| 27.27778 | 0.139285 | -0.16036 | 0.061559 | -0.12658 | 0.144033 | 0.214613 | -0.08624 | -0.02555 | -0.25072 | -0.145555 | 0 |
| 27.29167 | 0.14497 | -0.02056 | 0.030524 | 0.047897 | 0.111668 | 0.380269 | -0.01441 | 0.000967 | -0.20867 | -0.204222 | 0 |
| 27.30556 | 0.150729 | 0.016252 | 0.035957 | 0.149253 | 0.127616 | 0.340577 | -0.00251 | 0.019903 | -0.17685 | -0.091048 | 0 |
| 27.31944 | 0.141285 | -0.06748 | 0.146836 | 0.163262 | 0.19876 | 0.235722 | -0.02739 | 0.087548 | -0.12479 | 0.174845 | 0 |
| 27.33333 | 0.135504 | -0.13121 | 0.352563 | 0.272637 | 0.252914 | 0.3241 | 0.024067 | 0.303668 | -0.05901 | 0.350145 | 0 |
| 27.34722 | 0.112468 | 0.022166 | 0.5465 | 0.603524 | 0.198813 | 0.661694 | 0.125048 | 0.609425 | 0.032924 | 0.183694 | 0 |
| 27.36111 | 0.11888 | 0.261737 | 0.638737 | 0.873158 | 0.101022 | 0.981562 | 0.193576 | 0.755407 | 0.175879 | 0.035889 | 0 |
| 27.375 | 0.251263 | 0.346528 | 0.652714 | 0.898553 | 0.121665 | 1.109065 | 0.209789 | 0.668393 | 0.331513 | 0.169027 | 0 |
| 27.38889 | 0.486058 | 0.341032 | 0.670125 | 0.867189 | 0.132677 | 1.192372 | 0.213464 | 0.60444 | 0.400357 | 0.264063 | 0 |
| 27.40278 | 0.692214 | 0.376174 | 0.789981 | 0.839496 | 0.096733 | 1.191578 | 0.206676 | 0.615295 | 0.370131 | 0.305914 | 0 |
| 27.41667 | 0.901781 | 0.458538 | 0.92185 | 0.689965 | 0.037135 | 0.91468 | 0.136307 | 0.585852 | 0.30209 | 0.444526 | 0 |
| 27.43056 | 1.095452 | 0.670785 | 0.996245 | 0.529848 | -0.04072 | 0.603455 | 0.079671 | 0.610022 | 0.262402 | 0.607886 | 0 |
| 27.44444 | 1.284841 | 0.892934 | 0.949942 | 0.527453 | -0.1299 | 0.597689 | 0.226018 | 0.746012 | 0.334739 | 0.693814 | 0 |
| 27.45833 | 1.286569 | 0.948265 | 0.716711 | 0.614002 | -0.12681 | 0.776551 | 0.534744 | 0.868673 | 0.501226 | 0.684325 | 0 |

| | | | | | | | | | | | |
|---|---|---|---|---|---|---|---|---|---|---|---|
| 27.47222 | 1.097195 | 0.666458 | 0.28251 | 0.668134 | 0.040587 | 0.900969 | 0.72126 | 0.78223 | 0.60266 | 0.499485 | 0 |
| 27.48611 | 0.904308 | 0.288781 | -0.04066 | 0.977418 | 0.411246 | 0.841003 | 0.778985 | 0.692915 | 0.706434 | 0.603825 | 0 |
| 27.5 | 1.031379 | 0.151887 | 0.152265 | 1.488065 | 1.443598 | 0.732032 | 1.039478 | 0.740753 | 1.160803 | 1.335129 | 0 |
| 27.51389 | 1.30919 | 0.399862 | 0.954779 | 1.386509 | 1.860468 | 0.611208 | 1.338612 | 0.615704 | 1.903028 | 2.053001 | 0 |
| 27.52778 | 1.402971 | 0.92516 | 1.860197 | 0.949433 | 0.912851 | 0.377153 | 1.623359 | 0.524497 | 2.49456 | 2.402837 | 0 |
| 27.54167 | 1.108387 | 1.280232 | 1.628411 | 0.603071 | 0.598531 | -0.01618 | 1.754937 | 0.473227 | 2.268211 | 2.019796 | 0 |
| 27.55556 | 0.455197 | 1.165189 | 0.576552 | 0.233699 | 0.538766 | -0.3168 | 1.275174 | 0.148211 | 1.317184 | 1.085336 | 0 |
| 27.56944 | -0.07262 | 0.722789 | -0.5156 | 0.105707 | 0.561478 | -0.2731 | 0.573323 | -0.02174 | 0.456469 | 0.202196 | 0 |
| 27.58333 | -0.23593 | 0.325704 | -1.19887 | 0.165023 | 0.455369 | -0.14628 | -0.11826 | 0.208995 | -0.00707 | -0.181206 | 0 |
| 27.59722 | -0.12057 | 0.408096 | -1.2192 | 0.265576 | 0.224303 | -0.0821 | -0.72499 | 0.61222 | -0.07526 | -0.018686 | 0 |
| 27.61111 | 0.25909 | 0.803558 | -0.41821 | 0.450797 | 0.026718 | 0.351548 | -0.84214 | 1.063336 | 0.053854 | 0.415575 | 0 |
| 27.625 | 0.570333 | 1.216473 | 0.824568 | 0.517211 | -0.01118 | 0.820706 | -0.30473 | 1.477737 | 0.211141 | 0.827382 | 0 |
| 27.63889 | 0.584281 | 1.321825 | 1.760783 | 0.222951 | -0.05122 | 1.121406 | 0.412363 | 1.693521 | 0.402459 | 0.977371 | 0 |
| 27.65278 | 0.394035 | 1.343365 | 1.950441 | -0.091 | -0.20379 | 1.133746 | 1.015281 | 1.893051 | 0.470858 | 0.926515 | 0 |
| 27.66667 | 0.548766 | 0.887062 | 1.369189 | -0.08892 | -0.38635 | 0.714012 | 1.513124 | 1.919483 | 0.461155 | 0.850812 | 0 |
| 27.68056 | 1.141966 | 0.525473 | 0.215621 | 0.254926 | -0.4802 | 0.063842 | 0.954176 | 1.483171 | 0.095388 | 0.551224 | 0 |
| 27.69444 | 1.70668 | 0.071707 | -0.78856 | 0.500573 | -0.28319 | -0.52166 | 0.105088 | 1.015502 | -0.24101 | 0.030577 | 0 |
| 27.70833 | 1.226286 | -0.77326 | -1.14958 | 0.548145 | 0.092421 | -1.02481 | -0.88513 | 0.473386 | -0.05741 | -0.391689 | 0 |
| 27.72222 | 0.419384 | -1.15239 | -1.13748 | 0.397317 | 0.45056 | -1.05698 | -1.51125 | -0.08561 | 0.345723 | -0.414002 | 0 |
| 27.73611 | -0.46719 | -0.87468 | -0.82349 | 0.125294 | 0.737778 | -0.28524 | -1.7841 | -0.66089 | 0.508022 | -0.272808 | 0 |
| 27.75 | -1.01629 | -0.33374 | -0.63715 | -0.33184 | 0.59609 | 0.222165 | -1.50471 | -1.30241 | 0.255836 | -0.521474 | 0 |
| 27.76389 | -1.44606 | -0.16157 | -0.69173 | -0.72945 | 0.35829 | 0.202848 | -1.06958 | -1.65014 | -0.20073 | -0.809797 | 0 |
| 27.77778 | -1.57447 | -0.34961 | -0.89246 | -0.59037 | 0.276803 | -0.10454 | -0.58336 | -1.29306 | -0.60078 | -0.96293 | 0 |
| 27.79167 | -1.73832 | -0.91534 | -1.10663 | -0.24273 | 0.167081 | -0.34679 | -0.29677 | -0.90661 | -1.06676 | -1.099228 | 0 |
| 27.80556 | -1.71769 | -1.38763 | -0.83503 | -0.09735 | -0.06225 | -0.49385 | -0.06585 | -0.83856 | -1.40785 | -1.275726 | 0 |
| 27.81944 | -1.2177 | -1.18197 | -0.21086 | -0.2943 | -0.30631 | -0.72672 | 0.265545 | -0.84144 | -1.21112 | -1.179417 | 0 |
| 27.83333 | -0.57576 | -0.74949 | 0.25074 | -0.80956 | -0.56654 | -1.15207 | 0.477439 | -0.92419 | -0.80514 | -0.942228 | 0 |
| 27.84722 | -0.1954 | -0.43389 | 0.143665 | -1.29103 | -0.71208 | -1.50176 | 0.264266 | -1.17478 | -0.5698 | -0.91099 | 0 |
| 27.86111 | -0.07252 | -0.32419 | -0.43612 | -1.30636 | -0.80381 | -1.62766 | -0.28797 | -1.53843 | -0.70418 | -1.102615 | 0 |
| 27.875 | -0.37359 | -0.50439 | -1.10825 | -0.74056 | -0.82986 | -1.49934 | -0.75039 | -1.75623 | -1.16523 | -1.252673 | 0 |
| 27.88889 | -0.37909 | -0.97443 | -1.54379 | -0.2267 | -0.66625 | -1.31058 | -0.80772 | -1.51394 | -1.49842 | -1.335745 | 0 |
| 27.90278 | -0.15697 | -1.55072 | -1.46101 | -0.22822 | -0.53931 | -1.23204 | -0.58476 | -0.95037 | -1.35485 | -1.344033 | 0 |
| 27.91667 | 0.192184 | -2.06351 | -0.91311 | -0.80576 | -0.44808 | -1.09881 | -0.42038 | -0.46761 | -1.07467 | -1.140055 | 0 |
| 27.93056 | 0.457808 | -1.78181 | -0.3504 | -1.40777 | -0.20655 | -0.72874 | -0.31705 | -0.35049 | -0.66008 | -0.755532 | 0 |
| 27.94444 | 0.326351 | -1.09488 | -0.059 | -1.85421 | -0.00524 | -0.32124 | -0.26263 | -0.56466 | -0.2915 | -0.416965 | 0 |
| 27.95833 | -0.06963 | -0.59548 | -0.08498 | -1.54733 | 0.070139 | -0.11633 | -0.22923 | -0.81186 | -0.09381 | -0.266345 | 0 |
| 27.97222 | -0.29532 | -0.39723 | -0.19779 | -0.84267 | 0.097908 | -0.01293 | -0.19779 | -0.78489 | -0.14179 | -0.246307 | 0 |
| 27.98611 | -0.47757 | -0.45882 | -0.27472 | -0.32555 | 0.104166 | -0.00073 | -0.10841 | -0.53899 | -0.27691 | -0.264171 | 0 |
| 28 | -0.67599 | -1.13559 | -0.41767 | -0.25207 | 0.125471 | -0.22794 | 0.104341 | -0.10032 | -0.4897 | -0.308583 | 0 |
| 28.01389 | -1.03644 | -2.54196 | -0.78699 | -0.55384 | 0.070409 | -0.66253 | 0.20159 | 0.093367 | -0.91615 | -0.503564 | 0 |
| 28.02778 | -1.62775 | -3.42004 | -0.77106 | -0.56627 | 0.002489 | -0.87525 | 0.170211 | -0.37922 | -1.05715 | -0.744886 | 0 |
| 28.04167 | -2.06713 | -2.23618 | -0.03075 | -0.1985 | 0.00681 | -0.71195 | 0.17184 | -1.06634 | -0.52361 | -0.748351 | 0 |
| 28.05556 | -2.02304 | -0.40147 | 0.595921 | -0.11039 | 0.183888 | -0.27079 | 0.185189 | -1.47861 | 0.293832 | -0.389757 | 0 |
| 28.06944 | -1.61694 | 0.89733 | 0.816178 | -0.52713 | 0.428734 | 0.376826 | 0.110425 | -1.29889 | 0.563653 | -0.191066 | 0 |
| 28.08333 | -1.1219 | 1.28964 | 0.543371 | -0.60928 | 0.616366 | 0.996573 | -0.00118 | -0.6859 | 0.371233 | -0.13045 | 0 |
| 28.09722 | -0.54564 | 1.462477 | 0.417323 | -0.05183 | 0.768764 | 1.527493 | 0.043176 | -0.01121 | 0.316452 | 0.216989 | 0 |
| 28.11111 | -0.16606 | 1.657853 | 0.640072 | 0.624073 | 0.86621 | 1.992194 | 0.194094 | 0.466545 | 0.511862 | 0.729927 | 0 |
| 28.125 | 0.05173 | 1.672413 | 0.936529 | 0.804499 | 0.858574 | 2.168236 | 0.286534 | 0.568373 | 0.789167 | 1.073052 | 0 |
| 28.13889 | 0.095103 | 1.292966 | 0.986328 | 0.677362 | 0.705627 | 2.120947 | 0.308196 | 0.479011 | 0.94048 | 1.067213 | 0 |
| 28.15278 | 0.147964 | 0.926073 | 0.727545 | 0.443643 | 0.437361 | 1.912695 | 0.248975 | 0.432694 | 0.910252 | 0.760053 | 0 |
| 28.16667 | 0.317732 | 0.588165 | 0.312174 | 0.100626 | 0.28791 | 1.059555 | 0.140262 | 0.397782 | 0.669775 | 0.474011 | 0 |
| 28.18056 | 0.49014 | 0.290883 | 0.069858 | -0.24211 | 0.1606 | 0.495621 | 0.01963 | 0.28167 | 0.313148 | 0.211638 | 0 |
| 28.19444 | 0.549164 | 0.14452 | 0.013884 | -0.23719 | 0.076223 | 0.346254 | -0.00735 | 0.238844 | -0.00856 | 0.123235 | 0 |
| 28.20833 | 0.52424 | 0.22255 | 0.142093 | 0.078058 | 0.093229 | 0.490779 | 0.084581 | 0.180595 | -0.09937 | 0.240461 | 0 |
| 28.22222 | 0.51311 | 0.357015 | 0.481664 | 0.210065 | 0.212711 | 0.698602 | 0.203158 | 0.094815 | 0.072917 | 0.394038 | 0 |
| 28.23611 | 0.517365 | 0.410958 | 0.777973 | -0.01761 | 0.338963 | 0.757711 | 0.273988 | 0.057383 | 0.289469 | 0.367713 | 0 |
| 28.25 | 0.486928 | 0.338267 | 0.705494 | -0.16355 | 0.395727 | 0.699309 | 0.278732 | 0.090486 | 0.346838 | 0.260709 | 0 |

| | | | | | | | | | | | |
|---|---|---|---|---|---|---|---|---|---|---|---|
| 28.26389 | 0.414151 | 0.208189 | 0.391419 | 0.134631 | 0.347112 | 0.540898 | 0.23641 | 0.132699 | 0.237462 | 0.179086 | 0 |
| 28.27778 | 0.361452 | 0.112742 | 0.131655 | 0.537481 | 0.232515 | 0.325515 | 0.165665 | 0.109058 | 0.109604 | 0.119245 | 0 |
| 28.29167 | 0.415873 | 0.091502 | 0.081591 | 0.604126 | 0.110923 | 0.306556 | 0.119613 | 0.073025 | 0.081202 | 0.085147 | 0 |
| 28.30556 | 0.5009 | 0.121959 | 0.262515 | 0.298615 | 0.03267 | 0.528519 | 0.087067 | 0.098515 | 0.151433 | 0.092879 | 0 |
| 28.31944 | 0.501019 | 0.099972 | 0.527265 | 0.035056 | -0.00044 | 0.817039 | 0.02658 | 0.170162 | 0.213427 | 0.131547 | 0 |
| 28.33333 | 0.467529 | 0.053048 | 0.634534 | 0.132797 | -0.03618 | 0.935419 | -0.06387 | 0.317308 | 0.197072 | 0.170224 | 0 |
| 28.34722 | 0.46099 | 0.048378 | 0.538307 | 0.496102 | -0.02334 | 0.801537 | -0.06214 | 0.591301 | 0.158615 | 0.228058 | 0 |
| 28.36111 | 0.47797 | 0.097516 | 0.426335 | 0.784404 | 0.033676 | 0.64762 | 0.069774 | 0.897555 | 0.182766 | 0.373498 | 0 |
| 28.375 | 0.467706 | 0.162489 | 0.486828 | 0.841026 | 0.111377 | 0.591614 | 0.205494 | 1.098558 | 0.303172 | 0.545833 | 0 |
| 28.38889 | 0.466136 | 0.269275 | 0.778943 | 0.798769 | 0.179356 | 0.630315 | 0.236094 | 1.111938 | 0.535088 | 0.659963 | 0 |
| 28.40278 | 0.595325 | 0.4627 | 1.169532 | 0.757395 | 0.173354 | 0.720893 | 0.158699 | 1.056436 | 0.750828 | 0.746633 | 0 |
| 28.41667 | 0.833041 | 0.628741 | 1.299555 | 0.744436 | 0.042469 | 0.846812 | 0.018472 | 1.021924 | 0.882344 | 0.777216 | 0 |
| 28.43056 | 1.034365 | 0.708227 | 1.098847 | 0.715372 | -0.01168 | 1.069199 | 0.000559 | 1.073296 | 0.951002 | 0.78854 | 0 |
| 28.44444 | 1.051786 | 0.669491 | 0.820072 | 0.713639 | 0.125415 | 1.359419 | 0.166885 | 1.024423 | 0.948172 | 0.824176 | 0 |
| 28.45833 | 0.83331 | 0.495937 | 0.538886 | 0.641111 | 0.244976 | 1.284387 | 0.266132 | 0.787311 | 0.722839 | 0.678343 | 0 |
| 28.47222 | 0.499142 | 0.102686 | 0.245684 | 0.581215 | 0.218381 | 0.713807 | 0.196155 | 0.482947 | 0.265821 | 0.419397 | 0 |
| 28.48611 | 0.507109 | -0.10772 | 0.509661 | 1.124484 | 0.192441 | 0.289328 | 0.124213 | 0.460667 | 0.073258 | 0.628363 | 0 |
| 28.5 | 1.33969 | 0.54716 | 1.565893 | 2.683784 | 0.393391 | 0.454467 | 0.253159 | 0.936379 | 0.563388 | 1.751381 | 0 |
| 28.51389 | 2.460283 | 1.514364 | 2.64271 | 3.326367 | 0.881149 | 0.780364 | 0.580177 | 1.738884 | 1.5399 | 3.264363 | 0 |
| 28.52778 | 2.851985 | 2.01322 | 3.04773 | 2.712072 | 0.773521 | 1.07569 | 0.868177 | 2.003942 | 2.491583 | 3.964115 | 0 |
| 28.54167 | 2.09728 | 1.388569 | 2.16372 | 1.595309 | 0.417388 | 1.093683 | 1.008649 | 1.358611 | 3.10044 | 3.444466 | 0 |
| 28.55556 | 1.02953 | 0.773782 | 0.5821 | 0.299468 | 0.242823 | 0.313973 | 0.882704 | 0.535658 | 2.199483 | 1.687753 | 0 |
| 28.56944 | -0.20788 | 0.128393 | -0.88276 | -0.66056 | 0.279117 | -0.24818 | 0.596411 | -0.25951 | 0.466589 | 0.023071 | 0 |
| 28.58333 | -0.69582 | -0.72355 | -1.82011 | -0.86051 | 0.380346 | -0.77675 | 0.251645 | -0.73056 | -0.7946 | -1.114684 | 0 |
| 28.59722 | -0.78406 | -0.81999 | -1.89084 | 0.030063 | 0.422405 | -1.48111 | 0.137334 | -0.77222 | -1.36432 | -1.316191 | 0 |
| 28.61111 | -0.42617 | -0.80411 | -0.99839 | 1.093151 | 0.30062 | -0.91634 | 0.251269 | -0.31299 | -1.09029 | -0.726849 | 0 |
| 28.625 | 0.199112 | 0.291114 | 0.397792 | 1.685928 | 0.237894 | 0.37522 | 0.434432 | 0.338824 | -0.22079 | 0.062039 | 0 |
| 28.63889 | 0.380498 | 1.441579 | 1.553122 | 1.018407 | 0.396579 | 1.660651 | 0.381802 | 0.877814 | 0.913937 | 0.718431 | 0 |
| 28.65278 | 0.250092 | 1.2394 | 1.475442 | 0.27898 | 0.420549 | 2.396312 | 0.11975 | 1.003897 | 1.732138 | 0.834399 | 0 |
| 28.66667 | 0.154297 | 1.116279 | 1.059252 | -0.45863 | 0.288194 | 2.321295 | -0.20841 | 0.838581 | 1.265723 | 0.596576 | 0 |
| 28.68056 | 0.401188 | 0.615052 | 0.898372 | -1.0846 | 0.220189 | 1.228307 | -0.3817 | 0.81349 | 0.692422 | 0.289027 | 0 |
| 28.69444 | 0.525568 | 0.466059 | 0.244506 | -1.08962 | 0.273798 | 0.229221 | -0.3889 | 1.045194 | -0.06501 | 0.100014 | 0 |
| 28.70833 | 1.126183 | 0.22722 | -0.44894 | 0.036216 | 0.434558 | -0.90477 | -0.15836 | 0.952915 | -0.80967 | 0.216326 | 0 |
| 28.72222 | 2.02756 | -0.35259 | -0.37604 | 1.180659 | 0.551338 | -1.57928 | 0.167792 | 0.482704 | -0.895 | 0.243011 | 0 |
| 28.73611 | 1.584619 | -1.09124 | -0.49116 | 1.636827 | 0.485528 | -1.62558 | 0.32304 | 0.416092 | -0.261 | 0.30559 | 0 |
| 28.75 | 0.442888 | -1.33289 | -0.59521 | 0.822142 | 0.338852 | -1.12891 | 0.274884 | 0.827198 | 0.395435 | 0.301989 | 0 |
| 28.76389 | -0.74502 | 0.040346 | -0.67019 | -0.11475 | 0.256873 | -0.11861 | 0.120109 | 1.217061 | 0.826205 | 0.166454 | 0 |
| 28.77778 | -1.83534 | 0.53549 | -1.47525 | -0.67355 | 0.288129 | 0.661157 | -0.12408 | 1.047318 | 0.443191 | -0.355939 | 0 |
| 28.79167 | -2.14248 | -0.22898 | -2.4225 | -1.12327 | 0.205725 | 0.652075 | -0.24528 | 0.298157 | -0.61317 | -1.093576 | 0 |
| 28.80556 | -2.1859 | -1.78364 | -3.25902 | -1.16795 | -0.00972 | -0.29311 | -0.21919 | -0.94579 | -1.82665 | -1.965998 | 0 |
| 28.81944 | -2.06324 | -3.5719 | -2.61961 | -0.8017 | -0.2549 | -1.7452 | -0.20214 | -2.3189 | -2.62855 | -2.892261 | 0 |
| 28.83333 | -1.71859 | -4.97105 | -1.5465 | -0.53096 | -0.53002 | -3.05083 | -0.21466 | -3.12976 | -2.49325 | -3.289455 | 0 |
| 28.84722 | -1.64566 | -4.60052 | -0.91049 | -0.52804 | -0.76528 | -3.12449 | -0.33006 | -3.04298 | -2.02849 | -2.957798 | 0 |
| 28.86111 | -1.52425 | -3.34721 | -0.92453 | -0.97317 | -0.805 | -2.58228 | -0.4096 | -2.43112 | -1.82992 | -2.261737 | 0 |
| 28.875 | -1.6546 | -2.14442 | -1.32903 | -1.75978 | -0.72655 | -2.12917 | -0.42488 | -2.07356 | -1.82467 | -1.795328 | 0 |
| 28.88889 | -1.74324 | -1.25229 | -1.62155 | -2.89374 | -0.48372 | -1.72868 | -0.36417 | -1.99418 | -1.68204 | -1.467884 | 0 |
| 28.90278 | -1.52888 | -0.54644 | -1.5452 | -3.48958 | -0.2713 | -1.29105 | -0.15936 | -1.8893 | -1.46216 | -1.086038 | 0 |
| 28.91667 | -1.08464 | -0.22667 | -1.13462 | -2.5093 | -0.18876 | -1.05882 | 0.015581 | -1.66679 | -1.27721 | -0.824909 | 0 |
| 28.93056 | -0.69911 | -0.30387 | -0.68073 | -1.08441 | -0.28653 | -0.79372 | 0.055446 | -1.31278 | -1.04536 | -0.666415 | 0 |
| 28.94444 | -0.3334 | -0.68358 | -0.42933 | -0.15011 | -0.2746 | -0.44152 | -0.02871 | -0.7255 | -0.72285 | -0.541791 | 0 |
| 28.95833 | -0.08077 | -0.9278 | -0.27768 | 0.057169 | -0.16282 | -0.07638 | -0.09425 | -0.11304 | -0.29684 | -0.343619 | 0 |
| 28.97222 | 0.051833 | -0.85752 | -0.20682 | -0.14135 | 0.005148 | 0.178291 | -0.16836 | 0.224092 | 0.044644 | -0.100114 | 0 |
| 28.98611 | -0.14694 | -1.03717 | -0.32142 | -0.70956 | 0.047492 | -0.01561 | -0.35616 | -0.00468 | -0.05748 | -0.237186 | 0 |
| 29 | -0.70688 | -1.79051 | -0.95351 | -1.73423 | -0.16604 | -0.84776 | -0.59328 | -0.63585 | -0.70608 | -0.989638 | 0 |
| 29.01389 | -1.01697 | -2.39626 | -1.55407 | -2.32066 | -0.46736 | -1.74079 | -0.66994 | -1.06841 | -1.42343 | -1.525936 | 0 |
| 29.02778 | -1.10815 | -2.38571 | -1.8846 | -2.2053 | -0.9279 | -2.59983 | -0.62511 | -1.28977 | -2.01945 | -1.656234 | 0 |
| 29.04167 | -1.57587 | -2.86116 | -2.64645 | -2.07635 | -1.85942 | -4.05532 | -0.96532 | -2.02793 | -2.91321 | -2.121578 | 0 |

| | | | | | | | | | | | |
|---|---|---|---|---|---|---|---|---|---|---|---|
| 29.05556 | -2.1031 | -3.23015 | -3.41564 | -1.69237 | -2.98424 | -5.48643 | -1.29116 | -2.50627 | -3.30293 | -2.573245 | 0 |
| 29.06944 | -1.89848 | -2.44527 | -2.79627 | -1.19339 | -3.25442 | -5.09093 | -1.155 | -2.3244 | -2.30212 | -2.236802 | 0 |
| 29.08333 | -1.21352 | -1.11908 | -1.4528 | -0.94516 | -2.07593 | -2.81134 | -0.73862 | -1.71654 | -1.00309 | -1.35838 | 0 |
| 29.09722 | -0.6816 | -0.41907 | -0.42886 | -1.02033 | -0.85627 | -1.23015 | -0.60371 | -1.20987 | -0.31393 | -0.494331 | 0 |
| 29.11111 | -0.49406 | -0.12741 | 0.182266 | -0.99221 | -0.16252 | -0.29158 | -0.73631 | -0.88203 | -0.0944 | -0.032869 | 0 |
| 29.125 | -0.40444 | 0.256331 | 0.675323 | -0.54944 | 0.38113 | 0.443608 | -0.81242 | -0.60746 | 0.051688 | 0.252185 | 0 |
| 29.13889 | -0.13017 | 0.862217 | 1.129909 | 0.21998 | 1.032197 | 1.64984 | -0.61655 | -0.29854 | 0.396284 | 0.692076 | 0 |
| 29.15278 | 0.136207 | 1.518772 | 1.626462 | 0.753337 | 1.459743 | 3.016364 | -0.26295 | 0.102853 | 0.92164 | 1.056787 | 0 |
| 29.16667 | 0.354461 | 2.058047 | 2.050493 | 0.673136 | 1.489994 | 4.071974 | 0.023235 | 0.44794 | 1.696116 | 1.337981 | 0 |
| 29.18056 | 0.390817 | 2.126911 | 2.184461 | 0.239822 | 1.474518 | 3.65013 | 0.08063 | 0.639898 | 2.30853 | 1.508919 | 0 |
| 29.19444 | 0.090983 | 1.948771 | 1.700218 | -0.18151 | 1.303346 | 3.183893 | -0.11329 | 0.557024 | 2.204255 | 1.448191 | 0 |
| 29.20833 | -0.33843 | 1.65204 | 1.066466 | -0.43927 | 0.893555 | 2.749584 | -0.40178 | 0.299759 | 1.823478 | 0.92777 | 0 |
| 29.22222 | -0.64132 | 1.062674 | 0.546884 | -0.63299 | 0.381242 | 2.201138 | -0.5982 | -0.10865 | 1.095435 | 0.39544 | 0 |
| 29.23611 | -0.74373 | 0.544872 | 0.134869 | -0.67373 | -0.06806 | 1.360412 | -0.61211 | -0.32528 | 0.38373 | 0.022635 | 0 |
| 29.25 | -0.60884 | 0.32817 | -0.06118 | -0.51499 | -0.28019 | 0.627787 | -0.52075 | -0.36744 | -0.16393 | -0.14968 | 0 |
| 29.26389 | -0.47162 | 0.011149 | -0.10483 | -0.34256 | -0.3015 | -0.03711 | -0.48116 | -0.33691 | -0.55901 | -0.24335 | 0 |
| 29.27778 | -0.38897 | -0.35537 | -0.10465 | -0.28745 | -0.30488 | -0.40038 | -0.52367 | -0.22244 | -0.71829 | -0.33013 | 0 |
| 29.29167 | -0.32334 | -0.74577 | -0.2422 | -0.41413 | -0.36997 | -0.52082 | -0.58032 | -0.16922 | -0.76213 | -0.490533 | 0 |
| 29.30556 | -0.23603 | -1.10068 | -0.38266 | -0.57664 | -0.42269 | -0.53175 | -0.51697 | -0.15275 | -0.64791 | -0.62028 | 0 |
| 29.31944 | -0.1936 | -1.24776 | -0.31653 | -0.46413 | -0.46285 | -0.35061 | -0.37519 | -0.09325 | -0.49076 | -0.63376 | 0 |
| 29.33333 | -0.29206 | -1.33579 | -0.15013 | -0.13514 | -0.58059 | -0.14736 | -0.39588 | -0.18151 | -0.43435 | -0.679558 | 0 |
| 29.34722 | -0.48502 | -1.26798 | 0.012891 | 0.141808 | -0.75218 | -0.13533 | -0.62311 | -0.42277 | -0.49583 | -0.751596 | 0 |
| 29.36111 | -0.4747 | -0.82476 | 0.211378 | 0.265974 | -0.70023 | -0.04394 | -0.74068 | -0.49254 | -0.34177 | -0.579429 | 0 |
| 29.375 | -0.1383 | 0.081479 | 0.486215 | 0.303264 | -0.35192 | 0.223113 | -0.59587 | -0.30638 | 0.182902 | -0.071497 | 0 |
| 29.38889 | 0.126899 | 0.811264 | 0.643738 | 0.463105 | -0.05788 | 0.445606 | -0.30966 | -0.03255 | 0.712567 | 0.388067 | 0 |
| 29.40278 | 0.076363 | 1.066049 | 0.589921 | 0.861918 | 0.087238 | 0.50854 | -0.01654 | 0.250939 | 1.046187 | 0.518549 | 0 |
| 29.41667 | -0.00702 | 1.164588 | 0.389899 | 1.254469 | 0.10179 | 0.549084 | 0.127964 | 0.357304 | 1.266155 | 0.495744 | 0 |
| 29.43056 | 0.094064 | 0.685388 | 0.235382 | 1.095935 | -0.00653 | 0.633701 | 0.017758 | 0.212575 | 1.341637 | 0.38332 | 0 |
| 29.44444 | 0.290232 | 0.000752 | 0.313091 | 0.618059 | -0.13744 | 0.693732 | -0.21194 | 0.006883 | 1.286432 | 0.198074 | 0 |
| 29.45833 | 0.354583 | -0.57928 | 0.475359 | 0.140995 | -0.16389 | 0.725219 | -0.34808 | 0.008153 | 0.820623 | 0.029385 | 0 |
| 29.47222 | 0.236629 | -0.75966 | 0.518144 | 0.004519 | -0.12366 | 0.765667 | -0.29727 | 0.273621 | 0.273877 | -0.024295 | 0 |
| 29.48611 | 0.413336 | -0.34402 | 0.680751 | 0.82283 | -0.16235 | 0.84783 | -0.16608 | 0.713325 | 0.027748 | 0.401189 | 0 |
| 29.5 | 1.512487 | 0.604811 | 1.312471 | 2.734483 | -0.30282 | 1.150671 | 0.037732 | 1.404261 | 0.33 | 1.619146 | 0 |
| 29.51389 | 2.645603 | 1.368745 | 1.901533 | 4.061823 | -0.35935 | 1.18981 | 0.412 | 2.014649 | 1.111954 | 2.952471 | 0 |
| 29.52778 | 2.600019 | 1.858182 | 2.348161 | 3.75112 | -0.08213 | 1.127992 | 0.747214 | 2.03915 | 2.310951 | 3.456261 | 0 |
| 29.54167 | 2.168465 | 1.971843 | 2.145121 | 2.604857 | 0.240016 | 1.405605 | 0.832618 | 1.503138 | 3.026787 | 2.981714 | 0 |
| 29.55556 | 1.464031 | 1.537854 | 1.434402 | 1.033412 | 0.175232 | 0.953471 | 0.630166 | 0.756197 | 1.906054 | 1.793974 | 0 |
| 29.56944 | 0.615147 | 0.554764 | 0.532775 | -0.28901 | -0.00488 | 0.234548 | 0.179182 | 0.038013 | 0.462947 | 0.370335 | 0 |
| 29.58333 | 0.000375 | -0.25198 | -0.29249 | -0.98365 | -0.28501 | -0.32556 | -0.34462 | -0.50794 | -0.6037 | -0.689878 | 0 |
| 29.59722 | 0.038549 | -0.56141 | -0.93446 | -0.83214 | -0.48361 | -0.36545 | -0.71304 | -0.80222 | -1.05401 | -1.133537 | 0 |
| 29.61111 | 0.280129 | -0.57714 | -1.04818 | 0.112136 | -0.25722 | -0.14685 | -0.49724 | -0.63366 | -1.06206 | -0.933648 | 0 |
| 29.625 | 0.729684 | -0.34865 | -0.41731 | 1.325125 | 0.097315 | 0.195326 | 0.129338 | -0.10597 | -0.49755 | -0.216395 | 0 |
| 29.63889 | 1.194479 | 0.353529 | 0.569407 | 2.19672 | 0.657798 | 0.729113 | 1.002953 | 0.431253 | 0.523758 | 0.578469 | 0 |
| 29.65278 | 1.436691 | 1.289462 | 1.448212 | 1.639003 | 1.114401 | 1.345517 | 1.43914 | 0.894481 | 1.754548 | 1.292298 | 0 |
| 29.66667 | 1.280068 | 1.740404 | 1.381338 | 0.475465 | 1.113504 | 1.833345 | 1.294653 | 1.149321 | 2.614611 | 1.690136 | 0 |
| 29.68056 | 0.609847 | 1.462629 | 0.897343 | -0.34886 | 0.755012 | 1.496459 | 0.82413 | 0.81861 | 2.298027 | 1.470567 | 0 |
| 29.69444 | -0.20845 | -0.15882 | 0.22057 | -0.84989 | 0.312028 | 0.982749 | 0.149647 | 0.426297 | 1.054115 | 0.428454 | 0 |
| 29.70833 | -0.85837 | 0.031481 | 0.052973 | -1.19344 | 0.054458 | 0.527943 | -0.59581 | 0.201511 | -0.31668 | -0.776061 | 0 |
| 29.72222 | -0.88652 | 0.126226 | -0.21741 | -0.34702 | 0.054075 | 0.242879 | -0.51214 | -0.05177 | -0.97373 | -1.299049 | 0 |
| 29.73611 | 0.203657 | 0.093656 | -0.08677 | 1.653327 | 0.297792 | -0.03714 | -0.02387 | -0.28724 | -1.30126 | -0.9522 | 0 |
| 29.75 | 1.616416 | 0.015699 | 0.770192 | 2.811332 | 0.396235 | -0.42762 | 0.60101 | -0.17601 | -1.65126 | -0.190994 | 0 |
| 29.76389 | 1.945923 | -1.15481 | 1.056778 | 1.690651 | 0.26128 | -0.88646 | 1.077255 | -0.06873 | -1.43617 | 0.14267 | 0 |
| 29.77778 | 1.492646 | -2.45052 | 0.330334 | 0.099392 | -0.05965 | -1.08981 | 1.054163 | 0.047794 | -0.77306 | 0.042563 | 0 |
| 29.79167 | 0.643445 | -0.51285 | -1.18484 | -1.66338 | -0.43038 | -1.12661 | 0.409616 | 0.109979 | -0.1426 | -0.334374 | 0 |
| 29.80556 | -0.53664 | 0.81486 | -2.65811 | -4.25844 | -0.72686 | -1.12356 | -0.29723 | 0.22418 | 0.338721 | -0.962924 | 0 |
| 29.81944 | -1.42743 | 0.255615 | -3.68331 | -4.8845 | -0.87629 | -1.00379 | -1.21674 | 0.007157 | 0.125216 | -1.670579 | 0 |
| 29.83333 | -2.02893 | -0.86462 | -4.00701 | -2.36326 | -0.87603 | -1.05985 | -1.03674 | -0.56486 | -0.80907 | -2.040255 | 0 |

| | | | | | | | | | | | |
|---|---|---|---|---|---|---|---|---|---|---|---|
| 29.84722 | -2.32975 | -2.09941 | -2.87516 | -0.22292 | -0.73261 | -1.50366 | -0.56684 | -1.05839 | -1.83325 | -1.947849 | 0 |
| 29.86111 | -2.17292 | -3.54696 | -1.27677 | 0.561173 | -0.59833 | -2.19232 | 0.081859 | -1.43491 | -2.62353 | -1.757713 | 0 |
| 29.875 | -2.25433 | -5.07289 | -0.42645 | -0.33471 | -0.45415 | -2.20618 | 0.351723 | -1.76159 | -2.77479 | -1.838101 | 0 |
| 29.88889 | -2.13024 | -5.5205 | -0.1151 | -1.77908 | -0.35512 | -1.84629 | 0.148819 | -1.78154 | -2.49947 | -1.863336 | 0 |
| 29.90278 | -1.84783 | -4.13465 | -0.07164 | -2.86998 | -0.41811 | -1.38605 | -0.0323 | -1.48194 | -1.99456 | -1.821477 | 0 |
| 29.91667 | -1.58212 | -2.18449 | -0.18166 | -3.2168 | -0.47253 | -0.91996 | -0.38276 | -1.07802 | -1.44241 | -1.700712 | 0 |
| 29.93056 | -1.32502 | -0.58045 | -0.17085 | -2.67473 | -0.41425 | -0.57974 | -0.51893 | -0.78499 | -0.89575 | -1.311222 | 0 |
| 29.94444 | -0.9391 | 0.46398 | -0.06089 | -1.38636 | -0.2193 | -0.44832 | -0.15204 | -0.61957 | -0.54846 | -0.635789 | 0 |
| 29.95833 | -0.3712 | 0.911911 | 0.048049 | -0.46272 | 0.043738 | -0.43062 | 0.458873 | -0.43993 | -0.45675 | 0.224325 | 0 |
| 29.97222 | 0.067316 | 1.089322 | 0.014798 | -0.01355 | 0.194949 | -0.46413 | 1.060236 | -0.22765 | -0.35695 | 0.867096 | 0 |
| 29.98611 | 0.090637 | 0.605758 | -0.25874 | -0.1618 | 0.180816 | -0.71873 | 1.152858 | -0.22385 | -0.49686 | 0.543195 | 0 |
| 30 | -0.46116 | -0.66883 | -0.95677 | -1.15745 | -0.1667 | -1.41045 | 0.036155 | -0.69866 | -1.22042 | -1.064489 | 0 |
| 30.01389 | -0.9393 | -1.95458 | -1.80453 | -1.94343 | -0.96753 | -2.6236 | -1.06912 | -1.28864 | -2.2906 | -2.882481 | 0 |
| 30.02778 | -0.92176 | -2.47668 | -2.01983 | -1.44199 | -1.7985 | -3.15017 | -2.20023 | -1.54819 | -2.7452 | -3.564673 | 0 |
| 30.04167 | -0.66519 | -2.33591 | -1.2985 | -0.79 | -1.99248 | -3.09885 | -1.81443 | -1.40576 | -2.1257 | -3.001301 | 0 |
| 30.05556 | -0.22061 | -1.80724 | -0.57006 | -0.67899 | -1.66338 | -2.36823 | -0.64481 | -1.21278 | -1.12105 | -1.799436 | 0 |
| 30.06944 | 0.248495 | -0.99348 | 0.108698 | -0.74966 | -0.7934 | -0.87057 | 0.381967 | -0.92698 | -0.20469 | -0.449988 | 0 |
| 30.08333 | 0.506865 | -0.28351 | 0.716452 | -0.64442 | 0.123113 | 0.539468 | 1.009847 | -0.49919 | 0.225261 | 0.521457 | 0 |
| 30.09722 | 0.557725 | 0.097655 | 0.896234 | -0.41988 | 0.495923 | 1.14653 | 0.805735 | -0.03705 | 0.345377 | 0.95215 | 0 |
| 30.11111 | 0.438738 | 0.163761 | 0.653801 | -0.18423 | 0.587784 | 1.01489 | 0.423033 | 0.205472 | 0.362539 | 0.970913 | 0 |
| 30.125 | 0.176003 | 0.022665 | 0.305922 | -0.07915 | 0.684695 | 0.755271 | 0.085767 | 0.234967 | 0.37614 | 0.837513 | 0 |
| 30.13889 | -0.19069 | -0.16725 | 0.047292 | 0.205683 | 0.75596 | 0.71476 | -0.02686 | 0.203316 | 0.425015 | 0.719259 | 0 |
| 30.15278 | -0.40552 | -0.05164 | -0.01783 | 0.776905 | 0.852221 | 0.914266 | 0.152406 | 0.252939 | 0.512307 | 0.763279 | 0 |
| 30.16667 | -0.33719 | 0.472458 | 0.151108 | 1.249016 | 0.968191 | 1.328329 | 0.483884 | 0.319957 | 0.754521 | 1.008406 | 0 |
| 30.18056 | 0.045311 | 1.263991 | 0.444538 | 1.131903 | 0.90119 | 1.203714 | 0.54062 | 0.245512 | 1.047429 | 1.28468 | 0 |
| 30.19444 | 0.441225 | 1.939364 | 0.799364 | 0.776447 | 0.877968 | 1.35443 | 0.401246 | 0.183776 | 1.367177 | 1.254469 | 0 |
| 30.20833 | 0.581219 | 2.274925 | 1.07613 | 0.266163 | 0.873218 | 1.602295 | 0.093063 | 0.126803 | 1.591162 | 1.041873 | 0 |
| 30.22222 | 0.536921 | 2.195897 | 1.217908 | -0.12787 | 0.866411 | 1.543754 | -0.32265 | -0.019 | 1.663531 | 0.681567 | 0 |
| 30.23611 | 0.447551 | 1.880933 | 1.18172 | -0.27878 | 0.721153 | 1.266398 | -0.45813 | -0.13591 | 1.361479 | 0.405613 | 0 |
| 30.25 | 0.273444 | 1.338946 | 0.925901 | -0.22632 | 0.56779 | 1.018659 | -0.18533 | -0.09025 | 0.885025 | 0.284618 | 0 |
| 30.26389 | 0.067942 | 0.807554 | 0.619804 | -0.08434 | 0.455303 | 0.74075 | 0.151063 | 0.046639 | 0.461123 | 0.233973 | 0 |
| 30.27778 | -0.1015 | 0.355929 | 0.254875 | -0.25798 | 0.209979 | 0.344239 | 0.011734 | -0.03888 | 0.050492 | -0.058395 | 0 |
| 30.29167 | -0.191 | -0.0707 | -0.0049 | -0.71202 | -0.13433 | -0.12736 | -0.4637 | -0.35235 | -0.29839 | -0.478106 | 0 |
| 30.30556 | -0.1013 | -0.29421 | 0.010891 | -0.94044 | -0.31998 | -0.34071 | -0.73022 | -0.46547 | -0.34239 | -0.667536 | 0 |
| 30.31944 | 0.071188 | -0.4089 | 0.16254 | -0.82248 | -0.29119 | -0.26088 | -0.75058 | -0.3461 | -0.28111 | -0.671628 | 0 |
| 30.33333 | 0.106789 | -0.58499 | 0.107843 | -0.4872 | -0.27437 | -0.05347 | -0.74646 | -0.27228 | -0.37031 | -0.692997 | 0 |
| 30.34722 | 0.067125 | -0.6791 | -0.15241 | -0.14843 | -0.43373 | 0.147448 | -0.74293 | -0.34197 | -0.5678 | -0.73307 | 0 |
| 30.36111 | 0.08167 | -0.60264 | -0.36617 | 0.112587 | -0.60399 | 0.156282 | -0.63508 | -0.36487 | -0.62095 | -0.6454 | 0 |
| 30.375 | 0.223352 | -0.2781 | -0.36131 | 0.416793 | -0.56498 | 0.050237 | -0.36133 | -0.14761 | -0.35669 | -0.235214 | 0 |
| 30.38889 | 0.262825 | -0.01384 | -0.20764 | 0.650449 | -0.45385 | -0.09434 | -0.15909 | 0.079915 | -0.01598 | 0.152303 | 0 |
| 30.40278 | 0.030976 | -0.10978 | -0.02901 | 0.604973 | -0.44126 | -0.24341 | -0.30557 | 0.006046 | 0.183232 | 0.119905 | 0 |
| 30.41667 | -0.22773 | -0.38855 | 0.197881 | 0.424988 | -0.46708 | -0.21001 | -0.66598 | -0.33548 | 0.259657 | -0.078505 | 0 |
| 30.43056 | -0.45144 | -0.56453 | 0.393428 | 0.497834 | -0.50532 | -0.10956 | -0.91797 | -0.64943 | 0.303556 | -0.375593 | 0 |
| 30.44444 | -0.64995 | -0.43184 | 0.416181 | 0.60334 | -0.60522 | 0.068806 | -0.9616 | -0.80092 | 0.312096 | -0.702638 | 0 |
| 30.45833 | -0.47377 | -0.0037 | 0.286637 | 0.36896 | -0.79269 | 0.160661 | -0.89567 | -0.83088 | 0.276885 | -0.709003 | 0 |
| 30.47222 | -0.03383 | 0.626047 | 0.254367 | -0.10968 | -0.79467 | 0.18529 | -0.72229 | -0.66701 | 0.451057 | -0.312906 | 0 |
| 30.48611 | 0.480303 | 1.427685 | 0.512976 | -0.00853 | -0.55799 | 0.42639 | -0.39898 | -0.16201 | 0.907352 | 0.512138 | 0 |
| 30.5 | 1.292871 | 2.405422 | 1.247718 | 1.203453 | -0.21181 | 1.042467 | 0.025458 | 0.794497 | 1.710354 | 2.13305 | 0 |
| 30.51389 | 2.24684 | 2.526983 | 2.086399 | 2.9965 | 0.137933 | 1.444091 | 0.525017 | 1.583721 | 2.434921 | 3.922744 | 0 |
| 30.52778 | 2.319861 | 2.478968 | 2.10898 | 4.184352 | 0.513921 | 1.669274 | 1.433928 | 1.795265 | 2.744636 | 4.984092 | 0 |
| 30.54167 | 1.827268 | 2.166713 | 1.880096 | 3.788383 | 0.393903 | 1.550849 | 1.640468 | 1.44992 | 2.615143 | 4.442854 | 0 |
| 30.55556 | 1.020678 | 1.345702 | 1.073047 | 1.890196 | 0.246042 | 0.775529 | 0.806609 | 0.633783 | 1.624791 | 2.32362 | 0 |
| 30.56944 | 0.104435 | 0.154005 | 0.113039 | 0.111336 | 0.135276 | -0.21372 | -0.01673 | -0.14973 | 0.401445 | -0.002414 | 0 |
| 30.58333 | -0.5545 | -0.86806 | -0.77342 | -1.0433 | -0.11534 | -0.79286 | -0.69402 | -0.50302 | -0.69223 | -1.679291 | 0 |
| 30.59722 | -0.33769 | -1.48178 | -1.47119 | -1.47203 | -0.70586 | -0.746 | -1.07958 | -0.35009 | -1.45762 | -2.347934 | 0 |
| 30.61111 | 0.438282 | -1.80408 | -1.19825 | -0.76347 | -0.93244 | -0.23224 | -0.91255 | 0.182188 | -1.63489 | -2.02248 | 0 |
| 30.625 | 1.31365 | -2.06281 | -0.28443 | 0.645617 | -0.5228 | 0.62025 | -0.36808 | 0.805667 | -1.43175 | -0.941444 | 0 |

| | | | | | | | | | | | |
|---|---|---|---|---|---|---|---|---|---|---|---|
| 30.63889 | 1.731197 | -2.54902 | 0.828028 | 2.912346 | 0.01977 | 1.188155 | 0.292772 | 1.154987 | -0.89936 | 0.536699 | 0 |
| 30.65278 | 1.6199 | -1.56698 | 1.594734 | 4.497041 | 0.636699 | 1.462711 | 0.874252 | 1.185793 | -0.06279 | 1.881383 | 0 |
| 30.66667 | 1.328999 | 1.655313 | 2.007171 | 2.441663 | 0.815685 | 1.499778 | 1.245653 | 0.873204 | 1.389364 | 2.430951 | 0 |
| 30.68056 | 0.681166 | 5.032744 | 2.108945 | 0.417608 | 0.554154 | 1.022232 | 0.955029 | 0.033516 | 2.858534 | 1.705708 | 0 |
| 30.69444 | -0.12953 | 3.857834 | 0.961899 | -1.12316 | 0.01402 | 0.309993 | 0.208898 | -0.76396 | 2.623872 | 0.212733 | 0 |
| 30.70833 | -0.8034 | 2.358818 | -0.67983 | -2.46669 | -0.81076 | -0.60727 | -0.60523 | -1.08944 | 1.259983 | -1.362634 | 0 |
| 30.72222 | -1.10871 | 0.21719 | -2.01218 | -3.95993 | -1.62945 | -1.59044 | -0.83159 | -1.08267 | 0.061862 | -2.494534 | 0 |
| 30.73611 | -1.04197 | -0.83831 | -2.19047 | -4.45107 | -1.44686 | -1.77179 | -0.3895 | -0.78531 | -0.9823 | -2.562669 | 0 |
| 30.75 | -0.88523 | -1.02452 | -2.01496 | -1.17975 | -0.713 | -0.44987 | -0.17793 | -0.58954 | -1.98175 | -1.982242 | 0 |
| 30.76389 | -0.88149 | -2.16452 | -1.48936 | 2.071562 | -0.09746 | 0.3798 | -0.24897 | -0.73955 | -2.43807 | -1.097748 | 0 |
| 30.77778 | -0.72796 | -2.76543 | -0.52905 | 3.887039 | 0.362601 | 0.641831 | -0.28049 | -1.09907 | -2.31411 | -0.367675 | 0 |
| 30.79167 | -0.33724 | -3.47983 | 0.19474 | 1.789089 | 0.436315 | 0.003997 | -0.229 | -0.87736 | -1.97193 | -0.148375 | 0 |
| 30.80556 | -0.05798 | -4.66114 | 0.007878 | 0.121678 | -0.04008 | -0.91273 | -0.27589 | -0.53088 | -2.02105 | -0.653369 | 0 |
| 30.81944 | -0.16736 | -5.18557 | -0.94626 | -1.94371 | -1.05133 | -1.8805 | -0.70057 | -0.65552 | -2.00182 | -1.741862 | 0 |
| 30.83333 | -0.52786 | -4.35739 | -2.28085 | -4.21374 | -2.11423 | -2.66194 | -1.26766 | -1.09751 | -1.96834 | -2.979834 | 0 |
| 30.84722 | -0.8348 | -2.87174 | -2.29413 | -5.70928 | -2.29034 | -2.17017 | -1.26415 | -1.37509 | -1.19165 | -3.235181 | 0 |
| 30.86111 | -1.21641 | -1.6048 | -1.72591 | -5.0372 | -1.65552 | -1.48139 | -0.77278 | -1.47542 | -0.66886 | -2.642043 | 0 |
| 30.875 | -1.72185 | -0.68335 | -1.32343 | -3.51671 | -0.81485 | -1.00375 | -0.35693 | -1.64672 | -0.53485 | -1.689713 | 0 |
| 30.88889 | -1.76101 | 0.030667 | -0.95435 | -1.65052 | -0.07098 | -0.89897 | -0.47933 | -1.49552 | -1.17099 | -0.815961 | 0 |
| 30.90278 | -1.4275 | 0.31127 | -0.8591 | -0.52672 | 0.34178 | -1.1624 | -0.85356 | -1.07804 | -1.90314 | -0.359413 | 0 |
| 30.91667 | -1.02604 | 0.428255 | -0.9681 | -0.28346 | 0.24262 | -1.66697 | -0.68373 | -0.71614 | -1.85911 | -0.537743 | 0 |
| 30.93056 | -0.65448 | 0.594735 | -1.04379 | -0.48517 | -0.07246 | -1.69422 | -0.62718 | -0.37685 | -1.58178 | -0.884251 | 0 |
| 30.94444 | -0.37622 | 0.893398 | -0.99917 | -0.74933 | -0.32425 | -1.14723 | -0.34024 | -0.12441 | -1.04599 | -0.861675 | 0 |
| 30.95833 | -0.17795 | 1.224772 | -0.59139 | -0.81871 | -0.15839 | -0.22372 | 0.308208 | 0.024859 | -0.34974 | -0.179673 | 0 |
| 30.97222 | 0.092366 | 1.422027 | 0.097642 | -0.52252 | 0.425152 | 0.717105 | 1.078618 | 0.204432 | 0.253016 | 0.691148 | 0 |
| 30.98611 | 0.467296 | 1.288817 | 0.699564 | 0.023587 | 1.086617 | 1.319085 | 1.667132 | 0.399967 | 0.774877 | 1.025139 | 0 |
| 31 | 0.400983 | 0.766163 | 0.324403 | 0.558783 | 0.99429 | 0.459992 | 1.653077 | 0.590758 | 0.643589 | 0.067904 | 0 |
| 31.01389 | -0.55715 | -0.46611 | -1.31405 | 0.599975 | -0.4165 | -1.60355 | 0.48514 | 0.489916 | -0.47916 | -1.908653 | 0 |
| 31.02778 | -1.63137 | -1.88579 | -2.80027 | 0.366682 | -2.50221 | -3.47924 | -1.1654 | 0.108535 | -1.75892 | -3.348268 | 0 |
| 31.04167 | -1.14007 | -1.93201 | -2.48499 | -0.08384 | -2.75351 | -3.62107 | -1.34093 | -0.35554 | -2.14451 | -3.222564 | 0 |
| 31.05556 | -0.21779 | -1.42129 | -1.30647 | -0.67921 | -1.56316 | -2.88464 | -0.26845 | -0.83818 | -1.8278 | -2.003732 | 0 |
| 31.06944 | 0.814086 | -0.7477 | 0.027516 | -1.08101 | -0.26884 | -1.11374 | 0.614346 | -1.10358 | -1.10044 | -0.493664 | 0 |
| 31.08333 | 1.433273 | -0.15194 | 1.020789 | -1.07104 | 0.927647 | 0.330836 | 0.98787 | -1.02769 | -0.28158 | 0.410464 | 0 |
| 31.09722 | 1.601712 | 0.226032 | 1.684336 | -0.64879 | 1.297578 | 1.239212 | 0.934259 | -0.66078 | 0.277083 | 0.949636 | 0 |
| 31.11111 | 1.768612 | 0.725485 | 2.223479 | -0.16731 | 1.625449 | 1.906766 | 0.74312 | -0.25416 | 0.673849 | 1.651213 | 0 |
| 31.125 | 1.783424 | 1.12954 | 2.278494 | 0.164706 | 1.862368 | 2.360294 | 0.689667 | 0.099126 | 1.052803 | 2.279334 | 0 |
| 31.13889 | 2.007111 | 1.367201 | 2.134849 | 0.410165 | 1.74128 | 2.672231 | 0.644934 | 0.309114 | 1.264558 | 2.419699 | 0 |
| 31.15278 | 1.707092 | 1.142213 | 1.66273 | 0.596038 | 1.478638 | 2.822924 | 0.686542 | 0.469306 | 1.19216 | 2.399987 | 0 |
| 31.16667 | 1.118803 | 0.892001 | 1.107115 | 0.582577 | 1.388337 | 2.315962 | 0.836155 | 0.590959 | 1.136725 | 2.195022 | 0 |
| 31.18056 | 0.783189 | 0.868393 | 0.770675 | 0.435358 | 1.45994 | 1.772021 | 0.975884 | 0.61821 | 1.155252 | 1.803379 | 0 |
| 31.19444 | 0.400708 | 0.954177 | 0.498212 | 0.317105 | 1.488567 | 1.358243 | 0.891165 | 0.475201 | 1.061673 | 1.338033 | 0 |
| 31.20833 | 0.03714 | 0.953993 | 0.375882 | 0.210817 | 1.30383 | 0.798917 | 0.645309 | 0.252156 | 0.849529 | 1.032888 | 0 |
| 31.22222 | -0.20882 | 1.020797 | 0.505657 | 0.208199 | 0.914652 | 0.379916 | 0.402339 | 0.088513 | 0.743636 | 0.731822 | 0 |
| 31.23611 | -0.23776 | 1.02324 | 0.711768 | 0.290323 | 0.428724 | 0.172314 | 0.243413 | 0.009066 | 0.704742 | 0.141053 | 0 |
| 31.25 | 0.035503 | 1.077921 | 0.801686 | 0.207662 | -0.11781 | 0.153432 | 0.041727 | -0.12959 | 0.685419 | -0.18752 | 0 |
| 31.26389 | 0.66923 | 1.159063 | 0.87803 | 0.054893 | -0.17953 | 0.393058 | -0.13258 | -0.24325 | 0.671116 | 0.05641 | 0 |
| 31.27778 | 1.548547 | 1.30703 | 1.000743 | 0.04757 | 0.389337 | 0.915856 | -0.03236 | -0.12743 | 0.648418 | 0.606707 | 0 |
| 31.29167 | 1.897571 | 1.237626 | 0.979717 | 0.069381 | 1.041577 | 1.328621 | 0.229666 | 0.073238 | 0.585935 | 1.094888 | 0 |
| 31.30556 | 1.10823 | 0.711953 | 0.733945 | 0.047302 | 1.16046 | 1.269848 | 0.297702 | 0.177024 | 0.468295 | 1.090066 | 0 |
| 31.31944 | 0.070783 | 0.232087 | 0.386098 | 0.05626 | 0.666728 | 0.760511 | 0.283418 | 0.19565 | 0.32832 | 0.500878 | 0 |
| 31.33333 | -1.33769 | -0.31823 | 0.0002 | 0.146409 | 0.047011 | 0.187359 | 0.224891 | 0.232066 | 0.152844 | -0.357155 | 0 |
| 31.34722 | -1.78211 | -0.49381 | -0.17799 | 0.245704 | -0.60741 | -0.14437 | 0.120244 | 0.346031 | 0.051777 | -0.933137 | 0 |
| 31.36111 | -0.46252 | -0.00108 | 0.122773 | 0.244401 | -1.03207 | -0.04448 | -0.02178 | 0.460964 | 0.122894 | -0.632802 | 0 |
| 31.375 | 1.143318 | 0.566969 | 0.471287 | 0.179437 | -0.62646 | 0.343947 | -0.03441 | 0.556098 | 0.310149 | 0.315834 | 0 |
| 31.38889 | 2.251677 | 0.936097 | 0.739647 | 0.168428 | -0.10163 | 0.682759 | 0.221168 | 0.672317 | 0.51505 | 1.08226 | 0 |
| 31.40278 | 2.031231 | 0.81387 | 0.959187 | 0.207366 | 0.44073 | 0.795617 | 0.531541 | 0.71396 | 0.698386 | 1.293943 | 0 |
| 31.41667 | 0.693829 | 0.442687 | 1.019027 | 0.23831 | 0.556637 | 0.618606 | 0.591881 | 0.602318 | 0.799965 | 0.948683 | 0 |

| | | | | | | | | | | | |
|---|---|---|---|---|---|---|---|---|---|---|---|
| 31.43056 | -0.45522 | 0.124128 | 0.930476 | 0.296634 | 0.138594 | 0.31235 | 0.449788 | 0.431581 | 0.840172 | 0.248542 | 0 |
| 31.44444 | -0.58926 | 0.021435 | 0.882382 | 0.641244 | -0.50548 | 0.01006 | 0.137593 | 0.239193 | 0.871739 | -0.21167 | 0 |
| 31.45833 | 0.261817 | 0.29531 | 0.851455 | 1.55464 | -0.9959 | -0.07706 | -0.18357 | 0.079235 | 0.98226 | -0.043722 | 0 |
| 31.47222 | 1.121929 | 0.737031 | 0.908747 | 2.007296 | -0.79992 | 0.137905 | -0.21185 | 0.108299 | 1.121899 | 0.490468 | 0 |
| 31.48611 | 1.493622 | 1.177631 | 1.156629 | 2.364431 | -0.25934 | 0.579584 | 0.087488 | 0.470454 | 1.550468 | 1.341795 | 0 |
| 31.5 | 1.564069 | 1.933015 | 1.929832 | 4.005736 | 0.503525 | 1.233322 | 0.517032 | 1.221213 | 2.769475 | 2.779775 | 0 |
| 31.51389 | 1.791395 | 2.055121 | 2.920497 | 3.798619 | 0.944728 | 2.034809 | 0.593169 | 1.740722 | 3.446493 | 4.239624 | 0 |
| 31.52778 | 2.175253 | 2.338667 | 2.732599 | 2.490898 | 1.830524 | 2.421737 | 0.706513 | 1.46009 | 3.428771 | 5.216385 | 0 |
| 31.54167 | 2.020648 | 2.729974 | 2.603681 | 1.729227 | 2.167382 | 2.80324 | 0.837386 | 1.212483 | 3.570179 | 5.074759 | 0 |
| 31.55556 | 1.032962 | 2.150482 | 1.619196 | 1.001309 | 1.647949 | 1.835342 | 0.688463 | 0.846742 | 2.985622 | 3.251729 | 0 |
| 31.56944 | 0.077989 | 1.039558 | 0.611559 | 0.179224 | 0.791094 | 0.482931 | 0.385687 | 0.399436 | 1.760626 | 0.950466 | 0 |
| 31.58333 | -0.60241 | 0.301087 | -0.40359 | -0.68657 | -0.05078 | -0.70104 | 0.232103 | 0.117846 | 0.668526 | -0.64379 | 0 |
| 31.59722 | -1.02683 | -0.23494 | -1.36926 | -1.28269 | -0.7041 | -1.54571 | 0.206002 | 0.113698 | -0.29549 | -1.690168 | 0 |
| 31.61111 | -0.97455 | -0.6279 | -2.02307 | -1.63139 | -1.17988 | -2.1606 | 0.006331 | 0.335304 | -0.8575 | -2.12461 | 0 |
| 31.625 | -0.29774 | -0.88585 | -1.46739 | -0.30225 | -1.14716 | -2.4603 | -0.19261 | 0.538527 | -1.00831 | -2.094553 | 0 |
| 31.63889 | 0.665702 | -1.60553 | -0.56197 | 2.406061 | -0.9621 | -1.84228 | -0.36914 | 0.614466 | -1.31006 | -1.156941 | 0 |
| 31.65278 | 1.52422 | -2.46057 | 0.055471 | 4.241475 | -0.32482 | -0.60068 | -0.46382 | 0.496712 | -1.85721 | 0.084016 | 0 |
| 31.66667 | 1.499053 | -2.82638 | 0.543861 | 3.804752 | 0.368231 | 0.942571 | -0.28051 | 0.295184 | -1.66736 | 1.180254 | 0 |
| 31.68056 | 0.979474 | -2.73865 | 0.709104 | 2.244932 | 0.678147 | 2.021377 | 0.096949 | 0.365396 | -0.52795 | 1.593237 | 0 |
| 31.69444 | 0.60774 | -0.32832 | 0.965497 | 0.748816 | 0.52029 | 2.306738 | 0.547517 | 0.494847 | 1.18337 | 1.24086 | 0 |
| 31.70833 | 0.317787 | 2.772338 | 1.396859 | -0.85771 | -0.14791 | 2.033787 | 0.690253 | 0.216823 | 2.800846 | 0.25867 | 0 |
| 31.72222 | -0.16431 | 4.678275 | 0.869408 | -2.23654 | -0.97496 | 1.153282 | 0.272013 | 0.000727 | 2.610759 | -0.915586 | 0 |
| 31.73611 | -0.80931 | 3.899335 | -0.5915 | -3.64319 | -1.41699 | -0.40132 | -0.52977 | -0.26534 | 1.914942 | -1.683073 | 0 |
| 31.75 | -0.39298 | 2.827695 | -1.3541 | -4.02312 | -1.52709 | -1.29296 | -1.23336 | -0.34272 | 0.915722 | -2.162851 | 0 |
| 31.76389 | 0.046538 | 1.599473 | -1.64214 | -2.18395 | -2.18973 | -1.69799 | -1.78789 | -0.58837 | -0.35628 | -2.265087 | 0 |
| 31.77778 | 0.376014 | 0.337362 | -1.58342 | 0.381422 | -1.51644 | -1.97611 | -1.94042 | -0.96751 | -1.48286 | -2.163103 | 0 |
| 31.79167 | 0.031982 | -1.26099 | -1.83786 | 2.812643 | -0.58321 | -1.87041 | -1.84916 | -1.36194 | -2.46191 | -2.051889 | 0 |
| 31.80556 | -0.5629 | -2.95504 | -1.99227 | 4.544496 | -0.1337 | -0.92862 | -1.4057 | -1.57719 | -3.06044 | -1.766726 | 0 |
| 31.81944 | -1.12348 | -4.47223 | -2.25625 | 3.087027 | -0.0916 | -1.27221 | -0.86699 | -1.41936 | -3.0113 | -1.998803 | 0 |
| 31.83333 | -1.83713 | -4.43506 | -2.69761 | 0.447525 | -0.73545 | -2.25462 | -0.37827 | -1.20099 | -2.44366 | -2.219024 | 0 |
| 31.84722 | -2.0852 | -4.31425 | -3.307 | -2.03515 | -1.56174 | -2.78484 | -0.09741 | -1.06869 | -2.0087 | -2.462151 | 0 |
| 31.86111 | -1.50329 | -3.57228 | -3.67481 | -3.87188 | -2.27848 | -3.84035 | 0.019734 | -1.01482 | -1.93849 | -2.926661 | 0 |
| 31.875 | -1.07016 | -2.47363 | -3.04383 | -5.43086 | -2.5376 | -4.83541 | -0.08002 | -1.21201 | -1.94987 | -2.974798 | 0 |
| 31.88889 | -0.72442 | -1.67541 | -2.02143 | -4.99078 | -2.13389 | -3.6513 | -0.09662 | -1.45044 | -1.64303 | -2.701325 | 0 |
| 31.90278 | -0.33078 | -1.09303 | -0.96942 | -4.12489 | -1.5058 | -2.07457 | -0.08249 | -1.48537 | -1.26706 | -2.034261 | 0 |
| 31.91667 | -0.26333 | -0.69856 | -0.18928 | -2.07728 | -0.57202 | -0.59322 | -0.04716 | -1.30127 | -0.72529 | -1.144919 | 0 |
| 31.93056 | -0.21773 | -0.48326 | 0.081158 | -0.89865 | 0.404471 | 0.636691 | -0.03767 | -0.96152 | -0.06665 | -0.257912 | 0 |
| 31.94444 | -0.24987 | -0.35195 | 0.073691 | -0.32475 | 1.174993 | 1.134968 | -0.00212 | -0.65293 | 0.504526 | 0.373232 | 0 |
| 31.95833 | -0.2579 | -0.13427 | 0.13705 | -0.19047 | 1.590036 | 1.042159 | 0.058005 | -0.34073 | 0.813843 | 0.702591 | 0 |
| 31.97222 | -0.23338 | 0.130055 | 0.404917 | -0.14338 | 1.558709 | 0.873922 | 0.168511 | -0.13575 | 0.857714 | 0.751032 | 0 |
| 31.98611 | -0.34665 | 0.316218 | 0.448982 | -0.21855 | 1.069575 | 0.650748 | 0.197501 | -0.06757 | 0.405548 | 0.131217 | 0 |
| 32 | -0.82767 | 0.13658 | -0.50577 | -0.56858 | -0.18244 | -0.63094 | 0.042722 | -0.18143 | -0.58526 | -1.481668 | 0 |
| 32.01389 | -1.80883 | -0.65202 | -2.21945 | -1.04068 | -1.74553 | -2.64606 | -0.28809 | -0.4307 | -1.82992 | -3.000987 | 0 |
| 32.02778 | -2.36284 | -1.3332 | -3.32884 | -1.16841 | -2.47807 | -4.02798 | -0.79324 | -0.59461 | -2.54995 | -3.284648 | 0 |
| 32.04167 | -2.75467 | -1.67812 | -3.29922 | -0.55988 | -2.08014 | -4.49289 | -0.63783 | -0.58812 | -2.53225 | -2.731074 | 0 |
| 32.05556 | -2.91599 | -1.30083 | -2.7148 | -0.11538 | -1.11659 | -3.84017 | -0.20419 | -0.39616 | -2.26987 | -1.749605 | 0 |
| 32.06944 | -1.90221 | -0.4902 | -1.62787 | 0.080175 | -0.26306 | -2.73664 | 0.196957 | -0.27039 | -1.73949 | -0.828477 | 0 |
| 32.08333 | -0.54454 | 0.136247 | -0.3213 | -0.03093 | 0.177 | -1.44578 | 0.389728 | -0.2593 | -1.02369 | -0.217211 | 0 |
| 32.09722 | 0.602144 | 0.537274 | 1.191933 | -0.07538 | 0.384202 | -0.28021 | 0.466513 | -0.0741 | -0.17572 | 0.400364 | 0 |
| 32.11111 | 1.681228 | 0.832325 | 2.477886 | 0.077703 | 0.592161 | 0.751897 | 0.558348 | 0.312003 | 0.362218 | 0.848162 | 0 |
| 32.125 | 2.481594 | 1.041181 | 3.102578 | 0.250732 | 0.894704 | 1.848913 | 0.633493 | 0.650457 | 0.578241 | 1.295168 | 0 |
| 32.13889 | 2.758671 | 1.265621 | 3.329381 | 0.312839 | 1.145368 | 2.774939 | 0.764033 | 0.819828 | 0.798019 | 1.773561 | 0 |
| 32.15278 | 2.47977 | 1.212784 | 2.735928 | 0.160727 | 1.154244 | 3.61424 | 1.038375 | 0.767539 | 0.998114 | 1.95028 | 0 |
| 32.16667 | 1.988684 | 0.739261 | 1.826161 | -0.13337 | 1.287643 | 3.58269 | 1.183731 | 0.545426 | 1.194919 | 2.001359 | 0 |
| 32.18056 | 2.216182 | 0.743269 | 1.391166 | -0.36566 | 1.479923 | 3.496801 | 1.002075 | 0.340803 | 1.310975 | 2.114218 | 0 |
| 32.19444 | 2.287243 | 0.808116 | 0.693818 | -0.44319 | 1.24692 | 3.174365 | 0.696198 | 0.223908 | 1.418758 | 2.11201 | 0 |
| 32.20833 | 1.807282 | 0.826134 | 0.298325 | -0.3464 | 0.830364 | 2.73316 | 0.342434 | 0.177294 | 1.510001 | 1.76545 | 0 |

| | | | | | | | | | | | |
|---|---|---|---|---|---|---|---|---|---|---|---|
| 32.22222 | 1.854328 | 1.138129 | 0.57624 | -0.05994 | 0.435634 | 2.043245 | 0.271268 | 0.279719 | 0.983049 | 1.523146 | 0 |
| 32.23611 | 1.807732 | 1.53127 | 1.085419 | 0.098398 | 0.115159 | 1.251632 | 0.341418 | 0.431446 | 0.920088 | 1.293066 | 0 |
| 32.25 | 1.188943 | 1.339923 | 1.018387 | 0.05132 | -0.20971 | 0.36289 | 0.166155 | 0.441929 | 0.587434 | 0.667865 | 0 |
| 32.26389 | 0.317831 | 0.688893 | 0.653243 | -0.13006 | -0.606 | -0.48835 | -0.28117 | 0.277758 | 0.358719 | -0.147609 | 0 |
| 32.27778 | -0.3856 | 0.162292 | 0.339406 | -0.26567 | -0.84108 | -0.94552 | -0.52336 | 0.095297 | 0.319404 | -0.70099 | 0 |
| 32.29167 | -0.67115 | -0.11646 | 0.19755 | -0.26299 | -0.61994 | -0.73316 | -0.41271 | -0.05537 | 0.263576 | -0.79828 | 0 |
| 32.30556 | -0.50048 | -0.02759 | 0.317537 | -0.1593 | -0.06981 | 0.002845 | -0.10529 | -0.09089 | 0.234621 | -0.309796 | 0 |
| 32.31944 | 0.041432 | 0.193031 | 0.631622 | -0.00599 | 0.282422 | 0.719757 | 0.106125 | 0.017079 | 0.297632 | 0.315012 | 0 |
| 32.33333 | 0.609016 | 0.449836 | 0.839095 | 0.131991 | 0.378768 | 1.049808 | 0.212675 | 0.249965 | 0.350776 | 0.765301 | 0 |
| 32.34722 | 0.995244 | 0.655654 | 0.842966 | 0.24534 | 0.285137 | 0.943991 | 0.23768 | 0.559721 | 0.365231 | 0.909063 | 0 |
| 32.36111 | 0.848309 | 0.534393 | 0.625236 | 0.20715 | 0.052487 | 0.523779 | 0.184889 | 0.701166 | 0.325965 | 0.607574 | 0 |
| 32.375 | 0.297994 | 0.131263 | 0.325753 | -0.02359 | -0.25975 | -0.03394 | 0.072334 | 0.49492 | 0.263434 | -0.033735 | 0 |
| 32.38889 | -0.09526 | -0.19552 | 0.310933 | -0.12303 | -0.41122 | -0.22602 | 0.075997 | 0.269179 | 0.309085 | -0.423874 | 0 |
| 32.40278 | 0.095827 | -0.05444 | 0.739398 | 0.143078 | -0.25648 | 0.355118 | 0.245407 | 0.359571 | 0.533149 | -0.076426 | 0 |
| 32.41667 | 0.618336 | 0.351541 | 1.216878 | 0.401524 | 0.094455 | 1.218277 | 0.396473 | 0.523488 | 0.726905 | 0.536134 | 0 |
| 32.43056 | 0.896392 | 0.692284 | 1.233558 | 0.370758 | 0.312012 | 1.772251 | 0.404883 | 0.520505 | 0.649145 | 0.938763 | 0 |
| 32.44444 | 0.827418 | 0.976585 | 0.961859 | 0.23099 | 0.349261 | 1.91155 | 0.368796 | 0.552581 | 0.487624 | 1.118269 | 0 |
| 32.45833 | 0.758288 | 1.118939 | 0.637401 | 0.132678 | 0.24878 | 1.656364 | 0.292855 | 0.578472 | 0.393474 | 0.93918 | 0 |
| 32.47222 | 0.576922 | 0.976106 | 0.5094 | 0.170513 | 0.051459 | 1.013031 | 0.176375 | 0.476055 | 0.44099 | 0.495838 | 0 |
| 32.48611 | 0.615477 | 0.911949 | 0.887746 | 0.861381 | -0.12835 | 0.641175 | 0.031723 | 0.422776 | 0.844169 | 0.581099 | 0 |
| 32.5 | 0.866274 | 1.319987 | 2.163217 | 3.873482 | -0.11062 | 1.062948 | -0.2067 | 0.727731 | 1.750016 | 1.555997 | 0 |
| 32.51389 | 1.26509 | 1.87474 | 3.321562 | 5.488972 | 0.383006 | 2.36103 | -0.47562 | 1.288898 | 2.418339 | 3.056127 | 0 |
| 32.52778 | 2.152765 | 2.661284 | 3.279893 | 6.133499 | 1.469377 | 3.055023 | -0.29289 | 1.113214 | 2.2173 | 4.412885 | 0 |
| 32.54167 | 2.685287 | 3.64915 | 3.124302 | 4.751802 | 2.400989 | 3.207234 | 0.443559 | 0.933108 | 1.78749 | 4.79188 | 0 |
| 32.55556 | 1.559889 | 3.026231 | 2.249964 | 2.768278 | 2.345474 | 2.426065 | 0.813571 | 0.70814 | 1.603103 | 3.465166 | 0 |
| 32.56944 | 0.461201 | 2.013594 | 1.052212 | 1.629855 | 1.384981 | 1.119234 | 0.699094 | 0.515967 | 1.547766 | 1.473562 | 0 |
| 32.58333 | -0.12175 | 1.271685 | 0.188471 | 0.701759 | 0.412634 | 0.045029 | 0.542852 | 0.432897 | 1.549408 | 0.048315 | 0 |
| 32.59722 | -0.32691 | 0.665212 | -0.27182 | -0.24549 | -0.27429 | -0.92012 | 0.431063 | 0.423277 | 1.461302 | -0.888122 | 0 |
| 32.61111 | -0.48138 | 0.270323 | -0.50142 | -1.35723 | -0.68053 | -1.66309 | 0.271069 | 0.270137 | 1.13013 | -1.581719 | 0 |
| 32.625 | -0.5438 | -0.1542 | -0.63517 | -2.08554 | -0.8967 | -2.44729 | 0.017469 | 0.151145 | 0.797059 | -1.976131 | 0 |
| 32.63889 | -0.37585 | -0.96512 | -0.65532 | -3.51201 | -0.6313 | -2.48258 | -0.29014 | 0.393719 | 0.258595 | -2.028502 | 0 |
| 32.65278 | 0.042482 | -1.73798 | -0.70028 | -3.18977 | -0.55522 | -1.93601 | -0.37758 | 0.607579 | -0.42787 | -1.356774 | 0 |
| 32.66667 | 0.684111 | -2.26146 | -0.44771 | -0.50171 | -0.33924 | -1.0996 | -0.32518 | 0.763843 | -0.92561 | -0.328978 | 0 |
| 32.68056 | 1.259082 | -2.66109 | -0.18673 | 2.25941 | -0.21655 | -0.09388 | -0.32487 | 0.80382 | -1.01181 | 0.785363 | 0 |
| 32.69444 | 0.69248 | -3.46302 | -0.14024 | 4.937961 | -0.49257 | 0.114408 | -0.4546 | 0.602926 | -1.69211 | 1.786043 | 0 |
| 32.70833 | -0.12572 | -2.36845 | 0.235225 | 5.062573 | -0.95973 | 0.629711 | -0.55389 | 0.420074 | -1.56431 | 1.456912 | 0 |
| 32.72222 | 0.189701 | -0.793 | 0.289972 | 2.927076 | -0.78721 | 1.512684 | -0.56576 | 0.365263 | -0.46014 | 0.283988 | 0 |
| 32.73611 | 0.632799 | 1.035005 | 0.511458 | 1.33528 | -0.47585 | 1.631361 | -0.54205 | 0.198319 | 0.568074 | -0.965058 | 0 |
| 32.75 | 0.272491 | 2.971259 | 0.612776 | -0.09567 | -0.13336 | 1.472872 | -0.16124 | 0.097837 | 1.12617 | -1.57092 | 0 |
| 32.76389 | -0.16269 | 3.832131 | 0.27552 | -1.6256 | 0.017498 | -0.28197 | 0.328902 | -0.01362 | 1.106914 | -1.741631 | 0 |
| 32.77778 | -0.31131 | 3.100224 | -0.50525 | -2.8773 | -0.33418 | -2.21517 | 0.499041 | -0.23525 | 0.343407 | -1.608599 | 0 |
| 32.79167 | -0.38374 | 2.083403 | -1.24248 | -4.76346 | -0.72544 | -2.80126 | 0.298317 | -0.52196 | -0.32638 | -1.4512 | 0 |
| 32.80556 | -0.40766 | 0.500889 | -1.61276 | -6.17447 | -0.86867 | -2.28537 | -0.0688 | -0.80407 | -0.71711 | -1.355574 | 0 |
| 32.81944 | -0.30042 | -1.31756 | -2.06823 | -4.96408 | -0.82275 | -1.94338 | -0.50349 | -0.87274 | -0.86948 | -1.620433 | 0 |
| 32.83333 | -0.30024 | -2.7728 | -2.99511 | -1.95706 | -0.69619 | -1.85119 | -0.74039 | -0.8496 | -0.90192 | -2.465669 | 0 |
| 32.84722 | -0.43112 | -3.41214 | -3.4505 | -0.84671 | -0.81354 | -2.97163 | -0.72251 | -0.83892 | -0.97448 | -3.437713 | 0 |
| 32.86111 | -1.06794 | -3.6685 | -3.35273 | -0.62039 | -1.08961 | -4.32499 | -0.68456 | -0.82582 | -1.32093 | -4.059917 | 0 |
| 32.875 | -1.6942 | -3.46984 | -3.04944 | -0.70743 | -1.35691 | -4.73022 | -0.64087 | -0.76127 | -1.66542 | -4.17753 | 0 |
| 32.88889 | -2.02128 | -2.87515 | -2.44767 | -0.78828 | -1.54525 | -4.02121 | -0.476 | -0.83684 | -1.98289 | -3.530191 | 0 |
| 32.90278 | -1.63282 | -2.07249 | -1.6294 | -0.91113 | -1.31084 | -2.9071 | -0.10802 | -0.96482 | -2.02186 | -2.396046 | 0 |
| 32.91667 | -0.60392 | -1.16748 | -0.84606 | -1.08704 | -0.63256 | -1.63231 | 0.217394 | -1.05691 | -1.74568 | -1.161022 | 0 |
| 32.93056 | 0.397987 | -0.50965 | -0.17992 | -1.05294 | 0.112907 | -0.39335 | 0.434489 | -0.86789 | -1.17025 | -0.203681 | 0 |
| 32.94444 | 0.998854 | -0.06299 | 0.266697 | -0.71478 | 0.45204 | 0.58962 | 0.591531 | -0.47381 | -0.58864 | 0.47481 | 0 |
| 32.95833 | 1.285377 | 0.307688 | 0.455416 | -0.32891 | 0.610507 | 1.1788 | 0.724568 | -0.06817 | 0.027295 | 0.928228 | 0 |
| 32.97222 | 1.240933 | 0.475557 | 0.308936 | 0.041389 | 0.638466 | 1.014434 | 0.787151 | 0.170673 | 0.473218 | 0.894738 | 0 |
| 32.98611 | 0.798518 | 0.327222 | -0.08764 | 0.049655 | 0.297459 | 0.035734 | 0.666166 | 0.164247 | 0.527523 | 0.357731 | 0 |
| 33 | -0.42407 | -0.00608 | -0.97702 | -0.42664 | -0.55247 | -1.31102 | 0.353362 | -0.10291 | -0.06221 | -0.943487 | 0 |

| | | | | | | | | | | | |
|---|---|---|---|---|---|---|---|---|---|---|---|
| 33.01389 | -1.89955 | -0.54994 | -2.27516 | -0.82336 | -1.10571 | -2.03064 | -0.11469 | -0.407 | -0.91753 | -2.067775 | 0 |
| 33.02778 | -2.76558 | -1.03245 | -3.17639 | -0.40467 | -0.74403 | -2.01146 | -0.27793 | -0.27735 | -1.45956 | -2.161616 | 0 |
| 33.04167 | -2.9753 | -1.19986 | -3.05282 | 0.052546 | -0.50459 | -1.74398 | -0.00024 | -0.10401 | -1.05477 | -1.710826 | 0 |
| 33.05556 | -2.40224 | -0.90521 | -2.98643 | 0.159217 | -0.39373 | -1.27665 | 0.308396 | -0.13535 | -0.7172 | -1.415423 | 0 |
| 33.06944 | -1.12947 | -0.26963 | -2.20003 | 0.06413 | -0.25015 | -0.68987 | 0.446349 | -0.30452 | -0.45009 | -0.780728 | 0 |
| 33.08333 | -0.12745 | 0.236497 | -1.19702 | -0.05478 | -0.015 | -0.15838 | 0.574554 | -0.28572 | -0.1178 | -0.139323 | 0 |
| 33.09722 | 0.551664 | 0.496225 | -0.33149 | -0.11508 | 0.145816 | 0.125493 | 0.625783 | -0.15179 | 0.257192 | 0.353477 | 0 |
| 33.11111 | 0.859492 | 0.642371 | 0.320659 | -0.05427 | 0.004469 | 0.245887 | 0.587995 | 0.044883 | 0.631311 | 0.811374 | 0 |
| 33.125 | 0.761232 | 0.683356 | 0.863498 | 0.203812 | -0.08863 | 0.393814 | 0.598152 | 0.237103 | 0.705942 | 1.152571 | 0 |
| 33.13889 | 0.715667 | 0.717815 | 1.420483 | 0.442081 | 0.39367 | 0.711875 | 0.63387 | 0.312578 | 0.569252 | 1.164764 | 0 |
| 33.15278 | 0.822002 | 0.724197 | 2.033605 | 0.589279 | 0.749543 | 1.009975 | 0.710101 | 0.343425 | 0.5057 | 1.250622 | 0 |
| 33.16667 | 1.09063 | 0.923464 | 2.712011 | 0.676901 | 0.829716 | 0.992872 | 0.89792 | 0.41552 | 0.611848 | 1.288668 | 0 |
| 33.18056 | 1.502751 | 1.143838 | 3.066124 | 0.550299 | 0.738412 | 0.755081 | 0.926498 | 0.442579 | 0.900038 | 1.245535 | 0 |
| 33.19444 | 1.852721 | 1.166886 | 2.9077 | 0.274097 | 0.444645 | 0.453715 | 0.683849 | 0.419243 | 1.124294 | 1.17269 | 0 |
| 33.20833 | 2.133232 | 1.084275 | 2.491388 | 0.069761 | 0.099093 | 0.349782 | 0.34604 | 0.294769 | 0.959848 | 1.174105 | 0 |
| 33.22222 | 2.154917 | 0.987912 | 2.201212 | -0.05286 | -0.11787 | 0.695803 | 0.090564 | 0.211424 | 0.620304 | 1.288462 | 0 |
| 33.23611 | 2.045208 | 0.759466 | 1.754567 | -0.12357 | -0.27244 | 1.046774 | -0.07215 | 0.1286 | 0.301963 | 1.148664 | 0 |
| 33.25 | 1.814055 | 0.468515 | 1.272671 | -0.19329 | -0.21486 | 1.106718 | -0.03126 | 0.043338 | 0.163152 | 0.91717 | 0 |
| 33.26389 | 1.524805 | 0.250079 | 0.950151 | -0.23727 | -0.00671 | 0.990985 | 0.141449 | 0.081805 | 0.293374 | 0.822794 | 0 |
| 33.27778 | 1.070698 | 0.100536 | 0.673843 | -0.28878 | 0.123687 | 0.788296 | 0.269359 | 0.154497 | 0.468562 | 0.807305 | 0 |
| 33.29167 | 0.645247 | 0.025832 | 0.629076 | -0.2784 | 0.131916 | 0.449452 | 0.246879 | 0.162172 | 0.460234 | 0.642907 | 0 |
| 33.30556 | 0.385879 | 0.009566 | 0.622561 | -0.14371 | 0.047348 | 0.217232 | 0.190567 | 0.156156 | 0.343683 | 0.456703 | 0 |
| 33.31944 | 0.243177 | 0.050504 | 0.704385 | 0.090506 | 0.010875 | 0.130376 | 0.147059 | 0.17501 | 0.227924 | 0.358108 | 0 |
| 33.33333 | 0.185314 | 0.129747 | 0.83221 | 0.293622 | -0.02008 | 0.088328 | 0.093356 | 0.2142 | 0.136756 | 0.408373 | 0 |
| 33.34722 | 0.254947 | 0.217366 | 0.901132 | 0.418344 | -0.01414 | 0.108542 | 0.115345 | 0.26408 | 0.145826 | 0.524169 | 0 |
| 33.36111 | 0.347365 | 0.319846 | 0.862553 | 0.467917 | 0.115785 | 0.185717 | 0.25428 | 0.39941 | 0.342737 | 0.676626 | 0 |
| 33.375 | 0.418534 | 0.443082 | 0.71104 | 0.570415 | 0.342123 | 0.459686 | 0.501062 | 0.629053 | 0.611881 | 0.827044 | 0 |
| 33.38889 | 0.333805 | 0.508742 | 0.509191 | 0.698064 | 0.524741 | 0.789225 | 0.664564 | 0.762703 | 0.706904 | 0.876319 | 0 |
| 33.40278 | 0.219501 | 0.385281 | 0.466114 | 0.673193 | 0.462367 | 0.887304 | 0.531672 | 0.628096 | 0.620328 | 0.72745 | 0 |
| 33.41667 | 0.191005 | 0.194118 | 0.688829 | 0.492628 | 0.355209 | 0.909364 | 0.267465 | 0.425566 | 0.564834 | 0.646938 | 0 |
| 33.43056 | 0.311678 | 0.130576 | 0.971456 | 0.324797 | 0.421933 | 1.024504 | 0.166008 | 0.422328 | 0.651955 | 0.78563 | 0 |
| 33.44444 | 0.534571 | 0.162597 | 1.105279 | 0.219118 | 0.46594 | 1.170601 | 0.164018 | 0.479979 | 0.726277 | 0.941506 | 0 |
| 33.45833 | 0.774856 | 0.221453 | 1.045067 | 0.126425 | 0.420684 | 1.278454 | 0.204792 | 0.52334 | 0.650874 | 0.962099 | 0 |
| 33.47222 | 0.916814 | 0.267982 | 0.840141 | 0.075956 | 0.321754 | 1.431649 | 0.218227 | 0.466113 | 0.521769 | 0.87501 | 0 |
| 33.48611 | 1.114667 | 0.362219 | 0.752798 | 0.099411 | 0.437124 | 2.009833 | 0.065499 | 0.376267 | 0.646094 | 1.040085 | 0 |
| 33.5 | 1.604215 | 0.732245 | 1.189541 | 0.390809 | 0.962642 | 3.949269 | -0.25197 | 0.505616 | 1.079253 | 1.727015 | 0 |
| 33.51389 | 1.60025 | 1.227338 | 1.780175 | 0.968312 | 1.357914 | 5.563062 | -0.38695 | 0.732962 | 1.438441 | 2.510405 | 0 |
| 33.52778 | 1.250828 | 1.52988 | 2.733708 | 1.534876 | 1.616929 | 5.575518 | -0.15785 | 0.827936 | 2.021545 | 3.143178 | 0 |
| 33.54167 | 1.423082 | 1.673678 | 3.131377 | 2.37384 | 1.563109 | 5.791703 | 0.27325 | 0.637652 | 3.349519 | 3.467428 | 0 |
| 33.55556 | 1.324761 | 1.935482 | 2.426227 | 3.012656 | 1.464173 | 4.333972 | 0.470382 | 0.786218 | 2.887786 | 2.954834 | 0 |
| 33.56944 | 0.737548 | 2.02189 | 1.42103 | 2.935129 | 0.977835 | 2.306802 | 0.335107 | 0.567953 | 1.554934 | 1.78166 | 0 |
| 33.58333 | 0.236932 | 2.33097 | 0.639319 | 2.574534 | 0.689864 | 0.902197 | 0.120907 | 0.353569 | -0.06961 | 0.860488 | 0 |
| 33.59722 | -0.11006 | 3.084155 | -0.08693 | 2.651163 | 1.080896 | 0.620566 | 0.07381 | 0.247255 | -1.36162 | 0.248067 | 0 |
| 33.61111 | -0.40432 | 3.045304 | -0.62232 | 2.897356 | 1.276455 | 0.520467 | 0.129049 | 0.191793 | -1.34435 | -0.204029 | 0 |
| 33.625 | -0.33425 | 2.553208 | -1.01843 | 2.180003 | 0.786462 | -0.15575 | -0.02264 | 0.024114 | -0.5608 | -0.376796 | 0 |
| 33.63889 | -0.00206 | 2.104332 | -0.87655 | 0.705093 | 0.324906 | -0.5121 | -0.10382 | -0.23097 | -0.30644 | -0.447148 | 0 |
| 33.65278 | 0.263388 | 1.657893 | -0.51159 | -1.03552 | 0.1657 | -0.74711 | -0.07598 | -0.07271 | 0.315556 | -0.336158 | 0 |
| 33.66667 | 0.394066 | 0.804805 | -0.09664 | -3.28944 | 0.064536 | -1.06604 | 0.013061 | 0.235948 | 0.713922 | -0.196742 | 0 |
| 33.68056 | 0.26441 | -0.48555 | -0.03927 | -4.19607 | -0.01536 | -1.09969 | 0.096503 | 0.266749 | 0.801272 | -0.239857 | 0 |
| 33.69444 | 0.096795 | -1.90481 | 0.156804 | -4.05727 | 0.068649 | -1.49689 | 0.040999 | 0.173966 | 0.660525 | -0.248701 | 0 |
| 33.70833 | -0.23957 | -2.92334 | 0.293753 | -3.66352 | -0.16542 | -1.42074 | -0.08543 | 0.055455 | 0.235496 | -0.29661 | 0 |
| 33.72222 | -0.6262 | -3.85746 | 0.238595 | -2.04268 | -0.28249 | -0.99653 | -0.22479 | -0.11294 | -0.17367 | -0.633661 | 0 |
| 33.73611 | -0.13604 | -4.2122 | -0.21743 | -0.62643 | -0.45941 | -0.39666 | -0.43499 | -0.34523 | -0.68278 | -0.771818 | 0 |
| 33.75 | 0.259304 | -3.70241 | -0.90615 | -0.11282 | -0.80418 | 0.132953 | -0.72864 | -0.64094 | -1.1096 | -0.855571 | 0 |
| 33.76389 | 0.421318 | -2.15969 | -1.27265 | 0.007001 | -1.16918 | 0.686236 | -0.95971 | -0.9603 | -1.36058 | -0.760301 | 0 |
| 33.77778 | 0.228996 | -1.03668 | -1.15906 | -0.08219 | -1.31085 | 0.988302 | -1.15735 | -1.04198 | -1.23811 | -0.901551 | 0 |
| 33.79167 | -0.3212 | -0.61389 | -0.88182 | 0.199828 | -1.43311 | 0.598469 | -1.15864 | -1.04131 | -0.93497 | -1.119916 | 0 |

| | | | | | | | | | | | |
|---|---|---|---|---|---|---|---|---|---|---|---|
| 33.80556 | -0.96942 | -0.52661 | -0.85803 | 1.151756 | -1.61093 | -0.51024 | -0.98517 | -0.83482 | -0.7248 | -1.366917 | 0 |
| 33.81944 | -1.67315 | -0.47127 | -0.91307 | 1.894318 | -1.7312 | -2.11433 | -0.77577 | -0.62407 | -0.92093 | -1.476939 | 0 |
| 33.83333 | -2.11969 | -0.42293 | -1.12397 | 1.163617 | -1.81534 | -3.98435 | -0.66461 | -0.53797 | -1.3363 | -1.618759 | 0 |
| 33.84722 | -1.79732 | -0.37453 | -1.79576 | -0.35119 | -1.81056 | -5.89026 | -0.78 | -0.79475 | -1.81312 | -1.914614 | 0 |
| 33.86111 | -1.26611 | -0.26825 | -2.2271 | -2.00756 | -1.51679 | -5.53175 | -0.87151 | -1.10264 | -1.96702 | -2.225505 | 0 |
| 33.875 | -1.10302 | -0.35042 | -2.54374 | -3.14134 | -1.17411 | -4.3857 | -0.76328 | -1.34915 | -1.83932 | -2.675405 | 0 |
| 33.88889 | -1.02681 | -0.65662 | -2.91132 | -3.52669 | -1.19115 | -3.52637 | -0.74316 | -1.53345 | -1.70211 | -3.185104 | 0 |
| 33.90278 | -0.90059 | -0.95171 | -2.91135 | -2.63653 | -1.50534 | -2.98576 | -0.67408 | -1.45544 | -1.32574 | -3.244089 | 0 |
| 33.91667 | -0.83853 | -1.20694 | -2.43518 | -1.54275 | -1.65855 | -2.32847 | -0.5304 | -1.18843 | -0.89479 | -2.514487 | 0 |
| 33.93056 | -1.03884 | -1.41562 | -1.51539 | -0.54862 | -1.35698 | -1.66613 | -0.37012 | -0.78719 | -0.566 | -1.471861 | 0 |
| 33.94444 | -0.97653 | -1.26276 | -0.66937 | 0.146894 | -0.73392 | -1.28449 | -0.08997 | -0.26253 | -0.26822 | -0.650266 | 0 |
| 33.95833 | -0.6872 | -0.8428 | 0.00095 | 0.301563 | -0.1404 | -1.23324 | 0.129166 | 0.011711 | -0.00087 | -0.163117 | 0 |
| 33.97222 | -0.39909 | -0.59409 | 0.407916 | -0.23094 | 0.184838 | -1.06278 | 0.091782 | -0.04698 | 0.189753 | -0.128346 | 0 |
| 33.98611 | -0.59284 | -0.84429 | 0.247076 | -1.26133 | -0.72385 | -1.58708 | -0.19763 | -0.64432 | -0.23207 | -0.741225 | 0 |
| 34 | -1.9083 | -1.80594 | -0.70394 | -2.54952 | -2.97181 | -3.00687 | -0.66745 | -1.62293 | -1.48054 | -2.148216 | 0 |
| 34.01389 | -3.54458 | -2.58605 | -1.62642 | -2.73972 | -3.74217 | -3.74158 | -0.82112 | -2.14465 | -2.32003 | -3.05426 | 0 |
| 34.02778 | -4.57625 | -2.5583 | -1.76304 | -1.40649 | -1.63567 | -2.87531 | -0.38344 | -1.4777 | -1.8921 | -2.390335 | 0 |
| 34.04167 | -4.46086 | -2.5189 | -1.71056 | -0.07325 | -0.57432 | -2.27357 | -0.04447 | -0.57508 | -1.11828 | -1.291899 | 0 |
| 34.05556 | -3.8251 | -1.98007 | -1.45899 | 0.767853 | -0.16902 | -1.68059 | 0.072075 | 0.260034 | -0.80205 | -0.43485 | 0 |
| 34.06944 | -2.2528 | -0.94154 | -1.10849 | 0.885629 | -0.09752 | -0.87356 | 0.364258 | 0.741901 | -0.63927 | 0.116078 | 0 |
| 34.08333 | -0.60677 | 0.243676 | -0.68671 | 0.547173 | 0.145142 | -0.01902 | 0.650818 | 0.703273 | -0.2897 | 0.33937 | 0 |
| 34.09722 | 0.876531 | 1.136412 | -0.33327 | 0.196461 | 0.220991 | 0.570175 | 0.750762 | 0.608345 | 0.18515 | 0.213763 | 0 |
| 34.11111 | 2.230722 | 1.480462 | -0.10153 | 0.085237 | -0.04184 | 0.712481 | 0.690774 | 0.564354 | 0.550834 | 0.006619 | 0 |
| 34.125 | 3.05737 | 1.597483 | 0.11284 | 0.276431 | -0.01259 | 0.842434 | 0.610031 | 0.742719 | 0.981616 | 0.111943 | 0 |
| 34.13889 | 2.878635 | 1.453043 | 0.335953 | 0.588643 | 0.436487 | 0.934248 | 0.684248 | 0.934039 | 1.560682 | 0.499344 | 0 |
| 34.15278 | 2.154392 | 1.165815 | 0.663505 | 0.885567 | 0.735897 | 0.916385 | 1.017289 | 0.970222 | 2.179418 | 0.673253 | 0 |
| 34.16667 | 1.53809 | 0.982358 | 1.119974 | 0.968652 | 1.063811 | 1.036519 | 1.231663 | 0.885837 | 2.90323 | 0.988497 | 0 |
| 34.18056 | 1.063531 | 0.743386 | 1.458235 | 0.694347 | 1.306652 | 1.358964 | 1.024237 | 0.766752 | 2.576468 | 1.044206 | 0 |
| 34.19444 | 0.853662 | 0.491346 | 1.601721 | 0.337574 | 1.090663 | 1.227687 | 0.557685 | 0.561639 | 1.941495 | 0.915663 | 0 |
| 34.20833 | 0.632827 | 0.351086 | 1.39048 | 0.07167 | 0.679778 | 0.944545 | 0.297204 | 0.341868 | 1.151665 | 0.677459 | 0 |
| 34.22222 | 0.390288 | 0.203701 | 1.061097 | -0.0066 | 0.309358 | 0.393411 | 0.162225 | 0.175151 | 0.354122 | 0.435903 | 0 |
| 34.23611 | 0.105509 | 0.132736 | 0.821051 | 0.146605 | -0.01196 | -0.1212 | 0.009277 | 0.074378 | -0.35678 | 0.321735 | 0 |
| 34.25 | -0.01301 | 0.158814 | 0.703087 | 0.269179 | -0.24323 | -0.40587 | -0.0819 | -0.03045 | -0.94749 | 0.473586 | 0 |
| 34.26389 | 0.2714 | 0.150398 | 0.624164 | 0.053397 | -0.42493 | -0.50538 | -0.19777 | -0.2798 | -1.31972 | 0.513304 | 0 |
| 34.27778 | 0.700654 | 0.084574 | 0.621595 | -0.36578 | -0.57373 | -0.37432 | -0.27208 | -0.47547 | -0.86799 | 0.322159 | 0 |
| 34.29167 | 0.965923 | 0.068068 | 0.713557 | -0.58402 | -0.50312 | -0.26819 | -0.22457 | -0.35256 | -0.03292 | 0.212185 | 0 |
| 34.30556 | 0.862666 | 0.099363 | 0.741545 | -0.44514 | -0.25484 | -0.06873 | -0.01372 | -0.03368 | 0.750892 | 0.429444 | 0 |
| 34.31944 | 0.577408 | 0.121002 | 0.643536 | -0.1279 | 0.020405 | 0.239812 | 0.171307 | 0.223649 | 1.128327 | 0.543417 | 0 |
| 34.33333 | 0.339258 | 0.154123 | 0.538123 | 0.089045 | 0.210301 | 0.592812 | 0.218187 | 0.302699 | 1.156779 | 0.370415 | 0 |
| 34.34722 | 0.157773 | 0.162697 | 0.459447 | 0.157988 | 0.277841 | 0.779371 | 0.260546 | 0.228939 | 0.784713 | 0.34597 | 0 |
| 34.36111 | 0.007918 | 0.108611 | 0.397004 | 0.071662 | 0.254587 | 0.793238 | 0.269082 | 0.142607 | 0.30584 | 0.400975 | 0 |
| 34.375 | 0.070639 | 0.078763 | 0.469438 | -0.10458 | 0.211235 | 0.878924 | 0.209261 | 0.127233 | 0.174398 | 0.276584 | 0 |
| 34.38889 | 0.308344 | 0.116342 | 0.714194 | -0.2639 | 0.250279 | 1.085297 | 0.162101 | 0.193987 | 0.409676 | 0.148681 | 0 |
| 34.40278 | 0.45113 | 0.188479 | 0.925824 | -0.25302 | 0.369433 | 1.197114 | 0.14805 | 0.33397 | 0.763947 | 0.309033 | 0 |
| 34.41667 | 0.424104 | 0.247791 | 0.927087 | -0.06279 | 0.413152 | 0.8613 | 0.167286 | 0.514334 | 0.876998 | 0.650228 | 0 |
| 34.43056 | 0.335828 | 0.224796 | 0.777508 | 0.21746 | 0.291904 | 0.247977 | 0.133038 | 0.609193 | 0.668231 | 0.832039 | 0 |
| 34.44444 | 0.138914 | 0.054502 | 0.599699 | 0.420806 | 0.104357 | -0.20335 | 0.029694 | 0.569348 | 0.286611 | 0.747417 | 0 |
| 34.45833 | -0.19728 | -0.15006 | 0.458056 | 0.248979 | -0.09338 | -0.48878 | -0.11703 | 0.301237 | -0.04182 | 0.562497 | 0 |
| 34.47222 | -0.44862 | -0.22214 | 0.549366 | 0.031175 | -0.08017 | -0.3234 | -0.1679 | 0.063752 | -0.12926 | 0.488467 | 0 |
| 34.48611 | -0.15492 | -0.03759 | 1.093169 | 0.000806 | 0.413293 | 0.951923 | -0.14578 | 0.032954 | 0.26119 | 0.770538 | 0 |
| 34.5 | 0.475082 | 0.26202 | 1.898354 | -0.01265 | 1.089709 | 3.329741 | -0.19275 | 0.200692 | 1.189703 | 1.016196 | 0 |
| 34.51389 | 1.069847 | 0.318336 | 2.185648 | 0.139129 | 1.304057 | 4.941929 | -0.20889 | 0.535348 | 2.091028 | 1.153878 | 0 |
| 34.52778 | 1.455393 | 0.379938 | 2.100345 | 0.877185 | 1.457893 | 4.351204 | 0.230268 | 0.99442 | 3.048871 | 1.266469 | 0 |
| 34.54167 | 1.85223 | 0.654214 | 2.557204 | 1.182955 | 1.922882 | 3.58596 | 0.476201 | 1.303527 | 3.263225 | 1.694681 | 0 |
| 34.55556 | 1.712939 | 0.815332 | 2.508659 | 1.114423 | 2.227584 | 3.770781 | 0.673809 | 1.207191 | 2.972627 | 1.702231 | 0 |
| 34.56944 | 1.373989 | 0.846535 | 1.915212 | 1.050493 | 1.807892 | 2.665899 | 0.612762 | 0.855806 | 1.949896 | 1.378797 | 0 |
| 34.58333 | 1.076059 | 0.772372 | 1.330418 | 0.997122 | 1.180856 | 1.202645 | 0.528235 | 0.440158 | 0.552397 | 0.924107 | 0 |

| | | | | | | | | | | | |
|---|---|---|---|---|---|---|---|---|---|---|---|
| 34.59722 | 0.798001 | 0.778453 | 0.89008 | 1.188362 | 0.707759 | 0.17473 | 0.438528 | -0.08671 | -0.74315 | 0.504876 | 0 |
| 34.61111 | 0.195915 | 0.700079 | 0.54751 | 1.974919 | 0.508811 | 0.036906 | 0.486135 | -0.31968 | -1.03809 | 0.163129 | 0 |
| 34.625 | -0.87395 | 0.580137 | 0.17953 | 2.272256 | 0.406949 | -0.25948 | 0.52537 | -0.23364 | -1.24716 | 0.065915 | 0 |
| 34.63889 | -1.13739 | 0.553707 | -0.15239 | 1.864233 | 0.195688 | 0.13515 | 0.324481 | 0.04511 | -1.11382 | 0.381802 | 0 |
| 34.65278 | -0.16499 | 0.704926 | -0.46449 | 0.872949 | 0.441501 | -0.12599 | 0.141038 | 0.569397 | -0.68841 | 0.977687 | 0 |
| 34.66667 | 1.076995 | 0.893793 | -0.37655 | -0.61702 | 0.190487 | -0.13592 | 0.005514 | 0.790977 | -0.34494 | 1.347391 | 0 |
| 34.68056 | 1.667858 | 0.77945 | 0.024868 | -2.0439 | -0.0054 | -0.04807 | 0.110001 | 0.564959 | 0.173034 | 1.066293 | 0 |
| 34.69444 | 1.562948 | 0.601338 | 0.289128 | -2.55717 | -0.00778 | -0.57836 | 0.261743 | 0.355581 | 0.832785 | 0.494122 | 0 |
| 34.70833 | 1.03563 | 0.417409 | 0.651784 | -2.37318 | 0.372444 | -0.65955 | 0.470323 | 0.219033 | 1.44055 | 0.084295 | 0 |
| 34.72222 | 0.198984 | 0.059047 | 1.065656 | -1.3128 | 0.600435 | -0.54465 | 0.611539 | 0.080613 | 1.682205 | -0.498554 | 0 |
| 34.73611 | -0.82287 | -0.30862 | 0.796686 | -1.15121 | 0.499593 | -0.36514 | 0.644684 | -0.39493 | 1.516487 | -1.30281 | 0 |
| 34.75 | -1.12813 | -0.51686 | -0.01454 | -0.72166 | 0.496447 | -0.52445 | 0.500268 | -0.33998 | 1.190783 | -1.528969 | 0 |
| 34.76389 | -0.54352 | -0.62483 | -0.83355 | -0.198 | 0.724882 | -0.95355 | 0.263648 | 0.131607 | 0.728001 | -1.170257 | 0 |
| 34.77778 | -0.34994 | -0.61816 | -1.57432 | 0.160787 | 0.879485 | -1.42387 | 0.000438 | 0.48141 | 0.290067 | -0.910952 | 0 |
| 34.79167 | -0.6392 | -0.27657 | -3.04333 | 0.436663 | 0.694276 | -1.63397 | -0.15201 | 0.526465 | -0.25886 | -0.68154 | 0 |
| 34.80556 | -0.07795 | 0.221725 | -3.00581 | 0.521476 | 0.153106 | -1.46296 | -0.10113 | 0.169708 | -0.76074 | -0.414891 | 0 |
| 34.81944 | 0.470363 | 0.353528 | -1.45481 | 0.498616 | -0.29639 | -1.3554 | 0.015912 | -0.32273 | -1.10242 | -0.277244 | 0 |
| 34.83333 | 0.430524 | 0.139556 | -0.3657 | 0.274269 | -0.632 | -1.595 | 0.035195 | -0.84818 | -1.21763 | -0.793712 | 0 |
| 34.84722 | 0.49058 | 0.129431 | 0.090297 | -0.13082 | -0.86189 | -2.06081 | 0.088253 | -1.47555 | -1.16182 | -1.365014 | 0 |
| 34.86111 | 0.038956 | 0.132204 | -0.2896 | -0.52755 | -0.83917 | -2.29606 | 0.251177 | -1.72525 | -1.11182 | -1.720657 | 0 |
| 34.875 | -0.87013 | 0.050269 | -1.17729 | -0.45082 | -0.95892 | -2.1542 | 0.356999 | -1.56362 | -1.36182 | -1.663142 | 0 |
| 34.88889 | -1.16672 | 0.085779 | -2.09075 | -0.14737 | -1.03457 | -1.93116 | 0.428175 | -0.89638 | -1.28411 | -1.721788 | 0 |
| 34.90278 | -1.48739 | 0.299996 | -2.40554 | -0.03386 | -1.17992 | -1.76571 | 0.487913 | -0.05114 | -1.01013 | -1.516207 | 0 |
| 34.91667 | -1.51788 | 0.456858 | -2.19063 | -0.11517 | -1.16256 | -1.69135 | 0.535605 | 0.367414 | -0.73504 | -1.054296 | 0 |
| 34.93056 | -1.09601 | 0.453246 | -1.81984 | -0.11967 | -0.91039 | -0.92373 | 0.64812 | 0.562351 | -0.38388 | -0.572937 | 0 |
| 34.94444 | -0.53535 | 0.219481 | -1.35136 | -0.03665 | -0.38831 | -0.0488 | 0.729548 | 0.554853 | -0.02478 | -0.363383 | 0 |
| 34.95833 | 0.114749 | -0.01029 | -0.58482 | -0.05529 | 0.088288 | 0.638537 | 0.871157 | 0.371271 | 0.432526 | -0.345043 | 0 |
| 34.97222 | 0.670407 | -0.30262 | 0.29069 | -0.19832 | 0.472451 | 1.119718 | 0.94958 | 0.177697 | 0.999809 | -0.3696 | 0 |
| 34.98611 | 0.727422 | -0.82832 | 0.935759 | -0.43852 | 0.472289 | 1.165143 | 1.122033 | 0.306105 | 1.598412 | -0.521427 | 0 |
| 35 | -0.18235 | -1.52417 | 0.770079 | -0.82076 | -0.29727 | 0.398478 | 1.403004 | 0.141867 | 1.637646 | -0.929891 | 0 |
| 35.01389 | -1.85714 | -3.01527 | -0.26112 | -1.83019 | -1.16991 | -0.30077 | 1.116094 | -1.18762 | 0.902585 | -1.576084 | 0 |
| 35.02778 | -3.06314 | -3.67235 | -1.25869 | -1.39812 | -1.46332 | -0.70779 | 0.485274 | -1.17298 | 0.301551 | -2.202355 | 0 |
| 35.04167 | -2.09801 | -3.17702 | -1.71736 | -0.71156 | -1.46622 | -0.33825 | -0.03656 | -0.56171 | -0.03459 | -2.008963 | 0 |
| 35.05556 | -1.90833 | -1.93676 | -1.83361 | -0.46046 | -1.46965 | 0.090293 | 0.0772 | -0.31759 | 0.320601 | -1.17154 | 0 |
| 35.06944 | -1.34796 | -1.07275 | -1.5317 | -0.27071 | -0.85445 | 0.462027 | 0.89106 | 0.007189 | 1.066954 | 0.494619 | 0 |
| 35.08333 | -0.45276 | -0.06338 | -0.90656 | 0.533171 | 0.110804 | 0.714097 | 1.28942 | 0.718765 | -0.05336 | 1.823884 | 0 |
| 35.09722 | 0.552933 | 1.241646 | -0.24909 | 1.2773 | -0.38567 | 0.764338 | 0.539644 | 1.750017 | -0.30617 | 2.026597 | 0 |
| 35.11111 | 1.362088 | 2.249809 | 0.209728 | 0.960478 | -0.64709 | 0.729499 | -0.00616 | 1.498882 | 0.573123 | 1.351978 | 0 |
| 35.125 | 1.843205 | 2.297511 | 0.531647 | 0.387503 | -0.44847 | 0.65355 | 0.034667 | 1.036232 | 1.000608 | 0.909174 | 0 |
| 35.13889 | 2.111004 | 2.58206 | 0.85085 | 0.164116 | -0.0429 | 0.592993 | 0.418425 | 1.084465 | 1.272379 | 0.912484 | 0 |
| 35.15278 | 2.162989 | 2.71448 | 1.075736 | 0.100125 | 0.307276 | 0.479882 | 0.711462 | 1.355578 | 1.846981 | 0.936895 | 0 |
| 35.16667 | 1.911181 | 2.876076 | 1.253653 | 0.061568 | 0.395524 | 0.33518 | 0.699932 | 1.652472 | 1.25982 | 0.981693 | 0 |
| 35.18056 | 2.449785 | 2.787114 | 1.58182 | 0.334151 | 0.35912 | 0.399605 | 0.661542 | 2.124272 | 0.674636 | 1.02584 | 0 |
| 35.19444 | 2.830241 | 2.420144 | 1.684378 | 0.691579 | 0.421534 | 0.648182 | 0.631517 | 2.177902 | 1.917556 | 1.242842 | 0 |
| 35.20833 | 3.153942 | 1.910396 | 1.536879 | 0.775808 | 0.619868 | 0.860842 | 0.593388 | 1.57963 | 2.257644 | 1.245984 | 0 |
| 35.22222 | 2.243805 | 1.35882 | 1.115471 | 0.587438 | 0.880188 | 1.055231 | 0.468208 | 0.845294 | 1.80459 | 0.92275 | 0 |
| 35.23611 | 1.068373 | 0.527699 | 0.700742 | 0.225502 | 0.560949 | 1.07642 | 0.364191 | 0.181436 | 0.822496 | 0.323959 | 0 |
| 35.25 | 0.352573 | -0.11109 | 0.531657 | -0.19369 | 0.115228 | 0.690004 | 0.170418 | -0.3053 | -0.31725 | -0.186482 | 0 |
| 35.26389 | 0.210364 | -0.26389 | 0.627482 | -0.39965 | -0.06483 | 0.595396 | 0.065291 | -0.4865 | -1.02366 | -0.34257 | 0 |
| 35.27778 | 0.287009 | -0.0429 | 0.752586 | -0.30532 | -0.09302 | 0.784109 | 0.057407 | -0.39191 | -1.08051 | -0.263814 | 0 |
| 35.29167 | 0.30122 | 0.116385 | 0.746025 | -0.25884 | -0.09595 | 0.986067 | -0.01229 | -0.37739 | -0.75422 | -0.338043 | 0 |
| 35.30556 | 0.053878 | -0.01861 | 0.653893 | -0.38193 | -0.07645 | 1.063121 | -0.142 | -0.65629 | -0.25658 | -0.529863 | 0 |
| 35.31944 | -0.3477 | -0.30088 | 0.562506 | -0.56533 | 0.012582 | 0.772714 | -0.12088 | -0.95821 | 0.213548 | -0.550727 | 0 |
| 35.33333 | -0.50592 | -0.54951 | 0.447996 | -0.76373 | 0.063032 | 0.17469 | -0.01663 | -1.08717 | 0.341001 | -0.557397 | 0 |
| 35.34722 | -0.35449 | -0.72896 | 0.256152 | -0.86151 | 0.039712 | -0.14419 | -0.03943 | -1.0866 | 0.170909 | -0.6745 | 0 |
| 35.36111 | 0.207941 | -0.69588 | 0.057402 | -0.62553 | 0.02486 | -0.02645 | -0.06531 | -0.77332 | -0.04885 | -0.602808 | 0 |
| 35.375 | 0.868062 | -0.32618 | 0.089829 | -0.09095 | 0.084282 | 0.333547 | 0.060666 | -0.03923 | 0.0403 | -0.147275 | 0 |

| | | | | | | | | | | | |
|---|---|---|---|---|---|---|---|---|---|---|---|
| 35.38889 | 1.389452 | 0.118875 | 0.392032 | 0.115719 | 0.169898 | 0.592457 | 0.248987 | 0.542625 | 0.551813 | 0.261043 | 0 |
| 35.40278 | 1.524392 | 0.313852 | 0.5694 | -0.21467 | 0.185978 | 0.558477 | 0.290246 | 0.496204 | 1.144032 | 0.344468 | 0 |
| 35.41667 | 1.323452 | 0.454967 | 0.582454 | -0.57725 | 0.134568 | 0.38108 | 0.183273 | 0.309058 | 1.679728 | 0.294122 | 0 |
| 35.43056 | 1.107877 | 0.77935 | 0.637836 | -0.55324 | 0.090478 | 0.408731 | 0.167312 | 0.321609 | 2.002907 | 0.395903 | 0 |
| 35.44444 | 0.9045 | 1.018416 | 0.8526 | -0.31504 | 0.116256 | 0.64535 | 0.226841 | 0.449472 | 1.649927 | 0.487593 | 0 |
| 35.45833 | 0.366529 | 0.96004 | 1.003531 | -0.06648 | 0.218599 | 0.962749 | 0.253396 | 0.517856 | 1.083987 | 0.495056 | 0 |
| 35.47222 | -0.16409 | 0.682846 | 0.91235 | 0.234374 | 0.273417 | 1.03019 | 0.227031 | 0.607861 | 0.504083 | 0.467009 | 0 |
| 35.48611 | 0.118997 | 0.451777 | 0.82232 | 0.719265 | 0.289842 | 1.191097 | 0.019901 | 0.728803 | 0.052312 | 0.48052 | 0 |
| 35.5 | 1.036667 | 0.323652 | 1.107057 | 1.450854 | 0.335075 | 1.862379 | -0.37614 | 0.991497 | 0.080067 | 0.888453 | 0 |
| 35.51389 | 2.06752 | 0.19853 | 1.556955 | 1.505597 | 0.341711 | 2.813564 | -0.63613 | 1.618028 | 0.711072 | 1.865612 | 0 |
| 35.52778 | 3.162028 | 0.201195 | 1.486974 | 1.672523 | 0.106421 | 2.552997 | -0.37861 | 3.233843 | 1.751171 | 2.852236 | 0 |
| 35.54167 | 3.941124 | 0.64135 | 1.709506 | 1.940782 | 0.084902 | 2.116387 | 0.047309 | 4.390333 | 2.83441 | 3.315088 | 0 |
| 35.55556 | 2.879259 | 0.545458 | 2.363813 | 0.95807 | 0.429358 | 1.804166 | 0.230335 | 2.373661 | 3.036672 | 2.394877 | 0 |
| 35.56944 | 1.061274 | 0.334804 | 1.745583 | 0.412276 | 0.793009 | 1.46269 | 0.267477 | 0.841645 | 2.227136 | 1.106196 | 0 |
| 35.58333 | -0.27692 | 0.08266 | 0.955281 | 0.037743 | 0.953352 | 1.053393 | 0.114711 | -0.05904 | 1.320265 | 0.128048 | 0 |
| 35.59722 | -1.29995 | -0.24505 | 0.326238 | -0.19789 | 1.064495 | 0.378808 | 0.025401 | -0.75401 | 0.483671 | -0.485549 | 0 |
| 35.61111 | -1.69035 | -0.27058 | 0.029524 | -0.34274 | 1.274517 | -0.69508 | -0.0517 | -1.18809 | -0.26951 | -0.896599 | 0 |
| 35.625 | -1.633 | -0.6048 | -0.15725 | -0.65432 | 1.458649 | -1.47191 | 0.002836 | -1.69789 | -0.94802 | -1.076438 | 0 |
| 35.63889 | -1.71054 | -0.50995 | -0.36739 | -0.94125 | 1.596948 | -1.30407 | 0.058416 | -2.03212 | -0.89608 | -0.816468 | 0 |
| 35.65278 | -0.64776 | 0.049523 | -0.51827 | -1.30959 | 1.611175 | -0.67542 | 0.104759 | -1.98229 | -0.54984 | -0.575867 | 0 |
| 35.66667 | 0.788288 | 0.778002 | -0.5478 | -1.16715 | 1.390969 | -0.29931 | 0.038218 | -1.01909 | -0.31113 | -0.221774 | 0 |
| 35.68056 | 1.690826 | 1.110172 | -0.43395 | -0.88411 | 1.003965 | -0.02153 | -0.06069 | -0.01856 | -0.39654 | -0.049827 | 0 |
| 35.69444 | 2.311233 | 1.183168 | 0.001583 | -0.90622 | 0.673984 | 0.113212 | -0.13657 | 0.4287 | -0.36763 | 0.188243 | 0 |
| 35.70833 | 2.233239 | 0.891868 | 0.621019 | -0.72128 | 0.524006 | 0.441124 | -0.07775 | 0.482276 | 0.031244 | 0.370962 | 0 |
| 35.72222 | 1.015903 | 0.34939 | 0.890094 | -0.48004 | 0.578969 | 0.833131 | 0.071436 | 0.324894 | 0.645686 | 0.089787 | 0 |
| 35.73611 | 0.010585 | -0.28252 | 1.285903 | -0.39045 | 0.616367 | 1.170796 | 0.219066 | -0.10756 | 1.157609 | -0.655054 | 0 |
| 35.75 | -1.36262 | -0.94155 | 1.275385 | -0.784 | 0.676861 | 1.186172 | 0.264991 | -1.60322 | 1.505062 | -1.273496 | 0 |
| 35.76389 | -2.24444 | -1.46637 | 0.710658 | -1.7462 | 0.722607 | 1.176722 | 0.278479 | -3.84422 | 1.587373 | -1.398382 | 0 |
| 35.77778 | -2.79005 | -1.71199 | 0.255692 | -1.43719 | 0.71069 | 1.044719 | 0.25228 | -2.22791 | 1.299538 | -1.113897 | 0 |
| 35.79167 | -3.15303 | -1.63658 | -0.33548 | -0.80743 | 0.675322 | 0.690406 | 0.124814 | -2.18205 | 0.757538 | -1.282542 | 0 |
| 35.80556 | -2.5477 | -1.5962 | -0.93049 | -1.31492 | 0.669149 | 0.415621 | -0.02993 | -2.97883 | -0.05409 | -1.831897 | 0 |
| 35.81944 | -2.02638 | -1.08567 | -1.21952 | -1.82673 | 0.637526 | 0.342881 | -0.1246 | -3.349 | -0.7335 | -1.901974 | 0 |
| 35.83333 | -1.03937 | -0.43458 | -1.47438 | -1.9444 | 0.482257 | -0.0474 | -0.11491 | -2.59389 | -1.31371 | -1.867657 | 0 |
| 35.84722 | -0.59729 | -0.41582 | -1.49772 | -1.93825 | 0.20473 | -0.38541 | -0.07195 | -2.06749 | -1.69566 | -1.876794 | 0 |
| 35.86111 | -0.55141 | -0.88887 | -1.70438 | -1.96516 | -0.09753 | -0.60738 | -0.04457 | -2.07273 | -1.73913 | -2.224907 | 0 |
| 35.875 | -0.64754 | -1.46051 | -1.63503 | -1.90295 | -0.36115 | -0.73133 | 0.000237 | -2.42253 | -1.55043 | -2.664951 | 0 |
| 35.88889 | -0.81881 | -1.62017 | -1.43689 | -1.42754 | -0.61609 | -0.74584 | 0.125972 | -3.09075 | -1.349 | -2.858758 | 0 |
| 35.90278 | -0.45494 | -0.81611 | -1.18715 | -0.72007 | -0.94766 | -1.00292 | 0.204364 | -3.46642 | -1.07103 | -2.946997 | 0 |
| 35.91667 | 0.206471 | 0.206003 | -0.80655 | -0.12674 | -1.0242 | -1.11806 | 0.217954 | -3.96047 | -0.88241 | -2.423056 | 0 |
| 35.93056 | 0.735618 | 1.080128 | -0.76915 | 0.269614 | -0.81321 | -0.99486 | 0.325077 | -4.30602 | -0.47304 | -1.490526 | 0 |
| 35.94444 | 1.001753 | 1.67691 | -0.5948 | 0.558068 | -0.43758 | -0.59198 | 0.512423 | -2.09316 | 0.16479 | -0.297132 | 0 |
| 35.95833 | 0.995646 | 1.892573 | -0.40936 | 0.699585 | -0.07242 | -0.16063 | 0.681417 | -0.14595 | 0.750188 | 0.759923 | 0 |
| 35.97222 | 0.423767 | 1.564602 | -0.18346 | 0.992301 | 0.260791 | 0.219305 | 0.864409 | 1.7952 | 1.289825 | 1.594874 | 0 |
| 35.98611 | -0.75301 | 0.599655 | -0.2255 | 1.14551 | 0.180496 | 0.120423 | 0.933811 | 3.714652 | 1.530793 | 1.386634 | 0 |
| 36 | -2.40077 | -0.95682 | -1.12099 | 0.472588 | -0.4022 | -0.48246 | 0.747989 | 2.4092 | 0.979869 | -0.3272 | 0 |
| 36.01389 | -3.59712 | -2.08748 | -2.09604 | -0.68477 | -0.92591 | -0.85539 | 0.279978 | -0.62625 | 0.065758 | -2.534529 | 0 |
| 36.02778 | -3.14034 | -2.08855 | -1.77985 | -0.48023 | -0.80124 | -0.79105 | -0.11691 | -2.94669 | -0.33957 | -3.787527 | 0 |
| 36.04167 | -2.60813 | -2.09098 | -1.01437 | 0.283221 | -0.70804 | -0.68244 | -0.24841 | -3.60607 | -0.3269 | -2.634399 | 0 |
| 36.05556 | -1.13484 | -1.17232 | -0.39004 | 0.518171 | -0.69307 | -0.34922 | -0.09268 | -2.60549 | -0.40665 | -0.913416 | 0 |
| 36.06944 | -0.00435 | -0.0425 | -0.02104 | 0.701586 | -0.75881 | -0.00717 | 0.207932 | -1.36058 | -0.39706 | 0.066839 | 0 |
| 36.08333 | 0.732455 | 0.436253 | 0.156636 | 0.710819 | -0.60845 | 0.13616 | 0.320655 | -0.38457 | -0.35566 | 0.546515 | 0 |
| 36.09722 | 0.910966 | 0.395616 | 0.282254 | 0.690846 | -0.19186 | 0.073033 | 0.228964 | 0.421803 | -0.15501 | 1.407438 | 0 |
| 36.11111 | 0.898207 | 0.629839 | 0.351456 | 0.357276 | 0.066419 | 0.518056 | 0.087906 | 1.185189 | 0.05762 | 2.334361 | 0 |
| 36.125 | 1.375307 | 0.879066 | 0.331769 | -0.14735 | 0.065108 | 0.637643 | -0.0505 | 1.630017 | 0.301331 | 2.183513 | 0 |
| 36.13889 | 2.028155 | 1.035495 | 0.405001 | 0.062293 | -0.03286 | 0.565567 | -0.08215 | 1.830201 | 0.644246 | 1.609896 | 0 |
| 36.15278 | 2.459653 | 1.261411 | 0.476727 | 0.290191 | -0.03592 | 0.766147 | 0.236019 | 1.995683 | 1.199863 | 1.763155 | 0 |
| 36.16667 | 2.547062 | 1.510738 | 0.529972 | 0.37447 | 0.050964 | 0.694332 | 0.492139 | 2.111951 | 1.936013 | 2.058073 | 0 |

| | | | | | | | | | | | |
|---|---|---|---|---|---|---|---|---|---|---|---|
| 36.18056 | 2.64199 | 1.83682 | 0.64172 | 0.278295 | 0.128694 | 1.089275 | 0.655373 | 1.976624 | 2.381217 | 2.184804 | 0 |
| 36.19444 | 2.682128 | 1.92173 | 0.776808 | 0.022438 | 0.186298 | 1.200197 | 0.667908 | 1.595232 | 2.556515 | 2.058665 | 0 |
| 36.20833 | 2.387362 | 1.862151 | 0.887969 | -0.11427 | 0.228957 | 0.941554 | 0.609638 | 1.493175 | 2.555199 | 1.82994 | 0 |
| 36.22222 | 2.340802 | 1.842397 | 1.238626 | -0.05773 | 0.217885 | 0.494694 | 0.453453 | 1.38123 | 2.241214 | 1.665414 | 0 |
| 36.23611 | 1.957074 | 1.807882 | 1.186418 | -0.00622 | 0.205627 | 0.100095 | 0.122478 | 1.037988 | 1.336031 | 1.336402 | 0 |
| 36.25 | 1.911549 | 1.651059 | 1.131461 | 0.137641 | 0.421954 | 0.339193 | 0.13783 | 0.531989 | 0.655196 | 1.11109 | 0 |
| 36.26389 | 2.114399 | 1.649631 | 1.268532 | 0.564654 | 0.903707 | 1.075642 | 0.652431 | 0.31207 | 0.55355 | 1.164217 | 0 |
| 36.27778 | 2.023004 | 1.412749 | 1.132577 | 0.733372 | 1.010055 | 1.364905 | 0.910179 | 0.173128 | 0.545025 | 0.947173 | 0 |
| 36.29167 | 1.35931 | 0.70608 | 0.577749 | 0.278666 | 0.611002 | 1.010086 | 0.562777 | -0.13701 | 0.164755 | 0.299015 | 0 |
| 36.30556 | 0.50614 | -0.15483 | 0.27325 | -0.18041 | 0.089009 | 0.744207 | 0.261294 | -0.2252 | 0.039622 | -0.165739 | 0 |
| 36.31944 | -0.38972 | -0.73597 | 0.3977 | -0.27192 | -0.22771 | 0.843994 | 0.277288 | -0.09637 | 0.385114 | -0.082871 | 0 |
| 36.33333 | -0.42183 | -0.55238 | 0.381009 | -0.10898 | -0.31187 | 0.7431 | 0.3417 | 0.035207 | 0.36997 | 0.188851 | 0 |
| 36.34722 | 0.200228 | -0.13516 | 0.089105 | -0.07025 | -0.1893 | 0.435075 | 0.040962 | -0.19144 | -0.00277 | 0.026314 | 0 |
| 36.36111 | 0.497163 | -0.10939 | -0.12938 | -0.1177 | -0.2204 | 0.268905 | -0.36332 | -0.3648 | -0.12741 | -0.298764 | 0 |
| 36.375 | 0.416637 | -0.20679 | -0.11035 | -0.02633 | -0.29382 | 0.407266 | -0.38284 | -0.2548 | -0.03705 | -0.205514 | 0 |
| 36.38889 | 0.320954 | -0.20693 | 0.062085 | 0.082812 | -0.19722 | 0.811236 | -0.21467 | 0.108833 | 0.25777 | 0.053076 | 0 |
| 36.40278 | 0.33678 | -0.30392 | 0.318 | 0.118064 | -0.11254 | 1.297584 | -0.23657 | 0.588371 | 0.608614 | 0.092503 | 0 |
| 36.41667 | 0.391295 | -0.34968 | 0.678292 | 0.183192 | -0.07833 | 1.284893 | -0.31225 | 1.047744 | 0.908114 | 0.224565 | 0 |
| 36.43056 | 0.562624 | -0.22256 | 1.084979 | 0.243092 | -0.04058 | 0.89448 | -0.22823 | 1.337143 | 1.117482 | 0.462953 | 0 |
| 36.44444 | 0.731213 | -0.02593 | 0.98942 | 0.275887 | -0.03816 | 0.388969 | -0.07427 | 1.464641 | 1.17571 | 0.613285 | 0 |
| 36.45833 | 0.765712 | 0.054106 | 0.611611 | 0.282608 | 0.037094 | -0.25046 | -0.1243 | 1.427477 | 0.996461 | 0.55133 | 0 |
| 36.47222 | 0.544876 | 0.065212 | 0.348855 | 0.285902 | 0.035799 | -1.01441 | -0.26432 | 1.151421 | 0.805118 | 0.429447 | 0 |
| 36.48611 | 0.615619 | 0.07335 | 0.21514 | 0.349053 | 0.09491 | -0.76049 | -0.448 | 1.193728 | 0.608677 | 0.354122 | 0 |
| 36.5 | 1.447356 | 0.07148 | 1.504484 | 0.390651 | 0.101548 | -0.14959 | -0.80111 | 1.955648 | 0.458712 | 0.476275 | 0 |
| 36.51389 | 2.647155 | 0.22614 | 2.80249 | 0.341162 | 0.139623 | 0.548824 | -0.86228 | 3.233441 | 0.528396 | 0.79051 | 0 |
| 36.52778 | 3.460941 | 0.626318 | 3.260396 | 0.589804 | 0.537138 | 1.533252 | -0.37845 | 5.238788 | 1.331245 | 1.382819 | 0 |
| 36.54167 | 3.341962 | 1.020535 | 2.914668 | 1.006354 | 0.934349 | 1.685286 | 0.305804 | 7.639046 | 2.42071 | 2.181074 | 0 |
| 36.55556 | 2.242532 | 0.982285 | 2.128921 | 1.069361 | 1.229629 | 1.408251 | 0.525061 | 6.327948 | 3.218451 | 2.056436 | 0 |
| 36.56944 | 0.849439 | 0.719913 | 1.583877 | 0.869987 | 1.492897 | 1.123469 | 0.449908 | 3.707429 | 2.607713 | 1.486039 | 0 |
| 36.58333 | -0.52896 | 0.542235 | 1.140891 | 0.632654 | 1.432145 | 0.844568 | 0.176212 | 1.762384 | 2.042118 | 0.86136 | 0 |
| 36.59722 | -1.2396 | 0.569547 | 0.681806 | 0.430722 | 1.434137 | 0.485037 | -0.03248 | 0.828655 | 1.908893 | 0.407059 | 0 |
| 36.61111 | -1.79587 | 0.588251 | 0.214775 | 0.245846 | 1.599951 | 0.241578 | -0.11503 | 0.468646 | 1.937392 | 0.08748 | 0 |
| 36.625 | -1.80438 | 0.497648 | -0.29873 | 0.083154 | 1.768476 | 0.546929 | -0.06296 | -1.8E-05 | 1.621363 | -0.210411 | 0 |
| 36.63889 | -1.77636 | 0.332447 | -0.71351 | -0.00262 | 1.685658 | 1.218562 | 0.10669 | -0.66672 | 0.832146 | -0.482633 | 0 |
| 36.65278 | -1.84042 | -0.08551 | -0.8137 | -0.04146 | 1.498963 | 1.837003 | 0.121303 | -1.48153 | -0.1393 | -0.686363 | 0 |
| 36.66667 | -1.99834 | -0.58587 | -1.20169 | -0.01378 | 1.079901 | 1.968074 | -0.08356 | -2.32205 | -1.03217 | -1.036867 | 0 |
| 36.68056 | -1.58208 | -0.73299 | -1.89311 | -0.30211 | 0.961802 | 1.460403 | -0.33374 | -2.39899 | -1.72487 | -1.230085 | 0 |
| 36.69444 | 0.22623 | -0.49258 | -1.36096 | -0.68478 | 0.940119 | 0.84634 | -0.47108 | -1.87646 | -2.0509 | -1.162295 | 0 |
| 36.70833 | 1.636089 | -0.28686 | -0.57558 | -1.04024 | 0.845345 | 0.197572 | -0.4791 | -1.57972 | -1.84391 | -1.074796 | 0 |
| 36.72222 | 2.623871 | -0.29216 | 0.131435 | -1.36584 | 0.753121 | -0.10022 | -0.36011 | -1.55375 | -1.25828 | -1.046956 | 0 |
| 36.73611 | 2.116459 | -0.0108 | 1.210101 | -1.41482 | 0.829332 | 0.454716 | -0.19308 | -1.48023 | -0.38643 | -0.760211 | 0 |
| 36.75 | 2.163862 | 0.26038 | 2.17464 | -1.14737 | 0.864617 | 1.192283 | -0.00347 | -0.75597 | 0.310833 | -0.36835 | 0 |
| 36.76389 | 1.415862 | 0.555861 | 1.806786 | -0.79114 | 0.857488 | 1.385519 | 0.189331 | -0.52207 | 0.854203 | -0.158069 | 0 |
| 36.77778 | 0.197606 | 0.639244 | 1.057087 | -0.423 | 0.728259 | 1.28186 | 0.34996 | -0.89307 | 1.552211 | -0.207703 | 0 |
| 36.79167 | -0.80994 | 0.629205 | 0.618454 | 0.073947 | 0.59437 | 1.226375 | 0.410652 | -1.41705 | 2.026177 | -0.424702 | 0 |
| 36.80556 | -0.7985 | 0.52487 | 0.269952 | 0.216977 | 0.479439 | 1.149629 | 0.345355 | -1.62361 | 1.76476 | -0.798726 | 0 |
| 36.81944 | -1.35286 | 0.331579 | -0.28552 | 0.049013 | 0.40598 | 0.902948 | 0.187117 | -2.01402 | 0.677538 | -1.532233 | 0 |
| 36.83333 | -0.96609 | -0.11089 | -1.17708 | -0.40974 | 0.303959 | 0.638351 | 0.00024 | -2.90185 | -0.46074 | -1.985804 | 0 |
| 36.84722 | -1.28958 | -0.74746 | -1.91906 | -0.85027 | 0.089854 | 0.325011 | -0.10443 | -2.99918 | -1.4436 | -2.273492 | 0 |
| 36.86111 | -2.32281 | -1.22303 | -1.89615 | -1.17601 | -0.23433 | -0.20924 | -0.11469 | -2.66395 | -1.74405 | -2.408761 | 0 |
| 36.875 | -2.50998 | -1.32296 | -0.66298 | -1.53763 | -0.54897 | -0.55014 | -0.12643 | -2.01238 | -1.84272 | -2.873284 | 0 |
| 36.88889 | -2.13614 | -0.80396 | -0.6215 | -1.88268 | -0.78743 | -0.8213 | -0.28148 | -1.5439 | -1.78005 | -3.4523 | 0 |
| 36.90278 | -1.02935 | -0.38839 | -0.57593 | -1.48368 | -0.92465 | -0.89143 | -0.39709 | -1.94078 | -2.00321 | -3.263259 | 0 |
| 36.91667 | -0.22167 | -0.03843 | -0.1478 | -0.87498 | -0.99467 | -0.61442 | -0.45746 | -2.14901 | -1.83458 | -2.483349 | 0 |
| 36.93056 | 0.50022 | 0.152547 | 0.178721 | -0.73826 | -1.05876 | -0.22915 | -0.31003 | -2.26817 | -1.37894 | -1.756098 | 0 |
| 36.94444 | 0.98572 | 0.487584 | 0.338355 | -0.66474 | -1.00786 | 0.189072 | -0.13026 | -2.55883 | -0.92982 | -1.171859 | 0 |
| 36.95833 | 1.365132 | 0.830538 | 0.310263 | -0.5466 | -0.70163 | 0.326558 | 0.024979 | -2.82156 | -0.59313 | -0.856933 | 0 |

| | | | | | | | | | | | |
|---|---|---|---|---|---|---|---|---|---|---|---|
| 36.97222 | 1.302942 | 1.097887 | 0.258634 | -0.49904 | -0.35641 | 0.303095 | 0.141053 | -2.69477 | -0.22423 | -0.317533 | 0 |
| 36.98611 | 1.361536 | 1.093311 | 0.220642 | -0.4049 | -0.57143 | 0.195802 | 0.157521 | -2.47957 | 0.071549 | -0.087117 | 0 |
| 37 | 0.115827 | 0.147058 | 0.012842 | -0.50726 | -1.3037 | -0.04676 | -0.16293 | -2.95961 | 0.260012 | -0.636579 | 0 |
| 37.01389 | -1.61 | -0.88722 | -0.3531 | -1.16051 | -1.57088 | -0.18492 | -0.7435 | -3.09502 | 0.477823 | -1.756434 | 0 |
| 37.02778 | -2.36384 | -1.38957 | -0.68916 | -1.24332 | -1.29708 | -0.0771 | -0.825 | -2.42968 | 0.495557 | -2.556472 | 0 |
| 37.04167 | -1.9018 | -1.37199 | -0.76678 | -0.46567 | -1.23363 | -0.52661 | -0.83116 | -2.16723 | 0.458768 | -2.297158 | 0 |
| 37.05556 | -1.00931 | -1.20243 | -0.80965 | 0.243029 | -1.20388 | -0.75495 | -0.86339 | -2.40759 | 0.278372 | -1.678754 | 0 |
| 37.06944 | -0.46869 | -1.03032 | -0.85239 | 0.775002 | -0.92259 | -0.43881 | -0.45691 | -2.39525 | -0.08759 | -1.152616 | 0 |
| 37.08333 | -0.13453 | -0.85739 | -0.83596 | 0.743734 | -0.39408 | 0.084143 | -0.04882 | -1.99839 | -0.41702 | -0.670237 | 0 |
| 37.09722 | 0.242513 | -0.6296 | -0.67359 | 0.128932 | 0.108732 | 0.35155 | 0.062966 | -1.5402 | -0.53234 | -0.622931 | 0 |
| 37.11111 | 0.411343 | -0.49975 | -0.27493 | -0.29341 | 0.460253 | 0.528331 | 0.030287 | -1.15758 | -0.48757 | -0.589126 | 0 |
| 37.125 | 0.645433 | -0.23001 | 0.094616 | -0.32885 | 0.641239 | 0.954169 | 0.202957 | -0.4727 | -0.19693 | -0.194534 | 0 |
| 37.13889 | 1.129867 | 0.366509 | 0.598191 | 0.059866 | 0.966297 | 1.550286 | 0.697969 | 0.659433 | 0.355989 | 0.690901 | 0 |
| 37.15278 | 1.623542 | 0.928055 | 0.820541 | 0.391929 | 0.950146 | 1.749559 | 1.047413 | 1.855953 | 0.813622 | 1.47076 | 0 |
| 37.16667 | 2.162956 | 1.432388 | 0.764253 | 0.453096 | 0.628078 | 1.409822 | 1.116708 | 2.705533 | 0.909485 | 2.171726 | 0 |
| 37.18056 | 3.276209 | 2.17151 | 0.789361 | 0.503272 | 0.45899 | 1.214942 | 1.184044 | 3.214359 | 0.972488 | 2.905879 | 0 |
| 37.19444 | 4.26973 | 2.342293 | 0.851703 | 0.632208 | 0.35744 | 1.110411 | 1.198888 | 3.202031 | 0.924248 | 3.243886 | 0 |
| 37.20833 | 4.402905 | 2.341115 | 0.898335 | 0.695909 | 0.263251 | 1.253545 | 1.030247 | 3.16306 | 0.833872 | 3.222211 | 0 |
| 37.22222 | 3.929524 | 2.117343 | 0.944729 | 0.660855 | 0.233826 | 1.343984 | 0.834274 | 2.896352 | 0.895306 | 2.978522 | 0 |
| 37.23611 | 3.154801 | 1.783944 | 0.881998 | 0.456812 | 0.164771 | 1.254378 | 0.450801 | 2.40545 | 0.921182 | 2.2578 | 0 |
| 37.25 | 2.190776 | 1.444952 | 0.577097 | 0.117232 | 0.044258 | 0.981669 | -0.00231 | 1.551354 | 0.866385 | 1.252894 | 0 |
| 37.26389 | 0.923481 | 0.810769 | 0.222352 | -0.18527 | -0.11256 | 0.736726 | -0.22764 | 0.533484 | 0.796852 | 0.210517 | 0 |
| 37.27778 | 0.040865 | 0.374057 | 0.100344 | -0.32257 | -0.17984 | 0.613921 | -0.27636 | -0.19449 | 0.791498 | -0.405577 | 0 |
| 37.29167 | -0.12947 | 0.241813 | 0.116451 | -0.27189 | -0.11557 | 0.558082 | -0.23689 | -0.51554 | 0.784725 | -0.485253 | 0 |
| 37.30556 | 0.185563 | 0.220167 | 0.070669 | -0.16988 | -0.05736 | 0.519709 | -0.13422 | -0.44738 | 0.734226 | -0.291654 | 0 |
| 37.31944 | 0.425208 | 0.229607 | 0.012789 | -0.0875 | -0.0042 | 0.489624 | 0.023121 | -0.23203 | 0.679042 | -0.155445 | 0 |
| 37.33333 | 0.401604 | 0.161466 | 0.102952 | -0.14689 | 0.109856 | 0.553438 | 0.201612 | -0.10022 | 0.640719 | -0.212884 | 0 |
| 37.34722 | 0.40936 | 0.024248 | 0.267399 | -0.3095 | 0.168888 | 0.598349 | 0.244126 | -0.0657 | 0.606671 | -0.21014 | 0 |
| 37.36111 | 0.468432 | -0.14627 | 0.311783 | -0.43938 | 0.091169 | 0.523704 | 0.095367 | -0.02067 | 0.511015 | -0.193951 | 0 |
| 37.375 | 0.371895 | -0.29404 | 0.311556 | -0.46965 | -0.039 | 0.469292 | -0.16448 | 0.002715 | 0.463003 | -0.220468 | 0 |
| 37.38889 | 0.360415 | -0.31329 | 0.369989 | -0.43603 | -0.0194 | 0.533208 | -0.375 | 0.097907 | 0.410918 | -0.18802 | 0 |
| 37.40278 | 0.609215 | -0.19897 | 0.399546 | -0.31994 | 0.097604 | 0.516472 | -0.40656 | 0.2952 | 0.35815 | -0.055183 | 0 |
| 37.41667 | 0.83448 | -0.07682 | 0.538689 | -0.16318 | 0.253352 | 0.571486 | -0.23732 | 0.584589 | 0.330341 | 0.227533 | 0 |
| 37.43056 | 0.879887 | 0.01036 | 0.733808 | -0.00414 | 0.388501 | 0.669875 | 0.012778 | 0.813366 | 0.377718 | 0.513423 | 0 |
| 37.44444 | 0.755391 | -0.03235 | 0.831884 | 0.046016 | 0.457201 | 0.667517 | 0.026602 | 0.878727 | 0.347816 | 0.650333 | 0 |
| 37.45833 | 0.533911 | -0.10697 | 0.80267 | 0.067019 | 0.302786 | 0.59193 | -0.14271 | 0.880581 | 0.42086 | 0.483461 | 0 |
| 37.47222 | 0.21896 | -0.19723 | 0.788301 | -0.00218 | 0.121083 | 0.531785 | -0.32126 | 0.779479 | 0.480082 | 0.179774 | 0 |
| 37.48611 | 0.0502 | -0.32368 | 0.727392 | -0.18345 | 0.164554 | 0.288881 | -0.61872 | 0.802197 | 0.546728 | 0.008253 | 0 |
| 37.5 | 0.202222 | -0.26854 | 0.728077 | -0.18589 | 0.55983 | -0.00677 | -0.74055 | 1.205866 | 0.62292 | 0.266483 | 0 |
| 37.51389 | 0.842272 | 0.070132 | 0.888148 | 0.187477 | 0.912627 | -0.07482 | -0.31591 | 1.751291 | 0.800977 | 1.058644 | 0 |
| 37.52778 | 2.294302 | 0.80949 | 1.064229 | 0.784354 | 0.94873 | 0.16471 | -0.03086 | 2.445738 | 1.142938 | 2.135051 | 0 |
| 37.54167 | 4.465625 | 1.66589 | 1.277434 | 0.86187 | 0.745 | 0.423753 | 0.241247 | 3.01785 | 1.50521 | 2.846879 | 0 |
| 37.55556 | 3.049631 | 1.031909 | 1.395603 | 0.828901 | 1.026185 | 0.610655 | 0.513541 | 3.224923 | 1.901792 | 2.711903 | 0 |
| 37.56944 | 1.869108 | 0.698328 | 1.121252 | 0.414858 | 1.232546 | 0.701524 | 0.503286 | 3.035878 | 1.975623 | 2.106036 | 0 |
| 37.58333 | 0.633228 | 0.467513 | 0.846054 | -0.14141 | 1.070201 | 0.530978 | 0.208224 | 2.594433 | 1.725095 | 1.626349 | 0 |
| 37.59722 | -0.49256 | 0.11806 | 0.99106 | -0.54784 | 0.779234 | 0.37434 | -0.02847 | 2.352313 | 1.333311 | 1.110996 | 0 |
| 37.61111 | -1.24315 | -0.07755 | 1.296935 | -0.64139 | 0.548813 | 0.522419 | -0.22387 | 2.36645 | 1.068647 | 0.307329 | 0 |
| 37.625 | -1.51386 | -0.23935 | 1.145204 | -0.70627 | 0.672732 | 0.572316 | -0.34387 | 2.343554 | 0.969387 | -0.209184 | 0 |
| 37.63889 | -1.78932 | -0.36408 | 0.763885 | -0.857 | 0.738152 | 0.648664 | -0.28708 | 1.705395 | 0.993021 | -0.635567 | 0 |
| 37.65278 | -2.15753 | -0.68755 | 0.343746 | -0.92284 | 0.66775 | 0.700669 | -0.26307 | 0.322631 | 0.683764 | -1.188811 | 0 |
| 37.66667 | -1.91907 | -0.97857 | 0.901237 | -0.94905 | 0.510918 | 0.624077 | -0.5021 | -1.54273 | -0.31308 | -1.875168 | 0 |
| 37.68056 | -1.35776 | -0.86012 | 1.026573 | -0.60444 | 0.318421 | 0.52814 | -0.76516 | -3.54325 | -1.54573 | -2.615804 | 0 |
| 37.69444 | -0.74448 | -0.67477 | 0.264615 | -0.42003 | 0.193528 | 0.477977 | -0.86071 | -4.7145 | -1.7732 | -2.702355 | 0 |
| 37.70833 | -0.13723 | -0.46314 | -0.62161 | -0.28428 | 0.22585 | 0.431622 | -0.78015 | -4.50301 | -1.46824 | -2.408952 | 0 |
| 37.72222 | 0.431985 | 0.139982 | -0.7209 | -0.08111 | 0.334996 | 0.399299 | -0.72824 | -3.57269 | -1.28113 | -1.896686 | 0 |
| 37.73611 | 0.668073 | 0.780926 | 0.087369 | 0.089142 | 0.306924 | 0.440197 | -0.59802 | -2.43415 | -1.27614 | -1.12691 | 0 |
| 37.75 | 0.804092 | 0.70459 | -0.01327 | 0.091421 | 0.10268 | 0.461592 | -0.44685 | -1.56988 | -1.00245 | -0.393734 | 0 |

| | | | | | | | | | | | |
|---|---|---|---|---|---|---|---|---|---|---|---|
| 37.76389 | 0.568723 | 0.423952 | -0.18472 | -0.13905 | -0.17769 | 0.538878 | -0.28964 | -0.93218 | -0.54045 | 0.050864 | 0 |
| 37.77778 | 0.652026 | 0.09083 | -0.0488 | -0.44413 | -0.40449 | 0.679767 | -0.08528 | -0.21082 | -0.09188 | -0.204448 | 0 |
| 37.79167 | 0.350253 | -0.32625 | 0.267215 | -0.67323 | -0.44688 | 0.764995 | 0.005027 | 0.298244 | 0.037487 | -0.518644 | 0 |
| 37.80556 | -0.08605 | -0.53328 | 0.35466 | -0.73308 | -0.32435 | 0.66798 | -0.10543 | -0.2138 | -0.18109 | -0.891169 | 0 |
| 37.81944 | -0.56596 | -0.62996 | 0.281565 | -0.94097 | -0.33701 | 0.421864 | -0.3861 | -1.42558 | -0.24239 | -1.303639 | 0 |
| 37.83333 | -0.71345 | -0.63115 | 0.258344 | -1.26502 | -0.46022 | -0.00514 | -0.92483 | -2.93382 | -0.3328 | -1.859378 | 0 |
| 37.84722 | -0.91892 | -0.70371 | 0.501493 | -1.59701 | -1.03068 | -0.43653 | -1.69891 | -4.63012 | -0.48952 | -2.324792 | 0 |
| 37.86111 | -1.11555 | -0.54961 | 0.733976 | -1.85101 | -1.89142 | -0.70107 | -2.16573 | -5.69501 | -0.67755 | -2.740102 | 0 |
| 37.875 | -0.66365 | -0.43493 | 0.382561 | -1.79093 | -1.77748 | -0.62077 | -2.10809 | -5.1502 | -0.50891 | -2.929937 | 0 |
| 37.88889 | -0.39091 | -0.72263 | -0.40936 | -1.60231 | -1.45321 | -0.62304 | -2.00825 | -4.97861 | 0.141816 | -2.812897 | 0 |
| 37.90278 | -0.45347 | -1.18704 | -1.01327 | -1.50193 | -1.48134 | -0.49318 | -1.87506 | -5.01001 | 0.092116 | -2.857148 | 0 |
| 37.91667 | -1.14825 | -1.60117 | -1.15099 | -1.33545 | -1.03418 | -0.20113 | -1.43617 | -4.32217 | 0.015492 | -2.49088 | 0 |
| 37.93056 | -1.08045 | -1.40878 | -0.95118 | -1.11387 | -0.72843 | 0.073241 | -0.94432 | -3.52233 | 0.046617 | -2.00005 | 0 |
| 37.94444 | -0.17448 | -0.76841 | -0.6496 | -0.80942 | -0.39138 | 0.236204 | -0.43064 | -2.61505 | 0.190387 | -1.442804 | 0 |
| 37.95833 | 0.560237 | -0.24609 | -0.26212 | -0.35434 | -0.11751 | 0.461747 | 0.156585 | -1.62338 | 0.429278 | -0.956581 | 0 |
| 37.97222 | 0.41166 | -0.21371 | 0.217242 | 0.158914 | 0.174478 | 0.712813 | 0.817531 | -0.568 | 0.653944 | -0.534188 | 0 |
| 37.98611 | 0.526343 | -0.28495 | 0.665334 | 0.710487 | -0.11457 | 0.86861 | 0.943128 | -0.22619 | 0.76819 | -1.004833 | 0 |

**S1b Area under the motion curve (AUC) for control experiment**

| Day | Plant1 | Plant2 | Plant3 | Plant4 | Plant5 | Plant6 | Plant7 | Plant8 | Plant9 | Whole Tank ROI | Empty ROI |
|---|---|---|---|---|---|---|---|---|---|---|---|
| 18 | 0.450083 | 0.291443 | 0.200107 | 0.351355 | 0.030443 | 0.129136 | 2.748928 | 0.016816 | 0.001097 | 0.405243 | 0 |
| 18.01389 | 1.852455 | 1.740269 | 0.078379 | 0.089679 | 1.198368 | 0.092218 | 4.751277 | 0.41286 | 0.791296 | 1.599851 | 0 |
| 18.02778 | 2.642481 | 2.585331 | 0.552324 | 0.447297 | 2.516194 | 0.244314 | 2.99979 | 0.642641 | 1.212873 | 3.015904 | 0 |
| 18.04167 | 1.30252 | 1.281082 | 0.60062 | 0.184391 | 1.391648 | 0.440583 | 1.408413 | 0.334264 | 0.55164 | 2.143187 | 0 |
| 18.05556 | 0.103106 | 0.62603 | 0.370327 | 0.92628 | 0.076915 | 0.368268 | 0.042565 | 0.292843 | 0.200896 | 0.199701 | 0 |
| 18.06944 | 0.872961 | 2.621784 | 0.185485 | 2.646077 | 0.697789 | 0.04493 | 1.330566 | 0.988953 | 0.060511 | 1.649717 | 0 |
| 18.08333 | 0.451911 | 2.263823 | 0.050214 | 2.884997 | 0.669681 | 0.17018 | 1.835847 | 0.805798 | 0.059097 | 1.045999 | 0 |
| 18.09722 | 0.254443 | 0.607823 | 0.305041 | 0.951926 | 0.053135 | 0.085974 | 1.348679 | 0.012386 | 0.198906 | 0.353299 | 0 |
| 18.11111 | 0.51426 | 0.283509 | 0.394718 | 0.132834 | 0.319323 | 0.334738 | 0.769237 | 0.428334 | 0.308892 | 0.536401 | 0 |
| 18.125 | 0.586124 | 0.421887 | 0.350188 | 0.225591 | 0.395236 | 0.468771 | 0.304266 | 0.504036 | 0.420282 | 0.156197 | 0 |
| 18.13889 | 0.614704 | 0.449133 | 0.334477 | 0.265409 | 0.440221 | 0.57275 | 0.10134 | 0.507438 | 0.545897 | 0.149032 | 0 |
| 18.15278 | 0.552741 | 0.394598 | 0.269642 | 0.291163 | 0.40215 | 0.544969 | 0.334579 | 0.43025 | 0.515507 | 0.352378 | 0 |
| 18.16667 | 0.484487 | 0.315204 | 0.19548 | 0.277434 | 0.314374 | 0.482292 | 0.450543 | 0.35946 | 0.398543 | 0.35719 | 0 |
| 18.18056 | 0.519592 | 0.363085 | 0.243271 | 0.325782 | 0.333856 | 0.567138 | 0.534493 | 0.417827 | 0.360175 | 0.298919 | 0 |
| 18.19444 | 0.569875 | 0.464882 | 0.328587 | 0.397558 | 0.411239 | 0.789152 | 0.571803 | 0.525506 | 0.415985 | 0.344947 | 0 |
| 18.20833 | 0.52957 | 0.491375 | 0.317426 | 0.390513 | 0.381744 | 1.082185 | 0.463725 | 0.590586 | 0.395312 | 0.32645 | 0 |
| 18.22222 | 0.459583 | 0.435572 | 0.256901 | 0.369169 | 0.27941 | 1.399286 | 0.321775 | 0.71782 | 0.342847 | 0.379536 | 0 |
| 18.23611 | 0.510769 | 0.465375 | 0.284437 | 0.480562 | 0.222945 | 1.359487 | 0.34307 | 1.007999 | 0.393135 | 0.592202 | 0 |
| 18.25 | 0.661018 | 0.6068 | 0.418126 | 0.635033 | 0.140377 | 1.211758 | 0.506144 | 1.243165 | 0.536347 | 0.758685 | 0 |
| 18.26389 | 0.776581 | 0.713954 | 0.540901 | 0.670105 | 0.052297 | 1.101343 | 0.619379 | 1.171623 | 0.650931 | 0.649808 | 0 |
| 18.27778 | 0.805069 | 0.753668 | 0.596764 | 0.644982 | 0.070767 | 1.055067 | 0.641159 | 1.002113 | 0.676382 | 0.555865 | 0 |
| 18.29167 | 0.81841 | 0.805503 | 0.608509 | 0.679851 | 0.114747 | 1.01986 | 0.648312 | 0.885599 | 0.680153 | 0.588805 | 0 |
| 18.30556 | 0.846304 | 0.870449 | 0.612271 | 0.753381 | 0.251358 | 1.004732 | 0.641761 | 0.843981 | 0.687534 | 0.662296 | 0 |
| 18.31944 | 0.800407 | 0.853274 | 0.573174 | 0.721603 | 0.318617 | 0.912923 | 0.582839 | 0.763863 | 0.633341 | 0.668323 | 0 |
| 18.33333 | 0.632558 | 0.710302 | 0.452061 | 0.54258 | 0.37259 | 0.709578 | 0.372719 | 0.612167 | 0.489755 | 0.551497 | 0 |
| 18.34722 | 0.434733 | 0.542336 | 0.280458 | 0.329946 | 0.477231 | 0.507681 | 0.156217 | 0.451416 | 0.312119 | 0.370946 | 0 |
| 18.36111 | 0.251295 | 0.425702 | 0.111797 | 0.153578 | 0.662499 | 0.33669 | 0.010306 | 0.309625 | 0.147508 | 0.267325 | 0 |
| 18.375 | 0.104787 | 0.392656 | 0.025774 | 0.01021 | 0.999534 | 0.1949 | 0.085753 | 0.179719 | 0.015982 | 0.291913 | 0 |
| 18.38889 | 0.054184 | 0.482726 | 0.077156 | 0.037297 | 1.208066 | 0.176706 | 0.135281 | 0.084203 | 0.027759 | 0.484505 | 0 |
| 18.40278 | 0.128385 | 0.700908 | 0.000589 | 0.05949 | 1.348564 | 0.280907 | 0.040963 | 0.097423 | 0.05219 | 0.651095 | 0 |
| 18.41667 | 0.2588 | 0.866338 | 0.128802 | 0.215609 | 1.489346 | 0.423298 | 0.107357 | 0.187859 | 0.188876 | 0.834594 | 0 |
| 18.43056 | 0.343727 | 0.875512 | 0.212807 | 0.325553 | 1.554765 | 0.51273 | 0.204931 | 0.265524 | 0.289257 | 0.966138 | 0 |
| 18.44444 | 0.383146 | 0.775098 | 0.257908 | 0.394455 | 1.565662 | 0.576272 | 0.240301 | 0.308833 | 0.34804 | 1.01644 | 0 |
| 18.45833 | 0.469657 | 0.724362 | 0.353547 | 0.498114 | 1.6544 | 0.668658 | 0.347596 | 0.383239 | 0.438702 | 1.026604 | 0 |
| 18.47222 | 0.656283 | 0.844707 | 0.538375 | 0.688849 | 1.699778 | 0.833087 | 0.480853 | 0.515214 | 0.607193 | 1.068763 | 0 |
| 18.48611 | 1.00586 | 1.189838 | 0.827358 | 1.081267 | 1.807508 | 1.120264 | 0.412195 | 0.586667 | 0.856657 | 1.198163 | 0 |
| 18.5 | 1.425498 | 1.675218 | 1.15231 | 1.453276 | 1.958409 | 1.478379 | 0.234662 | 0.581421 | 1.116618 | 1.265365 | 0 |
| 18.51389 | 1.442717 | 1.739873 | 1.225942 | 1.070114 | 1.832037 | 1.450072 | 0.330398 | 0.668875 | 1.099914 | 1.351471 | 0 |
| 18.52778 | 0.932712 | 1.208169 | 0.907058 | 0.028367 | 1.290046 | 0.946098 | 0.486451 | 0.579612 | 0.748055 | 1.096237 | 0 |
| 18.54167 | 0.344999 | 0.544779 | 0.393849 | 0.631093 | 0.752692 | 0.381125 | 0.333248 | 0.350359 | 0.293522 | 0.697195 | 0 |
| 18.55556 | 0.003356 | 0.156486 | 0.060985 | 0.545835 | 0.424604 | 0.057763 | 0.110052 | 0.09565 | 0.027389 | 0.33619 | 0 |
| 18.56944 | 0.11948 | 0.015436 | 0.01425 | 0.323366 | 0.300571 | 0.042018 | 0.024472 | 0.045501 | 0.050723 | 0.189651 | 0 |
| 18.58333 | 0.169435 | 0.044972 | 0.021301 | 0.221573 | 0.216445 | 0.074608 | 0.103459 | 0.111091 | 0.05976 | 0.147609 | 0 |
| 18.59722 | 0.16233 | 0.027467 | 0.003658 | 0.100926 | 0.211147 | 0.038957 | 0.095838 | 0.082456 | 0.008346 | 0.236757 | 0 |
| 18.61111 | 0.158704 | 0.003666 | 0.031752 | 0.005068 | 0.262041 | 0.049654 | 0.040897 | 0.010025 | 0.057095 | 0.330407 | 0 |
| 18.625 | 0.148763 | 0.023542 | 0.064488 | 0.0758 | 0.307711 | 0.143485 | 0.052886 | 0.056496 | 0.115319 | 0.297954 | 0 |
| 18.63889 | 0.097338 | 0.088434 | 0.140365 | 0.187164 | 0.362616 | 0.204855 | 0.183951 | 0.123471 | 0.19086 | 0.276088 | 0 |
| 18.65278 | 0.075596 | 0.103758 | 0.165879 | 0.225568 | 0.36196 | 0.189866 | 0.266689 | 0.150438 | 0.203292 | 0.288482 | 0 |
| 18.66667 | 0.140047 | 0.078505 | 0.17722 | 0.21681 | 0.341606 | 0.179466 | 0.291193 | 0.176615 | 0.196279 | 0.289692 | 0 |
| 18.68056 | 0.032109 | 0.170111 | 0.255842 | 0.275211 | 0.410163 | 0.258868 | 0.354016 | 0.260677 | 0.277937 | 0.369119 | 0 |
| 18.69444 | 0.201742 | 0.377976 | 0.404064 | 0.450297 | 0.593013 | 0.451109 | 0.519475 | 0.424633 | 0.464543 | 0.595079 | 0 |
| 18.70833 | 0.365843 | 0.539679 | 0.550783 | 0.646188 | 0.783112 | 0.64215 | 0.69754 | 0.592759 | 0.650251 | 0.805553 | 0 |
| 18.72222 | 0.242648 | 0.448495 | 0.519257 | 0.623926 | 0.742198 | 0.599286 | 0.667717 | 0.554748 | 0.605509 | 0.75171 | 0 |
| 18.73611 | 0.062134 | 0.29124 | 0.45717 | 0.523293 | 0.624934 | 0.507034 | 0.573189 | 0.467889 | 0.490737 | 0.575352 | 0 |

| | | | | | | | | | | | |
|---|---|---|---|---|---|---|---|---|---|---|---|
| 18.75 | 0.093722 | 0.321209 | 0.554492 | 0.570288 | 0.674571 | 0.562827 | 0.627254 | 0.53249 | 0.545602 | 0.544007 | 0 |
| 18.76389 | 0.138201 | 0.369278 | 0.624564 | 0.619166 | 0.714761 | 0.579759 | 0.664788 | 0.584338 | 0.602765 | 0.571637 | 0 |
| 18.77778 | 0.121573 | 0.339012 | 0.596453 | 0.592738 | 0.667664 | 0.521441 | 0.623212 | 0.554536 | 0.583983 | 0.53219 | 0 |
| 18.79167 | 0.057968 | 0.290777 | 0.518336 | 0.536966 | 0.605769 | 0.458181 | 0.573489 | 0.488694 | 0.519541 | 0.458944 | 0 |
| 18.80556 | 0.021633 | 0.249824 | 0.451869 | 0.481032 | 0.538468 | 0.420605 | 0.531142 | 0.433387 | 0.451719 | 0.392232 | 0 |
| 18.81944 | 0.049714 | 0.262694 | 0.459455 | 0.447437 | 0.51738 | 0.419219 | 0.498354 | 0.412721 | 0.437139 | 0.42336 | 0 |
| 18.83333 | 0.114093 | 0.293477 | 0.514836 | 0.453218 | 0.546067 | 0.430813 | 0.507066 | 0.433683 | 0.465415 | 0.427211 | 0 |
| 18.84722 | 0.158137 | 0.318648 | 0.560335 | 0.457278 | 0.555707 | 0.420886 | 0.551031 | 0.449826 | 0.48539 | 0.402395 | 0 |
| 18.86111 | 0.108871 | 0.273727 | 0.527376 | 0.392112 | 0.468034 | 0.359103 | 0.549639 | 0.37897 | 0.435412 | 0.375104 | 0 |
| 18.875 | 0.057004 | 0.221058 | 0.48765 | 0.338105 | 0.396787 | 0.308183 | 0.531302 | 0.315398 | 0.392601 | 0.375438 | 0 |
| 18.88889 | 0.037172 | 0.190296 | 0.463112 | 0.316035 | 0.377393 | 0.285521 | 0.531178 | 0.289132 | 0.39176 | 0.374857 | 0 |
| 18.90278 | 0.022014 | 0.131994 | 0.417745 | 0.277066 | 0.332761 | 0.23838 | 0.529572 | 0.249826 | 0.362921 | 0.279438 | 0 |
| 18.91667 | 0.113485 | 0.0795 | 0.366216 | 0.240483 | 0.281122 | 0.200339 | 0.507163 | 0.212704 | 0.325921 | 0.155751 | 0 |
| 18.93056 | 0.211626 | 0.005942 | 0.279534 | 0.175385 | 0.20594 | 0.145496 | 0.444505 | 0.156872 | 0.260402 | 0.042532 | 0 |
| 18.94444 | 0.265533 | 0.050821 | 0.195388 | 0.107109 | 0.121363 | 0.063729 | 0.352163 | 0.098045 | 0.194456 | 0.010228 | 0 |
| 18.95833 | 0.259156 | 0.058638 | 0.162256 | 0.068737 | 0.072106 | 0.011902 | 0.308964 | 0.054816 | 0.165423 | 0.054756 | 0 |
| 18.97222 | 0.250736 | 0.08331 | 0.1424 | 0.060359 | 0.002044 | 0.087334 | 0.328923 | 0.01206 | 0.136696 | 0.088428 | 0 |
| 18.98611 | 0.392572 | 0.3124 | 0.07884 | 0.107438 | 0.274263 | 0.33505 | 0.569823 | 0.126899 | 0.033004 | 0.375584 | 0 |
| 19 | 0.502171 | 0.695654 | 0.012951 | 0.297646 | 0.760329 | 0.743491 | 1.672034 | 0.349166 | 0.387644 | 0.924012 | 0 |
| 19.01389 | 0.004634 | 0.565596 | 0.190845 | 0.676399 | 0.829387 | 0.71198 | 2.709803 | 0.339384 | 0.557476 | 0.506913 | 0 |
| 19.02778 | 0.847349 | 0.101186 | 0.425951 | 0.642632 | 0.36932 | 0.194903 | 0.985334 | 0.166441 | 0.354492 | 0.010083 | 0 |
| 19.04167 | 1.011076 | 0.456556 | 0.589008 | 0.12009 | 0.015229 | 0.182771 | 0.979658 | 0.129612 | 0.159455 | 0.075845 | 0 |
| 19.05556 | 0.77048 | 0.496331 | 0.488965 | 0.123146 | 0.089241 | 0.214903 | 1.827609 | 0.109066 | 0.090416 | 0.059132 | 0 |
| 19.06944 | 0.577116 | 0.434176 | 0.280815 | 0.073311 | 0.114445 | 0.185184 | 1.425666 | 0.047118 | 0.04739 | 0.032027 | 0 |
| 19.08333 | 0.386892 | 0.334849 | 0.112278 | 0.035238 | 0.141701 | 0.143587 | 0.67937 | 0.007354 | 0.01685 | 0.015533 | 0 |
| 19.09722 | 0.26881 | 0.271991 | 0.026835 | 0.071595 | 0.15883 | 0.107407 | 0.250602 | 0.032911 | 0.005148 | 0.027855 | 0 |
| 19.11111 | 0.208789 | 0.235613 | 0.012815 | 0.040725 | 0.148553 | 0.051144 | 0.140064 | 0.027298 | 0.014066 | 0.046595 | 0 |
| 19.125 | 0.151474 | 0.177732 | 0.049308 | 0.021446 | 0.101131 | 0.042398 | 0.177839 | 0.01979 | 0.065088 | 0.011432 | 0 |
| 19.13889 | 0.07047 | 0.0943 | 0.09931 | 0.091117 | 0.031343 | 0.130353 | 0.216914 | 0.072588 | 0.134994 | 0.08063 | 0 |
| 19.15278 | 0.0376 | 0.085712 | 0.092844 | 0.099114 | 0.016318 | 0.146002 | 0.18134 | 0.052387 | 0.134449 | 0.093322 | 0 |
| 19.16667 | 0.064027 | 0.122602 | 0.041976 | 0.036958 | 0.062456 | 0.111393 | 0.114181 | 0.012448 | 0.081299 | 0.042472 | 0 |
| 19.18056 | 0.066021 | 0.105026 | 0.028679 | 0.000297 | 0.087303 | 0.113895 | 0.081575 | 0.031736 | 0.063057 | 0.032615 | 0 |
| 19.19444 | 0.004055 | 0.031668 | 0.07655 | 0.053647 | 0.044621 | 0.157265 | 0.118411 | 0.012992 | 0.113128 | 0.044109 | 0 |
| 19.20833 | 0.113348 | 0.070516 | 0.174266 | 0.15971 | 0.04913 | 0.235112 | 0.216807 | 0.112916 | 0.212172 | 0.097384 | 0 |
| 19.22222 | 0.220317 | 0.160937 | 0.271716 | 0.229017 | 0.121022 | 0.317618 | 0.299509 | 0.20356 | 0.292928 | 0.139985 | 0 |
| 19.23611 | 0.19155 | 0.126538 | 0.244798 | 0.171101 | 0.082531 | 0.298341 | 0.265722 | 0.163711 | 0.244841 | 0.12762 | 0 |
| 19.25 | 0.005681 | 0.060828 | 0.044118 | 0.018016 | 0.082832 | 0.142562 | 0.108664 | 0.020888 | 0.048755 | 0.022415 | 0 |
| 19.26389 | 0.168409 | 0.200777 | 0.126036 | 0.141139 | 0.210149 | 0.003932 | 0.004378 | 0.150667 | 0.104639 | 0.170467 | 0 |
| 19.27778 | 0.138704 | 0.165315 | 0.101177 | 0.112608 | 0.156209 | 0.012793 | 0.00516 | 0.106175 | 0.083838 | 0.154382 | 0 |
| 19.29167 | 0.047494 | 0.086544 | 0.003207 | 0.077664 | 0.033249 | 0.076709 | 0.049459 | 0.011015 | 0.002699 | 0.053554 | 0 |
| 19.30556 | 0.02066 | 0.062285 | 0.03697 | 0.104453 | 0.007107 | 0.096007 | 0.077686 | 0.01992 | 0.036218 | 0.031293 | 0 |
| 19.31944 | 0.072107 | 0.110772 | 0.00528 | 0.177885 | 0.079963 | 0.045731 | 0.058724 | 0.028016 | 0.013774 | 0.048043 | 0 |
| 19.33333 | 0.202027 | 0.243967 | 0.124883 | 0.320155 | 0.187252 | 0.054943 | 0.020034 | 0.131358 | 0.127169 | 0.062074 | 0 |
| 19.34722 | 0.351939 | 0.395283 | 0.261041 | 0.553208 | 0.238426 | 0.090893 | 0.096852 | 0.231623 | 0.239656 | 0.083652 | 0 |
| 19.36111 | 0.399423 | 0.4533 | 0.296445 | 0.706161 | 0.162791 | 0.015105 | 0.066561 | 0.245039 | 0.25159 | 0.074819 | 0 |
| 19.375 | 0.182885 | 0.24855 | 0.064544 | 0.524441 | 0.127321 | 0.297728 | 0.184299 | 0.010188 | 0.022834 | 0.184969 | 0 |
| 19.38889 | 0.260224 | 0.197852 | 0.386382 | 0.033246 | 0.560763 | 0.703866 | 0.60489 | 0.428224 | 0.427215 | 0.670932 | 0 |
| 19.40278 | 0.605611 | 0.552504 | 0.717714 | 0.38703 | 0.866528 | 0.983404 | 0.915986 | 0.767606 | 0.751357 | 0.998921 | 0 |
| 19.41667 | 0.76934 | 0.724289 | 0.853156 | 0.636877 | 0.958876 | 1.091256 | 1.027731 | 0.92258 | 0.897029 | 1.098043 | 0 |
| 19.43056 | 0.843225 | 0.808757 | 0.902014 | 0.742909 | 0.912216 | 1.113134 | 1.025071 | 0.972281 | 0.940476 | 1.033501 | 0 |
| 19.44444 | 0.77835 | 0.74848 | 0.841716 | 0.668836 | 0.75471 | 1.003389 | 0.896636 | 0.868987 | 0.837233 | 0.854813 | 0 |
| 19.45833 | 0.612276 | 0.581755 | 0.699404 | 0.510811 | 0.557215 | 0.817097 | 0.724248 | 0.677272 | 0.653549 | 0.617294 | 0 |
| 19.47222 | 0.43929 | 0.406565 | 0.539315 | 0.347386 | 0.365063 | 0.607616 | 0.569611 | 0.501896 | 0.480467 | 0.392509 | 0 |
| 19.48611 | 0.239129 | 0.184114 | 0.323746 | 0.113787 | 0.109985 | 0.392541 | 0.425387 | 0.328024 | 0.28235 | 0.197264 | 0 |
| 19.5 | 0.000898 | 0.143447 | 0.02005 | 0.200158 | 0.241536 | 0.154819 | 0.190247 | 0.060501 | 0.033913 | 0.053812 | 0 |
| 19.51389 | 0.042708 | 0.306836 | 0.339052 | 0.320914 | 0.466784 | 0.068258 | 0.071984 | 0.211393 | 0.326897 | 0.279445 | 0 |

| | | | | | | | | | | | |
|---|---|---|---|---|---|---|---|---|---|---|---|
| 19.52778 | 0.25761 | 0.081709 | 0.312125 | 0.070685 | 0.321678 | 0.531118 | 0.031577 | 0.189853 | 0.285368 | 0.211784 | 0 |
| 19.54167 | 0.639425 | 0.297608 | 0.058753 | 0.251427 | 0.00872 | 1.539607 | 0.230739 | 0.088394 | 0.04468 | 0.062512 | 0 |
| 19.55556 | 0.849341 | 0.523941 | 0.140159 | 0.420379 | 0.21006 | 2.089206 | 0.389583 | 0.330723 | 0.136261 | 0.204931 | 0 |
| 19.56944 | 0.994014 | 0.658643 | 0.347074 | 0.46817 | 0.373842 | 1.705497 | 0.602935 | 0.496313 | 0.280149 | 0.331483 | 0 |
| 19.58333 | 1.061109 | 0.669198 | 0.5365 | 0.43842 | 0.454682 | 1.182557 | 0.960503 | 0.588153 | 0.368031 | 0.571507 | 0 |
| 19.59722 | 0.997398 | 0.550153 | 0.585866 | 0.375082 | 0.400537 | 0.741838 | 1.27931 | 0.599281 | 0.368504 | 0.522139 | 0 |
| 19.61111 | 0.97771 | 0.515193 | 0.590077 | 0.385949 | 0.410385 | 0.559961 | 1.495238 | 0.685778 | 0.435694 | 0.720429 | 0 |
| 19.625 | 0.943753 | 0.51065 | 0.575157 | 0.420019 | 0.474994 | 0.602486 | 1.46793 | 0.737005 | 0.505659 | 0.735538 | 0 |
| 19.63889 | 0.832832 | 0.429099 | 0.424336 | 0.358068 | 0.423173 | 0.560413 | 1.322349 | 0.638282 | 0.425979 | 0.768307 | 0 |
| 19.65278 | 0.703271 | 0.321955 | 0.162066 | 0.247216 | 0.294494 | 0.393887 | 0.998653 | 0.436386 | 0.255665 | 0.657544 | 0 |
| 19.66667 | 0.544897 | 0.239279 | 0.054989 | 0.15505 | 0.168994 | 0.186504 | 0.594101 | 0.242962 | 0.099115 | 0.465328 | 0 |
| 19.68056 | 0.430457 | 0.227323 | 0.150815 | 0.141752 | 0.113456 | 0.038667 | 0.295244 | 0.170085 | 0.019263 | 0.334843 | 0 |
| 19.69444 | 0.442147 | 0.292432 | 0.117796 | 0.220222 | 0.191528 | 0.024208 | 0.122846 | 0.220261 | 0.064933 | 0.324403 | 0 |
| 19.70833 | 0.526579 | 0.360072 | 0.041375 | 0.317224 | 0.323506 | 0.083582 | 0.006605 | 0.290846 | 0.175978 | 0.383095 | 0 |
| 19.72222 | 0.481143 | 0.326917 | 0.033306 | 0.334224 | 0.322009 | 0.040524 | 0.196399 | 0.288238 | 0.207966 | 0.374548 | 0 |
| 19.73611 | 0.265583 | 0.176964 | 0.005822 | 0.216598 | 0.140554 | 0.084479 | 0.398695 | 0.141298 | 0.091601 | 0.168714 | 0 |
| 19.75 | 0.062165 | 0.048673 | 0.04757 | 0.228369 | 0.015866 | 0.080044 | 0.445653 | 0.038252 | 0.020384 | 0.103855 | 0 |
| 19.76389 | 0.062199 | 0.111547 | 0.03518 | 0.788335 | 0.055292 | 0.159513 | 0.26981 | 0.004493 | 0.058518 | 0.178702 | 0 |
| 19.77778 | 0.121544 | 0.216643 | 0.090209 | 1.562426 | 0.186808 | 0.331264 | 0.123951 | 0.130346 | 0.148239 | 0.173791 | 0 |
| 19.79167 | 0.064714 | 0.132549 | 0.052012 | 1.451657 | 0.150156 | 0.205197 | 0.198363 | 0.074029 | 0.060897 | 0.050895 | 0 |
| 19.80556 | 0.042489 | 0.017822 | 0.214855 | 0.691346 | 0.045215 | 0.0014 | 0.341852 | 0.090621 | 0.089148 | 0.040537 | 0 |
| 19.81944 | 0.12094 | 0.070327 | 0.241571 | 0.238604 | 0.002364 | 0.120571 | 0.552207 | 0.202893 | 0.190644 | 0.087573 | 0 |
| 19.83333 | 0.12514 | 0.008236 | 0.190053 | 0.155149 | 0.021008 | 0.166051 | 0.713568 | 0.183003 | 0.178144 | 0.103802 | 0 |
| 19.84722 | 0.099364 | 0.087152 | 0.16435 | 0.120231 | 0.040388 | 0.159419 | 0.807083 | 0.124009 | 0.117363 | 0.116419 | 0 |
| 19.86111 | 0.062776 | 0.132653 | 0.164237 | 0.067852 | 0.066388 | 0.056093 | 0.713007 | 0.064921 | 0.066158 | 0.135564 | 0 |
| 19.875 | 0.073546 | 0.25174 | 0.060233 | 0.169262 | 0.203488 | 0.187029 | 0.498668 | 0.074331 | 0.074322 | 0.000551 | 0 |
| 19.88889 | 0.228383 | 0.360896 | 0.048208 | 0.37179 | 0.352894 | 0.372209 | 0.352712 | 0.324029 | 0.210591 | 0.187238 | 0 |
| 19.90278 | 0.27788 | 0.396761 | 0.086184 | 0.608244 | 0.423504 | 0.319722 | 0.373574 | 0.70421 | 0.264607 | 0.251713 | 0 |
| 19.91667 | 0.18275 | 0.374955 | 0.102159 | 0.927769 | 0.453044 | 0.129694 | 0.59283 | 0.981338 | 0.266195 | 0.216343 | 0 |
| 19.93056 | 0.042434 | 0.257474 | 0.096214 | 1.087491 | 0.410535 | 0.074305 | 0.780891 | 0.817294 | 0.217912 | 0.122652 | 0 |
| 19.94444 | 0.341411 | 0.045726 | 0.009304 | 0.754034 | 0.277536 | 0.262242 | 0.938617 | 0.48366 | 0.093888 | 0.0842 | 0 |
| 19.95833 | 0.606448 | 0.15214 | 0.1294 | 0.404934 | 0.108018 | 0.4157 | 0.979915 | 0.16372 | 0.059345 | 0.27419 | 0 |
| 19.97222 | 0.577652 | 0.133992 | 0.098825 | 0.331974 | 0.117315 | 0.324998 | 0.703071 | 0.108527 | 0.02253 | 0.20829 | 0 |
| 19.98611 | 0.084434 | 0.274522 | 0.286059 | 0.429176 | 0.492199 | 0.106868 | 0.035142 | 0.36563 | 0.372789 | 0.143339 | 0 |
| 20 | 0.535604 | 0.943924 | 0.997806 | 0.56228 | 1.147367 | 0.74406 | 0.847783 | 0.826011 | 1.068178 | 0.714364 | 0 |
| 20.01389 | 0.712157 | 1.300856 | 1.492397 | 0.454274 | 1.490426 | 1.124365 | 1.404994 | 0.947112 | 1.515951 | 1.039005 | 0 |
| 20.02778 | 0.359975 | 1.030767 | 1.28889 | 0.209716 | 1.183985 | 0.965811 | 1.224369 | 0.528722 | 1.276119 | 0.862711 | 0 |
| 20.04167 | 0.090383 | 0.600515 | 0.83312 | 0.282383 | 0.680885 | 0.632793 | 0.769629 | 0.179299 | 0.80926 | 0.493728 | 0 |
| 20.05556 | 0.001251 | 0.292647 | 0.525521 | 0.376193 | 0.311168 | 0.396767 | 0.444112 | 0.128237 | 0.475281 | 0.185156 | 0 |
| 20.06944 | 0.065879 | 0.078606 | 0.289784 | 0.308843 | 0.041718 | 0.186917 | 0.192769 | 0.090516 | 0.234474 | 0.063101 | 0 |
| 20.08333 | 0.087101 | 0.036134 | 0.131453 | 0.214716 | 0.078978 | 0.041764 | 0.038685 | 0.027876 | 0.105413 | 0.108002 | 0 |
| 20.09722 | 0.066252 | 0.051471 | 0.085523 | 0.20117 | 0.074348 | 0.006352 | 0.021447 | 0.035446 | 0.097261 | 0.014543 | 0 |
| 20.11111 | 0.054902 | 0.049269 | 0.078155 | 0.193631 | 0.057928 | 0.022117 | 0.031923 | 0.055537 | 0.093993 | 0.046879 | 0 |
| 20.125 | 0.095026 | 0.088545 | 0.036177 | 0.129448 | 0.101439 | 0.119424 | 0.000809 | 0.026338 | 0.036995 | 0.018394 | 0 |
| 20.13889 | 0.171443 | 0.154111 | 0.044398 | 0.035398 | 0.178793 | 0.244742 | 0.072633 | 0.052695 | 0.049217 | 0.07458 | 0 |
| 20.15278 | 0.232937 | 0.213949 | 0.109643 | 0.05445 | 0.241773 | 0.320229 | 0.146083 | 0.126244 | 0.128939 | 0.189013 | 0 |
| 20.16667 | 0.27678 | 0.259801 | 0.155829 | 0.130412 | 0.280085 | 0.377934 | 0.195006 | 0.176873 | 0.177363 | 0.259968 | 0 |
| 20.18056 | 0.331485 | 0.319512 | 0.233253 | 0.22991 | 0.323249 | 0.466917 | 0.252763 | 0.251844 | 0.230064 | 0.305447 | 0 |
| 20.19444 | 0.350409 | 0.337986 | 0.281984 | 0.275736 | 0.331756 | 0.488292 | 0.268409 | 0.289144 | 0.253566 | 0.332322 | 0 |
| 20.20833 | 0.299106 | 0.280931 | 0.243464 | 0.235208 | 0.273949 | 0.389638 | 0.20668 | 0.224955 | 0.206528 | 0.291241 | 0 |
| 20.22222 | 0.293492 | 0.247797 | 0.222584 | 0.232894 | 0.252785 | 0.343562 | 0.195745 | 0.193407 | 0.192595 | 0.257473 | 0 |
| 20.23611 | 0.349016 | 0.271688 | 0.271489 | 0.271167 | 0.292163 | 0.40962 | 0.266809 | 0.258083 | 0.244892 | 0.288144 | 0 |
| 20.25 | 0.362765 | 0.282308 | 0.297867 | 0.243906 | 0.304836 | 0.445956 | 0.293474 | 0.304762 | 0.264723 | 0.320842 | 0 |
| 20.26389 | 0.30492 | 0.2552 | 0.253935 | 0.16159 | 0.267755 | 0.404012 | 0.241121 | 0.280771 | 0.218998 | 0.323488 | 0 |
| 20.27778 | 0.26169 | 0.246492 | 0.222908 | 0.142729 | 0.268138 | 0.438775 | 0.22011 | 0.287791 | 0.20197 | 0.333442 | 0 |
| 20.29167 | 0.258221 | 0.256003 | 0.231951 | 0.203856 | 0.321067 | 0.609987 | 0.249525 | 0.335307 | 0.231413 | 0.321503 | 0 |

| | | | | | | | | | | | |
|---|---|---|---|---|---|---|---|---|---|---|---|
| 20.30556 | 0.271844 | 0.265616 | 0.242353 | 0.261788 | 0.353088 | 0.704168 | 0.272211 | 0.35917 | 0.253338 | 0.311794 | 0 |
| 20.31944 | 0.317259 | 0.304835 | 0.267842 | 0.309087 | 0.365261 | 0.684401 | 0.29617 | 0.372531 | 0.272397 | 0.328286 | 0 |
| 20.33333 | 0.331287 | 0.315614 | 0.268772 | 0.276687 | 0.334684 | 0.592763 | 0.281107 | 0.352885 | 0.258046 | 0.31755 | 0 |
| 20.34722 | 0.2729 | 0.2635 | 0.209761 | 0.144567 | 0.255496 | 0.442016 | 0.206387 | 0.281936 | 0.189083 | 0.221782 | 0 |
| 20.36111 | 0.230925 | 0.234402 | 0.165772 | 0.044587 | 0.206297 | 0.29972 | 0.154398 | 0.242533 | 0.140172 | 0.140126 | 0 |
| 20.375 | 0.270487 | 0.282943 | 0.203799 | 0.058724 | 0.247848 | 0.269392 | 0.183388 | 0.290125 | 0.176725 | 0.187233 | 0 |
| 20.38889 | 0.330903 | 0.343882 | 0.286099 | 0.122503 | 0.324515 | 0.31644 | 0.225022 | 0.360774 | 0.254182 | 0.295562 | 0 |
| 20.40278 | 0.303762 | 0.32976 | 0.297151 | 0.112325 | 0.319624 | 0.304331 | 0.169556 | 0.354716 | 0.255135 | 0.30317 | 0 |
| 20.41667 | 0.209505 | 0.264589 | 0.233342 | 0.061267 | 0.213533 | 0.234025 | 0.063058 | 0.290751 | 0.173355 | 0.205087 | 0 |
| 20.43056 | 0.140503 | 0.232831 | 0.197195 | 0.057471 | 0.035408 | 0.185832 | 0.016572 | 0.252873 | 0.118548 | 0.156748 | 0 |
| 20.44444 | 0.101202 | 0.212864 | 0.198086 | 0.041886 | 0.072826 | 0.159698 | 0.033044 | 0.235386 | 0.104742 | 0.184077 | 0 |
| 20.45833 | 0.066004 | 0.180283 | 0.18975 | 0.004327 | 0.089405 | 0.128028 | 0.092561 | 0.205447 | 0.087696 | 0.126292 | 0 |
| 20.47222 | 0.002669 | 0.147423 | 0.164848 | 0.067619 | 0.377387 | 0.052063 | 0.140893 | 0.168612 | 0.077806 | 0.074393 | 0 |
| 20.48611 | 0.153977 | 0.175948 | 0.227126 | 0.27456 | 0.532483 | 0.167949 | 0.158009 | 0.184097 | 0.138348 | 0.001336 | 0 |
| 20.5 | 0.41575 | 0.335058 | 0.509183 | 0.465608 | 0.739961 | 0.539758 | 0.214475 | 0.347406 | 0.34262 | 0.184055 | 0 |
| 20.51389 | 0.703618 | 0.483558 | 0.756644 | 0.149773 | 0.948621 | 0.670797 | 0.406034 | 0.578784 | 0.552106 | 0.169049 | 0 |
| 20.52778 | 0.75231 | 0.411846 | 0.659139 | 0.577762 | 0.83589 | 0.547326 | 0.580407 | 0.540408 | 0.536095 | 0.002922 | 0 |
| 20.54167 | 0.82657 | 0.02222 | 0.324668 | 0.486004 | 0.505568 | 0.5586 | 0.512817 | 0.101414 | 0.37898 | 0.126323 | 0 |
| 20.55556 | 0.831223 | 0.299048 | 0.045438 | 0.241247 | 0.201001 | 0.609362 | 0.32796 | 0.442177 | 0.221888 | 0.224482 | 0 |
| 20.56944 | 0.852664 | 0.477717 | 0.101988 | 0.057837 | 0.020289 | 0.530923 | 0.265131 | 0.843749 | 0.063613 | 0.379604 | 0 |
| 20.58333 | 0.837783 | 0.534732 | 0.188923 | 0.058708 | 0.043438 | 0.385257 | 0.478093 | 0.974777 | 0.120458 | 0.572002 | 0 |
| 20.59722 | 0.966928 | 0.549091 | 0.192074 | 0.029103 | 0.022507 | 0.30017 | 0.711811 | 0.92121 | 0.280836 | 0.707825 | 0 |
| 20.61111 | 1.033653 | 0.54194 | 0.137438 | 0.127898 | 0.026532 | 0.276473 | 0.589521 | 0.747646 | 0.369492 | 0.651543 | 0 |
| 20.625 | 0.966464 | 0.501138 | 0.102271 | 0.184738 | 0.04488 | 0.362676 | 0.177342 | 0.578375 | 0.344492 | 0.412015 | 0 |
| 20.63889 | 0.738054 | 0.406199 | 0.093397 | 0.05292 | 0.017597 | 0.461087 | 0.319161 | 0.447706 | 0.210778 | 0.323772 | 0 |
| 20.65278 | 0.507387 | 0.387848 | 0.043156 | 0.136217 | 0.075171 | 0.525667 | 0.704833 | 0.393801 | 0.136368 | 0.231378 | 0 |
| 20.66667 | 0.273708 | 0.328485 | 0.075739 | 0.218382 | 0.125507 | 0.491636 | 1.083224 | 0.333101 | 0.067119 | 0.048458 | 0 |
| 20.68056 | 0.048995 | 0.168181 | 0.220257 | 0.129764 | 0.050497 | 0.31385 | 1.404223 | 0.206522 | 0.102894 | 0.202486 | 0 |
| 20.69444 | 0.067229 | 0.016998 | 0.263084 | 0.047916 | 0.036479 | 0.204805 | 1.283609 | 0.099848 | 0.228786 | 0.304453 | 0 |
| 20.70833 | 0.125718 | 0.060294 | 0.164604 | 0.027485 | 0.056416 | 0.218592 | 0.692174 | 0.051604 | 0.177346 | 0.188952 | 0 |
| 20.72222 | 0.137368 | 0.026428 | 0.02353 | 0.059182 | 0.003515 | 0.342167 | 0.175603 | 0.066465 | 0.010219 | 0.013191 | 0 |
| 20.73611 | 0.158258 | 0.007346 | 0.123091 | 0.093093 | 0.038016 | 0.360688 | 0.882078 | 0.065049 | 0.084844 | 0.090307 | 0 |
| 20.75 | 0.242612 | 0.027213 | 0.044618 | 0.110065 | 0.007257 | 0.174251 | 1.136871 | 0.047188 | 0.033921 | 0.00821 | 0 |
| 20.76389 | 0.322987 | 0.054061 | 0.075296 | 0.122224 | 0.002829 | 0.012731 | 0.91507 | 0.10375 | 0.042908 | 0.082162 | 0 |
| 20.77778 | 0.346093 | 0.034457 | 0.111887 | 0.12851 | 0.049664 | 0.04314 | 0.336506 | 0.204986 | 0.081461 | 0.111211 | 0 |
| 20.79167 | 0.318912 | 0.023147 | 0.111693 | 0.070018 | 0.095667 | 0.158365 | 0.522268 | 0.2073 | 0.070882 | 0.219027 | 0 |
| 20.80556 | 0.301781 | 0.073095 | 0.179296 | 0.029395 | 0.053719 | 0.129618 | 1.285849 | 0.000134 | 0.103713 | 0.297724 | 0 |
| 20.81944 | 0.274217 | 0.061741 | 0.193799 | 0.008836 | 0.019569 | 0.033702 | 1.402272 | 0.226596 | 0.097658 | 0.284846 | 0 |
| 20.83333 | 0.267911 | 0.003562 | 0.117884 | 0.106101 | 0.055746 | 0.00618 | 0.806188 | 0.309891 | 0.008471 | 0.196761 | 0 |
| 20.84722 | 0.316948 | 0.026792 | 0.037912 | 0.209457 | 0.114945 | 0.04197 | 0.320943 | 0.281888 | 0.073446 | 0.184971 | 0 |
| 20.86111 | 0.365966 | 0.004552 | 0.022868 | 0.19411 | 0.154947 | 0.057432 | 0.174245 | 0.206385 | 0.076047 | 0.196992 | 0 |
| 20.875 | 0.451319 | 0.097734 | 0.014136 | 0.062962 | 0.093899 | 0.070107 | 0.140141 | 0.096092 | 0.045238 | 0.24656 | 0 |
| 20.88889 | 0.59573 | 0.197858 | 0.043275 | 0.160003 | 0.003958 | 0.2773 | 0.183202 | 0.078178 | 0.183154 | 0.344207 | 0 |
| 20.90278 | 0.759136 | 0.275535 | 0.117914 | 0.441778 | 0.059516 | 0.467124 | 0.341734 | 0.212468 | 0.281432 | 0.434023 | 0 |
| 20.91667 | 0.851984 | 0.314473 | 0.158335 | 0.522862 | 0.066197 | 0.52601 | 0.484978 | 0.15453 | 0.301281 | 0.420144 | 0 |
| 20.93056 | 0.813288 | 0.274437 | 0.118727 | 0.359564 | 0.028846 | 0.438678 | 0.527927 | 0.026009 | 0.217809 | 0.360617 | 0 |
| 20.94444 | 0.73262 | 0.225318 | 0.034715 | 0.198068 | 0.03748 | 0.381204 | 0.466135 | 0.198282 | 0.103363 | 0.297882 | 0 |
| 20.95833 | 0.64477 | 0.183159 | 0.031247 | 0.23433 | 0.087917 | 0.407684 | 0.34728 | 0.4095 | 0.045441 | 0.320778 | 0 |
| 20.97222 | 0.47861 | 0.126796 | 0.086514 | 0.202692 | 0.134423 | 0.460463 | 0.118362 | 0.471856 | 0.036717 | 0.125908 | 0 |
| 20.98611 | 0.09056 | 0.110064 | 0.270793 | 0.281096 | 0.33761 | 0.607817 | 0.343586 | 0.119124 | 0.283069 | 0.648508 | 0 |
| 21 | 0.365355 | 0.485755 | 0.582041 | 1.040232 | 0.705367 | 0.977024 | 0.958662 | 0.435066 | 0.646476 | 1.764691 | 0 |
| 21.01389 | 0.534744 | 0.642985 | 0.797044 | 1.255609 | 1.093487 | 0.869406 | 1.225858 | 0.666154 | 0.844969 | 1.984743 | 0 |
| 21.02778 | 0.347314 | 0.496643 | 0.721861 | 0.741767 | 0.805745 | 0.064008 | 0.83082 | 0.515134 | 0.774985 | 1.364087 | 0 |
| 21.04167 | 0.166478 | 0.275079 | 0.415007 | 0.243097 | 0.401573 | 0.348948 | 0.354496 | 0.123854 | 0.44982 | 0.34195 | 0 |
| 21.05556 | 0.081047 | 0.058569 | 0.155669 | 0.107282 | 0.128675 | 0.242743 | 0.087318 | 0.195597 | 0.18275 | 0.028081 | 0 |
| 21.06944 | 0.045292 | 0.039747 | 0.048907 | 0.087196 | 0.034358 | 0.086132 | 0.002597 | 0.253939 | 0.078753 | 0.002087 | 0 |

| | | | | | | | | | | | |
|---|---|---|---|---|---|---|---|---|---|---|---|
| 21.08333 | 0.069347 | 0.009198 | 0.081241 | 0.055206 | 0.063111 | 0.127186 | 0.044428 | 0.04391 | 0.106948 | 0.071114 | 0 |
| 21.09722 | 0.104844 | 0.095677 | 0.155999 | 0.00235 | 0.107102 | 0.304408 | 0.110123 | 0.110513 | 0.167644 | 0.308192 | 0 |
| 21.11111 | 0.097972 | 0.101866 | 0.179526 | 0.017989 | 0.114678 | 0.287969 | 0.135068 | 0.113249 | 0.176009 | 0.349529 | 0 |
| 21.125 | 0.088952 | 0.072236 | 0.158388 | 0.01906 | 0.101114 | 0.107248 | 0.145049 | 0.055654 | 0.154585 | 0.084459 | 0 |
| 21.13889 | 0.070099 | 0.04233 | 0.118629 | 0.001596 | 0.083692 | 0.00484 | 0.133334 | 0.02567 | 0.122456 | 0.05993 | 0 |
| 21.15278 | 0.059269 | 0.051604 | 0.111864 | 0.089086 | 0.054809 | 0.115708 | 0.099834 | 0.071746 | 0.121222 | 0.163514 | 0 |
| 21.16667 | 0.114295 | 0.12731 | 0.180992 | 0.101991 | 0.075737 | 0.402149 | 0.113889 | 0.191667 | 0.183899 | 0.421724 | 0 |
| 21.18056 | 0.181456 | 0.183909 | 0.23072 | 0.011374 | 0.115642 | 0.410213 | 0.174904 | 0.239476 | 0.233984 | 0.226424 | 0 |
| 21.19444 | 0.213356 | 0.183265 | 0.215831 | 0.142294 | 0.188937 | 0.141434 | 0.267606 | 0.15593 | 0.236585 | 0.231311 | 0 |
| 21.20833 | 0.193677 | 0.14313 | 0.175044 | 0.238482 | 0.188582 | 0.041403 | 0.317779 | 0.074476 | 0.205982 | 0.479485 | 0 |
| 21.22222 | 0.093383 | 0.087007 | 0.124152 | 0.157191 | 0.084929 | 0.064975 | 0.233847 | 0.047124 | 0.114386 | 0.019297 | 0 |
| 21.23611 | 0.007052 | 0.014217 | 0.051904 | 0.057552 | 0.034854 | 0.197733 | 0.121356 | 0.005444 | 0.006899 | 0.483597 | 0 |
| 21.25 | 0.010664 | 0.033454 | 0.018185 | 0.115067 | 0.053182 | 0.093708 | 0.125369 | 0.043631 | 0.017399 | 0.18604 | 0 |
| 21.26389 | 0.026654 | 0.019945 | 0.034699 | 0.185271 | 0.015809 | 0.030438 | 0.165049 | 0.038777 | 0.054816 | 0.084791 | 0 |
| 21.27778 | 0.023028 | 0.004663 | 0.047567 | 0.160394 | 0.001528 | 0.060383 | 0.162561 | 0.025539 | 0.090081 | 0.029807 | 0 |
| 21.29167 | 0.019069 | 0.029696 | 0.038537 | 0.145824 | 0.024147 | 0.050051 | 0.133404 | 0.058775 | 0.054033 | 0.108127 | 0 |
| 21.30556 | 0.06691 | 0.094647 | 0.028801 | 0.143704 | 0.086969 | 0.022284 | 0.076258 | 0.133922 | 0.01337 | 0.110055 | 0 |
| 21.31944 | 0.071709 | 0.123655 | 0.071041 | 0.17461 | 0.104902 | 0.049006 | 0.056338 | 0.169897 | 0.024279 | 0.038957 | 0 |
| 21.33333 | 0.057761 | 0.090811 | 0.146654 | 0.199692 | 0.074023 | 0.014032 | 0.073544 | 0.144293 | 0.068504 | 0.029651 | 0 |
| 21.34722 | 0.052124 | 0.051517 | 0.208608 | 0.171162 | 0.059095 | 0.012406 | 0.073012 | 0.105385 | 0.100132 | 0.014978 | 0 |
| 21.36111 | 0.009254 | 0.01696 | 0.299241 | 0.208134 | 0.031898 | 0.064605 | 0.111638 | 0.050965 | 0.141712 | 0.029499 | 0 |
| 21.375 | 0.049074 | 0.019615 | 0.382033 | 0.350034 | 0.005038 | 0.141543 | 0.207088 | 0.006747 | 0.22255 | 0.114015 | 0 |
| 21.38889 | 0.064922 | 0.038589 | 0.393554 | 0.472384 | 0.015758 | 0.181956 | 0.325335 | 0.03557 | 0.282299 | 0.231202 | 0 |
| 21.40278 | 0.066352 | 0.023414 | 0.323284 | 0.552509 | 0.005774 | 0.097398 | 0.467378 | 0.024519 | 0.265836 | 0.385526 | 0 |
| 21.41667 | 0.11456 | 0.009107 | 0.236745 | 0.733313 | 0.004393 | 0.077017 | 0.564142 | 0.027401 | 0.278006 | 0.646136 | 0 |
| 21.43056 | 0.17321 | 0.012065 | 0.175066 | 0.751955 | 0.021015 | 0.103791 | 0.608741 | 0.077895 | 0.285177 | 0.520558 | 0 |
| 21.44444 | 0.176698 | 0.01265 | 0.107501 | 0.537588 | 0.037412 | 0.015722 | 0.607361 | 0.133578 | 0.193024 | 0.239797 | 0 |
| 21.45833 | 0.146325 | 0.042026 | 0.00531 | 0.277353 | 0.128118 | 0.102086 | 0.41456 | 0.173974 | 0.069899 | 0.001699 | 0 |
| 21.47222 | 0.123098 | 0.049659 | 0.025008 | 0.074148 | 0.159233 | 0.349701 | 0.231226 | 0.294325 | 0.036613 | 0.111739 | 0 |
| 21.48611 | 0.190656 | 0.024374 | 0.060642 | 0.240942 | 0.250639 | 0.901862 | 0.266913 | 0.419591 | 0.095053 | 0.086027 | 0 |
| 21.5 | 0.39268 | 0.008146 | 0.29419 | 0.787873 | 0.613248 | 1.030215 | 0.339916 | 0.309955 | 0.048125 | 0.20787 | 0 |
| 21.51389 | 0.631683 | 0.025624 | 0.618705 | 1.11996 | 0.876034 | 0.476506 | 0.19367 | 0.023411 | 0.381906 | 0.208112 | 0 |
| 21.52778 | 0.478323 | 0.036214 | 0.630089 | 1.017395 | 0.887163 | 0.234687 | 0.177986 | 0.187324 | 0.635624 | 0.007562 | 0 |
| 21.54167 | 0.331625 | 0.046866 | 0.504451 | 0.609007 | 0.630181 | 0.400481 | 0.339012 | 0.097408 | 0.660849 | 0.146154 | 0 |
| 21.55556 | 0.275703 | 0.1947 | 0.393323 | 0.321563 | 0.314694 | 0.272278 | 0.296899 | 0.420017 | 0.617781 | 0.1873 | 0 |
| 21.56944 | 0.326305 | 0.282516 | 0.360109 | 0.288126 | 0.137291 | 0.140968 | 0.13129 | 0.642184 | 0.483765 | 0.063496 | 0 |
| 21.58333 | 0.525663 | 0.338796 | 0.2459 | 0.272038 | 0.067975 | 0.012301 | 0.336939 | 0.820711 | 0.230597 | 0.218763 | 0 |
| 21.59722 | 0.739402 | 0.332046 | 0.029603 | 0.105103 | 0.035268 | 0.171395 | 0.916107 | 0.963343 | 0.018059 | 0.514879 | 0 |
| 21.61111 | 0.674507 | 0.216401 | 0.246925 | 0.058685 | 0.049324 | 0.357698 | 1.417768 | 0.888905 | 0.181786 | 0.605551 | 0 |
| 21.625 | 0.393694 | 0.084988 | 0.443212 | 0.057184 | 0.04484 | 0.612319 | 1.606454 | 0.651869 | 0.29638 | 0.493139 | 0 |
| 21.63889 | 0.112125 | 0.019227 | 0.346304 | 0.066131 | 0.020118 | 0.855857 | 1.273113 | 0.416311 | 0.31243 | 0.253947 | 0 |
| 21.65278 | 0.213579 | 0.055374 | 0.015078 | 0.067509 | 0.029131 | 0.922745 | 0.416062 | 0.237187 | 0.179385 | 0.088617 | 0 |
| 21.66667 | 0.531645 | 0.046653 | 0.322922 | 0.019999 | 0.054933 | 0.727462 | 0.74435 | 0.089045 | 0.062085 | 0.525038 | 0 |
| 21.68056 | 0.703845 | 0.066739 | 0.482045 | 0.105983 | 0.012481 | 0.396397 | 1.760793 | 0.041862 | 0.294646 | 0.770765 | 0 |
| 21.69444 | 0.653432 | 0.092713 | 0.457885 | 0.250055 | 0.049477 | 0.107842 | 1.984772 | 0.131266 | 0.350271 | 0.714444 | 0 |
| 21.70833 | 0.388307 | 0.059616 | 0.310758 | 0.268183 | 0.067449 | 0.080037 | 1.461011 | 0.11666 | 0.218943 | 0.445184 | 0 |
| 21.72222 | 0.073046 | 0.098758 | 0.033737 | 0.091221 | 0.02187 | 0.112109 | 0.637185 | 0.028242 | 0.043186 | 0.077149 | 0 |
| 21.73611 | 0.432909 | 0.251122 | 0.185564 | 0.054251 | 0.095195 | 0.09766 | 0.156812 | 0.079983 | 0.013585 | 0.122198 | 0 |
| 21.75 | 0.428684 | 0.250606 | 0.242385 | 0.010353 | 0.068089 | 0.12026 | 0.787352 | 0.041362 | 0.03626 | 0.089822 | 0 |
| 21.76389 | 0.173146 | 0.171541 | 0.236007 | 0.065189 | 0.050183 | 0.088063 | 1.099608 | 0.2054 | 0.065899 | 0.000806 | 0 |
| 21.77778 | 0.169034 | 0.086008 | 0.132501 | 0.074408 | 0.021944 | 0.029917 | 0.90573 | 0.406041 | 0.109428 | 0.064728 | 0 |
| 21.79167 | 0.464921 | 0.023328 | 0.10019 | 0.051128 | 0.016392 | 0.018184 | 0.424949 | 0.584495 | 0.177796 | 0.231313 | 0 |
| 21.80556 | 0.641378 | 0.12093 | 0.335394 | 0.061229 | 0.005122 | 0.044264 | 0.01887 | 0.689535 | 0.201202 | 0.419313 | 0 |
| 21.81944 | 0.693237 | 0.415244 | 0.386824 | 0.102336 | 0.057435 | 0.052904 | 0.117009 | 0.666048 | 0.146519 | 0.516158 | 0 |
| 21.83333 | 0.671316 | 0.540937 | 0.29805 | 0.247905 | 0.071226 | 0.043191 | 0.101205 | 0.558202 | 0.089435 | 0.487919 | 0 |
| 21.84722 | 0.713013 | 0.329041 | 0.185368 | 0.368535 | 0.008996 | 0.016647 | 0.295505 | 0.486531 | 0.108472 | 0.451899 | 0 |

| | | | | | | | | | | | |
|---|---|---|---|---|---|---|---|---|---|---|---|
| 21.86111 | 0.755841 | 0.133714 | 0.124741 | 0.29324 | 0.042919 | 0.13197 | 0.681505 | 0.572253 | 0.188952 | 0.514813 | 0 |
| 21.875 | 0.78523 | 0.042139 | 0.125816 | 0.208637 | 0.051174 | 0.23085 | 0.924888 | 0.843617 | 0.274206 | 0.613015 | 0 |
| 21.88889 | 0.788583 | 0.003663 | 0.15961 | 0.32594 | 0.006649 | 0.25064 | 0.819101 | 1.166138 | 0.297869 | 0.669141 | 0 |
| 21.90278 | 0.752091 | 0.041694 | 0.209874 | 0.579345 | 0.042242 | 0.238957 | 0.656127 | 1.372857 | 0.275673 | 0.676569 | 0 |
| 21.91667 | 0.658993 | 0.048301 | 0.209202 | 0.752538 | 0.07169 | 0.188509 | 0.601835 | 1.308339 | 0.230595 | 0.725985 | 0 |
| 21.93056 | 0.568264 | 0.062622 | 0.195999 | 0.594297 | 0.055126 | 0.155318 | 0.778452 | 1.1532 | 0.205015 | 0.729219 | 0 |
| 21.94444 | 0.598225 | 0.179891 | 0.23985 | 0.408107 | 0.023383 | 0.195638 | 1.145437 | 1.082589 | 0.240218 | 0.795773 | 0 |
| 21.95833 | 0.657123 | 0.271603 | 0.239779 | 0.309933 | 0.051475 | 0.222754 | 1.214607 | 1.068416 | 0.250128 | 0.745052 | 0 |
| 21.97222 | 0.546745 | 0.255349 | 0.147279 | 0.205206 | 0.041699 | 0.210556 | 0.810561 | 0.982918 | 0.16046 | 0.450438 | 0 |
| 21.98611 | 0.264721 | 0.073805 | 0.138807 | 0.155021 | 0.305833 | 0.127778 | 0.086773 | 0.614012 | 0.126198 | 0.013979 | 0 |
| 22 | 0.089369 | 0.222922 | 0.566339 | 0.602217 | 0.691444 | 0.001847 | 0.654311 | 0.016933 | 0.538496 | 0.512719 | 0 |
| 22.01389 | 0.243359 | 0.288064 | 0.737689 | 0.684566 | 0.735636 | 0.070437 | 0.864784 | 0.407427 | 0.703457 | 0.755762 | 0 |
| 22.02778 | 0.185923 | 0.151244 | 0.512232 | 0.378668 | 0.569184 | 0.031017 | 0.539475 | 0.396468 | 0.471961 | 0.551748 | 0 |
| 22.04167 | 0.184895 | 0.072775 | 0.278079 | 0.218084 | 0.377317 | 0.179202 | 0.254293 | 0.25763 | 0.249967 | 0.352008 | 0 |
| 22.05556 | 0.252881 | 0.112461 | 0.205297 | 0.381842 | 0.281091 | 0.302857 | 0.213326 | 0.152013 | 0.207978 | 0.320937 | 0 |
| 22.06944 | 0.261862 | 0.105928 | 0.179847 | 0.429125 | 0.213954 | 0.23998 | 0.212214 | 0.054374 | 0.209302 | 0.265478 | 0 |
| 22.08333 | 0.243456 | 0.12496 | 0.196307 | 0.274442 | 0.173673 | 0.22834 | 0.205355 | 0.060437 | 0.228693 | 0.175525 | 0 |
| 22.09722 | 0.230214 | 0.160907 | 0.232463 | 0.158209 | 0.192916 | 0.242155 | 0.206943 | 0.124897 | 0.25103 | 0.140619 | 0 |
| 22.11111 | 0.173628 | 0.122335 | 0.194754 | 0.10233 | 0.16398 | 0.146659 | 0.157176 | 0.12949 | 0.197365 | 0.152609 | 0 |
| 22.125 | 0.089195 | 0.046513 | 0.11614 | 0.022053 | 0.087453 | 0.034917 | 0.072044 | 0.065331 | 0.099316 | 0.112664 | 0 |
| 22.13889 | 0.00253 | 0.024929 | 0.048637 | 0.069982 | 0.032747 | 0.004599 | 0.00045 | 0.017996 | 0.020746 | 0.059115 | 0 |
| 22.15278 | 0.073442 | 0.084064 | 0.000827 | 0.150269 | 0.003719 | 0.008506 | 0.047534 | 0.070182 | 0.022515 | 0.018955 | 0 |
| 22.16667 | 0.123579 | 0.116712 | 0.024956 | 0.191549 | 0.021117 | 0.000637 | 0.076064 | 0.068928 | 0.035661 | 0.046276 | 0 |
| 22.18056 | 0.126448 | 0.107923 | 0.017238 | 0.150943 | 0.029862 | 0.01218 | 0.073107 | 0.029011 | 0.019905 | 0.083708 | 0 |
| 22.19444 | 0.064773 | 0.049934 | 0.021839 | 0.011696 | 0.005575 | 0.052721 | 0.013032 | 0.032064 | 0.028081 | 0.026231 | 0 |
| 22.20833 | 0.002654 | 0.008662 | 0.051178 | 0.127529 | 0.015301 | 0.06932 | 0.051514 | 0.063364 | 0.058185 | 0.018645 | 0 |
| 22.22222 | 0.045964 | 0.034456 | 0.060365 | 0.251934 | 0.027018 | 0.032577 | 0.097664 | 0.04666 | 0.063432 | 0.052307 | 0 |
| 22.23611 | 0.102878 | 0.064727 | 0.091295 | 0.379796 | 0.053466 | 0.117298 | 0.151721 | 0.056841 | 0.102234 | 0.115014 | 0 |
| 22.25 | 0.175724 | 0.162382 | 0.175041 | 0.432668 | 0.106755 | 0.078747 | 0.203709 | 0.163118 | 0.208089 | 0.164 | 0 |
| 22.26389 | 0.228359 | 0.250066 | 0.254075 | 0.487875 | 0.170943 | 0.225427 | 0.234542 | 0.257966 | 0.304086 | 0.178437 | 0 |
| 22.27778 | 0.225755 | 0.228924 | 0.251032 | 0.564395 | 0.163239 | 0.071206 | 0.221495 | 0.252979 | 0.308726 | 0.207023 | 0 |
| 22.29167 | 0.186662 | 0.191504 | 0.20718 | 0.557832 | 0.085855 | 0.011508 | 0.163583 | 0.253538 | 0.275889 | 0.180472 | 0 |
| 22.30556 | 0.146111 | 0.153274 | 0.152066 | 0.555144 | 0.01268 | 0.06091 | 0.109645 | 0.262355 | 0.240566 | 0.140551 | 0 |
| 22.31944 | 0.121163 | 0.072661 | 0.101579 | 0.602076 | 0.004914 | 0.11156 | 0.118508 | 0.232935 | 0.206461 | 0.19926 | 0 |
| 22.33333 | 0.105229 | 0.034191 | 0.117514 | 0.617891 | 0.022814 | 0.1836 | 0.245331 | 0.262642 | 0.205819 | 0.236504 | 0 |
| 22.34722 | 0.068472 | 0.07264 | 0.182858 | 0.445563 | 0.001244 | 0.019429 | 0.503753 | 0.39318 | 0.216474 | 0.216281 | 0 |
| 22.36111 | 0.014192 | 0.076896 | 0.206507 | 0.371205 | 0.041174 | 0.230348 | 0.722619 | 0.473837 | 0.212832 | 0.151852 | 0 |
| 22.375 | 0.043995 | 0.008394 | 0.169295 | 0.4394 | 0.080116 | 0.101489 | 0.684553 | 0.467088 | 0.177808 | 0.240058 | 0 |
| 22.38889 | 0.122563 | 0.114623 | 0.109316 | 0.54881 | 0.13316 | 0.047423 | 0.446497 | 0.468497 | 0.134402 | 0.248796 | 0 |
| 22.40278 | 0.235304 | 0.155894 | 0.002398 | 0.52045 | 0.18731 | 0.087709 | 0.16634 | 0.49278 | 0.055641 | 0.09863 | 0 |
| 22.41667 | 0.441237 | 0.070632 | 0.134719 | 0.041776 | 0.326465 | 0.101771 | 0.072723 | 0.266577 | 0.161248 | 0.58737 | 0 |
| 22.43056 | 0.484228 | 1.042086 | 0.163672 | 0.543667 | 0.193863 | 0.227224 | 0.475709 | 0.036584 | 0.190806 | 0.392831 | 0 |
| 22.44444 | 0.129972 | 0.252017 | 0.027122 | 0.34145 | 0.581928 | 1.481288 | 0.786514 | 0.123558 | 0.069086 | 0.370165 | 0 |
| 22.45833 | 0.126013 | 0.07924 | 0.235778 | 0.407756 | 0.444159 | 0.648864 | 0.317518 | 0.600241 | 0.31941 | 0.119679 | 0 |
| 22.47222 | 0.275227 | 0.278349 | 0.25681 | 0.593156 | 0.189552 | 0.28591 | 0.000717 | 0.643553 | 0.277955 | 0.270301 | 0 |
| 22.48611 | 0.500505 | 0.449605 | 0.215403 | 0.246205 | 0.070407 | 0.27806 | 0.416567 | 0.462098 | 0.159979 | 0.402603 | 0 |
| 22.5 | 0.809124 | 0.629289 | 0.171957 | 0.167929 | 0.005341 | 0.54428 | 1.008841 | 0.536137 | 0.238647 | 0.678045 | 0 |
| 22.51389 | 0.804517 | 0.48097 | 0.074804 | 0.277458 | 0.10309 | 0.536098 | 1.191736 | 0.759019 | 0.322797 | 0.914917 | 0 |
| 22.52778 | 0.641588 | 0.225282 | 0.020368 | 0.05442 | 0.161721 | 0.246905 | 1.026643 | 0.908265 | 0.296685 | 0.58087 | 0 |
| 22.54167 | 0.553522 | 0.17363 | 0.022631 | 0.381187 | 0.125976 | 0.061164 | 0.887834 | 0.932327 | 0.189098 | 0.300851 | 0 |
| 22.55556 | 0.52099 | 0.203441 | 0.032266 | 0.146341 | 0.110661 | 0.045341 | 0.488039 | 0.657998 | 0.033344 | 0.002783 | 0 |
| 22.56944 | 0.462143 | 0.171523 | 0.111948 | 0.243057 | 0.148725 | 0.183733 | 0.083054 | 0.388065 | 0.199251 | 0.252033 | 0 |
| 22.58333 | 0.537702 | 0.157778 | 0.116337 | 0.400165 | 0.11163 | 0.210382 | 0.473974 | 0.329202 | 0.308985 | 0.180761 | 0 |
| 22.59722 | 0.656648 | 0.15874 | 0.011258 | 0.325739 | 0.065585 | 0.087199 | 0.58341 | 0.409636 | 0.222852 | 0.023035 | 0 |
| 22.61111 | 0.672723 | 0.074032 | 0.1898 | 0.200842 | 0.267513 | 0.025485 | 0.274637 | 0.587918 | 0.056939 | 0.117213 | 0 |
| 22.625 | 0.59876 | 0.0338 | 0.419663 | 0.016321 | 0.289172 | 0.085869 | 0.505921 | 0.836894 | 0.107665 | 0.179282 | 0 |

| | | | | | | | | | | | | |
|---|---|---|---|---|---|---|---|---|---|---|---|---|
| 22.63889 | 0.499476 | 0.103258 | 0.535527 | 0.32862 | 0.091715 | 0.128057 | 1.52756 | 1.079682 | 0.22583 | | 0.30234 | 0 |
| 22.65278 | 0.412969 | 0.262906 | 0.383791 | 0.424711 | 0.000813 | 0.067192 | 1.963006 | 1.039163 | 0.272166 | | 0.291237 | 0 |
| 22.66667 | 0.264133 | 0.558044 | 0.021648 | 0.058876 | 0.067086 | 0.105369 | 1.507299 | 0.718058 | 0.227268 | | 0.087016 | 0 |
| 22.68056 | 0.027314 | 0.460498 | 0.369134 | 0.474903 | 0.165312 | 0.359449 | 0.525299 | 0.292942 | 0.10135 | | 0.013168 | 0 |
| 22.69444 | 0.167192 | 0.223123 | 0.492595 | 0.715765 | 0.292991 | 0.533333 | 0.640664 | 0.15286 | 0.054712 | | 0.002193 | 0 |
| 22.70833 | 0.243195 | 0.856379 | 0.469385 | 0.71812 | 0.316174 | 0.567325 | 1.634788 | 0.507737 | 0.127097 | | 0.043312 | 0 |
| 22.72222 | 0.285277 | 1.098018 | 0.304511 | 0.711606 | 0.271216 | 0.493764 | 2.085243 | 0.648678 | 0.111484 | | 0.076637 | 0 |
| 22.73611 | 0.377294 | 1.081132 | 0.11781 | 0.567326 | 0.318837 | 0.468156 | 2.019529 | 0.51634 | 0.103978 | | 0.050997 | 0 |
| 22.75 | 0.380916 | 1.076711 | 0.116972 | 0.096849 | 0.618692 | 0.603537 | 1.349634 | 0.209026 | 0.065454 | | 0.034443 | 0 |
| 22.76389 | 0.218544 | 1.07535 | 0.300135 | 0.346469 | 1.071719 | 0.80726 | 0.342271 | 0.028873 | 0.010685 | | 0.157337 | 0 |
| 22.77778 | 0.031365 | 0.987633 | 0.288986 | 0.429843 | 1.187337 | 0.889989 | 0.450711 | 0.061359 | 0.02333 | | 0.335402 | 0 |
| 22.79167 | 0.008264 | 0.710588 | 0.103809 | 0.198043 | 1.032278 | 0.916312 | 0.721774 | 0.473404 | 0.030206 | | 0.33522 | 0 |
| 22.80556 | 0.155341 | 0.318278 | 0.098545 | 0.083817 | 0.832828 | 0.895839 | 0.557682 | 0.971823 | 0.069976 | | 0.074028 | 0 |
| 22.81944 | 0.328575 | 0.12895 | 0.230997 | 0.302739 | 0.539199 | 0.820576 | 0.07027 | 1.356628 | 0.255899 | | 0.359398 | 0 |
| 22.83333 | 0.464464 | 0.574596 | 0.332398 | 0.483307 | 0.423842 | 0.814709 | 0.525755 | 1.541432 | 0.418922 | | 0.897504 | 0 |
| 22.84722 | 0.561155 | 0.915971 | 0.424818 | 0.500585 | 0.662406 | 1.055169 | 1.0326 | 1.631563 | 0.512594 | | 1.244551 | 0 |
| 22.86111 | 0.597562 | 1.071185 | 0.469177 | 0.425313 | 1.135158 | 1.284019 | 1.273328 | 1.588985 | 0.593813 | | 1.362499 | 0 |
| 22.875 | 0.569019 | 0.950122 | 0.438145 | 0.443759 | 1.571865 | 1.37068 | 1.088801 | 1.453414 | 0.579202 | | 1.154226 | 0 |
| 22.88889 | 0.521083 | 0.537679 | 0.385399 | 0.575742 | 1.840749 | 1.357741 | 0.564941 | 1.361946 | 0.518218 | | 0.860854 | 0 |
| 22.90278 | 0.45513 | 0.052649 | 0.310294 | 0.603569 | 1.79879 | 1.159989 | 0.028949 | 1.39429 | 0.434113 | | 0.686762 | 0 |
| 22.91667 | 0.377841 | 0.250535 | 0.259416 | 0.597222 | 1.50567 | 0.98145 | 0.0777 | 1.380843 | 0.380835 | | 0.569372 | 0 |
| 22.93056 | 0.323451 | 0.172142 | 0.253708 | 0.542825 | 1.102084 | 0.861051 | 0.259659 | 1.216792 | 0.414991 | | 0.546004 | 0 |
| 22.94444 | 0.340669 | 0.219423 | 0.20934 | 0.434431 | 0.747669 | 0.726004 | 0.732246 | 0.95259 | 0.464777 | | 0.509237 | 0 |
| 22.95833 | 0.406421 | 0.687665 | 0.10307 | 0.286847 | 0.50077 | 0.607131 | 1.111771 | 0.714821 | 0.415635 | | 0.579321 | 0 |
| 22.97222 | 0.419299 | 0.98941 | 0.001373 | 0.110789 | 0.392081 | 0.499259 | 1.313712 | 0.494569 | 0.328537 | | 0.645585 | 0 |
| 22.98611 | 0.249727 | 1.133246 | 0.134791 | 0.11571 | 0.837102 | 0.462238 | 1.043871 | 0.121438 | 0.33332 | | 0.585422 | 0 |
| 23 | 0.118059 | 1.065442 | 0.431939 | 0.427856 | 1.867553 | 0.413186 | 0.270175 | 0.360014 | 0.463991 | | 0.381269 | 0 |
| 23.01389 | 0.472208 | 0.340821 | 0.630198 | 0.529576 | 1.510007 | 0.250737 | 0.313624 | 0.566709 | 0.237513 | | 0.057337 | 0 |
| 23.02778 | 0.607771 | 0.456622 | 0.446346 | 0.2809 | 0.129588 | 0.257213 | 0.415389 | 0.267795 | 0.261717 | | 0.39085 | 0 |
| 23.04167 | 0.896375 | 0.42757 | 0.071703 | 0.170182 | 0.232959 | 1.054166 | 0.234001 | 0.382996 | 0.540766 | | 1.181424 | 0 |
| 23.05556 | 1.06331 | 0.096645 | 0.166457 | 0.131942 | 0.078936 | 1.921227 | 0.024352 | 0.737742 | 0.439714 | | 1.179763 | 5.557098 |
| 23.06944 | 0.37927 | 0.398327 | 0.164966 | 0.173267 | 0.15058 | 0.026964 | 0.027964 | 0.100575 | 0.172217 | | 0.089059 | 0.011822 |
| 23.08333 | 0.263555 | 0.911498 | 0.041848 | 0.301265 | 0.054913 | 2.069573 | 0.208575 | 0.636226 | 0.021878 | | 0.928268 | 5.550682 |
| 23.09722 | 0.337183 | 0.625108 | 0.081122 | 0.044738 | 0.040449 | 1.488533 | 0.09134 | 0.442255 | 0.124407 | | 0.780255 | 0 |
| 23.11111 | 0.24399 | 0.230545 | 0.217807 | 0.086436 | 0.113577 | 0.16575 | 0.103432 | 0.0323 | 0.229908 | | 0.167936 | 0 |
| 23.125 | 0.240156 | 0.140214 | 0.317754 | 0.017236 | 0.235872 | 0.256293 | 0.269277 | 0.179236 | 0.337231 | | 0.105386 | 0 |
| 23.13889 | 0.132458 | 0.185928 | 0.32274 | 0.106756 | 0.294001 | 0.607948 | 0.333504 | 0.170202 | 0.342166 | | 0.063474 | 0 |
| 23.15278 | 0.011133 | 0.303888 | 0.252905 | 0.107821 | 0.260923 | 0.88486 | 0.319261 | 0.123296 | 0.24705 | | 0.325497 | 0 |
| 23.16667 | 0.064049 | 0.421601 | 0.129167 | 0.029867 | 0.063066 | 0.803781 | 0.274954 | 0.137014 | 0.127805 | | 0.515599 | 0 |
| 23.18056 | 0.074154 | 0.38399 | 0.081952 | 0.029117 | 0.026899 | 0.41934 | 0.354658 | 0.183916 | 0.135814 | | 0.286412 | 0 |
| 23.19444 | 0.332455 | 0.174656 | 0.208216 | 0.162987 | 0.14758 | 0.179572 | 0.694801 | 0.253566 | 0.287712 | | 0.108598 | 0 |
| 23.20833 | 0.397382 | 0.039916 | 0.390046 | 0.259697 | 0.390765 | 0.427794 | 1.043083 | 0.370054 | 0.417018 | | 0.189001 | 0 |
| 23.22222 | 0.284522 | 0.09643 | 0.499043 | 0.279366 | 0.487369 | 0.658062 | 1.01805 | 0.485068 | 0.470263 | | 0.105778 | 0 |
| 23.23611 | 0.24709 | 0.072 | 0.530827 | 0.319484 | 0.487907 | 0.552689 | 0.813105 | 0.489501 | 0.459219 | | 0.219935 | 0 |
| 23.25 | 0.321504 | 0.115486 | 0.538792 | 0.387296 | 0.467988 | 0.453834 | 0.604428 | 0.385611 | 0.433389 | | 0.367097 | 0 |
| 23.26389 | 0.433414 | 0.264503 | 0.556693 | 0.426855 | 0.419734 | 0.471952 | 0.506925 | 0.315028 | 0.431744 | | 0.428515 | 0 |
| 23.27778 | 0.459068 | 0.367924 | 0.537299 | 0.357658 | 0.403502 | 0.48431 | 0.465632 | 0.315902 | 0.415476 | | 0.393156 | 0 |
| 23.29167 | 0.27062 | 0.278854 | 0.420358 | 0.105411 | 0.380409 | 0.498715 | 0.409804 | 0.319218 | 0.316466 | | 0.180329 | 0 |
| 23.30556 | 0.01299 | 0.122464 | 0.290583 | 0.206215 | 0.289248 | 0.617004 | 0.337367 | 0.289179 | 0.226126 | | 0.122278 | 0 |
| 23.31944 | 0.171551 | 0.054484 | 0.210784 | 0.303406 | 0.14771 | 0.624154 | 0.250402 | 0.194338 | 0.197049 | | 0.166488 | 0 |
| 23.33333 | 0.155378 | 0.037443 | 0.158798 | 0.177001 | 0.027655 | 0.439261 | 0.135249 | 0.091445 | 0.175016 | | 0.008962 | 0 |
| 23.34722 | 0.116236 | 0.005869 | 0.108936 | 0.044109 | 0.213703 | 0.293655 | 0.010646 | 0.05521 | 0.093166 | | 0.018399 | 0 |
| 23.36111 | 0.120537 | 0.061801 | 0.044767 | 0.07536 | 0.370112 | 0.202727 | 0.059355 | 0.086952 | 0.070118 | | 0.138703 | 0 |
| 23.375 | 0.222737 | 0.236609 | 0.097771 | 0.226252 | 0.45976 | 0.057294 | 0.093077 | 0.374749 | 0.264579 | | 0.332111 | 0 |
| 23.38889 | 0.383703 | 0.483584 | 0.234893 | 0.435948 | 0.509314 | 0.065908 | 0.116069 | 0.659357 | 0.356337 | | 0.486769 | 0 |
| 23.40278 | 0.489563 | 0.660831 | 0.252481 | 0.610072 | 0.503324 | 0.053191 | 0.131408 | 0.865002 | 0.292576 | | 0.564061 | 0 |

| | | | | | | | | | | | |
|---|---|---|---|---|---|---|---|---|---|---|---|
| 23.41667 | 0.516265 | 0.70295 | 0.220368 | 0.673662 | 0.540728 | 0.059189 | 0.181686 | 0.928265 | 0.173242 | 0.67776 | 0 |
| 23.43056 | 0.47075 | 0.636208 | 0.193443 | 0.617327 | 0.614923 | 0.184076 | 0.336891 | 1.040358 | 0.144006 | 0.833193 | 0 |
| 23.44444 | 0.348143 | 0.592356 | 0.215746 | 0.457893 | 0.592473 | 0.106641 | 0.621235 | 1.061908 | 0.211998 | 0.875656 | 0 |
| 23.45833 | 0.248899 | 0.556179 | 0.252331 | 0.339532 | 0.476698 | 0.012014 | 0.884163 | 0.904884 | 0.262251 | 0.775496 | 0 |
| 23.47222 | 0.179312 | 0.474397 | 0.226787 | 0.200458 | 0.356052 | 0.084349 | 0.934602 | 0.55105 | 0.230049 | 0.529341 | 0 |
| 23.48611 | 0.008842 | 0.422229 | 0.048386 | 0.028437 | 0.254622 | 0.472531 | 0.821058 | 0.22159 | 0.070827 | 0.024354 | 0 |
| 23.5 | 0.356356 | 0.394986 | 0.246652 | 0.146198 | 0.324331 | 1.146637 | 0.825702 | 0.030102 | 0.074634 | 0.35604 | 0 |
| 23.51389 | 0.511941 | 0.343308 | 0.290549 | 0.072432 | 0.701265 | 0.806721 | 0.646857 | 0.077716 | 0.015882 | 0.056751 | 0 |
| 23.52778 | 0.252086 | 0.095982 | 0.059786 | 0.398134 | 0.526293 | 0.11139 | 0.506921 | 0.103625 | 0.122923 | 0.316632 | 0 |
| 23.54167 | 0.058118 | 0.077391 | 0.099829 | 0.49803 | 0.275716 | 0.218842 | 0.345985 | 0.336336 | 0.054238 | 0.286945 | 0 |
| 23.55556 | 0.228244 | 0.110252 | 0.042266 | 0.253144 | 0.146111 | 0.172714 | 0.232973 | 0.579843 | 0.391695 | 0.00937 | 0 |
| 23.56944 | 0.381085 | 0.002972 | 0.047663 | 0.043221 | 0.497305 | 0.084162 | 0.11361 | 0.892098 | 0.543537 | 0.189573 | 0 |
| 23.58333 | 0.646887 | 0.238604 | 0.244825 | 0.035403 | 0.562706 | 0.015287 | 0.020247 | 1.171574 | 0.389461 | 0.021425 | 0 |
| 23.59722 | 0.86818 | 0.438051 | 0.465995 | 0.103944 | 0.28941 | 0.04402 | 0.001878 | 1.267003 | 0.110936 | 0.288877 | 0 |
| 23.61111 | 0.80433 | 0.402306 | 0.535724 | 0.096058 | 0.05487 | 0.008173 | 0.195835 | 1.237505 | 0.219285 | 0.473336 | 0 |
| 23.625 | 0.579465 | 0.397318 | 0.428423 | 0.017637 | 0.425202 | 0.047235 | 0.737333 | 1.252868 | 0.521073 | 0.61535 | 0 |
| 23.63889 | 0.441341 | 0.682414 | 0.191844 | 0.243547 | 0.657622 | 0.016967 | 1.293049 | 1.210805 | 0.599689 | 0.891692 | 0 |
| 23.65278 | 0.400389 | 1.157168 | 0.115129 | 0.581909 | 1.02233 | 0.09817 | 1.235231 | 1.016517 | 0.580429 | 1.074165 | 0 |
| 23.66667 | 0.354399 | 1.38432 | 0.320387 | 0.738271 | 0.961697 | 0.111257 | 0.828884 | 0.704041 | 0.426389 | 0.921822 | 0 |
| 23.68056 | 0.351233 | 1.162101 | 0.279629 | 0.636614 | 0.386509 | 0.030945 | 0.381843 | 0.440687 | 0.20479 | 0.509084 | 0 |
| 23.69444 | 0.393775 | 0.634957 | 0.047692 | 0.285008 | 0.071883 | 0.097815 | 0.091648 | 0.181382 | 0.036329 | 0.124684 | 0 |
| 23.70833 | 0.384489 | 0.066376 | 0.241589 | 0.11781 | 0.026473 | 0.111834 | 0.19598 | 0.251246 | 0.105403 | 0.219088 | 0 |
| 23.72222 | 0.114067 | 0.502492 | 0.391774 | 0.532524 | 0.177694 | 0.14723 | 0.582343 | 0.719267 | 0.252583 | 0.513226 | 0 |
| 23.73611 | 0.171679 | 0.739377 | 0.337785 | 0.756834 | 0.043391 | 0.127831 | 0.630276 | 1.071034 | 0.354756 | 0.727803 | 0 |
| 23.75 | 0.335427 | 0.563012 | 0.133735 | 0.636215 | 0.277175 | 0.017495 | 0.399763 | 1.023842 | 0.357806 | 0.620286 | 0 |
| 23.76389 | 0.453743 | 0.331328 | 0.13965 | 0.312186 | 0.469146 | 0.040794 | 0.164692 | 0.796597 | 0.312326 | 0.379218 | 0 |
| 23.77778 | 0.458595 | 0.184971 | 0.35068 | 0.077616 | 0.603617 | 0.012887 | 0.215925 | 0.44691 | 0.228128 | 0.1418 | 0 |
| 23.79167 | 0.241728 | 0.002188 | 0.44694 | 0.028968 | 0.754438 | 0.056823 | 0.319076 | 0.124888 | 0.041509 | 0.005468 | 0 |
| 23.80556 | 0.044831 | 0.035723 | 0.51557 | 0.117059 | 0.844393 | 0.088994 | 0.41873 | 0.139843 | 0.098189 | 0.019131 | 0 |
| 23.81944 | 0.094949 | 0.307635 | 0.607727 | 0.243982 | 1.010993 | 0.18999 | 0.755691 | 0.600546 | 0.032743 | 0.340063 | 0 |
| 23.83333 | 0.313629 | 0.828291 | 0.618388 | 0.368855 | 1.298179 | 0.283857 | 1.068126 | 1.203074 | 0.354687 | 0.85472 | 0 |
| 23.84722 | 0.521873 | 1.170812 | 0.544226 | 0.425181 | 1.589091 | 0.312811 | 1.211871 | 1.58133 | 0.582402 | 1.223974 | 0 |
| 23.86111 | 0.674899 | 1.327025 | 0.427704 | 0.486389 | 1.916555 | 0.306146 | 1.27102 | 1.783796 | 0.70832 | 1.449794 | 0 |
| 23.875 | 0.839346 | 1.329816 | 0.43733 | 0.622316 | 2.118814 | 0.356622 | 1.359241 | 1.800849 | 0.810384 | 1.575696 | 0 |
| 23.88889 | 0.978607 | 1.305397 | 0.479762 | 0.685762 | 1.956463 | 0.384969 | 1.186898 | 1.55999 | 0.81884 | 1.450974 | 0 |
| 23.90278 | 0.976588 | 1.213702 | 0.450941 | 0.545407 | 1.709212 | 0.3817 | 0.782497 | 1.308305 | 0.736384 | 1.147652 | 0 |
| 23.91667 | 0.877822 | 1.019909 | 0.399273 | 0.430385 | 1.52355 | 0.394165 | 0.751715 | 1.277608 | 0.678187 | 0.93218 | 0 |
| 23.93056 | 0.713841 | 0.827329 | 0.361784 | 0.43866 | 1.418277 | 0.45725 | 1.012421 | 1.257498 | 0.614046 | 0.925075 | 0 |
| 23.94444 | 0.538882 | 0.753613 | 0.312862 | 0.419089 | 1.403222 | 0.486876 | 1.054179 | 1.136306 | 0.507755 | 0.922852 | 0 |
| 23.95833 | 0.51043 | 0.868483 | 0.276342 | 0.383874 | 1.461932 | 0.512464 | 0.900946 | 0.986261 | 0.459409 | 0.867329 | 0 |
| 23.97222 | 0.599484 | 1.071527 | 0.259861 | 0.438082 | 1.347701 | 0.567687 | 0.854378 | 0.831958 | 0.512135 | 0.781171 | 0 |
| 23.98611 | 0.498319 | 1.285851 | 0.196539 | 0.51031 | 0.905212 | 0.904335 | 0.866545 | 0.547555 | 0.529694 | 0.138848 | 0 |
| 24 | 0.197411 | 1.392817 | 0.052047 | 0.446564 | 0.40041 | 1.565283 | 0.985746 | 0.267357 | 0.357698 | 1.049073 | 0 |
| 24.01389 | 0.08944 | 1.092444 | 0.008749 | 0.299118 | 0.149994 | 1.445728 | 1.102496 | 0.069601 | 0.106386 | 1.149911 | 0 |
| 24.02778 | 0.293052 | 0.679775 | 0.102472 | 0.145381 | 0.315999 | 0.296531 | 1.014284 | 0.016148 | 0.061215 | 0.038622 | 0 |
| 24.04167 | 0.217098 | 0.385485 | 0.170344 | 0.223037 | 0.294314 | 0.181849 | 0.730435 | 0.270226 | 0.166708 | 0.689533 | 0 |
| 24.05556 | 0.187441 | 0.01895 | 0.118747 | 0.48429 | 0.069591 | 0.017359 | 0.5073 | 0.495953 | 0.216053 | 0.396216 | 0 |
| 24.06944 | 0.482919 | 0.386938 | 0.078963 | 0.580587 | 0.003247 | 0.078158 | 0.361708 | 0.310732 | 0.179817 | 0.09119 | 0 |
| 24.08333 | 0.349913 | 0.3854 | 0.100978 | 0.453937 | 0.07663 | 0.0331 | 0.28801 | 0.054638 | 0.131136 | 0.119552 | 0 |
| 24.09722 | 0.035381 | 0.139608 | 0.254147 | 0.263644 | 0.200311 | 0.248997 | 0.295138 | 0.076835 | 0.170771 | 0.69329 | 0 |
| 24.11111 | 0.373108 | 0.005214 | 0.428197 | 0.119387 | 0.235847 | 0.723481 | 0.303338 | 0.000831 | 0.275934 | 1.617811 | 0 |
| 24.125 | 0.266242 | 0.073489 | 0.275885 | 0.071282 | 0.030371 | 0.875489 | 0.121588 | 0.045995 | 0.199015 | 1.518037 | 0 |
| 24.13889 | 0.09323 | 0.249927 | 0.006783 | 0.236784 | 0.358441 | 0.486247 | 0.122949 | 0.132114 | 0.013103 | 0.4469 | 0 |
| 24.15278 | 0.267468 | 0.320862 | 0.05602 | 0.215304 | 0.320002 | 0.133439 | 0.216302 | 0.063378 | 0.084949 | 0.106677 | 0 |
| 24.16667 | 0.200468 | 0.290821 | 0.258227 | 0.089904 | 0.137361 | 0.010679 | 0.196068 | 0.075087 | 0.010305 | 0.036723 | 0 |
| 24.18056 | 0.172631 | 0.237405 | 0.781541 | 0.431612 | 0.114495 | 0.115204 | 0.061854 | 0.390257 | 0.152262 | 0.354305 | 0 |

| | | | | | | | | | | | |
|---|---|---|---|---|---|---|---|---|---|---|---|
| 24.19444 | 0.747144 | 1.407316 | 0.919927 | 1.528963 | 0.040458 | 0.285106 | 0.692621 | 0.986661 | 0.22109 | 0.960373 | 0 |
| 24.20833 | 0.905636 | 1.695046 | 0.55947 | 1.869613 | 0.082838 | 0.270538 | 1.008061 | 1.158688 | 0.26298 | 1.103398 | 0 |
| 24.22222 | 0.606769 | 0.882234 | 0.229077 | 1.088171 | 0.177875 | 0.198329 | 0.788699 | 0.697813 | 0.236957 | 0.710082 | 0 |
| 24.23611 | 0.375299 | 0.364594 | 0.153699 | 0.502946 | 0.207548 | 0.271153 | 0.617429 | 0.389015 | 0.188703 | 0.353735 | 0 |
| 24.25 | 0.208857 | 0.222147 | 0.225472 | 0.319809 | 0.172487 | 0.397859 | 0.613257 | 0.368802 | 0.175433 | 0.184009 | 0 |
| 24.26389 | 0.131837 | 0.172078 | 0.398668 | 0.224958 | 0.153102 | 0.362052 | 0.679121 | 0.333947 | 0.292475 | 0.356447 | 0 |
| 24.27778 | 0.187547 | 0.218627 | 0.500875 | 0.101823 | 0.128173 | 0.295335 | 0.699453 | 0.176601 | 0.42356 | 0.55168 | 0 |
| 24.29167 | 0.145003 | 0.199472 | 0.372586 | 0.183072 | 0.029999 | 0.278289 | 0.54251 | 0.039917 | 0.339423 | 0.313653 | 0 |
| 24.30556 | 0.050237 | 0.04961 | 0.135913 | 0.518264 | 0.243166 | 0.275869 | 0.296387 | 0.147927 | 0.144738 | 0.094783 | 0 |
| 24.31944 | 0.361907 | 0.19445 | 0.091615 | 0.767736 | 0.435876 | 0.153665 | 0.038266 | 0.208898 | 0.046932 | 0.326891 | 0 |
| 24.33333 | 0.691259 | 0.442766 | 0.291628 | 0.884179 | 0.550832 | 0.098496 | 0.180217 | 0.318124 | 0.221251 | 0.287702 | 0 |
| 24.34722 | 0.797539 | 0.434107 | 0.315156 | 0.773159 | 0.456463 | 0.125438 | 0.194964 | 0.327084 | 0.24597 | 0.156768 | 0 |
| 24.36111 | 0.510162 | 0.083222 | 0.131871 | 0.533139 | 0.247806 | 0.174016 | 0.037525 | 0.230571 | 0.062765 | 0.021588 | 0 |
| 24.375 | 0.07544 | 0.292743 | 0.087186 | 0.363412 | 0.152569 | 0.514577 | 0.109831 | 0.163941 | 0.190239 | 0.08351 | 0 |
| 24.38889 | 0.25594 | 0.519734 | 0.354625 | 0.074271 | 0.142433 | 0.861305 | 0.229307 | 0.029108 | 0.437281 | 0.165681 | 0 |
| 24.40278 | 0.361016 | 0.52392 | 0.684369 | 0.29196 | 0.079686 | 1.065892 | 0.35693 | 0.420087 | 0.63276 | 0.313244 | 0 |
| 24.41667 | 0.003181 | 0.047336 | 0.634326 | 0.167633 | 0.152571 | 0.860692 | 0.169554 | 0.369516 | 0.462882 | 0.060032 | 0 |
| 24.43056 | 0.598901 | 0.654828 | 0.170493 | 0.452984 | 0.397692 | 0.334281 | 0.353826 | 0.113348 | 0.022884 | 0.548086 | 0 |
| 24.44444 | 0.513833 | 0.742605 | 0.173705 | 0.875672 | 0.610188 | 0.027371 | 0.659965 | 0.471749 | 0.3243 | 0.824993 | 0 |
| 24.45833 | 0.216434 | 0.609183 | 0.341978 | 0.905258 | 0.76223 | 0.244026 | 0.665712 | 0.667824 | 0.453836 | 0.881589 | 0 |
| 24.47222 | 0.46076 | 0.80862 | 0.60929 | 0.971545 | 0.940587 | 0.581587 | 0.85169 | 1.008417 | 0.669619 | 1.192181 | 0 |
| 24.48611 | 1.179072 | 1.524972 | 0.833962 | 1.048084 | 1.108262 | 0.90669 | 1.363496 | 1.33816 | 0.953064 | 1.843633 | 0 |
| 24.5 | 1.495018 | 2.164418 | 0.741755 | 0.8766 | 1.208825 | 1.117874 | 2.082601 | 1.437605 | 1.198608 | 2.439048 | 0 |
| 24.51389 | 1.03674 | 1.848269 | 0.695725 | 0.621901 | 1.416465 | 0.898886 | 2.515457 | 1.391499 | 1.085975 | 2.566885 | 0 |
| 24.52778 | 0.732955 | 1.161753 | 0.863493 | 0.568085 | 1.67351 | 0.483421 | 2.482348 | 1.620287 | 0.727807 | 2.119294 | 0 |
| 24.54167 | 0.590713 | 0.631908 | 0.728374 | 0.639394 | 0.963922 | 0.312279 | 1.797918 | 2.069714 | 0.510116 | 1.357435 | 0 |
| 24.55556 | 0.40267 | 0.230254 | 0.208943 | 0.650381 | 0.043996 | 0.349219 | 0.759351 | 2.175103 | 0.195103 | 0.484332 | 0 |
| 24.56944 | 0.167237 | 0.003591 | 0.244074 | 0.365779 | 0.818439 | 0.474028 | 0.12062 | 1.73609 | 0.107007 | 0.18136 | 0 |
| 24.58333 | 0.250836 | 0.023289 | 0.239333 | 0.089659 | 1.228017 | 0.577086 | 0.465235 | 1.03918 | 0.251929 | 0.402213 | 0 |
| 24.59722 | 0.530143 | 0.06378 | 0.176936 | 0.057738 | 0.968611 | 0.559092 | 0.04969 | 0.764415 | 0.226933 | 0.029086 | 0 |
| 24.61111 | 0.928888 | 0.330398 | 0.822488 | 0.255133 | 0.098935 | 0.654939 | 1.048061 | 0.968313 | 0.061504 | 0.828688 | 0 |
| 24.625 | 1.15316 | 0.789946 | 1.347562 | 0.424172 | 1.048785 | 0.816671 | 2.20907 | 1.24052 | 0.602891 | 1.59854 | 0 |
| 24.63889 | 1.104006 | 1.390924 | 1.181623 | 0.436956 | 1.871392 | 0.824614 | 2.072327 | 1.179744 | 0.983491 | 1.723753 | 0 |
| 24.65278 | 0.817657 | 1.850985 | 0.576893 | 0.3836 | 1.961315 | 0.723803 | 1.064555 | 1.135483 | 1.124677 | 1.340438 | 0 |
| 24.66667 | 0.487591 | 1.778282 | 0.017093 | 0.375906 | 1.219871 | 0.519177 | 0.045853 | 1.392076 | 1.135787 | 0.846891 | 0 |
| 24.68056 | 0.32781 | 1.159754 | 0.21021 | 0.586969 | 0.319012 | 0.254442 | 0.530694 | 1.686777 | 0.956226 | 0.483617 | 0 |
| 24.69444 | 0.387817 | 0.645456 | 0.155682 | 0.760407 | 0.448554 | 0.056988 | 0.772248 | 1.727168 | 0.673734 | 0.392424 | 0 |
| 24.70833 | 0.563812 | 0.34163 | 0.037764 | 0.750911 | 0.804083 | 0.043449 | 0.424508 | 1.376279 | 0.370174 | 0.428822 | 0 |
| 24.72222 | 0.771512 | 0.101622 | 0.312673 | 0.604035 | 0.853182 | 0.054547 | 0.182872 | 0.790192 | 0.122527 | 0.300853 | 0 |
| 24.73611 | 0.7205 | 0.188546 | 0.504671 | 0.201466 | 0.727525 | 0.047926 | 0.576916 | 0.04833 | 0.010028 | 0.029681 | 0 |
| 24.75 | 0.385542 | 0.552087 | 0.389745 | 0.367219 | 0.570343 | 0.095631 | 0.111739 | 0.975743 | 0.098649 | 0.53889 | 0 |
| 24.76389 | 0.062329 | 0.719927 | 0.059946 | 0.645977 | 0.242983 | 0.083196 | 0.898952 | 1.825515 | 0.169064 | 1.001842 | 0 |
| 24.77778 | 0.110173 | 0.480397 | 0.236086 | 0.382107 | 0.185576 | 0.046576 | 1.782672 | 2.00643 | 0.151542 | 1.057298 | 0 |
| 24.79167 | 0.305635 | 0.015502 | 0.473093 | 0.133074 | 0.515113 | 0.143872 | 2.002428 | 1.535899 | 0.117212 | 0.702661 | 0 |
| 24.80556 | 0.472587 | 0.517387 | 0.587651 | 0.270831 | 0.506967 | 0.120512 | 1.036666 | 0.958286 | 0.098567 | 0.167956 | 0 |
| 24.81944 | 0.346197 | 0.67795 | 0.597543 | 0.382333 | 0.221935 | 0.056714 | 0.294002 | 0.396044 | 0.019553 | 0.234377 | 0 |
| 24.83333 | 0.143625 | 0.340381 | 0.532521 | 0.294974 | 0.233716 | 0.056009 | 0.704828 | 0.151878 | 0.115981 | 0.11186 | 0 |
| 24.84722 | 0.22913 | 0.187476 | 0.581169 | 0.326377 | 0.848715 | 0.317469 | 0.012012 | 0.488943 | 0.010295 | 0.54429 | 0 |
| 24.86111 | 0.475613 | 0.618828 | 0.619244 | 0.490388 | 1.508769 | 0.627493 | 1.029209 | 1.019215 | 0.255705 | 1.237625 | 0 |
| 24.875 | 0.682427 | 0.835385 | 0.642056 | 0.66502 | 1.912474 | 0.785248 | 1.499971 | 1.427226 | 0.601123 | 1.646922 | 0 |
| 24.88889 | 0.783938 | 1.058734 | 0.73595 | 0.883771 | 1.955016 | 0.85475 | 1.583478 | 1.661215 | 0.941681 | 1.732569 | 0 |
| 24.90278 | 0.812573 | 1.238241 | 0.7352 | 0.912569 | 1.736672 | 0.803134 | 1.389774 | 1.772998 | 0.933683 | 1.487708 | 0 |
| 24.91667 | 0.68159 | 1.217146 | 0.47799 | 0.765607 | 1.185684 | 0.549805 | 0.856534 | 1.636005 | 0.645127 | 1.012076 | 0 |
| 24.93056 | 0.65733 | 1.198035 | 0.264818 | 0.61598 | 0.674066 | 0.420404 | 0.307437 | 1.51804 | 0.444309 | 0.758968 | 0 |
| 24.94444 | 0.8549 | 1.242818 | 0.33108 | 0.519665 | 0.381868 | 0.387755 | 0.215186 | 1.479941 | 0.42139 | 0.817887 | 0 |
| 24.95833 | 1.059875 | 1.124341 | 0.394944 | 0.452619 | 0.338816 | 0.308366 | 0.532743 | 1.13469 | 0.41063 | 0.939476 | 0 |

| | | | | | | | | | | | |
|---|---|---|---|---|---|---|---|---|---|---|---|
| 24.97222 | 1.080099 | 0.822919 | 0.314787 | 0.482802 | 0.479107 | 0.221326 | 1.029604 | 0.567751 | 0.38588 | 0.958325 | 0 |
| 24.98611 | 0.78882 | 0.394459 | 0.007911 | 0.281119 | 0.386088 | 0.040998 | 1.116594 | 0.115405 | 0.145788 | 0.66032 | 0 |
| 25 | 0.226549 | 0.083957 | 0.645382 | 0.345547 | 0.109188 | 0.726844 | 0.472638 | 0.809673 | 0.488545 | 0.095187 | 0 |
| 25.01389 | 0.051239 | 0.093039 | 1.061176 | 0.819911 | 0.420351 | 1.285939 | 0.26095 | 0.87632 | 0.91588 | 0.162071 | 0 |
| 25.02778 | 0.427912 | 0.393818 | 0.907942 | 0.831164 | 1.197914 | 1.183852 | 0.564026 | 0.304104 | 0.678133 | 0.028753 | 0 |
| 25.04167 | 0.473865 | 0.761394 | 0.619675 | 0.571897 | 1.372918 | 0.749994 | 0.550501 | 0.106799 | 0.177835 | 0.228326 | 0 |
| 25.05556 | 0.181943 | 0.688831 | 0.362719 | 0.153785 | 0.843006 | 0.302256 | 0.337391 | 0.159753 | 0.21148 | 0.321831 | 0 |
| 25.06944 | 0.336525 | 0.235475 | 0.147743 | 0.266142 | 0.436168 | 0.019998 | 0.046606 | 0.090961 | 0.362914 | 0.33652 | 0 |
| 25.08333 | 0.825495 | 0.331654 | 0.075068 | 0.352259 | 0.269262 | 0.096796 | 0.095268 | 0.086881 | 0.302503 | 0.190841 | 0 |
| 25.09722 | 1.069078 | 0.685317 | 0.149572 | 0.15259 | 0.180293 | 0.025169 | 0.066344 | 0.200018 | 0.180334 | 0.06306 | 0 |
| 25.11111 | 0.997489 | 0.729738 | 0.296217 | 0.047654 | 0.043845 | 0.004038 | 0.003467 | 0.166118 | 0.076737 | 0.254793 | 0 |
| 25.125 | 0.871048 | 0.620105 | 0.436948 | 0.157207 | 0.172692 | 0.057009 | 0.121171 | 0.108414 | 0.037046 | 0.382252 | 0 |
| 25.13889 | 0.776432 | 0.490492 | 0.468877 | 0.186112 | 0.268235 | 0.104325 | 0.291966 | 0.081372 | 0.133846 | 0.385255 | 0 |
| 25.15278 | 0.688813 | 0.409388 | 0.357227 | 0.174419 | 0.245126 | 0.095299 | 0.406165 | 0.090003 | 0.161498 | 0.288114 | 0 |
| 25.16667 | 0.621143 | 0.357562 | 0.19966 | 0.137418 | 0.260267 | 0.025899 | 0.387822 | 0.135624 | 0.161232 | 0.202395 | 0 |
| 25.18056 | 0.574274 | 0.325189 | 0.077437 | 0.130945 | 0.322085 | 0.047202 | 0.297292 | 0.19364 | 0.123346 | 0.179349 | 0 |
| 25.19444 | 0.514266 | 0.298506 | 0.052044 | 0.194628 | 0.278866 | 0.108162 | 0.220576 | 0.221008 | 0.073375 | 0.193758 | 0 |
| 25.20833 | 0.451001 | 0.249188 | 0.044236 | 0.255082 | 0.018068 | 0.148652 | 0.149884 | 0.168231 | 0.073012 | 0.155013 | 0 |
| 25.22222 | 0.368564 | 0.154045 | 0.014943 | 0.255976 | 0.268857 | 0.17265 | 0.087051 | 0.068393 | 0.154021 | 0.047927 | 0 |
| 25.23611 | 0.222248 | 0.016492 | 0.100468 | 0.215745 | 0.235045 | 0.180573 | 0.031948 | 0.017764 | 0.190454 | 0.080154 | 0 |
| 25.25 | 0.032411 | 0.154871 | 0.142466 | 0.188067 | 0.02476 | 0.15847 | 0.243462 | 0.038997 | 0.082461 | 0.195619 | 0 |
| 25.26389 | 0.059146 | 0.288319 | 0.093186 | 0.187812 | 0.244787 | 0.087315 | 0.343628 | 0.06393 | 0.02717 | 0.123006 | 0 |
| 25.27778 | 0.056791 | 0.301778 | 0.043942 | 0.216284 | 0.29397 | 0.068465 | 0.196398 | 0.223546 | 0.035532 | 0.26286 | 0 |
| 25.29167 | 0.321986 | 0.114591 | 0.170207 | 0.347811 | 0.169164 | 0.025264 | 0.054495 | 0.390643 | 0.198294 | 0.53818 | 0 |
| 25.30556 | 0.51167 | 0.234789 | 0.220918 | 0.680549 | 0.100867 | 0.461392 | 0.219858 | 0.619035 | 0.296445 | 0.292787 | 0.006288 |
| 25.31944 | 0.64744 | 0.495527 | 0.281069 | 1.008654 | 0.194957 | 0.852391 | 0.300578 | 0.79578 | 0.384718 | 0.076873 | 0 |
| 25.33333 | 0.802662 | 0.47611 | 0.364898 | 1.128582 | 0.028419 | 0.661716 | 0.325392 | 0.822279 | 0.468476 | 0.296986 | 0 |
| 25.34722 | 0.952654 | 0.446968 | 0.445099 | 1.035732 | 0.145179 | 0.378087 | 0.215323 | 0.70583 | 0.414388 | 0.565414 | 0 |
| 25.36111 | 1.177279 | 0.579764 | 0.548729 | 0.900816 | 0.394442 | 0.266332 | 0.085808 | 0.567748 | 0.339838 | 0.652747 | 0 |
| 25.375 | 1.466545 | 0.858426 | 0.610344 | 0.849719 | 0.824342 | 0.210163 | 0.108891 | 0.513141 | 0.407503 | 0.680581 | 0 |
| 25.38889 | 1.262452 | 0.794426 | 0.596997 | 0.775722 | 1.127716 | 0.238964 | 0.232145 | 0.573249 | 0.608502 | 0.711834 | 0 |
| 25.40278 | 0.918052 | 0.538638 | 0.595474 | 0.704319 | 1.270296 | 0.3114 | 0.327084 | 0.580705 | 0.752974 | 0.755304 | 0 |
| 25.41667 | 0.558276 | 0.305587 | 0.60555 | 0.741548 | 1.151709 | 0.339804 | 0.391561 | 0.534322 | 0.779317 | 0.723964 | 0 |
| 25.43056 | 0.202035 | 0.080283 | 0.644699 | 0.907114 | 0.806119 | 0.325661 | 0.503614 | 0.451829 | 0.68817 | 0.61377 | 0 |
| 25.44444 | 0.051388 | 0.006826 | 0.737914 | 1.094674 | 0.441509 | 0.320853 | 0.720457 | 0.487508 | 0.564583 | 0.607262 | 0 |
| 25.45833 | 0.185796 | 0.094564 | 0.747318 | 1.012771 | 0.377509 | 0.337977 | 0.934832 | 0.663647 | 0.472472 | 0.695424 | 0 |
| 25.47222 | 0.350919 | 0.226528 | 0.601731 | 0.701367 | 0.534751 | 0.349328 | 1.004855 | 0.858212 | 0.466486 | 0.752979 | 0 |
| 25.48611 | 0.465757 | 0.599283 | 0.510929 | 0.435331 | 0.68228 | 0.340276 | 1.060639 | 1.094685 | 0.688548 | 1.026274 | 0 |
| 25.5 | 0.574863 | 1.52586 | 0.76852 | 0.391611 | 0.322023 | 0.166144 | 1.365817 | 1.419396 | 1.395158 | 1.266678 | 0 |
| 25.51389 | 0.55173 | 2.177132 | 1.140066 | 0.438769 | 2.284849 | 0.302042 | 1.605193 | 1.360126 | 1.605578 | 0.546994 | 0 |
| 25.52778 | 0.361291 | 1.66872 | 1.204751 | 0.408761 | 4.630025 | 0.405744 | 1.728947 | 1.050375 | 0.971178 | 0.094298 | 0 |
| 25.54167 | 0.304469 | 1.233528 | 0.517408 | 0.30852 | 2.454934 | 0.066886 | 1.432247 | 0.887673 | 0.324875 | 0.161087 | 0 |
| 25.55556 | 0.359822 | 0.857425 | 0.454977 | 0.038871 | 1.474986 | 0.162327 | 0.445909 | 0.876592 | 0.153642 | 0.024546 | 0 |
| 25.56944 | 0.478195 | 0.562989 | 0.978182 | 0.129769 | 0.991329 | 0.315674 | 0.544203 | 0.813797 | 0.489778 | 0.346201 | 0 |
| 25.58333 | 0.634077 | 0.409782 | 0.592088 | 0.160686 | 0.990889 | 0.381355 | 1.009492 | 0.787841 | 0.541683 | 0.455165 | 0 |
| 25.59722 | 0.715143 | 0.278656 | 0.37253 | 0.089075 | 1.076677 | 0.319634 | 0.690972 | 0.8804 | 0.235317 | 0.149976 | 0 |
| 25.61111 | 0.630601 | 0.043081 | 1.345592 | 0.058244 | 0.92762 | 0.235809 | 0.330055 | 1.28448 | 0.206889 | 0.425247 | 0 |
| 25.625 | 0.436925 | 0.185712 | 1.779455 | 0.119231 | 0.603362 | 0.236531 | 1.451139 | 1.755431 | 0.396004 | 0.792907 | 0 |
| 25.63889 | 0.253223 | 0.40508 | 1.049745 | 0.065087 | 0.386324 | 0.191758 | 1.818618 | 1.823529 | 0.3029 | 0.702626 | 0 |
| 25.65278 | 0.018212 | 0.51188 | 0.08173 | 0.057865 | 0.293751 | 0.059005 | 1.194889 | 1.555669 | 0.041945 | 0.27822 | 0 |
| 25.66667 | 0.096211 | 0.256237 | 0.971755 | 0.32875 | 0.347735 | 0.015577 | 0.197164 | 1.394238 | 0.16789 | 0.176384 | 0 |
| 25.68056 | 0.082921 | 0.293202 | 1.213077 | 0.524783 | 0.618785 | 0.113288 | 1.456625 | 1.294175 | 0.020208 | 0.425304 | 0 |
| 25.69444 | 0.035018 | 0.877658 | 0.765002 | 0.364887 | 0.871803 | 0.177739 | 1.946456 | 0.877587 | 0.490197 | 0.282333 | 0 |
| 25.70833 | 0.054945 | 1.360043 | 0.068428 | 0.076432 | 0.75082 | 0.178181 | 1.408232 | 0.323237 | 0.741774 | 0.201393 | 0 |
| 25.72222 | 0.456805 | 1.564737 | 0.671173 | 0.094989 | 0.375748 | 0.239009 | 0.018475 | 0.163145 | 0.57394 | 0.792865 | 0 |
| 25.73611 | 0.833571 | 1.290497 | 0.875053 | 0.022888 | 0.128734 | 0.248398 | 1.246195 | 0.724693 | 0.183294 | 0.911533 | 0 |

| | | | | | | | | | | | |
|---|---|---|---|---|---|---|---|---|---|---|---|
| 25.75 | 0.790052 | 0.732157 | 0.23513 | 0.361054 | 0.054521 | 0.105788 | 1.26891 | 1.354597 | 0.231656 | 0.364819 | 0 |
| 25.76389 | 0.518575 | 0.261013 | 0.685029 | 0.535112 | 0.008063 | 0.023015 | 0.470017 | 1.678016 | 0.36503 | 0.411451 | 0 |
| 25.77778 | 0.100971 | 0.165855 | 1.457784 | 0.488422 | 0.078742 | 0.128258 | 0.487752 | 1.685534 | 0.341089 | 0.944029 | 0 |
| 25.79167 | 0.222307 | 0.583123 | 1.383164 | 0.4459 | 0.089719 | 0.321562 | 1.200371 | 1.403434 | 0.338181 | 1.134055 | 0 |
| 25.80556 | 0.164542 | 0.871196 | 0.781603 | 0.48357 | 0.024315 | 0.555407 | 1.457468 | 1.002385 | 0.304459 | 0.990991 | 0 |
| 25.81944 | 0.041775 | 1.121761 | 0.285725 | 0.559493 | 0.137438 | 0.685508 | 1.493682 | 0.697802 | 0.35167 | 0.933957 | 0 |
| 25.83333 | 0.172928 | 1.38258 | 0.377373 | 0.472208 | 0.346332 | 0.754712 | 1.499595 | 0.609587 | 0.443654 | 1.135925 | 0 |
| 25.84722 | 0.485404 | 1.338826 | 0.700229 | 0.262409 | 0.462335 | 0.835845 | 1.280976 | 0.837236 | 0.439746 | 1.359501 | 0 |
| 25.86111 | 1.058827 | 1.480134 | 0.917052 | 0.309213 | 0.530553 | 1.012993 | 1.07598 | 1.47688 | 0.494835 | 1.650178 | 0 |
| 25.875 | 2.019018 | 1.981307 | 1.026518 | 0.63661 | 0.693362 | 1.107795 | 1.143604 | 2.109401 | 0.555734 | 1.928642 | 0 |
| 25.88889 | 2.153812 | 1.865311 | 0.863243 | 0.987049 | 0.765214 | 0.949143 | 1.246783 | 2.310734 | 0.596377 | 2.047206 | 0 |
| 25.90278 | 1.597136 | 1.326754 | 0.681921 | 1.120964 | 0.754665 | 0.593637 | 0.936134 | 2.33251 | 0.7898 | 1.89384 | 0 |
| 25.91667 | 0.663683 | 0.827195 | 0.414223 | 1.061929 | 0.740304 | 0.315241 | 0.620693 | 2.164073 | 1.001376 | 1.442159 | 0 |
| 25.93056 | 0.119893 | 0.464245 | 0.176509 | 0.707865 | 0.731625 | 0.267339 | 0.502907 | 1.605598 | 1.067643 | 0.969278 | 0 |
| 25.94444 | 0.318737 | 0.375982 | 0.092015 | 0.521035 | 0.724292 | 0.430995 | 0.724474 | 1.108253 | 0.983731 | 0.772171 | 0 |
| 25.95833 | 0.045375 | 0.42831 | 0.131831 | 0.414214 | 0.575098 | 0.568861 | 0.888831 | 0.664352 | 0.614447 | 0.745757 | 0 |
| 25.97222 | 0.408272 | 0.471216 | 0.246368 | 0.361041 | 0.386212 | 0.527515 | 0.707912 | 0.449001 | 0.265276 | 0.78468 | 0 |
| 25.98611 | 0.818031 | 0.618317 | 0.312682 | 0.3342 | 0.202091 | 0.318827 | 0.287968 | 0.355307 | 0.147663 | 0.896626 | 0 |
| 26 | 1.088712 | 0.862101 | 0.172427 | 0.271097 | 0.120416 | 0.072316 | 0.095025 | 0.285099 | 0.06678 | 1.005784 | 0 |
| 26.01389 | 1.523909 | 1.086427 | 0.063716 | 0.399811 | 0.387097 | 0.228255 | 0.10521 | 0.466028 | 0.009764 | 1.145488 | 0 |
| 26.02778 | 2.064451 | 1.195944 | 0.09712 | 0.552141 | 0.356491 | 0.077452 | 0.161927 | 0.837194 | 0.126123 | 1.161832 | 0 |
| 26.04167 | 2.363062 | 1.189423 | 0.079629 | 0.23672 | 0.237128 | 0.352403 | 0.235804 | 0.919227 | 0.316721 | 0.964501 | 0 |
| 26.05556 | 1.936023 | 0.959558 | 0.078521 | 0.056606 | 0.150411 | 0.282485 | 0.13324 | 0.723519 | 0.39581 | 0.744729 | 0 |
| 26.06944 | 1.146989 | 0.651738 | 0.011085 | 0.11324 | 0.020747 | 0.137085 | 0.090122 | 0.550977 | 0.402486 | 0.525777 | 0 |
| 26.08333 | 0.37925 | 0.384766 | 0.028868 | 0.380083 | 0.188648 | 0.099127 | 0.08925 | 0.362347 | 0.379049 | 0.384577 | 0 |
| 26.09722 | 0.227925 | 0.125409 | 0.047658 | 0.368681 | 0.294728 | 0.124072 | 0.010837 | 0.090364 | 0.294967 | 0.253625 | 0 |
| 26.11111 | 0.654666 | 0.146218 | 0.221159 | 0.160099 | 0.305237 | 0.134754 | 0.223156 | 0.109433 | 0.203147 | 0.084667 | 0 |
| 26.125 | 0.839648 | 0.372308 | 0.381745 | 0.115307 | 0.231576 | 0.126739 | 0.532786 | 0.249519 | 0.090783 | 0.193328 | 0 |
| 26.13889 | 0.877325 | 0.501232 | 0.483324 | 0.404148 | 0.04578 | 0.104245 | 0.793049 | 0.386306 | 0.055749 | 0.500196 | 0 |
| 26.15278 | 0.940313 | 0.589677 | 0.602447 | 0.533354 | 0.191283 | 0.00364 | 0.882432 | 0.481956 | 0.236182 | 0.690208 | 0 |
| 26.16667 | 1.015259 | 0.684616 | 0.677935 | 0.381154 | 0.395552 | 0.154957 | 0.805359 | 0.480095 | 0.404773 | 0.760852 | 0 |
| 26.18056 | 1.132921 | 0.786666 | 0.682317 | 0.14214 | 0.499695 | 0.228871 | 0.596194 | 0.396448 | 0.470247 | 0.72857 | 0 |
| 26.19444 | 1.237608 | 0.840371 | 0.557409 | 0.008364 | 0.53624 | 0.17176 | 0.374469 | 0.312126 | 0.41189 | 0.609099 | 0 |
| 26.20833 | 1.253485 | 0.870486 | 0.375233 | 0.053381 | 0.523182 | 0.097063 | 0.134435 | 0.281388 | 0.288612 | 0.390548 | 0 |
| 26.22222 | 1.175749 | 0.789781 | 0.224913 | 0.263652 | 0.430571 | 0.114839 | 0.075016 | 0.227399 | 0.151715 | 0.175858 | 0 |
| 26.23611 | 1.004209 | 0.506086 | 0.114481 | 0.34047 | 0.247637 | 0.188314 | 0.142416 | 0.067918 | 0.00562 | 0.023722 | 0 |
| 26.25 | 0.745308 | 0.216533 | 0.012402 | 0.162266 | 0.076589 | 0.236046 | 0.115813 | 0.07111 | 0.098077 | 0.174707 | 0 |
| 26.26389 | 0.570833 | 0.142211 | 0.023666 | 0.019542 | 0.07603 | 0.270685 | 0.027429 | 0.049819 | 0.082499 | 0.161894 | 0 |
| 26.27778 | 0.510536 | 0.166789 | 0.006043 | 0.117861 | 0.197925 | 0.281492 | 0.045281 | 0.105506 | 0.000579 | 0.071116 | 0 |
| 26.29167 | 0.456277 | 0.136171 | 0.026795 | 0.533 | 0.264732 | 0.275905 | 0.079904 | 0.291264 | 0.068806 | 0.016304 | 0 |
| 26.30556 | 0.393963 | 0.121976 | 0.075712 | 0.894368 | 0.30081 | 0.37444 | 0.105851 | 0.423638 | 0.135373 | 0.133964 | 0 |
| 26.31944 | 0.36252 | 0.187954 | 0.207864 | 0.853304 | 0.283278 | 0.587637 | 0.151618 | 0.580555 | 0.211711 | 0.263024 | 0 |
| 26.33333 | 0.37921 | 0.273237 | 0.371412 | 0.620975 | 0.218635 | 0.781841 | 0.184025 | 0.857279 | 0.274349 | 0.34201 | 0 |
| 26.34722 | 0.546247 | 0.362357 | 0.536216 | 0.557964 | 0.162855 | 0.806034 | 0.236613 | 1.176387 | 0.336082 | 0.459564 | 0 |
| 26.36111 | 0.81858 | 0.495836 | 0.626812 | 0.739242 | 0.14874 | 0.861853 | 0.358756 | 1.350032 | 0.393117 | 0.66098 | 0 |
| 26.375 | 1.105066 | 0.645893 | 0.618567 | 0.917312 | 0.172691 | 0.914858 | 0.520293 | 1.397364 | 0.430314 | 0.8733 | 0 |
| 26.38889 | 1.28928 | 0.724682 | 0.559516 | 0.917571 | 0.144775 | 0.910574 | 0.601081 | 1.405316 | 0.407759 | 1.021072 | 0 |
| 26.40278 | 1.231564 | 0.706873 | 0.553039 | 0.816846 | 0.039635 | 0.831878 | 0.547185 | 1.296912 | 0.356037 | 1.027423 | 0 |
| 26.41667 | 1.029685 | 0.649057 | 0.642657 | 0.680007 | 0.016397 | 0.646299 | 0.339808 | 1.112154 | 0.30602 | 0.899713 | 0 |
| 26.43056 | 0.793391 | 0.566045 | 0.819101 | 0.637151 | 0.00058 | 0.495646 | 0.224962 | 0.874103 | 0.400258 | 0.833623 | 0 |
| 26.44444 | 0.674357 | 0.432802 | 0.879243 | 0.661072 | 0.039671 | 0.382534 | 0.307005 | 0.730312 | 0.651003 | 0.85618 | 0 |
| 26.45833 | 0.539506 | 0.256415 | 0.637123 | 0.634121 | 0.103827 | 0.278052 | 0.429065 | 0.488705 | 0.757099 | 0.734607 | 0 |
| 26.47222 | 0.334148 | 0.022124 | 0.198231 | 0.516584 | 0.283124 | 0.134244 | 0.498525 | 0.208063 | 0.662417 | 0.550468 | 0 |
| 26.48611 | 0.105473 | 0.284394 | 0.044318 | 0.535306 | 0.469774 | 0.140788 | 0.815321 | 0.084856 | 0.68775 | 0.749983 | 0 |
| 26.5 | 0.064633 | 0.616009 | 0.600446 | 0.638898 | 1.207062 | 0.395461 | 1.319524 | 0.240477 | 1.244076 | 1.431129 | 0 |
| 26.51389 | 0.070395 | 0.443725 | 1.529096 | 0.553423 | 1.047573 | 0.298281 | 1.730918 | 0.719488 | 1.813804 | 1.816415 | 0 |

| | | | | | | | | | | | |
|---|---|---|---|---|---|---|---|---|---|---|---|
| 26.52778 | 0.276553 | 0.626069 | 2.067539 | 0.620981 | 0.218673 | 0.093044 | 1.548873 | 1.1626 | 1.611272 | 1.704973 | 0 |
| 26.54167 | 0.639572 | 1.525046 | 1.49739 | 0.592076 | 0.036608 | 0.226852 | 0.917028 | 1.271489 | 1.179514 | 1.462607 | 0 |
| 26.55556 | 0.643397 | 1.339206 | 0.277686 | 0.448836 | 0.118394 | 0.150793 | 0.064271 | 1.064726 | 0.633677 | 0.860053 | 0 |
| 26.56944 | 0.487697 | 0.697654 | 0.75744 | 0.317367 | 0.171967 | 0.07162 | 0.622492 | 0.814561 | 0.030401 | 0.271627 | 0 |
| 26.58333 | 0.373714 | 0.640604 | 1.226544 | 0.27036 | 0.261839 | 0.243054 | 0.912816 | 0.851884 | 0.475989 | 0.096018 | 0 |
| 26.59722 | 0.595239 | 1.017214 | 0.959076 | 0.327605 | 0.304662 | 0.116427 | 0.581957 | 1.320871 | 0.576635 | 0.380651 | 0 |
| 26.61111 | 0.851933 | 1.518005 | 0.085403 | 0.364111 | 0.319436 | 0.328571 | 0.501805 | 1.851434 | 0.174544 | 0.986086 | 0 |
| 26.625 | 1.164446 | 1.668559 | 1.575245 | 0.355782 | 0.243982 | 0.772149 | 1.56066 | 2.423247 | 0.372329 | 1.613779 | 0 |
| 26.63889 | 1.31211 | 1.399559 | 2.731779 | 0.372396 | 0.047406 | 0.924054 | 1.758814 | 2.675568 | 0.86769 | 1.83355 | 0 |
| 26.65278 | 1.006684 | 0.505381 | 2.394605 | 0.390252 | 0.140692 | 0.783736 | 1.252739 | 2.051977 | 0.744428 | 1.317111 | 0 |
| 26.66667 | 0.432076 | 0.590356 | 0.937587 | 0.421946 | 0.226014 | 0.462989 | 0.320898 | 0.994542 | 0.163478 | 0.332038 | 0 |
| 26.68056 | 0.015246 | 1.341669 | 0.664381 | 0.602832 | 0.154523 | 0.174847 | 0.565768 | 0.10989 | 0.010852 | 0.624116 | 0 |
| 26.69444 | 0.452952 | 1.356593 | 1.746937 | 0.527982 | 0.496996 | 0.061245 | 1.827826 | 1.140165 | 0.047512 | 1.311532 | 0 |
| 26.70833 | 0.729807 | 0.871838 | 2.271576 | 0.528954 | 1.711509 | 0.025661 | 2.662689 | 1.358872 | 0.057034 | 1.617102 | 0 |
| 26.72222 | 0.86273 | 0.465437 | 3.126449 | 1.740315 | 1.242413 | 0.703595 | 0.91544 | 1.206847 | 0.893584 | 2.227811 | 0 |
| 26.73611 | 1.39967 | 2.123333 | 5.404732 | 3.813266 | 0.655664 | 0.014155 | 1.304045 | 2.073078 | 3.063627 | 3.458413 | 0 |
| 26.75 | 0 | 0 | 0 | 0 | 0 | 0 | 0 | 0 | 0 | 0 | 0 |
| 26.76389 | 0 | 0 | 0 | 0 | 0 | 0 | 0 | 0 | 0 | 0 | 0 |
| 26.77778 | 0 | 0 | 0 | 0 | 0 | 0 | 0 | 0 | 0 | 0 | 0 |
| 26.79167 | 1.550495 | 1.518437 | 4.342729 | 3.304698 | 0.785358 | 0.224181 | 2.643063 | 2.606838 | 2.678288 | 1.599619 | 0 |
| 26.80556 | 0.703733 | 0.258818 | 1.858359 | 1.33187 | 0.296094 | 0.452546 | 1.49587 | 1.469213 | 1.154355 | 1.911017 | 0 |
| 26.81944 | 0.798988 | 0.80386 | 0.786572 | 0.553663 | 0.180695 | 0.232681 | 1.806236 | 0.400797 | 1.261113 | 0.441451 | 0 |
| 26.83333 | 0.859118 | 0.641029 | 0.099939 | 0.802911 | 0.309333 | 0.285506 | 2.561096 | 0.777189 | 0.987896 | 0.067298 | 0 |
| 26.84722 | 0.570138 | 0.235545 | 0.658517 | 1.017286 | 0.267014 | 0.36904 | 1.515853 | 1.856463 | 0.07028 | 0.740845 | 0 |
| 26.86111 | 0.043015 | 0.153378 | 1.248121 | 1.000094 | 0.444035 | 0.610939 | 0.371403 | 2.416202 | 0.555643 | 0.906894 | 0 |
| 26.875 | 0.593454 | 0.698744 | 1.505185 | 0.801317 | 0.665077 | 0.621492 | 0.0412 | 2.33113 | 1.066213 | 1.005675 | 0 |
| 26.88889 | 1.07822 | 1.565898 | 1.522808 | 0.705569 | 0.87811 | 0.449621 | 0.098922 | 2.060501 | 1.364602 | 1.096384 | 0 |
| 26.90278 | 1.113241 | 2.419175 | 1.331705 | 0.833496 | 0.988798 | 0.506486 | 0.038304 | 2.29262 | 1.516132 | 1.24891 | 0 |
| 26.91667 | 0.804768 | 2.823539 | 0.926777 | 1.095693 | 0.797032 | 0.792254 | 0.227884 | 2.455987 | 1.459338 | 1.525489 | 0 |
| 26.93056 | 0.538086 | 2.660793 | 0.514816 | 1.342845 | 0.503289 | 1.019675 | 0.463039 | 2.55259 | 1.421384 | 1.872543 | 0 |
| 26.94444 | 0.61625 | 2.159713 | 0.208682 | 1.671643 | 0.208302 | 1.026956 | 0.594815 | 2.582335 | 1.321169 | 2.028179 | 0 |
| 26.95833 | 0.80797 | 1.445925 | 0.047899 | 1.558884 | 0.151546 | 0.779761 | 0.464151 | 2.111932 | 0.975926 | 1.677433 | 0 |
| 26.97222 | 1.001863 | 0.788556 | 0.088638 | 1.14082 | 0.358098 | 0.57311 | 0.30018 | 1.384371 | 0.601471 | 1.092878 | 0 |
| 26.98611 | 1.142654 | 0.619191 | 0.251993 | 0.813717 | 0.303868 | 0.442467 | 0.171992 | 0.870761 | 0.374795 | 0.742051 | 0 |
| 27 | 1.153022 | 1.076335 | 0.413086 | 0.586158 | 0.098364 | 0.414178 | 0.081546 | 0.530931 | 0.266095 | 0.610147 | 0 |
| 27.01389 | 1.257074 | 2.023769 | 0.544176 | 0.494463 | 0.003199 | 0.645375 | 0.348426 | 0.373562 | 0.266705 | 0.654134 | 0 |
| 27.02778 | 1.463357 | 2.244724 | 0.160319 | 0.379717 | 0.296963 | 0.438775 | 0.555785 | 0.475199 | 0.098575 | 0.470819 | 0 |
| 27.04167 | 1.682206 | 0.766417 | 0.488148 | 0.241294 | 0.646287 | 0.089385 | 0.674166 | 0.530734 | 0.373054 | 0.039827 | 0 |
| 27.05556 | 1.717705 | 0.402182 | 0.952504 | 0.219078 | 0.661419 | 0.488193 | 0.593062 | 0.378622 | 0.635848 | 0.194016 | 0 |
| 27.06944 | 1.261696 | 0.514125 | 0.820212 | 0.1798 | 0.505148 | 0.633489 | 0.374478 | 0.064342 | 0.445801 | 0.097827 | 0 |
| 27.08333 | 0.621594 | 0.615185 | 0.475376 | 0.113181 | 0.433021 | 0.722679 | 0.22225 | 0.265201 | 0.169035 | 0.086474 | 0 |
| 27.09722 | 0.155161 | 0.346895 | 0.335009 | 0.047255 | 0.405354 | 0.754953 | 0.238798 | 0.384743 | 0.093854 | 0.144677 | 0 |
| 27.11111 | 0.082229 | 0.181724 | 0.415318 | 0.185521 | 0.43384 | 0.68773 | 0.320804 | 0.336691 | 0.235259 | 0.103249 | 0 |
| 27.125 | 0.174042 | 0.206692 | 0.500634 | 0.195972 | 0.409726 | 0.47987 | 0.321597 | 0.314134 | 0.323273 | 0.320216 | 0 |
| 27.13889 | 0.188539 | 0.322817 | 0.461728 | 0.114054 | 0.299226 | 0.27307 | 0.240617 | 0.318992 | 0.136894 | 0.289956 | 0 |
| 27.15278 | 0.178308 | 0.435517 | 0.377808 | 0.034093 | 0.221942 | 0.149345 | 0.15279 | 0.260623 | 0.094844 | 0.168429 | 0 |
| 27.16667 | 0.166898 | 0.492105 | 0.350449 | 0.198846 | 0.182896 | 0.190777 | 0.158391 | 0.137785 | 0.187693 | 0.012583 | 0 |
| 27.18056 | 0.168629 | 0.460701 | 0.346463 | 0.247254 | 0.161053 | 0.421055 | 0.250461 | 0.024252 | 0.133784 | 0.121729 | 0 |
| 27.19444 | 0.216318 | 0.350924 | 0.291914 | 0.160726 | 0.14276 | 0.568173 | 0.312314 | 0.186055 | 0.024216 | 0.165413 | 0 |
| 27.20833 | 0.256425 | 0.274502 | 0.204917 | 0.109359 | 0.10399 | 0.480529 | 0.247816 | 0.264489 | 0.009517 | 0.173934 | 0 |
| 27.22222 | 0.249802 | 0.203605 | 0.128864 | 0.185133 | 0.070918 | 0.239031 | 0.097027 | 0.143734 | 0.121307 | 0.223145 | 0 |
| 27.23611 | 0.202129 | 0.064232 | 0.086631 | 0.296579 | 0.033953 | 0.020442 | 0.049099 | 0.081784 | 0.238355 | 0.252653 | 0 |
| 27.25 | 0.172932 | 0.116643 | 0.066287 | 0.3291 | 0.073792 | 0.103595 | 0.130638 | 0.139681 | 0.304288 | 0.220751 | 0 |
| 27.26389 | 0.158728 | 0.229706 | 0.064819 | 0.265708 | 0.148464 | 0.031417 | 0.147967 | 0.035241 | 0.296057 | 0.136862 | 0 |
| 27.27778 | 0.139285 | 0.160356 | 0.061559 | 0.126576 | 0.144033 | 0.214613 | 0.086243 | 0.025546 | 0.250724 | 0.145555 | 0 |
| 27.29167 | 0.14497 | 0.020564 | 0.030524 | 0.047897 | 0.111668 | 0.380269 | 0.014406 | 0.000967 | 0.208666 | 0.204222 | 0 |

| | | | | | | | | | | | |
|---|---|---|---|---|---|---|---|---|---|---|---|
| 27.30556 | 0.150729 | 0.016252 | 0.035957 | 0.149253 | 0.127616 | 0.340577 | 0.002506 | 0.019903 | 0.176848 | 0.091048 | 0 |
| 27.31944 | 0.141285 | 0.067481 | 0.146836 | 0.163262 | 0.19876 | 0.235722 | 0.027391 | 0.087548 | 0.124794 | 0.174845 | 0 |
| 27.33333 | 0.135504 | 0.131211 | 0.352563 | 0.272637 | 0.252914 | 0.3241 | 0.024067 | 0.303668 | 0.059012 | 0.350145 | 0 |
| 27.34722 | 0.112468 | 0.022166 | 0.5465 | 0.603524 | 0.198813 | 0.661694 | 0.125048 | 0.609425 | 0.032924 | 0.183694 | 0 |
| 27.36111 | 0.11888 | 0.261737 | 0.638737 | 0.873158 | 0.101022 | 0.981562 | 0.193576 | 0.755407 | 0.175879 | 0.035889 | 0 |
| 27.375 | 0.251263 | 0.346528 | 0.652714 | 0.898553 | 0.121665 | 1.109065 | 0.209789 | 0.668393 | 0.331513 | 0.169027 | 0 |
| 27.38889 | 0.486058 | 0.341032 | 0.670125 | 0.867189 | 0.132677 | 1.192372 | 0.213464 | 0.60444 | 0.400357 | 0.264063 | 0 |
| 27.40278 | 0.692214 | 0.376174 | 0.789981 | 0.839496 | 0.096733 | 1.191578 | 0.206676 | 0.615295 | 0.370131 | 0.305914 | 0 |
| 27.41667 | 0.901781 | 0.458538 | 0.92185 | 0.689965 | 0.037135 | 0.91468 | 0.136307 | 0.585852 | 0.30209 | 0.444526 | 0 |
| 27.43056 | 1.095452 | 0.670785 | 0.996245 | 0.529848 | 0.040715 | 0.603455 | 0.079671 | 0.610022 | 0.262402 | 0.607886 | 0 |
| 27.44444 | 1.284841 | 0.892934 | 0.949942 | 0.527453 | 0.129895 | 0.597689 | 0.226018 | 0.746012 | 0.334739 | 0.693814 | 0 |
| 27.45833 | 1.286569 | 0.948265 | 0.716711 | 0.614002 | 0.126809 | 0.776551 | 0.534744 | 0.868673 | 0.501226 | 0.684325 | 0 |
| 27.47222 | 1.097195 | 0.666458 | 0.28251 | 0.668134 | 0.040587 | 0.900969 | 0.72126 | 0.78223 | 0.60266 | 0.499485 | 0 |
| 27.48611 | 0.904308 | 0.288781 | 0.040663 | 0.977418 | 0.411246 | 0.841003 | 0.778985 | 0.692915 | 0.706434 | 0.603825 | 0 |
| 27.5 | 1.031379 | 0.151887 | 0.152265 | 1.488065 | 1.443598 | 0.732032 | 1.039478 | 0.740753 | 1.160803 | 1.335129 | 0 |
| 27.51389 | 1.30919 | 0.399862 | 0.954779 | 1.386509 | 1.860468 | 0.611208 | 1.338612 | 0.615704 | 1.903028 | 2.053001 | 0 |
| 27.52778 | 1.402971 | 0.92516 | 1.860197 | 0.949433 | 0.912851 | 0.377153 | 1.623359 | 0.524497 | 2.49456 | 2.402837 | 0 |
| 27.54167 | 1.108387 | 1.280232 | 1.628411 | 0.603071 | 0.598531 | 0.016178 | 1.754937 | 0.473227 | 2.268211 | 2.019796 | 0 |
| 27.55556 | 0.455197 | 1.165189 | 0.576552 | 0.233699 | 0.538766 | 0.316803 | 1.275174 | 0.148211 | 1.317184 | 1.085336 | 0 |
| 27.56944 | 0.072619 | 0.722789 | 0.515604 | 0.105707 | 0.561478 | 0.273099 | 0.573323 | 0.021739 | 0.456469 | 0.202196 | 0 |
| 27.58333 | 0.235928 | 0.325704 | 1.19887 | 0.165023 | 0.455369 | 0.146277 | 0.118257 | 0.208995 | 0.007065 | 0.181206 | 0 |
| 27.59722 | 0.120568 | 0.408096 | 1.219198 | 0.265576 | 0.224303 | 0.082104 | 0.724992 | 0.61222 | 0.075257 | 0.018686 | 0 |
| 27.61111 | 0.25909 | 0.803558 | 0.418205 | 0.450797 | 0.026718 | 0.351548 | 0.842144 | 1.063336 | 0.053854 | 0.415575 | 0 |
| 27.625 | 0.570333 | 1.216473 | 0.824568 | 0.517211 | 0.011183 | 0.820706 | 0.304726 | 1.477737 | 0.211141 | 0.827382 | 0 |
| 27.63889 | 0.584281 | 1.321825 | 1.760783 | 0.222951 | 0.051219 | 1.121406 | 0.412363 | 1.693521 | 0.402459 | 0.977371 | 0 |
| 27.65278 | 0.394035 | 1.343365 | 1.950441 | 0.090995 | 0.203791 | 1.133746 | 1.015281 | 1.893051 | 0.470858 | 0.926515 | 0 |
| 27.66667 | 0.548766 | 0.887062 | 1.369189 | 0.088923 | 0.38635 | 0.714012 | 1.513124 | 1.919483 | 0.461155 | 0.850812 | 0 |
| 27.68056 | 1.141966 | 0.525473 | 0.215621 | 0.254926 | 0.480203 | 0.063842 | 0.954176 | 1.483171 | 0.095388 | 0.551224 | 0 |
| 27.69444 | 1.70668 | 0.071707 | 0.788561 | 0.500573 | 0.283186 | 0.521655 | 0.105088 | 1.015502 | 0.241009 | 0.030577 | 0 |
| 27.70833 | 1.226286 | 0.773257 | 1.149578 | 0.548145 | 0.092421 | 1.024806 | 0.885134 | 0.473386 | 0.057412 | 0.391689 | 0 |
| 27.72222 | 0.419384 | 1.152394 | 1.137477 | 0.397317 | 0.45056 | 1.056983 | 1.511246 | 0.085606 | 0.345723 | 0.414002 | 0 |
| 27.73611 | 0.467194 | 0.874677 | 0.823489 | 0.125294 | 0.737778 | 0.285244 | 1.784097 | 0.660889 | 0.508022 | 0.272808 | 0 |
| 27.75 | 1.016293 | 0.333741 | 0.637154 | 0.331842 | 0.59609 | 0.222165 | 1.504711 | 1.302405 | 0.255836 | 0.521474 | 0 |
| 27.76389 | 1.446058 | 0.161573 | 0.691732 | 0.729447 | 0.35829 | 0.202848 | 1.069577 | 1.650139 | 0.200732 | 0.809797 | 0 |
| 27.77778 | 1.57447 | 0.34961 | 0.892457 | 0.590367 | 0.276803 | 0.104541 | 0.583361 | 1.29306 | 0.600784 | 0.96293 | 0 |
| 27.79167 | 1.738318 | 0.91534 | 1.106627 | 0.242725 | 0.167081 | 0.346786 | 0.296767 | 0.906605 | 1.066757 | 1.099228 | 0 |
| 27.80556 | 1.717694 | 1.387627 | 0.835028 | 0.097351 | 0.062254 | 0.49385 | 0.065852 | 0.838561 | 1.407848 | 1.275726 | 0 |
| 27.81944 | 1.217702 | 1.181974 | 0.210862 | 0.294296 | 0.306314 | 0.726722 | 0.265545 | 0.841441 | 1.21112 | 1.179417 | 0 |
| 27.83333 | 0.57576 | 0.749488 | 0.25074 | 0.809559 | 0.566539 | 1.152065 | 0.477439 | 0.92419 | 0.805144 | 0.942228 | 0 |
| 27.84722 | 0.195403 | 0.433891 | 0.143665 | 1.291025 | 0.712075 | 1.501757 | 0.264266 | 1.174778 | 0.569798 | 0.91099 | 0 |
| 27.86111 | 0.07252 | 0.324193 | 0.43612 | 1.306355 | 0.80381 | 1.627664 | 0.287967 | 1.538434 | 0.704182 | 1.102615 | 0 |
| 27.875 | 0.373591 | 0.504386 | 1.108246 | 0.740562 | 0.829859 | 1.499337 | 0.750394 | 1.756229 | 1.165225 | 1.252673 | 0 |
| 27.88889 | 0.37909 | 0.97443 | 1.543791 | 0.226695 | 0.66625 | 1.310583 | 0.807716 | 1.513938 | 1.498419 | 1.335745 | 0 |
| 27.90278 | 0.156969 | 1.550724 | 1.461007 | 0.228222 | 0.539312 | 1.232037 | 0.584758 | 0.950366 | 1.354852 | 1.344033 | 0 |
| 27.91667 | 0.192184 | 2.063511 | 0.913111 | 0.805755 | 0.448075 | 1.098806 | 0.420384 | 0.467609 | 1.074667 | 1.140055 | 0 |
| 27.93056 | 0.457808 | 1.781814 | 0.350402 | 1.407767 | 0.206547 | 0.728738 | 0.317053 | 0.35049 | 0.66008 | 0.755532 | 0 |
| 27.94444 | 0.326351 | 1.09488 | 0.058998 | 1.854211 | 0.005237 | 0.321239 | 0.262628 | 0.564659 | 0.291498 | 0.416965 | 0 |
| 27.95833 | 0.069632 | 0.595481 | 0.084977 | 1.547328 | 0.070139 | 0.11633 | 0.229227 | 0.811855 | 0.093814 | 0.266345 | 0 |
| 27.97222 | 0.295321 | 0.397232 | 0.19779 | 0.842667 | 0.097908 | 0.012934 | 0.19779 | 0.784887 | 0.141794 | 0.246307 | 0 |
| 27.98611 | 0.477565 | 0.458817 | 0.274719 | 0.325551 | 0.104166 | 0.000728 | 0.108414 | 0.538987 | 0.27691 | 0.264171 | 0 |
| 28 | 0.675986 | 1.135593 | 0.417672 | 0.252065 | 0.125471 | 0.227942 | 0.104341 | 0.100317 | 0.489698 | 0.308583 | 0 |
| 28.01389 | 1.036438 | 2.541959 | 0.78699 | 0.553838 | 0.070409 | 0.662528 | 0.20159 | 0.093367 | 0.916149 | 0.503564 | 0 |
| 28.02778 | 1.627754 | 3.420043 | 0.771059 | 0.566274 | 0.002489 | 0.875254 | 0.170211 | 0.379218 | 1.057147 | 0.744886 | 0 |
| 28.04167 | 2.06713 | 2.236177 | 0.030749 | 0.198495 | 0.00681 | 0.711946 | 0.17184 | 1.066338 | 0.523605 | 0.748351 | 0 |
| 28.05556 | 2.023039 | 0.40147 | 0.595921 | 0.110391 | 0.183888 | 0.270792 | 0.185189 | 1.478608 | 0.293832 | 0.389757 | 0 |
| 28.06944 | 1.616943 | 0.89733 | 0.816178 | 0.52713 | 0.428734 | 0.376826 | 0.110425 | 1.298889 | 0.563653 | 0.191066 | 0 |

| | | | | | | | | | | | |
|---|---|---|---|---|---|---|---|---|---|---|---|
| 28.08333 | 1.1219 | 1.28964 | 0.543371 | 0.609281 | 0.616366 | 0.996573 | 0.001184 | 0.685902 | 0.371233 | 0.13045 | 0 |
| 28.09722 | 0.545643 | 1.462477 | 0.417323 | 0.051827 | 0.768764 | 1.527493 | 0.043176 | 0.011209 | 0.316452 | 0.216989 | 0 |
| 28.11111 | 0.166059 | 1.657853 | 0.640072 | 0.624073 | 0.86621 | 1.992194 | 0.194094 | 0.466545 | 0.511862 | 0.729927 | 0 |
| 28.125 | 0.05173 | 1.672413 | 0.936529 | 0.804499 | 0.858574 | 2.168236 | 0.286534 | 0.568373 | 0.789167 | 1.073052 | 0 |
| 28.13889 | 0.095103 | 1.292966 | 0.986328 | 0.677362 | 0.705627 | 2.120947 | 0.308196 | 0.479011 | 0.94048 | 1.067213 | 0 |
| 28.15278 | 0.147964 | 0.926073 | 0.727545 | 0.443643 | 0.437361 | 1.912695 | 0.248975 | 0.432694 | 0.910252 | 0.760053 | 0 |
| 28.16667 | 0.317732 | 0.588165 | 0.312174 | 0.100626 | 0.28791 | 1.059555 | 0.140262 | 0.397782 | 0.669775 | 0.474011 | 0 |
| 28.18056 | 0.49014 | 0.290883 | 0.069858 | 0.242105 | 0.1606 | 0.495621 | 0.01963 | 0.28167 | 0.313148 | 0.211638 | 0 |
| 28.19444 | 0.549164 | 0.14452 | 0.013884 | 0.23719 | 0.076223 | 0.346254 | 0.007346 | 0.238844 | 0.008563 | 0.123235 | 0 |
| 28.20833 | 0.52424 | 0.22255 | 0.142093 | 0.078058 | 0.093229 | 0.490779 | 0.084581 | 0.180595 | 0.099371 | 0.240461 | 0 |
| 28.22222 | 0.51311 | 0.357015 | 0.481664 | 0.210065 | 0.212711 | 0.698602 | 0.203158 | 0.094815 | 0.072917 | 0.394038 | 0 |
| 28.23611 | 0.517365 | 0.410958 | 0.777973 | 0.017613 | 0.338963 | 0.757711 | 0.273988 | 0.057383 | 0.289469 | 0.367713 | 0 |
| 28.25 | 0.486928 | 0.338267 | 0.705494 | 0.16355 | 0.395727 | 0.699309 | 0.278732 | 0.090486 | 0.346838 | 0.260709 | 0 |
| 28.26389 | 0.414151 | 0.208189 | 0.391419 | 0.134631 | 0.347112 | 0.540898 | 0.23641 | 0.132699 | 0.237462 | 0.179086 | 0 |
| 28.27778 | 0.361452 | 0.112742 | 0.131655 | 0.537481 | 0.232515 | 0.325515 | 0.165665 | 0.109058 | 0.109604 | 0.119245 | 0 |
| 28.29167 | 0.415873 | 0.091502 | 0.081591 | 0.604126 | 0.110923 | 0.306556 | 0.119613 | 0.073025 | 0.081202 | 0.085147 | 0 |
| 28.30556 | 0.5009 | 0.121959 | 0.262515 | 0.298615 | 0.03267 | 0.528519 | 0.087067 | 0.098515 | 0.151433 | 0.092879 | 0 |
| 28.31944 | 0.501019 | 0.099972 | 0.527265 | 0.035056 | 0.000437 | 0.817039 | 0.02658 | 0.170162 | 0.213427 | 0.131547 | 0 |
| 28.33333 | 0.467529 | 0.053048 | 0.634534 | 0.132797 | 0.036181 | 0.935419 | 0.063866 | 0.317308 | 0.197072 | 0.170224 | 0 |
| 28.34722 | 0.46099 | 0.048378 | 0.538307 | 0.496102 | 0.023337 | 0.801537 | 0.062138 | 0.591301 | 0.158615 | 0.228058 | 0 |
| 28.36111 | 0.47797 | 0.097516 | 0.426335 | 0.784404 | 0.033676 | 0.64762 | 0.069774 | 0.897555 | 0.182766 | 0.373498 | 0 |
| 28.375 | 0.467706 | 0.162489 | 0.486828 | 0.841026 | 0.111377 | 0.591614 | 0.205494 | 1.098558 | 0.303172 | 0.545833 | 0 |
| 28.38889 | 0.466136 | 0.269275 | 0.778943 | 0.798769 | 0.179356 | 0.630315 | 0.236094 | 1.111938 | 0.535088 | 0.659963 | 0 |
| 28.40278 | 0.595325 | 0.4627 | 1.169532 | 0.757395 | 0.173354 | 0.720893 | 0.158699 | 1.056436 | 0.750828 | 0.746633 | 0 |
| 28.41667 | 0.833041 | 0.628741 | 1.299555 | 0.744436 | 0.042469 | 0.846812 | 0.018472 | 1.021924 | 0.882344 | 0.777216 | 0 |
| 28.43056 | 1.034365 | 0.708227 | 1.098847 | 0.715372 | 0.011678 | 1.069199 | 0.000559 | 1.073296 | 0.951002 | 0.78854 | 0 |
| 28.44444 | 1.051786 | 0.669491 | 0.820072 | 0.713639 | 0.125415 | 1.359419 | 0.166885 | 1.024423 | 0.948172 | 0.824176 | 0 |
| 28.45833 | 0.83331 | 0.495937 | 0.538886 | 0.641111 | 0.244976 | 1.284387 | 0.266132 | 0.787311 | 0.722839 | 0.678343 | 0 |
| 28.47222 | 0.499142 | 0.102686 | 0.245684 | 0.581215 | 0.218381 | 0.713807 | 0.196155 | 0.482947 | 0.265821 | 0.419397 | 0 |
| 28.48611 | 0.507109 | 0.107724 | 0.509661 | 1.124484 | 0.192441 | 0.289328 | 0.124213 | 0.460667 | 0.073258 | 0.628363 | 0 |
| 28.5 | 1.33969 | 0.54716 | 1.565893 | 2.683784 | 0.393391 | 0.454467 | 0.253159 | 0.936379 | 0.563388 | 1.751381 | 0 |
| 28.51389 | 2.460283 | 1.514364 | 2.64271 | 3.326367 | 0.881149 | 0.780364 | 0.580177 | 1.738884 | 1.5399 | 3.264363 | 0 |
| 28.52778 | 2.851985 | 2.01322 | 3.04773 | 2.712072 | 0.773521 | 1.07569 | 0.868177 | 2.003942 | 2.491583 | 3.964115 | 0 |
| 28.54167 | 2.09728 | 1.388569 | 2.16372 | 1.595309 | 0.417388 | 1.093683 | 1.008649 | 1.358611 | 3.10044 | 3.444466 | 0 |
| 28.55556 | 1.02953 | 0.773782 | 0.5821 | 0.299468 | 0.242823 | 0.313973 | 0.882704 | 0.535658 | 2.199483 | 1.687753 | 0 |
| 28.56944 | 0.207879 | 0.128393 | 0.882756 | 0.660562 | 0.279117 | 0.248182 | 0.596411 | 0.259511 | 0.466589 | 0.023071 | 0 |
| 28.58333 | 0.695822 | 0.723553 | 1.820113 | 0.860507 | 0.380346 | 0.776746 | 0.251645 | 0.730561 | 0.794599 | 1.114684 | 0 |
| 28.59722 | 0.784063 | 0.819986 | 1.890842 | 0.030063 | 0.422405 | 1.48111 | 0.137334 | 0.772223 | 1.36432 | 1.316191 | 0 |
| 28.61111 | 0.426174 | 0.804109 | 0.998385 | 1.093151 | 0.30062 | 0.916335 | 0.251269 | 0.312989 | 1.090288 | 0.726849 | 0 |
| 28.625 | 0.199112 | 0.291114 | 0.397792 | 1.685928 | 0.237894 | 0.37522 | 0.434432 | 0.338824 | 0.220787 | 0.062039 | 0 |
| 28.63889 | 0.380498 | 1.441579 | 1.553122 | 1.018407 | 0.396579 | 1.660651 | 0.381802 | 0.877814 | 0.913937 | 0.718431 | 0 |
| 28.65278 | 0.250092 | 1.2394 | 1.475442 | 0.27898 | 0.420549 | 2.396312 | 0.11975 | 1.003897 | 1.732138 | 0.834399 | 0 |
| 28.66667 | 0.154297 | 1.116279 | 1.059252 | 0.458625 | 0.288194 | 2.321295 | 0.208405 | 0.838581 | 1.265723 | 0.596576 | 0 |
| 28.68056 | 0.401188 | 0.615052 | 0.898372 | 1.084595 | 0.220189 | 1.228307 | 0.381697 | 0.81349 | 0.692422 | 0.289027 | 0 |
| 28.69444 | 0.525568 | 0.466059 | 0.244506 | 1.089622 | 0.273798 | 0.229221 | 0.388901 | 1.045194 | 0.065012 | 0.100014 | 0 |
| 28.70833 | 1.126183 | 0.22722 | 0.448944 | 0.036216 | 0.434558 | 0.90477 | 0.158363 | 0.952915 | 0.809671 | 0.216326 | 0 |
| 28.72222 | 2.02756 | 0.352592 | 0.376039 | 1.180659 | 0.551338 | 1.579277 | 0.167792 | 0.482704 | 0.895 | 0.243011 | 0 |
| 28.73611 | 1.584619 | 1.091242 | 0.49116 | 1.636827 | 0.485528 | 1.625582 | 0.32304 | 0.416092 | 0.261 | 0.30559 | 0 |
| 28.75 | 0.442888 | 1.332888 | 0.59521 | 0.822142 | 0.338852 | 1.128909 | 0.274884 | 0.827198 | 0.395435 | 0.301989 | 0 |
| 28.76389 | 0.745016 | 0.040346 | 0.670192 | 0.114746 | 0.256873 | 0.118613 | 0.120109 | 1.217061 | 0.826205 | 0.166454 | 0 |
| 28.77778 | 1.835335 | 0.53549 | 1.475252 | 0.673551 | 0.288129 | 0.661157 | 0.124075 | 1.047318 | 0.443191 | 0.355939 | 0 |
| 28.79167 | 2.142484 | 0.228975 | 2.422504 | 1.123273 | 0.205725 | 0.652075 | 0.245277 | 0.298157 | 0.613171 | 1.093576 | 0 |
| 28.80556 | 2.185899 | 1.783642 | 3.259024 | 1.167947 | 0.009721 | 0.293112 | 0.219193 | 0.945793 | 1.826647 | 1.965998 | 0 |
| 28.81944 | 2.06324 | 3.571898 | 2.619609 | 0.801702 | 0.254902 | 1.745201 | 0.20214 | 2.318896 | 2.628549 | 2.892261 | 0 |
| 28.83333 | 1.718588 | 4.97105 | 1.546495 | 0.530961 | 0.530019 | 3.050827 | 0.214659 | 3.129761 | 2.493251 | 3.289455 | 0 |
| 28.84722 | 1.645664 | 4.600518 | 0.910486 | 0.528043 | 0.765279 | 3.124491 | 0.330055 | 3.04298 | 2.028493 | 2.957798 | 0 |

| | | | | | | | | | | | |
|---|---|---|---|---|---|---|---|---|---|---|---|
| 28.86111 | 1.524251 | 3.34721 | 0.924531 | 0.973173 | 0.804997 | 2.58228 | 0.409601 | 2.431122 | 1.829918 | 2.261737 | 0 |
| 28.875 | 1.654604 | 2.144417 | 1.329026 | 1.75978 | 0.726554 | 2.129165 | 0.424882 | 2.073562 | 1.824672 | 1.795328 | 0 |
| 28.88889 | 1.743236 | 1.252287 | 1.621545 | 2.89374 | 0.483717 | 1.728684 | 0.364174 | 1.994183 | 1.682036 | 1.467884 | 0 |
| 28.90278 | 1.528881 | 0.546442 | 1.5452 | 3.489584 | 0.271303 | 1.291048 | 0.159363 | 1.889297 | 1.46216 | 1.086038 | 0 |
| 28.91667 | 1.084637 | 0.226669 | 1.134618 | 2.509302 | 0.188759 | 1.058818 | 0.015581 | 1.666793 | 1.277214 | 0.824909 | 0 |
| 28.93056 | 0.699111 | 0.303874 | 0.680725 | 1.084411 | 0.286525 | 0.793717 | 0.055446 | 1.312784 | 1.045361 | 0.666415 | 0 |
| 28.94444 | 0.333396 | 0.683581 | 0.429333 | 0.150106 | 0.274596 | 0.441518 | 0.028714 | 0.725497 | 0.722848 | 0.541791 | 0 |
| 28.95833 | 0.080769 | 0.927801 | 0.27768 | 0.057169 | 0.162817 | 0.076376 | 0.094253 | 0.113039 | 0.296839 | 0.343619 | 0 |
| 28.97222 | 0.051833 | 0.857518 | 0.206823 | 0.141351 | 0.005148 | 0.178291 | 0.168361 | 0.224092 | 0.044644 | 0.100114 | 0 |
| 28.98611 | 0.146944 | 1.037171 | 0.321424 | 0.709562 | 0.047492 | 0.015614 | 0.35616 | 0.004677 | 0.057478 | 0.237186 | 0 |
| 29 | 0.706876 | 1.790509 | 0.953508 | 1.734228 | 0.166039 | 0.847758 | 0.593276 | 0.635851 | 0.706078 | 0.989638 | 0 |
| 29.01389 | 1.016966 | 2.396258 | 1.554074 | 2.320664 | 0.467362 | 1.740793 | 0.669943 | 1.06841 | 1.423428 | 1.525936 | 0 |
| 29.02778 | 1.108149 | 2.385712 | 1.884598 | 2.205297 | 0.927902 | 2.599831 | 0.625108 | 1.289766 | 2.019452 | 1.656234 | 0 |
| 29.04167 | 1.57587 | 2.861164 | 2.646447 | 2.076345 | 1.859417 | 4.055317 | 0.965316 | 2.027934 | 2.913208 | 2.121578 | 0 |
| 29.05556 | 2.103103 | 3.230145 | 3.415638 | 1.69237 | 2.98424 | 5.486432 | 1.291159 | 2.506268 | 3.302934 | 2.573245 | 0 |
| 29.06944 | 1.898481 | 2.445273 | 2.796267 | 1.193392 | 3.25442 | 5.090928 | 1.155002 | 2.324396 | 2.302123 | 2.236802 | 0 |
| 29.08333 | 1.213518 | 1.119077 | 1.452796 | 0.945158 | 2.075929 | 2.811338 | 0.738621 | 1.716541 | 1.003091 | 1.35838 | 0 |
| 29.09722 | 0.681597 | 0.419067 | 0.428858 | 1.020328 | 0.856273 | 1.230146 | 0.603708 | 1.209874 | 0.31393 | 0.494331 | 0 |
| 29.11111 | 0.494058 | 0.127405 | 0.182266 | 0.992209 | 0.162522 | 0.291581 | 0.736307 | 0.882031 | 0.094395 | 0.032869 | 0 |
| 29.125 | 0.404438 | 0.256331 | 0.675323 | 0.549441 | 0.38113 | 0.443608 | 0.812421 | 0.607459 | 0.051688 | 0.252185 | 0 |
| 29.13889 | 0.130171 | 0.862217 | 1.129909 | 0.21998 | 1.032197 | 1.64984 | 0.616546 | 0.298538 | 0.396284 | 0.692076 | 0 |
| 29.15278 | 0.136207 | 1.518772 | 1.626462 | 0.753337 | 1.459743 | 3.016364 | 0.262952 | 0.102853 | 0.92164 | 1.056787 | 0 |
| 29.16667 | 0.354461 | 2.058047 | 2.050493 | 0.673136 | 1.489994 | 4.071974 | 0.023235 | 0.44794 | 1.696116 | 1.337981 | 0 |
| 29.18056 | 0.390817 | 2.126911 | 2.184461 | 0.239822 | 1.474518 | 3.65013 | 0.08063 | 0.639898 | 2.30853 | 1.508919 | 0 |
| 29.19444 | 0.090983 | 1.948771 | 1.700218 | 0.181509 | 1.303346 | 3.183893 | 0.113289 | 0.557024 | 2.204255 | 1.448191 | 0 |
| 29.20833 | 0.338429 | 1.65204 | 1.066466 | 0.439273 | 0.893555 | 2.749584 | 0.401781 | 0.299759 | 1.823478 | 0.92777 | 0 |
| 29.22222 | 0.641321 | 1.062674 | 0.546884 | 0.632988 | 0.381242 | 2.201138 | 0.598197 | 0.108651 | 1.095435 | 0.39544 | 0 |
| 29.23611 | 0.743727 | 0.544872 | 0.134869 | 0.673731 | 0.06806 | 1.360412 | 0.612111 | 0.325282 | 0.38373 | 0.022635 | 0 |
| 29.25 | 0.608844 | 0.32817 | 0.061176 | 0.514994 | 0.280189 | 0.627787 | 0.520751 | 0.367439 | 0.163926 | 0.14968 | 0 |
| 29.26389 | 0.471619 | 0.011149 | 0.104828 | 0.34256 | 0.301501 | 0.037107 | 0.481157 | 0.33691 | 0.559007 | 0.24335 | 0 |
| 29.27778 | 0.388967 | 0.355374 | 0.10465 | 0.287453 | 0.304882 | 0.400377 | 0.52367 | 0.222441 | 0.718285 | 0.33013 | 0 |
| 29.29167 | 0.323337 | 0.74577 | 0.242201 | 0.414133 | 0.369974 | 0.520823 | 0.580319 | 0.169223 | 0.762133 | 0.490533 | 0 |
| 29.30556 | 0.236028 | 1.100679 | 0.38266 | 0.576644 | 0.42269 | 0.53175 | 0.51697 | 0.152746 | 0.647906 | 0.62028 | 0 |
| 29.31944 | 0.1936 | 1.247758 | 0.316525 | 0.464132 | 0.46285 | 0.350611 | 0.375187 | 0.093251 | 0.490763 | 0.63376 | 0 |
| 29.33333 | 0.292058 | 1.335789 | 0.150133 | 0.135138 | 0.58059 | 0.147357 | 0.395884 | 0.181505 | 0.434353 | 0.679558 | 0 |
| 29.34722 | 0.485015 | 1.267978 | 0.012891 | 0.141808 | 0.752175 | 0.135333 | 0.623108 | 0.422768 | 0.495825 | 0.751596 | 0 |
| 29.36111 | 0.474702 | 0.824758 | 0.211378 | 0.265974 | 0.700225 | 0.043935 | 0.740676 | 0.492536 | 0.341765 | 0.579429 | 0 |
| 29.375 | 0.138301 | 0.081479 | 0.486215 | 0.303264 | 0.35192 | 0.223113 | 0.595869 | 0.306384 | 0.182902 | 0.071497 | 0 |
| 29.38889 | 0.126899 | 0.811264 | 0.643738 | 0.463105 | 0.057884 | 0.445606 | 0.309656 | 0.032546 | 0.712567 | 0.388067 | 0 |
| 29.40278 | 0.076363 | 1.066049 | 0.589921 | 0.861918 | 0.087238 | 0.50854 | 0.016536 | 0.250939 | 1.046187 | 0.518549 | 0 |
| 29.41667 | 0.007024 | 1.164588 | 0.389899 | 1.254469 | 0.10179 | 0.549084 | 0.127964 | 0.357304 | 1.266155 | 0.495744 | 0 |
| 29.43056 | 0.094064 | 0.685388 | 0.235382 | 1.095935 | 0.006528 | 0.633701 | 0.017758 | 0.212575 | 1.341637 | 0.38332 | 0 |
| 29.44444 | 0.290232 | 0.000752 | 0.313091 | 0.618059 | 0.137442 | 0.693732 | 0.211935 | 0.006883 | 1.286432 | 0.198074 | 0 |
| 29.45833 | 0.354583 | 0.579283 | 0.475359 | 0.140995 | 0.16389 | 0.725219 | 0.348082 | 0.008153 | 0.820623 | 0.029385 | 0 |
| 29.47222 | 0.236629 | 0.759659 | 0.518144 | 0.004519 | 0.123663 | 0.765667 | 0.297265 | 0.273621 | 0.273877 | 0.024295 | 0 |
| 29.48611 | 0.413336 | 0.344023 | 0.680751 | 0.82283 | 0.162348 | 0.84783 | 0.166076 | 0.713325 | 0.027748 | 0.401189 | 0 |
| 29.5 | 1.512487 | 0.604811 | 1.312471 | 2.734483 | 0.302816 | 1.150671 | 0.037732 | 1.404261 | 0.33 | 1.619146 | 0 |
| 29.51389 | 2.645603 | 1.368745 | 1.901533 | 4.061823 | 0.35935 | 1.18981 | 0.412 | 2.014649 | 1.111954 | 2.952471 | 0 |
| 29.52778 | 2.600019 | 1.858182 | 2.348161 | 3.75112 | 0.082134 | 1.127992 | 0.747214 | 2.03915 | 2.310951 | 3.456261 | 0 |
| 29.54167 | 2.168465 | 1.971843 | 2.145121 | 2.604857 | 0.240016 | 1.405605 | 0.832618 | 1.503138 | 3.026787 | 2.981714 | 0 |
| 29.55556 | 1.464031 | 1.537854 | 1.434402 | 1.033412 | 0.175232 | 0.953471 | 0.630166 | 0.756197 | 1.906054 | 1.793974 | 0 |
| 29.56944 | 0.615147 | 0.554764 | 0.532775 | 0.289013 | 0.004875 | 0.234548 | 0.179182 | 0.038013 | 0.462947 | 0.370335 | 0 |
| 29.58333 | 0.000375 | 0.251982 | 0.292487 | 0.98365 | 0.28501 | 0.325559 | 0.344622 | 0.507943 | 0.6037 | 0.689878 | 0 |
| 29.59722 | 0.038549 | 0.561411 | 0.934456 | 0.832144 | 0.48361 | 0.365448 | 0.713044 | 0.802222 | 1.054007 | 1.133537 | 0 |
| 29.61111 | 0.280129 | 0.577138 | 1.048176 | 0.112136 | 0.257218 | 0.146846 | 0.497243 | 0.63366 | 1.062057 | 0.933648 | 0 |
| 29.625 | 0.729684 | 0.348645 | 0.417314 | 1.325125 | 0.097315 | 0.195326 | 0.129338 | 0.105974 | 0.49755 | 0.216395 | 0 |

| | | | | | | | | | | | |
|---|---|---|---|---|---|---|---|---|---|---|---|
| 29.63889 | 1.194479 | 0.353529 | 0.569407 | 2.19672 | 0.657798 | 0.729113 | 1.002953 | 0.431253 | 0.523758 | 0.578469 | 0 |
| 29.65278 | 1.436691 | 1.289462 | 1.448212 | 1.639003 | 1.114401 | 1.345517 | 1.43914 | 0.894481 | 1.754548 | 1.292298 | 0 |
| 29.66667 | 1.280068 | 1.740404 | 1.381338 | 0.475465 | 1.113504 | 1.833345 | 1.294653 | 1.149321 | 2.614611 | 1.690136 | 0 |
| 29.68056 | 0.609847 | 1.462629 | 0.897343 | 0.348863 | 0.755012 | 1.496459 | 0.82413 | 0.81861 | 2.298027 | 1.470567 | 0 |
| 29.69444 | 0.208453 | 0.158815 | 0.22057 | 0.849892 | 0.312028 | 0.982749 | 0.149647 | 0.426297 | 1.054115 | 0.428454 | 0 |
| 29.70833 | 0.858368 | 0.031481 | 0.052973 | 1.193439 | 0.054458 | 0.527943 | 0.595813 | 0.201511 | 0.316675 | 0.776061 | 0 |
| 29.72222 | 0.886521 | 0.126226 | 0.217409 | 0.347023 | 0.054075 | 0.242879 | 0.512138 | 0.051772 | 0.973728 | 1.299049 | 0 |
| 29.73611 | 0.203657 | 0.093656 | 0.086767 | 1.653327 | 0.297792 | 0.037137 | 0.023872 | 0.287243 | 1.30126 | 0.9522 | 0 |
| 29.75 | 1.616416 | 0.015699 | 0.770192 | 2.811332 | 0.396235 | 0.427622 | 0.60101 | 0.176013 | 1.651262 | 0.190994 | 0 |
| 29.76389 | 1.945923 | 1.154813 | 1.056778 | 1.690651 | 0.26128 | 0.886455 | 1.077255 | 0.068728 | 1.436171 | 0.14267 | 0 |
| 29.77778 | 1.492646 | 2.450524 | 0.330334 | 0.099392 | 0.059654 | 1.089806 | 1.054163 | 0.047794 | 0.773059 | 0.042563 | 0 |
| 29.79167 | 0.643445 | 0.512853 | 1.184841 | 1.663377 | 0.430376 | 1.126605 | 0.409616 | 0.109979 | 0.142601 | 0.334374 | 0 |
| 29.80556 | 0.536636 | 0.81486 | 2.658111 | 4.258442 | 0.726861 | 1.123563 | 0.29723 | 0.22418 | 0.338721 | 0.962924 | 0 |
| 29.81944 | 1.427429 | 0.255615 | 3.683313 | 4.884496 | 0.876289 | 1.003793 | 1.216739 | 0.007157 | 0.125216 | 1.670579 | 0 |
| 29.83333 | 2.028933 | 0.864621 | 4.007011 | 2.363257 | 0.876029 | 1.059853 | 1.036738 | 0.564856 | 0.809065 | 2.040255 | 0 |
| 29.84722 | 2.329753 | 2.09941 | 2.875162 | 0.222923 | 0.732609 | 1.503656 | 0.566835 | 1.058394 | 1.833253 | 1.947849 | 0 |
| 29.86111 | 2.172924 | 3.546955 | 1.276774 | 0.561173 | 0.598333 | 2.192322 | 0.081859 | 1.434907 | 2.623526 | 1.757713 | 0 |
| 29.875 | 2.254326 | 5.072891 | 0.426452 | 0.334708 | 0.454153 | 2.206179 | 0.351723 | 1.761593 | 2.774793 | 1.838101 | 0 |
| 29.88889 | 2.130238 | 5.520499 | 0.115102 | 1.779079 | 0.355124 | 1.846288 | 0.148819 | 1.781539 | 2.499469 | 1.863336 | 0 |
| 29.90278 | 1.847825 | 4.134653 | 0.071644 | 2.869981 | 0.418114 | 1.38605 | 0.032298 | 1.481943 | 1.994557 | 1.821477 | 0 |
| 29.91667 | 1.582123 | 2.184494 | 0.181662 | 3.216797 | 0.472525 | 0.919958 | 0.38276 | 1.078021 | 1.442406 | 1.700712 | 0 |
| 29.93056 | 1.325024 | 0.580452 | 0.170845 | 2.67473 | 0.414245 | 0.579736 | 0.51893 | 0.784994 | 0.89575 | 1.311222 | 0 |
| 29.94444 | 0.939099 | 0.46398 | 0.060891 | 1.386355 | 0.219296 | 0.448318 | 0.152038 | 0.619573 | 0.548458 | 0.635789 | 0 |
| 29.95833 | 0.371203 | 0.911911 | 0.048049 | 0.462721 | 0.043738 | 0.430616 | 0.458873 | 0.439928 | 0.456751 | 0.224325 | 0 |
| 29.97222 | 0.067316 | 1.089322 | 0.014798 | 0.013546 | 0.194949 | 0.464128 | 1.060236 | 0.227648 | 0.356951 | 0.867096 | 0 |
| 29.98611 | 0.090637 | 0.605758 | 0.258736 | 0.161803 | 0.180816 | 0.718728 | 1.152858 | 0.223848 | 0.496859 | 0.543195 | 0 |
| 30 | 0.461163 | 0.668827 | 0.956766 | 1.157445 | 0.1667 | 1.410445 | 0.036155 | 0.698656 | 1.220424 | 1.064489 | 0 |
| 30.01389 | 0.939296 | 1.954579 | 1.804533 | 1.943426 | 0.967526 | 2.623597 | 1.069116 | 1.288636 | 2.290601 | 2.882481 | 0 |
| 30.02778 | 0.921764 | 2.476679 | 2.019832 | 1.441991 | 1.798495 | 3.150169 | 2.200226 | 1.548189 | 2.745198 | 3.564673 | 0 |
| 30.04167 | 0.665185 | 2.335909 | 1.298499 | 0.790001 | 1.992483 | 3.098848 | 1.814431 | 1.405759 | 2.125701 | 3.001301 | 0 |
| 30.05556 | 0.220611 | 1.807238 | 0.570061 | 0.67899 | 1.663377 | 2.368229 | 0.644811 | 1.212782 | 1.121054 | 1.799436 | 0 |
| 30.06944 | 0.248495 | 0.993475 | 0.108698 | 0.74966 | 0.793399 | 0.870569 | 0.381967 | 0.926981 | 0.204685 | 0.449988 | 0 |
| 30.08333 | 0.506865 | 0.283511 | 0.716452 | 0.644423 | 0.123113 | 0.539468 | 1.009847 | 0.49919 | 0.225261 | 0.521457 | 0 |
| 30.09722 | 0.557725 | 0.097655 | 0.896234 | 0.419882 | 0.495923 | 1.14653 | 0.805735 | 0.037053 | 0.345377 | 0.95215 | 0 |
| 30.11111 | 0.438738 | 0.163761 | 0.653801 | 0.184227 | 0.587784 | 1.01489 | 0.423033 | 0.205472 | 0.362539 | 0.970913 | 0 |
| 30.125 | 0.176003 | 0.022665 | 0.305922 | 0.079154 | 0.684695 | 0.755271 | 0.085767 | 0.234967 | 0.37614 | 0.837513 | 0 |
| 30.13889 | 0.190688 | 0.167248 | 0.047292 | 0.205683 | 0.75596 | 0.71476 | 0.026861 | 0.203316 | 0.425015 | 0.719259 | 0 |
| 30.15278 | 0.405515 | 0.05164 | 0.017825 | 0.776905 | 0.852221 | 0.914266 | 0.152406 | 0.252939 | 0.512307 | 0.763279 | 0 |
| 30.16667 | 0.33719 | 0.472458 | 0.151108 | 1.249016 | 0.968191 | 1.328329 | 0.483884 | 0.319957 | 0.754521 | 1.008406 | 0 |
| 30.18056 | 0.045311 | 1.263991 | 0.444538 | 1.131903 | 0.90119 | 1.203714 | 0.54062 | 0.245512 | 1.047429 | 1.28468 | 0 |
| 30.19444 | 0.441225 | 1.939364 | 0.799364 | 0.776447 | 0.877968 | 1.35443 | 0.401246 | 0.183776 | 1.367177 | 1.254469 | 0 |
| 30.20833 | 0.581219 | 2.274925 | 1.07613 | 0.266163 | 0.873218 | 1.602295 | 0.093063 | 0.126803 | 1.591162 | 1.041873 | 0 |
| 30.22222 | 0.536921 | 2.195897 | 1.217908 | 0.127869 | 0.866411 | 1.543754 | 0.322654 | 0.019002 | 1.663531 | 0.681567 | 0 |
| 30.23611 | 0.447551 | 1.880933 | 1.18172 | 0.278776 | 0.721153 | 1.266398 | 0.458128 | 0.135913 | 1.361479 | 0.405613 | 0 |
| 30.25 | 0.273444 | 1.338946 | 0.925901 | 0.226319 | 0.56779 | 1.018659 | 0.185327 | 0.090248 | 0.885025 | 0.284618 | 0 |
| 30.26389 | 0.067942 | 0.807554 | 0.619804 | 0.084335 | 0.455303 | 0.74075 | 0.151063 | 0.046639 | 0.461123 | 0.233973 | 0 |
| 30.27778 | 0.101503 | 0.355929 | 0.254875 | 0.257984 | 0.209979 | 0.344239 | 0.011734 | 0.038881 | 0.050492 | 0.058395 | 0 |
| 30.29167 | 0.191002 | 0.070698 | 0.004903 | 0.712015 | 0.134334 | 0.12736 | 0.463696 | 0.352351 | 0.298393 | 0.478106 | 0 |
| 30.30556 | 0.101303 | 0.294214 | 0.010891 | 0.940436 | 0.31998 | 0.340708 | 0.730217 | 0.465469 | 0.342393 | 0.667536 | 0 |
| 30.31944 | 0.071188 | 0.408898 | 0.16254 | 0.822482 | 0.291193 | 0.26088 | 0.750577 | 0.346098 | 0.281106 | 0.671628 | 0 |
| 30.33333 | 0.106789 | 0.584992 | 0.107843 | 0.487199 | 0.274374 | 0.053469 | 0.746459 | 0.272276 | 0.370305 | 0.692997 | 0 |
| 30.34722 | 0.067125 | 0.679102 | 0.152409 | 0.14843 | 0.433727 | 0.147448 | 0.74293 | 0.341968 | 0.567799 | 0.73307 | 0 |
| 30.36111 | 0.08167 | 0.602635 | 0.366168 | 0.112587 | 0.603986 | 0.156282 | 0.635076 | 0.364873 | 0.620954 | 0.6454 | 0 |
| 30.375 | 0.223352 | 0.278099 | 0.361307 | 0.416793 | 0.564976 | 0.050237 | 0.361327 | 0.14761 | 0.356686 | 0.235214 | 0 |
| 30.38889 | 0.262825 | 0.013843 | 0.207636 | 0.650449 | 0.453845 | 0.09434 | 0.159093 | 0.079915 | 0.015976 | 0.152303 | 0 |
| 30.40278 | 0.030976 | 0.109783 | 0.029013 | 0.604973 | 0.441258 | 0.243409 | 0.305572 | 0.006046 | 0.183232 | 0.119905 | 0 |

| | | | | | | | | | | | |
|---|---|---|---|---|---|---|---|---|---|---|---|
| 30.41667 | 0.227732 | 0.38855 | 0.197881 | 0.424988 | 0.467076 | 0.210011 | 0.665979 | 0.335481 | 0.259657 | 0.078505 | 0 |
| 30.43056 | 0.451438 | 0.564525 | 0.393428 | 0.497834 | 0.505322 | 0.109555 | 0.917965 | 0.649429 | 0.303556 | 0.375593 | 0 |
| 30.44444 | 0.649949 | 0.43184 | 0.416181 | 0.60334 | 0.605224 | 0.068806 | 0.961595 | 0.800921 | 0.312096 | 0.702638 | 0 |
| 30.45833 | 0.473767 | 0.003697 | 0.286637 | 0.36896 | 0.792687 | 0.160661 | 0.895673 | 0.830882 | 0.276885 | 0.709003 | 0 |
| 30.47222 | 0.033827 | 0.626047 | 0.254367 | 0.109675 | 0.794668 | 0.18529 | 0.722292 | 0.667008 | 0.451057 | 0.312906 | 0 |
| 30.48611 | 0.480303 | 1.427685 | 0.512976 | 0.008529 | 0.557992 | 0.42639 | 0.398979 | 0.16201 | 0.907352 | 0.512138 | 0 |
| 30.5 | 1.292871 | 2.405422 | 1.247718 | 1.203453 | 0.211805 | 1.042467 | 0.025458 | 0.794497 | 1.710354 | 2.13305 | 0 |
| 30.51389 | 2.24684 | 2.526983 | 2.086399 | 2.9965 | 0.137933 | 1.444091 | 0.525017 | 1.583721 | 2.434921 | 3.922744 | 0 |
| 30.52778 | 2.319861 | 2.478968 | 2.10898 | 4.184352 | 0.513921 | 1.669274 | 1.433928 | 1.795265 | 2.744636 | 4.984092 | 0 |
| 30.54167 | 1.827268 | 2.166713 | 1.880096 | 3.788383 | 0.393903 | 1.550849 | 1.640468 | 1.44992 | 2.615143 | 4.442854 | 0 |
| 30.55556 | 1.020678 | 1.345702 | 1.073047 | 1.890196 | 0.246042 | 0.775529 | 0.806609 | 0.633783 | 1.624791 | 2.32362 | 0 |
| 30.56944 | 0.104435 | 0.154005 | 0.113039 | 0.111336 | 0.135276 | 0.213721 | 0.016727 | 0.149731 | 0.401445 | 0.002414 | 0 |
| 30.58333 | 0.554497 | 0.868055 | 0.773417 | 1.043304 | 0.115341 | 0.792862 | 0.694023 | 0.503017 | 0.69223 | 1.679291 | 0 |
| 30.59722 | 0.337694 | 1.481775 | 1.471185 | 1.472029 | 0.705858 | 0.746004 | 1.079579 | 0.350087 | 1.457615 | 2.347934 | 0 |
| 30.61111 | 0.438282 | 1.804083 | 1.198254 | 0.763468 | 0.932436 | 0.232242 | 0.912549 | 0.182188 | 1.634891 | 2.02248 | 0 |
| 30.625 | 1.31365 | 2.062808 | 0.284429 | 0.645617 | 0.5228 | 0.62025 | 0.368081 | 0.805667 | 1.43175 | 0.941444 | 0 |
| 30.63889 | 1.731197 | 2.549019 | 0.828028 | 2.912346 | 0.01977 | 1.188155 | 0.292772 | 1.154987 | 0.899362 | 0.536699 | 0 |
| 30.65278 | 1.6199 | 1.566983 | 1.594734 | 4.497041 | 0.636699 | 1.462711 | 0.874252 | 1.185793 | 0.062793 | 1.881383 | 0 |
| 30.66667 | 1.328999 | 1.655313 | 2.007171 | 2.441663 | 0.815685 | 1.499778 | 1.245653 | 0.873204 | 1.389364 | 2.430951 | 0 |
| 30.68056 | 0.681166 | 5.032744 | 2.108945 | 0.417608 | 0.554154 | 1.022232 | 0.955029 | 0.033516 | 2.858534 | 1.705708 | 0 |
| 30.69444 | 0.129526 | 3.857834 | 0.961899 | 1.12316 | 0.01402 | 0.309993 | 0.208898 | 0.763955 | 2.623872 | 0.212733 | 0 |
| 30.70833 | 0.803403 | 2.358818 | 0.679829 | 2.466686 | 0.810755 | 0.607266 | 0.605233 | 1.089435 | 1.259983 | 1.362634 | 0 |
| 30.72222 | 1.108706 | 0.21719 | 2.012177 | 3.959933 | 1.629445 | 1.59044 | 0.831592 | 1.082666 | 0.061862 | 2.494534 | 0 |
| 30.73611 | 1.041969 | 0.838311 | 2.190468 | 4.451074 | 1.446862 | 1.771786 | 0.389502 | 0.785314 | 0.982302 | 2.562669 | 0 |
| 30.75 | 0.885227 | 1.024516 | 2.014957 | 1.179748 | 0.712999 | 0.449866 | 0.177925 | 0.58954 | 1.981745 | 1.982242 | 0 |
| 30.76389 | 0.881485 | 2.164518 | 1.489361 | 2.071562 | 0.097461 | 0.3798 | 0.248966 | 0.739548 | 2.438071 | 1.097748 | 0 |
| 30.77778 | 0.727962 | 2.765427 | 0.529051 | 3.887039 | 0.362601 | 0.641831 | 0.28049 | 1.099065 | 2.314108 | 0.367675 | 0 |
| 30.79167 | 0.337242 | 3.479829 | 0.19474 | 1.789089 | 0.436315 | 0.003997 | 0.228997 | 0.877358 | 1.971934 | 0.148375 | 0 |
| 30.80556 | 0.057984 | 4.661144 | 0.007878 | 0.121678 | 0.040075 | 0.91273 | 0.275894 | 0.530884 | 2.021046 | 0.653369 | 0 |
| 30.81944 | 0.167361 | 5.185568 | 0.946263 | 1.943714 | 1.051328 | 1.880504 | 0.700567 | 0.655523 | 2.001819 | 1.741862 | 0 |
| 30.83333 | 0.527862 | 4.357389 | 2.280851 | 4.213741 | 2.114234 | 2.661943 | 1.26766 | 1.097506 | 1.968343 | 2.979834 | 0 |
| 30.84722 | 0.8348 | 2.871737 | 2.294126 | 5.709275 | 2.290342 | 2.170172 | 1.264147 | 1.375087 | 1.191647 | 3.235181 | 0 |
| 30.86111 | 1.216408 | 1.604804 | 1.725908 | 5.037201 | 1.655524 | 1.481386 | 0.772784 | 1.475417 | 0.668863 | 2.642043 | 0 |
| 30.875 | 1.721846 | 0.683353 | 1.323425 | 3.516705 | 0.814847 | 1.003747 | 0.356934 | 1.646718 | 0.534849 | 1.689713 | 0 |
| 30.88889 | 1.76101 | 0.030667 | 0.954352 | 1.650521 | 0.070976 | 0.898972 | 0.479325 | 1.495515 | 1.170992 | 0.815961 | 0 |
| 30.90278 | 1.427495 | 0.31127 | 0.859098 | 0.526722 | 0.34178 | 1.162399 | 0.853559 | 1.078038 | 1.903135 | 0.359413 | 0 |
| 30.91667 | 1.026043 | 0.428255 | 0.968104 | 0.283456 | 0.24262 | 1.666968 | 0.683729 | 0.716135 | 1.859106 | 0.537743 | 0 |
| 30.93056 | 0.654479 | 0.594735 | 1.043785 | 0.485174 | 0.072456 | 1.694221 | 0.62718 | 0.376847 | 1.581784 | 0.884251 | 0 |
| 30.94444 | 0.37622 | 0.893398 | 0.999167 | 0.74933 | 0.324254 | 1.147225 | 0.340235 | 0.124408 | 1.045985 | 0.861675 | 0 |
| 30.95833 | 0.177949 | 1.224772 | 0.591387 | 0.81871 | 0.158389 | 0.223721 | 0.308208 | 0.024859 | 0.349739 | 0.179673 | 0 |
| 30.97222 | 0.092366 | 1.422027 | 0.097642 | 0.522524 | 0.425152 | 0.717105 | 1.078618 | 0.204432 | 0.253016 | 0.691148 | 0 |
| 30.98611 | 0.467296 | 1.288817 | 0.699564 | 0.023587 | 1.086617 | 1.319085 | 1.667132 | 0.399967 | 0.774877 | 1.025139 | 0 |
| 31 | 0.400983 | 0.766163 | 0.324403 | 0.558783 | 0.99429 | 0.459992 | 1.653077 | 0.590758 | 0.643589 | 0.067904 | 0 |
| 31.01389 | 0.557149 | 0.466114 | 1.314049 | 0.599975 | 0.416496 | 1.603552 | 0.48514 | 0.489916 | 0.479158 | 1.908653 | 0 |
| 31.02778 | 1.631368 | 1.885787 | 2.800273 | 0.366682 | 2.50221 | 3.479243 | 1.165397 | 0.108535 | 1.758924 | 3.348268 | 0 |
| 31.04167 | 1.140068 | 1.932008 | 2.484985 | 0.083837 | 2.753509 | 3.621073 | 1.340929 | 0.355541 | 2.144507 | 3.222564 | 0 |
| 31.05556 | 0.21779 | 1.421285 | 1.306465 | 0.679211 | 1.563155 | 2.884642 | 0.26845 | 0.838179 | 1.827795 | 2.003732 | 0 |
| 31.06944 | 0.814086 | 0.747702 | 0.027516 | 1.081006 | 0.268838 | 1.113743 | 0.614346 | 1.103577 | 1.100435 | 0.493664 | 0 |
| 31.08333 | 1.433273 | 0.151936 | 1.020789 | 1.071038 | 0.927647 | 0.330836 | 0.98787 | 1.027686 | 0.281579 | 0.410464 | 0 |
| 31.09722 | 1.601712 | 0.226032 | 1.684336 | 0.648792 | 1.297578 | 1.239212 | 0.934259 | 0.660777 | 0.277083 | 0.949636 | 0 |
| 31.11111 | 1.768612 | 0.725485 | 2.223479 | 0.167308 | 1.625449 | 1.906766 | 0.74312 | 0.254164 | 0.673849 | 1.651213 | 0 |
| 31.125 | 1.783424 | 1.12954 | 2.278494 | 0.164706 | 1.862368 | 2.360294 | 0.689667 | 0.099126 | 1.052803 | 2.279334 | 0 |
| 31.13889 | 2.007111 | 1.367201 | 2.134849 | 0.410165 | 1.74128 | 2.672231 | 0.644934 | 0.309114 | 1.264558 | 2.419699 | 0 |
| 31.15278 | 1.707092 | 1.142213 | 1.66273 | 0.596038 | 1.478638 | 2.822924 | 0.686542 | 0.469306 | 1.19216 | 2.399987 | 0 |
| 31.16667 | 1.118803 | 0.892001 | 1.107115 | 0.582577 | 1.388337 | 2.315962 | 0.836155 | 0.590959 | 1.136725 | 2.195022 | 0 |
| 31.18056 | 0.783189 | 0.868393 | 0.770675 | 0.435358 | 1.45994 | 1.772021 | 0.975884 | 0.61821 | 1.155252 | 1.803379 | 0 |

| | | | | | | | | | | | |
|---|---|---|---|---|---|---|---|---|---|---|---|
| 31.19444 | 0.400708 | 0.954177 | 0.498212 | 0.317105 | 1.488567 | 1.358243 | 0.891165 | 0.475201 | 1.061673 | 1.338033 | 0 |
| 31.20833 | 0.03714 | 0.953993 | 0.375882 | 0.210817 | 1.30383 | 0.798917 | 0.645309 | 0.252156 | 0.849529 | 1.032888 | 0 |
| 31.22222 | 0.208817 | 1.020797 | 0.505657 | 0.208199 | 0.914652 | 0.379916 | 0.402339 | 0.088513 | 0.743636 | 0.731822 | 0 |
| 31.23611 | 0.237759 | 1.02324 | 0.711768 | 0.290323 | 0.428724 | 0.172314 | 0.243413 | 0.009066 | 0.704742 | 0.141053 | 0 |
| 31.25 | 0.035503 | 1.077921 | 0.801686 | 0.207662 | 0.117812 | 0.153432 | 0.041727 | 0.129591 | 0.685419 | 0.18752 | 0 |
| 31.26389 | 0.66923 | 1.159063 | 0.87803 | 0.054893 | 0.179528 | 0.393058 | 0.132577 | 0.243249 | 0.671116 | 0.05641 | 0 |
| 31.27778 | 1.548547 | 1.30703 | 1.000743 | 0.04757 | 0.389337 | 0.915856 | 0.032356 | 0.127427 | 0.648418 | 0.606707 | 0 |
| 31.29167 | 1.897571 | 1.237626 | 0.979717 | 0.069381 | 1.041577 | 1.328621 | 0.229666 | 0.073238 | 0.585935 | 1.094888 | 0 |
| 31.30556 | 1.10823 | 0.711953 | 0.733945 | 0.047302 | 1.16046 | 1.269848 | 0.297702 | 0.177024 | 0.468295 | 1.090066 | 0 |
| 31.31944 | 0.070783 | 0.232087 | 0.386098 | 0.05626 | 0.666728 | 0.760511 | 0.283418 | 0.19565 | 0.32832 | 0.500878 | 0 |
| 31.33333 | 1.337688 | 0.318232 | 0.0002 | 0.146409 | 0.047011 | 0.187359 | 0.224891 | 0.232066 | 0.152844 | 0.357155 | 0 |
| 31.34722 | 1.782112 | 0.493814 | 0.177986 | 0.245704 | 0.607412 | 0.144366 | 0.120244 | 0.346031 | 0.051777 | 0.933137 | 0 |
| 31.36111 | 0.462521 | 0.00108 | 0.122773 | 0.244401 | 1.032072 | 0.044477 | 0.021778 | 0.460964 | 0.122894 | 0.632802 | 0 |
| 31.375 | 1.143318 | 0.566969 | 0.471287 | 0.179437 | 0.62646 | 0.343947 | 0.034412 | 0.556098 | 0.310149 | 0.315834 | 0 |
| 31.38889 | 2.251677 | 0.936097 | 0.739647 | 0.168428 | 0.101628 | 0.682759 | 0.221168 | 0.672317 | 0.51505 | 1.08226 | 0 |
| 31.40278 | 2.031231 | 0.81387 | 0.959187 | 0.207366 | 0.44073 | 0.795617 | 0.531541 | 0.71396 | 0.698386 | 1.293943 | 0 |
| 31.41667 | 0.693829 | 0.442687 | 1.019027 | 0.23831 | 0.556637 | 0.618606 | 0.591881 | 0.602318 | 0.799965 | 0.948683 | 0 |
| 31.43056 | 0.455218 | 0.124128 | 0.930476 | 0.296634 | 0.138594 | 0.31235 | 0.449788 | 0.431581 | 0.840172 | 0.248542 | 0 |
| 31.44444 | 0.589262 | 0.021435 | 0.882382 | 0.641244 | 0.505477 | 0.01006 | 0.137593 | 0.239193 | 0.871739 | 0.21167 | 0 |
| 31.45833 | 0.261817 | 0.29531 | 0.851455 | 1.55464 | 0.995901 | 0.077058 | 0.183568 | 0.079235 | 0.98226 | 0.043722 | 0 |
| 31.47222 | 1.121929 | 0.737031 | 0.908747 | 2.007296 | 0.799918 | 0.137905 | 0.211845 | 0.108299 | 1.121899 | 0.490468 | 0 |
| 31.48611 | 1.493622 | 1.177631 | 1.156629 | 2.364431 | 0.25934 | 0.579584 | 0.087488 | 0.470454 | 1.550468 | 1.341795 | 0 |
| 31.5 | 1.564069 | 1.933015 | 1.929832 | 4.005736 | 0.503525 | 1.233322 | 0.517032 | 1.221213 | 2.769475 | 2.779775 | 0 |
| 31.51389 | 1.791395 | 2.055121 | 2.920497 | 3.798619 | 0.944728 | 2.034809 | 0.593169 | 1.740722 | 3.446493 | 4.239624 | 0 |
| 31.52778 | 2.175253 | 2.338667 | 2.732599 | 2.490898 | 1.830524 | 2.421737 | 0.706513 | 1.46009 | 3.428771 | 5.216385 | 0 |
| 31.54167 | 2.020648 | 2.729974 | 2.603681 | 1.729227 | 2.167382 | 2.80324 | 0.837386 | 1.212483 | 3.570179 | 5.074759 | 0 |
| 31.55556 | 1.032962 | 2.150482 | 1.619196 | 1.001309 | 1.647949 | 1.835342 | 0.688463 | 0.846742 | 2.985622 | 3.251729 | 0 |
| 31.56944 | 0.077989 | 1.039558 | 0.611559 | 0.179224 | 0.791094 | 0.482931 | 0.385687 | 0.399436 | 1.760626 | 0.950466 | 0 |
| 31.58333 | 0.602408 | 0.301087 | 0.403591 | 0.686568 | 0.050784 | 0.701039 | 0.232103 | 0.117846 | 0.668526 | 0.64379 | 0 |
| 31.59722 | 1.026834 | 0.234939 | 1.369258 | 1.282687 | 0.704102 | 1.545708 | 0.206002 | 0.113698 | 0.295491 | 1.690168 | 0 |
| 31.61111 | 0.974545 | 0.627895 | 2.023067 | 1.631387 | 1.179878 | 2.160598 | 0.006331 | 0.335304 | 0.857498 | 2.12461 | 0 |
| 31.625 | 0.29774 | 0.885851 | 1.467388 | 0.302245 | 1.147163 | 2.460301 | 0.192607 | 0.538527 | 1.008305 | 2.094553 | 0 |
| 31.63889 | 0.665702 | 1.605532 | 0.561973 | 2.406061 | 0.962098 | 1.842278 | 0.369135 | 0.614466 | 1.31006 | 1.156941 | 0 |
| 31.65278 | 1.52422 | 2.460571 | 0.055471 | 4.241475 | 0.324818 | 0.600684 | 0.463819 | 0.496712 | 1.857207 | 0.084016 | 0 |
| 31.66667 | 1.499053 | 2.826375 | 0.543861 | 3.804752 | 0.368231 | 0.942571 | 0.280513 | 0.295184 | 1.667358 | 1.180254 | 0 |
| 31.68056 | 0.979474 | 2.738646 | 0.709104 | 2.244932 | 0.678147 | 2.021377 | 0.096949 | 0.365396 | 0.527946 | 1.593237 | 0 |
| 31.69444 | 0.60774 | 0.328319 | 0.965497 | 0.748816 | 0.52029 | 2.306738 | 0.547517 | 0.494847 | 1.18337 | 1.24086 | 0 |
| 31.70833 | 0.317787 | 2.772338 | 1.396859 | 0.857709 | 0.147911 | 2.033787 | 0.690253 | 0.216823 | 2.800846 | 0.25867 | 0 |
| 31.72222 | 0.164311 | 4.678275 | 0.869408 | 2.236537 | 0.974956 | 1.153282 | 0.272013 | 0.000727 | 2.610759 | 0.915586 | 0 |
| 31.73611 | 0.809311 | 3.899335 | 0.591497 | 3.643187 | 1.416987 | 0.401324 | 0.529768 | 0.265337 | 1.914942 | 1.683073 | 0 |
| 31.75 | 0.39298 | 2.827695 | 1.354095 | 4.023119 | 1.527094 | 1.292963 | 1.233364 | 0.342717 | 0.915722 | 2.162851 | 0 |
| 31.76389 | 0.046538 | 1.599473 | 1.642138 | 2.183951 | 2.189734 | 1.697986 | 1.787889 | 0.588372 | 0.356276 | 2.265087 | 0 |
| 31.77778 | 0.376014 | 0.337362 | 1.583418 | 0.381422 | 1.516438 | 1.976112 | 1.940417 | 0.967505 | 1.482858 | 2.163103 | 0 |
| 31.79167 | 0.031982 | 1.260985 | 1.837857 | 2.812643 | 0.583212 | 1.870414 | 1.849159 | 1.361937 | 2.461906 | 2.051889 | 0 |
| 31.80556 | 0.562904 | 2.955036 | 1.992268 | 4.544496 | 0.133698 | 0.92862 | 1.405701 | 1.577187 | 3.060444 | 1.766726 | 0 |
| 31.81944 | 1.123477 | 4.472226 | 2.256253 | 3.087027 | 0.091601 | 1.272207 | 0.866985 | 1.419364 | 3.011303 | 1.998803 | 0 |
| 31.83333 | 1.837131 | 4.435059 | 2.697608 | 0.447525 | 0.735446 | 2.254624 | 0.37827 | 1.200989 | 2.443655 | 2.219024 | 0 |
| 31.84722 | 2.0852 | 4.314248 | 3.306995 | 2.035146 | 1.56174 | 2.784838 | 0.097409 | 1.068689 | 2.008696 | 2.462151 | 0 |
| 31.86111 | 1.50329 | 3.572283 | 3.674808 | 3.871876 | 2.278477 | 3.840347 | 0.019734 | 1.014822 | 1.938485 | 2.926661 | 0 |
| 31.875 | 1.070157 | 2.473634 | 3.043829 | 5.43086 | 2.537597 | 4.835412 | 0.080018 | 1.212013 | 1.949868 | 2.974798 | 0 |
| 31.88889 | 0.724422 | 1.675414 | 2.021433 | 4.990776 | 2.133889 | 3.651298 | 0.09662 | 1.450444 | 1.643028 | 2.701325 | 0 |
| 31.90278 | 0.330775 | 1.093033 | 0.969417 | 4.124889 | 1.505803 | 2.074574 | 0.082489 | 1.485367 | 1.267059 | 2.034261 | 0 |
| 31.91667 | 0.26333 | 0.698555 | 0.189275 | 2.07728 | 0.572022 | 0.593221 | 0.047164 | 1.301272 | 0.72529 | 1.144919 | 0 |
| 31.93056 | 0.217731 | 0.483261 | 0.081158 | 0.898646 | 0.404471 | 0.636691 | 0.037667 | 0.961518 | 0.06665 | 0.257912 | 0 |
| 31.94444 | 0.249869 | 0.35195 | 0.073691 | 0.324747 | 1.174993 | 1.134968 | 0.00212 | 0.65293 | 0.504526 | 0.373232 | 0 |
| 31.95833 | 0.257895 | 0.13427 | 0.13705 | 0.190466 | 1.590036 | 1.042159 | 0.058005 | 0.340733 | 0.813843 | 0.702591 | 0 |

| | | | | | | | | | | | |
|---|---|---|---|---|---|---|---|---|---|---|---|
| 31.97222 | 0.233384 | 0.130055 | 0.404917 | 0.143375 | 1.558709 | 0.873922 | 0.168511 | 0.135749 | 0.857714 | 0.751032 | 0 |
| 31.98611 | 0.346646 | 0.316218 | 0.448982 | 0.218553 | 1.069575 | 0.650748 | 0.197501 | 0.067569 | 0.405548 | 0.131217 | 0 |
| 32 | 0.827667 | 0.13658 | 0.505765 | 0.568582 | 0.18244 | 0.630937 | 0.042722 | 0.181427 | 0.585255 | 1.481668 | 0 |
| 32.01389 | 1.808833 | 0.652019 | 2.21945 | 1.04068 | 1.745525 | 2.646062 | 0.288094 | 0.430697 | 1.829924 | 3.000987 | 0 |
| 32.02778 | 2.362842 | 1.333196 | 3.328844 | 1.168407 | 2.478073 | 4.027976 | 0.793238 | 0.594612 | 2.549951 | 3.284648 | 0 |
| 32.04167 | 2.754673 | 1.678121 | 3.299219 | 0.559875 | 2.080139 | 4.492888 | 0.637831 | 0.588118 | 2.532254 | 2.731074 | 0 |
| 32.05556 | 2.91599 | 1.300829 | 2.714804 | 0.115376 | 1.116593 | 3.840173 | 0.204187 | 0.396161 | 2.269872 | 1.749605 | 0 |
| 32.06944 | 1.902205 | 0.490203 | 1.627871 | 0.080175 | 0.263062 | 2.736635 | 0.196957 | 0.270394 | 1.739485 | 0.828477 | 0 |
| 32.08333 | 0.544538 | 0.136247 | 0.321302 | 0.03093 | 0.177 | 1.445784 | 0.389728 | 0.2593 | 1.023685 | 0.217211 | 0 |
| 32.09722 | 0.602144 | 0.537274 | 1.191933 | 0.075383 | 0.384202 | 0.280207 | 0.466513 | 0.074098 | 0.175719 | 0.400364 | 0 |
| 32.11111 | 1.681228 | 0.832325 | 2.477886 | 0.077703 | 0.592161 | 0.751897 | 0.558348 | 0.312003 | 0.362218 | 0.848162 | 0 |
| 32.125 | 2.481594 | 1.041181 | 3.102578 | 0.250732 | 0.894704 | 1.848913 | 0.633493 | 0.650457 | 0.578241 | 1.295168 | 0 |
| 32.13889 | 2.758671 | 1.265621 | 3.329381 | 0.312839 | 1.145368 | 2.774939 | 0.764033 | 0.819828 | 0.798019 | 1.773561 | 0 |
| 32.15278 | 2.47977 | 1.212784 | 2.735928 | 0.160727 | 1.154244 | 3.61424 | 1.038375 | 0.767539 | 0.998114 | 1.95028 | 0 |
| 32.16667 | 1.988684 | 0.739261 | 1.826161 | 0.133373 | 1.287643 | 3.58269 | 1.183731 | 0.545426 | 1.194919 | 2.001359 | 0 |
| 32.18056 | 2.216182 | 0.743269 | 1.391166 | 0.365659 | 1.479923 | 3.496801 | 1.002075 | 0.340803 | 1.310975 | 2.114218 | 0 |
| 32.19444 | 2.287243 | 0.808116 | 0.693818 | 0.443185 | 1.24692 | 3.174365 | 0.696198 | 0.223908 | 1.418758 | 2.11201 | 0 |
| 32.20833 | 1.807282 | 0.826134 | 0.298325 | 0.346399 | 0.830364 | 2.73316 | 0.342434 | 0.177294 | 1.510001 | 1.76545 | 0 |
| 32.22222 | 1.854328 | 1.138129 | 0.57624 | 0.059942 | 0.435634 | 2.043245 | 0.271268 | 0.279719 | 0.983049 | 1.523146 | 0 |
| 32.23611 | 1.807732 | 1.53127 | 1.085419 | 0.098398 | 0.115159 | 1.251632 | 0.341418 | 0.431446 | 0.920088 | 1.293066 | 0 |
| 32.25 | 1.188943 | 1.339923 | 1.018387 | 0.05132 | 0.209711 | 0.36289 | 0.166155 | 0.441929 | 0.587434 | 0.667865 | 0 |
| 32.26389 | 0.317831 | 0.688893 | 0.653243 | 0.130055 | 0.605999 | 0.488354 | 0.281174 | 0.277758 | 0.358719 | 0.147609 | 0 |
| 32.27778 | 0.385601 | 0.162292 | 0.339406 | 0.265668 | 0.841077 | 0.945518 | 0.523364 | 0.095297 | 0.319404 | 0.70099 | 0 |
| 32.29167 | 0.671154 | 0.116458 | 0.19755 | 0.262991 | 0.619944 | 0.733159 | 0.412705 | 0.055374 | 0.263576 | 0.79828 | 0 |
| 32.30556 | 0.500479 | 0.027586 | 0.317537 | 0.159299 | 0.069809 | 0.002845 | 0.105287 | 0.090887 | 0.234621 | 0.309796 | 0 |
| 32.31944 | 0.041432 | 0.193031 | 0.631622 | 0.005985 | 0.282422 | 0.719757 | 0.106125 | 0.017079 | 0.297632 | 0.315012 | 0 |
| 32.33333 | 0.609016 | 0.449836 | 0.839095 | 0.131991 | 0.378768 | 1.049808 | 0.212675 | 0.249965 | 0.350776 | 0.765301 | 0 |
| 32.34722 | 0.995244 | 0.655654 | 0.842966 | 0.24534 | 0.285137 | 0.943991 | 0.23768 | 0.559721 | 0.365231 | 0.909063 | 0 |
| 32.36111 | 0.848309 | 0.534393 | 0.625236 | 0.20715 | 0.052487 | 0.523779 | 0.184889 | 0.701166 | 0.325965 | 0.607574 | 0 |
| 32.375 | 0.297994 | 0.131263 | 0.325753 | 0.023592 | 0.259749 | 0.033937 | 0.072334 | 0.49492 | 0.263434 | 0.033735 | 0 |
| 32.38889 | 0.095261 | 0.195519 | 0.310933 | 0.123028 | 0.411222 | 0.22602 | 0.075997 | 0.269179 | 0.309085 | 0.423874 | 0 |
| 32.40278 | 0.095827 | 0.05444 | 0.739398 | 0.143078 | 0.256475 | 0.355118 | 0.245407 | 0.359571 | 0.533149 | 0.076426 | 0 |
| 32.41667 | 0.618336 | 0.351541 | 1.216878 | 0.401524 | 0.094455 | 1.218277 | 0.396473 | 0.523488 | 0.726905 | 0.536134 | 0 |
| 32.43056 | 0.896392 | 0.692284 | 1.233558 | 0.370758 | 0.312012 | 1.772251 | 0.404883 | 0.520505 | 0.649145 | 0.938763 | 0 |
| 32.44444 | 0.827418 | 0.976585 | 0.961859 | 0.23099 | 0.349261 | 1.91155 | 0.368796 | 0.552581 | 0.487624 | 1.118269 | 0 |
| 32.45833 | 0.758288 | 1.118939 | 0.637401 | 0.132678 | 0.24878 | 1.656364 | 0.292855 | 0.578472 | 0.393474 | 0.93918 | 0 |
| 32.47222 | 0.576922 | 0.976106 | 0.5094 | 0.170513 | 0.051459 | 1.013031 | 0.176375 | 0.476055 | 0.44099 | 0.495838 | 0 |
| 32.48611 | 0.615477 | 0.911949 | 0.887746 | 0.861381 | 0.128351 | 0.641175 | 0.031723 | 0.422776 | 0.844169 | 0.581099 | 0 |
| 32.5 | 0.866274 | 1.319987 | 2.163217 | 3.873482 | 0.110624 | 1.062948 | 0.206697 | 0.727731 | 1.750016 | 1.555997 | 0 |
| 32.51389 | 1.26509 | 1.87474 | 3.321562 | 5.488972 | 0.383006 | 2.36103 | 0.47562 | 1.288898 | 2.418339 | 3.056127 | 0 |
| 32.52778 | 2.152765 | 2.661284 | 3.279893 | 6.133499 | 1.469377 | 3.055023 | 0.292889 | 1.113214 | 2.2173 | 4.412885 | 0 |
| 32.54167 | 2.685287 | 3.64915 | 3.124302 | 4.751802 | 2.400989 | 3.207234 | 0.443559 | 0.933108 | 1.78749 | 4.79188 | 0 |
| 32.55556 | 1.559889 | 3.026231 | 2.249964 | 2.768278 | 2.345474 | 2.426065 | 0.813571 | 0.70814 | 1.603103 | 3.465166 | 0 |
| 32.56944 | 0.461201 | 2.013594 | 1.052212 | 1.629855 | 1.384981 | 1.119234 | 0.699094 | 0.515967 | 1.547766 | 1.473562 | 0 |
| 32.58333 | 0.121748 | 1.271685 | 0.188471 | 0.701759 | 0.412634 | 0.045029 | 0.542852 | 0.432897 | 1.549408 | 0.048315 | 0 |
| 32.59722 | 0.326914 | 0.665212 | 0.271816 | 0.24549 | 0.274286 | 0.920115 | 0.431063 | 0.423277 | 1.461302 | 0.888122 | 0 |
| 32.61111 | 0.48138 | 0.270323 | 0.501417 | 1.357225 | 0.680528 | 1.663091 | 0.271069 | 0.270137 | 1.13013 | 1.581719 | 0 |
| 32.625 | 0.543795 | 0.154198 | 0.635165 | 2.085537 | 0.896698 | 2.447285 | 0.017469 | 0.151145 | 0.797059 | 1.976131 | 0 |
| 32.63889 | 0.375851 | 0.965123 | 0.655323 | 3.512012 | 0.631296 | 2.482579 | 0.290139 | 0.393719 | 0.258595 | 2.028502 | 0 |
| 32.65278 | 0.042482 | 1.737977 | 0.700283 | 3.189772 | 0.555224 | 1.936008 | 0.377579 | 0.607579 | 0.427869 | 1.356774 | 0 |
| 32.66667 | 0.684111 | 2.261463 | 0.447705 | 0.501711 | 0.339244 | 1.099598 | 0.325178 | 0.763843 | 0.925608 | 0.328978 | 0 |
| 32.68056 | 1.259082 | 2.661086 | 0.186733 | 2.25941 | 0.216553 | 0.093876 | 0.324873 | 0.80382 | 1.011813 | 0.785363 | 0 |
| 32.69444 | 0.69248 | 3.463019 | 0.140238 | 4.937961 | 0.492572 | 0.114408 | 0.454601 | 0.602926 | 1.692112 | 1.786043 | 0 |
| 32.70833 | 0.125715 | 2.368446 | 0.235225 | 5.062573 | 0.959727 | 0.629711 | 0.553891 | 0.420074 | 1.564306 | 1.456912 | 0 |
| 32.72222 | 0.189701 | 0.793004 | 0.289972 | 2.927076 | 0.787212 | 1.512684 | 0.56576 | 0.365263 | 0.460141 | 0.283988 | 0 |
| 32.73611 | 0.632799 | 1.035005 | 0.511458 | 1.33528 | 0.475847 | 1.631361 | 0.542053 | 0.198319 | 0.568074 | 0.965058 | 0 |

| | | | | | | | | | | | |
|---|---|---|---|---|---|---|---|---|---|---|---|
| 32.75 | 0.272491 | 2.971259 | 0.612776 | 0.095669 | 0.133359 | 1.472872 | 0.161239 | 0.097837 | 1.12617 | 1.57092 | 0 |
| 32.76389 | 0.162686 | 3.832131 | 0.27552 | 1.625602 | 0.017498 | 0.281967 | 0.328902 | 0.013616 | 1.106914 | 1.741631 | 0 |
| 32.77778 | 0.311308 | 3.100224 | 0.505247 | 2.877299 | 0.334183 | 2.215167 | 0.499041 | 0.235247 | 0.343407 | 1.608599 | 0 |
| 32.79167 | 0.38374 | 2.083403 | 1.242481 | 4.763461 | 0.725441 | 2.801256 | 0.298317 | 0.521961 | 0.326376 | 1.4512 | 0 |
| 32.80556 | 0.407661 | 0.500889 | 1.612758 | 6.174468 | 0.868671 | 2.28537 | 0.068797 | 0.804067 | 0.717111 | 1.355574 | 0 |
| 32.81944 | 0.300419 | 1.317557 | 2.068229 | 4.964077 | 0.822751 | 1.943376 | 0.503485 | 0.872743 | 0.869476 | 1.620433 | 0 |
| 32.83333 | 0.300244 | 2.772796 | 2.99511 | 1.957057 | 0.696186 | 1.851187 | 0.740393 | 0.849596 | 0.90192 | 2.465669 | 0 |
| 32.84722 | 0.431116 | 3.412137 | 3.450498 | 0.846709 | 0.813538 | 2.971633 | 0.722508 | 0.838916 | 0.974481 | 3.437713 | 0 |
| 32.86111 | 1.067943 | 3.668498 | 3.352726 | 0.620386 | 1.089611 | 4.32499 | 0.684562 | 0.825821 | 1.320929 | 4.059917 | 0 |
| 32.875 | 1.6942 | 3.46984 | 3.049438 | 0.707434 | 1.356906 | 4.730219 | 0.640868 | 0.761271 | 1.665419 | 4.17753 | 0 |
| 32.88889 | 2.021282 | 2.87515 | 2.447668 | 0.788275 | 1.545246 | 4.021206 | 0.476004 | 0.83684 | 1.982887 | 3.530191 | 0 |
| 32.90278 | 1.632817 | 2.072488 | 1.629399 | 0.911133 | 1.310843 | 2.907103 | 0.108024 | 0.964817 | 2.021862 | 2.396046 | 0 |
| 32.91667 | 0.603917 | 1.167475 | 0.846064 | 1.087042 | 0.632562 | 1.632314 | 0.217394 | 1.056906 | 1.745676 | 1.161022 | 0 |
| 32.93056 | 0.397987 | 0.509646 | 0.17992 | 1.052941 | 0.112907 | 0.39335 | 0.434489 | 0.867894 | 1.170253 | 0.203681 | 0 |
| 32.94444 | 0.998854 | 0.062992 | 0.266697 | 0.714784 | 0.45204 | 0.58962 | 0.591531 | 0.473805 | 0.588638 | 0.47481 | 0 |
| 32.95833 | 1.285377 | 0.307688 | 0.455416 | 0.328907 | 0.610507 | 1.1788 | 0.724568 | 0.068168 | 0.027295 | 0.928228 | 0 |
| 32.97222 | 1.240933 | 0.475557 | 0.308936 | 0.041389 | 0.638466 | 1.014434 | 0.787151 | 0.170673 | 0.473218 | 0.894738 | 0 |
| 32.98611 | 0.798518 | 0.327222 | 0.087639 | 0.049655 | 0.297459 | 0.035734 | 0.666166 | 0.164247 | 0.527523 | 0.357731 | 0 |
| 33 | 0.424069 | 0.006075 | 0.977022 | 0.426644 | 0.552468 | 1.311022 | 0.353362 | 0.102912 | 0.06221 | 0.943487 | 0 |
| 33.01389 | 1.89955 | 0.549943 | 2.275159 | 0.823355 | 1.10571 | 2.030635 | 0.114689 | 0.406996 | 0.917532 | 2.067775 | 0 |
| 33.02778 | 2.765577 | 1.032453 | 3.176388 | 0.404668 | 0.744033 | 2.011457 | 0.277933 | 0.277351 | 1.459561 | 2.161616 | 0 |
| 33.04167 | 2.975295 | 1.199862 | 3.052823 | 0.052546 | 0.504589 | 1.743981 | 0.000235 | 0.104008 | 1.054772 | 1.710826 | 0 |
| 33.05556 | 2.402235 | 0.905212 | 2.986426 | 0.159217 | 0.393725 | 1.276645 | 0.308396 | 0.135348 | 0.717203 | 1.415423 | 0 |
| 33.06944 | 1.129471 | 0.269627 | 2.200028 | 0.06413 | 0.25015 | 0.689873 | 0.446349 | 0.304517 | 0.450087 | 0.780728 | 0 |
| 33.08333 | 0.127445 | 0.236497 | 1.197022 | 0.054782 | 0.014996 | 0.158375 | 0.574554 | 0.285715 | 0.117798 | 0.139323 | 0 |
| 33.09722 | 0.551664 | 0.496225 | 0.331492 | 0.115079 | 0.145816 | 0.125493 | 0.625783 | 0.151789 | 0.257192 | 0.353477 | 0 |
| 33.11111 | 0.859492 | 0.642371 | 0.320659 | 0.054269 | 0.004469 | 0.245887 | 0.587995 | 0.044883 | 0.631311 | 0.811374 | 0 |
| 33.125 | 0.761232 | 0.683356 | 0.863498 | 0.203812 | 0.088634 | 0.393814 | 0.598152 | 0.237103 | 0.705942 | 1.152571 | 0 |
| 33.13889 | 0.715667 | 0.717815 | 1.420483 | 0.442081 | 0.39367 | 0.711875 | 0.63387 | 0.312578 | 0.569252 | 1.164764 | 0 |
| 33.15278 | 0.822002 | 0.724197 | 2.033605 | 0.589279 | 0.749543 | 1.009975 | 0.710101 | 0.343425 | 0.5057 | 1.250622 | 0 |
| 33.16667 | 1.09063 | 0.923464 | 2.712011 | 0.676901 | 0.829716 | 0.992872 | 0.89792 | 0.41552 | 0.611848 | 1.288668 | 0 |
| 33.18056 | 1.502751 | 1.143838 | 3.066124 | 0.550299 | 0.738412 | 0.755081 | 0.926498 | 0.442579 | 0.900038 | 1.245535 | 0 |
| 33.19444 | 1.852721 | 1.166886 | 2.9077 | 0.274097 | 0.444645 | 0.453715 | 0.683849 | 0.419243 | 1.124294 | 1.17269 | 0 |
| 33.20833 | 2.133232 | 1.084275 | 2.491388 | 0.069761 | 0.099093 | 0.349782 | 0.34604 | 0.294769 | 0.959848 | 1.174105 | 0 |
| 33.22222 | 2.154917 | 0.987912 | 2.201212 | 0.052863 | 0.117874 | 0.695803 | 0.090564 | 0.211424 | 0.620304 | 1.288462 | 0 |
| 33.23611 | 2.045208 | 0.759466 | 1.754567 | 0.123572 | 0.272441 | 1.046774 | 0.072148 | 0.1286 | 0.301963 | 1.148664 | 0 |
| 33.25 | 1.814055 | 0.468515 | 1.272671 | 0.193288 | 0.214859 | 1.106718 | 0.031258 | 0.043338 | 0.163152 | 0.91717 | 0 |
| 33.26389 | 1.524805 | 0.250079 | 0.950151 | 0.237268 | 0.00671 | 0.990985 | 0.141449 | 0.081805 | 0.293374 | 0.822794 | 0 |
| 33.27778 | 1.070698 | 0.100536 | 0.673843 | 0.288778 | 0.123687 | 0.788296 | 0.269359 | 0.154497 | 0.468562 | 0.807305 | 0 |
| 33.29167 | 0.645247 | 0.025832 | 0.629076 | 0.278398 | 0.131916 | 0.449452 | 0.246879 | 0.162172 | 0.460234 | 0.642907 | 0 |
| 33.30556 | 0.385879 | 0.009566 | 0.622561 | 0.14371 | 0.047348 | 0.217232 | 0.190567 | 0.156156 | 0.343683 | 0.456703 | 0 |
| 33.31944 | 0.243177 | 0.050504 | 0.704385 | 0.090506 | 0.010875 | 0.130376 | 0.147059 | 0.17501 | 0.227924 | 0.358108 | 0 |
| 33.33333 | 0.185314 | 0.129747 | 0.83221 | 0.293622 | 0.020078 | 0.088328 | 0.093356 | 0.2142 | 0.136756 | 0.408373 | 0 |
| 33.34722 | 0.254947 | 0.217366 | 0.901132 | 0.418344 | 0.014138 | 0.108542 | 0.115345 | 0.26408 | 0.145826 | 0.524169 | 0 |
| 33.36111 | 0.347365 | 0.319846 | 0.862553 | 0.467917 | 0.115785 | 0.185717 | 0.25428 | 0.39941 | 0.342737 | 0.676626 | 0 |
| 33.375 | 0.418534 | 0.443082 | 0.71104 | 0.570415 | 0.342123 | 0.459686 | 0.501062 | 0.629053 | 0.611881 | 0.827044 | 0 |
| 33.38889 | 0.333805 | 0.508742 | 0.509191 | 0.698064 | 0.524741 | 0.789225 | 0.664564 | 0.762703 | 0.706904 | 0.876319 | 0 |
| 33.40278 | 0.219501 | 0.385281 | 0.466114 | 0.673193 | 0.462367 | 0.887304 | 0.531672 | 0.628096 | 0.620328 | 0.72745 | 0 |
| 33.41667 | 0.191005 | 0.194118 | 0.688829 | 0.492628 | 0.355209 | 0.909364 | 0.267465 | 0.425566 | 0.564834 | 0.646938 | 0 |
| 33.43056 | 0.311678 | 0.130576 | 0.971456 | 0.324797 | 0.421933 | 1.024504 | 0.166008 | 0.422328 | 0.651955 | 0.78563 | 0 |
| 33.44444 | 0.534571 | 0.162597 | 1.105279 | 0.219118 | 0.46594 | 1.170601 | 0.164018 | 0.479979 | 0.726277 | 0.941506 | 0 |
| 33.45833 | 0.774856 | 0.221453 | 1.045067 | 0.126425 | 0.420684 | 1.278454 | 0.204792 | 0.52334 | 0.650874 | 0.962099 | 0 |
| 33.47222 | 0.916814 | 0.267982 | 0.840141 | 0.075956 | 0.321754 | 1.431649 | 0.218227 | 0.466113 | 0.521769 | 0.87501 | 0 |
| 33.48611 | 1.114667 | 0.362219 | 0.752798 | 0.099411 | 0.437124 | 2.009833 | 0.065499 | 0.376267 | 0.646094 | 1.040085 | 0 |
| 33.5 | 1.604215 | 0.732245 | 1.189541 | 0.390809 | 0.962642 | 3.949269 | 0.251967 | 0.505616 | 1.079253 | 1.727015 | 0 |
| 33.51389 | 1.60025 | 1.227338 | 1.780175 | 0.968312 | 1.357914 | 5.563062 | 0.386947 | 0.732962 | 1.438441 | 2.510405 | 0 |

| | | | | | | | | | | | |
|---|---|---|---|---|---|---|---|---|---|---|---|
| 33.52778 | 1.250828 | 1.52988 | 2.733708 | 1.534876 | 1.616929 | 5.575518 | 0.157846 | 0.827936 | 2.021545 | 3.143178 | 0 |
| 33.54167 | 1.423082 | 1.673678 | 3.131377 | 2.37384 | 1.563109 | 5.791703 | 0.27325 | 0.637652 | 3.349519 | 3.467428 | 0 |
| 33.55556 | 1.324761 | 1.935482 | 2.426227 | 3.012656 | 1.464173 | 4.333972 | 0.470382 | 0.786218 | 2.887786 | 2.954834 | 0 |
| 33.56944 | 0.737548 | 2.02189 | 1.42103 | 2.935129 | 0.977835 | 2.306802 | 0.335107 | 0.567953 | 1.554934 | 1.78166 | 0 |
| 33.58333 | 0.236932 | 2.33097 | 0.639319 | 2.574534 | 0.689864 | 0.902197 | 0.120907 | 0.353569 | 0.069609 | 0.860488 | 0 |
| 33.59722 | 0.110059 | 3.084155 | 0.086932 | 2.651163 | 1.080896 | 0.620566 | 0.07381 | 0.247255 | 1.361623 | 0.248067 | 0 |
| 33.61111 | 0.40432 | 3.045304 | 0.622323 | 2.897356 | 1.276455 | 0.520467 | 0.129049 | 0.191793 | 1.344351 | 0.204029 | 0 |
| 33.625 | 0.334249 | 2.553208 | 1.018425 | 2.180003 | 0.786462 | 0.155746 | 0.022644 | 0.024114 | 0.560803 | 0.376796 | 0 |
| 33.63889 | 0.002058 | 2.104332 | 0.876547 | 0.705093 | 0.324906 | 0.512101 | 0.103819 | 0.230967 | 0.306436 | 0.447148 | 0 |
| 33.65278 | 0.263388 | 1.657893 | 0.51159 | 1.035522 | 0.1657 | 0.747106 | 0.075976 | 0.072711 | 0.315556 | 0.336158 | 0 |
| 33.66667 | 0.394066 | 0.804805 | 0.096638 | 3.289442 | 0.064536 | 1.066042 | 0.013061 | 0.235948 | 0.713922 | 0.196742 | 0 |
| 33.68056 | 0.26441 | 0.485549 | 0.039269 | 4.196072 | 0.015364 | 1.099689 | 0.096503 | 0.266749 | 0.801272 | 0.239857 | 0 |
| 33.69444 | 0.096795 | 1.904806 | 0.156804 | 4.057274 | 0.068649 | 1.496886 | 0.040999 | 0.173966 | 0.660525 | 0.248701 | 0 |
| 33.70833 | 0.239571 | 2.923341 | 0.293753 | 3.663516 | 0.165422 | 1.420741 | 0.085431 | 0.055455 | 0.235496 | 0.29661 | 0 |
| 33.72222 | 0.626201 | 3.857462 | 0.238595 | 2.042678 | 0.282491 | 0.996527 | 0.224785 | 0.112942 | 0.173673 | 0.633661 | 0 |
| 33.73611 | 0.136043 | 4.212204 | 0.217426 | 0.626432 | 0.459413 | 0.396655 | 0.434987 | 0.345227 | 0.682776 | 0.771818 | 0 |
| 33.75 | 0.259304 | 3.702414 | 0.906149 | 0.112817 | 0.804179 | 0.132953 | 0.728641 | 0.640941 | 1.109599 | 0.855571 | 0 |
| 33.76389 | 0.421318 | 2.15969 | 1.272647 | 0.007001 | 1.169178 | 0.686236 | 0.959705 | 0.960295 | 1.360584 | 0.760301 | 0 |
| 33.77778 | 0.228996 | 1.036677 | 1.159063 | 0.082193 | 1.310849 | 0.988302 | 1.157351 | 1.041975 | 1.238106 | 0.901551 | 0 |
| 33.79167 | 0.321197 | 0.613893 | 0.881821 | 0.199828 | 1.433111 | 0.598469 | 1.158643 | 1.041311 | 0.934969 | 1.119916 | 0 |
| 33.80556 | 0.96942 | 0.526611 | 0.858025 | 1.151756 | 1.610926 | 0.510241 | 0.985165 | 0.834823 | 0.724801 | 1.366917 | 0 |
| 33.81944 | 1.673147 | 0.471268 | 0.913073 | 1.894318 | 1.731199 | 2.114332 | 0.775765 | 0.624067 | 0.920929 | 1.476939 | 0 |
| 33.83333 | 2.119689 | 0.422932 | 1.123971 | 1.163617 | 1.815344 | 3.984349 | 0.664614 | 0.537965 | 1.3363 | 1.618759 | 0 |
| 33.84722 | 1.797319 | 0.374525 | 1.795759 | 0.351186 | 1.810559 | 5.890259 | 0.779995 | 0.794749 | 1.813121 | 1.914614 | 0 |
| 33.86111 | 1.266105 | 0.268245 | 2.227097 | 2.007556 | 1.516785 | 5.531746 | 0.871511 | 1.10264 | 1.967018 | 2.225505 | 0 |
| 33.875 | 1.103017 | 0.350423 | 2.54374 | 3.141343 | 1.174111 | 4.385703 | 0.763281 | 1.349147 | 1.839318 | 2.675405 | 0 |
| 33.88889 | 1.026808 | 0.656616 | 2.911316 | 3.526688 | 1.191148 | 3.526366 | 0.743155 | 1.533449 | 1.702108 | 3.185104 | 0 |
| 33.90278 | 0.90059 | 0.951707 | 2.911351 | 2.636525 | 1.505336 | 2.985762 | 0.674076 | 1.455436 | 1.325738 | 3.244089 | 0 |
| 33.91667 | 0.83853 | 1.20694 | 2.435178 | 1.542753 | 1.658546 | 2.328468 | 0.530402 | 1.188434 | 0.894787 | 2.514487 | 0 |
| 33.93056 | 1.038841 | 1.415621 | 1.515393 | 0.548623 | 1.356977 | 1.666129 | 0.370122 | 0.787192 | 0.565999 | 1.471861 | 0 |
| 33.94444 | 0.976528 | 1.262757 | 0.669372 | 0.146894 | 0.733918 | 1.284487 | 0.089974 | 0.26253 | 0.268215 | 0.650266 | 0 |
| 33.95833 | 0.687201 | 0.842795 | 0.00095 | 0.301563 | 0.140395 | 1.233244 | 0.129166 | 0.011711 | 0.000868 | 0.163117 | 0 |
| 33.97222 | 0.399091 | 0.594091 | 0.407916 | 0.23094 | 0.184838 | 1.062783 | 0.091782 | 0.046981 | 0.189753 | 0.128346 | 0 |
| 33.98611 | 0.592836 | 0.84429 | 0.247076 | 1.261328 | 0.72385 | 1.587076 | 0.197629 | 0.644317 | 0.232065 | 0.741225 | 0 |
| 34 | 1.908296 | 1.805939 | 0.703935 | 2.549524 | 2.971812 | 3.006865 | 0.667451 | 1.622933 | 1.480536 | 2.148216 | 0 |
| 34.01389 | 3.544583 | 2.586047 | 1.626419 | 2.739717 | 3.742171 | 3.741575 | 0.821118 | 2.144651 | 2.320025 | 3.05426 | 0 |
| 34.02778 | 4.576253 | 2.558296 | 1.763041 | 1.406492 | 1.635666 | 2.875305 | 0.383439 | 1.477698 | 1.892104 | 2.390335 | 0 |
| 34.04167 | 4.460861 | 2.518899 | 1.710557 | 0.073251 | 0.574322 | 2.273566 | 0.044472 | 0.575084 | 1.118283 | 1.291899 | 0 |
| 34.05556 | 3.825096 | 1.980072 | 1.458989 | 0.767853 | 0.169019 | 1.68059 | 0.072075 | 0.260034 | 0.80205 | 0.43485 | 0 |
| 34.06944 | 2.252802 | 0.941539 | 1.108487 | 0.885629 | 0.097518 | 0.873557 | 0.364258 | 0.741901 | 0.639267 | 0.116078 | 0 |
| 34.08333 | 0.606766 | 0.243676 | 0.68671 | 0.547173 | 0.145142 | 0.019018 | 0.650818 | 0.703273 | 0.289703 | 0.33937 | 0 |
| 34.09722 | 0.876531 | 1.136412 | 0.333265 | 0.196461 | 0.220991 | 0.570175 | 0.750762 | 0.608345 | 0.18515 | 0.213763 | 0 |
| 34.11111 | 2.230722 | 1.480462 | 0.10153 | 0.085237 | 0.041844 | 0.712481 | 0.690774 | 0.564354 | 0.550834 | 0.006619 | 0 |
| 34.125 | 3.05737 | 1.597483 | 0.11284 | 0.276431 | 0.012591 | 0.842434 | 0.610031 | 0.742719 | 0.981616 | 0.111943 | 0 |
| 34.13889 | 2.878635 | 1.453043 | 0.335953 | 0.588643 | 0.436487 | 0.934248 | 0.684248 | 0.934039 | 1.560682 | 0.499344 | 0 |
| 34.15278 | 2.154392 | 1.165815 | 0.663505 | 0.885567 | 0.735897 | 0.916385 | 1.017289 | 0.970222 | 2.179418 | 0.673253 | 0 |
| 34.16667 | 1.53809 | 0.982358 | 1.119974 | 0.968652 | 1.063811 | 1.036519 | 1.231663 | 0.885837 | 2.90323 | 0.988497 | 0 |
| 34.18056 | 1.063531 | 0.743386 | 1.458235 | 0.694347 | 1.306652 | 1.358964 | 1.024237 | 0.766752 | 2.576468 | 1.044206 | 0 |
| 34.19444 | 0.853662 | 0.491346 | 1.601721 | 0.337574 | 1.090663 | 1.227687 | 0.557685 | 0.561639 | 1.941495 | 0.915663 | 0 |
| 34.20833 | 0.632827 | 0.351086 | 1.39048 | 0.07167 | 0.679778 | 0.944545 | 0.297204 | 0.341868 | 1.151665 | 0.677459 | 0 |
| 34.22222 | 0.390288 | 0.203701 | 1.061097 | 0.006597 | 0.309358 | 0.393411 | 0.162225 | 0.175151 | 0.354122 | 0.435903 | 0 |
| 34.23611 | 0.105509 | 0.132736 | 0.821051 | 0.146605 | 0.011956 | 0.121198 | 0.009277 | 0.074378 | 0.356777 | 0.321735 | 0 |
| 34.25 | 0.013009 | 0.158814 | 0.703087 | 0.269179 | 0.243229 | 0.405874 | 0.081897 | 0.03045 | 0.947489 | 0.473586 | 0 |
| 34.26389 | 0.2714 | 0.150398 | 0.624164 | 0.053397 | 0.424927 | 0.505382 | 0.197765 | 0.279799 | 1.319717 | 0.513304 | 0 |
| 34.27778 | 0.700654 | 0.084574 | 0.621595 | 0.365779 | 0.573731 | 0.374323 | 0.272076 | 0.475467 | 0.867988 | 0.322159 | 0 |
| 34.29167 | 0.965923 | 0.068068 | 0.713557 | 0.584015 | 0.503122 | 0.268186 | 0.224569 | 0.352561 | 0.032917 | 0.212185 | 0 |

| | | | | | | | | | | | |
|---|---|---|---|---|---|---|---|---|---|---|---|
| 34.30556 | 0.862666 | 0.099363 | 0.741545 | 0.445141 | 0.254843 | 0.068732 | 0.013718 | 0.03368 | 0.750892 | 0.429444 | 0 |
| 34.31944 | 0.577408 | 0.121002 | 0.643536 | 0.127904 | 0.020405 | 0.239812 | 0.171307 | 0.223649 | 1.128327 | 0.543417 | 0 |
| 34.33333 | 0.339258 | 0.154123 | 0.538123 | 0.089045 | 0.210301 | 0.592812 | 0.218187 | 0.302699 | 1.156779 | 0.370415 | 0 |
| 34.34722 | 0.157773 | 0.162697 | 0.459447 | 0.157988 | 0.277841 | 0.779371 | 0.260546 | 0.228939 | 0.784713 | 0.34597 | 0 |
| 34.36111 | 0.007918 | 0.108611 | 0.397004 | 0.071662 | 0.254587 | 0.793238 | 0.269082 | 0.142607 | 0.30584 | 0.400975 | 0 |
| 34.375 | 0.070639 | 0.078763 | 0.469438 | 0.104576 | 0.211235 | 0.878924 | 0.209261 | 0.127233 | 0.174398 | 0.276584 | 0 |
| 34.38889 | 0.308344 | 0.116342 | 0.714194 | 0.263895 | 0.250279 | 1.085297 | 0.162101 | 0.193987 | 0.409676 | 0.148681 | 0 |
| 34.40278 | 0.45113 | 0.188479 | 0.925824 | 0.253018 | 0.369433 | 1.197114 | 0.14805 | 0.33397 | 0.763947 | 0.309033 | 0 |
| 34.41667 | 0.424104 | 0.247791 | 0.927087 | 0.062787 | 0.413152 | 0.8613 | 0.167286 | 0.514334 | 0.876998 | 0.650228 | 0 |
| 34.43056 | 0.335828 | 0.224796 | 0.777508 | 0.21746 | 0.291904 | 0.247977 | 0.133038 | 0.609193 | 0.668231 | 0.832039 | 0 |
| 34.44444 | 0.138914 | 0.054502 | 0.599699 | 0.420806 | 0.104357 | 0.203349 | 0.029694 | 0.569348 | 0.286611 | 0.747417 | 0 |
| 34.45833 | 0.197275 | 0.150061 | 0.458056 | 0.248979 | 0.093378 | 0.488775 | 0.117034 | 0.301237 | 0.041818 | 0.562497 | 0 |
| 34.47222 | 0.448624 | 0.222141 | 0.549366 | 0.031175 | 0.080166 | 0.323398 | 0.167899 | 0.063752 | 0.12926 | 0.488467 | 0 |
| 34.48611 | 0.154919 | 0.037592 | 1.093169 | 0.000806 | 0.413293 | 0.951923 | 0.14578 | 0.032954 | 0.26119 | 0.770538 | 0 |
| 34.5 | 0.475082 | 0.26202 | 1.898354 | 0.012648 | 1.089709 | 3.329741 | 0.192751 | 0.200692 | 1.189703 | 1.016196 | 0 |
| 34.51389 | 1.069847 | 0.318336 | 2.185648 | 0.139129 | 1.304057 | 4.941929 | 0.208887 | 0.535348 | 2.091028 | 1.153878 | 0 |
| 34.52778 | 1.455393 | 0.379938 | 2.100345 | 0.877185 | 1.457893 | 4.351204 | 0.230268 | 0.99442 | 3.048871 | 1.266469 | 0 |
| 34.54167 | 1.85223 | 0.654214 | 2.557204 | 1.182955 | 1.922882 | 3.58596 | 0.476201 | 1.303527 | 3.263225 | 1.694681 | 0 |
| 34.55556 | 1.712939 | 0.815332 | 2.508659 | 1.114423 | 2.227584 | 3.770781 | 0.673809 | 1.207191 | 2.972627 | 1.702231 | 0 |
| 34.56944 | 1.373989 | 0.846535 | 1.915212 | 1.050493 | 1.807892 | 2.665899 | 0.612762 | 0.855806 | 1.949896 | 1.378797 | 0 |
| 34.58333 | 1.076059 | 0.772372 | 1.330418 | 0.997122 | 1.180856 | 1.202645 | 0.528235 | 0.440158 | 0.552397 | 0.924107 | 0 |
| 34.59722 | 0.798001 | 0.778453 | 0.89008 | 1.188362 | 0.707759 | 0.17473 | 0.438528 | 0.086714 | 0.743147 | 0.504876 | 0 |
| 34.61111 | 0.195915 | 0.700079 | 0.54751 | 1.974919 | 0.508811 | 0.036906 | 0.486135 | 0.31968 | 1.038093 | 0.163129 | 0 |
| 34.625 | 0.873948 | 0.580137 | 0.17953 | 2.272256 | 0.406949 | 0.259482 | 0.52537 | 0.233641 | 1.247155 | 0.065915 | 0 |
| 34.63889 | 1.137393 | 0.553707 | 0.152386 | 1.864233 | 0.195688 | 0.13515 | 0.324481 | 0.04511 | 1.113821 | 0.381802 | 0 |
| 34.65278 | 0.164994 | 0.704926 | 0.464486 | 0.872949 | 0.441501 | 0.125994 | 0.141038 | 0.569397 | 0.688414 | 0.977687 | 0 |
| 34.66667 | 1.076995 | 0.893793 | 0.376553 | 0.61702 | 0.190487 | 0.135919 | 0.005514 | 0.790977 | 0.344943 | 1.347391 | 0 |
| 34.68056 | 1.667858 | 0.77945 | 0.024868 | 2.043896 | 0.005396 | 0.048073 | 0.110001 | 0.564959 | 0.173034 | 1.066293 | 0 |
| 34.69444 | 1.562948 | 0.601338 | 0.289128 | 2.557165 | 0.007778 | 0.578355 | 0.261743 | 0.355581 | 0.832785 | 0.494122 | 0 |
| 34.70833 | 1.03563 | 0.417409 | 0.651784 | 2.373178 | 0.372444 | 0.659545 | 0.470323 | 0.219033 | 1.44055 | 0.084295 | 0 |
| 34.72222 | 0.198984 | 0.059047 | 1.065656 | 1.312799 | 0.600435 | 0.54465 | 0.611539 | 0.080613 | 1.682205 | 0.498554 | 0 |
| 34.73611 | 0.822873 | 0.308615 | 0.796686 | 1.151208 | 0.499593 | 0.365141 | 0.644684 | 0.394925 | 1.516487 | 1.30281 | 0 |
| 34.75 | 1.128126 | 0.51686 | 0.014537 | 0.721663 | 0.496447 | 0.524447 | 0.500268 | 0.339977 | 1.190783 | 1.528969 | 0 |
| 34.76389 | 0.54352 | 0.624827 | 0.833545 | 0.198002 | 0.724882 | 0.953549 | 0.263648 | 0.131607 | 0.728001 | 1.170257 | 0 |
| 34.77778 | 0.349935 | 0.61816 | 1.574323 | 0.160787 | 0.879485 | 1.42387 | 0.000438 | 0.48141 | 0.290067 | 0.910952 | 0 |
| 34.79167 | 0.639204 | 0.276568 | 3.043327 | 0.436663 | 0.694276 | 1.633974 | 0.152009 | 0.526465 | 0.258861 | 0.68154 | 0 |
| 34.80556 | 0.077954 | 0.221725 | 3.005807 | 0.521476 | 0.153106 | 1.462956 | 0.101129 | 0.169708 | 0.760736 | 0.414891 | 0 |
| 34.81944 | 0.470363 | 0.353528 | 1.454805 | 0.498616 | 0.296392 | 1.355395 | 0.015912 | 0.322734 | 1.102416 | 0.277244 | 0 |
| 34.83333 | 0.430524 | 0.139556 | 0.365699 | 0.274269 | 0.632004 | 1.595 | 0.035195 | 0.848178 | 1.21763 | 0.793712 | 0 |
| 34.84722 | 0.49058 | 0.129431 | 0.090297 | 0.130817 | 0.861894 | 2.060809 | 0.088253 | 1.475545 | 1.161822 | 1.365014 | 0 |
| 34.86111 | 0.038956 | 0.132204 | 0.289597 | 0.52755 | 0.83917 | 2.296064 | 0.251177 | 1.72525 | 1.111822 | 1.720657 | 0 |
| 34.875 | 0.870133 | 0.050269 | 1.177294 | 0.450823 | 0.958923 | 2.154195 | 0.356999 | 1.563623 | 1.361815 | 1.663142 | 0 |
| 34.88889 | 1.166722 | 0.085779 | 2.090751 | 0.147366 | 1.03457 | 1.931155 | 0.428175 | 0.896376 | 1.284112 | 1.721788 | 0 |
| 34.90278 | 1.487388 | 0.299996 | 2.405544 | 0.03386 | 1.179918 | 1.765712 | 0.487913 | 0.051136 | 1.01013 | 1.516207 | 0 |
| 34.91667 | 1.517884 | 0.456858 | 2.19063 | 0.115166 | 1.16256 | 1.691346 | 0.535605 | 0.367414 | 0.735038 | 1.054296 | 0 |
| 34.93056 | 1.096013 | 0.453246 | 1.819838 | 0.119668 | 0.910393 | 0.923729 | 0.64812 | 0.562351 | 0.383876 | 0.572937 | 0 |
| 34.94444 | 0.535347 | 0.219481 | 1.351355 | 0.036654 | 0.388314 | 0.048798 | 0.729548 | 0.554853 | 0.024781 | 0.363383 | 0 |
| 34.95833 | 0.114749 | 0.010293 | 0.58482 | 0.055291 | 0.088288 | 0.638537 | 0.871157 | 0.371271 | 0.432526 | 0.345043 | 0 |
| 34.97222 | 0.670407 | 0.302618 | 0.29069 | 0.198319 | 0.472451 | 1.119718 | 0.94958 | 0.177697 | 0.999809 | 0.3696 | 0 |
| 34.98611 | 0.727422 | 0.828316 | 0.935759 | 0.438522 | 0.472289 | 1.165143 | 1.122033 | 0.306105 | 1.598412 | 0.521427 | 0 |
| 35 | 0.182353 | 1.52417 | 0.770079 | 0.820764 | 0.297273 | 0.398478 | 1.403004 | 0.141867 | 1.637646 | 0.929891 | 0 |
| 35.01389 | 1.85714 | 3.015266 | 0.261115 | 1.830185 | 1.169913 | 0.300774 | 1.116094 | 1.18762 | 0.902585 | 1.576084 | 0 |
| 35.02778 | 3.06314 | 3.672351 | 1.258694 | 1.398122 | 1.463322 | 0.70779 | 0.485274 | 1.172983 | 0.301551 | 2.202355 | 0 |
| 35.04167 | 2.098014 | 3.177018 | 1.717358 | 0.711561 | 1.466224 | 0.338251 | 0.036556 | 0.561712 | 0.03459 | 2.008963 | 0 |
| 35.05556 | 1.90833 | 1.936761 | 1.833612 | 0.460461 | 1.469649 | 0.090293 | 0.0772 | 0.317591 | 0.320601 | 1.17154 | 0 |
| 35.06944 | 1.347961 | 1.072751 | 1.531695 | 0.270706 | 0.854448 | 0.462027 | 0.89106 | 0.007189 | 1.066954 | 0.494619 | 0 |

| | | | | | | | | | | | |
|---|---|---|---|---|---|---|---|---|---|---|---|
| 35.08333 | 0.45276 | 0.063382 | 0.906561 | 0.533171 | 0.110804 | 0.714097 | 1.28942 | 0.718765 | 0.053356 | 1.823884 | 0 |
| 35.09722 | 0.552933 | 1.241646 | 0.24909 | 1.2773 | 0.385668 | 0.764338 | 0.539644 | 1.750017 | 0.306168 | 2.026597 | 0 |
| 35.11111 | 1.362088 | 2.249809 | 0.209728 | 0.960478 | 0.647094 | 0.729499 | 0.00616 | 1.498882 | 0.573123 | 1.351978 | 0 |
| 35.125 | 1.843205 | 2.297511 | 0.531647 | 0.387503 | 0.448471 | 0.65355 | 0.034667 | 1.036232 | 1.000608 | 0.909174 | 0 |
| 35.13889 | 2.111004 | 2.58206 | 0.85085 | 0.164116 | 0.042904 | 0.592993 | 0.418425 | 1.084465 | 1.272379 | 0.912484 | 0 |
| 35.15278 | 2.162989 | 2.71448 | 1.075736 | 0.100125 | 0.307276 | 0.479882 | 0.711462 | 1.355578 | 1.846981 | 0.936895 | 0 |
| 35.16667 | 1.911181 | 2.876076 | 1.253653 | 0.061568 | 0.395524 | 0.33518 | 0.699932 | 1.652472 | 1.25982 | 0.981693 | 0 |
| 35.18056 | 2.449785 | 2.787114 | 1.58182 | 0.334151 | 0.35912 | 0.399605 | 0.661542 | 2.124272 | 0.674636 | 1.02584 | 0 |
| 35.19444 | 2.830241 | 2.420144 | 1.684378 | 0.691579 | 0.421534 | 0.648182 | 0.631517 | 2.177902 | 1.917556 | 1.242842 | 0 |
| 35.20833 | 3.153942 | 1.910396 | 1.536879 | 0.775808 | 0.619868 | 0.860842 | 0.593388 | 1.57963 | 2.257644 | 1.245984 | 0 |
| 35.22222 | 2.243805 | 1.35882 | 1.115471 | 0.587438 | 0.880188 | 1.055231 | 0.468208 | 0.845294 | 1.80459 | 0.92275 | 0 |
| 35.23611 | 1.068373 | 0.527699 | 0.700742 | 0.225502 | 0.560949 | 1.07642 | 0.364191 | 0.181436 | 0.822496 | 0.323959 | 0 |
| 35.25 | 0.352573 | 0.111088 | 0.531657 | 0.193685 | 0.115228 | 0.690004 | 0.170418 | 0.305299 | 0.317245 | 0.186482 | 0 |
| 35.26389 | 0.210364 | 0.26389 | 0.627482 | 0.399646 | 0.064827 | 0.595396 | 0.065291 | 0.486501 | 1.023659 | 0.34257 | 0 |
| 35.27778 | 0.287009 | 0.042895 | 0.752586 | 0.305315 | 0.09302 | 0.784109 | 0.057407 | 0.39191 | 1.080514 | 0.263814 | 0 |
| 35.29167 | 0.30122 | 0.116385 | 0.746025 | 0.258838 | 0.095946 | 0.986067 | 0.012291 | 0.377394 | 0.754221 | 0.338043 | 0 |
| 35.30556 | 0.053878 | 0.018612 | 0.653893 | 0.381927 | 0.076447 | 1.063121 | 0.141999 | 0.656291 | 0.256582 | 0.529863 | 0 |
| 35.31944 | 0.3477 | 0.300882 | 0.562506 | 0.565327 | 0.012582 | 0.772714 | 0.120882 | 0.958205 | 0.213548 | 0.550727 | 0 |
| 35.33333 | 0.505924 | 0.54951 | 0.447996 | 0.763732 | 0.063032 | 0.17469 | 0.016625 | 1.087166 | 0.341001 | 0.557397 | 0 |
| 35.34722 | 0.354486 | 0.72896 | 0.256152 | 0.861513 | 0.039712 | 0.144185 | 0.039427 | 1.0866 | 0.170909 | 0.6745 | 0 |
| 35.36111 | 0.207941 | 0.695884 | 0.057402 | 0.625531 | 0.02486 | 0.026447 | 0.065308 | 0.773322 | 0.04885 | 0.602808 | 0 |
| 35.375 | 0.868062 | 0.326181 | 0.089829 | 0.090949 | 0.084282 | 0.333547 | 0.060666 | 0.03923 | 0.0403 | 0.147275 | 0 |
| 35.38889 | 1.389452 | 0.118875 | 0.392032 | 0.115719 | 0.169898 | 0.592457 | 0.248987 | 0.542625 | 0.551813 | 0.261043 | 0 |
| 35.40278 | 1.524392 | 0.313852 | 0.5694 | 0.214673 | 0.185978 | 0.558477 | 0.290246 | 0.496204 | 1.144032 | 0.344468 | 0 |
| 35.41667 | 1.323452 | 0.454967 | 0.582454 | 0.577245 | 0.134568 | 0.38108 | 0.183273 | 0.309058 | 1.679728 | 0.294122 | 0 |
| 35.43056 | 1.107877 | 0.77935 | 0.637836 | 0.553239 | 0.090478 | 0.408731 | 0.167312 | 0.321609 | 2.002907 | 0.395903 | 0 |
| 35.44444 | 0.9045 | 1.018416 | 0.8526 | 0.315035 | 0.116256 | 0.64535 | 0.226841 | 0.449472 | 1.649927 | 0.487593 | 0 |
| 35.45833 | 0.366529 | 0.96004 | 1.003531 | 0.066483 | 0.218599 | 0.962749 | 0.253396 | 0.517856 | 1.083987 | 0.495056 | 0 |
| 35.47222 | 0.164088 | 0.682846 | 0.91235 | 0.234374 | 0.273417 | 1.03019 | 0.227031 | 0.607861 | 0.504083 | 0.467009 | 0 |
| 35.48611 | 0.118997 | 0.451777 | 0.82232 | 0.719265 | 0.289842 | 1.191097 | 0.019901 | 0.728803 | 0.052312 | 0.48052 | 0 |
| 35.5 | 1.036667 | 0.323652 | 1.107057 | 1.450854 | 0.335075 | 1.862379 | 0.376135 | 0.991497 | 0.080067 | 0.888453 | 0 |
| 35.51389 | 2.06752 | 0.19853 | 1.556955 | 1.505597 | 0.341711 | 2.813564 | 0.636129 | 1.618028 | 0.711072 | 1.865612 | 0 |
| 35.52778 | 3.162028 | 0.201195 | 1.486974 | 1.672523 | 0.106421 | 2.552997 | 0.378609 | 3.233843 | 1.751171 | 2.852236 | 0 |
| 35.54167 | 3.941124 | 0.64135 | 1.709506 | 1.940782 | 0.084902 | 2.116387 | 0.047309 | 4.390333 | 2.83441 | 3.315088 | 0 |
| 35.55556 | 2.879259 | 0.545458 | 2.363813 | 0.95807 | 0.429358 | 1.804166 | 0.230335 | 2.373661 | 3.036672 | 2.394877 | 0 |
| 35.56944 | 1.061274 | 0.334804 | 1.745583 | 0.412276 | 0.793009 | 1.46269 | 0.267477 | 0.841645 | 2.227136 | 1.106196 | 0 |
| 35.58333 | 0.276923 | 0.08266 | 0.955281 | 0.037743 | 0.953352 | 1.053393 | 0.114711 | 0.059035 | 1.320265 | 0.128048 | 0 |
| 35.59722 | 1.299949 | 0.245054 | 0.326238 | 0.197888 | 1.064495 | 0.378808 | 0.025401 | 0.754013 | 0.483671 | 0.485549 | 0 |
| 35.61111 | 1.690349 | 0.270575 | 0.029524 | 0.34274 | 1.274517 | 0.695077 | 0.051702 | 1.188087 | 0.269508 | 0.896599 | 0 |
| 35.625 | 1.633004 | 0.604796 | 0.157248 | 0.654322 | 1.458649 | 1.471907 | 0.002836 | 1.697893 | 0.94802 | 1.076438 | 0 |
| 35.63889 | 1.710538 | 0.509954 | 0.367388 | 0.941248 | 1.596948 | 1.304066 | 0.058416 | 2.032121 | 0.896084 | 0.816468 | 0 |
| 35.65278 | 0.647761 | 0.049523 | 0.518267 | 1.309592 | 1.611175 | 0.675418 | 0.104759 | 1.982285 | 0.549837 | 0.575867 | 0 |
| 35.66667 | 0.788288 | 0.778002 | 0.547795 | 1.167153 | 1.390969 | 0.299311 | 0.038218 | 1.019087 | 0.311126 | 0.221774 | 0 |
| 35.68056 | 1.690826 | 1.110172 | 0.433952 | 0.884107 | 1.003965 | 0.02153 | 0.060692 | 0.018556 | 0.396537 | 0.049827 | 0 |
| 35.69444 | 2.311233 | 1.183168 | 0.001583 | 0.906223 | 0.673984 | 0.113212 | 0.136572 | 0.4287 | 0.367626 | 0.188243 | 0 |
| 35.70833 | 2.233239 | 0.891868 | 0.621019 | 0.721275 | 0.524006 | 0.441124 | 0.077745 | 0.482276 | 0.031244 | 0.370962 | 0 |
| 35.72222 | 1.015903 | 0.34939 | 0.890094 | 0.480041 | 0.578969 | 0.833131 | 0.071436 | 0.324894 | 0.645686 | 0.089787 | 0 |
| 35.73611 | 0.010585 | 0.282516 | 1.285903 | 0.390446 | 0.616367 | 1.170796 | 0.219066 | 0.107564 | 1.157609 | 0.655054 | 0 |
| 35.75 | 1.362617 | 0.941551 | 1.275385 | 0.784 | 0.676861 | 1.186172 | 0.264991 | 1.603221 | 1.505062 | 1.273496 | 0 |
| 35.76389 | 2.244438 | 1.466369 | 0.710658 | 1.746203 | 0.722607 | 1.176722 | 0.278479 | 3.844216 | 1.587373 | 1.398382 | 0 |
| 35.77778 | 2.790048 | 1.711993 | 0.255692 | 1.437186 | 0.71069 | 1.044719 | 0.25228 | 2.227907 | 1.299538 | 1.113897 | 0 |
| 35.79167 | 3.153027 | 1.63658 | 0.335478 | 0.807433 | 0.675322 | 0.690406 | 0.124814 | 2.182052 | 0.757538 | 1.282542 | 0 |
| 35.80556 | 2.547703 | 1.596198 | 0.93049 | 1.314922 | 0.669149 | 0.415621 | 0.029928 | 2.978828 | 0.054085 | 1.831897 | 0 |
| 35.81944 | 2.026379 | 1.085671 | 1.219519 | 1.826725 | 0.637526 | 0.342881 | 0.124604 | 3.348995 | 0.733497 | 1.901974 | 0 |
| 35.83333 | 1.039366 | 0.434582 | 1.474375 | 1.944402 | 0.482257 | 0.0474 | 0.114911 | 2.593892 | 1.313705 | 1.867657 | 0 |
| 35.84722 | 0.59729 | 0.415815 | 1.497719 | 1.938248 | 0.20473 | 0.38541 | 0.07195 | 2.067493 | 1.695656 | 1.876794 | 0 |

| | | | | | | | | | | | |
|---|---|---|---|---|---|---|---|---|---|---|---|
| 35.86111 | 0.55141 | 0.888872 | 1.704375 | 1.965158 | 0.097528 | 0.607376 | 0.044571 | 2.072725 | 1.73913 | 2.224907 | 0 |
| 35.875 | 0.647543 | 1.460509 | 1.635033 | 1.902948 | 0.361152 | 0.731326 | 0.000237 | 2.422527 | 1.550428 | 2.664951 | 0 |
| 35.88889 | 0.818812 | 1.620168 | 1.436893 | 1.42754 | 0.616085 | 0.745838 | 0.125972 | 3.090746 | 1.349001 | 2.858758 | 0 |
| 35.90278 | 0.454942 | 0.816106 | 1.187145 | 0.720069 | 0.947657 | 1.002915 | 0.204364 | 3.466424 | 1.071028 | 2.946997 | 0 |
| 35.91667 | 0.206471 | 0.206003 | 0.806548 | 0.126741 | 1.024195 | 1.118062 | 0.217954 | 3.960468 | 0.882413 | 2.423056 | 0 |
| 35.93056 | 0.735618 | 1.080128 | 0.769154 | 0.269614 | 0.813211 | 0.994857 | 0.325077 | 4.306018 | 0.473036 | 1.490526 | 0 |
| 35.94444 | 1.001753 | 1.67691 | 0.594799 | 0.558068 | 0.437584 | 0.591982 | 0.512423 | 2.093155 | 0.16479 | 0.297132 | 0 |
| 35.95833 | 0.995646 | 1.892573 | 0.409359 | 0.699585 | 0.072424 | 0.160625 | 0.681417 | 0.145953 | 0.750188 | 0.759923 | 0 |
| 35.97222 | 0.423767 | 1.564602 | 0.183462 | 0.992301 | 0.260791 | 0.219305 | 0.864409 | 1.7952 | 1.289825 | 1.594874 | 0 |
| 35.98611 | 0.753008 | 0.599655 | 0.225496 | 1.14551 | 0.180496 | 0.120423 | 0.933811 | 3.714652 | 1.530793 | 1.386634 | 0 |
| 36 | 2.400768 | 0.95682 | 1.120993 | 0.472588 | 0.402201 | 0.482455 | 0.747989 | 2.4092 | 0.979869 | 0.3272 | 0 |
| 36.01389 | 3.597121 | 2.087476 | 2.096036 | 0.684771 | 0.925914 | 0.85539 | 0.279978 | 0.626252 | 0.065758 | 2.534529 | 0 |
| 36.02778 | 3.140335 | 2.088551 | 1.779847 | 0.480226 | 0.801238 | 0.791051 | 0.116907 | 2.946687 | 0.339568 | 3.787527 | 0 |
| 36.04167 | 2.608126 | 2.090976 | 1.014365 | 0.283221 | 0.708037 | 0.682437 | 0.248412 | 3.606071 | 0.326901 | 2.634399 | 0 |
| 36.05556 | 1.13484 | 1.172316 | 0.390038 | 0.518171 | 0.693073 | 0.349219 | 0.092677 | 2.60549 | 0.406646 | 0.913416 | 0 |
| 36.06944 | 0.004352 | 0.042504 | 0.021037 | 0.701586 | 0.758807 | 0.007172 | 0.207932 | 1.360582 | 0.397062 | 0.066839 | 0 |
| 36.08333 | 0.732455 | 0.436253 | 0.156636 | 0.710819 | 0.608454 | 0.13616 | 0.320655 | 0.384572 | 0.355659 | 0.546515 | 0 |
| 36.09722 | 0.910966 | 0.395616 | 0.282254 | 0.690846 | 0.191856 | 0.073033 | 0.228964 | 0.421803 | 0.155008 | 1.407438 | 0 |
| 36.11111 | 0.898207 | 0.629839 | 0.351456 | 0.357276 | 0.066419 | 0.518056 | 0.087906 | 1.185189 | 0.05762 | 2.334361 | 0 |
| 36.125 | 1.375307 | 0.879066 | 0.331769 | 0.147348 | 0.065108 | 0.637643 | 0.050502 | 1.630017 | 0.301331 | 2.183513 | 0 |
| 36.13889 | 2.028155 | 1.035495 | 0.405001 | 0.062293 | 0.032857 | 0.565567 | 0.082147 | 1.830201 | 0.644246 | 1.609896 | 0 |
| 36.15278 | 2.459653 | 1.261411 | 0.476727 | 0.290191 | 0.035922 | 0.766147 | 0.236019 | 1.995683 | 1.199863 | 1.763155 | 0 |
| 36.16667 | 2.547062 | 1.510738 | 0.529972 | 0.37447 | 0.050964 | 0.694332 | 0.492139 | 2.111951 | 1.936013 | 2.058073 | 0 |
| 36.18056 | 2.64199 | 1.83682 | 0.64172 | 0.278295 | 0.128694 | 1.089275 | 0.655373 | 1.976624 | 2.381217 | 2.184804 | 0 |
| 36.19444 | 2.682128 | 1.92173 | 0.776808 | 0.022438 | 0.186298 | 1.200197 | 0.667908 | 1.595232 | 2.556515 | 2.058665 | 0 |
| 36.20833 | 2.387362 | 1.862151 | 0.887969 | 0.114266 | 0.228957 | 0.941554 | 0.609638 | 1.493175 | 2.555199 | 1.82994 | 0 |
| 36.22222 | 2.340802 | 1.842397 | 1.238626 | 0.057725 | 0.217885 | 0.494694 | 0.453453 | 1.38123 | 2.241214 | 1.665414 | 0 |
| 36.23611 | 1.957074 | 1.807882 | 1.186418 | 0.006219 | 0.205627 | 0.100095 | 0.122478 | 1.037988 | 1.336031 | 1.336402 | 0 |
| 36.25 | 1.911549 | 1.651059 | 1.131461 | 0.137641 | 0.421954 | 0.339193 | 0.13783 | 0.531989 | 0.655196 | 1.11109 | 0 |
| 36.26389 | 2.114399 | 1.649631 | 1.268532 | 0.564654 | 0.903707 | 1.075642 | 0.652431 | 0.31207 | 0.55355 | 1.164217 | 0 |
| 36.27778 | 2.023004 | 1.412749 | 1.132577 | 0.733372 | 1.010055 | 1.364905 | 0.910179 | 0.173128 | 0.545025 | 0.947173 | 0 |
| 36.29167 | 1.35931 | 0.70608 | 0.577749 | 0.278666 | 0.611002 | 1.010086 | 0.562777 | 0.137007 | 0.164755 | 0.299015 | 0 |
| 36.30556 | 0.50614 | 0.154827 | 0.27325 | 0.180412 | 0.089009 | 0.744207 | 0.261294 | 0.225204 | 0.039622 | 0.165739 | 0 |
| 36.31944 | 0.389722 | 0.735972 | 0.3977 | 0.271917 | 0.227706 | 0.843994 | 0.277288 | 0.096368 | 0.385114 | 0.082871 | 0 |
| 36.33333 | 0.421825 | 0.552376 | 0.381009 | 0.108976 | 0.311873 | 0.7431 | 0.3417 | 0.035207 | 0.36997 | 0.188851 | 0 |
| 36.34722 | 0.200228 | 0.135155 | 0.089105 | 0.070254 | 0.189303 | 0.435075 | 0.040962 | 0.191441 | 0.002766 | 0.026314 | 0 |
| 36.36111 | 0.497163 | 0.109389 | 0.129379 | 0.117699 | 0.220395 | 0.268905 | 0.363318 | 0.364803 | 0.127405 | 0.298764 | 0 |
| 36.375 | 0.416637 | 0.206793 | 0.110352 | 0.026327 | 0.293818 | 0.407266 | 0.382838 | 0.254803 | 0.037046 | 0.205514 | 0 |
| 36.38889 | 0.320954 | 0.206934 | 0.062085 | 0.082812 | 0.197216 | 0.811236 | 0.214665 | 0.108833 | 0.25777 | 0.053076 | 0 |
| 36.40278 | 0.33678 | 0.303922 | 0.318 | 0.118064 | 0.112537 | 1.297584 | 0.236573 | 0.588371 | 0.608614 | 0.092503 | 0 |
| 36.41667 | 0.391295 | 0.349677 | 0.678292 | 0.183192 | 0.078333 | 1.284893 | 0.312251 | 1.047744 | 0.908114 | 0.224565 | 0 |
| 36.43056 | 0.562624 | 0.22256 | 1.084979 | 0.243092 | 0.040577 | 0.89448 | 0.228231 | 1.337143 | 1.117482 | 0.462953 | 0 |
| 36.44444 | 0.731213 | 0.02593 | 0.98942 | 0.275887 | 0.038159 | 0.388969 | 0.074266 | 1.464641 | 1.17571 | 0.613285 | 0 |
| 36.45833 | 0.765712 | 0.054106 | 0.611611 | 0.282608 | 0.037094 | 0.250459 | 0.1243 | 1.427477 | 0.996461 | 0.55133 | 0 |
| 36.47222 | 0.544876 | 0.065212 | 0.348855 | 0.285902 | 0.035799 | 1.014412 | 0.264322 | 1.151421 | 0.805118 | 0.429447 | 0 |
| 36.48611 | 0.615619 | 0.07335 | 0.21514 | 0.349053 | 0.09491 | 0.760494 | 0.447998 | 1.193728 | 0.608677 | 0.354122 | 0 |
| 36.5 | 1.447356 | 0.07148 | 1.504484 | 0.390651 | 0.101548 | 0.149589 | 0.801113 | 1.955648 | 0.458712 | 0.476275 | 0 |
| 36.51389 | 2.647155 | 0.22614 | 2.80249 | 0.341162 | 0.139623 | 0.548824 | 0.862279 | 3.233441 | 0.528396 | 0.79051 | 0 |
| 36.52778 | 3.460941 | 0.626318 | 3.260396 | 0.589804 | 0.537138 | 1.533252 | 0.378449 | 5.238788 | 1.331245 | 1.382819 | 0 |
| 36.54167 | 3.341962 | 1.020535 | 2.914668 | 1.006354 | 0.934349 | 1.685286 | 0.305804 | 7.639046 | 2.42071 | 2.181074 | 0 |
| 36.55556 | 2.242532 | 0.982285 | 2.128921 | 1.069361 | 1.229629 | 1.408251 | 0.525061 | 6.327948 | 3.218451 | 2.056436 | 0 |
| 36.56944 | 0.849439 | 0.719913 | 1.583877 | 0.869987 | 1.492897 | 1.123469 | 0.449908 | 3.707429 | 2.607713 | 1.486039 | 0 |
| 36.58333 | 0.528964 | 0.542235 | 1.140891 | 0.632654 | 1.432145 | 0.844568 | 0.176212 | 1.762384 | 2.042118 | 0.86136 | 0 |
| 36.59722 | 1.239602 | 0.569547 | 0.681806 | 0.430722 | 1.434137 | 0.485037 | 0.032483 | 0.828655 | 1.908893 | 0.407059 | 0 |
| 36.61111 | 1.795874 | 0.588251 | 0.214775 | 0.245846 | 1.599951 | 0.241578 | 0.115029 | 0.468646 | 1.937392 | 0.08748 | 0 |
| 36.625 | 1.80438 | 0.497648 | 0.298727 | 0.083154 | 1.768476 | 0.546929 | 0.062955 | 0.000018 | 1.621363 | 0.210411 | 0 |

| | | | | | | | | | | | |
|---|---|---|---|---|---|---|---|---|---|---|---|
| 36.63889 | 1.776361 | 0.332447 | 0.713511 | 0.002618 | 1.685658 | 1.218562 | 0.10669 | 0.666718 | 0.832146 | 0.482633 | 0 |
| 36.65278 | 1.840422 | 0.085506 | 0.813702 | 0.041461 | 1.498963 | 1.837003 | 0.121303 | 1.481525 | 0.139304 | 0.686363 | 0 |
| 36.66667 | 1.99834 | 0.585868 | 1.201687 | 0.013782 | 1.079901 | 1.968074 | 0.083563 | 2.322049 | 1.032171 | 1.036867 | 0 |
| 36.68056 | 1.582084 | 0.732993 | 1.893107 | 0.30211 | 0.961802 | 1.460403 | 0.333742 | 2.398988 | 1.724865 | 1.230085 | 0 |
| 36.69444 | 0.22623 | 0.49258 | 1.360959 | 0.684782 | 0.940119 | 0.84634 | 0.471084 | 1.876455 | 2.050904 | 1.162295 | 0 |
| 36.70833 | 1.636089 | 0.28686 | 0.575583 | 1.040244 | 0.845345 | 0.197572 | 0.479104 | 1.579715 | 1.843907 | 1.074796 | 0 |
| 36.72222 | 2.623871 | 0.29216 | 0.131435 | 1.365835 | 0.753121 | 0.100222 | 0.360114 | 1.553748 | 1.258282 | 1.046956 | 0 |
| 36.73611 | 2.116459 | 0.010795 | 1.210101 | 1.414815 | 0.829332 | 0.454716 | 0.193082 | 1.48023 | 0.386433 | 0.760211 | 0 |
| 36.75 | 2.163862 | 0.26038 | 2.17464 | 1.147369 | 0.864617 | 1.192283 | 0.003466 | 0.755971 | 0.310833 | 0.36835 | 0 |
| 36.76389 | 1.415862 | 0.555861 | 1.806786 | 0.791135 | 0.857488 | 1.385519 | 0.189331 | 0.522067 | 0.854203 | 0.158069 | 0 |
| 36.77778 | 0.197606 | 0.639244 | 1.057087 | 0.423004 | 0.728259 | 1.28186 | 0.34996 | 0.893066 | 1.552211 | 0.207703 | 0 |
| 36.79167 | 0.809939 | 0.629205 | 0.618454 | 0.073947 | 0.59437 | 1.226375 | 0.410652 | 1.417047 | 2.026177 | 0.424702 | 0 |
| 36.80556 | 0.798502 | 0.52487 | 0.269952 | 0.216977 | 0.479439 | 1.149629 | 0.345355 | 1.623607 | 1.76476 | 0.798726 | 0 |
| 36.81944 | 1.352864 | 0.331579 | 0.285521 | 0.049013 | 0.40598 | 0.902948 | 0.187117 | 2.014023 | 0.677538 | 1.532233 | 0 |
| 36.83333 | 0.966094 | 0.110887 | 1.17708 | 0.409743 | 0.303959 | 0.638351 | 0.00024 | 2.901847 | 0.460744 | 1.985804 | 0 |
| 36.84722 | 1.289581 | 0.747461 | 1.919055 | 0.850267 | 0.089854 | 0.325011 | 0.10443 | 2.99918 | 1.4436 | 2.273492 | 0 |
| 36.86111 | 2.322812 | 1.223032 | 1.896145 | 1.176006 | 0.234329 | 0.209244 | 0.114688 | 2.663949 | 1.744052 | 2.408761 | 0 |
| 36.875 | 2.509983 | 1.322957 | 0.66298 | 1.537628 | 0.548972 | 0.550138 | 0.126433 | 2.012382 | 1.842715 | 2.873284 | 0 |
| 36.88889 | 2.136141 | 0.803964 | 0.621497 | 1.882679 | 0.787434 | 0.821301 | 0.281484 | 1.543902 | 1.780051 | 3.4523 | 0 |
| 36.90278 | 1.029348 | 0.388389 | 0.575932 | 1.483675 | 0.924652 | 0.89143 | 0.39709 | 1.940781 | 2.003213 | 3.263259 | 0 |
| 36.91667 | 0.221671 | 0.03843 | 0.147804 | 0.874984 | 0.994666 | 0.614415 | 0.457455 | 2.149011 | 1.834583 | 2.483349 | 0 |
| 36.93056 | 0.50022 | 0.152547 | 0.178721 | 0.738255 | 1.058761 | 0.229148 | 0.310034 | 2.268171 | 1.378937 | 1.756098 | 0 |
| 36.94444 | 0.98572 | 0.487584 | 0.338355 | 0.664741 | 1.007859 | 0.189072 | 0.130262 | 2.558833 | 0.929818 | 1.171859 | 0 |
| 36.95833 | 1.365132 | 0.830538 | 0.310263 | 0.546601 | 0.701629 | 0.326558 | 0.024979 | 2.821559 | 0.593128 | 0.856933 | 0 |
| 36.97222 | 1.302942 | 1.097887 | 0.258634 | 0.499043 | 0.356406 | 0.303095 | 0.141053 | 2.694771 | 0.224229 | 0.317533 | 0 |
| 36.98611 | 1.361536 | 1.093311 | 0.220642 | 0.404903 | 0.571433 | 0.195802 | 0.157521 | 2.479572 | 0.071549 | 0.087117 | 0 |
| 37 | 0.115827 | 0.147058 | 0.012842 | 0.507257 | 1.303697 | 0.046755 | 0.162934 | 2.959611 | 0.260012 | 0.636579 | 0 |
| 37.01389 | 1.61 | 0.887216 | 0.353102 | 1.160509 | 1.570876 | 0.184917 | 0.743498 | 3.095018 | 0.477823 | 1.756434 | 0 |
| 37.02778 | 2.363844 | 1.389566 | 0.689164 | 1.243317 | 1.29708 | 0.0771 | 0.824998 | 2.429679 | 0.495557 | 2.556472 | 0 |
| 37.04167 | 1.901803 | 1.371994 | 0.766781 | 0.465674 | 1.233631 | 0.526614 | 0.831161 | 2.167225 | 0.458768 | 2.297158 | 0 |
| 37.05556 | 1.009311 | 1.202434 | 0.809652 | 0.243029 | 1.203876 | 0.754953 | 0.863393 | 2.407593 | 0.278372 | 1.678754 | 0 |
| 37.06944 | 0.468692 | 1.030319 | 0.852391 | 0.775002 | 0.922592 | 0.43881 | 0.456912 | 2.395252 | 0.087586 | 1.152616 | 0 |
| 37.08333 | 0.134526 | 0.857393 | 0.835961 | 0.743734 | 0.394075 | 0.084143 | 0.048821 | 1.998392 | 0.417024 | 0.670237 | 0 |
| 37.09722 | 0.242513 | 0.629597 | 0.673591 | 0.128932 | 0.108732 | 0.35155 | 0.062966 | 1.540199 | 0.53234 | 0.622931 | 0 |
| 37.11111 | 0.411343 | 0.499746 | 0.274926 | 0.293413 | 0.460253 | 0.528331 | 0.030287 | 1.157579 | 0.487571 | 0.589126 | 0 |
| 37.125 | 0.645433 | 0.230011 | 0.094616 | 0.32885 | 0.641239 | 0.954169 | 0.202957 | 0.472703 | 0.196928 | 0.194534 | 0 |
| 37.13889 | 1.129867 | 0.366509 | 0.598191 | 0.059866 | 0.966297 | 1.550286 | 0.697969 | 0.659433 | 0.355989 | 0.690901 | 0 |
| 37.15278 | 1.623542 | 0.928055 | 0.820541 | 0.391929 | 0.950146 | 1.749559 | 1.047413 | 1.855953 | 0.813622 | 1.47076 | 0 |
| 37.16667 | 2.162956 | 1.432388 | 0.764253 | 0.453096 | 0.628078 | 1.409822 | 1.116708 | 2.705533 | 0.909485 | 2.171726 | 0 |
| 37.18056 | 3.276209 | 2.17151 | 0.789361 | 0.503272 | 0.45899 | 1.214942 | 1.184044 | 3.214359 | 0.972488 | 2.905879 | 0 |
| 37.19444 | 4.26973 | 2.342293 | 0.851703 | 0.632208 | 0.35744 | 1.110411 | 1.198888 | 3.202031 | 0.924248 | 3.243886 | 0 |
| 37.20833 | 4.402905 | 2.341115 | 0.898335 | 0.695909 | 0.263251 | 1.253545 | 1.030247 | 3.16306 | 0.833872 | 3.222211 | 0 |
| 37.22222 | 3.929524 | 2.117343 | 0.944729 | 0.660855 | 0.233826 | 1.343984 | 0.834274 | 2.896352 | 0.895306 | 2.978522 | 0 |
| 37.23611 | 3.154801 | 1.783944 | 0.881998 | 0.456812 | 0.164771 | 1.254378 | 0.450801 | 2.40545 | 0.921182 | 2.2578 | 0 |
| 37.25 | 2.190776 | 1.444952 | 0.577097 | 0.117232 | 0.044258 | 0.981669 | 0.002309 | 1.551354 | 0.866385 | 1.252894 | 0 |
| 37.26389 | 0.923481 | 0.810769 | 0.222352 | 0.185269 | 0.11256 | 0.736726 | 0.227638 | 0.533484 | 0.796852 | 0.210517 | 0 |
| 37.27778 | 0.040865 | 0.374057 | 0.100344 | 0.322567 | 0.17984 | 0.613921 | 0.276356 | 0.194486 | 0.791498 | 0.405577 | 0 |
| 37.29167 | 0.129473 | 0.241813 | 0.116451 | 0.271886 | 0.115569 | 0.558082 | 0.236893 | 0.515541 | 0.784725 | 0.485253 | 0 |
| 37.30556 | 0.185563 | 0.220167 | 0.070669 | 0.169876 | 0.057361 | 0.519709 | 0.134224 | 0.447377 | 0.734226 | 0.291654 | 0 |
| 37.31944 | 0.425208 | 0.229607 | 0.012789 | 0.087495 | 0.004197 | 0.489624 | 0.023121 | 0.232034 | 0.679042 | 0.155445 | 0 |
| 37.33333 | 0.401604 | 0.161466 | 0.102952 | 0.14689 | 0.109856 | 0.553438 | 0.201612 | 0.100224 | 0.640719 | 0.212884 | 0 |
| 37.34722 | 0.40936 | 0.024248 | 0.267399 | 0.309499 | 0.168888 | 0.598349 | 0.244126 | 0.065701 | 0.606671 | 0.21014 | 0 |
| 37.36111 | 0.468432 | 0.146274 | 0.311783 | 0.439384 | 0.091169 | 0.523704 | 0.095367 | 0.020665 | 0.511015 | 0.193951 | 0 |
| 37.375 | 0.371895 | 0.294038 | 0.311556 | 0.469647 | 0.038998 | 0.469292 | 0.164476 | 0.002715 | 0.463003 | 0.220468 | 0 |
| 37.38889 | 0.360415 | 0.313293 | 0.369989 | 0.436026 | 0.019395 | 0.533208 | 0.375003 | 0.097907 | 0.410918 | 0.18802 | 0 |
| 37.40278 | 0.609215 | 0.198965 | 0.399546 | 0.319936 | 0.097604 | 0.516472 | 0.40656 | 0.2952 | 0.35815 | 0.055183 | 0 |

| | | | | | | | | | | | |
|---|---|---|---|---|---|---|---|---|---|---|---|
| 37.41667 | 0.83448 | 0.076817 | 0.538689 | 0.163177 | 0.253352 | 0.571486 | 0.237318 | 0.584589 | 0.330341 | 0.227533 | 0 |
| 37.43056 | 0.879887 | 0.01036 | 0.733808 | 0.00414 | 0.388501 | 0.669875 | 0.012778 | 0.813366 | 0.377718 | 0.513423 | 0 |
| 37.44444 | 0.755391 | 0.032349 | 0.831884 | 0.046016 | 0.457201 | 0.667517 | 0.026602 | 0.878727 | 0.347816 | 0.650333 | 0 |
| 37.45833 | 0.533911 | 0.106971 | 0.80267 | 0.067019 | 0.302786 | 0.59193 | 0.142706 | 0.880581 | 0.42086 | 0.483461 | 0 |
| 37.47222 | 0.21896 | 0.197228 | 0.788301 | 0.002178 | 0.121083 | 0.531785 | 0.321263 | 0.779479 | 0.480082 | 0.179774 | 0 |
| 37.48611 | 0.0502 | 0.32368 | 0.727392 | 0.183446 | 0.164554 | 0.288881 | 0.618723 | 0.802197 | 0.546728 | 0.008253 | 0 |
| 37.5 | 0.202222 | 0.268537 | 0.728077 | 0.185886 | 0.55983 | 0.00677 | 0.740545 | 1.205866 | 0.62292 | 0.266483 | 0 |
| 37.51389 | 0.842272 | 0.070132 | 0.888148 | 0.187477 | 0.912627 | 0.074818 | 0.315908 | 1.751291 | 0.800977 | 1.058644 | 0 |
| 37.52778 | 2.294302 | 0.80949 | 1.064229 | 0.784354 | 0.94873 | 0.16471 | 0.030861 | 2.445738 | 1.142938 | 2.135051 | 0 |
| 37.54167 | 4.465625 | 1.66589 | 1.277434 | 0.86187 | 0.745 | 0.423753 | 0.241247 | 3.01785 | 1.50521 | 2.846879 | 0 |
| 37.55556 | 3.049631 | 1.031909 | 1.395603 | 0.828901 | 1.026185 | 0.610655 | 0.513541 | 3.224923 | 1.901792 | 2.711903 | 0 |
| 37.56944 | 1.869108 | 0.698328 | 1.121252 | 0.414858 | 1.232546 | 0.701524 | 0.503286 | 3.035878 | 1.975623 | 2.106036 | 0 |
| 37.58333 | 0.633228 | 0.467513 | 0.846054 | 0.141406 | 1.070201 | 0.530978 | 0.208224 | 2.594433 | 1.725095 | 1.626349 | 0 |
| 37.59722 | 0.492563 | 0.11806 | 0.99106 | 0.547837 | 0.779234 | 0.37434 | 0.028466 | 2.352313 | 1.333311 | 1.110996 | 0 |
| 37.61111 | 1.24315 | 0.077549 | 1.296935 | 0.641387 | 0.548813 | 0.522419 | 0.22387 | 2.36645 | 1.068647 | 0.307329 | 0 |
| 37.625 | 1.513857 | 0.239348 | 1.145204 | 0.706265 | 0.672732 | 0.572316 | 0.343874 | 2.343554 | 0.969387 | 0.209184 | 0 |
| 37.63889 | 1.789323 | 0.364084 | 0.763885 | 0.857 | 0.738152 | 0.648664 | 0.287077 | 1.705395 | 0.993021 | 0.635567 | 0 |
| 37.65278 | 2.15753 | 0.687548 | 0.343746 | 0.922836 | 0.66775 | 0.700669 | 0.263067 | 0.322631 | 0.683764 | 1.188811 | 0 |
| 37.66667 | 1.919067 | 0.978566 | 0.901237 | 0.949047 | 0.510918 | 0.624077 | 0.502102 | 1.542733 | 0.313079 | 1.875168 | 0 |
| 37.68056 | 1.357762 | 0.860116 | 1.026573 | 0.604441 | 0.318421 | 0.52814 | 0.765155 | 3.543251 | 1.545733 | 2.615804 | 0 |
| 37.69444 | 0.744476 | 0.674768 | 0.264615 | 0.420025 | 0.193528 | 0.477977 | 0.860708 | 4.714501 | 1.773204 | 2.702355 | 0 |
| 37.70833 | 0.137234 | 0.463137 | 0.621613 | 0.284284 | 0.22585 | 0.431622 | 0.780151 | 4.503006 | 1.468242 | 2.408952 | 0 |
| 37.72222 | 0.431985 | 0.139982 | 0.720899 | 0.081106 | 0.334996 | 0.399299 | 0.728239 | 3.572688 | 1.281133 | 1.896686 | 0 |
| 37.73611 | 0.668073 | 0.780926 | 0.087369 | 0.089142 | 0.306924 | 0.440197 | 0.598023 | 2.43415 | 1.276143 | 1.12691 | 0 |
| 37.75 | 0.804092 | 0.70459 | 0.013273 | 0.091421 | 0.10268 | 0.461592 | 0.446849 | 1.569875 | 1.002454 | 0.393734 | 0 |
| 37.76389 | 0.568723 | 0.423952 | 0.18472 | 0.139046 | 0.177686 | 0.538878 | 0.289641 | 0.932179 | 0.540446 | 0.050864 | 0 |
| 37.77778 | 0.652026 | 0.09083 | 0.0488 | 0.444127 | 0.40449 | 0.679767 | 0.085284 | 0.210822 | 0.091878 | 0.204448 | 0 |
| 37.79167 | 0.350253 | 0.326253 | 0.267215 | 0.673227 | 0.446884 | 0.764995 | 0.005027 | 0.298244 | 0.037487 | 0.518644 | 0 |
| 37.80556 | 0.086054 | 0.533277 | 0.35466 | 0.733081 | 0.324348 | 0.66798 | 0.105434 | 0.2138 | 0.181094 | 0.891169 | 0 |
| 37.81944 | 0.565962 | 0.629955 | 0.281565 | 0.94097 | 0.337008 | 0.421864 | 0.386102 | 1.425584 | 0.242388 | 1.303639 | 0 |
| 37.83333 | 0.713454 | 0.631147 | 0.258344 | 1.265024 | 0.460219 | 0.005143 | 0.924825 | 2.933818 | 0.332801 | 1.859378 | 0 |
| 37.84722 | 0.918917 | 0.703707 | 0.501493 | 1.597009 | 1.030681 | 0.436526 | 1.698912 | 4.63012 | 0.489515 | 2.324792 | 0 |
| 37.86111 | 1.115547 | 0.549611 | 0.733976 | 1.851009 | 1.89142 | 0.701072 | 2.165729 | 5.695012 | 0.677552 | 2.740102 | 0 |
| 37.875 | 0.663647 | 0.434931 | 0.382561 | 1.790934 | 1.777478 | 0.620766 | 2.108089 | 5.150202 | 0.508908 | 2.929937 | 0 |
| 37.88889 | 0.390912 | 0.722626 | 0.409358 | 1.602306 | 1.45321 | 0.623036 | 2.008247 | 4.978612 | 0.141816 | 2.812897 | 0 |
| 37.90278 | 0.453468 | 1.187037 | 1.013271 | 1.501934 | 1.481341 | 0.493183 | 1.875063 | 5.010008 | 0.092116 | 2.857148 | 0 |
| 37.91667 | 1.148252 | 1.601171 | 1.150986 | 1.335454 | 1.034183 | 0.20113 | 1.436171 | 4.322173 | 0.015492 | 2.49088 | 0 |
| 37.93056 | 1.080445 | 1.408779 | 0.951175 | 1.113874 | 0.728433 | 0.073241 | 0.94432 | 3.52233 | 0.046617 | 2.00005 | 0 |
| 37.94444 | 0.174476 | 0.768405 | 0.6496 | 0.809415 | 0.391378 | 0.236204 | 0.430644 | 2.615053 | 0.190387 | 1.442804 | 0 |
| 37.95833 | 0.560237 | 0.246094 | 0.26212 | 0.354344 | 0.117512 | 0.461747 | 0.156585 | 1.623378 | 0.429278 | 0.956581 | 0 |
| 37.97222 | 0.41166 | 0.213705 | 0.217242 | 0.158914 | 0.174478 | 0.712813 | 0.817531 | 0.567999 | 0.653944 | 0.534188 | 0 |
| 37.98611 | 0.526343 | 0.284947 | 0.665334 | 0.710487 | 0.114568 | 0.86861 | 0.943128 | 0.226188 | 0.76819 | 1.004833 | 0 |

**S1c 12 hour integrated motion for control experiment**

| Day | Control Plant1 | Control Plant2 | Control Plant3 | Control Plant4 | Control Plant5 | Control Plant6 | Control Plant7 | Control Plant8 | Control Plant9 | Control_avg | Control_SEM |
|---|---|---|---|---|---|---|---|---|---|---|---|
| 18.25 | 0.305206912 | 0.395773571 | 0.158614839 | 0.268939003 | 0.347275023 | 0.294418402 | 0.351588903 | 0.258143172 | 0.199760279 | 0.286635567 | 0.025076067 |
| 18.75 | 0.106800365 | 0.144397255 | 0.185159101 | 0.181450973 | 0.250155129 | 0.175511762 | 0.19713137 | 0.149485461 | 0.17877067 | 0.17431801 | 0.013192968 |
| 19.25 | 0.156420872 | 0.143875498 | 0.130259851 | 0.123806729 | 0.127199833 | 0.153728289 | 0.252154222 | 0.110745026 | 0.12278811 | 0.146775381 | 0.014098654 |
| 19.75 | 0.196173305 | 0.131892257 | 0.089706758 | 0.214104622 | 0.116612258 | 0.193459529 | 0.277266078 | 0.154694339 | 0.089360821 | 0.162585552 | 0.020975598 |
| 20.25 | 0.11415598 | 0.147125918 | 0.150938348 | 0.091234554 | 0.158229462 | 0.184234103 | 0.130199418 | 0.126024334 | 0.139020876 | 0.137906999 | 0.008942937 |
| 20.75 | 0.252305126 | 0.103466111 | 0.075077014 | 0.086282724 | 0.064684565 | 0.158608645 | 0.283097523 | 0.150846307 | 0.083362508 | 0.139747836 | 0.026580114 |
| 21.25 | 0.055527701 | 0.049337632 | 0.094249357 | 0.128095153 | 0.069212808 | 0.086417682 | 0.130751355 | 0.067528589 | 0.093477369 | 0.086066405 | 0.009714305 |
| 21.75 | 0.248337741 | 0.073138047 | 0.130721906 | 0.137530084 | 0.06303762 | 0.122171677 | 0.353420948 | 0.27063406 | 0.112868975 | 0.167984562 | 0.0330913 |
| 22.25 | 0.081011443 | 0.073008753 | 0.084819114 | 0.173310304 | 0.084876465 | 0.081774557 | 0.128415153 | 0.110056344 | 0.095616162 | 0.101432033 | 0.010648522 |
| 22.75 | 0.200193997 | 0.248926517 | 0.113519889 | 0.175686465 | 0.281841329 | 0.282761042 | 0.415927739 | 0.387860133 | 0.12173155 | 0.247605407 | 0.035679298 |
| 23.25 | 0.153210852 | 0.162521341 | 0.129917912 | 0.12365609 | 0.174164052 | 0.238550478 | 0.1828323 | 0.198440399 | 0.130741599 | 0.166003891 | 0.012460867 |
| 23.75 | 0.229773887 | 0.311341235 | 0.154517494 | 0.172205557 | 0.404777872 | 0.112883459 | 0.314085915 | 0.425860793 | 0.181733502 | 0.256353301 | 0.037533119 |
| 24.25 | 0.165574074 | 0.226247923 | 0.145360923 | 0.246668853 | 0.128132195 | 0.188307279 | 0.226130781 | 0.15878599 | 0.117765461 | 0.178108164 | 0.015442487 |
| 24.75 | 0.295271242 | 0.370380272 | 0.231558242 | 0.235314783 | 0.427234607 | 0.195948102 | 0.472285511 | 0.606576296 | 0.217201364 | 0.339085602 | 0.046791116 |
| 25.25 | 0.268819312 | 0.183053447 | 0.18612012 | 0.246873803 | 0.213322611 | 0.146318819 | 0.151297881 | 0.17941643 | 0.160965576 | 0.192909778 | 0.014112186 |
| 25.75 | 0.248346026 | 0.426741847 | 0.348296254 | 0.185813894 | 0.369072474 | 0.181598008 | 0.504969338 | 0.586524068 | 0.241711201 | 0.34367479 | 0.047614093 |
| 26.25 | 0.446532713 | 0.255012616 | 0.161532378 | 0.210996368 | 0.109269333 | 0.16392543 | 0.147417905 | 0.271468617 | 0.141561149 | 0.211968501 | 0.03434656 |
| 26.75 | 0.266448729 | 0.455750692 | 0.443701689 | 0.295518066 | 0.170877253 | 0.178782693 | 0.382698118 | 0.625809208 | 0.323946059 | 0.34928139 | 0.048466294 |
| 27.25 | 0.270012922 | 0.226604223 | 0.209600057 | 0.175692614 | 0.102080526 | 0.256532911 | 0.120387743 | 0.175155115 | 0.122341506 | 0.184267513 | 0.020356346 |
| 27.75 | 0.341423756 | 0.406973187 | 0.410107265 | 0.293876833 | 0.213356116 | 0.305299341 | 0.356880759 | 0.453885996 | 0.349863909 | 0.347963018 | 0.024023776 |
| 28.25 | 0.331536439 | 0.34868431 | 0.272928327 | 0.218339857 | 0.119352893 | 0.424187494 | 0.071152655 | 0.258731617 | 0.221753306 | 0.251851878 | 0.036922297 |
| 28.75 | 0.547573368 | 0.59935118 | 0.605068007 | 0.549514068 | 0.181671582 | 0.559667247 | 0.151332977 | 0.558471547 | 0.578527206 | 0.481241909 | 0.059919543 |
| 29.25 | 0.259453703 | 0.561776217 | 0.437935388 | 0.360730679 | 0.367242681 | 0.747511698 | 0.241233181 | 0.291311602 | 0.502291262 | 0.418831823 | 0.054649799 |
| 29.75 | 0.593512609 | 0.646744502 | 0.495083415 | 0.783862692 | 0.19604797 | 0.455130481 | 0.282919308 | 0.351982447 | 0.610946782 | 0.490692245 | 0.063008329 |
| 30.25 | 0.160373154 | 0.402632438 | 0.261063506 | 0.27494863 | 0.336127428 | 0.426750536 | 0.285249121 | 0.209898257 | 0.355830559 | 0.301430403 | 0.029101865 |
| 30.75 | 0.449470763 | 0.951608752 | 0.592581025 | 1.031727896 | 0.298492567 | 0.524646474 | 0.328769798 | 0.404532808 | 0.71769848 | 0.588836507 | 0.087703829 |
| 31.25 | 0.497998205 | 0.393835201 | 0.492933012 | 0.219273347 | 0.464657154 | 0.548993149 | 0.238491062 | 0.189845453 | 0.398001028 | 0.382669735 | 0.044856384 |
| 31.75 | 0.400414015 | 0.939057232 | 0.693057278 | 1.068972541 | 0.53803546 | 0.85347414 | 0.244541895 | 0.378338038 | 0.819289354 | 0.659464439 | 0.094518234 |
| 32.25 | 0.620832742 | 0.353541481 | 0.615518092 | 0.126260064 | 0.318206618 | 0.824087644 | 0.195948854 | 0.190664607 | 0.414127047 | 0.40657635 | 0.078538393 |
| 32.75 | 0.388134181 | 0.920762152 | 0.614111816 | 1.11675571 | 0.375977837 | 0.887618582 | 0.22006895 | 0.287479064 | 0.554461339 | 0.596152181 | 0.104872868 |
| 33.25 | 0.510148721 | 0.244352334 | 0.68946101 | 0.146752301 | 0.158228596 | 0.394012468 | 0.170998144 | 0.14921227 | 0.276331483 | 0.304388592 | 0.063587935 |
| 33.75 | 0.369030376 | 0.763857921 | 0.576962055 | 0.84188286 | 0.477093929 | 1.044219197 | 0.204772903 | 0.286833744 | 0.518418597 | 0.564785731 | 0.091396855 |
| 34.25 | 0.588199893 | 0.331647885 | 0.404383824 | 0.218331526 | 0.257545955 | 0.441873839 | 0.175301398 | 0.251927689 | 0.462768785 | 0.347997866 | 0.045212036 |
| 34.75 | 0.420903505 | 0.216670107 | 0.58383427 | 0.395025922 | 0.366556625 | 0.686236931 | 0.191972722 | 0.275223242 | 0.557596677 | 0.410446667 | 0.056961943 |
| 35.25 | 0.594958507 | 0.616304032 | 0.399596645 | 0.250875273 | 0.191050602 | 0.293792439 | 0.167827672 | 0.404083025 | 0.418387812 | 0.370764001 | 0.053818184 |
| 35.75 | 0.707103742 | 0.406045802 | 0.445687266 | 0.503907657 | 0.335699326 | 0.439730493 | 0.102982875 | 0.959790488 | 0.513324962 | 0.490474735 | 0.079381506 |
| 36.25 | 0.672882781 | 0.443870533 | 0.316931968 | 0.141006339 | 0.152405592 | 0.329137533 | 0.151893146 | 0.54774806 | 0.372220015 | 0.347566219 | 0.061925879 |
| 36.75 | 0.756714323 | 0.268323492 | 0.52900146 | 0.331910212 | 0.422746531 | 0.401516086 | 0.126530663 | 1.091160529 | 0.674586309 | 0.511387734 | 0.097368291 |
| 37.25 | 0.59109622 | 0.370835771 | 0.261495704 | 0.18249994 | 0.210443005 | 0.348362762 | 0.210340518 | 0.661668666 | 0.278632667 | 0.346152806 | 0.057195067 |
| 37.75 | 0.508772128 | 0.299724408 | 0.321290558 | 0.363507104 | 0.331989379 | 0.232830347 | 0.332223455 | 1.276194727 | 0.390653145 | 0.450798361 | 0.106141845 |

**S1d Day-to-day motion rate for control experiment**

| Day | Control Plant1 | Control Plant2 | Control Plant3 | Control Plant4 | Control Plant5 | Control Plant6 | Control Plant7 | Control Plant8 | Control Plant9 | Control_avg | Control_SEM |
|---|---|---|---|---|---|---|---|---|---|---|---|
| 18.5 | 0.354261031 | 0.278042484 | 0.222354081 | 0.340091525 | 0.246253209 | 0.350988929 | 0.533695537 | 0.268137455 | 0.214345202 | 0.312018828 | 0.093224162 |
| 19.5 | 0.370417544 | 0.254140683 | 0.23112862 | 0.182657333 | 0.231750445 | 0.34775738 | 0.415883635 | 0.280561634 | 0.22572344 | 0.282224524 | 0.073781367 |
| 20.5 | 0.307916386 | 0.122701513 | 0.227435374 | 0.272769786 | 0.138249643 | 0.222006561 | 0.488386394 | 0.343228941 | 0.207340637 | 0.258892804 | 0.105665302 |
| 21.5 | 0.290300087 | 0.329427591 | 0.201029003 | 0.3518727 | 0.367243823 | 0.370246294 | 0.554241559 | 0.504848666 | 0.220115948 | 0.354369519 | 0.11019235 |
| 22.5 | 0.385520836 | 0.479537681 | 0.286484282 | 0.29707439 | 0.582962431 | 0.36267816 | 0.508354049 | 0.626049053 | 0.313485247 | 0.426905125 | 0.119578967 |
| 23.5 | 0.479415387 | 0.622248961 | 0.387861645 | 0.495349498 | 0.571457686 | 0.398314855 | 0.722347523 | 0.784638433 | 0.349908993 | 0.534615887 | 0.143608911 |
| 24.5 | 0.524391866 | 0.624553233 | 0.543301438 | 0.43843035 | 0.589369412 | 0.331433633 | 0.673117609 | 0.783399395 | 0.417146959 | 0.547238211 | 0.131336012 |
| 25.5 | 0.714162734 | 0.717016107 | 0.609711595 | 0.514668629 | 0.291791281 | 0.346432075 | 0.544941337 | 0.899537082 | 0.478922667 | 0.568575945 | 0.180097925 |
| 26.5 | 0.62487895 | 0.636637606 | 0.6210471 | 0.486690858 | 0.328317504 | 0.572756106 | 0.489896719 | 0.638997139 | 0.48517234 | 0.54271048 | 0.098368313 |
| 27.5 | 0.891934801 | 0.952583296 | 0.892409905 | 0.794300233 | 0.305092753 | 0.989019985 | 0.225106272 | 0.826904873 | 0.804701666 | 0.74245042 | 0.262976272 |
| 28.5 | 0.866340084 | 1.215109844 | 0.946860624 | 1.169296936 | 0.566520957 | 1.21652066 | 0.525567823 | 0.657999509 | 1.115722405 | 0.919993205 | 0.264205277 |
| 29.5 | 0.622157627 | 1.380859991 | 0.865871574 | 1.315093068 | 0.639965808 | 0.961597407 | 0.616966399 | 0.621073474 | 1.091707554 | 0.901699211 | 0.288277561 |
| 30.5 | 0.919646187 | 1.354494142 | 1.207424048 | 1.332483163 | 1.007990288 | 1.415056915 | 0.487231013 | 0.57993118 | 1.247289989 | 1.061282992 | 0.320549953 |
| 31.5 | 1.019256861 | 1.28980319 | 1.250817152 | 1.275896769 | 0.695844004 | 1.723540415 | 0.417673499 | 0.486133304 | 0.98660356 | 1.016174306 | 0.400054818 |
| 32.5 | 0.898060224 | 1.015810699 | 1.279911531 | 0.992039467 | 0.645043123 | 1.479614321 | 0.377975673 | 0.442170203 | 0.806731657 | 0.881928544 | 0.342385909 |
| 33.5 | 1.013478406 | 0.550398631 | 1.00899256 | 0.613450879 | 0.634540095 | 1.157844551 | 0.36962503 | 0.528773473 | 1.030441109 | 0.76750497 | 0.267533642 |
| 34.5 | 1.310087694 | 1.027734758 | 0.858682364 | 0.769853203 | 0.53108963 | 0.754727627 | 0.273560797 | 1.375820042 | 0.932632073 | 0.870465354 | 0.328697315 |
| 35.5 | 1.443923321 | 0.713199789 | 0.857875263 | 0.478053385 | 0.576516414 | 0.736973641 | 0.287098191 | 1.660779257 | 1.054218748 | 0.867626445 | 0.423026696 |
| 36.5 | 1.101621279 | 0.674672797 | 0.592893687 | 0.548571849 | 0.547462828 | 0.583246242 | 0.552003335 | 1.951808275 | 0.677408368 | 0.80329874 | 0.438006098 |

**S1e 1 hour integrated motion for control experiment**

| IntervalMid | Control1 | Control2 | Control3 | Control4 | Control5 | Control6 | Control7 | Control8 | Control9 |
|---|---|---|---|---|---|---|---|---|---|
| 18.01388889 | 0.047205 | 0.044148 | 0.006314 | 0.006792 | 0.034329 | 0.003874 | 0.105912 | 0.010314 | 0.019421 |
| 18.02777778 | 0.058611 | 0.056889 | 0.012386 | 0.008116 | 0.052933 | 0.007093 | 0.084439 | 0.014114 | 0.026171 |
| 18.04166667 | 0.037157 | 0.040094 | 0.014749 | 0.0121 | 0.037336 | 0.010373 | 0.040689 | 0.011139 | 0.01748 |
| 18.05555556 | 0.01654 | 0.035798 | 0.010602 | 0.032521 | 0.015578 | 0.008486 | 0.019612 | 0.013256 | 0.007041 |
| 18.06944444 | 0.015979 | 0.056482 | 0.005497 | 0.063218 | 0.014876 | 0.004363 | 0.031525 | 0.021365 | 0.002646 |
| 18.08333333 | 0.014106 | 0.05387 | 0.004104 | 0.065056 | 0.014516 | 0.003273 | 0.044104 | 0.018145 | 0.002622 |
| 18.09722222 | 0.010243 | 0.026132 | 0.007326 | 0.034178 | 0.007606 | 0.0047 | 0.036823 | 0.008742 | 0.005318 |
| 18.11111111 | 0.01298 | 0.011088 | 0.010032 | 0.010022 | 0.007549 | 0.008502 | 0.022163 | 0.009535 | 0.00859 |
| 18.125 | 0.015981 | 0.010947 | 0.009928 | 0.005899 | 0.010764 | 0.012813 | 0.010272 | 0.013499 | 0.011773 |
| 18.13888889 | 0.016446 | 0.011908 | 0.00895 | 0.007275 | 0.011652 | 0.014995 | 0.005844 | 0.013536 | 0.01408 |
| 18.15277778 | 0.01531 | 0.010788 | 0.007425 | 0.007814 | 0.010826 | 0.014896 | 0.008479 | 0.011996 | 0.013718 |
| 18.16666667 | 0.014176 | 0.00964 | 0.006277 | 0.008138 | 0.009477 | 0.014421 | 0.012293 | 0.010882 | 0.011616 |
| 18.18055556 | 0.014539 | 0.01046 | 0.007018 | 0.009212 | 0.009676 | 0.016706 | 0.014523 | 0.011949 | 0.010659 |
| 18.19444444 | 0.015201 | 0.01239 | 0.008457 | 0.010496 | 0.010681 | 0.022414 | 0.014874 | 0.014302 | 0.011024 |
| 18.20833333 | 0.014504 | 0.013078 | 0.008475 | 0.010748 | 0.010098 | 0.030228 | 0.012646 | 0.016837 | 0.01076 |
| 18.22222222 | 0.013608 | 0.012694 | 0.007748 | 0.011176 | 0.00808 | 0.036391 | 0.010072 | 0.021071 | 0.010237 |
| 18.23611111 | 0.014876 | 0.013702 | 0.008638 | 0.013648 | 0.006012 | 0.037014 | 0.010514 | 0.027618 | 0.011566 |
| 18.25 | 0.018121 | 0.016618 | 0.011539 | 0.016811 | 0.003861 | 0.033919 | 0.013713 | 0.032402 | 0.0147 |
| 18.26388889 | 0.020967 | 0.019364 | 0.01456 | 0.018196 | 0.002193 | 0.031038 | 0.01657 | 0.031865 | 0.017462 |
| 18.27777778 | 0.022258 | 0.021019 | 0.01627 | 0.018333 | 0.002143 | 0.029384 | 0.017708 | 0.028205 | 0.018638 |
| 18.29166667 | 0.022835 | 0.022466 | 0.016848 | 0.019153 | 0.003831 | 0.028469 | 0.017914 | 0.02512 | 0.018918 |
| 18.30555556 | 0.022996 | 0.023609 | 0.01671 | 0.020196 | 0.006501 | 0.027377 | 0.017463 | 0.023177 | 0.018671 |
| 18.31944444 | 0.021387 | 0.022828 | 0.015352 | 0.019022 | 0.008758 | 0.024584 | 0.01514 | 0.020721 | 0.016972 |
| 18.33333333 | 0.017363 | 0.019557 | 0.012207 | 0.014838 | 0.010702 | 0.019721 | 0.010309 | 0.016942 | 0.013368 |
| 18.34722222 | 0.012176 | 0.015421 | 0.007811 | 0.009417 | 0.013816 | 0.014317 | 0.00483 | 0.012671 | 0.00876 |
| 18.36111111 | 0.007237 | 0.012406 | 0.003679 | 0.004495 | 0.019457 | 0.009555 | 0.001823 | 0.008683 | 0.004327 |
| 18.375 | 0.003577 | 0.011762 | 0.00167 | 0.001467 | 0.026872 | 0.006272 | 0.002202 | 0.005231 | 0.001439 |
| 18.38888889 | 0.002372 | 0.014299 | 0.001255 | 0.001002 | 0.033085 | 0.005758 | 0.002759 | 0.003094 | 0.000859 |
| 18.40277778 | 0.003957 | 0.019103 | 0.001438 | 0.002583 | 0.037462 | 0.008068 | 0.002254 | 0.003242 | 0.002229 |
| 18.41666667 | 0.006873 | 0.02298 | 0.003271 | 0.005668 | 0.040847 | 0.011391 | 0.003199 | 0.00513 | 0.004994 |
| 18.43055556 | 0.009232 | 0.023559 | 0.005641 | 0.008758 | 0.042809 | 0.014063 | 0.005261 | 0.007137 | 0.007746 |
| 18.44444444 | 0.01097 | 0.021875 | 0.007515 | 0.011198 | 0.044031 | 0.016208 | 0.007175 | 0.008795 | 0.009889 |
| 18.45833333 | 0.013741 | 0.021309 | 0.01044 | 0.014441 | 0.045654 | 0.019074 | 0.009836 | 0.011045 | 0.012727 |
| 18.47222222 | 0.019362 | 0.025025 | 0.015678 | 0.020535 | 0.047649 | 0.023994 | 0.011955 | 0.013891 | 0.017429 |
| 18.48611111 | 0.028427 | 0.034025 | 0.023232 | 0.029893 | 0.050508 | 0.031611 | 0.010694 | 0.015764 | 0.023869 |
| 18.5 | 0.036803 | 0.043612 | 0.030263 | 0.035125 | 0.052475 | 0.038383 | 0.008416 | 0.016794 | 0.029096 |
| 18.51388889 | 0.036414 | 0.044188 | 0.031328 | 0.025152 | 0.048004 | 0.036977 | 0.009597 | 0.017353 | 0.028226 |
| 18.52777778 | 0.025369 | 0.032646 | 0.023847 | 0.012208 | 0.035867 | 0.025857 | 0.011365 | 0.015128 | 0.020066 |
| 18.54166667 | 0.011292 | 0.017043 | 0.012193 | 0.012753 | 0.022361 | 0.012265 | 0.008771 | 0.009555 | 0.009462 |
| 18.55555556 | 0.003272 | 0.006064 | 0.003681 | 0.014209 | 0.013212 | 0.003741 | 0.004013 | 0.004077 | 0.002771 |
| 18.56944444 | 0.002859 | 0.001613 | 0.000769 | 0.00982 | 0.008626 | 0.001503 | 0.001823 | 0.002068 | 0.00131 |
| 18.58333333 | 0.00431 | 0.000923 | 0.00042 | 0.006024 | 0.00656 | 0.001599 | 0.002272 | 0.002432 | 0.00124 |

| | | | | | | | | | |
|---|---|---|---|---|---|---|---|---|---|
| 18.59722222 | 0.004533 | 0.000719 | 0.000419 | 0.002976 | 0.006255 | 0.001404 | 0.002334 | 0.001986 | 0.000927 |
| 18.61111111 | 0.004365 | 0.000405 | 0.000914 | 0.001298 | 0.007243 | 0.001957 | 0.001601 | 0.001104 | 0.001652 |
| 18.625 | 0.003844 | 0.000967 | 0.002091 | 0.002388 | 0.008612 | 0.00376 | 0.002296 | 0.001712 | 0.003324 |
| 18.63888889 | 0.00291 | 0.002112 | 0.003549 | 0.004692 | 0.009687 | 0.00516 | 0.004774 | 0.003152 | 0.004863 |
| 18.65277778 | 0.002698 | 0.0026 | 0.004509 | 0.005938 | 0.009918 | 0.005306 | 0.007004 | 0.004173 | 0.005512 |
| 18.66666667 | 0.002693 | 0.002992 | 0.00539 | 0.006489 | 0.010106 | 0.005609 | 0.008355 | 0.005308 | 0.006068 |
| 18.68055556 | 0.002819 | 0.005533 | 0.00759 | 0.008455 | 0.012187 | 0.007974 | 0.010547 | 0.007796 | 0.008449 |
| 18.69444444 | 0.005566 | 0.010179 | 0.011214 | 0.012653 | 0.016523 | 0.012522 | 0.014517 | 0.011824 | 0.012898 |
| 18.70833333 | 0.008167 | 0.013235 | 0.014062 | 0.016435 | 0.020149 | 0.016213 | 0.017932 | 0.015034 | 0.016462 |
| 18.72222222 | 0.006342 | 0.011999 | 0.014212 | 0.016787 | 0.020086 | 0.016304 | 0.018098 | 0.01507 | 0.016333 |
| 18.73611111 | 0.003199 | 0.00939 | 0.013806 | 0.015561 | 0.018518 | 0.015112 | 0.016954 | 0.014049 | 0.01481 |
| 18.75 | 0.002693 | 0.009048 | 0.015213 | 0.015854 | 0.018672 | 0.015364 | 0.017309 | 0.014703 | 0.015172 |
| 18.76388889 | 0.003415 | 0.009714 | 0.016667 | 0.016676 | 0.019248 | 0.015582 | 0.017917 | 0.015665 | 0.016216 |
| 18.77777778 | 0.003051 | 0.009292 | 0.016221 | 0.016261 | 0.018443 | 0.01445 | 0.017255 | 0.015154 | 0.015905 |
| 18.79166667 | 0.0018 | 0.008128 | 0.014479 | 0.014915 | 0.016789 | 0.012906 | 0.015981 | 0.013648 | 0.014408 |
| 18.80555556 | 0.001048 | 0.007313 | 0.013066 | 0.013517 | 0.015278 | 0.011935 | 0.01482 | 0.012279 | 0.012917 |
| 18.81944444 | 0.001633 | 0.007421 | 0.013095 | 0.012702 | 0.014717 | 0.011735 | 0.014131 | 0.011754 | 0.01244 |
| 18.83333333 | 0.003028 | 0.008113 | 0.014232 | 0.012577 | 0.015036 | 0.011818 | 0.01433 | 0.012013 | 0.012871 |
| 18.84722222 | 0.003745 | 0.008365 | 0.01502 | 0.012221 | 0.014761 | 0.011331 | 0.014991 | 0.011891 | 0.012997 |
| 18.86111111 | 0.003006 | 0.00755 | 0.014602 | 0.010969 | 0.013115 | 0.010051 | 0.01515 | 0.010578 | 0.012145 |
| 18.875 | 0.001806 | 0.006293 | 0.013651 | 0.009614 | 0.011382 | 0.008757 | 0.014885 | 0.00902 | 0.011197 |
| 18.88888889 | 0.001065 | 0.005095 | 0.01272 | 0.008661 | 0.010308 | 0.007761 | 0.014745 | 0.007941 | 0.010688 |
| 18.90277778 | 0.001352 | 0.003707 | 0.011561 | 0.007713 | 0.009195 | 0.006685 | 0.014566 | 0.006955 | 0.010024 |
| 18.91666667 | 0.003199 | 0.002062 | 0.009929 | 0.006482 | 0.007645 | 0.005448 | 0.013808 | 0.005779 | 0.008855 |
| 18.93055556 | 0.005571 | 0.000988 | 0.007782 | 0.00485 | 0.005655 | 0.003855 | 0.012141 | 0.004337 | 0.00723 |
| 18.94444444 | 0.006957 | 0.001154 | 0.005782 | 0.003183 | 0.003616 | 0.001978 | 0.010124 | 0.002832 | 0.005658 |
| 18.95833333 | 0.007185 | 0.001746 | 0.004599 | 0.002118 | 0.001858 | 0.001214 | 0.009021 | 0.001526 | 0.004597 |
| 18.97222222 | 0.008008 | 0.003734 | 0.003652 | 0.002062 | 0.002434 | 0.003622 | 0.010671 | 0.001429 | 0.003277 |
| 18.98611111 | 0.010681 | 0.009748 | 0.002174 | 0.003978 | 0.009103 | 0.010423 | 0.02181 | 0.004271 | 0.0041 |
| 19 | 0.009733 | 0.015759 | 0.002053 | 0.009577 | 0.018224 | 0.017597 | 0.045998 | 0.008088 | 0.009485 |
| 19.01388889 | 0.009436 | 0.013389 | 0.005699 | 0.015924 | 0.019364 | 0.016405 | 0.05609 | 0.008294 | 0.012896 |
| 19.02777778 | 0.018822 | 0.008504 | 0.011332 | 0.014457 | 0.010995 | 0.008921 | 0.039306 | 0.005569 | 0.009902 |
| 19.04166667 | 0.025278 | 0.01049 | 0.014534 | 0.006986 | 0.003396 | 0.005384 | 0.033141 | 0.003713 | 0.005304 |
| 19.05555556 | 0.02173 | 0.013079 | 0.012832 | 0.003053 | 0.00214 | 0.00554 | 0.042087 | 0.002742 | 0.002692 |
| 19.06944444 | 0.016053 | 0.011802 | 0.008076 | 0.002118 | 0.003193 | 0.005062 | 0.03721 | 0.001463 | 0.001403 |
| 19.08333333 | 0.011248 | 0.009555 | 0.003696 | 0.001496 | 0.003866 | 0.004026 | 0.021076 | 0.000658 | 0.000599 |
| 19.09722222 | 0.00787 | 0.007739 | 0.001241 | 0.001522 | 0.004222 | 0.002844 | 0.009171 | 0.000698 | 0.000286 |
| 19.11111111 | 0.005818 | 0.006395 | 0.000707 | 0.001212 | 0.003869 | 0.001751 | 0.004921 | 0.000745 | 0.000683 |
| 19.125 | 0.004043 | 0.00476 | 0.001463 | 0.001213 | 0.002654 | 0.001849 | 0.004949 | 0.000969 | 0.001939 |
| 19.13888889 | 0.002292 | 0.003139 | 0.002366 | 0.002103 | 0.001251 | 0.003119 | 0.005507 | 0.001509 | 0.003261 |
| 19.15277778 | 0.001456 | 0.002697 | 0.002271 | 0.002266 | 0.000878 | 0.003707 | 0.004818 | 0.001318 | 0.003369 |
| 19.16666667 | 0.001609 | 0.003027 | 0.001427 | 0.001204 | 0.001587 | 0.003352 | 0.003412 | 0.000757 | 0.002501 |
| 19.18055556 | 0.00139 | 0.00253 | 0.001221 | 0.000633 | 0.001956 | 0.003448 | 0.002748 | 0.000617 | 0.002226 |
| 19.19444444 | 0.001302 | 0.001659 | 0.002473 | 0.001856 | 0.001567 | 0.004608 | 0.003717 | 0.001185 | 0.003483 |

| | | | | | | | | | |
|---|---|---|---|---|---|---|---|---|---|
| 19.20833333 | 0.003132 | 0.002317 | 0.004839 | 0.004181 | 0.001833 | 0.006563 | 0.005913 | 0.003072 | 0.005767 |
| 19.22222222 | 0.005177 | 0.003604 | 0.006684 | 0.005478 | 0.002595 | 0.008116 | 0.007511 | 0.004748 | 0.007242 |
| 19.23611111 | 0.00423 | 0.003298 | 0.005593 | 0.004092 | 0.002562 | 0.007339 | 0.006525 | 0.003832 | 0.005773 |
| 19.25 | 0.002579 | 0.003118 | 0.003188 | 0.002419 | 0.003183 | 0.004079 | 0.003385 | 0.002473 | 0.003104 |
| 19.26388889 | 0.003342 | 0.004359 | 0.002759 | 0.002867 | 0.004579 | 0.001133 | 0.000851 | 0.002975 | 0.002374 |
| 19.27777778 | 0.003426 | 0.004291 | 0.002303 | 0.003083 | 0.00386 | 0.000738 | 0.000446 | 0.002597 | 0.00191 |
| 19.29166667 | 0.001766 | 0.002783 | 0.001004 | 0.002586 | 0.001596 | 0.001821 | 0.001262 | 0.001029 | 0.000871 |
| 19.30555556 | 0.001118 | 0.002235 | 0.000572 | 0.003225 | 0.000885 | 0.002184 | 0.00183 | 0.000548 | 0.000617 |
| 19.31944444 | 0.002548 | 0.003665 | 0.001197 | 0.005419 | 0.00246 | 0.001683 | 0.001494 | 0.00144 | 0.001326 |
| 19.33333333 | 0.005751 | 0.006903 | 0.003584 | 0.009524 | 0.004812 | 0.001712 | 0.001359 | 0.003627 | 0.003526 |
| 19.34722222 | 0.009065 | 0.010332 | 0.006551 | 0.014811 | 0.005742 | 0.001749 | 0.001947 | 0.005831 | 0.005959 |
| 19.36111111 | 0.009262 | 0.010767 | 0.006378 | 0.017291 | 0.004801 | 0.002909 | 0.002877 | 0.005083 | 0.005317 |
| 19.375 | 0.007121 | 0.007974 | 0.005638 | 0.012419 | 0.006793 | 0.009128 | 0.007223 | 0.004817 | 0.005031 |
| 19.38888889 | 0.00909 | 0.008311 | 0.010799 | 0.006791 | 0.01469 | 0.018673 | 0.016042 | 0.011349 | 0.01131 |
| 19.40277778 | 0.015561 | 0.014077 | 0.018576 | 0.010029 | 0.022588 | 0.026125 | 0.02406 | 0.020042 | 0.019632 |
| 19.41666667 | 0.020747 | 0.019513 | 0.023098 | 0.016692 | 0.02567 | 0.029716 | 0.027754 | 0.024896 | 0.024208 |
| 19.43055556 | 0.022459 | 0.02146 | 0.024298 | 0.019386 | 0.02457 | 0.030006 | 0.027601 | 0.025945 | 0.025106 |
| 19.44444444 | 0.020918 | 0.020052 | 0.022811 | 0.017996 | 0.020686 | 0.02734 | 0.024601 | 0.023524 | 0.022698 |
| 19.45833333 | 0.01696 | 0.016101 | 0.019304 | 0.014152 | 0.015515 | 0.022536 | 0.020241 | 0.018927 | 0.018228 |
| 19.47222222 | 0.012014 | 0.010965 | 0.014596 | 0.009162 | 0.009704 | 0.016839 | 0.015895 | 0.013952 | 0.013172 |
| 19.48611111 | 0.006378 | 0.006377 | 0.008381 | 0.005383 | 0.00574 | 0.010747 | 0.011185 | 0.008461 | 0.007494 |
| 19.5 | 0.00197 | 0.005402 | 0.004881 | 0.005799 | 0.00736 | 0.00535 | 0.006096 | 0.004586 | 0.004702 |
| 19.51388889 | 0.002388 | 0.005825 | 0.007016 | 0.006338 | 0.010394 | 0.005711 | 0.00254 | 0.004675 | 0.006757 |
| 19.52777778 | 0.008315 | 0.005332 | 0.007098 | 0.004956 | 0.00777 | 0.018542 | 0.002541 | 0.004719 | 0.006544 |
| 19.54166667 | 0.016568 | 0.008339 | 0.003957 | 0.006902 | 0.003814 | 0.03958 | 0.006129 | 0.004843 | 0.003549 |
| 19.55555556 | 0.02314 | 0.013918 | 0.004765 | 0.010836 | 0.005574 | 0.051552 | 0.0112 | 0.008654 | 0.004148 |
| 19.56944444 | 0.027073 | 0.017434 | 0.009519 | 0.012466 | 0.009809 | 0.046408 | 0.01775 | 0.013274 | 0.007393 |
| 19.58333333 | 0.028567 | 0.017689 | 0.01393 | 0.011945 | 0.011693 | 0.03342 | 0.026411 | 0.015777 | 0.009616 |
| 19.59722222 | 0.028011 | 0.015866 | 0.01596 | 0.010934 | 0.01157 | 0.022404 | 0.034822 | 0.01717 | 0.0107 |
| 19.61111111 | 0.02706 | 0.014522 | 0.016258 | 0.010882 | 0.01178 | 0.017113 | 0.039845 | 0.018804 | 0.012122 |
| 19.625 | 0.025681 | 0.01365 | 0.015033 | 0.011 | 0.012386 | 0.016148 | 0.039954 | 0.019431 | 0.013007 |
| 19.63888889 | 0.023005 | 0.011742 | 0.011013 | 0.009607 | 0.011221 | 0.014703 | 0.035495 | 0.017014 | 0.011203 |
| 19.65277778 | 0.019335 | 0.009113 | 0.00558 | 0.006997 | 0.008202 | 0.010658 | 0.027179 | 0.012181 | 0.007197 |
| 19.66666667 | 0.015441 | 0.007138 | 0.002937 | 0.004855 | 0.00518 | 0.005594 | 0.017237 | 0.007586 | 0.003286 |
| 19.68055556 | 0.012833 | 0.00685 | 0.003295 | 0.004575 | 0.004079 | 0.002 | 0.009079 | 0.005579 | 0.001407 |
| 19.69444444 | 0.012787 | 0.008141 | 0.002971 | 0.006246 | 0.005695 | 0.001185 | 0.003802 | 0.00626 | 0.002258 |
| 19.70833333 | 0.013725 | 0.009302 | 0.001624 | 0.008256 | 0.008059 | 0.00161 | 0.002309 | 0.007571 | 0.004339 |
| 19.72222222 | 0.012184 | 0.00827 | 0.00079 | 0.008349 | 0.007695 | 0.00173 | 0.005542 | 0.007004 | 0.004747 |
| 19.73611111 | 0.007462 | 0.005066 | 0.000643 | 0.006915 | 0.004298 | 0.002011 | 0.009996 | 0.00423 | 0.002858 |
| 19.75 | 0.00314 | 0.00268 | 0.000945 | 0.01015 | 0.00158 | 0.002806 | 0.010832 | 0.001544 | 0.001326 |
| 19.76388889 | 0.00214 | 0.003392 | 0.001445 | 0.023385 | 0.002175 | 0.005072 | 0.007703 | 0.001233 | 0.001984 |
| 19.77777778 | 0.002569 | 0.004704 | 0.001858 | 0.037256 | 0.004021 | 0.007134 | 0.004973 | 0.002356 | 0.002888 |
| 19.79166667 | 0.002038 | 0.003469 | 0.002841 | 0.035813 | 0.003697 | 0.00516 | 0.00599 | 0.002563 | 0.002494 |
| 19.80555556 | 0.001879 | 0.001656 | 0.005023 | 0.02134 | 0.001687 | 0.002282 | 0.00996 | 0.003182 | 0.002985 |

| | | | | | | | | | |
|---|---|---|---|---|---|---|---|---|---|
| 19.81944444 | 0.002844 | 0.001158 | 0.006167 | 0.009192 | 0.000493 | 0.002837 | 0.014999 | 0.004718 | 0.004504 |
| 19.83333333 | 0.003268 | 0.001208 | 0.005459 | 0.004647 | 0.000589 | 0.004251 | 0.01935 | 0.004812 | 0.004613 |
| 19.84722222 | 0.002685 | 0.002189 | 0.004743 | 0.003218 | 0.001168 | 0.003757 | 0.021116 | 0.003444 | 0.003327 |
| 19.86111111 | 0.002073 | 0.004196 | 0.003841 | 0.002953 | 0.002616 | 0.003185 | 0.018971 | 0.002279 | 0.00225 |
| 19.875 | 0.003043 | 0.006924 | 0.002312 | 0.005404 | 0.005738 | 0.005572 | 0.014327 | 0.003733 | 0.002954 |
| 19.88888889 | 0.005612 | 0.009516 | 0.001686 | 0.010563 | 0.009255 | 0.008689 | 0.010956 | 0.009907 | 0.005279 |
| 19.90277778 | 0.006715 | 0.010621 | 0.002241 | 0.017473 | 0.011479 | 0.007926 | 0.011755 | 0.018846 | 0.006986 |
| 19.91666667 | 0.004763 | 0.009751 | 0.002686 | 0.024662 | 0.012084 | 0.004538 | 0.016251 | 0.024196 | 0.007048 |
| 19.93055556 | 0.004229 | 0.006497 | 0.00211 | 0.026783 | 0.010775 | 0.003754 | 0.021481 | 0.021525 | 0.005527 |
| 19.94444444 | 0.009248 | 0.00348 | 0.001696 | 0.020837 | 0.007456 | 0.007045 | 0.025264 | 0.01353 | 0.003229 |
| 19.95833333 | 0.014805 | 0.003361 | 0.002548 | 0.013166 | 0.004242 | 0.009852 | 0.025011 | 0.006386 | 0.001633 |
| 19.97222222 | 0.012821 | 0.004824 | 0.004258 | 0.010403 | 0.005798 | 0.008143 | 0.016814 | 0.005183 | 0.003314 |
| 19.98611111 | 0.008904 | 0.011298 | 0.011589 | 0.012171 | 0.015619 | 0.008908 | 0.011258 | 0.011568 | 0.012752 |
| 20 | 0.012971 | 0.02405 | 0.026209 | 0.013945 | 0.029704 | 0.018884 | 0.021776 | 0.020589 | 0.027952 |
| 20.01388889 | 0.01611 | 0.031781 | 0.036608 | 0.01167 | 0.03689 | 0.02749 | 0.033904 | 0.022562 | 0.037335 |
| 20.02777778 | 0.010573 | 0.02752 | 0.034051 | 0.008028 | 0.031523 | 0.025617 | 0.032107 | 0.015166 | 0.033871 |
| 20.04166667 | 0.003764 | 0.017531 | 0.024171 | 0.007991 | 0.01984 | 0.018251 | 0.022276 | 0.007052 | 0.023402 |
| 20.05555556 | 0.001103 | 0.008781 | 0.015097 | 0.009331 | 0.00934 | 0.011203 | 0.012852 | 0.003655 | 0.013849 |
| 20.06944444 | 0.001529 | 0.003375 | 0.008587 | 0.008393 | 0.003289 | 0.005641 | 0.00603 | 0.002341 | 0.007289 |
| 20.08333333 | 0.002127 | 0.001405 | 0.004432 | 0.006524 | 0.001903 | 0.001922 | 0.002025 | 0.001262 | 0.003768 |
| 20.09722222 | 0.001906 | 0.001308 | 0.002643 | 0.00563 | 0.001983 | 0.000532 | 0.000788 | 0.001072 | 0.002736 |
| 20.11111111 | 0.001883 | 0.001657 | 0.001931 | 0.004985 | 0.002025 | 0.001181 | 0.000598 | 0.0012 | 0.002238 |
| 20.125 | 0.002892 | 0.002642 | 0.001354 | 0.003388 | 0.003053 | 0.003512 | 0.000737 | 0.001117 | 0.001508 |
| 20.13888889 | 0.004659 | 0.004241 | 0.001629 | 0.001769 | 0.004867 | 0.006452 | 0.002029 | 0.001791 | 0.001836 |
| 20.15277778 | 0.006348 | 0.005846 | 0.002913 | 0.001908 | 0.006545 | 0.008772 | 0.003888 | 0.003348 | 0.003364 |
| 20.16666667 | 0.007764 | 0.007313 | 0.004546 | 0.003786 | 0.007814 | 0.010715 | 0.005478 | 0.005082 | 0.004956 |
| 20.18055556 | 0.008959 | 0.008589 | 0.00628 | 0.006014 | 0.008738 | 0.0125 | 0.006729 | 0.006734 | 0.006188 |
| 20.19444444 | 0.009246 | 0.008864 | 0.007227 | 0.00706 | 0.008755 | 0.01273 | 0.006918 | 0.007327 | 0.006554 |
| 20.20833333 | 0.008626 | 0.00797 | 0.006885 | 0.006799 | 0.007864 | 0.011188 | 0.006094 | 0.006475 | 0.005967 |
| 20.22222222 | 0.008577 | 0.007279 | 0.006668 | 0.006751 | 0.007442 | 0.010322 | 0.006007 | 0.006041 | 0.00581 |
| 20.23611111 | 0.009405 | 0.007455 | 0.007385 | 0.007077 | 0.00793 | 0.011172 | 0.007103 | 0.007044 | 0.006577 |
| 20.25 | 0.00958 | 0.00758 | 0.007786 | 0.006393 | 0.008122 | 0.011844 | 0.007603 | 0.007975 | 0.006898 |
| 20.26388889 | 0.008571 | 0.007217 | 0.007143 | 0.004929 | 0.007698 | 0.011755 | 0.006915 | 0.008015 | 0.006283 |
| 20.27777778 | 0.007545 | 0.006974 | 0.00647 | 0.00452 | 0.007813 | 0.013136 | 0.006464 | 0.008275 | 0.005933 |
| 20.29166667 | 0.007292 | 0.007112 | 0.006453 | 0.00564 | 0.008773 | 0.016409 | 0.006885 | 0.00915 | 0.006376 |
| 20.30555556 | 0.007772 | 0.007584 | 0.006837 | 0.007198 | 0.00967 | 0.018769 | 0.00757 | 0.009904 | 0.007017 |
| 20.31944444 | 0.008595 | 0.00827 | 0.00727 | 0.008032 | 0.009849 | 0.018512 | 0.007956 | 0.010119 | 0.007335 |
| 20.33333333 | 0.0087 | 0.00833 | 0.00705 | 0.006993 | 0.008959 | 0.016055 | 0.007394 | 0.009446 | 0.006789 |
| 20.34722222 | 0.007695 | 0.007479 | 0.005931 | 0.004239 | 0.007305 | 0.012337 | 0.005891 | 0.008051 | 0.005392 |
| 20.36111111 | 0.006981 | 0.00705 | 0.005174 | 0.002031 | 0.006361 | 0.009103 | 0.004851 | 0.007341 | 0.004487 |
| 20.375 | 0.007658 | 0.007946 | 0.005969 | 0.001976 | 0.007129 | 0.00802 | 0.005182 | 0.008219 | 0.005193 |
| 20.38888889 | 0.008584 | 0.009031 | 0.007452 | 0.002889 | 0.008448 | 0.008379 | 0.005576 | 0.009489 | 0.006529 |
| 20.40277778 | 0.007972 | 0.008805 | 0.007734 | 0.002836 | 0.008176 | 0.008049 | 0.004356 | 0.009451 | 0.006513 |
| 20.41666667 | 0.005995 | 0.007582 | 0.006674 | 0.00203 | 0.005431 | 0.006654 | 0.002168 | 0.008258 | 0.005003 |

| | | | | | | | | | |
|---|---|---|---|---|---|---|---|---|---|
| 20.43055556 | 0.004109 | 0.006549 | 0.005735 | 0.001515 | 0.00248 | 0.005315 | 0.000898 | 0.007166 | 0.003578 |
| 20.44444444 | 0.00284 | 0.005825 | 0.005438 | 0.001011 | 0.001878 | 0.004398 | 0.001217 | 0.006452 | 0.002887 |
| 20.45833333 | 0.001638 | 0.005006 | 0.005156 | 0.000821 | 0.004368 | 0.003249 | 0.002493 | 0.005659 | 0.002486 |
| 20.47222222 | 0.001565 | 0.004521 | 0.005185 | 0.002876 | 0.00956 | 0.002778 | 0.003697 | 0.005047 | 0.00265 |
| 20.48611111 | 0.005044 | 0.005794 | 0.007835 | 0.007516 | 0.015155 | 0.006442 | 0.004662 | 0.00614 | 0.004841 |
| 20.5 | 0.01173 | 0.009233 | 0.013904 | 0.009414 | 0.020563 | 0.013321 | 0.006896 | 0.010123 | 0.009553 |
| 20.51388889 | 0.017884 | 0.011903 | 0.018622 | 0.009326 | 0.024119 | 0.016866 | 0.011159 | 0.014204 | 0.01377 |
| 20.52777778 | 0.021075 | 0.009232 | 0.016664 | 0.01244 | 0.021708 | 0.016139 | 0.014442 | 0.012229 | 0.013912 |
| 20.54166667 | 0.022477 | 0.005245 | 0.009402 | 0.012438 | 0.014222 | 0.015791 | 0.013431 | 0.008232 | 0.010527 |
| 20.55555556 | 0.023206 | 0.007625 | 0.003594 | 0.007127 | 0.006443 | 0.016029 | 0.009957 | 0.012705 | 0.006155 |
| 20.56944444 | 0.023433 | 0.012425 | 0.003044 | 0.002886 | 0.001979 | 0.014281 | 0.00928 | 0.021559 | 0.003261 |
| 20.58333333 | 0.024272 | 0.014557 | 0.004666 | 0.001419 | 0.000901 | 0.011122 | 0.013425 | 0.025795 | 0.004065 |
| 20.59722222 | 0.026426 | 0.015103 | 0.004934 | 0.0017 | 0.000799 | 0.008764 | 0.0173 | 0.024756 | 0.007303 |
| 20.61111111 | 0.027783 | 0.01482 | 0.003953 | 0.003261 | 0.000836 | 0.008443 | 0.014362 | 0.020798 | 0.009474 |
| 20.625 | 0.025727 | 0.013545 | 0.003023 | 0.003821 | 0.00093 | 0.010159 | 0.008773 | 0.016334 | 0.008814 |
| 20.63888889 | 0.020486 | 0.011815 | 0.002307 | 0.002964 | 0.001078 | 0.012573 | 0.010559 | 0.012969 | 0.006267 |
| 20.65277778 | 0.014073 | 0.010489 | 0.001774 | 0.003776 | 0.002038 | 0.013917 | 0.019528 | 0.010892 | 0.003824 |
| 20.66666667 | 0.007665 | 0.008424 | 0.002881 | 0.00488 | 0.002616 | 0.012658 | 0.029691 | 0.008795 | 0.002594 |
| 20.68055556 | 0.003048 | 0.004735 | 0.005412 | 0.003652 | 0.001826 | 0.009195 | 0.035939 | 0.005875 | 0.003484 |
| 20.69444444 | 0.002147 | 0.001823 | 0.006327 | 0.001758 | 0.001249 | 0.006542 | 0.032386 | 0.003179 | 0.005124 |
| 20.70833333 | 0.003167 | 0.001139 | 0.004277 | 0.001125 | 0.001061 | 0.006834 | 0.019747 | 0.001872 | 0.004123 |
| 20.72222222 | 0.00388 | 0.000837 | 0.002325 | 0.001659 | 0.000705 | 0.008775 | 0.013371 | 0.001733 | 0.001963 |
| 20.73611111 | 0.004837 | 0.000475 | 0.002183 | 0.002468 | 0.000603 | 0.008596 | 0.021365 | 0.001693 | 0.001485 |
| 20.75 | 0.006712 | 0.000804 | 0.001997 | 0.003024 | 0.000384 | 0.005013 | 0.02827 | 0.001828 | 0.001358 |
| 20.76388889 | 0.008574 | 0.001179 | 0.002133 | 0.003354 | 0.000435 | 0.001686 | 0.022941 | 0.003192 | 0.001397 |
| 20.77777778 | 0.009264 | 0.001015 | 0.002853 | 0.00312 | 0.001374 | 0.001787 | 0.014655 | 0.005007 | 0.001922 |
| 20.79166667 | 0.008928 | 0.001068 | 0.003573 | 0.002069 | 0.002047 | 0.003399 | 0.01852 | 0.004304 | 0.00227 |
| 20.80555556 | 0.00831 | 0.001605 | 0.004612 | 0.000956 | 0.001546 | 0.003134 | 0.031224 | 0.003015 | 0.002611 |
| 20.81944444 | 0.007765 | 0.00139 | 0.004755 | 0.001064 | 0.001032 | 0.001411 | 0.034004 | 0.0053 | 0.002135 |
| 20.83333333 | 0.007826 | 0.000664 | 0.003246 | 0.00299 | 0.001708 | 0.000611 | 0.023164 | 0.007835 | 0.001306 |
| 20.84722222 | 0.008804 | 0.000428 | 0.001504 | 0.004994 | 0.00306 | 0.001025 | 0.011266 | 0.0075 | 0.001607 |
| 20.86111111 | 0.010418 | 0.000928 | 0.000679 | 0.004588 | 0.003602 | 0.001576 | 0.005622 | 0.005491 | 0.00188 |
| 20.875 | 0.012947 | 0.002763 | 0.000656 | 0.003334 | 0.002408 | 0.003298 | 0.004429 | 0.003311 | 0.002428 |
| 20.88888889 | 0.01668 | 0.00534 | 0.001518 | 0.005727 | 0.00112 | 0.007582 | 0.005891 | 0.003229 | 0.004812 |
| 20.90277778 | 0.020597 | 0.007385 | 0.003038 | 0.010878 | 0.001314 | 0.012066 | 0.009386 | 0.004567 | 0.007273 |
| 20.91666667 | 0.022753 | 0.008187 | 0.003842 | 0.012827 | 0.001533 | 0.013596 | 0.012775 | 0.003802 | 0.007651 |
| 20.93055556 | 0.0223 | 0.00756 | 0.00299 | 0.01 | 0.001121 | 0.012393 | 0.013937 | 0.002811 | 0.005835 |
| 20.94444444 | 0.020301 | 0.006307 | 0.001524 | 0.006875 | 0.001331 | 0.011172 | 0.012552 | 0.005778 | 0.003264 |
| 20.95833333 | 0.017366 | 0.004989 | 0.001276 | 0.006038 | 0.002415 | 0.011507 | 0.008882 | 0.010341 | 0.001604 |
| 20.97222222 | 0.011754 | 0.003797 | 0.003299 | 0.006395 | 0.004822 | 0.013447 | 0.006442 | 0.010225 | 0.002791 |
| 20.98611111 | 0.007119 | 0.005782 | 0.008404 | 0.012536 | 0.010521 | 0.018424 | 0.012251 | 0.007953 | 0.008676 |
| 21 | 0.009417 | 0.011976 | 0.015499 | 0.025119 | 0.019735 | 0.023828 | 0.024214 | 0.011496 | 0.016812 |
| 21.01388889 | 0.012376 | 0.015753 | 0.020125 | 0.029814 | 0.025681 | 0.019304 | 0.029453 | 0.015851 | 0.021607 |
| 21.02777778 | 0.009693 | 0.013273 | 0.018443 | 0.02071 | 0.021573 | 0.00935 | 0.022514 | 0.012641 | 0.019755 |

| | | | | | | | | | |
|---|---|---|---|---|---|---|---|---|---|
| 21.04166667 | 0.005287 | 0.007676 | 0.011858 | 0.009273 | 0.012066 | 0.006977 | 0.0113 | 0.006656 | 0.012898 |
| 21.05555556 | 0.002596 | 0.003 | 0.005384 | 0.003784 | 0.004814 | 0.006393 | 0.003693 | 0.00534 | 0.006209 |
| 21.06944444 | 0.001673 | 0.001023 | 0.002324 | 0.002339 | 0.001809 | 0.003765 | 0.000951 | 0.00519 | 0.003106 |
| 21.08333333 | 0.002006 | 0.001068 | 0.002551 | 0.001389 | 0.001859 | 0.004479 | 0.0014 | 0.003141 | 0.003196 |
| 21.09722222 | 0.002618 | 0.0021 | 0.003978 | 0.000541 | 0.002722 | 0.007111 | 0.002776 | 0.002626 | 0.004293 |
| 21.11111111 | 0.002707 | 0.002581 | 0.004677 | 0.000399 | 0.003039 | 0.006858 | 0.003648 | 0.002727 | 0.004682 |
| 21.125 | 0.002403 | 0.002005 | 0.00427 | 0.000401 | 0.002782 | 0.003523 | 0.003878 | 0.001738 | 0.00422 |
| 21.13888889 | 0.002003 | 0.001448 | 0.003524 | 0.000773 | 0.002245 | 0.001616 | 0.003552 | 0.001241 | 0.003616 |
| 21.15277778 | 0.002104 | 0.001895 | 0.003634 | 0.001957 | 0.001868 | 0.004433 | 0.003103 | 0.002506 | 0.003811 |
| 21.16666667 | 0.003259 | 0.003404 | 0.004893 | 0.002114 | 0.002236 | 0.009238 | 0.00349 | 0.004823 | 0.005021 |
| 21.18055556 | 0.004796 | 0.004711 | 0.00596 | 0.001854 | 0.003444 | 0.009472 | 0.005078 | 0.00574 | 0.00617 |
| 21.19444444 | 0.005568 | 0.004816 | 0.005815 | 0.003711 | 0.004737 | 0.005101 | 0.007138 | 0.004346 | 0.006341 |
| 21.20833333 | 0.00482 | 0.003865 | 0.004792 | 0.005392 | 0.004521 | 0.002008 | 0.007896 | 0.002444 | 0.005298 |
| 21.22222222 | 0.002691 | 0.002301 | 0.0033 | 0.004239 | 0.002731 | 0.002563 | 0.006297 | 0.00121 | 0.003067 |
| 21.23611111 | 0.00082 | 0.001034 | 0.001709 | 0.00269 | 0.001443 | 0.003848 | 0.00418 | 0.000706 | 0.001011 |
| 21.25 | 0.000382 | 0.000702 | 0.000854 | 0.003284 | 0.00109 | 0.002886 | 0.00373 | 0.000913 | 0.00067 |
| 21.26388889 | 0.000604 | 0.000542 | 0.000939 | 0.004486 | 0.0006 | 0.001493 | 0.004292 | 0.001019 | 0.001508 |
| 21.27777778 | 0.000637 | 0.000409 | 0.001169 | 0.004527 | 0.000299 | 0.001398 | 0.00433 | 0.001032 | 0.002007 |
| 21.29166667 | 0.000889 | 0.001102 | 0.001066 | 0.004137 | 0.00095 | 0.001269 | 0.003511 | 0.001924 | 0.001469 |
| 21.30555556 | 0.00156 | 0.002379 | 0.001161 | 0.004221 | 0.002104 | 0.000997 | 0.002377 | 0.003448 | 0.00073 |
| 21.31944444 | 0.001862 | 0.003005 | 0.002205 | 0.00481 | 0.002575 | 0.000933 | 0.001823 | 0.004292 | 0.000906 |
| 21.33333333 | 0.001662 | 0.002478 | 0.003979 | 0.005175 | 0.002167 | 0.000621 | 0.00192 | 0.003916 | 0.001815 |
| 21.34722222 | 0.001189 | 0.001464 | 0.005994 | 0.005209 | 0.001556 | 0.000718 | 0.0023 | 0.00282 | 0.002851 |
| 21.36111111 | 0.000831 | 0.00073 | 0.008258 | 0.00651 | 0.000888 | 0.001966 | 0.003496 | 0.001487 | 0.004209 |
| 21.375 | 0.001197 | 0.000658 | 0.010117 | 0.009587 | 0.000401 | 0.003678 | 0.005911 | 0.000695 | 0.006035 |
| 21.38888889 | 0.001703 | 0.000835 | 0.010364 | 0.012829 | 0.000294 | 0.004186 | 0.009202 | 0.000711 | 0.007312 |
| 21.40277778 | 0.002168 | 0.000656 | 0.008867 | 0.016047 | 0.00022 | 0.003151 | 0.012668 | 0.000778 | 0.007583 |
| 21.41666667 | 0.003255 | 0.000373 | 0.006749 | 0.019244 | 0.000247 | 0.002467 | 0.015308 | 0.001092 | 0.007688 |
| 21.43055556 | 0.004428 | 0.000319 | 0.004822 | 0.01927 | 0.000582 | 0.002086 | 0.01659 | 0.0022 | 0.007232 |
| 21.44444444 | 0.004673 | 0.000551 | 0.002746 | 0.014614 | 0.001555 | 0.001648 | 0.015542 | 0.003604 | 0.005147 |
| 21.45833333 | 0.004114 | 0.001016 | 0.000994 | 0.0081 | 0.003145 | 0.003956 | 0.011581 | 0.005388 | 0.002566 |
| 21.47222222 | 0.00405 | 0.001151 | 0.000805 | 0.004629 | 0.004842 | 0.011829 | 0.007944 | 0.00821 | 0.001654 |
| 21.48611111 | 0.00623 | 0.00074 | 0.003059 | 0.009333 | 0.008846 | 0.022109 | 0.007673 | 0.010024 | 0.001909 |
| 21.5 | 0.011165 | 0.00046 | 0.008804 | 0.020393 | 0.016341 | 0.023881 | 0.00792 | 0.007381 | 0.003981 |
| 21.51388889 | 0.014822 | 0.000664 | 0.015012 | 0.028092 | 0.022587 | 0.015402 | 0.006286 | 0.003778 | 0.010053 |
| 21.52777778 | 0.013333 | 0.001006 | 0.016551 | 0.026137 | 0.022782 | 0.00935 | 0.006171 | 0.003441 | 0.016069 |
| 21.54166667 | 0.009842 | 0.002254 | 0.014113 | 0.017757 | 0.017099 | 0.009083 | 0.008006 | 0.005571 | 0.017883 |
| 21.55555556 | 0.008398 | 0.004992 | 0.011467 | 0.010696 | 0.0097 | 0.007542 | 0.00739 | 0.01097 | 0.016529 |
| 21.56944444 | 0.010097 | 0.007629 | 0.009441 | 0.008124 | 0.004564 | 0.003934 | 0.006225 | 0.017535 | 0.01261 |
| 21.58333333 | 0.014702 | 0.008973 | 0.006122 | 0.006509 | 0.002142 | 0.00234 | 0.011953 | 0.022548 | 0.006688 |
| 21.59722222 | 0.018604 | 0.008467 | 0.003834 | 0.003756 | 0.001304 | 0.00495 | 0.024909 | 0.025252 | 0.003115 |
| 21.61111111 | 0.017237 | 0.005902 | 0.006713 | 0.001942 | 0.001241 | 0.01041 | 0.037209 | 0.023563 | 0.004708 |
| 21.625 | 0.010931 | 0.002817 | 0.010275 | 0.001661 | 0.001105 | 0.016932 | 0.040999 | 0.018118 | 0.007548 |
| 21.63888889 | 0.005774 | 0.001242 | 0.007992 | 0.001784 | 0.000793 | 0.022547 | 0.031727 | 0.011956 | 0.007643 |

| | | | | | | | | | |
|---|---|---|---|---|---|---|---|---|---|
| 21.65277778 | 0.007437 | 0.001227 | 0.004857 | 0.001536 | 0.000926 | 0.023811 | 0.019789 | 0.006804 | 0.005092 |
| 21.66666667 | 0.013755 | 0.001496 | 0.007937 | 0.001483 | 0.001052 | 0.019264 | 0.025455 | 0.003175 | 0.004154 |
| 21.68055556 | 0.018005 | 0.001895 | 0.012117 | 0.003347 | 0.000898 | 0.011306 | 0.043408 | 0.002111 | 0.006956 |
| 21.69444444 | 0.01666 | 0.002165 | 0.011865 | 0.006071 | 0.001242 | 0.004806 | 0.04994 | 0.002924 | 0.008431 |
| 21.70833333 | 0.010438 | 0.002158 | 0.00773 | 0.006095 | 0.001432 | 0.002639 | 0.0385 | 0.002728 | 0.005773 |
| 21.72222222 | 0.006717 | 0.00353 | 0.003915 | 0.003506 | 0.001433 | 0.002791 | 0.020085 | 0.001758 | 0.002215 |
| 21.73611111 | 0.009497 | 0.005914 | 0.004495 | 0.001459 | 0.001947 | 0.00297 | 0.012071 | 0.001594 | 0.00074 |
| 21.75 | 0.010163 | 0.006416 | 0.006294 | 0.000973 | 0.001955 | 0.00296 | 0.019661 | 0.002556 | 0.001056 |
| 21.76388889 | 0.006556 | 0.00472 | 0.005881 | 0.001494 | 0.001322 | 0.002266 | 0.02703 | 0.00596 | 0.001927 |
| 21.77777778 | 0.006779 | 0.002548 | 0.004175 | 0.001841 | 0.000767 | 0.001153 | 0.023167 | 0.011125 | 0.003212 |
| 21.79166667 | 0.012085 | 0.001761 | 0.004641 | 0.001652 | 0.000416 | 0.000768 | 0.012323 | 0.015726 | 0.004627 |
| 21.80555556 | 0.016951 | 0.004725 | 0.00804 | 0.001916 | 0.000584 | 0.001108 | 0.004026 | 0.018261 | 0.005047 |
| 21.81944444 | 0.018744 | 0.010364 | 0.009771 | 0.003568 | 0.001328 | 0.001342 | 0.002459 | 0.017916 | 0.004053 |
| 21.83333333 | 0.019089 | 0.012682 | 0.008113 | 0.006713 | 0.001451 | 0.001083 | 0.00427 | 0.015757 | 0.003013 |
| 21.84722222 | 0.019814 | 0.009255 | 0.005511 | 0.008876 | 0.000918 | 0.001448 | 0.00954 | 0.014608 | 0.00344 |
| 21.86111111 | 0.020902 | 0.004435 | 0.003894 | 0.008081 | 0.001014 | 0.003552 | 0.01794 | 0.017185 | 0.005282 |
| 21.875 | 0.021631 | 0.001539 | 0.003722 | 0.007198 | 0.001055 | 0.005863 | 0.023267 | 0.023789 | 0.007189 |
| 21.88888889 | 0.021628 | 0.000633 | 0.004548 | 0.009999 | 0.000741 | 0.006744 | 0.022356 | 0.031589 | 0.007956 |
| 21.90277778 | 0.020498 | 0.00094 | 0.005476 | 0.015536 | 0.001131 | 0.006368 | 0.01898 | 0.036251 | 0.007499 |
| 21.91666667 | 0.018322 | 0.001395 | 0.005724 | 0.018602 | 0.001672 | 0.005356 | 0.018321 | 0.035713 | 0.006541 |
| 21.93055556 | 0.016623 | 0.002454 | 0.005841 | 0.016314 | 0.001426 | 0.004825 | 0.022946 | 0.03262 | 0.006117 |
| 21.94444444 | 0.016818 | 0.004819 | 0.006357 | 0.011948 | 0.001065 | 0.005343 | 0.02975 | 0.030464 | 0.006497 |
| 21.95833333 | 0.017078 | 0.006795 | 0.006019 | 0.008564 | 0.001167 | 0.005915 | 0.030453 | 0.029183 | 0.006256 |
| 21.97222222 | 0.013995 | 0.005945 | 0.004675 | 0.006079 | 0.00306 | 0.005359 | 0.020295 | 0.025335 | 0.004842 |
| 21.98611111 | 0.008094 | 0.004346 | 0.006884 | 0.00776 | 0.009339 | 0.00325 | 0.011378 | 0.015471 | 0.006607 |
| 22 | 0.00477 | 0.005609 | 0.013953 | 0.014195 | 0.016836 | 0.001402 | 0.015696 | 0.007329 | 0.013241 |
| 22.01388889 | 0.005292 | 0.006599 | 0.017736 | 0.01632 | 0.018972 | 0.001207 | 0.020301 | 0.00853 | 0.016787 |
| 22.02777778 | 0.005556 | 0.004606 | 0.014168 | 0.011528 | 0.015634 | 0.002164 | 0.015264 | 0.010125 | 0.013176 |
| 22.04166667 | 0.005615 | 0.002842 | 0.008845 | 0.00831 | 0.011145 | 0.004807 | 0.00876 | 0.007387 | 0.008194 |
| 22.05555556 | 0.006615 | 0.002803 | 0.006031 | 0.009798 | 0.00801 | 0.007117 | 0.006202 | 0.004278 | 0.006078 |
| 22.06944444 | 0.007084 | 0.00312 | 0.005287 | 0.010518 | 0.00613 | 0.007022 | 0.005855 | 0.002231 | 0.005939 |
| 22.08333333 | 0.006799 | 0.003589 | 0.00559 | 0.00789 | 0.005238 | 0.00652 | 0.005763 | 0.002084 | 0.006373 |
| 22.09722222 | 0.006094 | 0.003952 | 0.005944 | 0.004814 | 0.005024 | 0.005967 | 0.005392 | 0.003054 | 0.006445 |
| 22.11111111 | 0.00463 | 0.00314 | 0.005126 | 0.002673 | 0.004225 | 0.003961 | 0.00412 | 0.00312 | 0.005174 |
| 22.125 | 0.002462 | 0.001669 | 0.003303 | 0.001503 | 0.002581 | 0.001535 | 0.002095 | 0.001932 | 0.002894 |
| 22.13888889 | 0.001165 | 0.001253 | 0.001488 | 0.002169 | 0.001088 | 0.000365 | 0.000837 | 0.001191 | 0.001134 |
| 22.15277778 | 0.001896 | 0.002151 | 0.000523 | 0.003903 | 0.000426 | 0.000155 | 0.001192 | 0.001578 | 0.000704 |
| 22.16666667 | 0.003105 | 0.002954 | 0.000472 | 0.004752 | 0.000526 | 0.000153 | 0.001894 | 0.001646 | 0.00079 |
| 22.18055556 | 0.003064 | 0.002656 | 0.000564 | 0.003508 | 0.0006 | 0.00054 | 0.001634 | 0.001104 | 0.000719 |
| 22.19444444 | 0.001796 | 0.001503 | 0.000778 | 0.002096 | 0.000391 | 0.001298 | 0.001046 | 0.001087 | 0.000932 |
| 22.20833333 | 0.000806 | 0.000706 | 0.001282 | 0.003602 | 0.000439 | 0.001555 | 0.001484 | 0.001427 | 0.001444 |
| 22.22222222 | 0.001371 | 0.000988 | 0.001828 | 0.007022 | 0.000853 | 0.001748 | 0.002768 | 0.001483 | 0.001995 |
| 22.23611111 | 0.002968 | 0.002266 | 0.002903 | 0.010029 | 0.001672 | 0.002402 | 0.0042 | 0.002246 | 0.003305 |
| 22.25 | 0.004741 | 0.004441 | 0.00483 | 0.012035 | 0.003041 | 0.003474 | 0.005512 | 0.004452 | 0.005712 |

| | | | | | | | | | |
|---|---|---|---|---|---|---|---|---|---|
| 22.26388889 | 0.00596 | 0.006191 | 0.006488 | 0.0137 | 0.004249 | 0.004172 | 0.00621 | 0.006472 | 0.007812 |
| 22.27777778 | 0.006018 | 0.006246 | 0.00669 | 0.015101 | 0.004051 | 0.002634 | 0.005841 | 0.007066 | 0.008315 |
| 22.29166667 | 0.005175 | 0.005314 | 0.005677 | 0.015522 | 0.002414 | 0.001077 | 0.004572 | 0.0071 | 0.007646 |
| 22.30555556 | 0.004167 | 0.003963 | 0.004256 | 0.015765 | 0.000806 | 0.001701 | 0.003482 | 0.007022 | 0.006691 |
| 22.31944444 | 0.003428 | 0.002311 | 0.003283 | 0.016508 | 0.000315 | 0.003247 | 0.004111 | 0.006881 | 0.005967 |
| 22.33333333 | 0.002778 | 0.001484 | 0.003607 | 0.015857 | 0.00036 | 0.00346 | 0.007729 | 0.007996 | 0.005796 |
| 22.34722222 | 0.00178 | 0.00178 | 0.00479 | 0.013057 | 0.000462 | 0.003144 | 0.013718 | 0.010575 | 0.005914 |
| 22.36111111 | 0.000978 | 0.001631 | 0.005314 | 0.011301 | 0.001137 | 0.004039 | 0.018289 | 0.012555 | 0.005694 |
| 22.375 | 0.001561 | 0.001447 | 0.004545 | 0.012492 | 0.002323 | 0.003339 | 0.017627 | 0.013031 | 0.004881 |
| 22.38888889 | 0.003642 | 0.002733 | 0.002711 | 0.014288 | 0.003707 | 0.001973 | 0.01211 | 0.013173 | 0.003488 |
| 22.40277778 | 0.007183 | 0.003452 | 0.001728 | 0.01133 | 0.005793 | 0.002254 | 0.005916 | 0.011949 | 0.002826 |
| 22.41666667 | 0.011125 | 0.0093 | 0.003024 | 0.00797 | 0.007181 | 0.003601 | 0.005469 | 0.007379 | 0.003951 |
| 22.43055556 | 0.010692 | 0.016714 | 0.003397 | 0.010212 | 0.009001 | 0.014149 | 0.012574 | 0.003217 | 0.00425 |
| 22.44444444 | 0.006043 | 0.011287 | 0.003151 | 0.011349 | 0.012513 | 0.026657 | 0.016432 | 0.006138 | 0.004503 |
| 22.45833333 | 0.004564 | 0.004784 | 0.005246 | 0.012154 | 0.011526 | 0.021284 | 0.009877 | 0.013664 | 0.006846 |
| 22.47222222 | 0.008173 | 0.007538 | 0.0067 | 0.01278 | 0.006206 | 0.010408 | 0.005108 | 0.016316 | 0.00719 |
| 22.48611111 | 0.014482 | 0.012548 | 0.005969 | 0.008705 | 0.002331 | 0.009627 | 0.012796 | 0.01461 | 0.005809 |
| 22.5 | 0.0203 | 0.015202 | 0.004404 | 0.005969 | 0.001279 | 0.013213 | 0.02518 | 0.015926 | 0.006667 |
| 22.51388889 | 0.021248 | 0.012615 | 0.002375 | 0.005398 | 0.002592 | 0.01294 | 0.030687 | 0.020573 | 0.008201 |
| 22.52777778 | 0.018342 | 0.007675 | 0.00096 | 0.00533 | 0.003837 | 0.007577 | 0.0287 | 0.02436 | 0.007675 |
| 22.54166667 | 0.015761 | 0.005389 | 0.00068 | 0.006688 | 0.003641 | 0.002879 | 0.02285 | 0.023826 | 0.004918 |
| 22.55555556 | 0.014289 | 0.005222 | 0.001383 | 0.006368 | 0.003445 | 0.00233 | 0.013521 | 0.018308 | 0.00316 |
| 22.56944444 | 0.013771 | 0.004891 | 0.002587 | 0.007171 | 0.003609 | 0.004328 | 0.007834 | 0.012245 | 0.005145 |
| 22.58333333 | 0.015237 | 0.004485 | 0.002471 | 0.009508 | 0.003039 | 0.004803 | 0.011211 | 0.010112 | 0.007223 |
| 22.59722222 | 0.017526 | 0.003815 | 0.002282 | 0.008698 | 0.003544 | 0.002849 | 0.013302 | 0.012058 | 0.005636 |
| 22.61111111 | 0.018061 | 0.002365 | 0.005629 | 0.005165 | 0.006179 | 0.001556 | 0.011379 | 0.016822 | 0.003086 |
| 22.625 | 0.016456 | 0.001701 | 0.010866 | 0.003904 | 0.006511 | 0.002259 | 0.019542 | 0.023204 | 0.003459 |
| 22.63888889 | 0.013963 | 0.003495 | 0.013017 | 0.007627 | 0.003288 | 0.002841 | 0.038361 | 0.028024 | 0.005774 |
| 22.65277778 | 0.011039 | 0.008244 | 0.0092 | 0.00859 | 0.001114 | 0.002554 | 0.048339 | 0.026917 | 0.006927 |
| 22.66666667 | 0.006726 | 0.012774 | 0.005529 | 0.007065 | 0.002085 | 0.004426 | 0.038215 | 0.019224 | 0.00575 |
| 22.68055556 | 0.003375 | 0.011821 | 0.008698 | 0.011975 | 0.004797 | 0.009428 | 0.022212 | 0.010117 | 0.003366 |
| 22.69444444 | 0.004201 | 0.012244 | 0.012665 | 0.018226 | 0.007413 | 0.013843 | 0.023899 | 0.007683 | 0.002346 |
| 22.70833333 | 0.00652 | 0.021069 | 0.012055 | 0.019886 | 0.008309 | 0.015012 | 0.041635 | 0.012618 | 0.002919 |
| 22.72222222 | 0.008271 | 0.028705 | 0.008307 | 0.01881 | 0.008177 | 0.014049 | 0.054339 | 0.016121 | 0.003153 |
| 22.73611111 | 0.009867 | 0.030118 | 0.004563 | 0.013494 | 0.010608 | 0.014122 | 0.051902 | 0.013128 | 0.002673 |
| 22.75 | 0.009428 | 0.02993 | 0.004527 | 0.007691 | 0.01825 | 0.01724 | 0.035146 | 0.006689 | 0.001705 |
| 22.76388889 | 0.005898 | 0.029271 | 0.006988 | 0.00847 | 0.027427 | 0.021584 | 0.017256 | 0.002279 | 0.000765 |
| 22.77777778 | 0.002011 | 0.026119 | 0.006819 | 0.009751 | 0.031102 | 0.02433 | 0.013649 | 0.00434 | 0.000608 |
| 22.79166667 | 0.001411 | 0.018938 | 0.004133 | 0.006318 | 0.028366 | 0.025128 | 0.017027 | 0.01375 | 0.001067 |
| 22.80555556 | 0.004497 | 0.010251 | 0.003694 | 0.004642 | 0.02248 | 0.024504 | 0.013246 | 0.026206 | 0.002959 |
| 22.81944444 | 0.008868 | 0.007991 | 0.006201 | 0.008143 | 0.016216 | 0.023276 | 0.0085 | 0.036295 | 0.006949 |
| 22.83333333 | 0.01263 | 0.015237 | 0.009171 | 0.012291 | 0.014231 | 0.024341 | 0.014961 | 0.04216 | 0.011155 |
| 22.84722222 | 0.015169 | 0.024151 | 0.011467 | 0.013262 | 0.020026 | 0.02923 | 0.026835 | 0.0444 | 0.014152 |
| 22.86111111 | 0.016148 | 0.027837 | 0.012509 | 0.012465 | 0.031282 | 0.03468 | 0.032417 | 0.043493 | 0.015829 |

| | | | | | | | | | |
|---|---|---|---|---|---|---|---|---|---|
| 22.875 | 0.015671 | 0.024369 | 0.01202 | 0.013115 | 0.042497 | 0.037383 | 0.027888 | 0.040679 | 0.015767 |
| 22.88888889 | 0.014349 | 0.014431 | 0.01055 | 0.01527 | 0.048973 | 0.036432 | 0.015609 | 0.038692 | 0.014234 |
| 22.90277778 | 0.012564 | 0.006205 | 0.008788 | 0.016528 | 0.048222 | 0.032355 | 0.004865 | 0.038412 | 0.012273 |
| 22.91666667 | 0.010655 | 0.005041 | 0.00752 | 0.016256 | 0.041057 | 0.027666 | 0.003083 | 0.037311 | 0.011186 |
| 22.93055556 | 0.009482 | 0.005654 | 0.006779 | 0.014703 | 0.030955 | 0.023816 | 0.009231 | 0.033104 | 0.011636 |
| 22.94444444 | 0.0098 | 0.009018 | 0.005385 | 0.011795 | 0.021515 | 0.020279 | 0.019694 | 0.026644 | 0.012223 |
| 22.95833333 | 0.010922 | 0.017946 | 0.002895 | 0.00777 | 0.01487 | 0.016941 | 0.029649 | 0.019978 | 0.011282 |
| 22.97222222 | 0.01038 | 0.026387 | 0.001671 | 0.004334 | 0.014736 | 0.01436 | 0.033216 | 0.012676 | 0.009764 |
| 22.98611111 | 0.0072 | 0.030009 | 0.004881 | 0.005348 | 0.027318 | 0.012756 | 0.025497 | 0.007621 | 0.010133 |
| 23 | 0.006653 | 0.025034 | 0.011312 | 0.010424 | 0.042238 | 0.01069 | 0.013179 | 0.009779 | 0.010408 |
| 23.01388889 | 0.011599 | 0.015304 | 0.014852 | 0.012277 | 0.034841 | 0.008138 | 0.009117 | 0.012231 | 0.008338 |
| 23.02777778 | 0.017945 | 0.011678 | 0.011074 | 0.008761 | 0.013904 | 0.012634 | 0.009572 | 0.010315 | 0.00904 |
| 23.04166667 | 0.024054 | 0.009781 | 0.005251 | 0.005231 | 0.004684 | 0.029769 | 0.006304 | 0.012302 | 0.012382 |
| 23.05555556 | 0.023627 | 0.007078 | 0.003955 | 0.004218 | 0.00376 | 0.034192 | 0.002157 | 0.013605 | 0.011058 |
| 23.06944444 | 0.014482 | 0.012533 | 0.003738 | 0.005415 | 0.003021 | 0.028088 | 0.002006 | 0.010938 | 0.005597 |
| 23.08333333 | 0.008636 | 0.019767 | 0.00229 | 0.005698 | 0.002089 | 0.039268 | 0.003725 | 0.012606 | 0.002364 |
| 23.09722222 | 0.008208 | 0.016613 | 0.00293 | 0.003314 | 0.001732 | 0.036197 | 0.003435 | 0.010785 | 0.003476 |
| 23.11111111 | 0.007398 | 0.008517 | 0.005795 | 0.001631 | 0.003496 | 0.014419 | 0.003941 | 0.004765 | 0.006399 |
| 23.125 | 0.00595 | 0.00484 | 0.008167 | 0.001581 | 0.006106 | 0.008933 | 0.006774 | 0.003896 | 0.008657 |
| 23.13888889 | 0.003585 | 0.005666 | 0.008445 | 0.002351 | 0.007533 | 0.016368 | 0.008719 | 0.004465 | 0.00881 |
| 23.15277778 | 0.001519 | 0.00844 | 0.006651 | 0.002446 | 0.006104 | 0.022093 | 0.00866 | 0.003846 | 0.006695 |
| 23.16666667 | 0.001482 | 0.010633 | 0.004119 | 0.001366 | 0.002875 | 0.020221 | 0.008499 | 0.004036 | 0.004434 |
| 23.18055556 | 0.003783 | 0.009474 | 0.003481 | 0.001744 | 0.001836 | 0.012653 | 0.01166 | 0.005267 | 0.004772 |
| 23.19444444 | 0.007892 | 0.00537 | 0.00617 | 0.004269 | 0.00495 | 0.008377 | 0.019357 | 0.007369 | 0.007835 |
| 23.20833333 | 0.009804 | 0.002437 | 0.010329 | 0.006679 | 0.009837 | 0.011758 | 0.026382 | 0.010269 | 0.011056 |
| 23.22222222 | 0.008427 | 0.002117 | 0.013326 | 0.007902 | 0.012871 | 0.015949 | 0.02703 | 0.012706 | 0.012616 |
| 23.23611111 | 0.00764 | 0.002472 | 0.01458 | 0.009067 | 0.013411 | 0.015398 | 0.02256 | 0.012845 | 0.012653 |
| 23.25 | 0.009191 | 0.003941 | 0.015035 | 0.010562 | 0.012803 | 0.013419 | 0.017562 | 0.010943 | 0.012207 |
| 23.26388889 | 0.01144 | 0.007031 | 0.015205 | 0.011102 | 0.011882 | 0.01307 | 0.014472 | 0.009247 | 0.011891 |
| 23.27777778 | 0.011265 | 0.008883 | 0.014248 | 0.008664 | 0.011161 | 0.013467 | 0.012833 | 0.008792 | 0.010966 |
| 23.29166667 | 0.007037 | 0.007278 | 0.011587 | 0.00538 | 0.010094 | 0.014575 | 0.011268 | 0.008636 | 0.008851 |
| 23.30555556 | 0.003251 | 0.004016 | 0.008419 | 0.005703 | 0.007685 | 0.016367 | 0.00927 | 0.007583 | 0.006707 |
| 23.31944444 | 0.003552 | 0.001867 | 0.006048 | 0.006875 | 0.004252 | 0.016004 | 0.00676 | 0.005342 | 0.005522 |
| 23.33333333 | 0.004157 | 0.000939 | 0.004426 | 0.004872 | 0.002894 | 0.012475 | 0.003691 | 0.003003 | 0.004446 |
| 23.34722222 | 0.00353 | 0.000771 | 0.002927 | 0.002365 | 0.00573 | 0.008537 | 0.001499 | 0.002006 | 0.002996 |
| 23.36111111 | 0.004028 | 0.002542 | 0.002057 | 0.002924 | 0.009817 | 0.005253 | 0.001545 | 0.004193 | 0.003458 |
| 23.375 | 0.006595 | 0.007074 | 0.0033 | 0.006693 | 0.012493 | 0.002661 | 0.002511 | 0.010388 | 0.006636 |
| 23.38888889 | 0.010276 | 0.012949 | 0.005695 | 0.011863 | 0.013762 | 0.001683 | 0.003171 | 0.017767 | 0.008818 |
| 23.40277778 | 0.013049 | 0.017418 | 0.006668 | 0.016179 | 0.014283 | 0.001607 | 0.003893 | 0.023039 | 0.007741 |
| 23.41666667 | 0.013839 | 0.01877 | 0.006157 | 0.01788 | 0.015276 | 0.00247 | 0.005775 | 0.026124 | 0.005438 |
| 23.43055556 | 0.012541 | 0.017831 | 0.005715 | 0.016432 | 0.01641 | 0.003708 | 0.010255 | 0.02827 | 0.004675 |
| 23.44444444 | 0.009833 | 0.016508 | 0.006092 | 0.013004 | 0.015809 | 0.002843 | 0.017108 | 0.028257 | 0.005766 |
| 23.45833333 | 0.00712 | 0.015133 | 0.006578 | 0.009288 | 0.013208 | 0.001493 | 0.023084 | 0.023769 | 0.006712 |
| 23.47222222 | 0.00428 | 0.013383 | 0.005238 | 0.005339 | 0.010024 | 0.004536 | 0.024822 | 0.015476 | 0.005508 |

| | | | | | | | | | |
|---|---|---|---|---|---|---|---|---|---|
| 23.48611111 | 0.003843 | 0.011902 | 0.00396 | 0.002802 | 0.008261 | 0.015111 | 0.023628 | 0.007113 | 0.0031 |
| 23.5 | 0.008566 | 0.010802 | 0.005779 | 0.002731 | 0.011143 | 0.024809 | 0.021662 | 0.002497 | 0.001639 |
| 23.51388889 | 0.011336 | 0.008178 | 0.006163 | 0.004786 | 0.015647 | 0.019941 | 0.018238 | 0.002008 | 0.001593 |
| 23.52777778 | 0.00746 | 0.004255 | 0.003541 | 0.009491 | 0.014094 | 0.008669 | 0.013935 | 0.004315 | 0.002194 |
| 23.54166667 | 0.004143 | 0.002507 | 0.002095 | 0.01144 | 0.008499 | 0.005012 | 0.009944 | 0.009418 | 0.004327 |
| 23.55555556 | 0.00622 | 0.002089 | 0.001611 | 0.007275 | 0.007398 | 0.004503 | 0.006427 | 0.016584 | 0.009591 |
| 23.56944444 | 0.01137 | 0.002464 | 0.002656 | 0.002604 | 0.011829 | 0.002474 | 0.003336 | 0.024553 | 0.012974 |
| 23.58333333 | 0.01766 | 0.006377 | 0.006967 | 0.001514 | 0.013279 | 0.001102 | 0.001083 | 0.031266 | 0.009954 |
| 23.59722222 | 0.022136 | 0.010535 | 0.011893 | 0.002357 | 0.008308 | 0.000774 | 0.001527 | 0.034327 | 0.005768 |
| 23.61111111 | 0.021224 | 0.011389 | 0.013652 | 0.002178 | 0.005725 | 0.000747 | 0.007853 | 0.034687 | 0.007435 |
| 23.625 | 0.016699 | 0.013051 | 0.011003 | 0.002603 | 0.010853 | 0.000831 | 0.02058 | 0.034403 | 0.012924 |
| 23.63888889 | 0.012934 | 0.020273 | 0.006439 | 0.007546 | 0.019186 | 0.001245 | 0.031657 | 0.032576 | 0.015978 |
| 23.65277778 | 0.011087 | 0.030424 | 0.005156 | 0.0149 | 0.025444 | 0.002254 | 0.031892 | 0.027416 | 0.015187 |
| 23.66666667 | 0.010142 | 0.035333 | 0.007191 | 0.018716 | 0.023141 | 0.002442 | 0.022742 | 0.019898 | 0.011375 |
| 23.68055556 | 0.010074 | 0.030163 | 0.00644 | 0.015948 | 0.012546 | 0.001882 | 0.011696 | 0.012269 | 0.006058 |
| 23.69444444 | 0.010578 | 0.01735 | 0.004282 | 0.009197 | 0.003866 | 0.00235 | 0.005286 | 0.007324 | 0.002659 |
| 23.70833333 | 0.008867 | 0.008821 | 0.006407 | 0.007314 | 0.002101 | 0.003255 | 0.007402 | 0.009744 | 0.00347 |
| 23.72222222 | 0.005447 | 0.012575 | 0.009465 | 0.01347 | 0.002953 | 0.003709 | 0.013826 | 0.019172 | 0.006704 |
| 23.73611111 | 0.005506 | 0.017668 | 0.008341 | 0.018628 | 0.003761 | 0.002919 | 0.015574 | 0.02698 | 0.009166 |
| 23.75 | 0.009002 | 0.015255 | 0.005173 | 0.01626 | 0.007409 | 0.001414 | 0.011073 | 0.02719 | 0.009602 |
| 23.76388889 | 0.011816 | 0.009796 | 0.005304 | 0.009293 | 0.012633 | 0.000778 | 0.006563 | 0.021277 | 0.008407 |
| 23.77777778 | 0.011199 | 0.004885 | 0.008944 | 0.003447 | 0.016881 | 0.000857 | 0.006358 | 0.012606 | 0.005626 |
| 23.79166667 | 0.006853 | 0.001563 | 0.012223 | 0.001754 | 0.020534 | 0.001497 | 0.008839 | 0.005809 | 0.002843 |
| 23.80555556 | 0.002961 | 0.002648 | 0.014485 | 0.003521 | 0.023988 | 0.00295 | 0.013279 | 0.00698 | 0.001879 |
| 23.81944444 | 0.003808 | 0.010273 | 0.016315 | 0.006763 | 0.028921 | 0.005228 | 0.020821 | 0.017667 | 0.0036 |
| 23.83333333 | 0.008639 | 0.021771 | 0.016588 | 0.00977 | 0.036086 | 0.007434 | 0.028499 | 0.031861 | 0.009198 |
| 23.84722222 | 0.014113 | 0.031229 | 0.014823 | 0.011844 | 0.044395 | 0.008442 | 0.033076 | 0.042705 | 0.015471 |
| 23.86111111 | 0.018827 | 0.035796 | 0.012757 | 0.01403 | 0.052368 | 0.008901 | 0.035508 | 0.048262 | 0.01951 |
| 23.875 | 0.02314 | 0.03675 | 0.012376 | 0.016783 | 0.056324 | 0.009752 | 0.035947 | 0.048233 | 0.021861 |
| 23.88888889 | 0.026202 | 0.035794 | 0.012832 | 0.017634 | 0.053757 | 0.010474 | 0.031358 | 0.043258 | 0.022114 |
| 23.90277778 | 0.026456 | 0.033005 | 0.012367 | 0.015326 | 0.047906 | 0.010712 | 0.024331 | 0.037876 | 0.020624 |
| 23.91666667 | 0.023931 | 0.028339 | 0.011189 | 0.012811 | 0.042879 | 0.011301 | 0.022905 | 0.035563 | 0.018797 |
| 23.93055556 | 0.019753 | 0.023807 | 0.00997 | 0.011992 | 0.040023 | 0.012469 | 0.026602 | 0.034229 | 0.016764 |
| 23.94444444 | 0.015986 | 0.022243 | 0.008777 | 0.011533 | 0.039491 | 0.013496 | 0.027929 | 0.031364 | 0.014507 |
| 23.95833333 | 0.014995 | 0.024737 | 0.007815 | 0.011284 | 0.039408 | 0.014441 | 0.025767 | 0.027367 | 0.013463 |
| 23.97222222 | 0.015331 | 0.029843 | 0.006893 | 0.012294 | 0.035157 | 0.017723 | 0.024141 | 0.022206 | 0.013982 |
| 23.98611111 | 0.012455 | 0.034973 | 0.004896 | 0.013231 | 0.024712 | 0.027373 | 0.024814 | 0.015239 | 0.013397 |
| 24 | 0.006823 | 0.035861 | 0.002148 | 0.011823 | 0.012889 | 0.03806 | 0.027365 | 0.007999 | 0.009385 |
| 24.01388889 | 0.004648 | 0.029566 | 0.001195 | 0.008265 | 0.007058 | 0.033009 | 0.029202 | 0.002935 | 0.004387 |
| 24.02777778 | 0.006199 | 0.019705 | 0.002667 | 0.005645 | 0.007474 | 0.015421 | 0.026816 | 0.002584 | 0.002747 |
| 24.04166667 | 0.006352 | 0.010206 | 0.003902 | 0.00747 | 0.006765 | 0.004705 | 0.020711 | 0.007309 | 0.004241 |
| 24.05555556 | 0.007465 | 0.005627 | 0.003381 | 0.012307 | 0.003033 | 0.002047 | 0.01463 | 0.010923 | 0.005407 |
| 24.06944444 | 0.010439 | 0.008182 | 0.002623 | 0.014579 | 0.001061 | 0.001436 | 0.010547 | 0.008139 | 0.004908 |
| 24.08333333 | 0.008459 | 0.009009 | 0.003716 | 0.012167 | 0.002478 | 0.002732 | 0.008562 | 0.00345 | 0.004256 |

| | | | | | | | | | |
|---|---|---|---|---|---|---|---|---|---|
| 24.09722222 | 0.005512 | 0.004652 | 0.007205 | 0.007643 | 0.004952 | 0.008712 | 0.008206 | 0.001452 | 0.005199 |
| 24.11111111 | 0.007277 | 0.001552 | 0.009628 | 0.003984 | 0.004878 | 0.017857 | 0.007107 | 0.000865 | 0.0064 |
| 24.125 | 0.006936 | 0.002792 | 0.006852 | 0.003463 | 0.004549 | 0.02056 | 0.004649 | 0.001562 | 0.004771 |
| 24.13888889 | 0.005001 | 0.00621 | 0.002399 | 0.005279 | 0.007411 | 0.01376 | 0.004054 | 0.002594 | 0.002154 |
| 24.15277778 | 0.005754 | 0.008212 | 0.002618 | 0.005259 | 0.007888 | 0.005304 | 0.00522 | 0.002319 | 0.001342 |
| 24.16666667 | 0.005841 | 0.007916 | 0.009403 | 0.005741 | 0.004925 | 0.001875 | 0.004655 | 0.004193 | 0.00179 |
| 24.18055556 | 0.008978 | 0.01509 | 0.019036 | 0.017237 | 0.002825 | 0.003654 | 0.007031 | 0.012793 | 0.003722 |
| 24.19444444 | 0.017865 | 0.032966 | 0.022089 | 0.037216 | 0.001932 | 0.006639 | 0.01705 | 0.02446 | 0.005954 |
| 24.20833333 | 0.02198 | 0.039442 | 0.01575 | 0.044141 | 0.002667 | 0.007115 | 0.024288 | 0.027791 | 0.006833 |
| 24.22222222 | 0.017323 | 0.026556 | 0.008134 | 0.03159 | 0.004487 | 0.006516 | 0.022242 | 0.02044 | 0.006428 |
| 24.23611111 | 0.010877 | 0.012733 | 0.005291 | 0.016763 | 0.005316 | 0.007906 | 0.018311 | 0.01281 | 0.005485 |
| 24.25 | 0.006423 | 0.006812 | 0.006967 | 0.009497 | 0.0049 | 0.009923 | 0.017521 | 0.010143 | 0.005778 |
| 24.26388889 | 0.004584 | 0.005451 | 0.010581 | 0.006052 | 0.004214 | 0.009842 | 0.018548 | 0.008426 | 0.008222 |
| 24.27777778 | 0.004527 | 0.005617 | 0.012313 | 0.004248 | 0.003052 | 0.008549 | 0.018198 | 0.005049 | 0.010271 |
| 24.29166667 | 0.003665 | 0.004633 | 0.009597 | 0.006849 | 0.002995 | 0.007832 | 0.01445 | 0.002808 | 0.008661 |
| 24.30555556 | 0.004218 | 0.003425 | 0.005111 | 0.013801 | 0.006613 | 0.006831 | 0.00815 | 0.003782 | 0.004693 |
| 24.31944444 | 0.010176 | 0.00612 | 0.004241 | 0.020402 | 0.011568 | 0.004734 | 0.003841 | 0.006138 | 0.003193 |
| 24.33333333 | 0.017653 | 0.010515 | 0.006875 | 0.022981 | 0.013847 | 0.003306 | 0.004123 | 0.00814 | 0.005107 |
| 24.34722222 | 0.01942 | 0.009682 | 0.007318 | 0.020581 | 0.011886 | 0.003635 | 0.00422 | 0.008353 | 0.005389 |
| 24.36111111 | 0.013148 | 0.006203 | 0.004626 | 0.015298 | 0.007671 | 0.006861 | 0.002638 | 0.006612 | 0.003901 |
| 24.375 | 0.006368 | 0.008253 | 0.004589 | 0.009266 | 0.004829 | 0.014337 | 0.003378 | 0.00408 | 0.006115 |
| 24.38888889 | 0.006586 | 0.01289 | 0.010283 | 0.005583 | 0.003591 | 0.022938 | 0.006426 | 0.00446 | 0.011789 |
| 24.40277778 | 0.006814 | 0.011215 | 0.016373 | 0.005735 | 0.003155 | 0.026762 | 0.007727 | 0.008603 | 0.015039 |
| 24.41666667 | 0.00671 | 0.008843 | 0.014747 | 0.007501 | 0.005434 | 0.021677 | 0.007291 | 0.008837 | 0.010982 |
| 24.43055556 | 0.011908 | 0.014581 | 0.007979 | 0.013537 | 0.01082 | 0.01081 | 0.010675 | 0.007416 | 0.005784 |
| 24.44444444 | 0.012799 | 0.019092 | 0.005971 | 0.021594 | 0.01653 | 0.004396 | 0.016246 | 0.011977 | 0.007815 |
| 24.45833333 | 0.009774 | 0.019233 | 0.010187 | 0.025401 | 0.021356 | 0.007618 | 0.019744 | 0.019554 | 0.013205 |
| 24.47222222 | 0.01609 | 0.026051 | 0.016629 | 0.027059 | 0.026053 | 0.016069 | 0.025921 | 0.027936 | 0.01907 |
| 24.48611111 | 0.029958 | 0.041826 | 0.020965 | 0.027391 | 0.030319 | 0.024395 | 0.039314 | 0.035572 | 0.026211 |
| 24.5 | 0.036152 | 0.053487 | 0.020925 | 0.023772 | 0.034322 | 0.028065 | 0.055862 | 0.038923 | 0.030807 |
| 24.51388889 | 0.029871 | 0.048769 | 0.02081 | 0.01867 | 0.039689 | 0.023605 | 0.066638 | 0.040562 | 0.028461 |
| 24.52777778 | 0.021482 | 0.033359 | 0.021883 | 0.016649 | 0.039774 | 0.015125 | 0.064431 | 0.04654 | 0.021192 |
| 24.54166667 | 0.016091 | 0.018443 | 0.017564 | 0.017342 | 0.025315 | 0.010119 | 0.047483 | 0.055103 | 0.013494 |
| 24.55555556 | 0.010856 | 0.007611 | 0.009655 | 0.016013 | 0.012989 | 0.010311 | 0.02387 | 0.056639 | 0.006995 |
| 24.56944444 | 0.006861 | 0.001811 | 0.006503 | 0.010219 | 0.020201 | 0.013016 | 0.010179 | 0.046434 | 0.004591 |
| 24.58333333 | 0.008327 | 0.000791 | 0.006248 | 0.004186 | 0.029466 | 0.01519 | 0.007644 | 0.031798 | 0.005818 |
| 24.59722222 | 0.015556 | 0.003342 | 0.009831 | 0.003196 | 0.022668 | 0.016321 | 0.011199 | 0.024558 | 0.005328 |
| 24.61111111 | 0.024591 | 0.010518 | 0.02201 | 0.00689 | 0.015384 | 0.01865 | 0.030242 | 0.027372 | 0.006617 |
| 24.625 | 0.030133 | 0.022925 | 0.032634 | 0.010697 | 0.028249 | 0.021617 | 0.052351 | 0.032147 | 0.01563 |
| 24.63888889 | 0.02902 | 0.037658 | 0.029776 | 0.011678 | 0.046895 | 0.022151 | 0.051516 | 0.032885 | 0.025657 |
| 24.65277778 | 0.022409 | 0.047717 | 0.016337 | 0.010973 | 0.048708 | 0.019385 | 0.029495 | 0.03363 | 0.030338 |
| 24.66666667 | 0.014727 | 0.045606 | 0.005703 | 0.011961 | 0.032778 | 0.014004 | 0.011715 | 0.038933 | 0.030226 |
| 24.68055556 | 0.010632 | 0.032939 | 0.004119 | 0.016043 | 0.016017 | 0.007535 | 0.013052 | 0.045089 | 0.025847 |
| 24.69444444 | 0.011578 | 0.019391 | 0.003884 | 0.019852 | 0.014029 | 0.00286 | 0.017359 | 0.04526 | 0.018569 |

| | | | | | | | | | |
|---|---|---|---|---|---|---|---|---|---|
| 24.70833333 | 0.015882 | 0.009933 | 0.003777 | 0.019905 | 0.020208 | 0.001378 | 0.012529 | 0.036597 | 0.010671 |
| 24.72222222 | 0.019634 | 0.005093 | 0.00811 | 0.015003 | 0.022486 | 0.001392 | 0.009494 | 0.020868 | 0.004342 |
| 24.73611111 | 0.018042 | 0.007158 | 0.011887 | 0.009543 | 0.01999 | 0.001709 | 0.010059 | 0.012935 | 0.001675 |
| 24.75 | 0.010791 | 0.013977 | 0.009334 | 0.010985 | 0.014661 | 0.002239 | 0.011801 | 0.026565 | 0.002614 |
| 24.76388889 | 0.004308 | 0.017169 | 0.005179 | 0.014176 | 0.008624 | 0.002143 | 0.025641 | 0.046064 | 0.004086 |
| 24.77777778 | 0.004085 | 0.011779 | 0.006981 | 0.010717 | 0.007842 | 0.002224 | 0.044908 | 0.05121 | 0.004093 |
| 24.79166667 | 0.008292 | 0.007144 | 0.012291 | 0.006383 | 0.011964 | 0.003159 | 0.04739 | 0.04192 | 0.003365 |
| 24.80555556 | 0.01109 | 0.012002 | 0.015597 | 0.007341 | 0.01216 | 0.003067 | 0.030346 | 0.026726 | 0.002319 |
| 24.81944444 | 0.009088 | 0.015373 | 0.016078 | 0.009239 | 0.008226 | 0.002014 | 0.016177 | 0.01321 | 0.001761 |
| 24.83333333 | 0.00599 | 0.010737 | 0.015582 | 0.009018 | 0.010681 | 0.003376 | 0.011914 | 0.008255 | 0.001818 |
| 24.84722222 | 0.007483 | 0.009265 | 0.01607 | 0.009987 | 0.023888 | 0.009156 | 0.012209 | 0.014923 | 0.002724 |
| 24.86111111 | 0.012936 | 0.015698 | 0.017095 | 0.013696 | 0.04013 | 0.016373 | 0.024794 | 0.027462 | 0.007797 |
| 24.875 | 0.018225 | 0.023252 | 0.018329 | 0.018779 | 0.050616 | 0.0212 | 0.038977 | 0.038437 | 0.016664 |
| 24.88888889 | 0.02127 | 0.029105 | 0.019786 | 0.02323 | 0.052494 | 0.022902 | 0.04206 | 0.045296 | 0.023737 |
| 24.90277778 | 0.021463 | 0.033003 | 0.018641 | 0.024129 | 0.045931 | 0.020908 | 0.036247 | 0.047522 | 0.023987 |
| 24.91666667 | 0.019674 | 0.033823 | 0.013583 | 0.021248 | 0.033209 | 0.016133 | 0.023682 | 0.045577 | 0.018529 |
| 24.93055556 | 0.0198 | 0.033722 | 0.009297 | 0.017481 | 0.020248 | 0.01235 | 0.011712 | 0.042722 | 0.013577 |
| 24.94444444 | 0.023799 | 0.033389 | 0.00918 | 0.014638 | 0.012338 | 0.010446 | 0.008823 | 0.038976 | 0.01179 |
| 24.95833333 | 0.028158 | 0.029961 | 0.009971 | 0.013248 | 0.010685 | 0.008513 | 0.016044 | 0.02998 | 0.011309 |
| 24.97222222 | 0.02784 | 0.021977 | 0.00717 | 0.011801 | 0.011688 | 0.0055 | 0.025754 | 0.016567 | 0.009223 |
| 24.98611111 | 0.02003 | 0.011776 | 0.006778 | 0.009657 | 0.009448 | 0.007154 | 0.02594 | 0.011168 | 0.008097 |
| 25 | 0.00898 | 0.004551 | 0.016388 | 0.012445 | 0.007117 | 0.01931 | 0.016131 | 0.018132 | 0.014158 |
| 25.01388889 | 0.005257 | 0.00461 | 0.025526 | 0.019559 | 0.014915 | 0.031129 | 0.010823 | 0.019906 | 0.020822 |
| 25.02777778 | 0.00959 | 0.011403 | 0.024283 | 0.021209 | 0.029091 | 0.030581 | 0.013469 | 0.011051 | 0.017014 |
| 25.04166667 | 0.010817 | 0.018093 | 0.017431 | 0.014783 | 0.033241 | 0.020737 | 0.013906 | 0.004705 | 0.008648 |
| 25.05555556 | 0.008155 | 0.01649 | 0.010367 | 0.007956 | 0.024272 | 0.009545 | 0.008833 | 0.003592 | 0.006692 |
| 25.06944444 | 0.01167 | 0.010357 | 0.005092 | 0.007211 | 0.013782 | 0.003049 | 0.003652 | 0.002976 | 0.00861 |
| 25.08333333 | 0.021226 | 0.011001 | 0.003107 | 0.0078 | 0.008021 | 0.001658 | 0.002108 | 0.003227 | 0.007974 |
| 25.09722222 | 0.027508 | 0.016889 | 0.004656 | 0.004896 | 0.004678 | 0.00105 | 0.001607 | 0.004535 | 0.005138 |
| 25.11111111 | 0.027327 | 0.019201 | 0.008187 | 0.002813 | 0.00306 | 0.000627 | 0.00135 | 0.004449 | 0.002575 |
| 25.125 | 0.024417 | 0.017086 | 0.011382 | 0.003807 | 0.004566 | 0.001544 | 0.003735 | 0.003224 | 0.001977 |
| 25.13888889 | 0.021616 | 0.013962 | 0.012027 | 0.004888 | 0.006627 | 0.002507 | 0.007717 | 0.002508 | 0.003238 |
| 25.15277778 | 0.019272 | 0.011575 | 0.009604 | 0.004669 | 0.007075 | 0.002228 | 0.010362 | 0.002757 | 0.004292 |
| 25.16666667 | 0.017398 | 0.010067 | 0.005792 | 0.004029 | 0.007554 | 0.001349 | 0.010272 | 0.003853 | 0.004217 |
| 25.18055556 | 0.015861 | 0.009073 | 0.002823 | 0.004125 | 0.008217 | 0.001587 | 0.008354 | 0.005166 | 0.003342 |
| 25.19444444 | 0.014263 | 0.008135 | 0.001568 | 0.005384 | 0.006235 | 0.002862 | 0.006169 | 0.005583 | 0.002383 |
| 25.20833333 | 0.012395 | 0.006604 | 0.00108 | 0.006672 | 0.004055 | 0.004015 | 0.004218 | 0.004346 | 0.002593 |
| 25.22222222 | 0.009794 | 0.003985 | 0.001212 | 0.006825 | 0.005492 | 0.004684 | 0.002472 | 0.002242 | 0.003969 |
| 25.23611111 | 0.005871 | 0.002374 | 0.002489 | 0.00608 | 0.005304 | 0.004807 | 0.002739 | 0.000992 | 0.004287 |
| 25.25 | 0.002404 | 0.004268 | 0.003324 | 0.005415 | 0.003676 | 0.004061 | 0.00599 | 0.001109 | 0.002657 |
| 25.26388889 | 0.001441 | 0.007176 | 0.002589 | 0.005416 | 0.005613 | 0.002789 | 0.007827 | 0.002711 | 0.001197 |
| 25.27777778 | 0.003436 | 0.006989 | 0.002439 | 0.006724 | 0.006958 | 0.001733 | 0.005492 | 0.006262 | 0.002059 |
| 25.29166667 | 0.00842 | 0.005318 | 0.004203 | 0.011059 | 0.005091 | 0.00403 | 0.003648 | 0.011277 | 0.005059 |
| 25.30555556 | 0.013839 | 0.007498 | 0.006202 | 0.018872 | 0.00393 | 0.012503 | 0.005519 | 0.016837 | 0.008166 |

| | | | | | | | | | |
|---|---|---|---|---|---|---|---|---|---|
| 25.31944444 | 0.01812 | 0.011819 | 0.007972 | 0.026572 | 0.003606 | 0.019638 | 0.007961 | 0.021062 | 0.010655 |
| 25.33333333 | 0.02226 | 0.013158 | 0.010111 | 0.029872 | 0.002757 | 0.017735 | 0.008102 | 0.021848 | 0.012056 |
| 25.34722222 | 0.026981 | 0.01354 | 0.012527 | 0.028478 | 0.004953 | 0.011696 | 0.005846 | 0.019456 | 0.011369 |
| 25.36111111 | 0.033151 | 0.017118 | 0.014951 | 0.025605 | 0.012211 | 0.007784 | 0.003443 | 0.01635 | 0.010428 |
| 25.375 | 0.037311 | 0.021466 | 0.016433 | 0.023444 | 0.02202 | 0.006428 | 0.00372 | 0.015051 | 0.012245 |
| 25.38888889 | 0.034094 | 0.020736 | 0.016665 | 0.021566 | 0.030209 | 0.006941 | 0.006252 | 0.015558 | 0.01651 |
| 25.40277778 | 0.025395 | 0.01512 | 0.016621 | 0.020319 | 0.033472 | 0.008344 | 0.008874 | 0.015757 | 0.020096 |
| 25.41666667 | 0.015532 | 0.008542 | 0.017023 | 0.02149 | 0.030416 | 0.009144 | 0.011207 | 0.014592 | 0.020832 |
| 25.43055556 | 0.00704 | 0.003285 | 0.018284 | 0.02535 | 0.02226 | 0.009111 | 0.014717 | 0.013371 | 0.018891 |
| 25.44444444 | 0.003407 | 0.001309 | 0.019916 | 0.028536 | 0.014352 | 0.009065 | 0.019996 | 0.014517 | 0.015901 |
| 25.45833333 | 0.005374 | 0.002934 | 0.019683 | 0.026539 | 0.012023 | 0.009348 | 0.024965 | 0.018563 | 0.013722 |
| 25.47222222 | 0.009399 | 0.007965 | 0.017095 | 0.019797 | 0.014787 | 0.009562 | 0.027814 | 0.02413 | 0.014542 |
| 25.48611111 | 0.012898 | 0.020493 | 0.016612 | 0.013636 | 0.015426 | 0.008306 | 0.031194 | 0.031021 | 0.022491 |
| 25.5 | 0.01505 | 0.040473 | 0.022139 | 0.011509 | 0.025078 | 0.006768 | 0.037482 | 0.036761 | 0.035309 |
| 25.51388889 | 0.014164 | 0.052423 | 0.029538 | 0.011652 | 0.066123 | 0.008166 | 0.043786 | 0.036042 | 0.038733 |
| 25.52777778 | 0.010964 | 0.046862 | 0.028243 | 0.010867 | 0.097221 | 0.008197 | 0.045106 | 0.030198 | 0.026895 |
| 25.54166667 | 0.009236 | 0.034675 | 0.018712 | 0.007394 | 0.076492 | 0.004874 | 0.034995 | 0.025711 | 0.012323 |
| 25.55555556 | 0.010433 | 0.024384 | 0.016705 | 0.003584 | 0.044418 | 0.004911 | 0.019919 | 0.023991 | 0.007791 |
| 25.56944444 | 0.013544 | 0.016619 | 0.020857 | 0.003188 | 0.030893 | 0.00816 | 0.017665 | 0.022861 | 0.011631 |
| 25.58333333 | 0.017094 | 0.011536 | 0.017603 | 0.003752 | 0.028124 | 0.009708 | 0.022598 | 0.022707 | 0.012559 |
| 25.59722222 | 0.018715 | 0.007015 | 0.01863 | 0.002758 | 0.028277 | 0.008725 | 0.018899 | 0.026619 | 0.008467 |
| 25.61111111 | 0.016759 | 0.003823 | 0.033633 | 0.002256 | 0.024551 | 0.007137 | 0.01946 | 0.036144 | 0.007258 |
| 25.625 | 0.012206 | 0.005692 | 0.041349 | 0.002512 | 0.017505 | 0.006254 | 0.035076 | 0.045964 | 0.00904 |
| 25.63888889 | 0.006678 | 0.010471 | 0.027505 | 0.002134 | 0.011596 | 0.004716 | 0.043634 | 0.048321 | 0.007248 |
| 25.65277778 | 0.00268 | 0.011702 | 0.015173 | 0.003539 | 0.009178 | 0.002259 | 0.030594 | 0.043952 | 0.003852 |
| 25.66666667 | 0.002039 | 0.00915 | 0.022488 | 0.008612 | 0.011167 | 0.001413 | 0.021152 | 0.039155 | 0.002763 |
| 25.68055556 | 0.002063 | 0.011947 | 0.028909 | 0.012106 | 0.017063 | 0.002916 | 0.035117 | 0.033751 | 0.004851 |
| 25.69444444 | 0.001444 | 0.023671 | 0.019524 | 0.009243 | 0.02162 | 0.004493 | 0.046929 | 0.023421 | 0.0121 |
| 25.70833333 | 0.004179 | 0.035851 | 0.010924 | 0.004255 | 0.019092 | 0.005369 | 0.033204 | 0.011717 | 0.017692 |
| 25.72222222 | 0.012515 | 0.040139 | 0.015874 | 0.002009 | 0.011327 | 0.006282 | 0.01869 | 0.009543 | 0.014395 |
| 25.73611111 | 0.020236 | 0.033874 | 0.018447 | 0.003485 | 0.004776 | 0.005844 | 0.026248 | 0.020605 | 0.00814 |
| 25.75 | 0.020363 | 0.020943 | 0.0141 | 0.00889 | 0.001707 | 0.003354 | 0.029542 | 0.035499 | 0.007025 |
| 25.76388889 | 0.01339 | 0.009861 | 0.021271 | 0.013331 | 0.001037 | 0.001945 | 0.018727 | 0.044418 | 0.009047 |
| 25.77777778 | 0.006547 | 0.008166 | 0.034609 | 0.013596 | 0.001773 | 0.004174 | 0.018374 | 0.044809 | 0.009621 |
| 25.79166667 | 0.004931 | 0.015301 | 0.034762 | 0.012943 | 0.001962 | 0.009214 | 0.03018 | 0.038158 | 0.00918 |
| 25.80555556 | 0.004119 | 0.023939 | 0.022445 | 0.013698 | 0.001915 | 0.014708 | 0.038951 | 0.028514 | 0.009019 |
| 25.81944444 | 0.002924 | 0.031231 | 0.012017 | 0.014408 | 0.004483 | 0.018619 | 0.041281 | 0.020886 | 0.01008 |
| 25.83333333 | 0.006063 | 0.03629 | 0.012088 | 0.012266 | 0.008975 | 0.021047 | 0.040096 | 0.019126 | 0.011658 |
| 25.84722222 | 0.015296 | 0.038475 | 0.018714 | 0.009071 | 0.012511 | 0.023885 | 0.035677 | 0.026118 | 0.012625 |
| 25.86111111 | 0.032098 | 0.043614 | 0.024728 | 0.010538 | 0.015394 | 0.027567 | 0.031782 | 0.040975 | 0.013786 |
| 25.875 | 0.050352 | 0.05075 | 0.02662 | 0.017844 | 0.018628 | 0.029012 | 0.032014 | 0.0556 | 0.015296 |
| 25.88888889 | 0.055026 | 0.04888 | 0.023854 | 0.025914 | 0.020684 | 0.024998 | 0.031759 | 0.06294 | 0.017627 |
| 25.90277778 | 0.041748 | 0.037125 | 0.018342 | 0.029798 | 0.020936 | 0.017025 | 0.02597 | 0.063471 | 0.022065 |
| 25.91666667 | 0.021142 | 0.023926 | 0.011714 | 0.027449 | 0.020603 | 0.010357 | 0.018614 | 0.057405 | 0.026807 |

| | | | | | | | | | |
|---|---|---|---|---|---|---|---|---|---|
| 25.93055556 | 0.008488 | 0.014803 | 0.005967 | 0.020824 | 0.020332 | 0.008895 | 0.016326 | 0.045024 | 0.028614 |
| 25.94444444 | 0.005575 | 0.01142 | 0.003419 | 0.015029 | 0.019134 | 0.011793 | 0.019727 | 0.031156 | 0.025344 |
| 25.95833333 | 0.005679 | 0.011832 | 0.004181 | 0.011879 | 0.015699 | 0.014557 | 0.022292 | 0.020041 | 0.017208 |
| 25.97222222 | 0.011666 | 0.013813 | 0.006509 | 0.010212 | 0.010761 | 0.013491 | 0.018004 | 0.013317 | 0.008977 |
| 25.98611111 | 0.021757 | 0.017847 | 0.007251 | 0.009032 | 0.006325 | 0.008594 | 0.009576 | 0.010033 | 0.004357 |
| 26 | 0.031384 | 0.023812 | 0.005009 | 0.008863 | 0.005764 | 0.004804 | 0.00405 | 0.009663 | 0.002021 |
| 26.01388889 | 0.043062 | 0.029381 | 0.002757 | 0.01127 | 0.008688 | 0.00421 | 0.003246 | 0.014266 | 0.001475 |
| 26.02777778 | 0.055666 | 0.032415 | 0.002344 | 0.012089 | 0.009286 | 0.005108 | 0.004617 | 0.021248 | 0.004019 |
| 26.04166667 | 0.060601 | 0.031489 | 0.002326 | 0.007515 | 0.006814 | 0.007394 | 0.005325 | 0.023605 | 0.008023 |
| 26.05555556 | 0.051265 | 0.026113 | 0.001721 | 0.003216 | 0.00388 | 0.007323 | 0.004114 | 0.020259 | 0.010492 |
| 26.06944444 | 0.032009 | 0.018387 | 0.0009 | 0.004605 | 0.002643 | 0.004554 | 0.002797 | 0.015193 | 0.010971 |
| 26.08333333 | 0.014815 | 0.010741 | 0.000809 | 0.008626 | 0.004811 | 0.00319 | 0.001941 | 0.009486 | 0.010108 |
| 26.09722222 | 0.010346 | 0.005429 | 0.002398 | 0.008872 | 0.007523 | 0.003347 | 0.00232 | 0.004531 | 0.00814 |
| 26.11111111 | 0.016506 | 0.005487 | 0.006054 | 0.005585 | 0.007894 | 0.003613 | 0.006875 | 0.00388 | 0.0055 |
| 26.125 | 0.022301 | 0.009667 | 0.010194 | 0.00552 | 0.005654 | 0.00342 | 0.014457 | 0.006908 | 0.003059 |
| 26.13888889 | 0.024546 | 0.013642 | 0.013548 | 0.010118 | 0.003572 | 0.002353 | 0.020842 | 0.010445 | 0.003045 |
| 26.15277778 | 0.026203 | 0.016425 | 0.016432 | 0.012861 | 0.005722 | 0.001851 | 0.023356 | 0.012711 | 0.006478 |
| 26.16666667 | 0.028498 | 0.019066 | 0.018338 | 0.009985 | 0.010292 | 0.003767 | 0.021454 | 0.012768 | 0.010528 |
| 26.18055556 | 0.03138 | 0.021516 | 0.018055 | 0.004679 | 0.013411 | 0.005448 | 0.016474 | 0.011008 | 0.012202 |
| 26.19444444 | 0.033761 | 0.02318 | 0.015086 | 0.001474 | 0.014551 | 0.004649 | 0.010275 | 0.009042 | 0.010991 |
| 26.20833333 | 0.034169 | 0.023411 | 0.010644 | 0.00263 | 0.01398 | 0.003338 | 0.004989 | 0.007655 | 0.007922 |
| 26.22222222 | 0.032008 | 0.020529 | 0.006525 | 0.006397 | 0.011333 | 0.003577 | 0.002964 | 0.005584 | 0.00415 |
| 26.23611111 | 0.027288 | 0.014017 | 0.003238 | 0.007687 | 0.006961 | 0.005052 | 0.003303 | 0.003016 | 0.001813 |
| 26.25 | 0.021289 | 0.007509 | 0.001132 | 0.004754 | 0.003311 | 0.006466 | 0.002788 | 0.001805 | 0.001974 |
| 26.26388889 | 0.016649 | 0.004637 | 0.000457 | 0.002217 | 0.002962 | 0.007354 | 0.0015 | 0.001918 | 0.001831 |
| 26.27777778 | 0.014223 | 0.00425 | 0.000434 | 0.005474 | 0.005115 | 0.007705 | 0.001374 | 0.003834 | 0.001059 |
| 26.29166667 | 0.012618 | 0.003897 | 0.00094 | 0.014432 | 0.00714 | 0.008387 | 0.002159 | 0.00772 | 0.0019 |
| 26.30555556 | 0.011158 | 0.003945 | 0.002681 | 0.022049 | 0.007984 | 0.011197 | 0.003078 | 0.011938 | 0.003828 |
| 26.31944444 | 0.010404 | 0.005355 | 0.005992 | 0.022375 | 0.007542 | 0.016191 | 0.004119 | 0.016959 | 0.005786 |
| 26.33333333 | 0.011578 | 0.007617 | 0.010326 | 0.018425 | 0.006135 | 0.020537 | 0.005252 | 0.024108 | 0.007615 |
| 26.34722222 | 0.015905 | 0.010374 | 0.01438 | 0.017195 | 0.004813 | 0.022609 | 0.007056 | 0.031667 | 0.009303 |
| 26.36111111 | 0.022837 | 0.013888 | 0.016725 | 0.020512 | 0.004396 | 0.023921 | 0.010239 | 0.036624 | 0.010782 |
| 26.375 | 0.029986 | 0.017447 | 0.01683 | 0.024246 | 0.004437 | 0.025015 | 0.013892 | 0.038542 | 0.011538 |
| 26.38888889 | 0.034133 | 0.019459 | 0.015907 | 0.024787 | 0.003485 | 0.024777 | 0.015761 | 0.038229 | 0.011124 |
| 26.40277778 | 0.033209 | 0.019358 | 0.01603 | 0.022439 | 0.00167 | 0.022365 | 0.014134 | 0.035495 | 0.009902 |
| 26.41666667 | 0.028363 | 0.017854 | 0.018455 | 0.019542 | 0.000507 | 0.018195 | 0.010082 | 0.030523 | 0.009502 |
| 26.43055556 | 0.022853 | 0.015375 | 0.021945 | 0.018162 | 0.000397 | 0.014029 | 0.007616 | 0.024935 | 0.012205 |
| 26.44444444 | 0.018622 | 0.011723 | 0.022324 | 0.01801 | 0.001276 | 0.010686 | 0.008806 | 0.019607 | 0.017079 |
| 26.45833333 | 0.014497 | 0.006721 | 0.016331 | 0.016985 | 0.003684 | 0.007451 | 0.011553 | 0.013304 | 0.019636 |
| 26.47222222 | 0.00912 | 0.004063 | 0.007485 | 0.015296 | 0.007916 | 0.004773 | 0.015566 | 0.006873 | 0.019234 |
| 26.48611111 | 0.004234 | 0.008381 | 0.006162 | 0.015459 | 0.016873 | 0.005634 | 0.023949 | 0.004293 | 0.022792 |
| 26.5 | 0.002119 | 0.013612 | 0.019266 | 0.016434 | 0.027302 | 0.008542 | 0.036009 | 0.008926 | 0.034651 |
| 26.51388889 | 0.003347 | 0.014788 | 0.039765 | 0.016436 | 0.024451 | 0.007535 | 0.04396 | 0.019736 | 0.045021 |
| 26.52777778 | 0.008771 | 0.022367 | 0.049733 | 0.01658 | 0.010566 | 0.004939 | 0.039901 | 0.029973 | 0.043166 |

| | | | | | | | | | |
|---|---|---|---|---|---|---|---|---|---|
| 26.54166667 | 0.015271 | 0.034829 | 0.037083 | 0.015653 | 0.002849 | 0.004844 | 0.023939 | 0.033127 | 0.031972 |
| 26.55555556 | 0.016764 | 0.034035 | 0.019515 | 0.012549 | 0.003093 | 0.004167 | 0.011584 | 0.029274 | 0.017203 |
| 26.56944444 | 0.013837 | 0.023438 | 0.020966 | 0.009402 | 0.005029 | 0.00373 | 0.015431 | 0.024623 | 0.008128 |
| 26.58333333 | 0.012711 | 0.020806 | 0.028956 | 0.008234 | 0.006947 | 0.004682 | 0.021042 | 0.026661 | 0.010826 |
| 26.59722222 | 0.016779 | 0.029118 | 0.022431 | 0.008956 | 0.008268 | 0.005587 | 0.017906 | 0.037118 | 0.012526 |
| 26.61111111 | 0.024052 | 0.039735 | 0.018786 | 0.009803 | 0.008247 | 0.010734 | 0.021849 | 0.051715 | 0.009014 |
| 26.625 | 0.031201 | 0.043435 | 0.041442 | 0.010056 | 0.005936 | 0.019423 | 0.037375 | 0.065094 | 0.012409 |
| 26.63888889 | 0.033301 | 0.034535 | 0.06551 | 0.010353 | 0.00333 | 0.023639 | 0.043965 | 0.068239 | 0.019807 |
| 26.65277778 | 0.026094 | 0.020838 | 0.05874 | 0.010936 | 0.003853 | 0.020517 | 0.031842 | 0.053987 | 0.0175 |
| 26.66666667 | 0.013098 | 0.021026 | 0.034265 | 0.012757 | 0.005189 | 0.013087 | 0.017085 | 0.028826 | 0.007516 |
| 26.68055556 | 0.006358 | 0.032155 | 0.02787 | 0.014969 | 0.007167 | 0.006069 | 0.02278 | 0.016351 | 0.001616 |
| 26.69444444 | 0.011465 | 0.034213 | 0.044652 | 0.015193 | 0.019861 | 0.002243 | 0.047806 | 0.026035 | 0.001131 |
| 26.70833333 | 0.019273 | 0.024762 | 0.065393 | 0.023099 | 0.03585 | 0.005668 | 0.056032 | 0.035172 | 0.007328 |
| 26.72222222 | 0.02677 | 0.027264 | 0.096731 | 0.054325 | 0.033694 | 0.010049 | 0.040261 | 0.040595 | 0.034082 |
| 26.73611111 | 0.025431 | 0.032723 | 0.096777 | 0.065048 | 0.017734 | 0.005083 | 0.024469 | 0.037174 | 0.048756 |
| 26.75 | 0.00972 | 0.014745 | 0.037533 | 0.026481 | 0.004553 | 9.83E-05 | 0.009056 | 0.014396 | 0.021275 |
| 26.76388889 | 0 | 0 | 0 | 0 | 0 | 0 | 0 | 0 | 0 |
| 26.77777778 | 0.010767 | 0.010545 | 0.030158 | 0.022949 | 0.005454 | 0.001557 | 0.018355 | 0.018103 | 0.018599 |
| 26.79166667 | 0.026422 | 0.022887 | 0.073221 | 0.055148 | 0.012964 | 0.006256 | 0.047097 | 0.046409 | 0.045215 |
| 26.80555556 | 0.02609 | 0.019722 | 0.061431 | 0.045292 | 0.010821 | 0.009458 | 0.051674 | 0.041292 | 0.04339 |
| 26.81944444 | 0.02195 | 0.017414 | 0.024524 | 0.022515 | 0.006714 | 0.008357 | 0.05326 | 0.021167 | 0.032392 |
| 26.83333333 | 0.02144 | 0.016121 | 0.011423 | 0.022061 | 0.007405 | 0.008144 | 0.058641 | 0.02647 | 0.022967 |
| 26.84722222 | 0.014183 | 0.008788 | 0.018508 | 0.02665 | 0.00894 | 0.011351 | 0.041418 | 0.047961 | 0.011695 |
| 26.86111111 | 0.008678 | 0.008618 | 0.032361 | 0.026519 | 0.01264 | 0.015364 | 0.015971 | 0.062639 | 0.01561 |
| 26.875 | 0.016029 | 0.021644 | 0.040148 | 0.022974 | 0.018419 | 0.015997 | 0.003838 | 0.063465 | 0.028144 |
| 26.88888889 | 0.026827 | 0.043401 | 0.040851 | 0.021152 | 0.023681 | 0.014078 | 0.001926 | 0.060727 | 0.036886 |
| 26.90277778 | 0.028538 | 0.064082 | 0.035507 | 0.024085 | 0.025366 | 0.015659 | 0.002801 | 0.063206 | 0.040668 |
| 26.91666667 | 0.022645 | 0.074493 | 0.025695 | 0.030331 | 0.021432 | 0.021602 | 0.006647 | 0.067758 | 0.040668 |
| 26.93055556 | 0.017342 | 0.071561 | 0.015035 | 0.037868 | 0.013972 | 0.026796 | 0.012144 | 0.070441 | 0.039051 |
| 26.94444444 | 0.017907 | 0.058515 | 0.006806 | 0.043368 | 0.007441 | 0.026759 | 0.0147 | 0.068258 | 0.034998 |
| 26.95833333 | 0.022459 | 0.040556 | 0.00273 | 0.041182 | 0.006038 | 0.021942 | 0.012662 | 0.056879 | 0.026906 |
| 26.97222222 | 0.027461 | 0.025293 | 0.003314 | 0.032321 | 0.008136 | 0.016448 | 0.008587 | 0.039941 | 0.017734 |
| 26.98611111 | 0.030835 | 0.021551 | 0.006984 | 0.023295 | 0.00739 | 0.013002 | 0.00504 | 0.025395 | 0.01123 |
| 27 | 0.032679 | 0.033303 | 0.011266 | 0.017226 | 0.003499 | 0.013307 | 0.004747 | 0.016015 | 0.008151 |
| 27.01388889 | 0.035629 | 0.051171 | 0.01154 | 0.013575 | 0.00279 | 0.014887 | 0.009265 | 0.012175 | 0.006237 |
| 27.02777778 | 0.040736 | 0.050553 | 0.009396 | 0.010383 | 0.008635 | 0.011197 | 0.014821 | 0.01288 | 0.005812 |
| 27.04166667 | 0.045455 | 0.029026 | 0.014508 | 0.00751 | 0.015632 | 0.007679 | 0.017342 | 0.013301 | 0.010281 |
| 27.05555556 | 0.044301 | 0.014479 | 0.022315 | 0.005967 | 0.017182 | 0.0118 | 0.015519 | 0.009391 | 0.014518 |
| 27.06944444 | 0.033769 | 0.014206 | 0.021308 | 0.004805 | 0.014616 | 0.017207 | 0.010863 | 0.005365 | 0.011781 |
| 27.08333333 | 0.018473 | 0.014524 | 0.014625 | 0.003149 | 0.012337 | 0.019679 | 0.007346 | 0.006802 | 0.006095 |
| 27.09722222 | 0.007043 | 0.010352 | 0.010838 | 0.002731 | 0.01165 | 0.02028 | 0.007088 | 0.009523 | 0.004111 |
| 27.11111111 | 0.003428 | 0.006368 | 0.011571 | 0.004266 | 0.011686 | 0.018127 | 0.008347 | 0.00953 | 0.006164 |
| 27.125 | 0.004298 | 0.006374 | 0.013044 | 0.004802 | 0.010781 | 0.013337 | 0.008365 | 0.008916 | 0.007074 |
| 27.13888889 | 0.005065 | 0.008943 | 0.012513 | 0.003182 | 0.008543 | 0.008162 | 0.006636 | 0.008422 | 0.004805 |

| | | | | | | | | | |
|---|---|---|---|---|---|---|---|---|---|
| 27.15277778 | 0.004945 | 0.011708 | 0.010887 | 0.002646 | 0.006431 | 0.005295 | 0.004893 | 0.006792 | 0.003571 |
| 27.16666667 | 0.004727 | 0.013059 | 0.009897 | 0.004716 | 0.0052 | 0.006611 | 0.005 | 0.003892 | 0.004195 |
| 27.18055556 | 0.005003 | 0.012253 | 0.009273 | 0.005931 | 0.004498 | 0.011118 | 0.006747 | 0.002586 | 0.00333 |
| 27.19444444 | 0.005956 | 0.00998 | 0.007883 | 0.004709 | 0.003823 | 0.014152 | 0.007798 | 0.004589 | 0.001331 |
| 27.20833333 | 0.006798 | 0.007663 | 0.005768 | 0.003921 | 0.002928 | 0.01228 | 0.006285 | 0.005964 | 0.001143 |
| 27.22222222 | 0.006654 | 0.00518 | 0.003814 | 0.00539 | 0.001943 | 0.006799 | 0.00341 | 0.004401 | 0.003406 |
| 27.23611111 | 0.005743 | 0.003116 | 0.002558 | 0.00769 | 0.001477 | 0.002663 | 0.002263 | 0.003104 | 0.006266 |
| 27.25 | 0.004908 | 0.003661 | 0.001972 | 0.008476 | 0.002292 | 0.001799 | 0.003183 | 0.002753 | 0.007937 |
| 27.26388889 | 0.004373 | 0.005114 | 0.001788 | 0.006855 | 0.003575 | 0.002646 | 0.003561 | 0.001637 | 0.007966 |
| 27.27777778 | 0.004044 | 0.003965 | 0.001517 | 0.003936 | 0.003807 | 0.00584 | 0.002325 | 0.000606 | 0.006987 |
| 27.29166667 | 0.004027 | 0.001512 | 0.001101 | 0.002581 | 0.003437 | 0.009137 | 0.000816 | 0.000329 | 0.005867 |
| 27.30555556 | 0.004081 | 0.000837 | 0.001731 | 0.003539 | 0.003928 | 0.009008 | 0.000325 | 0.000891 | 0.004772 |
| 27.31944444 | 0.00395 | 0.001961 | 0.004737 | 0.005197 | 0.005403 | 0.00789 | 0.000565 | 0.003463 | 0.003371 |
| 27.33333333 | 0.003644 | 0.002445 | 0.009712 | 0.009112 | 0.006274 | 0.010733 | 0.001393 | 0.009058 | 0.001915 |
| 27.34722222 | 0.003329 | 0.003037 | 0.014474 | 0.016339 | 0.005219 | 0.018257 | 0.003248 | 0.015819 | 0.002088 |
| 27.36111111 | 0.004177 | 0.006196 | 0.017199 | 0.022558 | 0.003629 | 0.02593 | 0.005014 | 0.019366 | 0.004974 |
| 27.375 | 0.007691 | 0.008999 | 0.018155 | 0.024566 | 0.003313 | 0.0305 | 0.00574 | 0.018727 | 0.008606 |
| 27.38888889 | 0.013303 | 0.009755 | 0.019326 | 0.024114 | 0.003359 | 0.032537 | 0.005857 | 0.01731 | 0.010433 |
| 27.40277778 | 0.019252 | 0.010777 | 0.022027 | 0.022473 | 0.002523 | 0.031182 | 0.005299 | 0.016812 | 0.010019 |
| 27.41666667 | 0.024939 | 0.013639 | 0.025208 | 0.019092 | 0.00147 | 0.025169 | 0.003882 | 0.016646 | 0.008588 |
| 27.43055556 | 0.030399 | 0.018702 | 0.026835 | 0.015813 | 0.001725 | 0.018884 | 0.003623 | 0.017722 | 0.008067 |
| 27.44444444 | 0.034387 | 0.023645 | 0.025089 | 0.015269 | 0.002967 | 0.017885 | 0.007406 | 0.02063 | 0.009952 |
| 27.45833333 | 0.034411 | 0.023999 | 0.018513 | 0.01683 | 0.002945 | 0.021193 | 0.014005 | 0.022678 | 0.013471 |
| 27.47222222 | 0.030453 | 0.017847 | 0.009183 | 0.020331 | 0.0043 | 0.023746 | 0.019141 | 0.021709 | 0.016757 |
| 27.48611111 | 0.027342 | 0.009694 | 0.003584 | 0.028549 | 0.016019 | 0.023021 | 0.023047 | 0.0202 | 0.022058 |
| 27.5 | 0.029696 | 0.006892 | 0.009028 | 0.037084 | 0.035826 | 0.020252 | 0.029143 | 0.019376 | 0.034244 |
| 27.51388889 | 0.035088 | 0.013033 | 0.027236 | 0.036184 | 0.042204 | 0.016192 | 0.037084 | 0.017338 | 0.051815 |
| 27.52777778 | 0.036274 | 0.024517 | 0.043775 | 0.027003 | 0.029755 | 0.009595 | 0.04403 | 0.014847 | 0.063614 |
| 27.54166667 | 0.028298 | 0.032297 | 0.039539 | 0.016592 | 0.018394 | 0.005044 | 0.044503 | 0.011244 | 0.057973 |
| 27.55555556 | 0.014524 | 0.030093 | 0.022897 | 0.008168 | 0.015538 | 0.006409 | 0.033879 | 0.005496 | 0.037216 |
| 27.56944444 | 0.005808 | 0.020392 | 0.01949 | 0.004237 | 0.014702 | 0.007009 | 0.017639 | 0.002783 | 0.015536 |
| 27.58333333 | 0.004618 | 0.012377 | 0.028698 | 0.00487 | 0.011781 | 0.004498 | 0.010659 | 0.007305 | 0.003791 |
| 27.59722222 | 0.005112 | 0.01351 | 0.028163 | 0.007965 | 0.006463 | 0.004597 | 0.016739 | 0.017339 | 0.001468 |
| 27.61111111 | 0.008396 | 0.022442 | 0.020001 | 0.011697 | 0.002006 | 0.011152 | 0.018847 | 0.029282 | 0.002737 |
| 27.625 | 0.013778 | 0.031655 | 0.026584 | 0.011862 | 0.000697 | 0.021628 | 0.012944 | 0.039669 | 0.006101 |
| 27.63888889 | 0.014812 | 0.036135 | 0.043726 | 0.00732 | 0.002204 | 0.029148 | 0.014894 | 0.046929 | 0.010326 |
| 27.65277778 | 0.013341 | 0.033997 | 0.048825 | 0.00343 | 0.005869 | 0.028492 | 0.027473 | 0.051383 | 0.012537 |
| 27.66666667 | 0.018288 | 0.025298 | 0.034059 | 0.003637 | 0.010116 | 0.018233 | 0.034692 | 0.050105 | 0.010337 |
| 27.68055556 | 0.031523 | 0.013956 | 0.017979 | 0.007634 | 0.011319 | 0.009468 | 0.02449 | 0.040981 | 0.006201 |
| 27.69444444 | 0.04015 | 0.010015 | 0.020433 | 0.012529 | 0.00791 | 0.014805 | 0.014233 | 0.027691 | 0.004408 |
| 27.70833333 | 0.031796 | 0.01924 | 0.029342 | 0.013848 | 0.006379 | 0.025196 | 0.023518 | 0.014221 | 0.004872 |
| 27.72222222 | 0.017585 | 0.027449 | 0.0295 | 0.010195 | 0.012023 | 0.023778 | 0.039526 | 0.009066 | 0.008728 |
| 27.73611111 | 0.016459 | 0.022469 | 0.023761 | 0.006804 | 0.017515 | 0.012845 | 0.045723 | 0.018818 | 0.011233 |
| 27.75 | 0.027402 | 0.011831 | 0.019372 | 0.010545 | 0.015891 | 0.006475 | 0.040716 | 0.034138 | 0.008475 |

| | | | | | | | | | |
|---|---|---|---|---|---|---|---|---|---|
| 27.76388889 | 0.038076 | 0.00699 | 0.02023 | 0.016535 | 0.011038 | 0.005086 | 0.029356 | 0.040943 | 0.008737 |
| 27.77777778 | 0.043981 | 0.012334 | 0.024884 | 0.014951 | 0.007493 | 0.005269 | 0.017591 | 0.035714 | 0.017146 |
| 27.79166667 | 0.047006 | 0.024777 | 0.027366 | 0.008147 | 0.004675 | 0.008972 | 0.00863 | 0.027395 | 0.028765 |
| 27.80555556 | 0.044385 | 0.033837 | 0.020747 | 0.005081 | 0.004152 | 0.014314 | 0.00482 | 0.023786 | 0.035372 |
| 27.81944444 | 0.032839 | 0.031257 | 0.010469 | 0.010385 | 0.008621 | 0.021523 | 0.007461 | 0.023928 | 0.032189 |
| 27.83333333 | 0.01781 | 0.021631 | 0.005944 | 0.022253 | 0.014941 | 0.031476 | 0.01031 | 0.026837 | 0.02355 |
| 27.84722222 | 0.007216 | 0.013482 | 0.006765 | 0.032625 | 0.019406 | 0.040161 | 0.008986 | 0.033418 | 0.018395 |
| 27.86111111 | 0.004959 | 0.011018 | 0.014751 | 0.032252 | 0.021872 | 0.043447 | 0.011046 | 0.041721 | 0.021829 |
| 27.875 | 0.008325 | 0.016024 | 0.029142 | 0.020932 | 0.021735 | 0.041229 | 0.018031 | 0.045589 | 0.03148 |
| 27.88888889 | 0.00895 | 0.027805 | 0.039284 | 0.009876 | 0.018762 | 0.03717 | 0.02049 | 0.039823 | 0.038312 |
| 27.90277778 | 0.006147 | 0.042635 | 0.037354 | 0.01034 | 0.015229 | 0.033843 | 0.01665 | 0.02696 | 0.036686 |
| 27.91666667 | 0.006939 | 0.051803 | 0.025261 | 0.022552 | 0.011403 | 0.028878 | 0.012101 | 0.015528 | 0.028919 |
| 27.93055556 | 0.009959 | 0.046681 | 0.011617 | 0.038024 | 0.006017 | 0.019983 | 0.009147 | 0.012036 | 0.018655 |
| 27.94444444 | 0.008195 | 0.031716 | 0.003843 | 0.046274 | 0.001994 | 0.01033 | 0.007441 | 0.015914 | 0.009284 |
| 27.95833333 | 0.005284 | 0.018632 | 0.002963 | 0.040219 | 0.00169 | 0.003936 | 0.006381 | 0.020648 | 0.004312 |
| 27.97222222 | 0.007902 | 0.012839 | 0.005245 | 0.02471 | 0.00257 | 0.000993 | 0.005092 | 0.020282 | 0.004544 |
| 27.98611111 | 0.013378 | 0.017017 | 0.00809 | 0.012124 | 0.002998 | 0.001683 | 0.003604 | 0.013633 | 0.008231 |
| 28 | 0.019903 | 0.036611 | 0.013174 | 0.009608 | 0.002955 | 0.007772 | 0.003602 | 0.005785 | 0.015086 |
| 28.01388889 | 0.030393 | 0.066941 | 0.019185 | 0.013375 | 0.001867 | 0.016863 | 0.004706 | 0.004627 | 0.023466 |
| 28.02777778 | 0.04416 | 0.080682 | 0.016388 | 0.013089 | 0.000571 | 0.021701 | 0.004957 | 0.01332 | 0.024681 |
| 28.04166667 | 0.054063 | 0.057596 | 0.00992 | 0.007456 | 0.001389 | 0.017847 | 0.004855 | 0.027712 | 0.016654 |
| 28.05555556 | 0.053682 | 0.027336 | 0.014158 | 0.006572 | 0.005579 | 0.011322 | 0.004532 | 0.036961 | 0.011631 |
| 28.06944444 | 0.044297 | 0.024207 | 0.019248 | 0.012319 | 0.011512 | 0.014035 | 0.002828 | 0.033071 | 0.012447 |
| 28.08333333 | 0.0306 | 0.034299 | 0.016113 | 0.012483 | 0.016877 | 0.027066 | 0.001083 | 0.018624 | 0.011268 |
| 28.09722222 | 0.016523 | 0.040781 | 0.014015 | 0.009285 | 0.020973 | 0.041971 | 0.001956 | 0.008159 | 0.010528 |
| 28.11111111 | 0.006455 | 0.044796 | 0.018292 | 0.014614 | 0.023332 | 0.053334 | 0.004985 | 0.010505 | 0.014787 |
| 28.125 | 0.002532 | 0.04372 | 0.024302 | 0.020211 | 0.02284 | 0.058678 | 0.007468 | 0.01446 | 0.021046 |
| 28.13888889 | 0.002708 | 0.036003 | 0.025255 | 0.018075 | 0.0188 | 0.057797 | 0.007999 | 0.013605 | 0.024864 |
| 28.15277778 | 0.004922 | 0.025926 | 0.019122 | 0.011564 | 0.012974 | 0.048652 | 0.006572 | 0.012098 | 0.023825 |
| 28.16666667 | 0.008844 | 0.01662 | 0.009873 | 0.00616 | 0.008151 | 0.03144 | 0.003813 | 0.010486 | 0.017798 |
| 28.18055556 | 0.012828 | 0.009128 | 0.003235 | 0.005709 | 0.004759 | 0.016646 | 0.001298 | 0.008333 | 0.00906 |
| 28.19444444 | 0.014672 | 0.005573 | 0.001665 | 0.005518 | 0.002821 | 0.011659 | 0.000826 | 0.006527 | 0.002984 |
| 28.20833333 | 0.014658 | 0.006574 | 0.005415 | 0.00419 | 0.003301 | 0.014072 | 0.002637 | 0.004825 | 0.001946 |
| 28.22222222 | 0.01436 | 0.009358 | 0.013079 | 0.003582 | 0.005956 | 0.018373 | 0.005312 | 0.00297 | 0.003713 |
| 28.23611111 | 0.01413 | 0.010536 | 0.019049 | 0.002839 | 0.008933 | 0.020231 | 0.007152 | 0.002084 | 0.006935 |
| 28.25 | 0.013232 | 0.008998 | 0.017919 | 0.003329 | 0.010261 | 0.018731 | 0.007416 | 0.002577 | 0.008476 |
| 28.26388889 | 0.011644 | 0.006024 | 0.01125 | 0.006738 | 0.009184 | 0.014629 | 0.00637 | 0.003229 | 0.006468 |
| 28.27777778 | 0.010784 | 0.003647 | 0.005113 | 0.012595 | 0.00641 | 0.010406 | 0.004773 | 0.002943 | 0.003735 |
| 28.29166667 | 0.011765 | 0.002901 | 0.003871 | 0.014197 | 0.003382 | 0.010189 | 0.003416 | 0.002456 | 0.002941 |
| 28.30555556 | 0.013324 | 0.003024 | 0.007874 | 0.008586 | 0.001227 | 0.015143 | 0.002224 | 0.003057 | 0.004149 |
| 28.31944444 | 0.013684 | 0.002604 | 0.013553 | 0.003483 | 0.000484 | 0.021514 | 0.001417 | 0.005251 | 0.005384 |
| 28.33333333 | 0.013174 | 0.001767 | 0.016213 | 0.005533 | 0.000668 | 0.024232 | 0.001503 | 0.009695 | 0.005321 |
| 28.34722222 | 0.012969 | 0.001718 | 0.014844 | 0.01326 | 0.000809 | 0.022126 | 0.001791 | 0.016649 | 0.004841 |
| 28.36111111 | 0.013088 | 0.002819 | 0.01304 | 0.02018 | 0.001403 | 0.018669 | 0.002828 | 0.024201 | 0.005745 |

| | | | | | | | | | |
|---|---|---|---|---|---|---|---|---|---|
| 28.375 | 0.013052 | 0.004804 | 0.015131 | 0.022675 | 0.003026 | 0.017091 | 0.004978 | 0.029213 | 0.009196 |
| 28.38888889 | 0.013856 | 0.008082 | 0.022321 | 0.022194 | 0.004468 | 0.017869 | 0.005808 | 0.030409 | 0.014751 |
| 28.40277778 | 0.01729 | 0.012663 | 0.030678 | 0.021236 | 0.003948 | 0.02027 | 0.003972 | 0.029491 | 0.020271 |
| 28.41666667 | 0.022887 | 0.016864 | 0.033802 | 0.020567 | 0.001875 | 0.024192 | 0.001363 | 0.028983 | 0.024073 |
| 28.43055556 | 0.027455 | 0.018852 | 0.029981 | 0.020061 | 0.001328 | 0.030171 | 0.001295 | 0.029118 | 0.02592 |
| 28.44444444 | 0.027578 | 0.017661 | 0.022763 | 0.019332 | 0.003524 | 0.035225 | 0.00417 | 0.027149 | 0.024793 |
| 28.45833333 | 0.022344 | 0.01225 | 0.014886 | 0.017896 | 0.00579 | 0.032236 | 0.006217 | 0.021403 | 0.01847 |
| 28.47222222 | 0.016241 | 0.005618 | 0.010694 | 0.020334 | 0.006071 | 0.020843 | 0.005435 | 0.015374 | 0.00922 |
| 28.48611111 | 0.019813 | 0.006009 | 0.019659 | 0.038291 | 0.006921 | 0.012131 | 0.004845 | 0.016255 | 0.006776 |
| 28.5 | 0.039214 | 0.018864 | 0.04364 | 0.068183 | 0.012919 | 0.01374 | 0.008408 | 0.02828 | 0.019027 |
| 28.51388889 | 0.063279 | 0.038813 | 0.068743 | 0.083671 | 0.020342 | 0.021464 | 0.015845 | 0.04457 | 0.042603 |
| 28.52777778 | 0.071261 | 0.048121 | 0.075708 | 0.071846 | 0.019761 | 0.027954 | 0.023092 | 0.049343 | 0.06683 |
| 28.54166667 | 0.056084 | 0.03864 | 0.055259 | 0.043071 | 0.012855 | 0.02484 | 0.026168 | 0.036506 | 0.075639 |
| 28.55555556 | 0.030307 | 0.021281 | 0.029241 | 0.019825 | 0.008209 | 0.013679 | 0.023406 | 0.018677 | 0.055319 |
| 28.56944444 | 0.014869 | 0.012181 | 0.028943 | 0.01723 | 0.008204 | 0.011021 | 0.016161 | 0.012398 | 0.027273 |
| 28.58333333 | 0.016553 | 0.016635 | 0.04454 | 0.016747 | 0.010154 | 0.022797 | 0.008591 | 0.017312 | 0.023751 |
| 28.59722222 | 0.018681 | 0.021997 | 0.045835 | 0.013985 | 0.010596 | 0.032328 | 0.0054 | 0.017972 | 0.032038 |
| 28.61111111 | 0.012747 | 0.018884 | 0.02976 | 0.027099 | 0.008761 | 0.025618 | 0.00746 | 0.012063 | 0.026151 |
| 28.625 | 0.008367 | 0.019638 | 0.023244 | 0.038079 | 0.008146 | 0.023107 | 0.01043 | 0.012975 | 0.016985 |
| 28.63888889 | 0.008404 | 0.030651 | 0.03458 | 0.02779 | 0.010081 | 0.042311 | 0.009151 | 0.021516 | 0.026256 |
| 28.65277778 | 0.007187 | 0.034977 | 0.038634 | 0.014132 | 0.010596 | 0.060935 | 0.005762 | 0.025862 | 0.039194 |
| 28.66666667 | 0.006666 | 0.028382 | 0.031197 | 0.015839 | 0.008452 | 0.057411 | 0.006377 | 0.024268 | 0.034417 |
| 28.68055556 | 0.010293 | 0.019531 | 0.021531 | 0.025816 | 0.006961 | 0.034772 | 0.009449 | 0.02438 | 0.018858 |
| 28.69444444 | 0.017906 | 0.012322 | 0.012752 | 0.022917 | 0.00835 | 0.017997 | 0.009152 | 0.026783 | 0.011334 |
| 28.70833333 | 0.033371 | 0.008841 | 0.010545 | 0.016269 | 0.011766 | 0.025125 | 0.006065 | 0.023845 | 0.017912 |
| 28.72222222 | 0.046986 | 0.014053 | 0.011751 | 0.028016 | 0.014047 | 0.039506 | 0.005674 | 0.016211 | 0.019866 |
| 28.73611111 | 0.039164 | 0.026861 | 0.013566 | 0.036642 | 0.012925 | 0.041384 | 0.007561 | 0.014876 | 0.012586 |
| 28.75 | 0.022329 | 0.026371 | 0.016332 | 0.023582 | 0.009862 | 0.027792 | 0.006895 | 0.02283 | 0.013042 |
| 28.76388889 | 0.026168 | 0.013535 | 0.023686 | 0.01198 | 0.007922 | 0.014078 | 0.004439 | 0.029921 | 0.017299 |
| 28.77777778 | 0.045543 | 0.009308 | 0.041967 | 0.017952 | 0.007214 | 0.014535 | 0.004261 | 0.025068 | 0.016151 |
| 28.79166667 | 0.057682 | 0.019285 | 0.066523 | 0.028389 | 0.004926 | 0.015683 | 0.00579 | 0.017982 | 0.024279 |
| 28.80555556 | 0.059566 | 0.051168 | 0.080279 | 0.029589 | 0.003334 | 0.020719 | 0.006151 | 0.03131 | 0.047882 |
| 28.81944444 | 0.055771 | 0.096517 | 0.069755 | 0.022933 | 0.007288 | 0.047461 | 0.00582 | 0.060509 | 0.066507 |
| 28.83333333 | 0.049626 | 0.125795 | 0.045994 | 0.016609 | 0.014446 | 0.07619 | 0.006677 | 0.080704 | 0.066969 |
| 28.84722222 | 0.045376 | 0.121662 | 0.029806 | 0.017779 | 0.0199 | 0.082515 | 0.008919 | 0.080881 | 0.058196 |
| 28.86111111 | 0.044089 | 0.093329 | 0.028393 | 0.029404 | 0.02154 | 0.072349 | 0.010932 | 0.069297 | 0.052174 |
| 28.875 | 0.045671 | 0.061725 | 0.03614 | 0.051295 | 0.01904 | 0.059509 | 0.011275 | 0.059531 | 0.049731 |
| 28.88888889 | 0.046319 | 0.036079 | 0.042481 | 0.076645 | 0.013648 | 0.047761 | 0.009115 | 0.055217 | 0.046187 |
| 28.90277778 | 0.040872 | 0.01786 | 0.040601 | 0.085988 | 0.008438 | 0.037289 | 0.004851 | 0.051664 | 0.040858 |
| 28.91666667 | 0.030537 | 0.009053 | 0.031216 | 0.066615 | 0.006495 | 0.029183 | 0.001708 | 0.045387 | 0.035152 |
| 28.93055556 | 0.019557 | 0.010542 | 0.020315 | 0.033529 | 0.007197 | 0.021443 | 0.001078 | 0.034846 | 0.028408 |
| 28.94444444 | 0.010046 | 0.018047 | 0.012619 | 0.010012 | 0.006934 | 0.012175 | 0.001438 | 0.019978 | 0.01936 |
| 28.95833333 | 0.003797 | 0.023588 | 0.008274 | 0.002818 | 0.004204 | 0.005365 | 0.002678 | 0.008164 | 0.009453 |
| 28.97222222 | 0.002301 | 0.025556 | 0.007033 | 0.007288 | 0.001532 | 0.003115 | 0.005466 | 0.00393 | 0.003081 |

| | | | | | | | | | |
|---|---|---|---|---|---|---|---|---|---|
| 28.98611111 | 0.00731 | 0.032794 | 0.012522 | 0.02288 | 0.001848 | 0.007342 | 0.010236 | 0.006037 | 0.006012 |
| 29 | 0.0179 | 0.048711 | 0.026267 | 0.04513 | 0.005881 | 0.023972 | 0.015366 | 0.016283 | 0.020091 |
| 29.01388889 | 0.026729 | 0.062283 | 0.041293 | 0.059589 | 0.014088 | 0.048119 | 0.017766 | 0.028211 | 0.038697 |
| 29.02777778 | 0.033397 | 0.069645 | 0.055345 | 0.061164 | 0.029046 | 0.07636 | 0.020038 | 0.039416 | 0.058163 |
| 29.04166667 | 0.044187 | 0.078737 | 0.073563 | 0.055905 | 0.052993 | 0.112478 | 0.026715 | 0.054527 | 0.077422 |
| 29.05555556 | 0.053337 | 0.081713 | 0.085236 | 0.046212 | 0.076961 | 0.139716 | 0.032657 | 0.065034 | 0.082092 |
| 29.06944444 | 0.0494 | 0.064165 | 0.072646 | 0.034891 | 0.08034 | 0.128331 | 0.030137 | 0.061608 | 0.061877 |
| 29.08333333 | 0.034772 | 0.035434 | 0.042574 | 0.0285 | 0.057379 | 0.082943 | 0.022472 | 0.048384 | 0.032099 |
| 29.09722222 | 0.021325 | 0.014477 | 0.017311 | 0.027625 | 0.027437 | 0.038633 | 0.018627 | 0.034849 | 0.011982 |
| 29.11111111 | 0.014404 | 0.00646 | 0.010199 | 0.024682 | 0.01085 | 0.015673 | 0.020061 | 0.024871 | 0.00385 |
| 29.125 | 0.009952 | 0.010433 | 0.018492 | 0.016049 | 0.01359 | 0.019643 | 0.020678 | 0.016635 | 0.004125 |
| 29.13888889 | 0.005562 | 0.024302 | 0.031678 | 0.012102 | 0.02712 | 0.046942 | 0.016031 | 0.009079 | 0.012263 |
| 29.15277778 | 0.005257 | 0.041374 | 0.044676 | 0.016665 | 0.037789 | 0.081629 | 0.008095 | 0.006612 | 0.027331 |
| 29.16666667 | 0.008583 | 0.053901 | 0.054944 | 0.016246 | 0.041071 | 0.10285 | 0.002709 | 0.011379 | 0.045989 |
| 29.18055556 | 0.008521 | 0.057366 | 0.056386 | 0.009266 | 0.039878 | 0.101084 | 0.002068 | 0.015866 | 0.059149 |
| 29.19444444 | 0.006328 | 0.053309 | 0.04619 | 0.007237 | 0.034547 | 0.088663 | 0.004924 | 0.014262 | 0.059309 |
| 29.20833333 | 0.009786 | 0.043858 | 0.030417 | 0.011757 | 0.024109 | 0.075585 | 0.010521 | 0.008786 | 0.048241 |
| 29.22222222 | 0.016422 | 0.030016 | 0.015938 | 0.016521 | 0.011973 | 0.059113 | 0.015349 | 0.00585 | 0.030542 |
| 29.23611111 | 0.019011 | 0.017226 | 0.006096 | 0.017329 | 0.005539 | 0.03854 | 0.016272 | 0.007824 | 0.014075 |
| 29.25 | 0.016896 | 0.008419 | 0.002514 | 0.01421 | 0.006458 | 0.018424 | 0.014825 | 0.009702 | 0.008824 |
| 29.26388889 | 0.01348 | 0.004902 | 0.002608 | 0.01033 | 0.008251 | 0.007655 | 0.013936 | 0.008776 | 0.01389 |
| 29.27777778 | 0.010923 | 0.010192 | 0.003863 | 0.009247 | 0.008897 | 0.009435 | 0.014645 | 0.006604 | 0.019151 |
| 29.29166667 | 0.008831 | 0.020469 | 0.006748 | 0.011753 | 0.010191 | 0.013707 | 0.015287 | 0.004956 | 0.020073 |
| 29.30555556 | 0.006868 | 0.029131 | 0.009195 | 0.014108 | 0.011654 | 0.013437 | 0.013816 | 0.003944 | 0.017699 |
| 29.31944444 | 0.006356 | 0.03425 | 0.008096 | 0.011389 | 0.013396 | 0.009586 | 0.01155 | 0.003616 | 0.014332 |
| 29.33333333 | 0.008769 | 0.036023 | 0.004373 | 0.006085 | 0.016501 | 0.005421 | 0.012431 | 0.006104 | 0.012884 |
| 29.34722222 | 0.012061 | 0.032615 | 0.00269 | 0.004755 | 0.019341 | 0.003208 | 0.016547 | 0.010553 | 0.012276 |
| 29.36111111 | 0.010922 | 0.020826 | 0.006402 | 0.006785 | 0.017393 | 0.003099 | 0.018752 | 0.011904 | 0.00946 |
| 29.375 | 0.006099 | 0.012493 | 0.012691 | 0.009275 | 0.010152 | 0.006498 | 0.01557 | 0.007902 | 0.009862 |
| 29.38888889 | 0.003253 | 0.019237 | 0.016414 | 0.014524 | 0.003854 | 0.01127 | 0.008554 | 0.004322 | 0.018432 |
| 29.40277778 | 0.001991 | 0.028527 | 0.015371 | 0.023899 | 0.00232 | 0.013971 | 0.003269 | 0.006193 | 0.028272 |
| 29.41666667 | 0.001281 | 0.028338 | 0.011147 | 0.031019 | 0.002065 | 0.015558 | 0.002015 | 0.008181 | 0.034168 |
| 29.43055556 | 0.003371 | 0.017612 | 0.008151 | 0.028225 | 0.001752 | 0.017432 | 0.002607 | 0.005482 | 0.03636 |
| 29.44444444 | 0.007147 | 0.008793 | 0.009284 | 0.017174 | 0.003092 | 0.019072 | 0.005484 | 0.001628 | 0.032883 |
| 29.45833333 | 0.008584 | 0.013326 | 0.012375 | 0.006282 | 0.004089 | 0.020207 | 0.008371 | 0.002061 | 0.022233 |
| 29.47222222 | 0.008619 | 0.016963 | 0.015225 | 0.006756 | 0.003983 | 0.021558 | 0.007699 | 0.008811 | 0.009695 |
| 29.48611111 | 0.017887 | 0.014254 | 0.022167 | 0.030449 | 0.005216 | 0.025083 | 0.004633 | 0.021559 | 0.004579 |
| 29.5 | 0.042249 | 0.020294 | 0.036161 | 0.0719 | 0.007829 | 0.030132 | 0.004538 | 0.038448 | 0.012498 |
| 29.51388889 | 0.065304 | 0.036114 | 0.051831 | 0.101453 | 0.007664 | 0.032349 | 0.011173 | 0.051894 | 0.033784 |
| 29.52777778 | 0.069542 | 0.049007 | 0.060715 | 0.098395 | 0.005303 | 0.03369 | 0.019021 | 0.052751 | 0.060838 |
| 29.54166667 | 0.05834 | 0.05097 | 0.056061 | 0.069404 | 0.005121 | 0.033977 | 0.021129 | 0.040289 | 0.071323 |
| 29.55555556 | 0.039664 | 0.038905 | 0.038519 | 0.034449 | 0.004134 | 0.024633 | 0.015779 | 0.021205 | 0.050707 |
| 29.56944444 | 0.018713 | 0.020134 | 0.019392 | 0.018021 | 0.003264 | 0.01214 | 0.009258 | 0.009307 | 0.023859 |
| 29.58333333 | 0.004545 | 0.011251 | 0.014251 | 0.021448 | 0.007351 | 0.008688 | 0.010982 | 0.01289 | 0.018919 |

| | | | | | | | | | |
|---|---|---|---|---|---|---|---|---|---|
| 29.59722222 | 0.002483 | 0.013555 | 0.022289 | 0.019167 | 0.010482 | 0.008356 | 0.01575 | 0.01907 | 0.026207 |
| 29.61111111 | 0.009226 | 0.014336 | 0.023945 | 0.016538 | 0.007607 | 0.005934 | 0.012756 | 0.015108 | 0.025525 |
| 29.625 | 0.020375 | 0.011305 | 0.017029 | 0.034438 | 0.007706 | 0.008796 | 0.012214 | 0.008867 | 0.017923 |
| 29.63888889 | 0.031634 | 0.016286 | 0.020863 | 0.051094 | 0.017551 | 0.020827 | 0.024822 | 0.012937 | 0.022914 |
| 29.65277778 | 0.037138 | 0.03245 | 0.033661 | 0.041321 | 0.027779 | 0.036483 | 0.035944 | 0.0234 | 0.046163 |
| 29.66666667 | 0.031991 | 0.043284 | 0.035474 | 0.020408 | 0.028447 | 0.045199 | 0.033698 | 0.027859 | 0.064457 |
| 29.68055556 | 0.018807 | 0.033503 | 0.023587 | 0.014049 | 0.020386 | 0.04034 | 0.021476 | 0.022311 | 0.057394 |
| 29.69444444 | 0.013091 | 0.012582 | 0.009663 | 0.022514 | 0.009955 | 0.027708 | 0.011939 | 0.013005 | 0.032798 |
| 29.70833333 | 0.019526 | 0.002417 | 0.003777 | 0.024887 | 0.003299 | 0.015844 | 0.012871 | 0.006119 | 0.018481 |
| 29.72222222 | 0.019688 | 0.002622 | 0.00399 | 0.024589 | 0.003197 | 0.007297 | 0.011416 | 0.004113 | 0.02476 |
| 29.73611111 | 0.02021 | 0.002286 | 0.008063 | 0.044896 | 0.007263 | 0.005172 | 0.008062 | 0.005571 | 0.036302 |
| 29.75 | 0.037378 | 0.008888 | 0.018638 | 0.062268 | 0.009386 | 0.012353 | 0.015994 | 0.004917 | 0.041944 |
| 29.76388889 | 0.048617 | 0.033166 | 0.02232 | 0.043695 | 0.006795 | 0.02285 | 0.026456 | 0.002509 | 0.036782 |
| 29.77777778 | 0.038713 | 0.045616 | 0.020155 | 0.024672 | 0.005632 | 0.029116 | 0.024967 | 0.001905 | 0.021701 |
| 29.79166667 | 0.023029 | 0.029799 | 0.037209 | 0.053365 | 0.011439 | 0.031018 | 0.015074 | 0.003416 | 0.009701 |
| 29.80555556 | 0.021834 | 0.016654 | 0.070725 | 0.104616 | 0.019169 | 0.030399 | 0.015422 | 0.003927 | 0.006564 |
| 29.81944444 | 0.037642 | 0.015213 | 0.097443 | 0.113824 | 0.023302 | 0.029104 | 0.026163 | 0.005579 | 0.00971 |
| 29.83333333 | 0.054271 | 0.028363 | 0.101198 | 0.068291 | 0.02334 | 0.032133 | 0.026785 | 0.015245 | 0.024837 |
| 29.84722222 | 0.061537 | 0.059794 | 0.076626 | 0.023405 | 0.020414 | 0.043469 | 0.015641 | 0.028587 | 0.049299 |
| 29.86111111 | 0.062013 | 0.099071 | 0.040661 | 0.011667 | 0.016552 | 0.056212 | 0.007516 | 0.039513 | 0.068438 |
| 29.875 | 0.061193 | 0.133425 | 0.015589 | 0.0209 | 0.012929 | 0.058687 | 0.006487 | 0.046803 | 0.074115 |
| 29.88888889 | 0.058074 | 0.140615 | 0.005058 | 0.046964 | 0.01099 | 0.050589 | 0.004734 | 0.047268 | 0.067835 |
| 29.90277778 | 0.051445 | 0.110933 | 0.003056 | 0.074554 | 0.011555 | 0.038461 | 0.00414 | 0.040441 | 0.055076 |
| 29.91666667 | 0.044008 | 0.063084 | 0.004207 | 0.083183 | 0.012343 | 0.026428 | 0.009144 | 0.030715 | 0.040105 |
| 29.93055556 | 0.035912 | 0.026454 | 0.004057 | 0.069115 | 0.010558 | 0.017554 | 0.010921 | 0.022692 | 0.026266 |
| 29.94444444 | 0.024822 | 0.016808 | 0.002366 | 0.041043 | 0.006226 | 0.013243 | 0.008902 | 0.017112 | 0.01701 |
| 29.95833333 | 0.012145 | 0.023452 | 0.001193 | 0.016148 | 0.003484 | 0.012317 | 0.014792 | 0.011994 | 0.012631 |
| 29.97222222 | 0.004142 | 0.025669 | 0.002336 | 0.004525 | 0.004267 | 0.014428 | 0.025918 | 0.007771 | 0.01158 |
| 29.98611111 | 0.004929 | 0.020623 | 0.010341 | 0.010379 | 0.005023 | 0.023 | 0.023626 | 0.009542 | 0.017855 |
| 30 | 0.013557 | 0.027069 | 0.027617 | 0.030695 | 0.01029 | 0.0428 | 0.015933 | 0.020207 | 0.036308 |
| 30.01388889 | 0.022649 | 0.048991 | 0.045734 | 0.045044 | 0.027085 | 0.06811 | 0.030379 | 0.033501 | 0.059353 |
| 30.02777778 | 0.023945 | 0.064193 | 0.049602 | 0.03901 | 0.045535 | 0.083492 | 0.050583 | 0.040214 | 0.068797 |
| 30.04166667 | 0.017172 | 0.062193 | 0.03602 | 0.025701 | 0.051714 | 0.081362 | 0.044958 | 0.038698 | 0.056373 |
| 30.05555556 | 0.009409 | 0.048221 | 0.01769 | 0.020123 | 0.042449 | 0.060457 | 0.024208 | 0.033044 | 0.031753 |
| 30.06944444 | 0.008503 | 0.028317 | 0.010444 | 0.019602 | 0.023426 | 0.032284 | 0.016796 | 0.024763 | 0.012192 |
| 30.08333333 | 0.012639 | 0.011515 | 0.016929 | 0.017072 | 0.010664 | 0.0215 | 0.022274 | 0.013628 | 0.006949 |
| 30.09722222 | 0.014313 | 0.004462 | 0.021963 | 0.011586 | 0.011825 | 0.026718 | 0.021141 | 0.005408 | 0.008879 |
| 30.11111111 | 0.011189 | 0.00311 | 0.017429 | 0.006024 | 0.016362 | 0.027303 | 0.012066 | 0.004743 | 0.010046 |
| 30.125 | 0.006816 | 0.002613 | 0.009118 | 0.003807 | 0.018841 | 0.022501 | 0.004315 | 0.006102 | 0.010693 |
| 30.13888889 | 0.006687 | 0.002839 | 0.002905 | 0.008802 | 0.021172 | 0.021521 | 0.002027 | 0.006212 | 0.012073 |
| 30.15277778 | 0.009298 | 0.00516 | 0.001625 | 0.020892 | 0.02381 | 0.026886 | 0.005664 | 0.007147 | 0.015307 |
| 30.16666667 | 0.007814 | 0.015698 | 0.00531 | 0.030603 | 0.025624 | 0.033157 | 0.011533 | 0.007905 | 0.021311 |
| 30.18055556 | 0.006035 | 0.034304 | 0.012775 | 0.029787 | 0.025337 | 0.035349 | 0.013655 | 0.006908 | 0.029282 |
| 30.19444444 | 0.010479 | 0.051511 | 0.021662 | 0.020493 | 0.024516 | 0.038298 | 0.009973 | 0.005138 | 0.037312 |

| | | | | | | | | | |
|---|---|---|---|---|---|---|---|---|---|
| 30.20833333 | 0.014865 | 0.060313 | 0.028955 | 0.009977 | 0.024242 | 0.04238 | 0.00632 | 0.003169 | 0.043146 |
| 30.22222222 | 0.014601 | 0.059359 | 0.032595 | 0.00556 | 0.023106 | 0.041363 | 0.008309 | 0.002088 | 0.043609 |
| 30.23611111 | 0.011844 | 0.050672 | 0.0313 | 0.006332 | 0.019976 | 0.035383 | 0.009891 | 0.002646 | 0.036608 |
| 30.25 | 0.007378 | 0.037267 | 0.02537 | 0.005665 | 0.016056 | 0.028087 | 0.006804 | 0.002521 | 0.024949 |
| 30.26388889 | 0.003547 | 0.022986 | 0.016808 | 0.004535 | 0.011725 | 0.019753 | 0.003467 | 0.001544 | 0.012901 |
| 30.27777778 | 0.003208 | 0.011042 | 0.007878 | 0.009113 | 0.007011 | 0.01081 | 0.004432 | 0.003311 | 0.005976 |
| 30.29166667 | 0.004061 | 0.005497 | 0.001914 | 0.018211 | 0.005546 | 0.006525 | 0.011593 | 0.008396 | 0.006873 |
| 30.30555556 | 0.003228 | 0.007417 | 0.001314 | 0.023718 | 0.007399 | 0.007428 | 0.018574 | 0.011315 | 0.00878 |
| 30.31944444 | 0.002434 | 0.011785 | 0.003082 | 0.021337 | 0.008172 | 0.006361 | 0.020679 | 0.00993 | 0.008854 |
| 30.33333333 | 0.002444 | 0.01568 | 0.003685 | 0.013509 | 0.008845 | 0.003578 | 0.020739 | 0.00856 | 0.011038 |
| 30.34722222 | 0.002241 | 0.017679 | 0.005409 | 0.006227 | 0.012124 | 0.003504 | 0.019912 | 0.009174 | 0.01477 |
| 30.36111111 | 0.003152 | 0.015017 | 0.008653 | 0.005489 | 0.015324 | 0.003543 | 0.016489 | 0.008468 | 0.015044 |
| 30.375 | 0.005494 | 0.008144 | 0.009003 | 0.011088 | 0.015193 | 0.002438 | 0.010533 | 0.005139 | 0.009377 |
| 30.38888889 | 0.005417 | 0.002886 | 0.005594 | 0.01613 | 0.013291 | 0.003349 | 0.006841 | 0.002177 | 0.003971 |
| 30.40277778 | 0.003837 | 0.004319 | 0.003219 | 0.015871 | 0.012524 | 0.005494 | 0.009974 | 0.002969 | 0.004459 |
| 30.41666667 | 0.006513 | 0.010079 | 0.005682 | 0.013561 | 0.013061 | 0.005368 | 0.017746 | 0.009211 | 0.006987 |
| 30.43055556 | 0.012365 | 0.013538 | 0.009729 | 0.014056 | 0.014465 | 0.003458 | 0.024052 | 0.016912 | 0.008187 |
| 30.44444444 | 0.015452 | 0.009944 | 0.010503 | 0.014399 | 0.01742 | 0.002832 | 0.02595 | 0.021404 | 0.008366 |
| 30.45833333 | 0.011329 | 0.007398 | 0.008638 | 0.010076 | 0.020731 | 0.003996 | 0.024134 | 0.021734 | 0.009145 |
| 30.47222222 | 0.007095 | 0.018635 | 0.009086 | 0.004145 | 0.020417 | 0.00665 | 0.019022 | 0.016159 | 0.014489 |
| 30.48611111 | 0.015884 | 0.040881 | 0.017556 | 0.009237 | 0.014739 | 0.014448 | 0.010734 | 0.012399 | 0.027612 |
| 30.5 | 0.036895 | 0.060872 | 0.035381 | 0.037583 | 0.007775 | 0.027468 | 0.00677 | 0.023158 | 0.046965 |
| 30.51388889 | 0.056295 | 0.069016 | 0.052288 | 0.079033 | 0.006956 | 0.038888 | 0.017427 | 0.039981 | 0.064756 |
| 30.52777778 | 0.060513 | 0.067025 | 0.056836 | 0.105233 | 0.010831 | 0.043983 | 0.034954 | 0.046001 | 0.07319 |
| 30.54166667 | 0.048577 | 0.056653 | 0.04821 | 0.094801 | 0.010748 | 0.038517 | 0.038344 | 0.037006 | 0.066665 |
| 30.55555556 | 0.027591 | 0.034806 | 0.028745 | 0.053334 | 0.007092 | 0.023025 | 0.022711 | 0.019911 | 0.043515 |
| 30.56944444 | 0.012389 | 0.017512 | 0.014393 | 0.021918 | 0.004388 | 0.01386 | 0.010653 | 0.009974 | 0.021666 |
| 30.58333333 | 0.010772 | 0.023416 | 0.021743 | 0.025486 | 0.007443 | 0.017677 | 0.017252 | 0.010457 | 0.022524 |
| 30.59722222 | 0.011584 | 0.039137 | 0.034125 | 0.032992 | 0.01708 | 0.01748 | 0.026151 | 0.009621 | 0.036405 |
| 30.61111111 | 0.017555 | 0.049672 | 0.028834 | 0.02531 | 0.021483 | 0.012713 | 0.022727 | 0.010556 | 0.042772 |
| 30.625 | 0.033311 | 0.05888 | 0.018022 | 0.034493 | 0.013874 | 0.018478 | 0.013483 | 0.020476 | 0.037484 |
| 30.63888889 | 0.044416 | 0.06061 | 0.02455 | 0.076162 | 0.008327 | 0.030967 | 0.012694 | 0.029871 | 0.02287 |
| 30.65277778 | 0.04375 | 0.05096 | 0.041838 | 0.09964 | 0.014645 | 0.038982 | 0.022826 | 0.030554 | 0.016766 |
| 30.66666667 | 0.034438 | 0.068822 | 0.053597 | 0.068041 | 0.019599 | 0.038087 | 0.030004 | 0.020595 | 0.039584 |
| 30.68055556 | 0.019589 | 0.108185 | 0.049909 | 0.030556 | 0.013458 | 0.026766 | 0.023365 | 0.011835 | 0.067572 |
| 30.69444444 | 0.012108 | 0.104911 | 0.032726 | 0.035629 | 0.009673 | 0.015621 | 0.013737 | 0.018409 | 0.065043 |
| 30.70833333 | 0.019757 | 0.06106 | 0.030095 | 0.069559 | 0.022673 | 0.021632 | 0.015632 | 0.027955 | 0.036151 |
| 30.72222222 | 0.028214 | 0.025219 | 0.04788 | 0.103039 | 0.038309 | 0.038611 | 0.018458 | 0.028056 | 0.016431 |
| 30.73611111 | 0.028319 | 0.020266 | 0.058389 | 0.097513 | 0.036362 | 0.038777 | 0.01242 | 0.02252 | 0.027835 |
| 30.75 | 0.025652 | 0.035082 | 0.05354 | 0.061681 | 0.020627 | 0.02119 | 0.006905 | 0.018777 | 0.051277 |
| 30.76388889 | 0.023446 | 0.056382 | 0.038352 | 0.063958 | 0.008823 | 0.012856 | 0.006641 | 0.021998 | 0.063694 |
| 30.77777778 | 0.018574 | 0.077606 | 0.019043 | 0.080797 | 0.008743 | 0.01158 | 0.007215 | 0.026493 | 0.062765 |
| 30.79166667 | 0.010142 | 0.099904 | 0.006433 | 0.052687 | 0.008856 | 0.010851 | 0.007044 | 0.023505 | 0.057493 |
| 30.80555556 | 0.00431 | 0.124914 | 0.008033 | 0.027612 | 0.010887 | 0.025764 | 0.010287 | 0.018018 | 0.055666 |

| | | | | | | | | | |
|---|---|---|---|---|---|---|---|---|---|
| 30.81944444 | 0.006393 | 0.13465 | 0.029036 | 0.057103 | 0.029562 | 0.050942 | 0.020449 | 0.020413 | 0.055507 |
| 30.83333333 | 0.014291 | 0.116473 | 0.054181 | 0.11167 | 0.05257 | 0.065101 | 0.03125 | 0.029345 | 0.049515 |
| 30.84722222 | 0.023707 | 0.081289 | 0.059688 | 0.143538 | 0.057989 | 0.058914 | 0.031727 | 0.036966 | 0.034865 |
| 30.86111111 | 0.034649 | 0.046977 | 0.049093 | 0.13403 | 0.044557 | 0.042616 | 0.021991 | 0.041477 | 0.021279 |
| 30.875 | 0.044591 | 0.020848 | 0.036994 | 0.095286 | 0.023307 | 0.030471 | 0.013653 | 0.043503 | 0.020205 |
| 30.88888889 | 0.046329 | 0.007333 | 0.028411 | 0.051003 | 0.009018 | 0.027528 | 0.015063 | 0.039693 | 0.033194 |
| 30.90277778 | 0.039181 | 0.00751 | 0.025282 | 0.020746 | 0.006925 | 0.033963 | 0.019932 | 0.030331 | 0.047475 |
| 30.91666667 | 0.028709 | 0.01224 | 0.02666 | 0.010964 | 0.006246 | 0.04299 | 0.019779 | 0.02005 | 0.050022 |
| 30.93055556 | 0.018828 | 0.017438 | 0.028159 | 0.013911 | 0.004943 | 0.043074 | 0.015822 | 0.011071 | 0.042143 |
| 30.94444444 | 0.011006 | 0.025044 | 0.025233 | 0.019462 | 0.006107 | 0.029253 | 0.011221 | 0.004518 | 0.027941 |
| 30.95833333 | 0.005726 | 0.03309 | 0.01583 | 0.020203 | 0.007404 | 0.016054 | 0.014134 | 0.002629 | 0.013878 |
| 30.97222222 | 0.005764 | 0.037206 | 0.010321 | 0.013107 | 0.014551 | 0.020674 | 0.028698 | 0.00579 | 0.011324 |
| 30.98611111 | 0.009916 | 0.033096 | 0.012647 | 0.007837 | 0.024949 | 0.026495 | 0.042125 | 0.011077 | 0.016989 |
| 31 | 0.012683 | 0.022828 | 0.018489 | 0.012091 | 0.024248 | 0.026685 | 0.037906 | 0.014385 | 0.017647 |
| 31.01388889 | 0.021852 | 0.02489 | 0.03995 | 0.01476 | 0.030066 | 0.049627 | 0.026311 | 0.011661 | 0.023339 |
| 31.02777778 | 0.034444 | 0.042845 | 0.065275 | 0.009842 | 0.056767 | 0.084605 | 0.028867 | 0.007379 | 0.042649 |
| 31.04166667 | 0.028676 | 0.049799 | 0.063033 | 0.008428 | 0.066475 | 0.094486 | 0.028581 | 0.011512 | 0.054693 |
| 31.05555556 | 0.016595 | 0.038349 | 0.035593 | 0.017523 | 0.042699 | 0.072945 | 0.017307 | 0.021774 | 0.04792 |
| 31.06944444 | 0.022772 | 0.02131 | 0.016544 | 0.027168 | 0.021031 | 0.037798 | 0.017257 | 0.028285 | 0.029932 |
| 31.08333333 | 0.036683 | 0.008872 | 0.026065 | 0.026888 | 0.023762 | 0.020935 | 0.024475 | 0.026526 | 0.013477 |
| 31.09722222 | 0.044481 | 0.009233 | 0.045923 | 0.017611 | 0.035752 | 0.03275 | 0.024997 | 0.018079 | 0.010483 |
| 31.11111111 | 0.048072 | 0.01949 | 0.058401 | 0.007973 | 0.04452 | 0.051479 | 0.021598 | 0.008807 | 0.018594 |
| 31.125 | 0.05099 | 0.030221 | 0.061912 | 0.006298 | 0.049246 | 0.06458 | 0.019218 | 0.005288 | 0.028083 |
| 31.13888889 | 0.052116 | 0.034765 | 0.05702 | 0.01098 | 0.047386 | 0.073109 | 0.018514 | 0.008241 | 0.033153 |
| 31.15277778 | 0.045417 | 0.031553 | 0.045607 | 0.015172 | 0.04227 | 0.073848 | 0.019821 | 0.012769 | 0.033233 |
| 31.16666667 | 0.032833 | 0.026351 | 0.032275 | 0.015254 | 0.039689 | 0.064075 | 0.023158 | 0.01576 | 0.032089 |
| 31.18055556 | 0.02143 | 0.024882 | 0.021852 | 0.012294 | 0.040255 | 0.050127 | 0.025549 | 0.01599 | 0.031312 |
| 31.19444444 | 0.011262 | 0.025908 | 0.014882 | 0.008892 | 0.039867 | 0.036718 | 0.023636 | 0.012644 | 0.028668 |
| 31.20833333 | 0.004749 | 0.026965 | 0.012192 | 0.006576 | 0.034798 | 0.023167 | 0.017945 | 0.007417 | 0.024336 |
| 31.22222222 | 0.004809 | 0.027909 | 0.014576 | 0.006372 | 0.024735 | 0.012021 | 0.01176 | 0.003043 | 0.021122 |
| 31.23611111 | 0.004999 | 0.028786 | 0.018964 | 0.00692 | 0.013124 | 0.006097 | 0.006465 | 0.001641 | 0.019712 |
| 31.25 | 0.006792 | 0.030126 | 0.022175 | 0.005282 | 0.00586 | 0.006057 | 0.003191 | 0.003552 | 0.019074 |
| 31.26388889 | 0.020295 | 0.03266 | 0.024712 | 0.002535 | 0.006015 | 0.012885 | 0.002356 | 0.005163 | 0.018584 |
| 31.27777778 | 0.039333 | 0.034797 | 0.0268 | 0.001524 | 0.013887 | 0.024676 | 0.002965 | 0.003968 | 0.017735 |
| 31.29166667 | 0.044805 | 0.03121 | 0.025654 | 0.001622 | 0.025229 | 0.033632 | 0.005482 | 0.003131 | 0.015893 |
| 31.30555556 | 0.029061 | 0.020095 | 0.019679 | 0.001529 | 0.027981 | 0.032145 | 0.007698 | 0.004326 | 0.012853 |
| 31.31944444 | 0.017969 | 0.010377 | 0.010461 | 0.002127 | 0.017645 | 0.020682 | 0.007565 | 0.005558 | 0.008873 |
| 31.33333333 | 0.031446 | 0.009461 | 0.00392 | 0.00413 | 0.009501 | 0.008886 | 0.005927 | 0.006985 | 0.004762 |
| 31.34722222 | 0.037253 | 0.009076 | 0.003326 | 0.006127 | 0.01593 | 0.003615 | 0.003383 | 0.009619 | 0.002634 |
| 31.36111111 | 0.026739 | 0.007382 | 0.006214 | 0.006347 | 0.022903 | 0.004009 | 0.001376 | 0.012667 | 0.00422 |
| 31.375 | 0.034728 | 0.014383 | 0.012535 | 0.005359 | 0.016574 | 0.009827 | 0.002165 | 0.015594 | 0.008738 |
| 31.38888889 | 0.053319 | 0.022591 | 0.020207 | 0.005025 | 0.008823 | 0.017396 | 0.007002 | 0.018158 | 0.014157 |
| 31.40277778 | 0.048666 | 0.020879 | 0.025535 | 0.005705 | 0.010693 | 0.020087 | 0.013029 | 0.018768 | 0.018832 |
| 31.41666667 | 0.026904 | 0.012662 | 0.027276 | 0.00681 | 0.011754 | 0.016286 | 0.015035 | 0.016321 | 0.021795 |

| | | | | | | | | | |
|---|---|---|---|---|---|---|---|---|---|
| 31.43055556 | 0.015233 | 0.004947 | 0.026127 | 0.010228 | 0.009301 | 0.008704 | 0.011313 | 0.011838 | 0.023278 |
| 31.44444444 | 0.013164 | 0.00321 | 0.02463 | 0.021762 | 0.014899 | 0.002844 | 0.006309 | 0.006869 | 0.024763 |
| 31.45833333 | 0.01552 | 0.009369 | 0.024264 | 0.039985 | 0.022897 | 0.002098 | 0.004976 | 0.003514 | 0.027487 |
| 31.47222222 | 0.027773 | 0.020465 | 0.026567 | 0.055095 | 0.019827 | 0.006475 | 0.004825 | 0.005321 | 0.03317 |
| 31.48611111 | 0.039398 | 0.034898 | 0.035777 | 0.074596 | 0.012654 | 0.017572 | 0.006277 | 0.015767 | 0.048558 |
| 31.5 | 0.044536 | 0.049297 | 0.055117 | 0.098434 | 0.015355 | 0.035285 | 0.011908 | 0.032317 | 0.073166 |
| 31.51388889 | 0.050848 | 0.058208 | 0.07294 | 0.097874 | 0.02933 | 0.053644 | 0.016735 | 0.042797 | 0.090911 |
| 31.52777778 | 0.056684 | 0.065711 | 0.076315 | 0.072984 | 0.047036 | 0.067233 | 0.019747 | 0.040787 | 0.096349 |
| 31.54166667 | 0.050344 | 0.069091 | 0.066383 | 0.048268 | 0.054259 | 0.068497 | 0.021318 | 0.03286 | 0.09413 |
| 31.55555556 | 0.028921 | 0.056045 | 0.044817 | 0.02716 | 0.043433 | 0.048311 | 0.018056 | 0.022954 | 0.078486 |
| 31.56944444 | 0.01244 | 0.031463 | 0.022541 | 0.014211 | 0.022784 | 0.024321 | 0.01175 | 0.012246 | 0.049829 |
| 31.58333333 | 0.016039 | 0.013032 | 0.019361 | 0.019688 | 0.011089 | 0.023824 | 0.007333 | 0.0052 | 0.023564 |
| 31.59722222 | 0.025213 | 0.009714 | 0.035869 | 0.033912 | 0.018325 | 0.041341 | 0.004517 | 0.004726 | 0.014701 |
| 31.61111111 | 0.022734 | 0.016504 | 0.047797 | 0.033665 | 0.029243 | 0.057828 | 0.002856 | 0.009186 | 0.020964 |
| 31.625 | 0.015526 | 0.027813 | 0.038332 | 0.032236 | 0.030808 | 0.061969 | 0.005283 | 0.014075 | 0.029057 |
| 31.63888889 | 0.021898 | 0.045538 | 0.018381 | 0.064971 | 0.023585 | 0.046844 | 0.009685 | 0.015723 | 0.038095 |
| 31.65277778 | 0.036203 | 0.064952 | 0.00845 | 0.10204 | 0.01375 | 0.027682 | 0.010953 | 0.013216 | 0.046471 |
| 31.66666667 | 0.038207 | 0.075361 | 0.012863 | 0.097888 | 0.012079 | 0.0313 | 0.00779 | 0.010087 | 0.039721 |
| 31.68055556 | 0.028234 | 0.059944 | 0.02033 | 0.062802 | 0.015589 | 0.050639 | 0.007097 | 0.010561 | 0.027129 |
| 31.69444444 | 0.01745 | 0.042831 | 0.028034 | 0.031946 | 0.012963 | 0.060199 | 0.013071 | 0.010916 | 0.039552 |
| 31.70833333 | 0.009775 | 0.073273 | 0.032143 | 0.032644 | 0.012438 | 0.052275 | 0.015278 | 0.006453 | 0.065249 |
| 31.72222222 | 0.010109 | 0.111307 | 0.025883 | 0.062319 | 0.024408 | 0.032928 | 0.01225 | 0.003358 | 0.069009 |
| 31.73611111 | 0.015111 | 0.106282 | 0.023656 | 0.09407 | 0.037056 | 0.022562 | 0.017812 | 0.00607 | 0.051086 |
| 31.75 | 0.011401 | 0.07746 | 0.034318 | 0.096343 | 0.046256 | 0.032536 | 0.033225 | 0.010688 | 0.028491 |
| 31.76388889 | 0.005987 | 0.044194 | 0.043207 | 0.06092 | 0.051549 | 0.046285 | 0.046872 | 0.017271 | 0.021605 |
| 31.77777778 | 0.005768 | 0.02455 | 0.046159 | 0.039996 | 0.040318 | 0.052227 | 0.052208 | 0.026981 | 0.040166 |
| 31.79166667 | 0.006964 | 0.040378 | 0.050357 | 0.073272 | 0.019559 | 0.04615 | 0.04892 | 0.036587 | 0.065744 |
| 31.80555556 | 0.015842 | 0.080856 | 0.056102 | 0.104088 | 0.006543 | 0.034721 | 0.038386 | 0.04122 | 0.080515 |
| 31.81944444 | 0.032271 | 0.113434 | 0.063905 | 0.077542 | 0.007308 | 0.039775 | 0.02443 | 0.039006 | 0.080047 |
| 31.83333333 | 0.047798 | 0.122615 | 0.0761 | 0.041786 | 0.021696 | 0.059488 | 0.011951 | 0.033959 | 0.068801 |
| 31.84722222 | 0.052158 | 0.115527 | 0.090183 | 0.058262 | 0.042621 | 0.081004 | 0.004117 | 0.03023 | 0.05833 |
| 31.86111111 | 0.042791 | 0.096753 | 0.095142 | 0.105623 | 0.060113 | 0.106257 | 0.001506 | 0.029933 | 0.054413 |
| 31.875 | 0.030334 | 0.070798 | 0.081833 | 0.136975 | 0.065886 | 0.119184 | 0.001919 | 0.033953 | 0.051953 |
| 31.88888889 | 0.01979 | 0.048038 | 0.055945 | 0.135676 | 0.057717 | 0.098698 | 0.00247 | 0.038877 | 0.04516 |
| 31.90277778 | 0.011453 | 0.031667 | 0.028816 | 0.106374 | 0.039705 | 0.058289 | 0.002144 | 0.039739 | 0.034045 |
| 31.91666667 | 0.007466 | 0.020649 | 0.009924 | 0.063737 | 0.021211 | 0.027067 | 0.001489 | 0.035065 | 0.019335 |
| 31.93055556 | 0.006588 | 0.014007 | 0.002953 | 0.029162 | 0.01775 | 0.020844 | 0.000865 | 0.026925 | 0.009466 |
| 31.94444444 | 0.006773 | 0.009177 | 0.002539 | 0.012074 | 0.03017 | 0.027422 | 0.000694 | 0.018112 | 0.013122 |
| 31.95833333 | 0.006938 | 0.005212 | 0.005227 | 0.005896 | 0.041068 | 0.028425 | 0.001991 | 0.010209 | 0.020763 |
| 31.97222222 | 0.00744 | 0.004935 | 0.009694 | 0.004832 | 0.040118 | 0.023894 | 0.004115 | 0.004721 | 0.020381 |
| 31.98611111 | 0.012183 | 0.006244 | 0.01256 | 0.00798 | 0.026947 | 0.019489 | 0.00421 | 0.003141 | 0.015653 |
| 32 | 0.026464 | 0.008621 | 0.025555 | 0.016642 | 0.022083 | 0.031658 | 0.003966 | 0.00598 | 0.023653 |
| 32.01388889 | 0.047279 | 0.019263 | 0.057455 | 0.026516 | 0.042719 | 0.069104 | 0.009807 | 0.011371 | 0.047188 |
| 32.02777778 | 0.064508 | 0.034698 | 0.084558 | 0.027343 | 0.060985 | 0.10552 | 0.017447 | 0.015334 | 0.065709 |

| | | | | | | | | | |
|---|---|---|---|---|---|---|---|---|---|
| 32.04166667 | 0.074918 | 0.041599 | 0.087792 | 0.016691 | 0.053854 | 0.117041 | 0.015785 | 0.015049 | 0.068641 |
| 32.05555556 | 0.072839 | 0.033125 | 0.071921 | 0.006047 | 0.03178 | 0.103541 | 0.008633 | 0.011464 | 0.061191 |
| 32.06944444 | 0.050451 | 0.016788 | 0.043693 | 0.00213 | 0.012637 | 0.074717 | 0.00686 | 0.008307 | 0.047031 |
| 32.08333333 | 0.024954 | 0.009028 | 0.024045 | 0.00151 | 0.006953 | 0.041031 | 0.01002 | 0.005994 | 0.027518 |
| 32.09722222 | 0.02382 | 0.014188 | 0.035993 | 0.001801 | 0.010678 | 0.019153 | 0.013063 | 0.004997 | 0.012065 |
| 32.11111111 | 0.044765 | 0.022522 | 0.064238 | 0.003344 | 0.017106 | 0.025229 | 0.015394 | 0.009365 | 0.010267 |
| 32.125 | 0.065299 | 0.02903 | 0.08342 | 0.006194 | 0.024493 | 0.050171 | 0.017982 | 0.016894 | 0.016088 |
| 32.13888889 | 0.072769 | 0.033231 | 0.086787 | 0.007202 | 0.030137 | 0.076479 | 0.022222 | 0.021234 | 0.022031 |
| 32.15277778 | 0.067409 | 0.030767 | 0.073801 | 0.005331 | 0.032927 | 0.094348 | 0.027948 | 0.020141 | 0.027703 |
| 32.16666667 | 0.060231 | 0.023851 | 0.054024 | 0.005508 | 0.036177 | 0.099142 | 0.030611 | 0.015272 | 0.032631 |
| 32.18055556 | 0.060474 | 0.021069 | 0.036822 | 0.009082 | 0.038156 | 0.095491 | 0.026973 | 0.010076 | 0.036359 |
| 32.19444444 | 0.059708 | 0.022122 | 0.021369 | 0.0111 | 0.033362 | 0.087352 | 0.019006 | 0.006708 | 0.039295 |
| 32.20833333 | 0.053862 | 0.02499 | 0.012963 | 0.008305 | 0.023217 | 0.074194 | 0.011475 | 0.00596 | 0.037651 |
| 32.22222222 | 0.050859 | 0.032178 | 0.017613 | 0.003921 | 0.012617 | 0.056051 | 0.008517 | 0.008112 | 0.030529 |
| 32.23611111 | 0.046241 | 0.038476 | 0.026149 | 0.002139 | 0.006081 | 0.034093 | 0.00778 | 0.011004 | 0.023685 |
| 32.25 | 0.031274 | 0.034028 | 0.026218 | 0.002299 | 0.007921 | 0.017123 | 0.006631 | 0.011063 | 0.017039 |
| 32.26388889 | 0.015349 | 0.02 | 0.018502 | 0.004008 | 0.015714 | 0.015869 | 0.008694 | 0.007588 | 0.01128 |
| 32.27777778 | 0.012224 | 0.007847 | 0.010622 | 0.006419 | 0.020195 | 0.021615 | 0.012088 | 0.003637 | 0.008758 |
| 32.29166667 | 0.015475 | 0.002936 | 0.007306 | 0.006604 | 0.014936 | 0.016769 | 0.010098 | 0.002062 | 0.007508 |
| 32.30555556 | 0.0119 | 0.002532 | 0.010168 | 0.00408 | 0.007236 | 0.010129 | 0.005065 | 0.001765 | 0.007156 |
| 32.31944444 | 0.00828 | 0.005996 | 0.016805 | 0.002106 | 0.007038 | 0.017307 | 0.003682 | 0.002604 | 0.008199 |
| 32.33333333 | 0.015658 | 0.012141 | 0.021894 | 0.003579 | 0.009202 | 0.026134 | 0.005341 | 0.007477 | 0.009475 |
| 32.34722222 | 0.023943 | 0.015941 | 0.021877 | 0.005763 | 0.006955 | 0.024039 | 0.006062 | 0.014379 | 0.009772 |
| 32.36111111 | 0.020763 | 0.012887 | 0.0168 | 0.004745 | 0.004513 | 0.014066 | 0.004721 | 0.017062 | 0.008893 |
| 32.375 | 0.010691 | 0.006892 | 0.011026 | 0.002621 | 0.006828 | 0.005678 | 0.002816 | 0.013612 | 0.008069 |
| 32.38888889 | 0.004058 | 0.004005 | 0.011715 | 0.002866 | 0.009296 | 0.005841 | 0.003262 | 0.009673 | 0.009825 |
| 32.40277778 | 0.006286 | 0.004555 | 0.020879 | 0.00563 | 0.007074 | 0.014962 | 0.006689 | 0.010499 | 0.014599 |
| 32.41666667 | 0.015478 | 0.010068 | 0.030602 | 0.009145 | 0.00526 | 0.031694 | 0.010022 | 0.013382 | 0.018306 |
| 32.43055556 | 0.02249 | 0.018838 | 0.032263 | 0.009542 | 0.007415 | 0.046349 | 0.010938 | 0.014702 | 0.01745 |
| 32.44444444 | 0.022983 | 0.026142 | 0.026352 | 0.006704 | 0.008745 | 0.050359 | 0.009968 | 0.015307 | 0.014013 |
| 32.45833333 | 0.020284 | 0.029101 | 0.01907 | 0.004631 | 0.006238 | 0.043315 | 0.007853 | 0.015178 | 0.011914 |
| 32.47222222 | 0.017553 | 0.02766 | 0.017666 | 0.009271 | 0.003334 | 0.030025 | 0.004704 | 0.013565 | 0.01472 |
| 32.48611111 | 0.01857 | 0.028611 | 0.03089 | 0.040047 | 0.002908 | 0.023322 | 0.003101 | 0.014232 | 0.02694 |
| 32.5 | 0.025091 | 0.037685 | 0.059276 | 0.097898 | 0.005088 | 0.035612 | 0.006394 | 0.021994 | 0.046962 |
| 32.51388889 | 0.038536 | 0.053686 | 0.083932 | 0.145729 | 0.016292 | 0.061389 | 0.010075 | 0.030686 | 0.061139 |
| 32.52777778 | 0.057333 | 0.075323 | 0.090317 | 0.156304 | 0.039741 | 0.081099 | 0.010451 | 0.030892 | 0.060003 |
| 32.54166667 | 0.063078 | 0.090179 | 0.081795 | 0.127815 | 0.059839 | 0.082608 | 0.013844 | 0.025608 | 0.051357 |
| 32.55555556 | 0.043516 | 0.081356 | 0.060253 | 0.082765 | 0.058867 | 0.06374 | 0.019235 | 0.019898 | 0.045427 |
| 32.56944444 | 0.018084 | 0.057813 | 0.031548 | 0.046734 | 0.038389 | 0.032705 | 0.019129 | 0.01509 | 0.043389 |
| 32.58333333 | 0.007164 | 0.036265 | 0.011812 | 0.02277 | 0.017254 | 0.014788 | 0.015388 | 0.012535 | 0.042416 |
| 32.59722222 | 0.008729 | 0.019947 | 0.008566 | 0.017708 | 0.011401 | 0.024641 | 0.011639 | 0.010761 | 0.038904 |
| 32.61111111 | 0.012732 | 0.009445 | 0.013263 | 0.035038 | 0.017584 | 0.046483 | 0.00688 | 0.007741 | 0.031379 |
| 32.625 | 0.013506 | 0.010721 | 0.016855 | 0.06278 | 0.021564 | 0.062779 | 0.00414 | 0.006709 | 0.020714 |
| 32.63888889 | 0.009292 | 0.026545 | 0.018376 | 0.085412 | 0.018851 | 0.06492 | 0.006773 | 0.010737 | 0.012098 |

| | | | | | | | | | |
|---|---|---|---|---|---|---|---|---|---|
| 32.65277778 | 0.007951 | 0.046545 | 0.017386 | 0.072175 | 0.014451 | 0.051765 | 0.009517 | 0.016477 | 0.014166 |
| 32.66666667 | 0.01854 | 0.061958 | 0.012378 | 0.04481 | 0.010071 | 0.029369 | 0.009395 | 0.02041 | 0.022853 |
| 32.68055556 | 0.027047 | 0.076713 | 0.006676 | 0.069156 | 0.008784 | 0.009734 | 0.009927 | 0.020656 | 0.032232 |
| 32.69444444 | 0.019234 | 0.083025 | 0.004878 | 0.11943 | 0.01501 | 0.006614 | 0.012416 | 0.016873 | 0.041391 |
| 32.70833333 | 0.007872 | 0.062451 | 0.006255 | 0.124932 | 0.022217 | 0.020045 | 0.014779 | 0.012558 | 0.036673 |
| 32.72222222 | 0.007902 | 0.034649 | 0.009213 | 0.085083 | 0.020903 | 0.036711 | 0.015469 | 0.009367 | 0.021199 |
| 32.73611111 | 0.011999 | 0.040516 | 0.013373 | 0.039537 | 0.013002 | 0.043391 | 0.012577 | 0.00597 | 0.018906 |
| 32.75 | 0.009309 | 0.075067 | 0.013976 | 0.02189 | 0.005278 | 0.033744 | 0.008288 | 0.002831 | 0.027273 |
| 32.76388889 | 0.006314 | 0.095387 | 0.011591 | 0.043223 | 0.00349 | 0.029528 | 0.009153 | 0.002502 | 0.025579 |
| 32.77777778 | 0.008118 | 0.084139 | 0.017559 | 0.084331 | 0.009801 | 0.052177 | 0.011287 | 0.006987 | 0.014723 |
| 32.79166667 | 0.010323 | 0.053944 | 0.031965 | 0.129019 | 0.018429 | 0.07016 | 0.008087 | 0.014467 | 0.011898 |
| 32.80555556 | 0.010413 | 0.030575 | 0.04539 | 0.153309 | 0.022816 | 0.06469 | 0.006524 | 0.020853 | 0.018264 |
| 32.81944444 | 0.009088 | 0.041033 | 0.060724 | 0.125414 | 0.022294 | 0.055717 | 0.012612 | 0.023605 | 0.023319 |
| 32.83333333 | 0.00925 | 0.071356 | 0.079923 | 0.067534 | 0.021032 | 0.059843 | 0.018797 | 0.023686 | 0.025332 |
| 32.84722222 | 0.015489 | 0.092122 | 0.092006 | 0.029659 | 0.023701 | 0.084163 | 0.01993 | 0.023286 | 0.028971 |
| 32.86111111 | 0.029592 | 0.098743 | 0.091704 | 0.019409 | 0.030206 | 0.113554 | 0.018976 | 0.022582 | 0.036679 |
| 32.875 | 0.044984 | 0.093634 | 0.082634 | 0.019608 | 0.037144 | 0.123657 | 0.01696 | 0.022119 | 0.046074 |
| 32.88888889 | 0.051178 | 0.078421 | 0.066487 | 0.022188 | 0.039988 | 0.108887 | 0.011812 | 0.02361 | 0.053146 |
| 32.90277778 | 0.040909 | 0.056858 | 0.045504 | 0.025678 | 0.03333 | 0.079637 | 0.006316 | 0.026551 | 0.053974 |
| 32.91666667 | 0.022491 | 0.034146 | 0.024316 | 0.028737 | 0.018673 | 0.045591 | 0.006787 | 0.027406 | 0.046413 |
| 32.93055556 | 0.016658 | 0.015623 | 0.010226 | 0.027137 | 0.0091 | 0.020893 | 0.011652 | 0.022684 | 0.032464 |
| 32.94444444 | 0.025563 | 0.006551 | 0.008116 | 0.019524 | 0.011302 | 0.019107 | 0.016265 | 0.013081 | 0.016492 |
| 32.95833333 | 0.033407 | 0.008013 | 0.010323 | 0.009819 | 0.016052 | 0.027511 | 0.019638 | 0.005422 | 0.007753 |
| 32.97222222 | 0.031707 | 0.011014 | 0.008062 | 0.003204 | 0.015173 | 0.022524 | 0.020591 | 0.003984 | 0.010425 |
| 32.98611111 | 0.022653 | 0.007889 | 0.010147 | 0.00394 | 0.012402 | 0.016645 | 0.017173 | 0.004181 | 0.011045 |
| 33 | 0.024626 | 0.006176 | 0.029978 | 0.011988 | 0.017417 | 0.032558 | 0.01033 | 0.005396 | 0.010899 |
| 33.01388889 | 0.048533 | 0.01485 | 0.060443 | 0.017208 | 0.024361 | 0.051276 | 0.005977 | 0.008293 | 0.023311 |
| 33.02777778 | 0.072264 | 0.026491 | 0.081116 | 0.011703 | 0.021516 | 0.05415 | 0.004658 | 0.007401 | 0.033968 |
| 33.04166667 | 0.077211 | 0.030121 | 0.085198 | 0.004646 | 0.014909 | 0.047056 | 0.004075 | 0.004311 | 0.029766 |
| 33.05555556 | 0.06187 | 0.022777 | 0.077956 | 0.003022 | 0.01071 | 0.034633 | 0.007385 | 0.004717 | 0.020412 |
| 33.06944444 | 0.033254 | 0.011673 | 0.059608 | 0.002377 | 0.006313 | 0.019547 | 0.012331 | 0.007153 | 0.01205 |
| 33.08333333 | 0.013445 | 0.008603 | 0.034205 | 0.002005 | 0.002958 | 0.007862 | 0.015425 | 0.007137 | 0.006548 |
| 33.09722222 | 0.014516 | 0.012995 | 0.015144 | 0.002356 | 0.00216 | 0.00455 | 0.016765 | 0.004404 | 0.008774 |
| 33.11111111 | 0.021055 | 0.017113 | 0.012752 | 0.002968 | 0.00169 | 0.007021 | 0.016666 | 0.003324 | 0.015457 |
| 33.125 | 0.021511 | 0.018937 | 0.024084 | 0.006278 | 0.003996 | 0.012121 | 0.016793 | 0.005775 | 0.018142 |
| 33.13888889 | 0.020935 | 0.019744 | 0.039848 | 0.011648 | 0.011288 | 0.019636 | 0.017889 | 0.008373 | 0.01632 |
| 33.15277778 | 0.02396 | 0.021456 | 0.056942 | 0.015955 | 0.018906 | 0.025866 | 0.0205 | 0.009826 | 0.015226 |
| 33.16666667 | 0.031292 | 0.025798 | 0.073082 | 0.017315 | 0.021857 | 0.026047 | 0.023836 | 0.011229 | 0.01826 |
| 33.18055556 | 0.041311 | 0.030403 | 0.081611 | 0.014247 | 0.019105 | 0.020533 | 0.023853 | 0.011944 | 0.024557 |
| 33.19444444 | 0.050982 | 0.03168 | 0.078979 | 0.008113 | 0.011992 | 0.013974 | 0.018335 | 0.010943 | 0.028531 |
| 33.20833333 | 0.057459 | 0.030023 | 0.070081 | 0.003239 | 0.005283 | 0.012841 | 0.010184 | 0.008474 | 0.025446 |
| 33.22222222 | 0.058946 | 0.026525 | 0.060058 | 0.002077 | 0.004217 | 0.019362 | 0.004162 | 0.005877 | 0.017378 |
| 33.23611111 | 0.055968 | 0.020662 | 0.048493 | 0.003426 | 0.006095 | 0.027056 | 0.001848 | 0.003555 | 0.009635 |
| 33.25 | 0.049987 | 0.013518 | 0.036459 | 0.00519 | 0.004923 | 0.029522 | 0.001917 | 0.002063 | 0.0064 |

| | | | | | | | | | |
|---|---|---|---|---|---|---|---|---|---|
| 33.26388889 | 0.041211 | 0.007425 | 0.026714 | 0.006643 | 0.002444 | 0.026924 | 0.004052 | 0.00251 | 0.008462 |
| 33.27777778 | 0.029941 | 0.003312 | 0.020326 | 0.007592 | 0.002681 | 0.020952 | 0.006438 | 0.00384 | 0.011741 |
| 33.29166667 | 0.019077 | 0.001123 | 0.01774 | 0.00687 | 0.00302 | 0.013225 | 0.006623 | 0.00441 | 0.012033 |
| 33.30555556 | 0.011529 | 0.000663 | 0.017907 | 0.004558 | 0.001649 | 0.007044 | 0.005382 | 0.00451 | 0.009552 |
| 33.31944444 | 0.007344 | 0.001669 | 0.019886 | 0.004294 | 0.000619 | 0.003933 | 0.004014 | 0.005003 | 0.006502 |
| 33.33333333 | 0.006033 | 0.003662 | 0.022708 | 0.007612 | 0.000453 | 0.002886 | 0.003119 | 0.006024 | 0.004495 |
| 33.34722222 | 0.00724 | 0.006141 | 0.024285 | 0.011099 | 0.00114 | 0.003411 | 0.004016 | 0.007929 | 0.005355 |
| 33.36111111 | 0.009501 | 0.009029 | 0.023176 | 0.013365 | 0.004082 | 0.006525 | 0.007812 | 0.01175 | 0.010022 |
| 33.375 | 0.010543 | 0.011908 | 0.019402 | 0.01602 | 0.0092 | 0.013155 | 0.01334 | 0.016807 | 0.015788 |
| 33.38888889 | 0.009067 | 0.012818 | 0.015247 | 0.018332 | 0.012875 | 0.020316 | 0.016402 | 0.019323 | 0.018375 |
| 33.40277778 | 0.006693 | 0.010232 | 0.014793 | 0.017619 | 0.012533 | 0.024119 | 0.013857 | 0.016975 | 0.017447 |
| 33.41666667 | 0.006342 | 0.006278 | 0.01955 | 0.013773 | 0.011074 | 0.025907 | 0.00856 | 0.013205 | 0.01668 |
| 33.43055556 | 0.009368 | 0.004291 | 0.025952 | 0.009454 | 0.011563 | 0.028673 | 0.005302 | 0.012154 | 0.018021 |
| 33.44444444 | 0.01497 | 0.004703 | 0.029355 | 0.006177 | 0.012323 | 0.032251 | 0.004853 | 0.013234 | 0.019135 |
| 33.45833333 | 0.020841 | 0.006066 | 0.028025 | 0.003805 | 0.011313 | 0.035827 | 0.005499 | 0.013839 | 0.017707 |
| 33.47222222 | 0.025855 | 0.007775 | 0.024154 | 0.002623 | 0.010426 | 0.042719 | 0.004908 | 0.012721 | 0.016254 |
| 33.48611111 | 0.032989 | 0.011977 | 0.024551 | 0.004622 | 0.014991 | 0.065282 | 0.004175 | 0.011974 | 0.020092 |
| 33.5 | 0.041134 | 0.021209 | 0.034111 | 0.012843 | 0.025836 | 0.107441 | 0.006642 | 0.014725 | 0.029466 |
| 33.51388889 | 0.042052 | 0.032756 | 0.051969 | 0.026822 | 0.036774 | 0.143409 | 0.00822 | 0.019441 | 0.041512 |
| 33.52777778 | 0.038368 | 0.041394 | 0.072076 | 0.044527 | 0.042742 | 0.15629 | 0.006777 | 0.021017 | 0.061327 |
| 33.54166667 | 0.037651 | 0.047311 | 0.079324 | 0.06455 | 0.043106 | 0.149256 | 0.008158 | 0.020066 | 0.080614 |
| 33.55555556 | 0.033404 | 0.052545 | 0.065312 | 0.07871 | 0.037981 | 0.116434 | 0.010758 | 0.019292 | 0.074167 |
| 33.56944444 | 0.021089 | 0.05771 | 0.041025 | 0.079566 | 0.02854 | 0.068401 | 0.00876 | 0.015803 | 0.042134 |
| 33.58333333 | 0.009177 | 0.067833 | 0.019351 | 0.074551 | 0.023878 | 0.032859 | 0.004519 | 0.010572 | 0.021221 |
| 33.59722222 | 0.005982 | 0.080171 | 0.009969 | 0.074821 | 0.028667 | 0.018499 | 0.002761 | 0.007221 | 0.028731 |
| 33.61111111 | 0.008701 | 0.081444 | 0.016319 | 0.073791 | 0.030696 | 0.01262 | 0.002462 | 0.004548 | 0.032022 |
| 33.625 | 0.007464 | 0.071223 | 0.024554 | 0.055295 | 0.022044 | 0.009334 | 0.001932 | 0.003271 | 0.019253 |
| 33.63888889 | 0.004179 | 0.058471 | 0.022799 | 0.032123 | 0.011125 | 0.013382 | 0.002127 | 0.00388 | 0.010342 |
| 33.65277778 | 0.006409 | 0.043229 | 0.013864 | 0.042122 | 0.005006 | 0.021336 | 0.001867 | 0.004252 | 0.011469 |
| 33.66666667 | 0.009138 | 0.026063 | 0.005168 | 0.082017 | 0.002154 | 0.027631 | 0.001379 | 0.005634 | 0.017671 |
| 33.68055556 | 0.007081 | 0.02556 | 0.002305 | 0.109298 | 0.001138 | 0.033072 | 0.001716 | 0.006551 | 0.020674 |
| 33.69444444 | 0.004844 | 0.050128 | 0.00449 | 0.110931 | 0.002209 | 0.038293 | 0.001833 | 0.004654 | 0.016374 |
| 33.70833333 | 0.008348 | 0.080618 | 0.006826 | 0.093243 | 0.004736 | 0.037048 | 0.003032 | 0.002763 | 0.009064 |
| 33.72222222 | 0.011306 | 0.103128 | 0.006864 | 0.058162 | 0.008263 | 0.026461 | 0.006736 | 0.004351 | 0.008789 |
| 33.73611111 | 0.008039 | 0.111002 | 0.010969 | 0.023669 | 0.013927 | 0.013353 | 0.012663 | 0.01003 | 0.018395 |
| 33.75 | 0.007472 | 0.095672 | 0.022933 | 0.005966 | 0.022479 | 0.009367 | 0.019805 | 0.017968 | 0.029601 |
| 33.76388889 | 0.009243 | 0.062906 | 0.032017 | 0.001451 | 0.030926 | 0.017318 | 0.026426 | 0.025024 | 0.035201 |
| 33.77777778 | 0.008337 | 0.033659 | 0.03106 | 0.002578 | 0.036278 | 0.022648 | 0.030785 | 0.028372 | 0.033137 |
| 33.79166667 | 0.012783 | 0.019382 | 0.026255 | 0.011344 | 0.040194 | 0.018719 | 0.030971 | 0.027496 | 0.026617 |
| 33.80555556 | 0.027314 | 0.01485 | 0.024382 | 0.030539 | 0.044348 | 0.025926 | 0.027116 | 0.02316 | 0.022955 |
| 33.81944444 | 0.04469 | 0.013139 | 0.026445 | 0.042389 | 0.047838 | 0.060578 | 0.022231 | 0.018201 | 0.027104 |
| 33.83333333 | 0.053541 | 0.011748 | 0.034422 | 0.031755 | 0.049809 | 0.110926 | 0.020035 | 0.017325 | 0.037546 |
| 33.84722222 | 0.048475 | 0.010002 | 0.048212 | 0.0269 | 0.048286 | 0.147893 | 0.021501 | 0.022431 | 0.048122 |
| 33.86111111 | 0.037726 | 0.00876 | 0.061067 | 0.052136 | 0.041793 | 0.148191 | 0.022822 | 0.030203 | 0.052684 |

| | | | | | | | | | |
|---|---|---|---|---|---|---|---|---|---|
| 33.875 | 0.031243 | 0.01129 | 0.071013 | 0.082062 | 0.035112 | 0.123816 | 0.021814 | 0.037044 | 0.051026 |
| 33.88888889 | 0.028175 | 0.018162 | 0.078318 | 0.089106 | 0.035151 | 0.100168 | 0.020303 | 0.040774 | 0.04562 |
| 33.90277778 | 0.025462 | 0.02616 | 0.077564 | 0.071823 | 0.040697 | 0.082127 | 0.018206 | 0.039116 | 0.036447 |
| 33.91666667 | 0.025115 | 0.033203 | 0.064563 | 0.043546 | 0.042913 | 0.064645 | 0.014618 | 0.03208 | 0.025565 |
| 33.93055556 | 0.027033 | 0.036812 | 0.042606 | 0.019353 | 0.035461 | 0.048231 | 0.009449 | 0.021009 | 0.015937 |
| 33.94444444 | 0.025549 | 0.033222 | 0.019827 | 0.007944 | 0.020592 | 0.037975 | 0.004717 | 0.009194 | 0.007662 |
| 33.95833333 | 0.019097 | 0.0246 | 0.007494 | 0.006812 | 0.00833 | 0.033429 | 0.003056 | 0.002312 | 0.003192 |
| 33.97222222 | 0.014432 | 0.019967 | 0.007388 | 0.014061 | 0.008569 | 0.034346 | 0.003544 | 0.005208 | 0.004253 |
| 33.98611111 | 0.024257 | 0.028393 | 0.011153 | 0.036827 | 0.031975 | 0.050304 | 0.008017 | 0.020545 | 0.014822 |
| 34 | 0.055236 | 0.048904 | 0.022787 | 0.063195 | 0.072289 | 0.078767 | 0.016345 | 0.041909 | 0.038286 |
| 34.01388889 | 0.094262 | 0.066225 | 0.039721 | 0.065524 | 0.083971 | 0.092815 | 0.018702 | 0.051319 | 0.055644 |
| 34.02777778 | 0.119152 | 0.070983 | 0.04766 | 0.039069 | 0.052693 | 0.081707 | 0.011337 | 0.039411 | 0.050156 |
| 34.04166667 | 0.120299 | 0.066501 | 0.046133 | 0.016117 | 0.020509 | 0.063215 | 0.003781 | 0.020055 | 0.034241 |
| 34.05555556 | 0.099749 | 0.051532 | 0.03984 | 0.017324 | 0.007013 | 0.045197 | 0.003839 | 0.012757 | 0.023345 |
| 34.06944444 | 0.062066 | 0.02852 | 0.030296 | 0.021433 | 0.003536 | 0.023936 | 0.010079 | 0.016994 | 0.01646 |
| 34.08333333 | 0.030159 | 0.017815 | 0.01955 | 0.015114 | 0.004228 | 0.01029 | 0.016782 | 0.019144 | 0.009749 |
| 34.09722222 | 0.031879 | 0.027757 | 0.010103 | 0.00712 | 0.004368 | 0.012999 | 0.019744 | 0.017252 | 0.008409 |
| 34.11111111 | 0.058301 | 0.039547 | 0.004508 | 0.004468 | 0.002203 | 0.019705 | 0.019044 | 0.017221 | 0.015753 |
| 34.125 | 0.077945 | 0.042559 | 0.004605 | 0.008519 | 0.003497 | 0.023136 | 0.018021 | 0.020721 | 0.028297 |
| 34.13888889 | 0.076174 | 0.039371 | 0.010057 | 0.016245 | 0.01126 | 0.02519 | 0.020804 | 0.024868 | 0.043628 |
| 34.15277778 | 0.060594 | 0.033104 | 0.019326 | 0.023114 | 0.02064 | 0.026413 | 0.027434 | 0.026113 | 0.061269 |
| 34.16666667 | 0.043709 | 0.026902 | 0.03029 | 0.024425 | 0.02896 | 0.030197 | 0.031284 | 0.024366 | 0.07335 |
| 34.18055556 | 0.031381 | 0.020559 | 0.039154 | 0.018715 | 0.03311 | 0.034598 | 0.026652 | 0.020701 | 0.069428 |
| 34.19444444 | 0.023637 | 0.014425 | 0.042029 | 0.010008 | 0.028943 | 0.033048 | 0.016922 | 0.015499 | 0.052855 |
| 34.20833333 | 0.017428 | 0.009703 | 0.037804 | 0.003385 | 0.019164 | 0.024376 | 0.009127 | 0.009865 | 0.031937 |
| 34.22222222 | 0.010548 | 0.006189 | 0.030095 | 0.001607 | 0.0091 | 0.012865 | 0.004381 | 0.005323 | 0.015394 |
| 34.23611111 | 0.004266 | 0.004361 | 0.023655 | 0.003951 | 0.004003 | 0.007234 | 0.001824 | 0.002461 | 0.013994 |
| 34.25 | 0.002798 | 0.004172 | 0.019801 | 0.005128 | 0.006412 | 0.009988 | 0.002575 | 0.002882 | 0.024802 |
| 34.26388889 | 0.008725 | 0.003779 | 0.017868 | 0.005151 | 0.011575 | 0.012437 | 0.005205 | 0.007399 | 0.030937 |
| 34.27777778 | 0.018324 | 0.002692 | 0.017923 | 0.009507 | 0.014413 | 0.010571 | 0.006712 | 0.010995 | 0.021449 |
| 34.29166667 | 0.024272 | 0.002223 | 0.019377 | 0.013743 | 0.012742 | 0.006802 | 0.005104 | 0.008432 | 0.011699 |
| 34.30555556 | 0.022699 | 0.002693 | 0.019723 | 0.011126 | 0.007175 | 0.004482 | 0.00294 | 0.004469 | 0.018493 |
| 34.31944444 | 0.016366 | 0.003441 | 0.017825 | 0.005486 | 0.003514 | 0.007925 | 0.00399 | 0.005442 | 0.028919 |
| 34.33333333 | 0.009817 | 0.004111 | 0.015134 | 0.003222 | 0.004992 | 0.015311 | 0.006029 | 0.007347 | 0.029351 |
| 34.34722222 | 0.004602 | 0.004084 | 0.012875 | 0.00331 | 0.007087 | 0.02045 | 0.007003 | 0.006272 | 0.021056 |
| 34.36111111 | 0.001696 | 0.003185 | 0.011965 | 0.002819 | 0.006932 | 0.022533 | 0.007 | 0.004454 | 0.010908 |
| 34.375 | 0.003177 | 0.002656 | 0.014237 | 0.003783 | 0.00644 | 0.025253 | 0.005901 | 0.004105 | 0.007391 |
| 34.38888889 | 0.007906 | 0.003472 | 0.019609 | 0.006149 | 0.007509 | 0.029491 | 0.004733 | 0.005897 | 0.012206 |
| 34.40277778 | 0.011352 | 0.005146 | 0.024256 | 0.005783 | 0.009738 | 0.030145 | 0.004344 | 0.009557 | 0.019546 |
| 34.41666667 | 0.011355 | 0.006312 | 0.024705 | 0.004139 | 0.010331 | 0.021998 | 0.004275 | 0.013693 | 0.022126 |
| 34.43055556 | 0.008574 | 0.005221 | 0.021401 | 0.006379 | 0.007648 | 0.010838 | 0.003216 | 0.015987 | 0.017362 |
| 34.44444444 | 0.005631 | 0.00336 | 0.016909 | 0.009084 | 0.004125 | 0.007941 | 0.002149 | 0.01423 | 0.008912 |
| 34.45833333 | 0.00682 | 0.004005 | 0.014342 | 0.006597 | 0.002578 | 0.010447 | 0.002998 | 0.00858 | 0.003469 |
| 34.47222222 | 0.008677 | 0.004388 | 0.018402 | 0.002168 | 0.004632 | 0.014496 | 0.004157 | 0.003206 | 0.0039 |

| | | | | | | | | | |
|---|---|---|---|---|---|---|---|---|---|
| 34.48611111 | 0.008566 | 0.003884 | 0.032181 | 0.000316 | 0.013864 | 0.03859 | 0.004529 | 0.002294 | 0.012787 |
| 34.5 | 0.015104 | 0.006111 | 0.049136 | 0.001147 | 0.027061 | 0.087176 | 0.00514 | 0.006734 | 0.032859 |
| 34.51388889 | 0.028265 | 0.008879 | 0.058125 | 0.008112 | 0.035804 | 0.121978 | 0.005839 | 0.015735 | 0.058477 |
| 34.52777778 | 0.040506 | 0.012031 | 0.062108 | 0.021364 | 0.042658 | 0.119655 | 0.007956 | 0.026581 | 0.079528 |
| 34.54166667 | 0.047728 | 0.017387 | 0.067524 | 0.030261 | 0.0523 | 0.106208 | 0.012892 | 0.033394 | 0.087139 |
| 34.55555556 | 0.046195 | 0.021746 | 0.065901 | 0.030988 | 0.056847 | 0.095788 | 0.016921 | 0.031762 | 0.077489 |
| 34.56944444 | 0.038451 | 0.022783 | 0.05326 | 0.029254 | 0.048779 | 0.071564 | 0.016858 | 0.023326 | 0.051561 |
| 34.58333333 | 0.030029 | 0.022012 | 0.037959 | 0.029397 | 0.033871 | 0.03643 | 0.014637 | 0.012659 | 0.026374 |
| 34.59722222 | 0.019917 | 0.021037 | 0.025403 | 0.037144 | 0.021564 | 0.011035 | 0.013135 | 0.006481 | 0.021367 |
| 34.61111111 | 0.014332 | 0.019158 | 0.015032 | 0.051462 | 0.014808 | 0.003528 | 0.013446 | 0.006665 | 0.02824 |
| 34.625 | 0.021397 | 0.016764 | 0.007354 | 0.05822 | 0.010544 | 0.004799 | 0.012926 | 0.005778 | 0.032265 |
| 34.63888889 | 0.023012 | 0.016614 | 0.006589 | 0.047734 | 0.00861 | 0.004554 | 0.009135 | 0.006203 | 0.028911 |
| 34.65277778 | 0.017669 | 0.019843 | 0.010124 | 0.029355 | 0.008814 | 0.003632 | 0.00425 | 0.013714 | 0.019692 |
| 34.66666667 | 0.027686 | 0.022722 | 0.008628 | 0.028826 | 0.005749 | 0.003097 | 0.00182 | 0.018863 | 0.010773 |
| 34.68055556 | 0.041498 | 0.021209 | 0.004968 | 0.05043 | 0.001452 | 0.005628 | 0.003384 | 0.015809 | 0.010582 |
| 34.69444444 | 0.040482 | 0.016663 | 0.008715 | 0.06619 | 0.002732 | 0.012947 | 0.007665 | 0.010383 | 0.022772 |
| 34.70833333 | 0.026619 | 0.010383 | 0.018461 | 0.059836 | 0.009397 | 0.016959 | 0.012597 | 0.006071 | 0.037473 |
| 34.72222222 | 0.01567 | 0.005862 | 0.02486 | 0.042708 | 0.014395 | 0.01468 | 0.016237 | 0.005383 | 0.043899 |
| 34.73611111 | 0.020645 | 0.008286 | 0.018566 | 0.030117 | 0.014556 | 0.012496 | 0.016675 | 0.008406 | 0.041014 |
| 34.75 | 0.025157 | 0.013661 | 0.011523 | 0.019393 | 0.015398 | 0.016442 | 0.013256 | 0.008378 | 0.032125 |
| 34.76388889 | 0.017813 | 0.01656 | 0.022611 | 0.008878 | 0.019623 | 0.026774 | 0.007139 | 0.007532 | 0.020395 |
| 34.77777778 | 0.013074 | 0.014845 | 0.048788 | 0.006641 | 0.02207 | 0.037745 | 0.002893 | 0.011256 | 0.010882 |
| 34.79166667 | 0.011849 | 0.009674 | 0.074075 | 0.010803 | 0.016813 | 0.042741 | 0.002817 | 0.011834 | 0.010893 |
| 34.80555556 | 0.008788 | 0.007455 | 0.072984 | 0.013738 | 0.009006 | 0.041078 | 0.002571 | 0.008254 | 0.020019 |
| 34.81944444 | 0.010064 | 0.007419 | 0.043619 | 0.012451 | 0.009569 | 0.040061 | 0.001168 | 0.011551 | 0.02905 |
| 34.83333333 | 0.012653 | 0.005292 | 0.015809 | 0.00818 | 0.016821 | 0.045876 | 0.001212 | 0.024268 | 0.032635 |
| 34.84722222 | 0.010074 | 0.003685 | 0.005805 | 0.007385 | 0.022187 | 0.055644 | 0.003214 | 0.038365 | 0.032313 |
| 34.86111111 | 0.00999 | 0.003084 | 0.012825 | 0.011366 | 0.0243 | 0.061161 | 0.006581 | 0.045067 | 0.032967 |
| 34.875 | 0.020458 | 0.002212 | 0.032882 | 0.010948 | 0.02633 | 0.059275 | 0.009676 | 0.039923 | 0.035553 |
| 34.88888889 | 0.032576 | 0.003624 | 0.053919 | 0.005413 | 0.029222 | 0.054043 | 0.011814 | 0.023663 | 0.034307 |
| 34.90277778 | 0.039301 | 0.007935 | 0.063142 | 0.002293 | 0.031646 | 0.04968 | 0.013469 | 0.009487 | 0.028051 |
| 34.91666667 | 0.039022 | 0.011576 | 0.059768 | 0.002666 | 0.030663 | 0.042168 | 0.015328 | 0.009363 | 0.019889 |
| 34.93055556 | 0.029481 | 0.010992 | 0.049873 | 0.002716 | 0.023414 | 0.024914 | 0.017787 | 0.014215 | 0.010608 |
| 34.94444444 | 0.015843 | 0.006267 | 0.035468 | 0.001724 | 0.012329 | 0.011527 | 0.020683 | 0.01419 | 0.006014 |
| 34.95833333 | 0.009967 | 0.003769 | 0.019526 | 0.0024 | 0.007204 | 0.016983 | 0.02376 | 0.010244 | 0.013123 |
| 34.97222222 | 0.01516 | 0.010027 | 0.014597 | 0.006184 | 0.010455 | 0.028077 | 0.02703 | 0.007172 | 0.02799 |
| 34.98611111 | 0.016025 | 0.02419 | 0.020363 | 0.013168 | 0.011905 | 0.026726 | 0.031921 | 0.006471 | 0.040516 |
| 35 | 0.020481 | 0.047861 | 0.019007 | 0.027154 | 0.015533 | 0.015714 | 0.035029 | 0.012343 | 0.040113 |
| 35.01388889 | 0.048332 | 0.077966 | 0.017715 | 0.040828 | 0.028475 | 0.01186 | 0.028614 | 0.025626 | 0.026003 |
| 35.02777778 | 0.07001 | 0.094007 | 0.031221 | 0.037069 | 0.03863 | 0.014268 | 0.014744 | 0.02844 | 0.010696 |
| 35.04166667 | 0.063663 | 0.083077 | 0.045327 | 0.02279 | 0.040732 | 0.01024 | 0.004414 | 0.018153 | 0.004801 |
| 35.05555556 | 0.050435 | 0.056412 | 0.04803 | 0.013217 | 0.036528 | 0.006812 | 0.007514 | 0.008362 | 0.012102 |
| 35.06944444 | 0.035118 | 0.028789 | 0.040303 | 0.01066 | 0.022843 | 0.012003 | 0.021866 | 0.007297 | 0.017416 |
| 35.08333333 | 0.019489 | 0.016953 | 0.024958 | 0.018155 | 0.010151 | 0.018434 | 0.027844 | 0.022186 | 0.010277 |

| | | | | | | | | | |
|---|---|---|---|---|---|---|---|---|---|
| 35.09722222 | 0.020283 | 0.033309 | 0.011212 | 0.028113 | 0.01062 | 0.020641 | 0.016492 | 0.039706 | 0.008603 |
| 35.11111111 | 0.035558 | 0.055825 | 0.008335 | 0.024901 | 0.01478 | 0.019978 | 0.004074 | 0.040167 | 0.017035 |
| 35.125 | 0.049719 | 0.065465 | 0.014749 | 0.013192 | 0.01102 | 0.018261 | 0.00343 | 0.032332 | 0.026713 |
| 35.13888889 | 0.05714 | 0.070667 | 0.02298 | 0.005666 | 0.005844 | 0.016107 | 0.010993 | 0.031672 | 0.037447 |
| 35.15277778 | 0.057973 | 0.075605 | 0.029555 | 0.002958 | 0.007312 | 0.013111 | 0.017648 | 0.037834 | 0.043237 |
| 35.16666667 | 0.058577 | 0.078151 | 0.035867 | 0.003871 | 0.010121 | 0.010763 | 0.019256 | 0.047117 | 0.035009 |
| 35.18055556 | 0.066951 | 0.075489 | 0.042373 | 0.009871 | 0.010662 | 0.012379 | 0.018434 | 0.056104 | 0.031435 |
| 35.19444444 | 0.078224 | 0.066235 | 0.045052 | 0.017313 | 0.012653 | 0.017756 | 0.017486 | 0.05597 | 0.046996 |
| 35.20833333 | 0.079041 | 0.052776 | 0.040789 | 0.019657 | 0.017649 | 0.023785 | 0.015878 | 0.042934 | 0.057204 |
| 35.22222222 | 0.060486 | 0.035804 | 0.031032 | 0.015112 | 0.020425 | 0.028109 | 0.013153 | 0.02397 | 0.046454 |
| 35.23611111 | 0.032869 | 0.017537 | 0.021171 | 0.008556 | 0.014704 | 0.02707 | 0.009493 | 0.01051 | 0.026159 |
| 35.25 | 0.013777 | 0.00704 | 0.016608 | 0.007031 | 0.005946 | 0.021193 | 0.005349 | 0.008879 | 0.017227 |
| 35.26388889 | 0.007363 | 0.004734 | 0.017633 | 0.009016 | 0.002347 | 0.018506 | 0.002489 | 0.011599 | 0.023924 |
| 35.27777778 | 0.007539 | 0.003237 | 0.019991 | 0.008813 | 0.002408 | 0.021873 | 0.001336 | 0.011442 | 0.027354 |
| 35.29166667 | 0.006551 | 0.002044 | 0.020129 | 0.008367 | 0.002509 | 0.026523 | 0.001555 | 0.012521 | 0.019761 |
| 35.30555556 | 0.005255 | 0.003156 | 0.018169 | 0.011028 | 0.001815 | 0.026979 | 0.002897 | 0.01839 | 0.010284 |
| 35.31944444 | 0.008717 | 0.008124 | 0.015465 | 0.015808 | 0.001143 | 0.019328 | 0.00278 | 0.025416 | 0.007116 |
| 35.33333333 | 0.011903 | 0.014784 | 0.011907 | 0.020516 | 0.001239 | 0.008794 | 0.001344 | 0.0293 | 0.007406 |
| 35.34722222 | 0.009881 | 0.018773 | 0.007067 | 0.021613 | 0.001162 | 0.003399 | 0.001117 | 0.028012 | 0.005081 |
| 35.36111111 | 0.011378 | 0.016992 | 0.0032 | 0.015302 | 0.001206 | 0.003685 | 0.001602 | 0.018559 | 0.002145 |
| 35.375 | 0.023149 | 0.010188 | 0.004369 | 0.006411 | 0.002523 | 0.008931 | 0.003025 | 0.009683 | 0.004731 |
| 35.38888889 | 0.035912 | 0.006096 | 0.010023 | 0.00373 | 0.004237 | 0.014423 | 0.005895 | 0.011255 | 0.015889 |
| 35.40277778 | 0.040012 | 0.008344 | 0.014676 | 0.007794 | 0.004697 | 0.014517 | 0.007033 | 0.012806 | 0.031386 |
| 35.41666667 | 0.036661 | 0.013911 | 0.016473 | 0.01335 | 0.003789 | 0.01201 | 0.005723 | 0.009972 | 0.045183 |
| 35.43055556 | 0.030859 | 0.021056 | 0.018824 | 0.01388 | 0.002998 | 0.012805 | 0.005172 | 0.009734 | 0.050941 |
| 35.44444444 | 0.022801 | 0.026224 | 0.02324 | 0.008679 | 0.003761 | 0.018487 | 0.006072 | 0.012072 | 0.044352 |
| 35.45833333 | 0.012511 | 0.025148 | 0.026195 | 0.004739 | 0.005742 | 0.025007 | 0.006671 | 0.014535 | 0.030014 |
| 35.47222222 | 0.005651 | 0.019288 | 0.025351 | 0.008712 | 0.007328 | 0.029265 | 0.005051 | 0.0171 | 0.014892 |
| 35.48611111 | 0.009991 | 0.013264 | 0.025445 | 0.021693 | 0.008251 | 0.03663 | 0.004465 | 0.021229 | 0.004783 |
| 35.5 | 0.029582 | 0.009011 | 0.031899 | 0.035601 | 0.00904 | 0.053677 | 0.00978 | 0.030068 | 0.006413 |
| 35.51388889 | 0.057873 | 0.006402 | 0.039638 | 0.042601 | 0.007812 | 0.06974 | 0.014076 | 0.051815 | 0.022593 |
| 35.52777778 | 0.085644 | 0.008627 | 0.043336 | 0.047163 | 0.004441 | 0.069694 | 0.010005 | 0.086639 | 0.048943 |
| 35.54166667 | 0.096691 | 0.014093 | 0.050485 | 0.045223 | 0.0049 | 0.059652 | 0.004886 | 0.099918 | 0.072616 |
| 35.55555556 | 0.074729 | 0.014355 | 0.056824 | 0.029647 | 0.01206 | 0.049913 | 0.005385 | 0.069301 | 0.077326 |
| 35.56944444 | 0.036658 | 0.009012 | 0.047293 | 0.012641 | 0.020616 | 0.040159 | 0.006111 | 0.028583 | 0.061189 |
| 35.58333333 | 0.020244 | 0.005175 | 0.027655 | 0.004761 | 0.02614 | 0.027419 | 0.003627 | 0.011901 | 0.037162 |
| 35.59722222 | 0.031716 | 0.005857 | 0.01137 | 0.005391 | 0.030256 | 0.017403 | 0.001508 | 0.019133 | 0.017758 |
| 35.61111111 | 0.043845 | 0.00966 | 0.003768 | 0.010678 | 0.035223 | 0.022506 | 0.000914 | 0.033528 | 0.013685 |
| 35.625 | 0.046298 | 0.01382 | 0.00494 | 0.018004 | 0.0402 | 0.034326 | 0.000804 | 0.045944 | 0.021261 |
| 35.63888889 | 0.039596 | 0.011627 | 0.009794 | 0.026711 | 0.043498 | 0.033024 | 0.001559 | 0.053781 | 0.022847 |
| 35.65277778 | 0.02635 | 0.009632 | 0.013554 | 0.03283 | 0.043127 | 0.020515 | 0.002126 | 0.048721 | 0.01602 |
| 35.66666667 | 0.027189 | 0.018859 | 0.014221 | 0.031444 | 0.03748 | 0.008997 | 0.00168 | 0.028049 | 0.010893 |
| 35.68055556 | 0.045008 | 0.029038 | 0.009842 | 0.026678 | 0.028284 | 0.003164 | 0.002057 | 0.010312 | 0.010221 |
| 35.69444444 | 0.059351 | 0.030336 | 0.007348 | 0.023735 | 0.019972 | 0.004785 | 0.002858 | 0.009432 | 0.008077 |

| | | | | | | | | | |
|---|---|---|---|---|---|---|---|---|---|
| 35.70833333 | 0.054122 | 0.02303 | 0.014817 | 0.019645 | 0.015979 | 0.012699 | 0.002524 | 0.011932 | 0.007471 |
| 35.72222222 | 0.029692 | 0.013008 | 0.025605 | 0.014388 | 0.01596 | 0.022765 | 0.003053 | 0.008609 | 0.017224 |
| 35.73611111 | 0.016665 | 0.012889 | 0.032898 | 0.014201 | 0.017282 | 0.030284 | 0.005379 | 0.014884 | 0.031014 |
| 35.75 | 0.034585 | 0.025222 | 0.031579 | 0.025727 | 0.018699 | 0.032777 | 0.007136 | 0.04971 | 0.039966 |
| 35.76388889 | 0.060011 | 0.038794 | 0.020503 | 0.039678 | 0.019672 | 0.031836 | 0.00746 | 0.079997 | 0.041523 |
| 35.77777778 | 0.076233 | 0.045326 | 0.010816 | 0.037695 | 0.019579 | 0.027476 | 0.006305 | 0.072792 | 0.034333 |
| 35.79166667 | 0.08086 | 0.045704 | 0.012897 | 0.030326 | 0.018962 | 0.01973 | 0.003693 | 0.066464 | 0.019922 |
| 35.80555556 | 0.071353 | 0.041074 | 0.023722 | 0.036556 | 0.018411 | 0.012948 | 0.002148 | 0.079783 | 0.011106 |
| 35.81944444 | 0.053054 | 0.029181 | 0.033638 | 0.048005 | 0.01685 | 0.007978 | 0.002736 | 0.085213 | 0.019686 |
| 35.83333333 | 0.032656 | 0.016463 | 0.039347 | 0.053151 | 0.012547 | 0.005716 | 0.002961 | 0.073641 | 0.035115 |
| 35.84722222 | 0.019343 | 0.014966 | 0.042876 | 0.05407 | 0.00687 | 0.0099 | 0.002107 | 0.061122 | 0.044751 |
| 35.86111111 | 0.016303 | 0.025375 | 0.045427 | 0.053969 | 0.005284 | 0.016191 | 0.00112 | 0.059969 | 0.046697 |
| 35.875 | 0.018509 | 0.037709 | 0.044523 | 0.04999 | 0.009972 | 0.019555 | 0.001188 | 0.069504 | 0.042979 |
| 35.88888889 | 0.019029 | 0.038312 | 0.039555 | 0.038042 | 0.017646 | 0.022402 | 0.00317 | 0.083823 | 0.036941 |
| 35.90277778 | 0.013439 | 0.024017 | 0.032068 | 0.020795 | 0.024553 | 0.026873 | 0.005227 | 0.097112 | 0.030371 |
| 35.91666667 | 0.011135 | 0.016029 | 0.024787 | 0.008633 | 0.026453 | 0.029402 | 0.006704 | 0.108982 | 0.022978 |
| 35.93055556 | 0.018607 | 0.028078 | 0.020414 | 0.0085 | 0.021446 | 0.025693 | 0.009587 | 0.101845 | 0.013842 |
| 35.94444444 | 0.025936 | 0.043934 | 0.016445 | 0.014481 | 0.012228 | 0.016246 | 0.014107 | 0.059988 | 0.010783 |
| 35.95833333 | 0.023728 | 0.048796 | 0.01109 | 0.020483 | 0.005856 | 0.007865 | 0.019025 | 0.02903 | 0.020521 |
| 35.97222222 | 0.018029 | 0.039038 | 0.006957 | 0.026595 | 0.005378 | 0.004998 | 0.023223 | 0.051743 | 0.033754 |
| 35.98611111 | 0.030073 | 0.025838 | 0.012191 | 0.026083 | 0.007111 | 0.006546 | 0.024167 | 0.08079 | 0.037023 |
| 36 | 0.063553 | 0.03195 | 0.031691 | 0.019274 | 0.01327 | 0.013477 | 0.018818 | 0.063606 | 0.024696 |
| 36.01388889 | 0.08844 | 0.050141 | 0.049256 | 0.016127 | 0.021217 | 0.020724 | 0.009895 | 0.045892 | 0.010076 |
| 36.02777778 | 0.086708 | 0.058025 | 0.04632 | 0.013392 | 0.022475 | 0.021666 | 0.005293 | 0.070317 | 0.007443 |
| 36.04166667 | 0.065913 | 0.051686 | 0.029157 | 0.010867 | 0.020211 | 0.017397 | 0.004906 | 0.088641 | 0.009722 |
| 36.05555556 | 0.033904 | 0.031098 | 0.012607 | 0.014036 | 0.019812 | 0.009639 | 0.004456 | 0.070678 | 0.010675 |
| 36.06944444 | 0.013028 | 0.011761 | 0.004089 | 0.018279 | 0.019577 | 0.00347 | 0.005758 | 0.039661 | 0.010809 |
| 36.08333333 | 0.016529 | 0.009102 | 0.004282 | 0.019542 | 0.015053 | 0.002448 | 0.007488 | 0.017719 | 0.008774 |
| 36.09722222 | 0.023976 | 0.012898 | 0.007449 | 0.017012 | 0.007351 | 0.005558 | 0.006017 | 0.016759 | 0.005023 |
| 36.11111111 | 0.028352 | 0.0176 | 0.009145 | 0.010783 | 0.002707 | 0.01213 | 0.003162 | 0.03071 | 0.003969 |
| 36.125 | 0.039423 | 0.023774 | 0.009861 | 0.00496 | 0.001594 | 0.016381 | 0.001882 | 0.043579 | 0.009059 |
| 36.13888889 | 0.0548 | 0.029246 | 0.01124 | 0.003904 | 0.001158 | 0.017604 | 0.003131 | 0.050598 | 0.019373 |
| 36.15277778 | 0.065934 | 0.035202 | 0.013114 | 0.007064 | 0.001081 | 0.01939 | 0.007266 | 0.055094 | 0.034583 |
| 36.16666667 | 0.070804 | 0.042498 | 0.015128 | 0.009149 | 0.001851 | 0.022528 | 0.013025 | 0.056918 | 0.051758 |
| 36.18055556 | 0.073008 | 0.049348 | 0.017988 | 0.006622 | 0.003435 | 0.028285 | 0.017158 | 0.053197 | 0.064271 |
| 36.19444444 | 0.072178 | 0.052378 | 0.021412 | 0.003038 | 0.005071 | 0.030772 | 0.018061 | 0.046252 | 0.069788 |
| 36.20833333 | 0.068039 | 0.052003 | 0.026329 | 0.002144 | 0.005987 | 0.024847 | 0.016254 | 0.041408 | 0.068806 |
| 36.22222222 | 0.062681 | 0.051075 | 0.031609 | 0.001638 | 0.006044 | 0.014104 | 0.011382 | 0.036761 | 0.05815 |
| 36.23611111 | 0.056712 | 0.04937 | 0.032937 | 0.001443 | 0.007299 | 0.007181 | 0.005807 | 0.027703 | 0.03867 |
| 36.25 | 0.054823 | 0.046942 | 0.032763 | 0.005876 | 0.013564 | 0.012876 | 0.007296 | 0.016764 | 0.022222 |
| 36.26388889 | 0.05669 | 0.044188 | 0.033341 | 0.013891 | 0.022496 | 0.026773 | 0.016339 | 0.009231 | 0.016023 |
| 36.27777778 | 0.05222 | 0.035981 | 0.028552 | 0.016042 | 0.024547 | 0.033441 | 0.02108 | 0.005523 | 0.012558 |
| 36.29166667 | 0.036443 | 0.020693 | 0.017787 | 0.010216 | 0.016119 | 0.028676 | 0.015952 | 0.004669 | 0.006348 |
| 36.30555556 | 0.019176 | 0.012165 | 0.010569 | 0.006329 | 0.007061 | 0.023212 | 0.009463 | 0.004748 | 0.004369 |

| | | | | | | | | | |
|---|---|---|---|---|---|---|---|---|---|
| 36.31944444 | 0.011857 | 0.015133 | 0.010067 | 0.005786 | 0.005946 | 0.022051 | 0.008039 | 0.003147 | 0.008193 |
| 36.33333333 | 0.009956 | 0.013721 | 0.008672 | 0.00389 | 0.007227 | 0.019203 | 0.006956 | 0.002488 | 0.007832 |
| 36.34722222 | 0.009163 | 0.006473 | 0.004782 | 0.00255 | 0.006326 | 0.013071 | 0.005465 | 0.005437 | 0.003492 |
| 36.36111111 | 0.011189 | 0.003894 | 0.003182 | 0.002305 | 0.006416 | 0.009584 | 0.007989 | 0.008166 | 0.002046 |
| 36.375 | 0.011468 | 0.005069 | 0.002862 | 0.001758 | 0.006981 | 0.013157 | 0.009331 | 0.006828 | 0.003189 |
| 36.38888889 | 0.00969 | 0.006421 | 0.003837 | 0.002153 | 0.005561 | 0.023106 | 0.007283 | 0.007367 | 0.008064 |
| 36.40277778 | 0.009624 | 0.008086 | 0.009558 | 0.003487 | 0.003477 | 0.032578 | 0.006945 | 0.016204 | 0.016549 |
| 36.41666667 | 0.011681 | 0.008513 | 0.019164 | 0.005052 | 0.002151 | 0.033068 | 0.007565 | 0.027924 | 0.024599 |
| 36.43055556 | 0.015609 | 0.005699 | 0.02665 | 0.006564 | 0.001373 | 0.024047 | 0.005854 | 0.036019 | 0.029992 |
| 36.44444444 | 0.01938 | 0.002281 | 0.025524 | 0.007482 | 0.001069 | 0.013353 | 0.00348 | 0.039541 | 0.031009 |
| 36.45833333 | 0.019497 | 0.001384 | 0.017788 | 0.007826 | 0.001029 | 0.013224 | 0.004078 | 0.037993 | 0.027595 |
| 36.47222222 | 0.01716 | 0.001791 | 0.010587 | 0.008357 | 0.001414 | 0.02111 | 0.007645 | 0.034195 | 0.022329 |
| 36.48611111 | 0.022385 | 0.001968 | 0.015858 | 0.009546 | 0.002272 | 0.018646 | 0.013621 | 0.038156 | 0.01723 |
| 36.5 | 0.04276 | 0.003073 | 0.041851 | 0.010219 | 0.003039 | 0.01117 | 0.020226 | 0.057906 | 0.014267 |
| 36.51388889 | 0.070851 | 0.007987 | 0.072013 | 0.011547 | 0.006375 | 0.019309 | 0.020168 | 0.09487 | 0.019769 |
| 36.52777778 | 0.08966 | 0.017356 | 0.084986 | 0.017549 | 0.014918 | 0.03681 | 0.013368 | 0.148264 | 0.038969 |
| 36.54166667 | 0.086024 | 0.025345 | 0.077907 | 0.025499 | 0.025246 | 0.043834 | 0.010522 | 0.186422 | 0.065216 |
| 36.55555556 | 0.060253 | 0.025729 | 0.060808 | 0.027882 | 0.033934 | 0.039064 | 0.012541 | 0.166683 | 0.07962 |
| 36.56944444 | 0.031044 | 0.020586 | 0.044705 | 0.023903 | 0.039219 | 0.031248 | 0.011119 | 0.107675 | 0.07275 |
| 36.58333333 | 0.021854 | 0.016486 | 0.03158 | 0.01782 | 0.040218 | 0.0229 | 0.005797 | 0.055978 | 0.059728 |
| 36.59722222 | 0.033361 | 0.015761 | 0.018884 | 0.012083 | 0.040975 | 0.014279 | 0.002474 | 0.027002 | 0.054148 |
| 36.61111111 | 0.046081 | 0.015581 | 0.009792 | 0.006983 | 0.044462 | 0.010522 | 0.00226 | 0.012264 | 0.051424 |
| 36.625 | 0.049868 | 0.013306 | 0.010595 | 0.00288 | 0.047379 | 0.017736 | 0.002414 | 0.007885 | 0.041752 |
| 36.63888889 | 0.049983 | 0.008667 | 0.017635 | 0.000902 | 0.046102 | 0.03348 | 0.002761 | 0.019548 | 0.023784 |
| 36.65277778 | 0.051775 | 0.007565 | 0.024601 | 0.00069 | 0.040024 | 0.047643 | 0.003006 | 0.041332 | 0.014881 |
| 36.66666667 | 0.051522 | 0.013821 | 0.035487 | 0.002577 | 0.032087 | 0.050233 | 0.004321 | 0.059199 | 0.027281 |
| 36.68055556 | 0.037422 | 0.01767 | 0.044089 | 0.009047 | 0.027386 | 0.039828 | 0.008487 | 0.062476 | 0.045367 |
| 36.69444444 | 0.02549 | 0.013924 | 0.036046 | 0.018833 | 0.025607 | 0.023268 | 0.012188 | 0.053692 | 0.053268 |
| 36.70833333 | 0.042516 | 0.009434 | 0.018358 | 0.028688 | 0.0235 | 0.009317 | 0.012426 | 0.045761 | 0.04859 |
| 36.72222222 | 0.062502 | 0.006125 | 0.014226 | 0.036019 | 0.02209 | 0.005922 | 0.00967 | 0.042829 | 0.032965 |
| 36.73611111 | 0.062643 | 0.003987 | 0.032821 | 0.037103 | 0.022753 | 0.015291 | 0.005207 | 0.036598 | 0.016264 |
| 36.75 | 0.054584 | 0.007552 | 0.051154 | 0.031255 | 0.023723 | 0.029339 | 0.002704 | 0.024404 | 0.012933 |
| 36.76388889 | 0.036064 | 0.013968 | 0.047537 | 0.021893 | 0.022971 | 0.036425 | 0.005084 | 0.018703 | 0.024802 |
| 36.77777778 | 0.018201 | 0.017108 | 0.031524 | 0.011883 | 0.020197 | 0.035942 | 0.009027 | 0.02587 | 0.041561 |
| 36.79166667 | 0.018167 | 0.016823 | 0.017805 | 0.005471 | 0.016642 | 0.033918 | 0.010532 | 0.037158 | 0.051176 |
| 36.80555556 | 0.02611 | 0.013962 | 0.010027 | 0.003867 | 0.013606 | 0.030754 | 0.008948 | 0.046377 | 0.043286 |
| 36.81944444 | 0.031044 | 0.00902 | 0.014014 | 0.005033 | 0.011079 | 0.024957 | 0.004999 | 0.059399 | 0.024865 |
| 36.83333333 | 0.031768 | 0.009033 | 0.031658 | 0.011936 | 0.007665 | 0.017393 | 0.002028 | 0.075117 | 0.021129 |
| 36.84722222 | 0.04075 | 0.019645 | 0.047995 | 0.022821 | 0.004986 | 0.0104 | 0.002249 | 0.080307 | 0.035361 |
| 36.86111111 | 0.058647 | 0.031364 | 0.044266 | 0.032916 | 0.007691 | 0.008984 | 0.003196 | 0.071802 | 0.047045 |
| 36.875 | 0.065826 | 0.032451 | 0.026692 | 0.042597 | 0.01472 | 0.014797 | 0.004507 | 0.057171 | 0.050066 |
| 36.88888889 | 0.054247 | 0.023051 | 0.017235 | 0.04713 | 0.02117 | 0.021418 | 0.007545 | 0.048896 | 0.051431 |
| 36.90277778 | 0.03067 | 0.011244 | 0.013341 | 0.039757 | 0.025218 | 0.022351 | 0.010647 | 0.052601 | 0.052924 |
| 36.91666667 | 0.013701 | 0.00429 | 0.007293 | 0.027583 | 0.027589 | 0.016315 | 0.011264 | 0.059076 | 0.048967 |

| | | | | | | | | | |
|---|---|---|---|---|---|---|---|---|---|
| 36.93055556 | 0.015332 | 0.005772 | 0.005858 | 0.020946 | 0.028611 | 0.008762 | 0.008387 | 0.064196 | 0.038349 |
| 36.94444444 | 0.026644 | 0.013599 | 0.008095 | 0.018155 | 0.026223 | 0.006485 | 0.004136 | 0.070885 | 0.026609 |
| 36.95833333 | 0.034854 | 0.022545 | 0.008455 | 0.015674 | 0.019219 | 0.007953 | 0.002231 | 0.075672 | 0.016252 |
| 36.97222222 | 0.037032 | 0.028608 | 0.007279 | 0.013539 | 0.013791 | 0.007837 | 0.003226 | 0.074241 | 0.00773 |
| 36.98611111 | 0.028763 | 0.02383 | 0.00495 | 0.012612 | 0.019465 | 0.005149 | 0.004299 | 0.073705 | 0.004357 |
| 37 | 0.022244 | 0.015796 | 0.004163 | 0.017916 | 0.032984 | 0.003293 | 0.00852 | 0.079818 | 0.007426 |
| 37.01388889 | 0.039581 | 0.022993 | 0.009779 | 0.028275 | 0.039879 | 0.003428 | 0.017187 | 0.080412 | 0.011883 |
| 37.02777778 | 0.057219 | 0.034988 | 0.017349 | 0.028561 | 0.037491 | 0.006012 | 0.022393 | 0.070289 | 0.013387 |
| 37.04166667 | 0.049839 | 0.037055 | 0.021058 | 0.01679 | 0.034502 | 0.013092 | 0.023269 | 0.063693 | 0.011746 |
| 37.05555556 | 0.03048 | 0.033383 | 0.022489 | 0.011991 | 0.031694 | 0.01719 | 0.020937 | 0.065123 | 0.00766 |
| 37.06944444 | 0.014453 | 0.028614 | 0.023267 | 0.017616 | 0.023911 | 0.011922 | 0.012681 | 0.063864 | 0.006046 |
| 37.08333333 | 0.006807 | 0.023435 | 0.022208 | 0.016607 | 0.012635 | 0.006657 | 0.004288 | 0.055085 | 0.010097 |
| 37.09722222 | 0.007159 | 0.018169 | 0.01707 | 0.008993 | 0.007443 | 0.009136 | 0.001424 | 0.043308 | 0.013676 |
| 37.11111111 | 0.011879 | 0.01291 | 0.009153 | 0.007254 | 0.011601 | 0.016405 | 0.002267 | 0.030056 | 0.011836 |
| 37.125 | 0.019667 | 0.00921 | 0.007377 | 0.007021 | 0.018813 | 0.027687 | 0.007876 | 0.019183 | 0.008593 |
| 37.13888889 | 0.031449 | 0.013133 | 0.014663 | 0.005837 | 0.024472 | 0.040308 | 0.018377 | 0.02533 | 0.011962 |
| 37.15277778 | 0.045416 | 0.025382 | 0.020858 | 0.009006 | 0.024269 | 0.044856 | 0.027149 | 0.049145 | 0.020088 |
| 37.16666667 | 0.064067 | 0.041419 | 0.021795 | 0.01251 | 0.018509 | 0.040168 | 0.031006 | 0.072787 | 0.025035 |
| 37.18055556 | 0.090174 | 0.056373 | 0.022185 | 0.014527 | 0.013219 | 0.034376 | 0.032526 | 0.085669 | 0.026241 |
| 37.19444444 | 0.112629 | 0.06387 | 0.023549 | 0.017108 | 0.00998 | 0.032565 | 0.032028 | 0.08876 | 0.025381 |
| 37.20833333 | 0.118091 | 0.063485 | 0.024952 | 0.018645 | 0.007762 | 0.034455 | 0.028428 | 0.086281 | 0.024217 |
| 37.22222222 | 0.107061 | 0.058054 | 0.025485 | 0.017184 | 0.00622 | 0.036083 | 0.021872 | 0.078897 | 0.024623 |
| 37.23611111 | 0.086319 | 0.049515 | 0.022818 | 0.011748 | 0.00422 | 0.033572 | 0.012071 | 0.064296 | 0.025028 |
| 37.25 | 0.058749 | 0.038088 | 0.015684 | 0.006087 | 0.002541 | 0.027461 | 0.004743 | 0.041956 | 0.023964 |
| 37.26388889 | 0.028324 | 0.023893 | 0.007793 | 0.005627 | 0.00312 | 0.021313 | 0.005097 | 0.019533 | 0.02258 |
| 37.27777778 | 0.00788 | 0.012505 | 0.003746 | 0.007655 | 0.004082 | 0.017518 | 0.007064 | 0.009986 | 0.021976 |
| 37.29166667 | 0.003371 | 0.007485 | 0.002805 | 0.007196 | 0.003252 | 0.015624 | 0.006141 | 0.011618 | 0.021494 |
| 37.30555556 | 0.006429 | 0.006332 | 0.001879 | 0.004855 | 0.001628 | 0.014494 | 0.00367 | 0.011405 | 0.020363 |
| 37.31944444 | 0.009983 | 0.005839 | 0.001383 | 0.003415 | 0.00122 | 0.014253 | 0.002653 | 0.007025 | 0.018979 |
| 37.33333333 | 0.011373 | 0.004005 | 0.003376 | 0.004797 | 0.002728 | 0.015242 | 0.004656 | 0.00346 | 0.017827 |
| 37.34722222 | 0.011727 | 0.002474 | 0.006594 | 0.00837 | 0.003742 | 0.015791 | 0.005453 | 0.001752 | 0.016424 |
| 37.36111111 | 0.011931 | 0.004242 | 0.008351 | 0.011513 | 0.00271 | 0.014688 | 0.004162 | 0.000762 | 0.014526 |
| 37.375 | 0.010921 | 0.007275 | 0.009062 | 0.012602 | 0.001309 | 0.013858 | 0.005551 | 0.000861 | 0.012833 |
| 37.38888889 | 0.011819 | 0.007775 | 0.010077 | 0.011539 | 0.001218 | 0.014251 | 0.009174 | 0.003429 | 0.01141 |
| 37.40277778 | 0.016759 | 0.005473 | 0.01186 | 0.008605 | 0.00325 | 0.014845 | 0.009899 | 0.00884 | 0.010122 |
| 37.41666667 | 0.021931 | 0.002521 | 0.015352 | 0.004517 | 0.006895 | 0.016176 | 0.006208 | 0.015818 | 0.009698 |
| 37.43055556 | 0.023261 | 0.000902 | 0.01971 | 0.00151 | 0.01033 | 0.017908 | 0.00201 | 0.021459 | 0.009956 |
| 37.44444444 | 0.02031 | 0.001264 | 0.022224 | 0.001133 | 0.011151 | 0.018034 | 0.001449 | 0.023968 | 0.010376 |
| 37.45833333 | 0.014182 | 0.00308 | 0.022399 | 0.001266 | 0.008221 | 0.01655 | 0.004398 | 0.023746 | 0.011595 |
| 37.47222222 | 0.007097 | 0.00573 | 0.021574 | 0.00177 | 0.004927 | 0.013503 | 0.00975 | 0.022512 | 0.013387 |
| 37.48611111 | 0.003622 | 0.00773 | 0.020633 | 0.003854 | 0.007014 | 0.007752 | 0.015967 | 0.024929 | 0.015253 |
| 37.5 | 0.009006 | 0.006464 | 0.021331 | 0.005158 | 0.015256 | 0.00262 | 0.016776 | 0.034481 | 0.018011 |
| 37.51388889 | 0.029035 | 0.00846 | 0.024782 | 0.009342 | 0.023151 | 0.00223 | 0.009745 | 0.049682 | 0.023388 |
| 37.52777778 | 0.068726 | 0.023299 | 0.02982 | 0.018181 | 0.024688 | 0.00575 | 0.004298 | 0.067088 | 0.031889 |

| | | | | | | | | | |
|---|---|---|---|---|---|---|---|---|---|
| 37.54166667 | 0.099133 | 0.035925 | 0.034824 | 0.023174 | 0.024062 | 0.01127 | 0.007131 | 0.081294 | 0.04205 |
| 37.55555556 | 0.086347 | 0.03075 | 0.036041 | 0.020379 | 0.027986 | 0.016296 | 0.012303 | 0.08683 | 0.050586 |
| 37.56944444 | 0.051535 | 0.020112 | 0.03114 | 0.0125 | 0.031677 | 0.017671 | 0.012002 | 0.082577 | 0.052626 |
| 37.58333333 | 0.025195 | 0.012163 | 0.02642 | 0.008649 | 0.028835 | 0.014846 | 0.006585 | 0.073452 | 0.046938 |
| 37.59722222 | 0.019872 | 0.005425 | 0.028647 | 0.013045 | 0.022066 | 0.012514 | 0.003396 | 0.067122 | 0.037919 |
| 37.61111111 | 0.031199 | 0.003559 | 0.032848 | 0.017617 | 0.017706 | 0.01383 | 0.005695 | 0.065478 | 0.030833 |
| 37.625 | 0.042085 | 0.006391 | 0.030217 | 0.020215 | 0.018281 | 0.016081 | 0.008324 | 0.060826 | 0.027781 |
| 37.63888889 | 0.050347 | 0.011494 | 0.020949 | 0.023216 | 0.019561 | 0.017849 | 0.008202 | 0.042201 | 0.025272 |
| 37.65277778 | 0.055718 | 0.018873 | 0.016338 | 0.025359 | 0.017948 | 0.01857 | 0.009134 | 0.027037 | 0.018567 |
| 37.66666667 | 0.051065 | 0.024339 | 0.022033 | 0.023787 | 0.013944 | 0.017201 | 0.014114 | 0.048273 | 0.019831 |
| 37.68055556 | 0.037355 | 0.023428 | 0.022354 | 0.017902 | 0.009314 | 0.014988 | 0.020091 | 0.092665 | 0.035957 |
| 37.69444444 | 0.020722 | 0.018561 | 0.015121 | 0.012005 | 0.006468 | 0.013304 | 0.022686 | 0.121356 | 0.045558 |
| 37.70833333 | 0.010076 | 0.01209 | 0.015477 | 0.007428 | 0.006807 | 0.012087 | 0.02187 | 0.120092 | 0.041603 |
| 37.72222222 | 0.011592 | 0.010584 | 0.014936 | 0.00372 | 0.008353 | 0.0116 | 0.019685 | 0.097795 | 0.036852 |
| 37.73611111 | 0.017863 | 0.016711 | 0.006312 | 0.002436 | 0.007302 | 0.012092 | 0.016466 | 0.06952 | 0.033582 |
| 37.75 | 0.019757 | 0.018153 | 0.002074 | 0.002854 | 0.004791 | 0.01321 | 0.012371 | 0.045181 | 0.026538 |
| 37.76388889 | 0.018011 | 0.011412 | 0.002997 | 0.00565 | 0.00599 | 0.015411 | 0.007718 | 0.025313 | 0.015106 |
| 37.77777778 | 0.015438 | 0.006471 | 0.003816 | 0.011809 | 0.009955 | 0.018496 | 0.003231 | 0.011473 | 0.00529 |
| 37.79166667 | 0.00999 | 0.008865 | 0.006513 | 0.017525 | 0.011268 | 0.019984 | 0.001394 | 0.007091 | 0.002416 |
| 37.80555556 | 0.007558 | 0.014047 | 0.008737 | 0.021391 | 0.009949 | 0.01752 | 0.004181 | 0.01494 | 0.004459 |
| 37.81944444 | 0.013413 | 0.016836 | 0.008168 | 0.026945 | 0.010129 | 0.010534 | 0.012517 | 0.041658 | 0.006935 |
| 37.83333333 | 0.020221 | 0.018027 | 0.009026 | 0.035195 | 0.01589 | 0.006032 | 0.027324 | 0.082801 | 0.009705 |
| 37.84722222 | 0.025464 | 0.017973 | 0.013856 | 0.04382 | 0.030646 | 0.010967 | 0.045058 | 0.12423 | 0.013815 |
| 37.86111111 | 0.026484 | 0.015541 | 0.016333 | 0.049236 | 0.045771 | 0.017079 | 0.056517 | 0.147016 | 0.016344 |
| 37.875 | 0.019679 | 0.014876 | 0.013253 | 0.048855 | 0.047914 | 0.017817 | 0.058265 | 0.145653 | 0.012758 |
| 37.88888889 | 0.013187 | 0.0213 | 0.015379 | 0.045121 | 0.042814 | 0.016389 | 0.055553 | 0.139704 | 0.006143 |
| 37.90277778 | 0.016987 | 0.032624 | 0.024909 | 0.041261 | 0.037848 | 0.012573 | 0.049962 | 0.134172 | 0.002372 |
| 37.91666667 | 0.0266 | 0.040265 | 0.029628 | 0.036713 | 0.029709 | 0.006727 | 0.039526 | 0.119283 | 0.001179 |
| 37.93055556 | 0.024192 | 0.036022 | 0.025715 | 0.030365 | 0.020017 | 0.004054 | 0.02608 | 0.097096 | 0.002077 |
| 37.94444444 | 0.013817 | 0.022164 | 0.017448 | 0.021438 | 0.01131 | 0.006996 | 0.013626 | 0.072054 | 0.005949 |
| 37.95833333 | 0.011851 | 0.010238 | 0.00966 | 0.011646 | 0.005562 | 0.013004 | 0.010843 | 0.044651 | 0.011826 |
| 37.97222222 | 0.013263 | 0.006656 | 0.009458 | 0.009602 | 0.004035 | 0.019139 | 0.018991 | 0.020733 | 0.017398 |
