## Supplemental Table S2a for "Leaf movements as a quantitative metric for early stress detection"

**Supplementary Table S2.** Raw motion data from: **(a)** different stress treatments in lettuce and their respective controls

**Supplementary Table S2.** Raw motion data from: (a) 150 mM Gradual NaCl stress

| Time | Control1 | Control2 | Control3 | Control4 | Control5 | Control6 | Control7 | Control8 | Control9 | Control10 | Control11 | Control12 | Stress1 | Stress2 | Stress3 | Stress4 | Stress5 | Stress6 | Stress7 | Stress8 | Stress9 | Stress10 | Stress11 | Stress12 |
|---|---|---|---|---|---|---|---|---|---|---|---|---|---|---|---|---|---|---|---|---|---|---|---|---|
| 18.09722 | 1.704239 | 1.08016 | 0.925252 | -0.19446 | 1.028643 | -0.06545 | 1.057191 | 0.111242 | 1.508922 | 0.34051 | -0.33673 | 1.194786 | 1.233282 | -1.65741 | 0.182183 | -0.05595 | 0.243828 | 0.040067 | -0.08195 | 0.009816 | 1.646202 | 1.289581 | 1.21658 | 0.149476 |
| 18.11111 | 0.930757 | 0.747874 | 0.549492 | -0.12995 | 0.453548 | -0.05781 | 0.604172 | 0.066766 | 2.083702 | 0.291365 | -0.2096 | 0.590316 | 0.595183 | -1.53756 | 0.075028 | -0.05905 | 0.056271 | 0.060677 | -0.06167 | -0.00523 | 1.057959 | 1.048163 | 0.649739 | 0.083587 |
| 18.125 | 0.39559 | 0.361766 | 0.311262 | -0.09334 | 0.198664 | -0.05414 | 0.121145 | 0.027602 | 1.109271 | 0.172795 | -0.15927 | 0.271464 | 0.236371 | -0.77859 | 0.007556 | -0.10076 | -0.02206 | 0.079414 | -0.05605 | -0.03385 | 0.415885 | 0.578051 | 0.305182 | -0.09026 |
| 18.13889 | 0.16207 | 0.147618 | 0.268798 | -0.08175 | 0.064079 | -0.05901 | -0.01687 | -0.01657 | 0.65747 | 0.091338 | -0.14838 | 0.131999 | 0.074864 | -0.37089 | -0.0182 | -0.13685 | -0.03635 | 0.094941 | -0.04777 | -0.04858 | 0.086207 | 0.191424 | 0.116958 | -0.08994 |
| 18.15278 | 0.074911 | 0.050375 | 0.325899 | -0.09691 | -0.00537 | -0.09041 | -0.05505 | -0.04635 | 0.376281 | 0.023256 | -0.1755 | 0.062955 | 0.02815 | -0.21727 | -0.01006 | -0.11927 | -0.0282 | 0.109303 | -0.02471 | -0.05079 | -0.01792 | 0.010226 | 0.03572 | -0.08296 |
| 18.16667 | 0.055192 | 0.02494 | 0.203995 | -0.11126 | -0.02917 | -0.13727 | -0.04631 | -0.04882 | 0.246685 | -0.01027 | -0.21547 | 0.047379 | 0.043863 | -0.17285 | 0.00404 | -0.084 | -0.02233 | 0.105188 | -0.0157 | -0.05583 | -0.02261 | -0.01935 | 0.031949 | -0.08177 |
| 18.18056 | 0.056574 | 0.026593 | 0.089797 | -0.10228 | -0.01348 | -0.15745 | -0.02734 | -0.01986 | 0.222816 | -0.00541 | -0.23382 | 0.068756 | 0.072511 | -0.21067 | 0.010193 | -0.07061 | -0.02063 | 0.089894 | -0.02224 | -0.06298 | -0.01791 | -0.00547 | 0.062722 | -0.07486 |
| 18.19444 | 0.059824 | 0.045358 | 0.081557 | -0.07961 | 0.01527 | -0.12834 | -0.00423 | 0.0199 | 0.228278 | 0.018049 | -0.22577 | 0.080956 | 0.09105 | -0.24194 | 0.015565 | -0.06521 | -0.01667 | 0.08134 | -0.02335 | -0.07105 | -0.01034 | 0.01886 | 0.085958 | -0.07643 |
| 18.20833 | 0.045375 | 0.065659 | 0.073654 | -0.0652 | 0.023402 | -0.09071 | 0.013326 | 0.033386 | 0.192289 | 0.037053 | -0.19554 | 0.057783 | 0.104998 | -0.2613 | 0.023979 | -0.05824 | -0.01414 | 0.074303 | -0.01772 | -0.08806 | -0.00901 | 0.020313 | 0.073626 | -0.06731 |
| 18.22222 | 0.02139 | 0.079442 | 0.057402 | -0.06893 | 0.011471 | -0.07087 | 0.014878 | 0.019975 | 0.13742 | 0.037261 | -0.17174 | 0.018855 | 0.115397 | -0.26739 | 0.034658 | -0.0541 | -0.01515 | 0.064281 | -0.01131 | -0.11703 | -0.01348 | 0.008503 | 0.037458 | -0.05552 |
| 18.23611 | 0.008963 | 0.080796 | 0.04673 | -0.06972 | 0.009089 | -0.05966 | 0.009149 | 0.008193 | 0.103853 | 0.027927 | -0.17946 | 0.002514 | 0.112013 | -0.25765 | 0.046398 | -0.0534 | -0.01733 | 0.05658 | -0.01063 | -0.15175 | -0.00717 | 0.012944 | 0.01777 | -0.07677 |
| 18.25 | 0.003819 | 0.050688 | 0.036568 | -0.06972 | 0.015249 | -0.05984 | 0.008771 | -0.00022 | 0.092099 | 0.012812 | -0.21316 | 0.005837 | 0.107626 | -0.26081 | 0.055966 | -0.05332 | -0.01615 | 0.055336 | -0.01409 | -0.20187 | -0.00177 | 0.028885 | 0.023008 | -0.10365 |
| 18.26389 | -0.01321 | 0.018738 | 0.016211 | -0.07995 | -0.00025 | -0.07153 | -0.00928 | -0.00829 | 0.084718 | -0.01078 | -0.25411 | 0.0087 | 0.134241 | -0.2425 | 0.058293 | -0.04375 | -0.00912 | 0.058411 | -0.00806 | -0.27342 | -0.00552 | 0.025045 | 0.028608 | -0.10712 |
| 18.27778 | -0.01945 | 0.02721 | 0.009778 | -0.08048 | -0.01731 | -0.07479 | -0.02469 | -0.00469 | 0.083888 | -0.02815 | -0.3019 | 0.011218 | 0.155059 | -0.24441 | 0.0496 | -0.02651 | -0.00594 | 0.057041 | -0.00334 | -0.32948 | -0.00932 | 0.014451 | 0.035313 | -0.11549 |
| 18.29167 | -0.0026 | 0.048985 | 0.033163 | -0.05297 | -0.00273 | -0.06531 | -0.00646 | 0.008404 | 0.106537 | -0.01736 | -0.37425 | 0.020139 | 0.161653 | -0.23999 | 0.033957 | -0.0114 | -0.01073 | 0.047535 | -0.0073 | -0.33698 | -0.00938 | 0.020588 | 0.051387 | -0.12787 |
| 18.30556 | 0.009384 | 0.0657 | 0.056563 | -0.05153 | 0.018799 | -0.06772 | 0.014703 | 0.012447 | 0.125973 | 0.012725 | -0.43515 | 0.014467 | 0.147212 | -0.19951 | 0.020572 | -0.0053 | -0.00914 | 0.03943 | -0.00758 | -0.29296 | -0.00258 | 0.034131 | 0.055451 | -0.11794 |
| 18.31944 | 0.010003 | 0.087489 | 0.05072 | -0.07467 | 0.022139 | -0.08054 | 0.010396 | 0.008033 | 0.115391 | 0.043162 | -0.53806 | -0.00292 | 0.126726 | -0.14515 | 0.015337 | -0.01211 | 0.003646 | 0.040505 | -0.00017 | -0.2461 | 0.00615 | 0.043798 | 0.045881 | -0.09698 |
| 18.33333 | 0.015943 | 0.107935 | 0.052361 | -0.09622 | 0.028348 | -0.07553 | -0.00377 | 0.007596 | 0.103025 | 0.074895 | -0.72658 | -0.00053 | 0.108357 | -0.10867 | 0.014603 | -0.02679 | 0.014318 | 0.042479 | 0.004558 | -0.22843 | 0.002799 | 0.03858 | 0.040713 | -0.08743 |
| 18.34722 | 0.018439 | 0.082526 | 0.061451 | -0.08443 | 0.029605 | -0.06031 | -0.01678 | 0.003057 | 0.097998 | 0.077653 | -0.81523 | 0.013323 | 0.094068 | -0.0941 | 0.014964 | -0.04275 | 0.028915 | 0.035808 | 0.001837 | -0.22743 | -0.00631 | 0.030564 | 0.037915 | -0.09787 |
| 18.36111 | 0.008066 | 0.032642 | 0.075447 | -0.08553 | 0.017273 | -0.0584 | -0.01791 | -0.00575 | 0.100812 | 0.047143 | -0.77907 | 0.022876 | 0.081589 | -0.08126 | 0.012713 | -0.05222 | 0.122503 | 0.031094 | -0.00124 | -0.23773 | 0.00227 | 0.05106 | 0.034698 | -0.10078 |
| 18.375 | 0.011657 | 0.037668 | 0.108907 | -0.07355 | 0.032789 | -0.04907 | 0.014092 | 0.005989 | 0.137052 | 0.039083 | -0.61168 | 0.033643 | 0.072071 | -0.0727 | 0.007331 | -0.05163 | 0.270346 | 0.029998 | -0.00065 | -0.27027 | 0.014543 | 0.076925 | 0.037533 | -0.08647 |
| 18.38889 | 0.028091 | 0.084608 | 0.123246 | -0.0531 | 0.068287 | -0.03067 | 0.035364 | 0.027744 | 0.170515 | 0.052787 | -0.41025 | 0.035659 | 0.060307 | -0.07286 | 0.004247 | -0.04672 | 0.245297 | 0.026564 | -0.00422 | -0.30774 | 0.001434 | 0.071426 | 0.049687 | -0.08704 |
| 18.40278 | 0.031134 | 0.110849 | 0.115185 | -0.05139 | 0.081225 | -0.02738 | 0.013488 | 0.033861 | 0.164698 | 0.0584 | -0.30044 | 0.033294 | 0.052822 | -0.06899 | 0.011295 | -0.0435 | 0.132063 | 0.026597 | -0.01226 | -0.37347 | -0.02059 | 0.057972 | 0.067526 | -0.1088 |
| 18.41667 | 0.027134 | 0.11108 | 0.099806 | -0.0654 | 0.078546 | -0.03749 | -0.0064 | 0.025439 | 0.153645 | 0.065891 | -0.24994 | 0.033051 | 0.05455 | -0.06485 | 0.035634 | -0.05019 | 0.069534 | 0.029231 | -0.01535 | -0.44924 | -0.02331 | 0.059094 | 0.081676 | -0.12319 |
| 18.43056 | 0.021826 | 0.10635 | 0.09607 | -0.07811 | 0.065741 | -0.04778 | 0.003967 | 0.017745 | 0.166954 | 0.091546 | -0.23466 | 0.016475 | 0.060956 | -0.05751 | 0.077936 | -0.0656 | 0.066125 | 0.037335 | -0.00971 | -0.49898 | -0.00398 | 0.084422 | 0.081503 | -0.12538 |
| 18.44444 | 0.024823 | 0.121411 | 0.107827 | -0.07071 | 0.064261 | -0.04893 | 0.029658 | 0.020742 | 0.199853 | 0.122711 | -0.23156 | -0.01254 | NaN | NaN | NaN | NaN | NaN | NaN | NaN | NaN | NaN | NaN | NaN | NaN |
| 18.45833 | 0.027848 | 0.093635 | 0.102695 | -0.0634 | 0.03704 | -0.06406 | 0.050683 | 0.020906 | 0.218442 | 0.120188 | -0.24793 | -0.05814 | NaN | NaN | NaN | NaN | NaN | NaN | NaN | NaN | NaN | NaN | NaN | NaN |
| 18.47222 | 0.009145 | -0.019 | 0.050213 | -0.08972 | -0.04066 | -0.111 | 0.051974 | 0.004351 | 0.195087 | 0.065229 | -0.29653 | -0.12552 | NaN | NaN | NaN | NaN | NaN | NaN | NaN | NaN | NaN | NaN | NaN | NaN |
| 18.48611 | 0.001238 | -0.02401 | 0.014411 | -0.11804 | -0.03103 | -0.13048 | 0.08749 | -0.00246 | 0.217075 | 0.057546 | -0.31866 | -0.13185 | NaN | NaN | NaN | NaN | NaN | NaN | NaN | NaN | NaN | NaN | NaN | NaN |
| 18.5 | 0.120459 | 0.363693 | 0.108934 | -0.08995 | 0.286925 | -0.06405 | 0.49662 | 0.153459 | 0.574782 | 0.263373 | -0.22936 | -0.03143 | NaN | NaN | NaN | NaN | NaN | NaN | NaN | NaN | NaN | NaN | NaN | NaN |
| 18.51389 | 0.351209 | 1.098077 | 0.394108 | -0.01156 | 0.927119 | 0.064998 | 1.462515 | 0.865333 | 1.297152 | 0.703257 | -0.06302 | 0.129413 | NaN | NaN | NaN | NaN | NaN | NaN | NaN | NaN | NaN | NaN | NaN | NaN |
| 18.52778 | 0.415777 | 1.439728 | 0.413162 | 0.000497 | 1.29237 | 0.202305 | 1.742592 | 0.874584 | 1.700223 | 1.236928 | 0.077115 | 0.203991 | NaN | NaN | NaN | NaN | NaN | NaN | NaN | NaN | NaN | NaN | NaN | NaN |
| 18.54167 | 0.255432 | 0.961569 | 0.01294 | -0.06386 | 0.937187 | 0.263495 | 1.020748 | 0.243244 | 1.337319 | 1.235061 | 0.160468 | 0.159501 | NaN | NaN | NaN | NaN | NaN | NaN | NaN | NaN | NaN | NaN | NaN | NaN |
| 18.55556 | 0.164228 | 0.425619 | -0.04542 | -0.03347 | 0.629301 | 0.204488 | 0.532155 | 0.185583 | 0.920984 | 1.022121 | 0.19765 | 0.14869 | NaN | NaN | NaN | NaN | NaN | NaN | NaN | NaN | NaN | NaN | NaN | NaN |
| 18.56944 | 0.225682 | 0.217633 | -0.00254 | 0.079623 | 0.592518 | 0.197243 | 0.33099 | 0.197956 | 0.719661 | 0.881827 | 0.209381 | 0.109176 | NaN | NaN | NaN | NaN | NaN | NaN | NaN | NaN | NaN | NaN | NaN | NaN |
| 18.58333 | 0.297751 | 0.142827 | 0.017153 | 0.133536 | 0.634243 | 0.179981 | 0.278175 | 0.185198 | 0.688481 | 0.859321 | 0.215956 | 0.037671 | NaN | NaN | NaN | NaN | NaN | NaN | NaN | NaN | NaN | NaN | NaN | NaN |
| 18.59722 | 0.376033 | 0.136101 | 0.038503 | 0.136965 | 0.661974 | 0.165093 | 0.292784 | 0.140422 | 0.799074 | 0.797108 | 0.210865 | -0.01081 | NaN | NaN | NaN | NaN | NaN | NaN | NaN | NaN | NaN | NaN | NaN | NaN |
| 18.61111 | 0.443632 | 0.182972 | 0.081847 | 0.131852 | 0.686129 | 0.1601 | 0.264576 | 0.105498 | 0.955881 | 0.735573 | 0.200528 | -0.02315 | NaN | NaN | NaN | NaN | NaN | NaN | NaN | NaN | NaN | NaN | NaN | NaN |
| 18.625 | 0.474295 | 0.258897 | 0.133516 | 0.118312 | 0.627135 | 0.145488 | 0.182809 | 0.084239 | 1.097203 | 0.703533 | 0.224568 | 0.00391 | NaN | NaN | NaN | NaN | NaN | NaN | NaN | NaN | NaN | NaN | NaN | NaN |
| 18.63889 | 0.482366 | 0.34679 | 0.188311 | 0.113196 | 0.572142 | 0.121836 | 0.076998 | 0.128943 | 1.233704 | 0.618925 | 0.301194 | 0.053348 | NaN | NaN | NaN | NaN | NaN | NaN | NaN | NaN | NaN | NaN | NaN | NaN |
| 18.65278 | 0.4084 | 0.441963 | 0.214806 | 0.119872 | 0.471913 | 0.075726 | -0.01389 | 0.237108 | 1.325311 | 0.535766 | 0.349553 | 0.075222 | NaN | NaN | NaN | NaN | NaN | NaN | NaN | NaN | NaN | NaN | NaN | NaN |
| 18.66667 | 0.326109 | 0.476866 | 0.20698 | 0.12526 | 0.354829 | 0.038906 | -0.07083 | 0.289394 | 1.326709 | 0.482486 | 0.32883 | 0.070882 | NaN | NaN | NaN | NaN | NaN | NaN | NaN | NaN | NaN | NaN | NaN | NaN |
| 18.68056 | 0.230973 | 0.449906 | 0.149113 | 0.107812 | 0.215178 | 0.040432 | -0.08496 | 0.227968 | 1.197509 | 0.434837 | 0.302638 | 0.06279 | NaN | NaN | NaN | NaN | NaN | NaN | NaN | NaN | NaN | NaN | NaN | NaN |
| 18.69444 | 0.154345 | 0.385849 | 0.094942 | 0.07087 | 0.097247 | 0.051501 | -0.05843 | 0.130695 | 0.98657 | 0.358391 | 0.283471 | 0.060241 | NaN | NaN | NaN | NaN | NaN | NaN | NaN | NaN | NaN | NaN | NaN | NaN |
| 18.70833 | 0.087946 | 0.339553 | 0.055265 | 0.095685 | 0.050179 | 0.060334 | -0.03231 | 0.103064 | 0.807501 | 0.235847 | 0.270377 | 0.043008 | NaN | NaN | NaN | NaN | NaN | NaN | NaN | NaN | NaN | NaN | NaN | NaN |
| 18.72222 | 0.012852 | 0.31593 | 0.02827 | 0.104762 | -0.00469 | 0.058827 | -0.02945 | 0.09285 | 0.641135 | 0.117196 | 0.262236 | 0.009764 | NaN | NaN | NaN | NaN | NaN | NaN | NaN | NaN | NaN | NaN | NaN | NaN |
| 18.73611 | -0.04299 | 0.293938 | 0.015366 | 0.120803 | -0.1121 | 0.063135 | -0.02295 | 0.078835 | 0.485438 | 0.028446 | 0.259537 | 0.002697 | NaN | NaN | NaN | NaN | NaN | NaN | NaN | NaN | NaN | NaN | NaN | NaN |
| 18.75 | -0.0679 | 0.320742 | 0.014606 | 0.109993 | -0.2231 | 0.068005 | 0.001436 | 0.07064 | 0.353323 | -0.03992 | 0.263513 | 0.018529 | NaN | NaN | NaN | NaN | NaN | NaN | NaN | NaN | NaN | NaN | NaN | NaN |
| 18.76389 | -0.07833 | 0.387995 | 0.014908 | 0.08728 | -0.28059 | 0.069117 | 0.017446 | 0.07239 | 0.252746 | -0.10076 | 0.261789 | 0.025974 | NaN | NaN | NaN | NaN | NaN | NaN | NaN | NaN | NaN | NaN | NaN | NaN |
| 18.77778 | -0.07513 | 0.400967 | 0.017972 | 0.080586 | -0.22059 | 0.103588 | 0.017116 | 0.08791 | 0.224325 | -0.09077 | 0.24306 | 0.025924 | NaN | NaN | NaN | NaN | NaN | NaN | NaN | NaN | NaN | NaN | NaN | NaN |
| 18.79167 | -0.04494 | 0.3271 | 0.020191 | 0.082739 | -0.12171 | 0.255278 | 0.005182 | 0.10956 | 0.239561 | 0.042385 | 0.217565 | 0.030766 | NaN | NaN | NaN | NaN | NaN | NaN | NaN | NaN | NaN | NaN | NaN | NaN |
| 18.80556 | -0.01504 | 0.237097 | 0.020973 | 0.11243 | -0.0618 | 0.352355 | -0.00827 | 0.106997 | 0.231158 | 0.181501 | 0.219326 | 0.031665 | NaN | NaN | NaN | NaN | NaN | NaN | NaN | NaN | NaN | NaN | NaN | NaN |
| 18.81944 | -0.00319 | 0.196929 | 0.029002 | 0.174707 | -0.05267 | 0.319168 | -0.035 | 0.099107 | 0.200847 | 0.268508 | 0.274586 | 0.008618 | NaN | NaN | NaN | NaN | NaN | NaN | NaN | NaN | NaN | NaN | NaN | NaN |
| 18.83333 | -0.03042 | 0.182281 | 0.013552 | 0.184723 | -0.07553 | 0.288953 | -0.08025 | 0.0762 | 0.167409 | 0.249483 | 0.426193 | -0.00915 | NaN | NaN | NaN | NaN | NaN | NaN | NaN | NaN | NaN | NaN | NaN | NaN |
| 18.84722 | -0.0856 | 0.170477 | 0.004201 | 0.143438 | -0.15074 | 0.239653 | -0.13125 | 0.046087 | 0.11089 | 0.155082 | 0.627126 | -0.00271 | NaN | NaN | NaN | NaN | NaN | NaN | NaN | NaN | NaN | NaN | NaN | NaN |
| 18.86111 | -0.12848 | 0.204911 | 0.005339 | 0.115467 | -0.26518 | 0.208357 | -0.19078 | 0.012726 | 0.022004 | 0.000158 | 0.588528 | 0.018145 | NaN | NaN | NaN | NaN | NaN | NaN | NaN | NaN | NaN | NaN | NaN | NaN |
| 18.875 | -0.1759 | 0.245067 | 0.002054 | 0.10242 | -0.40705 | 0.190889 | -0.26024 | 0.008288 | -0.08086 | -0.19513 | 0.410174 | 0.02652 | NaN | NaN | NaN | NaN | NaN | NaN | NaN | NaN | NaN | NaN | NaN | NaN |
| 18.88889 | -0.21403 | 0.20966 | -0.00339 | 0.071741 | -0.47538 | 0.17274 | -0.3828 | 0.026786 | -0.14751 | -0.2911 | 0.283999 | 0.000519 | NaN | NaN | NaN | NaN | NaN | NaN | NaN | NaN | NaN | NaN | NaN | NaN |
| 18.90278 | -0.28869 | 0.151289 | -0.00785 | 0.05298 | -0.47102 | 0.132246 | -0.48382 | 0.043241 | -0.20537 | -0.18031 | 0.235331 | -0.03195 | NaN | NaN | NaN | NaN | NaN | NaN | NaN | NaN | NaN | NaN | NaN | NaN |
| 18.91667 | -0.39467 | 0.125886 | -0.03215 | 0.04817 | -0.42819 | 0.103804 | -0.56156 | 0.040806 | -0.288 | -0.01807 | 0.233902 | -0.04862 | NaN | NaN | NaN | NaN | NaN | NaN | NaN | NaN | NaN | NaN | NaN | NaN |
| 18.93056 | -0.50646 | 0.177268 | -0.05041 | 0.042781 | -0.36999 | 0.089841 | -0.64441 | 0.035213 | -0.35329 | 0.040056 | 0.246682 | -0.0519 | NaN | NaN | NaN | NaN | NaN | NaN | NaN | NaN | NaN | NaN | NaN | NaN |
| 18.94444 | -0.52786 | 0.253431 | -0.04259 | 0.033388 | -0.31113 | 0.084161 | -0.70172 | 0.027629 | -0.35942 | -0.02112 | 0.259618 | -0.06627 | NaN | NaN | NaN | NaN | NaN | NaN | NaN | NaN | NaN | NaN | NaN | NaN |
| 18.95833 | -0.54267 | 0.192233 | -0.04189 | 0.030185 | -0.25963 | 0.086589 | -0.75774 | 0.00392 | -0.32648 | -0.17611 | 0.26519 | -0.08632 | NaN | NaN | NaN | NaN | NaN | NaN | NaN | NaN | NaN | NaN | NaN | NaN |
| 18.97222 | -0.44727 | 0.053366 | -0.05828 | 0.034867 | -0.19478 | 0.081178 | -0.69593 | -0.02418 | -0.31038 | -0.36281 | 0.287744 | -0.09438 | NaN | NaN | NaN | NaN | NaN | NaN | NaN | NaN | NaN | NaN | NaN | NaN |
| 18.98611 | -0.32968 | 0.002927 | -0.06905 | 0.035678 | -0.16409 | 0.07996 | -0.5893 | -0.01856 | -0.32385 | -0.47423 | 0.322148 | -0.09313 | NaN | NaN | NaN | NaN | NaN | NaN | NaN | NaN | NaN | NaN | NaN | NaN |
| 19 | -0.25472 | -0.10622 | -0.13273 | 0.034926 | -0.20206 | 0.021574 | -0.55227 | 0.040161 | -0.47445 | -0.57316 | 0.330404 | -0.17731 | NaN | NaN | NaN | NaN | NaN | NaN | NaN | NaN | NaN | NaN | NaN | NaN |
| 19.01389 | -0.30498 | -0.61512 | -0.28182 | -0.06734 | -0.36369 | -0.13218 | -0.83013 | 0.107118 | -0.9897 | -1.0425 | 0.186828 | -0.53102 | NaN | NaN | NaN | NaN | NaN | NaN | NaN | NaN | NaN | NaN | NaN | NaN |
| 19.02778 | -0.36074 | -1.33183 | -0.53246 | -0.22942 | -0.66352 | -0.48929 | -1.37278 | -0.15054 | -1.95015 | -1.93024 | -0.22623 | -1.15338 | NaN | NaN | NaN | NaN | NaN | NaN | NaN | NaN | NaN | NaN | NaN | NaN |
| 19.04167 | -0.32752 | -1.25265 | -0.72979 | -0.28737 | -0.56158 | -0.78604 | -1.89993 | -1.01335 | -3.12412 | -2.24336 | -0.79047 | -1.7309 | NaN | NaN | NaN | NaN | NaN | NaN | NaN | NaN | NaN | NaN | NaN | NaN |
| 19.05556 | -0.266 | -0.21864 | -0.1967 | -0.28926 | 0.081553 | -0.51033 | -1.15101 | -0.51076 | -1.05821 | -1.51319 | -1.21428 | -0.8252 | NaN | NaN | NaN | NaN | NaN | NaN | NaN | NaN | NaN | NaN | NaN | NaN |
| 19.06944 | -0.20872 | 0.559562 | 0.395414 | -0.28332 | 0.339484 | -0.28888 | 0.158057 | 0.446896 | 1.598898 | -0.16104 | -1.34083 | 0.480384 | NaN | NaN | NaN | NaN | NaN | NaN | NaN | NaN | NaN | NaN | NaN | NaN |
| 19.08333 | -0.15415 | 0.48369 | 0.203698 | -0.2475 | 0.151059 | -0.22339 | 0.67152 | 0.498934 | 1.832796 | 0.602676 | -1.35437 | 0.64028 | NaN | NaN | NaN | NaN | NaN | NaN | NaN | NaN | NaN | NaN | NaN | NaN |
| 19.09722 | -0.11354 | 0.224621 | 0.080935 | -0.21974 | -0.03286 | -0.17114 | 0.445542 | 0.096213 | 0.750257 | 0.606167 | -1.18031 | 0.333425 | NaN | NaN | NaN | NaN | NaN | NaN | NaN | NaN | NaN | NaN | NaN | NaN |
| 19.11111 | -0.06921 | 0.1291 | 0.036103 | -0.16908 | -0.05668 | -0.14034 | 0.228866 | -0.02108 | 0.310711 | 0.369031 | -0.88476 | 0.128891 | NaN | NaN | NaN | NaN | NaN | NaN | NaN | NaN | NaN | NaN | NaN | NaN |
| 19.125 | -0.03169 | 0.090408 | 0.00532 | -0.10804 | -0.03074 | -0.11821 | 0.100098 | -0.06746 | 0.126418 | 0.189024 | -0.65478 | 0.02887 | NaN | NaN | NaN | NaN | NaN | NaN | NaN | NaN | NaN | NaN | NaN | NaN |
| 19.13889 | -0.01 | 0.070177 | -0.00508 | -0.04503 | -0.01542 | -0.09707 | 0.025356 | -0.04506 | 0.052326 | 0.08942 | -0.54027 | 0.001914 | NaN | NaN | NaN | NaN | NaN | NaN | NaN | NaN | NaN | NaN | NaN | NaN |
| 19.15278 | 0.003074 | 0.092617 | -0.00195 | -0.02565 | 0.000699 | -0.07573 | 0.011876 | -0.02194 | 0.043722 | 0.054203 | -0.46492 | 0.014483 | NaN | NaN | NaN | NaN | NaN | NaN | NaN | NaN | NaN | NaN | NaN | NaN |
| 19.16667 | 0.012038 | 0.103275 | 0.017252 | -0.03191 | 0.010185 | -0.05449 | -0.00372 | -0.00206 | 0.011706 | 0.064809 | -0.35497 | 0.051563 | NaN | NaN | NaN | NaN | NaN | NaN | NaN | NaN | NaN | NaN | NaN | NaN |
| 19.18056 | -0.00382 | 0.03029 | 0.062528 | -0.02413 | -0.03167 | -0.05511 | -0.18622 | 0.023062 | -0.26717 | 0.00273 | -0.26283 | 0.090636 | NaN | NaN | NaN | NaN | NaN | NaN | NaN | NaN | NaN | NaN | NaN | NaN |
| 19.19444 | -0.04078 | -0.0028 | 0.092838 | -0.02674 | -0.1794 | -0.09427 | -0.3959 | 0.016115 | -0.55545 | -0.10637 | -0.24375 | 0.026216 | NaN | NaN | NaN | NaN | NaN | NaN | NaN | NaN | NaN | NaN | NaN | NaN |
| 19.20833 | 0.022804 | 0.152977 | 0.036032 | -0.11205 | -0.02314 | -0.12129 | -0.12734 | -0.04098 | -0.07607 | 0.023907 | -0.19729 | -0.05993 | NaN | NaN | NaN | NaN | NaN | NaN | NaN | NaN | NaN | NaN | NaN | NaN |
| 19.22222 | 0.153222 | 0.396231 | -0.0502 | -0.16802 | 0.222 | -0.0555 | 0.363518 | -0.03214 | 0.636754 | 0.329453 | -0.07301 | 0.007594 | NaN | NaN | NaN | NaN | NaN | NaN | NaN | NaN | NaN | NaN | NaN | NaN |
| 19.23611 | 0.201054 | 0.434003 | -0.06721 | -0.0308 | 0.174648 | 0.023837 | 0.429892 | 0.04023 | 0.601729 | 0.409463 | 0.010027 | 0.104179 | NaN | NaN | NaN | NaN | NaN | NaN | NaN | NaN | NaN | NaN | NaN | NaN |
| 19.25 | 0.149823 | 0.329083 | -0.04695 | 0.010254 | 0.110433 | 0.018643 | 0.193336 | 0.0492 | 0.235493 | 0.15188 | 0.020868 | 0.104494 | NaN | NaN | NaN | NaN | NaN | NaN | NaN | NaN | NaN | NaN | NaN | NaN |
| 19.26389 | 0.081072 | 0.1965 | -0.0467 | -0.00881 | 0.034671 | -0.02404 | 0.007097 | 0.004999 | 0.030521 | -0.10857 | -0.00162 | 0.063059 | NaN | NaN | NaN | NaN | NaN | NaN | NaN | NaN | NaN | NaN | NaN | NaN |
| 19.27778 | 0.045443 | 0.044725 | -0.0487 | -0.02661 | 0.008478 | -0.04245 | -0.04852 | -0.01353 | -0.00202 | -0.15799 | -0.02197 | 0.029975 | 0.006351 | 0.03629 | -0.01197 | -0.02968 | 0.032819 | -0.14157 | -0.03295 | -0.02184 | -0.05135 | -0.02519 | -0.08463 | -0.2317 |
| 19.29167 | 0.059921 | 0.003323 | -0.01999 | -0.01136 | 0.038191 | -0.02308 | 0.001593 | 0.018973 | 0.052121 | -0.03005 | -0.02339 | 0.028103 | 0.003188 | 0.032059 | -0.02577 | -0.02177 | 0.038272 | -0.06661 | -0.03543 | -0.02015 | -0.02623 | -0.04185 | -0.04804 | -0.17613 |
| 19.30556 | 0.106914 | 0.103948 | 0.030523 | 0.027964 | 0.083422 | 0.013583 | 0.098393 | 0.074513 | 0.134505 | 0.135729 | 0.006035 | 0.070213 | 0.015556 | 0.038087 | -0.03138 | -0.00766 | 0.066244 | -0.00224 | -0.01648 | -0.00407 | 0.051151 | -0.0297 | 0.020001 | -0.07039 |
| 19.31944 | 0.144492 | 0.244255 | 0.050084 | 0.048433 | 0.099436 | 0.027931 | 0.165418 | 0.094588 | 0.187494 | 0.254779 | 0.027513 | 0.099488 | 0.033303 | 0.044291 | -0.03284 | 0.007731 | 0.103441 | 0.052165 | 0.002842 | 0.015211 | 0.109322 | 0.015926 | 0.079709 | 0.02621 |
| 19.33333 | 0.141209 | 0.277477 | 0.027105 | 0.033146 | 0.085292 | 0.014288 | 0.170434 | 0.077622 | 0.194984 | 0.270651 | 0.019122 | 0.0917 | 0.038953 | 0.037626 | -0.03637 | 0.013724 | 0.123899 | 0.091141 | 0.001446 | 0.025264 | 0.094358 | 0.053725 | 0.095946 | 0.095827 |
| 19.34722 | 0.127346 | 0.253696 | 0.003123 | 0.023015 | 0.081108 | 0.002884 | 0.158404 | 0.064858 | 0.191078 | 0.219604 | 0.018839 | 0.07221 | 0.032207 | 0.013353 | -0.03452 | 0.011118 | 0.117608 | 0.112805 | -0.00705 | 0.026044 | 0.04067 | 0.066627 | 0.084564 | 0.118559 |
| 19.36111 | 0.118757 | 0.216298 | -0.00264 | 0.027339 | 0.089199 | 0.002833 | 0.158556 | 0.063061 | 0.186585 | 0.158695 | 0.031067 | 0.049123 | 0.020571 | -0.00141 | -0.03584 | 0.009116 | 0.109603 | 0.129066 | -0.00983 | 0.024379 | 0.008582 | 0.070751 | 0.07166 | 0.064016 |
| 19.375 | 0.12258 | 0.184714 | -0.00407 | 0.02599 | 0.093269 | 0.005077 | 0.174587 | 0.058437 | 0.189872 | 0.127574 | 0.041633 | 0.066344 | 0.019539 | 0.019403 | -0.04243 | 0.00813 | 0.114056 | 0.181563 | -0.00868 | 0.021393 | 0.02478 | 0.058618 | 0.098656 | -0.01259 |
| 19.38889 | 0.144731 | 0.187909 | 0.003575 | 0.031153 | 0.107706 | 0.014744 | 0.214571 | 0.054314 | 0.235221 | 0.142921 | 0.0555 | 0.089806 | 0.030154 | 0.052756 | -0.04063 | 0.00835 | 0.137523 | 0.247133 | -0.00707 | 0.023919 | 0.069109 | 0.046568 | 0.189349 | -0.01948 |
| 19.40278 | 0.172787 | 0.202541 | 0.0119 | 0.040325 | 0.125829 | 0.024405 | 0.243588 | 0.048229 | 0.295961 | 0.182559 | 0.069598 | 0.085661 | 0.035144 | 0.074512 | -0.03052 | 0.009608 | 0.163316 | 0.302637 | -0.01103 | 0.033288 | 0.097283 | 0.053521 | 0.2919 | 0.034642 |
| 19.41667 | 0.203196 | 0.172013 | 0.0092 | 0.033075 | 0.124333 | 0.022628 | 0.233992 | 0.040559 | 0.315082 | 0.199803 | 0.080244 | 0.05813 | 0.033706 | 0.078274 | -0.01888 | 0.010998 | 0.183488 | 0.278727 | -0.01747 | 0.043041 | 0.087998 | 0.093807 | 0.354749 | 0.091452 |
| 19.43056 | 0.252105 | 0.141393 | 0.009067 | 0.022802 | 0.123323 | 0.024237 | 0.233689 | 0.047079 | 0.312637 | 0.209531 | 0.095351 | 0.050507 | 0.036016 | 0.073616 | -0.00591 | 0.016596 | 0.213231 | 0.273159 | -0.0097 | 0.047894 | 0.069515 | 0.128081 | 0.3798 | 0.151298 |
| 19.44444 | 0.316651 | 0.127687 | 0.016731 | 0.016528 | 0.119503 | 0.028277 | 0.258835 | 0.053756 | 0.335258 | 0.203503 | 0.101223 | 0.046714 | 0.04069 | 0.070822 | -0.00512 | 0.023702 | 0.251563 | 0.288174 | 0.008037 | 0.048482 | 0.050532 | 0.144856 | 0.38937 | 0.190363 |
| 19.45833 | 0.342293 | 0.053487 | 0.015428 | 0.003308 | 0.06298 | 0.002235 | 0.278513 | 0.038839 | 0.396951 | 0.163941 | 0.085146 | -0.02316 | 0.058471 | 0.076015 | -0.02584 | 0.039156 | 0.264009 | 0.345014 | 0.022835 | 0.05417 | 0.052152 | 0.130987 | 0.401778 | 0.1755 |
| 19.47222 | 0.321209 | -0.06243 | 0.004059 | -0.01227 | -0.01137 | -0.0438 | 0.286562 | 0.025665 | 0.471494 | 0.080632 | 0.055702 | -0.14976 | 0.093726 | 0.101005 | -0.04393 | 0.076366 | 0.267313 | 0.396257 | 0.045931 | 0.081281 | 0.08794 | 0.122926 | 0.42209 | 0.149846 |
| 19.48611 | 0.300354 | -0.02897 | 0.004833 | -0.01698 | 0.010188 | -0.04262 | 0.333191 | 0.010005 | 0.559708 | 0.053662 | 0.065733 | -0.10837 | 0.120165 | 0.129324 | -0.01992 | 0.111377 | 0.266925 | 0.41278 | 0.064237 | 0.119255 | 0.135489 | 0.121963 | 0.537152 | 0.137571 |
| 19.5 | 0.444997 | 0.308027 | 0.069956 | -0.0067 | 0.306457 | 0.057399 | 0.633991 | 0.023324 | 1.040211 | 0.267144 | 0.18581 | 0.444627 | 0.11468 | 0.112522 | 0.018115 | 0.089099 | 0.26124 | 0.330071 | 0.034546 | 0.136825 | 0.192579 | 0.090615 | 0.847223 | 0.047374 |
| 19.51389 | 0.924878 | 0.872826 | 0.218653 | 0.004616 | 0.78865 | 0.253474 | 1.559941 | 0.090838 | 2.299098 | 0.694685 | 0.531069 | 2.353106 | 0.076576 | 0.088991 | 0.005779 | 0.024173 | 0.23123 | 0.684405 | -0.04049 | 0.146086 | 0.298854 | 0.026307 | 1.690446 | -0.0198 |
| 19.52778 | 1.206838 | 1.168807 | 0.34443 | -0.05466 | 1.036047 | 0.452585 | 1.293581 | 0.106147 | 3.432819 | 1.050869 | 1.226247 | 6.328777 | 0.036128 | 0.10264 | -0.07108 | -0.01798 | 0.197205 | 1.01192 | -0.14775 | 0.177244 | 0.352911 | -0.04542 | 2.401745 | 0.214032 |

| | | | | | | | | | | | | | | | | | | | | | | | | |
|---|---|---|---|---|---|---|---|---|---|---|---|---|---|---|---|---|---|---|---|---|---|---|---|---|
| 19.54167 | 0.992885 | 0.923481 | 0.371931 | 0.111105 | 0.904315 | 0.568662 | 1.117371 | 0.114173 | 3.013266 | 1.222256 | 2.628079 | 7.904811 | -0.01084 | 0.109749 | -0.16631 | -0.01502 | 0.125727 | 0.433891 | -0.24228 | 0.23592 | 0.221549 | -0.04894 | 2.111192 | 0.386907 |
| 19.55556 | 0.828623 | 0.662423 | 0.418892 | 0.589909 | 0.76953 | 0.529019 | 0.642353 | 0.138567 | 2.176547 | 1.216293 | 2.840682 | 5.737824 | -0.0352 | 0.153353 | -0.15448 | 0.071152 | 0.059065 | 0.170863 | -0.18534 | 0.437424 | 0.131861 | 0.076856 | 1.456072 | 0.323188 |
| 19.56944 | 0.856467 | 0.593959 | 0.642453 | 1.137731 | 0.81937 | 0.753464 | 0.443386 | 0.158042 | 1.799822 | 1.157879 | 3.041516 | 3.922197 | 0.037798 | 0.20538 | -0.05459 | 0.198728 | 0.14154 | 0.144282 | -0.01384 | 0.748626 | 0.134463 | 0.207992 | 0.860779 | 0.234927 |
| 19.58333 | 0.695713 | 0.563343 | 0.867403 | 1.415374 | 0.833708 | 0.801143 | 0.352665 | 0.125661 | 1.440169 | 0.934811 | 6.051561 | 2.387977 | 0.12235 | 0.221453 | -0.01005 | 0.2651 | 0.156427 | 0.158095 | 0.061852 | 0.821887 | 0.141295 | 0.214734 | 0.502348 | 0.165328 |
| 19.59722 | 0.410099 | 0.56267 | 0.944978 | 1.254666 | 0.770179 | 0.797634 | 0.226809 | 0.070837 | 1.258631 | 0.553938 | 4.330519 | 1.57552 | 0.202674 | 0.17032 | -0.01684 | 0.492792 | 0.085415 | 0.113015 | 0.065498 | 0.704074 | 0.166383 | 0.195393 | 0.417698 | 0.159914 |
| 19.61111 | 0.128522 | 0.681324 | 1.023406 | 0.882364 | 0.586281 | 0.756432 | 0.143201 | 0.037302 | 1.111163 | 0.339993 | 2.121178 | 1.063103 | 0.288486 | 0.104734 | -0.02474 | 1.247876 | -0.003 | 0.058081 | 0.069813 | 0.617626 | 0.239738 | 0.225735 | 0.498189 | 0.302923 |
| 19.625 | 0.070243 | 0.856072 | 0.853389 | 0.698121 | 0.322409 | 0.612552 | 0.137599 | 0.037908 | 0.876345 | 0.314056 | 2.411204 | 0.600816 | 0.363142 | 0.049681 | -0.04931 | 2.085475 | -0.02582 | 0.036272 | 0.091815 | 0.514286 | 0.326713 | 0.277606 | 0.654412 | 0.538921 |
| 19.63889 | 0.38522 | 0.978109 | 0.682478 | 0.583077 | -0.01942 | 0.620123 | 0.149737 | 0.074347 | 0.537687 | 0.37344 | 3.175691 | 0.325952 | 0.383007 | 0.02974 | -0.12296 | 2.345043 | 0.05343 | 0.069937 | 0.126684 | 0.386836 | 0.357988 | 0.312027 | 0.698677 | 0.733723 |
| 19.65278 | 0.901945 | 1.004342 | 0.544952 | 0.422835 | -0.24027 | 0.690974 | 0.125195 | 0.100215 | 0.271195 | 0.501177 | 3.721768 | 0.154 | 0.350371 | 0.035617 | -0.23571 | 1.362501 | 0.168169 | 0.127949 | 0.131651 | 0.298739 | 0.276842 | 0.263125 | 0.652513 | 0.700333 |
| 19.66667 | 1.120045 | 0.957603 | 0.467113 | 0.35536 | -0.20403 | 0.687536 | 0.113256 | 0.099624 | 0.179384 | 0.612471 | 3.499789 | 0.003835 | 0.282289 | 0.05454 | -0.34099 | 0.811199 | 0.205942 | 0.173826 | 0.096484 | 0.217959 | 0.158404 | 0.199492 | 0.484906 | 0.488104 |
| 19.68056 | 0.715034 | 0.901023 | 0.398605 | 0.322606 | 0.034645 | 0.541212 | 0.137186 | 0.096857 | 0.184006 | 0.627754 | 2.714291 | -0.09754 | 0.226421 | 0.120394 | -0.41937 | 0.525869 | 0.178882 | 0.21214 | 0.059645 | 0.139901 | 0.072297 | 0.143921 | 0.232992 | 0.191862 |
| 19.69444 | 0.187395 | 0.915305 | 0.320325 | 0.281113 | 0.211253 | 0.46327 | 0.157966 | 0.089098 | 0.150026 | 0.409423 | 1.874594 | 0.003354 | 0.15994 | 0.11682 | -0.44788 | 0.355119 | 0.122447 | 0.261003 | 0.041913 | 0.082277 | 0.02026 | 0.11286 | -0.05628 | -0.03614 |
| 19.70833 | -0.23213 | 0.926856 | 0.219543 | 0.238529 | 0.221316 | 0.478517 | 0.139454 | 0.056338 | 0.064753 | 0.102038 | 1.318327 | 0.092854 | 0.105699 | 0.097897 | -0.49262 | 0.24766 | 0.059523 | 0.291259 | 0.031966 | 0.028329 | -0.00217 | 0.071677 | -0.25039 | -0.20526 |
| 19.72222 | -0.54542 | 0.865913 | 0.135117 | 0.197298 | 0.191526 | 0.496953 | 0.101865 | 0.022855 | -0.00828 | -0.18357 | 0.990363 | 0.111716 | 0.07641 | 0.042476 | -0.4862 | 0.208295 | -0.00632 | 0.311681 | 0.018933 | -0.00349 | 0.013352 | 0.037798 | -0.26867 | -0.33704 |
| 19.73611 | -0.75807 | 0.739263 | 0.092912 | 0.135512 | 0.208849 | 0.465702 | 0.066702 | 0.019423 | -0.03819 | -0.28874 | 0.799195 | 0.183521 | 0.068378 | 0.019571 | -0.44068 | 0.19424 | -0.07328 | 0.294688 | 0.005306 | 0.001095 | 0.052707 | 0.044687 | -0.16209 | -0.27312 |
| 19.75 | -0.55087 | 0.624429 | 0.089326 | 0.074054 | 0.190586 | 0.389714 | 0.040155 | 0.025177 | -0.02201 | -0.28556 | 0.631572 | 0.300381 | 0.066226 | 0.024385 | -0.38371 | 0.142612 | -0.13016 | 0.199523 | 0.001449 | 0.038604 | 0.089996 | 0.112022 | -0.01317 | 0.016651 |
| 19.76389 | -0.01964 | 0.551508 | 0.115903 | 0.033025 | 0.115274 | 0.298302 | 0.043316 | 0.033866 | -0.01272 | -0.05404 | 0.469377 | 0.311793 | 0.051158 | 0.037104 | -0.30739 | 0.009794 | -0.12878 | 0.095509 | -0.00143 | 0.073359 | 0.097763 | 0.177368 | 0.126672 | 0.270099 |
| 19.77778 | 0.523429 | 0.497625 | 0.150172 | 0.041102 | 0.035301 | 0.209459 | 0.088035 | 0.038522 | 0.010556 | 0.314131 | 0.369376 | 0.270312 | 0.012651 | 0.026553 | -0.2442 | -0.05013 | -0.03499 | 0.021883 | -0.00878 | 0.0756 | 0.053579 | 0.200017 | 0.197898 | 0.347002 |
| 19.79167 | 0.658103 | 0.429482 | 0.16529 | 0.0711 | -0.02227 | 0.146331 | 0.130017 | 0.037178 | 0.023771 | 0.525615 | 0.350494 | 0.237211 | -0.03738 | 0.006441 | -0.15776 | -0.04061 | 0.095564 | -0.01599 | -0.0136 | 0.046267 | -0.00351 | 0.210699 | 0.079191 | 0.293428 |
| 19.80556 | 0.410594 | 0.279112 | 0.17272 | 0.063811 | -0.05353 | 0.135863 | 0.131611 | 0.031534 | -0.04023 | 0.514819 | 0.314429 | 0.187716 | -0.0836 | -0.00118 | -0.0845 | -0.07234 | 0.184234 | -0.05645 | -0.02302 | 0.007064 | -0.04212 | 0.163765 | -0.1692 | 0.137193 |
| 19.81944 | 0.116885 | 0.178737 | 0.148113 | 0.059249 | -0.0525 | 0.143958 | 0.118308 | 0.016696 | -0.16234 | 0.349763 | 0.263734 | 0.068934 | -0.11068 | 0.025758 | -0.05314 | -0.06996 | 0.190079 | -0.07595 | -0.03466 | -0.03057 | -0.04813 | 0.069492 | -0.32394 | -0.09536 |
| 19.83333 | -0.18485 | 0.144902 | 0.103023 | 0.078682 | -0.03716 | 0.115891 | 0.103387 | 0.001437 | -0.30793 | 0.091489 | 0.223132 | -0.10247 | -0.11092 | 0.060341 | -0.03687 | -0.01112 | 0.156265 | -0.07555 | -0.03774 | -0.05663 | -0.0394 | -0.01578 | -0.30922 | -0.21491 |
| 19.84722 | -0.45785 | 0.073761 | 0.067493 | 0.071197 | -0.00784 | 0.010332 | 0.055907 | 0.0111 | -0.42289 | -0.25737 | 0.227932 | -0.30761 | -0.08566 | 0.059617 | -0.01376 | 0.073915 | 0.103241 | -0.06228 | -0.02988 | -0.04732 | -0.0432 | -0.0717 | -0.19529 | -0.1622 |
| 19.86111 | -0.47203 | -0.02666 | 0.044965 | 0.018576 | 0.051477 | -0.1313 | -0.01538 | 0.02666 | -0.49982 | -0.57458 | 0.180902 | -0.60902 | -0.07204 | 0.032111 | 0.01155 | 0.11196 | 0.026526 | -0.03116 | -0.02099 | 0.008422 | -0.05875 | -0.11019 | -0.12651 | -0.00728 |
| 19.875 | -0.22057 | -0.14096 | 0.048406 | -0.0336 | 0.088902 | -0.24591 | -0.09659 | 0.025142 | -0.5515 | -0.75438 | 0.110715 | -1.04238 | -0.07491 | -0.03951 | 0.031752 | 0.067061 | -0.05187 | -0.00153 | -0.00661 | 0.049866 | -0.06221 | -0.12242 | -0.15358 | 0.135179 |
| 19.88889 | 0.0213 | -0.21926 | 0.058848 | -0.04607 | 0.045583 | -0.30161 | -0.22167 | 0.013069 | -0.6087 | -0.68184 | 0.051676 | -1.349 | -0.09422 | -0.05099 | 0.047732 | -0.01216 | -0.1407 | 0.00807 | 0.007443 | 0.049445 | -0.07041 | -0.14489 | -0.26387 | 0.141787 |
| 19.90278 | -0.01035 | -0.37791 | -0.01131 | -0.03961 | -0.11311 | -0.26402 | -0.26623 | -0.00567 | -0.70218 | -0.3849 | -0.00041 | -1.3189 | -0.13951 | 0.003628 | 0.034563 | -0.06079 | -0.28496 | -0.02572 | -0.01195 | 0.018773 | -0.13759 | -0.21134 | -0.57535 | 0.036247 |
| 19.91667 | -0.33136 | -0.61103 | -0.1743 | -0.02767 | -0.34437 | -0.18514 | -0.25772 | -0.0224 | -0.87501 | -0.20059 | -0.05706 | -1.22455 | -0.20114 | 0.030306 | 0.004088 | -0.04871 | -0.4076 | -0.11168 | -0.06307 | -0.04314 | -0.23613 | -0.28652 | -1.03014 | -0.11779 |
| 19.93056 | -0.82456 | -0.83542 | -0.37451 | 0.006235 | -0.55253 | -0.13348 | -0.28861 | -0.0251 | -1.01726 | -0.20481 | -0.08465 | -1.2784 | -0.29501 | -0.00152 | -0.02456 | 0.006834 | -0.42136 | -0.2086 | -0.09961 | -0.12844 | -0.35124 | -0.35185 | -1.47944 | -0.25677 |
| 19.94444 | -1.28658 | -0.97754 | -0.56477 | 0.03625 | -0.75192 | -0.12899 | -0.46338 | -0.04485 | -0.93016 | -0.41186 | -0.07661 | -1.4661 | -0.42728 | -0.04829 | -0.03221 | 0.048451 | -0.35793 | -0.26818 | -0.12178 | -0.2084 | -0.37854 | -0.38761 | -1.63602 | -0.33192 |
| 19.95833 | -1.37562 | -1.08827 | -0.60021 | 0.027738 | -0.76293 | -0.12595 | -0.69997 | -0.07118 | -0.74861 | -0.58491 | -0.07707 | -1.822 | -0.5199 | -0.03564 | -0.022 | 0.077872 | -0.3816 | -0.27811 | -0.13587 | -0.20737 | -0.37672 | -0.41352 | -1.43133 | -0.53232 |
| 19.97222 | -1.16214 | -1.03643 | -0.71429 | -0.01688 | -0.46117 | -0.12358 | -0.83418 | -0.07388 | -0.63498 | -0.78558 | -0.1074 | -2.06409 | -0.51673 | 0.001511 | -0.01448 | 0.062702 | -0.29402 | -0.25487 | -0.14012 | -0.12979 | -0.35195 | -0.51074 | -0.99744 | -0.71432 |
| 19.98611 | -0.95927 | -0.90046 | -0.8058 | -0.06536 | -0.35796 | -0.1092 | -0.77736 | -0.06484 | -0.60503 | -0.89595 | -0.17565 | -1.83126 | -0.43278 | -0.01701 | -0.01368 | 0.030003 | -0.19192 | -0.20769 | -0.11644 | -0.0344 | -0.28624 | -0.51142 | -0.60901 | -0.80015 |
| 20 | -0.79598 | -0.71852 | -0.75831 | -0.08497 | -0.30663 | -0.15471 | -0.69181 | -0.04412 | -0.57444 | -0.96762 | -0.32905 | -1.38047 | -0.29574 | -0.02876 | 0.025979 | 0.054468 | -0.05897 | -0.13331 | -0.02608 | 0.023404 | -0.17137 | -0.3494 | -0.35779 | -0.63105 |
| 20.01389 | -0.67382 | -0.66615 | -0.72671 | -0.06812 | -0.23455 | -0.36194 | -0.69109 | -0.02537 | -0.52117 | -1.37707 | -0.61701 | -1.46824 | -0.14497 | 0.031158 | 0.111734 | 0.117499 | 0.078489 | -0.01147 | 0.140419 | -0.0311 | -0.04164 | -0.09137 | -0.38095 | -0.42231 |
| 20.02778 | -0.50966 | -0.61418 | -0.86283 | -0.01278 | -0.11856 | -1.11784 | -0.74762 | 0.053787 | -0.49604 | -2.3936 | -1.26457 | -2.10323 | -0.0301 | 0.057421 | 0.188673 | 0.023825 | 0.123523 | 0.070471 | 0.267038 | -0.45624 | -0.02944 | 0.099106 | -0.87429 | -0.54797 |
| 20.04167 | -0.45522 | -0.53496 | -1.04357 | -0.02823 | 0.008821 | -1.4641 | -0.81184 | 0.005618 | -0.51913 | -2.26743 | -3.05861 | -3.34587 | 0.017356 | 0.004076 | 0.219164 | -0.81656 | 0.100452 | -0.07535 | 0.268814 | -0.77338 | -0.10296 | 0.167461 | -1.62891 | -0.84892 |
| 20.05556 | -0.48899 | -0.43046 | -0.80182 | -0.36409 | -0.03654 | -0.72481 | -0.64194 | -0.33953 | -0.53668 | -0.99526 | -3.32166 | -2.39412 | 0.003958 | -0.09392 | 0.14508 | -1.81293 | 0.00736 | 0.027203 | 0.167953 | -0.4621 | -0.0969 | 0.132654 | -1.21919 | -0.55308 |
| 20.06944 | -0.46753 | -0.26081 | -0.51942 | -0.39293 | -0.15046 | -0.40823 | -0.15747 | -0.2558 | -0.45632 | 0.248308 | -1.47577 | 0.14164 | -0.07136 | -0.16355 | -0.00411 | -1.1149 | -0.14346 | 0.235746 | 0.069071 | -0.23987 | -0.02458 | 0.049262 | -0.16683 | 0.061429 |
| 20.08333 | -0.32274 | -0.12867 | -0.32313 | -0.24067 | -0.1564 | -0.21742 | 0.114711 | -0.17141 | -0.2909 | 0.83356 | -0.16577 | 1.074485 | -0.11085 | -0.15463 | -0.0895 | -0.33896 | -0.16578 | 0.657804 | 0.020438 | -0.17398 | -0.02034 | -0.0135 | 0.289226 | 0.209912 |
| 20.09722 | -0.07046 | -0.09948 | -0.2259 | -0.16012 | -0.1212 | -0.13876 | 0.181946 | -0.10275 | -0.0788 | 0.524651 | 0.185598 | 0.943597 | -0.09893 | -0.12771 | -0.09549 | -0.02199 | -0.07071 | 0.465607 | -0.03376 | -0.11721 | -0.04765 | -0.03001 | 0.360217 | 0.03563 |
| 20.11111 | 0.20603 | -0.08021 | -0.15861 | -0.111 | -0.10942 | -0.10282 | 0.143143 | -0.06637 | 0.218016 | 0.16634 | 0.141606 | 0.689477 | -0.07921 | -0.11448 | -0.07825 | 0.100115 | -0.01106 | 0.23447 | -0.06309 | -0.1064 | -0.05543 | -0.01869 | 0.402043 | -0.06679 |
| 20.125 | 0.422268 | -0.03194 | -0.1155 | -0.08944 | -0.12894 | -0.07746 | 0.014638 | -0.06151 | 0.497492 | -0.0119 | 0.028673 | 0.493264 | -0.06547 | -0.11374 | -0.07184 | 0.109862 | -0.00232 | 0.096972 | -0.06586 | -0.11658 | -0.04345 | -0.01336 | 0.41373 | -0.09986 |
| 20.13889 | 0.557633 | 0.007534 | -0.09089 | -0.07971 | -0.12531 | -0.05305 | -0.07604 | -0.07413 | 0.591681 | -0.10879 | -0.04369 | 0.318286 | -0.04862 | -0.10591 | -0.07165 | 0.096107 | 0.01875 | 0.020515 | -0.06048 | -0.12285 | -0.03709 | -0.02524 | 0.346306 | -0.15232 |
| 20.15278 | 0.563036 | 0.028137 | -0.06773 | -0.06499 | -0.07966 | -0.04131 | -0.05991 | -0.05907 | 0.456262 | -0.15291 | -0.05715 | 0.228203 | -0.02998 | -0.0839 | -0.07113 | 0.071775 | 0.066552 | -0.01312 | -0.05186 | -0.12541 | -0.04028 | -0.03585 | 0.264327 | -0.19273 |
| 20.16667 | 0.529216 | 0.02336 | -0.03921 | -0.04411 | -0.01554 | -0.03701 | 0.016114 | -0.01427 | 0.332575 | -0.18521 | -0.01816 | 0.23703 | -0.01148 | -0.07559 | -0.07672 | 0.036623 | 0.087823 | -0.01734 | -0.03953 | -0.12531 | -0.03124 | -0.0377 | 0.279834 | -0.0441 |
| 20.18056 | 0.479289 | 0.007553 | 0.013793 | -0.01184 | 0.030022 | -0.02904 | 0.028892 | 0.014554 | 0.372517 | -0.22425 | 0.034236 | 0.254956 | 0.034164 | -0.22326 | -0.12381 | -0.02873 | 0.055027 | -0.03914 | -0.04425 | -0.14158 | -0.1127 | -0.09388 | 0.318893 | 0.112877 |
| 20.19444 | 0.427086 | 0.006426 | 0.098637 | 0.010022 | 0.029843 | -0.01435 | -0.20114 | 0.015503 | 0.322149 | -0.28994 | -0.04973 | 0.012175 | 0.178504 | -0.87454 | -0.32624 | -0.29072 | -0.15476 | -0.1951 | -0.12964 | -0.28913 | -0.5838 | -0.37637 | 0.226604 | -0.13354 |
| 20.20833 | 0.40232 | 0.040322 | 0.157923 | 0.031002 | 0.010412 | -0.00815 | -0.37325 | 0.00482 | 0.222925 | -0.27672 | -0.10684 | -0.17465 | 0.254156 | -1.3746 | -0.43639 | -0.39535 | -0.38009 | -0.21187 | -0.17358 | -0.44738 | -0.84538 | -0.61736 | 0.102323 | -0.26499 |
| 20.22222 | 0.377496 | 0.113368 | 0.131951 | 0.092054 | 0.04895 | -0.02262 | 0.098152 | 0.010716 | 0.670415 | 0.020373 | 0.190318 | 0.372247 | -0.00356 | -0.05671 | -0.11469 | 0.058953 | 0.060761 | 0.008101 | -0.00071 | -0.11843 | 0.034907 | 0.080605 | 0.256105 | 0.054127 |
| 20.23611 | 0.341437 | 0.214732 | 0.082458 | 0.114449 | 0.094489 | -0.0326 | 0.51842 | 0.008972 | 1.071842 | 0.27737 | 0.54366 | 0.950066 | -0.25375 | 1.181929 | 0.215127 | 0.537744 | 0.507843 | 0.230333 | 0.163734 | 0.228767 | 0.913669 | 0.806317 | 0.491591 | 0.435895 |
| 20.25 | 0.232581 | 0.224525 | 0.092996 | 0.052699 | 0.098108 | -0.01523 | 0.266166 | -0.00869 | 0.8305 | 0.160026 | 0.524356 | 0.874378 | -0.13673 | 0.595369 | 0.144663 | 0.549129 | 0.372331 | 0.204461 | 0.103717 | 0.15766 | 0.587249 | 0.532881 | 0.4458 | 0.428372 |
| 20.26389 | 0.09098 | 0.17763 | 0.163522 | 0.053554 | 0.101921 | 0.007747 | 0.031163 | 0.045056 | 0.557382 | 0.004487 | 0.361026 | 0.666843 | 0.025235 | 0.037212 | -0.03832 | 0.280534 | 0.121235 | 0.065877 | 0.007197 | 0.034581 | 0.108388 | 0.169657 | 0.303788 | 0.066036 |
| 20.27778 | 0.019125 | 0.146561 | 0.220719 | 0.065765 | 0.131504 | 0.020264 | 0.049016 | 0.106542 | 0.525033 | 0.025435 | 0.231939 | 0.55248 | 0.074418 | -0.04418 | -0.07556 | 0.214446 | 0.078898 | 0.046798 | -0.00603 | 0.001474 | 0.041625 | 0.083498 | 0.292851 | 0.024257 |
| 20.29167 | 0.019084 | 0.163148 | 0.230682 | 0.055849 | 0.129071 | 0.018413 | 0.12311 | 0.102656 | 0.58115 | 0.073107 | 0.177918 | 0.413932 | 0.082569 | -0.01992 | -0.06358 | 0.136711 | 0.108211 | 0.048285 | 0.003033 | -0.01029 | 0.063377 | 0.050704 | 0.300208 | 0.075267 |
| 20.30556 | 0.11596 | 0.200546 | 0.209611 | 0.038113 | 0.133357 | 0.01233 | 0.215649 | 0.07166 | 0.767861 | 0.165687 | 0.220861 | 0.434357 | 0.088082 | -0.01644 | -0.02886 | 0.069949 | 0.133803 | 0.04735 | 0.014962 | -0.02052 | 0.083032 | 0.045945 | 0.320978 | 0.098187 |
| 20.31944 | 0.310746 | 0.224778 | 0.205036 | 0.028897 | 0.188816 | 0.012514 | 0.296676 | 0.066173 | 0.983976 | 0.273218 | 0.31408 | 0.572001 | 0.098285 | -0.02239 | -0.01143 | 0.036096 | 0.1282 | 0.050706 | 0.017431 | -0.05254 | 0.11406 | 0.087622 | 0.333047 | 0.0805 |
| 20.33333 | 0.524236 | 0.222613 | 0.223167 | 0.034532 | 0.252648 | 0.022062 | 0.363044 | 0.08727 | 1.00675 | 0.341126 | 0.621941 | 0.689375 | 0.109791 | -0.01257 | -0.01257 | 0.029301 | 0.121086 | 0.055711 | 0.020249 | -0.13368 | 0.175533 | 0.143449 | 0.333751 | 0.05889 |
| 20.34722 | 0.629139 | 0.20884 | 0.253831 | 0.04821 | 0.257029 | 0.032637 | 0.440002 | 0.106005 | 0.924381 | 0.371358 | 1.189307 | 0.731977 | 0.136757 | 0.025597 | -0.01148 | 0.06185 | 0.154203 | 0.055812 | 0.017612 | -0.17526 | 0.250513 | 0.172659 | 0.3667 | 0.078226 |
| 20.36111 | 0.606762 | 0.22747 | 0.280742 | 0.053865 | 0.216649 | 0.047502 | 0.577725 | 0.101255 | 0.769295 | 0.36419 | 1.450118 | 0.772145 | 0.181109 | 0.079541 | -0.00296 | 0.111561 | 0.240957 | 0.061512 | 0.006193 | -0.12075 | 0.282896 | 0.144759 | 0.406048 | 0.123344 |
| 20.375 | 0.577181 | 0.279999 | 0.305233 | 0.035012 | 0.178741 | 0.062278 | 0.685026 | 0.098601 | 0.674528 | 0.362032 | 1.317661 | 0.818953 | 0.221916 | 0.144849 | 0.002903 | 0.137909 | 0.310026 | 0.097059 | -0.00015 | -0.08243 | 0.28809 | 0.102498 | 0.469942 | 0.153467 |
| 20.38889 | 0.548993 | 0.35702 | 0.311917 | 0.033502 | 0.183985 | 0.089287 | 0.670904 | 0.095085 | 0.775157 | 0.386993 | 1.155173 | 0.907766 | 0.248877 | 0.204002 | 0.003833 | 0.149061 | 0.299757 | 0.146262 | 0.004229 | -0.06671 | 0.292056 | 0.086054 | 0.561672 | 0.145031 |
| 20.40278 | 0.531679 | 0.396342 | 0.322851 | 0.068475 | 0.236753 | 0.119104 | 0.646759 | 0.105998 | 1.028001 | 0.412542 | 1.074054 | 0.999651 | 0.289881 | 0.199177 | 0.011637 | 0.148046 | 0.254572 | 0.183924 | 0.01645 | -0.04566 | 0.301255 | 0.126002 | 0.613109 | 0.124143 |
| 20.41667 | 0.471385 | 0.37092 | 0.322152 | 0.136075 | 0.284783 | 0.121558 | 0.566308 | 0.092386 | 1.208227 | 0.463703 | 0.940184 | 1.071749 | 0.338975 | 0.171543 | 0.026055 | 0.140368 | 0.244527 | 0.193539 | 0.037565 | -0.03387 | 0.319558 | 0.213928 | 0.626456 | 0.147071 |
| 20.43056 | 0.382089 | 0.338805 | 0.397241 | 0.203905 | 0.29262 | 0.118695 | 0.580905 | 0.084788 | 1.247617 | 0.593501 | 0.926754 | 1.167621 | 0.357226 | 0.1744 | 0.022191 | 0.11095 | 0.319708 | 0.190862 | 0.041381 | -0.02054 | 0.322938 | 0.28591 | 0.641688 | 0.266373 |
| 20.44444 | 0.225303 | 0.37068 | 0.464051 | 0.248945 | 0.326035 | 0.158361 | 0.630349 | 0.091992 | 1.224699 | 0.743943 | 0.992536 | 1.275038 | 0.367066 | 0.385526 | -0.0008 | 0.125259 | 0.42922 | 0.202742 | 0.035378 | -0.01654 | 0.379998 | 0.308171 | 0.65874 | 0.3848 |
| 20.45833 | 0.091076 | 0.420583 | 0.459679 | 0.280554 | 0.312567 | 0.240367 | 0.685533 | 0.117588 | 1.115996 | 0.779168 | 0.9488 | 1.310263 | 0.366421 | 0.81447 | -0.02839 | 0.248189 | 0.544387 | 0.169896 | 0.069239 | -0.0242 | 0.518486 | 0.301787 | 0.646748 | 0.362204 |
| 20.47222 | -0.00155 | 0.390088 | 0.361359 | 0.209591 | 0.240249 | 0.334556 | 0.580142 | 0.112647 | 0.925135 | 0.661615 | 0.826742 | 1.252102 | 0.349709 | 0.867002 | -0.04786 | 0.404642 | 0.538633 | 0.12901 | 0.144357 | -0.03173 | 0.642544 | 0.335184 | 0.681872 | 0.276847 |
| 20.48611 | 0.030834 | 0.45316 | 0.311762 | 0.120933 | 0.268745 | 0.353384 | 0.335155 | 0.07895 | 0.736333 | 0.402777 | 0.863172 | 1.261079 | 0.357675 | 0.504653 | -0.03267 | 0.465566 | 0.453713 | 0.144858 | 0.175373 | -0.01521 | 0.674841 | 0.414572 | 0.796585 | 0.237485 |
| 20.5 | 0.291817 | 0.972345 | 0.486412 | 0.171823 | 0.705851 | 0.558854 | 0.138754 | 0.088475 | 0.749574 | 0.431131 | 1.136616 | 1.918885 | 0.42559 | 0.408882 | 0.008197 | 0.451768 | 0.304393 | 0.134497 | 0.112963 | 0.069158 | 0.565771 | 0.595831 | 1.305108 | 0.108014 |
| 20.51389 | 0.798111 | 1.973677 | 1.011831 | 0.321494 | 1.593638 | 1.034154 | 0.119227 | 0.157985 | 1.037074 | 1.428865 | 1.698576 | 3.341955 | 0.534708 | 0.600418 | -0.00303 | 0.650267 | 0.180554 | 0.088615 | 0.036355 | 0.292818 | 0.513591 | 0.761276 | 2.409322 | -0.09805 |
| 20.52778 | 1.692166 | 2.62391 | 1.437638 | 0.425133 | 1.605276 | 1.584704 | 0.2599 | 0.191298 | 1.145583 | 2.589799 | 2.110723 | 3.779788 | 0.472575 | 0.86785 | -0.08417 | 0.606723 | 0.046182 | 0.09176 | 0.037525 | 0.530928 | 0.53292 | 0.72081 | 3.003244 | -0.17439 |
| 20.54167 | 1.751072 | 2.446343 | 1.40754 | 0.405122 | 1.048661 | 1.583404 | 0.289885 | 0.168137 | 0.985261 | 2.901271 | 1.991536 | 2.564935 | 0.235302 | 1.115575 | -0.17428 | -0.00034 | -0.07186 | 0.044354 | 0.037311 | 0.659094 | 0.488636 | 0.550143 | 2.547136 | -0.08104 |
| 20.55556 | 0.982936 | 2.393358 | 1.267787 | 0.316774 | 0.880779 | 0.948767 | 0.08403 | 0.179775 | 0.895285 | 2.269593 | 1.708142 | 1.512333 | 0.117325 | 1.195879 | -0.1473 | -0.16266 | -0.06939 | 0.045716 | 0.149701 | 1.003756 | 0.332576 | 0.600151 | 1.517934 | -0.00349 |
| 20.56944 | 0.484232 | 2.71741 | 1.159931 | 0.250434 | 1.317171 | 0.616263 | -0.0883 | 0.168944 | 1.081729 | 1.43569 | 1.389185 | 0.860821 | 0.152345 | 1.001501 | -0.03868 | -0.03674 | 0.090872 | 0.129609 | 0.226013 | 1.314846 | 0.161245 | 0.672793 | 0.442841 | -0.01228 |
| 20.58333 | 0.000378 | 2.825467 | 0.879398 | 0.200664 | 2.805549 | 0.396035 | -0.27646 | 0.105995 | 0.895693 | 0.251179 | 0.966861 | 0.262326 | 0.171088 | 0.711903 | 0.003356 | 0.046659 | 0.164828 | 0.169717 | 0.239719 | 1.045755 | -0.01549 | 0.587333 | -0.48081 | 0.075988 |
| 20.59722 | -0.51288 | 2.246307 | 0.493527 | 0.286118 | 2.065981 | 0.219676 | -0.36331 | 0.053753 | 0.341131 | -0.43033 | 0.65371 | -0.65091 | 0.168892 | 0.498066 | -0.00939 | 0.050393 | 0.11388 | 0.132385 | 0.218476 | 0.572691 | -0.1257 | 0.440041 | -1.05853 | 0.206537 |
| 20.61111 | -0.97507 | 1.446074 | 0.239217 | 0.44501 | 0.785822 | 0.094968 | -0.28493 | 0.049424 | -0.07461 | -0.10879 | 0.46094 | -1.45343 | 0.162712 | 0.428758 | -0.0205 | 0.041755 | 0.018287 | 0.09124 | 0.172356 | 0.119061 | -0.18895 | 0.331554 | -1.1129 | 0.307272 |
| 20.625 | -0.88146 | 0.839947 | 0.08476 | 0.563204 | 0.207152 | 0.215847 | -0.00705 | 0.028979 | -0.36027 | 0.572735 | 0.294088 | -1.60671 | 0.11376 | 0.438782 | -0.03922 | 0.019511 | -0.04619 | 0.04471 | 0.102066 | -0.09328 | -0.20209 | 0.245126 | -0.5055 | 0.414132 |
| 20.63889 | -0.08909 | 0.505963 | 0.131101 | 0.562253 | -0.13552 | 0.480984 | 0.285899 | 0.039548 | -0.55096 | 1.226991 | 0.136555 | -1.03829 | 0.033601 | 0.531047 | -0.0554 | 0.015673 | -0.00271 | 0.00905 | 0.06632 | -0.04109 | -0.05324 | 0.207275 | 0.152949 | 0.446901 |
| 20.65278 | 0.891652 | 0.328093 | 0.33567 | 0.539433 | -0.40247 | 0.820139 | 0.420195 | 0.058003 | -0.60682 | 1.571946 | -0.06976 | -0.27518 | -0.05139 | 0.606784 | -0.04959 | 0.038613 | 0.115761 | -0.01524 | 0.092729 | 0.191277 | 0.258322 | 0.240802 | 0.646247 | 0.371544 |
| 20.66667 | 1.739997 | 0.198042 | 0.50319 | 0.415024 | -0.34544 | 0.845671 | 0.261191 | 0.078437 | -0.54579 | 1.416352 | -0.21397 | -0.04605 | -0.12745 | 0.565591 | -0.04362 | 0.054145 | 0.188482 | -0.04193 | 0.153742 | 0.383825 | 0.543878 | 0.352803 | 0.92362 | 0.268608 |
| 20.68056 | 1.593293 | 0.143932 | 0.492827 | 0.265786 | -0.07644 | 0.412687 | -0.0094 | 0.060673 | -0.29415 | 0.565229 | -0.23594 | -0.19311 | -0.20181 | 0.443881 | -0.05226 | 0.037178 | 0.154572 | -0.06574 | 0.157532 | 0.417583 | 0.546387 | 0.443776 | 0.92729 | 0.173673 |
| 20.69444 | 1.160418 | 0.102483 | 0.404499 | 0.174779 | 0.492899 | 0.007526 | -0.22023 | 0.037139 | 0.118516 | -0.39025 | -0.12996 | -0.24279 | -0.22735 | 0.294809 | -0.05496 | 0.02183 | 0.058932 | -0.07257 | 0.120786 | 0.076316 | 0.300524 | 0.34696 | 0.507386 | 0.006755 |
| 20.70833 | 0.482892 | 0.03505 | 0.317352 | 0.163264 | 1.1056 | -0.22681 | -0.40336 | 0.019089 | 0.514684 | -1.2042 | 0.000996 | -0.04448 | -0.18819 | 0.169415 | -0.04819 | 0.020839 | -0.00319 | -0.08699 | 0.07628 | -0.91953 | -0.02846 | 0.181255 | -0.06125 | -0.12673 |
| 20.72222 | -0.2188 | -0.13889 | 0.225503 | 0.23145 | 1.217806 | -0.28113 | -0.43147 | -0.00203 | 0.652381 | -1.10368 | 0.177071 | 0.023041 | -0.10705 | 0.071859 | -0.04619 | 0.030904 | -0.03785 | -0.12693 | 0.024023 | -1.70103 | -0.27751 | 0.00631 | -0.31762 | -0.0236 |
| 20.73611 | -0.77334 | -0.334 | 0.216299 | 0.315168 | 0.845188 | -0.29411 | -0.287 | -0.00994 | 0.510067 | -0.42223 | 0.375626 | 0.004921 | -0.05002 | 0.05655 | -0.04367 | 0.067825 | 0.014307 | -0.18124 | -0.02471 | -1.39173 | -0.38597 | -0.09569 | -0.12704 | 0.184211 |
| 20.75 | -1.18702 | -0.48246 | 0.196954 | 0.333106 | 0.269328 | -0.32798 | -0.19648 | 0.009697 | 0.148939 | 0.216694 | 0.735504 | 0.060921 | -0.03138 | 0.119773 | -0.04409 | 0.10184 | 0.017942 | -0.22277 | -0.05627 | -1.20661 | -0.36896 | -0.08319 | 0.324363 | 0.320275 |
| 20.76389 | -1.63626 | -0.54636 | 0.089444 | 0.288906 | -0.29656 | -0.15242 | -0.03041 | -0.00249 | -0.23083 | 0.530685 | 0.864224 | 0.290639 | -0.03582 | 0.202715 | -0.04264 | 0.115284 | -0.15565 | -0.26128 | -0.06895 | -1.13435 | -0.35069 | 0.003219 | 0.631764 | 0.245606 |
| 20.77778 | -1.20977 | -0.44913 | -0.05301 | 0.224201 | -0.50108 | 0.177938 | 0.208537 | -0.00861 | -0.50273 | 0.525298 | 0.658524 | 0.413184 | -0.04405 | 0.303878 | -0.03192 | 0.123661 | -0.39312 | -0.2129 | -0.01966 | -0.83776 | -0.23667 | 0.146692 | 0.750598 | 0.180898 |
| 20.79167 | -0.09319 | -0.27966 | -0.10534 | 0.200476 | -0.44093 | 0.682464 | 0.485399 | 0.028028 | -0.5183 | 0.246783 | 0.348746 | 0.162568 | -0.05878 | 0.329406 | -0.01892 | 0.116943 | -0.37783 | -0.08331 | 0.0417 | -0.52381 | 0.03296 | 0.2847 | 0.577742 | 0.09821 |
| 20.80556 | 0.815305 | -0.15 | -0.12393 | 0.193357 | -0.29934 | 0.882417 | 0.375197 | 0.039606 | -0.28179 | -0.43275 | 0.078986 | -0.26284 | -0.07517 | 0.216665 | -0.01555 | 0.084029 | -0.14068 | 0.071667 | 0.045283 | -0.33573 | 0.241201 | 0.275828 | 0.122982 | -0.0596 |
| 20.81944 | 1.670718 | -0.07675 | -0.14159 | 0.158262 | -0.35189 | 0.573125 | 0.070214 | 0.015797 | 0.067725 | -1.31697 | -0.0312 | -0.68659 | -0.10417 | 0.028191 | -0.01993 | 0.056458 | 0.070574 | 0.180462 | 0.028085 | -0.22458 | 0.239498 | 0.177585 | -0.43681 | -0.1708 |
| 20.83333 | 1.56841 | -0.03056 | -0.11676 | 0.100753 | -0.35792 | 0.251052 | -0.15485 | -0.00892 | 0.264491 | -1.48795 | -0.17928 | -0.98028 | -0.11401 | -0.15827 | -0.03042 | 0.052253 | 0.172219 | 0.206737 | 0.008903 | -0.10668 | 0.086137 | -0.04619 | -0.83904 | -0.16987 |
| 20.84722 | 0.866921 | -0.03694 | -0.09837 | 0.064226 | -0.31873 | -0.01764 | -0.26264 | -0.04465 | 0.188203 | -0.81526 | -0.42025 | -0.89215 | -0.1146 | -0.29631 | -0.04095 | 0.048813 | 0.210957 | 0.071506 | -0.0267 | -0.07456 | -0.10125 | -0.25169 | -0.80949 | -0.00982 |
| 20.86111 | 0.211162 | -0.08563 | -0.07592 | 0.058124 | -0.19018 | -0.22167 | -0.22492 | -0.07437 | -0.04377 | -0.09949 | -0.44011 | -0.46201 | -0.12193 | -0.34397 | -0.04803 | 0.034138 | 0.260163 | -0.10255 | -0.07 | -0.1516 | -0.27524 | -0.32885 | -0.38465 | 0.15245 |
| 20.875 | -0.42233 | -0.1707 | -0.10666 | 0.025674 | -0.08376 | -0.32566 | -0.18112 | -0.01692 | -0.1972 | 0.267422 | -0.34539 | -0.13898 | -0.1451 | -0.31476 | -0.04919 | 0.026142 | 0.22672 | -0.23287 | -0.12006 | -0.23967 | -0.39979 | -0.36197 | 0.102987 | 0.180063 |
| 20.88889 | -1.06872 | -0.32005 | -0.18816 | -0.07794 | -0.10332 | -0.41126 | -0.15315 | 0.020899 | -0.18182 | 0.099217 | -0.21765 | -0.25116 | -0.19475 | -0.25232 | -0.02988 | 0.024068 | 0.038959 | -0.28333 | -0.16525 | -0.24843 | -0.38921 | -0.43414 | 0.440795 | 0.07422 |
| 20.90278 | -1.72574 | -0.45045 | -0.3237 | -0.20179 | -0.21048 | -0.47442 | -0.2042 | 0.03301 | -0.02292 | -0.35157 | -0.17246 | -0.6391 | -0.25063 | -0.20583 | -0.00628 | 0.004391 | -0.19107 | -0.2758 | -0.21645 | -0.17585 | -0.3362 | -0.48991 | 0.462374 | -0.08576 |
| 20.91667 | -2.1388 | -0.52231 | -0.48371 | -0.22383 | -0.303 | -0.38242 | -0.27052 | 0.02128 | 0.091257 | -0.86755 | -0.2316 | -0.91844 | -0.29948 | -0.21975 | -0.00914 | -0.02225 | -0.34361 | -0.27119 | -0.1997 | -0.13234 | -0.33737 | -0.4591 | 0.257318 | -0.219 |
| 20.93056 | -1.85623 | -0.53259 | -0.52326 | -0.13515 | -0.30469 | -0.20813 | -0.3018 | -0.03082 | -0.05477 | -1.16515 | -0.33757 | -0.95071 | -0.3282 | -0.36551 | -0.01116 | -0.02871 | -0.4214 | -0.2639 | -0.15082 | -0.133 | -0.44671 | -0.33924 | -0.11221 | -0.24503 |
| 20.94444 | -1.56329 | -0.71891 | -0.50644 | -0.05419 | -0.32647 | -0.09813 | -0.32556 | -0.12936 | -0.44161 | -1.223 | -0.42003 | -0.8228 | -0.3751 | -0.68671 | -0.00442 | -0.02545 | -0.52248 | -0.20931 | -0.09736 | -0.178 | -0.59851 | -0.29424 | -0.51281 | -0.06936 |
| 20.95833 | -0.9185 | -1.01548 | -0.62251 | -0.0568 | -0.33332 | -0.05315 | -0.35303 | -0.25607 | -0.88538 | -0.65106 | -0.56396 | -0.68176 | -0.50843 | -0.99127 | -0.00299 | -0.01504 | -0.65652 | -0.11058 | -0.07443 | -0.24494 | -0.69515 | -0.352 | -0.75672 | 0.045149 |
| 20.97222 | -0.20408 | -1.23464 | -0.81223 | -0.09024 | -0.38497 | -0.05425 | -0.35864 | -0.32444 | -1.25796 | 0.134122 | -0.78335 | -0.61278 | -0.70417 | -1.15297 | -0.00944 | -0.03205 | -0.80876 | -0.06077 | -0.08238 | -0.25409 | -0.67711 | -0.45846 | -0.77886 | -0.0313 |
| 20.98611 | 0.397468 | -1.20938 | -0.99151 | -0.06454 | -0.34884 | -0.07263 | -0.30177 | -0.20921 | -1.36081 | 0.752935 | -0.90263 | -0.53667 | -0.75788 | -1.09785 | -0.01611 | -0.03933 | -0.63351 | -0.04816 | -0.09924 | -0.21542 | -0.55724 | -0.48894 | -0.58341 | -0.27043 |
| 21 | 0.585997 | -0.99177 | -1.02502 | -0.04549 | -0.30892 | -0.11429 | -0.2691 | -0.03944 | -1.10343 | 0.79834 | -0.94116 | -0.56914 | -0.58504 | -0.91931 | -0.00802 | -0.02646 | -0.2326 | 0.005372 | -0.07432 | -0.17639 | -0.35062 | -0.35197 | -0.45599 | -0.44298 |

| | | | | | | | | | | | | | | | | | | | | | | | | |
|---|---|---|---|---|---|---|---|---|---|---|---|---|---|---|---|---|---|---|---|---|---|---|---|---|
| 21.01389 | 0.121745 | -0.87736 | -1.10793 | -0.11268 | -0.28597 | -0.19332 | -0.4924 | 0.028268 | -1.07933 | -0.08457 | -0.95772 | -1.02679 | -0.37391 | -0.8118 | 0.000961 | -0.01845 | -0.05051 | 0.094007 | -0.02961 | -0.31639 | -0.20174 | -0.24779 | -0.74051 | -0.6154 |
| 21.02778 | -0.61831 | -0.90974 | -1.11401 | -0.13692 | -0.21376 | -0.34911 | -0.88656 | -0.06215 | -1.2611 | -1.4494 | -1.01839 | -1.82422 | -0.32672 | -0.6829 | -0.02004 | 0.035536 | -0.22807 | 0.135496 | -0.04759 | -0.63693 | -0.22184 | -0.27626 | -1.62282 | -0.93425 |
| 21.04167 | -1.36567 | -1.13395 | -1.10198 | -0.01907 | -0.10028 | -0.57164 | -1.16511 | -0.07811 | -1.39198 | -2.43568 | -1.1224 | -2.7591 | -0.31703 | -0.51943 | 0.058973 | 0.096951 | -0.47638 | 0.128106 | -0.06488 | -0.81579 | -0.30617 | -0.37731 | -2.80799 | -1.12043 |
| 21.05556 | -1.98995 | -1.09385 | -1.1564 | 0.048702 | -0.17475 | -0.57478 | -1.27594 | -0.04249 | -1.08143 | -1.8591 | -1.06397 | -2.67882 | -0.26701 | -0.40869 | 0.102703 | 0.018009 | -0.65409 | 0.107934 | -0.10224 | -0.7582 | -0.32656 | -0.39439 | -2.68077 | -1.01825 |
| 21.06944 | -1.5307 | -0.80354 | -0.95514 | 0.149971 | -0.32282 | -0.34663 | -0.94221 | -0.05666 | -0.76618 | -0.5368 | -0.66293 | -1.67552 | -0.28609 | -0.33449 | 0.013675 | -0.09087 | -0.72068 | 0.101916 | -0.10811 | -0.49733 | -0.29813 | -0.17485 | -1.14286 | -0.52911 |
| 21.08333 | -0.5967 | -0.41637 | -0.55395 | 0.159469 | -0.3402 | -0.15004 | -0.44205 | -0.10586 | -0.37127 | 0.441818 | -0.32614 | -0.64406 | -0.30706 | -0.20352 | -0.06591 | -0.08348 | -0.43908 | 0.095195 | -0.06277 | -0.26115 | -0.16963 | 0.085934 | -0.0347 | -0.04743 |
| 21.09722 | 0.083644 | -0.06751 | -0.23875 | -0.02228 | -0.23178 | -0.07515 | -0.08803 | -0.15663 | 0.04195 | 0.797255 | -0.22191 | -0.04327 | -0.2895 | -0.10659 | -0.05277 | -0.01936 | -0.0255 | 0.083058 | 0.005622 | -0.09574 | -0.00801 | 0.074433 | 0.481182 | 0.184324 |
| 21.11111 | 0.487767 | 0.061914 | -0.09318 | -0.21739 | -0.16276 | -0.04789 | -0.01767 | -0.17597 | 0.19914 | 0.606692 | -0.20387 | 0.040371 | -0.27254 | -0.06849 | -0.02861 | 0.007105 | 0.309801 | 0.071169 | 0.014762 | 0.038006 | 0.097249 | -0.11606 | 0.64501 | 0.140167 |
| 21.125 | 0.567391 | 0.067334 | 0.009397 | -0.40001 | -0.18879 | -0.02755 | -0.16127 | -0.18807 | 0.091078 | 0.139113 | -0.16148 | -0.06061 | -0.24043 | -0.1077 | -0.03032 | 0.004601 | 0.478523 | 0.00232 | -0.02237 | 0.163643 | 0.112721 | -0.23561 | 0.571446 | -0.06826 |
| 21.13889 | 0.470554 | -0.01517 | 0.022004 | -0.47702 | -0.23708 | -0.03632 | -0.23834 | -0.10076 | -0.04921 | -0.31263 | -0.17134 | -0.15448 | -0.17313 | -0.15394 | -0.02948 | -0.00212 | 0.447174 | -0.01821 | -0.06777 | 0.306549 | 0.087061 | -0.24999 | 0.363245 | -0.26 |
| 21.15278 | 0.358446 | -0.09134 | -0.04709 | -0.36467 | -0.18878 | -0.04786 | -0.14274 | 0.007815 | -0.12933 | -0.57637 | -0.217 | -0.20579 | -0.11448 | -0.1374 | -0.01832 | -0.00238 | 0.301877 | -0.03171 | -0.10078 | 0.404812 | 0.052556 | -0.22903 | 0.196171 | -0.30208 |
| 21.16667 | 0.313557 | -0.11714 | -0.09331 | -0.08867 | -0.03384 | -0.04135 | -0.06018 | 0.081607 | -0.13965 | -0.71339 | -0.25258 | -0.03543 | -0.0808 | -0.06803 | -0.01378 | -0.00607 | 0.154562 | -0.04932 | -0.11536 | 0.364294 | 0.021319 | -0.23996 | 0.108296 | 0.035719 |
| 21.18056 | 0.288073 | -0.14399 | -0.08907 | 0.117509 | 0.06409 | -0.03581 | -0.06714 | 0.021557 | -0.04288 | -0.69965 | -0.25151 | 0.207035 | -0.0537 | -0.00309 | -0.02028 | -0.01923 | 0.06127 | -0.05444 | -0.09751 | 0.258401 | -0.01977 | -0.28218 | 0.153711 | 0.416051 |
| 21.19444 | 0.228917 | -0.20373 | -0.16492 | -0.00211 | 0.008523 | -0.0614 | -0.08267 | -0.08658 | 0.107637 | -0.47897 | -0.30439 | 0.16265 | -0.05559 | -0.01434 | -0.02576 | -0.0451 | 0.065942 | -0.04075 | -0.07907 | 0.177314 | -0.05621 | -0.33026 | 0.235853 | 0.335836 |
| 21.20833 | 0.221186 | -0.19563 | -0.18839 | -0.19852 | -0.05573 | -0.07069 | -0.00889 | -0.1208 | 0.407726 | -0.06435 | -0.32214 | 0.027046 | -0.08718 | -0.08976 | -0.02685 | -0.06426 | 0.136637 | -0.01392 | -0.08086 | 0.121969 | -0.02261 | -0.32523 | 0.469231 | 0.002359 |
| 21.22222 | 0.394397 | 0.000076 | 0.138458 | -0.02283 | -0.0229 | 0.029168 | 0.163042 | -0.06421 | 0.868983 | 0.517672 | -0.05906 | 0.22023 | -0.06893 | -0.12717 | -0.04086 | -0.02159 | 0.330034 | -0.01389 | -0.07709 | 0.078658 | 0.090245 | -0.20588 | 1.05302 | -0.23174 |
| 21.23611 | 0.648353 | 0.185529 | 0.430045 | 0.216599 | 0.060906 | 0.11755 | 0.250871 | 0.044235 | 1.00063 | 0.718305 | 0.133609 | 0.420295 | 0.007088 | -0.10231 | -0.03828 | 0.034211 | 0.538845 | -0.05215 | -0.06393 | 0.076445 | 0.170076 | -0.11348 | 1.191178 | -0.21672 |
| 21.25 | 0.653531 | 0.180856 | 0.317822 | 0.188898 | 0.111671 | 0.094391 | 0.134005 | 0.140973 | 0.657358 | 0.257259 | -8.1E-05 | 0.326343 | 0.054464 | -0.0383 | -0.01488 | 0.033327 | 0.622011 | -0.09872 | -0.06329 | 0.087009 | 0.148824 | -0.13093 | 0.659685 | 0.028858 |
| 21.26389 | 0.434662 | 0.115695 | 0.194115 | 0.177301 | 0.179912 | 0.048045 | 0.024703 | 0.094106 | 0.361933 | -0.32178 | -0.14244 | 0.053208 | 0.024329 | 0.032175 | -0.00711 | 0.010881 | 0.56395 | -0.14005 | -0.06335 | 0.105423 | 0.124318 | -0.13077 | 0.311474 | 0.180059 |
| 21.27778 | 0.055874 | 0.014176 | 0.176755 | 0.205031 | 0.195444 | 0.043489 | 0.019067 | -0.09859 | 0.436227 | -0.68464 | -0.15284 | -0.07364 | -0.04385 | 0.031631 | -0.00274 | 0.002179 | 0.397857 | -0.10809 | -0.03315 | 0.117742 | 0.133646 | -0.05852 | 0.253789 | 0.094491 |
| 21.29167 | -0.28682 | -0.07938 | 0.199987 | 0.183742 | 0.144394 | 0.063287 | 0.161015 | -0.09767 | 0.862252 | -0.3957 | -0.05801 | 0.298782 | -0.05467 | 0.008441 | -0.0012 | -0.00366 | 0.284587 | -0.01026 | 0.01662 | 0.111964 | 0.154243 | 0.025433 | 0.313296 | -0.01192 |
| 21.30556 | -0.31274 | -0.01372 | 0.258428 | 0.193853 | 0.127836 | 0.089407 | 0.317569 | 0.041774 | 1.367742 | 0.446113 | 0.062885 | 0.862594 | 0.023513 | 0.075466 | -0.00649 | -0.01226 | 0.313793 | 0.104055 | 0.049799 | 0.082003 | 0.170478 | 0.071211 | 0.392601 | -0.05294 |
| 21.31944 | -0.19366 | 0.089803 | 0.318377 | 0.211045 | 0.120565 | 0.097659 | 0.401316 | 0.167259 | 1.434571 | 1.063277 | 0.109162 | 1.004317 | 0.121977 | 0.184761 | -0.01049 | -0.01809 | 0.397842 | 0.168443 | 0.065098 | 0.051711 | 0.188424 | 0.10366 | 0.37024 | -0.00397 |
| 21.33333 | -0.04294 | 0.212858 | 0.40021 | 0.208388 | 0.140189 | 0.108636 | 0.465052 | 0.316985 | 1.141865 | 0.877079 | 0.115298 | 0.667312 | 0.158901 | 0.251632 | -0.00398 | -0.00965 | 0.396167 | 0.192058 | 0.090386 | 0.05737 | 0.214203 | 0.193447 | 0.241506 | 0.074717 |
| 21.34722 | 0.092555 | 0.289496 | 0.538733 | 0.185248 | 0.170312 | 0.138049 | 0.540739 | 0.22894 | 0.942988 | 0.316248 | 0.123696 | 0.422513 | 0.155789 | 0.279819 | 0.002156 | 0.009551 | 0.370188 | 0.178767 | 0.120789 | 0.093158 | 0.249548 | 0.306844 | 0.131131 | 0.125761 |
| 21.36111 | 0.335963 | 0.296965 | 0.695712 | 0.190052 | 0.176788 | 0.194346 | 0.653164 | 0.085529 | 0.876013 | -0.16363 | 0.165745 | 0.686513 | 0.143135 | 0.253067 | -0.00722 | 0.016683 | 0.299635 | 0.152029 | 0.143613 | 0.136644 | 0.281136 | 0.328899 | 0.146043 | 0.201368 |
| 21.375 | 0.571216 | 0.275192 | 0.791541 | 0.266595 | 0.187209 | 0.22547 | 0.794289 | -0.00394 | 0.859484 | 0.078073 | 0.212645 | 1.312616 | 0.171696 | 0.210617 | -0.01862 | 0.026741 | 0.242173 | 0.122321 | 0.150009 | 0.142816 | 0.314014 | 0.232464 | 0.209109 | 0.259142 |
| 21.38889 | 0.738229 | 0.279877 | 0.739157 | 0.335575 | 0.196362 | 0.297764 | 0.868878 | 0.077872 | 0.831895 | 1.127016 | 0.243474 | 1.791458 | 0.272461 | 0.237007 | -0.01625 | 0.064607 | 0.270937 | 0.140347 | 0.152863 | 0.125115 | 0.337402 | 0.139194 | 0.251917 | 0.220473 |
| 21.40278 | 0.783668 | 0.319438 | 0.661101 | 0.376276 | 0.185907 | 0.345491 | 0.841157 | 0.235769 | 0.766607 | 2.087024 | 0.275041 | 1.591314 | 0.364259 | 0.327494 | -0.01295 | 0.111282 | 0.369545 | 0.179634 | 0.149393 | 0.097124 | 0.346079 | 0.146793 | 0.285975 | 0.118666 |
| 21.41667 | 0.811411 | 0.396408 | 0.584503 | 0.351152 | 0.17415 | 0.335394 | 0.587925 | 0.245097 | 0.599272 | 1.786323 | 0.319662 | 1.100039 | 0.401669 | 0.410194 | -0.01484 | 0.14091 | 0.47624 | 0.210061 | 0.153705 | 0.084779 | 0.378753 | 0.258187 | 0.374617 | 0.038543 |
| 21.43056 | 0.840941 | 0.504223 | 0.478458 | 0.326211 | 0.20323 | 0.342135 | 0.406951 | 0.203208 | 0.498932 | 1.068525 | 0.390624 | 0.884451 | 0.353801 | 0.427359 | -0.0125 | 0.146801 | 0.54999 | 0.228313 | 0.165048 | 0.115245 | 0.460268 | 0.407611 | 0.547057 | 0.12173 |
| 21.44444 | 0.980538 | 0.58661 | 0.475288 | 0.262684 | 0.240842 | 0.333858 | 0.279744 | 0.177149 | 0.497829 | 0.264032 | 0.469006 | 1.030539 | 0.326866 | 0.41776 | -0.00755 | 0.155516 | 0.561941 | 0.312241 | 0.164622 | 0.194261 | 0.531011 | 0.500509 | 0.627964 | 0.318674 |
| 21.45833 | 1.071996 | 0.581315 | 0.544106 | 0.145737 | 0.245658 | 0.285589 | 0.268886 | 0.164768 | 0.553924 | -0.10586 | 0.49601 | 1.406671 | 0.355762 | 0.451781 | -0.01922 | 0.184194 | 0.526278 | 0.406069 | 0.186102 | 0.252917 | 0.544146 | 0.567076 | 0.632645 | 0.469508 |
| 21.47222 | 1.047193 | 0.476248 | 0.487834 | 0.080612 | 0.199593 | 0.230767 | 0.164814 | 0.117549 | 0.518542 | 0.395524 | 0.44833 | 1.576275 | 0.427728 | 0.456051 | -0.04546 | 0.246109 | 0.545642 | 0.326364 | 0.210504 | 0.285061 | 0.565661 | 0.575466 | 0.659993 | 0.50407 |
| 21.48611 | 0.662073 | 0.535732 | 0.418392 | 0.102503 | 0.229754 | 0.270772 | -0.05146 | 0.098549 | 0.412523 | 1.172581 | 0.433458 | 1.186237 | 0.528424 | 0.393317 | -0.0252 | 0.263253 | 0.517995 | 0.094685 | 0.258977 | 0.333018 | 0.607849 | 0.520676 | 0.746849 | 0.453687 |
| 21.5 | 0.268844 | 0.998794 | 0.628591 | 0.342338 | 0.55027 | 0.48259 | -0.24109 | 0.165508 | 0.263832 | 1.646353 | 0.789196 | 1.272249 | 0.515052 | 0.414469 | 0.004963 | 0.238903 | 0.387088 | -0.11398 | 0.331279 | 0.536275 | 0.665823 | 0.555777 | 1.070203 | 0.305551 |
| 21.51389 | 0.116758 | 2.056285 | 1.403594 | 0.833462 | 1.419989 | 0.962816 | -0.33763 | 0.266933 | 0.533679 | 2.2683 | 1.842907 | 2.672009 | 0.493462 | 0.770694 | -0.00584 | 0.19594 | 0.200341 | -0.12986 | 0.433809 | 1.205265 | 0.774403 | 0.779103 | 1.936413 | 0.075101 |
| 21.52778 | 0.382047 | 2.939217 | 1.951033 | 0.964765 | 1.512081 | 1.051427 | -0.09577 | 0.279152 | 1.030087 | 3.28115 | 2.528413 | 3.870128 | 0.414057 | 1.374781 | 0.061149 | 0.13957 | 0.1745 | 0.034005 | 0.207055 | 1.414396 | 0.612321 | 0.928946 | 2.59332 | 0.114122 |
| 21.54167 | 1.040952 | 2.360632 | 1.910113 | 0.556398 | 1.112596 | 0.548409 | 0.413689 | 0.23839 | 1.605592 | 3.338106 | 1.632274 | 2.950477 | 0.304266 | 1.698669 | -0.06787 | 0.052943 | 0.093409 | 0.151243 | 0.222944 | 0.783184 | 0.345375 | 1.298826 | 2.478282 | 0.443794 |
| 21.55556 | 1.771283 | 1.968446 | 1.516152 | 0.238237 | 1.084536 | 0.203484 | 0.112181 | 0.198942 | 1.953301 | 2.240251 | 1.124001 | 1.914048 | 0.159316 | 1.438467 | -0.12197 | 0.101125 | -0.02691 | 0.146831 | 0.34429 | 0.423919 | 0.281222 | 1.322772 | 1.81384 | 0.544371 |
| 21.56944 | 1.666148 | 1.737681 | 1.115424 | 0.143667 | 1.070418 | -0.03004 | -0.22538 | 0.21827 | 1.532994 | 1.006406 | 0.78651 | 1.074729 | 0.128624 | 1.082958 | -0.04938 | 0.131556 | 0.036216 | 0.076213 | 0.31134 | 0.229005 | 0.113519 | 1.010696 | 0.799111 | 0.167359 |
| 21.58333 | 0.9227 | 1.379225 | 0.658752 | 0.062422 | 0.885601 | -0.25712 | -0.65491 | 0.188945 | 0.872542 | -0.59629 | 0.404767 | 0.290223 | 0.043626 | 0.644347 | -0.00669 | 0.006549 | 0.016071 | -0.06199 | 0.172369 | 0.052977 | -0.13102 | 0.604287 | -0.08562 | -0.20723 |
| 21.59722 | 0.368577 | 1.06062 | 0.346722 | -0.01008 | 0.634557 | -0.51461 | -0.89662 | 0.136565 | 0.460554 | -2.54222 | 0.009967 | -0.8456 | -0.08986 | 0.107214 | -0.01058 | -0.10058 | -0.14935 | -0.18094 | -0.02228 | -0.11911 | -0.30279 | 0.198543 | -0.76336 | -0.43894 |
| 21.61111 | 0.090178 | 0.552447 | 0.091552 | -0.0019 | 0.410076 | -0.68261 | -0.76689 | 0.066089 | -0.04313 | -2.99258 | -0.3239 | -1.50585 | -0.1748 | -0.207 | -0.01479 | -0.08201 | -0.20998 | -0.17254 | -0.12195 | -0.28196 | -0.36322 | -0.20851 | -1.30632 | -0.58548 |
| 21.625 | 1.082254 | 0.077842 | -0.13547 | 0.10395 | 0.183702 | -0.5308 | -0.17598 | -0.05117 | -0.57143 | -0.97033 | -0.45403 | -0.96716 | -0.23309 | 0.079852 | -0.01268 | -0.06654 | -0.19632 | -0.09011 | -0.11223 | -0.3493 | -0.29556 | -0.31507 | -1.50433 | -0.26952 |
| 21.63889 | 0.743395 | -0.49183 | -0.28248 | 0.255355 | -0.05868 | -0.13733 | 0.595401 | -0.12356 | -0.6221 | 1.139582 | -0.25806 | 0.172261 | -0.2841 | 0.609706 | -0.01728 | -0.03578 | -0.10713 | 0.100766 | -0.05386 | -0.21152 | -0.14199 | -0.07193 | -0.94122 | 0.390094 |
| 21.65278 | -0.68412 | -0.82205 | -0.28227 | 0.302736 | -0.25782 | 0.295479 | 0.881868 | -0.11804 | -0.3206 | 2.361255 | 0.161972 | 0.474679 | -0.24433 | 0.989713 | -0.02817 | 0.019742 | -0.01188 | 0.261317 | 0.026134 | 0.055848 | 0.093269 | 0.284109 | -0.0614 | 0.845673 |
| 21.66667 | -0.54458 | -0.98719 | -0.08906 | 0.241666 | -0.3781 | 0.665213 | 0.508744 | -0.06122 | 0.03065 | 1.773173 | 0.554357 | -0.03252 | -0.19029 | 0.948806 | -0.03172 | 0.080086 | 0.018058 | 0.320557 | 0.069888 | 0.301289 | 0.294695 | 0.594213 | 0.625924 | 0.750048 |
| 21.68056 | 0.481811 | -0.91693 | 0.157599 | 0.133617 | -0.39367 | 0.860938 | 0.006705 | -0.02495 | 0.110594 | 0.658244 | 0.742098 | -0.51235 | -0.12025 | 0.606482 | -0.03253 | 0.117265 | 0.027376 | 0.24862 | 0.064927 | 0.436994 | 0.367514 | 0.665496 | 0.746114 | 0.288212 |
| 21.69444 | 1.773771 | -0.71377 | 0.272955 | 0.048996 | -0.22861 | 0.711976 | -0.48242 | -0.0303 | -0.10106 | -0.26434 | 0.581935 | -0.96276 | -0.06631 | 0.214584 | -0.03491 | 0.109106 | 0.105424 | 0.15224 | 0.024508 | 0.393911 | 0.231327 | 0.439156 | 0.700495 | -0.23235 |
| 21.70833 | 2.266485 | -0.49871 | 0.219654 | -0.00985 | -0.03691 | 0.352543 | -0.94948 | -0.03177 | -0.21713 | -1.2748 | 0.377333 | -1.21476 | -0.04845 | -0.22947 | -0.0416 | 0.099999 | 0.161078 | -0.01658 | -0.02064 | 0.286517 | 0.023525 | 0.126523 | 0.608248 | -0.63976 |
| 21.72222 | 1.354878 | -0.25307 | 0.201396 | -0.06515 | 0.137519 | 0.089463 | -0.96667 | -0.05384 | 0.118742 | -2.12705 | 0.158478 | -0.91105 | -0.0795 | -0.64624 | -0.03841 | 0.067746 | 0.07872 | -0.17313 | 0.000938 | 0.130011 | -0.13861 | -0.17626 | 0.586406 | -0.55761 |
| 21.73611 | 0.355248 | -0.1337 | 0.156739 | -0.05353 | 0.225425 | -0.18161 | -0.40033 | -0.04087 | 0.365029 | -2.0063 | -0.06508 | -0.07278 | -0.10749 | -0.88513 | -0.027 | 0.032952 | -0.10959 | -0.24566 | 0.01887 | -0.06139 | -0.24631 | -0.32661 | 0.40461 | -0.04828 |
| 21.75 | -0.09968 | -0.17057 | 0.035996 | 0.017471 | 0.257074 | -0.4803 | 0.162058 | -0.00858 | 0.38325 | -0.61074 | -0.20944 | 0.719952 | -0.07563 | -0.73342 | -0.02141 | 0.002907 | -0.21661 | -0.25567 | 0.016649 | -0.23896 | -0.29422 | -0.41497 | 0.038257 | 0.393905 |
| 21.76389 | -0.55656 | -0.23837 | -0.12758 | 0.120858 | 0.252599 | -0.70019 | 0.195846 | -0.00453 | 0.218263 | 0.749598 | -0.33615 | 1.225038 | -0.02262 | -0.3663 | -0.02487 | -0.04507 | -0.14408 | -0.29927 | -0.03051 | -0.37326 | -0.29749 | -0.44344 | -0.25706 | 0.566991 |
| 21.77778 | -0.84772 | -0.47762 | -0.31702 | 0.172302 | 0.158306 | -0.62401 | -0.03424 | 0.031071 | 0.105487 | 1.571492 | -0.40241 | 1.124618 | -0.03817 | -0.11096 | -0.02643 | -0.0953 | -0.03191 | -0.28126 | -0.08586 | -0.32122 | -0.22572 | -0.33018 | -0.24919 | 0.107954 |
| 21.79167 | -1.174 | -0.709 | -0.50048 | 0.155568 | 0.011898 | -0.33166 | -0.24491 | 0.022176 | 0.038199 | 1.219266 | -0.31966 | 0.371882 | -0.15792 | -0.02425 | -0.0202 | -0.10837 | 0.03776 | -0.09953 | -0.14008 | -0.15392 | -0.14532 | -0.08202 | -0.03688 | -0.23825 |
| 21.80556 | -1.56047 | -0.97015 | -0.60156 | 0.120048 | -0.13466 | -0.15813 | -0.45925 | -0.03029 | -0.19115 | 0.263675 | -0.06479 | -0.53461 | -0.31078 | -0.16065 | -0.01786 | -0.07142 | 0.013156 | 0.090205 | -0.18486 | 0.039137 | -0.02989 | 0.217711 | 0.192884 | -0.52245 |
| 21.81944 | -1.67074 | -1.25464 | -0.55785 | 0.062276 | -0.25455 | -0.12833 | -0.81399 | -0.10659 | -0.71492 | -0.71256 | 0.250398 | -1.28301 | -0.43329 | -0.52047 | -0.02064 | -0.04282 | -0.11517 | 0.086056 | -0.21641 | 0.176828 | 0.082289 | 0.353554 | 0.251913 | -0.67142 |
| 21.83333 | -1.5189 | -1.44679 | -0.50521 | -0.04313 | -0.3522 | 0.02244 | -0.84163 | -0.21474 | -1.28108 | -1.8016 | 0.335726 | -1.29477 | -0.52703 | -0.9156 | -0.01844 | 0.01343 | -0.29466 | -0.12078 | -0.23317 | 0.197758 | 0.138707 | 0.184384 | 0.102159 | -0.44154 |
| 21.84722 | -0.98884 | -1.17378 | -0.49844 | -0.09229 | -0.45105 | 0.206841 | -0.51371 | -0.37135 | -1.58859 | -2.57209 | 0.158009 | -0.86508 | -0.58582 | -1.09858 | -0.01547 | 0.007604 | -0.42752 | -0.3023 | -0.22775 | 0.115669 | 0.113065 | -0.17014 | -0.18 | -0.11596 |
| 21.86111 | -0.5367 | -1.06213 | -0.56774 | -0.08887 | -0.53048 | 0.103726 | -0.42301 | -0.49921 | -1.35388 | -2.14154 | -0.1505 | -0.6284 | -0.52042 | -0.98399 | -0.0271 | -0.08785 | -0.51666 | -0.34018 | -0.29596 | -0.01343 | 0.018911 | -0.51506 | -0.40249 | 0.088466 |
| 21.875 | -0.40029 | -1.26938 | -0.71735 | -0.0405 | -0.65074 | -0.25546 | -0.69802 | -0.46627 | -1.066 | -1.35352 | -0.51163 | -0.79136 | -0.46403 | -0.87592 | -0.0426 | -0.17848 | -0.61863 | -0.44342 | -0.41533 | -0.19278 | -0.10049 | -0.75581 | -0.58141 | -0.11879 |
| 21.88889 | -0.78284 | -1.62558 | -0.87574 | -0.01028 | -0.72234 | -0.58439 | -1.08914 | -0.35572 | -0.94489 | -0.44058 | -0.82374 | -1.08257 | -0.53069 | -0.742 | -0.04515 | -0.2915 | -0.6465 | -0.66615 | -0.46319 | -0.42808 | -0.24604 | -0.94613 | -0.65494 | -0.73572 |
| 21.90278 | -1.21415 | -1.73757 | -1.04535 | -0.02833 | -0.78265 | -0.64345 | -1.40141 | -0.20111 | -1.13079 | 0.36993 | -1.02068 | -1.35763 | -0.65558 | -0.51861 | -0.03788 | -0.40088 | -0.65597 | -0.78384 | -0.49209 | -0.63363 | -0.38168 | -1.11 | -0.36286 | -1.46956 |
| 21.91667 | -1.49332 | -1.71602 | -1.09991 | -0.08465 | -0.75051 | -0.47705 | -1.44372 | -0.11907 | -1.41788 | 0.357383 | -1.14846 | -1.58491 | -0.78638 | -0.44581 | -0.03125 | -0.48178 | -0.62354 | -0.75416 | -0.50125 | -0.68159 | -0.48042 | -1.1517 | -0.04067 | -1.50426 |
| 21.93056 | -1.32926 | -1.63925 | -1.0045 | -0.14088 | -0.52146 | -0.29531 | -1.1047 | -0.20132 | -1.53525 | -0.44208 | -1.21069 | -1.77379 | -0.90019 | -0.59444 | -0.03271 | -0.43182 | -0.69513 | -0.66836 | -0.58539 | -0.47186 | -0.44262 | -1.13357 | 0.266205 | -0.90578 |
| 21.94444 | -1.073 | -1.27917 | -0.78548 | -0.1805 | -0.30417 | -0.21216 | -0.46606 | -0.40085 | -1.36038 | -1.53318 | -1.05442 | -1.66713 | -0.96879 | -0.93627 | -0.01671 | -0.31365 | -0.6913 | -0.55353 | -0.56083 | -0.19604 | -0.30786 | -0.88018 | 0.414166 | -0.12177 |
| 21.95833 | -0.92076 | -0.88981 | -0.61182 | -0.21547 | -0.13787 | -0.13831 | 0.111759 | -0.50578 | -0.89753 | -2.60367 | -0.78112 | -1.09741 | -0.92178 | -1.37887 | 0.000466 | -0.18518 | -0.63252 | -0.44779 | -0.48058 | 0.005219 | -0.1492 | -0.48839 | 0.462614 | 0.42014 |
| 21.97222 | -1.00029 | -0.66717 | -0.50038 | -0.18873 | -0.12105 | -0.11132 | 0.239973 | -0.39472 | -0.53603 | -3.03471 | -0.55438 | -0.74031 | -0.88862 | -1.62427 | -0.01702 | -0.14768 | -0.58652 | -0.40384 | -0.36055 | 0.086565 | -0.07064 | -0.21999 | 0.363101 | 0.359286 |
| 21.98611 | -0.94681 | -0.57276 | -0.41152 | -0.07565 | -0.13665 | -0.15574 | 0.134214 | -0.22926 | -0.30394 | -2.25963 | -0.42298 | -0.45091 | -0.72067 | -1.46448 | -0.04119 | -0.20611 | -0.52227 | -0.37113 | -0.25039 | 0.056696 | -0.0564 | -0.19779 | 0.189673 | -0.14622 |
| 22 | -0.73052 | -0.55792 | -0.40394 | -0.00357 | -0.12653 | -0.19513 | -0.03717 | -0.08214 | -0.30097 | -1.17827 | -0.38892 | -0.24145 | -0.50781 | -1.10868 | -0.01187 | -0.19986 | -0.41581 | -0.29592 | -0.18242 | -0.07457 | -0.08667 | -0.26412 | -0.04392 | -0.71471 |
| 22.01389 | -0.72496 | -0.5927 | -0.66742 | -0.15113 | -0.12434 | -0.26802 | -0.22851 | -0.04704 | -0.5724 | -0.66608 | -0.77105 | -0.29338 | -0.32484 | -0.71178 | 0.059372 | -0.10083 | -0.39873 | -0.23739 | -0.19063 | -0.47074 | -0.17944 | -0.39225 | -0.56356 | -1.12475 |
| 22.02778 | -1.05165 | -0.66403 | -1.15486 | -0.67131 | -0.08074 | -0.45426 | -0.36575 | -0.04899 | -0.89223 | -1.17941 | -1.82351 | -0.812 | -0.29172 | -0.58063 | 0.118355 | -0.01649 | -0.55765 | -0.26625 | -0.24999 | -1.02779 | -0.38322 | -0.60707 | -1.46224 | -1.42266 |
| 22.04167 | -1.34243 | -0.75323 | -1.52968 | -0.92758 | 0.010087 | -0.63594 | -0.47033 | -0.1462 | -0.85548 | -2.22766 | -2.82651 | -1.58374 | -0.38938 | -0.56961 | 0.18127 | -0.05923 | -1.04026 | -0.42683 | -0.25082 | -0.86433 | -0.59052 | -0.76991 | -2.40129 | -1.7193 |
| 22.05556 | -1.49029 | -1.14562 | -1.47537 | -0.6022 | -0.00466 | -0.54111 | -0.62692 | -0.38371 | -0.58528 | -1.93318 | -1.73771 | -1.91509 | -0.48499 | -0.54758 | 0.151338 | 0.063042 | -1.72214 | -0.38215 | -0.05984 | -0.43704 | -0.53233 | -0.59856 | -1.69741 | -1.33548 |
| 22.06944 | -1.13805 | -1.21934 | -0.87573 | 0.006224 | -0.07178 | -0.36966 | -0.75931 | -0.25019 | -0.39932 | -0.80841 | -0.08453 | -1.71686 | -0.42624 | -0.42522 | 0.028757 | 0.17043 | -1.57984 | -0.13888 | 0.019229 | -0.21032 | -0.4738 | -0.28685 | -0.58025 | -0.52823 |
| 22.08333 | -0.49858 | -0.87594 | -0.11511 | 0.514885 | -0.0874 | -0.21749 | -0.65812 | 0.167319 | -0.33006 | 0.21707 | 0.468362 | -0.89483 | -0.26831 | -0.21723 | -0.02373 | 0.129112 | -0.80569 | -0.05145 | 0.032653 | -0.12064 | -0.36825 | -0.1616 | -0.12496 | 0.07614 |
| 22.09722 | -0.07249 | -0.39167 | 0.307499 | 0.639428 | -0.0515 | -0.11193 | -0.39588 | 0.650811 | -0.39172 | 1.448546 | 0.367802 | 0.226435 | -0.17176 | 0.074364 | -0.01709 | 0.058868 | -0.07268 | 0.008404 | 0.080297 | -0.03098 | -0.13951 | -0.22629 | 0.126116 | 0.487521 |
| 22.11111 | 0.16022 | 0.050814 | 0.295825 | 0.502558 | -0.01744 | -0.0532 | -0.2165 | 0.743569 | -0.28288 | 2.479216 | 0.188916 | 0.827712 | -0.15674 | 0.349436 | -0.01762 | 0.011041 | 0.298759 | 0.064225 | 0.15257 | 0.027701 | 0.01956 | -0.31197 | 0.25958 | 0.436694 |
| 22.125 | 0.21432 | 0.224033 | 0.180126 | 0.298655 | 0.039536 | -0.03844 | -0.29015 | 0.675402 | -0.15352 | 3.013766 | 0.07039 | 0.770939 | -0.19895 | 0.540422 | -0.01865 | -0.02226 | 0.397953 | 0.078556 | 0.184264 | 0.062264 | 0.037309 | -0.3104 | 0.201456 | 0.138046 |
| 22.13889 | 0.057522 | 0.178715 | 0.07697 | 0.144293 | 0.062929 | -0.04661 | -0.48957 | 0.374245 | -0.0931 | 2.715307 | -0.02254 | 0.398984 | -0.27243 | 0.632217 | -0.01712 | -0.04029 | 0.358645 | 0.053176 | 0.173526 | 0.115937 | -0.01336 | -0.20337 | 0.062583 | -0.20302 |
| 22.15278 | -0.07728 | 0.087714 | 0.04679 | 0.059293 | 0.036378 | -0.0441 | -0.32681 | 0.022296 | -0.08048 | 1.975205 | -0.04421 | -0.01821 | -0.35874 | 0.61691 | -0.01591 | -0.05453 | 0.301305 | 0.01241 | 0.153708 | 0.197919 | -0.01985 | -0.06089 | -0.03647 | -0.40955 |
| 22.16667 | -0.12441 | 0.141977 | 0.038623 | 0.028201 | -0.01595 | -0.02505 | 0.174686 | -0.28615 | -0.09989 | 1.245206 | -0.03421 | -0.27463 | -0.37757 | 0.570013 | -0.00901 | -0.06827 | 0.282418 | -0.02335 | 0.103024 | 0.25024 | 0.001503 | 0.036618 | -0.07888 | -0.4167 |
| 22.18056 | 0.02192 | 0.210593 | 0.021394 | 0.006863 | -0.08734 | -0.02979 | 0.384493 | -0.41678 | -0.12158 | 0.377492 | -0.07378 | -0.19534 | -0.26735 | 0.410069 | -0.00086 | -0.07758 | 0.263117 | -0.0446 | 0.00638 | 0.215123 | -0.03081 | 0.042374 | -0.07805 | -0.17512 |
| 22.19444 | 0.315334 | 0.248157 | -0.00637 | 0.00348 | -0.1201 | -0.05974 | 0.256527 | -0.28886 | -0.13854 | -0.46841 | -0.14988 | 0.20912 | -0.11666 | 0.168079 | 0.000665 | -0.06449 | 0.258784 | -0.06637 | -0.1153 | 0.131253 | -0.06956 | 0.033493 | -0.04576 | 0.156482 |
| 22.20833 | 0.461314 | 0.22276 | -0.00608 | 0.036854 | -0.07093 | -0.06785 | 0.132327 | -0.04427 | -0.12804 | -1.10604 | -0.1802 | 0.519694 | -0.08237 | -0.08237 | -0.00361 | -0.03928 | 0.269406 | -0.10538 | -0.2113 | 0.040311 | -0.04756 | 0.062633 | -0.00823 | 0.236865 |
| 22.22222 | 0.456434 | 0.11955 | 0.002978 | 0.063699 | 0.014615 | -0.04514 | 0.000975 | 0.017981 | -0.12179 | -1.27978 | -0.15769 | 0.417329 | -0.19533 | -0.22047 | -0.00648 | -0.01517 | 0.265986 | -0.13516 | -0.23712 | -0.01805 | -0.01939 | 0.072254 | 0.016952 | -0.02322 |
| 22.23611 | 0.357377 | 0.010147 | 0.015629 | 0.044283 | 0.048357 | -0.02327 | -0.15273 | -0.10944 | -0.15011 | -0.63976 | -0.11574 | -0.05009 | -0.32012 | -0.24777 | -0.00239 | -0.01329 | 0.235922 | -0.1237 | -0.18041 | -0.05853 | 0.009972 | 0.019646 | 0.043728 | -0.30481 |
| 22.25 | 0.127411 | -0.06839 | 0.00807 | -0.00904 | 0.03394 | -0.01862 | -0.18381 | -0.26291 | -0.17482 | 0.250925 | -0.09085 | -0.4821 | -0.32372 | -0.26386 | 0.00016 | -0.04496 | 0.155708 | -0.08796 | -0.10925 | -0.09341 | 0.01765 | -0.06387 | 0.022919 | -0.33832 |
| 22.26389 | -0.08875 | -0.09087 | -0.00158 | -0.01176 | -0.00662 | -0.02379 | -0.12571 | -0.2952 | -0.15875 | 0.437779 | -0.09376 | -0.60272 | -0.14037 | -0.27258 | -0.0037 | -0.08748 | 0.083121 | -0.0752 | -0.06451 | -0.12135 | -0.02412 | -0.11643 | 0.002593 | -0.15215 |
| 22.27778 | -0.18007 | -0.07301 | 0.018564 | 0.048165 | -0.03663 | -0.01752 | -0.10172 | -0.08361 | -0.10355 | -0.05676 | -0.09298 | -0.34219 | 0.05562 | -0.18089 | -0.00828 | -0.08696 | 0.046507 | -0.07447 | -0.03327 | -0.12296 | -0.06289 | -0.11694 | 0.013043 | 0.053974 |
| 22.29167 | -0.18389 | -0.08297 | 0.062655 | 0.109475 | -0.05572 | -0.00989 | -0.01966 | 0.249264 | -0.04364 | -0.52301 | -0.07682 | 0.059426 | 0.086787 | 0.060534 | -0.01044 | -0.03029 | 0.065646 | -0.0501 | -0.01187 | -0.09565 | -0.1237 | -0.0459 | 0.041608 | 0.075875 |
| 22.30556 | -0.09887 | -0.07124 | 0.153288 | 0.137643 | -0.03547 | -0.00729 | 0.178764 | 0.34553 | 0.008395 | -0.60409 | -0.00637 | 0.282314 | -0.0729 | 0.312213 | -0.00444 | 0.044171 | 0.135032 | 0.006209 | -0.00243 | -0.06073 | -0.15293 | 0.033039 | 0.029801 | -0.12627 |
| 22.31944 | 0.139507 | -0.03585 | 0.237534 | 0.171303 | 0.012372 | 0.000625 | 0.366962 | 0.165114 | 0.079572 | -0.1623 | 0.142463 | 0.247504 | -0.23121 | 0.461657 | 0.00429 | 0.093402 | 0.221731 | 0.081473 | 0.018856 | -0.03292 | -0.06024 | 0.07621 | 0.058703 | -0.23576 |
| 22.33333 | 0.507505 | 0.061474 | 0.307397 | 0.216703 | 0.04877 | 0.019455 | 0.479959 | -0.06244 | 0.195008 | 0.716541 | 0.280446 | 0.221203 | -0.24087 | 0.456678 | 0.008762 | 0.100831 | 0.264716 | 0.159293 | 0.070656 | -0.00025 | 0.075998 | 0.090052 | 0.131593 | -0.06278 |
| 22.34722 | 0.888269 | 0.201523 | 0.384106 | 0.264952 | 0.093657 | 0.05802 | 0.561973 | -0.08007 | 0.358634 | 1.655479 | 0.293354 | 0.470348 | -0.0201 | 0.489867 | 0.010907 | 0.09008 | 0.27857 | 0.243766 | 0.158158 | 0.036079 | 0.203477 | 0.107348 | 0.247073 | 0.27345 |
| 22.36111 | 1.010153 | 0.283709 | 0.430671 | 0.302217 | 0.10431 | 0.089189 | 0.558724 | 0.265762 | 0.510228 | 1.672719 | 0.20682 | 0.773011 | 0.296607 | 0.670262 | 0.009779 | 0.066661 | 0.276229 | 0.307861 | 0.22656 | 0.050179 | 0.258069 | 0.133987 | 0.364057 | 0.544645 |
| 22.375 | 0.685516 | 0.219231 | 0.424164 | 0.300717 | 0.06525 | 0.084282 | 0.480718 | 0.649978 | 0.582609 | 0.780784 | 0.113071 | 1.050956 | 0.486594 | 0.883988 | 0.004832 | 0.053153 | 0.295724 | 0.351612 | 0.18837 | 0.050443 | 0.217321 | 0.145959 | 0.434528 | 0.608451 |
| 22.38889 | 0.402266 | 0.167352 | 0.490389 | 0.33948 | 0.095348 | 0.117705 | 0.49768 | 0.715701 | 0.632361 | -0.27162 | 0.15977 | 1.04555 | 0.452626 | 0.88813 | 0.009042 | 0.086231 | 0.362727 | 0.379915 | 0.148529 | 0.077453 | 0.170019 | 0.130248 | 0.458276 | 0.560302 |
| 22.40278 | 0.329847 | 0.154195 | 0.572035 | 0.382183 | 0.172046 | 0.191652 | 0.603404 | 0.456326 | 0.655325 | -0.19682 | 0.311631 | 0.830061 | 0.305794 | 0.847035 | 0.016828 | 0.139563 | 0.451404 | 0.452147 | 0.153523 | 0.124825 | 0.123704 | 0.073816 | 0.463612 | 0.493608 |
| 22.41667 | 0.474572 | 0.13618 | 0.575195 | 0.344901 | 0.162438 | 0.232259 | 0.654794 | 0.007648 | 0.673062 | 1.118382 | 0.463606 | 0.617319 | 0.151278 | 0.858563 | 0.01886 | 0.160021 | 0.467906 | 0.514493 | 0.197047 | 0.149733 | 0.100262 | 0.049608 | 0.455721 | 0.411663 |
| 22.43056 | 0.781566 | 0.16948 | 0.53539 | 0.221073 | 0.093244 | 0.24719 | 0.708415 | -0.11883 | 0.734401 | 2.287613 | 0.468361 | 0.620793 | 0.084974 | 0.859688 | 0.021771 | 0.15209 | 0.489104 | 0.616206 | 0.288043 | 0.16093 | 0.198439 | 0.191276 | 0.486435 | 0.332659 |
| 22.44444 | 1.11664 | 0.229435 | 0.471388 | 0.104721 | 0.023513 | 0.258972 | 0.74255 | 0.22978 | 0.900353 | 2.032843 | 0.383544 | 0.738065 | 0.17828 | 0.749085 | 0.015137 | 0.143436 | 0.504059 | 0.60938 | 0.352133 | 0.169911 | 0.362007 | 0.409913 | 0.561922 | 0.326555 |
| 22.45833 | 1.262941 | 0.264436 | 0.507744 | 0.042315 | 0.013313 | 0.265839 | 0.646696 | 0.707463 | 1.011851 | 1.05415 | 0.354876 | 0.810079 | 0.435773 | 0.533545 | -0.01052 | 0.183268 | 0.547773 | 0.531914 | 0.328884 | 0.182164 | 0.52334 | 0.576904 | 0.604144 | 0.378846 |
| 22.47222 | 1.084424 | 0.227098 | 0.605384 | 0.102484 | 0.046025 | 0.287279 | 0.509476 | 0.761451 | 1.049979 | 0.014776 | 0.402864 | 0.724848 | 0.674812 | 0.265935 | -0.02621 | 0.27352 | 0.645494 | 0.436701 | 0.305684 | 0.219209 | 0.646836 | 0.665537 | 0.671252 | 0.465154 |

| | | | | | | | | | | | | | | | | | | | | | | | | |
|---|---|---|---|---|---|---|---|---|---|---|---|---|---|---|---|---|---|---|---|---|---|---|---|---|
| 22.48611 | 0.710566 | 0.221249 | 0.670643 | 0.109376 | 0.067729 | 0.279234 | 0.461287 | 0.454287 | 1.16199 | -0.82565 | 0.54602 | 0.615685 | 0.719784 | 0.117192 | 0.003137 | 0.270303 | 0.693415 | 0.343057 | 0.339897 | 0.258153 | 0.704727 | 0.71762 | 0.784619 | 0.456394 |
| 22.5 | 0.48314 | 0.418231 | 0.833998 | 0.292013 | 0.111472 | 0.405157 | 0.454863 | 0.166287 | 2.021674 | 0.138396 | 0.961969 | 0.744063 | 0.717966 | 0.436157 | 0.021247 | 0.146746 | 0.719947 | 0.22737 | 0.390656 | 0.34009 | 0.73122 | 0.740264 | 0.95552 | 0.380531 |
| 22.51389 | 0.817734 | 0.939903 | 1.480043 | 0.87285 | 0.236813 | 0.738548 | 0.630638 | 0.225816 | 3.719094 | 2.577468 | 1.686513 | 1.442505 | 0.849271 | 1.301301 | -0.01017 | 0.095296 | 0.776099 | 0.151391 | 0.722467 | 0.873682 | 1.006609 | 0.92876 | 1.547347 | 0.451729 |
| 22.52778 | 1.45178 | 1.737731 | 2.154031 | 1.356069 | 0.248635 | 0.884539 | 1.040906 | 0.691352 | 3.174451 | 5.020569 | 1.618751 | 2.205899 | 0.890378 | 2.421958 | -0.03243 | 0.284475 | 0.842479 | 0.311786 | 0.961246 | 1.216331 | 1.032809 | 1.219286 | 1.990711 | 0.658101 |
| 22.54167 | 1.696098 | 1.613905 | 1.789835 | 1.072692 | 0.05148 | 0.66369 | 1.733618 | 0.903972 | 1.629741 | 4.525907 | 0.978919 | 2.507425 | 0.608958 | 2.932629 | -0.06531 | 0.345505 | 0.627807 | 0.689121 | 0.614877 | 0.692358 | 0.709698 | 1.420162 | 1.503851 | 0.908097 |
| 22.55556 | 1.695277 | 1.223246 | 1.619082 | 0.757119 | 0.007556 | 0.701703 | 0.948431 | 0.410325 | 1.181199 | 2.643174 | 0.619827 | 1.9143 | 0.399735 | 2.353737 | -0.00405 | 0.627065 | 0.462414 | 0.781651 | 0.360673 | 0.373646 | 0.657303 | 1.506412 | 1.049377 | 0.982153 |
| 22.56944 | 0.963981 | 0.945096 | 1.62666 | 0.494469 | 0.199753 | 0.658591 | -0.08166 | 0.230413 | 0.79382 | 1.19103 | 0.373852 | 0.859828 | 0.381661 | 1.454081 | 0.098104 | 0.451838 | 0.408709 | 0.303754 | 0.270033 | 0.364024 | 0.441428 | 1.432627 | 0.628779 | 0.262333 |
| 22.58333 | 0.161928 | 1.007558 | 1.275933 | 0.189718 | 0.268006 | 0.374447 | -0.98534 | 0.147167 | 0.231129 | 0.135177 | 0.205008 | 0.115233 | 0.36467 | 0.435746 | 0.061241 | 0.145305 | 0.309121 | -0.23858 | 0.107463 | 0.426853 | 0.121532 | 0.877003 | 0.225828 | -0.61275 |
| 22.59722 | -0.74836 | 0.95972 | 0.880715 | -0.12682 | 0.196384 | 0.150509 | -1.4952 | -0.02922 | 0.09211 | -1.32796 | 0.125044 | -0.73494 | 0.334 | -0.72369 | -0.06272 | -0.15361 | 0.148753 | -0.64196 | -0.14903 | 0.487547 | -0.25046 | 0.279149 | -0.17199 | -1.371 |
| 22.61111 | -1.46372 | 0.976319 | 0.468942 | -0.26045 | 0.193309 | -0.09031 | -1.56759 | -0.40934 | 0.327243 | -2.59354 | 0.159144 | -0.73305 | 0.355927 | -1.2938 | -0.12182 | -0.40572 | -0.00196 | -0.80192 | -0.21143 | 0.470207 | -0.33763 | 0.124855 | -0.68281 | -1.35456 |
| 22.625 | -1.55307 | 1.159334 | 0.150232 | -0.15588 | 0.286789 | -0.19196 | -0.97577 | -0.82932 | 0.621743 | -2.73842 | 0.089909 | 0.021923 | 0.452056 | -1.66475 | -0.06511 | -0.30482 | -0.03147 | -0.64441 | 0.012734 | 0.365259 | -0.12532 | 0.309943 | -0.74303 | -0.75441 |
| 22.63889 | -1.07492 | 1.350536 | -0.07228 | 0.068964 | 0.328992 | -0.15922 | -0.02741 | -0.24428 | 0.46034 | -0.06956 | 0.072047 | 0.876498 | 0.559831 | -0.76373 | 0.03892 | -0.03392 | -0.01495 | -0.28526 | 0.227754 | 0.253386 | 0.099357 | 0.550922 | -0.58265 | -0.10614 |
| 22.65278 | -0.03477 | 1.593274 | -0.28186 | 0.293562 | 0.3119 | -0.06474 | 1.18114 | 0.4064 | 0.096397 | 2.503619 | 0.078301 | 1.490961 | 0.579866 | 0.5467 | 0.13737 | 0.104692 | 0.037249 | 0.072717 | 0.243441 | 0.276678 | 0.200979 | 0.650106 | -0.34576 | 0.669497 |
| 22.66667 | 1.081795 | 1.640993 | -0.46009 | 0.384195 | 0.240238 | 0.004599 | 2.158325 | 0.967363 | 0.065199 | 3.182892 | 0.108001 | 1.67386 | 0.436734 | 1.809967 | 0.220749 | 0.16125 | 0.06298 | 0.247249 | 0.16197 | 0.310845 | 0.330625 | 0.825531 | -0.23925 | 1.256552 |
| 22.68056 | 1.83049 | 1.563426 | -0.60646 | 0.414755 | 0.114337 | -0.03909 | 0.929891 | 0.561323 | 0.172188 | 2.249587 | 0.239096 | 1.114332 | 0.196693 | 2.312716 | 0.275751 | 0.211356 | 0.050315 | 0.270744 | 0.140699 | 0.229698 | 0.251468 | 0.574491 | -0.13825 | 0.956569 |
| 22.69444 | 1.751551 | 1.312823 | -0.43995 | 0.42335 | 0.009907 | -0.04878 | 0.050542 | 0.205579 | 0.226561 | 0.791061 | 0.384446 | 0.11101 | -0.0271 | 1.754098 | 0.258313 | 0.229598 | 0.084209 | 0.30144 | 0.060877 | 0.065209 | 0.180791 | 0.246304 | 0.292035 | 0.37087 |
| 22.70833 | 1.499386 | 0.99862 | -0.06333 | 0.301727 | -0.03061 | -0.05771 | -0.83634 | 0.006819 | 0.096241 | -0.18036 | 0.479764 | -0.55771 | -0.1517 | 0.532379 | 0.131183 | 0.335936 | 0.174169 | 0.388068 | -0.0606 | -0.20046 | -0.03632 | 0.089325 | 0.775439 | -0.20996 |
| 22.72222 | 1.090209 | 0.508792 | 0.045345 | 0.089952 | 0.006859 | 0.03578 | -1.89394 | -0.10486 | -0.4001 | -0.67403 | 0.443514 | -0.81654 | -0.28733 | -0.57861 | -0.05634 | 0.358853 | 0.189766 | 0.338467 | -0.19557 | -0.45848 | -0.15736 | -0.20186 | 1.064062 | -0.71668 |
| 22.73611 | 0.307297 | 0.094613 | 0.132645 | -0.14039 | 0.078832 | 0.180901 | -2.31414 | -0.2424 | -0.67532 | -1.50597 | 0.31397 | -1.01805 | -0.41131 | -1.39407 | -0.2004 | 0.117085 | 0.127279 | 0.085606 | -0.17641 | -0.49963 | -0.02157 | -0.32037 | 0.882465 | -0.8753 |
| 22.75 | -0.66471 | -0.21449 | 0.170133 | -0.29595 | 0.10871 | 0.239077 | -1.34422 | -0.53255 | -0.65926 | -2.49653 | 0.106246 | -1.44587 | -0.32978 | -1.87868 | -0.22453 | -0.26187 | 0.046213 | -0.23363 | -0.00973 | -0.35091 | 0.271134 | -0.0094 | 0.449119 | -0.58569 |
| 22.76389 | -1.50186 | -0.42628 | 0.08611 | -0.28794 | 0.089264 | 0.223466 | 0.000736 | -0.86369 | -0.50539 | -3.08048 | -0.03509 | -1.11294 | 0.010401 | -1.89742 | -0.12101 | -0.54651 | -0.06194 | -0.44866 | 0.127876 | -0.11235 | 0.273624 | 0.126687 | 0.027919 | -0.10426 |
| 22.77778 | -1.72528 | -0.65346 | -0.03963 | -0.06656 | -0.01452 | 0.142338 | 1.121635 | -0.4615 | -0.30094 | -1.97522 | -0.15031 | -0.22023 | 0.326538 | -1.24275 | 0.025588 | -0.46673 | -0.12705 | -0.555 | 0.139776 | 0.082733 | -0.07069 | 0.000356 | -0.31797 | 0.29354 |
| 22.79167 | -1.55322 | -0.73205 | -0.15268 | 0.132447 | -0.09966 | 0.029379 | 1.082333 | -0.09844 | -0.20156 | -0.2671 | -0.26706 | 0.804239 | 0.41927 | -0.3391 | 0.118656 | -0.17739 | -0.1877 | -0.55504 | 0.050172 | 0.108696 | -0.42606 | -0.35708 | -0.53388 | 0.474612 |
| 22.80556 | -1.15833 | -0.7044 | -0.22427 | 0.276662 | -0.09296 | -0.18631 | 0.30767 | 0.324943 | -0.11631 | 1.354944 | -0.30748 | 1.448197 | 0.275914 | 0.28088 | 0.147174 | -0.02769 | -0.21987 | -0.47677 | -0.09149 | -0.00912 | -0.63534 | -0.67328 | -0.50647 | 0.3048 |
| 22.81944 | -0.23466 | -0.63953 | -0.32189 | 0.287263 | 0.01098 | -0.25188 | -0.54791 | 0.805933 | 0.118264 | 2.58754 | -0.24177 | 0.699112 | 0.005477 | 0.545973 | 0.088164 | 0.006084 | -0.2212 | -0.28313 | -0.20891 | -0.14168 | -0.6629 | -0.8915 | -0.45072 | -0.13659 |
| 22.83333 | 0.236989 | -0.59719 | -0.50082 | 0.135503 | 0.07592 | -0.32086 | -1.58766 | 0.504806 | 0.142686 | 1.692397 | -0.16406 | -0.05176 | -0.32291 | 0.507715 | -0.06844 | -0.03376 | -0.26644 | -0.10928 | -0.30814 | -0.31017 | -0.53403 | -0.96443 | -0.48818 | -0.66862 |
| 22.84722 | -0.01719 | -0.71196 | -0.66112 | -0.13108 | 0.040376 | -0.37875 | -2.35941 | -0.1364 | -0.18786 | 0.171351 | -0.16739 | -0.63136 | -0.67818 | 0.083346 | -0.23592 | -0.10746 | -0.29608 | -0.01485 | -0.38064 | -0.54188 | -0.36015 | -1.01701 | -0.51314 | -1.17864 |
| 22.86111 | -0.40443 | -0.95031 | -0.76576 | -0.36207 | -0.02551 | -0.36288 | -1.3989 | -0.72316 | -0.62264 | -1.01868 | -0.29696 | -1.22747 | -0.9922 | -0.71178 | -0.29991 | -0.15311 | -0.36907 | -0.0374 | -0.36425 | -0.62758 | -0.15496 | -0.97759 | -0.52342 | -0.97042 |
| 22.875 | -0.72461 | -1.23678 | -0.94665 | -0.38642 | -0.15752 | -0.29635 | -0.56304 | -1.09876 | -1.0462 | -2.378 | -0.55689 | -1.46745 | -1.10571 | -1.63341 | -0.22847 | -0.14595 | -0.43999 | -0.14816 | -0.32289 | -0.64653 | -0.13301 | -0.82179 | -0.56045 | -0.44451 |
| 22.88889 | -1.34047 | -1.40737 | -1.09942 | -0.24795 | -0.26671 | -0.28295 | 0.045205 | -0.77222 | -1.4991 | -4.22545 | -0.64474 | -1.2794 | -1.00744 | -2.42086 | -0.15453 | -0.22348 | -0.49146 | -0.26916 | -0.338 | -0.79679 | -0.41339 | -0.80687 | -0.62665 | -0.03191 |
| 22.90278 | -2.27395 | -1.33133 | -1.18828 | -0.04848 | -0.22858 | -0.27499 | 0.145355 | -0.32451 | -1.45787 | -4.00501 | -0.4738 | -0.97017 | -0.93493 | -2.73731 | -0.07894 | -0.2366 | -0.46072 | -0.38966 | -0.43408 | -0.85152 | -0.64351 | -0.99369 | -0.66594 | 0.056192 |
| 22.91667 | -1.60166 | -1.10276 | -1.24692 | 0.092758 | -0.15202 | -0.27391 | -0.20238 | -0.3706 | -1.1176 | -2.72034 | -0.30687 | -0.7074 | -0.8393 | -2.32537 | 0.007524 | -0.20422 | -0.44564 | -0.54821 | -0.48189 | -0.64186 | -0.65706 | -1.29397 | -0.69457 | -0.22572 |
| 22.93056 | -0.87501 | -0.7263 | -1.2918 | 0.112953 | -0.09127 | -0.29055 | -0.58836 | 0.011983 | -0.93625 | -1.26442 | -0.26385 | -0.48242 | -0.88289 | -1.31186 | 0.051903 | -0.13628 | -0.41333 | -0.57683 | -0.42998 | -0.33685 | -0.46923 | -1.61483 | -0.69115 | -0.68969 |
| 22.94444 | -0.29724 | -0.09339 | -1.14917 | 0.060554 | -0.06462 | -0.29207 | -0.9775 | 0.362317 | -0.8993 | -0.00638 | -0.28257 | -0.38419 | -1.06401 | -0.50121 | 0.000327 | -0.21813 | -0.40176 | -0.44429 | -0.36727 | -0.13229 | -0.21095 | -1.68811 | -0.62521 | -1.11545 |
| 22.95833 | 0.114629 | 0.268713 | -0.89225 | -0.01479 | -0.0678 | -0.26609 | -1.04818 | 0.28101 | -0.89726 | 0.569722 | -0.3544 | -0.49871 | -1.21054 | 0.039504 | -0.07778 | -0.37726 | -0.36962 | -0.37416 | -0.32019 | -0.14356 | -0.02279 | -1.46212 | -0.51612 | -1.2541 |
| 22.97222 | 0.119237 | 0.322652 | -0.62899 | -0.0691 | -0.08341 | -0.25817 | -0.64644 | 0.262575 | -0.77762 | 0.389279 | -0.48515 | -0.61658 | -1.18599 | 0.170472 | -0.10091 | -0.44993 | -0.30865 | -0.31456 | -0.29157 | -0.26498 | 0.067762 | -0.93401 | -0.39542 | -0.9821 |
| 22.98611 | -0.09923 | 0.224737 | -0.39143 | -0.08947 | -0.09521 | -0.25122 | 0.053673 | 0.143348 | -0.61706 | -0.54218 | -0.50975 | -0.61188 | -0.72052 | 0.062176 | -0.05827 | -0.3533 | -0.18185 | -0.17479 | -0.25449 | -0.30596 | 0.018976 | -0.53382 | -0.27069 | -0.45997 |
| 23 | -0.42875 | 0.03809 | -0.35856 | -0.18255 | -0.11853 | -0.23563 | 0.488459 | -0.12574 | -0.63837 | -1.7144 | -0.53114 | -0.57323 | -0.31023 | -0.10834 | 0.010473 | -0.18019 | -0.11351 | -0.00707 | -0.22707 | -0.31873 | -0.15955 | -0.37366 | -0.30565 | -0.15032 |
| 23.01389 | -0.81268 | -0.35807 | -0.58514 | -0.39915 | -0.16803 | -0.23341 | 0.340766 | -0.38427 | -0.78556 | -2.67742 | -0.75006 | -0.58411 | -0.20457 | -0.32484 | 0.079476 | -0.06185 | -0.19797 | 0.082262 | -0.2469 | -0.57284 | -0.39397 | -0.43057 | -0.62177 | -0.35294 |
| 23.02778 | -1.30044 | -0.64063 | -0.89499 | -0.37063 | -0.18412 | -0.32192 | -0.18472 | -0.43149 | -1.05245 | -3.48389 | -1.49287 | -1.17517 | -0.37916 | -0.67269 | 0.135583 | -0.0722 | -0.4551 | 0.045519 | -0.32382 | -1.15769 | -0.72416 | -0.55624 | -1.33796 | -0.99122 |
| 23.04167 | -1.85023 | -0.63727 | -1.41253 | -0.2492 | -0.16071 | -0.50476 | -0.81261 | -0.69419 | -0.9995 | -4.67437 | -2.2442 | -2.1012 | -0.71255 | -1.08036 | 0.184119 | -0.13181 | -0.95493 | -0.0589 | -0.36348 | -1.26269 | -1.03126 | -0.69441 | -2.14887 | -1.50111 |
| 23.05556 | -2.26775 | -0.74947 | -1.76267 | -0.27309 | -0.16082 | -0.49047 | -1.54558 | -1.59016 | -0.69217 | -5.22213 | -2.21941 | -2.85766 | -1.13825 | -1.38632 | 0.157035 | -0.09057 | -1.4722 | -0.19861 | -0.16534 | -0.77485 | -1.11901 | -0.74123 | -1.92427 | -1.2353 |
| 23.06944 | -1.80713 | -0.9522 | -1.22519 | -0.24479 | -0.13165 | -0.3006 | -1.56688 | -2.89466 | -0.44437 | -3.81172 | -1.48993 | -2.47369 | -1.02356 | -1.30726 | 0.054782 | 0.045797 | -1.39282 | -0.34007 | 0.024925 | -0.32561 | -0.96497 | -0.41714 | -1.00792 | -0.87062 |
| 23.08333 | -0.89627 | -0.84112 | -0.35286 | -0.05001 | -0.03664 | -0.19554 | -0.89829 | -2.60467 | -0.38479 | -1.50098 | -0.72251 | -1.46743 | -0.35539 | -0.70781 | 0.001296 | 0.084323 | -0.96291 | -0.34616 | 0.058246 | -0.04545 | -0.57523 | 0.160517 | -0.25513 | -0.53117 |
| 23.09722 | 0.01433 | -0.51888 | 0.399845 | 0.084471 | -0.05583 | -0.17903 | -0.03325 | -0.90597 | -0.37772 | 0.195898 | 0.145851 | -0.38972 | 0.266038 | -0.08993 | 0.000759 | 0.065051 | -0.57114 | -0.15633 | 0.048674 | 0.1332 | -0.16433 | 0.45623 | 0.201673 | 0.241237 |
| 23.11111 | 0.526551 | -0.24939 | 0.761142 | 0.087646 | -0.19307 | -0.13271 | 0.588936 | 0.497882 | -0.42901 | 1.431156 | 0.730648 | 0.421238 | 0.520741 | 0.424826 | 0.004397 | 0.057527 | -0.28484 | 0.016949 | 0.021583 | 0.200728 | 0.004994 | 0.512286 | 0.28782 | 0.712073 |
| 23.125 | 0.539859 | -0.08493 | 0.722391 | 0.069034 | -0.24893 | -0.09278 | 0.655495 | 1.304651 | -0.52078 | 2.214044 | 0.808481 | 0.781428 | 0.444712 | 0.783045 | 0.004586 | 0.054251 | -0.13094 | 0.107978 | -0.00327 | 0.162977 | -0.03227 | 0.490537 | 0.184939 | 0.723592 |
| 23.13889 | 0.409612 | -0.05974 | 0.533676 | 0.065492 | -0.12915 | -0.07124 | 0.473389 | 1.549647 | -0.49087 | 2.461587 | 0.750452 | 0.89928 | 0.277951 | 0.89387 | 0.002185 | 0.064349 | -0.07778 | 0.146722 | 0.005407 | 0.103545 | 0.010039 | 0.401528 | 0.071825 | 0.678129 |
| 23.15278 | 0.360291 | -0.05614 | 0.347477 | 0.037628 | 0.070312 | -0.05407 | 0.356386 | 1.586944 | -0.29262 | 2.617162 | 0.548447 | 0.707173 | 0.061647 | 0.769866 | 0.005744 | 0.070029 | -0.03008 | 0.152599 | 0.050515 | 0.034051 | 0.170168 | 0.21224 | 0.017938 | 0.636913 |
| 23.16667 | 0.346531 | 0.037974 | 0.144372 | 0.00897 | 0.199734 | -0.05049 | 0.463651 | 1.433843 | -0.14219 | 2.467291 | 0.526112 | 0.3857 | -0.10369 | 0.608189 | 0.013454 | 0.059622 | 0.045415 | 0.15589 | 0.082248 | -0.03872 | 0.270602 | -0.08857 | 0.021184 | 0.55924 |
| 23.18056 | 0.346929 | 0.153412 | 0.020322 | -0.0074 | 0.110575 | -0.057 | 0.688263 | 1.218651 | -0.09543 | 2.217962 | 0.472869 | 0.166215 | -0.20687 | 0.51077 | 0.01106 | 0.022679 | 0.080821 | 0.161271 | 0.074193 | -0.12074 | 0.229087 | -0.37842 | 0.073722 | 0.483135 |
| 23.19444 | 0.377097 | 0.230536 | -0.00464 | 0.003711 | -0.0934 | -0.04325 | 0.739951 | 1.134188 | -0.107 | 2.024919 | 0.444601 | 0.11751 | -0.20991 | 0.431879 | 0.000213 | -0.02016 | 0.09246 | 0.152283 | 0.036177 | -0.1811 | 0.183408 | -0.46112 | 0.127731 | 0.443938 |
| 23.20833 | 0.502768 | 0.210172 | 0.014853 | 0.01376 | -0.30947 | -0.01711 | 0.454132 | 0.991097 | -0.06274 | 1.773553 | 0.329765 | 0.064634 | -0.07949 | 0.389521 | -0.00808 | -0.06562 | 0.091947 | 0.132777 | -0.02504 | -0.20514 | 0.143389 | -0.36073 | 0.146548 | 0.482216 |
| 23.22222 | 0.612241 | 0.1543 | 0.01237 | 0.021348 | -0.26292 | -0.0126 | 0.022984 | 0.932404 | 0.021291 | 1.281597 | 0.222766 | -0.00217 | 0.062502 | 0.390293 | -0.0085 | -0.10697 | 0.079258 | 0.117869 | -0.09674 | -0.22542 | 0.075545 | -0.09193 | 0.118799 | 0.390743 |
| 23.23611 | 0.583723 | 0.096812 | 0.001834 | 0.024414 | -0.07586 | -0.01851 | -0.479 | 0.771967 | -0.01065 | 0.778242 | 0.149821 | -0.00378 | 0.053999 | 0.377663 | -0.00247 | -0.12219 | 0.07802 | 0.090127 | -0.17018 | -0.20799 | 0.019114 | 0.182379 | 0.066377 | 0.223687 |
| 23.25 | 0.46897 | 0.066033 | -0.01309 | 0.004735 | 0.054011 | -0.03298 | -0.92286 | 0.641303 | -0.12062 | 0.211808 | 0.059959 | 0.048115 | -0.12065 | 0.284467 | 0.000336 | -0.11035 | 0.101663 | 0.062244 | -0.22428 | -0.13948 | 0.010206 | 0.23793 | 0.031869 | 0.028036 |
| 23.26389 | 0.3115 | 0.054023 | -0.07809 | -0.05766 | 0.057861 | -0.06102 | -0.82776 | 0.31998 | -0.119 | -0.20623 | 0.012549 | 0.128632 | -0.33258 | 0.181298 | 0.005973 | -0.08471 | 0.141165 | 0.041755 | -0.22373 | -0.08681 | 0.037941 | 0.018117 | 0.031993 | -0.12259 |
| 23.27778 | 0.157588 | 0.045167 | -0.14085 | -0.11739 | -0.01018 | -0.05242 | -0.41736 | 0.047826 | -0.00951 | -0.43828 | -0.03576 | 0.277884 | -0.37554 | 0.114823 | 0.014335 | -0.04643 | 0.182661 | 0.021933 | -0.15944 | -0.08096 | 0.088732 | -0.25816 | 0.038374 | -0.23505 |
| 23.29167 | 0.053264 | 0.05138 | -0.11525 | -0.12242 | -0.03034 | 0.015275 | 0.007385 | -0.09672 | 0.012989 | -0.59026 | -0.07436 | 0.541492 | -0.07916 | 0.059759 | 0.016146 | -0.01321 | 0.232841 | 0.007264 | -0.09709 | -0.0917 | 0.127495 | -0.31106 | 0.046321 | -0.27373 |
| 23.30556 | 0.037532 | 0.037536 | -0.02739 | -0.0706 | -0.00298 | 0.062856 | 0.189143 | -0.18271 | 0.012968 | -0.59185 | -0.07784 | 0.75605 | 0.364383 | 0.036964 | 0.011514 | -0.00144 | 0.262955 | 0.010463 | -0.07294 | -0.10115 | 0.13336 | -0.0988 | 0.036513 | -0.11809 |
| 23.31944 | 0.071893 | 0.039345 | 0.038566 | -0.00541 | 0.026069 | 0.058678 | 0.106623 | -0.23814 | 0.077049 | -0.25675 | -0.06082 | 0.662685 | 0.487175 | 0.071545 | 0.011539 | 0.00655 | 0.254522 | 0.044071 | -0.06553 | -0.09631 | 0.105271 | 0.10345 | 0.006373 | 0.164547 |
| 23.33333 | 0.142529 | 0.037593 | 0.057728 | 0.037333 | 0.012753 | 0.024404 | 0.02459 | -0.15605 | 0.160635 | 0.257554 | -0.03804 | 0.542507 | 0.373391 | 0.19051 | 0.019626 | 0.033588 | 0.26809 | 0.101151 | -0.03517 | -0.06939 | 0.087715 | 0.176776 | -0.01315 | 0.288848 |
| 23.34722 | 0.249343 | 0.031468 | 0.080846 | 0.030945 | -0.05676 | -0.00253 | 0.135223 | 0.177991 | 0.237805 | 0.581357 | -1.5E-05 | 0.463196 | 0.138677 | 0.320087 | 0.036544 | 0.070534 | 0.323871 | 0.152952 | 0.035285 | -0.04062 | 0.086848 | 0.168856 | -0.00459 | 0.116918 |
| 23.36111 | 0.402495 | 0.025263 | 0.145362 | 0.021324 | -0.08071 | 0.007126 | 0.338291 | 0.52673 | 0.256224 | 0.694862 | 0.042351 | 0.506554 | -0.06227 | 0.419024 | 0.073141 | 0.097278 | 0.373429 | 0.210502 | 0.141687 | -0.02181 | 0.134464 | 0.161512 | 0.057057 | -0.12442 |
| 23.375 | 0.537207 | 0.051941 | 0.204118 | 0.061896 | 0.012012 | 0.025585 | 0.532076 | 0.607887 | 0.244991 | 0.706265 | 0.073342 | 0.65715 | -0.0618 | 0.63804 | 0.116667 | 0.143624 | 0.380873 | 0.296086 | 0.2172 | 0.002169 | 0.215526 | 0.206987 | 0.154916 | -0.16304 |
| 23.38889 | 0.660338 | 0.095671 | 0.252456 | 0.132286 | 0.144275 | -0.05058 | 0.666082 | 0.455248 | 0.259155 | 0.864349 | 0.073777 | 0.859315 | 0.15523 | 1.007658 | 0.137973 | 0.170538 | 0.37941 | 0.380124 | 0.197121 | 0.032273 | 0.255304 | 0.277003 | 0.252857 | 0.07124 |
| 23.40278 | 0.779941 | 0.119506 | 0.307602 | 0.182592 | 0.264464 | -0.2429 | 0.675664 | 0.286066 | 0.2893 | 1.185124 | 0.083757 | 0.964674 | 0.453784 | 1.127859 | 0.157925 | 0.177463 | 0.381697 | 0.452575 | 0.132985 | 0.046496 | 0.238786 | 0.268748 | 0.320785 | 0.465519 |
| 23.41667 | 0.837671 | 0.11834 | 0.335382 | 0.211482 | 0.212001 | -0.26433 | 0.497131 | 0.162089 | 0.281013 | 1.294556 | 0.143887 | 0.890739 | 0.588592 | 0.837717 | 0.158601 | 0.175493 | 0.397275 | 0.493458 | 0.075828 | 0.075102 | 0.217542 | 0.192899 | 0.373346 | 0.767373 |
| 23.43056 | 0.763777 | 0.084732 | 0.331136 | 0.189029 | 0.049366 | -0.01605 | 0.342099 | 0.12848 | 0.264107 | 1.120558 | 0.223049 | 0.646342 | 0.472879 | 0.473362 | 0.17798 | 0.187824 | 0.375475 | 0.459567 | 0.027235 | 0.097791 | 0.21517 | 0.124551 | 0.391497 | 0.609938 |
| 23.44444 | 0.768342 | 0.030934 | 0.404369 | 0.134213 | -0.12036 | 0.13803 | 0.347435 | 0.104494 | 0.322186 | 1.115296 | 0.321521 | 0.453151 | 0.237942 | 0.276056 | 0.176223 | 0.206719 | 0.366811 | 0.470677 | 0.016389 | 0.105653 | 0.294283 | 0.168837 | 0.378538 | 0.26446 |
| 23.45833 | 0.853465 | -0.0086 | 0.525562 | 0.125703 | -0.1692 | 0.215247 | 0.498416 | 0.097435 | 0.502659 | 1.335646 | 0.33097 | 0.400718 | 0.090346 | 0.261198 | 0.152303 | 0.302725 | 0.495297 | 0.52407 | 0.115821 | 0.133127 | 0.469562 | 0.462659 | 0.413618 | 0.076596 |
| 23.47222 | 0.966349 | 0.006756 | 0.545986 | 0.171221 | -0.04589 | 0.254317 | 0.545089 | 0.304528 | 0.660198 | 1.631315 | 0.277357 | 0.37824 | 0.126546 | 0.388087 | 0.127093 | 0.431851 | 0.70754 | 0.577793 | 0.256584 | 0.166302 | 0.595265 | 0.830806 | 0.506487 | 0.193807 |
| 23.48611 | 0.979709 | 0.132863 | 0.396968 | 0.193705 | 0.199684 | 0.243155 | 0.421157 | 0.399942 | 0.760905 | 1.65809 | 0.287279 | 0.450205 | 0.358664 | 0.579765 | 0.137749 | 0.476768 | 0.872157 | 0.606718 | 0.310035 | 0.226292 | 0.621128 | 1.008225 | 0.637235 | 0.46811 |
| 23.5 | 1.194756 | 0.466453 | 0.423661 | 0.274461 | 0.381756 | 0.314968 | 0.481425 | 0.412799 | 1.483954 | 1.768225 | 0.627537 | 1.005152 | 0.754959 | 0.954666 | 0.223919 | 0.437881 | 0.903143 | 0.644629 | 0.377919 | 0.556899 | 0.83627 | 1.168265 | 0.984465 | 0.80832 |
| 23.51389 | 1.918686 | 1.123048 | 0.942058 | 0.617571 | 0.361942 | 0.669998 | 0.952645 | 0.752701 | 2.922916 | 2.847547 | 1.328575 | 2.084174 | 1.283962 | 1.590941 | 0.331709 | 0.285271 | 1.083894 | 0.815061 | 0.691828 | 1.44125 | 1.797013 | 1.469763 | 2.119667 | 1.116195 |
| 23.52778 | 2.089753 | 1.691448 | 1.34891 | 0.770456 | 0.273211 | 0.817851 | 1.315396 | 1.149103 | 3.204423 | 5.135376 | 1.548011 | 2.609192 | 1.528133 | 2.354568 | 0.409997 | -0.02342 | 1.035373 | 0.957487 | 0.605771 | 2.287665 | 2.48321 | 1.677541 | 3.428444 | 1.020949 |
| 23.54167 | 2.146461 | 1.331873 | 1.045245 | 0.649709 | 0.103342 | 0.716171 | 1.757284 | 1.249734 | 2.525679 | 6.602774 | 1.219772 | 2.098051 | 1.186001 | 2.57054 | 0.492867 | -0.17787 | 0.519053 | 0.673174 | 0.370645 | 1.705942 | 2.137826 | 1.586 | 2.119441 | 0.64243 |
| 23.55556 | 2.057218 | 0.747484 | 0.75327 | 0.588864 | -0.01103 | 0.636799 | 0.874979 | 1.096812 | 2.30085 | 4.662307 | 0.939648 | 1.517677 | 0.736228 | 2.502888 | 0.305297 | 0.029457 | 0.412649 | 0.17292 | 0.139367 | 1.052959 | 1.603262 | 1.507245 | 1.377529 | 0.34361 |
| 23.56944 | 1.143233 | 0.872518 | 0.801903 | 0.501443 | -0.01732 | 0.375178 | -0.27177 | 0.669452 | 2.019442 | 2.109671 | 0.531851 | 1.005779 | 0.447499 | 1.955583 | 0.204923 | 0.190406 | 0.410248 | -0.34046 | 0.174766 | 0.681561 | 0.774709 | 1.08775 | 0.912455 | -0.01953 |
| 23.58333 | 0.262712 | 0.908913 | 1.135229 | 0.382233 | 0.001577 | -0.2493 | -1.42452 | 0.14695 | 1.562913 | 0.60784 | 0.236963 | 0.380822 | 0.207932 | 1.000992 | 0.203551 | 0.18098 | 0.320892 | -0.61877 | 0.108718 | 0.36357 | -0.09314 | 0.443466 | 0.614527 | -0.28242 |
| 23.59722 | -0.36206 | 0.606576 | 1.361333 | 0.14854 | 0.114561 | -0.82674 | -1.61835 | -0.18219 | 1.090438 | -1.2029 | 0.105131 | 0.053263 | 0.103546 | -0.12993 | 0.10612 | 0.035433 | 0.21747 | -0.63101 | 0.045372 | 0.272417 | -0.7202 | 0.032805 | 0.550468 | -0.154 |
| 23.61111 | -0.85853 | 0.651529 | 1.117553 | -0.04259 | 0.250586 | -0.57963 | -1.10305 | -0.357 | 1.436147 | -3.57738 | 0.169595 | 0.325808 | 0.128114 | -0.91881 | 0.05121 | -0.09925 | 0.200861 | -0.42516 | 0.052187 | 0.337537 | -0.27907 | -0.25524 | 0.604586 | 0.086892 |
| 23.625 | -1.21 | 0.759485 | 0.594575 | -0.0831 | 0.244392 | -0.21261 | -0.23745 | -0.29435 | 1.546268 | -2.71541 | 0.25531 | 0.864901 | 0.314699 | -0.88074 | 0.119355 | -0.17555 | 0.198213 | -0.12932 | 0.194164 | 0.393326 | 0.113863 | -0.11049 | 0.585168 | 0.336747 |
| 23.63889 | -1.28642 | 0.795227 | 0.336766 | -0.11358 | 0.142827 | 0.12979 | 0.912611 | -0.20038 | 1.256926 | -1.40666 | 0.337728 | 1.432622 | 0.574355 | -0.66526 | 0.334607 | -0.23847 | 0.234354 | 0.163988 | 0.282917 | 0.473923 | 0.367228 | 0.311815 | 0.494896 | 0.638039 |
| 23.65278 | -1.0669 | 0.699253 | 0.153347 | -0.14063 | -0.01414 | 0.278194 | 1.466377 | -0.07569 | 1.1679 | 0.721475 | 0.396995 | 1.708002 | 0.722516 | -0.23726 | 0.330778 | -0.16658 | 0.235211 | 0.384359 | 0.243352 | 0.535079 | 0.53607 | 0.844436 | 0.232918 | 0.721722 |
| 23.66667 | -0.34388 | 0.690243 | -0.00961 | -0.05547 | -0.11966 | 0.20484 | 1.514956 | 0.246843 | 1.085779 | 2.832031 | 0.450465 | 0.714956 | 0.640472 | 0.429499 | 0.302459 | -0.04272 | 0.089146 | 0.518424 | 0.124198 | 0.751396 | 0.550525 | 1.233032 | -0.15309 | 0.407634 |
| 23.68056 | 0.207989 | 0.833802 | 0.042902 | 0.143175 | -0.11419 | 0.087845 | 0.892811 | 0.581909 | 0.757827 | 3.443838 | 0.44941 | -0.0189 | 0.448622 | 1.027099 | 0.194074 | 0.122707 | -0.08391 | 0.621552 | 0.015261 | 0.682804 | 0.121688 | 1.138574 | -0.53178 | -0.18614 |
| 23.69444 | 0.671776 | 0.863882 | -0.00598 | 0.338945 | -0.06035 | -0.00321 | 0.21213 | 0.661274 | 0.600488 | 2.66809 | 0.418323 | -0.53931 | 0.194594 | 1.183052 | -0.04944 | 0.28055 | -0.25408 | 0.595104 | 0.014947 | 0.192501 | -0.47796 | 0.738453 | -0.72702 | -0.63757 |
| 23.70833 | 1.139985 | 0.570464 | -0.23122 | 0.401808 | 0.138455 | -0.07484 | -0.53008 | 0.593272 | -0.04377 | 0.393412 | 0.297013 | -0.66656 | -0.0543 | 0.990546 | -0.13393 | 0.384311 | -0.35586 | 0.372303 | 0.056795 | -0.11287 | -0.82258 | 0.073321 | -0.76667 | -0.68352 |
| 23.72222 | 0.881772 | 0.19124 | -0.33999 | 0.327535 | 0.324214 | -0.07694 | -1.41094 | 0.37233 | -0.65743 | -1.37068 | 0.113978 | -0.7309 | -0.25911 | 0.619095 | -0.09849 | 0.38036 | -0.31517 | -0.08486 | 0.047376 | -0.33062 | -0.47888 | -0.65396 | -0.50136 | -0.36374 |
| 23.73611 | 0.200492 | 0.023864 | -0.4884 | 0.221105 | 0.331465 | 0.064403 | -1.60453 | 0.08863 | -1.11939 | -3.06163 | -0.07823 | -0.92029 | -0.39307 | 0.091092 | -0.06467 | 0.364321 | -0.0798 | -0.40967 | 0.037127 | -0.32331 | -0.01136 | -1.32011 | -0.16478 | 0.052672 |
| 23.75 | -0.4678 | -0.13083 | -0.48123 | 0.141888 | 0.17441 | 0.30558 | -0.81453 | -0.27143 | -1.22031 | -4.32948 | -0.18326 | -0.59332 | -0.42631 | -0.50377 | -0.07965 | 0.078466 | 0.155047 | -0.46118 | 0.144134 | -0.19686 | 0.198458 | -1.24671 | 0.04026 | 0.593018 |
| 23.76389 | -0.96536 | -0.46943 | -0.43577 | 0.048508 | -0.11339 | 0.519911 | -0.05557 | -0.68319 | -0.74057 | -4.34708 | -0.27011 | 0.031175 | -0.25216 | -1.05364 | -0.13985 | -0.21585 | 0.250634 | -0.45647 | 0.250704 | -0.06673 | 0.378771 | -0.35895 | 0.045862 | 0.869459 |
| 23.77778 | -1.32872 | -0.94819 | -0.32633 | 0.030118 | -0.32485 | 0.570135 | 0.449575 | -0.91501 | -0.70506 | -3.74286 | -0.28744 | 0.681968 | 0.076686 | -1.37162 | -0.13481 | -0.39637 | 0.18286 | -0.45429 | 0.238537 | -0.14623 | 0.110995 | 0.497329 | -0.145 | 0.290762 |
| 23.79167 | -1.64739 | -1.19743 | -0.28648 | 0.001627 | -0.37353 | 0.367183 | 0.35953 | -0.78732 | -0.40379 | -1.55688 | -0.14515 | 0.192097 | 0.343875 | -1.34096 | -0.08938 | -0.44408 | -0.02603 | -0.56139 | 0.124431 | -0.26949 | -0.39681 | 0.931249 | -0.33864 | -0.40435 |
| 23.80556 | -1.24652 | -1.03819 | -0.42173 | 0.033432 | -0.19708 | 0.123514 | 0.048761 | -0.49934 | 0.037928 | 0.28935 | 0.054944 | -0.35748 | 0.453094 | -1.07256 | -0.05508 | -0.40164 | -0.22116 | -0.65925 | 0.02008 | -0.39926 | -0.95959 | 0.487326 | -0.31919 | -0.60919 |
| 23.81944 | -0.57257 | -0.7047 | -0.56245 | 0.124904 | 0.109134 | -0.10989 | -0.37324 | -0.22592 | 0.21301 | 0.759968 | 0.139792 | -0.51857 | 0.346238 | -0.85807 | -0.03875 | -0.29521 | -0.33727 | -0.37221 | -0.07823 | -0.56364 | -1.29095 | -0.4688 | -0.13958 | -0.55557 |
| 23.83333 | -0.08002 | -0.59015 | -0.61766 | 0.183222 | 0.385357 | -0.27079 | -1.00846 | -0.04602 | 0.034337 | 0.115769 | 0.064449 | -0.90948 | 0.031147 | -0.63103 | -0.02903 | -0.13879 | -0.36783 | -0.05254 | -0.18205 | -0.34646 | -0.72689 | -1.65178 | -0.07677 | -0.40839 |
| 23.84722 | -0.06135 | -0.65522 | -0.7337 | 0.17391 | 0.309223 | -0.43775 | -1.41553 | 0.072246 | -0.10378 | -0.32933 | 0.017519 | -1.7001 | -0.40637 | -0.40206 | -0.08653 | -0.00822 | -0.30944 | 0.151031 | -0.31063 | -0.19212 | -0.33493 | -2.44255 | -0.2347 | -0.28501 |
| 23.86111 | -0.29197 | -0.73647 | -0.84883 | 0.110615 | 0.034403 | -0.5047 | -1.12984 | 0.025477 | -0.31688 | -1.02117 | -0.11065 | -1.9339 | -0.94618 | -0.32472 | -0.22593 | 0.000787 | -0.29012 | 0.138102 | -0.39874 | -0.18014 | -0.16179 | -2.17538 | -0.59283 | -0.22329 |
| 23.875 | -0.73077 | -0.80422 | -0.89239 | 0.006108 | -0.35066 | -0.44696 | -0.62143 | -0.13172 | -0.7488 | -2.48554 | -0.31393 | -1.58487 | -1.48962 | -0.3639 | -0.40853 | -0.07041 | -0.38222 | 0.020028 | -0.47076 | -0.3173 | -0.23737 | -1.52853 | -1.0652 | -0.37195 |
| 23.88889 | -1.2615 | -0.9821 | -0.81719 | -0.1239 | -0.36936 | -0.42451 | -0.24757 | -0.1931 | -1.17027 | -3.70324 | -0.53965 | -0.8606 | -1.69382 | -0.52902 | -0.42783 | -0.15183 | -0.53571 | -0.08971 | -0.41907 | -0.56046 | -0.55182 | -1.03884 | -1.47044 | -0.68798 |
| 23.90278 | -1.52991 | -1.1721 | -0.69747 | -0.20387 | -0.33819 | -0.5776 | -0.00282 | -0.29438 | -1.18925 | -5.12884 | -0.6826 | -0.04215 | -1.42964 | -0.77394 | -0.35421 | -0.20943 | -0.78447 | -0.20145 | -0.30747 | -0.88363 | -1.05091 | -1.03588 | -1.58071 | -0.95302 |
| 23.91667 | -1.97329 | -1.09928 | -0.60115 | -0.12912 | -0.02199 | -0.65727 | 0.053907 | -0.77297 | -1.14364 | -5.87877 | -0.70186 | 0.11722 | -1.02761 | -1.0746 | -0.24869 | -0.19337 | -1.02105 | -0.30201 | -0.21732 | -1.05468 | -1.35175 | -1.10946 | -1.34729 | -0.96702 |
| 23.93056 | -2.18746 | -0.77535 | -0.55124 | 0.038804 | 0.423789 | -0.67493 | -0.10275 | -0.56759 | -1.22868 | -3.5714 | -0.6598 | -0.05367 | -0.68395 | -1.37878 | -0.18545 | -0.13021 | -1.07626 | -0.41566 | -0.17145 | -0.88913 | -1.1322 | -0.91335 | -0.97141 | -0.74447 |
| 23.94444 | -1.42082 | -0.48989 | -0.54879 | 0.169547 | 0.566631 | -0.56946 | -0.3024 | -0.08714 | -1.06627 | -1.28533 | -0.44565 | -0.49484 | -0.46324 | -1.23237 | -0.08724 | -0.04988 | -0.77451 | -0.54112 | -0.11837 | -0.5602 | -0.6211 | -0.59315 | -0.57959 | -0.39976 |

| | | | | | | | | | | | | | | | | | | | | | | | | |
|---|---|---|---|---|---|---|---|---|---|---|---|---|---|---|---|---|---|---|---|---|---|---|---|---|
| 23.95833 | -0.45614 | -0.22725 | -0.59816 | 0.159568 | 0.351804 | -0.34085 | -0.45769 | 0.187099 | -0.79263 | 0.545926 | -0.13769 | -0.8023 | -0.35523 | -0.84405 | 0.009128 | 0.004384 | -0.36606 | -0.57961 | -0.0494 | -0.24158 | -0.11044 | -0.24668 | -0.31066 | 0.050859 |
| 23.97222 | 0.216854 | -0.05126 | -0.49374 | 0.042007 | 0.059368 | -0.16323 | -0.36824 | 0.285672 | -0.60776 | 1.649761 | 0.13702 | -1.00625 | -0.3762 | -0.54579 | 0.034687 | -0.0335 | -0.07462 | -0.55888 | -0.03361 | -0.06637 | 0.169771 | -0.06612 | -0.05376 | 0.272837 |
| 23.98611 | 0.463208 | 0.099559 | -0.29265 | -0.08247 | -0.24539 | -0.06502 | -0.20814 | 0.115595 | -0.64375 | 1.998542 | 0.268355 | -1.41408 | -0.46276 | -0.32035 | -0.04524 | -0.15444 | 0.037515 | -0.50642 | -0.09274 | 0.000227 | 0.125613 | -0.2667 | 0.083451 | 0.207226 |
| 24 | 0.226406 | 0.207524 | -0.27771 | -0.17847 | -0.39623 | -0.07972 | -0.15215 | -0.269 | -0.89856 | 1.218216 | 0.165923 | -1.83702 | -0.5612 | -0.08622 | -0.09641 | -0.23088 | -0.03224 | -0.37546 | -0.184 | -0.11217 | -0.10207 | -0.71665 | 0.051905 | 0.004042 |
| 24.01389 | -0.37517 | 0.141529 | -0.60647 | -0.24867 | -0.38931 | -0.29444 | -0.41127 | -0.64459 | -1.16739 | -0.67874 | -0.2382 | -1.93884 | -0.63755 | -0.08097 | -0.06706 | -0.21457 | -0.29607 | -0.21398 | -0.25465 | -0.44332 | -0.50578 | -1.18881 | -0.34742 | -0.41926 |
| 24.02778 | -1.07402 | -0.06428 | -0.96556 | -0.24519 | -0.15366 | -0.71145 | -0.74623 | -0.66616 | -1.14031 | -2.53599 | -0.85473 | -1.88305 | -0.74366 | -0.41013 | -0.09623 | -0.26393 | -0.61677 | -0.18588 | -0.3032 | -1.11044 | -1.0915 | -1.54995 | -1.1415 | -1.1155 |
| 24.04167 | -1.86096 | -0.36612 | -1.18245 | -0.13862 | 0.49739 | -1.15087 | -1.07389 | -0.45709 | -0.75523 | -4.12303 | -1.52475 | -2.59199 | -1.04032 | -1.00323 | -0.22144 | -0.41423 | -0.76487 | -0.28862 | -0.30294 | -1.47754 | -1.42743 | -1.73351 | -1.93285 | -1.7147 |
| 24.05556 | -2.6598 | -0.6194 | -1.3509 | -0.26101 | 0.823839 | -1.02871 | -1.38854 | -1.002 | -0.43419 | -3.24494 | -1.6693 | -3.45806 | -1.41038 | -1.95618 | -0.3391 | -0.50043 | -0.72412 | -0.45145 | -0.2425 | -0.67853 | -1.00175 | -1.59348 | -1.67685 | -1.66442 |
| 24.06944 | -2.23868 | -0.75377 | -1.23577 | -0.484 | 0.105025 | -0.68695 | -1.15454 | -1.94678 | -0.44998 | -1.19866 | -1.77512 | -2.77535 | -1.1832 | -2.15361 | -0.35664 | -0.27636 | -0.66728 | -0.55054 | -0.15228 | -0.08643 | -0.7794 | -1.1083 | -0.94891 | -1.36269 |
| 24.08333 | -1.35252 | -0.59065 | -0.90504 | -0.36582 | -0.34564 | -0.46797 | -0.36346 | -1.52096 | -0.56451 | -0.20891 | -1.44362 | -2.27419 | -0.45084 | -1.12938 | -0.27259 | -0.08275 | -0.47715 | -0.44393 | -0.1409 | -0.02255 | -0.69484 | -0.29598 | -0.34528 | -0.66704 |
| 24.09722 | -0.4373 | -0.25315 | -0.35646 | -0.07327 | -0.45364 | -0.3168 | 0.38535 | -0.78669 | -0.63675 | 0.046526 | -0.6689 | -1.06257 | 0.286306 | -0.0018 | 0.017911 | 0.179532 | -0.20133 | -0.15445 | -0.063 | 0.01237 | -0.29103 | 0.514313 | 0.114843 | 0.089975 |
| 24.11111 | 0.149787 | 0.013207 | 0.163416 | 0.186796 | -0.14327 | -0.1759 | 0.858992 | 0.045782 | -0.57471 | 0.102542 | 0.112394 | -0.14038 | 0.669325 | 0.814061 | 0.301312 | 0.342409 | -0.07321 | 0.095333 | 0.001899 | 0.197116 | 0.345735 | 0.912207 | 0.386313 | 0.600842 |
| 24.125 | 0.295818 | 0.133451 | 0.334072 | 0.322945 | 0.276517 | -0.09252 | 0.81893 | 0.7125 | -0.36815 | 0.104969 | 0.409019 | 0.360774 | 0.740789 | 1.171519 | 0.424148 | 0.273392 | -0.08773 | 0.251291 | 0.008394 | 0.192485 | 0.499156 | 1.018963 | 0.315163 | 0.761971 |
| 24.13889 | 0.284965 | 0.10472 | 0.265303 | 0.258093 | 0.385341 | -0.07356 | 0.650531 | 1.015458 | -0.17723 | 0.063549 | 0.423909 | 0.580472 | 0.582954 | 1.141783 | 0.409774 | 0.144611 | -0.09619 | 0.278944 | 0.008917 | 0.029662 | 0.253853 | 1.077344 | 0.134972 | 0.701968 |
| 24.15278 | 0.311822 | 0.029204 | -0.20047 | 0.109653 | 0.050206 | -0.10035 | 0.43276 | 0.835137 | -0.19573 | 0.016183 | 0.351224 | 0.736027 | 0.332368 | 0.725295 | 0.324366 | -0.06622 | -0.07823 | 0.23168 | 0.045523 | -0.04799 | -0.10024 | 0.942286 | 0.044027 | 0.737407 |
| 24.16667 | 0.219731 | -0.05513 | -0.73097 | -0.0514 | -0.08749 | -0.07905 | 0.300743 | 0.747827 | -0.19917 | 0.184549 | 0.605129 | 0.811148 | 0.016889 | -0.45701 | 0.431183 | -0.58408 | -0.42076 | 0.070371 | 0.210556 | 0.096963 | -0.66002 | 0.928706 | 0.027913 | 1.309361 |
| 24.18056 | -0.29205 | -0.18433 | -0.04518 | -0.15003 | 0.214411 | 0.239421 | 1.34888 | 1.573724 | 0.363491 | 1.630979 | 1.234347 | 1.031523 | -0.25506 | -1.77426 | 0.545692 | -1.08679 | -1.02505 | -0.16727 | 0.049771 | 0.136555 | -1.2087 | 0.464833 | -0.2205 | 1.211024 |
| 24.19444 | -1.19442 | -0.30441 | 1.237064 | -0.75571 | 0.528278 | 0.392548 | 2.052542 | 1.856739 | 0.95377 | 3.294262 | 1.864247 | 2.152788 | -0.62109 | -1.59017 | -0.04514 | -0.76837 | -1.05356 | -0.09581 | -1.45231 | -0.39404 | -1.33155 | -0.52533 | -1.10554 | -1.11635 |
| 24.20833 | 0.092305 | 0.160409 | 0.173347 | -0.18564 | -0.36579 | 0.229621 | 0.816864 | 0.873143 | -0.43605 | 2.802308 | 0.568812 | 2.410812 | -0.97398 | -0.27797 | -0.76814 | -0.78487 | -1.05753 | 0.019617 | -1.77426 | -0.79725 | -1.11412 | -1.02944 | -1.72208 | -2.56999 |
| 24.22222 | 1.319169 | 0.192571 | -1.10603 | 0.394706 | -0.06587 | -0.22026 | -1.53894 | -1.07106 | -1.72847 | -0.11097 | -0.3934 | 1.145172 | -0.67357 | 0.04865 | -0.65711 | -1.30648 | -0.26822 | -0.10922 | -0.67685 | -0.64755 | -0.56036 | -1.04047 | -1.55476 | -1.48435 |
| 24.23611 | 1.165424 | 0.206509 | -0.5253 | 0.107756 | -0.2153 | -0.0595 | -1.52933 | -1.26499 | -1.73475 | -0.19241 | -0.26109 | 1.05863 | -0.48295 | -0.51139 | -0.36594 | -1.69392 | 0.022472 | -0.17218 | -0.11989 | -0.45699 | -0.14103 | -0.49981 | -1.28382 | -0.54011 |
| 24.25 | 1.1587 | 0.231914 | 0.038551 | 0.030308 | 0.087384 | 0.128432 | -0.52516 | 0.109222 | -0.37099 | 0.947527 | 0.299376 | 1.671863 | -0.28147 | -0.55594 | -0.13484 | -1.05843 | -0.06207 | -0.20569 | -0.08828 | -0.41168 | 0.248789 | -0.28952 | -0.67044 | -0.0269 |
| 24.26389 | 1.359277 | 0.199264 | 0.147152 | -0.05414 | 0.065802 | 0.140351 | -0.58338 | 0.249443 | -0.11699 | 0.798261 | 0.25649 | 1.536457 | -0.10071 | -0.22872 | -0.0086 | -0.13617 | -0.13046 | -0.2212 | -0.17364 | -0.37734 | 0.425482 | -0.24938 | -0.20396 | 0.209045 |
| 24.27778 | 1.413376 | 0.142402 | 0.12003 | -0.15665 | 0.043751 | 0.102415 | -0.63448 | 0.092915 | -0.14961 | 0.376924 | 0.152535 | 1.133815 | -0.01259 | -0.01467 | 0.036457 | 0.283702 | -0.15097 | -0.21384 | -0.17169 | -0.33976 | 0.344269 | -0.23185 | -0.00853 | 0.203543 |
| 24.29167 | 1.266076 | 0.104762 | 0.171194 | -0.20103 | 0.069713 | 0.141127 | -0.39792 | -0.00367 | -0.14763 | 0.140063 | 0.178995 | 0.805555 | 0.017374 | 0.049668 | 0.035284 | 0.367087 | -0.13581 | -0.1692 | -0.05391 | -0.32884 | 0.175159 | -0.24503 | 0.056586 | 0.220495 |
| 24.30556 | 1.304809 | 0.161605 | 0.296944 | -0.14653 | 0.141578 | 0.29919 | -0.02364 | 0.09573 | -0.0701 | 0.294708 | 0.351298 | 0.593631 | 0.041014 | 0.117762 | 0.017119 | 0.651108 | -0.10784 | -0.07826 | -0.02182 | -0.32136 | 0.102468 | -0.27458 | 0.174883 | 0.430704 |
| 24.31944 | 1.331785 | 0.213219 | 0.332195 | -0.01049 | 0.250689 | 0.365887 | 0.310981 | 0.257227 | 0.058041 | 0.511227 | 0.478682 | 0.465311 | 0.044847 | 0.112984 | -0.00624 | 0.635239 | -0.03049 | -0.02487 | -0.05421 | -0.25526 | 0.094464 | -0.25681 | 0.188194 | 0.448932 |
| 24.33333 | 1.020599 | 0.185151 | 0.251914 | 0.129352 | 0.297623 | 0.259286 | 0.417546 | 0.247663 | 0.04116 | 0.506921 | 0.416998 | 0.172949 | 0.02896 | 0.032932 | -0.02017 | 0.140127 | 0.054671 | -0.03862 | -0.0216 | -0.18248 | 0.061031 | -0.18969 | 0.048788 | 0.198455 |
| 24.34722 | 0.655116 | 0.107139 | 0.147536 | 0.158384 | 0.225956 | 0.145014 | 0.292236 | 0.084159 | -0.02932 | 0.339368 | 0.253164 | -0.14474 | 0.033893 | 0.020058 | -0.01736 | -0.06473 | 0.092301 | -0.02666 | 0.030963 | -0.13843 | 0.04968 | -0.13085 | 0.000957 | 0.09677 |
| 24.36111 | 0.382781 | 0.06371 | 0.079303 | 0.048479 | 0.06757 | 0.088512 | 0.045728 | -0.02008 | -0.05254 | -0.01691 | 0.111476 | -0.35179 | 0.053243 | 0.049731 | -0.00651 | -0.01751 | 0.075496 | 0.006167 | 0.04869 | -0.09125 | 0.08606 | -0.08116 | 0.005872 | 0.13818 |
| 24.375 | 0.205004 | 0.028801 | 0.07023 | -0.06101 | -0.05317 | 0.114423 | -0.01488 | 0.148154 | 0.02349 | -0.36199 | 0.150552 | -0.32417 | 0.07265 | 0.072346 | 0.003178 | 0.119866 | 0.044495 | 0.022413 | 0.061037 | 0.000336 | 0.16082 | -0.00716 | 0.023605 | 0.2309 |
| 24.38889 | 0.150198 | -0.01504 | 0.148089 | -0.06642 | -0.07465 | 0.234742 | 0.19287 | 0.47891 | 0.276555 | -0.36214 | 0.375208 | 0.008545 | 0.097082 | 0.111495 | 0.009273 | 0.314237 | 0.048074 | 0.027738 | 0.085309 | 0.127793 | 0.248391 | 0.112951 | 0.115952 | 0.337508 |
| 24.40278 | 0.286105 | -0.03244 | 0.26861 | 0.100725 | 0.042154 | 0.421472 | 0.553064 | 0.925251 | 0.406993 | 0.087818 | 0.615837 | 0.376244 | 0.120721 | 0.175337 | 0.021725 | 0.506403 | 0.059376 | 0.036913 | 0.149189 | 0.223069 | 0.299368 | 0.252532 | 0.250981 | 0.398237 |
| 24.41667 | 0.581694 | 0.034971 | 0.381507 | 0.30242 | 0.201588 | 0.591095 | 0.907673 | 1.140836 | 0.328428 | 0.825847 | 0.685818 | 0.571379 | 0.128214 | 0.226889 | 0.037374 | 0.62783 | 0.05754 | 0.057296 | 0.214237 | 0.241805 | 0.252013 | 0.342727 | 0.317472 | 0.362331 |
| 24.43056 | 0.910576 | 0.115695 | 0.461494 | 0.366284 | 0.295968 | 0.684926 | 0.947336 | 0.952872 | 0.203835 | 1.540299 | 0.644673 | 0.580966 | 0.104775 | 0.219174 | 0.038275 | 0.474887 | 0.05908 | 0.066129 | 0.200848 | 0.187127 | 0.167937 | 0.323238 | 0.239315 | 0.244948 |
| 24.44444 | 1.11944 | 0.162312 | 0.466822 | 0.318323 | 0.209992 | 0.615639 | 0.730788 | 0.645118 | 0.157403 | 1.960063 | 0.547758 | 0.595417 | 0.091406 | 0.193 | 0.027567 | 0.202174 | 0.057155 | 0.072782 | 0.138291 | 0.143512 | 0.124439 | 0.223571 | 0.139799 | 0.166504 |
| 24.45833 | 1.175742 | 0.181536 | 0.443803 | 0.228614 | 0.050476 | 0.490124 | 0.468735 | 0.289693 | 0.241926 | 2.144347 | 0.424595 | 0.664367 | 0.124222 | 0.207722 | 0.011058 | 0.188654 | 0.088216 | 0.092707 | 0.135952 | 0.176031 | 0.177384 | 0.127776 | 0.129838 | 0.225631 |
| 24.47222 | 1.046648 | 0.157094 | 0.363607 | 0.132276 | -0.11843 | 0.441705 | 0.251609 | 0.021923 | 0.434189 | 2.078679 | 0.325221 | 0.703199 | 0.184303 | 0.251095 | -0.00842 | 0.32885 | 0.149609 | 0.113543 | 0.166593 | 0.273228 | 0.278346 | 0.054735 | 0.210956 | 0.328073 |
| 24.48611 | 0.80278 | 0.130718 | 0.332376 | 0.114989 | -0.14826 | 0.478697 | 0.214588 | -0.06024 | 0.655251 | 1.583245 | 0.311437 | 0.727736 | 0.230671 | 0.295872 | 0.024586 | 0.474883 | 0.212644 | 0.144376 | 0.199397 | 0.404675 | 0.355309 | 0.027332 | 0.355465 | 0.365892 |
| 24.5 | 0.829656 | 0.272046 | 0.613608 | 0.328788 | 0.006955 | 0.811642 | 0.548276 | 0.075514 | 1.03527 | 1.191698 | 0.536885 | 0.795932 | 0.29988 | 0.461923 | 0.141786 | 0.729427 | 0.271547 | 0.218597 | 0.272544 | 0.580586 | 0.370449 | 0.093529 | 0.556162 | 0.362994 |
| 24.51389 | 1.432688 | 0.628938 | 1.575907 | 0.613628 | 0.250351 | 1.535203 | 1.224081 | 0.338795 | 1.697574 | 1.71851 | 1.067748 | 1.324256 | 0.512714 | 0.908613 | 0.296909 | 1.123588 | 0.393393 | 0.395362 | 0.377891 | 1.019957 | 0.411445 | 0.295786 | 0.954995 | 0.472689 |
| 24.52778 | 1.908404 | 0.862623 | 2.115819 | 0.738864 | 0.379969 | 2.020338 | 1.708021 | 0.742866 | 1.833807 | 3.196292 | 1.621826 | 1.828036 | 0.830079 | 1.36332 | 0.396718 | 1.072197 | 0.549743 | 0.696352 | 0.422947 | 1.90073 | 0.564696 | 0.553814 | 1.500439 | 0.85829 |
| 24.54167 | 2.447802 | 0.592291 | 1.53395 | 0.639403 | 0.443893 | 1.529465 | 1.685207 | 0.846346 | 1.409967 | 4.837334 | 1.788548 | 1.6739 | 1.02689 | 1.358911 | 0.245324 | 0.60393 | 0.664292 | 0.681922 | 0.368288 | 2.425431 | 0.693286 | 0.82814 | 1.57832 | 1.043242 |
| 24.55556 | 3.476647 | 0.224421 | 1.135528 | 0.608373 | 0.353663 | 1.130986 | 1.060419 | 0.875565 | 2.601795 | 4.896615 | 1.496559 | 1.160962 | 0.954213 | 1.221796 | 0.097026 | 0.107648 | 0.480711 | 0.479889 | 0.292456 | 1.939485 | 0.675395 | 0.775472 | 0.847438 | 0.658804 |
| 24.56944 | 2.626234 | 0.335156 | 1.236786 | 0.578887 | 0.21582 | 0.845992 | 0.288682 | 0.969099 | 3.275782 | 4.194284 | 0.908786 | 0.815821 | 0.664349 | 1.066821 | 0.144731 | 0.072643 | 0.439214 | 0.330271 | 0.210721 | 1.089841 | 0.475574 | 0.757569 | 0.395505 | 0.275975 |
| 24.58333 | 1.129779 | 0.489323 | 1.392978 | 0.497652 | 0.000737 | 0.629134 | -0.48443 | 0.691912 | 2.263445 | 2.311646 | 0.227031 | 0.476822 | 0.387921 | 0.800091 | 0.186829 | 0.040989 | 0.356603 | 0.331763 | 0.062849 | 0.476246 | 0.255366 | 0.632936 | 0.021712 | 0.084773 |
| 24.59722 | 0.1721 | 0.494769 | 1.420584 | 0.425593 | -0.13019 | 0.41693 | -1.14066 | 0.257917 | 1.172731 | 0.225637 | -0.2837 | 0.165281 | 0.221504 | 0.550888 | 0.177533 | 0.012036 | 0.137656 | 0.380296 | -0.05529 | -0.07078 | 0.108415 | 0.326381 | -0.32337 | 0.180917 |
| 24.61111 | -0.57238 | 0.415783 | 1.036634 | 0.504726 | -0.05468 | 0.309938 | -1.24679 | 0.092954 | 0.321434 | -1.66067 | -0.09865 | -0.15412 | 0.304095 | 0.383802 | 0.173982 | 0.063214 | 0.029305 | 0.311029 | -0.03092 | -0.46307 | 0.019414 | -0.02452 | -0.10458 | 0.363861 |
| 24.625 | -1.18159 | 0.384572 | 0.901898 | 0.870106 | 0.019259 | 0.220087 | -1.08384 | -0.0378 | 1.221304 | -2.5405 | 0.462426 | -0.3109 | 0.411065 | 0.30721 | 0.200811 | 0.146658 | 0.093911 | 0.227996 | 0.070977 | -0.4029 | 0.074631 | -0.37018 | 0.46474 | 0.41521 |
| 24.63889 | -1.44971 | 0.341427 | 0.904389 | 0.848817 | 0.360034 | 0.070674 | -0.36761 | 0.047711 | 1.634529 | -3.51677 | 0.949781 | -0.34225 | 0.366118 | 0.294882 | 0.217039 | 0.163095 | 0.313628 | 0.12442 | 0.154341 | -0.02054 | 0.376299 | -0.69728 | 0.85156 | 0.302404 |
| 24.65278 | -1.31568 | 0.328403 | 0.757464 | 0.638584 | 0.364945 | -0.05237 | 0.469872 | -0.05534 | 1.4649 | -3.79609 | 0.876803 | 0.081065 | 0.210823 | 0.262077 | 0.187223 | 0.077668 | 0.496139 | -0.00177 | 0.200734 | 0.403742 | 0.716651 | -1.01954 | 0.506615 | 0.157867 |
| 24.66667 | -1.21582 | 0.34468 | 0.611796 | 0.610112 | 0.251869 | -0.01256 | 1.425617 | -0.34869 | 1.190615 | -2.79969 | 0.732547 | 0.703497 | 0.108449 | 0.17849 | 0.161549 | -0.00246 | 0.440676 | -0.12095 | 0.169186 | 0.563938 | 0.812444 | -1.04145 | 0.14279 | 0.065503 |
| 24.68056 | -0.78745 | 0.448049 | 0.541623 | 0.288933 | 0.110317 | 0.013963 | 1.603128 | -0.28939 | 0.901651 | 0.334633 | 0.582578 | 1.089992 | 0.191914 | 0.085336 | 0.176196 | -0.02712 | 0.242367 | -0.16458 | 0.137795 | 0.488629 | 0.567212 | -0.58486 | -0.15554 | -0.04162 |
| 24.69444 | 0.357096 | 0.617382 | 0.238805 | -0.12437 | -0.05913 | -0.09188 | 0.968163 | 0.140863 | 0.385158 | 3.844193 | 0.158393 | 1.019086 | 0.3648 | 0.081534 | 0.209904 | -0.06313 | 0.028373 | -0.06793 | 0.154042 | 0.527397 | 0.428204 | 0.257549 | -0.48957 | -0.1791 |
| 24.70833 | 1.122127 | 0.860986 | -0.11461 | -0.36435 | -0.25274 | 0.005113 | -0.0174 | 0.54777 | -0.03403 | 5.234736 | -0.41091 | 0.556872 | 0.48303 | 0.270925 | 0.220682 | -0.06899 | -0.06044 | 0.157349 | 0.196973 | 0.337589 | 0.232532 | 1.062771 | -0.44518 | -0.14704 |
| 24.72222 | 1.673456 | 1.224517 | -0.37744 | -0.47428 | -0.19551 | 0.063404 | -0.9696 | 0.630098 | 0.212724 | 4.108068 | -0.27165 | -0.02135 | 0.51923 | 0.624408 | 0.225815 | -0.00749 | 0.011436 | 0.342494 | 0.248013 | 0.044758 | 0.060671 | 1.698612 | -0.06798 | 0.058537 |
| 24.73611 | 1.954033 | 1.375502 | -0.56412 | -0.40892 | -0.04331 | 0.084264 | -1.69229 | 0.543262 | 1.483529 | 3.206125 | 0.129942 | -0.1314 | 0.50633 | 0.961773 | 0.214447 | 0.021604 | 0.228083 | 0.449048 | 0.208068 | -0.38582 | -0.39686 | 1.377813 | 0.370935 | 0.248992 |
| 24.75 | 1.94572 | 1.089924 | -0.64102 | -0.3147 | 0.158306 | 0.120892 | -1.69405 | 0.222462 | 1.004972 | 1.489718 | 0.381941 | -0.03294 | 0.375598 | 1.136052 | 0.217673 | -0.0006 | 0.494927 | 0.435861 | 0.150701 | -0.72942 | -1.08783 | 0.490592 | 0.65455 | 0.302843 |
| 24.76389 | 0.944422 | 0.643675 | -0.6481 | -0.3526 | 0.236753 | 0.075965 | -1.5445 | -0.01074 | -0.0003 | -0.48234 | 0.441229 | -0.29345 | 0.158722 | 1.035835 | 0.197933 | -0.02616 | 0.555305 | 0.339811 | 0.199391 | -0.47341 | -0.62169 | -0.30556 | 0.616084 | 0.259607 |
| 24.77778 | -0.14452 | 0.127044 | -0.66398 | -0.33052 | 0.075035 | 0.207737 | -0.84038 | -0.13976 | -0.93922 | -2.12416 | 0.301896 | -0.76775 | -0.0261 | 0.682566 | 0.161392 | -0.04831 | 0.370016 | 0.240291 | 0.187135 | 0.038047 | 0.150752 | -0.78866 | 0.326976 | 0.05573 |
| 24.79167 | -1.07017 | -0.19103 | -0.58441 | -0.44782 | -0.15726 | 0.222691 | -0.0302 | -0.26649 | -1.80303 | -3.81804 | -0.14419 | -1.18957 | -0.17616 | 0.312492 | 0.112351 | -0.09599 | 0.002071 | 0.23149 | 0.100871 | 0.492977 | 0.799212 | -1.00043 | -0.05617 | -0.22076 |
| 24.80556 | -1.6383 | -0.34231 | -0.52086 | -0.65813 | -0.42875 | 0.113415 | 0.496134 | -0.3771 | -1.54873 | -4.56839 | -0.54676 | -1.24517 | -0.28484 | -0.04875 | 0.052696 | -0.22469 | -0.3703 | 0.202899 | 0.005621 | 0.488845 | 0.85614 | -1.10035 | -0.42521 | -0.25135 |
| 24.81944 | -1.48516 | -0.45328 | -0.41856 | -0.88053 | -0.29482 | -0.01683 | 0.39317 | -0.4311 | -0.92324 | -4.60776 | -0.6818 | -0.78641 | -0.36803 | -0.47529 | 0.010323 | -0.24983 | -0.60411 | 0.124316 | -0.09936 | 0.168293 | 0.324099 | -1.15898 | -0.50077 | -0.04977 |
| 24.83333 | -1.12482 | -0.53453 | -0.32103 | -0.51926 | -0.03445 | -0.13087 | -0.03238 | -0.47514 | -0.69061 | -4.97824 | -0.26486 | -0.4044 | -0.41645 | -0.91781 | -0.00983 | -0.11567 | -0.77023 | -0.00873 | -0.20559 | -0.2062 | -0.27354 | -1.05282 | -0.29365 | 0.06706 |
| 24.84722 | -1.04756 | -0.54211 | -0.31496 | -0.24339 | 0.260879 | -0.22117 | -0.51931 | -0.40375 | -0.70978 | -4.50008 | 0.017338 | 0.171555 | -0.46188 | -1.21915 | -0.0273 | -0.08049 | -0.4683 | -0.15259 | -0.29669 | -0.61824 | -1.21289 | -0.26252 | 0.083598 | -0.03086 |
| 24.86111 | -1.29526 | -0.51248 | -0.32411 | -0.50993 | 0.304434 | -0.29874 | -1.00342 | -0.21673 | -0.58329 | -3.4025 | 0.042704 | 0.779335 | -0.59032 | -1.31053 | -0.06207 | -0.02029 | -0.17501 | -0.25655 | -0.16474 | -0.95595 | -2.17245 | 0.559484 | 0.222928 | -0.2635 |
| 24.875 | -1.9349 | -0.53362 | -0.48821 | -1.03567 | 0.134166 | -0.34211 | -1.35788 | -0.0429 | -0.44098 | -2.09339 | -0.0693 | 1.020318 | -0.70849 | -1.2696 | -0.11143 | 0.031494 | -0.02719 | -0.36351 | -0.00961 | -0.6545 | -1.64137 | 0.999643 | 0.022968 | -0.57975 |
| 24.88889 | -2.26147 | -0.64037 | -0.7858 | -1.4004 | -0.00724 | -0.33321 | -1.18948 | -0.13555 | -0.41693 | -0.97653 | -0.3153 | 0.288205 | -0.6402 | -1.11483 | -0.17383 | -0.01933 | -0.13695 | -0.49632 | 0.073027 | -0.2946 | -0.66119 | 0.761482 | -0.31067 | -0.9571 |
| 24.90278 | -2.34125 | -0.74188 | -1.17305 | -1.5399 | -0.09101 | -0.38646 | -0.96114 | -0.42327 | -0.61241 | -0.26553 | -0.58921 | -0.69515 | -0.45634 | -0.99365 | -0.24733 | -0.34868 | -0.36072 | -0.54223 | 0.012561 | -0.1911 | -0.3548 | 0.134654 | -0.57775 | -1.22033 |
| 24.91667 | -2.41665 | -0.76414 | -1.17349 | -1.30244 | -0.0592 | -0.42084 | -0.493 | -0.5387 | -0.90338 | -0.01694 | -0.84601 | -1.46272 | -0.38697 | -0.87426 | -0.27135 | -0.25373 | -0.59512 | -0.42233 | -0.14813 | -0.337 | -0.51681 | -0.44538 | -0.63377 | -1.11431 |
| 24.93056 | -2.07111 | -0.73252 | -1.04327 | -1.1034 | 0.006474 | -0.51782 | -0.0855 | -0.65342 | -1.03267 | 0.040103 | -1.13458 | -1.6549 | -0.47458 | -0.72226 | -0.23441 | 0.040875 | -0.64824 | -0.30048 | -0.27738 | -0.52647 | -0.78446 | -0.67292 | -0.38781 | -0.62459 |
| 24.94444 | -1.33899 | -0.5628 | -0.89321 | -0.4171 | 0.005954 | -0.58558 | 0.172743 | -1.00669 | -0.96531 | -0.01784 | -1.19008 | -1.75537 | -0.59972 | -0.55776 | -0.20896 | 0.185476 | -0.48497 | -0.25742 | -0.34906 | -0.73499 | -1.15562 | -0.77502 | -0.07993 | -0.04604 |
| 24.95833 | -0.35434 | -0.24857 | -0.70923 | -0.11004 | -0.03126 | -0.64699 | 0.227577 | -1.49499 | -0.79259 | -0.30717 | -0.71989 | -1.74397 | -0.72458 | -0.49888 | -0.14453 | 0.029427 | -0.26016 | -0.31121 | -0.40248 | -0.7314 | -1.21844 | -0.75126 | 0.077368 | 0.23721 |
| 24.97222 | 0.330334 | -0.08788 | -0.53758 | -0.16543 | -0.05034 | -0.63608 | 0.072159 | -1.54674 | -0.54819 | -0.87069 | -0.10879 | -1.44263 | -0.60557 | -0.50828 | -0.07059 | -0.13887 | 0.021763 | -0.36461 | -0.32153 | -0.45315 | -0.63497 | -0.61101 | 0.021471 | 0.181999 |
| 24.98611 | 0.600913 | 0.069605 | -0.54083 | -0.21275 | -0.01284 | -0.44049 | -0.16254 | -1.15261 | -0.29776 | -1.4637 | 0.424094 | -0.73962 | -0.34411 | -0.51689 | -0.00597 | -0.17877 | 0.152981 | -0.39126 | -0.20991 | -0.20075 | 0.057818 | -0.30075 | -0.09311 | 0.071821 |
| 25 | 0.375999 | 0.138864 | -0.60295 | -0.30925 | -0.00564 | -0.23664 | -0.44568 | -0.67572 | -0.13428 | -1.79843 | 0.498789 | 0.128108 | -0.19608 | -0.52826 | 0.066271 | -0.23135 | 0.073536 | -0.3162 | -0.04545 | -0.13029 | 0.205764 | 0.148131 | -0.32498 | -0.00525 |
| 25.01389 | -0.35369 | -0.01412 | -0.81471 | -0.54035 | -0.12501 | -0.32648 | -0.80542 | -0.4743 | -0.26674 | -1.92232 | -0.15861 | 0.386074 | -0.37178 | -0.54682 | 0.094448 | -0.51917 | -0.21854 | -0.23709 | -0.01551 | -0.37174 | -0.2104 | 0.299367 | -0.75126 | -0.26731 |
| 25.02778 | -1.24676 | -0.20547 | -1.04493 | -0.70314 | -0.25792 | -0.64394 | -1.21936 | -0.50525 | -0.66045 | -1.97517 | -1.07991 | -0.1674 | -0.78616 | -0.67967 | 0.026662 | -1.01051 | -0.38484 | -0.23453 | -0.18516 | -0.96113 | -0.85432 | 0.078871 | -1.35342 | -0.82517 |
| 25.04167 | -2.08346 | -0.42026 | -1.13601 | -0.78951 | -0.14403 | -1.05558 | -1.57646 | -0.695 | -1.30521 | -2.04951 | -1.87224 | -1.00411 | -1.17517 | -1.08628 | -0.09742 | -1.33963 | -0.56815 | -0.25464 | -0.37347 | -1.1691 | -1.48102 | -0.19385 | -1.87176 | -1.25688 |
| 25.05556 | -2.96992 | -0.70648 | -1.08391 | -1.04715 | -0.02579 | -1.30278 | -1.80131 | -1.03762 | -0.6013 | -1.79675 | -2.23945 | -1.7626 | -1.26842 | -1.84921 | -0.20163 | -0.7272 | -1.04399 | -0.19141 | -0.43221 | -0.87476 | -1.64477 | -0.49994 | -1.45661 | -0.86315 |
| 25.06944 | -2.75665 | -0.95684 | -1.04529 | -0.90321 | -0.00587 | -1.18946 | -1.3154 | -0.97435 | 0.391141 | -1.29863 | -1.98408 | -1.81672 | -0.97345 | -2.41957 | -0.20914 | 0.412431 | -0.401 | -0.16311 | -0.28184 | -0.38353 | -0.64103 | -0.54996 | -0.3777 | -0.16677 |
| 25.08333 | -1.97804 | -0.67824 | -0.76487 | -0.41425 | 0.042428 | -0.84124 | -0.38877 | -0.28642 | 1.214484 | -0.50611 | -1.35914 | -1.49089 | -0.33817 | -1.95679 | -0.19564 | 0.867547 | 0.728116 | -0.18888 | 0.028713 | 0.044964 | 0.930305 | 0.030222 | 0.384754 | 0.394202 |
| 25.09722 | -0.75289 | -0.21603 | -0.38503 | 0.0427 | 0.100098 | -0.37993 | 0.291377 | -0.02437 | 1.470995 | 0.283464 | -0.62732 | -0.85585 | 0.235521 | -0.89863 | -0.13507 | 0.612302 | 1.272285 | -0.12398 | 0.247407 | 0.423557 | 1.939316 | 0.64383 | 0.477464 | 0.887844 |
| 25.11111 | 0.191818 | 0.119038 | -0.06266 | 0.330739 | 0.208925 | -0.04911 | 0.640376 | -0.01974 | 1.061463 | 0.829093 | 0.376908 | -0.25249 | 0.312498 | -0.01631 | -0.04349 | 0.338266 | 0.971036 | -0.01269 | 0.203206 | 0.710779 | 2.277023 | 0.885011 | 0.452709 | 0.879213 |
| 25.125 | 0.713628 | 0.226093 | 0.12641 | 0.478193 | 0.337856 | 0.120783 | 0.635145 | -0.03584 | 0.425162 | 0.954557 | 1.004614 | 0.136682 | 0.193368 | 0.314264 | 0.037253 | 0.161412 | 0.527012 | 0.048833 | 0.072039 | 0.783898 | 1.983223 | 0.853664 | 0.296445 | 0.389266 |
| 25.13889 | 0.785638 | 0.157789 | 0.189211 | 0.392731 | 0.275459 | 0.177269 | 0.485138 | -0.06392 | -0.02157 | 0.652892 | 0.993842 | 0.305042 | 0.133452 | 0.277081 | 0.064434 | -0.05438 | 0.190771 | 0.047501 | -0.02773 | 0.415071 | 1.142406 | 0.691375 | 0.045116 | -0.12317 |
| 25.15278 | 0.589656 | 0.085378 | 0.133145 | 0.211607 | 0.128072 | 0.124611 | 0.344144 | -0.0608 | -0.26703 | 0.214394 | 0.644944 | 0.31538 | 0.120475 | 0.127111 | 0.044333 | -0.2585 | -0.02214 | 0.035986 | -0.06869 | 0.033682 | 0.416584 | 0.510812 | -0.13072 | -0.4743 |
| 25.16667 | 0.306388 | 0.049697 | 0.11069 | 0.018562 | -0.06029 | 0.036065 | 0.285198 | -0.02499 | -0.38124 | 0.002557 | 0.425582 | 0.287483 | 0.05382 | 0.029201 | 0.007658 | -0.33569 | -0.08869 | 0.036333 | -0.08537 | -0.26289 | -0.21441 | 0.335127 | -0.22697 | -0.52655 |
| 25.18056 | 0.048555 | 0.048122 | 0.094891 | -0.14463 | -0.23942 | -0.04612 | 0.253256 | 0.025616 | -0.29265 | 0.006018 | 0.416888 | 0.323346 | -0.04677 | -0.01485 | -0.02035 | -0.25691 | -0.06424 | 0.023301 | -0.08487 | -0.51659 | -0.73598 | 0.167047 | -0.2712 | -0.34689 |
| 25.19444 | -0.04622 | 0.064165 | 0.095227 | -0.184 | -0.27619 | -0.0683 | 0.212629 | 0.098861 | 0.033885 | 0.08528 | 0.550781 | 0.481198 | -0.10736 | -0.01971 | -0.03967 | -0.10999 | -0.03669 | 0.02025 | -0.05904 | -0.48357 | -0.93624 | 0.021113 | -0.22153 | -0.00993 |
| 25.20833 | 0.14555 | 0.066187 | 0.107924 | -0.08528 | -0.04622 | -0.02018 | 0.182896 | 0.236259 | 0.264619 | 0.190357 | 0.66702 | 0.670544 | -0.10106 | 0.009 | -0.0459 | -0.00645 | -0.06653 | 0.042703 | 0.000184 | -0.1903 | -0.38157 | -0.16853 | -0.12021 | 0.276038 |
| 25.22222 | 0.463545 | 0.05967 | 0.118768 | 0.014952 | 0.201215 | 0.101505 | 0.112232 | 0.343786 | 0.260123 | 0.258867 | 0.658504 | 0.901106 | -0.05201 | 0.068849 | -0.03878 | 0.009868 | -0.0995 | 0.073433 | 0.056005 | 0.067573 | 0.268505 | -0.31809 | 0.003216 | 0.403658 |
| 25.23611 | 0.677666 | 0.048414 | 0.099456 | -0.00599 | 0.199734 | 0.233756 | 0.012797 | 0.421793 | 0.171649 | 0.255916 | 0.615497 | 1.139229 | 0.01396 | 0.137728 | -0.03637 | -0.03174 | -0.0986 | 0.090199 | 0.047383 | 0.144397 | 0.380762 | -0.36406 | 0.114243 | 0.283952 |
| 25.25 | 0.717813 | 0.047007 | 0.063925 | -0.14098 | 0.023723 | 0.284047 | -0.09894 | 0.547396 | 0.115564 | 0.254751 | 0.523887 | 1.303506 | 0.05798 | 0.188918 | -0.03177 | -0.07808 | -0.07432 | 0.123974 | 0.011791 | 0.014037 | 0.083216 | -0.32608 | 0.237437 | 0.067222 |
| 25.26389 | 0.553508 | 0.024557 | 0.024575 | -0.27249 | -0.16949 | 0.16422 | -0.22721 | 0.689188 | 0.10308 | 0.251753 | 0.451612 | 1.366587 | 0.085709 | 0.22566 | -0.02916 | -0.0985 | -0.05441 | 0.133328 | -0.0149 | -0.13139 | -0.22217 | -0.28942 | 0.261027 | -0.06971 |
| 25.27778 | 0.334508 | -0.00624 | 0.021469 | -0.27257 | -0.19754 | 0.038134 | -0.30147 | 0.681752 | 0.184254 | 0.197434 | 0.380418 | 1.236062 | 0.133401 | 0.282202 | -0.03582 | -0.07508 | -0.0362 | 0.109788 | -0.01357 | -0.14273 | -0.27523 | -0.27922 | 0.144584 | 0.002386 |
| 25.29167 | 0.138654 | -0.03186 | 0.03574 | -0.15718 | 0.054037 | -0.05404 | -0.23139 | 0.596865 | 0.2887 | 0.155114 | 0.330488 | 0.937655 | 0.226113 | 0.372678 | -0.0401 | -0.01513 | -0.01097 | 0.077049 | 0.035238 | -0.03433 | -0.07963 | -0.17043 | -0.01965 | 0.315475 |
| 25.30556 | -0.0355 | -0.06171 | 0.035833 | 0.055233 | 0.257982 | -0.02161 | -0.09709 | 0.459466 | 0.223343 | 0.144403 | 0.316399 | 0.711315 | 0.31422 | 0.455085 | -0.05797 | 0.031305 | -0.01853 | 0.047762 | 0.084451 | 0.068049 | 0.068431 | -0.05567 | -0.10228 | 0.474504 |
| 25.31944 | -0.19351 | -0.0698 | 0.037038 | 0.211746 | 0.241472 | 0.08567 | 0.014572 | 0.479371 | 0.209237 | 0.182195 | 0.278088 | 0.493556 | 0.341703 | 0.511144 | -0.07092 | 0.053065 | -0.04132 | 0.032814 | 0.077471 | 0.090329 | 0.022332 | -0.00516 | -0.05034 | 0.351938 |
| 25.33333 | -0.30318 | -0.06641 | 0.042864 | 0.309714 | 0.13919 | 0.242831 | 0.134538 | 0.543234 | 0.401141 | 0.228486 | 0.229582 | 0.293924 | 0.281636 | 0.542408 | -0.05786 | 0.066614 | -0.03552 | 0.040912 | 0.066457 | 0.084279 | -0.05383 | 0.027084 | -0.04122 | 0.127158 |
| 25.34722 | -0.33853 | -0.07332 | 0.063833 | 0.362752 | 0.093171 | 0.366802 | 0.209566 | 0.622552 | 0.56061 | 0.218638 | 0.222823 | 0.071604 | 0.207419 | 0.515965 | -0.02803 | 0.048219 | 0.04921 | 0.056011 | 0.079158 | 0.115953 | 0.050677 | 0.021093 | 0.017872 | 0.03042 |
| 25.36111 | -0.30513 | -0.05676 | 0.093536 | 0.3504 | 0.120109 | 0.439542 | 0.241384 | 0.346134 | 0.447739 | 0.183054 | 0.219107 | -0.19711 | 0.236646 | 0.469086 | -0.00497 | 0.038131 | 0.251214 | 0.070224 | 0.074587 | 0.221163 | 0.422849 | 0.018287 | 0.123004 | 0.156288 |
| 25.375 | -0.24896 | -0.04889 | 0.150014 | 0.325203 | 0.175608 | 0.413385 | 0.277726 | 0.074446 | 0.202299 | 0.158832 | 0.243858 | -0.35905 | 0.390858 | 0.481841 | 0.020677 | 0.060825 | 0.418712 | 0.08872 | 0.058115 | 0.392063 | 0.93075 | 0.096983 | 0.197058 | 0.434103 |
| 25.38889 | -0.29627 | -0.07609 | 0.214924 | 0.300828 | 0.198498 | 0.281993 | 0.369024 | -0.03944 | 0.050407 | 0.145552 | 0.293578 | -0.31581 | 0.519365 | 0.522259 | 0.042648 | 0.086128 | 0.370709 | 0.116734 | 0.040167 | 0.508431 | 1.217894 | 0.225343 | 0.227012 | 0.697126 |
| 25.40278 | -0.28985 | -0.08525 | 0.254249 | 0.284907 | 0.188396 | 0.174561 | 0.453085 | -0.00877 | 0.144152 | 0.233799 | 0.398659 | -0.11839 | 0.570803 | 0.568489 | 0.095808 | 0.119212 | 0.215259 | 0.157454 | 0.051865 | 0.456098 | 1.021872 | 0.306806 | 0.292871 | 0.771931 |
| 25.41667 | -0.17151 | -0.0421 | 0.275466 | 0.356855 | 0.19059 | 0.267549 | 0.557647 | 0.106569 | 0.34779 | 0.462761 | 0.569551 | 0.144152 | 0.556475 | 0.564672 | 0.133359 | 0.159776 | 0.141174 | 0.211059 | 0.085066 | 0.339085 | 0.58684 | 0.274631 | 0.47533 | 0.666109 |

| | | | | | | | | | | | | | | | | | | | | | | | | |
|---|---|---|---|---|---|---|---|---|---|---|---|---|---|---|---|---|---|---|---|---|---|---|---|---|
| 25.43056 | 0.080509 | 0.043176 | 0.284633 | 0.485702 | 0.184508 | 0.435223 | 0.540415 | 0.371534 | 0.62005 | 0.570154 | 0.586653 | 0.342816 | 0.510076 | 0.486658 | 0.158096 | 0.190566 | 0.173674 | 0.292976 | 0.110066 | 0.285349 | 0.364501 | 0.207164 | 0.635547 | 0.468115 |
| 25.44444 | 0.446744 | 0.09724 | 0.296077 | 0.607557 | 0.146063 | 0.601096 | 0.43714 | 0.515871 | 0.639638 | 0.47992 | 0.409161 | 0.449644 | 0.465678 | 0.361278 | 0.123694 | 0.190572 | 0.255041 | 0.271544 | 0.096512 | 0.382257 | 0.512605 | 0.161099 | 0.543074 | 0.334683 |
| 25.45833 | 0.748191 | 0.096338 | 0.313559 | 0.64893 | 0.133854 | 0.536174 | 0.370992 | 0.561413 | 0.464982 | 0.467171 | 0.225433 | 0.473049 | 0.523817 | 0.291854 | 0.06547 | 0.215635 | 0.300235 | 0.204197 | 0.106495 | 0.604155 | 0.918461 | 0.191324 | 0.230857 | 0.302599 |
| 25.47222 | 0.897376 | 0.072734 | 0.286293 | 0.59084 | 0.135925 | 0.644577 | 0.355678 | 0.542738 | 0.143544 | 0.517036 | 0.215981 | 0.578932 | 0.612991 | 0.334596 | 0.017087 | 0.268688 | 0.257542 | 0.169919 | 0.166297 | 0.734353 | 1.117301 | 0.381101 | -0.04416 | 0.359925 |
| 25.48611 | 0.880958 | 0.077973 | 0.263633 | 0.559754 | 0.171292 | 0.649395 | 0.442576 | 0.371694 | -0.57739 | 0.663018 | 0.410006 | 0.766326 | 0.700449 | 0.4687 | 0.048488 | 0.288016 | 0.218905 | 0.223431 | 0.228755 | 0.702174 | 0.753239 | 0.681009 | -0.16078 | 0.457836 |
| 25.5 | 1.100002 | 0.169285 | 0.331267 | 0.756253 | 0.278997 | 0.613779 | 0.727837 | 0.256723 | -1.85095 | 1.246838 | 0.743676 | 1.034611 | 0.729126 | 0.749142 | 0.180453 | 0.442907 | 0.169128 | 0.375801 | 0.262001 | 0.707308 | 0.1845 | 0.637598 | 0.02409 | 0.526565 |
| 25.51389 | 1.769752 | 0.364935 | 0.567594 | 1.299043 | 0.448711 | 1.216781 | 1.385648 | 0.062509 | -0.54056 | 2.500315 | 1.387962 | 1.762072 | 0.936207 | 1.243933 | 0.386392 | 0.849614 | 0.164699 | 0.63583 | 0.228408 | 1.317169 | 0.004748 | 0.617656 | 0.57705 | 0.668075 |
| 25.52778 | 2.69387 | 0.476934 | 0.751652 | 1.599321 | 0.493259 | 2.011729 | 1.760657 | -0.2294 | 1.389712 | 3.210859 | 2.145499 | 2.331864 | 1.415214 | 1.842689 | 0.40794 | 1.08549 | 0.33521 | 1.096793 | 0.238899 | 1.8552 | 0.618361 | 1.305283 | 1.360227 | 1.274758 |
| 25.54167 | 3.163647 | 0.374466 | 0.789483 | 1.182015 | 0.589617 | 1.560661 | 1.607753 | -0.03056 | 1.502404 | 3.012042 | 2.380005 | 2.150476 | 1.712732 | 2.31347 | 0.2534 | 0.711798 | 0.62231 | 1.298354 | 0.124167 | 1.86035 | 1.499054 | 2.307545 | 1.739584 | 1.41698 |
| 25.55556 | 3.743152 | 0.288304 | 0.564324 | 0.882489 | 0.495456 | 1.170844 | 1.345945 | 0.217767 | 1.696757 | 3.342059 | 1.909686 | 1.47914 | 1.511073 | 2.469943 | 0.340585 | 0.273013 | 0.731005 | 1.074651 | 0.154024 | 1.868559 | 1.536135 | 2.629492 | 1.643325 | 1.170708 |
| 25.56944 | 4.560006 | 0.219396 | 0.63888 | 0.775586 | 0.324971 | 1.541396 | 0.795675 | 0.409015 | 3.087447 | 3.585485 | 1.16603 | 1.085045 | 1.137853 | 2.665076 | 0.426401 | 0.136195 | 0.725093 | 0.836293 | 0.177562 | 1.34252 | 1.402643 | 1.700054 | 1.31418 | 0.778058 |
| 25.58333 | 3.53513 | 0.149225 | 0.471395 | 0.574113 | 0.161793 | 0.907005 | 0.275248 | 0.388974 | 3.496019 | 3.49411 | 0.438113 | 0.81811 | 0.640592 | 2.817351 | 0.360353 | 0.079866 | 0.702532 | 0.445202 | 0.104443 | 0.476394 | 0.605678 | 0.45741 | 0.739191 | 0.110666 |
| 25.59722 | 2.218831 | 0.127148 | 0.250671 | 0.413287 | 0.016814 | 0.51654 | -0.13034 | 0.154213 | 2.26696 | 2.33968 | -0.47179 | 0.613844 | -0.01174 | 2.784027 | 0.181955 | -0.04404 | 0.138495 | 0.102892 | -0.03247 | -0.47686 | -0.33235 | -0.80058 | 0.175024 | -0.61568 |
| 25.61111 | 1.026638 | 0.09802 | 0.261407 | 0.18355 | -0.07263 | 0.320764 | -0.54206 | -0.17331 | 0.792653 | 0.697651 | -1.28521 | 0.339451 | -0.7047 | 2.142606 | 0.01694 | -0.12498 | -0.26583 | -0.08419 | -0.18149 | -1.43823 | -1.00923 | -1.95853 | -0.32899 | -1.04295 |
| 25.625 | -0.17455 | 0.057892 | 0.586089 | -0.44162 | -0.01792 | 0.336822 | -0.8873 | -0.33831 | -0.78346 | -0.86904 | -1.14699 | -0.16805 | -0.79937 | 0.938149 | -0.10638 | -0.10824 | -0.14145 | -0.02933 | -0.09668 | -1.26073 | -2.12257 | -2.89205 | -0.59425 | -0.6825 |
| 25.63889 | -0.62846 | -0.00297 | 0.70309 | -0.69026 | 0.072746 | 0.240118 | -0.89457 | -0.33572 | -0.99573 | -0.99017 | -0.16997 | -0.76572 | -0.48243 | -0.07424 | -0.1374 | -0.06135 | -0.0713 | 0.082167 | 0.10345 | -1.14612 | -2.52351 | -2.14555 | -0.14899 | 0.115256 |
| 25.65278 | -1.1518 | 0.018407 | 0.839555 | -0.56447 | 0.288589 | -0.01621 | -0.38874 | -0.35522 | 0.50602 | -1.4714 | 1.084007 | -1.08306 | -0.01476 | -0.68321 | -0.27803 | -0.08831 | -0.0149 | 0.06142 | 0.292362 | -0.28016 | -0.62877 | -1.27478 | 0.398882 | 0.657791 |
| 25.66667 | -1.4505 | 0.114917 | 0.749678 | -0.08117 | 0.522568 | -0.17657 | 0.399441 | -0.5898 | 0.752795 | -2.37851 | 1.87061 | -0.94945 | 0.67596 | -1.33674 | -0.2486 | -0.17086 | 0.315586 | 0.130085 | 0.350856 | 0.369992 | 1.32454 | 0.128867 | 0.680828 | 0.774656 |
| 25.68056 | -2.01066 | 0.117629 | 0.680889 | 0.182877 | 0.558484 | -0.25115 | 1.253113 | -0.88272 | 0.446267 | -1.81931 | 1.331404 | -0.25862 | 1.073986 | -2.02931 | 0.039558 | -0.10324 | 0.56981 | -0.07078 | 0.011963 | 1.076524 | 2.67095 | 1.314467 | 0.618366 | 0.439051 |
| 25.69444 | -2.48177 | 0.049516 | 0.76931 | 0.292522 | 0.433501 | -0.47105 | 1.19084 | -0.1906 | 0.20305 | -0.5776 | 0.422728 | 0.447369 | 1.004626 | -2.54604 | 0.478992 | 0.039502 | 0.82324 | -0.14909 | -0.10107 | 1.283984 | 2.482454 | 2.032341 | 0.335411 | -0.13009 |
| 25.70833 | -1.6911 | -0.05305 | 0.591354 | 0.391493 | 0.194465 | -0.47105 | 0.74761 | 0.419495 | 0.44639 | 0.466209 | -0.14463 | 0.834039 | 0.710805 | -2.83533 | 0.639908 | 0.101465 | 0.516645 | -0.15648 | -0.09224 | 1.898907 | 1.925256 | 3.174925 | 0.034563 | -0.49704 |
| 25.72222 | -0.3981 | -0.10263 | 0.365808 | 0.475192 | -0.04289 | -0.17197 | 0.150994 | 0.51542 | 0.85675 | 1.783605 | -0.52662 | 0.798394 | 0.23358 | -2.78275 | 0.533552 | 0.047272 | 0.010766 | -0.14126 | -0.01093 | 1.094675 | 0.732084 | 2.066851 | -0.14086 | -0.58322 |
| 25.73611 | -0.01963 | -0.03974 | 0.294315 | 0.520099 | -0.07003 | 0.189182 | -0.38313 | 0.464123 | 1.097209 | 2.99092 | -1.50731 | 0.600377 | -0.08004 | -2.14714 | 0.549169 | 0.012283 | -0.44024 | -0.15773 | -0.06004 | 0.063088 | -0.39154 | 0.666369 | -0.28227 | 0.059455 |
| 25.75 | 0.412682 | 0.022061 | 0.117601 | 0.400523 | 0.106487 | 0.401068 | -0.71525 | 0.868402 | 0.772991 | 3.181629 | -1.46832 | 0.251161 | 0.027641 | -0.94907 | 0.309355 | 0.090822 | -0.57522 | -0.26288 | -0.07913 | -0.87146 | -1.62616 | -0.62359 | -0.4281 | 0.740928 |
| 25.76389 | 0.678321 | 0.062602 | -0.14794 | 0.092418 | 0.236081 | 0.356477 | -0.92319 | 0.463573 | 0.365581 | 2.569468 | -0.63976 | -0.17782 | 0.229233 | -0.05167 | 0.020621 | 0.007938 | -0.45644 | -0.31311 | 0.031265 | -1.04176 | -2.67976 | -1.61938 | -0.2419 | 1.051542 |
| 25.77778 | 0.9759 | 0.011998 | -0.30744 | -0.327 | 0.240189 | 0.195363 | -1.01214 | 0.015109 | 0.153878 | 1.582295 | 0.282143 | -0.32754 | 0.069791 | 0.323968 | -0.24752 | -0.09226 | -0.15058 | -0.36861 | 0.082766 | -1.218 | -2.61546 | -2.06975 | 0.147647 | 1.010875 |
| 25.79167 | 1.181142 | -0.05458 | -0.46292 | -0.77653 | 0.008174 | -0.0274 | -1.04865 | -0.22817 | 0.138812 | -0.15885 | 0.964738 | -0.40844 | -0.47083 | 0.291353 | -0.36201 | -0.17638 | 0.180336 | -0.34509 | 0.087233 | -0.8383 | -2.1201 | -2.06947 | 0.518062 | 0.328107 |
| 25.80556 | 0.927865 | -0.15213 | -0.56095 | -0.98442 | -0.09629 | -0.14243 | -0.81282 | -0.22845 | 0.116864 | -1.81946 | 0.896456 | -0.52575 | -0.95203 | 0.097352 | -0.30959 | -0.26206 | 0.232635 | -0.31468 | 0.009096 | -0.3345 | -0.61163 | -1.83761 | 0.534659 | -0.5833 |
| 25.81944 | 0.224706 | -0.19515 | -0.58044 | -0.97088 | -0.18947 | 0.048271 | -0.58637 | -0.09468 | 0.019216 | -2.82426 | 0.274924 | -0.83776 | -1.33669 | -0.1582 | -0.1945 | -0.37392 | -0.0807 | -0.2682 | -0.04429 | -0.07885 | 0.829642 | -1.22656 | 0.441742 | -1.26667 |
| 25.83333 | -0.80676 | -0.18336 | -0.59242 | -0.8762 | -0.34276 | 0.238531 | -0.32467 | 0.01529 | 0.100781 | -3.38975 | -0.44521 | -1.19914 | -1.58987 | -0.49709 | -0.20413 | -0.47771 | -0.48796 | -0.23492 | -0.09212 | 0.159357 | 1.796655 | -0.73495 | 0.244445 | -1.28688 |
| 25.84722 | -1.82161 | -0.18714 | -0.4846 | -0.83364 | -0.22632 | 0.155765 | -0.13686 | 0.056714 | -0.08122 | -3.51743 | -0.909 | -1.55597 | -1.54862 | -0.79475 | -0.31741 | -0.51093 | -0.66073 | -0.32527 | -0.20157 | 0.350791 | 2.123005 | -0.39815 | -0.06318 | -1.0699 |
| 25.86111 | -2.18764 | -0.27568 | -0.28707 | -0.95869 | -0.02212 | -0.29606 | -0.09372 | -0.18026 | -0.29284 | -2.58726 | -1.11989 | -1.70702 | -1.34403 | -1.07506 | -0.37231 | -0.4781 | -0.63686 | -0.48508 | -0.40146 | 0.012863 | 1.188195 | -0.16286 | -0.45035 | -1.01041 |
| 25.875 | -1.99956 | -0.39535 | -0.27429 | -1.18213 | 0.240492 | -0.77558 | -0.09404 | -0.55744 | -0.80666 | -1.33248 | -1.00076 | -1.48723 | -0.99574 | -1.2243 | -0.31765 | -0.42726 | -0.48813 | -0.61895 | -0.54663 | -0.3127 | 0.108148 | 0.098355 | -0.8811 | -1.23704 |
| 25.88889 | -1.64613 | -0.40628 | -0.39171 | -1.15385 | 0.350982 | -1.18568 | -0.02137 | -1.01975 | -1.33812 | -0.64767 | -0.61082 | -1.01767 | -0.62714 | -1.04109 | -0.16321 | -0.31291 | -0.32163 | -0.54379 | -0.64095 | -0.56773 | -0.65561 | 0.4601 | -1.08682 | -1.4171 |
| 25.90278 | -1.4144 | -0.28376 | -0.57812 | -0.88506 | 0.495203 | -1.05788 | 0.065367 | -1.39049 | -1.81735 | -0.5676 | -0.2841 | -0.75886 | -0.43755 | -0.81149 | -0.02317 | -0.19032 | -0.23147 | -0.37792 | -0.58384 | -0.67361 | -1.01202 | 0.707609 | -1.00857 | -1.28299 |
| 25.91667 | -1.25575 | -0.13116 | -0.93458 | -0.87024 | 0.428529 | -1.06956 | -0.02911 | -0.95628 | -2.06152 | -0.77543 | -0.28192 | -0.66273 | -0.43149 | -0.67646 | 0.056469 | -0.15339 | -0.24227 | -0.24429 | -0.80127 | -0.86035 | -1.42525 | 0.700166 | -0.79405 | -0.94417 |
| 25.93056 | -1.4092 | -0.01261 | -1.10662 | -0.87893 | 0.239277 | -1.06011 | -0.29266 | -0.53526 | -1.76698 | -0.88467 | -0.66156 | -0.55281 | -0.46039 | -0.65871 | 0.067241 | -0.16046 | -0.20254 | -0.24365 | -0.67787 | -1.11202 | -2.26027 | 0.356771 | -0.55851 | -0.21466 |
| 25.94444 | -0.83665 | 0.070356 | -0.7314 | -0.81678 | 0.044693 | -0.22906 | -0.58394 | -0.33577 | -1.00636 | -0.99976 | -0.92605 | -0.31078 | -0.34957 | -0.60798 | 0.050216 | -0.15921 | -0.05434 | -0.24868 | -0.27476 | -1.23683 | -2.62116 | -0.13197 | -0.32013 | 0.311978 |
| 25.95833 | -0.36208 | 0.075881 | -0.38589 | -0.71017 | -0.0144 | 0.033476 | -0.7626 | -0.08153 | -0.33251 | -1.29678 | -0.95655 | 0.008769 | -0.09257 | -0.52442 | 0.005344 | -0.15071 | -0.09514 | -0.27547 | 0.085621 | -1.00368 | -2.40208 | -0.55167 | -0.1672 | 0.420133 |
| 25.97222 | -0.16936 | 0.054452 | -0.18195 | -0.54396 | 0.048302 | -0.4471 | -0.66153 | 0.057389 | -0.34508 | -1.21974 | -0.74383 | 0.327842 | 0.187403 | -0.43681 | -0.04407 | -0.13599 | -0.42115 | -0.28534 | 0.174705 | -0.76623 | -1.64255 | -0.64338 | -0.12787 | 0.219472 |
| 25.98611 | -0.10522 | 0.130348 | -0.0319 | -0.38227 | 0.085284 | -0.4914 | -0.27562 | -0.10509 | -0.55455 | -0.69354 | -0.34397 | 0.582114 | 0.318877 | -0.33174 | -0.06772 | -0.10069 | -0.73683 | -0.20083 | 0.127414 | -0.6392 | -0.92568 | -0.44049 | -0.17277 | 0.194192 |
| 26 | -0.28306 | 0.05897 | -0.16818 | -0.33701 | -0.11992 | -0.57257 | -0.00766 | -0.30329 | -1.10679 | -0.52732 | -0.18026 | 0.619383 | 0.216446 | -0.25288 | -0.10163 | -0.11691 | -0.81528 | -0.07279 | 0.027642 | -0.58251 | -0.62307 | -0.14695 | -0.3638 | 0.07849 |
| 26.01389 | -0.80657 | -0.11207 | -0.4798 | -0.53146 | -0.46315 | -0.66618 | -0.06089 | -0.35995 | -1.52923 | -0.68314 | -0.42336 | 0.314297 | -0.13553 | -0.17834 | -0.14968 | -0.30917 | -0.89232 | -0.02274 | -0.07921 | -0.5591 | -0.69959 | 0.098725 | -0.67883 | -0.20321 |
| 26.02778 | -1.60018 | -0.19719 | -0.70716 | -0.70629 | -0.69714 | -0.71266 | -0.16442 | -0.463 | -1.98231 | -0.69656 | -0.89239 | -0.11843 | -0.65792 | -0.05064 | -0.11381 | -0.59268 | -0.9388 | -0.11515 | -0.15345 | -0.45339 | -1.11412 | 0.255637 | -0.84603 | -0.5794 |
| 26.04167 | -2.54174 | -0.24264 | -0.58185 | -0.87053 | -0.73616 | -0.90636 | -0.34251 | -0.39189 | -2.32284 | -0.68843 | -0.92937 | -0.70703 | -1.28069 | -0.16368 | -0.05004 | -0.75445 | -0.92648 | -0.28216 | -0.32261 | -0.42275 | -1.73324 | 0.316866 | -0.84445 | -0.84074 |
| 26.05556 | -3.34794 | -0.52886 | -0.3201 | -1.23016 | -0.60236 | -1.37497 | -0.84934 | -0.17582 | -2.28016 | -0.81272 | -0.99793 | -1.50573 | -1.72655 | -0.78512 | -0.28028 | -0.76098 | -0.92683 | -0.47742 | -0.6461 | -0.628 | -2.73915 | 0.174765 | -0.82234 | -0.80465 |
| 26.06944 | -3.09449 | -0.84347 | -0.30452 | -1.35625 | -0.2554 | -1.47313 | -1.10778 | -0.11981 | -1.64592 | -0.84139 | -1.50721 | -1.92805 | -1.44112 | -1.43758 | -0.63398 | -0.59743 | -0.63203 | -0.59137 | -0.77992 | -0.80163 | -3.58428 | -0.01237 | -0.92597 | -0.57034 |
| 26.08333 | -1.85346 | -0.83304 | -0.38885 | -0.9043 | 0.252883 | -1.19617 | -0.71716 | -0.04698 | -1.31559 | -0.76244 | -1.31967 | -1.66792 | -0.5711 | -1.26046 | -0.47981 | -0.23639 | -0.10252 | -0.52807 | -0.49373 | -0.56657 | -3.15376 | -0.03451 | -0.85334 | -0.35168 |
| 26.09722 | -0.68153 | -0.58718 | -0.32472 | -0.25422 | 0.599483 | -0.65977 | -0.3033 | 0.074313 | -0.81874 | -0.927 | -0.06714 | -0.8343 | 0.281602 | -0.58417 | -0.28417 | 0.090978 | 0.439666 | -0.37702 | -0.09827 | -0.20523 | -1.47099 | 0.013252 | -0.54378 | 0.044173 |
| 26.11111 | 0.089505 | -0.31116 | -0.14723 | 0.199393 | 0.564847 | -0.0749 | -0.06383 | 0.259988 | -0.19032 | -1.52227 | 1.26444 | -0.30896 | 0.785241 | -0.09947 | -0.14136 | 0.126932 | 0.894329 | -0.11351 | 0.300475 | 0.022689 | 0.385698 | 0.178566 | -0.11206 | 0.537553 |
| 26.125 | 0.441443 | -0.05536 | 0.053385 | 0.395932 | 0.407928 | 0.259876 | 0.025043 | 0.250909 | 0.502658 | -1.11748 | 2.340541 | -0.02855 | 0.809262 | 0.045246 | -0.05938 | 0.033333 | 0.781646 | 0.175591 | 0.65709 | 0.083354 | 2.225354 | 0.619747 | 0.311692 | 0.989312 |
| 26.13889 | 0.526906 | 0.125541 | 0.204513 | 0.318539 | 0.262151 | 0.144795 | 0.119237 | 0.243442 | 0.905118 | -0.1103 | 2.230622 | 0.069215 | 0.654426 | -0.02928 | -0.0102 | -0.02665 | 0.428874 | 0.251265 | 0.73143 | 0.033625 | 3.590438 | 1.125443 | 0.684497 | 0.963301 |
| 26.15278 | 0.428397 | 0.199325 | 0.189687 | 0.231375 | 0.076131 | -0.16267 | 0.225518 | 0.107924 | 1.188124 | 0.338026 | 1.565494 | 0.106555 | 0.527836 | -0.08434 | 0.004638 | -0.03494 | 0.225204 | 0.148145 | 0.64504 | 0.025546 | 4.504057 | 1.481005 | 0.855375 | 0.744397 |
| 26.16667 | 0.211663 | 0.166736 | 0.043208 | 0.120149 | -0.0952 | -0.51787 | 0.271512 | 0.123356 | 1.087062 | 0.304247 | 0.687132 | 0.085056 | 0.423553 | -0.03881 | -0.00855 | -0.02805 | 0.108105 | -0.02605 | 0.50544 | 0.258433 | 4.269715 | 1.725725 | 0.5995 | 0.442497 |
| 26.18056 | 0.026806 | 0.121188 | -0.12037 | 0.017031 | -0.21852 | -0.74694 | 0.231459 | 0.038656 | 0.472681 | 0.239633 | -0.02865 | 0.038805 | 0.256275 | 0.087806 | -0.01201 | -0.02588 | 0.022146 | -0.19811 | 0.365656 | 0.360828 | 3.679318 | 1.629706 | 0.122672 | 0.011725 |
| 26.19444 | -0.07137 | 0.099431 | -0.1625 | -0.01794 | -0.21426 | -0.50496 | 0.185084 | -0.14859 | 0.065176 | 0.200586 | -0.30097 | 0.038188 | 0.075557 | 0.280992 | 0.006095 | -0.0027 | -0.00285 | -0.29393 | 0.255274 | 0.386385 | 2.857279 | 0.873444 | -0.17934 | -0.20302 |
| 26.20833 | -0.03403 | 0.087316 | -0.0572 | 0.085666 | -0.14355 | -0.18994 | 0.181233 | -0.21017 | -0.19799 | 0.135753 | -0.07806 | 0.093439 | -0.12219 | 0.416307 | 0.015635 | 0.029775 | 0.016226 | -0.20961 | 0.172254 | 0.296394 | 1.836622 | -0.37818 | -0.15498 | -0.01458 |
| 26.22222 | 0.078412 | 0.107296 | 0.112174 | 0.196206 | -0.02239 | 0.079907 | 0.196451 | -0.15953 | -0.31324 | 0.032928 | 0.431125 | 0.133688 | -0.31373 | 0.516239 | 0.006565 | 0.07164 | 0.055236 | -0.01108 | 0.082139 | 0.147585 | 0.973681 | -1.41732 | 0.023208 | 0.369495 |
| 26.23611 | 0.241173 | 0.115674 | 0.216697 | 0.249823 | 0.064055 | 0.1947 | 0.160393 | -0.01019 | -0.23264 | -0.02574 | 0.858561 | 0.159501 | -0.34206 | 0.634082 | -0.00419 | 0.12281 | 0.097461 | 0.172096 | -0.00473 | 0.025545 | 0.162473 | -1.67178 | 0.247896 | 0.705812 |
| 26.25 | 0.183716 | 0.101198 | 0.301305 | 0.233885 | 0.093361 | 0.181264 | 0.088808 | 0.176407 | -0.02595 | -0.01319 | 0.76777 | 0.20099 | -0.17431 | 0.824078 | -0.0099 | 0.167885 | 0.104294 | 0.240727 | -0.07572 | -0.0461 | -0.64204 | -1.32107 | 0.468462 | 0.765413 |
| 26.26389 | -0.08239 | 0.088523 | 0.3039 | 0.17487 | 0.086464 | 0.129949 | -0.00789 | 0.065785 | 0.075247 | -0.01515 | 0.069233 | 0.218261 | 0.018929 | 0.958394 | -0.02223 | 0.148415 | 0.045833 | 0.227899 | -0.10711 | -0.04895 | -1.13476 | -0.91676 | 0.574469 | 0.56704 |
| 26.27778 | -0.32174 | 0.070023 | 0.287992 | 0.11397 | 0.082007 | 0.15618 | -0.08713 | -0.05633 | 0.009983 | -0.00996 | -0.53679 | 0.224718 | 0.068787 | 1.141703 | -0.02918 | 0.093847 | -0.02493 | 0.16978 | -0.10195 | 0.038704 | -0.93179 | -0.37627 | 0.416242 | 0.264715 |
| 26.29167 | -0.38544 | 0.059031 | 0.25724 | 0.077558 | 0.062161 | 0.243932 | -0.12321 | -0.30326 | -0.20919 | 0.020666 | -0.56127 | 0.232089 | -0.0888 | 1.209201 | -0.02666 | 0.069407 | -0.04642 | 0.102912 | -0.00876 | 0.143918 | -0.28335 | 0.077023 | 0.09285 | -0.02272 |
| 26.30556 | -0.2605 | 0.051046 | 0.268353 | 0.021775 | 0.039544 | 0.320055 | -0.12148 | -0.35169 | -0.40137 | 0.050355 | -0.11631 | 0.253811 | -0.25996 | 1.164754 | -0.02541 | 0.115237 | -0.0081 | 0.089439 | 0.128708 | 0.272095 | 0.371792 | 0.404161 | 0.016501 | -0.04032 |
| 26.31944 | -0.08348 | 0.032828 | 0.242391 | -0.05062 | 0.003597 | 0.356439 | -0.11403 | -0.22455 | -0.36021 | 0.045876 | 0.352637 | 0.264491 | -0.28415 | 1.060109 | -0.01777 | 0.195451 | 0.058746 | 0.177978 | 0.264028 | 0.366891 | 0.568395 | 0.498283 | 0.133594 | 0.184578 |
| 26.33333 | 0.022279 | 0.014474 | 0.201775 | -0.07512 | -0.07244 | 0.352256 | -0.10141 | 0.000254 | 0.060295 | 0.060843 | 0.524935 | 0.281402 | -0.08586 | 0.964037 | -0.00745 | 0.221752 | 0.125342 | 0.352168 | 0.300079 | 0.380304 | 0.175888 | 0.47049 | 0.044356 | 0.311796 |
| 26.34722 | 0.147361 | 0.038949 | 0.144884 | 0.001589 | -0.16236 | 0.398299 | -0.07217 | 0.267036 | 0.464415 | 0.128176 | 0.420708 | 0.322857 | 0.207355 | 0.880628 | -0.00925 | 0.196127 | 0.167486 | 0.52349 | 0.261533 | 0.350646 | -0.33768 | 0.401224 | 0.108435 | 0.36497 |
| 26.36111 | 0.233091 | 0.091337 | 0.102993 | 0.140427 | -0.16169 | 0.518465 | -0.07928 | 0.357058 | 0.552284 | 0.293073 | 0.279293 | 0.395061 | 0.386564 | 0.820306 | -0.02596 | 0.155899 | 0.178021 | 0.605674 | 0.214698 | 0.350695 | -0.31945 | 0.354463 | 0.161124 | 0.322149 |
| 26.375 | 0.226679 | 0.115261 | 0.087058 | 0.276689 | -0.01081 | 0.590426 | -0.08356 | 0.447872 | 0.436277 | 0.50019 | 0.145552 | 0.44366 | 0.368102 | 0.724158 | -0.0347 | 0.159727 | 0.180406 | 0.542236 | 0.20753 | 0.451257 | 0.391427 | 0.484478 | 0.211219 | 0.241783 |
| 26.38889 | 0.242046 | 0.125883 | 0.12637 | 0.36567 | 0.125686 | 0.546806 | -0.01442 | 0.27309 | 0.237267 | 0.505872 | 0.095178 | 0.360638 | 0.233776 | 0.608957 | -0.01862 | 0.212096 | 0.137422 | 0.477896 | 0.365352 | 0.649913 | 1.372806 | 0.721473 | 0.284862 | 0.178098 |
| 26.40278 | 0.387496 | 0.136737 | 0.209999 | 0.499438 | 0.168159 | 0.440662 | 0.132696 | 0.02924 | -0.01462 | 0.373909 | 0.219188 | 0.317446 | 0.196562 | 0.524709 | 0.001662 | 0.276999 | 0.069573 | 0.514758 | 0.357795 | 0.83787 | 1.808135 | 1.015598 | 0.427292 | 0.174268 |
| 26.41667 | 0.695136 | 0.135684 | 0.21258 | 0.677865 | 0.173063 | 0.391743 | 0.296132 | -0.16728 | -0.25826 | 0.324894 | 0.483347 | 0.35438 | 0.315231 | 0.489433 | 0.002521 | 0.298686 | 0.027054 | 0.582111 | 0.206031 | 0.802686 | 1.200754 | 1.08838 | 0.542244 | 0.276839 |
| 26.43056 | 0.963557 | 0.118777 | 0.202052 | 0.814901 | 0.213077 | 0.465172 | 0.347034 | -0.14193 | -0.40034 | 0.412099 | 0.58806 | 0.371412 | 0.508217 | 0.4846 | -0.01228 | 0.233138 | 0.035497 | 0.510561 | 0.1127 | 0.674121 | 0.631982 | 0.937709 | 0.496633 | 0.454979 |
| 26.44444 | 1.06588 | 0.094459 | 0.188669 | 0.785419 | 0.252942 | 0.626912 | 0.374127 | 0.076303 | -0.48221 | 0.609019 | 0.706669 | 0.350473 | 0.667138 | 0.437686 | -0.00272 | 0.15478 | 0.07732 | 0.403647 | 0.039024 | 0.644839 | 0.676797 | 0.590098 | 0.40156 | 0.669516 |
| 26.45833 | 0.920668 | 0.05557 | 0.182569 | 0.786968 | 0.203279 | 0.720883 | 0.346891 | 0.145511 | -0.85952 | 0.807496 | 0.753629 | 0.370261 | 0.736052 | 0.423112 | 0.101672 | 0.30212 | 0.121952 | 0.339855 | 0.004978 | 0.785922 | 1.414461 | 0.22536 | 0.326659 | 0.771722 |
| 26.47222 | 0.711815 | 0.020347 | 0.181225 | 0.830323 | 0.084337 | 0.663476 | 0.350724 | 0.187228 | -0.90129 | 0.724911 | 0.4632 | 0.406987 | 0.766832 | 0.486949 | 0.284475 | 0.660226 | 0.17635 | 0.365879 | 0.073104 | 0.924865 | 1.782583 | 0.079646 | 0.275271 | 0.683304 |
| 26.48611 | 0.632496 | 0.042042 | 0.218795 | 0.915956 | 0.038516 | 0.567253 | 0.351753 | 0.384167 | -0.16216 | 0.628138 | 0.030821 | 0.442984 | 0.855183 | 0.595772 | 0.32886 | 0.65851 | 0.220602 | 0.521653 | 0.231333 | 0.859738 | 1.242291 | 0.213042 | 0.277997 | 0.570153 |
| 26.5 | 0.787897 | 0.117547 | 0.345139 | 1.183033 | 0.151603 | 0.573098 | 0.464651 | 0.445578 | 0.7489 | 0.87139 | 0.238673 | 0.681245 | 1.065884 | 0.805388 | 0.396921 | 0.618452 | 0.238345 | 0.740823 | 0.350971 | 0.86062 | 0.676546 | 0.374846 | 0.404672 | 0.508159 |
| 26.51389 | 1.510437 | 0.262214 | 0.543556 | 2.026772 | 0.312906 | 0.80458 | 0.80092 | 0.637318 | 1.290256 | 1.724971 | 1.347709 | 1.250484 | 1.613934 | 1.372086 | 0.622052 | 0.928868 | 0.214681 | 1.030565 | 0.253711 | 1.189544 | 0.710054 | 0.454552 | 0.762642 | 0.715364 |
| 26.52778 | 2.675583 | 0.491775 | 0.674926 | 2.463683 | 0.399617 | 1.011378 | 1.062132 | 1.21637 | 0.911948 | 3.01234 | 2.829419 | 1.573864 | 2.484311 | 1.90426 | 0.64808 | 1.112596 | 0.180678 | 1.492182 | 0.265543 | 1.637952 | 1.400648 | 0.807838 | 1.403132 | 1.073309 |
| 26.54167 | 3.18704 | 0.327213 | 0.574365 | 1.833673 | 0.415643 | 0.813405 | 1.003812 | 2.162159 | 0.220167 | 4.067387 | 3.64699 | 1.520148 | 2.714777 | 1.948339 | 0.451268 | 0.891012 | 0.125743 | 1.196887 | 0.147928 | 1.945772 | 2.027664 | 1.151551 | 1.915354 | 1.18116 |
| 26.55556 | 3.257361 | 0.337048 | 0.406587 | 1.256519 | 0.486068 | 0.641009 | 0.818095 | 1.828455 | -0.53342 | 5.254834 | 2.846583 | 1.062281 | 2.127965 | 1.77158 | 0.479383 | 0.500641 | 0.195186 | 0.800746 | -0.0229 | 1.784921 | 1.845158 | 1.190489 | 1.625388 | 0.922224 |
| 26.56944 | 3.287322 | 0.431794 | 0.409867 | 1.132731 | 0.36823 | 0.37999 | 0.702197 | 1.256945 | -0.66352 | 5.07466 | 1.634323 | 0.657501 | 1.439855 | 1.838799 | 0.583154 | 0.334105 | 0.358586 | 0.788976 | 0.00027 | 0.808002 | 1.092586 | 1.285524 | 1.215137 | 0.547346 |
| 26.58333 | 2.869089 | 0.423237 | 0.471213 | 0.810615 | 0.169314 | -0.01969 | 0.507767 | 0.760287 | 1.20131 | 4.571307 | 0.411999 | 0.367284 | 0.726469 | 2.31778 | 0.671566 | 0.215928 | 0.37793 | 0.567104 | -0.00496 | -0.28964 | 0.306619 | 0.954429 | 0.782361 | 0.118983 |
| 26.59722 | 2.498638 | 0.344591 | 0.539955 | 0.314958 | 0.046664 | -0.31512 | 0.307383 | 0.174595 | 1.128422 | 3.449971 | -0.68398 | 0.172313 | -0.06795 | 2.886724 | 0.499135 | 0.016557 | 0.281073 | 0.116547 | -0.10548 | -1.30271 | -0.79761 | 0.520403 | 0.234599 | -0.22062 |
| 26.61111 | 1.891807 | 0.22632 | 0.431053 | -0.11894 | -0.00853 | -0.39828 | 0.059577 | -0.40137 | 0.69696 | 1.552009 | -1.57706 | 0.076689 | -0.87235 | 2.905416 | 0.214003 | -0.05129 | 0.156809 | 0.030788 | -0.04901 | -1.66956 | -2.66913 | 0.01723 | -0.36876 | -0.62632 |
| 26.625 | 1.234003 | 0.111326 | 0.380067 | -0.2368 | 0.000372 | -0.31967 | -0.17403 | -0.5367 | 0.200424 | -0.30411 | -2.11853 | -0.1928 | -1.21334 | 2.045647 | -0.13419 | -0.03434 | 0.086643 | 0.335003 | 0.023254 | -1.26593 | -3.26903 | -0.47742 | -0.99595 | -0.96337 |
| 26.63889 | 0.558009 | 0.027035 | 0.045398 | -0.20273 | 0.022781 | -0.10774 | -0.27795 | -0.81168 | -0.41192 | -1.90189 | -1.40314 | -0.55657 | -1.17309 | 0.838919 | -0.40964 | 0.029869 | 0.06296 | 0.64282 | 0.133991 | -0.78434 | -2.45194 | -0.78829 | -0.77872 | -0.66383 |
| 26.65278 | -0.26507 | -0.0558 | -0.00667 | -0.28236 | 0.094969 | 0.067553 | -0.02626 | -1.31636 | -0.73192 | -2.72216 | 0.120886 | -0.99545 | -0.89622 | -0.5461 | -0.59205 | 0.082443 | 0.095485 | 0.600835 | 0.284469 | 0.005841 | -0.29703 | -0.93039 | 0.330703 | 0.261866 |
| 26.66667 | -0.98077 | -0.14729 | -0.00554 | -0.30111 | 0.220377 | 0.074677 | 0.361304 | -0.65015 | 0.193789 | -3.46343 | 1.644941 | -1.28019 | -0.46143 | -2.2878 | -0.55751 | 0.061089 | 0.162616 | 0.238506 | 0.383184 | 0.879824 | 2.092857 | -0.86332 | 1.115958 | 0.289715 |
| 26.68056 | -1.73082 | -0.18606 | -0.09297 | 0.134492 | 0.430259 | 0.099922 | 0.688656 | -0.01393 | 0.554742 | -4.1139 | 2.452215 | -1.33769 | 0.796308 | -3.90783 | -0.3063 | 0.021372 | 0.296351 | 0.148682 | 0.322475 | 1.579333 | 4.142543 | -0.66738 | 1.542398 | 0.122165 |
| 26.69444 | -1.98507 | -0.21451 | 0.09926 | 0.732671 | 0.602199 | 0.037 | 0.723441 | -0.12693 | 0.046829 | -2.8555 | 1.777481 | -0.93474 | 2.251519 | -4.92904 | 0.075209 | -0.0211 | 0.464264 | 0.119661 | 0.199776 | 2.055829 | 3.094035 | -0.48085 | 1.345564 | 0.040714 |
| 26.70833 | -1.77244 | -0.23663 | 0.34771 | 1.089715 | 0.527666 | -0.58101 | 0.396041 | -0.23566 | -0.60013 | -1.22281 | 0.330543 | -0.32635 | 2.552369 | -4.67401 | 0.436613 | -0.01956 | 0.543273 | 0.064974 | 0.044203 | 1.100716 | 0.664163 | -0.32619 | 0.795429 | 0.12136 |
| 26.72222 | -1.35111 | -0.29353 | 0.525035 | 1.202873 | 0.282948 | -1.0442 | 0.011771 | -0.2045 | -0.42133 | -0.34542 | -0.8523 | 0.23252 | 1.793548 | -3.08096 | 0.670571 | 0.020811 | 0.505085 | -0.2322 | -0.05963 | -0.03871 | -1.05786 | -0.29145 | 0.03393 | 0.205764 |
| 26.73611 | -0.75024 | -0.30389 | 0.225609 | 0.952516 | -0.03592 | -0.91804 | -0.41772 | 0.087232 | 0.014509 | -0.08812 | -2.09073 | 0.650604 | 0.942436 | -1.3975 | 0.750404 | 0.059779 | 0.477413 | -0.61917 | -0.08756 | -0.99754 | -3.08773 | -0.31272 | -0.60737 | 0.333756 |
| 26.75 | -1.05877 | -0.29148 | 0.292423 | 0.494514 | -0.4364 | -0.77757 | -0.70478 | 0.121192 | 0.36056 | 0.014897 | -3.18472 | 1.055178 | 0.096145 | -0.00307 | 0.635982 | 0.10779 | 0.265854 | -0.89825 | -0.11649 | -1.09126 | -3.48263 | -0.36288 | -1.06324 | 0.325535 |
| 26.76389 | -1.06508 | -0.24182 | 0.081242 | 0.088565 | -0.51255 | -0.46522 | -0.8888 | 0.152142 | 0.383533 | 0.021893 | -3.35949 | 1.01346 | -0.55073 | 1.022159 | 0.429455 | 0.186922 | 0.03395 | -0.91447 | -0.14621 | -0.99178 | -2.87454 | -0.40349 | -1.13689 | 0.226189 |
| 26.77778 | -0.46247 | -0.20396 | -0.0332 | -0.53979 | -0.40968 | -0.10899 | -0.89539 | 0.053508 | -0.09433 | -0.03937 | -2.70522 | 0.764278 | -0.9373 | 1.371544 | 0.283831 | 0.224389 | -0.34733 | -0.8975 | -0.10254 | -0.69671 | -1.0135 | -0.40751 | -0.6484 | 0.085037 |
| 26.79167 | 0.048502 | -0.18357 | 0.012245 | -0.96638 | -0.22499 | 0.117024 | -0.70259 | -0.23555 | -0.51463 | -0.11073 | -1.1613 | 0.365959 | -1.24865 | 0.899492 | 0.085051 | 0.105188 | -0.44478 | -0.87622 | -0.03443 | -0.52506 | 0.647797 | -0.42037 | -0.24635 | -0.12087 |
| 26.80556 | 0.236306 | -0.14387 | -0.09249 | -1.02772 | -0.0241 | 0.057408 | -0.3572 | -0.34234 | -0.56008 | -0.12228 | 0.310685 | -0.05105 | -1.36797 | 0.277049 | -0.08613 | -0.0794 | -0.6718 | -0.68033 | -0.05665 | -0.59872 | 1.336873 | -0.3803 | 0.017517 | -0.24167 |
| 26.81944 | 0.073229 | -0.07685 | -0.32726 | -0.71415 | 0.097901 | -0.09809 | -0.15528 | 0.235417 | -0.33463 | -0.1415 | 1.02906 | -0.38513 | -1.46513 | -0.03539 | -0.21259 | -0.26258 | -0.82888 | -0.51566 | -0.14058 | -1.04198 | 0.232431 | -0.36032 | -0.01026 | -0.4272 |
| 26.83333 | -0.50539 | -0.03056 | -0.48757 | -0.33241 | 0.198234 | -0.51091 | -0.05821 | 0.409069 | 0.162537 | -0.20372 | 1.001756 | -0.63996 | -1.57702 | -0.25573 | -0.31988 | -0.40603 | -0.68591 | -0.49413 | -0.21463 | -1.22241 | -1.45155 | -0.40362 | -0.19768 | -0.78944 |
| 26.84722 | -1.12026 | 0.027479 | -0.63475 | -0.26309 | 0.196773 | -0.94992 | -0.10582 | -0.09286 | 0.168591 | -0.28262 | 0.664072 | -0.96709 | -1.64083 | -0.45129 | -0.505 | -0.52076 | -0.37117 | -0.63609 | -0.30342 | -1.26937 | -3.58253 | -0.4335 | -0.44643 | -1.23456 |
| 26.86111 | -1.38179 | 0.072739 | -0.79804 | -0.69147 | -0.02649 | -1.15855 | -0.27695 | -0.90466 | -0.57914 | -0.34243 | -0.00412 | -0.95707 | -1.76659 | -0.70931 | -0.78105 | -0.70173 | -0.11133 | -0.98522 | -0.40127 | -0.86429 | -3.95173 | -0.51115 | -0.54561 | -1.11076 |
| 26.875 | -2.09695 | 0.088721 | -0.91473 | -1.34647 | -0.34315 | -0.90377 | -0.48776 | -1.33861 | -1.09741 | -0.47754 | -1.04679 | -0.99432 | -1.7655 | -1.01869 | -0.98488 | -0.82211 | -0.07681 | -1.28937 | -0.37974 | -0.68015 | -3.42025 | -0.49422 | -0.51861 | -0.73304 |
| 26.88889 | -2.1643 | 0.045496 | -0.87533 | -1.8272 | -0.56254 | -0.54058 | -0.58292 | -1.38016 | -1.44575 | -0.63218 | -2.07646 | -0.95959 | -1.49955 | -1.36753 | -0.92981 | -0.7695 | -0.14268 | -1.24881 | -0.22801 | -0.70508 | -2.02547 | -0.37428 | -0.61019 | -0.61782 |

| | | | | | | | | | | | | | | | | | | | | | | | | |
|---|---|---|---|---|---|---|---|---|---|---|---|---|---|---|---|---|---|---|---|---|---|---|---|---|
| 26.90278 | -1.7809 | -0.01442 | -0.65548 | -1.90226 | -0.51551 | -0.54954 | -0.51472 | -1.12547 | -1.56974 | -0.84389 | -2.5108 | -1.07617 | -1.16629 | -1.82727 | -0.83344 | -0.67359 | -0.27531 | -1.0573 | -0.2639 | -0.90234 | -1.56017 | -0.24477 | -0.65367 | -0.63135 |
| 26.91667 | -1.13162 | -0.06789 | -0.4892 | -1.76578 | -0.23389 | -0.80166 | -0.37918 | -0.64611 | -1.47818 | -0.77399 | -1.68561 | -0.83683 | -0.91732 | -2.15552 | -0.71578 | -0.62661 | -0.48352 | -0.90247 | -0.30102 | -1.13666 | -0.87084 | -0.16656 | -0.62163 | -0.54064 |
| 26.93056 | -0.75101 | -0.0654 | -0.3026 | -1.39382 | 0.144444 | -0.91534 | -0.15584 | -0.26307 | -1.17254 | -0.4428 | -0.25354 | -0.41599 | -0.57245 | -2.19125 | -0.54386 | -0.57794 | -0.49786 | -0.83738 | -0.14166 | -1.00502 | -0.54221 | -0.26572 | -0.48602 | -0.3223 |
| 26.94444 | -0.61205 | -0.07676 | -0.20517 | -0.83056 | 0.522492 | -0.79891 | 0.001135 | -0.06275 | -0.49706 | -0.1568 | 0.754194 | -0.05297 | -0.12882 | -2.01516 | -0.31113 | -0.44226 | -0.19192 | -0.64473 | -0.07066 | -0.61289 | -0.47148 | -0.41847 | -0.28339 | -0.15216 |
| 26.95833 | -0.57159 | -0.15409 | -0.07477 | -0.26774 | 0.587428 | -0.48204 | 0.020852 | -0.06845 | 0.147742 | -0.17904 | 0.880909 | 0.085254 | 0.272511 | -1.5994 | -0.19597 | -0.31823 | 0.219808 | -0.36894 | -0.09246 | -0.34596 | -0.59324 | -0.56446 | -0.15346 | -0.01236 |
| 26.97222 | -0.43313 | -0.17631 | 0.166563 | 0.038181 | 0.169195 | -0.15992 | 0.080496 | 0.045013 | 0.659061 | -0.33582 | 0.430033 | 0.137975 | 0.468905 | -1.13661 | -0.15781 | -0.23004 | 0.61066 | -0.12859 | -0.11515 | -0.18118 | -0.74603 | -0.57757 | -0.03681 | 0.082895 |
| 26.98611 | -0.3413 | -0.06003 | 0.393211 | 0.155518 | -0.25253 | 0.063646 | 0.108962 | 0.333948 | 0.869517 | -0.46583 | -0.08119 | 0.208225 | 0.477606 | -0.81381 | -0.14963 | -0.10919 | 0.688796 | 0.050989 | -0.11555 | -0.11849 | -0.70107 | -0.43368 | 0.048833 | 0.162622 |
| 27 | -0.30029 | 0.006274 | 0.340154 | -0.03951 | -0.50501 | -0.05353 | -0.01059 | 0.469312 | 0.510507 | -0.52024 | -0.53106 | 0.197359 | 0.234477 | -0.67056 | -0.16143 | 0.010405 | 0.32065 | 0.024926 | -0.14391 | -0.13507 | -0.88793 | -0.24431 | 0.023111 | 0.09601 |
| 27.01389 | -0.53789 | 0.006072 | -0.03278 | -0.47701 | -0.661 | -0.58332 | -0.21016 | 0.026917 | -0.32251 | -0.48532 | -1.08364 | -0.00774 | -0.16708 | -0.78743 | -0.20738 | -0.17294 | -0.12069 | -0.23463 | -0.21402 | -0.23262 | -1.35526 | -0.02786 | -0.16565 | -0.17259 |
| 27.02778 | -1.16339 | 0.025363 | -0.45056 | -0.77785 | -0.72479 | -1.13108 | -0.29005 | -0.50337 | -1.49792 | -0.37979 | -1.76387 | -0.27017 | -0.59432 | -0.97554 | -0.20163 | -0.58788 | -0.74314 | -0.55456 | -0.24752 | -0.44542 | -1.98744 | 0.203616 | -0.42299 | -0.53814 |
| 27.04167 | -1.50555 | 0.085636 | -0.45316 | -0.93158 | -0.6103 | -1.17548 | -0.28569 | -0.34589 | -2.74853 | -0.35019 | -2.13756 | -0.41017 | -1.07316 | -1.09126 | -0.18363 | -0.90128 | -1.42693 | -0.82324 | -0.27793 | -0.62675 | -2.58182 | 0.214076 | -0.62598 | -0.74682 |
| 27.05556 | -1.89983 | -0.19567 | -0.19678 | -1.28926 | -0.58901 | -1.09209 | -0.47394 | -0.47429 | -4.1189 | -0.55164 | -1.80794 | -0.41554 | -1.48235 | -1.17886 | -0.33164 | -0.82517 | -1.49114 | -1.20378 | -0.38716 | -0.59209 | -3.05406 | 0.0428 | -0.86384 | -0.87357 |
| 27.06944 | -2.35923 | -0.71011 | -0.10703 | -2.09729 | -0.63583 | -0.97853 | -0.54282 | -0.33904 | -5.35997 | -0.69255 | -1.3021 | -0.41587 | -1.42594 | -1.54922 | -0.5688 | -0.58707 | -1.58048 | -1.53767 | -0.5711 | -0.7739 | -2.57487 | -0.05707 | -0.91945 | -0.72484 |
| 27.08333 | -2.25554 | -0.8061 | -0.03521 | -2.1408 | -0.31801 | -0.60743 | -0.38478 | -0.20694 | -3.35934 | -0.57937 | -0.79752 | -0.32916 | -0.85243 | -1.76205 | -0.60098 | -0.25466 | 0.000489 | -0.91266 | -0.59725 | -0.76902 | -1.47904 | -0.07568 | -0.75171 | -0.30453 |
| 27.09722 | -1.31499 | -0.62737 | 0.099136 | -1.46239 | 0.117601 | -0.12713 | -0.16807 | 0.198583 | 0.161622 | -0.3117 | -0.25007 | -0.12918 | -0.11542 | -1.29198 | -0.4815 | 0.182167 | 1.621707 | -0.20264 | -0.54264 | -0.35113 | -0.48061 | -0.09947 | -0.46764 | 0.034843 |
| 27.11111 | -0.45751 | -0.3921 | 0.212113 | -0.75496 | 0.358982 | 0.275503 | 0.001558 | 0.650308 | 1.809328 | -0.12894 | 0.316096 | 0.052944 | 0.450614 | -0.7527 | -0.30745 | 0.485296 | 1.806976 | 0.273552 | -0.33452 | -0.00826 | 0.206744 | -0.14594 | -0.22328 | 0.233992 |
| 27.125 | 0.159168 | -0.1854 | 0.318415 | -0.37102 | 0.358498 | 0.50717 | 0.095323 | 0.724402 | 3.075585 | -0.05446 | 0.901444 | 0.054911 | 0.697318 | -0.48359 | -0.16432 | 0.587141 | 1.202424 | 0.555834 | -0.14205 | 0.120923 | 0.535275 | -0.17123 | -0.00989 | 0.383095 |
| 27.13889 | 0.469732 | -0.02286 | 0.455172 | -0.23311 | 0.307609 | 0.5386 | 0.083275 | 0.387425 | 2.973966 | 0.019892 | 1.54241 | -0.01428 | 0.71747 | -0.52019 | -0.08057 | 0.307902 | 0.524816 | 0.673753 | 0.018318 | 0.050371 | 0.877003 | -0.18122 | 0.180779 | 0.538668 |
| 27.15278 | 0.459282 | 0.101863 | 0.654577 | -0.23382 | 0.287542 | 0.39166 | 0.036048 | 0.211218 | 2.588103 | 0.045604 | 1.700305 | -0.09273 | 0.598817 | -0.60667 | -0.05148 | 0.04968 | 0.195948 | 0.671042 | 0.131016 | -0.06332 | 1.346577 | -0.15092 | 0.39883 | 0.620758 |
| 27.16667 | 0.219345 | 0.190915 | 0.534143 | -0.25483 | 0.262914 | 0.228252 | 0.00268 | 0.177116 | 2.010005 | -0.01381 | 1.619773 | -0.11323 | 0.503509 | -0.46313 | -0.04093 | -0.17546 | 0.29126 | 0.669904 | 0.222034 | 0.025708 | 1.67838 | -0.0657 | 0.587503 | 0.694967 |
| 27.18056 | -0.12155 | 0.244518 | 0.191017 | -0.21732 | 0.236584 | 0.017353 | 0.002473 | 0.489319 | 1.262672 | -0.07466 | 1.38105 | -0.04376 | 0.432584 | -0.28953 | -0.01608 | -0.37324 | 0.606329 | 0.631673 | 0.286436 | 0.120929 | 1.768462 | 0.039005 | 0.809168 | 0.611869 |
| 27.19444 | -0.26203 | 0.312041 | 0.126707 | -0.23155 | 0.177114 | -0.07108 | 0.149656 | 0.190366 | 1.125244 | 0.138193 | 1.018193 | 0.150717 | 0.272067 | -0.13106 | 0.001968 | -0.44762 | 0.846727 | 0.500852 | 0.231732 | 0.130829 | 1.504007 | 0.064921 | 0.693179 | 0.334446 |
| 27.20833 | 0.15497 | 0.50969 | 0.368359 | -0.47888 | 0.271579 | 0.237853 | 0.827064 | -1.22422 | 4.107879 | 1.385965 | 0.430392 | 0.829235 | -0.18177 | -0.11901 | -0.0019 | -0.58556 | 0.453499 | 0.160752 | -0.16213 | -0.05252 | 0.263496 | -0.07308 | -0.31854 | -0.39632 |
| 27.22222 | -0.5941 | 0.201245 | 0.375747 | -0.77815 | -0.15655 | 0.125727 | 0.94316 | -1.71171 | 2.989558 | 1.172731 | 0.114027 | 0.810339 | -1.03172 | -0.67585 | -0.04931 | -1.08074 | -0.807 | -0.52198 | -1.01086 | -0.44429 | -2.15326 | -0.30888 | -2.19633 | -1.93768 |
| 27.23611 | 0.893534 | -0.29321 | 0.260853 | 0.794875 | -0.06859 | -0.43476 | -1.5512 | 1.00078 | 0.139297 | -1.06638 | -0.37072 | -1.21639 | -1.94359 | -1.74693 | -0.16873 | -1.59682 | -2.17202 | -1.00369 | -2.06509 | -0.72499 | -4.10302 | -0.35578 | -3.21504 | -3.01762 |
| 27.25 | 2.560261 | -0.91696 | 0.058887 | 2.641187 | 0.078218 | -0.55222 | -3.10159 | 3.227167 | -0.81445 | -3.09839 | -0.89013 | -3.07313 | -1.44866 | -1.43057 | -0.27469 | -1.21652 | -1.21549 | -0.6121 | -1.31847 | -0.43197 | -2.58258 | -0.06241 | -2.13434 | -1.71023 |
| 27.26389 | 0.672471 | -0.20137 | -0.38147 | 1.331002 | -0.32515 | -0.49565 | -1.60313 | 2.113082 | -3.17997 | -2.671 | -0.30497 | -1.55657 | -0.60373 | 0.037762 | -0.22848 | -0.41836 | -0.19031 | -0.19112 | -0.0361 | -0.21277 | -0.69759 | 0.202223 | -0.9867 | -0.50315 |
| 27.27778 | 0.521845 | 0.243108 | 0.435577 | 0.514073 | -0.19864 | 0.132843 | 0.374679 | 0.630721 | 0.760993 | 0.316703 | 0.461969 | 0.5292 | -0.18126 | 0.494777 | -0.08287 | -0.16372 | 0.074138 | -0.15319 | 0.162933 | -0.07866 | -0.35592 | 0.134487 | -0.65723 | -0.2233 |
| 27.29167 | 0.238553 | 0.29857 | 0.492491 | 0.066676 | -0.01209 | 0.326999 | 0.872329 | 0.490189 | 1.811485 | 1.032535 | 0.501814 | 1.01723 | -0.0965 | 0.222016 | -0.01247 | -0.15407 | -0.01069 | -0.15991 | 0.061855 | -0.04037 | -0.09771 | 0.038919 | -0.38065 | -0.11303 |
| 27.30556 | -0.11567 | 0.194919 | 0.150717 | -0.06976 | 0.014547 | 0.247235 | 0.332043 | 0.64819 | 1.072961 | 0.030401 | 0.19333 | 0.453266 | -0.10217 | -0.00187 | -0.0082 | -0.20205 | -0.01985 | -0.12996 | 0.055233 | -0.04041 | 0.212128 | 0.023649 | -0.04762 | 0.004128 |
| 27.31944 | -0.19319 | 0.108693 | -0.10988 | -0.03126 | -0.07069 | 0.08459 | -0.39509 | 0.445363 | -0.27412 | -1.05067 | -0.34484 | -0.37404 | -0.09287 | -0.09038 | -0.01018 | -0.26619 | -0.00551 | -0.10941 | 0.094099 | -0.04992 | 0.304725 | 0.037021 | 0.131673 | 0.057578 |
| 27.33333 | -0.06696 | 0.016812 | -0.19168 | 0.17722 | -0.1162 | -0.01198 | -0.93386 | 0.080259 | -1.46483 | -1.57563 | -0.79039 | -0.98749 | -0.0634 | -0.09169 | -0.00392 | -0.26164 | 0.00615 | -0.07491 | 0.137048 | -0.04383 | 0.290162 | 0.051034 | 0.172257 | 0.07478 |
| 27.34722 | 0.174446 | -0.07186 | 0.061556 | 0.276616 | -0.16211 | -0.02646 | -0.68774 | -0.02979 | -1.46083 | -1.01647 | -0.88043 | -0.7207 | -0.03801 | -0.06485 | -0.00249 | -0.20742 | 0.012551 | -0.03575 | 0.1509 | -0.03629 | 0.243766 | 0.05311 | 0.135402 | 0.040675 |
| 27.36111 | 0.296342 | 0.053751 | 0.548144 | 0.192282 | -0.14543 | 0.107098 | 0.14186 | 0.011019 | -0.31461 | 0.114296 | -0.50832 | 0.28927 | -0.00059 | -0.03286 | 0.0017 | -0.12563 | 0.017355 | -0.00887 | 0.164083 | -0.02345 | 0.230663 | 0.051732 | 0.102621 | -0.01907 |
| 27.375 | 0.269017 | 0.157432 | 1.021046 | 0.120566 | -0.09353 | 0.194382 | 0.597242 | 0.115474 | 0.602895 | 0.727857 | 0.133679 | 0.695716 | 0.025352 | -0.01384 | 0.01111 | -0.06913 | 0.021937 | -0.00261 | 0.171206 | -0.03229 | 0.188747 | 0.043382 | 0.08197 | -0.10755 |
| 27.38889 | 0.338435 | 0.08152 | 0.77898 | 0.104106 | -0.1602 | 0.225662 | 0.105161 | 0.488664 | 0.627984 | 0.074578 | 0.608484 | 0.070535 | 0.012161 | -0.02826 | 0.008531 | -0.06883 | -0.00068 | -0.01402 | 0.148563 | -0.0535 | 0.116379 | 0.007127 | 0.04771 | -0.19907 |
| 27.40278 | 0.458712 | -0.03025 | 0.306063 | 0.160253 | -0.19075 | 0.310382 | -0.53621 | 0.614636 | -0.05063 | -0.78981 | 0.687719 | -0.92333 | 0.081254 | 0.055491 | 0.013841 | -0.07006 | -0.01356 | 0.018654 | 0.228088 | -0.04165 | 0.242131 | -0.01156 | 0.105552 | -0.13272 |
| 27.41667 | 0.431319 | 0.034518 | 0.198868 | 0.116767 | -0.03795 | 0.42487 | -0.06597 | 0.293424 | 0.013491 | -0.35962 | 0.419973 | -0.61926 | 0.25024 | 0.351253 | 0.047269 | 0.016216 | 0.032648 | 0.129472 | 0.511869 | 0.026557 | 0.611747 | 0.026008 | 0.348542 | 0.152235 |
| 27.43056 | 0.026862 | 0.265973 | 0.221693 | 0.200745 | 0.125809 | 0.565558 | 0.929922 | -0.21574 | 0.784473 | 0.80958 | -0.18299 | 0.695027 | 0.216927 | 0.593105 | 0.058853 | 0.093747 | 0.067654 | 0.203893 | 0.731378 | 0.08177 | 0.783462 | 0.042418 | 0.495273 | 0.293199 |
| 27.44444 | 0.11132 | 0.180156 | 0.352607 | 0.595233 | 0.293575 | 0.615226 | 1.156335 | -0.13895 | 0.692131 | 1.104132 | -0.53427 | 1.053837 | -0.04205 | 0.422576 | 0.038803 | 0.076556 | 0.025299 | 0.136953 | 0.487222 | 0.072907 | 0.383307 | 0.024228 | 0.263553 | 0.033685 |
| 27.45833 | 0.561501 | 0.080836 | 0.304203 | 0.924566 | 0.31121 | 0.49456 | 0.471198 | 0.356421 | -0.47475 | 0.2939 | -0.53547 | 0.291601 | -0.25537 | 0.050973 | -0.00376 | 0.02653 | -0.04586 | 0.018624 | 0.068105 | 0.048523 | -0.14674 | 0.016015 | -0.08462 | -0.3597 |
| 27.47222 | 0.821391 | -0.00951 | 0.301014 | 1.025104 | 0.203924 | 0.250699 | -0.00547 | 0.617382 | -1.1044 | -0.2181 | -0.20336 | -0.35469 | -0.29938 | -0.13775 | -0.04076 | 0.034507 | -0.07377 | -0.02976 | -0.13212 | 0.052724 | -0.35616 | 0.030935 | -0.21335 | -0.53411 |
| 27.48611 | 0.416368 | 0.078241 | 0.376148 | 0.566144 | 0.17138 | 0.381987 | 0.668675 | 0.005597 | 0.578143 | 1.342109 | -0.07944 | 0.704141 | -0.16699 | -0.06752 | 0.004304 | 0.062493 | -0.05094 | 0.007301 | -0.01962 | 0.085416 | -0.15686 | 0.03149 | -0.08899 | -0.29378 |
| 27.5 | 0.010081 | 0.266458 | 0.448187 | 0.524761 | 0.206236 | 0.798694 | 1.801108 | -0.75934 | 2.807936 | 3.531173 | 0.168893 | 2.287891 | 0.097795 | 0.190707 | 0.25289 | 0.078581 | 0.008805 | 0.099012 | 0.279629 | 0.155287 | 0.313072 | -0.0258 | 0.185056 | 0.143099 |
| 27.51389 | 0.776092 | 0.266032 | 0.495361 | 2.039187 | 0.310493 | 1.236148 | 1.970363 | -0.17047 | 3.215963 | 4.22356 | 0.867952 | 2.254511 | 0.287631 | 0.384008 | 0.859182 | 0.037564 | -0.00697 | 0.160163 | 0.442394 | 0.195017 | 0.558396 | -0.15874 | 0.393528 | 0.473681 |
| 27.52778 | 2.3459 | 0.151526 | 0.661263 | 2.953964 | 0.420654 | 1.681478 | 1.066223 | 1.165103 | 1.621002 | 3.685701 | 1.759285 | 1.566733 | 0.125422 | 0.232332 | 0.245967 | -0.14648 | -0.1165 | 0.09717 | 0.0558 | 0.104919 | -0.00867 | -0.267 | 0.242527 | 0.111066 |
| 27.54167 | 2.818525 | 0.11599 | 0.778809 | 2.300898 | 0.219579 | 1.36882 | 0.771171 | 1.848731 | 0.07631 | 3.033775 | 2.167439 | 1.184661 | -0.1584 | -0.03011 | -0.07002 | -0.31135 | -0.17065 | -0.00517 | -0.47157 | 0.008384 | -0.69128 | -0.24907 | -0.0758 | -0.36464 |
| 27.55556 | 3.436883 | 0.172076 | 0.817965 | 1.973254 | 0.137475 | 1.141687 | 0.721148 | 1.908776 | -0.16356 | 2.877616 | 1.845732 | 0.709658 | -0.20177 | -0.07436 | -0.11924 | -0.24786 | -0.0867 | -0.00058 | -0.46338 | 0.058603 | -0.52135 | -0.09015 | -0.0577 | -0.33043 |
| 27.56944 | 3.228105 | 0.23857 | 0.671972 | 1.704706 | 0.144053 | 0.821035 | 0.683742 | 1.368723 | 0.329304 | 3.011626 | 0.887052 | 0.395509 | -0.14882 | 0.054918 | -0.08538 | -0.04447 | 0.014293 | 0.047205 | -0.16863 | 0.167975 | -0.11456 | 0.061578 | 0.096872 | -0.03919 |
| 27.58333 | 2.115895 | 0.213686 | 0.432451 | 1.185815 | 0.095966 | 0.371275 | 0.588056 | 0.50537 | 0.734215 | 2.326038 | -0.20094 | 0.499165 | -0.19084 | 0.090313 | -0.02427 | 0.093094 | 0.038433 | 0.03657 | -0.03252 | 0.201494 | 0.00326 | 0.111266 | 0.126216 | 0.05894 |
| 27.59722 | 1.385106 | 0.226883 | 0.236232 | 1.03404 | 0.119344 | 0.532406 | 0.679831 | -0.348 | 0.825423 | 2.040158 | -0.95715 | 0.71074 | -0.27458 | 0.061306 | 0.006052 | 0.158857 | 0.013292 | 0.009968 | 0.001384 | 0.174956 | -0.04287 | 0.079497 | 0.107776 | 0.028348 |
| 27.61111 | 0.44591 | 0.169298 | -0.02113 | 0.853377 | 0.253361 | 0.602254 | 0.700268 | -1.06822 | 0.917417 | 1.684462 | -1.48033 | 0.749074 | -0.34498 | 0.02387 | 0.005206 | 0.18911 | -0.03157 | -0.01006 | 0.015398 | 0.13641 | -0.08569 | 0.002441 | 0.087625 | -0.00554 |
| 27.625 | -0.20234 | 0.062178 | -0.03122 | 0.33658 | 0.100136 | 0.344994 | 0.519816 | -1.26529 | 1.032583 | 1.412325 | -1.55945 | 0.553475 | -0.38931 | -0.04276 | -0.00507 | 0.201248 | -0.07214 | -0.0214 | 0.006437 | 0.097765 | -0.14443 | -0.09542 | 0.020464 | -0.04971 |
| 27.63889 | -0.06959 | 0.016378 | -0.30313 | -0.094 | -0.09514 | 0.239436 | 0.248809 | -0.81281 | 1.22119 | 1.645152 | -0.91398 | 0.594861 | -0.37746 | -0.10957 | -0.00307 | 0.177737 | -0.08965 | -0.03029 | -0.012 | 0.080969 | -0.18077 | -0.12988 | -0.13111 | -0.0864 |
| 27.65278 | -0.15576 | 0.014619 | -0.34369 | -0.43841 | -0.23546 | 0.229575 | -0.06197 | 0.256649 | 1.02447 | 1.434208 | 0.01647 | 0.334951 | -0.31657 | -0.14389 | -0.00102 | 0.166397 | -0.08785 | -0.04379 | -0.0122 | 0.078551 | -0.17676 | -0.10919 | -0.31365 | -0.09426 |
| 27.66667 | 0.178036 | 0.040105 | 0.088431 | -0.58077 | -0.18615 | 0.011401 | -0.2874 | 1.357866 | 0.434758 | 0.627574 | 0.998743 | -0.18307 | -0.24041 | -0.16015 | -0.00489 | 0.171666 | -0.08552 | -0.05945 | 0.016381 | 0.059899 | -0.14 | -0.08655 | -0.337 | -0.0836 |
| 27.68056 | 0.246867 | 0.029188 | 0.386811 | -0.35808 | 0.02844 | -0.0406 | -0.36046 | 1.514764 | 0.144428 | -0.4363 | 1.915086 | -0.62604 | -0.16219 | -0.16159 | -0.00968 | 0.143348 | -0.08422 | -0.07051 | 0.044345 | 0.032081 | -0.08807 | -0.07695 | -0.21891 | -0.07894 |
| 27.69444 | -0.55673 | -0.05195 | 0.287602 | -0.10484 | 0.336442 | 0.123164 | -0.20794 | 1.235658 | -0.11172 | -1.72447 | 1.605096 | -1.13284 | -0.11339 | -0.1215 | -0.01472 | 0.111634 | -0.07907 | -0.06918 | 0.057364 | 0.022217 | -0.05331 | -0.04713 | -0.12075 | -0.07516 |
| 27.70833 | -0.95623 | -0.11165 | 0.068215 | 0.262385 | 0.513467 | 0.179536 | 0.140116 | 0.442261 | -0.7379 | -3.01646 | 0.537059 | -1.24562 | -0.08649 | -0.0706 | -0.02315 | 0.088605 | -0.06215 | -0.04991 | 0.083771 | 0.023275 | -0.02352 | -0.02694 | -0.0748 | -0.06207 |
| 27.72222 | -0.1239 | -0.09271 | -0.19303 | 0.899897 | 0.645383 | -0.19774 | 0.528735 | -0.57887 | -1.42901 | -2.62398 | -0.41711 | -0.91722 | -0.05483 | -0.03013 | -0.02969 | 0.058259 | -0.04109 | -0.02143 | 0.100301 | 0.025895 | 0.012296 | -0.01656 | -0.02936 | -0.06353 |
| 27.73611 | 0.423945 | -0.09398 | -0.34481 | 0.853077 | 0.547005 | -0.61407 | 0.712612 | -1.48025 | -1.39625 | -1.81458 | -1.24506 | -0.68331 | -0.02573 | -0.01357 | -0.03506 | 0.044256 | -0.02758 | 0.00273 | 0.103083 | 0.032691 | 0.051743 | -0.00841 | -0.01943 | -0.06668 |
| 27.75 | 0.691102 | -0.11984 | -0.37508 | 0.983631 | 0.205353 | -0.68869 | 0.677058 | -1.45401 | -1.25406 | -1.10057 | -1.70321 | -0.46056 | -0.0047 | -0.01783 | -0.03918 | 0.047591 | -0.01373 | 0.016106 | 0.102567 | 0.046599 | 0.095245 | -0.00267 | -0.01503 | -0.05104 |
| 27.76389 | 0.784558 | -0.1358 | -0.48117 | 1.063258 | -0.09485 | -0.29505 | 0.476686 | 0.069371 | -0.20849 | -0.39493 | -2.02126 | -0.10397 | 0.015406 | -0.02838 | -0.04574 | 0.065817 | -0.00224 | 0.023265 | 0.096126 | 0.066558 | 0.125921 | -0.01286 | 0.009737 | -0.04895 |
| 27.77778 | 0.646049 | -0.14545 | -0.71629 | 0.656868 | -0.20754 | -0.05898 | 0.270969 | 1.037424 | 0.582311 | 0.098917 | -1.89676 | 0.185177 | 0.035945 | -0.04918 | -0.05042 | 0.082338 | -0.01052 | 0.026512 | 0.09147 | 0.083305 | 0.151816 | -0.03552 | 0.008912 | -0.04839 |
| 27.79167 | 0.195465 | -0.18291 | -0.65678 | 0.159782 | -0.24442 | 0.046869 | -0.07335 | 0.959921 | 0.857002 | 0.150859 | -1.24514 | 0.005223 | 0.062094 | -0.06752 | -0.04643 | 0.101313 | -0.02261 | 0.021629 | 0.0987 | 0.090589 | 0.173424 | -0.04653 | -0.014 | -0.0211 |
| 27.80556 | -0.15627 | -0.16587 | -0.49341 | -0.37553 | -0.16907 | 0.013833 | -0.48452 | -0.2599 | 0.486467 | -0.14147 | -0.82148 | -0.4344 | 0.081973 | -0.07376 | -0.04014 | 0.117967 | -0.02225 | 0.008705 | 0.097247 | 0.094374 | 0.175092 | -0.04874 | -0.02763 | -0.00427 |
| 27.81944 | -0.56329 | -0.12647 | -0.57344 | -0.86082 | -0.21059 | 0.077295 | -0.93729 | -0.77205 | 0.256138 | -0.32196 | -0.5294 | -0.6853 | 0.087138 | -0.09125 | -0.02858 | 0.122883 | -0.02405 | 0.008545 | 0.08855 | 0.101665 | 0.190349 | -0.0554 | -0.03082 | 0.003049 |
| 27.83333 | -0.88717 | -0.11896 | -0.72557 | -1.21012 | -0.45822 | -0.07548 | -1.08146 | -0.84118 | -0.0755 | -0.20976 | -0.28519 | -0.41373 | 0.076053 | -0.11791 | -0.02051 | 0.10519 | -0.02055 | 0.024627 | 0.091933 | 0.122655 | 0.218337 | -0.07166 | -0.01953 | 0.015119 |
| 27.84722 | -1.13446 | -0.1234 | -0.84092 | -1.2987 | -0.59265 | -0.508 | -1.1573 | -1.53355 | -0.53407 | -0.10119 | -0.12984 | -0.09164 | 0.070801 | -0.13241 | -0.02649 | 0.066101 | 0.005055 | 0.055059 | 0.09545 | 0.13716 | 0.272089 | -0.08379 | 0.029485 | 0.05596 |
| 27.86111 | -1.38702 | -0.21252 | -0.87432 | -1.13093 | -0.43073 | -0.78397 | -1.09388 | -1.58696 | -0.89015 | -0.39788 | 0.025748 | -0.16148 | 0.067754 | -0.10381 | -0.04581 | 0.035445 | 0.016851 | 0.081873 | 0.095752 | 0.146876 | 0.319263 | -0.09498 | 0.075006 | 0.061547 |
| 27.875 | -1.71525 | -0.27806 | -0.76997 | -0.89624 | -0.25522 | -0.87581 | -0.78247 | -0.81743 | -0.9083 | -0.72023 | 0.146087 | -0.54376 | 0.044244 | -0.09719 | -0.07897 | 0.022142 | -0.00118 | 0.09083 | 0.087179 | 0.137273 | 0.350404 | -0.10277 | 0.086652 | 0.052631 |
| 27.88889 | -2.12299 | -0.28952 | -0.62317 | -0.92547 | -0.28501 | -0.85448 | -0.70351 | -0.23851 | -1.01124 | -0.7438 | 0.192302 | -0.7513 | 0.022243 | -0.10493 | -0.1076 | 0.02068 | -0.00927 | 0.082186 | 0.079008 | 0.114537 | 0.343531 | -0.11034 | 0.081054 | 0.03149 |
| 27.90278 | -2.25526 | -0.24413 | -0.45136 | -1.18924 | -0.39621 | -0.65927 | -0.62632 | 0.116849 | -0.7486 | -0.68599 | 0.094964 | -0.90271 | 0.038124 | -0.09118 | -0.08932 | 0.037145 | 0.007696 | 0.07091 | 0.082707 | 0.113755 | 0.292944 | -0.11998 | 0.082154 | 0.043499 |
| 27.91667 | -2.21831 | -0.23368 | -0.37165 | -1.24382 | -0.46904 | -0.63062 | -0.49352 | -0.29812 | -0.5315 | -0.70496 | -0.0632 | -0.70789 | 0.082705 | -0.07089 | -0.05248 | 0.058377 | 0.024625 | 0.084366 | 0.108051 | 0.132077 | 0.3555 | -0.12374 | 0.109144 | 0.067069 |
| 27.93056 | -2.04488 | -0.14862 | -0.23502 | -0.95342 | -0.4679 | -0.55808 | -0.25583 | -0.58034 | -0.42223 | -0.63008 | -0.06654 | -0.24668 | 0.125973 | -0.01679 | -0.02144 | 0.06832 | 0.027487 | 0.096228 | 0.121781 | 0.16378 | 0.499259 | -0.11133 | 0.139664 | 0.093215 |
| 27.94444 | -1.56346 | 0.041013 | -0.02967 | -0.56022 | -0.27516 | -0.30214 | 0.126438 | -0.6745 | -0.50586 | -0.27402 | 0.00003 | 0.424101 | 0.156008 | 0.039425 | -0.01411 | 0.074518 | 0.037328 | 0.115313 | 0.131597 | 0.189125 | 0.623495 | -0.09667 | 0.158749 | 0.132998 |
| 27.95833 | -0.94911 | 0.167231 | 0.113767 | -0.25708 | -0.03812 | 0.00312 | 0.312289 | -0.68238 | -0.41359 | 0.061302 | 0.284044 | 0.855696 | 0.162987 | 0.066171 | -0.03116 | 0.078936 | 0.053291 | 0.143165 | 0.123069 | 0.218067 | 0.704825 | -0.08126 | 0.17428 | 0.141188 |
| 27.97222 | -0.3169 | 0.219095 | 0.015154 | -0.16472 | 0.276478 | -0.01491 | 0.070545 | -0.5148 | 0.047518 | -0.45284 | 0.497205 | 0.452489 | 0.143028 | 0.069665 | -0.04126 | 0.061899 | 0.055676 | 0.152972 | 0.123271 | 0.270162 | 0.775996 | -0.07219 | 0.200704 | 0.147363 |
| 27.98611 | 0.617905 | -0.02476 | -0.26688 | -0.461 | 0.493069 | -0.3516 | -0.87502 | 0.629292 | 0.800444 | -2.47726 | 0.733964 | -1.35434 | 0.09511 | 0.047194 | -0.01403 | 0.032093 | 0.043613 | 0.141078 | 0.131517 | 0.317864 | 0.817315 | -0.07295 | 0.223019 | 0.14817 |
| 28 | 1.172375 | -0.34997 | -0.62217 | -0.47715 | 0.307276 | -0.74081 | -2.1586 | 1.643012 | 1.566483 | -5.17574 | 0.721166 | -3.35939 | -0.10594 | -0.19819 | 0.053616 | -0.05765 | -0.05668 | 0.065301 | 0.014808 | 0.302676 | 0.445533 | -0.13769 | 0.060797 | -0.07457 |
| 28.01389 | 0.633051 | -0.54659 | -0.72953 | -0.63935 | -0.01546 | -0.97452 | -2.10024 | 1.392519 | 0.479641 | -4.96095 | 0.118434 | -3.3335 | -0.71295 | -0.98065 | 0.161391 | -0.4131 | -0.51525 | -0.30475 | -0.56517 | 0.195446 | -1.021 | -0.50122 | -0.3872 | -0.71319 |
| 28.02778 | -0.80294 | -0.09589 | -0.69363 | -0.86861 | -0.27288 | -0.91088 | -0.65553 | -0.01159 | -0.66654 | -2.42744 | -0.83498 | -1.0719 | -1.7974 | -2.41705 | 0.28366 | -1.50835 | -1.35898 | -1.57673 | -1.69558 | -0.02041 | -3.60009 | -1.42525 | -1.51723 | -1.46683 |
| 28.04167 | -1.43198 | -0.02878 | -0.72336 | -0.92981 | -0.23744 | -0.77912 | 0.025545 | -0.80455 | -0.58844 | -1.6565 | -1.07842 | -0.35046 | -2.18193 | -2.66801 | 0.322025 | -2.90119 | -1.54431 | -2.12956 | -2.03422 | -0.16755 | -5.03194 | -1.89275 | -0.83398 | -0.97549 |
| 28.05556 | -1.28235 | -0.6523 | -0.70388 | -1.49812 | -0.46906 | -0.62586 | -0.46612 | -0.78525 | 0.062189 | -2.32757 | -0.68144 | -0.96138 | -1.73142 | -1.42626 | 0.197988 | -2.45081 | -1.21238 | -0.58958 | -0.57818 | -0.28901 | -3.76587 | -1.39135 | -0.1382 | -0.54706 |
| 28.06944 | -1.10546 | -0.95451 | -0.46148 | -1.07208 | -0.60619 | -0.51561 | -0.73478 | -0.79514 | 0.158304 | -1.6469 | -0.3028 | -0.98741 | -0.87411 | -0.34674 | 0.053416 | -0.7762 | -0.66063 | 0.285578 | -0.26349 | -0.29303 | -1.63374 | -0.61547 | -0.24413 | -0.1785 |
| 28.08333 | -0.67718 | -0.83957 | -0.22207 | -0.40235 | -0.41486 | -0.32787 | -0.44176 | -0.70142 | -0.04402 | 0.119252 | -0.51467 | -0.24855 | -0.0833 | 0.194253 | -0.00841 | 0.178454 | -0.19757 | 0.450714 | -0.06151 | -0.17754 | -0.22555 | -0.21662 | -0.22212 | 0.069535 |
| 28.09722 | 0.00823 | -0.43644 | -0.00593 | 0.395943 | -0.16599 | -0.05948 | -0.14154 | -0.18278 | -0.18454 | 1.276761 | -0.57992 | 0.270212 | 0.356614 | 0.348038 | -0.00677 | 0.517718 | 0.128741 | 0.374077 | 0.113054 | -0.05993 | 0.502749 | -0.02265 | -0.09264 | 0.180951 |
| 28.11111 | 0.266254 | -0.03501 | 0.149078 | 0.51978 | 0.113023 | 0.103527 | 0.280269 | 0.326747 | 0.354471 | 2.248222 | -0.37907 | 0.664017 | 0.498564 | 0.43202 | 0.010112 | 0.525216 | 0.270667 | 0.321564 | 0.202263 | 0.028016 | 0.771087 | 0.077065 | -0.00841 | 0.205862 |
| 28.125 | 0.241601 | 0.164235 | 0.499013 | 0.388105 | 0.172507 | 0.215735 | 0.456635 | 0.633431 | 0.753713 | 1.932914 | 0.044489 | 0.479161 | 0.510775 | 0.492157 | 0.026766 | 0.457379 | 0.339068 | 0.313949 | 0.170157 | 0.081271 | 0.781251 | 0.168837 | 0.010675 | 0.192356 |
| 28.13889 | 0.482611 | 0.121696 | 0.860369 | 0.336176 | -0.04915 | 0.35579 | -0.00225 | 0.852389 | 0.620134 | 0.110272 | 0.472642 | -0.21016 | 0.414895 | 0.445957 | 0.017109 | 0.406046 | 0.346151 | 0.290699 | -0.00482 | 0.07638 | 0.738931 | 0.202486 | 0.014177 | 0.165273 |
| 28.15278 | 0.46532 | 0.182167 | 1.083626 | 0.153203 | -0.11227 | 0.440463 | 0.13472 | 0.555238 | 0.680442 | -0.60757 | 0.514422 | -0.3113 | 0.306718 | 0.28956 | -0.00285 | 0.356557 | 0.323722 | 0.271253 | -0.09519 | 0.04422 | 0.726127 | 0.24066 | 0.004488 | 0.141585 |
| 28.16667 | 0.410134 | 0.243352 | 1.053268 | -0.06399 | -0.12719 | 0.415468 | 0.309498 | 0.060037 | 0.583669 | -0.54491 | 0.407264 | -0.02384 | 0.26045 | 0.111267 | -0.00576 | 0.304074 | 0.354575 | 0.289368 | 0.029432 | 0.029223 | 0.677507 | 0.255329 | 0.008123 | 0.161823 |
| 28.18056 | 0.248369 | 0.177281 | 0.679122 | -0.09997 | -0.14217 | 0.399452 | 0.235235 | -0.16391 | 0.504151 | -0.58742 | 0.358334 | 0.040806 | 0.241288 | 0.088135 | 0.009065 | 0.262873 | 0.39079 | 0.342953 | 0.134914 | 0.041562 | 0.646558 | 0.281982 | 0.004245 | 0.173905 |
| 28.19444 | 0.004641 | 0.12353 | 0.408388 | -0.15242 | -0.07483 | 0.278339 | 0.235867 | -0.31388 | 0.314924 | -0.26015 | 0.309529 | 0.263499 | 0.228135 | 0.159874 | 0.022419 | 0.227768 | 0.379557 | 0.37618 | 0.191725 | 0.068548 | 0.617979 | 0.280089 | -0.00748 | 0.159295 |
| 28.20833 | -0.00362 | -0.04369 | 0.257282 | -0.01984 | -0.08526 | 0.183402 | -0.05084 | -0.0254 | 0.258602 | -0.56234 | 0.285474 | 0.071938 | 0.190163 | 0.202204 | 0.019147 | 0.197292 | 0.346588 | 0.369354 | 0.207457 | 0.09787 | 0.572013 | 0.240194 | -0.02113 | 0.131175 |
| 28.22222 | 0.168702 | -0.14966 | 0.326646 | 0.157843 | -0.06302 | 0.16563 | -0.31291 | 0.223916 | 0.226174 | -0.94076 | 0.362337 | -0.16871 | 0.135131 | 0.212174 | 0.002878 | 0.192643 | 0.28754 | 0.355256 | 0.19484 | 0.114585 | 0.523203 | 0.188622 | -0.03093 | 0.105042 |
| 28.23611 | 0.230751 | -0.06477 | 0.530105 | 0.256343 | 0.028194 | 0.158363 | -0.28575 | 0.43676 | 0.203254 | -0.68922 | 0.402714 | -0.16601 | 0.116792 | 0.189329 | -0.01176 | 0.254058 | 0.257581 | 0.351514 | 0.206603 | 0.105481 | 0.485228 | 0.166822 | -0.03721 | 0.100477 |
| 28.25 | 0.171304 | 0.11243 | 0.708339 | 0.193131 | 0.140228 | 0.13925 | -0.06626 | 0.442687 | 0.238704 | 0.081396 | 0.365428 | 0.045579 | 0.137163 | 0.171403 | -0.01695 | 0.254761 | 0.280881 | 0.365774 | 0.193213 | 0.087424 | 0.457131 | 0.151757 | -0.04446 | 0.09912 |
| 28.26389 | -0.22662 | 0.493439 | 0.629139 | -0.18043 | 0.309657 | 0.083808 | 0.64901 | 0.049784 | 0.548324 | 2.069702 | 0.003839 | 0.880314 | 0.142946 | 0.16368 | -0.01504 | 0.207381 | 0.34287 | 0.364417 | 0.172836 | 0.07126 | 0.389732 | 0.161625 | -0.05171 | 0.090369 |
| 28.27778 | -0.44835 | 0.616861 | 0.233007 | -0.43826 | 0.365601 | 0.074966 | 0.988231 | -0.32321 | 0.874427 | 2.96787 | -0.25954 | 1.231471 | 0.110424 | 0.15939 | -0.01267 | 0.171534 | 0.36295 | 0.359121 | 0.15547 | 0.056608 | 0.334941 | 0.151051 | -0.06671 | 0.076915 |
| 28.29167 | -0.09541 | 0.362125 | 0.267529 | -0.06084 | 0.073275 | 0.082779 | 0.198433 | -0.17131 | 0.327025 | 0.823874 | -0.00246 | 0.272159 | 0.06061 | 0.148166 | -0.00873 | 0.141427 | 0.244398 | 0.296285 | 0.128462 | 0.041046 | 0.289777 | 0.112065 | -0.07676 | 0.046537 |
| 28.30556 | 0.409687 | -0.11372 | 0.535985 | 0.593627 | -0.23042 | 0.117618 | -0.90499 | 0.325341 | -0.53715 | -2.17521 | 0.372023 | -1.13622 | 0.049305 | 0.129821 | -0.00386 | 0.114531 | 0.144989 | 0.252856 | 0.095702 | 0.036032 | 0.220336 | 0.063581 | -0.08917 | 0.001922 |
| 28.31944 | 0.487479 | -0.16527 | 0.456237 | 0.958594 | -0.16671 | 0.059037 | -1.09698 | 0.577538 | -0.20285 | -2.2861 | 0.294887 | -1.2637 | 0.053327 | 0.112731 | 0.003578 | 0.086352 | 0.053207 | 0.206162 | 0.068283 | 0.054915 | 0.136348 | 0.030828 | -0.11154 | -0.03251 |
| 28.33333 | -0.14071 | 0.201897 | 0.117969 | 0.388566 | 0.2934 | -0.09841 | -0.03868 | 0.399195 | -0.16017 | 0.498828 | -0.15391 | 0.181982 | 0.041515 | 0.113995 | 0.010774 | 0.058017 | 0.016765 | 0.184596 | 0.046109 | 0.075714 | 0.0709 | 0.0066 | -0.10729 | -0.05053 |
| 28.34722 | -0.56525 | 0.485455 | -0.00476 | -0.21764 | 0.55342 | -0.20447 | 0.754305 | 0.037637 | -0.15549 | 2.460685 | -0.50524 | 1.26132 | 0.01949 | 0.130374 | 0.011352 | 0.0368 | 0.01172 | 0.174557 | 0.026142 | 0.081795 | 0.028292 | -0.00886 | -0.08483 | -0.06487 |
| 28.36111 | -0.26224 | 0.353102 | 0.189742 | -0.1763 | 0.382998 | -0.05278 | 0.511819 | -0.03004 | -0.20305 | 1.625258 | -0.2404 | 0.700474 | 0.007906 | 0.125023 | 0.004594 | 0.021412 | -0.00503 | 0.174135 | 0.025692 | 0.073732 | 0.017413 | -0.03109 | -0.07611 | -0.06178 |

| | | | | | | | | | | | | | | | | | | | | | | | | |
|---|---|---|---|---|---|---|---|---|---|---|---|---|---|---|---|---|---|---|---|---|---|---|---|---|
| 28.375 | 0.269596 | 0.045783 | 0.334444 | 0.326879 | 0.027221 | 0.129463 | -0.26293 | 0.347804 | -0.04347 | -0.41351 | 0.194875 | -0.38477 | 0.018166 | 0.087367 | -0.00227 | 0.016669 | -0.02146 | 0.181682 | 0.049984 | 0.067868 | 0.042583 | -0.04096 | -0.07167 | -0.03276 |
| 28.38889 | 0.458657 | 0.024836 | 0.379467 | 0.533839 | -0.07253 | 0.227734 | -0.38583 | 0.588013 | 0.116597 | -0.75682 | 0.406607 | -0.50766 | 0.047422 | 0.072722 | 0.000784 | 0.027142 | -0.00914 | 0.190006 | 0.07828 | 0.083243 | 0.082017 | -0.03022 | -0.05378 | -0.01156 |
| 28.40278 | 0.163046 | 0.207215 | 0.357241 | 0.389552 | 0.152529 | 0.249973 | 0.213033 | 0.465651 | -0.08044 | 0.768005 | 0.110933 | 0.171931 | 0.079416 | 0.106901 | 0.007667 | 0.034447 | 0.003981 | 0.184545 | 0.08862 | 0.094732 | 0.104958 | -0.04438 | -0.05141 | -0.03159 |
| 28.41667 | 0.186479 | 0.268439 | 0.269712 | 0.605872 | 0.186306 | 0.266548 | 0.288544 | 0.398079 | -0.22667 | 0.884626 | -0.02699 | 0.261706 | 0.130644 | 0.168488 | 0.017164 | 0.045645 | 0.001412 | 0.160667 | 0.084683 | 0.087831 | 0.11366 | -0.09404 | -0.07836 | -0.07562 |
| 28.43056 | 0.441646 | 0.221065 | 0.285027 | 1.02679 | 0.124783 | 0.370903 | 0.074091 | 0.455887 | -0.26796 | 0.103407 | 0.127436 | -0.05163 | 0.179979 | 0.202904 | 0.028097 | 0.08568 | -0.01052 | 0.155479 | 0.111922 | 0.094349 | 0.134469 | -0.1302 | -0.0863 | -0.06728 |
| 28.44444 | 0.349964 | 0.308214 | 0.195293 | 0.643547 | 0.243195 | 0.473623 | 0.488797 | 0.358967 | -0.46389 | 0.838567 | -0.0885 | 0.376751 | 0.180057 | 0.178081 | 0.025772 | 0.140923 | -0.01809 | 0.176887 | 0.167599 | 0.127498 | 0.180841 | -0.12297 | -0.05619 | -0.01595 |
| 28.45833 | 0.238914 | 0.297253 | 0.203267 | 0.165788 | 0.253167 | 0.52078 | 0.717278 | 0.342644 | -0.52686 | 1.393542 | -0.31788 | 0.69867 | 0.14391 | 0.160826 | 0.014076 | 0.233375 | -1.5E-05 | 0.20798 | 0.207004 | 0.156058 | 0.215361 | -0.12769 | -0.02941 | 0.002343 |
| 28.47222 | 0.258065 | 0.153568 | 0.317989 | 0.033396 | 0.045787 | 0.460784 | 0.41158 | 0.364674 | -0.23908 | 0.628614 | -0.08826 | 0.265651 | 0.133761 | 0.183489 | 0.006596 | 0.354675 | 0.027104 | 0.224679 | 0.214939 | 0.178225 | 0.240461 | -0.13382 | -0.01225 | -0.00147 |
| 28.48611 | 0.239426 | 0.107961 | 0.260171 | 0.120689 | -0.01093 | 0.392154 | 0.405396 | 0.367179 | -0.30979 | 0.831338 | -0.03318 | 0.245088 | 0.162081 | 0.245792 | 0.018743 | 0.383701 | 0.05836 | 0.217409 | 0.199859 | 0.213369 | 0.338579 | -0.10288 | 0.023858 | 0.028931 |
| 28.5 | 0.228927 | 0.326348 | 0.119708 | 0.478042 | 0.421887 | 0.3688 | 1.16288 | 0.20371 | -0.964 | 2.416353 | -0.13975 | 0.982539 | 0.331351 | 0.441469 | 0.021862 | 0.405221 | 0.210786 | 0.271129 | 0.162163 | 0.365916 | 0.880652 | 0.025917 | 0.113137 | 0.140296 |
| 28.51389 | 0.688069 | 0.615031 | 0.382528 | 1.475146 | 0.688926 | 0.687752 | 1.63748 | 0.510203 | -1.39824 | 3.369761 | 0.244982 | 1.27996 | 0.794473 | 0.880681 | -0.01244 | 0.645575 | 0.642135 | 0.43141 | 0.115953 | 0.700369 | 2.270714 | 0.259267 | 0.167145 | 0.347711 |
| 28.52778 | 1.879638 | 0.468545 | 1.111752 | 3.161492 | 0.751759 | 1.363168 | 1.294515 | 1.215258 | 0.061273 | 2.681335 | 1.132576 | 0.8222 | 1.196303 | 1.316566 | -0.05967 | 0.844114 | 1.117281 | 0.682276 | 0.175188 | 0.962302 | 3.737439 | 0.257392 | 0.204881 | 0.759215 |
| 28.54167 | 2.864091 | 0.094937 | 1.796799 | 2.805484 | 0.558999 | 2.01767 | 0.650906 | 2.371751 | 2.139019 | 1.86813 | 1.084323 | 0.309947 | 0.973254 | 1.378911 | -0.08054 | 0.745723 | 1.171772 | 0.579524 | 0.188343 | 0.918258 | 4.047299 | 0.124327 | 0.461514 | 1.045796 |
| 28.55556 | 2.639962 | 0.033074 | 1.40772 | 0.863192 | 0.491988 | 1.578059 | 0.729599 | 2.500864 | 2.42598 | 1.190584 | 0.380256 | 0.173189 | 0.661902 | 1.375828 | -0.11915 | 0.511322 | 1.107924 | 0.375144 | 0.37069 | 0.918843 | 3.963602 | 0.046193 | 0.805233 | 1.202207 |
| 28.56944 | 2.053564 | 0.165794 | 0.439506 | 0.537019 | 0.541478 | 1.025444 | 0.934491 | 1.745869 | 1.065706 | 0.697056 | -0.0637 | 0.214237 | 0.532416 | 1.201913 | -0.0673 | 0.413476 | 1.095592 | 0.310358 | 0.504807 | 1.047119 | 3.690074 | 0.132795 | 0.958749 | 1.115492 |
| 28.58333 | 1.553881 | 0.180233 | -0.28333 | 0.887197 | 0.526916 | 0.719962 | 0.96999 | 0.906966 | -0.36004 | -0.08098 | -0.16007 | 0.356358 | 0.506572 | 1.032243 | 0.015341 | 0.40028 | 1.115593 | 0.317992 | 0.501045 | 1.112723 | 3.106349 | 0.252601 | 1.011432 | 0.943411 |
| 28.59722 | 0.902235 | 0.13499 | -0.58471 | 0.775802 | 0.442696 | 0.509421 | 0.743599 | 0.144788 | -1.12821 | -0.98621 | -0.12862 | 0.41898 | 0.423417 | 0.802657 | 0.053497 | 0.359601 | 0.984264 | 0.311711 | 0.444413 | 1.024767 | 2.546438 | 0.315751 | 1.326543 | 0.885178 |
| 28.61111 | 0.481312 | 0.108929 | -0.45673 | 0.274059 | 0.327291 | 0.473792 | 0.473021 | -0.67395 | -0.86099 | -1.43644 | -0.16719 | 0.344331 | 0.323419 | 0.590248 | 0.073696 | 0.32616 | 0.834282 | 0.29206 | 0.394585 | 0.880606 | 1.982702 | 0.323226 | 1.291926 | 0.810302 |
| 28.625 | 0.247622 | 0.109036 | -0.33219 | 0.378735 | 0.082409 | 0.286804 | 0.243855 | -1.05501 | -0.08802 | -0.99816 | -0.32584 | 0.306126 | 0.212821 | 0.445591 | 0.089517 | 0.318778 | 0.670115 | 0.257882 | 0.441596 | 0.72396 | 1.570049 | 0.266008 | 1.211263 | 0.689982 |
| 28.63889 | 0.163395 | 0.101468 | -0.11161 | 0.530743 | -0.0748 | 0.166491 | -0.07064 | -0.75719 | 0.64506 | -0.13257 | -0.5516 | 0.340205 | 0.12729 | 0.375911 | 0.073226 | 0.282365 | 0.474453 | 0.217204 | 0.458326 | 0.658172 | 1.209369 | 0.175958 | 1.090863 | 0.500896 |
| 28.65278 | 0.165094 | 0.069488 | 0.053487 | 0.752405 | -0.11607 | 0.160708 | -0.42691 | -0.13898 | 0.765365 | 0.577297 | -0.52745 | 0.290101 | 0.09915 | 0.370722 | 0.028307 | 0.226312 | 0.364474 | 0.211953 | 0.447979 | 0.5843 | 1.055019 | 0.110966 | 0.891298 | 0.405884 |
| 28.66667 | 0.336084 | 0.046138 | 0.018038 | 0.841615 | -0.08226 | 0.277536 | -0.79794 | 0.4477 | 0.566161 | 0.876399 | 0.014184 | 0.199361 | 0.158411 | 0.335139 | 0.005872 | 0.209522 | 0.277731 | 0.224295 | 0.462952 | 0.562193 | 0.910882 | 0.052474 | 0.78484 | 0.377107 |
| 28.68056 | 0.451938 | 0.042247 | 0.110181 | 0.762159 | -0.14053 | 0.358315 | -0.75298 | 1.37356 | 0.075822 | 0.699492 | 0.535372 | 0.038085 | 0.263176 | 0.30871 | 0.013962 | 0.215908 | 0.253213 | 0.232682 | 0.37165 | 0.515476 | 0.94179 | -0.01545 | 0.692831 | 0.385145 |
| 28.69444 | 0.459778 | 0.062434 | 0.256857 | 0.515215 | -0.14287 | 0.433398 | -0.62861 | 2.42952 | -0.43597 | 0.185729 | 0.640857 | -0.18055 | 0.338031 | 0.671824 | 0.049215 | 0.203154 | 0.282683 | 0.216364 | 0.296314 | 0.495504 | 0.878168 | -0.05519 | 0.651635 | 0.397511 |
| 28.70833 | 0.104702 | 0.047058 | 0.525421 | 0.434061 | -0.10319 | 0.482089 | -0.42813 | 2.787816 | -0.57641 | -0.3556 | 0.36941 | -0.37271 | 0.345117 | 1.192335 | 0.127849 | 0.181541 | 0.288215 | 0.190467 | 0.242801 | 0.463049 | 0.79546 | -0.07422 | 0.600024 | 0.3773 |
| 28.72222 | -0.28819 | -0.01835 | 0.071894 | 0.135752 | 0.006138 | 0.48328 | 0.053691 | 2.692435 | -0.4801 | -0.76222 | 0.058031 | -0.51285 | 0.294192 | 1.028596 | 0.171967 | 0.153983 | 0.248378 | 0.143873 | 0.191085 | 0.461453 | 0.723375 | -0.08841 | 0.522824 | 0.318362 |
| 28.73611 | -0.62242 | -0.0677 | -0.54585 | -0.29513 | 0.142996 | 0.385381 | 0.608202 | 1.652564 | -0.19389 | -1.04783 | -0.00547 | -0.48544 | 0.209247 | 0.455726 | 0.17696 | 0.107819 | 0.223886 | 0.080841 | 0.10921 | 0.43905 | 0.662534 | -0.09675 | 0.484267 | 0.270253 |
| 28.75 | -0.40474 | -0.0517 | -0.58522 | -0.37348 | 0.185706 | 0.266108 | 0.818951 | 0.024744 | -0.01141 | -1.34641 | 0.057858 | -0.26812 | 0.097966 | 0.170036 | 0.097694 | 0.060073 | 0.205493 | 0.021675 | 0.064523 | 0.418246 | 0.617208 | -0.10696 | 0.453696 | 0.235736 |
| 28.76389 | -0.05871 | 0.007148 | -0.6121 | 0.170029 | 0.258061 | 0.181609 | 0.854054 | -1.41838 | -0.03632 | -1.47705 | 0.057418 | -0.06672 | 0.004112 | 0.097837 | -0.05611 | 0.023259 | 0.22164 | -0.02058 | 0.068823 | 0.345624 | 0.597867 | -0.08668 | 0.397188 | 0.219077 |
| 28.77778 | 0.554624 | 0.01729 | -0.75397 | 0.761467 | 0.436744 | -0.02696 | 0.794759 | -2.31568 | -0.40333 | -1.37932 | 0.205886 | 0.042212 | -0.06778 | 0.06746 | -0.17096 | 0.021386 | 0.226476 | -0.03035 | 0.08816 | 0.282715 | 0.531381 | -0.02662 | 0.348766 | 0.209881 |
| 28.79167 | 0.735833 | -0.02253 | -0.52804 | 0.955354 | 0.553445 | -0.24637 | 0.649336 | -2.32343 | -0.70433 | -0.96037 | 0.329521 | 0.098875 | -0.10268 | 0.05141 | -0.20306 | 0.066704 | 0.206154 | 0.013133 | 0.078934 | 0.218904 | 0.445383 | 0.016082 | 0.324758 | 0.208382 |
| 28.80556 | 0.596018 | -0.05541 | -0.3299 | 0.644652 | 0.451506 | -0.45258 | 0.328429 | -1.72632 | -0.62767 | -0.72179 | -0.03677 | 0.089648 | -0.10678 | 0.073022 | -0.13471 | 0.102431 | 0.171944 | 0.045572 | 0.038784 | 0.163488 | 0.333523 | 0.0012 | 0.297112 | 0.198759 |
| 28.81944 | 0.080333 | -0.09321 | -0.34471 | -0.04301 | 0.164089 | -0.46892 | 0.056831 | -1.42025 | -0.21568 | -0.75666 | -0.35725 | 0.041725 | -0.04341 | 0.062709 | -0.08305 | 0.05789 | 0.137805 | 0.017959 | 0.005903 | 0.1321 | 0.223686 | -0.04697 | 0.282328 | 0.154803 |
| 28.83333 | -0.63349 | -0.13728 | -0.33028 | -0.55016 | -0.1777 | -0.39128 | -0.06219 | -1.15876 | 0.178578 | -1.12767 | -0.56892 | -0.0655 | 0.024262 | -0.01345 | -0.04763 | -0.0655 | 0.122691 | -0.05943 | -0.00961 | 0.102699 | 0.121643 | -0.09242 | 0.269459 | 0.137639 |
| 28.84722 | -1.22991 | -0.17566 | -0.20594 | -0.65443 | -0.39522 | -0.37528 | -0.17073 | -1.05559 | 0.40789 | -1.77803 | -0.567 | -0.22485 | 0.066975 | -0.11855 | -0.02226 | -0.2279 | 0.12547 | -0.17053 | -0.03388 | 0.090689 | 0.07551 | -0.09418 | 0.262992 | 0.146381 |
| 28.86111 | -1.53213 | -0.24939 | -0.21576 | -0.57296 | -0.75133 | -0.39859 | -0.37199 | -1.12351 | 0.547494 | -2.41869 | -0.3766 | -0.34097 | 0.060829 | -0.1952 | -0.02644 | -0.40378 | 0.136867 | -0.27123 | -0.07679 | 0.106508 | 0.085388 | -0.07506 | 0.227006 | 0.162627 |
| 28.875 | -1.53354 | -0.37231 | -0.35844 | -0.22367 | -0.74485 | -0.39732 | -0.51617 | -1.27638 | 0.174838 | -2.36186 | -0.21965 | -0.30423 | 0.024259 | -0.25243 | -0.06427 | -0.57982 | 0.138128 | -0.34079 | -0.10797 | 0.113618 | 0.101584 | -0.04398 | 0.15507 | 0.172528 |
| 28.88889 | -1.17831 | -0.41892 | -0.32549 | -0.45608 | -0.63937 | -0.34745 | -0.57811 | -1.25934 | -0.39216 | -1.92054 | -0.22074 | -0.15753 | -0.00855 | -0.25395 | -0.08797 | -0.69565 | 0.132932 | -0.32992 | -0.09159 | 0.094337 | 0.179764 | -0.01282 | 0.087008 | 0.162177 |
| 28.90278 | -0.82383 | -0.57297 | -0.17744 | -0.70233 | -0.49155 | -0.3371 | -0.53276 | -1.10434 | -0.51134 | -1.08649 | -0.08449 | -0.00273 | -0.03605 | -0.22945 | -0.08188 | -0.61409 | 0.136881 | -0.29204 | -0.05638 | 0.10058 | 0.266234 | 0.004913 | 0.083532 | 0.148512 |
| 28.91667 | -0.87776 | -0.40155 | 0.036367 | -0.93129 | -0.32097 | -0.28921 | -0.46428 | -0.83478 | -0.07811 | -0.28258 | 0.134896 | 0.108487 | -0.04679 | -0.25547 | -0.06133 | -0.37463 | 0.143462 | -0.2042 | -0.02745 | 0.107524 | 0.373458 | 0.005096 | 0.121515 | 0.138574 |
| 28.93056 | -0.93711 | -0.06353 | 0.0116 | -0.97154 | -0.11245 | -0.16662 | -0.38057 | -0.53299 | 0.59063 | 0.360227 | 0.377233 | 0.179716 | -0.08812 | -0.39192 | -0.04391 | -0.15306 | 0.140394 | -0.17225 | -0.01214 | 0.091539 | 0.4009 | -0.00747 | 0.156256 | 0.11209 |
| 28.94444 | -0.84082 | 0.182595 | 0.045299 | -0.81296 | -0.06176 | -0.05802 | -0.2973 | -0.20497 | 1.203646 | 0.830011 | 0.610171 | 0.268047 | -0.14123 | -0.41764 | -0.0301 | -0.00225 | 0.123931 | -0.2082 | -0.03009 | 0.046391 | 0.384304 | -0.02926 | 0.176217 | 0.079753 |
| 28.95833 | -0.58358 | 0.274201 | 0.046005 | -0.57575 | -0.15399 | 0.000771 | -0.2423 | -0.04581 | 1.479565 | 0.995486 | 0.732861 | 0.341384 | -0.14481 | -0.32934 | -0.01559 | 0.056452 | 0.104594 | -0.26805 | -0.03258 | 0.026652 | 0.365666 | -0.03067 | 0.174343 | 0.062268 |
| 28.97222 | -0.2356 | 0.316361 | 0.027945 | -0.48143 | -0.26295 | 0.034703 | -0.1946 | 0.060293 | 1.364492 | 0.653271 | 0.569094 | 0.408575 | -0.10036 | -0.25474 | -0.00576 | -0.00207 | 0.089822 | -0.32561 | -0.04457 | -0.00523 | 0.302012 | -0.03738 | 0.148518 | 0.056573 |
| 28.98611 | -0.28636 | 0.254626 | -0.00099 | -0.77496 | -0.28434 | -0.02843 | -0.40289 | -0.0047 | 0.884979 | -0.32008 | 0.3471 | 0.318759 | -0.09513 | -0.24219 | -0.00267 | -0.10775 | 0.07604 | -0.35159 | -0.0505 | -0.03915 | 0.198667 | -0.04587 | 0.111668 | 0.052323 |
| 29 | -0.9613 | -0.07996 | -0.06956 | -1.17472 | -0.16212 | -0.28311 | -0.61714 | -0.23458 | 0.022638 | -1.9181 | -0.01065 | -0.10366 | -0.08382 | -0.23813 | 0.042077 | -0.14333 | 0.080415 | -0.32242 | -0.02746 | -0.05232 | 0.126135 | -0.0558 | 0.093394 | 0.068329 |
| 29.01389 | -1.43175 | -0.28273 | -0.29794 | -1.24362 | 0.002243 | -0.51417 | -0.90534 | -0.61 | -1.03566 | -3.9605 | -0.40317 | -0.76125 | -0.14757 | -0.39789 | 0.166048 | -0.0906 | 0.008413 | -0.32242 | 0.041843 | -0.13741 | -0.02584 | -0.21888 | -0.07177 | 0.019119 |
| 29.02778 | -1.45217 | -0.18608 | -0.55151 | -1.75947 | 0.215086 | -0.54375 | -0.90584 | -0.67612 | -1.77693 | -5.30073 | -0.85369 | -1.45331 | -0.37808 | -0.74223 | 0.260581 | -0.03254 | -0.31762 | -0.51445 | 0.169529 | -0.52659 | -0.64754 | -0.55609 | -0.65838 | -0.08489 |
| 29.04167 | -1.58411 | -0.19287 | -0.48561 | -1.4623 | 0.189991 | -0.41274 | -0.71132 | -1.06874 | -1.71403 | -4.47663 | -0.9961 | -1.96043 | -0.41019 | -1.03734 | 0.204877 | 0.00998 | -1.08962 | -0.67217 | -0.02118 | -1.13588 | -1.5865 | -0.89422 | -1.27442 | -0.14177 |
| 29.05556 | -1.65642 | -0.42998 | -0.30009 | -0.79662 | 0.054362 | -0.65966 | -0.49373 | -1.31859 | -1.4872 | -1.54731 | -0.70975 | -1.23533 | -0.43555 | -0.6978 | 0.176936 | -0.04991 | -0.98739 | -0.37812 | -0.33528 | -0.55915 | -1.21685 | -0.85484 | -0.81935 | -0.44389 |
| 29.06944 | -1.27996 | -0.32237 | -0.01988 | -0.15865 | 0.08752 | -0.59114 | -0.17303 | -1.1607 | -0.72183 | 1.378207 | -0.55522 | -0.05006 | -0.28434 | -0.14608 | 0.107597 | -0.10996 | -0.05066 | 0.096566 | -0.19292 | 0.180657 | 0.12351 | -0.56528 | -0.13485 | -0.31529 |
| 29.08333 | -0.83488 | -0.07449 | 0.335334 | 0.439066 | 0.238408 | -0.3061 | 0.056824 | -0.25079 | 0.288264 | 3.201602 | -0.50959 | 0.961671 | -0.01577 | 0.325377 | 0.072026 | 0.154654 | 0.324294 | 0.271642 | 0.091936 | 0.280697 | 0.504268 | -0.29144 | 0.378595 | 0.021213 |
| 29.09722 | -0.29659 | 0.014986 | 0.497176 | 0.709576 | 0.327068 | -0.02401 | 0.186277 | 0.650438 | 1.33802 | 3.726008 | -0.46175 | 1.536023 | 0.248853 | 0.484157 | 0.024466 | 0.451258 | 0.301127 | 0.287817 | 0.162384 | 0.225485 | 0.464451 | -0.15819 | 0.522944 | 0.152159 |
| 29.11111 | 0.162499 | 0.155955 | 0.432052 | 0.604216 | 0.289881 | 0.177254 | 0.207344 | 0.739886 | 1.676208 | 3.356419 | -0.32595 | 1.501565 | 0.37248 | 0.504931 | 0.014357 | 0.457933 | 0.259009 | 0.308789 | 0.101648 | 0.168072 | 0.376549 | -0.07493 | 0.355289 | 0.157965 |
| 29.125 | 0.394031 | 0.330922 | 0.372835 | 0.542035 | 0.188874 | 0.295807 | 0.227918 | 1.258169 | 1.183882 | 2.987166 | -0.17272 | 1.223024 | 0.331993 | 0.502498 | 0.011619 | 0.300371 | 0.261119 | 0.323589 | 0.034337 | 0.127598 | 0.292673 | 0.011707 | 0.197257 | 0.15053 |
| 29.13889 | 0.677139 | 0.424098 | 0.530843 | 0.572305 | 0.078243 | 0.352047 | 0.246094 | 1.116368 | 0.175535 | 2.519779 | -0.07884 | 0.825693 | 0.188795 | 0.449062 | 0.035319 | 0.184981 | 0.261388 | 0.298441 | 0.01416 | 0.113259 | 0.261092 | 0.080397 | 0.189377 | 0.151345 |
| 29.15278 | 0.807477 | 0.43888 | 0.670923 | 0.557696 | 0.036674 | 0.339858 | 0.249023 | 0.873261 | -0.24877 | 1.740524 | 0.013875 | 0.440414 | 0.059386 | 0.385838 | 0.055094 | 0.091563 | 0.233545 | 0.192404 | 0.021301 | 0.120658 | 0.287596 | 0.106951 | 0.186397 | 0.15142 |
| 29.16667 | 0.738076 | 0.497449 | 0.76353 | 0.580328 | 0.04758 | 0.290464 | 0.231608 | 0.430426 | -0.02148 | 0.959094 | 0.096794 | 0.137124 | -0.03567 | 0.313788 | 0.055021 | 0.033372 | 0.209631 | 0.076419 | 0.014429 | 0.136616 | 0.332044 | 0.129502 | 0.086037 | 0.142241 |
| 29.18056 | 0.811221 | 0.414057 | 0.607306 | 0.606094 | 0.061054 | 0.300895 | 0.247728 | 0.232274 | 0.240866 | 0.124372 | 0.168221 | -0.05922 | -0.12498 | 0.219137 | 0.041148 | -0.01779 | 0.184038 | 0.021945 | 0.007168 | 0.1602 | 0.380986 | 0.139292 | 0.043359 | 0.125799 |
| 29.19444 | 0.591648 | 0.293054 | 0.466345 | 0.678745 | 0.088484 | 0.339346 | 0.280256 | 0.113941 | 0.47056 | -0.40859 | 0.251701 | -0.14293 | -0.16607 | 0.141262 | 0.037668 | -0.06298 | 0.165579 | 0.060802 | 0.008811 | 0.193274 | 0.453649 | 0.122226 | 0.131272 | 0.124113 |
| 29.20833 | 0.381815 | 0.158017 | 0.456902 | 0.760129 | 0.09104 | 0.382366 | 0.312146 | -0.01075 | 0.625978 | -0.68856 | 0.358334 | -0.11834 | -0.16698 | 0.108837 | 0.031059 | -0.09176 | 0.161807 | 0.12944 | 0.016651 | 0.209319 | 0.50029 | 0.073849 | 0.252343 | 0.118496 |
| 29.22222 | 0.212715 | 0.048673 | 0.423832 | 0.763484 | 0.048568 | 0.386814 | 0.287624 | 0.05646 | 0.748528 | -0.77665 | 0.437133 | -0.01309 | -0.16346 | 0.106 | 0.022721 | -0.10353 | 0.16201 | 0.145405 | 0.040206 | 0.232836 | 0.502703 | 0.039047 | 0.299308 | 0.107062 |
| 29.23611 | -0.01177 | -0.03758 | 0.345139 | 0.666369 | 0.013564 | 0.358507 | 0.21403 | 0.246101 | 0.987574 | -0.4737 | 0.458776 | 0.096915 | -0.15679 | 0.108522 | 0.019839 | -0.11696 | 0.163683 | 0.120686 | 0.063503 | 0.264496 | 0.469434 | 0.033243 | 0.24495 | 0.110699 |
| 29.25 | -0.16108 | -0.09058 | 0.307707 | 0.461145 | 0.011834 | 0.34015 | 0.104655 | 0.400904 | 0.9992 | -0.17978 | 0.375475 | 0.154903 | -0.12804 | 0.106028 | 0.013501 | -0.10749 | 0.173113 | 0.113584 | 0.06753 | 0.269452 | 0.454671 | 0.016675 | 0.139527 | 0.097707 |
| 29.26389 | -0.31521 | -0.08066 | 0.251978 | 0.258292 | -0.00711 | 0.309251 | 0.008877 | 0.336748 | 0.892411 | -0.0773 | 0.29996 | 0.18832 | -0.09882 | 0.108562 | 0.012183 | -0.08241 | 0.192623 | 0.132904 | 0.066587 | 0.278599 | 0.479547 | -0.01291 | 0.059204 | 0.102959 |
| 29.27778 | -0.41964 | -0.03389 | 0.17912 | 0.038131 | -0.08557 | 0.215177 | -0.09177 | 0.140065 | 0.629911 | -0.27677 | 0.232446 | 0.163658 | -0.06064 | 0.068291 | 0.020606 | -0.01301 | 0.167333 | 0.123616 | 0.072163 | 0.346386 | 0.658046 | -0.04369 | 0.045539 | 0.115223 |
| 29.29167 | -0.38689 | -0.11579 | 0.114824 | -0.0139 | -0.17332 | 0.106086 | -0.11792 | -0.22423 | 0.258538 | -0.58349 | 0.18102 | 0.142139 | -0.01418 | -0.04245 | 0.011912 | 0.030566 | 0.075406 | 0.031728 | 0.040446 | 0.387145 | 0.803525 | -0.08293 | 0.051335 | 0.082742 |
| 29.30556 | -0.35938 | -0.08228 | 0.12887 | 0.130666 | -0.16283 | 0.046613 | -0.0084 | -0.28343 | 0.065044 | -0.69906 | 0.175855 | 0.22216 | 0.013457 | -0.12261 | -0.00957 | 0.041299 | 0.007559 | -0.05611 | -0.01569 | 0.395851 | 0.773645 | -0.11696 | 0.079681 | 0.039581 |
| 29.31944 | -0.3259 | -0.09088 | 0.098839 | 0.419251 | -0.06949 | 0.018129 | 0.144858 | 0.081628 | 0.21425 | -0.47241 | 0.227501 | 0.386436 | 0.03801 | -0.06813 | -0.00982 | 0.03644 | 0.014315 | -0.04613 | -0.04675 | 0.365613 | 0.595594 | -0.10176 | 0.147141 | 0.04913 |
| 29.33333 | -0.31797 | -0.12689 | 0.091706 | 0.698868 | 0.006133 | 0.002404 | 0.239036 | 0.543618 | 0.520767 | 0.063365 | 0.298275 | 0.503283 | 0.076513 | 0.051525 | 0.002117 | 0.021926 | 0.04007 | 0.019646 | -0.04418 | 0.332153 | 0.508183 | -0.06005 | 0.165928 | 0.061964 |
| 29.34722 | -0.14995 | -0.19499 | 0.134454 | 0.749381 | 0.007741 | 0.058928 | 0.261469 | 0.873571 | 0.814174 | 0.528598 | 0.32412 | 0.482421 | 0.123748 | 0.193134 | 0.007359 | 0.005535 | 0.061786 | 0.074925 | -0.01191 | 0.313143 | 0.454264 | -0.02648 | 0.141038 | 0.054084 |
| 29.36111 | 0.043455 | -0.13437 | 0.213948 | 0.540137 | -0.00694 | 0.180209 | 0.240003 | 0.823215 | 0.909659 | 0.722602 | 0.305708 | 0.391541 | 0.180405 | 0.31167 | 0.008614 | -0.00306 | 0.076269 | 0.091726 | 0.026188 | 0.332638 | 0.444385 | 0.010201 | 0.102951 | 0.050596 |
| 29.375 | 0.294726 | 0.01848 | 0.265125 | 0.392596 | -0.0588 | 0.230121 | 0.188096 | 0.523167 | 0.73558 | 0.581083 | 0.281294 | 0.223924 | 0.209367 | 0.358823 | 0.006539 | 0.01304 | 0.054711 | 0.073339 | 0.038354 | 0.343315 | 0.424463 | 0.027305 | 0.053119 | 0.053779 |
| 29.38889 | 0.441429 | 0.120579 | 0.258936 | 0.269449 | -0.06426 | 0.185931 | 0.134248 | 0.294372 | 0.508543 | 0.225482 | 0.267742 | 0.044798 | 0.236661 | 0.283898 | 0.003193 | 0.061375 | 0.022946 | 0.060114 | 0.012756 | 0.309532 | 0.416344 | 0.02581 | 0.019531 | 0.033651 |
| 29.40278 | 0.464955 | 0.142731 | 0.235152 | 0.333233 | -0.0364 | 0.145973 | 0.143911 | 0.188905 | 0.342808 | -0.08266 | 0.253439 | -0.06784 | 0.248293 | 0.144652 | 0.010164 | 0.128079 | 0.014072 | 0.073015 | -0.00045 | 0.277232 | 0.408515 | 0.022433 | 0.023298 | 0.021964 |
| 29.41667 | 0.482644 | 0.161367 | 0.209182 | 0.521086 | -0.03004 | 0.140463 | 0.243572 | 0.302617 | 0.264057 | -0.16005 | 0.266216 | -0.00963 | 0.238401 | 0.036318 | 0.024459 | 0.16243 | 0.005034 | 0.08283 | -0.00203 | 0.252504 | 0.386368 | 0.040152 | 0.031482 | 0.017005 |
| 29.43056 | 0.580046 | 0.141557 | 0.108568 | 0.680715 | -0.08357 | 0.194614 | 0.418447 | 0.557764 | 0.185283 | -0.13605 | 0.298989 | 0.133929 | 0.180368 | -0.01403 | 0.025307 | 0.122604 | -0.01951 | 0.076306 | -0.00008 | 0.220312 | 0.325809 | 0.04446 | 0.03184 | 0.010245 |
| 29.44444 | 0.692435 | 0.11179 | -0.08088 | 0.780254 | -0.17313 | 0.307331 | 0.61317 | 0.790495 | 0.195701 | -0.03341 | 0.285587 | 0.256039 | 0.123498 | -0.00916 | 0.009864 | 0.067871 | -0.02173 | 0.071882 | 0.000108 | 0.190802 | 0.237246 | 0.01353 | 0.015219 | 0.011025 |
| 29.45833 | 0.730005 | 0.04066 | -0.01869 | 0.656943 | -0.2703 | 0.385863 | 0.701889 | 0.964316 | 0.309587 | 0.119198 | 0.232638 | 0.226218 | 0.098117 | 0.01409 | -0.01407 | 0.079145 | 0.009295 | 0.080478 | 0.033633 | 0.151192 | 0.202998 | -0.03702 | -0.00376 | 0.0229 |
| 29.47222 | 0.746774 | 0.057064 | 0.073338 | 0.482696 | -0.30541 | 0.536456 | 0.744713 | 1.05313 | 0.525255 | 0.262168 | 0.220449 | 0.022128 | 0.103669 | 0.037646 | -0.02705 | 0.171408 | 0.034339 | 0.096371 | 0.090256 | 0.129394 | 0.209419 | -0.0831 | -0.00791 | 0.042762 |
| 29.48611 | 0.907214 | 0.22245 | 0.151142 | 0.756885 | -0.11152 | 0.866413 | 0.864465 | 1.445024 | 0.628061 | 0.579811 | 0.340632 | -0.14061 | 0.118452 | 0.104458 | -0.0056 | 0.250948 | 0.028179 | 0.106971 | 0.108336 | 0.148129 | 0.22462 | -0.05422 | 0.006007 | 0.041181 |
| 29.5 | 1.266326 | 0.491294 | 0.309898 | 1.585021 | 0.359913 | 1.57627 | 0.952671 | 1.971444 | 1.262756 | 1.468696 | 0.574582 | -0.04717 | 0.193261 | 0.237664 | 0.001928 | 0.248706 | 0.03372 | 0.102167 | 0.084333 | 0.253915 | 0.274481 | 0.082569 | 0.065576 | -0.00599 |
| 29.51389 | 1.857868 | 0.400029 | 0.468068 | 1.926303 | 0.70739 | 1.436638 | 0.641617 | 2.676482 | 2.322134 | 2.737201 | 0.68604 | 0.209645 | 0.377845 | 0.489359 | -0.03749 | 0.227957 | 0.104827 | 0.111786 | 0.079937 | 0.468319 | 0.443933 | 0.278768 | 0.219598 | -0.0619 |
| 29.52778 | 2.004985 | 0.216881 | 0.606665 | 1.362478 | 0.929379 | 0.781931 | -0.11988 | 3.038035 | 3.035846 | 3.123491 | 0.962652 | 0.397106 | 0.502355 | 0.848722 | -0.09501 | 0.299861 | 0.136433 | 0.074919 | 0.123976 | 0.566525 | 0.543773 | 0.272297 | 0.349869 | 0.062104 |
| 29.54167 | 2.231319 | 0.215095 | 0.784462 | 0.528301 | 0.923324 | 0.448989 | -0.63522 | 3.484162 | 4.619332 | 3.190135 | 1.134231 | 0.398439 | 0.43933 | 1.076518 | -0.11227 | 0.357724 | 0.029558 | -0.0524 | 0.16983 | 0.437925 | 0.416312 | 0.018974 | 0.321824 | 0.176011 |
| 29.55556 | 2.68872 | 0.308101 | 1.292142 | 0.608608 | 0.771375 | 0.582084 | -0.43986 | 3.262922 | 3.777184 | 2.188365 | 0.967004 | 0.269309 | 0.302096 | 1.086114 | -0.05662 | 0.306492 | -0.00482 | -0.1216 | 0.184012 | 0.311603 | 0.312671 | -0.16545 | 0.299396 | 0.292961 |
| 29.56944 | 2.335526 | 0.275047 | 0.953493 | 0.974865 | 0.644095 | 0.74767 | 0.112032 | 2.216105 | 1.316455 | 1.185629 | 0.831683 | 0.233567 | 0.235966 | 0.984994 | 0.077136 | 0.31369 | 0.115256 | -0.05094 | 0.195599 | 0.37046 | 0.309716 | -0.18393 | 0.387735 | 0.310292 |
| 29.58333 | 1.857673 | 0.235669 | 0.622774 | 1.001594 | 0.438768 | 0.790625 | 0.518721 | 1.41797 | -0.99143 | 0.80456 | 0.692715 | 0.186311 | 0.156852 | 0.811983 | 0.082854 | 0.184502 | 0.182558 | 0.014834 | 0.143669 | 0.414189 | 0.312785 | -0.12242 | 0.372216 | 0.264839 |
| 29.59722 | 1.098784 | 0.189925 | 0.346563 | 1.039374 | 0.139813 | 0.667401 | 0.662171 | 0.654289 | -3.12853 | 0.304736 | 0.472594 | 0.022306 | 0.013596 | 0.441563 | -0.00901 | -0.02264 | 0.163084 | 0.040466 | 0.053186 | 0.41298 | 0.336453 | 0.021874 | 0.252242 | 0.201678 |
| 29.61111 | 0.434775 | 0.103921 | 0.026637 | 1.238084 | -0.25878 | 0.402599 | 0.655204 | -0.23385 | -2.76127 | 0.488896 | 0.364893 | -0.46107 | -0.10322 | 0.019699 | -0.11915 | -0.26272 | 0.105301 | 0.004648 | -0.00268 | 0.362572 | 0.348943 | 0.112053 | 0.07275 | 0.129972 |
| 29.625 | -0.01554 | 0.015682 | -0.22307 | 1.180331 | -0.61439 | 0.172765 | 0.580245 | -1.21633 | -1.16667 | 0.30778 | 0.430082 | -0.77298 | -0.1629 | -0.25976 | -0.22127 | -0.495 | 0.013303 | -0.09114 | -0.04888 | 0.224392 | 0.367116 | 0.062267 | -0.07209 | 0.060981 |
| 29.63889 | 0.454127 | -0.01597 | -0.15751 | 1.07454 | -0.61565 | 0.077716 | 0.384705 | -1.95098 | 0.144624 | 0.558494 | 0.467867 | -0.72179 | -0.16892 | -0.38446 | -0.2207 | -0.62884 | -0.00171 | -0.17187 | -0.06388 | 0.16366 | 0.364277 | -0.03421 | -0.11549 | 0.024009 |
| 29.65278 | 0.598653 | -0.03563 | 0.002773 | 0.912818 | -0.38855 | 0.024395 | 0.107246 | -2.42513 | 0.953079 | 1.414943 | 0.370824 | -0.51105 | -0.13171 | -0.45899 | -0.14846 | -0.66197 | 0.027805 | -0.16542 | -0.05248 | 0.152198 | 0.41403 | -0.15282 | -0.09332 | 0.010064 |
| 29.66667 | 0.58172 | -0.08636 | 0.131667 | 0.736336 | -0.23845 | -0.08871 | -0.18282 | -2.66158 | 1.395161 | 1.572878 | 0.165588 | -0.16908 | -0.10924 | -0.48072 | -0.09373 | -0.72956 | 0.060616 | -0.13385 | -0.0392 | 0.141788 | 0.460564 | -0.18676 | -0.03958 | -0.00536 |
| 29.68056 | 0.471796 | -0.15979 | 0.187131 | 0.618522 | -0.13527 | -0.14959 | -0.23963 | -0.87734 | 1.254133 | 1.092971 | -0.05708 | 0.241285 | -0.08026 | -0.43997 | -0.07334 | -0.74678 | 0.063513 | -0.09936 | -0.0212 | 0.171304 | 0.425984 | -0.13402 | 0.062196 | 0.005704 |
| 29.69444 | 0.364422 | -0.15378 | 0.24192 | 0.510063 | -0.08883 | -0.06036 | -0.13 | 0.954224 | 0.469019 | 0.327934 | -0.19073 | 0.512829 | -0.037 | -0.32923 | -0.02885 | -0.56894 | 0.083178 | -0.03456 | -0.0062 | 0.2263 | 0.406152 | -0.03968 | 0.174786 | 0.064132 |
| 29.70833 | 0.002562 | -0.15956 | 0.303138 | 0.310846 | -0.14025 | 0.106005 | -0.02402 | 3.383515 | -0.58891 | -0.59597 | -0.2276 | 0.52078 | -0.02627 | -0.25579 | 0.010428 | -0.34356 | 0.117445 | 0.041988 | -0.00881 | 0.232963 | 0.373581 | 0.016851 | 0.225736 | 0.124926 |
| 29.72222 | -0.2508 | -0.32888 | 0.321116 | 0.181315 | -0.24271 | 0.12309 | 0.083778 | 4.657341 | -1.32763 | -1.12887 | -0.10764 | 0.183049 | -0.06551 | -0.28507 | -2.7E-05 | -0.26057 | 0.064741 | 0.04927 | -0.08042 | 0.079438 | 0.190184 | -0.04409 | 0.087572 | 0.110152 |
| 29.73611 | -0.2979 | -0.38582 | 0.354059 | 0.171409 | -0.16076 | 0.301757 | 0.19132 | 4.806477 | -1.23256 | -1.06534 | 0.119927 | -0.30494 | -0.06009 | -0.35645 | -0.01655 | -0.22515 | -0.01469 | 0.010668 | -0.14843 | -0.11311 | -0.011 | -0.13133 | -0.12933 | 0.061643 |
| 29.75 | -0.18923 | -0.20337 | 0.291012 | 0.252238 | -0.11928 | 0.53234 | 0.203263 | 4.448106 | -1.20875 | -0.87511 | 0.45038 | -0.82235 | 0.024223 | -0.38086 | 0.01141 | -0.10059 | -0.01271 | -0.00868 | -0.12334 | -0.19435 | -0.0928 | -0.12122 | -0.29033 | 0.064079 |
| 29.76389 | -0.00958 | 0.034667 | 0.19866 | 0.318726 | -0.11377 | 0.591348 | 0.087662 | 3.144885 | -1.09537 | -0.78859 | 0.73686 | -1.23982 | 0.110319 | -0.33993 | 0.060983 | -0.00143 | 0.020157 | -0.01452 | -0.07776 | -0.21362 | -0.11455 | -0.09128 | -0.35412 | 0.078528 |
| 29.77778 | 0.331882 | 0.231962 | -0.18916 | 0.381349 | -0.09745 | 0.580845 | -0.09334 | 1.444195 | -0.94859 | -0.61093 | 0.784003 | -1.28838 | 0.146139 | -0.25465 | 0.119187 | 0.015239 | 0.025375 | -0.08597 | -0.06636 | -0.21077 | -0.10838 | -0.06746 | -0.31957 | 0.085851 |
| 29.79167 | 0.574198 | 0.296577 | -0.60101 | 0.423578 | -0.05119 | 0.29247 | -0.30519 | 0.18774 | -0.47529 | -0.59825 | 0.687274 | -1.01363 | 0.121522 | -0.22819 | 0.127918 | 0.018438 | 0.020858 | -0.1399 | -0.05462 | -0.17616 | -0.08707 | -0.06117 | -0.2157 | 0.055561 |
| 29.80556 | 0.551441 | 0.219039 | -0.62605 | 0.322228 | -0.00715 | 0.064032 | -0.42356 | -0.89006 | -0.03592 | -0.73968 | 0.574727 | -0.57518 | 0.058405 | -0.32187 | 0.0738 | 0.029574 | 0.015081 | -0.19148 | -0.06096 | -0.16488 | -0.07183 | -0.06983 | -0.15692 | 0.007569 |
| 29.81944 | 0.414415 | 0.149845 | -0.44086 | 0.120844 | 0.014002 | -0.12474 | -0.27422 | -2.50418 | -0.09232 | -1.14143 | 0.400613 | -0.45873 | -0.01148 | -0.46886 | -0.0063 | 0.029794 | 0.009084 | -0.21129 | -0.06769 | -0.1394 | -0.05363 | -0.07238 | -0.11199 | -0.01707 |
| 29.83333 | -0.21495 | -0.14425 | -0.39094 | -0.09922 | 0.016796 | -0.25067 | -0.28753 | -3.18548 | -0.73139 | -1.81077 | 0.231971 | -0.67863 | -0.07628 | -0.57052 | -0.05692 | 0.033249 | 0.014286 | -0.1702 | -0.08014 | -0.11219 | -0.00014 | -0.05315 | -0.09494 | 0.005946 |

| | | | | | | | | | | | | | | | | | | | | | | | | |
|---|---|---|---|---|---|---|---|---|---|---|---|---|---|---|---|---|---|---|---|---|---|---|---|---|
| 29.84722 | -0.82346 | -0.26905 | -0.33665 | -0.27919 | 0.006641 | -0.50162 | -0.34396 | -3.0765 | -0.78382 | -2.76062 | 0.094412 | -0.9414 | -0.14003 | -0.5832 | -0.06472 | 0.030126 | 0.018543 | -0.06059 | -0.11879 | -0.11604 | -0.016 | -0.02097 | -0.15798 | 0.01907 |
| 29.86111 | -1.18385 | -0.22714 | -0.34672 | -0.39836 | -0.16379 | -0.70653 | -0.48831 | -2.72786 | -0.34564 | -2.8701 | 0.023544 | -0.97574 | -0.17839 | -0.52564 | -0.05484 | 0.024253 | -0.01007 | 0.017966 | -0.16098 | -0.21056 | -0.17204 | 0.005748 | -0.32589 | 0.001083 |
| 29.875 | -1.14476 | -0.12139 | -0.48631 | -0.49662 | -0.34917 | -0.95009 | -0.64145 | -2.42225 | 0.183267 | -2.71467 | -0.12367 | -0.81373 | -0.13321 | -0.44344 | -0.00802 | 0.055896 | -0.02092 | 0.044471 | -0.15365 | -0.37759 | -0.42906 | 0.030119 | -0.47957 | -0.0088 |
| 29.88889 | -1.16428 | -0.15115 | -0.70527 | -0.71774 | -0.57959 | -1.01507 | -0.32321 | -2.67377 | 0.522983 | -2.22298 | -0.43087 | -0.62354 | -0.1001 | -0.30402 | 0.055719 | 0.141111 | -0.01505 | -0.00502 | -0.10755 | -0.5965 | -0.72842 | 0.025705 | -0.58929 | -0.00802 |
| 29.90278 | -0.8745 | -0.19396 | -0.91911 | -1.07944 | -0.5612 | -0.97875 | -0.40969 | -2.17574 | 0.360108 | -1.44732 | -0.86122 | -0.11132 | -0.13625 | -0.22189 | 0.066415 | 0.182778 | -0.06903 | -0.10212 | -0.12749 | -0.86699 | -1.02042 | -0.04534 | -0.68305 | -0.03957 |
| 29.91667 | -0.60239 | -0.13669 | -0.52568 | -1.37642 | -0.41411 | -0.71146 | -0.81673 | -1.92726 | 0.342432 | -0.56265 | -1.20133 | 0.733468 | -0.1787 | -0.20805 | 0.04766 | 0.217228 | -0.09054 | -0.19202 | -0.12451 | -1.03694 | -1.34961 | -0.1124 | -0.81336 | -0.0987 |
| 29.93056 | -0.42699 | -0.15168 | 0.026995 | -1.52939 | -0.1941 | -0.34805 | -1.24787 | -2.05987 | 0.313274 | 0.106971 | -1.22064 | 1.183875 | -0.11703 | -0.14964 | 0.05076 | 0.295772 | -0.02025 | -0.20132 | -0.09297 | -1.00479 | -1.3504 | -0.12219 | -0.82079 | -0.14095 |
| 29.94444 | -0.26203 | -0.22554 | 0.56786 | -0.74941 | 0.112404 | -0.01395 | -1.1866 | -1.35939 | 0.076627 | 0.558517 | -0.35995 | 0.880466 | -0.07725 | -0.06614 | 0.055476 | 0.339779 | 0.041827 | -0.1607 | -0.07024 | -0.92876 | -1.26408 | -0.10758 | -0.74657 | -0.17102 |
| 29.95833 | -0.12638 | -0.13672 | 0.897274 | -0.83654 | 0.307972 | 0.353119 | -0.39358 | -0.66435 | 0.195709 | 0.880725 | -0.05562 | 0.4883 | -0.05348 | -0.01033 | 0.042112 | 0.342246 | 0.057653 | -0.15065 | -0.05327 | -0.79198 | -1.05571 | -0.05784 | -0.64462 | -0.20414 |
| 29.97222 | 0.091503 | 0.050616 | 1.01284 | -0.7461 | 0.419706 | 0.592171 | 0.484766 | 0.153815 | 0.461425 | 0.787815 | 0.151144 | 0.334473 | -0.04054 | 0.033177 | 0.025945 | 0.36138 | 0.083415 | -0.1415 | -0.05864 | -0.68833 | -0.89056 | 0.030207 | -0.45415 | -0.14979 |
| 29.98611 | 0.171989 | 0.098342 | 1.16128 | -0.77712 | 0.299985 | 0.604327 | 0.494116 | 0.395149 | 0.617603 | 0.074532 | 0.264469 | 0.123284 | -0.05438 | 0.03589 | -0.00233 | 0.274403 | 0.058726 | -0.14523 | -0.08462 | -0.57681 | -0.76884 | 0.110087 | -0.27687 | -0.12252 |
| 30 | -0.41423 | -0.11419 | 0.804145 | -0.5198 | -0.00106 | 0.198089 | -0.20362 | -0.1989 | 0.274453 | -1.34659 | -0.13089 | -0.45565 | -0.06062 | -0.02212 | 0.035429 | 0.133726 | -0.05513 | -0.10898 | -0.16933 | -0.50178 | -0.75825 | 0.126118 | -0.21881 | -0.11872 |
| 30.01389 | -1.1224 | -0.29123 | -0.04044 | -0.53357 | -0.27515 | -0.35402 | -0.44954 | -1.33569 | -0.56663 | -3.20028 | -0.7439 | -1.29524 | -0.08628 | -0.21143 | 0.171516 | 0.089234 | -0.18759 | -0.06433 | -0.16934 | -0.3571 | -0.61697 | -0.01427 | -0.26019 | -0.18394 |
| 30.02778 | -1.5067 | -0.55 | -0.65466 | -1.04508 | -0.38307 | -0.82941 | -0.21895 | -2.16066 | -1.28575 | -4.34627 | -1.08005 | -1.74344 | -0.18758 | -0.49513 | 0.254626 | 0.201676 | -0.45887 | -0.15421 | -0.18554 | -0.17321 | -0.31903 | -0.42032 | -0.4751 | -0.16792 |
| 30.04167 | -1.70026 | -0.79902 | -0.71101 | -1.37345 | -0.55007 | -1.1285 | -0.53025 | -1.87862 | -1.03525 | -3.07297 | -0.99134 | -1.63086 | -0.39881 | -0.71652 | 0.052271 | 0.243978 | -0.92184 | -0.3109 | -0.13925 | -0.14001 | -0.07119 | -0.79383 | -0.71792 | -0.0324 |
| 30.05556 | -1.96498 | -1.1657 | -0.47645 | -1.63542 | -0.64418 | -1.18175 | -0.6567 | -1.58702 | -0.73512 | -1.52247 | -0.74922 | -1.21471 | -0.60149 | -0.54314 | -0.01346 | 0.075793 | -0.60625 | -0.26562 | -0.31616 | -0.43124 | -0.28331 | -0.58231 | -0.63227 | -0.06704 |
| 30.06944 | -1.8678 | -1.07076 | -0.2005 | -1.55105 | -0.51591 | -0.88356 | -0.29199 | -1.67325 | -0.46339 | -0.625 | -0.6178 | -0.34484 | -0.55565 | -0.27717 | 0.037636 | -0.25835 | 0.095478 | -0.06992 | -0.14826 | -0.58745 | -0.50592 | -0.1762 | -0.3765 | -0.23343 |
| 30.08333 | -1.35301 | -0.46468 | 0.084831 | -1.18921 | -0.3306 | -0.70577 | -0.03935 | -1.56738 | -0.21234 | -0.24382 | -0.26381 | 0.492856 | -0.08622 | 0.084836 | -0.00542 | -0.20778 | 0.180665 | 0.010851 | -0.07251 | -0.46423 | -0.31028 | 0.092137 | -0.17155 | 0.007064 |
| 30.09722 | -0.50312 | 0.218825 | 0.314273 | -0.78029 | -0.20185 | -0.53017 | 0.052745 | -1.03777 | 0.089394 | 0.074445 | 0.01974 | 1.131385 | 0.333838 | 0.486041 | 0.030538 | -0.04355 | 0.022751 | 0.030316 | -0.05601 | -0.27468 | -0.10764 | 0.193384 | -0.00725 | 0.099296 |
| 30.11111 | 0.157743 | 0.841846 | 0.413725 | -0.452 | -0.04172 | -0.32256 | 0.031003 | -0.4504 | 0.376172 | 0.661328 | 0.161192 | 1.572758 | 0.328788 | 0.698326 | 0.124456 | 0.126893 | -0.03519 | 0.073741 | -0.06903 | -0.13827 | -0.06856 | 0.261506 | 0.131537 | -0.07125 |
| 30.125 | 0.631879 | 0.476394 | 0.48347 | -0.10973 | 0.161513 | -0.10027 | -0.0046 | -0.17521 | 0.52325 | 1.322986 | 0.239163 | 1.499088 | 0.160975 | 0.60912 | 0.120241 | 0.232943 | -0.01472 | 0.083894 | -0.06261 | -0.06831 | -0.10195 | 0.288517 | 0.193383 | -0.10897 |
| 30.13889 | 0.925787 | 0.615461 | 0.576654 | 0.374849 | 0.300537 | 0.069122 | 0.206927 | -0.18507 | 0.526344 | 2.947699 | 0.238847 | 1.648805 | 0.031942 | 0.346307 | 0.054561 | 0.227579 | -0.0128 | 0.038218 | -0.02199 | -0.01381 | -0.07199 | 0.202536 | 0.141824 | -0.11186 |
| 30.15278 | 0.851035 | 0.537673 | 0.591385 | 0.806355 | 0.359953 | 0.141226 | 0.539557 | -0.18239 | 0.592308 | 2.456326 | 0.186096 | 1.447768 | -0.04067 | 0.065808 | 0.015297 | 0.208006 | -0.00415 | -0.03904 | 0.005667 | 0.024177 | -0.08608 | 0.080552 | -0.0048 | -0.08364 |
| 30.16667 | 0.591615 | 0.378503 | 0.577902 | 0.990732 | 0.356157 | 0.155767 | 0.657487 | -0.04253 | 0.683573 | 1.852918 | 0.2735 | 1.199112 | -0.04821 | -0.28312 | 0.005249 | 0.175779 | -0.00462 | -0.1272 | 0.017754 | 0.028382 | -0.18601 | -0.00288 | -0.15528 | -0.03519 |
| 30.18056 | 0.317523 | 0.523408 | 0.563927 | 0.995383 | 0.340973 | 0.137801 | 0.448184 | 0.163986 | 0.831568 | 1.932213 | 0.520508 | 0.490335 | -0.05838 | -0.49017 | -0.00213 | 0.101279 | -0.01986 | -0.19833 | 0.030207 | -0.00619 | -0.26697 | -0.00856 | -0.20492 | 0.010949 |
| 30.19444 | 0.154078 | 0.55241 | 0.433322 | 1.036196 | 0.282239 | 0.153058 | 0.321114 | 0.35846 | 1.071448 | 1.394701 | 0.784637 | -0.05438 | -0.10103 | -0.43959 | -0.01361 | 0.077628 | -0.03367 | -0.22516 | 0.034529 | -0.04662 | -0.27134 | 0.025111 | -0.17164 | 0.02229 |
| 30.20833 | 0.576741 | 0.330071 | 0.416821 | 0.602963 | 0.231895 | 0.151641 | 0.66671 | 0.678106 | 1.079762 | 0.62518 | 0.656472 | -0.40085 | -0.17878 | -0.39595 | -0.03444 | 0.070175 | -0.05819 | -0.19703 | 0.029302 | -0.07307 | -0.25946 | 0.01561 | -0.11637 | 0.005169 |
| 30.22222 | 1.064791 | 0.304644 | 0.654264 | -0.04321 | 0.187548 | 0.171718 | 0.471848 | 1.08476 | 1.030472 | 0.030722 | 0.701322 | -0.42763 | -0.24608 | -0.4289 | -0.04889 | 0.077981 | -0.06946 | -0.12573 | 0.032478 | -0.0675 | -0.24882 | -0.03635 | -0.06278 | -0.00026 |
| 30.23611 | 1.300399 | 0.24397 | 0.88446 | -0.22625 | 0.153024 | 0.298318 | 0.248516 | 1.573877 | 1.090713 | -0.06896 | 0.826646 | -0.28955 | -0.23513 | -0.46231 | -0.03253 | 0.091938 | -0.04372 | -0.06115 | 0.035966 | -0.02493 | -0.21278 | -0.04654 | -0.03625 | -0.00435 |
| 30.25 | 1.086489 | 0.174956 | 1.098141 | -0.25425 | 0.132093 | 0.49701 | 0.149672 | 1.939932 | 1.150512 | 0.208071 | 0.63412 | -0.07083 | -0.13276 | -0.47603 | 0.005708 | 0.089138 | -0.01858 | -0.03399 | 0.031888 | 0.023384 | -0.14001 | 0.017051 | -0.04647 | -0.01807 |
| 30.26389 | 0.628007 | 0.05899 | 1.065807 | -0.12494 | 0.096197 | 0.589338 | 0.138312 | 1.716899 | 0.960161 | 0.53767 | 0.589673 | 0.121245 | 0.023242 | -0.441 | 0.041023 | 0.060986 | -0.01738 | -0.02505 | 0.036075 | 0.046749 | -0.04394 | 0.11526 | -0.05031 | -0.0159 |
| 30.27778 | 0.278987 | -0.11371 | 0.842375 | -0.01035 | 0.043605 | 0.505868 | 0.108169 | 1.227419 | 0.856314 | 0.640332 | 0.499511 | 0.140633 | 0.188839 | -0.34747 | 0.072371 | 0.012709 | -0.02394 | -0.02509 | 0.043322 | 0.047822 | 0.011673 | 0.169307 | -0.01939 | 0.015895 |
| 30.29167 | -0.00362 | -0.35026 | 0.505619 | 0.113832 | 0.017515 | 0.315934 | 0.05868 | 0.806404 | 0.664925 | 0.336991 | 0.407279 | -0.01694 | 0.279449 | -0.17672 | 0.098534 | -0.06321 | -0.03303 | -0.02256 | 0.045906 | 0.033838 | 0.030809 | 0.164479 | 0.022657 | 0.047119 |
| 30.30556 | -0.21807 | -0.55212 | 0.212122 | 0.21927 | 0.007027 | 0.028751 | -0.03875 | 0.415826 | 0.392968 | -0.33824 | 0.247855 | -0.22246 | 0.310775 | -0.01353 | 0.085879 | -0.13768 | -0.04333 | -0.00198 | 0.045621 | 0.010998 | 0.024539 | 0.116842 | 0.050195 | 0.03541 |
| 30.31944 | -0.13038 | -1.03742 | -0.04035 | 0.255346 | -0.02342 | -0.27754 | -0.16884 | -0.00253 | 0.142356 | -0.90633 | 0.139146 | -0.42892 | 0.196695 | 0.107644 | 0.044332 | -0.19805 | -0.03853 | 0.038751 | 0.029097 | -0.01603 | 0.027968 | 0.037504 | 0.051028 | -0.00757 |
| 30.33333 | 0.010213 | -1.42749 | -0.14182 | 0.245322 | -0.02985 | -0.35184 | -0.26972 | -0.23322 | 0.068828 | -0.96077 | 0.173004 | -0.66907 | 0.05605 | 0.135613 | 0.007769 | -0.19244 | -0.00838 | 0.085035 | 0.001606 | -0.01944 | 0.021667 | -0.0253 | 0.034453 | -0.03677 |
| 30.34722 | 0.125967 | -1.17624 | -0.10272 | 0.226892 | 0.005213 | -0.14241 | -0.14333 | -0.07985 | 0.250886 | -0.56261 | 0.365195 | -0.52787 | -0.09691 | 0.123803 | 0.004108 | -0.18269 | 0.026057 | 0.138572 | -0.01093 | 0.008146 | 0.006925 | -0.02369 | 0.020413 | -0.04037 |
| 30.36111 | 0.39405 | -1.15521 | -0.01093 | 0.154251 | 0.082557 | 0.188261 | 0.160006 | 0.372263 | 0.579131 | 0.121752 | 0.521699 | -0.2035 | -0.18309 | 0.103705 | 0.021993 | -0.10472 | 0.024709 | 0.169904 | 0.001009 | 0.055046 | 0.016042 | 0.03206 | -0.00765 | -0.02254 |
| 30.375 | 0.574639 | -1.12935 | 0.126581 | 0.111546 | 0.121781 | 0.387041 | 0.361857 | 0.818854 | 0.514811 | 0.618696 | 0.607285 | 0.080654 | -0.16691 | 0.152143 | 0.039203 | -0.04009 | 0.020261 | 0.181348 | 0.03962 | 0.119432 | 0.059593 | 0.115197 | -0.01745 | -0.00704 |
| 30.38889 | 0.371846 | -1.1403 | 0.157421 | 0.076732 | 0.149642 | 0.446585 | 0.487661 | 1.103715 | 0.753915 | 0.823577 | 0.500929 | 0.146016 | -0.03502 | 0.292313 | 0.051348 | 0.026051 | 0.005586 | 0.168814 | 0.0787 | 0.179795 | 0.118698 | 0.16152 | -0.00461 | 0.026751 |
| 30.40278 | 0.259399 | -0.75455 | 0.168247 | 0.079812 | 0.170464 | 0.421582 | 0.534564 | 1.18031 | 1.020594 | 0.896815 | 0.527133 | -0.03377 | 0.117567 | 0.440745 | 0.075385 | 0.088733 | 0.013681 | 0.146848 | 0.097034 | 0.217702 | 0.117977 | 0.143554 | 0.01606 | 0.045964 |
| 30.41667 | 0.17559 | -0.29725 | 0.132699 | 0.118224 | 0.173034 | 0.402507 | 0.377956 | 1.265314 | 1.168395 | 0.750217 | 0.556571 | -0.31423 | 0.258415 | 0.487022 | 0.105262 | 0.152332 | 0.034917 | 0.126195 | 0.107942 | 0.226495 | 0.098145 | 0.137124 | 0.032992 | 0.047282 |
| 30.43056 | 0.161372 | 0.062617 | 0.166037 | 0.224747 | 0.124381 | 0.398938 | -0.10516 | 1.371165 | 1.141077 | 0.180258 | 0.572914 | -0.4431 | 0.30481 | 0.472511 | 0.124266 | 0.181762 | 0.050399 | 0.124696 | 0.113789 | 0.22013 | 0.091522 | 0.192508 | 0.03573 | 0.028446 |
| 30.44444 | 0.37499 | 0.258553 | 0.1806 | 0.31336 | 0.055112 | 0.377873 | -0.67812 | 1.291003 | 0.998712 | -0.56376 | 0.480845 | -0.36723 | 0.264626 | 0.422145 | 0.121058 | 0.170626 | 0.07019 | 0.126309 | 0.122578 | 0.199313 | 0.118209 | 0.189593 | 0.02374 | 0.024015 |
| 30.45833 | 0.559226 | 0.314627 | 0.173353 | 0.346199 | -0.04817 | 0.290674 | -0.90927 | 1.236674 | 0.847145 | -0.7297 | 0.475951 | -0.10446 | 0.144349 | 0.383916 | 0.020769 | 0.259839 | 0.108516 | 0.137653 | 0.155008 | 0.180447 | 0.24962 | 0.084963 | 0.032128 | 0.046461 |
| 30.47222 | 0.751641 | 0.446816 | 0.234846 | 0.23942 | -0.10325 | 0.296847 | -0.51443 | 1.315955 | 0.87752 | -0.18256 | 0.53114 | 0.143231 | 0.025271 | 0.34227 | -0.07872 | 0.390988 | 0.150203 | 0.153725 | 0.219021 | 0.19228 | 0.4448 | -0.07781 | 0.046596 | 0.096253 |
| 30.48611 | 0.947884 | 0.800628 | 0.39184 | 0.297341 | -0.00132 | 0.583827 | 0.479306 | 1.738543 | 1.231257 | 0.883961 | 0.694981 | 0.255793 | -0.01967 | 0.27224 | -0.01107 | 0.408455 | 0.160427 | 0.153521 | 0.249468 | 0.234562 | 0.438761 | -0.06377 | 0.058813 | 0.109273 |
| 30.5 | 1.54634 | 1.496135 | 0.819973 | 0.752871 | 0.214355 | 1.444509 | 1.791353 | 2.537932 | 1.988701 | 2.103579 | 1.027766 | 0.524486 | 0.030906 | 0.269772 | 0.151437 | 0.322184 | 0.135403 | 0.133531 | 0.206921 | 0.324523 | 0.267053 | 0.087854 | 0.08376 | 0.054176 |
| 30.51389 | 1.932936 | 2.57282 | 1.243731 | 1.69141 | 0.279567 | 2.450588 | 2.938969 | 2.439629 | 2.497802 | 2.392081 | 1.010823 | 1.27264 | 0.105787 | 0.349256 | 0.133639 | 0.258686 | 0.179589 | 0.135285 | 0.188136 | 0.379371 | 0.155732 | 0.221794 | 0.210383 | 0.013443 |
| 30.52778 | 0.763588 | 3.472127 | 1.58864 | 1.22428 | 0.171877 | 1.750602 | 2.382901 | 2.395327 | 4.09507 | 2.464847 | 0.809902 | 2.125829 | 0.181099 | 0.562525 | 0.138598 | 0.158191 | 0.227274 | 0.105511 | 0.141432 | 0.334134 | 0.028348 | 0.242511 | 0.401723 | -0.00415 |
| 30.54167 | 2.154908 | 3.13085 | 1.339907 | 0.451694 | 0.240316 | 1.8127 | 2.037445 | 2.492028 | 5.255082 | 1.662699 | 0.455259 | 1.93519 | 0.305942 | 0.725666 | 0.349636 | 0.055117 | 0.154243 | -0.04657 | 0.000556 | 0.206821 | -0.08784 | 0.098183 | 0.54698 | 0.039808 |
| 30.55556 | 1.920394 | 2.762545 | 1.119098 | 0.765192 | 0.284137 | 1.573782 | 1.485304 | 3.204777 | 5.155907 | 1.331071 | 0.653645 | 1.107675 | 0.328318 | 0.672793 | 0.349588 | 0.095901 | 0.121317 | -0.22976 | -0.01256 | 0.187825 | -0.02814 | -0.02168 | 0.612291 | 0.17336 |
| 30.56944 | 1.630856 | 2.031318 | 1.031299 | 1.128659 | 0.26878 | 1.046803 | 0.92509 | 3.126364 | 2.668378 | 1.031734 | 0.9054 | 0.433829 | 0.30769 | 0.520565 | 0.227988 | 0.216521 | 0.191372 | -0.11164 | 0.103286 | 0.280796 | 0.094932 | -0.01075 | 0.563819 | 0.254143 |
| 30.58333 | 1.435743 | 1.412449 | 0.588147 | 1.257765 | 0.191053 | 0.300534 | 0.227804 | 2.860044 | -0.14498 | 0.622197 | 0.946891 | -0.03294 | 0.177221 | 0.309434 | 0.052318 | 0.210558 | 0.186098 | -0.02809 | 0.141409 | 0.285375 | 0.09155 | -0.01146 | 0.356372 | 0.168265 |
| 30.59722 | 0.79654 | 0.936661 | 0.063691 | 1.27251 | 0.201346 | -0.14918 | -0.23748 | 2.751298 | -1.87529 | 0.410383 | 0.809648 | -0.40015 | 0.00349 | 0.033227 | -0.15905 | 0.103698 | 0.117223 | -0.05646 | 0.106344 | 0.273689 | -0.03124 | -0.04524 | 0.11901 | 0.02246 |
| 30.61111 | -0.25987 | 0.493341 | -0.33626 | 1.180996 | 0.231443 | -0.17262 | -0.37935 | 2.328284 | -2.58956 | 0.179773 | 0.450572 | -0.76915 | -0.13286 | -0.26587 | -0.32473 | -0.04464 | 0.019983 | -0.11006 | 0.092395 | 0.273991 | -0.08472 | -0.05347 | -0.05447 | -0.07982 |
| 30.625 | -1.00135 | 0.067563 | -0.61954 | 1.085017 | 0.221247 | 0.14206 | -0.59503 | 2.018558 | -2.90043 | 0.085586 | 0.182845 | -0.87753 | -0.1985 | -0.4975 | -0.45085 | -0.24059 | -0.06733 | -0.07434 | 0.094612 | 0.276902 | -0.10856 | -0.03877 | -0.14251 | -0.10797 |
| 30.63889 | -0.82512 | -0.231 | -0.6064 | 0.846862 | 0.202106 | 0.386295 | -0.68125 | 1.373837 | -2.22527 | 0.113744 | 0.579362 | -0.77307 | -0.23621 | -0.63944 | -0.45207 | -0.37481 | -0.09131 | -0.02365 | 0.068565 | 0.259747 | -0.11109 | -0.04487 | -0.15409 | -0.10462 |
| 30.65278 | 0.151876 | -0.5068 | -0.93732 | 0.728568 | 0.206743 | 0.131224 | -1.27891 | 0.40828 | -1.12373 | 0.083979 | 0.867295 | -0.73027 | -0.27798 | -0.6756 | -0.35102 | -0.46677 | -0.07343 | -0.03095 | 0.026245 | 0.219522 | -0.08424 | -0.06775 | -0.09611 | -0.07381 |
| 30.66667 | 0.399728 | -0.6632 | -1.12702 | 0.618254 | 0.185885 | -0.0632 | -1.40981 | -0.48899 | -0.48816 | 0.017326 | 0.850335 | -0.37873 | -0.28718 | -0.61957 | -0.29291 | -0.49946 | -0.01698 | -0.08096 | 0.012084 | 0.20928 | 0.004061 | -0.10134 | -5.7E-05 | -0.01484 |
| 30.68056 | -0.44113 | -0.55133 | -0.57253 | 0.511215 | 0.211825 | -0.07932 | -0.53922 | -0.98547 | -0.32922 | -0.0779 | 0.392555 | -0.00898 | -0.28799 | -0.53605 | -0.2563 | -0.46632 | -0.01321 | -0.11102 | 0.021203 | 0.245085 | 0.123128 | -0.11021 | 0.073392 | 0.042725 |
| 30.69444 | -0.89267 | -0.66941 | 0.17864 | 0.435981 | 0.286175 | -0.24192 | 1.189687 | -0.83598 | -0.12219 | -0.20459 | -0.10756 | 0.420319 | -0.28095 | -0.51969 | -0.22633 | -0.36928 | -0.01742 | -0.09875 | 0.043268 | 0.321923 | 0.319031 | -0.1089 | 0.140678 | 0.104682 |
| 30.70833 | -1.04412 | -0.58952 | 0.860031 | 0.43767 | 0.346364 | -0.39336 | 1.672121 | -1.03331 | 0.301205 | -0.29531 | -0.19118 | 0.711411 | -0.25334 | -0.52902 | -0.17922 | -0.2561 | 0.035036 | -0.08569 | 0.068916 | 0.400737 | 0.575421 | -0.14731 | 0.198185 | 0.153671 |
| 30.72222 | 0.259172 | -0.52624 | 1.21228 | 0.531321 | 0.302801 | -0.37858 | 0.821552 | -1.38604 | 0.039091 | -0.43573 | -0.22039 | 0.859123 | -0.23221 | -0.59347 | -0.11415 | -0.21427 | 0.062195 | -0.11156 | 0.066166 | 0.450272 | 0.745142 | -0.28319 | 0.18778 | 0.148029 |
| 30.73611 | 0.767438 | -0.30176 | 0.870385 | 0.487637 | 0.213515 | 0.055859 | -0.06122 | -0.79932 | -0.7229 | -0.63084 | -0.43182 | 0.771162 | -0.21462 | -0.672 | -0.11292 | -0.24088 | 0.025706 | -0.13504 | 0.030472 | 0.423967 | 0.782181 | -0.41039 | 0.174887 | 0.132874 |
| 30.75 | 0.815959 | -0.19728 | 0.576136 | 0.49172 | 0.119723 | 0.72434 | -0.50081 | 0.534533 | -1.02479 | -0.71825 | -0.29359 | 0.446412 | -0.19642 | -0.73036 | -0.12523 | -0.30318 | -0.02272 | -0.14011 | -0.04819 | 0.390643 | 0.69051 | -0.47003 | 0.183939 | 0.116226 |
| 30.76389 | 0.789945 | -0.18558 | 0.281986 | 0.426389 | 0.078595 | 0.567602 | -0.4402 | 1.650685 | -0.76481 | -0.64565 | 0.097725 | -0.157 | -0.17548 | -0.80252 | -0.1242 | -0.44245 | -0.07407 | -0.13922 | -0.13615 | 0.285572 | 0.516153 | -0.50338 | 0.15003 | 0.083034 |
| 30.77778 | 1.058301 | -0.0707 | -0.22859 | 0.311009 | 0.039729 | -0.11605 | -0.29231 | 2.129599 | -0.42538 | -0.54563 | 0.374983 | -0.91596 | -0.1826 | -0.91266 | -0.14431 | -0.60167 | -0.15129 | -0.13504 | -0.19163 | 0.127479 | 0.260867 | -0.51931 | 0.053421 | 0.04657 |
| 30.79167 | 1.661654 | 0.121218 | -0.89336 | 0.263716 | 0.018064 | -0.39829 | -0.30267 | 1.763951 | -0.2177 | -0.66281 | 0.631213 | -1.5219 | -0.21942 | -0.97957 | -0.12376 | -0.68069 | -0.20969 | -0.12045 | -0.22429 | -0.07735 | 0.03744 | -0.5162 | -0.01843 | 0.015089 |
| 30.80556 | 1.862623 | 0.182413 | -0.89938 | 0.270872 | -0.00758 | -0.38029 | -0.96124 | 0.893637 | -0.13427 | -0.58828 | 0.661891 | -1.84733 | -0.27543 | -0.99794 | -0.11176 | -0.5567 | -0.26072 | -0.08573 | -0.18052 | -0.22326 | -0.1197 | -0.45116 | -0.02389 | 0.012121 |
| 30.81944 | 1.054897 | 0.062646 | -0.76938 | 0.293771 | -0.114 | -0.38313 | -2.19338 | 0.224473 | -0.33358 | -0.75363 | 0.49114 | -1.97099 | -0.32197 | -0.97358 | -0.09944 | -0.43406 | -0.20994 | -0.03305 | -0.12928 | -0.29936 | -0.24347 | -0.33116 | -0.05663 | 0.026094 |
| 30.83333 | 0.29326 | -0.13899 | -0.12948 | 0.196426 | -0.15241 | -0.36836 | -2.17036 | -0.20849 | -0.52218 | -1.07943 | 0.132866 | -1.75513 | -0.37601 | -0.9205 | -0.09298 | -0.38529 | -0.15512 | -0.00284 | -0.1375 | -0.31185 | -0.36494 | -0.20154 | -0.12294 | -0.00642 |
| 30.84722 | -0.63324 | -0.26852 | -0.04509 | 0.093987 | -0.22741 | -0.28318 | -1.98488 | -0.76999 | -0.79943 | -1.52305 | -0.26926 | -1.48877 | -0.38169 | -0.82163 | -0.05581 | -0.40141 | -0.11104 | 0.007882 | -0.17254 | -0.35543 | -0.50251 | -0.07799 | -0.26146 | -0.08016 |
| 30.86111 | -1.46546 | -0.33349 | -0.0493 | 0.009677 | -0.29671 | -0.33619 | -1.80115 | -1.61452 | -1.07458 | -2.02895 | -0.51864 | -1.26648 | -0.33522 | -0.69244 | 0.005556 | -0.35205 | -0.09261 | 0.027943 | -0.20961 | -0.43718 | -0.67852 | 0.063429 | -0.398 | -0.15361 |
| 30.875 | -2.24093 | -0.47928 | -0.1074 | -0.15128 | -0.37705 | -0.51782 | -1.59217 | -2.46088 | -1.20238 | -2.35791 | -0.55857 | -1.12417 | -0.30581 | -0.54978 | 0.046811 | -0.25331 | -0.11517 | 0.040672 | -0.23752 | -0.5801 | -1.03546 | 0.191089 | -0.47509 | -0.18372 |
| 30.88889 | -3.1081 | -0.90968 | -0.10533 | -0.24519 | -0.31429 | -0.65384 | -0.8337 | -2.43457 | -0.96477 | -2.44781 | -0.50298 | -0.86145 | -0.29481 | -0.41689 | 0.037601 | -0.1773 | -0.16065 | 0.013289 | -0.25004 | -0.71586 | -1.24685 | 0.305124 | -0.50947 | -0.18559 |
| 30.90278 | -2.3212 | -1.21481 | -0.02572 | -0.21558 | -0.14564 | -0.6476 | -0.46813 | -2.1053 | -0.86993 | -2.17938 | -0.51198 | -0.64995 | -0.27826 | -0.25582 | 0.018596 | -0.09373 | -0.166 | -0.00708 | -0.19923 | -0.80112 | -1.30267 | 0.431287 | -0.50834 | -0.16254 |
| 30.91667 | -1.7236 | -1.62627 | 0.070985 | -0.21878 | 0.002784 | -0.55774 | -0.4052 | -2.41556 | -0.65496 | -1.78953 | -0.38757 | -0.30166 | -0.21018 | -0.0842 | 0.014965 | 0.014105 | -0.1152 | -0.00454 | -0.14422 | -0.75701 | -1.02403 | 0.542889 | -0.51055 | -0.11593 |
| 30.93056 | -0.86795 | -1.74285 | 0.137071 | -0.16509 | 0.094326 | -0.34956 | -0.43069 | -2.75225 | -0.291 | -0.95336 | -0.53519 | 0.158657 | -0.13492 | 0.069106 | 0.031897 | 0.094882 | -0.06311 | 0.025382 | -0.09976 | -0.73734 | -0.60627 | 0.663472 | -0.46851 | -0.07759 |
| 30.94444 | -0.45396 | -1.66758 | 0.225764 | -0.16763 | 0.199484 | -0.2137 | -0.31862 | -2.58812 | 0.30427 | -0.07134 | -0.52155 | 0.677079 | -0.0673 | 0.179802 | 0.042723 | 0.139618 | -0.03506 | 0.040253 | -0.0269 | -0.68656 | -0.3994 | 0.682681 | -0.38686 | -0.06057 |
| 30.95833 | -0.14735 | -1.55029 | 0.330284 | -0.16481 | 0.284401 | -0.15789 | -0.09695 | -1.96943 | 1.021147 | 0.715908 | 0.11721 | 1.081929 | -0.02956 | 0.244327 | 0.057696 | 0.13731 | -0.02955 | 0.047203 | -0.00646 | -0.66113 | -0.2628 | 0.636547 | -0.33623 | -0.04499 |
| 30.97222 | 0.285706 | -1.11461 | 0.39933 | -0.09323 | 0.319417 | -0.09416 | 0.119508 | -1.24833 | 1.588237 | 1.322623 | -0.01775 | 1.269515 | 0.017938 | 0.316085 | 0.064776 | 0.153231 | -0.00835 | 0.067968 | 0.008294 | -0.61488 | -0.25527 | 0.576913 | -0.30841 | -0.01332 |
| 30.98611 | 0.508256 | -0.56667 | 0.359634 | -0.06965 | 0.248844 | -0.02509 | 0.17829 | -0.7758 | 1.396082 | 1.463446 | -0.09057 | 1.030704 | 0.107796 | 0.389147 | 0.111937 | 0.185844 | -0.01695 | 0.110076 | -0.00566 | -0.52641 | -0.25074 | 0.49998 | -0.19661 | 0.012745 |
| 31 | 0.107175 | -0.26874 | 0.181901 | -0.23091 | 0.081025 | -0.17164 | -0.11243 | -0.97029 | 0.315795 | 0.689535 | -0.39477 | 0.267925 | 0.257567 | 0.458479 | 0.272916 | 0.233452 | -0.10891 | 0.117967 | -0.13363 | -0.42759 | -0.34164 | 0.41477 | -0.07799 | 0.054979 |
| 31.01389 | -0.73521 | -0.34898 | -0.05456 | -0.40517 | -0.05206 | -0.49699 | -0.72318 | -1.78919 | -1.54777 | -1.26583 | -0.9151 | -0.80816 | 0.435594 | 0.905327 | 0.543311 | 0.498892 | -0.31456 | 0.058193 | -0.29118 | -0.31904 | -0.50969 | 0.258346 | -0.01492 | 0.093308 |
| 31.02778 | -1.29901 | -0.17922 | -0.16868 | -0.40417 | -0.15401 | -0.7611 | -1.30862 | -2.70794 | -3.17341 | -2.93288 | -1.62586 | -2.10634 | 0.660922 | 2.069775 | 0.673936 | 1.237316 | -0.57709 | -0.01298 | -0.31383 | -0.27487 | -0.57226 | -0.11752 | -0.18167 | 0.065523 |
| 31.04167 | -1.54904 | -0.30884 | -0.26772 | -0.40005 | -0.29713 | -1.05843 | -1.34098 | -2.99326 | -3.60399 | -3.05889 | -2.49745 | -2.26937 | 0.704005 | 2.703417 | 0.459788 | 1.110374 | -0.75173 | -0.01053 | -0.28824 | -0.35156 | -0.40918 | -0.76396 | -0.49832 | -0.03489 |
| 31.05556 | -1.70709 | -0.7576 | -0.22622 | -0.4902 | -0.48397 | -1.33073 | -1.07844 | -2.91678 | -2.13972 | -1.93857 | -2.25923 | -1.56174 | 0.405638 | 1.880524 | 0.15439 | 0.313209 | -0.64722 | 0.026914 | -0.29408 | -0.51927 | -0.28005 | -1.10609 | -0.57979 | -0.08882 |
| 31.06944 | -1.38819 | -1.56925 | 0.04249 | -0.54797 | -0.56468 | -0.80717 | -1.50946 | -2.39594 | -1.03011 | -1.31231 | -2.31587 | -1.28686 | 0.108477 | 0.667725 | -0.05412 | -0.29778 | -0.40851 | -0.02895 | -0.18845 | -0.69373 | -0.34152 | -0.8451 | -0.41336 | -0.15256 |
| 31.08333 | -1.03438 | -2.2457 | 0.152975 | -0.60268 | -0.58351 | -0.55385 | -1.15819 | -1.90938 | -0.54472 | -1.09799 | -0.63038 | -1.07282 | -0.0438 | 0.085054 | -0.10212 | -0.26587 | -0.19469 | -0.11779 | -0.14113 | -0.63245 | -0.36413 | -0.26828 | -0.2073 | -0.09298 |
| 31.09722 | -0.63328 | -2.28894 | 0.161324 | -0.55381 | -0.55481 | -0.44998 | -0.57198 | -1.10478 | 0.04714 | -0.83662 | 0.616878 | -0.61613 | -0.10433 | -0.10043 | 0.043486 | -0.13837 | 0.004422 | -0.17298 | -0.04465 | -0.46923 | -0.26098 | 0.016388 | -0.03565 | 0.020538 |
| 31.11111 | -0.25148 | -1.90123 | 0.13349 | -0.53202 | -0.47718 | -0.33409 | -0.33418 | -0.56401 | 0.564882 | -0.52376 | 1.526657 | -0.06861 | -0.21928 | -0.18935 | 0.090891 | -0.01395 | 0.138759 | -0.17639 | 0.051048 | -0.22086 | 0.054145 | 0.102398 | 0.050431 | 0.096114 |
| 31.125 | -0.1143 | -1.30476 | 0.188314 | -0.49559 | -0.34144 | -0.28775 | -0.47836 | -0.23774 | 1.200134 | -0.25146 | 2.158658 | 0.38393 | -0.34321 | -0.04772 | 0.021205 | 0.057402 | 0.182806 | -0.1512 | 0.070895 | -0.07918 | 0.23228 | 0.099229 | 0.060774 | 0.082552 |
| 31.13889 | -0.01164 | -0.95174 | 0.246776 | -0.46173 | -0.23187 | -0.32128 | -0.72666 | -0.06002 | 1.147165 | -0.0469 | 1.319652 | 0.704692 | -0.33351 | 0.285016 | -0.02683 | 0.067339 | 0.039146 | -0.11966 | 0.021194 | -0.04421 | 0.180425 | 0.051632 | 0.050936 | 0.035099 |
| 31.15278 | 0.047219 | -0.79884 | 0.301979 | -0.41033 | -0.12718 | -0.32401 | -0.69672 | -0.11809 | 0.741544 | 0.009912 | 0.439244 | 0.897648 | -0.1946 | 0.470541 | -0.00576 | 0.035386 | -0.12655 | -0.10342 | -0.04214 | -0.06874 | 0.09038 | 0.005087 | 0.042526 | -0.01201 |
| 31.16667 | 0.301009 | -0.50199 | 0.349614 | -0.20641 | -0.01883 | -0.29336 | -0.29677 | -0.28742 | 0.214012 | 0.036449 | -0.05654 | 1.106106 | -0.02674 | 0.202884 | 0.082428 | -0.00247 | -0.23543 | -0.11193 | -0.06125 | -0.09899 | 0.06238 | 0.028997 | 0.009147 | -0.05179 |
| 31.18056 | 0.623538 | 0.148744 | 0.345428 | 0.133897 | 0.107631 | -0.13566 | 0.258961 | -0.39808 | -0.29295 | 0.344648 | -0.22339 | 1.163759 | 0.075643 | -0.16745 | 0.185848 | 0.014825 | -0.2576 | -0.14083 | -0.01882 | -0.08648 | -0.01471 | -0.03409 | -0.01941 | -0.07389 |
| 31.19444 | 0.988774 | 0.925397 | 0.346415 | 0.268213 | 0.236927 | 0.019737 | 0.905609 | -0.39342 | -0.45718 | 1.013158 | -1.23014 | 1.217455 | 0.086352 | -0.53724 | 0.169858 | 0.06857 | -0.24801 | -0.19206 | 0.018988 | -0.03569 | -0.2064 | -0.27404 | -0.01285 | -0.09026 |
| 31.20833 | 1.160535 | 1.8459 | 0.410983 | 0.203041 | 0.294764 | 0.150118 | 1.296634 | -0.17789 | -0.08864 | 1.714647 | -1.50988 | 1.747696 | 0.032798 | -0.78271 | 0.068824 | 0.099727 | -0.24916 | -0.25507 | 0.013478 | 0.000866 | -0.35006 | -0.58132 | 0.008594 | -0.08258 |
| 31.22222 | 1.333418 | 2.621965 | 0.564222 | 0.175778 | 0.288026 | 0.229257 | 1.287223 | 0.029787 | 0.429717 | 2.382778 | -0.40491 | 1.81395 | -0.02855 | -0.805 | -0.03639 | 0.101704 | -0.26291 | -0.26861 | -0.01461 | 0.001362 | -0.29655 | -0.61688 | -0.00477 | -0.06665 |
| 31.23611 | 1.307988 | 1.599959 | 0.60604 | 0.249006 | 0.301384 | 0.294709 | 1.238796 | 0.23295 | 1.395188 | 2.880528 | 0.938497 | 2.048599 | -0.10758 | -0.41184 | -0.07991 | 0.095246 | -0.25829 | -0.15591 | -0.04512 | -0.00872 | -0.11741 | -0.26867 | -0.04352 | -0.04797 |
| 31.25 | 1.434784 | 1.770644 | 0.608396 | 0.242414 | 0.305196 | 0.327023 | 1.241848 | 0.46308 | 3.09498 | 2.401952 | 2.456195 | 1.919481 | -0.15275 | 0.036177 | -0.05182 | 0.074336 | -0.25212 | -0.05545 | -0.05483 | 0.004442 | -0.01922 | 0.13207 | -0.07135 | -0.02637 |
| 31.26389 | 1.455647 | 1.386649 | 0.572184 | 0.250452 | 0.258788 | 0.284742 | 1.355957 | 0.625607 | 2.862078 | 1.624608 | 2.245497 | 1.347222 | -0.138 | 0.3776 | 0.030048 | 0.040389 | -0.24739 | -0.01384 | -0.02732 | 0.003774 | -0.03911 | 0.30967 | -0.03888 | 0.009157 |
| 31.27778 | 1.308013 | 1.338855 | 0.529019 | 0.296551 | 0.134167 | 0.208161 | 1.159542 | 0.583872 | 1.407005 | 0.549797 | 1.261605 | 0.783907 | -0.05798 | 0.598127 | 0.087697 | -0.01498 | -0.26341 | -0.00779 | -0.01302 | -0.01882 | -0.06766 | 0.339146 | 0.035652 | 0.023448 |
| 31.29167 | 1.17695 | 0.915478 | 0.469707 | 0.303119 | -0.02917 | 0.077902 | 0.588764 | 0.207292 | -0.24743 | -1.18155 | 0.402097 | 0.098378 | 0.014586 | 0.573043 | -0.06097 | 0.013064 | -0.297 | 0.035371 | 0.05077 | -0.01484 | -0.06962 | 0.219146 | 0.136255 | -0.16712 |
| 31.30556 | 1.01299 | 0.614589 | 0.434928 | 0.146127 | -0.06724 | 0.074323 | 0.054138 | -0.16528 | -1.69481 | -1.5791 | -0.10797 | -0.35418 | -0.05535 | 0.319249 | -0.83683 | 0.55539 | -0.17762 | 0.373676 | 0.503874 | 0.005413 | -0.47157 | -0.21709 | 0.40352 | -1.22702 |

| | | | | | | | | | | | | | | | | | | | | | | | | |
|---|---|---|---|---|---|---|---|---|---|---|---|---|---|---|---|---|---|---|---|---|---|---|---|---|
| 31.31944 | 0.763856 | 0.600437 | 0.383218 | 0.002653 | 0.082553 | 0.302684 | 0.052101 | -0.14227 | -1.69337 | -0.40474 | 0.090391 | -0.3387 | 0.178139 | 0.749338 | -1.56438 | 0.446157 | -1.89963 | 0.145868 | 0.484163 | -0.22146 | -2.58393 | -0.73431 | 0.69153 | -2.43986 |
| 31.33333 | 0.646458 | 0.404862 | 0.401266 | -0.05844 | 0.185644 | 0.451116 | 0.005824 | 0.011605 | -1.18292 | 0.372381 | 0.276542 | -0.16185 | -0.09787 | 1.030279 | -0.34077 | -0.18356 | -0.61604 | -0.23201 | -0.00938 | -0.01937 | -0.24417 | -0.38532 | 0.174256 | -0.53475 |
| 31.34722 | 0.670814 | -0.3766 | 0.414158 | -0.10622 | 0.143029 | 0.44931 | -0.26973 | 0.143017 | -0.05895 | 0.031131 | 0.629025 | -0.10049 | -0.12453 | 0.27246 | 1.122083 | -0.52078 | 1.454537 | -0.28244 | -0.33747 | -0.10875 | 1.800333 | 0.463978 | -0.5213 | 2.242999 |
| 31.36111 | 0.877632 | -1.01725 | 0.52645 | -0.13435 | 0.091971 | 0.328855 | -0.44629 | 0.232033 | 0.771826 | -0.08697 | 0.61081 | 0.080104 | 0.185865 | 0.118762 | 0.820571 | -0.56765 | 0.186477 | -0.1561 | -0.32006 | -0.25807 | 0.246512 | 0.588122 | -0.18292 | 1.217141 |
| 31.375 | 0.871816 | -1.06847 | 0.524051 | -0.12721 | 0.064796 | 0.202315 | -0.31481 | 0.196692 | 1.074099 | -0.02291 | 0.466637 | 0.313522 | 0.063885 | 0.362402 | 0.067177 | -0.12357 | 0.064445 | 0.057477 | 0.053094 | 0.005913 | -0.0252 | 0.491172 | -0.04241 | 0.142494 |
| 31.38889 | 0.784478 | -1.19389 | 0.465538 | -0.11099 | 0.011718 | 0.0637 | -0.18703 | 0.112504 | 0.609291 | -0.43183 | 0.169591 | 0.241447 | 0.046437 | 0.232212 | -0.04481 | -0.01462 | 0.08095 | 0.082705 | 0.102515 | 0.06625 | -0.04134 | 0.412101 | 0.000728 | -0.04668 |
| 31.40278 | 0.509566 | -1.4941 | 0.351075 | -0.11362 | -0.02094 | 0.03285 | -0.18408 | 0.00632 | -0.01571 | -1.07391 | -0.05241 | -0.08709 | 0.076286 | 0.02585 | 0.044501 | 0.01573 | 0.113592 | 0.068417 | 0.092602 | 0.056833 | -0.03115 | 0.357792 | 0.060037 | -0.07677 |
| 31.41667 | 0.3172 | -1.58157 | 0.253516 | -0.07016 | -0.04157 | 0.132503 | -0.26801 | -0.08016 | -0.52085 | -1.59036 | -0.30581 | -0.3286 | 0.115312 | -0.0984 | 0.195268 | 0.042983 | 0.164051 | 0.095949 | 0.110961 | 0.095562 | 0.008403 | 0.264921 | 0.126222 | -0.08396 |
| 31.43056 | 0.157193 | -1.72646 | 0.204584 | -0.04909 | -0.04453 | 0.253181 | -0.41373 | -0.1222 | -0.74111 | -1.76281 | -0.4588 | -0.35209 | 0.146332 | -0.03926 | 0.356687 | 0.089874 | 0.194367 | 0.167576 | 0.145073 | 0.173521 | 0.051404 | 0.204275 | 0.177106 | -0.06225 |
| 31.44444 | 0.161804 | -1.86645 | 0.28852 | -0.06341 | -0.03285 | 0.270465 | -0.36164 | -0.17465 | -0.65214 | -1.22937 | -0.61075 | -0.14907 | 0.126536 | 0.192817 | 0.3763 | 0.125379 | 0.185524 | 0.21408 | 0.142032 | 0.224799 | 0.046214 | 0.182671 | 0.173204 | -0.04625 |
| 31.45833 | 0.215923 | -2.16381 | 0.362 | -0.0824 | -0.03467 | 0.231449 | -0.23268 | -0.20522 | -0.32207 | -0.5802 | -0.24328 | 0.07439 | 0.067595 | 0.425576 | 0.280483 | 0.139659 | 0.112325 | 0.216926 | 0.107989 | 0.200588 | 0.018258 | 0.154154 | 0.132426 | -0.07298 |
| 31.47222 | 0.31383 | -2.06319 | 0.322747 | -0.07312 | -0.01839 | 0.221603 | -0.07543 | -0.08784 | 0.170238 | -0.17634 | 0.005369 | 0.226944 | 0.043872 | 0.528613 | 0.162561 | 0.143777 | 0.037367 | 0.224386 | 0.100058 | 0.192825 | -0.00818 | 0.113373 | 0.089315 | -0.12533 |
| 31.48611 | 0.740222 | -1.03358 | 0.436527 | 0.158286 | 0.113732 | 0.500484 | 0.541237 | 0.446873 | 0.646097 | 0.623502 | 0.09838 | 0.342875 | 0.060533 | 0.496746 | 0.135237 | 0.169093 | 0.063191 | 0.221687 | 0.147312 | 0.200869 | -0.00723 | 0.137977 | 0.08769 | -0.11687 |
| 31.5 | 1.50391 | 1.282507 | 0.684122 | 0.796287 | 0.414861 | 1.202926 | 1.935935 | 1.542772 | 1.06231 | 2.100396 | 0.414122 | 0.709839 | 0.085114 | 0.457162 | 0.159264 | 0.185368 | 0.214456 | 0.201423 | 0.193608 | 0.245046 | 0.111255 | 0.220099 | 0.156026 | -0.04745 |
| 31.51389 | 2.243783 | 3.090779 | 1.113415 | 1.544356 | 0.659845 | 1.581962 | 3.550787 | 2.800185 | 1.225083 | 3.256242 | 1.209283 | 1.231979 | 0.10619 | 0.455776 | 0.199994 | 0.14019 | 0.433938 | 0.164131 | 0.274817 | 0.342968 | 0.362186 | 0.349223 | 0.333452 | 0.042233 |
| 31.52778 | 2.471671 | 2.929799 | 1.470061 | 1.348663 | 0.614697 | 1.237294 | 3.207731 | 2.675593 | 0.862959 | 2.905206 | 2.063814 | 1.83745 | 0.132329 | 0.611083 | 0.328237 | 0.119641 | 0.56428 | 0.128324 | 0.34862 | 0.346705 | 0.546937 | 0.33493 | 0.532418 | 0.117648 |
| 31.54167 | 1.844458 | 2.831003 | 1.373615 | 0.607781 | 0.58033 | 0.594773 | 2.272138 | 2.174293 | 1.046487 | 2.71004 | 1.646178 | 2.344992 | 0.166528 | 0.865991 | 0.387535 | 0.090414 | 0.544595 | 0.056189 | 0.315609 | 0.170661 | 0.454839 | 0.193986 | 0.659458 | 0.141961 |
| 31.55556 | 0.848438 | 2.524474 | 1.014218 | 0.35939 | 0.647008 | 0.849167 | 2.143663 | 2.386084 | 2.807348 | 2.708975 | 1.024261 | 2.094461 | 0.18527 | 0.83273 | 0.49043 | 0.220207 | 0.286422 | 0.033552 | 0.233876 | 0.141306 | 0.340367 | 0.305255 | 0.6731 | 0.17995 |
| 31.56944 | 0.360169 | 2.784349 | 0.692535 | 0.296519 | 0.575601 | 1.446715 | 3.582966 | 2.633604 | 2.876276 | 1.764274 | 0.331003 | 1.13199 | 0.18395 | 0.589087 | 0.46059 | 0.370138 | 0.212877 | 0.08531 | 0.284235 | 0.238025 | 0.362093 | 0.482406 | 0.568519 | 0.191438 |
| 31.58333 | 1.540043 | 3.308035 | 0.338927 | 0.282734 | 0.440329 | 1.28385 | 3.491104 | 2.688444 | 2.562787 | 1.189398 | -0.50411 | 0.431021 | 0.187248 | 0.393581 | 0.436673 | 0.418451 | 0.20967 | 0.11659 | 0.331236 | 0.286006 | 0.337826 | 0.345637 | 0.508686 | 0.144638 |
| 31.59722 | 1.638678 | 2.990862 | 0.016606 | 0.328033 | 0.280179 | 1.43867 | 1.826174 | 2.591413 | 1.838791 | 0.628405 | -0.52594 | -0.03338 | 0.15382 | 0.255605 | 0.213486 | 0.356455 | 0.126811 | 0.080196 | 0.292256 | 0.328415 | 0.268461 | 0.1139 | 0.492901 | 0.070878 |
| 31.61111 | 1.430182 | 2.174972 | -0.29677 | 0.409178 | 0.178511 | 0.739647 | 0.342033 | 1.952767 | 0.902984 | 0.28756 | 0.207824 | -0.52051 | 0.093523 | 0.094702 | -0.03888 | 0.345645 | 0.031798 | 0.031981 | 0.236171 | 0.371538 | 0.228959 | -0.12979 | 0.41667 | -0.01979 |
| 31.625 | 1.242442 | 1.523212 | -0.54903 | 0.50415 | 0.060772 | 0.002322 | -1.02939 | 1.399914 | -0.31946 | -0.09355 | 0.02 | -0.91197 | 0.021275 | -0.08781 | -0.263 | 0.297636 | -0.02531 | 0.01493 | 0.171905 | 0.361812 | 0.259607 | -0.2841 | 0.140829 | -0.08555 |
| 31.63889 | 0.532938 | 0.881271 | -0.90484 | 0.52632 | -0.06427 | -0.64785 | -2.2785 | 1.01187 | -1.18109 | -0.52957 | -0.34298 | -0.74302 | -0.01614 | -0.27291 | -0.41727 | 0.269128 | -0.04993 | 0.035958 | 0.088563 | 0.330818 | 0.291958 | -0.20879 | 0.02035 | -0.11251 |
| 31.65278 | -0.34689 | 0.226532 | -1.10538 | 0.407106 | -0.14538 | -0.47131 | -3.18916 | 0.817882 | -1.35227 | -0.75092 | 0.27089 | -0.9468 | -0.0171 | -0.43186 | -0.42738 | 0.219173 | -0.13021 | 0.060682 | 0.021684 | 0.309143 | 0.325729 | -0.17214 | 0.130298 | -0.09708 |
| 31.66667 | -0.00765 | -0.45812 | -1.28256 | 0.234854 | -0.13361 | -0.35053 | -2.62896 | 0.610015 | -1.15526 | -1.08158 | 0.80384 | -0.91009 | 0.001461 | -0.53692 | -0.51342 | 0.180732 | -0.20188 | 0.099088 | 0.029982 | 0.332472 | 0.311018 | -0.23662 | 0.186414 | -0.05283 |
| 31.68056 | -0.56853 | -1.1064 | -0.85315 | 0.031331 | -0.11288 | -0.12083 | -2.12844 | 0.488764 | -0.69608 | -1.32812 | 1.126724 | -0.89878 | 0.01544 | -0.55026 | -0.48064 | 0.155105 | -0.21856 | 0.134452 | 0.071534 | 0.371843 | 0.26973 | -0.31806 | 0.237254 | -0.00422 |
| 31.69444 | -1.29877 | -1.58826 | -0.20454 | -0.16663 | -0.09026 | 0.027116 | -2.56988 | 0.450173 | -0.46803 | -1.55629 | 1.000754 | -0.49629 | -0.01854 | -0.43568 | -0.50914 | 0.123345 | -0.21568 | 0.170553 | 0.064595 | 0.432567 | 0.240859 | -0.21128 | 0.204994 | 0.032101 |
| 31.70833 | -2.30043 | -1.65518 | 0.134636 | -0.14625 | -0.00775 | 0.081068 | -2.82173 | 0.577269 | -0.38487 | -1.68141 | 1.11404 | -0.05105 | -0.10741 | -0.22684 | -0.43554 | 0.090436 | 0.029216 | 0.180866 | 0.008543 | 0.491667 | 0.20157 | 0.007308 | 0.197953 | 0.04435 |
| 31.72222 | -0.21182 | -1.68253 | 0.112427 | 0.141563 | 0.085832 | 0.173046 | -2.5622 | 0.859827 | -0.1694 | -1.90983 | 1.322855 | 0.161042 | -0.18304 | -0.04332 | -0.30144 | 0.03132 | 0.177132 | 0.17037 | -0.06056 | 0.522475 | 0.150736 | 0.125045 | 0.149272 | 0.033406 |
| 31.73611 | -0.50843 | -2.49068 | 0.00714 | 0.418379 | 0.119463 | 0.22628 | -1.98419 | 0.830362 | -0.38951 | -2.13057 | 1.018229 | 0.140551 | -0.24875 | 0.015616 | -0.17318 | -0.07707 | 0.275077 | 0.124216 | -0.10125 | 0.516293 | 0.086071 | 0.169739 | 0.07714 | 0.012324 |
| 31.75 | -0.52138 | -2.853 | 0.000883 | 0.478477 | 0.032955 | 0.207418 | -1.45048 | 0.697496 | -0.68603 | -2.26047 | -0.28923 | 0.090533 | -0.28487 | -0.05585 | -0.07259 | -0.16938 | 0.323187 | 0.066243 | -0.12798 | 0.457331 | -0.05208 | 0.125712 | 0.087467 | -0.02004 |
| 31.76389 | -0.92199 | -3.06437 | -0.00507 | 0.304431 | -0.03416 | 0.213642 | -1.04104 | 0.512716 | -1.1396 | -2.5002 | -0.88288 | 0.118402 | -0.28226 | -0.19968 | 0.000467 | -0.23612 | 0.285946 | -0.00967 | -0.15569 | 0.361088 | -0.23944 | 0.055126 | 0.092609 | -0.05971 |
| 31.77778 | -0.21608 | -2.81606 | -0.09069 | 0.061527 | -0.03745 | 0.235253 | -0.75419 | 0.281011 | -1.11387 | -2.25982 | 0.357211 | -0.09984 | -0.28527 | -0.40099 | 0.004193 | -0.30529 | 0.206371 | -0.08755 | -0.17694 | 0.30119 | -0.35342 | -0.1073 | 0.069957 | -0.0927 |
| 31.79167 | 0.374451 | -2.92848 | -0.14228 | -0.16158 | -0.00383 | 0.226723 | -0.6826 | 0.152573 | -1.0845 | -1.91681 | -0.06593 | -0.53412 | -0.28245 | -0.67287 | -0.09438 | -0.33291 | 0.038883 | -0.15288 | -0.20043 | 0.240783 | -0.35761 | -0.2811 | 0.060699 | -0.10143 |
| 31.80556 | 0.314442 | -2.66058 | -0.15501 | -0.30071 | -0.02105 | 0.170447 | -0.82281 | -0.03555 | -0.75906 | -2.0501 | -0.02213 | -1.03027 | -0.28998 | -0.89754 | -0.22935 | -0.34293 | -0.07254 | -0.16607 | -0.20965 | 0.169653 | -0.28872 | -0.46248 | 0.036475 | -0.08231 |
| 31.81944 | 0.131392 | -1.97691 | -0.17557 | -0.19711 | -0.02149 | 0.01262 | -0.90079 | -0.1044 | -0.46936 | -2.43801 | -0.24707 | -1.50516 | -0.30255 | -1.03826 | -0.30813 | -0.37869 | -0.13196 | -0.15667 | -0.21593 | 0.08921 | -0.26656 | -0.78371 | -0.04866 | -0.0575 |
| 31.83333 | 0.171841 | -1.29613 | -0.47808 | 0.022046 | 0.053608 | -0.07475 | -1.10562 | -0.09055 | -0.10509 | -2.84905 | 0.079253 | -2.02197 | -0.33002 | -0.97999 | -0.32818 | -0.41134 | -0.19798 | -0.15346 | -0.21479 | -0.00755 | -0.25628 | -0.98448 | -0.17503 | -0.00011 |
| 31.84722 | 0.276957 | -0.86377 | -0.81984 | 0.114832 | 0.074811 | -0.15886 | -1.43655 | -0.13387 | -0.16454 | -2.57248 | 0.522457 | -2.69261 | -0.33156 | -0.89022 | -0.33916 | -0.40487 | -0.25944 | -0.16672 | -0.18711 | -0.10558 | -0.31493 | -0.84399 | -0.24961 | 0.043145 |
| 31.86111 | 0.267474 | -0.78349 | -0.76743 | 0.033356 | 0.023954 | -0.30363 | -1.95423 | -0.26708 | -0.59565 | -2.22416 | 0.492999 | -2.46559 | -0.32707 | -0.7174 | -0.24912 | -0.33714 | -0.31727 | -0.23938 | -0.21841 | -0.19475 | -0.39196 | -0.46017 | -0.23934 | 0.027475 |
| 31.875 | 0.280729 | -0.79347 | -0.62572 | -0.12764 | 0.008062 | -0.45231 | -2.1893 | -0.52278 | -0.46859 | -1.70496 | 0.074997 | -2.0526 | -0.34922 | -0.59604 | -0.14429 | -0.29611 | -0.33346 | -0.36958 | -0.27281 | -0.24593 | -0.44965 | -0.1609 | -0.20025 | -0.02285 |
| 31.88889 | 0.275845 | -1.01121 | -0.62561 | -0.25335 | -0.00273 | -0.49053 | -1.96179 | -0.82947 | -0.08441 | -0.93879 | -0.01001 | -1.72959 | -0.34906 | -0.46507 | -0.06506 | -0.30802 | -0.42297 | -0.48608 | -0.28692 | -0.28698 | -0.48486 | -0.01407 | -0.18918 | -0.05173 |
| 31.90278 | 0.307851 | -1.00392 | -0.56716 | -0.29933 | -0.06669 | -0.43843 | -1.72557 | -1.05134 | 0.132226 | -0.41186 | 0.237741 | -1.26962 | -0.34906 | -0.40867 | -0.03618 | -0.31077 | -0.43027 | -0.49117 | -0.29313 | -0.34513 | -0.53299 | 0.000533 | -0.20417 | -0.07328 |
| 31.91667 | 0.261786 | -0.72205 | -0.47766 | -0.28673 | -0.11035 | -0.35271 | -1.33222 | -1.01624 | 0.214929 | -0.17436 | 0.26532 | -0.80603 | -0.29251 | -0.36093 | -0.02175 | -0.2575 | -0.41156 | -0.40476 | -0.2428 | -0.33265 | -0.58951 | -0.10653 | -0.25703 | -0.10331 |
| 31.93056 | 0.426995 | -0.5092 | -0.23643 | -0.19583 | -0.13569 | -0.15676 | -1.04665 | -0.791 | 0.250811 | -0.01865 | 0.436244 | -0.46948 | -0.2024 | -0.31649 | 0.011306 | -0.1541 | -0.29406 | -0.31522 | -0.20349 | -0.25117 | -0.56669 | -0.24635 | -0.32356 | -0.10151 |
| 31.94444 | 0.698969 | -0.16097 | 0.094283 | -0.07742 | -0.14614 | 0.083065 | -0.75015 | -0.50029 | 0.277912 | 0.119642 | 0.47584 | -0.12525 | -0.12092 | -0.31587 | 0.034105 | -0.05251 | -0.20232 | -0.26356 | -0.15315 | -0.19181 | -0.57447 | -0.48149 | -0.38169 | -0.06619 |
| 31.95833 | 0.936451 | 0.521743 | 0.367346 | 0.023666 | -0.16479 | 0.287839 | -0.54929 | -0.20504 | 0.356978 | 0.257128 | 0.633176 | 0.195828 | -0.05448 | -0.27823 | 0.023823 | 0.037936 | -0.13114 | -0.24746 | -0.12512 | -0.23881 | -0.5434 | -0.68414 | -0.3628 | -0.03195 |
| 31.97222 | 1.113076 | 1.118336 | 0.605391 | 0.035568 | -0.167 | 0.431553 | -0.39648 | -0.01442 | 0.451025 | 0.186256 | 0.590164 | 0.469743 | 0.026485 | -0.13654 | 0.003803 | 0.131499 | -0.05138 | -0.24386 | -0.10469 | -0.26803 | -0.50039 | -0.64883 | -0.28311 | -0.02593 |
| 31.98611 | 1.170725 | 1.117122 | 0.580904 | -0.23166 | -0.15784 | 0.391852 | -0.49924 | -0.15776 | 0.315805 | -0.33092 | 0.102106 | 0.460978 | 0.074157 | -0.00734 | -0.00403 | 0.170796 | -0.02527 | -0.24541 | -0.09658 | -0.2494 | -0.41841 | -0.43295 | -0.20136 | -0.03379 |
| 32 | 0.624911 | 0.540967 | 0.230262 | -1.07279 | -0.22148 | 0.073738 | -0.99219 | -0.6323 | -0.19853 | -1.49935 | -0.58892 | 0.086695 | 0.110602 | -0.02468 | 0.013895 | 0.194164 | -0.02725 | -0.25731 | -0.14208 | -0.20642 | -0.428 | -0.25728 | -0.10928 | -0.05256 |
| 32.01389 | -0.15405 | -0.5371 | -0.27098 | -1.58624 | -0.24237 | -0.29356 | -1.24483 | -0.98148 | -0.91941 | -2.96741 | -1.43878 | -0.65018 | 0.105366 | -0.29658 | 0.098618 | 0.223724 | -0.26297 | -0.3248 | -0.28841 | -0.14518 | -0.68477 | -0.28508 | -0.16603 | -0.1205 |
| 32.02778 | -0.50811 | -1.11611 | -0.58216 | -1.35473 | -0.14501 | -0.42118 | -1.32383 | -1.04466 | -1.32473 | -4.17767 | -1.89922 | -1.58837 | 0.018516 | -0.72434 | 0.213949 | 0.258234 | -0.87112 | -0.39773 | -0.44944 | -0.10417 | -1.00377 | -0.68297 | -0.36766 | -0.17989 |
| 32.04167 | -0.76356 | -1.34899 | -0.66304 | -1.10793 | -0.15517 | -0.47358 | -1.08767 | -1.0206 | -1.69776 | -2.64268 | -1.4626 | -1.75742 | -0.14416 | -0.92366 | 0.26411 | 0.104948 | -1.2936 | -0.25355 | -0.53698 | -0.17346 | -0.85436 | -1.12847 | -0.40507 | -0.10141 |
| 32.05556 | -1.01309 | -1.49276 | -0.58793 | -1.03685 | -0.35325 | -0.43128 | -0.96853 | -0.88018 | -1.41964 | 0.337057 | -1.15818 | -1.48871 | -0.42078 | -0.86391 | 0.050549 | -0.28024 | -1.20756 | -0.13992 | -0.4143 | -0.42646 | -0.46772 | -1.19651 | -0.31139 | -0.01422 |
| 32.06944 | -0.46847 | -1.58713 | -0.42255 | -0.57637 | -0.44817 | -0.2559 | -0.92716 | -0.7351 | -0.94871 | 1.243998 | -1.15016 | -0.77893 | -0.58336 | -0.70455 | -0.0955 | -0.55329 | -0.95875 | -0.24155 | -0.33662 | -0.58979 | -0.40747 | -1.08479 | -0.34954 | -0.06115 |
| 32.08333 | 0.308374 | -1.59922 | -0.15898 | 0.058183 | -0.33479 | -0.03158 | -0.49628 | -0.43739 | -0.48075 | 0.453046 | -1.6487 | -0.18444 | -0.54752 | -0.38468 | -0.03039 | -0.47194 | -0.50104 | -0.20252 | -0.17973 | -0.41187 | -0.33623 | -0.8058 | -0.3076 | -0.08935 |
| 32.09722 | 0.877437 | -1.62019 | 0.13719 | 0.521168 | -0.18462 | 0.183944 | -0.15944 | -0.05921 | -0.17316 | -0.1778 | -1.60406 | 0.215215 | -0.3835 | -0.06135 | 0.1041 | -0.22397 | 0.117997 | -0.0682 | 0.026292 | -0.12416 | -0.09554 | -0.33836 | -0.1573 | -0.02978 |
| 32.11111 | 1.056469 | -1.33967 | 0.325441 | 0.627737 | -0.13537 | 0.347961 | 0.158865 | 0.297558 | 0.226023 | -0.35363 | -0.57328 | 0.367303 | -0.19453 | 0.053698 | 0.140297 | -0.03249 | 0.682235 | -0.0232 | 0.146699 | 0.039213 | -0.00365 | 0.040413 | -0.03231 | 0.039368 |
| 32.125 | 1.102958 | -1.01068 | 0.324492 | 0.58278 | -0.13872 | 0.444251 | 0.243667 | 0.472737 | 0.393963 | -0.29816 | -0.16569 | 0.302006 | -0.05825 | 0.049085 | 0.088239 | -0.01272 | 0.84199 | -0.0271 | 0.145275 | 0.073026 | -0.02589 | 0.250478 | 0.010725 | 0.069241 |
| 32.13889 | 1.175863 | -0.82117 | 0.369633 | 0.462479 | -0.1445 | 0.423684 | 0.128412 | 0.497662 | 0.372713 | -0.22297 | 0.305693 | 0.274901 | -0.02185 | -0.01805 | 0.039857 | -0.03774 | 0.614314 | -0.02714 | 0.076775 | 0.035476 | -0.05708 | 0.332648 | 0.00714 | 0.027403 |
| 32.15278 | 1.427666 | -0.63482 | 0.604398 | 0.387887 | -0.11095 | 0.240544 | 0.015118 | 0.445839 | 0.31444 | -0.08304 | 0.305921 | 0.299251 | -0.04199 | -0.15149 | 0.03423 | -0.07263 | 0.203601 | -0.01643 | 0.00051 | 0.001479 | -0.05262 | 0.309393 | 0.018206 | -0.04522 |
| 32.16667 | 1.334359 | -0.29123 | 0.746997 | 0.352742 | -0.06939 | 0.017796 | 0.029872 | 0.322739 | 0.168046 | 0.152822 | 0.09584 | 0.386432 | -0.06124 | -0.27698 | 0.043132 | -0.06337 | -0.14288 | 0.013271 | -0.02009 | -0.00915 | -0.09213 | 0.266335 | 0.063925 | -0.0734 |
| 32.18056 | 1.423355 | 0.138582 | 0.709507 | 0.37111 | -0.01052 | -0.09175 | 0.206881 | 0.178628 | 0.075766 | 0.389084 | 0.230637 | 0.460858 | -0.05562 | -0.28493 | 0.041312 | -0.01449 | -0.28561 | 0.057913 | 0.033931 | -0.01002 | -0.12803 | 0.142329 | 0.087486 | -0.04119 |
| 32.19444 | 1.527539 | 0.606396 | 0.646538 | 0.431485 | 0.090256 | -0.13264 | 0.470598 | 0.070711 | 0.307375 | 0.499168 | 0.528333 | 0.702853 | -0.0392 | -0.17301 | 0.031251 | 0.03363 | -0.30683 | 0.073773 | 0.122568 | 0.012956 | -0.05143 | -0.04209 | 0.049252 | 0.00964 |
| 32.20833 | 1.707747 | 0.939583 | 0.638765 | 0.505662 | 0.193854 | -0.10431 | 0.683654 | 0.06252 | 0.38312 | 0.538348 | 0.728674 | 0.975118 | -0.04346 | -0.05009 | 0.034539 | 0.056812 | -0.32687 | 0.063091 | 0.171062 | 0.055687 | 0.041596 | -0.17801 | 0.005595 | 0.030164 |
| 32.22222 | 1.249418 | 1.117297 | 0.569029 | 0.543998 | 0.255576 | -0.00317 | 0.755598 | 0.140208 | 0.511702 | 0.445881 | 0.731651 | 1.13948 | -0.07683 | 0.028372 | 0.028941 | 0.043354 | -0.37976 | 0.037647 | 0.116911 | 0.080661 | -0.01175 | -0.18581 | 0.013897 | 0.018877 |
| 32.23611 | 0.886478 | 1.184928 | 0.529911 | 0.509779 | 0.286307 | 0.186438 | 0.698634 | 0.276245 | 0.597582 | 0.386161 | 1.182275 | 1.028713 | -0.12028 | 0.044348 | 0.013147 | 0.011299 | -0.40697 | 0.012501 | -0.04068 | 0.087883 | -0.13532 | -0.10493 | 0.050858 | 0.014435 |
| 32.25 | 0.695336 | 1.083211 | 0.676269 | 0.458066 | 0.258395 | 0.363483 | 0.741989 | 0.457885 | 0.620145 | 0.43338 | 1.365032 | 1.039737 | -0.15292 | 0.015223 | 0.005965 | -0.02596 | -0.36205 | 0.012646 | -0.20979 | 0.103621 | -0.1222 | -0.04856 | 0.0806 | 0.029439 |
| 32.26389 | 0.247755 | 1.001874 | 0.776853 | 0.393051 | 0.195868 | 0.396532 | 0.705587 | 0.534904 | 0.45445 | 0.516209 | 1.119843 | 0.986252 | -0.15534 | 0.001072 | 0.009269 | -0.01952 | -0.27471 | 0.038831 | -0.30808 | 0.137924 | -0.02931 | -0.0634 | 0.101347 | 0.03066 |
| 32.27778 | -0.13789 | 1.031428 | 0.722369 | 0.333589 | 0.142269 | 0.321518 | 0.689905 | 0.449633 | 0.249937 | 0.647341 | 0.634574 | 0.877257 | -0.11731 | 0.037235 | 0.007297 | 0.045192 | -0.15576 | 0.06799 | -0.28464 | 0.189159 | -0.01181 | -0.08195 | 0.123969 | 0.054585 |
| 32.29167 | -0.01644 | 0.89672 | 0.513868 | 0.298756 | 0.096915 | 0.271111 | 0.628486 | 0.400216 | 0.300195 | 0.747055 | 0.02509 | 0.788506 | -0.06584 | 0.085386 | -0.00386 | 0.107297 | -0.03171 | 0.06766 | -0.18661 | 0.250508 | 0.043521 | -0.06836 | 0.114542 | 0.080532 |
| 32.30556 | 0.300637 | 0.826429 | 0.290575 | 0.266055 | 0.074261 | 0.343833 | 0.571942 | 0.221676 | 0.301675 | 0.7849 | -0.42506 | 0.640194 | -0.02443 | 0.1437 | -0.01476 | 0.129486 | 0.036223 | 0.046946 | -0.08411 | 0.21535 | 0.151935 | -0.06168 | 0.115611 | 0.08648 |
| 32.31944 | 0.442853 | 0.702083 | 0.159668 | 0.285833 | 0.064724 | 0.506261 | 0.646996 | -0.09991 | 0.307839 | 0.772865 | -0.47327 | 0.468221 | 0.036664 | 0.228426 | -0.02031 | 0.105791 | 0.068631 | 0.047519 | -0.004 | 0.138221 | 0.140717 | -0.05119 | 0.08661 | 0.09142 |
| 32.33333 | 0.625718 | 0.515074 | 0.195227 | 0.295145 | 0.028472 | 0.728861 | 0.778641 | -0.44992 | 0.270651 | 0.705506 | 0.103801 | 0.253149 | 0.126285 | 0.308679 | -0.02766 | 0.046404 | 0.066827 | 0.067835 | 0.0142 | 0.040595 | 0.108743 | 0.051868 | 0.034978 | 0.079443 |
| 32.34722 | 0.656664 | 0.345489 | 0.310777 | 0.314263 | 0.035014 | 0.938957 | 0.760551 | -0.50122 | 0.144297 | 0.588882 | 0.414379 | 0.074228 | 0.215063 | 0.343657 | -0.03454 | -0.02339 | 0.034667 | 0.074115 | -0.01794 | -0.02294 | 0.149979 | 0.216495 | -0.00466 | 0.049575 |
| 32.36111 | 0.609568 | 0.228016 | 0.364534 | 0.285437 | 0.048053 | 0.996509 | 0.536325 | -0.34956 | 0.108694 | 0.484268 | 0.509051 | 0.078077 | 0.271366 | 0.340149 | -0.02247 | -0.0712 | -0.04865 | 0.058758 | -0.04761 | -0.08498 | 0.107703 | 0.284491 | 0.005208 | 0.015474 |
| 32.375 | 0.545739 | 0.109341 | 0.294177 | 0.170544 | 0.041735 | 0.909127 | 0.29771 | -0.27698 | 0.079698 | 0.35682 | 0.565113 | 0.155505 | 0.26841 | 0.354331 | 0.008975 | -0.10493 | -0.15779 | 0.051629 | -0.03848 | -0.09719 | 0.048156 | 0.291089 | 0.052879 | 0.001206 |
| 32.38889 | 0.331513 | -0.12607 | 0.210373 | 0.002604 | 0.002791 | 0.670897 | 0.066193 | -0.39072 | 0.118716 | 0.193866 | 0.475553 | 0.159007 | 0.231128 | 0.320071 | 0.042404 | -0.12276 | -0.21482 | 0.070011 | 0.010419 | -0.07207 | 0.083246 | 0.177904 | 0.118904 | 0.007374 |
| 32.40278 | 0.242214 | -0.18223 | 0.138151 | -0.06535 | -0.01442 | 0.475742 | -0.14319 | -0.50458 | 0.180089 | 0.03335 | 0.422376 | 0.082875 | 0.192279 | 0.22405 | 0.057254 | -0.10817 | -0.20493 | 0.082697 | 0.044916 | -0.01745 | 0.086282 | 0.016852 | 0.164547 | 0.010739 |
| 32.41667 | 0.315022 | -0.53231 | 0.294259 | -0.07343 | -0.0133 | 0.510763 | -0.12623 | -0.17907 | 0.363598 | -0.04064 | 0.778306 | 0.018621 | 0.150224 | 0.106361 | 0.058913 | -0.07765 | -0.15361 | 0.06324 | 0.027103 | 0.019649 | 0.052177 | -0.14027 | 0.10912 | -0.00125 |
| 32.43056 | 0.606681 | -0.79915 | 0.494743 | 0.019744 | 0.005376 | 0.557665 | 0.03892 | 0.415275 | 0.576896 | -0.00028 | 1.251292 | 0.048066 | 0.133302 | 0.078056 | 0.059561 | -0.06359 | -0.04801 | 0.044732 | 0.022574 | 0.080339 | 0.095726 | -0.22402 | 0.036205 | -0.01159 |
| 32.44444 | 0.857061 | -0.89761 | 0.644913 | 0.062363 | -0.05098 | 0.556014 | 0.235195 | 0.64731 | 0.845777 | 0.057718 | 1.63312 | 0.074828 | 0.157273 | 0.167154 | 0.066246 | -0.02023 | 0.105979 | 0.053865 | 0.096983 | 0.151485 | 0.152891 | -0.12707 | 0.046496 | -0.00141 |
| 32.45833 | 1.159222 | -0.8571 | 0.79671 | 0.012503 | -0.16387 | 0.42051 | 0.34894 | 0.603716 | 0.621005 | 0.071064 | 0.570554 | -0.00916 | 0.196769 | 0.323361 | 0.044582 | 0.054939 | 0.260454 | 0.08956 | 0.208031 | 0.165758 | 0.177186 | 0.121384 | 0.08878 | -0.02799 |
| 32.47222 | 1.261654 | -0.71149 | 0.89289 | 0.002726 | -0.20014 | 0.322045 | 0.443937 | 0.360509 | 0.30277 | 0.11189 | 0.447082 | -0.10602 | 0.227631 | 0.469905 | 0.049555 | 0.115963 | 0.349553 | 0.134433 | 0.266431 | 0.20054 | 0.209997 | 0.36048 | 0.113397 | -0.07956 |
| 32.48611 | 1.484447 | -0.36437 | 0.919267 | 0.188957 | -0.13243 | 0.409345 | 0.856885 | 0.258863 | 0.438422 | 0.473191 | 0.667697 | -0.07517 | 0.223243 | 0.573593 | 0.087194 | 0.119338 | 0.352142 | 0.161477 | 0.27276 | 0.181637 | 0.205699 | 0.494699 | 0.150395 | -0.07837 |
| 32.5 | 1.607807 | 0.399307 | 1.015395 | 0.686224 | 0.06865 | 0.682104 | 1.545396 | 0.64436 | 1.160474 | 1.083029 | 0.747451 | 0.237365 | 0.182896 | 0.576237 | 0.132659 | 0.085677 | 0.343624 | 0.165589 | 0.267278 | 0.186664 | 0.285241 | 0.550912 | 0.213954 | 0.01296 |
| 32.51389 | 1.881773 | 1.205985 | 0.954691 | 1.231904 | 0.239923 | 0.784769 | 2.893702 | 1.504705 | 1.605916 | 1.138353 | 1.337004 | 0.930883 | 0.169326 | 0.511055 | 0.126303 | 0.058613 | 0.421853 | 0.165307 | 0.342926 | 0.105066 | 0.557675 | 0.603019 | 0.30426 | 0.120343 |
| 32.52778 | 2.26236 | 1.841709 | 0.918419 | 1.262786 | 0.093026 | 0.499214 | 3.514357 | 2.210706 | 1.113118 | 1.109856 | 2.969458 | 1.841208 | 0.275942 | 0.58894 | 0.06626 | 0.087578 | 0.579396 | 0.134863 | 0.496829 | -0.03919 | 0.719259 | 0.743398 | 0.415633 | 0.12069 |
| 32.54167 | 2.457976 | 2.204717 | 1.239252 | 1.123049 | -0.04769 | -0.1419 | 3.515461 | 2.820729 | 0.592963 | 1.421433 | 3.400468 | 2.805136 | 0.401007 | 0.937784 | 0.021187 | 0.063824 | 0.713899 | 0.062621 | 0.7203 | -0.12022 | 0.529234 | 0.829206 | 0.416616 | 0.040732 |
| 32.55556 | 2.208668 | 3.590088 | 1.26601 | 1.129403 | 0.107141 | 0.052951 | 3.940451 | 2.856214 | 0.49175 | 2.190078 | 2.310354 | 2.351241 | 0.414423 | 1.347054 | 0.101109 | 0.112309 | 0.692178 | 0.049646 | 0.895672 | -0.02445 | 0.323877 | 0.825679 | 0.349828 | 0.043263 |
| 32.56944 | 1.67682 | 4.966351 | 0.859779 | 0.908514 | 0.160007 | 0.203313 | 4.114769 | 2.419366 | 1.327146 | 3.38479 | 0.496007 | 1.967862 | 0.384294 | 1.366064 | 0.252918 | 0.247185 | 0.615067 | 0.088712 | 0.949391 | 0.167316 | 0.288473 | 0.711901 | 0.316953 | 0.14381 |
| 32.58333 | 1.569293 | 3.340864 | 0.475178 | 0.52785 | 0.089179 | 0.169407 | 4.177933 | 2.087917 | 1.756066 | 3.156547 | 0.105984 | 1.68298 | 0.316428 | 1.061246 | 0.252423 | 0.321899 | 0.489472 | 0.026383 | 0.826403 | 0.337504 | 0.273874 | 0.403819 | 0.28856 | 0.157398 |
| 32.59722 | 0.556613 | 2.628826 | -0.12552 | 0.544252 | 0.174269 | 0.26879 | 3.480008 | 1.226536 | 1.906148 | 2.874205 | -1.26903 | 1.143075 | 0.226633 | 0.82822 | 0.158566 | 0.306448 | 0.300801 | -0.10926 | 0.654386 | 0.460018 | 0.1821 | 0.06455 | 0.165434 | 0.098074 |
| 32.61111 | 0.69095 | 2.717835 | -0.6324 | 0.757102 | 0.087191 | 0.313768 | 5.500778 | 0.901013 | 1.916796 | 2.776227 | -1.59331 | 0.826209 | 0.155582 | 0.543336 | 0.073904 | 0.262153 | 0.142808 | -0.17477 | 0.430367 | 0.546216 | 0.091079 | -0.20321 | 0.021062 | 0.038709 |
| 32.625 | 2.567266 | 2.362728 | 0.126664 | 0.704308 | 0.027185 | 0.74297 | 5.335098 | 1.637398 | 1.375383 | 2.025644 | -0.47178 | 0.48984 | 0.142863 | 0.177728 | -0.0165 | 0.155076 | -0.01728 | -0.10951 | 0.13727 | 0.502157 | 0.008102 | -0.35796 | -0.03265 | 0.01494 |
| 32.63889 | 2.752712 | 2.040451 | 0.126438 | 0.802494 | 0.10546 | 0.742073 | 2.783523 | 2.157082 | 0.849658 | 0.853343 | -0.5172 | 0.18707 | 0.162719 | -0.17926 | -0.07893 | 0.13028 | -0.13497 | -0.05287 | -0.20962 | 0.442081 | -0.02269 | -0.51668 | 0.014957 | 0.01537 |
| 32.65278 | 2.381825 | 1.721678 | -0.00583 | 0.758547 | 0.13497 | 0.620278 | 1.298347 | 1.430594 | 0.362486 | -0.59748 | -0.47225 | -0.04195 | 0.135491 | -0.49144 | -0.14124 | 0.088495 | -0.22146 | -0.00462 | -0.53617 | 0.412246 | -0.03673 | -0.5201 | 0.030635 | -0.01924 |
| 32.66667 | 1.824133 | 1.400127 | -0.1214 | 0.472059 | 0.089159 | 0.499784 | -0.70152 | 0.703416 | 0.048221 | -1.88339 | -0.85248 | -0.26697 | 0.084792 | -0.81209 | -0.17621 | -0.0008 | -0.34316 | -0.00264 | -0.70107 | 0.354641 | 0.032495 | -0.49967 | -0.03901 | -0.06608 |
| 32.68056 | 1.156841 | 1.153821 | -0.11245 | 0.111955 | -0.0453 | 0.281645 | -3.26851 | 0.288218 | -0.10408 | -2.65447 | 1.296418 | -0.62908 | 0.088507 | -0.80929 | -0.17796 | -0.00234 | -0.43527 | 0.090425 | -0.46601 | 0.328673 | 0.169591 | -0.43125 | -0.1363 | -0.05382 |
| 32.69444 | 0.582749 | 1.037562 | -0.17448 | -0.1946 | -0.1634 | 0.098043 | -3.52789 | 0.218291 | -0.3497 | -2.43177 | 2.234107 | -1.00216 | 0.159637 | -0.85515 | -0.15857 | 0.035883 | -0.4543 | 0.166274 | -0.08048 | 0.320823 | 0.320487 | -0.31611 | -0.20114 | -0.01011 |
| 32.70833 | -0.56139 | 0.652384 | -0.18729 | -0.44397 | -0.25505 | 0.070793 | -3.60064 | 0.33161 | -0.60531 | -1.99495 | 1.98706 | -1.18263 | 0.178478 | -0.72962 | -0.1378 | 0.063541 | -0.37923 | 0.17287 | 0.194269 | 0.349801 | 0.386752 | -0.18771 | -0.16416 | -0.00195 |
| 32.72222 | -1.41906 | 0.100323 | -0.20958 | -0.47425 | -0.27103 | 0.127288 | -2.81984 | 0.590953 | -0.61271 | -2.17169 | 1.885568 | -0.95777 | 0.13084 | -0.49364 | -0.12904 | 0.103877 | -0.21689 | 0.182006 | 0.214645 | 0.378919 | 0.379386 | -0.00656 | -0.02621 | -0.05091 |
| 32.73611 | -1.25256 | -0.19892 | -0.17451 | -0.25599 | -0.23485 | 0.141918 | -2.04306 | 0.894998 | -0.45158 | -1.77192 | 0.687548 | -0.64715 | 0.069131 | -0.06526 | -0.10825 | 0.13819 | 0.029142 | 0.165508 | 0.102406 | 0.387036 | 0.311169 | 0.219555 | 0.086329 | -0.15208 |
| 32.75 | -1.27942 | -0.52745 | -0.16504 | -0.16373 | -0.23062 | 0.017535 | -1.94769 | 1.015595 | -0.10168 | -1.72949 | 0.484409 | -0.50526 | 0.021374 | 0.388508 | -0.06457 | 0.14449 | 0.247935 | 0.14625 | 0.044245 | 0.36297 | 0.201674 | 0.393922 | 0.119301 | -0.26303 |
| 32.76389 | -1.17395 | -1.0463 | -0.1665 | -0.3177 | -0.32415 | -0.11035 | -2.35868 | 0.944269 | 0.090492 | -1.68775 | 0.808777 | -0.63545 | -0.00281 | 0.669422 | 0.021028 | 0.090715 | 0.375356 | 0.088387 | 0.056788 | 0.332399 | 0.10592 | 0.471493 | 0.13869 | -0.39205 |
| 32.77778 | -1.12678 | -1.88523 | -0.25731 | -0.4643 | -0.4069 | -0.04797 | -2.77137 | 0.917879 | 0.137763 | -1.84225 | 0.349671 | -0.85597 | -0.01871 | 0.687535 | 0.097429 | -0.03364 | 0.304315 | 0.018476 | 0.00847 | 0.30977 | 0.069606 | 0.421279 | 0.121333 | -0.44346 |

| | | | | | | | | | | | | | | | | | | | | | | | | |
|---|---|---|---|---|---|---|---|---|---|---|---|---|---|---|---|---|---|---|---|---|---|---|---|---|
| 32.79167 | -1.1686 | -2.12248 | -0.20878 | -0.57657 | -0.48731 | 0.12286 | -3.33304 | 0.782419 | 0.183919 | -1.9787 | -1.00424 | -0.98664 | -0.03693 | 0.112519 | 0.115462 | -0.20591 | 0.099362 | -0.06023 | -0.20122 | 0.272753 | 0.067146 | 0.171637 | 0.070648 | -0.43557 |
| 32.80556 | -1.04202 | -2.12121 | -0.23587 | -0.64271 | -0.65979 | 0.229382 | -3.50972 | 0.480528 | -0.06246 | -2.00337 | -1.34346 | -0.87522 | -0.04413 | -0.53774 | 0.082367 | -0.33757 | -0.11597 | -0.09267 | -0.45722 | 0.246912 | 0.07605 | -0.18706 | 0.012336 | -0.45586 |
| 32.81944 | -0.84096 | -1.99478 | -0.34296 | -0.76012 | -0.9382 | 0.138353 | -4.34305 | 0.352614 | 0.096166 | -1.66429 | -1.27078 | -0.64854 | -0.07851 | -1.01582 | 0.034664 | -0.39169 | -0.25235 | -0.1281 | -0.67381 | 0.214395 | 0.040672 | -0.55505 | -0.02747 | -0.50003 |
| 32.83333 | -1.06038 | -1.72727 | -0.53654 | -0.81742 | -1.02217 | 0.00734 | -5.25805 | 0.582362 | 0.151561 | -1.40141 | -1.26864 | -0.37999 | -0.1856 | -1.21793 | 0.002878 | -0.45556 | -0.15201 | -0.17205 | -0.64101 | 0.151991 | -0.05203 | -0.6623 | -0.07013 | -0.48373 |
| 32.84722 | -1.03829 | -1.45381 | -0.98894 | -0.80032 | -0.70736 | -0.14601 | -5.07662 | 0.766275 | 0.053491 | -1.28724 | -1.29823 | -0.4055 | -0.28767 | -1.05724 | -0.01042 | -0.44878 | 0.103563 | -0.19358 | -0.4863 | 0.076557 | -0.13293 | -0.56371 | -0.06772 | -0.39573 |
| 32.86111 | -1.19678 | -1.92687 | -1.35235 | -0.79274 | -0.47312 | -0.28113 | -5.31071 | 0.663437 | -0.22704 | -1.37896 | -1.61742 | -0.67955 | -0.32189 | -0.79761 | -0.01256 | -0.40276 | 0.367923 | -0.14705 | -0.30138 | 0.016313 | -0.17114 | -0.32979 | -0.03915 | -0.29994 |
| 32.875 | -0.88162 | -2.75914 | -1.61047 | -0.73791 | -0.334 | -0.40727 | -6.03712 | 0.411713 | -0.64488 | -1.6368 | -1.6289 | -0.99166 | -0.31823 | -0.55166 | -0.05255 | -0.33086 | 0.377505 | -0.13094 | -0.21458 | -0.09574 | -0.2444 | -0.18286 | -0.07275 | -0.27584 |
| 32.88889 | -0.36816 | -3.16036 | -1.7235 | -0.81154 | -0.30705 | -0.52962 | -6.33191 | 0.091512 | -0.71971 | -1.80108 | -1.37663 | -1.15124 | -0.31938 | -0.47167 | -0.14423 | -0.23946 | 0.154655 | -0.17142 | -0.25069 | -0.22631 | -0.37404 | -0.11979 | -0.1969 | -0.27154 |
| 32.90278 | -0.18081 | -2.60206 | -1.362 | -0.74176 | -0.27114 | -0.51652 | -6.81849 | -0.04924 | -0.31687 | -1.32354 | -1.10167 | -1.15421 | -0.32296 | -0.48571 | -0.19921 | -0.153 | -0.07303 | -0.22203 | -0.37266 | -0.36718 | -0.51662 | -0.09503 | -0.34503 | -0.31982 |
| 32.91667 | 0.134303 | -1.90572 | -1.04345 | -0.56831 | -0.1777 | -0.45985 | -6.10298 | -0.16226 | 0.081557 | -0.81489 | -0.27318 | -1.06525 | -0.3032 | -0.43155 | -0.1908 | -0.12972 | -0.23117 | -0.22616 | -0.47466 | -0.46264 | -0.64053 | -0.09633 | -0.44444 | -0.36454 |
| 32.93056 | 0.214047 | -1.481 | -0.37569 | -0.4476 | -0.03379 | -0.36542 | -3.54655 | -0.27566 | 0.132156 | -0.34359 | 0.375687 | -0.8294 | -0.24841 | -0.30002 | -0.18878 | -0.12213 | -0.23853 | -0.16925 | -0.39917 | -0.50238 | -0.7192 | -0.09781 | -0.46598 | -0.41332 |
| 32.94444 | 0.481768 | -1.09734 | 0.096364 | -0.39816 | 0.112788 | -0.22356 | -2.10463 | -0.22909 | 0.021379 | -0.1318 | 1.361179 | -0.55984 | -0.16394 | -0.09268 | -0.14263 | -0.11137 | -0.14377 | -0.11154 | -0.21592 | -0.44546 | -0.70597 | -0.11716 | -0.43761 | -0.34644 |
| 32.95833 | 0.656329 | -0.49912 | 0.205172 | -0.25259 | 0.144241 | -0.09633 | -1.4852 | 0.007984 | 0.174601 | -0.00569 | 1.56606 | -0.24049 | -0.06383 | 0.078826 | -0.07304 | -0.08022 | -0.02735 | -0.06957 | -0.04991 | -0.34319 | -0.66162 | -0.14395 | -0.36565 | -0.23306 |
| 32.97222 | 0.788544 | 0.098795 | 0.063354 | 0.057884 | 0.038223 | -0.13519 | -0.53134 | 0.273569 | 0.495204 | 0.088208 | 1.900913 | -0.03849 | 0.029236 | 0.165962 | -0.01649 | 0.011464 | -0.00645 | -0.0329 | -0.00243 | -0.22611 | -0.57163 | -0.14158 | -0.2904 | -0.13162 |
| 32.98611 | 0.894111 | 0.180454 | -0.38579 | 0.070242 | -0.22982 | -0.20514 | -0.07778 | 0.218956 | 0.476877 | -0.09049 | 1.37774 | -0.08375 | 0.107102 | 0.206146 | 0.061715 | 0.110279 | -0.02107 | -0.01208 | -0.04623 | -0.12333 | -0.45778 | -0.05461 | -0.1974 | -0.04732 |
| 33 | 0.169699 | -0.24296 | -0.94304 | -0.54275 | -0.59938 | -0.40636 | 0.015209 | -0.58154 | 0.016033 | -0.95946 | -0.07115 | -0.34213 | 0.192336 | 0.174581 | 0.203743 | 0.17016 | -0.05076 | -0.03814 | -0.14717 | -0.02654 | -0.49612 | 0.017846 | -0.14372 | 0.013839 |
| 33.01389 | -0.45205 | -0.70717 | -1.55716 | -1.02685 | -0.6309 | -0.24252 | -0.45323 | -1.60262 | -0.80461 | -1.97194 | -1.62461 | -0.60179 | 0.201472 | -0.11446 | 0.244654 | 0.09682 | -0.23758 | -0.19 | -0.29184 | 0.004634 | -1.00818 | -0.15912 | -0.17181 | -0.09176 |
| 33.02778 | -0.31816 | -0.95435 | -1.81943 | -0.79726 | -0.71564 | -0.00045 | 1.142604 | -2.4797 | -1.8204 | -2.16893 | -1.07806 | -0.96084 | 0.030699 | -0.50168 | 0.186358 | -0.01575 | -0.74913 | -0.40179 | -0.4985 | -0.0229 | -1.84979 | -0.6539 | -0.31581 | -0.29622 |
| 33.04167 | -0.54277 | -1.13364 | -2.16772 | -0.39573 | -0.77294 | -0.11472 | 0.630631 | -2.67811 | -2.10128 | -1.26508 | -10.4985 | -1.35607 | -0.21787 | -0.70163 | 0.152632 | -0.06882 | -1.10673 | -0.41649 | -0.76296 | -0.21969 | -1.86357 | -1.03093 | -0.44598 | -0.39659 |
| 33.05556 | -0.87304 | -1.5588 | -2.33325 | -0.05726 | -0.67675 | -0.05245 | 0.106556 | -1.97147 | -1.94745 | -0.23296 | -5.90276 | -1.09635 | -0.46744 | -0.61086 | 0.02887 | -0.19233 | -1.10432 | -0.20844 | -0.87242 | -0.61198 | -1.05971 | -0.9405 | -0.48391 | -0.38213 |
| 33.06944 | -0.96922 | -1.92132 | -1.57253 | 0.278104 | -0.2092 | 0.154799 | 0.152304 | -0.80814 | -1.10068 | 0.198642 | -2.70151 | -0.46818 | -0.61428 | -0.49318 | -0.07639 | -0.31871 | -0.89135 | -0.11297 | -0.7005 | -0.85228 | -0.43471 | -0.69461 | -0.50916 | -0.3544 |
| 33.08333 | -0.8727 | -2.13661 | -0.59621 | 0.405548 | 0.442026 | 0.459882 | 0.301054 | 0.246156 | -0.30484 | 0.140969 | -1.13657 | 0.057758 | -0.47145 | -0.29095 | -0.07339 | -0.2383 | -0.58736 | -0.07636 | -0.3382 | -0.67953 | -0.22169 | -0.44203 | -0.31776 | -0.18476 |
| 33.09722 | -0.55051 | -1.88855 | 0.405302 | 0.369634 | 0.688583 | 0.543037 | 0.270051 | 0.762679 | 0.12637 | -0.01617 | -0.2843 | 0.48301 | -0.1625 | -0.06311 | 0.146281 | 0.014583 | -0.19771 | -0.03965 | -0.03058 | -0.25256 | 0.00974 | -0.07996 | 0.089925 | 0.032175 |
| 33.11111 | -0.19423 | -1.66626 | 0.941203 | 0.285135 | 0.378249 | 0.423254 | 0.244479 | 0.8903 | 0.287224 | -0.20133 | 0.370554 | 0.780333 | 0.020529 | 0.025623 | 0.215519 | 0.22985 | 0.061819 | -0.00915 | 0.102902 | 0.050183 | 0.245366 | 0.105018 | 0.352081 | 0.101297 |
| 33.125 | 0.003954 | -1.42992 | 0.758923 | 0.244378 | -0.10263 | 0.352766 | 0.157185 | 0.822208 | 0.1754 | -0.35552 | 0.498162 | 1.064095 | 0.077738 | -0.01195 | 0.137323 | 0.282807 | 0.20857 | 0.002061 | 0.205036 | 0.118434 | 0.360337 | 0.143379 | 0.288211 | 0.073677 |
| 33.13889 | -0.15341 | -1.37498 | 0.153803 | 0.257778 | -0.35514 | 0.318703 | 0.088441 | 0.541742 | 0.045294 | -0.43036 | 0.422604 | 1.225452 | 0.067826 | -0.09099 | 0.080783 | 0.196216 | 0.375433 | 0.01038 | 0.338316 | 0.075777 | 0.339247 | 0.062521 | 0.202003 | -0.01446 |
| 33.15278 | -0.15297 | -1.28108 | 0.015632 | 0.329824 | -0.32679 | 0.36219 | 0.03973 | 0.252183 | 0.036664 | -0.38719 | 0.588461 | 0.754408 | 0.039558 | -0.18086 | 0.088604 | 0.048129 | 0.391979 | 0.002834 | 0.392966 | -0.01081 | 0.248708 | -0.04401 | 0.099367 | -0.14824 |
| 33.16667 | -0.48489 | -1.1258 | 0.379951 | 0.399604 | -0.19555 | 0.510581 | 0.021393 | -0.05528 | 0.050367 | -0.26779 | 0.694816 | 0.443704 | -0.00157 | -0.24618 | 0.13503 | -0.04398 | 0.346562 | -0.004 | 0.32151 | -0.08544 | 0.119936 | -0.07936 | -0.01178 | -0.21384 |
| 33.18056 | -0.40513 | -0.88867 | 0.774597 | 0.447843 | -0.08839 | 0.665326 | 0.044678 | -0.37072 | 0.088392 | -0.16911 | 1.31689 | 0.335371 | -0.06194 | -0.24357 | 0.15659 | -0.05219 | 0.176553 | -0.00058 | 0.242568 | -0.0979 | -0.0015 | -0.04178 | -0.06618 | -0.15403 |
| 33.19444 | -0.15506 | -0.7562 | 0.906374 | 0.465367 | 0.09627 | 0.732545 | 0.126986 | -0.44238 | 0.205856 | -0.07937 | 1.33117 | 0.297709 | -0.13198 | -0.14713 | 0.119705 | -0.00751 | 0.123429 | 0.00697 | 0.180805 | -0.05926 | -0.08978 | 0.011934 | -0.09247 | -0.07445 |
| 33.20833 | -0.00103 | -0.56428 | 0.96857 | 0.446207 | 0.338273 | 0.638496 | 0.212828 | -0.23518 | 0.275483 | 0.021118 | 0.486754 | 0.269013 | -0.18144 | -0.03775 | 0.053903 | 0.053256 | 0.176985 | -0.02706 | 0.111238 | -0.0241 | -0.14458 | 0.050537 | -0.18043 | -0.04478 |
| 33.22222 | 0.317351 | -0.41512 | 0.982119 | 0.40794 | 0.51231 | 0.361552 | 0.275992 | 0.177176 | 0.321792 | 0.070032 | 0.421734 | 0.255741 | -0.15785 | 0.018439 | -0.0111 | 0.076815 | 0.131468 | -0.09125 | 0.037359 | -0.03175 | -0.16943 | 0.066474 | -0.29165 | -0.03218 |
| 33.23611 | 0.675948 | -0.28861 | 0.859493 | 0.330032 | 0.44922 | -0.06052 | 0.169492 | 0.507066 | 0.224184 | 0.029374 | 0.293689 | 0.232935 | -0.06473 | 0.027485 | -0.05182 | 0.078887 | -0.00859 | -0.10529 | 0.006937 | -0.01908 | -0.16571 | 0.064431 | -0.29097 | -0.02961 |
| 33.25 | 0.699934 | -0.25132 | 0.621281 | 0.256801 | 0.166854 | -0.36679 | -0.08378 | 0.750091 | 0.138439 | -0.00608 | 0.178618 | 0.234698 | 0.053636 | 0.002809 | -0.04718 | 0.068313 | -0.12939 | -0.06811 | 0.010529 | 0.004167 | -0.14191 | 0.043791 | -0.18306 | -0.03988 |
| 33.26389 | 0.618219 | -0.24384 | 0.113417 | 0.21973 | -0.20544 | -0.17825 | -0.24323 | 0.854114 | 0.136897 | 0.013523 | 0.206805 | 0.263295 | 0.127927 | -0.03966 | -0.02927 | 0.023101 | -0.203 | -0.0366 | -0.0406 | 0.031356 | -0.13164 | -0.0285 | -0.01526 | -0.03424 |
| 33.27778 | 0.541099 | -0.31251 | -0.57499 | 0.195148 | -0.4193 | 0.091025 | -0.03361 | 0.769094 | 0.253148 | -0.00942 | 0.134722 | 0.295962 | 0.134488 | -0.11331 | -0.02312 | -0.02707 | -0.24849 | -0.0241 | -0.17271 | 0.019498 | -0.14516 | -0.13836 | 0.148777 | -0.01069 |
| 33.29167 | 0.56736 | -0.36915 | -1.04081 | 0.170756 | -0.3437 | 0.253918 | 0.233207 | 0.680706 | 0.382106 | -0.15152 | 0.028818 | 0.321865 | 0.111448 | -0.20751 | -0.03461 | -0.02784 | -0.27207 | -0.02908 | -0.29781 | -0.05516 | -0.18544 | -0.20921 | 0.235423 | 0.011222 |
| 33.30556 | 0.49034 | -0.37581 | -0.75408 | 0.161988 | -0.10797 | 0.27093 | 0.359809 | 0.638568 | 0.336136 | -0.24513 | -0.04022 | 0.269401 | 0.102537 | -0.28276 | -0.08845 | -0.00239 | -0.2755 | -0.03378 | -0.3596 | -0.12477 | -0.19178 | -0.19715 | 0.198984 | 0.019641 |
| 33.31944 | 0.4142 | -0.32855 | -0.00151 | 0.141359 | 0.074424 | 0.246736 | 0.303356 | 0.537212 | 0.335118 | -0.20862 | -0.0468 | 0.178527 | 0.119831 | -0.28263 | -0.17103 | -0.03658 | -0.28953 | -0.04237 | -0.3756 | -0.17871 | -0.14302 | -0.12987 | 0.090923 | 0.00756 |
| 33.33333 | 0.376484 | -0.32273 | 0.591136 | 0.117097 | 0.07373 | 0.267104 | 0.207817 | 0.505716 | 0.324091 | -0.11393 | -0.0088 | 0.040059 | 0.110166 | -0.19285 | -0.23867 | -0.12305 | -0.29954 | -0.04384 | -0.3067 | -0.20659 | -0.08028 | -0.10374 | -0.03241 | -0.00455 |
| 33.34722 | 0.277893 | -0.40265 | 0.66449 | 0.07209 | -0.08705 | 0.290786 | 0.222228 | 0.515591 | 0.16344 | -0.11566 | 0.033457 | -0.07728 | 0.095934 | -0.12272 | -0.27214 | -0.1821 | -0.23918 | -0.02493 | -0.19365 | -0.17558 | 0.001557 | -0.13892 | -0.05054 | 0.025177 |
| 33.36111 | 0.130236 | -0.46211 | 0.10991 | -0.02218 | -0.30964 | 0.272257 | 0.216142 | 0.486513 | -0.03235 | -0.16388 | -0.00214 | -0.16685 | 0.135843 | -0.09133 | -0.16305 | -0.22502 | -0.15127 | 0.018014 | -0.07306 | -0.10182 | 0.078923 | -0.15827 | 0.057901 | 0.075141 |
| 33.375 | 0.099606 | -0.40301 | -0.31222 | -0.05039 | -0.40651 | 0.28176 | 0.241044 | 0.471103 | -0.09579 | -0.14259 | 0.041517 | -0.16237 | 0.163346 | -0.04862 | -0.01102 | -0.26181 | -0.15623 | 0.047743 | 0.006052 | -0.08601 | 0.054998 | -0.16886 | 0.116719 | 0.030823 |
| 33.38889 | 0.221349 | -0.35163 | -0.0704 | 0.043909 | -0.33829 | 0.41964 | 0.500276 | 0.556672 | -0.00902 | 0.051501 | 0.1088 | -0.0605 | 0.149526 | 0.000014 | 0.114605 | -0.28142 | -0.20659 | 0.037907 | 0.068863 | -0.1323 | 0.000232 | -0.14963 | 0.077684 | -0.02563 |
| 33.40278 | 0.305127 | -0.368 | 0.41318 | 0.164072 | -0.15502 | 0.575424 | 0.690967 | 0.658294 | 0.100563 | 0.238382 | 0.160115 | 0.06797 | 0.140865 | 0.084179 | 0.183109 | -0.2121 | -0.23593 | 0.04925 | 0.154185 | -0.13278 | 0.018344 | -0.01335 | 0.057949 | 0.010079 |
| 33.41667 | 0.282903 | -0.39899 | 0.782257 | 0.254089 | 0.195567 | 0.639115 | 0.767576 | 0.828005 | 0.260456 | 0.313489 | 0.26455 | 0.194774 | 0.157059 | 0.096943 | 0.162896 | -0.14031 | -0.23757 | 0.058721 | 0.174307 | -0.10597 | 0.032913 | 0.123265 | 0.046144 | 0.027418 |
| 33.43056 | 0.189869 | -0.38084 | 0.914593 | 0.296654 | 0.527097 | 0.63584 | 0.596501 | 1.0044 | 0.353703 | 0.287686 | 0.476821 | 0.266914 | 0.138253 | 0.04301 | 0.08363 | -0.10385 | -0.20534 | 0.037241 | 0.145815 | -0.07817 | 0.026825 | 0.18169 | 0.018418 | -0.01733 |
| 33.44444 | 0.16143 | -0.48345 | 0.852603 | 0.301721 | 0.548529 | 0.539746 | 0.360295 | 1.065457 | 0.303307 | 0.178589 | 0.657746 | 0.21497 | 0.121976 | -0.03412 | 0.009888 | -0.03647 | -0.14315 | 0.021074 | 0.166731 | -0.0422 | 0.082426 | 0.234134 | 0.057498 | -0.00492 |
| 33.45833 | 0.059962 | -0.69396 | 0.68612 | 0.240686 | 0.204582 | 0.424395 | 0.212507 | 1.002037 | 0.138348 | -0.01055 | 0.770594 | 0.049736 | 0.137637 | -0.10976 | -0.0451 | 0.043439 | -0.07919 | 0.025766 | 0.202607 | -0.01835 | 0.150775 | 0.253701 | 0.122468 | -0.00504 |
| 33.47222 | 0.047204 | -0.83727 | 0.595058 | 0.184412 | -0.2193 | 0.378917 | 0.186828 | 0.900203 | 0.065959 | -0.07199 | 0.913116 | -0.10981 | 0.163564 | -0.17435 | -0.05235 | 0.086691 | -0.0135 | 0.055306 | 0.219711 | -0.00157 | 0.172538 | 0.166439 | 0.132003 | -0.01571 |
| 33.48611 | 0.234069 | -0.62028 | 0.523782 | 0.221602 | -0.41805 | 0.475869 | 0.384888 | 0.795319 | 0.202319 | 0.24018 | 1.422687 | -0.01823 | 0.180075 | -0.17833 | 0.054628 | 0.125423 | 0.066593 | 0.105786 | 0.261614 | 0.019994 | 0.167699 | 0.126659 | 0.074607 | -0.01243 |
| 33.5 | 0.905066 | 0.242812 | 0.531758 | 0.445201 | -0.11845 | 0.646254 | 0.592721 | 0.600428 | 0.482692 | 0.717034 | 2.125716 | 0.383991 | 0.162108 | -0.16624 | 0.239277 | 0.199032 | 0.101441 | 0.168646 | 0.338552 | -0.00033 | 0.29549 | 0.160061 | 0.037575 | 0.041192 |
| 33.51389 | 1.512987 | 1.263452 | 0.782124 | 0.56191 | 0.381009 | 0.684191 | 0.690724 | 0.518278 | 0.589162 | 0.742387 | 2.355566 | 0.824319 | 0.152348 | -0.17662 | 0.348575 | 0.313017 | 0.088187 | 0.223807 | 0.430407 | -0.08996 | 0.597797 | 0.265493 | 0.111653 | 0.113374 |
| 33.52778 | 1.835534 | 1.526294 | 1.383981 | 0.628279 | 0.261398 | 0.484563 | 0.546439 | 0.91205 | 0.600859 | 0.649148 | 2.775666 | 1.307358 | 0.240595 | -0.12318 | 0.247622 | 0.288161 | 0.154174 | 0.178077 | 0.56175 | -0.12137 | 0.721048 | 0.40358 | 0.351399 | 0.175486 |
| 33.54167 | 1.800832 | 1.076664 | 2.252635 | 0.821124 | 0.12423 | 0.523608 | 0.379992 | 1.470293 | 0.828265 | 0.991874 | 3.00204 | 1.917607 | 0.323175 | 0.104447 | 0.103706 | 0.227884 | 0.458581 | 0.038202 | 0.774755 | -0.06401 | 0.517893 | 0.300431 | 0.540156 | 0.193523 |
| 33.55556 | 1.89801 | 0.938491 | 2.748839 | 1.23824 | 0.342326 | 0.616858 | 0.17674 | 1.78184 | 0.722303 | 1.272206 | 4.224037 | 2.275812 | 0.322028 | 0.476417 | 0.09722 | 0.368949 | 0.856405 | -0.00807 | 1.049881 | 0.044612 | 0.457085 | 0.344145 | 0.459416 | 0.211146 |
| 33.56944 | 1.473471 | 1.400333 | 2.463714 | 1.277141 | 0.175884 | 0.656691 | -0.1341 | 1.844711 | 0.391529 | 0.812658 | 7.492945 | 2.243568 | 0.282624 | 0.728586 | 0.141217 | 0.452545 | 1.080049 | 0.044773 | 1.054447 | 0.323163 | 0.555453 | 0.505038 | 0.309749 | 0.269834 |
| 33.58333 | 0.837838 | 2.2237 | 2.310175 | 0.843501 | -0.34612 | 0.624017 | 0.291215 | 1.478854 | 0.287314 | 0.399507 | 8.492417 | 2.119178 | 0.224729 | 0.594751 | 0.054838 | 0.358096 | 1.017692 | 0.0052 | 1.00839 | 0.536701 | 0.588232 | 0.351369 | 0.19122 | 0.28577 |
| 33.59722 | 0.471633 | 2.965882 | 2.128931 | 0.498096 | -0.39151 | 0.575377 | 1.265179 | 0.717342 | 0.382103 | 0.360177 | 6.03851 | 2.001596 | 0.094897 | 0.198904 | -0.04444 | 0.231239 | 0.784322 | -0.05379 | 0.803056 | 0.668966 | 0.563721 | 0.098882 | 0.11464 | 0.245033 |
| 33.61111 | 0.406787 | 3.121435 | 1.814756 | 0.194439 | -0.24102 | 0.637999 | 1.770123 | 0.172314 | 0.38449 | 0.304242 | 2.66192 | 1.594645 | -0.01481 | -0.07054 | -0.09286 | 0.185503 | 0.575998 | -0.05712 | 0.561025 | 0.691657 | 0.52738 | -0.06227 | 0.002424 | 0.216379 |
| 33.625 | 0.567482 | 3.569086 | 1.418744 | 0.259138 | -0.37909 | 0.749773 | 1.744256 | 0.087661 | 0.54475 | 0.182139 | 0.223245 | 0.988208 | -0.07244 | -0.22986 | -0.13444 | 0.201209 | 0.409355 | -0.06669 | 0.18085 | 0.602187 | 0.390396 | -0.20436 | -0.17338 | 0.092671 |
| 33.63889 | 0.678815 | 3.844658 | 1.082276 | 0.338566 | -0.01617 | 0.712151 | 1.380507 | 0.169394 | 0.723895 | 0.173818 | -1.60617 | 0.25882 | -0.08494 | -0.3202 | -0.13807 | 0.211986 | 0.234665 | -0.06693 | -0.20425 | 0.527102 | 0.180462 | -0.3465 | -0.27007 | -0.0675 |
| 33.65278 | 0.790675 | 3.686841 | 0.866225 | 0.431372 | 0.160147 | 0.636645 | 1.429888 | -0.06945 | 0.783078 | 0.258893 | -7.38107 | -0.29481 | -0.04307 | -0.2666 | -0.05637 | 0.329151 | 0.105079 | 0.011325 | -0.53383 | 0.476845 | 0.048248 | -0.37669 | -0.19607 | -0.18053 |
| 33.66667 | 0.928488 | 3.420133 | 0.70462 | 0.297807 | -0.07054 | 0.562077 | 1.598287 | -0.32135 | 0.708753 | 0.389437 | -6.8045 | -0.7908 | 0.028082 | -0.22248 | 0.062088 | 0.51929 | -0.01923 | 0.131162 | -0.74675 | 0.377892 | -0.00906 | -0.29697 | -0.124 | -0.26376 |
| 33.68056 | 1.021964 | 3.186421 | 0.417911 | 0.045314 | -0.37513 | 0.459651 | 1.821938 | -0.46628 | 0.272089 | 0.230189 | -3.56546 | -1.30219 | 0.079997 | -0.10689 | 0.150969 | 0.654455 | -0.17064 | 0.220411 | -0.61768 | 0.271594 | -0.05046 | -0.1899 | -0.11197 | -0.34007 |
| 33.69444 | 0.978471 | 3.054436 | 0.241317 | -0.11105 | -0.61613 | 0.431745 | 2.029358 | -0.27158 | -0.34996 | -0.10635 | -1.81495 | -1.50136 | 0.104408 | -0.06836 | 0.23748 | 0.684703 | -0.18982 | 0.245399 | -0.42899 | 0.215667 | -0.02454 | -0.02123 | -0.00955 | -0.278 |
| 33.70833 | 0.751592 | 2.381209 | 0.307057 | -0.15516 | -0.87165 | 0.45832 | 1.944587 | 0.043317 | -1.09528 | -0.26134 | -1.47736 | -1.40113 | 0.090002 | -0.1343 | 0.283313 | 0.576936 | -0.10965 | 0.204041 | -0.21219 | 0.238532 | 0.104768 | 0.095694 | 0.083293 | -0.20231 |
| 33.72222 | 0.258707 | 1.038438 | 0.224858 | -0.15564 | -0.96464 | 0.405713 | 1.505635 | 0.169322 | -0.74984 | -0.32626 | -1.01082 | -1.34036 | 0.058992 | -0.20061 | 0.240803 | 0.424094 | -0.04251 | 0.122302 | -0.0007 | 0.308742 | 0.230521 | 0.124503 | 0.095306 | -0.12958 |
| 33.73611 | -0.05808 | -0.07667 | 0.017149 | -0.08089 | -1.00941 | 0.376115 | 0.955566 | 0.204413 | -0.80401 | -0.38749 | 1.283199 | -1.17079 | 0.053707 | -0.2305 | 0.142109 | 0.260777 | -0.1299 | 0.040218 | -0.01175 | 0.396155 | 0.279821 | 0.04371 | 0.074933 | -0.11526 |
| 33.75 | -0.40567 | -0.82804 | -0.2343 | -0.03549 | -1.0034 | 0.338253 | 0.759154 | 0.266726 | -0.54178 | -0.47079 | 2.872747 | -0.94582 | 0.060423 | -0.24443 | 0.040535 | 0.100993 | -0.23666 | 0.002578 | -0.19204 | 0.551184 | 0.313706 | -0.05721 | 0.126192 | -0.16772 |
| 33.76389 | -0.79854 | -1.2932 | -0.51858 | -0.09536 | -1.20048 | 0.3045 | 0.62026 | 0.326067 | -0.55834 | -0.49514 | 3.800821 | -1.03039 | 0.031436 | -0.29046 | -0.0352 | -0.0403 | -0.20576 | -0.00822 | -0.45496 | 0.695761 | 0.333404 | -0.11144 | 0.155981 | -0.26159 |
| 33.77778 | -0.8882 | -1.03649 | -0.8275 | -0.1754 | -1.37613 | 0.229033 | 0.39976 | 0.432375 | -0.6687 | -0.56364 | 5.201402 | -1.39075 | -0.02398 | -0.33587 | -0.09621 | -0.11528 | -0.04056 | -0.00377 | -0.61569 | 0.717887 | 0.276583 | -0.119 | 0.128446 | -0.32255 |
| 33.79167 | -0.90614 | -0.70928 | -1.47878 | -0.32648 | -1.5799 | 0.196864 | 0.120744 | 0.375698 | -0.46451 | -0.60349 | 4.548028 | -1.69842 | -0.04635 | -0.27918 | -0.13239 | -0.10732 | 0.141027 | 0.015262 | -0.59502 | 0.621751 | 0.12117 | -0.09861 | 0.092632 | -0.37085 |
| 33.80556 | -1.3211 | -0.8938 | -2.16283 | -0.39768 | -1.98632 | 0.125451 | -0.21202 | 0.26478 | -0.495 | -0.66999 | 10.95313 | -1.82762 | -0.02757 | -0.1711 | -0.1365 | -0.07092 | 0.2261 | 0.021039 | -0.36943 | 0.450558 | -0.06029 | -0.08265 | 0.067841 | -0.26084 |
| 33.81944 | -2.44718 | -0.98384 | -2.80832 | -0.34019 | -2.40762 | -0.00856 | -0.47231 | 0.11835 | -0.50023 | -0.71003 | 7.923407 | -1.77535 | -0.03027 | -0.09317 | -0.10222 | -0.11984 | 0.129676 | -0.01044 | -0.17075 | 0.202793 | -0.19465 | -0.10634 | -0.01388 | -0.2819 |
| 33.83333 | -3.30184 | -1.13622 | -3.04823 | -0.29105 | -2.63059 | -0.24629 | -0.73051 | 0.176794 | -0.3113 | -0.66602 | 4.394439 | -1.5948 | -0.08888 | -0.03628 | -0.04509 | -0.27554 | -0.07816 | -0.0417 | -0.06149 | -0.03912 | -0.21949 | -0.14879 | -0.09899 | -0.29933 |
| 33.84722 | -3.17826 | -1.23489 | -3.06158 | -0.29193 | -2.14763 | -0.32873 | -1.02194 | 0.332412 | -0.02097 | -0.59691 | 1.325046 | -1.23733 | -0.17326 | -0.00952 | 0.014681 | -0.40721 | -0.30333 | -0.06823 | -0.00648 | -0.18644 | -0.19584 | -0.12966 | -0.25535 |
| 33.86111 | -2.8375 | -1.61821 | -3.04415 | -0.20603 | -1.5042 | -0.33209 | -1.21707 | 0.502403 | 0.370172 | -0.54421 | -2.23672 | -0.93705 | -0.24797 | -0.04421 | 0.03052 | -0.50809 | -0.56056 | -0.09614 | 0.00825 | -0.26419 | -0.17332 | -0.22459 | -0.19609 | -0.21347 |
| 33.875 | -1.94398 | -1.91932 | -2.79778 | -0.14372 | -1.57352 | -0.33914 | -1.24798 | 0.581978 | 0.660604 | -0.50768 | -7.11458 | -0.58762 | -0.31428 | -0.1579 | -0.00954 | -0.57709 | -0.7309 | -0.11497 | -0.06321 | -0.35252 | -0.17243 | -0.21731 | -0.26346 | -0.22335 |
| 33.88889 | -2.12932 | -2.28334 | -2.4114 | -0.16068 | -1.75487 | -0.47336 | -1.22593 | 0.638909 | 0.646057 | -0.46708 | -8.30276 | -0.1766 | -0.36111 | -0.30297 | -0.07928 | -0.59453 | -0.92282 | -0.11869 | -0.17267 | -0.43325 | -0.21953 | -0.22468 | -0.3311 | -0.24136 |
| 33.90278 | -1.86063 | -2.22155 | -2.31704 | -0.1762 | -1.23565 | -0.56881 | -1.79335 | 0.643971 | 0.719256 | -0.45508 | -5.15253 | 0.241415 | -0.37425 | -0.47299 | -0.17185 | -0.63287 | -0.99743 | -0.12569 | -0.23753 | -0.44954 | -0.28796 | -0.17563 | -0.33523 | -0.23672 |
| 33.91667 | -1.6378 | -1.92521 | -2.4528 | -0.14749 | -1.33082 | -0.72641 | -2.51578 | 0.62408 | 0.890989 | -0.45493 | -2.97978 | 0.532357 | -0.3379 | -0.57448 | -0.23666 | -0.56161 | -0.8451 | -0.10787 | -0.19331 | -0.36762 | -0.38082 | -0.08895 | -0.30888 | -0.24734 |
| 33.93056 | -1.57457 | -1.31136 | -2.87778 | -0.13618 | 0.296712 | -0.83354 | -2.93657 | 0.550171 | 1.346308 | -0.31311 | -2.84318 | 0.652725 | -0.2308 | -0.55449 | -0.2393 | -0.46426 | -0.54823 | -0.07036 | -0.12172 | -0.24964 | -0.44024 | 0.020046 | -0.26815 | -0.22632 |
| 33.94444 | -1.01694 | -0.56698 | -2.84374 | -0.08539 | 0.288168 | -0.93633 | -2.71818 | 0.502611 | 1.705582 | -0.13454 | -1.98525 | 0.910438 | -0.09955 | -0.47731 | -0.21702 | -0.39532 | -0.41965 | -0.04436 | -0.10253 | -0.17904 | -0.45005 | 0.102104 | -0.2406 | -0.23289 |
| 33.95833 | -0.73212 | 0.273159 | -2.25091 | 0.040061 | 0.04137 | -1.00713 | -2.56371 | 0.465099 | 1.851659 | 0.03532 | -0.81798 | 0.996882 | -0.01822 | -0.36071 | -0.15347 | -0.30342 | -0.38707 | -0.05281 | -0.13697 | -0.14548 | -0.42937 | 0.107089 | -0.30062 | -0.2329 |
| 33.97222 | -0.55168 | 0.575565 | -1.23214 | 0.411653 | 0.1164 | -0.84221 | -1.52684 | 0.415828 | 1.95994 | 0.347413 | -0.39333 | 0.938516 | -0.01822 | -0.21055 | -0.06761 | -0.18722 | -0.31891 | -0.0478 | -0.12163 | -0.12002 | -0.32182 | 0.156559 | -0.26812 | -0.22435 |
| 33.98611 | -0.52899 | 0.014083 | -0.87714 | 0.856686 | 0.206277 | -0.95841 | -0.85331 | 0.24113 | 1.373636 | 0.193585 | -0.91292 | 0.64547 | -0.02329 | -0.00899 | 0.081307 | 0.060172 | -0.18341 | -0.01993 | -0.06359 | -0.07627 | -0.20286 | 0.249894 | -0.2145 | -0.16369 |
| 34 | -0.95027 | -0.57127 | -1.29245 | 0.769834 | -0.0618 | -1.03292 | -1.26542 | -0.04049 | 0.045417 | -0.67441 | -1.73444 | 0.073899 | -0.04422 | 0.210095 | 0.271953 | 0.289706 | -0.10098 | -0.04544 | -0.05866 | -0.03644 | -0.34712 | 0.287866 | -0.13137 | -0.09204 |
| 34.01389 | -1.61108 | -1.22556 | -1.68206 | 0.186944 | -1.0633 | -0.66701 | -2.28023 | -0.09651 | -1.12873 | -1.64343 | -2.17076 | -0.47424 | -0.01553 | 0.359112 | 0.358187 | 0.287596 | -0.21583 | -0.17057 | -0.09624 | -0.05419 | -0.99742 | 0.150213 | -0.13259 | -0.14904 |
| 34.02778 | -1.70187 | -1.69401 | -1.68618 | -0.07663 | -1.72585 | -0.36831 | -2.66462 | -0.05588 | -1.92875 | -2.13046 | -1.9247 | -0.90735 | -0.00903 | 0.434654 | 0.321006 | 0.101735 | -0.60001 | -0.34033 | -0.22645 | -0.1544 | -2.10199 | -0.18796 | -0.31674 | -0.35599 |
| 34.04167 | -1.47562 | -2.26084 | -1.84823 | -0.0929 | -1.62857 | -0.20362 | -2.26153 | 0.188326 | -2.13555 | -1.63348 | -1.10566 | -0.91679 | -0.08429 | 0.314935 | 0.21867 | -0.11652 | -0.96605 | -0.39747 | -0.62941 | -0.32767 | -2.37664 | -0.50186 | -0.56661 | -0.53792 |
| 34.05556 | -1.54462 | -2.88591 | -2.24674 | -0.20442 | -0.28142 | -0.51095 | -1.64669 | 0.3955 | -2.11016 | -0.97609 | -0.34416 | -0.61854 | -0.24593 | 0.037703 | 0.018445 | -0.35176 | -0.97447 | -0.19024 | -0.8962 | -0.64938 | -1.2811 | -0.61063 | -0.64038 | -0.54419 |
| 34.06944 | -1.74114 | -3.25922 | -2.44768 | -0.19794 | 1.296937 | -0.43317 | -1.21318 | 0.120373 | -1.89007 | -0.40879 | 0.307435 | -0.34245 | -0.32565 | -0.14724 | -0.19373 | -0.58965 | -0.7893 | -0.01502 | -0.89381 | -0.73129 | -0.38039 | -0.49308 | -0.54159 | -0.39525 |
| 34.08333 | -1.52719 | -3.02414 | -1.98729 | -0.15165 | 2.182928 | -0.36464 | -0.9441 | -0.53281 | -1.15015 | -0.17521 | 0.397881 | -0.16511 | -0.21034 | -0.13204 | -0.11347 | -0.5571 | -0.58837 | 0.000106 | -0.38183 | -0.54219 | -0.17858 | -0.33641 | -0.36557 | -0.01767 |
| 34.09722 | -1.64882 | -2.67919 | -1.3383 | -0.05183 | 1.833795 | -0.07976 | -0.63228 | -0.80837 | -0.68463 | -0.05036 | -0.05396 | -0.06167 | -0.0695 | -0.01858 | 0.017568 | -0.28426 | -0.15276 | -0.03238 | 0.056274 | -0.31732 | -0.00827 | -0.12927 | 0.035226 | 0.369707 |
| 34.11111 | -1.33264 | -2.36514 | -0.91751 | -0.01224 | 1.12699 | -0.01831 | -0.45387 | -0.55807 | -0.31153 | -0.0361 | -0.28721 | 0.072094 | 0.011268 | 0.023927 | 0.058606 | -0.01015 | 0.163629 | -0.04276 | 0.280984 | -0.17249 | 0.281182 | 0.01884 | 0.313037 | 0.465225 |
| 34.125 | -1.04871 | -2.09151 | -0.70571 | 0.041619 | 0.714336 | 0.050705 | -0.31554 | -0.16586 | 0.006762 | -0.14059 | -0.05383 | 0.214895 | 0.060235 | -0.00883 | 0.050198 | 0.16626 | 0.231429 | 0.007113 | 0.343219 | -0.0664 | 0.509187 | 0.08443 | 0.369275 | 0.469899 |
| 34.13889 | -0.90025 | -1.69712 | -0.8089 | 0.101901 | 0.57049 | 0.014672 | -0.26445 | 0.066325 | 0.21222 | -0.21197 | 0.254588 | 0.262798 | 0.084814 | -0.07669 | 0.034902 | 0.129898 | 0.193409 | 0.077547 | 0.387179 | 0.015023 | 0.565921 | 0.085718 | 0.464358 | 0.349056 |
| 34.15278 | -0.63701 | -1.36418 | -1.16336 | 0.136052 | 0.325463 | 0.030864 | -0.34248 | 0.180645 | 0.251942 | -0.19428 | 0.637076 | 0.256134 | 0.1112 | -0.13447 | 0.023547 | -0.09621 | 0.09872 | 0.109356 | 0.45977 | 0.04528 | 0.437084 | 0.051033 | 0.434645 | 0.08051 |
| 34.16667 | -0.19523 | -1.18589 | -1.00412 | 0.212497 | -0.25991 | 0.08853 | -0.27726 | 0.243291 | 0.127213 | -0.07795 | 1.247478 | 0.262639 | 0.121883 | -0.14483 | 0.03265 | -0.24812 | -0.0402 | 0.115175 | 0.479227 | -0.00331 | 0.227078 | 0.056211 | 0.314726 | -0.07753 |
| 34.18056 | 0.00996 | -1.14572 | -0.36992 | 0.376088 | -0.68551 | 0.24036 | -0.04747 | 0.287019 | 0.108759 | 0.031053 | 1.778474 | 0.287505 | 0.094605 | -0.1374 | 0.056735 | -0.20328 | -0.11096 | 0.112928 | 0.525524 | -0.06057 | -0.02573 | 0.094335 | 0.218001 | -0.09156 |
| 34.19444 | 0.266444 | -1.16714 | 0.066491 | 0.474413 | -0.42954 | 0.300274 | 0.09845 | 0.277944 | 0.255325 | 0.017007 | 2.032505 | 0.281047 | 0.035222 | -0.08306 | 0.09215 | -0.06264 | -0.09991 | 0.093886 | 0.507118 | -0.07956 | -0.23455 | 0.091864 | 0.172494 | -0.02922 |
| 34.20833 | 0.279869 | -1.19374 | 0.254305 | 0.509951 | 0.237208 | 0.385565 | 0.045744 | 0.247216 | 0.488994 | -0.03902 | 2.014163 | 0.274293 | -0.02109 | 0.016517 | 0.133768 | 0.027774 | -0.08681 | 0.067382 | 0.431977 | -0.04891 | -0.34487 | 0.041509 | 0.113415 | 0.021603 |
| 34.22222 | 0.293012 | -1.20218 | 0.35135 | 0.499012 | 0.776051 | 0.465452 | -0.14588 | 0.285043 | 0.512071 | -0.01138 | 2.017458 | 0.304651 | -0.04858 | 0.089928 | 0.15124 | 0.043395 | -0.14189 | 0.040352 | 0.28562 | -0.05222 | -0.3568 | -0.00365 | 0.028538 | 0.005826 |
| 34.23611 | 0.376671 | -1.06966 | 0.338546 | 0.491225 | 1.09323 | 0.523096 | -0.20208 | 0.383407 | 0.308503 | 0.057116 | 2.000583 | 0.334842 | -0.02251 | 0.1182 | 0.114149 | 0.026175 | -0.20961 | 0.004122 | 0.220275 | -0.08509 | -0.33276 | -0.0152 | -0.01289 | -0.03149 |
| 34.25 | 0.375536 | -0.88686 | 0.207396 | 0.528951 | 1.188733 | 0.504828 | -0.11388 | 0.406392 | -0.01625 | 0.070725 | 1.914228 | 0.289997 | 0.069474 | 0.145057 | 0.055126 | -0.05678 | -0.23776 | -0.01158 | 0.243528 | -0.09062 | -0.4202 | -0.01491 | -0.03352 | -0.07937 |

| | | | | | | | | | | | | | | | | | | | | | | | | |
|---|---|---|---|---|---|---|---|---|---|---|---|---|---|---|---|---|---|---|---|---|---|---|---|---|
| 34.26389 | 0.394615 | -0.6949 | 0.065564 | 0.453934 | 1.066823 | 0.433929 | -0.12004 | 0.317426 | -0.20405 | 0.065929 | 1.885432 | 0.237538 | 0.167458 | 0.166991 | 0.02261 | -0.18062 | -0.26748 | -0.01458 | 0.243497 | -0.064 | -0.62593 | -0.02191 | -0.04354 | -0.13567 |
| 34.27778 | 0.381056 | -0.56394 | 0.0414 | 0.282605 | 0.363455 | 0.385523 | -0.13623 | 0.219368 | -0.07909 | 0.080089 | 1.842569 | 0.207913 | 0.213442 | 0.189996 | 0.036352 | -0.22899 | -0.26539 | 0.001836 | 0.193971 | -0.02578 | -0.81177 | 0.0172 | 0.052524 | -0.13674 |
| 34.29167 | 0.505724 | -0.58968 | 0.123939 | 0.114435 | -0.62868 | 0.361359 | -0.09554 | 0.214478 | 0.04998 | 0.058362 | 1.754857 | 0.162296 | 0.233242 | 0.220039 | 0.090123 | -0.16941 | -0.19689 | 0.039824 | 0.108534 | 0.017891 | -0.80734 | 0.080752 | 0.186508 | -0.05552 |
| 34.30556 | 0.727606 | -0.75731 | 0.15129 | -0.02672 | -1.43357 | 0.348783 | -0.06892 | 0.289013 | 0.170769 | 0.007315 | 1.599225 | 0.0334 | 0.220056 | 0.221502 | 0.136178 | -0.06669 | -0.13985 | 0.054529 | -0.021 | 0.046435 | -0.55582 | 0.079054 | 0.211281 | 0.030105 |
| 34.31944 | 0.811056 | -0.99525 | 0.036318 | -0.1304 | -1.76214 | 0.36815 | -0.03077 | 0.39467 | 0.266092 | -0.06515 | 1.367032 | -0.10312 | 0.190966 | 0.157421 | 0.133514 | -0.01239 | -0.12584 | 0.040218 | -0.18548 | 0.045732 | -0.33649 | 0.03014 | 0.143339 | 0.063714 |
| 34.33333 | 0.803893 | -1.22873 | -0.04543 | -0.18828 | -1.65351 | 0.366236 | 0.005316 | 0.448079 | 0.225661 | -0.13103 | 1.110341 | -0.21133 | 0.145355 | 0.119366 | 0.062772 | -0.00297 | -0.13181 | 0.027212 | -0.30211 | 0.01936 | -0.26199 | -0.04109 | 0.025915 | 0.062979 |
| 34.34722 | 0.76736 | -1.40796 | -0.03274 | -0.20946 | -1.18774 | 0.415096 | -0.01677 | 0.464594 | 0.078498 | -0.16697 | 0.925491 | -0.30797 | 0.090896 | 0.107545 | -0.06788 | -0.02535 | -0.15963 | -0.00227 | -0.27704 | -0.01967 | -0.29607 | -0.13625 | -0.14655 | 0.023942 |
| 34.36111 | 0.519119 | -1.45625 | 0.041411 | -0.09423 | -0.4755 | 0.437967 | -0.0292 | 0.513127 | -0.06084 | -0.19108 | 0.804097 | -0.37915 | 0.068719 | 0.091239 | -0.17038 | -0.01445 | -0.18491 | -0.03683 | -0.1498 | -0.06564 | -0.30979 | -0.20662 | -0.23715 | -0.0263 |
| 34.375 | 0.302801 | -1.43864 | 0.163993 | 0.009182 | 0.030599 | 0.43693 | -0.03691 | 0.624708 | -0.21641 | -0.15587 | 0.87371 | -0.44307 | 0.104788 | 0.101865 | -0.17454 | 0.041622 | -0.12289 | -0.02991 | 0.073518 | -0.08857 | -0.26302 | -0.20925 | -0.17435 | -0.01238 |
| 34.38889 | 0.157489 | -1.4766 | 0.272107 | 0.003042 | -0.00964 | 0.428884 | -0.06639 | 0.743599 | -0.37081 | -0.15215 | 0.526742 | -0.50293 | 0.145017 | 0.112191 | -0.07787 | 0.132439 | -0.00956 | -0.00206 | 0.237154 | -0.095 | -0.18428 | -0.10945 | -0.07078 | 0.041396 |
| 34.40278 | -0.10034 | -1.46186 | 0.304349 | -0.08373 | -0.49848 | 0.42059 | -0.13852 | 0.815407 | -0.41134 | -0.14472 | 0.585317 | -0.46062 | 0.098618 | 0.089502 | 0.034883 | 0.181546 | 0.06626 | -0.00177 | 0.287023 | -0.11186 | -0.14221 | 0.026844 | -0.01936 | 0.049809 |
| 34.41667 | -0.14466 | -1.4545 | 0.269256 | -0.20953 | -0.80828 | 0.482888 | -0.2828 | 0.746096 | -0.27421 | -0.11788 | 0.434173 | -0.39626 | 0.038837 | 0.06533 | 0.143668 | 0.217304 | 0.144268 | -0.00237 | 0.223035 | -0.11226 | -0.11133 | 0.166708 | 0.012572 | 0.056724 |
| 34.43056 | -0.09197 | -1.39437 | 0.246139 | -0.15305 | -0.39849 | 0.557926 | -0.43465 | 0.717569 | -0.29496 | -0.05714 | 0.645022 | -0.38057 | 0.063735 | 0.08509 | 0.234865 | 0.265309 | 0.255672 | 0.042584 | 0.187213 | -0.07189 | -0.03205 | 0.259138 | 0.028679 | 0.111739 |
| 34.44444 | -0.13814 | -1.34494 | 0.238354 | -0.05 | 0.282195 | 0.501442 | -0.56899 | 0.710309 | -0.5403 | -0.04222 | 0.659789 | -0.47219 | 0.119352 | 0.121546 | 0.242568 | 0.293685 | 0.274683 | 0.079379 | 0.093118 | -0.04452 | 0.027559 | 0.24875 | 0.002492 | 0.132319 |
| 34.45833 | -0.19489 | -1.34106 | 0.349702 | -0.02388 | 0.758144 | 0.410923 | -0.69605 | 0.72022 | -0.78593 | -0.08482 | 0.900476 | -0.59025 | 0.141492 | 0.114154 | 0.155745 | 0.230246 | 0.210469 | 0.083726 | -0.00872 | -0.05704 | -0.00123 | 0.152989 | -0.03878 | 0.06792 |
| 34.47222 | -0.02332 | -1.3151 | 0.494692 | 0.021245 | 0.974093 | 0.356766 | -0.77525 | 0.673988 | -0.94009 | -0.04215 | 1.392425 | -0.58099 | 0.156123 | 0.100133 | 0.058239 | 0.156848 | 0.174678 | 0.0864 | -0.0398 | -0.07168 | -0.04545 | 0.055537 | 0.00323 | 0.031213 |
| 34.48611 | 0.205395 | -1.09954 | 0.348697 | 0.032119 | 0.998327 | 0.318474 | -0.80706 | 0.657783 | -0.63808 | 0.194298 | 1.956162 | -0.50214 | 0.197909 | 0.127471 | 0.043608 | 0.157686 | 0.254521 | 0.110558 | 0.016805 | -0.02431 | 0.007737 | 0.071668 | 0.09871 | 0.078582 |
| 34.5 | 0.824437 | -0.57238 | 0.122452 | 0.105108 | 1.217348 | 0.403776 | -0.66111 | 0.737441 | 0.079633 | 0.651064 | 2.773957 | -0.21315 | 0.185609 | 0.100343 | 0.111881 | 0.263232 | 0.374215 | 0.167676 | 0.129533 | 0.031961 | 0.190366 | 0.187653 | 0.125826 | 0.173704 |
| 34.51389 | 1.324324 | 0.036943 | 0.075131 | 0.375658 | 1.979602 | 0.368666 | -0.27844 | 0.66942 | 1.525257 | 1.070821 | 2.812068 | 0.279941 | 0.113182 | 0.012734 | 0.14044 | 0.421284 | 0.468606 | 0.270382 | 0.285108 | 0.05836 | 0.430881 | 0.327653 | 0.114858 | 0.2429 |
| 34.52778 | 1.21138 | 0.494467 | 0.115064 | 0.670974 | 3.249323 | 0.394081 | 0.359845 | 0.693749 | 2.983718 | 1.080157 | 2.950896 | 0.776436 | 0.061624 | -0.03281 | 0.080693 | 0.46396 | 0.567592 | 0.301502 | 0.496137 | 0.024411 | 0.428643 | 0.409167 | 0.344828 | 0.302321 |
| 34.54167 | 1.015417 | 0.916917 | 0.528494 | 0.976624 | 3.782494 | 0.499202 | 0.959096 | 1.277139 | 2.394653 | 0.79104 | 3.584525 | 1.227971 | 0.047005 | 0.057243 | 0.033686 | 0.335461 | 0.672618 | 0.160041 | 0.69907 | 0.0157 | 0.297571 | 0.421636 | 0.769139 | 0.396204 |
| 34.55556 | 0.97267 | 1.451359 | 1.17972 | 0.95701 | 2.031739 | 0.750281 | 1.058923 | 1.85535 | 1.805416 | 0.504679 | 3.293586 | 1.351213 | 0.02825 | 0.324137 | 0.214077 | 0.399845 | 0.796022 | 0.067263 | 1.092203 | 0.075169 | 0.451275 | 0.345489 | 1.035249 | 0.506092 |
| 34.56944 | 1.046102 | 1.700906 | 1.597098 | 0.638954 | -0.02632 | 0.746566 | 0.666028 | 1.964869 | 0.288362 | 0.366642 | 2.733958 | 1.485718 | 0.043227 | 0.548187 | 0.369856 | 0.605896 | 1.089149 | 0.121448 | 1.065556 | 0.236641 | 0.765275 | 0.312535 | 0.803549 | 0.447663 |
| 34.58333 | 1.189387 | 1.703049 | 1.657161 | 0.304753 | -1.66682 | 0.692867 | 0.141257 | 2.00707 | -1.11306 | 0.413591 | 2.158373 | 1.373505 | 0.026415 | 0.584603 | 0.389958 | 0.653576 | 1.159261 | 0.118187 | 0.963797 | 0.328841 | 0.886108 | 0.272952 | 0.472648 | 0.262896 |
| 34.59722 | 1.273769 | 1.654445 | 1.677043 | 0.156098 | -2.45082 | 0.645227 | -0.12547 | 1.921028 | -1.43436 | 0.719045 | 1.440319 | 1.390443 | -0.02539 | 0.399672 | 0.2795 | 0.533017 | 1.055931 | 0.038123 | 0.794715 | 0.378535 | 0.80212 | 0.292001 | -0.0196 | 0.05919 |
| 34.61111 | 1.226501 | 1.750296 | 1.875932 | 0.199681 | -1.91369 | 0.568683 | -0.10219 | 1.968462 | -0.89565 | 0.827235 | 0.82491 | 1.308992 | -0.06205 | 0.217878 | 0.182679 | 0.440805 | 0.936698 | -0.03137 | 0.602871 | 0.371241 | 0.714803 | 0.194351 | -0.43889 | -0.03138 |
| 34.625 | 1.251004 | 1.850882 | 2.260019 | 0.4103 | -0.68323 | 0.534849 | 0.126041 | 2.052037 | -0.27422 | 0.732935 | 0.415266 | 1.249744 | -0.0689 | 0.098403 | 0.061488 | 0.429759 | 0.811776 | -0.09046 | 0.325295 | 0.384319 | 0.554203 | 0.112 | -0.60426 | -0.00249 |
| 34.63889 | 1.073843 | 1.807847 | 2.276486 | 0.55925 | 0.234357 | 0.506671 | 0.359686 | 1.892639 | 0.016935 | 0.488219 | 0.103194 | 0.94333 | -0.05536 | 0.053542 | 0.00358 | 0.493193 | 0.609666 | -0.12588 | -0.02542 | 0.372942 | 0.367233 | 0.032872 | -0.46494 | 0.122558 |
| 34.65278 | 1.063555 | 1.585814 | 1.811726 | 0.432913 | 0.471522 | 0.539908 | 0.481802 | 1.442488 | -0.28492 | 0.342916 | -0.08791 | 0.634735 | -0.03445 | 0.043471 | 0.025537 | 0.469612 | 0.226745 | -0.10428 | -0.34323 | 0.397929 | 0.185926 | 0.024365 | -0.25852 | 0.225665 |
| 34.66667 | 1.051266 | 1.667951 | 1.455455 | 0.274247 | 0.253648 | 0.496404 | 0.464419 | 0.97958 | -1.01574 | 0.272123 | -0.23635 | 0.255782 | -0.03452 | -0.00376 | 0.067859 | 0.403925 | -0.25507 | -0.04386 | -0.70679 | 0.453668 | -0.00143 | -0.00533 | -0.04771 | 0.230018 |
| 34.68056 | 1.154544 | 1.807738 | 1.212126 | 0.146058 | -0.30118 | 0.481616 | 0.407066 | 0.479041 | -1.61474 | 0.174017 | -0.36769 | -0.07783 | -0.06807 | -0.1143 | 0.082563 | 0.310485 | -0.59512 | 0.02374 | -0.7683 | 0.489172 | -0.12597 | -0.0277 | -0.01128 | 0.149078 |
| 34.69444 | 1.180674 | 1.721794 | 1.068152 | -0.02958 | -1.46092 | 0.527732 | 0.447796 | 0.009097 | -2.16034 | -0.10017 | -0.36505 | -0.35688 | -0.07572 | -0.24603 | 0.058893 | 0.158863 | -0.73029 | 0.06358 | -0.61198 | 0.488644 | -0.10948 | -0.04179 | -0.09255 | 0.040673 |
| 34.70833 | 1.183567 | 1.37267 | 0.961969 | -0.35617 | -3.01314 | 0.510394 | 0.491347 | -0.19071 | -2.04751 | -0.25267 | -0.31278 | -0.54806 | -0.06833 | -0.35904 | 0.003326 | 0.012774 | -0.67675 | 0.060256 | -0.22253 | 0.394559 | -0.01995 | -0.06449 | -0.15584 | -0.03438 |
| 34.72222 | 1.231034 | 0.958526 | 0.834494 | -0.65268 | -3.62482 | 0.5107 | 0.386834 | -0.1724 | -1.66339 | -0.4482 | -0.08414 | -0.6938 | -0.07913 | -0.40992 | -0.07419 | -0.13357 | -0.53856 | -0.00321 | 0.213027 | 0.299986 | 0.102766 | -0.09713 | -0.12462 | -0.12596 |
| 34.73611 | 1.303339 | 0.618358 | 0.723456 | -0.67765 | -3.09069 | 0.462092 | 0.033939 | -0.06001 | -1.30753 | -0.53126 | 0.530011 | -0.95814 | -0.09579 | -0.42231 | -0.15117 | -0.28143 | -0.32433 | -0.12833 | 0.441094 | 0.238911 | 0.190684 | -0.2053 | -0.07495 | -0.27584 |
| 34.75 | 1.488438 | 0.482001 | 0.447107 | -0.60738 | -2.06121 | 0.389909 | -0.4937 | 0.075049 | -1.36333 | -0.55866 | 1.614986 | -1.32969 | -0.07827 | -0.43073 | -0.23593 | -0.46289 | -0.16715 | -0.26593 | 0.296495 | 0.261317 | 0.112613 | -0.28686 | 0.013238 | -0.30228 |
| 34.76389 | 1.652817 | 0.457417 | 0.217327 | -0.53055 | -1.05328 | 0.358839 | -0.85805 | 0.189193 | -1.42154 | -0.57067 | 1.788956 | -1.40237 | -0.05097 | -0.42543 | -0.33 | -0.56818 | -0.17828 | -0.32369 | -0.15836 | 0.346358 | -0.07707 | -0.3616 | 0.127645 | -0.3777 |
| 34.77778 | 1.72437 | 0.308343 | 0.146517 | -0.55548 | -0.7014 | 0.398739 | -0.8999 | 0.225666 | -1.22955 | -0.77725 | 1.223945 | -1.51314 | -0.05676 | -0.53385 | -0.35909 | -0.61545 | -0.37631 | -0.29323 | -0.64743 | 0.403321 | -0.22002 | -0.37492 | 0.228161 | -0.39635 |
| 34.79167 | 1.639688 | -0.02809 | 0.239056 | -0.66737 | -0.96551 | 0.501477 | -0.90949 | 0.14905 | -0.76808 | -0.90866 | 0.895406 | -1.50723 | -0.06656 | -0.58574 | -0.37291 | -0.5841 | -0.70032 | -0.22088 | -1.05775 | 0.452607 | -0.26163 | -0.28711 | 0.31758 | -0.36832 |
| 34.80556 | 1.393124 | -0.52913 | 0.180237 | -0.72812 | -1.69177 | 0.472246 | -1.02798 | 0.048135 | -0.55128 | -0.92497 | 1.31019 | -1.4032 | -0.03698 | -0.52495 | -0.37149 | -0.46253 | -0.96304 | -0.08072 | -1.15005 | 0.45488 | -0.15031 | -0.20921 | 0.243275 | -0.32501 |
| 34.81944 | 0.98382 | -1.00035 | 0.173959 | -0.7967 | -2.09452 | 0.496059 | -1.10104 | -0.13645 | -0.54995 | -0.81747 | 1.406693 | -1.34105 | 0.030311 | -0.46885 | -0.3586 | -0.21064 | -1.04428 | 0.043973 | -1.03285 | 0.349445 | -0.00159 | -0.11259 | 0.157929 | -0.33963 |
| 34.83333 | 0.705945 | -1.35364 | 0.218625 | -0.8224 | -1.61991 | 0.373738 | -0.99831 | -0.36715 | -0.21899 | -0.76299 | 1.404058 | -1.27295 | 0.069305 | -0.30733 | -0.34316 | 0.083973 | -0.89902 | 0.131478 | -0.73801 | 0.248132 | -0.00805 | -0.06381 | 0.032748 | -0.36176 |
| 34.84722 | 0.427569 | -1.5699 | 0.290813 | -0.84232 | -1.68432 | 0.178884 | -0.86849 | -0.7546 | 0.143625 | -0.79324 | 1.195195 | -1.16159 | 0.102883 | -0.06428 | -0.28353 | 0.251207 | -0.70175 | 0.172917 | -0.48798 | 0.155165 | -0.09564 | 0.008128 | -0.05862 | -0.37566 |
| 34.86111 | 0.097376 | -1.5525 | 0.232057 | -0.89059 | -2.20686 | 0.142075 | -0.7759 | -1.18141 | 0.344727 | -0.80725 | 0.749858 | -0.85666 | 0.078034 | 0.186617 | -0.20947 | 0.360708 | -0.60821 | 0.153279 | -0.33029 | 0.043383 | -0.23002 | 0.065263 | -0.10782 | -0.30792 |
| 34.875 | -0.3071 | -1.59853 | 0.184781 | -0.88618 | -2.93188 | 0.112645 | -0.80567 | -0.89183 | 0.617646 | -0.75416 | 0.364561 | -0.50174 | -0.00083 | 0.356562 | -0.13821 | 0.301075 | -0.50293 | 0.057193 | -0.31559 | -0.0549 | -0.32735 | 0.060486 | -0.18081 | -0.23191 |
| 34.88889 | -0.69611 | -1.78134 | 0.200025 | -0.80813 | -3.81808 | 0.114178 | -0.74333 | -0.33699 | 0.883557 | -0.6634 | 0.042648 | -0.06271 | -0.10319 | 0.309983 | -0.12099 | 0.234079 | -0.34408 | -0.05968 | -0.31068 | -0.16396 | -0.37489 | 0.017666 | -0.1562 | -0.14988 |
| 34.90278 | -0.91159 | -1.6319 | 0.34116 | -0.60711 | -3.83092 | 0.126356 | -0.57079 | -0.04804 | 1.22077 | -0.67549 | -0.44574 | 0.274095 | -0.15549 | 0.238327 | -0.112 | 0.137771 | -0.15835 | -0.12506 | -0.25692 | -0.21083 | -0.34701 | -0.04621 | -0.05686 | -0.09902 |
| 34.91667 | -0.92413 | -1.28022 | 0.439577 | -0.3366 | -3.41813 | 0.200052 | -0.25364 | 0.062037 | 1.382128 | -0.76304 | -0.77812 | 0.464669 | -0.1743 | 0.125731 | -0.10844 | 0.024265 | 0.000747 | -0.11792 | -0.1387 | -0.24547 | -0.23365 | -0.04566 | -0.01254 | -0.07635 |
| 34.93056 | -0.79611 | -1.05521 | 0.514551 | -0.08046 | -2.64787 | 0.348868 | 0.03843 | 0.102398 | 1.508302 | -0.56685 | -0.49221 | 0.527773 | -0.16641 | 0.097305 | -0.12294 | -0.07749 | 0.080304 | -0.09483 | 0.024869 | -0.28752 | -0.11882 | -0.01591 | 0.022735 | -0.09278 |
| 34.94444 | -0.54544 | -0.81528 | 0.33757 | 0.038183 | -2.16563 | 0.529001 | 0.248759 | 0.244411 | 1.671399 | -0.12195 | -0.41025 | 0.572641 | -0.15766 | 0.064224 | -0.126 | -0.09236 | 0.149229 | -0.05244 | 0.199419 | -0.29175 | -0.04392 | -0.02414 | 0.039402 | -0.10467 |
| 34.95833 | -0.44318 | -0.50575 | 0.220012 | 0.165838 | -1.50701 | 0.448819 | 0.401788 | 0.415817 | 1.620958 | 0.25426 | -0.52026 | 0.504656 | -0.1305 | 0.053323 | -0.10189 | -0.04755 | 0.143314 | -0.00453 | 0.349523 | -0.30447 | -0.15824 | -0.04311 | -0.00726 | -0.13442 |
| 34.97222 | -0.35921 | 0.279046 | 0.233626 | 0.339439 | -1.03587 | 0.207054 | 0.452511 | 0.407214 | 1.64724 | 0.241193 | -1.17205 | 0.279685 | -0.08428 | 0.012009 | -0.05714 | 0.061499 | 0.163122 | 0.037956 | 0.43779 | -0.23677 | -0.28787 | -0.03183 | -0.03129 | -0.09867 |
| 34.98611 | -0.73594 | 1.305088 | 0.338286 | 0.341969 | -0.94101 | -0.1311 | 0.471966 | -0.16075 | 1.390783 | -0.4338 | -2.50147 | -0.32608 | -0.02986 | 0.09773 | 0.083293 | 0.253998 | 0.166646 | 0.059026 | 0.471873 | -0.11117 | -0.36536 | 0.059972 | 0.063053 | -0.07679 |
| 35 | -0.91133 | 1.405917 | 0.177742 | 0.035573 | -1.62969 | -0.38538 | 0.134103 | -0.7926 | 0.476885 | -1.56144 | -3.44494 | -0.82552 | 0.072838 | 0.173798 | 0.329631 | 0.401253 | 0.196903 | 0.076974 | 0.378734 | -0.02315 | -0.54695 | 0.159168 | 0.251085 | -0.1064 |
| 35.01389 | -0.78334 | 0.386968 | 0.242234 | -0.15119 | -2.65505 | -0.36922 | -0.19294 | -0.89207 | -0.41975 | -2.90109 | -2.98277 | -0.81658 | 0.181676 | 0.18235 | 0.442503 | 0.261606 | 0.035599 | -0.00625 | 0.126671 | -0.19759 | -1.00256 | 0.064094 | 0.106266 | -0.29466 |
| 35.02778 | -0.34795 | -0.5213 | 0.344449 | 0.115365 | -2.76366 | 0.014304 | -0.45462 | -0.57175 | -0.56918 | -3.26065 | -1.91415 | -0.4744 | 0.186499 | 0.056622 | 0.343756 | -0.11395 | -0.34169 | -0.24139 | -0.29564 | -0.555 | -1.78379 | -0.3073 | -0.25302 | -0.43794 |
| 35.04167 | -0.03888 | -0.95856 | 0.250143 | 0.45567 | -2.12479 | 0.356491 | -0.62923 | -0.25057 | -0.49283 | -3.39047 | -0.82048 | -0.11744 | 0.087096 | -0.12929 | 0.245441 | -0.59762 | -0.8503 | -0.2655 | -0.83461 | -0.60893 | -1.89505 | -0.55206 | -0.47935 | -0.43641 |
| 35.05556 | 0.011494 | -1.20204 | 0.041824 | 0.52933 | -1.3093 | 0.459971 | -0.79618 | -0.10376 | -0.90838 | -3.54053 | 0.150428 | 0.046765 | -0.09726 | -0.24172 | 0.145497 | -1.12101 | -0.94467 | 0.034643 | -0.94035 | -0.55414 | -1.1997 | -0.39875 | -0.22028 | -0.34042 |
| 35.06944 | -0.04627 | -1.05663 | 0.011088 | 0.394192 | -0.38497 | 0.488646 | -0.72397 | -0.08907 | -0.92177 | -2.92234 | 0.764099 | -0.02618 | -0.27022 | -0.2478 | 0.008445 | -1.28474 | -0.86298 | 0.238787 | -0.94394 | -0.60575 | -0.66573 | -0.33206 | -0.24085 | -0.23289 |
| 35.08333 | 0.060778 | -0.95136 | 0.027415 | 0.334275 | 0.068557 | 0.461379 | -0.58338 | -0.11694 | -0.69019 | -1.90756 | 0.868424 | -0.18107 | -0.31749 | -0.13525 | -0.04261 | -0.94645 | -0.62213 | 0.23446 | -0.59854 | -0.53168 | -0.23981 | -0.32351 | -0.27097 | -0.08944 |
| 35.09722 | 0.31632 | -0.94587 | 0.077697 | 0.39571 | 0.029057 | 0.465926 | -0.49488 | -0.08358 | -0.4481 | -0.92734 | 0.808744 | -0.18995 | -0.23505 | 0.00588 | -0.00245 | -0.43483 | -0.26081 | 0.2046 | -0.10437 | -0.3314 | 0.278769 | -0.15991 | 0.143463 | 0.162534 |
| 35.11111 | 0.502853 | -1.15227 | 0.23466 | 0.344947 | 0.033552 | 0.495946 | -0.3917 | 0.065626 | -0.32597 | -0.26248 | 0.652526 | -0.07379 | -0.13356 | 0.129939 | 0.038118 | -0.02254 | 0.15762 | 0.205442 | 0.259105 | -0.13583 | 0.717716 | 0.017174 | 0.34125 | 0.374123 |
| 35.125 | 0.580055 | -1.29092 | 0.322851 | 0.314379 | 0.315665 | 0.470468 | -0.20863 | 0.252666 | -0.29797 | -0.05984 | 0.517303 | 0.088704 | -0.08098 | 0.179098 | 0.058827 | 0.18514 | 0.429126 | 0.263323 | 0.396726 | -0.00391 | 0.743372 | 0.112243 | 0.322699 | 0.480752 |
| 35.13889 | 0.565168 | -1.17784 | 0.314046 | 0.237705 | 0.734324 | 0.401199 | -0.00071 | 0.428424 | -0.29164 | -0.32461 | 0.422984 | 0.154419 | -0.06188 | 0.12677 | 0.034632 | 0.232624 | 0.458759 | 0.307044 | 0.413446 | 0.057449 | 0.559138 | 0.093454 | 0.324764 | 0.526867 |
| 35.15278 | 0.478679 | -0.93558 | 0.25417 | 0.226565 | 1.17996 | 0.330345 | 0.118758 | 0.550277 | -0.22963 | -0.83519 | 0.506162 | 0.215769 | -0.05668 | 0.045137 | -0.01334 | 0.220191 | 0.332168 | 0.314849 | 0.40838 | 0.014326 | 0.412966 | 0.024481 | 0.298717 | 0.442707 |
| 35.16667 | 0.451104 | -0.8842 | 0.179468 | 0.311145 | 1.520734 | 0.348331 | 0.158776 | 0.568823 | -0.15139 | -1.41515 | 0.689881 | 0.247877 | -0.03191 | -0.00682 | -0.03823 | 0.22877 | 0.163296 | 0.240827 | 0.451145 | -0.03734 | 0.366391 | 0.015108 | 0.312648 | 0.352674 |
| 35.18056 | 0.431033 | -0.93665 | 0.174979 | 0.351574 | 1.992329 | 0.343813 | 0.107975 | 0.53892 | 0.16111 | -1.66076 | 0.887326 | 0.236602 | 0.010599 | 0.00583 | -0.00326 | 0.182845 | 0.024843 | 0.182778 | 0.493646 | -0.05134 | 0.285731 | 0.054766 | 0.395681 | 0.235878 |
| 35.19444 | 0.495401 | -0.98253 | 0.255928 | 0.483909 | 2.227696 | 0.318899 | 0.015867 | 0.474382 | 0.262592 | -1.57745 | 1.025833 | 0.214135 | 0.032081 | 0.009884 | 0.05914 | 0.036465 | -0.06675 | 0.151405 | 0.427964 | -0.04819 | 0.174828 | 0.082456 | 0.458778 | 0.184552 |
| 35.20833 | 0.539955 | -0.95351 | 0.38647 | 0.608294 | 2.498339 | 0.351957 | -0.10295 | 0.325731 | 0.437487 | -1.32157 | 0.991384 | 0.197894 | 0.039047 | -0.0023 | 0.105996 | -0.05388 | -0.12376 | 0.16544 | 0.267943 | -0.06849 | 0.118077 | 0.101248 | 0.493209 | 0.196678 |
| 35.22222 | 0.531646 | -0.94566 | 0.496569 | 0.623201 | 2.863101 | 0.453378 | -0.13333 | 0.233626 | 0.368337 | -1.05024 | 0.961541 | 0.214162 | 0.057067 | -0.01764 | 0.116405 | -0.07477 | -0.17421 | 0.150642 | 0.148498 | -0.08368 | 0.149501 | 0.108052 | 0.439664 | 0.227317 |
| 35.23611 | 0.417493 | -0.72265 | 0.569697 | 0.748693 | 2.814825 | 0.519775 | -0.07391 | 0.297438 | 0.292481 | -0.84896 | 0.796232 | 0.161999 | 0.090715 | -0.01843 | 0.11816 | -0.11564 | -0.189 | 0.089963 | 0.099798 | -0.09296 | 0.139774 | 0.108886 | 0.26349 | 0.263393 |
| 35.25 | 0.375415 | -0.41516 | 0.607124 | 0.792921 | 2.738465 | 0.550628 | -0.02861 | 0.379555 | 0.308593 | -0.78215 | 0.739803 | 0.148752 | 0.121806 | -0.01676 | 0.107619 | -0.12521 | -0.20996 | 0.005608 | 0.118221 | -0.11461 | 0.047396 | 0.123451 | 0.057218 | 0.275328 |
| 35.26389 | 0.356458 | -0.14375 | 0.563879 | 0.777803 | 1.912547 | 0.498361 | -0.04955 | 0.354759 | 0.382214 | -0.8352 | 0.670338 | 0.087918 | 0.128048 | -0.03476 | 0.115204 | -0.10395 | -0.25039 | -0.05544 | 0.181114 | -0.15517 | -0.05374 | 0.121299 | -0.12663 | 0.3275 |
| 35.27778 | 0.292936 | -0.07617 | 0.387726 | 0.756549 | 1.096376 | 0.445167 | -0.12498 | 0.228466 | 0.395651 | -1.00779 | 0.632738 | -0.02621 | 0.115935 | -0.03957 | 0.142981 | -0.12122 | -0.30495 | -0.05289 | 0.281099 | -0.16454 | -0.18264 | 0.105692 | -0.18353 | 0.350813 |
| 35.29167 | 0.221548 | -0.02958 | 0.202919 | 0.759897 | 0.796699 | 0.358646 | -0.2225 | 0.118539 | 0.313509 | -1.38462 | 0.753129 | -0.07533 | 0.135858 | 0.003925 | 0.154169 | -0.15403 | -0.34032 | -0.01248 | 0.302367 | -0.10794 | -0.33319 | 0.119032 | -0.19816 | 0.299933 |
| 35.30556 | 0.285138 | -0.04159 | 0.11184 | 0.735927 | 0.649626 | 0.317225 | -0.27134 | 0.098129 | 0.364067 | -1.72996 | 0.981517 | -0.03845 | 0.162515 | 0.071108 | 0.117728 | -0.17157 | -0.38995 | 0.036126 | 0.148727 | -0.0519 | -0.43341 | 0.181073 | -0.20441 | 0.246729 |
| 35.31944 | 0.372378 | -0.07488 | 0.052473 | 0.421261 | 0.01902 | 0.25697 | -0.24748 | 0.105578 | 0.44266 | -1.82535 | 1.14568 | 0.004191 | 0.180194 | 0.101629 | 0.07983 | -0.15731 | -0.39969 | 0.088823 | -0.03156 | -0.00498 | -0.3723 | 0.250296 | -0.13214 | 0.22956 |
| 35.33333 | 0.413898 | -0.18374 | -0.04698 | 0.196262 | -0.5143 | 0.275768 | -0.21453 | 0.132566 | 0.508275 | -1.57928 | 1.251629 | -0.03567 | 0.184736 | 0.102631 | 0.047133 | -0.05758 | -0.36171 | 0.137174 | -0.12427 | 0.038993 | -0.19568 | 0.249488 | -0.00746 | 0.193963 |
| 35.34722 | 0.449183 | -0.30636 | -0.1298 | 0.195143 | -0.36139 | 0.28526 | -0.17214 | 0.178208 | 0.54105 | -1.25567 | 1.225756 | -0.09321 | 0.181855 | 0.049457 | -0.00095 | 0.069844 | -0.38446 | 0.16825 | -0.11349 | 0.074232 | -0.04035 | 0.179288 | 0.094615 | 0.098101 |
| 35.36111 | 0.467124 | -0.37794 | -0.21154 | 0.135757 | -0.24036 | 0.289421 | -0.17326 | 0.163849 | 0.578773 | -1.01518 | 1.094636 | -0.12342 | 0.163744 | -0.04226 | -0.00626 | 0.143456 | -0.36648 | 0.183758 | -0.0463 | 0.099234 | 0.110581 | 0.105935 | 0.156285 | -0.00826 |
| 35.375 | 0.459465 | -0.46567 | -0.2501 | -0.00465 | -0.56526 | 0.294225 | -0.17068 | 0.133969 | 0.74911 | -0.85204 | 1.052954 | -0.13224 | 0.159303 | -0.07775 | 0.002301 | 0.138819 | -0.30928 | 0.180919 | 0.049436 | 0.099157 | 0.240416 | 0.066623 | 0.174677 | -0.07802 |
| 35.38889 | 0.483035 | -0.5418 | -0.19487 | -0.0537 | -0.62893 | 0.383982 | -0.09841 | 0.212904 | 0.914097 | -0.87259 | 1.046366 | -0.13733 | 0.14946 | -0.04469 | 0.036369 | 0.144754 | -0.20627 | 0.180241 | 0.12571 | 0.081856 | 0.307446 | 0.084781 | 0.204989 | -0.07022 |
| 35.40278 | 0.534965 | -0.64052 | -0.14532 | 0.025745 | -0.28728 | 0.431338 | -0.02137 | 0.429914 | 0.843043 | -0.90775 | 1.156251 | -0.1501 | 0.116931 | 0.027097 | 0.074582 | 0.192929 | -0.08775 | 0.146617 | 0.161348 | 0.051114 | 0.311989 | 0.121957 | 0.226937 | 0.006853 |
| 35.41667 | 0.660922 | -0.74284 | -0.1603 | 0.039758 | 0.039322 | 0.437393 | 0.019261 | 0.637513 | 0.852152 | -0.90475 | 1.285766 | -0.14216 | 0.09827 | 0.068596 | 0.087508 | 0.235721 | 0.035166 | 0.092607 | 0.155518 | 0.035832 | 0.283153 | 0.148321 | 0.218623 | 0.089333 |
| 35.43056 | 0.751514 | -0.8408 | -0.14271 | 0.021609 | 0.214108 | 0.391404 | 0.070467 | 0.664121 | 0.556775 | -0.79089 | 1.510095 | -0.07216 | 0.134027 | 0.094073 | 0.08981 | 0.285062 | 0.136685 | 0.093955 | 0.196359 | 0.064275 | 0.281775 | 0.15794 | 0.190146 | 0.175578 |
| 35.44444 | 0.803278 | -0.93414 | -0.04012 | -0.04873 | 0.677846 | 0.238946 | 0.066872 | 0.681818 | 0.088417 | -0.69302 | 1.918177 | 0.258924 | 0.170637 | 0.129497 | 0.072419 | 0.337954 | 0.176469 | 0.12179 | 0.241093 | 0.067715 | 0.280466 | 0.159163 | 0.174135 | 0.238156 |
| 35.45833 | 0.896773 | -0.91746 | 0.09384 | 0.091106 | 1.490777 | 0.189973 | -0.02209 | 0.663242 | -0.16609 | -0.69721 | 2.112981 | 0.241497 | 0.135832 | 0.118354 | -0.00773 | 0.335248 | 0.114316 | 0.075918 | 0.206699 | 0.024284 | 0.255635 | 0.102811 | 0.139457 | 0.222973 |
| 35.47222 | 1.070725 | -0.37689 | 0.173465 | 0.543839 | 2.035504 | 0.414587 | -0.1019 | 0.379851 | 0.877057 | -0.79688 | 1.261961 | -0.74466 | 0.099456 | 0.11224 | -0.08211 | 0.315934 | 0.037467 | 0.035531 | 0.216343 | -0.0059 | 0.248854 | 0.047793 | 0.113706 | 0.182819 |
| 35.48611 | 1.488426 | -0.02514 | 0.525713 | 0.905818 | 2.709357 | 0.456742 | -0.12464 | 0.567486 | 1.035673 | -0.54864 | 1.326901 | -0.98137 | 0.120339 | 0.145383 | -0.0384 | 0.330781 | 0.038281 | 0.048436 | 0.313328 | 0.033003 | 0.2994 | 0.084696 | 0.10844 | 0.196076 |
| 35.5 | 1.732163 | -0.6976 | 1.356705 | 0.488669 | 4.099556 | -0.07939 | 0.014389 | 1.745144 | -1.4635 | 0.50785 | 3.357545 | 0.587265 | 0.103902 | 0.112034 | 0.087535 | 0.353683 | 0.134444 | 0.08481 | 0.47562 | 0.097226 | 0.474587 | 0.166007 | 0.05593 | 0.2106 |
| 35.51389 | 1.437403 | -3.68711 | 2.645642 | -1.58089 | 6.399976 | -2.73188 | -0.03841 | 3.260598 | -7.16245 | 0.448233 | 8.192095 | 4.060876 | 0.072244 | 0.029412 | 0.179108 | 0.441971 | 0.291836 | 0.190631 | 0.670543 | 0.204433 | 0.773134 | 0.248339 | 0.149128 | 0.223173 |
| 35.52778 | 1.045684 | -13.6441 | 4.926499 | -4.59951 | 7.701188 | -2.83442 | 1.176752 | 1.061395 | -6.39806 | 0.967454 | 11.39029 | 5.169963 | 0.206579 | 0.164728 | 0.02148 | 0.763848 | 0.37861 | 0.36444 | 0.832802 | 0.344507 | 0.945705 | 0.170301 | 0.67142 | 0.404622 |
| 35.54167 | 1.630467 | 0 | 0.837862 | 0.230724 | 1.841161 | 0.041726 | 2.338915 | 3.368873 | 0.997261 | 1.154025 | 3.943658 | -1.67412 | 0.380794 | 0.268977 | -1.30908 | 1.617909 | 0.007244 | 0.318119 | 0.857711 | 0.09882 | 0.656681 | -0.92548 | 0.906642 | 1.245883 |
| 35.55556 | 0.99057 | 11.47655 | 0.179505 | 2.48764 | -4.46391 | 5.349113 | 1.738646 | -0.23299 | 7.804894 | 5.21948 | -5.7902 | -5.72255 | -0.31391 | -1.07739 | -2.40487 | 1.417779 | -1.59452 | -0.24851 | 0.842352 | -0.75863 | -1.76252 | -2.70843 | 2.146179 | 2.712416 |
| 35.56944 | 0.962738 | 3.431101 | -0.78831 | 4.135095 | -1.16477 | 2.156323 | 1.386404 | -1.35783 | 7.516228 | 2.806571 | -3.01479 | -3.00421 | 0.178334 | 0.146155 | -0.0374 | 0.852869 | 0.833758 | -0.25733 | 0.321859 | -0.13027 | -0.29893 | -0.34243 | 0.937542 | 0.277453 |
| 35.58333 | 1.167242 | 1.354448 | 1.198343 | 2.554108 | 2.369986 | 0.816487 | 1.232464 | 0.697119 | 1.502416 | 1.040994 | 1.803437 | -0.51423 | 0.852493 | 1.423324 | 2.743869 | -0.59812 | 2.57305 | 0.200018 | 0.619206 | 1.584814 | 1.956684 | 2.564846 | -0.15793 | 0.219772 |
| 35.59722 | 1.31354 | 0.802598 | 1.687031 | 1.281019 | 1.938523 | 0.937521 | 0.992012 | 1.487133 | -0.14917 | 0.341395 | 2.861892 | 0.249818 | -0.06643 | 0.053321 | 1.596377 | -0.88368 | 1.081134 | -0.14292 | 0.525582 | 0.808174 | 0.523844 | 1.098296 | -0.35819 | -0.75371 |
| 35.61111 | 1.382085 | 1.070974 | 1.592906 | 0.322624 | -0.55283 | 1.075978 | 0.851046 | 1.530529 | -0.19997 | 0.238777 | 2.377264 | 0.362788 | -0.00358 | 0.172337 | 0.393309 | 0.035433 | 1.249277 | -0.31845 | 0.315531 | 0.537928 | 0.473852 | 0.257393 | -0.40303 | -0.12997 |
| 35.625 | 1.376215 | 1.432321 | 1.125602 | -0.28369 | -2.72378 | 0.931419 | 0.685963 | 1.298687 | 0.20769 | 0.337327 | 1.375909 | 0.074119 | 0.009955 | 0.138081 | 0.132872 | 0.12774 | 1.376732 | -0.29373 | 0.076146 | 0.458014 | 0.500001 | 0.053791 | -0.66354 | -0.06263 |
| 35.63889 | 1.326782 | 1.686527 | 0.786995 | -0.73213 | -3.83058 | 0.674735 | 0.619552 | 1.208255 | 0.653628 | 0.201136 | 0.749969 | -0.18441 | 0.001466 | 0.067771 | 0.099532 | 0.152327 | 1.309545 | -0.2381 | -0.24977 | 0.403188 | 0.382612 | -0.01962 | -0.71457 | 0.020517 |
| 35.65278 | 1.23152 | 1.554323 | 0.67621 | -1.06989 | -4.68969 | 0.663432 | 0.378788 | 0.895583 | 0.436613 | -0.05499 | 0.601225 | -0.15451 | 0.00399 | 0.02371 | 0.103253 | 0.172057 | 1.217039 | -0.24281 | -0.4877 | 0.340586 | 0.187312 | -0.0496 | -0.81602 | 0.115436 |
| 35.66667 | 1.19879 | 1.426388 | 0.606352 | -1.30408 | -3.72295 | 0.758929 | 0.197176 | 1.008558 | 0.31368 | -0.43639 | 0.987466 | 0.110369 | 0.009101 | -0.07909 | 0.1175 | 0.054738 | 0.860363 | -0.27429 | -0.78642 | 0.299743 | -0.09886 | -0.14184 | -0.76393 | 0.136263 |
| 35.68056 | 1.14176 | 2.202012 | 0.496353 | -1.34408 | -3.44329 | 0.901346 | 0.261704 | 0.988547 | 0.690832 | -0.83465 | 1.209659 | -0.10299 | 0.001003 | -0.25034 | 0.119954 | -0.07046 | 0.410784 | -0.23643 | -0.99706 | 0.312707 | -0.25824 | -0.23475 | -0.66531 | 0.117119 |
| 35.69444 | 0.956179 | 3.828664 | 0.375797 | -1.27798 | -2.88621 | 0.893788 | 0.271322 | 0.715476 | 0.487008 | -1.17851 | 1.352824 | -0.30918 | -0.02474 | -0.45256 | 0.105198 | -0.15561 | -0.03132 | -0.13609 | -0.87931 | 0.369058 | -0.30227 | -0.31383 | -0.31721 | 0.122659 |
| 35.70833 | 0.829079 | 4.885248 | 0.357872 | -0.88721 | -1.15982 | 0.924768 | 0.1418 | 0.705989 | -0.32063 | -1.2996 | 1.793362 | -0.09626 | -0.07714 | -0.55704 | 0.04004 | -0.22975 | -0.31245 | -0.0664 | -0.56646 | 0.383762 | -0.20529 | -0.38935 | 0.131411 | 0.123475 |
| 35.72222 | 0.79903 | 4.179708 | 0.342693 | -0.11016 | 0.230074 | 0.922049 | 0.148176 | 0.742966 | -0.45828 | -1.12604 | 1.765806 | -0.19515 | -0.16691 | -0.58789 | -0.05499 | -0.26969 | -0.3503 | -0.03276 | -0.19914 | 0.354261 | -0.07151 | -0.36941 | 0.417052 | 0.071364 |

| | | | | | | | | | | | | | | | | | | | | | | | | |
|---|---|---|---|---|---|---|---|---|---|---|---|---|---|---|---|---|---|---|---|---|---|---|---|---|
| 35.73611 | 1.070432 | 3.524758 | 0.405698 | 0.543723 | 1.125734 | 0.922087 | 0.154714 | 0.767643 | -0.40353 | -0.80665 | 1.606486 | -0.45233 | -0.2319 | -0.60641 | -0.14959 | -0.3046 | -0.23876 | -0.02953 | 0.087681 | 0.33765 | 0.014206 | -0.42581 | 0.47544 | -0.00241 |
| 35.75 | 1.185795 | 2.885529 | 0.497651 | 0.938029 | 1.682038 | 1.035675 | 0.181629 | 0.751275 | -0.17286 | -0.53816 | 1.522201 | -0.73206 | -0.25294 | -0.6928 | -0.22204 | -0.28987 | -0.25275 | -0.05318 | 0.289139 | 0.322845 | 0.088856 | -0.39895 | 0.42922 | -0.07574 |
| 35.76389 | 1.344128 | 2.777301 | 0.484958 | 1.224289 | 1.176344 | 0.889034 | 0.340665 | 0.317029 | 0.367377 | -0.23544 | 0.846601 | -1.1213 | -0.23897 | -0.73205 | -0.2762 | -0.28555 | -0.34219 | -0.10232 | 0.371638 | 0.373231 | 0.094375 | -0.38432 | 0.247518 | -0.11481 |
| 35.77778 | 1.309296 | 2.091105 | 0.472542 | 0.808736 | 0.191694 | 0.906791 | 0.329783 | 0.11673 | 0.484726 | -0.25577 | 0.672685 | -1.00599 | -0.19478 | -0.84753 | -0.30409 | -0.3334 | -0.49794 | -0.17964 | 0.217605 | 0.426269 | -0.04584 | -0.41216 | 0.039183 | -0.11459 |
| 35.79167 | 1.136159 | 1.149703 | 0.524583 | -0.02012 | -0.95413 | 1.085642 | 0.186317 | 0.173939 | 0.355031 | -0.54329 | 0.936798 | -0.76726 | -0.16998 | -0.96818 | -0.30472 | -0.45693 | -0.83376 | -0.2594 | -0.15695 | 0.433875 | -0.28403 | -0.41989 | -0.17205 | -0.17024 |
| 35.80556 | 1.235076 | 0.92844 | 0.515857 | -0.61645 | -2.09842 | 1.089517 | 0.127587 | -0.17203 | 0.519848 | -0.84749 | 0.896984 | -0.70413 | -0.1684 | -0.98678 | -0.27564 | -0.55665 | -1.23042 | -0.24234 | -0.55985 | 0.373149 | -0.49252 | -0.41178 | -0.31331 | -0.2428 |
| 35.81944 | 1.55428 | 1.08988 | 0.488633 | -0.76891 | -2.21699 | 0.652988 | 0.031029 | -0.76726 | 0.589945 | -1.05458 | 0.618297 | -0.59706 | -0.19294 | -0.92366 | -0.25716 | -0.63074 | -1.39407 | -0.15511 | -0.75657 | 0.25914 | -0.6268 | -0.36051 | -0.28803 | -0.30705 |
| 35.83333 | 1.792134 | 1.458745 | 0.491946 | -0.67393 | -2.53596 | 0.238041 | -0.1772 | -0.81358 | 0.551069 | -1.01612 | 0.534453 | -0.53807 | -0.26287 | -0.88735 | -0.24111 | -0.63594 | -1.27733 | -0.04936 | -0.81275 | 0.090563 | -0.80909 | -0.33211 | -0.28959 | -0.27385 |
| 35.84722 | 1.684789 | 1.473142 | 0.643566 | -0.70547 | -3.08211 | -0.0657 | -0.38679 | -0.79332 | 0.646041 | -0.9218 | 0.544791 | -0.5288 | -0.30491 | -0.75881 | -0.25596 | -0.57292 | -0.9631 | 0.078328 | -0.87926 | -0.08017 | -0.88939 | -0.27563 | -0.15637 | -0.30085 |
| 35.86111 | 1.340531 | 0.926808 | 0.737316 | -1.02192 | -3.68087 | -0.11722 | -0.61045 | -0.76157 | 0.731755 | -0.84156 | 0.552903 | -0.2912 | -0.28804 | -0.66271 | -0.22003 | -0.36679 | -0.70712 | 0.198303 | -0.84135 | -0.16745 | -0.87815 | -0.17491 | -0.03829 | -0.26719 |
| 35.875 | 1.130909 | 0.245645 | 0.78049 | -1.11013 | -3.87354 | -0.07811 | -0.69311 | -0.70026 | 0.455849 | -0.80953 | 0.518057 | 0.047125 | -0.21578 | -0.51639 | -0.22335 | -0.06928 | -0.41245 | 0.226463 | -0.81415 | -0.21156 | -0.83518 | -0.09709 | -0.012 | -0.25754 |
| 35.88889 | 1.035571 | 0.03878 | 0.722215 | -0.68426 | -3.88806 | -0.07605 | -0.52081 | -0.82382 | 0.442878 | -0.85267 | 0.273514 | 0.146721 | -0.12681 | -0.43765 | -0.22094 | 0.207651 | -0.16495 | 0.163142 | -0.6549 | -0.29523 | -0.81973 | 0.007785 | -0.0615 | -0.19647 |
| 35.90278 | 0.991267 | 0.04722 | 0.578072 | -0.56142 | -3.53967 | -0.09695 | -0.3147 | -0.72575 | 0.60399 | -0.76786 | 0.008228 | 0.129668 | -0.0188 | -0.30486 | -0.15012 | 0.399571 | 0.054258 | 0.092723 | -0.4262 | -0.41485 | -0.69692 | 0.14474 | -0.01115 | -0.06916 |
| 35.91667 | 0.829214 | -0.01117 | 0.479703 | -0.67007 | -3.3308 | 0.049382 | -0.20336 | -0.62936 | 0.278682 | -0.6617 | -0.05252 | 0.167865 | 0.019514 | -0.19281 | -0.09781 | 0.494127 | 0.108026 | -0.05335 | -0.26766 | -0.53598 | -0.50316 | 0.212736 | 0.044521 | 0.008913 |
| 35.93056 | 0.694256 | -0.04755 | 0.428357 | -0.53528 | -2.70598 | 0.225167 | -0.01097 | -0.64211 | -0.19557 | -0.55834 | -0.07473 | 0.240511 | -0.00275 | -0.16059 | -0.09967 | 0.574902 | 0.021964 | -0.23519 | -0.14943 | -0.6425 | -0.36834 | 0.127549 | 0.097183 | -0.00742 |
| 35.94444 | 0.661599 | -0.00047 | 0.561294 | -0.19725 | -1.61515 | 0.39776 | 0.295101 | -0.66737 | -0.19187 | -0.42076 | -0.03156 | 0.272472 | -0.06093 | -0.12222 | -0.114 | 0.463322 | 0.007046 | -0.30277 | 0.019304 | -0.64398 | -0.20945 | 0.021567 | 0.160525 | -0.07321 |
| 35.95833 | 0.7266 | 0.176107 | 0.703526 | 0.126613 | -0.95435 | 0.700902 | 0.534373 | -0.4898 | 0.056133 | -0.4848 | 0.018041 | 0.306734 | -0.09102 | -0.08274 | -0.08474 | 0.4173 | 0.093112 | -0.21228 | 0.265941 | -0.49113 | -0.00334 | -0.02266 | 0.241073 | -0.10286 |
| 35.97222 | 0.700178 | 0.376343 | 0.71196 | 0.227545 | -0.94648 | 0.759108 | 0.665057 | -0.37385 | 0.537517 | -0.68844 | -0.15507 | -0.06794 | -0.09488 | -0.02107 | -0.0479 | 0.508348 | 0.21769 | -0.04173 | 0.510309 | -0.31575 | 0.155479 | -0.05744 | 0.290455 | -0.10716 |
| 35.98611 | 0.47719 | 0.455309 | 0.156611 | 0.34396 | -1.70035 | 0.398865 | 0.673141 | -0.72454 | 1.245086 | -1.33827 | -0.73192 | -1.55053 | -0.08972 | -0.01719 | -0.02786 | 0.634272 | 0.292758 | 0.082353 | 0.659967 | -0.20681 | 0.179261 | -0.08076 | 0.289209 | -0.14355 |
| 36 | -0.40565 | -0.01151 | -0.44965 | -0.01695 | -2.49039 | -0.18624 | 0.421239 | -1.26914 | 1.368053 | -2.44416 | -1.21476 | -2.43786 | -0.03772 | 0.013256 | -0.0033 | 0.634569 | 0.210009 | 0.14142 | 0.63942 | -0.20397 | -2.7E-05 | -0.07465 | 0.223878 | -0.24137 |
| 36.01389 | -1.25151 | -1.53541 | -0.10017 | -1.11471 | -2.80856 | 0.464286 | -0.16322 | -1.00968 | 0.097023 | -2.68292 | -0.0529 | 0.11471 | 0.093772 | 0.376395 | 0.115231 | 0.629924 | -0.18426 | 0.126785 | 0.263829 | -0.3493 | -0.56919 | -0.03276 | 0.033309 | -0.40746 |
| 36.02778 | -1.59409 | -2.77602 | 0.462215 | -0.44447 | -2.9687 | 0.258409 | -0.57151 | -0.7741 | -0.57739 | -2.47305 | 1.10197 | 2.203588 | 0.233894 | 1.069807 | 0.355963 | 0.673563 | -0.90414 | 0.05179 | -0.34742 | -0.60654 | -1.43305 | -0.09498 | -0.31175 | -0.60634 |
| 36.04167 | -1.46507 | -1.91238 | 0.016197 | 0.602561 | -4.40409 | 0.468182 | -0.4355 | -1.26956 | 0.356641 | -4.17018 | 0.425814 | -0.75525 | 0.177466 | 1.232021 | 0.450774 | 0.72434 | -1.31695 | 0.047715 | -0.77722 | -0.70245 | -1.90351 | -0.12642 | -0.58975 | -0.65498 |
| 36.05556 | -1.13896 | -0.65407 | -0.3021 | 0.792017 | -5.08012 | -1.04138 | -0.31692 | -1.55586 | 1.405218 | -5.47204 | -0.2749 | -3.10015 | -0.08194 | 0.812208 | 0.238738 | 0.499433 | -1.13923 | 0.186961 | -0.82469 | -0.63817 | -1.42065 | -0.04859 | -0.58813 | -0.26524 |
| 36.06944 | -1.26814 | -1.50035 | 0.031317 | 0.724117 | -3.76041 | -0.22942 | -0.32103 | -0.62897 | 0.333527 | -4.77034 | 0.976943 | -0.50077 | -0.24529 | 0.582219 | 0.010202 | 0.230045 | -0.63886 | 0.292346 | -0.58933 | -0.51291 | -0.50164 | 0.111171 | -0.46516 | -0.02325 |
| 36.08333 | -1.71875 | -2.79934 | 0.464388 | 0.638436 | -4.04987 | -0.08681 | -0.4088 | 0.417408 | -1.5207 | -2.92005 | 2.323389 | 2.700784 | -0.24027 | 0.473468 | -0.0226 | 0.206822 | -0.25589 | 0.285644 | -0.18285 | -0.31219 | 0.270834 | 0.20193 | -0.2976 | 0.110686 |
| 36.09722 | -1.35795 | -1.89264 | 0.032361 | 0.700277 | -2.20601 | 0.451974 | -0.21298 | 0.313496 | -0.31224 | -2.27774 | 1.256906 | 1.187132 | -0.12228 | 0.345491 | 0.047365 | 0.136543 | -0.02182 | 0.285152 | 0.128179 | -0.04061 | 0.972638 | 0.195504 | 0.152929 | 0.209505 |
| 36.11111 | -0.42046 | -0.13674 | -0.40009 | 0.732925 | -0.81025 | 1.850163 | 0.059256 | -0.30317 | 0.871592 | -2.35957 | -0.16008 | -2.39599 | -0.03467 | 0.145961 | 0.081907 | -0.08473 | 0.15821 | 0.338227 | 0.292011 | 0.110333 | 1.514018 | 0.124459 | 0.655977 | 0.289914 |
| 36.125 | -0.27659 | 0.39941 | -0.1946 | 0.84841 | -0.48874 | 1.72198 | 0.057445 | -0.00278 | 1.315195 | -2.11629 | -0.09245 | -2.11046 | -0.02074 | 0.056425 | 0.069871 | -0.27219 | 0.283825 | 0.353952 | 0.319503 | 0.189332 | 1.686884 | 0.06674 | 0.910039 | 0.347802 |
| 36.13889 | -0.3983 | -0.17859 | -0.11552 | 0.691059 | -0.14625 | 0.861502 | 0.087568 | 0.345381 | 0.80556 | -1.77361 | 0.08779 | -1.11726 | -0.03085 | 0.067853 | 0.04988 | -0.38808 | 0.307571 | 0.273429 | 0.280614 | 0.185157 | 1.373899 | 0.062329 | 0.796896 | 0.443368 |
| 36.15278 | -0.59257 | -0.98472 | -0.12197 | 0.542447 | -0.05428 | 0.41463 | 0.066452 | 0.687895 | 0.283576 | -1.8897 | 0.582884 | 0.080881 | -0.04802 | 0.055225 | 0.031233 | -0.52415 | 0.212832 | 0.152588 | 0.272508 | 0.096826 | 0.905756 | 0.014176 | 0.534531 | 0.34043 |
| 36.16667 | -0.65409 | -3.97834 | 0.050507 | 0.603806 | -0.8553 | -0.71948 | 0.057962 | 1.6531 | -1.36274 | -0.97433 | 2.05202 | 3.714728 | -0.06167 | 0.018149 | -0.00164 | -0.71101 | 0.054013 | 0.06773 | 0.314619 | -0.03684 | 0.409371 | -0.0345 | 0.360393 | 0.259506 |
| 36.18056 | -0.33214 | -4.44428 | 0.38357 | 0.674894 | -1.47702 | -0.00891 | 0.479955 | 2.278427 | -1.13049 | -0.55951 | 2.005138 | 4.647708 | -0.04521 | -0.00752 | -0.0199 | -0.69331 | -0.09868 | 0.028402 | 0.424927 | -0.12926 | 0.052959 | -0.00619 | 0.414256 | 0.175593 |
| 36.19444 | 0.078913 | -2.40038 | 0.478233 | 0.819871 | -0.074 | 0.880888 | 0.906307 | 1.914063 | 0.428643 | -1.04442 | 1.331929 | 1.866597 | -0.00626 | 0.056066 | -0.006 | -0.58992 | -0.12394 | 0.037055 | 0.518287 | -0.13457 | -0.06926 | 0.102163 | 0.489548 | 0.214582 |
| 36.20833 | 0.326907 | -0.83835 | 0.447816 | 0.824447 | 1.258692 | 1.548923 | 0.88128 | 1.380241 | 1.197874 | -1.38565 | 0.681187 | -0.27646 | 0.024354 | 0.1755 | -0.0038 | -0.35614 | -0.07694 | 0.111766 | 0.513706 | -0.10021 | 0.024286 | 0.201184 | 0.505118 | 0.233064 |
| 36.22222 | 0.578633 | 0.312035 | 0.497764 | 0.770995 | 1.896898 | 1.271447 | 0.696921 | 0.911402 | 1.515482 | -1.01668 | 0.235787 | -1.60217 | 0.060611 | 0.255184 | -0.03428 | -0.11133 | -0.05962 | 0.213988 | 0.409417 | -0.07807 | 0.203275 | 0.262562 | 0.353502 | 0.175113 |
| 36.23611 | 1.130857 | 2.165003 | 0.557375 | 0.433946 | 2.497675 | 1.240951 | 0.749942 | 0.226307 | 1.71355 | -0.06076 | -0.32993 | -3.63223 | 0.068958 | 0.235395 | -0.08622 | 0.13894 | -0.10339 | 0.269881 | 0.247678 | -0.09895 | 0.230844 | 0.26725 | 0.160608 | 0.08153 |
| 36.25 | 1.036533 | 1.957655 | 0.69504 | 0.4668 | 2.368619 | 1.433795 | 0.630487 | 0.447921 | 1.15912 | 0.324797 | -0.38726 | -3.01695 | 0.057118 | 0.215402 | -0.09899 | 0.379239 | -0.15953 | 0.225893 | 0.073536 | -0.14705 | 0.142038 | 0.233163 | 0.074001 | 0.030699 |
| 36.26389 | 0.533124 | -0.91009 | 0.943298 | 0.902665 | 1.385865 | 0.716461 | 0.277393 | 1.660889 | 1.46666 | -0.00896 | 1.185514 | 1.182994 | 0.068974 | 0.210757 | -0.04328 | 0.5125 | -0.15656 | 0.165087 | 0.00988 | -0.14929 | 0.140184 | 0.251008 | 0.157902 | 0.155458 |
| 36.27778 | 0.329009 | -1.99601 | 0.924618 | 0.87843 | 0.555956 | 0.033269 | -0.0068 | 2.028703 | 1.641581 | -0.31791 | 1.738624 | 2.916334 | 0.148726 | 0.263262 | 0.038721 | 0.529702 | -0.06509 | 0.118039 | 0.047235 | -0.09566 | 0.205539 | 0.318785 | 0.297834 | 0.306927 |
| 36.29167 | 0.546128 | 0.39696 | 0.502161 | 0.713963 | 1.648512 | 0.802227 | -0.04763 | 1.070165 | 1.179101 | -0.6145 | 0.527403 | -0.40567 | 0.228563 | 0.353773 | 0.072257 | 0.361856 | 0.005441 | 0.069387 | 0.025278 | -0.05852 | 0.235195 | 0.312418 | 0.283655 | 0.341296 |
| 36.30556 | 0.856915 | 2.697713 | 0.11224 | 0.664465 | 3.008434 | 0.568985 | -0.0599 | 0.152225 | 0.447878 | -0.95048 | -0.27411 | -3.36255 | 0.24693 | 0.390854 | 0.045368 | 0.021828 | 0.015149 | 0.017046 | -0.12082 | -0.07902 | 0.186583 | 0.272372 | 0.125205 | 0.268719 |
| 36.31944 | 0.5215 | 1.773694 | 0.115837 | 0.811528 | 3.402003 | 1.275682 | -0.14518 | 0.395532 | 0.282589 | -1.52629 | -0.19573 | -2.56322 | 0.225002 | 0.400813 | 0.01315 | -0.22692 | 0.046191 | 0.041516 | -0.24653 | -0.06886 | 0.202731 | 0.240283 | -0.00857 | 0.176457 |
| 36.33333 | -0.03155 | 0.268051 | 0.089339 | 0.694005 | 2.402089 | 0.716299 | -0.19918 | 0.919504 | 0.51074 | -2.46759 | 0.560297 | -0.59965 | 0.228949 | 0.460742 | 0.011332 | -0.2499 | 0.121454 | 0.128695 | -0.2556 | -0.0142 | 0.219039 | 0.231448 | -0.09402 | -0.02493 |
| 36.34722 | -0.25605 | -0.03521 | -0.15075 | 0.576484 | 1.349951 | 0.406571 | -0.1688 | 1.000607 | 0.435847 | -2.80914 | 0.596493 | -0.21996 | 0.226485 | 0.429599 | 0.011684 | -0.14762 | 0.177855 | 0.202816 | -0.21364 | 0.007905 | 0.235202 | 0.216458 | -0.15077 | -0.22947 |
| 36.36111 | -0.65707 | -1.08411 | -0.47776 | 0.286665 | 0.310036 | -0.02278 | -0.16753 | 1.399826 | 0.665564 | -3.37096 | 1.010924 | 1.114981 | 0.222554 | 0.324826 | 0.020207 | 0.071903 | 0.200763 | 0.254572 | -0.14089 | -0.00252 | 0.306832 | 0.16059 | -0.15563 | -0.31499 |
| 36.375 | -1.11704 | -3.10286 | -0.90359 | -0.62323 | -1.69617 | -0.45215 | -0.18308 | 2.160065 | 1.622604 | -4.57389 | 1.698241 | 4.064064 | 0.20999 | 0.201636 | 0.047292 | 0.248858 | 0.220145 | 0.292455 | -0.0696 | -0.0003 | 0.374675 | 0.064763 | -0.12682 | -0.29236 |
| 36.38889 | -0.65134 | -2.24781 | -0.64262 | -0.6829 | -2.33425 | -0.64474 | -0.05361 | 2.094914 | 1.489125 | -3.75267 | 1.31925 | 3.039317 | 0.185702 | 0.058029 | 0.069258 | 0.345795 | 0.249112 | 0.308633 | -0.00455 | 0.027779 | 0.456791 | 0.01366 | -0.03971 | -0.28861 |
| 36.40278 | -0.04479 | 0.525326 | -0.06377 | 0.516598 | -0.41124 | 1.05713 | 0.094581 | 0.704355 | -0.24254 | -1.61895 | -0.1563 | -1.50351 | 0.161477 | -0.01063 | 0.088711 | 0.403174 | 0.278487 | 0.284903 | 0.029433 | 0.053452 | 0.493613 | 0.034866 | 0.127256 | -0.24496 |
| 36.41667 | 0.208668 | 1.391184 | 0.061803 | 1.084956 | 0.567866 | 1.870677 | 0.182667 | 0.267081 | -0.89576 | -0.76497 | -0.60454 | -3.31783 | 0.140864 | -0.00887 | 0.103186 | 0.437799 | 0.300459 | 0.209087 | 0.027387 | 0.074512 | 0.464789 | 0.097808 | 0.288255 | -0.17036 |
| 36.43056 | -0.11828 | 0.012029 | -0.19492 | 0.616953 | 0.486582 | 1.175738 | 0.13661 | 0.662423 | -0.23255 | -0.88532 | -0.37656 | -1.87318 | 0.140888 | 0.01886 | 0.108249 | 0.410257 | 0.343068 | 0.151366 | 0.018773 | 0.064808 | 0.393048 | 0.17926 | 0.358732 | -0.10982 |
| 36.44444 | -0.57346 | -1.36247 | -0.50795 | 0.175704 | 0.176112 | 0.730869 | 0.101936 | 1.028796 | 0.333136 | -1.30474 | 0.304192 | -0.2437 | 0.159395 | 0.030684 | 0.091409 | 0.372191 | 0.335978 | 0.117917 | 0.0304 | 0.046186 | 0.319482 | 0.223056 | 0.290488 | -0.05668 |
| 36.45833 | -0.60213 | -2.06568 | -0.67312 | -0.05671 | 0.141581 | 0.400234 | 0.084977 | 1.022524 | 0.531326 | -0.99709 | 0.691629 | 0.446128 | 0.174297 | 0.029038 | 0.020317 | 0.35219 | 0.297491 | 0.114282 | 0.044768 | 0.019033 | 0.292797 | 0.175798 | 0.147157 | 0.001402 |
| 36.47222 | -0.54561 | -2.43588 | -0.43804 | 0.033505 | 0.284718 | 0.534029 | 0.242578 | 0.77851 | 0.827275 | -0.60549 | 1.077523 | 1.025305 | 0.186725 | 0.014956 | -0.05809 | 0.347314 | 0.234919 | 0.164633 | 0.063891 | -0.0121 | 0.309323 | 0.119049 | -0.00082 | 0.051056 |
| 36.48611 | -0.71296 | -3.49024 | 0.089319 | -0.20656 | 0.131249 | -0.10493 | 0.521484 | 0.615048 | 1.503968 | -0.69679 | 2.0487 | 2.447256 | 0.190356 | 0.011077 | -0.03888 | 0.296898 | 0.244638 | 0.278682 | 0.103688 | -0.00083 | 0.351337 | 0.109187 | -0.11749 | 0.105537 |
| 36.5 | -0.58083 | -3.4672 | 0.520341 | -0.22284 | 0.563552 | 0.241615 | 0.537923 | 0.551463 | 1.702924 | 0.159144 | 2.802364 | 2.60818 | 0.191511 | 0.070737 | 0.093797 | 0.207636 | 0.398089 | 0.449803 | 0.213944 | 0.072129 | 0.499098 | 0.18042 | -0.0797 | 0.138617 |
| 36.51389 | -0.56559 | -2.88011 | 0.708187 | 0.024308 | 1.417742 | -0.15847 | 0.550423 | 0.817988 | 1.358826 | 2.104274 | 2.751607 | 1.1041 | 0.127023 | 0.079224 | 0.204062 | 0.170883 | 0.663209 | 0.571216 | 0.393234 | 0.15882 | 0.773597 | 0.268014 | 0.157113 | 0.145767 |
| 36.52778 | -0.61213 | -2.60157 | 0.514085 | 0.126399 | 1.250334 | 0.533152 | 0.798482 | 1.524681 | 0.604237 | 6.889932 | 2.561696 | -0.02093 | 0.050161 | -0.03447 | 0.145379 | 0.239836 | 0.839547 | 0.498764 | 0.421998 | 0.211658 | 0.892549 | 0.22558 | 0.698538 | 0.151719 |
| 36.54167 | -0.1727 | -1.16167 | 0.254501 | 0.649822 | 2.446749 | 1.023705 | 1.496175 | 2.354582 | 0.026383 | 12.15802 | 2.373704 | -0.60805 | 0.061397 | -0.12317 | -0.03886 | 0.412483 | 0.998897 | 0.336591 | 0.332332 | 0.237899 | 0.802942 | 0.140972 | 1.197338 | 0.140731 |
| 36.55556 | 0.542508 | 0.037452 | 0.421401 | 1.798174 | 4.366897 | 1.673172 | 1.574761 | 2.322945 | -0.44801 | 7.985423 | 2.285608 | -0.90882 | 0.124035 | -0.13829 | -0.04845 | 0.508385 | 1.219057 | 0.212503 | 0.460133 | 0.338842 | 0.755956 | 0.162402 | 1.304405 | 0.246434 |
| 36.56944 | 0.575953 | -0.29086 | 0.584633 | 1.698364 | 6.156508 | 1.404627 | 0.810215 | 1.779416 | -0.62164 | 3.599544 | 1.71108 | -0.62371 | 0.113192 | -0.11694 | 0.038091 | 0.402891 | 1.370463 | 0.120669 | 0.733032 | 0.463723 | 0.797804 | 0.218278 | 1.101148 | 0.310637 |
| 36.58333 | 0.417816 | -0.8676 | 0.724269 | 1.290887 | 5.109328 | 0.813361 | 0.272438 | 1.535471 | -0.51563 | 0.565418 | 1.999868 | 0.017757 | 0.049231 | -0.15541 | 0.014876 | 0.152377 | 1.247705 | -0.05134 | 0.788214 | 0.564719 | 0.756112 | 0.177875 | 0.765553 | 0.232609 |
| 36.59722 | 0.75234 | -0.41261 | 0.960561 | 0.991382 | 3.396128 | 0.760315 | 0.298818 | 1.563673 | -0.38629 | -1.59103 | 1.628923 | 0.149333 | -0.00684 | -0.24341 | -0.05001 | -0.01274 | 1.03551 | -0.21144 | 0.771812 | 0.604856 | 0.639603 | 0.102422 | 0.458684 | 0.160042 |
| 36.61111 | 1.415123 | 0.831853 | 1.142289 | 0.956941 | 1.420875 | 1.053734 | 0.266657 | 1.441821 | -0.16562 | -2.89492 | 0.897927 | -0.32233 | -0.01936 | -0.28349 | -0.08413 | -0.05899 | 0.785231 | -0.29472 | 0.837262 | 0.610209 | 0.472684 | -0.00476 | 0.063837 | 0.11195 |
| 36.625 | 1.570212 | 1.879108 | 1.02713 | 0.721378 | 0.035921 | 1.127131 | 0.202957 | 1.276991 | 0.044134 | -3.02339 | 0.681869 | -0.57515 | -0.00801 | -0.28558 | -0.08756 | -0.06319 | 0.829504 | -0.32321 | 0.923574 | 0.59064 | 0.31868 | -0.06204 | -0.30897 | 0.103076 |
| 36.63889 | 1.343475 | 2.620406 | 0.73661 | 0.332448 | -0.76184 | 1.187438 | 0.13562 | 0.984932 | 0.206344 | -2.12044 | 0.54223 | -0.58594 | 0.026083 | -0.28336 | -0.06866 | -0.02421 | 0.827349 | -0.29092 | 1.012127 | 0.560959 | 0.182854 | -0.07639 | -0.62107 | 0.156007 |
| 36.65278 | 0.964659 | 2.970081 | 0.537169 | -0.04225 | -0.83224 | 0.952246 | 0.112377 | 0.814424 | 0.322156 | -1.69588 | 0.71444 | -0.38922 | 0.061173 | -0.18682 | -0.01777 | 0.086374 | 0.818907 | -0.20435 | 1.031967 | 0.520027 | 0.075745 | -0.05874 | -0.73279 | 0.241989 |
| 36.66667 | 0.484188 | 2.53563 | 0.403086 | -0.67528 | -1.36228 | 0.666638 | 0.16861 | 0.768736 | 0.404206 | -2.22834 | 0.900473 | -0.22343 | 0.087932 | -0.0251 | 0.04959 | 0.209744 | 0.521124 | -0.1527 | 0.961452 | 0.438326 | -0.16313 | -0.04051 | -0.87469 | 0.310127 |
| 36.68056 | 0.193502 | 2.804268 | 0.310743 | -1.2491 | -2.63496 | 0.516429 | 0.057452 | 0.542426 | 0.466271 | -3.75921 | 0.741684 | -0.38119 | 0.110367 | 0.075695 | 0.103325 | 0.258467 | 0.027539 | -0.14979 | 0.74067 | 0.331885 | -0.51089 | -0.09024 | -0.95629 | 0.240616 |
| 36.69444 | 0.353999 | 3.257666 | 0.288252 | -1.33725 | -3.99755 | 0.4468 | -0.10717 | 0.181392 | 0.495283 | -5.65833 | 0.424714 | -0.72912 | 0.093547 | 0.090468 | 0.14694 | 0.222491 | -0.60243 | -0.14387 | 0.490642 | 0.276504 | -0.76338 | -0.15789 | -0.84874 | 0.143774 |
| 36.70833 | 0.642156 | 4.330093 | 0.25701 | -0.82328 | -4.20987 | 0.450777 | -0.34128 | -0.16158 | 0.467372 | -6.91284 | 0.186158 | -1.15314 | 0.032634 | -0.00794 | 0.149469 | 0.099428 | -1.12992 | -0.12118 | 0.208424 | 0.377229 | -0.88437 | -0.21458 | -0.61165 | 0.072262 |
| 36.72222 | 0.941045 | 4.687203 | 0.182063 | -0.10989 | -3.5617 | 0.358018 | -0.50515 | -0.30923 | 0.381854 | -7.1689 | 0.159712 | -1.22023 | -0.06304 | -0.20154 | 0.051511 | -0.08429 | -1.51206 | -0.16214 | -0.11817 | 0.458742 | -0.84738 | -0.29786 | -0.42601 | -0.02258 |
| 36.73611 | 0.878403 | 4.144474 | 0.384993 | -0.07571 | -2.75467 | 0.509013 | -0.46787 | -0.18808 | 0.319854 | -5.46454 | 0.539673 | -0.82342 | -0.1403 | -0.41735 | -0.0924 | -0.23916 | -1.70075 | -0.23599 | -0.37501 | 0.477808 | -0.82992 | -0.37356 | -0.22884 | -0.13921 |
| 36.75 | 0.982851 | 3.102971 | 0.475891 | 0.195784 | -1.08304 | 0.647598 | -0.22784 | 0.007378 | 0.340104 | -4.98888 | 0.73298 | -0.61587 | -0.16664 | -0.71843 | -0.2055 | -0.34251 | -1.52974 | -0.29471 | -0.52611 | 0.403873 | -0.83021 | -0.45157 | 0.040981 | -0.18656 |
| 36.76389 | 1.188243 | 2.299185 | 0.416044 | 0.3793 | -1.22089 | 0.804011 | -0.05666 | 0.183988 | 0.412512 | -3.10656 | 0.610632 | -0.64936 | -0.17769 | -0.9821 | -0.27524 | -0.45526 | -1.33808 | -0.35213 | -0.50563 | 0.2705 | -0.78226 | -0.57658 | 0.247739 | -0.25733 |
| 36.77778 | 1.104802 | 1.449292 | 0.508649 | 0.358624 | -1.09195 | 0.943322 | 0.132371 | 0.425485 | 0.37922 | -3.8871 | 0.775571 | -0.39716 | -0.18284 | -1.1417 | -0.29119 | -0.57377 | -0.94198 | -0.35968 | -0.55901 | 0.158975 | -0.78945 | -0.61582 | 0.290753 | -0.28921 |
| 36.79167 | 0.839655 | 0.839223 | 0.62548 | 0.258733 | -1.21715 | 0.63246 | 0.182737 | 0.705982 | 0.409915 | -4.58308 | 1.178717 | -0.10967 | -0.22454 | -1.3057 | -0.29557 | -0.65735 | -0.72897 | -0.2893 | -0.68759 | 0.054113 | -0.91801 | -0.62764 | 0.103398 | -0.29043 |
| 36.80556 | 0.745454 | 0.807052 | 0.591762 | 0.271606 | -1.13739 | 0.558424 | 0.081775 | 0.953227 | 0.557843 | -3.46674 | 1.182458 | -0.13232 | -0.25799 | -1.22136 | -0.26448 | -0.71452 | -0.54569 | -0.18903 | -0.78927 | -0.06789 | -1.07934 | -0.60694 | -0.22272 | -0.37721 |
| 36.81944 | 0.709399 | 0.649147 | 0.510132 | 0.22985 | -0.8331 | 0.504819 | -0.10794 | 1.092094 | 0.690328 | -1.27314 | 0.739054 | -0.31691 | -0.27562 | -1.23374 | -0.18422 | -0.69704 | -0.35886 | -0.08525 | -1.0203 | -0.20717 | -1.13747 | -0.4982 | -0.4613 | -0.42094 |
| 36.83333 | 0.612503 | 0.449679 | 0.567325 | 0.208424 | -0.561 | 0.38788 | -0.31414 | 1.091809 | 0.687684 | -0.41063 | 0.241443 | -0.42028 | -0.29085 | -1.1042 | -0.09848 | -0.67769 | -0.31006 | -0.00543 | -1.1347 | -0.42932 | -1.00592 | -0.38246 | -0.46811 | -0.32946 |
| 36.84722 | 0.550482 | 0.37938 | 0.639062 | 0.337278 | -0.47579 | 0.23239 | -0.52997 | 0.969583 | 0.819684 | -0.00988 | -0.0526 | -0.48325 | -0.30479 | -0.90762 | -0.06719 | -0.68556 | -0.35579 | 0.051743 | -1.39937 | -0.72085 | -0.83344 | -0.29782 | -0.34382 | -0.23681 |
| 36.86111 | 0.327488 | 0.293634 | 0.502548 | 0.371681 | -0.37084 | 0.255012 | -0.46349 | 0.773744 | 0.81968 | 0.639111 | -0.3355 | -0.52347 | -0.28379 | -0.6489 | -0.05692 | -0.61402 | -0.30207 | 0.085065 | -1.29977 | -0.9566 | -0.66315 | -0.20828 | -0.13825 | -0.32317 |
| 36.875 | -0.23885 | 0.05593 | 0.212037 | 0.343209 | -0.53311 | 0.305621 | -0.28799 | 0.725846 | 0.812149 | 0.509336 | -0.48629 | -0.44612 | -0.21723 | -0.38498 | -0.02887 | -0.52215 | -0.13163 | 0.179396 | -1.03981 | -1.09255 | -0.49112 | -0.10418 | 0.132621 | -0.30215 |
| 36.88889 | -0.67305 | -0.35915 | 0.433834 | 0.195631 | -1.01754 | 0.31394 | -0.1719 | 0.776699 | 0.867244 | 0.187714 | -0.47381 | -0.25419 | -0.17405 | -0.22169 | -0.01883 | -0.4376 | 0.064991 | 0.225693 | -0.69604 | -1.10049 | -0.27768 | -0.06764 | 0.254174 | -0.29277 |
| 36.90278 | -0.91018 | -0.74523 | 0.676498 | -0.04492 | -1.36612 | 0.163964 | -0.22225 | 0.824748 | 0.967269 | -0.12697 | -0.73161 | -0.11434 | -0.1657 | -0.165 | -0.03271 | -0.29465 | 0.232028 | 0.139202 | -0.51336 | -1.00702 | -0.1439 | -0.08161 | 0.27166 | -0.29187 |
| 36.91667 | -0.95171 | -1.1696 | 0.825235 | -0.17766 | -1.32544 | 0.049492 | -0.31988 | 0.989189 | 0.977863 | -0.39103 | -1.1601 | -0.03005 | -0.15348 | -0.20732 | -0.05466 | -0.12666 | 0.302971 | -0.02205 | -0.35652 | -0.81569 | -0.09392 | -0.10208 | 0.20772 | -0.23955 |
| 36.93056 | -0.76176 | -1.34814 | 0.903289 | -0.15455 | -1.30951 | 0.1391 | -0.2653 | 1.074987 | 1.089484 | -0.98534 | -1.3816 | 0.0382 | -0.14121 | -0.19912 | -0.0744 | 0.00803 | 0.378083 | -0.18301 | -0.1993 | -0.58031 | -0.1611 | -0.11831 | 0.146966 | -0.217 |
| 36.94444 | -0.62 | -1.318 | 0.938209 | 0.030701 | -1.34868 | 0.199159 | -0.1203 | 1.038121 | 1.290246 | -1.13523 | -1.22138 | 0.226879 | -0.12109 | -0.13853 | -0.07445 | 0.088669 | 0.431282 | -0.24571 | -0.0811 | -0.35445 | -0.23293 | -0.08318 | 0.133004 | -0.21481 |
| 36.95833 | -0.53398 | -1.14875 | 0.889015 | 0.312201 | -1.00799 | 0.425624 | 0.073313 | 0.85966 | 1.301651 | -0.77753 | -1.06021 | 0.200895 | -0.08133 | -0.0802 | -0.05537 | 0.145132 | 0.499946 | -0.17217 | -0.03035 | -0.23519 | -0.19692 | -0.02934 | 0.130022 | -0.16623 |
| 36.97222 | -0.56066 | -0.52908 | 0.653912 | 0.604176 | -0.19495 | 0.874681 | 0.150882 | 0.496286 | 1.085875 | 0.382684 | -1.17638 | -0.63659 | -0.05961 | -0.06248 | -0.04502 | 0.193099 | 0.541983 | -0.0631 | -0.0119 | -0.17894 | -0.09684 | -0.009 | 0.15431 | -0.14362 |
| 36.98611 | -0.7481 | -0.01422 | 0.315452 | 0.681534 | 0.128068 | 1.103116 | 0.116419 | 0.309436 | 0.819653 | 0.747216 | -1.20714 | -1.48211 | -0.03638 | -0.0515 | -0.03863 | 0.245549 | 0.578437 | 0.053382 | 0.052088 | -0.12858 | -0.0279 | 0.019416 | 0.235945 | -0.10597 |
| 37 | -1.02216 | -0.43731 | 0.258396 | 0.310518 | -0.85584 | 0.237535 | 0.143057 | 0.68187 | 0.920687 | -0.37776 | -0.50558 | -0.79458 | 0.08991 | 0.108649 | 0.05067 | 0.335797 | 0.542818 | 0.186139 | 0.131512 | -0.12733 | -0.01865 | 0.097546 | 0.383289 | -0.13415 |
| 37.01389 | -1.22193 | -1.29172 | 0.361162 | -0.09329 | -2.36125 | -0.4858 | -0.02676 | 1.040135 | 1.203943 | -2.51801 | 0.021953 | -0.12359 | 0.448954 | 0.83969 | 0.29561 | 0.588709 | 0.036031 | 0.25204 | -0.03339 | -0.27649 | -0.28829 | 0.162312 | 0.336291 | -0.35251 |
| 37.02778 | -1.46913 | -1.18388 | 0.382753 | 0.162853 | -2.6376 | -0.70476 | -0.47972 | 0.685833 | 0.957496 | -2.96159 | -0.39536 | -0.87156 | 0.879012 | 2.301451 | 0.607168 | 0.742663 | -0.6999 | 0.186853 | -0.5214 | -0.54749 | -0.89758 | 0.116404 | -0.22 | -0.68756 |
| 37.04167 | -1.61718 | -0.03413 | 0.261695 | 0.848728 | -2.87567 | -1.29941 | -0.9375 | -0.16156 | 0.3421 | -2.41448 | -0.92292 | -2.3563 | 0.802051 | 2.437605 | 0.675125 | 0.722908 | -0.92368 | 0.107476 | -0.79191 | -0.60148 | -1.25704 | 0.03056 | -0.88982 | -0.76164 |
| 37.05556 | -1.37262 | -0.66758 | 0.758422 | 0.688375 | -5.81317 | -1.18323 | -0.63058 | 0.374076 | 0.396611 | -2.45047 | -0.04805 | -1.1745 | 0.432018 | 1.260428 | 0.419131 | 0.713262 | -0.63616 | 0.009606 | -0.71025 | -0.39724 | -0.82472 | 0.095363 | -0.83287 | -0.49757 |
| 37.06944 | -0.543 | -3.73877 | 1.536562 | -0.57117 | -3.91194 | -1.79134 | 0.095426 | 2.207147 | 1.19335 | -5.27305 | 3.190052 | 2.706303 | 0.195437 | 0.46927 | 0.134614 | 0.522346 | -0.30476 | 0.077347 | -0.34541 | -0.19424 | -0.2225 | 0.178511 | -0.35985 | -0.20367 |
| 37.08333 | -0.31561 | -3.05603 | 1.327384 | -0.41843 | -2.69934 | -2.58555 | 0.13329 | 1.611977 | 0.866601 | -4.00188 | 3.004455 | 2.047423 | 0.139257 | 0.373894 | 0.018144 | 0.311403 | -0.02202 | 0.185011 | -0.02282 | -0.03204 | 0.287415 | 0.218938 | 0.108088 | -0.01915 |
| 37.09722 | -0.5356 | 0.374639 | 0.800352 | 1.147852 | -0.67673 | -0.18826 | -0.4744 | -0.91753 | -0.53622 | 0.526611 | 0.066089 | -1.87893 | 0.07933 | 0.303147 | -0.044 | 0.132428 | 0.141597 | 0.242737 | 0.15554 | 0.120701 | 0.602294 | 0.19015 | 0.671435 | 0.048317 |
| 37.11111 | -0.51599 | 0.674184 | 0.574904 | 1.272214 | -0.05227 | 0.239023 | -0.88237 | -1.27568 | -0.41432 | 1.265158 | -0.11609 | -2.20151 | -0.00637 | 0.162751 | -0.12704 | -0.09992 | 0.214834 | 0.294064 | 0.165847 | 0.18717 | 0.836722 | 0.072036 | 1.287088 | 0.100141 |
| 37.125 | -0.24456 | -0.95415 | 0.605326 | 0.548148 | -0.55279 | -0.79235 | -0.75675 | -0.34764 | -0.15492 | -0.33511 | 0.864336 | -0.45146 | -0.05873 | 0.100591 | -0.1486 | -0.0067 | 0.232275 | 0.467594 | 0.129627 | 0.221756 | 0.998136 | 0.011376 | 1.366385 | 0.110685 |
| 37.13889 | -0.42768 | -0.86228 | 0.282567 | 0.535605 | -0.27483 | -0.50827 | -0.83823 | -0.24889 | -0.32606 | 0.082149 | -0.09669 | -0.69907 | -0.06671 | 0.191033 | -0.11607 | 0.07646 | 0.221885 | 0.492015 | 0.109664 | 0.161905 | 1.037826 | 0.059544 | 1.067742 | 0.08707 |
| 37.15278 | -0.44753 | -1.37388 | 0.245585 | 0.33449 | -0.29461 | -1.0813 | -0.46511 | 0.243119 | 0.1536 | -0.24383 | 0.005532 | -0.21545 | -0.07621 | 0.31678 | -0.08662 | -0.01322 | 0.227477 | 0.325049 | 0.081949 | 0.134595 | 0.925564 | 0.084832 | 0.589222 | 0.033511 |
| 37.16667 | -0.12164 | -3.06022 | 0.879801 | -0.41037 | -1.04072 | -1.29366 | 0.214461 | 1.334145 | 0.740881 | -2.3005 | 1.936078 | 1.862897 | -0.08687 | 0.220354 | -0.05914 | -0.28316 | 0.210201 | 0.116676 | 0.064747 | 0.115654 | 0.612353 | 0.047881 | 0.148117 | -0.02392 |
| 37.18056 | 0.12017 | -3.25554 | 1.112068 | -0.52771 | -0.96883 | -0.94908 | 0.399853 | 1.65745 | 1.196426 | -2.5522 | 2.65945 | 2.50348 | -0.06054 | 0.082249 | -0.0179 | -0.45686 | 0.167429 | 0.058852 | 0.096705 | 0.100565 | 0.324021 | 0.049624 | -0.05797 | -0.0425 |
| 37.19444 | 0.019651 | -2.9323 | 1.066883 | -0.43763 | -0.71994 | -0.64425 | 0.687508 | 1.625101 | 1.218214 | -1.93961 | 2.683388 | 2.102989 | 0.00022 | -0.06394 | 0.00346 | -0.38152 | 0.104744 | 0.175535 | 0.194903 | 0.107561 | 0.168516 | 0.052738 | 0.035585 | -0.04633 |

| | | | | | | | | | | | | | | | | | | | | | | | | |
|---|---|---|---|---|---|---|---|---|---|---|---|---|---|---|---|---|---|---|---|---|---|---|---|---|
| 37.20833 | -0.25559 | -1.90545 | 0.73964 | -0.02418 | 0.204722 | -0.11372 | 0.680671 | 1.120629 | 0.8956 | -0.72793 | 1.868687 | 0.916348 | 0.066719 | -0.09897 | -0.05763 | -0.17026 | 0.057297 | 0.302006 | 0.29478 | 0.070511 | 0.091301 | 0.0317 | 0.210264 | -0.05414 |
| 37.22222 | -0.72019 | -0.4887 | 0.251141 | 0.836205 | 1.924669 | -0.00295 | 0.326161 | 0.310957 | 0.388829 | 1.412648 | 0.072405 | -1.23721 | 0.110539 | -0.06144 | -0.16899 | 0.026622 | 0.011826 | 0.261501 | 0.308976 | -0.00683 | 0.056254 | 0.010116 | 0.264144 | -0.07263 |
| 37.23611 | -0.47785 | -0.67879 | 0.485218 | 0.601782 | 1.607901 | 0.16617 | 0.31633 | 0.520628 | 0.36737 | 0.819979 | 0.499213 | -0.67224 | 0.109615 | 0.034285 | -0.23257 | 0.210463 | -0.05138 | 0.153244 | 0.246597 | -0.06581 | 0.116597 | 0.017897 | 0.10506 | -0.08713 |
| 37.25 | 0.222155 | -1.97602 | 1.207895 | -0.23333 | 0.074436 | -0.29995 | 0.710802 | 1.301418 | 1.1559 | -1.52916 | 2.592621 | 1.646257 | 0.08811 | 0.218737 | -0.19665 | 0.440245 | -0.09469 | 0.180907 | 0.196341 | -0.1185 | 0.195244 | 0.01244 | -0.13084 | -0.07938 |
| 37.26389 | 0.168917 | -1.3346 | 1.051092 | -0.11362 | 0.388201 | -0.4119 | 0.472853 | 1.125717 | 0.894629 | -0.8612 | 2.155691 | 1.025827 | 0.055548 | 0.405311 | -0.13007 | 0.618805 | -0.11604 | 0.306713 | 0.17327 | -0.17207 | 0.187712 | -0.02283 | -0.28293 | -0.05511 |
| 37.27778 | -0.04377 | -0.0653 | 0.648606 | 0.3436 | 1.67712 | 0.020003 | -0.04129 | 0.731365 | 0.182893 | 0.5808 | 1.110504 | -0.46199 | 0.043581 | 0.5349 | -0.06756 | 0.617498 | -0.09178 | 0.316746 | 0.182373 | -0.20309 | 0.091839 | -0.03656 | -0.27625 | -0.01157 |
| 37.29167 | 0.272745 | -1.02943 | 1.011666 | 0.010973 | 1.177917 | -1.2752 | 0.206413 | 0.940674 | 0.668439 | -0.95591 | 2.43326 | 0.824525 | 0.050224 | 0.513643 | -0.00378 | 0.746234 | -0.01796 | 0.216903 | 0.214471 | -0.17745 | 0.064099 | -0.0007 | -0.11703 | 0.037611 |
| 37.30556 | 0.223767 | -1.23146 | 0.962126 | -0.0978 | 0.786413 | -1.0945 | 0.070691 | 0.825441 | 0.644204 | -1.20227 | 2.616018 | 0.990747 | 0.080808 | 0.418035 | 0.0698 | 0.82042 | 0.049105 | 0.135269 | 0.229129 | -0.13739 | 0.077905 | 0.015745 | 0.058715 | 0.056657 |
| 37.31944 | -0.38911 | 0.109055 | 0.260945 | 0.502328 | 1.642226 | -1.63868 | -0.52507 | 0.135416 | -0.25421 | 0.735596 | 1.042667 | -0.96344 | 0.117197 | 0.325444 | 0.109844 | 0.70566 | 0.067753 | 0.087171 | 0.231295 | -0.11563 | 0.110491 | 0.047901 | 0.167581 | 0.042008 |
| 37.33333 | -0.45331 | -0.17025 | 0.2351 | 0.518844 | 0.7686 | -1.78067 | -0.89633 | 0.067085 | -0.40503 | 0.484156 | 1.335929 | -0.83773 | 0.153684 | 0.275192 | 0.098532 | 0.519326 | 0.028131 | 0.104487 | 0.209793 | -0.07965 | 0.136095 | 0.189138 | 0.167822 | 0.005108 |
| 37.34722 | -0.22281 | -1.10902 | 0.478809 | 0.164918 | -0.56235 | -1.79517 | -2.31224 | 0.343575 | 0.028092 | -0.72903 | 2.163393 | 0.252493 | 0.186374 | 0.266672 | 0.069012 | 0.2637 | -0.01663 | 0.162268 | 0.167338 | -0.03062 | 0.143513 | 0.314117 | 0.090357 | -0.0366 |
| 37.36111 | -0.30267 | -1.16545 | 0.423411 | 0.172039 | -0.77611 | -1.85247 | -3.59271 | 0.499027 | -0.09863 | -0.54418 | 1.917117 | 0.133309 | 0.173557 | 0.299068 | 0.031017 | 0.064685 | -0.03282 | 0.185001 | 0.098182 | -0.01752 | 0.133147 | 0.272731 | 0.037626 | -0.06114 |
| 37.375 | -0.4107 | -1.06186 | 0.449917 | 0.242273 | -0.74997 | -1.63739 | -2.23951 | 0.839807 | -0.08694 | -0.22949 | 1.861776 | -0.00736 | 0.133343 | 0.351878 | -0.00167 | -0.0489 | -0.02922 | 0.198342 | 0.077926 | -0.0309 | 0.108954 | 0.168602 | 0.065202 | -0.05661 |
| 37.38889 | -0.10526 | -1.6515 | 0.854028 | -0.12306 | -1.68837 | 0.498379 | -0.76951 | 1.243306 | 0.551042 | -1.33435 | 2.979979 | 1.322388 | 0.096953 | 0.389845 | 0.003762 | 0.011355 | 0.027143 | 0.273425 | 0.181278 | -0.00032 | 0.173975 | 0.132939 | 0.190585 | -0.00786 |
| 37.40278 | 0.348304 | -2.12781 | 1.315128 | -0.4403 | -2.53132 | 0.342755 | -0.49018 | 1.253283 | 1.046679 | -2.054 | 3.757837 | 2.402293 | 0.072562 | 0.405632 | 0.049792 | 0.132013 | 0.120692 | 0.361839 | 0.289179 | 0.055816 | 0.300025 | 0.146043 | 0.337353 | 0.036238 |
| 37.41667 | 0.114002 | -1.15384 | 0.862788 | -0.21464 | -1.86557 | -0.19894 | 0.034124 | 1.006745 | 0.573652 | -1.00081 | 2.68957 | 1.213195 | 0.04977 | 0.378435 | 0.093836 | 0.268017 | 0.190561 | 0.445574 | 0.282744 | 0.080703 | 0.387597 | 0.098802 | 0.404551 | 0.014033 |
| 37.43056 | -0.35964 | 0.167311 | 0.086898 | 0.518342 | -0.02007 | -0.07629 | 0.192578 | 0.407901 | -0.07445 | 0.721677 | 0.63766 | -1.17759 | 0.035909 | 0.339333 | 0.116594 | 0.329646 | 0.19708 | 0.468786 | 0.213251 | 0.080313 | 0.427522 | 0.022862 | 0.374726 | -0.02432 |
| 37.44444 | -0.69327 | 1.172942 | -0.45558 | 1.023002 | 1.580452 | -1.57073 | 0.47901 | -0.07115 | -0.96829 | 1.888511 | -0.99194 | -3.00643 | 0.032897 | 0.265747 | 0.124096 | 0.28949 | 0.172677 | 0.428117 | 0.183759 | 0.090987 | 0.442961 | 0.00942 | 0.320493 | -0.01614 |
| 37.45833 | -0.36535 | 0.90362 | -0.27332 | 0.937976 | 1.438233 | -1.77458 | 0.201029 | 0.117388 | -0.87013 | 1.31503 | -0.42115 | -2.58381 | 0.060661 | 0.168279 | 0.110064 | 0.250105 | 0.174099 | 0.393569 | 0.219747 | 0.100889 | 0.425465 | 0.048374 | 0.258122 | -0.00792 |
| 37.47222 | 0.525362 | -0.81756 | 0.779711 | -0.1464 | -0.58208 | 1.296623 | -0.04789 | 0.953975 | 0.372926 | -0.70085 | 2.41686 | 0.357041 | 0.103725 | 0.096801 | 0.066731 | 0.3241 | 0.179708 | 0.362986 | 0.265285 | 0.108889 | 0.33947 | 0.077275 | 0.242335 | -0.00943 |
| 37.48611 | 1.651413 | -2.12052 | 1.60511 | -0.93585 | -1.68082 | 4.648595 | -0.28211 | 1.92802 | 1.452186 | -2.10039 | 4.782524 | 2.850316 | 0.149462 | 0.085356 | 0.067962 | 0.498759 | 0.159535 | 0.373512 | 0.270874 | 0.112181 | 0.291344 | 0.087267 | 0.318309 | 0.031807 |
| 37.5 | 2.276862 | -1.35086 | 1.308967 | -0.61766 | -0.07355 | 3.733809 | 0.103693 | 2.225151 | 1.922383 | -1.69201 | 4.212451 | 2.456162 | 0.153936 | 0.07498 | 0.178954 | 0.719505 | 0.232469 | 0.433229 | 0.322144 | 0.128014 | 0.41525 | 0.088677 | 0.504887 | 0.136759 |
| 37.51389 | 2.35067 | -0.68322 | 0.816974 | 0.387615 | 3.014169 | 0.775656 | 0.167112 | 1.993321 | 1.010069 | -0.44735 | 1.783441 | 0.898072 | 0.105337 | 0.02857 | 0.33288 | 0.887965 | 0.371824 | 0.503853 | 0.475172 | 0.117388 | 0.747258 | 0.041329 | 0.809241 | 0.199199 |
| 37.52778 | 2.139578 | -0.56097 | 0.637988 | 1.039316 | 3.796007 | 0.35023 | 0.362093 | 1.505292 | 0.69573 | -0.12684 | 0.573816 | 0.134216 | 0.011182 | -0.01061 | 0.290513 | 0.817647 | 0.374296 | 0.542315 | 0.53161 | 0.016436 | 0.904099 | -0.07195 | 1.227531 | 0.166512 |
| 37.54167 | 1.57232 | -0.93697 | 0.662761 | 1.216001 | 3.921316 | 0.477116 | 0.7431 | 1.451571 | 0.862297 | -0.15126 | 0.309353 | 0.094345 | -0.08301 | -0.07663 | 0.104346 | 0.590739 | 0.281214 | 0.563887 | 0.402281 | -0.06228 | 0.755402 | -0.13329 | 1.50137 | 0.042315 |
| 37.55556 | 0.976409 | -0.84076 | 0.703469 | 1.177571 | 4.17722 | 1.309823 | 1.268651 | 1.738861 | 1.271802 | -0.1689 | 0.344925 | 0.048267 | -0.07533 | -0.08775 | 0.015828 | 0.346319 | 0.374185 | 0.470577 | 0.41988 | 0.062058 | 0.666276 | 0.000795 | 1.156084 | 0.035741 |
| 37.56944 | 0.72824 | -0.54644 | 0.656892 | 0.882922 | 3.323217 | 0.982001 | 1.691359 | 2.119553 | 1.545697 | -0.19545 | 0.715027 | 0.042564 | -0.0197 | -0.02868 | 0.0814 | 0.116919 | 0.636914 | 0.325483 | 0.577571 | 0.286951 | 0.722459 | 0.16653 | 0.738072 | 0.09286 |
| 37.58333 | 0.565412 | 0.26083 | 0.445572 | 0.904415 | 3.076016 | 0.846534 | 1.700336 | 2.128944 | 1.233141 | 0.083368 | 0.860078 | -0.28022 | 0.004052 | -0.03548 | 0.09058 | -0.09356 | 0.719024 | 0.045179 | 0.642796 | 0.351691 | 0.66338 | 0.23103 | 0.535748 | 0.100891 |
| 37.59722 | 0.40268 | 0.945187 | 0.423626 | 1.04599 | 2.639802 | 0.640285 | 1.480448 | 1.912021 | 0.827593 | 0.274478 | 1.15992 | -0.43335 | -0.0331 | -0.0541 | 0.033536 | -0.17494 | 0.714192 | -0.27066 | 0.627021 | 0.320013 | 0.492117 | 0.209668 | 0.313634 | 0.08105 |
| 37.61111 | 0.465705 | 0.892048 | 0.621597 | 0.692966 | 1.533752 | 0.90337 | 1.482268 | 1.750696 | 0.779161 | 0.099366 | 1.594116 | -0.15467 | -0.06334 | -0.02593 | -0.01734 | -0.20442 | 0.6847 | -0.49535 | 0.651074 | 0.304001 | 0.35614 | 0.177326 | 0.138567 | 0.065126 |
| 37.625 | 0.37996 | 0.506307 | 0.766992 | 0.557016 | 0.903938 | 1.14188 | 1.324569 | 1.71263 | 0.933987 | -0.02544 | 1.656319 | 0.0318 | -0.05921 | -0.06903 | -0.03977 | -0.12231 | 0.686458 | -0.63828 | 0.800452 | 0.293897 | 0.293026 | 0.104196 | 0.098872 | 0.085671 |
| 37.63889 | -0.03733 | 0.447604 | 0.634308 | 0.805557 | 1.09582 | 0.876884 | 1.091832 | 1.45669 | 0.820988 | 0.137024 | 1.35196 | -0.09399 | -0.03947 | -0.13362 | -0.05411 | -0.02445 | 0.654044 | -0.60856 | 0.937652 | 0.273842 | 0.305057 | 0.011737 | 0.136638 | 0.149003 |
| 37.65278 | -0.18337 | 0.173483 | 0.700365 | 0.787853 | 0.806278 | 0.895436 | 0.903889 | 1.381428 | 0.853904 | 0.109463 | 1.495099 | -0.04244 | -0.03539 | -0.20605 | -0.06735 | -0.04764 | 0.649956 | -0.49219 | 1.007693 | 0.181354 | 0.294416 | -0.0541 | 0.170712 | 0.200853 |
| 37.66667 | 0.161535 | -0.1416 | 0.971199 | 0.225385 | -0.46661 | 1.596316 | 0.944851 | 1.716716 | 1.208657 | -0.25744 | 2.028198 | 0.293025 | -0.03923 | -0.26602 | -0.05184 | -0.24555 | 0.681152 | -0.34678 | 0.948086 | 0.079623 | 0.245237 | -0.10344 | 0.055718 | 0.249949 |
| 37.68056 | 0.294079 | 0.061289 | 0.979195 | -0.14511 | -1.33782 | 1.384955 | 1.001251 | 1.742974 | 1.185984 | -0.17541 | 2.354085 | 0.260834 | -0.02635 | -0.25167 | 0.002394 | -0.54483 | 0.76162 | -0.22886 | 0.905982 | -0.00501 | 0.118066 | -0.13724 | -0.11282 | 0.288075 |
| 37.69444 | 0.132604 | 0.498097 | 0.705345 | -0.03526 | -1.43711 | 0.693723 | 0.873948 | 1.346449 | 0.724959 | 0.180007 | 2.179765 | -0.07246 | 0.001179 | -0.29819 | 0.054492 | -0.67585 | 0.725269 | -0.1376 | 0.862342 | -0.02137 | -0.02863 | -0.16747 | -0.23005 | 0.337446 |
| 37.70833 | 0.089943 | 0.686248 | 0.688124 | -0.12198 | -1.91727 | 0.744689 | 0.804181 | 1.220129 | 0.647499 | 0.086388 | 2.131027 | -0.10687 | -0.0093 | -0.44482 | 0.061288 | -0.56236 | 0.47551 | -0.12941 | 0.565159 | 0.008493 | -0.21534 | -0.22085 | -0.3021 | 0.360404 |
| 37.72222 | 0.003886 | 0.781739 | 0.708674 | -0.03641 | -2.27564 | 0.668676 | 0.804904 | 1.209588 | 0.628386 | -0.02046 | 2.083349 | -0.19751 | -0.03016 | -0.53894 | 0.04323 | -0.32351 | 0.282316 | -0.1578 | 0.224469 | 0.033746 | -0.36456 | -0.26919 | -0.34042 | 0.290618 |
| 37.73611 | -0.15471 | 0.940839 | 0.619052 | 0.267491 | -2.35937 | 0.372214 | 0.806318 | 1.009596 | 0.453508 | 0.043361 | 1.767313 | -0.34682 | -0.03917 | -0.63671 | 0.023753 | -0.10269 | 0.140263 | -0.20353 | -0.09321 | 0.116742 | -0.43877 | -0.2904 | -0.5032 | 0.183506 |
| 37.75 | -0.17551 | 0.455545 | 0.747867 | 0.347046 | -2.59908 | 0.567023 | 0.69977 | 1.223325 | 0.692755 | -0.25472 | 1.802102 | -0.27129 | -0.06906 | -0.64243 | -0.0038 | -0.05904 | -0.09842 | -0.26108 | -0.33198 | 0.231618 | -0.47619 | -0.34099 | -0.69234 | 0.052273 |
| 37.76389 | -0.11521 | -0.12084 | 0.878654 | 0.354349 | -2.72945 | 0.919411 | 0.669233 | 1.574337 | 0.946666 | -0.47958 | 1.807374 | -0.12439 | -0.11032 | -0.79139 | -0.04234 | -0.131 | -0.38363 | -0.31605 | -0.51746 | 0.410608 | -0.52435 | -0.41777 | -0.88333 | -0.05925 |
| 37.77778 | -0.03831 | -0.2505 | 0.919059 | 0.355059 | -1.95075 | 0.692044 | 0.598863 | 1.643933 | 0.837271 | -0.35157 | 1.499543 | -0.16222 | -0.15414 | -0.93072 | -0.08281 | -0.22716 | -0.60483 | -0.33992 | -0.66018 | 0.534681 | -0.59962 | -0.41516 | -0.85743 | -0.13675 |
| 37.79167 | -0.08028 | -0.34795 | 0.935987 | 0.361521 | -0.94512 | 0.493643 | 0.56891 | 1.554214 | 0.607351 | -0.12571 | 1.150101 | -0.22098 | -0.16587 | -0.87694 | -0.11079 | -0.36577 | -0.77839 | -0.25572 | -0.72405 | 0.455811 | -0.61825 | -0.42917 | -0.69531 | -0.18019 |
| 37.80556 | -0.14023 | -0.37801 | 0.896884 | 0.386896 | -0.17845 | 0.530329 | 0.603904 | 1.563171 | 0.560445 | 0.003204 | 0.983518 | -0.24987 | -0.12414 | -0.81109 | -0.10904 | -0.46836 | -0.83344 | -0.12429 | -0.76801 | 0.257483 | -0.51234 | -0.44541 | -0.56061 | -0.1689 |
| 37.81944 | -0.17527 | -0.33949 | 0.840835 | 0.438162 | 0.248736 | 0.604085 | 0.584976 | 1.527439 | 0.478688 | 0.106689 | 0.867441 | -0.23438 | -0.07449 | -0.70884 | -0.08203 | -0.4326 | -0.72875 | 0.006951 | -0.75609 | 0.13815 | -0.4514 | -0.42151 | -0.50565 | -0.16499 |
| 37.83333 | -0.1597 | -0.52114 | 0.814692 | 0.476692 | 0.283611 | 0.724584 | 0.63852 | 1.536659 | 0.514693 | 0.107214 | 0.636249 | -0.13089 | -0.06494 | -0.65762 | -0.06376 | -0.28109 | -0.68838 | 0.053314 | -0.74752 | 0.034621 | -0.33251 | -0.39908 | -0.4353 | -0.18669 |
| 37.84722 | -0.10054 | -0.74234 | 0.833227 | 0.536728 | 0.045661 | 0.745725 | 0.74287 | 1.629349 | 0.643868 | 0.103012 | 0.510777 | -0.02993 | -0.09553 | -0.59278 | -0.05444 | -0.1753 | -0.64947 | 0.017558 | -0.71074 | -0.08969 | -0.21856 | -0.37974 | -0.2736 | -0.20052 |
| 37.86111 | -0.02473 | -0.85045 | 0.875901 | 0.547629 | -0.3564 | 0.732687 | 0.800447 | 1.628006 | 0.76379 | 0.137014 | 0.136553 | 0.014331 | -0.11856 | -0.61261 | -0.02497 | 0.006795 | -0.68855 | -0.07753 | -0.66303 | -0.09775 | -0.19334 | -0.3384 | -0.13868 | -0.1886 |
| 37.875 | 0.112876 | -0.79792 | 0.859174 | 0.482444 | -0.61042 | 0.494849 | 0.679193 | 1.498413 | 0.749282 | 0.148076 | -0.15002 | 0.013053 | -0.14055 | -0.74532 | 0.018318 | 0.188878 | -0.65226 | -0.17571 | -0.54434 | -0.04313 | -0.16456 | -0.30597 | 0.018865 | -0.18284 |
| 37.88889 | 0.128856 | -0.76622 | 0.802599 | 0.451945 | -0.72705 | 0.337422 | 0.484367 | 1.330172 | 0.574613 | 0.124364 | -0.30356 | 0.01472 | -0.17744 | -0.79112 | 0.021885 | 0.289786 | -0.6182 | -0.28525 | -0.42157 | -0.02926 | -0.11966 | -0.28373 | 0.184766 | -0.11311 |
| 37.90278 | 0.27295 | -0.70495 | 0.781324 | 0.437028 | -0.78043 | 0.388842 | 0.30078 | 1.359603 | 0.483305 | 0.092048 | -0.27267 | -0.04722 | -0.21947 | -0.76346 | -0.01313 | 0.286435 | -0.31952 | -0.34461 | -0.23424 | -0.07017 | -0.03585 | -0.23302 | 0.325109 | -0.06791 |
| 37.91667 | 0.332651 | -0.71108 | 0.78594 | 0.428857 | -0.90594 | 0.23446 | 0.140406 | 1.188905 | 0.503022 | 0.066135 | -0.35119 | -0.09908 | -0.22715 | -0.71531 | -0.05988 | 0.249227 | -0.02546 | -0.36811 | -0.10272 | -0.10333 | 0.032657 | -0.16012 | 0.400357 | -0.05502 |
| 37.93056 | 0.390128 | -0.64372 | 0.765232 | 0.455663 | -1.15233 | 0.096576 | 0.093212 | 1.046241 | 0.441324 | 0.038697 | -0.47335 | -0.10429 | -0.23356 | -0.60595 | -0.11535 | 0.200908 | 0.177382 | -0.26893 | -0.03638 | -0.13304 | 0.051606 | -0.14735 | 0.387697 | -0.05809 |
| 37.94444 | 0.37655 | -0.49833 | 0.784184 | 0.512573 | -1.18252 | 0.011238 | 0.217394 | 0.922503 | 0.53763 | 0.066168 | -0.51294 | -0.03841 | -0.23835 | -0.49927 | -0.18592 | 0.124901 | 0.244067 | -0.15979 | -0.00562 | -0.16687 | 0.048353 | -0.11166 | 0.316194 | -0.05751 |
| 37.95833 | 0.387477 | -0.34198 | 0.800693 | 0.698066 | -0.64832 | 0.130395 | 0.435192 | 0.798737 | 0.852771 | 0.122071 | -0.5624 | 0.012713 | -0.22038 | -0.39248 | -0.22558 | 0.055614 | 0.272831 | -0.0354 | 0.057769 | -0.15561 | 0.028794 | -0.07725 | 0.209373 | -0.05048 |
| 37.97222 | 0.451881 | -0.10201 | 0.818272 | 0.859468 | 0.08852 | 0.418855 | 0.656676 | 0.693663 | 1.014567 | 0.163094 | -0.59266 | -0.00654 | -0.17231 | -0.2793 | -0.20662 | 0.070417 | 0.281339 | 0.093004 | 0.111345 | -0.11203 | 0.022467 | -0.04924 | 0.133986 | -0.0374 |
| 37.98611 | 0.49183 | 0.148501 | 0.787297 | 0.880335 | 0.339122 | 0.690836 | 0.729611 | 0.558117 | 0.985511 | 0.169715 | -0.72852 | -0.0181 | -0.12551 | -0.25649 | -0.09692 | 0.180916 | 0.263556 | 0.171151 | 0.117386 | -0.05069 | 0.024525 | -0.03898 | 0.157613 | 0.013463 |
| 38 | 0.401632 | 0.132592 | 0.649722 | 0.583638 | -0.09583 | 0.515356 | 0.544952 | 0.630397 | 0.67595 | 0.251677 | -0.94584 | -0.17503 | -0.04332 | -0.12469 | 0.05087 | 0.259801 | 0.166373 | 0.208604 | 0.06654 | -0.05744 | -0.02558 | -0.02308 | 0.229362 | 0.056492 |

**Supplementary Table S2.** Raw motion data from: (a) 100 mM NaCl stress

| Time | Control1 | Control2 | Control3 | Control4 | Control5 | Control6 | Control7 | Control8 | Control9 | Stress1 | Stress2 | Stress3 | Stress4 | Stress5 | Stress6 | Stress7 | Stress8 | Stress9 | Stress10 |
|---|---|---|---|---|---|---|---|---|---|---|---|---|---|---|---|---|---|---|---|
| 18 | -0.45008 | -0.29144 | -0.20011 | 0.351355 | 0.030443 | 0.129136 | -2.74893 | -0.01682 | -0.0011 | -0.98856 | -2.25749 | -0.12453 | 0.040678 | 0.277007 | 0.127137 | -0.21298 | -0.26966 | -0.80866 | -1.41879 |
| 18.01389 | -1.85246 | -1.74027 | 0.078379 | 0.089679 | -1.19837 | 0.092218 | -4.75128 | -0.41286 | -0.7913 | -2.30241 | -3.97933 | -0.41882 | 0.140722 | 0.379627 | 0.048173 | -0.22195 | -1.70562 | -1.84023 | -2.32701 |
| 18.02778 | -2.64248 | -2.58533 | 0.552324 | -0.4473 | -2.51619 | -0.24431 | -2.99979 | -0.64264 | -1.21287 | -2.32628 | -3.15407 | -0.49107 | 0.197972 | 0.233928 | -0.16997 | 0.028689 | -3.55014 | -2.05778 | -1.66786 |
| 18.04167 | -1.30252 | -1.28108 | 0.60062 | -0.18439 | -1.39165 | -0.44058 | -1.40841 | -0.33426 | -0.55164 | -0.8529 | -1.19828 | -0.32425 | 0.081461 | -0.06655 | -0.20061 | 0.051249 | -1.71013 | -0.89893 | -0.49095 |
| 18.05556 | -0.10311 | 0.62603 | 0.370327 | 0.92628 | -0.07692 | -0.36827 | -0.04257 | 0.292843 | -0.2009 | -0.29311 | 0.66679 | -0.20652 | -0.09641 | -0.2137 | -0.13428 | -0.0799 | -0.71171 | -0.34115 | -0.06736 |
| 18.06944 | 0.872961 | 2.621784 | 0.185485 | 2.646077 | 0.697789 | 0.04493 | 1.330566 | 0.988953 | -0.06051 | -0.28395 | 3.287262 | -0.10268 | -0.17006 | -0.21994 | -0.0951 | -0.13544 | -0.36645 | -0.17847 | -0.01167 |
| 18.08333 | 0.451911 | 2.263823 | -0.05021 | 2.884997 | 0.669681 | 0.17018 | 1.835847 | 0.805798 | -0.0591 | -0.18461 | 2.277652 | -0.06949 | -0.15185 | -0.21636 | -0.08576 | -0.1257 | -0.19726 | -0.15041 | -0.00238 |
| 18.09722 | -0.25444 | 0.607823 | -0.30504 | 0.951926 | 0.053135 | -0.08597 | 1.348679 | 0.012386 | -0.19891 | -0.01707 | 0.886802 | -0.09671 | -0.13046 | -0.27694 | -0.10332 | -0.10562 | -0.1541 | -0.16857 | -0.04443 |
| 18.11111 | -0.51426 | -0.28351 | -0.39472 | -0.13283 | -0.31932 | -0.33474 | 0.769237 | -0.42833 | -0.30889 | 0.033119 | 0.448366 | -0.13628 | -0.11811 | -0.25019 | -0.12104 | -0.08034 | -0.13748 | -0.16796 | -0.06755 |
| 18.125 | -0.58612 | -0.42189 | -0.35019 | -0.22559 | -0.39524 | -0.46877 | 0.304266 | -0.50404 | -0.42028 | 0.045772 | 0.317548 | -0.16129 | -0.12262 | -0.18684 | -0.13811 | -0.07163 | -0.12961 | -0.14844 | -0.05275 |
| 18.13889 | -0.6147 | -0.44913 | -0.33448 | -0.26541 | -0.44022 | -0.57275 | -0.10134 | -0.50744 | -0.5459 | 0.051768 | 0.274435 | -0.16396 | -0.1301 | -0.18408 | -0.14606 | -0.09035 | -0.12826 | -0.12735 | -0.0374 |
| 18.15278 | -0.55274 | -0.3946 | -0.26964 | -0.29116 | -0.40215 | -0.54497 | -0.33458 | -0.43025 | -0.51551 | 0.062763 | 0.287456 | -0.13561 | -0.11635 | -0.19528 | -0.1304 | -0.08215 | -0.09391 | -0.07244 | 0.001573 |
| 18.16667 | -0.48449 | -0.3152 | -0.19548 | -0.27743 | -0.31437 | -0.48229 | -0.45054 | -0.35946 | -0.39854 | 0.086857 | 0.301765 | -0.11288 | -0.12157 | -0.21326 | -0.13379 | -0.06865 | -0.07981 | -0.02458 | 0.041745 |
| 18.18056 | -0.51959 | -0.36309 | -0.24327 | -0.32578 | -0.33386 | -0.56714 | -0.53449 | -0.41783 | -0.36018 | 0.135395 | 0.272697 | -0.11793 | -0.14797 | -0.21548 | -0.14413 | -0.08111 | -0.11043 | -0.02815 | 0.052456 |
| 18.19444 | -0.56988 | -0.46488 | -0.32859 | -0.39756 | -0.41124 | -0.78915 | -0.5718 | -0.52551 | -0.41599 | 0.198316 | 0.211134 | -0.1386 | -0.15988 | -0.17211 | -0.13839 | -0.10245 | -0.13146 | -0.06602 | 0.035731 |
| 18.20833 | -0.52957 | -0.49138 | -0.31743 | -0.39051 | -0.38174 | -1.08219 | -0.46373 | -0.59059 | -0.39531 | 0.214807 | 0.10789 | -0.14963 | -0.16502 | -0.14191 | -0.13222 | -0.11959 | -0.10908 | -0.11598 | 0.014617 |
| 18.22222 | -0.45958 | -0.43557 | -0.2569 | -0.36917 | -0.27941 | -1.39929 | -0.32178 | -0.71782 | -0.34285 | 0.15698 | 0.012594 | -0.10586 | -0.14142 | -0.15633 | -0.11052 | -0.09151 | -0.03555 | -0.10671 | 0.020346 |
| 18.23611 | -0.51077 | -0.46538 | -0.28444 | -0.48056 | -0.22295 | -1.35949 | -0.34307 | -1.008 | -0.39314 | 0.110061 | -0.08866 | -0.05887 | -0.1004 | -0.16715 | -0.09 | -0.04882 | 0.009334 | -0.03199 | 0.018883 |
| 18.25 | -0.66102 | -0.6068 | -0.41813 | -0.63503 | -0.14038 | -1.21176 | -0.50614 | -1.24317 | -0.53635 | 0.046106 | -0.17887 | -0.07209 | -0.10028 | -0.16087 | -0.09744 | -0.05985 | -0.02714 | 0.022642 | -0.02932 |
| 18.26389 | -0.77658 | -0.71395 | -0.5409 | -0.67011 | -0.0523 | -1.10134 | -0.61938 | -1.17162 | -0.65093 | -0.03793 | -0.21519 | -0.11039 | -0.12915 | -0.15449 | -0.11074 | -0.0983 | -0.0954 | 0.017367 | -0.08789 |
| 18.27778 | -0.80507 | -0.75367 | -0.59676 | -0.64498 | -0.07077 | -1.05507 | -0.64116 | -1.00211 | -0.67638 | -0.09991 | -0.19312 | -0.12082 | -0.13005 | -0.1378 | -0.10285 | -0.11165 | -0.13679 | 0.001585 | -0.11203 |
| 18.29167 | -0.81841 | -0.8055 | -0.60851 | -0.67985 | -0.11475 | -1.01986 | -0.64831 | -0.8856 | -0.68015 | -0.11151 | -0.16455 | -0.10003 | -0.1095 | -0.13207 | -0.07731 | -0.10446 | -0.15295 | -0.00413 | -0.1087 |
| 18.30556 | -0.8463 | -0.87045 | -0.61227 | -0.75338 | -0.25136 | -1.00473 | -0.64176 | -0.84398 | -0.68753 | -0.09959 | -0.13245 | -0.07099 | -0.0987 | -0.19523 | -0.07858 | -0.09927 | -0.16552 | -0.02924 | -0.11882 |
| 18.31944 | -0.80041 | -0.85327 | -0.57317 | -0.7216 | -0.31862 | -0.91292 | -0.58284 | -0.76386 | -0.63334 | -0.08546 | -0.09329 | -0.01511 | -0.09397 | -0.2913 | -0.10416 | -0.09265 | -0.1696 | -0.05423 | -0.11859 |
| 18.33333 | -0.63256 | -0.7103 | -0.45206 | -0.54258 | -0.37259 | -0.70958 | -0.37272 | -0.61217 | -0.48976 | -0.07322 | -0.03381 | 0.07648 | -0.0896 | -0.38766 | -0.12852 | -0.08516 | -0.17276 | -0.06131 | -0.09847 |
| 18.34722 | -0.43473 | -0.54234 | -0.28046 | -0.32995 | -0.47723 | -0.50768 | -0.15622 | -0.45142 | -0.31212 | -0.06657 | -0.00107 | 0.090255 | -0.08048 | -0.43429 | -0.19093 | -0.07185 | -0.18903 | -0.06313 | -0.0841 |
| 18.36111 | -0.2513 | -0.4257 | -0.1118 | -0.15358 | -0.6625 | -0.33669 | -0.01031 | -0.30963 | -0.14751 | -0.06648 | -0.06194 | 0.046649 | -0.05618 | -0.59368 | -0.28211 | -0.05436 | -0.21916 | -0.07854 | -0.07531 |
| 18.375 | -0.10479 | -0.39266 | 0.025774 | -0.01021 | -0.99953 | -0.1949 | 0.085753 | -0.17972 | -0.01598 | -0.06311 | -0.15591 | 0.012695 | -0.04812 | -0.32834 | -0.29248 | -0.05163 | -0.27065 | -0.11258 | -0.0623 |
| 18.38889 | -0.05418 | -0.48273 | 0.077156 | 0.037297 | -1.20807 | -0.17671 | 0.135281 | -0.0842 | 0.027759 | -0.05177 | -0.19799 | 0.008488 | -0.05136 | -0.57641 | -0.2135 | -0.05192 | -0.31565 | -0.12679 | -0.04773 |
| 18.40278 | -0.12839 | -0.70091 | -0.00059 | -0.05949 | -1.34856 | -0.28091 | 0.040963 | -0.09742 | -0.05219 | -0.04988 | -0.19786 | -0.00419 | -0.06392 | -0.6467 | -0.16847 | -0.05223 | -0.34437 | -0.11524 | -0.04332 |
| 18.41667 | -0.2588 | -0.86634 | -0.1288 | -0.21561 | -1.48935 | -0.4233 | -0.10736 | -0.18786 | -0.18888 | -0.05585 | -0.16603 | -0.02412 | -0.0761 | -0.43185 | -0.17002 | -0.0611 | -0.34832 | -0.10597 | -0.0474 |
| 18.43056 | -0.34373 | -0.87551 | -0.21281 | -0.32555 | -1.55477 | -0.51273 | -0.20493 | -0.26552 | -0.28926 | -0.05019 | -0.1233 | -0.04665 | -0.08378 | -0.33853 | -0.17593 | -0.06685 | -0.3297 | -0.11209 | -0.03997 |
| 18.44444 | -0.38315 | -0.7751 | -0.25791 | -0.39446 | -1.56566 | -0.57627 | -0.2403 | -0.30883 | -0.34804 | -0.04501 | -0.13565 | -0.07887 | -0.09655 | -0.28783 | -0.19284 | -0.07403 | -0.30625 | -0.1377 | -0.03574 |
| 18.45833 | -0.46966 | -0.72436 | -0.35355 | -0.49811 | -1.6544 | -0.66866 | -0.3476 | -0.38324 | -0.4387 | -0.05291 | -0.1696 | -0.11929 | -0.11256 | -0.1545 | -0.2294 | -0.08993 | -0.27748 | -0.17697 | -0.05346 |
| 18.47222 | -0.65628 | -0.84471 | -0.53838 | -0.68885 | -1.69978 | -0.83309 | -0.48085 | -0.51521 | -0.60719 | -0.03835 | -0.11855 | -0.15559 | -0.12816 | -0.01733 | -0.25024 | -0.10032 | -0.26 | -0.19794 | -0.03423 |
| 18.48611 | -1.00586 | -1.18984 | -0.82736 | -1.08127 | -1.80751 | -1.12026 | -0.4122 | -0.58667 | -0.85666 | 0.17655 | 0.249733 | -0.18481 | -0.1748 | -0.24124 | -0.23788 | -0.09825 | -0.28747 | -0.17854 | 0.264874 |
| 18.5 | -1.4255 | -1.67522 | -1.15231 | -1.45328 | -1.95841 | -1.47838 | -0.23466 | -0.58142 | -1.11662 | 0.6737 | 1.209881 | -0.20061 | -0.26057 | -0.79409 | -0.24463 | -0.10217 | -0.345 | -0.12634 | 1.021621 |
| 18.51389 | -1.44272 | -1.73987 | -1.22594 | -1.07011 | -1.83204 | -1.45007 | -0.3304 | -0.66888 | -1.09991 | 0.982794 | 3.934183 | -0.20596 | -0.28857 | -0.15433 | -0.25314 | -0.13918 | -0.35605 | -0.11508 | 1.348902 |
| 18.52778 | -0.93271 | -1.20817 | -0.90706 | 0.028367 | -1.29005 | -0.9461 | -0.48645 | -0.57961 | -0.74806 | 0.642552 | 3.567908 | -0.19101 | -0.2081 | -0.17433 | -0.1848 | -0.16284 | -0.28777 | -0.1262 | 0.755819 |
| 18.54167 | -0.345 | -0.54478 | -0.39385 | 0.631093 | -0.75269 | -0.38113 | -0.33325 | -0.35036 | -0.29352 | 0.268074 | 1.971922 | -0.11447 | -0.07974 | -0.09463 | -0.06854 | -0.10312 | -0.17487 | -0.09045 | 0.365777 |
| 18.55556 | -0.00336 | -0.15649 | -0.06099 | 0.545835 | -0.4246 | -0.05776 | -0.11005 | -0.09565 | -0.02739 | 0.080199 | 1.101988 | -0.07046 | -0.01028 | -0.0285 | -0.00783 | -0.05711 | -0.10986 | -0.0724 | 0.187332 |
| 18.56944 | 0.11948 | -0.01544 | 0.01425 | 0.323366 | -0.30057 | 0.042018 | 0.024472 | 0.045501 | 0.050723 | 0.043845 | 0.554901 | -0.08751 | -0.01619 | 0.019907 | 0.00167 | -0.07053 | -0.10133 | -0.10922 | 0.087172 |
| 18.58333 | 0.169435 | 0.044972 | 0.021301 | 0.221573 | -0.21645 | 0.074608 | 0.103459 | 0.111091 | 0.05976 | 0.104376 | 0.362307 | -0.08317 | -0.02604 | 0.055441 | 0.005616 | -0.06652 | -0.09094 | -0.11907 | 0.022939 |
| 18.59722 | 0.16233 | 0.027467 | -0.00366 | 0.100926 | -0.21115 | 0.038957 | 0.095838 | 0.082456 | 0.008346 | 0.18693 | 0.312255 | -0.07182 | -0.04042 | 0.028979 | -0.00813 | -0.0523 | -0.06271 | -0.11635 | 0.006812 |
| 18.61111 | 0.158704 | -0.00367 | -0.03175 | 0.005068 | -0.26204 | -0.04965 | 0.040897 | 0.010025 | -0.0571 | 0.232812 | 0.234664 | -0.06926 | -0.05549 | 0.014229 | -0.01824 | -0.04769 | -0.03118 | -0.1327 | -0.00855 |
| 18.625 | 0.148763 | -0.02354 | -0.06449 | -0.0758 | -0.30771 | -0.14349 | -0.05289 | -0.0565 | -0.11532 | 0.231931 | 0.064112 | -0.10212 | -0.08277 | 0.099232 | -0.03455 | -0.06598 | -0.06267 | -0.15775 | -0.0309 |
| 18.63889 | 0.097338 | -0.08843 | -0.14037 | -0.18716 | -0.36262 | -0.20486 | -0.18395 | -0.12347 | -0.19086 | 0.160068 | -0.06152 | -0.15465 | -0.09918 | 0.449852 | -0.05256 | -0.08607 | -0.12437 | -0.1934 | -0.0553 |
| 18.65278 | 0.075596 | -0.10376 | -0.16588 | -0.22557 | -0.36196 | -0.18987 | -0.26669 | -0.15044 | -0.20329 | 0.040065 | -0.04854 | -0.14595 | -0.07611 | 0.501742 | -0.05932 | -0.0577 | -0.12125 | -0.1896 | -0.03169 |
| 18.66667 | 0.140047 | -0.07851 | -0.17722 | -0.21681 | -0.34161 | -0.17947 | -0.29119 | -0.17662 | -0.19628 | -0.09977 | -0.03156 | -0.12553 | -0.0713 | 0.223984 | -0.07462 | -0.05319 | -0.10215 | -0.17288 | -0.01382 |
| 18.68056 | 0.032109 | -0.17011 | -0.25584 | -0.27521 | -0.41016 | -0.25887 | -0.35402 | -0.26068 | -0.27794 | -0.22178 | -0.06178 | -0.12117 | -0.07826 | 0.004981 | -0.0712 | -0.0809 | -0.09171 | -0.17238 | -0.02298 |
| 18.69444 | -0.20174 | -0.37798 | -0.40406 | -0.4503 | -0.59301 | -0.45111 | -0.51948 | -0.42463 | -0.46454 | -0.27582 | -0.08393 | -0.11374 | -0.0751 | -0.03397 | -0.05485 | -0.0908 | -0.0852 | -0.17424 | -0.02285 |
| 18.70833 | -0.36584 | -0.53968 | -0.55078 | -0.64619 | -0.78311 | -0.64215 | -0.69754 | -0.59276 | -0.65025 | -0.28707 | -0.08418 | -0.11607 | -0.08075 | -0.04709 | -0.05513 | -0.0977 | -0.10538 | -0.17396 | -0.02042 |
| 18.72222 | -0.24265 | -0.4485 | -0.51926 | -0.62393 | -0.7422 | -0.59929 | -0.66772 | -0.55475 | -0.60551 | -0.22276 | -0.06328 | -0.09513 | -0.08354 | -0.08688 | -0.04004 | -0.07666 | -0.11362 | -0.14468 | -0.01102 |
| 18.73611 | -0.06213 | -0.29124 | -0.45717 | -0.52329 | -0.62493 | -0.50703 | -0.57319 | -0.46789 | -0.49074 | -0.13071 | -0.06547 | -0.06867 | -0.09159 | -0.15677 | -0.01246 | -0.04873 | -0.11111 | -0.11295 | 0.003145 |
| 18.75 | -0.09372 | -0.32121 | -0.55449 | -0.57029 | -0.67457 | -0.56283 | -0.62725 | -0.53249 | -0.5456 | -0.07235 | -0.07995 | -0.06928 | -0.08648 | -0.15405 | -0.00016 | -0.05237 | -0.11355 | -0.12097 | 0.008991 |
| 18.76389 | -0.1382 | -0.36928 | -0.62456 | -0.61917 | -0.71476 | -0.57976 | -0.66479 | -0.58434 | -0.60277 | -0.07203 | -0.09673 | -0.09107 | -0.07576 | -0.12347 | -0.01546 | -0.07872 | -0.13039 | -0.14645 | -0.00012 |
| 18.77778 | -0.12157 | -0.33901 | -0.59645 | -0.59274 | -0.66766 | -0.52144 | -0.62321 | -0.55454 | -0.58398 | -0.10081 | -0.14403 | -0.12423 | -0.07314 | -0.13898 | -0.03377 | -0.11274 | -0.1457 | -0.16957 | -0.03253 |
| 18.79167 | -0.05797 | -0.29078 | -0.51834 | -0.53697 | -0.60577 | -0.45818 | -0.57349 | -0.48869 | -0.51954 | -0.08868 | -0.14434 | -0.11001 | -0.05286 | -0.16205 | -0.02913 | -0.09254 | -0.11453 | -0.14721 | -0.04547 |
| 18.80556 | -0.02163 | -0.24982 | -0.45187 | -0.48103 | -0.53847 | -0.42061 | -0.53114 | -0.43339 | -0.45172 | -0.05768 | -0.1078 | -0.07655 | -0.02827 | -0.16341 | -0.01185 | -0.05037 | -0.07792 | -0.10782 | -0.02663 |
| 18.81944 | -0.04971 | -0.26269 | -0.45946 | -0.44744 | -0.51738 | -0.41922 | -0.49835 | -0.41272 | -0.43714 | -0.05544 | -0.08806 | -0.07198 | -0.00487 | -0.13338 | 0.015235 | -0.03933 | -0.06313 | -0.0999 | -0.01555 |
| 18.83333 | -0.11409 | -0.29348 | -0.51484 | -0.45322 | -0.54607 | -0.43081 | -0.50707 | -0.43368 | -0.46542 | -0.06414 | -0.09432 | -0.08261 | -0.00118 | -0.13222 | 0.010025 | -0.05063 | -0.07621 | -0.10804 | -0.04802 |
| 18.84722 | -0.15814 | -0.31865 | -0.56034 | -0.45728 | -0.55571 | -0.42089 | -0.55103 | -0.44983 | -0.48539 | -0.06939 | -0.13451 | -0.09484 | -0.01758 | -0.16689 | -0.02113 | -0.06779 | -0.10667 | -0.11879 | -0.10344 |
| 18.86111 | -0.10887 | -0.27373 | -0.52738 | -0.39211 | -0.46803 | -0.3591 | -0.54964 | -0.37897 | -0.43541 | -0.01716 | -0.12875 | -0.05821 | 0.002753 | -0.13582 | -0.00232 | -0.03075 | -0.0859 | -0.07725 | -0.09348 |
| 18.875 | -0.057 | -0.22106 | -0.48765 | -0.33811 | -0.39679 | -0.30818 | -0.5313 | -0.3154 | -0.3926 | 0.040612 | -0.10915 | -0.01823 | 0.034398 | -0.08035 | 0.024826 | 0.011893 | -0.05143 | -0.03071 | -0.06306 |
| 18.88889 | -0.03717 | -0.1903 | -0.46311 | -0.31604 | -0.37739 | -0.28552 | -0.53118 | -0.28913 | -0.39176 | 0.030812 | -0.14374 | -0.0379 | 0.029988 | -0.07066 | 0.012572 | -0.00585 | -0.05595 | -0.05338 | -0.09168 |
| 18.90278 | 0.022014 | -0.13199 | -0.41775 | -0.27707 | -0.33276 | -0.23838 | -0.52957 | -0.24983 | -0.36292 | 0.005796 | -0.19944 | -0.06943 | 0.00768 | -0.12346 | -0.00452 | -0.0332 | -0.07061 | -0.08192 | -0.14249 |
| 18.91667 | 0.113485 | -0.0795 | -0.36622 | -0.24048 | -0.28112 | -0.20034 | -0.50716 | -0.2127 | -0.32592 | -0.01836 | -0.26818 | -0.08147 | -0.0153 | -0.25784 | -0.00897 | -0.04458 | -0.06534 | -0.08739 | -0.1695 |
| 18.93056 | 0.211626 | -0.00594 | -0.27953 | -0.17539 | -0.20594 | -0.1455 | -0.44451 | -0.15687 | -0.2604 | -0.04403 | -0.32083 | -0.06758 | -0.02768 | -0.49243 | -0.00246 | -0.0325 | -0.04831 | -0.07153 | -0.16366 |
| 18.94444 | 0.265533 | 0.050821 | -0.19539 | -0.10711 | -0.12136 | -0.06373 | -0.35216 | -0.09805 | -0.19446 | -0.05707 | -0.3737 | -0.04706 | -0.03665 | -0.46344 | -0.00148 | -0.01588 | -0.04847 | -0.0518 | -0.14883 |
| 18.95833 | 0.259156 | 0.058638 | -0.16226 | -0.06874 | -0.07211 | 0.011902 | -0.30896 | -0.05482 | -0.16542 | -0.0968 | -0.34635 | -0.03595 | -0.018 | -0.18405 | 0.006504 | -0.00031 | -0.04491 | -0.05709 | -0.1294 |
| 18.97222 | 0.250736 | 0.08331 | -0.1424 | -0.06036 | -0.00204 | 0.087334 | -0.32892 | -0.01206 | -0.1367 | -0.14222 | -0.33196 | -0.03497 | 0.013961 | -0.10129 | 0.021448 | 0.011032 | -0.03049 | -0.07737 | -0.1452 |
| 18.98611 | 0.392572 | 0.3124 | -0.07884 | -0.10744 | 0.274263 | 0.33505 | -0.56982 | 0.126899 | 0.033004 | -0.24809 | -0.68723 | -0.02397 | 0.060257 | -0.05063 | 0.043836 | 0.016056 | 0.006741 | -0.13215 | -0.47434 |
| 19 | 0.502171 | 0.695654 | 0.012951 | -0.29765 | 0.760329 | 0.743491 | -1.67203 | 0.349166 | 0.387644 | -0.53712 | -1.49896 | 0.056235 | 0.140351 | 0.062439 | 0.086309 | 0.084975 | 0.074332 | -0.2004 | -1.33048 |
| 19.01389 | -0.00463 | 0.565596 | 0.190845 | -0.6764 | 0.829387 | 0.71198 | -2.7098 | 0.339384 | 0.557476 | -1.04889 | -3.09687 | 0.121564 | 0.112833 | 0.110978 | 0.057655 | 0.179663 | 0.082223 | -0.27576 | -1.77952 |
| 19.02778 | -0.84735 | -0.10119 | 0.425951 | -0.64263 | 0.36932 | 0.194903 | -0.98533 | 0.166441 | 0.354492 | -1.21358 | -3.04554 | 0.085664 | 0.01568 | -0.00813 | -0.08619 | 0.15377 | -0.00557 | -0.3888 | -1.14117 |
| 19.04167 | -1.01108 | -0.45656 | 0.589008 | -0.12009 | 0.015229 | -0.18277 | 0.979658 | 0.129612 | 0.159455 | -0.58986 | -1.45607 | 0.029487 | -0.02462 | -0.10133 | -0.20497 | 0.052548 | -0.0845 | -0.45191 | -0.5281 |
| 19.05556 | -0.77048 | -0.49633 | 0.488965 | 0.123146 | -0.08924 | -0.2149 | 1.827609 | 0.109066 | 0.090416 | -0.0939 | -0.67428 | -0.0048 | -0.02527 | -0.13213 | -0.20784 | -0.00703 | -0.11518 | -0.34079 | -0.29252 |
| 19.06944 | -0.57712 | -0.43418 | 0.280815 | 0.073311 | -0.11445 | -0.18518 | 1.425666 | 0.047118 | 0.04739 | 0.079693 | -0.33078 | -0.03386 | -0.04286 | -0.15336 | -0.15805 | -0.02773 | -0.13275 | -0.24309 | -0.23332 |
| 19.08333 | -0.38689 | -0.33485 | 0.112278 | -0.03524 | -0.1417 | -0.14359 | 0.67937 | -0.00735 | 0.01685 | 0.118998 | -0.18398 | -0.05489 | -0.06174 | -0.17097 | -0.13649 | -0.03889 | -0.13376 | -0.15572 | -0.19673 |
| 19.09722 | -0.26881 | -0.27199 | 0.026835 | -0.0716 | -0.15883 | -0.10741 | 0.250602 | -0.03291 | 0.005148 | 0.128671 | -0.10111 | -0.06865 | -0.05199 | -0.13354 | -0.12344 | -0.04338 | -0.11436 | -0.13217 | -0.14892 |
| 19.11111 | -0.20879 | -0.23561 | 0.012815 | -0.04073 | -0.14855 | -0.05114 | 0.140064 | -0.0273 | 0.014066 | 0.136778 | -0.0518 | -0.06919 | -0.03196 | -0.11052 | -0.08276 | -0.03593 | -0.09178 | -0.10844 | -0.09904 |
| 19.125 | -0.15147 | -0.17773 | 0.049308 | 0.021446 | -0.10113 | 0.042398 | 0.177839 | 0.01979 | 0.065088 | 0.131281 | -0.01226 | -0.05592 | -0.01846 | -0.09523 | -0.04555 | -0.02494 | -0.07903 | -0.08907 | -0.07658 |
| 19.13889 | -0.07047 | -0.0943 | 0.09931 | 0.091117 | -0.03134 | 0.130353 | 0.216914 | 0.072588 | 0.134994 | 0.11172 | 0.007861 | -0.05714 | -0.02291 | -0.08157 | -0.04234 | -0.02392 | -0.08502 | -0.08866 | -0.08324 |
| 19.15278 | -0.0376 | -0.08571 | 0.092844 | 0.099114 | -0.01632 | 0.146002 | 0.18134 | 0.052387 | 0.134449 | 0.085813 | 0.005549 | -0.06958 | -0.04173 | -0.08177 | -0.05346 | -0.03193 | -0.1022 | -0.10281 | -0.09155 |

| | | | | | | | | | | | | | | | | | | | |
|---|---|---|---|---|---|---|---|---|---|---|---|---|---|---|---|---|---|---|---|
| 19.16667 | -0.06403 | -0.1226 | 0.041976 | 0.036958 | -0.06246 | 0.111393 | 0.114181 | -0.01245 | 0.081299 | 0.079125 | 0.02061 | -0.08014 | -0.03681 | -0.07992 | -0.0472 | -0.03436 | -0.09584 | -0.11033 | -0.08907 |
| 19.18056 | -0.06602 | -0.10503 | 0.028679 | -0.0003 | -0.0873 | 0.113895 | 0.081575 | -0.03174 | 0.063057 | 0.083982 | 0.068733 | -0.09006 | -0.01803 | -0.06849 | -0.03454 | -0.03447 | -0.07034 | -0.10787 | -0.09141 |
| 19.19444 | -0.00406 | -0.03167 | 0.07655 | 0.053647 | -0.04462 | 0.157265 | 0.118411 | 0.012992 | 0.113128 | 0.067121 | 0.076664 | -0.05417 | -0.03198 | -0.06957 | -0.03061 | -0.03931 | -0.06923 | -0.09838 | -0.09816 |
| 19.20833 | 0.113348 | 0.070516 | 0.174266 | 0.15971 | 0.04913 | 0.235112 | 0.216807 | 0.112916 | 0.212172 | 0.042023 | 0.022202 | -0.02838 | -0.05306 | -0.09259 | -0.02872 | -0.0308 | -0.08147 | -0.08492 | -0.08499 |
| 19.22222 | 0.220317 | 0.160937 | 0.271716 | 0.229017 | 0.121022 | 0.317618 | 0.299509 | 0.20356 | 0.292928 | 0.06904 | 0.027141 | 0.011784 | -0.02224 | -0.14946 | -0.01212 | 0.025232 | -0.05873 | -0.03593 | -0.04216 |
| 19.23611 | 0.19155 | 0.126538 | 0.244798 | 0.171101 | 0.082531 | 0.298341 | 0.265722 | 0.163711 | 0.244841 | 0.088531 | 0.05526 | 0.047423 | -0.00369 | -0.3286 | -0.02453 | 0.063973 | -0.06411 | 0.002534 | -0.02959 |
| 19.25 | -0.00568 | -0.06083 | 0.044118 | -0.01802 | -0.08283 | 0.142562 | 0.108664 | -0.02089 | 0.048755 | -0.00413 | -0.00213 | -0.03744 | -0.07305 | -0.4571 | -0.10079 | -0.02859 | -0.13979 | -0.07958 | -0.10326 |
| 19.26389 | -0.16841 | -0.20078 | -0.12604 | -0.14114 | -0.21015 | 0.003932 | -0.00438 | -0.15067 | -0.10464 | -0.09939 | -0.0211 | -0.11987 | -0.10788 | -0.37206 | -0.11863 | -0.12553 | -0.15542 | -0.16021 | -0.14861 |
| 19.27778 | -0.1387 | -0.16532 | -0.10118 | -0.11261 | -0.15621 | 0.012793 | 0.00516 | -0.10618 | -0.08384 | -0.08847 | 0.042315 | -0.0882 | -0.04167 | -0.20961 | -0.04674 | -0.09325 | -0.08163 | -0.13038 | -0.10021 |
| 19.29167 | -0.04749 | -0.08654 | -0.00321 | -0.07766 | -0.03325 | 0.076709 | 0.049459 | -0.01102 | 0.002699 | -0.05268 | 0.048331 | -0.03715 | 0.005798 | -0.15372 | -0.00352 | -0.03262 | -0.04235 | -0.07321 | -0.05589 |
| 19.30556 | -0.02066 | -0.06229 | 0.03697 | -0.10445 | -0.00711 | 0.096007 | 0.077686 | 0.01992 | 0.036218 | -0.02628 | 0.034752 | -0.01244 | 0.010067 | -0.20027 | -0.01905 | 0.003838 | -0.05297 | -0.04211 | -0.04301 |
| 19.31944 | -0.07211 | -0.11077 | -0.00528 | -0.17789 | -0.07996 | 0.045731 | 0.058724 | -0.02802 | -0.01377 | -0.00734 | 0.073161 | -0.00925 | 0.012594 | -0.1868 | -0.05597 | 0.019258 | -0.07327 | -0.03322 | -0.04487 |
| 19.33333 | -0.20203 | -0.24397 | -0.12488 | -0.32016 | -0.18725 | -0.05494 | -0.02003 | -0.13136 | -0.12717 | -0.0133 | 0.119076 | -0.02896 | 0.023406 | -0.04096 | -0.06366 | -0.0078 | -0.0804 | -0.06355 | -0.05056 |
| 19.34722 | -0.35194 | -0.39528 | -0.26104 | -0.55321 | -0.23843 | -0.09089 | -0.09685 | -0.23162 | -0.23966 | -0.00547 | 0.170923 | -0.03164 | 0.039701 | 0.019683 | -0.0202 | -0.03287 | -0.06054 | -0.09708 | -0.03749 |
| 19.36111 | -0.39942 | -0.4533 | -0.29645 | -0.70616 | -0.16279 | 0.015105 | -0.06656 | -0.24504 | -0.25159 | 0.045546 | 0.25869 | -0.00222 | 0.033214 | -0.08402 | 0.030249 | -0.02123 | -0.05615 | -0.09487 | -0.02365 |
| 19.375 | -0.18289 | -0.24855 | -0.06454 | -0.52444 | 0.127321 | 0.297728 | 0.184299 | -0.01019 | -0.02283 | 0.130785 | 0.397674 | 0.051552 | 0.012232 | -0.09647 | 0.063955 | 0.02832 | -0.03356 | -0.04056 | 0.016352 |
| 19.38889 | 0.260224 | 0.197852 | 0.386382 | -0.03325 | 0.560763 | 0.703866 | 0.60489 | 0.428224 | 0.427215 | 0.213864 | 0.659173 | 0.096877 | 0.03447 | -0.05003 | 0.073235 | 0.082902 | 0.011937 | 0.033091 | 0.088656 |
| 19.40278 | 0.605611 | 0.552504 | 0.717714 | 0.38703 | 0.866528 | 0.983404 | 0.915986 | 0.767606 | 0.751357 | 0.206807 | 1.043407 | 0.06266 | 0.04307 | -0.04441 | 0.042474 | 0.054516 | 0.005475 | 0.032269 | 0.101648 |
| 19.41667 | 0.76934 | 0.724289 | 0.853156 | 0.636877 | 0.958876 | 1.091256 | 1.027731 | 0.92258 | 0.897029 | 0.145277 | 1.392875 | -0.00436 | 0.022972 | -0.15338 | 0.006342 | -0.01249 | -0.02967 | -0.01827 | 0.068604 |
| 19.43056 | 0.843225 | 0.808757 | 0.902014 | 0.742909 | 0.912216 | 1.113134 | 1.025071 | 0.972281 | 0.940476 | 0.143432 | 1.350462 | -0.00394 | 0.035382 | -0.44614 | 0.01182 | -0.0093 | -0.01445 | -0.02249 | 0.065882 |
| 19.44444 | 0.77835 | 0.74848 | 0.841716 | 0.668836 | 0.75471 | 1.003389 | 0.896636 | 0.868987 | 0.837233 | 0.173432 | 1.293016 | 0.042008 | 0.064773 | -0.76433 | 0.041303 | 0.036127 | 0.026646 | 0.011345 | 0.092785 |
| 19.45833 | 0.612276 | 0.581755 | 0.699404 | 0.510811 | 0.557215 | 0.817097 | 0.724248 | 0.677272 | 0.653549 | 0.182956 | 1.197626 | 0.073135 | 0.065106 | -0.16323 | 0.064765 | 0.064984 | 0.046243 | 0.040245 | 0.125078 |
| 19.47222 | 0.43929 | 0.406565 | 0.539315 | 0.347386 | 0.365063 | 0.607616 | 0.569611 | 0.501896 | 0.480467 | 0.17965 | 1.011704 | 0.056842 | 0.01924 | 0.141245 | 0.045674 | 0.057509 | 0.024605 | 0.043434 | 0.140525 |
| 19.48611 | 0.239129 | 0.184114 | 0.323746 | 0.113787 | 0.109985 | 0.392541 | 0.425387 | 0.328024 | 0.28235 | 0.288814 | 1.081541 | -0.00733 | -0.06286 | 0.098871 | -0.01664 | 0.02815 | -0.03594 | 0.065119 | 0.179207 |
| 19.5 | -0.0009 | -0.14345 | -0.02005 | -0.20016 | -0.24154 | 0.154819 | 0.190247 | 0.060501 | -0.03391 | 0.527136 | 1.56843 | -0.08566 | -0.12645 | -0.06356 | -0.08171 | -0.00468 | -0.12709 | 0.122032 | 0.340675 |
| 19.51389 | -0.04271 | -0.30684 | -0.33905 | -0.32091 | -0.46678 | 0.068258 | -0.07198 | -0.21139 | -0.3269 | 0.59089 | 1.745268 | -0.11192 | -0.12147 | -0.19863 | -0.09787 | -0.02793 | -0.1847 | 0.12639 | 0.701286 |
| 19.52778 | 0.25761 | -0.08171 | -0.31213 | -0.07069 | -0.32168 | 0.531118 | -0.03158 | -0.18985 | -0.28537 | 0.381444 | 1.715542 | -0.08574 | -0.09763 | -0.23255 | -0.04184 | -0.03345 | -0.14367 | 0.071808 | 1.13598 |
| 19.54167 | 0.639425 | 0.297608 | -0.05875 | 0.251427 | -0.00872 | 1.539607 | 0.230739 | 0.088394 | -0.04468 | 0.17432 | 1.448874 | -0.04947 | -0.05918 | -0.20898 | 0.040664 | -0.03693 | -0.05969 | 0.01753 | 1.164467 |
| 19.55556 | 0.849341 | 0.523941 | 0.140159 | 0.420379 | 0.21006 | 2.089206 | 0.389583 | 0.330723 | 0.136261 | 0.091906 | 0.906526 | -0.01642 | -0.00613 | -0.1848 | 0.113704 | -0.0291 | 0.01086 | -0.00353 | 0.855835 |
| 19.56944 | 0.994014 | 0.658643 | 0.347074 | 0.46817 | 0.373842 | 1.705497 | 0.602935 | 0.496313 | 0.280149 | 0.093418 | 0.64521 | 0.020408 | 0.023421 | -0.02201 | 0.123721 | -0.00053 | 0.039204 | 0.013781 | 0.709064 |
| 19.58333 | 1.061109 | 0.669198 | 0.5365 | 0.43842 | 0.454682 | 1.182557 | 0.960503 | 0.588153 | 0.368031 | 0.094215 | 0.535112 | 0.026172 | 0.018841 | 0.094927 | 0.091058 | 0.015422 | 0.030408 | 0.016402 | 0.663494 |
| 19.59722 | 0.997398 | 0.550153 | 0.585866 | 0.375082 | 0.400537 | 0.741838 | 1.27931 | 0.599281 | 0.368504 | 0.079214 | 0.439736 | 0.014991 | 0.056605 | 0.02387 | 0.085294 | 0.020226 | 0.034702 | -0.00337 | 0.602136 |
| 19.61111 | 0.97771 | 0.515193 | 0.590077 | 0.385949 | 0.410385 | 0.559961 | 1.495238 | 0.685778 | 0.435694 | 0.105705 | 0.408751 | 0.04211 | 0.113004 | -0.1012 | 0.114275 | 0.055881 | 0.067639 | 0.027933 | 0.545725 |
| 19.625 | 0.943753 | 0.51065 | 0.575157 | 0.420019 | 0.474994 | 0.602486 | 1.46793 | 0.737005 | 0.505659 | 0.117197 | 0.380959 | 0.053294 | 0.092342 | -0.09359 | 0.121994 | 0.050904 | 0.095033 | 0.048708 | 0.494732 |
| 19.63889 | 0.832832 | 0.429099 | 0.424336 | 0.358068 | 0.423173 | 0.560413 | 1.322349 | 0.638282 | 0.425979 | 0.070968 | 0.180713 | 0.040317 | 0.054565 | 0.055933 | 0.112017 | -0.00146 | 0.10667 | 0.020632 | 0.439142 |
| 19.65278 | 0.703271 | 0.321955 | 0.162066 | 0.247216 | 0.294494 | 0.393887 | 0.998653 | 0.436386 | 0.255665 | 0.051745 | -0.08213 | 0.059052 | 0.058148 | 0.146497 | 0.096909 | 0.011062 | 0.107321 | 0.03084 | 0.401742 |
| 19.66667 | 0.544897 | 0.239279 | -0.05499 | 0.15505 | 0.168994 | 0.186504 | 0.594101 | 0.242962 | 0.099115 | 0.046023 | -0.2618 | 0.049183 | 0.041713 | 0.07276 | 0.055492 | 0.055832 | 0.086637 | 0.03378 | 0.366757 |
| 19.68056 | 0.430457 | 0.227323 | -0.15082 | 0.141752 | 0.113456 | 0.038667 | 0.295244 | 0.170085 | 0.019263 | 0.050123 | -0.23067 | 0.029228 | 0.019602 | -0.00251 | 0.042042 | 0.089116 | 0.098455 | 0.011303 | 0.351645 |
| 19.69444 | 0.442147 | 0.292432 | -0.1178 | 0.220222 | 0.191528 | 0.024808 | 0.122846 | 0.220261 | 0.064933 | 0.087381 | -0.09772 | 0.060193 | 0.010204 | -0.01188 | 0.05133 | 0.129953 | 0.122206 | 0.021384 | 0.327928 |
| 19.70833 | 0.526579 | 0.360072 | -0.04138 | 0.317224 | 0.323506 | 0.083582 | 0.006605 | 0.290846 | 0.175978 | 0.114046 | 0.097827 | 0.067352 | -0.00265 | 0.031776 | 0.069514 | 0.119837 | 0.125219 | 0.024985 | 0.255637 |
| 19.72222 | 0.481143 | 0.326917 | 0.033306 | 0.334224 | 0.322009 | 0.040524 | -0.1964 | 0.288238 | 0.207966 | 0.102694 | 0.230769 | 0.030968 | 0.025667 | 0.105294 | 0.105335 | 0.103309 | 0.173613 | 0.02539 | 0.1954 |
| 19.73611 | 0.265583 | 0.176964 | -0.00582 | 0.216598 | 0.140554 | -0.08448 | -0.3987 | 0.141298 | 0.091601 | 0.057573 | 0.206638 | -0.00233 | 0.039022 | 0.098461 | 0.09637 | 0.086962 | 0.233535 | 0.03301 | 0.144329 |
| 19.75 | 0.062165 | 0.048673 | -0.04757 | 0.228369 | -0.01587 | -0.08004 | -0.44565 | -0.03825 | -0.02038 | -0.00055 | 0.097463 | -0.00354 | 0.011499 | 0.023795 | 0.050192 | 0.066692 | 0.250959 | 0.01518 | 0.097235 |
| 19.76389 | 0.062199 | 0.111547 | 0.03518 | 0.788335 | 0.055292 | 0.159513 | -0.26981 | 0.004493 | 0.058518 | -0.01897 | 0.0043 | 0.042674 | 0.035283 | -0.00163 | 0.054281 | 0.088504 | 0.263096 | 0.028551 | 0.103652 |
| 19.77778 | 0.121544 | 0.216643 | 0.090209 | 1.562426 | 0.186808 | 0.331264 | -0.12395 | 0.130346 | 0.148239 | -0.0396 | -0.07001 | 0.059495 | 0.065925 | -0.00987 | 0.060602 | 0.095 | 0.30474 | 0.055476 | 0.108611 |
| 19.79167 | 0.064714 | 0.132549 | -0.05201 | 1.451657 | 0.150156 | 0.205197 | -0.19836 | 0.074029 | 0.060897 | -0.01804 | -0.14777 | 0.01984 | 0.034805 | -0.01676 | 0.030618 | 0.045949 | 0.366048 | 0.028154 | 0.069293 |
| 19.80556 | -0.04249 | -0.01782 | -0.21486 | 0.691346 | 0.045215 | 0.0014 | -0.34185 | -0.09062 | -0.08915 | 0.102199 | -0.21944 | 0.001485 | -0.00464 | 0.102618 | 0.003498 | 0.016408 | 0.413068 | 0.012105 | 0.031449 |
| 19.81944 | -0.12094 | -0.07033 | -0.24157 | 0.238604 | -0.00236 | -0.12057 | -0.55221 | -0.20289 | -0.19064 | 0.177779 | -0.33521 | 0.004185 | -0.01404 | 0.158695 | 0.000832 | 0.01483 | 0.417068 | 0.019072 | 0.005005 |
| 19.83333 | -0.12514 | 0.008236 | -0.19005 | 0.155149 | 0.021008 | -0.16605 | -0.71357 | -0.183 | -0.17814 | 0.103979 | -0.40596 | 0.022974 | 0.015192 | 0.134803 | 0.024388 | 0.030318 | 0.516102 | 0.029456 | 0.002477 |
| 19.84722 | -0.09936 | 0.087152 | -0.16435 | 0.120231 | 0.040388 | -0.15942 | -0.80708 | -0.12401 | -0.11736 | -0.04962 | -0.38614 | 0.040323 | 0.03264 | 0.11821 | 0.04791 | 0.042618 | 0.763685 | 0.043229 | 0.017279 |
| 19.86111 | -0.06278 | 0.132653 | -0.16424 | 0.067852 | 0.066388 | -0.05609 | -0.71301 | -0.06492 | -0.06616 | -0.17129 | -0.23871 | 0.019941 | 0.008481 | 0.124357 | 0.030306 | 0.024634 | 0.971734 | 0.030326 | 0.000789 |
| 19.875 | 0.073546 | 0.25174 | -0.06023 | 0.169262 | 0.203488 | 0.187029 | -0.49867 | 0.074331 | 0.074322 | -0.21725 | -0.08963 | 0.008507 | -0.00324 | 0.147105 | -0.02291 | 0.015751 | 0.947754 | 0.032207 | -0.03898 |
| 19.88889 | 0.228383 | 0.360896 | 0.048208 | 0.37179 | 0.352894 | 0.372209 | -0.35271 | 0.324029 | 0.210591 | -0.25537 | -0.15007 | 0.009057 | -0.00199 | 0.154302 | -0.09043 | 0.005059 | 0.954028 | 0.039597 | -0.09972 |
| 19.90278 | 0.27788 | 0.396761 | 0.086184 | 0.608244 | 0.423504 | 0.319722 | -0.37357 | 0.70421 | 0.264607 | -0.25236 | -0.415 | 0.007053 | -0.01281 | 0.150984 | -0.12327 | -0.01089 | 0.946473 | 0.019503 | -0.12923 |
| 19.91667 | 0.18275 | 0.374955 | 0.102159 | 0.927769 | 0.453044 | 0.129694 | -0.59283 | 0.981338 | 0.266195 | -0.19345 | -0.7032 | -0.00754 | -0.04036 | 0.114583 | -0.11909 | -0.03427 | 0.904173 | -0.00222 | -0.13481 |
| 19.93056 | -0.04243 | 0.257474 | 0.096214 | 1.087491 | 0.410535 | -0.07431 | -0.78089 | 0.817294 | 0.217912 | -0.17317 | -0.89252 | -0.03421 | -0.0501 | 0.053799 | -0.09716 | -0.06057 | 0.842318 | -0.01549 | -0.16002 |
| 19.94444 | -0.34141 | 0.045726 | 0.009304 | 0.754034 | 0.277536 | -0.26224 | -0.93862 | 0.48366 | 0.093888 | -0.23743 | -1.0775 | -0.02179 | -0.01236 | 0.047861 | -0.04774 | -0.0395 | 0.913507 | 0.000106 | -0.20681 |
| 19.95833 | -0.60645 | -0.15214 | -0.1294 | 0.404934 | 0.108018 | -0.4157 | -0.97992 | 0.16372 | -0.05935 | -0.34099 | -1.14644 | 0.000342 | 0.005476 | 0.09473 | -0.02329 | -0.02043 | 1.242595 | 0.007899 | -0.2912 |
| 19.97222 | -0.57765 | -0.13399 | -0.09883 | 0.331974 | 0.117315 | -0.325 | -0.70307 | 0.108527 | -0.02253 | -0.40497 | -1.07721 | 0.012111 | 0.007029 | 0.151996 | -0.02616 | -0.02478 | 1.562518 | -0.00577 | -0.38461 |
| 19.98611 | -0.08443 | 0.274522 | 0.286059 | 0.429176 | 0.492199 | 0.106868 | -0.03514 | 0.36563 | 0.372789 | -0.43307 | -0.97113 | 0.038177 | 0.043947 | 0.272657 | -0.02138 | -0.04042 | 1.053811 | -0.03609 | -0.43256 |
| 20 | 0.535604 | 0.943924 | 0.997806 | 0.56228 | 1.147367 | 0.74406 | 0.847783 | 0.826011 | 1.068178 | -0.55542 | -1.0113 | 0.088658 | 0.123007 | 0.413677 | 0.002568 | -0.04281 | 0.054662 | -0.09688 | -0.47978 |
| 20.01389 | 0.712157 | 1.300856 | 1.492397 | 0.454274 | 1.490426 | 1.124365 | 1.404994 | 0.947112 | 1.515951 | -0.73003 | -1.12601 | 0.137413 | 0.160538 | 0.276616 | 0.004358 | 0.011975 | -1.07489 | -0.10039 | -0.56045 |
| 20.02778 | 0.359975 | 1.030767 | 1.28889 | 0.209716 | 1.183985 | 0.965811 | 1.224369 | 0.528722 | 1.276119 | -0.61445 | -0.77059 | 0.115516 | 0.080323 | -0.09177 | -0.06112 | 0.037543 | -1.82966 | -0.04415 | -0.61643 |
| 20.04167 | 0.090383 | 0.600515 | 0.83312 | 0.282383 | 0.680885 | 0.671328 | 0.769629 | 0.179299 | 0.80926 | -0.09821 | 0.071605 | 0.040094 | -0.02996 | -0.35543 | -0.12573 | -0.0356 | -1.52198 | -0.04487 | -0.50348 |
| 20.05556 | -0.00125 | 0.292647 | 0.525521 | 0.376193 | 0.311168 | 0.396767 | 0.444112 | 0.128237 | 0.475281 | 0.285054 | 0.55378 | -0.02428 | -0.07708 | -0.33538 | -0.13806 | -0.11064 | -1.06824 | -0.09343 | -0.27571 |
| 20.06944 | -0.06588 | 0.078606 | 0.289784 | 0.308843 | 0.041718 | 0.186917 | 0.192769 | 0.090516 | 0.234474 | 0.305973 | 0.477832 | -0.05933 | -0.09225 | -0.12587 | -0.13531 | -0.15383 | -0.77523 | -0.11394 | -0.11026 |
| 20.08333 | -0.0871 | -0.03613 | 0.131453 | 0.214716 | -0.07898 | 0.041764 | 0.038685 | 0.027876 | 0.105413 | 0.156693 | 0.258034 | -0.0641 | -0.09292 | 0.03606 | -0.12617 | -0.16381 | -0.59625 | -0.10053 | -0.0581 |
| 20.09722 | -0.06625 | -0.05147 | 0.085523 | 0.20117 | -0.07435 | 0.006352 | 0.021447 | 0.035446 | 0.097261 | 0.040982 | 0.077018 | -0.04854 | -0.07292 | 0.057278 | -0.10966 | -0.13726 | -0.43505 | -0.06925 | -0.04149 |
| 20.11111 | -0.0549 | -0.04927 | 0.078155 | 0.193631 | -0.05793 | -0.02212 | 0.031923 | 0.055537 | 0.093993 | -0.00956 | -0.04147 | -0.03384 | -0.0551 | -0.01221 | -0.10151 | -0.09291 | -0.3062 | -0.05651 | -0.02851 |
| 20.125 | -0.09503 | -0.08855 | 0.036177 | 0.129448 | -0.10144 | -0.11942 | 0.000809 | 0.026338 | 0.036995 | -0.02802 | -0.11286 | -0.03861 | -0.06672 | -0.07493 | -0.10133 | -0.05201 | -0.21817 | -0.06185 | -0.03483 |
| 20.13889 | -0.17144 | -0.15411 | -0.0444 | 0.035398 | -0.17879 | -0.24474 | -0.07263 | -0.0527 | -0.04922 | -0.0486 | -0.17553 | -0.06234 | -0.09548 | -0.05764 | -0.10779 | -0.04085 | -0.16035 | -0.06983 | -0.05707 |
| 20.15278 | -0.23294 | -0.21395 | -0.10964 | -0.05445 | -0.24177 | -0.32023 | -0.14608 | -0.12624 | -0.12894 | -0.07729 | -0.22199 | -0.08544 | -0.11929 | -0.06772 | -0.12028 | -0.04382 | -0.13227 | -0.08496 | -0.07226 |
| 20.16667 | -0.27678 | -0.2598 | -0.15583 | -0.13041 | -0.28009 | -0.37793 | -0.19501 | -0.17687 | -0.17736 | -0.08903 | -0.218 | -0.08873 | -0.11942 | -0.13898 | -0.11797 | -0.04572 | -0.12024 | -0.09757 | -0.0672 |
| 20.18056 | -0.33149 | -0.31951 | -0.23325 | -0.22991 | -0.32325 | -0.46692 | -0.25276 | -0.25184 | -0.23006 | -0.08143 | -0.19792 | -0.07593 | -0.10709 | -0.15493 | -0.09793 | -0.03681 | -0.09255 | -0.09062 | -0.05554 |
| 20.19444 | -0.35041 | -0.33799 | -0.28198 | -0.27574 | -0.33176 | -0.48829 | -0.26841 | -0.28914 | -0.25357 | -0.0754 | -0.1905 | -0.06869 | -0.10397 | -0.10034 | -0.08143 | -0.04492 | -0.06161 | -0.08329 | -0.05857 |
| 20.20833 | -0.29911 | -0.28093 | -0.24346 | -0.23521 | -0.27395 | -0.38964 | -0.20668 | -0.22496 | -0.20653 | -0.07305 | -0.18816 | -0.06926 | -0.10577 | 0.064853 | -0.07635 | -0.05514 | -0.03688 | -0.08488 | -0.07342 |
| 20.22222 | -0.29349 | -0.2478 | -0.22258 | -0.23289 | -0.25279 | -0.34356 | -0.19575 | -0.19341 | -0.1926 | -0.07695 | -0.18074 | -0.07128 | -0.09824 | 0.31877 | -0.08026 | -0.05839 | -0.02418 | -0.09138 | -0.08485 |
| 20.23611 | -0.34902 | -0.27169 | -0.27149 | -0.27117 | -0.29216 | -0.40962 | -0.26681 | -0.25808 | -0.24489 | -0.07676 | -0.16463 | -0.06064 | -0.07684 | 0.275626 | -0.07474 | -0.05119 | -0.01492 | -0.08695 | -0.07083 |
| 20.25 | -0.36277 | -0.28231 | -0.29787 | -0.24391 | -0.30484 | -0.44596 | -0.29347 | -0.30476 | -0.26472 | -0.07421 | -0.1566 | -0.05012 | -0.06046 | 0.052387 | -0.06516 | -0.03997 | -0.00966 | -0.06833 | -0.05341 |
| 20.26389 | -0.30492 | -0.2552 | -0.25394 | -0.16159 | -0.26776 | -0.40401 | -0.24112 | -0.28077 | -0.219 | -0.07446 | -0.15807 | -0.05214 | -0.06497 | 0.10866 | -0.07064 | -0.03597 | -0.01209 | -0.05401 | -0.05972 |
| 20.27778 | -0.26169 | -0.24649 | -0.22291 | -0.14273 | -0.26814 | -0.43878 | -0.22011 | -0.28779 | -0.20197 | -0.06318 | -0.15133 | -0.05233 | -0.07176 | 0.187048 | -0.07749 | -0.03267 | -0.01795 | -0.05655 | -0.07245 |
| 20.29167 | -0.25822 | -0.256 | -0.23195 | -0.20386 | -0.32107 | -0.60999 | -0.24953 | -0.33531 | -0.23141 | -0.0643 | -0.15737 | -0.06551 | -0.08098 | -0.2338 | -0.08671 | -0.03921 | -0.04878 | -0.06986 | -0.08428 |
| 20.30556 | -0.27184 | -0.26562 | -0.24235 | -0.26179 | -0.35309 | -0.70417 | -0.27221 | -0.35917 | -0.25334 | -0.07445 | -0.15503 | -0.07618 | -0.08267 | -0.43565 | -0.08885 | -0.0474 | -0.08366 | -0.06734 | -0.08056 |
| 20.31944 | -0.31726 | -0.30484 | -0.26784 | -0.30909 | -0.36526 | -0.6844 | -0.29617 | -0.37253 | -0.2724 | -0.06395 | -0.1117 | -0.06043 | -0.05934 | -0.28167 | -0.07199 | -0.0436 | -0.08647 | -0.05483 | -0.05811 |
| 20.33333 | -0.33129 | -0.31561 | -0.26877 | -0.27669 | -0.33468 | -0.59276 | -0.28111 | -0.35289 | -0.25805 | -0.05785 | -0.08012 | -0.0558 | -0.05234 | -0.04004 | -0.0624 | -0.04717 | -0.08408 | -0.05743 | -0.04689 |
| 20.34722 | -0.2729 | -0.2635 | -0.20976 | -0.14457 | -0.2555 | -0.44202 | -0.20639 | -0.28194 | -0.18908 | -0.05534 | -0.07705 | -0.06379 | -0.06071 | 0.36256 | -0.05821 | -0.05224 | -0.08153 | -0.06393 | -0.03456 |

| | | | | | | | | | | | | | | | | | | | |
|---|---|---|---|---|---|---|---|---|---|---|---|---|---|---|---|---|---|---|---|
| 20.36111 | -0.23093 | -0.2344 | -0.16577 | -0.04459 | -0.2063 | -0.29972 | -0.1544 | -0.24253 | -0.14017 | -0.03714 | -0.06415 | -0.05635 | -0.05199 | 0.361508 | -0.05083 | -0.04604 | -0.05848 | -0.06256 | -0.01439 |
| 20.375 | -0.27049 | -0.28294 | -0.2038 | -0.05872 | -0.24785 | -0.26939 | -0.18339 | -0.29013 | -0.17673 | -0.01989 | -0.06088 | -0.04545 | -0.03242 | -0.02016 | -0.05302 | -0.04036 | -0.03499 | -0.05822 | -0.0057 |
| 20.38889 | -0.3309 | -0.34388 | -0.2861 | -0.1225 | -0.32452 | -0.31644 | -0.22502 | -0.36077 | -0.25418 | -0.00985 | -0.08386 | -0.04395 | -0.024 | -0.21103 | -0.05614 | -0.043 | -0.02466 | -0.05361 | 0.0082 |
| 20.40278 | -0.30376 | -0.32976 | -0.29715 | -0.11233 | -0.31962 | -0.30433 | -0.16956 | -0.35472 | -0.25514 | 0.008113 | -0.07885 | -0.03837 | -0.01983 | -0.09554 | -0.04542 | -0.04166 | -0.00498 | -0.04837 | 0.046329 |
| 20.41667 | -0.20951 | -0.26459 | -0.23334 | -0.06127 | -0.21353 | -0.23403 | -0.06306 | -0.29075 | -0.17336 | 0.042402 | -0.00948 | -0.02104 | -0.00943 | 0.16065 | -0.02834 | -0.02725 | 0.028407 | -0.04053 | 0.081691 |
| 20.43056 | -0.1405 | -0.23283 | -0.1972 | -0.05747 | -0.03541 | -0.18583 | -0.01657 | -0.25287 | -0.11855 | 0.086188 | 0.121127 | -0.01581 | -0.01364 | 0.250414 | -0.03187 | -0.02297 | 0.051836 | -0.04349 | 0.090923 |
| 20.44444 | -0.1012 | -0.21286 | -0.19809 | -0.04189 | 0.072826 | -0.1597 | 0.033044 | -0.23539 | -0.10474 | 0.116239 | 0.275487 | -0.03086 | -0.04351 | 0.089481 | -0.05642 | -0.03855 | 0.067615 | -0.05472 | 0.100456 |
| 20.45833 | -0.066 | -0.18028 | -0.18975 | -0.00433 | -0.08941 | -0.12803 | 0.092561 | -0.20545 | -0.0877 | 0.178184 | 0.440528 | -0.03454 | -0.05303 | -0.17255 | -0.062 | -0.04731 | 0.10073 | -0.04985 | 0.154011 |
| 20.47222 | -0.00267 | -0.14742 | -0.16485 | 0.067619 | -0.37739 | -0.05206 | 0.140893 | -0.16861 | -0.07781 | 0.281727 | 0.647734 | -0.0316 | -0.02642 | -0.71632 | -0.04728 | -0.04483 | 0.182165 | -0.03476 | 0.202421 |
| 20.48611 | 0.153977 | -0.17595 | -0.22713 | 0.27456 | -0.53248 | 0.167949 | 0.158009 | -0.1841 | -0.13835 | 0.483968 | 1.146784 | -0.0694 | -0.03236 | -0.697 | -0.05517 | -0.07018 | 0.442856 | -0.00064 | 0.319179 |
| 20.5 | 0.41575 | -0.33506 | -0.50918 | 0.465608 | -0.73996 | 0.539758 | 0.214475 | -0.34741 | -0.34262 | 0.828786 | 2.039202 | -0.13609 | -0.09668 | -0.41334 | -0.09703 | -0.15492 | 1.199259 | 0.063482 | 0.61332 |
| 20.51389 | 0.703618 | -0.48356 | -0.75664 | -0.14977 | -0.94862 | 0.670797 | 0.406034 | -0.57878 | -0.55211 | 0.893423 | 1.787074 | -0.16267 | -0.16573 | -0.30672 | -0.12042 | -0.25753 | 2.107923 | 0.076894 | 0.841103 |
| 20.52778 | 0.75231 | -0.41185 | -0.65914 | -0.57776 | -0.83589 | 0.547326 | 0.580407 | -0.54041 | -0.5361 | 0.536732 | 0.692976 | -0.145 | -0.17988 | -0.23537 | -0.07554 | -0.21068 | 2.863172 | 0.005234 | 0.83843 |
| 20.54167 | 0.82657 | -0.02222 | -0.32467 | -0.486 | -0.50557 | 0.5586 | 0.512817 | -0.10141 | -0.37898 | 0.32603 | -0.22677 | -0.09683 | -0.10285 | -0.3135 | 0.019877 | -0.06698 | 3.368075 | -0.04692 | 0.82965 |
| 20.55556 | 0.831223 | 0.299048 | -0.04544 | -0.24125 | -0.201 | 0.609362 | 0.32796 | 0.442177 | -0.22189 | 0.326095 | -0.79478 | -0.03746 | -0.01235 | -0.40359 | 0.076834 | 0.010691 | 3.487566 | -0.02376 | 0.853903 |
| 20.56944 | 0.852664 | 0.477717 | 0.101988 | -0.05784 | -0.02029 | 0.530923 | 0.265131 | 0.843749 | -0.06361 | 0.371328 | -0.52856 | -0.02496 | 0.029985 | -0.30439 | 0.073422 | 0.016724 | 2.131293 | -0.00349 | 0.814222 |
| 20.58333 | 0.837783 | 0.534732 | 0.188923 | 0.058708 | 0.043438 | 0.385257 | 0.478093 | 0.974777 | 0.120458 | 0.438015 | 0.517324 | -0.01666 | 0.060071 | -0.09983 | 0.113818 | 0.023795 | 1.13539 | -0.00489 | 0.791603 |
| 20.59722 | 0.966928 | 0.549091 | 0.192074 | 0.029103 | 0.022507 | 0.30017 | 0.711811 | 0.92121 | 0.280836 | 0.562795 | 1.530352 | -0.00515 | 0.080923 | -0.0174 | 0.1671 | 0.041698 | 0.596326 | -0.01309 | 0.706398 |
| 20.61111 | 1.033653 | 0.54194 | 0.137438 | -0.1279 | -0.02653 | 0.276473 | 0.589521 | 0.747646 | 0.369492 | 0.726013 | 1.348995 | -0.03646 | 0.049363 | -0.00823 | 0.16372 | 0.013441 | 0.18406 | -0.0461 | 0.604578 |
| 20.625 | 0.966464 | 0.501138 | 0.102271 | -0.18474 | -0.04488 | 0.362676 | 0.177342 | 0.578375 | 0.344492 | 0.829756 | 0.625383 | -0.04106 | 0.008643 | 0.186233 | 0.115702 | 0.006207 | -0.29681 | -0.04044 | 0.463688 |
| 20.63889 | 0.738054 | 0.406199 | 0.093397 | -0.05292 | -0.0176 | 0.461087 | -0.31916 | 0.447706 | 0.210778 | 0.673155 | 0.050098 | -0.01369 | -0.01774 | 0.243909 | 0.060175 | 0.021638 | -0.81857 | -0.00293 | 0.288676 |
| 20.65278 | 0.507387 | 0.387848 | 0.043156 | 0.136217 | 0.075171 | 0.525667 | -0.70483 | 0.393801 | 0.136368 | 0.40998 | -0.53134 | 0.010802 | -0.03916 | 0.028758 | 0.004256 | 0.029952 | -1.08237 | 0.018285 | 0.087645 |
| 20.66667 | 0.273708 | 0.328485 | -0.07574 | 0.218382 | 0.125507 | 0.491636 | -1.08322 | 0.333101 | 0.067119 | 0.208544 | -1.31972 | 0.020118 | -0.04355 | -0.02896 | -0.02583 | 0.024863 | -0.98634 | 0.02253 | -0.10635 |
| 20.68056 | 0.048995 | 0.168181 | -0.22026 | 0.129764 | 0.050497 | 0.31385 | -1.40422 | 0.206522 | -0.10289 | 0.023609 | -1.85106 | -0.00914 | -0.03893 | 0.142674 | 0.037816 | -0.00917 | -0.8472 | -0.00665 | -0.23499 |
| 20.69444 | -0.06723 | 0.016998 | -0.26308 | 0.047916 | -0.03648 | 0.204805 | -1.28361 | 0.099848 | -0.22879 | -0.05904 | -1.40103 | -0.03359 | -0.01944 | 0.024467 | 0.096454 | -0.02801 | -0.59033 | -0.0272 | -0.26184 |
| 20.70833 | -0.12572 | -0.06029 | -0.1646 | 0.027485 | -0.05642 | 0.218592 | -0.69217 | 0.051604 | -0.17735 | 0.008877 | -0.46623 | -0.03403 | -0.00288 | -0.09201 | 0.097687 | -0.02506 | -0.27278 | -0.02496 | -0.19773 |
| 20.72222 | -0.13737 | -0.02643 | 0.02353 | 0.059182 | 0.003515 | 0.342167 | 0.175603 | 0.066465 | -0.01022 | 0.151309 | 0.274729 | -0.01565 | -0.00442 | -0.03692 | 0.036874 | -0.0031 | 0.09694 | -0.00883 | -0.04835 |
| 20.73611 | -0.15826 | 0.007346 | 0.123091 | 0.093093 | 0.038016 | 0.360688 | 0.882078 | 0.065049 | 0.084844 | 0.25556 | 0.818281 | 0.013166 | 0.000888 | -0.12127 | -0.01242 | 0.034698 | 0.415227 | 0.015602 | 0.165393 |
| 20.75 | -0.24261 | -0.02721 | 0.044618 | 0.110065 | 0.007257 | 0.174251 | 1.136871 | 0.047188 | 0.033921 | 0.259816 | 1.083109 | 0.028494 | 0.021442 | -0.2669 | -0.02414 | 0.055639 | 0.529286 | 0.019936 | 0.354398 |
| 20.76389 | -0.32299 | -0.05406 | -0.0753 | 0.122224 | 0.002829 | 0.012731 | 0.91507 | 0.10375 | -0.04291 | 0.179318 | 0.803843 | 0.013437 | 0.025419 | -0.15172 | -0.04903 | 0.037412 | 0.427918 | 0.004017 | 0.341824 |
| 20.77778 | -0.34609 | -0.03446 | -0.11189 | 0.12851 | 0.049664 | 0.04314 | 0.336506 | 0.204986 | -0.08146 | 0.012981 | 0.052138 | -0.02132 | -0.00096 | -0.03466 | -0.07302 | -0.00601 | 0.179906 | -0.02184 | 0.138969 |
| 20.79167 | -0.31891 | -0.02315 | -0.11169 | 0.070018 | 0.095667 | 0.158365 | -0.52227 | 0.2073 | -0.07088 | -0.17281 | -0.89795 | -0.0299 | -0.01455 | 0.013294 | -0.03672 | -0.02362 | -0.04917 | -0.0267 | -0.10721 |
| 20.80556 | -0.30178 | -0.0731 | -0.1793 | -0.0294 | 0.053719 | 0.129618 | -1.28585 | -0.00013 | -0.10371 | -0.37282 | -1.76358 | -0.01727 | -0.01603 | 0.012687 | 0.015356 | -0.01842 | -0.25114 | -0.01115 | -0.32367 |
| 20.81944 | -0.27422 | -0.06174 | -0.1938 | -0.00884 | 0.019569 | 0.033702 | -1.40227 | -0.2266 | -0.09766 | -0.42646 | -1.66056 | 0.013589 | -0.01329 | 0.006226 | 0.045241 | 0.002883 | -0.47127 | 0.026533 | -0.52877 |
| 20.83333 | -0.26791 | -0.00356 | -0.11788 | 0.106101 | 0.055746 | 0.00618 | -0.80619 | -0.30989 | -0.00847 | -0.32849 | -0.64082 | 0.038676 | -0.0054 | 0.021514 | 0.048305 | 0.015985 | -0.72074 | 0.053215 | -0.63355 |
| 20.84722 | -0.31695 | 0.026792 | -0.03791 | 0.209457 | 0.114945 | 0.04197 | -0.32094 | -0.28189 | 0.073446 | -0.21877 | 0.301535 | 0.041014 | 0.003277 | 0.041177 | 0.027118 | 0.00461 | -0.85448 | 0.031815 | -0.63599 |
| 20.86111 | -0.36597 | 0.004552 | 0.022868 | 0.19411 | 0.154947 | 0.057432 | -0.17425 | -0.20639 | 0.076047 | -0.16641 | 0.517212 | 0.04261 | -0.01307 | 0.03606 | -0.01238 | -0.01531 | -0.68923 | 0.007936 | -0.60759 |
| 20.875 | -0.45132 | -0.09773 | 0.014136 | 0.062962 | 0.093899 | -0.07011 | -0.14014 | -0.09609 | -0.04524 | -0.26036 | -0.04852 | 0.019637 | -0.05866 | 0.029164 | -0.05577 | -0.02967 | -0.40623 | 0.003873 | -0.6633 |
| 20.88889 | -0.59573 | -0.19786 | -0.04328 | -0.16 | -0.00396 | -0.2773 | -0.1832 | 0.078178 | -0.18315 | -0.44163 | -0.84831 | -0.01441 | -0.10479 | 0.046346 | -0.06815 | -0.02677 | -0.06901 | 0.012975 | -0.73284 |
| 20.90278 | -0.75914 | -0.27554 | -0.11791 | -0.44178 | -0.05952 | -0.46712 | -0.34173 | 0.212468 | -0.28143 | -0.65136 | -1.45931 | -0.02311 | -0.13179 | 0.053824 | -0.07158 | -0.02469 | 0.051565 | 0.018813 | -0.79307 |
| 20.91667 | -0.85198 | -0.31447 | -0.15834 | -0.52286 | -0.0662 | -0.52601 | -0.48498 | 0.15453 | -0.30128 | -0.70742 | -1.58158 | -0.00817 | -0.11173 | 0.043705 | -0.10327 | -0.03577 | -0.15151 | 0.022656 | -0.78466 |
| 20.93056 | -0.81329 | -0.27444 | -0.11873 | -0.35956 | -0.02885 | -0.43868 | -0.52793 | -0.02601 | -0.21781 | -0.64142 | -0.99594 | -0.00075 | -0.04401 | 0.053446 | -0.15292 | -0.04466 | -0.41509 | 0.027848 | -0.70754 |
| 20.94444 | -0.73262 | -0.22532 | -0.03472 | -0.19807 | 0.03748 | -0.3812 | -0.46614 | -0.19828 | -0.10336 | -0.40838 | -0.42343 | 0.014304 | -0.00849 | 0.069371 | -0.16611 | -0.03106 | -0.53058 | 0.048654 | -0.57033 |
| 20.95833 | -0.64477 | -0.18316 | 0.031247 | -0.23433 | 0.087917 | -0.40768 | -0.34728 | -0.4095 | -0.04544 | -0.18478 | -0.14232 | 0.036536 | 0.010074 | 0.065465 | -0.15883 | -0.0074 | -0.59207 | 0.078153 | -0.50731 |
| 20.97222 | -0.47861 | -0.1268 | 0.086514 | -0.20269 | 0.134423 | -0.46046 | -0.11836 | -0.47186 | 0.036717 | -0.09001 | -0.09708 | 0.036946 | 0.022614 | 0.059571 | -0.12885 | -0.01209 | -0.76491 | 0.067121 | -0.48664 |
| 20.98611 | -0.09056 | 0.110064 | 0.270793 | 0.281096 | 0.33761 | -0.60782 | 0.343586 | -0.11912 | 0.283069 | -0.10185 | -0.07287 | 0.052941 | 0.065865 | 0.065478 | -0.08062 | -0.0186 | -0.95396 | 0.017363 | -0.47213 |
| 21 | 0.365355 | 0.485755 | 0.582041 | 1.040232 | 0.705367 | -0.97702 | 0.958662 | 0.435066 | 0.646476 | -0.16115 | -0.06587 | 0.095961 | 0.135329 | 0.096924 | 0.004707 | 0.00645 | -1.14462 | -0.05563 | -0.46459 |
| 21.01389 | 0.534744 | 0.642985 | 0.797044 | 1.255609 | 1.093487 | -0.86941 | 1.225858 | 0.666154 | 0.844969 | -0.17824 | -0.04931 | 0.134352 | 0.15883 | 0.120374 | 0.087889 | 0.068824 | -1.41605 | -0.07775 | -0.35295 |
| 21.02778 | 0.347314 | 0.496643 | 0.721861 | 0.741767 | 0.805745 | -0.06401 | 0.83082 | 0.515134 | 0.774985 | -0.15375 | -0.00706 | 0.121805 | 0.098003 | 0.060564 | 0.073296 | 0.083464 | -1.09165 | -0.04634 | -0.2123 |
| 21.04167 | 0.166478 | 0.275079 | 0.415007 | 0.243097 | 0.401573 | 0.348948 | 0.354496 | 0.123854 | 0.44982 | -0.14752 | 0.001765 | 0.06245 | 0.01744 | -0.0493 | -0.01574 | 0.02341 | -0.32163 | -0.05286 | -0.12904 |
| 21.05556 | 0.081047 | 0.058569 | 0.155669 | 0.107282 | 0.128675 | 0.242743 | 0.087318 | -0.1956 | 0.18275 | -0.11726 | -0.00245 | 0.021128 | -0.01132 | -0.09351 | -0.07674 | -0.01304 | 0.120139 | -0.07296 | -0.08371 |
| 21.06944 | 0.045292 | -0.03975 | 0.048907 | 0.087196 | 0.034358 | 0.086132 | 0.002597 | -0.25394 | 0.078753 | -0.05315 | 0.005574 | 0.012222 | -0.00016 | -0.06774 | -0.08167 | -0.01045 | 0.162529 | -0.06947 | -0.06655 |
| 21.08333 | 0.069347 | 0.009198 | 0.081241 | 0.055206 | 0.063111 | 0.127186 | 0.044428 | -0.04391 | 0.106948 | 0.006361 | 0.011693 | 0.002564 | -0.00129 | -0.04558 | -0.06521 | -0.01363 | 0.038006 | -0.06564 | -0.07396 |
| 21.09722 | 0.104844 | 0.095677 | 0.155999 | 0.00235 | 0.107102 | 0.304408 | 0.110123 | 0.110513 | 0.167644 | 0.01312 | -0.00658 | -0.02038 | -0.02444 | -0.06456 | -0.05615 | -0.03113 | -0.11128 | -0.07347 | -0.09401 |
| 21.11111 | 0.097972 | 0.101866 | 0.179526 | -0.01799 | 0.114678 | 0.287969 | 0.135068 | 0.113249 | 0.176009 | -0.01842 | -0.02974 | -0.03817 | -0.02905 | -0.08245 | -0.05315 | -0.038 | -0.20446 | -0.07887 | -0.10914 |
| 21.125 | 0.088952 | 0.072236 | 0.158388 | 0.01906 | 0.101114 | 0.107248 | 0.145049 | 0.055654 | 0.154585 | -0.04004 | -0.03318 | -0.03977 | -0.02237 | -0.06135 | -0.05754 | -0.0307 | -0.21802 | -0.06654 | -0.11776 |
| 21.13889 | 0.070099 | 0.04233 | 0.118629 | -0.0016 | 0.083692 | -0.00484 | 0.133334 | 0.02567 | 0.122456 | -0.04428 | -0.02161 | -0.02378 | -0.01794 | -0.02548 | -0.05649 | -0.01833 | -0.16908 | -0.03993 | -0.11611 |
| 21.15278 | 0.059269 | 0.051604 | 0.111864 | -0.08909 | 0.054809 | 0.115708 | 0.099834 | 0.071746 | 0.121222 | -0.05468 | -0.01527 | -0.00806 | 0.000104 | -0.0259 | -0.05974 | -0.01292 | -0.11002 | -0.03084 | -0.11512 |
| 21.16667 | 0.114295 | 0.12731 | 0.180992 | -0.10199 | 0.075737 | 0.402149 | 0.113889 | 0.191667 | 0.183899 | -0.0929 | -0.03345 | -0.02151 | -0.00065 | -0.06053 | -0.08391 | -0.02975 | -0.08955 | -0.05412 | -0.13575 |
| 21.18056 | 0.181456 | 0.183909 | 0.23072 | -0.01137 | 0.115642 | 0.410213 | 0.174904 | 0.239476 | 0.233984 | -0.1313 | -0.04614 | -0.04255 | -0.03406 | -0.07136 | -0.0941 | -0.04886 | -0.09507 | -0.07059 | -0.14948 |
| 21.19444 | 0.213356 | 0.183265 | 0.215831 | 0.142294 | 0.188937 | 0.141434 | 0.267606 | 0.15593 | 0.236585 | -0.13355 | -0.04119 | -0.04482 | -0.05743 | -0.06427 | -0.08089 | -0.05426 | -0.09158 | -0.06408 | -0.13877 |
| 21.20833 | 0.193677 | 0.14313 | 0.175044 | 0.238482 | 0.188582 | -0.0414 | 0.317779 | 0.074476 | 0.205982 | -0.1041 | -0.02757 | -0.03627 | -0.04571 | -0.06267 | -0.06799 | -0.04821 | -0.08115 | -0.05646 | -0.11514 |
| 21.22222 | 0.093383 | 0.087007 | 0.124152 | 0.157191 | 0.084929 | 0.064975 | 0.233847 | 0.047124 | 0.114386 | -0.06128 | -0.00225 | -0.02515 | -0.01163 | -0.0219 | -0.03984 | -0.02884 | -0.07642 | -0.04201 | -0.08475 |
| 21.23611 | -0.00705 | 0.014217 | 0.051904 | 0.057552 | -0.03485 | 0.197733 | 0.121356 | 0.005444 | -0.0069 | -0.03061 | 0.000571 | -0.03491 | -0.01092 | -0.01001 | -0.00898 | -0.0311 | -0.08902 | -0.03597 | -0.08446 |
| 21.25 | -0.01066 | -0.03345 | 0.018185 | 0.115067 | -0.05318 | 0.093708 | 0.125369 | -0.04363 | -0.0174 | -0.01592 | -0.0235 | -0.05745 | -0.04096 | -0.07574 | -0.00082 | -0.05454 | -0.10394 | -0.0503 | -0.10852 |
| 21.26389 | 0.026654 | -0.01995 | 0.034699 | 0.185271 | -0.01581 | 0.030438 | 0.165049 | -0.03878 | 0.054816 | 0.027288 | -0.01149 | -0.05033 | -0.04456 | -0.0938 | 0.015969 | -0.04663 | -0.09554 | -0.04885 | -0.09966 |
| 21.27778 | 0.023028 | -0.00466 | 0.047567 | 0.160394 | 0.001528 | 0.060383 | 0.162561 | -0.02554 | 0.090081 | 0.092574 | 0.033587 | -0.03689 | -0.03433 | -0.0427 | 0.040573 | -0.02098 | -0.07516 | -0.03423 | -0.0604 |
| 21.29167 | -0.01907 | -0.0297 | 0.038537 | 0.145824 | -0.02415 | 0.050051 | 0.133404 | -0.05878 | 0.054033 | 0.146043 | 0.064725 | -0.04297 | -0.03715 | 0.013206 | 0.05952 | -0.01202 | -0.05652 | -0.03225 | -0.02806 |
| 21.30556 | -0.06691 | -0.09465 | 0.028801 | 0.143704 | -0.08697 | -0.02228 | 0.076258 | -0.13392 | 0.01337 | 0.192367 | 0.068965 | -0.04586 | -0.02962 | 0.040526 | 0.112071 | -0.0141 | -0.03491 | -0.03293 | -0.00342 |
| 21.31944 | -0.07171 | -0.12366 | 0.071041 | 0.17461 | -0.1049 | -0.04901 | 0.056338 | -0.1699 | 0.024279 | 0.231804 | 0.057785 | -0.05002 | -0.0226 | -0.00209 | 0.204883 | -0.02585 | -0.02761 | -0.03547 | 0.009635 |
| 21.33333 | -0.05776 | -0.09081 | 0.146654 | 0.199692 | -0.07402 | -0.01403 | 0.073544 | -0.14429 | 0.068504 | 0.292256 | 0.037897 | -0.04109 | -0.02048 | -0.05132 | 0.218332 | -0.02555 | -0.02481 | -0.02481 | 0.033431 |
| 21.34722 | -0.05212 | -0.05152 | 0.208608 | 0.171162 | -0.0591 | 0.012406 | 0.073012 | -0.10539 | 0.100132 | 0.364773 | -0.01277 | -0.01127 | -0.00377 | -0.03166 | 0.129722 | -0.00177 | -0.00519 | 0.002891 | 0.072128 |
| 21.36111 | -0.00925 | -0.01696 | 0.299241 | 0.208134 | -0.0319 | 0.064605 | 0.111638 | -0.05097 | 0.141712 | 0.402286 | -0.10931 | -0.00063 | 0.006831 | 0.042959 | 0.078028 | 0.008631 | 0.012678 | 0.012094 | 0.091551 |
| 21.375 | 0.049074 | 0.019615 | 0.382033 | 0.350034 | 0.005038 | 0.141543 | 0.207088 | 0.006747 | 0.22255 | 0.398828 | -0.11132 | -0.02093 | -0.00756 | 0.122534 | 0.039161 | -0.00545 | 0.007844 | -0.00287 | 0.074517 |
| 21.38889 | 0.064922 | 0.038589 | 0.393554 | 0.472384 | 0.015758 | 0.181956 | 0.325335 | 0.03557 | 0.282299 | 0.39479 | 0.028484 | -0.03993 | -0.02427 | 0.074898 | -0.00712 | -0.02058 | -0.00836 | -0.01976 | 0.064594 |
| 21.40278 | 0.066352 | 0.023414 | 0.323284 | 0.552509 | 0.005774 | 0.097398 | 0.467378 | 0.024519 | 0.265836 | 0.36953 | 0.164599 | -0.04119 | -0.01424 | -0.07068 | -0.0144 | -0.02392 | -0.01199 | -0.02598 | 0.084717 |
| 21.41667 | 0.11456 | 0.009107 | 0.236745 | 0.733313 | 0.004393 | -0.07702 | 0.564142 | 0.027401 | 0.278006 | 0.344012 | 0.205784 | -0.0359 | 0.00069 | -0.14401 | 0.016183 | -0.02538 | -0.0059 | -0.01832 | 0.123221 |
| 21.43056 | 0.17321 | 0.012065 | 0.175066 | 0.751955 | 0.021015 | -0.10379 | 0.608741 | 0.077895 | 0.285177 | 0.343911 | 0.157513 | -0.02301 | -0.00981 | -0.12929 | 0.039231 | -0.03618 | 0.002711 | -0.0139 | 0.149958 |
| 21.44444 | 0.176698 | 0.01265 | 0.107501 | 0.537588 | -0.03741 | -0.01572 | 0.607361 | 0.133578 | 0.193024 | 0.405212 | 0.119965 | 0.051887 | -0.0369 | -0.16056 | 0.016767 | -0.068 | 0.00764 | -0.04492 | 0.133805 |
| 21.45833 | 0.146325 | -0.04203 | -0.00531 | 0.277353 | -0.12812 | 0.102086 | 0.41456 | 0.173974 | 0.069899 | 0.411868 | 0.280792 | 0.161072 | -0.06072 | -0.14487 | -0.04585 | -0.10992 | 0.019547 | -0.10152 | 0.081423 |
| 21.47222 | 0.123098 | -0.04966 | -0.02501 | 0.074148 | -0.15923 | 0.349701 | 0.231226 | 0.294325 | 0.036613 | 0.354553 | 0.752731 | 0.122658 | -0.05935 | -0.02137 | -0.08897 | -0.11718 | 0.132089 | -0.12853 | 0.18422 |
| 21.48611 | 0.190656 | 0.024374 | -0.06064 | -0.24094 | -0.25064 | 0.901862 | 0.266913 | 0.419591 | 0.095053 | 0.298071 | 1.531029 | -0.018 | -0.06228 | -0.04439 | -0.1043 | -0.1095 | 0.708563 | -0.05173 | 0.82815 |
| 21.5 | 0.39268 | -0.00815 | -0.29419 | -0.78787 | -0.61325 | 1.030215 | 0.339916 | 0.309955 | -0.04813 | 0.295207 | 3.02743 | -0.12263 | -0.12348 | -0.20506 | -0.08792 | -0.14032 | 2.06788 | 0.036582 | 2.153621 |
| 21.51389 | 0.631683 | -0.02562 | -0.61871 | -1.11996 | -0.87603 | 0.476506 | 0.19367 | -0.02341 | -0.38191 | 0.226313 | 2.609005 | -0.18003 | -0.18415 | -0.31042 | -0.02885 | -0.15666 | 2.743787 | 0.039969 | 2.557191 |
| 21.52778 | 0.478323 | -0.03621 | -0.63009 | -1.0174 | -0.88716 | -0.23469 | 0.177986 | -0.18732 | -0.63562 | 0.027917 | 1.696926 | -0.17884 | -0.14975 | -0.30336 | 0.059596 | -0.09038 | 1.693661 | -0.02264 | 1.659489 |
| 21.54167 | 0.331625 | 0.046866 | -0.50445 | -0.60901 | -0.63018 | -0.40048 | 0.339012 | 0.097408 | -0.66085 | -0.07692 | 0.850355 | -0.11254 | -0.06546 | -0.11863 | 0.090887 | 0.022941 | 0.907633 | -0.04901 | 0.729391 |

| | | | | | | | | | | | | | | | | | | | |
|---|---|---|---|---|---|---|---|---|---|---|---|---|---|---|---|---|---|---|---|
| 21.55556 | 0.275703 | 0.1947 | -0.39332 | -0.32156 | -0.31469 | -0.27228 | 0.296899 | 0.420017 | -0.61778 | -0.09969 | 0.057542 | -0.01853 | -0.03275 | 0.068525 | 0.087766 | 0.090399 | 0.427512 | -0.00584 | 0.23387 |
| 21.56944 | 0.326305 | 0.282516 | -0.36011 | -0.28813 | -0.13729 | -0.14097 | 0.13129 | 0.642184 | -0.48377 | -0.07305 | -0.71703 | 0.019246 | -0.04286 | 0.060035 | 0.061507 | 0.103215 | -0.06396 | 0.026114 | 0.101821 |
| 21.58333 | 0.525663 | 0.338796 | -0.2459 | -0.27204 | -0.06798 | 0.012301 | 0.336939 | 0.820711 | -0.2306 | 0.037896 | -1.47097 | -0.00166 | -0.02993 | -0.0343 | 0.08728 | 0.090395 | -0.55378 | 0.019513 | 0.203594 |
| 21.59722 | 0.739402 | 0.332046 | -0.0296 | -0.1051 | -0.03527 | 0.171395 | 0.916107 | 0.963343 | 0.018059 | 0.194279 | -1.5611 | -0.02494 | -0.01446 | -0.13381 | 0.054146 | 0.080781 | -0.69233 | -0.00235 | 0.279428 |
| 21.61111 | 0.674507 | 0.216401 | 0.246925 | 0.058685 | -0.04932 | 0.357698 | 1.417768 | 0.888905 | 0.181786 | 0.32373 | -0.27283 | -0.00775 | -0.01298 | -0.09531 | -0.04782 | 0.066319 | -0.41921 | -0.02629 | 0.360422 |
| 21.625 | 0.393694 | 0.084988 | 0.443212 | 0.057184 | -0.04484 | 0.612319 | 1.606454 | 0.651869 | 0.29638 | 0.375405 | 1.367352 | 0.0416 | 0.014648 | 0.079997 | 0.072656 | 0.073077 | 0.145354 | -0.00571 | 0.383935 |
| 21.63889 | 0.112125 | -0.01923 | 0.346304 | -0.06613 | -0.02012 | 0.855857 | 1.273113 | 0.416311 | 0.31243 | 0.375191 | 1.749305 | 0.044907 | 0.038944 | 0.055206 | 0.147345 | 0.069429 | 0.626673 | 0.029773 | 0.00053 |
| 21.65278 | -0.21358 | -0.05537 | 0.015078 | -0.06751 | 0.029131 | 0.922745 | 0.416062 | 0.237187 | 0.179385 | 0.346394 | 1.056228 | 0.04573 | 0.048 | -0.0745 | 0.108721 | 0.053874 | 0.937955 | 0.053335 | -0.2885 |
| 21.66667 | -0.53165 | -0.04665 | -0.32292 | -0.02 | 0.054933 | 0.727462 | -0.74435 | 0.089045 | -0.06209 | 0.297859 | 0.694362 | 0.026589 | 0.024635 | -0.05024 | 0.012725 | 0.017952 | 0.965185 | 0.063154 | -0.15685 |
| 21.68056 | -0.70385 | -0.06674 | -0.48205 | -0.10598 | 0.012481 | 0.396397 | -1.76079 | -0.04186 | -0.29465 | 0.216651 | 0.368442 | -0.07402 | -0.02502 | -0.07497 | -0.10366 | -0.05074 | 0.633164 | 0.044834 | 0.10955 |
| 21.69444 | -0.65343 | -0.09271 | -0.45789 | -0.25006 | -0.04948 | 0.107842 | -1.98477 | -0.13127 | -0.35027 | 0.095679 | -0.21908 | -0.10768 | -0.04279 | -0.13553 | -0.15538 | -0.06743 | 0.171174 | 0.038262 | 0.317145 |
| 21.70833 | -0.38831 | -0.05962 | -0.31076 | -0.26818 | -0.06745 | -0.08004 | -1.46101 | -0.11666 | -0.21894 | 0.023598 | -0.92286 | -0.0452 | -0.02201 | 0.050707 | -0.11408 | -0.01661 | -0.18306 | 0.053707 | 0.336801 |
| 21.72222 | 0.073046 | 0.098758 | -0.03374 | -0.09122 | 0.02187 | -0.11211 | -0.63719 | 0.028242 | -0.04319 | 0.093932 | -1.17733 | 0.0097 | 0.005214 | 0.244322 | -0.00403 | 0.059281 | -0.34709 | 0.073556 | 0.317587 |
| 21.73611 | 0.432909 | 0.251122 | 0.185564 | 0.054251 | 0.095195 | -0.09766 | 0.156812 | 0.079983 | 0.013585 | 0.210822 | -0.84245 | 0.016129 | 0.002002 | 0.175028 | 0.098759 | 0.091882 | -0.43377 | 0.079437 | 0.264724 |
| 21.75 | 0.428684 | 0.250606 | 0.242385 | 0.010353 | 0.068089 | -0.12026 | 0.787352 | -0.04136 | -0.03626 | 0.297455 | -0.28359 | -0.00296 | -0.01024 | 0.056447 | 0.150442 | 0.073448 | -0.55356 | 0.061234 | 0.169335 |
| 21.76389 | 0.173146 | 0.171541 | 0.236007 | -0.06519 | 0.050183 | -0.08806 | 1.099608 | -0.2054 | -0.0659 | 0.248257 | 0.278698 | -0.01157 | -0.00179 | -0.04901 | 0.135084 | 0.024355 | -0.78943 | 0.050207 | -0.10646 |
| 21.77778 | -0.16903 | 0.086008 | 0.132501 | -0.07441 | 0.021944 | -0.02992 | 0.90573 | -0.40604 | -0.10943 | 0.127116 | 0.703708 | 0.000884 | 0.013095 | -0.20782 | 0.068051 | -0.01982 | -1.03761 | 0.058375 | -0.56916 |
| 21.79167 | -0.46492 | 0.023328 | -0.10019 | -0.05113 | -0.01639 | 0.018184 | 0.424949 | -0.5845 | -0.1778 | 0.004688 | 0.736448 | 0.028663 | 0.004203 | -0.30087 | -0.01849 | -0.05039 | -1.01466 | 0.075789 | -0.93298 |
| 21.80556 | -0.64138 | 0.12093 | -0.33539 | -0.06123 | -0.00512 | 0.044264 | -0.01887 | -0.68954 | -0.2012 | -0.17597 | 0.240651 | 0.006977 | -0.0506 | -0.18148 | -0.12517 | -0.08044 | -0.77174 | 0.078612 | -1.0798 |
| 21.81944 | -0.69324 | 0.415244 | -0.38682 | -0.10234 | 0.057435 | 0.052904 | -0.11701 | -0.66605 | -0.14652 | -0.38005 | -0.44433 | -0.01531 | -0.075 | 0.098651 | -0.13188 | -0.0794 | -0.43587 | 0.084811 | -0.80831 |
| 21.83333 | -0.67132 | 0.540937 | -0.29805 | -0.24791 | 0.071226 | 0.043191 | -0.10121 | -0.5582 | -0.08944 | -0.59487 | -1.14953 | 0.013068 | -0.04765 | 0.199433 | 0.007965 | -0.05082 | -0.29641 | 0.10514 | -0.29637 |
| 21.84722 | -0.71301 | 0.329041 | -0.18537 | -0.36854 | 0.008996 | -0.01665 | -0.29551 | -0.48653 | -0.10847 | -0.61262 | -1.67106 | 0.016037 | -0.03364 | 0.076292 | 0.074348 | -0.04486 | -0.48169 | 0.106725 | 0.064831 |
| 21.86111 | -0.75584 | 0.133714 | -0.12474 | -0.29324 | -0.04292 | -0.13197 | -0.68151 | -0.57225 | -0.18895 | -0.3662 | -1.81999 | -0.00715 | -0.0803 | 0.02157 | 0.010834 | -0.04212 | -0.84599 | 0.104686 | 0.027545 |
| 21.875 | -0.78523 | 0.042139 | -0.12582 | -0.20864 | -0.05117 | -0.23085 | -0.92489 | -0.84362 | -0.27421 | -0.12003 | -1.44994 | -0.00569 | -0.06932 | 0.019077 | -0.13895 | -0.02416 | -1.11829 | 0.107295 | -0.35169 |
| 21.88889 | -0.78858 | 0.003663 | -0.15961 | -0.32594 | -0.00665 | -0.25064 | -0.8191 | -1.16614 | -0.29787 | -0.1057 | -0.73774 | 0.023725 | -0.01903 | -0.00595 | -0.33332 | -0.0056 | -1.24952 | 0.141759 | -0.75688 |
| 21.90278 | -0.75209 | -0.04169 | -0.20987 | -0.57935 | 0.042242 | -0.23896 | -0.65613 | -1.37286 | -0.27567 | -0.31052 | -0.17089 | 0.023446 | -0.05104 | -0.05924 | -0.50216 | -0.02571 | -1.2661 | 0.145991 | -1.02325 |
| 21.91667 | -0.65899 | -0.0483 | -0.2092 | -0.75254 | 0.07169 | -0.18851 | -0.60184 | -1.30834 | -0.2306 | -0.52369 | -0.10576 | 0.012951 | -0.13822 | -0.09915 | -0.49728 | -0.06465 | -1.14583 | 0.118976 | -1.08808 |
| 21.93056 | -0.56826 | -0.06262 | -0.196 | -0.5943 | 0.055126 | -0.15532 | -0.77845 | -1.1532 | -0.20502 | -0.69582 | -0.46853 | 0.01585 | -0.1684 | -0.09365 | -0.39377 | -0.10349 | -0.79974 | 0.11248 | -1.04219 |
| 21.94444 | -0.59823 | -0.17989 | -0.23985 | -0.40811 | -0.02338 | -0.19564 | -1.14544 | -1.08259 | -0.24022 | -0.77173 | -0.81129 | 0.005957 | -0.12534 | -0.069 | -0.25189 | -0.0971 | -0.43429 | 0.124625 | -0.97385 |
| 21.95833 | -0.65712 | -0.2716 | -0.23978 | -0.30993 | -0.05148 | -0.22275 | -1.21461 | -1.06842 | -0.25013 | -0.76457 | -0.83406 | -0.01362 | -0.06126 | -0.03994 | -0.1635 | -0.06341 | -0.27042 | 0.138896 | -0.77827 |
| 21.97222 | -0.54675 | -0.25535 | -0.14728 | -0.20521 | 0.041699 | -0.21056 | -0.81056 | -0.98292 | -0.16046 | -0.56009 | -0.53347 | -0.02918 | -0.02269 | -0.02233 | -0.1046 | -0.0326 | -0.22193 | 0.127392 | -0.48346 |
| 21.98611 | -0.26472 | -0.07381 | 0.138807 | 0.155021 | 0.305833 | -0.12778 | -0.08677 | -0.61401 | 0.126198 | -0.2965 | -0.23829 | 0.033747 | 0.046105 | -0.00229 | -0.00748 | 0.038974 | -0.27605 | 0.110041 | -0.20838 |
| 22 | 0.089369 | 0.222922 | 0.566339 | 0.602217 | 0.691444 | -0.00185 | 0.654311 | -0.01693 | 0.538496 | -0.09994 | -0.17407 | 0.156052 | 0.111151 | 0.065003 | 0.091347 | 0.12536 | -0.52561 | 0.064565 | -0.13098 |
| 22.01389 | 0.243359 | 0.288064 | 0.737689 | 0.684566 | 0.735636 | 0.070437 | 0.864784 | 0.407427 | 0.703457 | -0.07844 | -0.10664 | 0.187296 | 0.081846 | 0.17862 | 0.087936 | 0.134674 | -0.80163 | -0.01914 | -0.17385 |
| 22.02778 | 0.185923 | 0.151244 | 0.512232 | 0.378668 | 0.569184 | 0.031017 | 0.539475 | 0.396468 | 0.471961 | -0.10126 | 0.160756 | 0.117401 | 0.023414 | 0.236348 | 0.023554 | 0.064667 | -0.56141 | -0.1052 | -0.15964 |
| 22.04167 | 0.184895 | 0.072775 | 0.278079 | 0.218084 | 0.377317 | 0.179202 | 0.254293 | 0.25763 | 0.249967 | -0.10667 | 0.330721 | 0.056346 | 0.015793 | 0.137604 | -0.01689 | -0.03326 | 0.065695 | -0.1615 | -0.09577 |
| 22.05556 | 0.252881 | 0.112461 | 0.205297 | 0.381842 | 0.281091 | 0.302857 | 0.213326 | 0.152013 | 0.207978 | -0.11623 | 0.219095 | 0.015604 | 0.013442 | -0.01513 | -0.03658 | -0.10914 | 0.326412 | -0.19407 | -0.05994 |
| 22.06944 | 0.261862 | 0.105928 | 0.179847 | 0.429125 | 0.213954 | 0.23998 | 0.212214 | 0.054374 | 0.209302 | -0.08404 | 0.040187 | -0.00921 | 0.001208 | -0.02562 | -0.05789 | -0.11491 | 0.15461 | -0.20191 | -0.05361 |
| 22.08333 | 0.243456 | 0.12496 | 0.196307 | 0.274442 | 0.173673 | 0.22834 | 0.205355 | 0.060437 | 0.228693 | 0.012896 | -0.03402 | -0.01805 | 0.007034 | -0.0215 | -0.06359 | -0.09395 | -0.02249 | -0.19847 | -0.02491 |
| 22.09722 | 0.230214 | 0.160907 | 0.232463 | 0.158209 | 0.192916 | 0.242155 | 0.206943 | 0.124897 | 0.25103 | 0.093481 | -0.00691 | -0.02746 | 0.020987 | -0.07662 | -0.05315 | -0.07769 | -0.06092 | -0.21011 | 0.018159 |
| 22.11111 | 0.173628 | 0.122335 | 0.194754 | 0.10233 | 0.16398 | 0.146659 | 0.157176 | 0.12949 | 0.197365 | 0.094769 | 0.046759 | -0.03191 | 0.015402 | -0.35743 | -0.04169 | -0.0617 | -0.07992 | -0.18831 | 0.002964 |
| 22.125 | 0.089195 | 0.046513 | 0.11614 | 0.022053 | 0.087453 | 0.034917 | 0.072044 | 0.065331 | 0.099316 | 0.036958 | -0.00147 | -0.02062 | -0.02683 | -0.18125 | -0.03937 | -0.0498 | -0.14725 | -0.12564 | -0.102 |
| 22.13889 | 0.00253 | -0.02493 | 0.048637 | -0.06998 | 0.032747 | -0.0046 | 0.00045 | -0.018 | 0.020746 | -0.0072 | -0.05223 | -0.01904 | -0.07758 | -0.02158 | -0.04651 | -0.04334 | -0.18679 | -0.09998 | -0.18884 |
| 22.15278 | -0.07344 | -0.08406 | -0.00083 | -0.15027 | 0.003719 | -0.00851 | -0.04753 | -0.07018 | -0.02252 | 0.007204 | -0.0346 | -0.02951 | -0.07749 | -0.08574 | -0.04707 | -0.0402 | -0.18058 | -0.09125 | -0.1781 |
| 22.16667 | -0.12358 | -0.11671 | -0.02496 | -0.19155 | -0.02112 | -0.00064 | -0.07606 | -0.06893 | -0.03566 | 0.023876 | -0.07649 | -0.01685 | -0.04777 | -0.15505 | -0.03797 | -0.02555 | -0.18482 | -0.06599 | -0.14419 |
| 22.18056 | -0.12645 | -0.10792 | -0.01724 | -0.15094 | -0.02986 | 0.01218 | -0.07311 | -0.02901 | -0.01991 | -0.00487 | -0.26028 | -0.00469 | -0.03606 | -0.11943 | -0.03569 | -0.01795 | -0.18901 | -0.04073 | -0.10496 |
| 22.19444 | -0.06477 | -0.04993 | 0.021839 | -0.0117 | -0.00558 | 0.052721 | -0.01303 | 0.032064 | 0.028081 | -0.03723 | -0.44045 | -0.00636 | -0.01982 | -0.09178 | -0.02874 | -0.02578 | -0.14352 | -0.04909 | -0.00543 |
| 22.20833 | 0.002654 | 0.008662 | 0.051178 | 0.127529 | 0.015301 | 0.06932 | 0.051514 | 0.063364 | 0.058185 | -0.0601 | -0.32458 | -0.01595 | 0.001219 | -0.05242 | -0.0212 | -0.03908 | -0.07321 | -0.07917 | 0.106927 |
| 22.22222 | 0.045964 | 0.034456 | 0.060365 | 0.251934 | 0.027018 | -0.03258 | 0.097664 | 0.04666 | 0.063432 | -0.06086 | 0.025835 | -0.05635 | 0.01574 | 0.008707 | -0.02904 | -0.06528 | -0.01165 | -0.12014 | 0.171226 |
| 22.23611 | 0.102878 | 0.064727 | 0.091295 | 0.379796 | 0.053466 | -0.1173 | 0.151721 | 0.056841 | 0.102234 | -0.07005 | 0.219374 | -0.07959 | 0.016093 | 0.029795 | -0.03575 | -0.07439 | 0.009873 | -0.12871 | 0.142318 |
| 22.25 | 0.175724 | 0.162382 | 0.175041 | 0.432668 | 0.106755 | 0.078747 | 0.203709 | 0.163118 | 0.208089 | -0.08523 | 0.128811 | -0.04164 | -0.01494 | -0.05418 | -0.02521 | -0.0458 | -0.03104 | -0.0883 | 0.001731 |
| 22.26389 | 0.228359 | 0.250066 | 0.254075 | 0.487875 | 0.170943 | 0.225427 | 0.234542 | 0.257966 | 0.304086 | -0.08355 | 0.034819 | -0.00266 | -0.03916 | -0.12433 | -0.01199 | -0.01656 | -0.05568 | -0.04978 | -0.08587 |
| 22.27778 | 0.225755 | 0.228924 | 0.251032 | 0.564395 | 0.163239 | 0.071206 | 0.221495 | 0.252979 | 0.308726 | -0.02315 | 0.108598 | -0.00451 | -0.01764 | -0.09467 | -0.01371 | -0.01085 | 0.011706 | -0.04811 | -0.03944 |
| 22.29167 | 0.186662 | 0.191504 | 0.20718 | 0.557832 | 0.085855 | 0.011508 | 0.163583 | 0.253538 | 0.275889 | 0.060824 | 0.212597 | -0.01766 | 0.011973 | -0.10044 | -0.02166 | -0.00924 | 0.070613 | -0.06165 | 0.032273 |
| 22.30556 | 0.146111 | 0.153274 | 0.152066 | 0.555144 | 0.01268 | 0.06091 | 0.109645 | 0.262355 | 0.240566 | 0.148963 | 0.17584 | -0.02117 | 0.000035 | -0.10878 | -0.01855 | -0.00102 | 0.045444 | -0.076 | 0.0145 |
| 22.31944 | 0.121163 | 0.072661 | 0.101579 | 0.602076 | 0.004914 | -0.11156 | 0.118508 | 0.232935 | 0.206461 | 0.266847 | 0.124927 | -0.03417 | -0.02524 | -0.04844 | -0.02452 | -0.00322 | 0.010714 | -0.06766 | 0.017274 |
| 22.33333 | 0.105229 | 0.034191 | 0.117514 | 0.617891 | 0.022814 | -0.1836 | 0.245331 | 0.262642 | 0.205819 | 0.400328 | 0.209461 | -0.06123 | -0.02132 | -0.01935 | -0.03507 | -0.01977 | 0.042182 | -0.06356 | 0.132113 |
| 22.34722 | 0.068472 | 0.07264 | 0.182858 | 0.445563 | 0.001244 | 0.019429 | 0.503753 | 0.39318 | 0.216474 | 0.465258 | 0.249856 | -0.04798 | -0.01144 | -0.01818 | -0.01709 | -0.01649 | 0.069956 | -0.05922 | 0.244008 |
| 22.36111 | 0.014192 | 0.076896 | 0.206507 | 0.371205 | -0.04117 | 0.230348 | 0.722619 | 0.473837 | 0.212832 | 0.468929 | 0.210071 | -0.01237 | -0.02255 | -0.01896 | 0.004278 | -0.00027 | 0.035628 | -0.05516 | 0.283472 |
| 22.375 | -0.044 | -0.00839 | 0.169295 | 0.4394 | -0.08012 | 0.101489 | 0.684553 | 0.467088 | 0.177808 | 0.461878 | 0.068997 | -0.05946 | -0.06181 | -0.04047 | -0.00992 | -0.01384 | -0.03694 | -0.07904 | 0.267154 |
| 22.38889 | -0.12256 | -0.11462 | 0.109316 | 0.54881 | -0.13316 | -0.04742 | 0.446497 | 0.468497 | 0.134402 | 0.450277 | -0.11674 | -0.15432 | -0.14131 | -0.14718 | -0.0244 | -0.0625 | -0.11409 | -0.11502 | 0.280024 |
| 22.40278 | -0.2353 | -0.15589 | -0.0024 | 0.52045 | -0.18731 | -0.08771 | 0.16634 | 0.49278 | 0.055641 | 0.589364 | -0.33995 | -0.13857 | -0.1979 | -0.28812 | 0.010834 | -0.07935 | -0.15569 | -0.06617 | 0.320962 |
| 22.41667 | -0.44124 | 0.070632 | -0.13472 | 0.041776 | -0.32647 | -0.10177 | -0.07272 | 0.266577 | -0.16125 | 0.775881 | -0.49343 | 0.036992 | -0.15812 | -0.31933 | 0.050266 | -0.03996 | -0.16212 | 0.138198 | 0.323224 |
| 22.43056 | -0.48423 | 1.042086 | -0.16367 | -0.54367 | -0.19386 | 0.227224 | -0.47571 | -0.03658 | -0.19081 | 0.701586 | -0.39815 | 0.108181 | -0.12155 | -0.36182 | 0.095796 | 0.010895 | -0.1277 | 1.465411 | 0.224083 |
| 22.44444 | -0.12997 | 0.252017 | 0.027122 | -0.34145 | 0.581928 | 1.481288 | -0.78651 | 0.123558 | -0.06909 | 0.470625 | -0.16144 | 0.034234 | -0.11182 | -0.08983 | 0.099706 | 0.030334 | -0.08392 | 0.163906 | 0.166603 |
| 22.45833 | 0.126013 | 0.07924 | 0.235778 | 0.407756 | 0.444159 | 0.648864 | -0.31752 | 0.600241 | 0.31941 | 0.254578 | -0.09183 | 0.310947 | -0.10312 | 0.094947 | -0.01045 | 0.068065 | -0.13793 | -0.16245 | 0.04002 |
| 22.47222 | 0.275227 | 0.278349 | 0.25681 | 0.593156 | 0.189552 | 0.28591 | 0.000717 | 0.643553 | 0.277955 | 0.135023 | -0.06644 | 0.435927 | -0.11389 | 0.032576 | -0.16405 | 0.064565 | -0.19623 | -0.13105 | -0.11472 |
| 22.48611 | 0.500505 | 0.449605 | 0.215403 | 0.246205 | 0.070407 | 0.27806 | 0.416567 | 0.462098 | 0.159979 | 0.155685 | 0.611188 | 0.375277 | -0.086 | -0.11383 | -0.15295 | 0.035958 | 0.275683 | -0.00752 | 0.286236 |
| 22.5 | 0.809124 | 0.629289 | 0.171957 | -0.16793 | 0.005341 | 0.54428 | 1.008841 | 0.536137 | 0.238647 | 0.284478 | 3.020268 | 0.210475 | -0.00046 | -0.17326 | 0.048733 | 0.005433 | 1.401616 | 0.141452 | 1.401095 |
| 22.51389 | 0.804517 | 0.48097 | 0.074804 | -0.27746 | -0.10309 | 0.536098 | 1.191736 | 0.759019 | 0.322797 | 0.289766 | 4.472651 | 0.06485 | -0.00821 | -0.00454 | 0.207209 | -0.06407 | 1.609012 | 0.121864 | 1.908319 |
| 22.52778 | 0.641588 | 0.225282 | -0.02037 | 0.05442 | -0.16172 | 0.246905 | 1.026643 | 0.908265 | 0.296685 | 0.175145 | 3.342262 | -0.00437 | -0.07523 | 0.17742 | 0.257214 | -0.08343 | 1.07008 | -0.04986 | 1.750458 |
| 22.54167 | 0.553522 | 0.17363 | -0.02263 | 0.381187 | -0.12598 | 0.061164 | 0.887834 | 0.932327 | 0.189098 | 0.114851 | 1.632885 | 0.006502 | -0.04369 | 0.197494 | 0.187301 | 0.011916 | 0.689279 | -0.1278 | 1.302773 |
| 22.55556 | 0.52099 | 0.203441 | -0.03227 | 0.146341 | -0.11066 | -0.04534 | 0.488039 | 0.657998 | 0.033344 | 0.159733 | 0.479296 | -0.01398 | -0.01019 | 0.063174 | 0.02906 | 0.046886 | 0.335071 | -0.08875 | 0.695518 |
| 22.56944 | 0.462143 | 0.171523 | -0.11195 | -0.24306 | -0.14873 | -0.18373 | -0.08305 | 0.388065 | -0.19925 | 0.310664 | -0.37749 | -0.06701 | -0.02019 | -0.066 | -0.06955 | 0.018865 | -0.06462 | -0.04962 | 0.274751 |
| 22.58333 | 0.537702 | 0.157778 | -0.11634 | -0.40017 | -0.11163 | -0.21038 | -0.47397 | 0.329202 | -0.30899 | 0.470115 | -0.96521 | -0.05522 | 0.004673 | -0.08136 | -0.02792 | 0.028587 | -0.30592 | -0.01724 | 0.225645 |
| 22.59722 | 0.656648 | 0.15874 | 0.011258 | -0.32574 | 0.065585 | -0.0872 | -0.58341 | 0.409636 | -0.22285 | 0.406545 | -1.35551 | -0.01614 | 0.010513 | -0.06727 | 0.063416 | 0.060024 | -0.10199 | -0.03396 | 0.266703 |
| 22.61111 | 0.672723 | 0.074032 | 0.1898 | -0.20084 | 0.267513 | 0.025485 | -0.27464 | 0.587918 | -0.05694 | 0.115709 | -1.43597 | -0.01409 | -0.00386 | -0.06564 | 0.196258 | 0.089568 | 0.522103 | -0.05149 | 0.281828 |
| 22.625 | 0.59876 | -0.0338 | 0.419663 | 0.016321 | 0.289172 | 0.085869 | 0.505921 | 0.836894 | 0.107665 | -0.10822 | -1.30758 | -0.00812 | -0.01776 | -0.03119 | 0.320152 | 0.116048 | 0.880275 | 0.046303 | 0.297539 |
| 22.63889 | 0.499476 | -0.10326 | 0.535527 | 0.32862 | 0.091715 | 0.128057 | 1.52756 | 1.079682 | 0.22583 | -0.25578 | -1.19966 | -0.00869 | -0.05968 | 0.025397 | 0.335419 | 0.103703 | 0.600517 | 0.224102 | 0.26766 |
| 22.65278 | 0.412969 | -0.26291 | 0.383791 | 0.424711 | -0.00081 | 0.067192 | 1.963006 | 1.039163 | 0.272166 | -0.25297 | -0.92272 | 0.008361 | -0.06583 | 0.062784 | 0.263027 | 0.085316 | 0.186377 | 0.42394 | 0.158941 |
| 22.66667 | 0.264133 | -0.55804 | -0.02165 | 0.058876 | -0.06709 | -0.10537 | 1.507299 | 0.718058 | 0.227268 | -0.15122 | -0.2575 | 0.009351 | -0.03995 | 0.018705 | 0.134982 | 0.045004 | -0.17002 | 0.591841 | -0.06517 |
| 22.68056 | 0.027314 | -0.4605 | -0.36913 | -0.4749 | -0.16531 | -0.35945 | 0.525299 | 0.292942 | 0.10135 | -0.05774 | 0.626057 | -0.0371 | -0.03556 | -0.0513 | -0.04066 | -0.02919 | -0.57742 | 0.66363 | -0.22612 |
| 22.69444 | -0.16719 | 0.223123 | -0.4926 | -0.71577 | -0.29299 | -0.53333 | -0.64066 | -0.15286 | -0.05471 | 0.214899 | 1.310898 | -0.09107 | -0.03417 | -0.06464 | -0.2076 | -0.07753 | -0.89564 | 0.581809 | -0.26099 |
| 22.70833 | -0.2432 | 0.856379 | -0.46939 | -0.71812 | -0.31617 | -0.56733 | -1.63479 | -0.50774 | -0.1271 | 0.525114 | 1.372286 | -0.10406 | 0.021626 | -0.024 | -0.26535 | -0.1159 | -0.85441 | 0.36047 | -0.25506 |
| 22.72222 | -0.28528 | 1.098018 | -0.30451 | -0.71161 | -0.27122 | -0.49376 | -2.08524 | -0.64868 | -0.11148 | 0.555649 | 0.703149 | -0.0558 | 0.097572 | 0.012839 | -0.18694 | -0.13002 | -0.46298 | 0.047949 | -0.16857 |
| 22.73611 | -0.37729 | 1.081132 | -0.11781 | -0.56733 | -0.31884 | -0.46816 | -2.01953 | -0.51634 | -0.10398 | 0.181209 | -0.23273 | -0.01787 | 0.097972 | 0.004174 | -0.06661 | -0.11621 | -0.0242 | -0.24637 | -0.076 |

| | | | | | | | | | | | | | | | | | | | |
|---|---|---|---|---|---|---|---|---|---|---|---|---|---|---|---|---|---|---|---|
| 22.75 | -0.38092 | 1.076711 | 0.116972 | -0.09685 | -0.61869 | -0.60354 | -1.34963 | -0.20903 | -0.06545 | -0.13351 | -0.9559 | -0.00051 | 0.047531 | 0.019697 | 0.037866 | -0.08273 | 0.340061 | -0.41728 | 0.007193 |
| 22.76389 | -0.21854 | 1.07535 | 0.300135 | 0.346469 | -1.07172 | -0.80726 | -0.34227 | 0.028873 | -0.01069 | -0.31334 | -1.29907 | 0.011125 | 0.000248 | -0.00313 | 0.100103 | -0.03481 | 0.838766 | -0.45438 | 0.097864 |
| 22.77778 | -0.03137 | 0.987633 | 0.288986 | 0.429843 | -1.18734 | -0.88999 | 0.450711 | -0.06136 | 0.02333 | -0.46684 | -1.39214 | -0.00302 | -0.07098 | -0.08267 | 0.106057 | -0.0002 | 0.865351 | -0.35621 | 0.059845 |
| 22.79167 | -0.00826 | 0.710588 | 0.103809 | 0.198043 | -1.03228 | -0.91631 | 0.721774 | -0.4734 | 0.030206 | -0.57845 | -1.11797 | -0.02383 | -0.16088 | -0.12764 | 0.014246 | 0.006488 | 0.187974 | -0.25906 | -0.27158 |
| 22.80556 | -0.15534 | 0.318278 | -0.09855 | -0.08382 | -0.83283 | -0.89584 | 0.557682 | -0.97182 | -0.06998 | -0.61757 | -0.94486 | -0.05458 | -0.22181 | -0.13195 | -0.1555 | -0.02627 | -0.50864 | -0.20334 | -0.80695 |
| 22.81944 | -0.32858 | -0.12895 | -0.231 | -0.30274 | -0.5392 | -0.82058 | 0.07027 | -1.35663 | -0.2559 | -0.48986 | -0.8961 | -0.09412 | -0.19003 | -0.11329 | -0.33448 | -0.07988 | -1.03088 | -0.14928 | -1.30794 |
| 22.83333 | -0.46446 | -0.5746 | -0.3324 | -0.48331 | -0.42384 | -0.81471 | -0.52576 | -1.54143 | -0.41892 | -0.26318 | -0.93364 | -0.13432 | -0.13649 | -0.0699 | -0.44087 | -0.13634 | -1.3776 | -0.15 | -1.37397 |
| 22.84722 | -0.56116 | -0.91597 | -0.42482 | -0.50059 | -0.66241 | -1.05517 | -1.0326 | -1.63156 | -0.51259 | -0.0162 | -0.93269 | -0.18866 | -0.09832 | -0.03666 | -0.39209 | -0.18875 | -1.36874 | -0.20785 | -1.14262 |
| 22.86111 | -0.59756 | -1.07119 | -0.46918 | -0.42531 | -1.13516 | -1.28402 | -1.27333 | -1.58899 | -0.59381 | 0.07142 | -1.02268 | -0.2488 | -0.07765 | -0.06567 | -0.39188 | -0.23861 | -1.21011 | -0.33894 | -0.81972 |
| 22.875 | -0.56902 | -0.95012 | -0.43815 | -0.44376 | -1.57187 | -1.37068 | -1.0888 | -1.45341 | -0.5792 | -0.0843 | -1.1886 | -0.24077 | -0.03893 | -0.10751 | -0.41016 | -0.26165 | -1.12092 | -0.50126 | -0.64183 |
| 22.88889 | -0.52108 | -0.53768 | -0.3854 | -0.57574 | -1.84075 | -1.35774 | -0.56494 | -1.36195 | -0.51822 | -0.35077 | -1.24221 | -0.19674 | -0.0063 | -0.11255 | -0.41823 | -0.24906 | -1.06207 | -0.55414 | -0.64448 |
| 22.90278 | -0.45513 | -0.05265 | -0.31029 | -0.60357 | -1.79879 | -1.15999 | -0.02895 | -1.39429 | -0.43411 | -0.63286 | -0.89701 | -0.17172 | 0.002265 | -0.10119 | -0.46392 | -0.17286 | -0.94131 | -0.4222 | -0.81033 |
| 22.91667 | -0.37784 | 0.250535 | -0.25942 | -0.59722 | -1.50567 | -0.98145 | 0.0777 | -1.38084 | -0.38084 | -0.71037 | -0.39746 | -0.15638 | -0.00875 | -0.07141 | -0.48647 | -0.03019 | -0.69089 | -0.22316 | -0.96622 |
| 22.93056 | -0.32345 | 0.172142 | -0.25371 | -0.54283 | -1.10208 | -0.86105 | -0.25966 | -1.21679 | -0.41499 | -0.8147 | -0.11109 | -0.16209 | -0.01755 | 0.006692 | -0.44503 | 0.066446 | -0.39373 | -0.09008 | -0.97402 |
| 22.94444 | -0.34067 | -0.21942 | -0.20934 | -0.43443 | -0.74767 | -0.726 | -0.73225 | -0.95259 | -0.46478 | -0.80691 | -0.00402 | -0.15694 | 0.002338 | 0.053102 | -0.35177 | 0.043437 | -0.17016 | -0.01004 | -0.77446 |
| 22.95833 | -0.40642 | -0.68767 | -0.10307 | -0.28685 | -0.50077 | -0.60713 | -1.11177 | -0.71482 | -0.41564 | -0.68476 | -0.05984 | -0.1489 | 0.002137 | -0.00524 | -0.25065 | -0.05214 | -0.08955 | -0.00573 | -0.59943 |
| 22.97222 | -0.4193 | -0.98941 | -0.00137 | -0.11079 | -0.39208 | -0.49926 | -1.31371 | -0.49457 | -0.32854 | -0.42331 | -0.21662 | -0.1107 | -0.00393 | -0.1334 | -0.14929 | -0.14157 | -0.1292 | -0.06108 | -0.4408 |
| 22.98611 | -0.24973 | -1.13325 | 0.134791 | 0.11571 | -0.8371 | -0.46224 | -1.04387 | -0.12144 | -0.33332 | -0.15125 | -0.53365 | -0.05396 | 0.002762 | -0.28144 | -0.03751 | -0.07503 | -0.36135 | -0.45661 | -0.35973 |
| 23 | 0.118059 | -1.06544 | 0.431939 | 0.427856 | -1.86755 | -0.41319 | -0.27018 | 0.360014 | -0.46399 | -0.04834 | -0.94778 | 0.005943 | 0.046658 | -0.2173 | 0.132047 | 0.131694 | -0.64876 | -1.37687 | -0.32588 |
| 23.01389 | 0.472208 | -0.34082 | 0.630198 | 0.529576 | -1.51001 | 0.250737 | 0.313624 | 0.566709 | -0.23751 | -0.07916 | -0.90613 | 0.018051 | 0.087146 | 0.009531 | 0.197827 | 0.267808 | -0.54552 | -1.60619 | -0.33131 |
| 23.02778 | 0.607771 | 0.456622 | 0.446346 | 0.2809 | -0.12959 | 0.257213 | 0.415389 | 0.267795 | 0.261717 | -0.10371 | -0.2496 | -0.00702 | 0.060893 | 0.073009 | 0.074579 | 0.152825 | -0.21209 | -0.77948 | -0.28817 |
| 23.04167 | 0.896375 | 0.42757 | 0.071703 | 0.170182 | 0.232959 | -1.05417 | 0.234001 | -0.383 | 0.540766 | -0.27539 | 0.22371 | -0.00486 | -0.06177 | -0.07559 | 0.005345 | 0.033796 | -0.19286 | -0.23799 | -0.15287 |
| 23.05556 | 1.06331 | 0.096645 | -0.16646 | 0.131942 | -0.07894 | -1.92123 | -0.02435 | -0.73774 | 0.439714 | -0.95839 | 0.240931 | -0.00695 | -0.16277 | -0.26281 | -0.03071 | -0.00372 | -0.31468 | -0.18571 | -0.0357 |
| 23.06944 | 0.37927 | 0.398327 | -0.16497 | -0.17327 | -0.15058 | -0.02696 | 0.027964 | -0.10058 | 0.172217 | -0.13703 | 0.100847 | -0.08106 | -0.08982 | -0.26299 | -0.14881 | -0.0876 | -0.03989 | -0.15868 | -0.09243 |
| 23.08333 | -0.26356 | 0.911498 | -0.04185 | -0.30127 | 0.054913 | 2.069573 | 0.208575 | 0.636226 | -0.02188 | 0.758351 | 0.163573 | -0.10082 | 0.045003 | -0.10121 | -0.17895 | -0.11449 | 0.303619 | 0.019968 | -0.10498 |
| 23.09722 | -0.33718 | 0.625108 | -0.08112 | -0.04474 | 0.040449 | 1.488533 | 0.09134 | 0.442255 | -0.12441 | 0.176219 | 0.36671 | -0.06795 | 0.070672 | -0.01962 | -0.10429 | -0.059 | 0.250689 | 0.184518 | -0.02083 |
| 23.11111 | -0.24399 | 0.230545 | -0.21781 | 0.086436 | -0.11358 | 0.16575 | -0.10343 | -0.0323 | -0.22991 | -0.07107 | 0.428593 | -0.09452 | 0.012889 | -0.05113 | -0.05695 | -0.02924 | 0.03964 | 0.288489 | -0.00087 |
| 23.125 | -0.24016 | 0.140214 | -0.31775 | -0.01724 | -0.23587 | -0.25629 | -0.26928 | -0.17924 | -0.33723 | -0.18127 | 0.436189 | -0.12478 | -0.03635 | -0.08639 | -0.04604 | -0.03524 | -0.12926 | 0.3707 | -0.06366 |
| 23.13889 | -0.13246 | 0.185928 | -0.32274 | -0.10676 | -0.294 | -0.60795 | -0.3335 | -0.1702 | -0.34217 | -0.22644 | 0.107006 | -0.14258 | -0.09959 | -0.11317 | -0.10252 | -0.0703 | -0.25659 | 0.44164 | -0.15107 |
| 23.15278 | 0.011133 | 0.303888 | -0.25291 | -0.10782 | -0.26092 | -0.88486 | -0.31926 | -0.1233 | -0.24705 | -0.16661 | -0.32476 | -0.15958 | -0.13024 | -0.142 | -0.15083 | -0.09792 | -0.29041 | 0.618666 | -0.19274 |
| 23.16667 | 0.064049 | 0.421601 | -0.12917 | -0.02987 | -0.06307 | -0.80378 | -0.27495 | -0.13701 | -0.12781 | -0.02677 | -0.2915 | -0.13703 | -0.0451 | -0.11387 | -0.05823 | -0.0371 | -0.14774 | 0.92607 | -0.0943 |
| 23.18056 | -0.07415 | 0.38399 | -0.08195 | -0.02912 | 0.026899 | -0.41934 | -0.35466 | -0.18392 | -0.13581 | 0.032287 | 0.095421 | -0.13507 | 0.028453 | -0.07801 | 0.019242 | 0.030338 | -0.03071 | 1.010156 | -0.01543 |
| 23.19444 | -0.33246 | 0.174656 | -0.20822 | -0.16299 | -0.14758 | -0.17957 | -0.6948 | -0.25357 | -0.28771 | -0.06467 | 0.276246 | -0.1566 | -0.00255 | -0.08539 | -0.01264 | 0.022144 | -0.07813 | 0.770884 | -0.03672 |
| 23.20833 | -0.39738 | -0.03992 | -0.39005 | -0.2597 | -0.39077 | -0.42779 | -1.04308 | -0.37005 | -0.41702 | -0.23811 | 0.19002 | -0.14736 | -0.04063 | -0.09178 | -0.03544 | -0.00939 | -0.14302 | 0.5518 | -0.05638 |
| 23.22222 | -0.28452 | -0.09643 | -0.49904 | -0.27937 | -0.48737 | -0.65806 | -1.01805 | -0.48507 | -0.47026 | -0.32718 | 0.056976 | -0.11166 | -0.04903 | -0.08185 | -0.03841 | -0.02722 | -0.13057 | 0.447764 | -0.05841 |
| 23.23611 | -0.24709 | -0.072 | -0.53083 | -0.31948 | -0.48791 | -0.55269 | -0.81311 | -0.4895 | -0.45922 | -0.15906 | -0.0258 | -0.03789 | -0.0417 | -0.05725 | -0.03342 | -0.01293 | -0.06724 | 0.430187 | -0.05133 |
| 23.25 | -0.3215 | -0.11549 | -0.53879 | -0.3873 | -0.46799 | -0.45383 | -0.60443 | -0.38561 | -0.43339 | 0.074286 | -0.06424 | 0.006303 | -0.05586 | -0.07104 | -0.03503 | -0.01759 | -0.03242 | 0.433687 | -0.05582 |
| 23.26389 | -0.43341 | -0.2645 | -0.55669 | -0.42686 | -0.41973 | -0.47195 | -0.50693 | -0.31503 | -0.43174 | 0.077276 | -0.03145 | -0.06904 | -0.11999 | -0.14157 | -0.10027 | -0.11468 | -0.05656 | 0.365096 | -0.10306 |
| 23.27778 | -0.45907 | -0.36792 | -0.5373 | -0.35766 | -0.4035 | -0.48431 | -0.46563 | -0.3159 | -0.41548 | -0.0235 | 0.112602 | -0.12974 | -0.12675 | -0.1313 | -0.13384 | -0.17529 | -0.02588 | 0.281439 | -0.10201 |
| 23.29167 | -0.27062 | -0.27885 | -0.42036 | -0.10541 | -0.38041 | -0.49872 | -0.4098 | -0.31922 | -0.31647 | 0.005106 | 0.228489 | -0.06128 | -0.0537 | -0.02926 | -0.06759 | -0.10092 | 0.041867 | 0.226794 | -0.03467 |
| 23.30556 | 0.01299 | -0.12246 | -0.29058 | 0.206215 | -0.28925 | -0.617 | -0.33737 | -0.28918 | -0.22613 | 0.129639 | 0.161506 | 0.018078 | -0.0189 | 0.005873 | -0.01531 | -0.02461 | 0.035956 | 0.186686 | 0.007115 |
| 23.31944 | 0.171551 | -0.05448 | -0.21078 | 0.303406 | -0.14771 | -0.62415 | -0.2504 | -0.19434 | -0.19705 | 0.235092 | 0.153857 | 0.035386 | -0.04368 | -0.02435 | -0.04311 | -0.03813 | 0.010209 | 0.203258 | 0.03713 |
| 23.33333 | 0.155378 | -0.03744 | -0.1588 | 0.177001 | 0.027655 | -0.43926 | -0.13525 | -0.09145 | -0.17502 | 0.277935 | 0.330418 | 0.016626 | -0.07235 | -0.04402 | -0.0713 | -0.09229 | 0.027222 | 0.290601 | 0.133009 |
| 23.34722 | 0.116236 | -0.00587 | -0.10894 | 0.044109 | 0.213703 | -0.29366 | -0.01065 | -0.05521 | -0.09317 | 0.297067 | 0.499714 | 0.01216 | -0.06534 | -0.01565 | -0.05345 | -0.10102 | 0.087269 | 0.309084 | 0.303247 |
| 23.36111 | 0.120537 | 0.061801 | -0.04477 | 0.07536 | 0.370112 | -0.20273 | 0.059355 | 0.086952 | 0.070118 | 0.313734 | 0.476372 | 0.050441 | -0.0206 | 0.051476 | -0.00314 | -0.03774 | 0.146584 | 0.255722 | 0.468553 |
| 23.375 | 0.222737 | 0.236609 | 0.097771 | 0.226252 | 0.45976 | -0.05729 | 0.093077 | 0.374749 | 0.264579 | 0.385145 | 0.380038 | 0.104801 | 0.024866 | 0.100401 | 0.054277 | 0.05807 | 0.158081 | 0.285952 | 0.586124 |
| 23.38889 | 0.383703 | 0.483584 | 0.234893 | 0.435948 | 0.509314 | 0.065908 | 0.116069 | 0.659357 | 0.356337 | 0.46287 | 0.220148 | 0.09689 | 0.011663 | 0.088772 | 0.051736 | 0.095116 | 0.101802 | 0.393214 | 0.576441 |
| 23.40278 | 0.489563 | 0.660831 | 0.252481 | 0.610072 | 0.503324 | 0.053191 | 0.131408 | 0.865002 | 0.292576 | 0.419581 | 0.057077 | -0.00292 | -0.07801 | 0.013507 | -0.01911 | 0.042004 | 0.01817 | 0.453772 | 0.4231 |
| 23.41667 | 0.516265 | 0.70295 | 0.220368 | 0.673662 | 0.540728 | -0.05919 | 0.181686 | 0.928265 | 0.173242 | 0.367545 | 0.020577 | -0.05943 | -0.11975 | -0.02197 | -0.04611 | 0.020392 | 0.032419 | 0.457599 | 0.300151 |
| 23.43056 | 0.47075 | 0.636208 | 0.193443 | 0.617327 | 0.614923 | -0.18408 | 0.336891 | 1.040358 | 0.144006 | 0.420736 | 0.072904 | -0.00807 | -0.06194 | 0.022254 | 0.017008 | 0.086274 | 0.124446 | 0.47032 | 0.319799 |
| 23.44444 | 0.348143 | 0.592356 | 0.215746 | 0.457893 | 0.592473 | -0.10664 | 0.621235 | 1.061908 | 0.211998 | 0.461343 | -0.03453 | 0.04858 | -0.02588 | 0.043576 | 0.079257 | 0.13315 | 0.129927 | 0.490099 | 0.368926 |
| 23.45833 | 0.248899 | 0.556179 | 0.252331 | 0.339532 | 0.476698 | -0.01201 | 0.884163 | 0.904884 | 0.262251 | 0.475027 | -0.18176 | 0.071737 | -0.04701 | 0.025448 | 0.084659 | 0.110875 | 0.067306 | 0.461874 | 0.431599 |
| 23.47222 | 0.179312 | 0.474397 | 0.226787 | 0.200458 | 0.356052 | 0.084349 | 0.934602 | 0.55105 | 0.230049 | 0.497806 | -0.15411 | 0.092289 | -0.05227 | -0.00494 | 0.058705 | 0.085725 | 0.059416 | 0.468091 | 0.510259 |
| 23.48611 | -0.00884 | 0.422229 | 0.048386 | 0.028437 | 0.254622 | 0.472531 | 0.821058 | 0.22159 | 0.070827 | 0.743819 | 0.0812 | 0.100133 | -0.00434 | 0.019316 | 0.05953 | 0.069651 | 0.293946 | 0.779393 | 0.687671 |
| 23.5 | -0.35636 | 0.394986 | -0.24665 | -0.1462 | 0.324331 | 1.146637 | 0.825702 | -0.0301 | -0.07463 | 1.214621 | 0.37745 | 0.071995 | 0.043715 | 0.090231 | 0.133869 | 0.051899 | 0.809075 | 1.491895 | 1.090846 |
| 23.51389 | -0.51194 | 0.343308 | -0.29055 | 0.072432 | 0.701265 | 0.806721 | 0.646857 | -0.07772 | -0.01588 | 1.156289 | 0.586437 | 0.152662 | 0.078037 | 0.061277 | 0.090248 | 0.037502 | 1.18892 | 1.96295 | 1.35975 |
| 23.52778 | -0.25209 | 0.095982 | -0.05979 | 0.398134 | 0.526293 | 0.11139 | 0.506921 | 0.103625 | 0.122923 | 0.610802 | 0.495818 | 0.274573 | 0.065507 | 0.008134 | 0.049373 | 0.063823 | 1.285016 | 2.086995 | 1.101467 |
| 23.54167 | 0.058118 | -0.07739 | 0.099829 | 0.49803 | 0.275716 | -0.21884 | 0.345985 | 0.336336 | -0.05424 | 0.365359 | 0.637309 | 0.280038 | 0.02004 | 0.01198 | 0.056213 | 0.199824 | 1.121107 | 1.495705 | 0.836898 |
| 23.55556 | 0.228244 | -0.11025 | 0.042266 | 0.253144 | -0.14611 | -0.17271 | 0.232973 | 0.579843 | -0.3917 | 0.128535 | 0.894104 | 0.097855 | -0.01651 | -0.00139 | 0.028256 | 0.222217 | 0.836829 | 0.492531 | 0.461477 |
| 23.56944 | 0.381085 | -0.00297 | 0.047663 | 0.043221 | -0.49731 | -0.08416 | 0.11361 | 0.892098 | -0.54354 | -0.06089 | 1.052509 | -0.00695 | -0.01794 | -0.00792 | 0.034088 | 0.209534 | 0.633418 | -0.05032 | 0.232258 |
| 23.58333 | 0.646887 | 0.238604 | 0.244825 | -0.0354 | -0.56271 | 0.015287 | 0.020247 | 1.171574 | -0.38946 | -0.13741 | 1.637446 | 0.009436 | 0.033132 | 0.051167 | 0.112631 | 0.156253 | 0.535446 | -0.32029 | 0.338868 |
| 23.59722 | 0.86818 | 0.438051 | 0.465995 | -0.10394 | -0.28941 | 0.04402 | -0.00188 | 1.267003 | -0.11094 | -0.12942 | 2.772092 | 0.038058 | 0.093579 | 0.071718 | 0.18354 | 0.080276 | 0.371555 | -0.50352 | 0.732114 |
| 23.61111 | 0.80433 | 0.402306 | 0.535724 | -0.09606 | 0.05487 | 0.008173 | 0.195835 | 1.237505 | 0.219285 | -0.02943 | 3.78733 | 0.007363 | 0.081951 | 0.022842 | 0.189762 | -0.02754 | 0.399148 | -0.71286 | 1.149692 |
| 23.625 | 0.579465 | 0.397318 | 0.428423 | 0.017637 | 0.425202 | -0.04724 | 0.737333 | 1.252868 | 0.521073 | 0.170578 | 3.487742 | 0.040185 | 0.05884 | 0.040701 | 0.18972 | -0.07627 | 0.513369 | -0.78195 | 1.350844 |
| 23.63889 | 0.441341 | 0.682414 | 0.191844 | 0.243547 | 0.657622 | -0.01697 | 1.293049 | 1.210805 | 0.599689 | 0.419415 | 2.306375 | 0.094131 | 0.146096 | 0.23374 | 0.230976 | -0.05202 | 0.427822 | -0.47714 | 1.332805 |
| 23.65278 | 0.400389 | 1.157168 | -0.11513 | 0.581909 | 1.02233 | 0.09817 | 1.235231 | 1.016517 | 0.580429 | 0.445887 | 0.841132 | 0.094494 | 0.216537 | 0.354713 | 0.243149 | -0.02205 | 0.081554 | 0.297647 | 0.974878 |
| 23.66667 | 0.354399 | 1.38432 | -0.32039 | 0.738271 | 0.961697 | 0.111257 | 0.828884 | 0.704041 | 0.426389 | 0.232666 | -0.72864 | 0.01463 | 0.178992 | 0.173898 | 0.170718 | 0.017997 | -0.09367 | 1.05524 | 0.478327 |
| 23.68056 | 0.351233 | 1.162101 | -0.27963 | 0.636614 | 0.386509 | -0.03095 | 0.381843 | 0.440687 | 0.20479 | -0.06282 | -1.97988 | -0.06451 | 0.114145 | -0.00604 | 0.05498 | 0.125621 | 0.042865 | 1.080401 | 0.068791 |
| 23.69444 | 0.393775 | 0.634957 | -0.04769 | 0.285008 | -0.07188 | -0.09782 | 0.091648 | 0.181382 | 0.036329 | -0.27081 | -2.9055 | -0.0575 | 0.047979 | -0.06631 | 0.004355 | 0.209044 | 0.127432 | 0.465841 | -0.19644 |
| 23.70833 | 0.384489 | 0.066376 | 0.241589 | -0.11781 | 0.026473 | -0.11183 | -0.19598 | -0.25125 | -0.1054 | -0.14233 | -3.17886 | -0.02415 | -0.0519 | -0.1137 | -0.01197 | 0.19177 | -0.0405 | -0.16819 | -0.35428 |
| 23.72222 | 0.114067 | -0.50249 | 0.391774 | -0.53252 | 0.177694 | -0.14723 | -0.58234 | -0.71927 | -0.25258 | 0.082997 | -2.53337 | -0.03788 | -0.14469 | -0.16861 | -0.05135 | 0.101333 | -0.25266 | -0.71269 | -0.36677 |
| 23.73611 | -0.17168 | -0.73938 | 0.337785 | -0.75683 | 0.043391 | -0.12783 | -0.63028 | -1.07103 | -0.35476 | 0.240736 | -1.88922 | -0.06209 | -0.15645 | -0.11357 | -0.09192 | 0.019084 | -0.36143 | -0.92201 | -0.23383 |
| 23.75 | -0.33543 | -0.56301 | 0.133735 | -0.63622 | -0.27718 | -0.0175 | -0.39976 | -1.02384 | -0.35781 | 0.074615 | -1.69151 | -0.04538 | -0.04698 | 0.032524 | -0.06327 | -0.04148 | -0.41957 | -0.63183 | -0.04815 |
| 23.76389 | -0.45374 | -0.33133 | -0.13965 | -0.31219 | -0.46915 | 0.040794 | -0.16469 | -0.7966 | -0.31233 | -0.2656 | -1.41293 | -0.03232 | -0.00041 | 0.090104 | -0.04784 | -0.12668 | -0.46833 | -0.05324 | 0.03288 |
| 23.77778 | -0.4586 | -0.18497 | -0.35068 | -0.07762 | -0.60362 | -0.01289 | -0.21593 | -0.44691 | -0.22813 | -0.81979 | -1.16364 | -0.07506 | -0.08972 | 0.006764 | -0.13367 | -0.22767 | -0.61241 | 0.164562 | -0.05654 |
| 23.79167 | -0.24173 | -0.00219 | -0.44694 | -0.02897 | -0.75444 | -0.05682 | -0.31908 | -0.12489 | -0.04151 | -0.88946 | -1.00772 | -0.09927 | -0.11929 | -0.03462 | -0.15429 | -0.25589 | -0.88228 | -0.08426 | -0.27278 |
| 23.80556 | -0.04483 | 0.035723 | -0.51557 | -0.11706 | -0.84439 | -0.08899 | -0.41873 | -0.13984 | 0.098189 | -0.41714 | -0.65214 | -0.07854 | -0.05733 | -0.01724 | -0.1057 | -0.17544 | -1.04719 | -0.41564 | -0.64262 |
| 23.81944 | -0.09495 | -0.30764 | -0.60773 | -0.24398 | -1.01099 | -0.18999 | -0.75569 | -0.60055 | -0.03274 | 0.071206 | -0.15852 | -0.11884 | -0.06006 | -0.08392 | -0.16335 | -0.10242 | -0.99134 | -0.65386 | -1.0596 |
| 23.83333 | -0.31363 | -0.82829 | -0.61839 | -0.36886 | -1.29818 | -0.28386 | -1.06813 | -1.20307 | -0.35469 | 0.20777 | 0.069364 | -0.21356 | -0.10627 | -0.19251 | -0.32695 | -0.11692 | -0.91894 | -0.76079 | -1.26239 |
| 23.84722 | -0.52187 | -1.17081 | -0.54423 | -0.42518 | -1.58909 | -0.31281 | -1.21187 | -1.58133 | -0.5824 | -0.11248 | -0.13775 | -0.25819 | -0.0928 | -0.24702 | -0.43666 | -0.18231 | -0.94253 | -0.78487 | -1.22646 |
| 23.86111 | -0.6749 | -1.32703 | -0.4277 | -0.48639 | -1.91656 | -0.30615 | -1.27102 | -1.7838 | -0.70832 | -0.66408 | -0.52755 | -0.27657 | 0.01846 | -0.22911 | -0.50919 | -0.26535 | -1.05796 | -0.93351 | -1.10841 |
| 23.875 | -0.83935 | -1.32982 | -0.43733 | -0.62232 | -2.11881 | -0.35662 | -1.35924 | -1.80085 | -0.81038 | -1.40911 | -0.99637 | -0.28823 | -0.01322 | -0.20771 | -0.54932 | -0.32969 | -1.04865 | -1.13425 | -0.95638 |
| 23.88889 | -0.97861 | -1.3054 | -0.47976 | -0.68576 | -1.95646 | -0.38497 | -1.1869 | -1.55999 | -0.81884 | -1.35466 | -1.2094 | -0.22303 | -0.12277 | -0.13445 | -0.48272 | -0.33254 | -0.85985 | -1.13068 | -0.87633 |
| 23.90278 | -0.97659 | -1.2137 | -0.45094 | -0.54541 | -1.70921 | -0.3817 | -0.7825 | -1.30831 | -0.73638 | -0.62176 | -1.07509 | -0.15557 | -0.13706 | -0.07809 | -0.32834 | -0.22058 | -0.50438 | -0.94559 | -0.94668 |
| 23.91667 | -0.87782 | -1.01991 | -0.39927 | -0.43039 | -1.52355 | -0.39417 | -0.75172 | -1.27761 | -0.67819 | 0.09448 | -0.45669 | -0.27184 | -0.12321 | -0.13491 | -0.23037 | -0.08334 | -0.24116 | -0.76835 | -1.12896 |
| 23.93056 | -0.71384 | -0.82733 | -0.36178 | -0.43866 | -1.41828 | -0.45725 | -1.01242 | -1.2575 | -0.61405 | 0.372031 | 0.189402 | -0.33784 | -0.09878 | -0.20254 | -0.20652 | -0.0311 | -0.13411 | -0.62196 | -1.12801 |

| | | | | | | | | | | | | | | | | | | | |
|---|---|---|---|---|---|---|---|---|---|---|---|---|---|---|---|---|---|---|---|
| 23.94444 | -0.53888 | -0.75361 | -0.31286 | -0.41909 | -1.40322 | -0.48688 | -1.05418 | -1.13631 | -0.50776 | -0.10355 | 0.463057 | -0.2454 | -0.00905 | -0.17193 | -0.14633 | -0.08767 | -0.0646 | -0.47993 | -0.98209 |
| 23.95833 | -0.51043 | -0.86848 | -0.27634 | -0.38387 | -1.46193 | -0.51246 | -0.90095 | -0.98626 | -0.45941 | -0.88031 | 0.287292 | -0.13983 | 0.047816 | -0.13178 | -0.09995 | -0.11282 | -0.01942 | -0.36375 | -0.77389 |
| 23.97222 | -0.59948 | -1.07153 | -0.25986 | -0.43808 | -1.3477 | -0.56769 | -0.85438 | -0.83196 | -0.51214 | -1.40615 | -0.08087 | -0.12396 | 0.015241 | -0.10886 | -0.12665 | -0.12357 | -0.11203 | -0.37593 | -0.61554 |
| 23.98611 | -0.49832 | -1.28585 | -0.19654 | -0.51031 | -0.90521 | -0.90434 | -0.86655 | -0.54756 | -0.52969 | -1.17938 | -0.42014 | -0.0736 | -0.03078 | -0.0506 | -0.08875 | -0.047 | -0.47845 | -0.71501 | -0.51989 |
| 24 | -0.19741 | -1.39282 | -0.05205 | -0.44656 | -0.40041 | -1.56528 | -0.98575 | -0.26736 | -0.3577 | -0.69375 | -0.74113 | 0.093611 | 0.003204 | 0.052876 | 0.024979 | 0.152038 | -0.90279 | -1.3085 | -0.49255 |
| 24.01389 | -0.08944 | -1.09244 | 0.008749 | -0.29912 | -0.14999 | -1.44573 | -1.1025 | -0.0696 | -0.10639 | -0.36762 | -0.97657 | 0.150539 | 0.117705 | 0.111949 | 0.117581 | 0.287072 | -0.84746 | -1.23124 | -0.41784 |
| 24.02778 | -0.29305 | -0.67978 | -0.10247 | -0.14538 | -0.316 | -0.29653 | -1.01428 | -0.01615 | -0.06122 | -0.24048 | -1.14594 | 0.019578 | 0.141793 | 0.037169 | 0.122526 | 0.162699 | -0.42822 | -0.6673 | -0.31703 |
| 24.04167 | -0.2171 | -0.38549 | -0.17034 | -0.22304 | -0.29431 | 0.181849 | -0.73044 | -0.27023 | -0.16671 | -0.21196 | -1.04158 | -0.07805 | 0.07119 | -0.07209 | 0.083507 | 0.025053 | -0.28047 | -0.45287 | -0.30726 |
| 24.05556 | 0.187441 | 0.01895 | -0.11875 | -0.48429 | -0.06959 | 0.017359 | -0.5073 | -0.49595 | -0.21605 | -0.09648 | -0.56182 | -0.02339 | 0.013009 | -0.13546 | 0.066488 | -0.02247 | -0.32332 | -0.41106 | -0.24221 |
| 24.06944 | 0.482919 | 0.386938 | -0.07896 | -0.58059 | 0.003247 | -0.07816 | -0.36171 | -0.31073 | -0.17982 | 0.16179 | 0.037959 | 0.068997 | -0.0128 | -0.14881 | 0.059622 | -0.03597 | -0.23791 | -0.3263 | -0.14133 |
| 24.08333 | 0.349913 | 0.3854 | -0.10098 | -0.45394 | -0.07663 | -0.0331 | -0.28801 | -0.05464 | -0.13114 | 0.379126 | 0.556255 | 0.073751 | -0.04621 | -0.13918 | 0.044149 | -0.06273 | -0.0012 | -0.2782 | -0.09785 |
| 24.09722 | -0.03538 | 0.139608 | -0.25415 | -0.26364 | -0.20031 | 0.248997 | -0.29514 | 0.076835 | -0.17077 | 0.471768 | 0.806517 | -0.03606 | -0.0689 | -0.13439 | -0.01285 | -0.12139 | 0.265532 | -0.24695 | -0.05621 |
| 24.11111 | -0.37311 | -0.00521 | -0.4282 | -0.11939 | -0.23585 | 0.723481 | -0.30334 | 0.000831 | -0.27593 | 0.540623 | 0.766515 | -0.13072 | -0.03046 | -0.12197 | -0.07726 | -0.17162 | 0.499423 | -0.04814 | 0.05914 |
| 24.125 | -0.26624 | 0.073489 | -0.27589 | 0.071282 | 0.030371 | 0.875489 | -0.12159 | 0.045995 | -0.19902 | 0.955105 | 0.892411 | -0.06511 | 0.071618 | -0.0019 | 0.020808 | -0.10974 | 0.637899 | 0.220224 | 0.19728 |
| 24.13889 | 0.09323 | 0.249927 | 0.006783 | 0.236784 | 0.358441 | 0.486247 | 0.122949 | 0.132114 | 0.013103 | 0.045987 | 1.062181 | 0.064566 | 0.122936 | 0.086101 | 0.135683 | -0.00189 | 0.38779 | 0.327391 | 0.148298 |
| 24.15278 | 0.267468 | 0.320862 | 0.05602 | 0.215304 | 0.320002 | 0.133439 | 0.216302 | 0.063378 | 0.084949 | -0.27493 | 0.925728 | 0.029862 | 0.028826 | -0.01684 | 0.062494 | -0.01921 | -0.07611 | 0.172819 | -0.06341 |
| 24.16667 | 0.200468 | 0.290821 | -0.25823 | 0.089904 | 0.137361 | 0.010679 | 0.196068 | -0.07509 | -0.01031 | -0.71642 | -0.41077 | -0.21713 | -0.20104 | -0.24501 | -0.2097 | -0.20591 | -0.9882 | -0.32236 | -0.74328 |
| 24.18056 | -0.17263 | -0.23741 | -0.78154 | -0.43161 | 0.114495 | -0.1152 | -0.06185 | -0.39026 | -0.15226 | -1.53879 | -3.26914 | -0.68928 | -0.5201 | -0.58143 | -0.72737 | -0.32413 | -3.01189 | -1.49826 | -2.35611 |
| 24.19444 | -0.74714 | -1.40732 | -0.91993 | -1.52896 | 0.040458 | -0.28511 | -0.69262 | -0.98666 | -0.22109 | -3.52419 | -4.55307 | -0.74303 | -0.56418 | -0.73989 | -1.40314 | -0.23221 | -4.24015 | -3.96996 | -3.18549 |
| 24.20833 | -0.90564 | -1.69505 | -0.55947 | -1.86961 | -0.08284 | -0.27054 | -1.00806 | -1.15869 | -0.26298 | -3.411 | -1.83543 | -0.28694 | -0.16478 | -0.53678 | -0.74372 | 0.24072 | -2.11768 | -3.41912 | -1.66527 |
| 24.22222 | -0.60677 | -0.88223 | -0.22908 | -1.08817 | -0.17788 | -0.19833 | -0.7887 | -0.69781 | -0.23696 | -0.61511 | 0.053145 | 0.014299 | 0.042523 | -0.24671 | -0.20346 | 0.195473 | -0.21945 | -0.74941 | -0.00543 |
| 24.23611 | -0.3753 | -0.36459 | -0.1537 | -0.50295 | -0.20755 | -0.27115 | -0.61743 | -0.38902 | -0.1887 | 0.287791 | 0.760965 | 0.144998 | 0.058594 | -0.11783 | -0.04523 | 0.064662 | 0.434353 | 0.148627 | 0.550572 |
| 24.25 | -0.20886 | -0.22215 | -0.22547 | -0.31981 | -0.17249 | -0.39786 | -0.61326 | -0.3688 | -0.17543 | 0.326549 | 0.542842 | 0.181059 | 0.005534 | -0.09626 | -0.06653 | 0.012388 | 0.412343 | 0.243269 | 0.347456 |
| 24.26389 | -0.13184 | -0.17208 | -0.39867 | -0.22496 | -0.1531 | -0.36205 | -0.67912 | -0.33395 | -0.29248 | 0.208214 | 0.223617 | 0.066139 | -0.13018 | -0.17411 | -0.18232 | -0.06641 | 0.164024 | 0.221547 | 0.015062 |
| 24.27778 | -0.18755 | -0.21863 | -0.50088 | -0.10182 | -0.12817 | -0.29534 | -0.69945 | -0.1766 | -0.42356 | 0.124022 | 0.043683 | -0.04517 | -0.18964 | -0.16873 | -0.1975 | -0.09374 | -0.05569 | 0.19119 | -0.08255 |
| 24.29167 | -0.145 | -0.19947 | -0.37259 | 0.183072 | 0.029999 | -0.27829 | -0.54251 | 0.039917 | -0.33942 | 0.137007 | 0.029512 | -0.03669 | -0.16544 | -0.0687 | -0.07904 | -0.03762 | -0.18989 | 0.099414 | -0.04808 |
| 24.30556 | 0.050237 | -0.04961 | -0.13591 | 0.518264 | 0.243166 | -0.27587 | -0.29639 | 0.147927 | -0.14474 | 0.140273 | 0.019253 | -0.00109 | -0.14699 | -0.0249 | -0.02923 | -0.00797 | -0.27914 | 0.016355 | -0.0544 |
| 24.31944 | 0.361907 | 0.19445 | 0.091615 | 0.767736 | 0.435876 | -0.15367 | -0.03827 | 0.208898 | 0.046932 | 0.011718 | -0.03491 | -0.01337 | -0.09867 | -0.01618 | -0.04082 | -0.03969 | -0.28268 | 0.005104 | -0.12168 |
| 24.33333 | 0.691259 | 0.442766 | 0.291628 | 0.884179 | 0.550832 | 0.098496 | 0.180217 | 0.318124 | 0.221251 | -0.10089 | -0.06252 | 0.010258 | -0.05968 | 0.019207 | -0.02799 | -0.04055 | -0.1992 | 0.045435 | -0.16287 |
| 24.34722 | 0.797539 | 0.434107 | 0.315156 | 0.773159 | 0.456463 | 0.125438 | 0.194964 | 0.327084 | 0.24597 | -0.09382 | -0.05932 | 0.070386 | -0.04933 | 0.023813 | -0.01153 | -0.01125 | -0.13435 | 0.111477 | -0.16377 |
| 24.36111 | 0.510162 | 0.083222 | 0.131871 | 0.533139 | 0.247806 | -0.17402 | 0.037525 | 0.230571 | 0.062765 | -0.06986 | -0.08063 | 0.027055 | -0.05078 | -0.0339 | -0.06663 | -0.05089 | -0.13959 | 0.142817 | -0.18056 |
| 24.375 | 0.07544 | -0.29274 | -0.08719 | 0.363412 | 0.152569 | -0.51458 | -0.10983 | 0.163941 | -0.19024 | 0.027737 | -0.04667 | -0.00836 | -0.01382 | -0.01577 | -0.08043 | -0.06693 | -0.07973 | 0.170978 | -0.13734 |
| 24.38889 | -0.25594 | -0.51973 | -0.35463 | 0.074271 | 0.142433 | -0.86131 | -0.22931 | -0.02911 | -0.43728 | 0.347669 | 0.056547 | 0.02774 | 0.069991 | 0.071344 | -0.0307 | -0.01518 | 0.061434 | 0.213745 | -0.01332 |
| 24.40278 | -0.36102 | -0.52392 | -0.68437 | -0.29196 | 0.079686 | -1.06589 | -0.35693 | -0.42009 | -0.63276 | 0.757375 | 0.074949 | -0.0441 | 0.045365 | 0.036951 | -0.05522 | -0.04554 | 0.095061 | 0.197659 | 0.028518 |
| 24.41667 | 0.003181 | -0.04734 | -0.63433 | -0.16763 | 0.152571 | -0.86069 | -0.16955 | -0.36952 | -0.46288 | 1.157664 | 0.082359 | -0.07977 | 0.037886 | 0.015368 | -0.02217 | -0.05751 | 0.110185 | 0.189405 | 0.049704 |
| 24.43056 | 0.598901 | 0.654828 | -0.17049 | 0.452984 | 0.397692 | -0.33428 | 0.353826 | 0.113348 | 0.022884 | 1.476076 | 0.214965 | 0.019104 | 0.160489 | 0.053703 | 0.060555 | 0.009951 | 0.197247 | 0.235057 | 0.124921 |
| 24.44444 | 0.513833 | 0.742605 | 0.173705 | 0.875672 | 0.610188 | 0.027371 | 0.659965 | 0.471749 | 0.3243 | 1.563203 | 0.657863 | 0.037608 | 0.243977 | -0.04331 | 0.026947 | -0.00762 | 0.201435 | 0.219378 | 0.109264 |
| 24.45833 | 0.216434 | 0.609183 | 0.341978 | 0.905258 | 0.76223 | 0.244026 | 0.665712 | 0.667824 | 0.453836 | 1.44759 | 1.947844 | 0.007393 | 0.176973 | -0.13029 | -0.00508 | -0.02257 | 0.212816 | 0.18887 | 0.058375 |
| 24.47222 | 0.46076 | 0.80862 | 0.60929 | 0.971545 | 0.940587 | 0.581587 | 0.85169 | 1.008417 | 0.669619 | 1.522712 | 2.060657 | 0.078786 | 0.137689 | -0.11553 | 0.089037 | 0.079044 | 0.361099 | 0.284102 | 0.137238 |
| 24.48611 | 1.179072 | 1.524972 | 0.833962 | 1.048084 | 1.108262 | 0.90669 | 1.363496 | 1.33816 | 0.953064 | 2.15523 | 2.204787 | 0.149805 | 0.09758 | -0.17451 | 0.144277 | 0.190238 | 0.644992 | 0.562509 | 0.354796 |
| 24.5 | 1.495018 | 2.164418 | 0.741755 | 0.8766 | 1.208825 | 1.117874 | 2.082601 | 1.437605 | 1.198608 | 3.32002 | 2.961823 | 0.199758 | -0.02815 | -0.17953 | 0.108873 | 0.230141 | 1.182418 | 0.895675 | 0.604297 |
| 24.51389 | 1.03674 | 1.848269 | 0.695725 | 0.621901 | 1.416465 | 0.898886 | 2.515457 | 1.391499 | 1.085975 | 3.474076 | 2.814567 | 0.203222 | -0.10983 | -0.14156 | 0.071258 | 0.206352 | 1.450537 | 0.955175 | 0.578433 |
| 24.52778 | 0.732955 | 1.161753 | 0.863493 | 0.568085 | 1.67351 | 0.483421 | 2.482348 | 1.620287 | 0.727807 | 1.907287 | 1.584336 | 0.259403 | -0.0732 | -0.05941 | 0.07317 | 0.15223 | 1.008622 | 0.772869 | 0.399674 |
| 24.54167 | 0.590713 | 0.631908 | 0.728374 | 0.639394 | 0.963922 | 0.312279 | 1.797918 | 2.069714 | 0.510116 | 0.808408 | 0.795968 | 0.243367 | -0.02439 | -0.02226 | 0.126662 | 0.108862 | 0.495449 | 0.587118 | 0.272387 |
| 24.55556 | 0.40267 | 0.230254 | 0.208943 | 0.650381 | -0.044 | 0.349219 | 0.759351 | 2.175103 | 0.195103 | 0.332722 | 0.425208 | 0.147028 | 0.001347 | -0.01517 | 0.153883 | 0.090008 | 0.287905 | 0.313334 | 0.208881 |
| 24.56944 | 0.167237 | 0.003591 | -0.24407 | 0.365779 | -0.81844 | 0.474028 | -0.12062 | 1.73609 | -0.10701 | 0.18652 | 0.273221 | 0.10125 | 0.032332 | 0.036556 | 0.154543 | 0.091021 | 0.459181 | 0.194704 | 0.199201 |
| 24.58333 | 0.250836 | -0.02329 | -0.23933 | 0.089659 | -1.22802 | 0.577086 | -0.46524 | 1.03918 | -0.25193 | 0.149745 | 0.199078 | 0.097926 | 0.054427 | 0.108746 | 0.132192 | 0.091601 | 1.052844 | 0.172616 | 0.229709 |
| 24.59722 | 0.530143 | 0.06378 | 0.176936 | 0.057738 | -0.96861 | 0.559092 | -0.04969 | 0.764415 | -0.22693 | 0.148549 | 0.163709 | 0.093499 | 0.068001 | 0.148668 | 0.098044 | 0.077619 | 0.836124 | 0.155541 | 0.279726 |
| 24.61111 | 0.928888 | 0.330398 | 0.822488 | 0.255133 | -0.09894 | 0.654939 | 1.048061 | 0.968313 | 0.061504 | 0.140367 | 0.127994 | 0.087697 | 0.064245 | 0.154778 | 0.054396 | 0.053096 | 0.402583 | 0.119895 | 0.283118 |
| 24.625 | 1.15316 | 0.789946 | 1.347562 | 0.424172 | 1.048785 | 0.816671 | 2.20907 | 1.24052 | 0.602891 | 0.11264 | 0.05762 | 0.087462 | 0.046974 | 0.196057 | 0.028767 | 0.038534 | 0.232817 | 0.093886 | 0.250597 |
| 24.63889 | 1.104006 | 1.390924 | 1.181623 | 0.436956 | 1.871392 | 0.824614 | 2.072327 | 1.179744 | 0.983491 | 0.089786 | -0.03681 | 0.095507 | 0.046077 | 0.262048 | 0.049308 | 0.045545 | 0.203982 | 0.144939 | 0.250399 |
| 24.65278 | 0.817657 | 1.850985 | 0.576893 | 0.3836 | 1.961315 | 0.723803 | 1.064555 | 1.135483 | 1.124677 | 0.094319 | -0.09126 | 0.104186 | 0.063022 | 0.317485 | 0.084726 | 0.055623 | 0.200343 | 0.266323 | 0.241078 |
| 24.66667 | 0.487591 | 1.778282 | 0.017093 | 0.375906 | 1.219871 | 0.519177 | 0.045853 | 1.392076 | 1.135787 | 0.116022 | -0.04524 | 0.114578 | 0.069883 | 0.319846 | 0.089794 | 0.060304 | 0.154259 | 0.314339 | 0.228245 |
| 24.68056 | 0.32781 | 1.159754 | -0.21021 | 0.586969 | 0.319012 | 0.254442 | -0.53069 | 1.686777 | 0.956226 | 0.139657 | 0.039829 | 0.114312 | 0.060445 | 0.268343 | 0.06427 | 0.059689 | 0.105894 | 0.270173 | 0.234709 |
| 24.69444 | 0.387817 | 0.645456 | -0.15568 | 0.760407 | -0.44855 | 0.056988 | -0.77225 | 1.727168 | 0.673734 | 0.146127 | 0.07269 | 0.089337 | 0.043289 | 0.1806 | 0.031815 | 0.048233 | 0.094578 | 0.218168 | 0.235363 |
| 24.70833 | 0.563812 | 0.34163 | 0.037764 | 0.750911 | -0.80408 | -0.04345 | -0.42451 | 1.376279 | 0.370174 | 0.194811 | 0.067331 | 0.0745 | 0.03882 | 0.12341 | 0.029799 | 0.050068 | 0.134005 | 0.195686 | 0.216504 |
| 24.72222 | 0.771512 | 0.101622 | 0.312673 | 0.604035 | -0.85318 | -0.05455 | 0.182872 | 0.790192 | 0.122527 | 0.279436 | 0.086364 | 0.093084 | 0.065556 | 0.119717 | 0.070677 | 0.080848 | 0.185286 | 0.20512 | 0.221783 |
| 24.73611 | 0.7205 | -0.18855 | 0.504671 | 0.201466 | -0.72753 | -0.04793 | 0.576916 | -0.04833 | -0.01003 | 0.248321 | 0.107256 | 0.0791 | 0.074943 | 0.114732 | 0.089398 | 0.097337 | 0.182318 | 0.215232 | 0.220925 |
| 24.75 | 0.385542 | -0.55209 | 0.389745 | -0.36722 | -0.57034 | -0.09563 | 0.111739 | -0.97574 | -0.09865 | 0.14865 | 0.07467 | -0.0094 | 0.033185 | 0.092834 | 0.043545 | 0.060773 | 0.109893 | 0.179066 | 0.165593 |
| 24.76389 | 0.062329 | -0.71993 | 0.059946 | -0.64598 | -0.24298 | -0.0832 | -0.89895 | -1.82552 | -0.16906 | 0.071603 | 0.033721 | -0.03739 | 0.007455 | 0.121661 | 0.010269 | 0.01899 | 0.052483 | 0.137109 | 0.105277 |
| 24.77778 | -0.11017 | -0.4804 | -0.23609 | -0.38211 | 0.185576 | 0.046576 | -1.78267 | -2.00643 | -0.15154 | 0.076443 | 0.054428 | -0.0043 | 0.046817 | 0.18272 | 0.044626 | 0.036802 | 0.073691 | 0.154468 | 0.109129 |
| 24.79167 | -0.30564 | 0.015502 | -0.47309 | -0.13307 | 0.515113 | 0.143872 | -2.00243 | -1.5359 | -0.11721 | 0.09036 | 0.075818 | 0.042704 | 0.067002 | 0.165384 | 0.063671 | 0.058941 | 0.091699 | 0.183194 | 0.117093 |
| 24.80556 | -0.47259 | 0.517387 | -0.58765 | -0.27083 | 0.506967 | 0.120512 | -1.03667 | -0.95829 | -0.09857 | 0.055984 | 0.049195 | 0.035225 | 0.036863 | 0.107038 | 0.035163 | 0.035865 | 0.06953 | 0.168939 | 0.080595 |
| 24.81944 | -0.3462 | 0.67795 | -0.59754 | -0.38233 | 0.221935 | 0.056714 | 0.294002 | -0.39604 | 0.019553 | 0.033514 | 0.048769 | 0.033891 | 0.03556 | 0.099705 | 0.034169 | 0.02511 | 0.071622 | 0.145938 | 0.059619 |
| 24.83333 | -0.14363 | 0.340381 | -0.53252 | -0.29497 | -0.23372 | -0.05601 | 0.704828 | -0.15188 | 0.115981 | 0.048755 | 0.071114 | 0.053355 | 0.048368 | 0.076717 | 0.047993 | 0.041064 | 0.079675 | 0.141575 | 0.058891 |
| 24.84722 | -0.22913 | -0.18748 | -0.58117 | -0.32638 | -0.84872 | -0.31747 | -0.01201 | -0.48894 | 0.010295 | 0.036198 | 0.065143 | 0.039843 | 0.02401 | 0.017076 | 0.032828 | 0.03323 | 0.055882 | 0.127496 | 0.036382 |
| 24.86111 | -0.47561 | -0.61883 | -0.61924 | -0.49039 | -1.50877 | -0.62749 | -1.02921 | -1.01922 | -0.25571 | 0.00623 | 0.061293 | 0.022311 | 0.013015 | 0.015166 | 0.034695 | 0.016841 | 0.044022 | 0.116005 | 0.022442 |
| 24.875 | -0.68243 | -0.83539 | -0.64206 | -0.66502 | -1.91247 | -0.78525 | -1.49997 | -1.42723 | -0.60112 | 0.034563 | 0.095861 | 0.056796 | 0.048667 | 0.0481 | 0.087778 | 0.041886 | 0.070883 | 0.159265 | 0.075322 |
| 24.88889 | -0.78394 | -1.05873 | -0.73595 | -0.88377 | -1.95502 | -0.85475 | -1.58348 | -1.66122 | -0.94168 | 0.035982 | 0.077804 | 0.055899 | 0.041957 | 0.024124 | 0.0899 | 0.035625 | 0.052154 | 0.166176 | 0.093894 |
| 24.90278 | -0.81257 | -1.23824 | -0.7352 | -0.91257 | -1.73667 | -0.80313 | -1.38977 | -1.773 | -0.93368 | -0.01768 | 0.019336 | 0.001634 | -0.00175 | 0.005889 | 0.03964 | -0.01506 | 0.004187 | 0.106394 | 0.076597 |
| 24.91667 | -0.68159 | -1.21715 | -0.47799 | -0.76561 | -1.18568 | -0.54981 | -0.85653 | -1.63601 | -0.64513 | 0.008821 | 0.051864 | 0.024224 | 0.030434 | 0.057265 | 0.061045 | -0.00224 | 0.033802 | 0.102364 | 0.127031 |
| 24.93056 | -0.65733 | -1.19804 | -0.26482 | -0.61598 | -0.67407 | -0.4204 | -0.30744 | -1.51804 | -0.44431 | 0.045075 | 0.085039 | 0.059619 | 0.058695 | 0.09965 | 0.088102 | 0.018203 | 0.061063 | 0.115052 | 0.172832 |
| 24.94444 | -0.8549 | -1.24282 | -0.33108 | -0.51967 | -0.38187 | -0.38776 | -0.21519 | -1.47994 | -0.42139 | 0.009598 | 0.035318 | 0.030031 | 0.024012 | 0.113583 | 0.06232 | -0.00748 | 0.029688 | 0.066931 | 0.143936 |
| 24.95833 | -1.05988 | -1.12434 | -0.39494 | -0.45262 | -0.33882 | -0.30837 | -0.53274 | -1.13469 | -0.41063 | 0.007049 | 0.012728 | 0.029094 | 0.022297 | 0.129109 | 0.066736 | 0.004418 | 0.020848 | 0.044957 | 0.129399 |
| 24.97222 | -1.0801 | -0.82292 | -0.31479 | -0.4828 | -0.47911 | -0.22133 | -1.0296 | -0.56775 | -0.38588 | 0.00518 | -0.01316 | 0.027658 | 0.037439 | 0.126711 | 0.080171 | 0.025527 | 0.003109 | 0.032195 | 0.104549 |
| 24.98611 | -0.78882 | -0.39446 | 0.007911 | -0.28112 | -0.38609 | 0.040998 | -1.11659 | 0.115405 | -0.14579 | -0.1236 | -0.24757 | -0.01019 | 0.085244 | 0.173447 | 0.05777 | 0.025108 | -0.121 | -0.11748 | -0.07026 |
| 25 | -0.22655 | 0.083957 | 0.645382 | 0.345547 | -0.10919 | 0.726844 | -0.47264 | 0.809673 | 0.488545 | -0.40802 | -0.94898 | 0.019111 | 0.219864 | 0.338112 | -0.02811 | 0.056911 | -0.53429 | -0.45138 | -0.45289 |
| 25.01389 | -0.05124 | 0.093039 | 1.061176 | 0.819911 | -0.42035 | 1.285939 | 0.26095 | 0.87632 | 0.91588 | -0.75987 | -1.83027 | 0.085718 | 0.272135 | 0.390181 | -0.13505 | 0.051432 | -1.31029 | -1.08242 | -0.76221 |
| 25.02778 | -0.42791 | -0.39382 | 0.907942 | 0.831164 | -1.19791 | 1.183852 | 0.564026 | 0.304104 | 0.678133 | -0.63261 | -1.40653 | 0.087626 | 0.133547 | 0.190622 | -0.09589 | 0.04894 | -1.05128 | -1.06808 | -0.50113 |
| 25.04167 | -0.47387 | -0.76139 | 0.619675 | 0.571897 | -1.37292 | 0.749994 | 0.550501 | -0.1068 | 0.177835 | -0.08524 | 0.061045 | 0.063314 | 0.025113 | 0.049796 | -0.01825 | 0.0844 | 0.12715 | 0.014352 | 0.094818 |
| 25.05556 | -0.18194 | -0.68863 | 0.362719 | 0.153785 | -0.84301 | 0.302256 | 0.337391 | -0.15975 | -0.21148 | 0.317743 | 0.624146 | 0.055184 | -0.00578 | 0.017006 | -0.00623 | 0.089907 | 0.60695 | 0.518165 | 0.338281 |
| 25.06944 | 0.336525 | -0.23548 | 0.147743 | -0.26614 | -0.43617 | -0.02 | 0.046606 | -0.09096 | -0.36291 | 0.479518 | 0.49752 | 0.033889 | -0.01882 | 0.004638 | -0.02493 | 0.043041 | 0.430913 | 0.430767 | 0.247221 |
| 25.08333 | 0.825495 | 0.331654 | 0.075068 | -0.35226 | -0.26926 | -0.0968 | -0.09527 | 0.086881 | -0.3025 | 0.520525 | 0.367778 | 0.013691 | -0.02748 | 0.012983 | -0.04546 | 0.005896 | 0.238268 | 0.234035 | 0.130513 |
| 25.09722 | 1.069078 | 0.685317 | 0.149572 | -0.15259 | -0.18029 | -0.02517 | -0.06634 | 0.200018 | -0.18033 | 0.549524 | 0.325662 | 0.021072 | -0.01241 | 0.039606 | -0.028 | 0.010972 | 0.16606 | 0.126934 | 0.090327 |
| 25.11111 | 0.997489 | 0.729738 | 0.296217 | 0.047654 | -0.04385 | -0.00404 | 0.003467 | 0.166118 | -0.07674 | 0.574159 | 0.334333 | 0.037214 | 0.004061 | 0.029278 | -0.00879 | 0.023962 | 0.135984 | 0.126406 | 0.089382 |
| 25.125 | 0.871048 | 0.620105 | 0.436948 | 0.157207 | 0.172692 | -0.05701 | 0.121171 | 0.108414 | 0.037046 | 0.606547 | 0.407399 | 0.052615 | 0.011968 | 0.015056 | -0.01983 | 0.022562 | 0.154665 | 0.182045 | 0.102401 |

| | | | | | | | | | | | | | | | | | | | |
|---|---|---|---|---|---|---|---|---|---|---|---|---|---|---|---|---|---|---|---|
| 25.13889 | 0.776432 | 0.490492 | 0.468877 | 0.186112 | 0.268235 | -0.10433 | 0.291966 | 0.081372 | 0.133846 | 0.716891 | 0.540324 | 0.078079 | 0.029608 | 0.037447 | -0.04563 | 0.024624 | 0.249178 | 0.235451 | 0.136433 |
| 25.15278 | 0.688813 | 0.409388 | 0.357227 | 0.174419 | 0.245126 | -0.0953 | 0.406165 | 0.090003 | 0.161498 | 0.692239 | 0.606815 | 0.086719 | 0.038195 | 0.058805 | -0.08451 | 0.027238 | 0.351002 | 0.201384 | 0.161711 |
| 25.16667 | 0.621143 | 0.357562 | 0.19966 | 0.137418 | 0.260267 | -0.0259 | 0.387822 | 0.135624 | 0.161232 | 0.581869 | 0.530993 | 0.048496 | 0.027911 | 0.066453 | -0.07873 | 0.010787 | 0.340115 | 0.113551 | 0.156319 |
| 25.18056 | 0.574274 | 0.325189 | 0.077437 | 0.130945 | 0.322085 | 0.047202 | 0.297292 | 0.19364 | 0.123346 | 0.359717 | 0.427698 | -0.00021 | 0.004934 | 0.062106 | -0.06428 | -0.00828 | 0.24153 | 0.039151 | 0.114238 |
| 25.19444 | 0.514266 | 0.298506 | 0.052044 | 0.194628 | 0.278866 | 0.108162 | 0.220576 | 0.221008 | 0.073375 | 0.162028 | 0.328161 | -0.06006 | -0.06164 | 0.029342 | -0.09846 | -0.04409 | 0.117859 | -0.02248 | 0.015996 |
| 25.20833 | 0.451001 | 0.249188 | 0.044236 | 0.255082 | -0.01807 | 0.148652 | 0.149884 | 0.168231 | 0.073012 | 0.016295 | 0.315856 | -0.11993 | -0.14526 | -0.00289 | -0.17743 | -0.0881 | 0.068286 | -0.05402 | -0.05115 |
| 25.22222 | 0.368564 | 0.154045 | -0.01494 | 0.255976 | -0.26886 | 0.17265 | 0.087051 | 0.068393 | 0.154021 | 0.024534 | 0.492337 | -0.08357 | -0.10184 | 0.055124 | -0.14625 | -0.0459 | 0.172088 | 0.004806 | 0.050562 |
| 25.23611 | 0.222248 | 0.016492 | -0.10047 | 0.215745 | -0.23505 | 0.180573 | -0.03195 | -0.01776 | 0.190454 | 0.105873 | 0.543715 | 0.019259 | 0.018073 | 0.135259 | -0.04125 | 0.042368 | 0.205495 | 0.049421 | 0.174626 |
| 25.25 | 0.032411 | -0.15487 | -0.14247 | 0.188067 | 0.02476 | 0.15847 | -0.24346 | -0.039 | 0.082461 | 0.078589 | 0.354756 | 0.039714 | 0.026797 | 0.100008 | -0.03775 | 0.039388 | 0.02257 | -0.03291 | 0.11921 |
| 25.26389 | -0.05915 | -0.28832 | -0.09319 | 0.187812 | 0.244787 | 0.087315 | -0.34363 | 0.06393 | -0.02717 | -0.05475 | 0.173881 | -0.03043 | -0.06322 | 0.012073 | -0.0836 | -0.0394 | -0.17605 | -0.1607 | -0.01043 |
| 25.27778 | 0.056791 | -0.30178 | 0.043942 | 0.216284 | 0.29397 | -0.06847 | -0.1964 | 0.223546 | 0.035532 | -0.14571 | 0.351201 | -0.02914 | -0.11842 | 0.013753 | -0.04509 | -0.05671 | -0.19733 | -0.19244 | -0.03746 |
| 25.29167 | 0.321986 | -0.11459 | 0.170207 | 0.347811 | 0.169164 | -0.02526 | 0.054495 | 0.390643 | 0.198294 | -0.36062 | 0.696285 | 0.098389 | -0.10802 | 0.086429 | 0.002674 | 0.000939 | -0.04056 | -0.03141 | 0.051846 |
| 25.30556 | 0.51167 | 0.234789 | 0.220918 | 0.680549 | -0.10087 | 0.461392 | 0.219858 | 0.619035 | 0.296445 | -0.62744 | 0.134817 | -0.24975 | -0.04122 | 0.154277 | -0.07101 | 0.015629 | 0.159735 | 0.231367 | 0.123561 |
| 25.31944 | 0.64744 | 0.495527 | 0.281069 | 1.008654 | -0.19496 | 0.852391 | 0.300578 | 0.79578 | 0.384718 | -0.25506 | -0.4564 | -0.17772 | 0.010806 | 0.216801 | -0.13257 | -0.0133 | 0.222019 | 0.301342 | 0.112323 |
| 25.33333 | 0.802662 | 0.47611 | 0.364898 | 1.128582 | -0.02842 | 0.661716 | 0.325392 | 0.822279 | 0.468476 | -0.19183 | -0.21583 | -0.05117 | -0.00442 | 0.234277 | -0.07611 | -0.01378 | 0.124506 | 0.142791 | 0.089722 |
| 25.34722 | 0.952654 | 0.446968 | 0.445099 | 1.035732 | 0.145179 | 0.378087 | 0.215323 | 0.70583 | 0.414388 | -0.03804 | 0.037607 | 0.007099 | -0.03725 | 0.104367 | -0.02485 | 0.000688 | 0.031697 | 0.041057 | 0.106252 |
| 25.36111 | 1.177279 | 0.579764 | 0.548729 | 0.900816 | 0.394442 | 0.266332 | 0.085808 | 0.567748 | 0.339838 | 0.019407 | 0.074048 | 0.008278 | -0.06782 | -0.02688 | -0.04502 | -0.01075 | -0.008 | 0.054404 | 0.112072 |
| 25.375 | 1.466545 | 0.858426 | 0.610344 | 0.849719 | 0.824342 | 0.210163 | 0.108891 | 0.513141 | 0.407503 | 0.008385 | 0.042099 | -0.00565 | -0.07266 | -0.08112 | -0.07394 | -0.03212 | 0.010545 | 0.072286 | 0.095036 |
| 25.38889 | 1.262452 | 0.794426 | 0.596997 | 0.775722 | 1.127716 | 0.238964 | 0.232145 | 0.573249 | 0.608502 | 0.01124 | 0.040709 | -0.00012 | -0.02513 | -0.04391 | -0.06109 | -0.01728 | 0.093285 | 0.096447 | 0.121086 |
| 25.40278 | 0.918052 | 0.538638 | 0.595474 | 0.704319 | 1.270296 | 0.3114 | 0.327084 | 0.580705 | 0.752974 | 0.056687 | 0.072024 | 0.022357 | 0.031175 | 0.017894 | -0.02516 | 0.020885 | 0.190539 | 0.150133 | 0.215363 |
| 25.41667 | 0.558276 | 0.305587 | 0.60555 | 0.741548 | 1.151709 | 0.339804 | 0.391561 | 0.534322 | 0.779317 | 0.103929 | 0.099082 | 0.031577 | 0.040643 | 0.030714 | -0.01795 | 0.035719 | 0.234032 | 0.190321 | 0.302537 |
| 25.43056 | 0.202035 | 0.080283 | 0.644699 | 0.907114 | 0.806119 | 0.325661 | 0.503614 | 0.451829 | 0.68817 | 0.126248 | 0.118754 | 0.031671 | 0.013156 | -0.00366 | -0.03744 | 0.035682 | 0.211032 | 0.189574 | 0.302427 |
| 25.44444 | 0.051388 | -0.00683 | 0.737914 | 1.094674 | 0.441509 | 0.320853 | 0.720457 | 0.487508 | 0.564583 | 0.10957 | 0.117525 | 0.00644 | -0.01418 | -0.0393 | -0.06764 | 0.017194 | 0.158141 | 0.167396 | 0.251136 |
| 25.45833 | 0.185796 | 0.094564 | 0.747318 | 1.012771 | 0.377509 | 0.337977 | 0.934832 | 0.663647 | 0.472472 | 0.065168 | 0.086431 | -0.02839 | -0.02654 | -0.03929 | -0.0735 | -0.00658 | 0.121364 | 0.128255 | 0.148985 |
| 25.47222 | 0.350919 | 0.226528 | 0.601731 | 0.701367 | 0.534751 | 0.349328 | 1.004855 | 0.858212 | 0.466486 | 0.070136 | 0.097936 | -0.02263 | -0.00627 | 0.005852 | -0.02751 | 0.004795 | 0.133856 | 0.161944 | 0.097591 |
| 25.48611 | 0.465757 | 0.599283 | 0.510929 | 0.435331 | 0.68228 | 0.340276 | 1.060639 | 1.094685 | 0.688548 | 0.198718 | 0.229784 | -0.0071 | -0.03038 | 0.025433 | -0.00448 | 0.019135 | 0.181551 | 0.473605 | 0.299344 |
| 25.5 | 0.574863 | 1.52586 | 0.76852 | 0.391611 | 0.322023 | 0.166144 | 1.365817 | 1.419396 | 1.395158 | 0.414623 | 0.447189 | -0.01778 | -0.15468 | -0.01801 | -0.05261 | -0.01138 | 0.264637 | 1.202133 | 0.738855 |
| 25.51389 | 0.55173 | 2.177132 | 1.140066 | 0.438769 | -2.28485 | -0.30204 | 1.605193 | 1.360126 | 1.605578 | 0.538578 | 0.524149 | -0.01572 | -0.24207 | -0.0547 | -0.09057 | -0.04579 | 0.317331 | 1.796214 | 1.019217 |
| 25.52778 | 0.361291 | 1.66872 | 1.204751 | 0.408761 | -4.63003 | -0.40574 | 1.728947 | 1.050375 | 0.971178 | 0.465884 | 0.363847 | 0.008644 | -0.20084 | -0.06073 | -0.04235 | -0.05071 | 0.260307 | 1.883595 | 0.726628 |
| 25.54167 | 0.304469 | 1.233528 | 0.517408 | 0.30852 | -2.45493 | -0.06689 | 1.432247 | 0.887673 | 0.324875 | 0.318266 | 0.139172 | 0.012158 | -0.12883 | -0.06248 | 0.006409 | -0.03929 | 0.145847 | 1.705972 | 0.348052 |
| 25.55556 | 0.359822 | 0.857425 | -0.45498 | 0.038871 | -1.47499 | 0.162327 | 0.445909 | 0.876592 | -0.15364 | 0.193678 | -0.00907 | -0.03002 | -0.097 | -0.05276 | 0.020426 | -0.03781 | 0.08927 | 1.171241 | 0.062534 |
| 25.56944 | 0.478195 | 0.562989 | -0.97818 | -0.12977 | -0.99133 | 0.315674 | -0.5442 | 0.813797 | -0.48978 | 0.107633 | -0.06653 | -0.04851 | -0.0442 | 0.012078 | 0.06537 | -0.01801 | 0.098041 | 0.605123 | -0.1531 |
| 25.58333 | 0.634077 | 0.409782 | -0.59209 | -0.16069 | -0.99089 | 0.381355 | -1.00949 | 0.787841 | -0.54168 | 0.046191 | -0.01911 | 0.025857 | 0.037419 | 0.1009 | 0.149899 | 0.027104 | 0.124639 | 0.274138 | -0.12652 |
| 25.59722 | 0.715143 | 0.278656 | 0.37253 | -0.08908 | -1.07668 | 0.319634 | -0.69097 | 0.8804 | -0.23532 | -0.06865 | 0.053325 | 0.090027 | 0.037319 | 0.105817 | 0.17416 | 0.026043 | 0.090969 | 0.10405 | 0.145101 |
| 25.61111 | 0.630601 | 0.043081 | 1.345592 | 0.058244 | -0.92762 | 0.235809 | 0.330055 | 1.28448 | 0.206889 | -0.20242 | 0.018566 | 0.051377 | -0.00953 | 0.050279 | 0.123228 | -0.00922 | 0.012341 | -0.04368 | 0.33541 |
| 25.625 | 0.436925 | -0.18571 | 1.779455 | 0.119231 | -0.60336 | 0.236531 | 1.451139 | 1.755431 | 0.396004 | -0.22445 | -0.11307 | 0.008768 | -0.00314 | 0.022987 | 0.09038 | 0.020936 | 0.010852 | -0.16437 | 0.260284 |
| 25.63889 | 0.253223 | -0.40508 | 1.049745 | 0.065087 | -0.38632 | 0.191758 | 1.818618 | 1.823529 | 0.3029 | -0.03317 | -0.26825 | 0.024196 | 0.014695 | -0.00267 | 0.055863 | 0.107218 | 0.116079 | -0.21352 | 0.08332 |
| 25.65278 | 0.018212 | -0.51188 | -0.08173 | 0.057865 | -0.29375 | 0.059005 | 1.194889 | 1.555669 | 0.041945 | 0.236365 | -0.22046 | 0.050276 | -0.02016 | -0.04522 | -0.02475 | 0.130418 | 0.288776 | -0.27557 | -0.02678 |
| 25.66667 | -0.09621 | -0.25624 | -0.97176 | 0.32875 | -0.34774 | 0.015577 | -0.19716 | 1.394238 | -0.16789 | 0.598697 | -0.04258 | 0.04029 | -0.05208 | -0.06479 | -0.0998 | 0.060771 | 0.405225 | -0.36099 | -0.06675 |
| 25.68056 | -0.08292 | 0.293202 | -1.21308 | 0.524783 | -0.61879 | 0.113288 | -1.45663 | 1.294175 | 0.020208 | 0.857202 | 0.161778 | 0.033815 | -0.02939 | -0.05759 | -0.11706 | 0.01349 | 0.284049 | -0.40696 | -0.08688 |
| 25.69444 | -0.03502 | 0.877658 | -0.765 | 0.364887 | -0.8718 | 0.177739 | -1.94646 | 0.877587 | 0.490197 | 0.737481 | 0.273893 | 0.025186 | -0.02469 | -0.00127 | -0.10419 | -0.04092 | 0.060682 | -0.37494 | -0.02076 |
| 25.70833 | 0.054945 | 1.360043 | -0.06843 | 0.076432 | -0.75082 | 0.178181 | -1.40823 | 0.323237 | 0.741774 | 0.466483 | 0.279064 | 0.003424 | -0.0492 | 0.056975 | -0.0872 | -0.09387 | -0.05936 | -0.30379 | 0.162244 |
| 25.72222 | 0.456805 | 1.564737 | 0.671173 | 0.094989 | -0.37575 | 0.239009 | -0.01848 | -0.16315 | 0.57394 | 0.157361 | 0.227407 | 0.035276 | -0.02974 | 0.081874 | -0.04019 | -0.05091 | -0.03029 | -0.1339 | 0.291549 |
| 25.73611 | 0.833571 | 1.290497 | 0.875053 | 0.022888 | -0.12873 | 0.248398 | 1.246195 | -0.72469 | 0.183294 | -0.224 | 0.085722 | 0.040684 | -0.01625 | 0.025331 | -0.02326 | 0.001889 | -0.03033 | 0.057481 | 0.399845 |
| 25.75 | 0.790052 | 0.732157 | 0.23513 | -0.36105 | -0.05452 | 0.105788 | 1.26891 | -1.3546 | -0.23166 | -0.7554 | -0.11409 | -0.00992 | -0.01156 | -0.04996 | -0.02311 | 0.021792 | -0.10721 | 0.115073 | 0.294771 |
| 25.76389 | 0.518575 | 0.261013 | -0.68503 | -0.53511 | 0.008063 | -0.02302 | 0.470017 | -1.67802 | -0.36503 | -1.00353 | -0.19311 | -0.00519 | 0.043155 | -0.05317 | 0.038801 | 0.0756 | -0.13725 | 0.049654 | -0.00505 |
| 25.77778 | 0.100971 | -0.16586 | -1.45778 | -0.48842 | 0.078742 | -0.12826 | -0.48775 | -1.68553 | -0.34109 | -0.87849 | -0.17459 | -0.01722 | 0.064425 | -0.06004 | 0.065413 | 0.080802 | -0.13655 | -0.14619 | -0.42759 |
| 25.79167 | -0.22231 | -0.58312 | -1.38316 | -0.4459 | 0.089719 | -0.32156 | -1.20037 | -1.40343 | -0.33818 | -0.38622 | -0.15481 | -0.06754 | 0.042724 | -0.08165 | 0.029828 | 0.002743 | -0.09287 | -0.41227 | -0.87279 |
| 25.80556 | -0.16454 | -0.8712 | -0.7816 | -0.48357 | 0.024315 | -0.55541 | -1.45747 | -1.00239 | -0.30446 | 0.217179 | -0.17452 | -0.07283 | 0.060201 | -0.07307 | 0.042607 | -0.05415 | -0.02246 | -0.55355 | -1.01808 |
| 25.81944 | -0.04178 | -1.12176 | -0.28573 | -0.55949 | -0.13744 | -0.68551 | -1.49368 | -0.6978 | -0.35167 | 0.744205 | -0.42961 | -0.09114 | 0.063106 | -0.09902 | 0.043621 | -0.11173 | -0.05216 | -0.56905 | -0.89209 |
| 25.83333 | -0.17293 | -1.38258 | -0.37737 | -0.47221 | -0.34633 | -0.75471 | -1.4996 | -0.60959 | -0.44365 | 0.832869 | -0.81003 | -0.19056 | 0.030313 | -0.11017 | 0.015451 | -0.164 | -0.16428 | -0.57064 | -0.7514 |
| 25.84722 | -0.4854 | -1.33883 | -0.70023 | -0.26241 | -0.46234 | -0.83585 | -1.28098 | -0.83724 | -0.43975 | 0.382783 | -0.79619 | -0.17808 | 0.036404 | -0.05856 | 0.022818 | -0.10464 | -0.27148 | -0.49832 | -0.51351 |
| 25.86111 | -1.05883 | -1.48013 | -0.91705 | -0.30921 | -0.53055 | -1.01299 | -1.07598 | -1.47688 | -0.49484 | -0.28085 | -0.43506 | -0.17613 | 0.029513 | -0.01944 | 0.003416 | -0.04181 | -0.43954 | -0.43721 | -0.35877 |
| 25.875 | -2.01902 | -1.98131 | -1.02652 | -0.63661 | -0.69336 | -1.1078 | -1.1436 | -2.1094 | -0.55573 | -1.06531 | -0.11214 | -0.23301 | 0.011384 | -0.01135 | -0.05011 | -0.04076 | -0.66788 | -0.45473 | -0.32013 |
| 25.88889 | -2.15381 | -1.86531 | -0.86324 | -0.98705 | -0.76521 | -0.94914 | -1.24678 | -2.31073 | -0.59638 | -1.5071 | 0.094397 | -0.21384 | 0.042988 | -0.003 | -0.03522 | -0.03172 | -0.90567 | -0.48918 | -0.34673 |
| 25.90278 | -1.59714 | -1.32675 | -0.68192 | -1.12096 | -0.75467 | -0.59364 | -0.93613 | -2.33251 | -0.7898 | -1.5343 | 0.064525 | -0.14366 | 0.062977 | -0.01738 | 0.044231 | -0.02948 | -0.7778 | -0.49344 | -0.39369 |
| 25.91667 | -0.66368 | -0.8272 | -0.41422 | -1.06193 | -0.7403 | -0.31524 | -0.62069 | -2.16407 | -1.00138 | -1.40373 | -0.13711 | -0.08546 | 0.030591 | -0.04758 | 0.10665 | -0.08721 | -0.3933 | -0.49377 | -0.33705 |
| 25.93056 | 0.119893 | -0.46425 | -0.17651 | -0.70787 | -0.73163 | -0.26734 | -0.50291 | -1.6056 | -1.06764 | -0.88766 | -0.35632 | -0.04977 | 0.01631 | -0.09753 | 0.106884 | -0.11735 | -0.03671 | -0.44821 | -0.30881 |
| 25.94444 | 0.318737 | -0.37598 | -0.09202 | -0.52104 | -0.72429 | -0.431 | -0.72447 | -1.10825 | -0.98373 | -0.17935 | -0.40989 | -0.04093 | 0.018324 | -0.15085 | 0.034083 | -0.10762 | 0.099233 | -0.3526 | -0.29276 |
| 25.95833 | 0.045375 | -0.42831 | -0.13183 | -0.41421 | -0.5751 | -0.56886 | -0.88883 | -0.66435 | -0.61445 | 0.304786 | -0.39381 | -0.04647 | 0.034531 | -0.12137 | -0.00524 | -0.06017 | -0.01711 | -0.21872 | -0.28132 |
| 25.97222 | -0.40827 | -0.47122 | -0.24637 | -0.36104 | -0.38621 | -0.52752 | -0.70791 | -0.449 | -0.26528 | 0.399024 | -0.35108 | -0.05031 | 0.039705 | -0.0668 | -0.017 | 0.000549 | -0.2249 | -0.15515 | -0.31061 |
| 25.98611 | -0.81803 | -0.61832 | -0.31268 | -0.3342 | -0.20209 | -0.31883 | -0.28797 | -0.35531 | -0.14766 | 0.196391 | -0.46971 | -0.03536 | 0.082618 | 0.0322 | 0.012985 | 0.081731 | -0.45267 | -0.27182 | -0.43761 |
| 26 | -1.08871 | -0.8621 | -0.17243 | -0.2711 | 0.120416 | 0.072316 | 0.095025 | -0.2851 | -0.06678 | -0.19576 | -0.80247 | 0.022558 | 0.213975 | 0.229765 | 0.146823 | 0.214524 | -0.65667 | -0.64913 | -0.66514 |
| 26.01389 | -1.52391 | -1.08643 | 0.063716 | -0.39981 | 0.387097 | 0.228255 | 0.10521 | -0.46603 | 0.009764 | -0.60134 | -1.06447 | 0.061205 | 0.266556 | 0.302216 | 0.214212 | 0.220488 | -0.73171 | -1.0118 | -0.85189 |
| 26.02778 | -2.06445 | -1.19594 | 0.09712 | -0.55214 | 0.356491 | -0.07745 | -0.16193 | -0.83719 | -0.12612 | -0.70887 | -0.91709 | 0.045332 | 0.168664 | 0.148504 | 0.19046 | 0.06734 | -0.41623 | -0.9546 | -0.84131 |
| 26.04167 | -2.36306 | -1.18942 | 0.079629 | -0.23672 | 0.237128 | -0.3524 | -0.2358 | -0.91923 | -0.31672 | -0.6052 | -0.61897 | 0.005226 | 0.074857 | -0.01093 | 0.133301 | -0.03293 | 0.004314 | -0.70486 | -0.60837 |
| 26.05556 | -1.93602 | -0.95956 | 0.078521 | 0.056606 | 0.150411 | -0.28249 | -0.13324 | -0.72352 | -0.39581 | -0.4569 | -0.45706 | -0.0176 | 0.045596 | -0.06831 | 0.054315 | -0.08631 | 0.16422 | -0.35104 | -0.1781 |
| 26.06944 | -1.14699 | -0.65174 | 0.011085 | -0.11324 | 0.020747 | -0.13709 | -0.09012 | -0.55098 | -0.40249 | -0.22987 | -0.22899 | -0.01277 | 0.034404 | -0.06092 | -0.03764 | -0.11312 | 0.113027 | -0.01454 | 0.160002 |
| 26.08333 | -0.37925 | -0.38477 | -0.02887 | -0.38008 | -0.18865 | -0.09913 | -0.08925 | -0.36235 | -0.37905 | 0.031887 | 0.006114 | -0.00467 | 0.019488 | -0.04216 | -0.0992 | -0.08623 | 0.065972 | 0.274529 | 0.17281 |
| 26.09722 | 0.227925 | -0.12541 | 0.047658 | -0.36868 | -0.29473 | -0.12407 | -0.01084 | -0.09036 | -0.29497 | 0.201171 | 0.129685 | -0.00156 | 0.012652 | -0.02403 | -0.07346 | -0.03837 | 0.061359 | 0.432586 | 0.050986 |
| 26.11111 | 0.654666 | 0.146218 | 0.221159 | -0.1601 | -0.30524 | -0.13475 | 0.223156 | 0.109433 | -0.20315 | 0.200849 | 0.154089 | -0.00598 | 0.009375 | -0.00844 | -0.02583 | -0.00159 | 0.048308 | 0.411997 | 0.018852 |
| 26.125 | 0.839648 | 0.372308 | 0.381745 | 0.115307 | -0.23158 | -0.12674 | 0.532786 | 0.249519 | -0.09078 | 0.106465 | 0.086632 | -0.01958 | -0.00374 | -0.01477 | -0.00501 | 0.000637 | -0.01273 | 0.336086 | 0.058206 |
| 26.13889 | 0.877325 | 0.501232 | 0.483324 | 0.404148 | -0.04578 | -0.10425 | 0.793049 | 0.386306 | 0.055749 | 0.028658 | 0.019606 | -0.01716 | -0.00041 | -0.01168 | 0.008335 | 0.010902 | -0.06516 | 0.286449 | 0.029528 |
| 26.15278 | 0.940313 | 0.589677 | 0.602447 | 0.533354 | 0.191283 | -0.00364 | 0.882432 | 0.481956 | 0.236182 | 0.003308 | 0.010788 | 0.008456 | 0.023006 | 0.012631 | 0.033987 | 0.064336 | -0.03584 | 0.240913 | -0.10767 |
| 26.16667 | 1.015259 | 0.684616 | 0.677935 | 0.381154 | 0.395552 | 0.154957 | 0.805359 | 0.480095 | 0.404773 | -0.00553 | 0.054561 | 0.015797 | 0.015107 | 0.006877 | 0.036117 | 0.094209 | 0.023465 | 0.14253 | -0.26685 |
| 26.18056 | 1.132921 | 0.786666 | 0.682317 | 0.14214 | 0.499695 | 0.228871 | 0.596194 | 0.396448 | 0.470247 | -0.01018 | 0.076012 | -0.01142 | -0.02043 | -0.02477 | 0.016035 | 0.056722 | 0.03887 | 0.028011 | -0.29635 |
| 26.19444 | 1.237608 | 0.840371 | 0.557409 | -0.00836 | 0.53624 | 0.17176 | 0.374469 | 0.312126 | 0.41189 | -0.0032 | 0.037895 | -0.03147 | -0.03678 | -0.03036 | 0.004065 | -0.00377 | 0.001078 | -0.02012 | -0.22243 |
| 26.20833 | 1.253485 | 0.870486 | 0.375233 | 0.053381 | 0.523182 | 0.097063 | 0.134435 | 0.281388 | 0.288612 | -0.00553 | -0.02979 | -0.04055 | -0.03793 | -0.02705 | -0.00365 | -0.05392 | -0.09522 | 0.006325 | -0.12006 |
| 26.22222 | 1.175749 | 0.789781 | 0.224913 | 0.263652 | 0.430571 | 0.114839 | -0.07502 | 0.227399 | 0.151715 | 0.003518 | -0.05157 | -0.03752 | -0.04229 | -0.04179 | -0.01732 | -0.09668 | -0.18324 | 0.028524 | -0.05456 |
| 26.23611 | 1.004209 | 0.506086 | 0.114481 | 0.34047 | 0.247637 | 0.188314 | -0.14242 | 0.067918 | 0.00562 | 0.083714 | -0.0028 | -0.00906 | -0.03977 | -0.03446 | -0.01356 | -0.11183 | -0.18939 | -0.01519 | 0.016645 |
| 26.25 | 0.745308 | 0.216533 | 0.012402 | 0.162266 | 0.076589 | 0.236046 | -0.11581 | -0.07111 | -0.09808 | 0.189912 | 0.030555 | 0.038462 | -0.02971 | -0.0168 | 0.004304 | -0.09075 | -0.1322 | -0.11121 | 0.041279 |
| 26.26389 | 0.570833 | 0.142211 | -0.02367 | -0.01954 | 0.07603 | 0.270685 | -0.02743 | -0.04982 | -0.0825 | 0.198505 | 0.009439 | 0.033899 | -0.02691 | -0.01481 | -0.00151 | -0.06081 | -0.09987 | -0.20356 | -0.06487 |
| 26.27778 | 0.510536 | 0.166789 | 0.006043 | 0.117861 | 0.197925 | 0.281492 | 0.045281 | 0.105506 | -0.00058 | 0.120789 | 0.016744 | 0.004122 | -0.03385 | -0.02809 | -0.02882 | -0.04918 | -0.09128 | -0.22669 | -0.19743 |
| 26.29167 | 0.456277 | 0.136171 | 0.026795 | 0.533 | 0.264732 | 0.275905 | 0.079904 | 0.291264 | 0.068806 | 0.069117 | 0.123079 | -0.00422 | -0.02864 | -0.01668 | -0.03365 | -0.04432 | -0.04671 | -0.13146 | -0.18801 |
| 26.30556 | 0.393963 | 0.121976 | 0.075712 | 0.894368 | 0.30081 | 0.37444 | 0.105851 | 0.423638 | 0.135373 | 0.11321 | 0.288633 | 0.007424 | -0.00341 | 0.005274 | -0.00806 | -0.03363 | 0.032719 | 0.091355 | -0.00042 |
| 26.31944 | 0.36252 | 0.187954 | 0.207864 | 0.853304 | 0.283278 | 0.587637 | 0.151618 | 0.580555 | 0.211711 | 0.205105 | 0.413797 | 0.018636 | 0.008452 | 0.002668 | 0.00375 | -0.02501 | 0.090255 | 0.305438 | 0.189812 |

| | | | | | | | | | | | | | | | | | | | |
|---|---|---|---|---|---|---|---|---|---|---|---|---|---|---|---|---|---|---|---|
| 26.33333 | 0.37921 | 0.273237 | 0.371412 | 0.620975 | 0.218635 | 0.781841 | 0.184025 | 0.857279 | 0.274349 | 0.286065 | 0.386595 | 0.005692 | -0.02148 | -0.02494 | -0.02318 | -0.02395 | 0.085183 | 0.386425 | 0.238806 |
| 26.34722 | 0.546247 | 0.362357 | 0.536216 | 0.557964 | 0.162855 | 0.806034 | 0.236613 | 1.176387 | 0.336082 | 0.344638 | 0.314036 | -0.01749 | -0.05669 | -0.04845 | -0.05992 | -0.01816 | 0.088645 | 0.305091 | 0.220115 |
| 26.36111 | 0.81858 | 0.495836 | 0.626812 | 0.739242 | 0.14874 | 0.861853 | 0.358756 | 1.350032 | 0.393117 | 0.377272 | 0.345481 | -0.02681 | -0.07555 | -0.05291 | -0.07662 | -0.00155 | 0.142278 | 0.17627 | 0.229561 |
| 26.375 | 1.105066 | 0.645893 | 0.618567 | 0.917312 | 0.172691 | 0.914858 | 0.520293 | 1.397364 | 0.430314 | 0.396658 | 0.445305 | -0.0134 | -0.11318 | -0.03231 | -0.06232 | 0.026506 | 0.207034 | 0.090437 | 0.276795 |
| 26.38889 | 1.28928 | 0.724682 | 0.559516 | 0.917571 | 0.144775 | 0.910574 | 0.601081 | 1.405316 | 0.407759 | 0.370301 | 0.533618 | 0.010853 | -0.14832 | -0.0052 | -0.03961 | 0.064333 | 0.266307 | 0.086845 | 0.24947 |
| 26.40278 | 1.231564 | 0.706873 | 0.553039 | 0.816846 | 0.039635 | 0.831878 | 0.547185 | 1.296912 | 0.356037 | 0.304595 | 0.566258 | -0.00695 | -0.1464 | -0.00276 | -0.05067 | 0.07993 | 0.34335 | 0.111489 | 0.174602 |
| 26.41667 | 1.029685 | 0.649057 | 0.642657 | 0.680007 | -0.0164 | 0.646299 | 0.339808 | 1.112154 | 0.30602 | 0.258993 | 0.495062 | -0.02697 | -0.1039 | -0.0279 | -0.07715 | 0.070063 | 0.444297 | 0.136782 | 0.116609 |
| 26.43056 | 0.793391 | 0.566045 | 0.819101 | 0.637151 | -0.00058 | 0.495646 | 0.224962 | 0.874103 | 0.400258 | 0.302274 | 0.407648 | 0.01183 | -0.07222 | -0.05515 | -0.07236 | 0.061912 | 0.533908 | 0.168393 | 0.094627 |
| 26.44444 | 0.674357 | 0.432802 | 0.879243 | 0.661072 | 0.039671 | 0.382534 | 0.307005 | 0.730312 | 0.651003 | 0.364663 | 0.382941 | 0.04809 | -0.07256 | -0.06783 | -0.05076 | 0.051721 | 0.571238 | 0.186717 | 0.071554 |
| 26.45833 | 0.539506 | 0.256415 | 0.637123 | 0.634121 | 0.103827 | 0.278052 | 0.429065 | 0.488705 | 0.757099 | 0.354459 | 0.381027 | 0.01106 | -0.07713 | -0.1086 | -0.06973 | 0.012112 | 0.49362 | 0.158476 | 0.060529 |
| 26.47222 | 0.334148 | 0.022124 | 0.198231 | 0.516584 | 0.283124 | 0.134244 | 0.498525 | 0.208063 | 0.662417 | 0.297752 | 0.372638 | -0.02109 | -0.09014 | -0.14044 | -0.11911 | -0.02117 | 0.484228 | 0.185242 | 0.119824 |
| 26.48611 | 0.105473 | -0.28439 | 0.044318 | 0.535306 | 0.469774 | -0.14079 | 0.815321 | 0.084856 | 0.68775 | 0.379489 | 0.452228 | 0.006324 | -0.13337 | -0.17276 | -0.15258 | -0.05716 | 0.585812 | 0.436302 | 0.327266 |
| 26.5 | -0.06463 | -0.61601 | 0.600446 | 0.638898 | 1.207062 | -0.39546 | 1.319524 | 0.240477 | 1.244076 | 0.61568 | 0.68328 | 0.071602 | -0.22986 | -0.21515 | -0.19375 | -0.11091 | 0.873147 | 0.981982 | 0.708033 |
| 26.51389 | -0.0704 | -0.44373 | 1.529096 | 0.553423 | 1.047573 | -0.29828 | 1.730918 | 0.719488 | 1.813804 | 0.924349 | 0.92163 | 0.174431 | -0.23899 | -0.20788 | -0.2292 | -0.09075 | 0.864537 | 1.405262 | 1.07689 |
| 26.52778 | 0.276553 | 0.626069 | 2.067539 | 0.620981 | 0.218673 | 0.093044 | 1.548873 | 1.1626 | 1.611272 | 0.941735 | 0.806791 | 0.318561 | -0.18554 | -0.09454 | -0.19907 | -0.03585 | 0.618468 | 1.141444 | 1.152566 |
| 26.54167 | 0.639572 | 1.525046 | 1.49739 | 0.592076 | -0.03661 | 0.226852 | 0.917028 | 1.271489 | 1.179514 | 0.736693 | 0.324302 | 0.347831 | -0.13704 | 0.027859 | -0.09012 | 0.006932 | 0.460286 | 0.919684 | 0.901154 |
| 26.55556 | 0.643397 | 1.339206 | 0.277686 | 0.448836 | -0.11839 | 0.150793 | 0.064271 | 1.064726 | 0.633677 | 0.644324 | -0.08489 | 0.19228 | -0.01588 | 0.159313 | 0.065691 | 0.078987 | 0.341644 | 0.700614 | 0.578278 |
| 26.56944 | 0.487697 | 0.697654 | -0.75744 | 0.317367 | -0.17197 | -0.07162 | -0.62249 | 0.814561 | 0.030401 | 0.358826 | -0.39677 | 0.123316 | 0.028698 | 0.236256 | 0.185578 | 0.087041 | 0.126725 | 0.531002 | 0.220188 |
| 26.58333 | 0.373714 | 0.640604 | -1.22654 | 0.27036 | -0.26184 | -0.24305 | -0.91282 | 0.851884 | -0.47599 | 0.027497 | -0.52134 | 0.09746 | -0.00139 | 0.275257 | 0.237892 | 0.05897 | -0.09565 | 0.3309 | 0.024624 |
| 26.59722 | 0.595239 | 1.017214 | -0.95908 | 0.327605 | -0.30466 | -0.11643 | -0.58196 | 1.320871 | -0.57664 | 0.034274 | -0.29656 | 0.132814 | -0.02775 | 0.356139 | 0.287048 | 0.056611 | -0.13704 | 0.10987 | 0.223936 |
| 26.61111 | 0.851933 | 1.518005 | 0.085403 | 0.364111 | -0.31944 | 0.328571 | 0.501805 | 1.851434 | -0.17454 | 0.332973 | 0.009287 | 0.12784 | -0.03906 | 0.33256 | 0.259364 | 0.038977 | 0.006974 | -0.07204 | 0.409037 |
| 26.625 | 1.164446 | 1.668559 | 1.575245 | 0.355782 | -0.24398 | 0.772149 | 1.56066 | 2.423247 | 0.372329 | 0.547715 | 0.156718 | 0.056311 | -0.01221 | 0.230286 | 0.15686 | 0.010484 | 0.212252 | -0.15008 | 0.471707 |
| 26.63889 | 1.31211 | 1.399559 | 2.731779 | 0.372396 | -0.04741 | 0.924054 | 1.758814 | 2.675568 | 0.86769 | 0.579734 | 0.236094 | 0.013648 | 0.057558 | 0.167152 | 0.087387 | 0.051892 | 0.476052 | 0.01575 | 0.366998 |
| 26.65278 | 1.006684 | 0.505381 | 2.394605 | 0.390252 | -0.14069 | 0.783736 | 1.252739 | 2.051977 | 0.744428 | 0.338325 | 0.165149 | -0.00741 | 0.059251 | 0.068058 | -6.9E-05 | 0.089144 | 0.557468 | 0.229037 | -0.07257 |
| 26.66667 | 0.432076 | -0.59036 | 0.937587 | 0.421946 | -0.22601 | 0.462989 | 0.320898 | 0.994542 | 0.163478 | -0.14647 | -0.05627 | -0.0298 | -0.01668 | -0.08343 | -0.11963 | 0.082342 | 0.315135 | 0.219631 | -0.56644 |
| 26.68056 | -0.01525 | -1.34167 | -0.66438 | 0.602832 | -0.15452 | 0.174847 | -0.56577 | -0.10989 | -0.01085 | -0.34949 | -0.20892 | -0.00679 | -0.04707 | -0.15459 | -0.15645 | 0.055138 | 0.008362 | -0.00888 | -0.71637 |
| 26.69444 | -0.45295 | -1.35659 | -1.74694 | 0.527982 | 0.496996 | -0.06125 | -1.82783 | -1.14017 | 0.047512 | -0.32732 | -0.25137 | -0.00047 | -0.07938 | -0.16114 | -0.16292 | 0.028566 | -0.22914 | -0.26135 | -0.57369 |
| 26.70833 | -0.72981 | -0.87184 | -2.27158 | 0.528954 | 1.711509 | 0.025661 | -2.66269 | -1.35887 | 0.057034 | -0.39591 | -0.22303 | -0.05032 | -0.07866 | -0.11682 | -0.23428 | -0.04628 | -0.3127 | -0.26546 | -0.30824 |
| 26.72222 | 0 | 0 | 0 | 0 | 0 | 0 | 0 | 0 | 0 | 0 | 0 | 0 | 0 | 0 | 0 | 0 | 0 | 0 | 0 |
| 26.73611 | 0 | 0 | 0 | 0 | 0 | 0 | 0 | 0 | 0 | 0 | 0 | 0 | 0 | 0 | 0 | 0 | 0 | 0 | 0 |
| 26.75 | 0 | 0 | 0 | 0 | 0 | 0 | 0 | 0 | 0 | 0 | 0 | 0 | 0 | 0 | 0 | 0 | 0 | 0 | 0 |
| 26.76389 | 0 | 0 | 0 | 0 | 0 | 0 | 0 | 0 | 0 | 0 | 0 | 0 | 0 | 0 | 0 | 0 | 0 | 0 | 0 |
| 26.77778 | 0 | 0 | 0 | 0 | 0 | 0 | 0 | 0 | 0 | 0 | 0 | 0 | 0 | 0 | 0 | 0 | 0 | 0 | 0 |
| 26.79167 | 0 | 0 | 0 | 0 | 0 | 0 | 0 | 0 | 0 | 0 | 0 | 0 | 0 | 0 | 0 | 0 | 0 | 0 | 0 |
| 26.80556 | 0.703733 | -0.25882 | 1.858359 | -1.33187 | 0.296094 | -0.45255 | -1.49587 | 1.469213 | 1.154355 | -0.21062 | -0.51395 | 0.110598 | -0.32993 | 0.07445 | 0.30548 | 0.178314 | -0.26143 | -0.43446 | 0.760683 |
| 26.81944 | 0.798988 | 0.80386 | 0.786572 | -0.55366 | -0.1807 | -0.23268 | -1.80624 | 0.400797 | 1.261113 | 0.303217 | -0.74825 | -0.05719 | 0.000616 | 0.082567 | 0.117461 | 0.118721 | 0.176557 | -0.00737 | 0.369264 |
| 26.83333 | 0.859118 | 0.641029 | 0.099939 | -0.80291 | -0.30933 | -0.28551 | -2.5611 | -0.77719 | 0.987896 | 0.632925 | -0.7687 | -0.27437 | 0.023476 | 0.026776 | 0.08396 | 0.066439 | 0.185695 | -0.34701 | -0.14582 |
| 26.84722 | 0.570138 | 0.235545 | -0.65852 | -1.01729 | -0.26701 | -0.36904 | -1.51585 | -1.85646 | 0.07028 | 0.404287 | -0.48032 | -0.37517 | 0.049523 | 0.027117 | 0.102993 | 0.027701 | 0.061281 | -0.71117 | -0.61948 |
| 26.86111 | 0.043015 | -0.15338 | -1.24812 | -1.00009 | -0.44404 | -0.61094 | -0.3714 | -2.4162 | -0.55564 | -0.18673 | -0.16963 | -0.29847 | 0.027988 | -0.00622 | 0.092787 | -0.04677 | -0.19076 | -0.87635 | -0.85284 |
| 26.875 | -0.59345 | -0.69874 | -1.50519 | -0.80132 | -0.66508 | -0.62149 | 0.0412 | -2.33113 | -1.06621 | -0.91819 | -0.1097 | -0.2203 | 0.008924 | -0.05648 | 0.053225 | -0.13769 | -0.45433 | -0.91413 | -0.84047 |
| 26.88889 | -1.07822 | -1.5659 | -1.52281 | -0.70557 | -0.87811 | -0.44962 | 0.098922 | -2.0605 | -1.3646 | -1.493 | -0.18995 | -0.16097 | 0.032685 | -0.09301 | 0.028767 | -0.20313 | -0.6217 | -0.84251 | -0.62636 |
| 26.90278 | -1.11324 | -2.41918 | -1.33171 | -0.8335 | -0.9888 | -0.50649 | -0.0383 | -2.29262 | -1.51613 | -1.65857 | -0.26437 | -0.2055 | 0.071437 | -0.12136 | 0.0135 | -0.27406 | -0.75406 | -0.73811 | -0.57079 |
| 26.91667 | -0.80477 | -2.82354 | -0.92678 | -1.09569 | -0.79703 | -0.79225 | -0.22788 | -2.45599 | -1.45934 | -1.33743 | -0.31806 | -0.22939 | 0.126003 | -0.0956 | 0.035742 | -0.27005 | -0.78795 | -0.73952 | -0.71807 |
| 26.93056 | -0.53809 | -2.66079 | -0.51482 | -1.34285 | -0.50329 | -1.01968 | -0.46304 | -2.55259 | -1.42138 | -1.03071 | -0.50586 | -0.17417 | 0.146603 | -0.0622 | 0.049394 | -0.20226 | -0.85203 | -0.72038 | -0.9297 |
| 26.94444 | -0.61625 | -2.15971 | -0.20868 | -1.67164 | -0.2083 | -1.02696 | -0.59482 | -2.58234 | -1.32117 | -0.88051 | -0.75908 | -0.12142 | 0.113612 | -0.03919 | -0.00246 | -0.12874 | -0.82048 | -0.67053 | -0.92964 |
| 26.95833 | -0.80797 | -1.44593 | -0.0479 | -1.55888 | 0.151546 | -0.77976 | -0.46415 | -2.11193 | -0.97593 | -0.8009 | -0.83826 | -0.04448 | 0.107278 | 0.004555 | -0.02474 | -0.07648 | -0.63029 | -0.56518 | -0.64433 |
| 26.97222 | -1.00186 | -0.78856 | -0.08864 | -1.14082 | 0.358098 | -0.57311 | -0.30018 | -1.38437 | -0.60147 | -0.77518 | -0.77928 | 0.00705 | 0.085654 | 0.023396 | -0.015 | -0.04151 | -0.50239 | -0.46837 | -0.42351 |
| 26.98611 | -1.14265 | -0.61919 | -0.25199 | -0.81372 | 0.303868 | -0.44247 | -0.17199 | -0.87076 | -0.3748 | -0.83569 | -0.86996 | -0.01468 | 0.086539 | 0.066798 | -0.03013 | -0.02066 | -0.47748 | -0.52297 | -0.53661 |
| 27 | -1.15302 | -1.07634 | -0.41309 | -0.58616 | 0.098364 | -0.41418 | 0.081546 | -0.53093 | -0.2661 | -1.01494 | -1.26946 | -0.00665 | 0.203953 | 0.245543 | 0.024029 | 0.098379 | -0.50066 | -0.76171 | -0.80801 |
| 27.01389 | -1.25707 | -2.02377 | -0.54418 | -0.49446 | 0.003199 | -0.64538 | 0.348426 | -0.37356 | -0.26671 | -1.09252 | -1.62799 | -0.00131 | 0.289033 | 0.319869 | 0.082978 | 0.142856 | -0.52793 | -0.92529 | -0.82797 |
| 27.02778 | -1.46336 | -2.24472 | -0.16032 | -0.37972 | 0.296963 | -0.43878 | 0.555785 | -0.4752 | -0.09858 | -0.75765 | -1.62096 | -0.03468 | 0.209628 | 0.150588 | 0.077399 | 0.025745 | -0.50798 | -0.83375 | -0.5192 |
| 27.04167 | -1.68221 | -0.76642 | 0.488148 | -0.24129 | 0.646287 | 0.089385 | 0.674166 | -0.53073 | 0.373054 | -0.3335 | -1.17426 | 0.003637 | 0.111639 | -0.02686 | 0.016538 | -0.06983 | -0.29389 | -0.55184 | -0.26169 |
| 27.05556 | -1.71771 | 0.402182 | 0.952504 | -0.21908 | 0.661419 | 0.488193 | 0.593062 | -0.37862 | 0.635848 | -0.12624 | -0.51154 | 0.060915 | 0.0646 | -0.08027 | -0.03209 | -0.04452 | -0.00475 | -0.26073 | -0.10266 |
| 27.06944 | -1.2617 | 0.514125 | 0.820212 | -0.1798 | 0.505148 | 0.633489 | 0.374478 | -0.06434 | 0.445801 | -0.15947 | -0.00801 | 0.06193 | 0.03073 | -0.07316 | -0.09214 | -0.00231 | 0.101579 | -0.05351 | -0.02796 |
| 27.08333 | -0.62159 | 0.615185 | 0.475376 | -0.11318 | 0.433021 | 0.722679 | 0.22225 | 0.265201 | 0.169035 | -0.19609 | 0.225128 | 0.01241 | -0.00427 | -0.06358 | -0.12355 | -0.01422 | 0.061307 | 0.025614 | -0.01712 |
| 27.09722 | -0.15516 | 0.346895 | 0.335009 | 0.047255 | 0.405354 | 0.754953 | 0.238798 | 0.384743 | 0.093854 | -0.19769 | 0.247128 | -0.02992 | -0.02171 | -0.06168 | -0.11349 | -0.04863 | 0.017289 | 0.01172 | -0.02439 |
| 27.11111 | 0.082229 | 0.181724 | 0.415318 | 0.185521 | 0.43384 | 0.68773 | 0.320804 | 0.336691 | 0.235259 | -0.18818 | 0.119165 | -0.03853 | -0.02871 | -0.06222 | -0.08849 | -0.06839 | 0.042339 | -0.03111 | -0.01798 |
| 27.125 | 0.174042 | 0.206692 | 0.500634 | 0.195972 | 0.409726 | 0.47987 | 0.321597 | 0.314134 | 0.323273 | -0.14408 | 0.054378 | -0.02491 | -0.02131 | -0.05759 | -0.06507 | -0.07 | 0.103384 | -0.0655 | -0.0129 |
| 27.13889 | 0.188539 | 0.322817 | 0.461728 | 0.114054 | 0.299226 | 0.27307 | 0.240617 | 0.318992 | 0.136894 | -0.05809 | 0.076457 | -0.023 | -0.01067 | -0.0493 | -0.05076 | -0.05708 | 0.114464 | -0.07017 | -0.00386 |
| 27.15278 | 0.178308 | 0.435517 | 0.377808 | -0.03409 | 0.221942 | 0.149345 | 0.15279 | 0.260623 | -0.09484 | -0.04704 | 0.111841 | -0.06103 | -0.02824 | -0.05805 | -0.06394 | -0.06275 | 0.047702 | -0.06794 | -0.00227 |
| 27.16667 | 0.166898 | 0.492105 | 0.350449 | -0.19885 | 0.182896 | 0.190777 | 0.158391 | 0.137785 | -0.18769 | -0.1112 | 0.087284 | -0.09321 | -0.04882 | -0.06865 | -0.08595 | -0.07971 | -0.00276 | -0.0874 | -0.01794 |
| 27.18056 | 0.168629 | 0.460701 | 0.346463 | -0.24725 | 0.161053 | 0.421055 | 0.250461 | -0.02425 | -0.13378 | -0.11963 | -0.01324 | -0.08327 | -0.03657 | -0.0593 | -0.08411 | -0.09388 | -0.00582 | -0.1207 | -0.0345 |
| 27.19444 | 0.216318 | 0.350924 | 0.291914 | -0.16073 | 0.14276 | 0.568173 | 0.312314 | -0.18606 | -0.02422 | -0.086 | -0.14026 | -0.08431 | -0.00931 | -0.04974 | -0.06967 | -0.11409 | -0.04797 | -0.12357 | -0.03877 |
| 27.20833 | 0.256425 | 0.274502 | 0.204917 | -0.10936 | 0.10399 | 0.480529 | 0.247816 | -0.26449 | -0.00952 | -0.10179 | -0.25755 | -0.13876 | 0.003441 | -0.04864 | -0.05283 | -0.13792 | -0.17932 | -0.11246 | -0.03492 |
| 27.22222 | 0.249802 | 0.203605 | 0.128864 | -0.18513 | 0.070918 | 0.239031 | 0.097027 | -0.14373 | -0.12131 | -0.2046 | -0.35772 | -0.1825 | -0.00983 | -0.04972 | -0.04153 | -0.10554 | -0.35133 | -0.1449 | -0.03381 |
| 27.23611 | 0.202129 | 0.064232 | 0.086631 | -0.29658 | 0.033953 | 0.020442 | -0.0491 | 0.081784 | -0.23836 | -0.21059 | -0.43103 | -0.17678 | -0.0313 | -0.0552 | -0.0538 | -0.05963 | -0.42924 | -0.14917 | -0.02909 |
| 27.25 | 0.172932 | -0.11664 | 0.066287 | -0.3291 | 0.073792 | -0.1036 | -0.13064 | 0.139681 | -0.30429 | -0.00574 | -0.44492 | -0.12806 | -0.03091 | -0.05065 | -0.06014 | -0.01837 | -0.33128 | -0.0613 | -0.01953 |
| 27.26389 | 0.158728 | -0.22971 | 0.064819 | -0.26571 | 0.148464 | -0.03142 | -0.14797 | 0.035241 | -0.29606 | 0.254274 | -0.43018 | -0.07059 | -0.01468 | -0.03315 | -0.03818 | -0.00066 | -0.17197 | 0.034106 | -0.00358 |
| 27.27778 | 0.139285 | -0.16036 | 0.061559 | -0.12658 | 0.144033 | 0.214613 | -0.08624 | -0.02555 | -0.25072 | 0.304543 | -0.40963 | -0.03913 | -0.01373 | -0.02504 | -0.01155 | 0.004569 | -0.1303 | 0.061835 | 0.009115 |
| 27.29167 | 0.14497 | -0.02056 | 0.030524 | 0.047897 | 0.111668 | 0.380269 | -0.01441 | 0.000967 | -0.20867 | 0.217869 | -0.36005 | -0.02576 | -0.0151 | -0.02027 | -0.00804 | 0.00591 | -0.13517 | 0.089269 | 0.019357 |
| 27.30556 | 0.150729 | 0.016252 | 0.035957 | 0.149253 | 0.127616 | 0.340577 | -0.00251 | 0.019903 | -0.17685 | 0.176914 | -0.22235 | 0.005789 | -0.00032 | -0.00684 | -0.00305 | 0.013732 | -0.07011 | 0.144477 | 0.035903 |
| 27.31944 | 0.141285 | -0.06748 | 0.146836 | 0.163262 | 0.19876 | 0.235722 | -0.02739 | 0.087548 | -0.12479 | 0.234568 | -0.08371 | 0.032047 | 0.011492 | 0.007716 | 0.008319 | 0.01598 | 0.029388 | 0.17757 | 0.042687 |
| 27.33333 | 0.135504 | -0.13121 | 0.352563 | 0.272637 | 0.252914 | 0.3241 | 0.024067 | 0.303668 | -0.05901 | 0.350056 | 0.055021 | 0.036527 | 0.004541 | 0.014489 | 0.015442 | 0.00772 | 0.094656 | 0.175978 | 0.049068 |
| 27.34722 | 0.112468 | 0.022166 | 0.5465 | 0.603524 | 0.198813 | 0.661694 | 0.125048 | 0.609425 | 0.032924 | 0.441407 | 0.211229 | 0.017946 | -0.01349 | -0.00153 | 0.020896 | -0.00597 | 0.108572 | 0.176758 | 0.059309 |
| 27.36111 | 0.11888 | 0.261737 | 0.638737 | 0.873158 | 0.101022 | 0.981562 | 0.193576 | 0.755407 | 0.175879 | 0.535657 | 0.371991 | 0.003954 | -0.01325 | -0.02534 | 0.023746 | -0.01767 | 0.129625 | 0.192405 | 0.070375 |
| 27.375 | 0.251263 | 0.346528 | 0.652714 | 0.898553 | 0.121665 | 1.109065 | 0.209789 | 0.668393 | 0.331513 | 0.595931 | 0.523764 | 0.045416 | 0.014728 | -0.02485 | 0.043292 | -0.00872 | 0.202702 | 0.240678 | 0.082311 |
| 27.38889 | 0.486058 | 0.341032 | 0.670125 | 0.867189 | 0.132677 | 1.192372 | 0.213464 | 0.60444 | 0.400357 | 0.587327 | 0.639161 | 0.122939 | 0.027183 | -0.02064 | 0.069862 | 0.005269 | 0.276025 | 0.305086 | 0.039535 |
| 27.40278 | 0.692214 | 0.376174 | 0.789981 | 0.839496 | 0.096733 | 1.191578 | 0.206676 | 0.615295 | 0.370131 | 0.545164 | 0.683384 | 0.159708 | 0.000019 | -0.03721 | 0.056226 | 0.005546 | 0.34029 | 0.329826 | -0.05597 |
| 27.41667 | 0.901781 | 0.458538 | 0.92185 | 0.689965 | 0.037135 | 0.91468 | 0.136307 | 0.585852 | 0.30209 | 0.625929 | 0.674002 | 0.167555 | -0.02422 | -0.05847 | 0.028617 | 0.000385 | 0.351695 | 0.375419 | -0.07569 |
| 27.43056 | 1.095452 | 0.670785 | 0.996245 | 0.529848 | -0.04072 | 0.603455 | 0.079671 | 0.610022 | 0.262402 | 0.704295 | 0.640125 | 0.15477 | -0.04264 | -0.08049 | -0.0093 | -0.0277 | 0.310378 | 0.36606 | 0.039909 |
| 27.44444 | 1.284841 | 0.892934 | 0.949942 | 0.527453 | -0.1299 | 0.597689 | 0.226018 | 0.746012 | 0.334739 | 0.600543 | 0.614119 | 0.107885 | -0.07635 | -0.09923 | -0.03961 | -0.07604 | 0.237087 | 0.266914 | 0.139053 |
| 27.45833 | 1.286569 | 0.948265 | 0.716711 | 0.614002 | -0.12681 | 0.776551 | 0.534744 | 0.868673 | 0.501226 | 0.397185 | 0.611666 | 0.087898 | -0.06735 | -0.05187 | -0.02206 | -0.04691 | 0.197419 | 0.198138 | 0.171974 |
| 27.47222 | 1.097195 | 0.666458 | 0.28251 | 0.668134 | 0.040587 | 0.900969 | 0.72126 | 0.78223 | 0.60266 | 0.34774 | 0.698289 | 0.113149 | -0.04017 | 0.020382 | 0.010762 | 0.042903 | 0.227717 | 0.25623 | 0.238742 |
| 27.48611 | 0.904308 | 0.288781 | -0.04066 | 0.977418 | 0.411246 | 0.841003 | 0.778985 | 0.692915 | 0.706434 | 0.52741 | 0.868016 | 0.112594 | -0.14952 | -0.0039 | -0.06603 | 0.094431 | 0.253292 | 0.540969 | 0.322946 |
| 27.5 | 1.031379 | 0.151887 | 0.152265 | 1.488065 | 1.443598 | 0.732032 | 1.039478 | 0.740753 | 1.160803 | 1.006601 | 1.335803 | 0.120359 | -0.35982 | -0.10737 | -0.23589 | 0.126643 | 0.330477 | 1.116395 | 0.540839 |
| 27.51389 | 1.30919 | 0.399862 | 0.954779 | 1.386509 | 1.860468 | 0.611208 | 1.338612 | 0.615704 | 1.903028 | 1.653682 | 1.841947 | 0.29453 | -0.42447 | -0.16524 | -0.25878 | 0.113464 | 0.631997 | 1.452512 | 0.990393 |

| | | | | | | | | | | | | | | | | | | | |
|---|---|---|---|---|---|---|---|---|---|---|---|---|---|---|---|---|---|---|---|
| 27.52778 | 1.402971 | 0.92516 | 1.860197 | 0.949433 | 0.912851 | 0.377153 | 1.623359 | 0.524497 | 2.49456 | 1.387403 | 1.958771 | 0.507475 | -0.28974 | -0.06692 | -0.17314 | 0.09559 | 1.0673 | 1.248324 | 1.39898 |
| 27.54167 | 1.108387 | 1.280232 | 1.628411 | 0.603071 | 0.598531 | -0.01618 | 1.754937 | 0.473227 | 2.268211 | 0.455613 | 1.614503 | 0.575323 | -0.15365 | 0.14789 | -0.05349 | 0.062754 | 1.318363 | 0.915445 | 1.429431 |
| 27.55556 | 0.455197 | 1.165189 | 0.576552 | 0.233699 | 0.538766 | -0.3168 | 1.275174 | 0.148211 | 1.317184 | -0.0738 | 0.891967 | 0.492373 | -0.08964 | 0.303037 | 0.10425 | 0.018055 | 1.043484 | 0.644507 | 1.161288 |
| 27.56944 | -0.07262 | 0.722789 | -0.5156 | 0.105707 | 0.561478 | -0.2731 | 0.573323 | -0.02174 | 0.456469 | -0.5496 | 0.196545 | 0.365904 | 0.026737 | 0.347652 | 0.232503 | -0.00222 | 0.646443 | 0.468571 | 0.905137 |
| 27.58333 | -0.23593 | 0.325704 | -1.19887 | 0.165023 | 0.455369 | -0.14628 | -0.11826 | 0.208995 | -0.00707 | -0.4577 | -0.31434 | 0.262668 | 0.088407 | 0.324617 | 0.287287 | 0.01889 | 0.293038 | 0.457541 | 0.671169 |
| 27.59722 | -0.12057 | 0.408096 | -1.2192 | 0.265576 | 0.224303 | -0.0821 | -0.72499 | 0.61222 | -0.07526 | 0.332289 | -0.36362 | 0.135116 | 0.069103 | 0.247184 | 0.254613 | 0.021918 | 0.195848 | 0.575629 | 0.305823 |
| 27.61111 | 0.25909 | 0.803558 | -0.41821 | 0.450797 | 0.026718 | 0.351548 | -0.84214 | 1.063336 | 0.053854 | 1.201252 | -0.10545 | 0.161557 | 0.058695 | 0.245413 | 0.246964 | 0.030286 | 0.328533 | 0.620791 | 0.100133 |
| 27.625 | 0.570333 | 1.216473 | 0.824568 | 0.517211 | -0.01118 | 0.820706 | -0.30473 | 1.477737 | 0.211141 | 1.582924 | 0.099373 | 0.22961 | 0.147695 | 0.226459 | 0.20631 | 0.047125 | 0.362689 | 0.671462 | 0.183456 |
| 27.63889 | 0.584281 | 1.321825 | 1.760783 | 0.222951 | -0.05122 | 1.121406 | 0.412363 | 1.693521 | 0.402459 | 1.25748 | 0.047006 | 0.167878 | 0.147902 | 0.144393 | 0.119427 | 0.027385 | 0.156883 | 0.613442 | 0.345215 |
| 27.65278 | 0.394035 | 1.343365 | 1.950441 | -0.091 | -0.20379 | 1.133746 | 1.015281 | 1.893051 | 0.470858 | 0.65567 | -0.02877 | 0.018902 | 0.059856 | 0.042781 | 0.042813 | -0.00162 | -0.12098 | 0.364619 | 0.267779 |
| 27.66667 | 0.548766 | 0.887062 | 1.369189 | -0.08892 | -0.38635 | 0.714012 | 1.513124 | 1.919483 | 0.461155 | -0.45676 | -0.12569 | -0.0917 | -0.00266 | -0.04808 | 0.033342 | 0.032551 | -0.314 | 0.15245 | 0.048785 |
| 27.68056 | 1.141966 | 0.525473 | 0.215621 | 0.254926 | -0.4802 | 0.063842 | 0.954176 | 1.483171 | 0.095388 | -0.71995 | -0.38254 | -0.21889 | -0.06378 | -0.12021 | 0.053592 | 0.063064 | -0.38387 | 0.045533 | -0.14375 |
| 27.69444 | 1.70668 | 0.071707 | -0.78856 | 0.500573 | -0.28319 | -0.52166 | 0.105088 | 1.015502 | -0.24101 | -0.33504 | -0.63853 | -0.40578 | -0.1012 | -0.12208 | 0.036635 | 0.06388 | -0.33968 | -0.02526 | -0.13034 |
| 27.70833 | 1.226286 | -0.77326 | -1.14958 | 0.548145 | 0.092421 | -1.02481 | -0.88513 | 0.473386 | -0.05741 | 0.023696 | -0.64859 | -0.43668 | -0.09317 | -0.03174 | -0.04192 | 0.096786 | -0.21679 | -0.06615 | 0.014828 |
| 27.72222 | 0.419384 | -1.15239 | -1.13748 | 0.397317 | 0.45056 | -1.05698 | -1.51125 | -0.08561 | 0.345723 | 0.056852 | -0.2917 | -0.23081 | -0.01097 | 0.07833 | -0.14765 | 0.109035 | -0.09141 | -0.08375 | 0.095273 |
| 27.73611 | -0.46719 | -0.87468 | -0.82349 | 0.125294 | 0.737778 | -0.28524 | -1.7841 | -0.66089 | 0.508022 | -0.07872 | 0.145744 | 0.035619 | 0.072169 | 0.219299 | -0.16275 | 0.127799 | 0.090444 | -0.02919 | 0.124608 |
| 27.75 | -1.01629 | -0.33374 | -0.63715 | -0.33184 | 0.59609 | 0.222165 | -1.50471 | -1.30241 | 0.255836 | -0.29449 | 0.301728 | 0.169932 | 0.048062 | 0.270271 | -0.13008 | 0.11145 | 0.33885 | 0.057111 | 0.056626 |
| 27.76389 | -1.44606 | -0.16157 | -0.69173 | -0.72945 | 0.35829 | 0.202848 | -1.06958 | -1.65014 | -0.20073 | -0.67492 | 0.115156 | 0.141345 | -0.07177 | 0.224334 | -0.11664 | 0.04538 | 0.628172 | 0.03597 | -0.05758 |
| 27.77778 | -1.57447 | -0.34961 | -0.89246 | -0.59037 | 0.276803 | -0.10454 | -0.58336 | -1.29306 | -0.60078 | -1.05068 | -0.02712 | 0.025318 | -0.09272 | 0.202649 | -0.06188 | 0.017605 | 0.725343 | -0.15087 | -0.11702 |
| 27.79167 | -1.73832 | -0.91534 | -1.10663 | -0.24273 | 0.167081 | -0.34679 | -0.29677 | -0.90661 | -1.06676 | -1.26545 | -0.02587 | -0.13493 | 0.006586 | 0.124389 | -0.01672 | -0.01818 | 0.487582 | -0.46032 | -0.24801 |
| 27.80556 | -1.71769 | -1.38763 | -0.83503 | -0.09735 | -0.06225 | -0.49385 | -0.06585 | -0.83856 | -1.40785 | -0.90094 | 0.028445 | -0.28152 | 0.084489 | -0.00155 | 0.004251 | -0.08418 | 0.115915 | -0.72746 | -0.52364 |
| 27.81944 | -1.2177 | -1.18197 | -0.21086 | -0.2943 | -0.30631 | -0.72672 | 0.265545 | -0.84144 | -1.21112 | -0.25917 | 0.064815 | -0.30959 | 0.157672 | -0.10381 | 0.102604 | -0.07377 | -0.21935 | -0.81213 | -0.66758 |
| 27.83333 | -0.57576 | -0.74949 | 0.25074 | -0.80956 | -0.56654 | -1.15207 | 0.477439 | -0.92419 | -0.80514 | 0.131893 | -0.09941 | -0.31752 | 0.181614 | -0.1907 | 0.164243 | -0.05208 | -0.43729 | -0.79381 | -0.52994 |
| 27.84722 | -0.1954 | -0.43389 | 0.143665 | -1.29103 | -0.71208 | -1.50176 | 0.264266 | -1.17478 | -0.5698 | -0.10169 | -0.4541 | -0.38919 | 0.153493 | -0.2265 | 0.076849 | -0.05692 | -0.52159 | -0.77899 | -0.25755 |
| 27.86111 | -0.07252 | -0.32419 | -0.43612 | -1.30636 | -0.80381 | -1.62766 | -0.28797 | -1.53843 | -0.70418 | -0.58462 | -0.84363 | -0.43338 | 0.100614 | -0.18906 | -0.03344 | -0.03882 | -0.56404 | -0.82222 | -0.2601 |
| 27.875 | -0.37359 | -0.50439 | -1.10825 | -0.74056 | -0.82986 | -1.49934 | -0.75039 | -1.75623 | -1.16523 | -0.94251 | -1.00339 | -0.36056 | 0.0103 | -0.16935 | -0.12449 | -0.072 | -0.6367 | -0.91534 | -0.7146 |
| 27.88889 | -0.37909 | -0.97443 | -1.54379 | -0.2267 | -0.66625 | -1.31058 | -0.80772 | -1.51394 | -1.49842 | -1.09994 | -0.74224 | -0.22465 | -0.01078 | -0.07398 | -0.13202 | -0.17665 | -0.71225 | -0.96924 | -1.18505 |
| 27.90278 | -0.15697 | -1.55072 | -1.46101 | -0.22822 | -0.53931 | -1.23204 | -0.58476 | -0.95037 | -1.35485 | -1.12099 | -0.30445 | -0.17097 | -0.01679 | -0.00485 | -0.10804 | -0.23009 | -0.65686 | -0.89772 | -1.24542 |
| 27.91667 | 0.192184 | -2.06351 | -0.91311 | -0.80576 | -0.44808 | -1.09881 | -0.42038 | -0.46761 | -1.07467 | -1.06795 | -0.11049 | -0.22026 | -0.02048 | 0.013486 | -0.10094 | -0.21392 | -0.48742 | -0.77681 | -1.04482 |
| 27.93056 | 0.457808 | -1.78181 | -0.3504 | -1.40777 | -0.20655 | -0.72874 | -0.31705 | -0.35049 | -0.66008 | -0.87086 | -0.24113 | -0.23549 | 0.01747 | 0.022115 | -0.06935 | -0.14045 | -0.35331 | -0.63593 | -0.84028 |
| 27.94444 | 0.326351 | -1.09488 | -0.059 | -1.85421 | -0.00524 | -0.32124 | -0.26263 | -0.56466 | -0.2915 | -0.59318 | -0.39964 | -0.18035 | 0.032852 | -0.01449 | -0.07144 | -0.09778 | -0.36276 | -0.4571 | -0.65039 |
| 27.95833 | -0.06963 | -0.59548 | -0.08498 | -1.54733 | 0.070139 | -0.11633 | -0.22923 | -0.81186 | -0.09381 | -0.35124 | -0.34081 | -0.13315 | 0.015687 | -0.05461 | -0.07892 | -0.09333 | -0.43893 | -0.29437 | -0.52646 |
| 27.97222 | -0.29532 | -0.39723 | -0.19779 | -0.84267 | 0.097908 | -0.01293 | -0.19779 | -0.78489 | -0.14179 | -0.18092 | -0.06774 | -0.05461 | 0.008031 | -0.0288 | -0.07039 | -0.04503 | -0.40379 | -0.20802 | -0.49457 |
| 27.98611 | -0.47757 | -0.45882 | -0.27472 | -0.32555 | 0.104166 | -0.00073 | -0.10841 | -0.53899 | -0.27691 | -0.28813 | -0.0491 | 0.026127 | 0.016237 | 0.090661 | -0.04002 | 0.025943 | -0.33338 | -0.34214 | -0.66648 |
| 28 | -0.67599 | -1.13559 | -0.41767 | -0.25207 | 0.125471 | -0.22794 | 0.104341 | -0.10032 | -0.4897 | -0.88776 | -0.54935 | 0.028312 | 0.049065 | 0.302053 | 0.007236 | 0.083732 | -0.4607 | -0.81078 | -1.03565 |
| 28.01389 | -1.03644 | -2.54196 | -0.78699 | -0.55384 | 0.070409 | -0.66253 | 0.20159 | 0.093367 | -0.91615 | -1.45015 | -0.99941 | -0.08252 | 0.085206 | 0.372592 | 0.061567 | 0.12166 | -0.74248 | -1.25133 | -1.136 |
| 28.02778 | -1.62775 | -3.42004 | -0.77106 | -0.56627 | 0.002489 | -0.87525 | 0.170211 | -0.37922 | -1.05715 | -1.30017 | -0.81466 | -0.23954 | 0.073948 | 0.178294 | 0.006539 | 0.108646 | -0.87034 | -1.09351 | -0.82992 |
| 28.04167 | -2.06713 | -2.23618 | -0.03075 | -0.1985 | 0.00681 | -0.71195 | 0.17184 | -1.06634 | -0.52361 | -0.77498 | -0.37474 | -0.23285 | 0.050971 | -0.06714 | -0.11066 | 0.066872 | -0.6825 | -0.72323 | -0.43722 |
| 28.05556 | -2.02304 | -0.40147 | 0.595921 | -0.11039 | 0.183888 | -0.27079 | 0.185189 | -1.47861 | 0.293832 | -0.29939 | -0.19453 | -0.08301 | 0.022644 | -0.16916 | -0.1516 | 0.018367 | -0.37154 | -0.4567 | -0.25335 |
| 28.06944 | -1.61694 | 0.89733 | 0.816178 | -0.52713 | 0.428734 | 0.376826 | 0.110425 | -1.29889 | 0.563653 | 0.175505 | -0.18301 | 0.034229 | -0.01806 | -0.13836 | -0.10027 | -0.00307 | -0.19471 | -0.27874 | -0.21927 |
| 28.08333 | -1.1219 | 1.28964 | 0.543371 | -0.60928 | 0.616366 | 0.996573 | -0.00118 | -0.6859 | 0.371233 | 0.332153 | -0.14231 | 0.069546 | -0.03991 | -0.07263 | -0.06352 | -0.00395 | -0.07236 | -0.14178 | -0.14813 |
| 28.09722 | -0.54564 | 1.462477 | 0.417323 | -0.05183 | 0.768764 | 1.527493 | 0.043176 | -0.01121 | 0.316452 | 0.096464 | -0.0736 | 0.05957 | -0.04431 | -0.03409 | -0.03706 | -0.00146 | 0.008119 | -0.01164 | -0.03351 |
| 28.11111 | -0.16606 | 1.657853 | 0.640072 | 0.624073 | 0.86621 | 1.992194 | 0.194094 | 0.466545 | 0.511862 | -0.13549 | -0.01968 | 0.065185 | -0.0401 | -0.03029 | -0.0145 | -0.0062 | 0.094966 | 0.061384 | -0.00775 |
| 28.125 | 0.05173 | 1.672413 | 0.936529 | 0.804499 | 0.858574 | 2.168236 | 0.286534 | 0.568373 | 0.789167 | -0.2535 | 0.111262 | 0.124337 | -0.02316 | -0.02452 | 0.003119 | 0.003787 | 0.267985 | 0.014465 | -0.09365 |
| 28.13889 | 0.095103 | 1.292966 | 0.986328 | 0.677362 | 0.705627 | 2.120947 | 0.308196 | 0.479011 | 0.94048 | -0.33906 | 0.339481 | 0.166974 | -0.01954 | -0.02437 | -0.01601 | 0.02104 | 0.439371 | -0.14258 | -0.21561 |
| 28.15278 | 0.147964 | 0.926073 | 0.727545 | 0.443643 | 0.437361 | 1.912695 | 0.248975 | 0.432694 | 0.910252 | -0.34787 | 0.493336 | 0.117746 | -0.04438 | -0.03589 | -0.04947 | 0.008083 | 0.351448 | -0.25081 | -0.24357 |
| 28.16667 | 0.317732 | 0.588165 | 0.312174 | 0.100626 | 0.28791 | 1.059555 | 0.140262 | 0.397782 | 0.669775 | -0.23692 | 0.430292 | 0.058987 | -0.0615 | -0.04069 | -0.04971 | -0.00791 | 0.216786 | -0.19452 | -0.17209 |
| 28.18056 | 0.49014 | 0.290883 | 0.069858 | -0.24211 | 0.1606 | 0.495621 | 0.01963 | 0.28167 | 0.313148 | -0.11345 | 0.28687 | 0.048673 | -0.06446 | -0.04079 | -0.0461 | -0.01315 | 0.17015 | -0.0956 | -0.14626 |
| 28.19444 | 0.549164 | 0.14452 | 0.013884 | -0.23719 | 0.076223 | 0.346254 | -0.00735 | 0.238844 | -0.00856 | -0.0721 | 0.173786 | 0.042435 | -0.07036 | -0.0343 | -0.06546 | -0.01142 | 0.280405 | -0.1172 | -0.22288 |
| 28.20833 | 0.52424 | 0.22255 | 0.142093 | 0.078058 | 0.093229 | 0.490779 | 0.084581 | 0.180595 | -0.09937 | -0.10248 | 0.054426 | -0.00519 | -0.06934 | -0.02454 | -0.09164 | -0.007 | 0.379881 | -0.23415 | -0.29665 |
| 28.22222 | 0.51311 | 0.357015 | 0.481664 | 0.210065 | 0.212711 | 0.698602 | 0.203158 | 0.094815 | 0.072917 | -0.19803 | -0.15252 | -0.08723 | -0.05565 | -0.03466 | -0.11423 | -0.01313 | 0.285277 | -0.30905 | -0.32089 |
| 28.23611 | 0.517365 | 0.410958 | 0.777973 | -0.01761 | 0.338963 | 0.757711 | 0.273988 | 0.057383 | 0.289469 | -0.35334 | -0.4061 | -0.1337 | -0.04999 | -0.04892 | -0.12324 | -0.02047 | 0.018703 | -0.27468 | -0.29581 |
| 28.25 | 0.486928 | 0.338267 | 0.705494 | -0.16355 | 0.395727 | 0.699309 | 0.278732 | 0.090486 | 0.346838 | -0.50396 | -0.56926 | -0.07128 | -0.05444 | -0.0504 | -0.08364 | -0.01122 | -0.21725 | -0.2376 | -0.21922 |
| 28.26389 | 0.414151 | 0.208189 | 0.391419 | 0.134631 | 0.347112 | 0.540898 | 0.23641 | 0.132699 | 0.237462 | -0.5214 | -0.52209 | 0.048301 | -0.06237 | -0.04767 | -0.04564 | -2.4E-05 | -0.2657 | -0.25278 | -0.12901 |
| 28.27778 | 0.361452 | 0.112742 | 0.131655 | 0.537481 | 0.232515 | 0.325515 | 0.165665 | 0.109058 | 0.109604 | -0.33276 | -0.30793 | 0.137289 | -0.05389 | -0.03608 | -0.0411 | -0.00546 | -0.08844 | -0.19097 | -0.05384 |
| 28.29167 | 0.415873 | 0.091502 | 0.081591 | 0.604126 | 0.110923 | 0.306556 | 0.119613 | 0.073025 | 0.081202 | -0.01261 | -0.09876 | 0.176325 | -0.03507 | -0.02194 | -0.05793 | -0.02584 | 0.09 | -0.06892 | -0.01777 |
| 28.30556 | 0.5009 | 0.121959 | 0.262515 | 0.298615 | 0.03267 | 0.528519 | 0.087067 | 0.098515 | 0.151433 | 0.281144 | -0.03789 | 0.163907 | -0.02907 | -0.01465 | -0.04344 | -0.03573 | 0.126868 | 0.046846 | -0.04931 |
| 28.31944 | 0.501019 | 0.099972 | 0.527265 | 0.035056 | -0.00044 | 0.817039 | 0.02658 | 0.170162 | 0.213427 | 0.451442 | -0.02413 | 0.142122 | -0.01589 | -0.00499 | 0.016843 | -0.0083 | 0.111372 | 0.148819 | -0.09388 |
| 28.33333 | 0.467529 | 0.053048 | 0.634534 | 0.132797 | -0.03618 | 0.935419 | -0.06387 | 0.317308 | 0.197072 | 0.394367 | 0.015868 | 0.15649 | -0.01337 | -0.02263 | 0.041691 | 0.00439 | 0.107492 | 0.209355 | -0.08054 |
| 28.34722 | 0.46099 | 0.048378 | 0.538307 | 0.496102 | -0.02334 | 0.801537 | -0.06214 | 0.591301 | 0.158615 | 0.138643 | 0.028494 | 0.151899 | -0.05441 | -0.07397 | -0.01377 | -0.05063 | 0.081927 | 0.170388 | 0.008039 |
| 28.36111 | 0.47797 | 0.097516 | 0.426335 | 0.784404 | 0.033676 | 0.64762 | 0.069774 | 0.897555 | 0.182766 | -0.06074 | 0.078746 | 0.155785 | -0.06818 | -0.07292 | -0.03063 | -0.08629 | 0.084073 | 0.107559 | 0.13249 |
| 28.375 | 0.467706 | 0.162489 | 0.486828 | 0.841026 | 0.111377 | 0.591614 | 0.205494 | 1.098558 | 0.303172 | 0.013271 | 0.221756 | 0.205967 | -0.02086 | -0.01469 | 0.049721 | -0.04321 | 0.133049 | 0.131697 | 0.293482 |
| 28.38889 | 0.466136 | 0.269275 | 0.778943 | 0.798769 | 0.179356 | 0.630315 | 0.236094 | 1.111938 | 0.535088 | 0.289595 | 0.361135 | 0.269242 | 0.015833 | 0.007239 | 0.123111 | 0.011977 | 0.176278 | 0.209764 | 0.450467 |
| 28.40278 | 0.595325 | 0.4627 | 1.169532 | 0.757395 | 0.173354 | 0.720893 | 0.158699 | 1.056436 | 0.750828 | 0.627841 | 0.45786 | 0.310908 | 0.007523 | -0.03831 | 0.113668 | 0.036934 | 0.23704 | 0.2956 | 0.563119 |
| 28.41667 | 0.833041 | 0.628741 | 1.299555 | 0.744436 | 0.042469 | 0.846812 | 0.018472 | 1.021924 | 0.882344 | 0.888132 | 0.512455 | 0.286618 | -0.04359 | -0.12091 | 0.035713 | 0.012309 | 0.288975 | 0.329702 | 0.587368 |
| 28.43056 | 1.034365 | 0.708227 | 1.098847 | 0.715372 | -0.01168 | 1.069199 | 0.000559 | 1.073296 | 0.951002 | 0.831566 | 0.481576 | 0.195468 | -0.05906 | -0.14175 | -0.00804 | -0.02059 | 0.304131 | 0.283369 | 0.571923 |
| 28.44444 | 1.051786 | 0.669491 | 0.820072 | 0.713639 | 0.125415 | 1.359419 | 0.166885 | 1.024423 | 0.948172 | 0.548526 | 0.413783 | 0.113072 | -0.00635 | -0.08055 | 0.037554 | -0.0025 | 0.263843 | 0.199686 | 0.603567 |
| 28.45833 | 0.83331 | 0.495937 | 0.538886 | 0.641111 | 0.244976 | 1.284387 | 0.266132 | 0.787311 | 0.722839 | 0.348129 | 0.349144 | 0.095474 | 0.021727 | -0.05984 | 0.052934 | 0.01521 | 0.181334 | 0.155137 | 0.545668 |
| 28.47222 | 0.499142 | 0.102686 | 0.245684 | 0.581215 | 0.218381 | 0.713807 | 0.196155 | 0.482947 | 0.265821 | 0.381224 | 0.324996 | 0.065573 | -0.01014 | -0.08313 | 0.00722 | 0.005918 | 0.11771 | 0.217685 | 0.420735 |
| 28.48611 | 0.507109 | -0.10772 | 0.509661 | 1.124484 | 0.192441 | 0.289328 | 0.124213 | 0.460667 | 0.073258 | 0.798432 | 0.616268 | 0.072428 | -0.07159 | -0.13903 | -0.0535 | -0.01059 | 0.172907 | 0.520571 | 0.532875 |
| 28.5 | 1.33969 | 0.54716 | 1.565893 | 2.683784 | 0.393391 | 0.454467 | 0.253159 | 0.936379 | 0.563388 | 1.808829 | 1.389917 | 0.093991 | -0.1547 | -0.22127 | -0.10487 | -0.02634 | 0.325697 | 1.166892 | 0.985196 |
| 28.51389 | 2.460283 | 1.514364 | 2.64271 | 3.326367 | 0.881149 | 0.780364 | 0.580177 | 1.738884 | 1.5399 | 2.675374 | 1.929694 | 0.149699 | -0.16233 | -0.15194 | -0.22808 | -0.02374 | 0.41919 | 1.577656 | 1.630699 |
| 28.52778 | 2.851985 | 2.01322 | 3.04773 | 2.712072 | 0.773521 | 1.07569 | 0.868177 | 2.003942 | 2.491583 | 2.897526 | 1.908793 | 0.242195 | -0.10209 | 0.068271 | -0.00073 | 0.033479 | 0.684369 | 1.541955 | 1.804077 |
| 28.54167 | 2.09728 | 1.388569 | 2.16372 | 1.595309 | 0.417388 | 1.093683 | 1.008649 | 1.358611 | 3.10044 | 1.886086 | 1.501259 | 0.392756 | -0.06475 | 0.278742 | 0.142307 | 0.114425 | 1.111541 | 1.365596 | 1.576441 |
| 28.55556 | 1.02953 | 0.773782 | 0.5821 | 0.299468 | 0.242823 | 0.313973 | 0.882704 | 0.535658 | 2.199483 | 0.494527 | 0.967251 | 0.551249 | -0.09235 | 0.398612 | 0.091726 | 0.114278 | 1.13986 | 0.909043 | 0.939139 |
| 28.56944 | -0.20788 | 0.128393 | -0.88276 | -0.66056 | 0.279117 | -0.24818 | 0.596411 | -0.25951 | 0.466589 | -0.99372 | 0.429852 | 0.417511 | -0.138 | 0.395242 | 0.018679 | 0.074997 | 0.900171 | 0.44841 | 0.340813 |
| 28.58333 | -0.69582 | -0.72355 | -1.82011 | -0.86051 | 0.380346 | -0.77675 | 0.251645 | -0.73056 | -0.7946 | -2.13772 | -0.0094 | 0.20628 | -0.09261 | 0.342897 | -0.01919 | 0.080044 | 0.607235 | 0.272494 | 0.002094 |
| 28.59722 | -0.78406 | -0.81999 | -1.89084 | 0.030063 | 0.422405 | -1.48111 | 0.137334 | -0.77222 | -1.36432 | -1.21406 | -0.39106 | 0.099567 | 0.062611 | 0.319099 | 0.042832 | 0.082547 | 0.398673 | 0.479965 | -0.03834 |
| 28.61111 | -0.42617 | -0.80411 | -0.99839 | 1.093151 | 0.30062 | -0.91634 | 0.251269 | -0.31299 | -1.09029 | 0.075643 | -0.64161 | 0.015889 | 0.102707 | 0.235162 | 0.168217 | 0.033591 | 0.196129 | 0.950488 | 0.000714 |
| 28.625 | 0.199112 | 0.291114 | 0.397792 | 1.685928 | 0.237894 | 0.37522 | 0.434432 | 0.338824 | -0.22079 | 1.13762 | -0.64648 | -0.00577 | 0.000541 | 0.12371 | 0.260921 | -0.04214 | 0.088941 | 1.095908 | -0.00753 |
| 28.63889 | 0.380498 | 1.441579 | 1.553122 | 1.018407 | 0.396579 | 1.660651 | 0.381802 | 0.877814 | 0.913937 | 1.36682 | -0.36969 | 0.049845 | -0.03651 | 0.045179 | 0.27533 | -0.06651 | 0.002952 | 0.783931 | 0.020981 |
| 28.65278 | 0.250092 | 1.2394 | 1.475442 | 0.27898 | 0.420549 | 2.396312 | 0.11975 | 1.003897 | 1.732138 | 0.7736 | -0.22234 | 0.028727 | 0.03336 | 0.037388 | 0.210769 | -0.07038 | -0.14706 | 0.285175 | 0.16554 |
| 28.66667 | 0.154297 | 1.116279 | 1.059252 | -0.45863 | 0.288194 | 2.321295 | -0.20841 | 0.838581 | 1.265723 | 0.26295 | -0.17905 | -0.12297 | 0.104852 | 0.042341 | 0.150287 | -0.06222 | -0.23232 | -0.11504 | 0.324105 |
| 28.68056 | 0.401188 | 0.615052 | 0.898372 | -1.0846 | 0.220189 | 1.228307 | -0.3817 | 0.81349 | 0.692422 | -0.05597 | -0.40526 | -0.29705 | 0.088783 | -0.01299 | 0.117791 | -0.01009 | -0.18076 | -0.3403 | 0.211642 |
| 28.69444 | 0.525568 | 0.466059 | 0.244506 | -1.08962 | 0.273798 | 0.229221 | -0.3889 | 1.045194 | -0.06501 | -0.2322 | -0.64149 | -0.40041 | 0.009719 | -0.06879 | 0.04709 | 0.044255 | -0.09474 | -0.40813 | -0.20915 |
| 28.70833 | 1.126183 | 0.22722 | -0.44894 | 0.036216 | 0.434558 | -0.90477 | -0.15836 | 0.952915 | -0.80967 | -0.11489 | -0.52926 | -0.34931 | -0.00146 | -0.00409 | 0.051348 | 0.126499 | 0.045921 | -0.40668 | -0.49896 |

| | | | | | | | | | | | | | | | | | | | |
|---|---|---|---|---|---|---|---|---|---|---|---|---|---|---|---|---|---|---|---|
| 28.72222 | 2.02756 | -0.35259 | -0.37604 | 1.180659 | 0.551338 | -1.57928 | 0.167792 | 0.482704 | -0.895 | 0.128562 | -0.29051 | -0.1415 | 0.022622 | 0.078252 | 0.098022 | 0.173023 | 0.108755 | -0.30428 | -0.45917 |
| 28.73611 | 1.584619 | -1.09124 | -0.49116 | 1.636827 | 0.485528 | -1.62558 | 0.32304 | 0.416092 | -0.261 | 0.218396 | -0.17259 | 0.081971 | -0.02238 | -0.00915 | 0.128053 | 0.153756 | 0.079463 | -0.07602 | -0.3865 |
| 28.75 | 0.442888 | -1.33289 | -0.59521 | 0.822142 | 0.338852 | -1.12891 | 0.274884 | 0.827198 | 0.395435 | 0.066823 | -0.07173 | 0.216992 | -0.07992 | -0.07304 | 0.111258 | 0.127005 | 0.05792 | 0.195601 | -0.31999 |
| 28.76389 | -0.74502 | 0.040346 | -0.67019 | -0.11475 | 0.256873 | -0.11861 | 0.120109 | 1.217061 | 0.826205 | -0.19116 | -0.02211 | 0.197409 | -0.11164 | -0.02969 | 0.002683 | 0.106096 | 0.127356 | 0.257442 | -0.21787 |
| 28.77778 | -1.83534 | 0.53549 | -1.47525 | -0.67355 | 0.288129 | 0.661157 | -0.12408 | 1.047318 | 0.443191 | -0.38513 | -0.04003 | 0.061359 | -0.12512 | 0.094592 | -0.11512 | 0.061212 | 0.24126 | 0.023966 | -0.21671 |
| 28.79167 | -2.14248 | -0.22898 | -2.4225 | -1.12327 | 0.205725 | 0.652075 | -0.24528 | 0.298157 | -0.61317 | -0.5359 | -0.11174 | -0.12861 | -0.19892 | 0.185485 | -0.19312 | 0.01402 | 0.171493 | -0.25272 | -0.50904 |
| 28.80556 | -2.1859 | -1.78364 | -3.25902 | -1.16795 | -0.00972 | -0.29311 | -0.21919 | -0.94579 | -1.82665 | -0.87715 | -0.2853 | -0.35067 | -0.31253 | 0.050771 | -0.23475 | -0.06631 | -0.04833 | -0.40843 | -1.02696 |
| 28.81944 | -2.06324 | -3.5719 | -2.61961 | -0.8017 | -0.2549 | -1.7452 | -0.20214 | -2.3189 | -2.62855 | -1.32415 | -0.4791 | -0.50239 | -0.31679 | -0.13572 | -0.27287 | -0.14305 | -0.34266 | -0.56924 | -1.3826 |
| 28.83333 | -1.71859 | -4.97105 | -1.5465 | -0.53096 | -0.53002 | -3.05083 | -0.21466 | -3.12976 | -2.49325 | -1.60423 | -0.61262 | -0.613 | -0.25195 | -0.18492 | -0.31871 | -0.23488 | -0.56844 | -0.91452 | -1.13195 |
| 28.84722 | -1.64566 | -4.60052 | -0.91049 | -0.52804 | -0.76528 | -3.12449 | -0.33006 | -3.04298 | -2.02849 | -1.65514 | -0.64524 | -0.78123 | -0.1721 | -0.20649 | -0.28133 | -0.33137 | -0.69677 | -1.26584 | -0.8347 |
| 28.86111 | -1.52425 | -3.34721 | -0.92453 | -0.97317 | -0.805 | -2.58228 | -0.4096 | -2.43112 | -1.82992 | -1.46728 | -0.59256 | -0.86629 | -0.14772 | -0.22302 | -0.15425 | -0.32433 | -0.7942 | -1.34726 | -0.85813 |
| 28.875 | -1.6546 | -2.14442 | -1.32903 | -1.75978 | -0.72655 | -2.12917 | -0.42488 | -2.07356 | -1.82467 | -1.13316 | -0.502 | -0.71573 | -0.15328 | -0.2684 | -0.1646 | -0.27361 | -0.93397 | -1.1072 | -1.04161 |
| 28.88889 | -1.74324 | -1.25229 | -1.62155 | -2.89374 | -0.48372 | -1.72868 | -0.36417 | -1.99418 | -1.68204 | -0.70855 | -0.4149 | -0.52801 | -0.21324 | -0.31939 | -0.23792 | -0.23817 | -1.00771 | -0.88519 | -1.17976 |
| 28.90278 | -1.52888 | -0.54644 | -1.5452 | -3.48958 | -0.2713 | -1.29105 | -0.15936 | -1.8893 | -1.46216 | -0.41194 | -0.31649 | -0.37131 | -0.25937 | -0.33335 | -0.28894 | -0.19997 | -1.00725 | -0.76422 | -1.21188 |
| 28.91667 | -1.08464 | -0.22667 | -1.13462 | -2.5093 | -0.18876 | -1.05882 | 0.015581 | -1.66679 | -1.27721 | -0.37535 | -0.20661 | -0.2896 | -0.22643 | -0.33958 | -0.27533 | -0.14597 | -0.97434 | -0.77722 | -1.28674 |
| 28.93056 | -0.69911 | -0.30387 | -0.68073 | -1.08441 | -0.28653 | -0.79372 | 0.055446 | -1.31278 | -1.04536 | -0.62015 | -0.12884 | -0.2839 | -0.2027 | -0.30527 | -0.28357 | -0.11687 | -0.85536 | -0.87868 | -1.25196 |
| 28.94444 | -0.3334 | -0.68358 | -0.42933 | -0.15011 | -0.2746 | -0.44152 | -0.02871 | -0.7255 | -0.72285 | -0.86845 | -0.07425 | -0.22088 | -0.16027 | -0.29759 | -0.27097 | -0.10988 | -0.54327 | -0.8686 | -0.97122 |
| 28.95833 | -0.08077 | -0.9278 | -0.27768 | 0.057169 | -0.16282 | -0.07638 | -0.09425 | -0.11304 | -0.29684 | -0.73427 | -0.05932 | -0.08425 | -0.11099 | -0.24339 | -0.23029 | -0.09258 | -0.23313 | -0.72298 | -0.53888 |
| 28.97222 | 0.051833 | -0.85752 | -0.20682 | -0.14135 | 0.005148 | 0.178291 | -0.16836 | 0.224092 | 0.044644 | -0.32459 | -0.11844 | 0.022565 | -0.09582 | -0.15004 | -0.18571 | -0.08012 | -0.10286 | -0.54919 | -0.28354 |
| 28.98611 | -0.14694 | -1.03717 | -0.32142 | -0.70956 | 0.047492 | -0.01561 | -0.35616 | -0.00468 | -0.05748 | -0.41844 | -0.50051 | 0.061687 | -0.10594 | 0.032421 | -0.0989 | -0.08615 | -0.24971 | -0.64188 | -0.42723 |
| 29 | -0.70688 | -1.79051 | -0.95351 | -1.73423 | -0.16604 | -0.84776 | -0.59328 | -0.63585 | -0.70608 | -1.46222 | -1.28361 | 0.061562 | -0.13158 | 0.274026 | -0.00104 | -0.10866 | -0.55393 | -1.10304 | -0.90197 |
| 29.01389 | -1.01697 | -2.39626 | -1.55407 | -2.32066 | -0.46736 | -1.74079 | -0.66994 | -1.06841 | -1.42343 | -2.19531 | -1.48233 | 0.021 | -0.10258 | 0.267658 | 0.039732 | -0.07416 | -0.53494 | -1.28599 | -0.96174 |
| 29.02778 | -1.10815 | -2.38571 | -1.8846 | -2.2053 | -0.9279 | -2.59983 | -0.62511 | -1.28977 | -2.01945 | -1.4847 | -0.77448 | -0.12154 | -0.0729 | -0.09275 | -0.03542 | -0.06811 | -0.31457 | -0.87639 | -0.44756 |
| 29.04167 | -1.57587 | -2.86116 | -2.64645 | -2.07635 | -1.85942 | -4.05532 | -0.96532 | -2.02793 | -2.91321 | -0.65433 | -0.23492 | -0.29646 | -0.1736 | -0.52582 | -0.22145 | -0.14956 | -0.25865 | -0.54901 | 0.020802 |
| 29.05556 | -2.1031 | -3.23015 | -3.41564 | -1.69237 | -2.98424 | -5.48643 | -1.29116 | -2.50627 | -3.30293 | -0.3997 | -0.18934 | -0.29092 | -0.19078 | -0.69371 | -0.26539 | -0.16551 | -0.20885 | -0.31014 | 0.147586 |
| 29.06944 | -1.89848 | -2.44527 | -2.79627 | -1.19339 | -3.25442 | -5.09093 | -1.155 | -2.3244 | -2.30212 | -0.35058 | -0.32487 | -0.14181 | -0.09304 | -0.39376 | -0.15568 | -0.09823 | -0.09554 | -0.00819 | 0.056107 |
| 29.08333 | -1.21352 | -1.11908 | -1.4528 | -0.94516 | -2.07593 | -2.81134 | -0.73862 | -1.71654 | -1.00309 | -0.30413 | -0.41163 | -0.05982 | -0.04934 | -0.20445 | -0.04386 | -0.05892 | -0.04212 | 0.159205 | -0.05004 |
| 29.09722 | -0.6816 | -0.41907 | -0.42886 | -1.02033 | -0.85627 | -1.23015 | -0.60371 | -1.20987 | -0.31393 | -0.2631 | -0.33901 | -0.06619 | -0.06115 | -0.17421 | -0.02697 | -0.05536 | -0.03062 | 0.149678 | -0.07496 |
| 29.11111 | -0.49406 | -0.12741 | 0.182266 | -0.99221 | -0.16252 | -0.29158 | -0.73631 | -0.88203 | -0.0944 | -0.22362 | -0.19815 | -0.07976 | -0.05701 | -0.18496 | -0.03792 | -0.04187 | -0.02085 | 0.136884 | -0.04984 |
| 29.125 | -0.40444 | 0.256331 | 0.675323 | -0.54944 | 0.38113 | 0.443608 | -0.81242 | -0.60746 | 0.051688 | -0.15904 | -0.08636 | -0.08309 | -0.04626 | -0.1919 | -0.02874 | -0.0319 | -0.01404 | 0.219935 | -0.01909 |
| 29.13889 | -0.13017 | 0.862217 | 1.129909 | 0.21998 | 1.032197 | 1.64984 | -0.61655 | -0.29854 | 0.396284 | -0.05075 | -0.02198 | -0.09428 | -0.05464 | -0.2226 | -0.00396 | -0.048 | -0.00319 | 0.328609 | -0.00901 |
| 29.15278 | 0.136207 | 1.518772 | 1.626462 | 0.753337 | 1.459743 | 3.016364 | -0.26295 | 0.102853 | 0.92164 | 0.131444 | 0.063696 | -0.07649 | -0.03734 | -0.21149 | 0.028044 | -0.04814 | 0.058576 | 0.347449 | 0.014442 |
| 29.16667 | 0.354461 | 2.058047 | 2.050493 | 0.673136 | 1.489994 | 4.071974 | 0.023235 | 0.44794 | 1.696116 | 0.299189 | 0.123208 | -0.03224 | -0.00742 | -0.14109 | 0.033835 | -0.02182 | 0.103966 | 0.295849 | 0.070867 |
| 29.18056 | 0.390817 | 2.126911 | 2.184461 | 0.239822 | 1.474518 | 3.65013 | 0.08063 | 0.639898 | 2.30853 | 0.367373 | 0.10961 | -0.01907 | -0.01337 | -0.09521 | -0.00514 | -0.01518 | 0.066751 | 0.14701 | 0.093274 |
| 29.19444 | 0.090983 | 1.948771 | 1.700218 | -0.18151 | 1.303346 | 3.183893 | -0.11329 | 0.557024 | 2.204255 | 0.325964 | 0.067531 | -0.0303 | -0.02474 | -0.10527 | -0.02647 | -0.02433 | 0.011906 | -0.00109 | 0.057707 |
| 29.20833 | -0.33843 | 1.65204 | 1.066466 | -0.43927 | 0.893555 | 2.749584 | -0.40178 | 0.299759 | 1.823478 | 0.255985 | 0.021352 | -0.04242 | -0.02917 | -0.1481 | -0.01502 | -0.03204 | -0.00876 | -0.14591 | 0.00563 |
| 29.22222 | -0.64132 | 1.062674 | 0.546884 | -0.63299 | 0.381242 | 2.201138 | -0.5982 | -0.10865 | 1.095435 | 0.14559 | -0.02447 | -0.06033 | -0.04463 | -0.19984 | -0.0115 | -0.0421 | -0.01689 | -0.30202 | -0.04807 |
| 29.23611 | -0.74373 | 0.544872 | 0.134869 | -0.67373 | -0.06806 | 1.360412 | -0.61211 | -0.32528 | 0.38373 | 0.024828 | -0.04166 | -0.0717 | -0.05967 | -0.21425 | -0.02634 | -0.04571 | -0.02697 | -0.38064 | -0.09519 |
| 29.25 | -0.60884 | 0.32817 | -0.06118 | -0.51499 | -0.28019 | 0.627787 | -0.52075 | -0.36744 | -0.16393 | -0.05979 | -0.01612 | -0.05604 | -0.05164 | -0.18519 | -0.04659 | -0.0305 | -0.0321 | -0.33956 | -0.10487 |
| 29.26389 | -0.47162 | 0.011149 | -0.10483 | -0.34256 | -0.3015 | -0.03711 | -0.48116 | -0.33691 | -0.55901 | -0.08378 | 0.030124 | -0.02489 | -0.02532 | -0.12793 | -0.05865 | -0.01112 | -0.02518 | -0.22624 | -0.0688 |
| 29.27778 | -0.38897 | -0.35537 | -0.10465 | -0.28745 | -0.30488 | -0.40038 | -0.52367 | -0.22244 | -0.71829 | -0.12213 | 0.029065 | -0.02733 | -0.01651 | -0.10174 | -0.06521 | -0.02815 | -0.04498 | -0.12842 | -0.03243 |
| 29.29167 | -0.32334 | -0.74577 | -0.2422 | -0.41413 | -0.36997 | -0.52082 | -0.58032 | -0.16922 | -0.76213 | -0.16833 | 0.009166 | -0.05562 | -0.00984 | -0.10491 | -0.05087 | -0.0616 | -0.0707 | -0.09639 | -0.03336 |
| 29.30556 | -0.23603 | -1.10068 | -0.38266 | -0.57664 | -0.42269 | -0.53175 | -0.51697 | -0.15275 | -0.64791 | -0.13729 | 0.04524 | -0.05321 | 0.013165 | -0.06261 | -0.0019 | -0.05661 | -0.03463 | -0.06278 | -0.04472 |
| 29.31944 | -0.1936 | -1.24776 | -0.31653 | -0.46413 | -0.46285 | -0.35061 | -0.37519 | -0.09325 | -0.49076 | -0.02483 | 0.087102 | -0.03193 | 0.027081 | -0.00908 | 0.04533 | -0.02834 | 0.033095 | -0.02293 | -0.04629 |
| 29.33333 | -0.29206 | -1.33579 | -0.15013 | -0.13514 | -0.58059 | -0.14736 | -0.39588 | -0.18151 | -0.43435 | 0.14949 | 0.113889 | -0.00399 | 0.056467 | 0.015246 | 0.074324 | -0.00216 | 0.104354 | 0.020143 | -0.04177 |
| 29.34722 | -0.48502 | -1.26798 | 0.012891 | 0.141808 | -0.75218 | -0.13533 | -0.62311 | -0.42277 | -0.49583 | 0.314855 | 0.103291 | 0.07213 | 0.105124 | 0.029903 | 0.08541 | 0.024011 | 0.179413 | 0.043061 | -0.01307 |
| 29.36111 | -0.4747 | -0.82476 | 0.211378 | 0.265974 | -0.70023 | -0.04394 | -0.74068 | -0.49254 | -0.34177 | 0.369872 | 0.060899 | 0.191379 | 0.138351 | 0.037167 | 0.0727 | 0.021595 | 0.214327 | 0.023643 | 0.055354 |
| 29.375 | -0.1383 | 0.081479 | 0.486215 | 0.303264 | -0.35192 | 0.223113 | -0.59587 | -0.30638 | 0.182902 | 0.382919 | 0.085112 | 0.241754 | 0.141286 | 0.028429 | 0.033944 | -0.01071 | 0.213498 | 0.033926 | 0.226407 |
| 29.38889 | 0.126899 | 0.811264 | 0.643738 | 0.463105 | -0.05788 | 0.445606 | -0.30966 | -0.03255 | 0.712567 | 0.438208 | 0.152173 | 0.208384 | 0.160866 | 0.034164 | 0.018346 | -0.01251 | 0.236017 | 0.126332 | 0.599619 |
| 29.40278 | 0.076363 | 1.066049 | 0.589921 | 0.861918 | 0.087238 | 0.50854 | -0.01654 | 0.250939 | 1.046187 | 0.500068 | 0.189394 | 0.195074 | 0.179201 | 0.055796 | 0.052964 | 0.026419 | 0.268108 | 0.265403 | 0.911518 |
| 29.41667 | -0.00702 | 1.164588 | 0.389899 | 1.254469 | 0.10179 | 0.549084 | 0.127964 | 0.357304 | 1.266155 | 0.575454 | 0.209397 | 0.230583 | 0.158971 | 0.058518 | 0.080055 | 0.053933 | 0.271222 | 0.361896 | 0.754546 |
| 29.43056 | 0.094064 | 0.685388 | 0.235382 | 1.095935 | -0.00653 | 0.633701 | 0.017758 | 0.212575 | 1.341637 | 0.658276 | 0.232641 | 0.259171 | 0.122928 | 0.047422 | 0.081203 | 0.058996 | 0.286198 | 0.382419 | 0.527169 |
| 29.44444 | 0.290232 | 0.000752 | 0.313091 | 0.618059 | -0.13744 | 0.693732 | -0.21194 | 0.006883 | 1.286432 | 0.705364 | 0.258502 | 0.211338 | 0.076502 | 0.04587 | 0.040065 | 0.040787 | 0.299531 | 0.348728 | 0.482046 |
| 29.45833 | 0.354583 | -0.57928 | 0.475359 | 0.140995 | -0.16389 | 0.725219 | -0.34808 | 0.008153 | 0.820623 | 0.732224 | 0.275921 | 0.197208 | 0.061443 | 0.070022 | 0.002648 | 0.027033 | 0.269635 | 0.345446 | 0.638267 |
| 29.47222 | 0.236629 | -0.75966 | 0.518144 | 0.004519 | -0.12366 | 0.765667 | -0.29727 | 0.273621 | 0.273877 | 0.792206 | 0.367206 | 0.210159 | 0.088837 | 0.098574 | 0.00404 | 0.041521 | 0.285134 | 0.39687 | 0.791191 |
| 29.48611 | 0.413336 | -0.34402 | 0.680751 | 0.82283 | -0.16235 | 0.84783 | -0.16608 | 0.713325 | 0.027748 | 1.100779 | 0.817885 | 0.270343 | 0.090546 | 0.028877 | -0.01606 | 0.050495 | 0.29351 | 0.591622 | 0.924635 |
| 29.5 | 1.512487 | 0.604811 | 1.312471 | 2.734483 | -0.30282 | 1.150671 | 0.037732 | 1.404261 | 0.33 | 2.063546 | 1.655035 | 0.450091 | 0.023362 | -0.0863 | -0.07723 | 0.075955 | 0.440527 | 1.373776 | 1.391447 |
| 29.51389 | 2.645603 | 1.368745 | 1.901533 | 4.061823 | -0.35935 | 1.18981 | 0.412 | 2.014649 | 1.111954 | 2.65348 | 2.368352 | 0.63922 | 0.016633 | -0.01696 | -0.13418 | 0.137414 | 0.522143 | 2.007531 | 2.04231 |
| 29.52778 | 2.600019 | 1.858182 | 2.348161 | 3.75112 | -0.08213 | 1.127992 | 0.747214 | 2.03915 | 2.310951 | 2.461522 | 2.093885 | 0.909408 | 0.101502 | 0.205228 | 0.127534 | 0.192958 | 0.515428 | 2.052695 | 2.323853 |
| 29.54167 | 2.168465 | 1.971843 | 2.145121 | 2.604857 | 0.240016 | 1.405605 | 0.832618 | 1.503138 | 3.026787 | 2.170505 | 1.542164 | 0.917534 | 0.19426 | 0.398886 | 0.182781 | 0.293478 | 0.765203 | 1.880251 | 2.018759 |
| 29.55556 | 1.464031 | 1.537854 | 1.434402 | 1.033412 | 0.175232 | 0.953471 | 0.630166 | 0.756197 | 1.906054 | 1.096732 | 1.029632 | 0.635115 | 0.223062 | 0.459982 | 0.191117 | 0.333479 | 1.035444 | 1.401386 | 1.470131 |
| 29.56944 | 0.615147 | 0.554764 | 0.532775 | -0.28901 | -0.00488 | 0.234548 | 0.179182 | 0.038013 | 0.462947 | 0.195449 | 0.60444 | 0.308842 | 0.138061 | 0.398409 | 0.067955 | 0.159542 | 0.929411 | 0.705789 | 0.588557 |
| 29.58333 | 0.000375 | -0.25198 | -0.29249 | -0.98365 | -0.28501 | -0.32556 | -0.34462 | -0.50794 | -0.6037 | -0.9077 | 0.325994 | 0.122669 | 0.08909 | 0.37619 | -0.06876 | 0.016608 | 0.794821 | 0.131686 | -0.06448 |
| 29.59722 | 0.038549 | -0.56141 | -0.93446 | -0.83214 | -0.48361 | -0.36545 | -0.71304 | -0.80222 | -1.05401 | -1.57172 | 0.119186 | 0.06737 | 0.151334 | 0.434306 | -0.07219 | 0.021218 | 0.686633 | -0.20908 | -0.14921 |
| 29.61111 | 0.280129 | -0.57714 | -1.04818 | 0.112136 | -0.25722 | -0.14685 | -0.49724 | -0.63366 | -1.06206 | -1.19804 | -0.26444 | 0.006716 | 0.279957 | 0.423949 | 0.143614 | 0.084405 | 0.579161 | -0.34395 | 0.051117 |
| 29.625 | 0.729684 | -0.34865 | -0.41731 | 1.325125 | 0.097315 | 0.195326 | 0.129338 | -0.10597 | -0.49755 | -0.56812 | -0.69459 | -0.05886 | 0.363662 | 0.326632 | 0.396929 | 0.083957 | 0.426423 | -0.44622 | 0.083759 |
| 29.63889 | 1.194479 | 0.353529 | 0.569407 | 2.19672 | 0.657798 | 0.729113 | 1.002953 | 0.431253 | 0.523758 | 0.313885 | -0.78544 | -0.07028 | 0.39089 | 0.064463 | 0.533409 | 0.046563 | 0.260154 | -0.33228 | -0.12408 |
| 29.65278 | 1.436691 | 1.289462 | 1.448212 | 1.639003 | 1.114401 | 1.345517 | 1.43914 | 0.894481 | 1.754548 | 1.169053 | -0.67587 | -0.08883 | 0.172331 | -0.04872 | 0.483531 | -0.0005 | 0.132023 | -0.03964 | -0.43632 |
| 29.66667 | 1.280068 | 1.740404 | 1.381338 | 0.475465 | 1.113504 | 1.833345 | 1.294653 | 1.149321 | 2.614611 | 1.275449 | -0.66792 | 0.001742 | -0.06886 | 0.163489 | 0.42153 | -0.00923 | 0.047159 | 0.29304 | -0.64125 |
| 29.68056 | 0.609847 | 1.462629 | 0.897343 | -0.34886 | 0.755012 | 1.496459 | 0.82413 | 0.81861 | 2.298027 | 0.904417 | -0.73326 | 0.110491 | -0.2159 | 0.289495 | 0.367082 | 0.019504 | -0.01829 | 0.548576 | -0.7756 |
| 29.69444 | -0.20845 | -0.15882 | 0.22057 | -0.84989 | 0.312028 | 0.982749 | 0.149647 | 0.426297 | 1.054115 | 0.62891 | -0.60189 | 0.116723 | -0.27149 | 0.366478 | 0.378535 | 0.067749 | -0.05327 | 0.658799 | -0.77958 |
| 29.70833 | -0.85837 | 0.031481 | 0.052973 | -1.19344 | 0.054458 | 0.527943 | -0.59581 | 0.201511 | -0.31668 | 0.356207 | -0.14876 | 0.062161 | -0.26303 | 0.405885 | 0.400479 | 0.156096 | -0.02946 | 0.684862 | -0.43614 |
| 29.72222 | -0.88652 | 0.126226 | -0.21741 | -0.34702 | 0.054075 | 0.242879 | -0.51214 | -0.05177 | -0.97373 | -0.02584 | 0.172539 | -0.04617 | -0.26117 | 0.191753 | 0.363373 | 0.197457 | 0.041145 | 0.664022 | 0.217411 |
| 29.73611 | 0.203657 | 0.093656 | -0.08677 | 1.653327 | 0.297792 | -0.03714 | -0.02387 | -0.28724 | -1.30126 | -0.44169 | 0.253659 | -0.15285 | -0.24031 | 0.14097 | 0.290885 | 0.203685 | 0.103095 | 0.547856 | 0.629077 |
| 29.75 | 1.616416 | 0.015699 | 0.770192 | 2.811332 | 0.396235 | -0.42762 | 0.60101 | -0.17601 | -1.65126 | -0.42325 | 0.254254 | -0.23994 | -0.15057 | 0.055913 | 0.186988 | 0.168454 | 0.159905 | 0.34915 | 0.439706 |
| 29.76389 | 1.945923 | -1.15481 | 1.056778 | 1.690651 | 0.26128 | -0.88646 | 1.077255 | -0.06873 | -1.43617 | 0.010067 | 0.342694 | -0.2745 | -0.04026 | -0.09308 | 0.123932 | 0.169109 | 0.152536 | 0.19214 | -0.1387 |
| 29.77778 | 1.492646 | -2.45052 | 0.330334 | 0.099392 | -0.05965 | -1.08981 | 1.054163 | 0.047794 | -0.77306 | 0.532191 | 0.325544 | -0.26322 | 0.032626 | -0.17214 | 0.067552 | 0.160742 | 0.11118 | 0.098934 | -0.76225 |
| 29.79167 | 0.643445 | -0.51285 | -1.18484 | -1.66338 | -0.43038 | -1.12661 | 0.409616 | 0.109979 | -0.1426 | 0.628138 | 0.157523 | -0.30558 | -0.05305 | -0.18632 | -0.01981 | 0.116032 | 0.042458 | -0.14869 | -1.27013 |
| 29.80556 | -0.53664 | 0.81486 | -2.65811 | -4.25844 | -0.72686 | -1.12356 | -0.29723 | 0.22418 | 0.338721 | 0.145517 | -0.09946 | -0.40388 | -0.31443 | -0.22074 | -0.10812 | 0.036806 | -0.15633 | -0.62194 | -1.57931 |
| 29.81944 | -1.42743 | 0.255615 | -3.68331 | -4.8845 | -0.87629 | -1.00379 | -1.21674 | 0.007157 | 0.125216 | -0.66746 | -0.23504 | -0.5335 | -0.63168 | -0.2359 | -0.20009 | -0.05894 | -0.54753 | -1.12179 | -1.70677 |
| 29.83333 | -2.02893 | -0.86462 | -4.00701 | -2.36326 | -0.87603 | -1.05985 | -1.03674 | -0.56486 | -0.80907 | -1.48162 | -0.15737 | -0.56386 | -0.77381 | -0.26388 | -0.30736 | -0.15714 | -0.74336 | -1.42965 | -1.42046 |
| 29.84722 | -2.32975 | -2.09941 | -2.87516 | -0.22292 | -0.73261 | -1.50366 | -0.56684 | -1.05839 | -1.83325 | -1.77124 | -0.09579 | -0.59837 | -0.73602 | -0.31436 | -0.45498 | -0.25541 | -0.81911 | -1.53087 | -0.80988 |
| 29.86111 | -2.17292 | -3.54696 | -1.27677 | 0.561173 | -0.59833 | -2.19232 | 0.081859 | -1.43491 | -2.62353 | -1.55089 | -0.26012 | -0.61077 | -0.5765 | -0.34228 | -0.49204 | -0.36839 | -0.7449 | -1.36112 | -0.40977 |
| 29.875 | -2.25433 | -5.07289 | -0.42645 | -0.33471 | -0.45415 | -2.20618 | 0.351723 | -1.76159 | -2.77479 | -1.07868 | -0.46427 | -0.55117 | -0.22865 | -0.3084 | -0.39134 | -0.47062 | -0.52786 | -1.01003 | -0.27258 |
| 29.88889 | -2.13024 | -5.5205 | -0.1151 | -1.77908 | -0.35512 | -1.84629 | 0.148819 | -1.78154 | -2.49947 | -0.43355 | -0.58128 | -0.41223 | 0.016491 | -0.22744 | -0.29858 | -0.50911 | -0.29023 | -0.71104 | -0.44013 |
| 29.90278 | -1.84783 | -4.13465 | -0.07164 | -2.86998 | -0.41811 | -1.38605 | -0.0323 | -1.48194 | -1.99456 | 0.012762 | -0.60185 | -0.30243 | 0.072532 | -0.19295 | -0.26636 | -0.44881 | -0.17668 | -0.53051 | -0.7003 |

| | | | | | | | | | | | | | | | | | | | |
|---|---|---|---|---|---|---|---|---|---|---|---|---|---|---|---|---|---|---|---|
| 29.91667 | -1.58212 | -2.18449 | -0.18166 | -3.2168 | -0.47253 | -0.91996 | -0.38276 | -1.07802 | -1.44241 | 0.094529 | -0.58512 | -0.28583 | -0.04993 | -0.19675 | -0.30174 | -0.37582 | -0.20798 | -0.46905 | -0.71832 |
| 29.93056 | -1.32502 | -0.58045 | -0.17085 | -2.67473 | -0.41425 | -0.57974 | -0.51893 | -0.78499 | -0.89575 | -0.2179 | -0.43105 | -0.26856 | -0.23412 | -0.18645 | -0.28013 | -0.29425 | -0.34394 | -0.49405 | -0.45992 |
| 29.94444 | -0.9391 | 0.46398 | -0.06089 | -1.38636 | -0.2193 | -0.44832 | -0.15204 | -0.61957 | -0.54846 | -0.58584 | -0.27163 | -0.22115 | -0.34435 | -0.16111 | -0.22784 | -0.22554 | -0.47246 | -0.50374 | -0.23369 |
| 29.95833 | -0.3712 | 0.911911 | 0.048049 | -0.46272 | 0.043738 | -0.43062 | 0.458873 | -0.43993 | -0.45675 | -0.81704 | -0.115 | -0.14592 | -0.3439 | -0.12939 | -0.19263 | -0.16014 | -0.53736 | -0.50514 | -0.14034 |
| 29.97222 | 0.067316 | 1.089322 | 0.014798 | -0.01355 | 0.194949 | -0.46413 | 1.060236 | -0.22765 | -0.35695 | -0.94717 | -0.13086 | -0.07712 | -0.25268 | -0.09894 | -0.15931 | -0.12669 | -0.53268 | -0.54391 | -0.16104 |
| 29.98611 | 0.090637 | 0.605758 | -0.25874 | -0.1618 | 0.180816 | -0.71873 | 1.152858 | -0.22385 | -0.49686 | -1.28708 | -0.67019 | -0.08772 | -0.23633 | -0.02486 | -0.08993 | -0.15984 | -0.70393 | -0.90449 | -0.4106 |
| 30 | -0.46116 | -0.66883 | -0.95677 | -1.15745 | -0.1667 | -1.41045 | 0.036155 | -0.69866 | -1.22042 | -2.14857 | -1.76397 | -0.20928 | -0.36402 | 0.053802 | -0.03077 | -0.23998 | -1.19557 | -1.75859 | -0.89911 |
| 30.01389 | -0.9393 | -1.95458 | -1.80453 | -1.94343 | -0.96753 | -2.6236 | -1.06912 | -1.28864 | -2.2906 | -2.52178 | -2.30368 | -0.28945 | -0.34614 | 0.034651 | -0.03144 | -0.23306 | -1.3954 | -2.05595 | -1.00581 |
| 30.02778 | -0.92176 | -2.47668 | -2.01983 | -1.44199 | -1.7985 | -3.15017 | -2.20023 | -1.54819 | -2.7452 | -1.70073 | -1.48376 | -0.24289 | -0.15759 | -0.19792 | -0.11526 | -0.1911 | -0.85653 | -1.18567 | -0.5318 |
| 30.04167 | -0.66519 | -2.33591 | -1.2985 | -0.79 | -1.99248 | -3.09885 | -1.81443 | -1.40576 | -2.1257 | -0.8943 | -0.52793 | -0.17888 | -0.08129 | -0.44139 | -0.26199 | -0.19356 | -0.3784 | -0.53771 | -0.14851 |
| 30.05556 | -0.22061 | -1.80724 | -0.57006 | -0.67899 | -1.66338 | -2.36823 | -0.64481 | -1.21278 | -1.12105 | -0.52634 | -0.04118 | -0.09141 | -0.06548 | -0.38141 | -0.29909 | -0.14237 | -0.16167 | -0.34942 | -0.02976 |
| 30.06944 | 0.248495 | -0.99348 | 0.108698 | -0.74966 | -0.7934 | -0.87057 | 0.381967 | -0.92698 | -0.20469 | -0.39699 | 0.292272 | 0.021087 | -0.0282 | -0.2118 | -0.13987 | -0.06175 | -0.02704 | -0.24017 | 0.015575 |
| 30.08333 | 0.506865 | -0.28351 | 0.716452 | -0.64442 | 0.123113 | 0.539468 | 1.009847 | -0.49919 | 0.225261 | -0.18717 | 0.423748 | 0.099261 | -0.00468 | -0.09939 | 0.069542 | -0.02896 | 0.026172 | -0.12999 | 0.034718 |
| 30.09722 | 0.557725 | 0.097655 | 0.896234 | -0.41988 | 0.495923 | 1.14653 | 0.805735 | -0.03705 | 0.345377 | -0.00247 | 0.25367 | 0.087016 | -0.01285 | -0.07045 | 0.165795 | -0.04446 | 0.015135 | -0.05873 | 0.03818 |
| 30.11111 | 0.438738 | 0.163761 | 0.653801 | -0.18423 | 0.587784 | 1.01489 | 0.423033 | 0.205472 | 0.362539 | 0.072514 | 0.002527 | 0.016145 | -0.02218 | -0.07182 | 0.164705 | -0.06031 | -0.00208 | -0.04793 | 0.04357 |
| 30.125 | 0.176003 | 0.022665 | 0.305922 | -0.07915 | 0.684695 | 0.755271 | 0.085767 | 0.234967 | 0.37614 | 0.071595 | -0.14468 | -0.03255 | -0.00891 | -0.06274 | 0.120267 | -0.06642 | 0.008118 | -0.0081 | 0.050056 |
| 30.13889 | -0.19069 | -0.16725 | 0.047292 | 0.205683 | 0.75596 | 0.71476 | -0.02686 | 0.203316 | 0.425015 | 0.092094 | -0.03986 | -0.02952 | 0.006981 | -0.02263 | 0.061886 | -0.05431 | 0.047905 | 0.156804 | 0.035625 |
| 30.15278 | -0.40552 | -0.05164 | -0.01783 | 0.776905 | 0.852221 | 0.914266 | 0.152406 | 0.252939 | 0.512307 | 0.17997 | 0.154312 | 0.007414 | 0.033107 | 0.006582 | 0.066616 | -0.01569 | 0.086647 | 0.360559 | 0.04003 |
| 30.16667 | -0.33719 | 0.472458 | 0.151108 | 1.249016 | 0.968191 | 1.328329 | 0.483884 | 0.319957 | 0.754521 | 0.326324 | 0.205221 | 0.058571 | 0.069158 | 0.021144 | 0.120267 | 0.032891 | 0.116844 | 0.498892 | 0.086837 |
| 30.18056 | 0.045311 | 1.263991 | 0.444538 | 1.131903 | 0.90119 | 1.203714 | 0.54062 | 0.245512 | 1.047429 | 0.497025 | 0.09289 | 0.077473 | 0.068754 | 0.023913 | 0.127788 | 0.040072 | 0.185661 | 0.51364 | 0.090454 |
| 30.19444 | 0.441225 | 1.939364 | 0.799364 | 0.776447 | 0.877968 | 1.35443 | 0.401246 | 0.183776 | 1.367177 | 0.560393 | -0.0369 | 0.056519 | 0.026063 | 0.024244 | 0.072112 | -0.00236 | 0.215455 | 0.458832 | 0.059955 |
| 30.20833 | 0.581219 | 2.274925 | 1.07613 | 0.266163 | 0.873218 | 1.602295 | 0.093063 | 0.126803 | 1.591162 | 0.459543 | 0.022484 | 0.038319 | 0.001686 | 0.015365 | 0.02269 | -0.04688 | 0.155248 | 0.395804 | 0.041158 |
| 30.22222 | 0.536921 | 2.195897 | 1.217908 | -0.12787 | 0.866411 | 1.543754 | -0.32265 | -0.019 | 1.663531 | 0.321388 | 0.115604 | 0.030206 | 0.016019 | -0.00069 | 0.019065 | -0.06313 | 0.07705 | 0.324505 | 0.026546 |
| 30.23611 | 0.447551 | 1.880933 | 1.18172 | -0.27878 | 0.721153 | 1.266398 | -0.45813 | -0.13591 | 1.361479 | 0.181488 | 0.001612 | 0.016848 | 0.053294 | -0.02852 | 0.020791 | -0.05875 | 0.040481 | 0.18243 | 0.00881 |
| 30.25 | 0.273444 | 1.338946 | 0.925901 | -0.22632 | 0.56779 | 1.018659 | -0.18533 | -0.09025 | 0.885025 | 0.036066 | -0.17037 | -0.00851 | 0.081246 | -0.03677 | 0.020245 | -0.06794 | 0.049049 | 0.037706 | -0.00638 |
| 30.26389 | 0.067942 | 0.807554 | 0.619804 | -0.08434 | 0.455303 | 0.74075 | 0.151063 | 0.046639 | 0.461123 | -0.11739 | -0.27548 | -0.07787 | 0.091756 | -0.03587 | -0.02358 | -0.07722 | 0.016384 | -0.06572 | -0.01261 |
| 30.27778 | -0.1015 | 0.355929 | 0.254875 | -0.25798 | 0.209979 | 0.344239 | 0.011734 | -0.03888 | 0.050492 | -0.14317 | -0.27741 | -0.13321 | 0.104626 | -0.05166 | -0.0751 | -0.08356 | -0.04951 | -0.06454 | 0.045676 |
| 30.29167 | -0.191 | -0.0707 | -0.0049 | -0.71202 | -0.13433 | -0.12736 | -0.4637 | -0.35235 | -0.29839 | 0.02821 | -0.04439 | -0.09247 | 0.136636 | -0.03983 | -0.03018 | -0.05212 | 0.005592 | 0.097767 | 0.166857 |
| 30.30556 | -0.1013 | -0.29421 | 0.010891 | -0.94044 | -0.31998 | -0.34071 | -0.73022 | -0.46547 | -0.34239 | 0.148997 | 0.209109 | -0.02309 | 0.097662 | 0.0159 | -0.01089 | -0.01622 | 0.079744 | 0.173491 | 0.162777 |
| 30.31944 | 0.071188 | -0.4089 | 0.16254 | -0.82248 | -0.29119 | -0.26088 | -0.75058 | -0.3461 | -0.28111 | 0.106539 | 0.175791 | 0.013404 | 0.011727 | 0.028933 | -0.09376 | -0.00276 | 0.036972 | 0.001763 | 0.031576 |
| 30.33333 | 0.106789 | -0.58499 | 0.107843 | -0.4872 | -0.27437 | -0.05347 | -0.74646 | -0.27228 | -0.37031 | 0.01989 | 0.032386 | 0.000667 | -0.0124 | 0.029206 | -0.12677 | 0.013338 | 0.006844 | -0.15226 | -0.04091 |
| 30.34722 | 0.067125 | -0.6791 | -0.15241 | -0.14843 | -0.43373 | 0.147448 | -0.74293 | -0.34197 | -0.5678 | -0.00831 | 0.015073 | -0.01312 | 0.018329 | 0.019183 | -0.07062 | 0.029457 | 0.03257 | -0.09958 | -0.08356 |
| 30.36111 | 0.08167 | -0.60264 | -0.36617 | 0.112587 | -0.60399 | 0.156282 | -0.63508 | -0.36487 | -0.62095 | 0.079544 | 0.249572 | 0.038979 | 0.112125 | 0.063931 | 0.043367 | 0.040393 | 0.112102 | 0.105261 | -0.09753 |
| 30.375 | 0.223352 | -0.2781 | -0.36131 | 0.416793 | -0.56498 | 0.050237 | -0.36133 | -0.14761 | -0.35669 | 0.241436 | 0.598931 | 0.181007 | 0.275021 | 0.203861 | 0.196181 | 0.091605 | 0.229158 | 0.292279 | 0.036774 |
| 30.38889 | 0.262825 | -0.01384 | -0.20764 | 0.650449 | -0.45385 | -0.09434 | -0.15909 | 0.079915 | -0.01598 | 0.333504 | 0.673095 | 0.28187 | 0.334697 | 0.292316 | 0.261973 | 0.104573 | 0.267081 | 0.31786 | 0.131619 |
| 30.40278 | 0.030976 | -0.10978 | -0.02901 | 0.604973 | -0.44126 | -0.24341 | -0.30557 | 0.006046 | 0.183232 | 0.261704 | 0.366425 | 0.263846 | 0.221549 | 0.210589 | 0.152831 | 0.064517 | 0.199657 | 0.275725 | 0.078146 |
| 30.41667 | -0.22773 | -0.38855 | 0.197881 | 0.424988 | -0.46708 | -0.21001 | -0.66598 | -0.33548 | 0.259657 | 0.205387 | -0.04843 | 0.217018 | 0.097118 | 0.111989 | 0.007175 | 0.029536 | 0.089538 | 0.240303 | 0.035004 |
| 30.43056 | -0.45144 | -0.56453 | 0.393428 | 0.497834 | -0.50532 | -0.10956 | -0.91797 | -0.64943 | 0.303556 | 0.180156 | -0.21871 | 0.236133 | 0.070298 | 0.051734 | -0.06042 | 0.037312 | 0.034468 | 0.273298 | 0.071023 |
| 30.44444 | -0.64995 | -0.43184 | 0.416181 | 0.60334 | -0.60522 | 0.068806 | -0.9616 | -0.80092 | 0.312096 | 0.237544 | 0.032402 | 0.264825 | 0.136512 | 0.031556 | -0.06357 | 0.062065 | 0.15311 | 0.340756 | 0.092129 |
| 30.45833 | -0.47377 | -0.0037 | 0.286637 | 0.36896 | -0.79269 | 0.160661 | -0.89567 | -0.83088 | 0.276885 | 0.357011 | 0.399873 | 0.308945 | 0.198771 | 0.054424 | -0.03992 | 0.070541 | 0.397268 | 0.372447 | 0.107926 |
| 30.47222 | -0.03383 | 0.626047 | 0.254367 | -0.10968 | -0.79467 | 0.18529 | -0.72229 | -0.66701 | 0.451057 | 0.516297 | 0.592109 | 0.337752 | 0.192482 | 0.133868 | -0.01148 | 0.102004 | 0.482426 | 0.392068 | 0.172766 |
| 30.48611 | 0.480303 | 1.427685 | 0.512976 | -0.00853 | -0.55799 | 0.42639 | -0.39898 | -0.16201 | 0.907352 | 0.999974 | 0.724368 | 0.46047 | 0.077381 | 0.188561 | -0.02175 | 0.146497 | 0.493532 | 0.641228 | 0.399516 |
| 30.5 | 1.292871 | 2.405422 | 1.247718 | 1.203453 | -0.21181 | 1.042467 | 0.025458 | 0.794497 | 1.710354 | 1.968363 | 1.150457 | 0.672389 | -0.00162 | 0.158685 | -0.10583 | 0.249011 | 0.623328 | 1.484977 | 0.949983 |
| 30.51389 | 2.24684 | 2.526983 | 2.086399 | 2.9965 | 0.137933 | 1.444091 | 0.525017 | 1.583721 | 2.434921 | 2.728324 | 1.597962 | 0.894817 | 0.154704 | 0.248259 | -0.05424 | 0.490248 | 0.859282 | 1.888464 | 1.78489 |
| 30.52778 | 2.319861 | 2.478968 | 2.10898 | 4.184352 | 0.513921 | 1.669274 | 1.433928 | 1.795265 | 2.744636 | 2.322323 | 1.669281 | 0.969318 | 0.459171 | 0.465351 | 0.061496 | 0.714794 | 0.7311 | 1.514625 | 2.575767 |
| 30.54167 | 1.827268 | 2.166713 | 1.880096 | 3.788383 | 0.393903 | 1.550849 | 1.640468 | 1.44992 | 2.615143 | 1.784973 | 1.179489 | 0.884381 | 0.480679 | 0.481336 | 0.147788 | 0.613443 | 0.513708 | 1.248886 | 2.566799 |
| 30.55556 | 1.020678 | 1.345702 | 1.073047 | 1.890196 | 0.246042 | 0.775529 | 0.806609 | 0.633783 | 1.624791 | 1.221475 | 0.471776 | 0.466382 | 0.389119 | 0.327293 | 0.050788 | 0.363762 | 0.273439 | 0.850676 | 1.93596 |
| 30.56944 | 0.104435 | 0.154005 | 0.113039 | 0.111336 | 0.135276 | -0.21372 | -0.01673 | -0.14973 | 0.401445 | 0.793367 | -0.28485 | 0.090352 | 0.295156 | 0.132991 | -0.0107 | 0.047744 | 0.211587 | 0.63356 | 1.09722 |
| 30.58333 | -0.5545 | -0.86806 | -0.77342 | -1.0433 | -0.11534 | -0.79286 | -0.69402 | -0.50302 | -0.69223 | 0.392859 | -0.65621 | -0.0361 | 0.289633 | 0.135207 | -0.05419 | -0.0466 | 0.308717 | 0.553039 | 0.575765 |
| 30.59722 | -0.33769 | -1.48178 | -1.47119 | -1.47203 | -0.70586 | -0.746 | -1.07958 | -0.35009 | -1.45762 | 0.017746 | -0.27828 | 0.014759 | 0.388446 | 0.257904 | -0.085 | -0.03749 | 0.414418 | 0.539925 | 0.371036 |
| 30.61111 | 0.438282 | -1.80408 | -1.19825 | -0.76347 | -0.93244 | -0.23224 | -0.91255 | 0.182188 | -1.63489 | -0.48591 | 0.089887 | 0.047539 | 0.413153 | 0.328074 | -0.10618 | -0.04994 | 0.358016 | 0.441911 | 0.241803 |
| 30.625 | 1.31365 | -2.06281 | -0.28443 | 0.645617 | -0.5228 | 0.62025 | -0.36808 | 0.805667 | -1.43175 | -0.76632 | 0.234829 | -0.04004 | 0.31946 | 0.314089 | 0.047902 | -0.04241 | 0.290791 | 0.391874 | 0.184678 |
| 30.63889 | 1.731197 | -2.54902 | 0.828028 | 2.912346 | 0.01977 | 1.188155 | 0.292772 | 1.154987 | -0.89936 | -0.54945 | 0.053268 | -0.21846 | 0.258397 | 0.247889 | 0.181059 | -0.01028 | 0.147149 | 0.347572 | 0.166036 |
| 30.65278 | 1.6199 | -1.56698 | 1.594734 | 4.497041 | 0.636699 | 1.462711 | 0.874252 | 1.185793 | -0.06279 | 0.150995 | -0.30718 | -0.28365 | 0.228572 | 0.190765 | 0.230304 | 0.017271 | 0.057977 | 0.363895 | 0.190746 |
| 30.66667 | 1.328999 | 1.655313 | 2.007171 | 2.441663 | 0.815685 | 1.499778 | 1.245653 | 0.873204 | 1.389364 | 0.884225 | -0.86236 | -0.24357 | 0.268487 | 0.16416 | 0.222976 | 0.088552 | 0.021332 | 0.520977 | -0.06334 |
| 30.68056 | 0.681166 | 5.032744 | 2.108945 | 0.417608 | 0.554154 | 1.022232 | 0.955029 | 0.033516 | 2.858534 | 1.235127 | -1.22549 | 0.029365 | 0.282871 | 0.184192 | 0.241125 | 0.117913 | 0.087538 | 0.630799 | -0.58025 |
| 30.69444 | -0.12953 | 3.857834 | 0.961899 | -1.12316 | 0.01402 | 0.309993 | 0.208898 | -0.76396 | 2.623872 | 1.028626 | -0.79585 | 0.371467 | 0.211765 | 0.260996 | 0.383371 | 0.101093 | 0.260413 | 0.376637 | -0.90506 |
| 30.70833 | -0.8034 | 2.358818 | -0.67983 | -2.46669 | -0.81076 | -0.60727 | -0.60523 | -1.08944 | 1.259983 | 0.563388 | -0.36433 | 0.481193 | 0.03183 | 0.271223 | 0.464372 | 0.034777 | 0.304144 | -0.05317 | -0.79477 |
| 30.72222 | -1.10871 | 0.21719 | -2.01218 | -3.95993 | -1.62945 | -1.59044 | -0.83159 | -1.08267 | 0.061862 | 0.147299 | -0.325 | 0.331553 | -0.14759 | 0.202974 | 0.397592 | -0.06863 | 0.176994 | -0.42099 | -0.38337 |
| 30.73611 | -1.04197 | -0.83831 | -2.19047 | -4.45107 | -1.44686 | -1.77179 | -0.3895 | -0.78531 | -0.9823 | -0.10904 | -0.41287 | 0.118273 | -0.2097 | 0.226169 | 0.310066 | -0.07737 | 0.056045 | -0.48367 | 0.092107 |
| 30.75 | -0.88523 | -1.02452 | -2.01496 | -1.17975 | -0.713 | -0.44987 | -0.17793 | -0.58954 | -1.98175 | -0.25377 | -0.50354 | -0.04253 | -0.18951 | 0.22862 | 0.234105 | -0.0031 | 0.003806 | -0.36667 | 0.356099 |
| 30.76389 | -0.88149 | -2.16452 | -1.48936 | 2.071562 | -0.09746 | 0.3798 | -0.24897 | -0.73955 | -2.43807 | -0.39285 | -0.48474 | -0.22663 | -0.16825 | 0.056411 | 0.107547 | 0.041681 | -0.0533 | -0.22909 | 0.232594 |
| 30.77778 | -0.72796 | -2.76543 | -0.52905 | 3.887039 | 0.362601 | 0.641831 | -0.28049 | -1.09907 | -2.31411 | -0.54755 | -0.21715 | -0.46826 | -0.31805 | -0.20042 | -0.00937 | 0.00831 | -0.22899 | -0.02625 | -0.22721 |
| 30.79167 | -0.33724 | -3.47983 | 0.19474 | 1.789089 | 0.436315 | 0.003997 | -0.229 | -0.87736 | -1.97193 | -0.59626 | 0.118572 | -0.70819 | -0.41873 | -0.3736 | -0.08535 | -0.0924 | -0.45871 | 0.132618 | -0.34553 |
| 30.80556 | -0.05798 | -4.66114 | 0.007878 | 0.121678 | -0.04008 | -0.91273 | -0.27589 | -0.53088 | -2.02105 | -0.48381 | 0.278597 | -0.7741 | -0.39692 | -0.33533 | -0.09022 | -0.21195 | -0.69249 | 0.120603 | -0.24853 |
| 30.81944 | -0.16736 | -5.18557 | -0.94626 | -1.94371 | -1.05133 | -1.8805 | -0.70057 | -0.65552 | -2.00182 | -0.37185 | 0.06884 | -0.56429 | -0.47568 | -0.26261 | -0.06931 | -0.36841 | -0.98049 | -0.13085 | -0.33688 |
| 30.83333 | -0.52786 | -4.35739 | -2.28085 | -4.21374 | -2.11423 | -2.66194 | -1.26766 | -1.09751 | -1.96834 | -0.35832 | -0.23664 | -0.35902 | -0.61364 | -0.204 | -0.09559 | -0.4681 | -1.1105 | -0.37964 | -0.67768 |
| 30.84722 | -0.8348 | -2.87174 | -2.29413 | -5.70928 | -2.29034 | -2.17017 | -1.26415 | -1.37509 | -1.19165 | -0.44406 | -0.38948 | -0.23762 | -0.8926 | -0.26384 | -0.131 | -0.38934 | -0.96829 | -0.47133 | -1.01681 |
| 30.86111 | -1.21641 | -1.6048 | -1.72591 | -5.0372 | -1.65552 | -1.48139 | -0.77278 | -1.47542 | -0.66886 | -0.56628 | -0.35537 | -0.1724 | -0.93737 | -0.4314 | -0.16229 | -0.2692 | -0.68146 | -0.49 | -1.26109 |
| 30.875 | -1.72185 | -0.68335 | -1.32343 | -3.51671 | -0.81485 | -1.00375 | -0.35693 | -1.64672 | -0.53485 | -0.54272 | -0.21247 | -0.12541 | -0.65351 | -0.49609 | -0.14058 | -0.11878 | -0.31588 | -0.53221 | -1.29104 |
| 30.88889 | -1.76101 | 0.030667 | -0.95435 | -1.65052 | -0.07098 | -0.89897 | -0.47933 | -1.49552 | -1.17099 | -0.47709 | -0.13134 | -0.1039 | -0.37394 | -0.421 | -0.13183 | -0.04853 | -0.11499 | -0.65133 | -0.99523 |
| 30.90278 | -1.4275 | 0.31127 | -0.8591 | -0.52672 | 0.34178 | -1.1624 | -0.85356 | -1.07804 | -1.90314 | -0.38856 | -0.18072 | -0.13057 | -0.14221 | -0.28373 | -0.22714 | -0.08078 | -0.10114 | -0.84658 | -0.72437 |
| 30.91667 | -1.02604 | 0.428255 | -0.9681 | -0.28346 | 0.24262 | -1.66697 | -0.68373 | -0.71614 | -1.85911 | -0.23503 | -0.1163 | -0.13891 | -0.01039 | -0.20007 | -0.35354 | -0.12307 | -0.12818 | -1.02532 | -0.53757 |
| 30.93056 | -0.65448 | 0.594735 | -1.04379 | -0.48517 | -0.07246 | -1.69422 | -0.62718 | -0.37685 | -1.58178 | -0.09787 | 0.00621 | -0.20893 | 0.004163 | -0.18034 | -0.4254 | -0.15362 | -0.14604 | -1.13947 | -0.40365 |
| 30.94444 | -0.37622 | 0.893398 | -0.99917 | -0.74933 | -0.32425 | -1.14723 | -0.34024 | -0.12441 | -1.04599 | -0.07321 | 0.011258 | -0.21146 | -0.01998 | -0.17696 | -0.4092 | -0.16004 | -0.11826 | -1.18697 | -0.29835 |
| 30.95833 | -0.17795 | 1.224772 | -0.59139 | -0.81871 | -0.15839 | -0.22372 | 0.308208 | 0.024859 | -0.34974 | -0.10704 | -0.04471 | -0.03507 | -0.01932 | -0.15262 | -0.34309 | -0.11596 | -0.06524 | -1.01048 | -0.1844 |
| 30.97222 | 0.092366 | 1.422027 | 0.097642 | -0.52252 | 0.425152 | 0.717105 | 1.078618 | 0.204432 | 0.253016 | -0.17037 | -0.09712 | 0.14143 | 0.00174 | -0.08908 | -0.21126 | -0.02733 | 0.025225 | -0.68885 | -0.1197 |
| 30.98611 | 0.467296 | 1.288817 | 0.699564 | 0.023587 | 1.086617 | 1.319085 | 1.667132 | 0.399967 | 0.774877 | -0.49217 | -0.2391 | 0.174368 | 0.005446 | -0.00179 | -0.09197 | 0.006952 | 0.059849 | -0.67905 | -0.25467 |
| 31 | 0.400983 | 0.766163 | 0.324403 | 0.558783 | 0.99429 | 0.459992 | 1.653077 | 0.590758 | 0.643589 | -1.18746 | -0.76632 | 0.006235 | -0.0104 | -0.00795 | -0.0706 | -0.08633 | -0.03971 | -1.40971 | -0.59052 |
| 31.01389 | -0.55715 | -0.46611 | -1.31405 | 0.599975 | -0.4165 | -1.60355 | 0.48514 | 0.489916 | -0.47916 | -1.5863 | -1.3809 | -0.20439 | -0.0175 | -0.21595 | -0.16006 | -0.21419 | -0.22214 | -1.97619 | -0.68602 |
| 31.02778 | -1.63137 | -1.88579 | -2.80027 | 0.366682 | -2.50221 | -3.47924 | -1.1654 | 0.108535 | -1.75892 | -1.24753 | -1.31227 | -0.25169 | -0.01987 | -0.52227 | -0.29864 | -0.22644 | -0.38524 | -1.30256 | -0.42249 |
| 31.04167 | -1.14007 | -1.93201 | -2.48499 | -0.08384 | -2.75351 | -3.62107 | -1.34093 | -0.35554 | -2.14451 | -0.7464 | -0.70283 | -0.15825 | 0.010013 | -0.46662 | -0.37492 | -0.18634 | -0.37817 | -0.60878 | -0.18536 |
| 31.05556 | -0.21779 | -1.42129 | -1.30647 | -0.67921 | -1.56316 | -2.88464 | -0.26845 | -0.83818 | -1.8278 | -0.4433 | 0.084914 | -0.07597 | 0.060443 | -0.33369 | -0.34114 | -0.12808 | -0.16617 | -0.23772 | -0.04219 |
| 31.06944 | 0.814086 | -0.7477 | 0.027516 | -1.08101 | -0.26884 | -1.11374 | 0.614346 | -1.10358 | -1.10044 | -0.30833 | 0.508716 | 0.004134 | 0.090071 | -0.23993 | -0.21922 | -0.0468 | 0.009403 | -0.05799 | 0.058373 |
| 31.08333 | 1.433273 | -0.15194 | 1.020789 | -1.07104 | 0.927647 | 0.330836 | 0.98787 | -1.02769 | -0.28158 | -0.26807 | 0.77851 | 0.064289 | 0.058441 | -0.17364 | -0.04111 | 0.02522 | 0.109639 | 0.036171 | 0.068327 |
| 31.09722 | 1.601712 | 0.226032 | 1.684336 | -0.64879 | 1.297578 | 1.239212 | 0.934259 | -0.66078 | 0.277083 | -0.21956 | 0.666102 | 0.09146 | 0.008474 | -0.13614 | 0.085792 | 0.056334 | 0.140242 | 0.088213 | 0.022416 |

| | | | | | | | | | | | | | | | | | | | |
|---|---|---|---|---|---|---|---|---|---|---|---|---|---|---|---|---|---|---|---|
| 31.11111 | 1.768612 | 0.725485 | 2.223479 | -0.16731 | 1.625449 | 1.906766 | 0.74312 | -0.25416 | 0.673849 | -0.18105 | 0.527307 | 0.042744 | 0.001048 | -0.1085 | 0.139331 | 0.048688 | 0.159827 | 0.098718 | -0.02631 |
| 31.125 | 1.783424 | 1.12954 | 2.278494 | 0.164706 | 1.862368 | 2.360294 | 0.689667 | 0.099126 | 1.052803 | -0.15041 | 0.273804 | 0.012553 | 0.021065 | -0.09444 | 0.118112 | 0.031276 | 0.213649 | 0.051134 | -0.03781 |
| 31.13889 | 2.007111 | 1.367201 | 2.134849 | 0.410165 | 1.74128 | 2.672231 | 0.644934 | 0.309114 | 1.264558 | -0.18518 | -0.05452 | 0.025427 | 0.0253 | -0.09104 | 0.078296 | 0.019182 | 0.211418 | -0.00648 | 0.013513 |
| 31.15278 | 1.707092 | 1.142213 | 1.66273 | 0.596038 | 1.478638 | 2.822924 | 0.686542 | 0.469306 | 1.19216 | -0.19376 | -0.23595 | 0.060877 | 0.03064 | -0.09932 | 0.08125 | 0.022473 | 0.225495 | 0.018685 | 0.072168 |
| 31.16667 | 1.118803 | 0.892001 | 1.107115 | 0.582577 | 1.388337 | 2.315962 | 0.836155 | 0.590959 | 1.136725 | -0.08426 | -0.11354 | 0.073128 | 0.041073 | -0.10232 | 0.135212 | 0.04357 | 0.263978 | 0.096306 | 0.094029 |
| 31.18056 | 0.783189 | 0.868393 | 0.770675 | 0.435358 | 1.45994 | 1.772021 | 0.975884 | 0.61821 | 1.155252 | 0.041996 | 0.170479 | 0.003275 | 0.025043 | -0.11652 | 0.147724 | 0.056834 | 0.24256 | 0.134673 | 0.093313 |
| 31.19444 | 0.400708 | 0.954177 | 0.498212 | 0.317105 | 1.488567 | 1.358243 | 0.891165 | 0.475201 | 1.061673 | 0.089649 | 0.392375 | -0.09965 | -0.00478 | -0.14651 | 0.103201 | 0.044363 | 0.132305 | 0.164066 | 0.055959 |
| 31.20833 | 0.03714 | 0.953993 | 0.375882 | 0.210817 | 1.30383 | 0.798917 | 0.645309 | 0.252156 | 0.849529 | 0.047716 | 0.236348 | -0.10132 | 0.002553 | -0.14127 | 0.074759 | 0.023465 | 0.085334 | 0.277168 | -0.00254 |
| 31.22222 | -0.20882 | 1.020797 | 0.505657 | 0.208199 | 0.914652 | 0.379916 | 0.402339 | 0.088513 | 0.743636 | -0.08093 | -0.21116 | -0.04615 | 0.022408 | -0.09377 | 0.042679 | -0.00758 | 0.027806 | 0.310336 | -0.02737 |
| 31.23611 | -0.23776 | 1.02324 | 0.711768 | 0.290323 | 0.428724 | 0.172314 | 0.243413 | 0.009066 | 0.704742 | -0.33523 | -0.87529 | -0.1081 | -0.01024 | -0.05036 | -0.01704 | -0.09425 | -0.1985 | 0.166274 | -0.02086 |
| 31.25 | 0.035503 | 1.077921 | 0.801686 | 0.207662 | -0.11781 | 0.153432 | 0.041727 | -0.12959 | 0.685419 | -0.3511 | -0.97294 | -0.16071 | -0.02837 | -0.03124 | -0.06003 | -0.14868 | -0.33816 | -0.04858 | -0.03548 |
| 31.26389 | 0.66923 | 1.159063 | 0.87803 | 0.054893 | -0.17953 | 0.393058 | -0.13258 | -0.24325 | 0.671116 | -0.04379 | -0.2025 | -0.0592 | 0.035143 | -0.02795 | -0.04366 | -0.08766 | -0.14809 | -0.04561 | -0.08625 |
| 31.27778 | 1.548547 | 1.30703 | 1.000743 | 0.04757 | 0.389337 | 0.915856 | -0.03236 | -0.12743 | 0.648418 | 0.28158 | 0.503965 | 0.027821 | 0.085119 | -0.04168 | 0.028776 | -0.01512 | 0.035665 | 0.091575 | -0.09198 |
| 31.29167 | 1.897571 | 1.237626 | 0.979717 | 0.069381 | 1.041577 | 1.328621 | 0.229666 | 0.073238 | 0.585935 | 0.249885 | 0.637092 | 0.005243 | 0.06315 | -0.04586 | 0.055369 | 0.002703 | -0.04685 | 0.170649 | -0.01531 |
| 31.30556 | 1.10823 | 0.711953 | 0.733945 | 0.047302 | 1.16046 | 1.269848 | 0.297702 | 0.177024 | 0.468295 | 0.028754 | 0.368295 | -0.0408 | 0.023758 | -0.02152 | 0.044776 | 0.00121 | -0.11865 | 0.160567 | 0.0498 |
| 31.31944 | 0.070783 | 0.232087 | 0.386098 | 0.05626 | 0.666728 | 0.760511 | 0.283418 | 0.19565 | 0.32832 | -0.03177 | 0.012599 | -0.03275 | 0.032742 | 0.024125 | 0.038819 | 0.009084 | -0.00245 | 0.099034 | 0.054582 |
| 31.33333 | -1.33769 | -0.31823 | 0.0002 | 0.146409 | 0.047011 | 0.187359 | 0.224891 | 0.232066 | 0.152844 | 0.092003 | -0.23761 | 0.002369 | 0.067657 | 0.094789 | 0.066909 | 0.03351 | 0.180119 | 0.022941 | 0.062861 |
| 31.34722 | -1.78211 | -0.49381 | -0.17799 | 0.245704 | -0.60741 | -0.14437 | 0.120244 | 0.346031 | 0.051777 | 0.141716 | -0.31133 | -0.00238 | 0.064975 | 0.149476 | 0.079483 | 0.042671 | 0.196517 | -0.07776 | 0.083696 |
| 31.36111 | -0.46252 | -0.00108 | 0.122773 | 0.244401 | -1.03207 | -0.04448 | -0.02178 | 0.460964 | 0.122894 | 0.134687 | -0.07031 | -0.01752 | 0.044341 | 0.14892 | 0.067134 | 0.034622 | 0.13311 | -0.13656 | 0.088267 |
| 31.375 | 1.143318 | 0.566969 | 0.471287 | 0.179437 | -0.62646 | 0.343947 | -0.03441 | 0.556098 | 0.310149 | 0.110818 | 0.280182 | -0.00319 | 0.036095 | 0.133876 | 0.057566 | 0.04988 | 0.103811 | -0.0888 | 0.116296 |
| 31.38889 | 2.251677 | 0.936097 | 0.739647 | 0.168428 | -0.10163 | 0.682759 | 0.221168 | 0.672317 | 0.51505 | 0.060606 | 0.53234 | 0.008243 | 0.025661 | 0.100474 | 0.062145 | 0.074003 | 0.073584 | 0.039363 | 0.14196 |
| 31.40278 | 2.031231 | 0.81387 | 0.959187 | 0.207366 | 0.44073 | 0.795617 | 0.531541 | 0.71396 | 0.698386 | -0.03052 | 0.40065 | -0.00594 | -0.00742 | 0.03238 | 0.0823 | 0.062824 | 0.012282 | 0.152535 | 0.145146 |
| 31.41667 | 0.693829 | 0.442687 | 1.019027 | 0.23831 | 0.556637 | 0.618606 | 0.591881 | 0.602318 | 0.799965 | 0.001353 | 0.101542 | 0.007971 | -0.01477 | 0.016979 | 0.115172 | 0.052645 | 0.007994 | 0.275501 | 0.134555 |
| 31.43056 | -0.45522 | 0.124128 | 0.930476 | 0.296634 | 0.138594 | 0.31235 | 0.449788 | 0.431581 | 0.840172 | 0.19873 | -0.06072 | 0.135762 | 0.032948 | 0.111444 | 0.146582 | 0.072768 | 0.128661 | 0.436106 | 0.11338 |
| 31.44444 | -0.58926 | 0.021435 | 0.882382 | 0.641244 | -0.50548 | 0.01006 | 0.137593 | 0.239193 | 0.871739 | 0.362826 | 0.000749 | 0.238108 | 0.061161 | 0.194663 | 0.138211 | 0.075505 | 0.260865 | 0.502057 | 0.077619 |
| 31.45833 | 0.261817 | 0.29531 | 0.851455 | 1.55464 | -0.9959 | -0.07706 | -0.18357 | 0.079235 | 0.98226 | 0.474754 | 0.383781 | 0.26825 | 0.079353 | 0.234818 | 0.132737 | 0.071197 | 0.329461 | 0.520567 | 0.037618 |
| 31.47222 | 1.121929 | 0.737031 | 0.908747 | 2.007296 | -0.79992 | 0.137905 | -0.21185 | 0.108299 | 1.121899 | 0.536761 | 0.799479 | 0.276628 | 0.106822 | 0.256217 | 0.100661 | 0.099227 | 0.333264 | 0.563319 | 0.037676 |
| 31.48611 | 1.493622 | 1.177631 | 1.156629 | 2.364431 | -0.25934 | 0.579584 | 0.087488 | 0.470454 | 1.550468 | 0.730165 | 1.13172 | 0.247341 | 0.121198 | 0.239126 | 0.023215 | 0.145897 | 0.283777 | 0.795914 | 0.184594 |
| 31.5 | 1.564069 | 1.933015 | 1.929832 | 4.005736 | 0.503525 | 1.233322 | 0.517032 | 1.221213 | 2.769475 | 1.431623 | 1.340574 | 0.302289 | 0.179765 | 0.291391 | -0.07377 | 0.257785 | 0.355949 | 1.454138 | 0.446809 |
| 31.51389 | 1.791395 | 2.055121 | 2.920497 | 3.798619 | 0.944728 | 2.034809 | 0.593169 | 1.740722 | 3.446493 | 1.885251 | 1.414242 | 0.469333 | 0.304206 | 0.690783 | -0.07916 | 0.423734 | 0.63009 | 1.89208 | 0.777727 |
| 31.52778 | 2.175253 | 2.338667 | 2.732599 | 2.490898 | 1.830524 | 2.421737 | 0.706513 | 1.46009 | 3.428771 | 1.84232 | 1.49018 | 0.650902 | 0.498798 | 1.140985 | -0.021 | 0.627557 | 1.209011 | 1.73192 | 1.261387 |
| 31.54167 | 2.020648 | 2.729974 | 2.603681 | 1.729227 | 2.167382 | 2.80324 | 0.837386 | 1.212483 | 3.570179 | 1.608128 | 1.386411 | 0.612797 | 0.453605 | 1.105929 | 0.042996 | 0.765379 | 1.207449 | 1.496546 | 1.644603 |
| 31.55556 | 1.032962 | 2.150482 | 1.619196 | 1.001309 | 1.647949 | 1.835342 | 0.688463 | 0.846742 | 2.985622 | 1.338151 | 0.810462 | 0.381109 | 0.36426 | 0.720081 | 0.035816 | 0.619749 | 0.6801 | 1.095417 | 1.644538 |
| 31.56944 | 0.077989 | 1.039558 | 0.611559 | 0.179224 | 0.791094 | 0.482931 | 0.385687 | 0.399436 | 1.760626 | 1.077572 | 0.025173 | 0.191403 | 0.278095 | 0.296993 | 0.025231 | 0.425148 | 0.043211 | 0.854661 | 1.460966 |
| 31.58333 | -0.60241 | 0.301087 | -0.40359 | -0.68657 | -0.05078 | -0.70104 | 0.232103 | 0.117846 | 0.668526 | 0.848551 | -0.6485 | 0.059502 | 0.217973 | 0.044645 | -0.00975 | 0.219448 | -0.23988 | 0.805865 | 1.488812 |
| 31.59722 | -1.02683 | -0.23494 | -1.36926 | -1.28269 | -0.7041 | -1.54571 | 0.206002 | 0.113698 | -0.29549 | 0.615203 | -0.98654 | -0.00176 | 0.226239 | -0.06027 | -0.07692 | 0.109901 | -0.33256 | 0.716283 | 1.626249 |
| 31.61111 | -0.97455 | -0.6279 | -2.02307 | -1.63139 | -1.17988 | -2.1606 | 0.006331 | 0.335304 | -0.8575 | 0.336155 | -0.99179 | -0.03331 | 0.441103 | -0.07429 | -0.17366 | 0.076159 | -0.09534 | 0.518142 | 1.506467 |
| 31.625 | -0.29774 | -0.88585 | -1.46739 | -0.30225 | -1.14716 | -2.4603 | -0.19261 | 0.538527 | -1.00831 | 0.176904 | -0.35741 | -0.02154 | 0.624513 | 0.05427 | -0.2309 | 0.048277 | 0.240671 | 0.231924 | 1.166799 |
| 31.63889 | 0.665702 | -1.60553 | -0.56197 | 2.406061 | -0.9621 | -1.84228 | -0.36914 | 0.614466 | -1.31006 | 0.070426 | 0.192554 | -0.01242 | 0.332249 | 0.248508 | -0.17518 | 0.009861 | 0.415725 | 0.056548 | 0.630614 |
| 31.65278 | 1.52422 | -2.46057 | 0.055471 | 4.241475 | -0.32482 | -0.60068 | -0.46382 | 0.496712 | -1.85721 | 0.008231 | 0.441439 | -0.02466 | 0.106433 | 0.393017 | -0.01744 | -0.02266 | 0.365272 | 0.091348 | 0.065602 |
| 31.66667 | 1.499053 | -2.82638 | 0.543861 | 3.804752 | 0.368231 | 0.942571 | -0.28051 | 0.295184 | -1.66736 | 0.055151 | 0.104582 | -0.03393 | 0.034889 | 0.391544 | 0.127189 | 0.006513 | 0.200976 | 0.218844 | -0.42926 |
| 31.68056 | 0.979474 | -2.73865 | 0.709104 | 2.244932 | 0.678147 | 2.021377 | 0.096949 | 0.365396 | -0.52795 | 0.116559 | -0.61617 | -0.0784 | 0.0956 | 0.344896 | 0.172469 | 0.040519 | -0.03926 | 0.240236 | -0.73051 |
| 31.69444 | 0.60774 | -0.32832 | 0.965497 | 0.748816 | 0.52029 | 2.306738 | 0.547517 | 0.494847 | 1.18337 | 0.190748 | -1.2449 | -0.18283 | 0.222995 | 0.327028 | 0.116633 | 0.013858 | -0.31938 | 0.085194 | -0.83245 |
| 31.70833 | 0.317787 | 2.772338 | 1.396859 | -0.85771 | -0.14791 | 2.033787 | 0.690253 | 0.216823 | 2.800846 | 0.262305 | -1.25288 | -0.27298 | 0.370775 | 0.272026 | 0.036025 | 0.049037 | -0.37573 | -0.19955 | -0.88275 |
| 31.72222 | -0.16431 | 4.678275 | 0.869408 | -2.23654 | -0.97496 | 1.153282 | 0.272013 | 0.000727 | 2.610759 | 0.303071 | -0.76787 | -0.1969 | 0.424126 | 0.164116 | -0.02257 | 0.13966 | -0.24694 | -0.38353 | -0.76033 |
| 31.73611 | -0.80931 | 3.899335 | -0.5915 | -3.64319 | -1.41699 | -0.40132 | -0.52977 | -0.26534 | 1.914942 | 0.289716 | -0.54482 | -0.02097 | 0.401533 | 0.009147 | -0.06739 | 0.165258 | -0.09023 | -0.38965 | -0.52903 |
| 31.75 | -0.39298 | 2.827695 | -1.3541 | -4.02312 | -1.52709 | -1.29296 | -1.23336 | -0.34272 | 0.915722 | 0.233313 | -0.70427 | 0.147283 | 0.338443 | -0.10356 | -0.11321 | 0.125593 | -0.08264 | -0.29789 | -0.16594 |
| 31.76389 | 0.046538 | 1.599473 | -1.64214 | -2.18395 | -2.18973 | -1.69799 | -1.78789 | -0.58837 | -0.35628 | 0.108967 | -1.04494 | 0.172185 | 0.227315 | -0.18022 | -0.13733 | 0.043587 | -0.17321 | -0.19779 | 0.155257 |
| 31.77778 | 0.376014 | 0.337362 | -1.58342 | 0.381422 | -1.51644 | -1.97611 | -1.94042 | -0.96751 | -1.48286 | -0.10096 | -1.14375 | 0.045814 | -0.03308 | -0.20541 | -0.14576 | -0.08715 | -0.19205 | -0.23797 | 0.350533 |
| 31.79167 | 0.031982 | -1.26099 | -1.83786 | 2.812643 | -0.58321 | -1.87041 | -1.84916 | -1.36194 | -2.46191 | -0.35788 | -0.99909 | -0.11056 | -0.45794 | -0.11011 | -0.12359 | -0.21036 | -0.25286 | -0.43947 | 0.281989 |
| 31.80556 | -0.5629 | -2.95504 | -1.99227 | 4.544496 | -0.1337 | -0.92862 | -1.4057 | -1.57719 | -3.06044 | -0.64546 | -0.67732 | -0.1957 | -0.8337 | 0.016453 | -0.09148 | -0.27538 | -0.46301 | -0.65525 | -0.00102 |
| 31.81944 | -1.12348 | -4.47223 | -2.25625 | 3.087027 | -0.0916 | -1.27221 | -0.86699 | -1.41936 | -3.0113 | -0.89457 | -0.41061 | -0.17221 | -1.01236 | 0.010239 | 0.011113 | -0.30339 | -0.6989 | -0.86648 | -0.4342 |
| 31.83333 | -1.83713 | -4.43506 | -2.69761 | 0.447525 | -0.73545 | -2.25462 | -0.37827 | -1.20099 | -2.44366 | -0.92603 | -0.45196 | -0.0869 | -0.95431 | -0.11553 | 0.09471 | -0.35798 | -0.86703 | -0.90364 | -0.81356 |
| 31.84722 | -2.0852 | -4.31425 | -3.307 | -2.03515 | -1.56174 | -2.78484 | -0.09741 | -1.06869 | -2.0087 | -0.89888 | -0.84898 | -0.04815 | -0.80358 | -0.26911 | 0.11234 | -0.38073 | -0.85604 | -0.79196 | -0.93057 |
| 31.86111 | -1.50329 | -3.57228 | -3.67481 | -3.87188 | -2.27848 | -3.84035 | 0.019734 | -1.01482 | -1.93849 | -0.86038 | -1.19208 | -0.12114 | -0.56744 | -0.53057 | 0.102734 | -0.2531 | -0.83171 | -0.5944 | -0.90661 |
| 31.875 | -1.07016 | -2.47363 | -3.04383 | -5.43086 | -2.5376 | -4.83541 | -0.08002 | -1.21201 | -1.94987 | -0.78895 | -1.08741 | -0.15505 | -0.342 | -0.72732 | 0.081845 | -0.15602 | -0.80402 | -0.50142 | -0.90656 |
| 31.88889 | -0.72442 | -1.67541 | -2.02143 | -4.99078 | -2.13389 | -3.6513 | -0.09662 | -1.45044 | -1.64303 | -0.63571 | -0.49042 | -0.2388 | -0.2014 | -0.83305 | 0.017975 | -0.13593 | -0.76681 | -0.37197 | -0.78129 |
| 31.90278 | -0.33078 | -1.09303 | -0.96942 | -4.12489 | -1.5058 | -2.07457 | -0.08249 | -1.48537 | -1.26706 | -0.46702 | 0.081002 | -0.27881 | -0.16517 | -0.77656 | -0.03945 | -0.12836 | -0.65666 | -0.2539 | -0.64406 |
| 31.91667 | -0.26333 | -0.69856 | -0.18928 | -2.07728 | -0.57202 | -0.59322 | -0.04716 | -1.30127 | -0.72529 | -0.44479 | 0.3233 | -0.28165 | -0.24567 | -0.6682 | -0.11536 | -0.11869 | -0.44433 | -0.25382 | -0.48609 |
| 31.93056 | -0.21773 | -0.48326 | 0.081158 | -0.89865 | 0.404471 | 0.636691 | -0.03767 | -0.96152 | -0.06665 | -0.46944 | 0.179141 | -0.21794 | -0.48728 | -0.48332 | -0.19496 | -0.12257 | -0.26577 | -0.34914 | -0.3968 |
| 31.94444 | -0.24987 | -0.35195 | 0.073691 | -0.32475 | 1.174993 | 1.134968 | -0.00212 | -0.65293 | 0.504526 | -0.4908 | -0.21392 | -0.11297 | -0.49722 | -0.26529 | -0.23232 | -0.10198 | -0.16688 | -0.4191 | -0.30323 |
| 31.95833 | -0.2579 | -0.13427 | 0.13705 | -0.19047 | 1.590036 | 1.042159 | 0.058005 | -0.34073 | 0.813843 | -0.53206 | -0.53179 | 0.02719 | -0.16223 | -0.12459 | -0.22881 | -0.03204 | -0.1754 | -0.37651 | -0.21596 |
| 31.97222 | -0.23338 | 0.130055 | 0.404917 | -0.14338 | 1.558709 | 0.873922 | 0.168511 | -0.13575 | 0.857714 | -0.49048 | -0.38311 | 0.090051 | -0.04495 | -0.16351 | -0.20324 | 0.015133 | -0.18257 | -0.40046 | -0.21073 |
| 31.98611 | -0.34665 | 0.316218 | 0.448982 | -0.21855 | 1.069575 | 0.650748 | 0.197501 | -0.06757 | 0.405548 | -0.58452 | -0.36123 | -0.00164 | -0.06816 | -0.30781 | -0.20165 | -0.06762 | -0.18759 | -0.83771 | -0.48698 |
| 32 | -0.82767 | 0.13658 | -0.50577 | -0.56858 | -0.18244 | -0.63094 | 0.042722 | -0.18143 | -0.58526 | -1.16341 | -1.29691 | -0.14544 | -0.16535 | -0.38871 | -0.26309 | -0.31739 | -0.3839 | -1.78649 | -1.11075 |
| 32.01389 | -1.80883 | -0.65202 | -2.21945 | -1.04068 | -1.74553 | -2.64606 | -0.28809 | -0.4307 | -1.82992 | -1.45209 | -2.51627 | -0.24378 | -0.24276 | -0.4384 | -0.38582 | -0.50739 | -0.68942 | -2.21087 | -1.42647 |
| 32.02778 | -2.36284 | -1.3332 | -3.32884 | -1.16841 | -2.47807 | -4.02798 | -0.79324 | -0.59461 | -2.54995 | -0.96025 | -2.4333 | -0.23723 | -0.21447 | -0.51819 | -0.35416 | -0.4804 | -0.81957 | -1.4097 | -1.20297 |
| 32.04167 | -2.75467 | -1.67812 | -3.29922 | -0.55988 | -2.08014 | -4.49289 | -0.63783 | -0.58812 | -2.53225 | -0.36066 | -1.53411 | -0.20142 | -0.15242 | -0.48914 | -0.35316 | -0.31248 | -0.81423 | -0.67406 | -0.85053 |
| 32.05556 | -2.91599 | -1.30083 | -2.7148 | -0.11538 | -1.11659 | -3.84017 | -0.20419 | -0.39616 | -2.26987 | 0.105526 | -0.47842 | -0.09417 | -0.06409 | -0.36462 | -0.24632 | -0.14477 | -0.62528 | -0.09193 | -0.32619 |
| 32.06944 | -1.90221 | -0.4902 | -1.62787 | 0.080175 | -0.26306 | -2.73664 | 0.196957 | -0.27039 | -1.73949 | 0.375963 | 0.377002 | -0.01339 | 0.010153 | -0.23002 | -0.13966 | -0.05514 | -0.27315 | 0.283661 | -0.01207 |
| 32.08333 | -0.54454 | 0.136247 | -0.3213 | -0.03093 | 0.177 | -1.44578 | 0.389728 | -0.2593 | -1.02369 | 0.365283 | 0.973674 | -0.01718 | 0.010021 | -0.12284 | -0.11292 | -0.00698 | 0.050776 | 0.417646 | 0.164041 |
| 32.09722 | 0.602144 | 0.537274 | 1.191933 | -0.07538 | 0.384202 | -0.28021 | 0.466513 | -0.0741 | -0.17572 | 0.176657 | 1.180943 | -0.02182 | -0.00323 | -0.01674 | -0.04989 | 0.077736 | 0.332327 | 0.391096 | 0.205075 |
| 32.11111 | 1.681228 | 0.832325 | 2.477886 | 0.077703 | 0.592161 | 0.751897 | 0.558348 | 0.312003 | 0.362218 | 0.302166 | 1.31253 | 0.01497 | 0.000821 | 0.053409 | 0.014655 | 0.151515 | 0.532452 | 0.318002 | 0.156252 |
| 32.125 | 2.481594 | 1.041181 | 3.102578 | 0.250732 | 0.894704 | 1.848913 | 0.633493 | 0.650457 | 0.578241 | 0.279239 | 1.286839 | 0.025757 | 0.014591 | 0.027516 | 0.034404 | 0.17965 | 0.600652 | 0.239919 | 0.143807 |
| 32.13889 | 2.758671 | 1.265621 | 3.329381 | 0.312839 | 1.145368 | 2.774939 | 0.764033 | 0.819828 | 0.798019 | 0.110583 | 1.148613 | 0.006335 | 0.022857 | -0.06818 | 0.039051 | 0.219761 | 0.524421 | 0.17441 | 0.170922 |
| 32.15278 | 2.47977 | 1.212784 | 2.735928 | 0.160727 | 1.154244 | 3.61424 | 1.038375 | 0.767539 | 0.998114 | -0.06884 | 0.866118 | -0.03885 | 0.019974 | -0.15988 | 0.069745 | 0.247503 | 0.350078 | 0.155627 | 0.196233 |
| 32.16667 | 1.988684 | 0.739261 | 1.826161 | -0.13337 | 1.287643 | 3.58269 | 1.183731 | 0.545426 | 1.194919 | -0.20536 | 0.417552 | -0.09285 | -0.01026 | -0.1908 | 0.127942 | 0.231905 | 0.233665 | 0.193017 | 0.159503 |
| 32.18056 | 2.216182 | 0.743269 | 1.391166 | -0.36566 | 1.479923 | 3.496801 | 1.002075 | 0.340803 | 1.310975 | -0.17137 | 0.454521 | -0.07459 | -0.00933 | -0.18434 | 0.219851 | 0.235803 | 0.363499 | 0.271456 | 0.01824 |
| 32.19444 | 2.287243 | 0.808116 | 0.693818 | -0.44319 | 1.24692 | 3.174365 | 0.696198 | 0.223908 | 1.418758 | -0.04299 | 0.620869 | 0.021929 | 0.018149 | -0.21733 | 0.260281 | 0.250511 | 0.473801 | 0.169266 | -0.15498 |
| 32.20833 | 1.807282 | 0.826134 | 0.298325 | -0.3464 | 0.830364 | 2.73316 | 0.342434 | 0.177294 | 1.510001 | 0.107022 | 0.683876 | 0.108208 | 0.031809 | -0.22375 | 0.22275 | 0.214172 | 0.437356 | -0.01183 | -0.35165 |
| 32.22222 | 1.854328 | 1.138129 | 0.57624 | -0.05994 | 0.435634 | 2.043245 | 0.271268 | 0.279719 | 0.983049 | 0.203662 | 0.704771 | 0.12863 | 0.025233 | -0.14067 | 0.127949 | 0.117679 | 0.280045 | -0.11084 | -0.50658 |
| 32.23611 | 1.807732 | 1.53127 | 1.085419 | 0.098398 | 0.115159 | 1.251632 | 0.341418 | 0.431446 | 0.920088 | 0.14117 | 0.474759 | 0.027476 | -0.01294 | -0.10958 | 0.009258 | -0.04677 | -0.02042 | -0.10876 | -0.52254 |
| 32.25 | 1.188943 | 1.339923 | 1.018387 | 0.05132 | -0.20971 | 0.36289 | 0.166155 | 0.441929 | 0.587434 | -0.03848 | 0.019829 | -0.09554 | -0.0661 | -0.10183 | -0.05228 | -0.18933 | -0.27085 | -0.05241 | -0.449 |
| 32.26389 | 0.317831 | 0.688893 | 0.653243 | -0.13006 | -0.606 | -0.48835 | -0.28117 | 0.277758 | 0.358719 | -0.29378 | -0.47775 | -0.13113 | -0.09459 | -0.07742 | -0.03345 | -0.17563 | -0.37462 | -0.0023 | -0.31842 |
| 32.27778 | -0.3856 | 0.162292 | 0.339406 | -0.26567 | -0.84108 | -0.94552 | -0.52336 | 0.095297 | 0.319404 | -0.42824 | -0.74979 | -0.08073 | -0.08995 | -0.04928 | 0.024708 | -0.00305 | -0.33895 | -0.01284 | -0.23757 |
| 32.29167 | -0.67115 | -0.11646 | 0.19755 | -0.26299 | -0.61994 | -0.73316 | -0.41271 | -0.05537 | 0.263576 | -0.29896 | -0.57726 | -0.04803 | -0.06276 | -0.00117 | 0.077861 | 0.192678 | -0.20169 | -0.02607 | -0.19279 |

| | | | | | | | | | | | | | | | | | | | |
|---|---|---|---|---|---|---|---|---|---|---|---|---|---|---|---|---|---|---|---|
| 32.30556 | -0.50048 | -0.02759 | 0.317537 | -0.1593 | -0.06981 | 0.002845 | -0.10529 | -0.09089 | 0.234621 | 0.058498 | -0.09912 | -0.0334 | -0.02106 | 0.046179 | 0.079415 | 0.282143 | -0.09625 | 0.0223 | -0.16813 |
| 32.31944 | 0.041432 | 0.193031 | 0.631622 | -0.00599 | 0.282422 | 0.719757 | 0.106125 | 0.017079 | 0.297632 | 0.376469 | 0.391157 | -0.0166 | 0.010617 | 0.058875 | 0.024504 | 0.255072 | -0.02765 | 0.093606 | -0.15098 |
| 32.33333 | 0.609016 | 0.449836 | 0.839095 | 0.131991 | 0.378768 | 1.049808 | 0.212675 | 0.249965 | 0.350776 | 0.393648 | 0.621916 | -0.00223 | 0.035798 | 0.063449 | -0.02813 | 0.197937 | 0.024336 | 0.140423 | -0.07381 |
| 32.34722 | 0.995244 | 0.655654 | 0.842966 | 0.24534 | 0.285137 | 0.943991 | 0.23768 | 0.559721 | 0.365231 | 0.143212 | 0.383524 | -0.01761 | 0.044269 | 0.092413 | -0.05219 | 0.155609 | 0.00513 | 0.113886 | 0.066064 |
| 32.36111 | 0.848309 | 0.534393 | 0.625236 | 0.20715 | 0.052487 | 0.523779 | 0.184889 | 0.701166 | 0.325965 | -0.08131 | 0.050632 | -0.04097 | 0.031775 | 0.120141 | -0.03727 | 0.154407 | -0.01605 | 0.061751 | 0.149617 |
| 32.375 | 0.297994 | 0.131263 | 0.325753 | -0.02359 | -0.25975 | -0.03394 | 0.072334 | 0.49492 | 0.263434 | -0.20791 | 0.059259 | 0.021065 | 0.02551 | 0.153947 | 0.014698 | 0.189204 | 0.130109 | 0.099554 | 0.087287 |
| 32.38889 | -0.09526 | -0.19552 | 0.310933 | -0.12303 | -0.41122 | -0.22602 | 0.075997 | 0.269179 | 0.309085 | -0.13788 | 0.283583 | 0.119957 | 0.049719 | 0.201806 | 0.094157 | 0.254381 | 0.369613 | 0.218454 | 0.059087 |
| 32.40278 | 0.095827 | -0.05444 | 0.739398 | 0.143078 | -0.25648 | 0.355118 | 0.245407 | 0.359571 | 0.533149 | 0.10806 | 0.40012 | 0.175337 | 0.093898 | 0.267465 | 0.164387 | 0.275886 | 0.498782 | 0.36939 | 0.152691 |
| 32.41667 | 0.618336 | 0.351541 | 1.216878 | 0.401524 | 0.094455 | 1.218277 | 0.396473 | 0.523488 | 0.726905 | 0.349613 | 0.488518 | 0.185384 | 0.087161 | 0.305358 | 0.182873 | 0.236412 | 0.497802 | 0.439298 | 0.205214 |
| 32.43056 | 0.896392 | 0.692284 | 1.233558 | 0.370758 | 0.312012 | 1.772251 | 0.404883 | 0.520505 | 0.649145 | 0.340688 | 0.622963 | 0.187487 | 0.020033 | 0.288335 | 0.159272 | 0.170837 | 0.446394 | 0.401547 | 0.137775 |
| 32.44444 | 0.827418 | 0.976585 | 0.961859 | 0.23099 | 0.349261 | 1.91155 | 0.368796 | 0.552581 | 0.487624 | 0.155716 | 0.659668 | 0.118009 | -0.0285 | 0.193126 | 0.107689 | 0.098625 | 0.341765 | 0.288833 | 0.077967 |
| 32.45833 | 0.758288 | 1.118939 | 0.637401 | 0.132678 | 0.24878 | 1.656364 | 0.292855 | 0.578472 | 0.393474 | 0.056757 | 0.455381 | 0.074146 | -0.01347 | 0.165114 | 0.073219 | 0.04665 | 0.229011 | 0.198381 | 0.113767 |
| 32.47222 | 0.576922 | 0.976106 | 0.5094 | 0.170513 | 0.051459 | 1.013031 | 0.176375 | 0.476055 | 0.44099 | 0.082113 | 0.263715 | 0.144168 | 0.080597 | 0.235723 | 0.068232 | 0.067226 | 0.242916 | 0.275376 | 0.178689 |
| 32.48611 | 0.615477 | 0.911949 | 0.887746 | 0.861381 | -0.12835 | 0.641175 | 0.031723 | 0.422776 | 0.844169 | 0.315655 | 0.336746 | 0.212069 | 0.14444 | 0.300138 | 0.040859 | 0.177738 | 0.421994 | 0.545176 | 0.345537 |
| 32.5 | 0.866274 | 1.319987 | 2.163217 | 3.873482 | -0.11062 | 1.062948 | -0.2067 | 0.727731 | 1.750016 | 0.826859 | 0.803605 | 0.245185 | 0.15498 | 0.406913 | -0.03554 | 0.34739 | 0.711877 | 0.994907 | 0.601794 |
| 32.51389 | 1.26509 | 1.87474 | 3.321562 | 5.488972 | 0.383006 | 2.36103 | -0.47562 | 1.288898 | 2.418339 | 1.157261 | 1.505035 | 0.304718 | 0.112214 | 0.730436 | -0.12367 | 0.469884 | 1.211344 | 1.490018 | 0.851006 |
| 32.52778 | 2.152765 | 2.661284 | 3.279893 | 6.133499 | 1.469377 | 3.055023 | -0.29289 | 1.113214 | 2.2173 | 1.173601 | 2.389299 | 0.451617 | 0.149825 | 1.504266 | 0.084142 | 0.57489 | 1.88679 | 1.638422 | 0.889257 |
| 32.54167 | 2.685287 | 3.64915 | 3.124302 | 4.751802 | 2.400989 | 3.207234 | 0.443559 | 0.933108 | 1.78749 | 1.282514 | 2.380089 | 0.527233 | 0.20859 | 1.740922 | 0.240301 | 0.565598 | 1.572161 | 1.55517 | 0.681401 |
| 32.55556 | 1.559889 | 3.026231 | 2.249964 | 2.768278 | 2.345474 | 2.426065 | 0.813571 | 0.70814 | 1.603103 | 1.341116 | 1.798672 | 0.393946 | 0.192216 | 1.248174 | 0.188296 | 0.417693 | 0.812241 | 1.255284 | 0.369363 |
| 32.56944 | 0.461201 | 2.013594 | 1.052212 | 1.629855 | 1.384981 | 1.119234 | 0.699094 | 0.515967 | 1.547766 | 1.402996 | 0.359012 | 0.202811 | 0.263556 | 0.698409 | 0.106559 | 0.232766 | 0.067823 | 0.810659 | 0.208747 |
| 32.58333 | -0.12175 | 1.271685 | 0.188471 | 0.701759 | 0.412634 | 0.045029 | 0.542852 | 0.432897 | 1.549408 | 1.241035 | -0.91518 | 0.061234 | 0.404815 | 0.332826 | 0.015672 | 0.059553 | -0.60339 | 0.475887 | 0.288835 |
| 32.59722 | -0.32691 | 0.665212 | -0.27182 | -0.24549 | -0.27429 | -0.92012 | 0.431063 | 0.423277 | 1.461302 | 0.934648 | -1.72992 | -0.00896 | 0.489032 | 0.127905 | -0.05223 | -0.09344 | -1.10685 | 0.336725 | 0.559547 |
| 32.61111 | -0.48138 | 0.270323 | -0.50142 | -1.35723 | -0.68053 | -1.66309 | 0.271069 | 0.270137 | 1.13013 | 0.695042 | -0.90126 | -0.08265 | 0.471057 | 0.02859 | -0.11626 | -0.18356 | -1.14841 | 0.291729 | 0.868169 |
| 32.625 | -0.5438 | -0.1542 | -0.63517 | -2.08554 | -0.8967 | -2.44729 | 0.017469 | 0.151145 | 0.797059 | 0.571491 | -0.57172 | -0.15563 | 0.384526 | -0.02933 | -0.19036 | -0.15794 | -0.64541 | 0.268139 | 0.801427 |
| 32.63889 | -0.37585 | -0.96512 | -0.65532 | -3.51201 | -0.6313 | -2.48258 | -0.29014 | 0.393719 | 0.258595 | 0.645637 | -1.02904 | -0.16353 | 0.234084 | -0.01088 | -0.23132 | 0.006607 | -0.06636 | 0.192249 | 0.592501 |
| 32.65278 | 0.042482 | -1.73798 | -0.70028 | -3.18977 | -0.55522 | -1.93601 | -0.37758 | 0.607579 | -0.42787 | 0.689161 | -0.16308 | -0.11303 | 0.087421 | 0.020909 | -0.24889 | 0.079873 | 0.158849 | 0.085024 | 0.428447 |
| 32.66667 | 0.684111 | -2.26146 | -0.44771 | -0.50171 | -0.33924 | -1.0996 | -0.32518 | 0.763843 | -0.92561 | 0.50197 | 0.745407 | -0.07537 | -0.01646 | 0.079807 | -0.19164 | 0.119614 | 0.15091 | -0.01884 | 0.258278 |
| 32.68056 | 1.259082 | -2.66109 | -0.18673 | 2.25941 | -0.21655 | -0.09388 | -0.32487 | 0.80382 | -1.01181 | 0.272621 | 0.778536 | -0.06715 | -0.09148 | 0.169437 | -0.10787 | 0.188499 | 0.193104 | -0.03452 | 0.03819 |
| 32.69444 | 0.69248 | -3.46302 | -0.14024 | 4.937961 | -0.49257 | 0.114408 | -0.4546 | 0.602926 | -1.69211 | 0.156139 | 0.181013 | -0.12709 | -0.17571 | 0.270853 | 0.032503 | 0.297919 | 0.352892 | 0.03061 | -0.24152 |
| 32.70833 | -0.12572 | -2.36845 | 0.235225 | 5.062573 | -0.95973 | 0.629711 | -0.55389 | 0.420074 | -1.56431 | 0.058124 | -0.2023 | -0.21991 | -0.22351 | 0.254342 | 0.118979 | 0.386318 | 0.452577 | 0.202922 | -0.37914 |
| 32.72222 | 0.189701 | -0.793 | 0.289972 | 2.927076 | -0.78721 | 1.512684 | -0.56576 | 0.365263 | -0.46014 | -0.09936 | -0.36631 | -0.27403 | -0.21596 | 0.174489 | 0.128588 | 0.389954 | 0.18779 | 0.244113 | -0.38746 |
| 32.73611 | 0.632799 | 1.035005 | 0.511458 | 1.33528 | -0.47585 | 1.631361 | -0.54205 | 0.198319 | 0.568074 | -0.24422 | -0.59779 | -0.22459 | -0.11376 | 0.123871 | 0.103081 | 0.189865 | -0.30042 | 0.024181 | -0.33098 |
| 32.75 | 0.272491 | 2.971259 | 0.612776 | -0.09567 | -0.13336 | 1.472872 | -0.16124 | 0.097837 | 1.12617 | -0.33835 | -1.15802 | -0.10458 | -0.02832 | 0.031885 | 0.11589 | -0.08029 | -0.72462 | -0.3081 | -0.24094 |
| 32.76389 | -0.16269 | 3.832131 | 0.27552 | -1.6256 | 0.017498 | -0.28197 | 0.328902 | -0.01362 | 1.106914 | -0.3564 | -1.71349 | -0.04143 | -0.00755 | -0.16753 | 0.094046 | -0.2546 | -0.8991 | -0.49568 | -0.06759 |
| 32.77778 | -0.31131 | 3.100224 | -0.50525 | -2.8773 | -0.33418 | -2.21517 | 0.499041 | -0.23525 | 0.343407 | -0.16492 | -2.11287 | -0.00899 | 0.015306 | -0.36613 | 0.017632 | -0.30439 | -0.83339 | -0.51748 | 0.122825 |
| 32.79167 | -0.38374 | 2.083403 | -1.24248 | -4.76346 | -0.72544 | -2.80126 | 0.298317 | -0.52196 | -0.32638 | 0.089514 | -2.22262 | -0.01319 | 0.032782 | -0.49998 | -0.09718 | -0.29968 | -0.6702 | -0.48461 | 0.283397 |
| 32.80556 | -0.40766 | 0.500889 | -1.61276 | -6.17447 | -0.86867 | -2.28537 | -0.0688 | -0.80407 | -0.71711 | 0.145716 | -2.06341 | -0.03657 | 0.042722 | -0.584 | -0.15053 | -0.34355 | -0.56734 | -0.5631 | 0.324164 |
| 32.81944 | -0.30042 | -1.31756 | -2.06823 | -4.96408 | -0.82275 | -1.94338 | -0.50349 | -0.87274 | -0.86948 | -0.0456 | -1.54163 | -0.05197 | 0.001482 | -0.5306 | -0.20977 | -0.56111 | -0.62618 | -0.64512 | 0.230899 |
| 32.83333 | -0.30024 | -2.7728 | -2.99511 | -1.95706 | -0.69619 | -1.85119 | -0.74039 | -0.8496 | -0.90192 | -0.30245 | -1.07874 | -0.05418 | -0.10083 | -0.31374 | -0.26873 | -0.77015 | -0.81415 | -0.65196 | 0.027447 |
| 32.84722 | -0.43112 | -3.41214 | -3.4505 | -0.84671 | -0.81354 | -2.97163 | -0.72251 | -0.83892 | -0.97448 | -0.46739 | -1.00945 | -0.0654 | -0.15509 | -0.13208 | -0.21818 | -0.79447 | -0.97023 | -0.58627 | -0.30178 |
| 32.86111 | -1.06794 | -3.6685 | -3.35273 | -0.62039 | -1.08961 | -4.32499 | -0.68456 | -0.82582 | -1.32093 | -0.57547 | -1.03549 | -0.07376 | -0.07765 | -0.10691 | -0.12005 | -0.59363 | -0.92515 | -0.57981 | -0.63738 |
| 32.875 | -1.6942 | -3.46984 | -3.04944 | -0.70743 | -1.35691 | -4.73022 | -0.64087 | -0.76127 | -1.66542 | -0.6769 | -0.66007 | -0.04987 | 0.010678 | -0.2014 | 0.026836 | -0.39151 | -0.72788 | -0.50633 | -0.82953 |
| 32.88889 | -2.02128 | -2.87515 | -2.44767 | -0.78828 | -1.54525 | -4.02121 | -0.476 | -0.83684 | -1.98289 | -0.69879 | -0.02609 | -0.08464 | 0.053288 | -0.31167 | 0.135278 | -0.32495 | -0.49472 | -0.40909 | -0.75904 |
| 32.90278 | -1.63282 | -2.07249 | -1.6294 | -0.91113 | -1.31084 | -2.9071 | -0.10802 | -0.96482 | -2.02186 | -0.69922 | 0.396995 | -0.1363 | 0.050608 | -0.38844 | 0.185899 | -0.33308 | -0.39078 | -0.27658 | -0.57703 |
| 32.91667 | -0.60392 | -1.16748 | -0.84606 | -1.08704 | -0.63256 | -1.63231 | 0.217394 | -1.05691 | -1.74568 | -0.63123 | 0.42554 | -0.14566 | 0.031333 | -0.43654 | 0.156843 | -0.33726 | -0.46914 | -0.22385 | -0.41699 |
| 32.93056 | 0.397987 | -0.50965 | -0.17992 | -1.05294 | 0.112907 | -0.39335 | 0.434489 | -0.86789 | -1.17025 | -0.51623 | 0.257479 | -0.103 | -0.00468 | -0.48154 | 0.096792 | -0.28295 | -0.59947 | -0.29823 | -0.26909 |
| 32.94444 | 0.998854 | -0.06299 | 0.266697 | -0.71478 | 0.45204 | 0.58962 | 0.591531 | -0.47381 | -0.58864 | -0.35914 | -0.06206 | -0.03726 | 0.036991 | -0.46096 | 0.071039 | -0.18734 | -0.62561 | -0.41403 | -0.11736 |
| 32.95833 | 1.285377 | 0.307688 | 0.455416 | -0.32891 | 0.610507 | 1.1788 | 0.724568 | -0.06817 | 0.027295 | -0.24427 | -0.48139 | 0.022466 | 0.075907 | -0.33795 | 0.055369 | -0.07497 | -0.47263 | -0.41911 | 0.025863 |
| 32.97222 | 1.240933 | 0.475557 | 0.308936 | 0.041389 | 0.638466 | 1.014434 | 0.787151 | 0.170673 | 0.473218 | -0.25641 | -0.96459 | 0.018931 | 0.038941 | -0.24511 | 0.006509 | -0.04018 | -0.28662 | -0.29706 | 0.042065 |
| 32.98611 | 0.798518 | 0.327222 | -0.08764 | 0.049655 | 0.297459 | 0.035734 | 0.666166 | 0.164247 | 0.527523 | -0.62797 | -1.62773 | -0.04243 | -0.07046 | -0.17572 | -0.04104 | -0.1296 | -0.2894 | -0.47076 | -0.15597 |
| 33 | -0.42407 | -0.00608 | -0.97702 | -0.42664 | -0.55247 | -1.31102 | 0.353362 | -0.10291 | -0.06221 | -1.41404 | -2.61599 | -0.11343 | -0.20008 | -0.18005 | -0.03893 | -0.41733 | -0.64125 | -1.39574 | -0.57892 |
| 33.01389 | -1.89955 | -0.54994 | -2.27516 | -0.82336 | -1.10571 | -2.03064 | -0.11469 | -0.407 | -0.91753 | -1.86461 | -3.4113 | -0.14698 | -0.26866 | -0.26129 | -0.06195 | -0.61336 | -0.94954 | -2.1413 | -0.77728 |
| 33.02778 | -2.76558 | -1.03245 | -3.17639 | -0.40467 | -0.74403 | -2.01146 | -0.27793 | -0.27735 | -1.45956 | -1.25602 | -2.41978 | -0.14597 | -0.2235 | -0.43293 | -0.17661 | -0.49327 | -0.89392 | -1.65093 | -0.46034 |
| 33.04167 | -2.9753 | -1.19986 | -3.05282 | 0.052546 | -0.50459 | -1.74398 | -0.00024 | -0.10401 | -1.05477 | -0.78152 | -1.48823 | -0.18362 | -0.19999 | -0.42771 | -0.24272 | -0.28418 | -0.64054 | -0.97684 | -0.14194 |
| 33.05556 | -2.40224 | -0.90521 | -2.98643 | 0.159217 | -0.39373 | -1.27665 | 0.308396 | -0.13535 | -0.7172 | -0.65441 | -0.96278 | -0.17866 | -0.14823 | -0.2219 | -0.17816 | -0.15733 | -0.38401 | -0.46172 | -0.02857 |
| 33.06944 | -1.12947 | -0.26963 | -2.20003 | 0.06413 | -0.25015 | -0.68987 | 0.446349 | -0.30452 | -0.45009 | -0.32924 | -0.23751 | -0.042 | -0.09194 | -0.0758 | -0.03432 | -0.04379 | -0.0977 | 0.141769 | -0.05638 |
| 33.08333 | -0.12745 | 0.236497 | -1.19702 | -0.05478 | -0.015 | -0.15838 | 0.574554 | -0.28572 | -0.1178 | 0.162278 | 0.406845 | 0.097349 | -0.01551 | 0.012724 | 0.132767 | 0.054656 | 0.147504 | 0.733714 | -0.12915 |
| 33.09722 | 0.551664 | 0.496225 | -0.33149 | -0.11508 | 0.145816 | 0.125493 | 0.625783 | -0.15179 | 0.257192 | 0.40866 | 0.696143 | 0.095922 | 0.010635 | 0.030335 | 0.186642 | 0.10491 | 0.197704 | 1.023372 | -0.12412 |
| 33.11111 | 0.859492 | 0.642371 | 0.320659 | -0.05427 | 0.004469 | 0.245887 | 0.587995 | 0.044883 | 0.631311 | 0.383712 | 0.716327 | 0.045868 | 0.00429 | 0.01834 | 0.176646 | 0.089695 | 0.117493 | 1.066416 | -0.12983 |
| 33.125 | 0.761232 | 0.683356 | 0.863498 | 0.203812 | -0.08863 | 0.393814 | 0.598152 | 0.237103 | 0.705942 | 0.30171 | 0.713625 | 0.017135 | 0.005681 | 0.049924 | 0.126435 | 0.061537 | 0.082123 | 0.846081 | -0.1257 |
| 33.13889 | 0.715667 | 0.717815 | 1.420483 | 0.442081 | 0.39367 | 0.711875 | 0.63387 | 0.312578 | 0.569252 | 0.227902 | 0.8758 | 0.013113 | 0.002969 | 0.099017 | 0.039634 | 0.031604 | 0.140365 | 0.676868 | -0.05816 |
| 33.15278 | 0.822002 | 0.724197 | 2.033605 | 0.589279 | 0.749543 | 1.009975 | 0.710101 | 0.343425 | 0.5057 | 0.174679 | 0.92207 | 0.018745 | 0.0157 | 0.095611 | -0.00572 | 0.031079 | 0.237108 | 0.643225 | 0.060114 |
| 33.16667 | 1.09063 | 0.923464 | 2.712011 | 0.676901 | 0.829176 | 0.992872 | 0.89792 | 0.41552 | 0.611848 | 0.244867 | 0.908395 | 0.006221 | 0.039754 | 0.03278 | -0.01612 | 0.09096 | 0.314718 | 0.60099 | 0.153581 |
| 33.18056 | 1.502751 | 1.143838 | 3.066124 | 0.550299 | 0.738412 | 0.755081 | 0.926498 | 0.442579 | 0.900038 | 0.442407 | 0.81024 | -0.0214 | 0.031611 | -0.05528 | 0.009429 | 0.122948 | 0.329662 | 0.519224 | 0.142208 |
| 33.19444 | 1.852721 | 1.166886 | 2.9077 | 0.274097 | 0.444645 | 0.453715 | 0.683849 | 0.419243 | 1.124294 | 0.469781 | 0.840306 | -0.04513 | -0.00724 | -0.139 | 0.031767 | 0.093797 | 0.3159 | 0.310383 | 0.058667 |
| 33.20833 | 2.133232 | 1.084275 | 2.491388 | 0.069761 | 0.099093 | 0.349782 | 0.34604 | 0.294769 | 0.959848 | 0.265581 | 0.946493 | -0.05751 | -0.05716 | -0.18859 | 0.008411 | 0.031446 | 0.238168 | 0.011633 | -0.01759 |
| 33.22222 | 2.154917 | 0.987912 | 2.201212 | -0.05286 | -0.11787 | 0.695803 | 0.090564 | 0.211424 | 0.620304 | 0.046857 | 1.023732 | -0.062 | -0.1059 | -0.22579 | -0.07753 | -0.04378 | 0.158597 | -0.33557 | -0.10796 |
| 33.23611 | 2.045208 | 0.759466 | 1.754567 | -0.12357 | -0.27244 | 1.046774 | -0.07215 | 0.1286 | 0.301963 | 0.040889 | 1.122887 | -0.056 | -0.10688 | -0.19281 | -0.08858 | -0.09268 | 0.130142 | -0.55226 | -0.19077 |
| 33.25 | 1.814055 | 0.468515 | 1.272671 | -0.19329 | -0.21486 | 1.106718 | -0.03126 | 0.043338 | 0.163152 | 0.104244 | 1.107856 | -0.03927 | -0.08137 | -0.16396 | -0.02941 | -0.09418 | 0.085155 | -0.48236 | -0.17053 |
| 33.26389 | 1.524805 | 0.250079 | 0.950151 | -0.23727 | -0.00671 | 0.990985 | 0.141449 | 0.081805 | 0.293374 | 0.051642 | 0.92711 | -0.03519 | -0.03737 | -0.09384 | 0.03841 | -0.03345 | 0.040239 | -0.13883 | -0.06569 |
| 33.27778 | 1.070698 | 0.100536 | 0.673843 | -0.28878 | 0.123687 | 0.788296 | 0.269359 | 0.154497 | 0.468562 | -0.00124 | 0.871363 | -0.02617 | 0.002395 | -0.08095 | 0.08813 | 0.053639 | 0.112425 | 0.210679 | -0.03437 |
| 33.29167 | 0.645247 | 0.025832 | 0.629076 | -0.2784 | 0.131916 | 0.449452 | 0.246879 | 0.162172 | 0.460234 | 0.020189 | 0.937556 | 0.00519 | 0.062169 | -0.09359 | 0.14617 | 0.132259 | 0.246103 | 0.344933 | 0.008835 |
| 33.30556 | 0.385879 | 0.009566 | 0.622561 | -0.14371 | 0.047348 | 0.217232 | 0.190567 | 0.156156 | 0.343683 | 0.059335 | 0.890369 | 0.022966 | 0.086723 | -0.10352 | 0.191527 | 0.178546 | 0.311715 | 0.269083 | 0.090999 |
| 33.31944 | 0.243177 | 0.050504 | 0.704385 | 0.090506 | 0.010875 | 0.130376 | 0.147059 | 0.17501 | 0.227924 | 0.187726 | 0.824132 | 0.027255 | 0.047414 | -0.08514 | 0.246206 | 0.218402 | 0.359875 | 0.090394 | 0.02633 |
| 33.33333 | 0.185314 | 0.129747 | 0.83221 | 0.293622 | -0.02008 | 0.088328 | 0.093356 | 0.2142 | 0.136756 | 0.279663 | 0.733341 | 0.039134 | 0.021251 | -0.01656 | 0.244959 | 0.257556 | 0.340299 | -0.06244 | -0.03886 |
| 33.34722 | 0.254947 | 0.217366 | 0.901132 | 0.418344 | -0.01414 | 0.108542 | 0.115345 | 0.26408 | 0.145826 | 0.226647 | 0.250822 | 0.048378 | 0.028085 | 0.059941 | 0.132474 | 0.25408 | 0.187675 | -0.16742 | 0.035788 |
| 33.36111 | 0.347365 | 0.319846 | 0.862553 | 0.467917 | 0.115785 | 0.185717 | 0.25428 | 0.39941 | 0.342737 | 0.088446 | -0.23802 | 0.040976 | 0.032807 | 0.068748 | 0.020568 | 0.232654 | 0.039539 | -0.0924 | 0.070579 |
| 33.375 | 0.418534 | 0.443082 | 0.71104 | 0.570415 | 0.342123 | 0.459686 | 0.501062 | 0.629053 | 0.611881 | 0.038927 | -0.37475 | 0.05136 | 0.043223 | 0.06894 | 0.00144 | 0.290546 | 0.009319 | 0.122893 | 0.03438 |
| 33.38889 | 0.333805 | 0.508742 | 0.509191 | 0.698064 | 0.524741 | 0.789225 | 0.664564 | 0.762703 | 0.706904 | 0.047533 | -0.25515 | 0.03347 | 0.055439 | 0.101572 | 0.025781 | 0.380035 | 0.053386 | 0.302437 | 0.099693 |
| 33.40278 | 0.219501 | 0.385281 | 0.466114 | 0.673193 | 0.462367 | 0.887304 | 0.531672 | 0.628096 | 0.620328 | 0.072126 | 0.019086 | -0.02454 | 0.045827 | 0.138669 | 0.02259 | 0.429405 | 0.100584 | 0.345739 | 0.210524 |
| 33.41667 | 0.191005 | 0.194118 | 0.688829 | 0.492628 | 0.355209 | 0.909364 | 0.267465 | 0.425566 | 0.564834 | 0.166694 | 0.472372 | -0.0455 | 0.010078 | 0.170482 | 0.037167 | 0.447482 | 0.198188 | 0.291489 | 0.177956 |
| 33.43056 | 0.311678 | 0.130576 | 0.971456 | 0.324797 | 0.421933 | 1.024504 | 0.166008 | 0.422328 | 0.651955 | 0.262269 | 0.813895 | -0.02564 | -0.01408 | 0.172902 | 0.049676 | 0.408811 | 0.339077 | 0.265696 | 0.077891 |
| 33.44444 | 0.534571 | 0.162597 | 1.105279 | 0.219118 | 0.46594 | 1.170601 | 0.164018 | 0.479979 | 0.726277 | 0.317209 | 0.875345 | -0.00212 | 0.006488 | 0.171397 | 0.028033 | 0.356282 | 0.392008 | 0.173721 | 0.093819 |
| 33.45833 | 0.774856 | 0.221453 | 1.045067 | 0.126425 | 0.420684 | 1.278454 | 0.204792 | 0.52334 | 0.650874 | 0.230516 | 0.619759 | 0.001645 | 0.014005 | 0.137061 | -0.01505 | 0.308631 | 0.363428 | 0.059435 | 0.16939 |
| 33.47222 | 0.916814 | 0.267982 | 0.840141 | 0.075956 | 0.321754 | 1.431649 | 0.218227 | 0.466113 | 0.521769 | 0.147024 | 0.484122 | -0.02397 | -0.00415 | 0.107945 | -0.05262 | 0.311086 | 0.336779 | 0.039297 | 0.185733 |
| 33.48611 | 1.114667 | 0.362219 | 0.752798 | 0.099411 | 0.437124 | 2.009833 | 0.065499 | 0.376267 | 0.646094 | 0.200985 | 0.854554 | -0.02807 | 0.010751 | 0.090941 | -0.06497 | 0.435054 | 0.545706 | 0.249054 | 0.316019 |

| | | | | | | | | | | | | | | | | | | | |
|---|---|---|---|---|---|---|---|---|---|---|---|---|---|---|---|---|---|---|---|
| 33.5 | 1.604215 | 0.732245 | 1.189541 | 0.390809 | 0.962642 | 3.949269 | -0.25197 | 0.505616 | 1.079253 | 0.438571 | 1.804173 | 0.027572 | 0.105565 | 0.12251 | -0.11532 | 0.670967 | 1.00119 | 0.725204 | 0.660894 |
| 33.51389 | 1.60025 | 1.227338 | 1.780175 | 0.968312 | 1.357914 | 5.563062 | -0.38695 | 0.732962 | 1.438441 | 0.625994 | 2.414542 | 0.163729 | 0.260116 | 0.432199 | -0.22351 | 0.811252 | 1.245182 | 1.092195 | 0.897033 |
| 33.52778 | 1.250828 | 1.52988 | 2.733708 | 1.534876 | 1.616929 | 5.575518 | -0.15785 | 0.827936 | 2.021545 | 0.613757 | 2.293802 | 0.381622 | 0.388775 | 0.801233 | -0.14949 | 0.975267 | 1.53498 | 1.091361 | 0.991014 |
| 33.54167 | 1.423082 | 1.673678 | 3.131377 | 2.37384 | 1.563109 | 5.791703 | 0.27325 | 0.637652 | 3.349519 | 0.483643 | 2.451622 | 0.500096 | 0.30663 | 1.10235 | -0.02059 | 0.866152 | 2.134003 | 1.097934 | 0.838244 |
| 33.55556 | 1.324761 | 1.935482 | 2.426227 | 3.012656 | 1.464173 | 4.333972 | 0.470382 | 0.786218 | 2.887786 | 0.282879 | 1.997081 | 0.437295 | 0.0655 | 0.863964 | -0.01151 | 0.49768 | 1.426855 | 1.024699 | 0.554783 |
| 33.56944 | 0.737548 | 2.02189 | 1.42103 | 2.935129 | 0.977835 | 2.306802 | 0.335107 | 0.567953 | 1.554934 | 0.034485 | 1.049125 | 0.179122 | -0.09262 | 0.646971 | -0.06922 | 0.055361 | 0.670888 | 0.869792 | 0.267103 |
| 33.58333 | 0.236932 | 2.33097 | 0.639319 | 2.574534 | 0.689864 | 0.902197 | 0.120907 | 0.353569 | -0.06961 | -0.05563 | 0.004657 | -0.00987 | -0.11517 | 0.508578 | -0.09972 | -0.13316 | 0.0164 | 0.809512 | 0.193956 |
| 33.59722 | -0.11006 | 3.084155 | -0.08693 | 2.651163 | 1.080896 | 0.620566 | 0.07381 | 0.247255 | -1.36162 | 0.130711 | -0.88099 | -0.07426 | -0.02627 | 0.47137 | -0.11752 | -0.09085 | -0.47771 | 0.719465 | 0.175253 |
| 33.61111 | -0.40432 | 3.045304 | -0.62232 | 2.897356 | 1.276455 | 0.520467 | 0.129049 | 0.191793 | -1.34435 | 0.421884 | -1.56906 | -0.07299 | 0.224271 | 0.500497 | -0.155 | -0.02233 | -0.78354 | 0.570797 | 0.118841 |
| 33.625 | -0.33425 | 2.553208 | -1.01843 | 2.180003 | 0.786462 | -0.15575 | -0.02264 | 0.024114 | -0.5608 | 0.53385 | -1.30278 | -0.08154 | 0.327661 | 0.418131 | -0.15583 | -0.02422 | -1.11307 | 0.39177 | 0.03835 |
| 33.63889 | -0.00206 | 2.104332 | -0.87655 | 0.705093 | 0.324906 | -0.5121 | -0.10382 | -0.23097 | -0.30644 | 0.397179 | -1.64964 | -0.13981 | 0.218805 | 0.16551 | -0.19506 | -0.11697 | -1.1886 | 0.148733 | 0.015698 |
| 33.65278 | 0.263388 | 1.657893 | -0.51159 | -1.03552 | 0.1657 | -0.74711 | -0.07598 | -0.07271 | 0.315556 | 0.204519 | -1.39928 | -0.12832 | 0.138367 | 0.047579 | -0.14569 | -0.20082 | -1.07435 | -0.10127 | 0.101636 |
| 33.66667 | 0.394066 | 0.804805 | -0.09664 | -3.28944 | 0.064536 | -1.06604 | 0.013061 | 0.235948 | 0.713922 | 0.066245 | -0.67339 | -0.09038 | 0.126193 | 0.070333 | -0.0453 | -0.25471 | -0.91708 | -0.29928 | 0.248332 |
| 33.68056 | 0.26441 | -0.48555 | -0.03927 | -4.19607 | -0.01536 | -1.09969 | 0.096503 | 0.266749 | 0.801272 | -0.0048 | 0.023753 | -0.09622 | 0.134427 | 0.052794 | 0.025557 | -0.12844 | -0.44647 | -0.45959 | 0.358114 |
| 33.69444 | 0.096795 | -1.90481 | 0.156804 | -4.05727 | 0.068649 | -1.49689 | 0.040999 | 0.173966 | 0.660525 | 0.313272 | 0.529406 | -0.10353 | 0.081172 | -0.03135 | 0.045501 | -0.02438 | 0.08814 | -0.53141 | 0.343342 |
| 33.70833 | -0.23957 | -2.92334 | 0.293753 | -3.66352 | -0.16542 | -1.42074 | -0.08543 | 0.055455 | 0.235496 | 0.612519 | 0.511962 | -0.09072 | 0.010759 | 0.024265 | 0.037986 | 0.028496 | 0.590084 | -0.52167 | 0.280162 |
| 33.72222 | -0.6262 | -3.85746 | 0.238595 | -2.04268 | -0.28249 | -0.99653 | -0.22479 | -0.11294 | -0.17367 | 0.450045 | 0.24474 | -0.0636 | -0.01204 | 0.287564 | 0.043808 | 0.022761 | 0.991054 | -0.55317 | 0.262833 |
| 33.73611 | -0.13604 | -4.2122 | -0.21743 | -0.62643 | -0.45941 | -0.39666 | -0.43499 | -0.34523 | -0.68278 | 0.011271 | 0.022576 | -0.10159 | 0.012786 | 0.469609 | 0.023181 | 0.020272 | 0.961788 | -0.52555 | 0.172792 |
| 33.75 | 0.259304 | -3.70241 | -0.90615 | -0.11282 | -0.80418 | 0.132953 | -0.72864 | -0.64094 | -1.1096 | -0.37468 | -0.22459 | -0.16457 | 0.05174 | 0.45281 | 0.011913 | 0.006164 | 0.602113 | -0.39093 | -0.00582 |
| 33.76389 | 0.421318 | -2.15969 | -1.27265 | 0.007001 | -1.16918 | 0.686236 | -0.95971 | -0.9603 | -1.36058 | -0.66892 | -0.78868 | -0.23136 | 0.049274 | 0.231782 | -0.01958 | -0.07256 | 0.089504 | -0.17831 | -0.16401 |
| 33.77778 | 0.228996 | -1.03668 | -1.15906 | -0.08219 | -1.31085 | 0.988302 | -1.15735 | -1.04198 | -1.23811 | -0.83753 | -1.53429 | -0.29282 | -0.00416 | -0.1331 | -0.06372 | -0.20488 | -0.49855 | -0.01855 | -0.3013 |
| 33.79167 | -0.3212 | -0.61389 | -0.88182 | 0.199828 | -1.43311 | 0.598469 | -1.15864 | -1.04131 | -0.93497 | -0.79057 | -2.18146 | -0.2537 | -0.04748 | -0.61287 | -0.06343 | -0.25743 | -1.11285 | 0.017332 | -0.41471 |
| 33.80556 | -0.96942 | -0.52661 | -0.85803 | 1.151756 | -1.61093 | -0.51024 | -0.98517 | -0.83482 | -0.7248 | -0.5079 | -2.52429 | -0.11669 | -0.06848 | -1.15235 | -0.03775 | -0.30019 | -1.37958 | -0.06916 | -0.47696 |
| 33.81944 | -1.67315 | -0.47127 | -0.91307 | 1.894318 | -1.7312 | -2.11433 | -0.77577 | -0.62407 | -0.92093 | -0.3692 | -2.76803 | -0.04899 | -0.06289 | -1.5312 | -0.01938 | -0.44193 | -1.44598 | -0.24068 | -0.45709 |
| 33.83333 | -2.11969 | -0.42293 | -1.12397 | 1.163617 | -1.81534 | -3.98435 | -0.66461 | -0.53797 | -1.3363 | -0.42477 | -2.63222 | -0.06249 | -0.06214 | -1.23735 | -0.03023 | -0.53158 | -1.33712 | -0.46044 | -0.36093 |
| 33.84722 | -1.79732 | -0.37453 | -1.79576 | -0.35119 | -1.81056 | -5.89026 | -0.78 | -0.79475 | -1.81312 | -0.48322 | -1.99054 | -0.0877 | -0.09822 | -0.66464 | -0.06107 | -0.50837 | -0.99154 | -0.60025 | -0.25331 |
| 33.86111 | -1.26611 | -0.26825 | -2.2271 | -2.00756 | -1.51679 | -5.53175 | -0.87151 | -1.10264 | -1.96702 | -0.4276 | -1.02481 | -0.05983 | -0.05331 | -0.23071 | -0.06295 | -0.35116 | -0.58422 | -0.57852 | -0.18357 |
| 33.875 | -1.10302 | -0.35042 | -2.54374 | -3.14134 | -1.17411 | -4.3857 | -0.76328 | -1.34915 | -1.83932 | -0.36021 | -0.15605 | -0.02763 | -0.02324 | -0.02663 | -0.05917 | -0.22592 | -0.27395 | -0.46014 | -0.20469 |
| 33.88889 | -1.02681 | -0.65662 | -2.91132 | -3.52669 | -1.19115 | -3.52637 | -0.74316 | -1.53345 | -1.70211 | -0.16927 | 0.345342 | -0.01033 | -0.03304 | -0.09247 | -0.039 | -0.19488 | -0.19353 | -0.43057 | -0.28475 |
| 33.90278 | -0.90059 | -0.95171 | -2.91135 | -2.63653 | -1.50534 | -2.98576 | -0.67408 | -1.45544 | -1.32574 | 0.020066 | 0.458834 | 0.002591 | -0.06627 | -0.23144 | -0.01423 | -0.17521 | -0.25187 | -0.43671 | -0.33113 |
| 33.91667 | -0.83853 | -1.20694 | -2.43518 | -1.54275 | -1.65855 | -2.32847 | -0.5304 | -1.18843 | -0.89479 | 0.07511 | 0.44162 | 0.007076 | -0.08383 | -0.27863 | 0.01405 | -0.15542 | -0.2935 | -0.4085 | -0.2768 |
| 33.93056 | -1.03884 | -1.41562 | -1.51539 | -0.54862 | -1.35698 | -1.66613 | -0.37012 | -0.78719 | -0.566 | 0.021011 | 0.5381 | 0.049448 | -0.03921 | -0.2114 | 0.074005 | -0.09397 | -0.27409 | -0.43742 | -0.11354 |
| 33.94444 | -0.97653 | -1.26276 | -0.66937 | 0.146894 | -0.73392 | -1.28449 | -0.08997 | -0.26253 | -0.26822 | -0.0468 | 0.621659 | 0.102048 | -0.02034 | -0.14641 | 0.104441 | -0.02242 | -0.30504 | -0.48003 | 0.033225 |
| 33.95833 | -0.6872 | -0.8428 | 0.00095 | 0.301563 | -0.1404 | -1.23324 | 0.129166 | 0.011711 | -0.00087 | -0.08483 | 0.578838 | 0.110393 | -0.02725 | -0.15484 | 0.093859 | 0.018113 | -0.3353 | -0.4666 | 0.07562 |
| 33.97222 | -0.39909 | -0.59409 | 0.407916 | -0.23094 | 0.184838 | -1.06278 | 0.091782 | -0.04698 | 0.189753 | -0.0356 | 0.246823 | 0.099903 | -0.04321 | -0.18689 | 0.064689 | 0.030968 | -0.31998 | -0.32931 | 0.027184 |
| 33.98611 | -0.59284 | -0.84429 | 0.247076 | -1.26133 | -0.72385 | -1.58708 | -0.19763 | -0.64432 | -0.23207 | -0.38641 | -0.69842 | 0.033397 | -0.18856 | -0.29641 | 0.001381 | -0.12641 | -0.58622 | -0.47221 | -0.22746 |
| 34 | -1.9083 | -1.80594 | -0.70394 | -2.54952 | -2.97181 | -3.00687 | -0.66745 | -1.62293 | -1.48054 | -1.40607 | -2.48009 | -0.11255 | -0.39064 | -0.45923 | -0.07988 | -0.46802 | -1.22219 | -1.24917 | -0.69706 |
| 34.01389 | -3.54458 | -2.58605 | -1.62642 | -2.73972 | -3.74217 | -3.74158 | -0.82112 | -2.14465 | -2.32003 | -2.32336 | -3.38096 | -0.24499 | -0.37576 | -0.61477 | -0.02524 | -0.6558 | -1.38994 | -1.88127 | -0.9236 |
| 34.02778 | -4.57625 | -2.5583 | -1.76304 | -1.40649 | -1.63567 | -2.87531 | -0.38344 | -1.4777 | -1.8921 | -1.89091 | -1.99049 | -0.26786 | -0.21363 | -0.79084 | 0.043302 | -0.51144 | -0.88479 | -1.83017 | -0.69752 |
| 34.04167 | -4.46086 | -2.5189 | -1.71056 | -0.07325 | -0.57432 | -2.27357 | -0.04447 | -0.57508 | -1.11828 | -0.86412 | -1.1161 | -0.11978 | -0.10602 | -0.5629 | 0.065893 | -0.26131 | -0.43706 | -1.4854 | -0.43175 |
| 34.05556 | -3.8251 | -1.98007 | -1.45899 | 0.767853 | -0.16902 | -1.68059 | 0.072075 | 0.260034 | -0.80205 | -0.01533 | -0.66249 | 0.047246 | -0.01539 | -0.19351 | 0.016922 | -0.08199 | -0.13182 | -0.81579 | -0.14846 |
| 34.06944 | -2.2528 | -0.94154 | -1.10849 | 0.885629 | -0.09752 | -0.87356 | 0.364258 | 0.741901 | -0.63927 | 0.520359 | -0.36017 | 0.201382 | 0.036653 | -0.02727 | -0.05782 | 0.052033 | 0.124339 | -0.11169 | 0.021068 |
| 34.08333 | -0.60677 | 0.243676 | -0.68671 | 0.547173 | 0.145142 | -0.01902 | 0.650818 | 0.703273 | -0.2897 | 0.853277 | -0.18771 | 0.282451 | 0.063761 | 0.039685 | -0.09575 | 0.140933 | 0.35139 | 0.449583 | 0.068234 |
| 34.09722 | 0.876531 | 1.136412 | -0.33327 | 0.196461 | 0.220991 | 0.570175 | 0.750762 | 0.608345 | 0.18515 | 0.705082 | -0.09741 | 0.314254 | 0.089048 | 0.102138 | -0.0763 | 0.175889 | 0.386212 | 0.792717 | 0.08022 |
| 34.11111 | 2.230722 | 1.480462 | -0.10153 | 0.085237 | -0.04184 | 0.712481 | 0.690774 | 0.564354 | 0.550834 | 0.402253 | -0.0287 | 0.247103 | 0.111775 | 0.175612 | -0.0735 | 0.166394 | 0.210051 | 0.939568 | 0.15349 |
| 34.125 | 3.05737 | 1.597483 | 0.11284 | 0.276431 | -0.01259 | 0.842434 | 0.610031 | 0.742719 | 0.981616 | 0.036533 | 0.008102 | 0.16582 | 0.102947 | 0.253524 | -0.08281 | 0.155022 | 0.114747 | 0.967287 | 0.201535 |
| 34.13889 | 2.878635 | 1.453043 | 0.335953 | 0.588643 | 0.436487 | 0.934248 | 0.684248 | 0.934039 | 1.560682 | -0.17604 | 0.103776 | 0.102938 | 0.057859 | 0.342007 | -0.06852 | 0.124354 | 0.158173 | 1.016546 | 0.097368 |
| 34.15278 | 2.154392 | 1.165815 | 0.663505 | 0.885627 | 0.735897 | 0.916355 | 1.017289 | 0.970222 | 2.179418 | -0.15723 | 0.220211 | 0.057953 | -0.00536 | 0.388448 | -0.04169 | 0.108716 | 0.204552 | 1.110388 | -0.0035 |
| 34.16667 | 1.53809 | 0.982358 | 1.119974 | 0.968652 | 1.063811 | 1.036519 | 1.231663 | 0.885837 | 2.90323 | 0.06799 | 0.287926 | 0.012769 | -0.03863 | 0.383402 | -0.05667 | 0.120338 | 0.163182 | 1.10733 | 0.063761 |
| 34.18056 | 1.063531 | 0.743386 | 1.458235 | 0.694347 | 1.306652 | 1.358964 | 1.024237 | 0.766752 | 2.576468 | 0.345463 | 0.364297 | -0.02105 | -0.00596 | 0.277913 | -0.07273 | 0.115621 | 0.112412 | 0.978279 | 0.13982 |
| 34.19444 | 0.853662 | 0.491346 | 1.601721 | 0.337574 | 1.090663 | 1.227687 | 0.557685 | 0.561639 | 1.941495 | 0.522037 | 0.52778 | -0.04202 | 0.029249 | 0.147095 | -0.03978 | 0.081362 | 0.132224 | 0.659177 | 0.075454 |
| 34.20833 | 0.632827 | 0.351086 | 1.39048 | 0.07167 | 0.679778 | 0.944545 | 0.297204 | 0.341868 | 1.151665 | 0.42672 | 0.682333 | -0.09839 | -0.01316 | -0.0162 | 0.00351 | -0.00074 | 0.166706 | 0.39377 | -0.06083 |
| 34.22222 | 0.390288 | 0.203701 | 1.061097 | -0.0066 | 0.309358 | 0.393411 | 0.162225 | 0.175151 | 0.354122 | 0.244848 | 0.634445 | -0.14369 | -0.07211 | -0.12379 | 0.001925 | -0.10358 | 0.199565 | 0.196446 | -0.13714 |
| 34.23611 | 0.105509 | 0.132736 | 0.821051 | 0.146605 | -0.01196 | -0.1212 | 0.009277 | 0.074378 | -0.35678 | 0.055788 | 0.536871 | -0.11746 | -0.02888 | -0.189 | -0.0183 | -0.13811 | 0.177768 | 0.042409 | -0.1423 |
| 34.25 | -0.01301 | 0.158814 | 0.703087 | 0.269179 | -0.24323 | -0.40587 | -0.0819 | -0.03045 | -0.94749 | -0.01984 | 0.402168 | -0.06945 | -0.03065 | -0.18806 | -0.03601 | -0.10517 | 0.12569 | -0.12145 | -0.19585 |
| 34.26389 | 0.2714 | 0.150398 | 0.624164 | 0.053397 | -0.42493 | -0.50538 | -0.19777 | -0.2798 | -1.31972 | 0.217711 | 0.438026 | -0.0369 | -0.06452 | -0.11444 | -0.00858 | -0.07665 | 0.160634 | -0.28556 | -0.31836 |
| 34.27778 | 0.700654 | 0.084574 | 0.621595 | -0.36578 | -0.57373 | -0.37432 | -0.27208 | -0.47547 | -0.86799 | 0.639186 | 0.66008 | 0.013953 | 0.014703 | 0.046831 | 0.070341 | -0.01386 | 0.338841 | -0.25869 | -0.29269 |
| 34.29167 | 0.965923 | 0.068068 | 0.713557 | -0.58402 | -0.50312 | -0.26819 | -0.22457 | -0.35256 | -0.03292 | 0.733126 | 0.881079 | 0.028528 | 0.123697 | 0.137258 | 0.121773 | 0.043965 | 0.487617 | 0.007074 | -0.10099 |
| 34.30556 | 0.862666 | 0.099363 | 0.741545 | -0.44514 | -0.25484 | -0.06873 | -0.01372 | -0.03368 | 0.750892 | 0.404214 | 0.831664 | -0.02626 | 0.095878 | 0.074199 | 0.080441 | 0.053906 | 0.424687 | 0.200242 | -0.02213 |
| 34.31944 | 0.577408 | 0.121002 | 0.643536 | -0.1279 | 0.020405 | 0.239812 | 0.171307 | 0.223649 | 1.128327 | 0.090987 | 0.70134 | -0.03322 | 0.040812 | 0.002426 | 0.038837 | 0.048828 | 0.269096 | 0.135024 | -0.07119 |
| 34.33333 | 0.339258 | 0.154123 | 0.538123 | 0.089045 | 0.210301 | 0.592812 | 0.218187 | 0.302699 | 1.156779 | 0.066446 | 0.515957 | 0.056022 | 0.068378 | -0.00659 | 0.06036 | 0.066591 | 0.116957 | -0.05348 | -0.0633 |
| 34.34722 | 0.157773 | 0.162697 | 0.459447 | 0.157988 | 0.277841 | 0.779371 | 0.260546 | 0.228939 | 0.784713 | 0.270252 | 0.445435 | 0.129704 | 0.112813 | -0.02223 | 0.064922 | 0.036302 | 0.028837 | -0.12547 | 0.004023 |
| 34.36111 | 0.007918 | 0.108611 | 0.397004 | 0.071662 | 0.254587 | 0.793238 | 0.269082 | 0.142607 | 0.30584 | 0.464232 | 0.443631 | 0.133089 | 0.114982 | -0.00974 | 0.064352 | 0.007314 | 0.006154 | -0.09996 | 0.027396 |
| 34.375 | 0.070639 | 0.078763 | 0.469438 | -0.10458 | 0.211235 | 0.878924 | 0.209261 | 0.127233 | 0.174398 | 0.43744 | 0.449779 | 0.073146 | 0.083237 | 0.077382 | 0.085573 | 0.053357 | 0.036519 | -0.08838 | 0.022179 |
| 34.38889 | 0.308344 | 0.116342 | 0.714194 | -0.2639 | 0.250279 | 1.085297 | 0.162101 | 0.193987 | 0.409676 | 0.269773 | 0.55078 | -0.01373 | 0.037115 | 0.129156 | 0.072275 | 0.114305 | 0.043646 | -0.07786 | 0.022888 |
| 34.40278 | 0.45113 | 0.188479 | 0.925824 | -0.25302 | 0.369433 | 1.197114 | 0.14805 | 0.33397 | 0.763947 | 0.159633 | 0.782492 | -0.06659 | 0.015831 | 0.081857 | 0.029299 | 0.086486 | 0.096707 | 0.013116 | 0.026424 |
| 34.41667 | 0.424104 | 0.247791 | 0.927087 | -0.06279 | 0.413152 | 0.8613 | 0.167286 | 0.514334 | 0.876998 | 0.144854 | 1.009228 | -0.03019 | 0.066252 | 0.065687 | 0.017071 | 0.056497 | 0.270402 | 0.175661 | 0.123336 |
| 34.43056 | 0.335828 | 0.224796 | 0.777508 | 0.21746 | 0.291904 | 0.247977 | 0.133038 | 0.609193 | 0.668231 | 0.259353 | 1.064504 | 0.077025 | 0.140028 | 0.119201 | 0.059361 | 0.072925 | 0.494707 | 0.290993 | 0.269218 |
| 34.44444 | 0.138914 | 0.054502 | 0.599699 | 0.420806 | 0.104357 | -0.20335 | 0.029694 | 0.569348 | 0.286611 | 0.480906 | 1.265591 | 0.138774 | 0.141877 | 0.117939 | 0.078372 | 0.069662 | 0.580966 | 0.289046 | 0.283096 |
| 34.45833 | -0.19728 | -0.15006 | 0.458056 | 0.248979 | -0.09338 | -0.48878 | -0.11703 | 0.301237 | -0.04182 | 0.599811 | 1.301826 | 0.089495 | 0.090532 | 0.059348 | 0.051197 | 0.032772 | 0.52064 | 0.181431 | 0.189771 |
| 34.47222 | -0.44862 | -0.22214 | 0.549366 | 0.031175 | -0.08017 | -0.3234 | -0.1679 | 0.063752 | -0.12926 | 0.611509 | 1.260318 | 0.0095 | 0.065429 | 0.011929 | 0.004501 | 0.048972 | 0.410296 | 0.098044 | 0.127014 |
| 34.48611 | -0.15492 | -0.03759 | 1.093169 | 0.000806 | 0.413293 | 0.951923 | -0.14578 | 0.032954 | 0.26119 | 0.587997 | 1.541505 | -0.02101 | 0.093621 | -0.03368 | -0.05347 | 0.187043 | 0.557786 | 0.264653 | 0.281994 |
| 34.5 | 0.475082 | 0.26202 | 1.898354 | -0.01265 | 1.089709 | 3.329741 | -0.19275 | 0.200692 | 1.189703 | 0.678976 | 2.447086 | 0.035798 | 0.176437 | -0.0308 | -0.11265 | 0.430179 | 0.951028 | 0.733951 | 0.722335 |
| 34.51389 | 1.069847 | 0.318336 | 2.185648 | 0.139129 | 1.304057 | 4.941929 | -0.20889 | 0.535348 | 2.091028 | 0.977283 | 2.99405 | 0.292213 | 0.352213 | 0.082763 | -0.12786 | 0.775167 | 1.432389 | 1.133624 | 1.205493 |
| 34.52778 | 1.455393 | 0.379938 | 2.100345 | 0.877185 | 1.457893 | 4.351204 | 0.230268 | 0.99442 | 3.048871 | 1.365222 | 2.278818 | 0.59232 | 0.474921 | 0.520188 | -0.05644 | 1.246139 | 1.561949 | 1.237147 | 1.325962 |
| 34.54167 | 1.85223 | 0.654214 | 2.557204 | 1.182955 | 1.922882 | 3.58596 | 0.476201 | 1.303527 | 3.263225 | 1.138931 | 1.879479 | 0.596218 | 0.496509 | 1.14618 | -0.01142 | 1.273707 | 1.897093 | 1.091284 | 1.056928 |
| 34.55556 | 1.712939 | 0.815332 | 2.508659 | 1.114423 | 2.227584 | 3.770781 | 0.673809 | 1.207191 | 2.972627 | 0.463234 | 1.657821 | 0.296105 | 0.356412 | 1.017009 | -0.00643 | 0.914125 | 1.820991 | 0.833485 | 0.532709 |
| 34.56944 | 1.373989 | 0.846535 | 1.915212 | 1.050493 | 1.807892 | 2.665899 | 0.612762 | 0.855806 | 1.949896 | -0.1445 | 1.319146 | -0.02777 | 0.130391 | 0.851887 | -0.02597 | 0.584133 | 1.329627 | 0.626685 | 0.264606 |
| 34.58333 | 1.076059 | 0.772372 | 1.330418 | 0.997122 | 1.180856 | 1.202645 | 0.528235 | 0.440158 | 0.552397 | -0.58672 | 0.937317 | -0.19449 | -0.03022 | 0.654469 | -0.03485 | 0.405936 | 0.762468 | 0.553749 | 0.250744 |
| 34.59722 | 0.798001 | 0.778453 | 0.89008 | 1.188362 | 0.707759 | 0.17473 | 0.438528 | -0.08671 | -0.74315 | -0.85614 | 0.244206 | -0.27182 | -0.02129 | 0.517121 | -0.03583 | 0.268781 | 0.159271 | 0.478812 | 0.272295 |
| 34.61111 | 0.195915 | 0.700079 | 0.54751 | 1.974919 | 0.508811 | 0.036906 | 0.486135 | -0.31968 | -1.03809 | -0.78657 | -0.61279 | -0.263 | 0.055276 | 0.442077 | -0.01312 | 0.219776 | -0.38635 | 0.38518 | 0.286883 |
| 34.625 | -0.87395 | 0.580137 | 0.17953 | 2.272256 | 0.406949 | -0.25948 | 0.52537 | -0.23364 | -1.24716 | -0.35218 | -1.34323 | -0.18484 | 0.134516 | 0.374256 | 0.028434 | 0.13273 | -0.87139 | 0.265486 | 0.201089 |
| 34.63889 | -1.13739 | 0.553707 | -0.15239 | 1.864233 | 0.195688 | 0.13515 | 0.324481 | 0.04511 | -1.11382 | 0.107015 | -2.01298 | -0.12877 | 0.118927 | 0.296327 | 0.033036 | 0.024789 | -1.22415 | 0.122142 | 0.08668 |
| 34.65278 | -0.16499 | 0.704926 | -0.46449 | 0.872949 | 0.441501 | -0.12599 | 0.141038 | 0.569397 | -0.68841 | 0.466649 | -1.92789 | -0.08436 | 0.019425 | 0.243879 | 0.02338 | -0.08676 | -1.37088 | -0.12586 | 0.053881 |
| 34.66667 | 1.076995 | 0.893793 | -0.37655 | -0.61702 | 0.190487 | -0.13592 | 0.005514 | 0.790977 | -0.34494 | 0.633873 | -2.08431 | -0.0768 | -0.08146 | 0.218429 | 0.0158 | -0.19411 | -1.40529 | -0.44408 | 0.078368 |
| 34.68056 | 1.667858 | 0.77945 | 0.024868 | -2.0439 | -0.0054 | -0.04807 | 0.110001 | 0.564959 | 0.173034 | 0.560058 | -1.53569 | -0.0728 | -0.0213 | 0.281149 | 0.012914 | -0.22974 | -0.87929 | -0.64973 | 0.186578 |

| | | | | | | | | | | | | | | | | | | | |
|---|---|---|---|---|---|---|---|---|---|---|---|---|---|---|---|---|---|---|---|
| 34.69444 | 1.562948 | 0.601338 | 0.289128 | -2.55717 | -0.00778 | -0.57836 | 0.261743 | 0.355581 | 0.832785 | 0.354722 | -0.49204 | -0.03802 | 0.127851 | 0.467359 | 0.03714 | -0.11636 | -0.17943 | -0.68228 | 0.288293 |
| 34.70833 | 1.03563 | 0.417409 | 0.651784 | -2.37318 | 0.372444 | -0.65955 | 0.470323 | 0.219033 | 1.44055 | 0.112316 | 0.44134 | -0.04651 | 0.171035 | 0.688538 | 0.043109 | -0.01045 | 0.293165 | -0.51582 | 0.207779 |
| 34.72222 | 0.198984 | 0.059047 | 1.065656 | -1.3128 | 0.600435 | -0.54465 | 0.611539 | 0.080613 | 1.682205 | 0.008993 | 1.105769 | -0.09348 | 0.084269 | 0.768833 | 0.035391 | 0.054265 | 0.543326 | -0.39576 | 0.031545 |
| 34.73611 | -0.82287 | -0.30862 | 0.796686 | -1.15121 | 0.499593 | -0.36514 | 0.644684 | -0.39493 | 1.516487 | -0.14561 | 1.473729 | -0.12932 | -0.05184 | 0.627398 | 0.03488 | 0.093581 | 0.732594 | -0.2573 | -0.05027 |
| 34.75 | -1.12813 | -0.51686 | -0.01454 | -0.72166 | 0.496447 | -0.52445 | 0.500268 | -0.33998 | 1.190783 | -0.30226 | 1.411993 | -0.16361 | -0.11181 | 0.429779 | 0.037355 | 0.083872 | 0.873635 | -0.14516 | -0.06472 |
| 34.76389 | -0.54352 | -0.62483 | -0.83355 | -0.198 | 0.724882 | -0.95355 | 0.263648 | 0.131607 | 0.728001 | -0.4174 | 0.896604 | -0.1856 | -0.03862 | 0.301828 | 0.045058 | 0.141391 | 0.848586 | -0.00993 | -0.04395 |
| 34.77778 | -0.34994 | -0.61816 | -1.57432 | 0.160787 | 0.879485 | -1.42387 | 0.000438 | 0.48141 | 0.290067 | -0.49435 | -0.08767 | -0.17629 | 0.06476 | 0.226244 | 0.047527 | 0.201118 | 0.587821 | 0.056148 | -0.03076 |
| 34.79167 | -0.6392 | -0.27657 | -3.04333 | 0.436663 | 0.694276 | -1.63397 | -0.15201 | 0.526465 | -0.25886 | -0.50515 | -1.25938 | -0.14951 | 0.06256 | 0.119605 | 0.046438 | 0.254907 | 0.000595 | 0.119027 | -0.10592 |
| 34.80556 | -0.07795 | 0.221725 | -3.00581 | 0.521476 | 0.153106 | -1.46296 | -0.10113 | 0.169708 | -0.76074 | -0.409 | -2.34312 | -0.09497 | 0.020865 | -0.1139 | 0.083044 | 0.235539 | -0.78817 | 0.213996 | -0.19269 |
| 34.81944 | 0.470363 | 0.353528 | -1.45481 | 0.498616 | -0.29639 | -1.3554 | 0.015912 | -0.32273 | -1.10242 | -0.29818 | -2.71951 | -0.03151 | -0.03105 | -0.4827 | 0.093222 | 0.032835 | -1.62467 | 0.401541 | -0.20582 |
| 34.83333 | 0.430524 | 0.139556 | -0.3657 | 0.274269 | -0.632 | -1.595 | 0.035195 | -0.84818 | -1.21763 | -0.25885 | -2.37307 | 0.004138 | -0.05563 | -0.95875 | 0.087806 | -0.21337 | -2.20357 | 0.509567 | -0.17945 |
| 34.84722 | 0.49058 | 0.129431 | 0.090297 | -0.13082 | -0.86189 | -2.06081 | 0.088253 | -1.47555 | -1.16182 | -0.15197 | -2.12436 | 0.021792 | -0.05099 | -1.35007 | 0.065514 | -0.37818 | -2.14956 | 0.443799 | -0.16583 |
| 34.86111 | 0.038956 | 0.132204 | -0.2896 | -0.52755 | -0.83917 | -2.29606 | 0.251177 | -1.72525 | -1.11182 | -0.09158 | -1.74166 | 0.029458 | 0.038823 | -1.50371 | 0.065935 | -0.41379 | -1.86141 | 0.137623 | -0.09871 |
| 34.875 | -0.87013 | 0.050269 | -1.17729 | -0.45082 | -0.95892 | -2.1542 | 0.356999 | -1.56362 | -1.36182 | -0.18665 | -0.96107 | 0.070174 | 0.145249 | -1.3005 | 0.111168 | -0.34888 | -1.35947 | -0.22571 | 0.010336 |
| 34.88889 | -1.16672 | 0.085779 | -2.09075 | -0.14737 | -1.03457 | -1.93116 | 0.428175 | -0.89638 | -1.28411 | -0.20951 | -0.37387 | 0.120968 | 0.171666 | -0.67902 | 0.101824 | -0.37254 | -0.83206 | -0.44735 | 0.087813 |
| 34.90278 | -1.48739 | 0.299996 | -2.40554 | -0.03386 | -1.17992 | -1.76571 | 0.487913 | -0.05114 | -1.01013 | -0.27898 | 0.114152 | 0.132675 | 0.125479 | -0.30368 | 0.030522 | -0.37718 | -0.46303 | -0.61253 | 0.024179 |
| 34.91667 | -1.51788 | 0.456858 | -2.19063 | -0.11517 | -1.16256 | -1.69135 | 0.535605 | 0.367414 | -0.73504 | -0.20569 | 0.447656 | 0.130981 | 0.084131 | -0.09742 | 0.002425 | -0.23958 | -0.22134 | -0.70895 | -0.0393 |
| 34.93056 | -1.09601 | 0.453246 | -1.81984 | -0.11967 | -0.91039 | -0.92373 | 0.64812 | 0.562351 | -0.38388 | -0.09756 | 0.586983 | 0.123551 | 0.087754 | 0.050186 | -0.0026 | -0.03031 | 0.044 | -0.64038 | -0.04933 |
| 34.94444 | -0.53535 | 0.219481 | -1.35136 | -0.03665 | -0.38831 | -0.0488 | 0.729548 | 0.554853 | -0.02478 | 0.052092 | 0.64128 | 0.172052 | 0.130924 | 0.100579 | 0.004853 | 0.104168 | 0.27952 | -0.44773 | -0.00757 |
| 34.95833 | 0.114749 | -0.01029 | -0.58482 | -0.05529 | 0.088288 | 0.638537 | 0.871157 | 0.371271 | 0.432526 | 0.27713 | 0.639531 | 0.238562 | 0.159511 | 0.063521 | 0.043761 | 0.171866 | 0.456879 | -0.29647 | 0.094266 |
| 34.97222 | 0.670407 | -0.30262 | 0.29069 | -0.19832 | 0.472451 | 1.119718 | 0.94958 | 0.177697 | 0.999809 | 0.390732 | 0.514764 | 0.227693 | 0.124685 | 0.009452 | 0.083406 | 0.184614 | 0.481191 | -0.33392 | 0.151845 |
| 34.98611 | 0.727422 | -0.82832 | 0.935759 | -0.43852 | 0.472289 | 1.165143 | 1.122033 | 0.306105 | 1.598412 | 0.088116 | -0.25914 | 0.12318 | 0.01319 | -0.06358 | 0.103593 | 0.011918 | 0.066737 | -0.73081 | -0.06192 |
| 35 | -0.18235 | -1.52417 | 0.770079 | -0.82076 | -0.29727 | 0.398478 | 1.403004 | 0.141867 | 1.637646 | -1.0071 | -1.93166 | -0.07407 | -0.18372 | -0.20011 | 0.141805 | -0.36533 | -0.88877 | -1.51829 | -0.64829 |
| 35.01389 | -1.85714 | -3.01527 | -0.26112 | -1.83019 | -1.16991 | -0.30077 | 1.116094 | -1.18762 | 0.902585 | -2.09947 | -2.56193 | -0.32787 | -0.46989 | -0.2998 | 0.206772 | -0.64901 | -1.64819 | -1.90193 | -1.07473 |
| 35.02778 | -3.06314 | -3.67235 | -1.25869 | -1.39812 | -1.46332 | -0.70779 | 0.485274 | -1.17298 | 0.301551 | -1.82239 | -1.62592 | -0.40923 | -0.69092 | -0.39095 | 0.231914 | -0.62326 | -1.45694 | -1.59571 | -1.17558 |
| 35.04167 | -2.09801 | -3.17702 | -1.71736 | -0.71156 | -1.46622 | -0.33825 | -0.03656 | -0.56171 | -0.03459 | -0.79626 | -1.06299 | -0.26156 | -0.8109 | -0.42427 | 0.216572 | -0.44271 | -0.83534 | -1.17369 | -0.69834 |
| 35.05556 | -1.90833 | -1.93676 | -1.83361 | -0.46046 | -1.46965 | 0.090293 | 0.0772 | -0.31759 | 0.320601 | 0.209913 | -0.86964 | 0.070346 | -0.38703 | -0.24631 | 0.080297 | -0.19204 | -0.32389 | -0.5535 | -0.15734 |
| 35.06944 | -1.34796 | -1.07275 | -1.5317 | -0.27071 | -0.85445 | 0.462027 | 0.89106 | 0.007189 | 1.066954 | 1.04425 | -0.35987 | 0.414375 | 0.271121 | -0.03151 | -0.13367 | 0.099212 | 0.116719 | 0.183622 | 0.40933 |
| 35.08333 | -0.45276 | -0.06338 | -0.90656 | 0.533171 | 0.110804 | 0.714097 | 1.28942 | 0.718765 | -0.05336 | 1.597578 | 0.196538 | 0.662199 | 0.752706 | 0.089043 | -0.1802 | 0.318519 | 0.483608 | 0.756377 | 0.689979 |
| 35.09722 | 0.552933 | 1.241646 | -0.24909 | 1.2773 | -0.38567 | 0.764338 | 0.539644 | 1.750017 | -0.30617 | 1.328411 | 0.423565 | 0.755714 | 0.745413 | 0.095615 | -0.14947 | 0.412748 | 0.624076 | 1.012163 | 0.492701 |
| 35.11111 | 1.362088 | 2.249809 | 0.209728 | 0.960478 | -0.64709 | 0.729499 | -0.00616 | 1.498882 | 0.573123 | 0.845536 | 0.344761 | 0.717025 | 0.513586 | 0.074862 | -0.06752 | 0.389721 | 0.5114 | 1.050578 | 0.203072 |
| 35.125 | 1.843205 | 2.297511 | 0.531647 | 0.387503 | -0.44847 | 0.65355 | 0.034667 | 1.036232 | 1.000608 | 0.520758 | 0.203528 | 0.515282 | 0.34725 | 0.049635 | -0.07159 | 0.30851 | 0.266802 | 0.962763 | 0.071847 |
| 35.13889 | 2.111004 | 2.58206 | 0.85085 | 0.164116 | -0.0429 | 0.592993 | 0.418425 | 1.084465 | 1.272379 | 0.081516 | 0.118316 | 0.274832 | 0.209078 | 0.049343 | -0.11522 | 0.215215 | 0.132081 | 0.733005 | 0.079288 |
| 35.15278 | 2.162989 | 2.71448 | 1.075736 | 0.100125 | 0.307276 | 0.479882 | 0.711462 | 1.355578 | 1.846981 | -0.29582 | 0.125286 | 0.106838 | 0.18061 | 0.074106 | -0.10755 | 0.160765 | 0.166944 | 0.618255 | 0.131937 |
| 35.16667 | 1.911181 | 2.876076 | 1.253653 | 0.061568 | 0.395524 | 0.33518 | 0.699932 | 1.652472 | 1.25982 | -0.36104 | 0.143508 | 0.102774 | 0.229716 | 0.133587 | -0.08399 | 0.185127 | 0.248423 | 0.632079 | 0.167838 |
| 35.18056 | 2.449785 | 2.787114 | 1.58182 | 0.334151 | 0.35912 | 0.399605 | 0.661542 | 2.124272 | 0.674636 | -0.07826 | 0.057887 | 0.178308 | 0.187527 | 0.12828 | -0.08624 | 0.188937 | 0.208123 | 0.576027 | 0.13229 |
| 35.19444 | 2.830241 | 2.420144 | 1.684378 | 0.691579 | 0.421534 | 0.648182 | 0.631517 | 2.177902 | 1.917556 | 0.263652 | -0.07382 | 0.227453 | 0.030299 | 0.065905 | -0.07366 | 0.122947 | 0.148067 | 0.477967 | 0.058204 |
| 35.20833 | 3.153942 | 1.910396 | 1.536879 | 0.775808 | 0.619868 | 0.860842 | 0.593388 | 1.57963 | 2.257644 | 0.556786 | -0.13842 | 0.181238 | -0.07139 | 0.016553 | -0.01226 | 0.019264 | 0.146029 | 0.445207 | -0.00163 |
| 35.22222 | 2.243805 | 1.35882 | 1.115471 | 0.587438 | 0.880188 | 1.055231 | 0.468208 | 0.845294 | 1.80459 | 0.43904 | -0.10934 | -0.01482 | -0.08691 | -0.006 | 0.014259 | -0.12219 | 0.089175 | 0.332705 | -0.15865 |
| 35.23611 | 1.068373 | 0.527699 | 0.700742 | 0.225502 | 0.560949 | 1.07642 | 0.364191 | 0.181436 | 0.822496 | 0.02425 | 0.018343 | -0.26894 | -0.11574 | -0.05322 | -0.02977 | -0.18817 | 0.016127 | 0.14166 | -0.33672 |
| 35.25 | 0.352573 | -0.11109 | 0.531657 | -0.19369 | 0.115228 | 0.690004 | 0.170418 | -0.3053 | -0.31725 | -0.3893 | 0.183625 | -0.37979 | -0.18611 | -0.11552 | -0.06449 | -0.17869 | 0.014264 | -0.07488 | -0.33156 |
| 35.26389 | 0.210364 | -0.26389 | 0.627482 | -0.39965 | -0.06483 | 0.595396 | 0.065291 | -0.4865 | -1.02366 | -0.46714 | 0.294923 | -0.31592 | -0.20581 | -0.15766 | -0.0543 | -0.0565 | 0.087961 | -0.1243 | -0.2317 |
| 35.27778 | 0.287009 | -0.0429 | 0.752586 | -0.30532 | -0.09302 | 0.784109 | 0.057407 | -0.39191 | -1.08051 | 0.095194 | 0.355655 | -0.14952 | -0.18253 | -0.12656 | 0.014144 | 0.074608 | 0.285962 | -0.02986 | -0.17838 |
| 35.29167 | 0.30122 | 0.116385 | 0.746025 | -0.25884 | -0.09595 | 0.986067 | -0.01229 | -0.37739 | -0.75422 | 0.793114 | 0.371104 | -0.0171 | -0.12786 | -0.04851 | 0.061968 | 0.08699 | 0.509111 | 0.086266 | -0.05929 |
| 35.30556 | 0.053878 | -0.01861 | 0.653893 | -0.38193 | -0.07645 | 1.063121 | -0.142 | -0.65629 | -0.25658 | 1.177652 | 0.376115 | 0.088052 | -0.07408 | -0.00265 | 0.057378 | 0.016251 | 0.508292 | 0.133589 | 0.049904 |
| 35.31944 | -0.3477 | -0.30088 | 0.562506 | -0.56533 | 0.012582 | 0.772714 | -0.12088 | -0.95821 | 0.213548 | 1.052937 | 0.484415 | 0.133501 | 0.002522 | -0.02028 | 0.02566 | -0.05288 | 0.381414 | 0.153559 | 0.074968 |
| 35.33333 | -0.50592 | -0.54951 | 0.447996 | -0.76373 | 0.063032 | 0.17469 | -0.01663 | -1.08717 | 0.341001 | 0.560158 | 0.514498 | 0.096746 | 0.085023 | -0.06739 | -0.03667 | -0.11129 | 0.18011 | 0.127302 | 0.064203 |
| 35.34722 | -0.35449 | -0.72896 | 0.256152 | -0.86151 | 0.039712 | -0.14419 | -0.03943 | -1.0866 | 0.170909 | 0.019658 | 0.448738 | 0.014363 | 0.125436 | -0.08533 | -0.07073 | -0.09732 | -0.01613 | 0.04968 | 0.106004 |
| 35.36111 | 0.207941 | -0.69588 | 0.057402 | -0.62553 | 0.02486 | -0.02645 | -0.06531 | -0.77332 | -0.04885 | -0.30629 | 0.474787 | -0.0098 | 0.133012 | -0.03419 | 0.021578 | 0.017903 | -0.02608 | -0.0058 | 0.237617 |
| 35.375 | 0.868062 | -0.32618 | 0.089829 | -0.09095 | 0.084282 | 0.333547 | 0.060666 | -0.03923 | 0.0403 | -0.13775 | 0.466488 | 0.043832 | 0.140225 | 0.013906 | 0.110246 | 0.12242 | 0.002946 | -0.01032 | 0.361619 |
| 35.38889 | 1.389452 | 0.118875 | 0.392032 | 0.115719 | 0.169898 | 0.592457 | 0.248987 | 0.542625 | 0.551813 | 0.334298 | 0.529906 | 0.176909 | 0.179488 | 0.082672 | 0.114497 | 0.163041 | -0.0123 | 0.02933 | 0.406847 |
| 35.40278 | 1.524392 | 0.313852 | 0.5694 | -0.21467 | 0.185978 | 0.558477 | 0.290246 | 0.496204 | 1.144032 | 0.888475 | 0.669012 | 0.31036 | 0.244477 | 0.134718 | 0.122771 | 0.163704 | 0.028381 | 0.118168 | 0.336613 |
| 35.41667 | 1.323452 | 0.454967 | 0.582454 | -0.57725 | 0.134568 | 0.38108 | 0.183273 | 0.309058 | 1.679728 | 1.043577 | 0.606358 | 0.265996 | 0.247449 | 0.109388 | 0.049062 | 0.102471 | 0.054155 | 0.148163 | 0.148114 |
| 35.43056 | 1.107877 | 0.77935 | 0.637836 | -0.55324 | 0.090478 | 0.408731 | 0.167312 | 0.321609 | 2.002907 | 0.585714 | 0.531359 | 0.126925 | 0.099995 | 0.092062 | -0.06209 | 0.020889 | 0.057469 | 0.124397 | 0.003002 |
| 35.44444 | 0.9045 | 1.018416 | 0.8526 | -0.31504 | 0.116256 | 0.64535 | 0.226841 | 0.449472 | 1.649927 | 0.103105 | 0.539754 | 0.052474 | 0.015071 | 0.129244 | -0.04192 | 0.019086 | 0.09538 | 0.140532 | 0.002368 |
| 35.45833 | 0.366529 | 0.96004 | 1.003531 | -0.06648 | 0.218599 | 0.962749 | 0.253396 | 0.517856 | 1.083987 | -0.11684 | 0.724286 | 0.032841 | 0.106462 | 0.180732 | 0.066762 | 0.097839 | 0.16242 | 0.19121 | 0.074653 |
| 35.47222 | -0.16409 | 0.682846 | 0.91235 | 0.234374 | 0.273417 | 1.03019 | 0.227031 | 0.607861 | 0.504083 | -0.12369 | 0.97231 | 0.035122 | 0.203699 | 0.183705 | 0.093838 | 0.151071 | 0.249271 | 0.268336 | 0.108173 |
| 35.48611 | 0.118997 | 0.451777 | 0.82232 | 0.719265 | 0.289842 | 1.191097 | 0.019901 | 0.728803 | 0.052312 | 0.291505 | 1.206258 | 0.07495 | 0.19667 | 0.16832 | -0.00202 | 0.167117 | 0.53325 | 0.443074 | 0.150392 |
| 35.5 | 1.036667 | 0.323652 | 1.107057 | 1.450854 | 0.335075 | 1.862379 | -0.37614 | 0.991497 | 0.080067 | 1.087793 | 1.713946 | 0.259437 | 0.247391 | 0.191815 | -0.09255 | 0.266404 | 1.202335 | 0.863981 | 0.376263 |
| 35.51389 | 2.06752 | 0.19853 | 1.556955 | 1.505597 | 0.341711 | 2.813564 | -0.63613 | 1.618028 | 0.711072 | 1.469779 | 2.185461 | 0.660932 | 0.398896 | 0.273424 | -0.069 | 0.56093 | 1.624502 | 1.199762 | 0.707502 |
| 35.52778 | 3.162028 | 0.201195 | 1.486974 | 1.672523 | 0.106421 | 2.552997 | -0.37861 | 3.233843 | 1.751171 | 1.349714 | 2.950457 | 0.962332 | 0.363125 | 0.47011 | 0.091653 | 0.917217 | 1.72556 | 1.11943 | 0.934884 |
| 35.54167 | 3.941124 | 0.64135 | 1.709506 | 1.940782 | 0.084902 | 2.116387 | 0.047309 | 4.390333 | 2.83441 | 1.099794 | 2.800261 | 0.765608 | 0.176563 | 0.581845 | 0.085525 | 1.024864 | 2.021813 | 0.973129 | 0.922064 |
| 35.55556 | 2.879259 | 0.545458 | 2.363813 | 0.95807 | 0.429358 | 1.804166 | 0.230335 | 2.373661 | 3.036672 | 0.651364 | 2.434668 | 0.378318 | 0.097279 | 0.493895 | 0.017294 | 0.716163 | 1.841201 | 0.818491 | 0.52027 |
| 35.56944 | 1.061274 | 0.334804 | 1.745583 | 0.412276 | 0.793009 | 1.46269 | 0.267477 | 0.841645 | 2.227136 | 0.174213 | 2.019323 | -0.08111 | 0.021874 | 0.359928 | 0.021871 | 0.379963 | 1.587044 | 0.745051 | 0.044932 |
| 35.58333 | -0.27692 | 0.08266 | 0.955281 | 0.037743 | 0.953352 | 1.053393 | 0.114711 | -0.05904 | 1.320265 | -0.35413 | 1.536868 | -0.48246 | 0.004735 | 0.312892 | 0.123241 | 0.157826 | 1.451544 | 0.820731 | -0.3164 |
| 35.59722 | -1.29995 | -0.24505 | 0.326238 | -0.19789 | 1.064495 | 0.378808 | 0.025401 | -0.75401 | 0.483671 | -0.71438 | 0.822336 | -0.65123 | -0.14173 | 0.286173 | 0.236725 | 0.038515 | 1.229449 | 0.920015 | -0.36215 |
| 35.61111 | -1.69035 | -0.27058 | 0.029524 | -0.34274 | 1.274517 | -0.69508 | -0.0517 | -1.18809 | -0.26951 | -0.90122 | 0.063397 | -0.55665 | -0.20022 | 0.259932 | 0.170319 | 0.011252 | 0.813236 | 0.768347 | -0.10877 |
| 35.625 | -1.633 | -0.6048 | -0.15725 | -0.65432 | 1.458649 | -1.47191 | 0.002836 | -1.69789 | -0.94802 | -0.82517 | -0.79448 | -0.42363 | -0.14299 | 0.185072 | 0.003111 | -0.00867 | 0.259843 | 0.534614 | 0.039446 |
| 35.63889 | -1.71054 | -0.50995 | -0.36739 | -0.94125 | 1.596948 | -1.30407 | 0.058416 | -2.03212 | -0.89608 | -0.67581 | -1.81335 | -0.31263 | 0.026462 | 0.082256 | -0.06858 | -0.03343 | -0.44802 | 0.296093 | 0.136284 |
| 35.65278 | -0.64776 | 0.049523 | -0.51827 | -1.30959 | 1.611175 | -0.67542 | 0.104759 | -1.98229 | -0.54984 | -0.24779 | -2.46677 | -0.23613 | 0.013223 | 0.088586 | 0.013977 | 0.112437 | -1.16555 | 0.216346 | 0.10225 |
| 35.66667 | 0.788288 | 0.778002 | -0.5478 | -1.16715 | 1.390969 | -0.29931 | 0.038218 | -1.01909 | -0.31113 | 0.198274 | -2.64473 | -0.22018 | -0.2133 | 0.171896 | 0.137209 | 0.244216 | -1.64889 | 0.19854 | -0.04224 |
| 35.68056 | 1.690826 | 1.110172 | -0.43395 | -0.88411 | 1.003965 | -0.02153 | -0.06069 | -0.01856 | -0.39654 | 0.469728 | -2.8226 | -0.1904 | -0.42325 | 0.287636 | 0.16169 | 0.247289 | -1.7042 | 0.171915 | -0.21038 |
| 35.69444 | 2.311233 | 1.183168 | 0.001583 | -0.90622 | 0.673984 | 0.113212 | -0.13657 | 0.4287 | -0.36763 | 0.639223 | -2.24584 | -0.06237 | -0.39144 | 0.385163 | 0.105799 | 0.173012 | -1.60787 | 0.114189 | -0.2761 |
| 35.70833 | 2.233239 | 0.891868 | 0.621019 | -0.72128 | 0.524006 | 0.441124 | -0.07775 | 0.482276 | 0.031244 | 0.627034 | -1.76839 | 0.086509 | -0.18319 | 0.427362 | 0.043308 | 0.058367 | -1.48272 | -0.00906 | -0.16508 |
| 35.72222 | 1.015903 | 0.34939 | 0.890094 | -0.48004 | 0.578969 | 0.833131 | 0.071436 | 0.324894 | 0.645686 | 0.41602 | -1.37206 | 0.079705 | -0.02985 | 0.426113 | 0.036209 | -0.04841 | -1.1524 | -0.19069 | -0.14133 |
| 35.73611 | 0.010585 | -0.28252 | 1.285903 | -0.39045 | 0.616367 | 1.170796 | 0.219066 | -0.10756 | 1.157609 | 0.188519 | -0.87941 | -0.0625 | -0.04817 | 0.351883 | 0.062453 | -0.13315 | -0.7676 | -0.36055 | -0.27654 |
| 35.75 | -1.36262 | -0.94155 | 1.275385 | -0.784 | 0.676861 | 1.186172 | 0.264991 | -1.60322 | 1.505062 | -0.04592 | -0.30955 | -0.19748 | -0.14071 | 0.221375 | 0.098438 | -0.17006 | -0.53101 | -0.49658 | -0.38041 |
| 35.76389 | -2.24444 | -1.46637 | 0.710658 | -1.7462 | 0.722607 | 1.176722 | 0.278479 | -3.84422 | 1.587373 | -0.38401 | 0.385015 | -0.25968 | -0.25622 | 0.097154 | 0.137118 | -0.16219 | -0.28328 | -0.57315 | -0.38806 |
| 35.77778 | -2.79005 | -1.71199 | 0.255692 | -1.43719 | 0.71069 | 1.044719 | 0.25228 | -2.22791 | 1.299538 | -0.61826 | 0.904348 | -0.21756 | -0.28804 | -0.02686 | 0.170521 | -0.10711 | -0.00065 | -0.69782 | -0.1279 |
| 35.79167 | -3.15303 | -1.63658 | -0.33548 | -0.80743 | 0.675322 | 0.690406 | 0.124814 | -2.18205 | 0.757538 | -0.77762 | 0.864415 | -0.17602 | -0.23369 | -0.08083 | 0.142883 | -0.06951 | 0.198294 | -0.68792 | 0.048493 |
| 35.80556 | -2.5477 | -1.5962 | -0.93049 | -1.31492 | 0.669149 | 0.415621 | -0.02993 | -2.97883 | -0.05409 | -0.8228 | 0.337295 | -0.16106 | -0.15074 | -0.08212 | 0.041371 | -0.12186 | 0.20053 | -0.62464 | -0.04483 |
| 35.81944 | -2.02638 | -1.08567 | -1.21952 | -1.82673 | 0.637526 | 0.342881 | -0.1246 | -3.349 | -0.7335 | -0.65901 | -0.36868 | -0.15693 | -0.09421 | -0.01535 | 0.012571 | -0.25113 | -0.00402 | -0.56302 | -0.21189 |
| 35.83333 | -1.03937 | -0.43458 | -1.47438 | -1.9444 | 0.482257 | -0.0474 | -0.11491 | -2.59389 | -1.31371 | -0.4508 | -1.07731 | -0.12516 | -0.07342 | 0.088899 | 0.087461 | -0.37983 | -0.41479 | -0.50129 | -0.35848 |
| 35.84722 | -0.59729 | -0.41582 | -1.49772 | -1.93825 | 0.20473 | -0.38541 | -0.07195 | -2.06749 | -1.69566 | -0.37206 | -1.63038 | -0.04057 | -0.06814 | 0.189943 | 0.195179 | -0.41296 | -0.87148 | -0.3695 | -0.39847 |
| 35.86111 | -0.55141 | -0.88887 | -1.70438 | -1.96516 | -0.09753 | -0.60738 | -0.04457 | -2.07273 | -1.73913 | -0.34542 | -1.79305 | 0.01587 | -0.00697 | 0.243353 | 0.197769 | -0.28735 | -1.16462 | -0.25716 | -0.33465 |
| 35.875 | -0.64754 | -1.46051 | -1.63503 | -1.90295 | -0.36115 | -0.73133 | 0.000237 | -2.42253 | -1.55043 | -0.19982 | -1.62393 | 0.039998 | 0.144086 | 0.188027 | 0.138534 | -0.06667 | -1.26852 | -0.17716 | -0.15925 |

| | | | | | | | | | | | | | | | | | | | |
|---|---|---|---|---|---|---|---|---|---|---|---|---|---|---|---|---|---|---|---|
| 35.88889 | -0.81881 | -1.62017 | -1.43689 | -1.42754 | -0.61609 | -0.74584 | 0.125972 | -3.09075 | -1.349 | 0.117808 | -1.25832 | 0.089695 | 0.202581 | 0.067518 | 0.14361 | 0.133833 | -1.11827 | -0.05652 | -0.05066 |
| 35.90278 | -0.45494 | -0.81611 | -1.18715 | -0.72007 | -0.94766 | -1.00292 | 0.204364 | -3.46642 | -1.07103 | 0.47678 | -0.83469 | 0.154511 | 0.113432 | -0.05267 | 0.15497 | 0.21911 | -1.01633 | 0.000333 | -0.0247 |
| 35.91667 | 0.206471 | 0.206003 | -0.80655 | -0.12674 | -1.0242 | -1.11806 | 0.217954 | -3.96047 | -0.88241 | 0.739767 | -0.5735 | 0.210816 | 0.024271 | -0.1244 | 0.128541 | 0.231919 | -0.89711 | 0.026431 | -0.07107 |
| 35.93056 | 0.735618 | 1.080128 | -0.76915 | 0.269614 | -0.81321 | -0.99486 | 0.325077 | -4.30602 | -0.47304 | 0.819702 | -0.50056 | 0.242144 | -0.02201 | -0.12895 | 0.118198 | 0.243018 | -0.73329 | 0.061234 | 0.019909 |
| 35.94444 | 1.001753 | 1.67691 | -0.5948 | 0.558068 | -0.43758 | -0.59198 | 0.512423 | -2.09316 | 0.16479 | 0.602131 | -0.44867 | 0.245102 | -0.06396 | -0.09702 | 0.100223 | 0.195129 | -0.53197 | 0.035157 | 0.159071 |
| 35.95833 | 0.995646 | 1.892573 | -0.40936 | 0.699585 | -0.07242 | -0.16063 | 0.681417 | -0.14595 | 0.750188 | 0.23018 | -0.27296 | 0.255923 | -0.05634 | -0.04281 | 0.063757 | 0.128273 | -0.36057 | -0.11088 | 0.252913 |
| 35.97222 | 0.423767 | 1.564602 | -0.18346 | 0.992301 | 0.260791 | 0.219305 | 0.864409 | 1.7952 | 1.289825 | -0.09778 | -0.02152 | 0.304498 | 0.079325 | 0.033205 | 0.078303 | 0.111375 | -0.29915 | -0.27523 | 0.364993 |
| 35.98611 | -0.75301 | 0.599655 | -0.2255 | 1.14551 | 0.180496 | 0.120423 | 0.933811 | 3.714652 | 1.530793 | -0.63666 | -0.3574 | 0.225228 | 0.090615 | -0.02656 | 0.10771 | 0.0178 | -0.63335 | -0.58806 | 0.144218 |
| 36 | -2.40077 | -0.95682 | -1.12099 | 0.472588 | -0.4022 | -0.48246 | 0.747989 | 2.4092 | 0.979869 | -1.76219 | -1.53524 | -0.14168 | -0.3067 | -0.25276 | 0.11915 | -0.3346 | -1.36543 | -1.2644 | -0.64869 |
| 36.01389 | -3.59712 | -2.08748 | -2.09604 | -0.68477 | -0.92591 | -0.85539 | 0.279978 | -0.62625 | 0.065758 | -2.70886 | -2.17696 | -0.58723 | -0.76573 | -0.33069 | 0.178866 | -0.69577 | -1.50421 | -1.64708 | -1.38049 |
| 36.02778 | -3.14034 | -2.08855 | -1.77985 | -0.48023 | -0.80124 | -0.79105 | -0.11691 | -2.94669 | -0.33957 | -2.65356 | -1.58606 | -0.59077 | -0.99116 | -0.16395 | 0.268591 | -0.64409 | -0.7807 | -1.40753 | -1.27631 |
| 36.04167 | -2.60813 | -2.09098 | -1.01437 | 0.283221 | -0.70804 | -0.68244 | -0.24841 | -3.60607 | -0.3269 | -1.84124 | -0.80224 | -0.43072 | -0.82831 | -0.05521 | 0.113527 | -0.33511 | -0.23667 | -1.03084 | -0.75763 |
| 36.05556 | -1.13484 | -1.17232 | -0.39004 | 0.518171 | -0.69307 | -0.34922 | -0.09268 | -2.60549 | -0.40665 | -0.81385 | -0.19541 | -0.09216 | -0.29903 | 0.049018 | -0.10979 | -0.09115 | 0.062325 | -0.61926 | -0.1262 |
| 36.06944 | -0.00435 | -0.0425 | -0.02104 | 0.701586 | -0.75881 | -0.00717 | 0.207932 | -1.36058 | -0.39706 | 0.16383 | 0.420862 | 0.273638 | 0.330316 | 0.16006 | -0.24297 | 0.145989 | 0.288463 | -0.25446 | 0.519015 |
| 36.08333 | 0.732455 | 0.436253 | 0.156636 | 0.710819 | -0.60845 | 0.13616 | 0.320655 | -0.38457 | -0.35566 | 0.690442 | 0.957852 | 0.583626 | 0.721009 | 0.213761 | -0.25305 | 0.45878 | 0.242313 | 0.121336 | 0.925102 |
| 36.09722 | 0.910966 | 0.395616 | 0.282254 | 0.690846 | -0.19186 | 0.073033 | 0.228964 | 0.421803 | -0.15501 | 0.988884 | 0.963966 | 0.59208 | 0.693508 | 0.148982 | -0.03769 | 0.583627 | 0.071961 | 0.363391 | 1.03287 |
| 36.11111 | 0.898207 | 0.629839 | 0.351456 | 0.357276 | 0.066419 | 0.518056 | 0.087906 | 1.185189 | 0.05762 | 0.956444 | 0.456256 | 0.433844 | 0.531379 | 0.084252 | -0.02115 | 0.490048 | 0.013099 | 0.404899 | 0.792714 |
| 36.125 | 1.375307 | 0.879066 | 0.331769 | -0.14735 | 0.065108 | 0.637643 | -0.0505 | 1.630017 | 0.301331 | 0.900116 | -0.10094 | 0.368776 | 0.409897 | 0.032696 | -0.0537 | 0.288628 | 0.034736 | 0.35072 | 0.585315 |
| 36.13889 | 2.028155 | 1.035495 | 0.405001 | 0.062293 | -0.03286 | 0.565567 | -0.08215 | 1.830201 | 0.644246 | 0.722682 | -0.46546 | 0.411969 | 0.381407 | 0.003647 | -0.04116 | 0.131908 | 0.071439 | 0.389196 | 0.430105 |
| 36.15278 | 2.459653 | 1.261411 | 0.476727 | 0.290191 | -0.03592 | 0.766147 | 0.236019 | 1.995683 | 1.199863 | 0.658027 | -0.44902 | 0.541141 | 0.474832 | 0.027541 | -0.04873 | 0.176226 | 0.093595 | 0.521098 | 0.421979 |
| 36.16667 | 2.547062 | 1.510738 | 0.529972 | 0.37447 | 0.050964 | 0.694332 | 0.492139 | 2.111951 | 1.936013 | 0.50591 | -0.11832 | 0.66271 | 0.580883 | 0.068086 | -0.10438 | 0.343862 | 0.099062 | 0.792206 | 0.574832 |
| 36.18056 | 2.64199 | 1.83682 | 0.64172 | 0.278295 | 0.128694 | 1.089275 | 0.655373 | 1.976624 | 2.381217 | 0.322722 | 0.138991 | 0.705376 | 0.566817 | 0.062574 | -0.17446 | 0.399699 | 0.092256 | 1.000918 | 0.617057 |
| 36.19444 | 2.682128 | 1.92173 | 0.776808 | 0.022438 | 0.186298 | 1.200197 | 0.667908 | 1.595232 | 2.556515 | 0.164334 | 0.071052 | 0.539054 | 0.359768 | 0.052129 | -0.13806 | 0.289966 | 0.085864 | 0.850229 | 0.532924 |
| 36.20833 | 2.387362 | 1.862151 | 0.887969 | -0.11427 | 0.228957 | 0.941554 | 0.609638 | 1.493175 | 2.555199 | 0.130032 | -0.08064 | 0.279175 | 0.283718 | 0.095301 | -0.0579 | 0.10427 | 0.116731 | 0.576499 | 0.328372 |
| 36.22222 | 2.340802 | 1.842397 | 1.238626 | -0.05773 | 0.217885 | 0.494694 | 0.453453 | 1.38123 | 2.241214 | 0.21206 | -0.09474 | 0.009761 | 0.182509 | 0.096088 | 0.067284 | -0.03668 | 0.13401 | 0.42154 | -0.01151 |
| 36.23611 | 1.957074 | 1.807882 | 1.186418 | -0.00622 | 0.205627 | 0.100095 | 0.122478 | 1.037988 | 1.336031 | 0.173853 | -0.07223 | -0.22958 | -0.03326 | 0.027919 | 0.072863 | -0.17036 | 0.103725 | 0.245302 | -0.44183 |
| 36.25 | 1.911549 | 1.651059 | 1.131461 | 0.137641 | 0.421954 | 0.339193 | 0.13783 | 0.531989 | 0.655196 | 0.010558 | -0.13225 | -0.38413 | -0.30976 | -0.08608 | -0.02234 | -0.20637 | -0.0062 | 0.109671 | -0.79162 |
| 36.26389 | 2.114399 | 1.649631 | 1.268532 | 0.564654 | 0.903707 | 1.075642 | 0.652431 | 0.31207 | 0.55355 | -0.1066 | -0.10395 | -0.26756 | -0.31707 | -0.0929 | -0.1203 | -0.07122 | -0.06709 | 0.071576 | -0.85469 |
| 36.27778 | 2.023004 | 1.412749 | 1.132577 | 0.733372 | 1.010055 | 1.364905 | 0.910179 | 0.173128 | 0.545025 | 0.016643 | 0.101008 | 0.090799 | 0.082264 | 0.03493 | -0.06969 | 0.16616 | -0.00737 | 0.181217 | -0.48928 |
| 36.29167 | 1.35931 | 0.70608 | 0.577749 | 0.278666 | 0.611002 | 1.010086 | 0.562777 | -0.13701 | 0.164755 | 0.384614 | 0.313539 | 0.351432 | 0.379255 | 0.147836 | 0.008422 | 0.380607 | 0.047794 | 0.312623 | -0.01852 |
| 36.30556 | 0.50614 | -0.15483 | 0.27325 | -0.18041 | 0.089009 | 0.744207 | 0.261294 | -0.2252 | 0.039622 | 0.826915 | 0.314766 | 0.287962 | 0.299361 | 0.153938 | 0.096727 | 0.446395 | 0.106004 | 0.449054 | 0.305081 |
| 36.31944 | -0.38972 | -0.73597 | 0.3977 | -0.27192 | -0.22771 | 0.843994 | 0.277288 | -0.09637 | 0.385114 | 1.113647 | 0.294666 | 0.139204 | 0.095327 | 0.150283 | 0.277179 | 0.444151 | 0.229295 | 0.541591 | 0.455649 |
| 36.33333 | -0.42183 | -0.55238 | 0.381009 | -0.10898 | -0.31187 | 0.7431 | 0.3417 | 0.035207 | 0.36997 | 1.24973 | 0.430546 | 0.00576 | -0.04152 | 0.132384 | 0.335861 | 0.354974 | 0.260798 | 0.454396 | 0.363354 |
| 36.34722 | 0.200228 | -0.13516 | 0.089105 | -0.07025 | -0.1893 | 0.435075 | 0.040962 | -0.19144 | -0.00277 | 1.153053 | 0.649739 | -0.07048 | -0.03721 | 0.095381 | 0.178279 | 0.191436 | 0.144322 | 0.22176 | 0.198648 |
| 36.36111 | 0.497163 | -0.10939 | -0.12938 | -0.1177 | -0.2204 | 0.268905 | -0.36332 | -0.3648 | -0.12741 | 0.823678 | 0.679059 | -0.03595 | 0.105336 | 0.062317 | 0.030184 | 0.047537 | 0.042018 | 0.046698 | 0.2024 |
| 36.375 | 0.416637 | -0.20679 | -0.11035 | -0.02633 | -0.29382 | 0.407266 | -0.38284 | -0.2548 | -0.03705 | 0.292209 | 0.529239 | 0.077663 | 0.28994 | 0.062297 | -0.00918 | -0.01511 | 0.020777 | -0.06577 | 0.282797 |
| 36.38889 | 0.320954 | -0.20693 | 0.062085 | 0.082812 | -0.19722 | 0.811236 | -0.21467 | 0.108833 | 0.25777 | -0.16246 | 0.313094 | 0.338955 | 0.386774 | 0.05461 | 0.033987 | 0.0437 | 0.043709 | -0.07478 | 0.404266 |
| 36.40278 | 0.33678 | -0.30392 | 0.318 | 0.118064 | -0.11254 | 1.297584 | -0.23657 | 0.588371 | 0.608614 | -0.11761 | 0.178853 | 0.51132 | 0.360251 | 0.020052 | 0.086282 | 0.112557 | 0.104651 | 0.02728 | 0.447379 |
| 36.41667 | 0.391295 | -0.34968 | 0.678292 | 0.183192 | -0.07833 | 1.284893 | -0.31225 | 1.047744 | 0.908114 | 0.310701 | 0.158726 | 0.461132 | 0.25787 | 0.009483 | 0.103238 | 0.217965 | 0.204785 | 0.13418 | 0.369655 |
| 36.43056 | 0.562624 | -0.22256 | 1.084979 | 0.243092 | -0.04058 | 0.89448 | -0.22823 | 1.337143 | 1.117482 | 0.775092 | 0.263522 | 0.335451 | 0.191447 | 0.062058 | 0.111902 | 0.378989 | 0.296262 | 0.196312 | 0.242051 |
| 36.44444 | 0.731213 | -0.02593 | 0.98942 | 0.275887 | -0.03816 | 0.388969 | -0.07427 | 1.464641 | 1.17571 | 1.036426 | 0.514076 | 0.255947 | 0.221815 | 0.073436 | 0.068849 | 0.410962 | 0.270886 | 0.207026 | 0.192246 |
| 36.45833 | 0.765712 | 0.054106 | 0.611611 | 0.282608 | 0.037094 | -0.25046 | -0.1243 | 1.427477 | 0.996461 | 0.859695 | 0.525665 | 0.182395 | 0.175649 | -0.00447 | -0.02047 | 0.290502 | 0.166991 | 0.12728 | 0.138238 |
| 36.47222 | 0.544876 | 0.065212 | 0.348855 | 0.285902 | 0.035799 | -1.01441 | -0.26432 | 1.151421 | 0.805118 | 0.510228 | 0.407567 | 0.193663 | 0.170129 | -0.0306 | -0.05635 | 0.226019 | 0.174821 | 0.072767 | 0.119431 |
| 36.48611 | 0.615619 | 0.07335 | 0.21514 | 0.349053 | 0.09491 | -0.76049 | -0.448 | 1.193728 | 0.608677 | 0.361865 | 0.631862 | 0.35462 | 0.283258 | 0.008384 | -0.00047 | 0.373586 | 0.545296 | 0.247342 | 0.227686 |
| 36.5 | 1.447356 | 0.07148 | 1.504484 | 0.390651 | 0.101548 | -0.14959 | -0.80111 | 1.955648 | 0.458712 | 0.78331 | 1.21794 | 0.61985 | 0.415528 | -0.07167 | -0.0025 | 0.658759 | 1.281068 | 0.61712 | 0.423337 |
| 36.51389 | 2.647155 | 0.22614 | 2.80249 | 0.341162 | 0.139623 | 0.548824 | -0.86228 | 3.233441 | 0.528396 | 1.27086 | 2.190769 | 0.891612 | 0.471564 | -0.0968 | -0.03862 | 0.909636 | 2.031625 | 0.930749 | 0.601677 |
| 36.52778 | 3.460941 | 0.626318 | 3.260396 | 0.589804 | 0.537138 | 1.533252 | -0.37845 | 5.238788 | 1.331245 | 1.203188 | 2.510266 | 1.158783 | 0.481479 | 0.226693 | 0.003567 | 0.96532 | 2.056436 | 0.984098 | 0.740859 |
| 36.54167 | 3.341962 | 1.020535 | 2.914668 | 1.006354 | 0.934349 | 1.685286 | 0.305804 | 7.639046 | 2.42071 | 0.882228 | 2.66265 | 1.062621 | 0.451534 | 0.653637 | -0.00606 | 0.836039 | 2.216598 | 0.959204 | 0.795141 |
| 36.55556 | 2.242532 | 0.982285 | 2.128921 | 1.069361 | 1.229629 | 1.408251 | 0.525061 | 6.327948 | 3.218451 | 0.724857 | 2.989082 | 0.611155 | 0.349543 | 0.787964 | -0.05182 | 0.623929 | 1.98341 | 0.855387 | 0.553465 |
| 36.56944 | 0.849439 | 0.719913 | 1.583877 | 0.869987 | 1.492897 | 1.123469 | 0.449908 | 3.707429 | 2.607713 | 0.457167 | 3.170266 | 0.122014 | 0.070286 | 0.661369 | -0.07603 | 0.301502 | 1.891705 | 0.820944 | 0.224848 |
| 36.58333 | -0.52896 | 0.542235 | 1.140891 | 0.632654 | 1.432145 | 0.844568 | 0.176212 | 1.762384 | 2.042118 | 0.05292 | 3.189656 | -0.301 | -0.09179 | 0.495096 | -0.03989 | 0.052045 | 2.08166 | 0.92374 | -0.11198 |
| 36.59722 | -1.2396 | 0.569547 | 0.681806 | 0.430722 | 1.434137 | 0.485037 | -0.03248 | 0.828655 | 1.908893 | -0.41061 | 2.726715 | -0.57314 | -0.13523 | 0.380465 | -0.00911 | -0.08335 | 2.121154 | 1.118609 | -0.35638 |
| 36.61111 | -1.79587 | 0.588251 | 0.214775 | 0.245846 | 1.599951 | 0.241578 | -0.11503 | 0.468646 | 1.937392 | -0.73027 | 1.889163 | -0.73026 | -0.07669 | 0.26283 | 0.01227 | -0.14119 | 1.843484 | 1.097932 | -0.38043 |
| 36.625 | -1.80438 | 0.497648 | -0.29873 | 0.083154 | 1.768476 | 0.546929 | -0.06296 | -1.8E-05 | 1.621363 | -0.96027 | 0.638234 | -0.57963 | 0.049087 | 0.076568 | 0.039248 | -0.16391 | 1.291807 | 1.00912 | -0.30486 |
| 36.63889 | -1.77636 | 0.332447 | -0.71351 | -0.00262 | 1.685658 | 1.218562 | 0.10669 | -0.66672 | 0.832146 | -1.206 | -0.94155 | -0.32005 | 0.087898 | -0.04429 | 0.048817 | -0.1331 | 0.610274 | 0.764403 | -0.16742 |
| 36.65278 | -1.84042 | -0.08551 | -0.8137 | -0.04146 | 1.498963 | 1.837003 | 0.121303 | -1.48153 | -0.1393 | -0.73112 | -2.46407 | -0.17097 | 0.038004 | -0.04608 | 0.034001 | 0.052055 | -0.13965 | 0.471226 | 0.022884 |
| 36.66667 | -1.99834 | -0.58587 | -1.20169 | -0.01378 | 1.079901 | 1.968074 | -0.08356 | -2.32205 | -1.03217 | 0.002728 | -3.6447 | -0.15948 | -0.09358 | 0.027242 | 0.005362 | 0.233103 | -0.86522 | 0.331734 | 0.068313 |
| 36.68056 | -1.58208 | -0.73299 | -1.89311 | -0.30211 | 0.961802 | 1.460403 | -0.33374 | -2.39899 | -1.72487 | 0.608524 | -3.92575 | -0.14767 | -0.27145 | 0.100534 | -0.01445 | 0.325354 | -1.5681 | 0.252308 | -0.11805 |
| 36.69444 | 0.22623 | -0.49258 | -1.36096 | -0.68478 | 0.940119 | 0.84634 | -0.47108 | -1.87646 | -2.0509 | 1.048042 | -3.55836 | -0.00901 | -0.2414 | 0.2386 | -0.02009 | 0.3278 | -2.2511 | 0.177513 | -0.39471 |
| 36.70833 | 1.636089 | -0.28686 | -0.57558 | -1.04024 | 0.845345 | 0.197572 | -0.4791 | -1.57972 | -1.84391 | 1.175902 | -2.92542 | 0.203245 | -0.07155 | 0.414259 | -0.05399 | 0.250698 | -2.68892 | 0.053712 | -0.51873 |
| 36.72222 | 2.623871 | -0.29216 | 0.131435 | -1.36584 | 0.753121 | -0.10022 | -0.36011 | -1.55375 | -1.25828 | 0.909004 | -2.34585 | 0.321902 | 0.070112 | 0.543545 | -0.11295 | 0.135974 | -2.6804 | -0.08095 | -0.46643 |
| 36.73611 | 2.116459 | -0.0108 | 1.210101 | -1.41482 | 0.829332 | 0.454716 | -0.19308 | -1.48023 | -0.38643 | 0.726377 | -1.45562 | 0.231823 | 0.029415 | 0.524762 | -0.12884 | -0.06967 | -2.20768 | -0.29996 | -0.37752 |
| 36.75 | 2.163862 | 0.26038 | 2.17464 | -1.14737 | 0.864617 | 1.192283 | -0.00347 | -0.75597 | 0.310833 | 0.422831 | -0.71232 | 0.046951 | -0.11143 | 0.44398 | -0.09198 | -0.32976 | -1.58377 | -0.54342 | -0.37486 |
| 36.76389 | 1.415862 | 0.555861 | 1.806786 | -0.79114 | 0.857488 | 1.385519 | 0.189331 | -0.52207 | 0.854203 | 0.063006 | -0.27042 | -0.15228 | -0.13623 | 0.291121 | -0.04152 | -0.49677 | -1.0613 | -0.69836 | -0.43957 |
| 36.77778 | 0.197606 | 0.639244 | 1.057087 | -0.423 | 0.728259 | 1.28186 | 0.34996 | -0.89307 | 1.552211 | -0.28381 | -0.07946 | -0.30746 | -0.04168 | 0.086376 | -0.01365 | -0.50585 | -0.72095 | -0.77971 | -0.45914 |
| 36.79167 | -0.80994 | 0.629205 | 0.618454 | 0.073947 | 0.59437 | 1.226375 | 0.410652 | -1.41705 | 2.026177 | -0.71017 | 0.032725 | -0.46814 | 0.011761 | -0.19736 | -0.00812 | -0.50826 | -0.5582 | -0.75828 | -0.51408 |
| 36.80556 | -0.7985 | 0.52487 | 0.269952 | 0.216977 | 0.479439 | 1.149629 | 0.345355 | -1.62361 | 1.76476 | -1.30973 | 0.028963 | -0.62573 | -0.05714 | -0.42205 | -0.01303 | -0.58524 | -0.44614 | -0.74543 | -0.57152 |
| 36.81944 | -1.35286 | 0.331579 | -0.28552 | 0.049013 | 0.40598 | 0.902948 | 0.187117 | -2.01402 | 0.677538 | -1.38769 | -0.12386 | -0.723 | -0.17765 | -0.41735 | 0.01561 | -0.68951 | -0.42518 | -0.74225 | -0.48952 |
| 36.83333 | -0.96609 | -0.11089 | -1.17708 | -0.40974 | 0.303959 | 0.638351 | 0.00024 | -2.90185 | -0.46074 | -1.26636 | -0.25471 | -0.73484 | -0.24878 | -0.28679 | 0.057109 | -0.66968 | -0.44085 | -0.72833 | -0.39653 |
| 36.84722 | -1.28958 | -0.74746 | -1.91906 | -0.85027 | 0.089854 | 0.325011 | -0.10443 | -2.99918 | -1.4436 | -1.05761 | -0.24053 | -0.58164 | -0.18634 | -0.0552 | 0.076624 | -0.6253 | -0.46476 | -0.72058 | -0.31598 |
| 36.86111 | -2.32281 | -1.22303 | -1.89615 | -1.17601 | -0.23433 | -0.20924 | -0.11469 | -2.66395 | -1.74405 | -0.66779 | -0.15258 | -0.21561 | -0.09633 | 0.088726 | 0.072599 | -0.37748 | -0.505 | -0.76801 | -0.20189 |
| 36.875 | -2.50998 | -1.32296 | -0.66298 | -1.53763 | -0.54897 | -0.55014 | -0.12643 | -2.01238 | -1.84272 | -0.08645 | -0.00292 | 0.086402 | 0.05759 | 0.158002 | 0.047889 | -0.04039 | -0.53857 | -0.82031 | 0.046068 |
| 36.88889 | -2.13614 | -0.80396 | -0.6215 | -1.88268 | -0.78743 | -0.8213 | -0.28148 | -1.5439 | -1.78005 | 0.497887 | 0.195874 | 0.262318 | 0.171746 | 0.160047 | -0.00038 | 0.190567 | -0.6075 | -0.72802 | 0.336753 |
| 36.90278 | -1.02935 | -0.38839 | -0.57593 | -1.48368 | -0.92465 | -0.89143 | -0.39709 | -1.94078 | -2.00321 | 0.948073 | 0.299988 | 0.294343 | 0.197928 | 0.110095 | -0.06136 | 0.218372 | -0.69506 | -0.52425 | 0.447645 |
| 36.91667 | -0.22167 | -0.03843 | -0.1478 | -0.87498 | -0.99467 | -0.61442 | -0.45746 | -2.14901 | -1.83458 | 1.25554 | 0.285212 | 0.290174 | 0.224169 | 0.086346 | -0.07466 | 0.217887 | -0.73532 | -0.32607 | 0.372466 |
| 36.93056 | 0.50022 | 0.152547 | 0.178721 | -0.73826 | -1.05876 | -0.22915 | -0.31003 | -2.26817 | -1.37894 | 1.14244 | 0.203165 | 0.270355 | 0.270258 | 0.095948 | -0.05232 | 0.215606 | -0.75899 | -0.1692 | 0.098907 |
| 36.94444 | 0.98572 | 0.487584 | 0.338355 | -0.66474 | -1.00786 | 0.189072 | -0.13026 | -2.55883 | -0.92982 | 0.827031 | 0.112601 | 0.255987 | 0.303285 | 0.0806 | -0.03695 | 0.227264 | -0.71754 | -0.10109 | -0.19233 |
| 36.95833 | 1.365132 | 0.830538 | 0.310263 | -0.5466 | -0.70163 | 0.326558 | 0.024979 | -2.82156 | -0.59313 | 0.51241 | 0.068604 | 0.291837 | 0.382926 | 0.055782 | -0.01934 | 0.252953 | -0.50861 | -0.02991 | -0.31526 |
| 36.97222 | 1.302942 | 1.097887 | 0.258634 | -0.49904 | -0.35641 | 0.303095 | 0.141053 | -2.69477 | -0.22423 | 0.188761 | -0.01802 | 0.343329 | 0.364215 | 0.015686 | 0.022386 | 0.248159 | -0.29479 | 0.03926 | -0.26765 |
| 36.98611 | 1.361536 | 1.093311 | 0.220642 | -0.4049 | -0.57143 | 0.195802 | 0.157521 | -2.47957 | 0.071549 | -0.49142 | -0.48576 | 0.154485 | 0.27006 | -0.19697 | 0.067873 | 0.027454 | -0.51585 | -0.14562 | -0.38266 |
| 37 | 0.115827 | 0.147058 | 0.012842 | -0.50726 | -1.3037 | -0.04676 | -0.16293 | -2.95961 | 0.260012 | -2.04098 | -1.46123 | -0.45568 | -0.23764 | -0.52968 | 0.088512 | -0.48784 | -1.50224 | -0.67443 | -0.94602 |
| 37.01389 | -1.61 | -0.88722 | -0.3531 | -1.16051 | -1.57088 | -0.18492 | -0.7435 | -3.09502 | 0.477823 | -2.56413 | -1.76495 | -1.04625 | -0.76928 | -0.57936 | 0.052741 | -0.89837 | -1.89334 | -0.98171 | -1.39001 |
| 37.02778 | -2.36384 | -1.38957 | -0.68916 | -1.24332 | -1.29708 | -0.0771 | -0.825 | -2.42968 | 0.495557 | -1.83051 | -0.947 | -1.16089 | -0.74622 | -0.25204 | 0.073396 | -0.80129 | -1.15714 | -0.61203 | -1.13917 |
| 37.04167 | -1.9018 | -1.37199 | -0.76678 | -0.46567 | -1.23363 | -0.52661 | -0.83116 | -2.16723 | 0.458768 | -1.66874 | -0.23818 | -0.79337 | -0.44359 | -0.04 | 0.188951 | -0.41775 | -0.43345 | -0.21378 | -0.50486 |
| 37.05556 | -1.00931 | -1.20243 | -0.80965 | 0.243029 | -1.20388 | -0.75495 | -0.86339 | -2.40759 | 0.278372 | -1.00607 | 0.202763 | -0.26124 | 0.154415 | 0.149372 | 0.171141 | 0.035194 | -0.07089 | 0.050705 | -0.07718 |
| 37.06944 | -0.46869 | -1.03032 | -0.85239 | 0.775002 | -0.92259 | -0.43881 | -0.45691 | -2.39525 | -0.08759 | -0.28024 | 0.504729 | 0.236995 | 0.448696 | 0.206845 | 0.170559 | 0.50061 | 0.182842 | 0.245381 | 0.242255 |

| | | | | | | | | | | | | | | | | | | | |
|---|---|---|---|---|---|---|---|---|---|---|---|---|---|---|---|---|---|---|---|
| 37.08333 | -0.13453 | -0.85739 | -0.83596 | 0.743734 | -0.39408 | 0.084143 | -0.04882 | -1.99839 | -0.41702 | 0.371998 | 0.695705 | 0.603228 | 0.525301 | 0.192082 | 0.338558 | 0.900784 | 0.352577 | 0.318073 | 0.605733 |
| 37.09722 | 0.242513 | -0.6296 | -0.67359 | 0.128932 | 0.108732 | 0.35155 | 0.062966 | -1.5402 | -0.53234 | 0.749746 | 0.687124 | 0.747961 | 0.792917 | 0.135666 | 0.206827 | 0.969979 | 0.509285 | 0.266427 | 0.909963 |
| 37.11111 | 0.411343 | -0.49975 | -0.27493 | -0.29341 | 0.460253 | 0.528331 | 0.030287 | -1.15758 | -0.48757 | 1.083707 | 0.64337 | 0.651937 | 0.746816 | 0.090792 | 0.039572 | 0.814977 | 0.397426 | 0.205672 | 1.017716 |
| 37.125 | 0.645433 | -0.23001 | 0.094616 | -0.32885 | 0.641239 | 0.954169 | 0.202957 | -0.4727 | -0.19693 | 0.875825 | 0.429691 | 0.619511 | 0.641819 | 0.073584 | -0.09877 | 0.528919 | 0.34526 | 0.173954 | 0.779965 |
| 37.13889 | 1.129867 | 0.366509 | 0.598191 | 0.059866 | 0.966297 | 1.550286 | 0.697969 | 0.659433 | 0.355989 | 0.483823 | 0.241797 | 0.534599 | 0.439195 | 0.158782 | -0.10814 | 0.254434 | 0.272734 | 0.218334 | 0.703863 |
| 37.15278 | 1.623542 | 0.928055 | 0.820541 | 0.391929 | 0.950146 | 1.749559 | 1.047413 | 1.855953 | 0.813622 | 0.136729 | 0.09462 | 0.494319 | 0.391758 | 0.260764 | -0.06911 | 0.196288 | 0.304504 | 0.293832 | 0.661504 |
| 37.16667 | 2.162956 | 1.432388 | 0.764253 | 0.453096 | 0.628078 | 1.409822 | 1.116708 | 2.705533 | 0.909485 | -0.15435 | 0.04311 | 0.487892 | 0.44857 | 0.315966 | -0.10516 | 0.303354 | 0.297757 | 0.373997 | 0.580876 |
| 37.18056 | 3.276209 | 2.17151 | 0.789361 | 0.503272 | 0.45899 | 1.214942 | 1.184044 | 3.214359 | 0.972488 | 0.310705 | 0.096608 | 0.473029 | 0.465138 | 0.240346 | -0.18174 | 0.374676 | 0.24528 | 0.380872 | 0.631389 |
| 37.19444 | 4.26973 | 2.342293 | 0.851703 | 0.632208 | 0.35744 | 1.110411 | 1.198888 | 3.202031 | 0.924248 | 0.743134 | 0.208837 | 0.372694 | 0.303067 | 0.071045 | -0.26032 | 0.19226 | 0.1314 | 0.331395 | 0.622643 |
| 37.20833 | 4.402905 | 2.341115 | 0.898335 | 0.695909 | 0.263251 | 1.253545 | 1.030247 | 3.16306 | 0.833872 | 0.63685 | 0.125908 | 0.153368 | 0.00782 | -0.07272 | -0.24932 | -0.01251 | 0.028604 | 0.246494 | 0.552121 |
| 37.22222 | 3.929524 | 2.117343 | 0.944729 | 0.660855 | 0.233826 | 1.343984 | 0.834274 | 2.896352 | 0.895306 | 0.035949 | -0.03993 | -0.03497 | -0.14771 | -0.13053 | -0.143 | -0.10177 | 0.001991 | 0.132971 | 0.311946 |
| 37.23611 | 3.154801 | 1.783944 | 0.881998 | 0.456812 | 0.164771 | 1.254378 | 0.450801 | 2.40545 | 0.921182 | -0.59947 | -0.16125 | -0.10535 | -0.12077 | -0.2033 | -0.0504 | -0.09532 | -0.02501 | -0.03841 | -0.05363 |
| 37.25 | 2.190776 | 1.444952 | 0.577097 | 0.117232 | 0.044258 | 0.981669 | -0.00231 | 1.551354 | 0.866385 | -0.9675 | -0.33767 | -0.12462 | -0.09924 | -0.17572 | 0.017678 | -0.07392 | -0.05715 | -0.11542 | -0.39288 |
| 37.26389 | 0.923481 | 0.810769 | 0.222352 | -0.18527 | -0.11256 | 0.736726 | -0.22764 | 0.533484 | 0.796852 | -0.9641 | -0.33216 | -0.09599 | 0.026412 | -0.09474 | 0.049841 | 0.039122 | -0.02438 | -0.00576 | -0.53222 |
| 37.27778 | 0.040865 | 0.374057 | 0.100344 | -0.32257 | -0.17984 | 0.613921 | -0.27636 | -0.19449 | 0.791498 | -0.53157 | -0.076 | 0.101079 | 0.183377 | 0.0575 | 0.030755 | 0.192233 | 0.093518 | 0.125726 | -0.53397 |
| 37.29167 | -0.12947 | 0.241813 | 0.116451 | -0.27189 | -0.11557 | 0.558082 | -0.23689 | -0.51554 | 0.784725 | 0.041122 | 0.19811 | 0.278111 | 0.28456 | 0.116815 | -0.03414 | 0.319923 | 0.149142 | 0.235631 | -0.21537 |
| 37.30556 | 0.185563 | 0.220167 | 0.070669 | -0.16988 | -0.05736 | 0.519709 | -0.13422 | -0.44738 | 0.734226 | 0.857889 | 0.422807 | 0.413992 | 0.251799 | 0.043514 | -0.11569 | 0.353632 | 0.105052 | 0.308824 | 0.153818 |
| 37.31944 | 0.425208 | 0.229607 | 0.012789 | -0.0875 | -0.0042 | 0.489624 | 0.023121 | -0.23203 | 0.679042 | 1.36673 | 0.422304 | 0.544053 | 0.178113 | -0.05833 | -0.15331 | 0.306528 | 0.053016 | 0.395706 | 0.225159 |
| 37.33333 | 0.401604 | 0.161466 | 0.102952 | -0.14689 | 0.109856 | 0.553438 | 0.201612 | -0.10022 | 0.640719 | 1.141947 | 0.150547 | 0.52051 | 0.104292 | -0.04678 | -0.10777 | 0.251223 | 0.063367 | 0.415895 | 0.090957 |
| 37.34722 | 0.40936 | 0.024248 | 0.267399 | -0.3095 | 0.168888 | 0.598349 | 0.244126 | -0.0657 | 0.606671 | 0.855256 | -0.14797 | 0.288255 | 0.120817 | 0.005712 | -0.03505 | 0.198696 | 0.050931 | 0.317508 | -0.10713 |
| 37.36111 | 0.468432 | -0.14627 | 0.311783 | -0.43938 | 0.091169 | 0.523704 | 0.095367 | -0.02067 | 0.511015 | 0.579486 | -0.39374 | -0.09366 | 0.110933 | -0.00668 | 0.000405 | 0.128802 | -0.02614 | 0.135023 | -0.22652 |
| 37.375 | 0.371895 | -0.29404 | 0.311556 | -0.46965 | -0.039 | 0.469292 | -0.16448 | 0.002715 | 0.463003 | 0.464636 | -0.26346 | -0.24207 | 0.120735 | 0.020141 | 0.053372 | 0.145138 | -0.04294 | 0.118681 | -0.13489 |
| 37.38889 | 0.360415 | -0.31329 | 0.369989 | -0.43603 | -0.0194 | 0.533208 | -0.375 | 0.097907 | 0.410918 | 0.532875 | 0.12509 | -0.12422 | 0.287675 | 0.113538 | 0.092692 | 0.228249 | 0.017957 | 0.221825 | 0.169919 |
| 37.40278 | 0.609215 | -0.19897 | 0.399546 | -0.31994 | 0.097604 | 0.516472 | -0.40656 | 0.2952 | 0.35815 | 0.893288 | 0.3611 | 0.126245 | 0.40242 | 0.072288 | 0.012663 | 0.140819 | 0.023497 | 0.239673 | 0.447268 |
| 37.41667 | 0.83448 | -0.07682 | 0.538689 | -0.16318 | 0.253352 | 0.571486 | -0.23732 | 0.584589 | 0.330341 | 0.711132 | 0.282866 | 0.357718 | 0.359217 | 0.017383 | -0.0539 | 0.020007 | 0.022955 | 0.187179 | 0.578843 |
| 37.43056 | 0.879887 | 0.01036 | 0.733808 | -0.00414 | 0.388501 | 0.669875 | 0.012778 | 0.813366 | 0.377718 | 0.482248 | 0.056547 | 0.552397 | 0.286905 | 0.036316 | -0.01494 | 0.043134 | 0.03801 | 0.194335 | 0.547977 |
| 37.44444 | 0.755391 | -0.03235 | 0.831884 | 0.046016 | 0.457201 | 0.667517 | 0.026602 | 0.878727 | 0.347816 | 0.223098 | -0.26525 | 0.535411 | 0.051822 | -0.02648 | -0.01958 | 0.038981 | -0.04567 | 0.135269 | 0.365968 |
| 37.45833 | 0.533911 | -0.10697 | 0.80267 | 0.067019 | 0.302786 | 0.59193 | -0.14271 | 0.880581 | 0.42086 | -0.0877 | -0.49368 | 0.410045 | -0.07102 | -0.04605 | 0.009567 | 0.082437 | -0.10751 | 0.067024 | 0.073339 |
| 37.47222 | 0.21896 | -0.19723 | 0.788301 | -0.00218 | 0.121083 | 0.531785 | -0.32126 | 0.779479 | 0.480082 | -0.02904 | -0.40513 | 0.301835 | 0.040138 | 0.017039 | 0.08019 | 0.179808 | -0.07992 | 0.072207 | -0.05555 |
| 37.48611 | 0.0502 | -0.32368 | 0.727392 | -0.18345 | 0.164554 | 0.288881 | -0.61872 | 0.802197 | 0.546728 | 0.695452 | -0.14454 | 0.260093 | 0.139622 | -0.02246 | -0.01791 | 0.25791 | 0.092165 | 0.164081 | -0.02739 |
| 37.5 | 0.202222 | -0.26854 | 0.728077 | -0.18589 | 0.55983 | -0.00677 | -0.74055 | 1.205866 | 0.62292 | 1.743909 | 0.22369 | 0.594348 | 0.292919 | -0.12113 | -0.20001 | 0.419114 | 0.823326 | 0.4988 | 0.152458 |
| 37.51389 | 0.842272 | 0.070132 | 0.888148 | 0.187477 | 0.912627 | -0.07482 | -0.31591 | 1.751291 | 0.800977 | 2.414055 | 0.665015 | 1.267703 | 0.635399 | -0.03878 | -0.27751 | 0.655803 | 1.970826 | 0.906428 | 0.32347 |
| 37.52778 | 2.294302 | 0.80949 | 1.064229 | 0.784354 | 0.94873 | 0.16471 | -0.03086 | 2.445738 | 1.142938 | 2.024145 | 0.909067 | 1.634951 | 1.009922 | 0.131168 | -0.21693 | 0.727656 | 2.374088 | 1.026122 | 0.417151 |
| 37.54167 | 4.465625 | 1.66589 | 1.277434 | 0.86187 | 0.745 | 0.423753 | 0.241247 | 3.01785 | 1.50521 | 2.196534 | 0.686658 | 1.286529 | 0.875772 | 0.421156 | -0.12954 | 0.65957 | 2.567633 | 0.869784 | 0.494185 |
| 37.55556 | 3.049631 | 1.031909 | 1.395603 | 0.828901 | 1.026185 | 0.610655 | 0.513541 | 3.224923 | 1.901792 | 1.4576 | 0.233368 | 0.736278 | 0.431962 | 0.52906 | -0.04787 | 0.420596 | 2.196539 | 0.670189 | 0.407402 |
| 37.56944 | 1.869108 | 0.698328 | 1.121252 | 0.414858 | 1.232546 | 0.701524 | 0.503286 | 3.035878 | 1.975623 | 0.493238 | 0.166941 | 0.213257 | -0.03799 | 0.512676 | 0.004574 | 0.188575 | 2.017569 | 0.528893 | 0.213268 |
| 37.58333 | 0.633228 | 0.467513 | 0.846054 | -0.14141 | 1.070201 | 0.530978 | 0.208224 | 2.594433 | 1.725095 | -0.3386 | 0.55765 | -0.10842 | -0.2577 | 0.507805 | 0.01268 | 0.122354 | 1.933234 | 0.574234 | 0.067795 |
| 37.59722 | -0.49256 | 0.11806 | 0.99106 | -0.54784 | 0.779234 | 0.37434 | -0.02847 | 2.352313 | 1.333311 | -0.89667 | 1.066136 | -0.27968 | -0.11597 | 0.504539 | -0.04476 | 0.168355 | 2.006592 | 0.813367 | -0.06609 |
| 37.61111 | -1.24315 | -0.07755 | 1.296935 | -0.64139 | 0.548813 | 0.522419 | -0.22387 | 2.36645 | 1.068647 | -0.86525 | 1.221999 | -0.32402 | -0.00515 | 0.481386 | -0.19189 | 0.255946 | 1.819481 | 0.845983 | -0.17477 |
| 37.625 | -1.51386 | -0.23935 | 1.145204 | -0.70627 | 0.672732 | 0.572316 | -0.34387 | 2.343554 | 0.969387 | -0.9974 | 0.932374 | -0.27776 | 0.137754 | 0.410052 | -0.24862 | 0.29559 | 1.335806 | 0.827191 | -0.25237 |
| 37.63889 | -1.78932 | -0.36408 | 0.763885 | -0.857 | 0.738152 | 0.648664 | -0.28708 | 1.705395 | 0.993021 | -0.87701 | 0.25947 | -0.28786 | 0.202931 | 0.328804 | -0.20355 | 0.371691 | 0.594957 | 0.681611 | -0.34166 |
| 37.65278 | -2.15753 | -0.68755 | 0.343746 | -0.92284 | 0.66775 | 0.700669 | -0.26307 | 0.322631 | 0.683764 | -0.58079 | -0.55545 | -0.26686 | 0.231796 | 0.198418 | -0.18782 | 0.387329 | -0.26684 | 0.565027 | -0.25579 |
| 37.66667 | -1.91907 | -0.97857 | 0.901237 | -0.94905 | 0.510918 | 0.624077 | -0.5021 | -1.54273 | -0.31308 | -0.56685 | -1.29428 | -0.26846 | 0.127821 | -0.05801 | -0.24336 | 0.266445 | -1.01092 | 0.444636 | -0.06618 |
| 37.68056 | -1.35776 | -0.86012 | 1.026573 | -0.60444 | 0.318421 | 0.52814 | -0.76516 | -3.54325 | -1.54573 | -0.63837 | -1.60164 | -0.29755 | -0.00767 | -0.31292 | -0.30072 | 0.182567 | -1.68275 | 0.305904 | 0.004579 |
| 37.69444 | -0.74448 | -0.67477 | 0.264615 | -0.42003 | 0.193528 | 0.477977 | -0.86071 | -4.7145 | -1.7732 | -0.612 | -1.52992 | -0.23306 | -0.10428 | -0.43025 | -0.25558 | 0.188993 | -2.20375 | 0.20492 | -0.02853 |
| 37.70833 | -0.13723 | -0.46314 | -0.62161 | -0.28428 | 0.22585 | 0.431622 | -0.78015 | -4.50301 | -1.46824 | -0.20349 | -1.05655 | -0.14322 | -0.14474 | -0.36305 | -0.14891 | 0.175346 | -2.46022 | 0.166948 | -0.10684 |
| 37.72222 | 0.431985 | 0.139982 | -0.7209 | -0.08111 | 0.334996 | 0.399299 | -0.72824 | -3.57269 | -1.28113 | 0.298114 | -0.61285 | -0.01239 | -0.04517 | -0.07583 | -0.09456 | 0.102825 | -2.42208 | 0.095884 | -0.12586 |
| 37.73611 | 0.668073 | 0.780926 | 0.087369 | 0.089142 | 0.306924 | 0.440197 | -0.59802 | -2.43415 | -1.27614 | 0.586153 | -0.28568 | 0.078998 | 0.09269 | 0.319017 | -0.1291 | -0.0017 | -2.10382 | -0.04816 | -0.15248 |
| 37.75 | 0.804092 | 0.70459 | -0.01327 | 0.091421 | 0.10268 | 0.461592 | -0.44685 | -1.56988 | -1.00245 | 0.655136 | -0.16449 | 0.197099 | 0.186828 | 0.633244 | -0.22171 | -0.09982 | -1.77792 | -0.2345 | -0.15826 |
| 37.76389 | 0.568723 | 0.423952 | -0.18472 | -0.13905 | -0.17769 | 0.538878 | -0.28964 | -0.93218 | -0.54045 | 0.621166 | -0.11305 | 0.188842 | 0.175096 | 0.7453 | -0.30181 | -0.14836 | -1.45959 | -0.42837 | -0.21993 |
| 37.77778 | 0.652026 | 0.09083 | -0.0488 | -0.44413 | -0.40449 | 0.679767 | -0.08528 | -0.21082 | -0.09188 | 0.643217 | -0.07041 | 0.101601 | 0.112491 | 0.563636 | -0.35965 | -0.17142 | -1.22873 | -0.62859 | -0.37424 |
| 37.79167 | 0.350253 | -0.32625 | 0.267215 | -0.67323 | -0.44688 | 0.764995 | 0.005027 | 0.298244 | 0.037487 | 0.5328 | -0.00791 | -0.08988 | 0.113055 | 0.374768 | -0.34814 | -0.13286 | -0.81644 | -0.74398 | -0.53342 |
| 37.80556 | -0.08605 | -0.53328 | 0.35466 | -0.73308 | -0.32435 | 0.66798 | -0.10543 | -0.2138 | -0.18109 | 0.142103 | 0.036368 | -0.29215 | 0.144919 | 0.211708 | -0.21386 | -0.06569 | -0.38806 | -0.73668 | -0.6828 |
| 37.81944 | -0.56596 | -0.62996 | 0.281565 | -0.94097 | -0.33701 | 0.421864 | -0.3861 | -1.42558 | -0.24239 | -0.37224 | -0.03941 | -0.49364 | 0.139174 | 0.029209 | -0.13543 | -0.05804 | -0.23943 | -0.70869 | -0.71533 |
| 37.83333 | -0.71345 | -0.63115 | 0.258344 | -1.26502 | -0.46022 | -0.00514 | -0.92483 | -2.93382 | -0.3328 | -0.93814 | -0.15756 | -0.6108 | 0.060739 | -0.15638 | -0.19504 | -0.13668 | -0.39669 | -0.73481 | -0.71537 |
| 37.84722 | -0.91892 | -0.70371 | 0.501493 | -1.59701 | -1.03068 | -0.43653 | -1.69891 | -4.63012 | -0.48952 | -1.12011 | -0.25176 | -0.62013 | 0.031727 | -0.32194 | -0.3132 | -0.25009 | -0.62885 | -0.89516 | -0.60889 |
| 37.86111 | -1.11555 | -0.54961 | 0.733976 | -1.85101 | -1.89142 | -0.70107 | -2.16573 | -5.69501 | -0.67755 | -1.14192 | -0.21548 | -0.50689 | 0.093968 | -0.3717 | -0.2616 | -0.29675 | -0.74879 | -1.04449 | -0.33989 |
| 37.875 | -0.66365 | -0.43493 | 0.382561 | -1.79093 | -1.77748 | -0.62077 | -2.10809 | -5.1502 | -0.50891 | -1.12461 | -0.10106 | -0.28392 | 0.129384 | -0.3024 | -0.08841 | -0.19942 | -0.74304 | -1.11012 | -0.04428 |
| 37.88889 | -0.39091 | -0.72263 | -0.40936 | -1.60231 | -1.45321 | -0.62304 | -2.00825 | -4.97861 | 0.141816 | -1.14133 | 0.021591 | -0.06655 | 0.049715 | -0.21955 | 0.090066 | -0.07524 | -0.61398 | -0.98608 | 0.194972 |
| 37.90278 | -0.45347 | -1.18704 | -1.01327 | -1.50193 | -1.48134 | -0.49318 | -1.87506 | -5.01001 | 0.092116 | -0.86709 | 0.151352 | 0.174049 | 0.014827 | -0.118 | 0.17726 | 0.055016 | -0.43517 | -0.57764 | 0.359852 |
| 37.91667 | -1.14825 | -1.60117 | -1.15099 | -1.33545 | -1.03418 | -0.20113 | -1.43617 | -4.32217 | 0.015492 | -0.4971 | 0.221408 | 0.365771 | -0.00377 | -0.06493 | 0.167803 | 0.173256 | -0.35155 | -0.17066 | 0.428261 |
| 37.93056 | -1.08045 | -1.40878 | -0.95118 | -1.11387 | -0.72843 | 0.073241 | -0.94432 | -3.52233 | 0.046617 | -0.10086 | 0.168002 | 0.472007 | 0.014806 | -0.02025 | 0.123478 | 0.206679 | -0.38999 | 0.043291 | 0.40367 |
| 37.94444 | -0.17448 | -0.76841 | -0.6496 | -0.80942 | -0.39138 | 0.236204 | -0.43064 | -2.61505 | 0.190387 | 0.344185 | 0.119498 | 0.521619 | 0.091323 | 0.051637 | 0.046179 | 0.208251 | -0.42323 | 0.112272 | 0.318497 |
| 37.95833 | 0.560237 | -0.24609 | -0.26212 | -0.35434 | -0.11751 | 0.461747 | 0.156585 | -1.62338 | 0.429278 | 0.645637 | 0.09267 | 0.573109 | 0.261419 | 0.135056 | 0.004172 | 0.206619 | -0.41944 | 0.143946 | 0.222945 |
| 37.97222 | 0.41166 | -0.21371 | 0.217242 | 0.158914 | 0.174478 | 0.712813 | 0.817531 | -0.568 | 0.653944 | 0.826831 | 0.129169 | 0.556478 | 0.399776 | 0.157426 | 0.014438 | 0.199269 | -0.43306 | 0.169691 | 0.097289 |
| 37.98611 | 0.526343 | -0.28495 | 0.665334 | 0.710487 | -0.11457 | 0.86861 | 0.943128 | -0.22619 | 0.76819 | 0.08053 | -0.316 | 0.264413 | 0.311962 | 0.02294 | 0.189559 | 0.04912 | -0.5588 | -0.06943 | -0.20079 |

**Supplementary Table S2.** Raw motion data from: (a) 100 mM KCl stress

| Time | Control1 | Control2 | Control3 | Control4 | Control5 | Control6 | Control7 | Control8 | Control9 | Stress1 | Stress2 | Stress3 | Stress4 | Stress5 | Stress6 | Stress7 | Stress8 | Stress9 | Stress10 |
|---|---|---|---|---|---|---|---|---|---|---|---|---|---|---|---|---|---|---|---|
| 18 | -2.87263 | -3.06867 | -0.34082 | -0.68394 | -0.95362 | -1.14881 | -0.11342 | -1.47269 | -3.50278 | -2.14625 | -1.24194 | -0.48034 | -0.98765 | -1.09023 | -2.13874 | -1.35111 | -0.88972 | -0.49787 | -0.93979 |
| 18.01389 | -3.90168 | -2.6829 | -0.45645 | -1.23005 | -1.88639 | -1.54856 | -0.46749 | -1.64265 | -3.90748 | -4.09581 | -2.77177 | -1.16627 | -1.37469 | -1.56773 | -3.96434 | -2.57414 | -2.85845 | -0.78684 | -1.38208 |
| 18.02778 | -2.06005 | -0.70264 | -0.05799 | -0.59904 | -1.14819 | -0.85005 | -0.56269 | -0.35955 | -1.18808 | -2.86705 | -2.73388 | -0.91277 | 0.409252 | -0.98281 | -1.13819 | -1.69915 | -1.66785 | -0.18012 | -0.94783 |
| 18.04167 | -0.40481 | 0.614401 | 0.273322 | 0.349995 | 0.189829 | 0.010282 | -0.26136 | 0.940684 | 1.249857 | -0.61813 | -0.94356 | -0.22551 | 1.770624 | -0.08561 | 2.018785 | 0.070812 | -0.16205 | 0.486653 | -0.24736 |
| 18.05556 | 0.095372 | 1.092677 | 0.270688 | 0.343731 | 0.699447 | 0.523167 | -0.05376 | 1.238128 | 2.318539 | 0.157283 | -0.16745 | 0.205008 | 1.963162 | 0.222427 | 3.328587 | 0.891549 | 0.667362 | 0.67149 | -0.01214 |
| 18.06944 | 0.198141 | 0.835235 | 0.119688 | 0.140995 | 0.470448 | 0.551971 | -0.00093 | 0.7422 | 1.916237 | 0.161929 | 0.051567 | 0.367412 | 1.128853 | 0.227192 | 2.634786 | 0.888925 | 1.194722 | 0.502343 | 0.091688 |
| 18.08333 | 0.172325 | 0.308808 | 0.015436 | 0.032171 | 0.14148 | 0.2602 | 0.018977 | 0.326419 | 1.052364 | -0.00471 | 0.054865 | 0.313092 | 0.520835 | 0.122739 | 1.548316 | 0.547317 | 1.120245 | 0.285947 | 0.089354 |
| 18.09722 | 0.12599 | -0.05336 | -0.00249 | -0.00264 | 0.016463 | 0.1102 | 0.082684 | 0.094703 | 0.522671 | -0.13011 | 0.045093 | 0.149309 | 0.240257 | -0.00067 | 0.92673 | 0.316923 | 0.654544 | 0.160029 | 0.055082 |
| 18.11111 | 0.11089 | -0.10305 | 0.022783 | 0.00617 | 0.001174 | 0.078689 | 0.123513 | 0.033917 | 0.22584 | -0.22605 | 0.050512 | 0.046375 | 0.138333 | -0.08359 | 0.572554 | 0.189728 | 0.324442 | 0.14028 | 0.064428 |
| 18.125 | 0.129504 | -0.01588 | 0.030769 | 0.032987 | 0.007362 | 0.039216 | 0.130219 | 0.009672 | -0.02658 | -0.33299 | 0.013087 | -0.00921 | 0.043242 | -0.10722 | 0.364578 | 0.09236 | 0.158902 | 0.08919 | 0.038966 |
| 18.13889 | 0.207397 | 0.096659 | -0.08246 | 0.123841 | 0.031753 | -0.04188 | 0.049105 | -0.03798 | -0.19913 | -0.54091 | -0.11139 | -0.12254 | -0.12874 | -0.16458 | 0.024834 | -0.12583 | 0.0947 | -0.10085 | -0.11304 |
| 18.15278 | 0.25843 | 0.008798 | -0.0323 | 0.00575 | 0.068928 | -0.10788 | 0.543946 | -0.08638 | -0.22404 | -0.83042 | -0.23301 | -0.23084 | -0.22941 | -0.21089 | -0.20549 | -0.34812 | 0.123371 | -0.34252 | -0.32001 |
| 18.16667 | 0.176665 | -0.16384 | 0.75301 | 1.04578 | 0.027637 | -0.10217 | 0.900871 | -0.05746 | -0.06245 | -0.84497 | -0.09146 | -0.12686 | -0.24291 | -0.58294 | 0.102525 | -0.21348 | 0.120295 | -0.28242 | -0.27681 |
| 18.18056 | -0.00665 | -0.20418 | 0.123758 | 0.513595 | -0.04349 | -0.1074 | 0.078134 | -0.05256 | -0.05144 | -0.82609 | -0.0319 | -0.04868 | -0.11409 | -0.12053 | 0.163983 | -0.02918 | -0.16608 | -0.06974 | -0.07935 |
| 18.19444 | -0.16627 | -0.23104 | -0.04591 | -0.06918 | -0.03361 | -0.13099 | -0.08401 | -0.10521 | -0.2904 | -0.99327 | -0.30385 | -0.1519 | -0.18917 | -0.11401 | -0.26949 | -0.11065 | -0.52285 | -0.06151 | -0.0381 |
| 18.20833 | -0.08178 | -0.17175 | -0.05022 | -0.0983 | 0.005651 | -0.09492 | -0.07199 | -0.07841 | -0.37247 | -0.996 | -0.56593 | -0.22086 | -0.22334 | -0.19291 | -0.36279 | -0.17124 | -0.58403 | -0.08281 | -0.06657 |
| 18.22222 | 0.083028 | 0.010305 | -0.02152 | -0.03796 | -0.00894 | -0.04973 | -0.04599 | 0.008293 | -0.22049 | -0.85998 | -0.55516 | -0.21121 | -0.14117 | -0.2197 | -0.08721 | -0.12737 | -0.4085 | -0.04609 | -0.05739 |
| 18.23611 | 0.080131 | 0.151258 | -0.00906 | -0.01143 | -0.06903 | -0.05426 | -0.05488 | 0.03672 | -0.09384 | -0.75675 | -0.42192 | -0.18373 | -0.01694 | -0.17014 | 0.18867 | -0.06949 | -0.21003 | -0.00267 | -0.01767 |
| 18.25 | -0.08857 | 0.017564 | -0.03092 | -0.03492 | -0.1007 | -0.08662 | -0.08302 | -0.07248 | -0.25351 | -0.72687 | -0.34455 | -0.12209 | 0.002231 | -0.1145 | 0.218385 | -0.05072 | -0.06058 | 0.021986 | 0.016196 |
| 18.26389 | -0.21532 | -0.2237 | -0.03565 | -0.05599 | -0.06082 | -0.07744 | -0.06705 | -0.17992 | -0.562 | -0.57922 | -0.29735 | -0.06047 | -0.09012 | -0.06566 | 0.074914 | -0.06442 | -0.05842 | -0.00496 | -0.00502 |
| 18.27778 | -0.09824 | -0.14899 | 0.001418 | -0.03828 | -0.02195 | -0.04268 | -0.02661 | -0.11233 | -0.48925 | -0.44853 | -0.23974 | -0.08562 | 0.001435 | -0.07297 | 0.165458 | -0.04952 | -0.06945 | 0.021544 | -0.0482 |
| 18.29167 | 0.049278 | 0.080435 | 0.01655 | -0.01411 | -0.041 | -0.03375 | -0.04162 | 0.02749 | -0.16326 | -0.49272 | -0.27064 | -0.14607 | 0.366193 | -0.04773 | 0.536707 | 0.066874 | 0.064498 | 0.133985 | -0.18687 |
| 18.30556 | -0.07609 | -0.03495 | -0.00433 | -0.00824 | -0.06929 | -0.04141 | -0.0691 | -0.01215 | -0.08421 | -0.55342 | -0.36493 | -0.08138 | 0.567633 | 0.015621 | 0.716151 | 0.157322 | 0.176921 | 0.187716 | -0.35718 |
| 18.31944 | -0.17665 | -0.21735 | -0.00178 | -0.0086 | -0.02934 | -0.0079 | -0.03715 | -0.03586 | -0.11671 | -0.18815 | -0.00997 | 0.448749 | -0.06636 | -0.24158 | -0.02464 | -0.3749 | 0.124274 | -0.04024 | 0.218388 |
| 18.33333 | -0.01505 | 0.019968 | 0.009606 | -0.00116 | 0.015572 | 0.040379 | -0.00801 | 0.090761 | 0.035237 | 1.408992 | 1.478045 | 0.982687 | -2.28515 | -0.81033 | -2.15793 | -2.08086 | -0.10301 | -0.64922 | 1.674597 |
| 18.34722 | 0.188023 | 0.263958 | -0.01408 | -0.02847 | 0.007331 | 0.041418 | -0.03889 | 0.168839 | 0.189611 | 1.880334 | 0.696251 | 1.174361 | -3.29882 | -0.80275 | -2.42685 | -2.25387 | -0.66565 | -0.56517 | 0.621393 |
| 18.36111 | 0.180956 | 0.136105 | -0.02865 | -0.05326 | 0.006706 | 0.009495 | -0.06052 | 0.111787 | 0.091458 | 0.664685 | 0.20815 | 0.473557 | -1.35985 | -0.24162 | -0.507 | -0.52485 | -0.30094 | -0.07921 | 0.132988 |
| 18.375 | -0.02738 | -0.04607 | -0.00043 | -0.01258 | 0.017575 | 0.0009 | -0.04223 | 0.021821 | -0.05155 | -0.01833 | -0.0206 | 0.139681 | -0.1232 | -0.03233 | 0.034497 | -0.08613 | -0.035 | 0.001536 | 0.065168 |
| 18.38889 | -0.1536 | -0.12834 | 0.024507 | 0.04188 | 0.015373 | 0.026681 | -0.01813 | -0.01235 | -0.09145 | -0.08317 | -0.05302 | 0.100898 | 0.077396 | 0.011798 | 0.095071 | -0.05496 | 0.024794 | -0.04642 | 0.001154 |
| 18.40278 | -0.03189 | -0.02941 | 0.037008 | 0.039003 | 0.049557 | 0.051292 | -0.01075 | 0.065939 | -0.00927 | -0.09062 | -0.0172 | 0.063514 | 0.13107 | -0.02762 | 0.198677 | 0.008332 | 0.09764 | 0.034248 | 0.036068 |
| 18.41667 | 0.227033 | 0.21823 | 0.069479 | 0.0567 | 0.129866 | 0.124608 | 0.033198 | 0.216858 | 0.208982 | -0.08027 | 0.021326 | 0.041468 | 0.201079 | -0.05462 | 0.407904 | 0.128162 | 0.172606 | 0.190847 | 0.101115 |
| 18.43056 | 0.302994 | 0.329388 | 0.166894 | 0.1734 | 0.205735 | 0.269038 | 0.120008 | 0.34563 | 0.402642 | 0.024465 | 0.074356 | 0.162279 | 0.253456 | 0.028898 | 0.588797 | 0.220052 | 0.282824 | 0.290148 | 0.084424 |
| 18.44444 | 0.200525 | 0.267176 | 0.195079 | 0.242504 | 0.243617 | 0.34316 | 0.1663 | 0.343948 | 0.396234 | 0.120316 | 0.151404 | 0.310229 | 0.214184 | 0.206176 | 0.481489 | 0.206158 | 0.331539 | 0.217263 | 0.063169 |
| 18.45833 | 0.239765 | 0.269195 | 0.153128 | 0.176883 | 0.23031 | 0.2827 | 0.157907 | 0.292161 | 0.348998 | 0.171805 | 0.274276 | 0.33693 | 0.195178 | 0.305507 | 0.374846 | 0.200438 | 0.313115 | 0.170469 | 0.134257 |
| 18.47222 | 0.585867 | 0.705729 | 0.077229 | 0.078054 | 0.171719 | 0.186342 | 0.063746 | 0.386015 | 0.69915 | 0.207853 | 0.495599 | 0.36097 | 0.33865 | 0.360631 | 0.513825 | 0.299103 | 0.372349 | 0.249732 | 0.240552 |
| 18.48611 | 1.474137 | 1.703331 | 0.018403 | 0.009584 | 0.116902 | 0.154441 | -0.13413 | 0.695342 | 1.244987 | 0.09793 | 1.145233 | 0.559608 | 0.658537 | 0.621747 | 0.869817 | 0.456334 | 0.698032 | 0.446352 | 0.349311 |
| 18.5 | 3.748588 | 1.907584 | 0.148268 | 0.071535 | 0.315887 | 0.460745 | -0.0089 | 0.954663 | 2.302151 | 0.168019 | 3.334296 | 1.097046 | 1.023096 | 1.022937 | 1.463175 | 0.783925 | 1.411891 | 1.162168 | 0.520786 |
| 18.51389 | 4.31207 | 2.682216 | 0.213881 | 0.283989 | 0.463761 | 0.74355 | 0.31578 | 0.597103 | 2.385856 | 1.039236 | 3.643194 | 1.6262 | 1.146434 | 1.079797 | 1.430529 | 1.094646 | 1.590826 | 1.417049 | 0.652756 |
| 18.52778 | 2.229158 | 1.351436 | 0.096557 | 0.630933 | 0.725739 | 0.465506 | 0.242511 | 0.103998 | 0.98398 | 1.070919 | 1.789791 | 1.16544 | 1.020467 | 0.857593 | 0.889014 | 1.327922 | 0.876874 | 0.885243 | 0.684301 |
| 18.54167 | 1.379261 | 0.644207 | -0.01595 | 0.440173 | 0.624683 | 0.184074 | 0.097126 | 0.096478 | 0.584584 | 0.762898 | 0.831686 | 0.534386 | 0.766659 | 0.726802 | 0.670171 | 1.499723 | 0.500239 | 0.63713 | 0.367118 |
| 18.55556 | 0.750471 | 0.289146 | -0.0132 | 0.318121 | 0.421044 | 0.122693 | 0.04997 | 0.065273 | 0.331557 | 0.447686 | 0.330973 | 0.215126 | 0.438168 | 0.826416 | 0.395064 | 1.486617 | 0.295073 | 0.374642 | 0.275103 |
| 18.56944 | 0.287357 | -0.00935 | 0.0323 | 0.249444 | 0.241559 | 0.115161 | 0.038264 | -0.05421 | 0.029389 | 0.35713 | 0.224546 | 0.107302 | 0.317821 | 0.832383 | 0.319749 | 1.656183 | 0.196502 | 0.210246 | 0.170617 |
| 18.58333 | 0.361306 | 0.029864 | 0.032032 | 0.152125 | 0.187527 | 0.116962 | 0.005407 | 0.010511 | -0.20964 | 0.376979 | 0.247983 | 0.082725 | 0.330741 | 0.662774 | 0.245797 | 1.538924 | 0.173411 | 0.09446 | 0.113126 |
| 18.59722 | 0.52582 | 0.078079 | 0.014746 | 0.114094 | 0.215906 | 0.141275 | -0.0167 | 0.116483 | -0.48131 | 0.348172 | 0.174938 | 0.021525 | 0.298444 | 0.532046 | 0.179191 | 1.108303 | 0.160999 | -0.01302 | 0.04194 |
| 18.61111 | 0.379361 | -0.01513 | 0.025516 | 0.161567 | 0.240781 | 0.116355 | -0.01798 | 0.062999 | -0.63866 | 0.26838 | 0.063254 | -0.07886 | 0.211223 | 0.399566 | 0.207676 | 0.879277 | 0.142567 | -0.0413 | -0.04203 |
| 18.625 | -0.00317 | -0.16987 | 0.034287 | 0.204968 | 0.170726 | 0.004305 | -0.02599 | -0.04145 | -0.60742 | 0.156009 | -0.06723 | -0.16903 | 0.145007 | 0.295506 | 0.245513 | 0.845191 | 0.13107 | 0.055065 | -0.05891 |
| 18.63889 | -0.41077 | -0.25121 | 0.013388 | 0.196607 | 0.069025 | -0.08864 | -0.03197 | -0.09784 | -0.44813 | 0.01585 | -0.18112 | -0.15717 | 0.167045 | 0.215095 | 0.2595 | 0.630926 | 0.139622 | 0.210328 | -0.01669 |
| 18.65278 | -0.47061 | -0.22824 | -0.03762 | 0.144583 | -0.00871 | -0.10185 | -0.03224 | -0.0335 | -0.09414 | -0.09477 | -0.23386 | -0.15236 | 0.239307 | 0.143049 | 0.206741 | 0.440028 | 0.141009 | 0.332098 | 0.022844 |
| 18.66667 | -0.19718 | -0.05974 | -0.08098 | 0.072621 | -0.02889 | -0.05781 | -0.03131 | 0.109794 | 0.536514 | -0.1227 | -0.19925 | -0.1131 | 0.28871 | 0.089669 | 0.124209 | 0.238269 | 0.119321 | 0.370269 | 0.031931 |
| 18.68056 | -0.10352 | 0.023133 | -0.06693 | 0.095448 | -0.03001 | -0.02716 | -0.00755 | 0.045337 | 0.841924 | -0.09855 | -0.1259 | -0.06024 | 0.222543 | 0.070398 | 0.024629 | 0.02761 | 0.097987 | 0.384984 | 0.01859 |
| 18.69444 | -0.11195 | 0.077166 | -0.01668 | 0.122192 | -0.02946 | 0.014591 | 0.026163 | -0.07644 | 0.513971 | -0.04773 | -0.08483 | -0.02166 | 0.111856 | 0.073146 | -0.04583 | -0.21025 | 0.08745 | 0.380892 | -0.01219 |
| 18.70833 | 0.012407 | 0.251859 | 0.019836 | 0.056686 | -0.01119 | 0.068006 | 0.028473 | -0.05259 | 0.142296 | -0.01118 | -0.05738 | 0.007021 | 0.032971 | 0.106632 | -0.11357 | -0.39803 | 0.081249 | 0.30569 | -0.02485 |
| 18.72222 | 0.136613 | 0.375276 | 0.048473 | -0.03949 | 0.011654 | 0.089133 | 0.011615 | 0.0357 | -0.0091 | 0.00714 | -0.03182 | -0.00909 | -0.00953 | 0.141058 | -0.06969 | -0.40736 | 0.08025 | 0.165141 | -0.01576 |
| 18.73611 | 0.250166 | 0.403323 | 0.063615 | -0.12333 | 0.047206 | 0.095726 | -0.00767 | 0.130812 | -0.12009 | -0.00155 | -0.02038 | -0.06664 | -0.01088 | 0.131877 | 0.127633 | -0.23754 | 0.054298 | 0.048288 | -0.01175 |
| 18.75 | 0.139229 | 0.336443 | 0.048751 | -0.05971 | 0.064265 | 0.071295 | 0.014127 | 0.068853 | -0.44283 | -0.01907 | -0.02298 | -0.12101 | 0.010316 | 0.100428 | 0.323533 | -0.05758 | 0.002793 | -0.00233 | -0.01796 |
| 18.76389 | -0.19658 | 0.134145 | -0.00088 | 0.049703 | 0.046325 | 0.025751 | 0.046052 | -0.1218 | -0.77505 | -0.03148 | -0.02964 | -0.13145 | -0.03443 | 0.07884 | 0.367995 | 0.015613 | -0.03862 | -0.01016 | -0.02031 |
| 18.77778 | -0.37663 | 0.072194 | -0.03339 | 0.123491 | 0.012711 | 0.017708 | 0.079145 | -0.19356 | -0.73762 | -0.03029 | -0.02998 | -0.1138 | -0.08359 | 0.087506 | 0.239911 | 0.003679 | -0.01944 | 0.037183 | -0.01041 |
| 18.79167 | -0.31021 | 0.096854 | -0.02383 | 0.120574 | -0.00466 | 0.0514 | 0.109716 | -0.12084 | -0.29985 | -0.01903 | -0.02303 | -0.06123 | 0.035573 | 0.116385 | 0.109346 | -0.03352 | 0.006459 | 0.174502 | 0.002329 |
| 18.80556 | -0.09773 | 0.271017 | 0.001052 | 0.039285 | -0.01387 | 0.07811 | 0.085259 | 0.031637 | 0.233807 | -0.02845 | -0.03345 | -0.03526 | 0.206474 | 0.123282 | -0.0145 | -0.08924 | -0.01913 | 0.30905 | -0.01271 |
| 18.81944 | 0.10504 | 0.378885 | 0.041031 | -0.048 | -0.00703 | 0.082669 | 0.032438 | 0.15885 | 0.542274 | -0.04635 | -0.05776 | -0.07478 | 0.178004 | 0.111842 | -0.09429 | -0.13048 | -0.06122 | 0.219783 | -0.04068 |
| 18.83333 | 0.167769 | 0.437348 | 0.066453 | -0.07111 | 0.003527 | 0.058828 | -0.00449 | 0.172068 | 0.358599 | -0.00718 | -0.05637 | -0.10931 | -0.04026 | 0.092247 | -0.17348 | -0.18218 | -0.08994 | -0.04168 | -0.03726 |
| 18.84722 | -0.0253 | 0.402189 | 0.03018 | -0.06949 | -0.03267 | -0.00049 | -0.01872 | 0.047823 | -0.18097 | 0.0458 | -0.04733 | -0.10848 | -0.28576 | 0.073441 | -0.38042 | -0.29108 | -0.09699 | -0.33326 | -0.00393 |
| 18.86111 | -0.30433 | 0.38696 | -0.05866 | -0.09186 | -0.07791 | -0.04266 | -0.02051 | -0.08303 | -0.66294 | 0.003009 | -0.07215 | -0.12584 | -0.40852 | 0.057376 | -0.53744 | -0.36433 | -0.12706 | -0.5651 | 0.004945 |
| 18.875 | -0.56308 | 0.397744 | -0.11172 | -0.06301 | -0.07738 | -0.0334 | 0.029219 | -0.25602 | -0.80248 | -0.0881 | -0.09652 | -0.16233 | -0.48864 | 0.046563 | -0.47476 | -0.35574 | -0.19286 | -0.5601 | -0.03706 |
| 18.88889 | -0.5728 | 0.3642 | -0.07866 | -0.05676 | -0.07291 | -0.00701 | 0.047034 | -0.27019 | -0.74565 | -0.14123 | -0.08058 | -0.14772 | -0.58543 | 0.059898 | -0.35008 | -0.32135 | -0.21645 | -0.20296 | -0.06693 |
| 18.90278 | -0.40084 | 0.625938 | -0.03766 | -0.13516 | -0.06919 | -0.02069 | -0.02193 | -0.07895 | -0.66357 | -0.18386 | -0.05707 | -0.09109 | -0.75047 | 0.08646 | -0.35213 | -0.28981 | -0.19462 | 0.165358 | -0.05937 |
| 18.91667 | -0.52002 | 0.608974 | -0.08564 | -0.15697 | -0.13046 | -0.11373 | -0.06366 | -0.05295 | -0.8213 | -0.24504 | -0.05715 | -0.06672 | -0.76582 | 0.079932 | -0.39824 | -0.27489 | -0.21874 | 0.224875 | -0.03829 |
| 18.93056 | -0.68875 | 0.525431 | -0.21098 | -0.1762 | -0.22255 | -0.2209 | -0.10246 | -0.1384 | -0.90994 | -0.27797 | -0.0591 | -0.09971 | -0.4389 | 0.049417 | -0.35451 | -0.26582 | -0.28005 | -0.02148 | -0.00866 |
| 18.94444 | -0.54395 | 0.610414 | -0.26883 | -0.23139 | -0.21467 | -0.20017 | -0.12869 | -0.11551 | -0.6331 | -0.31419 | -0.06226 | -0.1437 | -0.05266 | 0.033324 | -0.24766 | -0.22444 | -0.32318 | -0.32479 | 0.013026 |
| 18.95833 | -0.57218 | 0.793595 | -0.24287 | -0.12805 | -0.13893 | -0.10576 | -0.04729 | -0.18448 | -0.32591 | -0.34301 | -0.07249 | -0.18363 | 0.012126 | 0.029427 | -0.21367 | -0.2038 | -0.328 | -0.56956 | 0.000405 |
| 18.97222 | -0.91292 | 0.554539 | -0.22424 | -0.02974 | -0.16071 | -0.0543 | 0.094116 | -0.47733 | -0.46728 | -0.36669 | -0.09794 | -0.23865 | -0.23707 | 0.004327 | -0.34625 | -0.25995 | -0.38825 | -0.65968 | -0.04441 |
| 18.98611 | -1.27381 | -0.28372 | -0.24443 | -0.20531 | -0.28317 | -0.15509 | 0.089472 | -0.79783 | -1.43017 | -0.60778 | -0.38225 | -0.46489 | -0.7781 | -0.14475 | -0.79759 | -0.63708 | -0.57507 | -0.92251 | -0.26222 |
| 19 | -2.05192 | -1.23252 | -0.31683 | -0.60427 | -0.46783 | -0.54275 | -0.15221 | -0.92074 | -2.62014 | -2.00797 | -1.44515 | -1.06478 | -1.8439 | -0.51184 | -2.19692 | -1.96091 | -1.31508 | -2.33091 | -0.98894 |
| 19.01389 | -2.02981 | -1.22722 | -0.34345 | -0.94941 | -0.64666 | -0.63224 | -0.38379 | -0.65792 | -2.08818 | -2.41112 | -3.4904 | -1.5374 | -0.61056 | -0.66192 | -2.47142 | -2.57271 | -1.2634 | -1.84732 | -1.95555 |
| 19.02778 | -0.67756 | 0.08064 | -0.12958 | -0.5842 | -0.62135 | -0.36651 | -0.31495 | -0.02236 | -0.56379 | -2.61779 | -5.26563 | -1.51105 | 2.440804 | 0.072427 | 0.187333 | -1.15725 | -0.2526 | 0.235265 | -2.44554 |
| 19.04167 | 0.088709 | 0.529106 | 0.058327 | 0.008171 | -0.1991 | -0.05701 | -0.10094 | 0.560325 | 0.706088 | -1.68569 | -3.18393 | -1.06043 | 2.591724 | 0.934396 | 1.447266 | 0.142565 | 0.298265 | 1.15158 | -2.15529 |
| 19.05556 | -0.04909 | 0.71313 | 0.153879 | 0.088299 | -0.14573 | 0.065965 | 0.08433 | 0.183307 | 0.801012 | 1.926789 | -0.17584 | -0.27897 | 0.990555 | 0.452853 | 0.948517 | 0.16734 | 0.190787 | 0.827312 | -1.06467 |
| 19.06944 | 0.102376 | 0.701521 | -0.00494 | 0.053689 | -0.16312 | -0.00225 | 0.004285 | 0.065871 | 0.009027 | 0.022609 | -0.28045 | -0.02951 | 0.719719 | 0.04806 | 0.446262 | -0.14243 | 0.048676 | 0.575328 | -0.05159 |
| 19.08333 | 0.230186 | 1.443287 | -0.2098 | -0.03141 | -0.11023 | -0.05665 | -0.10867 | -0.214 | -0.74836 | -1.9276 | -0.92763 | 0.001482 | 0.895475 | -0.11397 | 0.092155 | -0.34979 | 0.04304 | 0.515476 | 0.431044 |
| 19.09722 | 0.179507 | 0.944983 | -0.10819 | -0.07919 | -0.0449 | -0.04652 | -0.06889 | -0.23299 | -0.47534 | -0.061 | -0.3243 | 0.026985 | 0.566878 | -0.01096 | -0.01475 | -0.28221 | 0.009952 | 0.314524 | 0.22152 |
| 19.11111 | 0.0786 | 0.545695 | -0.032 | -0.03238 | -0.0582 | -0.03352 | -0.03167 | -0.01009 | 0.007771 | -0.04442 | -0.18241 | 0.007932 | 0.37791 | 0.037191 | 0.133938 | -0.12542 | -0.01947 | 0.259542 | -0.01045 |
| 19.125 | 0.050135 | 0.365446 | 0.002838 | -0.0339 | -0.05665 | -0.03892 | -0.04341 | 0.027088 | 0.088096 | -0.03814 | -0.08452 | -0.0044 | 0.337281 | 0.020841 | 0.225228 | -0.0257 | 0.014668 | 0.321103 | 0.004656 |
| 19.13889 | 0.044232 | 0.183481 | 0.006624 | -0.04388 | -0.02801 | -0.03176 | -0.04006 | 0.044026 | 0.068128 | -0.03972 | -0.05463 | -0.02453 | 0.270302 | 0.006457 | 0.175011 | 0.000229 | -0.01165 | 0.304124 | 0.050184 |
| 19.15278 | 0.072489 | 0.239531 | 0.010146 | -0.03066 | -0.03105 | -0.02368 | -0.03119 | 0.065077 | 0.075325 | -0.09308 | -0.03258 | -0.05325 | 0.255523 | -0.00956 | 0.132136 | -0.02542 | -0.0897 | 0.276956 | 0.01624 |

| | | | | | | | | | | | | | | | | | | | |
|---|---|---|---|---|---|---|---|---|---|---|---|---|---|---|---|---|---|---|---|
| 19.16667 | 0.04025 | 0.38588 | 0.005036 | 0.002233 | -0.03802 | -0.00866 | -0.02128 | 0.036228 | 0.07233 | -0.11883 | -0.02768 | -0.0359 | 0.28517 | -0.0065 | 0.114699 | -0.04799 | -0.05227 | 0.228045 | -0.02272 |
| 19.18056 | -0.04621 | 0.264884 | 0.008581 | 0.02689 | -0.03836 | -0.00176 | -0.00227 | -0.00935 | 0.021522 | -0.04961 | -0.0105 | -0.03337 | 0.290201 | -0.01497 | 0.115282 | -0.04356 | -0.05524 | 0.183089 | 0.009406 |
| 19.19444 | -0.12149 | 0.072347 | 0.024317 | 0.025957 | -0.02123 | -0.00044 | 0.01506 | -0.01732 | -0.00455 | -0.04996 | -0.01209 | -0.06935 | 0.242812 | -0.04284 | 0.062811 | -0.06919 | -0.18385 | 0.121916 | 0.014074 |
| 19.20833 | -0.18999 | -0.00437 | 0.051944 | 0.036374 | -0.00642 | 0.00959 | 0.040118 | -0.00795 | -0.05179 | -0.11725 | -0.03896 | -0.06464 | 0.166239 | -0.05697 | -0.01793 | -0.09807 | -0.17988 | 0.044208 | -0.02546 |
| 19.22222 | -0.19207 | -0.00087 | 0.045173 | 0.013715 | -0.03426 | 0.00517 | 0.036589 | -0.01606 | -0.08804 | -0.04205 | -0.06219 | -0.02467 | 0.132786 | -0.035 | -0.0253 | -0.07569 | -0.07648 | 0.073338 | -0.04712 |
| 19.23611 | -0.06412 | 0.265752 | 0.005444 | -0.05015 | -0.04966 | -0.01704 | -0.00974 | 0.013503 | -0.01312 | 0.051662 | -0.0728 | -0.00605 | 0.135663 | -0.01681 | -0.01151 | -0.05392 | -0.03882 | 0.152441 | -0.04236 |
| 19.25 | -0.04154 | 0.483585 | -0.0026 | -0.05743 | -0.05822 | -0.03589 | -0.04528 | 0.03701 | 0.104641 | 0.006228 | -0.07134 | -0.01234 | 0.149842 | -0.05023 | -0.02052 | -0.06905 | -0.00842 | 0.151219 | -0.04188 |
| 19.26389 | -0.0782 | 0.475512 | 0.010156 | -0.02681 | -0.0521 | -0.04609 | -0.05063 | 0.014593 | 0.108191 | -0.02637 | -0.07313 | -0.01904 | 0.119535 | -0.07039 | -0.03877 | -0.08168 | 0.016184 | 0.10334 | -0.07041 |
| 19.27778 | 0.002707 | 0.558113 | 0.049878 | -0.01445 | -0.00806 | -0.03045 | -0.03055 | 0.024291 | 0.109062 | -0.03254 | -0.06618 | -0.02678 | 0.023805 | -0.09179 | -0.00639 | -0.07175 | -0.0017 | 0.129617 | -0.07373 |
| 19.29167 | 0.06392 | 0.737414 | 0.084893 | -0.01111 | 0.015276 | -0.01233 | -0.02499 | 0.036062 | 0.163801 | -0.09228 | -0.03923 | -0.03723 | -0.03524 | -0.15491 | 0.078803 | -0.06301 | -0.00444 | 0.261575 | -0.02376 |
| 19.30556 | 0.13221 | 0.94171 | 0.090079 | -0.03277 | 0.018389 | -0.00189 | -0.0401 | 0.06348 | 0.269317 | -0.0464 | -0.03179 | 0.005032 | -0.05285 | -0.13121 | 0.144641 | -0.02186 | 0.050139 | 0.324512 | -0.01582 |
| 19.31944 | 0.294835 | 1.094678 | 0.096373 | -0.05783 | 0.045415 | 0.021084 | -0.05231 | 0.132702 | 0.409909 | 0.078248 | -0.05193 | 0.059631 | -0.04305 | -0.01509 | 0.215602 | 0.046828 | 0.10511 | 0.278485 | -0.06094 |
| 19.33333 | 0.318316 | 1.008715 | 0.104748 | -0.04665 | 0.052331 | 0.031755 | -0.05148 | 0.163043 | 0.449158 | 0.083024 | -0.04838 | 0.064838 | 0.062691 | 0.033424 | 0.339917 | 0.104508 | 0.109037 | 0.301011 | -0.08255 |
| 19.34722 | 0.134052 | 0.862524 | 0.096553 | -0.02908 | 0.025637 | 0.018146 | -0.04874 | 0.083465 | 0.361851 | 0.034537 | -0.05135 | 0.05191 | 0.228629 | 0.039339 | 0.45038 | 0.142239 | 0.091138 | 0.351025 | -0.0736 |
| 19.36111 | -0.0352 | 0.908461 | 0.07382 | -0.01021 | 0.030806 | 0.025089 | -0.04341 | 0.025638 | 0.336761 | -0.00345 | -0.05084 | 0.037387 | 0.392983 | 0.017163 | 0.476522 | 0.154626 | 0.069673 | 0.429183 | -0.01793 |
| 19.375 | -0.05777 | 1.058595 | 0.111804 | 0.084196 | 0.065433 | 0.064525 | 0.012538 | 0.048205 | 0.42651 | -0.03301 | 0.006638 | 0.019416 | 0.544878 | -0.03716 | 0.517504 | 0.160775 | 0.084268 | 0.560854 | 0.055126 |
| 19.38889 | -0.08198 | 0.976423 | 0.146051 | 0.178401 | 0.065288 | 0.095288 | 0.07149 | 0.068003 | 0.470508 | 0.016673 | 0.048504 | 0.048 | 0.54275 | -0.00011 | 0.54565 | 0.172215 | 0.119758 | 0.611795 | 0.065239 |
| 19.40278 | -0.0562 | 0.819231 | 0.130642 | 0.159773 | 0.076592 | 0.084765 | 0.048728 | 0.119751 | 0.445904 | 0.08864 | 0.037106 | 0.106868 | 0.414566 | 0.111453 | 0.433115 | 0.1657 | 0.127102 | 0.514108 | 0.028888 |
| 19.41667 | 0.155168 | 0.903109 | 0.148104 | 0.161332 | 0.150176 | 0.1402 | 0.064541 | 0.213043 | 0.505061 | 0.106248 | 0.039872 | 0.122254 | 0.300958 | 0.140063 | 0.372531 | 0.144169 | 0.133518 | 0.501029 | 0.022531 |
| 19.43056 | 0.302392 | 1.010805 | 0.213549 | 0.291057 | 0.269781 | 0.278712 | 0.169336 | 0.299763 | 0.551795 | 0.082115 | 0.057698 | 0.086143 | 0.286561 | 0.077215 | 0.371066 | 0.123657 | 0.125379 | 0.533753 | 0.052262 |
| 19.44444 | 0.320931 | 0.931577 | 0.225507 | 0.362565 | 0.325144 | 0.348074 | 0.22724 | 0.334537 | 0.582094 | 0.088811 | 0.063997 | 0.070597 | 0.345675 | 0.048605 | 0.372567 | 0.122217 | 0.124654 | 0.556412 | 0.089938 |
| 19.45833 | 0.318994 | 0.90933 | 0.193967 | 0.2764 | 0.269396 | 0.280851 | 0.183456 | 0.333067 | 0.63277 | 0.18541 | 0.095818 | 0.136308 | 0.413169 | 0.10814 | 0.449788 | 0.195936 | 0.176887 | 0.552198 | 0.132034 |
| 19.47222 | 0.555123 | 1.212055 | 0.147798 | 0.05182 | 0.159371 | 0.162629 | 0.03669 | 0.375858 | 0.842692 | 0.336797 | 0.142556 | 0.245165 | 0.478895 | 0.23843 | 0.540843 | 0.290332 | 0.283416 | 0.580726 | 0.173881 |
| 19.48611 | 1.194627 | 1.601033 | 0.23668 | -0.20812 | 0.166317 | 0.105518 | -0.13292 | 0.504876 | 1.495021 | 0.426993 | 0.256953 | 0.315296 | 0.645856 | 0.462916 | 0.507954 | 0.335506 | 0.541862 | 0.767849 | 0.25239 |
| 19.5 | 2.466492 | 1.736022 | 0.326623 | 0.033324 | 0.214049 | 0.311708 | -0.02727 | 0.494164 | 3.079907 | 0.955106 | 0.626195 | 0.588515 | 0.961222 | 0.584919 | 0.460785 | 0.388632 | 1.047465 | 1.576698 | 0.481473 |
| 19.51389 | 2.366604 | 0.301963 | 0.230444 | 0.506034 | 0.582491 | 0.651709 | 0.371406 | 0.345927 | 2.175165 | 0.96945 | 1.579927 | 0.355073 | 1.153854 | 1.215135 | 0.36427 | 0.535497 | 1.151476 | 1.502343 | 0.851665 |
| 19.52778 | 1.66808 | -0.4977 | 0.002761 | 0.419523 | 0.761375 | 0.725148 | 0.461039 | 0.218417 | 0.903455 | 0.461217 | 2.188496 | 0.46773 | 0.523559 | 0.799672 | -0.02625 | 0.67519 | 0.606887 | 0.519352 | 1.032794 |
| 19.54167 | 0.857613 | 0.748077 | -0.09637 | 0.355387 | 0.440118 | 0.502108 | 0.387661 | 0.223858 | 0.202371 | 0.374087 | 1.765616 | 0.182272 | -0.10718 | 0.490571 | -0.0902 | 0.932432 | 0.260284 | -0.05273 | 1.044741 |
| 19.55556 | 0.407737 | 1.641745 | -0.14892 | 0.272438 | 0.274474 | 0.277615 | 0.288212 | 0.276786 | 0.270263 | 0.413443 | 1.136003 | 0.042617 | -0.53096 | 0.46437 | -0.32444 | 1.053168 | 0.098617 | -0.30957 | 0.551855 |
| 19.56944 | 0.46308 | 1.835212 | -0.14887 | 0.114571 | 0.083701 | 0.16226 | 0.182887 | 0.189789 | 0.100414 | 0.451585 | 0.640982 | 0.071069 | -0.70207 | 0.411841 | -0.51136 | 0.744068 | 0.053963 | -0.36823 | 0.218674 |
| 19.58333 | 0.51768 | 1.634208 | -0.07628 | 0.033069 | -0.03525 | 0.118381 | 0.117331 | 0.086454 | -0.2615 | 0.469589 | 0.37542 | 0.130101 | -0.57655 | 0.246386 | -0.00589 | 0.62842 | 0.080108 | -0.28405 | 0.079445 |
| 19.59722 | 0.344072 | 0.986737 | 0.060511 | 0.039869 | -0.01811 | 0.104035 | 0.076454 | 0.160783 | -0.09779 | 0.536092 | 0.2869 | 0.140622 | -0.02278 | 0.123422 | 0.605469 | 0.601182 | 0.117224 | -0.02329 | 0.020543 |
| 19.61111 | 0.12273 | 0.348741 | 0.174331 | -0.00743 | 0.023206 | 0.073256 | 0.044569 | 0.326701 | 0.422836 | 0.646573 | 0.3165 | 0.076363 | 0.893013 | 0.102693 | 0.965148 | 0.645206 | 0.13404 | 0.360141 | -0.01987 |
| 19.625 | -0.05912 | 0.099279 | 0.113543 | -0.05438 | 0.080447 | -0.01324 | 0.01211 | 0.370945 | 0.969007 | 0.652176 | 0.342386 | -0.04386 | 1.307958 | 0.113131 | 0.321963 | 0.581645 | 0.196839 | 0.699225 | -0.03858 |
| 19.63889 | -0.12673 | 0.289587 | -0.04161 | -0.03774 | 0.181361 | -0.06013 | 0.003543 | 0.264935 | 1.502984 | 0.539444 | 0.316198 | -0.14874 | 0.666097 | 0.106592 | -0.17661 | 0.635749 | 0.229036 | 0.787176 | -0.04895 |
| 19.65278 | -0.17123 | 0.471828 | -0.19862 | -0.00152 | 0.14786 | -0.06268 | 0.017366 | 0.015489 | 1.770727 | 0.388353 | 0.293328 | -0.17127 | -0.24046 | 0.035068 | -0.3344 | 0.786901 | 0.11739 | 0.529714 | -0.05846 |
| 19.66667 | -0.3851 | 0.358194 | -0.28779 | 0.061623 | -0.007 | -0.09501 | 0.024159 | -0.13734 | 1.02425 | 0.457344 | 0.300444 | -0.11691 | -0.77313 | -0.05547 | -0.52082 | 0.771784 | -0.06268 | 0.208168 | -0.05304 |
| 19.68056 | -0.36544 | 0.416622 | -0.20097 | 0.07503 | -0.16946 | -0.03496 | 0.061085 | -0.11306 | 0.030391 | 0.746428 | 0.284608 | 0.00488 | -1.06244 | -0.11239 | -0.58377 | 0.598169 | -0.16922 | -0.08625 | -0.03685 |
| 19.69444 | 0.081483 | 0.874607 | -0.12606 | -0.02297 | -0.26916 | 0.060277 | 0.084397 | -0.0559 | -0.51936 | 0.927883 | 0.254985 | 0.16098 | -1.20445 | -0.12223 | -0.24043 | 0.459198 | -0.17816 | -0.33343 | -0.04227 |
| 19.70833 | 0.387381 | 0.966902 | -0.01244 | -0.1098 | -0.25535 | 0.064967 | 0.029195 | -0.03408 | -0.79191 | 0.74997 | 0.142489 | 0.260655 | -1.05295 | -0.09657 | 0.535491 | 0.410124 | -0.15268 | -0.60385 | -0.07415 |
| 19.72222 | 0.179558 | 0.502579 | 0.15155 | -0.05266 | -0.17985 | -0.01345 | -0.03361 | -0.05163 | -0.82039 | 0.425336 | 0.040895 | 0.179015 | 0.056717 | -0.08989 | 1.213311 | 0.387905 | -0.13417 | -0.70935 | -0.10936 |
| 19.73611 | -0.14986 | 0.218416 | 0.161833 | -0.00808 | -0.07295 | -0.08327 | -0.0653 | -0.05534 | -0.47614 | 0.105219 | -0.03029 | 0.001826 | 1.569927 | -0.09875 | 1.464141 | 0.347525 | -0.15746 | -0.32973 | -0.10342 |
| 19.75 | -0.22401 | 0.535703 | 0.015382 | -0.07444 | 0.083113 | -0.09996 | -0.07546 | -0.01663 | -0.0368 | -0.28589 | -0.1451 | -0.04705 | 1.758591 | -0.10883 | 0.925681 | 0.200949 | -0.13356 | 0.452864 | -0.05082 |
| 19.76389 | -0.24778 | 0.823847 | -0.10802 | -0.02977 | 0.164665 | -0.09263 | -0.0487 | -0.07177 | 0.269786 | -0.25213 | -0.16681 | 0.011827 | 0.738856 | -0.11 | 0.416063 | 0.077425 | 0.001185 | 0.968366 | -0.0356 |
| 19.77778 | -0.17549 | 0.782953 | -0.13424 | 0.024127 | 0.142526 | -0.06539 | 0.01027 | -0.19337 | 0.451262 | -0.18576 | -0.1194 | 0.060642 | -0.28052 | -0.09739 | -0.18189 | -0.02389 | -0.00727 | 0.513454 | -0.07885 |
| 19.79167 | 0.154055 | 0.906542 | -0.13399 | -0.13437 | 0.082485 | 0.00288 | 0.024951 | -0.1439 | 0.573758 | -0.14335 | -0.07941 | 0.068826 | -1.10408 | -0.06032 | -0.56636 | -0.0709 | -0.07324 | -0.19775 | -0.1007 |
| 19.80556 | 0.19228 | 0.83609 | -0.16435 | -0.27591 | 0.015834 | 0.055082 | -0.03002 | -0.00399 | 0.004375 | -0.00427 | 0.001139 | 0.049512 | -1.56884 | 0.015358 | -0.69178 | -0.07844 | -0.16333 | -0.80372 | -0.08391 |
| 19.81944 | -0.3346 | 0.113365 | -0.15573 | -0.21933 | -0.07589 | 0.051708 | -0.05453 | -0.11063 | -1.32142 | 0.120451 | 0.09757 | 0.005672 | -1.74927 | 0.062323 | -0.75756 | -0.12548 | -0.25199 | -1.29224 | -0.05075 |
| 19.83333 | -0.70594 | -0.62803 | -0.05085 | -0.06118 | -0.24088 | -0.03127 | -0.03363 | -0.28051 | -2.06438 | 0.186671 | 0.151079 | -0.04719 | -1.50218 | 0.053928 | -0.58076 | -0.13998 | -0.22008 | -1.19873 | -0.03715 |
| 19.84722 | -0.46717 | -0.84341 | 0.032707 | -0.19654 | -0.33861 | -0.10682 | -0.03982 | -0.25927 | -1.57049 | 0.214662 | 0.136268 | -0.04778 | -0.70408 | -0.02552 | -0.1869 | -0.11398 | -0.27689 | -0.74701 | -0.05305 |
| 19.86111 | -0.19417 | -1.02895 | 0.020009 | -0.467 | -0.40086 | -0.20045 | -0.11155 | -0.26832 | -0.93817 | 0.15245 | 0.09854 | 0.0432 | 0.163132 | -0.12023 | 0.248789 | -0.00835 | -0.29529 | -0.15299 | -0.0543 |
| 19.875 | -0.40779 | -1.42125 | -0.06602 | -0.46421 | -0.41501 | -0.2765 | -0.13496 | -0.45095 | -0.3951 | 0.050254 | 0.071993 | 0.102078 | 0.369324 | -0.19092 | 0.553682 | 0.127429 | -0.24959 | 0.461552 | -0.01187 |
| 19.88889 | -0.60695 | -1.27504 | -0.12954 | -0.30804 | -0.37187 | -0.21299 | -0.06471 | -0.69656 | -0.22893 | 0.004502 | 0.094101 | 0.080701 | 0.050346 | -0.22452 | 0.519047 | 0.165097 | -0.16515 | 0.59636 | 0.003698 |
| 19.90278 | -0.80675 | -0.8162 | -0.13451 | -0.22633 | -0.33814 | -0.1733 | -0.00667 | -0.82444 | -0.48337 | 0.033988 | 0.15747 | 0.037981 | -0.36286 | -0.22956 | 0.321735 | 0.11091 | -0.16528 | 0.247261 | -0.00653 |
| 19.91667 | -0.97785 | -0.54952 | -0.12587 | -0.27168 | -0.33259 | -0.27412 | -0.02231 | -0.68988 | -0.58728 | 0.059715 | 0.18617 | -0.00684 | -0.66071 | -0.23669 | 0.092251 | 0.022545 | -0.25698 | -0.15154 | -0.00993 |
| 19.93056 | -0.86916 | -0.43276 | -0.1516 | -0.53043 | -0.37161 | -0.41496 | -0.15442 | -0.54648 | -0.57399 | 0.110945 | 0.146112 | -0.03776 | -0.83765 | -0.28064 | -0.19371 | -0.08688 | -0.36317 | -0.48434 | 0.031376 |
| 19.94444 | -0.80336 | -0.46258 | -0.14707 | -0.64276 | -0.4 | -0.43869 | -0.23367 | -0.44059 | -0.57218 | 0.104646 | 0.074105 | -0.018 | -0.84604 | -0.36449 | -0.51468 | -0.21676 | -0.38836 | -0.69607 | 0.079476 |
| 19.95833 | -0.79846 | -0.6567 | -0.05622 | -0.41338 | -0.42077 | -0.33324 | -0.16903 | -0.39262 | -0.51083 | 0.037517 | 0.012037 | 0.034719 | -0.80369 | -0.41107 | -0.79304 | -0.39683 | -0.31917 | -0.70877 | 0.09024 |
| 19.97222 | -0.98356 | -1.10183 | -0.01353 | -0.1298 | -0.50004 | -0.33556 | -0.0599 | -0.57458 | -0.50175 | -0.1291 | -0.04864 | 0.019006 | -0.68272 | -0.43262 | -0.89781 | -0.57002 | -0.28546 | -0.5077 | 0.022302 |
| 19.98611 | -1.12218 | -1.99771 | -0.03213 | -0.06676 | -0.55064 | -0.467 | 0.003029 | -0.91794 | -1.07327 | -0.74577 | -0.34174 | -0.25016 | -0.48435 | -0.47599 | -1.05937 | -0.77206 | -0.35757 | -0.43381 | -0.19715 |
| 20 | -1.25541 | -2.89634 | -0.102 | -0.34283 | -0.63823 | -0.65478 | -0.13105 | -1.1063 | -2.06275 | -2.07344 | -1.41258 | -0.90223 | -0.5394 | -0.7078 | -1.71863 | -1.33008 | -1.23344 | -1.00139 | -0.75854 |
| 20.01389 | -1.58766 | -2.32155 | -0.17028 | -0.7355 | -0.77363 | -0.58688 | -0.24568 | -0.8573 | -2.13174 | -2.54998 | -2.59317 | -1.11849 | -0.77051 | -0.78401 | -2.34789 | -1.65412 | -1.92987 | -1.45571 | -1.36419 |
| 20.02778 | -0.44942 | -0.7294 | -0.13862 | -0.5616 | -0.41332 | -0.31009 | -0.14357 | -0.40776 | -1.44274 | -0.92313 | -1.46592 | -0.09295 | -0.9334 | -0.12717 | -2.2928 | -1.42428 | -0.28899 | -0.8866 | -0.5658 |
| 20.04167 | 0.054019 | 0.275743 | -0.14596 | -0.21485 | -0.00972 | -0.08832 | -0.03629 | -0.27136 | -0.88214 | 0.598599 | 0.516967 | 0.673386 | -0.99711 | 0.29067 | -1.898 | -0.85952 | 0.8718 | -0.19755 | 0.638733 |
| 20.05556 | 0.044159 | 0.68005 | -0.18371 | -0.04308 | 0.07844 | 0.001866 | -0.01179 | -0.26108 | -0.48637 | 0.600408 | 0.949743 | 0.719992 | -0.38619 | 0.604507 | -0.85193 | -0.0904 | 1.102238 | 0.065414 | 0.647673 |
| 20.06944 | 0.088681 | 0.605406 | -0.01668 | 0.123052 | 0.149842 | 0.152531 | 0.125492 | -0.0115 | 0.013455 | 0.09366 | 0.411483 | 0.456811 | 0.581989 | 0.851849 | 0.264941 | 0.494524 | 0.782181 | 0.150913 | 0.234409 |
| 20.08333 | 0.035856 | 0.12669 | 0.127031 | 0.167823 | 0.195984 | 0.229544 | 0.214008 | 0.170218 | 0.162824 | 0.196372 | 0.358132 | 0.176066 | 0.486508 | 0.417561 | 0.224123 | 0.425309 | 0.134764 | 0.041605 | 0.227362 |
| 20.09722 | -0.12831 | -0.22424 | 0.05367 | 0.072426 | 0.097125 | 0.11316 | 0.099328 | 0.065711 | -0.00064 | 0.202946 | 0.197721 | -0.09988 | 0.019432 | -0.00644 | -0.08848 | 0.14537 | -0.33797 | -0.09969 | 0.158652 |
| 20.11111 | -0.05152 | -0.05788 | -0.00444 | 0.025312 | 0.036625 | 0.042826 | 0.021005 | 0.013248 | 0.003084 | 0.080147 | -0.01803 | -0.10381 | -0.25279 | -0.10285 | -0.08899 | 0.003778 | -0.33012 | -0.12496 | -0.01048 |
| 20.125 | 0.194624 | 0.332912 | 0.053114 | 0.040407 | 0.085341 | 0.056111 | 0.035721 | 0.117678 | 0.159383 | 0.065131 | -0.03266 | 0.008736 | -0.20038 | -0.06452 | 0.076494 | -0.00042 | -0.15227 | -0.0182 | -0.04918 |
| 20.13889 | 0.122847 | 0.376945 | 0.102953 | 0.044273 | 0.077906 | 0.034483 | 0.047501 | 0.1188 | 0.124132 | 0.074955 | 0.009496 | 0.020238 | 0.078802 | -0.08106 | 0.222171 | 0.062565 | -0.03249 | 0.147816 | 0.013523 |
| 20.15278 | -0.10132 | 0.145965 | 0.059066 | 0.023125 | 0.02248 | -0.01183 | 0.025729 | 0.012521 | -0.00676 | 0.038076 | -0.00117 | -0.01179 | 0.293791 | -0.12444 | 0.203257 | 0.098747 | -0.00975 | 0.22531 | 0.046906 |
| 20.16667 | -0.11929 | 0.094618 | -0.01361 | 0.018215 | 0.024131 | -0.0286 | -0.04144 | 0.033951 | 0.037905 | -0.05121 | -0.06778 | -0.03416 | 0.316309 | -0.16544 | 0.081658 | 0.04298 | -0.06965 | 0.159423 | -0.0324 |
| 20.18056 | -0.05155 | 0.084161 | -0.02752 | 0.035964 | 0.014808 | -0.01936 | -0.04861 | 0.029103 | 0.081247 | -0.08542 | -0.09102 | -0.04152 | 0.301269 | -0.16193 | 0.08426 | 0.0038 | -0.09204 | 0.142522 | -0.10168 |
| 20.19444 | -0.15273 | -0.11707 | 0.005943 | 0.031959 | -0.03213 | 0.00674 | 0.050663 | -0.11574 | -0.06924 | -0.04858 | -0.05956 | -0.02908 | 0.32918 | -0.10994 | 0.205749 | 0.03242 | -0.0463 | 0.187276 | -0.07679 |
| 20.20833 | -0.1315 | -0.09224 | 0.034556 | 0.002104 | -0.00636 | 0.030223 | 0.088069 | -0.11396 | -0.06396 | -0.03204 | -0.03587 | -0.02903 | 0.339757 | -0.09085 | 0.33711 | 0.068205 | -0.01065 | 0.220695 | -0.02036 |
| 20.22222 | -0.01658 | 0.233624 | 0.041871 | -0.01242 | 0.055894 | 0.046661 | 0.054661 | 0.035666 | 0.162217 | -0.02507 | -0.0257 | -0.02366 | 0.336332 | -0.07545 | 0.431578 | 0.103265 | 0.016492 | 0.308174 | 0.015019 |
| 20.23611 | 0.067441 | 0.336606 | 0.05363 | 0.019855 | 0.081761 | 0.056134 | 0.034884 | 0.182333 | 0.335671 | -0.00996 | -0.02376 | -0.00616 | 0.318652 | -0.03756 | 0.441428 | 0.114285 | 0.053937 | 0.308946 | 0.004635 |
| 20.25 | 0.107717 | 0.295392 | 0.081595 | 0.075783 | 0.09663 | 0.077061 | 0.049952 | 0.277152 | 0.422373 | 0.022208 | 0.000038 | 0.012997 | 0.325117 | 0.002582 | 0.432623 | 0.10917 | 0.063666 | 0.221166 | -0.02028 |
| 20.26389 | 0.089343 | 0.179702 | 0.089926 | 0.113401 | 0.076648 | 0.105675 | 0.103784 | 0.239042 | 0.3789 | 0.042375 | -0.0461 | 0.029408 | 0.369087 | 0.102085 | 0.472964 | 0.142146 | 0.039764 | 0.112371 | -0.10892 |
| 20.27778 | 0.008085 | 0.10196 | -0.01097 | 0.033227 | -0.04025 | 0.009946 | 0.033671 | 0.033046 | 0.319508 | -0.40024 | -0.5167 | -0.24473 | 0.813607 | 0.269723 | 0.820828 | 0.307202 | -0.00985 | 0.06116 | -0.55341 |
| 20.29167 | -0.12029 | 0.095423 | -0.14197 | -0.08125 | -0.14628 | -0.1068 | -0.07886 | -0.15829 | 0.330926 | -1.51092 | -1.93955 | -0.87366 | 2.02188 | 0.469551 | 1.866424 | 0.775611 | -0.00174 | 0.274413 | -1.27219 |
| 20.30556 | -0.12686 | 0.203606 | -0.04315 | 0.027581 | -0.02528 | 0.042231 | 0.073724 | -0.04192 | 0.501895 | -2.07287 | -2.55776 | -0.75485 | 2.665311 | 0.801342 | 2.705864 | 1.35408 | 0.013509 | 0.822443 | -1.40651 |
| 20.31944 | -0.02948 | 0.480624 | 0.176784 | 0.216614 | 0.179763 | 0.276033 | 0.246409 | 0.246492 | 0.776731 | -0.26817 | -0.09433 | 0.001033 | 1.129472 | 0.457476 | 1.817321 | 1.031964 | 0.285809 | 0.845004 | -0.33359 |
| 20.33333 | 0.113495 | 0.643903 | 0.219842 | 0.240681 | 0.207712 | 0.271853 | 0.177281 | 0.346173 | 0.872795 | 0.0762 | 0.044413 | 0.117337 | 0.661937 | 0.124587 | 0.88291 | 0.438039 | 0.190889 | 0.547845 | 0.0074 |
| 20.34722 | 0.211475 | 0.60878 | 0.198821 | 0.180439 | 0.194674 | 0.220578 | 0.077539 | 0.294563 | 0.778811 | 0.021657 | 0.007314 | 0.056695 | 0.487444 | 0.051553 | 0.471163 | 0.151926 | 0.106409 | 0.359208 | 0.034841 |

| | | | | | | | | | | | | | | | | | | | |
|---|---|---|---|---|---|---|---|---|---|---|---|---|---|---|---|---|---|---|---|
| 20.36111 | 0.310062 | 0.641212 | 0.161056 | 0.228421 | 0.235544 | 0.262567 | 0.087682 | 0.304979 | 0.823575 | 0.129621 | 0.073438 | 0.093083 | 0.433854 | 0.145062 | 0.500731 | 0.162047 | 0.169876 | 0.362316 | 0.083774 |
| 20.375 | 0.430421 | 0.891538 | 0.172217 | 0.317893 | 0.258308 | 0.309417 | 0.122162 | 0.46353 | 1.109059 | 0.256691 | 0.131372 | 0.183701 | 0.564154 | 0.257912 | 0.763431 | 0.223385 | 0.245392 | 0.467416 | 0.125558 |
| 20.38889 | 0.450646 | 1.027046 | 0.190507 | 0.373718 | 0.263986 | 0.319747 | 0.115042 | 0.636111 | 1.288958 | 0.196831 | 0.128556 | 0.184256 | 0.769268 | 0.228491 | 0.889202 | 0.213079 | 0.274271 | 0.584522 | 0.134789 |
| 20.40278 | 0.307407 | 0.758852 | 0.19895 | 0.378551 | 0.284695 | 0.299512 | 0.114472 | 0.549804 | 1.101202 | 0.074821 | 0.139087 | 0.206245 | 1.028004 | 0.145017 | 0.774311 | 0.185119 | 0.333898 | 0.643767 | 0.147472 |
| 20.41667 | 0.326473 | 0.442792 | 0.215648 | 0.435959 | 0.303725 | 0.276747 | 0.13548 | 0.384205 | 0.800519 | 0.080105 | 0.182579 | 0.283644 | 1.209883 | 0.183034 | 0.721644 | 0.223773 | 0.415923 | 0.662772 | 0.199679 |
| 20.43056 | 0.506853 | 0.435184 | 0.220617 | 0.449867 | 0.3307 | 0.280717 | 0.161331 | 0.277322 | 0.709874 | 0.11176 | 0.226323 | 0.352108 | 1.212383 | 0.270736 | 0.7611 | 0.267036 | 0.500218 | 0.697343 | 0.231423 |
| 20.44444 | 0.558145 | 0.575293 | 0.215998 | 0.42519 | 0.370969 | 0.297333 | 0.187078 | 0.359677 | 0.614679 | 0.061299 | 0.225857 | 0.299492 | 0.956208 | 0.279516 | 0.724469 | 0.250209 | 0.500572 | 0.60173 | 0.217406 |
| 20.45833 | 0.531339 | 0.764461 | 0.248819 | 0.440001 | 0.402975 | 0.332659 | 0.225312 | 0.474664 | 0.43816 | 0.239049 | 0.212613 | 0.302292 | 0.601047 | 0.337712 | 0.585962 | 0.273694 | 0.473649 | 0.401892 | 0.228228 |
| 20.47222 | 0.71035 | 1.062544 | 0.290713 | 0.385819 | 0.334842 | 0.361656 | 0.201605 | 0.444651 | 0.518737 | 0.54363 | 0.250778 | 0.394077 | 0.313919 | 0.472585 | 0.544742 | 0.385919 | 0.522818 | 0.284097 | 0.297082 |
| 20.48611 | 1.519743 | 1.607579 | 0.266965 | 0.196728 | 0.319659 | 0.42922 | 0.177143 | 0.456537 | 1.234237 | 0.683847 | 0.472696 | 0.447988 | 0.329437 | 0.533148 | 0.896761 | 0.677889 | 0.836018 | 0.46277 | 0.451295 |
| 20.5 | 3.143524 | 2.714111 | 0.275987 | 0.563488 | 0.880447 | 0.70062 | 0.618938 | 0.713599 | 2.635812 | 1.209574 | 1.069271 | 0.597178 | 0.861576 | 0.883693 | 2.093654 | 1.489247 | 1.755418 | 0.942203 | 1.078955 |
| 20.51389 | 3.447853 | 2.943577 | 0.298279 | 1.222406 | 1.475221 | 1.174711 | 1.23981 | 0.895932 | 4.4164 | 1.929965 | 1.648252 | 0.74817 | 2.043514 | 0.736843 | 3.810949 | 2.416744 | 2.039082 | 1.138628 | 0.977134 |
| 20.52778 | 1.612542 | 1.680606 | 0.202783 | 0.887923 | 0.907893 | 0.815575 | 0.849387 | 0.791502 | 2.912127 | 2.148365 | 1.573841 | 0.615445 | 2.629505 | 0.37457 | 1.788602 | 1.22275 | 1.340336 | 0.794013 | 0.549475 |
| 20.54167 | 0.448295 | 0.558905 | -0.01546 | 0.532154 | 0.619371 | 0.512648 | 0.543411 | 0.286131 | 0.711897 | 1.655559 | 1.01982 | 0.299339 | 1.871386 | 0.351382 | 0.373982 | 0.504606 | 0.814885 | 0.254707 | 0.360097 |
| 20.55556 | -0.3099 | -0.22452 | -0.3452 | 0.313391 | 0.323634 | 0.289122 | 0.419414 | -0.24469 | -1.24214 | 0.981013 | 0.520098 | 0.04692 | 0.224234 | 0.250257 | -0.64131 | -0.027 | 0.370099 | -0.16009 | 0.241231 |
| 20.56944 | -0.68175 | -0.56133 | -0.53016 | 0.134649 | 0.057582 | 0.117027 | 0.287338 | -0.56378 | -2.36298 | 0.529939 | 0.254286 | 0.040017 | -0.97826 | 0.151385 | -1.55646 | -0.35176 | 0.110881 | -0.46694 | 0.211337 |
| 20.58333 | -0.77538 | -0.34482 | -0.37749 | 0.003696 | -0.01809 | 0.126418 | 0.206294 | -0.3491 | -2.65407 | 0.469855 | 0.285195 | 0.208592 | -1.65172 | 0.156739 | -1.88983 | -0.51297 | 0.067964 | -0.53723 | 0.246357 |
| 20.59722 | -0.55209 | 0.346517 | -0.0518 | -0.08622 | -0.04251 | 0.24059 | 0.157336 | 0.211511 | -1.72912 | 0.581178 | 0.366001 | 0.382296 | -1.47056 | 0.167455 | -1.50025 | -0.67632 | 0.12948 | -0.31704 | 0.24407 |
| 20.61111 | 0.258051 | 1.206937 | 0.128151 | -0.25058 | -0.08782 | 0.247138 | 0.023412 | 0.677972 | -0.37347 | 0.659942 | 0.323006 | 0.315825 | -0.81797 | 0.13945 | -0.92327 | -0.85021 | 0.22696 | 0.069718 | 0.098456 |
| 20.625 | 1.336084 | 1.720143 | 0.263134 | -0.51304 | -0.0808 | 0.044615 | -0.18503 | 0.861795 | 0.897225 | 0.467345 | 0.135993 | 0.111951 | -0.33686 | 0.021107 | -0.45467 | -0.90235 | 0.30358 | 0.464112 | -0.09995 |
| 20.63889 | 1.423555 | 1.243112 | 0.266985 | -0.4985 | -0.01088 | -0.18489 | -0.28561 | 0.724937 | 1.884322 | 0.062943 | -0.08753 | -0.01277 | 0.271339 | -0.10414 | 0.130345 | -0.80514 | 0.32778 | 0.632793 | -0.23016 |
| 20.65278 | 0.389328 | -0.00087 | 0.11794 | -0.31591 | 0.141173 | -0.22945 | -0.15136 | 0.317918 | 1.54329 | -0.27028 | -0.20046 | -0.04274 | 1.291462 | -0.09986 | 0.761087 | -0.55409 | 0.113189 | 0.367063 | -0.23079 |
| 20.66667 | -0.40591 | -0.80756 | -0.13296 | -0.20183 | 0.225944 | -0.13343 | 0.045095 | -0.20368 | 0.482138 | -0.33369 | -0.16036 | 0.11045 | 1.665477 | 0.062615 | 0.655876 | -0.1989 | -0.36594 | -0.22501 | -0.14355 |
| 20.68056 | -0.5019 | -0.75797 | -0.25016 | -0.06947 | 0.186293 | -0.0274 | 0.141022 | -0.52864 | -0.46635 | -0.07573 | -0.04831 | 0.298942 | 0.952081 | 0.173094 | 0.071063 | -0.12794 | -0.58786 | -0.70607 | -0.07007 |
| 20.69444 | -0.28311 | -0.13476 | -0.1759 | 0.031605 | 0.056452 | 0.099927 | 0.136445 | -0.4031 | -1.33409 | 0.242282 | 0.067779 | 0.356686 | -0.24747 | 0.101996 | -0.21168 | -0.09779 | -0.33843 | -0.80362 | -0.02513 |
| 20.70833 | -0.23711 | 0.550484 | 0.002009 | 0.076467 | -0.1099 | 0.236843 | 0.159933 | 0.163098 | -2.21221 | 0.579083 | 0.101108 | 0.133796 | -1.09803 | 0.007161 | -0.38653 | -0.00446 | 0.140962 | -0.36014 | -0.02065 |
| 20.72222 | -0.13838 | 0.912429 | 0.129002 | 0.052484 | -0.21623 | 0.275453 | 0.215686 | 0.487884 | -2.03598 | 0.62385 | 0.077365 | -0.07026 | -1.2986 | -0.03099 | -0.56712 | 0.143116 | 0.386659 | 0.300312 | -0.05897 |
| 20.73611 | -0.04567 | 0.891646 | 0.003483 | -0.19257 | -0.23757 | 0.101015 | 0.105885 | 0.180775 | -0.99365 | 0.338097 | 0.031812 | -0.23081 | -1.23559 | -0.09451 | -0.74445 | 0.202467 | 0.409947 | 0.796419 | -0.11949 |
| 20.75 | -0.02221 | 0.533528 | -0.21161 | -0.50684 | -0.21101 | -0.10439 | -0.06837 | -0.10731 | -0.10537 | 0.015779 | -0.02844 | -0.30372 | -1.03926 | -0.16179 | -0.75 | 0.149973 | 0.233462 | 0.636197 | -0.10564 |
| 20.76389 | -0.05842 | -0.0456 | -0.26614 | -0.57167 | -0.12489 | -0.14654 | -0.11013 | -0.39532 | 0.377996 | -0.17131 | -0.04778 | -0.22713 | -0.60194 | -0.14885 | -0.43223 | 0.087174 | -0.02157 | 0.282456 | -0.03479 |
| 20.77778 | -0.02874 | -0.62229 | -0.0037 | -0.32761 | -0.03113 | -0.03011 | 0.012488 | -0.39892 | 0.385569 | -0.24845 | -0.0271 | -0.08036 | -0.22709 | -0.10656 | -0.14522 | 0.061648 | -0.16887 | 0.091252 | 0.032993 |
| 20.79167 | 0.015434 | -0.76145 | 0.191 | -0.08095 | -0.01347 | 0.115779 | 0.182923 | -0.11693 | 0.206996 | -0.20287 | -0.0224 | 0.049223 | -0.03302 | -0.09994 | -0.06712 | 0.009014 | -0.21867 | 0.033295 | 0.091191 |
| 20.80556 | 0.062489 | -0.39507 | 0.241469 | -0.00816 | -0.04059 | 0.174328 | 0.20188 | 0.154782 | -0.0049 | -0.07644 | -0.02082 | 0.174592 | 0.10648 | -0.11476 | -0.09781 | -0.09835 | -0.1911 | -0.01636 | 0.13906 |
| 20.81944 | 0.182632 | 0.268843 | 0.028083 | -0.25105 | -0.11315 | 0.144764 | 0.050188 | 0.205799 | -0.29637 | 0.06249 | 0.004125 | 0.194059 | 0.232186 | -0.11881 | -0.12875 | -0.20594 | -0.12311 | -0.06613 | 0.167586 |
| 20.83333 | -0.09387 | 0.582521 | -0.28936 | -0.46244 | -0.25218 | 0.045844 | -0.08496 | -0.14558 | -0.93538 | 0.227924 | 0.04313 | 0.13889 | 0.346286 | -0.11683 | -0.12189 | -0.2551 | -0.03686 | -0.1238 | 0.158553 |
| 20.84722 | -0.84587 | -0.09766 | -0.49146 | -0.50526 | -0.36046 | -0.13268 | -0.10296 | -0.65636 | -1.71069 | 0.327237 | 0.041495 | 0.096135 | 0.497462 | -0.10143 | -0.01473 | -0.27299 | -0.00402 | -0.23279 | 0.110243 |
| 20.86111 | -1.39067 | -1.08458 | -0.47662 | -0.54384 | -0.3883 | -0.28795 | -0.15463 | -0.94283 | -1.93321 | 0.309777 | -0.00773 | 0.057279 | 0.681937 | -0.07282 | 0.236011 | -0.25282 | -0.03452 | -0.38664 | 0.023191 |
| 20.875 | -1.57714 | -1.93299 | -0.39358 | -0.55308 | -0.42855 | -0.36746 | -0.18027 | -1.03076 | -1.61759 | 0.308433 | -0.05841 | 0.003113 | 0.757383 | -0.04908 | 0.274208 | -0.28789 | -0.06565 | -0.54738 | -0.05099 |
| 20.88889 | -1.26729 | -2.26237 | -0.23319 | -0.48984 | -0.4738 | -0.30846 | -0.11825 | -0.86889 | -1.07748 | 0.31109 | -0.09059 | -0.0653 | 0.706185 | -0.0337 | 0.137846 | -0.36266 | -0.07255 | -0.64436 | -0.10448 |
| 20.90278 | -0.76242 | -1.82258 | 0.023128 | -0.38043 | -0.45967 | -0.14197 | -0.02726 | -0.61593 | -0.77782 | 0.262348 | -0.08777 | -0.12608 | 0.460436 | -0.02566 | -0.01936 | -0.33366 | -0.09193 | -0.62452 | -0.11643 |
| 20.91667 | -0.36044 | -0.95765 | 0.143502 | -0.30207 | -0.46793 | 0.013806 | 0.050488 | -0.42559 | -0.65577 | 0.174723 | -0.06761 | -0.1963 | 0.022231 | -0.02539 | -0.28678 | -0.32949 | -0.11324 | -0.50333 | -0.10714 |
| 20.93056 | -0.39492 | -0.29817 | -0.0349 | -0.22994 | -0.43608 | -0.02492 | 0.077771 | -0.365 | -0.54568 | 0.045813 | -0.05343 | -0.23255 | -0.38401 | -0.02875 | -0.42861 | -0.30072 | -0.16498 | -0.39585 | -0.14113 |
| 20.94444 | -0.95859 | -0.25331 | -0.28431 | -0.37928 | -0.4338 | -0.2371 | 0.023242 | -0.55923 | -0.63004 | -0.09322 | -0.05119 | -0.2153 | -0.72042 | -0.04667 | -0.47808 | -0.27852 | -0.20575 | -0.33912 | -0.18729 |
| 20.95833 | -1.46179 | -0.56436 | -0.38091 | -0.61583 | -0.45918 | -0.30917 | 0.048448 | -0.73373 | -0.78019 | -0.19563 | -0.06484 | -0.16501 | -0.9573 | -0.08727 | -0.51345 | -0.25255 | -0.24572 | -0.31489 | -0.18863 |
| 20.97222 | -1.58263 | -0.83653 | -0.40462 | -0.58385 | -0.36595 | -0.24943 | 0.20621 | -0.75597 | -0.94858 | -0.39061 | -0.1395 | -0.07071 | -1.33556 | -0.19243 | -0.68692 | -0.2863 | -0.3164 | -0.36886 | -0.1667 |
| 20.98611 | -1.58653 | -1.49305 | -0.4671 | -0.50529 | -0.387 | -0.36686 | 0.31284 | -0.95402 | -1.46609 | -0.6021 | -0.28674 | 0.028098 | -1.91374 | -0.36565 | -1.09403 | -0.44132 | -0.36065 | -0.43109 | -0.22126 |
| 21 | -1.78725 | -2.37836 | -0.60516 | -0.61248 | -0.68173 | -0.87904 | -0.20414 | -1.57113 | -2.4876 | -0.49094 | -0.39544 | 0.096022 | -2.04209 | -0.52305 | -1.60246 | -0.64366 | -0.31136 | -0.30205 | -0.31884 |
| 21.01389 | -1.9176 | -2.85652 | -0.73315 | -0.69423 | -0.83548 | -0.86719 | -0.56523 | -2.01542 | -2.95454 | -0.23651 | -0.42235 | 0.089764 | -1.66786 | -0.59285 | -1.73594 | -0.70171 | -0.27565 | 0.05912 | -0.23708 |
| 21.02778 | -1.10023 | -2.18907 | -0.59983 | -0.47631 | -0.55291 | -0.50078 | -0.4314 | -1.69696 | -1.79635 | 0.025754 | -0.36229 | 0.003711 | -0.82033 | -0.5751 | -1.33646 | -0.53071 | -0.36005 | 0.392892 | -0.01963 |
| 21.04167 | -0.65654 | -1.31786 | -0.22857 | -0.29723 | -0.27451 | -0.08543 | -0.16541 | -0.91645 | -0.47709 | 0.046454 | -0.33656 | -0.04889 | 0.073744 | -0.54701 | -0.67256 | -0.35175 | -0.49532 | 0.50276 | 0.065431 |
| 21.05556 | -0.60524 | -0.68729 | 0.097708 | -0.18932 | -0.19197 | 0.12678 | -0.00858 | -0.12907 | 0.261795 | -0.19165 | -0.3559 | -0.07975 | 0.545438 | -0.53296 | -0.23959 | -0.29556 | -0.52541 | 0.328983 | -0.03674 |
| 21.06944 | -0.42023 | -0.32259 | 0.315064 | -0.05906 | -0.17857 | 0.112132 | 0.075165 | 0.556148 | 0.521878 | -0.54254 | -0.36647 | -0.05079 | 0.535017 | -0.49474 | -0.25945 | -0.38556 | -0.43375 | 0.059153 | -0.20236 |
| 21.08333 | -0.09196 | -0.2597 | 0.292445 | 0.060522 | -0.10199 | 0.075918 | 0.207353 | 0.662437 | 0.565011 | -0.85757 | -0.3316 | -0.00822 | 0.409853 | -0.39712 | -0.60486 | -0.56215 | -0.39672 | -0.1551 | -0.33375 |
| 21.09722 | 0.15934 | -0.19278 | 0.186133 | 0.080254 | -0.04879 | 0.142239 | 0.357266 | 0.297555 | 0.154216 | -1.17685 | -0.44638 | -0.10811 | 0.241503 | -0.22794 | -1.4007 | -0.9176 | -0.56699 | -0.39003 | -0.58605 |
| 21.11111 | 0.258176 | 0.0054 | 0.122388 | 0.044194 | -0.06646 | 0.205561 | 0.372277 | -0.06073 | -0.1911 | -2.18494 | -0.9689 | -0.36778 | -0.47583 | -0.18185 | -3.13008 | -1.62185 | -1.18532 | -0.98672 | -1.30218 |
| 21.125 | 0.148936 | 0.055623 | -0.01307 | 0.003118 | -0.07964 | 0.203677 | 0.291195 | -0.29734 | -0.42759 | -1.84797 | -1.10985 | -0.40884 | -1.45642 | -0.24358 | -2.89959 | -1.29634 | -1.34263 | -1.50266 | -1.15721 |
| 21.13889 | -0.00265 | 0.059054 | -0.05719 | -0.04312 | -0.02829 | 0.214322 | 0.222698 | -0.31911 | -0.48732 | -0.59678 | -0.47016 | -0.24015 | -1.5045 | -0.21505 | -1.16864 | -0.49098 | -0.75335 | -1.32393 | -0.40004 |
| 21.15278 | -0.14174 | 0.055175 | 0.024661 | -0.03452 | 0.037446 | 0.229003 | 0.231529 | -0.12261 | -0.26256 | 0.040683 | 0.155337 | -0.10744 | -1.06069 | -0.00425 | 0.038499 | -0.05396 | -0.20916 | -0.97669 | 0.158767 |
| 21.16667 | -0.2615 | 0.092744 | 0.111195 | -0.01132 | 0.057873 | 0.200607 | 0.22258 | 0.152251 | 0.092728 | 0.182719 | 0.353093 | -0.0422 | -0.62873 | 0.048986 | 0.459161 | 0.076809 | 0.204359 | -0.65952 | 0.306538 |
| 21.18056 | -0.29331 | 0.234363 | 0.11985 | -0.05664 | 0.061882 | 0.161827 | 0.153351 | 0.342114 | 0.344669 | 0.072648 | 0.247423 | -0.00558 | -0.56397 | -0.01354 | 0.369186 | 0.055354 | 0.407443 | -0.37688 | 0.144162 |
| 21.19444 | -0.29055 | 0.201985 | 0.063397 | -0.0635 | 0.082735 | 0.127539 | 0.132158 | 0.355584 | 0.254311 | -0.01118 | 0.092369 | 0.103015 | -0.40776 | -0.06486 | 0.190687 | 0.03523 | 0.310649 | -0.28795 | -0.05222 |
| 21.20833 | -0.26575 | -0.01112 | 0.016333 | -0.00019 | 0.088087 | 0.101393 | 0.180243 | 0.262646 | -0.01666 | 0.024278 | 0.026589 | 0.230678 | -0.18498 | -0.00988 | 0.027068 | 0.055938 | 0.106815 | -0.28761 | -0.03982 |
| 21.22222 | -0.27935 | -0.10493 | -0.00374 | -0.00986 | 0.070399 | 0.083407 | 0.132078 | 0.13339 | -0.10445 | 0.022832 | 0.027055 | 0.195256 | -0.04486 | 0.061359 | -0.00706 | 0.051861 | -0.00696 | -0.2071 | 0.095385 |
| 21.23611 | -0.34211 | 0.061391 | -0.00033 | -0.08082 | 0.072996 | 0.119647 | 0.022348 | 0.02992 | 0.0571 | -0.06617 | -0.02033 | 0.063982 | -0.08608 | 0.01541 | -0.03194 | 0.006257 | -0.14567 | -0.16473 | 0.080339 |
| 21.25 | -0.24194 | 0.351618 | 0.096434 | -0.06599 | 0.117366 | 0.221043 | 0.009382 | 0.02788 | 0.137903 | -0.1073 | -0.10749 | 0.011654 | -0.20393 | -0.08425 | -0.18419 | -0.01317 | -0.23909 | -0.23645 | -0.11322 |
| 21.26389 | -0.10556 | 0.221885 | 0.125282 | 0.071306 | 0.146664 | 0.244108 | 0.142992 | 0.075415 | -0.07916 | -0.02368 | -0.11678 | 0.026419 | -0.22396 | -0.10629 | -0.27226 | 0.021508 | -0.13841 | -0.23372 | -0.25892 |
| 21.27778 | -0.34997 | -0.3266 | -0.02792 | 0.196908 | 0.13678 | 0.164472 | 0.300858 | 0.057198 | -0.26394 | 0.03297 | -0.04558 | -0.01729 | -0.05062 | -0.03035 | -0.08145 | 0.032485 | 0.032958 | -0.0257 | -0.18612 |
| 21.29167 | -0.28222 | -0.23379 | -0.0411 | 0.196448 | 0.179406 | 0.216642 | 0.366486 | 0.135819 | -0.22254 | -0.0089 | 0.017204 | -0.11885 | 0.235683 | 0.049242 | 0.218945 | 0.018676 | 0.096181 | 0.232583 | -0.03232 |
| 21.30556 | 0.200308 | 0.503466 | 0.063111 | 0.190106 | 0.254677 | 0.433283 | 0.430351 | 0.314545 | 0.148016 | -0.04218 | 0.034893 | -0.11249 | 0.406959 | 0.101039 | 0.325061 | 0.014372 | 0.0701 | 0.325267 | 0.068148 |
| 21.31944 | 0.491545 | 0.645216 | 0.191672 | 0.200927 | 0.282639 | 0.586908 | 0.50667 | 0.388865 | 0.292013 | 0.031485 | 0.056561 | 0.02373 | 0.341062 | 0.155459 | 0.287073 | 0.025417 | 0.08909 | 0.281173 | 0.052065 |
| 21.33333 | 0.487557 | 0.349043 | 0.205378 | 0.129888 | 0.220106 | 0.555506 | 0.50505 | 0.254699 | 0.130008 | 0.097818 | 0.066702 | 0.102734 | 0.122189 | 0.161163 | 0.279312 | 0.004391 | 0.125029 | 0.226692 | -0.00766 |
| 21.34722 | 0.44729 | 0.306074 | 0.105253 | 0.066239 | 0.160877 | 0.484059 | 0.412926 | 0.20593 | 0.168991 | 0.065974 | 0.010728 | 0.043006 | -0.09556 | 0.069108 | 0.277543 | -0.0489 | 0.079033 | 0.14576 | -0.07552 |
| 21.36111 | 0.576444 | 0.743848 | 0.075495 | 0.154313 | 0.193146 | 0.519966 | 0.401878 | 0.557853 | 0.573858 | 0.048448 | -0.02012 | 0.030906 | -0.14372 | 0.009911 | 0.232897 | -0.05123 | 0.046826 | 0.071561 | -0.1029 |
| 21.375 | 0.728506 | 1.294728 | 0.187873 | 0.405529 | 0.308811 | 0.618504 | 0.503069 | 1.086749 | 0.9421 | 0.086951 | 0.039374 | 0.118035 | -0.01143 | 0.038722 | 0.33213 | 0.008854 | 0.13294 | 0.115561 | -0.02171 |
| 21.38889 | 0.936768 | 1.663793 | 0.377406 | 0.557731 | 0.364934 | 0.646885 | 0.530675 | 1.183563 | 0.900593 | 0.099777 | 0.114809 | 0.131728 | 0.136306 | 0.07784 | 0.580421 | 0.065047 | 0.212928 | 0.298514 | 0.14597 |
| 21.40278 | 1.088686 | 1.578255 | 0.419045 | 0.411668 | 0.297979 | 0.643389 | 0.508815 | 0.795406 | 0.390345 | 0.072772 | 0.122335 | 0.061751 | 0.239403 | 0.069303 | 0.684737 | 0.108139 | 0.211709 | 0.519163 | 0.363082 |
| 21.41667 | 1.006917 | 1.043465 | 0.243214 | 0.243014 | 0.240168 | 0.594629 | 0.427465 | 0.170571 | -0.38745 | 0.05104 | 0.075145 | 0.077904 | 0.320957 | 0.082629 | 0.561241 | 0.152534 | 0.220533 | 0.583394 | 0.466621 |
| 21.43056 | 1.032307 | 0.713824 | 0.065307 | 0.181412 | 0.265972 | 0.498943 | 0.332749 | -0.19111 | -0.75255 | 0.105652 | 0.039082 | 0.187574 | 0.35819 | 0.158126 | 0.461865 | 0.173652 | 0.290631 | 0.444457 | 0.35216 |
| 21.44444 | 1.239777 | 0.747235 | 0.039703 | 0.239271 | 0.363443 | 0.477563 | 0.368059 | 0.100354 | -0.13031 | 0.144326 | 0.032733 | 0.223943 | 0.329828 | 0.17042 | 0.43656 | 0.111629 | 0.323426 | 0.310441 | 0.213334 |
| 21.45833 | 1.521175 | 1.181569 | 0.218955 | 0.352555 | 0.463798 | 0.5548 | 0.524216 | 0.830209 | 0.969048 | 0.107667 | 0.020082 | 0.178901 | 0.241721 | 0.12248 | 0.299621 | 0.046522 | 0.302967 | 0.224869 | 0.172225 |
| 21.47222 | 1.841102 | 1.805211 | 0.492152 | 0.394403 | 0.479746 | 0.618866 | 0.629969 | 1.521323 | 2.037957 | 0.08824 | 0.025632 | 0.226119 | 0.196987 | 0.131051 | 0.179368 | 0.059787 | 0.286672 | 0.169978 | 0.264783 |
| 21.48611 | 1.981071 | 2.23256 | 0.801458 | 0.426702 | 0.541193 | 0.638534 | 0.663365 | 1.255285 | 2.726939 | 0.119898 | 0.094006 | 0.339661 | 0.389329 | 0.235614 | 0.31504 | 0.108686 | 0.306029 | 0.212581 | 0.400745 |
| 21.5 | 3.168336 | 2.910693 | 0.727105 | 1.133281 | 1.135897 | 0.649528 | 0.736677 | 0.888693 | 4.38854 | 0.165303 | 0.210314 | 0.372047 | 0.699097 | 0.408003 | 0.636518 | 0.126932 | 0.35175 | 0.390721 | 0.489302 |
| 21.51389 | 3.560169 | 3.317632 | 0.400155 | 2.235372 | 1.786067 | 0.674614 | 0.951093 | 0.711903 | 4.677891 | 0.228639 | 0.257832 | 0.328449 | 0.790405 | 0.517535 | 0.836673 | 0.099984 | 0.389851 | 0.528604 | 0.498941 |
| 21.52778 | 2.237269 | 2.363909 | 0.20838 | 1.648789 | 1.08435 | 0.133703 | 0.280109 | 0.622129 | 3.178859 | 0.20695 | 0.21028 | 0.218175 | 0.567794 | 0.455639 | 0.688601 | 0.064773 | 0.356119 | 0.546304 | 0.461235 |
| 21.54167 | 0.894375 | 0.885848 | 0.003497 | 0.742964 | 0.723069 | -0.35438 | -0.35797 | 0.336731 | 1.574681 | 0.134091 | 0.133836 | 0.128031 | 0.318023 | 0.358406 | 0.309261 | 0.071723 | 0.284864 | 0.460803 | 0.362254 |

| | | | | | | | | | | | | | | | | | | | |
|---|---|---|---|---|---|---|---|---|---|---|---|---|---|---|---|---|---|---|---|
| 21.55556 | -0.71646 | -0.84812 | -0.31868 | 0.084516 | 0.433289 | -0.5982 | -0.72983 | -0.19095 | 0.085855 | 0.106013 | 0.174232 | 0.124827 | 0.334582 | 0.347944 | 0.177512 | 0.156368 | 0.303584 | 0.415155 | 0.284106 |
| 21.56944 | -1.76817 | -2.14676 | -0.66929 | -0.25182 | 0.281362 | -0.5961 | -0.69302 | -0.50753 | -1.43126 | 0.156781 | 0.331677 | 0.19329 | 0.784877 | 0.457363 | 0.53258 | 0.311131 | 0.451466 | 0.696204 | 0.315952 |
| 21.58333 | -1.97048 | -2.25562 | -0.54549 | -0.25549 | 0.141757 | -0.37687 | -0.33154 | -0.15613 | -2.45767 | 0.407329 | 0.438708 | 0.313811 | 1.24931 | 0.632072 | 0.874509 | 0.432485 | 0.581792 | 0.94256 | 0.355015 |
| 21.59722 | -1.14012 | -1.22481 | -0.06692 | -0.31607 | -0.05387 | -0.02429 | 0.102958 | 0.450027 | -2.64274 | 1.046045 | 0.579386 | 0.387498 | 1.329299 | 0.755127 | 0.956129 | 0.492729 | 0.562196 | 1.058166 | 0.302102 |
| 21.61111 | 0.678407 | 0.336238 | 0.61248 | -0.39843 | -0.22387 | 0.264757 | 0.349527 | 1.199007 | -1.92221 | 2.025538 | 0.941699 | 0.309112 | 1.491274 | 0.96993 | 1.424204 | 0.798422 | 1.185432 | 1.566381 | 0.352179 |
| 21.625 | 1.620076 | 2.206819 | 0.889257 | -0.27144 | -0.35302 | 0.342016 | 0.247105 | 1.332275 | -1.03745 | 3.220685 | 1.331477 | 0.379496 | 2.510868 | 1.116172 | 2.836735 | 0.845165 | 1.744806 | 0.847429 | 0.574517 |
| 21.63889 | 1.621056 | 2.831659 | 0.543963 | -0.17091 | -0.36346 | 0.175971 | 0.005849 | 0.685213 | -0.11661 | 2.193534 | 1.106051 | 0.568573 | 2.587235 | 0.679762 | 2.160865 | 0.386758 | 0.965816 | -0.50063 | 0.649911 |
| 21.65278 | 1.07896 | 2.333916 | 0.301435 | -0.19282 | -0.22132 | 0.106867 | -0.10563 | 0.012765 | 0.674085 | 0.954222 | 0.569805 | 0.359339 | 1.467543 | 0.211001 | 1.297918 | 0.135407 | 0.371176 | -0.60692 | 0.577095 |
| 21.66667 | -0.0605 | 0.729978 | -0.07435 | -0.04089 | 0.009176 | -0.09847 | -0.21752 | -0.61304 | 1.265561 | 0.217582 | 0.130714 | 0.02548 | 0.051577 | -0.27472 | 0.034596 | -0.01724 | 0.127646 | -0.22073 | 0.351951 |
| 21.68056 | -1.43626 | -1.07318 | -0.28391 | 0.064418 | 0.076904 | -0.17535 | -0.22296 | -0.90067 | 1.671327 | -0.03139 | -0.04053 | -0.13109 | -0.80939 | -0.59158 | -0.74815 | -0.13073 | 0.076146 | -0.08161 | 0.150543 |
| 21.69444 | -2.18314 | -2.06197 | -0.41594 | 0.005505 | -0.09069 | -0.11341 | -0.08401 | -0.78628 | 1.552846 | 0.118301 | 0.095533 | 0.045457 | -1.13889 | -0.5307 | -0.90122 | -0.22364 | 0.124607 | -0.03075 | 0.256947 |
| 21.70833 | -2.17369 | -2.10517 | -0.09417 | 0.068705 | -0.2322 | -0.04622 | 0.080151 | -0.2921 | 0.897005 | 0.423538 | 0.305288 | 0.311143 | -1.12469 | -0.18187 | -0.37013 | -0.33362 | 0.080898 | 0.086962 | 0.303358 |
| 21.72222 | -1.18356 | -1.37062 | 0.294395 | 0.209827 | -0.19522 | -0.04709 | 0.243479 | 0.529091 | 0.152589 | 0.463992 | 0.319892 | 0.461084 | -0.66673 | 0.193281 | 0.280281 | -0.31498 | -0.1524 | 0.12535 | 0.164249 |
| 21.73611 | 0.053088 | -0.2206 | 0.257336 | 0.165268 | -0.05995 | 0.124099 | 0.297455 | 1.13495 | -0.39456 | 0.2502 | 0.117196 | 0.435389 | -0.29074 | 0.482646 | 0.904346 | -0.20461 | -0.50832 | 0.096558 | -0.12069 |
| 21.75 | 0.555677 | 0.705057 | -0.19618 | -0.11145 | 0.029045 | 0.347012 | 0.295191 | 1.068051 | -1.07972 | -0.12215 | -0.13256 | 0.220086 | -0.36533 | 0.637374 | 0.497264 | -0.18669 | -0.68956 | 0.018291 | -0.37668 |
| 21.76389 | 0.055903 | 0.492675 | -0.38619 | -0.43464 | 0.081347 | 0.469797 | 0.191365 | 0.193508 | -1.99471 | -0.37878 | -0.23868 | -0.06989 | -0.57439 | 0.454011 | -0.58463 | -0.30873 | -0.70939 | -0.1145 | -0.50753 |
| 21.77778 | -0.98021 | -0.73497 | -0.39527 | -0.81307 | -0.13074 | 0.335928 | 0.034457 | -0.61126 | -2.33645 | -0.25851 | -0.19787 | -0.27655 | -0.3366 | 0.019658 | -1.64844 | -0.35868 | -0.65575 | -0.2888 | -0.58037 |
| 21.79167 | -1.94756 | -1.82763 | -0.325 | -0.81508 | -0.17621 | 0.073366 | -0.16469 | -1.03973 | -2.66414 | 0.10149 | -0.11709 | -0.21885 | 0.734169 | -0.33472 | -1.71005 | -0.25628 | -0.54815 | -0.4159 | -0.42572 |
| 21.80556 | -2.66353 | -2.26103 | -0.01081 | -0.60276 | -0.12817 | -0.06415 | -0.18578 | -0.85596 | -2.17315 | 0.368867 | -0.01477 | 0.171035 | 1.165745 | -0.50039 | -0.80841 | -0.06806 | -0.25227 | -0.32572 | -0.11192 |
| 21.81944 | -2.21406 | -1.70348 | 0.186503 | -0.42842 | -0.04414 | -0.25706 | -0.30611 | -0.19803 | -1.22389 | 0.429895 | 0.095708 | 0.507338 | 0.749747 | -0.38335 | 0.228016 | 0.234989 | -0.0456 | -0.12239 | 0.184773 |
| 21.83333 | -1.37786 | -1.02322 | 0.159006 | -0.20084 | 0.006918 | -0.43956 | -0.42221 | 0.398454 | -0.16148 | 0.243312 | 0.161333 | 0.639908 | 0.637735 | -0.11736 | 0.827625 | 0.46347 | 0.024931 | 0.054316 | 0.361265 |
| 21.84722 | -0.95584 | -0.53117 | -0.1011 | -0.01978 | -0.0608 | -0.55223 | -0.35833 | 0.009865 | 0.364231 | -0.03316 | 0.136378 | 0.628471 | 0.338935 | 0.06159 | 0.577963 | 0.511689 | -0.02622 | 0.062462 | 0.313515 |
| 21.86111 | -1.12747 | -0.45843 | -0.47613 | -0.20929 | -0.20326 | -0.53428 | -0.30674 | -0.86684 | 0.101649 | -0.25729 | -0.03048 | 0.28772 | -0.41947 | 0.069495 | -0.1281 | 0.382005 | -0.15583 | -0.1209 | 0.016809 |
| 21.875 | -1.64459 | -0.54855 | -0.669 | -0.50785 | -0.50636 | -0.45506 | -0.2647 | -1.39374 | -0.51276 | -0.23505 | -0.23447 | -0.32143 | -1.29645 | 0.003632 | -0.97974 | 0.272517 | -0.33504 | -0.32857 | -0.38521 |
| 21.88889 | -2.24938 | -0.85705 | -0.74109 | -0.69446 | -0.77685 | -0.41666 | -0.22259 | -1.55741 | -1.1516 | -0.1298 | -0.3351 | -0.65142 | -1.82846 | -0.10601 | -1.48817 | 0.136336 | -0.48544 | -0.43416 | -0.64317 |
| 21.90278 | -2.76108 | -1.48281 | -0.71651 | -0.57926 | -0.8698 | -0.42128 | -0.17561 | -1.3893 | -1.55747 | -0.07144 | -0.30311 | -0.41943 | -1.76646 | -0.26345 | -1.22624 | -0.04344 | -0.52416 | -0.39657 | -0.50413 |
| 21.91667 | -2.64667 | -1.8952 | -0.61754 | -0.35277 | -0.71591 | -0.37491 | -0.15247 | -0.94163 | -1.29601 | -0.09344 | -0.18961 | 0.032475 | -0.87445 | -0.38371 | -0.81438 | -0.31001 | -0.56037 | -0.25117 | -0.3024 |
| 21.93056 | -2.0882 | -1.99144 | -0.44727 | -0.31105 | -0.52445 | -0.29311 | -0.17977 | -0.37514 | -0.72684 | -0.24938 | -0.16171 | 0.43437 | 0.345084 | -0.44052 | -0.50843 | -0.54053 | -0.58092 | 0.116073 | -0.1514 |
| 21.94444 | -1.66352 | -1.99917 | -0.18175 | -0.29233 | -0.47748 | -0.19511 | -0.20962 | -0.0519 | -0.21903 | -0.53595 | -0.25837 | 0.671079 | 0.975333 | -0.48786 | -0.65677 | -0.64022 | -0.59037 | 0.457508 | -0.13876 |
| 21.95833 | -1.27 | -1.67012 | 0.089289 | -0.23057 | -0.3819 | -0.09023 | -0.14098 | -0.1559 | 0.19375 | -0.81027 | -0.3908 | 0.459778 | 0.621302 | -0.5588 | -0.87365 | -0.58793 | -0.66452 | 0.444449 | -0.33413 |
| 21.97222 | -1.18215 | -1.13161 | 0.104948 | -0.22631 | -0.31593 | -0.06933 | -0.04274 | -0.69363 | 0.316604 | -0.82142 | -0.50882 | -0.03422 | -0.22224 | -0.59651 | -1.27455 | -0.48956 | -0.70599 | 0.090934 | -0.64426 |
| 21.98611 | -1.7623 | -1.3185 | -0.20699 | -0.32037 | -0.33852 | -0.30604 | -0.03431 | -1.60283 | -0.16637 | -0.85496 | -0.5755 | -0.50117 | -1.10114 | -0.60194 | -1.81735 | -0.33606 | -0.69112 | -0.174 | -0.86002 |
| 22 | -2.04348 | -2.24061 | -0.744 | -0.51686 | -0.48534 | -0.6125 | -0.25949 | -2.57125 | -1.19461 | -1.35744 | -0.53613 | -0.69353 | -1.74618 | -0.5731 | -1.41435 | -0.21457 | -0.66299 | -0.20676 | -0.68168 |
| 22.01389 | -1.29763 | -2.57439 | -1.1273 | -0.58188 | -0.51724 | -0.72391 | -0.44873 | -3.30061 | -2.46222 | -1.16885 | -0.34941 | -0.49751 | -1.70156 | -0.5244 | -1.1987 | -0.23946 | -0.53358 | -0.07329 | -0.47394 |
| 22.02778 | -0.28964 | -2.44034 | -1.11726 | -0.45679 | -0.32099 | -0.52601 | -0.46727 | -2.67759 | -1.55154 | -0.71204 | -0.23339 | -0.23003 | -1.0218 | -0.47376 | -0.98485 | -0.17152 | -0.33751 | -0.00579 | -0.30415 |
| 22.04167 | 0.410531 | -1.76723 | -0.80125 | -0.33855 | -0.10414 | -0.24606 | -0.35392 | -1.22042 | -0.29432 | -0.32608 | -0.10572 | -0.08572 | -0.23904 | -0.43108 | -0.58771 | -0.0227 | -0.20701 | -0.06902 | -0.21445 |
| 22.05556 | 0.482029 | -0.84719 | -0.48987 | -0.25355 | -0.01249 | -0.11787 | -0.18811 | -0.41042 | 0.147007 | -0.09777 | -0.02686 | -0.04869 | 0.302181 | -0.46673 | -0.14211 | 0.079228 | -0.15082 | -0.07677 | -0.24133 |
| 22.06944 | -0.20275 | -0.41528 | -0.2954 | -0.10344 | -0.02409 | -0.0915 | 0.028534 | -0.07929 | -0.01804 | -0.26417 | -0.07951 | -0.09269 | 0.266451 | -0.52754 | 0.011222 | 0.107931 | -0.12468 | -0.11628 | -0.42095 |
| 22.08333 | -0.52514 | 0.086162 | -0.12091 | 0.015023 | 0.017452 | 0.008799 | 0.167229 | 0.363557 | 0.035339 | -0.56655 | -0.16311 | -0.18678 | -0.13613 | -0.51637 | -0.22246 | 0.082975 | -0.11133 | -0.18376 | -0.60359 |
| 22.09722 | -0.1341 | 0.860996 | -0.06563 | 0.107512 | 0.126072 | 0.188389 | 0.200747 | 1.09286 | 0.578189 | -0.95916 | -0.19688 | -0.35208 | -0.63209 | -0.48967 | -0.85589 | -0.07074 | -0.24099 | -0.1436 | -0.80199 |
| 22.11111 | 0.145714 | 1.120405 | 0.011107 | 0.190599 | 0.17188 | 0.209319 | 0.220791 | 1.58543 | 1.039082 | -1.37209 | -0.19431 | -0.67407 | -1.16996 | -0.56493 | -1.76207 | -0.38503 | -0.50045 | -0.04583 | -0.97797 |
| 22.125 | 0.31762 | 1.011559 | 0.259798 | 0.185487 | 0.148629 | 0.115306 | 0.198565 | 1.795563 | 1.255197 | -1.7698 | -0.08393 | -1.03843 | -1.77341 | -0.53648 | -2.37393 | -0.53214 | -0.51927 | 0.011574 | -1.02452 |
| 22.13889 | 0.727984 | 1.268475 | 0.506782 | 0.119953 | 0.157115 | 0.10283 | 0.11171 | 1.893763 | 1.434896 | -2.66789 | 0.065031 | -1.02235 | -1.59869 | -0.26727 | -2.17936 | -0.36698 | -0.16405 | -0.04016 | -0.79576 |
| 22.15278 | 1.204289 | 1.610577 | 0.589968 | 0.091878 | 0.138105 | 0.126312 | 0.030014 | 1.731933 | 1.39014 | -2.15034 | 0.196302 | -0.62772 | -0.65309 | -0.111 | -0.80801 | -0.14327 | 0.113481 | -0.11536 | -0.65541 |
| 22.16667 | 1.13968 | 1.571855 | 0.452361 | 0.125235 | 0.070117 | 0.116489 | 0.051857 | 1.187191 | 1.139675 | -1.11106 | 0.172611 | -0.33912 | 0.320405 | -0.06455 | 0.18347 | -0.05232 | 0.162409 | -0.11335 | -0.49414 |
| 22.18056 | 0.945068 | 1.409177 | 0.32029 | 0.157609 | 0.043543 | 0.122707 | 0.082311 | 0.60324 | 0.658837 | -0.20264 | 0.049811 | -0.14846 | 0.749128 | -0.06355 | 1.215008 | 0.016154 | 0.083512 | -0.07215 | -0.31703 |
| 22.19444 | 0.998093 | 1.622416 | 0.42436 | 0.210342 | 0.117278 | 0.214505 | 0.058672 | 0.441688 | 0.439248 | 0.168865 | -0.0255 | -0.02827 | 0.707825 | -0.04053 | 1.834045 | 0.102041 | 0.009162 | -0.01896 | -0.12193 |
| 22.20833 | 0.89776 | 1.428343 | 0.528246 | 0.229383 | 0.169947 | 0.298027 | 0.04308 | 0.470934 | 0.22312 | 0.514597 | 0.012433 | 0.039909 | 0.685938 | 0.00453 | 1.67197 | 0.130685 | 0.014821 | 0.112262 | 0.129901 |
| 22.22222 | 0.33466 | 0.344652 | 0.363492 | 0.179746 | 0.084683 | 0.17445 | 0.013393 | 0.401005 | -0.17859 | 0.92986 | 0.038705 | 0.107789 | 0.815369 | 0.040052 | 1.566264 | 0.147442 | 0.066397 | 0.334609 | 0.311806 |
| 22.23611 | -0.25961 | -0.95545 | -0.05819 | 0.147035 | -0.02736 | -0.04094 | -0.00763 | 0.095144 | -0.28632 | 1.157395 | -0.02152 | 0.167909 | 0.733038 | 0.026771 | 1.140304 | 0.134664 | 0.10286 | 0.517561 | 0.30678 |
| 22.25 | -0.45563 | -1.60113 | -0.40235 | 0.10457 | -0.0562 | -0.21117 | -0.00817 | -0.32179 | -0.30088 | 1.176033 | -0.06613 | 0.139976 | 0.356118 | -0.00392 | 0.596441 | 0.046845 | 0.274871 | 0.604512 | 0.248733 |
| 22.26389 | -0.13875 | -1.18668 | -0.4757 | -0.00219 | 0.037351 | -0.22975 | -0.00918 | -0.75993 | -0.39936 | 1.171313 | -0.01154 | 0.063802 | -0.00139 | -0.00039 | 0.244454 | 0.012852 | 0.411865 | 0.665157 | 0.313133 |
| 22.27778 | 0.295592 | -0.14776 | -0.28751 | -0.07078 | 0.15933 | -0.09075 | -0.02592 | -0.90582 | -0.64981 | 0.993203 | 0.078371 | -0.02987 | -0.239 | 0.059535 | 0.103425 | 0.047174 | 0.399637 | 0.667047 | 0.451925 |
| 22.29167 | 0.455215 | 0.487051 | -0.03507 | 0.010039 | 0.179554 | 0.036733 | 0.027528 | -0.65719 | -0.87634 | 0.729091 | 0.085882 | 0.057157 | -0.27827 | 0.125887 | 0.051939 | 0.117889 | 0.372184 | 0.636548 | 0.473767 |
| 22.30556 | 0.57282 | 0.600957 | 0.132213 | 0.111166 | 0.128581 | 0.118 | 0.045333 | -0.02476 | -0.52995 | 0.568457 | 0.051617 | 0.213788 | -0.07392 | 0.124579 | 0.024677 | 0.121121 | 0.396962 | 0.4807 | 0.265912 |
| 22.31944 | 0.861492 | 0.56728 | 0.197463 | 0.126065 | 0.131242 | 0.184779 | -0.00677 | 0.340007 | 0.273018 | 0.4238 | 0.020382 | 0.342248 | 0.156047 | 0.100084 | 0.12437 | 0.059235 | 0.338985 | 0.228511 | -0.02506 |
| 22.33333 | 1.076464 | 0.433784 | 0.190847 | 0.12037 | 0.172915 | 0.20474 | 0.000444 | 0.52353 | 1.184935 | 0.128444 | 0.007169 | 0.278219 | 0.29326 | 0.039631 | 0.414706 | 0.013601 | 0.182444 | 0.064757 | -0.12443 |
| 22.34722 | 1.055707 | 0.45144 | 0.22434 | 0.159692 | 0.19468 | 0.23058 | 0.060958 | 0.591038 | 1.659161 | -0.1712 | 0.038551 | 0.1371 | 0.274562 | -0.01839 | 0.55031 | -0.0266 | 0.010343 | 0.027172 | -0.06898 |
| 22.36111 | 1.041128 | 0.813618 | 0.300033 | 0.230431 | 0.14673 | 0.24674 | 0.104662 | 0.438057 | 0.963559 | -0.28823 | 0.004997 | 0.052144 | 0.054257 | -0.05989 | 0.20616 | -0.17693 | -0.0415 | -0.02074 | -0.1561 |
| 22.375 | 0.676919 | 0.823301 | 0.2587 | 0.353144 | 0.045678 | 0.172741 | 0.107927 | 0.573831 | -0.01418 | -0.15789 | -0.07942 | 0.064234 | -0.16033 | -0.07725 | -0.2489 | -0.31715 | 0.08691 | -0.03609 | -0.34004 |
| 22.38889 | 0.298582 | 0.586963 | 0.169392 | 0.383549 | 0.035107 | 0.145811 | 0.087757 | 0.725863 | -0.69645 | 0.079052 | -0.03405 | 0.10763 | -0.07289 | -0.03883 | -0.15503 | -0.19843 | 0.304458 | 0.12209 | -0.27994 |
| 22.40278 | 0.311078 | 0.867777 | 0.216081 | 0.348725 | 0.152103 | 0.253591 | 0.126492 | 0.727949 | -0.47094 | 0.234977 | 0.176112 | 0.103845 | 0.262108 | 0.052425 | 0.355089 | 0.065721 | 0.503368 | 0.336131 | 0.002661 |
| 22.41667 | 0.456677 | 1.236751 | 0.37168 | 0.300441 | 0.239175 | 0.345417 | 0.206497 | 0.746115 | 0.328399 | 0.172496 | 0.236839 | 0.079479 | 0.53444 | 0.093007 | 0.733524 | 0.229267 | 0.588992 | 0.358785 | 0.131903 |
| 22.43056 | 0.421737 | 1.021368 | 0.399796 | 0.265262 | 0.253138 | 0.287424 | 0.18881 | 0.939997 | 1.37338 | 0.057015 | 0.11862 | 0.087998 | 0.493754 | 0.0849 | 0.588814 | 0.234926 | 0.511546 | 0.283607 | 0.100326 |
| 22.44444 | 0.031637 | 0.220094 | 0.207692 | 0.294234 | 0.356037 | 0.178574 | 0.113505 | 0.705144 | 1.584044 | 0.10827 | 0.067025 | 0.20959 | 0.206931 | 0.124768 | 0.258659 | 0.116206 | 0.377475 | 0.190628 | 0.229703 |
| 22.45833 | 0.117126 | 0.086229 | 0.097646 | 0.234315 | 0.450331 | 0.223821 | 0.101734 | -0.07475 | 0.44282 | 0.38395 | 0.128576 | 0.426955 | -0.04742 | 0.210903 | 0.123411 | -0.05388 | 0.345207 | 0.154071 | 0.539007 |
| 22.47222 | 0.791915 | 1.233353 | 0.304841 | 0.028866 | 0.123055 | 0.387898 | 0.106815 | -0.28917 | -0.36388 | 0.677549 | 0.135496 | 0.593542 | -0.12463 | 0.249233 | 0.149965 | -0.1314 | 0.351854 | 0.093065 | 0.659891 |
| 22.48611 | 1.32324 | 2.570027 | 0.713299 | -0.1009 | 0.110293 | 0.466545 | 0.197861 | -0.10453 | -1.29593 | 0.684742 | 0.020213 | 0.633678 | 0.083483 | 0.199571 | 0.29879 | -0.13699 | 0.309766 | 0.011898 | 0.457702 |
| 22.5 | 1.355594 | 3.826669 | 1.442545 | 0.432283 | 0.687508 | 0.684424 | 0.746927 | 1.031687 | 0.201629 | 0.313622 | -0.02489 | 0.472793 | 0.46009 | 0.116115 | 0.440539 | -0.10706 | 0.164809 | 0.073019 | 0.234505 |
| 22.51389 | 2.094867 | 4.115823 | 1.88464 | 1.388691 | 1.269604 | 1.09621 | 1.317016 | 1.89765 | 3.098407 | 0.101233 | 0.030858 | 0.385766 | 0.648426 | 0.077264 | 0.31615 | 0.003939 | 0.015285 | 0.136111 | 0.074517 |
| 22.52778 | 1.513718 | 2.282567 | 1.439176 | 1.544832 | 1.069105 | 0.807934 | 0.947724 | 1.32269 | 3.499974 | 0.189736 | 0.062528 | 0.385794 | 0.373453 | 0.068898 | -0.0708 | 0.070796 | -0.0024 | 0.040719 | 0.003084 |
| 22.54167 | 0.711146 | 0.535585 | 0.589112 | 0.899099 | 0.649317 | 0.31055 | 0.325991 | 0.073387 | 1.942624 | 0.390571 | 0.059007 | 0.352265 | -0.05182 | 0.099816 | -0.26427 | 0.084053 | 0.180316 | 0.034568 | 0.178286 |
| 22.55556 | -0.04509 | -0.82772 | -0.24241 | 0.110494 | 0.500972 | -0.15038 | -0.21799 | -0.69432 | 0.721671 | 0.593393 | 0.084637 | 0.2077 | -0.28526 | 0.146743 | -0.06908 | 0.084828 | 0.456531 | 0.178155 | 0.543422 |
| 22.56944 | -0.7319 | -2.09724 | -0.9102 | -0.49069 | 0.403739 | -0.63769 | -0.73522 | -1.47297 | -0.46982 | 0.723586 | 0.121414 | 0.078587 | -0.22296 | 0.198315 | 0.276127 | 0.023349 | 0.480153 | 0.2667 | 0.60253 |
| 22.58333 | -1.03174 | -2.19575 | -1.08706 | -0.62689 | 0.264201 | -0.85926 | -0.86219 | -1.71441 | -1.53943 | 0.614725 | 0.221968 | 0.110298 | 0.17261 | 0.316622 | 0.586275 | 0.033337 | 0.311182 | 0.241181 | 0.312736 |
| 22.59722 | -0.95567 | -1.22046 | -0.75648 | -0.34652 | 0.133709 | -0.44927 | -0.57651 | -1.84831 | -2.84978 | 0.635879 | 0.554836 | 0.297514 | 0.805188 | 0.68118 | 0.898867 | 0.194681 | 0.30218 | 0.226038 | 0.227624 |
| 22.61111 | -0.3743 | 0.186509 | -0.11723 | 0.338682 | -0.04645 | 0.061904 | 0.044776 | 0.133152 | -1.54097 | 1.093084 | 1.146719 | 0.895695 | 1.569805 | 1.503197 | 1.561108 | 0.423696 | 0.699581 | 0.374321 | 0.465592 |
| 22.625 | 0.43374 | 1.548853 | 0.516311 | 0.822927 | -0.18662 | 0.485656 | 0.516079 | 2.183344 | 0.495395 | 1.854407 | 1.072909 | 1.039689 | 2.345647 | 2.289852 | 1.927971 | 0.131513 | 1.109359 | 0.530053 | 0.724674 |
| 22.63889 | 0.862237 | 1.915891 | 1.035406 | 0.47582 | -0.25684 | 0.2607 | 0.491279 | 2.911009 | 2.026649 | 1.796271 | 0.542315 | 0.806131 | 2.197239 | 1.3885 | 1.472464 | -0.08 | 0.868983 | 0.588048 | 0.907343 |
| 22.65278 | 0.645108 | 1.174453 | 0.702531 | 0.154197 | -0.23988 | -0.08679 | 0.178804 | 0.995817 | 1.025265 | 0.917889 | -0.02454 | 0.340294 | 1.800553 | 0.557391 | 1.554048 | 0.127774 | 0.368858 | 0.703494 | 0.754287 |
| 22.66667 | 0.053808 | -0.05179 | 0.216302 | -0.36856 | -0.16909 | -0.45696 | -0.13994 | -1.20101 | -0.49083 | 0.119915 | -0.56826 | -0.19746 | 0.84821 | -0.11546 | 1.371411 | 0.459057 | -0.18 | 0.493344 | -0.16739 |
| 22.68056 | -0.36878 | -1.19345 | -0.47875 | -0.93859 | -0.1776 | -0.82633 | -0.46598 | -2.44381 | -1.62755 | -0.86067 | -0.6463 | -0.83162 | -0.16104 | -0.60406 | 1.169478 | 0.830677 | -0.63865 | 0.179943 | -0.77533 |
| 22.69444 | -0.43281 | -1.98215 | -0.6429 | -0.91858 | -0.23493 | -0.68557 | -0.69625 | -2.51965 | -1.37597 | -1.50521 | 0.039646 | -1.18578 | -0.63735 | -0.71781 | 0.662517 | 1.137627 | -0.81551 | 0.085288 | -0.69972 |
| 22.70833 | -0.3905 | -2.13851 | -0.53403 | -0.45761 | -0.32451 | -0.25872 | -0.67923 | -1.25355 | -0.4379 | -0.98294 | 0.790398 | -0.11301 | -0.19809 | -0.48218 | 0.752505 | 0.933152 | -0.21708 | 0.143406 | -0.09784 |
| 22.72222 | -0.23786 | -1.09221 | -0.09971 | -0.04949 | -0.3709 | 0.11285 | -0.47516 | 0.359347 | 0.672918 | 0.546267 | 0.859218 | 1.037317 | 0.303939 | -0.1528 | 1.266445 | 0.210425 | 0.521848 | 0.259438 | 0.457757 |
| 22.73611 | -0.17254 | 0.251206 | 0.108681 | 0.450311 | -0.32257 | 0.181753 | -0.15744 | 1.552529 | 1.395272 | 1.20709 | 0.313443 | 1.298443 | 0.414494 | 0.110666 | 1.139265 | -0.46374 | 1.097131 | 0.292112 | 0.783169 |

| | | | | | | | | | | | | | | | | | | | |
|---|---|---|---|---|---|---|---|---|---|---|---|---|---|---|---|---|---|---|---|
| 22.75 | -0.21911 | 1.043336 | 0.113006 | 0.426514 | -0.1325 | -0.0659 | 0.09174 | 1.485247 | 1.079945 | 0.665808 | -0.12789 | 0.572747 | -0.58307 | 0.144991 | -0.14446 | -0.84659 | 0.682918 | -0.00033 | 0.704377 |
| 22.76389 | -0.24935 | 0.474181 | -0.41107 | 0.302455 | 0.024773 | -0.3499 | 0.13614 | 0.055094 | -0.44785 | 0.328447 | -0.304 | -0.12062 | -1.28703 | 0.10771 | -1.35661 | -1.12716 | 0.140161 | -0.29435 | 0.150871 |
| 22.77778 | -0.28875 | -0.6487 | -1.09044 | -0.10781 | 0.062989 | -0.54397 | 0.056649 | -1.73712 | -2.19478 | -0.07761 | -0.41538 | -0.39322 | -1.53625 | -0.01458 | -2.13276 | -1.32659 | -0.24462 | -0.30425 | -0.62529 |
| 22.79167 | -0.62528 | -1.93412 | -1.14748 | -0.5238 | 0.046171 | -0.54389 | -0.05187 | -2.99218 | -3.32014 | -1.04679 | -0.27516 | -0.71907 | -1.45786 | -0.19175 | -2.86285 | -1.3825 | -0.63824 | -0.18603 | -1.02769 |
| 22.80556 | -0.97062 | -3.03049 | -1.24474 | -0.7367 | -0.03919 | -0.45585 | -0.17356 | -2.16992 | -3.04354 | -1.42777 | 0.05999 | -1.04869 | -0.97445 | -0.26843 | -3.32576 | -1.16705 | -0.63469 | 0.102612 | -0.94467 |
| 22.81944 | -1.01183 | -3.15606 | -1.18181 | -0.72796 | -0.10392 | -0.35919 | -0.27036 | -0.92333 | -2.19626 | -0.29931 | 0.339217 | -0.41839 | -0.09508 | -0.19172 | -2.14395 | -0.68834 | 0.041074 | 0.497299 | -0.03594 |
| 22.83333 | -0.76458 | -2.45345 | -1.02526 | -0.55789 | -0.15884 | -0.3352 | -0.3398 | -0.54157 | -1.69617 | 1.008901 | 0.361706 | 0.66138 | 0.670706 | -0.06499 | -0.34037 | -0.25496 | 0.77198 | 0.894261 | 0.81429 |
| 22.84722 | -0.36549 | -1.73158 | -0.67309 | -0.44394 | -0.30275 | -0.43357 | -0.34233 | -0.85208 | -1.41605 | 1.075934 | 0.168076 | 1.224204 | 1.3156 | -0.04305 | 0.753801 | -0.1294 | 0.787307 | 0.96243 | 0.668836 |
| 22.86111 | -0.24087 | -1.08681 | -0.60815 | -0.56788 | -0.46362 | -0.53448 | -0.25894 | -1.37124 | -1.02884 | -0.01299 | -0.18364 | 0.725704 | 1.22449 | -0.14077 | 0.358611 | -0.20657 | -0.13058 | 0.636505 | 0.059718 |
| 22.875 | -0.22809 | -0.26861 | -0.77013 | -0.88695 | -0.552 | -0.61627 | -0.27788 | -1.91102 | -0.79559 | -0.79835 | -0.51795 | 0.25485 | 0.562804 | -0.24297 | 0.027866 | -0.25926 | -0.83601 | 0.288013 | -0.51468 |
| 22.88889 | -0.30757 | 0.371224 | -0.97654 | -1.13597 | -0.60339 | -0.79705 | -0.34159 | -2.14329 | -0.71681 | -1.77931 | -0.60419 | -0.28763 | -0.22719 | -0.32005 | -0.43365 | -0.20448 | -1.19543 | -0.04166 | -1.08717 |
| 22.90278 | -0.40736 | 0.39544 | -1.04577 | -1.06214 | -0.60903 | -0.78011 | -0.31808 | -1.58288 | -0.34668 | -2.27389 | -0.41424 | -0.89875 | -0.99593 | -0.46714 | -1.44136 | -0.20101 | -1.15944 | -0.26287 | -1.3094 |
| 22.91667 | -0.4055 | 0.295171 | -0.7405 | -0.70736 | -0.53209 | -0.52513 | -0.18351 | -0.56845 | 0.219413 | -1.85084 | -0.11992 | -0.83913 | -1.45086 | -0.63025 | -2.50394 | -0.22408 | -0.78316 | -0.39389 | -0.52166 |
| 22.93056 | -0.23456 | 0.32771 | -0.38196 | -0.37102 | -0.32738 | -0.26126 | -0.03491 | 0.29042 | 0.608562 | -0.73984 | -0.03919 | -0.47059 | -1.43962 | -0.67296 | -2.13421 | -0.03157 | -0.52068 | -0.44717 | 0.256643 |
| 22.94444 | 0.062783 | 0.30424 | -0.11047 | -0.05378 | -0.10145 | -0.07535 | 0.048389 | 0.687209 | 0.405035 | -0.02831 | -0.18276 | 0.076474 | -1.10361 | -0.50083 | -1.25714 | 0.152704 | -0.34647 | -0.39083 | 0.576838 |
| 22.95833 | 0.293718 | -0.03369 | 0.027423 | 0.201059 | 0.032751 | -0.09441 | 0.040177 | 0.498047 | -0.2104 | 0.083803 | -0.32917 | 0.307112 | -0.89112 | -0.41008 | -0.55522 | 0.243638 | -0.35327 | -0.27933 | 0.303446 |
| 22.97222 | 0.136742 | -0.796 | 0.098065 | 0.336779 | -0.00246 | -0.3175 | 0.018212 | -0.11828 | -0.73977 | -0.36709 | -0.46254 | 0.173475 | -0.57349 | -0.49175 | 0.142076 | 0.228686 | -0.45567 | -0.31004 | -0.16679 |
| 22.98611 | -0.48781 | -1.84104 | -0.18756 | 0.134652 | -0.02349 | -0.60333 | -0.08379 | -1.06286 | -1.04586 | -0.93261 | -0.56094 | -0.22827 | -0.33385 | -0.61489 | 0.840377 | 0.100632 | -0.49221 | -0.28392 | -0.81679 |
| 23 | -0.92655 | -2.78509 | -0.97311 | -0.30546 | -0.0784 | -0.84669 | -0.51565 | -1.82339 | -1.55902 | -1.20345 | -0.51573 | -0.69657 | -0.37457 | -0.60652 | 1.438921 | 0.052988 | -0.34546 | -0.1498 | -1.54103 |
| 23.01389 | -0.86491 | -3.09522 | -1.50058 | -0.67458 | -0.14256 | -0.95084 | -0.97197 | -2.25516 | -1.65187 | -1.54021 | -0.42159 | -0.83997 | -0.42112 | -0.52108 | 1.225617 | 0.011803 | -0.09835 | 0.010912 | -1.80415 |
| 23.02778 | -0.51321 | -2.39137 | -1.40907 | -0.73081 | -0.21077 | -0.82375 | -1.1982 | -1.60091 | -0.8563 | -1.67986 | -0.3082 | -0.54438 | -0.44057 | -0.42001 | 0.408998 | -0.17463 | 0.058981 | 0.095378 | -1.38692 |
| 23.04167 | -0.44564 | -1.65573 | -0.87669 | -0.72509 | -0.17556 | -0.19601 | -0.82385 | -0.71244 | 0.097439 | -0.97307 | -0.20578 | -0.06341 | -0.42512 | -0.31354 | -0.2531 | -0.36522 | 0.083758 | 0.068799 | -0.83088 |
| 23.05556 | -0.89366 | -1.47646 | -0.52714 | -0.73619 | -0.25278 | 0.082592 | -0.44605 | -0.52732 | 0.397522 | 0.026496 | -0.15887 | 0.260514 | -0.17754 | -0.18432 | -0.94025 | -0.57002 | 0.053834 | -0.01832 | -0.20163 |
| 23.06944 | -0.97833 | -1.30988 | -0.31733 | -0.57236 | -0.24935 | 0.038348 | -0.17101 | -0.74194 | 0.130587 | 0.771074 | -0.12925 | 0.249847 | 0.266003 | -0.05123 | -1.78427 | -0.91488 | -0.00045 | -0.08636 | 0.318306 |
| 23.08333 | -0.54198 | -0.95287 | -0.14201 | -0.23475 | -0.192 | 0.052383 | -0.09587 | -0.61723 | -0.02272 | 0.875224 | -0.10259 | -0.02397 | 0.74039 | -0.04177 | -1.80066 | -0.92653 | -0.14027 | -0.14022 | 0.414468 |
| 23.09722 | -0.07224 | -0.39138 | 0.016456 | 0.026309 | -0.05394 | 0.247422 | -0.03258 | -0.33837 | 0.011277 | 0.043662 | -0.19057 | -0.46651 | 0.811598 | -0.20918 | -1.44664 | -0.69538 | -0.33849 | -0.14578 | 0.040625 |
| 23.11111 | 0.082243 | -0.06243 | 0.100624 | 0.177228 | 0.118951 | 0.518534 | 0.036192 | -0.10707 | 0.132859 | -1.29732 | -0.47486 | -1.10852 | -0.05082 | -0.55236 | -1.15083 | -0.58545 | -0.55681 | -0.24017 | -0.70393 |
| 23.125 | 0.245092 | 0.245664 | 0.170649 | 0.270458 | 0.302485 | 0.719831 | 0.099921 | 0.152605 | 0.323555 | -3.31092 | -0.7326 | -1.2154 | -1.33975 | -0.73508 | -0.91313 | -0.72435 | -0.52216 | -0.38426 | -1.21926 |
| 23.13889 | 0.44093 | 0.41853 | 0.199623 | 0.30001 | 0.347229 | 0.607567 | 0.162001 | 0.469429 | 0.531299 | -4.55978 | -0.72261 | -0.81838 | -1.99929 | -0.50235 | -0.58891 | -0.77543 | -0.16322 | -0.39575 | -1.07221 |
| 23.15278 | 0.353024 | 0.074445 | 0.137252 | 0.235289 | 0.226002 | 0.287856 | 0.270215 | 0.763588 | 0.800744 | -3.38457 | -0.42672 | -0.56021 | -1.80205 | -0.20676 | -0.24407 | -0.66714 | 0.169278 | -0.2448 | -0.71902 |
| 23.16667 | 0.231014 | 0.021348 | 0.184686 | 0.278983 | 0.22536 | 0.029267 | 0.315889 | 1.29753 | 1.186032 | -2.96051 | -0.22364 | -0.34235 | -1.10483 | -0.06835 | 0.113037 | -0.42326 | 0.249264 | -0.07366 | -0.9131 |
| 23.18056 | 0.394676 | 1.2172 | 0.418483 | 0.440416 | 0.247601 | 0.029813 | 0.27867 | 1.771176 | 1.378887 | -2.21676 | -0.3104 | -0.09799 | -0.27555 | -0.07028 | 0.258037 | -0.27155 | 0.154335 | -0.09254 | -0.89283 |
| 23.19444 | 0.468181 | 1.963348 | 0.602403 | 0.512993 | 0.159171 | 0.117209 | 0.209924 | 1.749243 | 1.429222 | -1.05049 | -0.37035 | -0.03183 | 0.214572 | -0.09482 | 0.177168 | -0.07854 | 0.07909 | -0.23272 | -0.68585 |
| 23.20833 | 0.501405 | 1.949239 | 0.659501 | 0.482104 | 0.055373 | 0.224649 | 0.225952 | 1.647093 | 1.370369 | -0.21929 | -0.15167 | -0.00557 | 0.628316 | -0.05162 | 0.182579 | 0.146655 | 0.039573 | -0.25224 | -0.29841 |
| 23.22222 | 0.463849 | 1.928354 | 0.580011 | 0.415319 | -0.01784 | 0.288022 | 0.300928 | 1.539532 | 1.202461 | 0.493282 | 0.191335 | -0.05302 | 1.007944 | 0.005966 | 0.365879 | 0.424491 | 0.04211 | -0.11268 | 0.150823 |
| 23.23611 | 0.377233 | 1.466237 | 0.458561 | 0.304104 | -0.11155 | 0.318264 | 0.314814 | 1.187019 | 0.926012 | 1.096371 | 0.486604 | -0.06677 | 1.152271 | 0.061065 | 0.7688 | 0.703448 | 0.149374 | 0.098076 | 0.563565 |
| 23.25 | 0.366755 | 1.143006 | 0.402922 | 0.277316 | -0.11976 | 0.224853 | 0.253614 | 0.665078 | 0.814753 | 1.50783 | 0.561815 | -0.01423 | 1.231403 | 0.094533 | 1.379459 | 1.002743 | 0.324093 | 0.251727 | 0.803596 |
| 23.26389 | 0.216355 | 0.774246 | 0.331676 | 0.220022 | 0.009913 | 0.110074 | 0.202167 | 0.17808 | 0.525796 | 1.859877 | 0.514566 | 0.037988 | 1.078216 | 0.084103 | 1.830866 | 1.189171 | 0.457746 | 0.311617 | 0.882112 |
| 23.27778 | -0.01559 | 0.074448 | 0.083847 | 0.084303 | 0.126246 | -0.02017 | 0.179004 | -0.5231 | -0.60385 | 2.19686 | 0.607791 | 0.051987 | 0.855535 | 0.107181 | 1.568932 | 1.056774 | 0.571215 | 0.419457 | 1.098914 |
| 23.29167 | -0.03405 | -0.52657 | -0.13451 | 0.00305 | 0.143147 | -0.07915 | 0.108356 | -1.33192 | -1.30097 | 2.040743 | 0.55131 | 0.044622 | 0.681037 | 0.181071 | 1.123422 | 1.127875 | 0.627397 | 0.552537 | 1.07543 |
| 23.30556 | 0.099619 | -0.764 | -0.18741 | 0.018134 | 0.080766 | -0.03854 | 0.012788 | -1.67054 | -1.14563 | 1.437333 | 0.344356 | 0.097165 | 0.527198 | 0.232878 | 0.705247 | 1.016888 | 0.614908 | 0.536947 | 0.804957 |
| 23.31944 | 0.215686 | -0.57484 | -0.16257 | 0.070914 | -0.0293 | 0.067901 | -0.01058 | -1.43299 | -0.53999 | 0.943927 | 0.181862 | 0.176069 | 0.404688 | 0.276416 | 0.385821 | 0.677923 | 0.593394 | 0.469952 | 0.634941 |
| 23.33333 | 0.260594 | -0.00957 | -0.07806 | 0.160395 | -0.07225 | 0.172895 | 0.025632 | -0.86217 | 0.187376 | 0.474153 | -0.03776 | 0.185224 | 0.167448 | 0.239082 | 0.090424 | 0.169043 | 0.549257 | 0.337036 | 0.41955 |
| 23.34722 | 0.365399 | 0.74952 | 0.040947 | 0.282494 | 0.028338 | 0.24216 | 0.06884 | 0.054006 | 0.760486 | -0.35762 | -0.97489 | -0.31008 | -0.29755 | -0.13231 | -0.36465 | -0.5203 | -0.00376 | -0.30063 | -0.40801 |
| 23.36111 | 0.534213 | 1.404012 | 0.178947 | 0.385818 | 0.197433 | 0.279174 | 0.109162 | 0.989711 | 1.211268 | -1.50349 | -2.52492 | -1.2469 | -0.70946 | -0.72206 | -0.83958 | -1.30967 | -0.9908 | -1.21276 | -1.77106 |
| 23.375 | 0.636072 | 1.547703 | 0.325269 | 0.454186 | 0.273235 | 0.293859 | 0.182469 | 1.891736 | 1.357881 | -1.327 | -2.49421 | -1.22191 | -0.65244 | -0.75438 | -0.83725 | -1.41572 | -1.20708 | -1.27197 | -1.87314 |
| 23.38889 | 0.565553 | 0.936574 | 0.373643 | 0.408962 | 0.210546 | 0.283988 | 0.241722 | 2.178879 | 1.030197 | -0.40693 | -0.99785 | -0.26206 | -0.09536 | -0.30637 | -0.40588 | -0.73158 | -0.61586 | -0.51962 | -0.78476 |
| 23.40278 | 0.376799 | 0.083989 | 0.35673 | 0.316223 | 0.116771 | 0.243192 | 0.226423 | 1.931927 | 0.619457 | -0.39891 | -0.23038 | 0.214394 | 0.420825 | -0.14501 | -0.0732 | -0.18238 | -0.3158 | -0.10654 | -0.27037 |
| 23.41667 | 0.379701 | -0.291 | 0.389496 | 0.317957 | 0.070194 | 0.224317 | 0.24384 | 1.433698 | 0.491973 | -0.63538 | -0.22129 | 0.173987 | 0.514632 | -0.22867 | 0.097868 | -0.01206 | -0.20045 | -0.12973 | -0.25401 |
| 23.43056 | 0.633683 | -0.00458 | 0.461819 | 0.406748 | 0.04998 | 0.289257 | 0.316476 | 0.895119 | 0.454533 | -0.34865 | -0.23814 | 0.151214 | 0.15828 | -0.24448 | 0.325434 | 0.070941 | 0.012226 | -0.14506 | -0.1869 |
| 23.44444 | 0.844623 | 0.415706 | 0.405729 | 0.422656 | -0.01336 | 0.345616 | 0.361935 | 0.330761 | 0.238398 | 0.318508 | -0.04676 | 0.217622 | -0.21297 | -0.13722 | 0.490296 | 0.215704 | 0.301368 | 0.010808 | -0.02089 |
| 23.45833 | 0.692636 | 0.61446 | 0.26594 | 0.330308 | -0.02657 | 0.289872 | 0.299992 | -0.41205 | -0.20736 | 1.021873 | 0.245231 | 0.278262 | -0.25216 | 0.02431 | 0.449474 | 0.40037 | 0.690436 | 0.341179 | 0.187103 |
| 23.47222 | 0.549581 | 1.362367 | 0.094402 | 0.411529 | 0.164739 | 0.43971 | 0.15188 | -0.66063 | -0.45727 | 1.304193 | 0.459547 | 0.321717 | -0.09627 | 0.19505 | 0.269828 | 0.446076 | 0.899539 | 0.383631 | 0.308059 |
| 23.48611 | 0.669576 | 2.943256 | 0.314209 | 0.608085 | 0.458694 | 0.744733 | -0.00965 | 0.260445 | -0.55863 | 1.20632 | 0.471014 | 0.319428 | 0.207827 | 0.371464 | 0.101291 | 0.359838 | 0.667731 | 0.326811 | 0.352043 |
| 23.5 | 1.345119 | 4.931886 | 1.251467 | 0.697521 | 0.469342 | 0.891972 | 0.061455 | 2.150132 | 0.80673 | 0.916701 | 0.389678 | 0.281667 | 0.579026 | 0.446537 | 0.145111 | 0.182401 | 0.378321 | 0.275741 | 0.459825 |
| 23.51389 | 1.940335 | 5.607976 | 2.154514 | 1.158949 | 0.295128 | 0.935464 | 0.608105 | 3.744176 | 2.607241 | 0.475941 | 0.242293 | 0.31472 | 0.743656 | 0.462855 | 0.387535 | 0.168678 | 0.147145 | 0.156811 | 0.679841 |
| 23.52778 | 2.552049 | 4.087182 | 2.173351 | 1.415717 | 0.523077 | 0.743308 | 0.546008 | 3.455349 | 2.735596 | 0.254702 | 0.260051 | 0.408871 | 0.680273 | 0.445943 | 0.722418 | 0.299577 | 0.037681 | 0.123449 | 1.027032 |
| 23.54167 | 1.872206 | 2.509334 | 1.539262 | 1.045557 | 0.905874 | 0.275903 | 0.339355 | 2.092052 | 2.214891 | 0.595598 | 0.45049 | 0.549944 | 0.566755 | 0.504826 | 0.981023 | 0.41856 | 0.041802 | 0.187103 | 0.819112 |
| 23.55556 | 0.704418 | 0.654046 | 0.763905 | 0.593774 | 1.05973 | -0.17377 | 0.01061 | 0.622377 | 1.323472 | 0.874476 | 0.439281 | 0.595072 | 0.280865 | 0.44828 | 0.812616 | 0.356334 | 0.031354 | 0.208416 | 0.228642 |
| 23.56944 | -0.44782 | -1.23331 | -0.02174 | 0.002239 | 1.001623 | -0.69966 | -0.37218 | -1.1215 | 0.039119 | 0.490615 | 0.198009 | 0.434902 | -0.2503 | 0.337691 | 0.147091 | 0.123655 | -0.01527 | 0.166453 | -0.43832 |
| 23.58333 | -1.34263 | -2.38075 | -0.54922 | -0.58635 | 0.876279 | -1.04634 | -0.60394 | -2.7282 | -1.20547 | 0.190042 | 0.125398 | 0.228749 | -0.54975 | 0.259671 | -0.41392 | -0.0573 | -0.0287 | 0.106885 | -0.71395 |
| 23.59722 | -1.47755 | -2.61925 | -0.72007 | -0.87055 | 0.820085 | -0.82126 | -0.58726 | -3.50894 | -1.93281 | 0.702918 | 0.38742 | 0.261382 | -0.20517 | 0.343495 | -0.33612 | -0.02537 | 0.158249 | 0.159116 | -0.11821 |
| 23.61111 | -0.53006 | -2.07259 | -0.72383 | -0.71322 | 0.755394 | -0.08448 | -0.19751 | -4.21532 | -2.42521 | 1.900738 | 0.77714 | 0.715697 | 0.740645 | 0.670814 | 0.479072 | 0.138001 | 0.799308 | 0.538971 | 1.073621 |
| 23.625 | 0.884984 | -0.08231 | -0.27963 | -0.09102 | 0.552981 | 0.730136 | 0.088833 | -2.11781 | -0.93859 | 3.456572 | 1.055578 | 1.412406 | 1.914149 | 1.247961 | 1.664862 | 0.119403 | 1.678649 | 1.000675 | 1.707012 |
| 23.63889 | 1.74004 | 2.193477 | 0.203213 | 0.64934 | 0.184564 | 0.552588 | 0.280966 | 1.199239 | 0.466121 | 3.540584 | 1.281597 | 2.42272 | 2.294686 | 1.511323 | 2.244822 | -0.02393 | 1.932553 | 0.768725 | 1.785596 |
| 23.65278 | 1.664215 | 3.323087 | 0.778956 | 1.17745 | -0.12244 | 0.244808 | 0.387003 | 3.129516 | 1.266595 | 2.110651 | 1.09701 | 1.347968 | 1.81557 | 0.986849 | 1.402902 | -0.09305 | 1.404362 | 0.607887 | 0.768104 |
| 23.66667 | 1.012876 | 2.76596 | 1.110958 | 0.578467 | -0.42011 | 0.068719 | 0.350265 | 2.900781 | 1.462235 | 0.169786 | 0.316352 | 0.314916 | 1.356615 | 0.104023 | 0.627009 | -0.30569 | 1.050854 | 0.390669 | -0.56544 |
| 23.68056 | 0.323725 | 1.735973 | 1.312108 | 0.095556 | -0.75277 | -0.41657 | 0.238683 | 0.921458 | -0.14381 | -2.05413 | -0.53221 | -0.55529 | 0.563837 | -0.49202 | -0.45582 | -0.6734 | 0.744575 | 0.082096 | -1.51893 |
| 23.69444 | -0.38238 | 0.194724 | 0.964389 | -0.51904 | -0.82948 | -0.6742 | 0.034954 | -1.37532 | -1.58799 | -3.16082 | -0.87459 | -1.30077 | -0.04803 | -0.49773 | -0.90136 | -0.80712 | 0.482155 | -0.17666 | -2.02056 |
| 23.70833 | -0.6918 | -0.73008 | 0.306674 | -0.7408 | -0.62548 | -0.38731 | -0.07417 | -2.81135 | -2.02144 | -3.28889 | -0.5627 | -1.44006 | 0.109392 | -0.10045 | -0.957 | -0.8281 | 0.106986 | -0.22697 | -1.81852 |
| 23.72222 | -0.20132 | -1.17552 | -0.48701 | -0.7037 | -0.21691 | 0.015339 | -0.13661 | -3.99262 | -2.79869 | -0.9581 | 0.170727 | -1.02761 | 0.247208 | 0.230612 | -0.48596 | -0.55255 | -0.07528 | -0.01958 | -0.35054 |
| 23.73611 | 0.330003 | -1.66567 | -0.9294 | -0.20993 | 0.077701 | 0.13042 | -0.18672 | -4.22373 | -2.53727 | 0.943569 | 0.684825 | -0.03914 | 0.034237 | 0.33521 | 1.496676 | 0.486628 | -0.09137 | 0.156939 | 1.386268 |
| 23.75 | 0.609567 | -2.07436 | -0.85506 | 0.409361 | 0.207522 | 0.073915 | -0.17684 | -2.46688 | -1.05509 | 1.224787 | 0.753069 | 1.092976 | -0.06966 | 0.40453 | 1.700272 | 0.758107 | -0.12482 | 0.114721 | 1.965676 |
| 23.76389 | 0.498935 | -1.45534 | -0.38692 | 0.760083 | 0.218473 | -0.36959 | -0.40055 | 0.429073 | 0.264995 | 0.699169 | 0.493157 | 1.91633 | -0.71321 | 0.408706 | 0.959468 | 0.636741 | -0.28707 | -0.0409 | 1.252107 |
| 23.77778 | -0.06674 | -0.16463 | 0.147396 | 0.383259 | 0.104323 | -1.06596 | -0.42136 | 2.603735 | 1.098983 | -0.42404 | 0.037937 | 0.751517 | -2.06383 | 0.021931 | 0.065008 | 0.30202 | -0.34174 | -0.19815 | 0.118454 |
| 23.79167 | -0.50927 | 1.177546 | 0.596565 | -0.23918 | -0.08936 | -1.32121 | -0.24427 | 3.074147 | 0.94811 | -1.53785 | -0.46071 | -0.1353 | -2.46137 | -0.39839 | -0.79608 | -0.23087 | -0.34187 | -0.27614 | -0.88699 |
| 23.80556 | -0.85023 | 1.18859 | 0.783773 | -0.59041 | -0.35402 | -1.0537 | 0.008878 | 1.761403 | 0.18508 | -2.19715 | -0.70574 | -0.61555 | -2.05793 | -0.77794 | -1.48871 | -0.66945 | -0.19948 | -0.26117 | -1.53897 |
| 23.81944 | -0.72216 | 0.21761 | 0.291977 | -0.88642 | -0.59435 | -0.55273 | 0.075283 | -0.41246 | -0.96489 | -2.0742 | -0.32443 | -0.59531 | -1.51348 | -0.96529 | -1.72926 | -0.67583 | -0.13361 | -0.35103 | -1.9732 |
| 23.83333 | -0.34239 | -0.74857 | -0.74449 | -0.97016 | -0.60639 | -0.22876 | 0.027996 | -2.3749 | -1.85928 | -1.56586 | 0.22333 | -0.71116 | -0.49432 | -0.88851 | -2.26434 | -1.00019 | 0.002127 | -0.45484 | -1.61442 |
| 23.84722 | -0.18701 | -1.68215 | -1.55421 | -1.01097 | -0.5561 | -0.27326 | -0.05639 | -3.83371 | -2.31497 | 0.13622 | 0.35795 | -1.67811 | 0.005627 | -0.47728 | -1.29995 | -0.44988 | 0.031181 | -0.21613 | 0.083869 |
| 23.86111 | -0.03647 | -2.43437 | -1.72386 | -0.75584 | -0.54501 | -0.47191 | -0.19568 | -3.16258 | -1.51214 | 0.365535 | 0.058329 | -1.48528 | 0.414032 | -0.02491 | -0.38791 | 0.053686 | 0.086127 | 0.114123 | 1.542424 |
| 23.875 | -0.13017 | -2.88743 | -1.80004 | -0.53757 | -0.52898 | -0.56994 | -0.41624 | -2.21352 | -0.7742 | 0.318691 | -0.26454 | -0.34357 | 0.731465 | 0.230042 | 0.207536 | 0.255696 | 0.281039 | 0.441296 | 0.954364 |
| 23.88889 | -0.149 | -2.65508 | -1.55147 | -0.17367 | -0.45817 | -0.30922 | -0.36985 | -0.9076 | -0.31762 | 0.182483 | -0.55974 | 0.699986 | 1.094284 | 0.237383 | 0.229248 | 0.084671 | 0.300996 | 0.43353 | -0.01896 |
| 23.90278 | -0.11506 | -1.62173 | -1.1936 | 0.051687 | -0.33929 | 0.01168 | -0.31658 | -0.238 | -0.13314 | -1.45025 | -0.77436 | 1.525693 | 1.148809 | 0.089701 | -0.18994 | -0.26199 | 0.015609 | 0.27875 | -0.96588 |
| 23.91667 | -0.13935 | -0.71779 | -0.95128 | 0.00429 | -0.2825 | 0.03517 | -0.30979 | 0.199299 | 0.123123 | -1.65842 | -0.72879 | 1.168779 | 0.616799 | -0.1856 | -0.32731 | -0.29072 | -0.58098 | 0.031132 | -2.15943 |
| 23.93056 | -0.02124 | 0.161512 | -0.27289 | 0.017477 | -0.23773 | -0.21858 | -0.15336 | 0.613784 | 0.437424 | -2.00046 | -0.43886 | 0.389389 | -0.25696 | -0.48451 | -0.22549 | -0.14452 | -1.08252 | -0.29176 | -2.16483 |

| | | | | | | | | | | | | | | | | | | | |
|---|---|---|---|---|---|---|---|---|---|---|---|---|---|---|---|---|---|---|---|
| 23.94444 | -0.05251 | 0.874988 | 0.352747 | -0.23769 | -0.1648 | -0.51681 | 0.062326 | 0.906112 | 0.498318 | -2.32761 | -0.14897 | -0.34841 | -1.08097 | -0.64929 | -0.22596 | -0.14022 | -1.21737 | -0.58897 | -1.69495 |
| 23.95833 | -0.18107 | 1.109881 | 0.876419 | -0.50116 | -0.15853 | -0.67902 | 0.311648 | 1.196492 | 0.399497 | -1.47867 | -0.04406 | -0.9006 | -1.59088 | -0.78886 | -0.3127 | -0.33335 | -0.94496 | -0.61739 | -1.04053 |
| 23.97222 | -0.26499 | 0.59078 | 1.244792 | -0.50996 | -0.36988 | -0.96031 | 0.529573 | 1.224345 | 0.292205 | -1.16071 | -0.14114 | -1.41925 | -1.62898 | -0.87162 | -0.45373 | -0.60857 | -0.48224 | -0.23438 | -1.0995 |
| 23.98611 | -0.72927 | -1.05904 | 0.96016 | -0.311 | -0.55865 | -1.49528 | 0.360702 | 0.431825 | -0.12021 | -1.28312 | -0.27638 | -1.29844 | -1.37424 | -0.83058 | -0.60233 | -0.7949 | -0.30675 | -0.06943 | -0.86015 |
| 24 | -1.53834 | -2.33825 | -0.23048 | -0.78794 | -0.38459 | -1.78433 | -0.1477 | -1.04296 | -0.78626 | -1.26039 | -0.41384 | -1.00065 | -1.10631 | -0.55996 | -0.65277 | -0.82318 | -0.16605 | -0.12953 | -0.18036 |
| 24.01389 | -1.67357 | -2.55279 | -1.25806 | -1.45758 | -0.36055 | -1.76528 | -0.56151 | -1.43582 | -1.35537 | -0.99665 | -0.46339 | -0.35736 | -0.91538 | -0.38619 | -0.42267 | -0.4387 | -0.13661 | -0.21257 | 0.277331 |
| 24.02778 | -1.21636 | -2.20686 | -1.51788 | -1.5729 | -0.34871 | -1.93883 | -0.53582 | -0.82604 | -1.34114 | -0.61813 | -0.47327 | 0.167249 | -0.73923 | -0.32437 | -0.27408 | 0.051263 | -0.08411 | -0.19815 | 0.471493 |
| 24.04167 | -0.52169 | -1.88903 | -1.39272 | -1.09155 | -0.34277 | -1.57608 | -0.3745 | -0.09682 | -0.74104 | -0.49392 | -0.53305 | 0.452666 | -0.37829 | -0.28697 | -0.2158 | 0.165846 | -0.05408 | -0.08464 | -0.14559 |
| 24.05556 | -0.23886 | -1.35088 | -1.07638 | -0.6338 | -0.224 | -0.94048 | -0.27308 | 0.101338 | -0.3932 | -0.59713 | -0.63863 | 0.52412 | 0.250726 | -0.13462 | -0.23015 | -0.08836 | -0.01207 | 0.069664 | -0.48594 |
| 24.06944 | -0.07036 | -0.8412 | -0.5981 | -0.23181 | -0.10508 | -0.23765 | -0.06193 | 0.065978 | -0.10141 | -0.50897 | -0.57074 | 0.497144 | 0.764606 | 0.041595 | -0.1839 | -0.35606 | 0.056554 | 0.148828 | -0.60092 |
| 24.08333 | 0.27139 | -0.46857 | -0.17466 | 0.119035 | -0.03856 | 0.136617 | 0.153373 | -0.08534 | 0.08013 | -0.22845 | -0.36313 | 0.42976 | 0.925477 | 0.099399 | -0.10156 | -0.39494 | 0.101219 | 0.143654 | -0.6033 |
| 24.09722 | 0.839268 | 0.017055 | -0.14539 | 0.440502 | -0.03495 | 0.413334 | 0.170784 | -0.39287 | -0.03909 | -0.25297 | -0.3975 | 0.177329 | 0.492909 | 0.061162 | -0.31472 | -0.42671 | 0.005388 | 0.083291 | -0.6726 |
| 24.11111 | 1.78478 | 1.267036 | -0.29829 | 0.952348 | 0.407546 | 1.047033 | 0.158933 | -0.61725 | -0.35724 | -1.43744 | -0.80485 | -0.42699 | -0.42863 | -0.08166 | -0.93767 | -0.73559 | -0.22193 | -0.08175 | -1.02265 |
| 24.125 | 3.009239 | 3.074012 | -0.12232 | 1.649915 | 1.44551 | 2.009328 | 0.41883 | -0.28136 | -0.15996 | -2.56033 | -1.16477 | -1.25444 | -1.15994 | -0.27397 | -1.11157 | -1.05567 | -0.3499 | -0.12162 | -1.62627 |
| 24.13889 | 2.78679 | 3.0579 | 0.25126 | 1.905341 | 1.763352 | 2.123693 | 0.531716 | 0.413213 | 0.675821 | -3.05237 | -1.32327 | -1.32296 | -0.92852 | -0.33218 | -0.60869 | -1.04385 | -0.25966 | 0.110182 | -1.72588 |
| 24.15278 | 1.387258 | 1.350063 | 0.491065 | 1.506838 | 1.081911 | 2.218813 | 0.437322 | 0.807487 | 1.618184 | -3.28944 | -1.3197 | -0.75904 | -0.32099 | -0.26295 | -0.27609 | -0.79503 | -0.19741 | 0.151016 | -1.2791 |
| 24.16667 | 0.590878 | 0.480252 | 0.524411 | 0.929142 | 0.518562 | 1.603648 | 0.418497 | 0.638054 | 1.655934 | -1.98278 | -0.84552 | -0.32345 | -0.14966 | -0.17027 | -0.29073 | -0.42855 | -0.11467 | 0.004824 | -1.28345 |
| 24.18056 | 0.379665 | 0.346715 | 0.391261 | 0.324001 | 0.158618 | 0.970586 | 0.287381 | 0.262651 | 0.778298 | -0.49335 | -0.23695 | 0.051291 | -0.06916 | -0.03188 | -0.28248 | -0.16301 | 0.066746 | -0.1556 | -0.82158 |
| 24.19444 | 0.157645 | 0.093264 | 0.068146 | -0.13015 | 0.035675 | 0.549508 | 0.029214 | 0.018447 | 0.253908 | 0.020227 | -0.08623 | 0.370545 | 0.002025 | 0.029053 | -0.38042 | -0.18174 | 0.056393 | -0.34931 | -0.43207 |
| 24.20833 | 0.06533 | -0.05507 | -0.16247 | -0.34433 | 0.11879 | 0.424241 | -0.12871 | 0.022892 | 0.031546 | -0.66932 | -1.01 | -0.18068 | -0.64276 | -0.16882 | -1.60329 | -1.03682 | -0.80891 | -0.88677 | -0.93742 |
| 24.22222 | 0.349325 | 0.041217 | -0.05878 | -0.17409 | 0.218824 | 0.328955 | -0.0262 | 0.10235 | 0.07497 | -3.13045 | -3.16492 | -2.33174 | -2.85662 | -0.86034 | -5.12048 | -3.27887 | -2.80646 | -2.39939 | -2.70177 |
| 24.23611 | 0.736299 | 0.210027 | 0.173969 | 0.216591 | 0.210062 | 0.192857 | 0.168742 | 0.114195 | 0.243271 | -3.26364 | -2.94044 | -3.48514 | -4.23691 | -1.65577 | -4.1451 | -2.92239 | -3.03952 | -2.2864 | -3.08271 |
| 24.25 | 0.743418 | 0.222018 | 0.255856 | 0.369319 | 0.089152 | -0.13038 | 0.182989 | 0.022838 | 0.328037 | -0.45476 | -0.57144 | -1.1258 | -2.08442 | -0.9065 | -1.70429 | -1.13039 | -1.19443 | -1.04748 | -0.66587 |
| 24.26389 | 0.321633 | 0.082409 | 0.050102 | 0.168013 | 0.013977 | -0.41883 | 0.015934 | -0.02578 | 0.254087 | 1.176376 | 0.677241 | 0.016611 | 0.688188 | 0.119089 | 1.78854 | 1.102097 | 0.747808 | 0.561928 | 1.030888 |
| 24.27778 | -0.20516 | -0.06944 | -0.21548 | -0.15469 | -0.00226 | -0.65582 | -0.13447 | 0.00834 | 0.188967 | 0.900659 | 0.424404 | -0.05041 | 1.197227 | 0.451618 | 2.131942 | 1.170038 | 0.781214 | 0.590803 | 0.938333 |
| 24.29167 | -0.42363 | -0.06147 | -0.33403 | -0.37569 | -0.10873 | -0.60406 | -0.24796 | 0.061312 | 0.190902 | -0.13798 | -0.32309 | -0.70125 | 0.068703 | 0.073802 | -0.87897 | -0.56592 | -0.33645 | -0.54271 | 0.088782 |
| 24.30556 | -0.3338 | 0.003216 | -0.22823 | -0.37839 | -0.22827 | -0.58434 | -0.35202 | 0.057781 | 0.214466 | -0.65464 | -0.61081 | -0.70964 | -0.63638 | -0.16746 | -2.44341 | -1.503 | -0.69523 | -1.04608 | -0.48132 |
| 24.31944 | -0.05735 | 0.042645 | -0.01932 | -0.16465 | -0.19007 | -0.46038 | -0.25675 | 0.054729 | 0.231753 | -0.53315 | -0.43345 | -0.50343 | -0.44923 | -0.09931 | -1.58035 | -0.93479 | -0.4356 | -0.76004 | -0.48534 |
| 24.33333 | 0.387719 | 0.084829 | 0.264968 | 0.233527 | -0.08111 | -0.1549 | 0.005301 | 0.105965 | 0.246206 | -0.26581 | -0.21088 | -0.43382 | 0.21101 | -0.05629 | -0.28527 | -0.32606 | -0.26958 | -0.46553 | -0.34784 |
| 24.34722 | 0.889692 | 0.229923 | 0.374672 | 0.701216 | 0.013743 | 0.296511 | 0.229568 | 0.121556 | 0.221156 | -0.10839 | -0.11385 | -0.48529 | 0.677855 | -0.10091 | 0.187372 | -0.1958 | -0.26689 | -0.35945 | -0.32441 |
| 24.36111 | 0.947634 | 0.224996 | 0.219131 | 0.844938 | 0.061884 | 0.562895 | 0.281114 | 0.101761 | 0.110536 | -0.17443 | -0.18634 | -0.61178 | 0.638116 | -0.18149 | -0.03013 | -0.34944 | -0.37487 | -0.14865 | -0.42224 |
| 24.375 | 0.410034 | -0.10741 | 0.020044 | 0.597733 | 0.108397 | 0.541775 | 0.262073 | 0.226503 | 0.021276 | -0.25872 | -0.26916 | -0.61093 | 0.522317 | -0.22121 | -0.37605 | -0.42477 | -0.4334 | 0.294418 | -0.50091 |
| 24.38889 | 0.536114 | -0.14982 | 0.169644 | 0.633513 | 0.045757 | 0.673571 | 0.330931 | 0.749792 | 0.391433 | -0.18471 | -0.21824 | -0.48934 | 0.447367 | -0.1277 | -0.49881 | -0.35184 | -0.34492 | 0.750024 | -0.4574 |
| 24.40278 | 1.294981 | 0.301183 | 0.662627 | 0.696385 | 0.024942 | 0.4138 | 0.454692 | 1.247468 | 1.115971 | -0.06244 | -0.14499 | -0.16687 | 0.327591 | -0.05592 | -0.50629 | -0.28069 | -0.2487 | 0.673466 | -0.42893 |
| 24.41667 | 1.447591 | 0.636736 | 0.885245 | 0.55308 | 0.019995 | 0.142919 | 0.529859 | 0.977333 | 1.182314 | -0.17831 | -0.19193 | 0.025177 | 0.059103 | -0.10915 | -0.5471 | -0.29668 | -0.27171 | 0.408213 | -0.4777 |
| 24.43056 | 1.328212 | 0.571237 | 0.63992 | 0.666902 | 0.027009 | -0.17851 | 0.44735 | 0.437165 | 0.663797 | -0.34222 | -0.24849 | 0.007797 | -0.09862 | -0.20394 | -0.62782 | -0.32203 | -0.33639 | 0.147029 | -0.4133 |
| 24.44444 | 0.469889 | 0.143441 | 0.193111 | 0.557936 | 0.057492 | -0.6021 | 0.218454 | 0.089569 | 0.13773 | -0.29844 | -0.19448 | 0.078666 | -0.10255 | -0.15264 | -0.5412 | -0.25591 | -0.24419 | 0.148427 | -0.31253 |
| 24.45833 | -0.02693 | 0.077451 | -0.07328 | 0.176597 | 0.090022 | -0.53327 | 0.076472 | -0.01724 | -0.11365 | -0.13794 | -0.08016 | 0.283947 | 0.030655 | -0.02322 | -0.31564 | -0.12463 | -0.03462 | 0.28081 | -0.14841 |
| 24.47222 | 0.757167 | 1.034228 | -0.12801 | -0.25805 | 0.112483 | 0.208879 | 0.104091 | 0.079407 | 0.30796 | -0.12087 | -0.02236 | 0.300039 | 0.155973 | -0.02184 | -0.11789 | -0.03213 | 0.0358 | 0.415779 | -0.01415 |
| 24.48611 | 2.174009 | 2.322397 | 0.045715 | -0.34535 | 0.005506 | 1.0888 | 0.184791 | 0.738205 | 1.161313 | -0.06447 | 0.056658 | 0.304269 | 0.329595 | 0.004428 | 0.170827 | 0.105581 | 0.104036 | 0.537621 | 0.16777 |
| 24.5 | 2.803076 | 2.782147 | 1.001321 | 0.212431 | 0.080155 | 1.0824 | 0.519617 | 2.439106 | 1.90795 | 0.271116 | 0.265913 | 0.655471 | 0.691368 | 0.183016 | 0.711007 | 0.371525 | 0.328474 | 0.77359 | 0.456568 |
| 24.51389 | 2.653778 | 2.619539 | 2.489366 | 1.378386 | 0.359825 | 0.949884 | 1.112099 | 4.933286 | 2.729504 | 0.566185 | 0.433502 | 0.921658 | 0.888346 | 0.310756 | 1.047339 | 0.548114 | 0.537034 | 1.010608 | 0.598073 |
| 24.52778 | 2.015287 | 2.102757 | 2.412343 | 1.871836 | 0.462461 | 0.203807 | 0.912852 | 6.232429 | 3.461202 | 0.537281 | 0.426486 | 0.746508 | 0.664052 | 0.302204 | 0.913484 | 0.484887 | 0.594499 | 0.775359 | 0.465163 |
| 24.54167 | 1.525606 | 1.690997 | 1.525858 | 1.423272 | 0.475325 | -0.12154 | 0.555734 | 4.964117 | 2.774143 | 0.398178 | 0.357082 | 0.488441 | 0.40323 | 0.226075 | 0.624968 | 0.39671 | 0.420496 | 0.529931 | 0.328556 |
| 24.55556 | 0.537652 | 1.260037 | 0.966465 | 0.499618 | 0.466485 | -0.37731 | 0.360036 | 2.357789 | 1.433037 | 0.350918 | 0.319922 | 0.402203 | 0.277565 | 0.195586 | 0.478198 | 0.369495 | 0.315969 | 0.376504 | 0.267585 |
| 24.56944 | -0.88341 | 0.572073 | 0.372952 | -0.31683 | 0.446338 | -0.74316 | 0.102781 | 0.484741 | 0.35433 | 0.295642 | 0.279015 | 0.3595 | 0.208815 | 0.197008 | 0.38588 | 0.30426 | 0.280857 | 0.201709 | 0.208321 |
| 24.58333 | -1.94306 | 0.155805 | -0.12002 | -0.95455 | 0.356849 | -1.11955 | -0.21979 | -1.05149 | -0.69916 | 0.229761 | 0.287375 | 0.286065 | 0.190395 | 0.158551 | 0.365422 | 0.273138 | 0.243656 | 0.143638 | 0.18649 |
| 24.59722 | -1.93219 | 0.369407 | -0.48719 | -1.35426 | 0.393169 | -0.95328 | -0.36795 | -2.66888 | -1.51987 | 0.327289 | 0.516058 | 0.380979 | 0.398246 | 0.259991 | 0.825213 | 0.545633 | 0.529408 | 0.422688 | 0.394628 |
| 24.61111 | -0.92155 | 0.692274 | -0.48643 | -1.39464 | 0.578955 | -0.17894 | -0.28807 | -4.62757 | -2.31263 | 0.692759 | 0.872469 | 0.845971 | 1.059366 | 0.593258 | 2.500295 | 1.392541 | 1.196641 | 1.226733 | 0.908465 |
| 24.625 | 0.142799 | 0.776588 | -0.1605 | -0.55287 | 0.939982 | 0.550465 | -0.02792 | -5.50187 | -2.36356 | 0.90598 | 0.957011 | 1.224545 | 1.637707 | 0.741172 | 4.465517 | 2.588582 | 1.550034 | 1.96714 | 1.280044 |
| 24.63889 | 1.519901 | 0.790106 | 0.289722 | 0.463682 | 1.095528 | 0.613386 | 0.036542 | -4.51227 | -1.38781 | 0.759124 | 0.677523 | 1.333258 | 1.643135 | 0.614535 | 3.478757 | 2.236093 | 1.137325 | 1.215518 | 0.994337 |
| 24.65278 | 2.203613 | 0.875554 | 0.630331 | 0.577917 | 0.764265 | 0.2768 | 0.222334 | -1.43581 | -0.20158 | 0.520417 | 0.370409 | 1.02378 | 0.920104 | 0.428795 | 1.671408 | 1.242251 | 0.570816 | 0.139002 | 0.568375 |
| 24.66667 | 1.415069 | 0.9299 | 0.598957 | 0.051062 | 0.278078 | -0.19721 | 0.116637 | 1.613226 | 0.649089 | 0.357801 | 0.157898 | 0.610702 | 0.531319 | 0.27988 | 0.835207 | 0.730603 | 0.289143 | -0.067 | 0.250671 |
| 24.68056 | 0.43634 | 0.684975 | 0.69447 | -0.3849 | -0.04019 | -0.44857 | 0.126677 | 3.848174 | 1.277044 | 0.246174 | 0.043521 | 0.346581 | 0.294029 | 0.170319 | 0.487789 | 0.475872 | 0.181017 | 0.578478 | 0.079249 |
| 24.69444 | -0.51862 | 0.228651 | 1.032723 | -0.58803 | -0.18969 | -0.667 | 0.202312 | 4.569364 | 1.871749 | 0.166576 | 0.034681 | 0.218648 | 0.155559 | 0.096741 | 0.297234 | 0.321942 | 0.162905 | 1.001951 | 0.037514 |
| 24.70833 | -1.66663 | -0.14237 | 0.647083 | -0.66131 | -0.21301 | -1.09846 | -0.26703 | 4.14809 | 2.034364 | 0.115846 | 0.100504 | 0.180549 | 0.079757 | 0.067094 | 0.218008 | 0.269628 | 0.197664 | 0.898526 | 0.10188 |
| 24.72222 | -2.43343 | -0.1337 | 0.394991 | -0.75108 | -0.20912 | -0.85744 | -0.22818 | 1.453157 | 0.904435 | 0.107163 | 0.200601 | 0.169814 | 0.051481 | 0.077643 | 0.190393 | 0.281127 | 0.227879 | 0.717357 | 0.169923 |
| 24.73611 | -1.68532 | 0.581765 | 0.109847 | -0.78864 | 0.012598 | 0.059013 | 0.096581 | -1.28103 | -0.34693 | 0.157789 | 0.297447 | 0.142365 | 0.063538 | 0.090664 | 0.174106 | 0.297556 | 0.243452 | 0.570666 | 0.204502 |
| 24.75 | -0.91891 | 1.230609 | -0.11532 | -0.48713 | 0.076428 | 0.771863 | 0.449873 | -3.81869 | -1.68059 | 0.216488 | 0.303366 | 0.101559 | 0.081081 | 0.082486 | 0.162898 | 0.289601 | 0.238555 | 0.423541 | 0.191476 |
| 24.76389 | -0.07317 | 2.461897 | -0.41131 | -0.21927 | 0.067713 | 0.741326 | 0.390507 | -5.17067 | -2.19849 | 0.232435 | 0.182017 | 0.056672 | 0.075445 | 0.056256 | 0.151615 | 0.253277 | 0.171699 | 0.275314 | 0.127615 |
| 24.77778 | -0.06477 | 1.878733 | -0.63686 | -0.16511 | -0.01762 | 0.003066 | -0.20293 | -4.41195 | -1.64874 | 0.21173 | 0.052629 | 0.016867 | 0.057942 | 0.022117 | 0.146599 | 0.220675 | 0.095237 | 0.198775 | 0.062887 |
| 24.79167 | -0.0857 | 0.092789 | -0.82089 | 0.09325 | -0.12433 | -0.87859 | -0.58185 | -2.6198 | -0.91754 | 0.187796 | 0.024667 | 0.055497 | 0.052976 | 0.012391 | 0.114422 | 0.206655 | 0.046594 | 0.233142 | 0.068149 |
| 24.80556 | -0.13751 | -1.49894 | -0.65338 | 0.267575 | -0.25607 | -1.41302 | -0.8791 | -1.08132 | -0.61191 | 0.172221 | 0.124788 | 0.412889 | 0.049534 | 0.021217 | 0.051517 | 0.201927 | 0.029934 | 0.290813 | 0.124817 |
| 24.81944 | -0.55789 | -2.25254 | -0.21679 | -0.07162 | -0.33646 | -1.83887 | -0.92328 | -0.35213 | -0.38704 | 0.161405 | 0.240844 | 1.374875 | 0.047281 | 0.031551 | -0.00044 | 0.191305 | 0.036611 | 0.281318 | 0.174575 |
| 24.83333 | -1.40753 | -2.01885 | 0.273672 | -0.79269 | -0.39714 | -1.83311 | -0.66106 | 0.023969 | -0.06706 | 0.149166 | 0.253219 | 0.703858 | 0.050865 | 0.04831 | -0.01843 | 0.162035 | 0.061107 | 0.197801 | 0.175664 |
| 24.84722 | -2.92052 | -1.58198 | 0.417051 | -1.88029 | -0.54376 | -1.84744 | -0.2995 | 0.092056 | 0.123 | 0.159741 | 0.100108 | 0.277003 | 0.062944 | 0.085675 | 0.011756 | 0.139551 | 0.096353 | 0.090509 | 0.157341 |
| 24.86111 | -3.9675 | -1.53407 | 0.234395 | -2.58119 | -1.1374 | -1.55761 | -0.0762 | -0.02575 | 0.099772 | 0.193227 | -0.05374 | 0.311613 | 0.086064 | 0.109724 | 0.07921 | 0.130536 | 0.122306 | -0.00597 | 0.140067 |
| 24.875 | -3.57249 | -1.26682 | -0.34614 | -2.40029 | -1.5566 | -0.95251 | -0.03486 | -0.14188 | -0.01792 | 0.212145 | -0.07474 | 0.253518 | 0.088439 | 0.070633 | 0.094429 | 0.096444 | 0.103379 | -0.08984 | 0.114501 |
| 24.88889 | -2.78739 | -1.06799 | -0.70919 | -1.70392 | -1.77666 | -0.29913 | -0.16446 | -0.07632 | -0.15306 | 0.172122 | -0.00657 | 0.160751 | 0.057476 | -0.00836 | 0.065555 | 0.029529 | 0.037576 | -0.16056 | 0.078443 |
| 24.90278 | -1.69567 | -0.92658 | -0.73434 | -1.18123 | -1.6084 | 0.218354 | -0.23523 | 0.087324 | -0.24265 | 0.117627 | 0.129328 | 0.09948 | 0.021357 | -0.07506 | 0.022271 | -0.02925 | -0.04342 | -0.19478 | 0.059216 |
| 24.91667 | -0.76203 | -0.71263 | -0.60373 | -0.35157 | -0.91282 | 0.824537 | -0.15432 | 0.22897 | -0.22409 | 0.079585 | 0.292332 | 0.073151 | -0.0022 | -0.10732 | -0.00533 | -0.04422 | -0.09094 | -0.11327 | 0.077096 |
| 24.93056 | 0.208532 | -0.49672 | -0.51703 | 0.579906 | -0.18675 | 1.231493 | 0.010379 | 0.281416 | -0.13793 | 0.058171 | 0.245179 | 0.04964 | -0.00989 | -0.10976 | -0.02223 | -0.01634 | -0.10854 | -0.01208 | 0.108223 |
| 24.94444 | 0.539404 | -0.36956 | -0.54244 | 0.948274 | 0.257545 | 1.13879 | 0.11063 | 0.199117 | 0.01341 | 0.056152 | 0.02459 | 0.021338 | -0.01722 | -0.08842 | -0.05082 | 0.036341 | -0.10868 | 0.051765 | 0.122045 |
| 24.95833 | 0.511907 | -0.2045 | -0.65162 | 0.77023 | 0.275511 | 1.028415 | 0.143367 | 0.161891 | 0.148872 | 0.102475 | -0.12747 | -0.00181 | -0.02328 | -0.02811 | -0.06699 | 0.093205 | -0.09556 | 0.07822 | 0.106686 |
| 24.97222 | 0.085347 | -0.25583 | -0.66976 | 0.401338 | -0.11916 | 0.903686 | 0.137707 | 0.142056 | 0.232123 | 0.182132 | -0.12371 | 0.008049 | 0.003208 | 0.047446 | -0.05303 | 0.118902 | -0.05755 | 0.102858 | 0.097669 |
| 24.98611 | -1.23229 | -0.89791 | -0.73916 | -0.26686 | -0.80744 | -0.01672 | -0.08108 | -0.15751 | -0.0883 | 0.249186 | 0.048024 | 0.043479 | 0.057113 | 0.049528 | 0.000139 | 0.094617 | -0.02452 | 0.13054 | 0.089083 |
| 25 | -2.99854 | -1.44561 | -0.84955 | -1.86838 | -1.52522 | -1.28775 | -0.58704 | -0.54685 | -0.73956 | 0.256019 | 0.256175 | 0.04895 | 0.072009 | -0.02248 | 0.046615 | 0.054892 | -0.03849 | 0.146672 | 0.072638 |
| 25.01389 | -3.38441 | -1.01915 | -0.74576 | -2.83269 | -1.33493 | -2.09005 | -1.06059 | -0.5189 | -1.24651 | 0.172548 | 0.338854 | 0.017342 | 0.033257 | -0.11253 | 0.052873 | 0.033627 | -0.09147 | 0.13377 | 0.055703 |
| 25.02778 | -2.24594 | -0.56477 | -0.53526 | -1.76169 | -0.65 | -2.5941 | -1.13787 | -0.17519 | -1.29327 | 0.052347 | 0.222599 | -0.02778 | -0.0539 | -0.18338 | 0.014221 | 0.040719 | -0.15488 | 0.079357 | 0.076891 |
| 25.04167 | -1.3475 | -0.51147 | -0.37561 | -0.9951 | -0.36319 | -2.45486 | -0.84218 | 0.104075 | -0.92985 | 0.015566 | 0.097786 | -0.05242 | -0.14934 | -0.12182 | -0.0276 | 0.042805 | -0.19121 | 0.074503 | 0.125576 |
| 25.05556 | -0.61103 | -0.43521 | -0.1831 | -0.52603 | -0.25932 | -1.31174 | -0.51907 | 0.240716 | -0.46864 | 0.092015 | 0.00876 | -0.04603 | -0.17986 | -0.04469 | -0.04423 | 0.031705 | -0.13159 | 0.163634 | 0.167523 |
| 25.06944 | 0.223454 | -0.07633 | 0.04244 | -0.24124 | -0.08348 | -0.26586 | -0.1932 | 0.214375 | -0.11983 | 0.24442 | -0.06019 | -0.02635 | -0.10848 | 0.006959 | -0.02162 | 0.020268 | 0.013367 | 0.255501 | 0.166445 |
| 25.08333 | 1.191909 | 0.457019 | 0.297115 | 0.042764 | 0.068634 | 0.296351 | 0.161448 | 0.19201 | 0.195213 | 0.312383 | -0.10309 | -0.03378 | 0.045466 | 0.040771 | 0.025065 | -0.02044 | 0.084588 | 0.297459 | 0.099545 |
| 25.09722 | 1.834703 | 0.99722 | 0.520507 | 0.269906 | 0.125948 | 0.576384 | 0.441319 | 0.170393 | 0.495458 | 0.081747 | -0.17584 | -0.21809 | 0.008904 | -0.08954 | -0.22196 | -0.42358 | -0.12418 | 0.276262 | -0.30053 |
| 25.11111 | 1.618925 | 1.017006 | 0.568478 | 0.555797 | 0.233176 | 0.665144 | 0.513073 | 0.256411 | 0.756736 | -0.61886 | -0.57849 | -0.97371 | -0.70391 | -0.53632 | -1.84286 | -1.78783 | -0.86992 | -0.49877 | -1.35237 |
| 25.125 | 1.191832 | 0.577025 | 0.523207 | 0.897208 | 0.255193 | 0.575129 | 0.52935 | 0.257054 | 0.755948 | -1.94414 | -1.4706 | -2.43568 | -1.908 | -1.11507 | -4.30921 | -3.47687 | -1.68618 | -0.91579 | -2.33889 |

| | | | | | | | | | | | | | | | | | | | |
|---|---|---|---|---|---|---|---|---|---|---|---|---|---|---|---|---|---|---|---|
| 25.13889 | 0.700708 | 0.143768 | 0.361931 | 1.103176 | 0.195765 | 0.436579 | 0.398583 | 0.153522 | 0.374235 | -1.86774 | -1.45359 | -1.88533 | -1.07647 | -0.92756 | -2.82421 | -2.37543 | -1.22345 | -0.43019 | -1.51284 |
| 25.15278 | 0.200111 | 0.002693 | 0.065436 | 0.979472 | 0.139961 | 0.40735 | 0.174008 | 0.100713 | 0.007449 | -0.32554 | -0.15163 | -0.027 | 0.378828 | -0.09839 | 0.05371 | -0.07843 | -0.02257 | 0.27278 | -0.38531 |
| 25.16667 | 0.305461 | 0.088783 | -0.17302 | 0.855595 | 0.163157 | 0.655922 | 0.036249 | 0.128347 | 0.006528 | 0.454039 | 0.350745 | 0.552391 | 1.07817 | 0.425215 | 1.384654 | 0.928662 | 0.556067 | 0.544453 | 0.219981 |
| 25.18056 | 0.884684 | 0.169805 | -0.18875 | 0.795192 | 0.414995 | 1.035515 | 0.111137 | 0.136986 | 0.422315 | 0.625116 | 0.243108 | 0.647363 | 1.098299 | 0.555218 | 1.344301 | 0.912981 | 0.487571 | 0.567333 | 0.352075 |
| 25.19444 | 1.504437 | 0.19871 | 0.063818 | 0.860982 | 0.754407 | 1.262329 | 0.309973 | 0.140552 | 0.91974 | 0.657556 | 0.264982 | 0.754335 | 0.800278 | 0.383179 | 1.393557 | 0.911365 | 0.379416 | 0.430933 | 0.336737 |
| 25.20833 | 1.833442 | 0.331373 | 0.347443 | 0.887722 | 0.960305 | 1.453484 | 0.469518 | 0.166584 | 1.082495 | 0.541734 | 0.470132 | 0.753642 | 0.51272 | 0.222122 | 1.37319 | 0.984908 | 0.387554 | 0.313713 | 0.379346 |
| 25.22222 | 1.161316 | 0.340285 | 0.395452 | 0.707342 | 0.631261 | 1.19279 | 0.419949 | 0.177754 | 0.816303 | 0.518963 | 0.583746 | 0.694094 | 0.383377 | 0.205761 | 1.017788 | 0.810055 | 0.410015 | 0.340357 | 0.448781 |
| 25.23611 | 0.06531 | 0.298419 | 0.217358 | 0.168642 | 0.020961 | 0.880784 | 0.21692 | 0.169177 | 0.443721 | 0.49669 | 0.387653 | 0.68731 | 0.232526 | 0.257458 | 0.494029 | 0.471391 | 0.42873 | 0.196538 | 0.221763 |
| 25.25 | -0.55779 | 0.270571 | 0.008877 | -0.40692 | -0.43968 | 0.635398 | -0.01233 | 0.168415 | 0.18431 | 0.20454 | -0.11617 | 0.667147 | -0.03981 | 0.253613 | -0.05033 | 0.092323 | 0.381371 | -0.17725 | -0.25922 |
| 25.26389 | -0.37569 | 0.127017 | -0.14396 | -0.51035 | -0.43117 | 0.375962 | -0.127 | 0.147168 | 0.130918 | -0.02241 | -0.31991 | 0.474255 | -0.13253 | 0.130703 | -0.19858 | -0.00938 | 0.229489 | -0.21687 | -0.36165 |
| 25.27778 | 0.140035 | -0.09586 | -0.09192 | -0.11153 | -0.16346 | 0.193038 | -0.02356 | 0.116662 | 0.294995 | 0.064587 | 0.072534 | 0.179353 | 0.163544 | -0.04695 | 0.181611 | 0.246438 | 0.060978 | 0.277851 | 0.05902 |
| 25.29167 | 0.671013 | -0.16233 | 0.157887 | 0.411511 | 0.100436 | 0.084594 | 0.232222 | 0.105822 | 0.452661 | 0.274694 | 0.447531 | 0.066523 | 0.481268 | -0.10893 | 0.55393 | 0.511183 | -0.03704 | 0.636441 | 0.4117 |
| 25.30556 | 0.999952 | 0.074635 | 0.450919 | 0.677274 | 0.374939 | 0.006256 | 0.439228 | 0.167372 | 0.582736 | 0.354871 | 0.361837 | 0.108185 | 0.496186 | -0.05503 | 0.514692 | 0.515357 | -0.0435 | 0.472495 | 0.406058 |
| 25.31944 | 0.826159 | 0.309122 | 0.42158 | 0.499147 | 0.382438 | -0.14261 | 0.462839 | 0.214168 | 0.557308 | 0.369513 | 0.116008 | 0.14486 | 0.339446 | -0.00161 | 0.283893 | 0.369514 | 0.00762 | 0.185476 | 0.252291 |
| 25.33333 | 0.179379 | 0.229159 | 0.164666 | 0.12884 | 0.111486 | -0.41787 | 0.298148 | 0.191406 | 0.364324 | 0.387479 | 0.0444 | 0.243289 | 0.226339 | 0.028379 | 0.149028 | 0.268922 | 0.113981 | 0.084591 | 0.14245 |
| 25.34722 | -0.13624 | 0.090723 | -0.09752 | 0.022285 | -0.11201 | -0.47114 | 0.14028 | 0.145724 | 0.205963 | 0.444327 | 0.352923 | 0.215019 | 0.354052 | -0.01424 | 0.254834 | 0.326799 | 0.08689 | 0.341265 | 0.210284 |
| 25.36111 | 0.083954 | 0.050509 | -0.15281 | 0.228112 | -0.04724 | -0.26334 | 0.097863 | 0.140495 | 0.319285 | 0.480954 | 0.65255 | 0.00711 | 0.517704 | -0.10663 | 0.393196 | 0.418706 | -0.09898 | 0.571054 | 0.288554 |
| 25.375 | 0.448643 | 0.077826 | 0.068426 | 0.447443 | 0.134135 | -0.05251 | 0.16093 | 0.137175 | 0.602358 | 0.42344 | 0.470829 | -0.10008 | 0.434522 | -0.16601 | 0.247385 | 0.363186 | -0.20082 | 0.365285 | 0.176842 |
| 25.38889 | 0.51694 | 0.03276 | 0.298829 | 0.50762 | 0.318276 | 0.003393 | 0.245421 | 0.055083 | 0.678699 | 0.407361 | 0.239625 | -0.18213 | 0.228196 | -0.13554 | 0.106264 | 0.302867 | -0.1784 | 0.19477 | 0.109192 |
| 25.40278 | 0.285019 | -0.14063 | 0.44305 | 0.339851 | 0.476898 | -0.04447 | 0.258388 | -0.03834 | 0.361092 | 0.391108 | 0.208379 | -0.28191 | -0.03546 | 0.033715 | 0.099265 | 0.347318 | -0.03728 | 0.206571 | 0.14292 |
| 25.41667 | 0.038533 | -0.02342 | 0.384642 | 0.076961 | 0.502092 | 0.058071 | 0.214894 | -0.00128 | 0.080855 | 0.361547 | 0.22612 | -0.19978 | -0.32758 | 0.180278 | 0.1086 | 0.434992 | 0.135981 | 0.178449 | 0.10109 |
| 25.43056 | 0.017454 | 0.307116 | 0.137895 | -0.12492 | 0.34002 | 0.312018 | 0.128124 | 0.14107 | -0.07499 | 0.425982 | 0.194854 | 0.0866 | -0.46243 | 0.241086 | 0.096486 | 0.386375 | 0.267979 | 0.067321 | -0.02001 |
| 25.44444 | 0.166518 | 0.306117 | -0.13711 | -0.00763 | 0.120721 | 0.490799 | 0.055975 | 0.174025 | -0.11778 | 0.485843 | -0.12919 | 0.512486 | -0.24965 | 0.273885 | -0.09228 | 0.168583 | 0.368628 | -0.22529 | -0.21989 |
| 25.45833 | 0.400047 | 0.069859 | -0.17233 | 0.283285 | -0.03451 | 0.57681 | 0.080053 | 0.124829 | 0.144195 | 0.461642 | -0.58337 | 0.762513 | 0.047632 | 0.296023 | -0.35455 | -0.04486 | 0.386315 | -0.53496 | -0.32217 |
| 25.47222 | 0.665829 | 0.056698 | 0.071195 | 0.497138 | -0.03976 | 0.704889 | 0.239586 | 0.116901 | 0.659103 | 0.475325 | -0.27605 | 0.394846 | 0.339604 | 0.159848 | -0.13505 | 0.159305 | 0.189428 | -0.12508 | 0.063319 |
| 25.48611 | 1.036123 | 0.396124 | 0.475951 | 0.592923 | 0.267303 | 0.967611 | 0.466051 | 0.128615 | 0.925844 | 0.633091 | 0.363723 | -0.19767 | 0.340123 | 0.092198 | 0.266347 | 0.568498 | -0.04716 | 0.539229 | 0.490287 |
| 25.5 | 1.567776 | 0.925777 | 1.294122 | 0.794166 | 0.881738 | 1.35293 | 0.914344 | 0.176978 | 0.990545 | 0.537944 | 0.327741 | -0.31185 | -0.18764 | 0.155852 | 0.153471 | 0.523214 | 0.032874 | 0.422969 | 0.335272 |
| 25.51389 | 2.228082 | 1.319237 | 2.35248 | 1.531138 | 1.337287 | 1.481676 | 1.095663 | 0.17916 | 0.845689 | 0.572669 | 0.345202 | -0.16173 | -0.40614 | 0.236534 | 0.145875 | 0.448532 | 0.150335 | 0.354709 | 0.242421 |
| 25.52778 | 2.514291 | 1.877196 | 2.648444 | 1.895733 | 0.835454 | 1.131179 | 1.123283 | 0.661621 | 0.732336 | 0.864378 | 0.864841 | -0.08469 | 0.124978 | 0.222926 | 0.526308 | 0.681097 | 0.113957 | 0.813101 | 0.611971 |
| 25.54167 | 1.585759 | 2.04212 | 1.98371 | 1.450296 | 0.315823 | 0.731895 | 0.910804 | 1.3267 | 0.653234 | 0.996933 | 0.903166 | 0.171474 | 0.550025 | 0.227513 | 0.594387 | 0.6017 | 0.170685 | 0.821463 | 0.866094 |
| 25.55556 | 0.03259 | 1.378856 | 0.933613 | 0.211758 | -0.21142 | 0.017297 | 0.291966 | 1.60753 | 0.3065 | 0.939982 | 0.513418 | 0.4151 | 0.73775 | 0.266859 | 0.330612 | 0.292449 | 0.272731 | 0.411176 | 0.757513 |
| 25.56944 | -1.75796 | 0.79119 | -0.02063 | -1.20005 | -0.90959 | -0.75098 | -0.23019 | 1.632759 | -0.15226 | 0.793309 | 0.333569 | 0.443504 | 0.646067 | 0.269644 | 0.284235 | 0.190627 | 0.323177 | 0.28147 | 0.636421 |
| 25.58333 | -2.33721 | 0.476762 | -0.8547 | -1.74601 | -1.07868 | -1.18049 | -0.47432 | 1.710194 | -0.43518 | 0.646401 | 0.426926 | 0.284533 | 0.298701 | 0.21181 | 0.399346 | 0.24109 | 0.32971 | 0.418158 | 0.61629 |
| 25.59722 | -1.95056 | 0.127058 | -1.22655 | -1.70117 | -0.7581 | -0.93166 | -0.37663 | 1.714094 | -0.30586 | 0.980482 | 0.78531 | 0.362769 | 0.110894 | 0.288056 | 0.500041 | 0.425204 | 0.502285 | 0.94149 | 1.144964 |
| 25.61111 | -1.49554 | -0.50126 | -0.59484 | -1.71154 | -0.35843 | -0.88713 | -0.0811 | 1.360976 | 0.162845 | 1.939659 | 1.201301 | 1.700321 | 0.557718 | 0.700636 | 1.088027 | 0.918324 | 1.088828 | 2.204582 | 2.275772 |
| 25.625 | -0.17486 | -1.11107 | -0.01646 | -0.61111 | 0.079566 | -0.79826 | 0.041275 | 0.982618 | 0.897096 | 2.080222 | 0.918989 | 2.728383 | 1.207404 | 1.151816 | 1.96473 | 1.424289 | 1.512811 | 1.937636 | 2.429961 |
| 25.63889 | 1.382683 | -1.51184 | 0.149768 | 0.600003 | 0.465082 | -0.31095 | 0.104811 | -0.30546 | 1.262923 | 1.699035 | 0.543801 | 2.083377 | 1.543691 | 0.93019 | 1.824384 | 1.047059 | 1.417641 | 1.404193 | 1.743351 |
| 25.65278 | 2.226708 | -1.72326 | 0.477115 | 1.284704 | 0.734293 | -0.0532 | 0.050998 | -1.56758 | 0.856686 | 1.310821 | 0.206057 | 1.484083 | 1.613397 | 0.699552 | 1.858477 | 0.822038 | 1.048591 | 1.3119 | 1.236791 |
| 25.66667 | 1.356588 | -1.28419 | -0.07701 | 0.888052 | 0.667966 | 0.463153 | -0.03498 | -2.11427 | -0.00145 | 0.586237 | 0.004757 | 0.819862 | 0.9678 | 0.501815 | 1.74624 | 0.668569 | 0.475927 | 1.123429 | 0.755737 |
| 25.68056 | -0.08699 | -0.55151 | -0.89666 | -0.13523 | 0.297383 | 0.130657 | -0.10118 | -2.36923 | -0.68546 | -0.21057 | 0.071523 | 0.256802 | 0.34038 | 0.397764 | 1.164166 | 0.462972 | -0.18115 | 0.637511 | 0.389253 |
| 25.69444 | -1.15214 | 0.14423 | -1.14764 | -0.59336 | -0.03675 | 0.01042 | 0.007988 | -2.03951 | -1.07165 | -0.74334 | 0.259263 | -0.27441 | -0.20458 | 0.12651 | 0.597676 | 0.250871 | -0.38464 | 0.318597 | 0.158197 |
| 25.70833 | -2.06857 | 0.920606 | -0.57991 | -1.1073 | -0.27762 | -0.28015 | 0.082102 | -1.12289 | -1.18536 | -0.75397 | 0.467808 | -0.43425 | -0.64842 | -0.17441 | 0.064371 | -0.00418 | 0.033206 | 0.235857 | 0.015956 |
| 25.72222 | -2.68903 | 1.748553 | 0.083695 | -1.63112 | -0.94846 | -0.91069 | 0.060801 | -0.39362 | -0.86468 | -0.53571 | 0.379067 | -0.05542 | -0.97239 | 0.001993 | -0.56915 | -0.38822 | 0.625381 | 0.159291 | -0.25624 |
| 25.73611 | -2.32724 | 1.891364 | 0.582013 | -1.72116 | -1.2604 | -0.85205 | 0.28127 | 0.190357 | -0.07961 | -0.36672 | -0.17943 | 0.491722 | -0.76572 | 0.221107 | -1.13445 | -0.68887 | 0.835739 | -0.03759 | -0.57813 |
| 25.75 | -0.60671 | 1.571819 | 0.712489 | -1.03933 | -0.59769 | -1.5986 | 0.279464 | 0.613925 | 0.688542 | -0.25353 | -0.65117 | 0.671765 | -0.24449 | 0.006352 | -1.25219 | -0.82047 | 0.25763 | -0.26032 | -0.61575 |
| 25.76389 | 0.664145 | 0.832351 | 0.366444 | -0.58191 | 0.137907 | -1.70155 | 0.160744 | 0.694837 | 1.096537 | -0.36392 | -0.72266 | 0.617913 | 0.251544 | -0.28534 | -1.17462 | -0.88882 | -0.1818 | -0.36283 | -0.57006 |
| 25.77778 | 1.283243 | 0.041913 | -0.32626 | -0.27892 | 0.497679 | -1.25623 | -0.06787 | 0.375624 | 1.002178 | -0.71415 | -0.52295 | 0.014911 | 0.49541 | -0.57341 | -0.80845 | -0.55821 | -0.65656 | -0.34209 | -0.57415 |
| 25.79167 | 0.602953 | -0.79094 | -0.76627 | -0.34362 | 0.839484 | -0.78998 | -0.10407 | 0.008977 | 0.487922 | -0.90208 | -0.11225 | -0.43049 | 0.32399 | -0.79044 | -0.24302 | 0.092337 | -1.23939 | -0.18541 | -0.50409 |
| 25.80556 | -0.36961 | -1.57793 | -0.85386 | -0.70253 | 0.737481 | -0.74714 | -0.55642 | -0.38569 | -0.25385 | -0.72992 | 0.167553 | -0.9891 | -0.0644 | -0.8201 | 0.328138 | 0.604787 | -1.00914 | -0.09591 | -0.27295 |
| 25.81944 | -0.73838 | -1.97818 | -1.19061 | -0.74642 | 0.621102 | -1.09797 | -1.21958 | -0.65091 | -1.12756 | -0.08092 | 0.202136 | -0.99847 | -0.3879 | -0.36866 | 0.693621 | 0.815545 | -0.04826 | -0.03848 | 0.220862 |
| 25.83333 | -0.7643 | -1.33694 | -1.16327 | -0.68384 | 0.265709 | -0.99314 | -1.01139 | -0.65046 | -1.62508 | 0.510131 | 0.160369 | -0.70767 | -0.4536 | 0.158405 | 0.725717 | 0.828806 | 0.636266 | 0.004104 | 0.720147 |
| 25.84722 | -0.94765 | -0.54085 | -0.98834 | -0.55704 | -0.15447 | -0.71911 | -0.5555 | -0.3142 | -1.30261 | 0.923157 | -0.04575 | -0.08999 | -0.19857 | 0.566186 | 0.524837 | 0.563633 | 1.157441 | -0.02202 | 0.745702 |
| 25.86111 | -1.24688 | -0.15639 | -1.05229 | -0.51482 | -0.87142 | -0.7535 | -0.40131 | -0.0344 | -0.68777 | 1.025939 | -0.06134 | 0.580973 | 0.055003 | 0.587983 | 0.213557 | 0.157026 | 1.178944 | -0.04302 | 0.387819 |
| 25.875 | -1.2379 | 0.031406 | -0.94235 | -0.68749 | -1.88707 | -0.71913 | -0.39996 | 0.070931 | -0.08719 | 0.824749 | 0.173688 | 0.84225 | 0.293177 | 0.319732 | -0.11075 | -0.15236 | 0.336118 | -0.01235 | 0.097667 |
| 25.88889 | -0.90404 | 0.182859 | -0.40876 | -0.82207 | -1.2795 | -0.49219 | -0.45184 | 0.066272 | 0.30515 | 0.45715 | 0.330019 | 0.748363 | 0.334186 | -0.03533 | -0.37985 | -0.42953 | -0.20197 | -0.01013 | -0.07901 |
| 25.90278 | -0.70597 | 0.31591 | -0.04769 | -1.01778 | -0.49136 | -0.03162 | -0.40769 | 0.009618 | 0.367269 | -0.05175 | 0.1584 | 0.42197 | 0.153247 | -0.38408 | -0.67271 | -0.66713 | -0.58567 | -0.05165 | -0.16826 |
| 25.91667 | -0.67536 | 0.403899 | 0.284656 | -0.97883 | 0.310025 | 0.558551 | -0.18834 | -0.0755 | 0.244734 | -0.61759 | -0.19179 | -0.05743 | -0.12168 | -0.46007 | -1.00447 | -0.78223 | -0.76842 | -0.10003 | -0.22056 |
| 25.93056 | -0.47228 | 0.392175 | 0.37219 | -0.54558 | 0.792564 | 1.086675 | 0.050378 | -0.17835 | 0.154678 | -1.04485 | -0.50534 | -0.77846 | -0.34822 | -0.33637 | -1.28025 | -1.13317 | -0.37811 | -0.10267 | -0.31794 |
| 25.94444 | 0.014907 | 0.374775 | 0.226815 | -0.09673 | 0.442375 | 1.362036 | 0.198839 | -0.24369 | 0.208469 | -1.30623 | -0.68221 | -1.20574 | -0.45499 | -0.17763 | -1.3521 | -1.31734 | -0.02199 | -0.03973 | -0.48787 |
| 25.95833 | 0.580041 | 0.40174 | 0.097175 | 0.289732 | 0.329029 | 1.316728 | 0.236251 | -0.22423 | 0.344979 | -1.44483 | -0.54979 | -0.90187 | -0.38643 | -0.2245 | -1.18879 | -1.2459 | -0.08626 | 0.07808 | -0.59211 |
| 25.97222 | 0.957363 | 0.414046 | 0.174274 | 0.52191 | 0.148947 | 1.058191 | 0.199704 | -0.10967 | 0.447617 | -1.44586 | -0.31757 | -0.67686 | -0.26731 | -0.5786 | -0.99789 | -0.8026 | -0.26734 | 0.211434 | -0.59314 |
| 25.98611 | 0.469503 | 0.19812 | 0.166995 | 0.202443 | -0.20122 | 0.38053 | -0.00606 | -0.05755 | 0.111084 | -1.30044 | -0.28438 | -0.47304 | -0.17603 | -0.81157 | -0.79058 | -0.3454 | -0.41868 | 0.174965 | -0.60017 |
| 26 | -1.19222 | -0.33829 | -0.25381 | -0.87821 | -0.71668 | -0.7869 | -0.51619 | -0.2339 | -0.89156 | -1.03309 | -0.49133 | -0.28352 | -0.08782 | -0.76677 | -0.59077 | -0.02755 | -0.39164 | -0.01113 | -0.70849 |
| 26.01389 | -3.09519 | -0.82459 | -0.59701 | -1.98401 | -1.34393 | -2.38477 | -1.04381 | -0.59708 | -2.20545 | -0.62152 | -0.70659 | -0.17802 | -0.05197 | -0.5986 | -0.31244 | 0.124797 | -0.29662 | -0.19339 | -0.72058 |
| 26.02778 | -3.22818 | -0.78434 | -0.48659 | -1.97806 | -1.93374 | -2.16859 | -0.90999 | -0.77619 | -2.82273 | -0.19596 | -0.68849 | -0.08984 | -0.08364 | -0.44356 | -0.03484 | 0.009795 | -0.17509 | -0.33153 | -0.68786 |
| 26.04167 | -1.60725 | -0.52124 | -0.57545 | -1.56667 | -1.70016 | -1.26865 | -0.91487 | -0.48908 | -2.51006 | 0.164483 | -0.18709 | 0.017197 | -0.08742 | -0.32211 | 0.103422 | -0.18869 | -0.12194 | -0.29505 | -0.456 |
| 26.05556 | -0.50084 | -0.25227 | -1.01261 | -1.59353 | -0.75693 | -0.46246 | -1.71502 | -0.15186 | -1.429 | 0.320082 | 0.314183 | 0.119001 | -0.03242 | -0.20598 | 0.162479 | -0.32594 | -0.14032 | -0.1686 | -0.19149 |
| 26.06944 | 0.034309 | -0.09581 | -0.83543 | -0.32548 | 0.357107 | -0.00413 | 0.069755 | 0.221291 | -0.02295 | 0.28661 | 0.453234 | 0.108077 | -0.01642 | -0.05236 | 0.176203 | -0.33153 | -0.06581 | -0.08406 | 0.013777 |
| 26.08333 | 0.433752 | 0.129419 | -0.01342 | -0.20451 | 1.211424 | 0.203307 | -0.06517 | 0.397138 | 1.060041 | 0.233503 | 0.282995 | -0.00054 | -0.06118 | 0.037646 | 0.071037 | -0.32358 | 0.082813 | -0.08515 | 0.125631 |
| 26.09722 | 0.767043 | 0.203033 | 0.259237 | 0.203178 | 1.437633 | 0.298441 | 0.411692 | 0.339398 | 1.550095 | -0.06749 | -0.11181 | -0.31673 | -0.38174 | -0.19188 | -0.368 | -0.66844 | -0.02455 | -0.45841 | -0.20486 |
| 26.11111 | 1.083652 | 0.050417 | 0.356004 | 0.428666 | 1.108031 | 0.31944 | 0.593217 | 0.216858 | 1.599291 | -1.09063 | -0.71716 | -1.15285 | -1.55913 | -0.85327 | -1.3122 | -1.56337 | -0.6235 | -1.72219 | -1.2871 |
| 26.125 | 1.177437 | -0.13599 | 0.271443 | 0.698935 | 0.75187 | 0.255671 | 0.575514 | 0.129989 | 1.252194 | -1.82599 | -1.02553 | -1.91942 | -2.35405 | -1.15783 | -2.16014 | -2.22292 | -1.1585 | -2.6169 | -2.52763 |
| 26.13889 | 1.019922 | -0.08592 | 0.131319 | 0.768932 | 0.672365 | 0.135335 | 0.445798 | 0.144788 | 0.863329 | -2.00665 | -0.80303 | -1.23547 | -1.50295 | -0.70002 | -0.68415 | -1.10383 | -0.75681 | -1.82289 | -1.48198 |
| 26.15278 | 0.91781 | -0.02562 | 0.07903 | 0.680606 | 0.736919 | 0.066172 | 0.274769 | 0.125098 | 0.569074 | -0.13326 | -0.38679 | -0.07505 | -0.20292 | -0.13226 | 0.699323 | 0.074062 | 0.153688 | -0.93777 | -0.31534 |
| 26.16667 | 0.802784 | 0.065811 | 0.167332 | 0.546608 | 0.476337 | 0.097203 | 0.213998 | 0.101372 | 0.486534 | 0.544246 | 0.048685 | 0.343895 | 0.728094 | 0.134101 | 0.912006 | 0.340842 | 0.490342 | 0.030629 | 0.310688 |
| 26.18056 | 0.754952 | 0.257188 | 0.413396 | 0.463675 | 0.184607 | 0.289333 | 0.371336 | 0.11356 | 0.564615 | 0.576225 | 0.462464 | 0.318397 | 1.028457 | 0.163896 | 0.978334 | 0.373798 | 0.293723 | 0.755871 | 0.63647 |
| 26.19444 | 0.810469 | 0.474969 | 0.610212 | 0.473658 | 0.063532 | 0.649151 | 0.648532 | 0.166242 | 0.814046 | 0.181836 | 0.236053 | 0.558581 | 0.842851 | 0.152395 | 0.687662 | 0.206945 | 0.182892 | 0.514344 | 0.365767 |
| 26.20833 | 0.809855 | 0.478277 | 0.581586 | 0.503266 | -0.15531 | 0.793051 | 0.692911 | 0.192629 | 1.112536 | -0.37412 | -0.40872 | 0.636633 | 0.509295 | 0.167942 | -0.02508 | -0.10829 | 0.310913 | -0.35945 | -0.31443 |
| 26.22222 | 0.668148 | 0.294581 | 0.407755 | 0.558104 | -0.37023 | 0.627358 | 0.558344 | 0.202393 | 1.219479 | -0.54954 | -0.45026 | 0.353052 | 0.416616 | 0.109219 | -0.29327 | -0.18361 | 0.294332 | -0.59229 | -0.47253 |
| 26.23611 | 0.56848 | 0.216757 | 0.269047 | 0.603735 | -0.37655 | 0.54744 | 0.431849 | 0.210924 | 1.099291 | -0.11724 | 0.193114 | -0.06115 | 0.38424 | -0.00594 | 0.137272 | 0.025358 | -0.03085 | 0.120559 | 0.05059 |
| 26.25 | 0.521672 | 0.297031 | 0.17992 | 0.548217 | -0.1747 | 0.595992 | 0.257319 | 0.216471 | 0.818819 | 0.195295 | 0.421777 | -0.2636 | 0.130049 | -0.04409 | 0.419287 | 0.146579 | -0.27737 | 0.408443 | 0.305069 |
| 26.26389 | 0.447812 | 0.28282 | 0.114287 | 0.435613 | 0.03061 | 0.60817 | 0.089021 | 0.159991 | 0.47422 | -0.04233 | 0.005092 | -0.24485 | -0.31579 | -0.02507 | 0.191094 | 0.057022 | -0.28561 | -0.14467 | 0.047135 |
| 26.27778 | 0.419431 | 0.101858 | 0.156438 | 0.33768 | 0.148706 | 0.474866 | 0.053248 | 0.083339 | 0.296722 | -0.20823 | -0.08569 | -0.2125 | -0.37908 | -0.02428 | 0.057093 | -4.9E-05 | -0.17805 | -0.28033 | 0.036683 |
| 26.29167 | 0.351669 | -0.11105 | 0.245751 | 0.293718 | 0.17726 | 0.275708 | 0.161945 | 0.037083 | 0.498862 | -0.00681 | 0.050844 | -0.10261 | -0.17367 | -0.00438 | -0.01647 | 0.01256 | 0.083288 | -0.11066 | 0.13472 |
| 26.30556 | 0.037018 | -0.21676 | 0.14288 | 0.12784 | 0.12376 | 0.14126 | 0.203103 | 0.05978 | 0.818491 | 0.14363 | -0.01078 | 0.12309 | 0.014133 | 0.058052 | -0.17647 | 0.01307 | 0.277096 | -0.36237 | -0.07654 |
| 26.31944 | -0.35858 | -0.05109 | -0.09312 | -0.12231 | -0.04946 | 0.092057 | 0.088907 | 0.152503 | 0.606492 | 0.297889 | 0.150361 | 0.108612 | 0.105309 | 0.081993 | -0.047 | 0.090108 | 0.05232 | -0.34124 | -0.00132 |

| | | | | | | | | | | | | | | | | | | | |
|---|---|---|---|---|---|---|---|---|---|---|---|---|---|---|---|---|---|---|---|
| 26.33333 | -0.60528 | 0.17161 | -0.19332 | -0.32302 | -0.3348 | 0.03443 | -0.05939 | 0.215 | -0.06737 | 0.520113 | 0.535844 | -0.12979 | 0.154761 | 0.062241 | 0.331395 | 0.213556 | -0.24241 | 0.213169 | 0.430094 |
| 26.34722 | -0.36182 | 0.323354 | -0.01271 | -0.33039 | -0.46663 | -0.08802 | -0.06663 | 0.216706 | -0.52823 | 0.521524 | 0.511235 | -0.25411 | 0.087003 | 0.098524 | 0.430862 | 0.208234 | -0.17613 | 0.414171 | 0.564635 |
| 26.36111 | 0.089766 | 0.410664 | 0.341884 | -0.05641 | -0.32452 | -0.19344 | 0.124938 | 0.179008 | -0.11887 | 0.250733 | 0.121433 | -0.23846 | -0.05373 | 0.14029 | 0.275105 | 0.103619 | 0.025137 | 0.203352 | 0.399023 |
| 26.375 | 0.508246 | 0.303872 | 0.658347 | 0.245753 | -0.13108 | -0.31987 | 0.360541 | 0.096683 | 0.667916 | 0.01191 | -0.20393 | -0.15719 | -0.18321 | 0.117116 | 0.029061 | 0.002273 | 0.056284 | -0.09508 | 0.15859 |
| 26.38889 | 0.788775 | 0.020181 | 0.68616 | 0.525473 | 0.029735 | -0.44631 | 0.529902 | -0.00469 | 1.361425 | 0.090024 | -0.23993 | -0.01937 | -0.18322 | 0.054016 | -0.10554 | 0.031103 | -0.16109 | -0.34061 | 0.002032 |
| 26.40278 | 0.622314 | -0.18376 | 0.442177 | 0.423504 | 0.140529 | -0.46002 | 0.655823 | -0.05877 | 1.156891 | 0.476224 | 0.165675 | 0.055372 | 0.069126 | 0.014183 | 0.053217 | 0.245663 | -0.22667 | -0.19563 | 0.131986 |
| 26.41667 | 0.358777 | -0.04523 | 0.246777 | 0.300599 | 0.224493 | -0.18475 | 0.698045 | 0.013711 | 0.554694 | 0.630284 | 0.378298 | 0.191807 | 0.190046 | 0.068532 | 0.116918 | 0.350022 | 0.156406 | -0.13186 | 0.157262 |
| 26.43056 | 0.25112 | 0.327088 | 0.282357 | 0.165814 | 0.323658 | 0.287639 | 0.613113 | 0.187648 | 0.069452 | 0.313927 | 0.125822 | 0.287562 | 0.062776 | 0.155369 | -0.09916 | 0.177464 | 0.418927 | -0.35242 | -0.01913 |
| 26.44444 | 0.191888 | 0.412456 | 0.452771 | 0.067447 | 0.292228 | 0.499264 | 0.381884 | 0.248807 | -0.29308 | 0.096988 | 0.171044 | 0.086195 | -0.00302 | 0.15883 | -0.07141 | 0.071478 | 0.268883 | -0.17604 | 0.044266 |
| 26.45833 | 0.121096 | 0.189096 | 0.510146 | 0.045266 | 0.096941 | 0.456256 | 0.139655 | 0.150683 | -0.02075 | 0.129892 | 0.301668 | -0.1131 | -0.00186 | 0.126063 | -0.04088 | 0.034677 | 0.062989 | -0.13798 | 0.097603 |
| 26.47222 | 0.194128 | -0.05304 | 0.329854 | 0.181328 | -0.11638 | 0.405811 | 0.017162 | 0.02774 | 0.70833 | 0.495882 | 0.309188 | -0.09838 | 0.038385 | 0.103761 | -0.14245 | 0.063699 | -0.01698 | -0.34806 | -0.00915 |
| 26.48611 | 0.604031 | -0.10487 | 0.194625 | 0.634317 | -0.05465 | 0.622592 | 0.366318 | 0.114091 | 1.529848 | 0.973432 | 0.513549 | -0.04393 | 0.231712 | 0.059845 | 0.064538 | 0.246195 | 0.028888 | -0.06039 | 0.043137 |
| 26.5 | 1.662881 | 0.186909 | 0.580419 | 1.552276 | 0.285925 | 1.463497 | 1.213961 | 0.505922 | 1.721832 | 0.975732 | 0.496757 | 0.111658 | 0.325795 | 0.081926 | 0.266056 | 0.381029 | 0.373975 | 0.280159 | 0.122747 |
| 26.51389 | 2.74531 | 0.623391 | 1.686863 | 2.563618 | 0.434351 | 1.848912 | 2.072215 | 0.823903 | 1.634766 | 0.622877 | 0.444956 | 0.27531 | 0.30131 | 0.192148 | 0.389821 | 0.425082 | 0.72313 | 0.587483 | 0.241102 |
| 26.52778 | 2.75278 | 0.838397 | 2.475296 | 2.954784 | -0.00313 | 1.716162 | 2.647832 | 0.657633 | 1.580521 | 0.424782 | 0.426852 | 0.389194 | 0.229595 | 0.314047 | 0.359145 | 0.430015 | 0.783825 | 0.590408 | 0.303209 |
| 26.54167 | 2.621891 | 1.027748 | 2.568933 | 2.916647 | -0.62407 | 0.890152 | 2.514345 | 0.563083 | 1.102909 | 0.525969 | 0.255142 | 0.488124 | 0.157911 | 0.415613 | 0.167197 | 0.395035 | 0.734987 | 0.159717 | 0.154755 |
| 26.55556 | 1.480692 | 0.602299 | 1.44087 | 1.863209 | -1.19761 | 0.320856 | 1.239909 | 0.37312 | 0.435382 | 0.767438 | 0.473389 | 0.303614 | 0.275891 | 0.35439 | 0.434265 | 0.531411 | 0.469557 | 0.316863 | 0.338141 |
| 26.56944 | 0.622841 | 0.144621 | 0.155231 | 0.451531 | -1.4516 | -0.11367 | -0.07294 | 0.149653 | -0.32026 | 0.852675 | 0.677593 | -0.01391 | 0.426094 | 0.184558 | 0.684595 | 0.594532 | 0.235792 | 0.621377 | 0.578209 |
| 26.58333 | 0.126036 | -0.00048 | -0.94821 | -0.68964 | -1.34234 | -0.67031 | -1.3667 | -0.09465 | -1.04304 | 0.777944 | 0.378463 | -0.03338 | 0.388477 | 0.144085 | 0.514798 | 0.519634 | 0.289414 | 0.469591 | 0.486853 |
| 26.59722 | -0.43641 | 0.002485 | -1.63442 | -1.09048 | -1.32078 | -1.02241 | -2.17202 | -0.14614 | -1.58457 | 0.90815 | 0.298177 | 0.315261 | 0.700847 | 0.419492 | 0.703569 | 0.84217 | 0.640404 | 0.636031 | 0.833986 |
| 26.61111 | -0.76066 | 0.045114 | -1.40975 | -0.86889 | -0.92703 | -1.27474 | -2.06057 | -0.01671 | -1.11629 | 1.751632 | 0.590989 | 1.20073 | 1.685418 | 1.250322 | 1.823764 | 2.044109 | 1.143698 | 1.271241 | 1.733425 |
| 26.625 | -0.87774 | 0.080115 | -1.26522 | -0.33699 | -0.46833 | -1.97295 | -0.90443 | 0.124419 | -0.05435 | 2.121962 | 0.735972 | 1.77637 | 2.273043 | 1.704545 | 2.172659 | 2.31309 | 1.431494 | 1.343679 | 1.877073 |
| 26.63889 | -0.54386 | 0.036584 | -0.99957 | 0.120936 | -0.17852 | -1.92523 | 0.514502 | 0.044808 | 0.857143 | 1.681761 | 0.690595 | 1.537057 | 2.037769 | 1.727953 | 1.599422 | 1.733711 | 1.488205 | 0.909689 | 1.410646 |
| 26.65278 | 0.113476 | 0.0391 | -0.01487 | 0.439203 | 0.082822 | -1.80769 | 1.280531 | -0.13501 | 1.171181 | 0.985609 | 0.318 | 1.371808 | 1.822391 | 1.688726 | 1.528921 | 1.360776 | 1.473635 | 0.560084 | 0.783016 |
| 26.66667 | 1.210969 | 0.101535 | 1.262087 | 1.005504 | 0.690549 | -0.88551 | 2.1079 | -0.1954 | 0.461557 | 0.085203 | -0.10683 | 0.933974 | 1.005842 | 1.096602 | 1.585025 | 1.050982 | 0.928139 | 0.369849 | 0.362245 |
| 26.68056 | 1.856173 | 0.099757 | 1.352807 | 1.35233 | 1.02518 | 0.130308 | 1.68201 | -0.24516 | -0.07548 | -0.81303 | -0.46825 | 0.441318 | 0.195213 | 0.440582 | 1.316903 | 0.536463 | 0.25959 | 0.165967 | 0.026849 |
| 26.69444 | 1.443034 | -0.04342 | 0.246075 | 0.595274 | 0.866623 | 1.1338 | 0.032285 | -0.22529 | -0.41945 | -1.50603 | -0.72266 | 0.093742 | -0.6024 | -0.26901 | 0.807691 | -0.1531 | -0.51251 | 0.083517 | -0.30555 |
| 26.70833 | 0.292947 | -0.52866 | -0.09921 | -0.07063 | 0.568158 | 1.336539 | -1.04422 | -0.09944 | -0.66988 | -1.38687 | -0.43301 | -0.25052 | -0.973 | -0.82619 | 0.117236 | -0.89058 | -0.85685 | 0.136106 | -0.58173 |
| 26.72222 | -0.32312 | -0.77855 | -0.32957 | -0.24924 | 0.129974 | 1.256277 | -1.70243 | -0.00652 | -0.95515 | -0.8757 | 0.044738 | -0.3113 | -0.99259 | -0.96008 | -0.63531 | -1.37746 | -0.47937 | 0.18753 | -0.74111 |
| 26.73611 | -0.31224 | -0.43037 | -0.65695 | 0.059876 | -0.09903 | 1.116046 | -2.06406 | 0.015183 | -0.82371 | -0.03149 | 0.478976 | -0.0688 | -0.50759 | -0.65925 | -1.21111 | -1.26379 | 0.191729 | 0.062676 | -0.86762 |
| 26.75 | -0.40496 | 0.020083 | -0.5539 | -0.04109 | -0.32377 | 0.662118 | -1.36876 | -0.02927 | -0.62435 | 0.033845 | 0.242176 | 0.03792 | -0.00922 | -0.23018 | -1.42749 | -0.99237 | 0.634416 | -0.2637 | -1.01669 |
| 26.76389 | -0.92121 | 0.156002 | -0.35179 | -0.61964 | -0.89759 | -0.23551 | -0.48661 | -0.12245 | -0.35767 | -0.64276 | -0.2533 | 0.377097 | -0.06339 | -0.16622 | -1.40259 | -0.80022 | 0.358037 | -0.59613 | -0.94303 |
| 26.77778 | -2.11513 | 0.096796 | -0.48948 | -1.29725 | -0.80391 | -1.30656 | -0.00451 | -0.21685 | -0.0045 | -1.26047 | -0.75442 | 0.128341 | -0.62169 | -0.43896 | -1.17038 | -0.66855 | -0.36237 | -0.61825 | -0.74834 |
| 26.79167 | -1.40507 | -0.07784 | -0.71927 | -0.72142 | -0.42252 | -1.81116 | -0.02658 | -0.19362 | 0.334469 | -1.61818 | -1.03644 | -0.38119 | -1.06744 | -0.70166 | -0.83207 | -0.38115 | -1.0319 | -0.27384 | -0.46148 |
| 26.80556 | -0.53146 | -0.34601 | -0.53369 | -0.16996 | -0.17592 | -2.20606 | -0.04613 | -0.15448 | 0.456533 | -1.19674 | -0.65083 | -0.25788 | -0.81475 | -0.72281 | -0.55701 | -0.04596 | -1.2452 | -0.1309 | -0.11075 |
| 26.81944 | -0.49193 | -0.57013 | -0.01028 | -0.63769 | -0.19329 | -2.37856 | -0.06679 | -0.01784 | 0.229381 | 0.060451 | -0.09199 | -0.47087 | -0.14907 | -0.4302 | -0.25397 | 0.284941 | -0.93348 | -0.23101 | 0.274395 |
| 26.83333 | -0.60254 | -0.51952 | 0.237288 | -1.15927 | -0.01613 | -2.52954 | -0.24027 | 0.052717 | -0.41467 | 1.424753 | 0.424158 | -0.20659 | 0.342783 | -0.07811 | 0.04494 | 0.54647 | -0.13078 | -0.42879 | 0.867922 |
| 26.84722 | -0.55061 | -0.15716 | 0.183393 | -1.23298 | 0.700675 | -1.58135 | -0.6785 | -0.00437 | -0.89199 | 2.036859 | 0.530435 | 0.235229 | 0.452259 | 0.202454 | 0.192756 | 0.535573 | 0.581363 | -0.65546 | 1.396432 |
| 26.86111 | -0.41476 | 0.13293 | -0.15985 | -1.00713 | 0.979873 | -0.83937 | -1.09433 | -0.1107 | -1.14149 | 1.507923 | 0.108152 | 0.435389 | 0.235727 | 0.315641 | 0.138656 | 0.192233 | 0.810343 | -0.59448 | 1.348388 |
| 26.875 | -0.36547 | 0.221742 | -0.52107 | -0.82779 | 0.648707 | -0.60804 | -1.31435 | -0.19612 | -1.01606 | 0.462362 | -0.50248 | 0.142171 | -0.129 | 0.199194 | -0.03476 | -0.39253 | 0.354737 | -0.05677 | 0.896021 |
| 26.88889 | -0.54014 | 0.085255 | -0.6305 | -0.80832 | 0.234622 | -0.39861 | -1.1834 | -0.26968 | -0.7944 | -0.75238 | -0.83319 | -0.26859 | -0.28428 | -0.07691 | -0.35081 | -0.94495 | -0.29712 | 0.149853 | 0.315911 |
| 26.90278 | -0.83098 | -0.15772 | -0.4649 | -1.11508 | -0.15578 | -0.11108 | -0.72023 | -0.31796 | -0.67412 | -1.13234 | -0.54673 | -0.68229 | -0.36463 | -0.25434 | -0.72451 | -1.2863 | -0.85599 | 0.026985 | -0.33439 |
| 26.91667 | -0.79852 | -0.24697 | -0.14088 | -1.33109 | -0.4262 | 0.13606 | -0.27054 | -0.46359 | -0.57388 | -0.66257 | -0.23748 | -1.1214 | -0.84463 | -0.41405 | -1.01955 | -1.34294 | -1.24961 | -0.14869 | -0.94335 |
| 26.93056 | -0.33801 | -0.10121 | 0.226605 | -0.97944 | -0.73456 | 0.239277 | 0.19589 | -0.5224 | -0.28357 | -0.29855 | -0.28854 | -1.06714 | -1.41081 | -0.58059 | -1.0903 | -1.08297 | -0.87565 | -0.34157 | -1.33126 |
| 26.94444 | 0.166699 | 0.122373 | 0.380211 | -0.41001 | -0.6868 | 0.233763 | 0.368789 | -0.49881 | 0.142022 | -0.74209 | -0.6869 | -0.78136 | -1.18558 | -0.70245 | -0.8325 | -0.85251 | -0.45298 | -0.35561 | -1.47393 |
| 26.95833 | 0.485908 | 0.30819 | 0.183611 | 0.077387 | -0.67474 | 0.226194 | 0.394815 | -0.41833 | 0.577879 | -1.41757 | -1.05134 | -0.57866 | -0.8249 | -0.71079 | -0.41456 | -0.77404 | -0.21211 | -0.05991 | -1.22683 |
| 26.97222 | 0.559896 | 0.376988 | -0.0574 | 0.377877 | -0.54881 | 0.275934 | 0.321952 | -0.27387 | 0.842409 | -1.80858 | -1.01597 | -0.49725 | -0.59356 | -0.71669 | -0.15844 | -0.79617 | -0.35325 | 0.11657 | -0.8457 |
| 26.98611 | 0.296179 | 0.180501 | -0.24222 | 0.202173 | -0.59362 | 0.204061 | -0.12115 | -0.27172 | 0.418881 | -1.71833 | -0.61273 | -0.49616 | -0.48512 | -0.80887 | -0.04146 | -0.79962 | -0.72626 | -0.12421 | -0.58483 |
| 27 | -0.49315 | -0.40206 | -0.73327 | -0.67203 | -0.94933 | -0.31747 | -1.08675 | -0.62458 | -0.74126 | -1.02489 | -0.3203 | -0.32877 | -0.43003 | -0.74005 | 0.052276 | -0.64414 | -0.8755 | -0.48303 | -0.53017 |
| 27.01389 | -1.43136 | -1.14285 | -1.3229 | -1.57645 | -0.80501 | -0.81155 | -1.73078 | -0.85039 | -1.99283 | -0.4932 | -0.35098 | -0.20619 | -0.50771 | -0.56572 | 0.10921 | -0.38248 | -0.89891 | -0.71616 | -0.51579 |
| 27.02778 | -1.73471 | -1.39667 | -1.30192 | -1.97275 | -0.31905 | -0.77017 | -1.4117 | -0.58541 | -2.34393 | -0.39286 | -0.60155 | -0.05803 | -0.45974 | -0.35099 | 0.15509 | -0.20092 | -0.64881 | -0.72149 | -0.56311 |
| 27.04167 | -1.41453 | -1.24973 | -1.03047 | -1.24464 | -0.11826 | -0.51248 | -0.69929 | -0.15288 | -2.2673 | -0.54846 | -0.70414 | 0.03231 | -0.15861 | -0.20758 | 0.183371 | -0.14121 | -0.31112 | -0.52897 | -0.44536 |
| 27.05556 | -0.89793 | -0.99595 | -0.59091 | -0.5954 | 0.049161 | -0.2555 | -0.31291 | -0.24244 | -1.33346 | -0.80453 | -0.56259 | 0.044074 | 0.126065 | -0.16338 | 0.114512 | -0.11608 | 0.032582 | -0.21515 | -0.40427 |
| 27.06944 | -0.25026 | 0.061027 | -0.19444 | -0.30831 | 0.342635 | 0.160026 | -0.18867 | -0.23448 | -1.34016 | -0.93444 | -0.2509 | 0.045861 | 0.169264 | -0.15605 | 0.050892 | -0.02263 | 0.24478 | -0.06726 | -0.3739 |
| 27.08333 | 0.298263 | 1.134499 | 0.315788 | 0.01736 | 0.652572 | 0.676509 | 0.126118 | 0.095788 | -1.73815 | -0.74859 | 0.075739 | 0.002677 | -0.02645 | -0.14215 | -0.03323 | 0.060821 | 0.140728 | -0.2272 | -0.42537 |
| 27.09722 | 0.926587 | 1.931628 | 0.578995 | 0.251218 | 1.327597 | 1.344705 | 0.582999 | 0.75556 | -0.86068 | -0.54683 | -0.08864 | -0.25572 | -0.88405 | -0.30528 | -0.73453 | -0.42937 | -0.35735 | -0.94395 | -0.88573 |
| 27.11111 | 1.551851 | 2.589643 | 0.735731 | 0.247171 | 1.482757 | 0.474731 | 0.722897 | 1.472265 | -0.40656 | -1.04945 | -0.86642 | -0.91937 | -2.59836 | -0.7055 | -2.48548 | -1.83889 | -1.03116 | -1.77624 | -1.80856 |
| 27.125 | -0.06728 | -0.25636 | 2.406926 | 0.938483 | -0.81007 | -3.51857 | 0.47653 | -1.23108 | -0.00195 | -2.02188 | -1.24631 | -1.1338 | -3.41138 | -0.74943 | -2.31377 | -2.0053 | -1.31286 | -1.84441 | -1.77716 |
| 27.13889 | -1.82521 | -2.50762 | 0.029512 | 0.942081 | -0.52267 | -1.99815 | -0.18912 | -2.19044 | 0.895438 | -1.1947 | -0.66862 | -0.31376 | -2.32636 | -0.28457 | -0.66494 | -0.62566 | -0.83428 | -0.13378 | -0.48627 |
| 27.15278 | -0.82832 | -0.96281 | 0.076711 | 1.081985 | 0.442486 | 1.270206 | 0.467853 | -0.61391 | 2.02873 | 0.309336 | -0.2035 | 0.266902 | -0.65625 | 0.213121 | 0.485962 | 0.611803 | -0.07631 | 0.238077 | 0.06267 |
| 27.16667 | 0.30058 | 0.09067 | -0.58721 | 1.086531 | 1.070639 | 2.057738 | 1.056344 | 0.558146 | 2.279224 | 1.96284 | 0.922567 | 0.531197 | 0.222581 | 0.371596 | 0.755857 | 1.043243 | 0.357313 | -0.17111 | -0.1652 |
| 27.18056 | 0.435564 | 0.249954 | 0.260334 | 0.745531 | 0.938428 | 0.672257 | 0.743803 | 0.447845 | 2.034983 | 0.983945 | 0.726659 | 0.213976 | 0.806224 | 0.062497 | 0.925386 | 0.965786 | 0.38126 | 0.328828 | 0.184748 |
| 27.19444 | 0.088886 | -0.0512 | 0.612875 | 0.198593 | 0.494948 | 0.303085 | 0.656259 | 0.676987 | 2.018496 | -0.01676 | 0.233765 | -0.09031 | 0.964345 | -0.14066 | 0.95661 | 0.754899 | 0.312001 | 0.982422 | 0.669091 |
| 27.20833 | -0.66755 | -1.00122 | 1.323161 | -0.21912 | -0.38916 | 0.27502 | 0.818254 | 1.534068 | 3.4677 | 0.868952 | 0.824772 | -0.03706 | 0.902538 | -0.07607 | 0.764873 | 0.521802 | 0.266428 | 0.793559 | 0.287298 |
| 27.22222 | -0.88386 | -1.39517 | 1.74406 | 0.054756 | -1.15333 | 0.459748 | 0.969138 | 1.407148 | 3.591431 | 0.554261 | 0.195349 | -0.06579 | 0.746267 | -0.04221 | 0.533696 | 0.261279 | 0.459342 | 0.166039 | -0.24068 |
| 27.23611 | -0.02656 | -0.32969 | 1.486814 | 0.96525 | -0.03039 | 0.4073 | 0.942107 | 0.75996 | 2.14154 | 0.375406 | -0.18108 | -0.04943 | 0.758872 | 0.081789 | 0.442289 | 0.154916 | 1.024066 | -0.16915 | -0.51055 |
| 27.25 | 0.659447 | 0.526908 | 1.348972 | 1.439311 | 0.157215 | 0.259773 | 0.843286 | 0.316124 | 1.121003 | 0.118579 | 0.258188 | 0.18824 | 0.99458 | 0.312782 | 0.710914 | 0.339427 | 1.410165 | 0.210703 | -0.06815 |
| 27.26389 | 0.845438 | 0.798643 | 0.877769 | 0.96167 | -0.22031 | 0.091979 | 0.41117 | 0.059776 | 0.49619 | 0.557095 | 0.81159 | 0.403512 | 1.070588 | 0.498447 | 0.848072 | 0.540581 | 1.399742 | 0.53522 | 0.517681 |
| 27.27778 | 0.570583 | 0.574262 | 0.140903 | 0.014548 | -0.58261 | -0.07413 | -0.19009 | -0.03626 | -0.08616 | 0.346918 | 0.286871 | 0.291668 | 0.528308 | 0.395565 | 0.377393 | 0.348145 | 0.917247 | 0.000863 | 0.335251 |
| 27.29167 | 0.100665 | 0.02345 | -0.21492 | -0.55194 | -0.66549 | -0.16604 | -0.54798 | -0.04765 | -0.50726 | -0.73689 | -0.64516 | 0.027014 | -0.36127 | 0.136168 | -0.20794 | -0.06133 | 0.149122 | -0.60709 | -0.2086 |
| 27.30556 | -0.16195 | -0.34422 | -0.11141 | -0.32826 | -0.45895 | -0.12781 | -0.32429 | 0.032539 | -0.36891 | -0.70479 | -0.7787 | -0.06225 | -0.85192 | -0.0054 | -0.42888 | -0.27277 | -0.25687 | -0.45561 | -0.38508 |
| 27.31944 | -0.00657 | -0.29317 | 0.29411 | 0.375833 | -0.37238 | 0.017109 | 0.25789 | 0.188673 | 0.115505 | 0.161634 | 0.101575 | 0.196149 | -0.59989 | 0.138448 | -0.18658 | -0.07852 | -0.07774 | 0.168508 | -0.04916 |
| 27.33333 | 0.408725 | 0.189054 | 0.696631 | 1.048651 | -0.04986 | 0.317625 | 0.773525 | 0.334878 | 0.705698 | 1.387462 | 1.550456 | 0.678711 | 0.085844 | 0.551659 | 0.537866 | 0.51186 | 0.487176 | 0.923189 | 0.822876 |
| 27.34722 | 0.696441 | 0.583129 | 0.766528 | 1.185372 | 0.293353 | 0.454369 | 0.867815 | 0.3459 | 1.121613 | 1.486327 | 1.921098 | 0.734758 | 0.456729 | 0.559052 | 0.90123 | 0.80774 | 0.743765 | 0.916102 | 1.144853 |
| 27.36111 | 0.563829 | 0.423911 | 0.552565 | 0.818612 | 0.362915 | 0.307998 | 0.595686 | 0.228094 | 1.008249 | 0.305681 | 0.816959 | 0.357088 | 0.147616 | 0.214861 | 0.539451 | 0.506929 | 0.29488 | 0.065353 | 0.456602 |
| 27.375 | 0.242637 | 0.150695 | 0.318579 | 0.355909 | 0.423146 | 0.114528 | 0.247614 | 0.113865 | 0.6165 | -0.61431 | -0.0922 | 0.112812 | -0.25671 | -0.08504 | 0.088099 | 0.07295 | -0.36973 | -0.36936 | -0.07104 |
| 27.38889 | 0.02296 | -0.09929 | 0.216949 | -0.07604 | 0.604762 | -0.01331 | -0.05279 | 0.074777 | 0.367645 | -0.69068 | -0.24182 | 0.068639 | -0.48968 | -0.26244 | -0.19817 | -0.23318 | -0.79802 | -0.27697 | -0.06167 |
| 27.40278 | 0.026912 | -0.1846 | 0.228089 | -0.27747 | 0.756075 | -0.11521 | -0.215 | 0.042289 | 0.174167 | -0.08386 | 0.067706 | 0.074468 | -0.56054 | -0.29316 | -0.34152 | -0.40132 | -0.78662 | -0.07455 | 0.017197 |
| 27.41667 | 0.180694 | -0.11728 | 0.290363 | -0.11722 | 0.740367 | -0.18158 | -0.13755 | 0.024589 | 0.161724 | 0.793605 | 0.58973 | 0.102601 | -0.46119 | -0.12992 | -0.3098 | -0.25602 | -0.33535 | 0.073177 | -0.00922 |
| 27.43056 | 0.346595 | -0.0226 | 0.38127 | 0.2787 | 0.724196 | -0.15657 | 0.055262 | 0.056487 | 0.407776 | 1.200657 | 0.828833 | 0.103426 | -0.33537 | 0.044752 | -0.19433 | 0.000108 | 0.20839 | 0.123209 | -0.09047 |
| 27.44444 | 0.357703 | 0.073134 | 0.372549 | 0.49734 | 0.588612 | -0.02989 | 0.2543 | 0.108737 | 0.758191 | 0.744684 | 0.648825 | 0.07426 | -0.24447 | 0.147437 | -0.06266 | 0.246992 | 0.358215 | -0.02772 | -0.12865 |
| 27.45833 | 0.254562 | 0.054697 | 0.362376 | 0.474808 | 0.428832 | 0.046693 | 0.344191 | 0.119213 | 1.022621 | 0.233687 | 0.409125 | 0.08117 | -0.02419 | 0.190728 | 0.069387 | 0.175644 | 0.199858 | -0.13105 | -0.0494 |
| 27.47222 | 0.254422 | 0.033726 | 0.457484 | 0.3264 | 0.365904 | 0.133807 | 0.308196 | 0.195201 | 0.88623 | -0.04933 | 0.333069 | 0.150605 | 0.221385 | 0.1869 | 0.160541 | 0.140245 | 0.117027 | -0.05635 | 0.078922 |
| 27.48611 | 0.519157 | -0.03977 | 0.71043 | 0.477038 | 0.459731 | 0.39963 | 0.528714 | 0.456358 | 0.705454 | 0.123948 | 0.413867 | 0.253746 | 0.404493 | 0.140249 | 0.1999 | 0.046061 | 0.213501 | 0.150171 | 0.12166 |
| 27.5 | 1.046764 | -0.12473 | 1.227871 | 1.435561 | 0.746833 | 1.275328 | 1.493554 | 1.041204 | 1.01379 | 0.795294 | 0.571018 | 0.373385 | 0.513873 | 0.11708 | 0.132035 | -0.08538 | 0.498813 | 0.257979 | 0.021857 |
| 27.51389 | 1.782826 | 0.077607 | 1.890376 | 2.685944 | 1.013176 | 2.467346 | 2.797725 | 1.297592 | 1.49672 | 1.29807 | 0.58141 | 0.380909 | 0.5327 | 0.153211 | 0.139639 | -0.04022 | 0.670472 | 0.09356 | -0.11545 |

| | | | | | | | | | | | | | | | | | | | |
|---|---|---|---|---|---|---|---|---|---|---|---|---|---|---|---|---|---|---|---|
| 27.52778 | 1.966338 | 0.331635 | 2.398197 | 3.668155 | 0.601762 | 2.868521 | 3.824163 | 0.887002 | 1.915005 | 0.909642 | 0.318345 | 0.2406 | 0.446472 | 0.2168 | 0.228317 | 0.110663 | 0.615852 | -0.21581 | -0.16647 |
| 27.54167 | 1.818029 | 0.387084 | 2.366447 | 3.233058 | 0.453725 | 2.330406 | 3.601084 | 0.685061 | 1.602983 | 0.257723 | 0.09777 | 0.131014 | 0.440096 | 0.32713 | 0.398293 | 0.190115 | 0.569317 | -0.38493 | -0.09325 |
| 27.55556 | 1.265223 | 0.378706 | 1.701933 | 1.631131 | 0.1656 | 1.52361 | 2.145656 | 0.32957 | 0.731108 | -0.12398 | 0.154225 | 0.106769 | 0.603892 | 0.340383 | 0.494707 | 0.269551 | 0.571049 | -0.08111 | 0.251267 |
| 27.56944 | 0.636538 | 0.290488 | 0.838854 | 0.191568 | -0.16152 | 0.857089 | 0.707624 | 0.058291 | 0.047852 | -0.28674 | 0.307194 | 0.18634 | 0.74415 | 0.204803 | 0.480202 | 0.297114 | 0.560061 | 0.37126 | 0.536232 |
| 27.58333 | 0.126136 | 0.247051 | 0.028265 | -0.97282 | -0.41953 | 0.384025 | -0.29424 | -0.0708 | -0.66284 | -0.08005 | 0.287969 | 0.274506 | 0.595778 | 0.107769 | 0.246857 | 0.151044 | 0.53624 | 0.452718 | 0.385784 |
| 27.59722 | -0.21614 | 0.351062 | -0.60994 | -1.69804 | -0.49155 | 0.047048 | -0.78759 | -0.04749 | -1.10187 | 0.569087 | 0.276068 | 0.360636 | 0.692065 | 0.200693 | 0.308318 | 0.243899 | 0.625152 | 0.654494 | 0.347921 |
| 27.61111 | -0.32648 | 0.311534 | -0.83251 | -1.82455 | -0.48735 | -0.26118 | -1.12003 | 0.088745 | -1.20227 | 1.394085 | 0.545916 | 0.85223 | 1.305871 | 0.539452 | 0.934076 | 0.780543 | 0.999995 | 1.655126 | 0.781429 |
| 27.625 | -0.56352 | 0.40617 | -0.55764 | -1.42688 | -0.50505 | -0.62671 | -1.35137 | 0.176955 | -0.69777 | 2.041407 | 0.966074 | 1.851281 | 1.827193 | 0.847109 | 1.322458 | 1.120525 | 1.28016 | 1.966126 | 1.407281 |
| 27.63889 | -0.4254 | 0.453056 | -0.28652 | -0.7287 | -0.5117 | -1.30173 | -1.47403 | 0.210327 | 0.061023 | 2.229162 | 1.380313 | 0.998368 | 2.246303 | 0.912399 | 1.06495 | 0.925503 | 1.275567 | 1.401431 | 1.305837 |
| 27.65278 | -0.20555 | 0.181492 | -0.08174 | -0.03689 | -0.21596 | -2.11871 | -1.35193 | 0.117151 | 0.702799 | 1.619592 | 1.072873 | 0.355728 | 2.574965 | 0.832616 | 0.652561 | 0.615938 | 1.242071 | 1.077974 | 0.994414 |
| 27.66667 | -0.03528 | -0.4273 | 0.085682 | 0.490687 | 0.304073 | -2.56068 | -0.59557 | -0.02271 | 0.800528 | 0.60901 | 0.127175 | 0.147653 | 1.896283 | 0.631329 | 0.321816 | 0.462573 | 0.856272 | 0.718379 | 0.753907 |
| 27.68056 | 0.215837 | -0.59139 | 0.261392 | 0.531912 | 0.778874 | -2.16571 | 0.327361 | -0.14801 | 0.294413 | -0.40618 | -1.03567 | 0.265859 | 0.762108 | 0.346871 | -0.0166 | 0.338016 | 0.504863 | 0.39656 | 0.517843 |
| 27.69444 | 0.736393 | -0.32959 | 0.481647 | 0.465951 | 0.750956 | -1.23586 | 1.220292 | -0.28018 | -0.55271 | -1.62137 | -2.18278 | 0.468737 | -0.025 | 0.049647 | -0.39872 | 0.371021 | 0.172158 | 0.157648 | 0.444071 |
| 27.70833 | 1.21535 | -0.04344 | 0.51493 | 0.62373 | 1.02176 | -0.41936 | 1.964777 | -0.32127 | -0.85274 | -1.6086 | -1.30779 | 0.401322 | -0.31881 | -0.04745 | -0.56211 | 0.296394 | -0.15236 | -0.0497 | 0.43778 |
| 27.72222 | 0.869902 | 0.089887 | 0.222622 | 0.153315 | 0.86728 | 0.346375 | 2.089626 | -0.21594 | -0.47407 | -0.24281 | 0.159809 | 0.326999 | -0.35815 | 0.052776 | -0.49719 | 0.1606 | -0.50186 | -0.06188 | 0.520872 |
| 27.73611 | -0.00282 | 0.056512 | -0.06903 | -0.77495 | 0.048658 | 1.044748 | 1.652915 | 0.058252 | 0.08901 | 1.413481 | 1.559323 | 0.356587 | -0.37272 | 0.29006 | -0.25071 | 0.212627 | -0.45419 | 0.054041 | 0.576569 |
| 27.75 | -0.65234 | -0.18687 | -0.1426 | -1.34157 | -0.8056 | 1.722037 | 0.624095 | 0.197546 | 0.362323 | 2.463165 | 1.867462 | 0.348596 | -0.36836 | 0.552902 | 0.009324 | 0.372205 | -0.01132 | 0.107329 | 0.467123 |
| 27.76389 | -0.83346 | -0.37692 | 0.090989 | -1.43848 | -1.13521 | 1.69502 | -0.13066 | 0.209089 | 0.436942 | 1.750646 | 0.689925 | 0.253256 | -0.44795 | 0.665201 | 0.080915 | 0.299191 | 0.544333 | 0.067948 | 0.343245 |
| 27.77778 | -0.53545 | -0.45344 | 0.322314 | -1.08765 | -1.52244 | 1.463115 | -0.79553 | 0.135117 | 0.023131 | 0.381336 | -0.42093 | 0.097885 | -0.51826 | 0.71707 | -0.02003 | 0.027239 | 0.637853 | -0.03312 | 0.271663 |
| 27.79167 | 0.109533 | -0.31736 | -0.00358 | -0.57296 | -1.14318 | 0.8585 | -1.76788 | -0.00565 | -0.28407 | -0.74694 | -1.16957 | 0.035991 | -0.52208 | 0.571247 | -0.06758 | -0.23523 | 0.464658 | -0.106 | 0.274733 |
| 27.80556 | 0.663541 | 0.016763 | -0.70137 | -0.15334 | -0.88506 | -0.02147 | -2.37303 | -0.03467 | -0.7115 | -1.65921 | -1.28781 | 0.087946 | -0.34464 | 0.159336 | 0.01454 | -0.39215 | 0.316033 | -0.1175 | 0.334262 |
| 27.81944 | 0.569813 | 0.158119 | -0.91797 | -0.01315 | -0.30824 | -0.8674 | -1.76262 | 0.01521 | -0.87551 | -1.8558 | -0.76582 | 0.095553 | -0.03494 | -0.23839 | 0.084307 | -0.48319 | 0.108054 | -0.12741 | 0.387512 |
| 27.83333 | -0.37641 | -0.07797 | -0.99901 | -0.38341 | 0.292755 | -1.89286 | -0.71402 | 0.147324 | -0.53 | -1.10757 | -0.14109 | -0.13274 | 0.345606 | -0.53048 | 0.074168 | -0.37629 | -0.17158 | -0.29577 | 0.230639 |
| 27.84722 | -1.2358 | -0.49654 | -1.55531 | -1.21042 | 0.699414 | -2.59431 | -0.65129 | 0.1591 | -0.234 | -0.07319 | 0.287614 | -0.36601 | 0.711145 | -0.67689 | 0.070848 | -0.16022 | -0.57367 | -0.45841 | 0.033516 |
| 27.86111 | -1.4791 | -1.03461 | -1.97995 | -1.59742 | 0.930401 | -2.22523 | -1.09822 | -0.03525 | -0.20683 | 0.692561 | 0.356528 | -0.53795 | 0.788585 | -0.58914 | 0.075724 | -0.04162 | -0.69253 | -0.44098 | -0.04157 |
| 27.875 | -1.10058 | -1.35945 | -1.68089 | -1.57615 | 0.854024 | -1.38004 | -1.5551 | -0.23072 | -0.30464 | 0.594555 | 0.01919 | -0.50023 | 0.498791 | -0.45465 | 0.018081 | 0.004528 | -0.51344 | -0.37616 | -0.1384 |
| 27.88889 | -1.11716 | -1.32641 | -0.9242 | -1.69477 | 0.492169 | -1.101 | -1.67229 | -0.41109 | -0.45186 | -0.06429 | -0.57296 | -0.24675 | -0.18524 | -0.32053 | -0.09722 | 0.012304 | -0.20646 | -0.30587 | -0.32263 |
| 27.90278 | -0.98076 | -0.95413 | -0.04604 | -1.70774 | 0.07122 | -0.79123 | -1.11576 | -0.5898 | -0.8861 | -1.01924 | -1.20593 | 0.014842 | -0.99133 | -0.4124 | -0.17827 | -0.03665 | -0.09892 | -0.1104 | -0.52105 |
| 27.91667 | -0.49084 | -0.28006 | 0.673834 | -1.17181 | -0.37314 | -0.46611 | -0.42751 | -0.71879 | -0.60392 | -1.92628 | -1.51178 | -0.01968 | -1.8713 | -0.68425 | -0.21844 | -0.15672 | -0.27025 | 0.049127 | -0.77823 |
| 27.93056 | 0.105406 | 0.360838 | 1.156741 | -0.44069 | -0.63844 | -0.16911 | 0.16858 | -0.55472 | -0.17978 | -1.78529 | -1.15242 | -0.26936 | -2.09973 | -1.0105 | -0.29149 | -0.31692 | -0.74821 | 0.013503 | -1.03516 |
| 27.94444 | 0.501416 | 0.876551 | 1.043673 | 0.372127 | -0.85693 | 0.065173 | 0.736519 | -0.23958 | 0.275351 | -0.94014 | -0.4673 | -0.53741 | -1.65883 | -1.02564 | -0.27228 | -0.32749 | -1.12011 | -0.17444 | -1.11353 |
| 27.95833 | 0.680223 | 1.005556 | 0.793166 | 1.0318 | -0.82429 | 0.250266 | 1.249415 | -0.06619 | 0.666957 | -0.09147 | 0.032188 | -0.45588 | -0.83244 | -0.83931 | -0.14265 | -0.21293 | -1.28978 | -0.36629 | -1.01595 |
| 27.97222 | 0.587255 | 0.792676 | 0.571599 | 1.187366 | -0.56836 | 0.384195 | 1.500819 | -0.15475 | 1.180288 | 0.223455 | 0.040225 | -0.16354 | -0.18042 | -0.69108 | -0.04061 | -0.13714 | -1.23956 | -0.533 | -0.81545 |
| 27.98611 | 0.135828 | 0.13267 | -0.03876 | 0.359288 | -0.62698 | 0.251028 | 1.002116 | -0.37226 | 1.086955 | -0.1893 | -0.3761 | 0.027789 | 0.044216 | -0.56308 | -0.01133 | -0.18433 | -0.91755 | -0.71301 | -0.59121 |
| 28 | -1.04683 | -1.10374 | -1.44898 | -1.57871 | -1.13737 | -0.30044 | -0.58468 | -0.81915 | -0.48177 | -0.99631 | -0.78537 | -0.05938 | -0.06266 | -0.4737 | -0.07084 | -0.27688 | -0.45055 | -0.84079 | -0.489 |
| 28.01389 | -2.65321 | -2.88094 | -3.46719 | -3.45442 | -1.22459 | -0.86128 | -2.00999 | -0.98408 | -2.37436 | -1.68347 | -0.93458 | -0.22185 | -0.25995 | -0.32 | -0.19478 | -0.40876 | -0.12317 | -0.8187 | -0.48148 |
| 28.02778 | -3.28752 | -3.98202 | -4.83141 | -3.60682 | -0.50676 | -0.85238 | -1.83128 | -0.94658 | -2.62429 | -1.49285 | -0.64702 | -0.26718 | -0.3208 | -0.07312 | -0.31165 | -0.46137 | -0.05757 | -0.81623 | -0.45454 |
| 28.04167 | -2.25839 | -3.66661 | -3.19454 | -1.95028 | 0.024538 | -0.38756 | -0.81001 | -0.40411 | -2.04963 | -0.51805 | -0.14172 | -0.14342 | -0.28361 | 0.183848 | -0.31153 | -0.32139 | -0.12152 | -0.82103 | -0.31059 |
| 28.05556 | -1.09866 | -2.07864 | -1.18448 | -0.69368 | 0.335309 | -0.02244 | -0.28664 | 0.035837 | -1.98943 | 0.333535 | 0.111683 | 0.049463 | -0.24536 | 0.285955 | -0.23791 | -0.07657 | -0.20418 | -0.82133 | -0.15536 |
| 28.06944 | -0.1706 | -0.46402 | 0.108454 | 0.350821 | 0.465071 | 0.120129 | -0.08649 | 0.341065 | -1.5723 | 0.451473 | -0.05905 | 0.183886 | -0.26817 | 0.069627 | -0.16399 | 0.099313 | -0.23839 | -0.80706 | -0.05455 |
| 28.08333 | 0.423364 | 0.732271 | 1.139951 | 1.126588 | 0.345883 | 0.066507 | -0.04126 | 0.274198 | -0.77868 | 0.012405 | -0.32404 | 0.216115 | -0.29886 | 0.002432 | -0.27276 | -0.04278 | -0.17839 | -0.66061 | -0.04222 |
| 28.09722 | 0.88619 | 1.533517 | 1.826691 | 1.51655 | 0.178528 | -0.10084 | -0.09457 | 0.074248 | -0.14189 | -0.66744 | -0.47976 | -0.02633 | -0.73443 | -0.29642 | -1.10976 | -0.91028 | -0.28143 | -0.6978 | -0.3597 |
| 28.11111 | 1.269515 | 2.379079 | 2.329993 | 1.798095 | 0.20953 | -0.15025 | -0.1841 | 0.061717 | 0.315781 | -1.96734 | -0.91601 | -1.05895 | -2.25396 | -0.93717 | -2.95745 | -2.75514 | -0.83197 | -1.35829 | -1.09281 |
| 28.125 | 1.492178 | 3.025705 | 2.121893 | 1.862459 | 0.34387 | 0.034857 | -0.00435 | 0.05781 | 0.399734 | -3.55891 | -1.64276 | -2.00864 | -2.42919 | -1.36953 | -3.52734 | -3.34109 | -1.56881 | -1.44724 | -1.59634 |
| 28.13889 | 1.581638 | 2.910088 | 1.547105 | 1.638773 | 0.313346 | 0.26778 | 0.410094 | -0.07884 | 0.288958 | -2.77118 | -1.91119 | -0.79144 | -1.00989 | -0.79883 | -1.42305 | -1.8039 | -1.56622 | -0.4659 | -0.53721 |
| 28.15278 | 1.37912 | 2.312567 | 1.210423 | 1.115032 | 0.234042 | 0.447417 | 0.680919 | -0.04604 | 0.355839 | -0.75693 | -0.63671 | 0.323174 | 0.444085 | -0.11981 | 0.381318 | -0.17182 | -0.66115 | 0.119566 | 0.523159 |
| 28.16667 | 0.976575 | 1.547007 | 1.034427 | 0.689907 | 0.174972 | 0.594059 | 0.862434 | 0.143197 | 1.050843 | 0.529465 | 0.546013 | 0.715024 | 1.076766 | 0.286919 | 0.944899 | 0.631395 | -0.1198 | 0.451964 | 0.788509 |
| 28.18056 | 0.734069 | 1.118781 | 0.889994 | 0.403057 | 0.156735 | 0.513736 | 0.796665 | 0.266313 | 1.770608 | 1.044056 | 1.263849 | 0.595095 | 1.039459 | 0.38559 | 0.741272 | 0.805037 | 0.241941 | 0.55487 | 0.466599 |
| 28.19444 | 0.646677 | 0.78845 | 0.759968 | 0.298659 | 0.021106 | 0.385364 | 0.675683 | 0.252299 | 1.793856 | 1.048887 | 1.257636 | 0.347216 | 0.881052 | 0.376892 | 0.559714 | 0.790905 | 0.45327 | 0.543406 | 0.117413 |
| 28.20833 | 0.55817 | 0.446613 | 0.854171 | 0.2599 | 0.044852 | 0.345537 | 0.661689 | 0.197932 | 1.315816 | 0.899398 | 0.989692 | 0.052432 | 0.643183 | 0.259347 | 0.468199 | 0.636769 | 0.467657 | 0.35546 | -0.08701 |
| 28.22222 | 0.523005 | 0.258231 | 1.029716 | 0.363558 | 0.169232 | 0.361783 | 0.781681 | 0.064034 | 1.134323 | 0.954103 | 0.978086 | -0.1579 | 0.267061 | 0.143487 | 0.317396 | 0.51528 | 0.360605 | 0.128258 | -0.12526 |
| 28.23611 | 0.533597 | 0.218787 | 1.121845 | 0.435437 | 0.267834 | 0.356526 | 0.80632 | -0.0333 | 1.073524 | 1.194775 | 1.145775 | -0.13669 | -0.11618 | 0.11882 | 0.241308 | 0.515969 | 0.290853 | -0.13156 | -0.06105 |
| 28.25 | 0.574951 | 0.31474 | 1.077478 | 0.323254 | 0.274896 | 0.313519 | 0.707175 | 0.021414 | 1.12809 | 1.285015 | 1.187447 | 0.210466 | -0.13316 | 0.199453 | 0.140579 | 0.46991 | 0.474552 | -0.49547 | -0.20637 |
| 28.26389 | 0.63256 | 0.499361 | 0.8596 | 0.136614 | 0.210328 | 0.238744 | 0.510792 | 0.151667 | 0.982391 | 1.135073 | 0.78313 | 0.593647 | 0.013987 | 0.273822 | 0.200341 | 0.418826 | 0.808719 | -0.69916 | -0.47588 |
| 28.27778 | 0.613252 | 0.56242 | 0.534113 | -0.01291 | 0.168317 | 0.16294 | 0.348771 | 0.24677 | 0.478636 | 0.905722 | 0.683822 | 0.570712 | 0.164642 | 0.208405 | 0.228426 | 0.414949 | 0.837455 | -0.13434 | -0.35063 |
| 28.29167 | 0.374915 | 0.250772 | 0.278829 | -0.01883 | 0.170415 | 0.120352 | 0.274844 | 0.172484 | -0.20414 | 0.645844 | 0.730083 | 0.12917 | 0.349117 | 0.139936 | 0.246909 | 0.375835 | 0.577208 | 0.601057 | 0.111486 |
| 28.30556 | 0.030839 | -0.24071 | 0.065007 | 0.118963 | 0.135168 | 0.048872 | 0.18888 | -0.02393 | -0.81775 | 0.29921 | 0.300723 | -0.34729 | 0.131484 | 0.177651 | 0.087785 | 0.122236 | 0.321727 | 0.513864 | 0.348081 |
| 28.31944 | -0.09925 | -0.4466 | -0.06659 | 0.419587 | 0.136896 | -0.03878 | 0.112825 | -0.22424 | -1.05618 | 0.002819 | -0.19153 | -0.37158 | -0.24159 | 0.237434 | -0.07912 | -0.13976 | 0.080334 | -0.07664 | 0.110363 |
| 28.33333 | 0.136798 | -0.16708 | -0.0254 | 0.781979 | 0.190121 | -0.05232 | 0.142373 | -0.27644 | -0.67894 | -0.11645 | -0.34271 | 0.11952 | -0.43205 | 0.216387 | -0.1231 | -0.19223 | 0.224824 | -0.43251 | -0.26812 |
| 28.34722 | 0.458234 | 0.245271 | 0.267812 | 1.080932 | 0.246369 | 0.007842 | 0.3178 | -0.12293 | 0.112551 | -0.06808 | -0.2623 | 0.589069 | -0.39097 | 0.08865 | -0.1388 | -0.08643 | 0.475091 | -0.52741 | -0.71425 |
| 28.36111 | 0.804361 | 0.691055 | 0.753311 | 1.090073 | 0.241888 | 0.106522 | 0.518821 | 0.156682 | 0.719575 | 0.04661 | 0.192586 | 0.595278 | -0.15798 | -0.11981 | -0.03831 | 0.047255 | 0.39618 | -0.12113 | -0.52999 |
| 28.375 | 1.028117 | 1.084752 | 1.144264 | 0.723448 | 0.192938 | 0.142463 | 0.554829 | 0.272838 | 0.680219 | 0.014325 | 0.670259 | 0.169342 | 0.084821 | -0.28029 | 0.165663 | 0.120587 | -0.10773 | 0.584846 | 0.219981 |
| 28.38889 | 0.882763 | 1.011361 | 1.2055 | 0.184752 | 0.091575 | 0.105466 | 0.32889 | 0.041053 | 0.079498 | -0.15031 | 0.559015 | -0.17364 | 0.065634 | -0.21154 | 0.201289 | 0.044007 | -0.42961 | 0.684401 | 0.510728 |
| 28.40278 | 0.271682 | 0.263385 | 0.649486 | -0.31872 | -0.016 | 0.062689 | -0.03909 | -0.2477 | -0.71454 | -0.2514 | 0.089793 | -0.27261 | -0.06004 | 0.031901 | 0.062743 | -0.08872 | -0.49279 | 0.248815 | 0.276517 |
| 28.41667 | -0.44215 | -0.62985 | -0.04541 | -0.69137 | -0.07764 | 0.064238 | -0.24746 | -0.42317 | -1.34348 | -0.17797 | -0.11826 | -0.02677 | -0.06374 | 0.195977 | -0.07962 | -0.16023 | -0.3686 | -0.13598 | 0.041274 |
| 28.43056 | -0.82775 | -1.14173 | -0.63343 | -0.55564 | -0.08964 | 0.073438 | -0.24305 | -0.44188 | -1.27715 | 0.115955 | 0.020556 | 0.335374 | 0.012857 | 0.14022 | -0.16511 | -0.13789 | -0.12205 | -0.08152 | -0.13282 |
| 28.44444 | -0.60741 | -1.02328 | -0.75715 | -0.0446 | -0.04001 | 0.093947 | -0.01474 | -0.27333 | -0.6349 | 0.632428 | 0.429864 | 0.587013 | 0.137112 | 0.046541 | -0.1016 | 0.01604 | 0.094868 | 0.00267 | -0.19246 |
| 28.45833 | -0.15824 | -0.60237 | -0.20401 | 0.555281 | 0.058575 | 0.13192 | 0.169351 | -0.01464 | 0.09605 | 1.008428 | 0.833941 | 0.497864 | 0.29888 | 0.073535 | 0.119299 | 0.246236 | 0.141201 | 0.038454 | -0.04163 |
| 28.47222 | 0.238262 | -0.07595 | 0.586538 | 0.938 | 0.089853 | 0.128821 | 0.231973 | 0.166903 | 0.299086 | 0.963703 | 0.895063 | 0.296183 | 0.419109 | 0.138872 | 0.326303 | 0.381835 | 0.093061 | 0.041703 | 0.154604 |
| 28.48611 | 0.597534 | 0.411545 | 1.406496 | 1.325608 | 0.245473 | 0.255641 | 0.595218 | 0.455641 | 0.404722 | 0.60781 | 0.546878 | 0.120718 | 0.408671 | 0.139466 | 0.39537 | 0.408017 | 0.090629 | -0.01823 | 0.256839 |
| 28.5 | 1.087284 | 0.806529 | 2.03521 | 2.305265 | 0.849572 | 0.606681 | 1.597312 | 1.032751 | 1.392608 | 0.319083 | 0.099414 | 0.121913 | 0.246801 | 0.098536 | 0.317076 | 0.311368 | 0.176027 | -0.1088 | 0.246167 |
| 28.51389 | 1.176169 | 1.017711 | 2.523272 | 3.383499 | 1.631862 | 0.829012 | 2.325789 | 1.434806 | 2.370525 | 0.192023 | -0.13248 | 0.269105 | 0.103959 | 0.056145 | 0.212351 | 0.191449 | 0.300256 | 0.043055 | 0.234615 |
| 28.52778 | 0.968354 | 1.146848 | 3.175224 | 4.953343 | 1.771955 | 0.886435 | 2.701939 | 1.280572 | 3.00264 | 0.305736 | 0.035207 | 0.430697 | 0.025646 | 0.015699 | 0.154367 | 0.081411 | 0.437857 | 0.2922 | 0.262391 |
| 28.54167 | 1.271886 | 1.50301 | 3.376044 | 5.23976 | 1.956232 | 1.034128 | 2.683489 | 0.88246 | 3.044784 | 0.594063 | 0.507084 | 0.522491 | 0.09806 | 0.054815 | 0.160358 | 0.044094 | 0.626581 | 0.31295 | 0.220811 |
| 28.55556 | 1.25603 | 1.230316 | 2.264274 | 2.979341 | 1.605252 | 0.916453 | 2.089383 | 0.454177 | 1.655747 | 0.814666 | 1.014611 | 0.42143 | 0.228975 | 0.0959 | 0.18952 | 0.097299 | 0.509768 | 0.070107 | 0.137322 |
| 28.56944 | 0.983605 | 0.658843 | 1.344749 | 1.172588 | 0.794644 | 0.647626 | 1.475095 | -0.09349 | 0.298802 | 0.710923 | 1.109942 | 0.149802 | 0.281207 | 0.067433 | 0.206907 | 0.160617 | 0.285509 | -0.15609 | 0.060627 |
| 28.58333 | 0.801713 | 0.173982 | 0.470436 | -0.07003 | 0.09529 | 0.568019 | 1.026922 | -0.43663 | -0.63358 | 0.326001 | 0.5731 | -0.06581 | 0.265253 | 0.050479 | 0.296731 | 0.241878 | 0.132223 | -0.20508 | 0.168308 |
| 28.59722 | 0.884415 | -0.14418 | -0.53846 | -1.21785 | -0.30902 | 0.735866 | 0.577094 | -0.50928 | -0.66464 | 0.152819 | -0.06747 | 0.118396 | 0.38088 | 0.248712 | 0.619037 | 0.492791 | 0.448456 | -0.25266 | 0.579219 |
| 28.61111 | 0.809337 | -0.42739 | -1.27019 | -1.58375 | -0.52385 | 0.921849 | 0.525646 | -0.15879 | -0.52019 | 0.746505 | -0.30273 | 1.115409 | 0.746048 | 0.786362 | 1.230384 | 1.003338 | 1.502312 | -0.47964 | 0.910273 |
| 28.625 | 0.73749 | -0.669 | -1.14998 | -1.08548 | -0.76914 | 0.909113 | 0.552201 | -0.00658 | -0.62853 | 1.344331 | 0.347817 | 1.758937 | 1.164273 | 1.170656 | 1.290545 | 1.061153 | 1.507516 | -0.74019 | 1.019855 |
| 28.63889 | 0.737016 | -0.64254 | -0.31437 | -0.5021 | -0.92923 | 0.646202 | 0.448192 | 0.015803 | -0.70178 | 1.868558 | 1.360172 | 1.2533 | 1.273552 | 0.96684 | 0.829242 | 0.742477 | 0.912988 | -0.57619 | 1.224188 |
| 28.65278 | 0.526584 | -0.53053 | 0.353063 | -0.15349 | -0.99939 | 0.174528 | 0.139041 | -0.00804 | -0.73679 | 2.2343 | 1.826768 | 0.530224 | 1.34248 | 0.767199 | 0.508159 | 0.61546 | 0.745815 | -0.17041 | 0.994326 |
| 28.66667 | 0.039736 | -0.38495 | 0.781745 | -0.23807 | -0.81876 | -0.45969 | -0.41483 | 0.069768 | -1.02228 | 1.566425 | 0.923586 | 0.1441 | 1.099386 | 0.679483 | 0.336857 | 0.499825 | 0.518421 | -0.10534 | 0.691471 |
| 28.68056 | -0.58051 | -0.30805 | 0.775272 | -0.77957 | -0.40286 | -1.13283 | -1.20556 | 0.139529 | -1.22499 | 0.678259 | -0.41048 | 0.173674 | 0.643642 | 0.385033 | 0.255968 | 0.209269 | 0.18244 | -0.17607 | 0.249798 |
| 28.69444 | -1.38267 | -0.26154 | 0.603998 | -1.3922 | 0.046273 | -1.55338 | -1.88226 | 0.115206 | -0.97884 | -0.34149 | -1.81066 | 0.123133 | 0.171682 | 0.026037 | 0.178609 | -0.10277 | -0.22041 | -0.28262 | 0.171111 |
| 28.70833 | -1.33616 | -0.22851 | 0.145835 | -1.06961 | 0.433047 | -1.39823 | -1.72422 | 0.036816 | -0.45596 | -1.23149 | -2.72885 | -0.14763 | -0.27382 | -0.30004 | -0.04168 | -0.19133 | -0.41197 | -0.31724 | -0.17946 |

| | | | | | | | | | | | | | | | | | | | |
|---|---|---|---|---|---|---|---|---|---|---|---|---|---|---|---|---|---|---|---|
| 28.72222 | -0.69464 | -0.21677 | -0.38698 | -0.01047 | 1.091496 | -0.58296 | -0.91518 | -0.12493 | -0.03078 | -1.45131 | -1.89114 | -0.33545 | -0.36273 | -0.34131 | 0.017249 | -0.0003 | -0.31969 | -0.356 | -0.0256 |
| 28.73611 | -0.29063 | -0.25233 | -0.47756 | 0.623601 | 1.582717 | -0.03623 | -0.02581 | -0.18529 | -0.09897 | -0.83456 | -0.38764 | -0.25951 | -0.20742 | -0.21687 | 0.177137 | 0.251684 | -0.12869 | -0.42974 | 0.540135 |
| 28.75 | 0.068056 | -0.3016 | -0.80812 | 0.861159 | 1.408593 | 0.256584 | 0.703447 | -0.16291 | -0.77908 | 0.113069 | 1.058306 | 0.045823 | 0.000756 | 0.03307 | 0.288042 | 0.37465 | -0.04024 | -0.49097 | 0.784185 |
| 28.76389 | 0.353442 | -0.38622 | -1.77736 | 0.478917 | 1.094014 | 0.287477 | 1.182162 | -0.2577 | -0.85409 | 1.14702 | 1.996485 | 0.239003 | 0.097842 | 0.197185 | 0.297435 | 0.368559 | -0.15551 | -0.44191 | 0.469461 |
| 28.77778 | 0.574233 | -0.43892 | -2.3088 | -0.0259 | 0.512425 | 0.144386 | 1.421652 | -0.39469 | -0.67228 | 1.755629 | 1.224854 | 0.29755 | -0.0456 | 0.2812 | 0.244975 | 0.293206 | -0.07644 | -0.50704 | 0.069082 |
| 28.79167 | 0.578617 | -0.67585 | -0.55494 | -0.54742 | -0.22442 | -0.08013 | 1.221011 | -0.33594 | -0.36417 | 1.441262 | 0.111254 | 0.23803 | -0.21909 | 0.257881 | 0.19728 | 0.117074 | 0.016511 | -0.53803 | -0.18715 |
| 28.80556 | 0.668052 | -0.90984 | 0.607786 | -0.84022 | -0.98949 | -0.2039 | 0.88083 | -0.07519 | -0.24709 | 0.58417 | -0.79261 | 0.162873 | -0.22133 | 0.325639 | 0.196849 | -0.00404 | 0.149367 | -0.42647 | -0.16632 |
| 28.81944 | 0.427882 | -0.92526 | 1.066932 | -0.81902 | -1.51297 | -0.27237 | 0.265687 | 0.18621 | -0.47876 | -0.20416 | -1.21037 | 0.071854 | -0.12115 | 0.264926 | 0.174016 | 0.037136 | 0.283623 | -0.32708 | 0.060888 |
| 28.83333 | -0.20685 | -0.74696 | 1.356553 | -0.78161 | -1.36342 | -0.31674 | -0.36344 | 0.399745 | -0.78539 | -0.85804 | -1.08061 | -0.06496 | -0.04063 | 0.062061 | 0.237341 | 0.13875 | 0.358844 | -0.28391 | 0.25567 |
| 28.84722 | -0.94264 | -0.80604 | 1.25314 | -1.32687 | -1.2429 | -0.38072 | -1.39998 | 0.413728 | -1.41131 | -1.13598 | -0.80719 | -0.21527 | -0.07275 | -0.25467 | 0.273265 | 0.161447 | 0.152335 | -0.27654 | 0.342346 |
| 28.86111 | -1.87886 | -1.08932 | -0.1538 | -1.90611 | -1.32604 | -0.50239 | -3.02484 | 0.22697 | -1.82565 | -1.14325 | -0.71867 | -0.32297 | -0.22896 | -0.16287 | 0.230776 | 0.099637 | -0.30379 | -0.32264 | 0.266121 |
| 28.875 | -2.20595 | -1.25252 | -1.48492 | -2.18889 | -0.99784 | -0.55929 | -3.4307 | -0.14448 | -2.21511 | -0.91075 | -0.8425 | -0.31364 | -0.31614 | -0.00919 | 0.133812 | -0.01388 | -0.66614 | -0.49836 | -0.01351 |
| 28.88889 | -1.70707 | -1.17047 | -2.84181 | -1.83478 | -0.61019 | -0.47684 | -2.05251 | -0.78848 | -2.32166 | -0.66725 | -0.9366 | -0.24122 | -0.28225 | -0.01297 | -0.05284 | -0.15095 | -0.83072 | -0.79652 | -0.41691 |
| 28.90278 | -0.95366 | -0.91713 | -2.90972 | -1.34921 | -0.39525 | -0.48539 | -0.83682 | -1.02947 | -2.02534 | -0.61058 | -0.98904 | -0.24861 | -0.23862 | -0.24745 | -0.19417 | -0.24039 | -0.81253 | -0.95143 | -0.71903 |
| 28.91667 | -0.17287 | -0.52756 | -1.73334 | -1.03206 | -0.37491 | -0.384 | -0.13505 | -1.05591 | -1.35389 | -0.82429 | -1.14876 | -0.37977 | -0.42766 | -0.40821 | -0.26485 | -0.25496 | -0.82474 | -0.83091 | -0.87655 |
| 28.93056 | 0.440124 | -0.14199 | -0.26429 | -0.40049 | -0.26564 | -0.16733 | 0.24066 | -0.85552 | -0.641 | -1.16509 | -1.34085 | -0.69655 | -0.70052 | -0.48358 | -0.2511 | -0.20774 | -0.81724 | -0.5474 | -0.99613 |
| 28.94444 | 0.674729 | 0.12867 | 0.966608 | 0.282355 | 0.067307 | 0.014373 | 0.438662 | -0.4995 | 0.192544 | -1.34694 | -1.57902 | -0.97647 | -0.74232 | -0.56082 | -0.18207 | -0.20795 | -0.64696 | -0.39628 | -0.9509 |
| 28.95833 | 0.657234 | 0.152272 | 2.167837 | 0.691371 | 0.500955 | 0.197006 | 0.569068 | -0.12591 | 1.028338 | -1.11539 | -1.49636 | -1.12796 | -0.49648 | -0.90137 | -0.17362 | -0.27525 | -0.54429 | -0.45485 | -0.79436 |
| 28.97222 | 0.298546 | 0.038277 | 2.976806 | 0.790728 | 0.649507 | 0.320924 | 0.637979 | 0.155308 | 1.846581 | -0.66372 | -1.2562 | -0.77496 | -0.20696 | -1.07757 | -0.25277 | -0.37284 | -0.67443 | -0.58738 | -0.68998 |
| 28.98611 | -0.446 | -0.50887 | 2.50858 | 0.006429 | 0.196962 | 0.131673 | 0.318047 | -0.20904 | 1.526117 | -0.2334 | -1.06262 | -0.23509 | 0.001989 | -0.82598 | -0.25532 | -0.37086 | -0.7309 | -0.6438 | -0.56625 |
| 29 | -1.87541 | -2.09074 | -0.18431 | -1.93703 | -0.88685 | -0.38247 | -0.57923 | -1.50125 | -0.78032 | -0.02976 | -0.77505 | 0.160668 | 0.054024 | -0.4579 | -0.12753 | -0.18552 | -0.5458 | -0.43688 | -0.38798 |
| 29.01389 | -3.59599 | -4.46033 | -3.17533 | -3.82768 | -1.94166 | -0.7687 | -1.33116 | -3.06173 | -3.3319 | -0.09325 | -0.40144 | 0.236501 | -0.09899 | -0.05248 | -0.00183 | -0.01112 | -0.34067 | -0.23247 | -0.10566 |
| 29.02778 | -3.2711 | -5.27429 | -5.23101 | -3.72662 | -1.7358 | -0.784 | -1.16502 | -2.22012 | -3.78285 | -0.22228 | -0.17657 | 0.004406 | -0.38636 | 0.139059 | 0.063354 | 0.006322 | -0.33702 | -0.14568 | 0.099468 |
| 29.04167 | -2.42911 | -4.59652 | -4.37221 | -2.39469 | -0.7793 | -0.80625 | -0.69687 | -1.41065 | -2.94407 | -0.29131 | -0.20555 | -0.27929 | -0.63783 | 0.188911 | 0.102404 | -0.06159 | -0.39085 | -0.28675 | 0.146693 |
| 29.05556 | -1.35519 | -3.06177 | -3.66743 | -1.29813 | -0.09881 | -0.45328 | -0.41443 | -0.84256 | -2.5454 | -0.32847 | -0.33696 | -0.29808 | -0.61473 | -0.0653 | 0.101073 | -0.07677 | -0.32524 | -0.62576 | 0.116472 |
| 29.06944 | -0.07961 | -1.13019 | -2.69774 | -0.28693 | 0.346897 | -0.04772 | -0.18665 | -0.35717 | -2.14653 | -0.40325 | -0.34669 | -0.02575 | -0.28044 | -0.1883 | 0.082145 | -0.03896 | -0.14141 | -0.9253 | 0.110176 |
| 29.08333 | 0.893812 | 0.379397 | -1.69021 | 0.346228 | 0.619217 | 0.215417 | 0.057881 | -0.04046 | -1.41244 | -0.47006 | -0.29781 | 0.226727 | 0.102927 | -0.2668 | -0.02077 | -0.04523 | 0.061288 | -1.04414 | 0.091713 |
| 29.09722 | 1.442331 | 1.078486 | -0.64284 | 0.764061 | 0.684213 | 0.263789 | 0.360107 | 0.216279 | -0.71664 | -0.83388 | -0.5605 | -0.17924 | -0.09706 | -0.56193 | -0.58876 | -0.54021 | -0.07144 | -1.63732 | -0.2749 |
| 29.11111 | 1.837036 | 1.449651 | 0.281955 | 1.155366 | 0.80586 | 0.031302 | 0.518198 | 0.759018 | -0.26465 | -2.23401 | -1.46304 | -1.56289 | -1.58673 | -1.11226 | -1.81387 | -1.7356 | -0.97139 | -3.98941 | -1.35297 |
| 29.125 | 2.197283 | 1.824912 | 1.008554 | 1.036708 | 0.893014 | 0.035301 | 0.383148 | 1.079772 | 0.00737 | -2.81954 | -2.08248 | -2.78859 | -2.32363 | -1.52725 | -2.22046 | -2.17934 | -1.77004 | -5.89098 | -1.929 |
| 29.13889 | 2.073826 | 1.523135 | 1.887859 | 0.711332 | 0.776022 | 0.149115 | 0.177702 | 1.585791 | 0.203594 | -1.42182 | -1.58022 | -2.15492 | -1.2982 | -1.13608 | -0.77865 | -0.90108 | -1.03919 | -3.8613 | -0.852 |
| 29.15278 | 1.842325 | 1.312064 | 3.303465 | 0.832107 | 0.553878 | 0.281835 | 0.475816 | 1.58948 | 0.285314 | -0.10038 | -0.62577 | -0.35859 | 0.326836 | -0.2449 | 0.320313 | 0.271565 | -0.18069 | -1.40424 | 0.04197 |
| 29.16667 | 1.342191 | 1.35385 | 4.133137 | 0.812808 | 0.433013 | 0.316325 | 0.756374 | 1.365981 | 0.396706 | 0.28987 | -0.12326 | 0.438725 | 1.201232 | 0.309868 | 0.43993 | 0.539701 | 0.232851 | -0.16586 | 0.327006 |
| 29.18056 | 0.932786 | 1.861597 | 4.224241 | 0.557611 | 0.22711 | 0.294587 | 1.090691 | 1.098638 | 0.576718 | 0.084866 | 0.436725 | 0.654263 | 1.184571 | 0.380316 | 0.325355 | 0.536419 | 0.20491 | 0.331965 | 0.458829 |
| 29.19444 | 0.616137 | 2.758613 | 3.733651 | 0.329131 | 0.063968 | 0.262983 | 1.105106 | 0.717518 | 0.756641 | -0.10308 | 0.787527 | 0.879779 | 1.206041 | 0.481131 | 0.304791 | 0.476679 | 0.159415 | 0.677198 | 0.622297 |
| 29.20833 | 0.530947 | 3.203053 | 3.336333 | 0.14021 | 0.003186 | 0.303737 | 0.553261 | 0.44111 | 0.899472 | -0.05002 | 0.707161 | 1.068217 | 1.087806 | 0.477943 | 0.157278 | 0.172218 | 0.070421 | 0.763346 | 0.57536 |
| 29.22222 | 0.44008 | 2.618281 | 2.517059 | 0.00969 | -0.04617 | 0.317612 | 0.264653 | 0.170435 | 0.933498 | 0.066446 | 0.393983 | 0.911002 | 0.645359 | 0.437524 | -0.05358 | -0.13809 | -0.03418 | 0.604144 | 0.224167 |
| 29.23611 | 0.319897 | 1.786614 | 1.256342 | -0.09759 | -0.07159 | 0.314304 | 0.18013 | -0.36492 | 0.906514 | 0.111035 | 0.345707 | 0.584189 | 0.316625 | 0.355887 | -0.12813 | -0.20336 | 0.022872 | 0.206669 | 0.008045 |
| 29.25 | 0.462051 | 1.313799 | 0.577418 | 0.044338 | -0.00774 | 0.431442 | 0.110003 | -0.20208 | 1.256463 | 0.168112 | 1.10495 | 0.412985 | 0.326145 | 0.398163 | 0.088535 | 0.081892 | 0.127539 | 0.493655 | 0.450076 |
| 29.26389 | 0.682118 | 1.164822 | 0.454225 | 0.471306 | 0.16846 | 0.517295 | 0.061586 | 0.226929 | 1.576506 | 0.350333 | 1.90876 | 0.379977 | 0.458595 | 0.465489 | 0.357334 | 0.3187 | 0.211052 | 0.545507 | 0.888142 |
| 29.27778 | 0.741054 | 0.95113 | 0.273802 | 0.798248 | 0.190445 | 0.476928 | 0.182089 | 0.32396 | 1.630278 | 0.57313 | 1.786886 | 0.355439 | 0.410131 | 0.449627 | 0.305581 | 0.2188 | 0.166928 | -0.09443 | 0.682962 |
| 29.29167 | 0.711158 | 0.625938 | -0.0136 | 0.794464 | 0.121871 | 0.327554 | 0.201475 | 0.14259 | 1.438503 | 0.646318 | 1.220572 | 0.323311 | 0.238932 | 0.291492 | 0.176667 | 0.054364 | 0.003467 | -0.64089 | 0.145437 |
| 29.30556 | 0.613446 | 0.378155 | -0.06215 | 0.604392 | 0.086135 | 0.208887 | 0.251671 | -0.0047 | 1.261552 | 0.489656 | 0.703703 | 0.221594 | 0.058323 | 0.021478 | 0.056815 | 0.027889 | -0.13181 | -0.98777 | -0.33056 |
| 29.31944 | 0.519569 | 0.181485 | -0.05126 | 0.30745 | 0.058455 | 0.112161 | 0.220769 | -0.08969 | 1.069137 | 0.345963 | 0.223264 | 0.012749 | -0.0733 | -0.18954 | -0.02531 | 0.023807 | -0.09029 | -1.20528 | -0.7169 |
| 29.33333 | 0.506917 | 0.133243 | -0.01299 | 0.095091 | 0.053079 | 0.135612 | 0.176424 | -0.0285 | 1.021643 | 0.277883 | -0.11133 | -0.21983 | -0.21141 | -0.20034 | -0.03264 | 0.027908 | 0.117953 | -1.33372 | -0.78786 |
| 29.34722 | 0.654007 | 0.365114 | 0.050044 | 0.162273 | -0.00543 | 0.1923 | 0.163169 | 0.166689 | 1.146122 | 0.270776 | -0.13403 | -0.32239 | -0.23779 | -0.09992 | 0.037391 | 0.037423 | 0.189138 | -1.07043 | -0.39273 |
| 29.36111 | 0.771209 | 0.672979 | 0.078876 | 0.318518 | -0.23014 | 0.153731 | 0.126452 | 0.247783 | 1.154736 | 0.384767 | 0.050325 | -0.09262 | -0.11047 | 0.147085 | 0.135372 | 0.05243 | 0.10254 | -0.57714 | 0.120768 |
| 29.375 | 0.636582 | 0.720668 | 0.110273 | 0.384858 | -0.53225 | 0.060669 | 0.13083 | 0.159598 | 1.120133 | 0.586422 | 0.153739 | 0.33102 | 0.052012 | 0.337698 | 0.1961 | 0.112481 | 0.116499 | -0.22918 | 0.382167 |
| 29.38889 | 0.271924 | 0.575162 | 0.310449 | 0.472111 | -0.64848 | 0.021067 | 0.271396 | 0.099145 | 1.036342 | 0.512346 | -0.06781 | 0.44114 | 0.11219 | 0.286445 | 0.134238 | 0.137898 | 0.166746 | -0.25423 | 0.160161 |
| 29.40278 | -0.07522 | 0.366737 | 0.671261 | 0.594793 | -0.49844 | 0.036484 | 0.521542 | 0.077564 | 0.917754 | 0.20636 | -0.4988 | 0.150333 | 0.139992 | 0.094973 | 0.019712 | 0.098779 | 0.134568 | -0.32699 | -0.39658 |
| 29.41667 | -0.24229 | 0.18229 | 0.93937 | 0.623653 | -0.27475 | 0.016028 | 0.729056 | 0.007114 | 0.828483 | 0.099388 | -0.67713 | -0.17854 | 0.191185 | -0.03001 | -0.02245 | 0.067389 | 0.185938 | -0.26952 | -0.72893 |
| 29.43056 | -0.10458 | 0.096716 | 0.869901 | 0.65413 | -0.12989 | 0.023976 | 0.733782 | -0.10521 | 0.698079 | 0.178367 | -0.41419 | -0.37533 | 0.182097 | 0.013235 | 0.010955 | 0.070764 | 0.207254 | -0.32253 | -0.54057 |
| 29.44444 | 0.032252 | 0.039666 | 0.564595 | 0.576432 | -0.1286 | 0.056255 | 0.493622 | -0.14684 | 0.399637 | 0.230813 | 0.057912 | -0.26951 | 0.126809 | 0.184473 | 0.090406 | 0.095627 | 0.115433 | -0.37017 | -0.04418 |
| 29.45833 | 0.226734 | -0.01819 | 0.317929 | 0.471019 | -0.14337 | 0.00179 | 0.236121 | -0.08623 | 0.18311 | 0.241182 | 0.444887 | 0.132098 | -0.02274 | 0.342979 | 0.138652 | 0.083474 | -0.04608 | -0.26573 | 0.328824 |
| 29.47222 | 0.515156 | 0.019733 | 0.366935 | 0.539498 | 0.03138 | -0.08652 | 0.201234 | 0.046596 | 0.257623 | 0.280364 | 0.608218 | 0.49135 | -0.19228 | 0.326953 | 0.107388 | 0.042342 | -0.11896 | -0.11459 | 0.406074 |
| 29.48611 | 0.840657 | 0.257618 | 0.72408 | 1.042478 | 0.632373 | 0.052522 | 0.517785 | 0.263229 | 0.809175 | 0.465639 | 0.685708 | 0.651189 | -0.09725 | 0.157133 | 0.06616 | 0.060816 | 0.067718 | 0.045961 | 0.328104 |
| 29.5 | 1.217292 | 0.625351 | 1.436536 | 2.008341 | 1.610478 | 0.379029 | 1.254894 | 0.676692 | 1.964811 | 0.715835 | 0.691565 | 0.679216 | 0.261635 | 0.074196 | 0.079799 | 0.119027 | 0.404015 | 0.077078 | 0.245034 |
| 29.51389 | 1.541517 | 0.81887 | 2.346854 | 2.603294 | 2.34411 | 0.822861 | 2.093017 | 0.9893 | 2.549159 | 0.818565 | 0.563444 | 0.560663 | 0.610158 | 0.058761 | 0.118691 | 0.142886 | 0.632508 | 0.026562 | 0.240555 |
| 29.52778 | 1.664897 | 0.863652 | 2.871531 | 2.767526 | 2.053342 | 1.006824 | 2.517167 | 1.100368 | 2.404458 | 0.762337 | 0.418602 | 0.371022 | 0.662682 | 0.170055 | 0.146467 | 0.125452 | 0.615117 | 0.011475 | 0.33539 |
| 29.54167 | 1.803777 | 0.909079 | 3.632305 | 2.576469 | 1.662379 | 0.800344 | 2.330648 | 0.883006 | 2.834189 | 0.653885 | 0.351128 | 0.180299 | 0.431194 | 0.343712 | 0.109904 | 0.067244 | 0.476889 | -0.0681 | 0.440473 |
| 29.55556 | 1.223963 | 0.841222 | 2.737848 | 1.841104 | 1.221357 | 0.650534 | 1.5848 | 0.982696 | 2.516954 | 0.587668 | 0.409972 | 0.126255 | 0.12955 | 0.423042 | 0.074599 | 0.003563 | 0.354404 | -0.22396 | 0.496462 |
| 29.56944 | 0.533968 | 0.806669 | 0.929239 | 1.107657 | 1.200971 | 0.475678 | 1.015014 | 0.754731 | 1.32537 | 0.64515 | 0.55895 | 0.269128 | -0.00568 | 0.403733 | 0.078773 | 0.031243 | 0.431974 | -0.30572 | 0.434783 |
| 29.58333 | -0.1486 | 0.829932 | -0.12182 | 0.593861 | 1.364943 | 0.431797 | 0.747828 | 0.096343 | 0.354979 | 0.812608 | 0.686284 | 0.586148 | 0.161364 | 0.413883 | 0.158055 | 0.159527 | 0.670355 | -0.24782 | 0.315586 |
| 29.59722 | -0.46996 | 0.704367 | -1.26559 | 0.083327 | 1.350785 | 0.403147 | 0.532357 | -0.45439 | -0.2507 | 1.222438 | 0.93886 | 0.893662 | 0.535848 | 0.563634 | 0.289682 | 0.341883 | 1.005012 | -0.32385 | 0.43773 |
| 29.61111 | -0.64992 | 0.446809 | -2.36963 | -0.05771 | 0.979588 | 0.478768 | 0.648269 | -0.23145 | -0.29194 | 1.878548 | 1.3793 | 1.042964 | 0.862953 | 0.952923 | 0.3972 | 0.47797 | 1.483295 | -0.91501 | 0.992329 |
| 29.625 | -0.76857 | -0.04215 | -2.26569 | -0.08374 | 0.596874 | 0.614013 | 0.721476 | 0.114043 | -0.3357 | 2.207571 | 1.671519 | 1.043687 | 0.916862 | 1.202409 | 0.327358 | 0.413967 | 1.376527 | -2.04212 | 1.109539 |
| 29.63889 | -0.88232 | -0.74725 | -2.08867 | 0.176155 | -0.00775 | -0.26153 | 0.708244 | 0.214614 | -0.53408 | 1.575321 | 1.64953 | 1.21554 | 0.921349 | 1.236369 | 0.117223 | 0.23184 | 0.953467 | -2.4979 | 1.088481 |
| 29.65278 | -0.63939 | -1.28504 | -1.14552 | 0.357861 | -0.57331 | -0.76182 | 0.466274 | 0.259854 | -0.46547 | 1.309168 | 1.492456 | 1.484046 | 1.028607 | 1.150153 | 0.014934 | 0.239136 | 0.760643 | -1.64836 | 1.318964 |
| 29.66667 | 0.064631 | -1.35903 | 0.575346 | 0.573122 | -0.98796 | -0.31553 | 0.192431 | 0.265104 | 0.038314 | 1.114211 | 1.047502 | 1.041067 | 0.716482 | 0.695002 | 0.003214 | 0.058554 | 0.398126 | -0.86316 | 0.971363 |
| 29.68056 | 0.66499 | -1.22005 | 1.647015 | 0.56977 | -1.32813 | -0.06847 | -0.11295 | 0.231016 | 0.206578 | 0.755515 | -0.02764 | 0.013483 | 0.378485 | 0.102333 | -0.09471 | -0.13564 | -0.09311 | -0.75193 | 0.461071 |
| 29.69444 | 0.760187 | -1.04017 | 2.200291 | 0.625338 | -1.62009 | 0.288705 | -0.4214 | 0.186497 | 0.351042 | 0.27905 | -1.3794 | -0.90405 | 0.058261 | -0.45024 | -0.12762 | -0.24341 | -0.39386 | -0.98801 | -0.15612 |
| 29.70833 | 0.634908 | -1.00292 | 2.029582 | 0.641894 | -1.746 | 0.743019 | -0.60905 | 0.10972 | 0.642103 | -0.03678 | -2.74232 | -1.31306 | -0.20614 | -0.76573 | -0.19572 | -0.25075 | -0.49225 | -1.16311 | -0.6629 |
| 29.72222 | 0.391044 | -0.82003 | 1.866052 | 0.846305 | -1.0965 | 0.452048 | -0.88722 | 0.032995 | 0.768424 | -0.06844 | -2.79978 | -1.29568 | -0.25928 | -0.84518 | -0.165 | -0.17433 | -0.3903 | -1.04355 | -0.69822 |
| 29.73611 | 0.151977 | -0.52614 | 1.254861 | 0.87111 | -0.14718 | 0.001321 | -1.11851 | 0.0091 | 0.467437 | -0.04886 | -1.21631 | -0.93005 | -0.09057 | -0.66268 | -0.05717 | -0.02612 | -0.43474 | -0.75932 | -0.32801 |
| 29.75 | -0.24643 | -0.27991 | 0.602617 | 0.621048 | 0.681934 | -0.00443 | -1.09157 | -0.00929 | -0.23459 | 0.044512 | 0.195343 | -0.24592 | 0.10013 | -0.34752 | 0.043865 | 0.068184 | -0.67559 | -0.49984 | 0.103521 |
| 29.76389 | -0.70439 | -0.4095 | -0.2026 | 0.242795 | 1.364726 | 0.055661 | -0.8681 | -0.14787 | -0.4767 | 0.340684 | 1.303688 | 0.293165 | 0.131029 | 0.025477 | -0.02276 | -0.0491 | -0.91583 | -0.30923 | 0.246141 |
| 29.77778 | -1.20909 | -0.95464 | -1.38404 | -0.26198 | 1.447421 | -0.17761 | -0.66724 | -0.60857 | -0.78962 | 0.578282 | 1.534686 | 0.576447 | -0.00356 | 0.286487 | -0.17259 | -0.21085 | -0.78941 | -0.01913 | 0.01942 |
| 29.79167 | -1.83838 | -1.52319 | -2.00347 | -0.64636 | 1.245802 | -0.67276 | -0.25497 | -1.38573 | -0.921 | 0.583111 | 0.85256 | 0.669516 | -0.14571 | 0.375809 | -0.2022 | -0.25191 | -0.17813 | 0.170383 | -0.272 |
| 29.80556 | -1.90534 | -1.79973 | -2.75493 | -0.942 | 0.831013 | -0.1835 | -0.06822 | -1.78943 | -0.78908 | 0.394761 | -0.17212 | 0.596644 | -0.17535 | 0.34126 | -0.10326 | -0.05895 | 0.540325 | 0.13868 | -0.26961 |
| 29.81944 | -1.70587 | -1.74229 | -2.36102 | -1.06642 | 0.134493 | -0.1982 | 0.027709 | -1.48498 | -1.02262 | 0.250139 | -1.04445 | 0.452433 | -0.00856 | 0.317995 | 0.189926 | 0.181733 | 0.979545 | -0.08129 | 0.032224 |
| 29.83333 | -1.1248 | -1.4467 | -1.77372 | -1.00425 | -0.69366 | -0.21686 | 0.053583 | -1.1172 | -1.00109 | 0.161755 | -1.38196 | 0.219241 | 0.162652 | 0.21908 | 0.294234 | 0.270371 | 0.878682 | -0.37538 | 0.297024 |
| 29.84722 | -0.35182 | -1.10352 | -1.24334 | -0.87427 | -1.30049 | -0.06638 | -0.18344 | -0.78801 | -0.48711 | 0.074364 | -1.20484 | -0.04927 | 0.233598 | -0.04923 | 0.168244 | 0.212294 | 0.524008 | -0.57197 | 0.303034 |
| 29.86111 | 0.026275 | -0.9054 | -0.514 | -0.95063 | -1.6675 | 0.157649 | -0.53015 | -0.54881 | 0.044107 | -0.06862 | -0.97776 | -0.30237 | 0.191064 | -0.3645 | -0.03951 | 0.01318 | 0.175932 | -0.65295 | 0.067813 |
| 29.875 | 0.202441 | -0.68422 | -0.53015 | -1.08052 | -1.44362 | 0.108335 | -0.79382 | -0.37535 | 0.473378 | -0.27006 | -1.0376 | -0.53424 | 0.108077 | -0.37551 | -0.17461 | -0.25134 | -0.06309 | -0.65528 | -0.44042 |
| 29.88889 | 0.367035 | -0.39195 | -0.55608 | -1.06245 | -1.85435 | 0.085391 | -0.76302 | 0.003897 | 0.512056 | -0.65021 | -1.41716 | -0.65647 | -0.13823 | -0.13498 | -0.13129 | -0.34916 | -0.26866 | -0.49023 | -1.02947 |
| 29.90278 | 0.499873 | -0.27301 | 0.029519 | -0.7387 | -2.01203 | -0.03417 | -0.15994 | 0.490319 | 0.625692 | -1.03403 | -1.88767 | -0.57444 | -0.51403 | -0.01975 | -0.03069 | -0.28038 | -0.49461 | -0.18453 | -1.16092 |

| | | | | | | | | | | | | | | | | | | | |
|---|---|---|---|---|---|---|---|---|---|---|---|---|---|---|---|---|---|---|---|
| 29.91667 | 0.529806 | -0.09812 | 0.838149 | -0.17145 | -1.06883 | -0.29511 | 0.362793 | 0.644242 | 0.815734 | -1.21392 | -2.11935 | -0.42807 | -0.95181 | -0.1245 | -0.01669 | -0.18523 | -0.86302 | 0.078258 | -0.93283 |
| 29.93056 | 0.32313 | -0.03088 | 1.553535 | 0.11199 | -0.41404 | -0.24984 | 0.72623 | 0.559949 | 0.676518 | -1.14505 | -2.22663 | -0.40023 | -1.03411 | -0.32855 | -0.09355 | -0.07734 | -1.24559 | 0.258148 | -0.53562 |
| 29.94444 | 0.041769 | 0.158069 | 1.912976 | 0.306452 | 0.138593 | 0.05315 | 0.915106 | 0.386022 | 0.429989 | -0.96706 | -2.15349 | -0.57422 | -0.87449 | -0.60118 | -0.11749 | -0.0496 | -1.45927 | 0.351042 | -0.30468 |
| 29.95833 | -0.13643 | 0.293745 | 1.794273 | 0.425156 | 0.502523 | 0.131981 | 0.980231 | 0.189299 | 0.270568 | -0.84932 | -1.91353 | -0.85651 | -0.62595 | -0.94696 | -0.23085 | -0.11592 | -1.44281 | 0.414482 | -0.23489 |
| 29.97222 | -0.18011 | 0.22372 | 1.359935 | 0.35648 | 0.509939 | 0.219124 | 0.936771 | -0.17962 | -0.25277 | -0.71931 | -1.57046 | -0.88619 | -0.25883 | -1.16256 | -0.17334 | -0.13314 | -1.07169 | 0.38981 | -0.10359 |
| 29.98611 | -0.63489 | -0.53127 | 0.336424 | -0.42491 | -0.12706 | 0.014331 | 0.407658 | -0.96014 | -1.34004 | -0.52602 | -1.09808 | -0.61535 | 0.084023 | -1.0706 | 0.03011 | -0.00479 | -0.58092 | 0.3896 | 0.087034 |
| 30 | -2.04102 | -2.38446 | -1.58848 | -1.88313 | -1.49112 | -0.88142 | -0.80938 | -2.43368 | -3.90258 | -0.27921 | -0.69033 | -0.23836 | 0.240781 | -0.71532 | 0.242747 | 0.15403 | -0.28408 | 0.31897 | 0.209583 |
| 30.01389 | -3.50422 | -3.97927 | -4.13502 | -2.85302 | -2.45636 | -1.14642 | -2.0456 | -2.39902 | -5.32467 | 0.051523 | -0.33162 | 0.08168 | 0.275903 | -0.30583 | 0.33091 | 0.307195 | -0.203 | 0.161893 | 0.217257 |
| 30.02778 | -3.43169 | -3.82009 | -5.84428 | -3.31 | -1.68164 | -1.27819 | -3.24636 | -1.41055 | -4.52892 | 0.377735 | -0.07233 | 0.231253 | 0.236368 | 0.028282 | 0.281274 | 0.392204 | -0.16319 | -0.00947 | 0.154945 |
| 30.04167 | -2.62815 | -2.57974 | -4.8494 | -2.28091 | -1.22969 | -1.01172 | -2.94684 | -0.41892 | -3.07959 | 0.506212 | 0.030226 | 0.127539 | 0.145246 | 0.168899 | 0.163852 | 0.342736 | -0.12573 | -0.07303 | 0.060735 |
| 30.05556 | -2.052 | -2.15106 | -3.45758 | -0.87183 | -0.68438 | -0.55233 | -1.38273 | -0.00268 | -2.08023 | 0.402152 | -0.00458 | -0.07292 | 0.050984 | 0.084341 | 0.081632 | 0.19356 | -0.04881 | -0.01468 | -0.01671 |
| 30.06944 | -1.5028 | -1.82179 | -1.88076 | -0.41591 | -0.25988 | -0.22872 | 0.045209 | -0.06502 | -1.70645 | 0.167988 | -0.01596 | -0.22244 | -0.02592 | -0.02534 | 0.045165 | 0.06043 | 0.207088 | 0.113927 | -0.02919 |
| 30.08333 | -0.42637 | -0.77305 | -0.31092 | 0.434774 | 0.033501 | 0.097528 | 1.457753 | -0.17885 | -1.29397 | -0.03567 | 0.001301 | -0.24589 | -0.07624 | -0.03608 | 0.047663 | -0.04374 | 0.398456 | 0.199785 | -0.0134 |
| 30.09722 | 0.606507 | 0.295118 | 0.388174 | 1.177111 | 0.298983 | 0.382391 | 2.116616 | -0.18364 | -0.74544 | -0.40234 | -0.28043 | -0.72297 | -0.25049 | -0.27654 | 0.051842 | -0.40521 | -0.00322 | -0.32828 | -0.2247 |
| 30.11111 | 1.57638 | 1.169479 | 0.417054 | 1.609554 | 0.563469 | 0.599674 | 2.242152 | -0.03337 | -0.04074 | -1.4118 | -1.16077 | -2.08437 | -0.81448 | -1.22438 | -0.11549 | -1.27417 | -1.482 | -1.53209 | -0.90465 |
| 30.125 | 2.22207 | 1.828268 | 0.508948 | 1.818965 | 0.769755 | 0.682379 | 2.233018 | 0.109317 | 0.692229 | -2.14388 | -1.72356 | -3.24392 | -1.53194 | -2.28081 | -0.46293 | -1.80803 | -2.82083 | -1.99937 | -1.43579 |
| 30.13889 | 1.989351 | 1.794003 | 0.657163 | 1.507571 | 0.595532 | 0.600411 | 1.970211 | 0.07247 | 0.996667 | -1.12722 | -1.22449 | -2.4695 | -1.19296 | -1.79923 | -0.12086 | -1.20528 | -1.727 | -0.54989 | -0.56285 |
| 30.15278 | 1.855956 | 1.676969 | 0.84205 | 0.981277 | 0.833862 | 0.464613 | 1.362054 | -0.04952 | 0.709605 | 0.128116 | -0.39504 | -0.75332 | -0.249 | -0.55615 | 0.190192 | -0.33401 | -0.45008 | 0.894873 | 0.519371 |
| 30.16667 | 2.41273 | 1.378586 | 1.066193 | 0.451986 | 0.951397 | 0.733107 | 0.776675 | 0.102751 | 0.784873 | 0.571872 | 0.025047 | 0.296976 | 0.449646 | 0.267605 | 0.373877 | 0.382011 | 0.24622 | 1.269709 | 0.725419 |
| 30.18056 | 2.254738 | 0.396615 | 1.422982 | -0.50029 | 0.975056 | 1.085511 | 0.08221 | 0.545382 | 1.617136 | 0.413866 | 0.125216 | 0.887827 | 0.846853 | 0.660971 | 0.393065 | 0.763916 | 0.469269 | 0.816581 | 0.438078 |
| 30.19444 | 2.091375 | 0.039262 | 1.941568 | -0.487 | 0.857242 | 1.138308 | -0.08917 | 0.882316 | 1.869804 | 0.145811 | 0.173885 | 1.292866 | 1.013213 | 0.826888 | 0.482181 | 0.891145 | 0.631941 | 0.387829 | 0.219605 |
| 30.20833 | 2.319783 | 0.45738 | 2.665435 | 0.490121 | 0.643661 | 0.820091 | 0.18955 | 1.110518 | 1.899858 | -0.10171 | 0.170181 | 1.525409 | 0.888089 | 0.80655 | 0.59734 | 0.782498 | 0.715234 | 0.16824 | 0.131814 |
| 30.22222 | 2.041709 | 1.156042 | 3.159423 | 1.182381 | 0.544243 | 0.794201 | 0.459602 | 1.333636 | 1.563888 | -0.11206 | 0.127724 | 1.685368 | 0.658462 | 0.773721 | 0.531072 | 0.64358 | 0.678144 | 0.097934 | 0.119102 |
| 30.23611 | 1.795788 | 1.620494 | 3.214287 | 1.409302 | 0.411315 | 0.97342 | 0.880023 | 1.363061 | 1.333601 | 0.140599 | 0.254716 | 2.139221 | 0.638478 | 0.870344 | 0.450777 | 0.444845 | 0.676198 | 0.117414 | 0.109254 |
| 30.25 | 1.377258 | 1.900395 | 2.845568 | 1.227012 | 0.308154 | 0.88283 | 1.191352 | 1.280051 | 1.343577 | 0.319238 | 0.443278 | 2.627791 | 0.634831 | 0.913741 | 0.30424 | 0.257204 | 0.669781 | 0.115319 | 0.088214 |
| 30.26389 | 0.554736 | 1.901184 | 2.26019 | 0.762976 | 0.30323 | 0.763044 | 1.106939 | 1.090285 | 1.721339 | 0.143438 | 0.557773 | 2.475316 | 0.621971 | 0.900159 | 0.092873 | 0.092425 | 0.519414 | 0.087841 | 0.135599 |
| 30.27778 | -0.18755 | 1.842422 | 1.858614 | 0.258504 | 0.376677 | 0.702483 | 0.833453 | 0.805162 | 2.618805 | -0.13836 | 0.527415 | 1.999744 | 0.511166 | 0.828549 | -0.08793 | 0.124865 | 0.361969 | 0.101376 | 0.231481 |
| 30.29167 | -0.19299 | 1.440255 | 1.965897 | 0.25002 | 0.585349 | 0.678589 | 0.970463 | 0.655893 | 3.092043 | -0.19068 | 0.566592 | 1.170753 | 0.33564 | 0.706868 | -0.08893 | 0.289798 | 0.292287 | 0.189481 | 0.246821 |
| 30.30556 | 0.451132 | 1.013894 | 1.700633 | 0.638058 | 0.798048 | 0.633538 | 1.308338 | 0.933669 | 2.626886 | 0.043544 | 0.708398 | 0.40186 | 0.187197 | 0.469235 | -0.02659 | 0.3676 | 0.148992 | 0.288132 | 0.130009 |
| 30.31944 | 0.962266 | 0.860461 | 1.537675 | 0.919989 | 0.962328 | 0.643432 | 1.498462 | 1.325162 | 2.752783 | 0.436257 | 1.104662 | -0.15262 | 0.096727 | 0.202977 | -0.071 | 0.235781 | -0.05209 | 0.354504 | -0.03983 |
| 30.33333 | 1.077052 | 0.852981 | 0.835485 | 0.845517 | 1.066899 | 0.537615 | 1.038575 | 1.31979 | 2.163701 | 0.755481 | 1.393716 | -0.164 | 0.114344 | 0.006514 | -0.0677 | 0.110895 | -0.15526 | 0.382038 | -0.0636 |
| 30.34722 | 0.790903 | 0.759901 | -0.20315 | 0.621382 | 0.943688 | 0.180524 | 0.332895 | 0.632889 | 0.553615 | 0.780518 | 1.38618 | 0.16332 | 0.199177 | -0.01975 | 0.062121 | 0.113938 | -0.02965 | 0.303981 | -0.01934 |
| 30.36111 | 0.36617 | 0.411164 | -0.75112 | 0.394458 | 0.673375 | -0.04067 | -0.09259 | -0.14745 | -0.49698 | 0.555878 | 0.887212 | 0.562465 | 0.26282 | 0.060869 | 0.215219 | 0.265004 | 0.214235 | 0.167932 | -0.06881 |
| 30.375 | 0.380171 | 0.12296 | -0.52515 | 0.387061 | 0.352481 | 0.071379 | -0.03578 | -0.41118 | -0.28968 | 0.448893 | 0.491594 | 0.789225 | 0.31089 | 0.167811 | 0.246225 | 0.471951 | 0.484916 | 0.098557 | -0.09574 |
| 30.38889 | 1.023584 | 0.057434 | -0.1636 | 0.679111 | 0.142792 | 0.210272 | 0.331949 | -0.1328 | 0.41658 | 0.888725 | 0.602664 | 0.760878 | 0.437781 | 0.237361 | 0.247401 | 0.69276 | 0.878382 | 0.173117 | 0.065529 |
| 30.40278 | 1.773742 | 0.243778 | 0.116844 | 1.115963 | 0.1343 | 0.356728 | 0.641734 | 0.095942 | 1.069406 | 1.449094 | 0.830646 | 0.357171 | 0.409265 | 0.149395 | 0.313101 | 0.748182 | 0.990518 | 0.271915 | 0.305245 |
| 30.41667 | 2.234164 | 0.499216 | 0.304679 | 1.492219 | 0.184148 | 0.397225 | 0.727291 | 0.121636 | 1.593826 | 1.354284 | 0.772948 | 0.046583 | 0.460024 | 0.43273 | 0.373492 | 0.603377 | 0.656339 | 0.288689 | 0.470818 |
| 30.43056 | 2.059713 | 0.60695 | 0.235465 | 1.507456 | 0.168367 | 0.255676 | 0.596647 | -0.01109 | 1.277317 | 1.155281 | 0.402395 | 0.297164 | 0.586804 | 0.672366 | 0.298743 | 0.45481 | 0.455597 | 0.265118 | 0.554113 |
| 30.44444 | 1.575255 | 0.609082 | 0.094565 | 1.265986 | 0.04566 | 0.071732 | 0.39662 | -0.2845 | 0.236062 | 0.57404 | -0.35954 | 0.687621 | 0.696221 | 0.640431 | 0.088359 | 0.226121 | 0.38162 | 0.219511 | 0.427601 |
| 30.45833 | 1.082831 | 0.71364 | 0.095874 | 1.022655 | -0.19843 | 0.048931 | 0.268698 | -0.41682 | -0.77747 | 0.225701 | -0.72101 | 0.970815 | 0.571319 | 0.447765 | -0.06905 | 0.012458 | 0.279184 | 0.155382 | 0.190993 |
| 30.47222 | 0.893681 | 0.979219 | 0.207924 | 0.957427 | -0.28896 | 0.041782 | 0.27126 | -0.3502 | -1.35701 | 0.39926 | -0.41546 | 1.046744 | 0.313044 | 0.344257 | -0.00219 | -0.01818 | 0.167604 | 0.090221 | 0.074647 |
| 30.48611 | 0.867941 | 1.100717 | 0.46228 | 1.038251 | 0.119807 | -0.04469 | 0.429473 | -0.06761 | -0.85575 | 0.80866 | 0.385546 | 0.58439 | 0.061122 | 0.364438 | 0.145576 | 0.045248 | 0.115446 | 0.038455 | 0.074415 |
| 30.5 | 1.514239 | 1.212737 | 1.155202 | 1.721516 | 1.219365 | -0.02262 | 0.88131 | 0.409155 | 0.47926 | 1.037164 | 1.185636 | 0.16781 | -0.05247 | 0.402144 | 0.254405 | 0.188815 | 0.192998 | 0.030615 | 0.197599 |
| 30.51389 | 2.508045 | 1.285934 | 2.317647 | 2.643606 | 2.52922 | 0.252192 | 1.712915 | 1.012763 | 1.572271 | 0.738911 | 1.499589 | -0.18451 | 0.079259 | 0.408215 | 0.301974 | 0.335864 | 0.337152 | 0.074803 | 0.515135 |
| 30.52778 | 3.206381 | 1.517444 | 3.884331 | 3.061644 | 2.701469 | 0.477314 | 2.496759 | 1.643401 | 2.052809 | 0.422484 | 1.408404 | -0.2128 | 0.274871 | 0.365137 | 0.187651 | 0.297189 | 0.380025 | 0.094809 | 0.856255 |
| 30.54167 | 2.893307 | 2.021494 | 4.301991 | 2.67587 | 2.20816 | 0.295726 | 2.354091 | 2.333832 | 2.275819 | 0.183512 | 1.064741 | 0.161775 | 0.4543 | 0.34364 | 0.022756 | 0.124193 | 0.303765 | 0.04338 | 0.928687 |
| 30.55556 | 2.445659 | 1.838943 | 3.212917 | 1.673411 | 2.005351 | 0.594871 | 1.605754 | 1.842627 | 2.017377 | 0.24834 | 0.780765 | 0.721897 | 0.505587 | 0.383978 | 0.015067 | 0.03449 | 0.159882 | -0.00995 | 0.70164 |
| 30.56944 | 1.030323 | 1.397214 | 1.918875 | 0.995985 | 1.756557 | 0.653319 | 0.83351 | 1.47532 | 1.616886 | 0.524682 | 0.551215 | 1.163619 | 0.415782 | 0.505276 | 0.161175 | 0.107843 | 0.08007 | -0.0046 | 0.301967 |
| 30.58333 | -0.3839 | 1.088952 | 0.750314 | 0.382603 | 1.860512 | 0.645132 | 0.058995 | 1.311309 | 1.632887 | 0.626771 | 0.463821 | 1.080063 | 0.227386 | 0.550224 | 0.303534 | 0.2369 | 0.18399 | 0.020634 | -0.01321 |
| 30.59722 | -1.1438 | 0.885343 | -0.08231 | 0.045814 | 1.868294 | 0.628043 | -0.48618 | 0.890712 | 0.785192 | 0.639641 | 0.724813 | 0.755354 | 0.033158 | 0.476942 | 0.220087 | 0.287524 | 0.446672 | -0.0295 | -0.0846 |
| 30.61111 | -0.99398 | 0.771558 | -0.79672 | -0.11243 | 1.627098 | 0.549123 | -0.51618 | 0.362948 | -0.49626 | 0.861838 | 1.464017 | 0.579247 | -0.02702 | 0.422305 | 0.072262 | 0.296286 | 0.756952 | -0.15939 | 0.198223 |
| 30.625 | -1.22073 | 0.505127 | -1.52401 | -0.06081 | 1.185898 | 0.30724 | -0.18734 | -0.18115 | -1.06376 | 1.023591 | 1.852934 | 0.86673 | 0.245119 | 0.578926 | 0.186814 | 0.444246 | 1.20029 | -0.51512 | 0.870139 |
| 30.63889 | -1.28235 | -0.11186 | -1.99306 | -0.1852 | 0.520261 | 0.014561 | -0.02055 | -0.84934 | -1.17727 | 1.172773 | 1.616923 | 1.323571 | 0.753404 | 0.91364 | 0.510512 | 0.786664 | 1.338591 | -0.92769 | 1.591526 |
| 30.65278 | -0.96627 | -0.84354 | -1.76406 | -0.37242 | -0.19265 | -0.40483 | 0.064214 | -1.15581 | -1.16012 | 1.372556 | 1.191121 | 1.68636 | 1.102015 | 1.146104 | 0.656188 | 1.095959 | 1.277842 | -0.69896 | 1.841748 |
| 30.66667 | 0.673893 | -1.11258 | -2.32458 | -0.38857 | -0.86243 | -0.6117 | -0.0363 | -1.18175 | -1.45337 | 1.28453 | 0.550251 | 1.382365 | 1.016907 | 1.010961 | 0.527961 | 0.898313 | 0.990655 | -0.34144 | 1.25724 |
| 30.68056 | 1.359589 | -1.0503 | -2.04788 | -0.31342 | -1.39857 | -0.56815 | -0.17398 | -1.13653 | -0.97666 | 0.841306 | -0.11463 | 0.624544 | 0.619132 | 0.59486 | 0.058713 | 0.187029 | 0.533891 | -0.12624 | 0.193318 |
| 30.69444 | 1.081754 | -0.69345 | -0.10241 | -0.01296 | -1.39094 | -0.54221 | -0.21585 | -1.02908 | 0.198437 | 0.662297 | -0.74081 | -0.26212 | 0.256984 | 0.112278 | -0.42804 | -0.59524 | 0.081938 | -0.10591 | -0.8437 |
| 30.70833 | 1.025975 | -0.24223 | 1.116792 | 0.402948 | -1.50136 | -0.74191 | -0.16163 | -1.22835 | 1.082715 | 0.394502 | -1.33538 | -0.9858 | 0.06196 | -0.17833 | -0.55487 | -0.97117 | -0.29767 | -0.14693 | -1.36203 |
| 30.72222 | 0.867142 | -0.13767 | 1.643589 | 0.766201 | -1.86249 | -0.4761 | 0.010342 | -1.55013 | 1.572241 | -0.01162 | -1.95451 | -1.46988 | -0.01776 | -0.31025 | -0.50128 | -0.95099 | -0.38481 | -0.04446 | -1.68422 |
| 30.73611 | 0.409464 | -0.40467 | 1.242355 | 1.075551 | -1.88113 | -0.03015 | 0.178982 | -1.57125 | 2.115943 | 0.081459 | -2.05584 | -1.11179 | -0.08507 | -0.26131 | -0.22566 | -0.60453 | -0.26548 | 0.061546 | -1.66767 |
| 30.75 | -0.15437 | -0.70418 | 0.20293 | 1.382212 | -1.31318 | -0.05378 | 0.306772 | -1.01448 | 1.19082 | 0.577564 | -1.68908 | -0.28035 | -0.18601 | -0.37571 | 0.045515 | -0.14694 | -0.13921 | 0.23044 | -0.82607 |
| 30.76389 | -0.34166 | -0.82583 | -0.14377 | 1.209171 | -0.68838 | 0.041269 | 0.286942 | -0.74435 | 0.446189 | 0.808312 | -1.14441 | 0.442799 | -0.29414 | -0.50723 | 0.075979 | -0.00402 | -0.10741 | 0.4283 | 0.106231 |
| 30.77778 | -0.22334 | -0.80678 | -0.53965 | 0.771005 | -0.11147 | 0.069513 | 0.147881 | -0.59799 | -0.0917 | 0.521127 | -0.44747 | 0.958553 | -0.29815 | -0.48684 | -0.05236 | -0.18822 | -0.10173 | 0.557604 | 0.612054 |
| 30.79167 | 0.188454 | -0.68854 | -0.59809 | 0.105116 | 0.407455 | 0.066123 | -0.18684 | -0.44007 | -0.61878 | 0.001322 | 0.1396 | 1.026179 | -0.30178 | -0.31007 | -0.18383 | -0.40375 | -0.08125 | 0.50588 | 0.496189 |
| 30.80556 | 0.201074 | -0.49416 | -0.64217 | -0.59221 | 0.726612 | 0.229129 | -0.52445 | -0.25264 | -1.35649 | -0.30055 | 0.297784 | 0.790526 | -0.22845 | -0.07134 | -0.22257 | -0.36719 | -0.14953 | 0.395973 | 0.027925 |
| 30.81944 | -0.46449 | -0.42928 | -0.70733 | -0.93836 | 0.71488 | 0.662273 | -0.84535 | 0.135464 | -1.67982 | -0.3368 | 0.0958 | 0.521936 | -0.05792 | 0.18173 | -0.1527 | -0.21101 | -0.23462 | 0.256848 | -0.34383 |
| 30.83333 | -0.79533 | -0.7688 | -1.81319 | -1.31412 | 0.355645 | 0.756309 | -0.76944 | 0.166484 | -1.42304 | -0.07864 | 0.140996 | 0.166648 | 0.13228 | 0.335995 | -0.11225 | -0.12396 | -0.2651 | 0.149855 | -0.40786 |
| 30.84722 | -0.82779 | -1.3556 | -3.38776 | -1.5889 | -0.25352 | 0.28156 | -1.05472 | 0.060363 | -1.09174 | 0.417524 | 0.222873 | -0.32903 | 0.239886 | 0.313716 | -0.112 | -0.0974 | -0.21924 | 0.090906 | -0.3047 |
| 30.86111 | -0.91216 | -1.84699 | -4.04707 | -1.61835 | -0.96817 | 0.008663 | -1.29575 | -0.11293 | -0.90015 | 0.69439 | 0.068595 | -0.81634 | 0.194898 | 0.182397 | -0.1002 | -0.03462 | -0.10357 | 0.09405 | -0.17801 |
| 30.875 | -0.7271 | -1.65553 | -3.79227 | -1.52778 | -1.65122 | -0.02678 | -1.35909 | -0.19529 | -0.97697 | 0.712399 | -0.17456 | -0.92294 | 0.094255 | 0.01791 | -0.06763 | 0.048785 | 0.018314 | 0.142768 | -0.10214 |
| 30.88889 | -0.3488 | -1.2094 | -3.27238 | -1.3643 | -1.90405 | -0.20669 | -1.27575 | -0.45855 | -1.05595 | 0.241533 | -0.51676 | -0.9167 | -0.07386 | -0.19316 | -0.06213 | 0.01029 | 0.139124 | 0.167328 | -0.23371 |
| 30.90278 | 0.286297 | -0.6617 | -2.54589 | -1.07679 | -1.84824 | -0.45387 | -0.78795 | -0.8211 | -0.76872 | -0.28465 | -1.14036 | -0.96991 | -0.34879 | -0.43563 | -0.0865 | -0.16115 | 0.056393 | 0.172464 | -0.44032 |
| 30.91667 | 1.038441 | 0.168626 | -1.55852 | -0.24643 | -1.42726 | -0.45412 | 0.003793 | -0.89096 | -0.40511 | -0.65949 | -1.72449 | -1.01638 | -0.65377 | -0.7143 | -0.15543 | -0.38299 | -0.10875 | 0.173284 | -0.39132 |
| 30.93056 | 1.994213 | 0.264544 | -0.31552 | 0.451095 | -0.83316 | -0.35341 | 0.980509 | -0.63775 | 0.201499 | -0.58913 | -1.97272 | -0.85408 | -0.84822 | -0.75343 | -0.13486 | -0.43019 | -0.31203 | 0.225438 | -0.0696 |
| 30.94444 | 2.706017 | 0.195878 | 0.971204 | 0.899709 | -0.23253 | -0.27892 | 1.825737 | -0.38887 | 0.790779 | -0.4204 | -1.90028 | -0.75662 | -0.84144 | -0.60655 | -0.18199 | -0.40927 | -0.42806 | 0.275061 | 0.206219 |
| 30.95833 | 2.838253 | 0.357329 | 1.954419 | 0.881362 | 0.318197 | -0.23005 | 2.200109 | -0.4608 | 1.024024 | -0.51205 | -1.93362 | -0.89249 | -0.72043 | -0.44517 | -0.20778 | -0.36001 | -0.57923 | 0.363931 | 0.244068 |
| 30.97222 | 2.404517 | 0.494662 | 2.484465 | 0.83912 | 0.6667 | -0.21584 | 1.925898 | -0.56242 | 1.090984 | -0.72881 | -1.80848 | -0.85373 | -0.47553 | -0.32304 | -0.16404 | -0.21048 | -0.77522 | 0.443471 | 0.206806 |
| 30.98611 | 1.078559 | 0.228627 | 2.069025 | 0.711596 | 0.2235 | -0.5588 | 1.217976 | -0.57988 | 0.502032 | -1.11713 | -1.36387 | -0.57198 | -0.22651 | -0.24254 | 0.061349 | 0.091715 | -0.80175 | 0.439818 | 0.209405 |
| 31 | -1.60078 | -0.64537 | 0.122079 | -0.44411 | -1.1674 | -1.10723 | -0.26678 | -1.13793 | -0.77441 | -1.35257 | -0.96684 | -0.28845 | -0.07002 | -0.15552 | 0.290544 | 0.409965 | -0.79255 | 0.328967 | 0.283989 |
| 31.01389 | -3.71382 | -1.12834 | -2.43971 | -1.76097 | -2.21174 | -1.06923 | -1.87508 | -1.00686 | -1.86554 | -1.17482 | -0.70967 | -0.16593 | 0.001348 | -0.02334 | 0.448888 | 0.508732 | -0.72705 | 0.242379 | 0.350863 |
| 31.02778 | -3.42325 | -0.89762 | -3.51961 | -2.07442 | -2.0641 | -0.31758 | -3.26452 | -0.21935 | -1.98737 | -0.69148 | -0.42877 | -0.03768 | 0.034269 | 0.146741 | 0.432274 | 0.434076 | -0.56356 | 0.218882 | 0.299924 |
| 31.04167 | -2.54776 | -1.04821 | -3.30409 | -1.71635 | -1.36425 | 0.136014 | -4.18698 | 0.47461 | -1.8993 | -0.22186 | -0.25706 | 0.016618 | 0.024648 | 0.269212 | 0.268886 | 0.139361 | -0.359 | 0.211471 | 0.210402 |
| 31.05556 | -2.00405 | -0.85063 | -2.54522 | -1.47238 | -0.70668 | 0.362557 | -3.52619 | 0.449894 | -1.86448 | 0.018636 | -0.11234 | -0.06714 | 0.000427 | 0.273605 | 0.047906 | -0.1403 | -0.31258 | 0.184294 | 0.127542 |
| 31.06944 | -1.56443 | -1.13306 | -1.86589 | -0.91305 | -0.37198 | 0.310028 | -1.71394 | 0.411871 | -1.41131 | -0.00153 | 0.022678 | -0.22128 | -0.03471 | 0.118566 | -0.05917 | -0.1878 | -0.34236 | 0.156655 | 0.079301 |
| 31.08333 | -1.79495 | -1.55592 | -1.31647 | -0.42412 | -0.25677 | 0.371088 | 0.15298 | 0.363788 | -0.86132 | -0.19632 | 0.026408 | -0.33839 | -0.1178 | -0.09808 | -0.02187 | -0.07991 | -0.27881 | 0.0943 | 0.027087 |
| 31.09722 | -1.21186 | -1.05322 | -0.47118 | 0.074093 | -0.00104 | 0.575945 | 2.048875 | 0.438309 | -0.28905 | -0.4046 | -0.33394 | -0.58851 | -0.47522 | -0.63383 | -0.13947 | -0.43772 | -0.47412 | -0.27064 | -0.21176 |

| | | | | | | | | | | | | | | | | | | | |
|---|---|---|---|---|---|---|---|---|---|---|---|---|---|---|---|---|---|---|---|
| 31.11111 | -0.03859 | -0.02761 | 0.86632 | 0.916544 | 0.283462 | 1.005857 | 3.963056 | 0.59551 | 0.404122 | -0.74193 | -1.1962 | -1.30022 | -1.30357 | -1.66262 | -0.5569 | -1.57547 | -1.21149 | -1.01446 | -0.82579 |
| 31.125 | 1.195094 | 0.972756 | 1.350445 | 1.521197 | 0.387501 | 1.126302 | 4.64499 | 0.421275 | 1.004535 | -0.98343 | -1.83623 | -1.82541 | -2.05441 | -1.9323 | -0.85192 | -2.17396 | -1.74222 | -1.09477 | -1.21905 |
| 31.13889 | 1.717079 | 1.464085 | 1.240068 | 1.368769 | 0.374921 | 0.831793 | 4.067246 | 0.101993 | 1.066664 | -0.53165 | -1.07382 | -1.14964 | -1.28351 | -1.28187 | -0.43907 | -1.21609 | -1.33448 | -0.30418 | -0.42886 |
| 31.15278 | 1.422581 | 1.355521 | 0.371374 | 0.879778 | 0.312829 | 0.395469 | 2.76187 | -0.02465 | 0.631384 | 0.242443 | -0.24093 | -0.35816 | -0.27905 | -0.22067 | 0.004188 | -0.12922 | -0.77532 | 0.457226 | 0.484962 |
| 31.16667 | 0.91419 | 1.080211 | -0.22685 | 0.656622 | 0.196179 | -0.02697 | 1.489358 | -0.02835 | 0.050017 | 0.70492 | 0.003338 | 0.122876 | 0.353562 | 0.253959 | 0.112584 | 0.187951 | -0.33947 | 0.628732 | 0.67687 |
| 31.18056 | 0.818004 | 1.474597 | -0.20855 | 0.44851 | 0.019973 | -0.15802 | 0.688713 | 0.156718 | -0.32057 | 0.988383 | -0.04863 | 0.449501 | 0.485716 | 0.315541 | 0.125942 | 0.181875 | -0.03034 | 0.430017 | 0.403327 |
| 31.19444 | 1.081465 | 1.887086 | 0.218928 | 0.4642 | -0.02005 | 0.026519 | 0.375426 | 0.316441 | -0.41159 | 1.041023 | 0.133735 | 0.916815 | 0.58856 | 0.397631 | 0.176737 | 0.412544 | 0.416372 | 0.428583 | 0.217509 |
| 31.20833 | 1.400567 | 1.918654 | 0.828233 | 0.523564 | 0.077944 | 0.17384 | 0.372071 | 0.213599 | -0.15969 | 0.899235 | 0.490257 | 1.40873 | 0.749335 | 0.538794 | 0.346468 | 0.850182 | 1.01885 | 0.732317 | 0.302748 |
| 31.22222 | 1.912332 | 2.098978 | 1.434302 | 0.606583 | 0.208219 | 0.228194 | 0.546464 | 0.062013 | 0.214302 | 0.885012 | 0.762085 | 1.539339 | 0.765224 | 0.781072 | 0.462561 | 1.125988 | 1.311308 | 0.886795 | 0.415613 |
| 31.23611 | 2.540056 | 1.931454 | 1.650701 | 0.675518 | 0.344908 | 0.284388 | 0.845277 | -0.05701 | 0.68155 | 0.709469 | 0.64333 | 1.138335 | 0.625873 | 0.805251 | 0.619648 | 1.200032 | 1.09166 | 0.594438 | 0.243477 |
| 31.25 | 2.733412 | 1.529762 | 1.493953 | 0.621128 | 0.361958 | 0.279007 | 0.963825 | -0.17193 | 0.8956 | 0.262544 | 0.29134 | 0.643694 | 0.624762 | 0.749203 | 0.663433 | 1.023826 | 0.746735 | 0.325655 | -0.12334 |
| 31.26389 | 2.573783 | 1.032388 | 0.997631 | 0.44487 | 0.290388 | 0.250866 | 0.694713 | -0.22671 | 0.626894 | -0.25075 | -0.03283 | 0.200797 | 0.591438 | 0.608148 | 0.351271 | 0.308411 | 0.380267 | 0.191451 | -0.37286 |
| 31.27778 | 2.086786 | 0.266586 | 0.527027 | 0.283305 | 0.2249 | 0.198446 | 0.325923 | -0.15945 | 0.164004 | -0.45068 | -0.18338 | 0.002702 | 0.416896 | 0.271959 | -0.08822 | -0.43845 | 0.189947 | 0.083495 | -0.34324 |
| 31.29167 | 1.229656 | -0.42232 | 0.427741 | 0.147721 | 0.195759 | 0.096199 | 0.033463 | -0.03792 | -0.09298 | -0.15629 | -0.17723 | 0.30338 | 0.188483 | 0.064299 | -0.39617 | -0.84653 | 0.100172 | -0.02941 | -0.1733 |
| 31.30556 | 0.336387 | -0.62721 | 0.447495 | 0.014985 | 0.156354 | 0.100388 | -0.09162 | 0.095489 | -0.14833 | 0.202845 | -0.07689 | 0.670917 | 0.12873 | -0.07 | -0.24683 | -0.71876 | -0.02084 | -0.07795 | -0.0171 |
| 31.31944 | -0.19955 | -0.28175 | 0.651251 | -0.09682 | 0.070049 | 0.309961 | -0.04815 | 0.190806 | -0.09815 | 0.223485 | -0.04586 | 0.804411 | 0.083245 | -0.2065 | -0.12123 | -0.47056 | -0.17276 | -0.09691 | 0.028234 |
| 31.33333 | -0.38223 | 0.207855 | 0.89434 | -0.04719 | 0.066751 | 0.489632 | 0.138591 | 0.299893 | 0.032939 | -0.02857 | -0.09653 | 0.653395 | -0.05648 | -0.13272 | 0.059648 | -0.14188 | -0.20842 | -0.12622 | 0.001974 |
| 31.34722 | -0.27174 | 0.720608 | 1.043964 | 0.07487 | 0.052268 | 0.59826 | 0.424752 | 0.353009 | 0.200674 | -0.14742 | -0.10798 | 0.336307 | -0.21042 | 0.06909 | 0.137238 | 0.087235 | -0.11053 | -0.15303 | -0.00149 |
| 31.36111 | -0.28173 | 0.750472 | 1.136524 | 0.024937 | 0.001274 | 0.543385 | 0.395007 | 0.30542 | 0.311645 | -0.21708 | -0.16206 | -0.05547 | -0.31973 | 0.073724 | 0.015366 | -0.15253 | -0.07408 | -0.15636 | 0.009262 |
| 31.375 | -0.52155 | 0.1587 | 1.088161 | -0.03863 | -0.07506 | 0.269331 | -0.07447 | 0.214039 | 0.292097 | -0.23354 | -0.25597 | -0.36623 | -0.36016 | -0.17955 | -0.20586 | -0.58919 | -0.15796 | -0.15498 | -0.00376 |
| 31.38889 | -0.94369 | -0.45964 | 0.974087 | -0.19592 | -0.14202 | 0.050325 | -0.53563 | 0.195157 | 0.21874 | -0.07846 | -0.20109 | -0.26343 | -0.30156 | -0.42152 | -0.35381 | -0.7831 | -0.18693 | -0.13105 | 0.01153 |
| 31.40278 | -1.27022 | -0.67525 | 0.828817 | -0.33513 | -0.17895 | -0.01835 | -0.66979 | 0.26185 | 0.174329 | 0.035547 | -0.04259 | -0.05322 | -0.16043 | -0.37994 | -0.34819 | -0.65907 | -0.10853 | -0.09341 | 0.066446 |
| 31.41667 | -1.01009 | -0.27332 | 0.621769 | -0.26917 | -0.15262 | 0.207396 | -0.33852 | 0.360204 | 0.144047 | 0.060759 | 0.163711 | 0.221394 | -0.09109 | -0.16253 | -0.2252 | -0.37992 | 0.082078 | -0.04554 | 0.093493 |
| 31.43056 | -0.59202 | -0.04329 | 0.553373 | 0.002798 | -0.09417 | 0.329558 | -0.03202 | 0.392344 | 0.102946 | 0.037698 | 0.342208 | 0.348498 | -0.02231 | 0.103686 | -0.08219 | -0.08449 | 0.247437 | 0.021206 | 0.079884 |
| 31.44444 | -0.58341 | -0.59816 | 0.647764 | 0.310018 | -0.20567 | 0.038813 | -0.15977 | 0.278797 | 0.013442 | 0.199607 | 0.437918 | 0.372226 | 0.098892 | 0.263076 | 0.107992 | 0.254761 | 0.288628 | 0.107937 | 0.077782 |
| 31.45833 | -0.61167 | -0.98108 | 0.7462 | 0.439313 | -0.42764 | -0.23722 | -0.33764 | 0.13948 | -0.00331 | 0.348267 | 0.41982 | 0.301384 | 0.136705 | 0.328995 | 0.199438 | 0.423133 | 0.230749 | 0.148377 | 0.06346 |
| 31.47222 | 0.062767 | -0.2369 | 0.706171 | 0.47128 | -0.60393 | -0.00783 | -0.0662 | 0.09966 | 0.118904 | 0.403667 | 0.346395 | 0.180251 | 0.036578 | 0.288598 | 0.151793 | 0.331312 | 0.200605 | 0.114427 | 0.045782 |
| 31.48611 | 1.092318 | 0.884535 | 1.154295 | 0.725978 | -0.5945 | 0.527604 | 0.501013 | 0.23245 | 0.500413 | 0.35696 | 0.360097 | 0.128969 | -0.05589 | 0.241403 | 0.02051 | 0.113076 | 0.249755 | 0.058174 | 0.058257 |
| 31.5 | 1.912593 | 0.986447 | 2.710592 | 1.194621 | -0.42348 | 0.744437 | 1.038319 | 0.440406 | 1.179426 | 0.248989 | 0.466979 | 0.222513 | -0.07092 | 0.246788 | -0.11547 | -0.12162 | 0.297723 | 0.016662 | 0.085551 |
| 31.51389 | 2.537371 | 0.489285 | 3.710699 | 1.834001 | 0.186365 | 0.75537 | 1.980654 | 0.345846 | 1.523775 | 0.132952 | 0.543252 | 0.393959 | -0.02286 | 0.297403 | -0.13577 | -0.19435 | 0.299177 | 0.004177 | 0.100305 |
| 31.52778 | 3.252521 | 0.680736 | 3.308564 | 2.668562 | 1.064273 | 0.766436 | 3.279172 | -0.02379 | 1.334992 | 0.150879 | 0.567658 | 0.523176 | 0.054422 | 0.404275 | -0.05621 | -0.09479 | 0.371405 | 0.026871 | 0.124072 |
| 31.54167 | 3.662005 | 0.961074 | 2.78643 | 2.819789 | 1.425544 | 0.393447 | 3.209204 | -0.19227 | 1.480611 | 0.291869 | 0.540862 | 0.519034 | 0.151067 | 0.557076 | 0.040587 | 0.108121 | 0.544574 | 0.0248 | 0.14333 |
| 31.55556 | 3.411817 | 0.598663 | 2.043255 | 2.373578 | 1.176499 | 0.057788 | 1.803038 | -0.07667 | 0.981085 | 0.404995 | 0.501407 | 0.344435 | 0.190205 | 0.648322 | 0.058039 | 0.295831 | 0.694156 | -0.03899 | 0.104827 |
| 31.56944 | 2.454556 | 0.134493 | 1.510446 | 1.379017 | 0.961834 | -0.07898 | 0.570653 | 0.120785 | 0.716253 | 0.340063 | 0.44996 | 0.14432 | 0.200806 | 0.747431 | 0.018883 | 0.39079 | 0.697075 | -0.08594 | 0.022796 |
| 31.58333 | 1.515367 | 0.201343 | 1.036792 | 0.594438 | 1.360408 | -0.28237 | -0.23243 | 0.381083 | 1.199345 | 0.185152 | 0.441 | 0.058857 | 0.263331 | 0.849593 | 0.008183 | 0.404781 | 0.640739 | -0.0577 | 0.012654 |
| 31.59722 | 0.636262 | 0.334722 | 0.86763 | 0.033789 | 2.116308 | -0.41614 | -0.95415 | 0.639727 | 1.257909 | 0.331092 | 0.772104 | 0.360944 | 0.564192 | 1.144364 | 0.088206 | 0.596326 | 0.779938 | 0.082922 | 0.12421 |
| 31.61111 | -0.35779 | 0.311691 | 0.169199 | -0.12385 | 2.01314 | -0.35413 | -1.49079 | 0.755361 | 0.63276 | 0.872425 | 1.784367 | 0.86005 | 1.086579 | 1.763244 | 0.244571 | 1.027881 | 1.193769 | 0.311519 | 0.336913 |
| 31.625 | -0.95037 | 0.2963 | -0.10417 | -0.13002 | 1.491985 | -0.17064 | -1.36325 | 0.670607 | 0.007753 | 1.1119 | 2.535268 | 1.437948 | 1.229712 | 2.16372 | 0.315767 | 1.23635 | 1.356342 | 0.366171 | 0.547123 |
| 31.63889 | -1.6056 | 0.16699 | -0.16867 | -0.28532 | 1.401243 | 0.156583 | -1.17164 | 0.264641 | -1.04516 | 1.039803 | 1.754742 | 2.198972 | 0.939452 | 2.026213 | 0.386104 | 1.158573 | 1.207135 | 0.123242 | 0.802187 |
| 31.65278 | -1.49172 | -0.41049 | -0.20916 | -0.53395 | 1.165697 | -0.05967 | -1.18003 | -0.31013 | -2.0542 | 1.003731 | 0.887972 | 2.727781 | 0.776565 | 1.691755 | 0.192777 | 1.098888 | 1.092541 | -0.19627 | 1.174374 |
| 31.66667 | -1.33129 | -0.72625 | -0.36025 | -0.34146 | 0.727604 | -0.44417 | -1.3318 | -0.75095 | -2.00459 | 0.951933 | 0.139124 | 1.675313 | 0.675514 | 1.020037 | 0.059215 | 0.775871 | 0.679858 | -0.42263 | 0.885378 |
| 31.68056 | -0.40964 | -0.61351 | -0.17706 | 0.021465 | 0.172688 | -0.52962 | -1.28601 | -1.03751 | -1.53384 | 0.327586 | -0.52558 | 0.267633 | 0.417592 | 0.162498 | -0.04081 | 0.222926 | 0.130709 | -0.50931 | 0.265843 |
| 31.69444 | 0.279076 | -0.45681 | -0.17706 | 0.358041 | -0.58358 | -0.38505 | -0.95139 | -0.9951 | -0.96789 | -0.36218 | -1.09815 | -1.16896 | 0.221588 | -0.5663 | -0.28435 | -0.31695 | -0.3744 | -0.49356 | -0.32007 |
| 31.70833 | 0.994417 | -0.51932 | 0.006333 | 0.558642 | -1.59085 | -0.32452 | -0.57603 | -0.6038 | -0.32956 | -0.61082 | -1.47807 | -1.70686 | 0.120805 | -1.04036 | -0.4258 | -0.64292 | -0.50891 | -0.36677 | -0.64613 |
| 31.72222 | 1.683945 | -0.67642 | 0.091147 | 0.79726 | -2.21739 | -0.39924 | -0.1676 | -0.19318 | 0.723182 | -0.38055 | -1.65288 | -1.37437 | 0.091058 | -1.01193 | -0.57591 | -0.95301 | -0.35374 | -0.2171 | -0.72777 |
| 31.73611 | 1.745972 | -0.8017 | 0.165967 | 0.636257 | -2.2671 | -0.54708 | 0.227432 | -0.08856 | 1.575083 | -0.03635 | -1.83275 | -1.00835 | 0.020826 | -0.75431 | -0.48164 | -0.91555 | -0.53652 | -0.09966 | -0.73344 |
| 31.75 | 2.247867 | -0.7397 | 0.409129 | 0.672117 | -1.88998 | -1.05659 | 0.418554 | -0.43939 | 2.34012 | 0.113937 | -1.80188 | -0.71595 | -0.1224 | -0.80507 | -0.24286 | -0.57669 | -0.63555 | -0.02297 | -0.69928 |
| 31.76389 | 2.30544 | -0.37958 | 0.611995 | 0.149433 | -1.59983 | -1.34588 | 0.365336 | -1.07654 | 2.060138 | 0.131941 | -1.45358 | -0.48682 | -0.28945 | -0.92921 | 0.035803 | -0.23058 | -0.71223 | 0.043973 | -0.59379 |
| 31.77778 | 1.41479 | 0.279252 | 0.680078 | 0.012014 | -1.54036 | -1.1376 | 0.676277 | -0.92758 | 1.62439 | 0.037725 | -0.92076 | -0.38377 | -0.38528 | -0.96165 | 0.232838 | -0.06293 | -0.63735 | 0.037325 | -0.44331 |
| 31.79167 | 0.217723 | 0.689136 | 0.075923 | -0.36505 | -1.48224 | -1.20799 | 0.288525 | -0.83518 | 0.537427 | -0.07318 | -0.24347 | -0.26469 | -0.36214 | -0.88436 | 0.084908 | -0.09801 | -0.34171 | -0.01523 | -0.25591 |
| 31.80556 | -0.51995 | 0.446662 | -0.20186 | -0.69063 | -1.57375 | -1.62542 | -0.45084 | -0.97039 | -0.6051 | -0.16588 | 0.558969 | -0.03695 | -0.22472 | -0.6155 | -0.17899 | -0.23354 | 0.136509 | -0.03811 | -0.1215 |
| 31.81944 | -1.44829 | -0.56243 | -0.47291 | -1.054 | -1.505 | -2.03764 | -1.14495 | -0.93069 | -1.30349 | -0.20691 | 1.28271 | 0.279903 | -0.11451 | -0.09818 | -0.48034 | -0.35653 | 0.495516 | -0.02255 | 0.073347 |
| 31.83333 | -2.06507 | -1.35658 | -0.5933 | -0.87504 | -1.28728 | -1.99748 | -1.45975 | -0.66069 | -1.54614 | -0.34909 | 1.59274 | 0.367382 | -0.01636 | 0.434865 | -0.54321 | -0.28151 | 0.570691 | 0.034648 | 0.291152 |
| 31.84722 | -2.2689 | -1.82524 | -0.8575 | -0.85408 | -1.19583 | -1.58732 | -1.61138 | -0.61115 | -1.92994 | -0.54062 | 1.33508 | 0.410198 | 0.080008 | 0.780502 | -0.32398 | -0.07014 | 0.332213 | 0.117706 | 0.394934 |
| 31.86111 | -2.44246 | -1.85207 | -1.34991 | -0.90281 | -1.17064 | -1.49073 | -1.99382 | -0.66965 | -2.14779 | -0.57212 | 0.783031 | 0.399265 | 0.125871 | 0.822847 | -0.18922 | 0.091209 | -0.15572 | 0.172841 | 0.436877 |
| 31.875 | -2.39515 | -1.85165 | -1.88043 | -0.9862 | -1.03131 | -1.6438 | -2.59681 | -0.85133 | -2.24472 | -0.38809 | 0.128078 | 0.356669 | 0.186313 | 0.684317 | -0.06929 | 0.163886 | -0.58115 | 0.16329 | 0.159124 |
| 31.88889 | -2.20777 | -1.79894 | -2.26228 | -0.95993 | -0.85252 | -1.83549 | -2.70846 | -1.05624 | -1.97719 | -0.1857 | -0.37654 | 0.219475 | 0.259397 | 0.314005 | 0.009478 | 0.095016 | -0.73931 | 0.135549 | -0.25316 |
| 31.90278 | -1.96201 | -2.51848 | -1.96942 | -0.89307 | -0.59341 | -1.47421 | -2.01493 | -1.25858 | -1.50877 | 0.033106 | -0.68329 | -0.07655 | 0.285056 | -0.25526 | 0.04469 | 0.035059 | -0.60437 | 0.106293 | -0.61725 |
| 31.91667 | -1.10674 | -1.7144 | -1.19857 | -0.62894 | -0.34379 | -1.00661 | -1.39262 | -1.21054 | -1.46178 | 0.027197 | -0.96885 | -0.47051 | 0.245316 | -0.8154 | 0.082268 | 0.047855 | -0.35253 | 0.102663 | -0.8087 |
| 31.93056 | -0.09751 | -0.8894 | -0.52733 | -0.06057 | -0.08503 | -0.55374 | -0.76504 | -0.74777 | -0.93421 | -0.26521 | -1.32769 | -0.837 | 0.062391 | -1.23028 | 0.087848 | 0.083565 | -0.24192 | 0.084651 | -0.52232 |
| 31.94444 | 0.648721 | -0.23338 | 0.112249 | 0.53866 | 0.135399 | -0.11359 | -0.11305 | -0.40417 | -0.10581 | -0.62458 | -1.49722 | -1.03131 | -0.14273 | -1.50121 | 0.118999 | 0.105684 | -0.35852 | 0.057411 | -0.27433 |
| 31.95833 | 1.075597 | 0.610747 | 0.666171 | 1.018273 | 0.297367 | 0.093649 | 0.565248 | -0.23696 | 0.735631 | -0.77729 | -1.42843 | -0.71832 | -0.28838 | -1.32058 | 0.123157 | 0.11507 | -0.47928 | 0.016589 | 0.013607 |
| 31.97222 | 1.221614 | 1.15912 | 1.195453 | 1.241454 | 0.388497 | 0.151064 | 1.147015 | -0.20505 | 1.476814 | -0.67941 | -1.08258 | -0.45212 | -0.40044 | -1.04331 | 0.117894 | 0.046005 | -0.41377 | 0.021948 | 0.179513 |
| 31.98611 | 0.618379 | 0.897872 | 0.91745 | 0.590372 | 0.042908 | -0.26627 | 1.002835 | -0.36255 | 1.315099 | -0.41096 | -0.76377 | -0.36881 | -0.46181 | -0.66538 | 0.152587 | 0.038029 | -0.25537 | 0.04553 | 0.175273 |
| 32 | -0.87428 | -0.66587 | -1.00633 | -1.46464 | -0.73639 | -1.08147 | -0.37005 | -0.57561 | -0.62743 | -0.09575 | -0.47272 | -0.22923 | -0.52342 | -0.33748 | 0.279202 | 0.092725 | -0.08823 | 0.07769 | 0.074844 |
| 32.01389 | -2.47658 | -2.60806 | -3.08368 | -3.6143 | -1.46895 | -1.56004 | -2.33758 | -0.70504 | -2.86513 | 0.135467 | -0.49952 | -0.08572 | -0.53721 | -0.00705 | 0.347793 | 0.123631 | 0.103372 | 0.1382 | -0.00722 |
| 32.02778 | -3.24841 | -3.30825 | -2.9463 | -3.59276 | -1.364 | -1.26127 | -2.74619 | -0.73164 | -3.28998 | 0.141275 | -0.51471 | 0.009208 | -0.52033 | 0.097533 | 0.39138 | 0.124199 | 0.298785 | 0.207879 | -0.06734 |
| 32.04167 | -3.10062 | -3.31527 | -2.44988 | -2.90655 | -1.14977 | -1.61148 | -2.22608 | -0.60435 | -2.74327 | -0.14784 | -0.52569 | 0.004757 | -0.41389 | 0.000539 | 0.308752 | 0.013019 | 0.425478 | 0.278223 | -0.03302 |
| 32.05556 | -2.42711 | -3.48282 | -2.68988 | -2.18336 | -0.9657 | -1.82425 | -2.23938 | -0.34497 | -2.31061 | -0.37019 | -0.39296 | 0.005545 | -0.22505 | 0.083933 | 0.174745 | -0.01081 | 0.497129 | 0.322683 | 0.09381 |
| 32.06944 | -1.56575 | -3.17053 | -2.35559 | -1.09105 | -0.85209 | -1.47885 | -1.63565 | -0.22145 | -0.99269 | -0.33822 | -0.23415 | 0.065924 | -0.06029 | 0.329647 | 0.069199 | 0.040063 | 0.434052 | 0.31508 | 0.25982 |
| 32.08333 | -0.50521 | -2.20001 | -1.31646 | -0.1426 | -0.46548 | -0.62384 | -0.5194 | -0.07883 | 0.018891 | -0.12125 | -0.16803 | 0.03943 | -0.01992 | 0.577877 | 0.020115 | 0.080191 | 0.271751 | 0.160997 | 0.304609 |
| 32.09722 | 0.508571 | -0.9442 | -0.47366 | 0.467682 | 0.095069 | 0.470492 | 0.064461 | 0.194856 | 0.682492 | -0.15382 | -0.48924 | -0.4149 | -0.45745 | 0.020937 | -0.11742 | -0.48827 | -0.29652 | -0.47211 | -0.01861 |
| 32.11111 | 1.114929 | -0.24397 | 0.056566 | 0.787066 | 0.477819 | 1.375559 | 0.376294 | 0.37413 | 1.12169 | -0.98203 | -1.37635 | -1.38978 | -1.41881 | -1.64934 | -0.48643 | -1.83447 | -1.41434 | -1.85534 | -0.79323 |
| 32.125 | 1.406104 | 0.074953 | 0.368361 | 0.982631 | 0.690062 | 1.728056 | 0.709373 | 0.430955 | 1.294144 | -1.92809 | -1.78646 | -1.54021 | -1.47922 | -3.04424 | -0.62103 | -2.01483 | -2.05575 | -2.32613 | -1.1581 |
| 32.13889 | 1.50972 | 0.173519 | 0.578493 | 1.185448 | 0.84503 | 1.728922 | 1.005773 | 0.428695 | 1.402193 | -1.7252 | -0.67913 | -0.59808 | -0.54037 | -2.23102 | -0.15003 | -0.71124 | -1.38117 | -0.68928 | -0.48652 |
| 32.15278 | 1.734263 | 0.3962 | 0.616721 | 1.237271 | 0.9326 | 1.828237 | 1.090503 | 0.552625 | 1.464992 | -0.71109 | 0.28036 | -0.10174 | 0.341124 | -0.62073 | 0.310973 | 0.539709 | -0.46043 | 0.511028 | 0.289689 |
| 32.16667 | 2.05354 | 0.932578 | 0.626745 | 1.070934 | 0.868263 | 1.733265 | 0.953326 | 0.738371 | 1.635161 | 0.008121 | 0.285509 | -0.10538 | 0.520609 | 0.281503 | 0.321852 | 0.863912 | 0.138223 | 0.833722 | 0.596589 |
| 32.18056 | 2.023842 | 1.206203 | 0.688352 | 0.974678 | 0.76671 | 1.448257 | 0.922755 | 0.772576 | 2.097002 | 0.566769 | 0.000612 | -0.01851 | 0.432778 | 0.719369 | 0.240986 | 0.715582 | 0.40997 | 0.661462 | 0.529693 |
| 32.19444 | 1.827123 | 1.279406 | 0.733126 | 1.013099 | 0.603053 | 1.044159 | 1.035929 | 0.87468 | 2.547258 | 1.209557 | 0.020209 | 0.311195 | 0.419483 | 1.024108 | 0.224604 | 0.704086 | 0.58947 | 0.500135 | 0.636486 |
| 32.20833 | 1.760273 | 1.465675 | 0.695162 | 0.967734 | 0.541827 | 0.787082 | 1.179305 | 0.927673 | 2.513323 | 1.405292 | 0.038982 | 0.7856 | 0.360827 | 1.047166 | 0.202973 | 0.54165 | 0.683792 | 0.408273 | 0.649585 |
| 32.22222 | 1.523438 | 1.453677 | 0.67969 | 0.868655 | 0.429289 | 0.644545 | 1.687096 | 0.6494 | 2.509327 | 1.323898 | 0.085945 | 1.14797 | 0.253287 | 0.847951 | 0.170465 | 0.358263 | 0.574437 | 0.355104 | 0.586155 |
| 32.23611 | 1.268294 | 1.405586 | 0.601662 | 0.614702 | 0.385435 | 0.575003 | 2.136371 | 0.438377 | 2.2426 | 1.30146 | 0.273028 | 1.20671 | 0.082518 | 0.650142 | 0.141862 | 0.083698 | 0.368813 | 0.319172 | 0.574753 |
| 32.25 | 1.149327 | 1.678199 | 0.500682 | 0.252492 | 0.401193 | 0.536832 | 2.312487 | 0.412927 | 2.09586 | 1.08941 | 0.419433 | 1.184725 | 0.029343 | 0.562037 | 0.190174 | 0.03614 | 0.3001 | 0.243479 | 0.49101 |
| 32.26389 | 0.918925 | 1.853409 | 0.717352 | -0.10676 | 0.461806 | 0.380771 | 2.547019 | 0.411779 | 1.92116 | 0.710676 | 0.342497 | 1.262208 | 0.113855 | 0.514521 | 0.117566 | 0.085816 | 0.542171 | 0.22204 | 0.282356 |
| 32.27778 | 0.944308 | 1.925166 | 1.092215 | -0.11892 | 0.494547 | 0.194612 | 2.614276 | 0.379158 | 1.633756 | 0.271523 | 0.153698 | 1.022529 | 0.303914 | 0.64669 | 0.040832 | 0.144337 | 1.018878 | 0.18081 | 0.056352 |
| 32.29167 | 1.119192 | 1.782955 | 1.213453 | 0.124723 | 0.489242 | 0.214966 | 2.429473 | 0.301788 | 1.237695 | -0.05214 | -0.02595 | 0.546621 | 0.479611 | 0.951576 | -0.06346 | 0.25769 | 1.186021 | 0.100609 | -0.04168 |

| | | | | | | | | | | | | | | | | | | | |
|---|---|---|---|---|---|---|---|---|---|---|---|---|---|---|---|---|---|---|---|
| 32.30556 | 1.241807 | 1.544451 | 1.200194 | 0.322862 | 0.477859 | 0.545234 | 2.437708 | 0.219878 | 0.776796 | -0.15516 | -0.0931 | 0.084976 | 0.603864 | 1.048203 | -0.03782 | 0.258306 | 0.907608 | 0.025558 | 0.033962 |
| 32.31944 | 1.493524 | 1.644943 | 1.088752 | 0.361541 | 0.566907 | 0.874077 | 2.418913 | 0.316578 | 0.580718 | -0.20521 | -0.05138 | -0.16838 | 0.607376 | 0.88238 | 0.004777 | 0.146599 | 0.45531 | 0.001522 | 0.062627 |
| 32.33333 | 1.563756 | 1.788516 | 0.943617 | 0.243939 | 0.631983 | 0.880578 | 2.148538 | 0.386886 | 0.548389 | -0.28164 | -0.00873 | -0.16153 | 0.491038 | 0.514226 | 0.0161 | 0.053973 | 0.057968 | -0.02279 | -0.06916 |
| 32.34722 | 1.221607 | 1.776267 | 0.956046 | 0.094502 | 0.447685 | 0.613573 | 1.62753 | 0.264757 | 0.385642 | -0.39863 | -0.06627 | -0.14849 | 0.248 | 0.021114 | -0.04115 | -0.06826 | -0.21703 | -0.05776 | -0.25595 |
| 32.36111 | 1.025263 | 1.60298 | 0.969574 | 0.091756 | 0.225357 | 0.332673 | 1.003647 | 0.065049 | 0.091906 | -0.39459 | -0.13024 | -0.16182 | 0.094814 | -0.30419 | -0.03777 | -0.07326 | -0.25143 | -0.02142 | -0.2642 |
| 32.375 | 1.417994 | 1.638098 | 1.004817 | 0.315921 | 0.290689 | 0.375513 | 0.364885 | 0.132839 | 0.050265 | -0.20794 | -0.0555 | -0.09816 | 0.121232 | -0.30348 | 0.030591 | 0.067104 | -0.00883 | 0.095082 | -0.08132 |
| 32.38889 | 2.043661 | 1.817279 | 1.03796 | 0.570203 | 0.519564 | 0.666341 | 0.058462 | 0.411998 | 0.36552 | 0.08058 | 0.097122 | 0.080175 | 0.21556 | 0.094835 | 0.078625 | 0.214994 | 0.316048 | 0.17298 | 0.131843 |
| 32.40278 | 2.060826 | 1.745533 | 1.186707 | 0.641298 | 0.538178 | 0.735639 | 0.036193 | 0.39064 | 0.58927 | 0.293475 | 0.169338 | 0.282844 | 0.230481 | 0.55062 | 0.063753 | 0.25722 | 0.573764 | 0.155162 | 0.27938 |
| 32.41667 | 1.574095 | 1.474799 | 0.967434 | 0.415109 | 0.389437 | 0.411036 | -0.2045 | 0.140778 | 0.498467 | 0.336759 | 0.175233 | 0.409948 | 0.144155 | 0.743464 | 0.03895 | 0.241851 | 0.623693 | 0.112031 | 0.356144 |
| 32.43056 | 1.079627 | 1.112243 | 0.266225 | 0.008254 | 0.265144 | -0.12981 | -0.7895 | -0.02091 | 0.270176 | 0.364132 | 0.211499 | 0.48964 | 0.067719 | 0.699505 | 0.038432 | 0.235801 | 0.41854 | 0.127589 | 0.285678 |
| 32.44444 | 0.685645 | 0.690048 | -0.25447 | -0.34611 | 0.154522 | -0.68373 | -1.29018 | -0.08898 | 0.074226 | 0.46061 | 0.319144 | 0.610847 | 0.10914 | 0.51605 | 0.082226 | 0.286858 | 0.178046 | 0.206366 | 0.159849 |
| 32.45833 | 0.636967 | 0.631398 | -0.35448 | -0.35932 | 0.08426 | -1.01951 | -1.55279 | -0.09148 | -0.07298 | 0.511186 | 0.412406 | 0.785446 | 0.20678 | 0.330793 | 0.107629 | 0.345062 | 0.039469 | 0.252409 | 0.006929 |
| 32.47222 | 0.987631 | 1.104962 | -0.267 | -0.24264 | 0.124827 | -1.10445 | -1.4978 | 0.012868 | -0.22862 | 0.456231 | 0.394858 | 0.983764 | 0.226419 | 0.23422 | 0.068198 | 0.321472 | 0.098755 | 0.203006 | -0.10339 |
| 32.48611 | 1.273399 | 1.559976 | 0.403579 | 0.130676 | 0.254176 | -0.69372 | -1.00074 | 0.139252 | -0.01973 | 0.310162 | 0.387624 | 0.981457 | 0.181787 | 0.230569 | 0.061757 | 0.278384 | 0.290294 | 0.14762 | -0.08162 |
| 32.5 | 1.852898 | 2.02594 | 2.015691 | 1.090667 | 0.583177 | 0.253688 | 0.294205 | 0.309174 | 0.771169 | 0.107932 | 0.426146 | 0.726271 | 0.158445 | 0.266027 | 0.111398 | 0.248942 | 0.425023 | 0.105923 | 0.009573 |
| 32.51389 | 2.719033 | 1.899782 | 2.6062 | 1.926992 | 0.859052 | 1.344493 | 2.034777 | 0.494171 | 1.414304 | -0.00769 | 0.470962 | 0.308624 | 0.174364 | 0.318798 | 0.115922 | 0.183387 | 0.413138 | 0.045426 | 0.095429 |
| 32.52778 | 3.13328 | 1.485665 | 2.308598 | 2.254129 | 0.69902 | 1.779064 | 3.649319 | 0.79484 | 1.853037 | 0.062676 | 0.514836 | -0.09693 | 0.168048 | 0.35868 | 0.02972 | 0.028772 | 0.307392 | -0.00352 | 0.1141 |
| 32.54167 | 3.136932 | 1.236581 | 2.455046 | 3.370978 | 0.680896 | 1.752973 | 4.375818 | 1.158101 | 2.118336 | 0.286072 | 0.517159 | -0.27763 | 0.151407 | 0.315821 | -0.0149 | -0.0694 | 0.155687 | -0.01808 | 0.098258 |
| 32.55556 | 2.702285 | 0.954201 | 2.580285 | 3.749931 | 0.896608 | 1.537357 | 4.04504 | 1.204458 | 1.980531 | 0.504415 | 0.459113 | -0.06366 | 0.191785 | 0.221628 | 0.034024 | 0.000125 | 0.035899 | 0.018638 | 0.105457 |
| 32.56944 | 2.103603 | 0.813351 | 2.322117 | 3.120216 | 1.076733 | 1.173132 | 3.085385 | 1.011772 | 1.24362 | 0.579168 | 0.340613 | 0.332298 | 0.207585 | 0.284924 | 0.050003 | 0.126533 | 0.075869 | 0.029207 | 0.084494 |
| 32.58333 | 1.331536 | 0.605184 | 2.048359 | 1.74499 | 1.377138 | 0.332715 | 2.295281 | 0.886496 | 0.899906 | 0.585171 | 0.240376 | 0.639306 | 0.174983 | 0.474077 | -0.03552 | 0.085057 | 0.2424 | 0.002185 | 0.001569 |
| 32.59722 | 0.679105 | 0.299695 | 1.720206 | 0.815119 | 1.817332 | -0.1592 | 1.894048 | 0.628423 | 0.569937 | 0.773326 | 0.532071 | 0.962748 | 0.281726 | 1.04753 | -0.08392 | 0.287219 | 0.633331 | 0.183907 | 0.049348 |
| 32.61111 | 0.562096 | 0.240215 | 1.326358 | 0.086468 | 1.842403 | -0.39143 | 1.482954 | 0.498831 | 0.225302 | 1.317245 | 1.338942 | 1.73794 | 0.701425 | 2.05769 | 0.042146 | 0.82903 | 1.317968 | 0.618628 | 0.177373 |
| 32.625 | 0.473099 | 0.205769 | 1.136409 | -0.55605 | 1.653945 | -0.0248 | 1.226108 | 0.449065 | -0.06311 | 1.51551 | 2.333096 | 2.153143 | 0.954173 | 2.178574 | 0.180001 | 1.068253 | 1.663572 | 0.684883 | 0.083873 |
| 32.63889 | 0.276686 | 0.092099 | 0.737944 | -0.66971 | 1.209203 | 0.399766 | 0.92278 | 0.242558 | -0.64509 | 1.493448 | 2.319595 | 2.20003 | 0.757565 | 1.777659 | 0.232276 | 0.837497 | 1.444157 | 0.321931 | 0.10026 |
| 32.65278 | -0.05795 | -0.12619 | 0.53041 | -0.74148 | 0.803068 | 0.293887 | -0.02281 | 0.116889 | -1.12968 | 1.652728 | 2.199924 | 2.647393 | 0.453406 | 1.520972 | 0.267914 | 0.801209 | 1.160977 | 0.063765 | 0.461043 |
| 32.66667 | -0.32325 | -0.2614 | 0.269944 | -0.57517 | 0.518012 | -0.05691 | -1.41412 | 0.062122 | -1.59933 | 1.207561 | 1.618076 | 2.234248 | 0.202586 | 1.10407 | 0.32921 | 0.571769 | 0.874596 | -0.06661 | 0.81944 |
| 32.68056 | -0.55006 | -0.2724 | 0.102356 | -0.57754 | 0.247769 | -0.8632 | -2.97744 | -0.01464 | -2.05916 | 0.177006 | 0.857127 | 0.77156 | -0.05386 | 0.310603 | 0.372692 | 0.101721 | 0.018045 | -0.23358 | 1.09086 |
| 32.69444 | -0.65618 | -0.29129 | 0.232128 | -0.76629 | 0.129604 | -1.97984 | -3.76534 | -0.0523 | -2.21228 | -0.66367 | 0.212728 | -0.91192 | -0.24204 | -0.35575 | -0.03951 | -0.38309 | -0.87535 | -0.30853 | 1.175391 |
| 32.70833 | -0.63388 | -0.2761 | 0.413648 | -0.74179 | 0.181206 | -2.73768 | -3.76923 | 0.197385 | -1.62567 | -0.89309 | -0.19851 | -1.99244 | -0.43605 | -0.60409 | -0.16866 | -0.44849 | -1.04491 | -0.25406 | 1.311586 |
| 32.72222 | -0.39914 | -0.3253 | 0.581443 | -0.30469 | 0.318108 | -2.24588 | -2.16376 | 0.511429 | -0.37802 | -0.66973 | -0.43113 | -2.02246 | -0.24325 | -0.59492 | -0.26393 | -0.3176 | -0.5672 | -0.24402 | 0.867735 |
| 32.73611 | -0.08337 | -0.46981 | 0.727713 | 0.384147 | 0.330804 | -1.40558 | -0.79919 | 0.565931 | 0.711673 | -0.53071 | -0.62991 | -1.75235 | 0.034859 | -0.57234 | -0.21231 | -0.24639 | -0.27912 | -0.23027 | 0.194178 |
| 32.75 | 0.229923 | -0.53578 | 0.682376 | 1.028858 | 0.176045 | -0.64795 | -0.04911 | 0.33269 | 1.453877 | -0.23986 | -0.86534 | -1.44804 | 0.171868 | -0.68956 | -0.18845 | -0.27831 | -0.37585 | -0.05582 | -0.49892 |
| 32.76389 | 0.48372 | -0.43292 | 0.588383 | 1.661772 | -0.12465 | 0.068892 | 0.291861 | -0.05791 | 1.51522 | 0.061117 | -1.07918 | -1.083 | 0.081415 | -0.96824 | -0.16999 | -0.29836 | -0.85162 | 0.117951 | -1.07236 |
| 32.77778 | 0.336773 | -0.48322 | 0.482657 | 1.702174 | -0.63949 | 0.656859 | 0.334896 | -0.36149 | 2.000007 | 0.152942 | -1.25993 | -0.88825 | -0.11335 | -1.25284 | -0.06348 | -0.22082 | -1.1859 | 0.16728 | -1.5112 |
| 32.79167 | -0.25104 | -0.54139 | 0.385899 | 1.431746 | -1.17375 | 0.715541 | 0.320359 | -0.39158 | 2.149329 | -0.01697 | -1.42049 | -0.88136 | -0.33629 | -1.25337 | -0.02171 | -0.08154 | -1.1531 | 0.065592 | -1.44941 |
| 32.80556 | -0.74854 | -0.39799 | 0.253343 | 0.312814 | -1.5729 | 0.797633 | 0.360117 | -0.21891 | 1.626426 | -0.24568 | -1.11121 | -0.74 | -0.47165 | -1.00109 | 0.054228 | 0.171021 | -0.87635 | -0.06619 | -1.25757 |
| 32.81944 | -1.16943 | -0.28725 | 0.014634 | -1.12548 | -1.89111 | 0.525388 | 0.296513 | 0.022371 | 0.699331 | -0.40619 | -0.65844 | -0.30144 | -0.27673 | -0.65636 | 0.219686 | 0.396788 | -0.2456 | -0.07813 | -0.9643 |
| 32.83333 | -1.41796 | -0.28212 | -0.33729 | -2.0752 | -2.18709 | 0.013927 | 0.291306 | 0.207559 | -0.27736 | -0.36951 | -0.2104 | 0.290556 | 0.104537 | -0.25937 | 0.365736 | 0.499143 | 0.26395 | -0.04952 | -0.41832 |
| 32.84722 | -1.68566 | -0.36676 | -0.65983 | -2.40171 | -2.19318 | -0.56584 | 0.024413 | 0.290127 | -1.24347 | -0.3101 | 0.079836 | 0.648738 | 0.326416 | 0.079189 | 0.364401 | 0.371167 | 0.615725 | -0.01386 | -0.05718 |
| 32.86111 | -1.99972 | -0.53767 | -0.96362 | -2.32284 | -1.97382 | -1.05145 | -0.65065 | 0.15861 | -1.44044 | -0.41225 | 0.167826 | 0.713527 | 0.290329 | 0.312792 | 0.303249 | 0.16417 | 0.714439 | 0.047962 | 0.045075 |
| 32.875 | -2.16407 | -0.76214 | -1.02515 | -2.03739 | -1.5091 | -1.28515 | -1.34532 | -0.3185 | -2.04011 | -0.61746 | 0.123535 | 0.604965 | 0.170947 | 0.543184 | 0.153679 | 0.029933 | 0.566777 | 0.097816 | 0.110623 |
| 32.88889 | -2.13688 | -0.96178 | -0.89066 | -1.69404 | -0.85756 | -1.53981 | -1.83699 | -0.78343 | -2.85283 | -0.88519 | 0.047417 | 0.117029 | 0.116278 | 0.704684 | -0.02587 | -0.01221 | 0.147323 | 0.133104 | 0.143923 |
| 32.90278 | -1.74762 | -0.97066 | -0.68033 | -1.12204 | -0.15044 | -1.88933 | -1.93308 | -0.78121 | -2.80428 | -1.21689 | -0.15614 | -0.48082 | 0.053602 | 0.677506 | -0.13998 | -0.03016 | -0.396 | 0.073314 | 0.063644 |
| 32.91667 | -1.10667 | -0.72487 | -0.50029 | -0.70654 | 0.408212 | -1.77615 | -1.41665 | -0.77242 | -3.00531 | -1.36989 | -0.36286 | -0.85593 | -0.00911 | 0.525529 | -0.11678 | 0.149557 | -0.73069 | -0.04099 | -0.0637 |
| 32.93056 | -0.49008 | -0.47434 | -0.41496 | -0.28779 | 0.872113 | -1.23741 | -1.02977 | -0.69802 | -2.74708 | -1.14144 | -0.34481 | -0.75903 | 0.088626 | 0.262881 | 0.09406 | 0.275731 | -0.65623 | -0.02186 | -0.02203 |
| 32.94444 | 0.044889 | -0.27105 | -0.08026 | 0.310283 | 1.005949 | -0.35173 | -0.60334 | -0.40917 | -1.30558 | -0.51454 | -0.37983 | -0.54526 | 0.261533 | -0.06273 | 0.279192 | 0.343643 | -0.41017 | 0.124731 | -0.0017 |
| 32.95833 | 0.474746 | -0.05684 | 0.558627 | 1.157601 | 1.197392 | 0.527537 | 0.039398 | -0.12265 | 0.624616 | 0.13267 | -0.45433 | -0.31814 | 0.327799 | -0.47369 | 0.346831 | 0.287139 | -0.20901 | 0.269976 | -0.04549 |
| 32.97222 | 0.790839 | 0.065686 | 0.971373 | 2.006359 | 1.319215 | 1.118568 | 0.912422 | 0.102917 | 2.169722 | 0.511415 | -0.51525 | -0.12405 | 0.202655 | -0.89354 | 0.191742 | 0.181795 | -0.44568 | 0.308714 | -0.09462 |
| 32.98611 | 0.803541 | -0.20424 | 0.64073 | 1.871451 | 1.143337 | 0.810757 | 0.950279 | 0.142846 | 2.401631 | 0.656121 | -0.46702 | 0.071227 | 0.070271 | -1.00518 | 0.123968 | 0.155496 | -0.57672 | 0.306885 | -0.02615 |
| 33 | -0.19 | -0.86686 | -0.66373 | 0.150408 | 0.202414 | -0.89592 | -0.70327 | -0.11386 | 0.592463 | 0.673096 | -0.28627 | 0.266917 | -0.01332 | -0.85485 | 0.180323 | 0.213556 | -0.44569 | 0.293663 | 0.052644 |
| 33.01389 | -2.10524 | -2.52538 | -1.91415 | -2.14065 | -1.02237 | -2.89252 | -2.6947 | -0.66866 | -2.24051 | 0.547461 | -0.14023 | 0.475733 | -0.07975 | -0.58193 | 0.215908 | 0.200842 | -0.13363 | 0.232357 | 0.059088 |
| 33.02778 | -3.7434 | -4.27657 | -1.97896 | -3.51247 | -1.72678 | -3.28327 | -3.01068 | -1.00556 | -3.61313 | 0.258766 | -0.07268 | 0.723889 | -0.0916 | -0.31423 | 0.234599 | 0.14743 | 0.110053 | 0.163309 | 0.040069 |
| 33.04167 | -4.9145 | -4.55194 | -1.6301 | -3.33343 | -2.20736 | -2.9188 | -1.94246 | -0.98328 | -3.59506 | 0.004212 | -0.05782 | 0.898838 | 0.036617 | -0.06198 | 0.318832 | 0.177344 | 0.244221 | 0.188325 | 0.078529 |
| 33.05556 | -3.89419 | -4.36988 | -1.37468 | -2.4457 | -2.17355 | -2.746 | -1.53111 | -0.84804 | -4.33002 | -0.08243 | 0.033713 | 0.85171 | 0.223892 | 0.088677 | 0.513622 | 0.319565 | 0.33591 | 0.284534 | 0.162162 |
| 33.06944 | -1.85806 | -3.80791 | -1.07433 | -1.50288 | -1.47616 | -2.42614 | -1.08893 | -0.71275 | -3.66227 | -0.06207 | 0.099841 | 0.664017 | 0.350956 | 0.153409 | 0.694111 | 0.484777 | 0.36063 | 0.336218 | 0.218129 |
| 33.08333 | -0.65599 | -2.66475 | -0.91156 | -0.69818 | -0.80649 | -2.49133 | -0.69983 | -0.6414 | -2.74758 | -0.02092 | 0.06164 | 0.363661 | 0.240743 | 0.104357 | 0.661756 | 0.476646 | 0.298531 | 0.229021 | 0.179356 |
| 33.09722 | 0.14288 | -1.47377 | -0.69635 | -0.15752 | -0.08138 | -2.25503 | -0.34351 | -0.37267 | -1.86157 | -0.26641 | -0.32133 | -0.33371 | -0.21425 | -0.37453 | 0.365689 | 0.081186 | -0.38761 | -0.30165 | -0.0234 |
| 33.11111 | 0.692422 | -0.09159 | -0.42857 | 0.434842 | 0.479291 | -1.87476 | -0.0126 | -0.00935 | -1.15451 | -1.31347 | -1.2759 | -1.7893 | -0.97653 | -1.62716 | -0.16569 | -0.76285 | -2.20507 | -1.34022 | -0.40814 |
| 33.125 | 0.829974 | 0.505532 | -0.13205 | 0.920704 | 0.777633 | -1.26809 | 0.235926 | 0.062623 | -0.73774 | -2.51079 | -2.11697 | -2.66789 | -1.71272 | -2.67685 | -0.30661 | -0.96582 | -3.61743 | -1.86711 | -0.5314 |
| 33.13889 | 0.814555 | 0.485856 | 0.095884 | 1.202214 | 0.962173 | -0.65232 | 0.497419 | 0.008545 | -0.59482 | -2.11601 | -1.15584 | -1.50786 | -1.09694 | -1.66827 | -0.12437 | -0.2108 | -2.88486 | -0.56626 | -0.06838 |
| 33.15278 | 1.072757 | 0.646563 | 0.416095 | 1.540607 | 1.153054 | 0.454796 | 0.821979 | 0.120185 | -0.21609 | -0.85087 | -0.27879 | -0.19181 | -0.10003 | -0.62135 | 0.179193 | 0.267619 | -1.46095 | 0.405925 | 0.341061 |
| 33.16667 | 1.497898 | 0.856844 | 0.875465 | 2.040552 | 1.125727 | 1.593475 | 0.991123 | 0.328984 | 0.517987 | 0.299297 | 0.304308 | 0.433103 | 0.519724 | 0.051872 | 0.272738 | 0.143777 | -0.46698 | 0.429111 | 0.492808 |
| 33.18056 | 1.921559 | 0.761165 | 1.075493 | 1.738072 | 1.241598 | 2.358361 | 1.064097 | 0.486303 | 1.418678 | 1.047003 | 0.539045 | 0.604412 | 0.751542 | 0.489041 | 0.297518 | -0.14482 | 0.365369 | 0.252873 | 0.48086 |
| 33.19444 | 2.20875 | 0.680866 | 1.002052 | 1.276034 | 1.263901 | 2.728285 | 1.101216 | 0.642041 | 2.359369 | 1.337002 | 0.533983 | 0.650989 | 0.731564 | 0.720361 | 0.242982 | -0.24733 | 0.875793 | 0.282788 | 0.364053 |
| 33.20833 | 2.222959 | 0.428052 | 0.934735 | 0.96383 | 1.170588 | 2.753598 | 1.070535 | 0.760651 | 3.093558 | 1.083785 | 0.448622 | 0.736074 | 0.58767 | 0.833371 | 0.216682 | -0.17441 | 1.117884 | 0.319563 | 0.280747 |
| 33.22222 | 1.847826 | 0.033651 | 0.802522 | 0.763948 | 0.859099 | 2.500848 | 0.887121 | 0.831152 | 3.533142 | 0.725671 | 0.422632 | 1.009947 | 0.474534 | 0.842901 | 0.281366 | -0.09966 | 1.224441 | 0.229298 | 0.378057 |
| 33.23611 | 1.500434 | -0.2704 | 0.639664 | 0.506616 | 0.515695 | 2.003459 | 0.724817 | 0.930527 | 3.287387 | 0.664647 | 0.343356 | 1.245571 | 0.197952 | 0.717177 | 0.281737 | -0.1603 | 1.218923 | 0.105779 | 0.571687 |
| 33.25 | 1.143586 | -0.16255 | 0.417853 | 0.187822 | 0.261788 | 1.731035 | 0.563463 | 1.109032 | 2.711771 | 0.870332 | 0.167506 | 1.184856 | -0.00119 | 0.547592 | 0.185052 | -0.30465 | 1.082785 | 0.114587 | 0.325045 |
| 33.26389 | 0.611833 | -0.14947 | 0.136722 | -0.14832 | 0.001714 | 1.062423 | 0.279109 | 1.087587 | 2.547802 | 1.282421 | 0.293312 | 1.193044 | -0.00981 | 0.565084 | 0.146071 | -0.18657 | 1.054447 | 0.073922 | 0.114997 |
| 33.27778 | 0.027275 | -0.49789 | 0.005039 | -0.32124 | -0.12791 | 0.242606 | 0.036482 | 0.759217 | 2.732695 | 1.075829 | 0.110464 | 1.230943 | 0.100672 | 0.586391 | 0.167075 | -0.16593 | 1.146891 | -0.08844 | 0.045456 |
| 33.29167 | -0.15278 | -0.14897 | 0.058256 | -0.26376 | -0.09829 | -0.33391 | -0.18152 | 0.660754 | 2.685031 | 0.599507 | -0.18063 | 1.373142 | 0.180645 | 0.443311 | 0.074919 | -0.08458 | 1.249559 | -0.20852 | 0.049962 |
| 33.30556 | 0.092777 | 0.821187 | 0.270927 | -0.04151 | 0.081286 | -0.34258 | -0.14239 | 0.631763 | 2.313206 | 0.256195 | -0.30151 | 1.053353 | 0.274764 | 0.25445 | 0.05276 | 0.14859 | 1.323052 | -0.1628 | 0.13041 |
| 33.31944 | 0.287389 | 1.048121 | 0.381404 | 0.112118 | 0.23622 | -0.24494 | -0.0831 | 0.420077 | 1.321012 | 0.130106 | -0.3034 | 0.32238 | 0.329296 | 0.103088 | 0.038158 | 0.348139 | 1.120472 | -0.06922 | 0.220768 |
| 33.33333 | 0.518767 | 0.885372 | 0.433421 | 0.258591 | 0.426819 | -0.15389 | 0.175213 | 0.353818 | 1.095274 | 0.006334 | -0.30879 | -0.50213 | 0.240787 | -0.16077 | -0.03349 | 0.311623 | 0.439786 | -0.00633 | 0.210188 |
| 33.34722 | 0.941372 | 1.466648 | 0.455666 | 0.408759 | 0.652703 | -0.14471 | 0.293544 | 0.361442 | 1.150794 | -0.22748 | -0.34593 | -0.97908 | 0.127777 | -0.42099 | -0.11982 | 0.153975 | -0.17519 | -0.03806 | 0.193695 |
| 33.36111 | 1.01657 | 1.700539 | 0.404963 | 0.375565 | 0.692848 | -0.47985 | 0.131137 | 0.260479 | 0.848736 | -0.49916 | -0.30596 | -0.92812 | 0.102696 | -0.66917 | -0.06868 | 0.010511 | -0.44252 | -0.068 | 0.229696 |
| 33.375 | 0.822908 | 1.312223 | 0.5032 | 0.253216 | 0.524589 | -1.06902 | -0.09837 | 0.182514 | 0.226533 | -0.57031 | -0.32042 | -0.85512 | 0.135382 | -0.74055 | -0.02404 | -0.08675 | -0.58098 | -0.10335 | 0.194002 |
| 33.38889 | 0.780929 | 1.087222 | 0.728378 | 0.279859 | 0.381025 | -1.30409 | -0.08282 | 0.146571 | -0.26139 | -0.49806 | -0.30826 | -0.59444 | 0.163497 | -0.54827 | 0.008779 | -0.05963 | -0.33781 | -0.18431 | 0.175525 |
| 33.40278 | 0.905528 | 1.09953 | 0.90027 | 0.320005 | 0.301764 | -1.40868 | 0.059034 | 0.10336 | -0.45542 | -0.20867 | -0.09044 | -0.23918 | 0.250676 | -0.09783 | 0.073974 | 0.199598 | 0.253949 | -0.08794 | 0.204486 |
| 33.41667 | 0.926822 | 1.019394 | 0.968123 | 0.27133 | 0.191497 | -1.33321 | 0.2117 | 0.078037 | -0.41233 | 0.128721 | 0.210164 | 0.06252 | 0.306084 | 0.384065 | 0.097548 | 0.367862 | 0.806487 | 0.088161 | 0.223382 |
| 33.43056 | 0.99992 | 0.955273 | 0.974012 | 0.310089 | 0.173435 | -1.02626 | 0.346955 | 0.101971 | -0.31037 | 0.331857 | 0.377956 | 0.212432 | 0.319884 | 0.803629 | 0.04034 | 0.339353 | 1.062702 | 0.169716 | 0.209326 |
| 33.44444 | 1.029146 | 0.902236 | 0.818399 | 0.234088 | 0.134264 | -1.12099 | 0.253781 | 0.007555 | -0.27412 | 0.388018 | 0.466174 | 0.280126 | 0.325895 | 1.027821 | -0.05333 | 0.314017 | 0.936545 | 0.230045 | 0.243117 |
| 33.45833 | 0.82683 | 0.541677 | 0.576171 | 0.081022 | -0.0269 | -1.59304 | 0.011607 | -0.10149 | -0.38573 | 0.349806 | 0.438636 | 0.33165 | 0.293092 | 0.965684 | -0.17501 | 0.253651 | 0.512169 | 0.360997 | 0.209631 |
| 33.47222 | 0.506725 | 0.296415 | 0.511706 | 0.016345 | -0.13629 | -1.83649 | -0.16804 | -0.07991 | -0.51227 | 0.316157 | 0.147546 | 0.386154 | 0.193089 | 0.683932 | -0.23123 | 0.128508 | 0.083497 | 0.410663 | 0.150285 |
| 33.48611 | 0.399707 | 0.958991 | 0.763751 | 0.429395 | -0.07414 | -1.54082 | -0.00105 | 0.101389 | -0.19872 | 0.421444 | -0.12588 | 0.434686 | 0.155546 | 0.379365 | -0.16149 | 0.095479 | -0.19617 | 0.39396 | 0.138751 |

| | | | | | | | | | | | | | | | | | | | |
|---|---|---|---|---|---|---|---|---|---|---|---|---|---|---|---|---|---|---|---|
| 33.5 | 0.651487 | 2.424832 | 1.463057 | 1.243415 | 0.311875 | -0.29316 | 0.667346 | 0.449999 | 0.517337 | 0.570589 | -0.14476 | 0.512406 | 0.13549 | 0.219986 | -0.09018 | 0.147027 | -0.22401 | 0.337344 | 0.134506 |
| 33.51389 | 1.458478 | 3.564491 | 2.203797 | 2.122619 | 0.731209 | 0.588339 | 1.418797 | 0.76899 | 1.478971 | 0.619499 | -0.00232 | 0.48616 | 0.10103 | 0.218141 | -0.09286 | 0.148273 | -0.05241 | 0.21769 | 0.091991 |
| 33.52778 | 2.181524 | 2.745991 | 2.338331 | 2.615995 | 1.002854 | 1.200624 | 1.97421 | 0.874241 | 2.143286 | 0.546608 | 0.185441 | 0.467853 | 0.144123 | 0.373683 | -0.05157 | 0.139082 | 0.323674 | 0.122221 | 0.103217 |
| 33.54167 | 2.7069 | 2.268517 | 2.163577 | 2.800639 | 0.876984 | 1.537639 | 1.609849 | 0.855036 | 2.875505 | 0.434252 | 0.327056 | 0.378193 | 0.214065 | 0.559479 | 0.016488 | 0.145882 | 0.651764 | 0.106154 | 0.116532 |
| 33.55556 | 2.757392 | 2.38924 | 1.671492 | 2.276725 | 1.161302 | 1.472441 | 1.44966 | 0.831981 | 3.054784 | 0.311007 | 0.386443 | 0.256418 | 0.233713 | 0.580907 | -0.01732 | 0.143026 | 0.719472 | 0.089252 | 0.058355 |
| 33.56944 | 2.679938 | 2.623429 | 1.297065 | 1.949599 | 1.196041 | 1.724263 | 0.9402 | 0.763711 | 2.394697 | 0.266105 | 0.388142 | 0.290049 | 0.266086 | 0.508654 | -0.01195 | 0.170503 | 0.641907 | 0.097531 | 0.032547 |
| 33.58333 | 2.344456 | 2.546292 | 1.009988 | 1.586851 | 1.04755 | 1.501119 | 0.222239 | 0.65982 | 1.641607 | 0.349461 | 0.493241 | 0.437612 | 0.288931 | 0.443511 | 0.073181 | 0.237985 | 0.611616 | 0.141759 | 0.084841 |
| 33.59722 | 1.705866 | 2.549412 | 0.745991 | 1.183251 | 1.101846 | 1.206909 | -0.20657 | 0.646882 | 1.063344 | 0.527244 | 0.830486 | 0.726085 | 0.301583 | 0.745722 | 0.150352 | 0.406053 | 1.039029 | 0.282579 | 0.115338 |
| 33.61111 | 1.044556 | 2.251278 | 0.608802 | 0.765718 | 1.049198 | 1.135946 | -0.29305 | 0.705788 | 0.585109 | 0.981447 | 1.572687 | 1.267568 | 0.476762 | 1.656932 | 0.288597 | 0.725576 | 2.349395 | 0.662321 | 0.234588 |
| 33.625 | 0.372264 | 1.451283 | 0.435475 | 0.577186 | 0.786963 | 1.070773 | -0.02942 | 0.5117 | -0.06304 | 1.348512 | 1.850093 | 1.785655 | 0.76568 | 2.251846 | 0.31232 | 0.807282 | 2.646429 | 0.936042 | 0.339754 |
| 33.63889 | -0.18857 | 0.317246 | 0.411267 | 0.232495 | 0.302157 | 0.818529 | 0.391038 | -0.04691 | -0.79492 | 1.432054 | 1.441906 | 2.03507 | 0.932798 | 2.104326 | 0.195341 | 0.565206 | 2.594157 | 0.721111 | 0.206047 |
| 33.65278 | -0.90578 | -0.51651 | 0.417561 | -0.26274 | -0.35969 | 0.266512 | 0.563068 | -0.61191 | -1.4993 | 1.226198 | 1.181227 | 2.04017 | 0.688804 | 1.953994 | -0.02978 | 0.36157 | 2.814725 | 0.511348 | -0.09632 |
| 33.66667 | -1.22052 | -1.14821 | 0.377017 | -0.63544 | -0.93449 | -0.24686 | 0.534961 | -1.07086 | -2.00622 | 0.683819 | 1.059104 | 1.713129 | 0.424016 | 1.693267 | -0.13935 | -0.05504 | 2.401165 | 0.337478 | -0.19042 |
| 33.68056 | -1.21186 | -1.63783 | 0.271366 | -0.71724 | -1.1769 | -0.64846 | 0.456384 | -1.66394 | -2.69915 | -0.02209 | 1.007957 | 0.805391 | 0.097632 | 1.254141 | -0.20268 | -0.42568 | 1.462477 | -0.11844 | 0.067676 |
| 33.69444 | -0.98957 | -1.72175 | 0.260323 | -0.57796 | -1.09274 | -0.87727 | 0.314909 | -2.05714 | -3.20042 | -0.48443 | 1.033591 | -0.00186 | -0.14723 | 0.817219 | -0.27718 | -0.68081 | 0.665601 | -0.58051 | 0.589195 |
| 33.70833 | -0.91665 | -1.64089 | 0.318786 | -0.18818 | -0.88543 | -0.74665 | 0.230372 | -1.92349 | -3.13021 | -0.4107 | 1.016846 | -0.43617 | -0.44065 | 0.464682 | -0.21407 | -0.85866 | 0.199328 | -0.56114 | 0.8762 |
| 33.72222 | -0.7771 | -1.75956 | 0.532805 | 0.26678 | -0.57968 | -0.60226 | 0.391066 | -1.23794 | -2.59898 | -0.13282 | 0.89422 | -0.53913 | -0.28763 | 0.140951 | -0.01414 | -0.45481 | -0.13704 | -0.52543 | 1.139132 |
| 33.73611 | -0.42522 | -1.51189 | 0.859765 | 0.919807 | -0.28033 | -0.53804 | 0.586427 | -0.4905 | -1.4308 | 0.023591 | 0.736592 | -0.68944 | -0.05031 | -0.2455 | 0.127673 | -0.09047 | -0.42044 | -0.50837 | 0.969211 |
| 33.75 | -0.50502 | -1.18643 | 1.076116 | 1.304414 | -0.06991 | -0.35503 | 0.659241 | -0.08286 | 0.192793 | -0.08576 | 0.551452 | -0.71355 | 0.060364 | -0.59565 | 0.122449 | -0.0215 | -0.63982 | -0.37703 | 0.552406 |
| 33.76389 | -0.60008 | -0.80185 | 1.169193 | 1.30721 | -0.06596 | -0.09357 | 0.669082 | 0.025044 | 1.671384 | -0.30522 | 0.355342 | -0.57256 | 0.013011 | -0.80215 | 0.021049 | -0.01596 | -0.74546 | -0.1434 | 0.112996 |
| 33.77778 | -0.72403 | -0.65452 | 1.105764 | 1.0548 | -0.2545 | 0.282224 | 0.57917 | 0.075076 | 2.50818 | -0.51211 | 0.176734 | -0.45844 | -0.14123 | -0.95828 | -0.08131 | -0.07063 | -0.90679 | 0.012332 | -0.32009 |
| 33.79167 | -0.84193 | -0.31383 | 0.818015 | 0.627123 | -0.73123 | 0.649824 | 0.36587 | 0.180998 | 2.507263 | -0.56934 | 0.009549 | -0.38312 | -0.37974 | -1.14442 | -0.24854 | -0.18366 | -1.04665 | 0.000704 | -0.72031 |
| 33.80556 | -1.56781 | -0.60025 | 0.404818 | -0.09434 | -1.39576 | 0.842942 | -0.09156 | 0.293228 | 2.083381 | -0.28108 | -0.07839 | -0.15905 | -0.45245 | -1.20055 | -0.2045 | -0.17967 | -1.0699 | -0.02843 | -0.82937 |
| 33.81944 | -2.73902 | -1.72256 | -0.20679 | -1.05398 | -2.18249 | 0.872488 | -0.52665 | 0.244384 | 1.320492 | 0.138036 | -0.04314 | 0.234192 | -0.22954 | -0.88392 | 0.011242 | -0.02723 | -0.88653 | -0.00455 | -0.63591 |
| 33.83333 | -3.68632 | -3.08755 | -0.81405 | -1.75685 | -2.87542 | 0.752795 | -0.63147 | -0.01318 | 0.066866 | 0.421013 | -0.01717 | 0.639998 | 0.132747 | -0.53124 | 0.230715 | 0.09369 | -0.48742 | 0.015871 | -0.44322 |
| 33.84722 | -4.09546 | -3.41866 | -1.16775 | -2.18037 | -3.7108 | 0.650126 | -0.41965 | -0.34795 | -1.20191 | 0.503292 | -0.0502 | 0.897434 | 0.257136 | -0.22883 | 0.304788 | 0.140638 | -0.04317 | 0.01383 | -0.23485 |
| 33.86111 | -3.6255 | -3.30521 | -1.26228 | -2.50755 | -4.29012 | 0.47137 | -0.16228 | -0.61611 | -2.54417 | 0.403413 | -0.05438 | 0.938135 | 0.218265 | 0.011534 | 0.31869 | 0.222818 | 0.32335 | 0.072887 | 0.033765 |
| 33.875 | -3.47098 | -3.79904 | -1.31923 | -2.69744 | -4.09779 | 0.272725 | -0.04228 | -0.73545 | -3.86372 | 0.290473 | -0.00325 | 0.820418 | 0.13549 | 0.204599 | 0.384156 | 0.362532 | 0.493137 | 0.132325 | 0.184444 |
| 33.88889 | -2.99041 | -4.2018 | -1.39056 | -2.4764 | -3.44617 | 0.014067 | -0.06399 | -0.71076 | -4.63456 | 0.157842 | -0.00879 | 0.561017 | 0.065214 | 0.209755 | 0.367341 | 0.346872 | 0.306573 | 0.101394 | 0.149517 |
| 33.90278 | -2.08405 | -4.15982 | -1.11442 | -2.06681 | -2.37976 | -0.2417 | -0.11778 | -0.5812 | -4.64512 | -0.05227 | -0.12744 | 0.2791 | 0.02799 | 0.04299 | 0.341162 | 0.222558 | -0.03281 | -0.01414 | -0.0534 |
| 33.91667 | -1.39798 | -3.46221 | -0.728 | -1.44468 | -0.99676 | -0.45872 | -0.1816 | -0.53295 | -4.28265 | -0.33428 | -0.23323 | -0.05053 | 0.041774 | -0.07704 | 0.298867 | 0.101084 | -0.33102 | -0.1459 | -0.18096 |
| 33.93056 | -0.70057 | -2.86166 | -0.20974 | -0.63703 | 0.451682 | -0.32976 | -0.12943 | -0.42754 | -4.07281 | -0.60385 | -0.30802 | -0.45113 | 0.124376 | -0.02926 | 0.11582 | 0.062885 | -0.58074 | -0.20534 | -0.13743 |
| 33.94444 | 0.037537 | -2.35958 | 0.321057 | 0.324031 | 1.957594 | -0.01976 | 0.224659 | -0.21334 | -3.16079 | -0.91757 | -0.50815 | -0.87167 | 0.118282 | 0.13033 | 0.041901 | -0.02475 | -0.78052 | -0.23461 | -0.03566 |
| 33.95833 | 0.695221 | -1.66977 | 0.776042 | 1.267224 | 2.826437 | 0.626218 | 0.645816 | 0.069642 | -2.07806 | -1.11341 | -0.88397 | -1.20419 | -0.03935 | 0.139921 | -0.00099 | -0.09419 | -0.80219 | -0.28145 | 0.093019 |
| 33.97222 | 0.896783 | -1.17975 | 0.986965 | 1.89009 | 1.79378 | 1.246099 | 0.983374 | 0.3142 | -1.11411 | -1.10425 | -1.11779 | -1.53615 | -0.22846 | -0.02221 | -0.01145 | -0.05811 | -0.61244 | -0.24703 | 0.194933 |
| 33.98611 | 0.32007 | -1.36119 | 0.424526 | 1.708429 | 0.698739 | 0.650205 | 0.765451 | 0.013076 | -1.36916 | -0.80667 | -1.07158 | -1.10352 | -0.29596 | -0.27835 | 0.103435 | 0.08646 | -0.28658 | -0.07333 | 0.259138 |
| 34 | -1.48007 | -3.08387 | -1.21987 | -0.44681 | -0.3971 | -1.40988 | -0.2873 | -0.7922 | -3.68626 | -0.4258 | -0.8991 | -0.58512 | -0.24409 | -0.50176 | -0.0692 | 0.059906 | 0.013586 | 0.14566 | 0.217447 |
| 34.01389 | -3.92677 | -5.2103 | -2.2873 | -3.05727 | -2.0445 | -1.95074 | -1.1387 | -1.34375 | -4.83069 | -0.10144 | -0.76267 | -0.13683 | -0.1718 | -0.70309 | -0.24066 | 0.019385 | 0.190661 | 0.251072 | 0.100003 |
| 34.02778 | -4.27401 | -6.14139 | -2.46304 | -3.47646 | -2.87366 | -1.79283 | -0.9948 | -1.34511 | -4.79263 | 0.173814 | -0.62323 | 0.333015 | -0.06803 | -0.65386 | -0.26702 | 0.015369 | 0.335896 | 0.295703 | 0.01229 |
| 34.04167 | -3.85282 | -4.47115 | -2.4443 | -2.12684 | -2.66584 | -1.77038 | -0.77337 | -1.06165 | -4.88468 | 0.391071 | -0.44347 | 0.845908 | 0.137012 | -0.50395 | -0.04389 | 0.109946 | 0.431682 | 0.321639 | 0.036969 |
| 34.05556 | -3.73633 | -3.58232 | -1.84587 | -1.50224 | -2.35716 | -1.43604 | -0.46192 | -1.00235 | -4.14906 | 0.527604 | -0.36065 | 1.117493 | 0.264413 | -0.33157 | 0.123049 | 0.086196 | 0.490772 | 0.322752 | 0.174415 |
| 34.06944 | -2.95163 | -2.77106 | -1.24905 | -0.94364 | -1.48521 | -1.13494 | -0.15965 | -0.94306 | -3.34049 | 0.554439 | -0.21315 | 1.159644 | 0.209479 | -0.26162 | 0.164795 | -0.04612 | 0.440515 | 0.305713 | 0.314479 |
| 34.08333 | -1.69243 | -2.30849 | -0.69235 | -0.40177 | -0.64235 | -1.02357 | 0.085908 | -0.73663 | -2.26542 | 0.415897 | -0.19782 | 0.934994 | 0.081053 | -0.28507 | 0.038125 | -0.27069 | 0.239573 | 0.175968 | 0.298774 |
| 34.09722 | -0.68419 | -1.68347 | -0.38103 | 0.004623 | 0.027997 | -1.08339 | 0.472314 | -0.50025 | -1.45571 | -0.22109 | -0.65331 | 0.064038 | -0.08663 | -0.71355 | -0.3503 | -0.89956 | -0.57309 | -0.49977 | -0.08012 |
| 34.11111 | -0.23509 | -1.14625 | -0.17161 | 0.171081 | 0.454248 | -1.00778 | 0.651457 | -0.26028 | -0.82293 | -1.7347 | -1.96316 | -1.79107 | -0.90737 | -1.83895 | -0.93946 | -2.0828 | -2.24823 | -2.12958 | -0.90033 |
| 34.125 | 0.111659 | -0.74298 | 0.274611 | 0.418281 | 0.837559 | -0.7666 | 0.828741 | -0.118 | -0.41914 | -3.1251 | -3.25506 | -3.20196 | -1.6554 | -2.85256 | -1.15066 | -2.59945 | -3.40974 | -3.32991 | -1.41786 |
| 34.13889 | 0.69631 | -0.31019 | 0.639517 | 0.746439 | 1.031718 | -0.41255 | 0.872555 | -0.05623 | -0.00102 | -2.59084 | -1.64144 | -1.92744 | -0.82261 | -1.70852 | -0.49314 | -1.17917 | -1.98287 | -1.7754 | -0.68608 |
| 34.15278 | 1.407953 | -0.06574 | 0.793965 | 1.139652 | 0.498169 | 0.017109 | 0.705161 | -0.08771 | 0.261588 | -1.06771 | -0.2493 | 0.092708 | 0.302293 | -0.10106 | 0.277526 | 0.258299 | 0.271854 | 0.162069 | 0.310011 |
| 34.16667 | 1.880326 | 0.078542 | 1.019644 | 1.541047 | 0.55329 | 0.537935 | 0.701117 | -0.02274 | 0.524264 | 0.055335 | 0.509102 | 0.795299 | 0.764211 | 0.623487 | 0.431577 | 0.73531 | 1.275433 | 1.116193 | 0.72598 |
| 34.18056 | 1.805218 | 0.522518 | 1.270021 | 1.72953 | 0.905734 | 1.233563 | 1.00333 | 0.23359 | 0.963678 | 1.007367 | 0.822983 | 1.168976 | 0.773544 | 0.798244 | 0.522835 | 0.649378 | 1.445981 | 1.353411 | 0.732252 |
| 34.19444 | 1.62099 | 0.818901 | 1.308234 | 2.006021 | 1.292799 | 2.033253 | 1.364849 | 0.385483 | 1.37552 | 1.580123 | 0.869407 | 1.472206 | 1.159507 | 0.790336 | 0.724364 | 0.58212 | 1.456992 | 1.417583 | 0.667499 |
| 34.20833 | 1.207374 | 0.680942 | 1.054219 | 2.219623 | 1.351547 | 3.227984 | 1.338608 | 0.292929 | 1.333037 | 1.357628 | 0.715724 | 1.511588 | 1.748749 | 0.710397 | 0.80108 | 0.509311 | 1.293652 | 1.421728 | 0.437712 |
| 34.22222 | 0.779935 | 0.208274 | 0.783391 | 2.157902 | 1.030163 | 3.588087 | 1.116232 | 0.150001 | 1.002293 | 1.043194 | 0.591381 | 1.009113 | 1.17593 | 0.6813 | 0.623767 | 0.485556 | 1.079272 | 1.202362 | 0.255022 |
| 34.23611 | 0.236663 | -0.95617 | 0.62285 | 1.379716 | 0.2009 | 3.21992 | 0.816503 | 0.082734 | 1.065882 | 0.789721 | 0.73082 | 0.610517 | 0.683357 | 0.849712 | 0.635655 | 0.538323 | 1.089991 | 1.003336 | 0.23701 |
| 34.25 | -0.59046 | -3.15949 | 0.529375 | 0.857532 | -1.03103 | 2.146155 | 0.405541 | 0.125567 | 1.4406 | 0.78159 | 0.7902 | 0.359765 | 0.585179 | 0.945402 | 0.53517 | 0.552644 | 1.157362 | 0.912651 | 0.213299 |
| 34.26389 | -0.75074 | -3.79443 | 0.25257 | 0.212605 | -1.86377 | 1.507278 | 0.156558 | 0.237393 | 1.908014 | 0.948169 | 0.604208 | 0.125883 | 0.439666 | 1.070528 | 0.314676 | 0.366902 | 1.092062 | 0.731389 | 0.116559 |
| 34.27778 | -0.24646 | -2.27164 | -0.1147 | 0.279051 | -1.63959 | 1.031369 | 0.22656 | 0.334994 | 2.077788 | 1.290415 | 0.395097 | 0.230852 | 0.180953 | 1.171283 | 0.182578 | 0.246584 | 0.857605 | 0.389524 | 0.059301 |
| 34.29167 | 0.14416 | -1.59807 | -0.26729 | 0.159616 | -1.37939 | 0.597167 | 0.288194 | 0.361327 | 1.890145 | 1.72197 | 0.41323 | 0.880554 | 0.018541 | 1.023022 | 0.150439 | 0.292477 | 0.820662 | 0.056478 | 0.205919 |
| 34.30556 | 0.208639 | -1.6216 | -0.1903 | -0.14477 | -1.39041 | 0.044969 | 0.33532 | 0.336577 | 1.805903 | 1.841247 | 0.288196 | 1.125166 | -0.04002 | 0.733618 | 0.104588 | 0.295198 | 0.721367 | -0.2608 | 0.226598 |
| 34.31944 | 0.072986 | -1.62924 | -0.18841 | -0.40471 | -1.24304 | -0.76438 | 0.361414 | 0.298669 | 1.73888 | 1.365716 | -0.064 | 0.991022 | -0.01626 | 0.223028 | 0.008465 | 0.153137 | 0.509239 | -0.41861 | 0.165852 |
| 34.33333 | 0.004094 | -1.44468 | -0.15878 | -0.33891 | -1.0852 | -1.79475 | 0.282152 | 0.196912 | 1.822771 | 0.617561 | -0.19999 | 0.375713 | -0.01165 | -0.20378 | 0.027061 | -0.01482 | 0.261243 | -0.63532 | 0.097305 |
| 34.34722 | -0.04576 | -1.2556 | -0.14652 | -0.15061 | -1.00298 | -2.50674 | 0.149675 | 0.058067 | 2.039084 | 0.067412 | -0.23132 | 0.045469 | 0.098715 | -0.42709 | 0.077821 | -0.14805 | 0.061138 | -0.74317 | 0.035172 |
| 34.36111 | -0.42107 | -1.56213 | -0.0714 | -0.20926 | -1.05777 | -2.57053 | -0.13351 | -0.0562 | 2.297877 | -0.1515 | -0.17226 | 0.181079 | 0.236125 | -0.31219 | 0.093872 | -0.10332 | -0.03492 | -0.54788 | 0.15896 |
| 34.375 | -0.95487 | -2.40219 | 0.17907 | -0.63553 | -1.6226 | -2.11001 | -0.23092 | -0.15539 | 2.693628 | -0.14136 | -0.17943 | 0.269811 | 0.31081 | -0.1695 | 0.155612 | 0.062965 | -0.04449 | -0.29895 | 0.313894 |
| 34.38889 | -0.83869 | -2.26274 | 0.408277 | -0.67122 | -1.60053 | -1.14909 | -0.13326 | -0.21379 | 2.445923 | -0.08736 | -0.2235 | 0.121796 | 0.25461 | -0.15555 | 0.139036 | 0.12945 | -0.09497 | -0.10011 | 0.291986 |
| 34.40278 | -0.19068 | -1.22491 | 0.410711 | -0.1585 | -0.88839 | -0.34987 | 0.313451 | -0.07315 | 1.825676 | -0.18641 | -0.38267 | 0.01902 | 0.281251 | -0.07348 | 0.094393 | 0.033934 | -0.11093 | 0.03132 | 0.218227 |
| 34.41667 | -0.02728 | -0.6883 | 0.301123 | 0.05562 | -0.93576 | -0.1496 | 0.621806 | 0.012919 | 1.68554 | -0.31622 | -0.41995 | -0.05405 | 0.34086 | 0.086137 | 0.108096 | 0.043565 | -0.0047 | 0.155405 | 0.236025 |
| 34.43056 | -0.31428 | -0.67088 | 0.042903 | 0.105726 | -1.67029 | -0.45079 | 0.578586 | -0.0806 | 1.527927 | -0.3959 | -0.35509 | -0.1781 | 0.556903 | 0.303262 | 0.212501 | 0.201449 | 0.164552 | 0.238414 | 0.266195 |
| 34.44444 | -0.53702 | -0.51298 | -0.21391 | 0.288901 | -1.7741 | -0.82847 | 0.394191 | -0.25773 | 1.011985 | -0.32772 | -0.17566 | -0.07167 | 0.59935 | 0.485582 | 0.217012 | 0.294364 | 0.263391 | 0.295249 | 0.246802 |
| 34.45833 | -0.20612 | -0.47625 | -0.18935 | 0.219508 | -1.13549 | -0.90019 | 0.301432 | -0.27301 | 0.678225 | -0.08563 | 0.054168 | 0.2669 | 0.629012 | 0.567918 | 0.104056 | 0.27844 | 0.346059 | 0.403653 | 0.207893 |
| 34.47222 | 0.080905 | -0.09801 | 0.155783 | 0.276811 | -0.3127 | -0.53119 | 0.473563 | -0.17556 | 0.652877 | 0.307192 | 0.33744 | 0.661621 | 0.669737 | 0.592666 | 0.080036 | 0.276399 | 0.405729 | 0.564488 | 0.270254 |
| 34.48611 | 0.573324 | 0.618718 | 0.708319 | 0.592708 | 0.09297 | -0.12591 | 0.7854 | -0.09409 | 0.898793 | 0.674111 | 0.521058 | 0.88561 | 0.585539 | 0.514839 | 0.115017 | 0.267977 | 0.378001 | 0.659038 | 0.333171 |
| 34.5 | 1.256995 | 1.731054 | 1.603568 | 1.310777 | 0.545149 | 0.615703 | 1.254802 | 0.046628 | 1.857688 | 0.727471 | 0.444643 | 0.776833 | 0.176235 | 0.213898 | 0.028651 | 0.14267 | 0.242234 | 0.561195 | 0.257875 |
| 34.51389 | 2.007974 | 2.88103 | 2.533312 | 2.233331 | 1.319804 | 2.0657 | 1.79769 | 0.35015 | 3.094038 | 0.587825 | 0.310032 | 0.530442 | -0.28588 | -0.1265 | -0.04615 | 0.039603 | 0.148256 | 0.307531 | 0.170714 |
| 34.52778 | 2.831293 | 3.67295 | 3.387066 | 2.972525 | 2.000585 | 2.632243 | 2.248429 | 0.668236 | 3.675761 | 0.594832 | 0.309924 | 0.428274 | -0.3684 | -0.24928 | 0.027355 | 0.109445 | 0.16054 | 0.167255 | 0.179458 |
| 34.54167 | 3.329201 | 4.414964 | 3.412571 | 3.367244 | 2.232727 | 2.776269 | 2.364773 | 0.600496 | 4.138336 | 0.644303 | 0.367534 | 0.468069 | -0.17993 | -0.19354 | 0.092203 | 0.183588 | 0.158813 | 0.039782 | 0.133813 |
| 34.55556 | 3.078135 | 5.347975 | 2.601438 | 3.084925 | 2.065163 | 2.610835 | 1.96112 | 0.400751 | 4.213203 | 0.715438 | 0.341898 | 0.393368 | -0.01969 | -0.04903 | 0.092361 | 0.182641 | 0.235912 | -0.07908 | 0.048902 |
| 34.56944 | 2.266453 | 5.79653 | 1.625306 | 1.956765 | 1.831685 | 2.868307 | 1.307858 | 0.514119 | 4.508533 | 0.87303 | 0.308481 | 0.514936 | 0.140692 | 0.179158 | 0.133734 | 0.203141 | 0.362751 | -0.03353 | -0.00053 |
| 34.58333 | 1.429498 | 5.68836 | 0.699126 | 0.781481 | 1.287587 | 2.585379 | 0.66518 | 0.732643 | 4.413524 | 0.99131 | 0.289556 | 0.691382 | 0.21462 | 0.425155 | 0.257945 | 0.25469 | 0.550965 | 0.095527 | -0.06962 |
| 34.59722 | 1.241442 | 5.043516 | 0.013577 | -0.05763 | 0.811796 | 2.186733 | 0.435878 | 0.713596 | 4.579964 | 1.088627 | 0.503391 | 0.982136 | 0.1658 | 0.756852 | 0.270011 | 0.368288 | 0.703459 | 0.30906 | -0.02247 |
| 34.61111 | 1.251925 | 3.926664 | -0.27208 | -0.39549 | 0.75203 | 1.874384 | 0.638662 | 0.390273 | 3.731403 | 1.383384 | 1.251304 | 1.575676 | 0.15888 | 1.462183 | 0.270162 | 0.574111 | 1.290855 | 0.792962 | 0.238093 |
| 34.625 | 0.757873 | 2.712959 | -0.3508 | -0.42905 | 0.631115 | 1.33199 | 1.018093 | 0.028328 | 2.293724 | 1.560222 | 1.630055 | 1.812829 | 0.400301 | 2.233384 | 0.322194 | 0.732476 | 2.317463 | 1.234537 | 0.521899 |
| 34.63889 | 0.080645 | 0.805571 | -0.50799 | -1.0804 | -0.1063 | 0.124229 | 1.160093 | -0.32152 | 0.724533 | 1.389892 | 1.408492 | 1.574769 | 0.633489 | 2.237034 | 0.404818 | 0.627733 | 2.039772 | 1.288888 | 0.529269 |
| 34.65278 | -0.82041 | -0.95971 | -0.85359 | -1.55631 | -1.30547 | -2.04228 | 1.099188 | -0.57759 | -0.80796 | 1.193346 | 1.126557 | 1.393652 | 0.622984 | 1.957092 | 0.426068 | 0.372695 | 1.458712 | 0.950259 | 0.35191 |
| 34.66667 | -1.52461 | -2.39752 | -1.01631 | -0.96219 | -1.99044 | -2.9669 | 1.384252 | -0.79284 | -2.38214 | 0.865996 | 1.120026 | 1.220674 | 0.388417 | 1.701263 | 0.345155 | 0.198111 | 1.208857 | 0.685676 | 0.151954 |
| 34.68056 | -1.74519 | -3.20984 | -0.9611 | -0.4458 | -2.53624 | -3.38536 | 1.575168 | -0.99809 | -3.78118 | 0.408929 | 1.36848 | 0.961737 | 0.218045 | 1.308862 | -0.00331 | 0.011737 | 0.77998 | 0.185146 | -0.09349 |

| | | | | | | | | | | | | | | | | | | | |
|---|---|---|---|---|---|---|---|---|---|---|---|---|---|---|---|---|---|---|---|
| 34.69444 | -1.55344 | -3.39896 | -0.72369 | -0.22299 | -2.81879 | -2.94778 | 1.584038 | -1.00533 | -4.2025 | -0.10123 | 1.387853 | 0.504432 | 0.034067 | 1.080837 | -0.35968 | -0.21716 | 0.352381 | -0.54676 | -0.30385 |
| 34.70833 | -1.22119 | -2.94541 | -0.50639 | -0.04361 | -2.29293 | -2.05467 | 1.673703 | -0.77551 | -4.16858 | -0.4532 | 1.262868 | 0.062938 | -0.11337 | 1.073976 | -0.52349 | -0.476 | 0.197152 | -1.32848 | -0.06083 |
| 34.72222 | -0.71439 | -2.30283 | -0.1867 | 0.252741 | -1.38012 | -0.97308 | 1.715432 | -0.67919 | -4.18871 | -0.45119 | 1.009267 | -0.20371 | -0.07618 | 1.101112 | -0.45128 | -0.30231 | 0.422857 | -1.33334 | 0.20211 |
| 34.73611 | -0.37101 | -1.61746 | 0.349623 | 0.451192 | -0.47026 | -0.00759 | 1.711755 | -0.61311 | -3.91304 | -0.41658 | 0.92669 | -0.2569 | -0.00858 | 1.069879 | -0.26786 | -0.05989 | 0.439824 | -1.0594 | 0.302592 |
| 34.75 | -0.20911 | -1.08857 | 1.047628 | 0.85251 | 0.101137 | 0.745819 | 1.705951 | -0.43684 | -2.88053 | -0.51746 | 0.875451 | -0.44724 | 0.007559 | 0.683817 | -0.02575 | 0.119378 | 0.303758 | -0.89965 | 0.213702 |
| 34.76389 | -0.04641 | -0.6163 | 1.503602 | 0.963431 | 0.110558 | 0.951941 | 1.579379 | -0.22447 | -1.6844 | -0.64845 | 0.817397 | -0.6297 | -0.12898 | 0.166196 | 0.1256 | 0.067817 | 0.1258 | -0.82005 | -0.08707 |
| 34.77778 | -0.03629 | -0.2419 | 1.41991 | 0.779201 | -0.03199 | 0.548829 | 1.467305 | -0.26002 | -0.8423 | -0.60712 | 0.729907 | -0.67461 | -0.23866 | -0.20358 | 0.000849 | -0.01543 | -0.09136 | -0.76935 | -0.29592 |
| 34.79167 | -0.0096 | 0.07941 | 1.077753 | 0.611478 | -0.1911 | -0.04112 | 1.342538 | -0.49946 | -0.20595 | -0.27389 | 0.823203 | -0.531 | -0.07227 | -0.32456 | -0.27208 | -0.07793 | -0.29477 | -0.58566 | -0.17858 |
| 34.80556 | 0.068936 | 0.260292 | 0.762228 | 0.339797 | -0.46731 | -0.50416 | 1.081178 | -0.60266 | 0.27378 | 0.181311 | 0.898526 | -0.38565 | 0.147258 | -0.40637 | -0.57325 | -0.20088 | -0.43259 | -0.32809 | 0.079608 |
| 34.81944 | 0.115547 | 0.284131 | 0.284925 | -0.00132 | -0.88415 | -1.00818 | 0.747002 | -0.62214 | 0.566478 | 0.56617 | 0.763717 | -0.20868 | 0.2716 | -0.19403 | -0.69071 | -0.26417 | -0.42814 | 0.027668 | 0.182095 |
| 34.83333 | -0.0032 | 0.13773 | -0.19698 | -0.20438 | -1.37487 | -1.4574 | 0.389491 | -0.6676 | 0.61509 | 0.967177 | 0.51114 | -0.06573 | 0.253863 | 0.08922 | -0.48005 | -0.1471 | -0.26631 | 0.464208 | 0.105806 |
| 34.84722 | -0.26986 | -0.22993 | -0.78437 | -0.40397 | -1.4644 | -1.77779 | -0.01253 | -0.79741 | 0.14588 | 1.152039 | 0.223933 | 0.160093 | 0.386281 | 0.299869 | -0.17626 | -0.05109 | -0.11664 | 0.803188 | 0.01884 |
| 34.86111 | -0.71024 | -0.79739 | -1.20951 | -0.51272 | -1.53245 | -2.15947 | -0.19856 | -0.84768 | -0.72972 | 1.016264 | -0.02705 | 0.45328 | 0.540822 | 0.314295 | 0.129501 | 0.055722 | -0.07899 | 0.738581 | -0.01881 |
| 34.875 | -0.98774 | -1.37486 | -1.30842 | -0.61383 | -2.1314 | -2.72138 | -0.20347 | -0.78926 | -1.7001 | 0.713187 | -0.27845 | 0.555834 | 0.502499 | 0.16069 | 0.356471 | 0.084055 | -0.12968 | 0.170212 | -0.09886 |
| 34.88889 | -1.19621 | -1.43608 | -1.31376 | -0.47319 | -2.71506 | -3.18794 | -0.13451 | -0.62267 | -2.35879 | 0.149341 | -0.55406 | 0.499358 | 0.321039 | -0.0769 | 0.335806 | 0.111737 | -0.15224 | -0.47757 | -0.09436 |
| 34.90278 | 0 | 0 | 0 | 0 | 0 | 0 | 0 | 0 | 0 | -0.47675 | -0.74957 | 0.377047 | 0.185652 | -0.20026 | 0.233808 | 0.144143 | -0.17816 | -0.90222 | 0.085169 |
| 34.91667 | 0 | 0 | 0 | 0 | 0 | 0 | 0 | 0 | 0 | -0.78573 | -0.78769 | 0.301592 | 0.138832 | -0.13573 | 0.036364 | 0.170905 | -0.16785 | -0.87507 | 0.38561 |
| 34.93056 | 0 | 0 | 0 | 0 | 0 | 0 | 0 | 0 | 0 | -0.85633 | -0.87472 | 0.245969 | 0.1523 | 0.002339 | -0.12321 | 0.150612 | -0.06895 | -0.68218 | 0.649483 |
| 34.94444 | 0 | 0 | 0 | 0 | 0 | 0 | 0 | 0 | 0 | -0.99862 | -1.06424 | 0.070177 | 0.166088 | 0.132983 | -0.31831 | 0.081792 | 0.022562 | -0.43481 | 0.568318 |
| 34.95833 | 0 | 0 | 0 | 0 | 0 | 0 | 0 | 0 | 0 | -1.15055 | -1.45158 | -0.16128 | 0.18163 | 0.250124 | -0.41533 | 0.077233 | 0.018673 | -0.26672 | 0.293497 |
| 34.97222 | 0 | 0 | 0 | 0 | 0 | 0 | 0 | 0 | 0 | -1.13097 | -1.78517 | -0.23252 | 0.255038 | 0.303085 | -0.30302 | 0.134535 | -0.03718 | -0.01893 | 0.166155 |
| 34.98611 | 0 | 0 | 0 | 0 | 0 | 0 | 0 | 0 | 0 | -0.97595 | -1.98752 | -0.18251 | 0.278546 | 0.341783 | -0.119 | 0.212 | -0.0311 | 0.267281 | 0.200084 |
| 35 | 0 | 0 | 0 | 0 | 0 | 0 | 0 | 0 | 0 | -0.70015 | -1.93066 | 0.016886 | 0.335081 | 0.435128 | 0.048374 | 0.271593 | -0.08933 | 0.434342 | 0.297225 |
| 35.01389 | 0 | 0 | 0 | 0 | 0 | 0 | 0 | 0 | 0 | -0.50068 | -1.67158 | 0.170716 | 0.396798 | 0.47457 | 0.090065 | 0.263123 | -0.14695 | 0.22266 | 0.26559 |
| 35.02778 | 0 | 0 | 0 | 0 | 0 | 0 | 0 | 0 | 0 | -0.3892 | -1.53009 | 0.167968 | 0.419772 | 0.477378 | 0.060853 | 0.23522 | -0.20739 | -0.03608 | 0.177128 |
| 35.04167 | 0 | 0 | 0 | 0 | 0 | 0 | 0 | 0 | 0 | -0.27709 | -1.30507 | 0.145165 | 0.369222 | 0.408719 | 0.075835 | 0.219299 | -0.21735 | 0.047579 | 0.247342 |
| 35.05556 | 0 | 0 | 0 | 0 | 0 | 0 | 0 | 0 | 0 | -0.10461 | -1.17583 | 0.230831 | 0.3584 | 0.18593 | 0.061134 | 0.243213 | -0.16995 | 0.414839 | 0.263142 |
| 35.06944 | 0 | 0 | 0 | 0 | 0 | 0 | 0 | 0 | 0 | 0.050688 | -0.83636 | 0.358503 | 0.193494 | -0.18921 | 0.137844 | 0.22663 | -0.07935 | 0.741057 | 0.280295 |
| 35.08333 | 0 | 0 | 0 | 0 | 0 | 0 | 0 | 0 | 0 | -0.0104 | -0.75211 | 0.40868 | -0.03791 | -0.64226 | 0.2328 | 0.122893 | 0.003755 | 0.703227 | 0.181435 |
| 35.09722 | 0 | 0 | 0 | 0 | 0 | 0 | 0 | 0 | 0 | -0.56727 | -1.2157 | 0.033489 | -0.23483 | -0.938 | 0.11968 | -0.2176 | -0.29662 | 0.010111 | -0.06048 |
| 35.11111 | 0 | 0 | 0 | 0 | 0 | 0 | 0 | 0 | 0 | -1.77707 | -2.47017 | -1.13301 | -0.5489 | -2.31187 | -0.36506 | -1.05631 | -1.36859 | -1.77618 | -0.76968 |
| 35.125 | 0 | 0 | 0 | 0 | 0 | 0 | 0 | 0 | 0 | -2.9887 | -4.1262 | -2.51321 | -1.1799 | -4.00629 | -1.0507 | -1.78204 | -1.6273 | -3.42958 | -1.54627 |
| 35.13889 | 0 | 0 | 0 | 0 | 0 | 0 | 0 | 0 | 0 | -2.34427 | -2.68535 | -1.64621 | -1.18877 | -2.85494 | -0.70966 | -1.06428 | -1.39802 | -2.60362 | -1.2856 |
| 35.15278 | 0 | 0 | 0 | 0 | 0 | 0 | 0 | 0 | 0 | -0.72017 | -0.27356 | 0.052867 | -0.31835 | -0.94217 | -0.04064 | 0.153817 | -0.29729 | -0.78319 | -0.3388 |
| 35.16667 | 0 | 0 | 0 | 0 | 0 | 0 | 0 | 0 | 0 | 0.002337 | 0.557081 | 0.744144 | 0.390079 | -0.02775 | 0.126713 | 0.633625 | 0.34981 | 0.800657 | 0.242071 |
| 35.18056 | 0 | 0 | 0 | 0 | 0 | 0 | 0 | 0 | 0 | 0.342061 | 0.360688 | 0.70916 | 0.918285 | 0.463422 | 0.026659 | 0.594438 | 0.720405 | 1.895867 | 0.056783 |
| 35.19444 | 0 | 0 | 0 | 0 | 0 | 0 | 0 | 0 | 0 | 0.805552 | 0.089389 | 0.879239 | 1.378515 | 0.697859 | 0.111229 | 0.429339 | 0.85574 | 2.160145 | -0.00695 |
| 35.20833 | 0 | 0 | 0 | 0 | 0 | 0 | 0 | 0 | 0 | 1.043713 | -0.15115 | 1.442838 | 1.476008 | 0.744643 | 0.303882 | 0.338602 | 0.820199 | 2.251777 | 0.201203 |
| 35.22222 | 0 | 0 | 0 | 0 | 0 | 0 | 0 | 0 | 0 | 0.881317 | -0.13193 | 1.621868 | 1.271112 | 0.649075 | 0.404933 | 0.258318 | 0.517008 | 1.902413 | 0.315895 |
| 35.23611 | 0 | 0 | 0 | 0 | 0 | 0 | 0 | 0 | 0 | 0.61094 | -0.12655 | 1.137061 | 1.003717 | 0.578344 | 0.322181 | 0.071953 | 0.297306 | 1.80591 | 0.20668 |
| 35.25 | 0 | 0 | 0 | 0 | 0 | 0 | 0 | 0 | 0 | 0.342913 | -0.16175 | 0.586222 | 0.679266 | 0.442293 | 0.180409 | -0.11285 | 0.191206 | 0.952232 | -0.02021 |
| 35.26389 | 0 | 0 | 0 | 0 | 0 | 0 | 0 | 0 | 0 | 0.213527 | -0.03906 | 0.306975 | 0.394275 | 0.606258 | 0.182138 | -0.05521 | 0.160305 | 0.471167 | -0.07004 |
| 35.27778 | 0 | 0 | 0 | 0 | 0 | 0 | 0 | 0 | 0 | 0.287574 | 0.166915 | 0.27761 | 0.035861 | 1.188777 | 0.270891 | 0.103135 | 0.116083 | 0.033007 | 0.114756 |
| 35.29167 | 0 | 0 | 0 | 0 | 0 | 0 | 0 | 0 | 0 | 0.413077 | 0.323879 | 0.414474 | -0.25307 | 1.509737 | 0.329766 | 0.148367 | 0.228834 | -0.58063 | 0.278122 |
| 35.30556 | 0 | 0 | 0 | 0 | 0 | 0 | 0 | 0 | 0 | 0.470442 | 0.390127 | 0.37741 | -0.40408 | 1.346669 | 0.253172 | 0.141117 | 0.318106 | -1.09548 | 0.238814 |
| 35.31944 | 0 | 0 | 0 | 0 | 0 | 0 | 0 | 0 | 0 | 0.379471 | 0.303438 | 0.37377 | -0.3978 | 0.892506 | 0.135695 | 0.044396 | 0.378509 | -1.52518 | 0.156259 |
| 35.33333 | 0 | 0 | 0 | 0 | 0 | 0 | 0 | 0 | 0 | 0.24053 | 0.165718 | 0.224651 | -0.25494 | 0.168333 | 0.122917 | -0.03195 | 0.281773 | -1.93271 | 0.155824 |
| 35.34722 | 0 | 0 | 0 | 0 | 0 | 0 | 0 | 0 | 0 | 0.196362 | 0.080481 | 0.069145 | -0.09121 | -0.38747 | 0.202526 | 0.023783 | 0.182893 | -1.8233 | 0.175371 |
| 35.36111 | 0 | 0 | 0 | 0 | 0 | 0 | 0 | 0 | 0 | 0.153022 | 0.013922 | -0.17429 | -0.04558 | -0.71912 | 0.232036 | 0.103679 | 0.123694 | -1.45743 | 0.149111 |
| 35.375 | 0 | 0 | 0 | 0 | 0 | 0 | 0 | 0 | 0 | 0.052205 | -0.06337 | -0.3456 | -0.14736 | -0.84943 | 0.217873 | 0.096588 | 0.048612 | -1.11689 | 0.108945 |
| 35.38889 | 0 | 0 | 0 | 0 | 0 | 0 | 0 | 0 | 0 | 0.001423 | -0.09232 | -0.19034 | -0.16895 | -0.77454 | 0.283393 | 0.141893 | -0.03823 | -0.92173 | 0.086255 |
| 35.40278 | 0 | 0 | 0 | 0 | 0 | 0 | 0 | 0 | 0 | 0.079865 | -0.05269 | 0.084726 | -0.14273 | -0.4701 | 0.37232 | 0.209844 | -0.04938 | -0.70698 | 0.045891 |
| 35.41667 | 0 | 0 | 0 | 0 | 0 | 0 | 0 | 0 | 0 | 0.147187 | -0.01259 | 0.239802 | -0.01819 | -0.17279 | 0.358426 | 0.131711 | 0.03786 | -0.4382 | 0.042414 |
| 35.43056 | 0 | 0 | 0 | 0 | 0 | 0 | 0 | 0 | 0 | 0.12566 | -0.03346 | 0.241436 | 0.037061 | -0.06378 | 0.193737 | -0.0212 | 0.080314 | -0.13989 | 0.048288 |
| 35.44444 | 0 | 0 | 0 | 0 | 0 | 0 | 0 | 0 | 0 | 0.032642 | -0.13392 | 0.205009 | 0.009502 | -0.08307 | 0.053821 | -0.09775 | 0.043266 | 0.008225 | 0.004896 |
| 35.45833 | 0 | 0 | 0 | 0 | 0 | 0 | 0 | 0 | 0 | -0.0505 | -0.21916 | 0.186273 | -0.099 | -0.07429 | 0.015537 | -0.07994 | 0.009507 | 0.018701 | -0.05578 |
| 35.47222 | 0 | 0 | 0 | 0 | 0 | 0 | 0 | 0 | 0 | -0.03478 | -0.21732 | 0.2258 | -0.18203 | 0.015189 | -0.01325 | -0.03724 | 0.09688 | 0.08112 | -0.07785 |
| 35.48611 | 0 | 0 | 0 | 0 | 0 | 0 | 0 | 0 | 0 | 0.071751 | -0.10882 | 0.293251 | -0.20765 | 0.138106 | -0.04061 | -0.02247 | 0.249297 | 0.229095 | -0.06548 |
| 35.5 | 0 | 0 | 0 | 0 | 0 | 0 | 0 | 0 | 0 | 0.168902 | -0.0658 | 0.360064 | -0.15953 | 0.163264 | -0.03791 | -0.00926 | 0.29292 | 0.30803 | -0.04983 |
| 35.51389 | 0 | 0 | 0 | 0 | 0 | 0 | 0 | 0 | 0 | 0.243882 | -0.065 | 0.34934 | -0.05507 | 0.194335 | -0.00589 | 0.020343 | 0.255048 | 0.277151 | -0.01123 |
| 35.52778 | 0 | 0 | 0 | 0 | 0 | 0 | 0 | 0 | 0 | 0.23295 | -0.05487 | 0.278968 | 0.045803 | 0.284477 | -0.00982 | 0.018075 | 0.207161 | 0.21299 | 0.037369 |
| 35.54167 | 0 | 0 | 0 | 0 | 0 | 0 | 0 | 0 | 0 | 0.20028 | -0.03408 | 0.25605 | 0.083287 | 0.30824 | -0.11761 | -0.04007 | 0.211542 | 0.184598 | 0.031272 |
| 35.55556 | 0 | 0 | 0 | 0 | 0 | 0 | 0 | 0 | 0 | 0.236528 | -0.03865 | 0.29286 | 0.127606 | 0.232772 | -0.18144 | -0.11123 | 0.262097 | 0.192953 | -0.05194 |
| 35.56944 | 0 | 0 | 0 | 0 | 0 | 0 | 0 | 0 | 0 | 0.274476 | -0.04358 | 0.308707 | 0.187006 | 0.10308 | -0.05325 | -0.11463 | 0.303146 | 0.128489 | -0.10622 |
| 35.58333 | 0 | 0 | 0 | 0 | 0 | 0 | 0 | 0 | 0 | 0.33656 | 0.037824 | 0.412497 | 0.220421 | 0.067313 | 0.18597 | 0.015231 | 0.377249 | 0.0672 | -0.10952 |
| 35.59722 | 0 | 0 | 0 | 0 | 0 | 0 | 0 | 0 | 0 | 0.792235 | 0.524911 | 1.157378 | 0.238347 | 0.420808 | 0.400008 | 0.256208 | 0.842076 | 0.507038 | 0.006032 |
| 35.61111 | 0 | 0 | 0 | 0 | 0 | 0 | 0 | 0 | 0 | 2.051165 | 2.077045 | 2.676776 | 0.303453 | 1.364028 | 0.545646 | 0.646238 | 2.06507 | 1.551608 | 0.371689 |
| 35.625 | 0 | 0 | 0 | 0 | 0 | 0 | 0 | 0 | 0 | 2.777307 | 2.300729 | 3.521071 | 0.389401 | 2.068776 | 0.327046 | 0.924883 | 1.99721 | 2.064652 | 0.829588 |
| 35.63889 | 0 | 0 | 0 | 0 | 0 | 0 | 0 | 0 | 0 | 1.804826 | 1.294794 | 2.219727 | 0.571824 | 2.265165 | 0.402982 | 0.73698 | 1.38333 | 1.865875 | 0.949563 |
| 35.65278 | 0 | 0 | 0 | 0 | 0 | 0 | 0 | 0 | 0 | 1.1091 | 0.645863 | 1.338061 | 0.674232 | 2.069932 | 0.312143 | 0.511649 | 1.004667 | 1.72483 | 0.754008 |
| 35.66667 | 0 | 0 | 0 | 0 | 0 | 0 | 0 | 0 | 0 | 0.742802 | 0.389514 | 0.966497 | 0.438355 | 1.334067 | 0.172588 | 0.155873 | 0.711407 | 1.17265 | 0.440733 |
| 35.68056 | 0 | 0 | 0 | 0 | 0 | 0 | 0 | 0 | 0 | 0.423614 | 0.248389 | 0.799843 | 0.211603 | 0.174832 | -0.04855 | -0.35316 | 0.437854 | 0.355023 | 0.096357 |
| 35.69444 | 0 | 0 | 0 | 0 | 0 | 0 | 0 | 0 | 0 | 0.213212 | 0.241032 | 0.68637 | -0.03654 | -1.01673 | -0.26955 | -0.74158 | 0.277157 | -0.39531 | -0.27358 |
| 35.70833 | 0 | 0 | 0 | 0 | 0 | 0 | 0 | 0 | 0 | 0.338048 | 0.299274 | 0.682044 | -0.14045 | -1.7146 | -0.22115 | -0.78496 | 0.504259 | -1.11853 | -0.44175 |
| 35.72222 | 0 | 0 | 0 | 0 | 0 | 0 | 0 | 0 | 0 | 0.545999 | 0.242583 | 0.588966 | -0.08224 | -1.26139 | -0.12533 | -0.42757 | 0.355578 | -1.36764 | -0.49026 |
| 35.73611 | 0 | 0 | 0 | 0 | 0 | 0 | 0 | 0 | 0 | 0.596951 | 0.096644 | 0.325335 | 0.037561 | -0.4939 | 0.064256 | -0.06251 | 0.56575 | -1.37539 | -0.2086 |
| 35.75 | 0 | 0 | 0 | 0 | 0 | 0 | 0 | 0 | 0 | 0.557858 | -0.01327 | 0.178092 | 0.155363 | -0.06983 | 0.016089 | 0.116348 | 0.717354 | -1.49074 | 0.09886 |
| 35.76389 | 0 | 0 | 0 | 0 | 0 | 0 | 0 | 0 | 0 | 0.415364 | -0.00966 | 0.102133 | 0.184636 | -0.00205 | -0.0066 | 0.049162 | 0.695216 | -1.62252 | 0.249969 |
| 35.77778 | 0 | 0 | 0 | 0 | 0 | 0 | 0 | 0 | 0 | 0.134783 | 0.034092 | 0.000332 | 0.104428 | -0.15571 | -0.01265 | -0.13202 | 0.486023 | -1.92988 | 0.179944 |
| 35.79167 | 0 | 0 | 0 | 0 | 0 | 0 | 0 | 0 | 0 | -0.19268 | 0.142848 | -0.13303 | -0.16479 | -0.4539 | -0.09391 | -0.24218 | 0.033429 | -1.98011 | -0.08004 |
| 35.80556 | 0 | 0 | 0 | 0 | 0 | 0 | 0 | 0 | 0 | -0.48091 | 0.198781 | -0.25991 | -0.39486 | -0.60614 | -0.21311 | -0.24919 | -0.39266 | -1.83833 | -0.35516 |
| 35.81944 | 0 | 0 | 0 | 0 | 0 | 0 | 0 | 0 | 0 | -0.58937 | 0.200264 | -0.51745 | -0.47819 | -0.49196 | -0.31899 | -0.11048 | -0.58388 | -1.50608 | -0.48064 |
| 35.83333 | 0 | 0 | 0 | 0 | 0 | 0 | 0 | 0 | 0 | -0.51955 | 0.155018 | -0.81895 | -0.46697 | -0.14938 | -0.39644 | 0.097283 | -0.28523 | -0.7992 | -0.48463 |
| 35.84722 | 0 | 0 | 0 | 0 | 0 | 0 | 0 | 0 | 0 | -0.34431 | 0.11875 | -0.96692 | -0.2498 | 0.307374 | -0.47598 | 0.311802 | 0.093039 | -0.03672 | -0.32748 |
| 35.86111 | 0 | 0 | 0 | 0 | 0 | 0 | 0 | 0 | 0 | -0.17276 | 0.065235 | -0.8032 | 0.02599 | 0.664757 | -0.48718 | 0.378882 | 0.259411 | 0.652693 | -0.13084 |
| 35.875 | 0 | 0 | 0 | 0 | 0 | 0 | 0 | 0 | 0 | -0.18287 | 0.029475 | -0.56634 | 0.272965 | 1.050385 | -0.43658 | 0.377772 | 0.167015 | 1.273735 | -0.04832 |

| | | | | | | | | | | | | | | | | | | | |
|---|---|---|---|---|---|---|---|---|---|---|---|---|---|---|---|---|---|---|---|
| 35.88889 | 0 | 0 | 0 | 0 | 0 | 0 | 0 | 0 | 0 | -0.28919 | 0.043813 | -0.49447 | 0.451468 | 1.278733 | -0.43745 | 0.237357 | -0.04824 | 1.503323 | -0.05331 |
| 35.90278 | 0 | 0 | 0 | 0 | 0 | 0 | 0 | 0 | 0 | -0.43916 | 0.044549 | -0.40453 | 0.555503 | 1.328152 | -0.43663 | 0.101275 | -0.12096 | 1.40035 | -0.06594 |
| 35.91667 | 0 | 0 | 0 | 0 | 0 | 0 | 0 | 0 | 0 | -0.53393 | 0.064936 | -0.28993 | 0.437829 | 1.082013 | -0.4728 | 0.027091 | -0.16813 | 0.916826 | -0.06674 |
| 35.93056 | 0 | 0 | 0 | 0 | 0 | 0 | 0 | 0 | 0 | -0.51757 | 0.073823 | -0.16365 | 0.192192 | 0.636046 | -0.37303 | 0.068151 | -0.27392 | 0.096302 | -0.01192 |
| 35.94444 | 0 | 0 | 0 | 0 | 0 | 0 | 0 | 0 | 0 | -0.41073 | 0.093769 | -0.09311 | 0.068915 | 0.211424 | -0.2597 | 0.107647 | -0.43265 | -0.67088 | 0.05394 |
| 35.95833 | 0 | 0 | 0 | 0 | 0 | 0 | 0 | 0 | 0 | -0.31363 | 0.04754 | -0.08624 | 0.051623 | -0.0914 | -0.15019 | 0.176028 | -0.49577 | -0.81682 | 0.116166 |
| 35.97222 | 0 | 0 | 0 | 0 | 0 | 0 | 0 | 0 | 0 | -0.35743 | -0.07059 | -0.14535 | 0.082354 | -0.14097 | -0.05127 | 0.200317 | -0.47896 | -0.60425 | 0.175943 |
| 35.98611 | 0 | 0 | 0 | 0 | 0 | 0 | 0 | 0 | 0 | -0.52139 | -0.1905 | -0.17791 | 0.133865 | -0.21294 | -0.02184 | 0.15933 | -0.47319 | -0.35496 | 0.303741 |
| 36 | 0 | 0 | 0 | 0 | 0 | 0 | 0 | 0 | 0 | -0.67902 | -0.34495 | -0.22094 | 0.155942 | -0.25848 | -0.04453 | 0.01147 | -0.42177 | -0.10879 | 0.464472 |
| 36.01389 | 0 | 0 | 0 | 0 | 0 | 0 | 0 | 0 | 0 | -0.71246 | -0.51459 | -0.23506 | 0.136487 | -0.18613 | -0.08551 | -0.0821 | -0.38076 | 0.228359 | 0.552913 |
| 36.02778 | 0 | 0 | 0 | 0 | 0 | 0 | 0 | 0 | 0 | -0.68412 | -0.69403 | -0.20283 | 0.071262 | -0.14582 | -0.07975 | -0.01856 | -0.36237 | 0.59752 | 0.465093 |
| 36.04167 | 0 | 0 | 0 | 0 | 0 | 0 | 0 | 0 | 0 | -0.57785 | -0.81297 | -0.15053 | 0.022687 | 0.005557 | -0.00932 | 0.144871 | -0.30246 | 0.865043 | 0.28501 |
| 36.05556 | 0 | 0 | 0 | 0 | 0 | 0 | 0 | 0 | 0 | -0.42782 | -0.79847 | -0.07669 | 0.048264 | 0.213877 | 0.106901 | 0.319862 | -0.1843 | 1.113926 | 0.200023 |
| 36.06944 | 0 | 0 | 0 | 0 | 0 | 0 | 0 | 0 | 0 | -0.36493 | -0.75123 | -0.05765 | 0.193503 | 0.334909 | 0.122505 | 0.370973 | -0.04915 | 1.337363 | 0.287434 |
| 36.08333 | 0 | 0 | 0 | 0 | 0 | 0 | 0 | 0 | 0 | -0.44538 | -0.69118 | -0.14458 | 0.340351 | 0.287988 | 0.031979 | 0.278318 | -0.06435 | 1.12609 | 0.430968 |
| 36.09722 | 0 | 0 | 0 | 0 | 0 | 0 | 0 | 0 | 0 | -0.94367 | -0.94142 | -0.93389 | 0.0641 | -0.31354 | -0.27986 | -0.20699 | -0.66255 | -0.36023 | -0.0294 |
| 36.11111 | 0 | 0 | 0 | 0 | 0 | 0 | 0 | 0 | 0 | -2.46437 | -2.3809 | -3.31557 | -1.40405 | -2.05731 | -1.04625 | -1.41022 | -2.36146 | -3.34498 | -1.21937 |
| 36.125 | 0 | 0 | 0 | 0 | 0 | 0 | 0 | 0 | 0 | -3.66277 | -3.5905 | -4.85322 | -2.89833 | -3.04265 | -1.85301 | -2.54385 | -3.23701 | -4.69529 | -2.03106 |
| 36.13889 | 0 | 0 | 0 | 0 | 0 | 0 | 0 | 0 | 0 | -1.46748 | -1.75647 | -0.59847 | -1.70132 | -0.91558 | -0.92526 | -1.18677 | -0.66707 | -2.36692 | -1.10232 |
| 36.15278 | 0 | 0 | 0 | 0 | 0 | 0 | 0 | 0 | 0 | 1.058094 | 1.101182 | 1.856867 | 0.136217 | 1.148692 | 0.629956 | 0.975504 | 1.951808 | 0.058982 | 0.413043 |
| 36.16667 | 0 | 0 | 0 | 0 | 0 | 0 | 0 | 0 | 0 | 1.202599 | 1.668464 | 2.027425 | 1.421167 | 1.844774 | 1.353192 | 1.660749 | 2.32226 | 1.030706 | 1.215712 |
| 36.18056 | 0 | 0 | 0 | 0 | 0 | 0 | 0 | 0 | 0 | 0.427475 | 0.469311 | 1.007929 | 2.053676 | 1.457775 | 1.622572 | 1.19262 | 1.207348 | 1.068867 | 1.155967 |
| 36.19444 | 0 | 0 | 0 | 0 | 0 | 0 | 0 | 0 | 0 | 0.226628 | 0.148188 | 0.539886 | 1.947219 | 1.078636 | 1.399128 | 0.749759 | 0.430477 | 1.220991 | 1.025531 |
| 36.20833 | 0 | 0 | 0 | 0 | 0 | 0 | 0 | 0 | 0 | 0.298546 | 0.16455 | 0.564274 | 1.612813 | 0.614332 | 1.047468 | 0.465729 | 0.019537 | 1.184863 | 0.922453 |
| 36.22222 | 0 | 0 | 0 | 0 | 0 | 0 | 0 | 0 | 0 | 0.108095 | 0.141354 | 0.479922 | 1.185732 | 0.096746 | 0.715398 | 0.273274 | -0.09448 | 1.130439 | 0.538948 |
| 36.23611 | 0 | 0 | 0 | 0 | 0 | 0 | 0 | 0 | 0 | -0.17831 | 0.057385 | 0.270978 | 0.91306 | -0.00768 | 0.444576 | 0.119151 | -0.10061 | 1.303965 | 0.157469 |
| 36.25 | 0 | 0 | 0 | 0 | 0 | 0 | 0 | 0 | 0 | -0.14034 | -0.10148 | 0.010951 | 0.677888 | -0.02754 | 0.271797 | 0.074812 | -0.17459 | 1.498901 | -0.14716 |
| 36.26389 | 0 | 0 | 0 | 0 | 0 | 0 | 0 | 0 | 0 | 0.181149 | 0.037811 | -0.21923 | 0.436138 | -0.00391 | 0.06957 | 0.069392 | -0.34867 | 1.643494 | -0.39451 |
| 36.27778 | 0 | 0 | 0 | 0 | 0 | 0 | 0 | 0 | 0 | 0.405342 | 0.256517 | -0.12694 | 0.355067 | 0.095583 | -0.02259 | 0.052306 | -0.35167 | 1.690403 | -0.33603 |
| 36.29167 | 0 | 0 | 0 | 0 | 0 | 0 | 0 | 0 | 0 | 0.407364 | 0.273493 | -0.03415 | 0.31896 | 0.244085 | -0.01544 | 0.022889 | -0.22084 | 1.58112 | -0.02404 |
| 36.30556 | 0 | 0 | 0 | 0 | 0 | 0 | 0 | 0 | 0 | 0.351313 | 0.281959 | 0.31592 | 0.313414 | 0.346468 | 0.103634 | 0.039213 | -0.0771 | 1.11806 | 0.208559 |
| 36.31944 | 0 | 0 | 0 | 0 | 0 | 0 | 0 | 0 | 0 | 0.410995 | 0.373122 | 0.419918 | 0.244674 | 0.347159 | 0.217772 | 0.079053 | 0.103553 | 0.385324 | 0.206768 |
| 36.33333 | 0 | 0 | 0 | 0 | 0 | 0 | 0 | 0 | 0 | 0.482239 | 0.476472 | 0.435974 | 0.093179 | 0.342435 | 0.217689 | 0.133424 | 0.229306 | -0.36437 | 0.096665 |
| 36.34722 | 0 | 0 | 0 | 0 | 0 | 0 | 0 | 0 | 0 | 0.483747 | 0.494058 | 0.41466 | -0.14171 | 0.293907 | 0.081721 | 0.142521 | 0.290503 | -1.03332 | -0.09547 |
| 36.36111 | 0 | 0 | 0 | 0 | 0 | 0 | 0 | 0 | 0 | 0.418402 | 0.435274 | 0.237426 | -0.38438 | 0.30824 | -0.03787 | 0.176589 | 0.272925 | -1.40414 | -0.2414 |
| 36.375 | 0 | 0 | 0 | 0 | 0 | 0 | 0 | 0 | 0 | 0.427809 | 0.408878 | 0.023559 | -0.52372 | 0.469461 | -0.02528 | 0.245008 | 0.153166 | -1.45939 | -0.17882 |
| 36.38889 | 0 | 0 | 0 | 0 | 0 | 0 | 0 | 0 | 0 | 0.45259 | 0.41249 | -0.11909 | -0.51324 | 0.700939 | 0.094396 | 0.305569 | 0.010989 | -1.45527 | 0.094247 |
| 36.40278 | 0 | 0 | 0 | 0 | 0 | 0 | 0 | 0 | 0 | 0.568895 | 0.477968 | -0.01651 | -0.25399 | 0.827924 | 0.253182 | 0.373771 | 0.067986 | -1.26455 | 0.342465 |
| 36.41667 | 0 | 0 | 0 | 0 | 0 | 0 | 0 | 0 | 0 | 1.16809 | 0.611234 | 0.651898 | 0.173158 | 0.987381 | 0.394718 | 0.510654 | 0.483798 | 0.048239 | 0.314164 |
| 36.43056 | 0 | 0 | 0 | 0 | 0 | 0 | 0 | 0 | 0 | 3.04692 | 0.991189 | 2.03113 | 0.621684 | 1.428525 | 0.514544 | 0.620778 | 1.352014 | 2.350394 | 0.207418 |
| 36.44444 | 0 | 0 | 0 | 0 | 0 | 0 | 0 | 0 | 0 | 4.415386 | 2.470038 | 3.33812 | 0.928737 | 1.869244 | 0.660764 | 0.820497 | 2.249149 | 3.946914 | 0.492379 |
| 36.45833 | 0 | 0 | 0 | 0 | 0 | 0 | 0 | 0 | 0 | 3.900907 | 2.144689 | 3.233609 | 0.942747 | 1.971106 | 0.807119 | 0.885548 | 2.277495 | 3.641085 | 1.230395 |
| 36.47222 | 0 | 0 | 0 | 0 | 0 | 0 | 0 | 0 | 0 | 2.203604 | 1.092818 | 1.82326 | 1.137234 | 1.865028 | 0.952598 | 0.709235 | 1.486281 | 2.132035 | 1.933095 |
| 36.48611 | 0 | 0 | 0 | 0 | 0 | 0 | 0 | 0 | 0 | 1.162529 | 0.606984 | 1.020574 | 1.309633 | 1.565477 | 0.901802 | 0.603425 | 0.835344 | 1.733814 | 1.703216 |
| 36.5 | 2.682978 | 3.155329 | 1.743494 | 1.606142 | 0.853509 | 0.293732 | 1.524523 | -1.51307 | 0.315236 | 0.584886 | 0.719831 | 0.691862 | 1.119765 | 1.155501 | 0.706492 | 0.398026 | 0.547904 | 1.563925 | 0.942766 |
| 36.51389 | 3.901688 | 5.12186 | 1.932326 | 1.901729 | 2.206905 | 1.154379 | 2.274495 | 0.003234 | 1.583664 | 0.245621 | 0.555979 | 0.568745 | 0.525441 | 0.683394 | 0.412196 | -0.03286 | 0.463835 | 0.947292 | 0.144336 |
| 36.52778 | 2.908767 | 3.799983 | 1.387372 | 1.257387 | 2.734542 | 1.063013 | 2.020731 | 1.178815 | 2.111645 | 0.038802 | -0.12648 | 0.440178 | -0.04657 | 0.187965 | 0.162342 | -0.47272 | 0.386518 | -0.09684 | -0.8112 |
| 36.54167 | 1.619255 | 2.169008 | 0.859707 | 0.771951 | 3.714753 | 0.773039 | 1.622227 | 1.09343 | 1.308256 | 0.050785 | -0.05765 | 0.339852 | -0.34643 | -0.22544 | 0.084764 | -0.65687 | 0.319955 | -0.96913 | -1.57523 |
| 36.55556 | 1.508816 | 1.507085 | 0.578345 | 0.75542 | 3.712789 | 0.81043 | 1.33029 | 1.0474 | 1.241774 | 0.306427 | 0.104906 | 0.258417 | -0.19593 | -0.38601 | 0.239105 | -0.56986 | 0.256379 | -1.22835 | -1.66086 |
| 36.56944 | 1.874857 | 1.565283 | 0.487072 | 1.225283 | 3.805011 | 0.923302 | 1.259745 | 1.326943 | 1.65213 | 0.571163 | 0.169635 | 0.306654 | 0.347013 | -0.55457 | 0.489788 | -0.2736 | 0.273068 | -1.51192 | -1.28627 |
| 36.58333 | 2.234266 | 1.747949 | 0.538022 | 1.999187 | 4.285488 | 1.35442 | 1.692679 | 0.899489 | 1.498207 | 0.560545 | 0.195482 | 0.392725 | 0.711309 | -0.37344 | 0.742288 | 0.044772 | 0.238704 | -1.77045 | -0.5142 |
| 36.59722 | 2.30306 | 1.762622 | 0.596855 | 2.543786 | 3.494524 | 0.998845 | 2.037208 | 0.041655 | 0.704827 | 0.317344 | 0.093441 | 0.38391 | 0.656887 | -0.54584 | 0.663897 | 0.157682 | 0.218049 | -2.2289 | -0.39061 |
| 36.61111 | 2.105551 | 1.647022 | 0.719558 | 3.129033 | 3.260328 | 0.923087 | 1.88383 | -0.83811 | 0.012567 | -0.0331 | -0.05382 | 0.247752 | 0.095724 | -0.83132 | 0.387542 | 0.081737 | 0.146135 | -2.40617 | -0.833 |
| 36.625 | 1.843052 | 1.313859 | 0.964995 | 3.483381 | 2.714869 | 0.974576 | 2.229892 | -1.28131 | -0.80304 | -0.38425 | -0.19619 | 0.082264 | -0.59441 | -1.08157 | 0.188399 | -0.10076 | 0.141374 | -2.68956 | -1.10812 |
| 36.63889 | 1.474764 | 0.805673 | 1.302451 | 3.133095 | 1.789991 | 0.957196 | 2.154 | -0.97289 | -1.22848 | -0.69404 | 0.16401 | -0.07921 | -0.81867 | -1.15573 | 0.043432 | -0.33337 | 0.108436 | -2.19893 | -1.00883 |
| 36.65278 | 1.106113 | 0.28923 | 1.418309 | 1.605653 | 0.851874 | 0.254203 | 0.879024 | -0.72414 | -0.81337 | -0.85627 | 1.978341 | -0.16223 | -0.58855 | -1.18656 | 0.188812 | -0.3925 | 0.084963 | -1.4753 | -0.67574 |
| 36.66667 | 0.449986 | -0.19562 | 0.934637 | -0.0782 | -0.14787 | -0.69882 | -0.47848 | -0.85971 | -0.09143 | -0.81087 | 0.690823 | -0.09033 | -0.20243 | -0.92752 | 0.32299 | -0.26813 | 0.135658 | -1.23138 | 0.0333 |
| 36.68056 | 0.111561 | -0.67902 | 0.604203 | -1.44215 | -1.13161 | -1.27351 | -1.48405 | -1.82629 | 0.113057 | -0.68788 | -1.15398 | 0.01924 | 0.32011 | -0.58558 | 0.563227 | -0.01277 | 0.116751 | -0.93823 | 0.536798 |
| 36.69444 | -0.19224 | -1.07179 | 0.457796 | -1.79592 | -2.2515 | -1.75921 | -1.68004 | -3.12231 | -0.09703 | -0.50724 | -1.55147 | -0.06323 | 0.677198 | -0.25502 | 0.630202 | 0.224044 | -0.07314 | -0.48471 | 0.462726 |
| 36.70833 | -0.49547 | -1.40578 | 0.522219 | -1.92919 | -3.69976 | -1.15368 | -0.75117 | -4.18951 | -0.62411 | -0.3949 | -0.26106 | -0.21135 | 0.71398 | -0.11824 | 0.455353 | 0.191008 | -0.27515 | 0.13838 | -0.28321 |
| 36.72222 | -0.49715 | -1.48705 | 0.74733 | -1.48253 | -3.74098 | -1.43232 | -0.24917 | -3.52842 | -0.31441 | -0.30302 | -0.22339 | -0.30268 | 0.347396 | -0.11797 | 0.190343 | 0.097064 | -0.31865 | 0.855541 | -0.7889 |
| 36.73611 | -0.20663 | -1.35978 | 1.123471 | -0.96797 | -4.25322 | -1.20333 | 0.290178 | -1.59452 | -0.04542 | -0.2644 | -0.25923 | -0.32284 | -0.17657 | 0.163859 | -0.072 | 0.041916 | -0.28189 | 1.472358 | -0.78545 |
| 36.75 | 0.060778 | -1.05272 | 1.367128 | -0.53136 | -4.05609 | -1.84932 | 0.511471 | 0.939466 | 0.377729 | -0.33944 | -0.3303 | -0.27846 | -0.43345 | 0.621966 | -0.44178 | -0.01175 | -0.19343 | 1.818184 | -0.36594 |
| 36.76389 | 0.134628 | -0.77594 | 1.225017 | -0.1623 | -4.31025 | -1.84857 | 0.654118 | 2.883279 | 0.321352 | -0.59547 | -0.4157 | -0.27936 | -0.1098 | 0.91304 | -0.79208 | -0.09443 | -0.13086 | 2.011502 | -0.38846 |
| 36.77778 | 0.030556 | -0.67988 | 0.789911 | 0.361353 | -3.61241 | -1.74634 | 0.640631 | 3.910125 | 0.518976 | -0.82416 | 1.196895 | -0.28958 | 0.441408 | 1.020049 | -0.80991 | 0.032204 | -0.13306 | 1.737961 | -0.32372 |
| 36.79167 | -0.15193 | -0.66762 | 0.525265 | 0.976014 | -2.15567 | -1.51338 | 0.509581 | 3.557846 | 0.232447 | -0.89882 | 2.034466 | -0.30024 | 0.949864 | 0.935888 | -0.62329 | 0.313501 | -0.12629 | 1.367861 | -0.25301 |
| 36.80556 | -0.20461 | -0.65711 | 0.350175 | 1.541277 | -1.32467 | -1.39962 | 0.203563 | 3.06472 | -0.47574 | -0.82171 | 1.026473 | -0.28473 | 1.212197 | 0.829095 | -0.28933 | 0.610278 | -0.20005 | 0.816749 | 0.093051 |
| 36.81944 | 0.087254 | -0.65945 | -0.03086 | 1.94721 | -0.98078 | -1.50715 | -0.2071 | 2.61298 | -0.71808 | -0.69771 | -1.66774 | -0.27154 | 1.198286 | 0.648643 | -0.127 | 0.794421 | -0.2574 | 0.324364 | 0.247063 |
| 36.83333 | 0.326746 | -0.80711 | -0.84068 | 2.119653 | -0.5177 | -1.40168 | -0.50108 | 1.508887 | -1.74105 | -0.60421 | -1.54482 | -0.22808 | 1.146749 | 0.358172 | -0.08101 | 0.707768 | -0.30602 | -0.01908 | 0.37446 |
| 36.84722 | 0.232263 | -1.08077 | -0.95861 | 2.04076 | -0.10822 | -1.12738 | -0.8346 | -0.23152 | -3.79071 | -0.53042 | -0.36685 | -0.17833 | 0.968091 | 0.156982 | -0.17357 | 0.452049 | -0.31729 | -0.23073 | 0.229282 |
| 36.86111 | -0.22326 | -1.25883 | -0.91347 | 1.724325 | -0.24115 | -0.96535 | -1.3386 | -1.98245 | -3.56779 | -0.47203 | -0.16795 | -0.12204 | 0.582943 | 0.009239 | -0.18958 | 0.243851 | -0.28867 | -0.34827 | 0.263073 |
| 36.875 | -0.7791 | -1.2413 | -1.11517 | 1.235164 | -0.74311 | -1.04017 | -1.7453 | -3.50575 | -2.96658 | -0.35921 | -0.19603 | -0.3373 | 0.262424 | -0.03502 | -0.19702 | 0.145398 | -0.27441 | -0.47966 | 1.023534 |
| 36.88889 | -1.00906 | -0.92638 | -1.49176 | 0.867627 | -1.66015 | -1.75325 | -1.86423 | -4.10492 | -2.37964 | -0.25516 | -0.19221 | -0.68165 | 0.156421 | 0.042725 | -0.08084 | 0.251016 | -0.27291 | -0.62353 | 1.255689 |
| 36.90278 | -0.9237 | -0.56117 | -2.03781 | 0.758881 | -2.41932 | -2.27055 | -1.91387 | -5.10661 | -1.94924 | -0.15373 | -0.09721 | -0.58209 | 0.258064 | 0.204948 | 0.03881 | 0.439914 | -0.17668 | -0.41133 | 1.251602 |
| 36.91667 | -0.68807 | -0.1343 | -1.95104 | 0.792676 | -2.45771 | -1.88801 | -1.82128 | -6.03085 | -1.42725 | -0.12873 | -0.11313 | -0.28551 | 0.274239 | 0.343356 | 0.081274 | 0.583241 | -0.15526 | -0.19973 | 1.415987 |
| 36.93056 | -0.22211 | 0.304202 | -1.4755 | 0.723673 | -2.06887 | -1.33893 | -1.44512 | -6.05328 | -1.23944 | -0.43551 | 0.228402 | -0.5709 | 0.030166 | -0.06482 | -0.31912 | 0.342118 | -0.66525 | -0.86971 | 0.803859 |
| 36.94444 | 0.378597 | 0.735193 | -0.76678 | 0.652425 | -1.68834 | -0.84799 | -1.01074 | -4.50298 | -0.64269 | -1.34685 | 0.599138 | -1.79518 | -1.02242 | -1.86633 | -1.41038 | -0.69682 | -1.88064 | -2.88543 | -0.59405 |
| 36.95833 | 0.908105 | 1.260584 | -0.0011 | 0.526792 | -1.18139 | -0.47289 | -0.48268 | -1.44353 | 0.471752 | -2.27395 | 0.141559 | -2.45752 | -2.58849 | -2.97418 | -2.6594 | -1.95627 | -2.75803 | -4.27317 | -1.36076 |
| 36.97222 | 1.200148 | 1.474464 | 0.384996 | 0.347663 | -0.79004 | -0.27893 | -0.13233 | 1.679901 | 1.650765 | -1.21593 | -2.61483 | -0.72865 | -2.6446 | -1.05943 | -1.06691 | -1.31492 | -1.15872 | -2.42513 | -1.21695 |
| 36.98611 | 0.479115 | 0.1287 | 0.050896 | 0.12781 | -1.02908 | -0.57448 | -0.23091 | 2.810922 | 1.758696 | 0.563084 | -3.56938 | 1.000457 | -0.798 | 1.171217 | 0.996342 | 0.111621 | 1.117061 | 0.377772 | -0.24266 |
| 37 | -1.64533 | -2.89414 | -1.29928 | -0.14539 | -2.27466 | -1.59832 | -0.88359 | -0.27171 | -0.58656 | 0.893152 | -2.47071 | 1.240794 | 0.308832 | 1.746137 | 1.426784 | 0.747746 | 1.526671 | 1.401923 | 1.108473 |
| 37.01389 | -3.03451 | -3.66753 | -1.85195 | -0.02861 | -3.06542 | -2.18604 | -1.23728 | -4.26304 | -2.56215 | 0.452519 | -0.57419 | 0.60802 | 0.836708 | 1.526801 | 0.957646 | 0.848434 | 0.766601 | 1.117476 | 1.129909 |
| 37.02778 | -1.76592 | -1.42788 | -0.98134 | 0.662675 | -2.39821 | -1.60772 | -0.91772 | -5.40558 | -1.21185 | 0.292137 | 0.413145 | 0.3175 | 1.488804 | 1.691763 | 0.504521 | 0.962128 | 0.230022 | 1.146081 | 1.337547 |
| 37.04167 | -0.52179 | -0.02501 | -0.37325 | 1.346946 | -2.18123 | -0.96867 | -0.76317 | -4.21413 | 0.088772 | 0.14595 | 0.455822 | 0.238992 | 1.943292 | 1.212665 | 0.475142 | 0.990884 | 0.050751 | 1.353388 | 1.633247 |
| 37.05556 | -0.08575 | 0.059661 | -0.5802 | 1.620495 | -3.0125 | -0.37002 | -0.80511 | -3.09989 | -0.6551 | 0.032461 | 0.351105 | 0.109452 | 1.66617 | 0.759278 | 0.342645 | 0.759975 | -0.02299 | 1.388107 | 1.682744 |
| 37.06944 | -0.30878 | -0.33442 | -0.89242 | 1.185083 | -3.09097 | -0.10351 | -0.85054 | -2.52566 | -1.36359 | -0.11475 | 0.091655 | 0.160162 | 1.120692 | 0.370898 | 0.169863 | 0.408962 | -0.07659 | 1.164361 | 1.15717 |

| | | | | | | | | | | | | | | | | | | | |
|---|---|---|---|---|---|---|---|---|---|---|---|---|---|---|---|---|---|---|---|
| 37.08333 | -0.72997 | -0.73114 | -0.96569 | 1.012265 | -2.23145 | -0.08035 | -0.8119 | -1.84149 | -1.52227 | -0.12157 | -0.19014 | 0.283365 | 0.617188 | -0.12346 | 0.126686 | 0.100891 | -0.07537 | 0.900329 | 0.355249 |
| 37.09722 | -0.92919 | -1.09856 | -1.00177 | 0.868913 | -2.20122 | -0.09222 | -0.81648 | -1.29515 | -1.64691 | 0.003728 | -0.28146 | 0.421806 | 0.237197 | -0.33674 | 0.191662 | -0.04734 | -0.08078 | 0.813502 | -0.1495 |
| 37.11111 | -1.09704 | -1.30039 | -1.04117 | 0.777393 | -2.41204 | -0.15503 | -0.85192 | -0.7634 | -1.49398 | 0.149614 | -0.1898 | 0.49792 | 0.071671 | -0.36141 | 0.229818 | -0.03321 | -0.07688 | 0.552509 | -0.58459 |
| 37.125 | -0.85734 | -1.30869 | -1.18673 | 0.690286 | -2.40054 | -0.17145 | -0.71134 | -0.08407 | -1.06866 | 0.276327 | -0.1135 | 0.471756 | 0.027847 | -0.39251 | 0.213654 | 0.070152 | -0.02972 | 0.292245 | -1.02714 |
| 37.13889 | -0.55031 | -1.0485 | -0.93907 | 0.567581 | -1.94341 | -0.11823 | -0.52257 | 0.57646 | -0.2335 | 0.372589 | -0.10075 | 0.424534 | 0.072195 | -0.12772 | 0.174797 | 0.228238 | 0.10898 | 0.25126 | -1.09217 |
| 37.15278 | -0.14916 | -0.63624 | -0.53004 | 0.567523 | -1.11055 | -0.03271 | -0.34061 | 0.836415 | 0.8925 | 0.442066 | -0.13238 | 0.455059 | 0.174228 | 0.22438 | 0.142461 | 0.229368 | 0.246101 | 0.152923 | -0.83951 |
| 37.16667 | 0.229334 | -0.33554 | -0.18159 | 0.662678 | -0.36281 | 0.014089 | -0.11956 | 1.074549 | 1.811594 | 0.497364 | -0.01444 | 0.475636 | 0.27184 | 0.588999 | 0.116481 | 0.206037 | 0.319256 | -0.19789 | -0.56181 |
| 37.18056 | 0.419618 | -0.3583 | -0.01198 | 0.745838 | 0.042259 | -0.00102 | -0.06339 | 1.656241 | 2.495964 | 0.473927 | 0.05188 | 0.484469 | 0.300041 | 0.888228 | 0.111179 | 0.251855 | 0.336941 | -0.52297 | -0.31096 |
| 37.19444 | 0.634015 | -0.36184 | 0.163459 | 0.75132 | 0.352264 | 0.007129 | -0.0883 | 2.083616 | 3.120425 | 0.318867 | 0.05452 | 0.352954 | 0.052026 | 0.871115 | 0.007524 | 0.271351 | 0.216694 | -0.81231 | -0.09336 |
| 37.20833 | 0.951907 | -0.14123 | 0.33905 | 0.734538 | 0.65619 | 0.058184 | -0.14881 | 2.455087 | 3.383016 | 0.065966 | -0.0718 | 0.190914 | -0.38976 | 0.607207 | -0.15211 | 0.261131 | 0.023378 | -1.04481 | -0.03939 |
| 37.22222 | 1.081324 | 0.005097 | 0.433152 | 0.631778 | 0.83237 | 0.104442 | -0.10458 | 2.320398 | 3.871397 | -0.02323 | -0.16996 | 0.108575 | -0.79549 | 0.260151 | -0.24396 | 0.375694 | -0.1404 | -1.14355 | -0.0526 |
| 37.23611 | 1.015187 | 0.083049 | 0.457663 | 0.462147 | 0.886299 | 0.110091 | 0.055228 | 1.660957 | 3.661258 | 0.153621 | -0.05706 | 0.156114 | -0.93086 | 0.197703 | -0.16953 | 0.466802 | -0.07549 | -0.87211 | 0.05668 |
| 37.25 | 0.92194 | -0.01278 | 0.387699 | 0.252547 | 0.876806 | 0.068367 | 0.080348 | 2.022772 | 2.419148 | 0.42142 | 0.170937 | 0.214515 | -0.76271 | 0.425625 | -0.11353 | 0.292715 | 0.116631 | -0.37887 | 0.307886 |
| 37.26389 | 0.612448 | -0.34713 | 0.19871 | 0.156659 | 0.715955 | 0.005236 | -0.05364 | 2.361935 | 1.697127 | 0.569835 | 0.291887 | 0.161974 | -0.49537 | 0.824743 | -0.22887 | 0.004442 | 0.227946 | 0.027027 | 0.455648 |
| 37.27778 | 0.201237 | -0.69403 | 0.052585 | 0.374615 | 0.609981 | -0.02039 | -0.20562 | 3.260484 | 1.396098 | 0.542221 | 0.234826 | 0.147213 | -0.2439 | 1.140287 | -0.3739 | -0.25495 | 0.205158 | 0.307549 | 0.284335 |
| 37.29167 | -0.06138 | -0.69987 | 0.083971 | 0.728014 | 0.529111 | -0.00781 | -0.25783 | 3.51037 | 0.525095 | 0.415201 | 0.089778 | 0.16997 | -0.15758 | 1.056182 | -0.3913 | -0.27491 | 0.124135 | 0.462188 | -0.0792 |
| 37.30556 | 0.036464 | -0.38535 | 0.205479 | 0.992472 | 0.473752 | -0.0099 | -0.22814 | 3.180529 | 0.044398 | 0.338275 | 0.040146 | 0.178101 | -0.0422 | 0.835824 | -0.29627 | -0.08815 | 0.056167 | 0.424339 | -0.34683 |
| 37.31944 | 0.334403 | -0.09102 | 0.290467 | 1.149957 | 0.4173 | -0.08034 | -0.22352 | 2.408275 | -0.13073 | 0.372224 | 0.063301 | 0.165822 | 0.016913 | 0.634521 | -0.1816 | 0.115686 | 0.01326 | 0.282831 | -0.33795 |
| 37.33333 | 0.685324 | 0.164124 | 0.32541 | 1.279052 | 0.339454 | -0.16347 | -0.19316 | 1.631678 | 0.32468 | 0.444395 | 0.127626 | 0.173263 | 0.003545 | 0.446356 | -0.12916 | 0.158064 | 0.019963 | 0.229808 | -0.15607 |
| 37.34722 | 0.898161 | 0.296512 | 0.364049 | 1.396042 | 0.235043 | -0.16087 | -0.0554 | 1.441839 | 1.070308 | 0.455452 | 0.120485 | 0.189929 | -0.08546 | 0.344215 | -0.14881 | 0.011019 | 0.032997 | 0.405048 | 0.301267 |
| 37.36111 | 0.889115 | 0.353157 | 0.405112 | 1.425904 | 0.00436 | -0.05887 | 0.073499 | 1.628523 | 2.114842 | 0.345766 | 0.089102 | 0.217032 | -0.24624 | 0.359474 | -0.16883 | -0.17936 | 0.04782 | 0.619845 | 0.690342 |
| 37.375 | 0.821369 | 0.433642 | 0.419634 | 1.332153 | -0.43034 | -0.00141 | 0.208329 | 1.715263 | 2.597868 | 0.272859 | 0.11254 | 0.285332 | -0.30549 | 0.464532 | -0.20433 | -0.32571 | 0.090923 | 0.693422 | 0.920355 |
| 37.38889 | 0.778712 | 0.540652 | 0.470751 | 1.26579 | -0.78422 | -0.02367 | 0.389934 | 1.591171 | 2.562133 | 0.257206 | 0.064488 | 0.299208 | -0.12209 | 0.516323 | -0.2172 | -0.32717 | 0.089701 | 0.660968 | 0.715112 |
| 37.40278 | 0.714285 | 0.505992 | 0.572182 | 1.369666 | -0.94472 | -0.0398 | 0.616166 | 1.014378 | 1.731359 | 0.218299 | -0.06537 | 0.181526 | 0.104948 | 0.379463 | -0.25596 | -0.30785 | 0.042363 | 0.640934 | 0.196332 |
| 37.41667 | 0.510917 | 0.300218 | 0.641342 | 1.514348 | -1.0365 | -0.02472 | 0.664349 | -0.04689 | 0.470746 | 0.271388 | -0.09038 | 0.143778 | 0.256185 | 0.38314 | -0.22131 | -0.23012 | 0.081904 | 0.798269 | -0.27207 |
| 37.43056 | 0.302216 | 0.151649 | 0.587645 | 1.492041 | -1.23183 | 0.002991 | 0.469738 | -1.14775 | -0.84765 | 0.786519 | 0.311043 | 0.414291 | 0.582336 | 0.773736 | 0.183837 | 0.011436 | 0.523455 | 1.661849 | -0.28228 |
| 37.44444 | 0.19516 | 0.187685 | 0.481349 | 1.242333 | -1.37503 | -0.03114 | 0.188783 | -2.00044 | -1.76174 | 1.989074 | 1.290845 | 1.04887 | 1.195679 | 1.704359 | 0.856293 | 0.433868 | 1.279337 | 2.980338 | 0.775783 |
| 37.45833 | 0.095154 | 0.235869 | 0.381363 | 1.065893 | -1.33321 | -0.07752 | 0.13383 | -2.56458 | -2.33775 | 2.544826 | 1.563913 | 1.464106 | 1.482895 | 1.991068 | 1.047335 | 0.779323 | 1.605559 | 3.504247 | 2.954658 |
| 37.47222 | 0.093322 | 0.30167 | 0.406722 | 1.02036 | -1.20573 | -0.02171 | 0.363733 | -3.09471 | -2.83068 | 1.904751 | 1.240884 | 1.155404 | 1.495431 | 1.566472 | 0.845611 | 0.938837 | 1.151303 | 2.149202 | 4.016303 |
| 37.48611 | 0.672646 | 1.058479 | 0.868157 | 1.133489 | -0.53775 | 0.365526 | 1.210723 | -3.24139 | -2.99566 | 0.911428 | 1.480845 | 0.620948 | 1.419848 | 0.852845 | 0.604175 | 0.837773 | 0.527156 | 1.994918 | 3.20939 |
| 37.5 | 2.338724 | 3.021942 | 1.802187 | 1.556376 | 0.946011 | 1.050822 | 2.968888 | -1.82626 | -1.4163 | 0.369585 | 2.85869 | 0.469822 | 0.992126 | 0.657744 | 0.425619 | 0.690941 | 0.318781 | 2.780201 | 1.713615 |
| 37.51389 | 3.780563 | 4.525818 | 2.411731 | 1.835649 | 2.893224 | 1.276374 | 3.992285 | 1.032613 | 1.501813 | 0.04305 | 2.697882 | 0.624488 | 0.48956 | 0.813835 | 0.294711 | 0.235472 | 0.249128 | 2.587513 | 0.929148 |
| 37.52778 | 3.504687 | 4.174815 | 2.273274 | 1.506229 | 3.0094 | 0.797492 | 3.387257 | 2.881119 | 4.238117 | -0.05745 | 0.82018 | 0.781905 | 0.037803 | 0.712025 | 0.256138 | -0.42005 | 0.201647 | 1.53297 | -0.52853 |
| 37.54167 | 2.964335 | 3.294144 | 2.102916 | 0.878657 | 4.322257 | 0.393118 | 2.706727 | 2.984367 | 5.545267 | -0.04484 | -1.62033 | 0.816488 | -0.12511 | 0.524716 | 0.233121 | -0.69792 | 0.252367 | 0.85312 | -1.42813 |
| 37.55556 | 2.945872 | 2.847873 | 1.92132 | 0.560323 | 5.883757 | 0.311361 | 2.341501 | 2.376726 | 3.534031 | 0.145293 | -3.54116 | 0.573041 | -0.02261 | 0.104332 | 0.296626 | -0.72298 | 0.334037 | 0.53599 | -2.18318 |
| 37.56944 | 2.870122 | 2.71977 | 1.573406 | 0.784321 | 7.205378 | 0.514634 | 2.289656 | 1.222093 | 1.633988 | 0.375818 | -3.37225 | 0.450712 | 0.209843 | 0.059302 | 0.478875 | -0.51974 | 0.33669 | 0.441517 | -3.12686 |
| 37.58333 | 2.610145 | 2.636372 | 1.243285 | 1.147251 | 6.500893 | 1.197416 | 2.240893 | 0.291105 | 0.974833 | 0.414184 | -2.00208 | 0.688688 | 0.31065 | 0.380683 | 0.634429 | 0.007486 | 0.412711 | 0.365019 | -2.52637 |
| 37.59722 | 2.206359 | 2.282795 | 0.958595 | 1.514339 | 5.056666 | 1.786618 | 2.101442 | -0.93697 | -0.75455 | 0.290288 | -0.58181 | 0.894434 | 0.332543 | 0.625136 | 0.728197 | 0.280887 | 0.431384 | -0.05547 | -3.57625 |
| 37.61111 | 1.709867 | 1.883058 | 0.809282 | 1.527239 | 4.200567 | 2.151898 | 1.92055 | -1.73058 | -2.60959 | 0.062471 | -0.48666 | 0.91662 | 0.253994 | 0.681222 | 0.656405 | 0.059171 | 0.347204 | -0.82 | -4.30752 |
| 37.625 | 1.244374 | 1.580451 | 0.762643 | 1.355737 | 3.261911 | 2.215502 | 1.74644 | -1.90407 | -4.03693 | -0.31481 | -0.75818 | 0.642385 | -0.04565 | 0.61447 | 0.396241 | -0.47437 | 0.22599 | -1.7027 | -3.60666 |
| 37.63889 | 1.232268 | 1.360861 | 0.632806 | 1.143185 | 2.751481 | 1.82188 | 1.569966 | -2.16407 | -4.175 | -0.78806 | -1.0691 | 0.317535 | -0.3919 | 0.402914 | 0.049139 | -0.95041 | 0.150064 | -2.48157 | -0.49804 |
| 37.65278 | 0.900791 | 1.289209 | 0.422282 | 0.903871 | 2.457528 | 1.186523 | 1.314676 | -2.34683 | -3.90574 | -0.90751 | -1.10915 | 0.097405 | -0.48282 | 0.094328 | -0.09925 | -1.14206 | 0.188231 | -2.82351 | 1.574526 |
| 37.66667 | 0.776839 | 1.319603 | 0.416633 | 0.613946 | 1.825971 | 0.176002 | 1.0766 | -2.73511 | -3.27464 | -0.7431 | 0.521763 | 0.075379 | -0.23648 | -0.05434 | -0.04084 | -0.78056 | 0.283968 | -2.85878 | 1.974393 |
| 37.68056 | 0.528116 | 1.32139 | 0.498697 | 0.306398 | 0.609463 | -0.85602 | 0.928708 | -3.32278 | -2.9112 | -0.56226 | 1.870322 | 0.12736 | 0.131709 | -0.10295 | 0.033015 | -0.27342 | 0.359937 | -2.65715 | 0.31983 |
| 37.69444 | 0.148077 | 1.106166 | 0.538392 | -0.05548 | -0.93744 | -1.73663 | 0.855523 | -4.42621 | -2.86855 | -0.45725 | 3.615348 | 0.167354 | 0.326607 | -0.30161 | 0.045276 | 0.076284 | 0.343581 | -2.26932 | 0.682122 |
| 37.70833 | 0.036477 | 0.869814 | 0.39766 | -0.5544 | -2.24302 | -2.13145 | 0.62459 | -5.57298 | -2.29535 | -0.51695 | 1.575453 | 0.142308 | 0.126941 | -0.43972 | -0.03734 | -0.00291 | 0.240331 | -1.82359 | 0.937434 |
| 37.72222 | 0.128688 | 0.721925 | -0.09929 | -0.7427 | -3.26492 | -0.98158 | 0.089761 | -6.24966 | -1.91918 | -0.80816 | -2.0466 | 0.059375 | -0.23736 | -0.61819 | -0.16138 | -0.36966 | 0.067934 | -1.83543 | 1.217605 |
| 37.73611 | 0.144963 | 0.688501 | -0.87428 | -0.61271 | -3.68504 | -0.95846 | -0.68186 | -5.88514 | -0.95745 | -1.08886 | -3.18794 | -0.06736 | -0.82072 | -0.94292 | -0.32694 | -0.66335 | -0.11748 | -1.90162 | 0.538362 |
| 37.75 | 0.242213 | 0.613942 | -1.13145 | -0.46602 | -4.56638 | -0.87498 | -1.12211 | -4.44672 | -0.44121 | -1.33652 | -1.98083 | -0.17892 | -1.2971 | -0.90703 | -0.46067 | -0.6609 | -0.19196 | -1.71187 | -1.68898 |
| 37.76389 | 0.085588 | 0.44483 | -0.96527 | -0.52327 | -5.5674 | -0.26976 | -0.95571 | -2.90359 | 0.138274 | -1.28817 | -0.4864 | -0.19892 | -1.35835 | -0.58132 | -0.54098 | -0.37218 | -0.17011 | -1.39592 | -1.36457 |
| 37.77778 | -0.34408 | 0.263988 | -0.81296 | -0.50203 | -5.55601 | -0.17597 | -0.91073 | -1.62068 | 0.112223 | -1.12089 | -0.38658 | -0.14595 | -1.17741 | -0.18771 | -0.60681 | -0.02781 | -0.13562 | -1.06882 | 0.146246 |
| 37.79167 | -0.64458 | 0.103265 | -0.80695 | 0.043716 | -5.102 | -0.08791 | -0.69126 | -1.09513 | -0.52879 | -0.88712 | -0.10092 | -0.15816 | -0.68682 | -0.10344 | -0.60439 | 0.154697 | -0.11455 | -0.67629 | 1.725845 |
| 37.80556 | -0.56522 | -0.14665 | -0.85742 | 1.106788 | -4.61004 | -0.03225 | -0.06635 | -1.21181 | -1.6949 | 0 | 0 | 0 | 0 | 0 | 0 | 0 | 0 | 0 | 0 |
| 37.81944 | -0.71522 | -0.46525 | -0.80079 | 1.786566 | -3.68176 | -0.01925 | 0.326074 | -1.84671 | -3.10001 | 0 | 0 | 0 | 0 | 0 | 0 | 0 | 0 | 0 | 0 |
| 37.83333 | -0.83932 | -0.7932 | -0.82079 | 2.015351 | -2.70135 | -0.04976 | 0.284993 | -2.59014 | -4.1143 | 0 | 0 | 0 | 0 | 0 | 0 | 0 | 0 | 0 | 0 |
| 37.84722 | -1.0803 | -1.08814 | -0.933 | 2.07521 | -2.22255 | -0.12257 | -0.08294 | -3.58725 | -5.01839 | 0 | 0 | 0 | 0 | 0 | 0 | 0 | 0 | 0 | 0 |
| 37.86111 | -1.65596 | -1.45562 | -0.95162 | 1.997437 | -1.89497 | -0.19809 | -0.60741 | -4.47258 | -5.25725 | 0 | 0 | 0 | 0 | 0 | 0 | 0 | 0 | 0 | 0 |
| 37.875 | -1.86134 | -1.65318 | -0.80978 | 1.915954 | -1.67136 | -0.22525 | -1.04038 | -5.37802 | -5.0175 | 0 | 0 | 0 | 0 | 0 | 0 | 0 | 0 | 0 | 0 |
| 37.88889 | -1.26931 | -1.74003 | -0.56885 | 1.932886 | -1.43369 | -0.23934 | -1.36815 | -6.14924 | -5.0792 | 0 | 0 | 0 | 0 | 0 | 0 | 0 | 0 | 0 | 0 |
| 37.90278 | -1.08215 | -1.62789 | -0.33772 | 1.96947 | -1.20883 | -0.30184 | -1.57064 | -5.58451 | -4.5374 | 0 | 0 | 0 | 0 | 0 | 0 | 0 | 0 | 0 | 0 |
| 37.91667 | -0.2754 | -0.91694 | -0.16513 | 1.887389 | -0.99437 | -0.40626 | -1.59384 | -4.99504 | -4.11505 | 0 | 0 | 0 | 0 | 0 | 0 | 0 | 0 | 0 | 0 |
| 37.93056 | 0.19949 | -0.22919 | -0.01294 | 1.714016 | -0.60282 | -0.47793 | -1.48799 | -4.42633 | -4.06456 | 0 | 0 | 0 | 0 | 0 | 0 | 0 | 0 | 0 | 0 |
| 37.94444 | 0.585776 | 0.594678 | 0.198481 | 1.572525 | -0.00212 | -0.32085 | -0.95441 | -3.08245 | -3.2159 | 0 | 0 | 0 | 0 | 0 | 0 | 0 | 0 | 0 | 0 |

**Supplementary Table S2.** Raw motion data from: (a) 100 mM Boron stress

| Time | Control1 | Control2 | Control3 | Control4 | Control5 | Control6 | Control7 | Control8 | Control9 | Stress1 | Stress2 | Stress3 | Stress4 | Stress5 | Stress6 | Stress7 | Stress8 | Stress9 |
|---|---|---|---|---|---|---|---|---|---|---|---|---|---|---|---|---|---|---|
| 18 | -2.87263 | -3.06867 | -0.34082 | -0.68394 | -0.95362 | -1.14881 | -0.11342 | -1.47269 | -3.50278 | -4.12276 | -0.92714 | -13.2115 | -4.89392 | -4.24594 | -5.28422 | -4.73433 | -3.84629 | -0.05758 |
| 18.01389 | -3.90168 | -2.6829 | -0.45645 | -1.23005 | -1.88639 | -1.54856 | -0.46749 | -1.64265 | -3.90748 | -6.33423 | -0.98883 | -10.3332 | -6.433 | -3.45821 | -5.08873 | -3.72075 | -2.85943 | -0.53947 |
| 18.02778 | -2.06005 | -0.70264 | -0.05799 | -0.59904 | -1.14819 | -0.85005 | -0.56269 | -0.35955 | -1.18808 | -4.38308 | -0.12853 | -2.84706 | -3.99753 | -1.18927 | -1.93605 | -0.12757 | 0.242772 | 0.330558 |
| 18.04167 | -0.40481 | 0.614401 | 0.273322 | 0.349995 | 0.189829 | 0.010282 | -0.26136 | 0.940684 | 1.249857 | -0.43927 | 0.792103 | 1.215057 | -0.60642 | 0.384111 | 0.766056 | 1.593897 | 1.850557 | 1.465928 |
| 18.05556 | 0.095372 | 1.092677 | 0.270688 | 0.343731 | 0.699447 | 0.523167 | -0.05376 | 1.238128 | 2.318539 | 1.10777 | 0.725502 | 3.062842 | 1.394312 | 0.922219 | 1.609907 | 1.673636 | 1.700832 | 1.160047 |
| 18.06944 | 0.198141 | 0.835235 | 0.119688 | 0.140995 | 0.470448 | 0.551971 | -0.00093 | 0.7422 | 1.916237 | 1.509779 | 0.443846 | 3.534119 | 1.96333 | 0.720936 | 1.10346 | 1.148348 | 1.002423 | 0.053288 |
| 18.08333 | 0.172325 | 0.308808 | 0.015436 | 0.032171 | 0.14148 | 0.2602 | 0.018977 | 0.326419 | 1.052364 | 1.175948 | 0.12647 | 2.544301 | 1.507464 | 0.900092 | 0.910818 | 0.546773 | 0.434286 | -0.40145 |
| 18.09722 | 0.12599 | -0.05336 | -0.00249 | -0.00264 | 0.016463 | 0.1102 | 0.082684 | 0.094703 | 0.522671 | 0.575888 | -0.15436 | 0.810754 | 0.708952 | 1.034744 | 0.67721 | -0.01098 | -0.01907 | -0.36619 |
| 18.11111 | 0.11089 | -0.10305 | 0.022783 | 0.00617 | 0.001174 | 0.078689 | 0.123513 | 0.033917 | 0.22584 | 0.157414 | -0.25979 | -0.08111 | 0.159839 | 0.445971 | 0.223214 | -0.27123 | -0.18325 | -0.23541 |
| 18.125 | 0.129504 | -0.01588 | 0.030769 | 0.032987 | 0.007362 | 0.039216 | 0.130219 | 0.009672 | -0.02658 | -0.07463 | -0.27724 | -0.38356 | -0.04427 | -0.08952 | -0.10788 | -0.37227 | -0.26939 | -0.19455 |
| 18.13889 | 0.207397 | 0.096659 | -0.08246 | 0.123841 | 0.031753 | -0.04188 | 0.049105 | -0.03798 | -0.19913 | -0.34002 | -0.47886 | -0.72632 | -0.2288 | -0.40058 | -0.44457 | -0.80693 | -0.78136 | -0.55375 |
| 18.15278 | 0.25843 | 0.008798 | -0.0323 | 0.00575 | 0.068928 | -0.10788 | 0.543946 | -0.08638 | -0.22404 | -0.62472 | -0.89581 | -0.72332 | -0.37384 | -0.76961 | -0.86727 | -1.58779 | -1.75316 | -1.21376 |
| 18.16667 | 0.176665 | -0.16384 | 0.75301 | 1.04578 | 0.027637 | -0.10217 | 0.900871 | -0.05746 | -0.06245 | -0.59975 | -0.92529 | -0.10575 | -0.17977 | -0.6582 | -0.77869 | -1.6196 | -1.92283 | -1.17029 |
| 18.18056 | -0.00665 | -0.20418 | 0.123758 | 0.513595 | -0.04349 | -0.1074 | 0.078134 | -0.05256 | -0.05144 | -0.453 | -0.49215 | -0.45487 | -0.2052 | -0.3637 | -0.43069 | -1.02027 | -1.17723 | -0.4238 |
| 18.19444 | -0.16627 | -0.23104 | -0.04591 | -0.06918 | -0.03361 | -0.13099 | -0.08401 | -0.10521 | -0.2904 | -0.46629 | -0.197 | -1.49229 | -0.50859 | -0.48876 | -0.45297 | -0.77768 | -0.65652 | 0.119643 |
| 18.20833 | -0.08178 | -0.17175 | -0.05022 | -0.0983 | 0.005651 | -0.09492 | -0.07199 | -0.07841 | -0.37247 | -0.28684 | -0.04366 | -1.00868 | -0.37221 | -0.38428 | -0.37637 | -0.4624 | -0.29362 | 0.239387 |
| 18.22222 | 0.083028 | 0.010305 | -0.02152 | -0.03796 | -0.00894 | -0.04973 | -0.04599 | 0.008293 | -0.22049 | 0.065645 | 0.143166 | 0.295397 | 0.025699 | -0.03429 | -0.06872 | 0.073813 | 0.123088 | 0.213887 |
| 18.23611 | 0.080131 | 0.151258 | -0.00906 | -0.01143 | -0.06903 | -0.05426 | -0.05488 | 0.03672 | -0.09384 | 0.177102 | 0.235119 | 0.825819 | 0.254917 | 0.218887 | 0.117473 | 0.288675 | 0.311658 | 0.19475 |
| 18.25 | -0.08857 | 0.017564 | -0.03092 | -0.03492 | -0.1007 | -0.08662 | -0.08302 | -0.07248 | -0.25351 | 0.220673 | 0.215223 | 1.076072 | 0.295367 | 0.244441 | 0.17094 | 0.324156 | 0.267429 | 0.082274 |
| 18.26389 | -0.21532 | -0.2237 | -0.03565 | -0.05599 | -0.06082 | -0.07744 | -0.06705 | -0.17992 | -0.562 | 0.279979 | 0.148094 | 1.267528 | 0.223914 | 0.094314 | 0.152453 | 0.314064 | 0.129111 | -0.01619 |
| 18.27778 | -0.09824 | -0.14899 | 0.001418 | -0.03828 | -0.02195 | -0.04268 | -0.02661 | -0.11233 | -0.48925 | 0.055633 | 0.036922 | 0.591803 | 0.159007 | -0.02484 | -0.0337 | -0.02231 | -0.05562 | 0.097654 |
| 18.29167 | 0.049278 | 0.080435 | 0.01655 | -0.01411 | -0.041 | -0.03375 | -0.04162 | 0.02749 | -0.16326 | -0.19545 | -0.04397 | -0.04126 | 0.16939 | 0.024652 | -0.17333 | -0.35809 | -0.17088 | 0.162135 |
| 18.30556 | -0.07609 | -0.03495 | -0.00433 | -0.00824 | -0.06929 | -0.04141 | -0.0691 | -0.01215 | -0.08421 | -0.11627 | -0.0346 | 0.272778 | 0.143938 | 0.084594 | -0.09329 | -0.27854 | -0.18315 | 0.008895 |
| 18.31944 | -0.17665 | -0.21735 | -0.00178 | -0.0086 | -0.02934 | -0.0079 | -0.03715 | -0.03586 | -0.11671 | -0.00013 | -0.0167 | 0.43937 | 0.141971 | 0.088227 | -0.00384 | -0.17683 | -0.14503 | 0.013718 |
| 18.33333 | -0.01505 | 0.019968 | 0.009606 | -0.00116 | 0.015572 | 0.040379 | -0.00801 | 0.090761 | 0.035237 | -0.02358 | -0.00669 | 0.16987 | 0.346684 | 0.144683 | 0.020187 | -0.20185 | -0.02371 | 0.191295 |
| 18.34722 | 0.188023 | 0.263958 | -0.01408 | -0.02847 | 0.007331 | 0.041418 | -0.03889 | 0.168839 | 0.189611 | 0.088188 | 0.044304 | 0.766365 | 0.426881 | 0.251963 | 0.104929 | 0.061565 | 0.08571 | 0.05862 |
| 18.36111 | 0.180956 | 0.136105 | -0.02865 | -0.05326 | 0.006706 | 0.009495 | -0.06052 | 0.111787 | 0.091458 | 0.300694 | 0.093814 | 1.603349 | 0.22366 | 0.210755 | 0.217918 | 0.414433 | 0.128574 | -0.18002 |
| 18.375 | -0.02738 | -0.04607 | -0.00043 | -0.01258 | 0.017575 | 0.0009 | -0.04223 | 0.021821 | -0.05155 | 0.227874 | 0.121032 | 1.102693 | 0.117919 | 0.090609 | 0.132951 | 0.292449 | 0.143806 | -0.02529 |
| 18.38889 | -0.1536 | -0.12834 | 0.024507 | 0.04188 | 0.015373 | 0.026681 | -0.01813 | -0.01235 | -0.09145 | 0.00349 | 0.094656 | 0.190781 | 0.216035 | 0.10556 | 0.023429 | 0.025382 | 0.139529 | 0.16472 |
| 18.40278 | -0.03189 | -0.02941 | 0.037008 | 0.039003 | 0.049557 | 0.051292 | -0.01075 | 0.065939 | -0.00927 | -0.03751 | -0.00905 | 0.19016 | 0.195285 | 0.057695 | -0.03186 | -0.03031 | 0.062663 | 0.019524 |
| 18.41667 | 0.227033 | 0.21823 | 0.069479 | 0.0567 | 0.129866 | 0.124608 | 0.033198 | 0.216858 | 0.208982 | -0.01918 | -0.13878 | 0.711753 | 0.13126 | 0.029867 | -0.01575 | 0.049455 | -0.00727 | -0.21453 |
| 18.43056 | 0.302994 | 0.329388 | 0.166894 | 0.1734 | 0.205735 | 0.269038 | 0.120008 | 0.34563 | 0.402642 | 0.000495 | -0.17955 | 1.10245 | 0.219557 | 0.171012 | 0.10328 | 0.168569 | 0.02766 | -0.32162 |
| 18.44444 | 0.200525 | 0.267176 | 0.195079 | 0.242504 | 0.243617 | 0.34316 | 0.1663 | 0.343948 | 0.396234 | -0.05483 | -0.13527 | 0.706428 | 0.30717 | 0.205981 | 0.132214 | 0.107217 | 0.047305 | -0.2026 |
| 18.45833 | 0.239765 | 0.269195 | 0.153128 | 0.176883 | 0.23031 | 0.2827 | 0.157907 | 0.292161 | 0.348998 | -0.05188 | -0.03443 | 0.187269 | 0.365601 | 0.132638 | 0.122215 | 0.042425 | 0.077522 | 0.020734 |
| 18.47222 | 0.585867 | 0.705729 | 0.077229 | 0.078054 | 0.171719 | 0.186342 | 0.063746 | 0.386015 | 0.69915 | 0.595833 | 0.14446 | 2.082891 | 0.530308 | 0.415484 | 0.620051 | 0.819689 | 0.423923 | -0.09895 |
| 18.48611 | 1.474137 | 1.703331 | 0.018403 | 0.009584 | 0.116902 | 0.154441 | -0.13413 | 0.695342 | 1.244987 | 2.374769 | 0.316748 | 5.87303 | 1.04734 | 1.256804 | 1.730696 | 2.488723 | 1.132949 | -0.43653 |
| 18.5 | 3.748588 | 1.907584 | 0.148268 | 0.071535 | 0.315887 | 0.460745 | -0.0089 | 0.954663 | 2.302151 | 5.426234 | 0.412009 | 9.587125 | 3.087366 | 3.337046 | 3.195748 | 4.273596 | 1.901813 | -0.57134 |
| 18.51389 | 4.31207 | 2.682216 | 0.213881 | 0.283989 | 0.463761 | 0.74355 | 0.31578 | 0.597103 | 2.385856 | 7.228259 | 0.072767 | 9.229709 | 4.375506 | 3.616855 | 3.648931 | 3.276578 | 1.363094 | -0.71295 |
| 18.52778 | 2.229158 | 1.351436 | 0.096557 | 0.630933 | 0.725739 | 0.465506 | 0.242511 | 0.103998 | 0.98398 | 4.510508 | -0.2514 | 4.709502 | 3.202927 | 1.992988 | 2.642671 | 1.149115 | 0.234679 | -0.77027 |
| 18.54167 | 1.379261 | 0.644207 | -0.01595 | 0.440173 | 0.624683 | 0.184074 | 0.097126 | 0.096478 | 0.584584 | 2.042487 | -0.10723 | 2.673057 | 2.342561 | 0.799137 | 1.523673 | 0.218388 | -0.13127 | -0.46511 |
| 18.55556 | 0.750471 | 0.289146 | -0.0132 | 0.318121 | 0.421044 | 0.122693 | 0.04997 | 0.065273 | 0.331557 | 0.826953 | 0.20209 | 1.635758 | 1.633943 | 0.353537 | 0.720525 | -0.01154 | 0.065327 | 0.045932 |
| 18.56944 | 0.287357 | -0.00935 | 0.0323 | 0.249444 | 0.241559 | 0.115161 | 0.038264 | -0.05421 | 0.029389 | 0.47923 | 0.363614 | 0.659481 | 1.087217 | 0.265185 | 0.273829 | 0.217916 | 0.383297 | 0.460467 |
| 18.58333 | 0.361306 | 0.029864 | 0.032032 | 0.152125 | 0.187527 | 0.116962 | 0.005407 | 0.010511 | -0.20964 | 0.516603 | 0.301545 | 0.309157 | 0.563871 | 0.404437 | 0.349587 | 0.716619 | 0.590633 | 0.396241 |
| 18.59722 | 0.52582 | 0.078079 | 0.014746 | 0.114094 | 0.215906 | 0.141275 | -0.0167 | 0.116483 | -0.48131 | 0.695652 | 0.200362 | 0.63469 | 0.09877 | 0.652653 | 0.885419 | 1.244342 | 0.684851 | 0.046422 |
| 18.61111 | 0.379361 | -0.01513 | 0.025516 | 0.161567 | 0.240781 | 0.116355 | -0.01798 | 0.062999 | -0.63866 | 0.662634 | 0.162882 | 0.233155 | -0.30804 | 0.896056 | 1.065856 | 1.329789 | 0.753754 | -0.00175 |
| 18.625 | -0.00317 | -0.16987 | 0.034287 | 0.204968 | 0.170726 | 0.004305 | -0.02599 | -0.04145 | -0.60742 | 0.333038 | 0.162052 | -0.4166 | -0.53275 | 0.964433 | 0.705556 | 0.934163 | 0.692885 | 0.217692 |
| 18.63889 | -0.41077 | -0.25121 | 0.013388 | 0.196607 | 0.069025 | -0.08864 | -0.03197 | -0.09784 | -0.44813 | 0.042586 | 0.124684 | -0.00871 | -0.56055 | 1.019997 | 0.427631 | 0.689192 | 0.45879 | 0.105642 |
| 18.65278 | -0.47061 | -0.22824 | -0.03762 | 0.144583 | -0.00871 | -0.10185 | -0.03224 | -0.0335 | -0.09414 | -0.21522 | 0.118192 | 0.590552 | -0.27807 | 0.933436 | 0.09464 | 0.632709 | 0.267651 | 0.111363 |
| 18.66667 | -0.19718 | -0.05974 | -0.08098 | 0.072621 | -0.02889 | -0.05781 | -0.03131 | 0.109794 | 0.536514 | -0.37623 | 0.173793 | 0.349123 | 0.28757 | 0.547739 | -0.44514 | 0.250276 | 0.132264 | 0.492649 |
| 18.68056 | -0.10352 | 0.023133 | -0.06693 | 0.095448 | -0.03001 | -0.02716 | -0.00755 | 0.045337 | 0.841924 | -0.08759 | 0.158174 | -0.00681 | 0.446093 | 0.300391 | -0.44277 | 0.000808 | -0.02491 | 0.523357 |
| 18.69444 | -0.11195 | 0.077166 | -0.01668 | 0.122192 | -0.02946 | 0.014591 | 0.026163 | -0.07644 | 0.513971 | 0.241896 | 0.112913 | 0.673174 | 0.272977 | 0.317936 | 0.072878 | 0.149837 | -0.03276 | 0.219828 |
| 18.70833 | 0.012407 | 0.251859 | 0.019836 | 0.056686 | -0.01119 | 0.068006 | 0.028473 | -0.05259 | 0.142296 | 0.423101 | 0.107456 | 1.064689 | 0.112963 | 0.365592 | 0.397814 | 0.299831 | 0.026277 | 0.074278 |
| 18.72222 | 0.136613 | 0.375276 | 0.048473 | -0.03949 | 0.011654 | 0.089133 | 0.011615 | 0.0357 | -0.0091 | 0.354661 | 0.141894 | 0.677268 | 0.052949 | 0.370803 | 0.30314 | 0.176554 | 0.068385 | 0.164653 |
| 18.73611 | 0.250166 | 0.403323 | 0.063615 | -0.12333 | 0.047206 | 0.095726 | -0.00767 | 0.130812 | -0.12009 | 0.272177 | 0.164314 | 0.324655 | -0.10291 | 0.385343 | 0.191092 | 0.035923 | 0.083281 | 0.170175 |
| 18.75 | 0.139229 | 0.336443 | 0.048751 | -0.05971 | 0.064265 | 0.071295 | 0.014127 | 0.068853 | -0.44283 | 0.289121 | 0.159093 | 0.46363 | -0.27541 | 0.437621 | 0.270858 | 0.238852 | 0.101645 | 0.023307 |
| 18.76389 | -0.19658 | 0.134145 | -0.00088 | 0.049703 | 0.046325 | 0.025751 | 0.046052 | -0.1218 | -0.77505 | 0.269771 | 0.150954 | 0.739068 | -0.13215 | 0.427927 | 0.259851 | 0.628852 | 0.159938 | 0.034273 |
| 18.77778 | -0.37663 | 0.072194 | -0.03339 | 0.123491 | 0.012711 | 0.017708 | 0.079145 | -0.19356 | -0.73762 | 0.231863 | 0.138965 | 1.133431 | 0.333812 | 0.345478 | 0.10959 | 0.882638 | 0.219854 | 0.19318 |
| 18.79167 | -0.31021 | 0.096854 | -0.02383 | 0.120574 | -0.00466 | 0.0514 | 0.109716 | -0.12084 | -0.29985 | 0.217103 | 0.140438 | 1.669121 | 0.693097 | 0.223317 | -0.04118 | 0.865677 | 0.189068 | 0.246048 |
| 18.80556 | -0.09773 | 0.271017 | 0.001052 | 0.039285 | -0.01387 | 0.07811 | 0.085259 | 0.031637 | 0.233807 | 0.163655 | 0.128914 | 1.188787 | 0.532992 | 0.065266 | -0.19775 | 0.528601 | 0.029312 | 0.282736 |
| 18.81944 | 0.10504 | 0.378885 | 0.041031 | -0.048 | -0.00703 | 0.082669 | 0.032438 | 0.15885 | 0.542274 | 0.056572 | 0.163276 | -0.39939 | -0.05672 | 0.111025 | -0.39185 | -0.0418 | -0.20307 | 0.435779 |
| 18.83333 | 0.167769 | 0.437348 | 0.066453 | -0.07111 | 0.003527 | 0.058828 | -0.00449 | 0.172068 | 0.358599 | -0.01342 | 0.206609 | -1.5784 | -0.69392 | 0.335981 | -0.37748 | -0.32328 | -0.19931 | 0.407125 |
| 18.84722 | -0.0253 | 0.402189 | 0.03018 | -0.06949 | -0.03267 | -0.00049 | -0.01872 | 0.047823 | -0.18097 | -0.03403 | 0.157577 | -1.32065 | -1.23851 | 0.56834 | -0.01745 | -0.41763 | -0.31963 | 0.117651 |
| 18.86111 | -0.30433 | 0.38696 | -0.05866 | -0.09186 | -0.07791 | -0.04266 | -0.02051 | -0.08303 | -0.66294 | -0.14667 | 0.091064 | -0.78079 | -1.42799 | 0.612654 | 0.112775 | -0.76049 | -0.67252 | -0.13277 |
| 18.875 | -0.56308 | 0.397744 | -0.11172 | -0.06301 | -0.07738 | -0.0334 | 0.029219 | -0.25602 | -0.80248 | -0.26721 | 0.077721 | -0.57035 | -1.19856 | 0.412359 | -0.18862 | -1.23454 | -0.90969 | -0.10496 |
| 18.88889 | -0.5728 | 0.3642 | -0.07866 | -0.05676 | -0.07291 | -0.00701 | 0.047034 | -0.27019 | -0.74565 | -0.31625 | 0.038034 | -0.15572 | -0.8651 | 0.161795 | -0.45055 | -1.3592 | -0.83251 | -0.13195 |
| 18.90278 | -0.40084 | 0.625938 | -0.03766 | -0.13516 | -0.06919 | -0.02069 | -0.02193 | -0.07895 | -0.66357 | -0.41113 | -0.08164 | 0.623143 | -0.77804 | 0.060883 | -0.39413 | -1.12124 | -0.8529 | -0.30583 |
| 18.91667 | -0.52002 | 0.608974 | -0.08564 | -0.15697 | -0.13046 | -0.11373 | -0.06366 | -0.05295 | -0.8213 | -0.67179 | -0.19432 | 1.009742 | -0.85484 | 0.174756 | -0.40509 | -0.96128 | -0.94688 | -0.49648 |
| 18.93056 | -0.68875 | 0.525431 | -0.21098 | -0.1762 | -0.22255 | -0.2209 | -0.10246 | -0.1384 | -0.90994 | -0.9108 | -0.22243 | 0.784153 | -0.92537 | 0.290829 | -0.63343 | -1.0157 | -1.00882 | -0.56329 |
| 18.94444 | -0.54395 | 0.610414 | -0.26883 | -0.23139 | -0.21467 | -0.20017 | -0.12869 | -0.11551 | -0.6331 | -1.02161 | -0.10187 | -0.06441 | -0.87401 | 0.215829 | -1.03806 | -0.94468 | -0.78058 | -0.39877 |
| 18.95833 | -0.57218 | 0.793595 | -0.24287 | -0.12805 | -0.13893 | -0.10576 | -0.04729 | -0.18448 | -0.32591 | -1.08179 | 0.008347 | -1.70475 | -0.9358 | -0.05069 | -1.41596 | -0.63272 | -0.45473 | -0.08612 |
| 18.97222 | -0.91292 | 0.554539 | -0.22424 | -0.02974 | -0.16071 | -0.0543 | 0.094116 | -0.47733 | -0.46728 | -1.23597 | -0.18571 | -4.10153 | -1.01598 | -0.29289 | -1.90493 | -0.51603 | -0.52904 | -0.20962 |
| 18.98611 | -1.27381 | -0.28372 | -0.24443 | -0.20531 | -0.28317 | -0.15509 | 0.089472 | -0.79783 | -1.43017 | -2.33367 | -0.83812 | -6.81544 | -1.78496 | -0.94246 | -2.80605 | -1.64607 | -1.19533 | -0.66227 |
| 19 | -2.05192 | -1.23252 | -0.31683 | -0.60427 | -0.46783 | -0.54275 | -0.15221 | -0.92074 | -2.62014 | -5.13939 | -1.47512 | -9.93639 | -3.63999 | -2.62715 | -3.5687 | -3.74894 | -1.96816 | -1.14914 |
| 19.01389 | -2.02981 | -1.22722 | -0.34345 | -0.94941 | -0.64666 | -0.63224 | -0.38379 | -0.65792 | -2.08818 | -6.32903 | -1.33646 | -6.80774 | -3.26196 | -2.78676 | -2.7247 | -3.17777 | -1.33689 | -0.61291 |
| 19.02778 | -0.67756 | 0.08064 | -0.12958 | -0.5842 | -0.62135 | -0.36651 | -0.31495 | -0.02236 | -0.56379 | -1.62853 | -0.08808 | -0.91293 | -0.46408 | -0.39809 | -0.53809 | -0.12376 | 0.320656 | 0.450252 |
| 19.04167 | 0.088709 | 0.529106 | 0.058327 | 0.008171 | -0.1991 | -0.05701 | -0.10094 | 0.560325 | 0.706088 | 0.557387 | 1.115727 | 0.805863 | 0.973172 | 0.88599 | 1.020005 | 1.411565 | 1.591888 | 0.680293 |
| 19.05556 | -0.04909 | 0.71313 | 0.153879 | 0.088299 | -0.14573 | 0.065965 | 0.08433 | 0.183307 | 0.801012 | 0.426879 | 1.403746 | 0.11202 | 0.998474 | 0.434776 | 0.978946 | 1.097757 | 1.16978 | 0.167067 |
| 19.06944 | 0.102376 | 0.701521 | -0.00494 | 0.053689 | -0.16312 | -0.00225 | 0.004285 | 0.065871 | 0.009027 | 0.614738 | 0.527204 | 0.494241 | 0.273481 | -0.23357 | 0.408173 | 0.337186 | 0.459518 | 0.40457 |
| 19.08333 | 0.230186 | 1.443287 | -0.2098 | -0.03141 | -0.11023 | -0.05665 | -0.10867 | -0.214 | -0.74836 | 1.128181 | -0.12006 | 1.987498 | -0.18879 | -0.20645 | 0.294023 | -0.06239 | 0.190015 | 0.779595 |

| | | | | | | | | | | | | | | | | | | |
|---|---|---|---|---|---|---|---|---|---|---|---|---|---|---|---|---|---|---|
| 19.09722 | 0.179507 | 0.944983 | -0.10819 | -0.07919 | -0.0449 | -0.04652 | -0.06889 | -0.23299 | -0.47534 | 0.680978 | 0.063511 | 1.618495 | -0.08096 | -0.0319 | 0.213807 | -0.10283 | -0.41601 | 0.377815 |
| 19.11111 | 0.0786 | 0.545695 | -0.032 | -0.03238 | -0.0582 | -0.03352 | -0.03167 | -0.01009 | 0.007771 | -0.00454 | -0.02935 | 0.137618 | -0.05265 | -0.05094 | -0.08857 | 0.128938 | -0.13289 | 0.017256 |
| 19.125 | 0.050135 | 0.365446 | 0.002838 | -0.0339 | -0.05665 | -0.03892 | -0.04341 | 0.027088 | 0.088096 | -0.07289 | -0.03527 | 0.124226 | -0.08398 | -0.07288 | -0.08689 | 0.193507 | -0.08064 | -0.12129 |
| 19.13889 | 0.044232 | 0.183481 | 0.006624 | -0.04388 | -0.02801 | -0.03176 | -0.04006 | 0.044026 | 0.068128 | 0.06656 | 0.053121 | 0.639852 | -0.03564 | -0.00923 | 0.135718 | 0.337405 | 0.072431 | -0.0878 |
| 19.15278 | 0.072489 | 0.239531 | 0.010146 | -0.03066 | -0.03105 | -0.02368 | -0.03119 | 0.065077 | 0.075325 | 0.078653 | 0.107411 | 0.469695 | 0.108393 | 0.053531 | 0.145876 | 0.471931 | 0.222088 | 0.042261 |
| 19.16667 | 0.04025 | 0.38588 | 0.005036 | 0.002233 | -0.03802 | -0.00866 | -0.02128 | 0.036228 | 0.07233 | -0.03077 | 0.062611 | -0.1191 | 0.241459 | -0.00297 | -0.04938 | 0.391372 | 0.181085 | 0.130957 |
| 19.18056 | -0.04621 | 0.264884 | 0.008581 | 0.02689 | -0.03836 | -0.00176 | -0.00227 | -0.00935 | 0.021522 | -0.0486 | -0.00662 | 0.034084 | 0.123301 | -0.09714 | -0.09273 | 0.232556 | -0.02464 | 0.001953 |
| 19.19444 | -0.12149 | 0.072347 | 0.024317 | 0.025957 | -0.02123 | -0.00044 | 0.01506 | -0.01732 | -0.00455 | -0.09395 | -0.05546 | 0.304472 | -0.10047 | -0.1883 | -0.07792 | 0.104683 | -0.14963 | -0.14201 |
| 19.20833 | -0.18999 | -0.00437 | 0.051944 | 0.036374 | -0.00642 | 0.00959 | 0.040118 | -0.00795 | -0.05179 | -0.30853 | -0.1035 | -0.45439 | -0.09881 | -0.27746 | -0.28993 | -0.03431 | -0.09711 | -0.0493 |
| 19.22222 | -0.19207 | -0.00087 | 0.045173 | 0.013715 | -0.03426 | 0.00517 | 0.036589 | -0.01606 | -0.08804 | -0.37396 | -0.13622 | -0.54724 | -0.13531 | -0.27545 | -0.38876 | 0.035234 | -0.06225 | -0.10518 |
| 19.23611 | -0.06412 | 0.265752 | 0.005444 | -0.05015 | -0.04966 | -0.01704 | -0.00974 | 0.013503 | -0.01312 | -0.23161 | -0.2001 | 0.594664 | -0.43484 | -0.23031 | -0.1776 | 0.139868 | -0.18679 | -0.5631 |
| 19.25 | -0.04154 | 0.483585 | -0.0026 | -0.05743 | -0.05822 | -0.03589 | -0.04528 | 0.03701 | 0.104641 | -0.40716 | -0.40783 | 0.509779 | -0.60235 | -0.41318 | -0.35342 | -0.23671 | -0.41627 | -0.80294 |
| 19.26389 | -0.0782 | 0.475512 | 0.010156 | -0.02681 | -0.0521 | -0.04609 | -0.05063 | 0.014593 | 0.108191 | -0.68794 | -0.53643 | -0.66965 | -0.43169 | -0.62887 | -0.71118 | -0.73143 | -0.57008 | -0.61462 |
| 19.27778 | 0.002707 | 0.558113 | 0.049878 | -0.01445 | -0.00806 | -0.03045 | -0.03055 | 0.024291 | 0.109062 | -0.50144 | -0.38144 | -0.37181 | -0.20671 | -0.49895 | -0.55756 | -0.57559 | -0.52036 | -0.41343 |
| 19.29167 | 0.06392 | 0.737414 | 0.084893 | -0.01111 | 0.015276 | -0.01233 | -0.02499 | 0.036062 | 0.163801 | -0.17403 | -0.14508 | 0.550481 | -0.08435 | -0.1987 | -0.17933 | -0.17058 | -0.35068 | -0.31662 |
| 19.30556 | 0.13221 | 0.94171 | 0.090079 | -0.03277 | 0.018389 | -0.00189 | -0.0401 | 0.06348 | 0.269317 | -0.14016 | -0.08816 | 0.549156 | -0.02211 | -0.06007 | -0.11389 | -0.09117 | -0.2486 | -0.22018 |
| 19.31944 | 0.294835 | 1.094678 | 0.096373 | -0.05783 | 0.045415 | 0.021084 | -0.05231 | 0.132702 | 0.409909 | -0.26438 | -0.18451 | 0.146164 | -0.0415 | -0.11822 | -0.29124 | -0.159 | -0.2205 | -0.21615 |
| 19.33333 | 0.318316 | 1.008715 | 0.104748 | -0.04665 | 0.052331 | 0.031755 | -0.05148 | 0.163043 | 0.449158 | -0.3205 | -0.246 | -0.01171 | -0.08939 | -0.2017 | -0.42591 | 0.004022 | -0.18269 | -0.27179 |
| 19.34722 | 0.134052 | 0.862524 | 0.096553 | -0.02908 | 0.025637 | 0.018146 | -0.04874 | 0.083465 | 0.361851 | -0.26859 | -0.22127 | 0.101706 | -0.14813 | -0.21726 | -0.3842 | 0.330358 | -0.14939 | -0.31654 |
| 19.36111 | -0.0352 | 0.908461 | 0.07382 | -0.01021 | 0.030806 | 0.025089 | -0.04341 | 0.025638 | 0.336761 | -0.16517 | -0.11948 | 0.215283 | -0.0601 | -0.12244 | -0.26024 | 0.582289 | -0.10773 | -0.21273 |
| 19.375 | -0.05777 | 1.058595 | 0.111804 | 0.084196 | 0.065433 | 0.064525 | 0.012538 | 0.048205 | 0.42651 | -0.05787 | 0.031994 | 0.20328 | 0.282687 | 0.080352 | -0.1505 | 0.713898 | 0.047678 | 0.056123 |
| 19.38889 | -0.08198 | 0.976423 | 0.146051 | 0.178401 | 0.065288 | 0.095288 | 0.07149 | 0.068003 | 0.470508 | 0.098089 | 0.137671 | 0.570958 | 0.5059 | 0.2823 | 0.044725 | 0.820958 | 0.223748 | 0.190163 |
| 19.40278 | -0.0562 | 0.819231 | 0.130642 | 0.159773 | 0.076592 | 0.084765 | 0.048728 | 0.119751 | 0.445904 | 0.276846 | 0.168783 | 1.332886 | 0.468544 | 0.398514 | 0.282843 | 1.011828 | 0.372704 | 0.124892 |
| 19.41667 | 0.155168 | 0.903109 | 0.148104 | 0.161332 | 0.150176 | 0.1402 | 0.064541 | 0.213043 | 0.505061 | 0.256559 | 0.193448 | 1.157483 | 0.603864 | 0.428897 | 0.226441 | 0.963934 | 0.602165 | 0.242498 |
| 19.43056 | 0.302392 | 1.010805 | 0.213549 | 0.291057 | 0.269781 | 0.278712 | 0.169336 | 0.299763 | 0.551795 | 0.092379 | 0.28104 | 0.135721 | 0.883863 | 0.413759 | -0.00116 | 0.656211 | 0.818125 | 0.521373 |
| 19.44444 | 0.320931 | 0.931577 | 0.225507 | 0.362565 | 0.325144 | 0.348074 | 0.22724 | 0.334537 | 0.582094 | 0.131062 | 0.355523 | -0.01081 | 0.79152 | 0.450341 | 0.073468 | 0.548578 | 0.900172 | 0.545352 |
| 19.45833 | 0.318994 | 0.90933 | 0.193967 | 0.2764 | 0.269396 | 0.280851 | 0.183456 | 0.333067 | 0.63277 | 0.355426 | 0.368504 | 1.009835 | 0.511394 | 0.576134 | 0.412232 | 0.620939 | 0.774093 | 0.322811 |
| 19.47222 | 0.555123 | 1.212055 | 0.147798 | 0.05182 | 0.159371 | 0.162629 | 0.03669 | 0.375858 | 0.842692 | 0.817161 | 0.336173 | 2.178388 | 0.245578 | 0.617727 | 0.818053 | 0.842715 | 0.593185 | 0.100595 |
| 19.48611 | 1.194627 | 1.601033 | 0.23668 | -0.20812 | 0.166317 | 0.105518 | -0.13292 | 0.504876 | 1.495021 | 2.362297 | 0.165321 | 3.837758 | 0.305046 | 0.811874 | 1.265727 | 1.848271 | 0.587223 | -0.17227 |
| 19.5 | 2.466492 | 1.736022 | 0.326623 | 0.033324 | 0.214049 | 0.311708 | -0.02727 | 0.494164 | 3.079907 | 5.790143 | 0.116584 | 6.282307 | 1.828865 | 1.810309 | 1.872679 | 4.263909 | 1.379986 | -0.42825 |
| 19.51389 | 2.366604 | 0.301963 | 0.230444 | 0.506034 | 0.582491 | 0.651709 | 0.371406 | 0.345927 | 2.175165 | 7.158272 | 0.095587 | 4.671598 | 3.517297 | 2.057328 | 1.999835 | 2.447611 | 0.851555 | -0.54037 |
| 19.52778 | 1.66808 | -0.4977 | 0.002761 | 0.419523 | 0.761375 | 0.725148 | 0.461039 | 0.218417 | 0.903455 | 3.434566 | 0.262563 | 0.24331 | 1.855832 | 1.183239 | 2.112237 | 0.598313 | 0.638974 | -0.20819 |
| 19.54167 | 0.857613 | 0.748077 | -0.09637 | 0.355387 | 0.440118 | 0.502108 | 0.387661 | 0.223858 | 0.202371 | 1.355211 | 0.571505 | -1.29991 | 0.459746 | 0.651432 | 2.504141 | 0.017241 | 0.78201 | 0.201195 |
| 19.55556 | 0.407737 | 1.641745 | -0.14892 | 0.272438 | 0.274474 | 0.277615 | 0.288212 | 0.276786 | 0.270263 | 0.639091 | 0.705096 | -1.87466 | -0.26885 | 0.320825 | 1.895267 | -1.03268 | 0.556193 | 0.424525 |
| 19.56944 | 0.46308 | 1.835212 | -0.14887 | 0.114571 | 0.083701 | 0.16226 | 0.182887 | 0.189789 | 0.100414 | 0.601544 | 0.626083 | -2.25965 | -0.63942 | 0.233898 | 1.15176 | -2.00852 | -0.05361 | 0.42104 |
| 19.58333 | 0.51768 | 1.634208 | -0.07628 | 0.033069 | -0.03525 | 0.118381 | 0.117331 | 0.086454 | -0.2615 | 0.470796 | 0.504415 | -1.48063 | -0.04217 | 0.505762 | 0.827277 | -1.58642 | -0.25355 | 0.379447 |
| 19.59722 | 0.344072 | 0.986737 | 0.060511 | 0.039869 | -0.01811 | 0.104035 | 0.076454 | 0.160783 | -0.09779 | 0.402846 | 0.311862 | 0.611359 | 1.193744 | 0.688612 | 0.79331 | -0.00306 | -0.20574 | 0.365669 |
| 19.61111 | 0.12273 | 0.348741 | 0.174331 | -0.00743 | 0.023206 | 0.073256 | 0.044569 | 0.326701 | 0.422836 | 0.359971 | 0.105438 | 2.817074 | 2.245503 | 0.392711 | 0.514972 | 1.423078 | 0.017982 | 0.23576 |
| 19.625 | -0.05912 | 0.099279 | 0.113543 | -0.05438 | 0.080447 | -0.01324 | 0.01211 | 0.370945 | 0.969007 | 0.506321 | -0.05289 | 4.440379 | 3.029197 | -0.08442 | 0.014928 | 1.677964 | 0.340286 | 0.043147 |
| 19.63889 | -0.12673 | 0.289587 | -0.04161 | -0.03774 | 0.181361 | -0.06013 | 0.003543 | 0.264935 | 1.502984 | 0.821379 | -0.16891 | 2.292774 | 1.90304 | -0.20756 | -0.45732 | 1.061328 | 0.23687 | -0.09505 |
| 19.65278 | -0.17123 | 0.471828 | -0.19862 | -0.00152 | 0.14786 | -0.06268 | 0.017366 | 0.015489 | 1.770727 | -0.87127 | -0.17219 | -1.7107 | -0.90638 | -0.10094 | -0.86018 | 0.14368 | -0.08639 | -0.17478 |
| 19.66667 | -0.3851 | 0.358194 | -0.28779 | 0.061623 | -0.007 | -0.09501 | 0.024159 | -0.13734 | 1.02425 | -1.90904 | -0.09041 | -4.33712 | -2.59202 | -0.16682 | -1.10325 | -0.96032 | -0.64494 | -0.0788 |
| 19.68056 | -0.36544 | 0.416622 | -0.20097 | 0.07503 | -0.16946 | -0.03496 | 0.061085 | -0.11306 | 0.030391 | -1.1605 | -0.01493 | -3.89092 | -2.58547 | -0.28699 | -1.0495 | -1.93268 | -1.124 | 0.022349 |
| 19.69444 | 0.081483 | 0.874607 | -0.12606 | -0.02297 | -0.26916 | 0.060277 | 0.084397 | -0.0559 | -0.51936 | -0.43275 | -0.00854 | -1.76601 | -1.97147 | -0.15668 | -0.91336 | -1.81655 | -1.37019 | -0.15612 |
| 19.70833 | 0.387381 | 0.966902 | -0.01244 | -0.1098 | -0.25535 | 0.064967 | 0.029195 | -0.03408 | -0.79191 | -0.41129 | -0.06893 | 0.447315 | -1.20227 | -0.05168 | -0.82532 | -0.90342 | -1.02229 | -0.43532 |
| 19.72222 | 0.179558 | 0.502579 | 0.15155 | -0.05266 | -0.17985 | -0.01345 | -0.03361 | -0.05163 | -0.82039 | -0.65714 | -0.14314 | 1.911998 | -0.51339 | -0.29615 | -0.87663 | 0.261798 | -0.45108 | -0.63322 |
| 19.73611 | -0.14986 | 0.218416 | 0.161833 | -0.00808 | -0.07295 | -0.08327 | -0.0653 | -0.05534 | -0.47614 | -0.83466 | -0.1785 | 1.688979 | 0.02457 | -0.61602 | -0.95283 | 0.623929 | 0.092399 | -0.5246 |
| 19.75 | -0.22401 | 0.535703 | 0.015382 | -0.07444 | 0.083113 | -0.09996 | -0.07546 | -0.01663 | -0.0368 | -0.80653 | -0.10086 | 0.046108 | 0.3635 | -0.51398 | -0.69125 | 0.180429 | 0.356363 | -0.32633 |
| 19.76389 | -0.24778 | 0.823847 | -0.10802 | -0.02977 | 0.164665 | -0.09263 | -0.0487 | -0.07177 | 0.269786 | -0.49111 | 0.038048 | -1.07735 | 0.822484 | -0.09964 | -0.22075 | -0.42679 | 0.25233 | -0.26115 |
| 19.77778 | -0.17549 | 0.782953 | -0.13424 | 0.024127 | 0.142526 | -0.06539 | 0.01027 | -0.19337 | 0.451262 | -0.09839 | 0.139179 | -1.05041 | 1.496955 | 0.154411 | 0.072759 | -0.77999 | -0.13485 | -0.12195 |
| 19.79167 | 0.154055 | 0.906542 | -0.13399 | -0.13437 | 0.082485 | 0.00288 | 0.024951 | -0.1439 | 0.573758 | 0.026965 | 0.174962 | -0.33392 | 2.800013 | 0.220438 | 0.071231 | -0.71624 | -0.51474 | -0.00052 |
| 19.80556 | 0.19228 | 0.83609 | -0.16435 | -0.27591 | 0.015834 | 0.055082 | -0.03002 | -0.00399 | 0.004375 | -0.09183 | 0.156329 | -0.33583 | 1.824621 | 0.00926 | -0.21021 | -0.66563 | -0.6989 | 0.033789 |
| 19.81944 | -0.3346 | 0.113365 | -0.15573 | -0.21933 | -0.07589 | 0.051708 | -0.05453 | -0.11063 | -1.32142 | -0.33668 | 0.045511 | -1.25842 | -0.04125 | -0.31146 | -0.53712 | -0.85725 | -0.59127 | -0.09049 |
| 19.83333 | -0.70594 | -0.62803 | -0.05085 | -0.06118 | -0.24088 | -0.03127 | -0.03363 | -0.28051 | -2.06438 | -0.6283 | -0.15912 | -2.09272 | -2.32004 | -0.54715 | -0.78447 | -1.28024 | -0.56503 | -0.33515 |
| 19.84722 | -0.46717 | -0.84341 | 0.032707 | -0.19654 | -0.33861 | -0.10682 | -0.03982 | -0.25927 | -1.57049 | -0.74414 | -0.37351 | -2.82756 | -4.15117 | -0.89771 | -1.26198 | -1.73951 | -0.84485 | -0.75449 |
| 19.86111 | -0.19417 | -1.02895 | 0.020009 | -0.467 | -0.40086 | -0.20045 | -0.11155 | -0.26832 | -0.93817 | -0.86653 | -0.41873 | -2.05226 | -4.0626 | -1.10864 | -1.55199 | -1.72834 | -1.17886 | -1.09533 |
| 19.875 | -0.40779 | -1.42125 | -0.06602 | -0.46421 | -0.41501 | -0.2765 | -0.13496 | -0.45095 | -0.3951 | -0.9672 | -0.31068 | -1.12133 | -3.11812 | -0.6991 | -1.41042 | -1.34608 | -1.37369 | -1.06784 |
| 19.88889 | -0.60695 | -1.27504 | -0.12954 | -0.30804 | -0.37187 | -0.21299 | -0.06471 | -0.69656 | -0.22893 | -1.01738 | -0.14325 | -0.43554 | -1.94354 | -0.01638 | -1.0155 | -0.83846 | -1.14818 | -0.64071 |
| 19.90278 | -0.80675 | -0.8162 | -0.13451 | -0.22633 | -0.33814 | -0.1733 | -0.00667 | -0.82444 | -0.48337 | -0.88665 | 0.042772 | 0.578605 | -1.05337 | 0.207085 | -0.72034 | -0.41897 | -0.439 | -0.22118 |
| 19.91667 | -0.97785 | -0.54952 | -0.12587 | -0.27168 | -0.33259 | -0.27412 | -0.02231 | -0.68988 | -0.58728 | -0.90078 | 0.092737 | 1.358466 | 0.028444 | -0.11216 | -0.78716 | -0.38759 | 0.055942 | -0.10964 |
| 19.93056 | -0.86916 | -0.43276 | -0.1516 | -0.53043 | -0.37161 | -0.41496 | -0.15442 | -0.54648 | -0.57399 | -1.16635 | -0.02395 | 0.760779 | 0.596232 | -0.37847 | -0.9371 | -0.58974 | 0.100953 | -0.28687 |
| 19.94444 | -0.80336 | -0.46258 | -0.14707 | -0.64276 | -0.4 | -0.43869 | -0.23367 | -0.44059 | -0.57218 | -1.32097 | -0.04921 | -0.73715 | -0.08029 | -0.27425 | -0.75905 | -0.68073 | 0.085563 | -0.34135 |
| 19.95833 | -0.79846 | -0.6567 | -0.05622 | -0.41338 | -0.42077 | -0.33324 | -0.16903 | -0.39262 | -0.51083 | -1.19934 | 0.063759 | -2.37497 | -1.18635 | -0.24459 | -0.55064 | -0.49578 | 0.071372 | -0.20302 |
| 19.97222 | -0.98356 | -1.10183 | -0.01353 | -0.1298 | -0.50004 | -0.33556 | -0.0599 | -0.57458 | -0.50175 | -1.51156 | -0.04163 | -4.54386 | -2.39948 | -0.44161 | -1.00226 | -0.598 | -0.22793 | -0.20475 |
| 19.98611 | -1.12218 | -1.99771 | -0.03213 | -0.06676 | -0.55064 | -0.467 | 0.003029 | -0.91794 | -1.07327 | -2.73113 | -0.37552 | -6.31693 | -2.38637 | -0.55455 | -1.81522 | -0.7641 | -0.48667 | 0.09305 |
| 20 | -1.25541 | -2.89634 | -0.102 | -0.34283 | -0.63823 | -0.65478 | -0.13105 | -1.1063 | -2.06275 | -4.89705 | -0.41807 | -5.99657 | -2.40232 | -0.66249 | -1.84417 | -0.73249 | -0.00951 | 0.936182 |
| 20.01389 | -1.58766 | -2.32155 | -0.17028 | -0.7355 | -0.77363 | -0.58688 | -0.24568 | -0.8573 | -2.13174 | -5.25087 | -0.06274 | -3.73638 | -2.49272 | -0.24104 | -0.88189 | -0.14239 | 0.508039 | 0.96189 |
| 20.02778 | -0.44942 | -0.7294 | -0.13862 | -0.5616 | -0.41332 | -0.31009 | -0.14357 | -0.40776 | -1.44274 | -1.04539 | 0.114892 | -1.62035 | -1.45717 | -0.14271 | 0.201363 | 0.073195 | 0.654784 | 0.494703 |
| 20.04167 | 0.054019 | 0.275743 | -0.14596 | -0.21485 | -0.00972 | -0.08832 | -0.03629 | -0.27136 | -0.88214 | 0.69389 | 0.11506 | -1.38284 | -0.84182 | -0.28238 | 0.523632 | -0.13915 | 0.670873 | 0.165333 |
| 20.05556 | 0.044159 | 0.68005 | -0.18371 | -0.04308 | 0.07844 | 0.001866 | -0.01179 | -0.26108 | -0.48637 | 1.24185 | 0.076345 | -1.07298 | -0.48876 | -0.23788 | 0.280428 | -0.13364 | 0.578076 | 0.06366 |
| 20.06944 | 0.088681 | 0.605406 | -0.01668 | 0.123052 | 0.149842 | 0.152531 | 0.125492 | -0.0115 | 0.013455 | 0.993047 | 0.09346 | -0.34181 | 0.075181 | 0.055527 | 0.203993 | 0.134319 | 0.52027 | 0.229063 |
| 20.08333 | 0.035856 | 0.12669 | 0.127031 | 0.167823 | 0.195984 | 0.229544 | 0.214008 | 0.170218 | 0.162824 | 0.008608 | 0.077463 | 0.001787 | 0.413253 | 0.216441 | 0.237826 | 0.158013 | 0.396682 | 0.362424 |
| 20.09722 | -0.12831 | -0.22424 | 0.05367 | 0.072426 | 0.097125 | 0.11316 | 0.099328 | 0.065711 | -0.00064 | -0.4988 | 0.11108 | 0.454255 | 0.334887 | 0.246146 | 0.333053 | 0.057614 | 0.233667 | 0.191824 |
| 20.11111 | -0.05152 | -0.05788 | -0.00444 | 0.025312 | 0.036625 | 0.042826 | 0.021005 | 0.013248 | 0.003084 | -0.00459 | 0.28069 | 0.909326 | 0.321499 | 0.331602 | 0.475729 | 0.266051 | 0.355957 | 0.036105 |
| 20.125 | 0.194624 | 0.332912 | 0.053114 | 0.040407 | 0.085341 | 0.056111 | 0.035721 | 0.117678 | 0.159383 | 0.198033 | 0.37718 | 0.347193 | 0.536984 | 0.352825 | 0.31912 | 0.509977 | 0.599899 | 0.187088 |
| 20.13889 | 0.122847 | 0.376945 | 0.102953 | 0.044273 | 0.077906 | 0.034483 | 0.047501 | 0.1188 | 0.124132 | -0.22123 | 0.417034 | -0.0443 | 0.709254 | 0.329546 | 0.197745 | 0.524064 | 0.679272 | 0.380603 |
| 20.15278 | -0.10132 | 0.145965 | 0.059066 | 0.023125 | 0.02248 | -0.01183 | 0.025729 | 0.012521 | -0.00676 | -0.36388 | 0.474745 | 0.494347 | 0.703529 | 0.362225 | 0.43824 | 0.491623 | 0.700081 | 0.423516 |
| 20.16667 | -0.11929 | 0.094618 | -0.01361 | 0.018215 | 0.024131 | -0.0286 | -0.04144 | 0.033951 | 0.037905 | -0.22951 | 0.422423 | 0.762716 | 0.521279 | 0.317491 | 0.463305 | 0.456486 | 0.550972 | 0.333257 |
| 20.18056 | -0.05155 | 0.084161 | -0.02752 | 0.035964 | 0.014808 | -0.01936 | -0.04861 | 0.029103 | 0.081247 | 0.097827 | 0.32221 | 0.792867 | 0.264984 | 0.19513 | 0.311435 | 0.333801 | 0.187644 | 0.197407 |
| 20.19444 | -0.15273 | -0.11707 | 0.005943 | 0.031959 | -0.03213 | 0.00674 | 0.050663 | -0.11574 | -0.06924 | 0.136413 | 0.211187 | 0.401064 | 0.108279 | 0.035707 | 0.09786 | 0.085699 | -0.15804 | 0.083207 |
| 20.20833 | -0.1315 | -0.09224 | 0.034556 | 0.002104 | -0.00636 | 0.030223 | 0.088069 | -0.11396 | -0.06396 | -0.09803 | 0.130421 | -0.03446 | 0.154616 | -0.10749 | -0.09553 | -0.0288 | -0.15255 | -0.02126 |

| | | | | | | | | | | | | | | | | | | |
|---|---|---|---|---|---|---|---|---|---|---|---|---|---|---|---|---|---|---|
| 20.22222 | -0.01658 | 0.233624 | 0.041871 | -0.01242 | 0.055894 | 0.046166 | 0.054661 | 0.035666 | 0.162217 | 0.031422 | 0.200868 | 0.42495 | 0.348926 | -0.05583 | 0.077937 | 0.171136 | 0.207359 | -0.14045 |
| 20.23611 | 0.067441 | 0.336606 | 0.05363 | 0.019855 | 0.081761 | 0.056134 | 0.034884 | 0.182333 | 0.335671 | 0.028489 | 0.148922 | 0.503681 | 0.565814 | 0.000731 | 0.013942 | 0.184724 | 0.33499 | -0.21379 |
| 20.25 | 0.107717 | 0.295392 | 0.081595 | 0.075783 | 0.09663 | 0.077061 | 0.049952 | 0.277152 | 0.422373 | -0.19672 | -0.08323 | -0.19146 | 0.647121 | -0.10035 | -0.41867 | -0.0709 | 0.107719 | -0.20101 |
| 20.26389 | 0.089343 | 0.179702 | 0.089926 | 0.113401 | 0.076648 | 0.105675 | 0.103784 | 0.239042 | 0.3789 | -0.09763 | -0.13912 | -0.28573 | 0.526241 | -0.15054 | -0.50111 | -0.1091 | -0.00445 | -0.19463 |
| 20.27778 | 0.008085 | 0.10196 | -0.01097 | 0.033227 | -0.04025 | 0.009946 | 0.033671 | 0.033046 | 0.319508 | 0.314423 | -0.01399 | 0.167388 | 0.457231 | -0.06574 | -0.24687 | 0.092502 | 0.035453 | -0.23123 |
| 20.29167 | -0.12029 | 0.095423 | -0.14197 | -0.08125 | -0.14628 | -0.1068 | -0.07886 | -0.15829 | 0.330926 | 0.804661 | 0.132271 | 1.004372 | 0.503976 | 0.080039 | 0.133514 | 0.286817 | 0.054463 | -0.27231 |
| 20.30556 | -0.12686 | 0.203606 | -0.04315 | 0.027581 | -0.02528 | 0.042231 | 0.073724 | -0.04192 | 0.501895 | 1.004911 | 0.177584 | 1.505651 | 0.739585 | 0.183155 | 0.291266 | 0.377222 | 0.079908 | -0.15335 |
| 20.31944 | -0.02948 | 0.480624 | 0.176784 | 0.216614 | 0.179763 | 0.276033 | 0.246409 | 0.246492 | 0.776731 | 0.563033 | 0.090947 | 0.8786 | 0.915918 | 0.218855 | 0.042277 | 0.2853 | 0.211117 | 0.19878 |
| 20.33333 | 0.113495 | 0.643903 | 0.219842 | 0.240681 | 0.207712 | 0.271853 | 0.177281 | 0.346173 | 0.872795 | 0.2159 | 0.088213 | 0.311249 | 0.952521 | 0.294179 | -0.07923 | 0.303907 | 0.394416 | 0.462565 |
| 20.34722 | 0.211475 | 0.60878 | 0.198821 | 0.180439 | 0.194674 | 0.220578 | 0.077539 | 0.294563 | 0.778811 | 0.309981 | 0.174878 | 0.486362 | 0.728858 | 0.328564 | 0.09069 | 0.334394 | 0.409164 | 0.418883 |
| 20.36111 | 0.310062 | 0.641212 | 0.161056 | 0.228421 | 0.235544 | 0.262567 | 0.087682 | 0.304979 | 0.823575 | 0.443379 | 0.206354 | 0.733212 | 0.526397 | 0.2385 | 0.169166 | 0.322815 | 0.341249 | 0.298777 |
| 20.375 | 0.430421 | 0.891538 | 0.172217 | 0.317893 | 0.258308 | 0.309417 | 0.122162 | 0.46353 | 1.109059 | 0.686759 | 0.267554 | 0.913921 | 0.552203 | 0.194958 | 0.179786 | 0.648522 | 0.654884 | 0.370451 |
| 20.38889 | 0.450646 | 1.027046 | 0.190507 | 0.373718 | 0.263986 | 0.319747 | 0.115042 | 0.636111 | 1.288958 | 0.876729 | 0.355696 | 0.635895 | 0.66671 | 0.231834 | 0.159697 | 1.101968 | 1.152996 | 0.551524 |
| 20.40278 | 0.307407 | 0.758852 | 0.19895 | 0.378551 | 0.284695 | 0.299512 | 0.114472 | 0.549804 | 1.101202 | 0.868042 | 0.359087 | 0.355272 | 0.506312 | 0.320662 | 0.199108 | 1.181824 | 1.114662 | 0.614387 |
| 20.41667 | 0.326473 | 0.442792 | 0.215648 | 0.435959 | 0.303725 | 0.276747 | 0.13548 | 0.384205 | 0.800519 | 0.989063 | 0.350737 | 0.704706 | 0.332273 | 0.369455 | 0.317516 | 1.164631 | 0.866759 | 0.449551 |
| 20.43056 | 0.506853 | 0.435184 | 0.220617 | 0.449867 | 0.3307 | 0.280717 | 0.161331 | 0.277322 | 0.709874 | 1.014221 | 0.27739 | 0.763212 | 0.527123 | 0.336593 | 0.174996 | 1.232271 | 0.596979 | 0.338393 |
| 20.44444 | 0.558145 | 0.575293 | 0.215998 | 0.42519 | 0.370969 | 0.297333 | 0.187078 | 0.359677 | 0.614679 | 0.746915 | 0.180465 | 0.163469 | 0.818389 | 0.249598 | -0.05427 | 1.058754 | 0.401776 | 0.385885 |
| 20.45833 | 0.531339 | 0.764461 | 0.248819 | 0.440001 | 0.402975 | 0.332659 | 0.225312 | 0.474664 | 0.43816 | 0.773657 | 0.325897 | 0.321493 | 0.833758 | 0.291027 | 0.225137 | 1.01121 | 0.454131 | 0.509468 |
| 20.47222 | 0.71035 | 1.062544 | 0.290713 | 0.385819 | 0.334842 | 0.361656 | 0.201605 | 0.444651 | 0.518737 | 1.563686 | 0.829477 | 2.036511 | 0.613391 | 0.612111 | 1.275794 | 1.689535 | 0.945807 | 0.645482 |
| 20.48611 | 1.519743 | 1.607579 | 0.266965 | 0.196728 | 0.319659 | 0.42922 | 0.177143 | 0.456537 | 1.234237 | 3.061084 | 1.592923 | 5.154085 | 1.149906 | 0.956963 | 2.162167 | 2.824661 | 1.736234 | 0.878207 |
| 20.5 | 3.143524 | 2.714111 | 0.275987 | 0.563488 | 0.880447 | 0.70062 | 0.618938 | 0.713599 | 2.635812 | 5.437149 | 1.997298 | 7.703617 | 3.315354 | 1.840424 | 3.161962 | 3.967415 | 2.225192 | 1.111265 |
| 20.51389 | 3.447853 | 2.943577 | 0.298279 | 1.222406 | 1.475221 | 1.174711 | 1.23981 | 0.895932 | 4.4164 | 5.718787 | 1.757869 | 2.295605 | 3.542887 | 2.159383 | 3.387333 | 2.94205 | 1.828843 | 0.956717 |
| 20.52778 | 1.612542 | 1.680606 | 0.202783 | 0.887923 | 0.907893 | 0.815575 | 0.849387 | 0.791502 | 2.912127 | 2.569663 | 0.86635 | -7.37849 | 2.022128 | 1.362465 | 2.5342 | 1.360555 | 1.358372 | 0.492114 |
| 20.54167 | 0.448295 | 0.558905 | -0.01546 | 0.532154 | 0.619371 | 0.512648 | 0.543411 | 0.286131 | 0.711897 | 0.939554 | 0.359169 | -5.55883 | 0.912819 | 0.195175 | 1.370216 | 0.303464 | 0.905256 | 0.216547 |
| 20.55556 | -0.3099 | -0.22452 | -0.3452 | 0.313391 | 0.323634 | 0.289122 | 0.419414 | -0.24469 | -1.24214 | -0.18071 | 0.374815 | -2.71891 | -0.08538 | -0.58781 | 0.428072 | -1.26419 | 0.197308 | 0.073836 |
| 20.56944 | -0.68175 | -0.56133 | -0.53016 | 0.134649 | 0.057582 | 0.117027 | 0.287338 | -0.56378 | -2.36298 | -0.71315 | 0.572752 | -2.83705 | -0.5187 | -0.3206 | 0.512476 | -2.43364 | -0.19025 | 0.158976 |
| 20.58333 | -0.77538 | -0.34482 | -0.37749 | 0.003696 | -0.01809 | 0.126418 | 0.206294 | -0.3491 | -2.65407 | -1.10737 | 0.784864 | -2.70289 | -0.21946 | 0.76365 | 1.172387 | -2.32494 | -0.03303 | 0.553959 |
| 20.59722 | -0.55209 | 0.346517 | -0.0518 | -0.08622 | -0.04251 | 0.24059 | 0.157336 | 0.211511 | -1.72912 | -1.34314 | 0.818253 | -0.83641 | 0.402837 | 1.655248 | 1.187338 | -1.09023 | 0.294451 | 0.717543 |
| 20.61111 | 0.258051 | 1.206937 | 0.128151 | -0.25058 | -0.08782 | 0.247138 | 0.023412 | 0.677972 | -0.37347 | -0.74249 | 0.653104 | 1.15154 | 0.27385 | 1.892271 | 0.589343 | 0.417038 | 0.570138 | 0.840491 |
| 20.625 | 1.336084 | 1.720143 | 0.263134 | -0.51304 | -0.0808 | 0.044615 | -0.18503 | 0.861795 | 0.897225 | -0.12649 | 0.310599 | 2.304197 | 0.932774 | 1.190265 | -0.26128 | 2.229117 | 0.855726 | 0.979747 |
| 20.63889 | 1.423555 | 1.243112 | 0.266985 | -0.4985 | -0.01088 | -0.18489 | -0.28561 | 0.724937 | 1.884322 | 0.728968 | 0.076125 | 3.343262 | 1.277385 | 0.120061 | -0.71586 | 3.126977 | 1.062435 | 0.666663 |
| 20.65278 | 0.389328 | -0.00087 | 0.11794 | -0.31591 | 0.141173 | -0.22945 | -0.15136 | 0.317918 | 1.54329 | 0.810891 | 0.040192 | 4.793804 | 1.245364 | -0.35759 | -0.65919 | 2.512424 | 0.74149 | 0.129747 |
| 20.66667 | -0.40591 | -0.80756 | -0.13296 | -0.20183 | 0.225944 | -0.13343 | 0.045095 | -0.20368 | 0.482138 | 0.863938 | 0.101952 | 1.347413 | -1.04747 | -0.07914 | -0.29955 | 0.778652 | -0.39139 | -0.20385 |
| 20.68056 | -0.5019 | -0.75797 | -0.25016 | -0.06947 | 0.186293 | -0.0274 | 0.141022 | -0.52864 | -0.46635 | -1.07358 | 0.136952 | -3.00385 | -2.16193 | 0.417306 | 0.044913 | -0.88443 | -1.22336 | -0.03914 |
| 20.69444 | -0.28311 | -0.13476 | -0.1759 | 0.031605 | 0.056452 | 0.099927 | 0.136445 | -0.4031 | -1.33409 | -2.15432 | 0.067279 | -5.66906 | -2.66204 | 0.62869 | 0.133404 | -2.01874 | -1.40637 | -0.02571 |
| 20.70833 | -0.23711 | 0.550484 | 0.002009 | 0.076467 | -0.1099 | 0.236843 | 0.159933 | 0.163098 | -2.21221 | -1.44445 | -0.09269 | -5.03981 | -1.67328 | 0.315811 | 0.104604 | -2.68137 | -1.15824 | -0.0781 |
| 20.72222 | -0.13838 | 0.912429 | 0.129002 | 0.052484 | -0.21623 | 0.275453 | 0.215686 | 0.487884 | -2.03598 | -0.95947 | -0.25615 | -3.46789 | -1.49297 | -0.20678 | 0.015301 | -3.02171 | -1.07686 | -0.11767 |
| 20.73611 | -0.04567 | 0.891646 | 0.003483 | -0.19257 | -0.23757 | 0.101015 | 0.105885 | 0.180775 | -0.99365 | -0.0191 | -0.25011 | -0.93197 | -1.15756 | -0.50583 | -0.18336 | -2.28663 | -1.03854 | -0.49817 |
| 20.75 | -0.02221 | 0.533528 | -0.21161 | -0.50684 | -0.21101 | -0.10439 | -0.06837 | -0.10731 | -0.10537 | 0.255536 | -0.10321 | 2.340508 | -0.38138 | -0.63902 | -0.37259 | -1.32836 | -1.05815 | -0.84586 |
| 20.76389 | -0.05842 | -0.0456 | -0.26614 | -0.57167 | -0.12489 | -0.14654 | -0.11013 | -0.39532 | 0.377996 | 0.366267 | 0.106629 | 4.510866 | 0.873805 | -0.75117 | -0.60765 | -0.45028 | -0.76828 | -0.83705 |
| 20.77778 | -0.02874 | -0.62229 | -0.0037 | -0.32761 | -0.03113 | -0.03011 | 0.012488 | -0.39892 | 0.385569 | 1.09393 | 0.565712 | 4.269263 | 2.016965 | -0.70638 | -0.65087 | 0.563275 | -0.1097 | -0.41677 |
| 20.79167 | 0.015434 | -0.76145 | 0.191 | -0.08095 | -0.01347 | 0.115779 | 0.182923 | -0.11693 | 0.206996 | 1.515021 | 1.53634 | 2.489144 | 1.281379 | -0.6298 | -0.4046 | 1.564383 | 0.573399 | 0.047676 |
| 20.80556 | 0.062489 | -0.39507 | 0.241469 | -0.00816 | -0.04059 | 0.174328 | 0.20188 | 0.154782 | -0.0049 | 0.301199 | 2.045692 | 0.268144 | 0.111497 | -0.67497 | -0.38809 | 1.96436 | 0.700381 | 0.312368 |
| 20.81944 | 0.182632 | 0.268843 | 0.028083 | -0.25105 | -0.11315 | 0.144764 | 0.050188 | 0.205799 | -0.29637 | -1.38543 | 0.598635 | -2.21534 | -0.56975 | -0.72589 | -0.76817 | 1.288274 | 0.091069 | 0.437211 |
| 20.83333 | -0.09387 | 0.582521 | -0.28936 | -0.46244 | -0.25218 | 0.045844 | -0.08496 | -0.14558 | -0.93538 | -2.50469 | -0.92406 | -3.89486 | -1.50095 | -0.65427 | -1.19292 | -0.11667 | -0.76084 | 0.036947 |
| 20.84722 | -0.84587 | -0.09766 | -0.49146 | -0.50526 | -0.36046 | -0.13268 | -0.10296 | -0.65636 | -1.71069 | -2.39951 | -2.38819 | -4.45954 | -2.8174 | -0.63607 | -1.57022 | -1.60097 | -1.19319 | -0.53093 |
| 20.86111 | -1.39067 | -1.08458 | -0.47662 | -0.54384 | -0.3883 | -0.28795 | -0.15463 | -0.94283 | -1.93321 | -2.12504 | -3.1446 | -4.21438 | -3.98592 | -0.83865 | -1.6826 | -2.26655 | -0.98824 | -1.02246 |
| 20.875 | -1.57714 | -1.93299 | -0.39358 | -0.55308 | -0.42855 | -0.36746 | -0.18027 | -1.03076 | -1.61759 | -1.90755 | -2.76474 | -2.87297 | -3.98627 | -1.22609 | -1.39083 | -1.83347 | -0.68615 | -0.91821 |
| 20.88889 | -1.26729 | -2.26237 | -0.23319 | -0.48984 | -0.4738 | -0.30846 | -0.11825 | -0.86889 | -1.07748 | -2.00969 | -2.1265 | -1.78077 | -2.84371 | -1.62217 | -1.2347 | -1.2636 | -0.76599 | -0.67143 |
| 20.90278 | -0.76242 | -1.82258 | 0.023128 | -0.38043 | -0.45967 | -0.14197 | -0.02726 | -0.61593 | -0.77782 | -2.0601 | -1.20986 | -1.22922 | -1.41717 | -1.50403 | -1.34748 | -0.73289 | -1.14592 | -0.41759 |
| 20.91667 | -0.36044 | -0.95765 | 0.143502 | -0.30207 | -0.46793 | 0.013806 | 0.050488 | -0.42559 | -0.65577 | -1.44124 | -0.39608 | -0.6048 | 0.127671 | -0.80279 | -1.07359 | -0.4675 | -0.9937 | -0.17947 |
| 20.93056 | -0.39492 | -0.29817 | -0.0349 | -0.22994 | -0.43608 | -0.02492 | 0.077771 | -0.365 | -0.54568 | -0.49433 | 0.292024 | 0.167522 | 1.026456 | -0.16675 | -0.59254 | -0.16615 | -0.42058 | -0.01141 |
| 20.94444 | -0.95859 | -0.25331 | -0.28431 | -0.37928 | -0.4338 | -0.2371 | 0.023242 | -0.55923 | -0.63004 | -0.1365 | 0.582525 | 0.578618 | 0.881182 | -0.13715 | -0.59638 | -0.03596 | 0.321738 | -0.0371 |
| 20.95833 | -1.46179 | -0.56436 | -0.38091 | -0.61583 | -0.45918 | -0.30917 | 0.048448 | -0.73373 | -0.78019 | -0.56707 | 0.300541 | 0.615832 | 0.252788 | -0.47737 | -0.81586 | 0.021202 | 0.823731 | -0.18981 |
| 20.97222 | -1.58263 | -0.83653 | -0.40462 | -0.58385 | -0.36595 | -0.24943 | 0.20621 | -0.75597 | -0.94858 | -1.34145 | -0.25149 | -0.31407 | -0.36806 | -0.92844 | -1.4117 | -0.09304 | 0.669085 | -0.30166 |
| 20.98611 | -1.58653 | -1.49305 | -0.4671 | -0.50529 | -0.387 | -0.36686 | 0.31284 | -0.95402 | -1.46609 | -2.21327 | -1.24358 | -2.54143 | -1.30788 | -1.71153 | -2.48055 | -0.95949 | -0.43701 | -0.54151 |
| 21 | -1.78725 | -2.37836 | -0.60516 | -0.61248 | -0.68173 | -0.87904 | -0.20414 | -1.57113 | -2.4876 | -3.18369 | -2.36295 | -3.6806 | -2.50874 | -2.12653 | -2.86053 | -2.0264 | -1.59276 | -0.74385 |
| 21.01389 | -1.9176 | -2.85652 | -0.73315 | -0.69423 | -0.83548 | -0.86719 | -0.56523 | -2.01542 | -2.95454 | -3.08791 | -2.15395 | -3.04091 | -3.06107 | -1.69282 | -1.81652 | -2.32066 | -1.94571 | -0.98932 |
| 21.02778 | -1.10023 | -2.18907 | -0.59983 | -0.47631 | -0.55291 | -0.50078 | -0.4314 | -1.69696 | -1.79635 | -1.95387 | -1.19945 | -2.06402 | -2.84214 | -1.01296 | -0.83604 | -1.5248 | -1.41176 | -1.18148 |
| 21.04167 | -0.65654 | -1.31786 | -0.22857 | -0.29723 | -0.27451 | -0.08543 | -0.16541 | -0.91645 | -0.47709 | -1.06718 | -0.6687 | -1.65385 | -2.83417 | -0.72052 | -0.53478 | -0.35302 | -0.31297 | -0.76364 |
| 21.05556 | -0.60524 | -0.68729 | 0.097708 | -0.18932 | -0.19197 | 0.12678 | -0.00858 | -0.12907 | 0.261795 | -0.37742 | -0.48123 | -1.58549 | -2.06508 | -0.52367 | -0.34896 | 0.449583 | 0.750996 | 0.011736 |
| 21.06944 | -0.42023 | -0.32259 | 0.315064 | -0.05906 | -0.17857 | 0.112132 | 0.075165 | 0.556148 | 0.521878 | 0.034703 | -0.47447 | -1.18954 | -0.81166 | -0.3668 | -0.12704 | 0.495665 | 1.124025 | 0.545892 |
| 21.08333 | -0.09196 | -0.2597 | 0.292445 | 0.060522 | -0.10199 | 0.075918 | 0.207353 | 0.662437 | 0.565011 | 0.164077 | -0.46305 | -0.82549 | 0.040986 | -0.33645 | -0.05263 | 0.005796 | 0.680238 | 0.55304 |
| 21.09722 | 0.15934 | -0.19278 | 0.186133 | 0.080254 | -0.04879 | 0.142239 | 0.357266 | 0.297555 | 0.154216 | 0.200286 | -0.34917 | -0.25675 | 0.317395 | -0.24441 | 0.027995 | -0.44184 | 0.118778 | 0.286778 |
| 21.11111 | 0.258176 | 0.0054 | 0.122388 | 0.044194 | -0.06646 | 0.205561 | 0.372277 | -0.06073 | -0.1911 | 0.305249 | -0.16722 | 0.601239 | 0.34151 | -0.03063 | 0.210637 | -0.56529 | -0.10459 | 0.063768 |
| 21.125 | 0.148936 | 0.055623 | -0.01307 | 0.003118 | -0.07964 | 0.203677 | 0.291195 | -0.29734 | -0.42759 | 0.382935 | -0.0488 | 0.741376 | 0.357893 | 0.084596 | 0.234952 | -0.48905 | -0.18646 | -0.04677 |
| 21.13889 | -0.00265 | 0.059054 | -0.05719 | -0.04312 | -0.02829 | 0.214322 | 0.222698 | -0.31911 | -0.48732 | 0.31488 | 0.0103 | -0.04349 | 0.363064 | 0.02327 | 0.020424 | -0.37179 | -0.23078 | -0.07305 |
| 21.15278 | -0.14174 | 0.055175 | 0.024661 | -0.03452 | 0.037446 | 0.229003 | 0.231529 | -0.12261 | -0.26256 | 0.028691 | 0.074392 | -0.94798 | 0.420429 | -0.11539 | -0.25894 | -0.26585 | -0.1369 | -0.01868 |
| 21.16667 | -0.2615 | 0.092744 | 0.111195 | -0.01132 | 0.057873 | 0.200607 | 0.22258 | 0.152251 | 0.092728 | -0.14162 | 0.180119 | -0.74932 | 0.402379 | -0.10471 | -0.22074 | -0.05067 | 0.197269 | 0.006467 |
| 21.18056 | -0.29331 | 0.234363 | 0.11985 | -0.05664 | 0.061882 | 0.161827 | 0.153351 | 0.342114 | 0.344669 | -0.16703 | 0.305683 | 0.226647 | 0.202505 | 0.048789 | 0.156306 | 0.142263 | 0.436943 | -0.09259 |
| 21.19444 | -0.29055 | 0.201985 | 0.063397 | -0.0635 | 0.082735 | 0.127539 | 0.132158 | 0.355584 | 0.254311 | -0.28688 | 0.209457 | 0.682374 | -0.08333 | -0.00535 | 0.301225 | 0.052625 | 0.276351 | -0.22075 |
| 21.20833 | -0.26575 | -0.01112 | 0.016333 | -0.00019 | 0.088087 | 0.101393 | 0.180243 | 0.262646 | -0.01666 | -0.37223 | 0.008193 | 0.204533 | -0.1465 | -0.18967 | 0.141507 | -0.08327 | -0.01173 | -0.23359 |
| 21.22222 | -0.27935 | -0.10493 | -0.00374 | -0.00986 | 0.070399 | 0.083407 | 0.132078 | 0.13339 | -0.10445 | -0.30135 | -0.01559 | -0.13961 | -0.09877 | -0.20009 | 0.056514 | -0.05446 | -0.10202 | -0.2118 |
| 21.23611 | -0.34211 | 0.061391 | -0.00033 | -0.08082 | 0.072996 | 0.119647 | 0.022348 | 0.02992 | 0.0571 | -0.41636 | -0.09334 | -0.303 | -0.14296 | -0.28734 | -0.05743 | -0.25022 | -0.32046 | -0.45689 |
| 21.25 | -0.24194 | 0.351618 | 0.096434 | -0.06599 | 0.117366 | 0.221043 | 0.009382 | 0.02788 | 0.137903 | -0.65355 | -0.30323 | -0.24624 | -0.26016 | -0.50234 | -0.25483 | -0.64636 | -0.65894 | -0.84195 |
| 21.26389 | -0.10556 | 0.221885 | 0.125282 | 0.071306 | 0.146664 | 0.244108 | 0.142992 | 0.075415 | -0.07916 | -0.7188 | -0.41867 | -0.27755 | -0.28429 | -0.63811 | -0.51175 | -0.75675 | -0.63345 | -0.95434 |
| 21.27778 | -0.34997 | -0.3266 | -0.02792 | 0.196908 | 0.13678 | 0.164472 | 0.300858 | 0.057198 | -0.26394 | -0.75801 | -0.57284 | -1.09064 | -0.59326 | -0.93976 | -1.12922 | -0.73998 | -0.4476 | -1.05699 |
| 21.29167 | -0.28222 | -0.23379 | -0.0411 | 0.196448 | 0.179406 | 0.216642 | 0.366486 | 0.135819 | -0.22254 | -1.09687 | -1.0081 | -1.80664 | -1.25643 | -1.53019 | -1.82777 | -1.11329 | -0.77838 | -1.56082 |
| 21.30556 | 0.200308 | 0.503466 | 0.063111 | 0.190106 | 0.254677 | 0.433283 | 0.430351 | 0.314545 | 0.148016 | -1.45534 | -1.44045 | -1.80432 | -1.61396 | -1.88616 | -2.05509 | -1.63409 | -1.4218 | -2.03871 |
| 21.31944 | 0.491545 | 0.645216 | 0.191672 | 0.200927 | 0.282639 | 0.586908 | 0.50667 | 0.388865 | 0.292013 | -1.43926 | -1.52134 | -1.48036 | -1.33264 | -1.72008 | -1.90968 | -1.66027 | -1.54279 | -2.03431 |
| 21.33333 | 0.487557 | 0.349043 | 0.205378 | 0.129888 | 0.220106 | 0.555506 | 0.50505 | 0.254699 | 0.130008 | -1.10861 | -1.13691 | -1.1631 | -0.86612 | -1.13689 | -1.5066 | -1.1247 | -1.01527 | -1.56924 |

| | | | | | | | | | | | | | | | | | | |
|---|---|---|---|---|---|---|---|---|---|---|---|---|---|---|---|---|---|---|
| 21.34722 | 0.44729 | 0.306074 | 0.105253 | 0.066239 | 0.160877 | 0.484059 | 0.412926 | 0.20593 | 0.168991 | -0.48201 | -0.52558 | -0.44693 | -0.56961 | -0.48257 | -0.8276 | -0.42564 | -0.39208 | -0.90376 |
| 21.36111 | 0.576444 | 0.743848 | 0.075495 | 0.154313 | 0.193146 | 0.519966 | 0.401878 | 0.557853 | 0.573858 | 0.258954 | -0.09082 | 0.399568 | -0.21323 | -0.05691 | -0.14727 | 0.014885 | -0.10329 | -0.19134 |
| 21.375 | 0.728506 | 1.294728 | 0.187873 | 0.405529 | 0.308811 | 0.618504 | 0.503069 | 1.086749 | 0.9421 | 0.921927 | 0.173154 | 0.818497 | 0.2709 | 0.114033 | 0.385495 | 0.21422 | 0.075831 | 0.52071 |
| 21.38889 | 0.936768 | 1.663793 | 0.377406 | 0.557731 | 0.364934 | 0.646885 | 0.530675 | 1.183563 | 0.900593 | 1.263649 | 0.371913 | 0.890461 | 0.638467 | 0.201329 | 0.855008 | 0.444194 | 0.333689 | 0.967865 |
| 21.40278 | 1.088686 | 1.578255 | 0.419045 | 0.411668 | 0.297979 | 0.643389 | 0.508815 | 0.795406 | 0.390345 | 1.301776 | 0.598931 | 0.954683 | 0.480927 | 0.283763 | 1.259134 | 0.848042 | 0.723825 | 1.02682 |
| 21.41667 | 1.006917 | 1.043465 | 0.243214 | 0.243014 | 0.240168 | 0.594629 | 0.427465 | 0.170571 | -0.38745 | 1.232842 | 0.685413 | 0.592868 | -0.00554 | 0.268156 | 1.248518 | 0.973164 | 0.775743 | 0.821722 |
| 21.43056 | 1.032307 | 0.713824 | 0.065307 | 0.181412 | 0.265972 | 0.498943 | 0.332749 | -0.19111 | -0.75255 | 0.845274 | 0.504701 | -0.33387 | -0.33524 | 0.156034 | 0.650741 | 0.22845 | 0.019348 | 0.450601 |
| 21.44444 | 1.239777 | 0.747235 | 0.039703 | 0.239271 | 0.363443 | 0.477563 | 0.368059 | 0.100354 | -0.13031 | 0.561796 | 0.359199 | -0.62294 | 0.044114 | 0.018339 | 0.055703 | -0.33814 | -0.61498 | -0.00834 |
| 21.45833 | 1.521175 | 1.181569 | 0.218955 | 0.352555 | 0.463798 | 0.5548 | 0.524216 | 0.830209 | 0.969048 | 0.801976 | 0.550616 | 0.11121 | 0.699544 | -0.01505 | 0.084502 | -0.07432 | -0.29365 | -0.3161 |
| 21.47222 | 1.841102 | 1.805211 | 0.492152 | 0.394403 | 0.479746 | 0.618866 | 0.629969 | 1.521323 | 2.037957 | 1.463428 | 0.875842 | 1.098663 | 1.312493 | 0.228891 | 0.869611 | 0.800414 | 0.724762 | -0.22023 |
| 21.48611 | 1.981071 | 2.23256 | 0.801458 | 0.426702 | 0.541193 | 0.638534 | 0.663365 | 1.255285 | 2.726939 | 2.590551 | 1.457195 | 3.000049 | 2.301357 | 0.854464 | 2.444218 | 1.699674 | 1.657333 | 0.329702 |
| 21.5 | 3.168336 | 2.910693 | 0.727105 | 1.133281 | 1.135897 | 0.649528 | 0.736677 | 0.888693 | 4.38854 | 4.27331 | 2.799996 | 5.958674 | 4.882032 | 1.771487 | 4.222365 | 3.039204 | 1.616881 | 0.850041 |
| 21.51389 | 3.560169 | 3.317632 | 0.400155 | 2.235372 | 1.786067 | 0.674614 | 0.951093 | 0.711903 | 4.677891 | 3.948819 | 3.491019 | 7.515666 | 6.812062 | 2.053968 | 3.841435 | 3.668624 | 1.152209 | 0.7703 |
| 21.52778 | 2.237269 | 2.363909 | 0.20838 | 1.648789 | 1.08435 | 0.133703 | 0.280109 | 0.622129 | 3.178859 | 1.696744 | 2.233571 | 5.823851 | 4.726665 | 1.279001 | 1.161007 | 2.512559 | 0.150342 | 0.326961 |
| 21.54167 | 0.894375 | 0.885848 | 0.003497 | 0.742964 | 0.723069 | -0.35438 | -0.35797 | 0.336731 | 1.574681 | -0.17338 | 0.834189 | 2.795803 | 1.854879 | 0.162756 | -1.12864 | 0.907079 | -0.733 | -0.52136 |
| 21.55556 | -0.71646 | -0.84812 | -0.31868 | 0.084516 | 0.433289 | -0.5982 | -0.72983 | -0.19095 | 0.085855 | -1.41028 | 0.358634 | 0.362661 | -1.03814 | -0.71463 | -3.0974 | -0.43166 | -0.52355 | -1.1383 |
| 21.56944 | -1.76817 | -2.14676 | -0.66929 | -0.25182 | 0.281362 | -0.5961 | -0.69302 | -0.50753 | -1.43126 | -1.62279 | 0.009901 | -0.57018 | -2.86556 | -0.40674 | -2.74426 | -0.85356 | 0.378492 | -0.7739 |
| 21.58333 | -1.97048 | -2.25562 | -0.54549 | -0.25549 | 0.141757 | -0.37687 | -0.33154 | -0.15613 | -2.45767 | -0.73202 | -0.58079 | -0.45924 | -3.3165 | -0.22135 | -1.57921 | -1.21397 | 1.195247 | 0.095624 |
| 21.59722 | -1.14012 | -1.22481 | -0.06692 | -0.31607 | -0.05387 | -0.02429 | 0.102958 | 0.450027 | -2.64274 | 0.040949 | -1.35209 | 0.05455 | -2.7139 | 0.458405 | -0.81718 | -1.40791 | 1.551066 | 0.800721 |
| 21.61111 | 0.678407 | 0.336238 | 0.61248 | -0.39843 | -0.22387 | 0.264757 | 0.349527 | 1.199007 | -1.92221 | 0.000623 | -1.66295 | 0.745022 | -1.58042 | 1.468577 | -0.4172 | -1.38169 | 1.121482 | 1.268326 |
| 21.625 | 1.620076 | 2.206819 | 0.889257 | -0.27144 | -0.35302 | 0.342016 | 0.247105 | 1.332275 | -1.03745 | 0.361304 | -1.34065 | 1.38817 | -0.50721 | 1.84203 | -0.25944 | -1.39408 | 0.07174 | 0.580946 |
| 21.63889 | 1.621056 | 2.831659 | 0.543963 | -0.17091 | -0.36346 | 0.175971 | 0.005849 | 0.685213 | -0.11661 | 0.396066 | -0.71467 | 1.117804 | -0.14241 | 1.219434 | -0.24275 | -1.26767 | -1.0002 | -0.39758 |
| 21.65278 | 1.07896 | 2.333916 | 0.301435 | -0.19282 | -0.22132 | 0.106867 | -0.10563 | 0.012765 | 0.674085 | -0.08159 | -0.18107 | -0.27655 | -0.08878 | 0.329993 | -0.1102 | -0.90071 | -1.91372 | -1.00691 |
| 21.66667 | -0.0605 | 0.729978 | -0.07435 | -0.04089 | 0.009176 | -0.09847 | -0.21752 | -0.61304 | 1.265561 | -0.95304 | 0.196522 | -1.33336 | -0.24332 | -0.33118 | -0.06238 | 0.628016 | -1.19042 | -0.83357 |
| 21.68056 | -1.43626 | -1.07318 | -0.28391 | 0.064418 | 0.076904 | -0.17535 | -0.22296 | -0.90067 | 1.671327 | -1.24523 | 0.540717 | -1.06267 | -0.4042 | -0.69398 | -0.15588 | 2.175268 | 0.188924 | -0.38028 |
| 21.69444 | -2.18314 | -2.06197 | -0.41594 | 0.005505 | -0.09069 | -0.11341 | -0.08401 | -0.78628 | 1.552846 | -0.30658 | 0.665706 | 0.230093 | -0.50428 | -0.87912 | -0.02578 | 2.975318 | 1.192959 | 0.044217 |
| 21.70833 | -2.17369 | -2.10517 | -0.09417 | 0.068705 | -0.2322 | -0.04622 | 0.080151 | -0.2921 | 0.897005 | 1.00363 | 0.263752 | 0.86838 | -0.77626 | -0.71867 | 0.159691 | 2.632214 | 0.975258 | 0.458964 |
| 21.72222 | -1.18356 | -1.37062 | 0.294395 | 0.209827 | -0.19522 | -0.04709 | 0.243479 | 0.529091 | 0.152589 | 1.465866 | -0.31925 | 0.041638 | -0.79709 | -0.37521 | -0.01848 | 1.042526 | 0.275123 | 0.643792 |
| 21.73611 | 0.053088 | -0.2206 | 0.257336 | 0.165268 | -0.05995 | 0.124099 | 0.297455 | 1.13495 | -0.39456 | 0.479015 | -0.63945 | -1.37687 | -0.73315 | -0.08938 | -0.48994 | -0.85296 | -0.82391 | 0.558892 |
| 21.75 | 0.555677 | 0.705057 | -0.19618 | -0.11145 | 0.029045 | 0.347012 | 0.295191 | 1.068051 | -1.07972 | -0.79043 | -0.76362 | -2.57021 | -0.76412 | 0.176733 | -1.23678 | -2.59482 | -2.38281 | -0.3573 |
| 21.76389 | 0.055903 | 0.492675 | -0.38619 | -0.43464 | 0.081347 | 0.469797 | 0.191365 | 0.193508 | -1.99471 | -1.55808 | -0.70304 | -2.07955 | -0.70058 | 0.380935 | -1.87104 | -3.20973 | -3.02231 | -1.26989 |
| 21.77778 | -0.98021 | -0.73497 | -0.39527 | -0.81307 | -0.13074 | 0.335928 | 0.034457 | -0.61126 | -2.33645 | -2.03748 | -0.43578 | -1.08189 | -0.53904 | 0.320811 | -1.85701 | -2.41203 | -1.56703 | -1.2389 |
| 21.79167 | -1.94756 | -1.82763 | -0.325 | -0.81508 | -0.17621 | 0.073366 | -0.16469 | -1.03973 | -2.66414 | -2.35449 | -0.25996 | -0.97948 | -0.69625 | -0.06849 | -1.36441 | -1.47684 | 0.003671 | -0.75544 |
| 21.80556 | -2.66353 | -2.26103 | -0.01081 | -0.60276 | -0.12817 | -0.06415 | -0.18578 | -0.85596 | -2.17315 | -1.58962 | -0.42606 | -1.04762 | -0.82436 | -0.7306 | -0.75555 | -0.58889 | 0.857584 | -0.20634 |
| 21.81944 | -2.21406 | -1.70348 | 0.186503 | -0.42842 | -0.04414 | -0.25706 | -0.30611 | -0.19803 | -1.22389 | -1.09371 | -0.70775 | -1.41022 | -0.81723 | -1.20965 | -0.4881 | -0.02701 | 0.723974 | 0.079247 |
| 21.83333 | -1.37786 | -1.02322 | 0.159006 | -0.20084 | 0.006918 | -0.43956 | -0.42221 | 0.398454 | -0.16148 | -0.97351 | -0.81958 | -2.01522 | -1.01012 | -1.30249 | -0.90103 | -0.10062 | 0.051459 | -0.20129 |
| 21.84722 | -0.95584 | -0.53117 | -0.1011 | -0.01978 | -0.0608 | -0.55223 | -0.35833 | 0.009865 | 0.364231 | -1.16177 | -0.68788 | -2.42314 | -1.62876 | -1.05227 | -1.50935 | -0.56172 | -1.27113 | -0.76796 |
| 21.86111 | -1.12747 | -0.45843 | -0.47613 | -0.20929 | -0.20326 | -0.53428 | -0.30674 | -0.86684 | 0.101649 | -1.52862 | -0.70279 | -2.27577 | -2.01712 | -0.70634 | -1.79982 | -1.05591 | -2.5806 | -1.10884 |
| 21.875 | -1.64459 | -0.54855 | -0.669 | -0.50785 | -0.50636 | -0.45506 | -0.2647 | -1.39374 | -0.51276 | -1.79541 | -0.91545 | -1.85368 | -1.7111 | -0.4037 | -1.49592 | -1.20191 | -2.49038 | -1.10863 |
| 21.88889 | -2.24938 | -0.85705 | -0.74109 | -0.69446 | -0.77685 | -0.41666 | -0.22259 | -1.55741 | -1.1516 | -1.76961 | -0.74863 | -1.45076 | -1.13834 | -0.2436 | -1.291 | -0.78731 | -0.9894 | -0.95603 |
| 21.90278 | -2.76108 | -1.48281 | -0.71651 | -0.57926 | -0.8698 | -0.42128 | -0.17561 | -1.3893 | -1.55747 | -1.47283 | -0.8249 | -1.59337 | -0.63544 | -0.0731 | -1.18267 | -0.33903 | 0.508293 | -0.6007 |
| 21.91667 | -2.64667 | -1.8952 | -0.61754 | -0.35277 | -0.71591 | -0.37491 | -0.15247 | -0.94163 | -1.29601 | -0.9929 | -1.52081 | -1.74125 | -0.59882 | -0.10942 | -0.86131 | 0.182088 | 1.306063 | -0.04208 |
| 21.93056 | -2.0882 | -1.99144 | -0.44727 | -0.31105 | -0.52445 | -0.29311 | -0.17977 | -0.37514 | -0.72684 | -0.61648 | -1.7707 | -1.70753 | -0.87008 | -0.28595 | -0.52924 | 0.387608 | 0.979012 | 0.286861 |
| 21.94444 | -1.66352 | -1.99917 | -0.18175 | -0.29233 | -0.47748 | -0.19511 | -0.20962 | -0.0519 | -0.21903 | -0.54615 | 0.338247 | -0.81498 | -0.86014 | -0.0768 | -0.24333 | 0.322585 | -0.16836 | 0.163856 |
| 21.95833 | -1.27 | -1.67012 | 0.089289 | -0.23057 | -0.3819 | -0.09023 | -0.14098 | -0.1559 | 0.19375 | -0.75338 | 0.251449 | 0.007303 | -0.69186 | -0.04981 | -0.27783 | -0.05806 | -1.43113 | -0.30611 |
| 21.97222 | -1.18215 | -1.13161 | 0.104948 | -0.22631 | -0.31593 | -0.06933 | -0.04274 | -0.69363 | 0.316604 | -0.99668 | -0.13298 | -0.96325 | -0.77955 | -0.15877 | -0.84947 | -0.53548 | -1.91207 | -1.02362 |
| 21.98611 | -1.7623 | -1.3185 | -0.20699 | -0.32037 | -0.33852 | -0.30604 | -0.03431 | -1.60283 | -0.16637 | -1.47425 | -0.36217 | -2.92269 | -1.05377 | -0.46461 | -1.75044 | -1.37746 | -1.5829 | -1.49525 |
| 22 | -2.04348 | -2.24061 | -0.744 | -0.51686 | -0.48534 | -0.6125 | -0.25949 | -2.57125 | -1.19461 | -2.11669 | -0.48822 | -3.93307 | -1.76281 | -0.81812 | -2.76139 | -2.65101 | -2.36816 | -1.87332 |
| 22.01389 | -1.29763 | -2.57439 | -1.1273 | -0.58188 | -0.51724 | -0.72391 | -0.44873 | -3.30061 | -2.46222 | -2.40994 | -0.35281 | -4.79835 | -2.84401 | -0.72181 | -3.12211 | -4.01466 | -4.20648 | -2.55039 |
| 22.02778 | -0.28964 | -2.44034 | -1.11726 | -0.45679 | -0.32099 | -0.52601 | -0.46727 | -2.67759 | -1.55154 | -1.52607 | -0.12918 | -4.08631 | -1.89774 | -0.20399 | -2.27121 | -3.62899 | -5.1349 | -2.53318 |
| 22.04167 | 0.410531 | -1.76723 | -0.80125 | -0.33855 | -0.10414 | -0.24606 | -0.35392 | -1.22042 | -0.29432 | -0.29262 | 0.05287 | -1.00636 | -0.6675 | 0.106626 | -0.54746 | -1.53347 | -2.56163 | -0.7738 |
| 22.05556 | 0.482029 | -0.84719 | -0.48987 | -0.25355 | -0.01249 | -0.11787 | -0.18811 | -0.41042 | 0.147007 | 0.156349 | 0.148535 | 1.410613 | -0.03037 | -0.1336 | 0.272512 | 0.193811 | 0.336415 | 0.264206 |
| 22.06944 | -0.20275 | -0.41528 | -0.2954 | -0.10344 | -0.02409 | -0.0915 | 0.028534 | -0.07929 | -0.01804 | 0.013196 | 0.112672 | 2.161761 | 0.107523 | -0.43674 | 0.287483 | 1.601452 | 2.448468 | 0.772369 |
| 22.08333 | -0.52514 | 0.086162 | -0.12091 | 0.015023 | 0.017452 | 0.008799 | 0.167229 | 0.363557 | 0.035339 | -0.18718 | 0.160292 | 2.077519 | 0.107496 | -0.36615 | 0.077452 | 1.713262 | 3.325972 | 0.791352 |
| 22.09722 | -0.1341 | 0.860996 | -0.06563 | 0.107512 | 0.126072 | 0.188389 | 0.200747 | 1.09286 | 0.578189 | -0.19294 | 0.378908 | 1.736741 | 0.068875 | -0.19555 | 0.074214 | 1.463194 | 3.035219 | 0.727373 |
| 22.11111 | 0.145714 | 1.120405 | 0.011107 | 0.190599 | 0.17188 | 0.209319 | 0.220791 | 1.58543 | 1.039082 | -0.09168 | 0.480039 | 0.941706 | -0.01992 | -0.16019 | 0.173408 | 1.007015 | 2.42838 | 0.738928 |
| 22.125 | 0.31762 | 1.011559 | 0.259798 | 0.185487 | 0.148629 | 0.115306 | 0.198565 | 1.795563 | 1.255197 | 0.079629 | 0.483726 | -0.00593 | -0.04374 | -0.15042 | 0.354525 | 0.837411 | 2.405968 | 0.860488 |
| 22.13889 | 0.727984 | 1.268475 | 0.506782 | 0.119953 | 0.157115 | 0.10283 | 0.11171 | 1.893763 | 1.434896 | 0.503537 | 0.480934 | -0.94863 | 0.135264 | -0.0908 | 0.67339 | 0.674336 | 2.625348 | 1.238916 |
| 22.15278 | 1.204289 | 1.610577 | 0.589968 | 0.091878 | 0.138105 | 0.126312 | 0.030014 | 1.731933 | 1.39014 | 0.841295 | 0.536537 | -1.09764 | 0.441465 | -0.04361 | 0.884728 | 0.158825 | 2.030103 | 1.524128 |
| 22.16667 | 1.13968 | 1.571855 | 0.452361 | 0.125235 | 0.070117 | 0.116489 | 0.051857 | 1.187191 | 1.139675 | 0.968002 | 0.576392 | 0.052628 | 0.794561 | 0.029947 | 0.852527 | -0.47769 | 0.454785 | 1.401379 |
| 22.18056 | 0.945068 | 1.409177 | 0.32029 | 0.157609 | 0.043543 | 0.122707 | 0.082311 | 0.60324 | 0.658837 | 0.913047 | 0.57898 | 1.079869 | 1.094547 | 0.093664 | 0.679659 | -0.60367 | -0.62699 | 1.083128 |
| 22.19444 | 0.998093 | 1.622416 | 0.42436 | 0.210342 | 0.117278 | 0.214505 | 0.058672 | 0.441688 | 0.439248 | 0.62722 | 0.50568 | 0.851668 | 1.154571 | 0.001968 | 0.514607 | 0.13758 | -0.62441 | 0.826788 |
| 22.20833 | 0.89776 | 1.428343 | 0.528246 | 0.229383 | 0.169947 | 0.298027 | 0.04308 | 0.470934 | 0.22312 | 0.364138 | 0.411171 | 0.310388 | 0.887363 | -0.10985 | 0.393542 | 0.970564 | 0.374372 | 0.633989 |
| 22.22222 | 0.33466 | 0.344652 | 0.363492 | 0.179746 | 0.084683 | 0.17445 | 0.013393 | 0.401005 | -0.17859 | 0.143202 | 0.351703 | 0.256841 | 0.499991 | -0.12214 | 0.226461 | 1.012548 | 1.167584 | 0.337317 |
| 22.23611 | -0.25961 | -0.95545 | -0.05819 | 0.147035 | -0.02736 | -0.04094 | -0.00763 | 0.095144 | -0.28632 | -0.08976 | 0.277952 | -0.04765 | 0.208921 | -0.05338 | -0.20654 | 0.100912 | 0.627998 | -0.32991 |
| 22.25 | -0.45563 | -1.60113 | -0.40235 | 0.10457 | -0.0562 | -0.21117 | -0.00817 | -0.32179 | -0.30088 | -0.22225 | 0.247683 | -0.85774 | 0.012644 | 0.028125 | -0.61629 | -1.15296 | -0.6772 | -0.92493 |
| 22.26389 | -0.13875 | -1.18668 | -0.4757 | -0.00219 | 0.037351 | -0.22975 | -0.00918 | -0.75993 | -0.39936 | -0.21752 | 0.333767 | -1.56985 | -0.03442 | 0.079195 | -0.77146 | -1.29257 | -1.17076 | -1.09844 |
| 22.27778 | 0.295592 | -0.14776 | -0.28751 | -0.07078 | 0.15933 | -0.09075 | -0.02592 | -0.90582 | -0.64981 | -0.12224 | 0.496707 | -1.53155 | 0.045009 | 0.02749 | -0.6209 | -0.05664 | -0.28892 | -0.78437 |
| 22.29167 | 0.455215 | 0.487051 | -0.03507 | 0.010039 | 0.179554 | 0.036733 | 0.027528 | -0.65719 | -0.87634 | -0.07671 | 0.604732 | -0.75841 | 0.155751 | -0.02232 | -0.39293 | 0.917259 | 0.79691 | -0.3694 |
| 22.30556 | 0.57282 | 0.600957 | 0.132213 | 0.111166 | 0.128581 | 0.118 | 0.045333 | -0.02476 | -0.52995 | 0.006249 | 0.70372 | 0.036872 | 0.157565 | 0.000785 | -0.14455 | 0.776328 | 1.108451 | -0.09699 |
| 22.31944 | 0.861492 | 0.56728 | 0.197463 | 0.126065 | 0.131242 | 0.184779 | -0.00677 | 0.340007 | 0.273018 | 0.117616 | 0.806124 | 0.550547 | 0.072655 | 0.051855 | 0.055584 | 0.00355 | 0.601104 | 0.076728 |
| 22.33333 | 1.076464 | 0.433784 | 0.190847 | 0.12037 | 0.172915 | 0.20474 | 0.000444 | 0.52353 | 1.184935 | 0.170874 | 0.677841 | 0.484999 | 0.115738 | -0.02809 | 0.123934 | -0.58067 | -0.26352 | 0.16672 |
| 22.34722 | 1.055707 | 0.45144 | 0.22434 | 0.159692 | 0.19468 | 0.23058 | 0.060958 | 0.591038 | 1.659161 | 0.30867 | 0.515882 | 0.50025 | 0.404554 | -0.13659 | 0.206452 | -0.37087 | -0.40467 | 0.327337 |
| 22.36111 | 1.041128 | 0.813618 | 0.300033 | 0.230431 | 0.14673 | 0.24674 | 0.104662 | 0.438057 | 0.963559 | 0.775847 | 0.543277 | 1.384844 | 0.837416 | -0.03582 | 0.412159 | 0.467833 | 0.645381 | 0.630204 |
| 22.375 | 0.676919 | 0.823301 | 0.2587 | 0.353144 | 0.045678 | 0.172741 | 0.107927 | 0.573831 | -0.01418 | 1.398416 | 0.69964 | 2.444658 | 1.066046 | 0.204704 | 0.549019 | 1.286311 | 2.132876 | 0.823046 |
| 22.38889 | 0.298582 | 0.586963 | 0.169392 | 0.383549 | 0.035107 | 0.145811 | 0.087757 | 0.725863 | -0.69645 | 1.674155 | 0.90954 | 2.555475 | 0.992721 | 0.382006 | 0.498083 | 1.252435 | 2.476363 | 0.718758 |
| 22.40278 | 0.311078 | 0.867777 | 0.216081 | 0.348725 | 0.152103 | 0.253591 | 0.126492 | 0.727949 | -0.47094 | 1.293474 | 1.009932 | 1.083583 | 0.754644 | 0.385511 | 0.223163 | 0.296438 | 0.916066 | 0.329758 |
| 22.41667 | 0.456677 | 1.236751 | 0.37168 | 0.300441 | 0.239175 | 0.345417 | 0.206497 | 0.746115 | 0.328399 | 0.679776 | 0.94048 | -0.58672 | 0.573884 | 0.264229 | 0.027893 | -0.34308 | -0.52414 | -0.00018 |
| 22.43056 | 0.421737 | 1.021368 | 0.399796 | 0.265262 | 0.253138 | 0.287424 | 0.18881 | 0.939997 | 1.37338 | 0.288774 | 0.882332 | -0.62073 | 0.574399 | 0.243261 | 0.17933 | -0.02443 | -0.99659 | 0.08244 |
| 22.44444 | 0.031637 | 0.220094 | 0.207692 | 0.294234 | 0.356037 | 0.178574 | 0.113505 | 0.705144 | 1.584044 | 0.171649 | 0.950168 | 0.449555 | 0.594414 | 0.374071 | 0.579357 | 0.723884 | -0.19124 | 0.451846 |
| 22.45833 | 0.117126 | 0.086229 | 0.097646 | 0.234315 | 0.450331 | 0.223821 | 0.101734 | -0.07475 | 0.44282 | 0.303187 | 0.850313 | 1.466982 | 0.437099 | 0.429839 | 0.865476 | 1.457122 | 1.07861 | 0.715685 |

| | | | | | | | | | | | | | | | | | | |
|---|---|---|---|---|---|---|---|---|---|---|---|---|---|---|---|---|---|---|
| 22.47222 | 0.791915 | 1.233353 | 0.304841 | 0.028866 | 0.123055 | 0.387898 | 0.106815 | -0.28917 | -0.36388 | 0.94474 | 0.707126 | 2.147331 | 0.14735 | 0.462897 | 1.10447 | 1.729666 | 1.872574 | 0.788592 |
| 22.48611 | 1.32324 | 2.570027 | 0.713299 | -0.1009 | 0.110293 | 0.466545 | 0.197861 | -0.10453 | -1.29593 | 2.031075 | 0.586522 | 2.361728 | 0.201553 | 0.713846 | 1.885602 | 1.364714 | 1.768948 | 1.0223 |
| 22.5 | 1.355594 | 3.826669 | 1.442545 | 0.432283 | 0.687508 | 0.684424 | 0.746927 | 1.031687 | 0.201629 | 3.222975 | 0.281666 | 2.396412 | 0.752661 | 2.046297 | 4.269851 | 2.171339 | 1.46478 | 1.687728 |
| 22.51389 | 2.094867 | 4.115823 | 1.88464 | 1.388691 | 1.269604 | 1.09621 | 1.317016 | 1.89765 | 3.098407 | 3.306906 | -0.03192 | 3.229152 | 1.916423 | 3.83892 | 6.945529 | 2.906204 | 1.596192 | 2.491986 |
| 22.52778 | 1.513718 | 2.282567 | 1.439176 | 1.544832 | 1.069105 | 0.807934 | 0.947724 | 1.32269 | 3.499974 | 2.262561 | 0.350853 | 5.37322 | 2.812711 | 3.646789 | 5.300152 | 3.040342 | 1.70641 | 2.029553 |
| 22.54167 | 0.711146 | 0.535585 | 0.589112 | 0.899099 | 0.649317 | 0.31055 | 0.325991 | 0.073387 | 1.942624 | 0.577514 | 0.880586 | 4.536864 | 1.762042 | 1.852152 | 1.997011 | 2.418486 | 1.086612 | 0.908523 |
| 22.55556 | -0.04509 | -0.82772 | -0.24241 | 0.110494 | 0.500972 | -0.15038 | -0.21799 | -0.69432 | 0.721671 | -1.23875 | 0.714117 | 0.035537 | 0.197721 | -0.25884 | -1.00916 | 0.302222 | -0.68807 | -0.48703 |
| 22.56944 | -0.7319 | -2.09724 | -0.9102 | -0.49069 | 0.403739 | -0.63769 | -0.73522 | -1.47297 | -0.46982 | -2.37636 | 0.184564 | -3.94079 | -1.43877 | -1.75553 | -3.35372 | -1.62885 | -2.34541 | -1.74482 |
| 22.58333 | -1.03174 | -2.19575 | -1.08706 | -0.62689 | 0.264201 | -0.85926 | -0.86219 | -1.71441 | -1.53943 | -1.02448 | 0.063896 | -4.89183 | -1.85574 | -1.73788 | -4.36117 | -2.98547 | -2.02785 | -2.87408 |
| 22.59722 | -0.95567 | -1.22046 | -0.75648 | -0.34652 | 0.133709 | -0.44927 | -0.57651 | -1.84831 | -2.84978 | 2.634399 | 0.396935 | -1.47214 | -0.90266 | -1.01878 | -3.8052 | -3.02959 | -0.45349 | -2.40853 |
| 22.61111 | -0.3743 | 0.186509 | -0.11723 | 0.338682 | -0.04645 | 0.169904 | 0.044776 | 0.133152 | -1.54097 | 5.216999 | 0.359908 | 2.64674 | -0.06369 | -0.09419 | -2.06215 | -2.36025 | 1.863574 | -0.99452 |
| 22.625 | 0.43374 | 1.548853 | 0.516311 | 0.822927 | -0.18662 | 0.485656 | 0.516079 | 2.183344 | 0.495395 | 4.218028 | 0.333329 | 5.987578 | 0.437257 | 1.215795 | 0.265526 | -1.91152 | 2.26986 | 0.735707 |
| 22.63889 | 0.862237 | 1.915891 | 1.035406 | 0.47582 | -0.25684 | 0.2607 | 0.491279 | 2.911009 | 2.026649 | 1.977727 | 0.661776 | 3.729737 | 0.884072 | 1.538533 | 1.535232 | -1.912 | 0.746635 | 1.964727 |
| 22.65278 | 0.645108 | 1.174453 | 0.702531 | 0.154197 | -0.23988 | -0.08679 | 0.178804 | 0.995817 | 1.025265 | -0.68333 | 1.269906 | -0.54974 | -0.1768 | 0.848241 | 0.273128 | -1.98001 | -1.33388 | 1.576262 |
| 22.66667 | 0.053808 | -0.05179 | 0.216302 | -0.36856 | -0.16909 | -0.45696 | -0.13994 | -1.20101 | -0.49083 | -3.46438 | 1.160449 | -5.50954 | -0.8701 | -0.58887 | -1.22469 | -1.67869 | -2.9311 | 0.203565 |
| 22.68056 | -0.36878 | -1.19345 | -0.47875 | -0.93859 | -0.1776 | -0.82633 | -0.46598 | -2.44381 | -1.62755 | -3.78651 | 0.016427 | -6.79332 | -1.19541 | -1.32303 | -2.17396 | -1.27541 | -2.16285 | -0.68861 |
| 22.69444 | -0.43281 | -1.98215 | -0.6429 | -0.91858 | -0.23493 | -0.68557 | -0.69625 | -2.51965 | -1.37597 | -2.64069 | -0.64758 | -2.90638 | -0.81235 | -0.84287 | -2.3035 | -0.72771 | -0.53191 | -1.19181 |
| 22.70833 | -0.3905 | -2.13851 | -0.53403 | -0.45761 | -0.32451 | -0.25872 | -0.67923 | -1.25355 | -0.4379 | -1.04961 | -0.79033 | 0.838227 | 0.160855 | -0.12621 | -1.72213 | 0.610328 | 1.2902 | -1.68579 |
| 22.72222 | -0.23786 | -1.09221 | -0.09971 | -0.04949 | -0.3709 | 0.11285 | -0.47516 | 0.359347 | 0.672918 | 0.858269 | -0.30259 | 3.886696 | 1.151153 | 0.429308 | 0.064037 | 1.400999 | 0.943233 | -0.92743 |
| 22.73611 | -0.17254 | 0.251206 | 0.108681 | 0.450311 | -0.32257 | 0.181753 | -0.15744 | 1.552529 | 1.395272 | 0.979528 | -0.52873 | 3.812732 | 1.973195 | 0.71518 | 1.942666 | 1.738363 | 0.287337 | 0.514225 |
| 22.75 | -0.21911 | 1.043336 | 0.113006 | 0.426514 | -0.1325 | -0.0659 | 0.09174 | 1.485247 | 1.079945 | -0.34526 | -1.15294 | -1.11901 | 0.318307 | 0.593533 | 2.127758 | -0.05159 | -1.9069 | 1.279304 |
| 22.76389 | -0.24935 | 0.474181 | -0.41107 | 0.302455 | 0.024773 | -0.3499 | 0.13614 | 0.055094 | -0.44785 | -1.47157 | -0.55387 | -5.42652 | -1.15647 | -0.07068 | 0.545484 | -2.59045 | -3.49944 | 0.331679 |
| 22.77778 | -0.28875 | -0.6487 | -1.09044 | -0.10781 | 0.062989 | -0.54397 | 0.056649 | -1.73712 | -2.19478 | -1.46377 | 0.674484 | -7.45966 | -2.25419 | -0.68703 | -1.09253 | -4.65631 | -2.91482 | -1.23588 |
| 22.79167 | -0.62528 | -1.93412 | -1.14748 | -0.5238 | 0.046171 | -0.54389 | -0.05187 | -2.99218 | -3.32014 | -1.12438 | 1.205204 | -5.40947 | -2.69631 | -0.9464 | -2.52418 | -6.00453 | -1.89577 | -3.33721 |
| 22.80556 | -0.97062 | -3.03049 | -1.24474 | -0.7367 | -0.03919 | -0.45585 | -0.17356 | -2.16992 | -3.04354 | -0.94505 | 0.535225 | -1.62809 | -1.16148 | -0.77009 | -3.15186 | -5.0671 | -0.54879 | -4.06556 |
| 22.81944 | -1.01183 | -3.15606 | -1.18181 | -0.72796 | -0.10392 | -0.35919 | -0.27036 | -0.92333 | -2.19626 | -0.82339 | -0.40999 | 1.148768 | 0.147104 | -0.62882 | -2.29845 | -4.37964 | -0.15498 | -2.93597 |
| 22.83333 | -0.76458 | -2.45345 | -1.02526 | -0.55789 | -0.15884 | -0.3352 | -0.3398 | -0.54157 | -1.69617 | -0.89445 | -1.71943 | 0.73675 | 0.735641 | -0.78694 | -1.16951 | -2.8496 | 0.266893 | -1.91953 |
| 22.84722 | -0.36549 | -1.73158 | -0.67309 | -0.44394 | -0.30275 | -0.43357 | -0.34233 | -0.85208 | -1.41605 | -1.43184 | -3.01269 | -1.52938 | 0.248852 | -1.00955 | -0.54386 | -0.44279 | -0.39436 | -1.28674 |
| 22.86111 | -0.24087 | -1.08681 | -0.60815 | -0.56788 | -0.46362 | -0.53448 | -0.25894 | -1.37124 | -1.02884 | -2.09595 | -2.93122 | -3.60173 | -0.70356 | -1.21642 | -0.64716 | 0.212501 | -2.20406 | -1.09543 |
| 22.875 | -0.22809 | -0.26861 | -0.77013 | -0.88695 | -0.552 | -0.61627 | -0.27788 | -1.91102 | -0.79559 | -1.98969 | -1.75497 | -4.37555 | -1.96755 | -1.16667 | -0.99552 | -0.49972 | -1.01372 | -1.07503 |
| 22.88889 | -0.30757 | 0.371224 | -0.97654 | -1.13597 | -0.60339 | -0.79705 | -0.34159 | -2.14329 | -0.71681 | -1.50158 | -0.76557 | -4.29423 | -3.36464 | -0.87872 | -1.08109 | -1.77399 | -0.66392 | -1.10653 |
| 22.90278 | -0.40736 | 0.39544 | -1.04577 | -1.06214 | -0.60903 | -0.78011 | -0.31808 | -1.58288 | -0.34668 | -0.80901 | 0.07196 | -3.50488 | -3.72386 | -0.69109 | -0.66774 | -2.40955 | -0.42898 | -0.71926 |
| 22.91667 | -0.4055 | 0.295171 | -0.7405 | -0.70736 | -0.53209 | -0.52513 | -0.18351 | -0.56845 | 0.219413 | 0.012028 | 0.763795 | -2.39176 | -2.20351 | -0.60847 | 0.223195 | -1.63063 | -0.41578 | 0.031814 |
| 22.93056 | -0.23456 | 0.32771 | -0.38196 | -0.37102 | -0.32738 | -0.26126 | -0.03491 | 0.29042 | 0.608562 | 0.40198 | 1.019283 | -0.94694 | -0.70343 | -0.65213 | 0.846209 | -0.05069 | -0.35519 | 0.653871 |
| 22.94444 | 0.062783 | 0.30424 | -0.11047 | -0.05378 | -0.10145 | -0.07535 | 0.048389 | 0.687209 | 0.405035 | 0.278605 | 0.678625 | 1.113248 | -0.14726 | -0.84316 | 0.659161 | 1.177453 | -0.47616 | 0.72462 |
| 22.95833 | 0.293718 | -0.03369 | 0.027423 | 0.201059 | 0.032751 | -0.09441 | 0.040177 | 0.498047 | -0.2104 | 0.043927 | 0.222407 | 3.117193 | -0.39343 | -1.13469 | -0.23573 | 1.585238 | -0.57074 | 0.198926 |
| 22.97222 | 0.136742 | -0.796 | 0.098065 | 0.336779 | -0.00246 | -0.3175 | 0.018212 | -0.11828 | -0.73977 | -0.80701 | -0.44711 | 2.981998 | -0.78645 | -1.11694 | -1.56142 | 1.260877 | -0.17642 | -0.65538 |
| 22.98611 | -0.48781 | -1.84104 | -0.18756 | 0.134652 | -0.02349 | -0.60333 | -0.08379 | -1.06286 | -1.04586 | -2.40968 | -1.34603 | -1.10317 | -1.36908 | -0.90978 | -2.71332 | -0.18767 | -0.88524 | -1.41532 |
| 23 | -0.92655 | -2.78509 | -0.97311 | -0.30546 | -0.0784 | -0.84669 | -0.51565 | -1.82339 | -1.55902 | -3.86145 | -2.28475 | -5.36992 | -2.50127 | -1.23781 | -2.97238 | -2.65257 | -2.81131 | -1.81914 |
| 23.01389 | -0.86491 | -3.09522 | -1.50058 | -0.67458 | -0.14256 | -0.95084 | -0.97197 | -2.25516 | -1.65187 | -4.35793 | -3.10902 | -8.97963 | -3.11284 | -1.3386 | -2.20467 | -4.42599 | -4.98626 | -1.65309 |
| 23.02778 | -0.51321 | -2.39137 | -1.40907 | -0.73081 | -0.21077 | -0.82375 | -1.1982 | -1.60091 | -0.8563 | -3.22812 | -2.86935 | -8.53337 | -2.0157 | -0.54184 | -0.81161 | -3.04789 | -5.9778 | -0.90342 |
| 23.04167 | -0.44564 | -1.65573 | -0.87669 | -0.72509 | -0.17556 | -0.19601 | -0.82385 | -0.71244 | 0.097439 | -1.29011 | -1.53266 | -4.05725 | -0.5694 | 0.234603 | 0.190174 | -0.93534 | -5.23134 | -0.15356 |
| 23.05556 | -0.89366 | -1.47646 | -0.52714 | -0.73619 | -0.25278 | 0.082592 | -0.44605 | -0.52732 | 0.397522 | -0.24149 | -0.26518 | -1.08341 | 0.11084 | 0.279371 | 0.16856 | -0.25191 | -3.64655 | -0.11085 |
| 23.06944 | -0.97833 | -1.30988 | -0.31733 | -0.57236 | -0.24935 | 0.038348 | -0.17101 | -0.74194 | 0.130587 | 0.203697 | 0.653322 | 0.462826 | 0.105505 | -0.16692 | -0.21466 | -0.3538 | -2.08217 | -0.37346 |
| 23.08333 | -0.54198 | -0.95287 | -0.14201 | -0.23475 | -0.192 | 0.052383 | -0.09587 | -0.61723 | -0.02272 | 0.636297 | 1.206735 | 2.469152 | -0.01109 | -0.41431 | -0.17529 | -0.54329 | -0.57121 | -0.42544 |
| 23.09722 | -0.07224 | -0.39138 | 0.016456 | 0.026309 | -0.05394 | 0.247422 | -0.03258 | -0.33837 | 0.011277 | 0.856089 | 1.335776 | 5.621849 | -0.05564 | -0.2963 | 0.07166 | -0.60372 | 0.653071 | -0.34118 |
| 23.11111 | 0.082243 | -0.06243 | 0.100624 | 0.177228 | 0.118951 | 0.518534 | 0.036192 | -0.10707 | 0.132859 | 0.868156 | 1.319883 | 7.05875 | -0.0486 | 0.0055 | 0.083226 | -0.55613 | 1.31728 | -0.33206 |
| 23.125 | 0.245092 | 0.245664 | 0.170649 | 0.270458 | 0.302485 | 0.719831 | 0.099921 | 0.152605 | 0.323555 | 0.700877 | 1.12274 | 6.49601 | -0.12095 | 0.230005 | 0.0394 | -0.39826 | 1.776833 | -0.35614 |
| 23.13889 | 0.44093 | 0.41853 | 0.199623 | 0.30001 | 0.347229 | 0.607567 | 0.162001 | 0.469429 | 0.531299 | 0.316528 | 0.851788 | 5.579431 | -0.15295 | 0.130208 | -0.27483 | 0.112019 | 1.947315 | -0.38182 |
| 23.15278 | 0.353024 | 0.074445 | 0.137252 | 0.235289 | 0.226002 | 0.287856 | 0.270215 | 0.763588 | 0.800744 | -0.19474 | 0.512232 | 3.318713 | -0.19 | -0.08293 | -0.70888 | 0.879421 | 1.986197 | -0.27389 |
| 23.16667 | 0.231014 | 0.021348 | 0.184686 | 0.278983 | 0.22536 | 0.029267 | 0.315889 | 1.29753 | 1.186032 | -0.12077 | 0.477874 | 1.358776 | -0.17392 | -0.21219 | -0.3521 | 1.730271 | 2.263908 | 0.227428 |
| 23.18056 | 0.394676 | 1.2172 | 0.418483 | 0.440416 | 0.247601 | 0.029813 | 0.27867 | 1.771176 | 1.378887 | 0.396276 | 0.632238 | 0.936691 | -0.04879 | 0.088041 | 0.392987 | 2.18364 | 2.44862 | 0.571533 |
| 23.19444 | 0.468181 | 1.963348 | 0.602403 | 0.512993 | 0.159171 | 0.117209 | 0.209924 | 1.749243 | 1.429222 | 0.728797 | 0.69672 | 0.734362 | 0.199038 | 0.48918 | 0.581705 | 2.1673 | 2.187759 | 0.45949 |
| 23.20833 | 0.501405 | 1.949239 | 0.659501 | 0.482104 | 0.055373 | 0.224649 | 0.225952 | 1.647093 | 1.370369 | 0.867173 | 0.648725 | 0.15235 | 0.392288 | 0.813106 | 0.553019 | 2.11303 | 2.119557 | 0.403549 |
| 23.22222 | 0.463849 | 1.928354 | 0.580161 | 0.415319 | -0.01784 | 0.288022 | 0.300928 | 1.539532 | 1.202461 | 0.854158 | 0.530791 | -0.59275 | 0.600486 | 0.949065 | 0.673969 | 1.915176 | 2.1611 | 0.526347 |
| 23.23611 | 0.377233 | 1.466237 | 0.458561 | 0.304104 | -0.11155 | 0.318264 | 0.314814 | 1.187019 | 0.926012 | 0.89878 | 0.386898 | 0.265327 | 0.804757 | 0.983054 | 0.928913 | 1.46819 | 1.961883 | 0.674337 |
| 23.25 | 0.366755 | 1.143006 | 0.402922 | 0.277316 | -0.11976 | 0.224853 | 0.253614 | 0.665078 | 0.814753 | 0.889036 | 0.290351 | 2.37014 | 0.789727 | 1.02318 | 1.005447 | 0.655724 | 1.750197 | 0.580938 |
| 23.26389 | 0.216355 | 0.774246 | 0.331676 | 0.220022 | 0.009913 | 0.110074 | 0.202167 | 0.17808 | 0.525796 | 0.438307 | 0.150638 | 2.325751 | 0.408894 | 0.804512 | 0.570607 | -0.36733 | -0.79583 | 0.104955 |
| 23.27778 | -0.01559 | 0.074448 | 0.083847 | 0.084303 | 0.126246 | -0.02017 | 0.179004 | -0.5231 | -0.60385 | -0.24724 | 0.080035 | -0.21442 | -0.24667 | 0.355143 | -0.21116 | -1.48653 | -2.40343 | -0.59298 |
| 23.29167 | -0.03405 | -0.52657 | -0.13451 | 0.00305 | 0.143147 | -0.07915 | 0.108356 | -1.33192 | -1.30097 | -0.76896 | 0.185549 | -3.51156 | -0.829 | -0.04267 | -0.75836 | -2.07235 | -2.74069 | -0.96979 |
| 23.30556 | 0.099619 | -0.764 | -0.18741 | 0.018134 | 0.080766 | -0.03854 | 0.012788 | -1.67054 | -1.14563 | -0.9078 | 0.35475 | -4.6835 | -0.82101 | -0.13672 | -0.72787 | -1.23138 | -1.8867 | -0.77397 |
| 23.31944 | 0.215686 | -0.57484 | -0.16257 | 0.070914 | -0.0293 | 0.067901 | -0.01058 | -1.43299 | -0.53999 | -0.46714 | 0.417137 | -1.20366 | -0.40559 | 0.132326 | -0.4015 | 0.138779 | -0.89115 | -0.46045 |
| 23.33333 | 0.260594 | -0.00957 | -0.07806 | 0.160395 | -0.07225 | 0.172895 | 0.025632 | -0.86217 | 0.187376 | 0.069946 | 0.54385 | 2.832048 | 0.091614 | 0.43677 | -0.00605 | 1.186612 | -0.06651 | -0.28643 |
| 23.34722 | 0.365399 | 0.74952 | 0.040947 | 0.282494 | 0.028338 | 0.24216 | 0.06884 | 0.054006 | 0.760486 | 0.547918 | 0.793291 | 3.826892 | 0.531057 | 0.525954 | 0.436848 | 1.624288 | 0.402286 | -0.12754 |
| 23.36111 | 0.534213 | 1.404012 | 0.178947 | 0.385818 | 0.197433 | 0.279174 | 0.109162 | 0.989711 | 1.211268 | 0.647134 | 0.841723 | 0.563981 | 0.839413 | 0.426643 | 0.728162 | 1.200097 | 0.076783 | -0.02328 |
| 23.375 | 0.636072 | 1.547703 | 0.325269 | 0.454186 | 0.273235 | 0.293859 | 0.182469 | 1.891736 | 1.357881 | 0.556333 | 0.609503 | -3.37416 | 0.866106 | 0.41056 | 0.741226 | 0.489976 | -0.60061 | 0.069441 |
| 23.38889 | 0.565553 | 0.936574 | 0.373643 | 0.408962 | 0.210546 | 0.283988 | 0.241722 | 2.178879 | 1.030197 | 0.694022 | 0.350979 | -3.30112 | 0.790827 | 0.545432 | 0.591396 | -0.0071 | -0.96296 | 0.153524 |
| 23.40278 | 0.376799 | 0.083989 | 0.35673 | 0.316223 | 0.116771 | 0.243192 | 0.226423 | 1.931927 | 0.619457 | 0.85342 | 0.304622 | 1.115045 | 0.726432 | 0.656715 | 0.515589 | -0.12633 | -0.99587 | 0.214966 |
| 23.41667 | 0.379701 | -0.291 | 0.389496 | 0.317957 | 0.070194 | 0.224317 | 0.24384 | 1.433698 | 0.491973 | 1.284699 | 0.432638 | 5.763891 | 0.918319 | 0.569956 | 0.56594 | 0.32935 | -0.60137 | 0.302406 |
| 23.43056 | 0.633683 | -0.00458 | 0.461819 | 0.406748 | 0.04998 | 0.289257 | 0.316476 | 0.895119 | 0.454533 | 1.944565 | 0.726206 | 6.468486 | 1.138335 | 0.499924 | 0.856774 | 1.300487 | 0.225167 | 0.459967 |
| 23.44444 | 0.844623 | 0.415706 | 0.405729 | 0.422656 | -0.01336 | 0.345616 | 0.361935 | 0.330761 | 0.238398 | 1.374593 | 0.987847 | 2.201919 | 1.195242 | 0.440781 | 1.090645 | 2.208124 | 1.000802 | 0.573257 |
| 23.45833 | 0.692636 | 0.61446 | 0.26594 | 0.330308 | -0.02657 | 0.289872 | 0.299992 | -0.41205 | -0.20736 | -0.13479 | 0.81326 | -2.5758 | 0.729789 | 0.338843 | 0.878433 | 1.722255 | 0.923062 | 0.397936 |
| 23.47222 | 0.549581 | 1.362367 | 0.094402 | 0.411529 | 0.164739 | 0.43971 | 0.15188 | -0.66063 | -0.45727 | -0.73566 | 0.596038 | -5.73138 | -0.19678 | 0.367718 | 0.80765 | 0.439552 | 0.323904 | 0.165537 |
| 23.48611 | 0.669576 | 2.943256 | 0.314209 | 0.608085 | 0.458694 | 0.744733 | -0.00965 | 0.260445 | -0.55863 | 1.105458 | 0.763806 | -3.7917 | -0.47797 | 0.697534 | 1.446464 | -0.23705 | 0.014569 | 0.290637 |
| 23.5 | 1.345119 | 4.931886 | 1.251467 | 0.697521 | 0.469342 | 0.891972 | 0.061455 | 2.150132 | 0.80673 | 2.860274 | 1.229854 | 1.239114 | 1.333757 | 1.577511 | 2.629419 | 1.284557 | 0.581467 | 0.912627 |
| 23.51389 | 1.940335 | 5.607976 | 2.154514 | 1.158949 | 0.295128 | 0.935464 | 0.608105 | 3.744176 | 2.607241 | 4.10502 | 1.571382 | 5.997494 | 3.125888 | 3.133962 | 3.891457 | 3.59864 | 1.575465 | 1.535436 |
| 23.52778 | 2.552049 | 4.087182 | 2.173351 | 1.415717 | 0.523077 | 0.743308 | 0.546008 | 3.455349 | 2.735596 | 4.486002 | 1.490174 | 6.880723 | 4.172553 | 3.336567 | 4.398752 | 4.75502 | 2.275781 | 1.61956 |
| 23.54167 | 1.872206 | 2.509334 | 1.539262 | 1.045557 | 0.905874 | 0.275903 | 0.339355 | 2.092052 | 2.214891 | 2.886356 | 0.80867 | 2.475873 | 2.70088 | 2.048906 | 2.541028 | 3.223339 | 1.581028 | 0.778113 |
| 23.55556 | 0.704418 | 0.654046 | 0.763905 | 0.593774 | 1.05973 | -0.17377 | 0.01061 | 0.622377 | 1.323472 | 0.54327 | -0.04282 | -1.16499 | 0.288337 | 0.271139 | 0.690112 | 1.020041 | 0.33734 | 0.240185 |
| 23.56944 | -0.44782 | -1.23331 | -0.02174 | 0.002239 | 1.001623 | -0.69966 | -0.37218 | -1.1215 | 0.039119 | -1.58114 | -0.77381 | -4.36475 | -1.6745 | -1.0712 | -0.73932 | -0.56399 | -0.5599 | -0.07083 |
| 23.58333 | -1.34263 | -2.38075 | -0.54922 | -0.58635 | 0.876279 | -1.04634 | -0.60394 | -2.7282 | -1.20547 | -2.52226 | -1.25942 | -7.35958 | -2.9773 | -1.52496 | -1.66332 | -1.96865 | -1.42588 | -0.39095 |

| | | | | | | | | | | | | | | | | | | |
|---|---|---|---|---|---|---|---|---|---|---|---|---|---|---|---|---|---|---|
| 23.59722 | -1.47755 | -2.61925 | -0.72007 | -0.87055 | 0.820085 | -0.82126 | -0.58726 | -3.50894 | -1.93281 | -2.69225 | -1.33452 | -7.34381 | -3.54106 | -0.87753 | -2.23054 | -3.15677 | -1.74091 | -0.59928 |
| 23.61111 | -0.53006 | -2.07259 | -0.72383 | -0.71322 | 0.755394 | -0.08448 | -0.19751 | -4.21532 | -2.42521 | -2.07506 | -0.71878 | -4.51813 | -3.22578 | 0.259381 | -2.50359 | -3.80534 | -1.66536 | -0.99767 |
| 23.625 | 0.884984 | -0.08231 | -0.27963 | -0.09102 | 0.552981 | 0.730136 | 0.088833 | -2.11781 | -0.93859 | 0.170571 | 0.539536 | 1.261754 | -1.21799 | 1.665322 | -1.68037 | -1.57955 | -0.86912 | -0.83022 |
| 23.63889 | 1.74004 | 2.193477 | 0.203213 | 0.64934 | 0.184564 | 0.552588 | 0.280966 | 1.199239 | 0.466121 | 2.359737 | 2.060215 | 6.48499 | 1.877682 | 1.634099 | 0.039932 | 0.869819 | -0.19228 | -0.17823 |
| 23.65278 | 1.664215 | 3.323087 | 0.778956 | 1.17745 | -0.12244 | 0.244808 | 0.387003 | 3.129516 | 1.266595 | 3.441603 | 2.511076 | 7.91514 | 3.796801 | 0.773743 | 1.476104 | 1.890471 | 0.126456 | 0.921487 |
| 23.66667 | 1.012876 | 2.76596 | 1.110958 | 0.578467 | -0.42011 | 0.068719 | 0.350265 | 2.900781 | 1.462235 | 2.132944 | 1.211008 | 3.782334 | 3.189687 | 0.337613 | 2.584704 | 1.208175 | -0.02238 | 1.942472 |
| 23.68056 | 0.323725 | 1.735973 | 1.312108 | 0.095556 | -0.75277 | -0.41657 | 0.238683 | 0.921458 | -0.14381 | 0.208484 | -0.57432 | -1.92896 | 1.005475 | 0.200101 | 2.393906 | -0.10579 | -0.5238 | 1.047534 |
| 23.69444 | -0.38238 | 0.194724 | 0.964389 | -0.51904 | -0.82948 | -0.6742 | 0.034954 | -1.37532 | -1.58799 | -1.27621 | -1.90766 | -6.46849 | -0.92186 | -0.59824 | 0.847788 | -1.27118 | -1.04504 | 0.143571 |
| 23.70833 | -0.6918 | -0.73008 | 0.306674 | -0.7408 | -0.62548 | -0.38731 | -0.07417 | -2.81135 | -2.02144 | -1.90342 | -2.49537 | -6.9487 | -1.89475 | -0.83486 | -0.76118 | -1.8385 | -0.96209 | -0.36184 |
| 23.72222 | -0.20132 | -1.17552 | -0.48701 | -0.7037 | -0.21691 | 0.015339 | -0.13661 | -3.99262 | -2.79869 | -2.63136 | -2.34787 | -7.02255 | -3.14092 | -0.38398 | -2.38731 | -3.01833 | -1.12921 | -0.87553 |
| 23.73611 | 0.330003 | -1.66567 | -0.9294 | -0.20993 | 0.077701 | 0.13042 | -0.18672 | -4.22373 | -2.53727 | -2.33617 | -1.36537 | -3.61462 | -3.46575 | 0.103091 | -3.47109 | -1.70571 | -0.57524 | -1.41608 |
| 23.75 | 0.609567 | -2.07436 | -0.85506 | 0.409361 | 0.207522 | 0.073915 | -0.17684 | -2.46688 | -1.05509 | -1.39825 | -0.23556 | -0.45705 | -2.53075 | 0.286527 | -3.13032 | -0.04992 | -0.77557 | -1.3327 |
| 23.76389 | 0.498935 | -1.45534 | -0.38692 | 0.760083 | 0.218473 | -0.36959 | -0.40055 | 0.429073 | 0.264995 | 0.07171 | 0.568815 | -0.25486 | -0.68664 | -0.22495 | -1.70498 | 2.116287 | -0.17744 | -0.84117 |
| 23.77778 | -0.06674 | -0.16463 | 0.147396 | 0.383259 | 0.104323 | -1.06596 | -0.42136 | 2.603735 | 1.098983 | 1.34499 | 0.957533 | -1.98557 | 1.15799 | -1.07873 | 0.034228 | 2.254037 | 0.342128 | -0.18827 |
| 23.79167 | -0.50927 | 1.177546 | 0.596565 | -0.23918 | -0.08936 | -1.32121 | -0.24427 | 3.074147 | 0.94811 | 1.602043 | 0.641232 | -3.51644 | 2.175706 | -1.86863 | 1.108975 | 0.488104 | 0.059007 | 0.317776 |
| 23.80556 | -0.85023 | 1.18859 | 0.783773 | -0.59041 | -0.35402 | -1.0397 | 0.008878 | 1.761403 | 0.18508 | 0.763933 | -0.06473 | -3.84583 | 1.72288 | -2.61165 | 0.844066 | -1.50335 | -0.91743 | 0.467834 |
| 23.81944 | -0.72216 | 0.21761 | 0.291977 | -0.88642 | -0.59435 | -0.55273 | 0.075283 | -0.41246 | -0.96489 | -1.10176 | -1.23552 | -2.96324 | -0.27389 | -2.35031 | -0.32216 | -3.67908 | -2.26533 | 0.109459 |
| 23.83333 | -0.34239 | -0.74857 | -0.74449 | -0.97016 | -0.60639 | -0.22876 | 0.027996 | -2.3749 | -1.85928 | -2.73935 | -2.46445 | -2.05673 | -2.1642 | -1.36918 | -1.35828 | -4.24706 | -2.60451 | -0.65604 |
| 23.84722 | -0.18701 | -1.68215 | -1.55421 | -1.01097 | -0.5561 | -0.27326 | -0.05639 | -3.83371 | -2.31497 | -3.79763 | -2.66403 | -1.81785 | -3.4577 | -0.64286 | -1.83364 | -3.41912 | -2.35907 | -1.16097 |
| 23.86111 | -0.03647 | -2.43437 | -1.72386 | -0.75584 | -0.54501 | -0.47191 | -0.19568 | -3.16258 | -1.51214 | -3.95192 | -2.70965 | -1.60753 | -3.42138 | -0.32846 | -1.62599 | -2.34448 | -2.409 | -1.23713 |
| 23.875 | -0.13017 | -2.88743 | -1.80004 | -0.53757 | -0.52898 | -0.56994 | -0.41624 | -2.21352 | -0.7742 | -2.71838 | -1.77192 | -1.01919 | -3.17369 | -0.45081 | -1.45762 | -1.74809 | -1.92437 | -1.13031 |
| 23.88889 | -0.149 | -2.65508 | -1.55147 | -0.17367 | -0.45817 | -0.30922 | -0.36985 | -0.9076 | -0.31762 | -1.12481 | -0.55206 | 0.096345 | -2.10932 | -0.6875 | -0.90185 | -1.12643 | -0.90395 | -0.57423 |
| 23.90278 | -0.11506 | -1.62173 | -1.1936 | 0.051687 | -0.33929 | 0.01168 | -0.31658 | -0.238 | -0.13314 | 0.260058 | 0.49298 | 1.544223 | -0.48563 | -0.57896 | -0.15359 | -0.22932 | 0.011376 | 0.113215 |
| 23.91667 | -0.13935 | -0.71779 | -0.95128 | 0.00429 | -0.2825 | 0.03517 | -0.30979 | 0.199299 | 0.123123 | 1.275772 | 1.080846 | 2.533844 | 0.816127 | -0.38458 | 0.439016 | 0.579126 | 0.563321 | 0.606025 |
| 23.93056 | -0.02124 | 0.161512 | -0.27289 | 0.017477 | -0.23773 | -0.21858 | -0.15336 | 0.613784 | 0.437424 | 1.359822 | 0.899859 | 2.794316 | 1.510875 | -0.36718 | 0.743831 | 1.288251 | 1.144882 | 0.896095 |
| 23.94444 | -0.05251 | 0.874988 | 0.352747 | -0.23769 | -0.1648 | -0.51681 | 0.062326 | 0.906112 | 0.498318 | 0.705009 | 0.324487 | 2.146839 | 1.269026 | -0.33861 | 0.667198 | 2.01815 | 1.647675 | 0.828995 |
| 23.95833 | -0.18107 | 1.109881 | 0.876419 | -0.50116 | -0.15853 | -0.67902 | 0.311648 | 1.196492 | 0.399497 | -0.18399 | -0.33768 | 1.070927 | 0.55604 | -0.21774 | 0.352519 | 2.136991 | 1.740422 | 0.412155 |
| 23.97222 | -0.26499 | 0.59078 | 1.244792 | -0.50996 | -0.36988 | -0.96031 | 0.529573 | 1.224345 | 0.292205 | -1.58843 | -0.997 | 0.198617 | -0.35446 | -0.27615 | -0.40729 | 1.266359 | 0.896793 | -0.29037 |
| 23.98611 | -0.72927 | -1.05904 | 0.96016 | -0.311 | -0.55865 | -1.49528 | 0.360702 | 0.431825 | -0.12021 | -3.37019 | -1.43643 | -1.3714 | -1.51672 | -0.67435 | -1.62628 | -0.72302 | -1.13629 | -1.07309 |
| 24 | -1.53834 | -2.33825 | -0.23048 | -0.78794 | -0.38459 | -1.78433 | -0.1477 | -1.04296 | -0.78626 | -4.42495 | -1.92403 | -3.83138 | -2.8923 | -2.41518 | -2.5037 | -2.81795 | -3.00288 | -1.53135 |
| 24.01389 | -1.67357 | -2.55279 | -1.25806 | -1.45758 | -0.36055 | -1.76528 | -0.56151 | -1.43582 | -1.35537 | -4.18886 | -2.06328 | -4.8703 | -3.41413 | -4.69463 | -2.43846 | -3.91789 | -3.82698 | -1.54842 |
| 24.02778 | -1.21636 | -2.20686 | -1.51788 | -1.5729 | -0.34871 | -1.93883 | -0.53582 | -0.82604 | -1.34114 | -2.74983 | -1.38421 | -3.01346 | -2.19176 | -4.19804 | -1.88361 | -2.60788 | -3.39564 | -1.18135 |
| 24.04167 | -0.52169 | -1.88903 | -1.39272 | -1.09155 | -0.34277 | -1.57608 | -0.3745 | -0.09682 | -0.74104 | -1.06669 | -0.649 | -0.82885 | -0.82865 | -2.02412 | -1.2606 | -0.88416 | -2.65891 | -1.00136 |
| 24.05556 | -0.23886 | -1.35088 | -1.07638 | -0.6338 | -0.224 | -0.94048 | -0.27308 | 0.101338 | -0.3932 | -0.40482 | -0.30636 | 0.004789 | -0.10989 | -0.55731 | -0.85533 | -0.58612 | -2.34634 | -1.08087 |
| 24.06944 | -0.07036 | -0.8412 | -0.5981 | -0.23181 | -0.10508 | -0.23765 | -0.06193 | 0.065978 | -0.10141 | -0.24529 | 0.009101 | 0.164993 | 0.170396 | 0.388883 | -0.3198 | -0.71577 | -1.39555 | -0.789 |
| 24.08333 | 0.27139 | -0.46857 | -0.17466 | 0.119035 | -0.03856 | 0.136617 | 0.153373 | -0.08534 | 0.08013 | 0.129127 | 0.443319 | 0.283692 | 0.356138 | 1.086841 | 0.339336 | -0.39769 | -0.24614 | -0.04727 |
| 24.09722 | 0.839268 | 0.017055 | -0.14539 | 0.440502 | -0.03495 | 0.413334 | 0.170784 | -0.39287 | -0.03909 | 0.730909 | 0.800296 | 0.44256 | 0.496012 | 1.625613 | 0.927239 | -0.12901 | 0.315613 | 0.47455 |
| 24.11111 | 1.78478 | 1.267036 | -0.29829 | 0.952348 | 0.407546 | 1.047033 | 0.158933 | -0.61725 | -0.35724 | 0.982399 | 0.960416 | 0.785842 | 0.922261 | 2.103851 | 1.221556 | -0.55839 | -0.08858 | 0.324509 |
| 24.125 | 3.009239 | 3.074012 | -0.12232 | 1.649915 | 1.44551 | 2.009328 | 0.41883 | -0.28136 | -0.15996 | 0.668418 | 0.829822 | 1.333526 | 1.679871 | 2.982411 | 1.357267 | -0.94297 | -0.42348 | -0.224 |
| 24.13889 | 2.78679 | 3.0579 | 0.25126 | 1.905341 | 1.763352 | 2.123693 | 0.531716 | 0.413213 | 0.675821 | -0.00811 | 0.388862 | 1.88647 | 1.372185 | 1.670766 | 0.165344 | -0.57906 | -0.16114 | -0.22207 |
| 24.15278 | 1.387258 | 1.350063 | 0.491065 | 1.506838 | 1.081911 | 2.218813 | 0.437322 | 0.807487 | 1.618184 | -0.60151 | -0.10148 | 2.85605 | 0.580418 | 0.916855 | 0.555761 | 0.114809 | 0.395481 | 0.488006 |
| 24.16667 | 0.590878 | 0.480252 | 0.524411 | 0.929142 | 0.518562 | 1.603648 | 0.418497 | 0.638054 | 1.655934 | -0.14036 | 0.159103 | 3.548481 | 0.582151 | 1.152846 | 1.238687 | 0.700814 | 1.267026 | 1.256854 |
| 24.18056 | 0.379665 | 0.346715 | 0.391261 | 0.324001 | 0.158618 | 0.970586 | 0.287381 | 0.262651 | 0.778298 | 0.792375 | 0.815153 | 3.422907 | 0.994564 | 1.792929 | 1.44327 | 1.295227 | 2.073593 | 1.695418 |
| 24.19444 | 0.157645 | 0.093264 | 0.068146 | -0.13015 | 0.035675 | 0.549508 | 0.029214 | 0.018447 | 0.253908 | 0.997225 | 0.93646 | 3.001052 | 0.956274 | 1.817609 | 1.28489 | 1.455809 | 1.648587 | 1.56347 |
| 24.20833 | 0.06533 | -0.05507 | -0.16247 | -0.34433 | 0.11879 | 0.424241 | -0.12871 | 0.022892 | 0.031546 | 0.760163 | 0.728495 | 2.535492 | 0.754924 | 1.346219 | 1.11962 | 1.287098 | 0.789906 | 1.199165 |
| 24.22222 | 0.349325 | 0.041217 | -0.05878 | -0.17409 | 0.218824 | 0.328955 | -0.0262 | 0.10235 | 0.07497 | 0.834305 | 0.794008 | 2.326248 | 0.985281 | 1.126601 | 1.200065 | 1.21457 | 0.791413 | 1.1111 |
| 24.23611 | 0.736299 | 0.210027 | 0.173969 | 0.216591 | 0.210062 | 0.192857 | 0.168742 | 0.114195 | 0.243271 | 0.980256 | 0.913716 | 2.215973 | 1.238008 | 1.069487 | 1.232143 | 1.33981 | 1.386457 | 1.186976 |
| 24.25 | 0.743418 | 0.222018 | 0.255856 | 0.369319 | 0.089152 | -0.13038 | 0.182989 | 0.022838 | 0.328037 | 0.843112 | 0.66404 | 1.88307 | 0.888578 | 0.890502 | 0.939473 | 1.269743 | 1.680148 | 1.002467 |
| 24.26389 | 0.321633 | 0.082409 | 0.050102 | 0.168013 | 0.013977 | -0.41883 | 0.015934 | -0.02578 | 0.254087 | 0.361407 | 0.175346 | 1.232805 | -0.00021 | 0.421583 | 0.325289 | 0.841933 | 0.940533 | 0.436037 |
| 24.27778 | -0.20516 | -0.06944 | -0.21548 | -0.15469 | -0.00226 | -0.65582 | -0.13447 | 0.00834 | 0.188967 | -0.04045 | -0.02978 | 0.23412 | -0.59294 | -0.04072 | -0.11445 | 0.368326 | -0.21191 | -0.07394 |
| 24.29167 | -0.42363 | -0.06147 | -0.33403 | -0.37569 | -0.10873 | -0.60406 | -0.24796 | 0.061312 | 0.190902 | -0.06496 | 0.057233 | -0.58043 | -0.47836 | -0.28112 | -0.0566 | 0.147074 | -0.84769 | -0.24234 |
| 24.30556 | -0.3338 | 0.003216 | -0.22823 | -0.37839 | -0.22827 | -0.58434 | -0.35202 | 0.057781 | 0.214466 | -0.05403 | 0.147693 | -1.55969 | -0.05539 | -0.51856 | 0.083296 | 0.139006 | -0.79016 | -0.26864 |
| 24.31944 | -0.05735 | 0.042645 | -0.01932 | -0.16465 | -0.19007 | -0.46038 | -0.25675 | 0.054729 | 0.231753 | -0.11079 | 0.211961 | -2.59545 | 0.092428 | -0.85084 | 0.057093 | 0.005075 | -0.55265 | -0.28711 |
| 24.33333 | 0.387719 | 0.084829 | 0.264968 | 0.233527 | -0.08111 | -0.1549 | 0.005301 | 0.105965 | 0.246206 | 0.01631 | 0.360845 | -3.04394 | 0.198626 | -0.71355 | 0.060836 | -0.04324 | -0.20583 | -0.21212 |
| 24.34722 | 0.889692 | 0.229923 | 0.374672 | 0.701216 | 0.013743 | 0.296511 | 0.229568 | 0.121556 | 0.221156 | 0.284054 | 0.506172 | -2.01916 | 0.05172 | -0.07582 | 0.064188 | 0.037262 | 0.289849 | -0.14896 |
| 24.36111 | 0.947634 | 0.224996 | 0.219131 | 0.844938 | 0.061884 | 0.562895 | 0.281114 | 0.101761 | 0.110536 | 0.056133 | 0.145415 | -0.16384 | -0.37301 | 0.352986 | -0.29337 | -0.11841 | 0.105062 | -0.49698 |
| 24.375 | 0.410034 | -0.10741 | 0.020044 | 0.597733 | 0.108397 | 0.541775 | 0.262073 | 0.226503 | 0.021276 | -1.20124 | -1.09544 | 2.021268 | -1.3896 | -0.11915 | -1.36814 | -0.74947 | -1.19834 | -1.62795 |
| 24.38889 | 0.536114 | -0.14982 | 0.169644 | 0.633513 | 0.045757 | 0.673571 | 0.330931 | 0.749792 | 0.391433 | -2.69169 | -2.53019 | 4.090482 | -2.35763 | -1.09415 | -2.50341 | -1.32373 | -2.44203 | -2.81091 |
| 24.40278 | 1.294981 | 0.301183 | 0.662627 | 0.696385 | 0.024942 | 0.4138 | 0.454692 | 1.247468 | 1.115971 | -2.57715 | -2.43397 | 3.350465 | -1.94867 | -1.22711 | -2.29489 | -0.97597 | -1.94634 | -2.59776 |
| 24.41667 | 1.447591 | 0.636736 | 0.885245 | 0.55308 | 0.019995 | 0.142919 | 0.529859 | 0.977333 | 1.182314 | -1.14268 | -0.97949 | 0.801079 | -0.65411 | -0.45107 | -1.07952 | -0.42174 | -0.55853 | -1.44274 |
| 24.43056 | 1.328212 | 0.571237 | 0.63992 | 0.666902 | 0.027009 | -0.17851 | 0.44735 | 0.437165 | 0.663797 | 0.091817 | 0.154339 | -1.6145 | -0.14036 | -0.04635 | -0.17389 | -0.22007 | 0.118086 | -0.56473 |
| 24.44444 | 0.469889 | 0.143441 | 0.193111 | 0.557936 | 0.057492 | -0.6021 | 0.218454 | 0.089569 | 0.13773 | 0.556333 | 0.380983 | -4.13326 | -0.42618 | -0.31389 | 0.08362 | -0.14656 | -0.49297 | -0.07298 |
| 24.45833 | -0.02693 | 0.077451 | -0.07328 | 0.176597 | 0.090022 | -0.53327 | 0.076472 | -0.01724 | -0.11365 | 0.359656 | 0.290687 | -5.16106 | -0.59648 | -0.25818 | 0.287756 | -0.00122 | -1.02949 | 0.294332 |
| 24.47222 | 0.757167 | 1.034228 | -0.12801 | -0.25805 | 0.112483 | 0.208879 | 0.104091 | 0.079407 | 0.30796 | 0.330777 | 0.436566 | -2.52107 | 0.219669 | 0.606671 | 0.894433 | 0.441517 | -0.28199 | 0.701146 |
| 24.48611 | 2.174009 | 2.322397 | 0.045715 | -0.34535 | 0.005506 | 1.0888 | 0.184791 | 0.738205 | 1.161313 | 0.924163 | 0.999621 | 1.723408 | 1.32833 | 1.887203 | 1.840553 | 1.267019 | 1.467155 | 1.412092 |
| 24.5 | 2.803076 | 2.782147 | 1.001321 | 0.212431 | 0.080155 | 1.0824 | 0.519617 | 2.439106 | 1.90795 | 2.321274 | 1.881805 | 7.832901 | 2.434439 | 2.552777 | 2.374683 | 2.829471 | 2.855563 | 2.019426 |
| 24.51389 | 2.653778 | 2.619539 | 2.489366 | 1.378386 | 0.359825 | 0.949884 | 1.112099 | 4.933286 | 2.729504 | 3.528078 | 1.93396 | 9.812663 | 2.420956 | 1.739543 | 1.722113 | 5.380342 | 3.559551 | 1.727203 |
| 24.52778 | 2.015287 | 2.102757 | 2.412343 | 1.871836 | 0.462461 | 0.203807 | 0.912852 | 6.232429 | 3.461202 | 3.801985 | 1.167142 | 6.483045 | 2.655872 | 0.787515 | 1.174418 | 6.135345 | 4.030638 | 1.261556 |
| 24.54167 | 1.525606 | 1.690997 | 1.525858 | 1.423272 | 0.475325 | -0.12154 | 0.555734 | 4.964117 | 2.774143 | 2.70584 | 0.451753 | 2.02177 | 1.782076 | 0.110502 | 0.72737 | 5.835185 | 2.814928 | 0.512865 |
| 24.55556 | 0.537652 | 1.260037 | 0.966465 | 0.499618 | 0.466485 | -0.37731 | 0.360036 | 2.357789 | 1.433037 | 0.546864 | -0.39351 | -2.18554 | 0.312775 | -0.96671 | 0.065344 | 3.664457 | 0.610599 | -0.23958 |
| 24.56944 | -0.88341 | 0.572073 | 0.372952 | -0.31683 | 0.446338 | -0.74316 | 0.102781 | 0.484741 | 0.35433 | -1.13514 | -1.06452 | -6.02113 | -0.9619 | -1.9625 | -0.591 | 1.032973 | -0.99463 | -0.55977 |
| 24.58333 | -1.94306 | 0.155805 | -0.12002 | -0.95455 | 0.356849 | -1.11955 | -0.21979 | -1.05149 | -0.69916 | -2.06035 | -1.27165 | -8.31351 | -1.76223 | -1.75918 | -0.73385 | -1.0238 | -2.09987 | -0.59245 |
| 24.59722 | -1.93219 | 0.369407 | -0.48719 | -1.35426 | 0.393169 | -0.95328 | -0.36795 | -2.66888 | -1.51987 | -2.56985 | -0.88213 | -9.48806 | -2.1028 | -0.25043 | -0.55945 | -2.41359 | -2.54153 | -0.62778 |
| 24.61111 | -0.92155 | 0.692274 | -0.48643 | -1.39464 | 0.578955 | -0.17894 | -0.28807 | -4.62757 | -2.31263 | -2.68801 | -0.34027 | -7.32307 | -2.15451 | 1.653005 | -0.54524 | -3.39374 | -2.55893 | -0.73861 |
| 24.625 | 0.142799 | 0.776588 | -0.1605 | -0.55287 | 0.939982 | 0.550465 | -0.02792 | -5.50187 | -2.36356 | -2.08781 | 0.267917 | -1.68337 | -1.62152 | 2.612631 | -0.64237 | -3.0967 | -1.95292 | -0.47308 |
| 24.63889 | 1.519901 | 0.790106 | 0.289722 | 0.463682 | 1.095528 | 0.613386 | 0.036542 | -4.51227 | -1.38781 | -0.10766 | 0.545097 | 5.151437 | -0.24185 | 1.777665 | -0.39038 | -2.1874 | -1.14204 | -0.08654 |
| 24.65278 | 2.203613 | 0.875554 | 0.630331 | 0.577917 | 0.764265 | 0.2768 | 0.222334 | -1.43581 | -0.20158 | 2.453267 | 1.033729 | 5.776926 | 0.793692 | 0.206798 | 0.02986 | -1.00756 | -0.34235 | 0.177768 |
| 24.66667 | 1.415069 | 0.9299 | 0.598957 | 0.051062 | 0.278078 | -0.19721 | 0.116637 | 1.613226 | 0.649089 | 3.801992 | 1.44047 | 1.462573 | 1.498736 | -0.79988 | 0.634797 | 0.173357 | 0.145746 | 0.343264 |
| 24.68056 | 0.43634 | 0.684975 | 0.69447 | -0.3849 | -0.04019 | -0.44857 | 0.126677 | 3.848174 | 1.277044 | 2.779902 | 1.135411 | -3.27321 | 1.646284 | -1.36559 | 1.15177 | 1.361461 | 0.310548 | 0.632616 |
| 24.69444 | -0.51862 | 0.228651 | 1.032723 | -0.58803 | -0.18969 | -0.667 | 0.202312 | 4.569364 | 1.871749 | 0.800923 | -0.2927 | -6.39963 | 1.944441 | -1.7501 | 1.187042 | 1.699592 | 0.106104 | 1.038171 |
| 24.70833 | -1.66663 | -0.14237 | 0.647083 | -0.66131 | -0.21301 | -1.09846 | -0.26703 | 4.14809 | 2.034364 | -0.52913 | -0.63851 | -7.30993 | 1.451225 | -1.1112 | 0.891147 | 1.524127 | -0.01997 | 1.139833 |

| | | | | | | | | | | | | | | | | | | |
|---|---|---|---|---|---|---|---|---|---|---|---|---|---|---|---|---|---|---|
| 24.72222 | -2.43343 | -0.1337 | 0.394991 | -0.75108 | -0.20912 | -0.85744 | -0.22818 | 1.453157 | 0.904435 | -0.66062 | -0.12198 | -5.95788 | 0.609403 | 0.516233 | 0.621542 | 0.911329 | -0.29174 | 0.767604 |
| 24.73611 | -1.68532 | 0.581765 | 0.109847 | -0.78864 | 0.012598 | 0.059013 | 0.096581 | -1.28103 | -0.34693 | -0.99652 | 0.28301 | -4.16462 | -0.24724 | 1.800992 | 0.102519 | 0.501757 | -0.5512 | 0.078556 |
| 24.75 | -0.91891 | 1.230609 | -0.11532 | -0.48713 | 0.076428 | 0.771863 | 0.449873 | -3.81869 | -1.68059 | -1.67515 | 0.157598 | -2.73229 | -1.31082 | 1.257798 | -0.66127 | 0.039463 | -0.58233 | -0.34722 |
| 24.76389 | -0.07317 | 2.461897 | -0.41131 | -0.21927 | 0.067713 | 0.741326 | 0.390507 | -5.17067 | -2.19849 | -1.97034 | -0.56408 | -1.3334 | -1.9051 | -0.34652 | -1.1141 | -0.79724 | -0.63374 | -0.41983 |
| 24.77778 | -0.06477 | 1.878733 | -0.63686 | -0.16511 | -0.01762 | 0.003066 | -0.20293 | -4.41195 | -1.64874 | -1.96651 | -0.91019 | -0.585 | -1.57958 | -1.98459 | -0.97962 | -1.52882 | -0.82075 | -0.30571 |
| 24.79167 | -0.0857 | 0.092789 | -0.82089 | 0.09325 | -0.12433 | -0.87859 | -0.58185 | -2.6198 | -0.91754 | -1.31326 | -0.57564 | 0.066636 | -0.8189 | -2.76926 | -0.41767 | -1.02576 | -0.88894 | -0.18122 |
| 24.80556 | -0.13751 | -1.49894 | -0.65338 | 0.267575 | -0.25607 | -1.41302 | -0.8791 | -1.08132 | -0.61191 | -0.98135 | -0.1684 | 0.348956 | -0.26748 | -2.24201 | 0.000931 | -0.54283 | -0.74302 | -0.02079 |
| 24.81944 | -0.55789 | -2.25254 | -0.21679 | -0.07162 | -0.33646 | -1.83887 | -0.92328 | -0.35213 | -0.38704 | -0.43722 | 0.27551 | -0.05733 | 0.028716 | -1.43288 | 0.133832 | -0.04804 | -0.83888 | 0.107761 |
| 24.83333 | -1.40753 | -2.01885 | 0.273672 | -0.79269 | -0.39714 | -1.83311 | -0.66106 | 0.023969 | -0.06706 | 0.598847 | 0.201284 | -0.3205 | 0.172734 | -1.14798 | 0.208607 | 0.013429 | -0.95328 | 0.282676 |
| 24.84722 | -2.92052 | -1.58198 | 0.417051 | -1.88029 | -0.54376 | -1.84744 | -0.2995 | 0.092056 | 0.123 | 0.428328 | -0.94206 | -0.39004 | -0.02564 | -1.41706 | 0.073533 | 0.043289 | -0.76873 | 0.280452 |
| 24.86111 | -3.9675 | -1.53407 | 0.234395 | -2.58119 | -1.1374 | -1.55761 | -0.0762 | -0.02575 | 0.099772 | -0.17886 | -1.688 | -0.12637 | 0.046678 | -1.91303 | 0.095306 | -0.15621 | -0.33601 | 0.27724 |
| 24.875 | -3.57249 | -1.26682 | -0.34614 | -2.40029 | -1.5566 | -0.95251 | -0.03486 | -0.14188 | -0.01792 | -1.00409 | -1.19096 | 0.616291 | -0.22801 | -2.04736 | 0.341548 | -0.3581 | -0.01474 | 0.52715 |
| 24.88889 | -2.78739 | -1.06799 | -0.70919 | -1.70392 | -1.77666 | -0.29913 | -0.16446 | -0.07632 | -0.15306 | -1.1836 | -0.77331 | 0.82565 | -0.10085 | -1.22171 | 0.356103 | -0.3563 | 0.115316 | 0.593523 |
| 24.90278 | -1.69567 | -0.92658 | -0.73434 | -1.18123 | -1.6084 | 0.218354 | -0.23523 | 0.087324 | -0.24265 | -1.13154 | -0.09739 | 0.429709 | 0.3262 | 0.267945 | 0.239354 | -0.2226 | 0.060182 | 0.38371 |
| 24.91667 | -0.76203 | -0.71263 | -0.60373 | -0.35157 | -0.91282 | 0.824537 | -0.15432 | 0.22897 | -0.22409 | -0.79747 | 0.503693 | 0.391925 | 0.558644 | 1.473206 | 0.447319 | -0.04542 | 0.218559 | 0.415189 |
| 24.93056 | 0.208532 | -0.49672 | -0.51703 | 0.579906 | -0.18675 | 1.231493 | 0.010379 | 0.281416 | -0.13793 | -0.47368 | 1.002349 | 0.654384 | 0.500272 | 2.13748 | 0.65413 | 0.353768 | 0.49887 | 0.497701 |
| 24.94444 | 0.539404 | -0.36956 | -0.54244 | 0.948274 | 0.257545 | 1.13879 | 0.11063 | 0.199117 | 0.01341 | -0.31381 | 1.070219 | 0.821106 | 0.260157 | 2.099401 | 0.528538 | 0.8175 | 0.892397 | 0.4799 |
| 24.95833 | 0.511907 | -0.2045 | -0.65162 | 0.77023 | 0.275511 | 1.028415 | 0.143367 | 0.161891 | 0.148872 | -0.10346 | 0.514517 | 0.840262 | 0.078631 | 1.544064 | 0.136336 | 1.16358 | 0.927466 | 0.183279 |
| 24.97222 | 0.085347 | -0.25583 | -0.66976 | 0.401338 | -0.11916 | 0.903686 | 0.137707 | 0.142056 | 0.232123 | 0.240464 | -0.23592 | 0.425622 | -0.44348 | 0.808415 | -0.59615 | 1.138645 | 0.344263 | -0.59988 |
| 24.98611 | -1.23229 | -0.89791 | -0.73916 | -0.26686 | -0.80744 | -0.01672 | -0.08108 | -0.15751 | -0.0883 | -0.42558 | -1.25779 | -1.32282 | -1.24128 | -0.48413 | -1.92472 | -0.27855 | -1.10968 | -1.85809 |
| 25 | -2.99854 | -1.44561 | -0.84955 | -1.86838 | -1.52522 | -1.28775 | -0.58704 | -0.54685 | -0.73956 | -2.16912 | -2.31691 | -2.93545 | -2.36851 | -2.80792 | -3.41991 | -1.96272 | -3.30478 | -2.79637 |
| 25.01389 | -3.38441 | -1.01915 | -0.74576 | -2.83269 | -1.33493 | -2.09005 | -1.06059 | -0.5189 | -1.24651 | -3.25574 | -2.48071 | -2.624 | -2.52424 | -4.9802 | -3.61446 | -2.65798 | -4.46331 | -2.78376 |
| 25.02778 | -2.24594 | -0.56477 | -0.53526 | -1.76169 | -0.65 | -2.5941 | -1.13787 | -0.17519 | -1.29327 | -3.02996 | -1.39639 | -1.11139 | -1.71602 | -4.67765 | -2.80819 | -1.2099 | -3.51194 | -1.67465 |
| 25.04167 | -1.3475 | -0.51147 | -0.37561 | -0.9951 | -0.36319 | -2.45486 | -0.84218 | 0.104075 | -0.92985 | -2.11909 | -0.23203 | -0.21512 | -0.76822 | -2.65105 | -1.69837 | 0.260346 | -1.41692 | -0.6725 |
| 25.05556 | -0.61103 | -0.43521 | -0.1831 | -0.52603 | -0.25932 | -1.31174 | -0.51907 | 0.240716 | -0.46864 | -0.99198 | 0.248216 | 0.163892 | -0.10959 | -0.86657 | -0.66269 | 0.830292 | 0.085185 | 0.055367 |
| 25.06944 | 0.223454 | -0.07633 | 0.04244 | -0.24124 | -0.08348 | -0.26586 | -0.1932 | 0.214375 | -0.11983 | -0.02341 | 0.268643 | 0.453052 | 0.419476 | -0.15219 | 0.012531 | 0.869575 | 1.054066 | 0.599648 |
| 25.08333 | 1.191909 | 0.457019 | 0.297115 | 0.042764 | 0.068634 | 0.296351 | 0.161448 | 0.19201 | 0.195213 | 1.085302 | 0.129644 | 0.652254 | 0.658367 | -0.3046 | 0.557483 | 0.781294 | 1.719959 | 0.759549 |
| 25.09722 | 1.834703 | 0.99722 | 0.520507 | 0.269906 | 0.125948 | 0.576384 | 0.441319 | 0.170393 | 0.495458 | 1.792567 | -1.03958 | 0.884306 | 0.370125 | -0.58408 | 1.091044 | 0.175494 | 2.240109 | 0.677354 |
| 25.11111 | 1.618925 | 1.017006 | 0.568478 | 0.555797 | 0.233176 | 0.665144 | 0.513073 | 0.256411 | 0.756736 | 1.330868 | -0.65435 | 0.417176 | 0.085726 | -0.0848 | 1.205706 | -0.53994 | 1.942575 | 0.437345 |
| 25.125 | 1.191832 | 0.577025 | 0.523207 | 0.897208 | 0.255193 | 0.575129 | 0.52935 | 0.257054 | 0.755948 | 1.074775 | -0.28739 | 0.160069 | 0.010959 | 1.184778 | 1.301396 | -0.17223 | 1.644581 | 0.427281 |
| 25.13889 | 0.700708 | 0.143768 | 0.361931 | 1.103176 | 0.195765 | 0.436579 | 0.398583 | 0.153522 | 0.374235 | 1.32398 | 0.667357 | 0.536448 | 0.416528 | 2.019428 | 1.505436 | 0.473442 | 1.433859 | 0.817435 |
| 25.15278 | 0.200111 | 0.002693 | 0.065436 | 0.979472 | 0.139961 | 0.40735 | 0.174008 | 0.100713 | 0.007449 | 1.34709 | 0.942289 | 0.93774 | 0.655724 | 2.019898 | 1.54229 | 0.822866 | 0.714137 | 1.063563 |
| 25.16667 | 0.305461 | 0.088783 | -0.17302 | 0.855595 | 0.163157 | 0.655922 | 0.036249 | 0.128347 | 0.006528 | 1.110118 | 0.741607 | 0.93273 | 0.458747 | 1.775094 | 1.387706 | 0.344566 | -0.2428 | 0.825895 |
| 25.18056 | 0.884684 | 0.169805 | -0.18875 | 0.795192 | 0.414995 | 1.035515 | 0.111137 | 0.136986 | 0.422315 | 0.984203 | 0.691784 | 0.664916 | 0.299208 | 1.586996 | 0.986985 | 0.002955 | -0.44557 | 0.388287 |
| 25.19444 | 1.504437 | 0.19871 | 0.063818 | 0.860982 | 0.754407 | 1.262329 | 0.309973 | 0.140552 | 0.91974 | 1.092877 | 0.9295 | 0.498928 | 0.353053 | 1.532278 | 0.466596 | -0.14212 | 0.458067 | 0.148291 |
| 25.20833 | 1.833442 | 0.331373 | 0.347443 | 0.887722 | 0.960305 | 1.453484 | 0.469518 | 0.166584 | 1.082495 | 1.464412 | 0.981103 | 0.498012 | 0.598223 | 1.490513 | 0.19845 | 0.06169 | 1.419775 | 0.197577 |
| 25.22222 | 1.161316 | 0.340285 | 0.395452 | 0.707342 | 0.631261 | 1.19279 | 0.419949 | 0.177754 | 0.816303 | 1.211144 | 0.873144 | 0.635882 | 0.515118 | 1.112745 | 0.214673 | 0.630799 | 1.051312 | 0.35726 |
| 25.23611 | 0.06531 | 0.298419 | 0.217358 | 0.168642 | 0.020961 | 0.880784 | 0.21692 | 0.169177 | 0.443721 | 0.539795 | 0.637363 | 0.690803 | 0.277131 | 0.511964 | 0.2394 | 0.776903 | 0.058378 | 0.4587 |
| 25.25 | -0.55779 | 0.270571 | 0.008877 | -0.40692 | -0.43968 | 0.635398 | -0.01233 | 0.168415 | 0.18431 | -0.06899 | 0.392775 | 0.669896 | 0.059935 | 0.045097 | 0.191765 | 0.598685 | -0.81407 | 0.432268 |
| 25.26389 | -0.37569 | 0.127017 | -0.14396 | -0.51035 | -0.43117 | 0.375962 | -0.127 | 0.147168 | 0.130918 | -0.21359 | 0.049844 | 0.628864 | 0.176649 | -0.07014 | 0.201432 | 0.325201 | -0.80193 | 0.228526 |
| 25.27778 | 0.140035 | -0.09586 | -0.09192 | -0.11153 | -0.16346 | 0.193038 | -0.02356 | 0.116662 | 0.294995 | -0.11575 | -0.28505 | 0.450937 | 0.342196 | 0.170951 | 0.272923 | 0.024882 | 0.067082 | 0.03572 |
| 25.29167 | 0.671013 | -0.16233 | 0.157887 | 0.411511 | 0.100436 | 0.084594 | 0.232222 | 0.105822 | 0.452661 | 0.355245 | -0.33185 | 0.305432 | 0.475181 | 0.453955 | 0.294857 | -0.03152 | 1.092146 | -0.08997 |
| 25.30556 | 0.999952 | 0.074635 | 0.450919 | 0.677274 | 0.374939 | 0.006256 | 0.439228 | 0.167372 | 0.582736 | 0.712291 | -0.10976 | 0.263277 | 0.298853 | 0.464043 | 0.181527 | 0.311656 | 1.456513 | -0.1109 |
| 25.31944 | 0.826159 | 0.309122 | 0.42158 | 0.499147 | 0.382438 | -0.14261 | 0.462839 | 0.214168 | 0.557308 | 0.612996 | 0.160377 | 0.335219 | 0.126248 | 0.365145 | 0.122964 | 0.650702 | 0.938372 | 0.006072 |
| 25.33333 | 0.179379 | 0.229159 | 0.164666 | 0.12884 | 0.111486 | -0.41787 | 0.298148 | 0.191406 | 0.364324 | 0.151015 | 0.296897 | 0.407423 | 0.051852 | 0.127975 | 0.12538 | 0.778262 | 0.190886 | 0.133767 |
| 25.34722 | -0.13624 | 0.090723 | -0.09752 | 0.022285 | -0.11201 | -0.47114 | 0.14028 | 0.145724 | 0.205963 | -0.30294 | 0.231127 | 0.23634 | -0.05122 | -0.13415 | 0.013107 | 0.406272 | -0.12739 | -0.04013 |
| 25.36111 | 0.083954 | 0.050509 | -0.15281 | 0.228112 | -0.04724 | -0.26334 | 0.097863 | 0.140495 | 0.319285 | -0.13128 | 0.287279 | -0.04273 | -0.00259 | -0.09897 | -0.06676 | 0.114035 | 0.264006 | -0.27826 |
| 25.375 | 0.448643 | 0.077826 | 0.068426 | 0.447443 | 0.134135 | -0.05251 | 0.16093 | 0.137175 | 0.602358 | 0.702779 | 0.699689 | -0.12711 | 0.307147 | 0.15922 | -0.04933 | 0.187602 | 0.899876 | -0.26771 |
| 25.38889 | 0.51694 | 0.03276 | 0.298829 | 0.50762 | 0.318276 | 0.003393 | 0.245421 | 0.055083 | 0.678699 | 1.402191 | 1.12055 | -0.04119 | 0.509904 | 0.202115 | -0.09476 | 0.361887 | 0.995553 | -0.22381 |
| 25.40278 | 0.285019 | -0.14063 | 0.44305 | 0.339851 | 0.476898 | -0.04447 | 0.258388 | -0.03834 | 0.361092 | 1.495826 | 1.267611 | 0.080646 | 0.346546 | 0.071091 | -0.10018 | 0.531216 | 0.373991 | -0.18225 |
| 25.41667 | 0.038533 | -0.02342 | 0.384642 | 0.076961 | 0.502092 | 0.058071 | 0.214894 | -0.00128 | 0.080855 | 1.082657 | 1.044993 | 0.175148 | 0.153225 | -0.10057 | -0.06259 | 0.569982 | -0.31081 | -0.0378 |
| 25.43056 | 0.017454 | 0.307116 | 0.137895 | -0.12492 | 0.34002 | 0.312018 | 0.128124 | 0.14107 | -0.07499 | 0.282748 | 0.709369 | 0.280844 | 0.012484 | -0.12425 | -0.00137 | 0.542131 | -0.86751 | 0.135431 |
| 25.44444 | 0.166518 | 0.306117 | -0.13711 | -0.00763 | 0.120721 | 0.490799 | 0.055975 | 0.174025 | -0.11778 | -0.34473 | 0.401501 | 0.304188 | 0.097104 | 0.099773 | 0.119482 | 0.410677 | -1.20996 | 0.188719 |
| 25.45833 | 0.400047 | 0.069859 | -0.17233 | 0.283285 | -0.03451 | 0.57681 | 0.080053 | 0.124829 | 0.144195 | -0.23983 | 0.204922 | 0.31241 | 0.427293 | 0.613127 | 0.309547 | 0.25816 | -0.85949 | 0.164688 |
| 25.47222 | 0.665829 | 0.056698 | 0.071195 | 0.497138 | -0.03976 | 0.704889 | 0.239586 | 0.116901 | 0.659103 | 0.551206 | 0.24834 | 0.454091 | 0.806347 | 1.217527 | 0.477146 | 0.261431 | 0.249138 | 0.145123 |
| 25.48611 | 1.036123 | 0.396124 | 0.475951 | 0.592923 | 0.267303 | 0.967611 | 0.466051 | 0.128615 | 0.925844 | 1.418319 | 0.589557 | 1.055553 | 0.943738 | 2.058376 | 0.659513 | 0.410786 | 1.566094 | 0.163728 |
| 25.5 | 1.567776 | 0.925777 | 1.294122 | 0.794166 | 0.881738 | 1.35293 | 0.914344 | 0.176978 | 0.990545 | 1.997254 | 1.053872 | 2.430554 | 1.079113 | 3.398228 | 0.925985 | 0.760443 | 2.356941 | 0.3152 |
| 25.51389 | 2.228082 | 1.319237 | 2.35248 | 1.531138 | 1.337287 | 1.481676 | 1.095663 | 0.17916 | 0.845689 | 2.336558 | 1.46728 | 4.763895 | 1.038288 | 4.123533 | 1.136564 | 1.040189 | 2.517299 | 0.598537 |
| 25.52778 | 2.514291 | 1.877196 | 2.648444 | 1.895733 | 0.835454 | 1.131179 | 1.123283 | 0.661621 | 0.732336 | 2.284854 | 1.274793 | 6.113056 | 1.04516 | 2.723082 | 1.135648 | 1.004529 | 2.321254 | 0.639366 |
| 25.54167 | 1.585759 | 2.04212 | 1.98371 | 1.450296 | 0.315823 | 0.731895 | 0.910804 | 1.3267 | 0.653234 | 1.783897 | 0.451721 | 5.461313 | 0.989595 | 1.219423 | 0.763374 | 0.706777 | 1.266168 | 0.571549 |
| 25.55556 | 0.03259 | 1.378856 | 0.933613 | 0.211758 | -0.21142 | 0.017297 | 0.291966 | 1.60753 | 0.3065 | 0.671993 | 0.075993 | 3.043979 | 0.782014 | 0.045331 | 0.435901 | 0.662199 | -0.12291 | 0.766439 |
| 25.56944 | -1.75796 | 0.79119 | -0.02063 | -1.20005 | -0.90959 | -0.75098 | -0.23019 | 1.632759 | -0.15226 | -0.30379 | -0.18966 | 0.357676 | 0.383213 | -0.96661 | 0.275851 | 0.834104 | -1.15288 | 1.010223 |
| 25.58333 | -2.33721 | 0.476762 | -0.8547 | -1.74601 | -1.07868 | -1.18049 | -0.47432 | 1.710194 | -0.43518 | -0.47308 | -0.38904 | -1.65801 | 0.037522 | -1.64172 | 0.121912 | 1.182884 | -1.98511 | 1.266443 |
| 25.59722 | -1.95056 | 0.127058 | -1.22655 | -1.70117 | -0.7581 | -0.93166 | -0.37663 | 1.714094 | -0.30586 | 0.273505 | -0.49493 | -2.09461 | -0.14413 | -1.88986 | -0.10134 | 1.210283 | -2.37041 | 1.456377 |
| 25.61111 | -1.49554 | -0.50126 | -0.59484 | -1.71154 | -0.35843 | -0.88713 | -0.0811 | 1.360976 | 0.162845 | 0.778735 | -0.58671 | -2.53375 | -0.31387 | -1.73619 | -0.38046 | 1.127231 | -1.36523 | 1.341107 |
| 25.625 | -0.17486 | -1.11107 | -0.01646 | -0.61111 | 0.079566 | -0.79826 | 0.041275 | 0.982618 | 0.897096 | 1.148102 | -0.17924 | -3.81004 | -0.55907 | -0.70323 | -0.45076 | 1.13264 | 0.188893 | 0.797136 |
| 25.63889 | 1.382683 | -1.51184 | 0.149768 | 0.600003 | 0.465082 | -0.31095 | 0.104811 | -0.30546 | 1.262923 | 1.299465 | 0.512755 | -3.82291 | -0.5439 | 0.930427 | -0.27984 | 0.521714 | 1.128759 | 0.003355 |
| 25.65278 | 2.226708 | -1.72326 | 0.477115 | 1.284704 | 0.734293 | -0.0532 | 0.050998 | -1.56758 | 0.856686 | 0.533336 | 1.071983 | -2.97059 | -0.428 | 2.108568 | -0.10324 | -0.91082 | 0.790142 | -0.6084 |
| 25.66667 | 1.356588 | -1.28419 | -0.07701 | 0.888052 | 0.667966 | 0.463153 | -0.03498 | -2.11427 | -0.00145 | -1.20441 | 0.595738 | -1.33494 | -0.18589 | 2.904605 | -0.02411 | -0.48926 | 0.169234 | -0.469 |
| 25.68056 | -0.08699 | -0.55151 | -0.89666 | -0.13523 | 0.297383 | 0.130657 | -0.10118 | -2.36923 | -0.68546 | -2.47475 | -0.00025 | 1.246317 | 0.289707 | 1.766095 | 0.087936 | 0.153931 | -0.49844 | 0.228677 |
| 25.69444 | -1.15214 | 0.14423 | -1.14764 | -0.59336 | -0.03675 | 0.01042 | 0.007988 | -2.03951 | -1.07165 | -2.38278 | -0.30104 | 4.431362 | 0.691568 | -0.62074 | 0.164474 | 0.759984 | -1.16217 | 1.127228 |
| 25.70833 | -2.06857 | 0.920606 | -0.57991 | -1.1073 | -0.27762 | -0.28015 | 0.082102 | -1.12289 | -1.18536 | -1.83444 | -0.96206 | 3.085411 | 0.28437 | -1.94725 | 0.197716 | 0.91763 | -1.76212 | 1.981719 |
| 25.72222 | -2.68903 | 1.748553 | 0.083695 | -1.63112 | -0.94846 | -0.91069 | 0.060801 | -0.39362 | -0.86468 | -0.10597 | -1.38151 | 0.026812 | 0.530916 | -2.36302 | 0.102735 | 0.698928 | -1.99245 | 2.512657 |
| 25.73611 | -2.32724 | 1.891364 | 0.582013 | -1.72116 | -1.2604 | -0.85205 | 0.28127 | 0.190357 | -0.07961 | 1.770284 | -1.02051 | -2.60141 | 0.751153 | -1.81827 | -0.18055 | -0.43554 | -1.14813 | 2.197215 |
| 25.75 | -0.60671 | 1.571819 | 0.712489 | -1.03933 | -0.59769 | -1.5986 | 0.279464 | 0.613925 | 0.688542 | 3.800635 | 0.244103 | -3.61382 | 0.353979 | -1.0237 | -0.35827 | 0.009019 | -0.13303 | 0.17981 |
| 25.76389 | 0.664145 | 0.832351 | 0.366444 | -0.58191 | 0.137907 | -1.70155 | 0.160744 | 0.694837 | 1.096537 | 3.955823 | 1.308062 | -2.97053 | -0.43057 | -0.39119 | -0.22812 | 0.439434 | 0.485879 | -0.36549 |
| 25.77778 | 1.283243 | 0.041913 | -0.32626 | -0.27892 | 0.497679 | -1.25623 | -0.06787 | 0.375624 | 1.002178 | 1.505335 | 1.381345 | -3.33151 | -1.21265 | 0.287949 | 0.170803 | 0.136488 | 0.02872 | -0.24717 |
| 25.79167 | 0.602953 | -0.79094 | -0.76627 | -0.34362 | 0.839484 | -0.78998 | -0.10407 | 0.008977 | 0.487922 | -0.66299 | 0.679619 | -3.32089 | -1.32258 | 0.310274 | 0.238644 | -0.31204 | -0.57444 | -0.00581 |
| 25.80556 | -0.36961 | -1.57793 | -0.85386 | -0.70253 | 0.737481 | -0.74714 | -0.55642 | -0.38569 | -0.25385 | -2.52669 | -0.29966 | -2.75403 | -1.48462 | -0.64335 | -0.32893 | 0.00063 | -0.8408 | -0.07335 |
| 25.81944 | -0.73838 | -1.97818 | -1.19061 | -0.74642 | 0.621102 | -1.09797 | -1.21958 | -0.65091 | -1.12756 | -4.00797 | -1.55821 | -1.77694 | -1.52045 | -1.5879 | -1.02084 | 0.188252 | -1.31125 | -0.2683 |
| 25.83333 | -0.7643 | -1.33694 | -1.16327 | -0.68384 | 0.265709 | -0.99314 | -1.01139 | -0.65046 | -1.62508 | -3.52919 | -2.11933 | -0.83262 | -0.91664 | -1.82189 | -1.0732 | 0.182445 | -1.79024 | -0.35569 |

| | | | | | | | | | | | | | | | | | | |
|---|---|---|---|---|---|---|---|---|---|---|---|---|---|---|---|---|---|---|
| 25.84722 | -0.94765 | -0.54085 | -0.98834 | -0.55704 | -0.15447 | -0.71911 | -0.5555 | -0.3142 | -1.30261 | -3.11106 | -2.47111 | -0.23321 | -0.42481 | -1.69127 | -0.68073 | -0.28486 | -1.96113 | -0.19915 |
| 25.86111 | -1.24688 | -0.15639 | -1.05229 | -0.51482 | -0.87142 | -0.7535 | -0.40131 | -0.0344 | -0.68777 | -1.64749 | -2.34986 | 0.072316 | -0.07306 | -1.55335 | -0.15309 | -0.70517 | -1.68111 | 0.02123 |
| 25.875 | -1.2379 | 0.031406 | -0.94235 | -0.68749 | -1.88707 | -0.71913 | -0.39996 | 0.070931 | -0.08719 | -0.66825 | -1.68854 | 0.074176 | 0.124439 | -1.16115 | 0.32803 | -0.61921 | -0.86079 | 0.174998 |
| 25.88889 | -0.90404 | 0.182859 | -0.40876 | -0.82207 | -1.2795 | -0.49219 | -0.45184 | 0.066272 | 0.30515 | -0.25521 | -0.84272 | 0.139715 | 0.113983 | -0.6264 | 0.311777 | -0.0019 | 0.03524 | 0.226623 |
| 25.90278 | -0.70597 | 0.31591 | -0.04769 | -1.01778 | -0.49136 | -0.03162 | -0.40769 | 0.009618 | 0.367269 | -0.11851 | -0.1988 | 0.30133 | -0.078 | -0.2706 | 0.069734 | 0.560563 | 0.71156 | 0.228632 |
| 25.91667 | -0.67536 | 0.403899 | 0.284656 | -0.97883 | 0.310025 | 0.558551 | -0.18834 | -0.0755 | 0.244734 | -0.28602 | 0.349988 | 0.311651 | -0.34124 | 0.056865 | -0.19309 | 0.70277 | 1.02202 | 0.260743 |
| 25.93056 | -0.47228 | 0.392175 | 0.37219 | -0.54558 | 0.792564 | 1.086675 | 0.050378 | -0.17835 | 0.154678 | -0.23848 | 0.675408 | 0.131009 | -0.62888 | 0.490127 | -0.20604 | 0.599789 | 1.076645 | 0.359026 |
| 25.94444 | 0.014907 | 0.374775 | 0.226815 | -0.09673 | 0.442375 | 1.362036 | 0.198839 | -0.24369 | 0.208469 | -0.05394 | 0.814469 | -0.13621 | -0.552 | 0.850736 | 0.003578 | 0.39475 | 1.097656 | 0.527581 |
| 25.95833 | 0.580041 | 0.40174 | 0.097175 | 0.289732 | 0.329029 | 1.316728 | 0.236251 | -0.22423 | 0.344979 | -0.19538 | 0.787737 | 0.527684 | 0.218006 | 1.029847 | 0.251593 | 0.573529 | 1.466837 | 0.860831 |
| 25.97222 | 0.957363 | 0.414046 | 0.174274 | 0.52191 | 0.148947 | 1.058191 | 0.199704 | -0.10967 | 0.447617 | -0.32744 | 0.552986 | 1.422309 | 0.950735 | 0.924666 | 0.329643 | 0.644366 | 1.980373 | 1.094431 |
| 25.98611 | 0.469503 | 0.19812 | 0.166995 | 0.202443 | -0.20122 | 0.38053 | -0.00606 | -0.05755 | 0.111084 | -0.33345 | -0.12987 | 1.199639 | 0.798368 | 0.065471 | -0.16892 | -0.2915 | 1.049762 | 0.589879 |
| 26 | -1.19222 | -0.33829 | -0.25381 | -0.87821 | -0.71668 | -0.7869 | -0.51619 | -0.2339 | -0.89156 | -1.14673 | -1.32132 | -0.87452 | -0.51128 | -1.52065 | -1.41031 | -2.19798 | -1.47985 | -0.7548 |
| 26.01389 | -3.09519 | -0.82459 | -0.59701 | -1.98401 | -1.34393 | -2.38771 | -1.04381 | -0.59708 | -2.20545 | -2.32544 | -1.85818 | -2.35308 | -1.69055 | -2.1144 | -1.9673 | -3.43386 | -3.75554 | -1.69742 |
| 26.02778 | -3.22818 | -0.78434 | -0.48659 | -1.97806 | -1.93374 | -2.16859 | -0.90999 | -0.77619 | -2.82273 | -2.26065 | -1.42673 | -3.00925 | -3.17409 | -1.024 | -1.07922 | -1.7772 | -2.62059 | -1.61368 |
| 26.04167 | -1.60725 | -0.52124 | -0.57545 | -1.56667 | -1.70016 | -1.26865 | -0.91487 | -0.48908 | -2.51006 | -1.30074 | -0.92099 | -2.59931 | -2.75513 | -0.14691 | -0.3387 | -0.53926 | -0.86127 | -1.19525 |
| 26.05556 | -0.50084 | -0.25227 | -1.01261 | -1.59353 | -0.75693 | -0.46246 | -1.71502 | -0.15186 | -1.429 | -0.76685 | -0.38493 | -1.51205 | -1.65803 | 0.076811 | -0.19213 | 0.271736 | -0.10228 | -0.69174 |
| 26.06944 | 0.034309 | -0.09581 | -0.83543 | -0.32548 | 0.357107 | -0.00413 | 0.069755 | 0.221291 | -0.02295 | 0.023595 | 0.238407 | 0.047831 | -0.42295 | 0.077092 | -0.13467 | 1.248899 | 0.142759 | -0.3091 |
| 26.08333 | 0.433752 | 0.129419 | -0.01342 | -0.20451 | 1.211424 | 0.203307 | -0.06517 | 0.397138 | 1.060041 | 0.756072 | 0.597447 | 0.935756 | 0.191007 | 0.035158 | -0.01939 | 1.795362 | 0.367986 | -0.02013 |
| 26.09722 | 0.767043 | 0.203033 | 0.259237 | 0.203178 | 1.437633 | 0.298441 | 0.411692 | 0.339398 | 1.550095 | 1.162547 | 0.719054 | 1.131476 | 0.455165 | 0.168073 | 0.116417 | 1.531557 | 0.579056 | 0.193015 |
| 26.11111 | 1.083652 | 0.050417 | 0.356004 | 0.428666 | 1.108031 | 0.31944 | 0.593217 | 0.216858 | 1.599291 | 1.308603 | 0.717625 | 1.386064 | 0.741477 | 0.442427 | 0.14897 | 0.909542 | 0.881954 | 0.294176 |
| 26.125 | 1.177437 | -0.13599 | 0.271443 | 0.698935 | 0.75187 | 0.255671 | 0.575514 | 0.129989 | 1.252194 | 1.231587 | 0.670536 | 2.095926 | 1.23851 | 0.510561 | 0.03144 | 0.434 | 1.346741 | 0.255601 |
| 26.13889 | 1.019922 | -0.08592 | 0.131319 | 0.768932 | 0.672365 | 0.135335 | 0.445798 | 0.144788 | 0.863329 | 0.935501 | 0.510421 | 2.698962 | 1.855417 | 0.523769 | -0.00126 | 0.178689 | 2.397319 | 0.258576 |
| 26.15278 | 0.91781 | -0.02562 | 0.07903 | 0.680606 | 0.736919 | 0.066172 | 0.274769 | 0.125098 | 0.569074 | 0.733429 | 0.420428 | 2.600809 | 2.244563 | 0.605758 | 0.158757 | 0.183737 | 2.720767 | 0.379706 |
| 26.16667 | 0.802784 | 0.065811 | 0.167332 | 0.546608 | 0.476337 | 0.097203 | 0.213998 | 0.101372 | 0.486534 | 0.977269 | 0.363174 | 1.819915 | 2.044822 | 0.640972 | 0.222447 | 0.324583 | 2.845132 | 0.423397 |
| 26.18056 | 0.754952 | 0.257188 | 0.413396 | 0.463675 | 0.184607 | 0.289333 | 0.371336 | 0.11356 | 0.564615 | 0.977926 | 0.34285 | 2.491088 | 1.49678 | 0.566122 | 0.124496 | 0.444646 | 2.559186 | 0.458297 |
| 26.19444 | 0.810469 | 0.474969 | 0.610212 | 0.473658 | 0.063532 | 0.649151 | 0.648532 | 0.166242 | 0.814046 | 0.841815 | 0.358286 | 1.890188 | 0.770033 | 0.494907 | 0.113413 | 0.354023 | 1.842256 | 0.500088 |
| 26.20833 | 0.809855 | 0.478277 | 0.581586 | 0.503266 | -0.15531 | 0.793051 | 0.692911 | 0.192629 | 1.112536 | 0.664308 | 0.423555 | 1.222275 | 0.300383 | 0.600174 | 0.22166 | 0.231039 | 1.059303 | 0.538324 |
| 26.22222 | 0.668148 | 0.294581 | 0.407755 | 0.558104 | -0.37023 | 0.627358 | 0.558344 | 0.202393 | 1.219479 | 0.441142 | 0.537193 | 0.294886 | -0.0069 | 0.765836 | 0.315421 | 0.370297 | 0.212333 | 0.484238 |
| 26.23611 | 0.56848 | 0.216757 | 0.269047 | 0.603735 | -0.37655 | 0.54744 | 0.431849 | 0.210924 | 1.099291 | 0.286315 | 0.614856 | -0.54341 | -0.20367 | 0.70896 | 0.2042 | 0.43303 | -0.43704 | 0.335611 |
| 26.25 | 0.521672 | 0.297031 | 0.17992 | 0.548217 | -0.1747 | 0.595992 | 0.257319 | 0.216471 | 0.818819 | 0.272417 | 0.550426 | -0.68147 | -0.19007 | 0.251948 | -0.15405 | 0.346245 | -0.50042 | 0.124619 |
| 26.26389 | 0.447812 | 0.28282 | 0.114287 | 0.435613 | 0.03061 | 0.60817 | 0.089021 | 0.159991 | 0.47422 | 0.426644 | 0.408458 | 0.103789 | 0.190434 | -0.18485 | -0.55015 | 0.249558 | 0.019753 | 0.031379 |
| 26.27778 | 0.419431 | 0.101858 | 0.156438 | 0.33768 | 0.148706 | 0.474866 | 0.053248 | 0.083339 | 0.296722 | 0.684523 | 0.318238 | 1.293537 | 0.635035 | -0.30511 | -0.71817 | 0.263063 | 0.886746 | 0.058507 |
| 26.29167 | 0.351669 | -0.11105 | 0.245751 | 0.293718 | 0.17726 | 0.275708 | 0.161945 | 0.037083 | 0.498862 | 0.832438 | 0.305251 | 1.521856 | 0.693158 | -0.33203 | -0.69761 | 0.408434 | 1.730894 | 0.129365 |
| 26.30556 | 0.037018 | -0.21676 | 0.14288 | 0.12784 | 0.12376 | 0.14126 | 0.203103 | 0.05978 | 0.818491 | 0.683893 | 0.316068 | 0.76903 | 0.334901 | -0.3008 | -0.70258 | 0.595149 | 1.942904 | 0.151103 |
| 26.31944 | -0.35858 | -0.05109 | -0.09312 | -0.12231 | -0.04946 | 0.092057 | 0.088907 | 0.152503 | 0.606492 | 0.344936 | 0.296438 | -0.55917 | -0.11037 | -0.24684 | -0.71165 | 0.715228 | 1.07049 | 0.154991 |
| 26.33333 | -0.60528 | 0.17161 | -0.19332 | -0.32302 | -0.3348 | 0.03443 | -0.05939 | 0.215 | -0.06737 | 0.014443 | 0.238853 | -2.15598 | -0.66889 | -0.23961 | -0.70887 | 0.59828 | -0.2251 | 0.114085 |
| 26.34722 | -0.36182 | 0.323354 | -0.01271 | -0.33039 | -0.46663 | -0.08802 | -0.06663 | 0.216706 | -0.52823 | -0.01362 | 0.197622 | -2.45307 | -0.84983 | -0.1973 | -0.73659 | 0.31677 | -1.12757 | -0.04885 |
| 26.36111 | 0.089766 | 0.410664 | 0.341884 | -0.05641 | -0.32452 | -0.19344 | 0.124938 | 0.179008 | -0.11887 | 0.175059 | 0.229918 | -1.07771 | -0.42527 | -0.01832 | -0.58355 | 0.190645 | -0.58086 | -0.16279 |
| 26.375 | 0.508246 | 0.303872 | 0.658347 | 0.245753 | -0.13108 | -0.31987 | 0.360541 | 0.096683 | 0.667916 | 0.455587 | 0.285606 | 0.463616 | 0.286431 | 0.138847 | -0.24272 | 0.313644 | 0.620863 | -0.19535 |
| 26.38889 | 0.788775 | 0.020181 | 0.68616 | 0.525473 | 0.029735 | -0.44631 | 0.529902 | -0.00469 | 1.361425 | 0.838671 | 0.327886 | 1.528485 | 0.805948 | 0.190008 | 0.109487 | 0.554731 | 2.248161 | -0.19803 |
| 26.40278 | 0.622314 | -0.18376 | 0.442177 | 0.423504 | 0.140529 | -0.46002 | 0.655823 | -0.05877 | 1.156891 | 0.962016 | 0.450781 | 1.420928 | 0.632948 | 0.269687 | 0.362605 | 0.829488 | 2.900567 | -0.14974 |
| 26.41667 | 0.358777 | -0.04523 | 0.246777 | 0.300599 | 0.224493 | -0.18475 | 0.698045 | 0.013711 | 0.554694 | 0.865648 | 0.821231 | 0.303315 | 0.336489 | 0.421192 | 0.646115 | 1.054002 | 1.74562 | -0.07236 |
| 26.43056 | 0.25112 | 0.327088 | 0.282357 | 0.165814 | 0.323658 | 0.287639 | 0.613113 | 0.187648 | 0.069452 | 0.595643 | 1.231056 | -0.62936 | 0.03145 | 0.668509 | 0.794928 | 0.873667 | 0.395164 | 0.011666 |
| 26.44444 | 0.191888 | 0.412456 | 0.452771 | 0.067447 | 0.292228 | 0.499264 | 0.381884 | 0.248807 | -0.29308 | 0.390287 | 1.289502 | -1.05132 | -0.15823 | 0.746156 | 0.73688 | 0.541474 | -0.98061 | 0.102119 |
| 26.45833 | 0.121096 | 0.189096 | 0.510146 | 0.045266 | 0.096941 | 0.456256 | 0.139655 | 0.150683 | -0.02075 | 0.242654 | 1.064916 | -0.65151 | -0.2129 | 0.584054 | 0.496253 | 0.332829 | -1.78887 | 0.196007 |
| 26.47222 | 0.194128 | -0.05304 | 0.329854 | 0.181328 | -0.11638 | 0.405811 | 0.017162 | 0.02774 | 0.70833 | 0.100333 | 0.687333 | 0.042172 | -0.11835 | 0.299581 | 0.230624 | 0.140738 | -1.68777 | 0.137004 |
| 26.48611 | 0.604031 | -0.10487 | 0.194625 | 0.634317 | -0.05465 | 0.622592 | 0.366318 | 0.114091 | 1.529848 | -0.03888 | 0.482289 | 0.412204 | -0.00475 | 0.290676 | 0.049433 | -0.12176 | -0.71256 | 0.040323 |
| 26.5 | 1.662881 | 0.186909 | 0.580419 | 1.552276 | 0.285925 | 1.463497 | 1.213961 | 0.505922 | 1.721832 | 0.301177 | 1.033764 | 0.613436 | 0.190502 | 1.072164 | 0.065933 | 0.059959 | 1.016554 | -0.08229 |
| 26.51389 | 2.74531 | 0.623391 | 1.686863 | 2.563618 | 0.434351 | 1.848912 | 2.072215 | 0.823903 | 1.634766 | 1.130128 | 1.908527 | 1.110552 | 0.576365 | 2.270869 | 0.176913 | 0.730002 | 2.84693 | 0.139035 |
| 26.52778 | 2.75278 | 0.838397 | 2.475296 | 2.954784 | -0.00313 | 1.716162 | 2.647832 | 0.657633 | 1.580521 | 2.131836 | 1.901028 | 2.88705 | 1.026696 | 2.4457 | 0.524357 | 1.410367 | 3.628241 | 0.415642 |
| 26.54167 | 2.621891 | 1.027748 | 2.568933 | 2.916647 | -0.62407 | 0.890152 | 2.514345 | 0.563083 | 1.102909 | 2.273856 | 1.665434 | 4.103429 | 1.601161 | 2.111971 | 0.791205 | 1.966794 | 3.086871 | 0.425357 |
| 26.55556 | 1.480692 | 0.602299 | 1.44087 | 1.863209 | -1.19761 | 0.320856 | 1.239909 | 0.37312 | 0.435382 | 1.159266 | 0.526247 | 3.217178 | 1.216488 | 1.324973 | 0.680949 | 1.99398 | 2.385378 | 0.089172 |
| 26.56944 | 0.622841 | 0.144621 | 0.155231 | 0.451531 | -1.4516 | -0.11367 | -0.07294 | 0.149653 | -0.32026 | 0.048065 | -0.827 | 1.619738 | 0.548361 | 0.309381 | 0.291995 | 0.971585 | 0.818417 | -0.19194 |
| 26.58333 | 0.126036 | -0.00048 | -0.94821 | -0.68964 | -1.34234 | -0.67031 | -1.3667 | -0.09465 | -1.04304 | -1.24593 | -1.56716 | 0.359757 | 0.160063 | -0.43665 | 0.107536 | 0.655976 | -0.44325 | -0.1002 |
| 26.59722 | -0.43641 | 0.002485 | -1.63442 | -1.09048 | -1.32078 | -1.02241 | -2.17202 | -0.14614 | -1.58457 | -1.36655 | -1.46861 | -0.05551 | -0.17259 | -0.62353 | 0.019193 | 0.47415 | -0.6579 | 0.142101 |
| 26.61111 | -0.76066 | 0.045114 | -1.40975 | -0.86889 | -0.92703 | -1.27474 | -2.06057 | -0.01671 | -1.11629 | -0.88036 | -0.81621 | 0.33037 | -0.30926 | -0.38617 | 0.006424 | 1.47178 | -0.67438 | 0.409959 |
| 26.625 | -0.87774 | 0.080115 | -1.26522 | -0.33699 | -0.46833 | -1.97295 | -0.90443 | 0.124419 | -0.05435 | -0.08879 | 0.083233 | -0.04049 | -0.34562 | -0.29634 | -0.0748 | 2.119336 | -2.02991 | 0.462359 |
| 26.63889 | -0.54386 | 0.036584 | -0.99957 | 0.120936 | -0.17852 | -1.92523 | 0.514502 | 0.044808 | 0.857143 | 0.129711 | 0.599951 | -0.82698 | -0.46288 | -0.49649 | -0.07178 | 1.746784 | -2.26518 | 0.378175 |
| 26.65278 | 0.113476 | 0.0391 | -0.01487 | 0.439203 | 0.082822 | -1.80769 | 1.280531 | -0.13501 | 1.171181 | -0.05479 | 0.367374 | -0.57108 | -0.79611 | -0.8552 | -0.24966 | 0.054556 | -1.41003 | 0.323191 |
| 26.66667 | 1.210969 | 0.101535 | 1.262087 | 1.005504 | 0.690549 | -0.88551 | 2.1079 | -0.1954 | 0.461557 | -0.39326 | -0.0325 | -0.60115 | -1.02525 | -0.99397 | -0.46157 | -1.88349 | 0.091011 | 0.001823 |
| 26.68056 | 1.856173 | 0.099757 | 1.352807 | 1.35233 | 1.02518 | 0.130308 | 1.68201 | -0.24516 | -0.07548 | -0.03118 | -0.21114 | -1.01917 | -0.98608 | -0.76333 | -0.5628 | -1.59731 | 1.73546 | -0.33776 |
| 26.69444 | 1.443034 | -0.04342 | 0.246075 | 0.595274 | 0.866623 | 1.1338 | 0.032285 | -0.22529 | -0.41945 | 0.489673 | 0.114914 | -1.33094 | -0.46503 | -0.52881 | -0.52158 | -0.87012 | 1.799184 | -0.44335 |
| 26.70833 | 0.292947 | -0.52866 | -0.09921 | -0.07063 | 0.568158 | 1.336539 | -1.04422 | -0.09944 | -0.66988 | 0.697844 | 0.178327 | -1.389 | 0.231471 | 0.255077 | -0.27839 | -0.5377 | 0.822543 | -0.38663 |
| 26.72222 | -0.32312 | -0.77855 | -0.32957 | -0.24924 | 0.129974 | 1.256277 | -1.70243 | -0.00652 | -0.95515 | 0.017604 | 0.052004 | -1.28727 | 0.68553 | 1.340176 | -0.09361 | -0.77886 | -0.93336 | -0.35891 |
| 26.73611 | -0.31224 | -0.43037 | -0.65695 | 0.059876 | -0.09903 | 1.116046 | -2.06406 | 0.015183 | -0.82371 | -1.29978 | -0.65495 | -0.61418 | 0.47974 | 1.92092 | -0.0528 | -1.13695 | -1.46592 | -0.26382 |
| 26.75 | -0.40496 | 0.020083 | -0.5539 | -0.04109 | -0.32377 | 0.662118 | -1.36876 | -0.02927 | -0.62435 | -1.85654 | -1.02227 | 0.085204 | 0.215885 | 1.68962 | 0.213373 | -0.15887 | -1.05887 | -0.14301 |
| 26.76389 | -0.92121 | 0.156002 | -0.35179 | -0.61964 | -0.89759 | -0.23551 | -0.48661 | -0.12245 | -0.35767 | -0.87715 | -0.97987 | 0.555779 | 0.120154 | 0.924799 | 0.474465 | 0.713088 | -0.9952 | -0.19523 |
| 26.77778 | -2.11513 | 0.096796 | -0.48948 | -1.29725 | -0.80391 | -1.30656 | -0.00451 | -0.21685 | -0.0045 | -0.63228 | -0.48095 | 0.986042 | 0.172535 | 0.114889 | 0.563643 | 1.316903 | -0.45053 | -0.05246 |
| 26.79167 | -1.40507 | -0.07784 | -0.71927 | -0.72142 | -0.42252 | -1.81116 | -0.02658 | -0.19362 | 0.334469 | -0.20309 | 0.520419 | 0.543798 | 0.271082 | -0.64172 | 0.71316 | 0.940666 | -0.61792 | 0.456466 |
| 26.80556 | -0.53146 | -0.34601 | -0.53369 | -0.16996 | -0.17592 | -2.20606 | -0.04613 | -0.15448 | 0.456533 | 1.066696 | 1.447835 | -0.23014 | 0.219833 | -1.22745 | 0.920685 | -0.14328 | -1.24941 | 1.108349 |
| 26.81944 | -0.49193 | -0.57013 | -0.01028 | -0.63769 | -0.19329 | -2.37856 | -0.06679 | -0.01784 | 0.229381 | 1.747547 | 1.553156 | -0.4906 | 0.069555 | -1.45408 | 0.6831 | 0.017035 | -0.63261 | 1.210041 |
| 26.83333 | -0.60254 | -0.51952 | 0.237288 | -1.15927 | -0.01613 | -2.52954 | -0.24027 | 0.052717 | -0.41467 | 1.63779 | 0.312912 | -1.00058 | -0.01396 | -2.06609 | -0.42688 | 0.540226 | 0.078456 | 0.60926 |
| 26.84722 | -0.55061 | -0.15716 | 0.183393 | -1.23298 | 0.700675 | -1.58135 | -0.6785 | -0.00437 | -0.89199 | 0.45681 | -0.51617 | -1.44361 | -0.09688 | -1.89446 | -0.91962 | 0.767014 | 0.660978 | 0.287111 |
| 26.86111 | -0.41476 | 0.13293 | -0.15985 | -1.00713 | 0.979873 | -0.83937 | -1.09433 | -0.1107 | -1.14149 | -0.91426 | -1.54044 | -1.55957 | -0.36069 | -1.1621 | -0.90352 | 0.875674 | 0.820255 | 0.12516 |
| 26.875 | -0.36547 | 0.221742 | -0.52107 | -0.82779 | 0.648707 | -0.60804 | -1.31435 | -0.19612 | -1.01606 | -2.00295 | -1.86752 | -2.1167 | -0.92086 | -0.54484 | -0.63546 | 0.442609 | 0.650674 | 0.114306 |
| 26.88889 | -0.54014 | 0.085255 | -0.6305 | -0.80832 | 0.234622 | -0.39861 | -1.1834 | -0.26968 | -0.7944 | -1.42774 | -1.76264 | -3.68575 | -1.04338 | -0.10938 | -0.2818 | -0.21211 | -0.21669 | 0.292145 |
| 26.90278 | -0.83098 | -0.15772 | -0.4649 | -1.11508 | -0.15578 | -0.11108 | -0.72023 | -0.31796 | -0.67412 | -1.41606 | -1.4313 | -3.58245 | -0.43305 | 0.091566 | 0.001948 | -0.88797 | -1.42178 | 0.433414 |
| 26.91667 | -0.79852 | -0.24697 | -0.14088 | -1.33109 | -0.4262 | 0.13606 | -0.27054 | -0.46359 | -0.57388 | -1.61566 | -0.66287 | -2.86193 | -0.03006 | 0.172408 | 0.074901 | -0.9718 | -1.18086 | 0.377903 |
| 26.93056 | -0.33801 | -0.10121 | 0.226605 | -0.97944 | -0.73456 | 0.239277 | 0.19589 | -0.5224 | -0.28357 | -1.42758 | 0.027651 | -2.62232 | 0.237457 | 0.282825 | 0.148939 | 0.116461 | -0.23585 | 0.214561 |
| 26.94444 | 0.166699 | 0.122373 | 0.380211 | -0.41001 | -0.6868 | 0.233763 | 0.368789 | -0.49881 | 0.142022 | -0.3301 | 0.500724 | -2.42961 | 0.524946 | 0.383816 | 0.240705 | 1.229005 | 0.586544 | 0.153436 |
| 26.95833 | 0.485908 | 0.30819 | 0.183611 | 0.077387 | -0.67474 | 0.226194 | 0.394815 | -0.41833 | 0.577879 | 0.494087 | 0.765858 | -1.87579 | 0.89084 | 0.388798 | 0.284718 | 1.195656 | 1.000693 | 0.254696 |

| | | | | | | | | | | | | | | | | | | |
|---|---|---|---|---|---|---|---|---|---|---|---|---|---|---|---|---|---|---|
| 26.97222 | 0.559896 | 0.376988 | -0.0574 | 0.377877 | -0.54881 | 0.275934 | 0.321952 | -0.27387 | 0.842409 | 0.998961 | 0.768497 | -1.77738 | 0.860454 | 0.331785 | 0.167737 | 0.408517 | 1.172115 | 0.446959 |
| 26.98611 | 0.296179 | 0.180501 | -0.24222 | 0.202173 | -0.59362 | 0.204061 | -0.12115 | -0.27172 | 0.418881 | 0.803319 | 0.245214 | -2.90617 | -0.17503 | 0.007037 | -0.47919 | -0.74681 | 0.461345 | 0.203465 |
| 27 | -0.49315 | -0.40206 | -0.73327 | -0.67203 | -0.94933 | -0.31747 | -1.08675 | -0.62458 | -0.74126 | -0.59719 | -0.75688 | -6.97695 | -1.73997 | -0.81485 | -1.85817 | -2.18376 | -1.51917 | -0.99377 |
| 27.01389 | -1.43136 | -1.14285 | -1.3229 | -1.57645 | -0.80501 | -0.81155 | -1.73078 | -0.85039 | -1.99283 | -1.59884 | -1.3725 | -7.8223 | -2.15491 | -1.4159 | -2.13276 | -2.5359 | -2.61484 | -1.79029 |
| 27.02778 | -1.73471 | -1.39667 | -1.30192 | -1.97275 | -0.31905 | -0.77017 | -1.4117 | -0.58541 | -2.34393 | -1.42464 | -0.91474 | -3.72805 | -1.09939 | -1.25229 | -1.2255 | -1.2239 | -1.40743 | -1.56608 |
| 27.04167 | -1.41453 | -1.24973 | -1.03047 | -1.24464 | -0.11826 | -0.51248 | -0.69929 | -0.15288 | -2.2673 | -2.18238 | -0.62711 | -2.18347 | -0.49307 | -0.76101 | -0.93686 | -0.71205 | -0.02009 | -1.57275 |
| 27.05556 | -0.89793 | -0.99595 | -0.59091 | -0.5954 | 0.049161 | -0.2555 | -0.31291 | -0.24244 | -1.33346 | -2.95934 | -0.46422 | -1.95925 | -0.30459 | -0.26323 | -0.86761 | -0.69529 | -0.15472 | -1.6266 |
| 27.06944 | -0.25026 | 0.061027 | -0.19444 | -0.30831 | 0.342635 | 0.160026 | -0.18867 | -0.23448 | -1.34016 | -1.41557 | -0.32667 | -0.97997 | -0.06958 | 0.423993 | -0.13433 | -0.86612 | -0.98654 | -0.67983 |
| 27.08333 | 0.298263 | 1.134499 | 0.315788 | 0.01736 | 0.652572 | 0.676509 | 0.126118 | 0.095788 | -1.73815 | 0.263041 | -0.0202 | -0.18092 | 0.292343 | 0.834599 | 0.518579 | -0.52887 | -1.25023 | 0.393932 |
| 27.09722 | 0.926587 | 1.931628 | 0.578995 | 0.251218 | 1.327597 | 1.344705 | 0.582999 | 0.75556 | -0.86068 | 0.232347 | 0.608879 | -0.73394 | 0.117846 | 0.766528 | 0.282106 | -0.20259 | -0.67727 | 0.147605 |
| 27.11111 | 1.551851 | 2.589643 | 0.735731 | 0.247171 | 1.482757 | 0.474731 | 0.722897 | 1.472265 | -0.40656 | 0.12601 | 0.278392 | -1.95871 | -0.64698 | 0.245953 | -0.40642 | 0.20694 | -0.34683 | -0.60407 |
| 27.125 | -0.06728 | -0.25636 | 2.406926 | 0.938483 | -0.81007 | -3.51857 | 0.47653 | -1.23108 | -0.00195 | 0.006725 | -0.39677 | -2.73501 | -1.16473 | -0.30354 | -0.88096 | 0.415085 | -0.38471 | -0.89541 |
| 27.13889 | -1.82521 | -2.50762 | 0.029512 | 0.942081 | -0.52267 | -1.99815 | -0.18912 | -2.19044 | 0.895438 | 0.184273 | -0.50992 | -1.9072 | -0.72527 | -0.33209 | -0.63228 | 0.790625 | 0.39192 | -0.23512 |
| 27.15278 | -0.82832 | -0.96281 | 0.076711 | 1.081985 | 0.442486 | 1.270206 | 0.467853 | -0.61391 | 2.02873 | 0.650097 | -0.0506 | 0.372306 | 0.031673 | 0.052288 | -0.02678 | 1.423467 | 1.681112 | 0.630579 |
| 27.16667 | 0.30058 | 0.09067 | -0.58721 | 1.086531 | 1.070639 | 2.057738 | 1.056344 | 0.558146 | 2.279224 | 0.865979 | 0.317769 | 2.516576 | 0.324131 | 0.218244 | 0.184403 | 1.469015 | 2.411682 | 0.722127 |
| 27.18056 | 0.435564 | 0.249954 | 0.260334 | 0.745531 | 0.938428 | 0.672257 | 0.743803 | 0.447845 | 2.034983 | 0.591542 | 0.270836 | 3.24281 | -0.0014 | 0.125087 | -0.02299 | 1.221831 | 2.645277 | 0.234594 |
| 27.19444 | 0.088886 | -0.0512 | 0.612875 | 0.198593 | 0.494948 | 0.303085 | 0.656259 | 0.676987 | 2.018496 | -0.56461 | -0.67112 | 1.61378 | -1.1268 | -0.61772 | -0.79408 | 0.711929 | 2.436858 | -0.53319 |
| 27.20833 | -0.66755 | -1.00122 | 1.323161 | -0.21912 | -0.38916 | 0.27502 | 0.818254 | 1.534068 | 3.4677 | -2.89925 | -2.68897 | -1.11534 | -3.26319 | -2.39632 | -2.44915 | -0.55577 | 2.019764 | -1.89599 |
| 27.22222 | -0.88386 | -1.39517 | 1.74406 | 0.054756 | -1.15333 | 0.459748 | 0.969138 | 1.407148 | 3.591431 | -4.03044 | -3.43417 | -3.02335 | -4.2991 | -3.45683 | -3.50349 | -1.69755 | 1.026612 | -2.86109 |
| 27.23611 | -0.02656 | -0.32969 | 1.486814 | 0.96525 | -0.03039 | 0.4073 | 0.942107 | 0.75996 | 2.14154 | -2.13701 | -1.93586 | -2.08044 | -2.47851 | -1.91984 | -2.16615 | -0.74759 | 0.380375 | -1.59228 |
| 27.25 | 0.659447 | 0.526908 | 1.348972 | 1.439311 | 0.157215 | 0.259773 | 0.843286 | 0.316124 | 1.121003 | -0.03115 | -0.11876 | 0.248291 | -0.13648 | 0.311423 | -0.04179 | 0.846339 | 0.83888 | 0.516878 |
| 27.26389 | 0.845438 | 0.798643 | 0.877769 | 0.96167 | -0.22031 | 0.091979 | 0.41117 | 0.059776 | 0.49619 | 0.727643 | 0.67751 | 1.0244 | 0.766381 | 1.164214 | 0.882727 | 1.618483 | 1.09028 | 1.189328 |
| 27.27778 | 0.570583 | 0.574262 | 0.140903 | 0.014548 | -0.58261 | -0.07413 | -0.19009 | -0.03626 | -0.08616 | 0.651967 | 0.734386 | -0.51676 | 0.611562 | 0.876316 | 0.777027 | 1.570774 | 0.549201 | 0.707665 |
| 27.29167 | 0.100665 | 0.02345 | -0.21492 | -0.55194 | -0.66549 | -0.16604 | -0.54798 | -0.04765 | -0.50726 | 0.50475 | 0.882972 | -2.72154 | 0.587698 | 0.630578 | 0.709337 | 1.401843 | 0.109708 | 0.220249 |
| 27.30556 | -0.16195 | -0.34422 | -0.11141 | -0.32826 | -0.45895 | -0.12781 | -0.32429 | 0.032539 | -0.36891 | 0.469189 | 0.851312 | -3.72567 | 0.72754 | 0.740631 | 0.807042 | 1.4548 | 0.48422 | 0.42275 |
| 27.31944 | -0.00657 | -0.29317 | 0.29411 | 0.375833 | -0.37238 | 0.017109 | 0.25789 | 0.188673 | 0.115505 | 0.500955 | 0.833132 | -3.63509 | 0.813742 | 1.063445 | 0.914212 | 1.566337 | 1.491797 | 0.879936 |
| 27.33333 | 0.408725 | 0.189054 | 0.696631 | 1.048651 | -0.04986 | 0.317625 | 0.773525 | 0.334878 | 0.705698 | 0.586985 | 0.852814 | -3.25731 | 0.792312 | 1.260225 | 0.928362 | 1.517481 | 2.336517 | 1.052093 |
| 27.34722 | 0.696441 | 0.583129 | 0.766528 | 1.185372 | 0.293353 | 0.454369 | 0.867815 | 0.3459 | 1.121613 | 0.616416 | 0.653298 | -2.74345 | 0.455293 | 0.963623 | 0.742172 | 1.209201 | 2.177113 | 0.823375 |
| 27.36111 | 0.563829 | 0.423911 | 0.552565 | 0.818612 | 0.362915 | 0.307998 | 0.595686 | 0.228094 | 1.008249 | 0.521258 | 0.207816 | -2.69072 | 0.095144 | 0.504346 | 0.402947 | 0.746718 | 1.269892 | 0.356086 |
| 27.375 | 0.242637 | 0.150695 | 0.318579 | 0.355909 | 0.423146 | 0.114528 | 0.247614 | 0.113865 | 0.6165 | 0.38809 | 0.024675 | -2.2932 | -0.09122 | 0.224342 | 0.085043 | 0.474026 | 0.422304 | -0.07209 |
| 27.38889 | 0.02296 | -0.09929 | 0.216949 | -0.07604 | 0.604762 | -0.01331 | -0.05279 | 0.074777 | 0.367645 | 0.241724 | 0.046415 | -1.64247 | -0.13983 | 0.076724 | -0.12325 | 0.358419 | -0.15399 | -0.26726 |
| 27.40278 | 0.026912 | -0.1846 | 0.228089 | -0.27747 | 0.756075 | -0.11521 | -0.215 | 0.042289 | 0.174167 | 0.148303 | 0.165875 | -0.99651 | -0.17249 | 0.09497 | -0.1696 | 0.363255 | -0.23098 | -0.18086 |
| 27.41667 | 0.180694 | -0.11728 | 0.290363 | -0.11722 | 0.740367 | -0.18158 | -0.13755 | 0.024589 | 0.161724 | 0.269309 | 0.410297 | -0.2349 | 0.012746 | 0.327555 | 0.016739 | 0.611216 | 0.36163 | 0.077845 |
| 27.43056 | 0.346595 | -0.0226 | 0.38127 | 0.2787 | 0.724196 | -0.15657 | 0.055262 | 0.056487 | 0.407776 | 0.489866 | 0.670266 | 0.327285 | 0.170145 | 0.473168 | 0.282865 | 1.032803 | 1.373883 | 0.280169 |
| 27.44444 | 0.357703 | 0.073134 | 0.372549 | 0.49734 | 0.588612 | -0.02989 | 0.2543 | 0.108737 | 0.758191 | 0.58897 | 0.655376 | 0.204854 | 0.04604 | 0.293334 | 0.335642 | 1.212841 | 1.965059 | 0.288118 |
| 27.45833 | 0.254562 | 0.054697 | 0.362376 | 0.474808 | 0.428832 | 0.046693 | 0.344191 | 0.119213 | 1.022621 | 0.576435 | 0.589613 | -0.22635 | -0.07937 | -0.03672 | 0.298994 | 1.06519 | 1.612962 | 0.143239 |
| 27.47222 | 0.254422 | 0.033726 | 0.457484 | 0.3264 | 0.365904 | 0.133807 | 0.308196 | 0.195201 | 0.88623 | 0.488223 | 0.581383 | -0.48977 | -0.00651 | -0.07651 | 0.312638 | 1.113623 | 0.974792 | 0.135803 |
| 27.48611 | 0.519157 | -0.03977 | 0.71043 | 0.477038 | 0.459731 | 0.39963 | 0.528714 | 0.456358 | 0.705454 | 0.410811 | 0.678239 | -0.41082 | 0.287192 | 0.155862 | 0.572167 | 1.225711 | 1.138042 | 0.262359 |
| 27.5 | 1.046764 | -0.12473 | 1.227871 | 1.435561 | 0.746833 | 1.275328 | 1.493554 | 1.041204 | 1.01379 | 0.411655 | 1.165024 | 0.009309 | 0.502326 | 0.621763 | 1.141372 | 1.529265 | 2.693928 | 0.330063 |
| 27.51389 | 1.782826 | 0.077607 | 1.890376 | 2.685944 | 1.013176 | 2.467346 | 2.797725 | 1.297592 | 1.49672 | 0.536454 | 1.764936 | 0.257138 | 0.279019 | 1.004305 | 1.262675 | 1.49282 | 4.306514 | 0.430583 |
| 27.52778 | 1.966338 | 0.331635 | 2.398197 | 3.668155 | 0.601762 | 2.868521 | 3.824163 | 0.887002 | 1.915005 | 1.148943 | 2.456506 | 0.972344 | 0.029928 | 0.919417 | 1.139427 | 1.352377 | 5.19821 | 0.520018 |
| 27.54167 | 1.818029 | 0.387084 | 2.366447 | 3.233058 | 0.453725 | 2.330406 | 3.601084 | 0.685061 | 1.602983 | 1.400675 | 2.07617 | 1.882789 | 0.069688 | 0.713157 | 0.863907 | 1.198444 | 4.429129 | 0.490092 |
| 27.55556 | 1.265223 | 0.378706 | 1.701933 | 1.631131 | 0.1656 | 1.52361 | 2.145656 | 0.32957 | 0.731108 | 0.81852 | 0.917459 | 1.732263 | 0.100013 | 0.316962 | 0.370278 | 1.082717 | 3.709649 | -0.13677 |
| 27.56944 | 0.636538 | 0.290488 | 0.838854 | 0.191568 | -0.16152 | 0.857089 | 0.707624 | 0.058291 | 0.047852 | 0.217029 | -0.44881 | 0.218074 | 0.083983 | -0.14447 | -0.09623 | 0.784097 | 2.243598 | -0.55825 |
| 27.58333 | 0.126136 | 0.247051 | 0.028265 | -0.97282 | -0.41953 | 0.384025 | -0.29424 | -0.0708 | -0.66284 | -0.24389 | -1.58558 | -1.38871 | 0.174603 | -0.54435 | -0.57503 | 0.679511 | -0.0551 | -0.86119 |
| 27.59722 | -0.21614 | 0.351062 | -0.60994 | -1.69804 | -0.49155 | 0.047048 | -0.78759 | -0.04749 | -1.10187 | -0.53462 | -2.21377 | -2.55495 | 0.250018 | -0.77726 | -1.05197 | 0.049393 | -2.12132 | -0.68078 |
| 27.61111 | -0.32648 | 0.311534 | -0.83251 | -1.82455 | -0.48735 | -0.26118 | -1.12003 | 0.088745 | -1.20227 | -0.72454 | -2.08632 | -3.08511 | 0.277732 | -0.43001 | -0.94357 | -0.64241 | -4.02701 | -0.29005 |
| 27.625 | -0.56352 | 0.40617 | -0.55764 | -1.42688 | -0.50505 | -0.62671 | -1.35137 | 0.176955 | -0.69777 | -0.83509 | -1.6171 | -2.29853 | 0.42795 | 0.062098 | -0.45227 | -0.15428 | -4.86657 | 0.175326 |
| 27.63889 | -0.4254 | 0.453056 | -0.28652 | -0.7287 | -0.5117 | -1.30173 | -1.47403 | 0.210327 | 0.061023 | -0.59029 | 0.069211 | -1.2827 | 0.279971 | 0.392331 | 0.004542 | 0.522622 | -4.01406 | 0.475832 |
| 27.65278 | -0.20555 | 0.181492 | -0.08174 | -0.03689 | -0.21596 | -2.11871 | -1.35193 | 0.117151 | 0.702799 | -0.04825 | 1.527297 | -0.11487 | 0.093638 | 0.276134 | 0.217222 | 0.927866 | -1.62649 | 0.468544 |
| 27.66667 | -0.03528 | -0.4273 | 0.085682 | 0.490687 | 0.304073 | -2.56068 | -0.59557 | -0.02271 | 0.800528 | 0.783031 | 2.183911 | 1.461066 | 0.14907 | 0.036258 | 0.436739 | 0.235516 | 0.640297 | -0.22518 |
| 27.68056 | 0.215837 | -0.59139 | 0.261392 | 0.531912 | 0.778874 | -2.16571 | 0.327361 | -0.14801 | 0.294413 | 1.328305 | 1.99388 | 3.044457 | 0.165714 | -0.07931 | 0.393929 | -0.47859 | 2.256765 | -0.96726 |
| 27.69444 | 0.736393 | -0.32959 | 0.481647 | 0.465951 | 0.750956 | -1.23586 | 1.220292 | -0.28018 | -0.55271 | 1.592578 | 0.967214 | 3.269356 | -0.19269 | -0.20358 | -0.16418 | -1.27977 | 1.390428 | -1.08813 |
| 27.70833 | 1.21535 | -0.04344 | 0.51493 | 0.62373 | 1.02176 | -0.41936 | 1.964777 | -0.32127 | -0.85274 | 1.393883 | -0.1404 | 1.277209 | -0.42141 | -0.45833 | -0.80394 | -1.49401 | 0.904803 | -0.85539 |
| 27.72222 | 0.869902 | 0.089887 | 0.222622 | 0.153315 | 0.86728 | 0.346375 | 2.089626 | -0.21594 | -0.47407 | 0.421959 | -0.71393 | -0.94956 | -0.2255 | -0.55092 | -0.99129 | -0.5648 | -0.00619 | -0.65984 |
| 27.73611 | -0.00282 | 0.056512 | -0.06903 | -0.77495 | 0.048658 | 1.044748 | 1.652915 | 0.058252 | 0.08901 | -1.09934 | -1.33787 | -3.07214 | -0.04384 | -0.70152 | -1.16983 | 0.432404 | -0.67966 | -0.60258 |
| 27.75 | -0.65234 | -0.18687 | -0.1426 | -1.34157 | -0.8056 | 1.722037 | 0.624095 | 0.197546 | 0.362323 | -1.63848 | -1.40358 | -4.12964 | 0.07631 | -0.71667 | -1.123 | 1.029985 | -1.20112 | -0.38306 |
| 27.76389 | -0.83346 | -0.37692 | 0.090989 | -1.43848 | -1.13521 | 1.69502 | -0.13066 | 0.209089 | 0.436942 | -1.01016 | -1.25975 | -3.44992 | 0.082758 | -0.3163 | -0.73713 | 0.50789 | -2.00717 | 0.061539 |
| 27.77778 | -0.53545 | -0.45344 | 0.322314 | -1.08765 | -1.52244 | 1.463115 | -0.79553 | 0.135117 | 0.023131 | -0.62717 | -1.5006 | -1.70405 | -0.17797 | -0.05067 | -0.21179 | -0.37029 | -2.41557 | 0.166333 |
| 27.79167 | 0.109533 | -0.31736 | -0.00358 | -0.57296 | -1.14318 | 0.8585 | -1.76788 | -0.00565 | -0.28407 | -1.16431 | -1.9763 | -0.94262 | -0.82913 | -0.00549 | -0.04911 | -1.01136 | -1.55132 | -0.37908 |
| 27.80556 | 0.663541 | 0.016763 | -0.70137 | -0.15334 | -0.88506 | -0.02147 | -2.37303 | -0.03467 | -0.7115 | -0.8667 | -1.21893 | -0.73279 | -1.11288 | -0.21738 | -0.36091 | -0.79716 | -0.69897 | -0.87461 |
| 27.81944 | 0.569813 | 0.158119 | -0.91797 | -0.01315 | -0.30824 | -0.8674 | -1.76262 | 0.01521 | -0.87551 | -0.11634 | -0.33315 | -0.52085 | -0.84317 | -0.2105 | -0.56759 | -0.11443 | -0.34123 | -1.05466 |
| 27.83333 | -0.37641 | -0.07797 | -0.99901 | -0.38341 | 0.292755 | -1.89286 | -0.71402 | 0.147324 | -0.53 | 0.624388 | 0.802078 | -0.25268 | -0.22858 | 0.127343 | -0.66705 | 0.260439 | -0.14149 | -0.97448 |
| 27.84722 | -1.2358 | -0.49654 | -1.55531 | -1.21042 | 0.699414 | -2.59431 | -0.65129 | 0.1591 | -0.234 | 1.021705 | 1.775752 | -0.34475 | 0.016603 | 0.183893 | -0.92719 | 0.719626 | -0.12282 | -0.72122 |
| 27.86111 | -1.4791 | -1.03461 | -1.97995 | -1.59742 | 0.930401 | -2.22523 | -1.09822 | -0.03525 | -0.20683 | 0.631888 | 1.864476 | -0.65899 | -0.05682 | -0.03116 | -1.27741 | 0.797281 | -0.08613 | -0.36494 |
| 27.875 | -1.10058 | -1.35945 | -1.68089 | -1.57615 | 0.854024 | -1.38004 | -1.5551 | -0.23072 | -0.30464 | 0.300073 | 1.268688 | -0.96557 | -0.23277 | -0.2024 | -1.51344 | 0.439111 | -0.37028 | -0.11147 |
| 27.88889 | -1.11716 | -1.32641 | -0.9242 | -1.69477 | 0.492169 | -1.101 | -1.67229 | -0.41109 | -0.45186 | 0.099351 | 0.654664 | -1.10719 | -0.34162 | -0.2051 | -1.16277 | -0.12164 | -0.63311 | 0.160765 |
| 27.90278 | -0.98076 | -0.95413 | -0.04604 | -1.70774 | 0.07122 | -0.79123 | -1.11576 | -0.5898 | -0.8861 | -0.05923 | -0.0834 | -1.20742 | -0.32835 | -0.34303 | -0.4125 | -0.24116 | -0.57874 | 0.257751 |
| 27.91667 | -0.49084 | -0.28006 | 0.673834 | -1.17181 | -0.37314 | -0.46611 | -0.42751 | -0.71879 | -0.60392 | -0.17835 | -0.89352 | -1.24648 | -0.21331 | -0.57062 | -0.1183 | -0.17775 | -0.21896 | 0.188392 |
| 27.93056 | 0.105406 | 0.360838 | 1.156741 | -0.44069 | -0.63844 | -0.16911 | 0.16858 | -0.55472 | -0.17978 | -0.79598 | -1.35995 | -1.25654 | -0.11311 | -0.62095 | -0.0813 | -0.21214 | 0.320892 | 0.060473 |
| 27.94444 | 0.501416 | 0.876551 | 1.043673 | 0.372127 | -0.85693 | 0.065173 | 0.736519 | -0.23958 | 0.275351 | -1.0428 | -1.15683 | -1.01107 | -0.04928 | -0.30396 | -0.05811 | 0.262593 | 0.937896 | 0.039132 |
| 27.95833 | 0.680223 | 1.005556 | 0.793166 | 1.0318 | -0.82429 | 0.250266 | 1.249415 | -0.06619 | 0.666957 | -0.7823 | -0.68039 | -0.7222 | 0.048595 | -0.01222 | -0.0124 | 0.874503 | 1.594507 | 0.190922 |
| 27.97222 | 0.587255 | 0.792676 | 0.571599 | 1.187366 | -0.56836 | 0.384195 | 1.500819 | -0.15475 | 1.180288 | -0.3837 | -0.31026 | -0.59165 | 0.213916 | 0.145387 | 0.10678 | 1.506991 | 1.825707 | 0.427826 |
| 27.98611 | 0.135828 | 0.13267 | -0.03876 | 0.359288 | -0.62698 | 0.251028 | 1.002116 | -0.37226 | 1.086955 | -0.42856 | -0.45154 | -1.2618 | -0.05023 | -0.06735 | -0.26702 | 1.137106 | 0.652548 | 0.024675 |
| 28 | -1.04683 | -1.10374 | -1.44898 | -1.57871 | -1.13737 | -0.30044 | -0.58468 | -0.81915 | -0.48177 | -1.38313 | -1.49549 | -3.21018 | -1.06306 | -0.92298 | -1.69735 | -0.71503 | -1.60307 | -1.50664 |
| 28.01389 | -2.65321 | -2.88094 | -3.46719 | -3.45442 | -1.22459 | -0.86128 | -2.00999 | -0.98408 | -2.37436 | -2.71418 | -2.91681 | -5.02747 | -2.24201 | -2.04375 | -3.2853 | -2.86125 | -2.76747 | -3.20483 |
| 28.02778 | -3.28752 | -3.98202 | -4.83141 | -3.60682 | -0.50676 | -0.85238 | -1.83128 | -0.94658 | -2.62429 | -2.50363 | -2.82531 | -4.39676 | -2.18613 | -2.32927 | -3.59088 | -3.64458 | -2.39065 | -3.27993 |
| 28.04167 | -2.25839 | -3.66661 | -3.19454 | -1.95028 | 0.024538 | -0.38756 | -0.81001 | -0.40411 | -2.04963 | -1.40831 | -1.4017 | -1.82238 | -0.97755 | -1.06138 | -1.49657 | -2.72285 | -1.27056 | -1.60829 |
| 28.05556 | -1.09866 | -2.07864 | -1.18448 | -0.69368 | 0.335309 | -0.02244 | -0.28664 | 0.035837 | -1.98943 | -0.38793 | -0.31736 | -0.40993 | 0.549782 | 0.130338 | 0.167679 | -0.84572 | 0.021046 | 0.319375 |
| 28.06944 | -0.1706 | -0.46402 | 0.108454 | 0.350821 | 0.465071 | 0.120129 | -0.08649 | 0.341065 | -1.5723 | 0.76137 | 0.53896 | 0.503573 | 1.51701 | 0.955529 | 1.311113 | 1.413584 | 0.966004 | 1.828901 |
| 28.08333 | 0.423364 | 0.732271 | 1.139951 | 1.126588 | 0.345883 | 0.066507 | -0.04126 | 0.274198 | -0.77868 | 1.383627 | 1.045549 | 1.084007 | 1.251059 | 1.39283 | 2.147458 | 3.329502 | 1.237037 | 2.698228 |

| | | | | | | | | | | | | | | | | | | |
|---|---|---|---|---|---|---|---|---|---|---|---|---|---|---|---|---|---|---|
| 28.09722 | 0.88619 | 1.533517 | 1.826691 | 1.51655 | 0.178528 | -0.10084 | -0.09457 | 0.074248 | -0.14189 | 1.297187 | 0.955864 | 1.182279 | 0.797385 | 1.489472 | 2.288589 | 2.380288 | 0.240308 | 1.998454 |
| 28.11111 | 1.269515 | 2.379079 | 2.329993 | 1.798095 | 0.20953 | -0.15025 | -0.1841 | 0.061717 | 0.315781 | 0.894696 | 0.707712 | 1.102431 | 0.46367 | 1.453422 | 1.752546 | 1.652388 | -0.20722 | 1.385985 |
| 28.125 | 1.492178 | 3.025705 | 2.121893 | 1.862459 | 0.34387 | 0.034857 | -0.00435 | 0.05781 | 0.399734 | 0.39455 | 0.359417 | 1.085147 | 0.232374 | 1.546126 | 1.420515 | 0.952325 | -0.2026 | 0.980752 |
| 28.13889 | 1.581638 | 2.910088 | 1.547105 | 1.638773 | 0.313346 | 0.26778 | 0.410094 | -0.07884 | 0.288958 | 0.032831 | 0.103359 | 1.256757 | 0.128753 | 1.48418 | 1.147889 | 0.269563 | -0.21834 | 0.409993 |
| 28.15278 | 1.37912 | 2.312567 | 1.210423 | 1.115032 | 0.234042 | 0.447417 | 0.680919 | -0.04604 | 0.355839 | -0.11518 | 0.097607 | 1.683769 | 0.178983 | 1.08101 | 0.63383 | -0.20119 | -0.22747 | 0.083756 |
| 28.16667 | 0.976575 | 1.547007 | 1.034427 | 0.689907 | 0.174972 | 0.594059 | 0.862434 | 0.143197 | 1.050843 | -0.04618 | 0.177008 | 2.033469 | 0.34123 | 0.782305 | 0.396874 | 0.058694 | -0.01713 | 0.051833 |
| 28.18056 | 0.734069 | 1.118781 | 0.889994 | 0.403057 | 0.156735 | 0.513736 | 0.796665 | 0.266313 | 1.770608 | 0.125305 | 0.219842 | 2.486486 | 0.377026 | 0.425249 | 0.201808 | 0.69167 | 0.31626 | 0.230557 |
| 28.19444 | 0.646677 | 0.78845 | 0.759968 | 0.298659 | 0.021106 | 0.385364 | 0.675683 | 0.252299 | 1.793856 | 0.309834 | 0.181231 | 2.754456 | 0.296197 | 0.064029 | -0.00476 | 1.261181 | 0.44929 | 0.324364 |
| 28.20833 | 0.55817 | 0.446613 | 0.854171 | 0.2599 | 0.044852 | 0.345537 | 0.661689 | 0.197932 | 1.315816 | 0.384474 | 0.144194 | 2.859082 | 0.142304 | -0.18776 | -0.15756 | 1.232443 | 0.354263 | 0.187947 |
| 28.22222 | 0.523005 | 0.258231 | 1.029716 | 0.363558 | 0.169232 | 0.361783 | 0.781681 | 0.064034 | 1.134323 | 0.40287 | 0.060576 | 2.900474 | 0.065038 | -0.28652 | -0.19219 | 0.927367 | 0.094542 | -0.05042 |
| 28.23611 | 0.533597 | 0.218787 | 1.121845 | 0.435437 | 0.267834 | 0.356526 | 0.80632 | -0.0333 | 1.073524 | 0.526016 | 0.004537 | 2.841299 | 0.147431 | -0.1495 | -0.09414 | 0.85237 | -0.02179 | -0.10413 |
| 28.25 | 0.574951 | 0.31474 | 1.077478 | 0.323254 | 0.274896 | 0.313519 | 0.707175 | 0.021414 | 1.12809 | 0.54336 | 0.006234 | 2.290505 | 0.322016 | 0.159415 | 0.068145 | 0.895354 | 0.021044 | 0.00042 |
| 28.26389 | 0.63256 | 0.499361 | 0.8596 | 0.136614 | 0.210328 | 0.238744 | 0.510792 | 0.151667 | 0.982391 | 0.452344 | 0.121582 | 1.756622 | 0.496695 | 0.495543 | 0.2239 | 1.055204 | 0.301385 | 0.113133 |
| 28.27778 | 0.613252 | 0.56242 | 0.534113 | -0.01291 | 0.168317 | 0.16294 | 0.348771 | 0.24677 | 0.478636 | 0.507355 | 0.385933 | 1.151036 | 0.700579 | 0.709167 | 0.387143 | 1.131279 | 0.712711 | 0.153853 |
| 28.29167 | 0.374915 | 0.250772 | 0.278829 | -0.01883 | 0.170415 | 0.120352 | 0.274844 | 0.172484 | -0.20414 | 0.746403 | 0.752427 | 0.375271 | 0.772796 | 0.812182 | 0.601343 | 1.210642 | 0.993651 | 0.11182 |
| 28.30556 | 0.030839 | -0.24071 | 0.065007 | 0.118963 | 0.135168 | 0.048872 | 0.18888 | -0.02393 | -0.81775 | 1.048811 | 1.059968 | -0.3165 | 0.926495 | 0.970505 | 0.895373 | 1.431089 | 1.102357 | 0.350661 |
| 28.31944 | -0.09925 | -0.4466 | -0.06659 | 0.419587 | 0.136896 | -0.03878 | 0.112825 | -0.22424 | -1.05618 | 1.16825 | 1.163181 | -0.71874 | 1.140576 | 1.104436 | 1.156306 | 1.443778 | 1.005539 | 0.573552 |
| 28.33333 | 0.136798 | -0.16708 | -0.0254 | 0.781979 | 0.190121 | -0.05232 | 0.142373 | -0.27644 | -0.67894 | 0.899279 | 0.852512 | -0.77117 | 1.008832 | 0.901292 | 1.013269 | 1.021559 | 0.634764 | 0.411646 |
| 28.34722 | 0.458234 | 0.245271 | 0.267812 | 1.080932 | 0.246369 | 0.007842 | 0.3178 | -0.12293 | 0.112551 | 0.515757 | 0.387201 | -0.67554 | 0.473886 | 0.4916 | 0.565895 | 0.52946 | 0.192664 | 0.015152 |
| 28.36111 | 0.804361 | 0.691055 | 0.753311 | 1.090073 | 0.241888 | 0.106522 | 0.518821 | 0.156682 | 0.719575 | 0.286003 | 0.173938 | -0.48159 | 0.291772 | 0.229335 | 0.230537 | 0.425686 | 0.032601 | -0.18123 |
| 28.375 | 1.028117 | 1.084752 | 1.144264 | 0.723448 | 0.192938 | 0.142463 | 0.554829 | 0.272838 | 0.680219 | 0.268223 | 0.192396 | -0.29352 | 0.321343 | 0.163327 | 0.08714 | 0.629492 | 0.129781 | -0.08213 |
| 28.38889 | 0.882763 | 1.011361 | 1.2055 | 0.184752 | 0.091575 | 0.105466 | 0.32889 | 0.041053 | 0.079498 | 0.354578 | 0.228362 | -0.35563 | 0.383461 | 0.150306 | -0.02345 | 0.892479 | 0.252141 | 0.025914 |
| 28.40278 | 0.271682 | 0.263385 | 0.649486 | -0.31872 | -0.016 | 0.062689 | -0.03909 | -0.2477 | -0.71454 | 0.48606 | 0.162634 | -0.47247 | 0.459571 | 0.085178 | -0.20098 | 0.922908 | 0.310089 | -0.04365 |
| 28.41667 | -0.44215 | -0.62985 | -0.04541 | -0.69137 | -0.07764 | 0.064238 | -0.24746 | -0.42317 | -1.34348 | 0.64076 | 0.034283 | -0.33827 | 0.42507 | -0.1049 | -0.33959 | 0.756468 | 0.303071 | -0.24245 |
| 28.43056 | -0.82775 | -1.14173 | -0.63343 | -0.55564 | -0.08964 | 0.073438 | -0.24305 | -0.44188 | -1.27715 | 0.793465 | 0.042186 | 0.037205 | 0.370673 | -0.29699 | -0.32553 | 0.56623 | 0.342847 | -0.11246 |
| 28.44444 | -0.60741 | -1.02328 | -0.75715 | -0.0446 | -0.04001 | 0.093947 | -0.01474 | -0.27333 | -0.6349 | 0.897746 | 0.421782 | 0.540998 | 0.432597 | 0.101463 | 0.084568 | 0.896581 | 0.466289 | 0.321971 |
| 28.45833 | -0.15824 | -0.60237 | -0.20401 | 0.555281 | 0.058575 | 0.13192 | 0.169351 | -0.01464 | 0.09605 | 0.922018 | 0.885787 | 0.989044 | 0.590733 | 0.758838 | 0.634452 | 1.347436 | 0.522929 | 0.594728 |
| 28.47222 | 0.238262 | -0.07595 | 0.586538 | 0.938 | 0.089853 | 0.128821 | 0.231973 | 0.166903 | 0.299086 | 0.769418 | 1.193283 | 1.189616 | 0.657976 | 1.298736 | 0.975354 | 1.516947 | 0.406844 | 0.425951 |
| 28.48611 | 0.597534 | 0.411545 | 1.406496 | 1.325608 | 0.245473 | 0.255641 | 0.595218 | 0.455641 | 0.404722 | 0.61302 | 1.036271 | 1.593758 | 0.614619 | 1.121005 | 0.706221 | 1.464542 | 0.396465 | 0.313444 |
| 28.5 | 1.087284 | 0.806529 | 2.03521 | 2.305265 | 0.849572 | 0.606681 | 1.597312 | 1.032751 | 1.392608 | 0.38809 | 1.029149 | 2.59858 | 0.351464 | 0.422587 | 0.423471 | 1.430856 | 0.882425 | 0.537555 |
| 28.51389 | 1.176169 | 1.017711 | 2.523272 | 3.383499 | 1.631862 | 0.829012 | 2.325789 | 1.434806 | 2.370525 | 0.178421 | 1.501541 | 2.84878 | -0.27989 | 0.406241 | 0.934103 | 1.306955 | 1.278015 | 0.886761 |
| 28.52778 | 0.968354 | 1.146848 | 3.175224 | 4.953343 | 1.771955 | 0.886435 | 2.701939 | 1.280572 | 3.00264 | -0.27563 | 2.373604 | 2.867307 | -0.6847 | 0.384095 | 1.31836 | 1.392348 | 2.323577 | 0.783351 |
| 28.54167 | 1.271886 | 1.50301 | 3.376044 | 5.23976 | 1.956232 | 1.034128 | 2.683489 | 0.88246 | 3.044784 | 0.230662 | 1.941159 | 2.674079 | -0.20426 | 0.46981 | 1.181268 | 1.728167 | 2.555353 | 0.581127 |
| 28.55556 | 1.25603 | 1.230316 | 2.264274 | 2.979341 | 1.605252 | 0.916453 | 2.089383 | 0.454177 | 1.655747 | 0.323364 | 0.453036 | 2.94977 | 0.335432 | 0.113682 | 0.523762 | 1.405671 | 1.996286 | 0.214667 |
| 28.56944 | 0.983605 | 0.658843 | 1.344749 | 1.172588 | 0.794644 | 0.647626 | 1.475095 | -0.09349 | 0.298802 | 0.452374 | -0.55258 | 2.656384 | 0.598402 | -0.65101 | -0.19813 | 1.004062 | 2.40428 | -0.29506 |
| 28.58333 | 0.801713 | 0.173982 | 0.470436 | -0.07003 | 0.09529 | 0.568019 | 1.026922 | -0.43663 | -0.63358 | 0.586593 | -1.53357 | 1.528651 | 0.805189 | -1.25478 | -1.11608 | 0.774879 | 3.688144 | -0.75819 |
| 28.59722 | 0.884415 | -0.14418 | -0.53846 | -1.21785 | -0.30902 | 0.735866 | 0.577094 | -0.50928 | -0.66464 | 0.72871 | -2.05062 | -0.23338 | 0.993646 | -0.79276 | -1.45449 | 0.504352 | 3.691239 | -1.45809 |
| 28.61111 | 0.809337 | -0.42739 | -1.27019 | -1.58375 | -0.52385 | 0.921849 | 0.525646 | -0.15879 | -0.52019 | 0.349259 | -2.06131 | -0.93614 | 1.116661 | 0.145537 | -0.82145 | -0.03242 | 3.506415 | -1.26409 |
| 28.625 | 0.73749 | -0.669 | -1.14998 | -1.08548 | -0.76914 | 0.909113 | 0.552201 | -0.00658 | -0.62853 | -0.10554 | -1.36689 | -2.58035 | 1.015451 | 0.926351 | 0.165019 | -0.63973 | 1.839487 | -0.5446 |
| 28.63889 | 0.737016 | -0.64254 | -0.31437 | -0.5021 | -0.92923 | 0.646202 | 0.448192 | 0.015803 | -0.70178 | -0.11251 | -0.51324 | -2.42585 | 0.797097 | 1.419012 | 1.167323 | 0.004749 | -0.27935 | 0.133894 |
| 28.65278 | 0.526584 | -0.53053 | 0.353063 | -0.15349 | -0.99939 | 0.174528 | 0.139041 | -0.00804 | -0.73679 | -0.39831 | -0.01669 | -1.2197 | 0.839563 | 1.296101 | 1.532504 | 1.018119 | -2.12517 | 0.788347 |
| 28.66667 | 0.039736 | -0.38495 | 0.781745 | -0.23807 | -0.81876 | -0.45969 | -0.41483 | 0.069768 | -1.02228 | -0.19515 | 0.928477 | -0.01926 | 0.831937 | 0.890086 | 1.237254 | 1.834324 | -3.2595 | 0.984083 |
| 28.68056 | -0.58051 | -0.30805 | 0.775272 | -0.77957 | -0.40286 | -1.13283 | -1.20556 | 0.139529 | -1.22499 | 0.555749 | 1.276061 | 1.065524 | 0.602236 | 0.463798 | 0.11811 | 1.341605 | -2.66378 | 0.81568 |
| 28.69444 | -1.38267 | -0.26154 | 0.603998 | -1.3922 | 0.046273 | -1.55338 | -1.88226 | 0.115206 | -0.97884 | 1.231921 | 1.461128 | 1.916656 | 0.231168 | -0.35073 | -0.63921 | 0.073012 | -1.7813 | 0.344341 |
| 28.70833 | -1.33616 | -0.22851 | 0.145835 | -1.06961 | 0.433047 | -1.39823 | -1.72422 | 0.036816 | -0.45596 | 0.839188 | 1.229755 | 2.129078 | 0.188463 | -1.34917 | -1.4899 | -0.72632 | -0.7424 | -0.13013 |
| 28.72222 | -0.69464 | -0.21677 | -0.38698 | -0.01047 | 1.091496 | -0.58296 | -0.91518 | -0.12493 | -0.03078 | 0.788729 | 0.393155 | 2.38367 | 0.227501 | -2.22094 | -2.15937 | -0.77125 | -0.86287 | -0.83711 |
| 28.73611 | -0.29063 | -0.25233 | -0.47756 | 0.623601 | 1.582717 | -0.03623 | -0.02581 | -0.18529 | -0.09897 | 0.996188 | -0.43464 | 1.503272 | -0.09814 | -1.60169 | -2.05693 | -0.70831 | -0.49254 | -1.47947 |
| 28.75 | 0.068056 | -0.3016 | -0.80812 | 0.861159 | 1.408593 | 0.256584 | 0.703447 | -0.16291 | -0.77908 | 0.668589 | -0.66383 | 0.299212 | -0.2702 | -0.55579 | -1.65301 | -0.36437 | 0.280454 | -1.81821 |
| 28.76389 | 0.353442 | -0.38622 | -1.77736 | 0.478917 | 1.094014 | 0.287477 | 1.182162 | -0.2577 | -0.85409 | 0.373039 | -0.54076 | -0.89938 | -0.77797 | 0.657785 | -0.97574 | -1.00859 | 0.326763 | -1.62136 |
| 28.77778 | 0.574233 | -0.43892 | -2.3088 | -0.0259 | 0.512425 | 0.144386 | 1.421652 | -0.39469 | -0.67228 | 0.391508 | -0.5297 | -1.70145 | -1.57203 | 1.225188 | -0.10376 | -1.16355 | -0.22441 | -1.0249 |
| 28.79167 | 0.578617 | -0.67585 | -0.55494 | -0.54742 | -0.22442 | -0.08013 | 1.221011 | -0.33594 | -0.36417 | 0.220405 | -1.11613 | -2.43985 | -1.44044 | 0.768572 | 0.424691 | -0.55507 | -1.03865 | -0.24437 |
| 28.80556 | 0.668052 | -0.90984 | 0.607786 | -0.84022 | -0.98949 | -0.2039 | 0.88083 | -0.07519 | -0.24709 | -0.48232 | -1.68345 | -3.12101 | -0.65072 | -0.00639 | 0.723839 | 0.047714 | -1.84915 | 0.107078 |
| 28.81944 | 0.427882 | -0.92526 | 1.066932 | -0.81902 | -1.51297 | -0.27237 | 0.265687 | 0.18621 | -0.47876 | -0.96712 | -1.94115 | -3.4485 | 0.255327 | -0.86876 | -0.17443 | 0.057408 | -1.63432 | 0.249134 |
| 28.83333 | -0.20685 | -0.74696 | 1.356553 | -0.78161 | -1.36342 | -0.31674 | -0.36344 | 0.399745 | -0.78539 | -0.97916 | -1.61261 | -3.58351 | 0.469038 | -1.62884 | -1.4076 | 0.452235 | -0.7701 | 0.233872 |
| 28.84722 | -0.94264 | -0.80604 | 1.25314 | -1.32687 | -1.2429 | -0.38072 | -1.39998 | 0.413728 | -1.41131 | -0.97861 | -1.17575 | -3.48105 | 0.12963 | -1.75484 | -1.98382 | 0.76962 | 0.251515 | 0.187075 |
| 28.86111 | -1.87886 | -1.08932 | -0.1538 | -1.90611 | -1.32604 | -0.50239 | -3.02484 | 0.22697 | -1.82565 | -0.86512 | -1.08656 | -3.04566 | -0.08213 | -1.69992 | -2.55553 | 0.504883 | 2.197536 | -0.05462 |
| 28.875 | -2.20595 | -1.25252 | -1.48492 | -2.18889 | -0.99784 | -0.55929 | -3.4307 | -0.14448 | -2.21511 | -0.74459 | -1.07834 | -2.38836 | -0.3947 | -1.47085 | -2.80754 | -0.76045 | 2.870068 | -0.55228 |
| 28.88889 | -1.70707 | -1.17047 | -2.84181 | -1.83478 | -0.61019 | -0.47684 | -2.05251 | -0.78848 | -2.32166 | -0.07856 | -0.77831 | -1.68046 | -1.08469 | -0.934 | -2.6398 | -2.44692 | 1.998188 | -0.9189 |
| 28.90278 | -0.95366 | -0.91713 | -2.90972 | -1.34921 | -0.39525 | -0.48539 | -0.83682 | -1.02947 | -2.02534 | 0.168646 | -0.54372 | -0.94391 | -1.95976 | -0.19354 | -1.73205 | -3.33816 | 1.536091 | -0.77586 |
| 28.91667 | -0.17287 | -0.52756 | -1.73334 | -1.03206 | -0.37491 | -0.384 | -0.13505 | -1.05591 | -1.35389 | 0.27889 | -0.32582 | -0.14807 | -1.33051 | 0.51626 | -0.21462 | -2.81532 | 0.582946 | -0.42291 |
| 28.93056 | 0.440124 | -0.14199 | -0.26429 | -0.40049 | -0.26564 | -0.16733 | 0.24066 | -0.85552 | -0.641 | 0.38942 | 0.068856 | 0.453992 | 0.458698 | 1.127528 | 1.331482 | -1.59468 | 0.16114 | 0.008493 |
| 28.94444 | 0.674729 | 0.12867 | 0.966608 | 0.282355 | 0.067307 | 0.014373 | 0.438662 | -0.4995 | 0.192544 | 0.626917 | 0.708825 | 0.63794 | 1.696408 | 1.513185 | 2.085947 | -0.34504 | 0.337479 | 0.367499 |
| 28.95833 | 0.657234 | 0.152272 | 2.167837 | 0.691371 | 0.500955 | 0.197006 | 0.569068 | -0.12591 | 1.028338 | 0.988389 | 1.041524 | 0.42426 | 2.087485 | 1.441717 | 1.696228 | 0.793541 | 0.887646 | 0.824152 |
| 28.97222 | 0.298546 | 0.038277 | 2.976806 | 0.790728 | 0.649507 | 0.320924 | 0.637979 | 0.155308 | 1.846581 | 1.28665 | 1.052814 | -0.05374 | 1.523031 | 0.779608 | 0.916444 | 1.520717 | 1.305282 | 1.187562 |
| 28.98611 | -0.446 | -0.50887 | 2.50858 | 0.006429 | 0.196962 | 0.131673 | 0.318047 | -0.20904 | 1.526117 | 0.890388 | 0.307146 | -1.35569 | 0.223461 | -0.48106 | -0.52206 | 1.012229 | 0.446656 | 0.475547 |
| 29 | -1.87541 | -2.09074 | -0.18431 | -1.93703 | -0.88685 | -0.38247 | -0.57923 | -1.50125 | -0.78032 | -0.53999 | -1.18948 | -3.41544 | -1.51292 | -2.25279 | -2.75555 | -1.58576 | -1.79605 | -1.80575 |
| 29.01389 | -3.59599 | -4.46033 | -3.17533 | -3.82768 | -1.94166 | -0.7687 | -1.33116 | -3.06173 | -3.3319 | -2.05265 | -3.12019 | -3.76726 | -2.82766 | -3.4702 | -4.56594 | -5.61484 | -3.59517 | -2.42019 |
| 29.02778 | -3.2711 | -5.27429 | -5.23101 | -3.72662 | -1.7358 | -0.784 | -1.16502 | -2.22012 | -3.78285 | -2.8857 | -5.23744 | -2.88378 | -1.38144 | -2.62654 | -3.52775 | -9.06478 | -4.66724 | -1.86189 |
| 29.04167 | -2.42911 | -4.59652 | -4.37221 | -2.39469 | -0.7793 | -0.80625 | -0.69687 | -1.41065 | -2.94407 | -1.45297 | -4.26911 | -2.54117 | -0.20143 | -0.8331 | -1.74438 | -8.49244 | -4.47015 | -1.42235 |
| 29.05556 | -1.35519 | -3.06177 | -3.66743 | -1.29813 | -0.09881 | -0.45328 | -0.41443 | -0.84256 | -2.5454 | -0.25017 | -1.80696 | -2.02469 | 0.410104 | 0.153216 | -0.56256 | -5.58198 | -2.85096 | -0.54296 |
| 29.06944 | -0.07961 | -1.13019 | -2.69774 | -0.28693 | 0.346897 | -0.04772 | -0.18665 | -0.35717 | -2.14653 | 0.94636 | 0.066866 | -1.63465 | 0.685915 | 0.605617 | 0.162398 | -2.12952 | -0.77739 | 0.363264 |
| 29.08333 | 0.893812 | 0.379397 | -1.69021 | 0.346228 | 0.619217 | 0.215417 | 0.057881 | -0.04046 | -1.41244 | 1.863373 | 1.788348 | -1.41553 | 0.724702 | 1.037733 | 0.9044 | 0.615586 | 0.719594 | 0.958528 |
| 29.09722 | 1.442331 | 1.078486 | -0.64284 | 0.764061 | 0.684213 | 0.263789 | 0.360107 | 0.216279 | -0.71664 | 2.11425 | 3.104944 | -1.24067 | 0.749127 | 1.0635 | 1.536582 | 2.917431 | 1.869518 | 1.30703 |
| 29.11111 | 1.837036 | 1.449651 | 0.281955 | 1.155366 | 0.80586 | 0.031302 | 0.518198 | 0.759018 | -0.26465 | 1.93068 | 4.431814 | -0.86126 | 0.94709 | 1.089848 | 2.26621 | 4.82269 | 2.812239 | 1.612607 |
| 29.125 | 2.197283 | 1.824912 | 1.008554 | 1.036708 | 0.893014 | 0.035301 | 0.383148 | 1.079772 | 0.00737 | 1.750258 | 4.493139 | -0.37813 | 1.249844 | 0.849882 | 2.477903 | 5.527793 | 3.01762 | 1.990241 |
| 29.13889 | 2.073826 | 1.523135 | 1.887859 | 0.711332 | 0.776022 | 0.149115 | 0.177702 | 1.585791 | 0.203594 | 0.907491 | 3.454107 | 0.106225 | 1.634577 | 0.801749 | 2.356431 | 5.077092 | 2.213606 | 2.324294 |
| 29.15278 | 1.842325 | 1.312064 | 3.303465 | 0.832107 | 0.553878 | 0.281835 | 0.475816 | 1.58948 | 0.285314 | 0.66026 | 2.740178 | 0.522232 | 2.478535 | 0.986496 | 3.20896 | 3.481647 | 1.054603 | 2.392151 |
| 29.16667 | 1.342191 | 1.35385 | 4.133137 | 0.812808 | 0.433013 | 0.316325 | 0.756374 | 1.365981 | 0.396706 | 0.855036 | 2.23911 | 0.986294 | 2.508778 | 1.366452 | 3.098997 | 1.833377 | 0.733305 | 2.110358 |
| 29.18056 | 0.932786 | 1.861597 | 4.224241 | 0.557611 | 0.22711 | 0.294587 | 1.090691 | 1.098638 | 0.576718 | 0.759629 | 1.483258 | 1.531913 | 1.634643 | 2.109544 | 3.653611 | 0.536321 | 0.331181 | 1.836462 |
| 29.19444 | 0.616137 | 2.758613 | 3.733651 | 0.329131 | 0.063968 | 0.262983 | 1.105106 | 0.717518 | 0.756641 | 0.652942 | 0.797941 | 2.025672 | 0.780084 | 2.419643 | 3.265553 | -0.29515 | 0.163611 | 1.31923 |
| 29.20833 | 0.530947 | 3.203053 | 3.336333 | 0.14021 | 0.003186 | 0.303737 | 0.553261 | 0.44111 | 0.899472 | 0.3163 | 0.151299 | 2.396515 | 0.309786 | 2.09069 | 2.341886 | -0.14747 | 0.154778 | 0.754408 |

| | | | | | | | | | | | | | | | | | | |
|---|---|---|---|---|---|---|---|---|---|---|---|---|---|---|---|---|---|---|
| 29.22222 | 0.44008 | 2.618281 | 2.517059 | 0.00969 | -0.04617 | 0.317612 | 0.264653 | 0.170435 | 0.933498 | -0.08433 | -0.33802 | 2.668762 | 0.021277 | 1.748262 | 1.547265 | 0.191588 | 0.357905 | 0.297739 |
| 29.23611 | 0.319897 | 1.786614 | 1.256342 | -0.09759 | -0.07159 | 0.314304 | 0.18013 | -0.36492 | 0.906514 | -0.4351 | -0.52631 | 2.313802 | -0.19375 | 1.158021 | 0.698044 | 0.319626 | 0.434615 | -0.12294 |
| 29.25 | 0.462051 | 1.313799 | 0.577418 | 0.044338 | -0.00774 | 0.431442 | 0.110003 | -0.20208 | 1.256463 | -0.51927 | -0.37858 | 1.843745 | -0.27765 | 0.522103 | 0.023823 | 0.367337 | 0.372205 | -0.40699 |
| 29.26389 | 0.682118 | 1.164822 | 0.454225 | 0.471306 | 0.16846 | 0.517295 | 0.061586 | 0.226929 | 1.576506 | -0.11998 | 0.313284 | 1.665648 | -0.05086 | 0.399764 | -0.14761 | 0.638448 | 0.582736 | -0.19606 |
| 29.27778 | 0.741054 | 0.95113 | 0.273802 | 0.798248 | 0.190445 | 0.476928 | 0.182089 | 0.32396 | 1.630278 | 0.312534 | 0.840054 | 1.887062 | 0.11588 | 0.396241 | -0.13324 | 1.05213 | 0.717926 | 0.064162 |
| 29.29167 | 0.711158 | 0.625938 | -0.0136 | 0.794464 | 0.121871 | 0.327554 | 0.201475 | 0.14259 | 1.438503 | 0.336966 | 0.5961 | 1.95781 | -0.06659 | 0.13015 | -0.31147 | 0.706396 | 0.391921 | -0.15125 |
| 29.30556 | 0.613446 | 0.378155 | -0.06215 | 0.604392 | 0.086135 | 0.208887 | 0.251671 | -0.0047 | 1.261552 | 0.148594 | 0.021259 | 2.179934 | -0.39216 | -0.27992 | -0.547 | 0.527467 | -0.01847 | -0.5719 |
| 29.31944 | 0.519569 | 0.181485 | -0.05126 | 0.30745 | 0.058455 | 0.112161 | 0.220769 | -0.08969 | 1.069137 | 0.086571 | -0.38686 | 2.201871 | -0.73715 | -0.53539 | -0.6537 | 0.590906 | -0.14319 | -0.90568 |
| 29.33333 | 0.506917 | 0.133243 | -0.01299 | 0.095091 | 0.053079 | 0.135612 | 0.176424 | -0.0285 | 1.021643 | 0.2153 | -0.31691 | 1.83209 | -0.60555 | -0.37348 | -0.60692 | 0.792369 | 0.067014 | -0.87881 |
| 29.34722 | 0.654007 | 0.365114 | 0.050044 | 0.162273 | -0.00543 | 0.1923 | 0.163169 | 0.166689 | 1.146122 | 0.489322 | 0.213416 | 1.197781 | 0.025223 | 0.217289 | -0.22611 | 0.970543 | 0.422434 | -0.73553 |
| 29.36111 | 0.771209 | 0.672979 | 0.078876 | 0.318518 | -0.23014 | 0.153731 | 0.126452 | 0.247783 | 1.154736 | 0.648067 | 0.716934 | 0.364826 | 0.245656 | 0.830766 | 0.232353 | 1.021649 | 0.515111 | -0.63713 |
| 29.375 | 0.636582 | 0.720668 | 0.110273 | 0.384858 | -0.53225 | 0.060669 | 0.13083 | 0.159598 | 1.120133 | 0.58072 | 0.888356 | -0.4754 | 0.125787 | 1.060268 | 0.448108 | 0.998035 | 0.39882 | -0.46221 |
| 29.38889 | 0.271924 | 0.575162 | 0.310449 | 0.472111 | -0.64848 | 0.021067 | 0.271396 | 0.099145 | 1.036342 | 0.578137 | 0.793868 | -1.14581 | 0.492967 | 1.057451 | 0.605766 | 1.029078 | 0.310297 | -0.27701 |
| 29.40278 | -0.07522 | 0.366737 | 0.671261 | 0.594793 | -0.49844 | 0.036484 | 0.521542 | 0.077564 | 0.917754 | 0.426172 | 0.335196 | -1.66476 | 1.264699 | 0.872263 | 0.667931 | 0.957412 | 0.089921 | -0.37232 |
| 29.41667 | -0.24229 | 0.18229 | 0.93937 | 0.623653 | -0.27475 | 0.016028 | 0.729056 | 0.007114 | 0.828483 | 0.258344 | -0.28873 | -1.60781 | 1.149572 | 0.654278 | 0.613104 | 0.90242 | -0.03549 | -0.52624 |
| 29.43056 | -0.10458 | 0.096716 | 0.869901 | 0.65413 | -0.12989 | 0.023976 | 0.733782 | -0.10521 | 0.698079 | 0.204192 | -0.8339 | -1.27328 | 0.484611 | 0.492261 | 0.453471 | 0.853525 | 0.041912 | -0.49454 |
| 29.44444 | 0.032252 | 0.039666 | 0.564595 | 0.576432 | -0.1286 | 0.056255 | 0.493622 | -0.14684 | 0.399637 | 0.241251 | -0.89172 | -1.02979 | -0.23307 | 0.335837 | -0.02631 | 0.738253 | 0.216613 | -0.41695 |
| 29.45833 | 0.226734 | -0.01819 | 0.317929 | 0.471019 | -0.14337 | 0.00179 | 0.236121 | -0.08623 | 0.18311 | 0.377158 | -0.4971 | -0.89728 | -0.94554 | 0.026529 | -0.80858 | 0.510417 | 0.305934 | -0.29705 |
| 29.47222 | 0.515156 | 0.019733 | 0.366935 | 0.539498 | 0.03138 | -0.08652 | 0.201234 | 0.046596 | 0.257623 | 0.50542 | 0.19782 | -0.64385 | -1.18349 | -0.06448 | -1.12755 | 0.329408 | 0.375512 | -0.10235 |
| 29.48611 | 0.840657 | 0.257618 | 0.72408 | 1.042478 | 0.632373 | 0.052522 | 0.517785 | 0.263229 | 0.809175 | 0.884269 | 0.925832 | 0.525701 | 0.09913 | 0.171677 | -0.34018 | 1.001189 | 0.635044 | 0.248772 |
| 29.5 | 1.217292 | 0.625351 | 1.436536 | 2.008341 | 1.610478 | 0.379029 | 1.254894 | 0.676692 | 1.964811 | 1.356981 | 1.975646 | 2.838443 | 1.492834 | 0.642763 | 0.982048 | 1.659405 | 0.938315 | 0.901458 |
| 29.51389 | 1.541517 | 0.81887 | 2.346854 | 2.603294 | 2.34411 | 0.822861 | 2.093017 | 0.9893 | 2.549159 | 1.399189 | 3.457696 | 6.78777 | 2.61625 | 1.27122 | 2.570952 | 1.430743 | 0.110334 | 1.277902 |
| 29.52778 | 1.664897 | 0.863652 | 2.871531 | 2.767526 | 2.053342 | 1.006824 | 2.517167 | 1.100368 | 2.404458 | 1.091563 | 3.984446 | 9.654016 | 2.411175 | 1.983027 | 3.658356 | 0.942434 | -0.85434 | 1.278936 |
| 29.54167 | 1.803777 | 0.909079 | 3.632305 | 2.576469 | 1.662379 | 0.800344 | 2.330648 | 0.883006 | 2.834189 | 1.448851 | 3.453611 | 7.181112 | 1.016339 | 1.782467 | 2.643281 | 0.688125 | -1.67834 | 1.748375 |
| 29.55556 | 1.223963 | 0.841222 | 2.737848 | 1.841104 | 1.221357 | 0.650534 | 1.5848 | 0.982696 | 2.516954 | 1.699437 | 2.110816 | 5.671688 | 0.707362 | 1.13488 | 1.813005 | 0.045881 | -1.2834 | 1.299586 |
| 29.56944 | 0.533968 | 0.806669 | 0.929239 | 1.107657 | 1.200971 | 0.475678 | 1.015014 | 0.754731 | 1.32537 | 1.20173 | 1.057137 | 4.8547 | 0.49624 | 0.46482 | 1.023738 | -0.01804 | -0.40712 | 0.781544 |
| 29.58333 | -0.1486 | 0.829932 | -0.12182 | 0.593861 | 1.364943 | 0.431797 | 0.747828 | 0.096343 | 0.354979 | 0.598364 | -0.0608 | 4.659113 | 0.143606 | -0.65119 | 0.050114 | -0.03554 | 0.070063 | 0.09116 |
| 29.59722 | -0.46996 | 0.704367 | -1.26559 | 0.083327 | 1.350785 | 0.403147 | 0.532357 | -0.45439 | -0.2507 | 0.492651 | -0.99772 | 5.209395 | 0.002355 | -1.43968 | -1.02484 | -0.00402 | 0.340891 | -0.58407 |
| 29.61111 | -0.64992 | 0.446809 | -2.36963 | -0.05771 | 0.979588 | 0.478768 | 0.648269 | -0.23145 | -0.29194 | 0.133549 | -1.65063 | 4.655879 | -0.58776 | -1.54137 | -1.8145 | -0.19844 | 1.117049 | -0.92234 |
| 29.625 | -0.76857 | -0.04215 | -2.26569 | -0.08374 | 0.596874 | 0.614013 | 0.721476 | 0.114043 | -0.3357 | -0.11865 | -2.02222 | 3.638963 | -1.7915 | -0.65091 | -2.05358 | -0.9189 | 2.176482 | -1.31187 |
| 29.63889 | -0.88232 | -0.74725 | -2.08867 | 0.176155 | -0.00775 | -0.26153 | 0.708244 | 0.214614 | -0.53408 | -0.25858 | -2.21506 | 2.456001 | -1.564 | -0.00851 | -1.28727 | -2.50171 | 2.653149 | -1.42586 |
| 29.65278 | -0.63939 | -1.28504 | -1.14552 | 0.357861 | -0.57331 | -0.76182 | 0.466274 | 0.259854 | -0.46547 | -0.33477 | -1.30368 | 1.476285 | 0.339139 | 0.743929 | 0.41437 | -1.0521 | 2.736942 | -0.77035 |
| 29.66667 | 0.064631 | -1.35903 | 0.575346 | 0.573122 | -0.98796 | -0.31553 | 0.192431 | 0.265104 | 0.038314 | -0.49815 | -0.53409 | 0.756262 | 1.614031 | 1.491119 | 1.991137 | 1.898641 | 2.22054 | -0.13891 |
| 29.68056 | 0.66499 | -1.22005 | 1.647015 | 0.56977 | -1.32813 | -0.06847 | -0.11295 | 0.231016 | 0.206578 | -0.49933 | -0.11381 | 0.028381 | 1.661939 | 1.54955 | 2.670956 | 4.452819 | 1.363577 | 0.927447 |
| 29.69444 | 0.760187 | -1.04017 | 2.200291 | 0.625338 | -1.62009 | 0.288705 | -0.4214 | 0.186497 | 0.351042 | -0.6371 | 0.100243 | -0.90069 | 1.260013 | 1.428475 | 1.922835 | 4.514212 | 0.498869 | 1.135652 |
| 29.70833 | 0.634908 | -1.00292 | 2.029582 | 0.641894 | -1.746 | 0.743019 | -0.60905 | 0.10972 | 0.642103 | -0.46293 | 0.540866 | -1.41745 | 0.572492 | 0.609525 | 0.548981 | 2.461809 | -0.88141 | 1.10638 |
| 29.72222 | 0.391044 | -0.82003 | 1.866052 | 0.846305 | -1.0965 | 0.452048 | -0.88722 | 0.032995 | 0.768424 | -0.04298 | 1.090392 | -1.344 | 0.191571 | -0.2748 | -0.63164 | 0.273585 | -1.66319 | 0.413865 |
| 29.73611 | 0.151977 | -0.52614 | 1.254861 | 0.87111 | -0.14718 | 0.001321 | -1.11851 | 0.0091 | 0.467437 | 0.631495 | 1.336158 | -0.41106 | -0.13358 | -0.81532 | -1.34979 | -1.97299 | -1.57042 | -0.58211 |
| 29.75 | -0.24643 | -0.27991 | 0.602617 | 0.621048 | 0.681934 | -0.00443 | -1.09157 | -0.00929 | -0.23459 | 1.26342 | 1.00029 | 0.542889 | -0.80362 | -1.14338 | -2.16675 | -2.93557 | -0.81953 | -1.2122 |
| 29.76389 | -0.70439 | -0.4095 | -0.2026 | 0.242795 | 1.364726 | 0.055661 | -0.8681 | -0.14787 | -0.4767 | 1.622711 | -0.24837 | 0.949846 | -2.02141 | -1.28554 | -3.18006 | -2.75935 | -0.15436 | -2.02282 |
| 29.77778 | -1.20909 | -0.95464 | -1.38404 | -0.26198 | 1.447421 | -0.17761 | -0.66724 | -0.60857 | -0.78962 | 1.598755 | -1.37423 | 0.875454 | -2.48326 | -0.96815 | -3.60274 | -1.47708 | 0.171402 | -2.65794 |
| 29.79167 | -1.83838 | -1.52319 | -2.00347 | -0.64636 | 1.245802 | -0.67276 | -0.25497 | -1.38573 | -0.921 | 1.119581 | -2.0374 | 0.493968 | -2.83582 | -1.29691 | -3.5917 | -0.0385 | 0.344125 | -2.0848 |
| 29.80556 | -1.90534 | -1.79973 | -2.75493 | -0.942 | 0.831013 | -0.1835 | -0.06822 | -1.78943 | -0.78908 | 0.283944 | -2.56813 | -0.23193 | -1.98631 | -1.49861 | -2.96718 | 0.059937 | -0.41577 | -1.46901 |
| 29.81944 | -1.70587 | -1.74229 | -2.36102 | -1.06642 | 0.134493 | -0.1982 | 0.027709 | -1.48498 | -1.02262 | -0.87535 | -2.03612 | -1.26884 | -0.77592 | -1.282 | -2.00539 | -1.21233 | -1.52873 | -1.46024 |
| 29.83333 | -1.1248 | -1.4467 | -1.77372 | -1.00425 | -0.69366 | -0.21686 | 0.053583 | -1.1172 | -1.00109 | -1.71081 | -1.81724 | -2.08745 | -0.02183 | -0.83998 | -1.1632 | 0.090333 | -2.11604 | -0.92445 |
| 29.84722 | -0.35182 | -1.10352 | -1.24334 | -0.87427 | -1.30049 | -0.06638 | -0.18344 | -0.78801 | -0.48711 | -1.61406 | -1.57847 | -2.62057 | 0.201887 | -0.45223 | -0.63713 | -0.12302 | -2.94157 | -0.56698 |
| 29.86111 | 0.026275 | -0.9054 | -0.514 | -0.95063 | -1.6675 | 0.157649 | -0.53015 | -0.54881 | 0.044107 | -1.07097 | -1.37155 | -2.64551 | 0.333238 | -0.30871 | -0.3914 | -0.66586 | -1.44607 | 0.161073 |
| 29.875 | 0.202441 | -0.68422 | -0.53015 | -1.08052 | -1.44362 | 0.108335 | -0.79382 | -0.37535 | 0.473378 | -0.52707 | -1.62117 | -2.29814 | 0.477061 | -0.34165 | -0.25274 | -0.0063 | -0.09731 | 0.667416 |
| 29.88889 | 0.367035 | -0.39195 | -0.55608 | -1.06245 | -1.85435 | 0.085391 | -0.76302 | 0.003897 | 0.512056 | -0.19595 | -1.60938 | -1.28054 | 0.746037 | -0.52796 | -0.31846 | 0.990539 | 0.635835 | 0.665581 |
| 29.90278 | 0.499873 | -0.27301 | 0.029519 | -0.7387 | -2.01203 | -0.03417 | -0.15994 | 0.490319 | 0.625692 | 0.085476 | -0.75939 | -0.40562 | 0.965057 | -0.48061 | -0.44706 | 1.849311 | 1.195704 | 0.454504 |
| 29.91667 | 0.529806 | -0.09812 | 0.838149 | -0.17145 | -1.06883 | -0.29511 | 0.362793 | 0.644242 | 0.815734 | 0.254121 | 0.432817 | 0.0209 | 0.940023 | -0.33767 | -0.70315 | 2.563838 | 1.814145 | 0.300112 |
| 29.93056 | 0.32313 | -0.03088 | 1.553535 | 0.11199 | -0.41404 | -0.24984 | 0.72623 | 0.559949 | 0.676518 | 0.382283 | 1.628524 | 0.057347 | 0.785566 | -0.0494 | -0.76866 | 2.518878 | 2.163363 | 0.224741 |
| 29.94444 | 0.041769 | 0.158069 | 1.912976 | 0.306452 | 0.138593 | 0.05315 | 0.915106 | 0.386022 | 0.429989 | 0.760909 | 2.464741 | 0.062356 | 0.785047 | 0.365182 | -0.50584 | 2.665549 | 2.119552 | 0.271377 |
| 29.95833 | -0.13643 | 0.293745 | 1.794273 | 0.425156 | 0.502523 | 0.131981 | 0.980231 | 0.189299 | 0.270568 | 0.894268 | 2.328724 | 0.337127 | 0.848687 | 0.58415 | -0.15837 | 3.260539 | 1.748115 | 0.14888 |
| 29.97222 | -0.18011 | 0.22372 | 1.359935 | 0.35648 | 0.509939 | 0.219124 | 0.936771 | -0.17962 | -0.25277 | 0.643566 | 1.523397 | 0.650325 | 0.771053 | 0.552337 | -0.00456 | 3.020883 | 1.41049 | -0.05226 |
| 29.98611 | -0.63489 | -0.53127 | 0.336424 | -0.42491 | -0.12706 | 0.014331 | 0.407658 | -0.96014 | -1.34004 | -0.01209 | 0.398371 | 0.00285 | -0.2395 | -0.09839 | -1.35759 | 1.459552 | 0.014974 | -0.97335 |
| 30 | -2.04102 | -2.38446 | -1.58848 | -1.88313 | -1.49112 | -0.88142 | -0.80938 | -2.43368 | -3.90258 | -1.87895 | -1.36728 | -1.88096 | -2.67532 | -1.53289 | -4.95044 | -1.21911 | -3.57506 | -3.26263 |
| 30.01389 | -3.50422 | -3.97927 | -4.13502 | -2.85302 | -2.45636 | -1.14642 | -2.0456 | -2.39902 | -5.32467 | -2.95364 | -3.59579 | -3.03657 | -3.47149 | -2.00673 | -5.56957 | -2.90517 | -5.00298 | -2.88533 |
| 30.02778 | -3.43169 | -3.82009 | -5.84428 | -3.31 | -1.68164 | -1.27819 | -3.24636 | -1.41055 | -4.52892 | -2.83041 | -4.79294 | -0.67949 | -1.88368 | -1.04595 | -2.56534 | -2.18444 | -2.57716 | -0.91477 |
| 30.04167 | -2.62815 | -2.57974 | -4.8494 | -2.28091 | -1.22969 | -1.01172 | -2.94684 | -0.41892 | -3.07959 | -0.98359 | -3.28556 | 0.6984 | -0.34565 | 0.188528 | -0.22766 | -1.81698 | -0.89845 | 0.20321 |
| 30.05556 | -2.052 | -2.15106 | -3.45758 | -0.87183 | -0.68438 | -0.55233 | -1.38273 | -0.00268 | -2.08023 | -0.02136 | -2.76049 | 1.416513 | 0.384956 | 0.178375 | 0.189443 | -0.52078 | -0.07407 | 0.564822 |
| 30.06944 | -1.5028 | -1.82179 | -1.88076 | -0.41591 | -0.25988 | -0.22872 | 0.045209 | -0.06502 | -1.70645 | 0.693153 | -1.41816 | 1.865665 | 0.49366 | 0.279697 | -0.18327 | 0.372224 | -0.11515 | 0.516015 |
| 30.08333 | -0.42637 | -0.77305 | -0.31092 | 0.434774 | 0.033501 | 0.097528 | 1.457753 | -0.17885 | -1.29397 | 0.877348 | -0.23722 | 2.147605 | 0.285438 | 0.317692 | -0.26524 | 0.441731 | -0.38497 | 0.415165 |
| 30.09722 | 0.606507 | 0.295118 | 0.388174 | 1.177111 | 0.298983 | 0.382391 | 2.116616 | -0.18364 | -0.74544 | 0.802757 | 0.44021 | 2.292596 | 0.073567 | 0.377297 | -0.0896 | 0.28674 | -0.4623 | 0.379677 |
| 30.11111 | 1.57638 | 1.169479 | 1.417054 | 1.609554 | 0.563469 | 0.599674 | 2.242152 | -0.03337 | -0.04074 | 0.705812 | 0.854861 | 1.067297 | 0.068152 | 0.413147 | 0.135842 | 0.385402 | 0.099345 | 0.450299 |
| 30.125 | 2.22207 | 1.828268 | 0.508948 | 1.818965 | 0.769755 | 0.682379 | 2.233018 | 0.109317 | 0.692229 | 0.66702 | 1.001364 | 1.289563 | 0.139224 | 0.487502 | 0.307199 | 0.822855 | 0.737193 | 0.621276 |
| 30.13889 | 1.989351 | 1.794003 | 0.657163 | 1.507571 | 0.595532 | 0.600411 | 1.970211 | 0.07247 | 0.996667 | 0.697787 | 1.155787 | 0.402408 | 0.211848 | 0.591603 | 0.404782 | 1.185776 | 0.944027 | 0.672994 |
| 30.15278 | 1.855956 | 1.676969 | 0.84205 | 0.981277 | 0.833862 | 0.464613 | 1.362054 | -0.04952 | 0.709605 | 0.801569 | 1.59728 | -0.42777 | 0.232033 | 0.57235 | 0.497138 | 1.257728 | 0.787612 | 0.495606 |
| 30.16667 | 2.41273 | 1.378586 | 1.066193 | 0.451986 | 0.951397 | 0.733107 | 0.776675 | 0.102751 | 0.784873 | 0.509939 | 2.062993 | -0.46033 | 0.342164 | 0.296545 | 0.464145 | 1.160073 | 0.720038 | 0.233151 |
| 30.18056 | 2.254738 | 0.396615 | 1.422982 | -0.50029 | 0.975056 | 1.085511 | 0.08221 | 0.545382 | 1.617136 | 0.678408 | 2.324629 | -0.16493 | 0.644615 | 0.140219 | 1.037604 | 1.235639 | 1.192125 | 0.291773 |
| 30.19444 | 2.091375 | 0.039262 | 1.941568 | -0.487 | 0.857242 | 1.138308 | -0.08917 | 0.882316 | 1.869804 | 0.347165 | 2.793331 | 0.150073 | 0.924606 | 0.090843 | 1.729365 | 1.049094 | 1.85234 | 0.447732 |
| 30.20833 | 2.319783 | 0.45738 | 2.665435 | 0.490121 | 0.643661 | 0.820091 | 0.18955 | 1.110518 | 1.899858 | 0.108517 | 2.820895 | 0.094393 | 0.900604 | -0.16037 | 2.284317 | 0.598342 | 1.974967 | 0.325526 |
| 30.22222 | 2.041709 | 1.156042 | 3.159423 | 1.182381 | 0.544243 | 0.794201 | 0.459602 | 1.333636 | 1.563888 | -0.31113 | 2.150403 | -0.05933 | 0.878618 | -0.56472 | 2.728259 | 0.101171 | 0.66157 | -0.0042 |
| 30.23611 | 1.795788 | 1.620494 | 3.214287 | 1.409302 | 0.411315 | 0.97342 | 0.880023 | 1.363061 | 1.333601 | 0.038141 | 1.590203 | 0.127683 | 1.887129 | -0.37087 | 2.816596 | -0.26348 | -0.81444 | -0.14969 |
| 30.25 | 1.377258 | 1.900395 | 2.845568 | 1.227012 | 0.308154 | 0.88283 | 1.191352 | 1.280051 | 1.343577 | 0.374479 | 1.275074 | 0.44117 | 2.006197 | 0.16475 | 2.389662 | -0.56036 | -1.29047 | -0.2794 |
| 30.26389 | 0.554736 | 1.901184 | 2.26019 | 0.762976 | 0.30323 | 0.763044 | 1.106939 | 1.090285 | 1.721339 | -0.61937 | 0.363707 | -0.14145 | 0.241888 | 0.310784 | 0.895165 | -1.36803 | -1.80579 | -1.27755 |
| 30.27778 | -0.18755 | 1.842422 | 1.858614 | 0.258504 | 0.376677 | 0.702483 | 0.833453 | 0.805162 | 2.618805 | -2.90936 | -1.54789 | -2.07743 | -2.85681 | 0.01741 | -1.75845 | -3.2178 | -3.35919 | -3.75402 |
| 30.29167 | -0.19299 | 1.440255 | 1.965897 | 0.25002 | 0.585349 | 0.678589 | 0.970463 | 0.655893 | 3.092043 | -5.25708 | -4.12961 | -4.48338 | -5.72282 | -3.28388 | -4.5824 | -6.49948 | -5.49752 | -6.37209 |
| 30.30556 | 0.451132 | 1.013894 | 1.700633 | 0.638058 | 0.798048 | 0.633538 | 1.308338 | 0.933669 | 2.626886 | -6.1257 | -6.24474 | -6.11131 | -6.75502 | -5.24648 | -6.18745 | -7.29599 | -6.53981 | -7.3188 |
| 30.31944 | 0.962266 | 0.860461 | 1.537675 | 0.919989 | 0.962328 | 0.643432 | 1.498462 | 1.325162 | 2.752783 | -6.41533 | -7.00998 | -6.58688 | -6.90973 | -5.73098 | -6.64945 | -6.30011 | -6.79339 | -7.38422 |
| 30.33333 | 1.077052 | 0.852981 | 0.835485 | 0.845517 | 1.066899 | 0.537615 | 1.038575 | 1.31979 | 2.163701 | -6.57953 | -7.14138 | -6.00408 | -6.6768 | -5.64884 | -6.71308 | -5.43543 | -6.52972 | -7.13825 |

| | | | | | | | | | | | | | | | | | | |
|---|---|---|---|---|---|---|---|---|---|---|---|---|---|---|---|---|---|---|
| 30.34722 | 0.790903 | 0.759901 | -0.20315 | 0.621382 | 0.943688 | 0.180524 | 0.332895 | 0.632889 | 0.553615 | -6.60274 | -7.26861 | -4.97913 | -6.16095 | -5.46744 | -6.57696 | -5.09862 | -5.99659 | -6.63186 |
| 30.36111 | 0.36617 | 0.411164 | -0.75112 | 0.394458 | 0.673375 | -0.04067 | -0.09259 | -0.14745 | -0.49698 | -6.26042 | -6.93185 | -3.90987 | -5.46624 | -4.83376 | -5.98164 | -4.6788 | -5.33287 | -5.56684 |
| 30.375 | 0.380171 | 0.12296 | -0.52515 | 0.387061 | 0.352481 | 0.071379 | -0.03578 | -0.41118 | -0.28968 | -4.28425 | -4.83493 | -2.90456 | -4.34353 | -3.42983 | -4.31624 | -3.68422 | -4.07961 | -4.09157 |
| 30.38889 | 1.023584 | 0.057434 | -0.1636 | 0.679111 | 0.142792 | 0.210272 | 0.331949 | -0.1328 | 0.41658 | -1.66759 | -1.90736 | -1.7699 | -2.1025 | -1.47715 | -2.28784 | -2.18412 | -2.451 | -2.36185 |
| 30.40278 | 1.773742 | 0.243778 | 0.116844 | 1.115963 | 0.1343 | 0.356728 | 0.641734 | 0.095942 | 1.069406 | -0.02702 | 0.590874 | -0.60804 | -0.64147 | 0.178533 | -0.68171 | -0.41266 | -1.04007 | -1.06169 |
| 30.41667 | 2.234164 | 0.499216 | 0.304679 | 1.492219 | 0.184148 | 0.397225 | 0.727291 | 0.121636 | 1.593826 | 0.32312 | 2.158929 | 0.146944 | 0.105583 | 1.055969 | 0.120235 | 1.139854 | -0.20517 | -0.34491 |
| 30.43056 | 2.059713 | 0.60695 | 0.235465 | 1.507456 | 0.168367 | 0.255676 | 0.596647 | -0.01109 | 1.277317 | -0.14672 | 1.864996 | 0.516036 | 0.29616 | 1.09427 | 0.498237 | 2.319091 | 0.175886 | -0.08045 |
| 30.44444 | 1.575255 | 0.609082 | 0.094565 | 1.265986 | 0.04566 | 0.071732 | 0.39662 | -0.2845 | 0.236062 | -0.90373 | 0.51461 | 0.805641 | 0.200433 | 0.741276 | 0.670228 | 3.250048 | 0.274283 | -0.02679 |
| 30.45833 | 1.082831 | 0.71364 | 0.095874 | 1.022655 | -0.19843 | 0.048931 | 0.268698 | -0.41682 | -0.77747 | -1.06875 | -0.66794 | 1.16699 | 0.374485 | 0.482831 | 0.652273 | 2.861245 | 0.137907 | -0.07622 |
| 30.47222 | 0.893681 | 0.979219 | 0.207924 | 0.957427 | -0.28896 | 0.041782 | 0.27126 | -0.3502 | -1.35701 | -0.66628 | -0.95843 | 1.080858 | 0.430195 | 0.333812 | 0.512917 | 1.343604 | -0.03869 | -0.11046 |
| 30.48611 | 0.867941 | 1.100717 | 0.46228 | 1.038251 | 0.119807 | -0.04469 | 0.429473 | -0.06761 | -0.85575 | -0.27782 | -0.4296 | 0.39403 | -0.09874 | 0.382821 | 0.210164 | -0.0694 | 0.087428 | -0.20303 |
| 30.5 | 1.514239 | 1.212737 | 1.155202 | 1.721516 | 1.219365 | -0.02262 | 0.88131 | 0.409155 | 0.47926 | 0.106655 | 0.757078 | -0.514 | -0.49142 | 1.1171 | 0.311702 | -1.48543 | 1.031489 | 0.059188 |
| 30.51389 | 2.508045 | 1.285934 | 2.317647 | 2.643606 | 2.52922 | 0.252192 | 1.712915 | 1.012763 | 1.572271 | 1.099164 | 2.611404 | -0.71057 | 0.212412 | 1.898009 | 1.715494 | -0.73357 | 2.096577 | 1.227709 |
| 30.52778 | 3.206381 | 1.517444 | 3.884331 | 3.061644 | 2.701469 | 0.477314 | 2.496759 | 1.643401 | 2.052809 | 2.605842 | 5.306975 | -0.17529 | 2.154068 | 2.648277 | 3.5675 | 1.606688 | 2.092608 | 1.972461 |
| 30.54167 | 2.893307 | 2.021494 | 4.301991 | 2.67587 | 2.20816 | 0.295726 | 2.354091 | 2.333832 | 2.275819 | 3.412969 | 5.440295 | 0.258758 | 2.842539 | 3.00373 | 3.606127 | 4.581393 | 1.423127 | 2.320958 |
| 30.55556 | 2.445659 | 1.838943 | 3.212917 | 1.673411 | 2.005351 | 0.594871 | 1.605754 | 1.842627 | 2.017377 | 2.262343 | 3.086346 | 0.516789 | 2.207855 | 1.777661 | 2.160922 | 3.228927 | 0.65487 | 1.451864 |
| 30.56944 | 1.030323 | 1.397214 | 1.918875 | 0.995985 | 1.756557 | 0.653319 | 0.83351 | 1.47532 | 1.616886 | 0.515411 | 0.548465 | 0.911438 | 1.052978 | 0.333918 | 1.086044 | 0.584999 | 0.284967 | 0.485247 |
| 30.58333 | -0.3839 | 1.088952 | 0.750314 | 0.382603 | 1.860512 | 0.645132 | 0.058995 | 1.311309 | 1.632887 | -0.28022 | -0.94986 | 1.146513 | -0.31056 | -0.64903 | 0.070135 | 0.146563 | 0.45713 | -0.40056 |
| 30.59722 | -1.1438 | 0.885343 | -0.08231 | 0.045814 | 1.868294 | 0.628043 | -0.48618 | 0.890712 | 0.785192 | -0.50913 | -2.12745 | 1.511324 | -1.2543 | -1.5974 | -1.48583 | 0.002875 | 0.901142 | -1.28234 |
| 30.61111 | -0.99398 | 0.771558 | -0.79672 | -0.11243 | 1.627098 | 0.549123 | -0.51618 | 0.362948 | -0.49626 | -0.52641 | -3.21612 | 1.658905 | -1.19533 | -1.9487 | -2.50541 | -0.12214 | 1.675195 | -1.90321 |
| 30.625 | -1.22073 | 0.505127 | -1.52401 | -0.06081 | 1.185898 | 0.30724 | -0.18734 | -0.18115 | -1.06376 | 0.188792 | -4.50155 | 1.602087 | -2.14283 | -1.52434 | -2.83879 | -2.96832 | 1.387904 | -1.00533 |
| 30.63889 | -1.28235 | -0.11186 | -1.99306 | -0.1852 | 0.520261 | 0.014561 | -0.02055 | -0.84934 | -1.17727 | 0.77304 | -6.0217 | 1.492819 | -3.36894 | -1.72898 | -3.07971 | -3.88317 | 0.8949 | -0.31779 |
| 30.65278 | -0.96627 | -0.84354 | -1.76406 | -0.37242 | -0.19265 | -0.40483 | 0.064214 | -1.15581 | -1.16012 | 0.691443 | -5.75296 | 1.502421 | -2.60873 | -1.90269 | -3.38079 | -2.07484 | 1.308438 | 0.056734 |
| 30.66667 | 0.673893 | -1.11258 | -2.32458 | -0.38857 | -0.86243 | -0.6117 | -0.0363 | -1.18175 | -1.45337 | 0.580366 | -3.11926 | 1.286895 | -0.84094 | -1.61497 | -1.30524 | -0.29439 | 1.933894 | 0.304638 |
| 30.68056 | 1.359589 | -1.0503 | -2.04788 | -0.31342 | -1.39857 | -0.56815 | -0.17398 | -1.13653 | -0.97666 | 0.179312 | -0.82141 | 0.810918 | 0.661616 | -1.01201 | 0.705552 | 0.60979 | 1.751878 | 0.784873 |
| 30.69444 | 1.081754 | -0.69345 | -0.10241 | -0.01296 | -1.39094 | -0.54221 | -0.21585 | -1.02908 | 0.198437 | -0.31183 | 1.271733 | 0.262105 | 1.882449 | -0.9458 | 3.132905 | 1.206545 | 1.618024 | 1.665259 |
| 30.70833 | 1.025975 | -0.24223 | 1.116792 | 0.402948 | -1.50136 | -0.74191 | -0.16163 | -1.22835 | 1.082715 | -0.39305 | 2.334286 | 0.018696 | 2.935483 | -0.20599 | 4.60019 | 2.060891 | 1.854191 | 2.782625 |
| 30.72222 | 0.867142 | -0.13767 | 1.643589 | 0.766201 | -1.86249 | -0.4761 | 0.010342 | -1.55013 | 1.572241 | -0.54636 | 2.713778 | -0.16046 | 3.473689 | 0.26577 | 4.057173 | 1.926727 | 1.185325 | 2.278746 |
| 30.73611 | 0.409464 | -0.40467 | 1.242355 | 1.075551 | -1.88113 | -0.03015 | 0.178982 | -1.57125 | 2.115943 | -0.02453 | 2.961721 | -0.05651 | 2.819716 | 0.588975 | 2.639584 | 2.569247 | 0.907876 | 1.506241 |
| 30.75 | -0.15437 | -0.70418 | 0.20293 | 1.382212 | -1.31318 | -0.05378 | 0.306772 | -1.0148 | 1.19082 | 0.438943 | 1.994369 | -0.12948 | 1.981038 | 1.209961 | 2.268873 | 3.359531 | 0.481479 | 0.553228 |
| 30.76389 | -0.34166 | -0.82583 | -0.14377 | 1.209171 | -0.68838 | 0.041269 | 0.286942 | -0.74435 | 0.446189 | 1.057891 | 1.156679 | -0.25099 | 1.187077 | 1.188739 | 1.028275 | 2.318329 | -0.20277 | -0.35316 |
| 30.77778 | -0.22334 | -0.80678 | -0.53965 | 0.771005 | -0.11147 | 0.069513 | 0.147881 | -0.59799 | -0.0917 | 0.864516 | 0.034837 | 0.019633 | -0.08985 | 0.241489 | -0.6706 | 0.741953 | -0.57427 | -1.10166 |
| 30.79167 | 0.188454 | -0.68854 | -0.59809 | 0.105116 | 0.407455 | 0.066123 | -0.18684 | -0.44007 | -0.61878 | 0.456446 | -1.65211 | 0.51607 | -1.33275 | -0.73287 | -2.18428 | -0.54042 | -0.46809 | -1.72356 |
| 30.80556 | 0.201074 | -0.49416 | -0.64217 | -0.59221 | 0.726612 | 0.229129 | -0.52445 | -0.25264 | -1.35649 | 0.128835 | -3.58745 | 1.033697 | -2.42297 | -1.18485 | -3.11833 | -0.36893 | -0.05796 | -1.60924 |
| 30.81944 | -0.46449 | -0.42928 | -0.70733 | -0.93836 | 0.71488 | 0.662273 | -0.84535 | 0.135464 | -1.67982 | -0.23887 | -4.31761 | 1.209705 | -3.62118 | -0.76974 | -5.05591 | 0.001508 | -0.03511 | -0.38688 |
| 30.83333 | -0.79533 | -0.7688 | -1.81319 | -1.31412 | 0.355645 | 0.756309 | -0.76944 | 0.166484 | -1.42304 | -0.99717 | -5.37787 | 1.144472 | -4.46597 | -0.30899 | -6.25249 | -0.13819 | -0.39364 | 0.392902 |
| 30.84722 | -0.82779 | -1.3556 | -3.38776 | -1.5889 | -0.25352 | 0.28156 | -1.05472 | 0.060363 | -1.09174 | -1.54794 | -4.1427 | 0.667072 | -3.353 | -0.4406 | -4.2399 | -1.32315 | -0.75671 | 0.403557 |
| 30.86111 | -0.91216 | -1.84699 | -4.04707 | -1.61835 | -0.96817 | 0.008663 | -1.29575 | -0.11293 | -0.90015 | -2.20477 | -2.32556 | -0.03172 | -2.69816 | -0.53237 | -3.5253 | -4.13101 | -1.08414 | 0.052895 |
| 30.875 | -0.7271 | -1.65553 | -3.79227 | -1.52778 | -1.65122 | -0.02678 | -1.35909 | -0.19529 | -0.97697 | -2.41312 | -1.35875 | -0.52035 | -1.71251 | -0.38387 | -2.38771 | -4.51952 | -1.47002 | -0.2985 |
| 30.88889 | -0.3488 | -1.2094 | -3.27238 | -1.3643 | -1.90405 | -0.20669 | -1.27575 | -0.45855 | -1.05595 | -1.99447 | -0.77004 | -0.52395 | -0.91627 | -0.0423 | -1.58407 | -3.98085 | -1.49128 | -0.33698 |
| 30.90278 | 0.286297 | -0.6617 | -2.54589 | -1.07679 | -1.84824 | -0.45387 | -0.78795 | -0.8211 | -0.76872 | -1.46311 | -0.54048 | -0.2675 | -0.37867 | 0.406531 | -0.61813 | -4.63818 | -1.75983 | -0.1633 |
| 30.91667 | 1.038441 | 0.168626 | -1.55852 | -0.24643 | -1.42726 | -0.45412 | 0.003793 | -0.89096 | -0.40511 | -0.64953 | -0.37406 | 0.103476 | 0.304352 | 0.576947 | 0.028048 | -4.9765 | -1.66548 | 0.137278 |
| 30.93056 | 1.994213 | 0.264544 | -0.31552 | 0.451095 | -0.83316 | -0.35341 | 0.980509 | -0.63775 | 0.201499 | 0.299771 | -0.31688 | 0.707429 | 1.086196 | 0.074244 | 0.493195 | -2.96377 | -0.59047 | 0.509321 |
| 30.94444 | 2.706017 | 0.195878 | 0.971204 | 0.899709 | -0.23253 | -0.27892 | 1.825737 | -0.38887 | 0.790779 | 1.061936 | 0.395805 | 1.224754 | 1.905821 | -0.56268 | 1.48546 | -0.58476 | 0.606233 | 0.977614 |
| 30.95833 | 2.838253 | 0.357329 | 1.954419 | 0.881362 | 0.318197 | -0.23005 | 2.200109 | -0.4608 | 1.024024 | 1.442479 | 1.594442 | 1.625623 | 3.09471 | -0.19907 | 3.133394 | 1.313125 | 1.330978 | 1.258008 |
| 30.97222 | 2.404517 | 0.494662 | 2.484465 | 0.83912 | 0.6667 | -0.21584 | 1.925898 | -0.56242 | 1.090984 | 1.336902 | 2.507028 | 1.846149 | 3.703547 | 0.399991 | 4.47349 | 2.540479 | 1.432792 | 1.097092 |
| 30.98611 | 1.078559 | 0.228627 | 2.069025 | 0.711596 | 0.2235 | -0.5588 | 1.217976 | -0.57988 | 0.502032 | 0.525024 | 1.680826 | 1.304102 | 2.547259 | -0.28765 | 3.03349 | 1.913858 | 0.591572 | 0.047663 |
| 31 | -1.60078 | -0.64537 | 0.122079 | -0.44411 | -1.1674 | -1.10723 | -0.26678 | -1.13793 | -0.77441 | -1.08743 | -1.49491 | -0.19129 | 0.16803 | -1.98698 | -1.50087 | -0.99475 | -2.02821 | -1.98463 |
| 31.01389 | -3.71382 | -1.12834 | -2.43971 | -1.76097 | -2.21174 | -1.06923 | -1.87508 | -1.00686 | -1.86554 | -2.11108 | -5.40765 | -1.10949 | -1.78863 | -3.26345 | -5.34283 | -3.77418 | -4.2841 | -3.28903 |
| 31.02778 | -3.42325 | -0.89762 | -3.51961 | -2.07442 | -2.0641 | -0.31758 | -3.26452 | -1.21935 | -1.98737 | -1.81117 | -5.72169 | -0.66652 | -2.03941 | -2.46134 | -4.56004 | -3.78901 | -2.85956 | -2.88975 |
| 31.04167 | -2.54776 | -1.04821 | -3.30409 | -1.71635 | -1.36425 | 0.136014 | -4.18698 | 0.47461 | -1.8993 | -1.11302 | -4.7767 | -0.28658 | -1.88418 | -1.7736 | -2.73035 | -2.4641 | -0.96235 | -2.04237 |
| 31.05556 | -2.00405 | -0.85063 | -2.54522 | -1.47238 | -0.70668 | 0.362557 | -3.52619 | 0.449894 | -1.86448 | -0.60791 | -3.57252 | -0.23828 | -2.37701 | -1.49888 | -1.93734 | -1.68304 | -0.57263 | -1.84383 |
| 31.06944 | -1.56443 | -1.13306 | -1.86589 | -0.91305 | -0.37198 | 0.310028 | -1.71394 | 0.411871 | -1.41131 | -0.20027 | -2.14476 | -0.36775 | -2.36298 | -1.42303 | -1.88725 | -1.30489 | -1.05475 | -1.95509 |
| 31.08333 | -1.79495 | -1.55592 | -1.31647 | -0.42412 | -0.25677 | 0.371088 | 0.15298 | 0.363788 | -0.86132 | -0.55677 | -1.56215 | -1.04499 | -2.65487 | -1.69462 | -2.76103 | -2.01304 | -2.1748 | -2.47452 |
| 31.09722 | -1.21186 | -1.05322 | -0.47118 | 0.074093 | -0.00104 | 0.575945 | 2.048875 | 0.438309 | -0.28905 | -1.43878 | -1.70625 | -2.13191 | -3.78578 | -2.0369 | -3.89779 | -4.33223 | -3.92812 | -3.54039 |
| 31.11111 | -0.03859 | -0.02761 | 0.86632 | 0.916544 | 0.283462 | 1.005857 | 3.963056 | 0.59551 | 0.404122 | -1.36788 | -0.90367 | -2.26313 | -3.67232 | -1.13613 | -3.51629 | -5.59068 | -4.2551 | -3.53291 |
| 31.125 | 1.195094 | 0.972756 | 1.350445 | 1.521197 | 0.387501 | 1.126302 | 4.64499 | 0.421275 | 1.004535 | -0.09178 | 0.97432 | -1.24641 | -1.73091 | 1.15687 | -1.85189 | -3.80988 | -2.72748 | -1.87129 |
| 31.13889 | 1.717079 | 1.464085 | 1.240068 | 1.368769 | 0.374921 | 0.831793 | 4.067246 | 0.101993 | 1.066664 | 1.01777 | 2.046889 | -0.13303 | 0.051727 | 2.539949 | -0.18581 | -1.56152 | -0.60137 | -0.00485 |
| 31.15278 | 1.422581 | 1.355521 | 0.371374 | 0.879778 | 0.312829 | 0.395469 | 2.76187 | -0.02465 | 0.631384 | 1.360226 | 2.032489 | 0.540485 | 0.982829 | 2.30744 | 0.897921 | -0.78029 | 0.645757 | 0.969362 |
| 31.16667 | 0.91419 | 1.080211 | -0.22685 | 0.656622 | 0.196179 | -0.02697 | 1.489358 | -0.05236 | 0.050017 | 1.139595 | 1.961855 | 1.02602 | 1.400377 | 1.42744 | 1.417647 | -0.37282 | 0.917159 | 1.300199 |
| 31.18056 | 0.818004 | 1.474597 | -0.20855 | 0.44851 | 0.019973 | -0.15802 | 0.688713 | 0.156718 | -0.32057 | 0.73528 | 1.846851 | 1.44371 | 1.691581 | 0.788711 | 1.586903 | 0.256868 | 0.947837 | 1.320747 |
| 31.19444 | 1.081465 | 1.887086 | 0.218928 | 0.4642 | -0.02005 | 0.026519 | 0.375426 | 0.316441 | -0.41159 | 0.877347 | 1.439555 | 1.664512 | 1.976906 | 0.925692 | 1.67544 | 0.582028 | 1.295519 | 1.181898 |
| 31.20833 | 1.400567 | 1.918654 | 0.828233 | 0.523564 | 0.077944 | 0.17384 | 0.372071 | 0.213599 | -0.15969 | 1.561635 | 1.540434 | 1.642432 | 2.17105 | 1.521038 | 1.755663 | 0.844057 | 1.270322 | 1.046672 |
| 31.22222 | 1.912332 | 2.098978 | 1.434302 | 0.606583 | 0.208219 | 0.228194 | 0.546464 | 0.062013 | 0.214302 | 1.95179 | 2.221317 | 1.397695 | 2.149375 | 2.078384 | 1.805419 | 1.075335 | 0.864713 | 0.900699 |
| 31.23611 | 2.540056 | 1.931454 | 1.650701 | 0.675518 | 0.344908 | 0.284388 | 0.845277 | -0.05701 | 0.68155 | 1.721154 | 2.664479 | 0.987157 | 1.751885 | 2.127468 | 1.551188 | 1.111748 | 0.39428 | 0.696747 |
| 31.25 | 2.733412 | 1.529762 | 1.493953 | 0.621128 | 0.361958 | 0.279007 | 0.963825 | -0.17193 | 0.8956 | 1.131281 | 2.049844 | 0.582435 | 1.329766 | 1.500994 | 1.142172 | 0.897889 | -0.048 | 0.360229 |
| 31.26389 | 2.573783 | 1.032388 | 0.997631 | 0.44487 | 0.290388 | 0.250866 | 0.694713 | -0.22671 | 0.626894 | 0.405651 | 0.791663 | 0.313104 | 1.015943 | 0.580482 | 0.724711 | 0.731255 | 0.045082 | 0.045335 |
| 31.27778 | 2.086786 | 0.266586 | 0.527027 | 0.283305 | 0.2249 | 0.198446 | 0.325923 | -0.15945 | 0.164004 | -0.01872 | 0.201295 | 0.18091 | 1.000624 | -0.03431 | 0.511589 | 1.147569 | 0.447831 | 0.063914 |
| 31.29167 | 1.229656 | -0.42232 | 0.427741 | 0.147721 | 0.195759 | 0.096199 | 0.033463 | -0.03792 | -0.09298 | -0.00158 | 0.216447 | 0.153724 | 1.007872 | -0.07521 | 0.49909 | 1.514635 | 0.733435 | 0.315122 |
| 31.30556 | 0.336387 | -0.62721 | 0.447495 | 0.014985 | 0.156354 | 0.100388 | -0.09162 | 0.095489 | -0.14833 | 0.268692 | 0.734714 | 0.116939 | 0.957617 | -0.01208 | 0.610637 | 1.438354 | 0.739965 | 0.454282 |
| 31.31944 | -0.19955 | -0.28175 | 0.651251 | -0.09682 | 0.070049 | 0.309961 | -0.04815 | 0.190806 | -0.09815 | 0.468624 | 0.985059 | 0.009029 | 1.024668 | -0.12942 | 0.661636 | 1.606104 | 0.501169 | 0.415053 |
| 31.33333 | -0.38223 | 0.207855 | 0.89434 | -0.04719 | 0.066751 | 0.489632 | 0.138591 | 0.299893 | 0.032939 | 0.783216 | 1.249438 | -0.0293 | 1.193872 | -0.09108 | 0.868046 | 2.099603 | 0.250609 | 0.285333 |
| 31.34722 | -0.27174 | 0.720608 | 1.043964 | 0.07487 | 0.052268 | 0.59826 | 0.424752 | 0.353009 | 0.200674 | 1.055042 | 1.628416 | 0.043339 | 1.201495 | 0.052598 | 0.993705 | 2.117239 | 0.22329 | 0.155875 |
| 31.36111 | -0.28173 | 0.750472 | 1.136524 | 0.024937 | 0.001274 | 0.543385 | 0.395007 | 0.30542 | 0.311645 | 1.081253 | 1.869573 | 0.137609 | 1.02668 | 0.194044 | 1.006769 | 1.730066 | 0.2295 | 0.19163 |
| 31.375 | -0.52155 | 0.1587 | 1.088161 | -0.03863 | -0.07506 | 0.269331 | -0.07447 | 0.214039 | 0.292097 | 1.066833 | 1.95145 | 0.108738 | 0.734895 | 0.249771 | 0.770017 | 1.225065 | 0.17773 | 0.14936 |
| 31.38889 | -0.94369 | -0.45964 | 0.974087 | -0.19592 | -0.14202 | 0.050325 | -0.53563 | 0.195157 | 0.21874 | 0.916579 | 1.739901 | -0.01595 | 0.49431 | 0.068685 | 0.540981 | 0.672207 | 0.078237 | 0.107855 |
| 31.40278 | -1.27022 | -0.67525 | 0.828817 | -0.33513 | -0.17895 | -0.01835 | -0.66979 | 0.26185 | 0.174329 | 0.648362 | 1.208436 | -0.00341 | 0.337166 | -0.00999 | 0.407789 | 0.317845 | 0.068987 | 0.245881 |
| 31.41667 | -1.01009 | -0.27332 | 0.621769 | -0.26917 | -0.15262 | 0.207396 | -0.33852 | 0.360204 | 0.144047 | 0.598124 | 0.850197 | 0.238571 | 0.340495 | 0.062216 | 0.468512 | 0.022779 | 0.177937 | 0.415245 |
| 31.43056 | -0.59202 | -0.04329 | 0.553373 | 0.002798 | -0.09417 | 0.329558 | -0.03202 | 0.392344 | 0.102946 | 0.391468 | 0.683193 | 0.365336 | 0.318621 | 0.310954 | 0.506298 | -0.38385 | 0.067928 | 0.316846 |
| 31.44444 | -0.58341 | -0.59816 | 0.647764 | 0.310018 | -0.20567 | 0.038813 | -0.15977 | 0.278797 | 0.013442 | 0.188577 | 0.70622 | 0.063976 | 0.090437 | 0.422317 | 0.311999 | -1.04534 | -0.40406 | -0.13644 |
| 31.45833 | -0.61167 | -0.98108 | 0.7462 | 0.439313 | -0.42764 | -0.23722 | -0.33764 | 0.13948 | -0.00331 | 0.216862 | 0.839567 | -0.50723 | -0.30178 | 0.252279 | -0.20718 | -1.6545 | -1.01093 | -0.71115 |

| | | | | | | | | | | | | | | | | | | |
|---|---|---|---|---|---|---|---|---|---|---|---|---|---|---|---|---|---|---|
| 31.47222 | 0.062767 | -0.2369 | 0.706171 | 0.47128 | -0.60393 | -0.00783 | -0.0662 | 0.09966 | 0.118904 | 0.284295 | 1.027884 | -0.77233 | -0.50403 | 0.125597 | -0.57691 | -1.85968 | -1.22301 | -0.76762 |
| 31.48611 | 1.092318 | 0.884535 | 1.154295 | 0.725978 | -0.5945 | 0.527604 | 0.501013 | 0.23245 | 0.500413 | 0.879383 | 1.745589 | -0.29886 | -0.10243 | 0.532162 | 0.490288 | -0.86187 | -0.33848 | 0.233027 |
| 31.5 | 1.912593 | 0.986447 | 2.710592 | 1.194621 | -0.42348 | 0.744437 | 1.038319 | 0.440406 | 1.179426 | 2.209796 | 3.371629 | 1.069167 | 0.854318 | 1.518165 | 3.508276 | 0.961017 | 1.558085 | 2.013411 |
| 31.51389 | 2.537371 | 0.489285 | 3.710699 | 1.834001 | 0.186365 | 0.75537 | 1.980654 | 0.345846 | 1.523775 | 3.705844 | 4.568693 | 2.819568 | 1.733997 | 2.707957 | 6.874025 | 3.032619 | 2.395625 | 2.985219 |
| 31.52778 | 3.252521 | 0.680736 | 3.308564 | 2.668562 | 1.064273 | 0.766436 | 3.279172 | -0.02379 | 1.334992 | 4.221222 | 4.840998 | 3.475965 | 2.775405 | 3.736885 | 8.198602 | 4.090409 | 2.124512 | 2.693023 |
| 31.54167 | 3.662005 | 0.961074 | 2.78643 | 2.819789 | 1.425544 | 0.393447 | 3.209204 | -0.19227 | 1.480611 | 4.716321 | 5.532536 | 3.042793 | 3.512628 | 4.569601 | 9.245644 | 4.671327 | 2.28725 | 2.649176 |
| 31.55556 | 3.411817 | 0.598663 | 2.043255 | 2.373578 | 1.176499 | 0.057788 | 1.803038 | -0.07667 | 0.981085 | 3.325553 | 5.132698 | 2.167693 | 3.389115 | 5.105507 | 8.74202 | 5.091077 | 2.692316 | 2.533808 |
| 31.56944 | 2.454556 | 0.134493 | 1.510446 | 1.379017 | 0.961834 | -0.07898 | 0.570653 | 0.120785 | 0.716253 | 1.404866 | 3.670464 | 1.204891 | 2.321426 | 3.632984 | 6.43992 | 4.750809 | 2.592834 | 1.930325 |
| 31.58333 | 1.515367 | 0.201343 | 1.036792 | 0.594438 | 1.360408 | -0.28237 | -0.23243 | 0.381083 | 1.199345 | 0.033234 | 2.077751 | 0.352679 | 0.970924 | 2.095016 | 3.879954 | 3.629901 | 2.026101 | 1.158468 |
| 31.59722 | 0.636262 | 0.334722 | 0.86763 | 0.033789 | 2.116308 | -0.41614 | -0.95415 | 0.639727 | 1.257909 | -0.57107 | 0.570236 | -0.12663 | 0.061731 | 0.706635 | 2.171536 | 2.289479 | 1.460473 | 0.712217 |
| 31.61111 | -0.35779 | 0.311691 | 0.169199 | -0.12385 | 2.01314 | -0.35413 | -1.49079 | 0.755361 | 0.63276 | -0.66605 | -0.43954 | -0.01708 | -0.32943 | -0.04629 | 0.456906 | 2.32133 | 1.36023 | 1.00144 |
| 31.625 | -0.95037 | 0.2963 | -0.10417 | -0.13002 | 1.491985 | -0.17064 | -1.36325 | 0.670607 | 0.007753 | -0.06645 | -0.62578 | 0.347067 | -0.8191 | -0.64341 | -1.95246 | 2.884665 | 1.120712 | 0.972598 |
| 31.63889 | -1.6056 | 0.16699 | -0.16867 | -0.28532 | 1.401243 | 0.156583 | -1.17164 | 0.264641 | -1.04516 | 0.631502 | -0.22981 | 0.459331 | -1.24207 | -1.46319 | -4.59721 | 1.936291 | 0.675332 | 0.34422 |
| 31.65278 | -1.49172 | -0.41049 | -0.20916 | -0.53395 | 1.165697 | -0.05967 | -1.18003 | -0.31013 | -2.0542 | 0.62536 | -0.02191 | 0.43941 | -1.90007 | -1.5689 | -6.33831 | 0.655281 | 0.139976 | -0.47631 |
| 31.66667 | -1.33129 | -0.72625 | -0.36025 | -0.34146 | 0.727604 | -0.44417 | -1.3318 | -0.75095 | -2.00459 | 0.588248 | -0.00796 | 0.382888 | -2.04579 | -0.70196 | -5.69862 | -0.51592 | -0.43648 | -1.12299 |
| 31.68056 | -0.40964 | -0.61351 | -0.3813 | 0.021465 | 0.172688 | -0.52962 | -1.28601 | -1.03751 | -1.53384 | 0.239753 | -0.29813 | 0.290269 | -1.55887 | -0.32404 | -4.90207 | -1.62528 | -0.79253 | -1.60115 |
| 31.69444 | 0.279076 | -0.45681 | -0.17706 | 0.358041 | -0.58358 | -0.38505 | -0.95139 | -0.9951 | -0.96789 | -0.19761 | -0.99727 | 0.193654 | -1.34547 | -0.08589 | -4.25942 | -2.33425 | -1.09736 | -1.36673 |
| 31.70833 | 0.994417 | -0.51932 | 0.006333 | 0.558642 | -1.59085 | -0.32452 | -0.57603 | -0.6038 | -0.32956 | 0.119165 | -1.15668 | 0.125818 | -1.34235 | 0.298579 | -3.29063 | -2.16746 | -0.87102 | -0.77086 |
| 31.72222 | 1.683945 | -0.67642 | 0.091147 | 0.79726 | -2.21739 | -0.39924 | -0.1676 | -0.19318 | 0.723182 | 0.736501 | -0.41259 | 0.174456 | -0.89184 | 0.20546 | -1.42805 | -2.10794 | -0.29763 | -0.2386 |
| 31.73611 | 1.745972 | -0.8017 | 0.165967 | 0.636257 | -2.2671 | -0.54708 | 0.227432 | -0.08856 | 1.575083 | 0.982406 | 0.519959 | 0.306361 | -0.0905 | 0.157726 | 0.876017 | -1.60841 | 0.155299 | 0.377076 |
| 31.75 | 2.247867 | -0.7397 | 0.409129 | 0.672117 | -1.88998 | -1.05659 | 0.418554 | -0.43939 | 2.34012 | 0.509884 | 1.030539 | 0.391166 | 0.893015 | 0.593647 | 3.020807 | -1.14545 | 0.451783 | 1.005217 |
| 31.76389 | 2.30544 | -0.37958 | 0.611995 | 0.149433 | -1.59983 | -1.34588 | 0.365336 | -1.07654 | 2.060138 | -0.04388 | 1.044891 | 0.341722 | 1.488632 | 0.9478 | 4.180693 | -0.99816 | 0.517445 | 1.338622 |
| 31.77778 | 1.41479 | 0.279252 | 0.680078 | 0.012014 | -1.54036 | -1.1376 | 0.676277 | -0.92758 | 1.62439 | -0.45502 | 0.623983 | 0.066062 | 1.474486 | 0.594962 | 3.377638 | -0.18266 | 0.475335 | 1.084575 |
| 31.79167 | 0.217723 | 0.689136 | 0.075923 | -0.36505 | -1.48224 | -1.20799 | 0.288525 | -0.83518 | 0.537427 | -0.53391 | -0.29594 | -0.32802 | 1.000309 | 0.29264 | 1.243692 | 0.638279 | 0.93394 | 0.558051 |
| 31.80556 | -0.51995 | 0.446662 | -0.20186 | -0.69063 | -1.57375 | -1.62542 | -0.45084 | -0.97039 | -0.6051 | -0.87971 | -1.16285 | -0.67315 | 0.340734 | -0.30078 | -1.12144 | 0.599952 | 1.050256 | 0.328708 |
| 31.81944 | -1.44829 | -0.56243 | -0.47291 | -1.054 | -1.505 | -2.03764 | -1.14495 | -0.93069 | -1.30349 | -1.01267 | -1.81653 | -0.69607 | -0.23257 | -0.49593 | -3.27174 | 0.423583 | 0.815294 | 0.265241 |
| 31.83333 | -2.06507 | -1.35658 | -0.5933 | -0.87504 | -1.28728 | -1.99748 | -1.45975 | -0.66069 | -1.54614 | -1.33795 | -2.03238 | -0.36021 | -0.62497 | -0.76752 | -5.33318 | 0.380175 | 0.828393 | 0.098553 |
| 31.84722 | -2.2689 | -1.82524 | -0.8575 | -0.85408 | -1.19583 | -1.58732 | -1.61138 | -0.61115 | -1.92994 | -2.08781 | -2.67016 | -0.11471 | -0.98803 | -0.78934 | -6.26956 | 0.340406 | 0.882242 | -0.58835 |
| 31.86111 | -2.44246 | -1.85207 | -1.34991 | -0.90281 | -1.17064 | -1.49073 | -1.99382 | -0.66965 | -2.14779 | -2.7503 | -3.65865 | -0.09552 | -1.73974 | -0.99624 | -6.12614 | -0.4425 | 0.703212 | -1.51355 |
| 31.875 | -2.39515 | -1.85165 | -1.88043 | -0.9862 | -1.03131 | -1.6438 | -2.59681 | -0.85133 | -2.24472 | -2.94213 | -4.72671 | -0.36916 | -1.88387 | -1.36828 | -5.41609 | -1.23998 | 0.41449 | -2.36512 |
| 31.88889 | -2.20777 | -1.79894 | -2.26228 | -0.95993 | -0.85252 | -1.83549 | -2.70846 | -1.05624 | -1.97719 | -2.80763 | -4.32514 | -0.90603 | -1.88881 | -1.38046 | -4.08184 | -1.47596 | 0.061455 | -2.50247 |
| 31.90278 | -1.96201 | -2.51848 | -1.96942 | -0.89307 | -0.59341 | -1.47421 | -2.01493 | -1.25858 | -1.50877 | -2.57323 | -3.08377 | -0.97145 | -1.82325 | -1.39515 | -2.70519 | -1.76608 | -0.4001 | -2.32279 |
| 31.91667 | -1.10674 | -1.7144 | -1.19857 | -0.62894 | -0.34379 | -1.00661 | -1.39262 | -1.21054 | -1.46178 | -1.86718 | -2.34687 | -0.5292 | -1.27621 | -1.09272 | -1.71133 | -2.06897 | -0.70387 | -2.01855 |
| 31.93056 | -0.09751 | -0.8894 | -0.52733 | -0.06057 | -0.08503 | -0.55374 | -0.76504 | -0.74777 | -0.93421 | -0.90136 | -1.58086 | 0.065761 | -0.71522 | -0.57789 | -1.01367 | -2.36342 | -0.85424 | -1.72927 |
| 31.94444 | 0.648721 | -0.23338 | 0.112249 | 0.53866 | 0.135399 | -0.11359 | -0.11305 | -0.40417 | -0.10581 | 0.239877 | -0.45715 | 0.609025 | -0.4603 | -0.05801 | -0.67066 | -1.77426 | -0.61834 | -1.18297 |
| 31.95833 | 1.075597 | 0.610747 | 0.666171 | 1.018273 | 0.297367 | 0.093649 | 0.565248 | -0.23696 | 0.735631 | 1.39177 | 0.774081 | 1.06382 | -0.2496 | 0.662714 | -0.23121 | -1.1038 | -0.17343 | -0.38619 |
| 31.97222 | 1.221614 | 1.15912 | 1.195453 | 1.241454 | 0.388497 | 0.151064 | 1.147015 | -0.20505 | 1.476814 | 1.894128 | 1.569783 | 0.968113 | -0.06557 | 1.215236 | 0.292923 | -0.70068 | 0.016858 | 0.200782 |
| 31.98611 | 0.618379 | 0.897872 | 0.91745 | 0.590372 | 0.042908 | -0.26627 | 1.002835 | -0.36255 | 1.315099 | 1.377166 | 0.696297 | -0.04431 | -0.40108 | 0.634601 | -0.18939 | -1.9885 | -1.20326 | -0.74503 |
| 32 | -0.87428 | -0.66587 | -1.00633 | -1.46464 | -0.73639 | -1.08147 | -0.37005 | -0.57561 | -0.62743 | -1.37823 | -2.55256 | -2.10791 | -1.32985 | -1.52428 | -1.99485 | -5.21865 | -4.53897 | -3.3098 |
| 32.01389 | -2.47658 | -2.60806 | -3.08368 | -3.6143 | -1.46895 | -1.56004 | -2.33758 | -0.70504 | -2.86513 | -4.4017 | -5.50182 | -3.60069 | -1.69739 | -3.43934 | -2.82901 | -7.50411 | -6.42685 | -4.43371 |
| 32.02778 | -3.24841 | -3.30825 | -2.9463 | -3.59276 | -1.364 | -1.26127 | -2.74619 | -0.73164 | -3.28998 | -4.97669 | -4.55853 | -3.6136 | -0.95191 | -2.92084 | -1.71689 | -4.68371 | -4.0848 | -2.95551 |
| 32.04167 | -3.10062 | -3.31527 | -2.44988 | -2.90655 | -1.14977 | -1.61148 | -2.22608 | -0.60435 | -2.74327 | -4.75665 | -3.42634 | -2.80148 | -0.57935 | -1.98294 | -0.45623 | -2.4456 | -3.00106 | -1.26963 |
| 32.05556 | -2.42711 | -3.48282 | -2.68988 | -2.18336 | -0.9657 | -1.82425 | -2.23938 | -0.34497 | -2.31061 | -2.78515 | -2.6141 | -1.72159 | -0.00422 | -1.51625 | 0.297872 | -1.65045 | -2.33131 | -0.4327 |
| 32.06944 | -1.56575 | -3.17053 | -2.35559 | -1.09105 | -0.85209 | -1.47885 | -1.63565 | -0.22145 | -0.99269 | -1.42827 | -2.47339 | -0.3609 | 0.148534 | -1.30844 | 0.211321 | -1.15069 | -1.32553 | -0.36947 |
| 32.08333 | -0.50521 | -2.20001 | -1.31646 | -0.1426 | -0.46548 | -0.62384 | -0.5194 | -0.07883 | 0.018891 | -0.59986 | -1.68536 | 0.275181 | 0.445928 | -1.0665 | -0.30345 | -0.47817 | 0.575012 | -0.33646 |
| 32.09722 | 0.508571 | -0.9442 | -0.47366 | 0.467682 | 0.095069 | 0.470492 | 0.064461 | 0.194856 | 0.682492 | -0.25713 | -0.81571 | 0.234529 | 0.397497 | -0.86455 | -0.65679 | 0.102054 | 1.872113 | -0.2168 |
| 32.11111 | 1.114929 | -0.24397 | 0.056566 | 0.787066 | 0.477819 | 1.375559 | 0.376294 | 0.37413 | 1.12169 | -0.03449 | -0.03322 | 0.050488 | 0.234036 | -0.46915 | -0.74345 | 0.243747 | 1.668621 | -0.13184 |
| 32.125 | 1.406104 | 0.074953 | 0.368361 | 0.982631 | 0.690062 | 1.728056 | 0.709373 | 0.430955 | 1.294144 | 0.158242 | 0.787002 | -0.01844 | 0.163597 | 0.199464 | -0.6196 | 0.535429 | 1.69392 | -0.02286 |
| 32.13889 | 1.50972 | 0.173519 | 0.578493 | 1.185448 | 0.84503 | 1.728922 | 1.005773 | 0.428695 | 1.402193 | 0.495122 | 1.438426 | 0.173176 | 0.341749 | 0.967534 | -0.34904 | 0.918507 | 2.293871 | 0.148565 |
| 32.15278 | 1.734263 | 0.3962 | 0.616721 | 1.237271 | 0.9326 | 1.828237 | 1.090503 | 0.552625 | 1.464992 | 0.935134 | 1.551219 | 0.497176 | 0.656132 | 1.656605 | 0.020186 | 1.10861 | 2.194327 | 0.381365 |
| 32.16667 | 2.05354 | 0.932578 | 0.626745 | 1.070934 | 0.868263 | 1.733265 | 0.953326 | 0.738371 | 1.635161 | 1.557714 | 0.647229 | 0.715332 | 1.021141 | 2.277819 | 0.443169 | 1.184199 | 1.746897 | 0.721484 |
| 32.18056 | 2.023842 | 1.206203 | 0.688352 | 0.974678 | 0.76671 | 1.448257 | 0.922755 | 0.772576 | 2.097002 | 1.767154 | 0.7631 | 0.592495 | 1.14058 | 2.442478 | 0.760662 | 1.253497 | 1.537289 | 1.021352 |
| 32.19444 | 1.827123 | 1.279406 | 0.733126 | 1.013099 | 0.603053 | 1.044159 | 1.035929 | 0.87468 | 2.547258 | 1.444544 | 1.734775 | 0.357537 | 1.040051 | 2.565567 | 0.948942 | 1.339443 | 1.27539 | 1.069069 |
| 32.20833 | 1.760273 | 1.465675 | 0.695162 | 0.967734 | 0.541827 | 0.787082 | 1.179305 | 0.927673 | 2.513323 | 1.58423 | 3.090278 | 0.623421 | 0.803926 | 2.889449 | 1.147903 | 1.504564 | 1.105459 | 1.231027 |
| 32.22222 | 1.523438 | 1.453677 | 0.67969 | 0.868655 | 0.429289 | 0.644545 | 1.687096 | 0.6494 | 2.509327 | 1.805844 | 2.780667 | 1.166889 | 0.343291 | 2.804562 | 1.19531 | 1.48658 | 0.966104 | 1.290633 |
| 32.23611 | 1.268294 | 1.405586 | 0.601662 | 0.614702 | 0.385435 | 0.575003 | 2.136371 | 0.438377 | 2.2426 | 1.805367 | 2.170675 | 1.220516 | -0.10342 | 2.390917 | 1.0405 | 1.281584 | 0.841849 | 1.094518 |
| 32.25 | 1.149327 | 1.678199 | 0.500682 | 0.252492 | 0.401193 | 0.536832 | 2.312487 | 0.412927 | 2.09586 | 1.442958 | 1.308372 | 0.777733 | -0.48045 | 1.816976 | 0.853856 | 1.024012 | 0.550804 | 0.834716 |
| 32.26389 | 0.918925 | 1.853409 | 0.717352 | -0.10676 | 0.461806 | 0.380771 | 2.547019 | 0.411779 | 1.92116 | 0.729578 | 0.303296 | 0.188726 | -0.72169 | 1.377473 | 0.666556 | 0.860309 | 0.366762 | 0.464184 |
| 32.27778 | 0.944308 | 1.925166 | 1.092215 | -0.11892 | 0.494547 | 0.194612 | 2.614276 | 0.379158 | 1.633756 | 0.368074 | -0.25274 | -0.0685 | -0.75157 | 1.107194 | 0.494406 | 0.779235 | 0.273057 | 0.222354 |
| 32.29167 | 1.119192 | 1.782955 | 1.213453 | 0.124723 | 0.489242 | 0.214966 | 2.429473 | 0.301788 | 1.237695 | 0.527149 | -0.24886 | 0.229241 | -0.25329 | 1.125886 | 0.47293 | 0.854056 | 0.524272 | 0.300209 |
| 32.30556 | 1.241807 | 1.544451 | 1.200194 | 0.322862 | 0.477859 | 0.545234 | 2.437708 | 0.219878 | 0.776796 | 0.960556 | 0.217037 | 0.655624 | 0.506995 | 1.495712 | 0.52046 | 1.137273 | 0.880348 | 0.7132 |
| 32.31944 | 1.493524 | 1.644943 | 1.088752 | 0.361541 | 0.566907 | 0.874077 | 2.418913 | 0.316578 | 0.580718 | 1.017313 | 0.5591 | 0.769771 | 0.816953 | 1.705742 | 0.036156 | 1.487761 | 1.244023 | 1.100727 |
| 32.33333 | 1.563756 | 1.788516 | 0.943617 | 0.243939 | 0.631983 | 0.880578 | 2.148538 | 0.386886 | 0.548389 | 0.647845 | 0.253584 | 0.44303 | 0.328027 | 1.305939 | -1.26847 | 1.375934 | 1.283309 | 1.08177 |
| 32.34722 | 1.221607 | 1.776267 | 0.956046 | 0.094502 | 0.447685 | 0.613573 | 1.62753 | 0.264757 | 0.385642 | -0.16118 | -0.26247 | -0.19639 | -0.41921 | 0.01478 | -1.71124 | 0.817839 | 0.865997 | 0.532351 |
| 32.36111 | 1.025263 | 1.60298 | 0.969574 | 0.091756 | 0.225357 | 0.332673 | 1.003647 | 0.065049 | 0.091906 | -0.65618 | -0.26947 | -0.49823 | -0.70339 | -0.42961 | -1.38939 | 0.329284 | 0.50767 | 0.143313 |
| 32.375 | 1.417994 | 1.638098 | 1.004817 | 0.315921 | 0.290689 | 0.375513 | 0.364885 | 0.132839 | 0.050265 | 0.056673 | 0.459664 | -0.09446 | -0.40408 | -0.08945 | -0.77823 | 0.168427 | 0.49269 | 0.305636 |
| 32.38889 | 2.043661 | 1.817279 | 1.03796 | 0.570203 | 0.519564 | 0.666341 | 0.058462 | 0.411998 | 0.36552 | 1.099051 | 1.450746 | 0.39348 | 0.117485 | 0.622671 | -0.2493 | 0.225462 | 0.655277 | 0.79553 |
| 32.40278 | 2.060826 | 1.745533 | 1.186707 | 0.641298 | 0.538178 | 0.735639 | 0.036193 | 0.39064 | 0.58927 | 1.294047 | 2.252484 | 0.297551 | 0.363159 | 0.782382 | -1.5E-05 | 0.461007 | 0.723744 | 1.151885 |
| 32.41667 | 1.574095 | 1.474799 | 0.967434 | 0.415109 | 0.389437 | 0.411036 | -0.2045 | 0.140778 | 0.498467 | 1.04671 | 2.158095 | -0.07222 | 0.259008 | 0.496035 | -0.01423 | 0.657551 | 0.754188 | 1.207519 |
| 32.43056 | 1.079627 | 1.112243 | 0.266225 | 0.008254 | 0.265144 | -0.12981 | -0.7895 | -0.02091 | 0.270176 | 1.073679 | 1.682167 | 0.047676 | 0.382357 | 0.4772 | 0.260908 | 1.038545 | 1.087008 | 1.445977 |
| 32.44444 | 0.685645 | 0.690048 | -0.25447 | -0.34611 | 0.154522 | -0.68373 | -1.29018 | -0.08898 | 0.074226 | 1.453163 | 1.450812 | 0.721451 | 1.052835 | 0.973961 | 0.920145 | 1.761268 | 1.735757 | 2.07004 |
| 32.45833 | 0.636967 | 0.631398 | -0.35448 | -0.35932 | 0.08426 | -1.01951 | -1.55279 | -0.09148 | -0.07298 | 1.224166 | 1.322745 | 1.013892 | 1.437043 | 1.183112 | 1.067261 | 2.048502 | 1.876461 | 2.222915 |
| 32.47222 | 0.987631 | 1.104962 | -0.267 | -0.24264 | 0.124827 | -1.10445 | -1.4978 | 0.012868 | -0.22862 | 0.504775 | 0.93288 | 0.497502 | 0.896534 | 0.6605 | 0.478806 | 1.454814 | 1.134476 | 1.366601 |
| 32.48611 | 1.273399 | 1.559976 | 0.403579 | 0.130676 | 0.254176 | -0.69372 | -1.00074 | 0.139252 | -0.01973 | -0.18918 | 1.133483 | -0.24963 | 0.093892 | 0.287325 | 0.259653 | 1.171401 | 0.662114 | 0.706646 |
| 32.5 | 1.852898 | 2.02594 | 2.015691 | 1.090667 | 0.583177 | 0.253688 | 0.294205 | 0.309174 | 0.771169 | -0.04984 | 2.517026 | -0.32333 | 0.05493 | 0.700684 | 0.86904 | 2.115201 | 1.424934 | 1.112017 |
| 32.51389 | 2.719033 | 1.899782 | 2.6062 | 1.926992 | 0.859052 | 1.344493 | 2.034777 | 0.494171 | 1.414304 | 1.660892 | 3.248225 | 0.925917 | 1.258017 | 1.865986 | 1.459087 | 2.76785 | 1.704658 | 2.111192 |
| 32.52778 | 3.13328 | 1.485665 | 2.308598 | 2.254129 | 0.69902 | 1.779064 | 3.649319 | 0.79484 | 1.853037 | 4.843305 | 3.307893 | 2.215149 | 2.32629 | 3.44611 | 0.745578 | 2.958835 | 1.341491 | 1.604798 |
| 32.54167 | 3.136932 | 1.236581 | 2.455046 | 3.370978 | 0.680896 | 1.752973 | 4.375818 | 1.158101 | 2.118336 | 5.192293 | 4.65949 | 2.042296 | 2.144312 | 3.718625 | 0.427942 | 2.229854 | 0.448669 | 1.771728 |
| 32.55556 | 2.702285 | 0.954201 | 2.580285 | 3.749931 | 0.896608 | 1.537357 | 4.04504 | 1.204458 | 1.980531 | 3.931635 | 4.168595 | 0.688698 | 1.493264 | 3.41222 | 2.021342 | 1.354685 | 0.707682 | 2.129746 |
| 32.56944 | 2.103603 | 0.813351 | 2.322117 | 3.120216 | 1.076733 | 1.173132 | 3.085385 | 1.011772 | 1.24362 | 1.666947 | 2.896336 | -0.29845 | 1.306526 | 1.611282 | 3.803124 | 0.307985 | 0.263173 | 1.95138 |
| 32.58333 | 1.331536 | 0.605184 | 2.048359 | 1.74499 | 1.377138 | 0.332715 | 2.295281 | 0.886496 | 0.899906 | -1.1527 | 1.182346 | -0.8996 | 1.128558 | 0.245317 | 4.8637 | -0.88365 | -0.41312 | 0.877716 |

| | | | | | | | | | | | | | | | | | | |
|---|---|---|---|---|---|---|---|---|---|---|---|---|---|---|---|---|---|---|
| 32.59722 | 0.679105 | 0.299695 | 1.720206 | 0.815119 | 1.817332 | -0.1592 | 1.894048 | 0.628423 | 0.569937 | -4.25482 | -0.32805 | -1.53486 | 0.905217 | -0.20207 | 5.051175 | -1.30534 | -0.90513 | -0.17202 |
| 32.61111 | 0.562096 | 0.240215 | 1.326358 | 0.086468 | 1.842403 | -0.39143 | 1.482954 | 0.498831 | 0.225302 | -5.01167 | -1.62746 | -2.63295 | 0.355506 | -1.10635 | 4.719764 | -0.64561 | -1.05518 | -0.64933 |
| 32.625 | 0.473099 | 0.205769 | 1.136409 | -0.55605 | 1.653945 | -0.0248 | 1.226108 | 0.449065 | -0.06311 | -3.6553 | -2.68598 | -2.87787 | -0.4014 | -1.67346 | 3.113311 | -0.46467 | -1.50933 | -0.92258 |
| 32.63889 | 0.276686 | 0.092099 | 0.737944 | -0.66971 | 1.209203 | 0.399766 | 0.92278 | 0.242558 | -0.64509 | -3.10103 | -1.63602 | -2.10196 | -0.01628 | -1.46247 | 1.353503 | -0.43231 | -1.55367 | -1.29409 |
| 32.65278 | -0.05795 | -0.12619 | 0.53041 | -0.74148 | 0.803068 | 0.293887 | -0.02281 | 0.116889 | -1.12968 | -2.16139 | -1.03721 | -0.92734 | 0.253037 | -1.72355 | 0.667214 | -0.37891 | -0.95742 | -0.94154 |
| 32.66667 | -0.32325 | -0.2614 | 0.269944 | -0.57517 | 0.518012 | -0.05691 | -1.41412 | 0.062122 | -1.59933 | -1.13123 | -0.57107 | -0.10761 | 0.165455 | -0.90659 | 0.236367 | -0.37701 | -0.50152 | -0.43376 |
| 32.68056 | -0.55006 | -0.2724 | 0.102356 | -0.57754 | 0.247769 | -0.8632 | -2.97744 | -0.01464 | -2.05916 | 0.008192 | -0.33754 | 0.140505 | -0.53996 | -0.38471 | -0.7785 | -1.39774 | -0.76555 | -0.41783 |
| 32.69444 | -0.65618 | -0.29129 | 0.232128 | -0.76629 | 0.129604 | -1.97984 | -3.76534 | -0.0523 | -2.21228 | 0.758325 | 0.255286 | -0.00537 | -1.35052 | 0.316022 | -2.65063 | -1.1529 | -0.97185 | -0.33617 |
| 32.70833 | -0.63388 | -0.2761 | 0.413648 | -0.74179 | 0.181206 | -2.73768 | -3.76923 | 0.197385 | -1.62567 | 1.365164 | 1.101995 | -0.00661 | -1.53455 | 1.297281 | -4.06281 | -1.00284 | -0.90936 | -0.44517 |
| 32.72222 | -0.39914 | -0.3253 | 0.581443 | -0.30469 | 0.318108 | -2.24588 | -2.16376 | 0.511429 | -0.37802 | 1.899637 | 1.963115 | 0.305216 | -1.23754 | 2.256075 | -4.62063 | -1.11893 | -0.78945 | -0.6173 |
| 32.73611 | -0.08337 | -0.46981 | 0.727713 | 0.384147 | 0.330804 | -1.40558 | -0.79919 | 0.565931 | 0.711673 | 2.192137 | 3.069953 | 0.494527 | -0.79251 | 2.714066 | -4.28786 | -1.42918 | -0.65862 | -0.66094 |
| 32.75 | 0.229923 | -0.53578 | 0.682376 | 1.028858 | 0.176045 | -0.64795 | -0.04911 | 0.33269 | 1.453877 | 1.945529 | 3.737664 | 0.282509 | -0.83383 | 2.837951 | -3.3658 | -1.01123 | -0.7859 | -0.78407 |
| 32.76389 | 0.48372 | -0.43292 | 0.588383 | 1.661772 | -0.12465 | 0.068892 | 0.291861 | -0.05791 | 1.51522 | 1.155922 | 2.251047 | -0.14007 | -1.30523 | 3.100707 | -2.57225 | -0.56154 | -0.97832 | -0.81768 |
| 32.77778 | 0.336773 | -0.48322 | 0.482657 | 1.702174 | -0.63949 | 0.656859 | 0.334896 | -0.36149 | 2.000007 | -0.22423 | 0.635323 | -0.57277 | -1.88259 | 0.923176 | -2.94046 | -0.44782 | -1.14479 | -0.49932 |
| 32.79167 | -0.25104 | -0.54139 | 0.385899 | 1.431746 | -1.17375 | 0.715541 | 0.320359 | -0.39158 | 2.149329 | -1.69304 | -0.71206 | -0.85427 | -2.25611 | -2.22356 | -2.85004 | -0.28978 | -0.60156 | -0.02989 |
| 32.80556 | -0.74854 | -0.39799 | 0.253343 | 0.312814 | -1.5729 | 0.797633 | 0.360117 | -0.21891 | 1.626426 | -2.28047 | -1.74383 | -1.13318 | -2.14976 | -3.84456 | -2.51642 | 0.194069 | -0.23128 | 0.440866 |
| 32.81944 | -1.16943 | -0.28725 | 0.014634 | -1.12548 | -1.89111 | 0.525388 | 0.296513 | 0.022371 | 0.699331 | -2.85915 | -3.25644 | -1.21723 | -1.71312 | -3.00038 | -1.62421 | 0.364405 | -0.40401 | 0.999691 |
| 32.83333 | -1.41796 | -0.28212 | -0.33729 | -2.0752 | -2.18709 | 0.013927 | 0.291306 | 0.207559 | -0.27736 | -3.02125 | -5.22663 | -1.10783 | -1.16128 | -2.38403 | -0.74919 | 0.754765 | -0.30945 | 1.1438 |
| 32.84722 | -1.68566 | -0.36676 | -0.65983 | -2.40171 | -2.19318 | -0.56584 | 0.024413 | 0.290127 | -1.24347 | -2.41602 | -5.9137 | -0.75854 | -0.39973 | -2.26285 | -0.34065 | 0.551333 | -0.20565 | 1.047108 |
| 32.86111 | -1.99972 | -0.53767 | -0.96362 | -2.32284 | -1.97382 | -1.05145 | -0.65065 | 0.15861 | -1.44044 | -1.50094 | -4.43105 | -0.32916 | 0.118272 | -1.29811 | -0.43361 | -0.65856 | -0.60214 | 0.770228 |
| 32.875 | -2.16407 | -0.76214 | -1.02515 | -2.03739 | -1.5091 | -1.28515 | -1.34532 | -0.3185 | -2.04011 | -0.7686 | -3.76964 | -0.04934 | 0.172722 | -1.08732 | -0.57848 | -1.84264 | -1.18455 | 0.178232 |
| 32.88889 | -2.13688 | -0.96178 | -0.89066 | -1.69404 | -0.85756 | -1.53981 | -1.83699 | -0.78343 | -2.85283 | -0.66412 | -3.39014 | 0.142081 | 0.123356 | -1.79405 | -0.87413 | -2.04592 | -1.41984 | -0.32474 |
| 32.90278 | -1.74762 | -0.97066 | -0.68033 | -1.12204 | -0.15044 | -1.88933 | -1.93308 | -0.78121 | -2.80428 | -1.0886 | -2.23977 | 0.155661 | 0.141161 | -4.24208 | -1.4754 | -1.60234 | -1.1305 | -0.57803 |
| 32.91667 | -1.10667 | -0.72487 | -0.50029 | -0.70654 | 0.408212 | -1.77615 | -1.41665 | -0.77242 | -3.00531 | -1.30267 | -1.02563 | 0.448367 | 0.24808 | -3.42184 | -1.10087 | -0.64373 | -0.83709 | -0.76346 |
| 32.93056 | -0.49008 | -0.47434 | -0.41496 | -0.28779 | 0.872113 | -1.23741 | -1.02977 | -0.69802 | -2.74708 | -0.70735 | 0.091512 | 0.76305 | 0.354283 | -1.82599 | -0.62581 | 0.303286 | -0.59113 | -0.94307 |
| 32.94444 | 0.044889 | -0.27105 | -0.08026 | 0.310283 | 1.005949 | -0.35173 | -0.60334 | -0.40917 | -1.30558 | 0.434077 | 1.275139 | 1.136851 | 0.491649 | -0.06434 | -0.19231 | 1.028148 | 0.210534 | -0.80651 |
| 32.95833 | 0.474746 | -0.05684 | 0.558627 | 1.157601 | 1.197392 | 0.527537 | 0.039398 | -0.12265 | 0.624616 | 1.391696 | 2.45521 | 1.233849 | 0.596752 | 1.828279 | 0.275712 | 1.611355 | 1.376224 | -0.67999 |
| 32.97222 | 0.790839 | 0.065686 | 0.971373 | 2.006359 | 1.319215 | 1.118568 | 0.912422 | 0.102917 | 2.169722 | 1.986871 | 3.33385 | 1.022541 | 0.372811 | 3.31642 | 0.594291 | 1.95881 | 1.860824 | -0.33891 |
| 32.98611 | 0.803541 | -0.20424 | 0.64073 | 1.871451 | 1.143337 | 0.810757 | 0.950279 | 0.142846 | 2.401631 | 1.63345 | 2.803469 | 0.600211 | 0.041348 | 3.056892 | -0.00029 | 1.16676 | 1.062423 | -0.58531 |
| 33 | -0.19 | -0.86686 | -0.66373 | 0.150408 | 0.202414 | -0.89592 | -0.70327 | -0.11386 | 0.592463 | -0.22598 | 0.674167 | -0.46003 | -0.78306 | 0.223016 | -1.64847 | -0.69307 | -0.906 | -1.89409 |
| 33.01389 | -2.10524 | -2.52538 | -1.91415 | -2.14065 | -1.02237 | -2.89252 | -2.6947 | -0.66866 | -2.24051 | -2.43299 | -2.24692 | -1.44579 | -2.2131 | -3.75681 | -3.22099 | -2.19385 | -2.39079 | -2.89381 |
| 33.02778 | -3.7434 | -4.27657 | -1.97896 | -3.51247 | -1.72678 | -3.28327 | -3.01068 | -1.00556 | -3.61313 | -2.67082 | -3.5185 | -1.25554 | -2.18153 | -5.87819 | -3.84874 | -2.37591 | -2.91976 | -2.416 |
| 33.04167 | -4.9145 | -4.55194 | -1.6301 | -3.33343 | -2.20736 | -2.9188 | -1.94246 | -0.98328 | -3.59506 | -3.23266 | -4.30064 | -0.56054 | -2.84183 | -5.55297 | -3.77064 | -2.52312 | -3.21409 | -2.35107 |
| 33.05556 | -3.89419 | -4.36988 | -1.37468 | -2.4457 | -2.17355 | -2.746 | -1.53111 | -0.84804 | -4.33002 | -3.15612 | -5.36786 | -1.13674 | -1.9581 | -5.14953 | -3.23464 | -2.47323 | -4.0284 | -2.27948 |
| 33.06944 | -1.85806 | -3.80791 | -1.07433 | -1.50288 | -1.47616 | -2.42614 | -1.08893 | -0.71275 | -3.66227 | -1.8146 | -4.52333 | -0.49215 | -1.1682 | -3.48554 | -2.76801 | -0.95331 | -2.31966 | -1.87966 |
| 33.08333 | -0.65599 | -2.66475 | -0.91156 | -0.69818 | -0.80649 | -2.49133 | -0.69983 | -0.6414 | -2.74758 | -0.56944 | -3.03616 | 0.633105 | -0.24774 | -1.57446 | -1.47469 | -0.1265 | -0.55981 | -0.74085 |
| 33.09722 | 0.14288 | -1.47377 | -0.69635 | -0.15752 | -0.08138 | -2.25503 | -0.34351 | -0.37267 | -1.86157 | 0.160515 | -2.47865 | 0.885809 | 0.463292 | -0.41542 | -0.53921 | 0.60937 | 0.13377 | 0.614688 |
| 33.11111 | 0.692422 | -0.09159 | -0.42857 | 0.434842 | 0.479291 | -1.87476 | -0.0126 | -0.00935 | -1.15451 | 0.431849 | -1.47738 | 0.92607 | 0.806798 | -0.07053 | -0.2966 | 0.375025 | 0.138643 | 1.157446 |
| 33.125 | 0.829974 | 0.505532 | -0.13205 | 0.920704 | 0.777633 | -1.26809 | 0.235926 | 0.062623 | -0.73774 | 0.673214 | -0.60354 | 1.15531 | 1.026557 | 0.024774 | -0.4685 | 0.44672 | 0.338794 | 1.054504 |
| 33.13889 | 0.814555 | 0.485856 | 0.095884 | 1.202214 | 0.962173 | -0.65232 | 0.497419 | 0.008545 | -0.59482 | 0.980986 | 0.047724 | 1.380739 | 1.66535 | 0.321524 | -0.25022 | 0.37732 | 0.766074 | 0.980434 |
| 33.15278 | 1.072757 | 0.646563 | 0.416095 | 1.540607 | 1.153054 | 0.454796 | 0.821979 | 0.120185 | -0.21609 | 1.340996 | 0.68491 | 1.307034 | 1.899945 | 0.731758 | 0.423521 | 0.888212 | 1.316034 | 1.645378 |
| 33.16667 | 1.497898 | 0.856844 | 0.875465 | 2.040552 | 1.125727 | 1.593475 | 0.991123 | 0.328984 | 0.517987 | 1.640383 | 1.373477 | 0.848388 | 1.59387 | 1.189352 | 0.961424 | 1.252695 | 1.710099 | 2.233175 |
| 33.18056 | 1.921559 | 0.761165 | 1.075493 | 1.738072 | 1.241598 | 2.358361 | 1.064097 | 0.486303 | 1.418678 | 1.959622 | 2.083599 | 0.377945 | 1.003183 | 1.451877 | 0.933531 | 1.364324 | 2.162135 | 1.722042 |
| 33.19444 | 2.20875 | 0.680866 | 1.002052 | 1.276034 | 1.263901 | 2.728285 | 1.101216 | 0.642041 | 2.359369 | 2.191028 | 2.695454 | 0.289542 | 0.523674 | 1.424726 | 0.661896 | 0.988056 | 2.294395 | 0.908209 |
| 33.20833 | 2.222959 | 0.428052 | 0.934735 | 0.96383 | 1.170588 | 2.753598 | 1.070535 | 0.760651 | 3.093558 | 2.453632 | 3.160406 | 0.441386 | 0.442747 | 1.827484 | 0.480863 | 0.674389 | 2.061851 | 0.206375 |
| 33.22222 | 1.847826 | 0.033651 | 0.802522 | 0.763948 | 0.859099 | 2.500848 | 0.887121 | 0.831152 | 3.533142 | 2.698167 | 4.049293 | 1.10823 | 0.637172 | 2.662309 | 0.521508 | 1.019728 | 2.07501 | -0.02071 |
| 33.23611 | 1.500434 | -0.2704 | 0.639664 | 0.506616 | 0.515695 | 2.003459 | 0.724817 | 0.930527 | 3.287387 | 2.762648 | 3.896778 | 1.336669 | 0.78332 | 2.475175 | 0.638717 | 0.904078 | 1.999828 | -0.1014 |
| 33.25 | 1.143586 | -0.16255 | 0.417853 | 0.187822 | 0.261788 | 1.731035 | 0.563463 | 1.109032 | 2.711771 | 2.815043 | 2.836142 | 0.881385 | 0.694066 | 1.54809 | 0.758688 | 1.299785 | 2.032755 | 0.051539 |
| 33.26389 | 0.611833 | -0.14947 | 0.136722 | 0.001714 | 1.062423 | 0.279109 | 1.087587 | 2.547802 | 2.452055 | 1.786601 | 0.009311 | 0.253253 | 1.158006 | 0.739623 | 1.484517 | 1.283028 | -0.13264 | |
| 33.27778 | 0.027275 | -0.49789 | 0.005039 | -0.32124 | -0.12791 | 0.242606 | 0.036482 | 0.759217 | 2.732695 | 1.692521 | 1.022064 | -0.53874 | -0.21569 | 0.920575 | 0.593396 | 1.399769 | 0.962882 | -0.29172 |
| 33.29167 | -0.15278 | -0.14897 | 0.058256 | -0.26376 | -0.09829 | -0.33391 | -0.18152 | 0.660754 | 2.685031 | 1.500507 | 0.653995 | -0.76866 | -0.65848 | 1.339511 | 0.441492 | 1.65454 | 0.857177 | -0.45335 |
| 33.30556 | 0.092777 | 0.821187 | 0.270927 | -0.04151 | 0.081286 | -0.34258 | -0.14239 | 0.631763 | 2.313206 | 1.557902 | 0.421773 | -0.50723 | -0.58004 | 1.977434 | 0.547312 | 1.878249 | 0.984113 | -0.37703 |
| 33.31944 | 0.287389 | 1.048121 | 0.381404 | 0.112118 | 0.23622 | -0.24494 | -0.0831 | 0.420077 | 1.321012 | 1.611082 | 0.541567 | 0.127696 | 0.016853 | 2.063083 | 0.82703 | 1.529861 | 1.24893 | 0.102102 |
| 33.33333 | 0.518767 | 0.885372 | 0.433421 | 0.258591 | 0.426819 | -0.15389 | 0.175213 | 0.353818 | 1.095274 | 1.266399 | 0.929472 | 0.688638 | 0.628481 | 2.083717 | 0.841767 | 1.496173 | 1.751578 | 0.712389 |
| 33.34722 | 0.941372 | 1.466648 | 0.455666 | 0.408759 | 0.652703 | -0.14471 | 0.293544 | 0.361442 | 1.150794 | 0.735613 | 1.235479 | 0.659749 | 0.75892 | 2.133092 | 0.758929 | 1.333139 | 1.769299 | 1.202698 |
| 33.36111 | 1.01657 | 1.700539 | 0.404963 | 0.375565 | 0.692848 | -0.47985 | 0.131137 | 0.260479 | 0.848736 | 0.246458 | 1.190186 | 0.44578 | 0.55987 | 2.104201 | 0.470913 | 0.548298 | 1.103329 | 1.248494 |
| 33.375 | 0.822908 | 1.312223 | 0.5032 | 0.253216 | 0.524589 | -1.06902 | -0.09837 | 0.182514 | 0.226533 | 0.07307 | 0.957679 | 0.272617 | 0.282369 | 1.615755 | 0.294243 | -0.41489 | 0.1786 | 1.196452 |
| 33.38889 | 0.780929 | 1.087222 | 0.728378 | 0.279859 | 0.381025 | -1.30409 | -0.08282 | 0.146571 | -0.26139 | 0.175267 | 0.76117 | 0.234945 | 0.044088 | 1.178423 | 0.306437 | -0.91481 | -0.66128 | 0.785238 |
| 33.40278 | 0.905528 | 1.09953 | 0.90027 | 0.320005 | 0.301764 | -1.40868 | 0.059034 | 0.10336 | -0.45542 | 0.41265 | 0.655373 | 0.341474 | -0.12064 | 0.943709 | 0.321174 | -0.92965 | -1.30263 | 0.290368 |
| 33.41667 | 0.926822 | 1.019394 | 0.968123 | 0.27133 | 0.191497 | -1.33321 | 0.2117 | 0.078037 | -0.41233 | 0.66558 | 0.469402 | 0.386094 | -0.18398 | 0.983233 | 0.502241 | -0.13734 | -1.03939 | -0.17356 |
| 33.43056 | 0.99992 | 0.955273 | 0.974012 | 0.310089 | 0.173435 | -1.02626 | 0.346955 | 0.101971 | -0.31037 | 0.946155 | 0.431753 | 0.326175 | -0.20446 | 1.005603 | 0.753201 | 0.483593 | -0.58342 | -0.5834 |
| 33.44444 | 1.029146 | 0.902236 | 0.818399 | 0.234088 | 0.134264 | -1.12099 | 0.253781 | 0.007555 | -0.27412 | 1.244341 | 0.558697 | 0.129417 | -0.29742 | 1.140216 | 0.730808 | 0.976841 | 0.011616 | -0.60239 |
| 33.45833 | 0.82683 | 0.541677 | 0.576171 | 0.081022 | -0.0269 | -1.59304 | 0.011607 | -0.10149 | -0.38573 | 1.394785 | 0.73831 | -0.11767 | -0.4739 | 1.102451 | 0.045767 | 1.139998 | 0.600587 | -0.32073 |
| 33.47222 | 0.506725 | 0.296415 | 0.511706 | 0.016345 | -0.13629 | -1.83649 | -0.16804 | -0.07991 | -0.51227 | 1.394179 | 0.937771 | -0.21294 | -0.55273 | 1.08836 | -0.0667 | 1.120746 | 0.980001 | -0.18472 |
| 33.48611 | 0.399707 | 0.958991 | 0.763751 | 0.429395 | -0.07414 | -1.54082 | -0.00105 | 0.101389 | -0.19872 | 1.724329 | 1.501793 | 0.064619 | -0.36058 | 1.785451 | 0.476692 | 1.218667 | 1.346387 | 0.641925 |
| 33.5 | 0.651487 | 2.424832 | 1.463057 | 1.243415 | 0.311875 | -0.29316 | 0.667346 | 0.449999 | 0.517337 | 2.596542 | 2.613313 | 0.736152 | 0.271998 | 3.490057 | 1.677974 | 1.914442 | 1.792663 | 1.805694 |
| 33.51389 | 1.458478 | 3.564491 | 2.203797 | 2.122619 | 0.731209 | 0.588339 | 1.418797 | 0.76899 | 1.478971 | 3.182916 | 3.416136 | 1.458366 | 1.221054 | 5.070278 | 2.754104 | 2.087301 | 1.598565 | 2.823503 |
| 33.52778 | 2.181524 | 2.745991 | 2.338331 | 2.615995 | 1.002854 | 1.200624 | 1.97421 | 0.874241 | 2.143286 | 3.776782 | 4.03493 | 2.314055 | 2.319312 | 5.254165 | 2.365274 | 1.56357 | -0.37678 | 3.533706 |
| 33.54167 | 2.7069 | 2.268517 | 2.163577 | 2.800639 | 0.876984 | 1.537639 | 1.609849 | 0.855036 | 2.875505 | 4.952721 | 4.948502 | 2.848998 | 2.893206 | 4.783692 | 1.700712 | 2.089895 | -1.4572 | 4.320176 |
| 33.55556 | 2.757392 | 2.38924 | 1.671492 | 2.276725 | 1.161302 | 1.472441 | 1.44966 | 0.831981 | 3.054784 | 6.30159 | 4.696869 | 2.387801 | 2.639264 | 3.850704 | 1.092916 | 2.479982 | 0.625466 | 4.213815 |
| 33.56944 | 2.679938 | 2.623429 | 1.297065 | 1.949599 | 1.196041 | 1.724263 | 0.9402 | 0.763711 | 2.394697 | 4.630481 | 3.361464 | 1.719258 | 2.077328 | 1.995696 | 0.44428 | 1.574655 | 1.159634 | 3.545238 |
| 33.58333 | 2.344456 | 2.546292 | 1.009988 | 1.586851 | 1.04755 | 1.501119 | 0.222239 | 0.65982 | 1.641607 | 1.963008 | 2.520844 | 0.79408 | 1.288881 | 0.525171 | 0.492662 | 0.498363 | 0.159963 | 3.300589 |
| 33.59722 | 1.705866 | 2.549412 | 0.745991 | 1.183251 | 1.101846 | 1.206909 | -0.20657 | 0.646882 | 1.063344 | -0.02605 | 1.645756 | -0.38396 | 0.683114 | -0.11763 | 0.734691 | -0.80084 | -0.92273 | 2.226638 |
| 33.61111 | 1.044556 | 2.251278 | 0.608802 | 0.765718 | 1.049198 | 1.135946 | -0.29305 | 0.705788 | 0.585109 | -1.54246 | 0.298793 | -1.49155 | -0.12445 | -0.48127 | 1.61717 | -1.73109 | -2.38101 | 0.977735 |
| 33.625 | 0.372264 | 1.451283 | 0.435475 | 0.577186 | 0.786963 | 1.070773 | -0.02942 | 0.5117 | -0.06304 | -3.16011 | -1.02262 | -2.13013 | -0.71485 | -0.4619 | 2.88775 | -1.56007 | -2.85883 | -0.2318 |
| 33.63889 | -0.18857 | 0.317246 | 0.411267 | 0.232495 | 0.302157 | 0.818529 | 0.391038 | -0.04691 | -0.79492 | -3.95046 | -1.98941 | -2.43583 | -1.17312 | -0.66221 | 3.24292 | -1.29418 | -3.4306 | -1.81512 |
| 33.65278 | -0.90578 | -0.51651 | 0.417561 | -0.26274 | -0.35969 | 0.266512 | 0.563068 | -0.61191 | -1.4993 | -3.99122 | -1.77664 | -2.00022 | -1.02067 | -0.62348 | 2.572834 | -1.02913 | -3.45244 | -3.01397 |
| 33.66667 | -1.22052 | -1.14821 | 0.377017 | -0.63544 | -0.93449 | -0.24686 | 0.534961 | -1.07086 | -2.00622 | -3.75421 | -1.17697 | -1.15632 | -0.61704 | 0.316485 | 2.05591 | -0.32067 | -2.32349 | -3.09901 |
| 33.68056 | -1.21186 | -1.63783 | 0.271366 | -0.71724 | -1.1769 | -0.64846 | 0.456384 | -1.66394 | -2.69915 | -2.4206 | -0.57768 | -0.90573 | -0.47569 | 1.204642 | 1.710295 | 1.126864 | -0.32273 | -2.18743 |
| 33.69444 | -0.98957 | -1.72175 | 0.260323 | -0.57796 | -1.09274 | -0.87727 | 0.314909 | -2.05714 | -3.20042 | -0.82738 | -0.09344 | -0.7576 | -0.46499 | 1.978102 | 1.489084 | 1.854412 | 1.952135 | -1.58279 |
| 33.70833 | -0.91665 | -1.64089 | 0.318786 | -0.18818 | -0.88543 | -0.74665 | 0.230372 | -1.92349 | -3.13021 | 0.851783 | 0.107233 | -0.67333 | -0.43988 | 1.576562 | 0.645946 | 1.801221 | 3.375174 | -1.00503 |

| | | | | | | | | | | | | | | | | | | |
|---|---|---|---|---|---|---|---|---|---|---|---|---|---|---|---|---|---|---|
| 33.72222 | -0.7771 | -1.75956 | 0.532805 | 0.26678 | -0.57968 | -0.60226 | 0.391066 | -1.23794 | -2.59898 | 1.868067 | 0.268489 | -0.46629 | -0.46364 | 1.162436 | -0.23328 | 0.932072 | 3.80758 | -1.12537 |
| 33.73611 | -0.42522 | -1.51189 | 0.859765 | 0.919807 | -0.28033 | -0.53804 | 0.586427 | -0.4905 | -1.4308 | 2.921098 | 0.666092 | -0.01985 | -0.76171 | 1.031952 | -1.85763 | 0.48365 | 2.690321 | -1.60797 |
| 33.75 | -0.50502 | -1.18643 | 1.076116 | 1.304414 | -0.06991 | -0.35503 | 0.659241 | -0.08286 | 0.192793 | 3.6856 | 1.180192 | 0.336345 | -0.91389 | 1.002532 | -3.18124 | -0.57325 | 0.747234 | -2.09884 |
| 33.76389 | -0.60008 | -0.80185 | 1.169193 | 1.30721 | -0.06596 | -0.09357 | 0.669082 | 0.025044 | 1.671384 | 4.072614 | 1.935816 | 0.299819 | -0.45206 | 0.913198 | -3.55403 | -1.15965 | -0.86939 | -1.66877 |
| 33.77778 | -0.72403 | -0.65452 | 1.105764 | 1.0548 | -0.2545 | 0.282224 | 0.57917 | 0.075076 | 2.50818 | 2.816904 | 2.296721 | -0.09131 | -0.14801 | 0.292025 | -3.61256 | -1.83924 | -2.82655 | -0.77775 |
| 33.79167 | -0.84193 | -0.31383 | 0.818015 | 0.627123 | -0.73123 | 0.649824 | 0.36587 | 0.180998 | 2.507263 | 0.881029 | 1.519676 | -0.29225 | -0.61696 | -0.43741 | -3.92841 | -2.3818 | -4.35502 | -0.10589 |
| 33.80556 | -1.56781 | -0.60025 | 0.404818 | -0.09434 | -1.39576 | 0.842942 | -0.09156 | 0.293228 | 2.083381 | -0.4999 | 0.636679 | -0.32378 | -1.29786 | -1.05484 | -3.43839 | -2.73538 | -5.83798 | 0.290452 |
| 33.81944 | -2.73902 | -1.72256 | -0.20679 | -1.05398 | -2.18249 | 0.872488 | -0.52665 | 0.244384 | 1.320492 | -0.8096 | 0.42765 | -0.67198 | -1.04509 | -2.67658 | -2.38515 | -1.75015 | -5.66565 | -0.24561 |
| 33.83333 | -3.68632 | -3.08755 | -0.81405 | -1.75685 | -2.87542 | 0.752795 | -0.63147 | -0.01318 | 0.066866 | -1.67633 | 0.103556 | -2.14037 | -0.53392 | -2.53947 | -0.37238 | -0.17442 | -3.07967 | -0.00507 |
| 33.84722 | -4.09546 | -3.41866 | -1.16775 | -2.18037 | -3.7108 | 0.650126 | -0.41965 | -0.34795 | -1.20191 | -2.50705 | -0.59038 | -2.84666 | 0.090097 | -2.05687 | 1.665864 | 1.620057 | 0.895039 | 0.211979 |
| 33.86111 | -3.6255 | -3.30521 | -1.26228 | -2.50755 | -4.29012 | 0.47137 | -0.16228 | -0.61611 | -2.54417 | -2.08481 | -1.20531 | -2.70222 | 0.115869 | -2.50874 | 2.697819 | 3.082728 | 3.658948 | 0.431442 |
| 33.875 | -3.47098 | -3.79904 | -1.31923 | -2.69744 | -4.09779 | 0.272725 | -0.04228 | -0.73545 | -3.86372 | -2.43709 | -1.91237 | -1.74414 | -0.28448 | -3.82896 | 2.029565 | 2.64408 | 5.397013 | -0.23836 |
| 33.88889 | -2.99041 | -4.2018 | -1.39056 | -2.4764 | -3.44617 | 0.014067 | -0.06399 | -0.71076 | -4.63456 | -2.76986 | -2.36721 | -0.3484 | -0.66927 | -3.7839 | 0.542893 | 2.030883 | 5.4905 | -0.56268 |
| 33.90278 | -2.08405 | -4.15982 | -1.11442 | -2.06681 | -2.37976 | -0.2417 | -0.11778 | -0.5812 | -4.64512 | -2.24898 | -2.02287 | 0.848642 | -0.8317 | -3.59195 | -0.52294 | 1.312753 | 4.621843 | -0.45334 |
| 33.91667 | -1.39798 | -3.46221 | -0.728 | -1.44468 | -0.99676 | -0.45872 | -0.1816 | -0.53295 | -4.28265 | -1.9014 | -2.17875 | 1.409496 | -0.82062 | -3.36807 | -0.94996 | 0.964531 | 3.064064 | -1.12658 |
| 33.93056 | -0.70057 | -2.86166 | -0.20974 | -0.63703 | 0.451682 | -0.32976 | -0.12943 | -0.42754 | -4.07281 | -1.04031 | -3.34897 | 1.211139 | -0.91006 | -3.78604 | -1.1525 | 0.620883 | 0.06764 | -2.27114 |
| 33.94444 | 0.037537 | -2.35958 | 0.321057 | 0.324031 | 1.957594 | -0.01976 | 0.224659 | -0.21334 | -3.16079 | 0.09184 | -3.2075 | 1.12221 | -0.68945 | -3.00317 | -1.07837 | 0.292182 | -1.98409 | -2.33209 |
| 33.95833 | 0.695221 | -1.66977 | 0.776042 | 1.267224 | 2.826437 | 0.626218 | 0.645816 | 0.069642 | -2.07806 | 1.707584 | -1.27905 | 1.686743 | 0.129981 | -1.54097 | -0.34673 | 0.672491 | -2.06231 | -1.18811 |
| 33.97222 | 0.896783 | -1.17975 | 0.986965 | 1.89009 | 1.79378 | 1.246099 | 0.983374 | 0.3142 | -1.11411 | 3.207628 | 0.339777 | 2.021214 | 1.037077 | 0.648034 | 0.52791 | 1.133078 | -1.26251 | 0.165073 |
| 33.98611 | 0.32007 | -1.36119 | 0.424526 | 1.708429 | 0.698739 | 0.650205 | 0.765451 | 0.013076 | -1.36916 | 2.929816 | 0.735794 | 0.924398 | 0.914382 | 1.624633 | 0.196706 | 0.70457 | -1.16762 | 0.273855 |
| 34 | -1.48007 | -3.08387 | -1.21987 | -0.44681 | -0.3971 | -1.40988 | -0.2873 | -0.7922 | -3.68626 | -0.19966 | -1.12587 | -1.50561 | -1.06237 | -1.46784 | -2.07111 | -0.89398 | -3.29833 | -1.72546 |
| 34.01389 | -3.92677 | -5.2103 | -2.2873 | -3.05727 | -2.0445 | -1.95074 | -1.1387 | -1.34375 | -4.83069 | -2.70675 | -3.67598 | -3.69237 | -3.11773 | -6.06035 | -4.1024 | -2.93786 | -7.51859 | -4.18027 |
| 34.02778 | -4.27401 | -6.14139 | -2.46304 | -3.47646 | -2.87366 | -1.79283 | -0.9948 | -1.34511 | -4.79263 | -2.75928 | -5.86593 | -3.6159 | -2.73863 | -6.26062 | -3.6941 | -3.61666 | -10.3662 | -4.52453 |
| 34.04167 | -3.85282 | -4.47115 | -2.4443 | -2.12684 | -2.66584 | -1.77038 | -0.77337 | -1.06165 | -4.88468 | -2.95181 | -5.99615 | -2.77481 | -2.69732 | -4.44648 | -2.29405 | -2.88782 | -8.83264 | -2.97872 |
| 34.05556 | -3.73633 | -3.58232 | -1.84587 | -1.50224 | -2.35716 | -1.43604 | -0.46192 | -1.00235 | -4.14906 | -3.12031 | -4.04857 | -2.03337 | -2.21727 | -3.26858 | -1.53042 | -2.52178 | -7.55586 | -1.65303 |
| 34.06944 | -2.95163 | -2.77106 | -1.24905 | -0.94364 | -1.48521 | -1.13494 | -0.15965 | -0.94306 | -3.34049 | -3.03272 | -1.87032 | -1.91055 | -1.4261 | -2.47835 | -1.28782 | -1.95158 | -6.32478 | -0.94261 |
| 34.08333 | -1.69243 | -2.30849 | -0.69235 | -0.40177 | -0.64235 | -1.02357 | 0.085908 | -0.73663 | -2.26542 | -2.7068 | -0.50211 | -0.71566 | -0.66937 | -1.73208 | -0.84866 | -0.72222 | -5.1077 | -1.03457 |
| 34.09722 | -0.68419 | -1.68347 | -0.38103 | 0.004623 | 0.027997 | -1.08339 | 0.472314 | -0.50025 | -1.45571 | -1.85649 | 0.214186 | 0.743175 | -0.12234 | -1.02132 | -0.41562 | 0.606234 | -3.05737 | -1.37789 |
| 34.11111 | -0.23509 | -1.14625 | -0.17161 | 0.171081 | 0.454248 | -1.00778 | 0.651457 | -0.26028 | -0.82293 | -1.08488 | 0.972006 | 1.873277 | 0.193725 | -0.52805 | -0.22687 | 1.344283 | -0.84062 | -1.37537 |
| 34.125 | 0.111659 | -0.74298 | 0.274611 | 0.418281 | 0.837559 | -0.7666 | 0.828741 | -0.118 | -0.41914 | -0.21298 | 2.069146 | 3.372243 | 0.486489 | -0.22124 | -0.3029 | 2.058799 | 1.630766 | -0.99649 |
| 34.13889 | 0.69631 | -0.31019 | 0.639517 | 0.746439 | 1.031718 | -0.41255 | 0.872555 | -0.05623 | -0.00102 | 0.533213 | 3.143023 | 5.887616 | 0.397932 | -0.02238 | -0.52539 | 2.33082 | 3.496066 | -1.0125 |
| 34.15278 | 1.407953 | -0.06574 | 0.793965 | 1.139652 | 0.498169 | 0.017109 | 0.705161 | -0.08771 | 0.261588 | 0.992556 | 3.15602 | 3.729637 | 0.293791 | 0.092303 | -0.78743 | 1.630976 | 4.084153 | -1.46616 |
| 34.16667 | 1.880326 | 0.078542 | 1.019644 | 1.541047 | 0.55329 | 0.537935 | 0.701117 | -0.02274 | 0.524264 | 1.478367 | 2.027378 | 1.631365 | 0.743006 | 0.34052 | -0.95592 | 0.893522 | 4.558253 | -1.89366 |
| 34.18056 | 1.805218 | 0.522518 | 1.270021 | 1.72953 | 0.905734 | 1.233563 | 1.00333 | 0.23359 | 0.963678 | 2.365067 | 2.443799 | 1.834648 | 0.947234 | 0.938083 | -0.75897 | 1.233365 | 5.948432 | -1.79276 |
| 34.19444 | 1.62099 | 0.818901 | 1.308234 | 2.006021 | 1.292799 | 2.033253 | 1.364849 | 0.385483 | 1.37552 | 3.020527 | 2.346765 | 1.633198 | 1.028553 | 1.385153 | -0.48733 | 1.911423 | 5.594155 | -1.39113 |
| 34.20833 | 1.207374 | 0.680942 | 1.054219 | 2.219623 | 1.351547 | 3.227984 | 1.338608 | 0.292929 | 1.333037 | 3.273517 | 1.832468 | 1.000158 | 0.944277 | 1.986761 | -0.35965 | 1.505081 | 3.51687 | -0.82361 |
| 34.22222 | 0.779935 | 0.208274 | 0.783391 | 2.157902 | 1.030163 | 3.588087 | 1.116232 | 0.150001 | 1.002293 | 3.496304 | 1.919866 | 0.591906 | 0.828103 | 2.71229 | -0.11633 | 0.833942 | 2.068262 | -0.13168 |
| 34.23611 | 0.236663 | -0.95617 | 0.62285 | 1.379716 | 0.2009 | 3.21992 | 0.816503 | 0.082734 | 1.065882 | 4.244355 | 2.149658 | 0.526815 | 0.974345 | 3.234357 | 0.261444 | 1.071953 | 1.803399 | 0.376169 |
| 34.25 | -0.59046 | -3.15949 | 0.529375 | 0.857532 | -1.03103 | 2.146155 | 0.405541 | 0.125567 | 1.4406 | 4.237032 | 1.375696 | 0.381662 | 0.759844 | 2.819646 | 0.201377 | 0.959045 | 1.269117 | 0.231539 |
| 34.26389 | -0.75074 | -3.79443 | 0.25257 | 0.212605 | -1.86377 | 1.507278 | 0.156558 | 0.237393 | 1.908014 | 3.204552 | 0.7825 | -0.33453 | 0.07587 | 1.649205 | -0.31002 | 0.507311 | -0.14696 | -0.64989 |
| 34.27778 | -0.24646 | -2.27164 | -0.1147 | 0.279051 | -1.63959 | 1.031369 | 0.22656 | 0.334994 | 2.077788 | 2.402512 | 0.612746 | -2.02973 | -0.31624 | 0.703599 | -0.44853 | 0.550239 | -0.63588 | -0.91235 |
| 34.29167 | 0.14416 | -1.59807 | -0.26729 | 0.159616 | -1.37939 | 0.597167 | 0.288194 | 0.361327 | 1.890145 | 1.617242 | 0.868108 | -1.61834 | -0.18145 | 0.486038 | 0.051856 | 0.757931 | -0.41895 | -0.23596 |
| 34.30556 | 0.208639 | -1.6216 | -0.1903 | -0.14477 | -1.39041 | 0.044969 | 0.33532 | 0.336577 | 1.805903 | 0.611579 | 0.74291 | -0.36878 | -0.12306 | 0.54825 | 0.407903 | 0.924266 | -0.08382 | 0.161303 |
| 34.31944 | 0.072986 | -1.62924 | -0.18841 | -0.40471 | -1.24304 | -0.76438 | 0.361414 | 0.298669 | 1.73888 | -0.15171 | 0.584967 | -0.30634 | -0.25896 | 0.396954 | 0.337864 | 1.092022 | -0.00097 | 0.164175 |
| 34.33333 | 0.004094 | -1.44468 | -0.15878 | -0.33891 | -1.0852 | -1.79475 | 0.282152 | 0.196912 | 1.822771 | -0.30519 | 0.531187 | -0.0083 | -0.43372 | 0.221539 | 0.113754 | 0.680075 | -0.09494 | 0.241027 |
| 34.34722 | -0.04576 | -1.2556 | -0.14652 | -0.15061 | -1.00298 | -2.50674 | 0.149675 | 0.058067 | 2.039084 | -0.36637 | 0.128248 | 1.14587 | -0.57197 | -0.07426 | -0.21384 | 0.356397 | -0.43468 | 0.209317 |
| 34.36111 | -0.42107 | -1.56213 | -0.0714 | -0.20926 | -1.05777 | -2.57053 | -0.13351 | -0.0562 | 2.297877 | -0.65424 | -0.44572 | 2.002128 | -0.67197 | -0.53828 | -0.65688 | 0.100667 | -0.84301 | -0.29516 |
| 34.375 | -0.95487 | -2.40219 | 0.17907 | -0.63553 | -1.6226 | -2.11001 | -0.23092 | -0.15539 | 2.693628 | -0.45233 | -0.67046 | 1.780151 | -0.63487 | -0.65842 | -0.86437 | -0.0436 | -2.1137 | -0.5854 |
| 34.38889 | -0.83869 | -2.26274 | 0.408277 | -0.67122 | -1.60053 | -1.14909 | -0.13326 | -0.21379 | 2.445923 | -0.0961 | -0.47267 | 0.459301 | -0.39686 | -0.4068 | -0.73923 | -0.23497 | -4.22229 | -0.41831 |
| 34.40278 | -0.19068 | -1.22491 | 0.410711 | -0.1585 | -0.88839 | -0.34987 | 0.313451 | -0.07315 | 1.825676 | 0.285011 | -0.31825 | -0.7935 | -0.13944 | -0.24355 | -0.52895 | -0.20827 | -5.97658 | 0.011746 |
| 34.41667 | -0.02728 | -0.6883 | 0.301123 | 0.05562 | -0.93576 | -0.1496 | 0.621806 | 0.012919 | 1.68554 | 0.594369 | -0.26743 | -2.13447 | -0.11027 | -0.02414 | -0.38376 | 0.204009 | -4.26251 | 0.348356 |
| 34.43056 | -0.31428 | -0.67088 | 0.042903 | 0.105726 | -1.67029 | -0.45079 | 0.578586 | -0.0806 | 1.527927 | 1.2757 | -0.16562 | -4.59526 | -0.30716 | 0.161716 | -0.25257 | 0.321847 | -2.28433 | 0.390409 |
| 34.44444 | -0.53702 | -0.51298 | -0.21391 | 0.288901 | -1.7741 | -0.82847 | 0.394191 | -0.25773 | 1.011985 | 1.675221 | 0.038459 | -9.20368 | -0.36164 | 0.385121 | -0.06453 | 0.682423 | -2.03627 | 0.207781 |
| 34.45833 | -0.20612 | -0.47625 | -0.18935 | 0.219508 | -1.13549 | -0.90019 | 0.301432 | -0.27301 | 0.678225 | 1.437602 | 0.076883 | -10.8288 | -0.21027 | 0.519844 | 0.14069 | 0.806473 | -3.24084 | -0.23604 |
| 34.47222 | 0.080905 | -0.09801 | 0.155783 | 0.276811 | -0.3127 | -0.53119 | 0.473563 | -0.17556 | 0.652877 | 0.624349 | 0.219514 | -8.38939 | -0.26909 | 0.697241 | 0.337763 | 0.605872 | -3.21904 | -0.67972 |
| 34.48611 | 0.573324 | 0.618718 | 0.708319 | 0.592708 | 0.09297 | -0.12591 | 0.7854 | -0.09409 | 0.898793 | 0.387405 | 0.819193 | -4.6418 | 0.038642 | 1.290267 | 0.981557 | 0.333389 | -1.43258 | -0.04286 |
| 34.5 | 1.256995 | 1.731054 | 1.603568 | 1.310777 | 0.545149 | 0.615703 | 1.254802 | 0.046628 | 1.857688 | 0.78502 | 1.785454 | -0.95267 | 0.449191 | 2.454837 | 2.188695 | 1.213221 | 0.588559 | 1.971047 |
| 34.51389 | 2.007974 | 2.88103 | 2.533312 | 2.233331 | 1.319804 | 2.0657 | 1.79769 | 0.35015 | 3.094038 | 1.45629 | 2.378051 | 2.65719 | 1.101575 | 3.16783 | 2.864409 | 2.264324 | 2.758467 | 3.176279 |
| 34.52778 | 2.831293 | 3.67295 | 3.387066 | 2.972525 | 2.000585 | 2.632243 | 2.248429 | 0.668236 | 3.675761 | 2.45307 | 2.384404 | 3.403444 | 1.238513 | 3.744378 | 2.356648 | 2.077721 | 4.6957 | 2.14545 |
| 34.54167 | 3.329201 | 4.414964 | 3.412571 | 3.367244 | 2.232727 | 2.776269 | 2.364773 | 0.600496 | 4.138336 | 3.297597 | 1.922427 | -0.08661 | 1.236648 | 4.708848 | 2.320384 | 1.930271 | 4.465988 | 1.47779 |
| 34.55556 | 3.078135 | 5.347975 | 2.601438 | 3.084925 | 2.065163 | 2.610835 | 1.96112 | 0.400751 | 4.213203 | 4.28853 | 1.671008 | -1.14932 | 0.963337 | 5.919208 | 3.16479 | 2.395008 | 5.732436 | 1.887899 |
| 34.56944 | 2.266453 | 5.79653 | 1.625306 | 1.956765 | 1.831685 | 2.868307 | 1.307858 | 0.514119 | 4.508533 | 4.120399 | 1.725703 | -0.82723 | 1.119123 | 6.099535 | 3.960394 | 2.81794 | 8.285526 | 3.016427 |
| 34.58333 | 1.429498 | 5.68836 | 0.699126 | 0.781481 | 1.287587 | 2.585379 | 0.66518 | 0.732643 | 4.413524 | 3.836908 | 1.477762 | 0.171153 | 1.138388 | 5.516059 | 3.713199 | 1.781686 | 7.028538 | 4.062197 |
| 34.59722 | 1.241442 | 5.043516 | 0.013577 | -0.05763 | 0.811796 | 2.186733 | 0.435878 | 0.713596 | 4.579964 | 3.200006 | 1.687932 | 0.067009 | 1.396769 | 3.635409 | 2.056374 | 1.015426 | 3.325072 | 4.124727 |
| 34.61111 | 1.251925 | 3.926664 | -0.27208 | -0.39549 | 0.75203 | 1.874384 | 0.638662 | 0.390273 | 3.731403 | 2.404023 | 1.956025 | 0.037157 | 0.916938 | 1.596639 | 1.015546 | 0.894154 | -0.87255 | 3.376183 |
| 34.625 | 0.757873 | 2.712959 | -0.3508 | -0.42905 | 0.631115 | 1.33199 | 1.018093 | 0.028328 | 2.293724 | 1.173425 | 1.92172 | -1.1128 | 0.137356 | 0.085021 | 0.104483 | 0.985635 | -4.32194 | 2.223903 |
| 34.63889 | 0.080645 | 0.805571 | -0.50799 | -1.0804 | -0.1063 | 0.124229 | 1.160093 | -0.32152 | 0.724533 | -0.8861 | 1.198045 | -4.78934 | -0.38254 | -1.16969 | -1.00083 | 1.02974 | -4.51649 | 1.228895 |
| 34.65278 | -0.82041 | -0.95971 | -0.85359 | -1.55631 | -1.30547 | -2.04228 | 1.099188 | -0.57759 | -0.80796 | -1.54714 | 0.31948 | -6.30654 | -0.70789 | -1.5986 | -1.65944 | 0.891504 | -2.92834 | 0.877997 |
| 34.66667 | -1.52461 | -2.39752 | -1.01631 | -0.96219 | -1.99044 | -2.9669 | 1.384252 | -0.79284 | -2.38214 | -2.66554 | -0.50046 | -10.5509 | -0.92752 | -1.21306 | -0.99548 | 0.788489 | -1.25334 | 0.711774 |
| 34.68056 | -1.74519 | -3.20984 | -0.9611 | -0.4458 | -2.53624 | -3.38536 | 1.575168 | -0.99809 | -3.78118 | -3.1118 | -0.65395 | -7.75852 | -0.85413 | -1.0691 | -0.45115 | 0.801907 | 0.194007 | 0.221285 |
| 34.69444 | -1.55344 | -3.39896 | -0.72369 | -0.22299 | -2.81879 | -2.94778 | 1.584038 | -1.00533 | -4.2025 | -3.34081 | -0.49479 | -3.96907 | -0.82087 | -0.76999 | -0.20575 | 0.555581 | 0.071154 | -0.58456 |
| 34.70833 | -1.22119 | -2.94541 | -0.50639 | -0.04361 | -2.29293 | -2.05467 | 1.673703 | -0.77551 | -4.16858 | -2.12844 | -0.03434 | -1.60075 | -0.49289 | 0.188536 | 0.12605 | 0.527457 | -0.8851 | -1.47947 |
| 34.72222 | -0.71439 | -2.30283 | -0.1867 | 0.252741 | -1.38012 | -0.97308 | 1.715432 | -0.67919 | -4.18871 | 0.080953 | 0.761065 | 0.3818 | 0.007314 | 0.872125 | 0.630877 | 0.732274 | -0.09015 | -2.19517 |
| 34.73611 | -0.37101 | -1.61746 | 0.349623 | 0.451192 | -0.47026 | -0.00759 | 1.711755 | -0.61311 | -3.91304 | 1.961606 | 1.205638 | 4.210249 | 0.326553 | 1.236493 | 0.7905 | 0.707594 | 1.240286 | -2.40348 |
| 34.75 | -0.20911 | -1.08857 | 1.047628 | 0.85251 | 0.101137 | 0.745819 | 1.705951 | -0.43684 | -2.88053 | 2.670415 | 1.340821 | 7.440209 | 0.301237 | 1.275251 | 0.674238 | 0.827571 | 2.544408 | -2.6972 |
| 34.76389 | -0.04641 | -0.6163 | 1.503602 | 0.963431 | 0.110558 | 0.951941 | 1.579379 | -0.22447 | -1.6844 | 2.317293 | 1.140049 | 6.402171 | 0.219887 | 1.135972 | 0.516924 | 0.408206 | 3.166412 | -3.56228 |
| 34.77778 | -0.03629 | -0.2419 | 1.41991 | 0.779201 | -0.03199 | 0.548829 | 1.467305 | -0.26002 | -0.8423 | 1.600676 | 0.849999 | 4.108261 | 0.359709 | 0.919635 | 0.400764 | 0.687153 | 3.332635 | -4.07177 |
| 34.79167 | -0.0096 | 0.07941 | 1.077753 | 0.611478 | -0.1911 | -0.04112 | 1.342538 | -0.49946 | -0.20595 | 0.146359 | 0.485082 | 2.345481 | 0.409033 | 0.636026 | 0.22034 | 0.652451 | 1.519617 | -3.4364 |
| 34.80556 | 0.068936 | 0.260292 | 0.762228 | 0.339797 | -0.46731 | -0.50416 | 1.081178 | -0.60266 | 0.27378 | -1.43463 | 0.266767 | 0.416249 | 0.316739 | 0.461508 | -0.00906 | -0.18342 | -1.3954 | -2.34179 |
| 34.81944 | 0.115547 | 0.284131 | 0.284925 | -0.00132 | -0.88415 | -1.00818 | 0.747002 | -0.62214 | 0.566478 | -2.84998 | 0.296015 | 0.290219 | 0.262585 | 0.199012 | -0.25491 | -1.24703 | -3.69412 | -1.7306 |
| 34.83333 | -0.0032 | 0.13773 | -0.19698 | -0.20438 | -1.37487 | -1.4574 | 0.389491 | -0.6676 | 0.61509 | -2.65713 | 0.212282 | 0.327162 | 0.466682 | -0.32377 | -0.57574 | -0.92705 | -3.52451 | -1.34499 |

| | | | | | | | | | | | | | | | | | | |
|---|---|---|---|---|---|---|---|---|---|---|---|---|---|---|---|---|---|---|
| 34.84722 | -0.26986 | -0.22993 | -0.78437 | -0.40397 | -1.4644 | -1.77779 | -0.01253 | -0.79741 | 0.14588 | -2.02335 | 0.258514 | -1.36022 | -0.16352 | -0.73803 | -0.77676 | -0.61521 | -3.9772 | -1.15123 |
| 34.86111 | -0.71024 | -0.79739 | -1.20951 | -0.51272 | -1.53245 | -2.15947 | -0.19856 | -0.84768 | -0.72972 | -1.62414 | -0.02949 | -7.1002 | -0.63906 | -0.96303 | -0.78665 | -0.99827 | -3.98396 | -1.46701 |
| 34.875 | -0.98774 | -1.37486 | -1.30842 | -0.61383 | -2.1314 | -2.72138 | -0.20347 | -0.78926 | -1.7001 | -1.89183 | -0.59625 | -5.01772 | -0.79472 | -1.04716 | -0.71571 | -1.31038 | -5.33166 | -2.15044 |
| 34.88889 | -1.19621 | -1.43608 | -1.31376 | -0.47319 | -2.71506 | -3.18794 | -0.13451 | -0.62267 | -2.35879 | -2.42102 | -1.24573 | -5.63946 | -0.66006 | -1.11462 | -0.73075 | -1.11099 | -5.55192 | -2.43989 |
| 34.90278 | NaN | NaN | NaN | NaN | NaN | NaN | NaN | NaN | NaN | -2.99059 | -1.28767 | -7.54667 | -0.60348 | -1.19216 | -0.62412 | -0.24721 | -3.00153 | -2.03613 |
| 34.91667 | NaN | NaN | NaN | NaN | NaN | NaN | NaN | NaN | NaN | -3.74492 | -1.45087 | -12.3214 | -0.69204 | -1.12537 | -0.51453 | 0.264384 | -1.84892 | -1.55607 |
| 34.93056 | NaN | NaN | NaN | NaN | NaN | NaN | NaN | NaN | NaN | -3.89373 | -0.85476 | -8.0156 | -0.61682 | -0.71262 | -0.39935 | 0.517471 | -2.56091 | -1.66153 |
| 34.94444 | NaN | NaN | NaN | NaN | NaN | NaN | NaN | NaN | NaN | -3.33891 | -0.03571 | -3.83792 | -0.63833 | -0.05092 | -0.22725 | 0.397342 | -2.59558 | -1.92941 |
| 34.95833 | NaN | NaN | NaN | NaN | NaN | NaN | NaN | NaN | NaN | -1.85139 | 0.75099 | -2.10547 | -0.54455 | 0.406614 | -0.1037 | 0.220317 | -1.14966 | -1.77682 |
| 34.97222 | NaN | NaN | NaN | NaN | NaN | NaN | NaN | NaN | NaN | -0.76482 | 0.828775 | -2.64137 | -0.48038 | 0.427706 | -0.34461 | 0.035036 | 1.525056 | -1.52624 |
| 34.98611 | NaN | NaN | NaN | NaN | NaN | NaN | NaN | NaN | NaN | -0.97525 | 0.899928 | -2.34371 | -0.78618 | 0.120002 | -0.95884 | -0.29036 | 2.122478 | -1.8567 |
| 35 | NaN | NaN | NaN | NaN | NaN | NaN | NaN | NaN | NaN | -2.71181 | -0.0162 | -4.11411 | -0.78167 | -0.44225 | -1.34666 | -0.32123 | -0.65092 | -3.11139 |
| 35.01389 | NaN | NaN | NaN | NaN | NaN | NaN | NaN | NaN | NaN | -4.69844 | -1.00281 | -9.56828 | -0.73976 | -1.22823 | -1.38433 | -1.19867 | -4.31703 | -4.24201 |
| 35.02778 | NaN | NaN | NaN | NaN | NaN | NaN | NaN | NaN | NaN | -4.54635 | -1.78152 | -6.0897 | -1.10557 | -1.56303 | -1.11252 | -1.87512 | -4.96653 | -3.99221 |
| 35.04167 | NaN | NaN | NaN | NaN | NaN | NaN | NaN | NaN | NaN | -3.88894 | -1.63891 | -3.22019 | -1.03626 | -1.1888 | -0.72695 | -1.73461 | -4.18667 | -3.21815 |
| 35.05556 | NaN | NaN | NaN | NaN | NaN | NaN | NaN | NaN | NaN | -3.35622 | -1.17523 | -1.74029 | -0.86848 | -0.76773 | -0.97639 | -1.15233 | -4.42439 | -3.80258 |
| 35.06944 | NaN | NaN | NaN | NaN | NaN | NaN | NaN | NaN | NaN | -2.97171 | -0.88026 | -2.21255 | -0.86145 | -0.33235 | -0.99893 | -0.97258 | -4.68739 | -4.8867 |
| 35.08333 | NaN | NaN | NaN | NaN | NaN | NaN | NaN | NaN | NaN | -2.53029 | -0.68602 | -2.64697 | -0.86397 | -0.08593 | -0.86543 | -0.99025 | -6.46921 | -4.03648 |
| 35.09722 | NaN | NaN | NaN | NaN | NaN | NaN | NaN | NaN | NaN | -2.22637 | -0.37669 | -2.51344 | -0.84609 | -0.07364 | -0.50462 | -0.54913 | -6.16071 | -2.33673 |
| 35.11111 | NaN | NaN | NaN | NaN | NaN | NaN | NaN | NaN | NaN | -2.03112 | -0.26507 | -2.36924 | -0.42148 | -0.20814 | -0.34696 | -0.41296 | -4.54059 | -1.49093 |
| 35.125 | NaN | NaN | NaN | NaN | NaN | NaN | NaN | NaN | NaN | -1.83312 | -0.16061 | -2.64101 | -0.05544 | -0.43783 | -0.50811 | -0.03449 | -3.95933 | -1.42859 |
| 35.13889 | NaN | NaN | NaN | NaN | NaN | NaN | NaN | NaN | NaN | -1.73523 | -0.24053 | -3.02331 | 0.143598 | -0.55807 | -0.77405 | 0.119093 | -4.64148 | -1.81272 |
| 35.15278 | NaN | NaN | NaN | NaN | NaN | NaN | NaN | NaN | NaN | -1.60452 | -0.16362 | -2.88278 | 0.106921 | -0.49985 | -0.87588 | -0.00353 | -3.63502 | -1.83454 |
| 35.16667 | NaN | NaN | NaN | NaN | NaN | NaN | NaN | NaN | NaN | -1.21126 | 0.247789 | -2.56679 | -0.2469 | -0.27161 | -0.83968 | 0.140072 | -2.45369 | -1.48886 |
| 35.18056 | NaN | NaN | NaN | NaN | NaN | NaN | NaN | NaN | NaN | -0.49898 | 0.769687 | -2.01899 | -0.4841 | 0.091249 | -0.62549 | 0.576482 | -1.2473 | -1.10129 |
| 35.19444 | NaN | NaN | NaN | NaN | NaN | NaN | NaN | NaN | NaN | 0.1067 | 1.238567 | -1.36889 | -0.35038 | 0.447869 | -0.39709 | 1.467275 | 0.678273 | -0.60531 |
| 35.20833 | NaN | NaN | NaN | NaN | NaN | NaN | NaN | NaN | NaN | 0.325972 | 1.539055 | -1.06126 | -0.07005 | 0.640417 | -0.27645 | 2.171381 | 1.997323 | -0.34366 |
| 35.22222 | NaN | NaN | NaN | NaN | NaN | NaN | NaN | NaN | NaN | -0.46665 | 1.806091 | -1.33801 | 0.133406 | 0.8409 | -0.02299 | 0.730834 | 2.371494 | 0.059553 |
| 35.23611 | NaN | NaN | NaN | NaN | NaN | NaN | NaN | NaN | NaN | -2.09195 | 1.783522 | -1.72297 | 0.127463 | 1.082651 | 0.458948 | -0.02347 | 2.190749 | 0.597132 |
| 35.25 | NaN | NaN | NaN | NaN | NaN | NaN | NaN | NaN | NaN | -3.11597 | 1.360063 | -2.04168 | -0.06538 | 1.056242 | 0.907252 | -0.02763 | 2.495683 | 1.136433 |
| 35.26389 | NaN | NaN | NaN | NaN | NaN | NaN | NaN | NaN | NaN | -2.21367 | 0.969971 | -1.91292 | -0.19823 | 0.799496 | 1.08226 | 0.343803 | 0.993665 | 1.848582 |
| 35.27778 | NaN | NaN | NaN | NaN | NaN | NaN | NaN | NaN | NaN | -1.21536 | 0.424854 | -1.6005 | -0.18712 | 0.516185 | 0.729807 | 0.064117 | -0.45398 | 1.068886 |
| 35.29167 | NaN | NaN | NaN | NaN | NaN | NaN | NaN | NaN | NaN | -0.68465 | 0.083212 | -1.07691 | 0.000387 | 0.398088 | 0.640417 | -0.05253 | -0.44801 | 0.272588 |
| 35.30556 | NaN | NaN | NaN | NaN | NaN | NaN | NaN | NaN | NaN | -0.28032 | 0.296318 | -0.38359 | 0.22286 | 0.514056 | 0.451599 | -0.14107 | 0.002369 | -0.33912 |
| 35.31944 | NaN | NaN | NaN | NaN | NaN | NaN | NaN | NaN | NaN | -0.12719 | 0.335611 | 0.130136 | 0.225444 | 0.396227 | 0.203966 | -0.2393 | 0.055136 | -0.50329 |
| 35.33333 | NaN | NaN | NaN | NaN | NaN | NaN | NaN | NaN | NaN | -0.10118 | 0.269635 | 0.323506 | 0.124238 | 0.041076 | -0.1699 | -0.37775 | 0.015449 | -0.43661 |
| 35.34722 | NaN | NaN | NaN | NaN | NaN | NaN | NaN | NaN | NaN | -0.07447 | 0.199688 | 0.437418 | 0.005677 | -0.12698 | -0.35941 | -0.92528 | -0.46154 | -0.40159 |
| 35.36111 | NaN | NaN | NaN | NaN | NaN | NaN | NaN | NaN | NaN | -0.17968 | 0.193609 | 0.430673 | -0.31329 | -0.18591 | -0.37046 | -1.31 | -1.1922 | -0.78362 |
| 35.375 | NaN | NaN | NaN | NaN | NaN | NaN | NaN | NaN | NaN | -0.5909 | -0.0953 | -0.51103 | -0.65968 | -0.66881 | -0.688 | -1.13913 | -1.80934 | -1.65971 |
| 35.38889 | NaN | NaN | NaN | NaN | NaN | NaN | NaN | NaN | NaN | -0.92815 | -0.43196 | -2.90014 | -2.37765 | -1.25959 | -1.57273 | -0.67629 | -1.91609 | -2.39017 |
| 35.40278 | NaN | NaN | NaN | NaN | NaN | NaN | NaN | NaN | NaN | -0.82013 | 0.12215 | -3.24056 | -4.28022 | -0.88112 | -1.57749 | 0.048925 | -1.91454 | -2.49088 |
| 35.41667 | NaN | NaN | NaN | NaN | NaN | NaN | NaN | NaN | NaN | -0.24626 | 0.660004 | -1.05487 | -1.61122 | -0.03771 | -0.6029 | 0.935602 | -2.22441 | -1.72712 |
| 35.43056 | NaN | NaN | NaN | NaN | NaN | NaN | NaN | NaN | NaN | -0.17562 | 0.670088 | -0.40209 | -1.27877 | 0.178117 | 0.168978 | 1.004951 | -1.49377 | -0.64711 |
| 35.44444 | NaN | NaN | NaN | NaN | NaN | NaN | NaN | NaN | NaN | -0.45845 | 0.448383 | -0.33108 | -1.4448 | -0.01257 | 0.268229 | 0.713571 | -0.60785 | -0.12979 |
| 35.45833 | NaN | NaN | NaN | NaN | NaN | NaN | NaN | NaN | NaN | -0.58917 | 0.461144 | -0.54478 | -1.15272 | -0.13543 | 0.429671 | 0.253927 | -0.37251 | 0.059195 |
| 35.47222 | NaN | NaN | NaN | NaN | NaN | NaN | NaN | NaN | NaN | -0.62377 | 0.428702 | -0.33871 | -0.42178 | -0.17013 | 0.3559 | -0.90029 | -1.61 | -0.15828 |
| 35.48611 | NaN | NaN | NaN | NaN | NaN | NaN | NaN | NaN | NaN | -0.30364 | 0.408529 | 0.892847 | -0.18436 | -0.04141 | -0.47764 | -1.22622 | -4.45239 | -1.56797 |
| 35.5 | NaN | NaN | NaN | NaN | NaN | NaN | NaN | NaN | NaN | 0.47142 | 0.836587 | 2.952599 | 0.385187 | 0.497029 | -0.52602 | -1.66317 | -4.1321 | -2.9928 |
| 35.51389 | NaN | NaN | NaN | NaN | NaN | NaN | NaN | NaN | NaN | 1.842586 | 1.386592 | 4.07224 | 1.288102 | 1.362872 | 0.196804 | 0.522585 | -1.55541 | -1.52746 |
| 35.52778 | NaN | NaN | NaN | NaN | NaN | NaN | NaN | NaN | NaN | 3.300576 | 1.773845 | 4.005339 | 2.308381 | 1.888183 | 0.638722 | 2.101656 | 0.34416 | 0.10395 |
| 35.54167 | NaN | NaN | NaN | NaN | NaN | NaN | NaN | NaN | NaN | 4.928268 | 1.672535 | 4.382632 | 2.907861 | 2.10312 | 0.744046 | 2.517536 | 1.999598 | 1.077063 |
| 35.55556 | NaN | NaN | NaN | NaN | NaN | NaN | NaN | NaN | NaN | 6.044839 | 1.715196 | 4.737692 | 2.531897 | 2.336672 | 0.97463 | 1.695855 | 2.000392 | 1.088638 |
| 35.56944 | NaN | NaN | NaN | NaN | NaN | NaN | NaN | NaN | NaN | 6.310675 | 1.730258 | 4.364807 | 2.054349 | 2.2911 | 1.246184 | 0.936732 | 1.599781 | 0.947137 |
| 35.58333 | NaN | NaN | NaN | NaN | NaN | NaN | NaN | NaN | NaN | 6.52602 | 1.461076 | 4.008313 | 1.582483 | 2.014972 | 1.357739 | 0.262044 | 1.013508 | 0.271288 |
| 35.59722 | NaN | NaN | NaN | NaN | NaN | NaN | NaN | NaN | NaN | 6.443948 | 1.254757 | 3.631632 | 1.942475 | 1.768562 | 1.391554 | -0.22454 | 0.082368 | -0.53601 |
| 35.61111 | NaN | NaN | NaN | NaN | NaN | NaN | NaN | NaN | NaN | 6.235824 | 1.286855 | 3.340537 | 3.088261 | 1.849078 | 1.590557 | -0.58051 | -0.81902 | -1.0408 |
| 35.625 | NaN | NaN | NaN | NaN | NaN | NaN | NaN | NaN | NaN | 5.637225 | 1.317835 | 3.057593 | 3.259801 | 1.949535 | 1.849373 | -0.70396 | -1.45695 | -1.13699 |
| 35.63889 | NaN | NaN | NaN | NaN | NaN | NaN | NaN | NaN | NaN | 4.028594 | 1.429317 | 2.433484 | 1.465037 | 1.849128 | 1.696653 | -0.55704 | -1.95879 | -1.45685 |
| 35.65278 | NaN | NaN | NaN | NaN | NaN | NaN | NaN | NaN | NaN | 2.251689 | 1.321321 | 1.600107 | -0.75549 | 1.629508 | 1.276693 | -0.00233 | -2.45178 | -1.89182 |
| 35.66667 | NaN | NaN | NaN | NaN | NaN | NaN | NaN | NaN | NaN | 0.536839 | 1.333402 | 0.783754 | -2.4765 | 1.459763 | 0.684234 | 0.395154 | -2.43546 | -2.03109 |
| 35.68056 | NaN | NaN | NaN | NaN | NaN | NaN | NaN | NaN | NaN | 0.247481 | 1.188623 | -0.05657 | -3.04705 | 1.159752 | 0.282617 | 0.500265 | -1.9503 | -1.59973 |
| 35.69444 | NaN | NaN | NaN | NaN | NaN | NaN | NaN | NaN | NaN | 0.142537 | 1.009856 | -0.60614 | -2.8086 | 0.951762 | 0.141123 | 0.516376 | -1.41696 | -1.10274 |
| 35.70833 | NaN | NaN | NaN | NaN | NaN | NaN | NaN | NaN | NaN | -0.08847 | 0.82868 | -1.1171 | -1.70015 | 0.847849 | 0.169425 | 0.125424 | -1.05213 | -0.57307 |
| 35.72222 | NaN | NaN | NaN | NaN | NaN | NaN | NaN | NaN | NaN | -0.24735 | 0.743358 | -1.60826 | -2.04382 | 0.712365 | 0.321364 | 0.068303 | -0.41868 | -0.12917 |
| 35.73611 | NaN | NaN | NaN | NaN | NaN | NaN | NaN | NaN | NaN | -0.19369 | 0.737164 | -2.08265 | -1.36847 | 0.760382 | 0.455237 | 0.115484 | -0.25393 | -0.05354 |
| 35.75 | NaN | NaN | NaN | NaN | NaN | NaN | NaN | NaN | NaN | -0.11587 | 0.935006 | -2.3054 | -0.62682 | 0.792717 | 0.599041 | 0.245313 | -0.12948 | -0.24338 |
| 35.76389 | NaN | NaN | NaN | NaN | NaN | NaN | NaN | NaN | NaN | -0.02272 | 1.230055 | -1.90319 | 0.555443 | 1.017909 | 0.804383 | 0.246786 | -0.01003 | -0.31634 |
| 35.77778 | NaN | NaN | NaN | NaN | NaN | NaN | NaN | NaN | NaN | 0.218259 | 1.342746 | -1.26251 | 1.466689 | 1.126061 | 0.922867 | 0.234428 | 0.230686 | -0.22702 |
| 35.79167 | NaN | NaN | NaN | NaN | NaN | NaN | NaN | NaN | NaN | 0.364268 | 1.115172 | -0.74337 | 2.160216 | 1.035067 | 0.87592 | 0.374618 | 0.333547 | -0.09771 |
| 35.80556 | NaN | NaN | NaN | NaN | NaN | NaN | NaN | NaN | NaN | 0.25649 | 0.869018 | -0.60313 | 3.410026 | 0.801082 | 0.75343 | 0.474021 | 0.174133 | -0.0756 |
| 35.81944 | NaN | NaN | NaN | NaN | NaN | NaN | NaN | NaN | NaN | -0.33124 | 0.528929 | -0.64026 | 4.004775 | 0.530234 | 0.646198 | 0.3703 | -0.19833 | -0.25414 |
| 35.83333 | NaN | NaN | NaN | NaN | NaN | NaN | NaN | NaN | NaN | -1.57035 | 0.026893 | -0.50235 | 3.566799 | 0.348329 | 0.580552 | 0.127852 | -0.71575 | -0.48162 |
| 35.84722 | NaN | NaN | NaN | NaN | NaN | NaN | NaN | NaN | NaN | -3.26729 | -1.26519 | -0.40659 | 2.307048 | 0.101168 | 0.361084 | -0.15347 | -1.02383 | -0.92295 |
| 35.86111 | NaN | NaN | NaN | NaN | NaN | NaN | NaN | NaN | NaN | -4.64952 | -2.86945 | -0.42315 | 0.095582 | -0.37301 | -0.12522 | -0.32025 | -1.70306 | -1.37643 |
| 35.875 | NaN | NaN | NaN | NaN | NaN | NaN | NaN | NaN | NaN | -4.61887 | -3.44229 | -0.65658 | -2.48441 | -0.87546 | -0.62139 | -0.41583 | -2.22986 | -1.73717 |
| 35.88889 | NaN | NaN | NaN | NaN | NaN | NaN | NaN | NaN | NaN | -4.34425 | -3.41887 | -0.89043 | -3.87906 | -1.05766 | -0.86996 | -0.2246 | -2.08378 | -1.79866 |
| 35.90278 | NaN | NaN | NaN | NaN | NaN | NaN | NaN | NaN | NaN | -3.79938 | -3.26127 | -1.47659 | -4.14985 | -0.81982 | -0.80046 | -0.04034 | -1.61074 | -1.39027 |
| 35.91667 | NaN | NaN | NaN | NaN | NaN | NaN | NaN | NaN | NaN | -3.05755 | -1.91033 | -1.77597 | -3.78001 | -0.23592 | -0.38615 | 0.141505 | -0.80759 | -0.89426 |
| 35.93056 | NaN | NaN | NaN | NaN | NaN | NaN | NaN | NaN | NaN | -2.98462 | -0.33577 | -2.04797 | -2.25493 | 0.254647 | 0.136292 | 0.575277 | -0.3403 | -0.48447 |
| 35.94444 | NaN | NaN | NaN | NaN | NaN | NaN | NaN | NaN | NaN | -2.40065 | 0.68147 | -2.03843 | -2.7248 | 0.372555 | 0.271874 | 0.248901 | -0.1182 | -0.24899 |
| 35.95833 | NaN | NaN | NaN | NaN | NaN | NaN | NaN | NaN | NaN | -1.46747 | 1.375625 | -1.97629 | -3.43893 | 0.511077 | -0.09711 | -0.246 | -0.42158 | -0.39779 |

| | | | | | | | | | | | | | | | | | | |
|---|---|---|---|---|---|---|---|---|---|---|---|---|---|---|---|---|---|---|
| 35.97222 | NaN | NaN | NaN | NaN | NaN | NaN | NaN | NaN | NaN | -0.86096 | 1.604699 | -2.06797 | -2.37253 | 0.745649 | -0.4383 | -0.66014 | -0.81057 | -0.63979 |
| 35.98611 | NaN | NaN | NaN | NaN | NaN | NaN | NaN | NaN | NaN | -1.52343 | 0.059361 | -3.43155 | -2.40731 | 0.048613 | -1.18827 | -1.60305 | -1.92771 | -1.60157 |
| 36 | NaN | NaN | NaN | NaN | NaN | NaN | NaN | NaN | NaN | -4.23202 | -3.99603 | -9.04533 | -5.02386 | -2.513 | -2.9813 | -4.01046 | -3.89909 | -3.94094 |
| 36.01389 | NaN | NaN | NaN | NaN | NaN | NaN | NaN | NaN | NaN | -5.65133 | -5.72353 | -9.50822 | -7.65372 | -4.77116 | -3.61099 | -6.40925 | -5.43242 | -4.82191 |
| 36.02778 | NaN | NaN | NaN | NaN | NaN | NaN | NaN | NaN | NaN | -3.89684 | -3.15031 | -6.32002 | -6.75324 | -3.52484 | -2.0004 | -4.25973 | -4.13814 | -1.93506 |
| 36.04167 | NaN | NaN | NaN | NaN | NaN | NaN | NaN | NaN | NaN | -2.6359 | -1.983 | -3.89144 | -4.87098 | -2.16016 | -0.47639 | -1.72839 | -2.28456 | 0.556712 |
| 36.05556 | NaN | NaN | NaN | NaN | NaN | NaN | NaN | NaN | NaN | -2.752 | -1.28836 | -2.71069 | -4.48444 | -1.28938 | 0.410567 | -0.36117 | -1.11756 | 1.732277 |
| 36.06944 | NaN | NaN | NaN | NaN | NaN | NaN | NaN | NaN | NaN | -3.18163 | -1.28338 | -2.47582 | -4.3592 | -0.71387 | 0.116487 | 0.165109 | -0.59477 | 1.476957 |
| 36.08333 | NaN | NaN | NaN | NaN | NaN | NaN | NaN | NaN | NaN | -2.78106 | -1.67846 | -2.63656 | -4.01144 | -0.44638 | 0.035339 | 0.26408 | 0.025539 | 0.833605 |
| 36.09722 | NaN | NaN | NaN | NaN | NaN | NaN | NaN | NaN | NaN | -2.26051 | -1.2276 | -2.456 | -2.89698 | -0.40116 | 0.284258 | 0.403468 | 0.675792 | 0.445202 |
| 36.11111 | NaN | NaN | NaN | NaN | NaN | NaN | NaN | NaN | NaN | -1.86052 | -0.84627 | -2.00457 | -2.0041 | -0.57068 | 0.721216 | 0.406053 | 0.847499 | 0.302138 |
| 36.125 | NaN | NaN | NaN | NaN | NaN | NaN | NaN | NaN | NaN | -1.25321 | -0.6138 | -1.58436 | -1.26527 | -0.4936 | 0.507759 | 0.134863 | 0.488941 | -0.07678 |
| 36.13889 | NaN | NaN | NaN | NaN | NaN | NaN | NaN | NaN | NaN | -0.59011 | -0.32518 | -1.2142 | -0.62266 | -0.32109 | 0.475821 | -0.29344 | 0.072588 | -0.2019 |
| 36.15278 | NaN | NaN | NaN | NaN | NaN | NaN | NaN | NaN | NaN | -0.12994 | 0.102838 | -1.1019 | -0.15871 | -0.1512 | 0.481865 | -0.6786 | -0.02995 | -0.2461 |
| 36.16667 | NaN | NaN | NaN | NaN | NaN | NaN | NaN | NaN | NaN | 0.166608 | 0.478428 | -1.0786 | 0.406533 | -0.15347 | 0.467567 | -0.69304 | 0.128322 | -0.17061 |
| 36.18056 | NaN | NaN | NaN | NaN | NaN | NaN | NaN | NaN | NaN | 0.156955 | 0.696023 | -1.03819 | 1.216408 | -0.0587 | 0.382684 | -0.43559 | 0.370957 | 0.018899 |
| 36.19444 | NaN | NaN | NaN | NaN | NaN | NaN | NaN | NaN | NaN | 0.130811 | 1.041183 | -0.90003 | 1.925417 | 0.126 | 0.26276 | -0.07788 | 0.738276 | 0.152459 |
| 36.20833 | NaN | NaN | NaN | NaN | NaN | NaN | NaN | NaN | NaN | 0.309808 | 1.634692 | -0.52626 | 2.568182 | 0.444798 | 0.177728 | 0.269004 | 1.446376 | 0.159432 |
| 36.22222 | NaN | NaN | NaN | NaN | NaN | NaN | NaN | NaN | NaN | 0.536963 | 1.872094 | 0.022914 | 3.010006 | 0.665214 | 0.37353 | 0.764421 | 2.087729 | 0.378044 |
| 36.23611 | NaN | NaN | NaN | NaN | NaN | NaN | NaN | NaN | NaN | 0.762773 | 1.646039 | 0.372168 | 3.144682 | 0.885868 | 0.548567 | 0.975958 | 2.288343 | 0.595059 |
| 36.25 | NaN | NaN | NaN | NaN | NaN | NaN | NaN | NaN | NaN | 0.893616 | 2.039496 | 0.185613 | 2.570658 | 1.015275 | 0.628351 | 1.031055 | 2.140299 | 0.651504 |
| 36.26389 | NaN | NaN | NaN | NaN | NaN | NaN | NaN | NaN | NaN | 0.902821 | 2.272678 | -0.17268 | 1.914515 | 0.962542 | 0.522765 | 1.002843 | 2.17077 | 0.596097 |
| 36.27778 | NaN | NaN | NaN | NaN | NaN | NaN | NaN | NaN | NaN | NaN | NaN | NaN | NaN | NaN | NaN | NaN | NaN | NaN |
| 36.29167 | NaN | NaN | NaN | NaN | NaN | NaN | NaN | NaN | NaN | NaN | NaN | NaN | NaN | NaN | NaN | NaN | NaN | NaN |
| 36.30556 | NaN | NaN | NaN | NaN | NaN | NaN | NaN | NaN | NaN | NaN | NaN | NaN | NaN | NaN | NaN | NaN | NaN | NaN |
| 36.31944 | NaN | NaN | NaN | NaN | NaN | NaN | NaN | NaN | NaN | NaN | NaN | NaN | NaN | NaN | NaN | NaN | NaN | NaN |
| 36.33333 | NaN | NaN | NaN | NaN | NaN | NaN | NaN | NaN | NaN | NaN | NaN | NaN | NaN | NaN | NaN | NaN | NaN | NaN |
| 36.34722 | NaN | NaN | NaN | NaN | NaN | NaN | NaN | NaN | NaN | NaN | NaN | NaN | NaN | NaN | NaN | NaN | NaN | NaN |
| 36.36111 | NaN | NaN | NaN | NaN | NaN | NaN | NaN | NaN | NaN | NaN | NaN | NaN | NaN | NaN | NaN | NaN | NaN | NaN |
| 36.375 | NaN | NaN | NaN | NaN | NaN | NaN | NaN | NaN | NaN | NaN | NaN | NaN | NaN | NaN | NaN | NaN | NaN | NaN |
| 36.38889 | NaN | NaN | NaN | NaN | NaN | NaN | NaN | NaN | NaN | NaN | NaN | NaN | NaN | NaN | NaN | NaN | NaN | NaN |
| 36.40278 | NaN | NaN | NaN | NaN | NaN | NaN | NaN | NaN | NaN | NaN | NaN | NaN | NaN | NaN | NaN | NaN | NaN | NaN |
| 36.41667 | NaN | NaN | NaN | NaN | NaN | NaN | NaN | NaN | NaN | NaN | NaN | NaN | NaN | NaN | NaN | NaN | NaN | NaN |
| 36.43056 | NaN | NaN | NaN | NaN | NaN | NaN | NaN | NaN | NaN | NaN | NaN | NaN | NaN | NaN | NaN | NaN | NaN | NaN |
| 36.44444 | NaN | NaN | NaN | NaN | NaN | NaN | NaN | NaN | NaN | NaN | NaN | NaN | NaN | NaN | NaN | NaN | NaN | NaN |
| 36.45833 | NaN | NaN | NaN | NaN | NaN | NaN | NaN | NaN | NaN | NaN | NaN | NaN | NaN | NaN | NaN | NaN | NaN | NaN |
| 36.47222 | NaN | NaN | NaN | NaN | NaN | NaN | NaN | NaN | NaN | NaN | NaN | NaN | NaN | NaN | NaN | NaN | NaN | NaN |
| 36.48611 | NaN | NaN | NaN | NaN | NaN | NaN | NaN | NaN | NaN | NaN | NaN | NaN | NaN | NaN | NaN | NaN | NaN | NaN |
| 36.5 | 2.682978 | 3.155329 | 1.743494 | 1.606142 | 0.853509 | 0.293732 | 1.524523 | -1.51307 | 0.315236 | NaN | NaN | NaN | NaN | NaN | NaN | NaN | NaN | NaN |
| 36.51389 | 3.901688 | 5.12186 | 1.932326 | 1.901729 | 2.206905 | 1.154379 | 2.274495 | 0.003234 | 1.583664 | 3.529512 | 4.325847 | 3.296055 | 0.138816 | 1.969007 | 1.591528 | 4.214073 | 3.590922 | 2.482047 |
| 36.52778 | 2.908767 | 3.799983 | 1.387372 | 1.257387 | 2.734542 | 1.063013 | 2.020731 | 1.178815 | 2.111645 | 4.539392 | 4.243979 | 2.282846 | 1.542294 | 1.750712 | 1.464337 | 2.940454 | 1.988507 | 2.137987 |
| 36.54167 | 1.619255 | 2.169008 | 0.859707 | 0.771951 | 3.714753 | 0.773039 | 1.622227 | 1.09343 | 1.308256 | 4.389946 | 3.870024 | 1.344463 | 2.708937 | 1.554382 | 1.24759 | 1.995417 | 1.306259 | 1.678934 |
| 36.55556 | 1.508816 | 1.507085 | 0.578345 | 0.75542 | 3.712789 | 0.81043 | 1.33029 | 1.0474 | 1.241774 | 3.640202 | 3.601599 | 1.795904 | 3.847907 | 1.842141 | 1.440463 | 1.912513 | 1.906277 | 1.924822 |
| 36.56944 | 1.874857 | 1.565283 | 0.487072 | 1.225283 | 3.805011 | 0.923302 | 1.259745 | 1.326943 | 1.65213 | 3.156407 | 3.289183 | 2.894922 | 4.53315 | 1.946927 | 1.544765 | 2.296103 | 2.709569 | 1.962693 |
| 36.58333 | 2.234266 | 1.747949 | 0.538022 | 1.999187 | 4.285488 | 1.35442 | 1.692679 | 0.899489 | 1.498207 | 3.028873 | 2.24917 | 3.755459 | 5.979049 | 1.939777 | 1.413651 | 2.900421 | 3.359294 | 1.556388 |
| 36.59722 | 2.30306 | 1.762622 | 0.596855 | 2.543786 | 3.494524 | 0.998845 | 2.037208 | 0.041655 | 0.704827 | 3.326206 | 1.153519 | 3.922058 | 5.891806 | 1.818265 | 1.012656 | 3.219389 | 3.409049 | 0.852124 |
| 36.61111 | 2.105551 | 1.647022 | 0.719558 | 3.129033 | 3.260328 | 0.923087 | 1.88383 | -0.83811 | 0.012567 | 3.34781 | 0.277892 | 3.981763 | 5.563662 | 1.73354 | 0.428148 | 2.820142 | 2.65748 | 0.358836 |
| 36.625 | 1.843052 | 1.313859 | 0.964995 | 3.483381 | 2.714869 | 0.974576 | 2.229892 | -1.28131 | -0.80304 | 3.006438 | -0.47045 | 4.048723 | 4.142681 | 1.622746 | 0.615924 | 1.938905 | 1.588853 | 0.145041 |
| 36.63889 | 1.474764 | 0.805673 | 1.302451 | 3.133095 | 1.789991 | 0.957196 | 2.154 | -0.97289 | -1.22848 | 1.815318 | -1.21612 | 3.562481 | 2.232614 | 1.266677 | 1.266737 | 1.283209 | 0.785307 | 0.093401 |
| 36.65278 | 1.106113 | 0.28923 | 1.418309 | 1.605653 | 0.851874 | 0.254203 | 0.879024 | -0.72414 | -0.81337 | 0.732491 | -1.78528 | 3.168268 | -0.2964 | 0.950678 | 1.553362 | 1.104331 | 0.253818 | -0.14799 |
| 36.66667 | 0.449986 | -0.19562 | 0.934637 | -0.0782 | -0.14787 | -0.69882 | -0.47848 | -0.85971 | -0.09143 | -0.02798 | -2.25864 | 2.846678 | -2.08907 | 0.587683 | 1.528136 | 1.071441 | -0.03637 | -0.45115 |
| 36.68056 | 0.111561 | -0.67902 | 0.604203 | -1.44215 | -1.13161 | -1.27351 | -1.48405 | -1.82629 | 0.113057 | -0.27957 | -2.6002 | 2.186029 | -3.35166 | 0.240496 | 1.33197 | 0.949522 | -0.32008 | -0.62851 |
| 36.69444 | -0.19224 | -1.07179 | 0.457796 | -1.79592 | -2.2515 | -1.75921 | -1.68004 | -3.12231 | -0.09703 | -1.10296 | -2.43017 | 1.050471 | -4.96588 | 0.030528 | 1.005305 | 0.784712 | -0.48524 | -0.73239 |
| 36.70833 | -0.49547 | -1.40578 | 0.522219 | -1.92919 | -3.69976 | -1.15368 | -0.75117 | -4.18951 | -0.62411 | -2.24872 | -2.0639 | -0.3196 | -7.25072 | -0.03047 | 0.571363 | 0.652248 | -0.52068 | -0.69271 |
| 36.72222 | -0.49715 | -1.48705 | 0.74733 | -1.48253 | -3.74098 | -1.43232 | -0.24917 | -3.52842 | -0.31441 | -2.21596 | 0.102307 | -1.54041 | -8.31765 | 0.014938 | 0.32723 | 0.501576 | -0.50139 | -0.67511 |
| 36.73611 | -0.20663 | -1.35978 | 1.123471 | -0.96797 | -4.25322 | -1.20333 | 0.290178 | -1.59452 | -0.04542 | -1.5871 | 1.087723 | -2.27338 | -6.75931 | 0.392208 | 0.421542 | 0.318914 | -0.4786 | -0.57116 |
| 36.75 | 0.060778 | -1.05272 | 1.367128 | -0.53136 | -4.05609 | -1.84932 | 0.511471 | 0.939466 | 0.377729 | -1.2371 | 1.0891 | -2.51164 | -4.71669 | 0.444701 | 0.323803 | 0.076407 | -0.45821 | -0.34921 |
| 36.76389 | 0.134628 | -0.77594 | 1.225017 | -0.1623 | -4.31025 | -1.84857 | 0.654118 | 2.883279 | 0.321352 | -0.81166 | 0.871099 | -2.33755 | -2.85804 | 0.407825 | 0.273439 | -0.0297 | -0.34409 | -0.16147 |
| 36.77778 | 0.030556 | -0.67988 | 0.789911 | 0.361353 | -3.61241 | -1.74634 | 0.640631 | 3.910125 | 0.518976 | -0.49016 | 0.632131 | -1.85108 | -1.5356 | 0.10925 | 0.295382 | 0.072457 | -0.1448 | -0.19792 |
| 36.79167 | -0.15193 | -0.66762 | 0.525265 | 0.976014 | -2.15567 | -1.51338 | 0.509581 | 3.557846 | 0.232447 | -0.47715 | 0.432212 | -1.54129 | -0.51014 | -0.51781 | 0.297458 | 0.179684 | 0.042561 | -0.30345 |
| 36.80556 | -0.20461 | -0.65711 | 0.350175 | 1.541277 | -1.32467 | -1.39962 | 0.203563 | 3.06472 | -0.47574 | -0.47942 | 0.097543 | -1.37903 | 0.127549 | -0.91614 | 0.553816 | 0.388814 | 0.352071 | -0.23432 |
| 36.81944 | 0.087254 | -0.65945 | -0.03086 | 1.94721 | -0.98078 | -1.50715 | -0.2071 | 2.61298 | -0.71808 | -0.92835 | -0.48162 | -1.30805 | -0.65775 | -0.70503 | 0.860546 | 0.412674 | 0.516282 | -0.11277 |
| 36.83333 | 0.326746 | -0.80711 | -0.84068 | 2.119653 | -0.5177 | -1.40168 | -0.50108 | 1.508887 | -1.74105 | -1.51782 | -0.95306 | -1.58553 | -2.42429 | -0.72326 | 0.988746 | -0.03045 | 0.337825 | -0.25273 |
| 36.84722 | 0.232263 | -1.08077 | -0.95861 | 2.04076 | -0.10822 | -1.12738 | -0.8346 | -0.23152 | -3.79071 | -1.85461 | -1.37799 | -1.84404 | -3.72942 | -0.93585 | 0.978083 | -0.27615 | -0.14506 | -0.53954 |
| 36.86111 | -0.22326 | -1.25883 | -0.91347 | 1.724325 | -0.24115 | -0.96535 | -1.3386 | -1.98245 | -3.56779 | -1.95557 | -1.81513 | -1.91831 | -4.19544 | -1.08182 | 0.790762 | -0.48093 | -0.68118 | -0.82752 |
| 36.875 | -0.7791 | -1.2413 | -1.11517 | 1.235164 | -0.74311 | -1.04017 | -1.7453 | -3.50575 | -2.96658 | -2.04315 | -2.53812 | -1.69687 | -4.34325 | -1.05436 | 0.701635 | -0.62196 | -0.8844 | -0.96265 |
| 36.88889 | -1.00906 | -0.92638 | -1.49176 | 0.867627 | -1.66015 | -1.75325 | -1.86423 | -4.10492 | -2.37964 | -2.13172 | -2.35718 | -1.49308 | -4.40957 | -0.71446 | 0.85587 | -0.55939 | -1.11564 | -1.03608 |
| 36.90278 | -0.9237 | -0.56117 | -2.03781 | 0.758881 | -2.41932 | -2.27055 | -1.91387 | -5.10661 | -1.94924 | -2.52222 | -2.05244 | -1.43992 | -3.98147 | -0.35089 | 1.448153 | -0.38637 | -1.35437 | -0.941 |
| 36.91667 | -0.68807 | -0.1343 | -1.95104 | 0.792676 | -2.45771 | -1.88801 | -1.82128 | -6.03085 | -1.42725 | -2.64665 | -1.5385 | -1.3491 | -3.63719 | 0.157392 | 1.891397 | -0.30364 | -1.39065 | -0.74273 |
| 36.93056 | -0.22211 | 0.304202 | -1.4755 | 0.723673 | -2.06887 | -1.33893 | -1.44512 | -6.05328 | -1.23944 | -2.27372 | -1.15857 | -0.86177 | -2.71958 | 0.63401 | 1.86095 | -0.05399 | -1.12266 | -0.52464 |
| 36.94444 | 0.378597 | 0.735193 | -0.76678 | 0.652425 | -1.68834 | -0.84799 | -1.01074 | -4.50298 | -0.64269 | -1.23046 | -0.5703 | -0.30376 | -1.56948 | 0.973605 | 1.901027 | 0.257519 | -0.76095 | -0.22816 |
| 36.95833 | 0.908105 | 1.260584 | -0.0011 | 0.526792 | -1.18139 | -0.47289 | -0.48268 | -1.44353 | 0.471752 | -0.05204 | 0.294164 | -0.05353 | -0.35438 | 1.239046 | 2.050733 | 0.417177 | -0.35898 | 0.129404 |
| 36.97222 | 1.200148 | 1.474464 | 0.384996 | 0.347663 | -0.79004 | -0.27893 | -0.13233 | 1.679901 | 1.650765 | 0.713325 | 0.946537 | -0.2542 | 0.556021 | 1.523106 | 2.240528 | 0.389619 | -0.06953 | 0.224923 |
| 36.98611 | 0.479115 | 0.1287 | 0.050896 | 0.12781 | -1.02908 | -0.57448 | -0.23091 | 2.810922 | 1.758696 | -0.33457 | 0.26323 | -1.70815 | -0.56553 | 0.876687 | 1.318483 | -0.64168 | -1.12508 | -1.29147 |
| 37 | -1.64533 | -2.89414 | -1.29928 | -0.14539 | -2.27466 | -1.59832 | -0.88359 | -0.27171 | -0.58656 | -4.43458 | -2.37541 | -4.5482 | -5.23124 | -1.52749 | -1.73542 | -3.40612 | -3.81871 | -5.43128 |
| 37.01389 | -3.03451 | -3.66753 | -1.85195 | -0.02861 | -3.06542 | -2.18604 | -1.23728 | -4.26304 | -2.56215 | -6.27516 | -4.94367 | -5.16376 | -9.06247 | -3.55519 | -3.72627 | -5.18384 | -5.49665 | -8.31994 |
| 37.02778 | -1.76592 | -1.42788 | -0.98134 | 0.662675 | -2.39821 | -1.60772 | -0.91772 | -5.40558 | -1.21185 | -4.05046 | -4.09398 | -3.18686 | -7.71812 | -3.11802 | -3.01283 | -3.70202 | -4.29341 | -4.68041 |
| 37.04167 | -0.52179 | -0.02501 | -0.37325 | 1.346946 | -2.18123 | -0.96867 | -0.76317 | -4.21413 | 0.088772 | -1.62569 | -2.74398 | -1.59706 | -5.35681 | -1.75032 | -1.71092 | -1.79041 | -1.93198 | -1.55609 |
| 37.05556 | -0.08575 | 0.059661 | -0.5802 | 1.620495 | -3.0125 | -0.37002 | -0.80511 | -3.09989 | -0.6551 | -1.05473 | -2.50061 | -1.44517 | -4.68921 | -0.78847 | -0.63667 | -0.4993 | -0.45348 | -0.19226 |
| 37.06944 | -0.30878 | -0.33442 | -0.89242 | 1.185083 | -3.09097 | -0.10351 | -0.85054 | -2.52566 | -1.36359 | -2.3008 | -2.58756 | -1.47732 | -4.48213 | -0.20229 | 0.552146 | 0.169362 | 0.134838 | -0.07929 |
| 37.08333 | -0.72997 | -0.73114 | -0.96569 | 1.012265 | -2.23145 | -0.08035 | -0.8119 | -1.84149 | -1.52227 | -2.12032 | -2.39918 | -1.78613 | -3.46491 | 0.194148 | 1.534068 | 0.430248 | -0.40459 | -0.49881 |

| | | | | | | | | | | | | | | | | | | |
|---|---|---|---|---|---|---|---|---|---|---|---|---|---|---|---|---|---|---|
| 37.09722 | -0.92919 | -1.09856 | -1.00177 | 0.868913 | -2.20122 | -0.09222 | -0.81648 | -1.29515 | -1.64691 | -1.9363 | -2.09781 | -1.53047 | -2.54283 | 0.346432 | 1.925788 | 0.394479 | -0.74348 | -0.98856 |
| 37.11111 | -1.09704 | -1.30039 | -1.04117 | 0.777393 | -2.41204 | -0.15503 | -0.85192 | -0.7634 | -1.49398 | -1.46856 | -1.90657 | -1.36287 | -1.7989 | 0.425853 | 2.032692 | 0.364824 | -0.62128 | -1.37693 |
| 37.125 | -0.85734 | -1.30869 | -1.18673 | 0.690286 | -2.40054 | -0.17145 | -0.71134 | -0.08407 | -1.06866 | -1.0496 | -1.7023 | -1.15645 | -1.39166 | 0.614267 | 1.906367 | 0.332823 | -0.47957 | -1.69238 |
| 37.13889 | -0.55031 | -1.0485 | -0.93907 | 0.567581 | -1.94341 | -0.11823 | -0.52257 | 0.57646 | -0.2335 | -0.33417 | -1.20363 | -0.95877 | -0.79315 | 0.480173 | 1.279731 | 0.517392 | -0.30892 | -1.56597 |
| 37.15278 | -0.14916 | -0.63624 | -0.53004 | 0.567523 | -1.11055 | -0.03271 | -0.34061 | 0.836415 | 0.8925 | 0.342487 | -0.60673 | -0.68987 | 0.120527 | -0.00339 | 0.103943 | 0.872851 | 0.101774 | -1.2662 |
| 37.16667 | 0.229334 | -0.33554 | -0.18159 | 0.662678 | -0.36281 | 0.014089 | -0.11956 | 1.074549 | 1.811594 | 0.634881 | -0.13668 | -0.70547 | 0.764639 | -0.50958 | -1.21864 | 0.95718 | 0.468089 | -1.10482 |
| 37.18056 | 0.419618 | -0.3583 | -0.01198 | 0.745838 | 0.042259 | -0.00102 | -0.06339 | 1.656241 | 2.495964 | 0.739325 | 0.144737 | -0.86729 | 0.98177 | -0.74763 | -2.07715 | 1.06332 | 0.655201 | -0.9692 |
| 37.19444 | 0.634015 | -0.36184 | 0.163459 | 0.75132 | 0.352264 | 0.007129 | -0.0883 | 2.083616 | 3.120425 | 1.047476 | 0.377525 | -0.77425 | 1.224093 | -0.58609 | -2.2455 | 1.065431 | 0.935967 | -0.71748 |
| 37.20833 | 0.951907 | -0.14123 | 0.33905 | 0.734538 | 0.65619 | 0.058184 | -0.14881 | 2.455087 | 3.383016 | 1.500367 | 0.540248 | -0.42914 | 1.821842 | -0.27388 | -1.80488 | 1.007742 | 1.31961 | -0.30767 |
| 37.22222 | 1.081324 | 0.005097 | 0.433152 | 0.631778 | 0.83237 | 0.104442 | -0.10458 | 2.320398 | 3.871397 | 1.918168 | 0.727663 | 0.065381 | 2.552833 | 0.07768 | -1.20192 | 1.02612 | 1.686262 | 0.069243 |
| 37.23611 | 1.015187 | 0.083049 | 0.457663 | 0.462147 | 0.886299 | 0.110091 | 0.055228 | 1.660957 | 3.661258 | 2.324545 | 0.99459 | 0.475164 | 3.095134 | 0.56621 | -0.24311 | 1.201955 | 2.009981 | 0.483633 |
| 37.25 | 0.92194 | -0.01278 | 0.387699 | 0.252547 | 0.876806 | 0.068367 | 0.080348 | 2.022772 | 2.419148 | 2.295599 | 1.136715 | 0.603798 | 3.135856 | 0.936775 | 0.255992 | 1.186806 | 1.905427 | 0.856614 |
| 37.26389 | 0.612448 | -0.34713 | 0.19871 | 0.156659 | 0.715955 | 0.005236 | -0.05364 | 2.361935 | 1.697127 | 1.924193 | 0.855751 | 0.352688 | 2.597782 | 0.813332 | 0.004831 | 1.287116 | 1.655068 | 0.778442 |
| 37.27778 | 0.201237 | -0.69403 | 0.052585 | 0.374615 | 0.609981 | -0.02039 | -0.20562 | 3.260484 | 1.396098 | 1.212582 | 0.338989 | 0.083686 | 1.886423 | 0.675962 | -0.09616 | 1.329783 | 1.236723 | 0.376745 |
| 37.29167 | -0.06138 | -0.69987 | 0.083971 | 0.728014 | 0.529111 | -0.00781 | -0.25783 | 3.51037 | 0.525095 | 0.773521 | -0.05694 | 0.030458 | 1.488875 | 0.602102 | 0.066486 | 1.373245 | 1.126887 | 0.237253 |
| 37.30556 | 0.036464 | -0.38535 | 0.205479 | 0.992472 | 0.473752 | -0.0099 | -0.22814 | 3.180529 | 0.044398 | 0.931677 | -0.06129 | 0.146985 | 1.765044 | 0.890115 | 0.683559 | 1.184656 | 1.042271 | 0.305256 |
| 37.31944 | 0.334403 | -0.09102 | 0.290467 | 1.149957 | 0.4173 | -0.08034 | -0.22352 | 2.408275 | -0.13073 | 1.511459 | 0.10006 | 0.199745 | 2.23233 | 0.688964 | 1.281741 | 1.028163 | 0.990224 | 0.121582 |
| 37.33333 | 0.685324 | 0.164124 | 0.32541 | 1.279052 | 0.339454 | -0.16347 | -0.19316 | 1.631678 | 0.32468 | 1.722579 | 0.15255 | 0.069648 | 2.050079 | 0.903433 | 1.651986 | 0.951204 | 0.792081 | -0.33779 |
| 37.34722 | 0.898161 | 0.296512 | 0.364049 | 1.396042 | 0.235043 | -0.16087 | -0.0554 | 1.441839 | 1.070308 | 1.444464 | 0.184006 | 0.007818 | 1.179652 | 0.36394 | 1.40476 | 0.713069 | 0.549318 | -0.43892 |
| 37.36111 | 0.889115 | 0.353157 | 0.405112 | 1.425904 | 0.00436 | -0.05887 | 0.073499 | 1.628523 | 2.114842 | 1.200925 | 0.435256 | 0.097737 | -0.21526 | 0.141182 | 0.770998 | 0.527499 | 0.247562 | -0.29366 |
| 37.375 | 0.821369 | 0.433642 | 0.419634 | 1.332153 | -0.43034 | -0.00141 | 0.208329 | 1.715263 | 2.597868 | 1.118918 | 0.858423 | 0.233911 | -2.35603 | 0.095552 | -0.39005 | 0.598803 | 0.12386 | -0.06908 |
| 37.38889 | 0.778712 | 0.540652 | 0.470751 | 1.26579 | -0.78422 | -0.02367 | 0.389934 | 1.591171 | 2.562133 | 1.441769 | 1.183555 | 0.29745 | -2.60245 | 0.236765 | -0.88093 | 0.759109 | 0.130463 | 0.196591 |
| 37.40278 | 0.714285 | 0.505992 | 0.572182 | 1.369666 | -0.94472 | -0.0398 | 0.616166 | 1.014378 | 1.731359 | 1.694543 | 1.131794 | 0.257952 | -0.98135 | 0.609341 | -0.05174 | 0.8242 | 0.301143 | 0.391763 |
| 37.41667 | 0.510917 | 0.300218 | 0.641342 | 1.514348 | -1.0365 | -0.02472 | 0.664349 | -0.04689 | 0.470746 | 1.77938 | 0.958334 | 0.244645 | 0.315226 | 0.850198 | 0.894807 | 0.723204 | 0.578616 | 0.470157 |
| 37.43056 | 0.302216 | 0.151649 | 0.587645 | 1.492041 | -1.23183 | 0.002991 | 0.469738 | -1.14775 | -0.84765 | 1.721414 | 0.80114 | 0.339401 | 0.871005 | 1.024243 | 1.419873 | 0.686947 | 0.786739 | 0.545764 |
| 37.44444 | 0.19516 | 0.187685 | 0.481349 | 1.242333 | -1.37503 | -0.03114 | 0.188783 | -2.00044 | -1.76174 | 1.536308 | 0.61806 | 0.430415 | 0.845688 | 1.043675 | 1.455205 | 0.709848 | 0.891279 | 0.468246 |
| 37.45833 | 0.095154 | 0.235869 | 0.381363 | 1.065893 | -1.33321 | -0.07752 | 0.13383 | -2.56458 | -2.33775 | 1.195025 | 0.276086 | 0.25453 | 0.326469 | 0.865841 | 1.067944 | 0.664836 | 0.791119 | 0.022701 |
| 37.47222 | 0.093322 | 0.30167 | 0.406722 | 1.02036 | -1.20573 | -0.02171 | 0.363733 | -3.09471 | -2.83068 | 0.850664 | 0.002264 | 0.250192 | 0.126636 | 0.625848 | 0.673407 | 0.686871 | 0.766772 | -0.17879 |
| 37.48611 | 0.672646 | 1.058479 | 0.868157 | 1.133489 | -0.53775 | 0.365526 | 1.210723 | -3.24139 | -2.99566 | 1.67837 | 0.562673 | 1.487279 | 1.660462 | 0.591512 | 0.86595 | 1.345184 | 1.73627 | 0.698682 |
| 37.5 | 2.338724 | 3.021942 | 1.802187 | 1.556376 | 0.946011 | 1.050822 | 2.968888 | -1.82626 | -1.4163 | 4.323067 | 2.463046 | 4.41889 | 5.089079 | 0.891098 | 1.666062 | 2.559946 | 4.20609 | 2.575489 |
| 37.51389 | 3.780563 | 4.525818 | 2.411731 | 1.835649 | 2.893224 | 1.276374 | 3.992285 | 1.032613 | 1.501813 | 7.818107 | 4.548383 | 5.641072 | 7.093033 | 1.4221 | 3.041468 | 3.333013 | 4.822574 | 3.651512 |
| 37.52778 | 3.504687 | 4.174815 | 2.273274 | 1.506229 | 3.0094 | 0.797492 | 3.387257 | 2.881119 | 4.238117 | 7.513242 | 4.979956 | 3.861726 | 5.6732 | 1.648462 | 4.734476 | 2.43351 | 2.619595 | 3.113569 |
| 37.54167 | 2.964335 | 3.294144 | 2.102916 | 0.878657 | 4.322257 | 0.393118 | 2.706727 | 2.984367 | 5.545267 | 5.764399 | 4.684878 | 2.39953 | 4.266739 | 1.535973 | 5.434684 | 1.423979 | 1.32837 | 2.594628 |
| 37.55556 | 2.945872 | 2.847873 | 1.92132 | 0.560323 | 5.883757 | 0.311361 | 2.341501 | 2.376726 | 3.534031 | 5.170064 | 4.309114 | 1.788233 | 4.23547 | 1.863109 | 5.276759 | -0.44357 | 1.621695 | 2.942672 |
| 37.56944 | 2.870122 | 2.71977 | 1.573406 | 0.784321 | 7.205378 | 0.514634 | 2.289656 | 1.222093 | 1.633988 | 5.01031 | 3.602082 | 1.377195 | 4.011184 | 1.587762 | 2.893562 | 0.108671 | 2.16207 | 3.06728 |
| 37.58333 | 2.610145 | 2.636372 | 1.243285 | 1.147251 | 6.500893 | 1.197416 | 2.240893 | 0.291105 | 0.974833 | 3.967913 | 2.257931 | 1.073841 | 3.047777 | 0.527995 | 0.959174 | 0.144143 | 2.307329 | 2.720826 |
| 37.59722 | 2.206359 | 2.282795 | 0.958595 | 1.514339 | 5.056666 | 1.786618 | 2.101442 | -0.93697 | -0.75455 | 3.466287 | 0.948914 | 0.878941 | 2.473064 | -0.25363 | -0.20111 | 0.046313 | 1.99422 | 2.145302 |
| 37.61111 | 1.709867 | 1.883058 | 0.809282 | 1.527239 | 4.200567 | 2.151898 | 1.92055 | -1.73058 | -2.60959 | 3.15645 | 0.468284 | 0.881091 | 2.182551 | -0.46284 | -0.98951 | 0.30049 | 1.44145 | 1.800743 |
| 37.625 | 1.244374 | 1.580451 | 0.762643 | 1.355737 | 3.261911 | 2.215502 | 1.74644 | -1.90407 | -4.03693 | 2.539555 | 0.425725 | 0.843019 | 1.52638 | -0.11848 | -1.00491 | 0.285805 | 0.889471 | 1.703672 |
| 37.63889 | 1.232268 | 1.360861 | 0.632806 | 1.143185 | 2.751481 | 1.82188 | 1.569966 | -2.16407 | -4.175 | 1.904286 | 0.451985 | 0.731021 | 0.595853 | 0.423667 | -0.16249 | 0.1691 | 0.424651 | 1.377731 |
| 37.65278 | 0.900791 | 1.289209 | 0.422282 | 0.903871 | 2.457528 | 1.186523 | 1.314676 | -2.34683 | -3.90574 | 1.272375 | 0.383281 | 0.25127 | -0.47487 | 0.492095 | 0.27426 | 0.120658 | -0.00922 | 1.002016 |
| 37.66667 | 0.776839 | 1.319603 | 0.416633 | 0.613946 | 1.825971 | 0.176002 | 1.0766 | -2.73511 | -3.27464 | 0.676702 | 0.285149 | -0.30354 | -2.09384 | 0.42808 | 0.570993 | 0.091807 | -0.44021 | 0.652478 |
| 37.68056 | 0.528116 | 1.32139 | 0.498697 | 0.306398 | 0.609463 | -0.85602 | 0.928708 | -3.32278 | -2.9112 | 0.1053 | 0.354329 | -0.9574 | -4.98183 | 0.560068 | 1.050668 | 0.149298 | -0.58388 | 0.567248 |
| 37.69444 | 0.148077 | 1.106166 | 0.538392 | -0.05548 | -0.93744 | -1.73663 | 0.855523 | -4.42621 | -2.86855 | 0.140616 | 0.774178 | -2.00676 | -11.1974 | 0.822836 | 1.799371 | 0.245093 | -0.55147 | 0.837817 |
| 37.70833 | 0.036477 | 0.869814 | 0.39766 | -0.5544 | -2.24302 | -2.13145 | 0.62459 | -5.57298 | -2.29535 | 2.051095 | 1.124421 | -2.32763 | -14.8549 | 1.068182 | 2.148174 | 0.308174 | -0.29399 | 1.234787 |
| 37.72222 | 0.128688 | 0.721925 | -0.09929 | -0.7427 | -3.26492 | -0.98158 | 0.089761 | -6.24966 | -1.91918 | 2.397634 | 1.680243 | -0.59805 | -9.51078 | 1.159591 | 2.302311 | 0.489761 | -0.0381 | 0.889773 |
| 37.73611 | 0.144963 | 0.688501 | -0.87428 | -0.61271 | -3.68504 | -0.95846 | -0.68186 | -5.88514 | -0.95745 | 2.603882 | 2.096191 | -0.28946 | -5.05163 | 1.242093 | 2.200695 | 0.608173 | -0.34311 | -0.21105 |
| 37.75 | 0.242213 | 0.613942 | -1.13145 | -0.46602 | -4.56638 | -0.87498 | -1.12211 | -4.44672 | -0.44121 | 2.167852 | 1.857933 | -0.84486 | -2.83836 | 1.185341 | 1.476675 | 0.663481 | -0.89815 | -1.23672 |
| 37.76389 | 0.085588 | 0.44483 | -0.96527 | -0.52327 | -5.5674 | -0.26976 | -0.95571 | -2.90359 | 0.138274 | 0.550855 | 1.051326 | -3.00691 | -2.01644 | 0.858311 | 0.558402 | 0.732831 | -1.13446 | -1.62858 |
| 37.77778 | -0.34408 | 0.263988 | -0.81296 | -0.50203 | -5.55601 | -0.17597 | -0.91073 | -1.62068 | 0.112223 | -1.31119 | 0.031424 | -7.42568 | -2.73421 | 0.508986 | -0.23102 | 0.675372 | -1.33819 | -1.90266 |
| 37.79167 | -0.64458 | 0.103265 | -0.80695 | 0.043716 | -5.102 | -0.08791 | -0.69126 | -1.09513 | -0.52879 | -1.53547 | -0.40354 | -11.3525 | -2.5976 | 0.463986 | -0.78836 | 0.770766 | -1.69411 | -2.42758 |
| 37.80556 | -0.56522 | -0.14665 | -0.85742 | 1.106788 | -4.61004 | -0.03225 | -0.06635 | -1.21181 | -1.6949 | -0.41925 | -0.36722 | -5.59844 | -2.08684 | 0.292404 | -1.42253 | 0.83583 | -1.83458 | -3.0193 |
| 37.81944 | -0.71522 | -0.46525 | -0.80079 | 1.786566 | -3.68176 | -0.01925 | 0.326074 | -1.84671 | -3.10001 | 0.658644 | -0.41701 | -4.06553 | -2.78075 | -0.15048 | -1.75166 | 0.83096 | -1.94223 | -3.0347 |
| 37.83333 | -0.83932 | -0.7932 | -0.82079 | 2.015351 | -2.70135 | -0.04976 | 0.284993 | -2.59014 | -4.1143 | 1.06136 | -0.99557 | -3.74211 | -3.47515 | -0.40952 | -1.39717 | 0.73774 | -1.72838 | -2.85371 |
| 37.84722 | -1.0803 | -1.08814 | -0.933 | 2.07521 | -2.22255 | -0.12257 | -0.08294 | -3.58725 | -5.01839 | 1.024061 | -1.6438 | -4.10011 | -3.9688 | -0.56053 | -1.32903 | 0.284245 | -1.99175 | -2.62537 |
| 37.86111 | -1.65596 | -1.45562 | -0.95162 | 1.997437 | -1.89497 | -0.19809 | -0.60741 | -4.47258 | -5.25725 | 0.226782 | -2.13616 | -4.89437 | -4.37634 | -0.69404 | -1.40373 | -0.80643 | -2.47705 | -2.95912 |
| 37.875 | -1.86134 | -1.65318 | -0.80978 | 1.915954 | -1.67136 | -0.22525 | -1.04038 | -5.37802 | -5.0175 | -0.80401 | -2.49832 | -5.5453 | -4.66224 | -0.97838 | -1.26977 | -1.19058 | -3.45071 | -3.59853 |
| 37.88889 | -1.26931 | -1.74003 | -0.56885 | 1.932886 | -1.43369 | -0.23934 | -1.36815 | -6.14924 | -5.0792 | -1.65768 | -2.70769 | -5.76846 | -4.21592 | -1.13004 | -1.22369 | -0.9466 | -3.88061 | -3.88966 |
| 37.90278 | -1.08215 | -1.62789 | -0.33772 | 1.96947 | -1.20883 | -0.30184 | -1.57064 | -5.58451 | -4.5374 | -1.92792 | -2.7113 | -5.88507 | -3.44721 | -1.18174 | -1.13497 | -0.86018 | -3.38923 | -3.04383 |
| 37.91667 | -0.2754 | -0.91694 | -0.16513 | 1.887389 | -0.99437 | -0.40626 | -1.59384 | -4.99504 | -4.11505 | -1.81009 | -2.5784 | -5.02552 | -2.81 | -0.73176 | -1.03139 | -0.60291 | -3.02973 | -2.3199 |
| 37.93056 | 0.19949 | -0.22919 | -0.01294 | 1.714016 | -0.60282 | -0.47793 | -1.48799 | -4.42633 | -4.06456 | -1.97486 | -2.73055 | -3.90664 | -2.63139 | 0.009624 | -1.30338 | -0.36659 | -2.88664 | -2.0463 |
| 37.94444 | 0.585776 | 0.594678 | 0.198481 | 1.572525 | -0.00212 | -0.32085 | -0.95441 | -3.08245 | -3.2159 | -2.43389 | -2.7577 | -3.29035 | -3.17902 | 1.053952 | -2.85343 | -0.05808 | -2.98391 | -1.93448 |

**Supplementary Table S2.** Raw motion data from: (a) Nutrient withdrawal stress

| Time | Control1 | Control2 | Control3 | Control4 | Control5 | Control6 | Control7 | Control8 | Control9 | Control10 | Control11 | Control12 | Stress1 | Stress2 | Stress3 | Stress4 | Stress5 | Stress6 | Stress7 | Stress8 | Stress9 | Stress10 | Stress11 | Stress12 |
|---|---|---|---|---|---|---|---|---|---|---|---|---|---|---|---|---|---|---|---|---|---|---|---|---|
| 18 | 1.704239 | 1.08016 | 0.925252 | -0.19446 | 1.028643 | -0.06545 | 1.057191 | 0.111242 | 1.508922 | 0.34051 | -0.33673 | 1.194786 | 1.486662 | 2.092135 | -0.03221 | 0.107909 | 0.239621 | 0.003451 | 0.203964 | 0.141233 | 2.280024 | -0.34293 | 3.388659 | 0.719448 |
| 18.01389 | 0.930757 | 0.747874 | 0.549492 | -0.12995 | 0.453548 | -0.05781 | 0.604172 | 0.066766 | 2.083702 | 0.291365 | -0.2096 | 0.590316 | 1.316762 | 1.995332 | 0.009598 | 0.14583 | 0.116286 | -0.00532 | 0.03065 | 0.049555 | 2.074476 | -0.34028 | 2.092917 | 0.650311 |
| 18.02778 | 0.39559 | 0.361766 | 0.311262 | -0.09334 | 0.198664 | -0.05414 | 0.121145 | 0.027602 | 1.109271 | 0.172795 | -0.15927 | 0.271464 | 0.975335 | 1.312621 | 0.012921 | 0.121885 | 0.027031 | 0.011488 | -0.04624 | -0.01352 | 1.715955 | -0.5118 | 1.213808 | 0.362717 |
| 18.04167 | 0.16207 | 0.147618 | 0.268798 | -0.08175 | 0.064079 | -0.05901 | -0.01687 | -0.01657 | 0.65747 | 0.091338 | -0.14838 | 0.131999 | 0.540302 | 0.633458 | -0.00247 | 0.089218 | -0.02263 | 0.007558 | -0.07619 | -0.0402 | 1.112818 | -0.58811 | 0.652086 | 0.183307 |
| 18.05556 | 0.074911 | 0.050375 | 0.325899 | -0.09691 | -0.00537 | -0.09041 | -0.05505 | -0.04635 | 0.376281 | 0.023256 | -0.1755 | 0.062955 | 0.204705 | 0.264778 | -0.02028 | 0.052097 | -0.04171 | -0.00546 | -0.07805 | -0.05123 | 0.860957 | -0.50918 | 0.168056 | 0.078728 |
| 18.06944 | 0.055192 | 0.02494 | 0.203995 | -0.11126 | -0.02917 | -0.13727 | -0.04631 | -0.04882 | 0.246685 | -0.01027 | -0.21547 | 0.047379 | 0.067927 | 0.098647 | -0.02164 | 0.023809 | -0.02696 | -0.00819 | -0.05723 | -0.04108 | 0.790041 | -0.3914 | 0.033144 | 0.087441 |
| 18.08333 | 0.056574 | 0.026593 | 0.089797 | -0.10228 | -0.01348 | -0.15745 | -0.02734 | -0.01986 | 0.222816 | -0.00541 | -0.23382 | 0.068756 | 0.139309 | 0.061682 | -0.00517 | 0.039271 | 0.003675 | 0.005352 | -0.02602 | -0.01406 | 0.877319 | -0.27304 | -0.02006 | 0.183574 |
| 18.09722 | 0.059824 | 0.045358 | 0.081557 | -0.07961 | 0.01527 | -0.12834 | -0.00423 | 0.0199 | 0.228278 | 0.018049 | -0.22577 | 0.080956 | 0.277073 | 0.110415 | 0.002662 | 0.064959 | 0.023512 | 0.01263 | -0.0136 | -0.01836 | 0.901924 | -0.26473 | -0.29306 | 0.468306 |
| 18.11111 | 0.045375 | 0.065659 | 0.073654 | -0.0652 | 0.023402 | -0.09071 | 0.013326 | 0.033386 | 0.192289 | 0.037053 | -0.19554 | 0.057783 | 0.316625 | 0.179623 | -0.01117 | 0.063642 | 0.024282 | 0.006798 | -0.02554 | -0.05366 | 0.916373 | -0.42603 | -0.33637 | 0.830122 |
| 18.125 | 0.02139 | 0.079442 | 0.057402 | -0.06893 | 0.011471 | -0.07087 | 0.014878 | 0.019975 | 0.13742 | 0.037261 | -0.17174 | 0.018855 | 0.281255 | 0.211278 | -0.02681 | 0.057123 | 0.018133 | 0.006805 | -0.03431 | -0.06897 | 0.836833 | -0.47776 | 0.086719 | 0.52221 |
| 18.13889 | 0.008963 | 0.080796 | 0.04673 | -0.06972 | 0.009089 | -0.05966 | 0.009149 | 0.008193 | 0.103853 | 0.027927 | -0.17946 | 0.002514 | 0.263126 | 0.198438 | -0.02748 | 0.061776 | 0.012765 | 0.017828 | -0.03022 | -0.05742 | 0.753338 | -0.2895 | 0.434014 | 1.303784 |
| 18.15278 | 0.003819 | 0.050688 | 0.036568 | -0.06972 | 0.015249 | -0.05984 | 0.008771 | -0.00022 | 0.092099 | 0.012812 | -0.21316 | 0.005837 | 0.252378 | 0.175931 | -0.02327 | 0.058029 | 0.001551 | 0.025406 | -0.03046 | -0.05141 | 0.655407 | 0.017162 | 0.718173 | 1.762884 |
| 18.16667 | -0.01321 | 0.018738 | 0.016211 | -0.07995 | -0.00025 | -0.07153 | -0.00928 | -0.00829 | 0.084718 | -0.01078 | -0.25411 | 0.0087 | 0.202152 | 0.152494 | -0.01591 | 0.026321 | -0.00737 | 0.028181 | -0.03022 | -0.05844 | 0.64521 | 0.447203 | 0.737928 | 1.379905 |
| 18.18056 | -0.01945 | 0.02721 | 0.009778 | -0.08048 | -0.01731 | -0.07479 | -0.02469 | -0.00469 | 0.083888 | -0.02815 | -0.3019 | 0.011218 | 0.176392 | 0.120739 | -0.00585 | -0.0205 | -0.0185 | 0.027471 | -0.03179 | -0.07962 | 0.666825 | 0.267463 | 0.669404 | 0.918504 |
| 18.19444 | -0.0026 | 0.048985 | 0.033163 | -0.05297 | -0.00273 | -0.06531 | -0.00646 | 0.008404 | 0.106537 | -0.01736 | -0.37425 | 0.020139 | 0.218057 | 0.090956 | -0.00869 | -0.03371 | -0.03595 | 0.023301 | -0.03772 | -0.0997 | 0.65733 | 0.023173 | 0.590467 | 0.630731 |
| 18.20833 | 0.009384 | 0.0657 | 0.056563 | -0.05153 | 0.018799 | -0.06772 | 0.014703 | 0.012447 | 0.125973 | 0.012725 | -0.43515 | 0.014467 | 0.237558 | 0.082986 | -0.01024 | 0.003167 | -0.01592 | 0.02938 | -0.02581 | -0.086 | 0.601146 | -0.06369 | 0.501073 | 0.495642 |
| 18.22222 | 0.010003 | 0.087489 | 0.05072 | -0.07467 | 0.022139 | -0.08054 | 0.010396 | 0.008033 | 0.115391 | 0.043162 | -0.53806 | -0.00292 | 0.186473 | 0.09492 | -0.00559 | 0.033887 | 0.125237 | 0.043274 | 0.005382 | -0.05664 | 0.544023 | 0.017598 | 0.471234 | 0.456176 |
| 18.23611 | 0.015943 | 0.107935 | 0.052361 | -0.09622 | 0.028348 | -0.07553 | -0.00377 | 0.007596 | 0.103025 | 0.074895 | -0.72658 | -0.00053 | 0.142652 | 0.090417 | -0.01265 | 0.033287 | 0.492823 | 0.0352 | 0.021207 | -0.04373 | 0.478462 | 0.042153 | 0.455168 | 0.357878 |
| 18.25 | 0.018439 | 0.082526 | 0.061451 | -0.08443 | 0.029605 | -0.06031 | -0.01678 | 0.003057 | 0.097998 | 0.077653 | -0.81523 | 0.013323 | 0.134265 | 0.076402 | -0.02461 | 0.038628 | 0.760399 | 0.018029 | 0.017769 | -0.03181 | 0.405747 | -0.01594 | 0.43776 | 0.234164 |
| 18.26389 | 0.008066 | 0.032642 | 0.075447 | -0.08553 | 0.017273 | -0.0584 | -0.01791 | -0.00575 | 0.100812 | 0.047143 | -0.77907 | 0.022876 | 0.17918 | 0.079456 | -0.02545 | 0.083337 | 0.422282 | 0.026359 | 0.021703 | 0.006912 | 0.339089 | -0.10112 | 0.440071 | 0.142638 |
| 18.27778 | 0.011657 | 0.037668 | 0.108907 | -0.07355 | 0.032789 | -0.04907 | 0.014092 | 0.005989 | 0.137052 | 0.039083 | -0.61168 | 0.033643 | 0.284467 | 0.101484 | -0.01849 | 0.11572 | 0.232271 | 0.074247 | 0.034786 | 0.061094 | 0.285454 | -0.13015 | 0.455274 | 0.105888 |
| 18.29167 | 0.028091 | 0.084608 | 0.123246 | -0.0531 | 0.068287 | -0.03067 | 0.035364 | 0.027744 | 0.170515 | 0.052787 | -0.41025 | 0.035659 | 0.329962 | 0.099371 | -0.01834 | 0.046198 | 0.180201 | 0.129937 | 0.035096 | 0.085915 | 0.256202 | -0.11271 | 0.447235 | 0.072279 |
| 18.30556 | 0.031134 | 0.110849 | 0.115185 | -0.05139 | 0.081225 | -0.02738 | 0.013488 | 0.033861 | 0.164698 | 0.0584 | -0.30044 | 0.033294 | 0.264976 | 0.087236 | -0.02428 | 0.057354 | 0.146739 | 0.134176 | 0.016921 | 0.076541 | 0.247494 | -0.14999 | 0.422152 | 0.041324 |
| 18.31944 | 0.027134 | 0.11108 | 0.099806 | -0.0654 | 0.078546 | -0.03749 | -0.0064 | 0.025439 | 0.153645 | 0.065891 | -0.24994 | 0.033051 | 0.213687 | 0.087856 | -0.02461 | 0.093133 | 0.131974 | 0.081413 | 0.009863 | 0.064177 | 0.283682 | -0.15862 | 0.424288 | 0.026935 |
| 18.33333 | 0.021826 | 0.10635 | 0.09607 | -0.07811 | 0.065741 | -0.04778 | 0.003967 | 0.017745 | 0.166954 | 0.091546 | -0.23466 | 0.016475 | 0.262433 | 0.09231 | -0.01531 | 0.120183 | 0.136545 | 0.057629 | 0.027311 | 0.072281 | 0.317384 | -0.1677 | 0.459828 | 0.035032 |
| 18.34722 | 0.024823 | 0.121411 | 0.107827 | -0.07071 | 0.064261 | -0.04893 | 0.029658 | 0.020742 | 0.199853 | 0.122711 | -0.23156 | -0.01254 | 0.335252 | 0.1001 | -0.01482 | 0.123656 | 0.144927 | 0.052873 | 0.037499 | 0.115536 | 0.315194 | -0.19937 | 0.498293 | 0.016821 |
| 18.36111 | 0.027848 | 0.093635 | 0.102695 | -0.0634 | 0.03704 | -0.06406 | 0.050683 | 0.020906 | 0.218442 | 0.120188 | -0.24793 | -0.05814 | 0.332082 | 0.065445 | -0.08725 | 0.08365 | 0.082674 | -0.02081 | -0.03015 | 0.109851 | 0.212442 | -0.23236 | 0.450953 | -0.06254 |
| 18.375 | 0.009145 | -0.019 | 0.050213 | -0.08972 | -0.04066 | -0.111 | 0.051974 | 0.004351 | 0.195087 | 0.065229 | -0.29653 | -0.12552 | 0.234441 | -0.01513 | -0.21541 | 0.026364 | -0.0463 | -0.14499 | -0.1439 | 0.015569 | 0.037546 | -0.29147 | 0.320626 | -0.16617 |
| 18.38889 | 0.001238 | -0.02401 | 0.014411 | -0.11804 | -0.03103 | -0.13048 | 0.08749 | -0.00246 | 0.217075 | 0.057546 | -0.31866 | -0.13185 | 0.355739 | -0.003 | -0.22532 | 0.062237 | -0.02826 | -0.14554 | -0.15013 | -0.01863 | -0.021 | -0.3495 | 0.321233 | -0.15227 |
| 18.40278 | 0.120459 | 0.363693 | 0.108934 | -0.08995 | 0.286925 | -0.06405 | 0.49662 | 0.153459 | 0.574782 | 0.263373 | -0.22936 | -0.03143 | 1.407756 | 0.323594 | -0.08945 | 0.224414 | 0.295999 | 0.037718 | -0.0265 | 0.088398 | 0.215297 | -0.49242 | 0.64966 | 0.102255 |
| 18.41667 | 0.351209 | 1.098077 | 0.394108 | -0.01156 | 0.927119 | 0.064998 | 1.462515 | 0.865333 | 1.297152 | 0.703257 | -0.06302 | 0.129413 | 4.096512 | 0.730494 | -0.0019 | 0.46949 | 0.827374 | 0.281094 | 0.050657 | 0.227901 | 0.62916 | -1.58577 | 1.24322 | 0.651559 |
| 18.43056 | 0.415777 | 1.439728 | 0.413162 | 0.000497 | 1.29237 | 0.202305 | 1.742592 | 0.874584 | 1.700223 | 1.236928 | 0.077115 | 0.203991 | 6.063105 | 0.484659 | -0.1057 | 0.550624 | 0.817184 | 0.424824 | -0.12492 | 0.159342 | 0.711076 | -3.0716 | 1.43217 | 0.971782 |
| 18.44444 | 0.255432 | 0.961569 | 0.01294 | -0.06386 | 0.937187 | 0.263495 | 1.020748 | 0.243244 | 1.337319 | 1.235061 | 0.160468 | 0.159501 | 3.75881 | 0.274973 | -0.2868 | 0.114097 | 0.241614 | 0.212893 | -0.43435 | -0.07788 | 0.253086 | -2.69735 | 1.004885 | 0.18587 |
| 18.45833 | 0.164228 | 0.425619 | -0.04542 | -0.03347 | 0.629301 | 0.204488 | 0.532155 | 0.185583 | 0.920984 | 1.022121 | 0.19765 | 0.14869 | 2.091384 | 0.232187 | -0.20924 | -0.18059 | 0.001598 | 0.128254 | -0.36736 | -0.08864 | 0.04574 | -1.74668 | 0.752601 | -0.16443 |
| 18.47222 | 0.225682 | 0.217633 | -0.00254 | 0.079623 | 0.592518 | 0.197243 | 0.33099 | 0.197956 | 0.719661 | 0.881827 | 0.209381 | 0.109176 | 1.499124 | 0.468296 | 0.053669 | 0.06497 | 0.260521 | 0.246223 | -0.01874 | 0.147014 | 0.323715 | -1.09907 | 0.791085 | -0.11351 |
| 18.48611 | 0.297751 | 0.142827 | 0.017153 | 0.133536 | 0.634243 | 0.179981 | 0.278175 | 0.185198 | 0.688481 | 0.859321 | 0.215956 | 0.037671 | 1.231551 | 0.633818 | 0.161679 | 0.173707 | 0.505069 | 0.265498 | 0.156765 | 0.297491 | 0.417292 | -0.62818 | 0.827833 | -0.04108 |
| 18.5 | 0.376033 | 0.136101 | 0.038503 | 0.136965 | 0.661974 | 0.165093 | 0.292784 | 0.140422 | 0.799074 | 0.797108 | 0.210865 | -0.01081 | 1.200729 | 0.647503 | 0.140046 | 0.148863 | 0.523485 | 0.218109 | 0.181376 | 0.328974 | 0.301508 | 0.022203 | 0.771274 | 0.000257 |
| 18.51389 | 0.443632 | 0.182972 | 0.081847 | 0.131852 | 0.686129 | 0.1601 | 0.264576 | 0.105498 | 0.955881 | 0.735573 | 0.200528 | -0.02315 | 1.312279 | 0.635627 | 0.108083 | 0.126294 | 0.457816 | 0.216413 | 0.198092 | 0.329218 | 0.672757 | 0.578879 | 0.689087 | 0.069639 |
| 18.52778 | 0.474295 | 0.258897 | 0.133516 | 0.118312 | 0.627135 | 0.145488 | 0.182809 | 0.084239 | 1.097203 | 0.703533 | 0.224568 | 0.00391 | 1.427276 | 0.47404 | 0.08657 | 0.111451 | 0.364857 | 0.251859 | 0.230695 | 0.3488 | 2.024175 | 0.543924 | 0.578099 | 0.136061 |
| 18.54167 | 0.482366 | 0.34679 | 0.188311 | 0.113196 | 0.572142 | 0.121836 | 0.076998 | 0.128943 | 1.233704 | 0.618925 | 0.301194 | 0.053348 | 1.477324 | 0.331944 | 0.074265 | 0.101936 | 0.393151 | 0.30501 | 0.241177 | 0.398776 | 2.452532 | 0.092017 | 0.550421 | 0.159335 |
| 18.55556 | 0.4084 | 0.441963 | 0.214806 | 0.119872 | 0.471913 | 0.075726 | -0.01389 | 0.237108 | 1.325311 | 0.535766 | 0.349553 | 0.075222 | 1.418957 | 0.268013 | 0.075677 | 0.10873 | 0.496325 | 0.343089 | 0.245824 | 0.405541 | 2.223409 | -0.40498 | 0.610409 | 0.164535 |
| 18.56944 | 0.326109 | 0.476866 | 0.20698 | 0.12526 | 0.354829 | 0.038906 | -0.07083 | 0.289394 | 1.326709 | 0.482486 | 0.32883 | 0.070882 | 1.299516 | 0.287172 | 0.099717 | 0.147495 | 0.525597 | 0.316672 | 0.212592 | 0.350205 | 1.919296 | -0.61179 | 0.715723 | 0.167754 |
| 18.58333 | 0.230973 | 0.449906 | 0.149113 | 0.107812 | 0.215178 | 0.040432 | -0.08496 | 0.227968 | 1.197509 | 0.434837 | 0.302638 | 0.06279 | 1.080649 | 0.253022 | 0.077279 | 0.153771 | 0.442318 | 0.225494 | 0.131278 | 0.20256 | 1.639745 | -0.39343 | 0.704157 | 0.093978 |
| 18.59722 | 0.154345 | 0.385849 | 0.094942 | 0.07087 | 0.097247 | 0.051501 | -0.05843 | 0.130695 | 0.98657 | 0.358391 | 0.283471 | 0.060241 | 0.823666 | 0.259603 | 0.023425 | 0.138817 | 0.318406 | 0.117548 | 0.020081 | 0.042874 | 1.255291 | -0.01653 | 0.589983 | -0.02546 |
| 18.61111 | 0.087946 | 0.339553 | 0.055265 | 0.095685 | 0.050179 | 0.060334 | -0.03231 | 0.103064 | 0.807501 | 0.235847 | 0.270377 | 0.043008 | 0.63844 | 0.348457 | 0.015203 | 0.178126 | 0.26466 | 0.071406 | -0.00725 | -0.01181 | 0.895993 | 0.275785 | 0.483149 | -0.07838 |
| 18.625 | 0.012852 | 0.31593 | 0.02827 | 0.104762 | -0.00469 | 0.058827 | -0.02945 | 0.09285 | 0.641135 | 0.117196 | 0.262236 | 0.009764 | 0.477124 | 0.325449 | 0.02143 | 0.182772 | 0.23002 | 0.065838 | 0.020348 | -0.02834 | 0.715368 | 0.401885 | 0.379604 | -0.07934 |
| 18.63889 | -0.04299 | 0.293938 | 0.015366 | 0.120803 | -0.1121 | 0.063135 | -0.02295 | 0.078835 | 0.485438 | 0.028446 | 0.259537 | 0.002697 | 0.37554 | 0.33329 | 0.013328 | 0.159682 | 0.152729 | 0.078485 | 0.058849 | -0.03131 | 0.505129 | 0.397041 | 0.2884 | -0.06636 |
| 18.65278 | -0.0679 | 0.320742 | 0.014606 | 0.109993 | -0.2231 | 0.068005 | 0.001436 | 0.07064 | 0.353323 | -0.03992 | 0.263513 | 0.018529 | 0.377934 | 0.289714 | 0.013426 | 0.172627 | 0.105241 | 0.086221 | 0.094295 | 0.064325 | 0.322659 | 0.293941 | 0.247015 | -0.0549 |
| 18.66667 | -0.07833 | 0.387995 | 0.014908 | 0.08728 | -0.28059 | 0.069117 | 0.017446 | 0.07239 | 0.252746 | -0.10076 | 0.261789 | 0.025974 | 0.404986 | 0.172252 | 0.02338 | 0.216305 | 0.015831 | 0.113391 | 0.089374 | 0.22979 | 0.311794 | 0.138486 | 0.232874 | -0.04099 |
| 18.68056 | -0.07513 | 0.400967 | 0.017972 | 0.080586 | -0.22059 | 0.103588 | 0.017116 | 0.08791 | 0.224325 | -0.09077 | 0.24306 | 0.025924 | 0.390828 | 0.024607 | 0.02715 | 0.190413 | -0.01595 | 0.132606 | 0.049686 | 0.290844 | 0.399648 | -0.03973 | 0.174889 | -0.04812 |
| 18.69444 | -0.04494 | 0.3271 | 0.020191 | 0.082739 | -0.12171 | 0.255278 | 0.005182 | 0.10956 | 0.239561 | 0.042385 | 0.217565 | 0.030766 | 0.343909 | 0.006947 | 0.021559 | 0.143322 | -0.00459 | 0.126345 | 0.022062 | 0.185838 | 0.513586 | -0.24442 | 0.06906 | -0.06882 |
| 18.70833 | -0.01504 | 0.237097 | 0.020973 | 0.11243 | -0.0618 | 0.352355 | -0.00827 | 0.106997 | 0.231158 | 0.181501 | 0.219326 | 0.031665 | 0.322438 | 0.153883 | 0.011886 | 0.11059 | 0.038469 | 0.100421 | 0.02141 | 0.0288 | 0.564059 | -0.42021 | 0.043711 | -0.07579 |
| 18.72222 | -0.00319 | 0.196929 | 0.029002 | 0.174707 | -0.05267 | 0.319168 | -0.035 | 0.099107 | 0.200847 | 0.268508 | 0.274586 | 0.008618 | 0.323761 | 0.28019 | 0.002918 | 0.056867 | -0.01265 | 0.073216 | 0.013705 | -0.08095 | 0.485003 | -0.41756 | 0.109771 | -0.06777 |
| 18.73611 | -0.03042 | 0.182281 | 0.013552 | 0.184723 | -0.07553 | 0.288953 | -0.08025 | 0.0762 | 0.167409 | 0.249483 | 0.426193 | -0.00915 | 0.31442 | 0.223071 | 0.002935 | 0.017367 | -0.17369 | 0.057931 | -0.00179 | -0.1415 | 0.344393 | -0.15774 | 0.151581 | -0.05508 |
| 18.75 | -0.0856 | 0.170477 | 0.004201 | 0.143438 | -0.15074 | 0.239653 | -0.13125 | 0.046087 | 0.11089 | 0.155082 | 0.627126 | -0.00271 | 0.285901 | 0.216314 | 0.003852 | -0.00843 | -0.33858 | 0.052763 | -0.02551 | -0.1891 | 0.209683 | 0.14558 | 0.141828 | -0.06268 |
| 18.76389 | -0.12848 | 0.204911 | 0.005339 | 0.115467 | -0.26518 | 0.208357 | -0.19078 | 0.012726 | 0.022004 | 0.000158 | 0.588528 | 0.018145 | 0.223661 | 0.223201 | 0.006464 | -0.02201 | -0.42273 | 0.046046 | -0.04892 | -0.21685 | 0.095447 | 0.245203 | 0.14275 | -0.07886 |
| 18.77778 | -0.1759 | 0.245067 | 0.002054 | 0.10242 | -0.40705 | 0.190889 | -0.26024 | 0.008288 | -0.08086 | -0.19513 | 0.410174 | 0.02652 | 0.1163 | 0.17349 | 0.015395 | -0.02949 | -0.52854 | 0.034813 | -0.07273 | -0.22812 | 0.056786 | 0.123343 | 0.098803 | -0.09592 |
| 18.79167 | -0.21403 | 0.20966 | -0.00339 | 0.071741 | -0.47538 | 0.17274 | -0.3828 | 0.026786 | -0.14751 | -0.2911 | 0.283999 | 0.000519 | -0.05761 | 0.130387 | 0.010099 | -0.05123 | -0.70911 | 0.00736 | -0.12154 | -0.25271 | 0.062977 | -0.17216 | -0.05069 | -0.14639 |
| 18.80556 | -0.28869 | 0.151289 | -0.00785 | 0.05298 | -0.47102 | 0.132246 | -0.48382 | 0.043241 | -0.20537 | -0.18031 | 0.235331 | -0.03195 | -0.2552 | 0.030342 | -0.01396 | -0.0738 | -0.84757 | -0.01661 | -0.17934 | -0.28447 | 0.114167 | -0.64118 | -0.17399 | -0.21674 |
| 18.81944 | -0.39467 | 0.125886 | -0.03215 | 0.04817 | -0.42819 | 0.103804 | -0.56156 | 0.040806 | -0.288 | -0.01807 | 0.233902 | -0.04862 | -0.3973 | -0.12195 | -0.03714 | -0.08022 | -0.8356 | -0.04309 | -0.17642 | -0.31571 | 0.190303 | -1.09169 | -0.24722 | -0.30191 |
| 18.83333 | -0.50646 | 0.177268 | -0.05041 | 0.042781 | -0.36999 | 0.089841 | -0.64441 | 0.035213 | -0.35329 | 0.040056 | 0.246682 | -0.0519 | -0.44081 | -0.28839 | -0.05239 | -0.07691 | -0.80499 | -0.07289 | -0.14194 | -0.4113 | 0.152621 | -1.37617 | -0.33506 | -0.35302 |
| 18.84722 | -0.52786 | 0.253431 | -0.04259 | 0.033388 | -0.31113 | 0.084161 | -0.70172 | 0.027629 | -0.35942 | -0.02112 | 0.259618 | -0.06627 | -0.4 | -0.39839 | -0.0587 | -0.07243 | -0.75525 | -0.09739 | -0.1333 | -0.4967 | -0.08848 | -1.68065 | -0.39897 | -0.38368 |
| 18.86111 | -0.54267 | 0.192233 | -0.04189 | 0.030185 | -0.25963 | 0.086589 | -0.75774 | 0.00392 | -0.32648 | -0.17611 | 0.26519 | -0.08632 | -0.37892 | -0.35293 | -0.0632 | -0.07478 | -0.72953 | -0.11332 | -0.15457 | -0.51396 | -0.36168 | -2.02428 | -0.43428 | -0.36803 |
| 18.875 | -0.44727 | 0.053366 | -0.05828 | 0.034867 | -0.19478 | 0.081178 | -0.69593 | -0.02418 | -0.31038 | -0.36281 | 0.287744 | -0.09438 | -0.41071 | -0.14355 | -0.05768 | -0.06476 | -0.59629 | -0.1059 | -0.20901 | -0.50406 | -0.49802 | -2.37755 | -0.42729 | -0.3435 |
| 18.88889 | -0.32968 | 0.002927 | -0.06905 | 0.035678 | -0.16409 | 0.07996 | -0.5893 | -0.01856 | -0.32385 | -0.47423 | 0.322148 | -0.09313 | -0.53362 | 0.10339 | -0.0327 | -0.03704 | -0.37453 | -0.08102 | -0.12474 | -0.48581 | -0.54609 | -2.82906 | -0.35783 | -0.30876 |
| 18.90278 | -0.25472 | -0.10622 | -0.13273 | 0.034926 | -0.20206 | 0.021574 | -0.55227 | 0.040161 | -0.47445 | -0.57316 | 0.330404 | -0.17731 | -0.91796 | 0.169443 | 0.008118 | 0.037927 | -0.30613 | -0.1056 | -0.05635 | -0.424 | -0.7551 | -3.15624 | -0.36237 | -0.34923 |
| 18.91667 | -0.30498 | -0.61512 | -0.28182 | -0.06734 | -0.36369 | -0.13218 | -0.83013 | 0.107118 | -0.9897 | -1.0425 | 0.186828 | -0.53102 | -1.90242 | -0.08479 | 0.074688 | 0.204895 | -0.45416 | -0.24877 | 0.053221 | -0.28486 | -1.35664 | -3.02678 | -0.55619 | -0.59114 |
| 18.93056 | -0.36074 | -1.33183 | -0.53246 | -0.22942 | -0.66352 | -0.48929 | -1.37278 | -0.15054 | -1.95015 | -1.93024 | -0.22623 | -1.15338 | -4.69321 | -0.59387 | 0.228002 | 0.272122 | -0.57671 | -0.53249 | 0.28033 | -0.06954 | -2.43294 | -2.74725 | -0.92037 | -0.95799 |
| 18.94444 | -0.32752 | -1.25265 | -0.72979 | -0.28737 | -0.56158 | -0.78604 | -1.89993 | -1.01335 | -3.12412 | -2.24336 | -0.79047 | -1.7309 | -6.98869 | -1.27059 | 0.405119 | 0.082401 | -0.58326 | -0.52839 | 0.519504 | 0.068927 | -4.53544 | -3.60127 | -0.95728 | -0.89761 |
| 18.95833 | -0.266 | -0.21864 | -0.1967 | -0.28926 | 0.081553 | -0.51033 | -1.15101 | -0.51076 | -1.05821 | -1.51319 | -1.21428 | -0.8252 | -3.78328 | -0.69239 | 0.293639 | 0.270464 | -0.25099 | -0.14357 | 0.386536 | -0.10551 | -2.58057 | -2.75565 | -0.3473 | -0.18432 |
| 18.97222 | -0.20872 | 0.559562 | 0.395414 | -0.28332 | 0.339484 | -0.28888 | 0.158057 | 0.446896 | 1.598898 | -0.16104 | -1.34083 | 0.480384 | 1.157524 | -0.33799 | -0.01571 | 0.82045 | 0.05938 | 0.121889 | 0.002128 | -0.37281 | 0.816692 | 0.686773 | 0.291064 | 0.346162 |
| 18.98611 | -0.15415 | 0.48369 | 0.203698 | -0.2475 | 0.151059 | -0.22339 | 0.67152 | 0.498934 | 1.832796 | 0.602676 | -1.35437 | 0.64028 | 3.528725 | 0.213446 | -0.12877 | 1.381532 | 0.091468 | 0.326227 | -0.16158 | -0.33623 | 2.08383 | 0.991929 | 0.343674 | 0.287302 |
| 19 | -0.11354 | 0.224621 | 0.080935 | -0.21974 | -0.03286 | -0.17114 | 0.445542 | 0.096213 | 0.750257 | 0.606167 | -1.18031 | 0.333425 | 2.705853 | 0.619983 | -0.08414 | 0.819134 | 0.18393 | 0.216219 | -0.12479 | -0.1462 | 1.53 | 0.427678 | 0.1587 | 0.132199 |
| 19.01389 | -0.06921 | 0.1291 | 0.036103 | -0.16908 | -0.05668 | -0.14034 | 0.228866 | -0.02108 | 0.310711 | 0.369031 | -0.88476 | 0.128891 | 1.711204 | 0.381053 | -0.03737 | 0.397729 | 0.25879 | 0.130525 | -0.06391 | -0.01911 | 0.984118 | 0.160839 | 0.082771 | 0.059172 |
| 19.02778 | -0.03169 | 0.090408 | 0.00532 | -0.10804 | -0.03074 | -0.11821 | 0.100098 | -0.06746 | 0.126418 | 0.189024 | -0.65478 | 0.02887 | 0.906749 | 0.144546 | -0.01642 | 0.228297 | 0.274754 | 0.050051 | -0.02845 | 0.022106 | 0.410107 | 0.002214 | 0.077241 | 0.012591 |
| 19.04167 | -0.01 | 0.070177 | -0.00508 | -0.04503 | -0.01542 | -0.09707 | 0.025356 | -0.04506 | 0.052326 | 0.08942 | -0.54027 | 0.001914 | 0.316449 | 0.100713 | -0.01574 | 0.135926 | 0.271979 | 0.006776 | -0.00878 | -0.00032 | 0.09256 | -0.14115 | 0.08104 | -0.01612 |
| 19.05556 | 0.003074 | 0.092617 | -0.00195 | -0.02565 | 0.000699 | -0.07573 | 0.011876 | -0.02194 | 0.043722 | 0.054203 | -0.46492 | 0.014483 | 0.106537 | 0.101782 | -0.0176 | 0.072659 | 0.238134 | 0.001107 | -0.00298 | -0.01488 | 0.075969 | -0.1784 | 0.06486 | -0.03163 |
| 19.06944 | 0.012038 | 0.103275 | 0.017252 | -0.03191 | 0.010185 | -0.05449 | -0.00372 | -0.00206 | 0.011706 | 0.064809 | -0.35497 | 0.051563 | 0.018457 | -0.52344 | -0.02205 | -0.03753 | 0.078239 | 0.016282 | 0.016884 | 0.022952 | 0.129921 | -0.21215 | 0.011811 | -0.06987 |
| 19.08333 | -0.00382 | 0.03029 | 0.062528 | -0.02413 | -0.03167 | -0.05511 | -0.18622 | 0.023062 | -0.26717 | 0.00273 | -0.26283 | 0.090636 | -0.87463 | -4.0212 | -0.13942 | -0.59886 | -0.90887 | -0.02258 | 0.07248 | 0.121065 | -0.40096 | -0.77418 | -0.36812 | -0.35885 |
| 19.09722 | -0.04078 | -0.0028 | 0.092838 | -0.02674 | -0.1794 | -0.09427 | -0.3959 | 0.016115 | -0.55545 | -0.10637 | -0.24375 | 0.026216 | -2.2925 | -4.97282 | -0.24853 | -0.98169 | -1.44156 | -0.10264 | 0.010896 | 0.066341 | -1.31549 | -1.49781 | -0.76645 | -0.57642 |
| 19.11111 | 0.022804 | 0.152977 | 0.036032 | -0.11205 | -0.02314 | -0.12129 | -0.12734 | -0.04098 | -0.07607 | 0.023907 | -0.19729 | -0.05993 | -0.10825 | -0.12785 | -0.19978 | -0.16121 | -0.04583 | -0.2161 | -0.28109 | -0.22114 | 0.000679 | -0.61041 | -0.39175 | -0.38414 |
| 19.125 | 0.153222 | 0.396231 | -0.0502 | -0.16802 | 0.222 | -0.0555 | 0.363518 | -0.03214 | 0.636754 | 0.329453 | -0.07301 | 0.007594 | 2.691124 | 4.826686 | 0.082597 | 0.945323 | 1.910522 | -0.0133 | -0.28036 | -0.2487 | 1.888428 | 0.96249 | 0.470999 | 0.14892 |
| 19.13889 | 0.201054 | 0.434003 | -0.06721 | -0.0308 | 0.174648 | 0.023837 | 0.429892 | 0.04023 | 0.601729 | 0.409463 | 0.010027 | 0.104179 | 2.208527 | 4.586174 | 0.292442 | 0.77263 | 1.720998 | 0.254927 | 0.026741 | 0.071212 | 1.383725 | 1.084427 | 0.841167 | 0.419962 |
| 19.15278 | 0.149823 | 0.329083 | -0.04695 | 0.010254 | 0.110433 | 0.018643 | 0.193336 | 0.0492 | 0.235493 | 0.15188 | 0.020868 | 0.104494 | 0.621319 | 1.042584 | 0.182961 | 0.193091 | 0.699984 | 0.180176 | 0.121423 | 0.211052 | 0.386933 | 0.256556 | 0.490445 | 0.149827 |
| 19.16667 | 0.081072 | 0.1965 | -0.0467 | -0.00881 | 0.034671 | -0.02404 | 0.007097 | 0.004999 | 0.030521 | -0.10857 | -0.00162 | 0.063059 | -0.35726 | 0.122148 | 0.041956 | 0.02745 | 0.239825 | 0.025854 | 0.048037 | 0.150006 | -0.14823 | -0.23801 | 0.204353 | -0.05084 |
| 19.18056 | 0.045443 | 0.044725 | -0.0487 | -0.02661 | 0.008478 | -0.04245 | -0.04852 | -0.01353 | -0.00202 | -0.15799 | -0.02197 | 0.029975 | -0.71723 | -0.02462 | -0.02996 | -0.03169 | 0.116966 | -0.04657 | -0.01741 | 0.131799 | -0.35356 | -0.3268 | 0.084024 | -0.12856 |
| 19.19444 | 0.059921 | 0.003323 | -0.01999 | -0.01136 | 0.038191 | -0.02308 | 0.001593 | 0.018973 | 0.052121 | -0.03005 | -0.02339 | 0.028103 | -0.47452 | -0.02439 | -0.02137 | -0.03656 | 0.155809 | -0.0235 | -0.00215 | 0.235162 | -0.14874 | -0.1815 | 0.097189 | -0.09641 |
| 19.20833 | 0.106914 | 0.103948 | 0.030523 | 0.027964 | 0.083422 | 0.013583 | 0.098393 | 0.074513 | 0.134505 | 0.135729 | 0.006035 | 0.070213 | 0.106058 | 0.109678 | 0.040329 | 0.017187 | 0.327805 | 0.055972 | 0.093673 | 0.353068 | 0.336452 | 0.082461 | 0.205566 | 0.003608 |
| 19.22222 | 0.144492 | 0.244255 | 0.050084 | 0.048433 | 0.099436 | 0.027931 | 0.165418 | 0.094588 | 0.187494 | 0.254779 | 0.027513 | 0.099488 | 0.591058 | 0.282013 | 0.084755 | 0.057771 | 0.452082 | 0.114309 | 0.159773 | 0.38417 | 0.611067 | 0.278523 | 0.330677 | 0.06011 |
| 19.23611 | 0.141209 | 0.277477 | 0.027105 | 0.033146 | 0.085292 | 0.014288 | 0.170434 | 0.077622 | 0.194984 | 0.270651 | 0.019122 | 0.0917 | 0.72646 | 0.336353 | 0.069938 | 0.030056 | 0.409182 | 0.106409 | 0.118355 | 0.301185 | 0.600554 | 0.285925 | 0.377089 | 0.041496 |
| 19.25 | 0.127346 | 0.253696 | 0.003123 | 0.023015 | 0.081108 | 0.002884 | 0.158404 | 0.064858 | 0.191078 | 0.219604 | 0.018839 | 0.07221 | 0.651565 | 0.262753 | 0.040754 | -0.00718 | 0.29981 | 0.078551 | 0.089991 | 0.225106 | 0.523146 | 0.226103 | 0.356783 | 0.013546 |
| 19.26389 | 0.118757 | 0.216298 | -0.00264 | 0.027339 | 0.089199 | 0.002833 | 0.158556 | 0.063061 | 0.186585 | 0.158695 | 0.031067 | 0.049123 | 0.548862 | 0.182058 | 0.028989 | -0.0102 | 0.265494 | 0.074223 | 0.135751 | 0.335163 | 0.473567 | 0.21725 | 0.346738 | 0.002863 |
| 19.27778 | 0.12258 | 0.184714 | -0.00407 | 0.02599 | 0.093269 | 0.005077 | 0.174587 | 0.058437 | 0.189872 | 0.127574 | 0.041633 | 0.066344 | 0.557189 | 0.145655 | 0.028587 | -0.01061 | 0.270866 | 0.080118 | 0.171013 | 0.601827 | 0.56873 | 0.263603 | 0.344 | 0.001893 |
| 19.29167 | 0.144731 | 0.187909 | 0.003575 | 0.031153 | 0.107706 | 0.014744 | 0.214571 | 0.054314 | 0.235221 | 0.142921 | 0.0555 | 0.089806 | 0.73473 | 0.156109 | 0.035627 | -0.01658 | 0.318009 | 0.091996 | 0.14751 | 0.749362 | 0.758329 | 0.349507 | 0.363796 | 0.007801 |
| 19.30556 | 0.172787 | 0.202541 | 0.0119 | 0.040325 | 0.125829 | 0.024405 | 0.243588 | 0.048229 | 0.295961 | 0.182559 | 0.069598 | 0.085661 | 0.859632 | 0.179695 | 0.035988 | -0.02755 | 0.427534 | 0.091519 | 0.113334 | 0.566898 | 0.705556 | 0.380366 | 0.373172 | -0.00762 |
| 19.31944 | 0.203196 | 0.172013 | 0.0092 | 0.033075 | 0.124333 | 0.022628 | 0.233992 | 0.040559 | 0.315082 | 0.199803 | 0.080244 | 0.05813 | 0.741214 | 0.182754 | 0.02845 | -0.05382 | 0.493309 | 0.077111 | 0.105772 | 0.247898 | 0.50269 | 0.320394 | 0.400326 | -0.02295 |
| 19.33333 | 0.252105 | 0.141393 | 0.009067 | 0.022802 | 0.123323 | 0.024237 | 0.233689 | 0.047079 | 0.312637 | 0.209531 | 0.095351 | 0.050507 | 0.579461 | 0.171366 | 0.035073 | -0.05395 | 0.479664 | 0.068163 | 0.158792 | 0.076967 | 0.401232 | 0.298321 | 0.565409 | 0.002141 |
| 19.34722 | 0.316651 | 0.127687 | 0.016731 | 0.016528 | 0.119503 | 0.028277 | 0.258835 | 0.053756 | 0.335258 | 0.203503 | 0.101223 | 0.046714 | 0.524023 | 0.14814 | 0.019961 | -0.05572 | 0.493027 | 0.052005 | 0.19468 | 0.03453 | 0.407768 | 0.281775 | 0.688363 | 0.009977 |
| 19.36111 | 0.342293 | 0.053487 | 0.015428 | 0.003308 | 0.06298 | 0.002235 | 0.278513 | 0.038839 | 0.396951 | 0.163941 | 0.085146 | -0.02316 | 0.485707 | 0.089902 | -0.06322 | -0.10953 | 0.407957 | -0.01762 | 0.112678 | 0.020292 | 0.365624 | 0.211225 | 0.68313 | -0.05521 |
| 19.375 | 0.321209 | -0.06243 | 0.004059 | -0.01227 | -0.01137 | -0.0438 | 0.286562 | 0.025665 | 0.471494 | 0.080632 | 0.055702 | -0.14976 | 0.383093 | 0.064269 | -0.16769 | -0.17351 | 0.255963 | -0.10949 | -0.00735 | 0.068871 | 0.274002 | 0.141085 | 0.597215 | -0.13965 |
| 19.38889 | 0.300354 | -0.02897 | 0.004833 | -0.01698 | 0.010188 | -0.04262 | 0.333191 | 0.010005 | 0.559708 | 0.053662 | 0.065733 | -0.10837 | 0.425328 | 0.150589 | -0.17091 | -0.15799 | 0.255646 | -0.09066 | -0.01239 | 0.287139 | 0.380044 | 0.180093 | 0.681737 | -0.12562 |
| 19.40278 | 0.444997 | 0.308027 | 0.069956 | -0.0067 | 0.306457 | 0.057399 | 0.633991 | 0.023324 | 1.040211 | 0.267144 | 0.18581 | 0.444627 | 1.18757 | 0.470379 | -0.09325 | -0.09277 | 0.606828 | 0.080639 | 0.121576 | 0.519319 | 1.061094 | 0.546 | 0.855787 | 0.012212 |
| 19.41667 | 0.924878 | 0.872826 | 0.218653 | 0.004616 | 0.78865 | 0.253474 | 1.559941 | 0.090838 | 2.299098 | 0.694685 | 0.531069 | 2.353106 | 3.034641 | 0.947659 | -0.03452 | -0.09441 | 1.192954 | 0.299503 | 0.264616 | 0.543467 | 2.215458 | 1.207101 | 1.135784 | 0.145627 |
| 19.43056 | 1.206838 | 1.168807 | 0.34443 | -0.05466 | 1.036047 | 0.452585 | 1.293581 | 0.106147 | 3.432819 | 1.050869 | 1.226247 | 6.328777 | 4.16607 | 1.010487 | -0.08262 | -0.2666 | 1.307901 | 0.305249 | 0.229153 | 0.273524 | 2.253847 | 1.672575 | 1.08089 | 0.099257 |

| | | | | | | | | | | | | | | | | | | | | | | | | |
|---|---|---|---|---|---|---|---|---|---|---|---|---|---|---|---|---|---|---|---|---|---|---|---|---|
| 19.44444 | 0.992885 | 0.923481 | 0.371931 | 0.111105 | 0.904315 | 0.568662 | 1.117371 | 0.114173 | 3.013266 | 1.222256 | 2.628079 | 7.904811 | 2.40585 | 0.44828 | -0.24205 | -0.45605 | 0.72807 | 0.048213 | -0.07708 | 0.110914 | 1.39487 | 1.871463 | 0.48354 | -0.08543 |
| 19.45833 | 0.828623 | 0.662423 | 0.418892 | 0.589909 | 0.76953 | 0.529019 | 0.642353 | 0.138567 | 2.176547 | 1.216293 | 2.840682 | 5.737824 | 1.345424 | 0.209053 | -0.2046 | -0.36736 | 0.296301 | -0.01814 | -0.10422 | 0.393408 | 0.94055 | 1.914889 | 0.1986 | -0.09255 |
| 19.47222 | 0.856467 | 0.593959 | 0.642453 | 1.137731 | 0.81937 | 0.753464 | 0.443386 | 0.158042 | 1.799822 | 1.157879 | 3.041516 | 3.922197 | 1.034424 | 0.305529 | 0.00383 | -0.04359 | 0.226148 | 0.113247 | 0.134474 | 0.641911 | 0.866986 | 1.646271 | 0.144453 | 0.071365 |
| 19.48611 | 0.695713 | 0.563343 | 0.867403 | 1.415374 | 0.833708 | 0.801143 | 0.352665 | 0.125661 | 1.440169 | 0.934811 | 6.051561 | 2.387977 | 0.899553 | 0.371219 | 0.076786 | 0.103794 | 0.115626 | 0.168459 | 0.155732 | 0.348399 | 0.757376 | 1.231012 | 0.071681 | 0.16405 |
| 19.5 | 0.410099 | 0.56267 | 0.944978 | 1.254666 | 0.770179 | 0.797634 | 0.226809 | 0.070837 | 1.258631 | 0.553938 | 4.330519 | 1.57552 | 0.790105 | 0.386754 | 0.06089 | 0.063664 | -0.02294 | 0.125173 | 0.026803 | -0.10443 | 0.58271 | 1.119924 | 0.018938 | 0.204819 |
| 19.51389 | 0.128522 | 0.681324 | 1.023406 | 0.882364 | 0.586281 | 0.756432 | 0.143201 | 0.037302 | 1.111163 | 0.339993 | 2.121178 | 1.063103 | 0.816444 | 0.390396 | 0.065191 | 0.011398 | -0.06476 | 0.095733 | -0.0439 | -0.36716 | 0.523167 | 1.269235 | 0.028212 | 0.252613 |
| 19.52778 | 0.070243 | 0.856072 | 0.853389 | 0.698121 | 0.322409 | 0.612552 | 0.137599 | 0.037908 | 0.876345 | 0.314056 | 2.411204 | 0.600816 | 0.87447 | 0.199057 | 0.07434 | -0.00542 | -0.00787 | 0.093212 | -0.00244 | -0.32892 | 0.561671 | 1.293359 | 0.230963 | 0.277517 |
| 19.54167 | 0.38522 | 0.978109 | 0.682478 | 0.583077 | -0.01942 | 0.620123 | 0.149737 | 0.074347 | 0.537687 | 0.37344 | 3.175691 | 0.325952 | 0.887356 | -0.08004 | 0.063353 | 0.017407 | 0.144902 | 0.118847 | 0.136939 | -0.17553 | 0.651853 | 1.027949 | 0.581443 | 0.274095 |
| 19.55556 | 0.901945 | 1.004342 | 0.544952 | 0.422835 | -0.24027 | 0.690974 | 0.125195 | 0.100215 | 0.271195 | 0.501177 | 3.721768 | 0.154 | 0.763013 | -0.22107 | 0.029747 | 0.022939 | 0.301568 | 0.130787 | 0.224094 | -0.04128 | 0.651692 | 0.572554 | 0.812205 | 0.223661 |
| 19.56944 | 1.120045 | 0.957603 | 0.467113 | 0.35536 | -0.20403 | 0.687536 | 0.113256 | 0.099624 | 0.179384 | 0.612471 | 3.499789 | 0.003835 | 0.546757 | -0.09389 | -0.02468 | 0.007112 | 0.365138 | 0.125025 | 0.162561 | 0.143011 | 0.618419 | 0.14521 | 0.767819 | 0.135123 |
| 19.58333 | 0.715034 | 0.901023 | 0.398605 | 0.322606 | 0.034645 | 0.541212 | 0.137186 | 0.096857 | 0.184006 | 0.627754 | 2.714291 | -0.09754 | 0.316273 | 0.211133 | -0.03595 | 0.004523 | 0.302212 | 0.113134 | 0.13563 | 0.324771 | 0.557474 | -0.0679 | 0.503233 | 0.082613 |
| 19.59722 | 0.187395 | 0.915305 | 0.320325 | 0.281113 | 0.211253 | 0.46327 | 0.157966 | 0.089098 | 0.150026 | 0.409423 | 1.874594 | 0.003354 | 0.210066 | 0.358919 | -0.00687 | 0.016313 | 0.177758 | 0.094665 | 0.09131 | 0.382968 | 0.564459 | -0.11866 | 0.241332 | 0.064144 |
| 19.61111 | -0.23213 | 0.926856 | 0.219543 | 0.238529 | 0.221316 | 0.478517 | 0.139454 | 0.056338 | 0.064753 | 0.102038 | 1.318327 | 0.092854 | 0.238012 | 0.461224 | 0.011093 | 0.021102 | 0.017786 | 0.055784 | -0.01006 | 0.20992 | 0.535004 | -0.12232 | 0.028305 | 0.033224 |
| 19.625 | -0.54542 | 0.865913 | 0.135117 | 0.197298 | 0.191526 | 0.496953 | 0.101865 | 0.022855 | -0.00828 | -0.18357 | 0.990363 | 0.111716 | 0.258392 | 0.213601 | 0.006735 | 0.015274 | -0.11834 | 0.019325 | -0.08488 | -0.14098 | 0.449587 | -0.05266 | -0.15187 | 0.023558 |
| 19.63889 | -0.75807 | 0.739263 | 0.092912 | 0.135512 | 0.208849 | 0.465702 | 0.066702 | 0.019423 | -0.03819 | -0.28874 | 0.799195 | 0.183521 | 0.243326 | -0.07943 | 0.000318 | 0.01069 | -0.2059 | 0.019285 | -0.05747 | -0.52697 | 0.386725 | 0.17639 | -0.31578 | 0.032406 |
| 19.65278 | -0.55087 | 0.624429 | 0.089326 | 0.074054 | 0.190586 | 0.389714 | 0.040155 | 0.025177 | -0.02201 | -0.28556 | 0.631572 | 0.300381 | 0.210309 | -0.1941 | -0.00681 | 0.008126 | -0.23665 | 0.035814 | -0.09449 | -0.68255 | 0.374866 | 0.473991 | -0.42924 | 0.019429 |
| 19.66667 | -0.01964 | 0.551508 | 0.115903 | 0.033025 | 0.115274 | 0.298302 | 0.043316 | 0.033866 | -0.01272 | -0.05404 | 0.469377 | 0.311793 | 0.174227 | -0.21272 | -0.01691 | 0.004714 | -0.15239 | 0.0376 | 0.028763 | -0.55725 | 0.382699 | 0.662994 | -0.35673 | -0.00565 |
| 19.68056 | 0.523429 | 0.497625 | 0.150172 | 0.041102 | 0.035301 | 0.209459 | 0.088035 | 0.038522 | 0.010556 | 0.314131 | 0.369376 | 0.270312 | 0.148626 | -0.16528 | -0.01398 | 0.010934 | -0.01577 | 0.041422 | 0.154962 | -0.30156 | 0.375355 | 0.717908 | -0.18741 | -0.01624 |
| 19.69444 | 0.658103 | 0.429482 | 0.16529 | 0.0711 | -0.02227 | 0.146331 | 0.130017 | 0.037178 | 0.023771 | 0.525615 | 0.350494 | 0.237211 | 0.139487 | -0.09026 | -0.00107 | 0.012453 | 0.056108 | 0.031145 | 0.122832 | -0.00843 | 0.362744 | 0.722904 | -0.05059 | -0.02255 |
| 19.70833 | 0.410594 | 0.279112 | 0.17272 | 0.063811 | -0.05353 | 0.135863 | 0.131611 | 0.031534 | -0.04023 | 0.514819 | 0.314429 | 0.187716 | 0.114471 | 0.084942 | 0.009913 | -0.00255 | 0.059387 | -0.002 | 0.078014 | 0.24965 | 0.343072 | 0.541524 | 0.008192 | -0.04679 |
| 19.72222 | 0.116885 | 0.178737 | 0.148113 | 0.059249 | -0.0525 | 0.143958 | 0.118308 | 0.016696 | -0.16234 | 0.349763 | 0.263734 | 0.068934 | 0.082812 | 0.206388 | 0.018135 | -0.02773 | 0.035426 | -0.03306 | 0.08838 | 0.378835 | 0.304286 | 0.198298 | -0.03896 | -0.05723 |
| 19.73611 | -0.18485 | 0.144902 | 0.103023 | 0.078682 | -0.03716 | 0.115891 | 0.103387 | 0.001437 | -0.30793 | 0.091489 | 0.223132 | -0.10247 | 0.086914 | 0.129772 | 0.01598 | -0.04609 | -0.01032 | -0.05145 | 0.001899 | 0.321606 | 0.243586 | -0.04032 | -0.14538 | -0.04737 |
| 19.75 | -0.45785 | 0.073761 | 0.067493 | 0.071197 | -0.00784 | 0.010332 | 0.055907 | 0.0111 | -0.42289 | -0.25737 | 0.227932 | -0.30761 | 0.12228 | -0.02187 | 0.012786 | -0.03439 | -0.10342 | -0.05263 | -0.03929 | 0.09827 | 0.168029 | -0.09717 | -0.32655 | -0.04548 |
| 19.76389 | -0.47203 | -0.02666 | 0.044965 | 0.018576 | 0.051477 | -0.1313 | -0.01538 | 0.02666 | -0.49982 | -0.57458 | 0.180902 | -0.60902 | 0.124212 | -0.03418 | 0.00681 | -0.02072 | -0.18017 | -0.0487 | 0.067944 | -0.17781 | 0.073618 | -0.04571 | -0.48287 | -0.05829 |
| 19.77778 | -0.22057 | -0.14096 | 0.048406 | -0.0336 | 0.088902 | -0.24591 | -0.09659 | 0.025142 | -0.5515 | -0.75438 | 0.110715 | -1.04238 | 0.054584 | -0.11482 | -0.00829 | -0.02927 | -0.22016 | -0.04323 | -0.05989 | -0.41384 | 0.003968 | -0.08771 | -0.66059 | -0.09168 |
| 19.79167 | 0.0213 | -0.21926 | 0.058848 | -0.04607 | 0.045583 | -0.30161 | -0.22167 | 0.013069 | -0.6087 | -0.68184 | 0.051676 | -1.349 | -0.07888 | -0.20796 | -0.02859 | -0.04874 | -0.1943 | -0.064 | -0.27262 | -0.60858 | -0.0238 | -0.20843 | -0.76289 | -0.14902 |
| 19.80556 | -0.01035 | -0.37791 | -0.01131 | -0.03961 | -0.11311 | -0.26402 | -0.26623 | -0.00567 | -0.70218 | -0.3849 | -0.00041 | -1.3189 | -0.27889 | -0.3642 | -0.03828 | -0.05922 | -0.20513 | -0.09051 | -0.32322 | -0.71913 | -0.03921 | -0.33893 | -0.91554 | -0.22714 |
| 19.81944 | -0.33136 | -0.61103 | -0.1743 | -0.02767 | -0.34437 | -0.18514 | -0.25772 | -0.0224 | -0.87501 | -0.20059 | -0.05706 | -1.22455 | -0.53128 | -0.47775 | -0.05001 | -0.06962 | -0.39943 | -0.10752 | -0.20389 | -0.79588 | -0.09375 | -0.58105 | -0.94862 | -0.3432 |
| 19.83333 | -0.82456 | -0.83542 | -0.37451 | 0.006235 | -0.55253 | -0.13348 | -0.28861 | -0.0251 | -1.01726 | -0.20481 | -0.08465 | -1.2784 | -0.77098 | -0.28375 | -0.07586 | -0.0954 | -0.72867 | -0.12927 | -0.0147 | -0.74838 | -0.19267 | -1.10012 | -1.00706 | -0.48581 |
| 19.84722 | -1.28658 | -0.97754 | -0.56477 | 0.03625 | -0.75192 | -0.12899 | -0.46338 | -0.04485 | -0.93016 | -0.41186 | -0.07661 | -1.4661 | -0.96343 | -0.16541 | -0.09104 | -0.11276 | -1.11706 | -0.1506 | 0.127694 | -0.54624 | -0.30128 | -1.97249 | -1.17239 | -0.61103 |
| 19.86111 | -1.37562 | -1.08827 | -0.60021 | 0.027738 | -0.76293 | -0.12595 | -0.69997 | -0.07118 | -0.74861 | -0.58491 | -0.07707 | -1.822 | -1.06242 | -0.11578 | -0.06033 | -0.09705 | -1.36253 | -0.17096 | 0.197441 | -0.36131 | -0.3595 | -2.85124 | -1.27304 | -0.553 |
| 19.875 | -1.16214 | -1.03643 | -0.71429 | -0.01688 | -0.46117 | -0.12358 | -0.83418 | -0.07388 | -0.63498 | -0.78558 | -0.1074 | -2.06409 | -1.0708 | -0.13919 | -0.03263 | -0.07692 | -1.4916 | -0.18488 | 0.177065 | -0.20408 | -0.38743 | -3.17854 | -1.12016 | -0.50016 |
| 19.88889 | -0.95927 | -0.90046 | -0.8058 | -0.06536 | -0.35796 | -0.1092 | -0.77736 | -0.06484 | -0.60503 | -0.89595 | -0.17565 | -1.83126 | -1.03008 | -0.21876 | -0.04271 | -0.07618 | -1.32713 | -0.1945 | 0.032167 | -0.16439 | -0.4517 | -2.85095 | -0.92209 | -0.44793 |
| 19.90278 | -0.79598 | -0.71852 | -0.75831 | -0.08497 | -0.30663 | -0.15471 | -0.69181 | -0.04412 | -0.57444 | -0.96762 | -0.32905 | -1.38047 | -0.97542 | -0.20924 | -0.01796 | -0.01836 | -0.9832 | -0.17716 | -0.08181 | -0.16804 | -0.59432 | -2.31445 | -0.66686 | -0.39605 |
| 19.91667 | -0.67382 | -0.66615 | -0.72671 | -0.06812 | -0.23455 | -0.36194 | -0.69109 | -0.02537 | -0.52117 | -1.37707 | -0.61701 | -1.46824 | -1.04935 | -0.12646 | 0.098923 | 0.135368 | -0.55279 | -0.13159 | -0.0742 | -0.16216 | -1.07051 | -1.85971 | -0.44014 | -0.28787 |
| 19.93056 | -0.50966 | -0.61418 | -0.86283 | -0.01278 | -0.11856 | -1.11784 | -0.74762 | 0.053787 | -0.49604 | -2.3936 | -1.26457 | -2.10323 | -1.28783 | -0.00688 | 0.283704 | 0.354985 | -0.27236 | -0.1248 | 0.033409 | -0.17575 | -2.2387 | -1.38679 | -0.29725 | -0.23369 |
| 19.94444 | -0.45522 | -0.53496 | -1.04357 | -0.02823 | 0.008821 | -1.4641 | -0.81184 | 0.005618 | -0.51913 | -2.26743 | -3.05861 | -3.34587 | -1.88993 | 0.118803 | 0.452775 | 0.536835 | -0.09027 | -0.20289 | 0.146203 | 0.071456 | -2.66999 | -1.21868 | -0.29502 | -0.49491 |
| 19.95833 | -0.48899 | -0.43046 | -0.80182 | -0.36409 | -0.03654 | -0.72481 | -0.64194 | -0.33953 | -0.53668 | -0.99526 | -3.32166 | -2.39412 | -2.01602 | 0.024642 | 0.323026 | 0.388788 | -0.19141 | -0.05965 | 0.098046 | -0.43804 | -3.37679 | -1.15177 | -0.52474 | -1.44391 |
| 19.97222 | -0.46753 | -0.26081 | -0.51942 | -0.39293 | -0.15046 | -0.40823 | -0.15747 | -0.2558 | -0.45632 | 0.248308 | -1.47577 | 0.14164 | -1.06597 | -0.15768 | -0.01887 | 0.023317 | -0.28932 | -0.01427 | -0.12168 | -0.74242 | 0.374138 | -0.9827 | -0.78043 | -1.25128 |
| 19.98611 | -0.32274 | -0.12867 | -0.32313 | -0.24067 | -0.1564 | -0.21742 | 0.114711 | -0.17141 | -0.2909 | 0.83356 | -0.16577 | 1.074485 | -0.02251 | -0.11919 | -0.1682 | -0.1431 | -0.07309 | 0.057676 | -0.12908 | -0.24031 | 2.388235 | -0.83435 | -0.7287 | -0.65268 |
| 20 | -0.07046 | -0.09948 | -0.2259 | -0.16012 | -0.1212 | -0.13876 | 0.181946 | -0.10275 | -0.0788 | 0.524651 | 0.185598 | 0.943597 | 0.460569 | 0.01742 | -0.13968 | -0.13313 | 0.335035 | 0.098574 | 0.034893 | 0.367502 | 1.838343 | -0.65155 | -0.46193 | -0.28898 |
| 20.01389 | 0.20603 | -0.08021 | -0.15861 | -0.111 | -0.10942 | -0.10282 | 0.143143 | -0.06637 | 0.218016 | 0.16634 | 0.141606 | 0.689477 | 0.432466 | 0.037143 | -0.09751 | -0.11476 | 0.614245 | 0.024099 | 0.178211 | 0.514308 | 0.820768 | -0.45749 | -0.18552 | -0.15589 |
| 20.02778 | 0.422268 | -0.03194 | -0.1155 | -0.08944 | -0.12894 | -0.07746 | 0.014638 | -0.06151 | 0.497492 | -0.0119 | 0.028673 | 0.493264 | 0.282971 | -0.03802 | -0.07192 | -0.09168 | 0.737854 | -0.04179 | 0.191545 | 0.457034 | 0.291011 | -0.309 | -0.00625 | -0.11111 |
| 20.04167 | 0.557633 | 0.007534 | -0.09089 | -0.07971 | -0.12531 | -0.05305 | -0.07604 | -0.07413 | 0.591681 | -0.10879 | -0.04369 | 0.318286 | 0.16505 | -0.10805 | -0.05458 | -0.06678 | 0.769548 | -0.05692 | 0.115463 | 0.414937 | 0.003914 | -0.23897 | 0.065596 | -0.07652 |
| 20.05556 | 0.563036 | 0.028137 | -0.06773 | -0.06499 | -0.07966 | -0.04131 | -0.05991 | -0.05907 | 0.456262 | -0.15291 | -0.05715 | 0.228203 | 0.109905 | -0.1026 | -0.04111 | -0.04409 | 0.639742 | -0.04178 | 0.059361 | 0.4058 | -0.03608 | -0.17611 | 0.078511 | -0.05753 |
| 20.06944 | 0.529216 | 0.02336 | -0.03921 | -0.04411 | -0.01554 | -0.03701 | 0.016114 | -0.01427 | 0.332575 | -0.18521 | -0.01816 | 0.23703 | 0.06643 | -0.02697 | -0.03964 | -0.02908 | 0.423266 | -0.02049 | 0.07084 | 0.393631 | 0.029841 | -0.11828 | 0.145692 | -0.04036 |
| 20.08333 | 0.479289 | 0.007553 | 0.013793 | -0.01184 | 0.030022 | -0.02904 | 0.028892 | 0.014554 | 0.372517 | -0.22425 | 0.034236 | 0.254956 | -0.02734 | -0.70665 | -0.06592 | -0.14844 | 0.203728 | -0.00887 | 0.111544 | 0.362497 | 0.096816 | -0.0957 | 0.18054 | -0.1068 |
| 20.09722 | 0.427086 | 0.006426 | 0.098637 | 0.010022 | 0.029843 | -0.01435 | -0.20114 | 0.015503 | 0.322149 | -0.28994 | -0.04973 | 0.012175 | -0.34949 | -5.24834 | -0.16157 | -0.97384 | -0.48867 | 0.003262 | 0.150251 | 0.394163 | 0.195688 | -0.07626 | 0.101504 | -0.61868 |
| 20.11111 | 0.40232 | 0.040322 | 0.157923 | 0.031002 | 0.010412 | -0.00815 | -0.37325 | 0.00482 | 0.222925 | -0.27672 | -0.10684 | -0.17465 | -0.75932 | -7.28997 | 0.052703 | -1.62446 | -0.78533 | 0.096597 | 0.127922 | 0.321237 | 0.25083 | -0.36854 | 0.380941 | -0.7943 |
| 20.125 | 0.377496 | 0.113368 | 0.131951 | 0.092054 | 0.04895 | -0.02262 | 0.098152 | 0.010716 | 0.670415 | 0.020373 | 0.190318 | 0.372247 | 0.054595 | 0.092899 | 0.009953 | 0.010714 | 0.055682 | -0.02658 | 0.04836 | 0.078229 | 0.168743 | -0.01437 | 0.380315 | -0.03745 |
| 20.13889 | 0.341437 | 0.214732 | 0.082458 | 0.114449 | 0.094489 | -0.0326 | 0.51842 | 0.008972 | 1.071842 | 0.27737 | 0.54366 | 0.950066 | 1.013282 | 7.289759 | 0.022766 | 1.751746 | 1.047516 | -0.07725 | 0.03829 | -0.09132 | 0.229721 | 0.260269 | 0.458619 | 0.815217 |
| 20.15278 | 0.232581 | 0.224525 | 0.092996 | 0.052699 | 0.098108 | -0.01523 | 0.266166 | -0.00869 | 0.8305 | 0.160026 | 0.524356 | 0.874378 | 0.718209 | 5.400032 | 0.133508 | 1.043148 | 0.934466 | 0.07658 | 0.085555 | -0.00922 | 0.451823 | 0.003507 | 0.781592 | 0.595466 |
| 20.16667 | 0.09098 | 0.17763 | 0.163522 | 0.053554 | 0.101921 | 0.007747 | 0.031163 | 0.045056 | 0.557382 | 0.004487 | 0.361026 | 0.666843 | 0.438545 | 1.081706 | 0.085286 | 0.166979 | 0.472301 | 0.086842 | 0.005877 | 0.064081 | 0.391934 | 0.093935 | 0.545604 | 0.103554 |
| 20.18056 | 0.019125 | 0.146561 | 0.220719 | 0.065765 | 0.131504 | 0.020264 | 0.049016 | 0.106542 | 0.525033 | 0.025435 | 0.231939 | 0.55248 | 0.319359 | 0.404664 | 0.036375 | 0.034515 | 0.358254 | 0.068903 | -0.02247 | 0.02719 | 0.270722 | 0.07818 | 0.372106 | 0.007928 |
| 20.19444 | 0.019084 | 0.163148 | 0.230682 | 0.055849 | 0.129071 | 0.018413 | 0.12311 | 0.102656 | 0.58115 | 0.073107 | 0.177918 | 0.413932 | 0.252734 | 0.359776 | 0.031461 | 0.031481 | 0.401784 | 0.057225 | 0.018952 | 0.024627 | 0.135766 | 0.1082 | 0.319068 | 0.002597 |
| 20.20833 | 0.11596 | 0.200546 | 0.209611 | 0.038113 | 0.133357 | 0.01233 | 0.215649 | 0.07166 | 0.767861 | 0.165687 | 0.220861 | 0.434357 | 0.195506 | 0.157606 | 0.028012 | 0.015327 | 0.60012 | 0.039313 | 0.091023 | 0.105807 | 0.061416 | 0.190272 | 0.31743 | -0.00975 |
| 20.22222 | 0.310746 | 0.224778 | 0.205036 | 0.028897 | 0.188816 | 0.012514 | 0.296676 | 0.066173 | 0.983976 | 0.273218 | 0.31408 | 0.572001 | 0.161184 | -0.04975 | 0.026231 | -0.00614 | 0.759392 | 0.018999 | 0.154766 | 0.200915 | 0.043072 | 0.271782 | 0.315894 | -0.02114 |
| 20.23611 | 0.524236 | 0.222613 | 0.223167 | 0.034532 | 0.252648 | 0.022062 | 0.363044 | 0.08727 | 1.00675 | 0.341126 | 0.621941 | 0.689375 | 0.159533 | -0.05225 | 0.044509 | -0.00674 | 0.835273 | 0.019421 | 0.167332 | 0.237259 | 0.10283 | 0.328399 | 0.435104 | -0.00934 |
| 20.25 | 0.629139 | 0.20884 | 0.253831 | 0.04821 | 0.257029 | 0.032637 | 0.440002 | 0.106005 | 0.924381 | 0.371358 | 1.189307 | 0.731977 | 0.204676 | 0.271427 | 0.063029 | 0.014894 | 0.966056 | 0.044387 | 0.132429 | 0.22686 | 0.229489 | 0.320393 | 0.644719 | 0.006438 |
| 20.26389 | 0.606762 | 0.22747 | 0.280742 | 0.053865 | 0.216649 | 0.047502 | 0.577725 | 0.101255 | 0.769295 | 0.36419 | 1.450118 | 0.772145 | 0.267167 | 0.594833 | 0.053624 | 0.041385 | 1.157463 | 0.064339 | 0.125938 | 0.19793 | 0.329903 | 0.254406 | 0.850725 | -0.00406 |
| 20.27778 | 0.577181 | 0.279999 | 0.305233 | 0.035012 | 0.178741 | 0.062278 | 0.685026 | 0.098601 | 0.674528 | 0.362032 | 1.317661 | 0.818953 | 0.313169 | 0.438624 | 0.039565 | 0.052122 | 1.244487 | 0.06677 | 0.177188 | 0.195119 | 0.317288 | 0.227762 | 0.727063 | -0.01813 |
| 20.29167 | 0.548993 | 0.35702 | 0.311917 | 0.033502 | 0.183985 | 0.089287 | 0.670904 | 0.095085 | 0.775157 | 0.386993 | 1.155173 | 0.907766 | 0.341095 | 0.185131 | 0.030099 | 0.035095 | 1.119161 | 0.065792 | 0.249896 | 0.285318 | 0.260895 | 0.359422 | 0.543136 | -0.0128 |
| 20.30556 | 0.531679 | 0.396342 | 0.322851 | 0.068475 | 0.236753 | 0.119104 | 0.646759 | 0.105998 | 1.028001 | 0.412542 | 1.074054 | 0.999651 | 0.334118 | -0.01571 | 0.027393 | 0.016303 | 0.820323 | 0.065011 | 0.290456 | 0.442003 | 0.248808 | 0.565413 | 0.532512 | -0.00339 |
| 20.31944 | 0.471385 | 0.37092 | 0.322152 | 0.136075 | 0.284783 | 0.121558 | 0.566308 | 0.092386 | 1.208227 | 0.463703 | 0.940184 | 1.071749 | 0.300167 | -0.04812 | 0.028866 | 0.018361 | 0.508444 | 0.056927 | 0.299345 | 0.492899 | 0.259022 | 0.722832 | 0.867639 | 0.003296 |
| 20.33333 | 0.382089 | 0.338805 | 0.397241 | 0.203905 | 0.29262 | 0.118695 | 0.580905 | 0.084788 | 1.247617 | 0.593501 | 0.926754 | 1.167621 | 0.283125 | 0.158903 | 0.026015 | 0.034478 | 0.425927 | 0.050185 | 0.276826 | 0.435384 | 0.303953 | 0.825569 | 1.253886 | 0.015836 |
| 20.34722 | 0.225303 | 0.37068 | 0.464051 | 0.248945 | 0.326035 | 0.158361 | 0.630349 | 0.091992 | 1.224699 | 0.743943 | 0.992536 | 1.275038 | 0.354071 | 0.363511 | 0.025942 | 0.050605 | 0.607559 | 0.058856 | 0.228833 | 0.427398 | 0.378486 | 0.878637 | 1.340042 | 0.043189 |
| 20.36111 | 0.091076 | 0.420583 | 0.459679 | 0.280554 | 0.312567 | 0.240367 | 0.685533 | 0.117588 | 1.115996 | 0.779168 | 0.9488 | 1.310263 | 0.426975 | 0.30636 | -0.01022 | 0.033179 | 0.655936 | 0.035244 | 0.159984 | 0.417133 | 0.36351 | 0.840134 | 0.987245 | 0.044566 |
| 20.375 | -0.00155 | 0.390088 | 0.361359 | 0.209591 | 0.240249 | 0.334556 | 0.580142 | 0.112647 | 0.925135 | 0.661615 | 0.826742 | 1.252102 | 0.41227 | 0.13769 | -0.10096 | -0.03617 | 0.395596 | -0.03772 | 0.039094 | 0.325091 | 0.23945 | 0.779755 | 0.461018 | 0.013881 |
| 20.38889 | 0.030834 | 0.45316 | 0.311762 | 0.120933 | 0.268745 | 0.353384 | 0.335155 | 0.07895 | 0.736333 | 0.402777 | 0.863172 | 1.261079 | 0.536931 | 0.072084 | -0.10632 | -0.04489 | 0.143448 | -0.01047 | 0.015298 | 0.254487 | 0.275832 | 0.902583 | 0.123523 | 0.061144 |
| 20.40278 | 0.291817 | 0.972345 | 0.486412 | 0.171823 | 0.705851 | 0.558854 | 0.138754 | 0.088475 | 0.749574 | 0.431131 | 1.136616 | 1.918885 | 1.331276 | 0.066398 | 0.013307 | 0.018951 | 0.3074 | 0.162902 | 0.146352 | 0.213727 | 0.770113 | 1.472455 | 0.140052 | 0.227117 |
| 20.41667 | 0.798111 | 1.973677 | 1.011831 | 0.321494 | 1.593638 | 1.034154 | 0.119227 | 0.157985 | 1.037074 | 1.428865 | 1.698576 | 3.341955 | 3.20212 | -0.00421 | 0.105373 | 0.040244 | 0.751984 | 0.306789 | 0.27531 | 0.207556 | 1.5957 | 2.338161 | 0.529103 | 0.424856 |
| 20.43056 | 1.692166 | 2.62391 | 1.437638 | 0.425133 | 1.605276 | 1.584704 | 0.2599 | 0.191298 | 1.145583 | 2.589799 | 2.110723 | 3.779788 | 3.624666 | -0.11578 | 0.038002 | -0.07772 | 1.021265 | 0.224905 | 0.199375 | 0.274224 | 2.032792 | 2.93205 | 0.806798 | 0.513099 |
| 20.44444 | 1.751072 | 2.446343 | 1.40754 | 0.405122 | 1.048661 | 1.583404 | 0.289885 | 0.168137 | 0.985261 | 2.901271 | 1.991536 | 2.564935 | 1.721609 | -0.17379 | -0.17226 | -0.30002 | 0.842908 | 0.011918 | -0.1236 | 0.230191 | 1.153884 | 3.242165 | 0.439338 | 0.524939 |
| 20.45833 | 0.982936 | 2.393358 | 1.267787 | 0.316774 | 0.880779 | 0.948767 | 0.08403 | 0.179775 | 0.895285 | 2.269593 | 1.708142 | 1.512333 | 1.072411 | -0.09353 | -0.15149 | -0.23103 | 0.617335 | 0.032589 | -0.11549 | 0.477417 | 0.745588 | 3.520503 | 0.193287 | 0.564533 |
| 20.47222 | 0.484232 | 2.71741 | 1.159931 | 0.250434 | 1.317171 | 0.616263 | -0.0883 | 0.168944 | 1.081729 | 1.43569 | 1.389185 | 0.860821 | 1.281653 | 0.116314 | 0.075461 | 0.055574 | 0.604782 | 0.241991 | 0.124693 | 0.700509 | 0.801943 | 3.053651 | 0.368567 | 0.538659 |
| 20.48611 | 0.000378 | 2.825467 | 0.879398 | 0.200664 | 2.805549 | 0.396035 | -0.27646 | 0.105995 | 0.895693 | 0.251179 | 0.966861 | 0.262326 | 1.431796 | 0.104436 | 0.113285 | 0.139031 | 0.297224 | 0.244539 | 0.16854 | 0.493549 | 0.745966 | 2.203981 | 0.415726 | 0.297048 |
| 20.5 | -0.51288 | 2.246307 | 0.493527 | 0.286118 | 2.065981 | 0.219676 | -0.36331 | 0.053753 | 0.341131 | -0.43033 | 0.65371 | -0.65091 | 1.400734 | 0.028101 | 0.01589 | 0.041288 | -0.23364 | 0.118138 | 0.037866 | 0.162374 | 0.594822 | 1.428649 | 0.221684 | 0.089458 |
| 20.51389 | -0.97507 | 1.446074 | 0.239217 | 0.44501 | 0.785822 | 0.094968 | -0.28493 | 0.049424 | -0.07461 | -0.10879 | 0.46094 | -1.45343 | 1.330127 | 0.024463 | 0.033651 | -0.00159 | -0.77648 | 0.100399 | 0.072926 | -0.08206 | 0.606167 | 0.936623 | -0.08409 | 0.079233 |
| 20.52778 | -0.88146 | 0.839947 | 0.08476 | 0.563204 | 0.207152 | 0.215847 | -0.00705 | 0.028979 | -0.36027 | 0.572735 | 0.294088 | -1.60671 | 1.163174 | 0.088608 | 0.147316 | 0.039812 | -1.21071 | 0.219671 | 0.252299 | -0.26509 | 0.69011 | 0.612075 | -0.29256 | 0.163709 |
| 20.54167 | -0.08909 | 0.505963 | 0.131101 | 0.562253 | -0.13552 | 0.480984 | 0.285899 | 0.039548 | -0.55096 | 1.226991 | 0.136555 | -1.03829 | 0.760746 | 0.115701 | 0.183803 | 0.060458 | -1.1842 | 0.28686 | 0.204901 | -0.40514 | 0.690997 | 0.05025 | -0.31151 | 0.148013 |
| 20.55556 | 0.891652 | 0.328093 | 0.33567 | 0.539433 | -0.40247 | 0.820139 | 0.420195 | 0.058003 | -0.60682 | 1.571946 | -0.06976 | -0.27518 | 0.121557 | 0.093892 | 0.127305 | 0.057986 | -0.11742 | 0.26172 | 0.112767 | -0.30334 | 0.629979 | -0.87078 | -0.44575 | 0.11507 |
| 20.56944 | 1.739997 | 0.198042 | 0.50319 | 0.415024 | -0.34544 | 0.845671 | 0.261191 | 0.078437 | -0.54579 | 1.416352 | -0.21397 | -0.04605 | -0.55914 | 0.073167 | 0.10717 | 0.102018 | 1.065698 | 0.241733 | 0.16404 | 0.073285 | 0.623719 | -1.70888 | -0.47533 | 0.175592 |
| 20.58333 | 1.593293 | 0.143932 | 0.492827 | 0.265786 | -0.07644 | 0.412687 | -0.0094 | 0.060673 | -0.29415 | 0.565229 | -0.23594 | -0.19311 | -0.93342 | 0.06359 | 0.122985 | 0.144252 | 1.701769 | 0.227876 | 0.166214 | 0.758016 | 0.60159 | -2.00681 | 0.188302 | 0.216543 |
| 20.59722 | 1.160418 | 0.102483 | 0.404499 | 0.174779 | 0.492899 | 0.007526 | -0.22023 | 0.037139 | 0.118516 | -0.39025 | -0.12996 | -0.24279 | -0.74698 | 0.063106 | 0.124064 | 0.142666 | 1.869109 | 0.153914 | -0.01211 | 1.126912 | 0.561213 | -1.58064 | 1.086416 | 0.177376 |
| 20.61111 | 0.482892 | 0.03505 | 0.317352 | 0.163264 | 1.1056 | -0.22681 | -0.40336 | 0.019089 | 0.514684 | -1.2042 | 0.000996 | -0.04448 | -0.19116 | 0.025333 | 0.073985 | 0.115246 | 1.524222 | 0.047148 | -0.04408 | 0.993566 | 0.534061 | -0.85362 | 1.509587 | 0.03473 |
| 20.625 | -0.2188 | -0.13889 | 0.225503 | 0.23145 | 1.217806 | -0.28113 | -0.43147 | -0.00203 | 0.652381 | -1.10368 | 0.177071 | 0.023041 | 0.346859 | -0.05126 | -0.00789 | 0.040621 | 0.628729 | -0.00806 | -0.02226 | 0.61266 | 0.466109 | -0.18358 | 1.116467 | -0.15143 |
| 20.63889 | -0.77334 | -0.334 | 0.216299 | 0.315168 | 0.845188 | -0.29411 | -0.287 | -0.00994 | 0.510067 | -0.42223 | 0.375626 | 0.004921 | 0.637698 | -0.13104 | -0.03987 | -0.01522 | -0.27947 | 0.070186 | -0.05789 | 0.208437 | 0.375173 | 0.425851 | 0.416176 | -0.26811 |
| 20.65278 | -1.18702 | -0.48246 | 0.196954 | 0.333106 | 0.269328 | -0.32798 | -0.19648 | 0.009697 | 0.148939 | 0.216694 | 0.735504 | 0.060921 | 0.725672 | -0.13329 | -0.04482 | -0.0471 | -1.34288 | 0.14087 | -0.06222 | -0.16628 | 0.252678 | 0.822974 | -0.1363 | -0.28424 |
| 20.66667 | -1.63626 | -0.54636 | 0.089444 | 0.288906 | -0.29656 | -0.15242 | -0.03041 | -0.00249 | -0.23083 | 0.530685 | 0.864224 | 0.290639 | 0.568817 | -0.05373 | -0.02623 | -0.10482 | -2.16374 | 0.110377 | -0.07195 | -0.52649 | 0.13257 | 0.921352 | -0.471 | -0.29016 |
| 20.68056 | -1.20977 | -0.44913 | -0.05301 | 0.224201 | -0.50108 | 0.177938 | 0.208537 | -0.00861 | -0.50273 | 0.525298 | 0.658524 | 0.413184 | 0.08585 | 0.024646 | 0.02235 | -0.10097 | -2.00907 | 0.043222 | -0.02707 | -0.79913 | 0.095853 | 0.808361 | -0.66641 | -0.22564 |
| 20.69444 | -0.09319 | -0.27966 | -0.10534 | 0.200476 | -0.44093 | 0.682464 | 0.485399 | 0.028028 | -0.5183 | 0.246783 | 0.348746 | 0.162568 | -0.34837 | 0.043083 | 0.075569 | -0.00868 | -0.96818 | 0.011995 | -0.00552 | -0.8362 | 0.133802 | 0.528609 | -0.91545 | -0.16246 |
| 20.70833 | 0.815305 | -0.15 | -0.12393 | 0.193357 | -0.29934 | 0.882417 | 0.375197 | 0.039606 | -0.28179 | -0.43275 | 0.078986 | -0.26284 | -0.45578 | 0.05824 | 0.088781 | 0.097613 | 0.235327 | -0.03205 | -0.03037 | -0.48753 | 0.149316 | -0.14206 | -1.22299 | -0.11968 |
| 20.72222 | 1.670718 | -0.07675 | -0.14159 | 0.158262 | -0.35189 | 0.573125 | 0.070214 | 0.015797 | 0.067725 | -1.31697 | -0.0312 | -0.68659 | -0.33918 | -0.02616 | 0.048685 | 0.132571 | 1.327919 | -0.07258 | -0.00197 | -0.10716 | 0.063289 | -0.93549 | -1.16589 | -0.09071 |
| 20.73611 | 1.56841 | -0.03056 | -0.11676 | 0.100753 | -0.35792 | 0.251052 | -0.15485 | -0.00892 | 0.264491 | -1.48795 | -0.17928 | -0.98028 | 0.058833 | -0.14311 | -0.02699 | 0.084609 | 1.852423 | -0.05581 | 0.080384 | 0.22849 | -0.03269 | -1.55233 | -0.80031 | -0.06085 |
| 20.75 | 0.866921 | -0.03694 | -0.09837 | 0.064226 | -0.31873 | -0.01764 | -0.26264 | -0.04465 | 0.188203 | -0.81526 | -0.42025 | -0.89215 | 0.326163 | -0.31459 | -0.09687 | -0.02609 | 1.437296 | -0.00963 | 0.097912 | 0.482368 | -0.01921 | -1.74188 | -0.32889 | -0.08622 |
| 20.76389 | 0.211162 | -0.08563 | -0.07592 | 0.058124 | -0.19018 | -0.22167 | -0.22492 | -0.07437 | -0.04377 | -0.09949 | -0.44011 | -0.46201 | 0.209209 | -0.49773 | -0.10888 | -0.10965 | 0.728915 | 0.000861 | 0.051009 | 0.431962 | 0.005951 | -1.54623 | -0.13033 | -0.1576 |
| 20.77778 | -0.42233 | -0.1707 | -0.10666 | 0.025674 | -0.08376 | -0.32566 | -0.18112 | -0.01692 | -0.1972 | 0.267422 | -0.34539 | -0.13898 | -0.27543 | -0.41542 | -0.06059 | -0.1333 | -0.03015 | -0.04728 | -0.02727 | 0.214397 | -0.02162 | -1.33146 | -0.09631 | -0.25334 |
| 20.79167 | -1.06872 | -0.32005 | -0.18816 | -0.07794 | -0.10332 | -0.41126 | -0.15315 | 0.020899 | -0.18182 | 0.099217 | -0.21765 | -0.25116 | -0.65974 | -0.20296 | -0.01155 | -0.10451 | -0.82905 | -0.09655 | -0.07048 | -0.02979 | -0.07202 | -1.18951 | -0.17745 | -0.33289 |
| 20.80556 | -1.72574 | -0.45045 | -0.3237 | -0.20179 | -0.21048 | -0.47442 | -0.2042 | 0.03301 | -0.02292 | -0.35157 | -0.17246 | -0.6391 | -0.85137 | 0.013711 | -0.00138 | -0.0873 | -1.57825 | -0.14463 | -0.11584 | -0.29621 | -0.13625 | -1.26482 | -0.34296 | -0.39185 |
| 20.81944 | -2.1388 | -0.52231 | -0.48371 | -0.22383 | -0.303 | -0.38242 | -0.27052 | 0.02128 | 0.091257 | -0.86755 | -0.2316 | -0.91844 | -0.91587 | 0.050234 | -0.02285 | -0.08175 | -1.84253 | -0.1368 | -0.19717 | -0.49208 | -0.18508 | -1.62433 | -0.4367 | -0.43712 |
| 20.83333 | -1.85623 | -0.53259 | -0.52326 | -0.13515 | -0.30469 | -0.20813 | -0.3018 | -0.03082 | -0.05477 | -1.16515 | -0.33757 | -0.95071 | -1.09333 | -0.04911 | -0.04831 | -0.02453 | -1.54081 | -0.09266 | -0.26955 | -0.6001 | -0.20753 | -2.15039 | -0.34367 | -0.39217 |
| 20.84722 | -1.56329 | -0.71891 | -0.50644 | -0.05419 | -0.32647 | -0.09813 | -0.32556 | -0.12936 | -0.44161 | -1.223 | -0.42003 | -0.8228 | -1.12564 | -0.15231 | -0.06605 | 0.043918 | -1.10075 | -0.02833 | -0.38539 | -0.5975 | -0.18967 | -2.56444 | -0.23052 | -0.36684 |
| 20.86111 | -0.9185 | -1.01548 | -0.62251 | -0.0568 | -0.33332 | -0.05315 | -0.35303 | -0.25607 | -0.88538 | -0.65106 | -0.56396 | -0.68176 | -1.24898 | -0.25785 | -0.10536 | 0.065786 | -0.81332 | -0.01383 | -0.45715 | -0.59789 | -0.19256 | -2.87169 | -0.33791 | -0.37395 |
| 20.875 | -0.20408 | -1.23464 | -0.81223 | -0.09024 | -0.38497 | -0.05425 | -0.35864 | -0.32444 | -1.25796 | 0.134122 | -0.78335 | -0.61278 | -1.40591 | -0.39912 | -0.20634 | 0.014702 | -0.66056 | -0.07144 | -0.53932 | -0.53528 | -0.25723 | -2.98796 | -0.68697 | -0.41277 |
| 20.88889 | 0.397468 | -1.20938 | -0.99151 | -0.06454 | -0.34884 | -0.07263 | -0.30177 | -0.20921 | -1.36081 | 0.752935 | -0.90263 | -0.53667 | -1.39109 | -0.37115 | -0.28466 | -0.06793 | -0.65195 | -0.12872 | -0.5226 | -0.37158 | -0.36372 | -2.84249 | -1.03112 | -0.43318 |
| 20.90278 | 0.585997 | -0.99177 | -1.02502 | -0.04549 | -0.30892 | -0.11429 | -0.2691 | -0.03944 | -1.10343 | 0.79834 | -0.94116 | -0.56914 | -1.07002 | -0.16672 | -0.2601 | -0.06722 | -0.63396 | -0.09232 | -0.37891 | -0.17267 | -0.53087 | -2.02473 | -1.22043 | -0.28699 |

| | | | | | | | | | | | | | | | | | | | | | | | | |
|---|---|---|---|---|---|---|---|---|---|---|---|---|---|---|---|---|---|---|---|---|---|---|---|---|
| 20.91667 | 0.121745 | -0.87736 | -1.10793 | -0.11268 | -0.28597 | -0.19332 | -0.4924 | 0.028268 | -1.07933 | -0.08457 | -0.95772 | -1.02679 | -0.76986 | 0.147657 | -0.15742 | 0.037707 | -0.64735 | -0.02715 | -0.19266 | -0.13389 | -0.69752 | -1.21327 | -1.25387 | -0.03874 |
| 20.93056 | -0.61831 | -0.90974 | -1.11401 | -0.13692 | -0.21376 | -0.34911 | -0.88656 | -0.06215 | -1.2611 | -1.4494 | -1.01839 | -1.82422 | -0.59074 | 0.489733 | 0.013142 | 0.224452 | -0.64121 | 0.037306 | -0.04252 | -0.28697 | -0.96959 | -0.85547 | -1.22657 | 0.247981 |
| 20.94444 | -1.36567 | -1.13395 | -1.10198 | -0.01907 | -0.10028 | -0.57164 | -1.16511 | -0.07811 | -1.39198 | -2.43568 | -1.1224 | -2.7591 | -0.60048 | 0.774617 | 0.234298 | 0.4062 | -0.82829 | 0.063469 | 0.00601 | -0.52839 | -1.30539 | -0.89831 | -1.00353 | 0.420231 |
| 20.95833 | -1.98995 | -1.09385 | -1.1564 | 0.048702 | -0.17475 | -0.57478 | -1.27594 | -0.04249 | -1.08143 | -1.8591 | -1.06397 | -2.67882 | -0.99488 | 0.684251 | 0.243084 | 0.323327 | -0.77219 | -0.11425 | -0.30377 | -0.73448 | -1.38667 | -1.09049 | -0.85937 | 0.176113 |
| 20.97222 | -1.5307 | -0.80354 | -0.95514 | 0.149971 | -0.32282 | -0.34663 | -0.94221 | -0.05666 | -0.76618 | -0.5368 | -0.66293 | -1.67552 | -1.29995 | 0.158834 | 0.080068 | 0.056428 | -0.65351 | -0.27826 | -0.50859 | -1.03817 | -0.84224 | -1.11729 | -0.58244 | -0.07019 |
| 20.98611 | -0.5967 | -0.41637 | -0.55395 | 0.159469 | -0.3402 | -0.15004 | -0.44205 | -0.10586 | -0.37127 | 0.441818 | -0.32614 | -0.64406 | -1.11016 | -0.29727 | -0.02055 | -0.05967 | -0.432 | -0.23978 | -0.24232 | -0.68679 | -0.26429 | -0.83159 | -0.07707 | -0.1213 |
| 21 | 0.083644 | -0.06751 | -0.23875 | -0.02228 | -0.23178 | -0.07515 | -0.08803 | -0.15663 | 0.04195 | 0.797255 | -0.22191 | -0.04327 | -0.67689 | -0.48104 | -0.05937 | -0.04682 | -0.1513 | -0.14234 | 0.167662 | -0.23088 | 0.120814 | -0.54049 | 0.210791 | -0.08764 |
| 21.01389 | 0.487767 | 0.061914 | -0.09318 | -0.21739 | -0.16276 | -0.04789 | -0.01767 | -0.17597 | 0.19914 | 0.606692 | -0.20387 | 0.040371 | -0.40896 | -0.48919 | -0.12683 | -0.06829 | 0.090448 | -0.11108 | 0.25205 | -0.01642 | 0.23943 | -0.40262 | 0.336983 | -0.1183 |
| 21.02778 | 0.567391 | 0.067334 | 0.009397 | -0.40001 | -0.18879 | -0.02755 | -0.16127 | -0.18807 | 0.091078 | 0.139113 | -0.16148 | -0.06061 | -0.22759 | -0.34645 | -0.17016 | -0.10929 | 0.217014 | -0.09412 | 0.153449 | -0.02293 | 0.220972 | -0.37859 | 0.394498 | -0.17411 |
| 21.04167 | 0.470554 | -0.01517 | 0.022004 | -0.47702 | -0.23708 | -0.03632 | -0.23834 | -0.10076 | -0.04921 | -0.31263 | -0.17134 | -0.15448 | -0.10047 | -0.13572 | -0.16216 | -0.12877 | 0.317751 | -0.05843 | 0.093099 | -0.03351 | 0.154758 | -0.397 | 0.292064 | -0.17124 |
| 21.05556 | 0.358446 | -0.09134 | -0.04709 | -0.36467 | -0.18878 | -0.04786 | -0.14274 | 0.007815 | -0.12933 | -0.57637 | -0.217 | -0.20579 | -0.01783 | 0.099879 | -0.11997 | -0.11471 | 0.531204 | -0.02819 | 0.059072 | 0.055264 | 0.091166 | -0.37312 | 0.113022 | -0.1342 |
| 21.06944 | 0.313557 | -0.11714 | -0.09331 | -0.08867 | -0.03384 | -0.04135 | -0.06018 | 0.081607 | -0.13965 | -0.71339 | -0.25258 | -0.03543 | 0.01796 | 0.216628 | -0.07342 | -0.07686 | 0.592314 | -0.01438 | 0.044208 | 0.200791 | 0.063732 | -0.33288 | -0.01096 | -0.10916 |
| 21.08333 | 0.288073 | -0.14399 | -0.08907 | 0.117509 | 0.06409 | -0.03581 | -0.06714 | 0.021557 | -0.04288 | -0.69965 | -0.25151 | 0.207035 | -0.03456 | 0.146297 | -0.05164 | -0.04902 | 0.524393 | -0.02514 | 0.1324 | 0.330835 | 0.008545 | -0.26977 | -0.02265 | -0.03171 |
| 21.09722 | 0.228917 | -0.20373 | -0.16492 | -0.00211 | 0.008523 | -0.0614 | -0.08267 | -0.08658 | 0.107637 | -0.47897 | -0.30439 | 0.16265 | -0.20716 | -0.06104 | -0.09504 | -0.08559 | 0.347401 | -0.07408 | 0.242672 | 0.375852 | -0.17898 | -0.2061 | 0.080041 | -0.03431 |
| 21.11111 | 0.221186 | -0.19563 | -0.18839 | -0.19852 | -0.05573 | -0.07069 | -0.00889 | -0.1208 | 0.407726 | -0.06435 | -0.32214 | 0.027046 | -0.38405 | -0.24147 | -0.15643 | -0.16245 | 0.467312 | -0.12554 | 0.241357 | 0.385067 | -0.29377 | -0.06426 | 0.235654 | -0.11236 |
| 21.125 | 0.394397 | 0.000076 | 0.138458 | -0.02283 | -0.0229 | 0.029168 | 0.163042 | -0.06421 | 0.868983 | 0.517672 | -0.05906 | 0.22023 | -0.15598 | -0.1211 | -0.13884 | -0.18907 | 0.751127 | -0.0999 | 0.257073 | 0.507151 | 0.023334 | 0.210768 | 0.483776 | -0.15327 |
| 21.13889 | 0.648353 | 0.185529 | 0.430045 | 0.216599 | 0.060906 | 0.11755 | 0.250871 | 0.044235 | 1.00063 | 0.718305 | 0.133609 | 0.420295 | 0.139392 | 0.215126 | -0.04076 | -0.12924 | 0.610229 | -0.01172 | 0.285908 | 0.584562 | 0.280987 | 0.363931 | 0.571865 | -0.10418 |
| 21.15278 | 0.653531 | 0.180856 | 0.317822 | 0.188898 | 0.111671 | 0.094391 | 0.134005 | 0.140973 | 0.657358 | 0.257259 | -8.1E-05 | 0.326343 | 0.099668 | 0.446824 | 0.02772 | -0.04488 | 0.531384 | 0.032133 | 0.171043 | 0.432465 | 0.149895 | 0.275844 | 0.243823 | 0.000959 |
| 21.16667 | 0.434662 | 0.115695 | 0.194115 | 0.177301 | 0.179912 | 0.048045 | 0.024703 | 0.094106 | 0.361933 | -0.32178 | -0.14244 | 0.053208 | 0.006794 | 0.324204 | 0.036417 | 0.028465 | 0.16258 | 0.025653 | 0.003896 | 0.252773 | 0.006031 | 0.143692 | -0.11948 | 0.074238 |
| 21.18056 | 0.055874 | 0.014176 | 0.176755 | 0.205031 | 0.195444 | 0.043489 | 0.019067 | -0.09859 | 0.436227 | -0.68464 | -0.15284 | -0.07364 | 0.001993 | 0.221943 | 0.027862 | 0.068002 | 0.067255 | 0.025701 | -0.03216 | 0.20527 | 0.006385 | 0.111747 | -0.19508 | 0.054233 |
| 21.19444 | -0.28682 | -0.07938 | 0.199987 | 0.183742 | 0.144394 | 0.063287 | 0.161015 | -0.09767 | 0.862252 | -0.3957 | -0.05801 | 0.298782 | 0.094081 | 0.170554 | 0.024823 | 0.045364 | 0.05524 | 0.0288 | 0.073563 | 0.26356 | 0.040269 | 0.177766 | -0.08638 | 0.011101 |
| 21.20833 | -0.31274 | -0.01372 | 0.258428 | 0.193853 | 0.127836 | 0.089407 | 0.317569 | 0.041774 | 1.367742 | 0.446113 | 0.062885 | 0.862594 | 0.176111 | 0.177913 | 0.031601 | -0.0081 | 0.054571 | 0.017911 | 0.189954 | 0.308244 | 0.084985 | 0.269326 | 0.011506 | -0.01124 |
| 21.22222 | -0.19366 | 0.089803 | 0.318377 | 0.211045 | 0.120565 | 0.097659 | 0.401316 | 0.167259 | 1.434571 | 1.063277 | 0.109162 | 1.004317 | 0.153252 | 0.163391 | 0.049029 | -0.0241 | 0.197951 | 0.001342 | 0.206943 | 0.265224 | 0.114126 | 0.342794 | -0.01463 | -0.01268 |
| 21.23611 | -0.04294 | 0.212858 | 0.40021 | 0.208388 | 0.140189 | 0.108636 | 0.465052 | 0.316985 | 1.141865 | 0.877079 | 0.115298 | 0.667312 | 0.169732 | 0.202884 | 0.082516 | 0.02382 | 0.309086 | 0.006223 | 0.181731 | 0.209119 | 0.149285 | 0.49023 | -0.16902 | -0.00499 |
| 21.25 | 0.092555 | 0.289496 | 0.538733 | 0.185248 | 0.170312 | 0.138049 | 0.540739 | 0.22894 | 0.942988 | 0.316248 | 0.123696 | 0.422513 | 0.261935 | 0.390553 | 0.113474 | 0.080941 | 0.402168 | 0.032585 | 0.219413 | 0.219847 | 0.19041 | 0.601798 | -0.28657 | 0.035776 |
| 21.26389 | 0.335963 | 0.296965 | 0.695712 | 0.190052 | 0.176788 | 0.194346 | 0.653164 | 0.085529 | 0.876013 | -0.16363 | 0.165745 | 0.686513 | 0.171018 | 0.655341 | 0.0926 | 0.059479 | 0.41455 | 0.047942 | 0.323338 | 0.263376 | 0.225673 | 0.528315 | -0.10476 | 0.06379 |
| 21.27778 | 0.571216 | 0.275192 | 0.791541 | 0.266595 | 0.187209 | 0.22547 | 0.794289 | -0.00394 | 0.859484 | 0.078073 | 0.212645 | 1.312616 | 0.198358 | 0.618256 | 0.05123 | 0.006514 | 0.501846 | 0.065408 | 0.401054 | 0.391401 | 0.239907 | 0.447996 | 0.384999 | 0.056639 |
| 21.29167 | 0.738229 | 0.279877 | 0.739157 | 0.335575 | 0.196362 | 0.297764 | 0.868878 | 0.077872 | 0.831895 | 1.127016 | 0.243474 | 1.791458 | 0.291797 | 0.310594 | 0.054669 | 0.002547 | 0.693488 | 0.091949 | 0.41823 | 0.500197 | 0.247994 | 0.554329 | 1.143356 | 0.052052 |
| 21.30556 | 0.783668 | 0.319438 | 0.661101 | 0.376276 | 0.185907 | 0.345491 | 0.841157 | 0.235769 | 0.766607 | 2.087024 | 0.275041 | 1.591314 | 0.367646 | 0.128145 | 0.087218 | 0.032026 | 0.891912 | 0.086435 | 0.357713 | 0.526578 | 0.249179 | 0.739373 | 1.327833 | 0.055113 |
| 21.31944 | 0.811411 | 0.396408 | 0.584503 | 0.351152 | 0.17415 | 0.335394 | 0.587925 | 0.245097 | 0.599272 | 1.786323 | 0.319662 | 1.100039 | 0.442982 | 0.055612 | 0.177364 | 0.087542 | 0.850652 | 0.084948 | 0.299162 | 0.551513 | 0.319651 | 0.981479 | 0.79038 | 0.067781 |
| 21.33333 | 0.840941 | 0.504223 | 0.478458 | 0.326211 | 0.20323 | 0.342135 | 0.406951 | 0.203208 | 0.498932 | 1.068525 | 0.390624 | 0.884451 | 0.544288 | 0.047786 | 0.243614 | 0.08568 | 0.540603 | 0.111803 | 0.246324 | 0.587139 | 0.432367 | 1.217561 | 0.50243 | 0.111239 |
| 21.34722 | 0.980538 | 0.58661 | 0.475288 | 0.262684 | 0.240842 | 0.333858 | 0.279744 | 0.177149 | 0.497829 | 0.264032 | 0.469006 | 1.030539 | 0.591121 | 0.059323 | 0.211727 | 0.050115 | 0.215153 | 0.141693 | 0.181364 | 0.468204 | 0.412929 | 1.211463 | 0.457501 | 0.147509 |
| 21.36111 | 1.071996 | 0.581315 | 0.544106 | 0.145737 | 0.245658 | 0.285589 | 0.268886 | 0.164768 | 0.553924 | -0.10586 | 0.49601 | 1.406671 | 0.604211 | 0.063533 | 0.101628 | 0.025258 | 0.049492 | 0.117961 | 0.124808 | 0.231607 | 0.272789 | 1.135697 | 0.660311 | 0.117158 |
| 21.375 | 1.047193 | 0.476248 | 0.487834 | 0.080612 | 0.199593 | 0.230767 | 0.164814 | 0.117549 | 0.518542 | 0.395524 | 0.44833 | 1.576275 | 0.496311 | 0.048437 | -0.06072 | 0.006685 | 0.074644 | 0.073384 | 0.101149 | 0.068144 | 0.135806 | 1.138934 | 0.905491 | 0.088099 |
| 21.38889 | 0.662073 | 0.535732 | 0.418392 | 0.102503 | 0.229754 | 0.270772 | -0.05146 | 0.098549 | 0.412523 | 1.172581 | 0.433458 | 1.186237 | 0.486057 | 0.030212 | -0.08676 | 0.022581 | 0.291526 | 0.124667 | 0.160762 | 0.161012 | 0.235465 | 1.406455 | 1.414071 | 0.150865 |
| 21.40278 | 0.268844 | 0.998794 | 0.628591 | 0.342338 | 0.55027 | 0.48259 | -0.24109 | 0.165508 | 0.263832 | 1.646353 | 0.789196 | 1.272249 | 0.980325 | 0.00688 | 0.013623 | 0.033106 | 0.665688 | 0.296112 | 0.265665 | 0.53958 | 0.74618 | 2.343078 | 1.243919 | 0.326082 |
| 21.41667 | 0.116758 | 2.056285 | 1.403594 | 0.833462 | 1.419989 | 0.962816 | -0.33763 | 0.266933 | 0.533679 | 2.2683 | 1.842907 | 2.672009 | 2.754493 | 0.017437 | 0.156027 | -0.02919 | 1.164405 | 0.492923 | 0.335067 | 1.068633 | 1.610065 | 4.639056 | 1.041756 | 0.493219 |
| 21.43056 | 0.382047 | 2.939217 | 1.951033 | 0.964765 | 1.512081 | 1.051427 | -0.09577 | 0.279152 | 1.030087 | 3.28115 | 2.528413 | 3.870128 | 3.416691 | 0.109571 | 0.266303 | -0.14675 | 1.478729 | 0.682773 | 0.299443 | 1.247804 | 1.835784 | 5.847033 | 0.896213 | 0.512825 |
| 21.44444 | 1.040952 | 2.360632 | 1.910113 | 0.556398 | 1.112596 | 0.548409 | 0.413689 | 0.23839 | 1.605592 | 3.338106 | 1.632274 | 2.950477 | 1.974044 | 0.036866 | 0.195515 | -0.33846 | 1.313471 | 0.520001 | 0.054206 | 0.857372 | 1.204299 | 3.937358 | 0.576961 | 0.36818 |
| 21.45833 | 1.771283 | 1.968446 | 1.516152 | 0.238237 | 1.084536 | 0.203484 | 0.112181 | 0.198942 | 1.953301 | 2.240251 | 1.124001 | 1.914048 | 1.420514 | -0.03503 | 0.192869 | -0.32921 | 1.554989 | 0.459415 | -0.01033 | 0.489979 | 0.977463 | 2.188134 | 0.406991 | 0.339972 |
| 21.47222 | 1.666148 | 1.737681 | 1.115424 | 0.143667 | 1.070418 | -0.03004 | -0.22538 | 0.21827 | 1.532994 | 1.006406 | 0.78651 | 1.074729 | 1.1555 | -0.00299 | 0.211914 | -0.08712 | 1.443253 | 0.455653 | 0.15277 | 0.339644 | 1.182302 | 2.075049 | 0.382028 | 0.37254 |
| 21.48611 | 0.9227 | 1.379225 | 0.658752 | 0.062422 | 0.885601 | -0.25712 | -0.65491 | 0.188945 | 0.872542 | -0.59629 | 0.404767 | 0.290223 | 0.725809 | -0.02649 | 0.145663 | -0.0068 | 0.671223 | 0.286838 | 0.149962 | 0.077466 | 0.840832 | 2.025604 | 0.288618 | 0.186946 |
| 21.5 | 0.368577 | 1.06062 | 0.346722 | -0.01008 | 0.634557 | -0.51461 | -0.89662 | 0.136565 | 0.460554 | -2.54222 | 0.009967 | -0.8456 | 0.051277 | -0.1221 | 0.020653 | -0.02445 | -0.30615 | 0.14729 | 0.002267 | -0.24336 | 0.472244 | 1.513458 | -0.02022 | -0.05944 |
| 21.51389 | 0.090178 | 0.552447 | 0.091552 | -0.0019 | 0.410076 | -0.68261 | -0.76689 | 0.066089 | -0.04313 | -2.99258 | -0.3239 | -1.50585 | -0.79369 | -0.17425 | -0.07108 | 0.002869 | -1.59235 | 0.177457 | -0.05279 | -0.57129 | 0.303341 | 0.343148 | -0.40943 | -0.17986 |
| 21.52778 | 1.082254 | 0.077842 | -0.13547 | 0.10395 | 0.183702 | -0.5308 | -0.17598 | -0.05117 | -0.57143 | -0.97033 | -0.45403 | -0.96716 | -1.32905 | -0.18113 | -0.14252 | 0.037429 | -2.76922 | 0.223984 | -0.00406 | -0.86208 | 0.092845 | -0.98961 | -0.69654 | -0.11294 |
| 21.54167 | 0.743395 | -0.49183 | -0.28248 | 0.255355 | -0.05868 | -0.13733 | 0.595401 | -0.12356 | -0.6221 | 1.139582 | -0.25806 | 0.172261 | -1.75905 | -0.12596 | -0.17752 | 0.01267 | -2.81611 | 0.162843 | 0.060713 | -0.73259 | -0.01563 | -1.46139 | -0.6093 | 0.050723 |
| 21.55556 | -0.68412 | -0.82205 | -0.28227 | 0.302736 | -0.25782 | 0.295479 | 0.881868 | -0.11804 | -0.3206 | 2.361255 | 0.161972 | 0.474679 | -1.69412 | -0.07991 | -0.10489 | 0.005708 | -1.55237 | 0.011153 | 0.123713 | -0.18694 | -0.08758 | -1.13012 | -0.31591 | 0.23245 |
| 21.56944 | -0.54458 | -0.98719 | -0.08906 | 0.241666 | -0.3781 | 0.665213 | 0.508744 | -0.06122 | 0.03065 | 1.773173 | 0.554357 | -0.03252 | -1.1309 | -0.06082 | 0.016951 | 0.019494 | 0.142149 | -0.04542 | 0.229578 | 0.424896 | -0.25972 | -0.80537 | -0.21859 | 0.308709 |
| 21.58333 | 0.481811 | -0.91693 | 0.157599 | 0.133617 | -0.39367 | 0.860938 | 0.006705 | -0.02495 | 0.110594 | 0.658244 | 0.742098 | -0.51235 | -0.65015 | -0.04984 | 0.149585 | 0.006351 | 1.703131 | -0.00415 | 0.30221 | 0.95255 | -0.29999 | -0.82001 | -0.05047 | 0.242765 |
| 21.59722 | 1.773771 | -0.71377 | 0.272955 | 0.048996 | -0.22861 | 0.711976 | -0.48242 | -0.0303 | -0.10106 | -0.26434 | 0.581935 | -0.96276 | 0.208579 | -0.02865 | 0.206361 | 0.011361 | 2.440119 | 0.082502 | 0.284159 | 1.277925 | -0.20646 | -1.15145 | 0.56762 | 0.072751 |
| 21.61111 | 2.266485 | -0.49871 | 0.219654 | -0.00985 | -0.03691 | 0.352543 | -0.94948 | -0.03177 | -0.21713 | -1.2748 | 0.377333 | -1.21476 | 0.380997 | -0.00569 | 0.090129 | -0.0199 | 2.453532 | 0.240731 | 0.211739 | 1.111703 | -0.2622 | -1.49698 | 1.068701 | -0.12627 |
| 21.625 | 1.354878 | -0.25307 | 0.201396 | -0.06515 | 0.137519 | 0.089463 | -0.96667 | -0.05384 | 0.118742 | -2.12705 | 0.158478 | -0.91105 | 0.414456 | 0.043835 | -0.05187 | -0.07068 | 2.198354 | 0.291206 | 0.103086 | 0.577498 | -0.44538 | -1.4252 | 1.080653 | -0.23122 |
| 21.63889 | 0.355248 | -0.1337 | 0.156739 | -0.05353 | 0.225425 | -0.18161 | -0.40033 | -0.04087 | 0.365029 | -2.0063 | -0.06508 | -0.07278 | 0.107293 | 0.092213 | -0.1708 | -0.1202 | 1.067698 | 0.167102 | 0.064559 | 0.029955 | -0.65684 | -1.13466 | 0.676574 | -0.10596 |
| 21.65278 | -0.09968 | -0.17057 | 0.035996 | 0.017471 | 0.257074 | -0.4803 | 0.162058 | -0.00858 | 0.38325 | -0.61074 | -0.20944 | 0.719952 | -0.45585 | 0.096068 | -0.20025 | -0.05447 | -0.211 | -0.03772 | -0.00166 | -0.359 | -0.76847 | -0.74538 | 0.138354 | -0.00494 |
| 21.66667 | -0.55656 | -0.23837 | -0.12758 | 0.120858 | 0.252599 | -0.70019 | 0.195846 | -0.00453 | 0.218263 | 0.749598 | -0.33615 | 1.225038 | -0.61353 | 0.061563 | -0.16003 | -0.02086 | -1.5267 | -0.17833 | -0.10551 | -0.6073 | -0.69211 | -0.24669 | -0.13477 | 0.096211 |
| 21.68056 | -0.84772 | -0.47762 | -0.31702 | 0.172302 | 0.158306 | -0.62401 | -0.03424 | 0.031071 | 0.105487 | 1.571492 | -0.40241 | 1.124618 | -0.88887 | -0.02729 | -0.10659 | -0.03884 | -2.51193 | -0.17467 | -0.24846 | -0.82114 | -0.48361 | 0.185689 | -0.2592 | 0.286005 |
| 21.69444 | -1.174 | -0.709 | -0.50048 | 0.155568 | 0.011898 | -0.33166 | -0.24491 | 0.022176 | 0.038199 | 1.219266 | -0.31966 | 0.371882 | -0.84407 | -0.13013 | -0.03585 | -0.05025 | -2.86571 | -0.10948 | -0.35507 | -1.05772 | -0.3448 | 0.20031 | -0.39825 | 0.177513 |
| 21.70833 | -1.56047 | -0.97015 | -0.60156 | 0.120048 | -0.13466 | -0.15813 | -0.45925 | -0.03029 | -0.19115 | 0.263675 | -0.06479 | -0.53461 | -0.76044 | -0.19124 | 0.052459 | -0.07632 | -3.07273 | -0.14032 | -0.45174 | -1.06811 | -0.47004 | -0.36508 | -0.86817 | -0.05621 |
| 21.72222 | -1.67074 | -1.25464 | -0.55785 | 0.062276 | -0.25455 | -0.12833 | -0.81399 | -0.10659 | -0.71492 | -0.71256 | 0.250398 | -1.28301 | -0.76679 | -0.18163 | 0.089666 | -0.11901 | -2.41653 | -0.2057 | -0.41039 | -0.7579 | -0.7661 | -1.12772 | -1.11645 | -0.31051 |
| 21.73611 | -1.5189 | -1.44679 | -0.50521 | -0.04313 | -0.3522 | 0.02244 | -0.84163 | -0.21474 | -1.28108 | -1.8016 | 0.335726 | -1.29477 | -0.81257 | -0.11036 | 0.038552 | -0.1692 | -1.40974 | -0.2021 | -0.21013 | -0.3203 | -1.02243 | -1.57673 | -0.86922 | -0.49689 |
| 21.75 | -0.98884 | -1.17378 | -0.49844 | -0.09229 | -0.45105 | 0.206841 | -0.51371 | -0.37135 | -1.58859 | -2.57209 | 0.158009 | -0.86508 | -0.95623 | -0.05892 | -0.06175 | -0.26929 | -0.65146 | -0.16852 | 0.032486 | 0.004487 | -1.21894 | -1.61872 | -0.44761 | -0.34715 |
| 21.76389 | -0.5367 | -1.06213 | -0.56774 | -0.08887 | -0.53048 | 0.103726 | -0.42301 | -0.49921 | -1.35388 | -2.14154 | -0.1505 | -0.6284 | -1.1272 | -0.11086 | -0.16149 | -0.17367 | -0.47023 | -0.1663 | 0.102545 | 0.034524 | -1.21889 | -1.58301 | -0.24493 | -0.33257 |
| 21.77778 | -0.40029 | -1.26938 | -0.71735 | -0.0405 | -0.65074 | -0.25546 | -0.69802 | -0.46627 | -1.066 | -1.35352 | -0.51163 | -0.79136 | -1.05399 | -0.30217 | -0.22621 | -0.00927 | -0.70617 | -0.22687 | -0.00316 | -0.22365 | -1.05583 | -1.59166 | -0.78513 | -0.30009 |
| 21.79167 | -0.78284 | -1.62558 | -0.87574 | -0.01028 | -0.72234 | -0.58439 | -1.08914 | -0.35572 | -0.94489 | -0.44058 | -0.82374 | -1.08257 | -0.75275 | -0.52922 | -0.29197 | 0.136171 | -1.0329 | -0.32905 | -0.19103 | -0.56452 | -0.88513 | -1.71983 | -1.07064 | -0.08821 |
| 21.80556 | -1.21415 | -1.73757 | -1.04535 | -0.02833 | -0.78265 | -0.64345 | -1.40141 | -0.20111 | -1.13079 | 0.36993 | -1.02068 | -1.35763 | -0.64358 | -0.59268 | -0.3553 | 0.05951 | -1.23136 | -0.28617 | -0.47314 | -0.78169 | -0.75839 | -2.20651 | -1.13381 | -0.00847 |
| 21.81944 | -1.49332 | -1.71602 | -1.09991 | -0.08465 | -0.75051 | -0.47705 | -1.44372 | -0.11907 | -1.41788 | 0.357383 | -1.14846 | -1.58491 | -0.59602 | -0.4993 | -0.40249 | -0.20168 | -1.26019 | -0.28442 | -0.79039 | -0.85293 | -0.62465 | -2.70898 | -1.10189 | -0.01653 |
| 21.83333 | -1.32926 | -1.63925 | -1.0045 | -0.14088 | -0.52146 | -0.29531 | -1.1047 | -0.20132 | -1.53525 | -0.44208 | -1.21069 | -1.77379 | -0.5574 | -0.34754 | -0.33509 | -0.49534 | -1.08051 | -0.29006 | -0.99174 | -0.89959 | -0.65105 | -2.66153 | -0.94846 | -0.26893 |
| 21.84722 | -1.073 | -1.27917 | -0.78548 | -0.1805 | -0.30417 | -0.21216 | -0.46606 | -0.40085 | -1.36038 | -1.53318 | -1.05442 | -1.66713 | -0.5341 | -0.12426 | -0.17414 | -0.56612 | -0.95301 | -0.23316 | -0.95437 | -0.7162 | -0.5783 | -1.99023 | -0.58154 | -0.46873 |
| 21.86111 | -0.92076 | -0.88981 | -0.61182 | -0.21547 | -0.13787 | -0.13831 | 0.111759 | -0.50578 | -0.89753 | -2.60367 | -0.78112 | -1.09741 | -0.48451 | 0.00351 | -0.04465 | -0.51076 | -0.93713 | -0.17157 | -0.8092 | -0.42804 | -0.46509 | -1.34737 | -0.20809 | -0.49311 |
| 21.875 | -1.00029 | -0.66717 | -0.50038 | -0.18873 | -0.12105 | -0.11132 | 0.239973 | -0.39472 | -0.53603 | -3.03471 | -0.55438 | -0.74031 | -0.50036 | 0.036781 | -0.04678 | -0.38283 | -0.68406 | -0.15813 | -0.69739 | -0.32155 | -0.46034 | -0.88114 | -0.0209 | -0.46434 |
| 21.88889 | -0.94681 | -0.57276 | -0.41152 | -0.07565 | -0.13665 | -0.15574 | 0.134214 | -0.22926 | -0.30394 | -2.25963 | -0.42298 | -0.45091 | -0.51724 | 0.032012 | -0.11425 | -0.21666 | -0.44931 | -0.12461 | -0.60809 | -0.26122 | -0.47934 | -0.16092 | 0.04221 | -0.42066 |
| 21.90278 | -0.73052 | -0.55792 | -0.40394 | -0.00357 | -0.12653 | -0.19513 | -0.03717 | -0.08214 | -0.30097 | -1.17827 | -0.38892 | -0.24145 | -0.55323 | 0.073581 | -0.12162 | -0.01608 | -0.24027 | -0.02186 | -0.54912 | -0.22494 | -0.46759 | 0.596232 | -0.02577 | -0.29176 |
| 21.91667 | -0.72496 | -0.5927 | -0.66742 | -0.15113 | -0.12434 | -0.26802 | -0.22851 | -0.04704 | -0.5724 | -0.66608 | -0.77105 | -0.29338 | -0.61221 | 0.147785 | -0.03211 | 0.130241 | -0.19382 | -0.00152 | -0.4842 | -0.28054 | -0.61065 | 0.8285 | -0.29322 | -0.13463 |
| 21.93056 | -1.05165 | -0.66403 | -1.15486 | -0.67131 | -0.08074 | -0.45426 | -0.36575 | -0.04899 | -0.89223 | -1.17941 | -1.82351 | -0.812 | -0.64707 | 0.22545 | 0.161312 | 0.20493 | -0.27763 | -0.15397 | -0.44942 | -0.4412 | -0.76842 | 0.299625 | -0.67599 | -0.07986 |
| 21.94444 | -1.34243 | -0.75323 | -1.52968 | -0.92758 | 0.010087 | -0.63594 | -0.47033 | -0.1462 | -0.85548 | -2.22766 | -2.82651 | -1.58374 | -0.58266 | 0.240816 | 0.347136 | 0.261019 | -0.85354 | -0.28747 | -0.31185 | -0.71935 | -1.18084 | -0.47533 | -1.20086 | -0.11566 |
| 21.95833 | -1.49029 | -1.14562 | -1.47537 | -0.6022 | -0.00466 | -0.54111 | -0.62692 | -0.38371 | -0.58528 | -1.93318 | -1.73771 | -1.91509 | -0.55425 | 0.025113 | 0.28464 | 0.117245 | -1.17025 | -0.1629 | -0.43258 | -1.02306 | -0.81584 | -1.45514 | -1.89239 | -0.20614 |
| 21.97222 | -1.13805 | -1.21934 | -0.87573 | 0.006224 | -0.07178 | -0.36966 | -0.75931 | -0.25019 | -0.39932 | -0.80841 | -0.08453 | -1.71686 | -0.39855 | -0.36141 | 0.04228 | -0.08667 | -0.864 | 0.084824 | -0.60201 | -1.2382 | -0.06394 | -1.90739 | -1.55702 | -0.19049 |
| 21.98611 | -0.49858 | -0.87594 | -0.11511 | 0.514885 | -0.0874 | -0.21749 | -0.65812 | 0.167319 | -0.33006 | 0.21707 | 0.468362 | -0.89483 | -0.39478 | -0.6532 | -0.00633 | -0.00271 | -0.18227 | 0.31524 | -0.42388 | -0.99483 | 0.178732 | -1.36409 | -0.40006 | 0.048923 |
| 22 | -0.07249 | -0.39167 | 0.307499 | 0.639428 | -0.0515 | -0.11193 | -0.39588 | 0.650811 | -0.39172 | 1.448546 | 0.367802 | 0.226435 | -0.26322 | -0.50111 | 0.085114 | 0.265292 | 0.799734 | 0.305574 | -0.10118 | -0.43512 | 0.293706 | -0.68319 | 0.492732 | 0.220677 |
| 22.01389 | 0.16022 | 0.050814 | 0.295825 | 0.502558 | -0.01744 | -0.0532 | -0.2165 | 0.743569 | -0.28288 | 2.479216 | 0.188916 | 0.827712 | -0.22638 | -0.16619 | 0.197584 | 0.375021 | 0.181861 | 0.172367 | 0.064845 | 0.070824 | 0.198629 | -0.38225 | 0.888205 | 0.180871 |
| 22.02778 | 0.21432 | 0.224033 | 0.180126 | 0.298655 | 0.039536 | -0.03844 | -0.29015 | 0.675402 | -0.15352 | 3.013766 | 0.07039 | 0.770939 | -0.21852 | 0.374396 | 0.218968 | 0.343506 | -0.02422 | 0.184042 | 0.068675 | 0.369011 | 0.162535 | -0.21895 | 0.668167 | 0.096935 |
| 22.04167 | 0.057522 | 0.178715 | 0.07697 | 0.144293 | 0.062929 | -0.04661 | -0.48957 | 0.374245 | -0.0931 | 2.715307 | -0.02254 | 0.398984 | -0.1848 | 0.69458 | 0.189745 | 0.224707 | -0.09123 | 0.258519 | 0.080532 | 0.493944 | 0.194867 | -0.24098 | 0.432658 | 0.03907 |
| 22.05556 | -0.07728 | 0.087714 | 0.04679 | 0.059293 | 0.036378 | -0.0441 | -0.32681 | 0.022296 | -0.08048 | 1.975205 | -0.04421 | -0.01821 | -0.16589 | 0.879509 | 0.126322 | 0.042603 | -0.1158 | 0.229206 | 0.131298 | 0.554403 | 0.232151 | -0.26942 | 0.367948 | 0.015338 |
| 22.06944 | -0.12441 | 0.141977 | 0.038623 | 0.028201 | -0.01595 | -0.02505 | 0.174686 | -0.28615 | -0.09989 | 1.245206 | -0.03421 | -0.27463 | -0.079 | 0.586758 | 0.074271 | -0.14374 | -0.02499 | 0.190345 | 0.185551 | 0.648662 | 0.184356 | -0.23807 | 0.345477 | -0.0188 |
| 22.08333 | 0.02192 | 0.210593 | 0.021394 | 0.006863 | -0.08734 | -0.02979 | 0.384493 | -0.41678 | -0.12158 | 0.377492 | -0.07378 | -0.19534 | -0.02947 | 0.339556 | 0.029745 | -0.2784 | 0.18214 | 0.129699 | 0.198149 | 0.755691 | 0.099134 | -0.18963 | 0.315898 | -0.03193 |
| 22.09722 | 0.315334 | 0.248157 | -0.00637 | 0.00348 | -0.1201 | -0.05974 | 0.256527 | -0.28886 | -0.13854 | -0.46841 | -0.14988 | 0.20912 | -0.02351 | 0.092242 | -0.02175 | -0.29521 | 0.269968 | 0.031712 | 0.171073 | 0.714814 | 0.022407 | -0.18477 | 0.298313 | -0.04545 |
| 22.11111 | 0.461314 | 0.22276 | -0.00608 | 0.036854 | -0.07093 | -0.06785 | 0.132327 | -0.04427 | -0.12804 | -1.10604 | -0.1802 | 0.519694 | -0.04096 | -0.07878 | -0.0457 | -0.19575 | 0.168765 | -0.08246 | 0.15784 | 0.698686 | -0.02456 | -0.14936 | 0.301678 | -0.04445 |
| 22.125 | 0.456434 | 0.11955 | 0.002978 | 0.063699 | 0.014615 | -0.04514 | 0.000975 | 0.017981 | -0.12179 | -1.27978 | -0.15769 | 0.417329 | -0.06082 | -0.1717 | -0.04865 | 0.023839 | 0.081907 | -0.15428 | 0.216686 | 0.627813 | -0.01255 | -0.08837 | 0.272765 | -0.03057 |
| 22.13889 | 0.357377 | 0.010147 | 0.015629 | 0.044283 | 0.048357 | -0.02327 | -0.15273 | -0.10944 | -0.15011 | -0.63976 | -0.11574 | -0.05009 | -0.08787 | -0.17808 | -0.05036 | 0.1009 | 0.048033 | -0.12434 | 0.304485 | 0.521462 | 0.023496 | 0.007579 | 0.210793 | -0.03311 |
| 22.15278 | 0.127411 | -0.06839 | 0.00807 | -0.00904 | 0.03394 | -0.01862 | -0.18381 | -0.26291 | -0.17482 | 0.250925 | -0.09085 | -0.4821 | -0.11041 | -0.10729 | -0.04615 | -0.06802 | 0.085772 | -0.03901 | 0.31537 | 0.37371 | 0.063879 | 0.1211 | 0.117759 | -0.06881 |
| 22.16667 | -0.08875 | -0.09087 | -0.00158 | -0.01176 | -0.00662 | -0.02379 | -0.12571 | -0.2952 | -0.15875 | 0.437779 | -0.09376 | -0.60272 | -0.09918 | 0.047444 | -0.00747 | -0.34165 | 0.238362 | 0.020656 | 0.207868 | 0.227291 | 0.086213 | 0.177499 | 0.061134 | -0.07493 |
| 22.18056 | -0.18007 | -0.07301 | 0.018564 | 0.048165 | -0.03663 | -0.01752 | -0.10172 | -0.08361 | -0.10355 | -0.05676 | -0.09298 | -0.34219 | -0.07978 | 0.184638 | 0.048807 | -0.47756 | 0.378826 | 0.039815 | 0.099677 | 0.107801 | 0.066836 | 0.142997 | 0.128151 | 0.038625 |
| 22.19444 | -0.18389 | -0.08297 | 0.062655 | 0.109475 | -0.05572 | -0.00989 | -0.01966 | 0.249264 | -0.04364 | -0.52301 | -0.07682 | 0.059426 | -0.06745 | 0.198787 | 0.094422 | -0.21908 | 0.267932 | 0.003954 | 0.05816 | -0.02404 | 0.05835 | 0.097308 | 0.220756 | 0.122676 |
| 22.20833 | -0.09887 | -0.07124 | 0.153288 | 0.137643 | -0.03547 | -0.00729 | 0.178764 | 0.34553 | 0.008395 | -0.60409 | -0.00637 | 0.282314 | -0.03807 | 0.101412 | 0.12396 | 0.283927 | 0.080769 | -0.0387 | 0.04998 | -0.09972 | 0.106507 | 0.150087 | 0.298264 | 0.087645 |
| 22.22222 | 0.139507 | -0.03585 | 0.237534 | 0.171303 | 0.012372 | 0.000625 | 0.366962 | 0.165114 | 0.079572 | -0.1623 | 0.142463 | 0.247504 | 0.023633 | -0.0202 | 0.143967 | 0.590926 | -0.02663 | -0.02459 | 0.060447 | -0.00693 | 0.199425 | 0.276387 | 0.335643 | 0.056529 |
| 22.23611 | 0.507505 | 0.061474 | 0.307397 | 0.216703 | 0.04877 | 0.019455 | 0.479659 | -0.06244 | 0.195008 | 0.716541 | 0.280446 | 0.221203 | 0.170687 | -0.01806 | 0.153395 | 0.522578 | 0.188172 | 0.050441 | 0.121456 | 0.276738 | 0.267964 | 0.420138 | 0.338304 | 0.099031 |
| 22.25 | 0.888269 | 0.201523 | 0.384106 | 0.264952 | 0.093657 | 0.05802 | 0.561973 | -0.08007 | 0.358634 | 1.655479 | 0.293354 | 0.470348 | 0.305712 | 0.19291 | 0.14737 | 0.237788 | 0.566524 | 0.2088 | 0.260005 | 0.657816 | 0.254611 | 0.555833 | 0.350198 | 0.176169 |
| 22.26389 | 1.010153 | 0.283709 | 0.430671 | 0.302217 | 0.10431 | 0.089189 | 0.558724 | 0.265762 | 0.510228 | 1.672719 | 0.20682 | 0.773011 | 0.330582 | 0.395893 | 0.115464 | -0.02073 | 0.671345 | 0.385281 | 0.341762 | 0.945679 | 0.173469 | 0.629006 | 0.349992 | 0.178736 |
| 22.27778 | 0.685516 | 0.219231 | 0.424164 | 0.300717 | 0.06525 | 0.084282 | 0.480718 | 0.649978 | 0.582609 | 0.780784 | 0.113071 | 1.050956 | 0.300297 | 0.393771 | 0.10637 | -0.10915 | 0.4957 | 0.472723 | 0.321574 | 1.04857 | 0.106029 | 0.753217 | 0.397818 | 0.124471 |
| 22.29167 | 0.402266 | 0.167352 | 0.490389 | 0.33948 | 0.095348 | 0.117705 | 0.49768 | 0.715701 | 0.632361 | -0.27162 | 0.15977 | 1.04555 | 0.317107 | 0.348967 | 0.215179 | 0.046414 | 0.414074 | 0.508073 | 0.328087 | 1.046139 | 0.148883 | 1.038312 | 0.624098 | 0.203626 |
| 22.30556 | 0.329847 | 0.154195 | 0.572035 | 0.382183 | 0.172046 | 0.191652 | 0.603404 | 0.456326 | 0.655325 | -0.19682 | 0.311631 | 0.830061 | 0.380855 | 0.276652 | 0.355966 | 0.18991 | 0.440023 | 0.526169 | 0.416277 | 0.924794 | 0.243709 | 1.355487 | 0.84088 | 0.38537 |
| 22.31944 | 0.474572 | 0.13618 | 0.575195 | 0.344901 | 0.162438 | 0.232259 | 0.654794 | 0.007648 | 0.673062 | 1.118382 | 0.463606 | 0.617319 | 0.489537 | 0.209452 | 0.365155 | 0.201169 | 0.538652 | 0.46253 | 0.517739 | 0.712969 | 0.292533 | 1.613137 | 0.702258 | 0.613845 |
| 22.33333 | 0.781566 | 0.16948 | 0.53539 | 0.221073 | 0.093244 | 0.24719 | 0.708415 | -0.11883 | 0.734401 | 2.287613 | 0.468361 | 0.620793 | 0.523577 | 0.139483 | 0.242318 | 0.115625 | 0.627738 | 0.316505 | 0.634626 | 0.551946 | 0.30844 | 1.812573 | 0.380074 | 0.608739 |
| 22.34722 | 1.11664 | 0.229435 | 0.471388 | 0.104721 | 0.023513 | 0.258972 | 0.74255 | 0.22978 | 0.900353 | 2.032843 | 0.383544 | 0.738065 | 0.417947 | 0.104301 | 0.122993 | 0.02239 | 0.541854 | 0.120125 | 0.744934 | 0.572641 | 0.301745 | 1.88288 | 0.20573 | 0.558681 |
| 22.36111 | 1.262941 | 0.264436 | 0.507744 | 0.042315 | 0.013313 | 0.265839 | 0.646696 | 0.707463 | 1.011851 | 1.05415 | 0.354876 | 0.810079 | 0.301719 | 0.202775 | 0.02443 | -0.03651 | 0.474748 | 0.006792 | 0.609282 | 0.71471 | 0.226735 | 2.03557 | 0.074246 | 0.345884 |
| 22.375 | 1.084424 | 0.227098 | 0.605384 | 0.102484 | 0.046025 | 0.287279 | 0.509476 | 0.761451 | 1.049979 | 0.014776 | 0.402864 | 0.724848 | 0.261623 | 0.328419 | -0.02955 | -0.05174 | 0.462901 | 0.007597 | 0.340955 | 0.799113 | 0.105949 | 2.277657 | 0.209441 | 0.243472 |

| | | | | | | | | | | | | | | | | | | | | | | | | |
|---|---|---|---|---|---|---|---|---|---|---|---|---|---|---|---|---|---|---|---|---|---|---|---|---|
| 22.38889 | 0.710566 | 0.221249 | 0.670643 | 0.109376 | 0.067729 | 0.279234 | 0.461287 | 0.454287 | 1.16199 | -0.82565 | 0.54602 | 0.615685 | 0.417297 | 0.353718 | 0.051642 | -0.00955 | 0.602636 | 0.117543 | 0.190224 | 0.656007 | 0.137588 | 2.865386 | 0.569453 | 0.36263 |
| 22.40278 | 0.48314 | 0.418231 | 0.833998 | 0.292013 | 0.111472 | 0.405157 | 0.454863 | 0.166287 | 2.021674 | 0.138396 | 0.961969 | 0.744063 | 0.822577 | 0.11058 | 0.277377 | 0.093702 | 1.059931 | 0.32145 | 0.249944 | 0.557847 | 0.516204 | 3.844678 | 0.81604 | 0.597858 |
| 22.41667 | 0.817734 | 0.939903 | 1.480043 | 0.87285 | 0.236813 | 0.738548 | 0.630638 | 0.225816 | 3.719094 | 2.577468 | 1.686513 | 1.442505 | 1.619062 | -0.15403 | 0.431887 | 0.183139 | 1.840025 | 0.68767 | 0.416048 | 0.769365 | 1.33733 | 5.787532 | 0.728851 | 0.723458 |
| 22.43056 | 1.45178 | 1.737731 | 2.154031 | 1.356069 | 0.248635 | 0.884539 | 1.040906 | 0.691352 | 3.174451 | 5.020569 | 1.618751 | 2.205899 | 2.130142 | -0.32246 | 0.597087 | 0.152703 | 2.52913 | 1.183043 | 0.496008 | 0.976509 | 2.572776 | 7.038998 | 0.627741 | 0.580098 |
| 22.44444 | 1.696098 | 1.613905 | 1.789835 | 1.072692 | 0.05148 | 0.66369 | 1.733618 | 0.903972 | 1.629741 | 4.525907 | 0.978919 | 2.507425 | 1.867763 | -0.32778 | 0.358755 | 0.225783 | 2.48987 | 1.036965 | 0.29999 | 1.26643 | 2.188313 | 4.191146 | 0.518887 | 0.290134 |
| 22.45833 | 1.695277 | 1.223246 | 1.619082 | 0.757119 | 0.007556 | 0.701703 | 0.948431 | 0.410325 | 1.181199 | 2.643174 | 0.619827 | 1.9143 | 1.768494 | -0.09048 | 0.292448 | 0.251705 | 1.981696 | 0.814451 | 0.152879 | 1.31407 | 2.215807 | 1.470754 | 0.380984 | 0.087137 |
| 22.47222 | 0.963981 | 0.945096 | 1.62666 | 0.494469 | 0.199753 | 0.658591 | -0.08166 | 0.230413 | 0.79382 | 1.19103 | 0.373852 | 0.859828 | 1.945946 | 0.178615 | 0.398637 | 0.00943 | 1.31087 | 0.708548 | 0.19568 | 0.884133 | 2.506608 | -0.14897 | 0.49055 | -0.09862 |
| 22.48611 | 0.161928 | 1.007558 | 1.275933 | 0.189718 | 0.268006 | 0.374447 | -0.98534 | 0.147167 | 0.231129 | 0.135177 | 0.205008 | 0.115233 | 1.608972 | 0.235739 | 0.303533 | -0.33033 | 0.289662 | 0.324628 | 0.023359 | 0.708082 | 1.763008 | -1.09212 | 0.49219 | -0.24481 |
| 22.5 | -0.74836 | 0.95972 | 0.880715 | -0.12682 | 0.196384 | 0.150509 | -1.4952 | -0.02922 | 0.09211 | -1.32796 | 0.125044 | -0.73494 | 1.14193 | 0.095847 | 0.079173 | -0.39747 | -0.72274 | -0.14858 | -0.27415 | 0.382252 | 0.635504 | -1.84057 | 0.412564 | -0.24267 |
| 22.51389 | -1.46372 | 0.976319 | 0.468942 | -0.26045 | 0.193309 | -0.09031 | -1.56759 | -0.40934 | 0.327243 | -2.59354 | 0.159144 | -0.73305 | 0.329027 | -0.09042 | -0.10743 | -0.20535 | -1.48704 | -0.44223 | -0.32484 | -0.67103 | -0.11797 | -2.35277 | 0.240759 | 0.134251 |
| 22.52778 | -1.55307 | 1.159334 | 0.150232 | -0.15588 | 0.286789 | -0.19196 | -0.97577 | -0.82932 | 0.621743 | -2.73842 | 0.089909 | 0.021923 | 0.11751 | -0.23848 | -0.28204 | -0.09566 | -2.16245 | -0.35969 | -0.04816 | -1.61169 | -0.12439 | -2.22016 | 0.051758 | 0.489757 |
| 22.54167 | -1.07492 | 1.350536 | -0.07228 | 0.068964 | 0.328992 | -0.15922 | -0.02741 | -0.24428 | 0.46034 | -0.06956 | 0.072047 | 0.876498 | 0.284431 | -0.25584 | -0.31247 | 0.024901 | -1.65711 | -0.03463 | 0.373467 | -1.67667 | 0.201729 | -1.75518 | -0.21846 | 0.563979 |
| 22.55556 | -0.03477 | 1.593274 | -0.28186 | 0.293562 | 0.3119 | -0.06474 | 1.18114 | 0.4064 | 0.096397 | 2.503619 | 0.078301 | 1.490961 | 0.28092 | -0.1653 | -0.0817 | 0.11924 | -0.65469 | 0.523002 | 0.700825 | -0.83878 | 0.492017 | -1.34798 | -0.52986 | 0.515781 |
| 22.56944 | 1.081795 | 1.640993 | -0.46009 | 0.384195 | 0.240238 | 0.004599 | 2.158325 | 0.967363 | 0.065199 | 3.182892 | 0.108001 | 1.67386 | 0.430948 | 0.015338 | 0.208751 | 0.196384 | 0.072736 | 0.877003 | 0.640083 | 0.385974 | 0.557194 | -0.9475 | -0.91994 | 0.196706 |
| 22.58333 | 1.83049 | 1.563426 | -0.60646 | 0.414755 | 0.114337 | -0.03909 | 0.929891 | 0.561323 | 0.172188 | 2.249587 | 0.239096 | 1.114332 | 0.570915 | 0.157783 | 0.498583 | 0.340653 | 0.891668 | 0.605129 | 0.262154 | 1.285729 | 0.853606 | -0.07907 | -0.93749 | -0.18571 |
| 22.59722 | 1.751551 | 1.312823 | -0.43995 | 0.42335 | 0.009907 | -0.04878 | 0.050542 | 0.205579 | 0.226561 | 0.791061 | 0.384446 | 0.11101 | 0.133466 | 0.181353 | 0.536282 | 0.308471 | 1.735051 | 0.280694 | -0.11922 | 1.434814 | 1.084417 | 0.98837 | -0.51033 | -0.41008 |
| 22.61111 | 1.499386 | 0.99862 | -0.06333 | 0.301727 | -0.03061 | -0.05771 | -0.83634 | 0.006819 | 0.096241 | -0.18036 | 0.479764 | -0.55771 | -0.69187 | 0.167435 | 0.274703 | 0.19708 | 1.862499 | 0.017415 | -0.32189 | 1.263644 | 0.996366 | 2.105253 | -0.05027 | -0.59842 |
| 22.625 | 1.090209 | 0.508792 | 0.045345 | 0.089952 | 0.006859 | 0.03578 | -1.89394 | -0.10486 | -0.4001 | -0.67403 | 0.443514 | -0.81654 | -1.21151 | 0.117575 | 0.032699 | 0.085552 | 1.721982 | -0.22924 | -0.32342 | 0.981451 | 0.54698 | 2.242655 | 0.27408 | -0.52624 |
| 22.63889 | 0.307297 | 0.094613 | 0.132645 | -0.14039 | 0.078832 | 0.180901 | -2.31414 | -0.2424 | -0.67532 | -1.50597 | 0.31397 | -1.01805 | -0.96372 | 0.030974 | -0.17867 | -0.02888 | 1.320028 | -0.46431 | -0.36387 | 0.376171 | 0.232905 | 1.655274 | 0.500131 | 0.072053 |
| 22.65278 | -0.66471 | -0.21449 | 0.170133 | -0.29595 | 0.10871 | 0.239077 | -1.34422 | -0.53255 | -0.65926 | -2.49653 | 0.106246 | -1.44587 | -0.50997 | -0.01664 | -0.37539 | -0.12775 | 0.686407 | -0.60171 | -0.12729 | -0.24199 | -0.00754 | 0.97962 | 0.620465 | 0.471562 |
| 22.66667 | -1.50186 | -0.42628 | 0.08611 | -0.28794 | 0.089264 | 0.223466 | 0.000736 | -0.86369 | -0.50539 | -3.08048 | -0.03509 | -1.11294 | -0.1284 | -0.06053 | -0.39341 | -0.03453 | -0.23286 | -0.48469 | 0.177746 | -0.68847 | -0.30793 | 0.519265 | 0.496599 | 0.511144 |
| 22.68056 | -1.72528 | -0.65346 | -0.03963 | -0.06656 | -0.01452 | 0.142338 | 1.121635 | -0.4615 | -0.30094 | -1.97522 | -0.15031 | -0.22023 | 0.021885 | -0.12759 | -0.32127 | 0.057437 | -0.94612 | -0.13794 | 0.276106 | -1.0266 | -0.42238 | 0.191312 | 0.366887 | 0.463083 |
| 22.69444 | -1.55322 | -0.73205 | -0.15268 | 0.132447 | -0.09966 | 0.029379 | 1.082333 | -0.09844 | -0.20156 | -0.2671 | -0.26706 | 0.804239 | -0.07325 | -0.18452 | -0.18397 | 0.090712 | -1.46949 | 0.022022 | 0.123338 | -1.12819 | -0.39601 | 0.098268 | 0.222056 | 0.013734 |
| 22.70833 | -1.15833 | -0.7044 | -0.22427 | 0.276662 | -0.09296 | -0.18631 | 0.30767 | 0.324943 | -0.11631 | 1.354944 | -0.30748 | 1.448197 | -0.30381 | -0.15948 | -0.03933 | 0.017745 | -1.93551 | -0.03149 | -0.24627 | -0.92713 | -0.53155 | 0.009896 | -0.02943 | -0.50597 |
| 22.72222 | -0.23466 | -0.63953 | -0.32189 | 0.287263 | 0.01098 | -0.25188 | -0.54791 | 0.805933 | 0.118264 | 2.58754 | -0.24177 | 0.699112 | -0.5819 | -0.05349 | 0.100425 | -0.05036 | -2.15034 | -0.19837 | -0.59733 | -0.50784 | -0.50661 | -0.55016 | -0.74038 | -0.7625 |
| 22.73611 | 0.236989 | -0.59719 | -0.50082 | 0.135503 | 0.07592 | -0.32086 | -1.58766 | 0.504806 | 0.142686 | 1.692397 | -0.16406 | -0.05176 | -0.9273 | 0.02357 | 0.156406 | -0.13212 | -2.33105 | -0.37241 | -0.82894 | -0.31834 | -0.28229 | -1.58433 | -1.03842 | -0.91468 |
| 22.75 | -0.01719 | -0.71196 | -0.66112 | -0.13108 | 0.040376 | -0.37875 | -2.35941 | -0.1364 | -0.18786 | 0.171351 | -0.16739 | -0.63136 | -1.15045 | 0.102642 | 0.080022 | -0.23787 | -1.74715 | -0.46321 | -0.82551 | -0.41699 | -0.17286 | -2.64567 | -1.01372 | -1.10988 |
| 22.76389 | -0.40443 | -0.95031 | -0.76576 | -0.36207 | -0.02551 | -0.36288 | -1.3989 | -0.72316 | -0.62264 | -1.01868 | -0.29696 | -1.22747 | -1.0716 | 0.263046 | -0.07428 | -0.27905 | -1.08561 | -0.39353 | -0.76285 | -0.50426 | -0.31531 | -3.16532 | -0.67205 | -0.67346 |
| 22.77778 | -0.72461 | -1.23678 | -0.94665 | -0.38642 | -0.15752 | -0.29635 | -0.56304 | -1.09876 | -1.0462 | -2.378 | -0.55689 | -1.46745 | -1.06121 | 0.481177 | -0.25994 | -0.2292 | -0.996 | -0.27914 | -0.33954 | -0.62469 | -0.5159 | -3.08407 | -0.33807 | -0.08945 |
| 22.79167 | -1.34047 | -1.40737 | -1.09942 | -0.24795 | -0.26671 | -0.28295 | 0.045205 | -0.77222 | -1.4991 | -4.22545 | -0.64474 | -1.2794 | -1.07908 | 0.343021 | -0.50767 | -0.17696 | -0.99156 | -0.27016 | 0.093222 | -0.94565 | -0.70751 | -2.86854 | -0.19555 | 0.118074 |
| 22.80556 | -2.27395 | -1.33133 | -1.18828 | -0.04848 | -0.22858 | -0.27499 | 0.145355 | -0.32451 | -1.45787 | -4.00501 | -0.4738 | -0.97017 | -0.9867 | 0.035778 | -0.67592 | -0.25941 | -0.85555 | -0.31359 | 0.183341 | -1.18058 | -0.7331 | -2.4962 | -0.62709 | 0.097001 |
| 22.81944 | -1.60166 | -1.10276 | -1.24692 | 0.092758 | -0.15202 | -0.27391 | -0.20238 | -0.3706 | -1.1176 | -2.72034 | -0.30687 | -0.7074 | -0.77282 | -0.21362 | -0.62655 | -0.41981 | -0.84377 | -0.35607 | 0.094571 | -1.12793 | -0.67278 | -1.56649 | -1.03517 | -0.11416 |
| 22.83333 | -0.87501 | -0.7263 | -1.2918 | 0.112953 | -0.09127 | -0.29055 | -0.58836 | 0.011983 | -0.93625 | -1.26442 | -0.26305 | -0.48242 | -0.46211 | -0.24873 | -0.5292 | -0.6023 | -0.91412 | -0.38376 | -0.14147 | -0.81127 | -0.59478 | -0.62125 | -0.65602 | -0.5435 |
| 22.84722 | -0.29724 | -0.09339 | -1.14917 | 0.060554 | -0.06462 | -0.29207 | -0.9775 | 0.362317 | -0.8993 | -0.00638 | -0.28257 | -0.38419 | -0.21038 | -0.11457 | -0.54545 | -0.69201 | -0.6978 | -0.41618 | -0.53037 | -0.49808 | -0.45099 | 0.033577 | -0.28008 | -1.32561 |
| 22.86111 | 0.114629 | 0.268713 | -0.89225 | -0.01479 | -0.0678 | -0.26609 | -1.04818 | 0.28101 | -0.89726 | 0.569722 | -0.3544 | -0.49871 | -0.09836 | 0.240284 | -0.54968 | -0.69895 | -0.54271 | -0.40985 | -0.80055 | -0.34756 | -0.21814 | 0.228881 | -0.04441 | -1.78855 |
| 22.875 | 0.119237 | 0.322652 | -0.62899 | -0.0691 | -0.08341 | -0.25817 | -0.64644 | 0.262575 | -0.77762 | 0.389279 | -0.48515 | -0.61658 | -0.04292 | 0.396919 | -0.46951 | -0.35816 | -0.49102 | -0.36186 | -0.85583 | -0.02997 | 0.066232 | 0.051008 | 0.052238 | -1.51456 |
| 22.88889 | -0.09923 | 0.224737 | -0.39143 | -0.08947 | -0.09521 | -0.25122 | 0.053673 | 0.143348 | -0.61706 | -0.54218 | -0.50975 | -0.61188 | 0.039124 | 0.588597 | -0.37159 | -0.05688 | -0.48049 | -0.17319 | -0.6365 | 0.251109 | 0.237926 | -0.03078 | -0.01097 | -0.87644 |
| 22.90278 | -0.42875 | 0.03809 | -0.35856 | -0.18255 | -0.11853 | -0.23563 | 0.488459 | -0.12574 | -0.63837 | -1.7144 | -0.53114 | -0.57323 | -0.06547 | 0.558659 | -0.34399 | 0.069554 | -0.49066 | 0.086118 | -0.35151 | 0.097744 | 0.124641 | 0.161498 | -0.26414 | -0.42121 |
| 22.91667 | -0.81268 | -0.35807 | -0.58514 | -0.39915 | -0.16803 | -0.23341 | 0.340766 | -0.38427 | -0.78556 | -2.67742 | -0.75006 | -0.58411 | -0.36263 | 0.475303 | -0.24315 | 0.066998 | -0.66477 | -0.0066 | -0.33802 | -0.45544 | -0.25374 | 0.316941 | -0.45165 | -0.18458 |
| 22.93056 | -1.30044 | -0.64063 | -0.89499 | -0.37063 | -0.18412 | -0.32192 | -0.18472 | -0.43149 | -1.05245 | -3.48389 | -1.49287 | -1.17517 | -0.51852 | 0.408185 | -0.10046 | 0.038365 | -1.28222 | -0.23004 | -0.48184 | -1.15588 | -0.70862 | -0.0523 | -0.70244 | -0.14394 |
| 22.94444 | -1.85023 | -0.63727 | -1.41253 | -0.2492 | -0.16071 | -0.50476 | -0.81261 | -0.69419 | -0.9995 | -4.67437 | -2.2442 | -2.1012 | -0.47298 | 0.316343 | -0.00225 | -0.02269 | -2.18737 | -0.3913 | -0.45539 | -1.69049 | -1.00349 | -0.57667 | -1.01768 | -0.23443 |
| 22.95833 | -2.26775 | -0.74947 | -1.76267 | -0.27309 | -0.16082 | -0.49047 | -1.54558 | -1.59016 | -0.69217 | -5.22213 | -2.21941 | -2.85766 | -0.57369 | -0.1001 | -0.15446 | -0.28358 | -4.09252 | -0.7197 | -0.47781 | -2.1591 | -0.94408 | -1.14913 | -1.87767 | -0.53375 |
| 22.97222 | -1.80713 | -0.9522 | -1.22519 | -0.24479 | -0.13165 | -0.3006 | -1.56688 | -2.89466 | -0.44437 | -3.81172 | -1.48993 | -2.47369 | -0.64329 | -0.5999 | -0.36668 | -0.41918 | -3.84226 | -0.84622 | -0.35574 | -2.71362 | -0.71089 | -1.80667 | -2.60691 | -0.77005 |
| 22.98611 | -0.89627 | -0.84112 | -0.35286 | -0.05001 | -0.03664 | -0.19554 | -0.89829 | -2.60467 | -0.38479 | -1.50098 | -0.72251 | -1.46743 | -0.33027 | -0.74812 | -0.35756 | 0.079927 | -1.63258 | -0.20447 | -0.01935 | -1.99017 | -0.51422 | -1.6487 | -1.51102 | -0.50163 |
| 23 | 0.01433 | -0.51888 | 0.399845 | 0.084471 | -0.05583 | -0.17903 | -0.03325 | -0.90597 | -0.37772 | 0.195898 | 0.145851 | -0.38972 | -0.00737 | -0.55497 | -0.18545 | 0.613565 | 0.315854 | 0.545621 | 0.257807 | -0.8757 | -0.1488 | -1.13926 | -0.63627 | 0.027073 |
| 23.01389 | 0.526551 | -0.24939 | 0.761142 | 0.087646 | -0.19307 | -0.13271 | 0.588936 | 0.497882 | -0.42901 | 1.431156 | 0.730648 | 0.421238 | 0.048202 | -0.20838 | -0.00798 | 0.810087 | 2.092162 | 0.909359 | 0.367572 | -0.16345 | 0.105157 | -0.67553 | 0.067393 | 0.405415 |
| 23.02778 | 0.539859 | -0.08493 | 0.722391 | 0.069034 | -0.24893 | -0.09278 | 0.655495 | 1.304651 | -0.52078 | 2.214044 | 0.808481 | 0.781428 | 0.003124 | 0.120329 | 0.137584 | 0.576939 | 2.694146 | 0.970115 | 0.382022 | 0.243767 | 0.107286 | -0.35362 | 0.470137 | 0.584511 |
| 23.04167 | 0.409612 | -0.05974 | 0.533676 | 0.065492 | -0.12915 | -0.07124 | 0.473389 | 1.549647 | -0.49087 | 2.461587 | 0.750452 | 0.89928 | -0.05456 | 0.309549 | 0.127281 | 0.363416 | 2.154272 | 0.855102 | 0.331756 | 0.460569 | 0.037818 | -0.24332 | 0.445219 | 0.575973 |
| 23.05556 | 0.360291 | -0.05614 | 0.347477 | 0.037628 | 0.070312 | -0.05407 | 0.356386 | 1.586944 | -0.29262 | 2.617162 | 0.548447 | 0.707173 | -0.00429 | 0.372301 | 0.04543 | 0.151661 | 1.338622 | 0.689332 | 0.34247 | 0.600986 | 0.061997 | -0.16691 | 0.299977 | 0.362622 |
| 23.06944 | 0.346531 | 0.037974 | 0.144372 | 0.00897 | 0.199734 | -0.05049 | 0.463651 | 1.433843 | -0.14219 | 2.467291 | 0.526112 | 0.3857 | 0.048605 | 0.409642 | -0.03357 | 0.036989 | 0.898981 | 0.520982 | 0.338568 | 0.544652 | 0.097818 | -0.03072 | 0.188004 | -0.00475 |
| 23.08333 | 0.346929 | 0.153412 | 0.020322 | -0.0074 | 0.110575 | -0.057 | 0.688263 | 1.218651 | -0.09543 | 2.217962 | 0.472869 | 0.166215 | 0.062505 | 0.396771 | -0.06484 | -0.05568 | 0.586841 | 0.352215 | 0.270234 | 0.62066 | 0.018371 | 0.110108 | 0.118875 | -0.36618 |
| 23.09722 | 0.377097 | 0.230536 | -0.00464 | 0.003711 | -0.0934 | -0.04325 | 0.739951 | 1.134188 | -0.107 | 2.024919 | 0.444601 | 0.11751 | 0.044306 | 0.317695 | -0.01992 | -0.13001 | 0.505278 | 0.155105 | 0.102384 | 0.651259 | -0.08981 | 0.201051 | 0.139133 | -0.58243 |
| 23.11111 | 0.502768 | 0.210172 | 0.014853 | 0.01376 | -0.30947 | -0.01711 | 0.454132 | 0.991097 | -0.06274 | 1.773553 | 0.329765 | 0.064634 | 0.041585 | 0.227074 | 0.041211 | -0.08418 | 0.349707 | 0.023324 | 0.030605 | 0.716432 | -0.15913 | 0.210579 | 0.193118 | -0.66291 |
| 23.125 | 0.612241 | 0.1543 | 0.01237 | 0.021348 | -0.26292 | -0.0126 | 0.022984 | 0.932404 | 0.021291 | 1.281597 | 0.222766 | -0.00217 | 0.051206 | 0.117766 | 0.081703 | 0.000992 | 0.213646 | -0.0855 | -0.04108 | 0.691817 | -0.19179 | 0.166055 | 0.173019 | -0.45203 |
| 23.13889 | 0.583723 | 0.096812 | 0.001834 | 0.024414 | -0.07586 | -0.01851 | -0.479 | 0.771967 | -0.01065 | 0.778242 | 0.149821 | -0.00378 | 0.043523 | 0.072256 | 0.050749 | -0.02827 | 0.157652 | -0.12781 | -0.04276 | 0.661245 | -0.21602 | 0.150423 | 0.187341 | -0.04071 |
| 23.15278 | 0.46897 | 0.066033 | -0.01309 | 0.004735 | 0.054011 | -0.03298 | -0.92286 | 0.641303 | -0.12062 | 0.211808 | 0.059959 | 0.048115 | 0.062239 | 0.070248 | -0.0132 | -0.23125 | 0.144831 | -0.0767 | -0.00879 | 0.585356 | -0.20338 | 0.244881 | 0.184318 | 0.290186 |
| 23.16667 | 0.3115 | 0.054023 | -0.07809 | -0.05766 | 0.057861 | -0.06102 | -0.82776 | 0.31998 | -0.119 | -0.20623 | 0.012549 | 0.128632 | 0.098935 | 0.107957 | -0.04773 | -0.52434 | 0.163271 | 0.037579 | -0.00467 | 0.449536 | -0.15746 | 0.353255 | 0.173138 | 0.283429 |
| 23.18056 | 0.157588 | 0.045167 | -0.14085 | -0.11739 | -0.01018 | -0.05242 | -0.41736 | 0.047826 | -0.00951 | -0.43828 | -0.03576 | 0.277884 | 0.091659 | 0.136994 | -0.05778 | -0.69071 | 0.057907 | 0.105435 | 0.013163 | 0.300512 | -0.11475 | 0.419776 | 0.151867 | 0.012148 |
| 23.19444 | 0.053264 | 0.05138 | -0.11525 | -0.12242 | -0.03034 | 0.015275 | 0.007385 | -0.09672 | 0.012989 | -0.59026 | -0.07436 | 0.541492 | 0.07442 | 0.125602 | -0.07276 | -0.59341 | -0.10628 | 0.038689 | -0.01449 | 0.195321 | -0.10165 | 0.438376 | 0.085373 | -0.24487 |
| 23.20833 | 0.037532 | 0.037536 | -0.02739 | -0.0706 | -0.00298 | 0.062856 | 0.189143 | -0.18271 | 0.012968 | -0.59185 | -0.07784 | 0.75605 | 0.058542 | 0.097559 | -0.06346 | -0.20729 | 0.005033 | -0.06998 | -0.02155 | 0.125412 | -0.10274 | 0.418951 | -0.01673 | -0.21276 |
| 23.22222 | 0.071893 | 0.039345 | 0.038566 | -0.00541 | 0.026069 | 0.058678 | 0.106623 | -0.23814 | 0.077049 | -0.25675 | -0.06082 | 0.662685 | 0.035396 | 0.069638 | -0.0344 | 0.183853 | 0.204456 | -0.137 | 0.041256 | 0.06676 | -0.02765 | 0.416769 | -0.05605 | 0.117336 |
| 23.23611 | 0.142529 | 0.037593 | 0.057728 | 0.037333 | 0.012753 | 0.024404 | 0.02459 | -0.15605 | 0.160635 | 0.257554 | -0.03804 | 0.542507 | 0.03804 | 0.065139 | 0.006288 | 0.352913 | 0.196873 | -0.07868 | 0.088139 | 0.025646 | 0.118963 | 0.48612 | 0.054944 | 0.52264 |
| 23.25 | 0.249343 | 0.031468 | 0.080846 | 0.030945 | -0.05676 | -0.00253 | 0.135223 | 0.177991 | 0.237805 | 0.581357 | -1.5E-05 | 0.463196 | 0.064133 | 0.113692 | 0.042175 | 0.185066 | 0.177619 | 0.089068 | 0.018004 | 0.07704 | 0.275758 | 0.593716 | 0.201767 | 0.775989 |
| 23.26389 | 0.402495 | 0.025263 | 0.145362 | 0.021324 | -0.08071 | 0.007126 | 0.338291 | 0.52673 | 0.256224 | 0.694862 | 0.042351 | 0.506554 | 0.07668 | 0.187405 | 0.051459 | -0.06913 | 0.118557 | 0.31194 | -0.13834 | 0.282027 | 0.373679 | 0.677273 | 0.257534 | 0.627189 |
| 23.27778 | 0.537207 | 0.051941 | 0.204118 | 0.061896 | 0.012012 | 0.025585 | 0.532076 | 0.607887 | 0.244991 | 0.706265 | 0.073342 | 0.65715 | 0.098457 | 0.255555 | 0.042032 | -0.14391 | 0.163814 | 0.545015 | -0.11431 | 0.551727 | 0.365612 | 0.757286 | 0.261595 | 0.401287 |
| 23.29167 | 0.660338 | 0.095671 | 0.252456 | 0.132286 | 0.144275 | -0.05058 | 0.666082 | 0.455248 | 0.259155 | 0.864349 | 0.073777 | 0.859315 | 0.075971 | 0.1735 | 0.033259 | -0.03227 | 0.344816 | 0.544107 | 0.170119 | 0.790171 | 0.255161 | 0.815713 | 0.205532 | 0.284633 |
| 23.30556 | 0.779941 | 0.119506 | 0.307602 | 0.182592 | 0.264464 | -0.2429 | 0.675664 | 0.286066 | 0.2893 | 1.185124 | 0.083757 | 0.964674 | -0.01418 | 0.047039 | 0.064998 | 0.236636 | 0.50302 | 0.39214 | 0.464262 | 1.01961 | 0.158012 | 0.8876 | 0.12452 | 0.449072 |
| 23.31944 | 0.837671 | 0.11834 | 0.335382 | 0.211482 | 0.212001 | -0.26433 | 0.497131 | 0.162089 | 0.281013 | 1.294556 | 0.143887 | 0.890739 | -0.05354 | 0.014528 | 0.233576 | 0.343964 | 0.30207 | 0.173154 | 0.539882 | 1.124493 | 0.187967 | 0.95812 | 0.107027 | 0.940398 |
| 23.33333 | 0.763777 | 0.084732 | 0.331136 | 0.189029 | 0.049366 | -0.01605 | 0.342099 | 0.12848 | 0.264107 | 1.120558 | 0.223049 | 0.646342 | -0.0501 | 0.055241 | 0.40301 | 0.304261 | 0.308992 | -0.00737 | 0.433464 | 0.863976 | 0.253324 | 0.941204 | 0.159467 | 1.125112 |
| 23.34722 | 0.768342 | 0.030934 | 0.404369 | 0.134213 | -0.12036 | 0.13803 | 0.347435 | 0.104494 | 0.322186 | 1.115296 | 0.321521 | 0.453151 | 0.061177 | 0.120563 | 0.358765 | 0.167099 | 0.304236 | -0.05664 | 0.283167 | 0.613855 | 0.369704 | 1.014214 | 0.286865 | 0.73189 |
| 23.36111 | 0.853465 | -0.0086 | 0.525562 | 0.125703 | -0.1692 | 0.215247 | 0.498416 | 0.097435 | 0.502659 | 1.335646 | 0.33097 | 0.400718 | 0.122145 | 0.188306 | 0.249147 | -0.06243 | 0.262578 | 0.077237 | 0.265303 | 0.494537 | 0.451961 | 1.121961 | 0.434051 | 0.350784 |
| 23.375 | 0.966349 | 0.006756 | 0.545986 | 0.171221 | -0.04589 | 0.254317 | 0.545089 | 0.304528 | 0.660198 | 1.631315 | 0.277357 | 0.37824 | 0.09663 | 0.247076 | 0.112638 | -0.13472 | 0.325335 | 0.228948 | 0.370359 | 0.490407 | 0.453899 | 1.131321 | 0.552359 | 0.384891 |
| 23.38889 | 0.979709 | 0.132863 | 0.396968 | 0.193705 | 0.199684 | 0.243155 | 0.421157 | 0.399942 | 0.760905 | 1.65809 | 0.287279 | 0.450205 | 0.124324 | 0.189225 | 0.109455 | 0.058885 | 0.551671 | 0.343066 | 0.535593 | 0.648985 | 0.510634 | 1.27016 | 0.527392 | 0.984127 |
| 23.40278 | 1.194756 | 0.466453 | 0.423661 | 0.274461 | 0.381756 | 0.314968 | 0.481425 | 0.412799 | 1.483954 | 1.768225 | 0.627537 | 1.005152 | 0.453471 | 0.106028 | 0.327004 | 0.330569 | 0.869487 | 0.252176 | 0.715935 | 1.1264 | 1.110603 | 2.257154 | 0.713664 | 1.639512 |
| 23.41667 | 1.918686 | 1.123048 | 0.942058 | 0.617571 | 0.361942 | 0.669998 | 0.952645 | 0.752701 | 2.922916 | 2.847547 | 1.328575 | 2.084174 | 1.169637 | 0.046609 | 0.678473 | 0.412667 | 1.776898 | 0.282629 | 0.843302 | 2.02512 | 2.445591 | 4.525707 | 1.113352 | 1.724196 |
| 23.43056 | 2.089753 | 1.691448 | 1.34891 | 0.770456 | 0.273211 | 0.817851 | 1.315396 | 1.149103 | 3.204423 | 5.135376 | 1.548011 | 2.609192 | 1.88031 | -0.05972 | 0.799384 | 0.390135 | 1.828702 | 0.404953 | 0.655764 | 1.542585 | 3.806985 | 6.196884 | 1.657456 | 1.021214 |
| 23.44444 | 2.146461 | 1.331873 | 1.045245 | 0.649709 | 0.103342 | 0.716171 | 1.757284 | 1.249734 | 2.525679 | 6.602774 | 1.219772 | 2.098051 | 1.590869 | -0.27471 | 0.626282 | 1.151531 | 1.642421 | 0.592675 | 0.356959 | 0.723867 | 2.200637 | 3.903882 | 1.20458 | 0.185399 |
| 23.45833 | 2.057218 | 0.747484 | 0.75327 | 0.588864 | -0.01103 | 0.636799 | 0.874979 | 1.096812 | 2.30085 | 4.662307 | 0.939648 | 1.517677 | 1.159449 | -0.23099 | 0.506938 | 1.277541 | 1.084107 | 0.870343 | 0.205282 | 0.438443 | 1.827172 | 1.992448 | 0.523742 | -0.40341 |
| 23.47222 | 1.143233 | 0.872518 | 0.801903 | 0.501443 | -0.01732 | 0.375178 | -0.27177 | 0.669452 | 2.019442 | 2.109671 | 0.531851 | 1.005779 | 1.043888 | -0.00774 | 0.664124 | 0.338473 | 1.053372 | 0.987957 | 0.114974 | 0.499891 | 1.722485 | 1.44768 | 0.378932 | -0.83321 |
| 23.48611 | 0.262712 | 0.908913 | 1.135229 | 0.382233 | 0.001577 | -0.2493 | -1.42452 | 0.14695 | 1.562913 | 0.60784 | 0.236963 | 0.380822 | 0.997031 | 0.087015 | 0.814477 | -0.61089 | 0.904845 | 0.486878 | -0.1322 | 0.465745 | 1.445469 | 1.411084 | 0.385127 | -0.92054 |
| 23.5 | -0.36206 | 0.606576 | 1.361333 | 0.14854 | 0.114561 | -0.82674 | -1.61835 | -0.18219 | 1.090438 | -1.2029 | 0.105131 | 0.053263 | 0.729419 | 0.002686 | 0.885582 | -1.68029 | 0.325682 | 0.026521 | -0.39389 | 0.366413 | 1.073511 | 1.360788 | 0.440086 | -0.311 |
| 23.51389 | -0.85853 | 0.651529 | 1.117553 | -0.04259 | 0.250586 | -0.57963 | -1.10305 | -0.357 | 1.436147 | -3.57738 | 0.169595 | 0.325808 | 0.513474 | -0.10748 | 0.701253 | -1.71625 | -0.39298 | -0.43336 | -0.4217 | -0.17879 | 1.029618 | 0.970379 | 0.514492 | 0.975109 |
| 23.52778 | -1.21 | 0.759485 | 0.594575 | -0.0831 | 0.244392 | -0.21261 | -0.23745 | -0.29435 | 1.546268 | -2.71541 | 0.25531 | 0.864901 | 0.381363 | -0.02411 | 0.301206 | -0.62891 | -0.98087 | -0.67669 | -0.16054 | -0.37834 | 1.118945 | 0.379862 | 0.55216 | 1.500094 |
| 23.54167 | -1.28642 | 0.795227 | 0.336766 | -0.11358 | 0.142827 | 0.12979 | 0.912611 | -0.20038 | 1.256926 | -1.40666 | 0.337728 | 1.432622 | 0.564156 | 0.279265 | -0.06017 | -0.04834 | -0.66101 | -0.59663 | 0.395457 | -0.25629 | 0.975198 | -0.25308 | 0.505299 | 1.465215 |
| 23.55556 | -1.0669 | 0.699253 | 0.153347 | -0.14063 | -0.01414 | 0.278194 | 1.466377 | -0.07569 | 1.1679 | 0.721475 | 0.396995 | 1.708002 | 1.060965 | 0.328767 | -0.24036 | 0.205561 | -0.19959 | -0.43483 | 0.820732 | -0.05369 | 0.769001 | -0.80533 | 0.383258 | 0.773002 |
| 23.56944 | -0.34388 | 0.690243 | -0.00961 | -0.05547 | -0.11966 | 0.20484 | 1.514956 | 0.246843 | 1.085779 | 2.832031 | 0.450465 | 0.714956 | 1.012668 | 0.232014 | -0.17508 | 0.805115 | 0.209541 | -0.09712 | 0.907457 | 0.19433 | 0.606107 | -1.32339 | 0.189893 | -0.19995 |
| 23.58333 | 0.207989 | 0.833802 | 0.042902 | 0.143175 | -0.11419 | 0.087845 | 0.892811 | 0.581909 | 0.757827 | 3.443838 | 0.44941 | -0.0189 | 0.934467 | 0.181106 | -0.03415 | 1.492769 | 0.122749 | 0.404265 | 0.502408 | 0.418798 | 0.582994 | -1.77213 | -0.03093 | -1.14156 |
| 23.59722 | 0.671776 | 0.863882 | -0.00598 | 0.338945 | -0.06035 | -0.00321 | 0.21213 | 0.661274 | 0.600488 | 2.66809 | 0.418323 | -0.53931 | 1.057868 | 0.22485 | 0.053564 | 1.392839 | 0.009749 | 1.002385 | -0.02503 | 0.50653 | 0.917529 | -2.25085 | -0.27056 | -1.84813 |
| 23.61111 | 1.139985 | 0.570464 | -0.23122 | 0.401808 | 0.138455 | -0.07484 | -0.53008 | 0.593272 | -0.04377 | 0.393412 | 0.297013 | -0.66656 | 0.737599 | 0.203614 | 0.122034 | 0.513764 | 0.36346 | 1.174866 | -0.55006 | 0.68972 | 1.311889 | -2.63082 | -0.55926 | -2.41211 |
| 23.625 | 0.881772 | 0.19124 | -0.33999 | 0.327535 | 0.324214 | -0.07694 | -1.41094 | 0.37233 | -0.65935 | -1.37068 | 0.113978 | -0.7309 | 0.374664 | 0.088645 | 0.18668 | -0.21166 | 0.967997 | 0.701783 | -0.91803 | 0.634512 | 1.193188 | -2.08337 | -0.63974 | -1.46887 |
| 23.63889 | 0.200492 | 0.023864 | -0.4884 | 0.221105 | 0.331465 | 0.064403 | -1.60453 | 0.08863 | -1.11939 | -3.06163 | -0.07823 | -0.92029 | 0.060942 | 0.031871 | 0.155921 | -0.91713 | 1.332938 | 0.253353 | -1.00109 | 0.446684 | 0.770479 | -0.82352 | -0.43211 | 0.811137 |
| 23.65278 | -0.4678 | -0.13083 | -0.48123 | 0.141888 | 0.17441 | 0.30558 | -0.81453 | -0.27143 | -1.22031 | -4.32948 | -0.18326 | -0.59332 | -0.2188 | 0.071183 | -0.05602 | -1.42154 | 0.996135 | -0.28184 | -0.52846 | 0.226342 | 0.549158 | 0.313575 | 0.002333 | 1.850623 |
| 23.66667 | -0.96536 | -0.46943 | -0.43577 | 0.048508 | -0.11339 | 0.519911 | -0.05557 | -0.68319 | -0.74057 | -4.34708 | -0.27011 | 0.031175 | -0.4104 | 0.171084 | -0.28711 | -1.21589 | 0.137781 | -0.57612 | 0.279848 | 0.065353 | 0.412035 | 0.740627 | 0.334224 | 1.73101 |
| 23.68056 | -1.32872 | -0.94819 | -0.32633 | 0.030118 | -0.32485 | 0.570135 | 0.449575 | -0.91501 | -0.70506 | -3.74286 | -0.28744 | 0.681968 | -0.37147 | 0.160266 | -0.36868 | -0.3488 | -0.43204 | -0.66971 | 0.71143 | 0.018615 | 0.105889 | 0.535026 | 0.511505 | 0.927187 |
| 23.69444 | -1.64739 | -1.19743 | -0.28648 | 0.001627 | -0.37353 | 0.367183 | 0.35953 | -0.78732 | -0.40379 | -1.55688 | -0.14515 | 0.192097 | -0.14133 | -0.09125 | -0.37431 | -0.12634 | -0.56987 | -0.66579 | 0.514104 | 0.012759 | -0.37026 | 0.042486 | 0.578632 | -0.21573 |
| 23.70833 | -1.24652 | -1.03819 | -0.42173 | 0.033432 | -0.19708 | 0.123514 | 0.048761 | -0.49934 | 0.037928 | 0.28935 | 0.054944 | -0.35748 | -0.06823 | -0.32422 | -0.33295 | 0.324029 | -0.56563 | -0.59524 | 0.116428 | 0.016663 | -0.7169 | -0.58595 | 0.414333 | -1.39618 |
| 23.72222 | -0.57257 | -0.7047 | -0.56245 | 0.124904 | 0.109134 | -0.10989 | -0.37324 | -0.22592 | 0.21301 | 0.759968 | 0.139792 | -0.51857 | -0.0157 | -0.43178 | -0.30872 | 0.274773 | -0.93236 | -0.47852 | -0.42998 | -0.12032 | -0.77398 | -1.22219 | -0.02834 | -2.29056 |
| 23.73611 | -0.08002 | -0.59015 | -0.61766 | 0.183222 | 0.385357 | -0.27079 | -1.00846 | -0.04602 | 0.034337 | 0.115769 | 0.064449 | -0.90948 | -0.03562 | -0.53505 | -0.32247 | -0.05564 | -1.30077 | -0.21453 | -1.04395 | -0.28325 | -0.71727 | -1.58199 | -0.21039 | -2.56872 |
| 23.75 | -0.06135 | -0.65522 | -0.7337 | 0.17391 | 0.309223 | -0.43775 | -1.41553 | 0.072246 | -0.10378 | -0.32933 | 0.017519 | -1.7001 | -0.15826 | -0.68364 | -0.33567 | -0.37025 | -1.67998 | -0.11507 | -1.526 | -0.52527 | -0.53996 | -1.60314 | -0.28491 | -2.43211 |
| 23.76389 | -0.29197 | -0.73647 | -0.84883 | 0.110615 | 0.034403 | -0.5047 | -1.12984 | 0.025477 | -0.31688 | -1.02117 | -0.11065 | -1.9339 | -0.35614 | -0.69682 | -0.37065 | -0.58663 | -2.10335 | -0.20804 | -1.40425 | -0.88233 | -0.48085 | -2.0978 | -0.40837 | -1.59095 |
| 23.77778 | -0.73077 | -0.80422 | -0.89239 | 0.006108 | -0.35066 | -0.44696 | -0.62143 | -0.13172 | -0.7488 | -2.48554 | -0.31393 | -1.58487 | -0.68519 | -0.81412 | -0.43942 | -0.6148 | -1.81617 | -0.49981 | -0.90486 | -1.24034 | -0.67037 | -3.6089 | -0.68888 | -0.91424 |
| 23.79167 | -1.2615 | -0.9821 | -0.81719 | -0.1239 | -0.36936 | -0.42451 | -0.24757 | -0.1931 | -1.17027 | -3.70324 | -0.53965 | -0.8606 | -0.99349 | -0.83565 | -0.4978 | -0.71329 | -2.08732 | -0.82285 | -0.37675 | -1.3368 | -1.00234 | -4.34204 | -0.89334 | -0.54424 |
| 23.80556 | -1.52991 | -1.1721 | -0.69747 | -0.20387 | -0.33819 | -0.5776 | -0.00282 | -0.29438 | -1.18925 | -5.12884 | -0.6826 | -0.04215 | -1.14485 | -0.74576 | -0.5886 | -0.6835 | -2.02939 | -0.95588 | -0.0969 | -1.20599 | -1.45428 | -3.52235 | -0.86503 | -0.3461 |
| 23.81944 | -1.97329 | -1.09928 | -0.60115 | -0.12912 | -0.02199 | -0.65727 | 0.053907 | -0.77297 | -1.14364 | -5.87877 | -0.70186 | 0.11722 | -1.09802 | -0.52472 | -0.76827 | -0.54022 | -1.66106 | -0.63048 | -0.19607 | -0.99468 | -1.8605 | -2.14334 | -0.76014 | -0.24069 |
| 23.83333 | -2.18746 | -0.77535 | -0.55124 | 0.038804 | 0.423789 | -0.67493 | -0.10275 | -0.56759 | -1.22868 | -3.5714 | -0.6598 | -0.05367 | -0.92037 | -0.48509 | -0.75127 | -0.2842 | -1.16858 | -0.18714 | -0.40359 | -0.76025 | -2.15251 | -1.10528 | -0.64359 | -0.22394 |
| 23.84722 | -1.42082 | -0.48989 | -0.54879 | 0.169547 | 0.566631 | -0.56946 | -0.3024 | -0.08714 | -1.06627 | -1.28533 | -0.44565 | -0.49484 | -0.71042 | -0.48046 | -0.4628 | -0.12583 | -0.66289 | 0.139893 | -0.60245 | -0.34294 | -1.7929 | -0.73414 | -0.47979 | -0.50119 |

| | | | | | | | | | | | | | | | | | | | | | | | | |
|---|---|---|---|---|---|---|---|---|---|---|---|---|---|---|---|---|---|---|---|---|---|---|---|---|
| 23.86111 | -0.45614 | -0.22725 | -0.59816 | 0.159568 | 0.351804 | -0.34085 | -0.45769 | 0.187099 | -0.79263 | 0.545926 | -0.13769 | -0.8023 | -0.49979 | -0.45231 | -0.12624 | -0.07406 | -0.30331 | 0.277248 | -0.78226 | 0.118613 | -0.97509 | -0.56964 | -0.31687 | -0.97511 |
| 23.875 | 0.216854 | -0.05126 | -0.49374 | 0.042007 | 0.059368 | -0.16323 | -0.36824 | 0.285672 | -0.60776 | 1.649761 | 0.13702 | -1.00625 | -0.34648 | -0.3001 | -0.0132 | -0.20704 | -0.20967 | 0.222935 | -0.97992 | 0.281242 | -0.38413 | -0.45942 | -0.22021 | -1.21718 |
| 23.88889 | 0.463208 | 0.099559 | -0.29265 | -0.08247 | -0.24539 | -0.06502 | -0.20814 | 0.115595 | -0.64375 | 1.998542 | 0.268355 | -1.41408 | -0.22328 | -0.00586 | -0.05671 | -0.3554 | -0.31035 | 0.029828 | -0.89467 | 0.190304 | -0.14061 | -0.40373 | -0.09488 | -1.17091 |
| 23.90278 | 0.226406 | 0.207524 | -0.27771 | -0.17847 | -0.39623 | -0.07972 | -0.15215 | -0.269 | -0.89856 | 1.218216 | 0.165923 | -1.83702 | -0.07117 | 0.257357 | -0.10127 | -0.43133 | -0.51481 | -0.08114 | -0.62294 | 0.015715 | -0.15307 | -0.15601 | -0.08114 | -0.94536 |
| 23.91667 | -0.37517 | 0.141529 | -0.60647 | -0.24867 | -0.38931 | -0.29444 | -0.41127 | -0.64459 | -1.16739 | -0.67874 | -0.2382 | -1.93884 | -0.06599 | 0.352748 | -0.14925 | -0.69111 | -0.81665 | -0.0884 | -0.46999 | -0.29772 | -0.6445 | -0.04134 | -0.21518 | -0.65283 |
| 23.93056 | -1.07402 | -0.06428 | -0.96556 | -0.24519 | -0.15366 | -0.71145 | -0.74623 | -0.66616 | -1.14031 | -2.53599 | -0.85473 | -1.88305 | -0.10982 | 0.292496 | -0.28708 | -1.01762 | -1.51233 | -0.12529 | -0.4913 | -0.79036 | -1.35514 | -0.42944 | -0.36638 | -0.5213 |
| 23.94444 | -1.86096 | -0.36612 | -1.18245 | -0.13862 | 0.49739 | -1.15087 | -1.07389 | -0.45709 | -0.75523 | -4.12303 | -1.52475 | -2.59199 | 0.016423 | 0.379479 | -0.55554 | -1.32453 | -2.89469 | -0.2107 | -0.59849 | -1.35033 | -1.76057 | -0.96324 | -0.46188 | -0.67893 |
| 23.95833 | -2.6598 | -0.6194 | -1.3509 | -0.26101 | 0.823839 | -1.02871 | -1.38854 | -1.002 | -0.43419 | -3.24494 | -1.6693 | -3.45806 | -0.06182 | 0.288812 | -1.04769 | -1.81019 | -4.6216 | -0.47705 | -0.86232 | -1.99063 | -1.45495 | -1.2299 | -0.67532 | -1.13734 |
| 23.97222 | -2.23868 | -0.75377 | -1.23577 | -0.484 | 0.105025 | -0.68695 | -1.15454 | -1.94678 | -0.44998 | -1.19866 | -1.77512 | -2.77535 | -0.2506 | -0.13821 | -0.83539 | -1.71744 | -4.81556 | -0.69083 | -1.02382 | -2.52236 | -1.0288 | -1.29865 | -0.88617 | -1.57509 |
| 23.98611 | -1.35252 | -0.59065 | -0.90504 | -0.36582 | -0.34564 | -0.46797 | -0.36346 | -1.52096 | -0.56451 | -0.20891 | -1.44362 | -2.27419 | -0.21627 | -0.26834 | -0.49423 | 0.117414 | -2.40461 | -0.20338 | -0.38427 | -2.11907 | -0.72427 | -1.11964 | -0.78123 | -0.99217 |
| 24 | -0.4373 | -0.25315 | -0.35646 | -0.07327 | -0.45364 | -0.3168 | 0.38535 | -0.78669 | -0.63675 | 0.046526 | -0.6689 | -1.06257 | -0.02472 | -0.30459 | -0.20804 | 1.688389 | -0.15679 | 0.18797 | 0.390294 | -1.28458 | -0.47897 | -0.74425 | -0.31133 | 0.161375 |
| 24.01389 | 0.149787 | 0.013207 | 0.163416 | 0.186796 | -0.14327 | -0.1759 | 0.858992 | 0.045782 | -0.57471 | 0.102542 | 0.112394 | -0.14038 | 0.094198 | -0.12634 | -0.01287 | 2.02979 | 1.503877 | 0.405797 | 0.760196 | -0.60301 | -0.14146 | -0.35963 | 0.119321 | 1.436196 |
| 24.02778 | 0.295818 | 0.133451 | 0.334072 | 0.322945 | 0.276517 | -0.09252 | 0.81893 | 0.7125 | -0.36815 | 0.104969 | 0.409019 | 0.360774 | 0.168889 | 0.043494 | 0.128892 | 1.264569 | 2.281357 | 0.701236 | 0.753491 | -0.11684 | 0.08214 | -0.22664 | 0.27576 | 1.565418 |
| 24.04167 | 0.284965 | 0.10472 | 0.265303 | 0.258093 | 0.385341 | -0.07356 | 0.650531 | 1.015458 | -0.17723 | 0.063549 | 0.423909 | 0.580472 | 0.225127 | 0.014403 | 0.202122 | 0.49421 | 2.448614 | 0.86912 | 0.761346 | 0.085735 | 0.001276 | -0.25174 | 0.24509 | 0.973454 |
| 24.05556 | 0.311822 | 0.029204 | -0.20047 | 0.109653 | 0.050206 | -0.10035 | 0.43276 | 0.835137 | -0.19573 | 0.016183 | 0.351224 | 0.736027 | 0.269963 | 0.066379 | 0.181149 | 0.221786 | 2.16929 | 0.517185 | 0.707099 | 0.047774 | -0.23978 | -0.28747 | 0.215274 | 0.48776 |
| 24.06944 | 0.219731 | -0.05513 | -0.73097 | -0.0514 | -0.08749 | -0.07905 | 0.300743 | 0.747827 | -0.19917 | 0.184549 | 0.605129 | 0.811148 | 0.545779 | 1.724524 | 0.211218 | 0.744803 | 2.051007 | 0.660478 | 0.490491 | -0.0447 | -0.16586 | -0.25448 | 0.729872 | 0.843542 |
| 24.08333 | -0.29205 | -0.18433 | -0.04518 | -0.15003 | 0.214411 | 0.239421 | 1.34888 | 1.573724 | 0.363491 | 1.630979 | 1.234347 | 1.031523 | 0.377224 | 3.88662 | 0.758402 | 1.543561 | 2.428488 | 0.866275 | 0.537761 | 0.102713 | 0.453379 | 0.087395 | 1.382911 | 1.874174 |
| 24.09722 | -1.19442 | -0.30441 | 1.237064 | -0.75571 | 0.528278 | 0.392548 | 2.052542 | 1.856739 | 0.95377 | 3.294262 | 1.864247 | 2.152788 | -1.76351 | 0.993604 | 1.574056 | 0.210082 | -1.94661 | -0.15841 | -2.19244 | 1.844867 | -0.16673 | -0.54184 | 2.603409 | -0.6532 |
| 24.11111 | 0.092305 | 0.160409 | 0.173347 | -0.18564 | -0.36579 | 0.229621 | 0.816864 | 0.873143 | -0.43605 | 2.802308 | 0.568812 | 2.410812 | -0.12834 | 1.503164 | 0.381033 | 0.118666 | -3.70997 | -0.84782 | -2.04688 | 1.089081 | 1.050459 | -0.87347 | 0.128122 | 0.815269 |
| 24.125 | 1.319169 | 0.192571 | -1.10603 | 0.394706 | -0.06587 | -0.22026 | -1.53894 | -1.07106 | -1.72847 | -1.11097 | -0.3934 | 1.145172 | -0.28452 | -0.35738 | -0.16688 | -0.90252 | -0.28256 | -0.58716 | -0.16663 | 0.017597 | 0.645611 | 0.820061 | -0.27991 | 2.432102 |
| 24.13889 | 1.165424 | 0.206509 | -0.5253 | 0.107756 | -0.2153 | -0.0595 | -1.52933 | -1.26499 | -1.73475 | -0.19241 | -0.26109 | 1.05863 | -0.86971 | -5.19537 | -0.57999 | -3.63938 | -0.0395 | -0.85699 | -0.77224 | -0.04681 | 0.225882 | -0.18274 | -0.65356 | -1.55509 |
| 24.15278 | 1.1587 | 0.231914 | 0.038551 | 0.030308 | 0.087384 | 0.128432 | -0.52516 | 0.109222 | -0.37099 | 0.947527 | 0.299376 | 1.671863 | -0.32731 | -3.28478 | -0.5732 | -2.54238 | 0.144531 | -0.64841 | -0.52071 | -0.04969 | -0.51749 | -0.48391 | -0.42264 | -2.01021 |
| 24.16667 | 1.359277 | 0.199264 | 0.147152 | -0.05414 | 0.065802 | 0.140351 | -0.58338 | 0.249443 | -0.11699 | 0.798261 | 0.25649 | 1.536457 | 0.280407 | -0.30346 | -0.27081 | -0.95957 | 0.242006 | -0.2714 | 0.258856 | 0.227595 | -0.40025 | -0.11822 | 0.164289 | -1.57458 |
| 24.18056 | 1.413376 | 0.142402 | 0.12003 | -0.15665 | 0.043751 | 0.102415 | -0.63448 | 0.092915 | -0.14961 | 0.376924 | 0.152535 | 1.133815 | 0.269738 | 0.498812 | -0.13711 | -0.52616 | 0.161927 | -0.3038 | -0.10631 | 0.219177 | -0.14904 | -0.06215 | 0.200431 | -1.30379 |
| 24.19444 | 1.266076 | 0.104762 | 0.171194 | -0.20103 | 0.069713 | 0.141127 | -0.39792 | -0.00367 | -0.14763 | 0.140063 | 0.178995 | 0.805555 | 0.208676 | 0.811736 | -0.06342 | -0.37717 | 0.032933 | -0.36633 | -0.47908 | -0.03253 | 0.159153 | -0.0623 | 0.12783 | -0.66828 |
| 24.20833 | 1.304809 | 0.161605 | 0.296944 | -0.14653 | 0.141578 | 0.29919 | -0.02364 | 0.09573 | -0.0701 | 0.294708 | 0.351298 | 0.593631 | 0.307166 | 0.900982 | 0.016229 | -0.17232 | 0.023181 | -0.3386 | -0.31568 | -0.19944 | 0.349622 | 0.057105 | 0.217426 | 0.544272 |
| 24.22222 | 1.331785 | 0.213219 | 0.332195 | -0.01049 | 0.250689 | 0.365887 | 0.310981 | 0.257227 | 0.058041 | 0.511227 | 0.478682 | 0.465311 | 0.425411 | 0.646936 | 0.13684 | 0.030762 | 0.028963 | -0.21162 | 0.105725 | -0.2144 | 0.370562 | 0.181008 | 0.300429 | 1.522455 |
| 24.23611 | 1.020599 | 0.185151 | 0.251914 | 0.129352 | 0.297623 | 0.259286 | 0.417546 | 0.247663 | 0.04116 | 0.506921 | 0.416998 | 0.172949 | 0.387004 | 0.462138 | 0.248998 | 0.196362 | -0.08577 | -0.08622 | 0.449045 | -0.21988 | 0.29643 | 0.206864 | 0.228463 | 1.455765 |
| 24.25 | 0.655116 | 0.107139 | 0.147536 | 0.158384 | 0.225956 | 0.145014 | 0.292236 | 0.084159 | -0.02932 | 0.339368 | 0.253164 | -0.14474 | 0.288356 | 0.343858 | 0.360682 | 0.351386 | -0.20328 | -0.02465 | 0.671576 | -0.2125 | 0.258671 | 0.214495 | 0.108818 | 0.697512 |
| 24.26389 | 0.382781 | 0.06371 | 0.079303 | 0.048479 | 0.06757 | 0.088512 | 0.045728 | -0.02008 | -0.05254 | -0.01691 | 0.111476 | -0.35179 | 0.150856 | 0.26589 | 0.318282 | 0.35918 | -0.14597 | -0.02416 | 0.588843 | -0.07651 | 0.311391 | 0.223183 | 0.009689 | -0.14532 |
| 24.27778 | 0.205004 | 0.028801 | 0.07023 | -0.06101 | -0.05317 | 0.114423 | -0.01488 | 0.148154 | 0.02349 | -0.36199 | 0.150552 | -0.32417 | 0.168354 | 0.37878 | 0.206947 | 0.164038 | 0.064026 | -0.04387 | 0.397889 | 0.120184 | 0.402412 | 0.207349 | -0.03985 | -0.02805 |
| 24.29167 | 0.150198 | -0.01504 | 0.148089 | -0.06642 | -0.07465 | 0.234742 | 0.19287 | 0.47891 | 0.276555 | -0.36214 | 0.375208 | 0.008545 | 0.328476 | 0.535917 | 0.21405 | 0.160414 | 0.335677 | -0.00161 | 0.192477 | 0.450865 | 0.604068 | 0.220962 | 0.038992 | 1.527985 |
| 24.30556 | 0.286105 | -0.03244 | 0.26861 | 0.100725 | 0.042154 | 0.421472 | 0.553064 | 0.925251 | 0.406993 | 0.087818 | 0.615837 | 0.376244 | 0.498245 | 0.511964 | 0.411238 | 0.334452 | 0.539491 | 0.13987 | 0.247119 | 0.889856 | 0.973529 | 0.3598 | 0.252083 | 3.218329 |
| 24.31944 | 0.581694 | 0.034971 | 0.381507 | 0.30242 | 0.201588 | 0.591095 | 0.907673 | 1.140836 | 0.328428 | 0.825847 | 0.685818 | 0.571379 | 0.570739 | 0.495221 | 0.715513 | 0.501261 | 0.713716 | 0.335036 | 0.590396 | 1.364071 | 1.25625 | 0.657559 | 0.546492 | 2.689803 |
| 24.33333 | 0.910576 | 0.115695 | 0.461494 | 0.366284 | 0.295968 | 0.684926 | 0.947336 | 0.952872 | 0.203835 | 1.540299 | 0.644673 | 0.580966 | 0.59904 | 0.46513 | 1.047753 | 0.637283 | 0.914838 | 0.483473 | 0.864308 | 1.683538 | 1.353414 | 0.924205 | 0.785345 | 1.430662 |
| 24.34722 | 1.11944 | 0.162312 | 0.466822 | 0.318323 | 0.209992 | 0.615639 | 0.730788 | 0.645118 | 0.157403 | 1.960063 | 0.547758 | 0.595417 | 0.508828 | 0.44088 | 0.952481 | 0.555227 | 1.106805 | 0.4234 | 1.045355 | 1.82558 | 1.123829 | 0.97957 | 0.772238 | 0.350325 |
| 24.36111 | 1.175742 | 0.181536 | 0.443803 | 0.228614 | 0.050476 | 0.490124 | 0.468735 | 0.289693 | 0.241926 | 2.144347 | 0.424595 | 0.664367 | 0.469233 | 0.471905 | 0.612328 | 0.590876 | 1.108893 | 0.207903 | 1.235907 | 1.740861 | 0.837825 | 0.874794 | 0.582991 | -0.37419 |
| 24.375 | 1.046648 | 0.157094 | 0.363607 | 0.132276 | -0.11843 | 0.441705 | 0.251609 | 0.021923 | 0.434189 | 2.078679 | 0.325221 | 0.703199 | 0.37459 | 0.48483 | 0.216281 | 0.541367 | 0.87308 | -0.00906 | 0.824705 | 1.419313 | 0.6387 | 0.626275 | 0.360098 | -0.5851 |
| 24.38889 | 0.80278 | 0.130718 | 0.332376 | 0.114989 | -0.14826 | 0.478697 | 0.214588 | -0.06024 | 0.655251 | 1.583245 | 0.311437 | 0.727736 | 0.340683 | 0.338342 | 0.064688 | 0.258846 | 0.726636 | -0.1078 | 0.489506 | 0.986849 | 0.69002 | 0.561008 | 0.237397 | -0.17358 |
| 24.40278 | 0.829656 | 0.272046 | 0.613608 | 0.328788 | 0.006955 | 0.811642 | 0.548276 | 0.075514 | 1.03527 | 1.191698 | 0.536885 | 0.795932 | 0.492638 | 0.07775 | 0.28735 | 0.149414 | 0.838409 | -0.02216 | 0.593367 | 0.898899 | 1.179918 | 0.964816 | 0.254973 | 0.674287 |
| 24.41667 | 1.432688 | 0.628938 | 1.575907 | 0.613628 | 0.250351 | 1.535203 | 1.224081 | 0.338795 | 1.697574 | 1.71851 | 1.067748 | 1.324256 | 0.886267 | -0.1533 | 0.783086 | 0.394099 | 1.533808 | 0.234843 | 1.062618 | 1.365859 | 2.23336 | 1.855695 | 0.396086 | 1.444715 |
| 24.43056 | 1.908404 | 0.862623 | 2.115819 | 0.738864 | 0.379969 | 2.020338 | 1.708021 | 0.742866 | 1.833807 | 3.196292 | 1.621826 | 1.828036 | 1.354437 | -0.33927 | 1.165663 | 0.721703 | 2.195038 | 0.5939 | 1.216459 | 1.96028 | 3.277569 | 2.383736 | 0.543193 | 1.832219 |
| 24.44444 | 2.447802 | 0.592291 | 1.53395 | 0.639403 | 0.443893 | 1.529465 | 1.685207 | 0.846346 | 1.409967 | 4.837334 | 1.788548 | 1.6739 | 1.498742 | -0.47031 | 1.049864 | 1.055655 | 2.32083 | 0.787188 | 0.792819 | 2.278132 | 2.427655 | 1.56018 | 0.451135 | 1.257804 |
| 24.45833 | 3.476647 | 0.224421 | 1.135528 | 0.608373 | 0.353663 | 1.130986 | 1.060419 | 0.875565 | 2.601795 | 4.896615 | 1.496559 | 1.160962 | 1.284568 | -0.29283 | 1.013174 | 1.307591 | 2.227476 | 0.891629 | 0.556144 | 1.923287 | 1.589994 | 0.977808 | 0.341465 | 0.636925 |
| 24.47222 | 2.626234 | 0.335156 | 1.236786 | 0.578887 | 0.21582 | 0.845992 | 0.288682 | 0.969099 | 3.275782 | 4.194284 | 0.908786 | 0.815821 | 1.26843 | 0.08896 | 0.958645 | 1.043677 | 1.97262 | 0.80507 | 0.576254 | 1.42899 | 1.364067 | 0.764804 | 0.431255 | 0.34039 |
| 24.48611 | 1.129779 | 0.489323 | 1.392978 | 0.497652 | 0.000737 | 0.629134 | -0.48443 | 0.691912 | 2.263445 | 2.311646 | 0.227031 | 0.476822 | 1.200813 | 0.231262 | 0.743151 | 0.581544 | 1.521319 | 0.514185 | 0.354629 | 0.995202 | 1.153256 | 0.645647 | 0.478834 | -0.20278 |
| 24.5 | 0.1721 | 0.494769 | 1.420584 | 0.425593 | -0.13019 | 0.41693 | -1.14066 | 0.257917 | 1.172731 | 0.225637 | -0.2837 | 0.165281 | 0.989079 | 0.127079 | 0.336548 | 0.124982 | 1.042497 | 0.206407 | 0.055611 | 0.542028 | 1.0203 | 0.520605 | 0.464897 | -0.94937 |
| 24.51389 | -0.57238 | 0.415783 | 1.036634 | 0.504726 | -0.05468 | 0.309938 | -1.24679 | 0.092954 | 0.321434 | -1.66067 | -0.09865 | -0.15412 | 0.764042 | -0.01291 | -0.01043 | -0.24813 | 0.745673 | -0.06472 | -0.1575 | 0.099583 | 1.066335 | 0.618705 | 0.421962 | -1.51143 |
| 24.52778 | -1.18159 | 0.384572 | 0.901898 | 0.870106 | 0.019259 | 0.220087 | -1.08384 | -0.0378 | 1.221304 | -2.5405 | 0.462426 | -0.3109 | 0.552155 | -0.10874 | -0.18771 | -0.66498 | 0.445659 | -0.32908 | -0.26465 | -0.35316 | 1.136474 | 0.542113 | 0.472588 | -2.00326 |
| 24.54167 | -1.44971 | 0.341427 | 0.904389 | 0.848817 | 0.360034 | 0.070674 | -0.36761 | 0.047711 | 1.634529 | -3.51677 | 0.949781 | -0.34225 | 0.406193 | -0.12444 | -0.33068 | -1.15757 | 0.137442 | -0.44154 | -0.12743 | -0.77941 | 1.063226 | 0.424297 | 0.450199 | -1.6453 |
| 24.55556 | -1.31568 | 0.328403 | 0.757464 | 0.638584 | 0.364945 | -0.05237 | 0.469872 | -0.05534 | 1.4649 | -3.79609 | 0.876803 | 0.081065 | 0.438986 | -0.07018 | -0.28468 | -1.03441 | -0.09736 | -0.33093 | 0.274599 | -0.98259 | 0.959411 | 0.399144 | 0.351556 | -0.07013 |
| 24.56944 | -1.21582 | 0.34468 | 0.611796 | 0.610112 | 0.251869 | -0.01256 | 1.425617 | -0.34869 | 1.190615 | -2.79969 | 0.732547 | 0.703497 | 0.459028 | 0.017277 | 0.006601 | -0.29861 | -0.39549 | -0.04956 | 0.788791 | -0.48962 | 0.861041 | 0.372425 | 0.240272 | 1.747217 |
| 24.58333 | -0.78745 | 0.448049 | 0.541623 | 0.288933 | 0.110317 | 0.013963 | 1.603128 | -0.28939 | 0.901651 | 0.334633 | 0.582578 | 1.089992 | 0.435028 | 0.07391 | 0.309021 | 0.435781 | -0.65654 | 0.328773 | 1.281248 | 0.421182 | 0.808347 | 0.145587 | 0.035746 | 3.043112 |
| 24.59722 | 0.357096 | 0.617382 | 0.238805 | -0.12437 | -0.05913 | -0.09188 | 0.968163 | 0.140863 | 0.385158 | 3.844193 | 0.158393 | 1.019086 | 0.380951 | 0.052978 | 0.513676 | 1.451823 | -0.86929 | 0.697961 | 1.461267 | 1.412906 | 0.662975 | -0.22313 | -0.17032 | 3.714272 |
| 24.61111 | 1.122127 | 0.860986 | -0.11461 | -0.36435 | -0.25274 | 0.005113 | -0.0174 | 0.54777 | -0.03403 | 5.234736 | -0.41091 | 0.556872 | 0.342234 | -0.0235 | 0.413345 | 1.804092 | -1.03373 | 0.949764 | 1.037522 | 1.998944 | 0.435866 | -0.35483 | -0.33662 | 2.277627 |
| 24.625 | 1.673456 | 1.224517 | -0.37744 | -0.47428 | -0.19551 | 0.063404 | -0.9696 | 0.630098 | 0.212724 | 4.108068 | -0.27165 | -0.02135 | 0.32823 | -0.11441 | 0.26881 | 1.029115 | -1.03841 | 0.830766 | 0.441532 | 1.595326 | 0.266933 | -0.47693 | -0.41859 | 0.622618 |
| 24.63889 | 1.954033 | 1.375502 | -0.56412 | -0.40892 | -0.04331 | 0.084264 | -1.69229 | 0.543262 | 1.483529 | 3.206125 | 0.129942 | -0.1314 | 0.300878 | -0.09633 | 0.105453 | 0.561896 | -1.0223 | 0.561764 | -0.08362 | 0.888707 | 0.196847 | -0.48008 | -0.47454 | -0.49835 |
| 24.65278 | 1.94572 | 1.089924 | -0.64102 | -0.3147 | 0.158306 | 0.120892 | -1.69405 | 0.222462 | 1.004972 | 1.489718 | 0.381941 | -0.03294 | 0.217796 | -0.04056 | -0.04949 | 0.152728 | -0.68402 | 0.306494 | -0.36065 | 0.252736 | 0.206291 | -0.41508 | -0.28515 | -1.79192 |
| 24.66667 | 0.944422 | 0.643675 | -0.6481 | -0.3526 | 0.236753 | 0.075965 | -1.5445 | -0.01074 | -0.0003 | -0.48234 | 0.441229 | -0.29345 | 0.086879 | -0.12041 | -0.18004 | -0.24535 | -0.26927 | 0.067131 | -0.38747 | -0.30456 | 0.225825 | -0.31559 | -0.11502 | -3.14327 |
| 24.68056 | -0.14452 | 0.127044 | -0.66398 | -0.33052 | 0.075035 | 0.207737 | -0.84038 | -0.13976 | -0.93922 | -2.12416 | 0.301896 | -0.76775 | -0.05352 | -0.26269 | -0.28537 | -0.63291 | 0.067243 | -0.17136 | -0.40022 | -0.62965 | 0.170881 | -0.26479 | -0.00491 | -3.45564 |
| 24.69444 | -1.07017 | -0.19103 | -0.58441 | -0.44782 | -0.15726 | 0.222691 | -0.0302 | -0.26649 | -1.80303 | -3.81804 | -0.14419 | -1.18957 | -0.12184 | -0.38555 | -0.47236 | -1.04378 | 0.356943 | -0.43691 | -0.60197 | -0.84438 | -0.02449 | -0.18084 | 0.051814 | -2.80224 |
| 24.70833 | -1.6383 | -0.34231 | -0.52086 | -0.65813 | -0.42875 | 0.113415 | 0.496134 | -0.3771 | -1.54873 | -4.56839 | -0.54676 | -1.24517 | -0.18926 | -0.4891 | -0.65072 | -1.13536 | 0.393547 | -0.69437 | -0.94231 | -1.07994 | -0.38157 | -0.15232 | 0.051239 | -1.01693 |
| 24.72222 | -1.48516 | -0.45328 | -0.41856 | -0.88053 | -0.29482 | -0.01683 | 0.39317 | -0.4311 | -0.92324 | -4.60776 | -0.6818 | -0.78641 | -0.24299 | -0.53504 | -0.68924 | -0.71496 | 0.209971 | -0.79711 | -1.16133 | -1.30316 | -0.78365 | -0.16047 | 0.047418 | 0.242565 |
| 24.73611 | -1.12482 | -0.53453 | -0.32103 | -0.51926 | -0.03445 | -0.13087 | -0.03238 | -0.47514 | -0.69061 | -4.97824 | -0.26486 | -0.4044 | -0.29236 | -0.5833 | -0.58527 | -0.21292 | -0.09159 | -0.72916 | -1.03323 | -1.16589 | -1.10783 | -0.3502 | 0.137141 | 0.312068 |
| 24.75 | -1.04756 | -0.54211 | -0.31496 | -0.24339 | 0.260879 | -0.22117 | -0.51931 | -0.40375 | -0.70978 | -4.50008 | 0.017338 | 0.171555 | -0.43137 | -0.64377 | -0.46742 | 0.199767 | -0.34616 | -0.62475 | -0.86712 | -1.13915 | -1.22032 | -0.47582 | 0.157208 | -0.20943 |
| 24.76389 | -1.29526 | -0.51248 | -0.32411 | -0.50993 | 0.304434 | -0.29874 | -1.00342 | -0.21673 | -0.58329 | -3.4025 | 0.042704 | 0.779335 | -0.5705 | -0.73703 | -0.43686 | 0.159188 | -0.57465 | -0.29045 | -0.43818 | -0.87945 | -1.23259 | -0.38076 | -0.23115 | -0.57939 |
| 24.77778 | -1.9349 | -0.53362 | -0.48821 | -1.03567 | 0.134166 | -0.34211 | -1.35788 | -0.0429 | -0.44098 | -2.09339 | -0.0693 | 1.020318 | -0.75764 | -0.84451 | -0.53348 | -0.43807 | -0.74967 | -0.08372 | -0.28982 | -0.75582 | -1.37485 | -0.30015 | -0.47632 | -0.96605 |
| 24.79167 | -2.26147 | -0.64037 | -0.7858 | -1.4004 | -0.00724 | -0.33321 | -1.18948 | -0.13555 | -0.41693 | -0.97653 | -0.3153 | 0.288205 | -0.87317 | -1.03731 | -0.68817 | -1.37712 | -0.91935 | -0.13148 | -0.48019 | -0.74321 | -1.74553 | -0.37568 | -0.60231 | -1.75019 |
| 24.80556 | -2.34125 | -0.74188 | -1.17305 | -1.5399 | -0.09101 | -0.38646 | -0.96114 | -0.42327 | -0.61241 | -0.26553 | -0.58921 | -0.69515 | -1.01701 | -1.24909 | -0.87326 | -1.2859 | -1.11874 | -0.25121 | -0.816 | -0.93665 | -2.22547 | -0.52596 | -0.67269 | -2.79686 |
| 24.81944 | -2.41665 | -0.76414 | -1.17349 | -1.30244 | -0.0592 | -0.42084 | -0.493 | -0.5387 | -0.90338 | -0.01694 | -0.84601 | -1.46272 | -1.13616 | -1.34962 | -1.06272 | -0.93585 | -1.22918 | -0.3071 | -1.00895 | -1.07111 | -2.84396 | -0.74839 | -0.61717 | -2.94288 |
| 24.83333 | -2.07111 | -0.73252 | -1.04327 | -1.1034 | 0.006474 | -0.51782 | -0.0855 | -0.65342 | -1.03267 | 0.040103 | -1.13458 | -1.6549 | -1.07558 | -1.11971 | -0.99993 | -0.79429 | -1.05147 | -0.28684 | -0.76456 | -0.88551 | -2.62311 | -0.99311 | -0.5775 | -2.2314 |
| 24.84722 | -1.33899 | -0.5628 | -0.89321 | -0.4171 | 0.005954 | -0.58558 | 0.172743 | -1.00669 | -0.96531 | -0.01784 | -1.19008 | -1.75537 | -0.8913 | -0.74475 | -0.83718 | -0.68557 | -0.7695 | -0.33351 | -0.48808 | -0.65894 | -2.13544 | -1.28073 | -0.40902 | -1.34729 |
| 24.86111 | -0.35434 | -0.24857 | -0.70923 | -0.11004 | -0.03126 | -0.64699 | 0.227577 | -1.49499 | -0.79259 | -0.30717 | -0.71989 | -1.74397 | -0.76712 | -0.46461 | -0.75331 | -0.3666 | -0.6853 | -0.34354 | -0.49385 | -0.52841 | -1.61761 | -1.42449 | -0.14752 | -0.88508 |
| 24.875 | 0.330334 | -0.08788 | -0.53758 | -0.16543 | -0.05034 | -0.63608 | 0.072159 | -1.54674 | -0.54819 | -0.87069 | -0.10879 | -1.44263 | -0.74582 | -0.19612 | -0.58706 | -0.08032 | -0.68307 | -0.35988 | -0.71899 | -0.37819 | -1.21243 | -1.28703 | 0.006197 | -0.89393 |
| 24.88889 | 0.600913 | 0.069605 | -0.54083 | -0.21275 | -0.01284 | -0.44049 | -0.16254 | -1.15261 | -0.29776 | -1.4637 | 0.424094 | -0.73962 | -0.75685 | 0.233968 | -0.3154 | 0.078988 | -0.67495 | -0.21929 | -0.92714 | -0.14719 | -0.88024 | -0.95378 | -0.00678 | -1.05014 |
| 24.90278 | 0.375999 | 0.138864 | -0.60295 | -0.30925 | -0.00564 | -0.23664 | -0.44568 | -0.67572 | -0.13428 | -1.79843 | 0.498789 | 0.128108 | -0.73207 | 0.4359 | -0.09019 | 0.100196 | -0.6192 | -0.04669 | -0.93792 | 0.018661 | -0.8044 | -0.5909 | -0.13898 | -1.12334 |
| 24.91667 | -0.35369 | -0.01412 | -0.81471 | -0.54035 | -0.12501 | -0.32648 | -0.80542 | -0.4743 | -0.26674 | -1.92232 | -0.15861 | 0.386074 | -0.69519 | 0.348143 | -0.16113 | -0.13593 | -0.53136 | 0.003819 | -0.78314 | -0.11931 | -0.89649 | -0.38506 | -0.31503 | -1.15875 |
| 24.93056 | -1.24676 | -0.20547 | -1.04493 | -0.70314 | -0.25792 | -0.64394 | -1.21936 | -0.50525 | -0.66045 | -1.97517 | -1.07991 | -0.1674 | -0.60824 | 0.270176 | -0.54302 | -0.4568 | -0.44875 | -0.09603 | -0.71839 | -0.65417 | -1.15549 | -0.49802 | -0.53673 | -1.35159 |
| 24.94444 | -2.08346 | -0.42026 | -1.13601 | -0.78951 | -0.14403 | -1.05558 | -1.57646 | -0.695 | -1.30521 | -2.04951 | -1.87224 | -1.00411 | -0.32915 | 0.173525 | -1.182 | -0.5883 | -0.7993 | -0.25472 | -0.73029 | -1.51012 | -1.48452 | -0.73188 | -0.71172 | -1.88802 |
| 24.95833 | -2.96992 | -0.70648 | -1.08391 | -1.04715 | -0.02579 | -1.30278 | -1.80131 | -1.03762 | -0.6013 | -1.79675 | -2.23945 | -1.7626 | -0.16374 | -0.19935 | -1.71729 | -0.67469 | -1.5327 | -0.51093 | -0.54339 | -2.5833 | -1.91139 | -1.17401 | -1.10397 | -2.73737 |
| 24.97222 | -2.75665 | -0.95684 | -1.04529 | -0.90321 | -0.00587 | -1.18946 | -1.3154 | -0.97435 | 0.391141 | -1.29863 | -1.98408 | -1.81672 | -0.12841 | -0.68624 | -1.28862 | -0.35307 | -1.53247 | -0.74501 | -0.15058 | -2.5133 | -1.73774 | -1.56141 | -1.12148 | -1.93734 |
| 24.98611 | -1.97804 | -0.67824 | -0.76487 | -0.41425 | 0.042428 | -0.84124 | -0.38877 | -0.28642 | 1.214484 | -0.50611 | -1.35914 | -1.49089 | 0.051217 | -0.64769 | -0.60759 | -0.11167 | -0.72241 | -0.56089 | 0.537784 | -1.82427 | -0.91058 | -1.39925 | -0.53444 | -0.44513 |
| 25 | -0.75289 | -0.21603 | -0.38503 | 0.0427 | 0.100098 | -0.37993 | 0.291377 | -0.02437 | 1.470995 | 0.283464 | -0.62732 | -0.85585 | 0.203048 | -0.06015 | 0.012021 | 0.061346 | 0.072541 | -0.16076 | 1.204638 | -1.10675 | -0.16421 | -0.90105 | 0.136462 | 0.943752 |
| 25.01389 | 0.191818 | 0.119038 | -0.06266 | 0.330739 | 0.208925 | -0.04911 | 0.640376 | -0.01974 | 1.061463 | 0.829093 | 0.376908 | -0.25249 | 0.273259 | 0.424221 | 0.558205 | 0.270393 | 0.521446 | 0.239309 | 1.274473 | -0.51876 | 0.136715 | -0.37344 | 0.365838 | 2.066642 |
| 25.02778 | 0.713628 | 0.226093 | 0.12641 | 0.478193 | 0.337856 | 0.120783 | 0.635145 | -0.03584 | 0.425162 | 0.954557 | 1.004614 | 0.136682 | 0.267374 | 0.591118 | 0.808861 | 0.544641 | 0.61405 | 0.44225 | 1.268901 | -0.08289 | 0.167858 | -0.0296 | 0.339803 | 2.339888 |
| 25.04167 | 0.785638 | 0.157789 | 0.189211 | 0.392731 | 0.275459 | 0.177269 | 0.485138 | -0.06392 | -0.02157 | 0.652892 | 0.993842 | 0.305042 | 0.265679 | 0.384349 | 0.671124 | 0.51392 | 0.409536 | 0.490627 | 0.995523 | 0.140894 | 0.199887 | 0.093172 | 0.272656 | 2.086638 |
| 25.05556 | 0.589656 | 0.085378 | 0.133145 | 0.211607 | 0.128072 | 0.124611 | 0.344144 | -0.0608 | -0.26703 | 0.214394 | 0.644944 | 0.31538 | 0.209644 | 0.067926 | 0.387863 | 0.424251 | 0.157618 | 0.456218 | 0.712543 | 0.223626 | 0.150097 | 0.030039 | 0.112395 | 1.576444 |
| 25.06944 | 0.306388 | 0.049697 | 0.11069 | 0.018562 | -0.06029 | 0.036065 | 0.285198 | -0.02499 | -0.38124 | 0.002557 | 0.425582 | 0.287483 | 0.153928 | -0.16215 | 0.143229 | 0.282195 | -0.03017 | 0.384947 | 0.6646 | 0.180705 | 0.044316 | -0.10814 | -0.04466 | 0.971682 |
| 25.08333 | 0.048555 | 0.048122 | 0.094891 | -0.14463 | -0.23942 | -0.04612 | 0.253256 | 0.025616 | -0.29265 | 0.006018 | 0.416888 | 0.323346 | 0.083603 | -0.23004 | -0.03987 | 0.11973 | -0.08959 | 0.305286 | 0.679677 | 0.128238 | -0.03047 | -0.20556 | -0.16479 | 0.68039 |
| 25.09722 | -0.04622 | 0.064165 | 0.095227 | -0.184 | -0.27619 | -0.0683 | 0.212629 | 0.098861 | 0.033885 | 0.08528 | 0.550781 | 0.481198 | 0.05636 | -0.01147 | -0.13401 | 0.057947 | -0.02517 | 0.278902 | 0.633628 | 0.158232 | -0.03526 | -0.16479 | -0.12423 | 0.735506 |
| 25.11111 | 0.14555 | 0.066187 | 0.107924 | -0.08528 | -0.04622 | -0.02018 | 0.182896 | 0.236259 | 0.264619 | 0.190357 | 0.66702 | 0.670544 | 0.054443 | 0.304772 | -0.12468 | 0.124339 | 0.067603 | 0.298662 | 0.575899 | 0.249591 | 0.086021 | -0.07365 | 0.164712 | 1.010907 |
| 25.125 | 0.463545 | 0.05967 | 0.118768 | 0.014952 | 0.201215 | 0.101505 | 0.112232 | 0.343786 | 0.260123 | 0.258867 | 0.658504 | 0.901106 | 0.052651 | 0.465875 | -0.03488 | 0.239324 | 0.072539 | 0.292454 | 0.55321 | 0.342058 | 0.244545 | 0.077886 | 0.459098 | 1.182294 |
| 25.13889 | 0.677666 | 0.048414 | 0.099456 | -0.00599 | 0.199734 | 0.233756 | 0.012797 | 0.421793 | 0.171649 | 0.255916 | 0.615497 | 1.139229 | 0.08888 | 0.326043 | 0.049229 | 0.30791 | 0.024615 | 0.252235 | 0.565795 | 0.419427 | 0.320747 | 0.201087 | 0.558524 | 0.967593 |
| 25.15278 | 0.717813 | 0.047007 | 0.063925 | -0.14098 | 0.023723 | 0.284047 | -0.09894 | 0.547396 | 0.115564 | 0.254751 | 0.523887 | 1.303506 | 0.146633 | 0.164574 | 0.118077 | 0.273726 | -0.0114 | 0.193553 | 0.484382 | 0.500103 | 0.248138 | 0.248078 | 0.49854 | 0.599474 |
| 25.16667 | 0.553508 | 0.024557 | 0.024575 | -0.27249 | -0.16949 | 0.16422 | -0.22721 | 0.689188 | 0.10308 | 0.251753 | 0.451612 | 1.366587 | 0.182394 | 0.023428 | 0.123509 | 0.179876 | -0.04665 | 0.085114 | 0.283143 | 0.571872 | 0.08155 | 0.172391 | 0.403328 | 0.265136 |
| 25.18056 | 0.334508 | -0.00624 | 0.021469 | -0.27257 | -0.19754 | 0.038134 | -0.30147 | 0.681752 | 0.184254 | 0.197434 | 0.380418 | 1.236062 | 0.182268 | 0.004519 | 0.093408 | 0.073687 | -0.05199 | -0.0121 | 0.053471 | 0.615856 | -0.10793 | 0.078642 | 0.279176 | -0.05184 |
| 25.19444 | 0.138654 | -0.03186 | 0.03574 | -0.15718 | 0.054037 | -0.05404 | -0.23139 | 0.596865 | 0.2887 | 0.155114 | 0.330488 | 0.937655 | 0.17461 | 0.140493 | 0.060222 | 0.046207 | -0.01089 | -0.06248 | -0.17891 | 0.697136 | -0.16942 | 0.111899 | 0.201269 | -0.19013 |
| 25.20833 | -0.0355 | -0.06171 | 0.035833 | 0.055233 | 0.257982 | -0.02161 | -0.09709 | 0.459466 | 0.223343 | 0.144403 | 0.316399 | 0.711315 | 0.152276 | 0.255078 | 0.032925 | 0.027251 | 0.033367 | -0.09411 | -0.34366 | 0.69149 | -0.0669 | 0.144227 | 0.217498 | -0.17654 |
| 25.22222 | -0.19351 | -0.0698 | 0.037038 | 0.211746 | 0.241472 | 0.08567 | 0.014572 | 0.479371 | 0.209237 | 0.182195 | 0.278088 | 0.493556 | 0.138987 | 0.300751 | 0.043273 | 0.041705 | 0.05358 | -0.08509 | -0.25775 | 0.635094 | 0.18148 | 0.167108 | 0.327044 | -0.00404 |
| 25.23611 | -0.30318 | -0.06641 | 0.042864 | 0.309714 | 0.13919 | 0.242831 | 0.134538 | 0.543234 | 0.401141 | 0.228486 | 0.229582 | 0.293924 | 0.163531 | 0.325183 | 0.148733 | 0.093648 | 0.065402 | 0.041584 | 0.163846 | 0.620223 | 0.42642 | 0.288481 | 0.432772 | 0.21467 |
| 25.25 | -0.33853 | -0.07332 | 0.063833 | 0.362752 | 0.093171 | 0.366802 | 0.209566 | 0.622552 | 0.56061 | 0.218638 | 0.222823 | 0.071604 | 0.216851 | 0.2981 | 0.302676 | 0.128311 | 0.064261 | 0.180582 | 0.485906 | 0.619061 | 0.441417 | 0.385276 | 0.425759 | 0.429132 |
| 25.26389 | -0.30513 | -0.05676 | 0.093536 | 0.3504 | 0.120109 | 0.439542 | 0.241384 | 0.346134 | 0.447739 | 0.183054 | 0.219107 | -0.19711 | 0.304286 | 0.23633 | 0.430677 | 0.203685 | 0.068831 | 0.23098 | 0.433542 | 0.583911 | 0.291462 | 0.407094 | 0.311778 | 0.757332 |
| 25.27778 | -0.24896 | -0.04889 | 0.150014 | 0.325203 | 0.175608 | 0.413385 | 0.277726 | 0.074446 | 0.202299 | 0.158832 | 0.243858 | -0.35905 | 0.348035 | 0.179836 | 0.527885 | 0.303717 | 0.104648 | 0.228971 | 0.142857 | 0.558718 | 0.198199 | 0.451615 | 0.171258 | 1.270402 |
| 25.29167 | -0.29627 | -0.07609 | 0.214924 | 0.300828 | 0.198498 | 0.281993 | 0.369024 | -0.03944 | 0.050407 | 0.145552 | 0.293578 | -0.31581 | 0.332342 | 0.143025 | 0.517691 | 0.320751 | 0.114892 | 0.196112 | 0.040087 | 0.595461 | 0.114523 | 0.581677 | 0.084513 | 1.601115 |
| 25.30556 | -0.28985 | -0.08525 | 0.254249 | 0.284907 | 0.188396 | 0.174561 | 0.453085 | -0.00877 | 0.144152 | 0.233799 | 0.398659 | -0.11839 | 0.285498 | 0.209244 | 0.473662 | 0.194224 | 0.090907 | 0.199271 | 0.425893 | 0.691176 | 0.208899 | 0.690679 | 0.156322 | 1.581591 |
| 25.31944 | -0.17151 | -0.0421 | 0.275466 | 0.356855 | 0.19059 | 0.267549 | 0.557647 | 0.106569 | 0.34779 | 0.462761 | 0.569551 | 0.144152 | 0.312672 | 0.313297 | 0.432345 | 0.083534 | 0.060199 | 0.239262 | 0.972211 | 0.7936 | 0.607183 | 0.787887 | 0.391706 | 1.733228 |

| | | | | | | | | | | | | | | | | | | | | | | | | |
|---|---|---|---|---|---|---|---|---|---|---|---|---|---|---|---|---|---|---|---|---|---|---|---|---|
| 25.33333 | 0.080509 | 0.043176 | 0.284633 | 0.485702 | 0.184508 | 0.435223 | 0.540415 | 0.371534 | 0.62005 | 0.570154 | 0.586653 | 0.342816 | 0.335722 | 0.289182 | 0.458701 | 0.114114 | 0.071192 | 0.325669 | 1.31919 | 0.813462 | 1.009191 | 0.8699 | 0.573754 | 1.970113 |
| 25.34722 | 0.446744 | 0.09724 | 0.296077 | 0.607557 | 0.146063 | 0.601096 | 0.43714 | 0.515871 | 0.639638 | 0.47992 | 0.409161 | 0.449644 | 0.264925 | 0.241612 | 0.490623 | 0.180587 | 0.110152 | 0.349885 | 1.49699 | 0.804754 | 0.983148 | 0.915414 | 0.581562 | 1.611915 |
| 25.36111 | 0.748191 | 0.096338 | 0.313559 | 0.64893 | 0.133854 | 0.536174 | 0.370992 | 0.561413 | 0.464982 | 0.467171 | 0.225433 | 0.473049 | 0.154693 | 0.21105 | 0.399246 | 0.253262 | 0.098048 | 0.287653 | 1.385346 | 0.80124 | 0.83419 | 0.870374 | 0.492651 | 0.731105 |
| 25.375 | 0.897376 | 0.072734 | 0.286293 | 0.59084 | 0.135925 | 0.644577 | 0.355678 | 0.542738 | 0.143544 | 0.517036 | 0.215981 | 0.578932 | 0.121749 | 0.169179 | 0.262765 | 0.292596 | 0.047116 | 0.214704 | 1.231123 | 0.870076 | 0.757087 | 0.869264 | 0.432615 | 0.118307 |
| 25.38889 | 0.880958 | 0.077973 | 0.263633 | 0.559754 | 0.171292 | 0.649395 | 0.442576 | 0.371694 | -0.57739 | 0.663018 | 0.410006 | 0.766326 | 0.280007 | 0.261419 | 0.319014 | 0.433126 | 0.167471 | 0.231946 | 1.092581 | 1.051087 | 0.7607 | 1.072515 | 0.441098 | 0.11247 |
| 25.40278 | 1.100002 | 0.169285 | 0.331267 | 0.756253 | 0.278997 | 0.613779 | 0.727837 | 0.256723 | -1.85095 | 1.246838 | 0.743676 | 1.034611 | 0.520561 | 0.430613 | 0.584621 | 0.672752 | 0.629867 | 0.303666 | 0.942836 | 1.358112 | 0.880775 | 1.471337 | 0.54398 | 0.866543 |
| 25.41667 | 1.769752 | 0.364935 | 0.567594 | 1.299043 | 0.448711 | 1.216781 | 1.385648 | 0.062509 | -0.54056 | 2.500315 | 1.387962 | 1.762072 | 0.880997 | 0.448947 | 0.882717 | 1.022476 | 1.389214 | 0.487554 | 0.869435 | 1.893824 | 1.232126 | 2.488983 | 0.704944 | 2.264307 |
| 25.43056 | 2.69387 | 0.476934 | 0.751652 | 1.599321 | 0.493259 | 2.011729 | 1.760657 | -0.2294 | 1.389712 | 3.210859 | 2.145499 | 2.331864 | 1.263919 | 0.35261 | 1.053527 | 1.682735 | 2.009335 | 0.733253 | 0.839092 | 2.664411 | 1.438973 | 3.818268 | 0.788154 | 2.75234 |
| 25.44444 | 3.163647 | 0.374466 | 0.789483 | 1.182015 | 0.589617 | 1.560661 | 1.607753 | -0.03056 | 1.502404 | 3.012042 | 2.380005 | 2.150476 | 1.024449 | 0.018297 | 0.808857 | 2.047972 | 1.66075 | 0.971112 | 0.689879 | 2.573209 | 1.450377 | 3.237962 | 0.63794 | 1.500738 |
| 25.45833 | 3.743152 | 0.288304 | 0.564324 | 0.882489 | 0.495456 | 1.170844 | 1.345945 | 0.217767 | 1.696757 | 3.342059 | 1.909686 | 1.47914 | 0.705512 | -0.07539 | 0.658859 | 2.105189 | 1.388291 | 1.015971 | 0.241494 | 1.967518 | 1.529664 | 2.38346 | 0.576261 | 0.468104 |
| 25.47222 | 4.560006 | 0.219396 | 0.63888 | 0.775586 | 0.324971 | 1.541396 | 0.795675 | 0.409015 | 3.087447 | 3.585485 | 1.16603 | 1.085045 | 0.778775 | 0.047103 | 0.643718 | 1.270533 | 1.152781 | 0.790332 | 0.185902 | 1.690913 | 1.690862 | 2.158301 | 0.580581 | -0.25467 |
| 25.48611 | 3.53513 | 0.149225 | 0.471395 | 0.574113 | 0.161793 | 0.907005 | 0.275248 | 0.388974 | 3.496019 | 3.49411 | 0.438113 | 0.81811 | 0.68562 | 0.080781 | 0.480106 | 0.597704 | 0.876844 | 0.45123 | 0.254416 | 1.165784 | 1.570637 | 1.714472 | 0.351432 | -0.80377 |
| 25.5 | 2.218831 | 0.127148 | 0.250671 | 0.413287 | 0.016814 | 0.51654 | -0.13034 | 0.154213 | 2.26696 | 2.33968 | -0.47179 | 0.613844 | 0.478867 | 0.028034 | 0.317629 | 0.114191 | 0.603718 | 0.247292 | 0.481331 | 0.726998 | 1.329695 | 1.428503 | 0.120712 | -1.14253 |
| 25.51389 | 1.026638 | 0.09802 | 0.261407 | 0.18355 | -0.07263 | 0.320764 | -0.54206 | -0.17331 | 0.792653 | 0.697651 | -1.28521 | 0.339451 | 0.326693 | 0.004524 | 0.305719 | -0.19231 | 0.474322 | 0.12986 | 0.752129 | 0.417937 | 1.064664 | 1.245573 | 0.105927 | -1.05781 |
| 25.52778 | -0.17455 | 0.057892 | 0.586089 | -0.44162 | -0.01792 | 0.336822 | -0.8873 | -0.33831 | -0.78346 | -0.86904 | -1.14699 | -0.16805 | 0.285706 | 0.048683 | 0.325546 | -0.64131 | 0.429543 | -0.01244 | 0.852689 | 0.096019 | 0.813998 | 0.94721 | 0.06087 | -0.44803 |
| 25.54167 | -0.62846 | -0.00297 | 0.70309 | -0.69026 | 0.072746 | 0.240118 | -0.89457 | -0.33572 | -0.99573 | -0.99017 | -0.16997 | -0.76572 | 0.375533 | 0.133846 | 0.253975 | -1.58596 | 0.398921 | -0.27653 | 0.630298 | -0.32257 | 0.684346 | 0.615193 | 0.020143 | 0.589552 |
| 25.55556 | -1.1518 | 0.018407 | 0.839555 | -0.56447 | 0.288589 | -0.01621 | -0.38874 | -0.35522 | 0.50602 | -1.4714 | 1.084007 | -1.08306 | 0.536393 | 0.253094 | 0.392335 | -2.11998 | 0.326625 | -0.47692 | 0.287609 | -0.68284 | 0.888774 | 0.405838 | 0.249542 | 1.499306 |
| 25.56944 | -1.4505 | 0.114917 | 0.749678 | -0.08117 | 0.522568 | -0.17657 | 0.399441 | -0.5898 | 0.752795 | -2.37851 | 1.87061 | -0.94945 | 0.683199 | 0.31696 | 0.599844 | -0.99744 | 0.291798 | -0.22697 | -0.09948 | -0.50127 | 1.199106 | 0.357581 | 0.525217 | 2.253804 |
| 25.58333 | -2.01066 | 0.117629 | 0.680889 | 0.182877 | 0.558484 | -0.25115 | 1.253113 | -0.88272 | 0.446267 | -1.81931 | 1.331404 | -0.25862 | 0.692669 | 0.265675 | 0.609554 | -0.06943 | 0.269672 | 0.256595 | -0.08827 | -0.29977 | 1.451798 | 0.127726 | 0.291813 | 2.832708 |
| 25.59722 | -2.48177 | 0.049516 | 0.76931 | 0.292522 | 0.433501 | -0.47105 | 1.19084 | -0.1906 | 0.20305 | -0.5776 | 0.422728 | 0.447369 | 0.466285 | 0.133689 | 0.403524 | 0.776142 | 0.248381 | 0.846818 | 0.42478 | 0.15101 | 1.38863 | -0.41981 | 0.106568 | 1.854866 |
| 25.61111 | -1.6911 | -0.05305 | 0.591354 | 0.391493 | 0.194465 | -0.47105 | 0.74761 | 0.419495 | 0.44639 | 0.466209 | -0.14463 | 0.834039 | 0.248833 | -0.03039 | 0.041502 | 1.394878 | 0.186948 | 1.104893 | 0.681263 | 0.828681 | 1.146105 | -1.17045 | -0.06656 | 0.330857 |
| 25.625 | -0.3981 | -0.10263 | 0.365808 | 0.475192 | -0.04289 | -0.17197 | 0.150994 | 0.51542 | 0.85675 | 1.783605 | -0.52662 | 0.798394 | 0.237559 | -0.18951 | 0.055791 | 1.443996 | 0.003139 | 0.835432 | 0.90143 | 1.415787 | 0.770322 | -1.41976 | -0.23683 | -1.07944 |
| 25.63889 | -0.01963 | -0.03974 | 0.294315 | 0.520099 | -0.07003 | 0.189182 | -0.38313 | 0.464123 | 1.097209 | 2.99092 | -1.50731 | 0.600377 | 0.297898 | -0.27847 | 0.288596 | 0.746375 | -0.20057 | 0.554894 | 1.459558 | 1.530086 | 0.451628 | -1.25367 | -0.22237 | -2.57659 |
| 25.65278 | 0.412682 | 0.022061 | 0.117601 | 0.400523 | 0.106487 | 0.401068 | -0.71525 | 0.868402 | 0.772991 | 3.181629 | -1.46832 | 0.251161 | 0.286758 | -0.29074 | 0.539937 | 0.051604 | -0.3567 | 0.33666 | 0.933615 | 1.19814 | 0.249888 | -1.1284 | -0.38991 | -3.3819 |
| 25.66667 | 0.678321 | 0.062602 | -0.14794 | 0.092418 | 0.236081 | 0.356477 | -0.92319 | 0.463573 | 0.365581 | 2.569468 | -0.63976 | -0.17782 | 0.19051 | -0.20977 | 0.355498 | -0.21502 | -0.40454 | 0.178769 | 0.178403 | 0.616787 | 0.109705 | -1.12805 | -0.60992 | -3.38235 |
| 25.68056 | 0.9759 | 0.011998 | -0.30744 | -0.327 | 0.240189 | 0.195363 | -1.01214 | 0.015109 | 0.153878 | 1.582295 | 0.282143 | -0.32754 | 0.078677 | -0.15077 | 0.210817 | -0.53665 | -0.33029 | 0.034631 | -0.52724 | 0.151425 | -0.01908 | -1.03199 | -0.7951 | -1.02139 |
| 25.69444 | 1.181142 | -0.05458 | -0.46292 | -0.77653 | 0.008174 | -0.0274 | -1.04865 | -0.22817 | 0.138812 | -0.15885 | 0.964738 | -0.40844 | -0.05713 | -0.17141 | 0.126191 | -0.58811 | -0.3107 | -0.06671 | -1.05286 | -0.19697 | -0.03141 | -0.89043 | -0.84406 | 1.160277 |
| 25.70833 | 0.927865 | -0.15213 | -0.56095 | -0.98442 | -0.09629 | -0.14243 | -0.81282 | -0.22845 | 0.116864 | -1.81946 | 0.896456 | -0.52575 | -0.17117 | -0.25546 | -0.06411 | -0.55044 | -0.33782 | -0.20324 | -1.47768 | -0.51772 | -0.01678 | -0.72419 | -0.18419 | 1.369614 |
| 25.72222 | 0.224706 | -0.19515 | -0.58044 | -0.97088 | -0.18947 | 0.048271 | -0.58637 | -0.09468 | 0.019216 | -2.82426 | 0.274924 | -0.83776 | -0.21954 | -0.31649 | -0.10038 | -0.62238 | -0.38705 | -0.28321 | -1.7331 | -0.7592 | -0.05724 | -0.63324 | 0.202223 | 0.10844 |
| 25.73611 | -0.80676 | -0.18336 | -0.59242 | -0.8762 | -0.34276 | 0.238531 | -0.32467 | 0.01529 | 0.100781 | -3.38975 | -0.44521 | -1.19914 | -0.2146 | -0.4928 | -0.06423 | -0.42294 | -0.39761 | -0.39498 | -1.80485 | -0.95039 | -0.16714 | -0.62204 | 0.080924 | -1.31214 |
| 25.75 | -1.82161 | -0.18714 | -0.4846 | -0.83364 | -0.22632 | 0.155765 | -0.13686 | 0.056714 | -0.08122 | -3.51743 | -0.909 | -1.55597 | -0.23144 | -0.73689 | -0.08696 | -0.4505 | -0.39865 | -0.36668 | -1.6097 | -1.13597 | -0.31952 | -0.5509 | -0.0633 | -2.6536 |
| 25.76389 | -2.18764 | -0.27568 | -0.28707 | -0.95869 | -0.02212 | -0.29606 | -0.09372 | -0.18026 | -0.29284 | -2.58726 | -1.11989 | -1.70702 | -0.26621 | -0.93364 | -0.3558 | -0.98369 | -0.38183 | -0.04464 | -1.59142 | -1.44952 | -0.62588 | -0.36732 | -0.30393 | -3.48067 |
| 25.77778 | -1.99956 | -0.39535 | -0.27429 | -1.18213 | 0.240492 | -0.77558 | -0.09404 | -0.55744 | -0.80666 | -1.33248 | -1.00076 | -1.48723 | -0.30479 | -0.85129 | -1.13127 | -1.61331 | -0.44083 | 0.392797 | -1.32168 | -1.81151 | -1.02007 | -0.34335 | -0.62278 | -3.44287 |
| 25.79167 | -1.64613 | -0.40628 | -0.39171 | -1.15385 | 0.350982 | -1.18568 | -0.02137 | -1.01975 | -1.33812 | -0.64767 | -0.61082 | -1.01767 | -0.27266 | -0.85584 | -1.44866 | -1.62361 | -0.56982 | 0.068541 | -1.3586 | -2.13864 | -1.39014 | -0.54171 | -1.00942 | -2.98658 |
| 25.80556 | -1.4144 | -0.28376 | -0.57812 | -0.88506 | 0.495203 | -1.05788 | 0.065367 | -1.39049 | -1.81735 | -0.5676 | -0.2841 | -0.75886 | -0.31779 | -0.89845 | -0.91272 | -1.10765 | -0.74233 | -0.21749 | -1.69869 | -2.2313 | -1.67918 | -0.89272 | -0.79594 | -1.03072 |
| 25.81944 | -1.25575 | -0.13116 | -0.93458 | -0.87024 | 0.428529 | -1.06956 | -0.02911 | -0.95628 | -2.06152 | -0.77543 | -0.28192 | -0.66273 | -0.54403 | -1.27789 | -0.62962 | -0.83414 | -0.99234 | -0.45246 | -2.42855 | -2.71368 | -1.88661 | -1.56278 | -0.46568 | 0.698437 |
| 25.83333 | -1.4092 | -0.01261 | -1.10662 | -0.87893 | 0.239277 | -1.06011 | -0.29266 | -0.53526 | -1.76698 | -0.88467 | -0.66156 | -0.55281 | -0.66757 | -1.37713 | -0.89715 | -0.47406 | -1.23493 | -0.68471 | -2.41804 | -2.69036 | -1.85267 | -2.10208 | -0.67583 | 1.170701 |
| 25.84722 | -0.83665 | 0.070356 | -0.7314 | -0.81678 | 0.044693 | -0.22906 | -0.58394 | -0.33577 | -1.00636 | -0.99976 | -0.92605 | -0.31078 | -0.70177 | -0.92268 | -0.7334 | -0.07413 | -1.21401 | -0.58457 | -1.49683 | -2.09244 | -1.16347 | -1.87664 | -0.87353 | 0.742928 |
| 25.86111 | -0.36208 | 0.075881 | -0.38589 | -0.71017 | -0.0144 | 0.033476 | -0.7626 | -0.08153 | -0.33251 | -1.29678 | -0.95655 | 0.008769 | -0.68559 | -0.40699 | -0.54581 | 0.136123 | -0.99422 | -0.37096 | -0.66539 | -1.45181 | -0.7816 | -1.27527 | -0.47543 | -0.25374 |
| 25.875 | -0.16936 | 0.054452 | -0.18195 | -0.54396 | 0.048302 | -0.4471 | -0.66153 | 0.057389 | -0.34508 | -1.21974 | -0.74383 | 0.327842 | -0.56542 | -0.20227 | -0.42793 | 0.12721 | -0.73446 | -0.22579 | -0.18978 | -0.90309 | -0.65835 | -0.87425 | -0.55935 | -1.44555 |
| 25.88889 | -0.10522 | 0.130348 | -0.0319 | -0.38227 | 0.085284 | -0.4914 | -0.27562 | -0.10509 | -0.55455 | -0.69354 | -0.34397 | 0.582114 | -0.38719 | -0.17374 | -0.33085 | 0.085042 | -0.65092 | -0.11773 | -0.07111 | -0.27751 | -0.65875 | -0.64047 | -0.79051 | -2.81021 |
| 25.90278 | -0.28306 | 0.05897 | -0.16818 | -0.33701 | -0.11992 | -0.57257 | -0.00766 | -0.30329 | -1.10679 | -0.52732 | -0.18026 | 0.619383 | -0.32647 | -0.24432 | -0.35676 | -0.00786 | -0.69475 | -0.13502 | -0.25833 | 0.151077 | -0.86846 | -0.57045 | -1.13901 | -3.79669 |
| 25.91667 | -0.80657 | -0.11207 | -0.4798 | -0.53146 | -0.46315 | -0.66618 | -0.06089 | -0.35995 | -1.52923 | -0.68314 | -0.42336 | 0.314297 | -0.3786 | -0.26539 | -0.44676 | -0.16415 | -0.78276 | -0.26446 | -0.4345 | 0.075908 | -1.14319 | -0.63806 | -1.39088 | -2.93383 |
| 25.93056 | -1.60018 | -0.19719 | -0.70716 | -0.70629 | -0.69714 | -0.71266 | -0.16442 | -0.463 | -1.98231 | -0.69656 | -0.89239 | -0.11843 | -0.26561 | -0.29357 | -0.3896 | -0.15251 | -0.46746 | -0.24697 | -0.51295 | -0.12824 | -1.29734 | -0.7423 | -1.67713 | -1.84589 |
| 25.94444 | -2.54174 | -0.24264 | -0.58185 | -0.87053 | -0.73616 | -0.90636 | -0.34251 | -0.39189 | -2.32284 | -0.68843 | -0.92937 | -0.70703 | -0.00807 | -0.31524 | -0.3347 | -0.00921 | -0.19775 | -0.05118 | -0.6523 | -0.37146 | -1.28191 | -0.96751 | -1.74165 | -1.39463 |
| 25.95833 | -3.34794 | -0.52886 | -0.3201 | -1.23016 | -0.60236 | -1.37497 | -0.84934 | -0.17582 | -2.28016 | -0.81272 | -0.99793 | -1.50573 | -0.08339 | -0.43574 | -0.85258 | -0.30776 | -0.50751 | -0.27947 | -1.1808 | -0.95717 | -1.36939 | -1.15209 | -1.1455 | -1.63637 |
| 25.97222 | -3.09449 | -0.84347 | -0.30452 | -1.35625 | -0.2554 | -1.47313 | -1.10778 | -0.11981 | -1.64592 | -0.84139 | -1.50721 | -1.92805 | -0.33947 | -0.59449 | -1.3563 | -0.63481 | -0.95396 | -0.69041 | -1.28792 | -1.58388 | -1.30768 | -1.35537 | -0.49426 | -1.82209 |
| 25.98611 | -1.85346 | -0.83304 | -0.38885 | -0.9043 | 0.252883 | -1.19617 | -0.71716 | -0.04698 | -1.31559 | -0.76244 | -1.31967 | -1.66792 | -0.3571 | -0.34912 | -0.95953 | -0.32295 | -0.87621 | -0.74109 | -0.53424 | -1.49337 | -0.95601 | -0.83064 | 0.018735 | -0.91006 |
| 26 | -0.68153 | -0.58718 | -0.32472 | -0.25422 | 0.599483 | -0.65977 | -0.3033 | 0.074313 | -0.81874 | -0.927 | -0.06714 | -0.8343 | -0.19014 | 0.194448 | -0.27999 | 0.083579 | -0.53235 | -0.4169 | 0.372347 | -1.04504 | -0.39025 | -0.30199 | 0.18285 | 0.237768 |
| 26.01389 | 0.089505 | -0.31116 | -0.14723 | 0.199393 | 0.564847 | -0.0749 | -0.06383 | 0.259988 | -0.19032 | -1.52227 | 1.26444 | -0.30896 | -0.04593 | 0.54892 | 0.212439 | 0.400938 | -0.23662 | -0.15071 | 0.89269 | -0.66674 | -0.05978 | -0.00601 | -0.01122 | 1.467449 |
| 26.02778 | 0.441443 | -0.05536 | 0.053385 | 0.395932 | 0.407928 | 0.259876 | 0.025043 | 0.250909 | 0.502658 | -1.11748 | 2.340541 | -0.02855 | 0.038076 | 0.507881 | 0.360613 | 0.440871 | -0.05101 | -0.01629 | 0.944391 | -0.37877 | 0.136794 | 0.280867 | -0.00892 | 1.977279 |
| 26.04167 | 0.526906 | 0.125541 | 0.204513 | 0.318539 | 0.262151 | 0.144795 | 0.119237 | 0.243442 | 0.905118 | -0.1103 | 2.230622 | 0.069215 | 0.072365 | 0.42755 | 0.360452 | 0.313741 | 0.05248 | 0.086818 | 0.793761 | -0.15551 | 0.279084 | 0.442034 | 0.342071 | 1.884537 |
| 26.05556 | 0.428397 | 0.199325 | 0.189687 | 0.231375 | 0.076131 | -0.16267 | 0.225518 | 0.107924 | 1.188124 | 0.338026 | 1.565494 | 0.106555 | 0.076726 | 0.354753 | 0.276646 | 0.215713 | 0.086787 | 0.217232 | 0.689015 | -0.00783 | 0.302517 | 0.32925 | 0.701505 | 1.673315 |
| 26.06944 | 0.211663 | 0.166736 | 0.043208 | 0.120149 | -0.0952 | -0.51787 | 0.271512 | 0.123356 | 1.087062 | 0.304247 | 0.687132 | 0.085056 | 0.010816 | 0.36741 | 0.139578 | 0.176815 | 0.044866 | 0.316571 | 0.692923 | 0.056707 | 0.280265 | 0.202653 | 0.762476 | 1.530054 |
| 26.08333 | 0.026806 | 0.121188 | -0.12037 | 0.017031 | -0.21852 | -0.74694 | 0.231459 | 0.038656 | 0.472681 | 0.239633 | -0.02865 | 0.038805 | -0.0692 | 0.454945 | 0.033545 | 0.113532 | -0.01486 | 0.357088 | 0.802688 | 0.059092 | 0.134603 | 0.040199 | 0.461775 | 1.263045 |
| 26.09722 | -0.07137 | 0.099431 | -0.1625 | -0.01794 | -0.21426 | -0.50496 | 0.185084 | -0.14859 | 0.065176 | 0.200586 | -0.30097 | 0.038188 | -0.10504 | 0.501479 | -0.00181 | 0.062308 | -0.05565 | 0.401763 | 0.981502 | 0.066466 | 0.014406 | -0.06841 | 0.07836 | 1.11786 |
| 26.11111 | -0.03403 | 0.087316 | -0.0572 | 0.085666 | -0.14355 | -0.18994 | 0.181233 | -0.21017 | -0.19799 | 0.135753 | -0.07806 | 0.093439 | -0.09966 | 0.369534 | -0.005 | 0.048203 | -0.0815 | 0.478304 | 1.05826 | 0.110962 | -0.0864 | -0.08568 | -0.27207 | 0.955571 |
| 26.125 | 0.078412 | 0.107296 | 0.112174 | 0.196206 | -0.02239 | 0.079907 | 0.196451 | -0.15953 | -0.31324 | 0.032928 | 0.431125 | 0.133688 | -0.09719 | 0.215435 | 0.024296 | 0.095854 | -0.07708 | 0.509452 | 0.968174 | 0.203756 | -0.1244 | -0.05159 | -0.41137 | 0.803684 |
| 26.13889 | 0.241173 | 0.115674 | 0.216697 | 0.249823 | 0.064055 | 0.1947 | 0.160393 | -0.01019 | -0.23264 | -0.02574 | 0.858561 | 0.159501 | -0.07372 | 0.165857 | 0.102998 | 0.189881 | -0.01678 | 0.490584 | 0.659711 | 0.289055 | -0.10998 | 0.008681 | -0.27657 | 0.601954 |
| 26.15278 | 0.183716 | 0.101198 | 0.301305 | 0.233885 | 0.093361 | 0.181264 | 0.088808 | 0.176407 | -0.02595 | -0.01319 | 0.76777 | 0.20099 | -0.02219 | 0.164183 | 0.166531 | 0.210146 | 0.058473 | 0.432192 | 0.285834 | 0.303084 | -0.05648 | 0.045299 | -0.17382 | 0.416515 |
| 26.16667 | -0.08239 | 0.088523 | 0.3039 | 0.17487 | 0.086464 | 0.129949 | -0.00789 | 0.065785 | 0.075247 | -0.01515 | 0.069233 | 0.218261 | 0.00444 | 0.079998 | 0.140307 | 0.186234 | 0.104399 | 0.374217 | 0.068593 | 0.270634 | -0.04153 | 0.049549 | -0.15978 | 0.429563 |
| 26.18056 | -0.32174 | 0.070023 | 0.287992 | 0.11397 | 0.082007 | 0.15618 | -0.08713 | -0.05633 | 0.009983 | -0.00996 | -0.53679 | 0.224718 | -0.00732 | -0.05843 | 0.050315 | 0.111006 | 0.110784 | 0.339719 | 0.039795 | 0.228062 | -0.06382 | 0.026472 | -0.28162 | 0.660811 |
| 26.19444 | -0.38544 | 0.059031 | 0.25724 | 0.077558 | 0.062161 | 0.243932 | -0.12321 | -0.30326 | -0.20919 | 0.020666 | -0.56127 | 0.232089 | -0.01868 | -0.11613 | -0.02636 | 0.021988 | 0.110503 | 0.324308 | 0.233905 | 0.243995 | -0.08289 | 0.034086 | -0.44016 | 0.869843 |
| 26.20833 | -0.2605 | 0.051046 | 0.268353 | 0.021775 | 0.039544 | 0.320055 | -0.12148 | -0.35169 | -0.40137 | 0.050355 | -0.11631 | 0.253811 | 0.000232 | -0.04297 | -0.05331 | -0.00098 | 0.09354 | 0.255675 | 0.628626 | 0.275913 | -0.06081 | 0.055452 | -0.35956 | 0.809657 |
| 26.22222 | -0.08348 | 0.032828 | 0.242391 | -0.05062 | 0.003597 | 0.356439 | -0.11403 | -0.22455 | -0.36021 | 0.045876 | 0.352637 | 0.264491 | 0.021181 | 0.044722 | -0.01235 | 0.05937 | 0.07586 | 0.198217 | 0.966654 | 0.317019 | 0.011255 | 0.064292 | -0.11162 | 0.538917 |
| 26.23611 | 0.022279 | 0.014474 | 0.201775 | -0.07512 | -0.07244 | 0.352256 | -0.10141 | 0.000254 | 0.060295 | 0.060843 | 0.524935 | 0.281402 | 0.046544 | 0.07098 | 0.093103 | 0.124147 | 0.036167 | 0.183319 | 0.996177 | 0.3759 | 0.111892 | 0.034314 | 0.15451 | 0.369586 |
| 26.25 | 0.147361 | 0.038949 | 0.144884 | 0.001589 | -0.16236 | 0.398299 | -0.07217 | 0.267036 | 0.464415 | 0.128176 | 0.420708 | 0.322857 | 0.082144 | 0.063646 | 0.227562 | 0.197472 | 0.024347 | 0.22949 | 0.747058 | 0.482428 | 0.269412 | 0.043578 | 0.309765 | 0.565363 |
| 26.26389 | 0.233091 | 0.091337 | 0.102993 | 0.140427 | -0.16169 | 0.518465 | -0.07928 | 0.357058 | 0.552284 | 0.293073 | 0.279293 | 0.395061 | 0.105935 | 0.067902 | 0.304554 | 0.241797 | 0.049863 | 0.27786 | 0.474184 | 0.64718 | 0.432773 | 0.085718 | 0.379454 | 1.000454 |
| 26.27778 | 0.226679 | 0.115261 | 0.087058 | 0.276689 | -0.01081 | 0.590426 | -0.08356 | 0.447872 | 0.436277 | 0.50019 | 0.145552 | 0.44366 | 0.085352 | 0.074134 | 0.30951 | 0.237583 | 0.067343 | 0.260536 | 0.468555 | 0.72159 | 0.527741 | 0.119965 | 0.518343 | 1.470069 |
| 26.29167 | 0.242046 | 0.125883 | 0.12637 | 0.36567 | 0.125686 | 0.546806 | -0.01442 | 0.27309 | 0.237267 | 0.505872 | 0.095178 | 0.360638 | 0.084294 | 0.102121 | 0.303608 | 0.309101 | 0.102863 | 0.228142 | 0.768246 | 0.711111 | 0.614944 | 0.192917 | 0.738167 | 1.646261 |
| 26.30556 | 0.387496 | 0.136737 | 0.209999 | 0.499438 | 0.168159 | 0.440662 | 0.132696 | 0.02924 | -0.01462 | 0.373909 | 0.219188 | 0.317446 | 0.140135 | 0.170244 | 0.326066 | 0.388918 | 0.162474 | 0.241053 | 1.05017 | 0.662269 | 0.716827 | 0.318868 | 0.955095 | 1.438731 |
| 26.31944 | 0.695136 | 0.135684 | 0.21258 | 0.677865 | 0.173063 | 0.391743 | 0.296132 | -0.16728 | -0.25826 | 0.324894 | 0.483347 | 0.35438 | 0.186882 | 0.210961 | 0.319407 | 0.415553 | 0.236764 | 0.236314 | 1.215325 | 0.63826 | 0.815292 | 0.420113 | 0.989744 | 1.09287 |
| 26.33333 | 0.963557 | 0.118777 | 0.202052 | 0.814901 | 0.213077 | 0.465172 | 0.347034 | -0.14193 | -0.40034 | 0.412099 | 0.58806 | 0.371412 | 0.201433 | 0.208125 | 0.278387 | 0.45449 | 0.262057 | 0.218711 | 1.255941 | 0.621221 | 0.82837 | 0.414063 | 0.747318 | 0.739589 |
| 26.34722 | 1.06588 | 0.094459 | 0.188669 | 0.785419 | 0.252942 | 0.626912 | 0.374127 | 0.076303 | -0.48221 | 0.609019 | 0.706669 | 0.350473 | 0.188747 | 0.184771 | 0.225521 | 0.48242 | 0.243926 | 0.207444 | 1.064879 | 0.561687 | 0.729359 | 0.382846 | 0.286023 | 0.545384 |
| 26.36111 | 0.920668 | 0.05557 | 0.182569 | 0.786968 | 0.203279 | 0.720883 | 0.346891 | 0.145511 | -0.85952 | 0.807496 | 0.753629 | 0.370261 | 0.136179 | 0.155589 | 0.161168 | 0.395336 | 0.141674 | 0.212444 | 0.733432 | 0.527822 | 0.549269 | 0.372235 | -0.14571 | 0.754347 |
| 26.375 | 0.711815 | 0.020347 | 0.181225 | 0.830323 | 0.084337 | 0.663476 | 0.350724 | 0.187228 | -0.90129 | 0.724911 | 0.4632 | 0.406987 | 0.051455 | 0.12164 | 0.088036 | 0.322144 | 0.029605 | 0.195201 | 0.636093 | 0.500497 | 0.412277 | 0.359299 | -0.26044 | 1.281894 |
| 26.38889 | 0.632496 | 0.042042 | 0.218795 | 0.915956 | 0.038516 | 0.567253 | 0.351753 | 0.384167 | -0.16216 | 0.628138 | 0.030821 | 0.442984 | 0.039911 | 0.128596 | 0.149752 | 0.400369 | 0.05124 | 0.247105 | 0.965519 | 0.576797 | 0.468826 | 0.446053 | -0.02977 | 1.897798 |
| 26.40278 | 0.787897 | 0.117547 | 0.345139 | 1.183033 | 0.151603 | 0.573098 | 0.464651 | 0.445578 | 0.7489 | 0.87139 | 0.238673 | 0.681245 | 0.18377 | 0.170375 | 0.351626 | 0.688319 | 0.277786 | 0.402424 | 1.338082 | 0.838707 | 0.782809 | 0.770926 | 0.466524 | 2.24618 |
| 26.41667 | 1.510437 | 0.262214 | 0.543556 | 2.026772 | 0.312906 | 0.80458 | 0.80092 | 0.637318 | 1.290256 | 1.724971 | 1.347709 | 1.250484 | 0.459225 | 0.199148 | 0.544846 | 1.246435 | 0.698833 | 0.589655 | 1.336219 | 1.315728 | 1.373154 | 1.416816 | 1.126012 | 2.395978 |
| 26.43056 | 2.675583 | 0.491775 | 0.674926 | 2.463683 | 0.399617 | 1.011378 | 1.062132 | 1.21637 | 0.911948 | 3.01234 | 2.829419 | 1.573864 | 0.79367 | 0.160748 | 0.519207 | 2.029823 | 1.059601 | 0.733668 | 1.047556 | 1.771245 | 2.243062 | 1.999954 | 1.998132 | 2.417024 |
| 26.44444 | 3.18704 | 0.327213 | 0.574365 | 1.833673 | 0.415643 | 0.813405 | 1.003812 | 2.162159 | 0.220167 | 4.067387 | 3.64699 | 1.520148 | 0.787679 | 0.07056 | 0.344513 | 2.268469 | 0.853967 | 0.60967 | 0.719538 | 1.706538 | 2.167164 | 1.746638 | 2.104781 | 1.674713 |
| 26.45833 | 3.257361 | 0.337048 | 0.406587 | 1.256519 | 0.486068 | 0.641009 | 0.818095 | 1.828455 | -0.53342 | 5.254834 | 2.846583 | 1.062281 | 0.420628 | 0.056535 | 0.418647 | 1.858421 | 0.663689 | 0.554518 | 0.811018 | 0.995591 | 1.649181 | 1.305438 | 1.430702 | 0.81565 |
| 26.47222 | 3.287322 | 0.431794 | 0.409867 | 1.132731 | 0.36823 | 0.37999 | 0.702197 | 1.256945 | -0.66352 | 5.07466 | 1.634323 | 0.657501 | 0.330789 | 0.170504 | 0.505282 | 1.321469 | 0.71956 | 0.652381 | 1.009057 | 0.76205 | 1.358033 | 1.222448 | 1.221236 | 0.272637 |
| 26.48611 | 2.869089 | 0.423237 | 0.471213 | 0.810615 | 0.169314 | -0.01969 | 0.507767 | 0.760287 | 1.20131 | 4.571307 | 0.411999 | 0.367284 | 0.206879 | 0.127398 | 0.389027 | 0.722151 | 0.815532 | 0.595701 | 0.761353 | 0.552103 | 0.903417 | 1.104677 | 0.825203 | -0.42925 |
| 26.5 | 2.498638 | 0.344591 | 0.539955 | 0.314958 | 0.046664 | -0.31512 | 0.307383 | 0.174595 | 1.128422 | 3.449971 | -0.68398 | 0.172313 | 0.025723 | 0.030781 | 0.245027 | 0.1511 | 0.773761 | 0.422185 | 0.315075 | 0.497204 | 0.490469 | 0.992564 | 0.22796 | -1.1479 |
| 26.51389 | 1.891807 | 0.22632 | 0.431053 | -0.11894 | -0.00853 | -0.39828 | 0.059577 | -0.40137 | 0.69696 | 1.552009 | -1.57706 | 0.076689 | -0.09608 | 0.017616 | 0.183337 | -0.18068 | 0.760404 | 0.327694 | 0.074549 | 0.567978 | 0.267056 | 0.880486 | -0.285 | -1.58572 |
| 26.52778 | 1.234003 | 0.111326 | 0.380067 | -0.2368 | 0.000372 | -0.31967 | -0.17403 | -0.5367 | 0.200424 | -0.30411 | -2.11853 | -0.1928 | -0.09138 | 0.05623 | 0.228715 | -0.50419 | 0.659479 | 0.273066 | -0.10844 | 0.593586 | 0.335851 | 0.738154 | -0.43203 | -1.23408 |
| 26.54167 | 0.558009 | 0.027035 | 0.045398 | -0.20273 | 0.022781 | -0.10774 | -0.27795 | -0.81168 | -0.41192 | -1.90189 | -1.40314 | -0.55657 | 0.116515 | 0.179676 | 0.385045 | -0.66396 | 0.512945 | 0.235622 | -0.25878 | 0.699246 | 0.71927 | 0.69329 | -0.21353 | 0.314814 |
| 26.55556 | -0.26507 | -0.0558 | -0.00667 | -0.28236 | 0.094969 | 0.067553 | -0.02626 | -1.31636 | -0.73192 | -2.72216 | 0.120886 | -0.99545 | 0.382684 | 0.308171 | 0.725216 | -0.50478 | 0.466819 | 0.158444 | -0.18233 | 0.712596 | 1.10003 | 0.792959 | 0.334002 | 1.991516 |
| 26.56944 | -0.98077 | -0.14729 | -0.00554 | -0.30111 | 0.220377 | 0.074677 | 0.361304 | -0.65015 | 0.193789 | -3.46343 | 1.644941 | -1.28019 | 0.605329 | 0.345331 | 0.951913 | -0.08745 | 0.515333 | 0.088023 | 0.188339 | 0.633915 | 1.457994 | 0.859414 | 0.88303 | 2.907797 |
| 26.58333 | -1.73082 | -0.18606 | -0.09297 | 0.134492 | 0.430259 | 0.099922 | 0.688656 | -0.01393 | 0.554742 | -4.1139 | 2.452215 | -1.33769 | 0.76812 | 0.350974 | 0.954275 | 0.401524 | 0.456679 | 0.079652 | 1.06357 | 0.343893 | 1.846358 | 0.844902 | 1.075824 | 2.171933 |
| 26.59722 | -1.98507 | -0.21451 | 0.09926 | 0.732671 | 0.602199 | 0.037 | 0.723441 | -0.12693 | 0.046829 | -2.8555 | 1.777481 | -0.93474 | 0.818313 | 0.314023 | 0.822145 | 0.819614 | 0.387834 | 0.234902 | 2.106527 | 0.035671 | 2.524767 | 0.796924 | 0.856993 | 0.824652 |
| 26.61111 | -1.77244 | -0.23663 | 0.34771 | 1.089715 | 0.527666 | -0.58101 | 0.396041 | -0.23566 | -0.60013 | -1.22281 | 0.330543 | -0.32635 | 0.689041 | 0.247114 | 0.592964 | 0.938565 | 0.320682 | 0.395932 | 2.263984 | -0.19892 | 2.027927 | 0.556383 | 0.786718 | -0.02105 |
| 26.625 | -1.35111 | -0.29353 | 0.525035 | 1.202873 | 0.282948 | -1.0442 | 0.011771 | -0.2045 | -0.42133 | -0.34542 | -0.8523 | 0.23252 | 0.347194 | 0.123897 | 0.253339 | 0.69014 | 0.258984 | 0.431852 | 1.672822 | -0.31673 | 1.207121 | 0.399647 | 0.693547 | -0.59569 |
| 26.63889 | -0.75024 | -0.30389 | 0.225609 | 0.952516 | -0.03592 | -0.91804 | -0.41772 | 0.087232 | 0.014509 | -0.08812 | -2.09073 | 0.650604 | -0.01531 | -0.01192 | -0.11839 | 0.158415 | 0.178686 | 0.271174 | 0.9982 | -0.30189 | 0.547735 | 0.144007 | 0.124002 | -0.98434 |
| 26.65278 | -1.05877 | -0.29148 | 0.292423 | 0.494514 | -0.4364 | -0.77757 | -0.70478 | 0.121192 | 0.36056 | 0.014897 | -3.18472 | 1.055178 | -0.20894 | -0.06372 | -0.22749 | -0.43115 | 0.101071 | 0.118125 | 0.454925 | -0.16488 | 0.050723 | -0.31681 | -0.35775 | -1.32196 |
| 26.66667 | -1.06508 | -0.24182 | 0.081242 | 0.088565 | -0.51255 | -0.46522 | -0.8888 | 0.152142 | 0.383533 | 0.021893 | -3.35949 | 1.01346 | -0.22488 | -0.1452 | -0.12575 | -0.85441 | 0.059312 | 0.055904 | 0.111478 | -0.04014 | -0.59608 | -0.56794 | -0.71242 | -1.60476 |
| 26.68056 | -0.46247 | -0.20396 | -0.0332 | -0.53979 | -0.40968 | -0.10899 | -0.89539 | 0.053508 | -0.09433 | -0.03937 | -2.70522 | 0.764278 | -0.21936 | -0.18841 | 0.001744 | -1.17706 | 0.054313 | 0.049152 | -0.12044 | 0.04532 | -0.58294 | -0.55274 | -0.87563 | -2.12797 |
| 26.69444 | 0.048502 | -0.18357 | 0.012245 | -0.96638 | -0.22499 | 0.117024 | -0.70259 | -0.23555 | -0.51463 | -0.11073 | -1.1613 | 0.365959 | -0.07921 | -0.10796 | 0.053727 | -0.90099 | 0.038285 | 0.0681 | -0.29888 | 0.071336 | -0.56125 | -0.46932 | -0.88172 | -2.46451 |
| 26.70833 | 0.236306 | -0.14387 | -0.09249 | -1.02772 | -0.0241 | 0.057408 | -0.3572 | -0.34234 | -0.56008 | -0.12228 | 0.310685 | -0.05105 | 0.052761 | -0.04895 | 0.07087 | -0.4728 | -0.00917 | 0.108676 | -0.43854 | 0.062082 | -0.45721 | -0.34352 | -0.70356 | -2.24482 |
| 26.72222 | 0.073229 | -0.07685 | -0.32726 | -0.71415 | 0.097901 | -0.09809 | -0.15528 | 0.235417 | -0.33463 | -0.1415 | 1.02906 | -0.38513 | 0.073709 | -0.0279 | -0.05835 | -0.24345 | -0.09252 | 0.097633 | -0.62426 | 0.000071 | -0.25441 | -0.33419 | -0.51307 | -2.31214 |
| 26.73611 | -0.50539 | -0.03056 | -0.48757 | -0.33241 | 0.198234 | -0.51091 | -0.05821 | 0.409069 | 0.162537 | -0.20372 | 1.001756 | -0.63996 | -0.01644 | -0.03564 | -0.26372 | -0.28173 | -0.18211 | 0.048008 | -0.79771 | -0.15757 | -0.07822 | -0.46236 | -0.34661 | -2.41999 |
| 26.75 | -1.12026 | 0.027479 | -0.63475 | -0.26309 | 0.196773 | -0.94992 | -0.10582 | -0.09286 | 0.168591 | -0.28262 | 0.664072 | -0.96709 | -0.16975 | -0.10705 | -0.38865 | -0.3895 | -0.26199 | 0.012241 | -1.04608 | -0.36788 | 0.015096 | -0.56377 | -0.09529 | -2.22517 |
| 26.76389 | -1.38179 | 0.072739 | -0.79804 | -0.69147 | -0.02649 | -1.15855 | -0.27695 | -0.90466 | -0.57914 | -0.34243 | -0.00412 | -0.95707 | -0.35598 | -0.26652 | -0.40393 | -0.58987 | -0.34264 | -0.04479 | -1.38392 | -0.55423 | -0.17861 | -0.67604 | 0.037896 | -2.08845 |
| 26.77778 | -2.09695 | 0.088721 | -0.91473 | -1.34647 | -0.34315 | -0.90377 | -0.48776 | -1.33861 | -1.09741 | -0.47754 | -1.04679 | -0.99432 | -0.46895 | -0.70326 | -0.36468 | -0.79785 | -0.47215 | -0.10448 | -1.82037 | -0.67326 | -0.58576 | -0.80296 | -0.21824 | -2.28832 |
| 26.79167 | -2.1643 | 0.045496 | -0.87533 | -1.8272 | -0.56254 | -0.54058 | -0.58292 | -1.38016 | -1.44575 | -0.63218 | -2.07646 | -0.95959 | -0.22965 | -0.72369 | -0.40336 | -0.87199 | -0.58294 | -0.21419 | -2.21428 | -0.70743 | -1.26263 | -0.86975 | -0.63266 | -2.44374 |

| | | | | | | | | | | | | | | | | | | | | | | | | |
|---|---|---|---|---|---|---|---|---|---|---|---|---|---|---|---|---|---|---|---|---|---|---|---|---|
| 26.80556 | -1.7809 | -0.01442 | -0.65548 | -1.90226 | -0.51551 | -0.54954 | -0.51472 | -1.12547 | -1.56974 | -0.84389 | -2.5108 | -1.07617 | -0.24797 | -0.81926 | -0.45765 | -0.86454 | -0.75116 | -0.27277 | -2.61737 | -0.7809 | -1.47139 | -0.83549 | -0.91447 | -2.02926 |
| 26.81944 | -1.13162 | -0.06789 | -0.4892 | -1.76578 | -0.23389 | -0.80166 | -0.37918 | -0.64611 | -1.47818 | -0.77399 | -1.68561 | -0.83683 | -0.18078 | -0.69439 | -0.51733 | -0.94111 | -0.92935 | -0.32105 | -2.66548 | -0.77942 | -1.25831 | -0.8672 | -1.06739 | -1.15443 |
| 26.83333 | -0.75101 | -0.0654 | -0.3026 | -1.39382 | 0.144444 | -0.91534 | -0.15584 | -0.26307 | -1.17254 | -0.4428 | -0.25354 | -0.41599 | -0.15426 | -0.35169 | -0.48226 | -0.73526 | -0.98275 | -0.37663 | -2.46475 | -0.75017 | -1.22861 | -0.7936 | -1.12454 | -0.57276 |
| 26.84722 | -0.61205 | -0.07676 | -0.20517 | -0.83056 | 0.522492 | -0.79891 | 0.001135 | -0.06275 | -0.49706 | -0.1568 | 0.754194 | -0.05297 | -0.0365 | 0.018126 | -0.445 | -0.53766 | -1.09829 | -0.40873 | -2.25583 | -0.77152 | -1.23215 | -0.79676 | -1.12614 | -0.40303 |
| 26.86111 | -0.57159 | -0.15409 | -0.07477 | -0.26774 | 0.587428 | -0.48204 | 0.020852 | -0.06845 | 0.147742 | -0.17904 | 0.880909 | 0.085254 | 0.070946 | 0.023638 | -0.43522 | -0.32444 | -1.24398 | -0.41969 | -2.26318 | -0.92986 | -1.13736 | -0.97157 | -1.07506 | -0.48286 |
| 26.875 | -0.43313 | -0.17631 | 0.166563 | 0.038181 | 0.169195 | -0.15992 | 0.080496 | 0.045013 | 0.659061 | -0.33582 | 0.430033 | 0.137975 | 0.103389 | -0.06339 | -0.41828 | -0.17721 | -1.2175 | -0.3641 | -1.63621 | -0.88083 | -0.77651 | -0.92013 | -0.91581 | -0.54575 |
| 26.88889 | -0.3413 | -0.06003 | 0.393211 | 0.155518 | -0.25253 | 0.063646 | 0.108962 | 0.333948 | 0.869517 | -0.46583 | -0.08119 | 0.208225 | 0.150949 | -0.09072 | -0.26149 | -0.03108 | -0.98238 | -0.15183 | -0.31504 | -0.65877 | -0.26578 | -0.59208 | -0.65041 | -0.72886 |
| 26.90278 | -0.30029 | 0.006274 | 0.340154 | -0.03951 | -0.50501 | -0.05353 | -0.01059 | 0.469312 | 0.510507 | -0.52024 | -0.53106 | 0.197359 | 0.0749 | -0.13867 | -0.12743 | 0.055322 | -0.75131 | 0.001616 | 0.618437 | -0.43684 | -0.11253 | -0.32899 | -0.36622 | -1.03483 |
| 26.91667 | -0.53789 | 0.006072 | -0.03278 | -0.47701 | -0.661 | -0.58332 | -0.21016 | 0.026917 | -0.32251 | -0.48532 | -1.08364 | -0.00774 | -0.10485 | -0.17892 | -0.06541 | 0.057118 | -0.62668 | -0.04595 | 0.504933 | -0.50529 | -0.35994 | -0.29066 | -0.58165 | -1.36058 |
| 26.93056 | -1.16339 | 0.025363 | -0.45056 | -0.77785 | -0.72479 | -1.13108 | -0.29005 | -0.50337 | -1.49792 | -0.37979 | -1.76387 | -0.27017 | -0.11633 | -0.21524 | 0.07234 | 0.150684 | -0.40622 | -0.13542 | -0.03681 | -0.62808 | -0.91591 | -0.28925 | -1.09973 | -1.44149 |
| 26.94444 | -1.50555 | 0.085636 | -0.45316 | -0.93158 | -0.6103 | -1.17548 | -0.28569 | -0.34589 | -2.74853 | -0.35019 | -2.13756 | -0.41017 | -0.03309 | -0.41688 | 0.122693 | 0.373864 | -0.10823 | -0.03976 | -0.27944 | -0.60319 | -1.28658 | -0.20015 | -1.13559 | -1.59771 |
| 26.95833 | -1.89983 | -0.19567 | -0.19678 | -1.28926 | -0.58901 | -1.09209 | -0.47394 | -0.47429 | -4.1189 | -0.55164 | -1.80794 | -0.41554 | 0.122988 | -0.64835 | -0.14615 | 0.228721 | -0.18492 | -0.05334 | -0.37103 | -0.82421 | -1.53281 | -0.24416 | -0.86727 | -2.27115 |
| 26.97222 | -2.35923 | -0.71011 | -0.10703 | -2.09729 | -0.63583 | -0.97853 | -0.54282 | -0.33904 | -5.35997 | -0.69255 | -1.3021 | -0.41587 | 0.063096 | -0.77623 | -0.47827 | -0.14982 | -0.48382 | -0.20926 | -0.49832 | -1.27033 | -1.86934 | -0.57143 | -1.23356 | -2.53687 |
| 26.98611 | -2.25554 | -0.8061 | -0.03521 | -2.1408 | -0.31801 | -0.60743 | -0.38478 | -0.20694 | -3.35934 | -0.57937 | -0.79752 | -0.32916 | 0.007075 | -0.53836 | -0.5582 | -0.28765 | -0.56679 | -0.17033 | -0.5005 | -1.20624 | -1.47735 | -0.51535 | -0.96597 | -1.96727 |
| 27 | -1.31499 | -0.62737 | 0.099136 | -1.46239 | 0.117601 | -0.12713 | -0.16807 | 0.198583 | 0.161622 | -0.3117 | -0.25007 | -0.12918 | -0.02613 | -0.16744 | -0.42326 | -0.18192 | -0.46883 | -0.13655 | -0.22557 | -0.92969 | -0.73409 | -0.40648 | -0.55727 | -1.19826 |
| 27.01389 | -0.45751 | -0.3921 | 0.212113 | -0.75496 | 0.358982 | 0.275503 | 0.001558 | 0.650308 | 1.809328 | -0.12894 | 0.316096 | 0.052944 | -0.0521 | 0.079122 | -0.27952 | -0.11562 | -0.40579 | -0.10229 | 0.011314 | -0.66519 | -0.28375 | -0.32465 | -0.2785 | -0.61056 |
| 27.02778 | 0.159168 | -0.1854 | 0.318415 | -0.37102 | 0.358498 | 0.50717 | 0.095323 | 0.724402 | 3.075585 | -0.05446 | 0.901444 | 0.054911 | -0.11428 | 0.177902 | -0.19254 | -0.05057 | -0.33341 | -0.07402 | 0.274335 | -0.41001 | 0.179471 | -0.34585 | -0.07327 | -0.19483 |
| 27.04167 | 0.469732 | -0.02286 | 0.455172 | -0.23311 | 0.307609 | 0.5386 | 0.083275 | 0.387425 | 2.973966 | 0.019892 | 1.54241 | -0.01428 | -0.10226 | 0.185792 | -0.10136 | 0.023196 | -0.22523 | 0.010575 | 0.439629 | -0.16342 | 0.416224 | -0.34876 | 0.248093 | 0.13806 |
| 27.05556 | 0.459282 | 0.101863 | 0.654577 | -0.23382 | 0.287542 | 0.39166 | 0.036048 | 0.211218 | 2.588103 | 0.045604 | 1.700305 | -0.09273 | -0.02254 | 0.211837 | -0.01033 | 0.065483 | -0.1457 | 0.11228 | 0.495078 | 0.001537 | 0.48551 | -0.39714 | 0.510542 | 0.352268 |
| 27.06944 | 0.219345 | 0.190915 | 0.534143 | -0.25483 | 0.262914 | 0.228252 | 0.00268 | 0.177116 | 2.010005 | -0.01381 | 1.619773 | -0.11323 | 0.05046 | 0.29731 | 0.057138 | 0.055362 | -0.09745 | 0.180468 | 0.60022 | 0.07432 | 0.531258 | -0.41982 | 0.647095 | 0.518222 |
| 27.08333 | -0.12155 | 0.244518 | 0.191017 | -0.21732 | 0.236584 | 0.017353 | 0.002473 | 0.489319 | 1.262672 | -0.07466 | 1.38105 | -0.04376 | 0.080278 | 0.367251 | 0.078052 | 0.022346 | -0.0713 | 0.214664 | 0.774742 | 0.099815 | 0.58004 | -0.35044 | 0.530506 | 0.538278 |
| 27.09722 | -0.26203 | 0.312041 | 0.126707 | -0.23155 | 0.177114 | -0.07108 | 0.149656 | 0.190366 | 1.125244 | 0.138193 | 1.018193 | 0.150717 | 0.028846 | 0.326777 | 0.047754 | -0.1041 | -0.11406 | 0.166124 | 0.882883 | 0.036845 | 0.569936 | -0.27712 | 0.347363 | 0.361991 |
| 27.11111 | 0.15497 | 0.50969 | 0.368359 | -0.47888 | 0.271579 | 0.237853 | 0.827064 | -1.22422 | 4.107879 | 1.385965 | 0.430392 | 0.829235 | -0.10304 | -0.07429 | -0.00436 | -0.30564 | -0.28324 | 0.091889 | 0.695771 | -0.24552 | 0.313473 | -0.48156 | 0.11565 | 0.035024 |
| 27.125 | -0.5941 | 0.201245 | 0.375747 | -0.77815 | -0.15655 | 0.125727 | 0.94316 | -1.71171 | 2.989558 | 1.172731 | 0.114027 | 0.810339 | -0.31516 | -0.76244 | -0.05817 | -0.62571 | -0.47094 | 0.052525 | 0.187868 | -0.67461 | 0.045783 | -0.80372 | -0.19495 | -0.27103 |
| 27.13889 | 0.893534 | -0.29321 | 0.260853 | 0.794875 | -0.06859 | -0.43476 | -1.5512 | 1.00078 | 0.139297 | -1.06638 | -0.37072 | -1.21639 | -0.26514 | -0.6222 | -0.08037 | -0.38666 | -0.41775 | 0.045061 | 0.162929 | -0.44348 | 0.02186 | -0.60211 | -0.45314 | -0.13971 |
| 27.15278 | 2.560261 | -0.91696 | 0.058887 | 2.641187 | 0.078218 | -0.55222 | -3.10159 | 3.227167 | -0.81445 | -3.09839 | -0.89013 | -3.07313 | 0.031397 | 0.697037 | -0.02733 | 0.441636 | -0.10467 | 0.04898 | 0.995456 | 0.447941 | 0.19176 | 0.104399 | -0.20621 | 0.381492 |
| 27.16667 | 0.672471 | -0.20137 | -0.38147 | 1.331002 | -0.32515 | -0.49565 | -1.60313 | 2.113082 | -3.17997 | -2.671 | -0.30497 | -1.55657 | 0.249968 | 1.471681 | 0.114575 | 0.92039 | 0.205063 | 0.209712 | 1.535551 | 0.954793 | 0.416979 | 0.503134 | 0.082577 | 0.798428 |
| 27.18056 | 0.521845 | 0.243108 | 0.435577 | 0.514073 | -0.19864 | 0.132843 | 0.374679 | 0.630721 | 0.760993 | 0.316703 | 0.461969 | 0.5292 | 0.191721 | 1.029182 | 0.183225 | 0.647625 | 0.205814 | 0.359255 | 1.165707 | 0.776032 | 0.45414 | 0.371873 | 0.047284 | 0.698493 |
| 27.19444 | 0.238553 | 0.29857 | 0.492491 | 0.066676 | -0.01209 | 0.326999 | 0.872329 | 0.490189 | 1.811485 | 1.032535 | 0.501814 | 1.01723 | 0.13812 | 0.534926 | 0.101792 | 0.296513 | 0.103108 | 0.404273 | 0.620917 | 0.494872 | 0.336068 | 0.242979 | 0.155824 | 0.390557 |
| 27.20833 | -0.11567 | 0.194919 | 0.150717 | -0.06976 | 0.014547 | 0.247235 | 0.332043 | 0.64819 | 1.072961 | 0.030401 | 0.19333 | 0.453266 | 0.088831 | 0.34189 | -0.00895 | 0.215415 | 0.046627 | 0.312129 | 0.422493 | 0.245443 | 0.261791 | 0.139906 | 0.143082 | 0.167563 |
| 27.22222 | -0.19319 | 0.108693 | -0.10988 | -0.03126 | -0.07069 | 0.08459 | -0.39509 | 0.445363 | -0.27412 | -1.05067 | -0.34484 | -0.37404 | 0.047458 | 0.235204 | -0.01246 | 0.201917 | 0.016118 | 0.211459 | 0.442566 | 0.141245 | 0.210668 | 0.029297 | 0.11387 | 0.144254 |
| 27.23611 | -0.06696 | 0.016812 | -0.19168 | 0.17722 | -0.1162 | -0.01198 | -0.93386 | 0.080259 | -1.46483 | -1.57563 | -0.79039 | -0.98749 | 0.021747 | 0.206161 | 0.086481 | 0.176461 | -0.01689 | 0.165092 | 0.521652 | 0.180003 | 0.172198 | -0.05023 | 0.031588 | 0.184147 |
| 27.25 | 0.174446 | -0.07186 | 0.061556 | 0.276616 | -0.16211 | -0.02646 | -0.68774 | -0.02979 | -1.46083 | -1.01647 | -0.88043 | -0.7207 | 0.004651 | 0.194659 | 0.156622 | 0.127752 | -0.04439 | 0.167732 | 0.546196 | 0.235218 | 0.118683 | -0.07846 | -0.05284 | 0.144515 |
| 27.26389 | 0.296342 | 0.053751 | 0.548144 | 0.192282 | -0.14543 | 0.107098 | 0.14186 | 0.011019 | -0.31461 | 0.114296 | -0.50832 | 0.28927 | 0.019976 | 0.247336 | 0.150014 | 0.117978 | -0.01299 | 0.22307 | 0.647067 | 0.284787 | 0.138411 | -0.03339 | 0.039477 | 0.1671 |
| 27.27778 | 0.269017 | 0.157432 | 1.021046 | 0.120566 | -0.09353 | 0.194382 | 0.597242 | 0.115474 | 0.602895 | 0.727857 | 0.133679 | 0.695716 | 0.057918 | 0.317053 | 0.109706 | 0.146478 | 0.053779 | 0.247706 | 0.748198 | 0.300739 | 0.202885 | 0.005393 | 0.207241 | 0.312401 |
| 27.29167 | 0.338435 | 0.08152 | 0.77898 | 0.104106 | -0.1602 | 0.225662 | 0.105161 | 0.488664 | 0.627984 | 0.074578 | 0.608484 | 0.070535 | 0.077866 | 0.312957 | 0.054015 | 0.16248 | 0.063289 | 0.229796 | 0.717173 | 0.217249 | 0.188528 | -0.0438 | 0.267269 | 0.474325 |
| 27.30556 | 0.458712 | -0.03025 | 0.306063 | 0.160253 | -0.19075 | 0.310382 | -0.53621 | 0.614636 | -0.05063 | -0.78981 | 0.687719 | -0.92333 | 0.070509 | 0.264089 | -0.01205 | 0.120073 | 0.041121 | 0.202138 | 0.588482 | 0.117362 | 0.161364 | -0.11914 | 0.236442 | 0.538677 |
| 27.31944 | 0.431319 | 0.034518 | 0.198868 | 0.116767 | -0.03795 | 0.42487 | -0.06597 | 0.293424 | 0.013491 | -0.35962 | 0.419973 | -0.61926 | 0.064273 | 0.227809 | -0.01722 | 0.111392 | 0.052583 | 0.181421 | 0.482865 | 0.130652 | 0.20131 | -0.06543 | 0.219918 | 0.52097 |
| 27.33333 | 0.026862 | 0.265973 | 0.221693 | 0.200745 | 0.125809 | 0.565558 | 0.929922 | -0.21574 | 0.784473 | 0.80958 | -0.18299 | 0.695027 | 0.074463 | 0.212126 | 0.063296 | 0.146327 | 0.09973 | 0.174079 | 0.467569 | 0.284883 | 0.265841 | 0.065399 | 0.215727 | 0.516355 |
| 27.34722 | 0.11132 | 0.180156 | 0.352607 | 0.595233 | 0.293575 | 0.615226 | 1.156335 | -0.13895 | 0.692131 | 1.104132 | -0.53427 | 1.053837 | 0.074433 | 0.209123 | 0.152714 | 0.190923 | 0.117247 | 0.170204 | 0.544037 | 0.436353 | 0.366139 | 0.121342 | 0.221703 | 0.539558 |
| 27.36111 | 0.561501 | 0.080836 | 0.304203 | 0.924566 | 0.31121 | 0.49456 | 0.471198 | 0.356421 | -0.47475 | 0.2939 | -0.53547 | 0.291601 | 0.077286 | 0.222787 | 0.175006 | 0.235126 | 0.074427 | 0.129759 | 0.577347 | 0.405108 | 0.298235 | 0.10466 | 0.26291 | 0.481985 |
| 27.375 | 0.821391 | -0.00951 | 0.301014 | 1.025104 | 0.203924 | 0.250699 | -0.00547 | 0.617382 | -1.1044 | -0.2181 | -0.20336 | -0.35469 | 0.030997 | 0.20529 | 0.058761 | 0.226037 | -0.02005 | 0.028636 | 0.493761 | 0.22111 | 0.182858 | 0.057179 | 0.28619 | 0.243897 |
| 27.38889 | 0.416368 | 0.078241 | 0.376148 | 0.566144 | 0.17138 | 0.381987 | 0.668675 | 0.005597 | 0.578143 | 1.342109 | -0.07944 | 0.704141 | 0.011261 | 0.191738 | -0.04448 | 0.155591 | -0.01985 | -0.02837 | 0.465729 | 0.101677 | 0.136707 | 0.035842 | 0.346316 | 0.123954 |
| 27.40278 | 0.010081 | 0.266458 | 0.448187 | 0.524761 | 0.206236 | 0.798694 | 1.801108 | -0.75934 | 2.807936 | 3.531173 | 0.168893 | 2.287891 | 0.12298 | 0.22989 | 0.109069 | 0.173045 | 0.179712 | 0.083182 | 0.703665 | 0.326041 | 0.313824 | 0.104922 | 0.477994 | 0.473507 |
| 27.41667 | 0.776092 | 0.266032 | 0.495361 | 2.039187 | 0.310493 | 1.236148 | 1.970363 | -0.17047 | 3.215963 | 4.22356 | 0.867952 | 2.254511 | 0.252324 | 0.349825 | 0.487492 | 0.369987 | 0.423644 | 0.244167 | 1.190854 | 1.015531 | 0.635757 | 0.179186 | 0.663635 | 1.207941 |
| 27.43056 | 2.3459 | 0.151526 | 0.661263 | 2.953964 | 0.420654 | 1.681478 | 1.066223 | 1.165103 | 1.621002 | 3.685701 | 1.759285 | 1.566733 | 0.369127 | 0.484667 | 0.760273 | 0.872931 | 0.482319 | 0.214438 | 1.606081 | 1.977602 | 0.677598 | 0.093909 | 0.643215 | 1.876429 |
| 27.44444 | 2.818525 | 0.11599 | 0.778809 | 2.300898 | 0.219579 | 1.36882 | 0.771171 | 1.848731 | 0.07631 | 3.033775 | 2.167439 | 1.184661 | 0.525741 | 0.446799 | 0.639239 | 1.16871 | 0.345585 | -0.03923 | 1.543272 | 2.237651 | 0.668378 | -0.02561 | 0.552876 | 2.707631 |
| 27.45833 | 3.436883 | 0.172076 | 0.817965 | 1.973254 | 0.137475 | 1.141687 | 0.721148 | 1.908776 | -0.16356 | 2.877616 | 1.845732 | 0.709658 | 0.343353 | 0.380118 | 0.298244 | 1.01265 | 0.311945 | -0.05422 | 1.337792 | 1.526348 | 0.788192 | 0.035825 | 0.417028 | 1.724067 |
| 27.47222 | 3.228105 | 0.23857 | 0.671972 | 1.704706 | 0.144053 | 0.821035 | 0.683742 | 1.368723 | 0.329304 | 3.011626 | 0.887052 | 0.395509 | 0.318532 | 0.37534 | 0.233188 | 0.812844 | 0.435342 | 0.094459 | 1.227446 | 1.1298 | 0.922354 | 0.292718 | 0.368106 | 0.886908 |
| 27.48611 | 2.115895 | 0.213686 | 0.432451 | 1.185815 | 0.095966 | 0.371275 | 0.588056 | 0.50537 | 0.734215 | 2.326038 | -0.20094 | 0.499165 | 0.234459 | 0.360266 | 0.142224 | 0.496159 | 0.388325 | 0.131737 | 0.906054 | 0.72791 | 0.876391 | 0.483309 | 0.465393 | 0.349241 |
| 27.5 | 1.385106 | 0.226883 | 0.236232 | 1.03404 | 0.119344 | 0.532406 | 0.679831 | -0.348 | 0.825423 | 2.040158 | -0.95715 | 0.71074 | 0.139638 | 0.327845 | 0.01033 | 0.289644 | 0.227522 | 0.052878 | 0.612964 | 0.341933 | 0.73758 | 0.563791 | 0.637864 | 0.115697 |
| 27.51389 | 0.44591 | 0.169298 | -0.02113 | 0.853377 | 0.253361 | 0.602254 | 0.700268 | -1.06822 | 0.917417 | 1.684462 | -1.48033 | 0.749074 | 0.087259 | 0.31828 | -0.04137 | 0.244233 | 0.12199 | -0.00611 | 0.47391 | 0.162342 | 0.612151 | 0.615641 | 0.746451 | 0.112815 |
| 27.52778 | -0.20234 | 0.062178 | -0.03122 | 0.33658 | 0.100136 | 0.344994 | 0.519816 | -1.26529 | 1.032583 | 1.412325 | -1.55945 | 0.553475 | 0.026641 | 0.305894 | 0.060281 | 0.267782 | 0.081476 | -0.01543 | 0.431249 | 0.241431 | 0.646682 | 0.55731 | 0.790699 | 0.267162 |
| 27.54167 | -0.06959 | 0.016378 | -0.30313 | -0.094 | -0.09514 | 0.239436 | 0.248809 | -0.81281 | 1.22119 | 1.645152 | -0.91398 | 0.594861 | -0.02651 | 0.306654 | 0.253788 | 0.467192 | 0.082266 | -0.01959 | 0.386364 | 0.477917 | 0.796999 | 0.449699 | 0.790234 | 0.574061 |
| 27.55556 | -0.15576 | 0.014619 | -0.34369 | -0.43841 | -0.23546 | 0.229575 | -0.06197 | 0.256649 | 1.02447 | 1.434208 | 0.01647 | 0.334951 | -0.03706 | 0.318874 | 0.441496 | 0.658706 | 0.132629 | -0.01597 | 0.342452 | 0.736095 | 0.948658 | 0.380046 | 0.880944 | 0.986039 |
| 27.56944 | 0.178036 | 0.040105 | 0.088431 | -0.58077 | -0.18615 | 0.011401 | -0.2874 | 1.357866 | 0.434758 | 0.627574 | 0.998743 | -0.18307 | 0.066344 | 0.379414 | 0.565083 | 0.811462 | 0.232717 | 0.007929 | 0.317545 | 0.941392 | 1.139307 | 0.366218 | 1.035384 | 1.390582 |
| 27.58333 | 0.246867 | 0.029188 | 0.386811 | -0.35808 | 0.02844 | -0.0406 | -0.36046 | 1.514764 | 0.144428 | -0.4363 | 1.915086 | -0.62604 | 0.202544 | 0.444953 | 0.591491 | 0.891941 | 0.335282 | -0.00281 | 0.271299 | 0.950035 | 1.339031 | 0.36252 | 0.96704 | 1.385379 |
| 27.59722 | -0.55673 | -0.05195 | 0.287602 | -0.10484 | 0.336442 | 0.123164 | -0.20794 | 1.235658 | -0.11172 | -1.72447 | 1.605096 | -1.13284 | 0.277425 | 0.442871 | 0.483346 | 0.819946 | 0.350144 | -0.02562 | 0.16195 | 0.775703 | 1.307351 | 0.351369 | 0.665548 | 1.027116 |
| 27.61111 | -0.95623 | -0.11165 | 0.068215 | 0.262385 | 0.513467 | 0.179536 | 0.140116 | 0.442261 | -0.7379 | -3.01646 | 0.537059 | -1.24562 | 0.276349 | 0.417706 | 0.433855 | 0.727179 | 0.29528 | -0.02492 | 0.102219 | 0.485653 | 1.114048 | 0.347821 | 0.342404 | 0.556895 |
| 27.625 | -0.1239 | -0.09271 | -0.19303 | 0.899897 | 0.645383 | -0.19774 | 0.528735 | -0.57887 | -1.42901 | -2.62398 | -0.41711 | -0.91722 | 0.229771 | 0.359636 | 0.349673 | 0.548213 | 0.247204 | -0.03771 | 0.164339 | 0.179315 | 0.713137 | 0.268597 | 0.137319 | 0.212292 |
| 27.63889 | 0.423945 | -0.09398 | -0.34481 | 0.853077 | 0.547005 | -0.61407 | 0.712612 | -1.48025 | -1.39625 | -1.81458 | -1.24506 | -0.68331 | 0.153884 | 0.249484 | 0.189589 | 0.386765 | 0.17999 | -0.09121 | 0.236925 | -0.11432 | 0.311968 | 0.185483 | -0.04075 | 0.025266 |
| 27.65278 | 0.691102 | -0.11984 | -0.37508 | 0.983631 | 0.205353 | -0.68869 | 0.677058 | -1.45401 | -1.25406 | -1.10057 | -1.70321 | -0.46056 | 0.082468 | 0.16423 | -0.11116 | 0.265331 | 0.108397 | -0.13403 | 0.28498 | -0.25997 | -0.07375 | 0.12373 | -0.26401 | -0.10514 |
| 27.66667 | 0.784558 | -0.1358 | -0.48117 | 1.063258 | -0.09485 | -0.29505 | 0.476686 | 0.069371 | -0.20849 | -0.39493 | -2.02126 | -0.10397 | 0.005132 | 0.09488 | -0.21232 | 0.223726 | 0.078864 | -0.14 | 0.26553 | -0.21504 | -0.38739 | 0.081765 | -0.5195 | -0.27575 |
| 27.68056 | 0.646049 | -0.14545 | -0.71629 | 0.656868 | -0.20754 | -0.05898 | 0.270969 | 1.037424 | 0.582311 | 0.098917 | -1.89676 | 0.185177 | -0.11025 | -0.02043 | -0.21708 | 0.137916 | 0.091791 | -0.17127 | 0.116705 | -0.1762 | -0.56464 | -0.05135 | -0.78157 | -0.44766 |
| 27.69444 | 0.195465 | -0.18291 | -0.65678 | 0.159782 | -0.24442 | 0.046869 | -0.07335 | 0.959921 | 0.857002 | 0.150859 | -1.24514 | 0.005223 | -0.2973 | -0.14551 | -0.13261 | 0.007403 | 0.114975 | -0.1908 | -0.06877 | -0.14826 | -0.74112 | -0.36805 | -0.93881 | -0.6263 |
| 27.70833 | -0.15627 | -0.16587 | -0.49341 | -0.37553 | -0.16907 | 0.013833 | -0.48452 | -0.2599 | 0.486467 | -0.14147 | -0.82148 | -0.4344 | -0.42554 | -0.2608 | -0.05571 | -0.05709 | 0.117101 | -0.20608 | -0.2231 | -0.14688 | -0.92125 | -0.77641 | -0.7669 | -0.78534 |
| 27.72222 | -0.56329 | -0.12647 | -0.57344 | -0.86082 | -0.21059 | 0.077295 | -0.93729 | -0.77205 | 0.256138 | -0.32196 | -0.5294 | -0.6853 | -0.59873 | -0.41751 | -0.10453 | -0.14452 | 0.057295 | -0.23431 | -0.39153 | -0.21364 | -0.92606 | -0.7524 | -0.58869 | -0.88907 |
| 27.73611 | -0.88717 | -0.11896 | -0.72557 | -1.21012 | -0.45822 | -0.07548 | -1.08146 | -0.84118 | -0.0755 | -0.20976 | -0.28519 | -0.41373 | -0.68851 | -0.56245 | -0.31928 | -0.24321 | -0.04687 | -0.24246 | -0.56699 | -0.27496 | -0.83342 | -0.52896 | -0.53073 | -1.06531 |
| 27.75 | -1.13446 | -0.1234 | -0.84092 | -1.2987 | -0.59265 | -0.508 | -1.1573 | -1.53355 | -0.53407 | -0.10119 | -0.12984 | -0.09164 | -0.69729 | -0.738 | -0.38583 | -0.33287 | -0.14802 | -0.23217 | -0.65172 | -0.27546 | -0.73469 | -0.42448 | -0.55635 | -1.31696 |
| 27.76389 | -1.38702 | -0.21252 | -0.87432 | -1.13093 | -0.43073 | -0.78397 | -1.09388 | -1.58696 | -0.89015 | -0.39788 | 0.025748 | -0.16148 | -0.52665 | -0.87933 | -0.4147 | -0.39018 | -0.19224 | -0.25581 | -0.67639 | -0.28951 | -0.50062 | -0.49254 | -0.67906 | -1.48335 |
| 27.77778 | -1.71525 | -0.27806 | -0.76997 | -0.89624 | -0.25522 | -0.87581 | -0.78247 | -0.81743 | -0.9083 | -0.72023 | 0.146087 | -0.54376 | -0.5086 | -0.92199 | -0.28503 | -0.52367 | -0.20593 | -0.33376 | -0.79569 | -0.3831 | -0.36916 | -0.6143 | -0.70647 | -1.59327 |
| 27.79167 | -2.12299 | -0.28952 | -0.62317 | -0.92547 | -0.28501 | -0.85448 | -0.70351 | -0.23851 | -1.01124 | -0.7438 | 0.192302 | -0.7513 | -0.5123 | -1.04335 | -0.21514 | -0.65423 | -0.21783 | -0.3413 | -0.9746 | -0.48194 | -0.3009 | -0.57907 | -0.65675 | -1.87126 |
| 27.80556 | -2.25526 | -0.24413 | -0.45136 | -1.18924 | -0.39621 | -0.65927 | -0.62632 | 0.116849 | -0.7486 | -0.68599 | 0.094964 | -0.90271 | -0.36703 | -0.89692 | -0.1092 | -0.7389 | -0.17093 | -0.32612 | -1.23943 | -0.57691 | -0.32745 | -0.39218 | -0.67226 | -2.12277 |
| 27.81944 | -2.21831 | -0.23368 | -0.37165 | -1.24382 | -0.46904 | -0.63062 | -0.49352 | -0.29812 | -0.5315 | -0.70496 | -0.0632 | -0.70789 | -0.13154 | -0.80062 | -0.11579 | -0.92171 | -0.13577 | -0.28047 | -1.59657 | -0.60154 | -0.33888 | -0.19914 | -0.59393 | -2.35655 |
| 27.83333 | -2.04488 | -0.14862 | -0.23502 | -0.95342 | -0.4679 | -0.55808 | -0.25583 | -0.58034 | -0.42223 | -0.63008 | -0.06654 | -0.24668 | -0.05717 | -0.64996 | -0.16704 | -1.09811 | -0.18121 | -0.34149 | -1.97091 | -0.66251 | -0.35678 | -0.08369 | -0.38796 | -2.52514 |
| 27.84722 | -1.56346 | 0.041013 | -0.02967 | -0.56022 | -0.27516 | -0.30214 | 0.126438 | -0.6745 | -0.50586 | -0.27402 | 0.00003 | 0.424101 | -0.02384 | -0.48915 | -0.15971 | -1.09814 | -0.23456 | -0.34862 | -2.10348 | -0.6179 | -0.33938 | -0.01607 | -0.00774 | -2.17702 |
| 27.86111 | -0.94911 | 0.167231 | 0.113767 | -0.25708 | -0.03812 | 0.00312 | 0.312289 | -0.68238 | -0.41359 | 0.061302 | 0.284044 | 0.855696 | -0.00684 | -0.35529 | -0.12936 | -0.84539 | -0.30358 | -0.26394 | -1.95708 | -0.56419 | -0.28764 | -0.036 | 0.063273 | -1.53563 |
| 27.875 | -0.3169 | 0.219095 | 0.015154 | -0.16472 | 0.276478 | -0.01491 | 0.070545 | -0.5148 | 0.047518 | -0.45284 | 0.497205 | 0.452489 | -0.05187 | -0.29422 | -0.09948 | -0.63627 | -0.3402 | -0.2307 | -1.74143 | -0.40417 | -0.26439 | -0.19186 | 0.083213 | -1.3191 |
| 27.88889 | 0.617905 | -0.02476 | -0.26688 | -0.461 | 0.493069 | -0.3516 | -0.87502 | 0.629292 | 0.800444 | -2.47726 | 0.733964 | -1.35434 | -0.0655 | -0.23023 | -0.06445 | -0.4528 | -0.35669 | -0.20291 | -1.52269 | -0.2129 | -0.22001 | -0.32577 | 0.121087 | -1.25276 |
| 27.90278 | 1.172375 | -0.34997 | -0.62217 | -0.47715 | 0.307276 | -0.74081 | -2.1586 | 1.643012 | 1.566483 | -5.17574 | 0.721166 | -3.35939 | -0.01887 | -0.12385 | -0.01589 | -0.34838 | -0.3857 | -0.14008 | -1.34043 | -0.08496 | -0.29333 | -0.40512 | 0.090023 | -1.1491 |
| 27.91667 | 0.633051 | -0.54659 | -0.72953 | -0.63935 | -0.01546 | -0.97452 | -2.10024 | 1.392519 | 0.479641 | -4.96095 | 0.118434 | -3.3335 | -0.04 | 0.049871 | 0.012742 | -0.15882 | -0.44585 | -0.14119 | -1.12115 | -0.15363 | -0.44231 | -0.38237 | 0.026136 | -0.9706 |
| 27.93056 | -0.80294 | -0.09589 | -0.69363 | -0.86861 | -0.27288 | -0.91088 | -0.65553 | -0.01159 | -0.66654 | -2.42744 | -0.83498 | -1.0719 | -0.15332 | 0.170964 | 0.002766 | 0.128929 | -0.36492 | -0.23763 | -0.80531 | -0.46128 | -0.54 | -0.41803 | -0.0607 | -0.76265 |
| 27.94444 | -1.43198 | -0.02878 | -0.72336 | -0.92981 | -0.23744 | -0.77912 | 0.025545 | -0.80455 | -0.58844 | -1.6565 | -1.07842 | -0.35046 | -0.31116 | 0.122215 | 0.006247 | 0.407761 | -0.21385 | -0.22358 | -0.66301 | -0.67835 | -0.53622 | -0.57848 | -0.17167 | -0.64506 |
| 27.95833 | -1.28235 | -0.6523 | -0.70388 | -1.49812 | -0.46906 | -0.62586 | -0.46612 | -0.78525 | 0.062189 | -2.32757 | -0.68144 | -0.96138 | -0.36138 | -0.23407 | -0.05844 | 0.354662 | -0.10597 | -0.07225 | -0.96201 | -0.48058 | -0.60716 | -0.52793 | -0.44848 | -0.59028 |
| 27.97222 | -1.10546 | -0.95451 | -0.46148 | -1.07208 | -0.60619 | -0.51561 | -0.73478 | -0.79514 | 0.158304 | -1.6469 | -0.3028 | -0.98741 | -0.27646 | -0.76127 | -0.22621 | 0.038267 | -0.15497 | -0.05986 | -1.61735 | -0.19081 | -0.5376 | -0.47754 | -0.63495 | -0.54352 |
| 27.98611 | -0.67718 | -0.83957 | -0.22207 | -0.40235 | -0.41486 | -0.32787 | -0.44176 | -0.70142 | -0.04402 | 0.119252 | -0.51467 | -0.24855 | -0.19443 | -0.92686 | -0.27449 | -0.14555 | -0.2206 | -0.11946 | -2.00237 | -0.12945 | -0.51972 | -0.31111 | -0.56773 | -0.58281 |
| 28 | 0.00823 | -0.43644 | -0.00593 | 0.395943 | -0.16599 | -0.05948 | -0.14154 | -0.18278 | -0.18454 | 1.276761 | -0.57992 | 0.270212 | -0.14258 | -0.58989 | -0.22064 | -0.12998 | -0.23316 | -0.09073 | -1.73302 | -0.19317 | -0.63108 | -0.17058 | -0.42839 | -0.5966 |
| 28.01389 | 0.266254 | -0.03501 | 0.149078 | 0.51978 | 0.113023 | 0.103527 | 0.280269 | 0.326747 | 0.354471 | 2.248222 | -0.37907 | 0.664017 | -0.1057 | -0.29008 | -0.14261 | -0.05195 | -0.20172 | 0.038554 | -1.30975 | -0.209 | -0.51563 | -0.07265 | -0.23781 | -0.50001 |
| 28.02778 | 0.241601 | 0.164235 | 0.499013 | 0.388105 | 0.172507 | 0.215735 | 0.456635 | 0.633431 | 0.753713 | 1.932914 | 0.044489 | 0.479161 | -0.06403 | -0.06164 | -0.09342 | -0.00677 | -0.13596 | 0.137872 | -0.88992 | -0.19062 | -0.31558 | -0.00694 | -0.10883 | -0.30729 |
| 28.04167 | 0.482611 | 0.121696 | 0.860369 | 0.336176 | -0.04915 | 0.35579 | -0.00225 | 0.852389 | 0.620134 | 0.110272 | 0.472642 | -0.21016 | -0.04727 | 0.064947 | -0.07669 | -0.04162 | -0.10557 | 0.128567 | -0.62908 | -0.27629 | -0.22093 | 0.025301 | -0.06573 | -0.1459 |
| 28.05556 | 0.46532 | 0.182167 | 1.083626 | 0.153203 | -0.11227 | 0.440463 | 0.13472 | 0.555238 | 0.680442 | -0.60757 | 0.514422 | -0.3113 | 0.001777 | 0.150712 | -0.06148 | -0.06261 | -0.09967 | 0.087258 | -0.47171 | -0.36822 | -0.17306 | 0.088845 | -0.02291 | -0.08548 |
| 28.06944 | 0.410134 | 0.243352 | 1.053268 | -0.06399 | -0.12719 | 0.415468 | 0.309498 | 0.060037 | 0.583669 | -0.54491 | 0.407264 | -0.02384 | 0.093389 | 0.238746 | -0.0063 | -0.01157 | -0.07779 | 0.087841 | -0.35705 | -0.27441 | -0.11873 | 0.170396 | 0.030986 | -0.04038 |
| 28.08333 | 0.248369 | 0.177281 | 0.679122 | -0.09997 | -0.14217 | 0.399452 | 0.235235 | -0.16391 | 0.504151 | -0.58742 | 0.358334 | 0.040806 | 0.166112 | 0.306217 | 0.034942 | 0.034371 | -0.05713 | 0.052578 | -0.2421 | 0.039438 | -0.05627 | 0.132783 | 0.049366 | 0.007668 |
| 28.09722 | 0.004641 | 0.12353 | 0.408388 | -0.15242 | -0.07483 | 0.278339 | 0.235867 | -0.31388 | 0.314924 | -0.26015 | 0.309529 | 0.263499 | 0.206461 | 0.285194 | 0.022391 | 0.049056 | -0.05101 | -0.00571 | -0.15191 | 0.358416 | 0.024483 | 0.046866 | 0.041981 | 0.017735 |
| 28.11111 | -0.00362 | -0.04369 | 0.257282 | -0.01984 | -0.08526 | 0.183402 | -0.05084 | -0.0254 | 0.258602 | -0.56234 | 0.285474 | 0.071938 | 0.234534 | 0.264708 | -0.00625 | 0.061431 | -0.05705 | -0.02236 | -0.08667 | 0.520075 | 0.11831 | -0.02229 | 0.034886 | 0.007536 |
| 28.125 | 0.168702 | -0.14966 | 0.326646 | 0.157843 | -0.06302 | 0.16563 | -0.31291 | 0.223916 | 0.226174 | -0.94076 | 0.362337 | -0.16871 | 0.226682 | 0.253474 | -0.01101 | 0.0744 | -0.06212 | 0.019003 | -0.04185 | 0.451937 | 0.187918 | -0.02745 | 0.038341 | 0.000529 |
| 28.13889 | 0.230751 | -0.06477 | 0.530105 | 0.256343 | 0.028194 | 0.158363 | -0.28575 | 0.43676 | 0.203254 | -0.68922 | 0.402714 | -0.16601 | 0.215739 | 0.229236 | 0.001202 | 0.074471 | -0.06686 | 0.076836 | 0.00172 | 0.228899 | 0.223251 | 0.016692 | 0.050549 | -0.00728 |
| 28.15278 | 0.171304 | 0.11243 | 0.708339 | 0.193131 | 0.140228 | 0.13925 | -0.06626 | 0.442687 | 0.238704 | 0.081396 | 0.365428 | 0.045579 | 0.203089 | 0.227234 | 0.005649 | 0.055544 | -0.07482 | 0.127094 | 0.024886 | 0.040527 | 0.228277 | 0.059017 | 0.043467 | -0.01614 |
| 28.16667 | -0.22662 | 0.493439 | 0.629139 | -0.18043 | 0.309657 | 0.083808 | 0.64901 | 0.049784 | 0.548324 | 2.069702 | 0.003839 | 0.880314 | 0.222402 | 0.219217 | -0.00624 | 0.029207 | -0.08125 | 0.166735 | 0.014698 | -0.00027 | 0.218692 | 0.130621 | 0.02514 | -0.00856 |
| 28.18056 | -0.44835 | 0.616861 | 0.233007 | -0.43826 | 0.365601 | 0.074966 | 0.988231 | -0.32321 | 0.874427 | 2.96787 | -0.25954 | 1.231471 | 0.255404 | 0.236119 | 0.006551 | 0.03913 | -0.07292 | 0.213942 | 0.014875 | 0.091696 | 0.280426 | 0.324912 | 0.005139 | -0.00741 |
| 28.19444 | -0.09541 | 0.362125 | 0.267529 | -0.06084 | 0.073275 | 0.082779 | 0.198433 | -0.17131 | 0.327025 | 0.823874 | -0.00246 | 0.272159 | 0.263259 | 0.247382 | 0.016758 | 0.074089 | -0.06564 | 0.260306 | 0.007589 | 0.188742 | 0.293291 | 0.438904 | -0.01952 | -0.024 |
| 28.20833 | 0.409687 | -0.11372 | 0.535985 | 0.593627 | -0.23042 | 0.117618 | -0.90499 | 0.325341 | -0.53715 | -2.17521 | 0.372023 | -1.13622 | 0.238992 | 0.211765 | 0.009809 | 0.077956 | -0.05292 | 0.277802 | -0.01222 | 0.238222 | 0.219306 | 0.412179 | -0.02614 | -0.04091 |
| 28.22222 | 0.487479 | -0.16527 | 0.456237 | 0.958594 | -0.16671 | 0.059037 | -1.09698 | 0.577538 | -0.20285 | -2.2861 | 0.294887 | -1.2637 | 0.204062 | 0.172563 | -0.00162 | 0.073854 | -0.04599 | 0.280665 | -0.03706 | 0.223918 | 0.143756 | 0.238957 | -0.03579 | -0.07613 |
| 28.23611 | -0.14071 | 0.201897 | 0.117969 | 0.388566 | 0.2934 | -0.09841 | -0.03868 | 0.399195 | -0.16017 | 0.498828 | -0.15391 | 0.181982 | 0.147901 | 0.152799 | 0.003564 | 0.059322 | -0.0616 | 0.283224 | -0.06103 | 0.140382 | 0.08817 | 0.114671 | -0.03545 | -0.10521 |
| 28.25 | -0.56525 | 0.485455 | -0.00476 | -0.21764 | 0.55342 | -0.20447 | 0.754305 | 0.037637 | -0.15549 | 2.460685 | -0.50524 | 1.26132 | 0.130617 | 0.160486 | 0.02355 | 0.061116 | -0.05923 | 0.277879 | -0.04886 | 0.101412 | 0.083701 | 0.081271 | -0.00195 | -0.11153 |
| 28.26389 | -0.26224 | 0.353102 | 0.189742 | -0.1763 | 0.382998 | -0.05278 | 0.511819 | -0.03004 | -0.20305 | 1.625258 | -0.2404 | 0.700474 | 0.151088 | 0.177487 | 0.050926 | 0.065688 | -0.02636 | 0.211705 | 0.001599 | 0.138056 | 0.11232 | 0.118573 | 0.062521 | -0.11527 |

| | | | | | | | | | | | | | | | | | | | | | | | | |
|---|---|---|---|---|---|---|---|---|---|---|---|---|---|---|---|---|---|---|---|---|---|---|---|---|
| 28.27778 | 0.269596 | 0.045783 | 0.334444 | 0.326879 | 0.027221 | 0.129463 | -0.26293 | 0.347804 | -0.04347 | -0.41351 | 0.194875 | -0.38477 | 0.192857 | 0.190145 | 0.045294 | 0.069437 | 0.015796 | 0.185034 | 0.067664 | 0.180674 | 0.170725 | 0.207037 | 0.120265 | -0.07283 |
| 28.29167 | 0.458657 | 0.024836 | 0.379467 | 0.533839 | -0.07253 | 0.227734 | -0.38583 | 0.588013 | 0.116597 | -0.75682 | 0.406607 | -0.50766 | 0.232981 | 0.220936 | 0.025523 | 0.067366 | 0.043715 | 0.177727 | 0.117154 | 0.17924 | 0.221013 | 0.340535 | 0.163018 | 0.016029 |
| 28.30556 | 0.163046 | 0.207215 | 0.357241 | 0.389552 | 0.152529 | 0.249973 | 0.213033 | 0.465651 | -0.08044 | 0.768005 | 0.110933 | 0.171931 | 0.269918 | 0.229453 | -0.00685 | 0.048482 | 0.032545 | 0.153865 | 0.082553 | 0.128395 | 0.206429 | 0.408038 | 0.144223 | 0.044369 |
| 28.31944 | 0.186479 | 0.268439 | 0.269712 | 0.605872 | 0.186306 | 0.266548 | 0.288544 | 0.398079 | -0.22667 | 0.884626 | -0.02699 | 0.261706 | 0.30467 | 0.217702 | -0.02814 | 0.02542 | 0.021271 | 0.119919 | 0.014091 | 0.085895 | 0.16924 | 0.368421 | 0.083918 | 0.00235 |
| 28.33333 | 0.441646 | 0.221065 | 0.285027 | 1.02679 | 0.124783 | 0.370903 | 0.074091 | 0.455887 | -0.26796 | 0.103407 | 0.127436 | -0.05163 | 0.319161 | 0.226075 | -0.01198 | 0.037301 | 0.037166 | 0.139338 | 0.031013 | 0.087076 | 0.151602 | 0.330115 | 0.068163 | -0.01803 |
| 28.34722 | 0.349964 | 0.308214 | 0.195293 | 0.643547 | 0.243195 | 0.473623 | 0.488797 | 0.358967 | -0.46389 | 0.838567 | -0.0885 | 0.376751 | 0.278276 | 0.224464 | 0.009445 | 0.059942 | 0.047866 | 0.160447 | 0.107672 | 0.105973 | 0.143289 | 0.198363 | 0.101518 | 0.009252 |
| 28.36111 | 0.238914 | 0.297253 | 0.203267 | 0.165788 | 0.253167 | 0.52078 | 0.717278 | 0.342644 | -0.52686 | 1.393542 | -0.31788 | 0.69867 | 0.202569 | 0.23967 | -0.03277 | 0.075208 | 0.013987 | 0.103496 | 0.124428 | 0.081861 | 0.090627 | 0.106444 | 0.083185 | 0.006706 |
| 28.375 | 0.258065 | 0.153568 | 0.317989 | 0.033396 | 0.045787 | 0.460784 | 0.41158 | 0.364674 | -0.23908 | 0.628614 | -0.08826 | 0.265651 | 0.12218 | 0.241845 | -0.14439 | 0.070532 | -0.0642 | -0.0243 | 0.069328 | 0.00276 | -0.01868 | 0.014967 | -0.00346 | -0.07215 |
| 28.38889 | 0.239426 | 0.107961 | 0.260171 | 0.120689 | -0.01093 | 0.392154 | 0.405396 | 0.367179 | -0.30979 | 0.831338 | -0.03318 | 0.245088 | 0.104853 | 0.233087 | -0.16004 | 0.033445 | -0.0748 | -0.03993 | 0.074317 | -0.00472 | -0.037 | 0.00573 | 0.005958 | -0.08385 |
| 28.40278 | 0.228927 | 0.326348 | 0.119708 | 0.478042 | 0.421887 | 0.3688 | 1.16288 | 0.20371 | -0.964 | 2.416353 | -0.13975 | 0.982539 | 0.246909 | 0.208941 | 0.0793 | 0.007001 | 0.105062 | 0.082811 | 0.336103 | 0.207999 | 0.165803 | 0.137685 | 0.253741 | 0.267021 |
| 28.41667 | 0.688069 | 0.615031 | 0.382528 | 1.475146 | 0.688926 | 0.687752 | 1.63748 | 0.510203 | -1.39824 | 3.369761 | 0.244982 | 1.27996 | 0.414022 | 0.269 | 0.519944 | 0.065735 | 0.424 | 0.277275 | 0.923638 | 0.588282 | 0.537109 | 0.3336 | 0.652203 | 1.087886 |
| 28.43056 | 1.879638 | 0.468545 | 1.111752 | 3.161492 | 0.751759 | 1.363168 | 1.294515 | 1.215258 | 0.061273 | 2.681335 | 1.132576 | 0.8222 | 0.379425 | 0.553113 | 0.818461 | 0.150051 | 0.534582 | 0.400469 | 1.525161 | 0.680071 | 0.706096 | 0.35077 | 0.912905 | 2.091663 |
| 28.44444 | 2.864091 | 0.094937 | 1.796799 | 2.805484 | 0.558999 | 2.01767 | 0.650906 | 2.371751 | 2.139019 | 1.86813 | 1.084323 | 0.309947 | 0.137214 | 0.59705 | 0.566898 | 0.136059 | 0.146816 | 0.304799 | 1.552509 | 0.325198 | 0.579521 | -0.05284 | 0.750033 | 1.872711 |
| 28.45833 | 2.639962 | 0.033074 | 1.40772 | 0.863192 | 0.491988 | 1.578059 | 0.729599 | 2.500864 | 2.42598 | 1.190584 | 0.380256 | 0.173189 | -0.12111 | 0.407302 | 0.070783 | 0.078695 | -0.22164 | 0.060174 | 1.058794 | -0.09231 | 0.150476 | -0.4172 | 0.47266 | 0.838338 |
| 28.47222 | 2.053564 | 0.165794 | 0.439506 | 0.537019 | 0.541478 | 1.025444 | 0.934491 | 1.745869 | 1.065706 | 0.697056 | -0.0637 | 0.214237 | -0.01165 | 0.31939 | 0.147786 | 0.185316 | -0.15891 | 0.140427 | 0.846962 | -0.00875 | 0.016195 | -0.22522 | 0.270137 | 0.29212 |
| 28.48611 | 1.553881 | 0.180233 | -0.28333 | 0.887197 | 0.526916 | 0.719962 | 0.96999 | 0.906966 | -0.36004 | -0.08098 | -0.16007 | 0.356358 | 0.26907 | 0.48256 | 0.349545 | 0.345613 | 0.115908 | 0.354158 | 0.773826 | 0.214381 | 0.158113 | 0.128079 | 0.345693 | 0.266152 |
| 28.5 | 0.902235 | 0.13499 | -0.58471 | 0.775802 | 0.442696 | 0.509421 | 0.743599 | 0.144788 | -1.12821 | -0.98621 | -0.12862 | 0.41898 | 0.355271 | 0.496848 | 0.348978 | 0.334997 | 0.185115 | 0.330494 | 0.641726 | 0.20051 | 0.20703 | 0.224179 | 0.409324 | 0.22586 |
| 28.51389 | 0.481312 | 0.108929 | -0.45673 | 0.274059 | 0.327291 | 0.473792 | 0.473021 | -0.67395 | -0.86099 | -1.43644 | -0.16719 | 0.344331 | 0.281978 | 0.370531 | 0.214398 | 0.218644 | 0.104895 | 0.157136 | 0.464966 | 0.056709 | 0.159285 | 0.138611 | 0.360014 | 0.099486 |
| 28.52778 | 0.247622 | 0.109036 | -0.33219 | 0.378735 | 0.082409 | 0.286804 | 0.243855 | -1.05501 | -0.08802 | -0.99816 | -0.32584 | 0.306126 | 0.211913 | 0.254097 | 0.151802 | 0.13887 | 0.040765 | 0.003711 | 0.38367 | 0.013874 | 0.135808 | 0.13618 | 0.343844 | 0.129273 |
| 28.54167 | 0.163395 | 0.101468 | -0.11161 | 0.530743 | -0.0748 | 0.166491 | -0.07064 | -0.75719 | 0.64506 | -0.13257 | -0.5516 | 0.340205 | 0.1646 | 0.202283 | 0.132605 | 0.158413 | 0.022298 | -0.04876 | 0.400118 | 0.109016 | 0.136302 | 0.145925 | 0.458465 | 0.413993 |
| 28.55556 | 0.165094 | 0.069488 | 0.053487 | 0.752405 | -0.11607 | 0.160708 | -0.42691 | -0.13898 | 0.765365 | 0.577297 | -0.52745 | 0.290101 | 0.181148 | 0.237296 | 0.128594 | 0.281165 | 0.049623 | 0.020845 | 0.482198 | 0.135227 | 0.251006 | 0.221305 | 0.806718 | 0.815228 |
| 28.56944 | 0.336084 | 0.046138 | 0.018038 | 0.841615 | -0.08226 | 0.277536 | -0.79794 | 0.4477 | 0.566161 | 0.876399 | 0.014184 | 0.199361 | 0.254205 | 0.307746 | 0.210568 | 0.4107 | 0.091532 | 0.11832 | 0.630837 | 0.178673 | 0.374535 | 0.270909 | 1.244066 | 1.115363 |
| 28.58333 | 0.451938 | 0.042247 | 0.110181 | 0.762159 | -0.14053 | 0.358315 | -0.75298 | 1.37356 | 0.075822 | 0.699492 | 0.535372 | 0.038085 | 0.313377 | 0.361125 | 0.272251 | 0.512798 | 0.122165 | 0.1614 | 0.7203 | 0.198082 | 0.40941 | 0.273585 | 1.458533 | 1.151348 |
| 28.59722 | 0.459778 | 0.062434 | 0.256857 | 0.515215 | -0.14287 | 0.433398 | -0.62861 | 2.42952 | -0.43597 | 0.185729 | 0.640857 | -0.18055 | 0.30711 | 0.390487 | 0.357424 | 0.467277 | 0.119304 | 0.177325 | 0.725142 | 0.214935 | 0.400611 | 0.239959 | 1.178568 | 1.100823 |
| 28.61111 | 0.104702 | 0.047058 | 0.525421 | 0.434061 | -0.10319 | 0.482089 | -0.42813 | 2.787816 | -0.57641 | -0.3556 | 0.36941 | -0.37271 | 0.306081 | 0.366842 | 0.303562 | 0.323071 | 0.087904 | 0.127092 | 0.625061 | 0.109344 | 0.300045 | 0.169206 | 0.903729 | 0.973993 |
| 28.625 | -0.28819 | -0.01835 | 0.071894 | 0.135752 | 0.006138 | 0.48328 | 0.053691 | 2.692435 | -0.4801 | -0.76222 | 0.058031 | -0.51285 | 0.258781 | 0.294739 | 0.165309 | 0.132222 | 0.034394 | 0.035924 | 0.43174 | 0.005121 | 0.15278 | 0.101695 | 0.575507 | 0.737242 |
| 28.63889 | -0.62242 | -0.0677 | -0.54585 | -0.29513 | 0.142996 | 0.385381 | 0.608202 | 1.652564 | -0.19389 | -1.04783 | -0.00547 | -0.48544 | 0.202166 | 0.239974 | 0.027074 | -0.01145 | 0.007093 | -0.02689 | 0.30016 | -0.06289 | 0.026882 | 0.069776 | 0.343737 | 0.481816 |
| 28.65278 | -0.40474 | -0.0517 | -0.58522 | -0.37348 | 0.185706 | 0.266108 | 0.818951 | 0.024744 | -0.01141 | -1.34641 | 0.057858 | -0.26812 | 0.193403 | 0.256196 | -0.05006 | -0.09159 | -0.0011 | -0.05678 | 0.240821 | -0.11305 | -0.03388 | 0.032476 | 0.127234 | 0.250176 |
| 28.66667 | -0.05871 | 0.007148 | -0.6121 | 0.170029 | 0.258061 | 0.181609 | 0.854054 | -1.41838 | -0.03632 | -1.47705 | 0.057418 | -0.06672 | 0.181666 | 0.278634 | -0.06764 | -0.05079 | -0.00481 | -0.09759 | 0.227694 | -0.14063 | -0.0447 | 0.078973 | -0.01288 | 0.117443 |
| 28.68056 | 0.554624 | 0.01729 | -0.75397 | 0.761467 | 0.436744 | -0.02696 | 0.794759 | -2.31568 | -0.40333 | -1.37932 | 0.205886 | 0.042212 | 0.149671 | 0.259787 | -0.08984 | 0.060942 | -0.01311 | -0.13946 | 0.222105 | -0.18673 | -0.06748 | 0.168929 | -0.18372 | 0.056089 |
| 28.69444 | 0.735833 | -0.02253 | -0.52804 | 0.955354 | 0.553445 | -0.24637 | 0.649336 | -2.32343 | -0.70433 | -0.96037 | 0.329521 | 0.098875 | 0.081742 | 0.166612 | -0.17286 | 0.057085 | -0.05765 | -0.19728 | 0.213833 | -0.25663 | -0.10606 | 0.12013 | -0.36825 | -0.01303 |
| 28.70833 | 0.596018 | -0.05541 | -0.3299 | 0.644652 | 0.451506 | -0.45258 | 0.328429 | -1.72632 | -0.62767 | -0.72179 | -0.03677 | 0.089648 | -0.03973 | 0.058649 | -0.22485 | -0.00598 | -0.11753 | -0.26024 | 0.196037 | -0.34452 | -0.17966 | 0.03503 | -0.52157 | -0.08439 |
| 28.72222 | 0.080333 | -0.09321 | -0.34471 | -0.04301 | 0.164089 | -0.46892 | 0.056831 | -1.42025 | -0.21568 | -0.75666 | -0.35725 | 0.041725 | -0.13502 | -0.04488 | -0.20401 | -0.06959 | -0.13692 | -0.2689 | 0.161507 | -0.35611 | -0.22403 | -0.07268 | -0.66696 | -0.16236 |
| 28.73611 | -0.63349 | -0.13728 | -0.33028 | -0.55016 | -0.1777 | -0.39128 | -0.06219 | -1.15876 | 0.178578 | -1.12767 | -0.56892 | -0.0655 | -0.18534 | -0.08765 | -0.11778 | -0.09815 | -0.1169 | -0.24578 | 0.161404 | -0.3403 | -0.22673 | -0.14893 | -0.58902 | -0.21757 |
| 28.75 | -1.22991 | -0.17566 | -0.20594 | -0.65443 | -0.39522 | -0.37528 | -0.17073 | -1.05559 | 0.40789 | -1.77803 | -0.567 | -0.22485 | -0.1894 | -0.06524 | -0.049 | -0.05026 | -0.09676 | -0.23428 | 0.220658 | -0.33051 | -0.22703 | -0.08798 | -0.63454 | -0.20348 |
| 28.76389 | -1.53213 | -0.24939 | -0.21576 | -0.57296 | -0.75133 | -0.39859 | -0.37199 | -1.12351 | 0.547494 | -2.41869 | -0.3766 | -0.34097 | -0.25805 | -0.04364 | -0.0328 | -0.03931 | -0.09843 | -0.3303 | 0.22378 | -0.41659 | -0.1802 | 0.01881 | -0.63119 | -0.23058 |
| 28.77778 | -1.53354 | -0.37231 | -0.35844 | -0.22367 | -0.74485 | -0.39732 | -0.51617 | -1.27638 | 0.174838 | -2.36186 | -0.21965 | -0.30423 | -0.32137 | -0.08412 | -0.08175 | -0.09541 | -0.12543 | -0.33686 | 0.143113 | -0.52524 | -0.21282 | 0.023464 | -0.6724 | -0.30036 |
| 28.79167 | -1.17831 | -0.41892 | -0.32549 | -0.45608 | -0.63937 | -0.34745 | -0.57811 | -1.25934 | -0.39216 | -1.92054 | -0.22074 | -0.15753 | -0.34933 | -0.20239 | -0.08474 | -0.14481 | -0.10314 | -0.29051 | 0.077754 | -0.61788 | -0.26215 | -0.04722 | -0.65214 | -0.41653 |
| 28.80556 | -0.82383 | -0.57297 | -0.17744 | -0.70233 | -0.49155 | -0.3371 | -0.53276 | -1.10434 | -0.51134 | -1.08649 | -0.08449 | -0.00273 | -0.37647 | -0.26408 | -0.08623 | -0.13792 | -0.02499 | -0.2866 | -0.01495 | -0.56844 | -0.33148 | -0.12999 | -0.64367 | -0.43188 |
| 28.81944 | -0.87776 | -0.40155 | 0.036367 | -0.93129 | -0.32097 | -0.28921 | -0.46428 | -0.83478 | -0.07811 | -0.28258 | 0.134896 | 0.108487 | -0.34121 | -0.29839 | -0.07577 | -0.13204 | 0.011456 | -0.44829 | -0.10684 | -0.57491 | -0.40053 | -0.0688 | -0.63049 | -0.36289 |
| 28.83333 | -0.93711 | -0.06353 | 0.0116 | -0.97154 | -0.11245 | -0.16662 | -0.38057 | -0.53299 | 0.59063 | 0.360227 | 0.377233 | 0.179716 | -0.45465 | -0.37907 | -0.01378 | -0.14791 | -0.06283 | -0.58012 | -0.21184 | -0.53992 | -0.40517 | -0.09034 | -0.67234 | -0.24577 |
| 28.84722 | -0.84082 | 0.182595 | 0.045299 | -0.81296 | -0.06176 | -0.05802 | -0.2973 | -0.20497 | 1.203646 | 0.830011 | 0.610171 | 0.268047 | -0.63433 | -0.54935 | -0.02892 | -0.16956 | -0.1296 | -0.44954 | -0.23146 | -0.39287 | -0.35128 | -0.07689 | -0.59531 | -0.16395 |
| 28.86111 | -0.58358 | 0.274201 | 0.046005 | -0.57575 | -0.15399 | 0.000771 | -0.2423 | -0.04581 | 1.479565 | 0.995486 | 0.732861 | 0.341384 | -0.81417 | -0.58526 | -0.05729 | -0.13204 | -0.12204 | -0.30456 | -0.16508 | -0.21723 | -0.26253 | 0.054589 | -0.40201 | -0.1293 |
| 28.875 | -0.2356 | 0.316361 | 0.027945 | -0.48143 | -0.26295 | 0.034703 | -0.1946 | 0.060293 | 1.364492 | 0.653271 | 0.569094 | 0.408575 | -0.68466 | -0.58244 | -0.04027 | -0.02858 | -0.09843 | -0.14174 | -0.10946 | -0.07731 | -0.14521 | 0.177153 | -0.31382 | -0.12127 |
| 28.88889 | -0.28636 | 0.254626 | -0.00099 | -0.77496 | -0.28434 | -0.02843 | -0.40289 | -0.0047 | 0.884979 | -0.32008 | 0.3471 | 0.318759 | -0.47597 | -0.56364 | -0.02538 | 0.020079 | -0.09888 | -0.02095 | -0.11812 | -0.02294 | -0.18604 | 0.209002 | -0.14652 | -0.21257 |
| 28.90278 | -0.9613 | -0.07996 | -0.06956 | -1.17472 | -0.16212 | -0.28311 | -0.61714 | -0.23458 | 0.022638 | -1.9181 | -0.01065 | -0.10366 | -0.1792 | -0.46927 | -0.0112 | 0.075169 | -0.10398 | 0.093343 | -0.16882 | -0.08121 | -0.308 | 0.106873 | -0.08428 | -0.42961 |
| 28.91667 | -1.43175 | -0.28273 | -0.29794 | -1.24362 | 0.002243 | -0.51417 | -0.90534 | -0.61 | -1.03566 | -3.9605 | -0.40317 | -0.76125 | -0.04817 | -0.2636 | 0.050647 | 0.189087 | -0.08373 | 0.155024 | -0.21177 | -0.20956 | -0.36901 | -0.0419 | -0.28243 | -0.66722 |
| 28.93056 | -1.45217 | -0.18608 | -0.55151 | -1.75947 | 0.215086 | -0.54375 | -0.90584 | -0.67612 | -1.77693 | -5.30073 | -0.85369 | -1.45331 | 0.042438 | 0.007173 | 0.230752 | 0.425232 | 0.096749 | 0.180824 | -0.07513 | -0.35988 | -0.21567 | -0.11849 | -0.40024 | -0.553 |
| 28.94444 | -1.58411 | -0.19287 | -0.48561 | -1.4623 | 0.189991 | -0.41274 | -0.71132 | -1.06874 | -1.71403 | -4.47663 | -0.9961 | -1.96043 | 0.052739 | 0.107476 | 0.428957 | 0.627687 | 0.336017 | 0.231018 | 0.163611 | -0.33098 | -0.0314 | -0.17269 | -0.33526 | -0.27002 |
| 28.95833 | -1.65642 | -0.42998 | -0.30009 | -0.79662 | 0.054362 | -0.65966 | -0.49373 | -1.31859 | -1.4872 | -1.54731 | -0.70975 | -1.23533 | -0.21313 | -0.20105 | 0.295384 | 0.446451 | 0.259404 | 0.189969 | 0.032396 | -0.34828 | -0.18496 | -0.43783 | -0.29344 | -0.27196 |
| 28.97222 | -1.27996 | -0.32237 | -0.01988 | -0.15865 | 0.08752 | -0.59114 | -0.17303 | -1.1607 | -0.72183 | 1.378207 | -0.55522 | -0.05006 | -0.30589 | -0.69273 | -0.01007 | 0.052202 | -0.04086 | 0.042798 | -0.54713 | -0.3195 | -0.40339 | -0.55556 | -0.40596 | -0.41071 |
| 28.98611 | -0.83488 | -0.07449 | 0.335334 | 0.439066 | 0.238408 | -0.3061 | 0.056824 | -0.25079 | 0.288264 | 3.201602 | -0.50959 | 0.961671 | -0.30107 | -0.92006 | -0.12331 | -0.10307 | -0.11335 | 0.043509 | -0.88191 | -0.09879 | -0.32982 | -0.26557 | -0.34639 | -0.37795 |
| 29 | -0.29659 | 0.014986 | 0.497176 | 0.709576 | 0.327068 | -0.02401 | 0.186277 | 0.650438 | 1.33802 | 3.726008 | -0.46175 | 1.536023 | -0.22182 | -0.80358 | -0.10094 | -0.08514 | -0.04934 | 0.139798 | -0.77286 | 0.171608 | -0.09633 | 0.156963 | -0.24749 | -0.33635 |
| 29.01389 | 0.162499 | 0.155955 | 0.432052 | 0.604216 | 0.289881 | 0.177254 | 0.207344 | 0.739886 | 1.676208 | 3.356419 | -0.32595 | 1.501565 | -0.14254 | -0.57852 | -0.07624 | -0.06593 | -0.02277 | 0.191587 | -0.65869 | 0.282426 | -0.02048 | 0.470688 | -0.19536 | -0.28633 |
| 29.02778 | 0.394031 | 0.330922 | 0.372835 | 0.542035 | 0.188874 | 0.295807 | 0.227918 | 1.258169 | 1.183882 | 2.987166 | -0.17272 | 1.223024 | -0.1452 | -0.41339 | -0.05031 | -0.06659 | -0.0209 | 0.179386 | -0.51291 | 0.294325 | -0.02919 | 0.522869 | -0.17011 | -0.26268 |
| 29.04167 | 0.677139 | 0.424098 | 0.530843 | 0.572305 | 0.078243 | 0.352047 | 0.246094 | 1.116368 | 0.175535 | 2.519779 | -0.07884 | 0.825693 | -0.21006 | -0.25482 | -0.02845 | -0.06524 | -0.02517 | 0.122132 | -0.40868 | 0.167558 | -0.05492 | 0.412737 | -0.14598 | -0.24602 |
| 29.05556 | 0.807477 | 0.43888 | 0.670923 | 0.557696 | 0.036674 | 0.339858 | 0.249023 | 0.873261 | -0.24877 | 1.740524 | 0.013875 | 0.440414 | -0.22321 | -0.15935 | -0.04752 | -0.07392 | -0.03376 | 0.09093 | -0.38494 | 0.024419 | -0.10253 | 0.297256 | -0.13373 | -0.1878 |
| 29.06944 | 0.738076 | 0.497449 | 0.76353 | 0.580328 | 0.04758 | 0.290464 | 0.231608 | 0.430426 | -0.02148 | 0.959094 | 0.096794 | 0.137124 | -0.20665 | -0.12678 | -0.0624 | -0.07505 | -0.03138 | 0.091864 | -0.37825 | -0.02441 | -0.10441 | 0.220956 | -0.09098 | -0.11273 |
| 29.08333 | 0.811221 | 0.414057 | 0.607306 | 0.606094 | 0.061054 | 0.300895 | 0.247728 | 0.232274 | 0.240866 | 0.124372 | 0.168221 | -0.05922 | -0.12581 | -0.09083 | -0.01077 | -0.03961 | -0.01968 | 0.136854 | -0.37351 | 0.054519 | -0.06037 | 0.172829 | 0.006072 | -0.06673 |
| 29.09722 | 0.591648 | 0.293054 | 0.466345 | 0.678745 | 0.088484 | 0.339346 | 0.280256 | 0.113941 | 0.47056 | -0.40859 | 0.251701 | -0.14293 | -0.00289 | -0.06593 | 0.067717 | 0.000554 | -0.00662 | 0.169182 | -0.36003 | 0.238521 | -0.05242 | 0.09833 | 0.099743 | -0.0421 |
| 29.11111 | 0.381815 | 0.158017 | 0.456902 | 0.760129 | 0.09104 | 0.382366 | 0.312146 | -0.01075 | 0.625978 | -0.68856 | 0.358334 | -0.11834 | -0.04268 | -0.09203 | 0.100294 | 0.017988 | 0.008548 | 0.155372 | -0.3702 | 0.43694 | -0.078 | 0.001135 | 0.128707 | -0.04473 |
| 29.125 | 0.212715 | 0.048673 | 0.423832 | 0.763484 | 0.048568 | 0.386814 | 0.287624 | 0.05646 | 0.748528 | -0.77665 | 0.437133 | -0.01309 | -0.04259 | -0.08778 | 0.087967 | 0.031753 | 0.029763 | 0.11269 | -0.34567 | 0.55215 | -0.09053 | -0.03129 | 0.10054 | -0.05937 |
| 29.13889 | -0.01177 | -0.03758 | 0.345139 | 0.666369 | 0.013564 | 0.358507 | 0.21403 | 0.246101 | 0.987574 | -0.4737 | 0.458776 | 0.096915 | -0.02049 | -0.03279 | 0.064199 | 0.060113 | 0.023645 | 0.08988 | -0.3608 | 0.584398 | -0.12829 | 0.026108 | 0.057137 | -0.08198 |
| 29.15278 | -0.16108 | -0.09058 | 0.307707 | 0.461145 | 0.011834 | 0.34015 | 0.104655 | 0.400904 | 0.9992 | -0.17978 | 0.375475 | 0.154903 | -0.01189 | 0.00464 | 0.005518 | 0.069016 | -0.00152 | 0.054332 | -0.36163 | 0.520702 | -0.13853 | 0.153754 | -0.0131 | -0.11777 |
| 29.16667 | -0.31521 | -0.08066 | 0.251978 | 0.258292 | -0.00711 | 0.309251 | 0.008877 | 0.336748 | 0.892411 | -0.0773 | 0.29996 | 0.18832 | 0.020652 | -0.00452 | -0.0277 | 0.060983 | -0.01904 | 0.037308 | -0.35495 | 0.442229 | -0.10579 | 0.27847 | -0.05396 | -0.13389 |
| 29.18056 | -0.41964 | -0.03389 | 0.17912 | 0.038131 | -0.08557 | 0.215177 | -0.09177 | 0.140065 | 0.629911 | -0.27677 | 0.232446 | 0.163658 | 0.031732 | -0.03961 | -0.02276 | 0.053802 | -0.01441 | 0.054309 | -0.31666 | 0.333518 | -0.04505 | 0.30087 | -0.03936 | -0.11274 |
| 29.19444 | -0.38689 | -0.11579 | 0.114824 | -0.0139 | -0.17332 | 0.106086 | -0.11792 | -0.22423 | 0.258538 | -0.58349 | 0.18102 | 0.142139 | -0.01806 | -0.1038 | -0.04993 | 0.010922 | -0.03932 | 0.059946 | -0.31818 | 0.229524 | -0.06696 | 0.233018 | -0.0409 | -0.10162 |
| 29.20833 | -0.35938 | -0.08228 | 0.12887 | 0.130666 | -0.16283 | 0.046613 | -0.0084 | -0.28343 | 0.065044 | -0.69906 | 0.175855 | 0.22216 | -0.07817 | -0.15015 | -0.09149 | -0.02703 | -0.06891 | 0.064244 | -0.31737 | 0.164082 | -0.13929 | 0.123814 | -0.04447 | -0.08081 |
| 29.22222 | -0.3259 | -0.09088 | 0.098839 | 0.419251 | -0.06949 | 0.018129 | 0.144858 | 0.081628 | 0.21425 | -0.47241 | 0.227501 | 0.386436 | -0.06057 | -0.14282 | -0.06589 | 0.003719 | -0.0411 | 0.144982 | -0.26504 | 0.19863 | -0.08506 | 0.123041 | 0.03225 | -0.00905 |
| 29.23611 | -0.31797 | -0.12689 | 0.091706 | 0.698868 | 0.006133 | 0.002404 | 0.239036 | 0.543618 | 0.520767 | 0.063365 | 0.298275 | 0.503283 | 0.012499 | -0.09093 | 0.00567 | 0.064175 | 0.019708 | 0.269511 | -0.18711 | 0.259625 | 0.049395 | 0.135626 | 0.111054 | 0.046871 |
| 29.25 | -0.14995 | -0.19499 | 0.134454 | 0.749381 | 0.007741 | 0.058928 | 0.261469 | 0.873571 | 0.814174 | 0.528598 | 0.32412 | 0.482421 | 0.09564 | -0.00272 | 0.048644 | 0.109553 | 0.057779 | 0.320605 | -0.16097 | 0.246229 | 0.137289 | 0.128813 | 0.090691 | 0.01707 |
| 29.26389 | 0.043455 | -0.13437 | 0.213948 | 0.540137 | -0.00694 | 0.180209 | 0.240003 | 0.823215 | 0.909659 | 0.722602 | 0.305708 | 0.391541 | 0.143466 | 0.062385 | 0.055141 | 0.131587 | 0.049292 | 0.26421 | -0.17068 | 0.203687 | 0.136156 | 0.116245 | 0.036162 | -0.00296 |
| 29.27778 | 0.294726 | 0.01848 | 0.265125 | 0.392596 | -0.0588 | 0.230121 | 0.188096 | 0.523167 | 0.73558 | 0.581083 | 0.281294 | 0.223924 | 0.127606 | 0.056961 | 0.028302 | 0.154697 | 0.01302 | 0.197576 | -0.18105 | 0.148299 | 0.093945 | 0.144636 | 0.019858 | 0.001474 |
| 29.29167 | 0.441429 | 0.120579 | 0.258936 | 0.269449 | -0.06426 | 0.185931 | 0.134248 | 0.294372 | 0.508543 | 0.225482 | 0.267742 | 0.044798 | 0.100327 | 0.0094 | -0.00557 | 0.180901 | -0.01172 | 0.192893 | -0.19109 | 0.089432 | 0.048189 | 0.162553 | 0.001784 | -0.0149 |
| 29.30556 | 0.464955 | 0.142731 | 0.235152 | 0.333233 | -0.0364 | 0.145973 | 0.143911 | 0.188905 | 0.342808 | -0.08266 | 0.253439 | -0.06784 | 0.125332 | -0.03928 | -0.01057 | 0.228213 | -0.00529 | 0.173383 | -0.18566 | 0.097458 | 0.064873 | 0.225358 | -0.00118 | -0.04977 |
| 29.31944 | 0.482644 | 0.161367 | 0.209182 | 0.521086 | -0.03004 | 0.140463 | 0.243572 | 0.302617 | 0.264057 | -0.16005 | 0.266216 | -0.00963 | 0.160605 | -0.0906 | -0.00511 | 0.256592 | 0.010717 | 0.131801 | -0.17764 | 0.116364 | 0.073236 | 0.244719 | 0.014814 | -0.1008 |
| 29.33333 | 0.580046 | 0.141557 | 0.108568 | 0.680715 | -0.08357 | 0.194614 | 0.418447 | 0.557764 | 0.185283 | -0.13605 | 0.298989 | 0.133929 | 0.131909 | -0.12923 | -0.02081 | 0.183116 | -0.01355 | 0.094427 | -0.1955 | 0.108906 | 0.062166 | 0.217761 | -0.01236 | -0.14926 |
| 29.34722 | 0.692435 | 0.11179 | -0.08088 | 0.780254 | -0.17313 | 0.307331 | 0.61317 | 0.790495 | 0.195701 | -0.03341 | 0.285587 | 0.256039 | 0.080682 | -0.14057 | -0.04875 | 0.087384 | -0.06788 | 0.086344 | -0.21429 | 0.097104 | 0.036398 | 0.147643 | -0.07653 | -0.17052 |
| 29.36111 | 0.730005 | 0.04066 | -0.01869 | 0.656943 | -0.2703 | 0.385863 | 0.701889 | 0.964316 | 0.309587 | 0.119198 | 0.232638 | 0.226218 | 0.026737 | -0.102 | -0.1112 | 0.009305 | -0.11189 | 0.060239 | -0.23753 | 0.059304 | -0.01821 | 0.087921 | -0.13239 | -0.15632 |
| 29.375 | 0.746774 | 0.057064 | 0.073338 | 0.482696 | -0.30541 | 0.536456 | 0.744713 | 1.05313 | 0.525255 | 0.262168 | 0.220449 | 0.022128 | -0.03454 | -0.02639 | -0.20825 | -0.07299 | -0.11347 | -0.00706 | -0.24761 | -0.00961 | -0.09084 | 0.044405 | -0.12545 | -0.13674 |
| 29.38889 | 0.907214 | 0.22245 | 0.151142 | 0.756885 | -0.11152 | 0.866413 | 0.864465 | 1.445024 | 0.628061 | 0.579811 | 0.340632 | -0.14061 | -0.01508 | 0.021444 | -0.20638 | -0.1466 | -0.05382 | -0.01797 | -0.1925 | -0.05275 | -0.06476 | 0.094207 | -0.0207 | -0.07521 |
| 29.40278 | 1.266326 | 0.491294 | 0.309898 | 1.585021 | 0.359913 | 1.57627 | 0.952671 | 1.971444 | 1.262756 | 1.468696 | 0.574582 | -0.04717 | 0.137523 | -0.00705 | -0.03343 | -0.18508 | 0.071519 | 0.057566 | -0.05131 | 0.03961 | 0.09409 | 0.291589 | 0.235931 | 0.095417 |
| 29.41667 | 1.857868 | 0.400029 | 0.468068 | 1.926303 | 0.70739 | 1.436638 | 0.641617 | 2.676482 | 2.322134 | 2.737201 | 0.68604 | 0.209645 | 0.396821 | -0.01981 | 0.283023 | -0.05386 | 0.269196 | 0.186956 | 0.183539 | 0.277985 | 0.327056 | 0.493935 | 0.608629 | 0.427933 |
| 29.43056 | 2.004985 | 0.216881 | 0.606665 | 1.362478 | 0.929379 | 0.781931 | -0.11988 | 3.038035 | 3.035846 | 3.123491 | 0.962652 | 0.397106 | 0.547188 | 0.041362 | 0.47397 | 0.232074 | 0.310411 | 0.228714 | 0.194461 | 0.378382 | 0.376875 | 0.378311 | 0.589792 | 0.611879 |
| 29.44444 | 2.231319 | 0.215095 | 0.784462 | 0.528301 | 0.923324 | 0.448989 | -0.63522 | 3.484162 | 4.619332 | 3.190135 | 1.134231 | 0.398439 | 0.446314 | 0.120273 | 0.267072 | 0.220932 | 0.056106 | 0.095561 | -0.05521 | 0.054608 | 0.120373 | -0.06422 | 0.192233 | 0.405959 |
| 29.45833 | 2.68872 | 0.308101 | 1.292142 | 0.608608 | 0.771375 | 0.582084 | -0.43986 | 3.262922 | 3.777184 | 2.188365 | 0.967004 | 0.269309 | 0.397544 | 0.210848 | 0.097506 | 0.001047 | -0.08344 | 0.102742 | -0.14563 | -0.13135 | 0.011192 | -0.34206 | 0.159947 | 0.166298 |
| 29.47222 | 2.335526 | 0.275047 | 0.953493 | 0.974865 | 0.644095 | 0.74767 | 0.112032 | 2.216105 | 1.316455 | 1.185629 | 0.831683 | 0.233567 | 0.39281 | 0.270825 | 0.166604 | -0.02962 | 0.03479 | 0.211157 | 0.005388 | -0.00465 | 0.180018 | -0.26868 | 0.284654 | 0.181945 |
| 29.48611 | 1.857673 | 0.235669 | 0.622774 | 1.001594 | 0.438768 | 0.790625 | 0.518721 | 1.41797 | -0.99143 | 0.80456 | 0.692715 | 0.186311 | 0.263978 | 0.200412 | 0.154472 | -0.04462 | 0.080958 | 0.20126 | 0.054975 | 0.037264 | 0.232259 | -0.24337 | 0.253144 | 0.105156 |
| 29.5 | 1.098784 | 0.189925 | 0.346563 | 1.039374 | 0.139813 | 0.667401 | 0.662171 | 0.654289 | -3.12853 | 0.304736 | 0.472594 | 0.022306 | 0.193907 | 0.13138 | 0.103814 | -0.14426 | 0.041491 | 0.143297 | 0.072579 | -0.03551 | 0.205598 | -0.16299 | 0.156279 | -0.02522 |
| 29.51389 | 0.434775 | 0.103921 | 0.026637 | 1.238084 | -0.25878 | 0.402599 | 0.655204 | -0.23385 | -2.76127 | 0.488896 | 0.364893 | -0.46107 | 0.155832 | 0.066903 | 0.018786 | -0.25473 | -0.03275 | 0.092234 | 0.060207 | -0.09758 | 0.156134 | -0.03823 | 0.077858 | -0.17167 |
| 29.52778 | -0.01554 | 0.015682 | -0.22307 | 1.180331 | -0.61439 | 0.172765 | 0.580245 | -1.21633 | -1.16667 | 0.30778 | 0.430082 | -0.77298 | 0.092895 | -0.03377 | -0.07978 | -0.2532 | -0.11184 | 0.030527 | -0.04629 | -0.06593 | 0.077717 | 0.088662 | 0.008809 | -0.27443 |
| 29.54167 | 0.454127 | -0.01597 | -0.15751 | 1.07454 | -0.61565 | 0.077716 | 0.384705 | -1.95098 | 0.144624 | 0.558494 | 0.467867 | -0.72179 | 0.14031 | -0.00514 | -0.09246 | -0.09947 | -0.04998 | 0.046097 | -0.02945 | 0.098534 | 0.137216 | 0.264139 | 0.092892 | -0.1155 |
| 29.55556 | 0.598653 | -0.03563 | 0.002773 | 0.912818 | -0.38855 | 0.024395 | 0.107246 | -2.42513 | 0.953079 | 1.414943 | 0.370824 | -0.51105 | 0.235588 | 0.117417 | -0.03514 | 0.08115 | 0.041341 | 0.108388 | 0.005847 | 0.240125 | 0.28287 | 0.429172 | 0.288217 | 0.205562 |
| 29.56944 | 0.58172 | -0.08636 | 0.131667 | 0.736336 | -0.23845 | -0.08871 | -0.18282 | -2.66158 | 1.395161 | 1.572878 | 0.165588 | -0.16908 | 0.301572 | 0.151466 | -0.02361 | 0.126369 | 0.03881 | 0.084931 | -0.00963 | 0.177463 | 0.414291 | 0.378694 | 0.448989 | 0.413471 |
| 29.58333 | 0.471796 | -0.15979 | 0.187131 | 0.618522 | -0.13527 | -0.14959 | -0.23963 | -0.87734 | 1.254133 | 1.092971 | -0.05708 | 0.241285 | 0.353793 | 0.185835 | -0.00187 | 0.099488 | 0.032811 | 0.036216 | -0.02559 | 0.10433 | 0.502522 | 0.232468 | 0.550654 | 0.453912 |
| 29.59722 | 0.364422 | -0.15378 | 0.24192 | 0.510063 | -0.08883 | -0.06036 | -0.13 | 0.954224 | 0.469019 | 0.327934 | -0.19073 | 0.512829 | 0.470197 | 0.316335 | 0.117276 | 0.126758 | 0.11144 | 0.074945 | 0.098267 | 0.150315 | 0.557001 | 0.210119 | 0.58074 | 0.444858 |
| 29.61111 | 0.002562 | -0.15956 | 0.303138 | 0.310846 | -0.14025 | 0.106005 | -0.02402 | 3.383515 | -0.58891 | -0.59597 | -0.2276 | 0.52078 | 0.521919 | 0.388103 | 0.198355 | 0.152939 | 0.164569 | 0.106647 | 0.243026 | 0.207059 | 0.571493 | 0.239785 | 0.5267 | 0.358749 |
| 29.625 | -0.2508 | -0.32888 | 0.321116 | 0.181315 | -0.24271 | 0.12309 | 0.083778 | 4.657341 | -1.32763 | -1.12887 | -0.10764 | 0.183049 | 0.340521 | 0.24575 | 0.062458 | -0.04457 | 0.02075 | -0.03756 | 0.155807 | -0.02573 | 0.31897 | 0.092877 | 0.272727 | 0.087259 |
| 29.63889 | -0.2979 | -0.38582 | 0.354059 | 0.171409 | -0.16076 | 0.301757 | 0.19132 | 4.806477 | -1.23256 | -1.06534 | 0.119927 | -0.30494 | 0.158167 | 0.110514 | -0.09331 | -0.26526 | -0.13041 | -0.19639 | 0.041317 | -0.27804 | 0.077317 | -0.03255 | 0.029067 | -0.15497 |
| 29.65278 | -0.18923 | -0.20337 | 0.291012 | 0.252238 | -0.11928 | 0.53234 | 0.203263 | 4.448106 | -1.20875 | -0.87511 | 0.45038 | -0.82235 | 0.164392 | 0.13832 | -0.07192 | -0.20531 | -0.10156 | -0.19314 | 0.146592 | -0.30361 | 0.056602 | 0.012476 | 0.036074 | -0.17712 |
| 29.66667 | -0.00958 | 0.034667 | 0.19866 | 0.318726 | -0.11377 | 0.591348 | 0.087662 | 3.144885 | -1.09537 | -0.78859 | 0.73686 | -1.23982 | 0.197118 | 0.167568 | -0.02811 | -0.05229 | -0.02797 | -0.14884 | 0.322913 | -0.34525 | 0.104042 | 0.084878 | 0.093761 | -0.1096 |
| 29.68056 | 0.331882 | 0.231962 | -0.18916 | 0.381349 | -0.09745 | 0.580845 | -0.09334 | 1.444195 | -0.94859 | -0.61093 | 0.784003 | -1.28838 | 0.209381 | 0.144757 | -0.03642 | 0.038821 | 0.009911 | -0.09237 | 0.113876 | -0.37329 | 0.140709 | 0.189372 | 0.055881 | 0.005576 |
| 29.69444 | 0.574198 | 0.296577 | -0.60101 | 0.423578 | -0.05119 | 0.29247 | -0.30519 | 0.18774 | -0.47529 | -0.59825 | 0.687274 | -1.01363 | 0.231479 | 0.106877 | -0.09469 | 0.014692 | 0.0213 | -0.05178 | 0.005513 | -0.31095 | 0.09954 | 0.245713 | 0.042752 | 0.022833 |
| 29.70833 | 0.551441 | 0.219039 | -0.62605 | 0.322228 | -0.00715 | 0.064032 | -0.42356 | -0.89006 | -0.03592 | -0.73968 | 0.574727 | -0.57518 | 0.213645 | 0.131844 | -0.22257 | -0.12919 | -0.03525 | -0.07572 | -0.03536 | -0.32587 | 0.073718 | 0.229828 | -0.00455 | -0.00123 |
| 29.72222 | 0.414415 | 0.149845 | -0.44086 | 0.120844 | 0.014002 | -0.12474 | -0.27422 | -2.50418 | -0.09232 | -1.14143 | 0.400613 | -0.45873 | 0.159467 | 0.174979 | -0.34816 | -0.28506 | -0.13197 | -0.17705 | -0.1055 | -0.37807 | -0.01376 | 0.082065 | -0.02619 | -0.03439 |
| 29.73611 | -0.21495 | -0.14425 | -0.39094 | -0.09922 | 0.016796 | -0.25067 | -0.28753 | -3.18548 | -0.73139 | -1.81077 | 0.231971 | -0.67863 | 0.106897 | 0.197494 | -0.39604 | -0.21678 | -0.2036 | -0.26072 | -0.11066 | -0.39545 | -0.04011 | -0.06545 | -0.05866 | -0.13462 |

| | | | | | | | | | | | | | | | | | | | | | | | | |
|---|---|---|---|---|---|---|---|---|---|---|---|---|---|---|---|---|---|---|---|---|---|---|---|---|
| 29.75 | -0.82346 | -0.26905 | -0.33665 | -0.27919 | 0.006641 | -0.50162 | -0.34396 | -3.0765 | -0.78382 | -2.76062 | 0.094412 | -0.9414 | 0.050375 | 0.223626 | -0.36185 | -0.09165 | -0.21034 | -0.36614 | -0.09067 | -0.40399 | -0.09153 | -0.18417 | -0.10938 | -0.30378 |
| 29.76389 | -1.18385 | -0.22714 | -0.34672 | -0.39836 | -0.16379 | -0.70653 | -0.48831 | -2.72786 | -0.34564 | -2.8701 | 0.023544 | -0.97574 | -0.01815 | 0.240987 | -0.33015 | -0.08501 | -0.22992 | -0.45423 | -0.0807 | -0.52378 | -0.19076 | -0.3401 | -0.13606 | -0.40374 |
| 29.77778 | -1.14476 | -0.12139 | -0.48631 | -0.49662 | -0.34917 | -0.95009 | -0.64145 | -2.42225 | 0.183267 | -2.71467 | -0.12367 | -0.81373 | -0.0158 | 0.26518 | -0.19945 | 0.026373 | -0.21639 | -0.51092 | -0.07716 | -0.58339 | -0.23057 | -0.43573 | -0.19415 | -0.35388 |
| 29.79167 | -1.16428 | -0.15115 | -0.70527 | -0.71774 | -0.57959 | -1.01507 | -0.32321 | -2.67377 | 0.522983 | -2.22298 | -0.43087 | -0.62354 | -0.04217 | 0.259097 | -0.0318 | 0.117888 | -0.15958 | -0.46305 | -0.08051 | -0.80112 | -0.18922 | -0.342 | -0.19963 | -0.28758 |
| 29.80556 | -0.8745 | -0.19396 | -0.91911 | -1.07944 | -0.5612 | -0.97875 | -0.40969 | -2.17574 | 0.360108 | -1.44732 | -0.86122 | -0.11132 | -0.22058 | 0.10761 | -0.00032 | 0.040567 | -0.17699 | -0.46441 | -0.22516 | -1.0751 | -0.24061 | -0.39125 | -0.30731 | -0.36477 |
| 29.81944 | -0.60239 | -0.13669 | -0.52568 | -1.37642 | -0.41411 | -0.71146 | -0.81673 | -1.92726 | 0.342432 | -0.56265 | -1.20133 | 0.733468 | -0.38098 | -0.06769 | -0.0338 | -0.10402 | -0.17172 | -0.55537 | -0.33718 | -1.21306 | -0.28296 | -0.52152 | -0.37659 | -0.39962 |
| 29.83333 | -0.42699 | -0.15168 | 0.026995 | -1.52939 | -0.1941 | -0.34805 | -1.24787 | -2.05987 | 0.313274 | 0.106971 | -1.22064 | 1.183875 | -0.37341 | -0.08548 | 0.006112 | -0.11483 | -0.0868 | -0.47238 | -0.33306 | -1.03926 | -0.2004 | -0.44119 | -0.3225 | -0.29109 |
| 29.84722 | -0.26203 | -0.22554 | 0.56786 | -0.74941 | 0.112404 | -0.01395 | -1.1866 | -1.35939 | 0.076627 | 0.558517 | -0.35995 | 0.880466 | -0.39348 | -0.05099 | 0.034006 | -0.09144 | -0.03297 | -0.38833 | -0.32275 | -0.66321 | -0.08306 | -0.33799 | -0.27051 | -0.21138 |
| 29.86111 | -0.12638 | -0.13672 | 0.897274 | -0.83654 | 0.307972 | 0.353119 | -0.39358 | -0.66435 | 0.195709 | 0.880725 | -0.05562 | 0.4883 | -0.42172 | -0.0001 | 0.017625 | -0.08812 | 0.024862 | -0.41118 | -0.33302 | -0.3403 | -0.00607 | -0.24508 | -0.17521 | -0.12022 |
| 29.875 | 0.091503 | 0.050616 | 1.01284 | -0.7461 | 0.419706 | 0.592171 | 0.484766 | 0.153815 | 0.461425 | 0.787815 | 0.151144 | 0.334473 | -0.32217 | 0.159913 | 0.020835 | -0.06185 | 0.106411 | -0.3806 | -0.29346 | -0.03541 | 0.06696 | -0.1216 | -0.06842 | 0.013486 |
| 29.88889 | 0.171989 | 0.098342 | 1.16128 | -0.77712 | 0.299985 | 0.604327 | 0.494116 | 0.395149 | 0.617603 | 0.074532 | 0.264469 | 0.123284 | -0.15137 | 0.257647 | -0.00646 | -0.03807 | 0.11116 | -0.30457 | -0.26062 | 0.24438 | 0.055821 | -0.03251 | 0.008596 | 0.083987 |
| 29.90278 | -0.41423 | -0.11419 | 0.804145 | -0.5198 | -0.00106 | 0.198089 | -0.20362 | -0.1989 | 0.274453 | -1.34659 | -0.13089 | -0.45565 | -0.07241 | 0.18562 | -0.0755 | -0.00454 | 0.036386 | -0.19569 | -0.29874 | 0.363234 | -0.10823 | -0.05135 | -0.11461 | -0.09148 |
| 29.91667 | -1.1224 | -0.29123 | -0.04044 | -0.53357 | -0.27515 | -0.35402 | -0.44954 | -1.33569 | -0.56663 | -3.20028 | -0.7439 | -1.29524 | -0.07025 | 0.099392 | -0.1002 | 0.096669 | -0.01402 | -0.10062 | -0.36606 | 0.179123 | -0.27536 | -0.07935 | -0.29216 | -0.35785 |
| 29.93056 | -1.5067 | -0.55 | -0.65466 | -1.04508 | -0.38307 | -0.82941 | -0.21895 | -2.16066 | -1.28575 | -4.34627 | -1.08005 | -1.74344 | 0.023443 | 0.256009 | 0.040207 | 0.296838 | 0.169411 | -0.02474 | -0.21131 | 0.015105 | -0.1266 | 0.072498 | -0.20129 | -0.36111 |
| 29.94444 | -1.70026 | -0.79902 | -0.71101 | -1.37345 | -0.55007 | -1.1285 | -0.53025 | -1.87862 | -1.03525 | -3.07297 | -0.99134 | -1.63086 | 0.185646 | 0.532979 | 0.297641 | 0.485793 | 0.448981 | 0.109975 | 0.180038 | 0.019095 | 0.18989 | 0.232037 | 0.041971 | -0.19305 |
| 29.95833 | -1.96498 | -1.1657 | -0.47645 | -1.63542 | -0.64418 | -1.18175 | -0.6567 | -1.58702 | -0.73512 | -1.52247 | -0.74922 | -1.21471 | 0.111027 | 0.514277 | 0.280411 | 0.389598 | 0.425899 | 0.135328 | 0.345651 | 0.053801 | 0.206174 | 0.100377 | 0.01075 | -0.17735 |
| 29.97222 | -1.8678 | -1.07076 | -0.2005 | -1.55105 | -0.51591 | -0.88356 | -0.29199 | -1.67325 | -0.46339 | -0.625 | -0.6178 | -0.34484 | -0.1291 | 0.248266 | 0.084778 | 0.11279 | 0.160689 | 0.049662 | 0.107237 | 0.068086 | -0.00778 | -0.20058 | -0.04973 | -0.12686 |
| 29.98611 | -1.35301 | -0.46468 | 0.084831 | -1.18921 | -0.3306 | -0.70577 | -0.03935 | -1.56738 | -0.21234 | -0.24382 | -0.26381 | 0.492856 | -0.15332 | 0.067685 | 0.007927 | -0.00144 | 0.014352 | 0.021121 | -0.11764 | 0.121614 | -0.05568 | -0.25423 | -0.0717 | -0.12954 |
| 30 | -0.50312 | 0.218825 | 0.314273 | -0.78029 | -0.20185 | -0.53017 | 0.052745 | -1.03777 | 0.089394 | 0.074445 | 0.01974 | 1.131385 | -0.12662 | -0.04084 | 0.014406 | -0.02153 | -0.00057 | 0.106301 | -0.19656 | 0.185294 | 0.023111 | -0.13267 | -0.02375 | -0.06335 |
| 30.01389 | 0.157743 | 0.841846 | 0.413725 | -0.452 | -0.04172 | -0.32256 | 0.031003 | -0.4504 | 0.376172 | 0.661328 | 0.161192 | 1.572758 | -0.06703 | -0.08949 | 0.028027 | -0.03003 | 0.023602 | 0.206305 | -0.21604 | 0.178495 | 0.050699 | -0.06606 | 0.001654 | -0.03608 |
| 30.02778 | 0.631879 | 0.476394 | 0.48347 | -0.10973 | 0.161513 | -0.10027 | -0.0046 | -0.17521 | 0.52325 | 1.322986 | 0.239163 | 1.499088 | -0.07421 | -0.04291 | 0.014219 | -0.01521 | 0.041528 | 0.221632 | -0.19201 | 0.150475 | 0.045578 | -0.03367 | 0.024741 | -0.07889 |
| 30.04167 | 0.925787 | 0.615461 | 0.576654 | 0.374849 | 0.300537 | 0.069122 | 0.206927 | -0.18507 | 0.526344 | 2.947699 | 0.238847 | 1.648805 | -0.05148 | -0.0069 | 0.001704 | 0.008475 | 0.027386 | 0.21872 | -0.14102 | 0.122549 | 0.035307 | 0.014021 | 0.024423 | -0.13899 |
| 30.05556 | 0.851035 | 0.537673 | 0.591385 | 0.806355 | 0.359953 | 0.141226 | 0.539557 | -0.18239 | 0.592308 | 2.456326 | 0.186096 | 1.447768 | -0.04265 | -0.01415 | -0.00632 | 0.034075 | 0.006839 | 0.205342 | -0.08963 | 0.098878 | 0.033625 | 0.076352 | 0.007924 | -0.11014 |
| 30.06944 | 0.591615 | 0.378503 | 0.577902 | 0.990732 | 0.356157 | 0.155767 | 0.657487 | -0.04253 | 0.683573 | 1.852918 | 0.2735 | 1.199112 | -0.03976 | -0.01387 | -0.01902 | 0.047483 | 0.005098 | 0.254925 | -0.06199 | 0.111365 | 0.078837 | 0.125633 | -0.00016 | 0.013065 |
| 30.08333 | 0.317523 | 0.523408 | 0.563927 | 0.995383 | 0.340973 | 0.137801 | 0.448184 | 0.163986 | 0.831568 | 1.932213 | 0.520508 | 0.490335 | -0.04478 | 0.025466 | -0.04445 | 0.041686 | 0.01287 | 0.30519 | -0.05108 | 0.215029 | 0.149215 | 0.152778 | 0.010115 | 0.137102 |
| 30.09722 | 0.154078 | 0.55241 | 0.433322 | 1.036196 | 0.282239 | 0.153058 | 0.321114 | 0.35846 | 1.071448 | 1.394701 | 0.784637 | -0.05438 | -0.05156 | 0.042761 | -0.04525 | 0.023158 | 0.013292 | 0.370601 | -0.05725 | 0.351326 | 0.172166 | 0.162717 | 0.007367 | 0.190201 |
| 30.11111 | 0.576741 | 0.330071 | 0.416821 | 0.602963 | 0.231895 | 0.151641 | 0.66671 | 0.678106 | 1.079762 | 0.62518 | 0.656472 | -0.40085 | -0.05012 | -0.00361 | -0.04832 | 0.006249 | -0.00815 | 0.357784 | -0.07812 | 0.446481 | 0.128475 | 0.17566 | -0.0135 | 0.172442 |
| 30.125 | 1.064791 | 0.304644 | 0.654264 | -0.04321 | 0.187548 | 0.171718 | 0.471848 | 1.08476 | 1.030472 | 0.030722 | 0.701322 | -0.42763 | -0.0388 | -0.05009 | -0.01725 | 0.005345 | -0.01632 | 0.227548 | -0.02699 | 0.49335 | 0.103111 | 0.21724 | -0.02463 | 0.090646 |
| 30.13889 | 1.300399 | 0.24397 | 0.88446 | -0.22625 | 0.153024 | 0.298318 | 0.248516 | 1.573877 | 1.090713 | -0.06896 | 0.826646 | -0.28955 | -0.0133 | -0.04886 | 0.038384 | 0.030698 | -0.03422 | 0.119931 | 0.052565 | 0.475795 | 0.108793 | 0.271344 | 0.006515 | 0.075204 |
| 30.15278 | 1.086489 | 0.174956 | 1.098141 | -0.25425 | 0.132093 | 0.49701 | 0.149672 | 1.939932 | 1.150512 | 0.208071 | 0.63412 | -0.07083 | 0.012548 | -0.01867 | 0.06673 | 0.054533 | -0.0292 | 0.081539 | 0.057374 | 0.372965 | 0.120325 | 0.289059 | 0.050386 | 0.133648 |
| 30.16667 | 0.628007 | 0.05899 | 1.065807 | -0.12494 | 0.096197 | 0.589338 | 0.138312 | 1.716899 | 0.960161 | 0.53767 | 0.589673 | 0.121245 | 0.027745 | -0.01252 | 0.065633 | 0.076576 | -0.01236 | 0.0661 | 0.018662 | 0.265433 | 0.139545 | 0.256611 | 0.087375 | 0.215071 |
| 30.18056 | 0.278987 | -0.11371 | 0.842375 | -0.01035 | 0.043605 | 0.505868 | 0.108169 | 1.227419 | 0.856314 | 0.640332 | 0.499511 | 0.140633 | 0.026982 | -0.01826 | 0.060302 | 0.096423 | 0.000844 | 0.051017 | 0.027518 | 0.21187 | 0.139425 | 0.188742 | 0.089915 | 0.236931 |
| 30.19444 | -0.00362 | -0.35026 | 0.505619 | 0.113832 | 0.017515 | 0.315934 | 0.05868 | 0.806404 | 0.664925 | 0.336991 | 0.407279 | -0.01694 | 0.009636 | -0.0542 | 0.054923 | 0.082291 | -0.01357 | 0.023735 | 0.015599 | 0.184831 | 0.092419 | 0.117934 | 0.058692 | 0.161942 |
| 30.20833 | -0.21807 | -0.55212 | 0.212122 | 0.21927 | 0.007027 | 0.028751 | -0.03875 | 0.415826 | 0.392968 | -0.33824 | 0.247855 | -0.22246 | 0.000116 | -0.11413 | 0.038969 | 0.058995 | -0.03639 | 0.040241 | -0.06019 | 0.18432 | 0.03436 | 0.05381 | 0.012554 | 0.023413 |
| 30.22222 | -0.13038 | -1.03742 | -0.04035 | 0.255346 | -0.02342 | -0.27754 | -0.16884 | -0.00253 | 0.142356 | -0.90633 | 0.139146 | -0.42892 | -0.01674 | -0.16446 | -0.00543 | 0.053585 | -0.05202 | 0.053947 | -0.17102 | 0.175273 | 0.022457 | 0.015194 | -0.04501 | -0.1163 |
| 30.23611 | 0.010213 | -1.42749 | -0.14182 | 0.245322 | -0.02985 | -0.35184 | -0.26972 | -0.23322 | 0.068828 | -0.96077 | 0.173004 | -0.66907 | -0.06539 | -0.19358 | -0.05952 | 0.035504 | -0.07264 | 0.050867 | -0.24301 | 0.125464 | 0.022063 | -0.00712 | -0.10055 | -0.16134 |
| 30.25 | 0.125967 | -1.17624 | -0.10272 | 0.226892 | 0.005213 | -0.14241 | -0.14333 | -0.07985 | 0.250886 | -0.56261 | 0.365195 | -0.52787 | -0.07648 | -0.18276 | -0.0852 | 0.008364 | -0.06975 | 0.028092 | -0.21225 | 0.124529 | 0.064792 | 0.019711 | -0.07336 | -0.07721 |
| 30.26389 | 0.39405 | -1.15521 | -0.01093 | 0.154251 | 0.082557 | 0.188261 | 0.160006 | 0.372263 | 0.579131 | 0.121752 | 0.521699 | -0.2035 | -0.01158 | -0.11345 | -0.05234 | -0.00145 | -0.01545 | 0.038313 | -0.10446 | 0.218013 | 0.146957 | 0.110507 | 0.047495 | 0.033629 |
| 30.27778 | 0.574639 | -1.12935 | 0.126581 | 0.111546 | 0.121781 | 0.387041 | 0.361857 | 0.818854 | 0.514811 | 0.618696 | 0.607285 | 0.080654 | 0.086008 | -0.0359 | 0.008019 | 0.022391 | 0.058347 | 0.102488 | -0.00843 | 0.346659 | 0.223506 | 0.235256 | 0.161652 | 0.141666 |
| 30.29167 | 0.371846 | -1.1403 | 0.157421 | 0.076732 | 0.149642 | 0.446585 | 0.487661 | 1.103715 | 0.753915 | 0.823577 | 0.500929 | 0.146016 | 0.145556 | -0.00738 | 0.019959 | 0.039715 | 0.095313 | 0.153825 | 0.034783 | 0.394912 | 0.232175 | 0.30535 | 0.187081 | 0.198898 |
| 30.30556 | 0.259399 | -0.75455 | 0.168247 | 0.079812 | 0.170464 | 0.421582 | 0.534564 | 1.18031 | 1.020594 | 0.896815 | 0.527133 | -0.03377 | 0.123101 | -0.06957 | -0.00382 | 0.038712 | 0.084287 | 0.158357 | 0.01739 | 0.350156 | 0.198216 | 0.284824 | 0.112864 | 0.202664 |
| 30.31944 | 0.17559 | -0.29725 | 0.132699 | 0.118224 | 0.173034 | 0.402507 | 0.377956 | 1.265314 | 1.168395 | 0.750217 | 0.556571 | -0.31423 | 0.069874 | -0.15177 | -0.0341 | 0.028054 | 0.0473 | 0.138725 | -0.03438 | 0.26189 | 0.167536 | 0.228582 | 0.044893 | 0.14189 |
| 30.33333 | 0.161372 | 0.062617 | 0.166037 | 0.224747 | 0.124381 | 0.398938 | -0.10516 | 1.371165 | 1.141077 | 0.180258 | 0.572914 | -0.4431 | 0.043847 | -0.17513 | -0.03782 | 0.022367 | -0.00456 | 0.099725 | -0.06864 | 0.206896 | 0.11861 | 0.178462 | 0.003184 | 0.056418 |
| 30.34722 | 0.37499 | 0.258553 | 0.1806 | 0.31336 | 0.055112 | 0.377873 | -0.67812 | 1.291003 | 0.998712 | -0.56376 | 0.480845 | -0.36723 | 0.014685 | -0.15882 | -0.0403 | 0.031245 | -0.02847 | 0.089371 | 0.013807 | 0.135075 | 0.068104 | 0.129715 | 0.023347 | -0.0421 |
| 30.36111 | 0.559226 | 0.314627 | 0.173353 | 0.346199 | -0.04817 | 0.290674 | -0.90927 | 1.236674 | 0.847145 | -0.7297 | 0.475951 | -0.10446 | -0.0739 | -0.09323 | -0.10333 | 0.019267 | -0.03357 | 0.023983 | 0.0632 | 0.028146 | -0.02697 | 0.060329 | 0.037956 | -0.13297 |
| 30.375 | 0.751641 | 0.446816 | 0.234846 | 0.23942 | -0.10325 | 0.296847 | -0.51443 | 1.315955 | 0.87752 | -0.18256 | 0.53114 | 0.143231 | -0.19149 | 0.008893 | -0.23059 | -0.0337 | -0.04998 | -0.07376 | -0.11063 | -0.06319 | -0.14397 | -0.03162 | 0.011999 | -0.21828 |
| 30.38889 | 0.947884 | 0.800628 | 0.39184 | 0.297341 | -0.00132 | 0.583827 | 0.479306 | 1.738543 | 1.231257 | 0.883961 | 0.694981 | 0.255793 | -0.1686 | 0.020092 | -0.251 | -0.04502 | -0.02745 | -0.06815 | -0.24375 | -0.03997 | -0.12079 | -0.00499 | -0.02145 | -0.16836 |
| 30.40278 | 1.54634 | 1.496135 | 0.819973 | 0.752871 | 0.214355 | 1.444509 | 1.791353 | 2.537932 | 1.988701 | 2.103579 | 1.027766 | 0.524486 | 0.007507 | -0.19699 | -0.10108 | -0.05632 | 0.009778 | 0.017213 | -0.04416 | 0.107379 | 0.046947 | 0.159057 | 0.16012 | 0.126467 |
| 30.41667 | 1.932936 | 2.57282 | 1.243731 | 1.69141 | 0.279567 | 2.450588 | 2.938969 | 2.439629 | 2.497802 | 2.392081 | 1.010823 | 1.27264 | 0.151722 | -0.48665 | 0.054168 | -0.12044 | 0.057319 | 0.041921 | 0.247638 | 0.353813 | 0.219565 | 0.320233 | 0.439692 | 0.595155 |
| 30.43056 | 0.763588 | 3.472127 | 1.58864 | 1.22428 | 0.171877 | 1.750602 | 2.382901 | 2.395327 | 4.09507 | 2.464847 | 0.809902 | 2.125829 | 0.137098 | -0.55013 | 0.022227 | -0.18853 | 0.075978 | -0.01056 | 0.279361 | 0.431377 | 0.24899 | 0.220967 | 0.495163 | 0.873995 |
| 30.44444 | 2.154908 | 3.13085 | 1.339907 | 0.451694 | 0.240316 | 1.8127 | 2.037445 | 2.492028 | 5.255082 | 1.662699 | 0.455259 | 1.93519 | -0.03749 | -0.39973 | -0.18296 | -0.20892 | -0.0726 | -0.08751 | 0.071352 | 0.195505 | -0.01418 | -0.1124 | 0.247008 | 0.620457 |
| 30.45833 | 1.920394 | 2.762545 | 1.119098 | 0.765192 | 0.284137 | 1.573782 | 1.485304 | 3.204777 | 5.155907 | 1.331071 | 0.653645 | 1.107675 | -0.07079 | -0.1715 | -0.21109 | 0.102391 | -0.07061 | 0.057383 | -0.1058 | 0.051079 | -0.20188 | -0.22258 | 0.116476 | 0.231415 |
| 30.47222 | 1.630856 | 2.031318 | 1.031299 | 1.128659 | 0.26878 | 1.046803 | 0.92509 | 3.126364 | 2.668378 | 1.031734 | 0.9054 | 0.433829 | 0.084732 | 0.053679 | -0.00821 | 0.233536 | 0.094704 | 0.281761 | 0.049752 | 0.175744 | -0.05007 | 0.03852 | 0.204807 | 0.082081 |
| 30.48611 | 1.435743 | 1.412449 | 0.588147 | 1.257765 | 0.191053 | 0.300534 | 0.227804 | 2.860044 | -0.14498 | 0.622197 | 0.946891 | -0.03294 | 0.155325 | 0.129568 | 0.073238 | 0.135956 | 0.153205 | 0.335344 | 0.139734 | 0.225383 | 0.151768 | 0.200358 | 0.203692 | -0.03885 |
| 30.5 | 0.79654 | 0.936661 | 0.063691 | 1.27251 | 0.201346 | -0.14918 | -0.23748 | 2.751298 | -1.87529 | 0.410383 | 0.809648 | -0.40015 | 0.103048 | 0.104066 | 0.00619 | -0.06561 | 0.105258 | 0.196921 | 0.087808 | 0.112323 | 0.187693 | 0.219419 | 0.118524 | -0.15347 |
| 30.51389 | -0.25987 | 0.493341 | -0.33626 | 1.180996 | 0.231443 | -0.17262 | -0.37935 | 2.328284 | -2.58956 | 0.179773 | 0.450572 | -0.76915 | 0.058071 | 0.075984 | -0.0543 | -0.20842 | 0.078392 | 0.076636 | 0.0526 | 0.073781 | 0.194843 | 0.241474 | 0.099649 | -0.15672 |
| 30.52778 | -1.00135 | 0.067563 | -0.61954 | 1.085017 | 0.221247 | 0.14206 | -0.59503 | 2.018558 | -2.90043 | 0.085586 | 0.182845 | -0.87753 | 0.071108 | 0.047415 | -0.03614 | -0.23033 | 0.1019 | 0.045792 | 0.06461 | 0.168836 | 0.192764 | 0.239779 | 0.120899 | -0.04014 |
| 30.54167 | -0.82512 | -0.231 | -0.6064 | 0.846862 | 0.202106 | 0.386295 | -0.68125 | 1.373837 | -2.22527 | 0.113744 | 0.579362 | -0.77307 | 0.104886 | 0.039675 | 0.009559 | -0.09298 | 0.120554 | 0.032018 | 0.078395 | 0.271397 | 0.27413 | 0.234942 | 0.112372 | 0.066264 |
| 30.55556 | 0.151876 | -0.5068 | -0.93732 | 0.728568 | 0.206743 | 0.131224 | -1.27891 | 0.40828 | -1.12373 | 0.083979 | 0.867295 | -0.73027 | 0.118159 | 0.091608 | 0.05429 | 0.034641 | 0.121599 | 0.044915 | 0.070019 | 0.323918 | 0.215198 | 0.240937 | 0.137231 | 0.097828 |
| 30.56944 | 0.399728 | -0.6632 | -1.12702 | 0.618254 | 0.185885 | -0.0632 | -1.40981 | -0.48899 | -0.48816 | 0.017326 | 0.850335 | -0.37873 | 0.139849 | 0.166874 | 0.118263 | 0.103446 | 0.155863 | 0.091383 | 0.099334 | 0.330862 | 0.243948 | 0.28445 | 0.192945 | 0.130301 |
| 30.58333 | -0.44113 | -0.55133 | -0.57253 | 0.511215 | 0.211825 | -0.07932 | -0.53922 | -0.98547 | -0.32922 | -0.0779 | 0.392555 | -0.00898 | 0.150264 | 0.193226 | 0.138381 | 0.072989 | 0.17971 | 0.15112 | 0.117496 | 0.306857 | 0.228539 | 0.296262 | 0.224077 | 0.121929 |
| 30.59722 | -0.89267 | -0.66941 | 0.17864 | 0.435981 | 0.286175 | -0.24192 | 1.189687 | -0.83598 | -0.12219 | -0.20459 | -0.10756 | 0.420319 | 0.151219 | 0.193586 | 0.110378 | 0.021854 | 0.166265 | 0.196259 | 0.116098 | 0.250702 | 0.209582 | 0.278689 | 0.223741 | 0.01149 |
| 30.61111 | -1.04412 | -0.58952 | 0.860031 | 0.43767 | 0.346364 | -0.39336 | 1.672121 | -1.03331 | 0.301205 | -0.29531 | -0.19118 | 0.711411 | 0.143698 | 0.19504 | 0.05624 | -0.01381 | 0.146532 | 0.181579 | 0.103114 | 0.190762 | 0.1579 | 0.234711 | 0.258983 | -0.13923 |
| 30.625 | 0.259172 | -0.52624 | 1.21228 | 0.531321 | 0.302801 | -0.37858 | 0.821552 | -1.38604 | 0.039091 | -0.43573 | -0.22039 | 0.859123 | 0.11559 | 0.169366 | -0.03628 | -0.10272 | 0.085734 | 0.091763 | 0.065499 | 0.0898 | 0.075051 | 0.138778 | 0.232258 | -0.27465 |
| 30.63889 | 0.767438 | -0.30176 | 0.870385 | 0.487637 | 0.213515 | 0.055859 | -0.06122 | -0.79932 | -0.7229 | -0.63084 | -0.43182 | 0.771162 | 0.070673 | 0.148124 | -0.11914 | -0.21063 | 0.016837 | 0.007176 | 0.028643 | 0.022634 | 0.055722 | 0.102343 | 0.165904 | -0.27486 |
| 30.65278 | 0.815959 | -0.19728 | 0.576136 | 0.49172 | 0.119723 | 0.72434 | -0.50081 | 0.534533 | -1.02479 | -0.71825 | -0.29359 | 0.446412 | 0.051945 | 0.160944 | -0.16602 | -0.21056 | 0.003394 | -0.00741 | 0.000864 | 0.019148 | 0.10132 | 0.132153 | 0.153179 | -0.12412 |
| 30.66667 | 0.789945 | -0.18558 | 0.281986 | 0.426389 | 0.078595 | 0.567602 | -0.4402 | 1.650685 | -0.76481 | -0.64565 | 0.097725 | -0.157 | 0.0399 | 0.170364 | -0.18601 | -0.18702 | 0.009076 | -0.01089 | -0.00941 | 0.008286 | 0.112627 | 0.159783 | 0.15788 | -0.05938 |
| 30.68056 | 1.058301 | -0.0707 | -0.22859 | 0.311009 | 0.039729 | -0.11605 | -0.29231 | 2.129599 | -0.42538 | -0.54563 | 0.374983 | -0.91596 | 0.020732 | 0.106771 | -0.19331 | -0.17473 | -0.00405 | -0.03397 | -0.04385 | -0.06604 | 0.013411 | 0.131483 | 0.12161 | -0.11989 |
| 30.69444 | 1.661654 | 0.121218 | -0.89336 | 0.263716 | 0.018064 | -0.39829 | -0.30267 | 1.763951 | -0.2177 | -0.66281 | 0.631213 | -1.5219 | -0.00084 | 0.016194 | -0.19804 | -0.16635 | -0.03117 | -0.06723 | -0.13258 | -0.1522 | -0.05487 | 0.107269 | 0.1601 | -0.14374 |
| 30.70833 | 1.862623 | 0.182413 | -0.89938 | 0.270872 | -0.00758 | -0.38029 | -0.96124 | 0.893637 | -0.13427 | -0.58828 | 0.661891 | -1.84733 | 0.022072 | 0.014797 | -0.19002 | -0.18083 | -0.02841 | -0.07567 | -0.17985 | -0.18379 | -0.08096 | 0.049691 | 0.189044 | -0.11663 |
| 30.72222 | 1.054897 | 0.062646 | -0.76938 | 0.293771 | -0.114 | -0.38313 | -2.19338 | 0.224473 | -0.33358 | -0.75363 | 0.49114 | -1.97099 | 0.04921 | 0.099131 | -0.19467 | -0.19683 | -0.01313 | -0.1329 | -0.18732 | -0.22352 | -0.12953 | 0.014882 | 0.186893 | -0.12499 |
| 30.73611 | 0.29326 | -0.13899 | -0.12948 | 0.196426 | -0.15241 | -0.36836 | -2.17036 | -0.20849 | -0.52218 | -1.07943 | 0.132866 | -1.75513 | 0.037513 | 0.190289 | -0.2265 | -0.21558 | -0.01386 | -0.16188 | -0.22619 | -0.30154 | -0.16787 | -0.04494 | 0.10938 | -0.16732 |
| 30.75 | -0.63324 | -0.26852 | -0.04509 | 0.093987 | -0.22741 | -0.28318 | -1.98488 | -0.76999 | -0.79943 | -1.52305 | -0.26926 | -1.48877 | 0.010894 | 0.233081 | -0.24448 | -0.19896 | -0.04692 | -0.18875 | -0.23924 | -0.36089 | -0.20979 | -0.10263 | 0.05171 | -0.20072 |
| 30.76389 | -1.46546 | -0.33349 | -0.0493 | 0.009677 | -0.29671 | -0.33619 | -1.80115 | -1.61452 | -1.07458 | -2.02895 | -0.51864 | -1.26648 | 0.001196 | 0.244551 | -0.18363 | -0.14899 | -0.05788 | -0.18587 | -0.20034 | -0.35368 | -0.22122 | -0.10596 | 0.013696 | -0.21938 |
| 30.77778 | -2.24093 | -0.47928 | -0.1074 | -0.15128 | -0.37705 | -0.51782 | -1.59217 | -2.46088 | -1.20238 | -2.35791 | -0.55857 | -1.12417 | -0.02746 | 0.228245 | -0.08562 | -0.09385 | -0.04991 | -0.16682 | -0.28557 | -0.33552 | -0.27641 | -0.1606 | -0.03466 | -0.25624 |
| 30.79167 | -3.1081 | -0.90968 | -0.10533 | -0.24519 | -0.31429 | -0.65384 | -0.8337 | -2.43457 | -0.96477 | -2.44781 | -0.50298 | -0.86145 | -0.10933 | 0.131384 | -0.03455 | -0.10564 | -0.08893 | -0.23881 | -0.44363 | -0.32288 | -0.34193 | -0.26808 | -0.12674 | -0.33442 |
| 30.80556 | -2.3212 | -1.21481 | -0.02572 | -0.21558 | -0.14564 | -0.6476 | -0.46813 | -2.1053 | -0.86993 | -2.17938 | -0.51198 | -0.64995 | -0.15166 | 0.017161 | -0.0655 | -0.16815 | -0.12302 | -0.3199 | -0.5525 | -0.25184 | -0.33676 | -0.32908 | -0.19809 | -0.31177 |
| 30.81944 | -1.7236 | -1.62627 | 0.070985 | -0.21878 | 0.002784 | -0.55774 | -0.4052 | -2.41556 | -0.65496 | -1.78953 | -0.38757 | -0.30166 | -0.17073 | -0.0407 | -0.06421 | -0.18452 | -0.16435 | -0.37141 | -0.54184 | -0.22231 | -0.31317 | -0.35197 | -0.25543 | -0.23563 |
| 30.83333 | -0.86795 | -1.74285 | 0.137071 | -0.16509 | 0.094326 | -0.34956 | -0.43069 | -2.75225 | -0.291 | -0.95336 | -0.53519 | 0.158657 | -0.16189 | -0.10171 | -0.06734 | -0.15672 | -0.19546 | -0.31271 | -0.47217 | -0.32172 | -0.27422 | -0.32325 | -0.26325 | -0.16871 |
| 30.84722 | -0.45396 | -1.66758 | 0.225764 | -0.16763 | 0.199484 | -0.2137 | -0.31862 | -2.58812 | 0.30427 | -0.07134 | -0.52155 | 0.677079 | -0.1497 | -0.10367 | -0.03389 | -0.10299 | -0.18628 | -0.22761 | -0.35655 | -0.41319 | -0.14774 | -0.28384 | -0.20211 | -0.14311 |
| 30.86111 | -0.14735 | -1.55029 | 0.330284 | -0.16481 | 0.284401 | -0.15789 | -0.09695 | -1.96943 | 1.021147 | 0.715908 | 0.11721 | 1.081929 | -0.16841 | -0.06668 | -0.06633 | -0.08682 | -0.1697 | -0.17419 | -0.2736 | -0.45616 | -0.05873 | -0.27355 | -0.2098 | -0.1876 |
| 30.875 | 0.285706 | -1.11461 | 0.39933 | -0.09323 | 0.319417 | -0.09416 | 0.119508 | -1.24833 | 1.588237 | 1.322623 | -0.01775 | 1.269515 | -0.18969 | 0.012587 | -0.06079 | -0.08094 | -0.17709 | -0.17461 | -0.22397 | -0.43691 | -0.0561 | -0.33376 | -0.19841 | -0.17866 |
| 30.88889 | 0.508256 | -0.56667 | 0.359634 | -0.06965 | 0.248844 | -0.02509 | 0.17829 | -0.7758 | 1.396082 | 1.463446 | -0.09057 | 1.030704 | -0.15961 | -0.009 | -0.01013 | -0.06005 | -0.15692 | -0.16034 | -0.18853 | -0.33955 | -0.10852 | -0.35975 | -0.13669 | -0.09767 |
| 30.90278 | 0.107175 | -0.26874 | 0.181901 | -0.23091 | 0.081025 | -0.17164 | -0.11243 | -0.97029 | 0.315795 | 0.689535 | -0.39477 | 0.267925 | -0.15473 | -0.62611 | 0.062479 | -0.15712 | -0.24649 | -0.13143 | -0.21717 | -0.30881 | -0.18995 | -0.33536 | -0.14013 | -0.17637 |
| 30.91667 | -0.73521 | -0.34898 | -0.05456 | -0.40517 | -0.05206 | -0.49699 | -0.72318 | -1.78919 | -1.54777 | -1.26583 | -0.9151 | -0.80816 | -0.17717 | -1.65866 | 0.142205 | -0.278 | -0.42639 | -0.07245 | -0.34429 | -0.41344 | -0.28089 | -0.27679 | -0.25009 | -0.50125 |
| 30.93056 | -1.29901 | -0.17922 | -0.16868 | -0.40417 | -0.15401 | -0.7611 | -1.30862 | -2.70794 | -3.17341 | -2.93288 | -1.62586 | -2.10634 | -0.08555 | -1.56236 | 0.253732 | -0.08533 | -0.31999 | 0.084278 | -0.33211 | -0.29346 | -0.24462 | -0.14233 | -0.21822 | -0.50876 |
| 30.94444 | -1.54904 | -0.30884 | -0.26772 | -0.40005 | -0.29713 | -1.05843 | -1.34098 | -2.99326 | -3.60399 | -3.05889 | -2.49745 | -2.26937 | 0.172655 | -0.38022 | 0.384825 | 0.38427 | 0.110676 | 0.219072 | -0.05538 | 0.015187 | 0.04434 | -0.0316 | 0.01209 | -0.08233 |
| 30.95833 | -1.70709 | -0.7576 | -0.22622 | -0.4902 | -0.48397 | -1.33073 | -1.07844 | -2.91678 | -2.13972 | -1.93857 | -2.25923 | -1.56174 | 0.159498 | 0.163579 | 0.273675 | 0.384829 | 0.256691 | 0.070189 | 0.096314 | 0.076877 | 0.141819 | -0.11361 | 0.048643 | 0.174408 |
| 30.97222 | -1.38819 | -1.56925 | 0.04249 | -0.54797 | -0.56468 | -0.80717 | -1.50946 | -2.39594 | -1.03011 | -1.31231 | -2.31587 | -1.28686 | -0.15382 | 0.027294 | -0.03249 | 0.073392 | 0.090788 | -0.17055 | 0.081753 | -0.10477 | 0.049238 | -0.30747 | -0.11897 | 0.126612 |
| 30.98611 | -1.03438 | -2.2457 | 0.152975 | -0.60268 | -0.58351 | -0.55385 | -1.15819 | -1.90938 | -0.54472 | -1.09799 | -0.63038 | -1.07282 | -0.24099 | -0.05341 | -0.15741 | -0.03565 | -0.0429 | -0.21922 | 0.047058 | -0.12988 | 0.103496 | -0.32239 | -0.12239 | 0.056167 |
| 31 | -0.63328 | -2.28894 | 0.161324 | -0.55381 | -0.55481 | -0.44998 | -0.57198 | -1.10478 | 0.04714 | -0.83662 | 0.616878 | -0.61613 | -0.15878 | -0.10083 | -0.105 | -0.01516 | -0.04809 | -0.18152 | 0.008058 | -0.03788 | 0.185475 | -0.21687 | -0.03365 | 0.079817 |
| 31.01389 | -0.25148 | -1.90123 | 0.13349 | -0.53202 | -0.47718 | -0.33409 | -0.33418 | -0.56401 | 0.564882 | -0.52376 | 1.526657 | -0.06861 | -0.09368 | -0.15181 | -0.09517 | -0.02631 | -0.08014 | -0.1558 | -0.07955 | 0.043389 | 0.142279 | -0.15743 | -0.02694 | 0.075299 |
| 31.02778 | -0.1143 | -1.30476 | 0.188314 | -0.49559 | -0.34144 | -0.28775 | -0.47836 | -0.23774 | 1.200134 | -0.25146 | 2.158658 | 0.38393 | -0.06269 | -0.06166 | -0.09633 | -0.01517 | -0.06835 | -0.1182 | -0.12115 | 0.09095 | 0.035616 | -0.12375 | -0.02469 | 0.002395 |
| 31.04167 | -0.01164 | -0.95174 | 0.246776 | -0.46173 | -0.23187 | -0.32128 | -0.72666 | -0.06002 | 1.147165 | -0.0469 | 1.319652 | 0.704692 | -0.02494 | 0.008463 | -0.08874 | 0.014115 | -0.03459 | -0.05489 | -0.08691 | 0.092217 | -0.07884 | -0.09222 | 0.006638 | -0.10627 |
| 31.05556 | 0.047219 | -0.79884 | 0.301979 | -0.41033 | -0.12718 | -0.32401 | -0.69672 | -0.11809 | 0.741544 | 0.009912 | 0.439244 | 0.897648 | -0.00621 | -0.0323 | -0.07293 | 0.006491 | -0.03277 | -0.01498 | -0.03389 | 0.028017 | -0.14647 | -0.06673 | 0.004833 | -0.18427 |
| 31.06944 | 0.301009 | -0.50199 | 0.349614 | -0.20641 | -0.01883 | -0.29336 | -0.29677 | -0.28742 | 0.214012 | 0.036449 | -0.05654 | 1.106106 | -0.02033 | -0.09184 | -0.07595 | -0.0204 | -0.0436 | 0.001472 | 0.027877 | -0.04693 | -0.15129 | -0.04866 | -0.01483 | -0.18321 |
| 31.08333 | 0.623538 | 0.148744 | 0.345428 | 0.133897 | 0.107631 | -0.13566 | 0.258961 | -0.39808 | -0.29295 | 0.344648 | -0.22339 | 1.163759 | -0.01089 | -0.04791 | -0.05054 | -0.01093 | -0.0184 | 0.020349 | 0.099878 | -0.04463 | -0.03423 | 0.004504 | 0.002421 | -0.06897 |
| 31.09722 | 0.988774 | 0.925397 | 0.346415 | 0.268213 | 0.236927 | 0.019737 | 0.905609 | -0.39342 | -0.45718 | 1.013158 | -1.23014 | 1.217455 | 0.025649 | 0.019924 | -0.00406 | 0.013276 | 0.019275 | 0.049369 | 0.159151 | 0.027751 | 0.141492 | 0.088557 | 0.067112 | 0.105634 |
| 31.11111 | 1.160535 | 1.8459 | 0.410983 | 0.203041 | 0.294764 | 0.150118 | 1.296634 | -0.17789 | -0.08864 | 1.714647 | -1.50988 | 1.747696 | 0.050626 | 0.014028 | 0.009681 | 0.023271 | 0.030927 | 0.056178 | 0.141662 | 0.078381 | 0.234818 | 0.159327 | 0.102418 | 0.21356 |
| 31.125 | 1.333418 | 2.621965 | 0.564222 | 0.175778 | 0.288026 | 0.229257 | 1.287223 | 0.029787 | 0.429717 | 2.382778 | -0.40491 | 1.81395 | 0.05773 | -0.05609 | -0.031 | 0.000885 | 0.026346 | 0.048452 | 0.124767 | 0.097783 | 0.189107 | 0.197113 | 0.048566 | 0.214588 |
| 31.13889 | 1.307988 | 1.599959 | 0.60604 | 0.249006 | 0.301384 | 0.294709 | 1.238796 | 0.23295 | 1.395188 | 2.880528 | 0.938497 | 2.048599 | 0.056387 | -0.09014 | -0.06939 | -0.02681 | 0.009001 | 0.031389 | 0.102443 | 0.110738 | 0.116788 | 0.226958 | -0.01564 | 0.113083 |
| 31.15278 | 1.434784 | 1.770644 | 0.608396 | 0.242414 | 0.305196 | 0.327023 | 1.241848 | 0.46308 | 3.09498 | 2.401952 | 2.456195 | 1.919481 | 0.058315 | -0.00197 | -0.06916 | 0.009621 | 0.037545 | 0.060394 | 0.114353 | 0.182122 | 0.097185 | 0.260584 | -0.04171 | 0.039793 |
| 31.16667 | 1.455647 | 1.386649 | 0.572184 | 0.250452 | 0.258788 | 0.284742 | 1.355957 | 0.625607 | 2.862078 | 1.624608 | 2.245497 | 1.347222 | 0.081179 | 0.070121 | -0.02039 | 0.069311 | 0.103376 | 0.104459 | 0.109772 | 0.269239 | 0.105466 | 0.300901 | -0.02328 | 0.053875 |
| 31.18056 | 1.308013 | 1.338855 | 0.529019 | 0.296551 | 0.134167 | 0.208161 | 1.159542 | 0.583872 | 1.407005 | 0.549797 | 1.261605 | 0.783907 | 0.053444 | -0.41731 | 0.017874 | 0.008739 | 0.073134 | 0.098488 | 0.036231 | 0.30514 | 0.063854 | 0.282047 | -0.04142 | 0.008409 |
| 31.19444 | 1.17695 | 0.915478 | 0.469707 | 0.303119 | -0.02917 | 0.077902 | 0.588764 | 0.207292 | -0.24743 | -1.18155 | 0.402097 | 0.098378 | -0.08745 | -3.45498 | -0.03821 | -0.42367 | -0.17477 | 0.004841 | -0.15738 | 0.190635 | -0.08246 | 0.133615 | -0.22048 | -0.16141 |
| 31.20833 | 1.01299 | 0.614589 | 0.434928 | 0.146127 | -0.06724 | 0.074323 | 0.054138 | -0.16528 | -1.69481 | -1.5791 | -0.10797 | -0.35418 | -0.17646 | -6.51369 | -0.23434 | -0.90858 | -0.2214 | -0.06054 | -0.4143 | 0.09378 | -0.29088 | -0.0983 | -0.31665 | -0.30839 |

| | | | | | | | | | | | | | | | | | | | | | | | | |
|---|---|---|---|---|---|---|---|---|---|---|---|---|---|---|---|---|---|---|---|---|---|---|---|---|
| 31.22222 | 0.763856 | 0.600437 | 0.383218 | 0.002653 | 0.082553 | 0.302684 | 0.052101 | -0.14227 | -1.69337 | -0.40474 | 0.090391 | -0.3387 | 0.216839 | -0.4258 | -0.48351 | 0.480752 | 0.634292 | 0.130489 | -0.04205 | 0.272663 | -0.28336 | -0.23323 | 0.790199 | 0.468314 |
| 31.23611 | 0.646458 | 0.404862 | 0.401266 | -0.05844 | 0.185644 | 0.451116 | 0.005824 | 0.011605 | -1.18292 | 0.372381 | 0.276542 | -0.16185 | 0.718308 | 11.69669 | -0.51236 | 3.717605 | 2.110506 | 0.241218 | 1.168642 | 0.794332 | -0.98118 | -0.53145 | 1.770473 | 1.59511 |
| 31.25 | 0.670814 | -0.3766 | 0.414158 | -0.10622 | 0.143029 | 0.44931 | -0.26973 | 0.143017 | -0.05895 | 0.031131 | 0.629025 | -0.10049 | -0.14575 | 10.1794 | -0.22666 | 1.384766 | 1.141352 | -0.01332 | 0.860851 | 0.225132 | -1.12492 | 0.279346 | 0.588016 | 0.352323 |
| 31.26389 | 0.877632 | -1.01725 | 0.52645 | -0.13435 | 0.091971 | 0.328855 | -0.44629 | 0.232033 | 0.771826 | -0.08697 | 0.61081 | 0.080104 | -0.20892 | -4.60609 | 0.917947 | -1.32553 | -2.35016 | -0.41708 | -0.82391 | -0.38622 | 1.041178 | 0.693334 | -1.38638 | -1.17117 |
| 31.27778 | 0.871816 | -1.06847 | 0.524051 | -0.12721 | 0.064796 | 0.202315 | -0.31481 | 0.196692 | 1.074099 | -0.02291 | 0.466637 | 0.313522 | 0.033454 | -4.61489 | 0.874651 | -0.9435 | -1.16524 | -0.25391 | -0.32365 | -0.45459 | 0.28834 | 0.176575 | -1.00231 | -0.81413 |
| 31.29167 | 0.784478 | -1.19389 | 0.465538 | -0.11099 | 0.011718 | 0.0637 | -0.18703 | 0.112504 | 0.609291 | -0.43183 | 0.169591 | 0.241447 | 0.017851 | -0.58869 | 0.046525 | -0.14547 | -0.19364 | -0.05202 | -0.08655 | -0.05075 | -0.02387 | 0.079233 | -0.15453 | -0.0977 |
| 31.30556 | 0.509566 | -1.4941 | 0.351075 | -0.11362 | -0.02094 | 0.03285 | -0.18408 | 0.00632 | -0.01571 | -1.07391 | -0.05241 | -0.08709 | -0.00845 | -0.15279 | 0.011266 | 0.011983 | -0.05279 | -0.05402 | -0.02484 | 0.03742 | -0.06956 | 0.019822 | -0.0194 | -0.00817 |
| 31.31944 | 0.3172 | -1.58157 | 0.253516 | -0.07016 | -0.04157 | 0.132503 | -0.26801 | -0.08016 | -0.52085 | -1.59036 | -0.30581 | -0.3286 | -0.01122 | -0.20729 | 0.006428 | 0.032336 | -0.03556 | -0.0626 | 0.004881 | 0.051249 | -0.08927 | 0.017596 | -0.00983 | -0.09621 |
| 31.33333 | 0.157193 | -1.72646 | 0.204584 | -0.04909 | -0.04453 | 0.253181 | -0.41373 | -0.1222 | -0.74111 | -1.76281 | -0.4588 | -0.35209 | 0.018785 | -0.15625 | 0.051013 | 0.08223 | 0.02591 | -0.03631 | 0.017738 | 0.111931 | -0.04441 | 0.064793 | 0.052605 | -0.11539 |
| 31.34722 | 0.161804 | -1.86645 | 0.28852 | -0.06341 | -0.03285 | 0.270465 | -0.36164 | -0.17465 | -0.65214 | -1.22937 | -0.61075 | -0.14907 | 0.0356 | -0.12848 | 0.073001 | 0.103248 | 0.056654 | -0.00561 | -0.00172 | 0.13736 | -0.01184 | 0.108878 | 0.118018 | -0.05691 |
| 31.36111 | 0.215923 | -2.16381 | 0.362 | -0.0824 | -0.03467 | 0.231449 | -0.23268 | -0.20522 | -0.32207 | -0.5802 | -0.24328 | 0.07439 | -0.0379 | -0.02845 | -0.00638 | 0.046053 | 0.01401 | -0.02998 | -0.06856 | 0.079967 | -0.07151 | 0.077521 | 0.101483 | -0.0412 |
| 31.375 | 0.31383 | -2.06319 | 0.322747 | -0.07312 | -0.01839 | 0.221603 | -0.07543 | -0.08784 | 0.170238 | -0.17634 | 0.005369 | 0.226944 | -0.13042 | 0.099454 | -0.11086 | -0.01217 | -0.01258 | -0.09341 | -0.13283 | 0.033474 | -0.13464 | 0.019162 | 0.063834 | -0.0707 |
| 31.38889 | 0.740222 | -1.03358 | 0.436527 | 0.158286 | 0.113732 | 0.500484 | 0.541237 | 0.446873 | 0.646097 | 0.623502 | 0.09838 | 0.342875 | -0.10928 | 0.185178 | -0.09948 | 0.018802 | 0.035561 | -0.06909 | -0.1015 | 0.092449 | -0.09298 | 0.053957 | 0.091575 | -0.02067 |
| 31.40278 | 1.50391 | 1.282507 | 0.684122 | 0.796287 | 0.414861 | 1.202926 | 1.935935 | 1.542772 | 1.06231 | 2.100396 | 0.414122 | 0.709839 | 0.000056 | -0.03457 | 0.000847 | 0.010397 | 0.095755 | -0.02807 | -0.03019 | 0.164317 | 0.058523 | 0.179748 | 0.211471 | 0.214578 |
| 31.41667 | 2.243783 | 3.090779 | 1.113415 | 1.544356 | 0.659845 | 1.581962 | 3.550787 | 2.800185 | 1.225083 | 3.256242 | 1.209283 | 1.231979 | 0.116635 | -0.31987 | 0.102761 | -0.02396 | 0.146579 | -0.0489 | 0.037627 | 0.226203 | 0.238398 | 0.284186 | 0.445327 | 0.592335 |
| 31.43056 | 2.471671 | 2.929799 | 1.470061 | 1.348663 | 0.614697 | 1.237294 | 3.207731 | 2.675593 | 0.862959 | 2.905206 | 2.063814 | 1.83745 | 0.110548 | -0.42664 | 0.108131 | -0.1391 | 0.119873 | -0.1118 | 0.006259 | 0.238265 | 0.269649 | 0.235574 | 0.653997 | 0.783871 |
| 31.44444 | 1.844458 | 2.831003 | 1.373615 | 0.607781 | 0.58033 | 0.594773 | 2.272138 | 2.174293 | 1.046487 | 2.71004 | 1.646178 | 2.344992 | -0.09571 | -0.42005 | -0.05113 | -0.26154 | -0.04665 | -0.19699 | -0.25526 | 0.103767 | 0.038402 | 0.007409 | 0.621081 | 0.493968 |
| 31.45833 | 0.848438 | 2.524474 | 1.014218 | 0.35939 | 0.647008 | 0.849167 | 2.143663 | 2.386084 | 2.807348 | 2.708975 | 1.024261 | 2.094461 | -0.15568 | -0.30192 | -0.09939 | -0.09577 | -0.06365 | -0.11583 | -0.36035 | 0.043987 | -0.07444 | -0.05631 | 0.570445 | 0.195374 |
| 31.47222 | 0.360169 | 2.784349 | 0.692535 | 0.296519 | 0.575601 | 1.446715 | 3.582966 | 2.633604 | 2.876276 | 1.764274 | 0.331003 | 1.13199 | 0.005166 | -0.09411 | 0.014771 | 0.325278 | 0.106054 | 0.095885 | -0.07805 | 0.14705 | 0.08441 | 0.130928 | 0.556737 | 0.032899 |
| 31.48611 | 1.540043 | 3.308035 | 0.338927 | 0.282734 | 0.440329 | 1.28385 | 3.491104 | 2.688444 | 2.562787 | 1.189398 | -0.50411 | 0.431021 | 0.072225 | 0.032922 | 0.06603 | 0.369921 | 0.142493 | 0.17813 | 0.162557 | 0.175828 | 0.058917 | 0.20453 | 0.356165 | -0.10304 |
| 31.5 | 1.638678 | 2.990862 | 0.016606 | 0.328033 | 0.280179 | 1.43867 | 1.826174 | 2.591413 | 1.838791 | 0.628405 | -0.52594 | -0.03338 | 0.021703 | 0.055589 | 0.046117 | 0.143463 | 0.069959 | 0.130235 | 0.189015 | 0.104031 | -0.03145 | 0.190629 | 0.118074 | -0.21686 |
| 31.51389 | 1.430182 | 2.174972 | -0.29677 | 0.409178 | 0.178511 | 0.739647 | 0.342033 | 1.952767 | 0.902984 | 0.28756 | 0.207824 | -0.52051 | -0.02905 | 0.040106 | 0.036678 | -0.0786 | 0.031712 | 0.073854 | 0.154559 | 0.034254 | -0.05471 | 0.15367 | 0.0352 | -0.30148 |
| 31.52778 | 1.242442 | 1.523212 | -0.54903 | 0.50415 | 0.060772 | 0.002322 | -1.02939 | 1.399914 | -0.31946 | -0.09355 | 0.02 | -0.91197 | -0.05625 | 0.008774 | 0.045006 | -0.1667 | 0.047323 | 0.033829 | 0.12074 | -0.02342 | -0.01004 | 0.123401 | -1.3E-05 | -0.32632 |
| 31.54167 | 0.532938 | 0.881271 | -0.90484 | 0.52632 | -0.06427 | -0.64785 | -2.2785 | 1.01187 | -1.18109 | -0.52957 | -0.34298 | -0.74302 | -0.08529 | -0.02081 | 0.037832 | -0.10577 | 0.055942 | 0.020188 | 0.095225 | -0.07218 | 0.009329 | 0.093808 | -0.04097 | -0.28898 |
| 31.55556 | -0.34689 | 0.226532 | -1.10538 | 0.407106 | -0.14538 | -0.47131 | -3.18916 | 0.817882 | -1.35227 | -0.75092 | 0.27089 | -0.9468 | -0.10578 | -0.06569 | -0.00138 | -0.03985 | 0.023122 | -0.01128 | 0.057166 | -0.08755 | 0.019089 | 0.029022 | -0.0506 | -0.23603 |
| 31.56944 | -0.00765 | -0.45812 | -1.28256 | 0.234854 | -0.13361 | -0.35053 | -2.62896 | 0.610015 | -1.15526 | -1.08158 | 0.80384 | -0.91009 | -0.0904 | -0.10097 | -0.03718 | 0.015075 | -0.01278 | -0.01117 | 0.039387 | -0.02096 | 0.044982 | -0.04304 | -0.01446 | -0.10767 |
| 31.58333 | -0.56853 | -1.1064 | -0.85315 | 0.031331 | -0.11288 | -0.12083 | -2.12844 | 0.488764 | -0.69608 | -1.32812 | 1.126724 | -0.89878 | -0.0511 | -0.09681 | -0.02833 | 0.05931 | -0.014 | 0.035391 | 0.038547 | 0.064313 | 0.076111 | -0.06048 | 0.0652 | 0.045594 |
| 31.59722 | -1.29877 | -1.58826 | -0.20454 | -0.16663 | -0.09026 | 0.027116 | -2.56988 | 0.450173 | -0.46803 | -1.55629 | 1.000754 | -0.49629 | -0.02485 | -0.09246 | -0.00953 | 0.109086 | -9.6E-05 | 0.08678 | 0.005989 | 0.081169 | 0.113905 | -0.03047 | 0.142834 | 0.096636 |
| 31.61111 | -2.30043 | -1.65518 | 0.134636 | -0.14625 | -0.00775 | 0.081068 | -2.82173 | 0.577269 | -0.38487 | -1.68141 | 1.11404 | -0.05105 | -0.00499 | -0.09307 | 0.002382 | 0.109245 | 0.026277 | 0.093569 | -0.05671 | 0.044375 | 0.119462 | 0.007854 | 0.166833 | 0.003872 |
| 31.625 | -0.21182 | -1.68253 | 0.112427 | 0.141563 | 0.085832 | 0.173046 | -2.5622 | 0.859827 | -0.1694 | -1.90983 | 1.322855 | 0.161042 | 0.010752 | -0.0716 | 0.011852 | 0.047382 | 0.068847 | 0.066815 | -0.09874 | 0.017961 | 0.083574 | 0.037166 | 0.144614 | -0.08156 |
| 31.63889 | -0.50843 | -2.49068 | 0.00714 | 0.418379 | 0.119463 | 0.22628 | -1.98419 | 0.830362 | -0.38951 | -2.13057 | 1.018229 | 0.140551 | 0.022651 | -0.02008 | 0.023818 | -0.02157 | 0.10741 | 0.035524 | -0.07632 | 0.008893 | 0.057485 | 0.067296 | 0.122732 | -0.08515 |
| 31.65278 | -0.52138 | -2.853 | 0.000883 | 0.478477 | 0.032955 | 0.207418 | -1.45048 | 0.697496 | -0.68603 | -2.26047 | -0.28923 | 0.090533 | 0.016487 | 0.029561 | 0.032167 | -0.08917 | 0.105833 | 0.00557 | -0.03354 | -0.01684 | 0.008979 | 0.082165 | 0.081459 | -0.08262 |
| 31.66667 | -0.92199 | -3.06437 | -0.00507 | 0.304431 | -0.03416 | 0.213642 | -1.04104 | 0.512716 | -1.1396 | -2.5002 | -0.88288 | 0.118402 | -0.01533 | 0.051029 | -0.00999 | -0.19944 | 0.070497 | -0.0084 | 0.004484 | -0.06682 | -0.08277 | 0.053802 | 0.024165 | -0.13653 |
| 31.68056 | -0.21608 | -2.81606 | -0.09069 | 0.061527 | -0.03745 | 0.235253 | -0.75419 | 0.281011 | -1.11387 | -2.25982 | 0.357211 | -0.09984 | -0.03153 | 0.0494 | -0.05662 | -0.34113 | 0.018034 | -0.02685 | 0.036913 | -0.06266 | -0.18463 | 0.009837 | -0.03994 | -0.17549 |
| 31.69444 | 0.374451 | -2.92848 | -0.14228 | -0.16158 | -0.00383 | 0.226723 | -0.6826 | 0.152573 | -1.0845 | -1.91681 | -0.06593 | -0.53412 | -0.0232 | 0.051803 | -0.07157 | -0.24337 | -0.00811 | -0.02444 | 0.069944 | -0.01003 | -0.24621 | -0.00142 | -0.08048 | -0.23574 |
| 31.70833 | 0.314442 | -2.66058 | -0.15501 | -0.30071 | -0.02105 | 0.170447 | -0.82281 | -0.03555 | -0.75906 | -2.0501 | -0.02213 | -1.03027 | -0.00188 | 0.084852 | -0.06961 | -0.12884 | 0.007664 | 0.008041 | 0.069861 | -0.02204 | -0.38118 | -0.00059 | -0.07626 | -0.1858 |
| 31.72222 | 0.131392 | -1.97691 | -0.17557 | -0.19711 | -0.02149 | 0.01262 | -0.90079 | -0.1044 | -0.46936 | -2.43801 | -0.24707 | -1.50516 | -0.02162 | 0.105705 | -0.08822 | -0.13763 | 0.010387 | 0.024987 | 0.051237 | -0.09417 | -0.45067 | -0.02558 | -0.06627 | -0.18413 |
| 31.73611 | 0.171841 | -1.29613 | -0.47808 | 0.022046 | 0.053608 | -0.07475 | -1.10562 | -0.09055 | -0.10509 | -2.84905 | 0.079253 | -2.02197 | -0.04124 | 0.101318 | -0.12924 | -0.1777 | -0.03268 | 0.010542 | 0.006571 | -0.15447 | -0.3816 | -0.06828 | -0.17251 | -0.21852 |
| 31.75 | 0.276957 | -0.86377 | -0.81984 | 0.114832 | 0.074811 | -0.15886 | -1.43655 | -0.13387 | -0.16454 | -2.57248 | 0.522457 | -2.69261 | -0.02311 | 0.12912 | -0.12935 | -0.21105 | -0.04902 | 0.023115 | 0.017011 | -0.14447 | -0.16664 | -0.07479 | -0.20044 | -0.33816 |
| 31.76389 | 0.267474 | -0.78349 | -0.76743 | 0.033356 | 0.023954 | -0.30363 | -1.95423 | -0.26708 | -0.59565 | -2.22416 | 0.492999 | -2.46559 | -0.00342 | 0.171136 | -0.10166 | -0.207 | -0.04702 | 0.023964 | -0.02134 | -0.10892 | -0.16191 | -0.08375 | -0.18952 | -0.45879 |
| 31.77778 | 0.280729 | -0.79347 | -0.62572 | -0.12764 | 0.008062 | -0.45231 | -2.1893 | -0.52278 | -0.46859 | -1.70496 | 0.074997 | -2.0526 | -0.01261 | 0.187057 | -0.09063 | -0.19948 | -0.05902 | -0.0207 | -0.11273 | -0.13128 | -0.17121 | -0.10123 | -0.11118 | -0.51268 |
| 31.79167 | 0.275845 | -1.01121 | -0.62561 | -0.25335 | -0.00273 | -0.49053 | -1.96179 | -0.82947 | -0.08441 | -0.93879 | -0.01001 | -1.72959 | -0.0219 | 0.184563 | -0.06923 | -0.14838 | -0.04512 | -0.11149 | -0.18768 | -0.16013 | -0.17168 | -0.12256 | -0.06718 | -0.51175 |
| 31.80556 | 0.307851 | -1.00392 | -0.56716 | -0.29933 | -0.06669 | -0.43843 | -1.72557 | -1.05134 | 0.132226 | -0.41186 | 0.237741 | -1.26962 | -0.02949 | 0.172394 | -0.01201 | -0.10243 | -0.05944 | -0.23225 | -0.27561 | -0.22742 | -0.17011 | -0.17475 | -0.00197 | -0.50286 |
| 31.81944 | 0.261786 | -0.72205 | -0.47766 | -0.28673 | -0.11035 | -0.35271 | -1.33222 | -1.01624 | 0.214929 | -0.17436 | 0.26532 | -0.80603 | -0.02197 | 0.197696 | 0.05394 | -0.05129 | -0.06714 | -0.28842 | -0.28349 | -0.23909 | -0.15058 | -0.23186 | 0.028531 | -0.50101 |
| 31.83333 | 0.426995 | -0.5092 | -0.23643 | -0.19583 | -0.13569 | -0.15676 | -1.04665 | -0.791 | 0.250811 | -0.01865 | 0.436244 | -0.46948 | -0.00574 | 0.241184 | 0.105264 | 0.014698 | -0.04658 | -0.22923 | -0.18439 | -0.20742 | -0.06885 | -0.21743 | 0.063214 | -0.38335 |
| 31.84722 | 0.698969 | -0.16097 | 0.094283 | -0.07742 | -0.14614 | 0.083065 | -0.75015 | -0.50029 | 0.277912 | 0.119642 | 0.47584 | -0.12525 | 0.002825 | 0.228509 | 0.082145 | 0.01676 | -0.15962 | -0.17664 | -0.12612 | -0.16786 | -0.03364 | -0.24906 | 0.024345 | -0.1996 |
| 31.86111 | 0.936451 | 0.521743 | 0.367346 | 0.023666 | -0.16479 | 0.287839 | -0.54929 | -0.20504 | 0.356978 | 0.257128 | 0.633176 | 0.195828 | -0.04891 | 0.172654 | 0.018747 | -0.01333 | -0.20163 | -0.15166 | -0.11699 | -0.16701 | -0.06946 | -0.2784 | -0.00939 | 0.034554 |
| 31.875 | 1.113076 | 1.118336 | 0.605391 | 0.035568 | -0.167 | 0.431553 | -0.39648 | -0.01442 | 0.451025 | 0.186256 | 0.590164 | 0.469743 | -0.05693 | 0.156322 | 0.008783 | -0.00213 | -0.10541 | -0.13617 | -0.08552 | -0.16346 | -0.04468 | -0.2396 | -0.01874 | 0.225957 |
| 31.88889 | 1.170725 | 1.117122 | 0.580904 | -0.23166 | -0.15784 | 0.391852 | -0.49924 | -0.15776 | 0.315805 | -0.33092 | 0.102106 | 0.460978 | -0.0128 | 0.141929 | 0.030608 | 0.021618 | -0.06526 | -0.09647 | -0.00533 | -0.19847 | -0.0194 | -0.20811 | 0.025759 | 0.23475 |
| 31.90278 | 0.624911 | 0.540967 | 0.230262 | -1.07279 | -0.22148 | 0.073738 | -0.99219 | -0.6323 | -0.19853 | -1.49935 | -0.58892 | 0.086695 | -0.0446 | 0.041831 | 0.001095 | 0.030949 | -0.03932 | -0.0401 | 0.024506 | -0.23028 | -0.09528 | -0.194 | -0.02041 | -0.02521 |
| 31.91667 | -0.15405 | -0.5371 | -0.27098 | -1.58624 | -0.24237 | -0.29356 | -1.24483 | -0.98148 | -0.91941 | -2.96741 | -1.43878 | -0.65018 | -0.09429 | 0.012116 | 0.008576 | 0.102269 | -0.01969 | 0.05951 | 0.032883 | -0.2741 | -0.1761 | -0.1627 | -0.08448 | -0.32419 |
| 31.93056 | -0.50811 | -1.11611 | -0.58216 | -1.35473 | -0.14501 | -0.42118 | -1.32383 | -1.04466 | -1.32473 | -4.17767 | -1.89922 | -1.58837 | 0.097159 | 0.34057 | 0.229827 | 0.314148 | 0.178577 | 0.26073 | 0.102817 | -0.0253 | 0.068047 | 0.073466 | 0.079707 | -0.19827 |
| 31.94444 | -0.76356 | -1.34899 | -0.66304 | -1.10793 | -0.15517 | -0.47358 | -1.08767 | -1.0206 | -1.69776 | -2.64268 | -1.4626 | -1.75742 | 0.409547 | 0.69421 | 0.519042 | 0.509016 | 0.382643 | 0.384149 | 0.286314 | 0.340965 | 0.438902 | 0.287682 | 0.3241 | 0.121276 |
| 31.95833 | -1.01309 | -1.49276 | -0.58793 | -1.03685 | -0.35325 | -0.43128 | -0.96853 | -0.88018 | -1.41964 | 0.337057 | -1.15818 | -1.47821 | 0.345229 | 0.533474 | 0.443111 | 0.357948 | 0.262625 | 0.14138 | 0.39123 | 0.31978 | 0.375761 | 0.102156 | 0.242926 | 0.127191 |
| 31.97222 | -0.46847 | -1.58713 | -0.42255 | -0.57637 | -0.44817 | -0.2559 | -0.92716 | -0.7351 | -0.94871 | 1.243998 | -1.15016 | -0.77893 | 0.040253 | 0.147293 | 0.127767 | 0.044298 | 0.052094 | -0.19536 | 0.331357 | 0.021372 | 0.07862 | -0.23426 | -0.00244 | 0.083408 |
| 31.98611 | 0.308374 | -1.59922 | -0.15898 | 0.058183 | -0.33479 | -0.03158 | -0.49628 | -0.43739 | -0.48075 | 0.453046 | -1.6487 | -0.18444 | -0.11572 | 0.000109 | -0.01941 | -0.06669 | 0.008701 | -0.30921 | 0.238849 | -0.11987 | -0.06677 | -0.32942 | -0.08908 | 0.007219 |
| 32 | 0.877437 | -1.62019 | 0.13719 | 0.521168 | -0.18462 | 0.183944 | -0.15944 | -0.05921 | -0.17316 | -0.1778 | -1.60406 | 0.215215 | -0.07463 | -0.00625 | -0.01271 | -0.02687 | 0.051154 | -0.26896 | 0.123384 | -0.04984 | -0.02463 | -0.28907 | -0.03711 | -0.0372 |
| 32.01389 | 1.056469 | -1.33967 | 0.325441 | 0.627737 | -0.13537 | 0.347961 | 0.158865 | 0.297558 | 0.226023 | -0.35363 | -0.57328 | 0.367303 | -0.07472 | -0.01216 | -0.00899 | 0.007506 | 0.01196 | -0.22034 | -0.00912 | 0.043454 | 0.022287 | -0.2233 | 0.015908 | 0.02096 |
| 32.02778 | 1.102958 | -1.01068 | 0.324492 | 0.58278 | -0.13872 | 0.444251 | 0.243667 | 0.472737 | 0.393963 | -0.29816 | -0.16569 | 0.302006 | -0.06555 | -0.01561 | -0.02665 | 0.0142 | -0.02964 | -0.13074 | -0.11452 | 0.06111 | 0.007417 | -0.15448 | 0.009462 | 0.079965 |
| 32.04167 | 1.175863 | -0.82117 | 0.369633 | 0.462479 | -0.1445 | 0.423684 | 0.128412 | 0.497662 | 0.372713 | -0.22297 | 0.305693 | 0.274901 | -0.05085 | -0.0085 | -0.02632 | 0.01486 | -0.03809 | -0.07874 | -0.17565 | 0.020227 | 0.008825 | -0.0948 | -0.0342 | 0.084109 |
| 32.05556 | 1.427666 | -0.63482 | 0.604398 | 0.387887 | -0.11095 | 0.240544 | 0.015118 | 0.445839 | 0.31444 | -0.08304 | 0.305921 | 0.299251 | -0.02624 | 0.000972 | -0.00653 | 0.023793 | -0.01806 | -0.05392 | -0.19853 | -0.0196 | 0.011074 | -0.06267 | -0.06609 | 0.03489 |
| 32.06944 | 1.334359 | -0.29123 | 0.746997 | 0.352742 | -0.06939 | 0.017796 | 0.029872 | 0.322739 | 0.168046 | 0.152822 | 0.09584 | 0.386432 | 0.000539 | 0.002084 | 0.014008 | 0.042595 | -0.01077 | -0.04004 | -0.18011 | -0.0287 | -0.0032 | -0.03037 | -0.05246 | 0.057915 |
| 32.08333 | 1.423355 | 0.138582 | 0.709507 | 0.37111 | -0.01052 | -0.09175 | 0.206881 | 0.178628 | 0.075766 | 0.389084 | 0.230637 | 0.460858 | 0.022168 | 0.006366 | 0.02384 | 0.064826 | -0.00424 | -0.01813 | -0.16414 | -0.01133 | -0.00954 | 0.009852 | -0.00122 | 0.175612 |
| 32.09722 | 1.527539 | 0.606396 | 0.646538 | 0.431485 | 0.090256 | -0.13264 | 0.470598 | 0.070711 | 0.307375 | 0.499168 | 0.528333 | 0.702853 | 0.026451 | -0.0026 | 0.017241 | 0.06826 | -0.00274 | 0.008735 | -0.15278 | 0.006124 | 0.006976 | 0.044308 | 0.061224 | 0.269659 |
| 32.11111 | 1.707747 | 0.939583 | 0.638765 | 0.505662 | 0.193854 | -0.10431 | 0.683654 | 0.06252 | 0.38312 | 0.538348 | 0.728674 | 0.975118 | 0.003213 | -0.0295 | 0.007401 | 0.064572 | -0.00447 | 0.045372 | -0.15022 | 0.012596 | 0.02167 | 0.059312 | 0.128041 | 0.282242 |
| 32.125 | 1.249418 | 1.117297 | 0.569029 | 0.543998 | 0.255576 | -0.00317 | 0.755598 | 0.140208 | 0.511702 | 0.445881 | 0.731651 | 1.13948 | -0.01692 | -0.0411 | 0.008724 | 0.076731 | -0.01829 | 0.088496 | -0.14459 | 0.015344 | 0.018981 | 0.052424 | 0.158618 | 0.27356 |
| 32.13889 | 0.886478 | 1.184928 | 0.529911 | 0.509779 | 0.286307 | 0.186438 | 0.698634 | 0.276245 | 0.597582 | 0.386161 | 1.182275 | 1.028713 | -0.00735 | -0.0394 | 0.011081 | 0.110576 | -0.03052 | 0.120952 | -0.13108 | 0.026934 | 0.024711 | 0.042617 | 0.147502 | 0.223154 |
| 32.15278 | 0.695336 | 1.083211 | 0.676269 | 0.458066 | 0.258395 | 0.363483 | 0.741989 | 0.457885 | 0.620145 | 0.43338 | 1.365032 | 1.039737 | 0.005469 | -0.04599 | 0.011473 | 0.121361 | -0.02911 | 0.130099 | -0.11023 | 0.033186 | 0.04702 | 0.044158 | 0.129795 | 0.142712 |
| 32.16667 | 0.247755 | 1.001874 | 0.776853 | 0.393051 | 0.195868 | 0.396532 | 0.705587 | 0.534904 | 0.45445 | 0.516209 | 1.119843 | 0.986252 | -0.00051 | -0.05246 | 0.006782 | 0.11483 | -0.01045 | 0.093951 | -0.07517 | 0.024892 | 0.024129 | 0.035957 | 0.101934 | 0.081508 |
| 32.18056 | -0.13789 | 1.031428 | 0.722369 | 0.333589 | 0.142269 | 0.321518 | 0.689905 | 0.449633 | 0.249937 | 0.647341 | 0.634574 | 0.877257 | -0.00303 | -0.03888 | -0.00065 | 0.092059 | 0.023463 | 0.058369 | -0.06426 | 0.011368 | 0.000042 | 0.010069 | 0.068477 | 0.079637 |
| 32.19444 | -0.01644 | 0.89672 | 0.513868 | 0.298756 | 0.096915 | 0.271111 | 0.628486 | 0.400216 | 0.300195 | 0.747055 | 0.02509 | 0.788506 | -0.00676 | -0.021 | -0.00893 | 0.070265 | 0.033959 | 0.048908 | -0.08937 | 0.013956 | -0.0075 | -0.01962 | 0.038671 | 0.124677 |
| 32.20833 | 0.300637 | 0.826429 | 0.290575 | 0.266055 | 0.074261 | 0.343833 | 0.571942 | 0.221676 | 0.301675 | 0.7849 | -0.42506 | 0.640194 | 0.000105 | -0.00858 | 0.0053 | 0.061947 | 0.037407 | 0.055134 | -0.09679 | 0.035146 | 0.012268 | -0.0222 | 0.02175 | 0.174663 |
| 32.22222 | 0.442853 | 0.702803 | 0.159668 | 0.285833 | 0.064724 | 0.506261 | 0.646996 | -0.09991 | 0.307839 | 0.772865 | -0.47327 | 0.468221 | 0.029142 | 0.011312 | 0.051242 | 0.092243 | 0.053266 | 0.070624 | -0.07105 | 0.064577 | 0.053339 | 0.022443 | 0.038077 | 0.196684 |
| 32.23611 | 0.625718 | 0.515074 | 0.195227 | 0.295145 | 0.028472 | 0.728861 | 0.778641 | -0.44992 | 0.270651 | 0.705506 | 0.103801 | 0.253149 | 0.058163 | 0.015359 | 0.095172 | 0.117785 | 0.089262 | 0.084789 | -0.05248 | 0.099328 | 0.088167 | 0.080819 | 0.05044 | 0.189702 |
| 32.25 | 0.656664 | 0.345489 | 0.310777 | 0.314263 | 0.035014 | 0.938957 | 0.760551 | -0.50122 | 0.144297 | 0.588882 | 0.414379 | 0.074228 | 0.070686 | -0.00623 | 0.101975 | 0.125783 | 0.082748 | 0.080902 | -0.04699 | 0.131405 | 0.097622 | 0.110014 | 0.060614 | 0.16008 |
| 32.26389 | 0.609568 | 0.228016 | 0.364534 | 0.285437 | 0.048053 | 0.996509 | 0.536325 | -0.34956 | 0.108694 | 0.484268 | 0.509051 | 0.078077 | 0.058382 | -0.02987 | 0.071386 | 0.127518 | 0.068742 | 0.056909 | -0.04013 | 0.119297 | 0.081661 | 0.109066 | 0.051677 | 0.139547 |
| 32.27778 | 0.545739 | 0.109341 | 0.294177 | 0.170544 | 0.041735 | 0.909127 | 0.29771 | -0.27698 | 0.079698 | 0.35682 | 0.565113 | 0.155505 | 0.04222 | -0.02605 | 0.047447 | 0.127992 | 0.056583 | 0.014313 | -0.03075 | 0.034929 | 0.08919 | 0.124928 | 0.056929 | 0.164054 |
| 32.29167 | 0.331513 | -0.12607 | 0.210373 | 0.002604 | 0.002791 | 0.670897 | 0.066193 | -0.39072 | 0.118716 | 0.193866 | 0.475553 | 0.159007 | 0.033937 | -0.01376 | 0.049967 | 0.128612 | 0.05727 | -0.0243 | -0.02449 | -0.07768 | 0.12715 | 0.165078 | 0.085816 | 0.212324 |
| 32.30556 | 0.242214 | -0.18223 | 0.138151 | -0.06535 | -0.01442 | 0.475742 | -0.14319 | -0.50458 | 0.180089 | 0.03335 | 0.422376 | 0.082875 | 0.048678 | -0.03754 | 0.05272 | 0.118147 | 0.058164 | -0.01875 | -0.01863 | -0.09189 | 0.148022 | 0.200078 | 0.084831 | 0.238756 |
| 32.31944 | 0.315022 | -0.53231 | 0.294259 | -0.07343 | -0.0133 | 0.510763 | -0.12623 | -0.17907 | 0.363598 | -0.04064 | 0.778306 | 0.018621 | 0.065619 | -0.06485 | 0.048421 | 0.106084 | 0.060023 | 0.033116 | -0.04523 | 0.006812 | 0.127919 | 0.199065 | 0.057248 | 0.251617 |
| 32.33333 | 0.606681 | -0.79915 | 0.494743 | 0.019744 | 0.005376 | 0.557665 | 0.03892 | 0.415275 | 0.576896 | -0.00028 | 1.251292 | 0.048066 | 0.052413 | -0.11279 | 0.045493 | 0.104624 | 0.059997 | 0.041375 | -0.07453 | 0.056186 | 0.102072 | 0.17169 | 0.043162 | 0.21025 |
| 32.34722 | 0.857061 | -0.89761 | 0.644913 | 0.062363 | -0.05098 | 0.556014 | 0.235195 | 0.64731 | 0.845777 | 0.057718 | 1.63312 | 0.074828 | 0.024761 | -0.12348 | 0.052231 | 0.105732 | 0.048865 | 0.02566 | -0.11266 | 0.049724 | 0.100797 | 0.142456 | 0.051917 | 0.155889 |
| 32.36111 | 1.159222 | -0.8571 | 0.79671 | 0.012503 | -0.16387 | 0.42051 | 0.34894 | 0.603716 | 0.621005 | 0.071064 | 0.570554 | -0.00916 | -0.02241 | -0.06366 | 0.038257 | 0.11044 | 0.052088 | -0.01242 | -0.154 | 0.004428 | 0.054689 | 0.075226 | 0.024852 | 0.104792 |
| 32.375 | 1.261654 | -0.71149 | 0.89289 | 0.002726 | -0.20014 | 0.322045 | 0.443937 | 0.360509 | 0.30277 | 0.11189 | 0.447082 | -0.10602 | -0.10865 | 0.003105 | -0.03727 | 0.067514 | 0.037637 | -0.08181 | -0.18151 | -0.07674 | -0.04075 | -0.01593 | -0.03566 | 0.060897 |
| 32.38889 | 1.484447 | -0.36437 | 0.919267 | 0.188957 | -0.13243 | 0.409345 | 0.856885 | 0.258863 | 0.438422 | 0.473191 | 0.667697 | -0.07517 | -0.13737 | -0.03126 | -0.07429 | 0.008764 | 0.015047 | -0.10755 | -0.18935 | -0.10786 | -0.05817 | -0.02206 | -0.05722 | 0.064607 |
| 32.40278 | 1.607807 | 0.399307 | 1.015395 | 0.686224 | 0.06865 | 0.682104 | 1.545396 | 0.64436 | 1.160474 | 1.083029 | 0.747451 | 0.237365 | -0.0368 | -0.15293 | 0.011351 | 0.020114 | 0.024434 | -0.05927 | -0.18075 | -0.02606 | 0.077804 | 0.09943 | 0.055578 | 0.240033 |
| 32.41667 | 1.881773 | 1.205985 | 0.954691 | 1.231904 | 0.239923 | 0.784769 | 2.893702 | 1.504705 | 1.605916 | 1.138353 | 1.337004 | 0.930883 | 0.093644 | -0.28148 | 0.127745 | 0.049439 | 0.081576 | -0.02601 | -0.16464 | 0.090612 | 0.298076 | 0.218776 | 0.272419 | 0.534312 |
| 32.43056 | 2.26236 | 1.841709 | 0.918419 | 1.262786 | 0.093026 | 0.499214 | 3.514357 | 2.210706 | 1.113118 | 1.109856 | 2.969458 | 1.841208 | 0.083188 | -0.32487 | 0.138086 | 0.015041 | 0.073471 | -0.02647 | -0.1048 | 0.085367 | 0.284111 | 0.125326 | 0.306061 | 0.585681 |
| 32.44444 | 2.457976 | 2.204717 | 1.239252 | 1.123049 | -0.04769 | -0.1419 | 3.515461 | 2.820729 | 0.592963 | 1.421433 | 3.400468 | 2.805136 | -0.09578 | -0.33996 | -0.0328 | -0.1043 | -0.0703 | -0.12524 | -0.20987 | -0.10041 | 0.034233 | -0.11966 | 0.13372 | 0.28484 |
| 32.45833 | 2.208668 | 3.590088 | 1.26601 | 1.129403 | 0.107141 | 0.052951 | 3.940451 | 2.856214 | 0.49175 | 2.190078 | 2.310354 | 2.351241 | -0.13665 | -0.26014 | -0.14865 | -0.06088 | -0.09829 | -0.0708 | -0.28343 | -0.11579 | -0.08162 | -0.14688 | 0.075144 | 0.041565 |
| 32.47222 | 1.67682 | 4.966351 | 0.859779 | 0.908514 | 0.160007 | 0.203313 | 4.114769 | 2.419366 | 1.327146 | 3.38479 | 0.496007 | 1.967862 | 0.037878 | -0.05918 | -0.03197 | 0.145347 | 0.0697 | 0.157776 | -0.10514 | 0.098887 | 0.084565 | 0.036356 | 0.143104 | 0.068256 |
| 32.48611 | 1.569293 | 3.340864 | 0.475178 | 0.52785 | 0.089179 | 0.169407 | 4.177933 | 2.087917 | 1.756066 | 3.156547 | 0.105984 | 1.68298 | 0.130442 | 0.029444 | 0.065705 | 0.239442 | 0.154081 | 0.216836 | 0.019165 | 0.184949 | 0.116328 | 0.150231 | 0.145983 | 0.084004 |
| 32.5 | 0.556613 | 2.628826 | -0.12552 | 0.544252 | 0.174269 | 0.26879 | 3.480008 | 1.226536 | 1.906148 | 2.874205 | -1.26903 | 1.143075 | 0.105244 | -0.01779 | 0.068399 | 0.195602 | 0.140377 | 0.172618 | 0.032972 | 0.125957 | 0.03126 | 0.164997 | 0.08038 | 0.062036 |
| 32.51389 | 0.69095 | 2.717835 | -0.6324 | 0.757102 | 0.087191 | 0.313768 | 5.500778 | 0.901013 | 1.916796 | 2.776227 | -1.59331 | 0.826209 | 0.083366 | -0.05672 | 0.045706 | 0.141898 | 0.115701 | 0.125186 | 0.047476 | 0.076036 | -0.00873 | 0.146391 | 0.039221 | 0.046307 |
| 32.52778 | 2.567266 | 2.362728 | 0.126664 | 0.704308 | 0.027185 | 0.74297 | 5.335098 | 1.637398 | 1.375383 | 2.025644 | -0.47178 | 0.48984 | 0.08173 | -8.3E-05 | 0.032295 | 0.114173 | 0.095164 | 0.09806 | 0.056787 | 0.074294 | 0.012598 | 0.129596 | 0.037597 | 0.070589 |
| 32.54167 | 2.752712 | 2.040451 | 0.126438 | 0.802494 | 0.10546 | 0.742073 | 2.783523 | 2.157082 | 0.849658 | 0.853343 | -0.5172 | 0.18707 | 0.106054 | 0.11881 | 0.067074 | 0.122481 | 0.102088 | 0.109419 | 0.076633 | 0.116446 | 0.11975 | 0.129925 | 0.084966 | 0.145627 |
| 32.55556 | 2.381825 | 1.721678 | -0.00583 | 0.758547 | 0.13497 | 0.620278 | 1.298347 | 1.430594 | 0.362486 | -0.59748 | -0.47225 | -0.04195 | 0.100104 | 0.175915 | 0.082714 | 0.092595 | 0.087339 | 0.09212 | 0.06874 | 0.112227 | 0.211616 | 0.091789 | 0.143001 | 0.150889 |
| 32.56944 | 1.824133 | 1.400127 | -0.1214 | 0.472059 | 0.089159 | 0.499784 | -0.70152 | 0.703416 | 0.048221 | -1.88339 | -0.85248 | -0.26697 | 0.041252 | 0.107838 | 0.028794 | 0.007774 | 0.033696 | 0.006651 | 0.016428 | 0.05423 | 0.209188 | 0.002921 | 0.139666 | 0.080941 |
| 32.58333 | 1.156841 | 1.153821 | -0.11245 | 0.111955 | -0.0453 | 0.281645 | -3.26851 | 0.288218 | -0.10408 | -2.65447 | 1.296418 | -0.62908 | 0.022457 | 0.062579 | 0.011739 | 0.006173 | 0.047799 | -0.01946 | 0.043726 | 0.064411 | 0.198804 | -0.01713 | 0.165521 | 0.046007 |
| 32.59722 | 0.582749 | 1.037562 | -0.17448 | -0.1946 | -0.1634 | 0.098043 | -3.52789 | 0.218291 | -0.3497 | -2.43177 | 2.234107 | -1.00216 | 0.038172 | 0.071491 | 0.035613 | 0.077965 | 0.093009 | 0.004029 | 0.124416 | 0.117242 | 0.209399 | 0.004153 | 0.198882 | 0.042214 |
| 32.61111 | -0.56139 | 0.652384 | -0.18729 | -0.44397 | -0.25505 | 0.070793 | -3.60064 | 0.33161 | -0.60531 | -1.99495 | 1.98706 | -1.18263 | 0.00979 | 0.009356 | 0.022021 | 0.090139 | 0.062128 | -0.00993 | 0.107366 | 0.083188 | 0.109595 | -0.01826 | 0.171615 | 0.006768 |
| 32.625 | -1.41906 | 0.100323 | -0.20958 | -0.47425 | -0.27103 | 0.127288 | -2.81984 | 0.590953 | -0.61271 | -2.17169 | 1.885568 | -0.95777 | -0.02972 | -0.08777 | 0.002013 | 0.075485 | 0.012217 | -0.03224 | 0.058557 | 0.040276 | -0.00485 | -0.03726 | 0.128249 | -0.03588 |
| 32.63889 | -1.25256 | -0.19892 | -0.17451 | -0.25599 | -0.23485 | 0.141918 | -2.04306 | 0.894998 | -0.45158 | -1.77192 | 0.687548 | -0.64715 | -0.06658 | -0.15551 | -0.01754 | 0.066344 | -0.02354 | -0.03833 | 0.005844 | -0.0005 | -0.08539 | -0.0313 | 0.057884 | -0.08732 |
| 32.65278 | -1.27942 | -0.52745 | -0.16504 | -0.16373 | -0.23062 | 0.017535 | -1.94769 | 1.015595 | -0.10168 | -1.72949 | 0.484409 | -0.50526 | -0.10611 | -0.19892 | -0.06307 | 0.020723 | -0.07299 | -0.06502 | -0.09838 | -0.08314 | -0.16342 | -0.05457 | -0.06946 | -0.14852 |
| 32.66667 | -1.17395 | -1.0463 | -0.1665 | -0.3177 | -0.32415 | -0.11035 | -2.35868 | 0.944269 | 0.090492 | -1.68775 | 0.808777 | -0.63545 | -0.12464 | -0.18476 | -0.08861 | -0.02691 | -0.10348 | -0.06371 | -0.16638 | -0.12783 | -0.2203 | -0.07441 | -0.16143 | -0.18119 |
| 32.68056 | -1.12678 | -1.88523 | -0.25731 | -0.4643 | -0.4069 | -0.04797 | -2.77137 | 0.917879 | 0.137763 | -1.84225 | 0.349671 | -0.85597 | -0.09926 | -0.14164 | -0.07612 | -0.03854 | -0.09503 | -0.04923 | -0.153 | -0.11192 | -0.22742 | -0.05322 | -0.15797 | -0.17833 |

| | | | | | | | | | | | | | | | | | | | | | | | | |
|---|---|---|---|---|---|---|---|---|---|---|---|---|---|---|---|---|---|---|---|---|---|---|---|---|
| 32.69444 | -1.1686 | -2.12248 | -0.20878 | -0.57657 | -0.48731 | 0.12286 | -3.33304 | 0.782419 | 0.183919 | -1.9787 | -1.00424 | -0.98664 | -0.06122 | -0.11699 | -0.07254 | -0.08548 | -0.07014 | -0.03751 | -0.13839 | -0.08847 | -0.27066 | -0.03255 | -0.11302 | -0.15855 |
| 32.70833 | -1.04202 | -2.12121 | -0.23587 | -0.64271 | -0.65979 | 0.229382 | -3.50972 | 0.480528 | -0.06246 | -2.00337 | -1.34346 | -0.87522 | -0.02187 | -0.11384 | -0.06808 | -0.09679 | -0.03123 | -0.03904 | -0.15599 | -0.04202 | -0.30232 | -5E-06 | -0.02091 | -0.13431 |
| 32.72222 | -0.84096 | -1.99478 | -0.34296 | -0.76012 | -0.9382 | 0.138353 | -4.34305 | 0.352614 | 0.096166 | -1.66429 | -1.27078 | -0.64854 | -0.00018 | -0.07831 | -0.05869 | -0.0906 | -0.01052 | -0.03599 | -0.17249 | 0.010365 | -0.30785 | 0.029417 | 0.198357 | -0.13147 |
| 32.73611 | -1.06038 | -1.72727 | -0.53654 | -0.81742 | -1.02217 | 0.00734 | -5.25805 | 0.582362 | 0.151561 | -1.40141 | -1.26864 | -0.37999 | -0.00288 | -0.10012 | -0.07423 | -0.09225 | -0.02889 | -0.04665 | -0.17791 | 0.017159 | -0.36385 | 0.011265 | 0.717722 | -0.17093 |
| 32.75 | -1.03829 | -1.45381 | -0.98894 | -0.80032 | -0.70736 | -0.14601 | -5.07662 | 0.766275 | 0.053491 | -1.28724 | -1.29823 | -0.4055 | 0.005585 | -0.08751 | -0.05968 | -0.08533 | -0.02865 | -0.02316 | -0.14435 | 0.020736 | -0.29386 | 0.007056 | 0.618831 | -0.20404 |
| 32.76389 | -1.19678 | -1.92687 | -1.35235 | -0.79274 | -0.47312 | -0.28113 | -5.31071 | 0.663437 | -0.22704 | -1.37896 | -1.61742 | -0.67955 | 0.052652 | 0.012348 | 0.009065 | -0.04665 | 0.014015 | 0.0136 | -0.08228 | 0.029793 | -0.18812 | 0.057754 | -0.00902 | -0.2359 |
| 32.77778 | -0.88162 | -2.75914 | -1.61047 | -0.73791 | -0.334 | -0.40727 | -6.03712 | 0.411713 | -0.64488 | -1.6368 | -1.6289 | -0.99166 | 0.04987 | 0.010163 | -0.03041 | -0.00842 | 0.018433 | -0.01529 | -0.05122 | -0.01622 | -0.14885 | 0.073307 | -0.10897 | -0.26453 |
| 32.79167 | -0.36816 | -3.16036 | -1.7235 | -0.81154 | -0.30705 | -0.52962 | -6.33191 | 0.091512 | -0.71971 | -1.80108 | -1.37663 | -1.15124 | 0.010501 | -0.03246 | -0.11016 | -0.04932 | -0.02216 | -0.08824 | -0.05376 | -0.08126 | -0.20095 | 0.032253 | -0.17946 | -0.373 |
| 32.80556 | -0.18081 | -2.60206 | -1.362 | -0.74176 | -0.27114 | -0.51652 | -6.81849 | -0.04924 | -0.31687 | -1.32354 | -1.10167 | -1.15421 | -0.01023 | -0.01686 | -0.14503 | -0.08843 | -0.05322 | -0.12461 | -0.04441 | -0.09028 | -0.24849 | -0.00317 | -0.24126 | -0.44233 |
| 32.81944 | 0.134303 | -1.90572 | -1.04345 | -0.56831 | -0.1777 | -0.45985 | -6.10298 | -0.16226 | 0.081557 | -0.81489 | -0.27318 | -1.06525 | -0.01311 | 0.101824 | -0.1513 | -0.11788 | -0.04966 | -0.10834 | -0.03221 | -0.04593 | -0.2718 | -0.01293 | -0.73768 | -0.3687 |
| 32.83333 | 0.214047 | -1.481 | -0.37569 | -0.4476 | -0.03379 | -0.36542 | -3.54655 | -0.27566 | 0.132156 | -0.34359 | 0.375687 | -0.8294 | 0.007635 | 0.29948 | -0.12812 | -0.07503 | 0.019835 | -0.05201 | 0.018765 | 0.021574 | -0.23771 | -0.02544 | -1.04588 | -0.24912 |
| 32.84722 | 0.481768 | -1.09734 | 0.096364 | -0.39816 | 0.112788 | -0.22356 | -2.10463 | -0.22909 | 0.021379 | -0.1318 | 1.361179 | -0.55984 | 0.024129 | 0.39155 | -0.06584 | 0.030998 | 0.07022 | -0.03089 | 0.00779 | 0.068731 | -0.22037 | -0.03211 | -0.51944 | -0.13684 |
| 32.86111 | 0.656329 | -0.49912 | 0.205172 | -0.25259 | 0.144241 | -0.09633 | -1.4852 | 0.007984 | 0.174601 | -0.00569 | 1.56606 | -0.24049 | 0.016689 | 0.346162 | -0.05736 | 0.03957 | 0.068857 | -0.01337 | -0.05958 | 0.086446 | -0.21432 | -0.00997 | 0.002345 | -0.07205 |
| 32.875 | 0.788544 | 0.098795 | 0.063354 | 0.057884 | 0.038223 | -0.13519 | -0.53134 | 0.273569 | 0.495204 | 0.088208 | 1.900913 | -0.03849 | 0.015486 | 0.313382 | -0.0449 | 0.067437 | 0.052037 | 0.016893 | -0.07176 | 0.088075 | -0.15825 | 0.019278 | 0.443574 | 0.002664 |
| 32.88889 | 0.894111 | 0.180454 | -0.38579 | 0.070242 | -0.22982 | -0.20514 | -0.07778 | 0.218956 | 0.476877 | -0.09049 | 1.37774 | -0.08375 | 0.036138 | -0.01279 | -0.01134 | 0.050631 | -0.00955 | 0.023862 | -0.11443 | 0.06889 | -0.04549 | 0.056231 | 0.90319 | 0.022432 |
| 32.90278 | 0.169699 | -0.24296 | -0.94304 | -0.54275 | -0.59938 | -0.40636 | 0.015209 | -0.58154 | 0.016033 | -0.95946 | -0.07115 | -0.34213 | -0.02516 | -0.59863 | -0.01169 | -0.03057 | -0.11492 | 0.011459 | -0.16002 | -0.02295 | -0.04278 | 0.017786 | 0.883768 | -0.12338 |
| 32.91667 | -0.45205 | -0.70717 | -1.55716 | -1.02685 | -0.6309 | -0.24252 | -0.45323 | -1.60262 | -0.80461 | -1.97194 | -1.62461 | -0.60179 | -0.20198 | -0.71561 | -0.0249 | 0.002603 | -0.14177 | -0.02538 | -0.18155 | -0.23168 | -0.2076 | -0.05859 | 0.160546 | -0.29952 |
| 32.93056 | -0.31816 | -0.95435 | -1.81943 | -0.79726 | -0.71564 | -0.00045 | 1.142604 | -2.4797 | -1.8204 | -2.16893 | -1.07806 | -0.96084 | -0.05538 | -0.097 | 0.145026 | 0.236917 | 0.07798 | 0.035846 | -0.25685 | -0.18848 | -0.16899 | 0.128007 | -0.00851 | -0.13385 |
| 32.94444 | -0.54277 | -1.13364 | -2.16772 | -0.39573 | -0.77294 | -0.11472 | 0.630631 | -2.67811 | -2.10128 | -1.26508 | -10.4985 | -1.35607 | 0.447092 | 0.584018 | 0.455049 | 0.54088 | 0.418721 | 0.25059 | -0.2155 | 0.278157 | 0.260326 | 0.429822 | 0.28871 | 0.306279 |
| 32.95833 | -0.87304 | -1.5588 | -2.33325 | -0.05726 | -0.67675 | -0.05245 | 0.106556 | -1.97147 | -1.94745 | -0.23296 | -5.90276 | -1.09635 | 0.592414 | 0.677513 | 0.439355 | 0.490009 | 0.432704 | 0.281646 | 0.027307 | 0.522624 | 0.430062 | 0.370295 | 0.303271 | 0.442887 |
| 32.97222 | -0.96922 | -1.92132 | -1.57253 | 0.278104 | -0.2092 | 0.154799 | 0.152304 | -0.80814 | -1.10068 | 0.198642 | -2.70151 | -0.46818 | 0.24734 | 0.332252 | 0.133783 | 0.153114 | 0.144799 | 0.014044 | 0.055986 | 0.23437 | 0.168024 | 0.063533 | -0.16969 | 0.281914 |
| 32.98611 | -0.8727 | -2.13661 | -0.59621 | 0.405548 | 0.442026 | 0.459882 | 0.301054 | 0.246156 | -0.30484 | 0.140969 | -1.13657 | 0.057758 | 0.012611 | 0.157031 | -0.02563 | -0.0412 | -0.01128 | -0.15387 | 0.057494 | -0.01174 | 0.004357 | -0.06443 | -0.83844 | 0.128046 |
| 33 | -0.55051 | -1.88855 | 0.405302 | 0.369634 | 0.688583 | 0.543037 | 0.270051 | 0.762679 | 0.12637 | -0.01617 | -0.2843 | 0.48301 | -0.03865 | 0.112093 | -0.03214 | -0.02205 | -0.02502 | -0.14285 | 0.040658 | -0.07237 | -0.01293 | -0.04703 | -0.95077 | 0.080458 |
| 33.01389 | -0.19423 | -1.66626 | 0.941203 | 0.285135 | 0.378249 | 0.423254 | 0.244479 | 0.8903 | 0.287224 | -0.20133 | 0.370554 | 0.780333 | -0.08543 | 0.080836 | -0.05622 | -0.01381 | -0.06316 | -0.12213 | -0.07267 | -0.14826 | -0.05057 | -0.05131 | -0.65806 | 0.035152 |
| 33.02778 | 0.003954 | -1.42992 | 0.758923 | 0.244378 | -0.10263 | 0.352766 | 0.157185 | 0.822208 | 0.1754 | -0.35552 | 0.498162 | 1.064095 | -0.10345 | 0.05256 | -0.05657 | -0.01482 | -0.07389 | -0.11069 | -0.17212 | -0.20716 | -0.06126 | -0.0621 | -0.32261 | -0.0175 |
| 33.04167 | -0.15341 | -1.37498 | 0.153803 | 0.257778 | -0.35514 | 0.318703 | 0.088441 | 0.541742 | 0.045294 | -0.43036 | 0.422604 | 1.225452 | -0.07661 | 0.038064 | -0.02213 | 0.015635 | -0.03859 | -0.07672 | -0.22271 | -0.19056 | -0.03197 | -0.06381 | -0.17082 | -0.05869 |
| 33.05556 | -0.15297 | -1.28108 | 0.015632 | 0.329824 | -0.32679 | 0.36219 | 0.03973 | 0.252183 | 0.036664 | -0.38719 | 0.588461 | 0.754408 | -0.06566 | 0.007448 | 0.001806 | 0.02499 | -0.03649 | -0.10768 | -0.27703 | -0.15394 | -0.0239 | -0.08866 | -0.09817 | -0.06515 |
| 33.06944 | -0.48489 | -1.1258 | 0.379951 | 0.399604 | -0.19555 | 0.510581 | 0.021393 | -0.05528 | 0.050367 | -0.26779 | 0.694816 | 0.443704 | -0.05232 | 0.015318 | 0.028694 | 0.03468 | -0.03232 | -0.12229 | -0.285 | -0.11198 | -0.0232 | -0.08228 | -0.04766 | -0.02505 |
| 33.08333 | -0.40513 | -0.88867 | 0.774597 | 0.447843 | -0.08839 | 0.665326 | 0.044678 | -0.37072 | 0.088392 | -0.16911 | 1.31689 | 0.335371 | -0.00855 | 0.057598 | 0.059397 | 0.073017 | 0.00502 | -0.08931 | -0.22007 | -0.07144 | -0.00559 | -0.0275 | 0.022027 | 0.038922 |
| 33.09722 | -0.15506 | -0.7562 | 0.906374 | 0.465367 | 0.09627 | 0.732545 | 0.126986 | -0.44238 | 0.205856 | -0.07937 | 1.33117 | 0.297709 | 0.038905 | 0.100971 | 0.117317 | 0.121117 | 0.020096 | -0.02971 | -0.14117 | -0.04441 | 0.030919 | 0.019344 | 0.090078 | 0.101863 |
| 33.11111 | -0.00103 | -0.56428 | 0.96857 | 0.446207 | 0.338273 | 0.638496 | 0.212828 | -0.23518 | 0.275483 | 0.021118 | 0.486754 | 0.269013 | 0.026306 | 0.093586 | 0.127323 | 0.119123 | -0.02538 | -0.02894 | -0.16469 | -0.0724 | 0.027611 | 0.006521 | 0.080938 | 0.10281 |
| 33.125 | 0.317351 | -0.41512 | 0.982119 | 0.40794 | 0.51231 | 0.361552 | 0.275992 | 0.177176 | 0.321792 | 0.070032 | 0.421734 | 0.255741 | -0.03663 | 0.065142 | 0.011937 | 0.078807 | -0.07932 | -0.07965 | -0.25124 | -0.13242 | -0.01578 | -0.03745 | 0.038059 | 0.085471 |
| 33.13889 | 0.675948 | -0.28861 | 0.859493 | 0.330032 | 0.44922 | -0.06052 | 0.169492 | 0.507066 | 0.224184 | 0.029374 | 0.293689 | 0.232935 | -0.02589 | 0.07545 | -0.01178 | 0.078686 | -0.05465 | -0.06771 | -0.25094 | -0.11123 | -0.01138 | -0.03704 | 0.063329 | 0.11421 |
| 33.15278 | 0.699934 | -0.25132 | 0.621281 | 0.256801 | 0.166854 | -0.36679 | -0.08378 | 0.750091 | 0.138439 | -0.00608 | 0.178618 | 0.234698 | 0.057057 | 0.120077 | 0.054828 | 0.130163 | -0.00024 | -0.00593 | -0.16067 | -0.02284 | 0.044755 | -0.00126 | 0.122729 | 0.136828 |
| 33.16667 | 0.618219 | -0.24384 | 0.113417 | 0.21973 | -0.20544 | -0.17825 | -0.24323 | 0.854114 | 0.136897 | 0.013523 | 0.206805 | 0.263295 | 0.057047 | 0.111937 | 0.065537 | 0.117715 | -0.01232 | -0.01283 | -0.13019 | -0.01942 | 0.047646 | -0.00012 | 0.104257 | 0.094672 |
| 33.18056 | 0.541099 | -0.31251 | -0.57499 | 0.195148 | -0.4193 | 0.091025 | -0.03361 | 0.769094 | 0.253148 | -0.00942 | 0.134722 | 0.295962 | -0.03462 | 0.045086 | -0.01498 | 0.046237 | -0.07182 | -0.08419 | -0.17621 | -0.10555 | -0.00614 | -0.0507 | 0.036501 | 0.005751 |
| 33.19444 | 0.56736 | -0.36915 | -1.04081 | 0.170756 | -0.3437 | 0.253918 | 0.233207 | 0.680706 | 0.382106 | -0.15152 | 0.028818 | 0.321865 | -0.07448 | 0.016394 | -0.04531 | 0.007764 | -0.07038 | -0.10354 | -0.16226 | -0.14271 | 0.004883 | -0.06459 | 0.018451 | -0.04875 |
| 33.20833 | 0.49034 | -0.37581 | -0.75408 | 0.161988 | -0.10797 | 0.27093 | 0.359809 | 0.638568 | 0.336136 | -0.24513 | -0.04022 | 0.269401 | -0.0384 | 0.02825 | -0.00312 | 0.041699 | -0.00542 | -0.06652 | -0.13699 | -0.12533 | 0.069984 | -0.0231 | 0.036116 | -0.01177 |
| 33.22222 | 0.4142 | -0.32855 | -0.00151 | 0.141359 | 0.074424 | 0.246736 | 0.303356 | 0.537212 | 0.335118 | -0.20862 | -0.0468 | 0.178527 | -0.01969 | 0.016712 | 0.030682 | 0.064417 | 0.023625 | -0.06669 | -0.20885 | -0.16411 | 0.109364 | 0.005322 | 0.041202 | 0.038345 |
| 33.23611 | 0.376484 | -0.32273 | 0.591136 | 0.117097 | 0.07373 | 0.267104 | 0.207817 | 0.505716 | 0.324091 | -0.11393 | -0.0088 | 0.040059 | -0.03533 | -0.02802 | 0.023429 | 0.049042 | -0.0078 | -0.11192 | -0.31593 | -0.23691 | 0.104991 | 0.007237 | 0.007194 | 0.051251 |
| 33.25 | 0.277893 | -0.40265 | 0.66449 | 0.07209 | -0.08705 | 0.290786 | 0.222228 | 0.515591 | 0.16344 | -0.11566 | 0.033457 | -0.07728 | -0.03129 | -0.0423 | 0.026289 | 0.050313 | -0.0224 | -0.1049 | -0.3485 | -0.24777 | 0.084271 | 0.008923 | -0.00056 | 0.049842 |
| 33.26389 | 0.130236 | -0.46211 | 0.10991 | -0.02218 | -0.30964 | 0.272257 | 0.216142 | 0.486513 | -0.03235 | -0.16388 | -0.00214 | -0.16685 | 0.027693 | 0.012613 | 0.073748 | 0.091382 | 0.029916 | -0.036 | -0.29035 | -0.15837 | 0.104434 | 0.029176 | 0.037384 | 0.065102 |
| 33.27778 | 0.099606 | -0.40301 | -0.31222 | -0.05039 | -0.40651 | 0.28176 | 0.241044 | 0.471103 | -0.09579 | -0.14259 | 0.041517 | -0.16237 | 0.048141 | 0.055788 | 0.068285 | 0.07524 | 0.036749 | -0.01421 | -0.2691 | -0.11566 | 0.074995 | 0.010664 | -0.00769 | 0.033103 |
| 33.29167 | 0.221349 | -0.35163 | -0.0704 | 0.043909 | -0.33829 | 0.41964 | 0.500276 | 0.556672 | -0.00902 | 0.051501 | 0.1088 | -0.0605 | -0.00892 | 0.041471 | 0.00241 | 0.002877 | -0.03246 | -0.06638 | -0.28959 | -0.16482 | 0.008397 | -0.03801 | -0.08535 | -0.03921 |
| 33.30556 | 0.305127 | -0.368 | 0.41318 | 0.164072 | -0.15502 | 0.575424 | 0.690967 | 0.658294 | 0.100563 | 0.238382 | 0.160115 | 0.06797 | -0.01828 | 0.060163 | 0.000355 | -0.0074 | -0.05077 | -0.07287 | -0.20615 | -0.15672 | 0.022778 | -0.01305 | -0.05231 | -0.01969 |
| 33.31944 | 0.282903 | -0.39899 | 0.782257 | 0.254089 | 0.195567 | 0.639115 | 0.767576 | 0.828005 | 0.260456 | 0.313489 | 0.26455 | 0.194774 | 0.014711 | 0.090064 | 0.027627 | 0.03176 | -0.02769 | -0.04255 | -0.11711 | -0.14576 | 0.048266 | 0.028345 | 0.011482 | 0.019161 |
| 33.33333 | 0.189869 | -0.38084 | 0.914593 | 0.296654 | 0.527097 | 0.63584 | 0.596501 | 1.0044 | 0.353703 | 0.287686 | 0.476821 | 0.266914 | -0.0042 | 0.075825 | 0.015572 | 0.040703 | -0.01573 | -0.04154 | -0.09246 | -0.1487 | 0.037008 | 0.03219 | 0.034121 | 0.028042 |
| 33.34722 | 0.16143 | -0.48345 | 0.852603 | 0.301721 | 0.548529 | 0.539746 | 0.360295 | 1.065457 | 0.303307 | 0.178589 | 0.657746 | 0.21497 | -0.03001 | 0.068631 | 0.002553 | 0.04281 | -0.00605 | -0.04534 | -0.10119 | -0.11967 | 0.045194 | 0.038562 | 0.047042 | 0.068118 |
| 33.36111 | 0.059962 | -0.69396 | 0.68612 | 0.240686 | 0.204582 | 0.424395 | 0.212507 | 1.002037 | 0.138348 | -0.01055 | 0.770594 | 0.049736 | -0.09249 | 0.134204 | -0.03346 | 0.012171 | -0.03128 | -0.09325 | -0.16107 | -0.15434 | 0.001561 | -0.00595 | -0.00152 | 0.067229 |
| 33.375 | 0.047204 | -0.83727 | 0.595058 | 0.184412 | -0.2193 | 0.378917 | 0.186828 | 0.900203 | 0.065959 | -0.07199 | 0.913116 | -0.10981 | -0.191 | 0.227701 | -0.10448 | -0.05932 | -0.06763 | -0.18778 | -0.27191 | -0.23823 | -0.0988 | -0.10638 | -0.07216 | 0.017395 |
| 33.38889 | 0.234069 | -0.62028 | 0.523782 | 0.221602 | -0.41805 | 0.475869 | 0.384888 | 0.795319 | 0.202319 | 0.24018 | 1.422687 | -0.01823 | -0.18937 | 0.165856 | -0.104 | -0.06156 | -0.0602 | -0.17738 | -0.34391 | -0.20663 | -0.06367 | -0.09408 | -0.05909 | 0.00879 |
| 33.40278 | 0.905066 | 0.242812 | 0.531758 | 0.445201 | -0.11845 | 0.646254 | 0.592721 | 0.600428 | 0.482692 | 0.717034 | 2.125716 | 0.383991 | -0.06545 | -0.04279 | 0.001917 | 0.005925 | -0.00086 | -0.0439 | -0.32402 | -0.00558 | 0.164087 | 0.06676 | 0.091129 | 0.107345 |
| 33.41667 | 1.512987 | 1.263452 | 0.782124 | 0.56191 | 0.381009 | 0.684191 | 0.690724 | 0.518278 | 0.589162 | 0.742387 | 2.355566 | 0.824319 | 0.03263 | -0.20057 | 0.076956 | 0.024519 | 0.049614 | 0.018155 | -0.21357 | 0.191663 | 0.351264 | 0.150078 | 0.235602 | 0.222465 |
| 33.43056 | 1.835534 | 1.526294 | 1.383981 | 0.628279 | 0.261398 | 0.484563 | 0.546439 | 0.91205 | 0.600859 | 0.649148 | 2.775666 | 1.307358 | -0.03557 | -0.27561 | -0.02205 | -0.09436 | -0.03514 | -0.11845 | -0.10974 | 0.101204 | 0.205666 | -0.03614 | 0.169301 | 0.176183 |
| 33.44444 | 1.800832 | 1.076664 | 2.252635 | 0.821124 | 0.12423 | 0.523608 | 0.379992 | 1.470293 | 0.828265 | 0.991874 | 3.00204 | 1.917607 | -0.23718 | -0.31371 | -0.23741 | -0.22846 | -0.18023 | -0.39567 | -0.10385 | -0.34496 | -0.11935 | -0.28445 | 0.002422 | -0.00563 |
| 33.45833 | 1.89801 | 0.938491 | 2.748839 | 1.23824 | 0.342326 | 0.616858 | 0.17674 | 1.78184 | 0.722303 | 1.272206 | 4.224037 | 2.275812 | -0.19844 | -0.14807 | -0.21482 | -0.13499 | -0.10982 | -0.66171 | 0.024772 | -0.66847 | -0.1466 | -0.21543 | 0.102398 | 0.004848 |
| 33.47222 | 1.473471 | 1.400333 | 2.463714 | 1.277141 | 0.175884 | 0.656691 | -0.1341 | 1.844711 | 0.391529 | 0.812658 | 7.492945 | 2.243568 | 0.039831 | 0.100421 | 0.025446 | 0.135735 | 0.123245 | -0.33866 | 0.281471 | -0.41577 | 0.115485 | 0.052132 | 0.329596 | 0.173961 |
| 33.48611 | 0.837838 | 2.2237 | 2.310175 | 0.843501 | -0.34612 | 0.624017 | 0.291215 | 1.478854 | 0.287314 | 0.399507 | 8.492417 | 2.119178 | 0.12389 | 0.145924 | 0.12727 | 0.189491 | 0.181345 | 0.070891 | 0.194914 | -0.07827 | 0.171264 | 0.144738 | 0.293758 | 0.186319 |
| 33.5 | 0.471633 | 2.965882 | 2.128931 | 0.498096 | -0.39151 | 0.575377 | 1.265179 | 0.717342 | 0.382103 | 0.360177 | 6.03851 | 2.001596 | 0.080292 | 0.070111 | 0.105785 | 0.15106 | 0.122418 | 0.105145 | 0.019499 | 0.015524 | 0.045754 | 0.107434 | 0.131247 | 0.132483 |
| 33.51389 | 0.406787 | 3.121435 | 1.814756 | 0.194439 | -0.24102 | 0.637999 | 1.770123 | 0.172314 | 0.38449 | 0.304242 | 2.66192 | 1.594645 | 0.03531 | 0.00593 | 0.075475 | 0.130521 | 0.073343 | 0.055912 | -0.06424 | -0.01241 | -0.05274 | 0.066207 | 0.046795 | 0.081813 |
| 33.52778 | 0.567482 | 3.569086 | 1.418744 | 0.259138 | -0.37909 | 0.749773 | 1.744256 | 0.087661 | 0.54475 | 0.182139 | 0.223245 | 0.988208 | -0.05868 | -0.10075 | -0.01521 | 0.041047 | -0.01456 | -0.04741 | -0.1653 | -0.12505 | -0.17971 | -0.02217 | -0.04531 | -0.06818 |
| 33.54167 | 0.678815 | 3.844658 | 1.082276 | 0.338566 | -0.01617 | 0.712151 | 1.380507 | 0.169394 | 0.723895 | 0.173818 | -1.60617 | 0.25882 | -0.14488 | -0.205 | -0.10446 | -0.04272 | -0.10182 | -0.13819 | -0.22328 | -0.22934 | -0.2686 | -0.10152 | -0.07507 | -0.19195 |
| 33.55556 | 0.790675 | 3.686841 | 0.866225 | 0.431372 | 0.160147 | 0.636645 | 1.429888 | -0.06945 | 0.783078 | 0.258893 | -7.38107 | -0.29481 | -0.13582 | -0.19979 | -0.09554 | -0.05132 | -0.10715 | -0.1332 | -0.19381 | -0.20374 | -0.21251 | -0.09531 | 0.035963 | -0.1625 |
| 33.56944 | 0.928488 | 3.420133 | 0.70462 | 0.297807 | -0.07054 | 0.562077 | 1.598287 | -0.32135 | 0.708753 | 0.389437 | -6.8045 | -0.7908 | -0.06206 | -0.09717 | -0.02816 | 0.012287 | -0.04902 | -0.08207 | -0.11587 | -0.11513 | -0.10142 | -0.05061 | 0.142439 | -0.05399 |
| 33.58333 | 1.021964 | 3.186421 | 0.417911 | 0.045314 | -0.37513 | 0.459651 | 1.821938 | -0.46628 | 0.272089 | 0.230189 | -3.56546 | -1.30219 | 0.008475 | 0.022693 | 0.030305 | 0.087148 | 0.018129 | -0.03281 | -0.01454 | -0.0062 | 0.007489 | 0.00932 | 0.199342 | 0.062895 |
| 33.59722 | 0.978471 | 3.054436 | 0.241317 | -0.11105 | -0.61613 | 0.431745 | 2.029358 | -0.27158 | -0.34996 | -0.10635 | -1.81495 | -1.50136 | 0.03141 | 0.075178 | 0.053566 | 0.148405 | 0.053756 | 0.001244 | 0.058172 | 0.077075 | 0.072934 | 0.040108 | 0.216545 | 0.113426 |
| 33.61111 | 0.751592 | 2.381209 | 0.307057 | -0.15516 | -0.87165 | 0.45832 | 1.944587 | 0.043317 | -1.09528 | -0.26134 | -1.47736 | -1.40113 | 0.012764 | 0.055363 | 0.040227 | 0.16315 | 0.045365 | 0.013131 | 0.001788 | 0.089708 | 0.073663 | 0.016598 | 0.220745 | 0.074851 |
| 33.625 | 0.258707 | 1.038438 | 0.224858 | -0.15564 | -0.96464 | 0.405713 | 1.505635 | 0.169322 | -0.74984 | -0.32626 | -0.01082 | -1.34036 | -0.0115 | 0.018291 | 0.015081 | 0.093314 | 0.02278 | 0.007929 | -0.05993 | 0.060746 | 0.042132 | -0.00927 | 0.176835 | 0.001161 |
| 33.63889 | -0.05808 | -0.07667 | 0.017149 | -0.08089 | -1.00941 | 0.376115 | 0.955566 | 0.204413 | -0.80401 | -0.38749 | 1.283199 | -1.17079 | -0.02159 | -0.01589 | -0.00426 | 0.045148 | 0.011643 | -0.01609 | -0.11359 | 0.022083 | 0.024812 | -0.01429 | 0.113375 | -0.03282 |
| 33.65278 | -0.40567 | -0.82804 | -0.2343 | -0.03549 | -1.0034 | 0.338253 | 0.759154 | 0.266726 | -0.54178 | -0.47079 | 2.872747 | -0.94582 | -0.0288 | -0.04628 | -0.01258 | 0.013416 | 0.013173 | -0.05323 | -0.14305 | -0.01046 | 0.013013 | -0.00762 | 0.07813 | -0.03218 |
| 33.66667 | -0.79854 | -1.2932 | -0.51858 | -0.09536 | -1.20048 | 0.3045 | 0.62026 | 0.326067 | -0.55834 | -0.49514 | 3.800821 | -1.03039 | -0.04085 | -0.03898 | -0.01433 | -0.00663 | 0.008494 | -0.08793 | -0.17144 | -0.03048 | -0.03121 | -0.00493 | 0.119689 | -0.05611 |
| 33.68056 | -0.8882 | -1.03649 | -0.8275 | -0.1754 | -1.37613 | 0.229033 | 0.39976 | 0.432375 | -0.6687 | -0.56364 | 5.201402 | -1.39075 | -0.04406 | -0.00715 | -0.01211 | -0.01798 | 0.002988 | -0.11361 | -0.17792 | -0.05329 | -0.08603 | -0.00258 | 0.217122 | -0.07974 |
| 33.69444 | -0.90614 | -0.70928 | -1.47878 | -0.32648 | -1.5799 | 0.196864 | 0.120744 | 0.375698 | -0.46451 | -0.60349 | 4.548028 | -1.69842 | -0.02815 | -0.02041 | -0.00954 | -0.01213 | 0.031448 | -0.11439 | -0.09994 | -0.05169 | -0.08001 | 0.013489 | 0.288017 | -0.05679 |
| 33.70833 | -1.3211 | -0.8938 | -2.16283 | -0.39768 | -1.98632 | 0.125451 | -0.21202 | 0.26478 | -0.495 | -0.66999 | 10.95313 | -1.82762 | -0.01942 | -0.0578 | 0.005524 | -0.01225 | 0.057239 | -0.10024 | 0.008146 | -0.04626 | -0.05335 | 0.015823 | 0.341705 | -0.06554 |
| 33.72222 | -2.44718 | -0.98384 | -2.80832 | -0.34019 | -2.40762 | -0.00856 | -0.47231 | 0.11835 | -0.50023 | -0.71003 | 7.923407 | -1.77535 | -0.06249 | -0.09455 | -0.05528 | -0.03302 | 0.009728 | -0.13236 | 0.012913 | -0.10705 | -0.12937 | -0.05304 | 0.327122 | -0.14807 |
| 33.73611 | -3.30184 | -1.13622 | -3.04823 | -0.29105 | -2.63059 | -0.24629 | -0.73051 | 0.176794 | -0.3113 | -0.66602 | 4.394439 | -1.5948 | -0.09278 | -0.13455 | -0.1169 | -0.03754 | -0.045 | -0.17617 | -0.05002 | -0.16139 | -0.20782 | -0.14739 | 0.260641 | -0.19907 |
| 33.75 | -3.17826 | -1.23489 | -3.06158 | -0.29193 | -2.14763 | -0.32873 | -1.02194 | 0.332412 | -0.02097 | -0.59691 | 1.325046 | -1.23733 | -0.06545 | -0.11847 | -0.09382 | 0.009195 | -0.02562 | -0.16288 | -0.10382 | -0.1258 | -0.22304 | -0.15977 | 0.168664 | -0.2012 |
| 33.76389 | -2.8375 | -1.61821 | -3.04415 | -0.20603 | -1.5042 | -0.33209 | -1.21707 | 0.502403 | 0.370172 | -0.54421 | -2.23672 | -0.93705 | -0.0161 | -0.08314 | -0.06079 | 0.04559 | 0.022012 | -0.11727 | -0.12705 | -0.06512 | -0.23478 | -0.14912 | 0.133801 | -0.18886 |
| 33.77778 | -1.94398 | -1.91932 | -2.79778 | -0.14372 | -1.57352 | -0.33914 | -1.24798 | 0.581978 | 0.660604 | -0.50768 | -7.11458 | -0.58762 | 0.012369 | 0.052501 | -0.05275 | 0.059393 | 0.075172 | -0.06047 | -0.13609 | -0.01912 | -0.27003 | -0.12208 | 0.120723 | -0.16923 |
| 33.79167 | -2.12932 | -2.28334 | -2.4114 | -0.16068 | -1.75487 | -0.47336 | -1.22593 | 0.638909 | 0.646057 | -0.46708 | -8.30276 | -0.1766 | 0.019195 | 0.229593 | -0.03936 | 0.049023 | 0.106531 | -0.00603 | -0.12702 | 0.016268 | -0.32847 | -0.06924 | 0.086718 | -0.11374 |
| 33.80556 | -1.86063 | -2.22155 | -2.31704 | -0.1762 | -1.23565 | -0.56881 | -1.79335 | 0.643971 | 0.719256 | -0.45508 | -5.15253 | 0.241415 | 0.007212 | 0.233442 | -0.01653 | -0.00362 | 0.103954 | 0.011335 | -0.17223 | 0.033799 | -0.33586 | -0.01161 | 0.084493 | -0.12152 |
| 33.81944 | -1.6378 | -1.92521 | -2.4528 | -0.14749 | -0.33082 | -0.72641 | -2.51578 | 0.62408 | 0.890989 | -0.45493 | -2.97978 | 0.532357 | 0.012603 | 0.226291 | -0.00203 | -0.07428 | 0.105658 | 0.013186 | -0.23895 | 0.061195 | -0.32871 | 0.037083 | 0.138285 | -0.13207 |
| 33.83333 | -1.57457 | -1.31136 | -2.87778 | -0.13618 | 0.296712 | -0.83354 | -2.93657 | 0.550171 | 1.346308 | -0.31311 | -2.84318 | 0.652725 | 0.035468 | 0.2978 | 0.006057 | -0.05565 | 0.123423 | 0.025298 | -0.23088 | 0.089415 | -0.32379 | 0.079936 | 0.180472 | -0.14449 |
| 33.84722 | -1.01694 | -0.56698 | -2.84374 | -0.08539 | 0.288168 | -0.93633 | -2.71818 | 0.502611 | 1.705582 | -0.13454 | -1.98525 | 0.910438 | 0.027887 | 0.311485 | -0.01705 | -0.06584 | 0.095972 | 0.0035 | -0.05132 | 0.058854 | -0.33645 | 0.087821 | 0.110109 | -0.19959 |
| 33.86111 | -0.73212 | 0.273159 | -2.25091 | 0.040061 | 0.04137 | -1.00713 | -2.56371 | 0.465099 | 1.851659 | 0.03532 | -0.81798 | 0.996882 | -0.03278 | 0.214545 | -0.08565 | -0.11968 | 0.024646 | -0.05449 | 0.280181 | -0.02063 | -0.38767 | 0.054117 | 0.021001 | -0.20311 |
| 33.875 | -0.55168 | 0.575565 | -1.23214 | 0.411653 | 0.1164 | -0.84221 | -1.52684 | 0.415828 | 1.95994 | 0.347413 | -0.39333 | 0.938516 | -0.01639 | 0.165803 | -0.09322 | -0.08592 | 0.016892 | -0.03627 | 0.450693 | 0.002779 | -0.33004 | 0.049138 | 0.016433 | -0.21404 |
| 33.88889 | -0.52899 | 0.014083 | -0.87714 | 0.856686 | 0.206277 | -0.95841 | -0.85331 | 0.24113 | 1.373636 | 0.193585 | -0.91292 | 0.64547 | 0.089666 | -0.21354 | -0.04918 | -0.04375 | 0.018595 | 0.012386 | 0.260472 | 0.060082 | -0.16424 | 0.079719 | -0.11236 | -0.27097 |
| 33.90278 | -0.95027 | -0.57127 | -1.29245 | 0.769834 | -0.0618 | -1.03292 | -1.26542 | -0.04049 | 0.045417 | -0.67441 | -1.73444 | 0.073899 | 0.113879 | -0.92264 | -0.08024 | -0.17561 | -0.10064 | -0.03214 | -0.00424 | -0.07343 | -0.12752 | -0.01179 | -0.58916 | -0.40988 |
| 33.91667 | -1.61108 | -1.22556 | -1.68206 | 0.186944 | -1.0633 | -0.66701 | -2.28023 | -0.09651 | -1.12873 | -1.64343 | -2.17076 | -0.47424 | -0.03157 | -1.00434 | -0.08385 | -0.1614 | -0.14957 | -0.04379 | -0.15292 | -0.32864 | -0.22963 | -0.17057 | -0.67772 | -0.42015 |
| 33.93056 | -1.70187 | -1.69401 | -1.68618 | -0.07663 | -1.72585 | -0.36831 | -2.66462 | -0.05588 | -1.92875 | -2.13046 | -1.9247 | -0.90735 | -0.04317 | -0.31049 | 0.108179 | 0.069057 | 0.047448 | 0.080923 | -0.30915 | -0.56357 | -0.33207 | -0.06259 | -0.09741 | -0.20234 |
| 33.94444 | -1.47562 | -2.26084 | -1.84823 | -0.0929 | -1.62857 | -0.20362 | -2.26153 | 0.188326 | -2.13555 | -1.63348 | -1.10566 | -0.91679 | 0.24779 | 0.318422 | 0.401083 | 0.287446 | 0.313298 | 0.191622 | -0.40781 | -0.68461 | -0.15231 | 0.216167 | 0.568055 | 0.159712 |
| 33.95833 | -1.54462 | -2.88591 | -2.24674 | -0.20442 | -0.28142 | -0.51095 | -1.64669 | 0.3955 | -2.11016 | -0.97609 | -0.34416 | -0.61854 | 0.379453 | 0.396314 | 0.414451 | 0.242481 | 0.316666 | 0.12559 | -0.23169 | -0.4019 | -0.12723 | 0.222485 | 0.598302 | 0.335703 |
| 33.97222 | -1.74114 | -3.25922 | -2.44768 | -0.19794 | 1.296937 | -0.43317 | -1.21318 | 0.120373 | -1.89007 | -0.40879 | 0.307435 | -0.34245 | 0.174625 | 0.18986 | 0.190462 | 0.065285 | 0.135288 | -0.00874 | -0.02697 | -0.23557 | -0.21035 | 0.083811 | -0.04166 | 0.245447 |
| 33.98611 | -1.52719 | -3.02414 | -1.98729 | -0.15165 | 2.182928 | -0.36464 | -0.9441 | -0.53281 | -1.15015 | -0.17521 | 0.397881 | -0.16511 | 0.044896 | 0.085851 | 0.075454 | 0.021332 | 0.035029 | -0.05889 | 0.094344 | -0.25588 | -0.16183 | 0.040275 | -0.16209 | 0.173465 |
| 34 | -1.64882 | -2.67919 | -1.3383 | -0.05183 | 1.833795 | -0.07976 | -0.63228 | -0.80837 | -0.68463 | -0.05036 | -0.05396 | -0.06167 | -0.0194 | 0.045532 | 0.018569 | 0.00139 | -0.0372 | -0.11984 | 0.08978 | -0.44063 | -0.11819 | 0.043691 | -0.41128 | 0.091118 |
| 34.01389 | -1.33264 | -2.36514 | -0.91751 | -0.01224 | 1.12699 | -0.01831 | -0.45387 | -0.55807 | -0.31153 | -0.0361 | -0.28721 | 0.072094 | -0.11646 | -0.01321 | -0.0735 | -0.07145 | -0.1217 | -0.21763 | 0.01458 | -0.66383 | -0.1428 | 0.019126 | -0.84216 | 0.002483 |
| 34.02778 | -1.04871 | -2.09151 | -0.70571 | 0.041619 | 0.714336 | 0.050705 | -0.31554 | -0.16586 | 0.006762 | -0.14059 | -0.05383 | 0.214895 | -0.15021 | 0.001243 | -0.07897 | -0.1126 | -0.13027 | -0.2588 | -0.07001 | -0.80246 | -0.10728 | 0.044208 | -0.70305 | -0.03997 |
| 34.04167 | -0.90025 | -1.69712 | -0.8089 | 0.101901 | 0.57049 | 0.014672 | -0.26445 | 0.066325 | 0.21222 | -0.21197 | 0.254588 | 0.262798 | -0.11496 | 0.094565 | -0.01017 | -0.08366 | -0.0878 | -0.33407 | -0.13352 | -0.9479 | -0.02569 | 0.098988 | -0.45901 | -0.02053 |
| 34.05556 | -0.63701 | -1.36418 | -1.16336 | 0.136052 | 0.325463 | 0.030864 | -0.34248 | 0.180645 | 0.251942 | -0.19428 | 0.637076 | 0.256134 | -0.08511 | 0.149544 | 0.020277 | -0.05593 | -0.07373 | -0.49708 | -0.17951 | -1.11471 | 0.000657 | 0.10668 | -0.15459 | -0.02036 |
| 34.06944 | -0.19523 | -1.18589 | -1.00412 | 0.212497 | -0.25991 | 0.08853 | -0.27726 | 0.243291 | 0.127213 | -0.07795 | 1.247478 | 0.262639 | -0.06643 | 0.175288 | 0.021764 | -0.04123 | -0.06913 | -0.7607 | -0.16149 | -1.18741 | -0.01002 | 0.07719 | 0.044849 | -0.02658 |
| 34.08333 | 0.00996 | -1.14572 | -0.36992 | 0.376088 | -0.68551 | 0.24036 | -0.04747 | 0.287019 | 0.108759 | 0.031053 | 1.778474 | 0.287505 | -0.05236 | 0.214079 | 0.020843 | -0.0239 | -0.04556 | -0.84816 | -0.07738 | -0.99647 | 0.003035 | 0.041093 | 0.142735 | 0.010982 |
| 34.09722 | 0.266444 | -1.16714 | 0.066491 | 0.474413 | -0.42954 | 0.300274 | 0.09845 | 0.277944 | 0.255325 | 0.017007 | 2.032505 | 0.281047 | -0.06672 | 0.237649 | 0.027817 | 0.000222 | -0.03263 | -0.91677 | -0.03066 | -0.4948 | 0.004517 | 0.00034 | 0.130719 | 0.068877 |
| 34.11111 | 0.279869 | -1.19374 | 0.254305 | 0.509951 | 0.237208 | 0.385565 | 0.045744 | 0.247216 | 0.488994 | -0.03902 | 2.014163 | 0.274293 | -0.10696 | 0.177163 | 0.048227 | 0.031567 | -0.03299 | -0.93546 | -0.07217 | -0.18035 | -0.00368 | -0.02935 | 0.074013 | 0.124361 |
| 34.125 | 0.293012 | -1.20218 | 0.35135 | 0.499012 | 0.776051 | 0.465452 | -0.14588 | 0.285043 | 0.512071 | -0.01138 | 2.017458 | 0.304651 | -0.17088 | 0.105231 | 0.048247 | 0.045248 | -0.04359 | -0.35453 | -0.19394 | -0.0145 | 0.020021 | -0.02137 | 0.023816 | 0.171552 |
| 34.13889 | 0.376671 | -1.06966 | 0.338546 | 0.491225 | 1.09323 | 0.523096 | -0.20208 | 0.383407 | 0.308503 | 0.057116 | 2.000583 | 0.334842 | -0.19157 | 0.071628 | 0.045076 | 0.030817 | -0.06545 | -0.47382 | -0.34159 | -0.06577 | 0.101189 | -0.00467 | 0.02036 | 0.225693 |
| 34.15278 | 0.375536 | -0.88686 | 0.207396 | 0.528951 | 1.188733 | 0.504828 | -0.11388 | 0.406392 | -0.01625 | 0.070725 | 1.914228 | 0.289997 | -0.16442 | 0.095887 | 0.038609 | 0.017506 | -0.07655 | -0.55147 | -0.44067 | -0.02702 | 0.18034 | 0.000444 | 0.03946 | 0.208642 |

| | | | | | | | | | | | | | | | | | | | | | | | | |
|---|---|---|---|---|---|---|---|---|---|---|---|---|---|---|---|---|---|---|---|---|---|---|---|---|
| 34.16667 | 0.394615 | -0.6949 | 0.065564 | 0.453934 | 1.066823 | 0.433929 | -0.12004 | 0.317426 | -0.20405 | 0.065929 | 1.885432 | 0.237538 | -0.15554 | 0.049374 | 0.004336 | 0.009332 | -0.09239 | -0.656 | -0.4192 | -0.05646 | 0.15939 | -0.0206 | 0.045296 | 0.08994 |
| 34.18056 | 0.381056 | -0.56394 | 0.0414 | 0.282605 | 0.363455 | 0.385523 | -0.13623 | 0.219368 | -0.07909 | 0.080089 | 1.842569 | 0.207913 | -0.13145 | 0.021291 | -0.0088 | 0.031076 | -0.09167 | -0.74283 | -0.38296 | -0.07063 | 0.17095 | -0.02968 | 0.090799 | 0.006973 |
| 34.19444 | 0.505724 | -0.58968 | 0.123939 | 0.114435 | -0.62868 | 0.361359 | -0.09554 | 0.214478 | 0.04998 | 0.058362 | 1.754857 | 0.162296 | -0.07863 | 0.055863 | 0.027191 | 0.079863 | -0.04854 | -0.92611 | -0.29583 | -0.01339 | 0.217605 | -0.02583 | 0.132246 | 0.022071 |
| 34.20833 | 0.727606 | -0.75731 | 0.15129 | -0.02672 | -1.43357 | 0.348783 | -0.06892 | 0.289013 | 0.170769 | 0.007315 | 1.599225 | 0.0334 | -0.07857 | 0.07704 | 0.034593 | 0.084101 | -0.03015 | -0.85863 | -0.25043 | 0.008054 | 0.196924 | -0.04599 | 0.072204 | 0.055378 |
| 34.22222 | 0.811056 | -0.99525 | 0.036318 | -0.1304 | -1.76214 | 0.36815 | -0.03077 | 0.39467 | 0.266092 | -0.06515 | 1.367032 | -0.10312 | -0.12548 | 0.025293 | 0.012738 | 0.051606 | -0.05548 | -0.57191 | -0.25628 | -0.01484 | 0.135278 | -0.06853 | 0.003539 | 0.060424 |
| 34.23611 | 0.803893 | -1.22873 | -0.04543 | -0.18828 | -1.65351 | 0.366236 | 0.005316 | 0.448079 | 0.225661 | -0.13103 | 1.110341 | -0.21133 | -0.13846 | -0.031 | -0.0097 | 0.018025 | -0.08848 | -0.46823 | -0.27953 | -0.04418 | 0.075822 | -0.07076 | -0.02054 | 0.04937 |
| 34.25 | 0.76736 | -1.40796 | -0.03274 | -0.20946 | -1.18774 | 0.415096 | -0.01677 | 0.464594 | 0.078498 | -0.16697 | 0.925491 | -0.30797 | -0.15618 | -0.09918 | -0.07173 | -0.02523 | -0.13287 | -0.6727 | -0.34331 | -0.17303 | 0.029748 | -0.09786 | -0.09177 | 0.010478 |
| 34.26389 | 0.519119 | -1.45625 | 0.041411 | -0.09423 | -0.4755 | 0.437967 | -0.0292 | 0.513127 | -0.06084 | -0.19108 | 0.804097 | -0.37915 | -0.18334 | -0.15606 | -0.13603 | -0.0583 | -0.15584 | -0.98428 | -0.39495 | -0.37179 | 0.02224 | -0.1337 | -0.18265 | -0.01573 |
| 34.27778 | 0.302801 | -1.43864 | 0.163993 | 0.009182 | 0.030599 | 0.43693 | -0.03691 | 0.624708 | -0.21641 | -0.15587 | 0.87371 | -0.44307 | -0.13285 | -0.13126 | -0.10949 | -0.02503 | -0.11137 | -0.99378 | -0.33615 | -0.36145 | 0.119703 | -0.11509 | -0.14392 | -0.001 |
| 34.29167 | 0.157489 | -1.4766 | 0.272107 | 0.003042 | -0.00964 | 0.428884 | -0.06639 | 0.743599 | -0.37081 | -0.15215 | 0.526742 | -0.50293 | -0.07039 | -0.08628 | -0.05963 | 0.024125 | -0.08021 | -0.71092 | -0.27567 | -0.19535 | 0.185129 | -0.07179 | -0.04117 | -0.0017 |
| 34.30556 | -0.10034 | -1.46186 | 0.304349 | -0.08373 | -0.49848 | 0.42059 | -0.13852 | 0.815407 | -0.41134 | -0.14472 | 0.585317 | -0.46062 | -0.11419 | -0.09954 | -0.09237 | -0.02024 | -0.13465 | -0.58078 | -0.35581 | -0.141 | 0.141087 | -0.07852 | 0.011185 | -0.03323 |
| 34.31944 | -0.14466 | -1.4545 | 0.269256 | -0.20953 | -0.80828 | 0.482888 | -0.2828 | 0.746096 | -0.27421 | -0.11788 | 0.434173 | -0.39626 | -0.13696 | -0.07463 | -0.11377 | -0.04957 | -0.16597 | -0.5428 | -0.3602 | -0.14447 | 0.125386 | -0.09328 | 0.035079 | -0.0497 |
| 34.33333 | -0.09197 | -1.39437 | 0.246139 | -0.15305 | -0.39849 | 0.557926 | -0.43465 | 0.717569 | -0.29496 | -0.05714 | 0.645022 | -0.38057 | -0.08374 | -0.00502 | -0.03785 | 0.017895 | -0.09816 | -0.48055 | -0.25952 | -0.07337 | 0.195108 | -0.03718 | 0.082334 | -0.01529 |
| 34.34722 | -0.13814 | -1.34494 | 0.238354 | -0.05 | 0.282195 | 0.501442 | -0.56899 | 0.710309 | -0.5403 | -0.04222 | 0.659789 | -0.47219 | -0.07891 | 0.021887 | 0.000838 | 0.063644 | -0.05481 | -0.52918 | -0.31689 | -0.03732 | 0.238692 | 0.002483 | 0.097782 | 0.007783 |
| 34.36111 | -0.19489 | -1.34106 | 0.349702 | -0.02388 | 0.758144 | 0.410923 | -0.69605 | 0.72022 | -0.78593 | -0.08482 | 0.900476 | -0.59025 | -0.17966 | 0.022389 | -0.10046 | 0.012746 | -0.13074 | -0.45053 | -0.72875 | -0.12576 | 0.137963 | -0.09004 | 0.059029 | -0.06486 |
| 34.375 | -0.02332 | -1.3151 | 0.494692 | 0.021245 | 0.974093 | 0.356766 | -0.77525 | 0.673988 | -0.94009 | -0.04215 | 1.392425 | -0.58099 | -0.2867 | 0.081006 | -0.20294 | -0.04561 | -0.21988 | -0.35327 | -0.75527 | -0.20241 | 0.031498 | -0.2043 | 0.051368 | -0.1219 |
| 34.38889 | 0.205395 | -1.09954 | 0.348697 | 0.032119 | 0.998327 | 0.318474 | -0.80706 | 0.657783 | -0.63808 | 0.194298 | 1.956162 | -0.50214 | -0.25527 | 0.1186 | -0.15188 | -0.0258 | -0.18292 | -0.19821 | -0.62603 | -0.1404 | 0.056819 | -0.14396 | 0.135155 | -0.04339 |
| 34.40278 | 0.824437 | -0.57238 | 0.122452 | 0.105108 | 1.217348 | 0.403776 | -0.66111 | 0.737441 | 0.079633 | 0.651064 | 2.773957 | -0.21315 | -0.14769 | 0.008514 | -0.016 | 0.007964 | -0.04279 | -0.03134 | -0.38651 | -0.00684 | 0.281217 | 0.063279 | 0.282048 | 0.10949 |
| 34.41667 | 1.324324 | 0.036943 | 0.075131 | 0.375658 | 1.979602 | 0.368666 | -0.27844 | 0.66942 | 1.525257 | 1.070821 | 2.812068 | 0.279941 | -0.04647 | -0.11881 | 0.076388 | 0.015036 | 0.07684 | 0.076707 | -0.05005 | 0.08364 | 0.582102 | 0.214693 | 0.377402 | 0.250779 |
| 34.43056 | 1.21138 | 0.494467 | 0.115064 | 0.670974 | 3.249323 | 0.394081 | 0.359845 | 0.693749 | 2.983718 | 1.080157 | 2.950896 | 0.776436 | -0.11664 | -0.19018 | 0.007858 | -0.03731 | -0.0061 | 0.028972 | 0.163497 | -0.0036 | 0.542353 | 0.103116 | 0.228778 | 0.173432 |
| 34.44444 | 1.015417 | 0.916917 | 0.528494 | 0.976624 | 3.782494 | 0.499202 | 0.959096 | 1.277139 | 2.394653 | 0.79104 | 3.584525 | 1.227971 | -0.37473 | -0.29391 | -0.22439 | -0.15849 | -0.22586 | -0.16238 | 0.082023 | -0.27613 | 0.084407 | -0.14009 | -0.11465 | -0.059 |
| 34.45833 | 0.97267 | 1.451359 | 1.17972 | 0.95701 | 2.031739 | 0.750281 | 1.058923 | 1.85535 | 1.805416 | 0.504679 | 3.293586 | 1.351213 | -0.35274 | -0.2136 | -0.22315 | -0.05843 | -0.19318 | -0.1313 | 0.111463 | -0.25525 | -0.08574 | -0.11623 | -0.1507 | -0.03702 |
| 34.47222 | 1.046102 | 1.700906 | 1.597098 | 0.638954 | -0.02632 | 0.746566 | 0.666028 | 1.964869 | 0.288362 | 0.366642 | 2.733958 | 1.485718 | -0.00295 | 0.02339 | 0.045505 | 0.236498 | 0.084115 | 0.122559 | 0.412881 | 0.063959 | 0.186158 | 0.120897 | 0.071916 | 0.168494 |
| 34.48611 | 1.189387 | 1.703049 | 1.657161 | 0.304753 | -1.66682 | 0.692867 | 0.141257 | 2.00707 | -1.11306 | 0.413591 | 2.158373 | 1.373505 | 0.220008 | 0.09669 | 0.160489 | 0.301342 | 0.199321 | 0.194849 | 0.44588 | 0.17678 | 0.268792 | 0.158614 | 0.218731 | 0.183042 |
| 34.5 | 1.273769 | 1.654445 | 1.677043 | 0.156098 | -2.45082 | 0.645227 | -0.12547 | 1.921028 | -1.43436 | 0.719045 | 1.440319 | 1.390443 | 0.179351 | 0.045394 | 0.089021 | 0.164512 | 0.124006 | 0.093764 | 0.188185 | 0.089765 | 0.130129 | 0.033286 | 0.222687 | 0.056656 |
| 34.51389 | 1.226501 | 1.750296 | 1.875932 | 0.199681 | -1.91369 | 0.568683 | -0.10219 | 1.968462 | -0.89565 | 0.827235 | 0.82491 | 1.308992 | 0.046775 | -0.00494 | -0.00863 | 0.022125 | 0.00992 | -0.02296 | -0.03409 | -0.00324 | -0.02828 | -0.06551 | 0.238471 | -0.03474 |
| 34.52778 | 1.251004 | 1.850882 | 2.260019 | 0.4103 | -0.68323 | 0.534849 | 0.126041 | 2.052037 | -0.27422 | 0.732935 | 0.415266 | 1.249744 | -0.06791 | -0.07361 | -0.09019 | -0.09789 | -0.08672 | -0.1275 | -0.18719 | -0.07788 | -0.12756 | -0.11404 | 0.247471 | -0.09393 |
| 34.54167 | 1.073843 | 1.807847 | 2.276486 | 0.55925 | 0.234357 | 0.506671 | 0.359686 | 1.892639 | 0.016935 | 0.488219 | 0.103194 | 0.94333 | -0.07359 | -0.0989 | -0.08728 | -0.11758 | -0.09666 | -0.13875 | -0.18204 | -0.05427 | -0.07432 | -0.09341 | 0.256916 | -0.08628 |
| 34.55556 | 1.063555 | 1.585814 | 1.811726 | 0.432913 | 0.471522 | 0.539908 | 0.481802 | 1.442488 | -0.28492 | 0.342916 | -0.08791 | 0.634735 | 0.006351 | -0.03923 | -0.012 | -0.02467 | -0.0219 | -0.07455 | -0.04559 | 0.055332 | 0.067211 | -0.02399 | 0.251559 | -0.00185 |
| 34.56944 | 1.051266 | 1.667951 | 1.455455 | 0.274247 | 0.253648 | 0.496404 | 0.464419 | 0.97958 | -1.01574 | 0.272123 | -0.23635 | 0.255782 | 0.057391 | 0.032946 | 0.051426 | 0.09213 | 0.052079 | -0.02205 | 0.073998 | 0.143106 | 0.169976 | 0.055744 | 0.27633 | 0.075005 |
| 34.58333 | 1.154544 | 1.807738 | 1.212126 | 0.146058 | -0.30118 | 0.481616 | 0.407066 | 0.479041 | -1.61474 | 0.174017 | -0.36769 | -0.07783 | 0.065925 | 0.095013 | 0.083715 | 0.181006 | 0.087569 | 0.00025 | 0.1429 | 0.184257 | 0.222624 | 0.110939 | 0.235952 | 0.128508 |
| 34.59722 | 1.180674 | 1.721794 | 1.068152 | -0.02958 | -1.46092 | 0.527732 | 0.447796 | 0.009097 | -2.16034 | -0.10017 | -0.36505 | -0.35688 | 0.046883 | 0.114592 | 0.071513 | 0.19879 | 0.07965 | -0.00497 | 0.172276 | 0.178272 | 0.218791 | 0.094958 | 0.221409 | 0.135509 |
| 34.61111 | 1.183567 | 1.37267 | 0.961969 | -0.35617 | -3.01314 | 0.510394 | 0.491347 | -0.19071 | -2.04751 | -0.25267 | -0.31278 | -0.54806 | 0.010251 | 0.088844 | 0.032669 | 0.157252 | 0.051705 | -0.0373 | 0.149774 | 0.138953 | 0.154632 | 0.021662 | 0.272248 | 0.08635 |
| 34.625 | 1.231034 | 0.958526 | 0.834494 | -0.65268 | -3.62482 | 0.5107 | 0.386834 | -0.1724 | -1.66339 | -0.4482 | -0.08414 | -0.6938 | -0.01031 | 0.056146 | 0.019383 | 0.093214 | 0.044312 | -0.08015 | 0.09452 | 0.093606 | 0.106111 | -0.02123 | 0.381312 | 0.03093 |
| 34.63889 | 1.303339 | 0.618358 | 0.723456 | -0.67765 | -3.09069 | 0.462092 | 0.033939 | -0.06001 | -1.30753 | -0.53126 | 0.530011 | -0.95814 | -0.01215 | 0.044841 | 0.022455 | 0.03677 | 0.039342 | -0.1344 | 0.041416 | 0.055531 | 0.084366 | -0.02549 | 0.17677 | -0.01325 |
| 34.65278 | 1.488438 | 0.482001 | 0.447107 | -0.60738 | -2.06121 | 0.389909 | -0.4937 | 0.075049 | -1.36333 | -0.55866 | 1.614986 | -1.32969 | -0.04092 | 0.04532 | -0.0037 | -0.03586 | -0.00478 | -0.17838 | -0.02971 | 0.019508 | 0.019922 | -0.0375 | -0.01203 | -0.06775 |
| 34.66667 | 1.652817 | 0.457417 | 0.217327 | -0.53055 | -1.05328 | 0.358839 | -0.85805 | 0.189193 | -1.42154 | -0.57067 | 1.788956 | -1.40237 | -0.08519 | 0.036173 | -0.03533 | -0.08825 | -0.04819 | -0.22618 | -0.09092 | -0.01189 | -0.07497 | -0.04833 | -0.10226 | -0.13705 |
| 34.68056 | 1.72437 | 0.308343 | 0.146517 | -0.55548 | -0.7014 | 0.398739 | -0.8999 | 0.225666 | -1.22955 | -0.77725 | 1.223945 | -1.51314 | -0.10964 | 0.015042 | -0.0534 | -0.10796 | -0.05672 | -0.26583 | -0.11167 | -0.04096 | -0.12969 | -0.05093 | -0.21802 | -0.19623 |
| 34.69444 | 1.639688 | -0.02809 | 0.239056 | -0.66737 | -0.96551 | 0.501477 | -0.90949 | 0.14905 | -0.76808 | -0.90866 | 0.895406 | -1.50723 | -0.1341 | -0.01276 | -0.07851 | -0.1069 | -0.07269 | -0.30262 | -0.14995 | -0.0612 | -0.18262 | -0.07945 | -0.32156 | -0.24841 |
| 34.70833 | 1.393124 | -0.52913 | 0.180237 | -0.72812 | -1.69177 | 0.472246 | -1.02798 | 0.048135 | -0.55128 | -0.92497 | 1.31019 | -1.4032 | -0.15717 | -0.03532 | -0.08398 | -0.10084 | -0.08978 | -0.3051 | -0.18555 | -0.06063 | -0.23552 | -0.12811 | -0.32036 | -0.23797 |
| 34.72222 | 0.98382 | -1.00035 | 0.173959 | -0.7967 | -2.09452 | 0.496059 | -1.10104 | -0.13645 | -0.54995 | -0.81747 | 1.406693 | -1.34105 | -0.16487 | -0.03156 | -0.07615 | -0.06999 | -0.07817 | -0.28776 | -0.17459 | -0.05808 | -0.28792 | -0.16466 | -0.25011 | -0.21988 |
| 34.73611 | 0.705945 | -1.35364 | 0.218625 | -0.8224 | -1.61991 | 0.373738 | -0.99831 | -0.36715 | -0.21899 | -0.76299 | 1.404058 | -1.27295 | -0.13436 | 0.100033 | -0.03113 | 0.029655 | -0.02013 | -0.22537 | -0.09794 | -0.01781 | -0.25633 | -0.13254 | -0.16306 | -0.16574 |
| 34.75 | 0.427569 | -1.5699 | 0.290813 | -0.84232 | -1.68432 | 0.178884 | -0.86849 | -0.7546 | 0.143625 | -0.79324 | 1.195195 | -1.16159 | -0.07767 | 0.238018 | 0.021362 | 0.109178 | 0.042084 | -0.16801 | -0.03585 | 0.035165 | -0.22154 | -0.08976 | -0.0973 | -0.09711 |
| 34.76389 | 0.097376 | -1.5525 | 0.232057 | -0.89059 | -2.20686 | 0.142075 | -0.7759 | -1.18141 | 0.344727 | -0.80725 | 0.749858 | -0.85666 | -0.08741 | 0.200816 | 0.011995 | 0.096287 | 0.033446 | -0.19536 | -0.09569 | 0.027112 | -0.2635 | -0.1009 | -0.11981 | -0.14481 |
| 34.77778 | -0.3071 | -1.59853 | 0.184781 | -0.88618 | -2.93188 | 0.112645 | -0.80567 | -0.89183 | 0.617646 | -0.75416 | 0.364561 | -0.50174 | -0.16019 | 0.018868 | -0.07107 | -0.01316 | -0.04816 | -0.29885 | -0.21936 | -0.0314 | -0.35516 | -0.13547 | -0.26359 | -0.27908 |
| 34.79167 | -0.69611 | -1.78134 | 0.200025 | -0.80813 | -3.81808 | 0.114178 | -0.74333 | -0.33699 | 0.883557 | -0.6634 | 0.042648 | -0.06271 | -0.21114 | -0.0732 | -0.15144 | -0.1131 | -0.11442 | -0.42109 | -0.33331 | -0.07511 | -0.41968 | -0.15742 | -0.37731 | -0.37457 |
| 34.80556 | -0.91159 | -1.6319 | 0.34116 | -0.60711 | -3.83092 | 0.126356 | -0.57079 | -0.04804 | 1.22077 | -0.67549 | -0.44574 | 0.274095 | -0.20705 | -0.04406 | -0.18048 | -0.11603 | -0.11354 | -0.44573 | -0.4116 | -0.09404 | -0.44512 | -0.14544 | -0.38218 | -0.44448 |
| 34.81944 | -0.92413 | -1.28022 | 0.439577 | -0.3366 | -3.41813 | 0.200052 | -0.25364 | 0.062037 | 1.382128 | -0.76304 | -0.77812 | 0.464669 | -0.13584 | 0.043327 | -0.15783 | -0.09531 | -0.06688 | -0.39253 | -0.42903 | -0.07222 | -0.43802 | -0.07465 | -0.32659 | -0.41584 |
| 34.83333 | -0.79611 | -1.05521 | 0.514551 | -0.08046 | -2.64787 | 0.348868 | 0.03843 | 0.102398 | 1.508302 | -0.56685 | -0.49221 | 0.527773 | -0.04473 | 0.120231 | -0.10884 | -0.0436 | 0.010023 | -0.30107 | -0.40708 | 0.002177 | -0.41249 | 0.003958 | -0.1978 | -0.32818 |
| 34.84722 | -0.54544 | -0.81528 | 0.33757 | 0.038183 | -2.16563 | 0.529001 | 0.248759 | 0.244411 | 1.671399 | -0.12195 | -0.41025 | 0.572641 | -0.00295 | 0.206586 | -0.05772 | -0.01413 | 0.058295 | -0.21861 | -0.31459 | 0.038299 | -0.38481 | 0.031764 | -0.08154 | -0.2167 |
| 34.86111 | -0.44318 | -0.50575 | 0.220012 | 0.165838 | -1.50701 | 0.448819 | 0.401788 | 0.415817 | 1.620958 | 0.25426 | -0.52026 | 0.504656 | -0.00958 | 0.193986 | -0.0459 | 0.006321 | 0.050151 | -0.15394 | -0.23152 | 0.033194 | -0.38531 | 0.013757 | -0.03292 | -0.1842 |
| 34.875 | -0.35921 | 0.279046 | 0.233626 | 0.339439 | -1.03587 | 0.207054 | 0.452511 | 0.407214 | 1.64724 | 0.241193 | -1.17205 | 0.279685 | -0.03516 | 0.062254 | -0.04225 | 0.009005 | 0.012551 | -0.12009 | -0.13328 | 0.005053 | -0.38722 | -0.02798 | -0.07063 | -0.19304 |
| 34.88889 | -0.73594 | 1.305088 | 0.338286 | 0.341969 | -0.94101 | -0.1311 | 0.471966 | -0.16075 | 1.390783 | -0.4338 | -2.50147 | -0.32608 | -0.05435 | -0.51457 | -0.08243 | -0.02258 | -0.0682 | -0.17694 | -0.03362 | -0.04744 | -0.34846 | -0.0452 | -0.21179 | -0.30483 |
| 34.90278 | -0.91133 | 1.405917 | 0.177742 | 0.035573 | -1.62969 | -0.38538 | 0.134103 | -0.7926 | 0.476885 | -1.56144 | -3.44494 | -0.82552 | -0.06113 | -1.47562 | -0.09826 | -0.03397 | -0.14891 | -0.18425 | 0.146301 | -0.08883 | -0.20814 | 0.007448 | -0.38344 | -0.41718 |
| 34.91667 | -0.78334 | 0.386968 | 0.242234 | -0.15119 | -2.65505 | -0.36922 | -0.19294 | -0.89207 | -0.41975 | -2.90109 | -2.98277 | -0.81658 | -0.20414 | -1.65776 | -0.073 | 0.059037 | -0.16468 | -0.08318 | 0.042562 | -0.23973 | -0.23346 | 0.04152 | -0.54757 | -0.49338 |
| 34.93056 | -0.34795 | -0.5213 | 0.344449 | 0.115365 | -2.76366 | 0.014304 | -0.45462 | -0.57175 | -0.56918 | -3.26065 | -1.91415 | -0.4744 | -0.24413 | -0.87627 | 0.033259 | 0.094899 | -0.0638 | 0.013154 | -0.51365 | -0.36293 | -0.28899 | 0.052456 | -0.6293 | -0.40036 |
| 34.94444 | -0.03888 | -0.95856 | 0.250143 | 0.45567 | -2.12479 | 0.356491 | -0.62923 | -0.25057 | -0.49283 | -3.39047 | -0.82048 | -0.11744 | -0.06315 | 0.009761 | 0.281437 | 0.12456 | 0.164468 | 0.146342 | -0.70426 | 0.086985 | -0.19128 | 0.253683 | -0.28221 | 0.05055 |
| 34.95833 | 0.011494 | -1.20204 | 0.041824 | 0.52933 | -1.3093 | 0.459971 | -0.79618 | -0.10376 | -0.90838 | -3.54053 | 0.150428 | 0.046765 | -0.0337 | 0.319369 | 0.367618 | 0.066169 | 0.288675 | 0.127133 | -0.33395 | 0.511337 | -0.19525 | 0.347804 | 0.283667 | 0.504435 |
| 34.97222 | -0.04627 | -1.05663 | 0.011088 | 0.394192 | -0.38497 | 0.488646 | -0.72397 | -0.08907 | -0.92177 | -2.92234 | 0.764099 | -0.02618 | -0.16477 | 0.198251 | 0.154097 | -0.1136 | 0.137908 | -0.04434 | -0.13407 | 0.307572 | -0.33718 | 0.17075 | 0.380107 | 0.469261 |
| 34.98611 | 0.060778 | -0.95136 | 0.027415 | 0.334275 | 0.068557 | 0.461379 | -0.58338 | -0.11694 | -0.69019 | -1.90756 | 0.868424 | -0.18107 | -0.24151 | 0.051918 | -0.01532 | -0.18074 | -0.04101 | -0.15498 | -0.12736 | 0.049504 | -0.38159 | 0.009411 | 0.239656 | 0.214459 |
| 35 | 0.31632 | -0.94587 | 0.077697 | 0.39571 | 0.029057 | 0.465926 | -0.49488 | -0.08358 | -0.4481 | -0.92734 | 0.808744 | -0.18995 | -0.29325 | 0.141046 | -0.07647 | -0.11482 | -0.11949 | -0.17638 | -0.13313 | -0.0695 | -0.31871 | -0.01765 | 0.178393 | 0.068745 |
| 35.01389 | 0.502853 | -1.15227 | 0.23466 | 0.344947 | 0.033552 | 0.495946 | -0.3917 | 0.065626 | -0.32597 | -0.26248 | 0.652526 | -0.07379 | -0.36033 | 0.266497 | -0.08945 | -0.00324 | -0.12899 | -0.15913 | -0.14734 | -0.11932 | -0.24123 | 0.020138 | 0.265166 | 0.047751 |
| 35.02778 | 0.580055 | -1.29092 | 0.322851 | 0.314379 | 0.315665 | 0.470468 | -0.20863 | 0.252666 | -0.29797 | -0.05984 | 0.517303 | 0.088704 | -0.41613 | 0.402785 | -0.0461 | 0.083759 | -0.07499 | -0.09085 | -0.16012 | -0.08369 | -0.15566 | 0.078632 | 0.308331 | 0.086074 |
| 35.04167 | 0.565168 | -1.17784 | 0.314046 | 0.237705 | 0.734324 | 0.401199 | -0.00071 | 0.428424 | -0.29164 | -0.32461 | 0.422984 | 0.154419 | -0.51327 | 0.484016 | -0.0447 | 0.082014 | -0.06389 | -0.05216 | -0.19917 | -0.05542 | -0.11613 | 0.075315 | 0.195861 | 0.099953 |
| 35.05556 | 0.478679 | -0.93558 | 0.25417 | 0.226565 | 1.17996 | 0.330345 | 0.118758 | 0.550277 | -0.22963 | -0.83519 | 0.506162 | 0.215769 | -0.58355 | 0.277781 | -0.12548 | -0.03341 | -0.13047 | -0.08075 | -0.2778 | -0.10242 | -0.15077 | 0.016385 | 0.034663 | 0.035676 |
| 35.06944 | 0.451104 | -0.8842 | 0.179468 | 0.311145 | 1.520734 | 0.348331 | 0.158776 | 0.568823 | -0.15139 | -1.41515 | 0.689881 | 0.247877 | -0.50644 | 0.104455 | -0.14396 | -0.07808 | -0.16426 | -0.07968 | -0.30269 | -0.11891 | -0.1429 | 0.010399 | -0.0242 | 0.019979 |
| 35.08333 | 0.431033 | -0.93665 | 0.174979 | 0.351574 | 1.992329 | 0.343813 | 0.107975 | 0.53892 | 0.16111 | -1.66076 | 0.887326 | 0.236602 | -0.35152 | 0.05297 | -0.05629 | -0.00959 | -0.13708 | -0.00289 | -0.32189 | -0.0755 | -0.03981 | 0.058677 | 0.046811 | 0.105112 |
| 35.09722 | 0.495401 | -0.98253 | 0.255928 | 0.483909 | 2.227696 | 0.318899 | 0.015867 | 0.474382 | 0.262592 | -1.57745 | 1.025833 | 0.214135 | -0.2755 | 0.076938 | 0.012667 | 0.048492 | -0.11768 | 0.044042 | -0.4169 | -0.05975 | 0.021854 | 0.084671 | 0.088841 | 0.170127 |
| 35.11111 | 0.539955 | -0.95351 | 0.38647 | 0.608294 | 2.498339 | 0.351957 | -0.10295 | 0.325731 | 0.437487 | -1.32157 | 0.991384 | 0.197894 | -0.29631 | 0.111051 | 0.014279 | 0.066066 | -0.12281 | 0.027449 | -0.51821 | -0.0672 | 0.009308 | 0.081403 | 0.091725 | 0.14237 |
| 35.125 | 0.531646 | -0.94566 | 0.496569 | 0.623201 | 2.863101 | 0.453378 | -0.13333 | 0.233626 | 0.368337 | -1.05024 | 0.961541 | 0.214162 | -0.3212 | 0.145317 | 0.003894 | 0.092894 | -0.11691 | 0.0041 | -0.56025 | -0.04863 | 0.01578 | 0.078653 | 0.103455 | 0.12164 |
| 35.13889 | 0.417493 | -0.72265 | 0.569697 | 0.748693 | 2.814825 | 0.519775 | -0.07391 | 0.297438 | 0.292481 | -0.84896 | 0.796232 | 0.161999 | -0.31121 | 0.143417 | 0.009473 | 0.126531 | -0.09599 | 0.003243 | -0.54298 | -0.01265 | 0.072827 | 0.078059 | 0.153441 | 0.145963 |
| 35.15278 | 0.375415 | -0.41516 | 0.607124 | 0.792921 | 2.738465 | 0.550628 | -0.02861 | 0.379555 | 0.308593 | -0.78215 | 0.739803 | 0.148752 | -0.32596 | 0.110762 | -0.0078 | 0.098173 | -0.09525 | -0.00026 | -0.51928 | -0.02366 | 0.106318 | 0.046072 | 0.128386 | 0.125727 |
| 35.16667 | 0.356458 | -0.14375 | 0.563879 | 0.777803 | 1.912547 | 0.498361 | -0.04955 | 0.354759 | 0.382214 | -0.8352 | 0.670338 | 0.087918 | -0.32346 | 0.060862 | -0.04922 | 0.034968 | -0.12702 | -0.02914 | -0.49958 | -0.08511 | 0.073631 | -0.02225 | 0.08485 | 0.067109 |
| 35.18056 | 0.292936 | -0.07617 | 0.387726 | 0.756549 | 1.096376 | 0.445167 | -0.12498 | 0.228466 | 0.395651 | -1.00779 | 0.632738 | -0.02621 | -0.295 | 0.010153 | -0.07272 | 0.00384 | -0.15305 | -0.04175 | -0.46503 | -0.11897 | 0.04391 | -0.09533 | 0.044144 | 0.008444 |
| 35.19444 | 0.221548 | -0.02958 | 0.202919 | 0.759897 | 0.796699 | 0.358646 | -0.2225 | 0.118539 | 0.313509 | -1.38462 | 0.753129 | -0.07533 | -0.26496 | -0.06481 | -0.04622 | 0.026815 | -0.14022 | -0.00326 | -0.54469 | -0.10628 | 0.087664 | -0.12109 | 0.028796 | -0.06402 |
| 35.20833 | 0.285138 | -0.04159 | 0.11184 | 0.735927 | 0.649626 | 0.317225 | -0.27134 | 0.098129 | 0.364067 | -1.72996 | 0.981517 | -0.03845 | -0.2571 | -0.06509 | -0.02065 | 0.054682 | -0.11429 | 0.029903 | -0.70869 | -0.07556 | 0.177256 | -0.11139 | 0.050376 | -0.11899 |
| 35.22222 | 0.372378 | -0.07488 | 0.052473 | 0.421261 | 0.01902 | 0.25697 | -0.24748 | 0.105578 | 0.44266 | -1.82535 | 1.14568 | 0.004191 | -0.29465 | 0.006713 | -0.01726 | 0.058804 | -0.10288 | 0.041921 | -0.74389 | -0.04865 | 0.245504 | -0.08353 | 0.103177 | -0.06878 |
| 35.23611 | 0.413898 | -0.18374 | -0.04698 | 0.196262 | -0.5143 | 0.275768 | -0.21453 | 0.132566 | 0.508275 | -1.57928 | 1.251629 | -0.03567 | -0.37122 | 0.036118 | -0.03278 | 0.038069 | -0.10564 | 0.039807 | -0.68349 | -0.04835 | 0.269232 | -0.04379 | 0.119295 | 0.011955 |
| 35.25 | 0.449183 | -0.30636 | -0.1298 | 0.195143 | -0.36139 | 0.28526 | -0.17214 | 0.178208 | 0.54105 | -1.25567 | 1.225756 | -0.09321 | -0.46529 | -0.00973 | -0.05912 | 0.010397 | -0.12371 | -0.00189 | -0.68631 | -0.06938 | 0.246969 | -0.02419 | 0.061481 | 0.004064 |
| 35.26389 | 0.467124 | -0.37794 | -0.21154 | 0.135757 | -0.24036 | 0.289421 | -0.17326 | 0.163849 | 0.578773 | -1.01518 | 1.094636 | -0.12342 | -0.57028 | -0.09296 | -0.08804 | -0.04245 | -0.16239 | -0.06566 | -0.7185 | -0.09748 | 0.18844 | -0.04388 | -0.01797 | -0.05404 |
| 35.27778 | 0.459465 | -0.46567 | -0.2501 | -0.00465 | -0.56526 | 0.294225 | -0.17068 | 0.133969 | 0.74911 | -0.85204 | 1.052954 | -0.13224 | -0.56715 | -0.12056 | -0.10939 | -0.07625 | -0.19861 | -0.08546 | -0.74468 | -0.12416 | 0.142857 | -0.08607 | -0.06145 | -0.08669 |
| 35.29167 | 0.483035 | -0.5418 | -0.19487 | -0.0537 | -0.62893 | 0.383982 | -0.09841 | 0.212904 | 0.914097 | -0.87259 | 1.046366 | -0.13733 | -0.52579 | -0.11151 | -0.11266 | -0.0771 | -0.22429 | -0.07499 | -0.73797 | -0.11804 | 0.141632 | -0.10341 | -0.05386 | -0.06719 |
| 35.30556 | 0.534965 | -0.64052 | -0.14532 | 0.025745 | -0.28728 | 0.431338 | -0.02137 | 0.429914 | 0.843043 | -0.90775 | 1.156251 | -0.1501 | -0.51783 | -0.08756 | -0.11947 | -0.08438 | -0.23185 | -0.08234 | -0.7332 | -0.0891 | 0.129465 | -0.10239 | -0.04157 | -0.04262 |
| 35.31944 | 0.660922 | -0.74284 | -0.1603 | 0.039758 | 0.039322 | 0.437393 | 0.019261 | 0.637513 | 0.852152 | -0.90475 | 1.285766 | -0.14216 | -0.50677 | -0.06032 | -0.11993 | -0.0968 | -0.21749 | -0.0892 | -0.70049 | -0.07192 | 0.146173 | -0.0862 | -0.03908 | -0.04021 |
| 35.33333 | 0.751514 | -0.8408 | -0.14271 | 0.021609 | 0.214108 | 0.391404 | 0.070467 | 0.664121 | 0.556775 | -0.79089 | 1.510095 | -0.07216 | -0.42144 | -0.00501 | -0.05587 | -0.03795 | -0.16175 | -0.02273 | -0.57031 | -0.02082 | 0.215615 | -0.02248 | 0.017844 | 0.014847 |
| 35.34722 | 0.803278 | -0.93414 | -0.04012 | -0.04873 | 0.677846 | 0.238946 | 0.066872 | 0.681818 | 0.088417 | -0.69302 | 1.918177 | 0.258924 | -0.39799 | 0.029735 | -0.02028 | -0.01095 | -0.14408 | 0.015834 | -0.57028 | -0.00567 | 0.243124 | 0.016324 | 0.048598 | 0.083822 |
| 35.36111 | 0.896773 | -0.91746 | 0.09384 | 0.091106 | 1.490777 | 0.189973 | -0.02209 | 0.663242 | -0.16609 | -0.69721 | 2.112981 | 0.241497 | -0.54201 | 0.006352 | -0.14143 | -0.13666 | -0.26676 | -0.12312 | -0.78954 | -0.13053 | 0.107396 | -0.07116 | -0.04489 | 0.034743 |
| 35.375 | 1.070725 | -0.37689 | 0.173465 | 0.543839 | 2.035504 | 0.414587 | -0.1019 | 0.379851 | 0.877057 | -0.79688 | 1.261961 | -0.74466 | -0.69255 | 0.006757 | -0.27384 | -0.24174 | -0.37594 | -0.26149 | -0.93358 | -0.23635 | -0.03609 | -0.18599 | -0.12506 | -0.0419 |
| 35.38889 | 1.488426 | -0.02514 | 0.525713 | 0.905818 | 2.709357 | 0.456742 | -0.12464 | 0.567486 | 1.035673 | -0.54864 | 1.326901 | -0.98137 | -0.63317 | 0.055746 | -0.24726 | -0.21863 | -0.32314 | -0.22496 | -0.91338 | -0.17337 | 0.011577 | -0.17937 | -0.07763 | -0.03453 |
| 35.40278 | 1.732163 | -0.6976 | 1.356705 | 0.488669 | 4.099556 | -0.07939 | 0.014389 | 1.745144 | -1.4635 | 0.50785 | 3.357545 | 0.587265 | -0.48 | 0.085401 | -0.11738 | -0.15929 | -0.11054 | -0.10479 | -0.67299 | -0.0305 | 0.252828 | -0.08009 | 0.043554 | -0.01344 |
| 35.41667 | 1.437403 | -3.68711 | 2.645642 | -1.58089 | 6.399976 | -2.73188 | -0.03841 | 3.260598 | -7.16245 | 0.448233 | 8.192095 | 4.060876 | -0.31015 | 0.202835 | 0.013534 | -0.12237 | 0.170442 | -0.01782 | -0.14979 | 0.079689 | 0.567947 | 0.02962 | 0.230275 | 0.028169 |
| 35.43056 | 1.045684 | -13.6441 | 4.926499 | -4.59951 | 7.701188 | -2.83442 | 1.176752 | 1.061395 | -6.39806 | 0.967454 | 11.39029 | 5.169963 | -0.1 | 0.232507 | 0.004326 | -0.07801 | 0.248271 | -0.05185 | 0.420676 | 0.032931 | 0.656619 | 0.062013 | 0.350228 | 0.071045 |
| 35.44444 | 1.630467 | NaN | 0.837862 | 0.230724 | 1.841161 | 0.041726 | 2.338915 | 3.368873 | 0.997261 | 1.154025 | 3.943658 | -1.67412 | -0.00376 | 0.06673 | -0.15843 | -0.04932 | 0.047409 | -0.16918 | 0.544851 | -0.21279 | 0.327974 | 0.000376 | 0.262332 | 0.079745 |
| 35.45833 | 0.99057 | 11.47655 | 0.179505 | 2.48764 | -4.46391 | 5.349113 | 1.738646 | -0.23299 | 7.804894 | 5.21948 | -5.7902 | -5.72255 | -0.09046 | -0.13006 | -0.29496 | -0.04712 | -0.19528 | -0.20167 | 0.353986 | -0.3833 | 0.076511 | -0.07287 | 0.02874 | -0.02773 |
| 35.47222 | 0.962738 | 3.431101 | -0.78831 | 4.135095 | -1.16477 | 2.156323 | 1.386404 | -1.35783 | 7.516228 | 2.806571 | -3.01479 | -3.00421 | -0.26187 | -0.26871 | -0.31452 | -0.04673 | -0.26835 | -0.15271 | 0.266903 | -0.29097 | 0.030547 | -0.09356 | -0.2257 | -0.08408 |
| 35.48611 | 1.167242 | 1.354448 | 1.198343 | 2.554108 | 2.369986 | 0.816487 | 1.232464 | 0.697119 | 1.502416 | 1.040994 | 1.803437 | -0.51423 | -0.26956 | -0.22104 | -0.19597 | 0.027346 | -0.16898 | -0.03924 | 0.281039 | -0.10055 | 0.110432 | -0.01189 | -0.2113 | 0.021821 |
| 35.5 | 1.31354 | 0.802598 | 1.687031 | 1.281019 | 1.938523 | 0.937521 | 0.992012 | 1.487133 | -0.14917 | 0.341395 | 2.861892 | 0.249818 | -0.12971 | -0.01668 | -0.01546 | 0.089254 | -0.02608 | 0.091644 | 0.262087 | 0.037963 | 0.267274 | 0.094001 | -0.03571 | 0.162533 |
| 35.51389 | 1.382085 | 1.070974 | 1.592906 | 0.322624 | -0.55283 | 1.075978 | 0.851046 | 1.530529 | -0.19997 | 0.238777 | 2.377264 | 0.362788 | -0.07876 | 0.104868 | 0.059904 | 0.10094 | 0.035571 | 0.112445 | 0.192312 | 0.082187 | 0.32431 | 0.105021 | 0.080749 | 0.150416 |
| 35.52778 | 1.376215 | 1.432321 | 1.125602 | -0.28369 | -2.72378 | 0.931419 | 0.685963 | 1.298687 | 0.20769 | 0.337327 | 1.375909 | 0.074119 | -0.07723 | 0.101294 | 0.054232 | 0.092375 | 0.013828 | 0.071346 | 0.105413 | 0.058893 | 0.286571 | 0.086584 | 0.050766 | 0.042787 |
| 35.54167 | 1.326782 | 1.686527 | 0.786995 | -0.73213 | -3.83058 | 0.674735 | 0.619552 | 1.208255 | 0.653628 | 0.201136 | 0.749969 | -0.18441 | -0.05734 | 0.04563 | 0.043658 | 0.090697 | -0.01572 | 0.046688 | 0.025297 | 0.039964 | 0.246617 | 0.084753 | 0.037203 | 0.035197 |
| 35.55556 | 1.23152 | 1.554323 | 0.67621 | -1.06989 | -4.68969 | 0.663432 | 0.378788 | 0.895583 | 0.436613 | -0.05499 | 0.601225 | -0.15451 | -0.03512 | -0.01218 | 0.050248 | 0.08075 | -0.0267 | 0.045994 | -0.01758 | 0.052295 | 0.23523 | 0.078018 | 0.110908 | 0.098753 |
| 35.56944 | 1.19879 | 1.426388 | 0.606352 | -1.30408 | -3.72295 | 0.758929 | 0.197176 | 1.008558 | 0.31368 | -0.43639 | 0.987466 | 0.110369 | -0.0175 | -0.03602 | 0.068582 | 0.074564 | -0.01715 | 0.053867 | -0.02239 | 0.076407 | 0.246443 | 0.055276 | 0.189696 | 0.120399 |
| 35.58333 | 1.14176 | 2.202012 | 0.496353 | -1.34408 | -3.44329 | 0.901346 | 0.261704 | 0.988547 | 0.690832 | -0.83465 | 1.209659 | -0.10299 | 0.016516 | 0.011956 | 0.093544 | 0.093501 | 0.029264 | 0.084512 | 0.018175 | 0.120851 | 0.253272 | 0.053946 | 0.239224 | 0.141506 |
| 35.59722 | 0.956179 | 3.828664 | 0.375797 | -1.27798 | -2.88621 | 0.893788 | 0.271322 | 0.715476 | 0.487008 | -1.17851 | 1.352824 | -0.30918 | 0.033782 | 0.093224 | 0.099772 | 0.121294 | 0.084097 | 0.102275 | 0.052692 | 0.16265 | 0.245793 | 0.058611 | 0.239357 | 0.171262 |
| 35.61111 | 0.829079 | 4.885248 | 0.357872 | -0.88721 | -1.15982 | 0.924768 | 0.1418 | 0.705989 | -0.32063 | -1.2996 | 1.793362 | -0.09626 | 0.009672 | 0.137017 | 0.077558 | 0.113839 | 0.109695 | 0.090552 | 0.050753 | 0.162171 | 0.209459 | 0.038297 | 0.195137 | 0.163893 |
| 35.625 | 0.79903 | 4.179708 | 0.342693 | -0.11016 | 0.230074 | 0.922049 | 0.148176 | 0.742966 | -0.45828 | -1.12604 | 1.765806 | -0.19515 | -0.01615 | 0.134071 | 0.045967 | 0.100311 | 0.110309 | 0.080776 | 0.072406 | 0.142183 | 0.187348 | 0.046674 | 0.154323 | 0.138067 |

| | | | | | | | | | | | | | | | | | | | | | | | | |
|---|---|---|---|---|---|---|---|---|---|---|---|---|---|---|---|---|---|---|---|---|---|---|---|---|
| 35.63889 | 1.070432 | 3.524758 | 0.405698 | 0.543723 | 1.125734 | 0.922087 | 0.154714 | 0.767643 | -0.40353 | -0.80665 | 1.606486 | -0.45233 | -0.03854 | 0.095921 | 0.00985 | 0.092301 | 0.077453 | 0.066225 | 0.093917 | 0.099483 | 0.162099 | 0.065246 | 0.117214 | 0.087153 |
| 35.65278 | 1.185795 | 2.885529 | 0.497651 | 0.938029 | 1.682038 | 1.035675 | 0.181629 | 0.751275 | -0.17286 | -0.53816 | 1.522201 | -0.73206 | -0.05318 | 0.064791 | -0.02654 | 0.057061 | 0.045269 | 0.04444 | 0.080554 | 0.044525 | 0.128832 | 0.040642 | 0.107232 | 0.043421 |
| 35.66667 | 1.344128 | 2.777301 | 0.484958 | 1.224289 | 1.176344 | 0.889034 | 0.340665 | 0.317029 | 0.367377 | -0.23544 | 0.846601 | -1.1213 | -0.0656 | 0.016825 | -0.03098 | 0.028269 | 0.033583 | 0.03742 | 0.059635 | 0.005149 | 0.104589 | -0.0046 | 0.116415 | 0.014837 |
| 35.68056 | 1.309296 | 2.091105 | 0.472542 | 0.808736 | 0.191694 | 0.906791 | 0.329783 | 0.11673 | 0.484726 | -0.25577 | 0.672685 | -1.00599 | -0.09573 | -0.01114 | -0.01672 | 0.027943 | 0.008424 | 0.008019 | 0.048566 | -0.01767 | 0.0765 | -0.03601 | 0.128709 | 0.032065 |
| 35.69444 | 1.136159 | 1.149703 | 0.524583 | -0.02012 | -0.95413 | 1.085642 | 0.186317 | 0.173939 | 0.355031 | -0.54329 | 0.936798 | -0.76726 | -0.13503 | 0.100701 | -0.00253 | 0.052731 | -0.01557 | -0.03228 | 0.004975 | -0.00855 | 0.018769 | -0.05437 | 0.119083 | 0.099368 |
| 35.70833 | 1.235076 | 0.92844 | 0.515857 | -0.61645 | -2.09842 | 1.089517 | 0.127587 | -0.17203 | 0.519848 | -0.84749 | 0.896984 | -0.70413 | -0.1613 | 0.160297 | 0.039684 | 0.094044 | -0.00195 | -0.0409 | -0.04806 | 0.035663 | -0.00719 | -0.05115 | 0.112488 | 0.126964 |
| 35.72222 | 1.55428 | 1.08988 | 0.488633 | -0.76891 | -2.21699 | 0.652988 | 0.031029 | -0.76726 | 0.589945 | -1.05458 | 0.618297 | -0.59706 | -0.18828 | 0.179314 | 0.091727 | 0.107051 | 0.005122 | -0.03002 | -0.1053 | 0.044576 | 0.000015 | -0.05265 | 0.090211 | 0.054695 |
| 35.73611 | 1.792134 | 1.458745 | 0.491946 | -0.67393 | -2.53596 | 0.238041 | -0.1772 | -0.81358 | 0.551069 | -1.01612 | 0.534453 | -0.53807 | -0.27158 | 0.089295 | 0.047045 | 0.014671 | -0.07994 | -0.08524 | -0.25199 | -0.04485 | -0.091 | -0.13695 | -0.03313 | -0.05001 |
| 35.75 | 1.684789 | 1.473142 | 0.643566 | -0.70547 | -3.08211 | -0.0657 | -0.38679 | -0.79332 | 0.646041 | -0.9218 | 0.544791 | -0.5288 | -0.32919 | 0.12132 | -0.01454 | -0.03324 | -0.12921 | -0.11714 | -0.39014 | -0.0988 | -0.1464 | -0.19387 | -0.09652 | -0.07658 |
| 35.76389 | 1.340531 | 0.926808 | 0.737316 | -1.02192 | -3.68087 | -0.11722 | -0.61045 | -0.76157 | 0.731755 | -0.84156 | 0.552903 | -0.2912 | -0.3224 | 0.312812 | -0.01581 | 0.014627 | -0.08246 | -0.07527 | -0.40849 | -0.05699 | -0.10301 | -0.14773 | -0.06907 | -0.08019 |
| 35.77778 | 1.130909 | 0.245645 | 0.78049 | -1.11013 | -3.87354 | -0.07811 | -0.69311 | -0.70026 | 0.455849 | -0.80953 | 0.518057 | 0.047125 | -0.3436 | 0.324918 | -0.04235 | -0.00516 | -0.08867 | -0.09073 | -0.45987 | -0.08502 | -0.11469 | -0.17611 | -0.09546 | -0.14888 |
| 35.79167 | 1.035571 | 0.03878 | 0.722215 | -0.68426 | -3.88806 | -0.07605 | -0.52081 | -0.82382 | 0.442878 | -0.85267 | 0.273514 | 0.146721 | -0.35943 | 0.150251 | -0.07945 | -0.04545 | -0.10322 | -0.116 | -0.47074 | -0.13469 | -0.16372 | -0.205 | -0.06098 | -0.1841 |
| 35.80556 | 0.991267 | 0.04722 | 0.578072 | -0.56142 | -3.53967 | -0.09695 | -0.3147 | -0.72575 | 0.60399 | -0.76786 | 0.008228 | 0.129668 | -0.3066 | 0.107713 | -0.03576 | 0.005852 | -0.04938 | -0.05759 | -0.34435 | -0.08185 | -0.12781 | -0.10901 | 0.013646 | -0.11879 |
| 35.81944 | 0.829214 | -0.01117 | 0.479703 | -0.67007 | -3.3308 | 0.049382 | -0.20336 | -0.62936 | 0.278682 | -0.6617 | -0.05252 | 0.167865 | -0.29915 | -0.07262 | -0.0058 | 0.015785 | -0.04067 | -0.0485 | -0.27013 | -0.04182 | -0.15745 | -0.05886 | -0.01051 | -0.11094 |
| 35.83333 | 0.694256 | -0.04755 | 0.428357 | -0.53528 | -2.70598 | 0.225167 | -0.01097 | -0.64211 | -0.19557 | -0.55834 | -0.07473 | 0.240511 | -0.35767 | -0.12131 | -0.05272 | -0.04179 | -0.1086 | -0.14548 | -0.34416 | -0.12436 | -0.28436 | -0.12696 | -0.14233 | -0.1901 |
| 35.84722 | 0.661599 | -0.00047 | 0.561294 | -0.19725 | -1.61515 | 0.39776 | 0.295101 | -0.66737 | -0.19187 | -0.42076 | -0.03156 | 0.272472 | -0.33696 | -0.19977 | -0.07848 | -0.07139 | -0.12626 | -0.18267 | -0.34144 | -0.16574 | -0.35511 | -0.11507 | -0.21565 | -0.21343 |
| 35.86111 | 0.7266 | 0.176107 | 0.703526 | 0.126613 | -0.95435 | 0.700902 | 0.534373 | -0.4898 | 0.056133 | -0.4848 | 0.018041 | 0.306734 | -0.2252 | -0.12271 | -0.0247 | -0.01865 | -0.05429 | -0.09753 | -0.24863 | -0.05808 | -0.2892 | 0.014301 | -0.12098 | -0.09981 |
| 35.875 | 0.700178 | 0.376343 | 0.71196 | 0.227545 | -0.94648 | 0.759108 | 0.665057 | -0.37385 | 0.537517 | -0.68844 | -0.15507 | -0.06794 | -0.11131 | -0.04835 | 0.010175 | 0.011159 | 0.011305 | -0.0309 | -0.10041 | 0.01913 | -0.19636 | 0.091976 | -0.06998 | -0.05678 |
| 35.88889 | 0.47719 | 0.455309 | 0.156611 | 0.34396 | -1.70035 | 0.398865 | 0.673141 | -0.72454 | 1.245086 | -1.33827 | -0.73192 | -1.55053 | -0.03631 | -0.34634 | -0.13223 | -0.1509 | -0.07015 | -0.11047 | -0.12181 | -0.14581 | -0.12473 | 0.007459 | -0.18993 | -0.24194 |
| 35.90278 | -0.40565 | -0.01151 | -0.44965 | -0.01695 | -2.49039 | -0.18624 | 0.421239 | -1.26914 | 1.368053 | -2.44416 | -1.21476 | -2.43786 | 0.034134 | -1.02034 | -0.34765 | -0.45699 | -0.26606 | -0.2255 | -0.15015 | -0.43562 | -0.13629 | -0.14341 | -0.42461 | -0.56801 |
| 35.91667 | -1.25151 | -1.53541 | -0.10017 | -1.11471 | -2.80856 | 0.464286 | -0.16322 | -1.00968 | 0.097023 | -2.68292 | -0.0529 | 0.11471 | -0.08503 | -1.58447 | -0.38032 | -0.45616 | -0.34088 | -0.15665 | -0.51447 | -0.69707 | -0.22217 | -0.24649 | -0.51288 | -0.73174 |
| 35.93056 | -1.59409 | -2.77602 | 0.462215 | -0.44447 | -2.9687 | 0.258409 | -0.57151 | -0.7741 | -0.57739 | -2.47305 | 1.10197 | 2.203588 | -0.34127 | -1.32374 | -0.09653 | -0.12199 | -0.17822 | 0.112828 | -1.25741 | -0.94279 | -0.16575 | -0.14447 | -0.36707 | -0.50838 |
| 35.94444 | -1.46507 | -1.91238 | 0.016197 | 0.602561 | -4.40409 | 0.468182 | -0.4355 | -1.26956 | 0.356641 | -4.17018 | 0.425814 | -0.75525 | -0.27851 | -0.37417 | 0.340671 | 0.282677 | 0.173893 | 0.305813 | -1.85022 | -0.48277 | 0.254571 | 0.121043 | -0.01749 | 0.125939 |
| 35.95833 | -1.13896 | -0.65407 | -0.3021 | 0.792017 | -5.08012 | -1.04138 | -0.31692 | -1.55586 | 1.405218 | -5.47204 | -0.2749 | -3.10015 | -0.03742 | 0.114164 | 0.435087 | 0.303818 | 0.244476 | 0.197514 | -1.42743 | 0.498211 | 0.30153 | 0.299326 | 0.120943 | 0.514214 |
| 35.97222 | -1.26814 | -1.50035 | 0.031317 | 0.724117 | -3.76041 | -0.22942 | -0.32103 | -0.62897 | 0.333527 | -4.77034 | 0.976943 | -0.50077 | 0.094809 | 0.063275 | 0.164328 | 0.025896 | 0.020257 | -0.04625 | -0.65121 | 0.378584 | -0.08394 | 0.238496 | 0.044772 | 0.362842 |
| 35.98611 | -1.71875 | -2.79934 | 0.464388 | 0.638436 | -4.04987 | -0.08681 | -0.4088 | 0.417408 | -1.5207 | -2.92005 | 2.323389 | 2.700784 | 0.17443 | -0.08673 | 0.026041 | -0.07889 | -0.11743 | -0.09579 | -0.18891 | 0.170593 | -0.21854 | 0.152448 | 0.138315 | 0.15766 |
| 36 | -1.35795 | -1.89264 | 0.032361 | 0.700277 | -2.20601 | 0.451974 | -0.21298 | 0.313496 | -0.31224 | -2.27774 | 1.256906 | 1.187132 | 0.199567 | 0.04861 | 0.053663 | -0.01549 | -0.08974 | -0.02863 | -0.01223 | 0.13532 | -0.12484 | 0.160752 | 0.314586 | 0.155099 |
| 36.01389 | -0.42046 | -0.13674 | -0.40009 | 0.732925 | -0.81025 | 1.850163 | 0.059256 | -0.30317 | 0.871592 | -2.35957 | -0.16088 | -2.39599 | 0.11062 | 0.162999 | 0.05464 | 0.036386 | -0.05542 | -0.00552 | -0.0414 | 0.045094 | -0.07615 | 0.151608 | 0.409409 | 0.13879 |
| 36.02778 | -0.27659 | 0.39941 | -0.1946 | 0.84841 | -0.48874 | 1.72198 | 0.057445 | -0.00278 | 1.315195 | -2.11629 | -0.09245 | -2.11046 | -0.00191 | 0.28993 | 0.027216 | 0.064648 | -0.06168 | -0.01536 | -0.15834 | -0.03584 | -0.07121 | 0.091298 | 0.347971 | 0.07687 |
| 36.04167 | -0.3983 | -0.17859 | -0.11552 | 0.691059 | -0.14625 | 0.861502 | 0.087568 | 0.345381 | 0.80556 | -1.77361 | 0.08779 | -1.11726 | -0.08815 | 0.320816 | 0.005358 | 0.072651 | -0.05465 | -0.00133 | -0.25166 | -0.04269 | -0.05539 | 0.063332 | 0.243093 | 0.042401 |
| 36.05556 | -0.59257 | -0.98472 | -0.12197 | 0.542447 | -0.05428 | 0.41463 | 0.066452 | 0.687895 | 0.283576 | -1.8897 | 0.582884 | 0.080881 | -0.1894 | 0.269147 | -0.02925 | 0.030572 | -0.05795 | -0.00936 | -0.3733 | -0.0607 | -0.06554 | 0.039092 | 0.147244 | 0.013861 |
| 36.06944 | -0.65409 | -3.97834 | 0.050507 | 0.603806 | -0.8553 | -0.71948 | 0.057962 | 1.6531 | -1.36274 | -0.97433 | 2.05202 | 3.714728 | -0.29134 | 0.045469 | -0.08593 | -0.05855 | -0.11032 | -0.07927 | -0.55085 | -0.13464 | -0.13296 | -0.00051 | 0.028247 | -0.05032 |
| 36.08333 | -0.33214 | -4.44428 | 0.38357 | 0.674894 | -1.47702 | -0.00891 | 0.479955 | 2.278427 | -1.13049 | -0.55951 | 2.005138 | 4.647708 | -0.31183 | -0.0835 | -0.0888 | -0.10553 | -0.14215 | -0.11625 | -0.64399 | -0.15158 | -0.18465 | 0.01986 | -0.0077 | -0.06546 |
| 36.09722 | 0.078913 | -2.40038 | 0.478233 | 0.819871 | -0.074 | 0.880888 | 0.906307 | 1.914063 | 0.428643 | -1.04442 | 1.331929 | 1.866597 | -0.25223 | -0.05892 | -0.02116 | -0.07463 | -0.11234 | -0.07935 | -0.62431 | -0.08594 | -0.15697 | 0.092852 | 0.058467 | -0.01711 |
| 36.11111 | 0.326907 | -0.83835 | 0.447816 | 0.824447 | 1.258692 | 1.548923 | 0.88128 | 1.380241 | 1.197874 | -1.38565 | 0.681187 | -0.27646 | -0.18261 | 0.01645 | 0.032549 | -0.01663 | -0.07167 | -0.03583 | -0.56148 | -0.03667 | -0.11406 | 0.126965 | 0.120494 | -0.01677 |
| 36.125 | 0.578633 | 0.312035 | 0.497764 | 0.770995 | 1.896898 | 1.271447 | 0.696921 | 0.911402 | 1.515482 | -1.01668 | 0.235787 | -1.60217 | -0.15649 | 0.098392 | 0.047444 | 0.027855 | -0.05091 | -0.01033 | -0.52308 | -0.01455 | -0.07687 | 0.13727 | 0.138144 | -0.01353 |
| 36.13889 | 1.130857 | 2.165003 | 0.557375 | 0.433946 | 2.497675 | 1.240951 | 0.749942 | 0.226307 | 1.71355 | -0.06076 | -0.32993 | -3.63223 | -0.19914 | 0.133922 | 0.024552 | 0.019823 | -0.07265 | -0.03721 | -0.55248 | -0.04297 | -0.08098 | 0.0951 | 0.089897 | -0.01432 |
| 36.15278 | 1.036533 | 1.957655 | 0.69504 | 0.4668 | 2.368619 | 1.433795 | 0.630487 | 0.447921 | 1.15912 | 0.324797 | -0.38726 | -3.01695 | -0.27675 | 0.138447 | -0.01053 | -0.0368 | -0.10859 | -0.09628 | -0.61632 | -0.10161 | -0.12785 | 0.020715 | 0.030509 | -0.05327 |
| 36.16667 | 0.533124 | -0.91009 | 0.943298 | 0.902665 | 1.385865 | 0.716461 | 0.277393 | 1.660889 | 1.46666 | -0.00896 | 1.185514 | 1.182994 | -0.28413 | 0.146926 | 0.019666 | -0.03742 | -0.08247 | -0.08258 | -0.58418 | -0.07353 | -0.08154 | 0.027323 | 0.069241 | 0.001353 |
| 36.18056 | 0.329009 | -1.99601 | 0.924618 | 0.87843 | 0.555956 | 0.033269 | -0.0068 | 2.028703 | 1.641581 | -0.31791 | 1.738624 | 2.916334 | -0.18411 | 0.168326 | 0.101404 | 0.037529 | -0.00234 | 0.011782 | -0.53683 | 0.021445 | 0.053378 | 0.113176 | 0.177075 | 0.135623 |
| 36.19444 | 0.546128 | 0.39696 | 0.502161 | 0.713963 | 1.648512 | 0.802227 | -0.04763 | 1.070165 | 1.179101 | -0.6145 | 0.527403 | -0.40567 | -0.14447 | 0.153642 | 0.118921 | 0.060286 | 0.017912 | 0.045791 | -0.5611 | 0.02508 | 0.088035 | 0.091675 | 0.183595 | 0.127148 |
| 36.20833 | 0.856915 | 2.697713 | 0.11224 | 0.664465 | 3.008434 | 0.568985 | -0.0599 | 0.152225 | 0.447878 | -0.95048 | -0.27411 | -3.36255 | -0.20896 | 0.029312 | 0.02687 | -0.02039 | -0.05358 | -0.04307 | -0.70118 | -0.07475 | -0.02237 | -0.03474 | 0.072607 | -0.01077 |
| 36.22222 | 0.5215 | 1.773694 | 0.115837 | 0.811528 | 3.402003 | 1.275682 | -0.14518 | 0.395532 | 0.282589 | -1.52629 | -0.19573 | -2.56322 | -0.23194 | -0.02718 | -0.02674 | -0.05338 | -0.0922 | -0.09277 | -0.71414 | -0.12043 | -0.08156 | -0.08578 | 0.021968 | -0.08069 |
| 36.23611 | -0.03155 | 0.268051 | 0.089339 | 0.694005 | 2.402089 | 0.716299 | -0.19918 | 0.919504 | 0.51074 | -2.46759 | 0.560297 | -0.59965 | -0.1619 | 0.046005 | 0.01653 | -0.00633 | -0.05026 | -0.02802 | -0.58462 | -0.06116 | -0.00948 | -0.03592 | 0.044005 | -0.04626 |
| 36.25 | -0.25605 | -0.03521 | -0.15075 | 0.576484 | 1.349951 | 0.406571 | -0.1688 | 1.000607 | 0.435847 | -2.80914 | 0.596493 | -0.21996 | -0.11292 | 0.061907 | 0.05261 | 0.028851 | -0.0215 | 0.029745 | -0.40864 | -0.00479 | 0.047744 | 0.004458 | 0.091879 | -0.01017 |
| 36.26389 | -0.65707 | -1.08411 | -0.47776 | 0.286665 | 0.310036 | -0.02278 | -0.16753 | 1.399826 | 0.665564 | -3.37096 | 1.010924 | 1.114981 | -0.13779 | 0.010068 | 0.032315 | 0.022563 | -0.03459 | 0.033491 | -0.31903 | -0.00922 | 0.041984 | 0.013286 | 0.092748 | -0.01954 |
| 36.27778 | -1.11704 | -3.10286 | -0.90359 | -0.62323 | -1.69617 | -0.45215 | -0.18308 | 2.160065 | 1.622604 | -4.57389 | 1.698241 | 4.064064 | -0.17372 | -0.02566 | -0.02155 | 0.004978 | -0.06401 | -0.0023 | -0.361 | -0.06482 | 0.015381 | -0.00404 | 0.033255 | -0.04272 |
| 36.29167 | -0.65134 | -2.24781 | -0.64262 | -0.6829 | -2.33425 | -0.64474 | -0.05361 | 2.094914 | 1.489125 | -3.75267 | 1.31925 | 3.039317 | -0.205 | -0.05131 | -0.06826 | -0.01928 | -0.0798 | -0.03836 | -0.58035 | -0.1174 | -0.00066 | -0.02504 | 0.000663 | -0.05166 |
| 36.30556 | -0.04479 | 0.525326 | -0.06377 | 0.516598 | -0.41124 | 1.05713 | 0.094581 | 0.704355 | -0.24254 | -1.61895 | -0.1563 | -1.50351 | -0.22531 | -0.05655 | -0.06106 | -0.02424 | -0.07534 | -0.04077 | -1.05454 | -0.11928 | -0.00133 | -0.04104 | 0.003815 | -0.0402 |
| 36.31944 | 0.208668 | 1.391184 | 0.061803 | 1.084956 | 0.567866 | 1.870677 | 0.182667 | 0.267081 | -0.89576 | -0.76497 | -0.60454 | -3.31783 | -0.24795 | -0.02103 | -0.02505 | -0.01356 | -0.07171 | -0.02996 | -1.31528 | -0.09918 | 0.012599 | -0.04241 | 0.032718 | -0.00952 |
| 36.33333 | -0.11828 | 0.012029 | -0.19492 | 0.616953 | 0.486582 | 1.175738 | 0.13661 | 0.662423 | -0.23255 | -0.88532 | -0.37656 | -1.87318 | -0.26027 | -0.06259 | 0.009788 | 0.0028 | -0.06786 | -0.01629 | -1.17053 | -0.07191 | 0.03925 | -0.01225 | 0.068266 | 0.027652 |
| 36.34722 | -0.57346 | -1.36247 | -0.50795 | 0.175704 | 0.176112 | 0.730869 | 0.101936 | 1.028796 | 0.333136 | -1.30474 | 0.304192 | -0.2437 | -0.28399 | -0.09235 | -0.00159 | 0.005294 | -0.08799 | -0.02413 | -1.02507 | -0.07559 | 0.032939 | -0.01715 | 0.054379 | 0.064067 |
| 36.36111 | -0.60213 | -2.06568 | -0.67312 | -0.05671 | 0.141581 | 0.400234 | 0.084977 | 1.022524 | 0.531326 | -0.99709 | 0.691629 | 0.446128 | -0.40477 | -0.13828 | -0.08068 | -0.04825 | -0.16899 | -0.10749 | -0.97133 | -0.14389 | -0.05855 | -0.12632 | -0.02145 | 0.030462 |
| 36.375 | -0.54561 | -2.43588 | -0.43804 | 0.033505 | 0.284718 | 0.534029 | 0.242578 | 0.77851 | 0.827275 | -0.60549 | 1.077523 | 1.025305 | -0.54543 | -0.12126 | -0.17743 | -0.12919 | -0.27175 | -0.23611 | -0.90497 | -0.2315 | -0.15842 | -0.23829 | -0.11926 | -0.07203 |
| 36.38889 | -0.71296 | -3.49024 | 0.089319 | -0.20656 | 0.131249 | -0.10493 | 0.521484 | 0.615048 | 1.503968 | -0.69679 | 2.0487 | 2.447256 | -0.54082 | -0.03129 | -0.17409 | -0.13407 | -0.24636 | -0.25557 | -0.74642 | -0.239 | -0.14197 | -0.21693 | -0.11247 | -0.11652 |
| 36.40278 | -0.58083 | -3.4672 | 0.520341 | -0.22284 | 0.563552 | 0.241615 | 0.537923 | 0.551463 | 1.702924 | 0.159144 | 2.802364 | 2.60818 | -0.36681 | 0.026192 | -0.06324 | -0.0483 | -0.01289 | -0.14485 | -0.31556 | -0.14093 | 0.048421 | 0.011902 | 0.027305 | -0.05208 |
| 36.41667 | -0.56559 | -2.88011 | 0.708187 | 0.024308 | 1.417742 | -0.15847 | 0.550423 | 0.817988 | 1.358826 | 2.104274 | 2.751607 | 1.1041 | -0.20411 | 0.031568 | -0.00902 | 0.038924 | 0.239126 | -0.05165 | 0.407051 | -0.02003 | 0.296235 | 0.247952 | 0.191665 | 0.057118 |
| 36.43056 | -0.61213 | -2.60157 | 0.514085 | 0.126399 | 1.250334 | 0.533152 | 0.798482 | 1.524681 | 0.604237 | 6.889932 | 2.561696 | -0.02093 | -0.10104 | 0.036606 | -0.13014 | -0.04641 | 0.169173 | -0.11137 | 1.016505 | -0.01001 | 0.290386 | 0.15621 | 0.181829 | 0.080899 |
| 36.44444 | -0.1727 | -1.16167 | 0.254501 | 0.649822 | 2.446749 | 1.023705 | 1.496175 | 2.354582 | 0.026383 | 12.15802 | 2.373704 | -0.60805 | -0.05199 | -0.02879 | -0.40786 | -0.1792 | -0.12271 | -0.2441 | 1.067754 | -0.14974 | 0.014154 | -0.05988 | 0.00892 | -0.02803 |
| 36.45833 | 0.542508 | 0.037452 | 0.421401 | 1.798174 | 4.366897 | 1.673172 | 1.574761 | 2.322945 | -0.44801 | 7.985423 | 2.285608 | -0.90882 | 0.038014 | -0.05861 | -0.41798 | -0.08983 | -0.16991 | -0.14192 | 0.858329 | -0.12617 | -0.007 | -0.0737 | 0.018322 | 0.019865 |
| 36.47222 | 0.575953 | -0.29086 | 0.584633 | 1.698364 | 6.156508 | 1.404627 | 0.810215 | 1.779416 | -0.62164 | 3.599544 | 1.71108 | -0.62371 | 0.161653 | 0.035874 | -0.14089 | 0.108998 | -0.00836 | 0.089445 | 0.693089 | 0.076587 | 0.210004 | 0.062873 | 0.152803 | 0.149695 |
| 36.48611 | 0.417816 | -0.8676 | 0.724269 | 1.290887 | 5.109328 | 0.813361 | 0.272438 | 1.535471 | -0.51563 | 0.565418 | 1.999868 | 0.017757 | 0.109687 | 0.055206 | -0.02139 | 0.132017 | 0.013415 | 0.15673 | 0.427904 | 0.093723 | 0.248028 | 0.062511 | 0.1557 | 0.11857 |
| 36.5 | 0.75234 | -0.41261 | 0.960561 | 0.991382 | 3.396128 | 0.760315 | 0.298818 | 1.563673 | -0.38629 | -1.59103 | 1.628923 | 0.149333 | -0.01087 | -0.03766 | -0.00381 | 0.05512 | -0.04235 | 0.099439 | 0.188533 | 0.008641 | 0.172375 | -0.01883 | 0.124115 | -0.01046 |
| 36.51389 | 1.415123 | 0.831853 | 1.142289 | 0.956941 | 1.420875 | 1.053734 | 0.266657 | 1.441821 | -0.16562 | -2.89492 | 0.897927 | -0.32233 | -0.06597 | -0.10644 | 0.06401 | 0.036811 | -0.05296 | 0.050313 | 0.063237 | -0.00536 | 0.129741 | -0.05856 | 0.130133 | -0.08893 |
| 36.52778 | 1.570212 | 1.879108 | 1.02713 | 0.721378 | 0.035921 | 1.127131 | 0.202957 | 1.276991 | 0.044134 | -3.02339 | 0.681869 | -0.57515 | -0.11984 | -0.1562 | 0.109418 | 0.040088 | -0.04549 | 0.012138 | -0.01866 | 0.001406 | 0.124047 | -0.04991 | 0.105152 | -0.0962 |
| 36.54167 | 1.343475 | 2.620406 | 0.73661 | 0.332448 | -0.76184 | 1.187438 | 0.13562 | 0.984932 | 0.206344 | -2.12044 | 0.54223 | -0.58594 | -0.14802 | -0.12979 | 0.11462 | 0.041953 | -0.03279 | -0.00711 | -0.05769 | 0.002006 | 0.152831 | 0.035339 | 0.074629 | -0.09855 |
| 36.55556 | 0.964659 | 2.970081 | 0.537169 | -0.04225 | -0.83224 | 0.952246 | 0.112377 | 0.814424 | 0.322156 | -1.69588 | 0.71444 | -0.38922 | -0.08048 | -0.08857 | 0.10658 | 0.070973 | -0.00459 | 0.018857 | 0.017319 | 0.042554 | 0.22428 | 0.172656 | 0.105529 | -0.06713 |
| 36.56944 | 0.484188 | 2.53563 | 0.403086 | -0.67528 | -1.36228 | 0.666638 | 0.16861 | 0.768736 | 0.404206 | -2.22834 | 0.900473 | -0.22343 | 0.031678 | -0.01336 | 0.110716 | 0.114748 | 0.047732 | 0.060471 | 0.161693 | 0.102404 | 0.300446 | 0.228209 | 0.177667 | 0.04701 |
| 36.58333 | 0.193502 | 2.804268 | 0.310743 | -1.2491 | -2.63496 | 0.516429 | 0.057452 | 0.542426 | 0.466271 | -3.75921 | 0.741684 | -0.38119 | 0.058797 | 0.021405 | 0.110574 | 0.119798 | 0.062638 | 0.047382 | 0.173035 | 0.106667 | 0.266331 | 0.200531 | 0.217978 | 0.121706 |
| 36.59722 | 0.353999 | 3.257666 | 0.288252 | -1.33725 | -3.99755 | 0.4468 | -0.10717 | 0.181392 | 0.495283 | -5.65833 | 0.424714 | -0.72912 | 0.065211 | 0.082431 | 0.14338 | 0.128102 | 0.081759 | 0.042056 | 0.122872 | 0.111809 | 0.227998 | 0.185717 | 0.270284 | 0.2154 |
| 36.61111 | 0.642156 | 4.330093 | 0.25701 | -0.82328 | -4.20987 | 0.450777 | -0.34128 | -0.16158 | 0.467372 | -6.91284 | 0.186158 | -1.15314 | 0.129365 | 0.160544 | 0.184348 | 0.167681 | 0.129203 | 0.089361 | 0.182736 | 0.17075 | 0.261937 | 0.202074 | 0.355542 | 0.278548 |
| 36.625 | 0.941045 | 4.687203 | 0.182063 | -0.10989 | -3.5617 | 0.358018 | -0.50515 | -0.30923 | 0.381854 | -7.1689 | 0.159712 | -1.22023 | 0.155452 | 0.178255 | 0.155323 | 0.163383 | 0.129498 | 0.107308 | 0.235911 | 0.177292 | 0.274481 | 0.152014 | 0.359686 | 0.231557 |
| 36.63889 | 0.878403 | 4.144474 | 0.384993 | -0.07571 | -2.75467 | 0.509013 | -0.46787 | -0.18808 | 0.319854 | -5.46454 | 0.539673 | -0.82342 | 0.124302 | 0.238072 | 0.128791 | 0.161041 | 0.100418 | 0.115092 | 0.267521 | 0.172032 | 0.260467 | 0.118116 | 0.334232 | 0.176119 |
| 36.65278 | 0.982851 | 3.102971 | 0.475891 | 0.195784 | -1.08304 | 0.647598 | -0.22784 | 0.007378 | 0.340104 | -4.98888 | 0.73298 | -0.61587 | 0.073394 | 0.365922 | 0.139139 | 0.197417 | 0.066211 | 0.120889 | 0.284613 | 0.178145 | 0.233041 | 0.089691 | 0.339226 | 0.136265 |
| 36.66667 | 1.188243 | 2.299185 | 0.416044 | 0.3793 | -1.22089 | 0.804011 | -0.05666 | 0.183988 | 0.412512 | -3.10656 | 0.610632 | -0.64936 | 0.022233 | 0.369366 | 0.125378 | 0.190197 | 0.045971 | 0.084183 | 0.275694 | 0.147103 | 0.162581 | 0.014206 | 0.284545 | 0.05493 |
| 36.68056 | 1.104802 | 1.449292 | 0.508649 | 0.358624 | -1.09195 | 0.943322 | 0.132371 | 0.425485 | 0.37922 | -3.8871 | 0.775571 | -0.39716 | -0.02751 | 0.234771 | 0.088557 | 0.156513 | 0.012995 | 0.043539 | 0.22975 | 0.094196 | 0.073047 | -0.07902 | 0.2162 | -0.04216 |
| 36.69444 | 0.839655 | 0.839223 | 0.62548 | 0.258733 | -1.21715 | 0.63246 | 0.182737 | 0.705982 | 0.409915 | -4.58308 | 1.178717 | -0.10967 | -0.06035 | 0.14613 | 0.068945 | 0.105767 | -0.0142 | 0.001754 | 0.147516 | 0.046298 | 0.001034 | -0.14958 | 0.169917 | -0.06825 |
| 36.70833 | 0.745454 | 0.807052 | 0.591762 | 0.271606 | -1.13739 | 0.558424 | 0.081775 | 0.953227 | 0.557843 | -3.46674 | 1.182458 | -0.13232 | -0.08428 | -0.00845 | 0.069646 | 0.095143 | -0.00071 | -0.01711 | 0.051251 | 0.005106 | -0.00398 | -0.14791 | 0.112024 | -0.07461 |
| 36.72222 | 0.709399 | 0.649147 | 0.510132 | 0.22985 | -0.8331 | 0.504819 | -0.10794 | 1.092094 | 0.690328 | -1.27314 | 0.739054 | -0.31691 | -0.07266 | -0.13273 | 0.095241 | 0.116578 | 0.043596 | -0.00357 | 0.005954 | 0.00433 | 0.047495 | -0.09227 | 0.085725 | -0.06709 |
| 36.73611 | 0.612503 | 0.449679 | 0.567325 | 0.208424 | -0.561 | 0.38788 | -0.31414 | 1.091809 | 0.687684 | -0.41063 | 0.241443 | -0.42028 | -0.09086 | -0.09346 | 0.091778 | 0.094522 | 0.010737 | -0.01184 | -0.07225 | -0.0139 | 0.020295 | -0.09935 | -0.04793 | -0.07947 |
| 36.75 | 0.550482 | 0.37938 | 0.639062 | 0.337278 | -0.47579 | 0.23239 | -0.52997 | 0.969583 | 0.819684 | -0.00988 | -0.0526 | -0.48325 | -0.21326 | -0.12866 | 0.011542 | -0.04823 | -0.09864 | -0.10762 | -0.22352 | -0.12104 | -0.10757 | -0.17419 | -0.17305 | -0.16665 |
| 36.76389 | 0.327488 | 0.293634 | 0.502548 | 0.371681 | -0.37084 | 0.255012 | -0.46349 | 0.773744 | 0.81968 | 0.639111 | -0.3355 | -0.52347 | -0.3062 | 0.00222 | 0.086716 | 0.071755 | -0.15602 | -0.14151 | -0.30291 | -0.09692 | -0.29185 | -0.1545 | -0.22652 | -0.13753 |
| 36.77778 | -0.23885 | 0.05593 | 0.212037 | 0.343209 | -0.53311 | 0.305621 | -0.28799 | 0.725846 | 0.812149 | 0.509336 | -0.48629 | -0.44612 | -0.47754 | 0.023392 | 0.672423 | 1.041885 | -0.09201 | 0.072301 | -0.30881 | 0.262997 | -1.3158 | 0.305099 | -0.0882 | 0.36006 |
| 36.79167 | -0.67305 | -0.35915 | 0.433834 | 0.195631 | -1.01754 | 0.31394 | -0.1719 | 0.776699 | 0.867244 | 0.187714 | -0.47381 | -0.25419 | -0.22643 | -6.53048 | 1.000834 | 1.547247 | 0.630425 | 0.47134 | -0.25634 | 0.00425 | -3.20218 | 0.104072 | -1.67534 | 0.146846 |
| 36.80556 | -0.91018 | -0.74523 | 0.676498 | -0.04492 | -1.36612 | 0.163964 | -0.22225 | 0.824748 | 0.967269 | -0.12697 | -0.73161 | -0.11134 | -0.15457 | -0.93263 | 0.24183 | 0.116749 | 0.065351 | 0.139088 | -0.38231 | -0.24626 | -0.62131 | -0.21305 | -0.39269 | 0.033403 |
| 36.81944 | -0.95171 | -1.1696 | 0.825235 | -0.17766 | -1.32544 | 0.049492 | -0.31988 | 0.989189 | 0.977863 | -0.39103 | -1.1601 | -0.03005 | -0.08794 | 2.548994 | -0.26699 | -0.40811 | -0.06279 | -0.33396 | -0.23936 | -0.11664 | 1.475208 | -0.00527 | 0.92795 | -0.16595 |
| 36.83333 | -0.76176 | -1.34814 | 0.903289 | -0.15455 | -1.30951 | 0.1391 | -0.2653 | 1.074987 | 1.089484 | -0.98534 | -1.3816 | 0.0382 | -0.20896 | -2.73074 | 0.403928 | 0.489906 | 0.073552 | 0.365529 | -0.20919 | -0.1507 | -1.58371 | -0.0573 | -1.01665 | -0.05623 |
| 36.84722 | -0.62 | -1.318 | 0.938209 | 0.030701 | -1.34868 | 0.199159 | -0.1203 | 1.038121 | 1.290246 | -1.13523 | -1.22138 | 0.226879 | -0.51154 | 0.464615 | -0.05373 | -0.0463 | -0.06303 | -0.13151 | -0.48465 | -0.2899 | 0.165627 | -0.02822 | 0.234202 | -0.35408 |
| 36.86111 | -0.53398 | -1.14875 | 0.889015 | 0.312201 | -1.00799 | 0.425624 | 0.073313 | 0.85966 | 1.301651 | -0.77753 | -1.06021 | 0.200895 | -0.30792 | 6.70363 | -1.08579 | -1.06434 | -0.14705 | -0.7734 | -0.47066 | -0.52212 | 3.155424 | -0.06506 | 2.003966 | -0.73675 |
| 36.875 | -0.56066 | -0.52908 | 0.653912 | 0.604176 | -0.19495 | 0.874681 | 0.150882 | 0.496286 | 1.085875 | 0.382684 | -1.17638 | -0.63659 | 0.016923 | 0.539117 | -0.7534 | -0.82065 | 0.00127 | -0.31823 | -0.20327 | -0.47044 | 1.202192 | -0.40459 | 0.239128 | -0.68641 |
| 36.88889 | -0.7481 | -0.01422 | 0.315452 | 0.681534 | 0.128068 | 1.103116 | 0.116419 | 0.309436 | 0.819653 | 0.747216 | -1.20714 | -1.48211 | -0.09959 | -0.1801 | -0.24113 | -0.20881 | -0.09277 | -0.17238 | -0.26547 | -0.23619 | -0.11152 | -0.22164 | -0.13501 | -0.2645 |
| 36.90278 | -1.02216 | -0.43731 | 0.258396 | 0.310518 | -0.85584 | 0.237535 | 0.143057 | 0.68187 | 0.920687 | -0.37776 | -0.50558 | -0.79458 | -0.25275 | -0.10008 | 0.424325 | 0.563596 | 0.043127 | 0.045377 | -0.02501 | 0.165894 | -1.50564 | 0.154177 | -0.19629 | 0.297368 |
| 36.91667 | -1.22193 | -1.29172 | 0.361162 | -0.09329 | -2.36125 | -0.4858 | -0.02676 | 1.040135 | 1.203943 | -2.51801 | 0.021953 | -0.12359 | -0.2431 | -10.5482 | 0.974222 | 1.533406 | 0.212134 | 1.53108 | 0.819745 | 0.294525 | -5.98524 | -0.34293 | -4.13526 | 0.879196 |
| 36.93056 | -1.46913 | -1.18388 | 0.382753 | 0.162853 | -2.6376 | -0.70476 | -0.47972 | 0.685833 | 0.957496 | -2.96159 | -0.39536 | -0.87156 | 3.323819 | NaN | -1.24691 | 0.962261 | 0.460926 | 1.026333 | -7.54909 | -4.68227 | NaN | NaN | NaN | -2.48259 |
| 36.94444 | -1.61718 | -0.03413 | 0.261695 | 0.848728 | -2.87567 | -1.29941 | -0.9375 | -0.16156 | 0.3421 | -2.41448 | -0.92292 | -2.3563 | -2.79121 | NaN | 3.80381 | 0.278205 | -0.41178 | -0.37712 | 9.450485 | -6.77718 | NaN | NaN | NaN | 1.232843 |
| 36.95833 | -1.37262 | -0.66758 | 0.758422 | 0.688375 | -5.81317 | -1.18323 | -0.63058 | 0.374076 | 0.396611 | -2.45047 | -0.04805 | -1.1745 | 1.087278 | 13.14227 | -1.48276 | -1.87301 | -0.33716 | -1.4439 | -3.8703 | -0.70353 | 6.832738 | -0.31335 | 3.456004 | -1.28531 |
| 36.97222 | -0.543 | -3.73877 | 1.536562 | -0.57117 | -3.91194 | -1.79134 | 0.095426 | 2.207147 | 1.19335 | -5.27305 | 3.190052 | 2.706303 | -0.14286 | -0.12763 | -0.09481 | -0.13658 | -0.08765 | -0.10758 | -0.22752 | -0.13941 | -0.19245 | -0.15469 | -0.09897 | -0.12109 |
| 36.98611 | -0.31561 | -3.05603 | 1.327384 | -0.41843 | -2.69934 | -2.58555 | 0.13329 | 1.611977 | 0.866601 | -4.00188 | 3.004455 | 2.047423 | -1.49765 | -13.9213 | 1.449671 | 1.270852 | -0.05218 | 1.408846 | 3.304838 | 0.633113 | -6.55798 | -0.05308 | -4.36759 | 1.016201 |
| 37 | -0.5356 | 0.374639 | 0.800352 | 1.147852 | -0.67673 | -0.18826 | -0.4744 | -0.91753 | -0.53622 | 0.526611 | 0.066089 | -1.87893 | 3.836985 | NaN | -3.21672 | 0.136913 | 0.335953 | 0.834371 | -12.2573 | -2.31747 | NaN | NaN | NaN | -1.80405 |
| 37.01389 | -0.51599 | 0.674184 | 0.574904 | 1.272214 | -0.05227 | 0.239023 | -0.88237 | -1.27568 | -0.41432 | 1.265158 | -0.11609 | -2.20151 | -3.54827 | NaN | 0.821304 | 0.084423 | -0.87408 | 0.32838 | 3.589213 | NaN | NaN | NaN | NaN | NaN |
| 37.02778 | -0.24456 | -0.95415 | 0.605326 | 0.548148 | -0.55279 | -0.79235 | -0.75675 | -0.34764 | -0.15492 | -0.33511 | 0.864336 | -0.45146 | -1.04597 | NaN | 1.050048 | -0.21885 | 0.003225 | -0.31921 | -1.79947 | 0.104774 | -2.86357 | 3.985831 | 5.471158 | 0.348099 |
| 37.04167 | -0.42768 | -0.86228 | 0.282567 | 0.535605 | -0.27483 | -0.50827 | -0.83823 | -0.24889 | -0.32606 | 0.082149 | -0.09669 | -0.69907 | 0.188955 | 7.292838 | -0.14693 | -0.77615 | 0.023876 | -1.44313 | -1.33576 | -0.28436 | 4.702074 | 0.817971 | 4.632129 | -3.61988 |
| 37.05556 | -0.44753 | -1.37388 | 0.245585 | 0.33449 | -0.29461 | -1.0813 | -0.46511 | 0.243119 | 0.1536 | -0.24383 | 0.005532 | -0.21545 | 0.836884 | 8.217107 | -1.14541 | -0.76564 | -0.37743 | -1.10588 | -0.62006 | -1.13363 | 4.428383 | -0.67167 | 1.624305 | -0.94188 |
| 37.06944 | -0.12164 | -3.06022 | 0.879801 | -0.41037 | -1.04072 | -1.29366 | 0.214461 | 1.334145 | 0.740881 | -2.3005 | 1.936078 | 1.862897 | 0.390554 | -1.82753 | -0.96496 | -0.85681 | -0.21182 | -0.35138 | -0.36283 | -0.73671 | 1.41747 | -0.19833 | -0.10829 | -0.82515 |
| 37.08333 | 0.12017 | -3.25554 | 1.112068 | -0.52771 | -0.96883 | -0.94908 | 0.399853 | 1.65745 | 1.196426 | -2.5522 | 2.65945 | 2.50348 | -0.01533 | -0.44345 | -0.22106 | -0.16352 | -0.07764 | -0.0731 | -0.3181 | -0.18722 | 0.016372 | 0.059616 | -0.09746 | -0.09991 |
| 37.09722 | 0.019651 | -2.9323 | 1.066883 | -0.43763 | -0.71994 | -0.64425 | 0.687508 | 1.625101 | 1.218214 | -1.93961 | 2.683388 | 2.102989 | -0.03519 | -0.15829 | -0.01216 | 0.023084 | -0.01383 | 0.041139 | -0.28453 | -0.00336 | -0.03392 | 0.166405 | -0.08771 | 0.066268 |

| | | | | | | | | | | | | | | | | | | | | | | | | |
|---|---|---|---|---|---|---|---|---|---|---|---|---|---|---|---|---|---|---|---|---|---|---|---|---|
| 37.11111 | -0.25559 | -1.90545 | 0.73964 | -0.02418 | 0.204722 | -0.11372 | 0.680671 | 1.120629 | 0.8956 | -0.72793 | 1.868687 | 0.916348 | -0.02281 | -0.10227 | 0.04516 | 0.068849 | -0.00462 | 0.061208 | -0.29137 | 0.042423 | -0.0235 | 0.158835 | 0.020047 | 0.086483 |
| 37.125 | -0.72019 | -0.4887 | 0.251141 | 0.836205 | 1.924669 | -0.00295 | 0.326161 | 0.310957 | 0.388829 | 1.412648 | 0.072405 | -1.23721 | -0.08348 | -0.09112 | 0.005238 | 0.033813 | -0.06962 | -0.00982 | -0.39626 | -0.0231 | -0.09706 | 0.11068 | -0.0337 | 0.035785 |
| 37.13889 | -0.47785 | -0.67879 | 0.485218 | 0.601782 | 1.607901 | 0.16617 | 0.31633 | 0.520628 | 0.36737 | 0.819979 | 0.499213 | -0.67224 | -0.11594 | -0.08633 | -0.04578 | -0.01072 | -0.11517 | -0.06771 | -0.4659 | -0.10412 | -0.13964 | 0.098471 | -0.02926 | -0.02467 |
| 37.15278 | 0.222155 | -1.97602 | 1.207895 | -0.23333 | 0.074436 | -0.29995 | 0.710802 | 1.301418 | 1.1559 | -1.52916 | 2.592621 | 1.646257 | -0.11399 | -0.05522 | -0.04511 | -0.02769 | -0.10312 | -0.06449 | -0.49563 | -0.13256 | -0.10957 | 0.118802 | 0.002414 | -0.03495 |
| 37.16667 | 0.168917 | -1.3346 | 1.051092 | -0.11362 | 0.388201 | -0.4119 | 0.472853 | 1.125717 | 0.894629 | -0.8612 | 2.155691 | 1.025827 | -0.13412 | -0.01463 | -0.03965 | -0.03115 | -0.09857 | -0.07024 | -0.53745 | -0.14586 | -0.10552 | 0.138001 | 0.002188 | -0.00148 |
| 37.18056 | -0.04377 | -0.0653 | 0.648606 | 0.3436 | 1.67712 | 0.020003 | -0.04129 | 0.731365 | 0.182893 | 0.5808 | 1.110504 | -0.46199 | -0.14359 | 0.039727 | -0.02926 | -0.03488 | -0.09916 | -0.08056 | -0.51532 | -0.1214 | -0.11043 | 0.147075 | -0.00728 | 0.057861 |
| 37.19444 | 0.272745 | -1.02943 | 1.011666 | 0.010973 | 1.177917 | -1.2752 | 0.206413 | 0.940674 | 0.668439 | -0.95591 | 2.43326 | 0.824525 | -0.0878 | 0.110119 | 0.023812 | 0.022311 | -0.0496 | -0.0291 | -0.44701 | -0.04849 | -0.03745 | 0.192566 | 0.041231 | 0.109638 |
| 37.20833 | 0.223767 | -1.23146 | 0.962126 | -0.0978 | 0.786413 | -1.0945 | 0.070691 | 0.825441 | 0.644204 | -1.20227 | 2.616018 | 0.990747 | 0.001444 | 0.162451 | 0.085079 | 0.082653 | 0.015222 | 0.038569 | -0.33398 | 0.00579 | 0.057167 | 0.233806 | 0.104948 | 0.095898 |
| 37.22222 | -0.38911 | 0.109055 | 0.260945 | 0.502328 | 1.642226 | -1.63868 | -0.52507 | 0.135416 | -0.25421 | 0.735596 | 1.042667 | -0.96344 | 0.010074 | 0.117991 | 0.077577 | 0.077991 | 0.008284 | 0.036549 | -0.27867 | -0.02321 | 0.060306 | 0.184062 | 0.080803 | 0.058333 |
| 37.23611 | -0.45331 | -0.17025 | 0.2351 | 0.518844 | 0.7686 | -1.78067 | -0.89633 | 0.067085 | -0.40503 | 0.484156 | 1.335929 | -0.83773 | -0.01928 | 0.054241 | 0.035071 | 0.036248 | -0.04313 | 0.000239 | -0.26042 | -0.09098 | 0.015155 | 0.11539 | 0.019146 | 0.036111 |
| 37.25 | -0.22281 | -1.10902 | 0.478809 | 0.164918 | -0.56235 | -1.79517 | -2.31224 | 0.343575 | 0.028092 | -0.72903 | 2.163393 | 0.252493 | -0.03805 | 0.0537 | 0.01231 | 0.0153 | -0.07905 | -0.00727 | -0.17489 | -0.1054 | 0.016663 | 0.121275 | 0.024646 | 0.025572 |
| 37.26389 | -0.30267 | -1.16545 | 0.423411 | 0.172039 | -0.77611 | -1.85247 | -3.59271 | 0.499027 | -0.09863 | -0.54418 | 1.917117 | 0.133309 | -0.04976 | 0.05651 | -0.01558 | -0.01291 | -0.11602 | -0.03283 | -0.10032 | -0.12893 | -0.00122 | 0.119373 | 0.016617 | -0.01589 |
| 37.27778 | -0.4107 | -1.06186 | 0.449917 | 0.242273 | -0.74997 | -1.63739 | -2.23951 | 0.839807 | -0.08694 | -0.22949 | 1.861776 | -0.00736 | -0.06231 | 0.057859 | -0.05033 | -0.05033 | -0.14551 | -0.08163 | -0.10606 | -0.18312 | -0.06572 | 0.039316 | -0.02899 | -0.07717 |
| 37.29167 | -0.10526 | -1.6515 | 0.854028 | -0.12306 | -1.68837 | 0.498379 | -0.76951 | 1.243306 | 0.551042 | -1.33435 | 2.979979 | 1.322388 | -0.05664 | 0.063861 | -0.04713 | -0.05054 | -0.12053 | -0.08169 | -0.09368 | -0.1508 | -0.04656 | -0.00065 | -0.00096 | -0.05115 |
| 37.30556 | 0.348304 | -2.12781 | 1.315128 | -0.4403 | -2.53132 | 0.342755 | -0.49018 | 1.253283 | 1.046679 | -2.054 | 3.757837 | 2.402293 | -0.03961 | 0.083727 | -0.02873 | -0.02143 | -0.07857 | -0.04333 | -0.04226 | -0.07924 | 0.027032 | 0.030268 | 0.029789 | -0.0106 |
| 37.31944 | 0.114002 | -1.15384 | 0.862788 | -0.21464 | -1.86557 | -0.19894 | 0.034124 | 1.006745 | 0.573652 | -1.00081 | 2.68957 | 1.213195 | -0.0369 | 0.092964 | -0.00653 | 0.008643 | -0.05679 | -0.0091 | -0.02004 | -0.06798 | 0.037478 | 0.07691 | -0.00204 | -0.01494 |
| 37.33333 | -0.35964 | 0.167311 | 0.086898 | 0.518342 | -0.02007 | -0.07629 | 0.192578 | 0.407901 | -0.07445 | 0.721677 | 0.63766 | -1.17759 | -0.02167 | 0.06001 | 0.006113 | 0.009097 | -0.05511 | -0.0007 | -0.03926 | -0.08244 | 0.004774 | 0.072106 | -0.0353 | -0.03385 |
| 37.34722 | -0.69327 | 1.172942 | -0.45558 | 1.023002 | 1.580452 | -1.57073 | 0.47901 | -0.07115 | -0.96829 | 1.888511 | -0.99194 | -3.00643 | -0.02948 | -0.04575 | 0.152458 | 0.123273 | -0.08669 | 0.016644 | -0.04506 | 0.047384 | -0.15125 | 0.117399 | 0.01876 | 0.038318 |
| 37.36111 | -0.36535 | 0.90362 | -0.27332 | 0.937976 | 1.438233 | -1.77458 | 0.201029 | 0.117388 | -0.87013 | 1.31503 | -0.42115 | -2.58381 | -0.14796 | -1.29169 | 1.016462 | 0.846976 | -0.16562 | 0.238944 | -0.00881 | 0.742129 | -1.56348 | 0.789519 | 0.321522 | 0.62815 |
| 37.375 | 0.525362 | -0.81756 | 0.779711 | -0.1464 | -0.58208 | 1.296623 | -0.04789 | 0.953975 | 0.372926 | -0.70085 | 2.41686 | 0.357041 | 0.267833 | -11.1315 | 1.273502 | 0.975534 | -0.47833 | 0.747477 | 0.671662 | 0.776647 | -4.25759 | 0.709427 | -0.85892 | 0.88983 |
| 37.38889 | 1.651413 | -2.12052 | 1.60511 | -0.93585 | -1.68082 | 4.648595 | -0.28211 | 1.92802 | 1.452186 | -2.10039 | 4.782524 | 2.850316 | -0.29938 | -1.14287 | -0.4424 | -0.15522 | -0.40011 | -0.13519 | 0.038909 | -0.39302 | -1.07695 | -0.14227 | -1.00457 | -0.24937 |
| 37.40278 | 2.276862 | -1.35086 | 1.308967 | -0.61766 | -0.07355 | 3.733809 | 0.103693 | 2.225151 | 1.922383 | -1.69201 | 4.212451 | 2.456162 | -0.42675 | 4.90742 | -1.42263 | -0.79067 | 0.163808 | -0.66722 | -0.52935 | -0.66868 | 3.112382 | -0.65585 | 0.180481 | -1.01143 |
| 37.41667 | 2.35067 | -0.68322 | 0.816974 | 0.387615 | 3.014169 | 0.775656 | 0.167112 | 1.993321 | 1.010069 | -0.44735 | 1.783441 | 0.898072 | 0.112705 | -0.00787 | -0.8079 | -0.68874 | 0.268958 | -0.11135 | 0.187889 | -0.42019 | 1.197219 | -0.25436 | -0.10809 | -0.43611 |
| 37.43056 | 2.139578 | -0.56097 | 0.637988 | 1.039316 | 3.796007 | 0.35023 | 0.362093 | 1.505292 | 0.69573 | -0.12684 | 0.573816 | 0.134216 | 0.065 | 0.065857 | 0.009533 | 0.145893 | 0.062993 | -0.02341 | 0.161679 | 0.201985 | 0.248079 | 0.263699 | 0.19086 | 0.051387 |
| 37.44444 | 1.57232 | -0.93697 | 0.662761 | 1.216001 | 3.921316 | 0.477116 | 0.7431 | 1.451571 | 0.862297 | -0.15126 | 0.309353 | 0.094345 | -0.16393 | 0.113434 | -0.24725 | -0.04863 | -0.20821 | -0.18125 | 0.033649 | -0.0383 | -0.13394 | -0.0146 | 0.013164 | -0.01052 |
| 37.45833 | 0.976409 | -0.84076 | 0.703469 | 1.177571 | 4.17722 | 1.309823 | 1.268651 | 1.738861 | 1.271802 | -0.1689 | 0.344925 | 0.048267 | -0.19107 | 0.133353 | -0.37698 | -0.1276 | -0.20419 | -0.1339 | 0.188037 | -0.17078 | -0.14227 | -0.11383 | -0.02211 | -0.00416 |
| 37.47222 | 0.72824 | -0.54644 | 0.656892 | 0.882922 | 3.323217 | 0.982001 | 1.691359 | 2.119553 | 1.545697 | -0.19545 | 0.715027 | 0.042564 | 0.022671 | 0.304782 | -0.16465 | 0.08652 | 0.036799 | 0.132557 | 0.573542 | 0.042226 | 0.173215 | 0.110206 | 0.145321 | 0.160198 |
| 37.48611 | 0.565412 | 0.26083 | 0.445572 | 0.904415 | 3.076016 | 0.846534 | 1.700336 | 2.128944 | 1.233141 | 0.083368 | 0.860078 | -0.28022 | 0.141424 | 0.281336 | -0.00089 | 0.180504 | 0.130812 | 0.246909 | 0.715414 | 0.150267 | 0.323119 | 0.18512 | 0.192923 | 0.16956 |
| 37.5 | 0.40268 | 0.945187 | 0.423626 | 1.04599 | 2.639802 | 0.640285 | 1.480448 | 1.912021 | 0.827593 | 0.274478 | 1.15992 | -0.43335 | 0.106381 | 0.111001 | 0.027015 | 0.119826 | 0.052236 | 0.17615 | 0.584713 | 0.082247 | 0.234775 | 0.088571 | 0.121458 | 0.081959 |
| 37.51389 | 0.465705 | 0.892048 | 0.621597 | 0.692966 | 1.533752 | 0.90337 | 1.482268 | 1.750696 | 0.779161 | 0.099366 | 1.594116 | -0.15467 | 0.06351 | -0.00754 | 0.009661 | 0.070012 | -0.01622 | 0.095374 | 0.424742 | 0.023783 | 0.16056 | 0.020972 | 0.078893 | 0.043194 |
| 37.52778 | 0.37996 | 0.506307 | 0.766992 | 0.557016 | 0.903938 | 1.14188 | 1.324569 | 1.71263 | 0.933987 | -0.02544 | 1.656319 | 0.0318 | 0.047285 | -0.0773 | 0.005408 | 0.067693 | -0.03261 | 0.045661 | 0.287912 | 0.016854 | 0.145958 | -0.00706 | 0.057746 | 0.009688 |
| 37.54167 | -0.03733 | 0.447604 | 0.634308 | 0.805557 | 1.09582 | 0.876884 | 1.091832 | 1.45669 | 0.820988 | 0.137024 | 1.35196 | -0.09399 | 0.014628 | -0.10621 | -0.00208 | 0.059123 | -0.03992 | 0.004736 | 0.193233 | 0.014277 | 0.126174 | -0.03361 | 0.005828 | -0.0549 |
| 37.55556 | -0.18337 | 0.173483 | 0.700365 | 0.787853 | 0.806278 | 0.895436 | 0.903889 | 1.381428 | 0.853904 | 0.109463 | 1.495099 | -0.04244 | -0.0307 | -0.11519 | -0.01688 | 0.031702 | -0.05374 | -0.01488 | 0.097089 | 0.02329 | 0.111158 | -0.05611 | -0.02735 | -0.10119 |
| 37.56944 | 0.161535 | -0.1416 | 0.971199 | 0.225385 | -0.46661 | 1.596316 | 0.944851 | 1.716716 | 1.208657 | -0.25744 | 2.028198 | 0.293025 | -0.03242 | -0.04817 | 0.005445 | 0.04218 | -0.03199 | 0.010741 | 0.068401 | 0.103164 | 0.142979 | -0.05228 | 0.031249 | -0.07138 |
| 37.58333 | 0.294079 | 0.061289 | 0.979195 | -0.14511 | -1.33782 | 1.384955 | 1.001251 | 1.742974 | 1.185984 | -0.17541 | 2.354085 | 0.260834 | 0.014844 | 0.086404 | 0.075081 | 0.095587 | 0.037015 | 0.065494 | 0.139914 | 0.219375 | 0.183957 | 0.034282 | 0.120086 | 0.017484 |
| 37.59722 | 0.132604 | 0.498097 | 0.705345 | -0.03526 | -1.43711 | 0.693723 | 0.873948 | 1.346449 | 0.724959 | 0.180007 | 2.179765 | -0.07246 | 0.031356 | 0.201376 | 0.142412 | 0.120971 | 0.084926 | 0.09346 | 0.234403 | 0.254994 | 0.179853 | 0.151725 | 0.215408 | 0.109087 |
| 37.61111 | 0.089943 | 0.686248 | 0.688124 | -0.12198 | -1.91727 | 0.744689 | 0.804181 | 1.220129 | 0.647499 | 0.086388 | 2.131027 | -0.10687 | 0.024175 | 0.202267 | 0.18637 | 0.114366 | 0.073094 | 0.086201 | 0.251192 | 0.202987 | 0.155472 | 0.176376 | 0.216807 | 0.145205 |
| 37.625 | 0.003886 | 0.781739 | 0.708674 | -0.03641 | -2.27564 | 0.668676 | 0.804904 | 1.209588 | 0.628386 | -0.02046 | 2.083349 | -0.19751 | 0.060821 | 0.153511 | 0.18957 | 0.140429 | 0.070015 | 0.104575 | 0.277544 | 0.184 | 0.16537 | 0.14969 | 0.185671 | 0.127776 |
| 37.63889 | -0.15471 | 0.940839 | 0.619052 | 0.267491 | -2.35937 | 0.372214 | 0.806318 | 1.009596 | 0.453508 | 0.043361 | 1.767313 | -0.34682 | 0.133522 | 0.141533 | 0.205534 | 0.213677 | 0.111697 | 0.17139 | 0.330641 | 0.239889 | 0.210694 | 0.141778 | 0.214125 | 0.124819 |
| 37.65278 | -0.17551 | 0.455545 | 0.747867 | 0.347046 | -2.59908 | 0.567023 | 0.69977 | 1.223325 | 0.692755 | -0.25472 | 1.802102 | -0.27129 | 0.144588 | 0.153306 | 0.228992 | 0.236128 | 0.12097 | 0.191435 | 0.299271 | 0.238933 | 0.185726 | 0.115416 | 0.2115 | 0.113152 |
| 37.66667 | -0.11521 | -0.12084 | 0.878654 | 0.354349 | -2.72945 | 0.919411 | 0.669233 | 1.574337 | 0.946666 | -0.47958 | 1.807374 | -0.12439 | 0.064372 | 0.128294 | 0.195168 | 0.170066 | 0.057476 | 0.115618 | 0.16843 | 0.140116 | 0.082297 | 0.054084 | 0.167293 | 0.032801 |
| 37.68056 | -0.03831 | -0.2505 | 0.919059 | 0.355059 | -1.95075 | 0.692044 | 0.598863 | 1.643933 | 0.837271 | -0.35157 | 1.499543 | -0.16222 | -0.05143 | -0.07438 | 0.078981 | 0.050717 | -0.04928 | -0.00571 | -0.0226 | 0.00938 | -0.04662 | -0.05342 | 0.024037 | -0.1228 |
| 37.69444 | -0.08028 | -0.34795 | 0.935987 | 0.361521 | -0.94512 | 0.493643 | 0.56891 | 1.554214 | 0.607351 | -0.12571 | 1.150101 | -0.22098 | -0.12986 | -0.19406 | -0.01925 | -0.04611 | -0.12869 | -0.09491 | -0.18654 | -0.09424 | -0.14463 | -0.13144 | -0.11667 | -0.26031 |
| 37.70833 | -0.14023 | -0.37801 | 0.896884 | 0.386896 | -0.17845 | 0.530329 | 0.603904 | 1.563171 | 0.560445 | 0.003204 | 0.983518 | -0.24987 | -0.07692 | -0.08981 | 0.032162 | -0.01262 | -0.07976 | -0.05575 | -0.17282 | -0.05474 | -0.0926 | -0.10382 | -0.04764 | -0.23574 |
| 37.72222 | -0.17527 | -0.33949 | 0.840835 | 0.438162 | 0.248736 | 0.604085 | 0.584976 | 1.527439 | 0.478688 | 0.106689 | 0.867441 | -0.23438 | -0.01123 | 0.380521 | 0.307191 | 0.206228 | 0.007673 | 0.024374 | -0.07511 | 0.137309 | -0.08873 | 0.06046 | 0.108518 | -0.06753 |
| 37.73611 | -0.1597 | -0.52114 | 0.814692 | 0.476692 | 0.283611 | 0.724584 | 0.63852 | 1.536659 | 0.514693 | 0.107214 | 0.636249 | -0.13089 | -0.28809 | 2.126654 | 1.242528 | 0.727303 | -0.19516 | 0.051937 | 0.103343 | 0.639471 | -0.68648 | 0.39796 | 0.133511 | 0.220827 |
| 37.75 | -0.10054 | -0.74234 | 0.833227 | 0.536728 | 0.045661 | 0.745725 | 0.74287 | 1.629349 | 0.643868 | 0.103012 | 0.510777 | -0.02993 | 0.34391 | 10.15792 | 1.62537 | 0.589065 | -0.83867 | 0.559068 | 0.287514 | 0.381517 | -2.31655 | 0.258896 | -1.70223 | 0.719872 |
| 37.76389 | -0.02473 | -0.85045 | 0.875901 | 0.547629 | -0.3564 | 0.732687 | 0.800447 | 1.628006 | 0.76379 | 0.137014 | 0.136553 | 0.014331 | 0.051617 | NaN | 0.233586 | 0.03891 | 0.000848 | 0.049607 | -0.22816 | -0.00827 | -0.13515 | -0.14391 | 0.357429 | -0.07348 |
| 37.77778 | 0.112876 | -0.79792 | 0.859174 | 0.482444 | -0.61042 | 0.494849 | 0.679193 | 1.498413 | 0.749282 | 0.148076 | -0.15002 | 0.013053 | -0.23384 | -9.77818 | -1.20337 | -0.333 | 1.192993 | -0.5516 | -0.20501 | -0.35762 | 2.802578 | -0.41575 | 2.350715 | -0.61166 |
| 37.79167 | 0.128856 | -0.76622 | 0.802599 | 0.451945 | -0.72705 | 0.337422 | 0.484367 | 1.330172 | 0.574613 | 0.124364 | -0.30356 | 0.01472 | 0.212582 | -2.5166 | -0.7283 | -0.63658 | 0.308814 | -0.04926 | -0.14177 | -0.38225 | 0.782493 | -0.17678 | -0.37371 | -0.24059 |
| 37.80556 | 0.27295 | -0.70495 | 0.781324 | 0.437028 | -0.78043 | 0.388842 | 0.30078 | 1.359603 | 0.483305 | 0.092048 | -0.27267 | -0.04722 | -0.18285 | 2.794586 | 0.948641 | 0.744861 | -0.2677 | 0.06717 | 0.133629 | 0.334288 | -0.71532 | 0.256879 | 0.282143 | 0.226854 |
| 37.81944 | 0.332651 | -0.71108 | 0.78594 | 0.428857 | -0.90594 | 0.23446 | 0.140406 | 1.188905 | 0.503022 | 0.066135 | -0.35119 | -0.09908 | -0.1062 | 9.938325 | 1.368985 | 0.270045 | -1.32792 | 0.721462 | -0.05038 | 0.419086 | -3.40652 | 0.170417 | -1.82161 | 0.767069 |
| 37.83333 | 0.390128 | -0.64372 | 0.765232 | 0.455663 | -1.15233 | 0.096576 | 0.093212 | 1.046241 | 0.441324 | 0.038697 | -0.47335 | -0.10429 | 0.007192 | NaN | 0.044476 | 0.020066 | -0.03821 | -9.7E-05 | -0.22381 | -0.07268 | -0.18254 | -0.21722 | 0.276776 | -0.2075 |
| 37.84722 | 0.37655 | -0.49833 | 0.784184 | 0.512573 | -1.18252 | 0.011238 | 0.217394 | 0.922503 | 0.53763 | 0.066168 | -0.51294 | -0.03841 | -0.05495 | -9.26239 | -1.10644 | -0.44055 | 1.010633 | -0.73988 | -0.60318 | -0.59449 | 3.086235 | 0.003297 | 1.735537 | -1.55887 |
| 37.86111 | 0.387477 | -0.34198 | 0.800693 | 0.698066 | -0.64832 | 0.130395 | 0.435192 | 0.798737 | 0.852771 | 0.122071 | -0.5624 | 0.012713 | -0.04327 | 0.038382 | -0.01157 | -0.00542 | -0.01267 | 0.001086 | -0.1393 | -0.05919 | -0.08122 | 0.007052 | 0.010977 | -0.04712 |
| 37.875 | 0.451881 | -0.10201 | 0.818272 | 0.859468 | 0.08852 | 0.418855 | 0.656676 | 0.693663 | 1.014567 | 0.163094 | -0.59266 | -0.00654 | -0.014 | 4.69115 | 1.158425 | 0.320157 | -1.21672 | 0.834571 | -0.13536 | 0.438337 | -3.51535 | -0.30127 | -1.65216 | 1.378024 |
| 37.88889 | 0.49183 | 0.148501 | 0.787297 | 0.880335 | 0.339122 | 0.690836 | 0.729611 | 0.558117 | 0.985511 | 0.169715 | -0.72852 | -0.0181 | -0.08938 | NaN | -0.07062 | -0.09519 | 0.03681 | -0.13505 | -0.1614 | -0.19918 | 0.326879 | 0.028212 | 0.596221 | -0.45842 |
| 37.90278 | 0.401632 | 0.132592 | 0.649722 | 0.583638 | -0.09583 | 0.515356 | 0.544952 | 0.630397 | 0.67595 | 0.251677 | -0.94584 | -0.17503 | 0.22357 | -12.5886 | -1.707 | -0.77316 | 1.13007 | -0.79468 | -1.1707 | -1.24632 | 3.737789 | -0.52907 | 1.857884 | -1.78218 |
| 37.91667 | -0.36085 | 0.155433 | 0.403661 | 0.308738 | -0.47703 | -0.22751 | 0.217718 | 0.52734 | 0.040208 | 0.668644 | -1.53627 | -0.82153 | 0.459632 | -3.60029 | -2.08238 | -1.91679 | 0.366501 | -0.15877 | -0.76526 | -1.39437 | 1.214668 | -0.75386 | -0.71286 | -0.61916 |
| 37.93056 | -1.24059 | 0.304496 | 0.354573 | 0.232505 | -0.50157 | -1.1019 | -0.06766 | 0.581232 | -0.66094 | 1.059267 | -2.85847 | -1.5527 | 0.229756 | -2.3423 | -0.82661 | -0.84287 | 0.103187 | 0.050809 | -0.49366 | -1.05411 | 0.479588 | -0.48961 | -0.49209 | -0.42212 |
| 37.94444 | -1.58707 | -0.35053 | 0.735889 | -0.15643 | -1.12581 | -0.81378 | -0.25008 | 0.375593 | -0.62728 | 0.456527 | -3.25169 | -1.04468 | 0.30049 | -0.94993 | 0.288768 | 0.209579 | 0.208299 | 0.257729 | -0.29628 | -0.27728 | 0.339892 | 0.17131 | 0.045032 | -0.03511 |
| 37.95833 | -1.29186 | -0.01054 | 1.054504 | -0.4631 | -2.21298 | 0.534004 | -0.53895 | 0.453101 | -0.30862 | -0.54654 | -2.30453 | -0.05303 | 0.231564 | 3.32061 | 1.772552 | 1.362902 | -0.19518 | 0.249088 | 0.039618 | 0.807545 | -0.69981 | 0.909647 | 0.745552 | 0.466403 |
| 37.97222 | -0.6 | -0.80659 | 1.218928 | -0.2148 | -2.91496 | 1.173486 | -0.96085 | 0.552524 | -0.78693 | -0.82149 | -1.10444 | 0.285023 | -0.05556 | 6.208479 | 2.021392 | 0.944488 | -0.82143 | 0.633719 | 0.295508 | 0.755484 | -3.4916 | 0.080439 | -0.05403 | 1.02499 |
| 37.98611 | 0.053418 | 0.182055 | 1.235968 | -0.13699 | -2.87191 | 1.207287 | -0.90691 | 0.473269 | -1.0407 | -0.71225 | -0.12148 | 0.381865 | -0.21245 | 0.119833 | -0.14126 | -0.21142 | -0.16361 | -0.2062 | -0.52813 | 0.086445 | 0.186981 | -0.30955 | 0.279967 | -0.57784 |
| 38 | 0.348931 | -0.5905 | 1.01755 | 0.08828 | -2.52967 | 0.402677 | -0.52206 | 0.210943 | -1.14029 | -0.0357 | 0.111744 | -0.15513 | -0.16615 | -5.67083 | -2.02296 | -1.50392 | 0.279938 | -0.67425 | -0.94212 | -0.8063 | 3.507398 | -0.93703 | 0.184666 | -1.06216 |

**Supplementary Table S2.** Raw motion data from: (a) Water withdrawal stress

| Time | Control1 | Control2 | Control3 | Control4 | Control5 | Control6 | Control7 | Control8 | Control9 | Control10 | Control11 | Control12 | Stress1 | Stress2 | Stress3 | Stress4 | Stress5 | Stress6 | Stress7 | Stress8 | Stress9 | Stress10 | Stress11 | Stress12 |
|---|---|---|---|---|---|---|---|---|---|---|---|---|---|---|---|---|---|---|---|---|---|---|---|---|
| 18 | 1.898552 | -0.0292 | -0.21706 | 0.110467 | 0.070267 | -0.26271 | 0.365834 | 0.091008 | 0.339499 | -0.02305 | 1.642956 | -0.06257 | -0.10899 | -0.2967 | -0.20771 | 0.018848 | -0.03309 | 0.524463 | 0.015627 | -0.08747 | -0.09428 | -0.07119 | -0.09821 | 0.00318 |
| 18.01389 | 3.860303 | 0.13985 | 0.194306 | 0.182694 | 0.281276 | 0.234489 | 1.435947 | 0.785706 | 0.448581 | 0.088049 | 2.531331 | 0.574616 | 0.384988 | 0.012542 | -0.08404 | 0.14315 | 0.072912 | 0.951013 | 0.588312 | 0.174395 | 0.529982 | 0.380615 | 0.632137 | 0.517762 |
| 18.02778 | 3.290776 | -0.01732 | -0.08976 | 0.216762 | 0.252192 | 0.464573 | 1.340772 | 0.810213 | 0.260338 | 0.09094 | 1.64565 | 0.725784 | 0.194649 | 0.040644 | -0.06647 | 0.050026 | 0.05753 | 0.782302 | 1.220936 | 0.133318 | 0.674648 | 0.306823 | 0.302283 | 0.4996 |
| 18.04167 | 2.320418 | -0.06138 | -0.16162 | 0.249235 | 0.084673 | 0.801213 | 0.730266 | 0.403187 | 0.137481 | 0.124882 | 1.457433 | 0.671732 | -0.02765 | 0.013309 | -0.03688 | 0.025714 | 0.075537 | 0.394316 | 0.517959 | 0.063932 | 0.691583 | 0.1667 | 0.073071 | 0.297946 |
| 18.05556 | 1.681401 | 0.000758 | -0.06203 | 0.18947 | -0.02461 | 0.923902 | 0.400955 | 0.194977 | 0.074652 | 0.189835 | 0.929431 | 0.596096 | 0.011458 | 0.064058 | 0.075148 | 0.110649 | 0.124539 | 0.196691 | -0.02577 | 0.076341 | 0.695374 | 0.117805 | 0.200972 | 0.144903 |
| 18.06944 | 1.162109 | 0.071768 | 0.045934 | 0.086687 | -0.08406 | 0.777137 | 0.245941 | 0.060532 | -0.00046 | 0.239299 | 0.521909 | 0.538748 | 0.119174 | 0.094599 | 0.119486 | 0.162763 | 0.101549 | 0.150181 | -0.04179 | 0.073587 | 0.570085 | 0.125555 | 0.340406 | 0.102839 |
| 18.08333 | 0.855372 | 0.13051 | 0.128874 | 0.041135 | -0.08533 | 0.548755 | 0.166867 | 0.011973 | -0.01127 | 0.275942 | 0.37682 | 0.5063 | 0.137942 | 0.025315 | 0.068776 | 0.106503 | 0.006053 | 0.077716 | -0.1084 | 0.008142 | 0.390934 | 0.11743 | 0.290172 | 0.068492 |
| 18.09722 | 0.704377 | 0.163001 | 0.190115 | 0.043693 | -0.07276 | 0.415393 | 0.151756 | 0.003682 | 0.055007 | 0.304734 | 0.438251 | 0.542564 | 0.168755 | -0.02677 | 0.059701 | 0.051175 | -0.05073 | 0.085374 | -0.10518 | -0.00163 | 0.292217 | 0.113042 | 0.213617 | 0.055961 |
| 18.11111 | 0.668588 | 0.189295 | 0.217447 | 0.040148 | -0.04875 | 0.334776 | 0.146043 | 0.004105 | 0.091806 | 0.309412 | 0.475607 | 0.610793 | 0.25268 | -0.00179 | 0.119174 | 0.056972 | -0.04644 | 0.133406 | -0.0322 | 0.06298 | 0.260469 | 0.15982 | 0.232412 | 0.096883 |
| 18.125 | 0.693811 | 0.237492 | 0.245386 | 0.043193 | 0.010639 | 0.289172 | 0.188714 | 0.030773 | 0.103767 | 0.326506 | 0.414294 | 0.719381 | 0.294267 | 0.025464 | 0.141735 | 0.046802 | -0.05463 | 0.12882 | 0.0102 | 0.113576 | 0.21824 | 0.169801 | 0.267056 | 0.11937 |
| 18.13889 | 0.782825 | 0.238133 | 0.270539 | 0.069998 | 0.049977 | 0.274924 | 0.236141 | 0.091235 | 0.11817 | 0.325289 | 0.316689 | 0.784132 | 0.310234 | 0.05236 | 0.142553 | 0.049588 | -0.05932 | 0.130944 | 0.036827 | 0.192799 | 0.222424 | 0.144934 | 0.269838 | 0.145298 |
| 18.15278 | 0.785928 | 0.145555 | 0.248283 | 0.081525 | 0.043001 | 0.224512 | 0.227836 | 0.119902 | 0.088213 | 0.30285 | 0.255086 | 0.742381 | 0.300442 | 0.071385 | 0.131052 | 0.063419 | -0.04851 | 0.118069 | 0.051417 | 0.272912 | 0.252687 | 0.120575 | 0.250054 | 0.182679 |
| 18.16667 | 0.664265 | 0.023271 | 0.19933 | 0.071691 | 0.036533 | 0.159044 | 0.193646 | 0.10858 | 0.048432 | 0.281546 | 0.291722 | 0.652607 | 0.186571 | 0.028365 | 0.072025 | 0.048435 | -0.07263 | 0.063429 | 0.006811 | 0.27564 | 0.221852 | 0.066806 | 0.169372 | 0.187423 |
| 18.18056 | 0.458667 | -0.08985 | 0.150661 | 0.052994 | 0.021122 | 0.147025 | 0.164061 | 0.103891 | 0.097603 | 0.256684 | 0.357866 | 0.596884 | 0.08252 | 0.006587 | 0.066109 | 0.075082 | -0.06708 | 0.094728 | 0.001945 | 0.285433 | 0.232051 | 0.049168 | 0.107954 | 0.203149 |
| 18.19444 | 0.327271 | -0.17642 | 0.099835 | 0.00881 | -0.01665 | 0.142295 | 0.123222 | 0.099915 | 0.158078 | 0.226878 | 0.32068 | 0.584671 | 0.04399 | 0.041358 | 0.119047 | 0.126996 | -0.02029 | 0.149931 | 0.060791 | 0.3061 | 0.291377 | 0.087832 | 0.095296 | 0.220275 |
| 18.20833 | 0.276047 | -0.1906 | 0.063655 | -0.01811 | -0.02546 | 0.119922 | 0.094249 | 0.079213 | 0.157855 | 0.191272 | 0.190743 | 0.531909 | -0.02553 | 0.029303 | 0.107286 | 0.089088 | -0.03007 | 0.093816 | 0.03075 | 0.232439 | 0.26436 | 0.086724 | 0.05051 | 0.154846 |
| 18.22222 | 0.305577 | -0.14144 | 0.039184 | -0.00361 | -0.01481 | 0.099662 | 0.078644 | 0.060118 | 0.142724 | 0.146876 | 0.065854 | 0.455399 | -0.05582 | -0.00923 | 0.076804 | 0.031786 | -0.06035 | 0.039665 | -0.03159 | 0.182444 | 0.195461 | 0.033231 | 0.013154 | 0.083802 |
| 18.23611 | 0.368419 | -0.08306 | 0.024962 | 0.010235 | -0.01048 | 0.090622 | 0.069121 | 0.046565 | 0.119422 | 0.109031 | 0.016703 | 0.40698 | 0.0109 | 0.000099 | 0.109994 | 0.047133 | -0.02711 | 0.074428 | 0.001207 | 0.246369 | 0.190944 | 0.029202 | 0.046452 | 0.108297 |
| 18.25 | 0.394222 | -0.03029 | 0.042087 | 0.02075 | 0.016861 | 0.09559 | 0.070991 | 0.045751 | 0.086315 | 0.098772 | 0.109321 | 0.395499 | 0.042224 | -0.00795 | 0.124109 | 0.045033 | -0.00263 | 0.08799 | 0.027859 | 0.29055 | 0.192456 | 0.040047 | 0.070421 | 0.146434 |
| 18.26389 | 0.347236 | 0.024226 | 0.076877 | 0.039544 | 0.041741 | 0.113007 | 0.085311 | 0.059612 | 0.109591 | 0.121656 | 0.281884 | 0.422898 | 0.017093 | -0.05107 | 0.102717 | 0.005151 | -0.02411 | 0.086308 | 0.004993 | 0.245724 | 0.176338 | 0.017499 | 0.067302 | 0.14394 |
| 18.27778 | 0.351573 | 0.077739 | 0.109232 | 0.034387 | 0.035448 | 0.094669 | 0.086065 | 0.073794 | 0.164267 | 0.141354 | 0.358805 | 0.468383 | 0.041997 | -0.06421 | 0.129577 | -0.00196 | -0.01775 | 0.11914 | 0.011893 | 0.218893 | 0.177333 | 0.024518 | 0.105053 | 0.176255 |
| 18.29167 | 0.417866 | 0.14699 | 0.152288 | 0.03352 | 0.04501 | 0.079045 | 0.085595 | 0.093828 | 0.226213 | 0.163322 | 0.310857 | 0.542346 | 0.091776 | -0.06609 | 0.179775 | 0.004454 | -0.01231 | 0.131713 | 0.020531 | 0.216367 | 0.152588 | 0.042133 | 0.150484 | 0.235885 |
| 18.30556 | 0.507172 | 0.215969 | 0.186272 | 0.06002 | 0.052984 | 0.106496 | 0.07981 | 0.115916 | 0.252165 | 0.179452 | 0.241375 | 0.630357 | 0.163791 | -0.05091 | 0.225731 | 0.024658 | -0.00953 | 0.146017 | 0.027461 | 0.223284 | 0.040145 | 0.045203 | 0.163211 | 0.309025 |
| 18.31944 | 0.622565 | 0.241284 | 0.18303 | 0.060306 | 0.024855 | 0.117217 | 0.039824 | 0.118086 | 0.196901 | 0.158907 | 0.185363 | 0.663957 | 0.250012 | 0.002363 | 0.272175 | 0.059523 | 0.009537 | 0.177062 | 0.052467 | 0.223918 | -0.18494 | 0.070637 | 0.152299 | 0.393852 |
| 18.33333 | 0.645382 | 0.247859 | 0.179665 | 0.06104 | 0.004393 | 0.112759 | 0.010622 | 0.108418 | 0.122592 | 0.128952 | 0.196759 | 0.691302 | 0.286444 | 0.051634 | 0.275467 | 0.071033 | 0.003018 | 0.173072 | 0.038911 | 0.185419 | -0.23138 | 0.011558 | 0.133241 | 0.388927 |
| 18.34722 | 0.587394 | 0.260236 | 0.175661 | 0.088735 | -0.00387 | 0.117693 | 0.017646 | 0.098342 | 0.12194 | 0.119778 | 0.326372 | 0.725614 | 0.294535 | 0.098838 | 0.272188 | 0.084959 | -0.02127 | 0.165093 | 0.016302 | 0.16181 | 0.010253 | 0.025384 | 0.143122 | 0.350039 |
| 18.36111 | 0.447437 | 0.252011 | 0.146356 | 0.099512 | -0.01379 | 0.103797 | 0.026267 | 0.080422 | 0.145073 | 0.130916 | 0.382179 | 0.802415 | 0.330213 | 0.124112 | 0.254917 | 0.099575 | -0.03351 | 0.161267 | 0.022652 | 0.186031 | 0.187629 | 0.07585 | 0.18217 | 0.344196 |
| 18.375 | 0.309514 | 0.212986 | 0.118626 | 0.090336 | -0.00731 | 0.089105 | 0.028271 | 0.065488 | 0.182465 | 0.168755 | 0.305102 | 0.832073 | 0.33751 | 0.101307 | 0.20624 | 0.09873 | -0.04181 | 0.147248 | 0.0249 | 0.209455 | 0.243534 | 0.067106 | 0.207815 | 0.345437 |
| 18.38889 | 0.316448 | 0.191725 | 0.122442 | 0.072008 | 0.009098 | 0.104037 | 0.053796 | 0.087894 | 0.207367 | 0.226575 | 0.207753 | 0.736038 | 0.359222 | 0.094391 | 0.193649 | 0.120868 | -0.03241 | 0.166138 | 0.057136 | 0.252736 | 0.291882 | 0.066649 | 0.238439 | 0.357392 |
| 18.40278 | 0.46588 | 0.279507 | 0.186281 | 0.078864 | 0.022161 | 0.178017 | 0.114484 | 0.159627 | 0.194255 | 0.294068 | 0.197311 | 0.718041 | 0.389528 | 0.125561 | 0.211854 | 0.151751 | -0.01957 | 0.193209 | 0.093157 | 0.294939 | 0.291528 | 0.084972 | 0.270343 | 0.381979 |
| 18.41667 | 0.612714 | 0.415524 | 0.252087 | 0.120434 | 0.008977 | 0.277144 | 0.164233 | 0.221473 | 0.188708 | 0.329743 | 0.253946 | 0.782122 | 0.440762 | 0.186586 | 0.280141 | 0.181891 | -0.00896 | 0.240723 | 0.108431 | 0.336537 | 0.268619 | 0.087455 | 0.30211 | 0.390232 |
| 18.43056 | 0.817604 | 0.498902 | 0.253103 | 0.112595 | 0.003797 | 0.293524 | 0.188798 | 0.285025 | 0.214337 | 0.382446 | 0.451481 | 0.9066 | 0.516076 | 0.276152 | 0.414572 | 0.213263 | 0.018721 | 0.386135 | 0.174809 | 0.369637 | 0.286828 | 0.101964 | 0.377584 | 0.406087 |
| 18.44444 | 1.697427 | 1.085258 | 0.455302 | 0.066549 | 0.15562 | 0.382089 | 0.503977 | 0.597741 | 0.492988 | 0.855119 | 1.225789 | 1.802617 | 0.780747 | 0.399341 | 0.727273 | 0.357541 | 0.092796 | 1.035638 | 0.560117 | 0.571938 | 0.428814 | 0.210323 | 0.962297 | 0.918006 |
| 18.45833 | 3.664285 | 2.78462 | 1.111174 | 0.082472 | 0.504982 | 0.703704 | 1.383838 | 1.281755 | 1.142803 | 1.952877 | 2.764259 | 4.259661 | 1.366621 | 0.663719 | 1.391963 | 0.738662 | 0.260091 | 2.529336 | 1.845832 | 1.180415 | 0.736712 | 0.468832 | 2.389254 | 2.35095 |
| 18.47222 | 4.409522 | 3.832309 | 1.651755 | 0.158016 | 0.761831 | 1.01525 | 2.08311 | 1.663817 | 1.373087 | 2.570121 | 3.469726 | 6.42202 | 1.178778 | 0.772858 | 1.701106 | 0.981408 | 0.352405 | 3.048516 | 1.990655 | 1.395926 | 0.870063 | 0.526827 | 3.0661 | 3.531822 |
| 18.48611 | 2.620376 | 2.271989 | 1.126431 | 0.161207 | 0.649085 | 0.501862 | 1.646997 | 1.055174 | 0.946174 | 1.704262 | 2.203979 | 4.483005 | 0.428375 | 0.45335 | 1.213602 | 0.647261 | 0.196419 | 1.858594 | 0.80199 | 0.737151 | 0.452115 | 0.328757 | 2.011096 | 2.663164 |
| 18.5 | 0.76345 | 1.001131 | 0.486214 | 0.083686 | 0.402838 | 0.196922 | 0.889453 | 0.463025 | 0.516443 | 0.886551 | 0.915985 | 2.213341 | 0.031353 | 0.067379 | 0.576554 | 0.333025 | 0.040645 | 0.809607 | 0.259118 | 0.273394 | 0.162072 | 0.214815 | 0.823915 | 1.303537 |
| 18.51389 | -0.09838 | 0.46397 | 0.155192 | 0.017601 | 0.216416 | 0.107602 | 0.558711 | 0.230066 | 0.330818 | 0.600301 | 0.291226 | 1.141576 | 0.002174 | -0.1584 | 0.240702 | 0.131053 | -0.00108 | 0.503179 | 0.198494 | 0.126201 | 0.124643 | 0.216328 | 0.399544 | 0.654317 |
| 18.52778 | -0.35999 | 0.369125 | 0.043041 | 0.010494 | 0.145173 | 0.090342 | 0.470641 | 0.238611 | 0.340044 | 0.531352 | 0.274233 | 0.551798 | 0.070356 | -0.28333 | 0.168 | -0.00761 | -0.01997 | 0.502014 | 0.268852 | 0.132706 | 0.114657 | 0.249886 | 0.362812 | 0.529141 |
| 18.54167 | -0.38448 | 0.468502 | 0.014205 | 0.055838 | 0.180865 | 0.13087 | 0.488803 | 0.344201 | 0.456819 | 0.564418 | 0.466311 | 0.37378 | 0.022474 | -0.22067 | 0.18725 | -0.07891 | -0.0487 | 0.535424 | 0.22351 | 0.211844 | 0.080733 | 0.269821 | 0.326518 | 0.500469 |
| 18.55556 | -0.25261 | 0.628623 | -0.05141 | 0.097296 | 0.237614 | 0.147727 | 0.476715 | 0.434121 | 0.565185 | 0.506975 | 0.632639 | 0.471303 | -0.18535 | 0.074247 | 0.191743 | -0.06182 | -0.04421 | 0.595312 | 0.115016 | 0.316643 | 0.035786 | 0.2987 | 0.079733 | 0.559432 |
| 18.56944 | -0.04939 | 0.901696 | -0.13642 | 0.139535 | 0.270759 | 0.122018 | 0.419571 | 0.514198 | 0.576244 | 0.327381 | 0.617518 | 0.687656 | -0.48536 | 0.358591 | 0.145914 | -0.01754 | -0.00536 | 0.62568 | 0.051777 | 0.350726 | -0.01213 | 0.334565 | -0.01409 | 0.644648 |
| 18.58333 | 0.256839 | 1.324707 | -0.15721 | 0.256349 | 0.267022 | 0.156069 | 0.401293 | 0.549467 | 0.543537 | 0.233279 | 0.50226 | 0.994245 | -0.57201 | 0.343128 | 0.114932 | 0.052583 | 0.042994 | 0.653357 | 0.096878 | 0.2756 | -0.05338 | 0.343318 | 0.271968 | 0.712386 |
| 18.59722 | 0.61943 | 1.553521 | -0.05421 | 0.30648 | 0.228734 | 0.222344 | 0.505986 | 0.51158 | 0.415384 | 0.320445 | 0.402286 | 1.338628 | -0.23891 | 0.078659 | 0.178278 | 0.149623 | 0.098125 | 0.669013 | 0.203564 | 0.194177 | -0.03208 | 0.31479 | 0.659126 | 0.757833 |
| 18.61111 | 0.887209 | 1.369895 | 0.158335 | 0.211221 | 0.177497 | 0.293244 | 0.611108 | 0.435564 | 0.212962 | 0.456084 | 0.286694 | 1.568454 | 0.366926 | -0.15695 | 0.290893 | 0.241666 | 0.127711 | 0.697424 | 0.278254 | 0.112961 | -0.00281 | 0.283137 | 0.716249 | 0.727201 |
| 18.625 | 1.054019 | 1.007607 | 0.401911 | 0.109953 | 0.109293 | 0.381655 | 0.702797 | 0.353209 | 0.070921 | 0.548045 | 0.335947 | 1.502599 | 0.909463 | -0.27601 | 0.366161 | 0.317352 | 0.136744 | 0.777842 | 0.288578 | 0.084672 | 0.064949 | 0.260705 | 0.551619 | 0.643079 |
| 18.63889 | 1.065612 | 0.615556 | 0.541851 | 0.026892 | 0.027068 | 0.328558 | 0.824136 | 0.269877 | 0.040122 | 0.55141 | 0.429373 | 1.104866 | 1.026202 | -0.25533 | 0.372239 | 0.365747 | 0.149473 | 0.848203 | 0.265556 | 0.091452 | 0.125751 | 0.280694 | 0.453417 | 0.560283 |
| 18.65278 | 0.943955 | 0.20435 | 0.414734 | -0.03571 | -0.00072 | 0.142832 | 0.87255 | 0.218172 | 0.08636 | 0.434503 | 0.343814 | 0.681389 | 0.627648 | -0.15982 | 0.302169 | 0.256549 | 0.121379 | 0.7479 | 0.198126 | 0.071566 | 0.104837 | 0.296175 | 0.265647 | 0.516167 |
| 18.66667 | 0.831328 | -0.14859 | 0.112121 | -0.0649 | 0.036344 | 0.022913 | 0.823604 | 0.209006 | 0.191266 | 0.234504 | 0.156889 | 0.496797 | 0.174949 | -0.06889 | 0.217382 | 0.068597 | 0.056418 | 0.615275 | 0.090393 | 0.009966 | 0.044257 | 0.239117 | 0.168294 | 0.437441 |
| 18.68056 | 0.761275 | -0.29443 | -0.1469 | -0.03181 | 0.091163 | 0.06007 | 0.716061 | 0.199312 | 0.361839 | 0.20627 | 0.021738 | 0.579487 | -0.17239 | -0.01481 | 0.155572 | -0.06897 | 0.009824 | 0.522638 | -5.1E-05 | -0.03001 | 0.013473 | 0.130176 | 0.288488 | 0.370027 |
| 18.69444 | 0.643331 | -0.1152 | -0.19624 | 0.094409 | 0.145632 | 0.16356 | 0.577597 | 0.190419 | 0.487474 | 0.398492 | -0.02595 | 0.802808 | -0.56782 | 0.012355 | 0.105602 | -0.13871 | -0.01967 | 0.553337 | -0.00844 | -0.02643 | 0.020618 | 0.059267 | 0.4306 | 0.435835 |
| 18.70833 | 0.554957 | 0.433123 | -0.14443 | 0.277765 | 0.182749 | 0.268255 | 0.403416 | 0.193953 | 0.459659 | 0.500296 | -0.033 | 1.055078 | -0.88334 | -0.0052 | 0.1419 | -0.1086 | -0.01226 | 0.669858 | 0.057447 | 0.02952 | 0.025755 | -0.00033 | 0.405914 | 0.523394 |
| 18.72222 | 0.580339 | 1.094902 | -0.07244 | 0.385507 | 0.184322 | 0.351577 | 0.258743 | 0.199312 | 0.381381 | 0.396842 | 0.161264 | 1.324258 | -1.02683 | -0.08671 | 0.296448 | -0.03535 | 0.025342 | 0.89189 | 0.147981 | 0.119509 | -0.0125 | 0.005337 | 0.329283 | 0.53893 |
| 18.73611 | 0.589689 | 1.470058 | -0.09026 | 0.317449 | 0.161815 | 0.372931 | 0.141496 | 0.194881 | 0.343383 | 0.343575 | 0.506722 | 1.590324 | -0.83879 | -0.16971 | 0.442136 | -0.05633 | 0.047183 | 1.115992 | 0.18232 | 0.21262 | -0.06862 | 0.080641 | 0.28305 | 0.56907 |
| 18.75 | 0.579951 | 1.383811 | -0.10413 | 0.143432 | 0.155487 | 0.30967 | 0.156929 | 0.208758 | 0.307595 | 0.340616 | 0.619204 | 1.826117 | -0.54715 | -0.2217 | 0.426776 | -0.14832 | 0.046446 | 1.230652 | 0.151314 | 0.274787 | -0.09708 | 0.204967 | 0.221513 | 0.601531 |
| 18.76389 | 0.637512 | 0.961352 | -0.17433 | -0.00095 | 0.164231 | 0.229004 | 0.267138 | 0.300479 | 0.246787 | 0.434315 | 0.47304 | 1.888432 | -0.15313 | -0.28735 | 0.232901 | -0.18954 | 0.037914 | 1.121369 | 0.08933 | 0.24562 | -0.0537 | 0.239923 | 0.109334 | 0.504224 |
| 18.77778 | 0.577723 | 0.347278 | -0.33218 | -0.09605 | 0.137527 | 0.16591 | 0.502393 | 0.426804 | 0.173022 | 0.604774 | 0.184804 | 1.775482 | -0.06105 | -0.27316 | -0.02988 | -0.20835 | 0.011769 | 0.945236 | 0.037771 | 0.155295 | 0.071398 | 0.169303 | 0.033381 | 0.341253 |
| 18.79167 | 0.156694 | -0.55751 | -0.51236 | -0.18138 | 0.065475 | 0.045865 | 0.698805 | 0.515758 | 0.08173 | 0.681731 | -0.12409 | 1.471543 | -0.63058 | -0.20269 | -0.163 | -0.29095 | -0.03507 | 0.742194 | -0.01754 | 0.065097 | 0.181196 | 0.120832 | -0.07186 | 0.1713 |
| 18.80556 | -0.45608 | -1.47385 | -0.65286 | -0.23351 | -0.01689 | -0.13555 | 0.570101 | 0.504064 | -0.01609 | 0.352174 | -0.34686 | 0.969661 | -1.62282 | -0.24639 | -0.09992 | -0.38309 | -0.07779 | 0.550155 | -0.06979 | 0.017664 | 0.205876 | 0.059371 | -0.22331 | 0.033725 |
| 18.81944 | -0.80041 | -1.64032 | -0.62719 | -0.17455 | -0.06904 | -0.27805 | 0.190827 | 0.403677 | -0.08299 | -0.1958 | -0.40863 | 0.433988 | -1.93883 | -0.34948 | 0.138319 | -0.41151 | -0.11177 | 0.471072 | -0.10145 | -0.00572 | 0.143026 | -0.01962 | -0.35257 | -0.05904 |
| 18.83333 | -0.69954 | -1.04023 | -0.60639 | -0.13065 | -0.07538 | -0.34554 | -0.20547 | 0.30876 | -0.05649 | -0.65521 | -0.44459 | 0.12357 | -1.43151 | -0.30415 | 0.312739 | -0.40623 | -0.13074 | 0.434647 | -0.10933 | -0.02468 | -0.0669 | -0.12835 | -0.45116 | -0.09452 |
| 18.84722 | -0.37846 | -0.43616 | -0.63114 | -0.20011 | -0.08258 | -0.33612 | -0.59034 | 0.216588 | -0.0209 | -0.83539 | -0.52605 | 0.192136 | -0.7104 | -0.27004 | 0.262918 | -0.43966 | -0.14366 | 0.39015 | -0.09007 | -0.06235 | -0.31977 | -0.24091 | -0.54183 | -0.13946 |
| 18.86111 | -0.04572 | -0.51688 | -0.75922 | -0.38078 | -0.12088 | -0.24078 | -0.84174 | 0.148878 | -0.13578 | -0.8646 | -0.60979 | 0.525465 | -0.29879 | -0.23653 | 0.034663 | -0.51519 | -0.14435 | 0.281082 | -0.09502 | -0.12018 | -0.48337 | -0.24078 | -0.62423 | -0.22758 |
| 18.875 | -0.03339 | -1.1036 | -0.92826 | -0.55836 | -0.17551 | -0.2058 | -0.85328 | 0.143838 | -0.33545 | -0.73783 | -0.66735 | 0.731187 | -0.48859 | -0.17931 | -0.20166 | -0.57774 | -0.11592 | 0.170702 | -0.14428 | -0.12338 | -0.45606 | -0.11999 | -0.63833 | -0.32128 |
| 18.88889 | -0.30691 | -1.65624 | -1.01505 | -0.58191 | -0.22264 | -0.29334 | -0.66233 | 0.227308 | -0.48406 | -0.54119 | -0.6721 | 0.619184 | -1.08617 | -0.13303 | -0.31554 | -0.56845 | -0.07281 | 0.158306 | -0.21228 | -0.06999 | -0.38705 | 0.005329 | -0.56303 | -0.36772 |
| 18.90278 | -0.55278 | -1.61159 | -0.93841 | -0.48569 | -0.23363 | -0.47763 | -0.32613 | 0.353879 | -0.51042 | -0.23097 | -0.47325 | 0.407062 | -1.54043 | -0.09754 | -0.28347 | -0.49905 | -0.03373 | 0.219593 | -0.26334 | -0.06123 | -0.36223 | 0.022607 | -0.42401 | -0.35633 |
| 18.91667 | -0.66793 | -1.063 | -0.70938 | -0.29463 | -0.18365 | -0.58555 | 0.115449 | 0.434348 | -0.4425 | 0.064188 | -0.1844 | 0.282152 | -1.47649 | -0.10435 | -0.13644 | -0.41416 | -0.01274 | 0.319147 | -0.25945 | -0.08009 | -0.32395 | -0.02697 | -0.31768 | -0.31718 |
| 18.93056 | -0.67752 | -0.46542 | -0.54593 | -0.14964 | -0.11984 | -0.60463 | 0.676514 | 0.382499 | -0.28806 | 0.347327 | -0.14968 | 0.118082 | -1.08794 | -0.1218 | -0.0069 | -0.37537 | -0.03998 | 0.257976 | -0.22024 | -0.14762 | -0.28266 | -0.08063 | -0.35583 | -0.333 |
| 18.94444 | -1.11483 | -0.54852 | -0.66893 | -0.19562 | -0.18795 | -0.77834 | 0.66038 | 0.129418 | -0.23222 | -0.04336 | -0.7803 | -0.39132 | -1.03782 | -0.21785 | -0.15016 | -0.44552 | -0.14096 | -0.42465 | -0.19807 | -0.41039 | -0.36997 | -0.20746 | -0.78301 | -0.70728 |
| 18.95833 | -2.93198 | -1.99176 | -1.30275 | -0.48946 | -0.45956 | -1.64419 | -0.22782 | -0.3685 | -0.53746 | -1.49145 | -1.92964 | -1.68722 | -1.39165 | -0.53706 | -0.56921 | -0.8504 | -0.48126 | -2.04911 | -0.39849 | -1.06417 | -0.84381 | -0.47696 | -1.72915 | -1.70744 |
| 18.97222 | -5.75673 | -4.81803 | -2.25147 | -0.75619 | -0.50698 | -4.17066 | -0.70004 | -1.16741 | -1.04514 | -3.24459 | -3.79868 | -3.21457 | -2.62341 | -1.00782 | -0.88783 | -1.55194 | -0.76397 | -3.59976 | -1.50745 | -2.55605 | -1.7225 | -0.8672 | -3.0494 | -3.45217 |
| 18.98611 | -4.13346 | -3.87822 | -1.90061 | -0.49217 | -0.12756 | -2.93021 | -1.29773 | -0.66618 | -0.76994 | -2.34388 | -2.90168 | -2.52378 | -3.57871 | -0.91032 | -0.53788 | -1.53172 | -0.56472 | -2.13741 | -0.59689 | -1.77847 | -1.75003 | -0.81562 | -3.69696 | -3.37793 |
| 19 | 1.80437 | 0.672604 | -0.39684 | -0.02301 | -0.18924 | 0.832026 | 0.751716 | 0.271869 | -0.0302 | -0.32375 | 2.313108 | -0.01997 | 1.71695 | -0.31939 | -0.05336 | -0.05406 | -0.06427 | 0.611089 | 0.346047 | 0.340994 | -0.21838 | -0.13607 | 1.532583 | -0.0518 |
| 19.01389 | 4.369748 | 2.3538 | 0.322067 | 0.099459 | 0.020878 | 2.254807 | 3.106723 | 0.79084 | 0.339781 | 0.543506 | 3.281732 | 1.772306 | 3.781994 | 0.077298 | 0.080819 | 0.813488 | 0.301451 | 1.763606 | 0.796687 | 1.332266 | 0.880691 | 0.242297 | 3.928377 | 1.664333 |
| 19.02778 | 2.730336 | 1.6693 | 0.31903 | 0.181232 | 0.564043 | 1.384952 | 2.684537 | 0.822179 | 0.389741 | 0.544286 | 2.080129 | 2.060014 | 1.683106 | 0.131499 | 0.057899 | 0.568295 | 0.19974 | 1.316792 | 0.764748 | 0.976952 | 0.845479 | 0.154915 | 2.376396 | 1.427593 |
| 19.04167 | 1.642081 | 0.958819 | 0.252263 | 0.225824 | 0.489967 | 0.671072 | 1.53526 | 0.513341 | 0.375006 | 0.385768 | 1.569924 | 1.616605 | 0.75894 | 0.036423 | 0.016325 | 0.232484 | 0.048014 | 0.786763 | 0.428695 | 0.527644 | 0.490532 | 0.034392 | 1.133408 | 0.756955 |
| 19.05556 | 1.069507 | 0.505211 | 0.240747 | 0.242229 | 0.264641 | 0.377939 | 0.857725 | 0.338487 | 0.28795 | 0.305366 | 0.758107 | 1.220411 | 0.310925 | -0.04151 | 0.02006 | 0.107204 | 0.0298 | 0.36919 | 0.015 | 0.245066 | 0.304585 | -0.02165 | 0.314491 | 0.267764 |
| 19.06944 | 0.604521 | 0.218801 | 0.152007 | 0.185155 | 0.085783 | 0.199246 | 0.49146 | 0.221374 | 0.124779 | 0.216116 | 0.081521 | 0.862594 | 0.071507 | -0.07148 | 0.066681 | 0.069483 | 0.003595 | 0.190672 | -0.0797 | 0.094223 | 0.235096 | -0.03585 | -0.10839 | -0.01793 |
| 19.08333 | 0.32447 | 0.018073 | -0.04314 | 0.06641 | -0.02209 | 0.056169 | 0.268234 | 0.098632 | -0.07057 | 0.122128 | -0.35892 | 0.547213 | -0.11114 | -0.0716 | 0.088207 | 0.015482 | -0.02799 | 0.044369 | -0.15092 | -0.05058 | 0.148732 | -0.04156 | -0.3553 | -0.0869 |
| 19.09722 | 0.216621 | -0.00045 | -0.17742 | -0.00393 | -0.07084 | 0.034628 | 0.184252 | 0.034946 | -0.15892 | 0.081143 | -0.35292 | 0.361644 | -0.20232 | -0.05247 | 0.115037 | -0.04163 | -0.04471 | 0.000097 | -0.1516 | -0.06869 | 0.073834 | -0.03057 | -0.34884 | 0.053709 |
| 19.11111 | 0.149186 | 0.113624 | -0.08688 | -0.01007 | -0.09563 | 0.123714 | 0.171945 | 0.052893 | -0.07259 | 0.094773 | 0.040819 | 0.324183 | -0.15713 | -0.03433 | 0.167101 | -0.05605 | -0.03223 | 0.072232 | -0.09535 | 0.040432 | 0.0894 | -0.01055 | -0.0859 | 0.18962 |
| 19.125 | 0.087605 | 0.17189 | 0.157573 | -0.01514 | -0.08562 | 0.194397 | 0.173483 | 0.056859 | 0.078724 | 0.111901 | 0.311629 | 0.312357 | -0.09152 | -0.02396 | 0.185621 | -0.0197 | -0.00881 | 0.109571 | -0.07283 | 0.134259 | 0.110948 | -0.00033 | 0.143392 | 0.140787 |
| 19.13889 | 0.073271 | 0.099257 | 0.287737 | -0.02295 | -0.04248 | 0.186041 | 0.164385 | 0.038234 | 0.139675 | 0.115297 | 0.174214 | 0.369636 | -0.11808 | -0.0306 | 0.165995 | 0.031405 | 0.000646 | 0.073557 | -0.09791 | 0.076461 | 0.082033 | -0.02047 | 0.126263 | -0.03232 |
| 19.15278 | 0.102549 | -0.032 | 0.185773 | -0.02604 | -0.01669 | 0.11897 | 0.156523 | 0.004265 | 0.060552 | 0.122152 | -0.18317 | 0.531034 | -0.20354 | -0.04656 | 0.15823 | 0.0513 | -0.00422 | 0.039384 | -0.11651 | -0.06749 | 0.032622 | -0.05035 | -0.07617 | -0.13538 |
| 19.16667 | 0.109406 | -0.10513 | 0.01863 | -0.03784 | -0.00998 | 0.061122 | 0.157176 | -0.03154 | -0.04284 | 0.147524 | -0.45344 | 0.676217 | -0.2366 | -0.04798 | 0.186628 | 0.010325 | -0.01461 | 0.029949 | -0.10754 | -0.17527 | -0.00629 | -0.05792 | -0.28215 | -0.0573 |
| 19.18056 | 0.06431 | -0.05351 | -0.01419 | -0.04381 | -0.0033 | 0.062771 | 0.16124 | -0.03516 | -0.04756 | 0.178973 | -0.52916 | 0.682353 | -0.17632 | -0.02165 | 0.312834 | -0.02027 | -0.01503 | 0.076102 | -0.0937 | -0.11594 | 0.012097 | -0.01914 | -0.34853 | 0.129678 |
| 19.19444 | 0.025902 | 0.067152 | 0.133199 | -0.04647 | -0.01009 | 0.121165 | 0.146246 | -0.02592 | 0.05705 | 0.20136 | -0.20048 | 0.617968 | -0.07211 | 0.006264 | 0.50315 | -0.01503 | -0.01597 | 0.158281 | -0.08071 | 0.087205 | 0.070191 | 0.041683 | -0.21872 | 0.230045 |
| 19.20833 | 0.063682 | 0.172766 | 0.330192 | -0.05124 | -0.03215 | 0.218052 | 0.136025 | -0.00363 | 0.202514 | 0.230026 | 0.278868 | 0.564666 | -0.00748 | 0.025176 | 0.549285 | 0.016976 | -0.01962 | 0.260025 | -0.04558 | 0.211442 | 0.0827 | 0.054198 | 0.053084 | 0.164202 |
| 19.22222 | 0.195191 | 0.23216 | 0.42472 | -0.05377 | -0.03682 | 0.310885 | 0.155351 | 0.015565 | 0.290426 | 0.255227 | 0.428686 | 0.4895 | 0.020717 | 0.046538 | 0.517169 | 0.066381 | -0.00411 | 0.382918 | 0.0168 | 0.150154 | 0.057059 | 0.012772 | 0.265708 | 0.054526 |
| 19.23611 | 0.254306 | 0.244232 | 0.366514 | -0.04187 | 0.000673 | 0.337177 | 0.178663 | 0.025996 | 0.289194 | 0.261582 | 0.20437 | 0.421669 | 0.016302 | 0.047832 | 0.518095 | 0.091257 | 0.009197 | 0.414561 | 0.026707 | 0.026248 | 0.032516 | -0.04164 | 0.246991 | 0.044889 |
| 19.25 | 0.174117 | 0.220086 | 0.245733 | -0.00692 | 0.056335 | 0.307372 | 0.188453 | 0.029589 | 0.264132 | 0.259616 | -0.07167 | 0.382639 | 0.031272 | 0.02439 | 0.601881 | 0.067835 | 0.007588 | 0.36745 | -0.01096 | -0.01859 | 0.023794 | -0.06329 | 0.106306 | 0.196032 |
| 19.26389 | 0.13057 | 0.292958 | 0.229217 | 0.049215 | 0.108175 | 0.280465 | 0.213311 | 0.03311 | 0.287369 | 0.269636 | -0.14668 | 0.410745 | 0.152282 | 0.00801 | 0.73494 | 0.058282 | -0.01009 | 0.418546 | 0.053549 | 0.182208 | 0.076601 | 0.058621 | 0.160728 | 0.469784 |
| 19.27778 | 0.322182 | 0.562112 | 0.306167 | -0.00132 | 0.022971 | 0.170302 | 0.188644 | -0.213 | 0.159162 | 0.237453 | -0.09873 | 0.300584 | 0.510503 | -0.06861 | 0.956989 | 0.125811 | -0.08878 | 0.87576 | 0.413437 | 0.791712 | 0.342018 | 0.660551 | 0.846928 | 1.053514 |
| 19.29167 | 0.500178 | 0.874674 | 0.264204 | -0.27162 | -0.23186 | -0.06316 | 0.08687 | -0.79009 | -0.21516 | 0.182956 | -0.15331 | -0.04151 | 0.974749 | -0.26878 | 1.184926 | 0.151615 | -0.22976 | 1.562451 | 0.920739 | 1.481843 | 0.735972 | 2.170454 | 1.905797 | 1.864748 |
| 19.30556 | 0.395681 | 0.872695 | 0.39487 | -0.27047 | -0.32265 | -0.03627 | 0.049657 | -0.81324 | -0.33598 | 0.288202 | -0.00077 | -0.00766 | 1.021196 | 0.098821 | 1.02011 | 0.169397 | -0.22769 | 1.571718 | 0.646417 | 1.373562 | 0.754245 | 1.923409 | 1.950627 | 1.758064 |
| 19.31944 | 0.360708 | 0.772353 | 0.487286 | -0.00489 | -0.12706 | 0.189074 | 0.084441 | -0.31368 | 0.090992 | 0.34375 | 0.266292 | 0.275956 | 0.563397 | 0.35484 | 0.660036 | 0.233946 | -0.07026 | 0.85848 | 0.134788 | 0.644939 | 0.396596 | 0.444165 | 0.920211 | 0.797683 |
| 19.33333 | 0.380485 | 0.569474 | 0.335239 | 0.169238 | 0.009364 | 0.238889 | 0.09523 | -0.07039 | 0.340449 | 0.307986 | 0.10616 | 0.265867 | 0.190102 | 0.138753 | 0.506662 | 0.152653 | 0.039437 | 0.340877 | -0.14304 | 0.210816 | 0.128928 | 0.008903 | 0.109842 | 0.311681 |
| 19.34722 | 0.374325 | 0.372088 | 0.214596 | 0.182293 | 0.058409 | 0.194933 | 0.101039 | -0.05779 | 0.259152 | 0.249739 | -0.17011 | 0.160651 | 0.054411 | 0.067353 | 0.536813 | 0.119984 | 0.099242 | 0.217106 | -0.18689 | 0.163329 | 0.068362 | 0.017276 | -0.01045 | 0.231681 |
| 19.36111 | 0.420215 | 0.331618 | 0.354563 | 0.164285 | 0.128254 | 0.189841 | 0.126094 | -0.06034 | 0.175955 | 0.281986 | 0.037372 | 0.247753 | 0.175694 | 0.078191 | 0.46874 | 0.12622 | 0.145671 | 0.308246 | -0.08541 | 0.147672 | 0.081311 | 0.087089 | 0.378698 | 0.246249 |
| 19.375 | 0.58488 | 0.527506 | 0.597564 | 0.221107 | 0.201696 | 0.254874 | 0.166613 | -0.0212 | 0.303455 | 0.3894 | 0.696848 | 0.479885 | 0.436694 | 0.132893 | 0.370893 | 0.132448 | 0.166379 | 0.45726 | 0.052258 | 0.148434 | 0.126764 | 0.099313 | 0.720383 | 0.24962 |
| 19.38889 | 0.796803 | 0.724381 | 0.708649 | 0.341852 | 0.228348 | 0.340438 | 0.201973 | 0.007868 | 0.505024 | 0.467734 | 1.102879 | 0.59797 | 0.564709 | 0.170288 | 0.323202 | 0.138748 | 0.156371 | 0.486286 | 0.112458 | 0.162634 | 0.189042 | 0.060571 | 0.528158 | 0.234587 |
| 19.40278 | 1.02449 | 0.760255 | 0.639531 | 0.423002 | 0.195124 | 0.431301 | 0.223211 | -0.00191 | 0.513467 | 0.485708 | 0.681849 | 0.029706 | 0.526956 | 0.15559 | 0.328418 | 0.170772 | 0.13185 | 0.43169 | 0.076802 | 0.144817 | 0.181869 | 0.024818 | 0.164689 | 0.202092 |
| 19.41667 | 1.138101 | 0.692898 | 0.515131 | 0.408917 | 0.124064 | 0.500536 | 0.239663 | 0.017922 | 0.317743 | 0.512078 | -0.03208 | -1.1684 | 0.406326 | 0.168027 | 0.35543 | 0.234675 | 0.128539 | 0.445783 | 0.043676 | 0.124218 | 0.185614 | 0.033449 | 0.064747 | 0.273084 |
| 19.43056 | 1.07429 | 0.656337 | 0.499084 | 0.298654 | 0.115074 | 0.41272 | 0.31493 | 0.11923 | 0.145371 | 0.600413 | -0.34421 | -0.93671 | 0.359217 | 0.255849 | 0.388846 | 0.262355 | 0.137354 | 0.556667 | 0.133413 | 0.173479 | 0.208707 | 0.084596 | 0.358232 | 0.45051 |

| | | | | | | | | | | | | | | | | | | | | | | | | |
|---|---|---|---|---|---|---|---|---|---|---|---|---|---|---|---|---|---|---|---|---|---|---|---|---|
| 19.44444 | 1.528328 | 1.343502 | 0.934955 | 0.310524 | 0.405174 | 0.297796 | 0.635304 | 0.426552 | 0.413443 | 1.129026 | 0.597665 | -0.13187 | 0.995438 | 0.487015 | 0.701476 | 0.366575 | 0.192512 | 1.14802 | 0.62321 | 0.413686 | 0.337285 | 0.202274 | 1.035248 | 0.97765 |
| 19.45833 | 2.959167 | 3.880041 | 2.08917 | 0.585683 | 1.040851 | 0.230607 | 1.351283 | 0.911946 | 1.180252 | 2.738073 | 2.6801 | 1.017585 | 3.073132 | 0.978334 | 1.322337 | 0.768643 | 0.320482 | 2.759684 | 1.905073 | 0.962681 | 0.626341 | 0.387105 | 1.685153 | 2.161536 |
| 19.47222 | 4.053065 | 6.364546 | 2.847912 | 0.903418 | 1.528683 | 0.067885 | 2.029237 | 0.977281 | 0.998708 | 3.891494 | 4.169688 | 3.314742 | 4.660681 | 1.4492 | 1.320181 | 1.179779 | 0.428889 | 4.43105 | 2.194785 | 1.368582 | 0.716755 | 0.408131 | 1.531162 | 2.542426 |
| 19.48611 | 2.953935 | 4.456599 | 1.970762 | 0.727865 | 1.294418 | -0.58363 | 1.5327 | 0.496725 | 0.476771 | 1.698244 | 3.180921 | 3.029342 | 2.651913 | 0.96909 | 0.983568 | 1.089312 | 0.370935 | 3.122002 | 1.275826 | 0.930013 | 0.428069 | 0.257666 | 0.641399 | 1.047574 |
| 19.5 | 1.047232 | 1.762459 | 0.880676 | 0.387825 | 0.680407 | -0.38833 | 0.702662 | 0.166846 | 0.339892 | 0.630859 | 1.400766 | 1.001642 | 1.179397 | 0.398836 | 0.575108 | 0.712421 | 0.249034 | 1.804547 | 0.645057 | 0.380207 | 0.209779 | 0.152663 | 0.103789 | 0.159058 |
| 19.51389 | -0.17392 | 0.383647 | 0.052519 | 0.108646 | 0.323862 | -0.04248 | 0.145922 | 0.016382 | 0.013652 | -0.01285 | 0.331715 | -0.13474 | 0.524058 | 0.101232 | 0.290999 | 0.3951 | 0.172988 | 1.224586 | 0.286743 | 0.156209 | 0.164131 | 0.155091 | -0.15414 | -1.08734 |
| 19.52778 | -0.88882 | -0.38243 | -0.60541 | -0.14913 | 0.113707 | -0.03697 | -0.21894 | -0.06447 | -0.36097 | -0.44179 | -0.07665 | -0.14767 | 0.226965 | -0.12272 | 0.09303 | 0.132523 | 0.094431 | 0.894682 | 0.152217 | 0.074548 | 0.132281 | 0.193674 | -0.0628 | -1.3354 |
| 19.54167 | -1.04029 | -0.94948 | -1.03329 | -0.33797 | -0.01661 | -0.25442 | -0.49775 | -0.01162 | -0.53716 | -0.75029 | 0.208393 | -1.95193 | 0.118523 | -0.23174 | -0.05196 | -0.10943 | 0.008502 | 0.62783 | 0.142186 | 0.072537 | 0.038705 | 0.193723 | 0.311618 | -0.10265 |
| 19.55556 | -0.25891 | -1.07954 | -1.47364 | -0.30623 | -0.06886 | -0.2965 | -0.7511 | 0.072745 | -0.2134 | -0.93625 | 0.789917 | -2.12035 | 0.214982 | -0.07367 | -0.15687 | -0.24287 | -0.008 | 0.469182 | 0.264218 | 0.15383 | -0.09272 | 0.152988 | 0.175937 | 0.790104 |
| 19.56944 | 0.870485 | -0.9263 | -1.74429 | -0.09686 | -0.01263 | -0.31632 | -0.69199 | 0.139372 | 0.323197 | -0.81945 | 1.182181 | -1.25185 | 0.482657 | 0.359724 | -0.18791 | -0.15367 | 0.069362 | 0.450003 | 0.425421 | 0.278152 | -0.20708 | 0.147311 | -0.1341 | 0.734896 |
| 19.58333 | 1.603081 | -0.443 | -1.12078 | 0.252858 | 0.121733 | 0.001173 | -0.26069 | 0.090337 | 0.543407 | -0.20064 | 1.087915 | 0.499592 | 0.533161 | 0.510998 | -0.22134 | 0.025042 | 0.16007 | 0.509337 | 0.433753 | 0.319903 | -0.18784 | 0.218413 | 0.159067 | 0.388496 |
| 19.59722 | 1.834338 | 0.327416 | 0.237282 | 0.477248 | 0.209992 | 0.551089 | 0.249545 | 0.015223 | 0.379739 | 0.549133 | 0.763176 | 2.615159 | 0.262591 | 0.247262 | -0.18354 | 0.159484 | 0.22432 | 0.573817 | 0.346715 | 0.262533 | -0.02382 | 0.286283 | 0.405536 | 0.135526 |
| 19.61111 | 1.618208 | 1.222232 | 1.82204 | 0.547125 | 0.211323 | 0.951396 | 0.771075 | 0.095864 | 0.280317 | 1.191002 | 0.2203 | 3.075461 | 0.028461 | -0.09123 | -0.02149 | 0.228456 | 0.195443 | 0.488293 | 0.270876 | 0.189815 | 0.150373 | 0.379552 | 0.321332 | -0.05959 |
| 19.625 | 1.148401 | 2.013015 | 2.618875 | 0.520411 | 0.132988 | 1.018852 | 1.458492 | 0.3705 | 0.33358 | 1.620767 | -0.31066 | 2.136671 | 0.025028 | -0.32555 | 0.263596 | 0.255367 | 0.149012 | 0.363348 | 0.242707 | 0.193114 | 0.195328 | 0.372842 | 0.383976 | -0.21844 |
| 19.63889 | 0.393868 | 2.115823 | 2.234854 | 0.366794 | 0.036525 | 0.667868 | 2.311077 | 0.623052 | 0.508601 | 1.378813 | -0.43561 | 1.392983 | 0.120864 | -0.36862 | 0.518914 | 0.215472 | 0.130682 | 0.343396 | 0.254469 | 0.202936 | 0.223157 | 0.232098 | 0.698249 | -0.38197 |
| 19.65278 | -0.53547 | 1.001951 | 1.422316 | 0.062837 | 0.09818 | 0.444092 | 2.562154 | 0.633825 | 0.726034 | 0.284821 | -0.18401 | 0.366138 | 0.121624 | -0.37734 | 0.529515 | 0.044415 | 0.11502 | 0.43193 | 0.26297 | 0.174498 | 0.164198 | 0.073845 | 0.764774 | -0.11626 |
| 19.66667 | -1.32094 | -0.18001 | 0.47727 | -0.05932 | 0.121295 | 0.29168 | 1.314851 | 0.390457 | 0.840784 | -0.03296 | 0.089451 | -0.68956 | 0.090977 | -0.00095 | 0.338153 | -0.09497 | 0.041615 | 0.441259 | 0.232437 | 0.127088 | 0.065181 | -0.04993 | 0.547148 | 0.392719 |
| 19.68056 | -1.34791 | -1.03527 | -0.30208 | -0.03167 | 0.13197 | 0.176541 | 0.336447 | 0.127146 | 0.760484 | -0.27309 | 0.09687 | -2.00204 | 0.133683 | 0.771497 | 0.192561 | -0.08557 | -0.02838 | 0.379639 | 0.22156 | 0.118948 | 0.003251 | -0.11741 | 0.401194 | 0.554985 |
| 19.69444 | -0.32919 | -1.62618 | -0.87337 | 0.018066 | 0.207931 | 0.073105 | -0.03884 | -0.06482 | 0.465407 | -0.50769 | -0.16326 | -2.92595 | 0.210603 | 1.121282 | 0.118026 | -0.1367 | -0.05796 | 0.30862 | 0.208073 | 0.15764 | -0.01975 | -0.13096 | 0.272217 | 0.35581 |
| 19.70833 | 0.738158 | -1.88379 | -1.22146 | 0.029549 | 0.306606 | 0.054921 | -0.6618 | -0.16245 | 0.081821 | -0.63921 | -0.43295 | -2.23104 | 0.209613 | 0.571235 | 0.067344 | -0.25088 | -0.09127 | 0.219821 | 0.200682 | 0.196115 | -0.02859 | -0.16086 | 0.027679 | 0.132479 |
| 19.72222 | 1.032351 | -1.65726 | -1.69373 | 0.045586 | 0.31301 | 0.089666 | -1.28601 | -0.17551 | -0.23315 | -0.98048 | -0.37085 | -0.26651 | 0.164349 | 0.121272 | 0.029924 | -0.236 | -0.10198 | 0.109732 | 0.174759 | 0.175904 | -0.02801 | -0.13678 | -0.10365 | -0.0768 |
| 19.73611 | 0.607082 | -1.03978 | -2.24038 | 0.024843 | 0.173992 | -0.08285 | -1.19683 | -0.14458 | -0.3786 | -1.24685 | 0.058375 | 1.185312 | 0.026797 | -0.01607 | -0.00507 | -0.05848 | -0.07972 | 0.036628 | 0.133519 | 0.095177 | -0.04938 | -0.07572 | 0.023936 | -0.323 |
| 19.75 | -0.136 | -0.4972 | -2.13704 | -0.06772 | -0.04734 | -0.14068 | -1.07383 | -0.1042 | -0.32671 | -0.96009 | 0.294406 | 1.812091 | -0.18411 | 0.018294 | -0.05556 | 0.128258 | -0.0229 | 0.095964 | 0.135318 | 0.000133 | -0.04956 | 0.022902 | 0.058497 | -0.2874 |
| 19.76389 | -0.77123 | -0.34289 | -1.00168 | -0.17525 | -0.20832 | -0.11477 | -1.09289 | -0.03634 | -0.22207 | -0.89549 | 0.078437 | 1.593309 | -0.36306 | 0.027232 | -0.0538 | 0.133603 | -0.00382 | 0.292627 | 0.171145 | -0.04946 | 0.020884 | 0.122364 | -0.16091 | 0.099448 |
| 19.77778 | -1.17252 | -0.51359 | 0.376534 | -0.28618 | -0.24663 | -0.12521 | -0.85134 | 0.184819 | -0.23526 | -0.77064 | -0.41448 | 0.933296 | -0.52986 | -0.04974 | -0.06972 | -0.03767 | 0.010969 | 0.376395 | 0.153261 | -0.0905 | 0.090303 | 0.167787 | -0.37345 | 0.500047 |
| 19.79167 | -1.29516 | -0.70342 | 1.402212 | -0.37683 | -0.19486 | -0.07247 | -0.53424 | 0.470279 | -0.40779 | -0.50924 | -0.83322 | -0.20556 | -0.72814 | -0.17746 | -0.21325 | -0.24171 | 0.029325 | 0.149972 | 0.070013 | -0.18222 | 0.111224 | 0.118841 | -0.4348 | 0.567622 |
| 19.80556 | -0.88624 | -0.80967 | 1.416955 | -0.40671 | -0.14817 | -0.03411 | -0.19729 | 0.491079 | -0.5269 | -0.22671 | -1.27612 | -1.60503 | -0.8707 | -0.28788 | -0.3845 | -0.41604 | 0.021962 | -0.23116 | -0.00768 | -0.34795 | 0.080387 | 0.001691 | -0.37465 | 0.237563 |
| 19.81944 | -0.27648 | -0.94139 | 0.747761 | -0.37967 | -0.17022 | -0.08261 | 0.228283 | 0.108097 | -0.52008 | 0.105444 | -1.80218 | -2.69577 | -0.93584 | -0.30869 | -0.42712 | -0.5799 | -0.0701 | -0.56462 | -0.05417 | -0.55722 | -0.0034 | -0.08906 | -0.29929 | -0.15328 |
| 19.83333 | -0.21187 | -1.11436 | 0.02913 | -0.37453 | -0.26426 | -0.10483 | 0.811357 | -0.3567 | -0.42747 | 0.710714 | -2.16182 | -2.81147 | -1.09472 | -0.36764 | -0.33134 | -0.7071 | -0.21255 | -0.79128 | -0.13551 | -0.72676 | -0.11691 | -0.1504 | -0.19537 | -0.45962 |
| 19.84722 | -0.69341 | -1.16708 | -0.67307 | -0.44861 | -0.42165 | -0.29018 | 1.357786 | -0.56446 | -0.22454 | 1.204087 | -2.08174 | -2.62415 | -1.31853 | -0.57321 | -0.18574 | -0.80191 | -0.26139 | -0.9369 | -0.23061 | -0.79718 | -0.26496 | -0.17056 | -0.17716 | -0.73295 |
| 19.86111 | -1.22859 | -1.24112 | -1.34862 | -0.61282 | -0.65084 | -0.65216 | 1.322297 | -0.43473 | -0.10383 | 0.92951 | -1.66723 | -2.36216 | -1.51887 | -0.81889 | -0.13094 | -0.85899 | -0.25974 | -1.02286 | -0.27799 | -0.81208 | -0.39821 | -0.12656 | -0.31673 | -0.94549 |
| 19.875 | -1.58224 | -1.25056 | -1.81857 | -0.73738 | -0.86105 | -0.94379 | 1.524743 | -0.25184 | -0.18435 | 0.879267 | -1.36356 | -1.9754 | -1.63517 | -1.00802 | -0.26961 | -0.91936 | -0.23317 | -1.00638 | -0.29034 | -0.80455 | -0.49519 | -0.10898 | -0.57178 | -1.14228 |
| 19.88889 | -1.66641 | -1.16359 | -1.88185 | -0.75904 | -0.89985 | -0.91129 | 0.86312 | -0.1597 | -0.30834 | -0.16574 | -1.27014 | -1.67499 | -1.67195 | -0.8084 | -0.41174 | -0.81656 | -0.17245 | -0.85063 | -0.25501 | -0.76851 | -0.57778 | -0.14177 | -0.80917 | -1.23398 |
| 19.90278 | -1.33339 | -0.96021 | -1.57716 | -0.64778 | -0.69627 | -0.62306 | 0.13033 | -0.21284 | -0.30355 | -0.37005 | -1.28902 | -1.3061 | -1.55459 | -0.35418 | -0.40858 | -0.52759 | -0.09792 | -0.63358 | -0.23221 | -0.66987 | -0.60352 | -0.18852 | -0.89592 | -0.88614 |
| 19.91667 | -0.81149 | -0.63622 | -1.30519 | -0.40933 | -0.3769 | -0.32811 | -0.31403 | -0.35033 | -0.22256 | -0.28831 | -1.14004 | -0.845 | -1.19415 | 0.174209 | -0.21363 | -0.22502 | -0.05641 | -0.4539 | -0.20765 | -0.50108 | -0.51393 | -0.17904 | -0.87448 | -0.40723 |
| 19.93056 | -0.50924 | -0.42899 | -1.16559 | -0.24524 | -0.21909 | -0.23461 | -0.45438 | -0.49537 | -0.22883 | -0.35885 | -1.0081 | -0.53468 | -0.66035 | 0.512407 | -0.14103 | -0.01284 | -0.04811 | -0.39165 | -0.28806 | -0.3515 | -0.33099 | -0.15036 | -0.84529 | -0.22546 |
| 19.94444 | -0.85476 | -0.62467 | -1.18614 | -0.28084 | -0.34805 | -0.44412 | -0.81384 | -0.70922 | -0.39746 | -0.69195 | -1.34859 | -0.63772 | -0.5712 | 0.226697 | -0.27938 | -0.09766 | -0.15957 | -0.73786 | -0.683 | -0.43618 | -0.2046 | -0.33799 | -1.04571 | -0.61419 |
| 19.95833 | -1.89783 | -1.25316 | -1.3901 | -0.45685 | -0.74235 | -0.8822 | -1.16488 | -0.98637 | -0.7174 | -1.04641 | -2.40908 | -1.29293 | -1.2625 | -0.56087 | -0.54833 | -0.5283 | -0.40287 | -1.62783 | -1.62665 | -0.8596 | -0.43723 | -0.75541 | -1.67489 | -1.56354 |
| 19.97222 | -2.84065 | -1.80944 | -1.61443 | -0.56495 | -1.06033 | -1.15389 | -1.30895 | -1.10326 | -1.11809 | -1.2206 | -3.35101 | -2.17773 | -2.11828 | -1.17322 | -0.78486 | -0.90276 | -0.57444 | -2.12957 | -2.2773 | -1.18825 | -0.92 | -1.04383 | -2.09488 | -2.63223 |
| 19.98611 | -2.31902 | -1.59382 | -1.40761 | -0.52155 | -0.91165 | -0.92274 | -0.93301 | -0.9271 | -1.28104 | -0.99041 | -2.86203 | -2.46767 | -1.84356 | -0.95383 | -0.64267 | -0.77728 | -0.45124 | -1.56514 | -1.46433 | -0.87084 | -1.04508 | -0.83748 | -1.81968 | -2.4209 |
| 20 | -0.93397 | -0.887 | -0.85025 | -0.41963 | -0.5875 | -0.49031 | -0.4869 | -0.47428 | -0.76014 | -0.63085 | -1.38819 | -1.71595 | -0.80909 | -0.49326 | -0.25863 | -0.33971 | -0.19831 | -0.71332 | -0.5897 | -0.25192 | -0.71975 | -0.44162 | -1.09471 | -0.72914 |
| 20.01389 | 0.134629 | -0.08796 | -0.34522 | -0.31357 | -0.13747 | -0.16495 | -0.10631 | 0.057725 | 0.16999 | -0.31002 | 0.018842 | -0.78566 | -0.01883 | -0.24114 | -0.04882 | -0.01471 | -0.03864 | -0.03877 | 0.006246 | 0.18265 | -0.2307 | -0.17251 | -0.10152 | 0.873666 |
| 20.02778 | 0.513842 | 0.37684 | -0.1072 | -0.11214 | 0.291953 | 0.001333 | 0.109723 | 0.347245 | 1.048196 | -0.1144 | 0.90723 | 0.064631 | 0.477349 | -0.16841 | -0.04003 | 0.064491 | 0.010963 | 0.375111 | 0.382989 | 0.347895 | 0.178356 | -0.09335 | 0.882993 | 1.895384 |
| 20.04167 | 0.362763 | 0.329891 | -0.01324 | 0.142185 | 0.409839 | -0.02838 | -0.02176 | 0.288366 | 1.076724 | -0.08735 | 0.794226 | 0.457127 | 0.840223 | -0.13764 | -0.07656 | -0.03656 | 0.013418 | 0.505368 | 0.387661 | 0.296028 | 0.33665 | -0.07684 | 1.190402 | 0.952087 |
| 20.05556 | 0.195563 | 0.035848 | 0.03391 | 0.248794 | 0.303011 | -0.19203 | -0.33184 | 0.051896 | 0.535637 | -0.20277 | 0.290957 | 0.200644 | 0.677161 | -0.14264 | -0.12504 | -0.16002 | -0.04547 | 0.350741 | 0.180832 | 0.042024 | 0.219529 | -0.11528 | 0.791287 | 0.016189 |
| 20.06944 | 0.010853 | -0.28825 | 0.066676 | 0.165202 | 0.164732 | -0.31143 | -0.54196 | -0.23044 | 0.058916 | -0.39159 | -0.14776 | -0.26144 | 0.202401 | -0.19376 | -0.13722 | -0.22776 | -0.09572 | 0.140518 | -0.03281 | -0.21271 | 0.035077 | -0.15504 | 0.227331 | -0.51584 |
| 20.08333 | -0.29509 | -0.76374 | 0.062322 | 0.043907 | 0.297371 | -0.45737 | -0.58471 | -0.47388 | -0.22845 | -0.5627 | -0.61471 | -1.03876 | -0.21645 | -0.26416 | -0.12166 | -0.26059 | -0.05868 | 0.008071 | -0.17753 | -0.38995 | -0.11668 | -0.17765 | -0.21252 | -0.83949 |
| 20.09722 | -0.30507 | -1.04321 | 0.003434 | -0.06665 | 0.334375 | -0.60008 | -0.61995 | -0.35514 | -0.39194 | -0.67524 | -0.90525 | -1.71683 | -0.48529 | -0.26474 | -0.13349 | -0.299 | 0.021747 | -0.07683 | -0.26685 | -0.49595 | -0.23655 | -0.2038 | -0.50129 | -0.85579 |
| 20.11111 | 0.183094 | -0.35121 | 0.067183 | -0.0397 | -0.21809 | -0.42297 | -0.60579 | -0.00565 | -0.38042 | -0.58135 | -0.60263 | -0.76947 | -0.47993 | 0.008908 | -0.1067 | -0.1988 | 0.035863 | 0.034856 | -0.19222 | -0.2931 | -0.179 | -0.13361 | -0.45431 | -0.25081 |
| 20.125 | 0.649449 | 0.519964 | 0.225584 | 0.054224 | -0.65251 | 0.027163 | -0.52957 | 0.200783 | -0.17632 | -0.40399 | -0.02967 | 0.426421 | -0.31402 | 0.192917 | -0.08084 | -0.01912 | -0.04892 | 0.205359 | -0.02425 | -0.03372 | -0.01255 | -0.01865 | -0.19468 | 0.485067 |
| 20.13889 | 0.525101 | 0.560171 | 0.295704 | 0.049136 | -0.33638 | 0.23336 | -0.45861 | -0.04537 | 0.093903 | -0.36299 | 0.116357 | 0.376163 | -0.20167 | 0.005049 | -0.08514 | 0.007225 | -0.13213 | 0.232642 | 0.032469 | -0.01307 | 0.047866 | 0.023573 | 0.021667 | 0.600171 |
| 20.15278 | 0.158174 | 0.217231 | 0.273964 | -0.02995 | -0.01638 | 0.115266 | -0.44851 | -0.2539 | 0.283046 | -0.3821 | 0.044601 | 0.05906 | -0.096 | -0.19938 | 0.006483 | -0.07113 | -0.14343 | 0.212542 | -0.00134 | -0.16818 | 0.028737 | 0.007485 | 0.171401 | 0.12961 |
| 20.16667 | -0.11714 | 0.0254 | 0.247874 | -0.08048 | -0.04709 | -0.07203 | -0.4919 | -0.16679 | 0.201836 | -0.37205 | -0.04936 | -0.12619 | -0.07106 | -0.31252 | 0.142171 | -0.11425 | -0.07114 | 0.191982 | -0.04278 | -0.29329 | -0.02822 | -0.01749 | 0.22487 | -0.38308 |
| 20.18056 | -0.26661 | -0.09627 | 0.316374 | -0.08198 | -0.1149 | -0.13945 | -0.42314 | -0.00308 | -0.11242 | -0.35357 | -0.05749 | -0.19044 | -0.12347 | -0.26628 | 0.163842 | -0.12602 | 0.071623 | 0.166324 | -0.03466 | -0.27374 | -0.08684 | -0.02275 | 0.126623 | -0.45931 |
| 20.19444 | -0.05233 | -0.10089 | 0.429985 | -0.06145 | -0.10781 | 0.017118 | -0.29963 | 0.063968 | -0.38391 | -0.36713 | -0.00845 | -0.12093 | -0.15269 | -0.1127 | 0.024649 | -0.08395 | 0.143032 | 0.189829 | -0.00017 | -0.13692 | -0.09942 | -0.02344 | -0.01833 | -0.27603 |
| 20.20833 | 0.357949 | 0.03609 | 0.444084 | -0.03329 | -0.03804 | 0.176216 | -0.24018 | -0.04311 | -0.3291 | -0.37597 | 0.011935 | 0.039938 | -0.10388 | -0.03928 | -0.11304 | -0.0532 | 0.05953 | 0.239002 | 0.020549 | 0.036725 | -0.04717 | -0.02831 | -0.05819 | -0.21346 |
| 20.22222 | 0.535359 | 0.1978 | 0.384056 | -0.01981 | 0.04149 | 0.173533 | -0.23581 | -0.11085 | 0.068707 | -0.31328 | 0.023352 | 0.327445 | 0.007159 | -0.12798 | -0.07166 | -0.08755 | -0.08291 | 0.258318 | 0.03174 | 0.137381 | 0.023921 | -0.02033 | 0.014037 | -0.30936 |
| 20.23611 | 0.280265 | 0.265042 | 0.346291 | -0.03084 | 0.041395 | 0.085327 | -0.24725 | -0.02837 | 0.445558 | -0.23378 | 0.093841 | 0.536709 | 0.063402 | -0.26062 | 0.09637 | -0.13111 | -0.15064 | 0.242454 | 0.020175 | 0.119571 | 0.024251 | -0.01921 | 0.113086 | -0.28423 |
| 20.25 | -0.07981 | 0.226616 | 0.384108 | -0.05084 | -0.00357 | -0.00363 | -0.2493 | 0.084224 | 0.454297 | -0.16247 | 0.207116 | 0.536096 | 0.069041 | -0.2309 | 0.227663 | -0.10914 | -0.05738 | 0.217955 | -0.00114 | -0.01978 | -0.01434 | -0.01142 | 0.182104 | 0.096129 |
| 20.26389 | -0.35345 | 0.122305 | 0.437999 | -0.04643 | 0.020346 | 0.048666 | -0.1747 | 0.15152 | 0.080135 | -0.08049 | 0.315776 | 0.328804 | 0.099138 | 0.018428 | 0.177343 | -0.00837 | 0.135936 | 0.248866 | 0.026367 | -0.17071 | -0.00802 | 0.002174 | 0.157024 | 0.471364 |
| 20.27778 | -0.40546 | 0.048067 | 0.452702 | -0.00816 | 0.113257 | 0.228512 | -0.04243 | 0.123395 | -0.30439 | 0.008093 | 0.348667 | 0.110818 | 0.124543 | 0.21163 | -0.0171 | 0.110211 | 0.225558 | 0.317515 | 0.104587 | -0.15114 | 0.052243 | -0.00856 | 0.088616 | 0.306182 |
| 20.29167 | -0.07168 | 0.111465 | 0.38455 | 0.04745 | 0.156419 | 0.323021 | 0.039254 | 0.071329 | -0.17876 | 0.103326 | 0.287017 | 0.240774 | 0.116421 | 0.141837 | -0.18826 | 0.123607 | 0.147717 | 0.341973 | 0.149659 | 0.044911 | 0.137153 | -0.02302 | 0.14626 | -0.21896 |
| 20.30556 | 0.420728 | 0.305397 | 0.299064 | 0.082782 | 0.107043 | 0.196097 | 0.048749 | 0.125036 | 0.466639 | 0.206189 | 0.241809 | 0.681091 | 0.115933 | -0.01151 | -0.11942 | 0.049378 | -0.00278 | 0.307924 | 0.104769 | 0.225071 | 0.220066 | -0.00401 | 0.361387 | -0.53272 |
| 20.31944 | 0.639076 | 0.482898 | 0.26496 | 0.077881 | 0.044354 | -0.10027 | 0.035459 | 0.254916 | 0.99175 | 0.294308 | 0.282294 | 1.106374 | 0.166993 | 0.039616 | 0.136889 | 0.021415 | -0.04298 | 0.25586 | 0.035841 | 0.206805 | 0.288188 | 0.03271 | 0.517372 | -0.0479 |
| 20.33333 | 0.44042 | 0.525674 | 0.301285 | 0.100844 | 0.042155 | -0.30987 | 0.05913 | 0.414764 | 0.83212 | 0.345936 | 0.360237 | 0.980575 | 0.284213 | 0.373666 | 0.371495 | 0.106873 | 0.059915 | 0.214765 | 0.004982 | 0.070955 | 0.311267 | 0.067617 | 0.443064 | 0.833667 |
| 20.34722 | 0.236683 | 0.455509 | 0.415628 | 0.182693 | 0.094881 | -0.25256 | 0.12413 | 0.407433 | 0.277932 | 0.398674 | 0.458462 | 0.668505 | 0.419633 | 0.680451 | 0.354299 | 0.224236 | 0.190998 | 0.263465 | 0.060808 | 0.020052 | 0.257184 | 0.063057 | 0.25188 | 0.974131 |
| 20.36111 | 0.110164 | 0.304877 | 0.527025 | 0.245861 | 0.13903 | -0.01818 | 0.171012 | 0.324963 | -0.18038 | 0.455573 | 0.506056 | 0.479454 | 0.50076 | 0.636413 | 0.161132 | 0.289785 | 0.230815 | 0.371541 | 0.196557 | 0.11498 | 0.193525 | 0.021259 | 0.10099 | 0.207461 |
| 20.375 | 0.152448 | 0.274011 | 0.599828 | 0.227269 | 0.145499 | 0.11526 | 0.169322 | 0.272216 | -0.32852 | 0.489713 | 0.518703 | 0.556665 | 0.510023 | 0.480445 | -0.00563 | 0.309878 | 0.193115 | 0.487502 | 0.305342 | 0.276272 | 0.203087 | 0.013782 | 0.288036 | -0.33314 |
| 20.38889 | 0.627892 | 0.50569 | 0.684728 | 0.194361 | 0.156811 | 0.153417 | 0.161342 | 0.351443 | 0.106234 | 0.516588 | 0.442629 | 0.77659 | 0.434877 | 0.367552 | 0.007176 | 0.298761 | 0.169209 | 0.503837 | 0.29315 | 0.346116 | 0.28474 | 0.048843 | 0.559463 | -0.45592 |
| 20.40278 | 1.151698 | 0.90007 | 0.802618 | 0.238448 | 0.169166 | 0.165875 | 0.195095 | 0.530119 | 0.826286 | 0.554547 | 0.375097 | 0.981466 | 0.359305 | 0.442967 | 0.223972 | 0.285702 | 0.148686 | 0.362025 | 0.176049 | 0.247783 | 0.335819 | 0.081305 | 0.583263 | 0.818069 |
| 20.41667 | 1.214118 | 0.986255 | 0.942909 | 0.345609 | 0.135493 | 0.214797 | 0.21841 | 0.60662 | 1.112721 | 0.602596 | 0.467477 | 1.119287 | 0.440242 | 0.598978 | 0.449937 | 0.370164 | 0.180916 | 0.309922 | 0.159151 | 0.127416 | 0.368375 | 0.113766 | 0.382494 | 1.49235 |
| 20.43056 | 1.009583 | 0.724083 | 1.023379 | 0.403487 | 0.130861 | 0.304946 | 0.212404 | 0.485482 | 0.659556 | 0.641021 | 0.805252 | 1.022589 | 0.740174 | 0.630993 | 0.51608 | 0.51724 | 0.282532 | 0.489977 | 0.320436 | 0.153508 | 0.376277 | 0.153762 | 0.284624 | 0.92674 |
| 20.44444 | 1.557507 | 1.027439 | 1.406705 | 0.585155 | 0.492887 | 0.453524 | 0.37266 | 0.569128 | 0.274996 | 1.072481 | 1.843982 | 1.56098 | 1.771957 | 1.020375 | 0.782947 | 0.79176 | 0.335615 | 1.183104 | 0.732145 | 0.469352 | 0.621007 | 0.212084 | 0.590748 | 0.335806 |
| 20.45833 | 3.614756 | 3.154807 | 3.099254 | 1.098506 | 1.407771 | 0.541183 | 0.810036 | 1.200149 | 0.307953 | 2.630415 | 3.572825 | 3.987945 | 4.107778 | 2.620993 | 1.190588 | 1.416901 | 0.352805 | 2.45709 | 1.515112 | 1.204048 | 1.421013 | 0.38649 | 1.333123 | 0.971527 |
| 20.47222 | 4.621606 | 6.438427 | 3.423155 | 1.212483 | 1.963985 | 0.657272 | 1.022584 | 1.614444 | 0.663222 | 4.174962 | 4.22842 | 5.735128 | 5.32702 | 2.608486 | 2.1186 | 1.889493 | 0.380328 | 2.92016 | 2.099449 | 2.007892 | 1.830214 | 0.530587 | 1.676046 | 2.734474 |
| 20.48611 | 2.805471 | 5.537513 | 2.246012 | 0.702171 | 1.469304 | -0.37813 | 0.580066 | 1.232408 | 0.995066 | 1.842469 | 2.634625 | 3.783765 | 2.649173 | 1.231306 | 1.488383 | 1.339354 | 0.287003 | 1.962216 | 1.494905 | 1.306178 | 1.142107 | 0.500584 | 1.277684 | 3.56077 |
| 20.5 | 0.996813 | 2.422993 | 1.183068 | 0.250915 | 0.627837 | -0.40639 | 0.238535 | 0.474208 | 1.426787 | 0.819136 | 1.114441 | 2.200806 | 0.863418 | 0.219189 | 0.784271 | 0.796701 | 0.24456 | 1.102107 | 0.696094 | 0.522857 | 0.535094 | 0.318575 | 0.531318 | 3.006761 |
| 20.51389 | -0.17163 | 0.691048 | 0.158782 | 0.023565 | 0.095986 | -0.39112 | 0.467841 | -0.07882 | 0.375519 | 0.536715 | -0.25395 | 0.139764 | 0.00145 | -0.44654 | 0.232475 | 0.527941 | 0.152193 | 0.539945 | 0.116946 | 0.145198 | 0.283887 | 0.294113 | 0.08899 | 0.748337 |
| 20.52778 | -0.83841 | -0.08712 | -0.73633 | -0.05502 | -0.21538 | -0.72719 | 0.773246 | -0.47898 | -0.45364 | 0.506705 | -0.98524 | -1.55592 | -0.42559 | -0.97162 | -0.3151 | 0.222495 | -0.01224 | -0.05623 | -0.22099 | -0.10401 | 0.141965 | 0.223113 | -0.38693 | -1.17589 |
| 20.54167 | -1.07343 | -0.46472 | -1.50372 | -0.03033 | -0.3802 | -1.10308 | 0.921514 | -0.75729 | -1.13269 | 0.533429 | -1.14847 | -3.11155 | -0.2838 | -1.03375 | -0.80108 | -0.09187 | -0.10786 | -0.286 | -0.19482 | -0.19037 | 0.020809 | 0.138987 | -0.74545 | -2.59374 |
| 20.55556 | -0.65576 | -0.20686 | -1.93583 | 0.081008 | -0.23234 | -0.68656 | 1.291292 | -0.79748 | -1.17922 | 0.831236 | -0.33845 | -3.50972 | 0.369364 | -0.2414 | -0.89822 | -0.00244 | -0.05258 | -0.12776 | 0.252965 | 0.031374 | 0.052964 | 0.151991 | -0.12899 | -2.21186 |
| 20.56944 | 0.244364 | 0.514047 | -1.5942 | 0.194601 | 0.043336 | 0.148548 | 1.704284 | -0.42442 | -0.34442 | 1.069881 | 0.956712 | -2.12287 | 1.018999 | 0.883848 | -0.39267 | 0.241036 | 0.061882 | 0.142851 | 0.696338 | 0.278793 | 0.100406 | 0.216858 | 0.750593 | -0.54137 |
| 20.58333 | 1.081277 | 1.015674 | -1.00758 | 0.221356 | 0.100145 | 0.478126 | 1.029451 | 0.126286 | 0.73799 | 0.606534 | 1.899267 | 0.016739 | 0.925299 | 1.321494 | -0.01277 | 0.115332 | 0.17841 | 0.092224 | 0.723742 | 0.182535 | 0.027594 | 0.244645 | 0.925272 | 1.473724 |
| 20.59722 | 1.571174 | 0.798166 | -0.51216 | 0.174193 | -0.03502 | 0.19152 | 0.466798 | 0.575923 | 1.433744 | 0.37792 | 1.648553 | 1.941014 | 0.247969 | 0.907073 | 0.165425 | -0.08206 | 0.162739 | -0.21577 | 0.568716 | -0.06156 | -0.0603 | 0.161657 | 0.439579 | 2.646627 |
| 20.61111 | 1.833152 | 0.230138 | 0.330073 | 0.102764 | -0.09595 | 0.171962 | 0.231647 | 0.964807 | 1.112529 | 0.344321 | 1.023028 | 3.290707 | -0.21642 | 0.213706 | 0.023855 | -0.161 | 0.194507 | -0.37761 | 0.396421 | -0.24327 | -0.11073 | 0.061273 | 0.298662 | 1.928277 |
| 20.625 | 1.988696 | -0.04429 | 1.410164 | 0.10367 | -0.06905 | 0.131898 | 0.028421 | 1.27681 | 0.621356 | 0.4644 | 0.293237 | 3.950328 | -0.51914 | -0.20286 | -0.12576 | -0.02627 | 0.187125 | -0.38471 | 0.231456 | -0.35772 | -0.20794 | -0.03223 | 0.293985 | 0.942164 |
| 20.63889 | 1.83468 | -0.06527 | 2.238956 | 0.079325 | -0.13589 | -0.08936 | -0.14984 | 1.191503 | 0.161829 | 0.47027 | -0.43097 | 3.275041 | -0.39821 | -0.38201 | -0.16736 | 0.244325 | 0.211241 | -0.15149 | 0.12293 | -0.30994 | -0.23505 | -0.0196 | 0.213808 | -0.29056 |
| 20.65278 | 1.499486 | -0.2538 | 2.539479 | -0.02288 | -0.25525 | -0.18212 | -0.48246 | 0.731134 | -0.36016 | 0.261411 | -0.8343 | 1.772118 | 0.031705 | -0.46007 | 0.056899 | 0.428249 | 0.321799 | 0.095209 | 0.146506 | -0.12864 | -0.13646 | 0.102685 | 0.245065 | -1.64425 |
| 20.66667 | 1.08435 | -0.54526 | 2.392397 | -0.17802 | -0.20385 | -0.00887 | -0.74761 | 0.36175 | -0.70928 | 0.055369 | -0.2713 | 0.386123 | 0.416157 | -0.44863 | 0.391429 | 0.526629 | 0.257697 | 0.258049 | 0.246564 | 0.144065 | -0.00224 | 0.182018 | 0.480145 | -2.77087 |
| 20.68056 | 0.308664 | -0.38679 | 1.904648 | -0.20705 | 0.051119 | 0.221221 | -0.96394 | 0.156993 | -0.41378 | 0.06609 | 0.666766 | -0.31912 | 0.65261 | -0.05931 | 0.615612 | 0.513859 | 0.142477 | 0.365083 | 0.310652 | 0.441509 | 0.173057 | 0.237537 | 0.670779 | -2.21053 |
| 20.69444 | -0.63373 | 0.328958 | 1.280404 | -0.05831 | 0.320704 | 0.355337 | -1.20494 | -0.0415 | 0.313421 | 0.105083 | 1.018648 | -0.80497 | 0.74056 | 0.414523 | 0.672788 | 0.424409 | 0.033819 | 0.400028 | 0.368999 | 0.614947 | 0.378209 | 0.287025 | 0.536128 | -0.80213 |
| 20.70833 | -1.25394 | 1.294603 | 0.361522 | 0.275928 | 0.531589 | 0.238408 | -0.92774 | -0.24557 | 0.850699 | 0.023109 | 0.66929 | -1.23799 | 0.445727 | 0.497297 | 0.594094 | 0.240702 | 0.014226 | 0.336429 | 0.348189 | 0.509544 | 0.412663 | 0.287204 | 0.215138 | 1.116291 |
| 20.72222 | -1.19254 | 1.758583 | -0.56709 | 0.585233 | 0.591783 | -0.00794 | -0.23556 | -0.39588 | 0.778683 | -0.00498 | -0.24582 | -1.85574 | -0.06084 | 0.279701 | 0.077488 | -0.04094 | -0.04614 | 0.114525 | 0.221294 | 0.29771 | 0.22396 | 0.17614 | -0.00843 | 2.803705 |
| 20.73611 | -0.69873 | 1.458495 | -1.2694 | 0.643206 | 0.485044 | -0.37808 | 0.52084 | -0.52373 | 0.373869 | 0.032497 | -1.12802 | -2.14001 | -0.37461 | 0.02819 | -0.54169 | -0.21422 | -0.05355 | -0.2065 | 0.039928 | 0.031103 | -0.00573 | 0.052703 | -0.1864 | 2.619427 |
| 20.75 | -0.17939 | 0.613148 | -2.11736 | 0.486358 | 0.2282 | -0.81708 | 1.183372 | -0.5345 | 0.054607 | 0.021138 | -1.55683 | -1.69387 | -0.36277 | -0.16724 | -1.05937 | -0.3643 | 0.021703 | -0.43464 | -0.03241 | -0.21694 | -0.03689 | 0.024727 | -0.24906 | 1.50519 |
| 20.76389 | 0.183694 | -0.33838 | -3.01346 | 0.159229 | -0.15346 | -1.18431 | 1.364493 | -0.39675 | -0.09231 | 0.173296 | -1.0916 | -0.72868 | -0.21325 | -0.41336 | -1.09804 | -0.50441 | 0.029935 | -0.56076 | 0.000421 | -0.38047 | 0.022319 | 0.033056 | -0.15512 | 0.300425 |
| 20.77778 | 0.439424 | -1.2579 | -3.42981 | -0.22308 | -0.54985 | -0.76525 | 1.003558 | -0.09731 | -0.09711 | 0.28855 | -0.1372 | 0.324461 | -0.16152 | -0.7597 | -0.8029 | -0.59896 | 0.112757 | -0.52106 | 0.027058 | -0.49363 | 0.000688 | 0.019101 | -0.0885 | -1.215 |
| 20.79167 | 0.529017 | -2.07839 | -2.85782 | -0.53453 | -0.77486 | -0.21438 | 0.516412 | 0.194497 | -0.14804 | 0.230306 | 0.695493 | 1.413324 | -0.19917 | -0.86992 | -0.35425 | -0.51024 | 0.084438 | -0.38675 | -0.02648 | -0.56684 | -0.10488 | 0.026537 | 0.003217 | -2.23695 |
| 20.80556 | 0.357834 | -2.54984 | -2.14805 | -0.77624 | -0.82087 | -0.24948 | -0.06706 | 0.336097 | -0.27311 | -0.06847 | 1.103389 | 2.053307 | -0.37117 | -0.51921 | -0.05976 | -0.28408 | 0.03966 | -0.33718 | -0.10219 | -0.59662 | -0.20033 | 0.057347 | 0.191094 | -1.76476 |
| 20.81944 | -0.09475 | -2.42159 | -1.23995 | -0.95594 | -0.81607 | -0.47509 | -0.10124 | 0.169002 | -0.55715 | -0.24994 | 0.865689 | 1.294939 | -0.69012 | -0.2189 | -0.14759 | -0.11376 | -0.00194 | -0.51193 | -0.219 | -0.65419 | -0.27196 | 0.069116 | 0.13652 | -1.09156 |
| 20.83333 | -0.82327 | -1.89873 | -0.36963 | -1.0394 | -0.80247 | -0.45571 | -0.18263 | -0.23571 | -0.91688 | -0.42281 | 0.160442 | -0.3756 | -1.06828 | -0.16361 | -0.39521 | -0.18113 | -0.05824 | -0.76035 | -0.3583 | -0.73724 | -0.30848 | 0.01253 | -0.31305 | -0.1516 |
| 20.84722 | -1.54665 | -1.41491 | 0.456225 | -0.97834 | -0.68059 | -0.27478 | -0.50448 | -0.66492 | -0.95839 | -0.56571 | -0.88572 | -1.59332 | -1.39842 | -0.27877 | -0.54451 | -0.45411 | -0.15587 | -0.92759 | -0.56892 | -0.75828 | -0.41518 | -0.08156 | -0.93623 | 0.79144 |
| 20.86111 | -1.91418 | -1.20938 | 1.07023 | -0.82014 | -0.57965 | -0.23469 | -1.23475 | -1.03004 | -0.61319 | -0.64206 | -2.06734 | -2.23381 | -1.79681 | -0.345 | -0.73219 | -0.70389 | -0.22758 | -1.0167 | -0.79848 | -0.80942 | -0.56822 | -0.18874 | -1.37366 | 1.073632 |
| 20.875 | -1.60837 | -1.1174 | 1.053402 | -0.59968 | -0.53243 | -0.33662 | -1.89603 | -1.26217 | -0.23856 | -0.44413 | -3.06037 | -2.31118 | -2.05364 | -0.44685 | -1.02631 | -0.8304 | -0.2603 | -1.09348 | -0.93031 | -0.93724 | -0.67595 | -0.27606 | -1.29493 | 0.627782 |
| 20.88889 | -1.00019 | -0.87474 | 0.506643 | -0.32958 | -0.53911 | -0.39948 | -2.04516 | -1.26395 | 0.049872 | -0.33205 | -3.2234 | -2.0182 | -1.85777 | -0.60273 | -1.13406 | -0.87258 | -0.35361 | -0.98689 | -0.87038 | -1.05482 | -0.68355 | -0.27105 | -0.96918 | -0.26211 |
| 20.90278 | -0.82587 | -0.52279 | -0.17765 | -0.06753 | -0.56769 | -0.42955 | -2.17814 | -1.11007 | 0.179341 | -0.55126 | -2.17028 | -1.62488 | -1.40313 | -0.69117 | -0.65302 | -0.92528 | -0.48041 | -0.78579 | -0.75424 | -0.82385 | -0.54433 | -0.16301 | -0.74458 | -1.30251 |

| | | | | | | | | | | | | | | | | | | | | | | | | |
|---|---|---|---|---|---|---|---|---|---|---|---|---|---|---|---|---|---|---|---|---|---|---|---|---|
| 20.91667 | -1.01179 | -0.25933 | -0.69896 | 0.112762 | -0.51967 | -0.55821 | -2.12711 | -0.82043 | 0.027062 | -0.73391 | -0.76224 | -1.57237 | -0.91933 | -0.64848 | -0.23246 | -0.8159 | -0.47261 | -0.67044 | -0.60349 | -0.37063 | -0.40706 | -0.09478 | -0.62984 | -2.07081 |
| 20.93056 | -1.12418 | -0.21538 | -0.87958 | 0.173526 | -0.3657 | -0.76841 | -1.62796 | -0.56164 | -0.39692 | -0.78143 | -0.00313 | -1.79728 | -0.54278 | -0.52535 | -0.22641 | -0.42461 | -0.36433 | -0.67913 | -0.56971 | -0.06246 | -0.37155 | -0.14932 | -0.72766 | -2.5626 |
| 20.94444 | -1.37112 | -0.60608 | -0.85808 | 0.077125 | -0.33013 | -1.00286 | -1.57088 | -0.56694 | -0.93064 | -1.08997 | -0.36937 | -2.19246 | -0.41524 | -0.51373 | -0.63104 | -0.13253 | -0.32729 | -0.91545 | -0.78113 | -0.2336 | -0.47508 | -0.39679 | -1.17469 | -2.76676 |
| 20.95833 | -2.14443 | -1.68732 | -1.11182 | -0.20169 | -0.63588 | -1.35953 | -2.2814 | -0.83845 | -1.5728 | -1.95525 | -1.76787 | -2.9364 | -0.79064 | -0.93552 | -1.22523 | -0.42681 | -0.478 | -1.60848 | -1.31853 | -0.81774 | -0.7966 | -0.82566 | -2.14486 | -3.2315 |
| 20.97222 | -2.99071 | -2.72585 | -1.75195 | -0.50893 | -1.13671 | -1.77213 | -2.74735 | -1.12042 | -2.16044 | -2.127 | -3.19257 | -4.043 | -1.36259 | -1.59431 | -1.62064 | -1.02637 | -0.78825 | -2.44095 | -1.76979 | -1.39911 | -1.23032 | -1.08187 | -2.84856 | -4.21308 |
| 20.98611 | -2.59654 | -2.70056 | -2.1808 | -0.62979 | -1.45339 | -1.89751 | -2.46065 | -0.98708 | -2.44392 | -1.64766 | -3.49892 | -4.52977 | -1.51764 | -1.58677 | -1.31328 | -1.27003 | -0.76263 | -2.31414 | -1.6642 | -1.47123 | -1.15885 | -0.83979 | -2.34397 | -4.38419 |
| 21 | -1.25948 | -1.55055 | -2.15026 | -0.55715 | -1.04271 | -1.4678 | -1.38916 | -0.49644 | -1.93442 | -1.14625 | -2.23535 | -3.35419 | -1.30449 | -0.98005 | -0.43296 | -1.02365 | -0.29963 | -1.37077 | -1.04427 | -1.01708 | -0.70199 | -0.50106 | -1.2506 | -1.93482 |
| 21.01389 | 0.010343 | -0.35814 | -1.74585 | -0.48845 | -0.47156 | -0.77236 | -0.30719 | 0.088802 | -0.99836 | -0.66625 | -0.57991 | -1.58956 | -1.10231 | -0.38082 | 0.499037 | -0.59783 | 0.039351 | -0.53054 | -0.31971 | -0.37091 | -0.28384 | -0.34407 | -0.1582 | 0.207652 |
| 21.02778 | 0.792531 | 0.209439 | -1.26401 | -0.47897 | 0.060195 | -0.29458 | 0.264072 | 0.372994 | 0.011546 | -0.34137 | 0.771124 | 0.038897 | -0.7702 | 0.00627 | 0.765808 | -0.32333 | 0.176776 | 0.043024 | 0.359222 | 0.272229 | 0.029724 | -0.25877 | 0.765216 | 2.185285 |
| 21.04167 | 0.697808 | -0.13643 | -0.67223 | -0.48977 | 0.432967 | -0.06117 | 0.292831 | 0.275047 | 0.890092 | -0.22461 | 1.351412 | 1.223697 | -0.12693 | -0.10609 | 0.068238 | -0.249 | 0.232 | 0.389955 | 0.765651 | 0.710344 | 0.194553 | -0.1678 | 1.278057 | 2.061346 |
| 21.05556 | 0.051025 | -0.59916 | -0.64769 | -0.5018 | 0.554829 | -0.23164 | 0.133024 | 0.006729 | 1.30596 | -0.24323 | 0.776581 | 1.529743 | 0.502902 | -0.38966 | -0.57553 | -0.31083 | 0.151501 | 0.433316 | 0.595129 | 0.530196 | 0.174691 | -0.0597 | 1.216526 | 0.77258 |
| 21.06944 | -0.41368 | -0.70719 | -0.82355 | -0.41014 | 0.404062 | -0.55484 | -0.04656 | -0.22388 | 1.137209 | -0.33115 | 0.09004 | 0.888731 | 0.614043 | -0.64878 | -0.86327 | -0.33504 | 0.069413 | 0.237405 | 0.228723 | 0.237225 | 0.07515 | -0.01653 | 0.751792 | 0.116876 |
| 21.08333 | -0.71921 | -0.63718 | -0.93676 | -0.26653 | 0.167965 | -0.71641 | -0.16657 | -0.24439 | 0.878378 | -0.42734 | -0.34637 | 0.329026 | 0.308044 | -0.8439 | -0.84666 | -0.31605 | -0.00119 | 0.09341 | -0.06221 | -0.06581 | -0.03645 | -0.05514 | 0.268153 | -0.1878 |
| 21.09722 | -0.8543 | -0.49909 | -0.97568 | -0.19243 | 0.024058 | -0.6175 | -0.15142 | -0.12815 | 0.618582 | -0.44241 | -0.58672 | -0.01565 | 0.031179 | -0.76734 | -0.64735 | -0.31063 | -0.08008 | 0.007276 | -0.24008 | -0.31317 | -0.13215 | -0.07785 | -0.03689 | -0.25354 |
| 21.11111 | -0.7889 | -0.3686 | -0.94402 | -0.16422 | -0.06646 | -0.35605 | -0.13434 | 0.0552 | 0.38833 | -0.37104 | -0.74197 | -0.27045 | -0.11882 | -0.32634 | -0.37802 | -0.2249 | -0.0262 | -0.00862 | -0.26849 | -0.44664 | -0.14398 | -0.03101 | -0.20155 | -0.1245 |
| 21.125 | -0.393 | -0.29238 | -0.74088 | -0.12751 | -0.10123 | -0.06583 | -0.12209 | 0.150021 | 0.237768 | -0.22372 | -0.70581 | -0.33945 | -0.24547 | 0.007816 | -0.3564 | -0.13742 | 0.056065 | 0.046563 | -0.15859 | -0.44517 | -0.11106 | -0.01701 | -0.29853 | 0.12235 |
| 21.13889 | 0.222828 | -0.24605 | -0.36761 | -0.08392 | -0.02886 | 0.121656 | -0.07635 | 0.068594 | 0.229597 | -0.07691 | -0.29338 | -0.15067 | -0.40604 | -0.12555 | -0.56345 | -0.15797 | -0.01075 | 0.084512 | 0.00628 | -0.29381 | -0.11137 | -0.07475 | -0.27651 | 0.306938 |
| 21.15278 | 0.472143 | -0.21974 | -0.09864 | -0.07878 | 0.072265 | 0.035853 | -0.00687 | -0.0981 | 0.279428 | -0.06322 | 0.09931 | 0.1332 | -0.42805 | -0.47093 | -0.5432 | -0.15712 | -0.06263 | 0.111231 | 0.060325 | -0.04708 | -0.07491 | -0.08024 | -0.06473 | 0.32564 |
| 21.16667 | 0.232432 | -0.21584 | -0.09371 | -0.09514 | 0.103821 | -0.16628 | 0.06747 | -0.15757 | 0.332682 | -0.12494 | 0.014929 | 0.329912 | -0.2708 | -0.6341 | -0.25203 | -0.13579 | -0.06344 | 0.161419 | -0.11107 | -0.01134 | -0.02597 | -0.04626 | 0.173306 | 0.19938 |
| 21.18056 | -0.04872 | -0.21225 | -0.20712 | -0.09572 | 0.065957 | -0.2467 | 0.095747 | 0.021884 | 0.306051 | -0.1232 | -0.30915 | 0.34741 | -0.14266 | -0.47929 | -0.11966 | -0.16133 | -0.08287 | 0.113061 | -0.36944 | -0.29168 | -0.06966 | -0.06147 | 0.180413 | 0.073641 |
| 21.19444 | -0.17218 | -0.17228 | -0.26418 | -0.0797 | -0.06823 | -0.18491 | -0.03406 | 0.179027 | 0.044855 | -0.11462 | -0.51928 | 0.04675 | -0.15922 | -0.24433 | -0.25975 | -0.14842 | -0.03416 | 0.011171 | -0.44939 | -0.6123 | -0.17366 | -0.10946 | -0.0628 | -0.00515 |
| 21.20833 | -0.04947 | -0.08899 | -0.10895 | -0.02343 | -0.27083 | -0.04618 | -0.2307 | 0.074023 | -0.3076 | -0.12084 | -0.57362 | -0.41631 | -0.23489 | -0.01858 | -0.45774 | -0.10434 | 0.061987 | 0.020922 | -0.24459 | -0.60415 | -0.23907 | -0.14491 | -0.24733 | -0.09036 |
| 21.22222 | 0.253634 | -0.06456 | 0.201173 | 0.013951 | -0.38085 | 0.061869 | -0.27015 | -0.12959 | -0.50408 | -0.08642 | -0.42464 | -0.7047 | -0.31801 | -0.00629 | -0.60179 | -0.11628 | 0.044535 | 0.1193 | 0.023885 | -0.31141 | -0.22539 | -0.20037 | -0.21816 | -0.20017 |
| 21.23611 | 0.521691 | -0.22578 | 0.346506 | -0.04638 | -0.29593 | -0.00248 | -0.15382 | -0.32284 | -0.48431 | -0.06462 | -0.0526 | -0.53444 | -0.36225 | -0.1617 | -0.61354 | -0.19144 | -0.12854 | 0.190025 | 0.123129 | -0.12355 | -0.17886 | -0.22597 | -0.02412 | -0.17011 |
| 21.25 | 0.602989 | -0.35225 | 0.238258 | -0.10777 | -0.15968 | -0.14004 | -0.02849 | -0.21203 | -0.19997 | -0.05527 | 0.242056 | -0.10055 | -0.23285 | -0.21629 | -0.36408 | -0.26555 | -0.2768 | 0.27095 | 0.089134 | -0.02066 | -0.05445 | -0.1607 | 0.274395 | 0.097669 |
| 21.26389 | 0.539161 | -0.1355 | 0.129479 | -0.0296 | -0.09524 | -0.15946 | -0.00713 | 0.197618 | 0.237087 | 0.027267 | 0.097037 | 0.381996 | -0.03772 | -0.10533 | 0.007155 | -0.22508 | -0.24402 | 0.253953 | 0.038818 | -0.01835 | 0.065118 | -0.0577 | 0.406459 | 0.462368 |
| 21.27778 | 0.443679 | 0.28921 | 0.130383 | 0.075645 | -0.05238 | -0.0515 | -0.02098 | 0.395942 | 0.487848 | 0.128115 | -0.25587 | 0.588381 | 0.053946 | 0.000946 | 0.333645 | -0.03003 | 0.038202 | 0.070739 | 0.014588 | -0.16051 | 0.072078 | -0.00273 | 0.173623 | 0.581394 |
| 21.29167 | 0.336765 | 0.550513 | 0.16783 | 0.050466 | 0.017097 | 0.073649 | 0.032257 | 0.214989 | 0.349023 | 0.122178 | -0.26605 | 0.395565 | 0.046451 | 0.056735 | 0.383466 | 0.182466 | 0.342396 | -0.08759 | 0.073712 | -0.29682 | 0.008674 | -0.00185 | -0.17012 | 0.411128 |
| 21.30556 | 0.260302 | 0.50622 | 0.263904 | -0.04978 | 0.100916 | 0.173674 | 0.17855 | -0.00961 | 0.12865 | 0.056533 | 0.209644 | 0.100118 | 0.105318 | 0.1054 | 0.149889 | 0.271606 | 0.393126 | -0.01581 | 0.209517 | -0.19386 | 0.023923 | -0.00963 | -0.23063 | 0.157041 |
| 21.31944 | 0.339034 | 0.223232 | 0.403897 | -0.10717 | 0.174212 | 0.225819 | 0.336848 | -0.04116 | 0.162987 | 0.031039 | 0.836181 | 0.139698 | 0.311245 | 0.20018 | -0.14622 | 0.210757 | 0.246443 | 0.223535 | 0.28295 | 0.148504 | 0.123208 | -0.02122 | 0.090916 | 0.113197 |
| 21.33333 | 0.571032 | -0.15343 | 0.424593 | -0.09074 | 0.23144 | 0.230815 | 0.433552 | 0.235177 | 0.522076 | 0.071483 | 1.094251 | 0.622594 | 0.54625 | 0.34228 | -0.22175 | 0.08697 | 0.030257 | 0.432173 | 0.264945 | 0.441263 | 0.201833 | -0.04206 | 0.532324 | 0.304076 |
| 21.34722 | 0.808118 | -0.26269 | 0.337978 | -0.00123 | 0.263372 | 0.191031 | 0.416222 | 0.557043 | 0.997538 | 0.135136 | 0.747447 | 1.324438 | 0.643809 | 0.46795 | 0.059822 | 0.017482 | -0.12077 | 0.410085 | 0.143112 | 0.44928 | 0.217252 | -0.04531 | 0.840189 | 0.661862 |
| 21.36111 | 0.853289 | 0.133355 | 0.296182 | 0.071391 | 0.266156 | 0.172958 | 0.35537 | 0.645016 | 1.423549 | 0.160443 | 0.094111 | 1.765772 | 0.623514 | 0.56082 | 0.503306 | 0.128261 | 0.01932 | 0.230399 | 0.048174 | 0.197969 | 0.216312 | 0.001454 | 0.844534 | 1.159167 |
| 21.375 | 0.748468 | 0.717329 | 0.329436 | 0.083195 | 0.25098 | 0.334139 | 0.298023 | 0.539589 | 1.396678 | 0.137676 | -0.54007 | 1.685672 | 0.526338 | 0.572169 | 0.672597 | 0.360628 | 0.342682 | 0.11188 | 0.073588 | -0.14118 | 0.229185 | 0.066531 | 0.521649 | 1.26134 |
| 21.38889 | 0.6572 | 0.971115 | 0.403204 | 0.050221 | 0.240016 | 0.553264 | 0.34174 | 0.368407 | 0.888093 | 0.116163 | -0.5714 | 1.063541 | 0.517286 | 0.490983 | 0.419817 | 0.519542 | 0.605037 | 0.21373 | 0.219088 | -0.36077 | 0.266061 | 0.106227 | 0.174723 | 0.91875 |
| 21.40278 | 0.734726 | 0.753742 | 0.499556 | 0.04732 | 0.257461 | 0.596046 | 0.461467 | 0.341861 | 0.399353 | 0.149416 | 0.321948 | 0.532641 | 0.600711 | 0.481608 | -0.00169 | 0.560009 | 0.545616 | 0.483147 | 0.364824 | -0.19568 | 0.359503 | 0.131961 | 0.097851 | 0.404925 |
| 21.41667 | 0.948716 | 0.448144 | 0.641018 | 0.118125 | 0.281572 | 0.464605 | 0.523367 | 0.596319 | -0.01155 | 0.231689 | 1.5669 | 0.213853 | 0.583606 | 0.599076 | -0.32206 | 0.455893 | 0.260069 | 0.653401 | 0.393622 | 0.212567 | 0.379178 | 0.10084 | 0.285814 | -0.03662 |
| 21.43056 | 1.120086 | 0.301042 | 0.750605 | 0.206545 | 0.31032 | 0.197365 | 0.436537 | 0.881793 | -0.29088 | 0.33413 | 1.95092 | 0.18928 | 0.498497 | 0.858457 | 0.143557 | 0.349635 | 0.044803 | 0.631667 | 0.356061 | 0.659613 | 0.360743 | 0.038934 | 0.620593 | 0.080848 |
| 21.44444 | 1.967199 | 1.096414 | 1.303801 | 0.632149 | 0.775876 | 0.37434 | 0.581932 | 1.12524 | 0.154273 | 0.802775 | 2.235812 | 1.548721 | 1.148372 | 1.521211 | 1.836853 | 0.702112 | 0.235303 | 1.037176 | 0.659135 | 1.106226 | 0.703136 | 0.133058 | 1.43637 | 1.361219 |
| 21.45833 | 5.382612 | 3.670473 | 2.578067 | 1.797168 | 2.263292 | 1.218091 | 1.373632 | 1.937288 | 1.660882 | 2.352165 | 4.349483 | 4.933225 | 3.259265 | 3.452199 | 3.79724 | 2.048561 | 0.754805 | 2.334759 | 1.491242 | 1.661004 | 1.779391 | 0.356614 | 2.886098 | 4.267125 |
| 21.47222 | 6.621602 | 7.277342 | 3.707422 | 2.387814 | 3.759895 | 2.739815 | 2.423881 | 2.560842 | 3.292488 | 3.956919 | 5.518585 | 6.583646 | 5.169095 | 3.878021 | 4.33385 | 3.06538 | 1.145912 | 3.319686 | 2.469028 | 2.893309 | 2.464495 | 0.430429 | 3.151098 | 6.062298 |
| 21.48611 | 2.622597 | 6.504154 | 2.931088 | 1.665948 | 3.099501 | 2.291594 | 1.772978 | 2.146022 | 2.888517 | 2.197008 | 4.172691 | 4.528762 | 3.70906 | 2.330736 | 2.974069 | 2.241942 | 0.880197 | 2.573215 | 1.909716 | 2.336373 | 1.629181 | 0.267032 | 1.647726 | 4.340841 |
| 21.5 | 0.635136 | 2.831507 | 1.677713 | 0.586078 | 1.427123 | 1.013579 | 0.54912 | 1.099165 | 1.544212 | 0.933964 | 2.489829 | 2.506615 | 1.771508 | 0.837941 | 0.822807 | 1.028239 | 0.579123 | 1.71849 | 1.014729 | 0.88166 | 0.729152 | 0.103957 | 0.522798 | 1.913488 |
| 21.51389 | -0.48784 | 0.668415 | 0.800197 | 0.082591 | 0.129708 | 0.086783 | -0.17275 | -0.00891 | 0.158962 | 0.143852 | 0.374647 | 0.509576 | 0.453027 | -0.75473 | -0.93 | 0.594959 | 0.119828 | 0.608228 | -0.25175 | 0.024494 | 0.324828 | 0.062498 | -0.34264 | -0.11784 |
| 21.52778 | -1.52476 | -0.8607 | 0.086763 | -0.26267 | -1.01618 | -0.78615 | -0.7739 | -1.03728 | -1.21708 | -0.35623 | -1.53832 | -2.08141 | -0.25098 | -1.50455 | -1.69896 | 0.15838 | -0.4209 | -0.36551 | -1.0857 | -0.60887 | 0.114361 | 0.072812 | -1.47442 | -2.24616 |
| 21.54167 | -2.04887 | -1.67448 | -0.62351 | -0.45069 | -1.09462 | -0.92499 | -0.70258 | -1.8292 | -1.73568 | -0.47156 | -3.12793 | -3.81466 | -0.25727 | -1.20425 | -0.93346 | -0.10273 | -0.78642 | -0.74578 | -0.83242 | -0.69253 | -0.04608 | 0.122603 | -0.99911 | -2.52127 |
| 21.55556 | -1.7249 | -0.68582 | -1.06073 | -0.2016 | -0.35476 | -0.90858 | -0.16435 | -1.39523 | -0.97111 | -0.27801 | -1.35195 | -3.76901 | 0.27434 | -1.47969 | 0.363299 | -0.46367 | -0.6999 | -0.50772 | -0.09882 | 0.027057 | -0.0145 | 0.20553 | 0.194039 | -1.3238 |
| 21.56944 | -0.64146 | 1.170731 | -1.08307 | 0.012564 | 0.410961 | -0.37323 | 0.646627 | -0.15165 | 0.398279 | 0.013446 | 2.34867 | -1.89258 | 0.976328 | -0.81711 | 1.42105 | 0.066195 | -0.18692 | 0.10108 | 0.713135 | 0.662615 | 0.122276 | 0.289084 | 1.72418 | 1.070235 |
| 21.58333 | 0.38313 | 2.685028 | -1.13127 | 0.079997 | 0.58486 | 0.324027 | 1.218366 | 0.349393 | 1.915119 | 0.276594 | 4.190076 | 0.546823 | 1.346537 | 0.307859 | 0.753822 | 0.746785 | 0.553597 | 0.772831 | 1.296731 | 0.438385 | 0.22277 | 0.31899 | 2.292821 | 3.187922 |
| 21.59722 | 0.8554 | 2.417226 | -1.34748 | 0.254828 | 0.21224 | 0.116174 | 1.416695 | 0.123791 | 2.473071 | 0.319136 | 1.778491 | 1.238677 | 1.149428 | 0.84574 | -0.28257 | 0.775811 | 0.604207 | 0.690248 | 1.240159 | -0.23102 | 0.143024 | 0.287342 | 0.959401 | 3.102504 |
| 21.61111 | 0.94264 | 1.119673 | -1.61652 | 0.199347 | -0.17912 | -0.15527 | 1.163079 | 0.064177 | 1.347259 | 0.49579 | 0.410676 | 0.2917 | 0.867566 | 0.014283 | -1.70634 | 0.520977 | 0.82274 | 0.362135 | 0.619065 | -0.88098 | 0.035649 | 0.261109 | 0.476847 | 2.134139 |
| 21.625 | 0.748702 | 0.224912 | -1.34763 | 0.167866 | -0.46212 | -0.29985 | 0.993573 | 0.396345 | 0.311889 | 0.966783 | -0.52156 | -0.19609 | 0.499491 | -0.99669 | -2.35541 | 0.187988 | 0.832981 | -0.23222 | -0.01027 | -1.37367 | -0.07742 | 0.206806 | 0.178141 | 1.295957 |
| 21.63889 | 0.422344 | -0.36825 | -0.79828 | 0.183071 | -0.7501 | -0.56837 | 0.82619 | 0.946502 | -0.29978 | 1.88488 | -1.51953 | -0.47741 | -0.01084 | -0.16631 | -1.47024 | -0.16265 | 0.189275 | -0.88265 | -0.62492 | -1.59434 | -0.22696 | 0.129426 | -0.30954 | 0.098888 |
| 21.65278 | 0.243557 | -1.00673 | -0.13373 | 0.186717 | -1.01327 | -0.83569 | 0.469366 | 1.775458 | -0.40642 | 2.87041 | -2.58923 | -0.77992 | -0.37665 | 0.711571 | 0.121581 | -0.09448 | 0.029027 | -1.31276 | -0.95781 | -1.30379 | -0.22934 | 0.139935 | -0.90417 | -1.48201 |
| 21.66667 | 0.5471 | -1.44527 | 0.512031 | 0.331471 | -0.03109 | -0.19408 | 1.735739 | 2.430704 | -0.1716 | 3.890282 | -2.28665 | -1.02247 | -0.29074 | 1.567987 | 1.448878 | 0.211746 | -0.03883 | -0.96099 | -0.47343 | -0.22448 | -0.13317 | 0.211376 | -0.15932 | -2.19715 |
| 21.68056 | 1.507416 | -0.75706 | 1.230879 | 0.509983 | 0.781556 | 0.654417 | 2.163748 | 2.226698 | 0.258628 | 3.21781 | 0.636352 | -0.82636 | 0.191861 | 1.105514 | 1.888945 | 0.393366 | -0.0616 | -0.01789 | 0.16961 | 0.842595 | -0.00517 | 0.262302 | 0.823848 | -1.95206 |
| 21.69444 | 2.049589 | 1.118301 | 1.709853 | 0.568795 | 1.351875 | 1.155924 | 2.669545 | 1.399598 | 0.935402 | 2.811662 | 3.611681 | 0.23145 | 0.686526 | -0.31622 | 1.551298 | 0.491305 | 0.345878 | 0.674332 | 0.56163 | 1.177475 | 0.04841 | 0.250828 | 1.741395 | -0.54249 |
| 21.70833 | 1.395054 | 2.436046 | 1.661022 | 0.470476 | 0.86827 | 1.023248 | 1.795159 | 0.72308 | 1.202545 | 1.950679 | 2.732656 | 1.191652 | 0.955664 | -1.20518 | 0.960994 | 0.353737 | 0.591856 | 0.694341 | 0.779635 | 1.005282 | 0.039612 | 0.19366 | 1.564331 | 0.05265 |
| 21.72222 | 0.175457 | 2.174947 | 1.177716 | 0.322236 | 0.258604 | 0.536134 | 0.918693 | 0.098899 | 0.951841 | 1.185854 | 0.861323 | 1.219732 | 0.749484 | -1.24144 | 0.354196 | 0.412084 | 0.456462 | 0.493246 | 0.616364 | 0.970285 | 0.039286 | 0.138796 | 0.423939 | 1.305417 |
| 21.73611 | -0.85262 | 1.047297 | 0.728873 | 0.136871 | -0.17598 | 0.001411 | 0.293573 | -0.4641 | 0.468932 | 0.456652 | -0.71195 | 0.53257 | 0.013791 | -1.40957 | -0.72927 | 0.519532 | 0.442811 | 0.378589 | 0.422606 | 0.882721 | 0.065332 | 0.072451 | -0.07027 | 2.302068 |
| 21.75 | -1.32283 | 0.103494 | 0.445924 | -0.05978 | -0.55575 | -0.48004 | -0.41753 | -1.09439 | 0.091722 | -0.29708 | -2.47278 | -0.39315 | -0.65349 | -0.81203 | -1.18977 | 0.075499 | 0.00251 | 0.036461 | 0.155058 | 0.232154 | 0.114496 | 0.027127 | -0.61775 | 1.499874 |
| 21.76389 | -1.36517 | -0.47845 | 0.17175 | -0.1212 | -1.10321 | -0.96962 | -1.29723 | -1.62655 | -0.19871 | -0.96326 | -4.26379 | -1.09981 | -1.10479 | -0.16191 | -0.89745 | -0.52836 | -0.47542 | -0.50221 | -0.22607 | -0.75044 | 0.062785 | 0.005089 | -0.83625 | 0.709861 |
| 21.77778 | -1.06027 | -0.84109 | -0.33718 | -0.12893 | -1.44429 | -0.98474 | -2.02931 | -1.60606 | -0.46941 | -1.60675 | -3.2906 | -1.50358 | -1.22605 | 0.258767 | -0.27012 | -1.00916 | -0.84999 | -1.11679 | -0.70084 | -0.74218 | -0.00711 | 0.0072 | -0.09084 | -0.59756 |
| 21.79167 | -0.58962 | -1.17875 | -1.15036 | -0.17698 | -0.87081 | -0.52377 | -1.85861 | -1.17725 | -0.58402 | -2.14864 | -0.88019 | -1.45294 | -0.90552 | 0.128836 | 0.629749 | -1.1759 | -0.99935 | -1.61017 | -0.89511 | -0.3137 | -0.14962 | 0.057755 | 0.606063 | -2.06437 |
| 21.80556 | -0.13457 | -0.98826 | -2.02695 | -0.34168 | -0.56403 | -0.3128 | -1.46849 | -0.84578 | -0.73116 | -2.382 | 1.421772 | -0.90879 | -0.47989 | -0.39341 | 0.998425 | -1.0012 | -0.4495 | -1.67105 | -0.69424 | -0.06503 | -0.35408 | 0.040896 | 0.712619 | -2.45748 |
| 21.81944 | -0.21661 | -0.85294 | -2.68131 | -0.68724 | -0.53471 | -0.74922 | -1.29091 | -0.80464 | -0.81902 | -2.30715 | 1.964887 | -0.3677 | -0.31056 | -0.85519 | 0.088732 | -0.91948 | 0.093446 | -1.48562 | -0.4967 | -0.29588 | -0.64384 | -0.07727 | 0.215554 | -2.97586 |
| 21.83333 | -0.71636 | -1.12989 | -3.18697 | -1.0289 | -0.62587 | -1.34807 | -1.12726 | -0.88173 | -1.28061 | -1.90938 | 0.130068 | -0.49528 | -0.34771 | -1.25105 | -1.47533 | -1.09387 | 0.092882 | -1.52077 | -0.37596 | -1.04304 | -0.81079 | -0.20513 | -0.43193 | -3.44382 |
| 21.84722 | -1.14767 | -1.79005 | -3.45461 | -1.25875 | -0.77148 | -1.48284 | -1.298 | -0.86674 | -1.93386 | -1.47018 | -2.14374 | -1.47457 | -0.63109 | -1.47095 | -2.71393 | -1.31339 | -0.15591 | -1.37311 | -0.25818 | -2.03159 | -0.8796 | -0.24003 | -0.93175 | -1.34543 |
| 21.86111 | -1.25473 | -2.29287 | -3.40434 | -1.26482 | -0.96 | -1.20585 | -1.57107 | -0.7656 | -2.3841 | -1.10069 | -4.73284 | -2.84544 | -1.01857 | -1.4756 | -3.17082 | -1.34384 | -0.25317 | -0.91474 | -0.21475 | -2.45059 | -0.79647 | -0.20785 | -1.15661 | 0.720646 |
| 21.875 | -1.04863 | -2.17635 | -3.05381 | -1.01475 | -1.06395 | -0.86051 | -1.67117 | -0.60562 | -2.20675 | -1.02856 | -5.19025 | -3.08227 | -1.34093 | -1.33413 | -2.81171 | -1.02096 | -0.1101 | -0.35074 | -0.38496 | -2.39737 | -0.72024 | -0.16303 | -1.17539 | 1.613922 |
| 21.88889 | -0.65808 | -1.7989 | -2.11384 | -0.68322 | -1.00654 | -0.53666 | -1.65107 | -0.47081 | -1.37538 | -0.87954 | -3.80702 | -1.97402 | -1.51103 | -1.1432 | -1.77585 | -0.71258 | -0.07216 | -0.05492 | -0.58203 | -2.36187 | -0.64666 | -0.10567 | -0.98675 | 1.19515 |
| 21.90278 | -0.25018 | -1.42623 | -0.81169 | -0.53201 | -0.91176 | -0.24234 | -1.4217 | -0.33934 | -0.42205 | -0.74293 | -1.96646 | -0.60287 | -1.58446 | -0.86735 | -0.613 | -0.6811 | -0.43114 | -0.3216 | -0.76225 | -2.24562 | -0.64595 | -0.10426 | -0.70448 | 0.069108 |
| 21.91667 | -0.12554 | -1.1869 | 0.32482 | -0.54847 | -0.71538 | -0.07822 | -0.91008 | -0.13317 | 0.317637 | -0.25056 | -0.26142 | 0.326731 | -1.52222 | -0.53353 | 0.413637 | -0.7014 | -0.89583 | -0.80286 | -0.8837 | -1.53484 | -0.66634 | -0.18177 | -0.45355 | -0.65462 |
| 21.93056 | -0.29598 | -1.15872 | 1.132185 | -0.56689 | -0.49301 | -0.15461 | -0.30478 | 0.02776 | 0.473619 | 0.216605 | 0.953684 | 0.520614 | -1.27192 | -0.11394 | 0.731464 | -0.5825 | -1.28561 | -1.01696 | -0.87835 | -0.59041 | -0.64122 | -0.23384 | -0.297 | -1.16168 |
| 21.94444 | -0.86559 | -1.55451 | 0.969196 | -0.62438 | -0.5734 | -0.47393 | -0.31396 | -0.0285 | -0.1212 | 0.294227 | 0.804592 | -0.19505 | -1.05886 | -0.00013 | 0.068638 | -0.48551 | -1.25771 | -1.16202 | -0.90833 | -0.03982 | -0.65495 | -0.34867 | -0.631 | -1.91175 |
| 21.95833 | -2.04365 | -2.4861 | -0.61904 | -0.82544 | -1.17191 | -1.28315 | -1.33205 | -0.52919 | -1.35404 | -0.30847 | -1.41314 | -1.95747 | -1.39977 | -0.73717 | -1.42678 | -1.37084 | -1.55156 | -1.89227 | -1.3013 | -0.59736 | -0.93729 | -0.7792 | -1.79798 | -3.2402 |
| 21.97222 | -3.35994 | -3.71821 | -2.6334 | -1.02945 | -2.23099 | -2.2512 | -2.97501 | -1.31897 | -2.56933 | -1.26877 | -4.9512 | -4.27448 | -2.19822 | -2.24712 | -3.12442 | -2.83325 | -1.68915 | -2.81158 | -1.86444 | -1.91386 | -1.42208 | -1.22434 | -3.07508 | -5.08713 |
| 21.98611 | -3.48247 | -4.1253 | -3.27211 | -0.94525 | -2.62491 | -2.27497 | -3.31617 | -1.99692 | -2.95781 | -1.73063 | -6.93373 | -5.16767 | -2.5413 | -3.14025 | -3.28915 | -3.25081 | -1.53417 | -2.76792 | -1.99043 | -2.79742 | -1.4603 | -1.1683 | -3.13394 | -5.28267 |
| 22 | -2.57321 | -2.51479 | -2.29766 | -0.42295 | -1.93564 | -1.65682 | -2.01723 | -1.93009 | -2.49047 | -1.54276 | -5.08952 | -3.99577 | -2.09889 | -2.20566 | -2.40776 | -2.52946 | -0.86475 | -2.0192 | -1.27853 | -2.31409 | -1.08445 | -0.89913 | -2.47423 | -4.20536 |
| 22.01389 | -1.45398 | -1.17322 | -1.37666 | 0.133615 | -1.19203 | -1.0607 | -0.7458 | -0.99948 | -1.90533 | -1.34684 | -2.86849 | -2.76963 | -1.35755 | -0.56084 | -0.93598 | -1.44419 | 0.036263 | -1.15475 | -0.50575 | -1.43881 | -0.74789 | -0.71896 | -1.55025 | -2.6794 |
| 22.02778 | -0.25572 | -0.15819 | -0.75456 | 0.454024 | -0.4532 | -0.97154 | 0.29027 | 0.135962 | -1.18058 | -1.00941 | -1.34136 | -1.59733 | -0.48151 | 0.876253 | 0.302545 | -0.42167 | 0.805503 | -0.28941 | 0.006071 | -0.28288 | -0.27154 | -0.5653 | -0.22547 | -0.92735 |
| 22.04167 | 0.582129 | 0.386282 | -0.15689 | 0.267119 | 0.114646 | -0.90259 | 0.59024 | 0.844232 | -0.52395 | -0.52148 | 0.009304 | -0.65823 | 0.290645 | 1.697685 | 1.639261 | 0.23718 | 0.860668 | 0.209108 | 0.395414 | 1.054247 | 0.122286 | -0.29773 | 0.932468 | 0.526142 |
| 22.05556 | 0.902492 | 0.792073 | 0.405546 | 0.036598 | 0.448364 | -0.73048 | 0.509961 | 0.68276 | -0.06564 | -0.21675 | 1.164394 | -0.03048 | 0.856172 | 0.585106 | 2.147455 | 0.537655 | 0.445275 | 0.625337 | 0.572942 | 2.116737 | 0.314224 | -0.08783 | 1.442606 | 1.284835 |
| 22.06944 | 0.578229 | 0.974706 | 0.704674 | -0.06674 | 0.591384 | -0.46506 | 0.491118 | 0.188902 | 0.131873 | -0.2288 | 1.799951 | 0.265848 | 1.026044 | 0.199079 | 1.560056 | 0.536485 | 0.149057 | 0.792235 | 0.530392 | 1.974493 | 0.458272 | 0.041443 | 1.312537 | 1.333626 |
| 22.08333 | 0.07435 | 0.985474 | 0.744457 | -0.08623 | 0.532808 | 0.150055 | 0.390611 | -0.13915 | 0.163878 | -0.36131 | 2.083435 | 0.265061 | 0.688344 | -0.18767 | 0.558042 | 0.336592 | -0.06926 | 0.616482 | 0.322235 | 1.142948 | 0.436938 | 0.091863 | 0.774278 | 1.046968 |
| 22.09722 | -0.37666 | 0.960928 | 0.515984 | -0.0613 | 0.327064 | 0.717067 | 0.267368 | -0.28421 | 0.186965 | -0.47826 | 2.047993 | 0.149044 | 0.288596 | -0.44343 | -0.20308 | 0.116909 | -0.15119 | 0.425766 | 0.156346 | 0.529113 | 0.363474 | 0.142188 | 0.330568 | 0.655147 |
| 22.11111 | -0.53838 | 0.834792 | 0.249828 | -0.03027 | 0.247276 | 0.845281 | 0.272375 | -0.30255 | 0.367519 | -0.44593 | 1.890638 | 0.099493 | 0.0717 | -0.52296 | -0.65009 | 0.068301 | -0.13935 | 0.366846 | 0.038163 | 0.233316 | 0.355948 | 0.167339 | 0.074126 | 0.448273 |
| 22.125 | -0.39322 | 0.488006 | 0.113881 | 0.048608 | 0.35182 | 0.513154 | 0.444815 | -0.17626 | 0.585532 | -0.25141 | 1.810572 | 0.185004 | -0.03719 | -0.4681 | -0.77162 | 0.15962 | -0.06545 | 0.360076 | -0.08754 | 0.116829 | 0.318719 | 0.141998 | -0.03043 | 0.442177 |
| 22.13889 | -0.15583 | 0.028218 | 0.066048 | 0.098468 | 0.439876 | -0.07913 | 0.537959 | 0.017159 | 0.726767 | -0.09621 | 1.431685 | 0.369727 | -0.09777 | -0.24391 | -0.44773 | 0.360888 | 0.100571 | 0.383346 | -0.15141 | 0.182729 | 0.254662 | 0.101263 | -0.13089 | 0.559272 |
| 22.15278 | -0.01265 | -0.25521 | 0.10955 | 0.061754 | 0.564879 | -0.1162 | 0.494084 | 0.122135 | 0.848612 | -0.06545 | 0.943177 | 0.722127 | -0.11833 | -0.06662 | -0.00189 | 0.597974 | 0.259888 | 0.508067 | -0.09501 | 0.422667 | 0.161256 | 0.09014 | -0.22271 | 0.634151 |
| 22.16667 | -0.138 | -0.16172 | 0.216513 | -0.09768 | 0.670836 | 0.292816 | 0.482978 | 0.034743 | 0.943364 | -0.16823 | 0.547393 | 1.072652 | -0.09669 | -0.21383 | -0.06667 | 0.661311 | 0.27849 | 0.733914 | 0.037343 | 0.676253 | 0.108695 | 0.080439 | -0.23282 | 0.584395 |
| 22.18056 | -0.45356 | 0.297873 | 0.262119 | -0.25466 | 0.7263 | 0.722462 | 0.54551 | -0.16258 | 1.034013 | -0.32701 | 0.201251 | 1.270336 | -0.02975 | -0.50079 | -0.6403 | 0.524368 | 0.257371 | 0.958364 | 0.096966 | 0.626724 | 0.094713 | 0.058424 | -0.12793 | 0.439295 |
| 22.19444 | -0.70446 | 0.610188 | 0.206699 | -0.22921 | 0.668402 | 0.442548 | 0.656306 | -0.30528 | 1.046817 | -0.38553 | -0.12857 | 1.232496 | -0.06249 | -0.65557 | -1.17413 | 0.112301 | 0.056984 | 0.804114 | -0.06349 | 0.104682 | -0.00212 | -0.0393 | -0.05761 | 0.191266 |
| 22.20833 | -0.76607 | 0.234407 | 0.038954 | -0.08559 | 0.430538 | -0.36244 | 0.559073 | -0.26538 | 0.854969 | -0.31178 | -0.49723 | 0.915346 | -0.19342 | -0.45885 | -1.17543 | -0.17562 | -0.19821 | 0.346786 | -0.32593 | -0.64504 | -0.1719 | -0.13828 | -0.09223 | -0.06904 |
| 22.22222 | -0.60478 | -0.4138 | -0.16496 | -0.00932 | 0.153536 | -0.93138 | 0.154176 | -0.03733 | 0.627851 | -0.15007 | -0.91033 | 0.469915 | -0.28805 | -0.06076 | -0.52224 | -0.1003 | -0.03284 | -0.01826 | -0.45319 | -1.29465 | -0.27667 | -0.18206 | -0.10507 | -0.20664 |
| 22.23611 | -0.24497 | -0.81283 | -0.21965 | -0.08113 | -0.03331 | -0.5795 | -0.32119 | 0.154811 | 0.527318 | 0.013507 | -1.16577 | 0.455576 | -0.40312 | 0.148671 | 0.392135 | 0.020402 | 0.208411 | -0.15842 | -0.39968 | -1.56423 | -0.38556 | -0.27195 | -0.11527 | -0.38263 |
| 22.25 | 0.041286 | -0.70314 | -0.07628 | -0.21655 | -0.10805 | 0.030018 | -0.38323 | 0.019322 | 0.404835 | 0.037702 | -1.0134 | 0.553082 | -0.52486 | -0.08539 | 0.850252 | -0.03214 | 0.187297 | -0.21232 | -0.24312 | -1.03707 | -0.42335 | -0.36149 | -0.16763 | -0.70447 |
| 22.26389 | 0.039778 | -0.1475 | 0.138918 | -0.29217 | -0.09247 | 0.128684 | -0.17806 | -0.45691 | 0.150431 | -0.07615 | -0.42804 | 0.49958 | -0.50541 | -0.54383 | 0.405538 | -0.15771 | 0.007527 | -0.21501 | -0.09489 | -0.21509 | -0.30836 | -0.32082 | -0.12005 | -0.74921 |
| 22.27778 | -0.07412 | 0.470361 | 0.268935 | -0.12116 | -3.1E-05 | -0.33437 | 0.165663 | -0.83088 | -0.05534 | -0.14708 | 0.161479 | 0.379941 | -0.3715 | -1.00503 | -0.4293 | -0.21061 | -0.21298 | -0.21577 | -0.01213 | 0.360981 | -0.09176 | -0.17526 | 0.046407 | -0.27215 |
| 22.29167 | -0.05422 | 0.583315 | 0.192984 | 0.177941 | 0.095677 | -0.75917 | 0.439643 | -0.3987 | -0.10525 | -0.05811 | 0.564638 | 0.271297 | -0.31513 | -1.04218 | -1.10909 | -0.32123 | -0.42161 | -0.24385 | -0.00391 | 0.123039 | 0.068433 | -0.09267 | 0.181344 | 0.415141 |
| 22.30556 | 0.103888 | 0.264449 | -0.0502 | 0.299383 | 0.071182 | -0.78731 | 0.049804 | 0.577326 | -0.08379 | 0.11419 | 0.643933 | 0.068612 | -0.27151 | -0.5426 | -1.23465 | -0.33409 | -0.3888 | -0.19988 | 0.058125 | -0.54372 | 0.118673 | -0.06483 | 0.255808 | 0.892981 |
| 22.31944 | 0.375936 | -0.13767 | -0.29914 | 0.134158 | -0.00283 | -0.36422 | -0.41797 | 1.279279 | -0.094 | 0.303245 | 0.338756 | -0.10467 | -0.17619 | 0.200324 | -0.56448 | -0.09842 | 0.002709 | -0.06188 | 0.209765 | -0.90766 | 0.107168 | -0.06079 | 0.377142 | 0.982018 |
| 22.33333 | 0.733096 | -0.30607 | -0.16636 | -0.11197 | -0.00507 | 0.221453 | -0.30619 | 1.105753 | -0.15084 | 0.408259 | 0.08294 | -0.03564 | -0.07001 | 0.787872 | 0.437701 | 0.187506 | 0.458693 | 0.158609 | 0.352868 | -0.58713 | 0.096377 | -0.06641 | 0.554358 | 0.887687 |
| 22.34722 | 0.966957 | 0.11238 | 0.221548 | -0.28578 | 0.093062 | 0.520552 | 0.046771 | 0.353552 | -0.1978 | 0.347792 | 0.453602 | 0.152329 | 0.086752 | 1.043508 | 1.500451 | 0.361444 | 0.537085 | 0.542319 | 0.492619 | 0.247134 | 0.169055 | -0.01944 | 0.723192 | 0.707483 |
| 22.36111 | 1.094068 | 0.854799 | 0.506646 | -0.1273 | 0.331211 | 0.561433 | 0.522441 | -0.35715 | -0.0531 | 0.25888 | 1.547478 | 0.452159 | 0.39765 | 0.926642 | 2.217497 | 0.428409 | 0.454904 | 1.054751 | 0.577802 | 1.23713 | 0.360531 | 0.079275 | 0.893042 | 0.73603 |
| 22.375 | 1.172098 | 1.515409 | 0.625932 | 0.222479 | 0.727649 | 0.605947 | 1.180591 | -0.39804 | 0.345464 | 0.226307 | 2.810713 | 0.906313 | 0.686549 | 0.580624 | 1.442935 | 0.493084 | 0.38915 | 1.418519 | 0.573742 | 1.99805 | 0.534237 | 0.127879 | 0.926435 | 0.639613 |

| | | | | | | | | | | | | | | | | | | | | | | | | |
|---|---|---|---|---|---|---|---|---|---|---|---|---|---|---|---|---|---|---|---|---|---|---|---|---|
| 22.38889 | 1.092687 | 1.619359 | 0.607009 | 0.543556 | 1.051429 | 0.845357 | 1.203357 | 0.40032 | 0.821433 | 0.286496 | 3.419359 | 1.295209 | 0.885634 | 0.327162 | 0.073446 | 0.445641 | 0.537552 | 1.429891 | 0.457711 | 2.058394 | 0.649894 | 0.121505 | 0.702095 | 0.459517 |
| 22.40278 | 0.945163 | 1.174637 | 0.491967 | 0.461739 | 1.137554 | 1.17364 | 0.925253 | 1.494902 | 1.34836 | 0.47939 | 2.362929 | 1.475238 | 1.081639 | 0.48326 | -1.01924 | 0.48658 | 0.53696 | 1.27371 | 0.29119 | 1.414049 | 0.829535 | 0.182401 | 0.411217 | 0.125526 |
| 22.41667 | 0.744027 | 0.687666 | 0.299168 | 0.201959 | 0.893702 | 1.373541 | 0.512243 | 1.710314 | 1.674902 | 0.695382 | 0.95701 | 1.704143 | 1.432252 | 0.878284 | -1.16077 | 0.675167 | 0.734373 | 1.130259 | 0.170142 | 0.545627 | 0.814956 | 0.281727 | 0.29308 | -0.1652 |
| 22.43056 | 0.54709 | 0.307323 | 0.437086 | -0.06964 | 0.528052 | 0.928684 | 0.368368 | 0.966252 | 1.483438 | 0.807253 | -0.82309 | 1.911018 | 1.70413 | 0.957889 | -0.04976 | 0.697423 | 0.701989 | 0.881262 | 0.145372 | -0.20872 | 0.6578 | 0.284194 | 0.40712 | -0.22563 |
| 22.44444 | 1.214924 | 0.653121 | 1.516289 | 0.166605 | 0.454812 | 0.750979 | 0.511161 | 0.517825 | 1.474433 | 1.216467 | -2.09405 | 2.80457 | 2.614175 | 0.748725 | 1.806848 | 0.766471 | 0.368678 | 0.908158 | 0.339511 | -0.3747 | 0.859736 | 0.333698 | 1.227386 | 0.695376 |
| 22.45833 | 3.826149 | 2.477436 | 3.337338 | 1.595833 | 1.286607 | 1.411133 | 1.079977 | 1.902187 | 2.456475 | 2.320998 | 0.065198 | 4.656149 | 4.921554 | 1.060691 | 4.608471 | 1.54275 | 0.14866 | 1.62286 | 0.959173 | 0.747208 | 2.020272 | 0.692521 | 3.90818 | 3.877584 |
| 22.47222 | 6.393862 | 6.146278 | 4.678803 | 3.02217 | 2.405569 | 2.406376 | 1.473439 | 4.255511 | 3.80963 | 3.679471 | 4.048858 | 4.68409 | 7.599892 | 2.659853 | 5.495503 | 3.418824 | 0.925214 | 2.181251 | 2.197748 | 2.424971 | 3.225939 | 0.984473 | 5.181697 | 6.375644 |
| 22.48611 | 4.248795 | 7.141656 | 3.205324 | 3.122467 | 2.674827 | 2.211881 | 0.365563 | 3.434403 | 2.913117 | 2.674993 | 6.855788 | 3.601254 | 4.321258 | 2.606609 | 3.347392 | 3.543967 | 1.69827 | 2.090971 | 2.346546 | 2.198751 | 3.186334 | 0.912699 | 2.695831 | 5.403114 |
| 22.5 | 1.202737 | 3.73651 | 2.150192 | 1.865173 | 1.728404 | 1.536987 | -0.28308 | 2.078225 | 1.873918 | 2.179266 | 5.716734 | 2.410999 | 1.675762 | 2.72044 | 0.904769 | 2.577003 | 1.692868 | 2.067286 | 1.415948 | 1.509307 | 1.991116 | 0.629962 | 1.176694 | 2.792809 |
| 22.51389 | -0.56059 | 0.601899 | 1.343699 | 0.52423 | 0.619368 | 0.278676 | -0.31183 | 0.505714 | 0.628594 | 1.146258 | 2.271881 | 0.482626 | 0.620634 | 0.763969 | -0.50354 | 1.341403 | 0.445877 | 0.947805 | 0.448474 | 0.408801 | 0.747941 | 0.174993 | -0.01024 | 0.513493 |
| 22.52778 | -2.1074 | -1.17464 | 0.378412 | -0.60613 | -0.41765 | -0.8906 | -0.67609 | -1.00607 | -0.66259 | -0.1866 | -0.98398 | -1.83299 | -0.51755 | -0.37893 | -0.95371 | 0.15296 | -0.72212 | -0.22246 | -0.60081 | -0.94428 | -0.468 | -0.26341 | -1.92745 | -1.93192 |
| 22.54167 | -2.44634 | -1.41789 | -0.25048 | -1.54074 | -1.5687 | -2.29518 | -1.44979 | -2.67562 | -1.51656 | -0.90951 | -4.13075 | -3.14175 | -1.04122 | -1.07792 | -0.81436 | -0.41628 | -1.06084 | -1.1665 | -0.92361 | -1.69418 | -1.0901 | -0.42122 | -2.52457 | -3.74225 |
| 22.55556 | -1.55881 | -1.21059 | -0.96656 | -1.70914 | -1.15206 | -2.50873 | -1.03044 | -3.43492 | -0.93005 | -0.51933 | -8.08483 | -2.68014 | -0.59225 | -1.11508 | -0.18611 | 0.068783 | -0.13647 | -1.03938 | -0.33098 | -1.98395 | -1.13829 | -0.07515 | -0.40933 | -3.19816 |
| 22.56944 | -0.14594 | 0.61643 | -1.28443 | -0.44747 | -0.0356 | -1.6974 | 0.136866 | -2.13611 | 0.689703 | -0.13013 | -7.27807 | -0.73572 | 0.140683 | -0.79324 | 0.114028 | 0.093927 | 0.388665 | 0.1867 | 0.249296 | -0.69983 | -0.89409 | 0.473512 | 1.703768 | -0.16455 |
| 22.58333 | 1.016207 | 2.26681 | -1.05586 | 1.064142 | 1.268299 | -0.66491 | 1.503754 | -0.40654 | 1.940781 | -0.02646 | -2.41655 | 1.075368 | 0.111007 | -0.90602 | 0.092052 | -0.21003 | 0.656833 | 1.056316 | 0.906187 | 0.549479 | -0.51554 | 0.717227 | 3.998313 | 3.779579 |
| 22.59722 | 0.829533 | 3.054756 | -1.06664 | 2.092953 | 1.262135 | 0.581713 | 2.07074 | 0.073679 | 2.005644 | -0.2762 | 2.166496 | 0.330853 | -0.68189 | -0.7328 | -0.35698 | -0.69666 | 0.802654 | 0.905822 | 1.37453 | 1.284365 | -0.31196 | 0.607757 | 2.301127 | 3.716806 |
| 22.61111 | -0.1041 | 1.39125 | -1.57764 | 1.40647 | 0.144825 | 0.640654 | 1.583619 | -0.26857 | 1.24265 | -0.38639 | 7.781604 | -0.09048 | -0.90222 | -0.96154 | -1.23531 | -0.14434 | 0.644486 | 0.364434 | 1.262067 | 1.176741 | 0.2494 | 0.688762 | 0.52906 | 2.260361 |
| 22.625 | -0.99082 | -0.67785 | -1.74943 | 0.650777 | -0.57239 | -0.13822 | 1.096728 | -0.66564 | 0.582727 | -0.34042 | 8.092411 | -0.52131 | -1.06137 | -1.54245 | -2.57606 | 0.148764 | -0.07856 | -0.17931 | 0.742253 | 0.250537 | 0.458536 | 0.509265 | 0.011199 | 1.777695 |
| 22.63889 | -1.42789 | -1.01147 | -1.46764 | -0.01203 | -1.15827 | -1.17218 | 0.086311 | -0.64924 | 0.087816 | -0.091 | 3.989643 | -0.99348 | -1.28109 | -1.13755 | -2.16373 | 0.147942 | -0.10708 | -0.77403 | 0.089567 | -0.79059 | 0.115517 | 0.351582 | -0.1408 | 0.813986 |
| 22.65278 | -1.01418 | -0.57219 | -1.03206 | -0.52195 | -1.71673 | -2.55247 | -1.13929 | 0.324068 | -0.48929 | 0.539542 | 1.589697 | -1.30935 | -1.54265 | -0.27171 | -1.25367 | 0.094381 | -0.24136 | -1.39867 | -0.35072 | -1.9663 | -0.24791 | 0.149168 | -0.60654 | -0.24255 |
| 22.66667 | 0.255378 | -0.32037 | -0.78209 | -0.83046 | -1.70724 | -2.25629 | -2.11492 | 1.795341 | -0.86974 | 1.50354 | -0.41075 | -1.32237 | -1.27656 | 1.328905 | -0.51463 | 0.022644 | -0.94381 | -1.434 | -0.63022 | -2.48829 | -0.2586 | 0.199259 | -1.97218 | -1.93133 |
| 22.68056 | 1.526978 | -1.18291 | -0.58081 | -0.4809 | -0.52956 | -0.9835 | -1.83348 | 2.704219 | -0.76056 | 2.027693 | -2.69159 | -0.88068 | -0.12161 | 2.230749 | 0.50379 | 0.236763 | -0.57407 | -0.88231 | -0.64459 | -2.36288 | 0.002477 | 0.370582 | -1.36064 | -2.70912 |
| 22.69444 | 1.814868 | -2.05565 | -0.45324 | 0.216615 | 0.728988 | 0.457761 | -0.43021 | 2.766736 | -0.1378 | 1.728333 | -5.74743 | 0.310875 | 1.120536 | 2.285315 | 1.633646 | 0.390099 | -0.23092 | -0.16761 | -0.2279 | -1.36547 | 0.303 | 0.384849 | 0.399854 | -2.93749 |
| 22.70833 | 1.201528 | -1.26113 | -0.34534 | 0.773494 | 1.718665 | 1.106932 | 1.062017 | 2.321143 | 0.574169 | 1.354328 | -8.86703 | 1.412067 | 2.16506 | 1.175312 | 2.832357 | 0.325319 | 0.163313 | 0.38611 | 0.189168 | 0.949707 | 0.592373 | 0.202832 | 1.845099 | -2.84577 |
| 22.72222 | 0.454034 | -0.15134 | 0.113582 | 1.089989 | 2.081969 | 0.569218 | 1.956521 | 1.177518 | 1.048593 | 0.734496 | -6.3582 | 2.316717 | 2.516963 | 0.144047 | 2.244253 | 0.57743 | 0.584628 | 0.472317 | 0.404323 | 2.530192 | 0.702672 | 0.073943 | 2.462317 | -0.09946 |
| 22.73611 | -0.03804 | 1.059317 | 1.025226 | 0.857639 | 0.95826 | -0.10837 | 1.375841 | -0.16603 | 1.103255 | 0.040209 | -1.8518 | 2.311774 | 1.861511 | -0.97331 | 0.880112 | 0.464301 | 0.886457 | -0.06948 | 0.453909 | 2.635873 | 0.496654 | -0.11528 | 1.804414 | 2.458969 |
| 22.75 | -0.25359 | 1.925205 | 2.103286 | 0.219687 | -0.07133 | -0.70816 | 0.643343 | -1.37263 | 0.635929 | -0.48149 | 1.600641 | 1.250757 | 0.617683 | -2.04413 | -0.44477 | 0.243261 | 1.274487 | -0.76476 | 0.446245 | 2.378311 | -0.02821 | -0.46005 | 0.833397 | 3.365889 |
| 22.76389 | -0.38924 | 2.002171 | 2.426719 | -0.42744 | -0.96931 | -1.22744 | -0.29458 | -2.40428 | 0.107366 | -0.69391 | 5.581485 | -0.20728 | -0.53564 | -2.70055 | -0.74097 | -0.35839 | 0.923497 | -0.98748 | 0.356016 | 1.246478 | -0.66563 | -0.7117 | 0.132427 | 2.603712 |
| 22.77778 | -0.61521 | 1.550536 | 1.670565 | -0.82427 | -1.69657 | -0.9346 | -1.47502 | -2.92118 | -0.27263 | -0.62465 | 8.476497 | -1.75804 | -1.29006 | -2.60925 | -0.74938 | -1.28853 | 0.087286 | -0.92213 | 0.200441 | -0.37791 | -0.56986 | -0.41851 | -0.51805 | 1.763917 |
| 22.79167 | -0.9409 | 0.125187 | 0.369768 | -0.5766 | -1.78824 | -0.26563 | -2.21639 | -2.48285 | -0.57654 | -0.6656 | 5.706321 | -3.03238 | -1.6727 | -2.17354 | -1.05423 | -1.93993 | -0.81712 | -0.90705 | -0.15365 | -2.28973 | -0.13384 | 0.011646 | -1.59759 | 0.195533 |
| 22.80556 | -1.20067 | -1.39994 | -0.90558 | -0.4278 | -1.46211 | 0.07818 | -2.62112 | -1.76526 | -0.92174 | -0.99549 | 1.311614 | -3.62905 | -1.725 | -2.10253 | -0.49266 | -2.08643 | -1.06245 | -1.12106 | -0.70627 | -3.585 | 0.009644 | 0.338287 | -1.81392 | -1.1286 |
| 22.81944 | -1.45221 | -2.68572 | -1.845 | -0.54826 | -1.061 | -0.70251 | -2.07481 | -1.43571 | -1.43619 | -1.55238 | -2.04368 | -3.28942 | -1.67539 | -2.34887 | 0.012618 | -2.04888 | -0.61605 | -1.68569 | -1.29308 | -4.05514 | -0.34049 | 0.246191 | -1.06295 | -2.09645 |
| 22.83333 | -1.86005 | -3.06804 | -2.2377 | -1.05671 | -0.97177 | -2.06282 | -0.94582 | -1.92184 | -1.96808 | -2.138 | -6.73097 | -2.48474 | -1.79885 | -2.65165 | 0.018155 | -2.29515 | -0.46642 | -2.33276 | -1.5701 | -2.93093 | -0.80912 | 0.01103 | -0.6671 | -3.57606 |
| 22.84722 | -2.05083 | -2.7164 | -2.38304 | -1.8956 | -1.18513 | -2.59912 | -0.28919 | -2.68346 | -2.42441 | -2.25199 | -10.5136 | -1.70067 | -1.92044 | -2.16673 | -1.26451 | -2.20927 | -0.35078 | -2.53758 | -1.42371 | -1.28105 | -1.3979 | -0.44442 | -0.92098 | -4.31011 |
| 22.86111 | -1.89514 | -2.26469 | -2.7777 | -2.25213 | -1.13179 | -2.10141 | 0.088617 | -3.10936 | -2.0874 | -1.75255 | -8.88954 | -0.77663 | -2.02834 | -0.89616 | -2.5095 | -1.7548 | -0.41069 | -2.09934 | -1.02296 | -0.55308 | -1.9422 | -0.89814 | -1.79656 | -4.66384 |
| 22.875 | -1.49223 | -1.69258 | -3.06292 | -1.83545 | -0.64795 | -1.4635 | 0.372669 | -1.98223 | -1.28832 | -0.97325 | -5.32187 | 0.200276 | -1.96433 | 0.333877 | -3.53575 | -1.0652 | -0.45016 | -1.25288 | -0.49496 | 0.187507 | -1.95907 | -0.99743 | -1.3748 | -2.36496 |
| 22.88889 | -1.32519 | -1.00536 | -2.93112 | -1.045 | 0.111562 | -1.16602 | 0.72463 | -0.2616 | -0.729 | -0.36789 | -2.05358 | 1.040043 | -1.73888 | 0.636424 | -3.72943 | -0.33327 | -0.18383 | -0.45782 | -0.01988 | 0.754572 | -1.70958 | -0.48096 | -0.1285 | 0.726695 |
| 22.90278 | -1.15985 | -0.5376 | -2.17538 | -0.44169 | 0.503414 | -1.21806 | 0.750307 | 1.031695 | -0.49926 | -0.07997 | 0.737276 | 1.26352 | -1.38205 | 0.350678 | -2.36337 | 0.479162 | 0.316991 | -0.19167 | 0.223114 | 0.504265 | -1.06569 | 0.036804 | 0.832443 | 3.030957 |
| 22.91667 | -0.89028 | -0.26971 | -1.33263 | -0.28256 | 0.36887 | -1.26677 | 0.396919 | 1.150857 | -0.50894 | -0.10077 | 2.530909 | 0.814286 | -1.19226 | -0.37814 | -0.98553 | 0.600209 | 0.373708 | -0.37747 | 0.135984 | -0.49011 | -0.38652 | 0.189456 | 0.627875 | 3.270497 |
| 22.93056 | -0.60075 | -0.26771 | -0.8035 | -0.39747 | -0.04747 | -1.12488 | -0.11889 | 0.151338 | -0.53253 | -0.2225 | 3.01663 | 0.214563 | -1.57113 | -2.08431 | 0.031658 | 0.522439 | 0.050407 | -0.53441 | -0.1564 | -1.91724 | 0.17425 | 0.116974 | -0.46231 | 1.954869 |
| 22.94444 | -0.75789 | -0.85802 | -0.88497 | -0.74414 | -0.65214 | -0.89174 | -0.75246 | -1.02137 | -0.91882 | -0.42948 | 2.193955 | -0.46946 | -2.20952 | -2.84953 | 0.254357 | 0.057104 | -0.48625 | -1.00282 | -0.67781 | -3.34557 | 0.157033 | -0.18732 | -1.42712 | -0.17019 |
| 22.95833 | -2.13341 | -2.45652 | -2.08246 | -1.36392 | -1.64022 | -1.84726 | -1.87974 | -1.88748 | -2.25111 | -0.874 | -0.23545 | -1.7626 | -2.94642 | -2.8961 | -0.81759 | -1.52681 | -1.47276 | -2.53891 | -1.50474 | -4.49873 | -0.73041 | -0.74073 | -2.90836 | -2.57693 |
| 22.97222 | -3.97516 | -4.81287 | -3.57409 | -2.05251 | -3.5285 | -3.85682 | -3.98065 | -2.87285 | -4.1128 | -1.43671 | -3.29928 | -3.65013 | -4.69702 | -3.62426 | -2.36049 | -3.24175 | -1.92403 | -4.77052 | -2.45206 | -5.5776 | -2.25604 | -1.37892 | -4.87141 | -4.86851 |
| 22.98611 | -3.02218 | -4.75631 | -3.43581 | -1.92666 | -4.14709 | -4.48507 | -4.44093 | -3.71336 | -4.57343 | -1.46851 | -4.86754 | -4.29184 | -6.17971 | -3.8106 | -2.90442 | -3.81216 | -1.97652 | -4.9814 | -2.24258 | -5.56811 | -2.95687 | -1.51326 | -5.04329 | -5.30641 |
| 23 | -0.60349 | -2.61534 | -1.35051 | -1.16436 | -2.44223 | -2.55031 | -2.51177 | -3.10221 | -3.01669 | -0.8834 | -4.40786 | -3.17324 | -4.5566 | -2.26211 | -2.76146 | -3.12798 | -1.49268 | -2.82439 | -1.4307 | -4.0553 | -2.45445 | -0.93844 | -2.9175 | -3.50819 |
| 23.01389 | 0.471144 | -0.29871 | 0.23183 | -0.6496 | -1.07077 | -0.36665 | -0.84617 | -1.94939 | -1.60116 | -0.61019 | -3.82185 | -2.40831 | -1.57979 | -0.92843 | -2.49406 | -1.8927 | -0.71836 | -0.75244 | -0.70794 | -1.94047 | -1.21206 | -0.60394 | -1.8236 | -2.48976 |
| 23.02778 | 0.406156 | 0.614173 | 0.60241 | -0.12844 | -0.21704 | 0.483507 | 0.193327 | -0.60747 | -0.57553 | -0.624 | -2.45753 | -1.59146 | 1.123784 | 0.369862 | -0.68061 | -0.53329 | 0.084267 | 0.603595 | -0.41633 | -0.21752 | -0.1502 | -0.36938 | -0.9285 | -1.63827 |
| 23.04167 | 0.075033 | 0.176667 | 0.252747 | 0.547618 | 0.094974 | -0.25142 | 0.546593 | 1.24127 | -0.04745 | -0.30015 | -0.68882 | -0.65745 | 2.973229 | 1.483395 | 1.400495 | 0.491507 | 0.79411 | 0.646005 | 0.004242 | 0.608297 | 0.637554 | -0.07179 | -0.05526 | -0.66425 |
| 23.05556 | -0.2768 | -0.18555 | -0.31706 | 0.874684 | 0.186839 | -1.10044 | 0.599744 | 1.935092 | 0.10949 | 0.118932 | 0.292718 | -0.05451 | 2.857894 | 1.765902 | 2.093631 | 0.741498 | 0.759329 | 0.417611 | 0.18705 | 0.817196 | 0.87663 | 0.036126 | -0.03227 | 0.002841 |
| 23.06944 | -0.84033 | -0.30053 | -0.88183 | 0.710624 | 0.193594 | -1.34428 | 0.475229 | 1.592965 | -0.00712 | 0.201336 | 0.660196 | 0.014697 | 1.82038 | 1.747493 | 1.521943 | 0.445083 | 0.386363 | 0.140392 | 0.349517 | 0.57503 | 0.6252 | 0.007472 | -0.05919 | 0.260229 |
| 23.08333 | -1.41638 | 0.018685 | -0.57779 | 0.41511 | 0.235273 | -0.81327 | 0.340913 | 1.14188 | -0.18276 | 0.076913 | 0.673793 | -0.1691 | 1.204205 | 1.788761 | 0.760003 | 0.167407 | 0.155654 | 0.107118 | 0.527855 | 0.377264 | 0.243196 | 0.021437 | 0.147647 | 0.24971 |
| 23.09722 | -1.0096 | 0.565027 | 0.184414 | 0.351431 | 0.261737 | 0.107925 | 0.316685 | 0.845734 | -0.10849 | 0.09373 | 0.252973 | -0.29574 | 0.924574 | 1.615805 | 0.37993 | 0.124605 | 0.151771 | 0.278923 | 0.382639 | 0.585009 | 0.092048 | 0.076642 | 0.498098 | 0.086716 |
| 23.11111 | 0.057392 | 1.41746 | 0.961697 | 0.448946 | 0.459824 | 0.940517 | 0.536891 | 0.66389 | 0.287304 | 0.231023 | 0.204411 | -0.04493 | 0.597306 | 1.397194 | 0.195541 | 0.241127 | 0.180221 | 0.227345 | 0.121425 | 0.772943 | 0.133793 | 0.071082 | 0.65218 | -0.11257 |
| 23.125 | 1.107775 | 1.956642 | 1.180837 | 0.505525 | 0.704738 | 1.083866 | 0.940036 | 0.709707 | 0.809997 | 0.354157 | 0.87174 | 0.505082 | 0.543488 | 1.64295 | 0.211495 | 0.465324 | 0.233104 | 0.226003 | -0.06883 | 1.072044 | 0.31246 | 0.079889 | 0.671049 | 0.107207 |
| 23.13889 | 1.043101 | 1.364356 | 0.469496 | 0.383746 | 0.637016 | 0.488666 | 1.075106 | 0.67795 | 0.991341 | 0.324692 | 1.394183 | 0.877748 | 0.30565 | 1.535142 | 0.280348 | 0.596445 | 0.222277 | 0.292563 | 0.041346 | 0.707697 | 0.560713 | 0.111681 | 0.611913 | 0.337951 |
| 23.15278 | 0.45571 | -0.00211 | -0.50623 | 0.144882 | 0.42755 | -0.52222 | 0.889033 | 0.396313 | 0.806151 | 0.200562 | 1.590589 | 0.928292 | -0.27489 | 0.89072 | 0.161062 | 0.557563 | 0.139349 | 0.346096 | 0.403602 | 0.8885 | 0.59164 | 0.108228 | 0.200032 | 0.381798 |
| 23.16667 | -0.06587 | -0.59857 | -0.81166 | -0.06778 | 0.446838 | -0.60556 | 0.810199 | 0.081699 | 0.659008 | 0.106725 | 1.836702 | 0.896747 | -0.27539 | 0.291984 | -0.19877 | 0.472993 | 0.044657 | 0.803368 | 0.529159 | 1.116281 | 0.564628 | 0.12456 | -0.19081 | 0.620984 |
| 23.18056 | -0.4169 | 0.035597 | -0.27284 | -0.18942 | 0.610047 | 0.606857 | 0.681196 | -0.32896 | 0.667218 | -4.6E-05 | 1.739 | 0.800652 | -0.18763 | -0.58703 | -0.77387 | 0.421158 | -0.0753 | 1.160604 | 0.222188 | 1.22195 | 0.413496 | 0.111364 | -0.15607 | 0.806763 |
| 23.19444 | -0.27394 | 1.057547 | 0.543173 | -0.23732 | 0.640367 | 1.489026 | 0.531236 | -0.78027 | 0.821232 | -0.10409 | 1.336608 | 0.749506 | -0.30539 | -1.54906 | -1.41524 | 0.29886 | -0.14313 | 0.906166 | -0.19814 | 1.295739 | 0.15662 | 0.059094 | 0.149857 | 0.593499 |
| 23.20833 | 0.271608 | 1.5776 | 0.991384 | -0.24655 | 0.442904 | 1.043953 | 0.378032 | -1.00227 | 1.009134 | -0.15155 | 1.145106 | 0.872852 | -0.70644 | -2.05203 | -1.72242 | 0.202654 | -0.02515 | 0.252896 | -0.54886 | 1.301702 | -0.15673 | 0.013389 | 0.519926 | 0.06574 |
| 23.22222 | 0.58554 | 0.728124 | 0.409932 | -0.31462 | 0.066922 | 0.041156 | 0.172807 | -0.8285 | 0.870954 | -0.16885 | 0.927179 | 0.946636 | -0.64604 | -1.54796 | -2.12482 | 0.121571 | 0.153028 | -0.267 | -0.48326 | 1.007705 | -0.43186 | -0.02626 | 0.640858 | -0.60712 |
| 23.23611 | 0.356682 | -0.56676 | -0.31197 | -0.31663 | -0.2824 | -0.87367 | -0.06988 | -0.20417 | 0.555551 | -0.22396 | 0.687955 | 0.893419 | -0.20897 | -0.61266 | -2.2372 | -0.07846 | 0.095996 | -0.41154 | -0.27687 | 0.578396 | -0.47132 | -0.05233 | 0.366285 | -0.89811 |
| 23.25 | 0.131636 | -1.5828 | -0.6664 | -0.11628 | -0.2602 | -0.68634 | -0.17233 | 0.36012 | 0.386777 | -0.27564 | 0.463023 | 0.803449 | -0.0591 | 0.02519 | -1.75229 | -0.28131 | -0.10041 | -0.06774 | -0.3039 | 0.226679 | -0.34864 | -0.05659 | -0.07746 | -0.27281 |
| 23.26389 | -0.22144 | -1.32629 | -0.45598 | 0.061633 | 0.014813 | 0.311836 | -0.14405 | 0.3714 | 0.226004 | -0.29511 | -0.30821 | 0.507598 | -0.05544 | 0.25766 | -0.96635 | -0.27965 | -0.23646 | 0.378209 | -0.47775 | -0.04954 | -0.26316 | -0.06786 | -0.42812 | 0.248001 |
| 23.27778 | -0.43308 | -0.27658 | 0.075956 | 0.09866 | 0.368406 | 1.363908 | -0.02605 | -0.16525 | 0.065034 | -0.2363 | -1.1297 | 0.311391 | 0.010165 | 0.225505 | -0.15367 | -0.21034 | -0.3443 | 0.318984 | -0.55495 | -0.11068 | -0.27671 | -0.10255 | -0.39999 | 0.386215 |
| 23.29167 | -0.03083 | 0.755235 | 0.657641 | 0.168982 | 0.686474 | 1.708392 | 0.209141 | -0.4863 | 0.161633 | 0.018621 | -1.37313 | 0.504919 | 0.166427 | 0.143969 | 0.637835 | -0.23902 | -0.46828 | -0.17247 | -0.55174 | 0.028678 | -0.32248 | -0.13039 | 0.023265 | 0.11934 |
| 23.30556 | 0.525313 | 0.644202 | 0.75412 | 0.239694 | 0.600306 | 0.612359 | 0.310196 | -0.32717 | 0.18402 | 0.296708 | -1.30647 | 0.664132 | 0.141197 | 0.094332 | 1.157647 | -0.32746 | -0.43468 | -0.71072 | -0.37991 | 0.018063 | -0.38856 | -0.19943 | 0.520495 | -0.14665 |
| 23.31944 | 0.803377 | 0.033119 | 0.170758 | 0.261311 | 0.200142 | -0.88344 | 0.167735 | 0.247384 | -0.03637 | 0.447027 | -0.65815 | 0.484061 | 0.20484 | 0.284397 | 1.370263 | -0.25903 | -0.2055 | -0.81294 | 0.000694 | -0.12956 | -0.36748 | -0.26522 | 0.713353 | -0.31909 |
| 23.33333 | 0.665599 | -0.6521 | -0.49019 | 0.221238 | -0.09908 | -1.81143 | 0.060626 | 0.919398 | -0.31201 | 0.503632 | 0.156264 | 0.23458 | 0.252638 | 0.76728 | 1.58659 | -0.14482 | -0.02386 | -0.27938 | 0.345191 | -0.24989 | -0.14558 | -0.25363 | 0.560742 | 0.129526 |
| 23.34722 | 0.240225 | -0.70191 | -0.62176 | 0.168872 | 0.026376 | -0.89643 | 0.027111 | 1.452762 | -0.53856 | 0.548698 | 0.188232 | -0.05365 | 0.412777 | 1.243466 | 1.583597 | -0.04549 | -0.01171 | 0.384792 | 0.460331 | -0.28124 | 0.207506 | -0.15854 | 0.18338 | 0.761115 |
| 23.36111 | -0.20701 | 0.33213 | -0.13038 | 0.165874 | 0.438983 | 0.90381 | 0.183869 | 1.620298 | -0.56425 | 0.694052 | 0.414292 | 0.022872 | 0.70462 | 1.696979 | 1.511133 | 0.125569 | 0.006416 | 0.807983 | 0.46972 | 0.017035 | 0.565391 | -0.00501 | -0.04299 | 1.470265 |
| 23.375 | -0.30642 | 1.42516 | 0.641192 | 0.159005 | 0.84624 | 2.860243 | 0.51339 | 1.604122 | -0.35034 | 0.846421 | 1.252055 | 0.424941 | 0.77076 | 1.434902 | 1.202867 | 0.388448 | 0.158268 | 0.729335 | 0.387846 | 0.716981 | 0.839529 | 0.115062 | 0.19982 | 1.686603 |
| 23.38889 | 0.074583 | 1.495959 | 1.136564 | 0.091071 | 0.61252 | 2.358863 | 0.792848 | 0.965823 | -0.11471 | 0.891974 | 2.119727 | 0.67747 | 0.651607 | 0.250773 | 0.477353 | 0.651003 | 0.511676 | 0.390011 | 0.440393 | 1.472227 | 0.859903 | 0.215116 | 0.766145 | 1.291776 |
| 23.40278 | 0.806721 | 0.807304 | 0.626076 | 0.148388 | 0.268485 | 0.44833 | 0.94872 | 0.176664 | 0.168535 | 0.794701 | 3.235022 | 0.718783 | 0.409428 | -0.92239 | -0.3672 | 1.007729 | 1.094617 | 0.44216 | 0.744233 | 2.097858 | 1.0612 | 0.257117 | 1.344326 | 0.747792 |
| 23.41667 | 1.345103 | 0.216198 | -0.10545 | 0.432153 | 0.095654 | -1.39183 | 1.085211 | -0.35047 | 0.545727 | 0.73308 | 3.867968 | 0.751965 | 0.070578 | -1.77861 | -0.9225 | 1.378368 | 1.616807 | 1.065663 | 1.074437 | 2.343968 | 1.401376 | 0.223314 | 1.873192 | 0.537965 |
| 23.43056 | 1.51611 | 0.161066 | -0.39741 | 0.916447 | 0.38389 | -1.37697 | 1.044018 | -0.26736 | 0.967375 | 0.715419 | 3.591429 | 0.876428 | -0.02701 | -1.67589 | -1.15347 | 1.747538 | 1.910317 | 1.799955 | 1.091977 | 2.301291 | 1.443512 | 0.225567 | 1.877923 | 0.815195 |
| 23.44444 | 2.292332 | 1.302321 | 1.048517 | 1.68811 | 1.241499 | 0.809359 | 1.184744 | 1.010628 | 1.963039 | 1.141087 | 3.797243 | 2.400328 | 1.893998 | -0.07189 | 0.222677 | 2.104212 | 2.095235 | 2.513024 | 1.098152 | 2.330583 | 1.348806 | 0.302534 | 1.569222 | 1.543867 |
| 23.45833 | 4.517755 | 3.636787 | 3.739367 | 3.697243 | 3.078149 | 3.307897 | 2.002774 | 4.122226 | 4.188687 | 2.343164 | 6.300717 | 5.562693 | 5.645519 | 2.961591 | 3.342161 | 3.111731 | 2.951485 | 3.970575 | 1.597514 | 2.379946 | 1.708629 | 0.475082 | 2.535013 | 3.319814 |
| 23.47222 | 6.258179 | 6.484507 | 4.37668 | 4.282842 | 4.151104 | 3.950901 | 2.685904 | 7.138013 | 6.162417 | 3.412224 | 8.393847 | 4.839828 | 6.387644 | 5.483597 | 5.232991 | 4.936103 | 2.839782 | 4.494772 | 3.137793 | 2.364874 | 3.02458 | 1.051049 | 4.026922 | 4.921586 |
| 23.48611 | 3.912739 | 5.701323 | 4.083776 | 2.976476 | 3.201519 | 2.723001 | 2.157309 | 3.96776 | 4.73593 | 2.335543 | 4.800218 | 1.667981 | 3.128293 | 4.061506 | 4.847663 | 4.351854 | 2.013446 | 3.695278 | 3.002827 | 0.827189 | 3.042158 | 1.276335 | 2.894795 | 4.559495 |
| 23.5 | 2.72845 | 2.561033 | 1.941887 | 1.028117 | 1.43792 | 1.258759 | 1.499555 | 1.1846 | 2.537297 | 1.314487 | 1.027578 | -0.02978 | 2.105769 | 2.416472 | 2.895429 | 2.736845 | 0.724518 | 3.062019 | 1.564419 | 0.098382 | 1.988597 | 0.917236 | 2.376288 | 3.366128 |
| 23.51389 | 0.712336 | 0.23794 | 0.579278 | -0.31648 | 0.242883 | -0.26143 | 0.549138 | -0.86121 | 0.967128 | 0.552308 | -2.7266 | -0.44978 | 1.095637 | 0.904546 | 0.932075 | 1.304423 | -0.59171 | 1.568393 | 0.289829 | -1.29741 | 0.869638 | 0.322578 | 1.705712 | 1.171906 |
| 23.52778 | -2.13772 | -1.90215 | -0.31952 | -1.95247 | -1.01567 | -1.55015 | -0.82043 | -2.72727 | -0.39216 | -0.42031 | -6.92021 | -1.17967 | -1.07399 | -0.40387 | -0.77037 | 0.110373 | -1.50766 | -0.15715 | -0.91007 | -2.49261 | -0.07601 | -0.11901 | -0.39345 | -1.71996 |
| 23.54167 | -3.66333 | -3.24063 | -0.6599 | -2.17328 | -1.70769 | -2.04568 | -1.97793 | -3.63492 | -1.53993 | -1.26434 | -7.91759 | -2.07623 | -2.96665 | -1.55591 | -1.53411 | 0.103151 | -1.5785 | -1.7606 | -1.64458 | -2.58115 | -0.79018 | -0.48462 | -3.24418 | -4.73477 |
| 23.55556 | -3.9942 | -1.95368 | -0.07668 | -1.17331 | -1.03148 | -0.95652 | -1.62957 | -2.80659 | -1.84499 | -0.72337 | -4.44846 | -2.16758 | -2.77158 | -1.85973 | -2.21755 | 0.015786 | -0.56869 | -2.59689 | -0.74467 | -0.54158 | -0.97541 | -0.5565 | -2.99018 | -3.67232 |
| 23.56944 | -2.90401 | 0.784124 | 0.649728 | 0.587452 | -0.0905 | 0.177605 | -0.44162 | -0.42329 | -0.90006 | 0.58725 | 0.627651 | -0.82225 | -1.163 | -0.41332 | 0.346862 | 0.580585 | 0.152064 | -1.3147 | 0.493599 | 2.079906 | -0.78909 | -0.04413 | -0.4636 | 0.205169 |
| 23.58333 | -0.89861 | 3.004447 | 0.436303 | 1.767243 | 0.552313 | 0.957248 | 0.975489 | 1.774897 | 0.625309 | 1.691014 | 5.492107 | 0.510309 | 0.644275 | 1.873102 | 3.13888 | 0.76392 | 0.5647 | 0.177336 | 1.351727 | 3.181484 | -1.03754 | 0.484401 | 1.427758 | 3.278144 |
| 23.59722 | 1.032228 | 3.09956 | -0.2932 | 2.389128 | 0.541555 | 1.267697 | 1.918276 | 2.494185 | 1.847159 | 1.359905 | 9.621236 | 1.65309 | 0.638754 | 4.125862 | 2.943816 | 1.094754 | 0.646849 | 1.346984 | 1.652308 | 2.328477 | -0.51124 | 0.926215 | 1.374983 | 4.690068 |
| 23.61111 | 1.756406 | 1.246686 | -1.02993 | 1.793403 | 0.251403 | 0.508179 | 1.871582 | 2.195789 | 2.27022 | 0.600438 | 7.765254 | 2.591354 | -0.75923 | 3.412907 | 1.396151 | 0.962113 | 0.974744 | 1.570799 | 1.60439 | 1.120087 | -0.0763 | 0.815296 | 0.22549 | 2.224547 |
| 23.625 | 1.165229 | 0.134928 | -1.68756 | 0.281326 | 0.002086 | -0.85151 | 1.478156 | 0.454328 | 1.924541 | -0.05947 | 4.801426 | 3.015276 | -1.60702 | 1.656475 | 0.795626 | 1.157763 | 0.935067 | 1.070105 | 1.549297 | -0.0486 | 0.047435 | 0.527644 | 0.343194 | 0.124314 |
| 23.63889 | -0.14008 | -1.36727 | -2.27036 | -0.62447 | 0.043294 | -2.6571 | 0.819495 | -1.1509 | 1.00531 | -0.64422 | 1.432396 | 2.325413 | -1.80139 | -0.01243 | -0.41703 | 0.50347 | 0.49325 | 0.176641 | 0.939174 | -1.44689 | -0.0837 | 0.421909 | -0.11495 | -1.00744 |
| 23.65278 | -1.39833 | -2.3359 | -1.87968 | -1.36824 | -0.07236 | -3.83387 | -0.20264 | -2.42592 | 0.038341 | -0.87284 | -1.28786 | 1.610425 | -1.80079 | -1.53963 | -1.84341 | -0.87719 | -0.2122 | -0.81597 | -0.06634 | -3.43982 | -0.25567 | 0.48833 | -0.68851 | -0.88068 |
| 23.66667 | -2.16617 | -1.68369 | -0.71613 | -1.03415 | -0.41969 | -3.21567 | -1.37909 | -2.85176 | -0.64457 | -0.60253 | -3.76485 | 0.918495 | -2.17743 | -3.55148 | -2.61338 | -1.90227 | -1.23316 | -1.25762 | -0.5836 | -4.17928 | 0.082378 | 0.506978 | -1.34699 | -1.37899 |
| 23.68056 | -2.17779 | -1.32772 | 0.168933 | -0.18451 | -0.73778 | -0.86425 | -1.78652 | -3.47628 | -0.79801 | 0.108468 | -7.1617 | -0.13396 | -2.91194 | -4.5374 | -2.99411 | -1.78048 | -1.37277 | -1.09934 | -0.6079 | -3.21777 | 0.848091 | 0.127447 | -3.30369 | -2.99553 |
| 23.69444 | -1.50648 | -0.85423 | 0.076672 | 0.4275 | -0.88948 | 0.817475 | -1.24113 | -1.61305 | -0.60844 | 0.858956 | -9.90521 | -1.18559 | -2.25314 | -2.48149 | -3.8662 | -0.74144 | -0.70441 | -0.97428 | -0.46972 | -1.90199 | 1.162992 | -0.14808 | -3.6065 | -4.46313 |
| 23.70833 | -0.59297 | -0.66227 | -0.48425 | 0.394477 | -0.584 | 0.703403 | 0.075778 | 1.337034 | -0.4063 | 1.156205 | -6.71817 | -1.83176 | 0.415314 | 1.004317 | -2.07195 | 0.205725 | -0.1123 | -1.45276 | -0.88382 | -0.61997 | 0.880228 | -0.15329 | -1.33172 | -3.30571 |
| 23.72222 | -0.01326 | -0.13847 | -0.88275 | 0.133877 | 0.084815 | 0.050085 | 1.368131 | 3.484931 | -0.41905 | 0.885596 | -2.30547 | -1.04585 | 2.888758 | 2.793998 | 0.311089 | 0.79102 | -0.16527 | -1.49643 | -0.76466 | 0.828076 | 0.435643 | 0.019648 | 1.576252 | -0.17591 |
| 23.73611 | 0.017375 | 0.743815 | -1.30745 | -0.42115 | 0.485017 | -0.51559 | 1.625429 | 3.851903 | -0.30467 | 0.335783 | 0.319712 | -0.66881 | 3.625112 | 1.921322 | 2.86882 | 0.551731 | 0.211338 | -1.26766 | -0.44638 | 1.995512 | 0.347975 | 0.181309 | 4.06977 | 2.992418 |
| 23.75 | -0.03533 | 1.371641 | -1.656 | -0.81757 | 0.482035 | -1.00697 | 1.217052 | 2.968335 | -0.24575 | -0.20452 | 2.53622 | -0.11166 | 2.699639 | -0.19838 | 4.739917 | 0.251736 | 0.246772 | -1.30642 | -0.15376 | 1.968761 | 0.249421 | 0.237687 | 4.399785 | 4.596649 |
| 23.76389 | 0.110224 | 0.724456 | -0.543 | -0.618 | 0.36372 | -1.6082 | 0.482186 | 0.873454 | -0.31441 | -0.46137 | 4.523698 | 0.126976 | 1.058711 | -1.97367 | 4.446768 | -0.02841 | 0.289751 | -1.50727 | -0.09238 | 1.079134 | -0.26301 | -0.07069 | 1.733926 | 4.360771 |
| 23.77778 | 0.143449 | -0.15818 | 0.887906 | 0.076084 | 0.149882 | -2.23872 | -0.27507 | -1.44599 | -0.48907 | -0.57594 | 5.873587 | -0.12387 | -0.33555 | -2.88046 | 1.716965 | -0.33454 | 0.684692 | -1.16907 | 0.10382 | -0.26234 | -0.80165 | -0.43179 | -0.91453 | 1.939235 |
| 23.79167 | 0.000171 | -0.0459 | 1.507224 | 0.573257 | -0.19304 | -2.07118 | -1.00748 | -3.69675 | -0.72729 | -0.79079 | 5.719545 | -0.57462 | -1.20741 | -3.30125 | -1.18118 | -0.36895 | 1.203373 | -0.40869 | 0.108479 | -1.87752 | -1.18497 | -0.71021 | -1.45683 | 0.287842 |
| 23.80556 | -0.37026 | 0.157434 | 0.423652 | 0.214085 | -0.65069 | -1.20582 | -1.62831 | -5.00043 | -0.9384 | -1.10424 | 3.22912 | -0.88851 | -1.70161 | -2.98595 | -3.71066 | -0.79389 | 0.333869 | 0.086269 | -0.18403 | -2.782 | -1.45947 | -0.76935 | -1.02034 | -1.0605 |
| 23.81944 | -0.97434 | 0.382441 | -1.04698 | -0.92063 | -1.26621 | -0.43119 | -1.91999 | -4.83464 | -1.17067 | -1.43488 | -0.24445 | -1.0397 | -1.96562 | -2.28038 | -4.9885 | -1.40081 | -0.88272 | 0.264823 | -0.3525 | -3.31071 | -1.75872 | -0.76057 | -1.41862 | -3.51366 |
| 23.83333 | -1.37447 | -0.20706 | -2.26429 | -2.0799 | -1.56909 | -0.62959 | -1.70872 | -3.27943 | -1.52124 | -1.76911 | -3.73401 | -1.09614 | -2.13181 | -1.43575 | -4.25362 | -1.58994 | -1.53424 | -0.11519 | -0.67087 | -3.67337 | -1.96293 | -0.62079 | -1.82284 | -5.66584 |
| 23.84722 | -1.28009 | -2.06424 | -2.78914 | -2.92745 | -1.50771 | -1.5063 | -1.17226 | -1.60725 | -1.78939 | -2.02983 | -6.08842 | -1.09761 | -1.99917 | -0.72684 | -2.36939 | -2.13928 | -1.59511 | -0.89162 | -1.19336 | -3.35427 | -2.08022 | -0.64019 | -2.21935 | -5.51366 |

| | | | | | | | | | | | | | | | | | | | | | | | | |
|---|---|---|---|---|---|---|---|---|---|---|---|---|---|---|---|---|---|---|---|---|---|---|---|---|
| 23.86111 | -1.12656 | -3.2519 | -2.50516 | -2.87307 | -1.23664 | -2.14933 | -0.53382 | -0.21984 | -1.60457 | -1.98007 | -6.27019 | -0.89965 | -1.68673 | -0.06939 | -1.0697 | -1.99027 | -0.79239 | -1.58852 | -1.32209 | -2.45015 | -2.16632 | -0.6235 | -3.08226 | -5.35892 |
| 23.875 | -1.15763 | -2.94193 | -1.5408 | -1.9964 | -0.84146 | -1.79102 | -0.06573 | 1.336936 | -0.91552 | -1.56571 | -5.21262 | -0.62553 | -1.38953 | 0.651876 | -0.44075 | -1.61506 | -0.22595 | -1.95549 | -1.0744 | -1.62296 | -1.81841 | -0.48528 | -3.26218 | -2.93001 |
| 23.88889 | -1.2787 | -1.8946 | -0.29548 | -0.70445 | -0.46828 | -0.89926 | 0.273843 | 1.80328 | -0.30944 | -1.09088 | -3.8894 | -0.34294 | -1.41317 | 0.696554 | -0.18211 | -1.26725 | -0.20198 | -1.69797 | -0.6184 | -0.87657 | -1.26622 | -0.22002 | -2.00257 | -0.64922 |
| 23.90278 | -1.31005 | -1.24443 | 0.412806 | 0.248744 | -0.19719 | -0.31987 | 0.619007 | 1.404115 | -0.10899 | -0.67159 | -2.42814 | -0.02973 | -1.10544 | 0.187461 | 0.470536 | -0.83634 | -0.15224 | -1.25281 | -0.33231 | 0.033957 | -0.77797 | -0.00264 | -0.33952 | 1.134325 |
| 23.91667 | -1.0754 | -0.74606 | -0.12669 | 0.238052 | -0.05843 | -0.31588 | 0.824688 | 0.141715 | -0.28939 | -0.45866 | -0.7668 | 0.285148 | -1.00316 | -0.53043 | 1.151997 | -0.34181 | -0.17614 | -0.9255 | -0.15672 | 0.970866 | -0.44174 | 0.089399 | 1.347354 | 2.724312 |
| 23.93056 | -0.72337 | 0.032841 | -1.34249 | -0.22656 | -0.11004 | -0.59539 | 0.630428 | -1.58346 | -0.65322 | -0.4286 | 0.4608 | 0.637971 | -1.25023 | -1.25651 | 0.749774 | -0.09818 | -0.2626 | -0.71852 | 0.134508 | 1.317723 | -0.25113 | 0.106777 | 1.834208 | 3.210632 |
| 23.94444 | -0.69028 | 0.328351 | -2.83118 | -0.88319 | -0.45712 | -1.26023 | -0.11122 | -2.89377 | -1.44286 | -0.60772 | 0.444778 | 0.422025 | -1.75336 | -1.71529 | -0.68808 | -0.53735 | -0.74028 | -1.24699 | 0.017208 | 0.42669 | -0.46283 | -0.1573 | 0.044069 | 1.637854 |
| 23.95833 | -1.71648 | -1.1022 | -4.47722 | -1.8783 | -1.32534 | -2.40259 | -1.45283 | -3.49169 | -3.3831 | -1.27786 | -1.45257 | -1.00782 | -2.56232 | -2.38501 | -2.54236 | -1.99705 | -1.78554 | -2.99302 | -0.9825 | -1.37999 | -1.23112 | -0.75174 | -2.66442 | -1.33627 |
| 23.97222 | -3.0463 | -3.70684 | -5.90831 | -3.01607 | -2.73638 | -4.00779 | -3.76811 | -4.05268 | -6.10281 | -2.19625 | -3.99611 | -3.13405 | -4.09015 | -3.4619 | -3.75351 | -3.81226 | -2.65269 | -5.13504 | -2.48994 | -3.11629 | -2.19862 | -1.05712 | -5.63881 | -4.60714 |
| 23.98611 | -1.95509 | -4.57723 | -4.69015 | -3.06241 | -3.37558 | -4.94533 | -4.25708 | -4.27587 | -6.66737 | -2.42033 | -4.38706 | -4.22381 | -4.97038 | -3.16526 | -4.15899 | -4.32962 | -2.5221 | -5.86125 | -2.92432 | -3.72807 | -2.33724 | -0.98954 | -5.7669 | -5.48135 |
| 24 | 0.210867 | -2.77363 | -1.46091 | -1.99689 | -2.73352 | -3.79678 | -3.13842 | -3.3937 | -5.26821 | -1.85 | -2.49144 | -3.03856 | -3.29998 | -1.56216 | -2.95066 | -3.08168 | -1.35573 | -3.90977 | -1.63166 | -2.5356 | -1.86342 | -0.88735 | -3.07906 | -4.07588 |
| 24.01389 | 0.512291 | -0.62343 | 1.628097 | -1.50471 | -1.57381 | -0.83602 | -1.76604 | -2.36152 | -3.28476 | -1.07413 | -1.42901 | -1.09213 | -2.06922 | -0.77407 | -1.62407 | -1.78705 | -0.63154 | -1.34853 | -0.47059 | -1.45095 | -1.14522 | -0.57062 | -0.64226 | -3.3046 |
| 24.02778 | -0.15142 | 0.558775 | 2.61225 | -1.22862 | -0.60996 | 1.768094 | -0.88137 | -1.67787 | -1.3831 | -0.46446 | -1.94295 | -0.96618 | -1.34674 | -0.36701 | -0.76949 | -1.61908 | -0.12731 | 0.978555 | 0.225992 | -0.78471 | -0.69029 | -0.40251 | 0.441219 | -2.67443 |
| 24.04167 | -0.69076 | -0.01417 | 1.144444 | -0.94282 | 0.096308 | 2.328786 | -0.6191 | -1.02285 | 0.078695 | -0.11981 | -2.03647 | -1.12794 | -0.75444 | -0.14906 | -0.54173 | 0.144585 | 0.058804 | 1.775098 | 0.286691 | -0.608 | -0.37339 | -0.28048 | -0.43762 | -1.85016 |
| 24.05556 | -0.99731 | -0.34165 | 0.220804 | -0.61025 | 0.544066 | 1.983469 | -0.46989 | -0.42263 | 0.841827 | 0.173181 | -1.79366 | -1.40715 | -0.32376 | -0.02401 | -0.38963 | 0.493989 | 0.177284 | 1.488308 | 0.452562 | -0.35983 | -0.0292 | -0.08915 | -0.9403 | -1.04194 |
| 24.06944 | -0.95449 | 0.061832 | -0.09301 | -0.32806 | 0.749711 | 1.938448 | -0.19473 | 0.038125 | 1.149066 | 0.340959 | -1.15405 | -1.2327 | 0.032147 | 0.079755 | -0.21494 | 0.636632 | 0.319531 | 1.717933 | 0.781554 | -0.02533 | 0.270878 | 0.020613 | -1.66257 | -0.33211 |
| 24.08333 | -0.43768 | 0.916528 | 0.125931 | -0.18704 | 0.72717 | 2.123386 | 0.020889 | 0.324441 | 1.268168 | 0.219563 | -0.26346 | -0.48407 | 0.225936 | 0.194344 | -0.02318 | 0.682037 | 0.399056 | 1.530804 | 1.183998 | 0.165776 | 0.321821 | 0.094154 | -2.93493 | 0.124913 |
| 24.09722 | 0.147672 | 1.454273 | 0.489729 | -0.09591 | 0.510256 | 2.071203 | 0.10935 | 0.449528 | 1.167686 | -0.06423 | 0.625592 | 0.248437 | 0.145187 | 0.263757 | 0.17939 | 0.547081 | 0.313024 | 1.396876 | 1.190959 | 0.005562 | -0.04526 | -0.0053 | -2.28669 | 0.173697 |
| 24.11111 | 0.309925 | 1.315338 | 0.627579 | 0.037007 | 0.342438 | 1.778996 | 0.101332 | 0.592747 | 0.961873 | -0.2648 | 0.957634 | 0.544069 | -0.15368 | 0.243957 | 0.302863 | 0.319333 | 0.210691 | 1.033201 | 0.623122 | -0.30036 | -0.39079 | -0.13582 | -0.0927 | 0.0322 |
| 24.125 | -0.00573 | 0.949478 | 0.673747 | 0.250706 | 0.47887 | 1.267466 | 0.154138 | 0.725169 | 0.699184 | -0.27808 | 0.234671 | -0.04261 | -0.13184 | 0.264155 | 0.334437 | 0.169229 | 0.227794 | 0.572743 | 0.229665 | -0.49624 | -0.28891 | -0.26397 | 2.094006 | -0.02226 |
| 24.13889 | -0.34016 | 0.743282 | 0.657227 | 0.637928 | 1.067503 | 0.91262 | 0.615983 | 1.046148 | 1.051067 | 0.02532 | 0.512131 | -0.07444 | 0.66006 | 0.633446 | 0.814475 | 0.500993 | 0.332911 | 0.591022 | 0.148479 | -0.39405 | 0.270991 | -0.61555 | 2.525321 | 0.383114 |
| 24.15278 | -0.25913 | 1.225677 | 0.697081 | 1.357776 | 1.459021 | 1.022831 | 1.399259 | 2.538294 | 2.885647 | 0.858186 | 1.402257 | 0.302505 | 2.440122 | 1.79056 | 2.004957 | 1.386762 | 0.56329 | 1.569899 | 0.509225 | 0.147604 | 1.148967 | 0.305308 | 2.660831 | 2.265866 |
| 24.16667 | 0.282802 | 2.337073 | 0.846327 | 2.046456 | 1.666184 | 1.274809 | 1.983647 | 2.998203 | 3.749355 | 1.198337 | 2.355281 | 1.18583 | 4.131885 | 3.107881 | 3.077796 | 1.887323 | 0.798421 | 2.443206 | 1.126417 | 0.942082 | 1.266726 | 1.23943 | 2.406219 | 3.784953 |
| 24.18056 | 0.861434 | 2.312296 | 0.73756 | 1.671003 | 1.190283 | 1.658772 | 1.589824 | 2.334431 | 2.228148 | 0.657871 | 2.330394 | 1.512161 | 2.739473 | 2.545354 | 3.469885 | 1.417738 | 0.835738 | 2.28286 | 1.507221 | 1.120002 | 0.748062 | 1.461197 | -0.09821 | 3.111195 |
| 24.19444 | 0.83755 | 2.093382 | 0.668284 | 1.004667 | 0.634548 | 1.9126 | 0.819079 | 1.692661 | 0.71024 | 0.078331 | 2.199665 | 1.421257 | 1.302459 | 1.987022 | 3.099427 | 0.803567 | 0.49163 | 1.576719 | 0.917623 | 0.499442 | 0.286363 | 0.723911 | -1.20693 | 2.026478 |
| 24.20833 | -0.01255 | 0.895513 | -0.12413 | 0.498205 | 0.268611 | 1.119929 | 0.340679 | 0.963084 | 0.000003 | -0.28987 | 1.536262 | 0.51697 | 0.136889 | 1.320125 | 2.118848 | 0.258588 | 0.111746 | 0.576724 | -0.20995 | 0.057788 | 0.108693 | 0.121486 | -0.17001 | 1.19782 |
| 24.22222 | -1.06958 | -0.54636 | -1.13115 | 0.284564 | 0.009019 | -0.13357 | 0.103542 | 0.299517 | -0.2078 | -0.46393 | 0.769432 | -0.3789 | -0.96723 | 0.776043 | 1.214279 | -0.12698 | -0.09617 | -0.44484 | -1.18181 | 0.105337 | 0.206944 | -0.31476 | 1.715734 | 0.596046 |
| 24.23611 | -1.46325 | -1.55429 | -1.13077 | 0.253287 | -0.02132 | -1.54479 | 0.081471 | -0.17757 | 0.18278 | -0.25914 | 0.313112 | -0.53574 | -0.96152 | 0.402364 | 0.731115 | -0.11019 | 0.085353 | -0.78429 | -0.60441 | 0.586654 | 0.271259 | 0.050705 | 2.22803 | 0.098733 |
| 24.25 | -0.32203 | -0.79825 | 0.172129 | 0.351869 | 0.201807 | -2.14131 | 0.252558 | -0.51968 | 1.1242 | 0.171421 | 0.596624 | 0.366431 | -0.2722 | 0.166445 | 0.66227 | 0.398958 | 0.478581 | -0.11297 | 0.298604 | 1.04256 | 0.252657 | 0.759351 | 0.997498 | 0.258527 |
| 24.26389 | 1.244949 | 0.92057 | 1.746881 | 0.441607 | 0.323583 | -0.45103 | 0.337161 | -0.79322 | 1.697302 | 0.45788 | 1.344013 | 1.580508 | 0.316142 | -0.1799 | 0.62252 | 0.859256 | 0.634162 | 0.841676 | 1.08726 | 1.149547 | 0.442468 | 1.156983 | -0.0461 | 1.147186 |
| 24.27778 | 1.547678 | 2.154237 | 1.752317 | 0.352613 | 0.192579 | 1.9113 | 0.136061 | -1.03031 | 0.891973 | 0.124355 | 1.808387 | 1.953569 | 0.222098 | -0.76558 | 0.291857 | 0.576173 | 0.273698 | 1.147336 | 0.426541 | 0.70526 | 0.371832 | 0.551583 | -2.08151 | 1.675386 |
| 24.29167 | 0.564781 | 0.603023 | 0.055738 | 0.165065 | 0.001741 | 2.858766 | -0.10791 | -0.95803 | -0.38329 | -0.44701 | 1.027026 | 0.956394 | -0.53093 | -1.37356 | -0.30236 | -0.07278 | -0.18212 | 0.04132 | -0.51793 | 0.237987 | 0.219088 | 0.027631 | -2.10438 | 1.208591 |
| 24.30556 | -0.39624 | -0.7969 | -1.42353 | 0.010978 | -0.08859 | 1.059248 | -0.24769 | -0.50543 | -1.35858 | -0.7285 | 0.268102 | -0.09436 | -1.07066 | -1.59282 | -0.73253 | -0.54253 | -0.45332 | -1.35864 | -1.69447 | 0.058462 | -0.05408 | -0.4396 | -0.41097 | 0.34927 |
| 24.31944 | -1.12518 | -1.89891 | -1.30225 | -0.16365 | -0.20712 | -0.57321 | -0.40302 | -0.14216 | -1.2726 | -0.51033 | -0.6831 | -0.69723 | -1.407 | -1.63993 | -1.03836 | -0.85164 | -0.44222 | -2.29877 | -1.81818 | 0.09435 | -0.18939 | -0.48229 | 1.186291 | -0.54729 |
| 24.33333 | -1.16994 | -1.77127 | 0.116916 | -0.29373 | -0.23529 | -2.89541 | -0.46002 | 0.035739 | -0.77911 | -0.11394 | -1.19663 | -0.27354 | -1.7725 | -1.6113 | -1.40083 | -0.61329 | -0.13415 | -1.64701 | -0.97303 | -0.00225 | -0.196 | 0.07109 | 1.816538 | -0.9939 |
| 24.34722 | -0.35041 | -0.63897 | 1.739843 | -0.20589 | -0.10978 | -3.36399 | -0.28633 | 0.109357 | -0.01538 | 0.306055 | -0.67827 | 0.983758 | -1.64096 | -1.28741 | -1.47532 | 0.025536 | 0.285032 | -0.35604 | 0.091554 | -0.13785 | 0.165644 | 0.602543 | 0.787754 | -0.69235 |
| 24.36111 | 0.935162 | 0.691928 | 1.90608 | 0.122849 | 0.026994 | -0.90231 | -0.06603 | 0.12639 | 0.764015 | 0.456998 | 0.135597 | 1.878875 | -0.96941 | -0.72349 | -1.13901 | 0.707703 | 0.527837 | 1.082719 | 1.190116 | -0.2376 | 0.353915 | 0.361995 | -0.08249 | 0.066474 |
| 24.375 | 2.356161 | 1.412199 | 0.513451 | 0.53991 | 0.088442 | 2.059332 | 0.042311 | 0.305178 | 1.150402 | 0.413781 | 0.404987 | 2.045493 | 0.03802 | -0.08912 | -0.48687 | 0.877844 | 0.648896 | 1.980744 | 1.159921 | -0.13525 | 0.356303 | 0.107035 | -1.12076 | 1.074487 |
| 24.38889 | 2.246843 | 0.881872 | -1.08865 | 1.011562 | 0.225096 | 4.159281 | 0.090373 | 0.80696 | 0.912921 | 0.389957 | -0.33314 | 0.655515 | 0.99431 | 0.502333 | 0.335441 | 0.839103 | 0.777647 | 1.098084 | 0.259361 | 0.269828 | 0.321227 | -0.08391 | -2.20618 | 1.513232 |
| 24.40278 | 0.632573 | 0.17695 | -1.82074 | 1.3485 | 0.211365 | 2.670058 | 0.205332 | 1.789825 | 0.380509 | 0.516958 | -0.37355 | -0.18295 | 1.478136 | 1.056585 | 1.227282 | 0.753311 | 0.875009 | 0.229606 | -0.33986 | 1.023259 | 0.567514 | -0.15711 | -0.47124 | 1.628822 |
| 24.41667 | -0.67786 | -0.24251 | -0.64235 | 1.407329 | 0.339482 | 0.669088 | 0.46345 | 3.323041 | -0.2077 | 0.8959 | -0.50865 | -0.55538 | 1.483377 | 1.520823 | 2.037068 | 0.670788 | 1.022932 | -0.42516 | -0.7124 | 1.897299 | 0.993012 | 0.526697 | 1.466038 | 1.677086 |
| 24.43056 | -1.74495 | 0.31225 | 1.415448 | 1.285102 | 0.563313 | -1.31055 | 0.801888 | 4.212905 | -0.19079 | 1.397993 | 0.425119 | 0.746509 | 1.31561 | 1.852885 | 3.199811 | 0.812693 | 1.030955 | 0.174138 | 0.037933 | 2.577063 | 1.368828 | 1.491341 | 2.848789 | 1.678738 |
| 24.44444 | -0.62039 | 1.413571 | 3.71289 | 1.442335 | 1.108654 | -3.35063 | 1.414598 | 4.239173 | 1.411988 | 2.139981 | 2.991239 | 3.0616 | 2.042691 | 2.510947 | 3.913992 | 1.681327 | 1.010754 | 1.935155 | 0.977264 | 3.192917 | 1.72354 | 1.885334 | 2.819606 | 2.57262 |
| 24.45833 | 1.899053 | 3.04965 | 5.371637 | 2.508662 | 2.696788 | -0.74208 | 3.29534 | 6.12412 | 5.096762 | 4.268213 | 9.364789 | 6.955558 | 4.658743 | 4.17819 | 5.011572 | 3.537464 | 1.80329 | 4.538033 | 2.914215 | 4.766544 | 2.610713 | 1.472403 | 2.480562 | 5.567743 |
| 24.47222 | 3.523248 | 3.755345 | 6.141273 | 4.339337 | 3.734571 | 4.406972 | 4.424606 | 7.735724 | 8.206272 | 5.372117 | 13.3823 | 10.95561 | 4.139969 | 5.528791 | 6.660638 | 6.31566 | 2.78257 | 6.052732 | 3.760377 | 6.614553 | 3.881154 | 1.358941 | 1.455273 | 8.259385 |
| 24.48611 | 4.277922 | 3.162429 | 3.845328 | 2.869299 | 2.692961 | 6.752346 | 3.16985 | 4.599406 | 6.703151 | 3.788608 | 8.985708 | 6.311132 | 2.370979 | 5.024886 | 4.839242 | 5.125732 | 2.653211 | 4.634517 | 3.170432 | 5.330588 | 3.54117 | 0.975392 | -0.81652 | 5.862843 |
| 24.5 | 3.278983 | 2.183013 | 1.688452 | 1.099624 | 1.96464 | 6.337686 | 2.255799 | 1.230417 | 4.484905 | 2.362782 | 4.097175 | 2.714783 | 1.809669 | 3.906126 | 2.304488 | 2.766011 | 1.460808 | 2.421402 | 1.937484 | 2.483111 | 1.994682 | 0.760256 | -1.6265 | 2.944687 |
| 24.51389 | 1.455951 | 0.528495 | 0.950076 | -0.26775 | 0.936392 | 4.293899 | 1.050052 | -0.78806 | 2.078481 | 0.845796 | 1.052613 | 0.237813 | 1.166581 | 2.084864 | -0.73277 | 1.764883 | 0.395425 | 0.66827 | 0.565612 | 0.392202 | 0.921803 | 0.221843 | -0.72301 | 0.847134 |
| 24.52778 | -0.29851 | -1.57637 | 0.158754 | -1.27148 | -0.2835 | 1.501559 | -0.25014 | -2.69823 | 0.061557 | -0.84068 | -2.21916 | -2.02508 | 0.386148 | -0.51982 | -3.60595 | 0.598703 | -0.97533 | -0.52657 | -0.82112 | -1.78667 | 0.155314 | -0.21231 | 1.096228 | -1.1905 |
| 24.54167 | -1.76003 | -2.7659 | -0.44206 | -1.9886 | -1.43448 | -0.90145 | -1.34087 | -4.13129 | -1.62119 | -2.08101 | -5.69838 | -2.76075 | -1.47377 | -2.83909 | -3.94611 | -0.56414 | -2.45918 | -1.5725 | -1.20626 | -3.89627 | -0.84382 | -0.2206 | 2.603547 | -3.88907 |
| 24.55556 | -2.73451 | -1.46352 | -0.41839 | -0.88472 | -1.63646 | -3.28851 | -1.65551 | -3.84049 | -2.4562 | -2.09917 | -8.81373 | -1.67085 | -3.60345 | -4.03937 | -4.75 | -0.89942 | -2.84811 | -1.51599 | -1.74735 | -3.77286 | -1.18778 | -0.27655 | 2.019399 | -5.55332 |
| 24.56944 | -1.61821 | 0.243114 | 0.015646 | 0.757521 | -0.85968 | -4.90573 | -1.47427 | -1.34331 | -1.45808 | -0.67235 | -7.70215 | 0.568691 | -4.31862 | -3.98353 | -2.99854 | -0.02029 | -0.94611 | -0.43534 | -0.94247 | -2.24256 | -0.94441 | -0.31259 | 0.037623 | -4.61117 |
| 24.58333 | 0.487461 | 2.499846 | 0.329715 | 2.17288 | 0.233852 | -4.63005 | -0.49009 | 1.997437 | 0.35053 | 0.852393 | -5.29463 | 1.320311 | -2.97111 | -2.31247 | -0.16158 | 0.704558 | 0.648539 | 0.816823 | 0.173291 | 0.035797 | -0.42962 | -0.4395 | -1.52128 | -1.99505 |
| 24.59722 | 1.716316 | 2.685829 | 0.144218 | 2.718569 | 0.843058 | -1.05829 | 0.380245 | 4.711281 | 1.630413 | 1.896787 | -1.70546 | 1.457699 | -1.22274 | 0.05958 | 1.990368 | 0.904826 | 1.141664 | 0.810029 | 0.84328 | 2.630901 | 0.103105 | -0.27959 | -1.44825 | 2.231471 |
| 24.61111 | 0.863905 | 2.021781 | -0.44043 | 2.722196 | 0.651261 | 1.773787 | 0.798398 | 3.183673 | 1.829101 | 1.913671 | 2.724212 | 1.290766 | -0.87828 | 1.508648 | 4.263669 | 0.417198 | 1.251565 | 0.761948 | 0.448276 | 3.922621 | 0.307023 | -0.35249 | -0.99121 | 5.64083 |
| 24.625 | -0.24092 | 1.220424 | -1.33734 | 1.704025 | 0.359312 | 3.24554 | 0.896521 | 1.600459 | 1.324623 | 1.048948 | 7.853604 | 0.541632 | -0.62196 | 2.260249 | 3.789072 | -0.01348 | 1.318277 | 0.561835 | -0.14095 | 3.529198 | 0.241026 | -0.28276 | 0.373873 | 5.51845 |
| 24.63889 | -0.90998 | -1.1197 | -1.78798 | 0.207572 | 0.089486 | 2.068154 | 0.883832 | 0.775105 | 0.78973 | -0.18373 | 9.647674 | -0.09948 | -1.04707 | 2.786209 | 3.061689 | 0.007934 | 1.156533 | 0.188912 | -0.03597 | 2.319484 | 0.000693 | 0.6506 | 0.823858 | 3.864077 |
| 24.65278 | -1.39122 | -2.81073 | -1.51471 | -0.83337 | -0.07598 | -0.1247 | 0.875292 | -0.1531 | 0.196232 | -1.14666 | 6.630951 | 0.375911 | -1.71058 | 2.668649 | 0.151026 | 0.415289 | 1.514458 | -0.13544 | 1.121628 | 0.439234 | -0.51696 | 1.472246 | 1.218477 | 1.847338 |
| 24.66667 | -1.4346 | -1.19451 | -1.07546 | -1.53437 | -0.21775 | -1.84147 | 0.80001 | -0.98845 | -0.3304 | -1.30186 | 3.453205 | 1.912421 | -1.48567 | 1.692164 | -2.49581 | 0.704278 | 1.092448 | -0.46562 | 1.291943 | -1.43613 | -0.71456 | 2.06747 | 0.617805 | 0.340899 |
| 24.68056 | -0.78934 | 0.411725 | -0.91681 | -1.75551 | -0.05693 | -3.64866 | 0.784351 | -1.8413 | -0.44931 | -0.69743 | 1.57984 | 2.554606 | -0.09812 | 0.401418 | -5.75413 | 0.563382 | 0.779452 | -0.53061 | 0.998333 | -2.69663 | -0.29146 | 1.114 | -0.40625 | -0.44645 |
| 24.69444 | 0.193564 | 2.716259 | -0.95417 | -1.36844 | 0.370947 | -4.55682 | 0.815794 | -2.9605 | -0.03562 | -0.07634 | -0.3039 | 1.830993 | 1.616924 | -0.91448 | -6.01151 | -0.22098 | 0.425175 | -0.31792 | 0.758166 | -2.92945 | 0.384066 | -0.17006 | -1.17578 | -0.64958 |
| 24.70833 | 0.646768 | 2.693512 | -1.14572 | -0.64823 | 0.74396 | -3.20728 | 0.799357 | -4.58424 | 0.493062 | 0.200981 | -2.73738 | 0.200078 | 2.354843 | -1.95064 | -5.19762 | -0.86919 | -0.35563 | -0.14605 | 0.328724 | -2.24087 | 0.91659 | -1.01296 | -1.33036 | -0.61462 |
| 24.72222 | 0.306207 | 0.508674 | -1.38739 | 0.210886 | 0.901556 | -0.64477 | 0.851801 | -3.94454 | 0.392656 | 0.065671 | -4.66123 | -0.82017 | 1.959276 | -2.30388 | -4.5378 | -1.0568 | -0.78991 | -0.24395 | -0.6165 | -1.82053 | 0.971078 | -1.48389 | 0.239378 | -0.31247 |
| 24.73611 | -0.30031 | -1.60234 | -1.6066 | 0.731422 | 0.588856 | 1.49264 | 0.563279 | -1.01165 | -0.56396 | -0.13456 | -6.25471 | -1.2829 | 0.873349 | -2.3196 | -2.72621 | -0.89897 | -0.95514 | -0.67317 | -1.80658 | -1.55732 | 0.717469 | -1.10441 | 0.858978 | -1.8424 |
| 24.75 | -0.69424 | -3.97493 | -1.74998 | 0.422288 | 0.049443 | 1.710391 | 0.489169 | 2.541904 | -1.54541 | -0.30056 | -6.85087 | -1.03269 | -0.11293 | -2.19317 | -0.34055 | -0.7391 | -0.92546 | -1.16784 | -2.43529 | -1.30536 | 0.20099 | -0.40464 | 1.833401 | -3.79293 |
| 24.76389 | -0.49962 | -4.43778 | -1.54351 | -0.00684 | -0.28825 | 0.271568 | 0.399663 | 5.302214 | -2.27771 | -0.21502 | -5.53828 | -0.50018 | -0.68295 | -1.88351 | 2.093598 | -0.5047 | -0.06283 | -1.36912 | -1.62144 | -0.09311 | -0.42092 | -0.11898 | 1.762028 | -4.0978 |
| 24.77778 | -0.28944 | -1.91829 | -0.69393 | -0.54632 | -0.666 | -1.23939 | 0.113157 | 4.082978 | -2.52889 | 0.184218 | -3.42766 | -0.21662 | 0.138428 | -1.90203 | 4.658374 | -0.53698 | 0.055976 | -1.50815 | -1.21869 | 1.553076 | -1.00985 | -0.50523 | 0.839636 | -2.55478 |
| 24.79167 | -0.53272 | -0.04138 | 0.338609 | -1.15092 | -1.08253 | -2.16029 | -0.5028 | 1.7191 | -2.29408 | 0.532935 | -0.91289 | -0.29681 | 0.574707 | -2.27331 | 4.49899 | -0.78752 | 0.163073 | -1.23263 | -0.81474 | 2.415889 | -1.23254 | -0.77652 | -0.63775 | -0.67641 |
| 24.80556 | -0.77508 | 1.828471 | 0.924811 | -1.53424 | -1.00545 | -2.26152 | -0.74658 | 0.421705 | -1.51175 | 0.647629 | 1.022793 | -0.94076 | 0.803515 | -1.59338 | 1.821295 | -1.14049 | 0.045877 | -0.93905 | -0.32349 | 2.568191 | -1.36009 | -1.08899 | -2.27439 | 2.07443 |
| 24.81944 | -0.93164 | 3.134702 | 0.406342 | -1.2948 | -0.6576 | -2.04716 | -0.69435 | -1.09948 | -0.84106 | 0.534699 | 2.046719 | -1.71428 | 0.835747 | 0.006333 | -0.87674 | -1.3938 | -0.28431 | -0.49018 | 0.127715 | 3.041038 | -1.29083 | -0.91512 | -1.32182 | 4.082495 |
| 24.83333 | -1.2282 | 2.247764 | -0.75889 | -0.77923 | -0.31397 | -1.15757 | -0.45809 | -2.98252 | -0.83023 | -0.12586 | 1.649585 | -2.11284 | -0.01167 | 1.074652 | -3.86802 | -1.47683 | -0.63287 | -0.29992 | 0.304051 | 2.27948 | -1.11815 | -0.38578 | -0.6266 | 3.595948 |
| 24.84722 | -1.37547 | 0.338212 | -1.84305 | -0.67909 | -0.30283 | -0.26813 | -0.33311 | -5.03685 | -1.45644 | -1.00943 | 1.054627 | -2.08107 | -1.4591 | 0.722302 | -6.20473 | -1.47633 | -0.71747 | -0.44601 | 0.360078 | 0.520501 | -0.93774 | 0.380111 | 0.759892 | 2.484227 |
| 24.86111 | -1.00346 | -1.78332 | -2.85733 | -1.03887 | -0.5502 | 0.135843 | -0.31014 | -5.99313 | -2.07769 | -1.88319 | 0.366732 | -1.94953 | -2.7113 | 0.02008 | -4.73574 | -1.38445 | -0.80397 | -0.90744 | 0.147965 | -1.46294 | -0.86328 | 0.863547 | 1.124294 | 1.075804 |
| 24.875 | -0.8386 | -3.14077 | -3.32138 | -1.3651 | -0.88366 | 0.165352 | -0.29215 | -6.38316 | -1.76195 | -2.51006 | -0.27528 | -1.65866 | -3.20263 | -0.58387 | -3.08194 | -1.29646 | -1.19857 | -1.48391 | -0.05449 | -3.69386 | -0.81971 | 0.620922 | -0.3732 | -0.56445 |
| 24.88889 | -1.32901 | -3.05688 | -2.71196 | -1.30034 | -1.04526 | -0.279 | -0.32492 | -6.12567 | -0.9105 | -2.76767 | -0.4814 | -1.1675 | -3.24999 | -1.27196 | -1.61745 | -1.51379 | -1.53197 | -2.15097 | 0.112753 | -4.33631 | -0.73565 | 0.038202 | -2.37951 | -1.22799 |
| 24.90278 | -1.76933 | -1.66063 | -1.449 | -1.15122 | -1.00747 | -1.01159 | -0.42613 | -3.27703 | 0.371301 | -2.09589 | -0.45723 | -0.5208 | -2.81047 | -1.87076 | 0.531195 | -1.42946 | -1.42832 | -2.5435 | 0.198448 | -4.06068 | -0.74091 | -0.5179 | -3.67803 | -1.43225 |
| 24.91667 | -1.47327 | -0.258 | -0.06783 | -0.94225 | -0.78538 | -1.67484 | -0.3822 | 0.059146 | 1.217796 | -1.26906 | -0.55285 | -0.06946 | -1.9536 | -2.21901 | 1.975085 | -0.92737 | -0.62744 | -2.24485 | 0.215604 | -3.40589 | -0.71113 | -0.94308 | -3.09069 | -1.49799 |
| 24.93056 | -0.84466 | 0.481064 | 0.673537 | -0.78513 | -0.41984 | -2.07354 | -0.13465 | 2.671499 | 1.40703 | -0.59095 | -0.96801 | 0.015516 | -1.91986 | -2.31276 | 1.638731 | -0.18434 | 0.368542 | -1.31578 | 0.157402 | -2.94752 | -0.6483 | -0.82096 | -0.75846 | -1.67802 |
| 24.94444 | -0.68029 | -0.35872 | -0.46933 | -1.08237 | -0.33882 | -2.32695 | 0.005228 | 2.537944 | 0.21063 | -0.42506 | -2.05961 | -0.56393 | -2.12642 | -2.37396 | 0.217188 | -0.01509 | 0.913622 | -1.00641 | -0.26597 | -2.98779 | -0.81989 | -0.60396 | 0.451833 | -2.85129 |
| 24.95833 | -1.59977 | -3.04789 | -3.51968 | -2.19817 | -1.24933 | -3.61594 | -0.57891 | -0.2296 | -2.60167 | -0.98992 | -4.405 | -2.65471 | -3.35455 | -3.08915 | -1.82189 | -1.35468 | -0.1858 | -2.45099 | -1.20613 | -4.10772 | -1.29922 | -0.75012 | -2.07603 | -5.99625 |
| 24.97222 | -3.44925 | -7.8421 | -7.24722 | -3.8813 | -3.33559 | -6.00868 | -2.034 | -2.58019 | -5.46128 | -2.14692 | -7.74849 | -5.5001 | -5.97675 | -4.70656 | -4.87649 | -3.86084 | -1.85587 | -5.09719 | -2.16846 | -5.38059 | -2.02505 | -1.32847 | -5.37094 | -9.34879 |
| 24.98611 | -3.27416 | -11.2212 | -9.98508 | -3.96859 | -4.60554 | -7.21802 | -2.54552 | -3.43119 | -6.14036 | -2.98802 | -9.59954 | -7.11867 | -7.53905 | -5.16225 | -4.09416 | -4.871 | -2.86534 | -6.02238 | -2.11661 | -4.34909 | -2.54592 | -1.80076 | -5.52595 | -9.66598 |
| 25 | -0.11567 | -10.1706 | -7.59618 | -2.8321 | -3.57174 | -6.11767 | -1.32571 | -2.19617 | -5.23546 | -2.29204 | -7.63963 | -5.61442 | -6.88049 | -3.98359 | -2.48062 | -4.19964 | -2.36568 | -5.79783 | -1.83692 | -2.35871 | -2.70317 | -1.21973 | -4.2111 | -6.66667 |
| 25.01389 | 2.726548 | -7.18476 | -3.18267 | -1.9333 | -2.07104 | -4.78956 | -0.35545 | -1.89293 | -4.71653 | -1.29612 | -5.61863 | -3.34641 | -4.81013 | -2.34944 | -1.61424 | -3.60983 | -2.38 | -5.2361 | -2.19086 | -0.83921 | -1.93299 | 0.038761 | -1.70378 | -3.36494 |
| 25.02778 | 3.490449 | -3.4049 | 0.254483 | -0.98782 | -0.46402 | -2.73148 | 0.014153 | -1.89137 | -2.82703 | -0.2023 | -2.73679 | -1.32687 | -2.50949 | -0.80809 | -1.12807 | -2.28467 | -1.69991 | -3.31337 | -1.65308 | 0.083521 | -0.71871 | 0.862912 | -0.43331 | -2.42858 |
| 25.04167 | 2.36506 | -0.21602 | 3.261391 | -0.23739 | 0.34458 | -0.81785 | -0.25459 | -1.77623 | -1.02478 | 0.802531 | 0.101283 | 0.886709 | -0.62529 | -0.20795 | -0.59029 | -0.71921 | -0.44449 | -0.87699 | -0.56885 | 0.367178 | 0.242733 | 0.283331 | 1.998194 | -0.8288 |
| 25.05556 | 1.183489 | 2.358533 | 6.066649 | 0.149843 | 0.825817 | 0.691408 | -0.38428 | -1.43135 | 0.147232 | 0.974314 | 1.564932 | 2.289436 | 0.739956 | -0.05744 | -0.05478 | 0.532725 | 0.301563 | 0.661218 | -0.19916 | 0.305443 | 0.646538 | -0.04605 | 1.018833 | 0.59377 |
| 25.06944 | 0.179167 | 4.905589 | 5.679526 | 0.308865 | 1.338757 | 2.388344 | -0.12275 | -0.96528 | 0.953796 | 0.818993 | 1.669479 | 2.801924 | 1.222204 | 0.077304 | 0.209865 | 1.176262 | 0.743704 | 1.822806 | 0.121349 | 0.31001 | 0.938079 | 0.044285 | 0.088394 | 1.244361 |
| 25.08333 | -0.87129 | 6.787747 | 3.749081 | 0.496485 | 1.994362 | 3.548318 | 0.196624 | -0.44986 | 1.632339 | 0.62628 | 1.345788 | 2.737109 | 1.084999 | 0.184442 | 0.257279 | 1.321596 | 0.822755 | 2.729201 | 0.620848 | 0.314383 | 0.936164 | 0.382417 | -0.8048 | 1.423189 |
| 25.09722 | -1.17964 | 6.733044 | 1.744545 | 0.557622 | 2.221202 | 4.062361 | 0.161784 | -0.3561 | 1.450532 | 0.384567 | 0.773901 | 1.910674 | 0.521016 | 0.110507 | 0.100484 | 0.977883 | 0.663677 | 3.020409 | 1.002458 | 0.143355 | 0.70371 | 0.613941 | -1.98065 | 1.058789 |
| 25.11111 | -1.21005 | 4.586787 | 0.830603 | 0.490194 | 1.873031 | 4.835757 | -0.17534 | -0.5056 | 0.907226 | 0.179616 | 0.409327 | 0.964363 | -0.05762 | -0.06496 | -0.09533 | 0.788132 | 0.621288 | 3.213238 | 1.153619 | 0.010741 | 0.47583 | 0.383893 | -2.36091 | 0.749916 |
| 25.125 | -0.73878 | 3.246066 | 0.51514 | 0.60368 | 1.47866 | 4.717683 | -0.71553 | -0.33684 | 1.086933 | 0.151035 | 0.245348 | 0.442146 | -0.20498 | -0.10804 | -0.02044 | 0.923563 | 0.744625 | 3.38865 | 1.50739 | 0.02733 | 0.524827 | -0.19582 | -1.28161 | 0.940652 |
| 25.13889 | 0.34214 | 2.384799 | 0.588377 | 0.873909 | 1.077939 | 4.107037 | -1.37896 | 0.129602 | 1.660265 | 0.197511 | 0.178996 | 0.216471 | 0.097874 | 0.029047 | 0.376364 | 1.076112 | 0.929666 | 2.991132 | 1.637415 | 0.130689 | 0.770973 | -0.94409 | 0.840446 | 1.151731 |
| 25.15278 | 1.301388 | 1.294261 | 0.889136 | 1.131911 | 0.397504 | 3.02641 | -1.94849 | 0.70087 | 2.075197 | -0.25594 | 0.042617 | 0.105199 | 0.601114 | 0.333616 | 0.983023 | 1.090592 | 0.988865 | 2.319364 | 1.278334 | 0.189822 | 0.801293 | -1.12893 | 3.290081 | 1.273342 |
| 25.16667 | 0.963781 | 0.41713 | 1.178346 | 1.231118 | 0.252222 | 1.680684 | -2.09396 | 1.085865 | 2.044033 | -0.99055 | -0.06082 | 0.153644 | 1.037981 | 0.664063 | 1.496899 | 1.020545 | 0.882194 | 1.621627 | 0.549708 | 0.221223 | 0.54001 | -0.0244 | 2.226422 | 0.972972 |
| 25.18056 | -0.51085 | -0.22091 | 1.141504 | 1.309238 | 0.439352 | 0.464113 | -1.33278 | 1.335103 | 1.497841 | -0.88298 | -0.00462 | 0.446934 | 1.438354 | 1.573656 | 1.722138 | 0.854892 | 0.761467 | 0.540559 | -0.04735 | 0.270055 | 0.595944 | 1.379695 | -0.51784 | 0.65753 |
| 25.19444 | -1.86122 | -0.41015 | 0.799169 | 1.516586 | 0.559491 | -0.45912 | -0.65224 | 1.514622 | 1.011238 | -0.43328 | 0.157856 | 0.856366 | 1.448576 | 1.222159 | 1.884517 | 0.552734 | 0.628527 | -0.04416 | -0.48689 | 0.089516 | 0.425707 | 1.751106 | -3.54099 | 0.58845 |
| 25.20833 | -1.86583 | -0.05666 | 0.156847 | 1.444738 | 0.267817 | -1.17642 | -0.91252 | 1.615752 | 0.477235 | -0.47849 | 0.212941 | 1.092679 | 1.164214 | 0.903556 | 1.71014 | 0.161868 | 0.335546 | -0.26025 | -0.70698 | -0.05247 | 0.167768 | 0.658653 | -4.8647 | 0.677992 |
| 25.22222 | 0.013622 | 0.640826 | -0.38917 | 1.142094 | -0.09538 | -1.27886 | -1.46134 | 1.484442 | -0.25163 | -0.72477 | 0.097629 | 1.04291 | 0.793607 | 0.520016 | 1.638448 | -0.04299 | 0.086438 | -0.6725 | -0.67273 | 0.003705 | 0.475815 | -0.2859 | -2.1118 | 0.559413 |
| 25.23611 | 1.873076 | 1.338591 | -0.4503 | 0.791256 | -0.16001 | -0.33625 | -1.43278 | 1.091361 | -0.74529 | -0.78757 | 0.009507 | 0.80871 | 0.628234 | 0.578142 | 1.543975 | -0.06191 | -0.02531 | -1.09669 | -0.4119 | 0.028723 | 0.453816 | -1.49241 | 2.090279 | 0.56438 |
| 25.25 | 2.349012 | 1.637905 | 0.168944 | 0.711587 | 0.106823 | 1.238404 | -0.91988 | 1.000988 | -0.44059 | -0.51885 | -0.08203 | 0.702202 | 1.087139 | 1.065082 | 1.883216 | -0.01368 | -0.00524 | -1.42769 | -0.03853 | 0.306224 | 0.171392 | -1.39017 | 5.719818 | 0.704842 |
| 25.26389 | 1.166413 | 1.454751 | 1.084803 | 0.735361 | 0.353871 | 2.467038 | -0.39696 | 1.04265 | 0.312997 | -0.11869 | -0.30036 | 0.60028 | 2.02449 | 1.759246 | 2.146258 | 0.222141 | 0.027158 | -0.82926 | 0.288424 | 0.770288 | -0.00824 | 0.356393 | 2.914264 | 0.69858 |
| 25.27778 | -0.36355 | 0.969864 | 1.650892 | 0.641963 | 0.45813 | 1.593377 | -0.19605 | 0.896848 | 0.978817 | 0.122656 | -0.27376 | 0.677799 | 2.20907 | 1.577097 | 1.635453 | 0.087755 | -0.10957 | 0.041285 | 0.249158 | 0.810595 | 0.067452 | 2.054394 | 0.254554 | 0.098877 |
| 25.29167 | -1.64498 | 0.314056 | 1.363646 | 0.368861 | 0.372601 | -0.54276 | -0.32037 | 0.42643 | 1.200945 | 0.052007 | 0.114411 | 0.936285 | 1.473154 | 0.600757 | 0.610692 | -0.21185 | -0.36747 | 0.942632 | -0.19087 | 0.368104 | 0.4046 | 2.881352 | -2.04542 | -0.55269 |
| 25.30556 | -2.00386 | -0.28461 | 0.203454 | 0.182717 | 0.017684 | -2.64135 | -0.67529 | 0.077194 | 1.277133 | -0.27561 | 0.64947 | 2.044469 | 0.793571 | -0.14027 | -0.20444 | -0.27984 | -0.54235 | 0.997573 | -0.66641 | 0.170987 | 0.9315 | 1.360693 | -4.03659 | -0.4401 |
| 25.31944 | -0.90687 | -0.87508 | -0.95711 | 0.141997 | -0.34798 | -4.30613 | -0.86665 | 0.053087 | 1.116876 | -0.52733 | 0.918305 | 1.691657 | 0.20633 | -0.26795 | -0.54105 | -0.23668 | -0.62232 | 0.466145 | -0.85754 | 0.377875 | 0.962002 | 0.222925 | -3.53921 | 0.331603 |

| | | | | | | | | | | | | | | | | | | | | | | | | |
|---|---|---|---|---|---|---|---|---|---|---|---|---|---|---|---|---|---|---|---|---|---|---|---|---|
| 25.33333 | 0.891458 | -0.5342 | -1.2396 | 0.085247 | -0.37026 | -3.30443 | -0.77154 | -0.0013 | 0.317835 | -0.4471 | 0.627545 | 0.724808 | -0.43555 | -0.03285 | -0.50667 | -0.32209 | -0.58926 | -0.16173 | -0.67865 | 0.462456 | 0.2329 | -0.71502 | -1.40105 | 0.598964 |
| 25.34722 | 2.307206 | 0.394161 | -0.30515 | 0.065907 | 0.043654 | 0.281993 | -0.48751 | -0.34625 | -0.85931 | -0.03638 | 0.255095 | 0.036086 | -1.04169 | 0.089828 | -0.46074 | -0.40392 | -0.30778 | -0.89191 | -0.31199 | 0.481991 | -0.15045 | -1.49787 | 1.550282 | 0.057527 |
| 25.36111 | 2.34014 | 1.342905 | 0.95407 | 0.319284 | 0.611543 | 3.537403 | -0.07916 | -0.59085 | -1.66469 | 0.584115 | -0.67471 | -1.5067 | -1.26756 | -0.06052 | -0.44423 | -0.04632 | 0.1227 | -1.25038 | 0.057009 | 0.635616 | 0.073133 | -0.87982 | 3.885965 | -0.77733 |
| 25.375 | 0.722945 | 1.992507 | 2.134004 | 0.798751 | 0.89541 | 4.754662 | 0.243272 | -0.29548 | -1.3628 | 0.882901 | -1.59346 | -2.08677 | -1.0395 | -0.24747 | -0.76922 | 0.44404 | 0.57836 | -0.68067 | 0.38579 | 0.789761 | 0.336112 | 0.598718 | 2.888905 | -1.08888 |
| 25.38889 | -0.39344 | 1.398234 | 1.93057 | 1.135041 | 0.549719 | 2.601965 | 0.145147 | 0.041499 | -0.59403 | 0.61978 | -0.54689 | -0.44964 | -0.62018 | -0.51022 | -0.91705 | 0.599885 | 0.601785 | 0.714324 | 0.561882 | 0.896058 | 0.258384 | 2.166652 | 1.152801 | -0.96612 |
| 25.40278 | -1.68425 | 0.163061 | 0.73626 | 1.338647 | 0.012005 | 0.084277 | -0.16043 | 0.280049 | 0.57975 | 0.110968 | 1.108904 | 0.939364 | -0.23554 | -0.66395 | -1.01189 | 0.60987 | 0.578698 | 1.786519 | 0.361868 | 0.934524 | 0.698549 | 2.495407 | -0.25404 | -0.47664 |
| 25.41667 | -2.55901 | -1.53505 | -0.24127 | 1.548261 | -0.26184 | -2.46774 | -0.22059 | 0.599836 | 2.042352 | -0.0331 | 2.506335 | 1.74027 | 0.311782 | -0.25581 | -0.96807 | 0.790748 | 0.802646 | 2.008937 | 0.113768 | 1.04948 | 0.476458 | 1.212307 | -1.88503 | 0.64713 |
| 25.43056 | -1.95495 | -3.22163 | -0.7739 | 1.486173 | -0.04358 | -4.06121 | 0.269064 | 0.902017 | 2.91268 | 0.428531 | 1.523211 | 1.227189 | 0.79751 | 0.509401 | -0.45997 | 0.97222 | 0.857839 | 1.352332 | -0.27321 | 1.439645 | 0.62861 | 0.084091 | -3.77076 | 1.645967 |
| 25.44444 | 1.215052 | -1.89834 | 0.034098 | 1.45389 | 0.737121 | -3.14815 | 1.152559 | 1.682736 | 3.290572 | 1.513122 | 2.326631 | 1.742286 | 1.590152 | 1.674848 | 1.098932 | 1.117696 | 0.560221 | 0.707143 | -0.33473 | 2.138489 | 0.984327 | -1.02725 | -2.37278 | 2.027086 |
| 25.45833 | 5.018763 | 1.484269 | 3.111962 | 2.667515 | 2.027314 | 1.330604 | 2.51869 | 4.320813 | 4.292345 | 3.713417 | 5.897292 | 3.308203 | 3.843256 | 4.425242 | 4.798703 | 2.125079 | 0.804788 | 1.146995 | 0.755179 | 4.468682 | 2.638627 | -1.6352 | 1.047083 | 2.938741 |
| 25.47222 | 7.428908 | 4.582789 | 6.070875 | 4.168596 | 3.085622 | 7.619638 | 3.069657 | 7.055271 | 5.299633 | 4.883299 | 8.338155 | 5.392853 | 7.175932 | 6.882644 | 8.170649 | 4.121963 | 2.012045 | 1.507943 | 1.564732 | 6.340792 | 4.296231 | 0.354504 | 4.453371 | 4.486607 |
| 25.48611 | 4.880995 | 5.509755 | 6.587114 | 3.238653 | 3.034626 | 9.033997 | 1.947787 | 5.75607 | 5.073686 | 2.828379 | 7.839167 | 5.094908 | 7.507718 | 5.646201 | 7.59142 | 4.01488 | 2.590478 | 2.108333 | 2.217424 | 5.107703 | 4.045033 | 2.387735 | 5.78277 | 3.987062 |
| 25.5 | 1.996366 | 5.499407 | 4.597534 | 1.845644 | 2.441136 | 6.915719 | 1.740464 | 3.228868 | 3.480848 | 1.161774 | 6.155956 | 4.207166 | 4.618153 | 4.403812 | 4.72779 | 2.86315 | 2.223639 | 2.456608 | 2.142362 | 3.318567 | 3.102132 | 3.07371 | 3.811527 | 2.8886 |
| 25.51389 | -0.04956 | 3.407221 | 2.660624 | 0.551315 | 1.402141 | 4.17955 | 1.356219 | 1.342611 | 1.5001 | 0.272052 | 4.577102 | 3.207648 | 2.507111 | 2.466725 | 1.629838 | 2.089765 | 1.695606 | 2.913504 | 1.915274 | 2.127151 | 1.813896 | 2.220925 | 2.193114 | 1.706313 |
| 25.52778 | -2.07909 | 1.271682 | 1.084882 | -0.55687 | 0.521486 | 1.466007 | 0.462831 | -0.1011 | -0.45808 | -0.35055 | 2.576875 | 1.730816 | 1.029237 | 0.595555 | -0.83121 | 1.572647 | 0.022914 | 3.308278 | 0.960581 | 0.789909 | 0.451319 | 1.129569 | 0.292539 | 0.625309 |
| 25.54167 | -2.85803 | -0.50473 | -0.11863 | -1.27122 | -0.38013 | -0.62133 | -0.893 | -1.30105 | -2.10865 | -1.02507 | 0.484297 | 0.167869 | -0.77752 | -0.858 | -4.49727 | 0.99276 | -1.11811 | 1.788576 | -0.32168 | -0.5896 | -0.64177 | 0.228514 | -2.08402 | 0.12585 |
| 25.55556 | -2.21601 | -2.13092 | -1.65943 | -0.90048 | -1.5074 | -2.45622 | -2.22668 | -1.78815 | -1.8159 | -1.52007 | -0.97567 | -1.21011 | -3.02873 | -1.92462 | -4.83962 | 0.3601 | -1.43921 | 0.577366 | -1.22808 | -2.28994 | -0.80233 | -1.01922 | -5.03098 | 0.090812 |
| 25.56944 | 1.244359 | -3.35975 | -1.91326 | 0.298508 | -1.02928 | -4.42903 | -1.57613 | -1.21776 | -0.18358 | -0.99437 | -1.38742 | -1.78963 | -4.77751 | -2.47495 | -4.45815 | 0.053909 | -1.35596 | -0.19208 | -1.13938 | -2.79149 | -0.32312 | -2.03039 | -4.23706 | 0.893823 |
| 25.58333 | 4.179334 | -3.51709 | -1.45542 | 1.971328 | 0.54488 | -5.57673 | 0.152469 | -0.58854 | 2.084794 | 0.332327 | -0.75445 | -0.57031 | -4.82113 | -2.46523 | -3.87465 | -0.04161 | -0.76976 | -0.81939 | -0.31409 | -2.23106 | 0.145491 | -2.17797 | -1.62709 | 2.201467 |
| 25.59722 | 3.618397 | -1.02708 | 0.397339 | 2.935461 | 2.189642 | -3.48826 | 1.381674 | -0.13858 | 3.894526 | 1.904814 | -0.39643 | 1.060496 | -3.69046 | -2.24395 | -4.29528 | 0.686903 | 0.011888 | -1.26149 | 0.020151 | -1.03286 | 1.035187 | -0.56592 | 1.362078 | 2.493467 |
| 25.61111 | 2.310801 | 0.904456 | 2.232536 | 1.867734 | 2.508029 | 1.378935 | 1.235911 | 0.943588 | 4.086167 | 2.840262 | -0.74697 | 1.887731 | -2.79852 | -1.7049 | -3.54575 | 0.873398 | 0.531287 | -1.27886 | 0.072468 | 0.445509 | 1.704459 | 1.147635 | 3.137202 | 1.256461 |
| 25.625 | 0.896529 | 2.825935 | 2.775547 | 0.602982 | 0.722189 | 5.162916 | -0.15795 | 2.84763 | 3.210604 | 2.582879 | -1.16916 | 1.104683 | -1.88413 | -0.9953 | -1.33098 | 0.877305 | 0.383995 | -0.29708 | 0.462019 | 1.471068 | 1.192402 | 1.709132 | 1.954229 | -0.18012 |
| 25.63889 | -0.67263 | 4.144394 | 1.710701 | -0.10892 | -1.49724 | 5.127843 | -1.91231 | 4.631648 | 1.433862 | 1.221238 | -0.579 | 0.022463 | -1.1159 | -0.24362 | 1.104416 | 0.445396 | 0.253417 | 0.424298 | 0.749448 | 2.072274 | 0.210206 | 0.537461 | -0.06125 | -0.76881 |
| 25.65278 | -2.31356 | 3.144935 | -0.21131 | -0.84068 | -1.65822 | 2.397061 | -1.89363 | 4.621659 | -0.43401 | -0.41716 | 0.901404 | -0.63895 | 0.361265 | 0.790402 | 3.808312 | -0.38167 | 0.412696 | 0.6329 | 0.397456 | 2.823458 | -0.80986 | 0.047225 | -1.4149 | -0.86412 |
| 25.66667 | -3.34736 | 0.632359 | -1.19667 | -1.05778 | -1.57696 | -0.28252 | -1.26471 | 3.023712 | -2.03709 | -1.92894 | 2.602146 | -0.32769 | -0.09286 | 2.149168 | 4.802307 | -0.41422 | 0.390014 | 0.501422 | -0.08934 | 2.793069 | -1.6979 | -0.00322 | -3.4284 | -0.65664 |
| 25.68056 | -2.94991 | -1.10617 | -1.94698 | -0.57549 | -1.11246 | -2.75163 | -0.7062 | 0.632172 | -2.68838 | -2.19779 | 3.698883 | 0.672194 | 1.240992 | 3.334387 | 3.956585 | -0.38907 | 0.124327 | -0.31283 | 0.095551 | 1.363244 | -2.06903 | -0.16312 | -4.77892 | -0.3071 |
| 25.69444 | -1.06476 | -3.32251 | -2.88132 | -0.94731 | -0.79441 | -4.97666 | -0.86974 | -1.7622 | -2.13638 | -1.54781 | 3.970909 | 1.675271 | 2.294229 | 3.264947 | 1.465484 | -0.29713 | -0.30794 | -1.34288 | -0.23431 | 0.428996 | -1.08116 | -0.43854 | -2.96518 | 0.441661 |
| 25.70833 | 0.628139 | -4.26269 | -3.50474 | -0.97363 | 0.027363 | -6.13679 | 0.009987 | -3.69496 | -1.22473 | -0.60703 | 3.952165 | 2.092782 | 2.684407 | 1.326132 | -0.19348 | -0.56697 | -0.73288 | -2.01158 | 0.048508 | -0.45373 | -0.62887 | -0.21109 | 0.880619 | 1.208241 |
| 25.72222 | 0.51004 | -4.1083 | -3.45666 | -0.18565 | 1.252239 | -6.11913 | 1.506923 | -4.69484 | 0.128574 | 0.028381 | 2.706979 | 0.912222 | 1.388834 | -0.10574 | -1.81901 | -0.47633 | -0.48142 | -2.00379 | 0.237291 | -1.17548 | -0.08833 | 0.548367 | 3.890199 | 1.605745 |
| 25.73611 | -0.51879 | -1.79961 | -1.76346 | 0.052019 | 1.703407 | -3.36176 | 2.278684 | -4.59199 | 1.309261 | 0.139729 | 0.355814 | -0.7741 | -0.27943 | -0.92172 | -4.31041 | -0.02253 | 0.29919 | -2.80707 | -0.049 | -1.90722 | 0.487782 | 1.163584 | 4.681983 | 2.196286 |
| 25.75 | -1.39384 | 0.58828 | 1.177101 | 0.349159 | 1.743086 | 1.193921 | 2.204888 | -3.8479 | 1.522101 | -0.46236 | -0.92324 | -0.97732 | -0.32611 | -1.79841 | -5.3913 | 0.042324 | 0.853239 | -1.72314 | -0.42442 | -2.99998 | 0.622773 | 0.829583 | 2.183669 | 1.79503 |
| 25.76389 | -1.59556 | 2.496283 | 2.880872 | -0.17803 | 1.19521 | 2.983812 | 1.354434 | -2.74079 | 0.604383 | -1.17608 | -1.32165 | -0.71933 | -0.8639 | -2.46302 | -4.99992 | -0.54352 | -0.28489 | 0.696481 | -0.54308 | -4.67285 | -0.11815 | -0.02662 | -0.26813 | 0.546364 |
| 25.77778 | -1.94349 | 2.879966 | 2.82754 | -1.47898 | 0.12875 | 1.930938 | 0.097271 | -0.87683 | -0.08622 | -1.60194 | -1.92366 | 0.049193 | -2.16491 | -2.60116 | -4.35342 | -0.73138 | -0.5248 | 2.572284 | -0.61562 | -4.91486 | -0.90653 | -1.06499 | -2.98554 | -0.27377 |
| 25.79167 | -2.26591 | 2.154917 | 1.659177 | -2.48967 | -1.25353 | -0.17415 | -1.18836 | 0.702666 | -0.84463 | -1.29092 | -2.97922 | -0.3229 | -2.49305 | -2.09206 | -3.22608 | -0.80846 | -0.35004 | 1.655355 | -0.73382 | -4.67675 | -1.45969 | -2.53231 | -5.11326 | -0.83522 |
| 25.80556 | -1.33026 | 0.06453 | -0.3483 | -3.9655 | -2.42837 | -2.68686 | -2.3924 | 1.516924 | -1.58634 | -0.33593 | -4.2867 | -1.18977 | -2.9094 | -1.50333 | -2.42112 | -1.12874 | -0.80603 | -0.10064 | -1.07455 | -3.64514 | -1.61559 | -3.35437 | -4.03878 | -1.80374 |
| 25.81944 | -0.06846 | -2.45447 | -3.52892 | -4.92181 | -3.03848 | -4.95773 | -3.34585 | 1.718373 | -2.44275 | 0.631315 | -5.40577 | -2.23768 | -2.85516 | -1.16325 | -2.26141 | -1.48433 | -0.82687 | -2.04047 | -1.08776 | -2.35392 | -1.66104 | -2.39019 | -1.21506 | -3.4327 |
| 25.83333 | 0.072827 | -6.35126 | -6.49538 | -4.7296 | -3.38566 | -7.2373 | -3.54233 | 1.221615 | -3.50331 | 0.996271 | -6.35031 | -3.77724 | -2.51773 | -1.14546 | -2.25602 | -2.05482 | -0.3015 | -4.2181 | -0.91838 | -1.03749 | -2.01175 | -1.0772 | 1.185712 | -5.09028 |
| 25.84722 | -1.31133 | -7.46529 | -7.69871 | -4.31248 | -3.3148 | -6.72071 | -3.42708 | 0.325226 | -4.12032 | 0.529458 | -6.86352 | -4.60084 | -2.287 | -1.40794 | -1.95315 | -2.32043 | 0.027687 | -5.49774 | -1.1661 | 0.070932 | -2.27626 | -0.22224 | 1.878924 | -5.60563 |
| 25.86111 | -2.26379 | -5.74364 | -5.06094 | -3.76791 | -2.90829 | -5.25377 | -3.11229 | -0.55684 | -4.84202 | -0.59991 | -5.39039 | -4.23352 | -2.03813 | -1.48006 | -1.3177 | -2.2165 | -0.50953 | -5.47235 | -1.03085 | 1.334727 | -1.96377 | -0.08777 | 0.826864 | -4.29036 |
| 25.875 | -2.2764 | -3.18262 | -2.0198 | -3.05172 | -2.11016 | -2.89831 | -2.41192 | -1.29007 | -4.28858 | -1.88798 | -2.35202 | -3.16562 | -1.65345 | -1.26354 | -0.70247 | -1.85301 | -1.24202 | -4.20686 | -0.67024 | 2.635705 | -1.13731 | -0.16961 | -0.74244 | -2.25778 |
| 25.88889 | -1.18861 | -0.06972 | 0.515346 | -2.09056 | -1.47455 | -0.6308 | -1.60787 | -1.50436 | -2.56319 | -2.71454 | 0.906412 | -1.65046 | -1.38214 | -0.80173 | -0.01763 | -1.36811 | -1.52622 | -2.83503 | -0.45056 | 2.913784 | -0.32761 | 0.004022 | -3.16798 | -0.54145 |
| 25.90278 | 0.208189 | 2.548874 | 1.850946 | -0.87681 | -0.94061 | 1.816765 | -0.63451 | -1.13846 | -1.30429 | -2.69024 | 3.924245 | 0.471609 | -0.78607 | -0.02006 | 0.682608 | -0.7751 | -1.23485 | -1.19402 | -0.30849 | 3.759856 | 0.384198 | 0.312377 | -5.68002 | 0.597982 |
| 25.91667 | 1.049875 | 3.452707 | 1.800231 | 0.282973 | -0.64509 | 3.811183 | 0.218548 | -0.71216 | -0.32827 | -1.7459 | 5.383015 | 2.208972 | -0.1753 | 1.136586 | 1.518463 | -0.22333 | -0.84393 | 0.731872 | -0.07168 | 4.140744 | 0.976418 | 0.71178 | -3.46916 | 1.330279 |
| 25.93056 | 1.398582 | 2.669995 | 0.538325 | 1.201353 | -0.41257 | 4.316332 | 1.10525 | -0.58622 | 0.28503 | -0.72733 | 4.631503 | 2.434435 | 0.014456 | 1.440382 | 2.152727 | -0.04626 | -0.56557 | 2.257721 | 0.143251 | 3.030009 | 0.981728 | 0.987394 | 0.617148 | 1.38577 |
| 25.94444 | 0.302989 | 0.937434 | -1.52231 | 0.951659 | -0.67674 | 2.995175 | 1.495479 | -1.20254 | -0.22844 | -0.37102 | 2.597944 | 1.218428 | 0.364657 | 0.738417 | 1.81649 | -0.80517 | -0.87454 | 2.030673 | -0.02321 | 0.945223 | 0.199614 | 0.57818 | 3.769455 | 0.317856 |
| 25.95833 | -2.28928 | -1.98202 | -4.33898 | -1.03034 | -2.08221 | -1.16668 | -0.3599 | -2.92073 | -3.01967 | -1.35441 | -1.07341 | -2.10722 | -1.70771 | -1.04205 | -0.22634 | -2.77225 | -1.74312 | -0.96273 | -0.70559 | -2.26401 | -1.27183 | -0.71197 | 1.089277 | -2.22043 |
| 25.97222 | -6.10827 | -6.17821 | -7.88005 | -4.05056 | -4.57262 | -6.42662 | -2.7552 | -5.22628 | -7.07181 | -2.93568 | -5.90025 | -6.34878 | -5.0771 | -3.74889 | -2.82948 | -5.46236 | -3.00357 | -4.5769 | -1.2377 | -5.69133 | -3.05065 | -3.02647 | -4.40692 | -4.95728 |
| 25.98611 | -9.90655 | -7.96169 | -9.06243 | -4.50153 | -5.10462 | -7.58034 | -3.61111 | -5.60724 | -7.61204 | -3.68407 | -7.83036 | -8.46888 | -5.95534 | -4.36121 | -3.42586 | -5.74601 | -3.81237 | -4.44299 | -0.92725 | -6.75921 | -3.70925 | -4.66852 | -6.78137 | -4.60841 |
| 26 | -9.11585 | -5.84769 | -5.7049 | -2.62263 | -2.88478 | -4.94624 | -2.57606 | -4.04189 | -4.49022 | -2.97487 | -5.68356 | -6.25903 | -3.35167 | -3.23667 | -2.04099 | -4.03719 | -2.97757 | -2.92328 | -0.25399 | -5.77029 | -2.92399 | -5.01091 | -6.15952 | -2.54708 |
| 26.01389 | -5.98762 | -3.50244 | -2.73203 | -1.63201 | -1.26353 | -3.64121 | -1.63721 | -2.95015 | -2.18433 | -2.35185 | -4.05987 | -4.40238 | -2.34704 | -2.29924 | -1.16834 | -2.625 | -2.05623 | -1.93198 | 0.076056 | -4.70939 | -1.73984 | -3.6302 | -3.51381 | -1.37325 |
| 26.02778 | -2.38303 | -2.15731 | -1.10061 | -1.0758 | -0.27637 | -2.52336 | -0.59283 | -1.97098 | -1.04652 | -1.59548 | -2.78432 | -2.8543 | -2.52801 | -1.85766 | -0.99767 | -1.50609 | -0.78258 | -1.5423 | -0.01005 | -3.77695 | -0.81969 | -2.18591 | -2.07239 | -1.06611 |
| 26.04167 | 0.481483 | -1.25423 | -0.13167 | -0.68715 | 0.197828 | -1.69588 | 0.166175 | -0.85693 | -0.45786 | -0.84096 | -1.43606 | -1.45515 | -2.54494 | -1.4679 | -0.70079 | -0.53399 | 0.263095 | -1.31242 | -0.26825 | -2.77277 | 0.221933 | -1.27583 | -0.97427 | -0.95974 |
| 26.05556 | 3.491491 | -0.6675 | 0.468814 | -0.36823 | 0.229868 | -1.16756 | 0.345718 | -0.2402 | -0.36231 | -0.3991 | -0.63643 | -0.48035 | -2.12661 | -1.20114 | -0.00402 | 0.170839 | 0.926129 | -0.95307 | -0.27027 | -1.1725 | 0.532439 | -0.22456 | 0.510593 | -0.6777 |
| 26.06944 | 4.947691 | -0.20392 | 0.620781 | -0.12355 | 0.207189 | -0.96334 | 0.404824 | -0.07637 | -0.27276 | -0.25686 | -0.61809 | -0.07639 | -2.10781 | -1.21717 | 0.497567 | 0.466883 | 1.040321 | -0.54697 | -0.01627 | -0.61332 | 0.803416 | 0.581661 | 1.404167 | -0.34525 |
| 26.08333 | 5.22059 | 0.366613 | 0.545787 | 0.208465 | 0.471021 | -0.84785 | 0.717963 | -0.05499 | -0.01344 | -0.19032 | -0.64988 | 0.209004 | -2.22585 | -1.19315 | 0.531577 | 0.464866 | 0.702676 | -0.21388 | 0.263945 | -0.48783 | 0.998187 | 1.078792 | 1.671243 | -0.04399 |
| 26.09722 | 6.057945 | 0.799995 | 0.814465 | 0.620761 | 0.844809 | -0.56314 | 1.03846 | -0.09081 | 0.413659 | -0.0827 | 0.081295 | 0.701117 | -2.20261 | -1.10407 | 0.526347 | 0.428273 | 0.431885 | 0.074441 | 0.232629 | -0.43336 | 0.895672 | 1.786337 | 2.350921 | -0.16148 |
| 26.11111 | 6.487203 | 0.907002 | 2.351399 | 0.746734 | 0.875566 | -0.20019 | 1.182261 | -0.09909 | 0.663839 | 0.080726 | 1.117219 | 1.396013 | -2.09552 | -1.00861 | 0.483808 | 0.363193 | 0.263695 | 0.279657 | 0.010217 | -0.30104 | 0.471505 | 2.453098 | 3.100632 | -0.58136 |
| 26.125 | 5.745153 | 1.179945 | 4.164402 | 0.660043 | 0.839331 | 0.177943 | 0.992975 | -0.07538 | 0.869385 | 0.426727 | 2.050482 | 2.244224 | -1.75531 | -0.93452 | 0.495028 | 0.320334 | 0.173348 | 0.486513 | -0.21356 | -0.03935 | 0.146522 | 2.879475 | 2.68159 | -0.78747 |
| 26.13889 | 4.590142 | 2.009482 | 6.582783 | 0.619084 | 1.055661 | 0.585957 | 1.082441 | 0.002276 | 1.213843 | 1.108443 | 2.409625 | 3.392989 | -1.08703 | -0.68859 | 0.633393 | 0.287021 | 0.150223 | 0.835065 | -0.21737 | 0.250475 | 0.170276 | 2.84676 | 2.064639 | -0.43202 |
| 26.15278 | 3.528813 | 3.952893 | 6.010524 | 0.573086 | 1.393594 | 1.011441 | 1.017672 | 0.068509 | 1.678107 | 1.653736 | 2.268731 | 4.42661 | -0.44071 | -0.36836 | 0.751797 | 0.47176 | 0.218273 | 1.377153 | -0.00205 | 0.446409 | 0.554417 | 2.326175 | 1.895573 | 0.214013 |
| 26.16667 | 2.669574 | 5.82133 | 4.104934 | 0.412808 | 1.49784 | 1.34653 | 0.667296 | 0.061637 | 1.968295 | 1.633318 | 1.231185 | 3.970974 | -0.14094 | -0.1227 | 0.610621 | 0.953485 | 0.297668 | 2.091101 | 0.220135 | 0.416401 | 0.939924 | 1.565676 | 1.568407 | 0.749069 |
| 26.18056 | 1.028287 | 5.451596 | 2.123908 | 0.21192 | 1.397949 | 1.537765 | 0.253249 | 0.103757 | 2.390784 | 1.157591 | 0.624307 | 2.93434 | -0.07763 | 0.125038 | 0.417735 | 1.167354 | 0.244932 | 2.388073 | 0.392482 | 0.235494 | 0.947431 | 0.729294 | 1.018259 | 0.904797 |
| 26.19444 | -0.64229 | 5.989465 | 0.338545 | 0.188855 | 1.369432 | 1.956648 | -0.04321 | 0.185895 | 3.196104 | 0.778593 | 0.641329 | 2.223867 | 0.095373 | 0.364984 | 0.431889 | 1.204783 | 0.241051 | 2.687463 | 0.395154 | 0.112662 | 0.757157 | 0.124924 | 0.577614 | 0.976561 |
| 26.20833 | -2.70644 | 5.914361 | -1.01524 | 0.361089 | 1.389594 | 3.011078 | -0.1835 | 0.159888 | 3.511206 | 0.516599 | 0.848013 | 2.080259 | 0.364766 | 0.484354 | 0.646366 | 1.159741 | 0.361371 | 2.772217 | 0.181619 | 0.128982 | 0.790037 | -0.34521 | 0.203254 | 1.048655 |
| 26.22222 | -4.43373 | 3.550213 | -1.9766 | 0.482586 | 1.057982 | 3.350372 | -0.24367 | 0.022151 | 2.808345 | 0.091077 | 0.641234 | 1.515903 | 0.42251 | 0.329011 | 0.624867 | 0.874158 | 0.408147 | 2.612526 | -0.1131 | 0.165419 | 0.887035 | -0.83363 | -0.30202 | 0.772985 |
| 26.23611 | -3.7603 | 1.751396 | -2.50515 | 0.607974 | 0.646019 | 3.32715 | -0.07135 | -0.00359 | 1.976239 | -0.23741 | 0.17418 | 0.759457 | 0.400963 | 0.117011 | 0.566453 | 0.526022 | 0.256348 | 2.802762 | -0.15533 | 0.15172 | 1.241909 | -0.91634 | -0.065 | 0.535381 |
| 26.25 | -1.26355 | 1.080786 | -2.28804 | 0.916924 | 0.831944 | 3.733051 | 0.337424 | 0.14851 | 2.023489 | -0.10105 | -0.03913 | 0.291709 | 0.542553 | 0.16215 | 0.769428 | 0.46928 | 0.162405 | 3.178164 | 0.114858 | 0.197316 | 1.317244 | -0.38632 | 1.112802 | 0.666715 |
| 26.26389 | 0.979261 | 0.615412 | -1.11247 | 0.963777 | 0.867078 | 3.635085 | 0.530212 | 0.262154 | 1.734857 | 0.142127 | -0.09448 | -0.13612 | 0.501735 | 0.313385 | 0.859981 | 0.443994 | 0.113617 | 2.863426 | 0.437754 | 0.178705 | 0.555766 | 0.423856 | 2.125092 | 0.680931 |
| 26.27778 | 2.521063 | -0.13004 | 0.330693 | 0.568123 | 0.405515 | 2.542836 | 0.256278 | 0.190118 | 1.220982 | 0.218657 | -0.1621 | -0.46416 | 0.141591 | 0.333162 | 0.521633 | 0.253932 | 0.053965 | 1.877331 | 0.458741 | 0.019787 | -0.50632 | 1.129699 | 1.971752 | 0.311428 |
| 26.29167 | 3.259222 | -0.96432 | 1.387835 | 0.240357 | -0.01891 | 1.622391 | -0.22087 | 0.04353 | 0.749345 | 0.334155 | 0.093206 | -0.28909 | 0.032804 | 0.273295 | 0.279849 | 0.064786 | 0.014601 | 0.774053 | 0.090087 | -0.09072 | -1.3988 | 1.197478 | 0.808579 | 0.02649 |
| 26.30556 | 2.931848 | -1.45201 | 1.937904 | 0.393035 | -0.15129 | 1.065037 | -0.52727 | 0.050712 | 0.308245 | 0.708869 | 0.523253 | 0.446721 | 0.39489 | 0.332646 | 0.389841 | -0.02065 | 0.030047 | 0.119457 | -0.1352 | -0.05642 | -0.59176 | 0.717126 | -0.54222 | 0.138433 |
| 26.31944 | 0.641923 | -1.26593 | 1.486048 | 0.589687 | -0.17536 | 0.254017 | -0.56591 | 0.023492 | -0.30475 | 0.87752 | 0.539639 | 1.061527 | 0.517045 | 0.096436 | 0.403741 | -0.09621 | 0.078942 | -0.23407 | -0.0765 | -0.02113 | 0.807995 | 0.120758 | -2.50208 | 0.231382 |
| 26.33333 | -2.24731 | -0.61394 | 0.344304 | 0.584116 | -0.16029 | -0.90492 | -0.46081 | -0.04154 | -1.11866 | 0.555493 | 0.238503 | 0.942425 | 0.122794 | -0.42573 | 0.130701 | -0.2321 | 0.073991 | -0.60695 | 0.029801 | -0.06703 | 1.403366 | -0.60539 | -4.13339 | 0.059396 |
| 26.34722 | -4.23148 | 0.709086 | -0.65646 | 0.614021 | 0.057816 | -1.44822 | -0.19103 | 0.039383 | -1.26191 | 0.313396 | 0.19006 | 0.892155 | -0.17609 | -0.56067 | 0.106728 | -0.21718 | 0.054515 | -0.8186 | 0.134916 | -0.05802 | 0.655093 | -1.3132 | -3.47721 | 0.025637 |
| 26.36111 | -4.41638 | 2.606928 | -0.98058 | 0.852779 | 0.406398 | -0.78093 | 0.181812 | 0.303083 | -0.33934 | 0.565777 | 0.552838 | 0.866761 | -0.09261 | -0.171 | 0.449417 | 0.143715 | 0.114317 | -0.70423 | 0.210477 | 0.095204 | -0.21731 | -1.403 | -0.50106 | 0.347503 |
| 26.375 | -1.54115 | 4.419409 | -0.34572 | 0.972584 | 0.493371 | 0.269844 | 0.336082 | 0.534619 | 0.757576 | 1.067721 | 0.67437 | 0.689947 | 0.110648 | 0.37297 | 0.704851 | 0.496024 | 0.217476 | -0.49525 | 0.267831 | 0.244253 | -1.00437 | -0.46404 | 2.103618 | 0.637351 |
| 26.38889 | 2.655343 | 3.971186 | 1.131487 | 0.764304 | 0.238099 | 0.984321 | 0.13399 | 0.548802 | 1.323611 | 1.316548 | 0.280707 | 0.164893 | 0.257016 | 0.517495 | 0.587555 | 0.535196 | 0.201834 | -0.06789 | 0.243961 | 0.255654 | -1.12104 | 0.678293 | 3.836101 | 0.795485 |
| 26.40278 | 6.1535 | 1.808911 | 3.135615 | 0.501461 | 0.063959 | 1.348205 | -0.17031 | 0.460132 | 1.430662 | 1.196111 | -0.3078 | -0.30317 | 0.456085 | 0.245304 | 0.391522 | 0.34985 | 0.101668 | 0.647919 | 0.054754 | 0.185985 | -0.18158 | 2.276946 | 2.777985 | 0.880661 |
| 26.41667 | 5.299711 | -0.15536 | 4.720809 | 0.640021 | 0.144362 | 1.88546 | -0.41585 | 0.594222 | 1.578314 | 0.900518 | -0.48531 | -0.34924 | 0.618027 | -0.00312 | 0.410209 | 0.025686 | 0.063267 | 1.609339 | -0.10681 | 0.202448 | 1.047836 | 2.230676 | 1.200704 | 0.882829 |
| 26.43056 | 1.954876 | -1.67764 | 3.992315 | 0.903995 | 0.400201 | 2.04597 | -0.44681 | 0.705052 | 1.521485 | 0.558471 | -0.00136 | 0.328302 | 0.670624 | -0.04565 | 0.495797 | -0.1163 | 0.106595 | 2.651793 | -0.16511 | 0.401681 | 2.068587 | 1.153006 | -0.53579 | 0.500108 |
| 26.44444 | -0.54688 | -2.20443 | 1.623126 | 1.708965 | 1.131299 | 2.170129 | 0.393617 | 0.878469 | 1.757221 | 0.539869 | 1.740522 | 1.662827 | 1.069665 | 0.923697 | 0.982818 | 0.32206 | 0.401612 | 3.249302 | 0.059209 | 1.000234 | 2.253109 | 0.273254 | -2.27356 | 1.00455 |
| 26.45833 | -2.92944 | -2.2126 | 0.413374 | 4.332078 | 2.609421 | 4.656704 | 2.493098 | 2.696981 | 3.617293 | 2.49323 | 5.279921 | 4.169798 | 4.527178 | 4.293664 | 3.526631 | 1.744574 | 1.049968 | 5.13087 | 0.688804 | 2.590895 | 2.674004 | -0.81523 | -3.15255 | 3.614624 |
| 26.47222 | -0.791 | 1.484152 | 1.411145 | 6.179684 | 4.157917 | 6.208451 | 4.736978 | 5.25225 | 4.787074 | 4.227024 | 7.763749 | 6.302662 | 7.126861 | 6.286083 | 7.214995 | 3.416216 | 1.479271 | 6.210527 | 1.427568 | 4.346276 | 3.497418 | -1.06518 | -3.35841 | 5.349184 |
| 26.48611 | 2.426582 | 3.732021 | 2.73254 | 4.778658 | 3.944525 | 6.677001 | 4.67644 | 5.876312 | 3.382606 | 3.872145 | 4.360681 | 2.969825 | 6.21378 | 4.107971 | 6.254809 | 3.310051 | 1.553491 | 4.120488 | 1.812836 | 5.95648 | 3.050148 | 0.496575 | -1.31739 | 5.449312 |
| 26.5 | 5.11523 | 6.063798 | 3.400236 | 2.629169 | 2.805777 | 5.541973 | 3.655773 | 5.786176 | 2.577766 | 2.792437 | 1.185058 | 2.035456 | 5.116776 | 2.216024 | 3.977166 | 2.223199 | 1.716901 | 1.119465 | 2.140026 | 5.486202 | 1.324102 | 2.270119 | 2.533531 | 5.295895 |
| 26.51389 | 4.910106 | 4.381932 | 3.455292 | 1.784553 | 1.879425 | 3.090039 | 2.235508 | 4.198088 | 1.313288 | 1.689504 | 1.165988 | 2.07348 | 4.027411 | 1.349943 | 2.052796 | 1.51798 | 1.732771 | -0.01384 | 1.875624 | 4.465862 | -0.01465 | 3.991931 | 5.36483 | 3.040324 |
| 26.52778 | 1.838702 | 1.631414 | 3.680076 | 1.211056 | 0.753893 | 0.663882 | 0.464515 | 2.737411 | -0.22092 | 0.535911 | 0.804302 | 1.547615 | 1.804384 | 0.735712 | 0.915508 | 1.202539 | 1.489105 | -0.71725 | 1.492844 | 3.701753 | -0.0308 | 3.531497 | 3.279634 | 0.981505 |
| 26.54167 | -0.24599 | -0.2645 | 2.81394 | 0.265854 | -0.44748 | -1.67861 | -1.50614 | 1.633079 | -1.25477 | -0.31118 | 0.455648 | 0.822604 | -0.15214 | 0.254334 | 0.164385 | 1.015212 | 1.190226 | -1.42752 | 1.274743 | 3.098327 | 1.025985 | 1.630277 | 1.237284 | -1.07348 |
| 26.55556 | -1.61967 | -1.64719 | 1.155528 | -0.56881 | -0.53272 | -3.52732 | -2.0925 | 1.066386 | -2.28068 | -0.73522 | 1.157296 | 0.533136 | -2.00209 | 0.117842 | -0.17001 | 0.824807 | 0.452393 | -2.70446 | 0.940911 | 2.559017 | 1.602461 | 0.420393 | -0.16331 | -2.6788 |
| 26.56944 | -2.22963 | -1.60146 | -0.68643 | -0.84439 | 0.283821 | -3.01984 | -0.85764 | 1.165264 | -2.95895 | -0.87145 | 2.202072 | 1.219222 | -0.84226 | 0.206834 | 0.110174 | 0.522802 | 0.266911 | -2.91406 | 0.6375 | 2.053465 | 0.924113 | -1.08748 | -2.05884 | -2.62924 |
| 26.58333 | -0.06144 | -1.30639 | -1.98343 | 0.189464 | 1.373802 | -1.30108 | 0.415842 | 1.420255 | -0.84063 | -0.28624 | 3.098095 | 2.167205 | 0.544647 | 0.267858 | 0.377802 | 0.531003 | 0.390333 | 0.406954 | 0.853497 | 1.952448 | 0.737605 | -2.1416 | -3.97416 | -1.15276 |
| 26.59722 | 2.341977 | -2.7548 | -2.47958 | 1.805549 | 2.221833 | 1.215484 | 1.091954 | 1.216172 | 1.768728 | 0.160223 | 2.917426 | 2.891162 | 1.599723 | 0.211713 | 0.04046 | 0.754578 | 0.597159 | 3.682979 | 1.139432 | 1.730984 | 0.405913 | -2.36056 | -4.82061 | 0.768502 |
| 26.61111 | 3.958713 | -4.53308 | -2.0276 | 2.491216 | 2.602503 | 3.225218 | 1.59539 | 0.104066 | 3.361575 | 0.421473 | 2.163718 | 3.008932 | 1.317646 | -0.09323 | -0.83413 | 0.638604 | 0.524776 | 4.113478 | 0.891222 | 0.402837 | 0.135168 | -0.86122 | -2.85776 | 1.388808 |
| 26.625 | 3.650052 | -4.19476 | -0.44265 | 2.055067 | 2.624938 | 3.16994 | 1.663152 | -1.33508 | 2.625453 | 0.559777 | -0.10085 | 2.403418 | -0.91225 | -0.41705 | -1.74257 | 0.40632 | 0.377604 | 3.666257 | 0.38405 | -2.03258 | -0.24835 | 1.063322 | 0.628622 | 0.710026 |
| 26.63889 | 2.824804 | -0.97545 | 2.008091 | 1.381015 | 1.71797 | 2.590813 | 0.872704 | -2.28734 | 1.73731 | 0.840798 | -2.3834 | 1.067009 | -2.42641 | -0.55345 | -2.12802 | 0.154331 | 0.051338 | 2.877278 | -0.12385 | -4.28379 | -0.31122 | 2.555697 | 2.871736 | -0.49842 |
| 26.65278 | 1.244863 | 2.725382 | 3.345452 | 1.181639 | -0.01588 | 2.104958 | -0.70021 | -2.26212 | 1.383242 | 1.181184 | -2.86633 | -0.35216 | -3.53705 | -0.14085 | -2.17759 | -0.52238 | -0.46643 | 0.634892 | -0.41852 | -4.52364 | -0.22009 | 2.143547 | 2.863631 | -1.45438 |
| 26.66667 | -2.61329 | 4.306525 | 2.625192 | 1.38858 | -1.43615 | 1.202909 | -1.94206 | -1.20571 | 1.0311 | 1.140043 | -0.69111 | -0.60959 | -3.54037 | 0.50416 | -1.69075 | -1.12561 | -0.63099 | -1.37013 | -0.60423 | -3.82532 | 0.034175 | 0.897698 | 4.099406 | -1.49738 |
| 26.68056 | -6.04555 | 4.661266 | 1.039084 | 1.079302 | -1.95603 | 0.019955 | -2.49657 | 0.27102 | 0.632708 | 1.062775 | 1.665458 | -0.2565 | -0.9501 | 1.109762 | -0.41059 | -1.81284 | -0.55601 | -3.71318 | -0.69465 | -2.25489 | -0.68479 | -0.04469 | 1.588699 | -0.86578 |
| 26.69444 | -6.31806 | 2.668283 | -0.33612 | -0.02799 | -1.99487 | -1.22137 | -1.6048 | 2.144 | 0.281934 | 0.818253 | 3.110459 | -0.22934 | 1.301636 | 1.585744 | 0.994141 | -1.31459 | -0.30842 | -4.48428 | -0.35207 | -0.04539 | -0.98915 | -0.54777 | -0.96999 | 0.399921 |
| 26.70833 | -3.21849 | 0.763832 | -1.68374 | -1.11143 | -1.49395 | -2.50417 | 0.107179 | 3.909505 | 0.375511 | 0.687612 | 3.096742 | -0.51086 | 2.970844 | 1.477326 | 2.529325 | -0.14317 | -0.23439 | -3.54699 | -0.26682 | 2.279071 | -1.22465 | -2.14325 | -3.34216 | 1.361223 |
| 26.72222 | 0.123533 | -1.85715 | -3.62516 | -1.26471 | -0.72052 | -3.92982 | 0.489859 | 4.966389 | -0.1386 | 0.594014 | 2.121777 | 0.175201 | 2.903382 | 1.163269 | 3.913529 | 0.786413 | -0.21256 | -3.25331 | 0.005962 | 4.692442 | -0.82844 | -3.09146 | -2.09765 | 2.025744 |
| 26.73611 | 0.019911 | -4.14344 | -5.05177 | -1.03504 | -0.30215 | -5.37066 | 0.512128 | 4.879848 | -0.64924 | 0.205858 | 0.781481 | 1.089585 | 1.463063 | 0.971752 | 3.634 | 1.180006 | -0.12479 | -4.21959 | 0.719 | 6.041987 | -0.69692 | -1.44736 | -2.60615 | 1.480685 |
| 26.75 | -1.33886 | -4.3201 | -4.51759 | -1.25365 | -0.19714 | -2.26648 | 0.423341 | 3.565519 | -0.18927 | -0.0963 | -0.15073 | 1.201044 | -0.47792 | 0.666048 | 1.554295 | 0.906302 | 0.233178 | -3.0726 | 0.904319 | 5.007304 | 0.133135 | 0.496038 | -1.27009 | 0.790598 |
| 26.76389 | -1.28457 | -4.281 | -1.40809 | -1.61718 | -0.39696 | 1.055291 | 0.060997 | 1.699309 | 0.912668 | -0.24877 | -0.33484 | 0.871076 | -2.49788 | 0.288964 | -0.19508 | 0.265089 | 0.467633 | 0.401571 | 0.530513 | 3.051759 | 1.247266 | 1.562426 | -0.03021 | 0.291156 |
| 26.77778 | -1.35166 | -3.15705 | 1.511641 | -1.23427 | -0.77107 | 2.084144 | -0.39664 | 0.007059 | 1.331074 | -0.39438 | -0.2795 | 0.650004 | -2.92353 | -0.07482 | -1.62613 | -0.11972 | 0.263455 | 3.417035 | 0.195254 | 0.70402 | 1.45049 | 1.369993 | 0.978135 | -0.07865 |
| 26.79167 | -1.63206 | -0.99075 | 3.43787 | -1.17214 | -1.08537 | 0.805696 | -0.92927 | -1.55402 | 0.803769 | -0.5882 | -1.43134 | -0.06281 | -2.28508 | -0.4177 | -2.97529 | -0.34053 | 0.189538 | 3.410727 | -0.05907 | -1.69639 | 0.870263 | 0.336425 | 1.105979 | -0.60223 |

| | | | | | | | | | | | | | | | | | | | | | | | | |
|---|---|---|---|---|---|---|---|---|---|---|---|---|---|---|---|---|---|---|---|---|---|---|---|---|
| 26.80556 | -3.82622 | 1.183656 | 3.205149 | -1.44653 | -1.54958 | -0.50359 | -1.71592 | -2.9338 | -0.39843 | -0.99674 | -2.80239 | -1.02469 | -2.42164 | -0.77034 | -4.15111 | -0.53384 | -0.17809 | 1.118022 | -0.54398 | -4.22069 | -0.48425 | -1.26663 | 0.37867 | -1.34572 |
| 26.81944 | -5.88772 | 2.130076 | 0.506443 | -2.03362 | -1.72268 | -2.0339 | -2.31229 | -4.33772 | -2.07959 | -1.34749 | -3.91317 | -2.11882 | -3.30415 | -1.20649 | -4.73799 | -0.83181 | -0.66436 | -0.31174 | -0.90159 | -6.37506 | -2.15343 | -3.2901 | -1.20954 | -2.25098 |
| 26.83333 | -5.08481 | 2.131237 | -2.33345 | -2.73885 | -1.81311 | -4.18211 | -2.82018 | -5.79679 | -3.29945 | -1.48556 | -4.91618 | -3.5573 | -3.95345 | -1.66551 | -4.52179 | -1.1651 | -1.041 | -1.58963 | -1.07567 | -7.7424 | -3.21502 | -3.97092 | -2.65777 | -2.83337 |
| 26.84722 | -3.53714 | 0.670468 | -6.00286 | -3.44271 | -1.84503 | -5.4774 | -2.54949 | -6.60225 | -3.86641 | -1.42492 | -5.26625 | -4.20511 | -4.64311 | -2.47457 | -3.36078 | -1.55287 | -1.26216 | -2.466 | -0.94264 | -6.74784 | -3.22542 | -3.18706 | -3.39215 | -3.30656 |
| 26.86111 | -1.85887 | -0.95831 | -9.16361 | -3.48465 | -1.72735 | -4.51965 | -2.10565 | -7.26442 | -3.53977 | -1.1398 | -3.90454 | -2.84784 | -5.12242 | -2.97775 | -2.59858 | -1.69094 | -1.24994 | -3.16832 | -0.76726 | -5.53568 | -1.89311 | -1.6875 | -3.8546 | -3.03935 |
| 26.875 | -1.29199 | -2.30514 | -8.48115 | -1.96748 | -1.4277 | -3.09895 | -1.78026 | -5.63962 | -2.8587 | -0.83937 | -1.79647 | -1.4866 | -4.76551 | -2.636 | -1.93583 | -1.50296 | -1.02161 | -3.5191 | -0.58894 | -4.19301 | -0.57553 | -0.51667 | -2.80985 | -1.84437 |
| 26.88889 | 0.331428 | -3.68338 | -5.47561 | -0.43386 | -1.02199 | -0.97195 | -1.3612 | -3.3834 | -2.40968 | -0.53185 | -0.35389 | -0.19511 | -3.86454 | -2.26148 | -1.15222 | -1.19972 | -0.83538 | -3.4579 | -0.32024 | -3.28271 | 0.030255 | 0.34782 | -1.43607 | -1.15342 |
| 26.90278 | 1.717969 | -3.73316 | -0.78589 | 0.32807 | -0.57902 | 1.442155 | -0.84503 | -1.31286 | -2.20169 | -0.26706 | 0.36017 | 1.50476 | -2.34213 | -1.70364 | -0.43475 | -0.82046 | -0.79654 | -2.96684 | -0.13903 | -1.83773 | -0.33497 | 0.770123 | -0.02107 | -1.08064 |
| 26.91667 | 3.212876 | -2.5065 | 3.402852 | 0.515102 | -0.23381 | 3.177548 | -0.31876 | 0.964281 | -1.71743 | -0.08964 | 0.059219 | 1.51295 | -0.76199 | -0.84405 | 0.286793 | -0.61289 | -0.79273 | -3.22212 | -0.0493 | -0.88029 | -0.99047 | 0.764877 | 1.272187 | -0.99481 |
| 26.93056 | 4.958067 | -0.81834 | 5.314338 | 0.424901 | -0.14297 | 3.088842 | 0.024795 | 4.295336 | -1.26145 | -0.02419 | -1.31063 | -0.41535 | 0.252124 | 0.008809 | 0.774389 | -0.77649 | -0.81784 | -4.07368 | -0.24737 | -0.15708 | -1.20863 | 0.330124 | 2.231185 | -0.89335 |
| 26.94444 | 3.710266 | 0.06608 | 2.814558 | -0.70354 | -0.83814 | 0.685835 | -0.36375 | 5.126528 | -1.72333 | -0.27531 | -2.32269 | -2.17317 | 0.005109 | 0.014296 | 0.070328 | -1.52635 | -1.17403 | -4.8046 | -0.67158 | -0.13545 | -1.68942 | -0.51863 | 1.373424 | -1.54451 |
| 26.95833 | -1.3912 | -1.27605 | -1.84896 | -3.10084 | -3.22693 | -3.80977 | -2.24 | 0.53527 | -3.78508 | -0.91422 | -4.48377 | -4.24462 | -1.35154 | -1.92702 | -2.11424 | -3.03453 | -1.77658 | -5.50689 | -1.10957 | -1.57721 | -3.17659 | -1.54011 | -1.63073 | -3.53119 |
| 26.97222 | -6.98935 | -5.01936 | -6.29168 | -6.05102 | -7.96093 | -8.19477 | -5.20995 | -4.50113 | -5.92954 | -1.5413 | -7.25681 | -7.27873 | -2.51818 | -5.69528 | -4.50796 | -4.77412 | -2.34179 | -6.29157 | -1.13966 | -3.54743 | -6.09813 | -2.63628 | -4.84814 | -5.62577 |
| 26.98611 | -10.5722 | -7.34204 | -7.8461 | -6.32649 | -7.88497 | -7.78228 | -4.7961 | -6.59326 | -4.6361 | -0.50406 | -5.8266 | -6.95145 | -2.6016 | -5.27628 | -4.17237 | -3.8188 | -2.60544 | -6.1489 | -0.62765 | -4.04411 | -6.53698 | -3.74884 | -4.93105 | -4.17147 |
| 27 | -9.0461 | -4.64094 | -4.60434 | -3.96611 | -4.75708 | -4.70612 | -2.12224 | -4.56215 | -2.00636 | -0.08221 | -1.58065 | -3.34681 | -0.96009 | -1.07903 | -1.63331 | -1.43174 | -1.9751 | -4.91454 | -0.38623 | -3.2461 | -4.10549 | -2.77097 | -3.30885 | -1.59024 |
| 27.01389 | -7.00363 | -1.80535 | -3.15485 | -2.65085 | -2.31227 | -3.46648 | -0.4312 | -3.25574 | -0.38347 | -0.11582 | 0.743016 | -1.1957 | 0.433375 | 0.095822 | -0.64294 | -0.58695 | -1.39047 | -3.12227 | -0.13169 | -1.64613 | -1.84433 | -1.63528 | -2.26989 | -0.563 |
| 27.02778 | -4.621 | -0.01053 | -1.7898 | -2.14009 | -0.51578 | -2.77416 | 0.230106 | -2.45036 | 0.276167 | -0.08442 | 0.510875 | -0.45787 | 0.964006 | 0.159518 | -0.684 | -0.70311 | -0.99481 | -1.79495 | -0.08871 | -0.76869 | -0.8951 | -0.62544 | -1.60051 | -0.55606 |
| 27.04167 | -2.6437 | 0.715853 | -1.38202 | -1.73775 | 0.31343 | -1.97535 | 0.391758 | -1.71466 | 0.565708 | 0.135931 | -0.2503 | -0.34883 | 0.418265 | 0.087337 | -0.65103 | -0.82573 | -0.36605 | -0.79994 | -0.02635 | -0.09577 | -0.34713 | -0.12271 | -0.96092 | -0.57613 |
| 27.05556 | -1.1164 | 0.742309 | -1.02572 | -1.29582 | 0.552579 | -1.10473 | 0.438127 | -1.16977 | 0.62749 | 0.390102 | -0.77718 | -0.23992 | -0.15486 | 0.039235 | -0.33047 | -0.64565 | 0.312451 | -0.04053 | 0.069359 | 0.742627 | 0.277143 | 0.217514 | -0.45089 | -0.22917 |
| 27.06944 | 0.092583 | 0.926666 | -0.56598 | -0.8431 | 0.532913 | -0.46005 | 0.513939 | -0.90001 | 0.603266 | 0.536423 | -0.46919 | -0.02579 | -0.13915 | -0.05964 | -0.14727 | -0.25549 | 0.751304 | 0.298352 | 0.132632 | 1.473594 | 1.032741 | 0.500957 | -0.16338 | 0.284203 |
| 27.08333 | 1.287335 | 1.028307 | -0.22151 | -0.37866 | 0.726953 | -0.15452 | 0.765553 | -0.75555 | 0.718007 | 0.581246 | 0.286305 | 0.410993 | 0.151311 | -0.16442 | -0.19776 | 0.287839 | 0.910831 | 0.433994 | 0.166768 | 1.955798 | 2.319654 | 0.648561 | -0.05218 | 1.156187 |
| 27.09722 | 1.988193 | 0.910901 | -0.06063 | -0.06309 | 0.905446 | -0.08713 | 0.930592 | -0.81634 | 0.796783 | 0.45995 | 0.948422 | 0.832348 | 0.49468 | -0.1477 | -0.0835 | 0.750556 | 0.820725 | 0.483337 | 0.337331 | 1.81939 | 2.883528 | 0.156099 | -0.02259 | 1.415784 |
| 27.11111 | 2.47608 | 0.733665 | 0.035406 | 0.338301 | 0.871173 | -0.11745 | 0.859201 | -0.91309 | 0.481089 | 0.278982 | 1.187226 | 0.942103 | 0.499122 | 0.028365 | 0.339677 | 0.896002 | 0.608545 | 0.394718 | 0.5511 | 1.281582 | 2.815357 | 0.200281 | 0.117756 | 0.449602 |
| 27.125 | 3.381647 | 0.469254 | 0.111612 | 1.154681 | 1.135076 | -0.20445 | 0.640306 | -0.91321 | -0.03635 | 0.121369 | 0.837615 | 0.891537 | -0.10012 | 0.228072 | 0.748803 | 0.557727 | 0.409203 | 0.215237 | 0.396151 | 0.725529 | 2.440526 | 0.804364 | 0.411793 | -0.47151 |
| 27.13889 | 4.688931 | 0.225672 | 0.266401 | 1.41747 | 1.096747 | -0.22251 | 0.396712 | -0.81884 | -0.0067 | 0.112574 | 0.206588 | 0.704041 | -0.91092 | 0.192392 | 0.702945 | 0.002434 | 0.181527 | 0.010679 | 0.133629 | 0.495545 | 2.351713 | 1.103919 | 0.627961 | -1.4338 |
| 27.15278 | 6.390033 | 0.312118 | 0.766439 | 1.058029 | 1.008652 | -0.01492 | 0.266569 | -0.53792 | 0.400511 | 0.331782 | -0.04311 | 0.761852 | -0.9022 | 0.18441 | 0.409656 | -0.41737 | 0.072974 | -0.05887 | -0.0609 | 0.552641 | 2.431569 | 1.36224 | 1.09331 | -1.2237 |
| 27.16667 | 8.148365 | 0.79289 | 1.408928 | 0.951447 | 1.134442 | 0.253043 | 0.572633 | -0.14344 | 0.841813 | 0.650897 | 0.729782 | 1.153329 | 0.075098 | 0.296333 | 0.388961 | -0.11774 | 0.185117 | 0.264419 | -0.02718 | 0.94252 | 2.262433 | 1.905833 | 1.8619 | -0.00379 |
| 27.18056 | 8.591256 | 1.37338 | 1.752785 | 0.864684 | 1.524517 | 0.368532 | 0.824337 | 0.160465 | 1.022293 | 0.768405 | 1.853943 | 1.813931 | 0.783515 | 0.236669 | 0.599438 | 0.568714 | 0.279908 | 0.726144 | 0.170247 | 1.055054 | 1.905348 | 1.691212 | 2.60358 | 1.023714 |
| 27.19444 | 7.218344 | 1.728297 | 2.022832 | 0.605029 | 1.729182 | 0.372964 | 0.883044 | 0.203243 | 0.965432 | 0.613059 | 2.433597 | 2.320727 | 1.092647 | 0.097372 | 0.684957 | 1.041107 | 0.239869 | 0.8335 | 0.340831 | 0.6554 | 0.96722 | 1.466869 | 2.514925 | 1.33312 |
| 27.20833 | 4.764995 | 1.804725 | 2.502781 | 0.335046 | 1.450747 | 0.493344 | 0.608384 | 0.077156 | 0.843831 | 0.377318 | 1.804014 | 2.531711 | 0.892424 | 0.176496 | 0.775708 | 1.205574 | 0.20459 | 0.936143 | 0.433925 | 0.134741 | -0.04991 | 1.190754 | 2.542902 | 0.528752 |
| 27.22222 | 2.231008 | 1.63383 | 2.917849 | 0.217613 | 0.985963 | 0.771261 | 0.231159 | 0.046344 | 0.75628 | 0.175478 | 1.089255 | 2.724203 | 0.508847 | 0.358647 | 0.92027 | 0.943445 | 0.214313 | 1.104752 | 0.389368 | -0.28867 | -0.69815 | 1.105331 | 2.413587 | -0.21681 |
| 27.23611 | -0.02453 | 0.872068 | 3.035159 | 0.204862 | 0.701783 | 0.95678 | -0.0675 | 0.095449 | 0.606043 | 0.049196 | 0.85638 | 2.971337 | 0.151675 | 0.364862 | 0.758828 | 0.330047 | 0.052033 | 1.167062 | 0.219139 | 0.01622 | -0.698 | 0.92382 | 2.479241 | -0.45952 |
| 27.25 | -1.70022 | 0.785015 | 2.594755 | 0.184963 | 0.485943 | 0.888242 | -0.15587 | 0.091753 | 0.459303 | 0.031448 | 1.045474 | 2.62395 | 0.161906 | 0.170198 | 0.382135 | -0.22513 | -0.08484 | 0.936454 | 0.058362 | 0.357693 | -0.08777 | 0.64628 | 2.064015 | 0.094759 |
| 27.26389 | -2.97985 | 1.107488 | 1.953171 | 0.171421 | 0.708476 | 0.798874 | 0.092767 | 0.075391 | 0.438233 | 0.12732 | 1.502339 | 2.226873 | 0.485952 | -0.00813 | 0.176432 | -0.24528 | -0.0728 | 0.792803 | 0.000101 | 0.380426 | 0.812193 | 0.363709 | 1.515878 | 0.71944 |
| 27.27778 | -4.14024 | 1.375185 | 1.442403 | 0.213729 | 1.27428 | 0.805472 | 0.511706 | 0.085159 | 0.563925 | 0.235554 | 1.672494 | 2.137883 | 0.909773 | -0.05549 | 0.237045 | 0.100774 | 0.065679 | 0.86629 | 0.145915 | 0.190545 | 1.656568 | 0.151448 | 0.889583 | 1.181591 |
| 27.29167 | -4.11343 | 1.25305 | 1.00425 | 0.270728 | 1.57256 | 0.852474 | 0.654735 | 0.083958 | 0.624178 | 0.312072 | 1.068643 | 1.754586 | 0.881316 | -0.01214 | 0.444422 | 0.487059 | 0.237904 | 0.972679 | 0.32669 | -0.03314 | 1.724584 | 0.028771 | 0.270192 | 1.037779 |
| 27.30556 | -3.20176 | 0.893589 | 0.687763 | 0.28221 | 1.285134 | 1.01371 | 0.510614 | 0.065468 | 0.581993 | 0.393616 | 0.277592 | 1.404742 | 0.415378 | 0.058515 | 0.691015 | 0.633014 | 0.327127 | 1.053373 | 0.406444 | -0.21054 | 1.127287 | 0.068689 | -0.09369 | 0.712149 |
| 27.31944 | -2.44134 | 0.595494 | 0.299331 | 0.208523 | 0.747196 | 1.078627 | 0.200155 | 0.095333 | 0.392006 | 0.466281 | -0.21174 | 1.016508 | -0.08281 | 0.095509 | 0.742668 | 0.514038 | 0.334726 | 0.956178 | 0.347203 | -0.27287 | 0.37223 | 0.298341 | -0.29963 | 0.446924 |
| 27.33333 | -1.5363 | 0.447884 | -0.11064 | 0.080971 | 0.304216 | 1.062022 | -0.07604 | 0.157729 | 0.131358 | 0.488819 | -0.30437 | 0.621829 | -0.15008 | 0.132375 | 0.696623 | 0.279637 | 0.247157 | 0.787679 | 0.189154 | -0.14416 | -0.35396 | 0.725481 | -0.2118 | 0.258177 |
| 27.34722 | -0.0617 | 0.623199 | -0.33998 | 0.048414 | 0.397591 | 1.058437 | -0.04136 | 0.173919 | 0.014831 | 0.506458 | -0.07509 | 0.290366 | 0.355652 | 0.146283 | 0.677307 | 0.116643 | 0.084225 | 0.81998 | 0.021953 | 0.066367 | -0.7967 | 1.178586 | 0.20896 | 0.178633 |
| 27.36111 | 1.394031 | 0.930807 | -0.39167 | 0.157536 | 0.773046 | 1.054006 | 0.297217 | 0.196707 | 0.0917 | 0.502743 | 0.292866 | -2.6E-05 | 0.835992 | 0.080125 | 0.665391 | -0.02811 | -0.0647 | 0.888914 | -0.042 | 0.184161 | -0.41989 | 1.242951 | 0.625162 | 0.171073 |
| 27.375 | 2.98738 | 1.277368 | -0.32104 | 0.370447 | 1.035411 | 1.103102 | 0.632666 | 0.308496 | 0.293555 | 0.482666 | 0.579073 | -0.02972 | 1.023262 | 0.029521 | 0.701953 | -0.00057 | -0.09435 | 1.160717 | 0.080538 | 0.245149 | 0.749426 | 1.147449 | 0.979162 | 0.344029 |
| 27.38889 | 4.358763 | 1.366483 | -0.06725 | 0.575953 | 1.102951 | 1.233409 | 0.770742 | 0.399334 | 0.441078 | 0.44416 | 0.561579 | 0.231647 | 0.888606 | 0.077927 | 0.85428 | 0.190927 | 0.030713 | 1.602721 | 0.286861 | 0.39527 | 2.371099 | 0.999593 | 1.309109 | 0.706149 |
| 27.40278 | 4.45522 | 1.093115 | 0.634283 | 0.744559 | 0.981735 | 1.35182 | 0.781832 | 0.438422 | 0.53367 | 0.417357 | 0.319217 | 0.55405 | 0.632699 | 0.164056 | 0.864213 | 0.396353 | 0.256948 | 1.864921 | 0.424639 | 0.551487 | 3.220739 | 0.875133 | 1.508766 | 1.010916 |
| 27.41667 | 4.073317 | 0.630131 | 1.846588 | 0.852546 | 0.755466 | 1.526955 | 0.811456 | 0.546753 | 0.675314 | 0.466389 | 0.01833 | 0.79495 | 0.324872 | 0.230807 | 0.751073 | 0.468405 | 0.481056 | 1.825761 | 0.332009 | 0.682428 | 2.34839 | 0.644746 | 1.707358 | 0.954769 |
| 27.43056 | 4.135852 | 0.27089 | 3.390979 | 0.81208 | 0.515624 | 1.685299 | 0.805406 | 0.670484 | 0.850754 | 0.559668 | -0.23481 | 0.98293 | 0.172714 | 0.365977 | 0.56206 | 0.543356 | 0.596413 | 1.619708 | 0.203248 | 0.691254 | 0.831936 | 0.284833 | 1.818287 | 0.594293 |
| 27.44444 | 4.215116 | 0.750726 | 5.104309 | 1.273691 | 0.913211 | 2.11752 | 1.24435 | 0.883925 | 1.727122 | 0.827078 | 0.483625 | 1.795544 | 0.843116 | 0.996928 | 0.97838 | 0.996125 | 0.696658 | 2.268524 | 0.192131 | 0.949218 | -0.33522 | 0.006393 | 2.10374 | 0.903532 |
| 27.45833 | 5.065655 | 2.776432 | 9.284978 | 4.136403 | 2.647606 | 4.483647 | 2.934461 | 2.961749 | 4.792623 | 1.428967 | 3.213519 | 4.233238 | 2.856187 | 2.746573 | 2.875038 | 1.981808 | 1.197523 | 5.138462 | 0.390382 | 1.888725 | -0.20541 | 0.336391 | 3.409143 | 3.34133 |
| 27.47222 | 6.566792 | 4.638421 | 9.912864 | 6.706439 | 4.775477 | 7.411894 | 5.142781 | 6.377607 | 7.495941 | 2.021082 | 7.978046 | 6.008058 | 5.173794 | 5.004815 | 5.743105 | 2.680721 | 1.681092 | 8.025508 | 0.679133 | 3.96024 | 1.708336 | 1.254885 | 6.281752 | 6.444831 |
| 27.48611 | 6.444355 | 3.490157 | 7.076255 | 5.692504 | 4.658325 | 7.662645 | 5.149386 | 7.775344 | 5.390938 | 2.289751 | 8.584498 | 4.803968 | 5.311864 | 5.090132 | 5.247627 | 2.883933 | 1.789285 | 7.980531 | 0.78215 | 5.972967 | 2.929851 | 2.202164 | 4.754685 | 5.757051 |
| 27.5 | 3.422268 | 2.850878 | 4.147043 | 4.299265 | 3.522104 | 5.577185 | 4.242857 | 7.688432 | 3.682384 | 2.397263 | 5.851647 | 4.806289 | 4.751505 | 4.029016 | 4.082348 | 2.531036 | 1.446045 | 6.092409 | 0.922999 | 7.234711 | 3.983979 | 2.836005 | 2.131896 | 4.96424 |
| 27.51389 | 0.214118 | 2.082502 | 1.778059 | 2.700681 | 2.580959 | 3.239267 | 3.519047 | 5.97456 | 2.376622 | 1.952623 | 3.556194 | 3.518874 | 2.863635 | 3.228094 | 2.817117 | 1.961119 | 1.223437 | 4.025314 | 0.7901 | 6.085789 | 4.208304 | 2.908681 | -0.2349 | 3.181557 |
| 27.52778 | -3.17698 | 1.104878 | -0.29788 | 1.234481 | 1.864436 | 1.551552 | 2.432164 | 3.695845 | 0.804411 | 1.100098 | 1.41652 | 1.128674 | 1.239016 | 2.159908 | 1.650643 | 1.320389 | 0.921019 | 2.246829 | 0.538157 | 4.55613 | 2.536094 | 1.760764 | -3.0573 | 1.637553 |
| 27.54167 | -6.21611 | 0.149044 | -2.91278 | -0.30096 | 1.06063 | 0.026673 | 1.110401 | 2.04605 | -0.83044 | 0.043788 | -0.93111 | -0.97747 | -0.48344 | 1.063625 | -0.14344 | 0.622516 | 0.487779 | 0.585289 | 0.322282 | 2.791264 | 0.368682 | 0.184091 | -4.59641 | 0.097233 |
| 27.55556 | -9.00482 | -0.7928 | -5.52304 | -1.29902 | 0.688256 | -0.66765 | -0.12882 | 1.365261 | -2.34929 | -0.7755 | -1.15062 | -1.81184 | -1.48384 | 0.3569 | -0.5409 | 0.230444 | 0.376596 | -0.34937 | 0.028607 | 1.156779 | -1.42605 | -0.30341 | -3.47482 | -1.60047 |
| 27.56944 | -7.21761 | -1.40146 | -5.40603 | -0.91849 | 0.345843 | -1.50496 | -1.13046 | 1.373997 | -3.20362 | -0.75773 | 0.115209 | -1.85127 | -1.47474 | 0.205167 | 0.454896 | 0.320463 | 0.496948 | -0.58113 | 0.004975 | 1.082168 | -2.38122 | -0.30604 | -1.40419 | -1.84232 |
| 27.58333 | -2.70998 | -1.03842 | -2.88955 | -0.01922 | 0.545047 | -2.16108 | 0.014791 | 1.510212 | -1.95507 | 0.024192 | 1.556655 | -0.90306 | -0.5187 | 0.350591 | 1.25019 | 1.024309 | 1.12418 | -0.63995 | 0.278788 | 1.557155 | -2.65481 | -0.72605 | 1.149063 | -0.7904 |
| 27.59722 | 2.984149 | 0.097915 | 1.142244 | 0.538801 | 0.742541 | -2.46583 | 1.190042 | 1.475959 | -0.48175 | 0.558003 | 2.571791 | 0.53413 | 0.813442 | 0.348437 | 1.982278 | 1.451149 | 1.680324 | -0.62881 | 0.537961 | 1.790245 | -1.41644 | -0.80568 | 2.9968 | 0.472899 |
| 27.61111 | 7.663557 | 1.198135 | 5.174531 | 0.629006 | 0.947244 | -1.96615 | 1.582124 | 0.711669 | 0.304358 | 0.873691 | 2.027647 | 1.841086 | 1.890845 | -0.00112 | 1.658148 | 1.358092 | 1.355511 | -1.06931 | 0.861588 | 0.922686 | 0.689543 | -0.48842 | 4.612189 | 1.725809 |
| 27.625 | 9.105258 | 2.005842 | 6.994123 | 0.831372 | 1.03409 | -1.6385 | 1.668742 | -0.8941 | 0.200413 | 0.947737 | 0.504085 | 2.046504 | 1.372058 | -0.56365 | 0.601018 | 0.935317 | 0.723595 | -2.14482 | 0.92186 | -0.74264 | 2.347565 | 0.271744 | 3.746377 | 1.820614 |
| 27.63889 | 6.609958 | 2.038449 | 6.625799 | 1.13255 | 1.169178 | -1.95718 | 1.330807 | -2.76064 | 0.352402 | 0.884439 | -1.28603 | 0.893764 | 0.242401 | -1.10055 | -0.59799 | 0.453266 | 0.1377 | -3.29884 | 0.954401 | -2.41487 | 3.39002 | 1.140162 | 1.915712 | 0.918643 |
| 27.65278 | 3.496818 | 1.069561 | 4.599331 | 1.102382 | 0.895607 | -1.04876 | 1.032257 | -4.72322 | 0.942508 | 0.659353 | -3.04485 | 0.099045 | -0.66535 | -1.37066 | -1.72422 | 0.154575 | -0.36663 | -4.0861 | 0.85218 | -3.73167 | 3.408834 | 1.558002 | 0.104732 | 0.46987 |
| 27.66667 | 0.442543 | 0.084827 | 2.577973 | 0.978525 | 0.135386 | 0.924836 | -0.4306 | -5.12924 | 1.655639 | 0.429259 | -3.98229 | -0.21324 | -1.60322 | -1.46377 | -2.29441 | -0.05182 | -0.63512 | -2.6188 | 0.465759 | -4.70584 | 1.854116 | 1.289402 | -1.15005 | 0.116636 |
| 27.68056 | -1.7129 | -0.59227 | 0.549883 | 0.897781 | -0.41691 | 2.933249 | -1.74824 | -4.52398 | 1.924772 | 0.33918 | -3.39681 | -0.99383 | -2.3231 | -1.2383 | -2.00998 | -0.34394 | -0.76762 | -0.63796 | 0.020222 | -5.0105 | 0.393822 | 0.561319 | -1.63988 | -0.69253 |
| 27.69444 | -4.64189 | -0.88688 | -1.78829 | 1.265711 | -0.42246 | 4.036581 | -2.152 | -2.90299 | 1.462983 | 0.376919 | -1.60858 | -1.67252 | -2.36864 | -0.73354 | -1.41436 | -0.54278 | -0.76547 | 1.54344 | -0.49387 | -3.88856 | -0.90758 | -0.1566 | -2.03021 | -1.55516 |
| 27.70833 | -8.43429 | -0.82292 | -4.1135 | 1.477013 | -0.77311 | 4.225063 | -2.15494 | -0.65157 | 1.106381 | 0.782428 | -0.07063 | -0.64664 | -1.54693 | -0.02679 | -1.1266 | -0.81214 | -0.55618 | 3.406267 | -0.79659 | -2.43595 | -2.58273 | -0.24729 | -1.94366 | -1.46381 |
| 27.72222 | -9.04838 | -0.28412 | -5.33081 | 0.718153 | -1.48585 | 3.301261 | -2.20008 | 2.064807 | 0.298015 | 0.910947 | 0.361226 | 0.350276 | -0.39188 | 0.639206 | -1.42492 | -0.84356 | -0.09932 | 4.366607 | -0.64068 | -0.9434 | -3.27336 | -0.23596 | -1.4131 | -0.46545 |
| 27.73611 | -8.75547 | 0.468361 | -5.68571 | 0.221889 | -1.05446 | 1.907796 | -1.3194 | 5.343117 | 0.679978 | 0.669128 | 0.137165 | 0.896134 | 0.357639 | 0.935054 | -1.66242 | -0.74317 | 0.21498 | 2.167477 | -0.61869 | 0.503855 | -1.99509 | -0.69673 | -0.79975 | 0.29764 |
| 27.75 | -5.54964 | 0.862012 | -4.53828 | 0.093713 | -0.5273 | 0.159777 | -0.86257 | 7.011381 | 0.832761 | 0.119764 | -0.27761 | 0.942187 | 0.535072 | 0.901784 | -1.40827 | -0.52452 | 0.26729 | 0.809168 | -0.58179 | 1.487 | -0.4574 | -1.25763 | 0.601088 | 0.720282 |
| 27.76389 | -2.41536 | 0.539222 | -1.89103 | -0.34111 | -0.33195 | -1.73087 | -0.62433 | 5.519401 | 0.724338 | -0.31482 | -1.10575 | 0.26487 | 0.357343 | 0.752727 | -0.85353 | -0.18757 | 0.195022 | -0.56545 | -0.35936 | 1.700115 | 0.704683 | -1.82788 | 1.408775 | 0.450156 |
| 27.77778 | -0.66247 | -0.21886 | 0.197815 | -1.14746 | -0.78119 | -3.99219 | -0.88688 | 2.576201 | -0.17681 | -0.49494 | -1.59843 | -0.74303 | -0.25726 | 0.514946 | -0.26576 | 0.13615 | 0.1568 | -1.48043 | -0.12882 | 1.193795 | 0.464934 | -0.94562 | 1.933726 | -0.55882 |
| 27.79167 | -0.40434 | -1.32748 | 1.247 | -2.09014 | -1.44299 | -6.47884 | -1.24311 | 0.303325 | -1.07873 | -0.52828 | -1.44756 | -1.93255 | -1.12853 | 0.21118 | -0.21811 | -0.0938 | -0.1824 | -1.75298 | 0.10479 | 0.203223 | -0.88909 | -0.10669 | 1.250427 | -1.44041 |
| 27.80556 | -1.24935 | -2.27075 | 1.074722 | -2.36592 | -1.88403 | -6.22484 | -1.20979 | -1.5795 | -2.1383 | -0.69891 | -0.79229 | -2.77558 | -1.63772 | -0.09932 | -0.59068 | -0.79292 | -0.94102 | -2.36555 | -0.11046 | -0.83692 | -2.32249 | 0.076942 | -0.10862 | -2.19796 |
| 27.81944 | -3.07963 | -2.81878 | 0.257505 | -2.59045 | -2.06838 | -5.1339 | -1.02182 | -3.28433 | -2.83157 | -0.91475 | -0.21974 | -2.21768 | -1.98121 | -0.44018 | -1.49352 | -1.65306 | -2.10239 | -3.48554 | -0.7157 | -1.63678 | -3.21255 | -0.31355 | -2.19664 | -2.4869 |
| 27.83333 | -5.8318 | -2.73801 | -1.08615 | -3.19971 | -2.28329 | -4.91894 | -0.91398 | -5.12508 | -2.60818 | -1.0952 | -0.0886 | -1.24244 | -2.53095 | -0.75201 | -3.11749 | -2.23372 | -2.34642 | -5.25825 | -1.05956 | -2.33268 | -3.08893 | -0.78409 | -2.87551 | -2.13739 |
| 27.84722 | -6.86261 | -2.04661 | -2.56494 | -3.77695 | -2.39363 | -4.47172 | -1.03907 | -6.61771 | -2.11466 | -1.25458 | -0.39724 | -0.48771 | -2.79184 | -1.10703 | -3.8614 | -2.43312 | -2.09973 | -7.38248 | -0.97849 | -2.83852 | -1.67757 | -1.11913 | -4.52965 | -1.55278 |
| 27.86111 | -5.90817 | -1.10012 | -3.60869 | -4.10384 | -2.0285 | -2.89038 | -1.13507 | -7.76932 | -1.74703 | -1.22996 | -0.89268 | -0.23349 | -1.96225 | -1.36326 | -3.64331 | -2.87575 | -2.31474 | -6.20454 | -0.69302 | -3.13602 | -0.65582 | -1.26955 | -4.62951 | -1.08835 |
| 27.875 | -3.7682 | -0.01696 | -3.67304 | -3.82673 | -1.2069 | -0.43942 | -1.08859 | -8.12091 | -1.61324 | -0.86521 | -1.05305 | -0.31902 | -1.89619 | -1.23326 | -3.57246 | -3.06534 | -2.28766 | -4.91392 | -0.48127 | -3.10493 | -0.21527 | -1.06942 | -3.42613 | -0.94936 |
| 27.88889 | -1.08609 | 0.708831 | -2.71993 | -2.86234 | -0.29612 | 2.124132 | -0.81666 | -5.9619 | -1.74265 | -0.35979 | -1.12107 | -0.5285 | -1.83324 | -0.87455 | -3.09465 | -2.4171 | -1.7224 | -4.35696 | -0.31472 | -2.7059 | -0.22529 | -0.65254 | -1.39737 | -0.84224 |
| 27.90278 | 1.492299 | 0.871051 | -1.03908 | -1.48297 | 0.397691 | 4.638933 | -0.43254 | -3.64475 | -1.5714 | -0.06052 | -1.06857 | -0.86067 | -1.52527 | -0.47583 | -2.2245 | -1.26434 | -0.79368 | -3.02782 | -0.10729 | -2.6524 | -0.3783 | -0.19901 | 0.942736 | -0.48015 |
| 27.91667 | 3.67671 | 0.706416 | 0.663568 | -0.2077 | 0.714338 | 5.424707 | -0.05707 | -1.60053 | -0.98617 | 0.048495 | -0.87004 | -0.76763 | -1.22846 | -0.31216 | -1.07026 | -0.41343 | -0.25319 | -1.17006 | 0.062512 | -2.51196 | -0.41786 | 0.082062 | 2.668844 | 0.086508 |
| 27.93056 | 5.026221 | 0.480553 | 1.575703 | 0.63995 | 0.639264 | 5.509958 | 0.182019 | -0.49874 | -0.51771 | 0.075918 | -0.55602 | -0.42296 | -0.97414 | -0.39647 | -0.17347 | -0.04407 | -0.09952 | 0.519398 | 0.100101 | -1.94595 | -0.16202 | 0.116193 | 3.263723 | 0.490275 |
| 27.94444 | 3.992614 | -0.22834 | 0.748177 | 0.194096 | -0.23281 | 4.536574 | -0.23008 | -0.36413 | -0.7834 | -0.21128 | -1.1216 | -0.58897 | -1.21392 | -0.89499 | -0.62148 | -0.43777 | -0.3928 | 1.229992 | -0.1194 | -1.68813 | -0.37212 | -0.35069 | 2.565697 | -0.10992 |
| 27.95833 | -0.53071 | -1.89528 | -1.8476 | -1.87338 | -2.43996 | 0.446083 | -2.02724 | -1.24237 | -2.46501 | -1.08903 | -3.22524 | -2.28725 | -2.61986 | -1.93947 | -2.77145 | -1.82745 | -1.36982 | -0.41843 | -0.65913 | -2.63033 | -1.9435 | -1.52677 | -0.63292 | -2.4482 |
| 27.97222 | -6.17792 | -3.87585 | -4.82569 | -4.24299 | -5.47931 | -4.10542 | -4.98322 | -2.17469 | -5.04356 | -2.1376 | -5.36384 | -5.56749 | -5.15645 | -3.44794 | -5.63963 | -3.21574 | -2.43821 | -3.22253 | -1.04986 | -3.67968 | -4.17844 | -2.90872 | -4.41488 | -5.72757 |
| 27.98611 | -8.55413 | -4.20467 | -6.30986 | -4.49637 | -6.6446 | -7.25312 | -5.74906 | -1.68298 | -4.68552 | -2.09713 | -4.59992 | -5.84681 | -5.69223 | -3.03478 | -6.15513 | -2.97833 | -2.51271 | -3.91052 | -0.86614 | -3.79932 | -5.149 | -3.1645 | -5.15427 | -6.37637 |
| 28 | -5.7693 | -3.1861 | -5.59896 | -2.72963 | -5.26732 | -6.13876 | -3.8878 | -0.48559 | -2.45293 | -1.37034 | -2.64708 | -3.995 | -3.47678 | -1.6765 | -3.78952 | -1.92336 | -2.18017 | -3.41808 | -0.66175 | -2.65612 | -4.01127 | -1.87632 | -3.2369 | -3.74152 |
| 28.01389 | -3.05768 | -2.02171 | -3.85597 | -1.39677 | -4.24937 | -5.87056 | -2.75376 | -0.01055 | -0.98782 | -0.78192 | -1.03187 | -2.79212 | -1.32115 | -0.61802 | -0.99721 | -1.09955 | -1.68875 | -2.82201 | -0.57052 | -1.46538 | -2.67269 | -0.79684 | -1.69332 | -1.06131 |
| 28.02778 | -1.05263 | -0.96954 | -2.56495 | -0.90463 | -3.47248 | -5.16055 | -2.08277 | 0.012891 | -0.07365 | -0.26077 | 0.14901 | -1.83626 | -0.14096 | 0.141952 | 0.749113 | -0.57142 | -1.67025 | -2.19249 | -0.29647 | -0.38503 | -1.67773 | -0.23097 | -0.91895 | 0.748392 |
| 28.04167 | -0.27902 | -0.28571 | -1.34247 | -0.40258 | -2.07741 | -3.93149 | -1.26319 | -0.11343 | 0.604728 | 0.155753 | 0.744554 | -0.75642 | 0.400583 | 0.54181 | 0.470607 | -0.2173 | -1.03386 | -1.58066 | -0.02824 | 0.12283 | -0.93375 | 0.00904 | -0.62441 | 1.428645 |
| 28.05556 | 0.028468 | -0.26243 | -0.462 | 0.247849 | -0.10156 | -2.21633 | 0.298246 | -0.1626 | 0.961671 | 0.551221 | 0.761967 | 0.204713 | 0.616145 | 0.576182 | -0.29548 | 0.017773 | -0.36924 | -1.04401 | 0.135481 | 0.445187 | -0.36708 | 0.124778 | -0.24482 | 1.566248 |
| 28.06944 | 0.150145 | -0.93829 | -0.19083 | 0.834683 | 1.44028 | -0.89584 | 1.865984 | 0.218052 | 0.986704 | 0.858867 | 1.042579 | 1.114989 | 0.843455 | 0.3244 | -0.32081 | 0.187875 | -0.05901 | -0.39463 | 0.139042 | 1.190194 | 0.1208 | 0.299054 | 0.443782 | 2.098556 |
| 28.08333 | 0.137533 | -0.31634 | -0.14873 | 1.20003 | 2.02966 | -0.19139 | 2.565051 | 0.638754 | 0.808797 | 1.138098 | 1.700315 | 1.916867 | 1.307644 | 0.028223 | 0.889336 | 0.397201 | -0.01516 | 0.207712 | -0.07821 | 2.110724 | 0.526167 | 0.497142 | 0.881381 | 2.609334 |
| 28.09722 | 0.0873 | 0.725839 | -0.14171 | 1.137917 | 2.084035 | -0.56556 | 2.231493 | 0.574245 | 0.644464 | 1.361356 | 2.007745 | 2.321638 | 1.421774 | -0.19229 | 1.784779 | 0.642403 | -0.00914 | 0.479294 | -0.00469 | 2.44477 | 0.672071 | 0.453272 | 0.721592 | 2.41703 |
| 28.11111 | -0.0015 | 0.951464 | -0.32269 | 0.710792 | 1.775107 | -0.88725 | 1.932343 | 0.324147 | 0.572135 | 1.135684 | 1.740147 | 2.425365 | 1.213485 | -0.0757 | 1.603062 | 0.741323 | -0.00907 | 0.537814 | 0.219818 | 1.832102 | 0.612287 | 0.325472 | 0.279996 | 1.884182 |
| 28.125 | -0.07927 | 1.091202 | -0.36743 | 0.511047 | 1.345459 | -0.8154 | 1.565184 | 0.062594 | 0.785711 | 0.635417 | 1.005086 | 1.752589 | 0.784128 | 0.161321 | 0.65232 | 0.747899 | -0.00228 | 0.432465 | 0.295762 | 0.730776 | 0.486873 | 0.083667 | -0.06639 | 0.793891 |
| 28.13889 | 0.019161 | 1.294257 | -0.22478 | 0.393694 | 0.975809 | -0.84361 | 1.159675 | -0.16226 | 1.148471 | 0.2704 | 0.259244 | 1.25509 | 0.362191 | 0.402726 | -0.47611 | 0.566857 | 0.059187 | 0.497868 | 0.327731 | -0.417 | 0.347346 | -0.05863 | -0.13685 | -0.28043 |
| 28.15278 | 0.440985 | 1.298471 | -0.01999 | 0.37619 | 0.904838 | -0.56752 | 0.798106 | -0.33532 | 1.41303 | 0.129532 | -0.35864 | 0.835772 | -0.0474 | 0.469591 | -1.02615 | 0.176057 | 0.079428 | 0.591936 | 0.511853 | -1.00541 | 0.532479 | 0.075416 | 0.144653 | -1.00416 |
| 28.16667 | 1.130152 | 1.036665 | 0.13928 | 0.517393 | 0.575374 | -0.21763 | 0.466305 | -0.3448 | 1.412952 | 0.255495 | -0.64323 | 0.392018 | -0.35349 | 0.318903 | -0.64131 | -0.14004 | 0.083615 | 0.821804 | 0.545671 | -0.55046 | 1.10398 | 0.429351 | 0.862102 | -0.91561 |
| 28.18056 | 1.927998 | 0.603172 | 0.270894 | 0.685056 | 0.312609 | 0.102887 | 0.290722 | -0.08134 | 1.301281 | 0.637352 | -0.25354 | 0.11573 | -0.44008 | 0.089963 | 0.439556 | -0.22827 | 0.184621 | 1.33109 | 0.529079 | 0.741368 | 1.641444 | 0.750647 | 1.7464 | 0.081994 |
| 28.19444 | 2.634189 | 0.209956 | 0.420325 | 0.782636 | 0.138443 | 0.329639 | 0.263546 | 0.337375 | 1.226819 | 0.978691 | 0.466888 | 0.026368 | -0.10342 | -0.05156 | 1.557688 | -0.09837 | 0.309421 | 1.871534 | 0.387483 | 1.821475 | 1.881978 | 1.146713 | 2.130398 | 1.033893 |
| 28.20833 | 3.181719 | -0.08706 | 0.605604 | 0.895622 | 0.26636 | 0.514879 | 0.452699 | 0.639265 | 1.220088 | 0.992576 | 0.992441 | 0.178148 | 0.585376 | -0.0153 | 1.845962 | 0.127651 | 0.409243 | 2.40805 | 0.190516 | 2.321235 | 1.955416 | 1.309157 | 2.140954 | 1.283357 |
| 28.22222 | 3.58775 | -0.01416 | 0.804662 | 1.049061 | 0.686353 | 0.577347 | 0.750238 | 0.727394 | 1.118216 | 0.718467 | 0.800568 | 0.438112 | 1.111874 | 0.095752 | 1.045597 | 0.405076 | 0.452917 | 2.840576 | -0.0379 | 1.881192 | 1.662826 | 1.29318 | 2.329893 | 0.706114 |
| 28.23611 | 3.152005 | 0.280679 | 0.809789 | 0.899715 | 1.035161 | 0.477257 | 0.877511 | 0.546738 | 0.837098 | 0.298883 | 0.263481 | 0.595656 | 0.858098 | 0.229423 | -0.22932 | 0.588304 | 0.441623 | 2.157683 | -0.13413 | 1.0012 | 1.177677 | 1.09228 | 2.09442 | -0.05625 |
| 28.25 | 2.302452 | 0.706575 | 0.801235 | 0.635021 | 1.023922 | 0.412487 | 0.720765 | 0.446273 | 0.58535 | 0.121647 | -0.30287 | 0.677655 | 0.291198 | 0.265022 | -1.10928 | 0.518299 | 0.383393 | 1.871341 | -0.05607 | 0.438903 | 0.897784 | 0.879508 | 1.932951 | -0.50857 |
| 28.26389 | 1.406812 | 1.149549 | 0.829451 | 0.489489 | 0.901414 | 0.384802 | 0.622225 | 0.476108 | 0.432134 | 0.271976 | -0.52432 | 0.607969 | -0.19824 | 0.137379 | -1.13212 | 0.268345 | 0.32186 | 1.874799 | 0.181862 | 0.360867 | 0.801745 | 0.724551 | 2.073731 | -0.46853 |

| | | | | | | | | | | | | | | | | | | | | | | | | |
|---|---|---|---|---|---|---|---|---|---|---|---|---|---|---|---|---|---|---|---|---|---|---|---|---|
| 28.27778 | 0.610322 | 1.375972 | 0.824797 | 0.465519 | 0.683833 | 0.36613 | 0.591466 | 0.51637 | 0.319959 | 0.453694 | -0.14446 | 0.41664 | -0.41731 | 0.020529 | -0.28789 | 0.039038 | 0.262773 | 1.590719 | 0.428748 | 0.70332 | 0.802621 | 0.582536 | 2.188403 | 0.050749 |
| 28.29167 | 0.052409 | 1.341975 | 0.994238 | 0.483454 | 0.492744 | 0.531565 | 0.574301 | 0.383041 | 0.332089 | 0.463422 | 0.432041 | 0.285831 | -0.18552 | 0.023714 | 0.752393 | -0.12457 | 0.133267 | 1.352583 | 0.486361 | 1.139957 | 0.819942 | 0.476073 | 2.009532 | 0.708883 |
| 28.30556 | -0.33104 | 0.990365 | 1.080898 | 0.473988 | 0.382164 | 0.671751 | 0.486249 | 0.138126 | 0.416101 | 0.339392 | 0.842448 | 0.333265 | 0.273393 | 0.103914 | 1.398966 | -0.16691 | 0.086089 | 0.990422 | 0.39705 | 1.261436 | 0.769192 | 0.379605 | 1.364032 | 1.075293 |
| 28.31944 | -0.83252 | 0.50069 | 1.100092 | 0.423706 | 0.350248 | 0.764585 | 0.361568 | -0.02062 | 0.490728 | 0.199798 | 0.690646 | 0.471295 | 0.904565 | 0.225754 | 0.806073 | 0.022992 | 0.419738 | 0.566788 | 0.351455 | 0.90939 | 0.606601 | 0.28236 | 0.376526 | 0.825866 |
| 28.33333 | -1.47205 | 0.185568 | 1.004 | 0.363454 | 0.437859 | 0.816365 | 0.305068 | -0.00921 | 0.562509 | 0.124338 | 0.143546 | 0.626834 | 1.318868 | 0.204637 | -0.38723 | 0.37888 | 1.093473 | 0.259972 | 0.388134 | 0.272016 | 0.402881 | 0.154054 | -0.5705 | 0.093902 |
| 28.34722 | -1.93531 | 0.341 | 0.941022 | 0.369272 | 0.677975 | 0.893301 | 0.353627 | 0.133685 | 0.624218 | 0.117698 | -0.40291 | 0.836195 | 0.86896 | -0.11164 | -1.34718 | 0.706311 | 1.236973 | 0.188664 | 0.437612 | -0.30746 | 0.26205 | 0.006287 | -0.94988 | -0.38399 |
| 28.36111 | -2.16679 | 0.770493 | 0.903903 | 0.405192 | 0.946662 | 0.870288 | 0.475713 | 0.294712 | 0.552845 | 0.229578 | -0.6113 | 0.880001 | 0.080209 | -0.40691 | -1.40059 | 0.781174 | 1.107472 | 0.285529 | 0.411509 | -0.43995 | 0.160349 | -0.11646 | -0.42849 | -0.24456 |
| 28.375 | -2.34639 | 1.133467 | 0.708508 | 0.337158 | 0.956762 | 0.643676 | 0.553451 | 0.355687 | 0.352169 | 0.38831 | -0.37166 | 0.662615 | -0.66841 | -0.37791 | -0.44512 | 0.719475 | 0.948002 | 0.403018 | 0.433522 | 0.041262 | -0.11568 | -0.13994 | 0.513219 | 0.268296 |
| 28.38889 | -2.1362 | 1.399186 | 0.525998 | 0.27064 | 0.855096 | 0.424137 | 0.692713 | 0.359845 | 0.298102 | 0.493915 | 0.228198 | 0.410848 | -0.63547 | -0.06424 | 1.028151 | 0.603015 | 0.813038 | 0.503521 | 0.700057 | 0.392574 | -0.47845 | -0.50883 | 1.308921 | 0.706767 |
| 28.40278 | -1.28287 | 1.557184 | 0.554729 | 0.367345 | 0.783624 | 0.447673 | 0.908337 | 0.404276 | 0.496245 | 0.558436 | 0.809418 | 0.33831 | -0.03766 | 0.236105 | 2.011535 | 0.56181 | 0.689522 | 0.676037 | 0.903177 | 0.39204 | 0.05385 | 0.183612 | 1.681092 | 1.115357 |
| 28.41667 | -0.36527 | 1.124777 | 0.665469 | 0.499934 | 0.518901 | 0.608267 | 0.921705 | 0.415824 | 0.683599 | 0.611786 | 0.784249 | 0.285049 | 0.533095 | 0.322764 | 1.588946 | 0.40682 | 0.541905 | 0.739542 | 0.905073 | 0.544854 | 0.246501 | 0.587245 | 1.425196 | 1.384099 |
| 28.43056 | 0.469446 | 0.450706 | 0.77707 | 0.600654 | 0.070039 | 0.733674 | 0.627531 | 0.375271 | 0.709629 | 0.667122 | 0.309951 | 0.381713 | 0.997868 | 0.195565 | 0.29819 | 0.295267 | 0.466828 | 0.713426 | 0.621389 | 0.758635 | 0.467074 | 0.685484 | 0.985026 | 0.928382 |
| 28.44444 | 2.219283 | 0.279232 | 1.30407 | 1.499804 | 0.256316 | 1.191787 | 0.496604 | 0.389447 | 1.144903 | 1.011703 | -0.13232 | 1.282518 | 1.61543 | 0.358023 | -0.66857 | 0.658876 | 0.751603 | 1.263355 | 0.455401 | 0.808209 | 1.265035 | 1.167007 | 1.504088 | 0.760189 |
| 28.45833 | 7.746285 | 1.620421 | 2.734846 | 3.841145 | 2.059744 | 2.510501 | 1.633537 | 0.819456 | 2.714537 | 1.836958 | 0.354841 | 3.400351 | 2.774798 | 1.410229 | 0.116455 | 1.81502 | 1.558143 | 3.349482 | 0.444015 | 1.12781 | 2.738356 | 2.537504 | 3.628122 | 1.881909 |
| 28.47222 | 13.51187 | 4.171704 | 4.734983 | 5.208681 | 4.65428 | 5.004855 | 3.786101 | 2.222821 | 4.177674 | 2.983736 | 2.207074 | 5.265921 | 3.881749 | 2.96692 | 2.139606 | 2.921515 | 2.412649 | 6.58866 | 0.428156 | 2.296264 | 4.446973 | 3.224792 | 5.596504 | 3.308827 |
| 28.48611 | 14.71082 | 4.76732 | 4.226861 | 5.750027 | 4.606892 | 7.279901 | 4.763978 | 3.80782 | 3.892525 | 3.453982 | 3.833134 | 5.117429 | 3.481065 | 3.430783 | 4.887112 | 2.851697 | 2.192144 | 6.417474 | 0.570651 | 3.717035 | 5.243539 | 2.739026 | 3.081189 | 3.384521 |
| 28.5 | 11.78072 | 4.048923 | 3.928325 | 4.91882 | 3.591638 | 7.398434 | 3.842747 | 5.138624 | 3.676291 | 3.027084 | 4.412125 | 5.628352 | 2.875002 | 2.187348 | 5.062726 | 1.964172 | 1.695687 | 4.881537 | 0.763507 | 5.021788 | 4.489651 | 2.2268 | 1.188283 | 3.881746 |
| 28.51389 | 8.49035 | 3.087573 | 3.346168 | 2.887402 | 2.689924 | 6.543659 | 2.620049 | 6.368428 | 2.990652 | 2.040084 | 2.940809 | 4.188562 | 2.591942 | 1.20435 | 3.474563 | 1.529468 | 1.62756 | 3.118571 | 0.870718 | 4.907737 | 2.711632 | 1.545882 | 0.073363 | 3.32539 |
| 28.52778 | 4.882999 | 2.061672 | 3.070473 | 1.732976 | 1.641203 | 5.588533 | 1.164184 | 6.08654 | 1.990033 | 1.135444 | 1.199825 | 2.227782 | 1.283215 | 0.856775 | 2.626868 | 1.221479 | 1.35646 | 1.594245 | 0.492718 | 4.103515 | 0.708556 | 0.583335 | -1.15546 | 0.629487 |
| 28.54167 | 2.072509 | 1.038539 | 2.087623 | 0.399405 | 0.47144 | 4.020413 | -0.03736 | 4.292567 | 1.338497 | 0.262586 | -0.57733 | 0.680582 | 0.07764 | 0.271246 | 1.724622 | 0.843512 | 0.755331 | 0.221539 | -0.03489 | 3.398157 | -1.0019 | -0.67015 | -2.08869 | -1.82863 |
| 28.55556 | -0.47184 | -0.00983 | 0.333367 | -0.87961 | -0.30224 | 3.566833 | -0.97347 | 2.948455 | 0.966665 | -0.47492 | -1.93281 | -0.61469 | -1.17394 | 0.229175 | 0.688698 | 0.533325 | 0.006756 | -1.14639 | -0.102 | 2.551271 | -2.16271 | -1.53974 | -3.34919 | -3.73502 |
| 28.56944 | -2.27482 | -0.69325 | -0.79461 | -2.08012 | -0.43737 | 3.019887 | -1.22544 | 3.321016 | 0.783405 | -0.85552 | -1.22543 | -0.64633 | -2.48445 | 0.848113 | -0.0744 | 0.523795 | -0.55615 | -1.42874 | 0.442942 | 2.630556 | -2.19786 | -1.55794 | -3.55403 | -4.41824 |
| 28.58333 | -3.37834 | -1.20924 | -0.47514 | -1.33591 | -0.13679 | 2.480681 | -0.50326 | 3.533762 | 0.650681 | -0.59099 | 0.001374 | -0.03215 | -2.25999 | 1.160282 | -0.22038 | 0.605255 | -0.30912 | -0.82227 | 0.67626 | 3.591117 | -1.75276 | -0.68927 | -1.08385 | -3.88459 |
| 28.59722 | -4.79942 | -1.86248 | 0.203176 | -0.65994 | 0.304524 | 0.983749 | 0.599149 | 3.876441 | 0.904189 | 0.463652 | 1.478096 | 0.667104 | -0.06991 | 1.367503 | 0.896456 | 0.590603 | 0.296667 | -0.04206 | 0.792594 | 3.969386 | -0.92134 | 0.500668 | -0.34372 | -1.41871 |
| 28.61111 | -7.44334 | -1.22053 | 0.211744 | -0.45771 | 0.638123 | -0.52984 | 1.259406 | 4.010866 | 1.106118 | 1.152938 | 2.714741 | 1.103372 | 1.440414 | 0.963638 | 2.413357 | 0.413435 | 0.418632 | 0.230996 | 0.933607 | 3.730619 | -0.17203 | 1.66342 | -0.01485 | 1.410776 |
| 28.625 | -8.50591 | 0.328229 | -0.54781 | -0.62406 | 0.743868 | -2.16664 | 1.190732 | 2.875941 | 1.049277 | 1.038711 | 2.645363 | 1.368886 | 2.407705 | 0.285997 | 3.069113 | 0.673599 | 0.497557 | -0.15826 | 0.80089 | 3.411488 | 0.940165 | 2.698177 | 0.575113 | 3.529852 |
| 28.63889 | -9.14647 | 1.441456 | -1.608 | -1.41237 | 0.835402 | -3.20053 | 0.955741 | 0.784271 | 0.679374 | 0.677292 | 1.695447 | 0.815634 | 2.461716 | 0.170419 | 1.921354 | 1.402993 | 0.441575 | -0.6265 | 0.723616 | 2.300038 | 1.646128 | 3.01753 | 1.005524 | 4.484798 |
| 28.65278 | -6.28179 | 2.298722 | -1.7025 | -2.44132 | 0.875962 | -3.84793 | 0.867626 | -1.45178 | 0.740028 | 0.348989 | 0.530042 | -0.32862 | 1.659462 | -0.16784 | 0.313236 | 1.782544 | 0.15193 | -1.17906 | 0.438349 | 0.701377 | 1.927134 | 2.646693 | 1.909505 | 3.754082 |
| 28.66667 | -1.61196 | 2.432594 | -0.9948 | -2.58319 | 1.036746 | -4.4253 | 1.255092 | -3.41874 | 0.386358 | 0.211085 | -1.10917 | -1.0978 | 0.826156 | -0.77092 | -1.26051 | 1.43489 | -0.40022 | -1.17039 | 0.158166 | -1.22435 | 1.76644 | 1.639908 | 1.807526 | 1.878369 |
| 28.68056 | 2.228227 | 1.41816 | 0.273849 | -1.98617 | 1.128893 | -6.15765 | 1.593786 | -4.63303 | -0.09696 | 0.329228 | -2.42386 | -0.99677 | -0.0663 | -1.03169 | -2.59643 | 0.797366 | -0.96493 | -1.21112 | 0.015306 | -3.85386 | 1.444239 | 0.712968 | 1.348138 | 0.146093 |
| 28.69444 | 4.857793 | 0.279685 | 1.75075 | -0.785 | 0.954123 | -5.15849 | 1.649326 | -4.94071 | 0.152907 | 0.470001 | -2.05587 | -0.80791 | -1.45301 | -0.64764 | -2.91487 | 0.329855 | -0.96719 | -1.46702 | -0.07603 | -6.37257 | 0.934583 | -0.21971 | 0.222786 | -1.23001 |
| 28.70833 | 5.578253 | 0.185297 | 3.916829 | 0.915789 | 0.647329 | -3.39853 | 1.572824 | -4.8392 | 0.389479 | 0.710657 | -1.45076 | -0.76747 | -2.9415 | -0.33425 | -2.52492 | 0.013033 | -0.72693 | -1.06009 | -0.10779 | -7.26925 | 0.035184 | -1.64225 | -0.33834 | -2.46057 |
| 28.72222 | 4.324428 | 0.21665 | 5.20875 | 2.440214 | 0.429735 | -1.2678 | 1.250312 | -4.16391 | 0.52326 | 0.950188 | -0.52063 | -0.56273 | -2.8308 | -0.00829 | -1.84943 | -0.15997 | -0.12024 | 0.164598 | 0.207268 | -7.14435 | -0.62304 | -2.0309 | -0.36673 | -3.01607 |
| 28.73611 | 3.012098 | -0.03589 | 4.948264 | 3.531192 | 0.148704 | 0.676401 | 0.507 | -3.10194 | 0.414889 | 0.890277 | 0.136824 | -0.1546 | -1.4214 | -0.42873 | -1.47634 | -0.24667 | 0.354215 | 1.159692 | 0.397981 | -6.38434 | -0.98787 | -2.15591 | -0.60412 | -2.04944 |
| 28.75 | 1.200266 | -0.17891 | 3.388284 | 2.653389 | -0.2025 | 2.605501 | -0.26043 | -1.83823 | -0.21113 | 0.581253 | -0.11362 | 0.131328 | -0.22914 | -0.81654 | -1.4531 | -0.402 | 0.413383 | 1.68129 | 0.40394 | -4.83006 | -1.35151 | -1.87298 | -0.43481 | -0.27335 |
| 28.76389 | -1.61641 | -0.13661 | 0.779784 | 1.065425 | -0.5059 | 3.814078 | -0.72576 | -0.47034 | -0.86457 | 0.169772 | -1.06069 | -0.13079 | 0.718243 | -0.72657 | -2.43662 | -0.5445 | 0.204621 | 2.047023 | 0.273743 | -3.26458 | -1.52266 | -0.81232 | 0.097088 | 0.797607 |
| 28.77778 | -5.24465 | 0.094945 | -1.8086 | 0.084495 | -0.49972 | 3.972625 | -0.90662 | 0.684327 | -1.32479 | -0.2765 | -2.20911 | -0.63323 | 0.727485 | -0.65418 | -3.50206 | -0.44724 | -0.12186 | 2.722032 | 0.086695 | -1.81557 | -1.43407 | -0.09749 | 0.449759 | 1.498087 |
| 28.79167 | -7.8803 | 0.200873 | -3.85943 | -0.64796 | -0.61351 | 2.122635 | -0.9604 | 1.566472 | -1.90091 | -0.74824 | -2.9499 | -1.08784 | -0.17398 | -1.05863 | -4.58871 | -0.46581 | -0.37087 | 2.645442 | -0.31193 | -0.83769 | -1.01934 | 0.018244 | 0.124911 | 1.353027 |
| 28.80556 | -8.53191 | -0.0412 | -4.94112 | -1.45908 | -1.00947 | 0.057935 | -0.7848 | 1.862624 | -2.11372 | -1.23295 | -3.13546 | -1.70356 | -1.38425 | -1.31619 | -5.16528 | -0.65957 | -0.52348 | 1.58244 | -0.53745 | -0.59711 | -0.72898 | 0.150398 | 0.054895 | 0.506824 |
| 28.81944 | -7.34003 | -0.43949 | -5.14458 | -2.28173 | -1.26879 | -1.958 | -0.6843 | 1.718803 | -2.19019 | -1.67268 | -2.69685 | -2.21988 | -2.54878 | -1.02273 | -4.7469 | -0.86836 | -0.59536 | -0.28546 | -0.50705 | -0.89518 | -0.63147 | -0.05436 | 0.069275 | -0.66185 |
| 28.83333 | -5.71457 | -1.02564 | -4.11432 | -2.78207 | -1.70959 | -3.78771 | -0.67017 | 1.5194 | -2.05135 | -1.80816 | -1.74261 | -2.52757 | -2.87775 | -0.75009 | -3.82424 | -1.03582 | -0.4883 | -2.17728 | -0.8095 | -1.12535 | -0.64273 | -0.05043 | -0.2158 | -2.08829 |
| 28.84722 | -4.45422 | -1.63321 | -2.89446 | -3.30644 | -1.80355 | -4.67438 | -0.98296 | 0.62995 | -1.47416 | -1.28776 | -0.79801 | -2.10982 | -2.25358 | -0.48986 | -2.63783 | -1.46461 | -0.55716 | -4.13426 | -0.89375 | -1.22799 | -0.81615 | -0.24055 | -0.93431 | -3.56403 |
| 28.86111 | -3.71743 | -1.88058 | -1.99742 | -3.39177 | -1.54235 | -4.8384 | -1.32984 | -0.72998 | -0.87044 | -0.63331 | -0.29607 | -1.1232 | -1.27244 | -0.50401 | -1.59221 | -1.91938 | -0.58314 | -5.24027 | -0.755 | -1.24248 | -0.93841 | -0.62028 | -1.24623 | -4.51795 |
| 28.875 | -2.58012 | -1.44579 | -1.22345 | -2.89234 | -1.2433 | -4.96205 | -1.54164 | -1.91393 | -0.64233 | -0.31675 | -0.18476 | -0.30855 | -0.32133 | -0.52573 | -0.73297 | -1.85369 | -0.55004 | -4.20994 | -0.41425 | -1.03795 | -0.76518 | -0.54803 | -1.4963 | -3.46589 |
| 28.88889 | -0.7826 | -1.03952 | -0.54439 | -2.37839 | -0.92802 | -5.49574 | -1.90783 | -2.62269 | -0.75279 | -0.25375 | -0.45135 | 0.275652 | 0.41007 | -0.64256 | -0.33411 | -1.72198 | -0.76894 | -3.10181 | -0.12409 | -0.78064 | -0.39481 | -0.02812 | -1.89752 | -1.8103 |
| 28.90278 | 1.317503 | -0.53714 | 0.20648 | -1.73769 | -0.50693 | -4.84033 | -1.8832 | -2.86906 | -0.77815 | -0.22267 | -0.88366 | 0.503473 | 0.580937 | -0.76964 | -0.18474 | -1.5981 | -0.91996 | -3.09848 | -0.01039 | -0.39389 | -0.12685 | 0.344914 | -1.92749 | -0.15882 |
| 28.91667 | 2.931324 | -0.15624 | 1.043456 | -1.23722 | -0.33153 | -2.99548 | -1.72603 | -2.75722 | -0.65375 | -0.1516 | -0.91229 | 0.431442 | 0.364018 | -0.7187 | 0.238913 | -1.35035 | -0.70441 | -3.45866 | 0.013528 | 0.21383 | -0.02888 | 0.204171 | -1.78075 | 0.924825 |
| 28.93056 | 3.773627 | -0.00646 | 1.640971 | -0.99242 | -0.53618 | -1.55861 | -1.56225 | -2.73086 | -0.43042 | -0.09255 | -0.69367 | -0.01448 | -0.10124 | -0.35054 | 0.776584 | -0.83279 | -0.43764 | -3.0485 | -0.04027 | 0.664888 | -0.00881 | -0.31683 | -1.65938 | 0.987695 |
| 28.94444 | 2.468175 | -0.53276 | 1.066505 | -1.53025 | -1.45525 | -1.95106 | -1.62474 | -3.23437 | -0.64632 | -0.32527 | -1.10046 | -1.06119 | -1.12013 | -0.37733 | -0.09971 | -0.96991 | -0.61427 | -2.82343 | -0.35869 | 0.184712 | -0.42848 | -1.01491 | -2.64821 | -0.68009 |
| 28.95833 | -1.94576 | -2.16539 | -1.50698 | -3.26294 | -3.38508 | -4.72191 | -2.55263 | -5.41046 | -1.73623 | -1.0157 | -2.98538 | -3.59444 | -3.26542 | -1.57268 | -3.4239 | -2.35494 | -1.5487 | -3.92631 | -1.15051 | -1.40723 | -1.59715 | -2.57459 | -5.96359 | -4.02241 |
| 28.97222 | -8.13422 | -3.47584 | -4.56595 | -4.60692 | -6.02479 | -8.96523 | -2.85165 | -7.49229 | -2.80712 | -1.93255 | -5.84713 | -6.98395 | -6.05718 | -3.05164 | -8.70951 | -3.77751 | -2.54518 | -5.43339 | -1.95064 | -3.40943 | -2.69443 | -3.76475 | -8.63467 | -7.13507 |
| 28.98611 | -9.31474 | -2.66609 | -6.07839 | -4.10695 | -5.43552 | -10.2214 | -1.78926 | -4.98601 | -2.20053 | -2.25528 | -6.64811 | -7.23545 | -5.90716 | -2.94189 | -9.73472 | -3.2034 | -2.31679 | -4.01878 | -1.97536 | -4.56386 | -3.1686 | -2.17424 | -7.61023 | -6.48647 |
| 29 | -4.63042 | -1.1518 | -5.57374 | -2.86998 | -4.16233 | -7.42558 | -0.79757 | -2.44992 | -0.83154 | -1.73003 | -4.68333 | -5.31145 | -4.19006 | -1.691 | -7.0952 | -1.83336 | -1.61916 | -1.96043 | -1.41704 | -3.1755 | -3.29926 | -1.08078 | -4.3924 | -3.80954 |
| 29.01389 | -1.99928 | -0.02375 | -4.31993 | -1.60133 | -3.15566 | -5.48657 | -0.73319 | -0.29028 | -0.03929 | -0.99714 | -1.59602 | -4.06189 | -3.03879 | -0.73443 | -4.14567 | -1.07432 | -1.25714 | -1.22108 | -0.73196 | -1.31365 | -2.99697 | -1.40119 | -2.53631 | -1.62467 |
| 29.02778 | -1.03946 | 0.500717 | -2.55761 | -0.51943 | -2.12711 | -2.8751 | -0.47729 | 0.945865 | 0.214253 | -0.20604 | 0.338641 | -3.25607 | -2.19942 | -0.14562 | -1.82353 | -0.55764 | -0.84246 | -0.86441 | -0.13811 | 0.10832 | -2.11301 | -1.24845 | -0.81463 | -0.0621 |
| 29.04167 | -0.44657 | 0.552858 | -1.117 | 0.131512 | -1.20411 | -0.53472 | -0.26397 | 1.287384 | 0.334934 | 0.479459 | 0.970651 | -2.18232 | -1.4013 | 0.284178 | -0.24168 | -0.26691 | -0.66495 | -0.80072 | 0.317574 | 1.525937 | -0.94425 | -0.42019 | 0.460891 | 1.051411 |
| 29.05556 | -0.10406 | 0.378024 | -0.17752 | 0.387734 | -0.54517 | 1.354612 | -0.11821 | 1.163637 | 0.366993 | 0.997427 | 1.661906 | -1.20842 | -0.76261 | 0.341208 | 0.922483 | -0.115 | -0.30539 | -0.61916 | 0.546412 | 2.084087 | 0.018661 | 0.29624 | 1.218067 | 1.783698 |
| 29.06944 | 0.124708 | 0.337672 | 0.300403 | 0.493643 | -0.06696 | 2.75637 | 0.066308 | 1.32766 | 0.206998 | 1.355636 | 2.002774 | -0.55686 | -1.00482 | 0.161933 | 1.898305 | 0.033067 | -0.07561 | -0.39629 | 0.655591 | 2.293659 | 0.676538 | 0.936189 | 1.648985 | 2.456365 |
| 29.08333 | 0.23644 | 0.394284 | 0.571116 | 0.634629 | 0.443523 | 3.702786 | 0.388094 | 1.712757 | -0.01766 | 1.558914 | 2.482789 | -0.07683 | -3.01657 | -0.13005 | 2.705287 | 0.285912 | 0.103944 | -0.21482 | 0.65181 | 2.352946 | 0.975163 | 1.39054 | 2.143264 | 3.362189 |
| 29.09722 | 0.199253 | 0.289834 | 1.554393 | 0.810451 | 1.404553 | 4.054615 | 1.083227 | 2.064771 | 0.582872 | 2.000201 | 3.369976 | 0.629344 | -3.0745 | -0.21186 | 3.611375 | 0.841317 | 0.317113 | 0.028421 | 0.450674 | 2.504419 | 1.208142 | 1.746114 | 2.796825 | 3.938632 |
| 29.11111 | 0.349786 | 0.718097 | 2.825601 | 1.484869 | 4.497576 | 6.247025 | 2.230021 | 2.089603 | 1.604639 | 2.384132 | 2.992504 | 1.130784 | 0.124376 | 0.862714 | 4.308108 | 1.987407 | 0.184436 | 0.364201 | -0.4689 | 3.183484 | 1.084445 | 1.112502 | 2.064407 | 3.36856 |
| 29.125 | 0.939622 | 1.707792 | 2.772516 | 2.296345 | 6.442049 | 6.41906 | 0.978183 | 1.53313 | 2.13099 | 1.781375 | 2.205688 | 2.137527 | -0.12396 | -0.07141 | 0.862678 | 1.554219 | -1.08541 | -0.05846 | -1.28156 | 2.682172 | -0.42093 | -0.83441 | 0.989407 | 0.857472 |
| 29.13889 | 1.145699 | 2.087019 | 1.798404 | 2.660031 | 3.607426 | 4.217969 | 0.816836 | 0.187751 | 1.801755 | 1.265464 | 1.705857 | 4.001743 | -3.93658 | -5.50322 | -6.25094 | -3.57444 | -0.2359 | -2.12047 | -0.13275 | -4.29351 | -4.02907 | 2.671044 | -1.20404 | -4.35282 |
| 29.15278 | 0.605059 | 1.532941 | 0.991575 | 1.893616 | 2.381621 | 1.685344 | 1.145984 | -0.76329 | 0.994167 | 0.923255 | 0.471463 | 3.490771 | -5.22874 | -6.42744 | -1.0327 | -4.01549 | -2.15572 | -3.22713 | -2.44823 | -12.3676 | -2.40389 | 1.105666 | -0.76932 | -5.71298 |
| 29.16667 | 0.021705 | 1.118881 | 0.28722 | 1.13132 | 2.062026 | 0.575872 | 1.202051 | -0.69948 | 0.408746 | 0.457274 | 0.234137 | 2.140623 | -5.65566 | -5.68953 | -5.61098 | -5.73037 | -3.27583 | -6.70719 | -4.1117 | -10.985 | -5.44665 | -2.32838 | -3.93634 | -5.97514 |
| 29.18056 | -0.18127 | 1.211628 | -0.00461 | 0.900146 | 1.963668 | 0.246023 | 1.189166 | -0.08126 | 0.245372 | 0.309784 | 0.100967 | 1.231984 | -4.40052 | -5.95584 | -8.58153 | -6.91188 | -3.80261 | -8.65157 | -5.24644 | -8.60247 | -7.22105 | -3.76317 | -8.79004 | -7.96009 |
| 29.19444 | -0.22709 | 1.094398 | -0.07283 | 0.671273 | 1.846475 | 0.129507 | 1.027293 | 0.428169 | 0.198365 | 0.276682 | -0.14796 | 0.530588 | -5.50314 | -4.9267 | -8.99151 | -5.91245 | -4.34298 | -8.31842 | -5.99557 | -8.01 | -6.82917 | -3.27003 | -8.11653 | -7.93683 |
| 29.20833 | -0.19421 | 0.702733 | 0.008684 | 0.432503 | 1.500193 | -0.00503 | 0.755334 | 0.386357 | 0.116809 | 0.187982 | -0.51107 | 0.264202 | -4.37149 | -4.2473 | -7.10917 | -5.19476 | -4.75823 | -6.96717 | -5.33378 | -7.7885 | -5.39962 | -2.95545 | -7.30136 | -7.58583 |
| 29.22222 | -0.07352 | 0.358234 | 0.385162 | 0.253921 | 0.870759 | -0.2545 | 0.434391 | 0.060312 | 0.347891 | 0.206439 | -0.325 | 0.305302 | -3.17905 | -4.07227 | -4.23759 | -4.17898 | -4.59947 | -5.31474 | -4.95293 | -5.11455 | -4.21909 | -2.46161 | -6.35761 | -8.27209 |
| 29.23611 | 0.116645 | 0.215292 | 0.857663 | 0.168507 | 0.729389 | -0.30263 | 0.090354 | -0.26008 | 0.769007 | 0.258965 | 0.354101 | 0.618705 | -2.90048 | -3.4848 | -2.15658 | -3.0152 | -3.70982 | -3.69352 | -3.96304 | -3.65018 | -3.15955 | -2.39939 | -5.27959 | -5.85335 |
| 29.25 | 0.328108 | 0.167302 | 1.306337 | 0.205802 | 0.612681 | 0.166364 | 0.10127 | -0.40602 | 1.247086 | 0.370648 | 1.064921 | 1.056121 | -2.49868 | -2.26317 | -1.27024 | -1.95472 | -2.40732 | -2.40798 | -2.64961 | -2.73628 | -2.53798 | -2.21007 | -3.38265 | -3.02048 |
| 29.26389 | 0.495321 | 0.124241 | 1.533688 | 0.321473 | 0.277858 | 0.710788 | 0.06184 | -0.17734 | 1.421459 | 0.486798 | 1.464047 | 1.18161 | -2.15593 | -1.4466 | -0.99909 | -1.31361 | -1.50983 | -1.70139 | -1.90349 | -1.86391 | -1.98582 | -1.98334 | -2.01551 | -1.88235 |
| 29.27778 | 0.420933 | 0.049131 | 1.243406 | 0.346463 | -0.17107 | 0.732705 | 0.011899 | 0.324522 | 1.141657 | 0.482645 | 1.487077 | 0.840926 | -2.1033 | -1.11373 | -0.93543 | -1.02486 | -1.09554 | -1.48357 | -1.51924 | -1.29361 | -1.49818 | -1.57944 | -1.40369 | -1.41091 |
| 29.29167 | 0.259686 | -0.00154 | 0.829041 | 0.297468 | -0.58797 | 0.494441 | -0.05398 | 0.706235 | 0.730781 | 0.419629 | 1.085845 | 0.098106 | -1.98165 | -0.99633 | -0.90311 | -0.90584 | -0.93501 | -1.31833 | -1.40634 | -1.03861 | -1.17851 | -1.04236 | -0.9876 | -1.09469 |
| 29.30556 | 0.124934 | 0.177992 | 0.596396 | 0.31649 | -0.68635 | 0.342799 | -0.00038 | 0.722293 | 0.379788 | 0.498317 | 0.121293 | -0.54205 | -1.55142 | -0.90558 | -0.89294 | -0.82369 | -0.86079 | -1.13425 | -1.60447 | -0.89445 | -0.98199 | -0.7804 | -0.72284 | -0.92239 |
| 29.31944 | -0.01033 | 0.304568 | 0.537878 | 0.307122 | -0.67392 | 0.159386 | 0.020121 | 0.346859 | 0.105942 | 0.619045 | -0.67649 | -0.79272 | -1.35746 | -0.81346 | -0.8705 | -0.78548 | -0.82913 | -0.99246 | -1.68468 | -0.78678 | -0.89077 | -0.6615 | -0.55942 | -0.88975 |
| 29.33333 | -0.11154 | 0.311114 | 0.48922 | 0.220526 | -0.63052 | -0.04096 | -0.09293 | -0.08133 | 0.137326 | 0.639461 | -1.00115 | -0.61801 | -1.21074 | -0.67952 | -0.69863 | -0.74316 | -0.78878 | -0.75816 | -1.33586 | -0.70201 | -0.85976 | -0.5753 | -0.47134 | -0.91758 |
| 29.34722 | -0.06745 | 0.350961 | 0.566742 | 0.227559 | -0.29595 | 0.059536 | 0.045077 | -0.35025 | 0.655015 | 0.690029 | -0.44972 | -0.07385 | -0.97915 | -0.55799 | -0.54064 | -0.72111 | -0.70191 | -0.54993 | -1.00654 | -0.56528 | -0.80001 | -0.54337 | -0.48403 | -0.92047 |
| 29.36111 | 0.088423 | 0.427601 | 0.743817 | 0.384012 | 0.13557 | 0.493761 | 0.276562 | -0.43907 | 1.406172 | 0.874394 | 0.565767 | 0.781923 | -0.79102 | -0.52326 | -0.50809 | -0.82334 | -0.60609 | -0.48286 | -0.77435 | -0.40278 | -0.73984 | -0.53706 | -0.53507 | -0.87194 |
| 29.375 | 0.195525 | 0.546161 | 0.814985 | 0.58225 | 0.457004 | 1.019811 | 0.419496 | -0.34362 | 1.863723 | 1.045435 | 1.404499 | 1.671599 | -0.68057 | -0.46424 | -0.5021 | -1.03469 | -0.51036 | -0.44443 | -0.62579 | -0.23264 | -0.69254 | -0.48518 | -0.53165 | -0.67745 |
| 29.38889 | 0.245709 | 0.712451 | 0.758254 | 0.767778 | 0.573621 | 1.281486 | 0.524567 | -0.10427 | 1.619945 | 1.044804 | 1.673625 | 1.622933 | -0.60617 | -0.34522 | -0.41217 | -0.98492 | -0.37723 | -0.34229 | -0.51107 | -0.07639 | -0.62876 | -0.37097 | -0.46674 | -0.43473 |
| 29.40278 | 0.342432 | 0.914614 | 0.82404 | 0.997048 | 0.839015 | 1.241998 | 0.715899 | 0.121866 | 1.217438 | 0.948795 | 1.288517 | 0.890166 | -0.57301 | -0.27443 | -0.33949 | -0.66447 | -0.28649 | -0.27934 | -0.46121 | -0.02185 | -0.58076 | -0.27465 | -0.39145 | -0.30145 |
| 29.41667 | 0.456646 | 1.044827 | 0.984484 | 1.104069 | 1.149583 | 0.889837 | 0.915545 | 0.231702 | 0.726327 | 0.840917 | 0.498306 | -0.07697 | -0.54424 | -0.20657 | -0.32729 | -0.35726 | -0.22633 | -0.24027 | -0.44138 | 0.008414 | -0.53382 | -0.2092 | -0.30137 | -0.19991 |
| 29.43056 | 0.630617 | 1.043157 | 1.081405 | 1.014439 | 1.155427 | 0.464649 | 0.921395 | 0.242975 | 0.17068 | 0.663121 | -0.49991 | -0.88469 | -0.47777 | -0.11109 | -0.36806 | -0.18501 | -0.17922 | -0.17779 | -0.3627 | -0.07718 | -0.39042 | -0.14071 | -0.22199 | -0.11566 |
| 29.44444 | 2.003829 | 1.398773 | 1.498141 | 1.184417 | 1.335863 | 0.421987 | 0.774207 | 0.146612 | 0.305004 | 0.543763 | -1.06231 | -0.34892 | -0.42724 | -0.08867 | -0.41387 | -0.1121 | -0.13469 | -0.07727 | -0.22937 | -0.10266 | -0.25836 | -0.04708 | -0.1376 | -0.04298 |
| 29.45833 | 7.978701 | 2.514641 | 2.850043 | 1.751539 | 2.248129 | 1.82545 | 0.925942 | 0.031111 | 1.595764 | 0.706452 | -0.4072 | 1.856958 | -0.39334 | -0.10527 | -0.35106 | -0.01016 | -0.01847 | 0.102928 | -0.03965 | 0.126731 | -0.01247 | 0.147157 | 0.023977 | 0.08159 |
| 29.47222 | 12.58513 | 3.668236 | 4.038858 | 2.827199 | 3.187017 | 4.681499 | 1.364408 | 0.48577 | 2.69189 | 1.070462 | 1.161496 | 3.681703 | -0.27742 | -0.09604 | -0.17156 | 0.082173 | 0.123348 | 0.195865 | 0.113079 | 0.348655 | 0.197824 | 0.342471 | 0.111435 | 0.155312 |
| 29.48611 | 14.08236 | 3.658099 | 3.520905 | 3.898688 | 3.378555 | 6.931525 | 1.313774 | 1.314628 | 2.979077 | 1.023367 | 3.323765 | 4.638695 | -0.16991 | -0.08461 | -0.04152 | 0.087252 | 0.184029 | 0.105874 | 0.149875 | 0.337809 | 0.217927 | 0.481949 | 0.035089 | 0.10899 |
| 29.5 | 12.9928 | 3.095078 | 3.22031 | 4.137168 | 3.077234 | 7.363173 | 1.175506 | 1.805524 | 2.811785 | 0.745533 | 5.610719 | 5.492703 | -0.13218 | -0.05832 | 0.025298 | 0.086692 | 0.215597 | 0.010948 | 0.163603 | 0.22449 | 0.181835 | 0.65948 | -0.05167 | 0.043053 |
| 29.51389 | 9.769433 | 2.126937 | 2.851128 | 3.381583 | 2.652109 | 8.064843 | 1.119149 | 1.77512 | 2.343893 | 0.657868 | 5.613711 | 5.737472 | -0.14428 | -0.04313 | 0.045948 | 0.093213 | 0.244217 | -0.03681 | 0.189759 | 0.134219 | 0.144122 | 0.822301 | -0.0858 | 0.003198 |
| 29.52778 | 6.096227 | 1.485091 | 1.816702 | 2.302338 | 1.751459 | 8.497226 | 1.128802 | 1.241095 | 1.547479 | 0.464248 | 1.894406 | 3.038945 | -0.12622 | -0.04921 | 0.040668 | 0.085076 | 0.273674 | -0.06941 | 0.175557 | 0.063259 | 0.109462 | 0.893224 | -0.08221 | -0.01363 |
| 29.54167 | 3.469041 | 1.07541 | 0.140569 | 1.928487 | 0.524896 | 7.377134 | 0.636083 | 0.547513 | 0.26082 | 0.315806 | -1.46906 | -0.02524 | -0.06418 | -0.02741 | 0.0729 | 0.109573 | 0.322808 | -0.05754 | 0.293445 | 0.050008 | 0.114589 | 0.849726 | -0.04918 | 0.007024 |
| 29.55556 | 1.237175 | 0.850546 | -1.52272 | 1.503375 | -0.69366 | 5.172714 | -0.20208 | 0.0262 | -0.86553 | -0.06158 | -5.04982 | -3.15404 | -0.00237 | 0.004189 | 0.124226 | 0.13741 | 0.363235 | -0.02989 | 0.490615 | 0.060246 | 0.132595 | 0.801519 | -0.01881 | 0.035632 |
| 29.56944 | -0.58706 | 1.00872 | -1.62548 | 0.936998 | -1.57551 | 4.088646 | -0.8351 | 0.37442 | -1.384 | -0.41623 | -5.85746 | -3.87807 | 0.017 | 0.000011 | 0.13048 | 0.13216 | 0.376823 | -0.02408 | 0.585194 | 0.051469 | 0.132626 | 0.756765 | -0.01166 | 0.043895 |
| 29.58333 | -0.55187 | 1.334898 | -0.56062 | 0.386338 | -1.55741 | 3.301253 | -0.94467 | 1.231274 | -1.06772 | -0.46069 | -4.23592 | -1.67248 | 0.003257 | -0.02855 | 0.099452 | 0.112117 | 0.350745 | -0.04324 | 0.584158 | 0.022649 | 0.120431 | 0.745288 | -0.01408 | 0.045782 |
| 29.59722 | -0.71414 | 1.674081 | 0.2723 | 0.461916 | -0.90167 | 2.722339 | -0.46569 | 2.581306 | 0.091578 | -0.40946 | -1.49206 | 0.444991 | -0.02599 | -0.03468 | 0.090771 | 0.112633 | 0.311098 | -0.05879 | 0.593122 | 0.010501 | 0.136415 | 0.825946 | 0.021147 | 0.149032 |
| 29.61111 | -2.37624 | 1.845695 | 0.508915 | 0.593097 | -0.39539 | 1.520925 | -0.12083 | 3.648503 | 1.333981 | -0.02221 | 1.205651 | 2.227541 | -0.05139 | 0.000523 | 0.122961 | 0.160236 | 0.299984 | -0.03512 | 0.688922 | 0.029485 | 0.20775 | 1.128002 | 0.129687 | 0.4205 |
| 29.625 | -4.67606 | 1.632385 | 0.269129 | 0.400307 | 0.094811 | -0.20587 | 0.510015 | 4.634583 | 1.866506 | 0.586859 | 3.534733 | 3.016066 | -0.0122 | 0.054469 | 0.178131 | 0.238751 | 0.355057 | 0.019741 | 0.887153 | 0.071499 | 0.282078 | 1.519924 | 0.185929 | 0.574323 |
| 29.63889 | -5.03802 | 1.154097 | -0.50305 | -0.27992 | 0.281756 | -1.44659 | 0.636734 | 5.045839 | 2.211022 | 1.166584 | 5.107175 | 3.525792 | 0.034751 | 0.051441 | 0.166229 | 0.270358 | 0.397599 | 0.020147 | 1.022465 | 0.06484 | 0.260042 | 1.59472 | 0.16008 | 0.457383 |
| 29.65278 | -5.87455 | 0.574893 | -1.51881 | -1.33024 | 0.278496 | -2.37635 | 0.797553 | 4.656643 | 1.356558 | 1.527671 | 4.484643 | 3.352173 | -0.01053 | -0.01626 | 0.091086 | 0.237989 | 0.396153 | -0.04595 | 0.717045 | 0.011326 | 0.211182 | 1.43005 | 0.099376 | 0.275976 |
| 29.66667 | -7.64241 | 0.054548 | -1.62289 | -1.44788 | 0.222176 | -3.96355 | 0.898173 | 3.56182 | 0.523886 | 1.691142 | 2.752665 | 2.02035 | -0.07422 | -0.06663 | 0.037522 | 0.21396 | 0.446935 | -0.08678 | 0.877273 | -0.02326 | 0.179194 | 1.376978 | 0.066282 | 0.184536 |
| 29.68056 | -7.08411 | -0.47843 | -0.95126 | -1.16513 | 0.292647 | -5.60224 | 0.977357 | 1.842162 | -0.2437 | 1.209402 | 0.993927 | 0.265162 | -0.05984 | -0.05898 | 0.04985 | 0.257253 | 0.511092 | -0.06637 | 1.992475 | -0.01105 | 0.20409 | 1.4232 | 0.051484 | 0.153962 |
| 29.69444 | -4.28024 | -0.52447 | -0.00817 | -0.91245 | 0.201983 | -6.99413 | 1.066568 | -0.0955 | -0.88959 | 0.630433 | -0.58771 | -1.28081 | 0.019186 | 0.001411 | 0.118048 | 0.334919 | 0.623388 | 0.005181 | 2.562148 | 0.040651 | 0.273763 | 1.404152 | 0.072383 | 0.174049 |
| 29.70833 | -1.19291 | -0.44394 | 1.197027 | -0.43004 | 0.17583 | -7.25604 | 1.253198 | -2.02899 | -1.1359 | 0.119326 | -2.62332 | -2.71447 | 0.043864 | 0.021863 | 0.138133 | 0.349252 | 0.673992 | 0.050217 | 1.961233 | 0.052149 | 0.287351 | 1.296998 | 0.065493 | 0.179836 |
| 29.72222 | 1.165698 | -0.25579 | 2.640599 | 0.314351 | 0.105129 | -6.0577 | 0.884456 | -3.32702 | -1.20641 | -0.30086 | -3.22006 | -2.3715 | 0.00752 | -0.01305 | 0.089403 | 0.32227 | 0.695066 | 0.03178 | 1.266667 | 0.016125 | 0.215084 | 1.138798 | 0.019417 | 0.154476 |
| 29.73611 | 2.571271 | 0.20032 | 3.902518 | 0.876944 | 0.161912 | -4.31747 | 0.294019 | -3.86837 | -0.5886 | -0.74269 | -2.38285 | -2.23186 | 0.021798 | -0.01503 | 0.077932 | 0.339018 | 0.781335 | 0.027189 | 1.056066 | 0.019598 | 0.169478 | 1.180041 | 0.016495 | 0.143917 |

| | | | | | | | | | | | | | | | | | | | | | | | | |
|---|---|---|---|---|---|---|---|---|---|---|---|---|---|---|---|---|---|---|---|---|---|---|---|---|
| 29.75 | 3.145812 | 0.669751 | 4.752402 | 0.866239 | -0.39331 | -2.12564 | -0.33012 | -3.8304 | -0.18781 | -0.93627 | -1.3609 | -1.91864 | 0.06191 | 0.001189 | 0.091685 | 0.390326 | 0.979132 | 0.031713 | 1.138685 | 0.046454 | 0.154956 | 1.294204 | 0.044231 | 0.130807 |
| 29.76389 | 3.030375 | 0.691537 | 3.945425 | 0.25107 | -0.87965 | 0.060039 | -0.65758 | -3.62408 | 0.227844 | -0.89615 | -0.67729 | -1.28472 | 0.072866 | 0.012727 | 0.094683 | 0.455438 | 1.349821 | 0.031023 | 1.239571 | 0.056504 | 0.159156 | 1.456192 | 0.061264 | 0.091826 |
| 29.77778 | 1.699899 | 0.366927 | 3.01864 | -0.53733 | -1.05329 | 2.756022 | -0.71432 | -3.2853 | 0.25339 | -0.45981 | -0.42506 | -0.47805 | 0.075905 | 0.032255 | 0.094001 | 0.513301 | 1.632994 | 0.03572 | 1.414473 | 0.061286 | 0.162568 | 1.442694 | 0.078908 | 0.06258 |
| 29.79167 | -0.15589 | -0.06752 | 1.171288 | -1.36188 | -0.97358 | 5.579223 | -0.94308 | -2.76265 | -0.51281 | 0.136888 | -0.58824 | -0.47381 | 0.051521 | 0.014698 | 0.066121 | 0.507357 | 1.589286 | 0.015864 | 1.683023 | 0.042644 | 0.16691 | 1.242239 | 0.069139 | 0.04016 |
| 29.80556 | -2.53942 | -0.39363 | -1.12921 | -1.91399 | -1.07732 | 5.910375 | -1.32305 | -1.93978 | -1.31059 | 0.680069 | -1.176 | -1.01597 | 0.016023 | -0.02516 | 0.042645 | 0.480088 | 1.504254 | -0.01143 | 1.690054 | 0.013111 | 0.288505 | 1.213048 | 0.05068 | 0.043401 |
| 29.81944 | -5.59771 | -0.73808 | -3.13212 | -2.27255 | -1.30867 | 5.515416 | -1.50322 | -1.57355 | -1.6997 | 0.928452 | -2.2197 | -1.81226 | 0.029134 | -0.01822 | 0.034119 | 0.503329 | 1.483559 | 0.001044 | 1.357049 | 0.023368 | 0.576744 | 1.556503 | 0.069286 | 0.102834 |
| 29.83333 | -7.57971 | -0.81761 | -4.26603 | -2.51107 | -1.26163 | 1.349435 | -1.26626 | -1.98235 | -1.86189 | 0.656222 | -3.60198 | -2.71387 | 0.057782 | 0.008275 | 0.058413 | 0.497808 | 1.535593 | 0.023544 | 1.292943 | 0.04725 | 0.524271 | 1.572752 | 0.082142 | 0.135439 |
| 29.84722 | -7.15823 | -0.65064 | -4.6169 | -2.76705 | -1.75728 | -2.66712 | -0.94363 | -2.52446 | -1.88799 | -0.17832 | -4.08825 | -2.99763 | 0.055447 | 0.011853 | 0.053546 | 0.480852 | 1.498136 | 0.024634 | 1.924435 | 0.03813 | 0.324801 | 1.324337 | 0.066896 | 0.082853 |
| 29.86111 | -5.50058 | -0.39238 | -4.71729 | -2.49598 | -2.04116 | -5.12263 | -0.80529 | -2.64491 | -1.88658 | -1.15946 | -3.61186 | -2.16327 | 0.044202 | 0.012188 | 0.047098 | 0.452443 | 1.414402 | 0.025488 | 2.821912 | 0.019787 | 0.199059 | 1.148097 | 0.055832 | 0.030221 |
| 29.875 | -3.97152 | -0.20829 | -4.02983 | -2.05002 | -1.68782 | -5.83271 | -0.77837 | -2.30292 | -1.74326 | -1.63129 | -3.09907 | -1.67855 | 0.054528 | 0.022907 | 0.06386 | 0.416314 | 1.198914 | 0.04582 | 3.473366 | 0.033944 | 0.169625 | 1.12915 | 0.063375 | 0.027199 |
| 29.88889 | -2.81212 | -0.09973 | -3.11839 | -1.63567 | -1.45538 | -6.83396 | -0.85454 | -1.40599 | -1.16457 | -1.89249 | -2.19949 | -0.95125 | 0.075712 | 0.027338 | 0.093086 | 0.486716 | 1.138976 | 0.05528 | 3.06095 | 0.067015 | 0.169089 | 1.236494 | 0.090732 | 0.06715 |
| 29.90278 | -1.5429 | 0.014916 | -2.66976 | -1.15585 | -1.17645 | -6.86786 | -0.65006 | -0.44418 | -0.31687 | -1.97829 | -0.69735 | 0.100961 | 0.083219 | 0.013204 | 0.090401 | 0.656407 | 1.32732 | 0.040156 | 2.320828 | 0.08775 | 0.156006 | 1.164653 | 0.102492 | 0.144438 |
| 29.91667 | -0.55319 | -0.02579 | -2.02315 | -0.62262 | -0.52264 | -4.72362 | 0.029612 | 0.207012 | 0.319894 | -0.66804 | 0.709998 | 1.230666 | 0.085678 | 0.000911 | 0.084974 | 0.870816 | 1.808531 | 0.025643 | 1.80951 | 0.114459 | 0.129749 | 0.896343 | 0.107784 | 0.233528 |
| 29.93056 | 0.015511 | -0.26478 | -1.4652 | -0.12205 | -0.02373 | -2.8815 | 0.440348 | 0.41269 | 0.746526 | 0.036708 | 1.494283 | 2.141234 | 0.124403 | 0.017756 | 0.099832 | 0.89207 | 2.307574 | 0.016698 | 1.555528 | 0.156971 | 0.105873 | 0.657212 | 0.106751 | 0.251112 |
| 29.94444 | -0.39265 | -0.72244 | -1.48245 | -0.43518 | -0.3355 | -4.12887 | 0.380132 | -0.04695 | 0.619175 | 0.364569 | 0.971844 | 1.588115 | 0.156977 | 0.016533 | 0.028935 | 0.536209 | 1.636762 | -0.10783 | 0.98805 | 0.069444 | -0.04934 | 0.331405 | 0.004425 | 0.089109 |
| 29.95833 | -2.08608 | -1.70573 | -2.62537 | -2.16376 | -1.67692 | -6.71724 | -0.78167 | -1.53994 | -0.71128 | -0.04982 | -1.64504 | -1.69936 | 0.131079 | -0.02301 | -0.16394 | -0.11071 | 0.154883 | -0.36711 | -0.09448 | -0.19189 | -0.45773 | -0.48644 | -0.22699 | -0.10998 |
| 29.97222 | -3.79648 | -2.93454 | -4.2341 | -4.21537 | -3.13538 | -10.7369 | -1.38982 | -3.46591 | -2.40862 | -0.67401 | -6.19759 | -6.79088 | 0.059974 | -0.03393 | -0.22177 | -0.94216 | -1.65094 | -0.45303 | -1.73541 | -0.36708 | -0.93439 | -2.31158 | -0.33632 | -0.08732 |
| 29.98611 | -3.38732 | -3.58398 | -4.89317 | -4.56212 | -3.55718 | -14.2148 | -1.27622 | -4.6047 | -3.25054 | -0.80368 | -8.64587 | -8.53405 | -0.07583 | -0.05021 | -0.15309 | -1.67309 | -3.3542 | -0.37617 | -3.3009 | -0.3909 | -1.29883 | -4.57727 | -0.26654 | -0.05836 |
| 30 | -1.83544 | -2.67259 | -4.43926 | -3.63946 | -3.06576 | -14.1641 | -1.20559 | -3.34644 | -1.8451 | -0.43399 | -5.99196 | -5.16213 | -0.21256 | -0.11159 | -0.1506 | -1.44586 | -3.42226 | -0.31417 | -3.01395 | -0.40555 | -1.37262 | -3.3461 | -0.20676 | -0.18442 |
| 30.01389 | -1.20467 | -1.50551 | -3.4271 | -2.64294 | -2.20215 | -11.5335 | -0.6636 | -2.1105 | -0.60171 | -0.01101 | -2.49688 | -2.71828 | -0.25655 | -0.15861 | -0.21566 | -1.09822 | -2.60916 | -0.27159 | -3.40401 | -0.37063 | -1.08156 | -2.49212 | -0.20693 | -0.30037 |
| 30.02778 | -0.80898 | -0.29894 | -2.27234 | -1.59519 | -1.25902 | -6.63736 | -0.30182 | -0.73814 | 0.192942 | 0.219076 | 0.182512 | -1.263 | -0.23238 | -0.17026 | -0.24068 | -1.08166 | -1.99538 | -0.22886 | -3.5382 | -0.28215 | -0.72113 | -1.85782 | -0.21629 | -0.33176 |
| 30.04167 | -0.26124 | 1.075095 | -0.96868 | -0.39404 | -0.44539 | -2.00387 | -0.13776 | 0.376396 | 0.665911 | 0.293555 | 2.03775 | -0.11987 | -0.19205 | -0.156 | -0.21294 | -1.04647 | -1.78625 | -0.18947 | -2.24589 | -0.20138 | -0.46353 | -1.27146 | -0.21343 | -0.29572 |
| 30.05556 | 0.054434 | 1.903263 | -0.21717 | 0.660588 | 0.130388 | 0.27332 | 0.035657 | 0.85188 | 0.935715 | 0.339609 | 2.844861 | 0.555174 | -0.156 | -0.1409 | -0.18189 | -0.74469 | -1.37925 | -0.16321 | -1.40626 | -0.16193 | -0.35146 | -0.84098 | -0.18578 | -0.24283 |
| 30.06944 | 0.235065 | 2.079782 | 0.159877 | 1.242135 | 0.530494 | 1.473379 | 0.227642 | 0.93585 | 1.240498 | 0.496064 | 2.91821 | 0.976375 | -0.14118 | -0.1542 | -0.15805 | -0.5079 | -0.91195 | -0.14965 | -1.04706 | -0.13927 | -0.31189 | -0.61812 | -0.15291 | -0.2022 |
| 30.08333 | 0.37753 | 2.349538 | 0.329241 | 1.441468 | 0.81225 | 2.675082 | 0.397497 | 1.243216 | 1.898047 | 0.760819 | 3.486444 | 1.505647 | -0.11899 | -0.05851 | -0.1159 | -0.34966 | -0.61771 | -0.13114 | -0.96381 | -0.1203 | -0.31811 | -0.5823 | -0.12375 | -0.17548 |
| 30.09722 | 0.315767 | 2.346548 | 0.405138 | 0.943869 | 0.880767 | 3.899054 | 0.412479 | 1.973361 | 2.602683 | 0.988455 | 4.22914 | 1.789325 | 0.012244 | -0.03643 | 0.047053 | -0.21964 | -0.4277 | -0.09497 | -1.00551 | -0.13258 | -0.36144 | -0.66979 | -0.17033 | -0.2571 |
| 30.11111 | 0.002773 | 1.537313 | 0.509785 | 0.558856 | 0.641116 | 4.999757 | 0.039683 | 2.635866 | 3.096709 | 0.773428 | 4.579689 | 1.725926 | 0.736915 | -0.02716 | 0.850824 | -0.0014 | -0.08569 | 0.016347 | -1.17935 | -0.33505 | -0.72596 | -1.11511 | -0.6575 | -0.98078 |
| 30.125 | -0.47512 | 0.675864 | 0.751181 | 0.014494 | -0.05286 | 6.379014 | -0.49052 | 2.321307 | 2.727718 | 0.231714 | 3.694147 | 1.210594 | 2.144702 | -0.19006 | 2.218902 | 0.488213 | 0.671439 | 0.175659 | -1.45069 | -0.87798 | -1.34892 | -2.13838 | -0.71084 | -2.17674 |
| 30.13889 | -0.56169 | -0.12954 | 1.112269 | 0.266786 | 0.002177 | 7.22269 | -0.12914 | 1.777712 | 2.373765 | 0.242082 | 2.295077 | 1.103302 | 2.362057 | -0.37605 | 2.30572 | 0.905687 | 1.93062 | 0.192809 | -0.1844 | -0.85382 | -1.13806 | -1.53995 | -0.58037 | -1.62984 |
| 30.15278 | -0.04667 | 0.03269 | 1.203938 | 0.599172 | 0.717651 | 7.121875 | 0.82747 | 0.922531 | 2.017727 | 0.874644 | 1.691545 | 1.905678 | 1.0954 | -0.07992 | 1.379917 | 0.797568 | 1.091071 | 0.06733 | 1.512921 | -0.16361 | -0.17581 | 0.212777 | -0.15527 | 0.417272 |
| 30.16667 | 0.183509 | 0.455917 | 1.026712 | 1.012119 | 1.057922 | 5.553643 | 1.034583 | 0.069789 | 1.698341 | 1.041331 | 0.650433 | 2.225791 | 0.693498 | 0.096646 | 0.75598 | 0.921603 | 1.119155 | 0.12313 | 2.971196 | 0.119943 | 0.430686 | 1.17919 | 0.196371 | 1.662936 |
| 30.18056 | 0.147472 | 0.315267 | 0.915819 | 1.344884 | 1.084491 | 4.275888 | 1.010489 | -0.83562 | 1.543306 | 0.78084 | -0.59517 | 2.147044 | 1.558062 | 0.31102 | 0.573885 | 1.272212 | 1.62753 | 0.377569 | 2.733124 | 0.526084 | 0.667676 | 1.641838 | 0.476279 | 2.216254 |
| 30.19444 | 0.211019 | -0.07263 | 0.730739 | 1.27912 | 1.127223 | 3.157381 | 1.092146 | -1.49513 | 1.305663 | 0.433568 | -1.54589 | 2.03096 | 2.694255 | 0.59998 | 0.429736 | 1.529289 | 2.014231 | 0.61768 | 2.128591 | 0.878074 | 0.902173 | 2.141723 | 0.648507 | 2.494607 |
| 30.20833 | 0.256695 | -0.73904 | 0.555069 | 0.966914 | 1.251058 | 1.968772 | 0.829008 | -1.52493 | 1.219772 | 0.082044 | -1.18142 | 1.884261 | 2.509323 | 1.048094 | 0.558097 | 1.657996 | 2.266177 | 0.869122 | 2.648722 | 0.790214 | 1.008197 | 2.716296 | 0.652608 | 2.227246 |
| 30.22222 | 0.260239 | -1.02858 | 0.331725 | 0.638884 | 1.407969 | 1.060984 | 0.596402 | -0.85342 | 1.015242 | -0.19145 | -0.19238 | 1.57727 | 1.610097 | 1.514936 | 0.810594 | 1.348689 | 2.413042 | 1.161042 | 2.628047 | 0.790326 | 1.031598 | 3.014897 | 0.634455 | 1.712617 |
| 30.23611 | 0.272339 | -0.53403 | 0.100401 | 0.329802 | 1.309546 | 0.306625 | 0.305598 | 0.096293 | 0.677587 | -0.35968 | 0.862706 | 0.785403 | 1.273626 | 1.737652 | 1.154563 | 1.053321 | 2.187432 | 1.178744 | 2.116115 | 1.123899 | 1.070142 | 2.780972 | 0.690769 | 1.546076 |
| 30.25 | 0.283281 | 0.195065 | -0.08102 | 0.2443 | 0.885388 | -0.47832 | 0.079767 | 0.856896 | 0.256722 | -0.30656 | 1.629459 | 0.047509 | 1.335829 | 1.769756 | 1.604588 | 1.451505 | 2.513613 | 1.0035 | 2.192075 | 1.572369 | 1.085911 | 2.232188 | 0.940958 | 1.573706 |
| 30.26389 | 0.314622 | 0.713456 | -0.21151 | 0.209199 | 0.420416 | -1.22397 | -0.0013 | 1.191084 | -0.04034 | -0.21195 | 1.756858 | -0.31949 | 1.613004 | 1.463274 | 1.864582 | 1.884806 | 3.456475 | 0.970607 | 2.449586 | 1.856848 | 1.326868 | 1.744337 | 1.561843 | 1.613725 |
| 30.27778 | 0.295902 | 0.820171 | -0.29479 | 0.100235 | -0.09535 | -1.81836 | -0.01747 | 1.034743 | -0.15167 | -0.08463 | 1.328067 | -0.38143 | 2.023526 | 2.287177 | 2.280714 | 1.968166 | 2.970119 | 1.076419 | 2.681708 | 1.827033 | 1.907101 | 1.590736 | 2.477773 | 1.852499 |
| 30.29167 | 0.269316 | 0.689609 | -0.27626 | -0.05502 | -0.59076 | -1.89298 | -0.20297 | 0.635751 | -0.00478 | 0.127911 | 0.679211 | -0.33035 | 2.406106 | 3.884682 | 3.044931 | 1.382259 | 1.91808 | 1.304796 | 2.726576 | 1.844421 | 2.033672 | 1.523743 | 3.27915 | 2.312377 |
| 30.30556 | 0.329304 | 0.202097 | -0.21543 | -0.18591 | -0.88694 | -1.53375 | -0.12738 | 0.206139 | 0.379217 | 0.355242 | -0.00819 | -0.21986 | 2.81207 | 2.908509 | 3.531103 | 1.195191 | 1.289007 | 1.601055 | 2.807111 | 2.200911 | 1.812613 | 1.372202 | 3.800808 | 2.578471 |
| 30.31944 | 0.340793 | -0.14444 | -0.09827 | -0.17808 | -0.78197 | -1.02675 | 0.028169 | -0.12991 | 0.740209 | 0.522393 | -0.42484 | -0.05874 | 2.688538 | 1.693218 | 3.643054 | 1.13727 | 0.873759 | 2.442627 | 2.230211 | 2.596164 | 2.592002 | 1.080865 | 3.39901 | 3.70509 |
| 30.33333 | 0.262813 | -0.13621 | -0.02297 | -0.05973 | -0.54135 | -0.51081 | 0.154899 | -0.21006 | 1.023234 | 0.629671 | -0.24849 | 0.080829 | 2.394047 | 1.664044 | 4.366074 | 0.97079 | 0.700068 | 2.869264 | 2.188068 | 2.595296 | 3.497926 | 0.834391 | 3.021444 | 3.946582 |
| 30.34722 | 0.116067 | 0.164297 | 0.079489 | 0.174108 | -0.15411 | 0.180674 | 0.339733 | 0.048584 | 1.212812 | 0.741561 | 0.572087 | 0.150749 | 1.893204 | 1.90732 | 4.265116 | 1.173513 | 0.652293 | 2.730161 | 2.572175 | 2.336994 | 4.375676 | 0.751437 | 2.459942 | 2.951972 |
| 30.36111 | 0.055812 | 0.528484 | 0.196826 | 0.424742 | 0.128998 | 0.68623 | 0.463042 | 0.365748 | 1.191938 | 0.848144 | 1.48255 | 0.0569 | 2.046924 | 1.894063 | 3.8116 | 1.712111 | 0.664752 | 2.349183 | 2.532228 | 2.567363 | 5.127658 | 0.815692 | 2.229914 | 2.557486 |
| 30.375 | 0.101077 | 1.104185 | 0.2477 | 0.580852 | 0.316753 | 0.968833 | 0.52305 | 0.348892 | 0.910121 | 0.928562 | 1.731178 | -0.09585 | 2.698152 | 2.076178 | 4.269256 | 2.074534 | 0.684578 | 2.26351 | 2.358556 | 3.265564 | 4.168512 | 0.937973 | 2.699416 | 2.929772 |
| 30.38889 | 0.313062 | 1.307714 | 0.286637 | 0.606127 | 0.471133 | 1.250616 | 0.604335 | 0.123244 | 0.665172 | 1.037283 | 1.346775 | -0.10935 | 2.700355 | 2.763009 | 4.439735 | 2.43525 | 0.851539 | 2.186191 | 2.070392 | 4.533285 | 2.729348 | 0.912549 | 3.716035 | 4.325191 |
| 30.40278 | 0.508575 | 0.853802 | 0.497965 | 0.615363 | 0.550712 | 1.536321 | 0.80444 | -0.11725 | 0.753064 | 1.156999 | 0.582966 | 0.071764 | 2.608737 | 2.730818 | 4.08561 | 2.912282 | 1.165209 | 2.09297 | 1.766133 | 4.812891 | 2.365064 | 0.764573 | 4.153379 | 5.235367 |
| 30.41667 | 0.545366 | 0.443355 | 0.672297 | 0.59177 | 0.560249 | 1.882778 | 1.044611 | -0.38983 | 1.064874 | 1.170445 | -0.25267 | 0.325665 | 2.037019 | 2.555296 | 3.384085 | 2.86282 | 1.078867 | 2.30472 | 1.563141 | 5.45209 | 2.788692 | 0.770408 | 4.227679 | 6.007563 |
| 30.43056 | 0.645787 | 0.205675 | 0.746658 | 0.532902 | 0.624477 | 2.038342 | 1.067276 | -0.72767 | 1.379297 | 1.083564 | -0.80393 | 0.605088 | 1.596424 | 2.579264 | 2.480816 | 2.609176 | 1.010365 | 2.726409 | 1.498864 | 6.207268 | 3.716916 | 0.811705 | 4.185921 | 5.286496 |
| 30.44444 | 0.999961 | -0.17568 | 1.105983 | 0.882768 | 1.176591 | 2.094829 | 1.073921 | -0.85148 | 1.681238 | 1.120521 | -0.70809 | 1.215749 | 2.017222 | 2.828768 | 2.518426 | 3.250678 | 1.377209 | 3.175061 | 1.868905 | 6.051674 | 4.267409 | 0.97597 | 3.967533 | 4.187719 |
| 30.45833 | 3.523582 | 0.220573 | 2.312172 | 1.908116 | 3.009449 | 2.938114 | 1.470903 | -0.66435 | 2.392583 | 1.507741 | 0.36463 | 2.221068 | 2.367497 | 3.168839 | 3.921704 | 4.868071 | 1.825579 | 3.795536 | 2.572028 | 6.043961 | 5.30294 | 1.563595 | 4.377609 | 5.007265 |
| 30.47222 | 9.115474 | 1.406499 | 4.166383 | 3.380765 | 4.14795 | 3.762168 | 1.554574 | -0.2373 | 2.997788 | 1.707357 | 1.526818 | 3.962298 | 2.585578 | 4.229924 | 4.912463 | 6.413154 | 1.785924 | 4.958623 | 2.793182 | 6.457649 | 6.58255 | 2.145382 | 4.821488 | 6.084271 |
| 30.48611 | 11.2568 | 2.010757 | 3.959398 | 3.772888 | 4.600649 | 4.333398 | 1.850592 | 0.671449 | 2.965739 | 1.650037 | 2.11679 | 4.382803 | 2.603544 | 4.229003 | 5.350622 | 6.414178 | 1.35267 | 5.558672 | 2.411267 | 7.682365 | 6.556542 | 2.276242 | 4.098751 | 6.190439 |
| 30.5 | 10.61654 | 2.13938 | 3.070617 | 4.019941 | 4.41655 | 4.397167 | 1.714374 | 2.088275 | 2.997111 | 1.680824 | 2.763237 | 3.894612 | 2.077826 | 3.491171 | 6.627707 | 6.313463 | 2.00234 | 5.055004 | 1.821969 | 7.993073 | 7.424262 | 2.589175 | 3.388036 | 5.359808 |
| 30.51389 | 9.601788 | 1.883099 | 2.918972 | 3.490789 | 3.304386 | 4.520748 | 1.402503 | 2.897549 | 3.346954 | 1.667406 | 4.870173 | 3.29794 | 1.450778 | 2.638453 | 6.496078 | 5.497046 | 1.86715 | 3.956466 | 1.491849 | 6.683971 | 7.002654 | 3.253953 | 2.343699 | 4.129279 |
| 30.52778 | 6.427459 | 1.593777 | 2.309335 | 2.055695 | 1.949411 | 4.278555 | 1.029241 | 2.518524 | 2.536487 | 1.414305 | 2.937529 | 2.329364 | 1.202432 | 2.509993 | 5.283322 | 4.241337 | 1.750973 | 2.850034 | 1.358429 | 4.794885 | 5.294992 | 3.138844 | 2.05647 | 3.264193 |
| 30.54167 | 3.78291 | 1.534584 | 1.319565 | 0.483762 | 0.923277 | 3.823676 | 0.495875 | 1.243338 | 1.012225 | 0.883083 | 0.254912 | 0.829273 | 1.539762 | 2.940582 | 4.838318 | 3.322873 | 1.612239 | 2.542196 | 1.13607 | 3.781825 | 4.091849 | 2.691801 | 1.956405 | 2.787478 |
| 30.55556 | 3.16101 | 1.407752 | 1.166107 | -0.43142 | 0.792953 | 3.958507 | 0.111443 | -0.2081 | -0.30832 | 0.237455 | -2.3766 | -0.60908 | 1.978961 | 2.784181 | 3.919578 | 2.821306 | 1.477411 | 2.240474 | 0.899616 | 3.185438 | 3.550977 | 2.328756 | 1.912873 | 2.287665 |
| 30.56944 | 3.934757 | 0.605017 | 1.297423 | -0.68647 | 0.485639 | 4.059454 | -0.18207 | -1.32578 | -0.97768 | -0.33712 | -4.45836 | -1.12593 | 1.946603 | 1.978861 | 2.873054 | 2.169611 | 1.233646 | 1.623977 | 0.733959 | 2.939938 | 2.957745 | 2.002937 | 1.977776 | 1.928244 |
| 30.58333 | 4.206022 | 0.623953 | 1.281632 | -0.18292 | -0.28665 | 3.530294 | -0.60262 | -1.81243 | -0.14233 | -0.74521 | -5.11353 | -0.94817 | 1.777927 | 1.500187 | 2.344235 | 1.546438 | 1.132233 | 1.185421 | 0.702426 | 2.742033 | 2.444805 | 1.700292 | 1.954986 | 1.810037 |
| 30.59722 | 4.06261 | 0.755937 | 1.059254 | 0.552649 | -1.08298 | 2.518042 | -0.98959 | -1.68547 | 0.337165 | -0.61042 | -4.29033 | -0.1419 | 1.620977 | 1.186439 | 2.166248 | 1.536753 | 1.036791 | 0.916884 | 0.737262 | 2.152318 | 1.990597 | 1.442638 | 1.611933 | 1.724221 |
| 30.61111 | 3.402088 | 0.611983 | 1.007875 | 1.086945 | -1.93458 | 1.721513 | -0.77129 | -1.24534 | 0.545247 | -0.28535 | -2.71242 | 0.897741 | 1.450884 | 1.030484 | 2.049914 | 1.820539 | 0.822427 | 0.780733 | 0.730567 | 1.481024 | 1.750581 | 1.429882 | 1.388409 | 1.740255 |
| 30.625 | 2.145828 | 0.605744 | 0.968487 | 1.148043 | -1.71519 | 1.186945 | -0.61453 | -0.30955 | 0.776274 | 0.002044 | -0.62002 | 1.315972 | 1.331102 | 1.357871 | 1.924611 | 2.032818 | 0.633723 | 0.762027 | 0.664438 | 0.905462 | 1.698837 | 1.746609 | 1.25928 | 1.80419 |
| 30.63889 | 0.464689 | 0.771824 | 0.809583 | 0.676201 | -2.15785 | 0.714287 | -0.46019 | 0.756755 | 0.66097 | 0.347695 | 1.240534 | 1.092731 | 1.255092 | 1.42408 | 1.655723 | 2.047479 | 0.619662 | 0.727582 | 0.544146 | 0.566986 | 1.592254 | 2.074353 | 1.185477 | 1.766134 |
| 30.65278 | -1.27297 | 0.726549 | 0.678229 | 0.377114 | -2.77981 | 0.297417 | -0.19763 | 2.06269 | 0.081184 | 0.768787 | 2.126815 | 0.735721 | 1.234801 | 1.407061 | 1.35293 | 1.741746 | 0.521157 | 0.640121 | 0.495791 | 0.472541 | 1.39454 | 2.135016 | 1.025355 | 1.611407 |
| 30.66667 | -2.39595 | 0.790378 | 0.639565 | 0.628371 | -1.31067 | -0.1818 | 0.210972 | 3.686363 | -0.33353 | 1.227245 | 4.003531 | 0.233802 | 1.206663 | 1.278241 | 1.30404 | 1.369265 | 0.433281 | 0.578296 | 0.516489 | 0.723162 | 1.304736 | 1.913653 | 0.944317 | 1.44991 |
| 30.68056 | -3.47747 | 1.211565 | 0.645932 | 0.576496 | 0.711983 | -0.98084 | 0.581907 | 4.471043 | -0.24663 | 1.47226 | 3.699413 | 0.200707 | 1.14544 | 1.116648 | 1.289721 | 1.158713 | 0.630587 | 0.624164 | 0.604432 | 0.82375 | 1.198945 | 1.596962 | 0.857827 | 1.238697 |
| 30.69444 | -5.68755 | 1.178239 | 0.452549 | 0.409765 | 2.308467 | -2.31757 | 1.022239 | 4.575071 | 0.290862 | 1.539287 | 1.596117 | -0.12952 | 0.975617 | 0.899976 | 1.091986 | 0.981523 | 0.824146 | 0.708079 | 0.664851 | 0.804623 | 0.960224 | 1.161425 | 0.728179 | 1.005725 |
| 30.70833 | -8.16333 | 1.226734 | 0.085445 | 0.251857 | 3.401067 | -2.68264 | 1.360325 | 3.974168 | 0.855397 | 1.501628 | -0.13921 | -0.30611 | 0.76377 | 0.702007 | 0.730241 | 0.792067 | 0.870853 | 0.698098 | 0.625397 | 0.76132 | 0.743378 | 0.761791 | 0.553036 | 0.832547 |
| 30.72222 | -8.30217 | 1.062591 | -0.33695 | 0.375711 | 2.494297 | -3.24116 | 1.35166 | 1.987526 | 1.363357 | 1.302975 | -1.70549 | 0.024287 | 0.579727 | 0.530927 | 0.45868 | 0.597115 | 0.798713 | 0.664835 | 0.628624 | 0.659009 | 0.609776 | 0.731599 | 0.432112 | 0.721581 |
| 30.73611 | -7.12857 | 0.57064 | -0.60359 | 0.356407 | 2.630767 | -3.49304 | 1.414886 | -0.45605 | 1.425079 | 0.946937 | -3.58614 | 0.299644 | 0.463917 | 0.397347 | 0.319994 | 0.415581 | 0.653709 | 0.673915 | 0.732649 | 0.555854 | 0.617773 | 1.018049 | 0.229569 | 0.6094 |
| 30.75 | -5.56815 | -0.24814 | -0.69987 | 0.353877 | 1.656264 | -3.46986 | 1.452067 | -2.75412 | 0.84848 | 0.474152 | -3.46974 | 0.138349 | 0.433666 | 0.242568 | 0.291447 | 0.20475 | 0.448741 | 0.546781 | 0.695426 | 0.323253 | 0.582099 | 1.296064 | -0.00769 | 0.518126 |
| 30.76389 | -3.47385 | -0.67729 | -0.83597 | -0.09648 | 0.841636 | -3.32139 | 0.731029 | -3.89251 | -0.09647 | -0.09987 | -3.60817 | -0.25208 | 0.382126 | -0.00602 | 0.245473 | -0.03488 | 0.155185 | 0.400809 | 0.489818 | -0.31646 | 0.473226 | 1.085378 | -0.11092 | 0.452551 |
| 30.77778 | -1.73764 | -0.35148 | -0.82234 | -0.43556 | 0.207868 | -2.40398 | -0.18146 | -4.77622 | -0.98219 | -0.78063 | -3.31584 | -0.7484 | 0.180773 | -0.38584 | 0.166478 | -0.28177 | -0.15365 | 0.415193 | 0.285347 | -1.0263 | 0.587977 | 0.852286 | -0.19848 | 0.491162 |
| 30.79167 | -0.76025 | 0.199232 | -0.77378 | -0.74818 | -0.35424 | -2.20844 | -1.00714 | -5.24805 | -1.57303 | -1.47148 | -2.75874 | -1.58915 | -0.07969 | -0.68756 | 0.091657 | -0.46804 | -0.34218 | 0.444577 | 0.158442 | -1.1238 | 0.619421 | 0.57359 | -0.18825 | 0.545931 |
| 30.80556 | -0.70917 | 0.476931 | -0.63307 | -0.73334 | -0.79969 | -2.18243 | -1.25464 | -4.98295 | -1.85483 | -2.01513 | -2.33385 | -2.33296 | -0.29081 | -0.99027 | -0.03629 | -0.61179 | -0.3169 | 0.291287 | 0.033252 | -0.60033 | 0.636622 | 0.255918 | -0.09189 | 0.519977 |
| 30.81944 | -1.16684 | 0.258102 | -0.61001 | -0.79343 | -1.11115 | -3.58228 | -1.53608 | -3.93116 | -2.06687 | -2.31551 | -2.0101 | -2.43676 | -0.34215 | -1.04531 | -0.26522 | -0.63994 | -0.29212 | 0.078089 | -0.04469 | 0.02954 | 0.563567 | -0.11376 | 0.04382 | 0.416712 |
| 30.83333 | -1.63432 | -0.27772 | -0.77557 | -1.13885 | -2.17582 | -4.45795 | -1.48547 | -2.78615 | -2.23094 | -2.63866 | -1.91496 | -2.14609 | -0.17778 | -0.80224 | -0.57903 | -0.53883 | -0.32757 | 0.046317 | -0.11784 | 0.495318 | 0.48503 | -0.4234 | 0.11356 | 0.216913 |
| 30.84722 | -1.87562 | -0.65131 | -1.05484 | -1.46334 | -3.75894 | -4.50197 | -1.57954 | -1.89977 | -2.56783 | -2.29406 | -1.97356 | -1.59718 | 0.100085 | -0.45501 | -0.89561 | -0.66431 | -0.30386 | 0.048096 | -0.10447 | 0.392 | 0.353925 | -0.60523 | -0.01545 | -0.04605 |
| 30.86111 | -1.84238 | -1.59019 | -1.09658 | -1.40171 | -3.94361 | -3.92344 | -1.43303 | -1.4845 | -2.86436 | -1.51022 | -1.87892 | -0.94955 | 0.297185 | -0.17088 | -1.17464 | -0.71866 | -0.20559 | 0.209616 | -0.03324 | -0.15388 | 0.111345 | -0.66033 | -0.37785 | -0.34068 |
| 30.875 | -1.12197 | -2.05366 | -0.60404 | -1.32694 | -4.10002 | -2.32651 | -0.78267 | -1.35751 | -2.75352 | -0.91113 | -1.4774 | -0.26696 | 0.243778 | 0.019467 | -1.54903 | -0.58175 | -0.16606 | 0.302813 | 0.004745 | -0.77003 | -0.09495 | -0.67992 | -0.72638 | -0.66166 |
| 30.88889 | 0.096919 | -0.99355 | 0.057503 | -0.95518 | -2.69773 | 0.133119 | -0.16581 | -1.23695 | -2.25942 | -0.30653 | -0.65804 | 0.264172 | 0.17403 | 0.072585 | -1.84996 | -0.45663 | -0.25571 | 0.396233 | -0.15382 | -1.31809 | -0.24675 | -0.78049 | -0.96886 | -0.93691 |
| 30.90278 | 1.503601 | 0.77734 | 0.602614 | -0.49797 | -1.49688 | 2.026548 | 0.400638 | -0.77608 | -1.55143 | 0.380739 | 0.23471 | 0.802961 | 0.129305 | -0.03391 | -1.70545 | -0.48859 | -0.35169 | 0.403394 | -0.33296 | -2.14798 | -0.49297 | -0.90177 | -1.10211 | -0.96311 |
| 30.91667 | 2.703154 | 1.909017 | 0.918587 | -0.16255 | -0.70063 | 3.412683 | 0.838744 | -0.24293 | -0.72292 | 0.887193 | 0.720145 | 1.167716 | 0.026804 | -0.14483 | -0.78408 | -0.67874 | -0.07792 | 0.227102 | -0.27576 | -2.23615 | -0.46899 | -0.9915 | -1.19211 | -0.81074 |
| 30.93056 | 2.924671 | 2.490983 | 0.956108 | 0.025828 | -0.08413 | 3.872389 | 0.939224 | 0.332305 | -0.07512 | 1.074778 | 0.88946 | 1.080674 | -0.19517 | -0.23566 | 0.2594 | -0.46218 | 0.021186 | -0.11599 | -0.27614 | -1.44221 | -0.28592 | -1.11461 | -1.01766 | -0.53383 |
| 30.94444 | 1.213686 | 2.095444 | 0.294949 | -0.18102 | 0.002647 | 2.554605 | 0.466869 | 0.448114 | 0.011538 | 0.887731 | 0.332691 | 0.289841 | -0.42148 | -0.25548 | 0.82963 | -0.47485 | -0.25812 | -0.29636 | -0.33423 | -0.52646 | -0.27213 | -1.36054 | -0.8189 | -0.54601 |
| 30.95833 | -2.91736 | 0.070581 | -1.59099 | -1.5557 | -0.95817 | -1.64158 | -0.74783 | -0.71283 | -1.02157 | 0.207385 | -1.70829 | -1.91344 | -0.63339 | -0.58716 | 0.395458 | -1.29855 | -1.02587 | -0.46253 | -0.68099 | -0.26778 | -0.95759 | -2.11711 | -1.11257 | -1.11769 |
| 30.97222 | -7.418 | -3.09607 | -4.08732 | -3.4035 | -2.38131 | -7.75375 | -1.74875 | -2.873 | -2.63843 | -0.44951 | -5.24337 | -5.10684 | -0.98945 | -1.4769 | -0.6957 | -2.94583 | -1.97181 | -0.86409 | -1.51496 | -0.29703 | -2.43626 | -3.89022 | -1.54988 | -1.75697 |
| 30.98611 | -8.17872 | -4.6021 | -4.5513 | -3.39755 | -2.31294 | -9.04655 | -0.95585 | -4.02713 | -3.09193 | -0.53509 | -6.9824 | -5.68744 | -1.22036 | -1.9699 | -1.26272 | -3.36256 | -1.79778 | -0.80262 | -1.74353 | 0.525682 | -3.53414 | -4.12838 | -1.63562 | -1.44654 |
| 31 | -4.99817 | -3.40105 | -3.98053 | -2.32105 | -1.39695 | -6.57695 | -0.58489 | -4.07521 | -2.25513 | -0.23195 | -5.37881 | -3.79866 | -0.20777 | -0.4545 | 0.03286 | -1.32376 | -0.2277 | -0.12381 | -0.5505 | 1.979433 | -1.37322 | -1.04785 | -0.78698 | -0.18609 |
| 31.01389 | -3.37999 | -1.94872 | -3.6822 | -1.40865 | -0.40264 | -7.48401 | -1.01785 | -3.80051 | -1.1526 | 0.020185 | -4.67515 | -2.70287 | 0.931254 | 1.157988 | 2.683442 | 0.695225 | 1.405834 | 0.624524 | 1.068315 | 2.41825 | 1.669433 | 1.452905 | 0.965231 | 1.108121 |
| 31.02778 | -2.319 | -0.68564 | -2.69051 | 0.075025 | 0.328478 | -6.81383 | -0.14737 | -3.14283 | -0.55704 | 0.184925 | -3.96407 | -2.06328 | 1.251971 | 1.369392 | 3.682215 | 1.440757 | 1.919863 | 0.890769 | 1.68586 | 1.671685 | 2.700371 | 2.040527 | 1.725112 | 1.312673 |
| 31.04167 | -1.25987 | 0.501023 | -1.12275 | 0.968152 | 0.785223 | -3.42445 | 0.13398 | -2.20776 | -0.45308 | 0.342367 | -1.91486 | -1.3628 | 0.853844 | 0.630463 | 2.763317 | 1.712094 | 1.800887 | 0.667444 | 1.552259 | 0.679748 | 2.007597 | 1.852228 | 1.822204 | 0.817984 |
| 31.05556 | -0.3443 | 1.332844 | 0.099054 | 1.093589 | 1.013118 | -0.85436 | 0.054169 | -1.39093 | -0.614 | 0.368229 | -1.63693 | -0.60273 | 0.650732 | 0.266285 | 2.241241 | 1.99437 | 1.617378 | 0.393268 | 1.332407 | 0.472804 | 1.40338 | 1.920154 | 1.718063 | 0.719151 |
| 31.06944 | 0.286045 | 2.033671 | 0.780625 | 1.101901 | 1.165236 | 0.407104 | 0.037564 | -0.81481 | -0.39716 | 0.337149 | -0.88407 | 0.762729 | 0.764454 | 0.254765 | 1.758439 | 2.295486 | 1.398399 | 0.145353 | 1.02757 | 0.936093 | 1.108742 | 1.787046 | 1.446162 | 0.950493 |
| 31.08333 | 0.863176 | 2.699399 | 0.971814 | 1.227628 | 1.357834 | 1.11036 | 0.103961 | -0.23855 | 0.278002 | 0.262129 | 0.987764 | 2.771104 | 1.05359 | 0.475323 | 0.951632 | 2.184694 | 1.121127 | 0.114029 | 0.756721 | 1.651655 | 0.937655 | 1.449093 | 1.265697 | 1.22874 |
| 31.09722 | 1.279629 | 3.067155 | 0.890563 | 1.410062 | 1.469447 | 1.489457 | 0.207987 | 0.717482 | 1.103215 | 0.201199 | 2.777609 | 3.619149 | 1.127785 | 0.662607 | 0.402426 | 2.00878 | 0.823256 | 0.223282 | 0.53299 | 2.044276 | 0.809446 | 1.388341 | 1.397019 | 1.521196 |
| 31.11111 | 1.691715 | 2.754675 | 0.808169 | 1.347242 | 1.486546 | 1.725491 | 0.370162 | 1.602505 | 1.868092 | 0.256751 | 3.552118 | 3.812591 | 0.930729 | 0.877407 | 0.445363 | 1.946409 | 0.544792 | 0.292759 | 0.385699 | 1.886013 | 0.913232 | 1.364109 | 1.199396 | 1.83292 |
| 31.125 | 1.870419 | 2.340318 | 0.695012 | 1.2375 | 1.419948 | 1.87162 | 0.584223 | 2.174979 | 2.715409 | 0.490562 | 4.048804 | 3.672867 | 1.191095 | 1.13374 | 0.981989 | 1.753975 | 0.338142 | 0.481473 | 0.402227 | 1.782737 | 1.367587 | 1.332808 | 1.033587 | 2.150441 |
| 31.13889 | 2.009236 | 2.970458 | 0.70142 | 1.29317 | 1.492909 | 2.026268 | 0.893239 | 2.414128 | 3.480171 | 0.815876 | 4.052444 | 3.325283 | 1.694954 | 1.473385 | 1.48903 | 1.979264 | 0.399819 | 0.825656 | 0.571418 | 1.519364 | 1.723663 | 1.144235 | 0.783897 | 2.16446 |
| 31.15278 | 1.72572 | 3.662291 | 0.525722 | 1.534373 | 1.621476 | 2.358787 | 1.229048 | 2.289118 | 4.042016 | 0.935183 | 2.743877 | 3.007596 | 1.675668 | 1.329672 | 1.13102 | 1.852818 | 0.28776 | 0.847863 | 0.714392 | 0.646926 | 1.987657 | 1.237934 | 0.348043 | 1.56841 |
| 31.16667 | 1.105692 | 2.467028 | 0.096905 | 1.649237 | 1.617654 | 2.56968 | 1.456656 | 2.169265 | 4.44804 | 0.515706 | 2.449021 | 2.484705 | 1.171119 | 0.976736 | 0.245434 | 1.480272 | 0.06352 | 0.580634 | 0.707046 | -0.23114 | 1.752461 | 1.405009 | -0.22956 | 0.706739 |
| 31.18056 | 0.908701 | 1.391354 | -0.10508 | 1.376279 | 1.639931 | 2.370641 | 1.593996 | 2.217453 | 4.070811 | 0.910778 | 4.323682 | 1.847317 | 0.822883 | 0.651409 | -0.36173 | 1.019333 | -0.05149 | 0.320639 | 0.648111 | -0.69114 | 1.530777 | 1.350048 | -0.73816 | 0.006483 |
| 31.19444 | 0.739138 | 1.098062 | -0.12539 | 1.05544 | 1.648264 | 1.913855 | 1.556871 | 2.520759 | 2.167715 | 1.287542 | 5.918103 | 1.191683 | 0.641753 | 0.219477 | -0.88456 | 0.522522 | -0.02039 | 0.292577 | 0.484736 | -0.68266 | 1.166897 | 1.136821 | -0.99661 | -0.2667 |
| 31.20833 | 0.572896 | 0.659518 | -0.06274 | 0.80848 | 1.648253 | 1.739124 | 1.389549 | 2.252989 | 1.234976 | 1.062629 | 5.198721 | 0.435601 | 0.463787 | -0.11328 | -1.34493 | -0.14482 | -0.07757 | 0.211099 | 0.192338 | -0.43686 | 0.753126 | 0.970365 | -0.785 | -0.15842 |

| | | | | | | | | | | | | | | | | | | | | | | | | |
|---|---|---|---|---|---|---|---|---|---|---|---|---|---|---|---|---|---|---|---|---|---|---|---|---|
| 31.22222 | 0.410633 | -0.0359 | 0.068361 | 0.571744 | 1.633584 | 1.877445 | 1.160987 | 1.814508 | 0.67581 | 0.796566 | 3.522063 | -0.1882 | 0.302065 | -0.23629 | -1.6943 | -0.60291 | 0.019787 | 0.132487 | -0.05661 | -0.1033 | 0.658944 | 0.857833 | -0.39742 | 0.323962 |
| 31.23611 | 0.234088 | -0.87034 | 0.353373 | 0.271131 | 1.548332 | 2.050409 | 0.98551 | 1.014066 | 0.3623 | 0.490739 | 1.066847 | -0.57791 | 0.054059 | -0.24378 | -1.38987 | -0.95901 | 0.201476 | 0.191613 | -0.14749 | 0.11582 | 0.384522 | 0.770141 | -0.12818 | 0.723565 |
| 31.25 | -0.14545 | -1.6021 | 0.633407 | 0.038475 | 1.326758 | 1.912767 | 0.848506 | 0.24396 | -0.03921 | 0.099572 | -1.83704 | -0.93179 | -0.16843 | -0.20581 | -0.93502 | -1.16332 | 0.36191 | 0.169747 | -0.08511 | 0.146048 | -0.02833 | 0.598886 | -0.12498 | 0.845946 |
| 31.26389 | -0.41943 | -1.40257 | 0.849213 | -0.04215 | 1.05135 | 1.442544 | 0.63849 | -0.37435 | -0.33852 | -0.2919 | -3.79301 | -1.20623 | -0.33107 | -0.05855 | -0.21944 | -1.15032 | 0.416952 | 0.173209 | 0.068532 | -0.06265 | -0.45642 | 0.41366 | -0.15233 | 0.558245 |
| 31.27778 | -0.35081 | -0.71964 | 0.955566 | 0.078053 | 0.719897 | 0.875833 | 0.562299 | -0.70179 | -0.20762 | -0.39085 | -4.31887 | -0.99519 | -0.42582 | 0.035626 | 0.370861 | -1.2403 | 0.43068 | 0.137056 | 0.172329 | -0.2118 | -0.85081 | 0.224158 | -0.24186 | 0.041771 |
| 31.29167 | -0.09144 | -0.2865 | 0.601327 | 0.298952 | 0.51232 | 0.384948 | 0.557411 | -0.86844 | 0.080983 | -0.27798 | -3.71418 | -1.11727 | -0.39792 | 0.0525 | 0.719993 | -1.41343 | 0.366042 | 0.101841 | 0.205036 | -0.1271 | -1.05764 | 0.053935 | -0.30606 | -0.12731 |
| 31.30556 | 0.134936 | 0.145924 | 0.120019 | 0.500446 | 0.347671 | 0.066154 | 0.48332 | -0.67093 | 0.256563 | -0.21145 | -3.25274 | -1.10683 | -0.21972 | -0.21735 | 0.876736 | -1.46104 | 0.259104 | 0.063921 | 0.145698 | 0.260062 | -0.96667 | -0.02727 | -0.1731 | -0.16627 |
| 31.31944 | 0.326513 | 0.410178 | -0.14052 | 0.615662 | 0.147569 | -0.1666 | 0.320986 | -0.39458 | 0.380779 | -0.12791 | -2.79598 | -0.71634 | -0.0041 | -0.58074 | 0.763077 | -1.34305 | 0.236325 | -0.03297 | 0.019447 | 0.969743 | -0.71079 | -0.05092 | 0.047279 | -0.09198 |
| 31.33333 | 0.451695 | 0.492786 | -0.15205 | 0.553784 | 0.004345 | -0.3808 | 0.138256 | -0.06788 | 0.580753 | -0.05447 | -1.76023 | 0.261324 | 0.132402 | -0.96527 | 0.503255 | -1.03023 | 0.282491 | -0.13759 | -0.10782 | 1.770546 | -0.46046 | -0.08805 | 0.559669 | -0.00287 |
| 31.34722 | 0.400137 | 0.414974 | -0.00929 | 0.455243 | -0.07761 | -0.51262 | 0.064094 | 0.178902 | 0.91582 | 0.084338 | -0.22936 | 1.500697 | 0.170528 | -0.89683 | 0.244053 | -0.34483 | 0.478642 | 0.011841 | -0.12887 | 2.544297 | -0.2112 | -0.14388 | 1.198512 | 0.439505 |
| 31.36111 | 0.236068 | 0.451774 | 0.075269 | 0.416642 | -0.0659 | -0.48588 | 0.09979 | 0.151957 | 1.254457 | 0.202375 | 1.121433 | 2.463216 | 0.189935 | -0.4253 | 0.354661 | 0.577952 | 0.85709 | 0.274438 | 0.009354 | 2.739634 | 0.202201 | -0.15959 | 1.529093 | 1.312426 |
| 31.375 | 0.227484 | 0.630549 | -0.05375 | 0.446232 | 0.062966 | -0.25784 | 0.142511 | 0.104077 | 1.335794 | 0.278517 | 1.538517 | 2.49624 | 0.273515 | 0.184375 | 0.672389 | 1.5865 | 1.328933 | 0.484033 | 0.280241 | 2.222252 | 0.571702 | -0.05931 | 1.311574 | 1.75683 |
| 31.38889 | 0.494104 | 0.986409 | -0.10185 | 0.697252 | 0.260868 | 0.027563 | 0.22187 | 0.054036 | 1.367326 | 0.366937 | 1.782461 | 2.160119 | 0.458967 | 1.056507 | 0.865631 | 2.417895 | 1.546368 | 0.570439 | 0.585821 | 1.363715 | 0.50265 | 0.044196 | 0.870703 | 1.53431 |
| 31.40278 | 0.852766 | 1.49165 | 0.15694 | 1.069671 | 0.402075 | 0.214821 | 0.441854 | -0.10373 | 1.612791 | 0.424229 | 2.428126 | 1.800752 | 0.61199 | 1.904071 | 0.792721 | 2.504307 | 1.394081 | 0.469509 | 0.699384 | 0.376173 | 0.302988 | 0.026494 | 0.421776 | 0.758445 |
| 31.41667 | 1.109404 | 1.874658 | 0.352703 | 1.291216 | 0.587251 | 0.385991 | 0.517481 | -0.052 | 1.569116 | 0.498853 | 2.645674 | 1.26841 | 0.705958 | 2.159483 | 0.493476 | 2.072787 | 1.026298 | 0.322735 | 0.641545 | -0.44232 | 0.263001 | 0.096444 | -0.16238 | 0.578884 |
| 31.43056 | 1.20059 | 2.054706 | 0.330389 | 1.423634 | 0.797647 | 0.60998 | 0.49145 | 0.125697 | 1.399183 | 0.589513 | 2.487759 | 0.661818 | 0.793177 | 1.982099 | 0.206016 | 1.702867 | 0.641316 | 0.125447 | 0.595735 | -0.97233 | 0.482671 | 0.449908 | -0.67345 | 0.474748 |
| 31.44444 | 1.22818 | 2.282939 | 0.478824 | 1.756052 | 1.173999 | 0.759084 | 0.57935 | 0.200905 | 1.404503 | 0.919837 | 2.502776 | 0.669855 | 0.868577 | 1.322532 | -0.12109 | 1.430826 | 0.409799 | 0.189649 | 0.832008 | -1.05605 | 1.198043 | 1.214781 | -0.51509 | 0.881122 |
| 31.45833 | 2.171358 | 2.75937 | 1.188302 | 2.304137 | 1.791457 | 0.815663 | 0.740661 | 0.395093 | 1.707714 | 1.543788 | 2.164653 | 1.602867 | 1.215086 | 0.982055 | -0.1018 | 1.762001 | 0.680717 | 0.673898 | 1.653393 | -0.6673 | 3.310353 | 3.24681 | 0.813917 | 1.786198 |
| 31.47222 | 5.989917 | 2.921406 | 3.025429 | 2.451227 | 1.934664 | 0.702561 | 0.818018 | 0.779706 | 2.212364 | 2.01064 | 2.021811 | 3.159997 | 1.594028 | 1.44031 | 0.711102 | 2.200754 | 1.343807 | 1.290957 | 2.422226 | 0.007618 | 5.417837 | 4.902282 | 3.065495 | 2.255092 |
| 31.48611 | 8.597651 | 2.530062 | 3.836535 | 2.23122 | 1.812419 | 0.783607 | 0.781751 | 1.504315 | 2.474507 | 2.140182 | 2.889361 | 4.068656 | 1.495857 | 1.940087 | 1.806322 | 2.295929 | 2.17768 | 1.427009 | 2.174246 | 1.038743 | 4.800569 | 3.484866 | 4.176125 | 2.561627 |
| 31.5 | 9.66847 | 2.25585 | 3.207657 | 1.812385 | 1.881407 | 1.576308 | 0.825314 | 2.519783 | 2.49575 | 1.924498 | 5.132809 | 4.470794 | 1.213756 | 2.170228 | 2.452907 | 2.189958 | 2.866587 | 1.26101 | 1.809903 | 1.426064 | 4.054933 | 2.176122 | 4.080783 | 2.687003 |
| 31.51389 | 8.389481 | 1.64185 | 3.130185 | 1.45103 | 2.768373 | 2.631635 | 0.897263 | 3.300166 | 2.036862 | 1.181799 | 7.919248 | 4.56045 | 0.933945 | 1.829532 | 2.235838 | 2.205174 | 3.294453 | 1.024081 | 1.525954 | 1.125203 | 3.892155 | 1.753268 | 4.154751 | 2.505011 |
| 31.52778 | 5.660899 | 1.360321 | 2.577982 | 1.230523 | 2.838107 | 2.898715 | 1.07146 | 2.584369 | 1.021781 | 0.568868 | 5.203156 | 3.078786 | 0.578066 | 0.763247 | 1.454799 | 1.572709 | 2.359589 | 1.060441 | 0.99221 | 0.705253 | 2.987277 | 1.505746 | 3.614801 | 1.908646 |
| 31.54167 | 3.862677 | 0.063749 | 1.661158 | 0.439986 | 2.028995 | 2.242727 | 1.338595 | 1.120689 | 0.092356 | 0.215718 | 1.480089 | 1.075235 | 0.137768 | -0.17587 | 0.19447 | 0.156852 | 0.266565 | 0.8483 | 0.401327 | 0.121396 | 1.718179 | 0.956365 | 2.312455 | 1.230628 |
| 31.55556 | 3.6919 | -1.59538 | 0.646284 | -0.315 | 1.705835 | 1.121633 | 1.604142 | -0.46071 | -0.60928 | -0.00163 | -2.01111 | -0.5734 | -0.23262 | -1.02002 | -1.28242 | -1.12621 | -1.52215 | 0.431511 | -0.12723 | -0.76345 | 0.396222 | 0.238126 | 1.141764 | 0.546781 |
| 31.56944 | 4.840603 | -2.17832 | 0.237013 | -0.57562 | 1.563594 | 0.100166 | 1.396945 | -1.16176 | -0.86254 | -0.22808 | -5.53159 | -1.44023 | -0.40439 | -1.41859 | -2.51888 | -1.97517 | -2.63374 | -0.04839 | -0.55754 | -1.83485 | -0.5867 | -0.3913 | -0.37799 | -0.35875 |
| 31.58333 | 6.203944 | -1.24172 | 0.098525 | -0.60125 | 1.32133 | 0.110879 | 1.064165 | -1.73526 | -0.21473 | -0.40351 | -7.95762 | -0.5946 | -0.49627 | -1.1317 | -3.46353 | -1.99529 | -2.02138 | -0.25488 | -0.66226 | -2.20232 | -1.05305 | -0.6731 | -1.26966 | -1.15587 |
| 31.59722 | 6.252973 | 0.163285 | 0.600966 | -0.20685 | 1.227391 | 0.432807 | 0.954037 | -1.99406 | 0.544076 | -0.43951 | -6.53756 | 0.846372 | -0.51274 | -0.80649 | -3.42488 | -2.46549 | -1.58115 | -0.40688 | -0.70374 | -1.74741 | -1.08481 | -0.55364 | -1.47686 | -1.75984 |
| 31.61111 | 5.969181 | 1.54465 | 1.159101 | 0.270167 | 1.14609 | 0.93804 | 0.839191 | -1.31278 | 1.074887 | -0.29292 | -4.06803 | 1.849806 | -0.40583 | -0.26467 | -2.77257 | -2.40254 | -0.74382 | -0.44382 | -0.62709 | -0.8975 | -0.98912 | -0.68759 | -1.52849 | -1.42188 |
| 31.625 | 5.70751 | 2.913669 | 1.428044 | 0.809351 | 0.996314 | 1.462455 | 0.569484 | -0.78415 | 1.501263 | -0.18749 | -2.44838 | 1.890573 | -0.42008 | -0.07589 | -1.3572 | -2.00412 | 0.156993 | -0.23788 | -0.31627 | -0.14311 | -1.21035 | -0.90398 | -1.42802 | -0.78984 |
| 31.63889 | 5.204648 | 2.940765 | 1.046852 | 1.024792 | 0.683136 | 1.588143 | 0.069907 | -0.22235 | 1.413901 | -0.0977 | -0.87833 | 1.013492 | -0.49922 | 0.295089 | 0.118991 | -1.73497 | 0.976028 | 0.050034 | 0.015752 | 0.201629 | -1.49455 | -0.99955 | -0.8386 | -0.18364 |
| 31.65278 | 3.949173 | 2.393727 | 0.497469 | 0.617268 | -0.06215 | 0.470108 | -0.59557 | 0.575088 | 0.558107 | 0.092115 | 1.327639 | -0.28831 | -0.47485 | 0.378509 | 1.255782 | -0.51929 | 1.562005 | 0.423856 | 0.305281 | 0.182031 | -1.80206 | -1.16037 | -0.09971 | 0.112451 |
| 31.66667 | 2.069733 | 1.012603 | -0.02411 | 0.001374 | -1.03916 | -0.9413 | -0.45551 | 1.533573 | -0.31326 | 0.156133 | 2.50044 | -1.50337 | -0.35501 | 0.138712 | 1.907207 | 0.215009 | 1.878059 | 0.585848 | 0.611541 | 0.125319 | -2.15681 | -0.96447 | 0.523385 | 0.200228 |
| 31.68056 | 0.375647 | 0.475946 | -0.32943 | -0.25185 | -1.25652 | -2.22743 | -0.36165 | 2.267208 | -0.94599 | 0.27376 | 3.172885 | -1.97145 | -0.17603 | -0.30697 | 2.40104 | 0.592789 | 1.63812 | 0.736023 | 0.672118 | -0.20998 | -2.24548 | -0.67725 | 0.697592 | 0.59836 |
| 31.69444 | -1.02851 | 0.778459 | -0.28579 | -0.19998 | -0.67283 | -2.72769 | 0.05486 | 2.322617 | -0.92765 | 0.423036 | 3.711562 | -2.39386 | 0.109031 | -0.42603 | 2.155918 | 0.835207 | 0.952721 | 0.422033 | 0.512827 | -0.61376 | -1.7686 | -0.14186 | 0.611371 | 0.968357 |
| 31.70833 | -2.20125 | 0.739883 | -0.11442 | 0.109999 | -0.11511 | -2.03489 | 0.298183 | 1.889735 | -0.31647 | 0.576635 | 2.759174 | -2.88834 | 0.46005 | -0.27848 | 1.449357 | 0.793711 | -0.08326 | 0.223836 | 0.339523 | -0.37526 | -0.98114 | 0.461767 | 0.630953 | 1.317672 |
| 31.72222 | -3.24995 | -0.42313 | -0.28539 | 0.315225 | 0.199598 | -0.71549 | 0.49645 | 1.441433 | 0.456346 | 0.552726 | 0.516917 | -2.39524 | 0.701147 | 0.200392 | 0.827853 | 0.605954 | -0.97875 | 0.108375 | 0.164191 | -0.08599 | -0.11613 | 0.984871 | 0.583753 | 1.419351 |
| 31.73611 | -4.21101 | -1.13543 | -0.81425 | 0.21702 | 0.255601 | 0.359034 | 0.510293 | 0.534729 | 0.869561 | 0.539036 | -2.38433 | -1.76331 | 0.755763 | 0.603416 | -0.03455 | 0.647641 | -1.26043 | -0.02292 | -0.07661 | 0.065445 | 0.487125 | 1.220752 | 0.150916 | 0.887876 |
| 31.75 | -4.54196 | -0.84459 | -1.17378 | -0.24381 | -0.07414 | 1.241562 | 0.470999 | -1.06276 | 0.965513 | 0.768747 | -4.93615 | -1.34516 | 0.61341 | 0.709479 | -0.92485 | 0.285319 | -1.16433 | -0.19829 | -0.3499 | -0.06468 | 0.594185 | 1.050903 | -0.4635 | 0.319739 |
| 31.76389 | -5.48786 | -0.78389 | -1.3163 | -0.91476 | -0.68704 | 2.098236 | 0.37912 | -2.62568 | 0.492603 | 0.616563 | -6.04938 | -1.45278 | 0.323455 | 0.695145 | -1.99994 | -0.03203 | -0.84727 | -0.33632 | -0.52532 | -0.31742 | 0.498092 | 0.537893 | -1.34488 | 0.040472 |
| 31.77778 | -7.19234 | -0.04981 | -1.14814 | -1.03725 | -0.74733 | 3.006695 | 0.501044 | -3.34578 | -0.23351 | 0.350878 | -5.82185 | -2.05242 | -0.01776 | 0.41266 | -3.37084 | -0.28638 | -0.31533 | -0.55762 | -0.62817 | -0.55424 | 0.311207 | -0.22931 | -1.83187 | 0.008812 |
| 31.79167 | -8.20802 | 0.642889 | -0.86871 | -0.71232 | -0.20296 | 2.53167 | 0.984687 | -3.82754 | -0.87895 | 0.024625 | -5.63522 | -2.54277 | -0.25871 | 0.035553 | -3.97611 | -0.5708 | 0.089962 | -0.70625 | -0.76254 | -0.90594 | -0.37197 | -1.04263 | -2.20326 | 0.046582 |
| 31.80556 | -8.81276 | 1.077091 | -0.89776 | -0.79247 | 0.177497 | 1.218701 | 1.322889 | -3.97418 | -1.23908 | -0.48549 | -5.08347 | -2.7586 | -0.39826 | -0.50088 | -3.88851 | -1.1051 | 0.158152 | -0.88023 | -0.75949 | -1.39516 | -1.397 | -1.6699 | -2.48627 | -0.09102 |
| 31.81944 | -8.3533 | 1.401247 | -1.16178 | -1.50613 | 0.188272 | 0.173107 | 1.513068 | -4.07655 | -1.58629 | -1.23368 | -5.60818 | -3.06961 | -0.51994 | -0.71648 | -3.6316 | -1.77639 | -0.02528 | -1.18491 | -0.89808 | -1.96382 | -2.023 | -2.09226 | -2.89319 | -0.46133 |
| 31.83333 | -6.54535 | 1.011054 | -1.71699 | -1.95835 | 0.113304 | -0.48837 | 1.15935 | -3.92585 | -1.81896 | -1.85796 | -5.22312 | -3.21446 | -0.65466 | -0.62769 | -3.31486 | -2.22058 | -0.47208 | -1.69682 | -0.93145 | -2.11544 | -1.82525 | -2.33919 | -3.40549 | -0.83737 |
| 31.84722 | -4.99368 | -0.30232 | -2.02685 | -1.40282 | -0.03107 | -0.58767 | 0.681474 | -3.32473 | -1.76964 | -1.95355 | -4.58501 | -2.60526 | -0.81444 | -0.71906 | -2.20371 | -2.68038 | -1.16955 | -1.45328 | -1.07746 | -2.07295 | -1.40982 | -2.48799 | -4.36055 | -0.96673 |
| 31.86111 | -3.67028 | -1.56055 | -1.7169 | -1.25612 | -0.12146 | -0.47487 | -0.07815 | -2.65422 | -1.5014 | -1.75263 | -3.98411 | -1.73997 | -0.96575 | -1.19354 | -3.0079 | -3.10039 | -1.73969 | -1.19539 | -1.30741 | -1.88988 | -1.19711 | -2.44191 | -5.28365 | -0.96461 |
| 31.875 | -2.37205 | -1.83692 | -0.85139 | -1.14985 | -0.08434 | -0.46862 | -0.84177 | -2.08898 | -1.21784 | -1.39677 | -3.08789 | -1.16872 | -1.02475 | -1.34098 | -2.67979 | -3.405 | -2.3039 | -0.82184 | -1.41595 | -1.49224 | -1.1691 | -2.49493 | -4.72503 | -1.18306 |
| 31.88889 | -1.00774 | -0.83831 | -0.13182 | -0.89917 | 0.082357 | -1.12124 | -1.26653 | -1.48118 | -1.04359 | -0.79993 | -1.63192 | -0.67789 | -0.96987 | -1.0233 | -2.42414 | -3.67839 | -2.72206 | -0.49005 | -1.41566 | -0.92917 | -1.28692 | -2.65188 | -4.15624 | -1.43715 |
| 31.90278 | 0.394236 | -0.07665 | 0.068459 | -0.81928 | 0.139157 | -1.98124 | -1.30807 | -0.66747 | -0.70088 | -0.24091 | 0.002953 | -0.19086 | -0.84692 | -0.8999 | -2.16708 | -3.82394 | -2.55932 | -0.76005 | -1.4145 | -0.57328 | -1.41326 | -2.23184 | -3.44741 | -1.56076 |
| 31.91667 | 1.806888 | 0.307505 | 0.063502 | -0.92285 | 0.051921 | -3.35231 | -1.29744 | 0.257831 | -0.27112 | 0.169744 | 1.454967 | 0.350658 | -0.69597 | -0.5535 | -1.81695 | -3.4446 | -1.73653 | -0.99051 | -1.21273 | -0.42608 | -1.44691 | -1.64289 | -2.82403 | -1.60521 |
| 31.93056 | 2.781563 | 0.299656 | 0.051274 | -0.80776 | -0.21665 | -3.71334 | -1.16103 | 1.094836 | 0.045583 | 0.53883 | 2.457232 | 0.841364 | -0.53892 | -0.16477 | -1.29472 | -2.98738 | -1.03478 | -0.94452 | -0.8151 | -0.41004 | -1.44736 | -1.18846 | -2.44213 | -1.81613 |
| 31.94444 | 1.713812 | -0.27423 | -0.64007 | -0.91043 | -0.72451 | -3.87664 | -1.17389 | 1.17193 | -0.18184 | 0.627293 | 1.98079 | 0.497519 | -0.60022 | -0.35498 | -1.26315 | -2.96675 | -1.00672 | -1.02647 | -0.69214 | -0.88115 | -2.10078 | -1.32159 | -2.50856 | -2.46043 |
| 31.95833 | -1.94789 | -2.15401 | -2.83699 | -2.06742 | -1.46873 | -6.12312 | -2.87854 | -0.09553 | -1.13256 | -0.08397 | -0.93783 | -1.36579 | -1.26717 | -1.85031 | -2.072 | -4.59279 | -2.10525 | -1.76765 | -1.35259 | -2.16527 | -4.00266 | -2.48678 | -3.17074 | -3.88127 |
| 31.97222 | -6.18489 | -5.49168 | -6.10186 | -4.10041 | -2.08038 | -8.78528 | -2.71251 | -2.1498 | -2.26892 | -1.19453 | -5.24116 | -4.12736 | -2.62987 | -4.25545 | -2.72089 | -6.99723 | -4.02188 | -2.60971 | -2.42046 | -3.58257 | -6.06706 | -4.00924 | -4.35537 | -5.52944 |
| 31.98611 | -6.74409 | -8.13966 | -6.94169 | -4.62198 | -2.05141 | -8.03934 | -1.88655 | -3.78277 | -2.5313 | -1.62603 | -7.99736 | -4.53 | -2.50256 | -4.51557 | -2.24533 | -5.86227 | -4.02196 | -1.82647 | -2.58626 | -2.84414 | -4.90014 | -3.87751 | -3.64814 | -4.22561 |
| 32 | -4.85954 | -7.82382 | -5.65472 | -3.20857 | -1.62727 | -4.56642 | -0.82489 | -3.92758 | -1.66618 | -1.05163 | -8.9787 | -2.93819 | -1.00813 | -2.86955 | -0.88563 | -3.16164 | -2.03164 | -0.13077 | -1.40947 | -1.69232 | -2.13238 | -2.04096 | -2.1874 | -1.53733 |
| 32.01389 | -5.57142 | -5.9302 | -3.24462 | -1.57511 | -1.18851 | -2.3793 | -0.23249 | -3.34373 | -0.63802 | -0.24648 | -8.8141 | -1.75189 | 0.19534 | -1.49536 | 0.013114 | -1.57038 | -0.26853 | 0.92272 | -0.48745 | -1.1011 | -0.67234 | -0.53144 | -0.64398 | 0.042963 |
| 32.02778 | -3.32146 | -3.11129 | -0.88717 | -0.3223 | -0.66709 | -1.45996 | -0.06397 | -2.55614 | -0.42894 | 0.352528 | -7.19062 | -1.17124 | 0.613241 | -0.32986 | 0.239709 | -0.87281 | 0.502163 | 0.999779 | 0.031393 | -0.75346 | -0.33594 | 0.299851 | -0.24832 | 0.532123 |
| 32.04167 | -2.1207 | -0.95793 | 1.039983 | 0.486164 | -0.15965 | -0.84387 | -0.03766 | -1.9457 | -0.91618 | 0.693084 | -5.60711 | -0.93781 | 0.596272 | 0.957137 | 0.303861 | -0.34206 | 0.805156 | 0.801429 | 0.508865 | -0.20027 | -0.13041 | 0.932705 | -0.00213 | 0.519589 |
| 32.05556 | -1.44053 | 0.339327 | 1.278932 | 0.963901 | 0.172041 | -0.09253 | 0.097152 | -1.87478 | -0.59972 | 0.873942 | -3.83539 | -0.51632 | 0.397439 | 2.051004 | 0.356744 | 0.068507 | 0.926914 | 0.762722 | 0.886241 | 0.214332 | 0.10506 | 1.290036 | 0.045623 | 0.452189 |
| 32.06944 | -0.95975 | 1.327951 | 0.803861 | 1.437658 | 0.553207 | 0.588887 | 0.288623 | -1.0476 | 0.41759 | 0.964132 | -1.32235 | 0.179817 | -0.00114 | 2.03769 | 0.325393 | 0.357447 | 1.177113 | 0.46026 | 1.090814 | 0.218565 | 0.201413 | 1.376576 | -0.27038 | 0.389381 |
| 32.08333 | -0.34752 | 2.219816 | 0.525348 | 2.05202 | 1.036588 | 0.925827 | 0.445784 | -0.74124 | 0.998839 | 0.787755 | 0.293355 | 0.701897 | -0.20592 | 1.052441 | 0.14445 | 0.653624 | 1.482091 | 0.292559 | 0.956797 | -0.10889 | 0.166326 | 1.120446 | -0.57901 | 0.348882 |
| 32.09722 | -0.1266 | 2.670868 | 0.744121 | 2.648679 | 1.283778 | 0.840202 | 0.312989 | -0.53079 | 0.78282 | 0.558874 | 1.224254 | 0.778508 | -0.10488 | 0.072768 | -0.03991 | 0.841154 | 1.649405 | 0.134086 | 0.601742 | -0.5788 | 0.096056 | 0.804393 | -0.58204 | 0.373584 |
| 32.11111 | 0.03786 | 2.849351 | 0.910815 | 2.744515 | 1.310436 | 0.813211 | 0.079437 | -0.2518 | 0.346987 | 0.349415 | 1.885547 | 0.674802 | 0.01085 | -0.58554 | -0.36991 | 0.946955 | 1.275833 | -0.26849 | 0.306195 | -0.94509 | 0.021261 | 0.138964 | -0.38318 | 0.45402 |
| 32.125 | 1.147318 | 2.946646 | 0.721391 | 2.579108 | 1.250218 | 1.139438 | 0.055447 | 0.079064 | 0.530998 | 0.387077 | 2.72372 | 0.853736 | 0.025439 | -0.63933 | -0.6471 | 0.941047 | 0.855204 | -0.54104 | 0.207697 | -0.76424 | -0.04056 | -0.49298 | -0.19132 | 0.427283 |
| 32.13889 | 2.720351 | 3.115904 | 0.171928 | 2.130068 | 1.071267 | 1.420464 | 0.130481 | 0.445318 | 1.220538 | 0.688044 | 3.609534 | 1.26579 | 0.033111 | -0.06163 | -0.50775 | 0.898719 | 0.560031 | -0.3896 | 0.25396 | -0.19412 | -0.0505 | -0.8949 | -0.03881 | 0.408643 |
| 32.15278 | 3.954774 | 4.089088 | -0.37591 | 1.659603 | 0.828343 | 1.40139 | 0.237418 | 0.794729 | 1.913295 | 1.030896 | 4.336642 | 1.860985 | 0.117728 | 0.991514 | -0.24586 | 1.004905 | 0.537242 | 0.001218 | 0.33532 | 0.466819 | 0.018127 | -0.60273 | 0.033759 | 0.419693 |
| 32.16667 | 4.467928 | 5.361956 | -0.66876 | 1.357612 | 0.745799 | 1.47913 | 0.394878 | 1.349013 | 2.536353 | 1.339189 | 4.97175 | 3.112471 | 0.169861 | 1.849994 | -0.01861 | 1.119363 | 0.549928 | 0.520223 | 0.349338 | 0.918845 | 0.142073 | -0.1582 | 0.026508 | 0.357514 |
| 32.18056 | 4.289377 | 6.213067 | -0.52435 | 1.282764 | 0.825457 | 1.712058 | 0.571568 | 2.022635 | 2.986053 | 1.358798 | 6.14843 | 5.114376 | 0.0906 | 1.914339 | -0.04506 | 0.995271 | 0.555397 | 0.731029 | 0.325871 | 0.959983 | 0.230424 | 0.39799 | 0.027127 | 0.2758 |
| 32.19444 | 2.845706 | 5.195939 | 0.113793 | 1.336357 | 0.974593 | 2.0153 | 0.668945 | 2.557363 | 3.168002 | 1.337863 | 6.68704 | 5.907993 | -0.08568 | 1.450695 | -0.31588 | 0.761158 | 0.519475 | 0.631647 | 0.228821 | 0.773438 | 0.274233 | 0.815036 | 0.364487 | 0.236158 |
| 32.20833 | 2.390818 | 3.98932 | 0.8285 | 1.413046 | 1.117465 | 2.232695 | 0.690029 | 2.852809 | 3.163005 | 1.349178 | 6.128376 | 5.764722 | -0.01245 | 1.142474 | -0.09966 | 0.556948 | 0.442517 | 0.431196 | 0.203781 | 0.717091 | 0.399367 | 0.780854 | 0.663948 | 0.211484 |
| 32.22222 | 2.371306 | 3.34997 | 1.273036 | 1.46537 | 1.23319 | 2.262457 | 0.693674 | 2.967203 | 3.001697 | 1.3307 | 5.136028 | 5.240706 | 0.338974 | 0.926739 | 0.483596 | 0.483341 | 0.385006 | 0.299854 | 0.344745 | 0.890786 | 0.680771 | 0.469632 | 0.658142 | 0.297852 |
| 32.23611 | 1.965588 | 3.712104 | 1.404548 | 1.399612 | 1.265492 | 2.229005 | 0.6649 | 3.032888 | 2.615569 | 1.194553 | 4.235332 | 4.261113 | 0.478201 | 0.632869 | 0.582327 | 0.442471 | 0.29957 | 0.169597 | 0.479705 | 0.911296 | 0.878597 | 0.304552 | 0.703098 | 0.403916 |
| 32.25 | 1.314235 | 2.130397 | 1.164485 | 1.06512 | 1.155342 | 2.057288 | 0.666536 | 2.822728 | 2.160734 | 0.990257 | 2.406615 | 3.259117 | 0.20587 | 0.29629 | 0.128127 | 0.208226 | 0.153054 | 0.028762 | 0.514622 | 0.721869 | 0.778038 | 0.301071 | 0.787243 | 0.493065 |
| 32.26389 | 1.173868 | 1.112482 | 0.591886 | 0.74236 | 1.015814 | 1.755924 | 0.696569 | 2.278879 | 1.554737 | 0.827385 | 2.431116 | 1.805725 | -0.08049 | 0.206805 | -0.22971 | 0.062204 | 0.046151 | 0.055719 | 0.513594 | 0.636811 | 0.580196 | 0.448577 | 0.666502 | 0.550929 |
| 32.27778 | 1.312754 | 0.307127 | 0.123185 | 0.275934 | 0.832252 | 1.4737 | 0.725931 | 1.797367 | 0.878645 | 0.701219 | 2.222949 | 0.102596 | -0.04283 | 0.418014 | -0.09144 | 0.425555 | 0.207354 | 0.351174 | 0.611547 | 0.955784 | 0.644234 | 0.678402 | 0.451093 | 0.683797 |
| 32.29167 | 1.354434 | -0.86328 | 0.171505 | -0.19123 | 0.591198 | 1.35925 | 0.742838 | 1.224947 | 0.305389 | 0.630609 | 1.060332 | -1.33776 | 0.289421 | 0.695842 | 0.431697 | 1.30411 | 0.517779 | 0.73893 | 0.7953 | 1.550189 | 0.92333 | 0.797203 | 0.306565 | 0.917543 |
| 32.30556 | 1.240552 | -2.04187 | 0.471251 | -0.49876 | 0.40892 | 1.252654 | 0.78089 | 0.655213 | -0.05764 | 0.630487 | -0.52695 | -2.42368 | 0.385795 | 0.223298 | 0.547042 | 1.645574 | 0.670488 | 0.558491 | 0.745689 | 1.638807 | 1.019235 | 0.837879 | 0.335833 | 0.959819 |
| 32.31944 | 1.086282 | -2.77439 | 0.587187 | -0.51925 | 0.38292 | 0.935573 | 0.770133 | 0.09545 | -0.26332 | 0.598164 | -1.21904 | -3.24683 | 0.228771 | -0.68133 | 0.255865 | 1.518473 | 0.692678 | 0.301066 | 0.575891 | 1.333771 | 0.974037 | 0.879777 | 0.40052 | 0.890696 |
| 32.33333 | 0.770258 | -3.41219 | 0.331434 | -0.49832 | 0.407115 | 0.547899 | 0.728596 | -0.44181 | -0.31734 | 0.508396 | -3.39452 | -4.08111 | 0.177216 | -0.96239 | 0.044032 | 1.603012 | 0.787207 | 0.266558 | 0.560823 | 1.329839 | 1.023953 | 0.935885 | 0.545164 | 0.975794 |
| 32.34722 | 0.275134 | -3.57038 | -0.02522 | -0.79844 | 0.304291 | 0.320176 | 0.666162 | -0.81024 | -0.09692 | 0.427607 | -3.78186 | -4.20608 | 0.088891 | -0.42656 | -0.16634 | 1.773839 | 0.802939 | 0.255087 | 0.503873 | 1.287148 | 0.969632 | 0.804069 | 0.610949 | 0.968095 |
| 32.36111 | -0.35945 | -3.32845 | -0.21454 | -1.06779 | 0.152388 | 0.252217 | 0.609117 | -1.03613 | 0.308181 | 0.288221 | -2.63683 | -2.13367 | 0.038705 | 0.566228 | -0.21307 | 1.804575 | 0.914123 | 0.385573 | 0.43734 | 1.180668 | 0.878556 | 0.545531 | 0.586778 | 0.937628 |
| 32.375 | -0.55671 | -2.63254 | -0.27412 | -0.88154 | 0.170072 | 0.250025 | 0.497241 | -1.01037 | 0.539843 | 0.128461 | -2.09395 | -1.02172 | 0.418584 | 1.85175 | 0.323609 | 2.282698 | 1.250273 | 0.821197 | 0.560864 | 1.620041 | 1.069385 | 0.420554 | 0.847278 | 1.23379 |
| 32.38889 | -0.57287 | -1.51024 | -0.06008 | -0.31891 | 0.378867 | 0.250115 | 0.43488 | -0.96565 | 0.526522 | 0.059667 | -2.25969 | -0.81558 | 0.686375 | 2.556098 | 0.950878 | 2.589837 | 1.378825 | 1.109399 | 0.692601 | 2.014892 | 1.329719 | 0.64678 | 1.388267 | 1.667322 |
| 32.40278 | -0.35883 | -0.57626 | 0.617857 | 0.242283 | 0.596687 | 0.256886 | 0.472923 | -0.77633 | 0.568733 | 0.160859 | -2.32438 | -0.8719 | 0.519283 | 2.059554 | 0.97937 | 2.072063 | 1.20199 | 0.872081 | 0.594991 | 1.698284 | 1.295978 | 0.960225 | 1.847616 | 1.744863 |
| 32.41667 | -0.02331 | 0.120338 | 1.317081 | 0.513011 | 0.699671 | 0.274092 | 0.593259 | -0.53549 | 0.735304 | 0.316492 | -2.01527 | -0.71101 | 0.406149 | 1.415801 | 0.937253 | 1.408064 | 0.844593 | 0.625841 | 0.507355 | 1.352387 | 1.201037 | 1.162611 | 1.879767 | 1.628915 |
| 32.43056 | 0.308211 | 0.65473 | 1.455506 | 0.693242 | 0.752224 | 0.290527 | 0.6339 | -0.25731 | 0.88526 | 0.427707 | -1.29092 | -0.16943 | 0.49327 | 1.177202 | 1.099792 | 1.035268 | 0.601995 | 0.385931 | 0.579507 | 1.186511 | 1.327866 | 1.301452 | 1.683566 | 1.742923 |
| 32.44444 | 0.848719 | 1.362898 | 1.037357 | 1.146771 | 0.966738 | 0.367181 | 0.560509 | 0.246727 | 1.178659 | 0.535481 | 0.216633 | 1.126374 | 0.898876 | 1.333549 | 1.666321 | 1.043936 | 0.653331 | 0.24322 | 0.88195 | 1.429056 | 2.149511 | 1.813947 | 1.875436 | 2.471392 |
| 32.45833 | 1.843718 | 2.59315 | 0.967999 | 2.134191 | 1.581292 | 0.482063 | 0.512479 | 0.876819 | 1.739619 | 0.864511 | 2.785004 | 3.32922 | 1.851888 | 2.47158 | 3.439354 | 1.862295 | 1.569994 | 0.340131 | 1.67778 | 2.225244 | 4.672146 | 3.2407 | 3.122011 | 4.546498 |
| 32.47222 | 3.461649 | 4.411011 | 2.342884 | 3.007526 | 2.145328 | 0.297506 | 0.541026 | 1.002946 | 2.126879 | 1.523484 | 4.602532 | 4.971972 | 2.7886 | 3.95856 | 6.55608 | 3.792905 | 3.168482 | 1.091848 | 2.656809 | 2.77764 | 6.108582 | 3.967909 | 4.504135 | 6.289265 |
| 32.48611 | 4.827167 | 5.139426 | 4.541486 | 3.291074 | 2.270018 | 0.132406 | 0.59688 | 0.879746 | 2.041711 | 2.079762 | 5.28235 | 4.735796 | 2.575791 | 4.525358 | 7.3139 | 5.36147 | 3.781884 | 2.058513 | 2.751738 | 2.794278 | 4.287368 | 3.418352 | 4.648209 | 4.675329 |
| 32.5 | 5.300335 | 5.191373 | 4.261723 | 3.30892 | 2.208825 | 0.921238 | 0.636745 | 1.241399 | 2.624179 | 2.209383 | 7.71415 | 5.27887 | 1.895013 | 4.979546 | 6.71331 | 5.625592 | 4.322366 | 2.548523 | 2.434684 | 3.083491 | 2.750653 | 3.132806 | 4.961284 | 3.225942 |
| 32.51389 | 4.799333 | 4.883435 | 3.455083 | 2.990253 | 1.990617 | 1.554928 | 0.608471 | 1.71408 | 2.850051 | 1.924651 | 10.40069 | 7.569042 | 1.46039 | 3.488468 | 3.89647 | 4.3897 | 4.064329 | 1.43696 | 2.385466 | 3.398168 | 2.415603 | 2.874921 | 4.981353 | 2.64644 |
| 32.52778 | 4.561929 | 4.98619 | 1.547632 | 1.851466 | 1.214748 | 1.506471 | 0.457254 | 1.78501 | 2.275387 | 1.55383 | 10.12215 | 6.994471 | 1.160721 | 1.201457 | 1.723685 | 3.769807 | 3.858829 | 0.909409 | 2.040164 | 2.205984 | 2.177176 | 2.50154 | 3.62087 | 2.033052 |
| 32.54167 | 4.031495 | 3.175288 | -0.73849 | 0.107048 | 0.305881 | 1.735624 | 0.367946 | 1.956831 | 1.498107 | 1.059805 | 8.717585 | 6.437244 | 0.985357 | -0.36208 | 0.116524 | 3.086508 | 2.767733 | 0.47936 | 1.977159 | 0.442564 | 1.910486 | 1.965118 | 1.256678 | 0.977179 |
| 32.55556 | 3.82195 | 1.444066 | -2.22314 | -0.89965 | -0.58257 | 1.773084 | 0.376943 | 1.998849 | 0.711171 | 0.59024 | 5.321986 | 4.707814 | 0.728275 | -1.43269 | -1.41409 | 0.725636 | 0.340708 | -0.21119 | 1.406946 | -1.0547 | 2.263617 | 1.801538 | -0.99725 | -0.15904 |
| 32.56944 | 3.958577 | 0.707386 | -2.16142 | -1.86005 | -1.05349 | 1.397746 | 0.665146 | 1.597371 | -0.07244 | 0.266978 | 2.037865 | 2.82227 | 0.514549 | -2.05881 | -2.75569 | -0.92215 | -1.47079 | -1.2285 | 0.282249 | -2.39368 | 2.878688 | 1.769655 | -2.81487 | -0.90618 |
| 32.58333 | 4.504905 | 0.314157 | -0.45718 | -2.48076 | -1.07332 | 0.877957 | 0.924813 | 0.945719 | -0.68655 | 0.428736 | -1.43917 | 1.12551 | 0.261873 | -1.86464 | -3.51389 | -1.2875 | -1.97225 | -2.59825 | -0.20311 | -2.38717 | 2.427897 | 1.917386 | -2.86091 | -0.83814 |
| 32.59722 | 4.544514 | -0.36598 | 1.632461 | -2.19742 | -0.76066 | 0.455309 | 1.273942 | 0.205733 | -1.01349 | 0.752538 | -5.04381 | -0.78192 | 0.228639 | -0.22596 | -3.57251 | -1.99516 | -2.04339 | -1.27908 | -0.46323 | -1.93985 | 1.578461 | 1.310016 | -2.15261 | -0.281 |
| 32.61111 | 4.479904 | -1.16982 | 3.503477 | -1.31592 | -0.13592 | 0.393936 | 1.396808 | -0.36989 | -0.74065 | 0.87729 | -8.42797 | -2.43211 | 0.480696 | 2.046968 | -3.83816 | -2.36893 | -1.71081 | 0.002764 | -0.60572 | -1.20382 | 0.665485 | 1.183939 | -1.83275 | 0.21791 |
| 32.625 | 5.102234 | -1.53836 | 3.80413 | -0.42025 | 0.588738 | 0.806064 | 1.321217 | -0.59028 | -0.08504 | 0.672905 | -9.89774 | -4.10986 | 0.58292 | 3.395801 | -3.94743 | -1.49981 | -1.05396 | 0.725254 | -0.56194 | -0.32064 | -0.54779 | 1.394714 | -0.16155 | 0.893647 |
| 32.63889 | 6.080918 | -0.56627 | 2.210858 | 0.148679 | 1.062538 | 1.273678 | 1.252652 | -0.94462 | 0.507751 | 0.359398 | -9.82125 | -5.02044 | 0.443921 | 2.920906 | -2.70191 | -0.00859 | 0.213871 | 1.615237 | -0.01636 | 0.765226 | -1.88622 | 1.261421 | 2.59816 | 1.417063 |
| 32.65278 | 7.219678 | -0.33374 | -0.09197 | 0.956761 | 1.283293 | 1.349839 | 1.080208 | -1.39537 | 0.898994 | 0.120993 | -7.34725 | -3.55164 | 0.182889 | 1.554805 | -1.65184 | 1.259216 | 1.441011 | 2.319501 | 0.417296 | 1.768911 | -2.67525 | 0.785282 | 4.997209 | 1.779672 |
| 32.66667 | 6.692066 | -1.3143 | -2.10038 | 1.509227 | 1.382496 | 1.007958 | 0.780818 | -1.57996 | 1.184294 | -0.03343 | -4.12991 | -1.45738 | 0.056303 | 0.238134 | -0.22942 | 2.050719 | 2.233812 | 2.315503 | 0.928401 | 2.255674 | -1.64481 | 0.127365 | 6.159698 | 2.242362 |
| 32.68056 | 5.831519 | -2.2033 | -2.81965 | 1.422032 | 1.192725 | 0.400665 | 0.435594 | -1.3003 | 1.320432 | -0.10163 | -2.97537 | -1.19992 | 0.114738 | -1.42694 | 0.966813 | 1.272296 | 2.119192 | 1.507847 | 1.261625 | 2.063908 | -0.31825 | -0.58176 | 6.223189 | 2.917056 |

| | | | | | | | | | | | | | | | | | | | | | | | | |
|---|---|---|---|---|---|---|---|---|---|---|---|---|---|---|---|---|---|---|---|---|---|---|---|---|
| 32.69444 | 4.431548 | -1.545 | -1.96173 | 0.958979 | 0.662804 | -0.45144 | 0.205634 | -0.65822 | 1.094032 | 0.041275 | 0.326675 | -1.3278 | 0.184057 | -2.68702 | 1.450662 | 0.369395 | 1.629777 | 0.386243 | 1.329148 | 1.214015 | 0.224249 | -0.94372 | 4.296848 | 3.173903 |
| 32.70833 | 2.686733 | 0.783075 | -0.34362 | 0.315322 | -0.10636 | -1.56556 | 0.237804 | 0.353556 | 0.659509 | 0.240653 | 4.464682 | -0.67315 | 0.142393 | -2.46065 | 1.168725 | 0.051162 | 1.009003 | -0.94119 | 1.10445 | -0.03721 | -0.04139 | -1.27173 | 1.305568 | 2.186166 |
| 32.72222 | 1.227832 | 3.165055 | 1.226241 | -0.01028 | -0.54268 | -2.17012 | 0.57187 | 1.208049 | 0.050181 | 0.327071 | 6.253441 | 0.177103 | 0.062299 | -1.2246 | 0.753202 | -0.09714 | 0.28673 | -1.64717 | 0.740754 | -1.13899 | -0.48047 | -1.12958 | -1.62618 | 0.661463 |
| 32.73611 | -0.14557 | 4.433306 | 1.993762 | -0.08652 | -0.47961 | -2.30901 | 0.894747 | 0.851992 | -0.60715 | 0.35374 | 4.979679 | 0.678271 | 0.04718 | 0.68772 | -0.03943 | -0.63813 | -0.10336 | -1.19926 | 0.580091 | -1.78634 | -0.62066 | -0.84674 | -3.619 | -0.6064 |
| 32.75 | -1.92726 | 4.19121 | 1.097419 | -0.20394 | -0.3822 | -2.35731 | 0.985027 | -0.39626 | -1.06749 | 0.291384 | 2.419538 | 0.792409 | 0.076851 | 2.287664 | -0.76238 | -1.25622 | -0.60325 | -0.64605 | 0.514586 | -2.25557 | -0.46674 | -0.45421 | -4.81834 | -1.19968 |
| 32.76389 | -3.53 | 3.617198 | -0.62723 | -0.17467 | -0.2917 | -2.5303 | 0.838945 | -1.65142 | -1.17135 | -0.19604 | 0.279042 | 0.588053 | 0.063685 | 3.305711 | -1.27389 | -0.76801 | -0.54715 | -0.122 | 0.453606 | -2.68529 | -0.32357 | -0.30036 | -4.83695 | -1.49818 |
| 32.77778 | -4.47904 | 3.111984 | -2.56719 | 0.009633 | -0.19185 | -2.84984 | 0.608832 | -2.79097 | -0.94006 | -0.93914 | -3.26789 | -0.49603 | -0.06072 | 2.383379 | -1.93793 | -0.55126 | -0.7487 | 0.313192 | 0.347008 | -2.6922 | -0.18308 | -0.28391 | -3.61917 | -1.57506 |
| 32.79167 | -5.5456 | 1.290389 | -4.21335 | -0.04344 | -0.11466 | -2.74216 | 0.368704 | -3.40409 | -0.92318 | -1.12961 | -6.30299 | -2.09868 | -0.35307 | 0.816121 | -3.57837 | -0.71608 | -0.97586 | 0.524315 | 0.113419 | -2.26024 | -0.05025 | -0.37549 | -3.02897 | -1.22496 |
| 32.80556 | -7.18196 | -0.27742 | -5.15553 | -0.2821 | -0.03895 | -1.85441 | 0.206455 | -3.35543 | -1.02773 | -0.95601 | -7.72326 | -3.24589 | -0.73561 | -0.91444 | -5.71348 | -1.31611 | -1.18591 | 0.486216 | 0.064444 | -1.73503 | 0.026338 | -0.54377 | -2.82715 | -1.04302 |
| 32.81944 | -9.08747 | -1.20587 | -4.47964 | -0.6298 | -0.01063 | -0.90621 | 0.071336 | -3.20054 | -1.05414 | -0.95391 | -7.51836 | -4.25154 | -0.86492 | -2.61195 | -7.12361 | -1.90612 | -1.35324 | 0.223928 | -0.03206 | -1.40997 | -0.07736 | -0.78789 | -2.76184 | -1.06869 |
| 32.83333 | -10.5674 | -2.66863 | -2.55377 | -0.98124 | -0.10144 | -0.17837 | -0.034 | -2.90659 | -1.14779 | -1.06384 | -7.36905 | -5.31606 | -0.59018 | -4.08821 | -7.82118 | -2.41765 | -1.33223 | -0.15215 | -0.02106 | -1.29347 | -0.32786 | -1.1116 | -2.74463 | -1.19833 |
| 32.84722 | -11.449 | -4.06839 | -0.81078 | -1.21907 | -0.11408 | 0.245175 | -0.10976 | -2.90017 | -1.32026 | -0.94553 | -7.30784 | -5.73076 | -0.40209 | -3.02226 | -6.44349 | -2.80423 | -1.22933 | -0.47596 | -0.02154 | -1.22647 | -0.57776 | -1.36218 | -2.76584 | -1.35249 |
| 32.86111 | -11.311 | -4.90453 | -0.0596 | -0.87609 | -0.02509 | 0.380392 | -0.1289 | -2.74511 | -1.25894 | -0.52775 | -8.09831 | -5.00374 | -0.44631 | -1.69921 | -4.43312 | -3.07655 | -1.36276 | -0.76203 | 0.049282 | -1.17837 | -0.708 | -1.39077 | -2.92085 | -1.37451 |
| 32.875 | -9.03072 | -3.95073 | -0.00354 | -0.31522 | -0.00241 | 0.352568 | -0.00886 | -2.37674 | -1.12845 | -0.13903 | -7.92062 | -3.50947 | -0.49809 | -1.04111 | -2.63077 | -3.44566 | -1.46496 | -0.98216 | 0.184208 | -0.98923 | -0.77732 | -1.26014 | -2.76056 | -1.33124 |
| 32.88889 | -5.31943 | -4.10218 | 0.270042 | 0.148455 | 0.060165 | 0.13982 | 0.17513 | -1.99002 | -0.88603 | 0.095935 | -6.20246 | -2.60644 | -0.38938 | -0.87507 | -1.20433 | -3.37106 | -1.43748 | -0.77063 | 0.303628 | -0.61004 | -0.67414 | -1.01954 | -1.99429 | -1.2694 |
| 32.90278 | -3.12831 | -3.88128 | 0.963244 | 0.489744 | 0.013507 | -0.19846 | 0.208266 | -1.74253 | -0.67991 | 0.095362 | -3.49834 | -1.67036 | -0.18543 | -0.40132 | -0.0054 | -2.60701 | -0.98011 | -0.33053 | 0.414876 | -0.20876 | -0.50721 | -0.61926 | -1.56111 | -1.1392 |
| 32.91667 | -1.28557 | -3.09869 | 1.510359 | 0.88522 | 0.040216 | -0.34492 | 0.214997 | -1.44694 | -0.46551 | 0.011446 | -0.84256 | -0.64777 | 0.023249 | 0.326765 | 1.061126 | -1.30667 | -0.38587 | 0.064393 | 0.426437 | 0.027893 | -0.42023 | -0.25147 | -1.37398 | -1.03087 |
| 32.93056 | -0.95521 | -1.77099 | 1.574407 | 1.008433 | 0.197518 | -0.18034 | 0.308041 | -0.93909 | -0.38438 | -0.02652 | 1.301637 | 0.002079 | -0.02095 | 0.502162 | 1.907903 | -0.294 | 0.014963 | 0.156166 | 0.249307 | 0.00177 | -0.51416 | -0.04915 | -1.45119 | -1.04219 |
| 32.94444 | -1.97862 | -0.86909 | 0.913573 | 0.538795 | 0.064488 | -0.4265 | 0.355287 | -0.55134 | -0.6604 | -0.1692 | 2.212096 | -0.21797 | -0.05078 | -0.25795 | 1.565269 | -0.56535 | -0.30925 | -0.34585 | -0.16169 | -0.56322 | -1.20174 | -0.26152 | -2.02637 | -1.80855 |
| 32.95833 | -5.46848 | -2.03018 | -0.92929 | -0.51116 | -0.58981 | -1.50713 | 0.184568 | -1.18276 | -1.25883 | -0.70764 | 0.339794 | -1.58266 | -1.59317 | -2.19171 | -1.19354 | -2.92193 | -1.84126 | -1.57715 | -1.05526 | -1.9216 | -3.01165 | -0.91198 | -3.32549 | -3.9735 |
| 32.97222 | -8.40822 | -6.07286 | -4.00124 | -1.60164 | -1.3743 | -2.55788 | -0.1567 | -2.60154 | -1.76648 | -1.30631 | -3.8402 | -3.39506 | -3.29103 | -4.27573 | -5.01916 | -6.8192 | -4.15277 | -3.00487 | -2.0292 | -3.93684 | -6.38279 | -2.46762 | -4.73693 | -7.27974 |
| 32.98611 | -7.5292 | -8.44067 | -5.87435 | -2.03897 | -1.6167 | -2.60409 | -0.41076 | -3.18517 | -1.7683 | -1.38025 | -8.11719 | -4.40783 | -3.83384 | -5.63599 | -6.73458 | -7.87847 | -4.01939 | -3.07924 | -1.72061 | -4.79647 | -8.6602 | -2.89654 | -4.40801 | -6.29397 |
| 33 | -6.23387 | -7.47049 | -6.87647 | -1.59166 | -1.0609 | -2.04934 | -0.30666 | -2.71335 | -1.40721 | -1.0448 | -8.95876 | -4.1884 | -3.36031 | -4.53797 | -4.41022 | -6.19336 | -2.1484 | -1.59069 | -0.55163 | -3.73379 | -4.10563 | -2.79546 | -2.69836 | -4.8287 |
| 33.01389 | -6.1256 | -5.20172 | -6.1889 | -0.95702 | -0.55103 | -1.37533 | 0.001231 | -2.00217 | -0.75481 | -0.20045 | -7.50602 | -2.95117 | -1.72381 | -1.19836 | -2.347 | -3.60781 | -0.8048 | 0.071329 | 0.393593 | -2.27686 | -2.22857 | -1.67757 | -0.95625 | -3.73203 |
| 33.02778 | -5.56386 | -2.75921 | -2.3882 | -0.20437 | 0.06042 | -0.63879 | 0.280081 | -1.26292 | 0.06474 | 0.664291 | -5.60487 | -1.59273 | 0.523608 | 1.457481 | -0.97309 | -1.2981 | 0.044307 | 1.11232 | 0.839346 | -1.27994 | -1.23177 | -0.32676 | 0.151599 | -2.67123 |
| 33.04167 | -4.86129 | -1.273 | 1.772985 | 0.300282 | 0.501765 | -0.13992 | 0.464892 | -0.62774 | 0.646583 | 1.387312 | -4.01958 | -0.58338 | 1.549057 | 3.419646 | -0.19009 | 0.16416 | 0.573926 | 1.783497 | 0.9638 | -0.27993 | -0.50856 | 0.777059 | 0.702271 | -1.04741 |
| 33.05556 | -3.3423 | -0.1728 | 4.496834 | 0.713047 | 0.743909 | 0.152559 | 0.460598 | -0.19078 | 0.675772 | 1.444529 | -2.58009 | 0.002332 | 1.750717 | 4.340607 | 0.295834 | 0.935728 | 0.828898 | 1.900474 | 1.026365 | 0.545979 | 0.080012 | 1.688364 | 0.859764 | 0.164279 |
| 33.06944 | -1.79791 | 0.538209 | 5.323069 | 1.01759 | 0.81597 | 0.038313 | 0.382306 | -0.03003 | 0.309016 | 1.283342 | -1.55877 | 0.301766 | 1.356049 | 4.477164 | 0.642578 | 1.224852 | 0.896316 | 1.979098 | 1.093566 | 0.747873 | 0.494323 | 1.989457 | 0.910998 | 0.458188 |
| 33.08333 | -0.84496 | 0.889561 | 4.505093 | 1.211603 | 0.969026 | -0.23209 | 0.370859 | -0.02396 | 0.081828 | 1.137556 | -0.84999 | 0.419105 | 0.326167 | 3.743553 | 0.636227 | 1.281585 | 1.010813 | 1.499272 | 1.094104 | 0.364138 | 0.565272 | 2.409686 | 0.780505 | 0.418784 |
| 33.09722 | -0.4818 | 1.041032 | 4.072348 | 1.433864 | 1.313665 | -0.3584 | 0.35526 | -0.04642 | 0.191103 | 1.162712 | -0.31076 | 0.452362 | -0.04299 | 2.709756 | 0.282698 | 1.418064 | 1.048706 | 0.79375 | 0.880097 | -0.09924 | 0.31482 | 2.21784 | 0.626743 | 0.023711 |
| 33.11111 | -0.4183 | 0.990465 | 3.066075 | 1.715714 | 1.414554 | -0.41124 | 0.316519 | -0.03071 | 0.378924 | 1.184714 | 0.103126 | 0.475669 | -0.42545 | 1.47974 | 0.033595 | 1.494753 | 1.086047 | 0.257329 | 0.554053 | -0.39039 | -0.08172 | 1.569748 | 0.367651 | -0.13913 |
| 33.125 | -0.63468 | 0.830989 | 0.523285 | 1.754437 | 1.343824 | -0.39096 | 0.273652 | 0.045648 | 0.495759 | 1.278339 | 0.533313 | 0.533076 | -0.6368 | 0.309727 | -0.14819 | 1.243696 | 1.049381 | -0.31746 | 0.202946 | -0.55932 | -0.42044 | 1.183779 | 0.036655 | -0.24009 |
| 33.13889 | -1.04709 | 0.826712 | -2.22318 | 1.646038 | 1.237548 | -0.18882 | 0.355566 | 0.186368 | 0.620451 | 1.460772 | 1.151405 | 0.705449 | -0.62252 | -0.3789 | -0.21766 | 0.86161 | 0.871916 | -0.92038 | -0.06404 | -0.66933 | -0.47647 | 0.514516 | -0.17846 | -0.21433 |
| 33.15278 | -1.54103 | 1.090123 | -4.40877 | 1.548602 | 1.203663 | 0.25171 | 0.611833 | 0.418433 | 0.826844 | 1.687567 | 2.096799 | 0.970129 | -0.38778 | -0.51859 | -0.2112 | 0.626949 | 0.670334 | -1.17494 | -0.10111 | -0.69825 | -0.24024 | -0.20638 | -0.26128 | -0.10854 |
| 33.16667 | -2.8799 | 1.210644 | -5.1407 | 1.415275 | 1.203852 | 0.768541 | 0.933739 | 0.667877 | 1.027416 | 1.729287 | 3.295574 | 1.221683 | -0.25012 | -0.3808 | -0.11085 | 0.453984 | 0.540039 | -0.90905 | 0.181355 | -0.61182 | -0.01602 | -0.62197 | -0.03873 | -0.01146 |
| 33.18056 | -6.10637 | 1.115771 | -3.72821 | 1.107329 | 1.126679 | 1.106599 | 1.067575 | 0.807535 | 1.008947 | 1.457913 | 4.339845 | 1.225302 | -0.16605 | 0.064258 | 0.022353 | 0.261422 | 0.480267 | -0.40011 | 0.430594 | -0.50411 | -0.00865 | -0.69322 | 0.47547 | -0.03054 |
| 33.19444 | -7.02206 | 1.568468 | -1.3225 | 0.905608 | 1.020691 | 1.342409 | 1.00601 | 0.911956 | 0.893325 | 1.166685 | 5.005058 | 1.06479 | 0.02839 | 0.94964 | 0.465907 | 0.467566 | 0.374795 | 0.127467 | 0.51919 | -0.03547 | 0.172754 | -0.50121 | 1.069664 | -0.09158 |
| 33.20833 | -5.01305 | 2.371302 | 1.000055 | 1.090497 | 0.907979 | 1.608809 | 0.835891 | 1.013006 | 0.936103 | 0.972592 | 5.983696 | 1.305984 | 0.168703 | 1.681117 | 0.86125 | 0.807716 | 0.22397 | 0.603858 | 0.442715 | 0.476308 | 0.375683 | -0.07084 | 1.444731 | -0.26221 |
| 33.22222 | -4.09992 | 2.907015 | 2.194264 | 1.217462 | 0.850407 | 1.747215 | 0.72045 | 1.056837 | 1.043419 | 0.835064 | 7.539338 | 2.096389 | 0.057764 | 1.642233 | 0.807379 | 0.724533 | 0.172793 | 0.600795 | 0.35667 | 0.389062 | 0.278825 | 0.266168 | 1.532076 | -0.28291 |
| 33.23611 | -2.89125 | 2.906743 | 1.652468 | 1.288085 | 0.920158 | 1.615415 | 0.76043 | 1.063799 | 1.03986 | 0.716567 | 8.762297 | 2.453563 | 0.106417 | 1.248348 | 0.693457 | 0.665719 | 0.198509 | 0.394219 | 0.358389 | 0.195516 | 0.162887 | 0.550865 | 1.521318 | -0.03871 |
| 33.25 | -2.02132 | 2.778341 | 0.075254 | 1.050193 | 0.919593 | 1.262538 | 0.881251 | 1.003838 | 0.989588 | 0.431097 | 8.355169 | 2.531682 | 0.356671 | 0.734206 | 0.718993 | 0.853873 | 0.20662 | 0.397904 | 0.300843 | 0.246488 | 0.186477 | 0.64074 | 1.22446 | 0.321944 |
| 33.26389 | -1.95434 | 2.417686 | -1.96807 | 0.521821 | 0.833748 | 1.006046 | 1.037016 | 1.05079 | 0.968315 | 0.074612 | 7.322134 | 2.863241 | 0.486949 | 0.215425 | 0.642817 | 0.892179 | 0.205481 | 0.488543 | 0.216198 | 0.353965 | 0.241546 | 0.610924 | 0.761445 | 0.562714 |
| 33.27778 | -2.07497 | 1.338053 | -3.74775 | 0.161397 | 0.745456 | 0.92189 | 1.16526 | 1.016352 | 1.165929 | -0.10286 | 5.634436 | 3.29004 | 0.602505 | 0.176295 | 0.686433 | 0.913128 | 0.297793 | 0.657195 | 0.275506 | 0.581287 | 0.431076 | 0.673834 | 0.453379 | 0.739724 |
| 33.29167 | -2.01336 | 0.962114 | -4.32012 | 0.045069 | 0.689416 | 0.741844 | 1.239121 | 0.770013 | 1.208932 | -0.02776 | 3.666291 | 2.926309 | 0.653573 | 0.835954 | 0.846618 | 1.01333 | 0.329019 | 0.869465 | 0.309225 | 0.905063 | 0.7186 | 0.771281 | 0.3737 | 0.735547 |
| 33.30556 | -1.61697 | 0.017314 | -3.51753 | 0.042207 | 0.603422 | 0.487823 | 1.241274 | 0.486777 | 1.177741 | 0.24204 | 1.911514 | 2.171371 | 0.469267 | 1.67794 | 0.937347 | 0.903385 | 0.18201 | 0.920571 | 0.23213 | 1.097596 | 0.885407 | 0.807142 | 0.486901 | 0.540398 |
| 33.31944 | -1.01438 | -0.22604 | -1.98622 | 0.195544 | 0.61121 | 0.310847 | 1.111598 | 0.220332 | 1.030277 | 0.471463 | 0.234922 | 0.944148 | 0.165301 | 1.944286 | 0.849533 | 0.462136 | 0.040945 | 0.808218 | 0.239198 | 0.991839 | 0.796464 | 0.763962 | 0.798896 | 0.312535 |
| 33.33333 | -0.41472 | -0.23357 | -0.999 | 0.49729 | 0.687915 | 0.173568 | 0.921095 | -0.03559 | 0.824402 | 0.590069 | -1.38523 | -0.45083 | 0.038246 | 1.510975 | 0.734686 | 0.161019 | -0.00292 | 0.646719 | 0.355819 | 0.811739 | 0.603128 | 0.686691 | 1.075509 | 0.169261 |
| 33.34722 | 0.12473 | -0.31664 | -1.04377 | 0.753401 | 0.805514 | 0.017525 | 0.764699 | -0.3067 | 0.565992 | 0.581828 | -2.78797 | -1.6952 | 0.063139 | 0.55278 | 0.693267 | 0.019063 | -0.00956 | 0.513391 | 0.442665 | 0.604124 | 0.401332 | 0.51689 | 1.168244 | 0.056746 |
| 33.36111 | 0.452351 | -0.4625 | -1.8736 | 0.865435 | 0.91043 | -0.18427 | 0.770712 | -0.46449 | 0.342496 | 0.483085 | -3.97153 | -2.72973 | -0.01517 | -0.52656 | 0.527828 | -0.16714 | -0.0037 | 0.232355 | 0.474152 | 0.313943 | 0.087549 | 0.177125 | 0.969966 | -0.01874 |
| 33.375 | 0.766086 | -0.52706 | -2.78164 | 0.888929 | 1.039806 | -0.34445 | 0.915259 | -0.52644 | 0.200349 | 0.323693 | -4.7286 | -3.43936 | -0.10526 | -1.13062 | 0.326395 | -0.25389 | 0.073615 | 0.081052 | 0.460054 | 0.195001 | -0.05108 | -0.0818 | 0.666947 | 0.035589 |
| 33.38889 | 0.803878 | -0.4064 | -2.66958 | 0.921159 | 1.111264 | -0.30983 | 0.979058 | -0.54625 | 0.273842 | 0.4183 | -4.59811 | -3.19784 | 0.121328 | -1.24035 | 0.235284 | -0.27813 | 0.153897 | 0.117311 | 0.451477 | 0.318604 | 0.070668 | -0.18112 | 0.543928 | 0.206703 |
| 33.40278 | 0.87361 | -0.10907 | -1.13889 | 1.046445 | 1.218253 | -0.01309 | 1.004945 | -0.47972 | 0.660447 | 0.615006 | -4.33914 | -2.82224 | 0.435036 | -0.44995 | 0.316246 | -0.41686 | 0.227686 | 0.124993 | 0.507423 | 0.610866 | 0.265231 | -0.06076 | 0.949962 | 0.423369 |
| 33.41667 | 0.851982 | 0.2445 | 1.097239 | 1.232299 | 1.291202 | 0.339922 | 0.985328 | -0.34089 | 1.076161 | 0.850768 | -3.27043 | -1.58137 | 0.368176 | 0.581871 | 0.313557 | -0.21649 | 0.399492 | 0.178705 | 0.576435 | 0.916142 | 0.4699 | 0.23051 | 1.254005 | 0.576609 |
| 33.43056 | 0.747961 | 0.564088 | 2.829236 | 1.235372 | 1.267558 | 0.511943 | 0.89973 | -0.21372 | 1.309954 | 0.925081 | -1.86004 | -0.43848 | 0.217282 | 1.132656 | 0.25681 | 0.464623 | 0.559565 | 0.356475 | 0.556443 | 1.185084 | 0.742733 | 0.496242 | 1.52148 | 0.679203 |
| 33.44444 | 1.019681 | 1.129195 | 3.821531 | 1.166935 | 1.20678 | 0.762821 | 0.857572 | -0.04295 | 1.732026 | 1.086965 | -0.42043 | 0.588733 | 0.653009 | 1.388705 | 0.531748 | 1.506983 | 0.802236 | 0.593527 | 0.503676 | 1.783926 | 1.565441 | 1.132542 | 2.154386 | 1.280882 |
| 33.45833 | 2.720562 | 2.359588 | 4.519815 | 1.449065 | 1.3065 | 1.301652 | 0.981471 | 0.331102 | 2.545782 | 1.44661 | 1.471766 | 1.933703 | 1.714467 | 1.640286 | 1.394134 | 2.702279 | 1.221472 | 0.474293 | 0.648381 | 2.891128 | 3.070701 | 2.475067 | 3.233736 | 2.803873 |
| 33.47222 | 4.638563 | 3.226719 | 6.113839 | 2.219518 | 1.584644 | 1.795478 | 1.357303 | 0.67946 | 3.131469 | 2.433137 | 3.223141 | 3.226576 | 3.44241 | 2.126585 | 2.857716 | 3.573857 | 1.719991 | 0.469739 | 0.954752 | 3.879661 | 4.6872 | 3.839814 | 6.344309 | 4.794806 |
| 33.48611 | 4.962736 | 2.932505 | 6.516288 | 2.94994 | 1.69363 | 1.92995 | 1.743496 | 0.865245 | 2.810117 | 3.352686 | 4.076222 | 3.228063 | 5.343591 | 2.911092 | 4.449893 | 3.947791 | 2.177439 | 1.574358 | 1.174985 | 4.053068 | 3.317991 | 3.666919 | 9.393755 | 4.920414 |
| 33.5 | 4.481664 | 2.770245 | 5.707883 | 3.54829 | 1.649737 | 2.009769 | 1.692846 | 1.123456 | 2.130759 | 2.528684 | 4.641802 | 2.804135 | 5.813754 | 4.409951 | 4.774016 | 4.87645 | 3.040952 | 2.81968 | 1.620996 | 4.541832 | 1.950451 | 3.195437 | 5.505974 | 4.340743 |
| 33.51389 | 3.279123 | 3.091586 | 4.34602 | 3.719068 | 1.52839 | 2.560684 | 1.549997 | 1.411876 | 1.802417 | 1.900942 | 5.199851 | 2.780386 | 4.926966 | 4.835066 | 4.235784 | 5.845164 | 3.786451 | 2.862599 | 1.998186 | 5.580059 | 2.065156 | 2.750509 | 3.499967 | 2.089379 |
| 33.52778 | 2.078314 | 3.72023 | 1.641792 | 2.200539 | 1.328298 | 2.289482 | 1.297605 | 1.4753 | 2.041655 | 1.255085 | 6.200655 | 2.803086 | 3.052599 | 3.940381 | 4.10158 | 5.027724 | 3.020164 | 2.452447 | 1.978895 | 4.874003 | 2.921681 | 2.494852 | 2.958317 | 2.010972 |
| 33.54167 | 2.495618 | 4.592678 | -0.76731 | 0.402085 | 1.134575 | 1.751232 | 0.92152 | 1.576045 | 2.070423 | 0.604229 | 6.742232 | 3.14618 | 1.169986 | 2.560083 | 2.112991 | 2.442655 | 1.341124 | 1.505664 | 2.716478 | 3.210517 | 2.375074 | 2.125132 | 2.021575 | 3.290822 |
| 33.55556 | 3.223853 | 5.016107 | -3.39365 | -0.50982 | 0.810344 | 1.055375 | 0.480948 | 1.805624 | 1.883965 | -0.26339 | 6.325563 | 3.446138 | -0.68797 | 1.019487 | -0.24743 | 0.95282 | 1.993792 | 0.402001 | 2.586651 | 2.215526 | 1.171962 | 1.440712 | 1.480232 | 3.544729 |
| 33.56944 | 3.327882 | 4.60837 | -4.76959 | -0.42474 | 0.374797 | 0.350635 | 0.133849 | 1.277803 | 1.661818 | -0.82674 | 5.309992 | 4.828447 | -1.60121 | 0.80958 | -1.67661 | 1.196501 | 1.369633 | -0.36083 | 1.841842 | 1.401303 | 0.881533 | 0.548566 | 0.587075 | 2.368503 |
| 33.58333 | 3.958124 | 3.365693 | -5.92818 | -0.28832 | 0.246026 | -0.03637 | -0.00662 | 0.750214 | 1.394205 | -0.7538 | 4.709912 | 5.518043 | -1.81279 | 2.946753 | -3.62078 | 1.599146 | 0.431668 | -1.35493 | 0.5488 | -0.05793 | 0.668999 | 0.041453 | -0.53667 | 0.74546 |
| 33.59722 | 5.388739 | 2.237955 | -4.17544 | -0.24823 | 0.424516 | 0.27448 | 0.024439 | 0.795118 | 1.154035 | -0.67722 | 4.080737 | 4.912612 | -0.81946 | 5.255953 | -3.9109 | 1.309286 | -0.74153 | -1.98347 | -1.11917 | -1.29388 | 0.08812 | 0.092036 | -1.75228 | -1.85531 |
| 33.61111 | 7.081505 | 1.797874 | -1.16429 | 0.012196 | 0.550399 | 0.54625 | 0.475851 | 0.596111 | 0.901818 | -0.05406 | 2.206809 | 4.040593 | 0.309197 | 5.15616 | -2.79523 | 0.882481 | -1.46233 | -3.02308 | -1.82655 | -2.59881 | -0.2378 | 0.131773 | -2.1216 | -2.43868 |
| 33.625 | 8.605715 | 1.16395 | 1.522321 | 0.389504 | 0.541726 | 0.664583 | 0.952704 | -0.00716 | 0.706312 | 0.534765 | 0.042326 | 2.953627 | 1.634473 | 3.25946 | -1.9111 | 0.479375 | -1.78978 | -2.42726 | -1.27876 | -2.94303 | -0.21636 | 0.258775 | -2.81261 | -2.08899 |
| 33.63889 | 11.37576 | 0.135041 | 4.001965 | 1.021544 | 0.688104 | 0.877504 | 1.290071 | -0.59961 | 0.627498 | 0.787119 | -2.28151 | 0.957889 | 2.091564 | 2.213005 | -0.2401 | -1.54158 | -2.60485 | -0.6614 | -0.53327 | -2.75676 | 0.289121 | 0.486677 | -3.20975 | -2.1524 |
| 33.65278 | 14.77935 | -0.94369 | 6.060391 | 2.069725 | 0.919271 | 1.215248 | 1.444359 | -0.83708 | 0.671777 | 0.75877 | -4.41232 | -1.00537 | 1.125899 | 1.21779 | 1.181291 | -1.84291 | -2.36109 | 0.963357 | 0.083934 | -1.80428 | 0.330154 | 0.873189 | -3.48916 | -1.42159 |
| 33.66667 | 14.22028 | -2.08912 | 6.754589 | 2.70924 | 1.043516 | 1.243543 | 1.586585 | -1.14533 | 0.614202 | 0.659184 | -6.58926 | -2.54457 | 0.30136 | -0.18739 | 2.557696 | -0.03835 | -0.35635 | 1.978828 | 0.976103 | -1.30924 | 0.293154 | 1.150657 | -2.31512 | -0.12943 |
| 33.68056 | 10.83107 | -2.88701 | 5.739126 | 2.657601 | 1.053922 | 0.839296 | 1.563704 | -1.21022 | 0.69557 | 0.533649 | -7.04107 | -2.81604 | -0.9979 | -2.22953 | 3.667333 | 1.848826 | 1.738242 | 2.576142 | 1.321371 | -0.85165 | 0.604104 | 1.257589 | -0.66635 | 0.759491 |
| 33.69444 | 7.633956 | -2.60718 | 3.809995 | 2.254807 | 1.128333 | 0.478333 | 1.261657 | -1.06208 | 0.933556 | 0.398315 | -5.62136 | -1.08011 | -1.97079 | -3.92667 | 3.470268 | 2.0866 | 3.372044 | 2.169694 | 1.910292 | -0.64254 | 0.765069 | 1.317902 | 0.75417 | 1.631491 |
| 33.70833 | 6.116124 | -2.0583 | 2.45918 | 1.734552 | 1.144065 | 0.276625 | 1.100001 | -0.74153 | 1.135572 | 0.393149 | -2.83829 | 0.565983 | -2.56896 | -2.76559 | 5.65121 | 0.855861 | 3.544298 | 1.146441 | 2.220515 | -0.50099 | 0.676646 | 1.219261 | 2.357826 | 2.333284 |
| 33.72222 | 3.491387 | -1.71735 | 0.978289 | 0.960353 | 0.97406 | 0.160062 | 1.073413 | -0.36386 | 1.289869 | 0.520431 | -0.87816 | 0.969772 | -2.8262 | -1.43609 | 4.062718 | -0.3805 | 2.805235 | -0.15978 | 1.66347 | -0.48133 | 0.446293 | 1.000708 | 3.513622 | 2.691607 |
| 33.73611 | 0.069884 | -1.41013 | -0.60939 | 0.005366 | 0.489787 | -0.04541 | 1.024059 | -0.10678 | 1.328058 | 0.607266 | 0.564553 | 0.711166 | -2.95232 | -0.92221 | 1.67771 | -1.21058 | 1.436737 | -1.33515 | 0.878909 | -0.5818 | 0.301015 | 0.696454 | 3.971349 | 2.388578 |
| 33.75 | -2.99278 | -1.1882 | -1.88166 | -0.77788 | 0.154569 | -0.48072 | 0.886252 | -0.07098 | 1.234109 | 0.497009 | 2.06581 | 0.350792 | -2.84949 | -1.27564 | 1.21543 | -2.66287 | -0.09045 | -2.89306 | 0.473709 | -0.66326 | 0.203724 | 0.456621 | 3.754717 | 2.000093 |
| 33.76389 | -5.76402 | -1.35255 | -2.87949 | -1.21927 | 0.046291 | -1.09325 | 0.690453 | -0.21567 | 0.994039 | 0.292129 | 3.762738 | -0.06923 | -2.03016 | -0.93453 | -2.87964 | -3.92947 | -1.59462 | -3.84519 | 0.489422 | -0.64259 | 0.002666 | 0.396265 | 3.725549 | 1.698266 |
| 33.77778 | -8.05724 | -0.49209 | -4.19441 | -1.64001 | -0.03082 | -2.06068 | 0.284798 | -0.53002 | 0.601589 | 0.013322 | 4.518725 | -0.29361 | -0.98136 | -0.47313 | -3.48752 | -3.6583 | -2.0127 | -3.97795 | 0.761457 | -0.3697 | -0.10667 | 0.450898 | 3.429329 | 1.334304 |
| 33.79167 | -10.7084 | 0.097304 | -6.00232 | -2.04825 | -0.16665 | -3.13706 | -0.20399 | -0.65528 | 0.180453 | -0.35749 | 3.891416 | -0.22534 | -0.09621 | -0.10671 | -2.08219 | -2.72393 | -1.69812 | -3.4657 | 0.801514 | -0.02284 | -0.09157 | 0.471225 | 2.109093 | 0.232142 |
| 33.80556 | -14.3541 | 0.348898 | -6.68454 | -2.32704 | -0.27495 | -3.5775 | -0.53226 | -1.24528 | -0.21851 | -0.45634 | 2.214356 | -0.02592 | 0.533641 | -0.35229 | -2.86272 | -2.14108 | -2.3118 | -2.88993 | 0.655257 | -0.04009 | -0.13946 | 0.352766 | 0.085804 | -1.08762 |
| 33.81944 | -16.9042 | 0.307258 | -5.60624 | -2.09352 | -0.5455 | -3.42588 | -0.84602 | -1.92413 | -0.62781 | -0.55566 | 0.404433 | 0.192663 | 0.259453 | -0.99508 | -1.40996 | -1.69448 | -2.6795 | -2.23338 | 0.589716 | -0.43671 | -0.0335 | 0.140028 | -2.01052 | -2.12994 |
| 33.83333 | -19.0251 | -0.42141 | -2.51272 | -1.64605 | -0.97052 | -2.38998 | -1.11585 | -2.25588 | -0.98094 | -0.74678 | -1.71254 | 0.401934 | -0.88765 | -1.59688 | -1.1041 | -1.63419 | -2.54845 | -1.84356 | 0.454835 | -1.02492 | 0.101479 | -0.05448 | -4.33412 | -2.83418 |
| 33.84722 | -18.0254 | -1.55325 | 1.316298 | -0.75257 | -1.20927 | -1.61327 | -1.38616 | -2.3717 | -1.11819 | -0.81429 | -3.90859 | 0.455941 | -1.98297 | -1.82089 | -1.06376 | -1.30899 | -2.57107 | -1.53838 | -0.10754 | -1.65157 | 0.224347 | -0.18024 | -6.78708 | -3.17921 |
| 33.86111 | -14.4557 | -2.25928 | 5.058458 | 0.003655 | -1.28423 | -1.03471 | -1.38685 | -2.54064 | -0.98772 | -0.74327 | -6.0523 | 0.474463 | -2.49466 | -1.85803 | -1.65993 | -0.73247 | -2.22122 | -1.15482 | -0.71881 | -2.24043 | 0.232543 | -0.36852 | -8.25672 | -3.38547 |
| 33.875 | -11.8708 | -2.43668 | 5.263953 | 0.496623 | -0.94464 | -0.82339 | -1.01049 | -2.26299 | -0.71834 | -0.38655 | -7.07249 | 0.542805 | -1.9998 | -1.66535 | -1.53853 | -0.47191 | -1.2694 | -0.77664 | -0.30826 | -3.00799 | 0.320367 | -0.57834 | -9.25311 | -3.18551 |
| 33.88889 | -8.33558 | -2.28577 | 3.492127 | 0.730176 | -0.59879 | -0.97987 | -0.3957 | -1.88191 | -0.42709 | 0.087676 | -6.76556 | 0.553046 | -1.30224 | -1.17842 | -1.11112 | -0.11504 | -0.47926 | -0.4591 | -0.06054 | -3.46866 | 0.03663 | -0.81191 | -8.35515 | -2.64387 |
| 33.90278 | -5.06564 | -2.14586 | 1.885501 | 1.012539 | -0.16031 | -0.78759 | 0.172962 | -1.57626 | -0.19193 | 0.443061 | -5.47775 | 0.402649 | -0.61132 | -0.62278 | -1.01544 | 0.387108 | 0.473859 | -0.13673 | 0.222773 | -3.26586 | 0.150203 | -0.91022 | -6.43676 | -1.55091 |
| 33.91667 | -1.81128 | -1.48593 | -0.15345 | 1.081084 | 0.242363 | -0.39041 | 0.550605 | -1.21635 | -0.07498 | 0.577764 | -4.6861 | 0.090812 | 0.363433 | -0.1312 | -0.45282 | 0.628249 | 1.366595 | 0.039532 | 0.363936 | -2.28571 | 0.203851 | -0.78051 | -4.76618 | -0.84441 |
| 33.93056 | 1.227498 | -0.68985 | -2.58014 | 0.855213 | 0.367544 | -0.21718 | 0.695693 | -1.01942 | -0.11836 | 0.546432 | -3.80367 | -0.26836 | 1.098593 | 0.242079 | -0.15813 | 0.95844 | 1.85111 | 0.291856 | 0.539739 | -1.53994 | 0.235251 | -0.5483 | -3.22811 | -0.73049 |
| 33.94444 | 3.155572 | -0.56638 | -4.86187 | 0.207397 | 0.219549 | -0.6793 | 0.555123 | -1.08175 | -0.34483 | 0.315647 | -3.6625 | -1.112 | 0.565481 | 0.118126 | -0.27505 | 0.105153 | 1.263637 | 0.231095 | 0.442796 | -0.93244 | 0.018758 | -0.56607 | -1.88239 | -1.28302 |
| 33.95833 | 0.893662 | -1.72766 | -7.00793 | -0.78499 | -0.1766 | -1.8515 | 0.017436 | -1.80347 | -0.63422 | -0.33478 | -5.08368 | -2.92334 | -2.59381 | -1.17631 | -1.01163 | -2.56227 | -0.79101 | -1.07306 | -0.17487 | -1.61008 | -1.22567 | -1.20089 | -1.75292 | -3.13545 |
| 33.97222 | -2.8352 | -3.50205 | -10.093 | -1.7102 | -0.83278 | -2.88287 | -0.74288 | -2.68902 | -1.07632 | -1.05605 | -7.03815 | -4.40962 | -8.07358 | -3.46847 | -2.0596 | -7.14732 | -3.93989 | -3.28195 | -0.90869 | -3.66841 | -3.26367 | -2.03788 | -2.90341 | -6.38871 |
| 33.98611 | -4.55175 | -3.93356 | -11.6052 | -1.80931 | -1.18015 | -2.47749 | -1.02765 | -2.87975 | -1.28207 | -1.16322 | -7.20215 | -4.97592 | -8.1205 | -4.06964 | -2.60271 | -8.87027 | -4.56394 | -3.99368 | -1.28745 | -4.60303 | -3.28681 | -1.9123 | -3.79085 | -7.03326 |
| 34 | -2.67053 | -3.34292 | -11.038 | -1.0485 | -0.83696 | -1.24008 | -0.71372 | -2.17326 | -0.88539 | -0.87114 | -5.63692 | -4.45985 | -4.51793 | -2.67453 | -3.09935 | -6.54888 | -3.04745 | -2.94822 | -1.19253 | -3.34383 | -1.48417 | -0.9629 | -3.1978 | -3.87899 |
| 34.01389 | -0.71505 | -2.3376 | -6.14185 | -0.0309 | -0.21813 | -0.06164 | -0.25424 | -1.4113 | -0.27056 | -0.47518 | -4.50988 | -4.24704 | -2.50106 | -0.87735 | -2.97419 | -4.19838 | -0.94165 | -1.40493 | -0.43595 | -2.06622 | -0.2909 | -0.23236 | -2.70356 | -1.83201 |
| 34.02778 | 0.050911 | -1.37233 | -5.02659 | 0.507248 | 0.249198 | 0.402251 | 0.126253 | -0.72899 | 0.257916 | -0.09767 | -3.71992 | -2.88058 | -2.18557 | 0.382563 | -1.89324 | -3.13964 | 0.534139 | -0.00399 | 0.825774 | -1.30033 | 0.056071 | 0.132767 | -2.49614 | -0.86043 |
| 34.04167 | -0.10105 | -0.60711 | -3.52294 | 0.692798 | 0.687425 | 0.546953 | 0.517216 | -0.23175 | 0.582314 | 0.284891 | -2.65339 | -1.70697 | -0.72493 | 1.094344 | -1.33391 | -2.08152 | 0.874516 | 0.702172 | 1.487937 | -0.84208 | 0.09477 | 0.327908 | -2.21141 | -0.60757 |
| 34.05556 | -0.40742 | -0.02486 | -1.70326 | 0.669775 | 0.877662 | 0.455157 | 0.754284 | 0.044149 | 0.627562 | 0.51035 | -2.04035 | -0.72109 | 0.973054 | 1.652554 | -0.71392 | -1.01648 | 0.882656 | 1.081011 | 1.438242 | -0.59129 | 0.19937 | 0.506416 | -1.46185 | -0.26647 |
| 34.06944 | -0.34481 | 0.372466 | 0.373487 | 0.573544 | 0.788604 | 0.34881 | 0.740705 | 0.071796 | 0.53655 | 0.555499 | -1.39363 | -0.22029 | 2.97783 | 1.985373 | 0.038449 | -0.0757 | 0.850041 | 1.374346 | 1.339021 | -0.32879 | 0.316507 | 0.646147 | -0.28673 | 0.302405 |
| 34.08333 | 0.095387 | 0.73896 | 2.599268 | 0.581804 | 0.671619 | 0.35746 | 0.615442 | -0.03141 | 0.548323 | 0.552991 | -0.69345 | 0.21582 | 4.54507 | 1.803551 | 0.636669 | 0.40455 | 0.821129 | 1.479247 | 1.297995 | -0.08158 | 0.185733 | 0.701283 | 0.735453 | 0.70078 |
| 34.09722 | 0.556131 | 1.14408 | 4.622553 | 0.823018 | 0.714182 | 0.529876 | 0.533905 | -0.07874 | 0.740117 | 0.528562 | -0.14391 | 0.47019 | 5.134857 | 1.763417 | 1.074424 | 0.544119 | 0.681158 | 1.682714 | 1.189801 | 0.044581 | -0.02298 | 0.599456 | 1.346856 | 0.84301 |
| 34.11111 | 0.604951 | 1.158637 | 6.449137 | 0.91066 | 0.779863 | 0.666881 | 0.499824 | -0.06456 | 0.812065 | 0.474826 | -0.02145 | 0.453178 | 3.179291 | 1.459723 | 1.227472 | 0.483344 | 0.391778 | 1.759702 | 0.97095 | 0.06661 | -0.12409 | 0.401269 | 1.388225 | 0.800853 |
| 34.125 | 0.329601 | 0.886357 | 8.14017 | 0.763176 | 0.818027 | 0.7973 | 0.492784 | 0.061967 | 0.770788 | 0.496624 | -0.10201 | 0.51855 | 0.637419 | 0.986725 | 1.148188 | 0.327864 | 0.183437 | 1.511905 | 0.695213 | 0.067136 | -0.04955 | 0.247738 | 1.277639 | 0.75686 |
| 34.13889 | 0.191383 | 0.805654 | 9.816031 | 0.738477 | 0.92902 | 0.827445 | 0.604011 | 0.282427 | 0.895898 | 0.763829 | 0.077319 | 0.732436 | -1.50031 | 0.547257 | 1.042259 | 0.130484 | 0.135981 | 1.099031 | 0.43189 | 0.108173 | 0.115485 | 0.115539 | 1.223136 | 0.885917 |
| 34.15278 | 0.340097 | 0.995836 | 8.903859 | 0.690817 | 1.218789 | 0.954104 | 0.874816 | 0.332081 | 1.078089 | 1.182482 | 0.376083 | 0.921846 | -2.70142 | 0.242227 | 0.873481 | -0.44929 | 0.221314 | 0.700014 | 0.305322 | 0.168342 | 0.246294 | -0.03427 | 1.13208 | 1.098413 |

| | | | | | | | | | | | | | | | | | | | | | | | | |
|---|---|---|---|---|---|---|---|---|---|---|---|---|---|---|---|---|---|---|---|---|---|---|---|---|
| 34.16667 | 0.468677 | 1.315564 | 6.597595 | 0.904374 | 1.646343 | 1.130895 | 1.194121 | 0.174129 | 1.145702 | 1.604247 | 0.63961 | 1.075228 | -1.69363 | 0.048394 | 1.0096 | -1.54422 | 0.432592 | 0.395214 | 0.373182 | 0.283063 | 0.35859 | 0.082094 | 0.995778 | 1.35879 |
| 34.18056 | 0.211128 | 1.744465 | 4.004124 | 1.600589 | 1.649288 | 1.27789 | 1.24517 | 0.0894 | 1.188067 | 1.77807 | 0.908161 | 1.227118 | 0.532045 | 0.153521 | 1.164568 | -1.66063 | 0.71889 | 0.056129 | 0.515128 | 0.246355 | 0.469359 | 0.257494 | 0.853032 | 1.505692 |
| 34.19444 | -0.4277 | 2.333748 | 3.035032 | 3.39257 | 0.97998 | 1.282477 | 0.62858 | 0.194447 | 1.300865 | 2.065066 | 1.22144 | 1.420873 | 2.970185 | 0.466225 | 1.176007 | -0.47902 | 0.88436 | -0.33544 | 0.481572 | -0.05355 | 0.632182 | 0.271303 | 0.68761 | 1.623881 |
| 34.20833 | -0.75941 | 3.001255 | 1.998579 | 3.627439 | 1.192067 | 1.213961 | 0.460585 | 0.279388 | 1.300995 | 2.296713 | 1.432056 | 1.440318 | 4.100606 | 0.867719 | 1.014798 | 0.540943 | 0.996029 | -0.38676 | 0.512767 | -0.27826 | 1.023381 | 0.388491 | 0.643802 | 1.85101 |
| 34.22222 | -0.53552 | 3.359303 | 0.764777 | 2.148585 | 2.01076 | 0.781639 | 1.254145 | -0.03615 | 0.966977 | 1.925767 | 1.230781 | 0.815975 | 4.020462 | 1.04645 | 1.027582 | 0.993668 | 1.090177 | -0.00399 | 0.68081 | -0.21542 | 1.209839 | 0.44847 | 0.802367 | 1.823281 |
| 34.23611 | 0.502365 | 3.749159 | -0.39306 | 0.002769 | 1.762034 | 0.385783 | 0.981739 | -0.61268 | 0.300533 | 0.840064 | 0.222863 | -0.9725 | 2.020826 | 0.715722 | 0.25435 | 0.828779 | 0.759478 | -0.26298 | 0.524772 | -0.65462 | 0.704 | -0.05093 | 0.265911 | 0.955974 |
| 34.25 | 2.592311 | 5.178985 | -0.70878 | 0.008317 | 1.888072 | 1.116851 | 0.957691 | -0.32586 | 0.091146 | 0.027166 | -0.6405 | -3.00366 | -1.02311 | 0.464351 | -1.90989 | 0.332177 | -1.02042 | -1.66061 | -0.28895 | -2.22202 | -0.5783 | -1.10978 | -1.1376 | -0.36766 |
| 34.26389 | 2.992804 | 6.094679 | -1.06832 | 0.956068 | 2.220208 | 2.580168 | 1.666467 | 0.865076 | 1.012942 | 0.805582 | 0.783696 | -1.66162 | -2.83942 | 0.901966 | -3.95028 | 0.892304 | -0.17765 | -2.71321 | -0.69587 | -2.59809 | -1.11213 | -1.45376 | -1.40666 | -0.77295 |
| 34.27778 | 0.832661 | 4.758631 | -2.98896 | 0.636081 | 1.692176 | 2.677464 | 1.685556 | 1.205076 | 1.567199 | 1.613989 | 2.666192 | 0.787103 | -1.25318 | 1.208928 | -1.08006 | 1.749702 | 0.70527 | -2.01477 | 0.084891 | -0.83744 | -0.07006 | -0.60528 | 0.13381 | 0.093926 |
| 34.29167 | -0.96118 | 2.560237 | -5.2976 | -0.04723 | 0.897541 | 1.675068 | 1.118278 | 0.674142 | 1.218157 | 1.488084 | 2.755127 | 1.331124 | 0.934724 | 1.26664 | 0.182474 | 1.449297 | 0.662692 | -0.80842 | 0.427311 | 0.416454 | 0.765771 | -0.14857 | 1.158864 | 0.53701 |
| 34.30556 | -1.40539 | 1.354113 | -5.47513 | 0.256571 | 0.739388 | 1.097095 | 0.948874 | 0.322573 | 0.919746 | 1.316412 | 2.607486 | 1.494558 | 1.862633 | 1.500922 | -0.09934 | 1.080566 | 0.452688 | 0.376751 | 0.635599 | 0.635326 | 0.99168 | -0.04486 | 1.500043 | 0.508823 |
| 34.31944 | -1.05264 | 0.461774 | -4.95116 | 0.708926 | 0.835536 | 0.770495 | 1.000786 | 0.136912 | 0.913497 | 1.050171 | 2.821246 | 1.601914 | 0.505623 | 1.665434 | -0.37201 | 0.990575 | 0.415519 | 1.058929 | 0.7663 | 0.599606 | 0.938381 | -0.04546 | 1.614378 | 0.563899 |
| 34.33333 | -0.45357 | -0.44642 | -5.22174 | 0.761694 | 0.813005 | 0.331004 | 0.957967 | -0.32641 | 0.87741 | 0.555159 | 3.745213 | 1.321366 | -1.67169 | 1.530584 | -0.25365 | 0.999061 | 0.30969 | 1.005797 | 0.829114 | 0.729674 | 0.907632 | -0.06684 | 1.473235 | 0.78851 |
| 34.34722 | 0.123236 | -0.95136 | -4.89556 | 0.818568 | 0.904581 | 0.255249 | 1.008468 | -0.77707 | 0.924726 | 0.274776 | 3.095737 | 0.734523 | -2.71633 | 1.317167 | 0.094022 | 1.13243 | 0.239853 | 0.555643 | 0.830312 | 1.092519 | 0.960705 | 0.059831 | 1.162 | 1.046936 |
| 34.36111 | 0.504135 | -1.07217 | -3.04823 | 1.022117 | 1.204006 | 0.526884 | 1.296449 | -0.78026 | 0.930849 | 0.346561 | 0.73747 | 0.34901 | -1.93028 | 1.01688 | 0.428368 | 1.094584 | 0.371815 | 0.290543 | 0.837392 | 1.468319 | 1.098474 | 0.31013 | 1.396157 | 1.170162 |
| 34.375 | 0.725565 | -0.90622 | -0.64514 | 1.264415 | 1.663292 | 0.706452 | 1.629148 | -0.48268 | 0.896445 | 0.494924 | 0.245654 | 0.140828 | -0.01952 | 0.741727 | 0.615243 | 0.786526 | 0.464799 | 0.401299 | 0.868599 | 1.387729 | 1.15799 | 0.420704 | 1.803294 | 0.980131 |
| 34.38889 | 0.826004 | -0.52997 | 2.216856 | 1.576754 | 2.253988 | 1.085919 | 1.967074 | -0.3038 | 0.925172 | 0.627539 | -0.2442 | 0.045989 | 2.355384 | 0.804303 | 0.652471 | 0.449026 | 0.456666 | 0.808244 | 0.791088 | 1.059842 | 1.04448 | 0.339522 | 1.469181 | 0.6563 |
| 34.40278 | 0.75292 | -0.19243 | 4.142203 | 1.917045 | 2.767196 | 1.777039 | 2.242788 | -0.23774 | 1.192671 | 0.836478 | -0.80604 | -0.07055 | 3.548631 | 1.030002 | 0.666303 | 0.337985 | 0.345868 | 1.265428 | 0.774773 | 0.848379 | 0.895072 | 0.263916 | 1.06235 | 0.420426 |
| 34.41667 | 0.4355 | 0.20787 | 4.144462 | 2.390272 | 3.066458 | 2.256331 | 2.318331 | -0.12063 | 1.675934 | 1.094444 | -1.25984 | -0.05553 | 2.55693 | 0.985779 | 0.355068 | 0.384436 | 0.186117 | 1.315198 | 0.82989 | 0.804373 | 0.776841 | 0.314772 | 1.104141 | 0.374261 |
| 34.43056 | 0.082837 | 0.531602 | 3.302561 | 2.494729 | 2.952455 | 2.124135 | 2.326836 | 0.009007 | 2.11159 | 1.376473 | -1.56228 | 0.183066 | -0.55891 | 0.577566 | 0.071596 | 0.592896 | 0.195894 | 0.845757 | 0.967619 | 0.870322 | 0.704861 | 0.489346 | 1.372758 | 0.57981 |
| 34.44444 | 0.139617 | 1.024917 | 2.16446 | 2.112429 | 2.743438 | 1.970328 | 2.496143 | 0.208934 | 2.293434 | 1.94174 | -1.80737 | 0.546919 | -3.09038 | 0.100713 | -0.07224 | 1.279995 | 0.560463 | 0.154037 | 1.169 | 1.391683 | 0.867143 | 0.875619 | 1.805919 | 1.181069 |
| 34.45833 | 0.956197 | 2.055039 | 2.703624 | 1.894312 | 3.002336 | 2.232391 | 3.239973 | 0.703654 | 2.469131 | 2.912388 | -1.75208 | 1.33376 | -3.24385 | 0.068777 | 0.200283 | 2.759702 | 1.371141 | -0.71073 | 1.384692 | 2.665083 | 1.414043 | 1.472122 | 2.343436 | 2.274412 |
| 34.47222 | 2.036219 | 3.084299 | 5.529518 | 2.919462 | 3.359981 | 2.538873 | 3.807364 | 1.237193 | 2.391843 | 3.557437 | -0.84124 | 2.053999 | -0.66097 | 0.686006 | 1.254313 | 3.714258 | 2.112043 | -0.60919 | 1.478615 | 3.830476 | 2.034253 | 1.726212 | 4.64026 | 3.939107 |
| 34.48611 | 2.669458 | 3.111667 | 7.797435 | 4.180475 | 3.452068 | 3.096546 | 3.84916 | 1.297393 | 1.895624 | 3.393624 | 0.04941 | 2.098398 | 3.22731 | 1.852722 | 1.988746 | 3.577492 | 2.240114 | 1.414351 | 1.495768 | 4.757454 | 2.141207 | 1.385547 | 7.306365 | 4.194666 |
| 34.5 | 2.32712 | 2.400398 | 6.497386 | 4.921196 | 3.283378 | 2.204707 | 3.839581 | 1.019463 | 1.259316 | 2.773468 | 0.248483 | 1.80923 | 5.458702 | 3.106251 | 2.060559 | 3.641668 | 2.813348 | 4.49384 | 1.660158 | 4.706864 | 2.15654 | 1.320862 | 7.886753 | 4.535667 |
| 34.51389 | 1.304035 | 1.873451 | 3.760805 | 5.207086 | 2.888205 | 0.8881 | 3.137435 | 0.686496 | 0.79793 | 2.294196 | 0.194259 | 1.610325 | 7.489429 | 3.443533 | 1.259714 | 3.910408 | 3.077095 | 5.551716 | 1.832665 | 4.423352 | 2.234539 | 1.601328 | 7.721435 | 4.145092 |
| 34.52778 | 0.253361 | 1.45949 | 1.671272 | 3.093606 | 1.825178 | 0.654107 | 2.015245 | 0.423477 | 0.736736 | 2.04245 | 0.231033 | 1.435594 | 7.635516 | 3.383337 | 0.792151 | 3.624247 | 2.882577 | 4.693692 | 1.71409 | 3.305988 | 2.281831 | 1.711325 | 6.630922 | 3.398335 |
| 34.54167 | -0.18954 | 1.269851 | -0.29008 | -0.30985 | 0.779986 | 0.451869 | 0.856492 | 0.111676 | 0.885303 | 1.795095 | 0.38646 | 1.208014 | 5.117114 | 2.534495 | 0.113062 | 3.115084 | 2.507897 | 3.15523 | 1.623142 | 2.203405 | 1.783698 | 1.780665 | 4.277499 | 2.680947 |
| 34.55556 | 0.187995 | 1.28233 | -2.27815 | -3.48009 | -0.05578 | 0.447453 | -0.0512 | -0.11549 | 1.074678 | 1.479189 | 0.600106 | 1.027684 | 1.245249 | 0.270607 | -0.73324 | 2.757318 | 2.277134 | 1.071847 | 1.520737 | 1.533894 | 0.786668 | 1.574084 | 1.975901 | 2.493602 |
| 34.56944 | 0.762741 | 1.087361 | -3.81936 | -3.86191 | -0.63173 | -0.08827 | -0.48629 | -0.28716 | 1.16146 | 1.218398 | 0.75609 | 0.846823 | -2.05972 | -1.58833 | -0.60381 | 2.324022 | 2.055562 | -0.40549 | 1.135267 | 1.123416 | -0.03884 | 1.241478 | 1.012543 | 3.090228 |
| 34.58333 | 0.771733 | 0.766918 | -6.68568 | -2.66315 | -0.68577 | -1.47905 | -0.13726 | -0.3378 | 1.046842 | 1.005768 | 0.791658 | 0.750128 | -4.97744 | -1.89498 | -0.3886 | 1.584539 | 1.625058 | -1.3089 | 0.755814 | 1.066519 | -0.10533 | 1.162837 | 1.248947 | 4.12007 |
| 34.59722 | 0.870369 | 0.650934 | -7.52626 | -2.07835 | -0.17876 | -1.72714 | 0.539254 | -0.04186 | 1.027017 | 1.084896 | 1.072398 | 0.909137 | -6.41438 | -1.17001 | -0.15786 | 1.28225 | 1.088454 | -2.14679 | 0.37731 | 0.80598 | 0.464639 | 1.241648 | 0.507427 | 2.863963 |
| 34.61111 | 1.408217 | 0.717786 | -6.98314 | -0.54576 | 0.656612 | -1.51597 | 1.217211 | 0.421737 | 1.150404 | 1.319645 | 1.633279 | 1.223008 | -5.48524 | -0.90183 | 0.038364 | 1.106364 | 0.689612 | -2.73304 | -0.09446 | 0.086497 | 0.819008 | 1.052611 | -1.13389 | 0.478206 |
| 34.625 | 1.831902 | 0.720707 | -5.48574 | 0.417922 | 1.365647 | -0.79607 | 1.638776 | 0.620109 | 1.158872 | 1.508619 | 1.826462 | 1.374248 | -3.08199 | -0.48142 | 0.489933 | 1.157016 | 0.432706 | -2.34861 | 0.005075 | -0.92276 | 1.117727 | 0.625901 | -3.3019 | -1.33013 |
| 34.63889 | 2.605486 | 0.688463 | -2.11742 | 0.978167 | 1.854074 | 0.21467 | 1.798085 | 0.576549 | 1.071103 | 1.558035 | 1.560201 | 1.271903 | -0.27841 | 0.204597 | 0.940493 | 1.324481 | 0.43834 | -0.90902 | 0.674151 | -2.12913 | 1.16593 | 0.243262 | -5.06726 | -1.93055 |
| 34.65278 | 4.198257 | 0.685651 | 2.224592 | 1.802048 | 2.167427 | 1.038943 | 1.766188 | 0.537174 | 0.977272 | 1.413507 | 0.952172 | 0.878968 | 2.391735 | 1.006932 | 0.926181 | 1.66951 | 0.371452 | 0.379561 | 1.201739 | -2.5927 | 0.982907 | 0.044252 | -4.8311 | -1.93857 |
| 34.66667 | 5.647563 | 0.84909 | 5.132593 | 2.855995 | 2.275947 | 1.996962 | 1.661886 | 0.537515 | 0.928237 | 1.220152 | 0.299772 | 0.387304 | 4.29499 | 1.692101 | 0.874924 | 2.009829 | 0.297252 | 0.983191 | 1.336877 | -2.2943 | 0.593582 | 0.085415 | -4.82895 | -1.91397 |
| 34.68056 | 6.006238 | 1.094616 | 6.288761 | 3.444224 | 2.320806 | 2.660648 | 1.676197 | 0.545746 | 0.970204 | 1.195403 | -0.22009 | 0.171356 | 5.060577 | 1.967977 | 1.296302 | 2.298542 | 0.340195 | 1.051463 | 1.300198 | -2.15774 | 0.125035 | 0.195433 | -4.90948 | -1.31469 |
| 34.69444 | 5.298078 | 1.338703 | 5.554824 | 3.044678 | 2.440757 | 2.714999 | 1.80122 | 0.474671 | 1.027196 | 1.347309 | -0.72816 | 0.003468 | 3.615026 | 1.9883 | 1.257773 | 2.414185 | 0.615315 | 0.924866 | 1.321403 | -1.96057 | 0.188421 | 0.187409 | -5.29618 | -0.33539 |
| 34.70833 | 3.837794 | 1.41322 | 5.105895 | 4.414983 | 2.572115 | 2.648828 | 1.982376 | 0.405031 | 1.019718 | 1.567383 | -1.34591 | -0.29546 | 1.974442 | 1.59605 | 0.315746 | 2.30052 | 0.86597 | 0.161535 | 1.329097 | -1.54449 | 0.091606 | 0.130915 | -5.34356 | 0.179141 |
| 34.72222 | 2.629891 | 1.398303 | 5.291497 | 5.914616 | 2.683414 | 2.634559 | 2.191584 | 0.342421 | 0.972636 | 1.834731 | -1.69276 | -0.55788 | -0.09243 | 1.011406 | -0.13185 | 1.947308 | 1.19558 | -0.57847 | 1.232649 | -1.09106 | 0.152431 | 0.106596 | -4.60257 | 0.194266 |
| 34.73611 | 1.655847 | 1.354057 | 4.158065 | 5.55282 | 2.759691 | 2.210453 | 2.299756 | 0.232816 | 0.99325 | 2.044646 | -1.53021 | -0.5278 | -2.69438 | 0.488635 | 0.024641 | 1.270767 | 1.528805 | -0.96662 | 1.174565 | -0.51171 | 0.288867 | 0.107274 | -2.68055 | 0.707019 |
| 34.75 | 0.574562 | 1.100876 | 1.980133 | 3.501552 | 2.616391 | 1.054849 | 2.188262 | 0.137964 | 0.954424 | 1.989874 | -1.07451 | -0.49734 | -4.77329 | 0.493008 | -0.32987 | 0.418436 | 1.611548 | -1.22164 | 1.239576 | -0.20523 | 0.468197 | 0.050338 | 0.688756 | 1.36981 |
| 34.76389 | -0.75437 | 0.765583 | -0.18124 | 0.536022 | 2.322188 | 0.069752 | 1.842394 | 0.056347 | 0.865046 | 1.638122 | -0.77805 | -0.44237 | -5.69313 | 0.406226 | -1.13985 | -0.3529 | 1.49994 | -1.27671 | 1.416501 | 0.103494 | 0.607 | -0.08191 | 4.613592 | 2.284563 |
| 34.77778 | -2.31036 | 0.550534 | -2.40784 | -1.94177 | 1.800333 | -0.91072 | 1.328598 | -0.08548 | 0.780458 | 1.139849 | -0.53626 | -0.2953 | -4.7497 | 0.523111 | -1.38836 | -0.65375 | 1.382549 | -1.22169 | 1.526959 | 0.432123 | 0.659394 | -0.12609 | 7.647781 | 3.332737 |
| 34.79167 | -4.09985 | 0.426746 | -4.63623 | -2.90165 | 1.161149 | -2.06157 | 0.73767 | -0.24203 | 0.706829 | 0.64886 | -0.16743 | -0.02314 | -4.58912 | 0.809478 | -1.73956 | -0.28914 | 1.474711 | -1.46013 | 1.337868 | 0.68657 | 0.582099 | -0.06179 | 6.87287 | 3.611788 |
| 34.80556 | -5.76113 | 0.352304 | -7.03991 | -1.74751 | 0.591685 | -3.33414 | 0.219631 | -0.34988 | 0.573889 | 0.151693 | 0.440887 | 0.140087 | -3.92516 | 0.955446 | -1.92989 | 0.487293 | 1.681602 | -2.29969 | 0.793091 | 0.753858 | 0.294723 | -0.06347 | 6.358687 | 3.262935 |
| 34.81944 | -7.40008 | 0.307571 | -8.14396 | -1.06838 | 0.179456 | -4.35785 | -0.09661 | -0.44758 | 0.280966 | -0.35091 | 1.049739 | 0.082769 | -3.30872 | 0.687041 | -1.73788 | 1.199757 | 1.569603 | -3.48136 | 0.249115 | 0.518568 | -0.13697 | 0.021807 | 4.752937 | 2.486612 |
| 34.83333 | -8.80809 | 0.4264 | -8.30872 | -1.26884 | -0.14034 | -4.59643 | -0.35711 | -0.71459 | -0.04645 | -0.82254 | 1.338728 | -0.14821 | -2.69575 | 0.151631 | -1.06479 | 1.650272 | 1.226851 | -4.02378 | -0.08059 | -0.00035 | -0.37141 | 0.137393 | 1.56116 | 0.954213 |
| 34.84722 | -9.61383 | 0.538471 | -9.05967 | -0.72239 | -0.43896 | -3.39653 | -0.51447 | -1.06169 | -0.28101 | -1.13863 | 1.292987 | -0.48955 | -1.98317 | -0.26947 | -0.38255 | 1.875955 | 0.96467 | -3.95731 | -0.10225 | -0.73405 | -0.34804 | 0.180023 | -2.59072 | -1.2617 |
| 34.86111 | -9.42879 | 0.556085 | -6.73933 | -0.63501 | -0.6263 | -2.56565 | -0.55814 | -1.1384 | -0.46246 | -1.21954 | 0.924998 | -0.8335 | -1.62503 | -0.38967 | 0.164728 | 1.888743 | 0.922084 | -3.41199 | 0.094255 | -1.87639 | -0.11807 | 0.223961 | -6.56944 | -3.02525 |
| 34.875 | -7.45733 | 0.458833 | -3.06201 | -0.48818 | -0.61587 | -2.03396 | -0.38101 | -1.01808 | -0.61945 | -1.07989 | 0.390533 | -1.0505 | -1.51064 | -0.33588 | 0.726374 | 1.696991 | 0.525679 | -2.16315 | 0.539626 | -2.79974 | 0.237712 | 0.411551 | -8.63459 | -4.09358 |
| 34.88889 | -4.89116 | 0.455678 | 1.460679 | -0.04482 | -0.36791 | -1.53502 | -0.07493 | -0.78794 | -0.70449 | -0.79069 | -0.15358 | -0.87447 | -1.42883 | -0.23249 | 1.379602 | 1.228107 | -0.19519 | -0.98999 | 1.059647 | -3.10551 | 0.572528 | 0.691584 | -10.0718 | -3.69171 |
| 34.90278 | -2.73753 | 0.555117 | 5.725939 | 0.51449 | -0.1562 | -0.95579 | 0.201606 | -0.61496 | -0.66756 | -0.4567 | -0.41258 | -0.39965 | -0.90777 | -0.26116 | 1.665757 | 0.861632 | -1.0992 | -0.24184 | 1.380322 | -2.60978 | 0.742021 | 0.892932 | -7.48514 | -1.48167 |
| 34.91667 | -0.94843 | 0.48582 | 8.421213 | 0.874564 | 0.04299 | 0.044557 | 0.404353 | -0.3497 | -0.57159 | -0.11795 | -0.3893 | 0.065559 | -0.20378 | -0.45956 | 1.383662 | 0.62677 | -1.62335 | 0.387241 | 1.450274 | -2.39213 | 0.704301 | 0.908543 | -6.15565 | 0.001369 |
| 34.93056 | 0.430425 | 0.26819 | 8.593512 | 0.866244 | 0.128365 | 0.402117 | 0.4434 | -0.31942 | -0.50331 | 0.039699 | -0.3918 | 0.494919 | 0.327148 | -1.18598 | 0.716172 | 0.32082 | -1.82624 | 1.18228 | 1.53919 | -2.12559 | 0.575502 | 0.741709 | -5.08068 | 0.21693 |
| 34.94444 | 0.449354 | 0.057217 | 6.743805 | 0.632973 | -0.09395 | -0.10552 | 0.199969 | -0.3863 | -0.66803 | -0.29834 | -1.06744 | 0.331577 | 0.255018 | -2.0459 | -0.19185 | -0.36476 | -2.17448 | 1.531182 | 1.60501 | -2.33428 | 0.225501 | 0.316163 | -4.38009 | -0.18921 |
| 34.95833 | -1.94335 | -0.64113 | 1.488166 | -0.62858 | -0.80958 | -1.46834 | -0.33298 | -1.43106 | -1.178 | -1.22088 | -3.52077 | -0.76153 | -1.37568 | -2.71725 | -2.10171 | -2.16461 | -3.24109 | -0.0447 | 1.362456 | -3.0964 | -0.42972 | -0.42408 | -5.37468 | -1.86995 |
| 34.97222 | -6.1763 | -1.57434 | -5.11126 | -1.90078 | -1.51458 | -2.73003 | -0.96579 | -2.831 | -1.88644 | -2.40489 | -6.96952 | -2.27035 | -4.25498 | -3.07752 | -4.62059 | -6.32627 | -4.94684 | -3.09506 | 0.289383 | -4.53009 | -1.21345 | -1.2043 | -6.3472 | -3.97527 |
| 34.98611 | -7.1055 | -1.78282 | -9.67696 | -1.89448 | -1.42942 | -2.90285 | -1.32138 | -3.34419 | -2.20387 | -2.80134 | -8.82308 | -3.64969 | -6.82493 | -2.55279 | -6.17454 | -7.60015 | -4.45907 | -6.36905 | -1.42915 | -4.63128 | -1.6069 | -1.41905 | -5.7252 | -4.3562 |
| 35 | -5.2923 | -1.1657 | -12.4173 | -0.80289 | -0.87278 | -1.56762 | -1.07173 | -2.7515 | -1.66121 | -2.38423 | -7.04803 | -3.7642 | -7.24356 | -0.98909 | -4.87306 | -6.07661 | -2.66045 | -7.8543 | -1.96954 | -2.95872 | -1.042 | -0.92821 | -3.67249 | -2.62547 |
| 35.01389 | -3.05483 | -0.52161 | -12.5935 | 0.204686 | -0.19283 | -0.78677 | -0.53242 | -1.84234 | -0.93089 | -1.80831 | -4.64202 | -3.60125 | -6.43646 | 0.156789 | -4.25535 | -3.37606 | -0.9882 | -6.07329 | -1.46048 | -1.64607 | -0.26764 | -0.41018 | -2.61835 | -0.53138 |
| 35.02778 | -1.91649 | -0.13386 | -9.83324 | 0.437871 | 0.412673 | -0.42238 | 0.033078 | -1.2049 | -0.45395 | -1.27065 | -3.44547 | -3.3913 | -2.39999 | 0.912124 | -2.59627 | -1.67163 | 0.164732 | -3.32414 | -0.60995 | -0.95649 | 0.187469 | -0.14856 | -1.74808 | 0.97702 |
| 35.04167 | -1.7059 | 0.127381 | -6.71044 | 0.579614 | 0.864469 | -0.19272 | 0.404636 | -0.48534 | -0.09391 | -1.01569 | -2.68931 | -2.32464 | 1.327961 | 1.212107 | -1.13288 | -1.21713 | 0.258053 | -1.4117 | 0.056111 | -0.50944 | 0.349957 | -0.01674 | -0.97727 | 1.589268 |
| 35.05556 | -1.51523 | 0.241264 | -5.47529 | 0.627386 | 1.001892 | 0.096469 | 0.529876 | 0.090221 | 0.223469 | -0.70836 | -2.33173 | -1.36872 | 4.376066 | 1.107145 | -0.72772 | -2.033 | -0.86003 | -0.15159 | 0.509348 | -0.22916 | 0.252935 | -0.04565 | -0.38956 | 1.406349 |
| 35.06944 | -1.16273 | 0.368364 | -4.79334 | 0.612594 | 0.938231 | 0.415321 | 0.596115 | 0.271795 | 0.473734 | -0.43071 | -1.73417 | -0.70855 | 4.1752 | 0.792287 | -0.45834 | -1.87696 | -1.05704 | 0.708084 | 0.922037 | -0.02969 | 0.160582 | -0.1536 | 0.068583 | 1.371465 |
| 35.08333 | -0.57628 | 0.552856 | -3.86342 | 0.6559 | 0.883563 | 0.585792 | 0.637748 | 0.132202 | 0.632606 | -0.13866 | -1.29206 | -0.10607 | 4.007368 | 0.610887 | -0.40438 | -0.6123 | -0.10389 | 1.592203 | 1.443326 | 0.129327 | 0.231218 | -0.11425 | 0.387328 | 1.317116 |
| 35.09722 | -0.0616 | 0.687565 | -2.99724 | 0.690925 | 0.848586 | 0.59533 | 0.610059 | -0.16165 | 0.69008 | 0.091976 | -1.06209 | 0.487847 | 4.045613 | 0.447338 | -0.28849 | 0.326675 | 0.704282 | 2.310916 | 2.122739 | 0.195753 | 0.378174 | 0.058356 | 0.394938 | 1.043522 |
| 35.11111 | 0.259899 | 0.661153 | -2.06318 | 0.656371 | 0.92406 | 0.501887 | 0.60036 | -0.35899 | 0.64497 | 0.153043 | -1.01964 | 0.972773 | 3.128361 | 0.47892 | -0.07028 | 0.709541 | 0.990009 | 2.731932 | 2.83576 | 0.170655 | 0.453283 | 0.233313 | 0.300068 | 0.682605 |
| 35.125 | 0.430964 | 0.57802 | -1.20996 | 0.694195 | 1.121161 | 0.341154 | 0.653899 | -0.25508 | 0.583212 | 0.177592 | -0.97337 | 1.377003 | 1.90433 | 0.645371 | 0.175987 | 0.634795 | 0.873737 | 2.641654 | 3.407194 | 0.216503 | 0.457923 | 0.386448 | 0.163439 | 0.320565 |
| 35.13889 | 0.502249 | 0.572975 | -0.43984 | 0.764931 | 1.421074 | 0.211073 | 0.841318 | -0.00456 | 0.565776 | 0.332791 | -0.83043 | 1.700131 | 0.185658 | 0.513465 | 0.204152 | 0.407619 | 0.550838 | 2.195018 | 3.319332 | 0.371824 | 0.444129 | 0.466397 | 0.028623 | 0.087552 |
| 35.15278 | 0.615 | 0.662837 | 0.311355 | 0.830728 | 1.76793 | 0.163525 | 1.103564 | 0.183144 | 0.516261 | 0.599223 | -0.59765 | 2.049398 | -1.32965 | 0.381594 | 0.102319 | 0.022703 | 0.114969 | 1.648102 | 2.846564 | 0.459258 | 0.351312 | 0.442553 | 0.112135 | -0.0066 |
| 35.16667 | 0.834453 | 0.785527 | 1.024194 | 0.894461 | 2.067047 | 0.189888 | 1.332285 | 0.146971 | 0.461409 | 0.814694 | -0.36137 | 2.296111 | -1.99002 | 0.278313 | 0.009721 | -0.37292 | -0.24704 | 1.108948 | 2.310088 | 0.452648 | 0.249184 | 0.329446 | 0.389698 | 0.144229 |
| 35.18056 | 1.118133 | 0.937608 | 1.56514 | 0.906977 | 2.255891 | 0.240528 | 1.461308 | -0.13632 | 0.452872 | 0.991654 | -0.29641 | 2.212351 | -1.64115 | 0.340031 | 0.052708 | -0.49 | -0.29074 | 0.618903 | 1.910816 | 0.355746 | 0.282571 | 0.290309 | 0.882017 | 0.606388 |
| 35.19444 | 1.29586 | 1.050125 | 2.295731 | 0.892879 | 2.316562 | 0.319059 | 1.479027 | -0.46052 | 0.490533 | 1.288323 | -0.44632 | 1.777668 | -0.52548 | 0.453533 | 0.354013 | -0.40533 | -0.18244 | 0.260378 | 1.566391 | 0.166467 | 0.430693 | 0.28604 | 1.271161 | 1.085854 |
| 35.20833 | 1.296521 | 1.102348 | 3.461445 | 0.865865 | 2.234518 | 0.471039 | 1.431838 | -0.56527 | 0.530564 | 1.607094 | -0.64291 | 1.226024 | 1.763254 | 0.59242 | 0.850862 | -0.0752 | -0.0206 | -0.01433 | 1.242044 | -0.06377 | 0.540108 | 0.188678 | 1.398464 | 1.343031 |
| 35.22222 | 1.152583 | 1.110758 | 6.034279 | 0.839611 | 2.222675 | 0.73235 | 1.538776 | -0.55739 | 0.50402 | 1.834882 | -0.74838 | 0.951991 | 3.906995 | 0.8113 | 1.300767 | 0.614632 | 0.449407 | -0.1647 | 0.898402 | -0.1693 | 0.578501 | 0.061645 | 1.335191 | 1.405381 |
| 35.23611 | 1.107745 | 1.034161 | 6.667294 | 0.832726 | 2.240176 | 1.086868 | 1.796297 | -0.44537 | 0.541202 | 1.919564 | -0.78821 | 0.886983 | 5.273202 | 1.044797 | 1.530253 | 1.360827 | 1.061238 | -0.12138 | 0.686141 | -0.08437 | 0.597905 | -0.01402 | 1.220584 | 1.439529 |
| 35.25 | 1.317925 | 0.982931 | 8.792404 | 0.872507 | 2.188663 | 1.133294 | 2.016194 | -0.20234 | 0.670251 | 2.011022 | -0.77748 | 0.831105 | 5.730038 | 1.175671 | 1.680061 | 1.971613 | 1.562784 | 0.0407 | 0.700599 | 0.065618 | 0.659808 | 0.041533 | 1.117947 | 1.479078 |
| 35.26389 | 1.550762 | 0.992423 | 9.999644 | 0.826518 | 2.046905 | 0.888219 | 2.028143 | -0.03211 | 0.824276 | 2.024327 | -0.73165 | 0.769164 | 5.05483 | 1.202947 | 1.859403 | 2.060622 | 1.502885 | 0.20676 | 0.828969 | 0.145385 | 0.785262 | 0.162128 | 1.121159 | 1.535847 |
| 35.27778 | 2.208374 | 0.97678 | 12.11214 | 0.74292 | 1.874288 | 0.749126 | 1.837206 | -0.03673 | 0.871978 | 1.882291 | -0.69596 | 0.598501 | 4.310772 | 1.157036 | 1.829486 | 1.740668 | 1.143232 | 0.084923 | 0.861327 | 0.127702 | 0.861814 | 0.243378 | 1.168656 | 1.502061 |
| 35.29167 | 2.789616 | 0.924728 | 13.64261 | 0.975524 | 1.689058 | 1.144886 | 1.720446 | -0.02751 | 0.892852 | 1.720548 | -0.68048 | 0.440945 | 3.592855 | 1.202258 | 1.712272 | 1.522652 | 0.885415 | -0.20709 | 0.762743 | 0.098852 | 0.889161 | 0.249742 | 1.324809 | 1.382773 |
| 35.30556 | 2.642942 | 0.936556 | 11.54475 | 1.731687 | 1.489171 | 1.985829 | 1.696786 | 0.120557 | 0.984581 | 1.698584 | -0.63737 | 0.312086 | 3.469228 | 1.255152 | 1.552388 | 1.525829 | 0.861516 | -0.37124 | 0.742657 | 0.019276 | 0.873363 | 0.197639 | 1.384312 | 1.30358 |
| 35.31944 | 3.404218 | 0.975469 | 7.902891 | 3.47028 | 1.387451 | 2.236963 | 1.7773 | 0.21798 | 1.152137 | 1.678672 | -0.59594 | 0.25267 | 3.232824 | 1.196715 | 1.185988 | 1.504473 | 0.832818 | -0.16802 | 0.846464 | -0.12555 | 0.809554 | 0.070483 | 1.189342 | 1.263725 |
| 35.33333 | 2.035053 | 0.972332 | 3.402203 | 2.933008 | 1.408964 | 1.685926 | 1.909484 | 0.129981 | 1.234554 | 1.59819 | -0.59701 | 0.219842 | 2.587849 | 1.154392 | 0.805527 | 1.162553 | 0.561811 | 0.167056 | 0.987397 | -0.14906 | 0.788478 | -0.0386 | 0.923443 | 1.271789 |
| 35.34722 | 0.094473 | 0.971501 | 0.331813 | 0.234674 | 1.545038 | 1.613211 | 2.044363 | -0.02306 | 1.216115 | 1.493153 | -0.52637 | 0.054359 | 1.708219 | 1.094508 | 0.424949 | 0.812404 | 0.379081 | 0.496052 | 1.109046 | -0.01975 | 0.858665 | -0.05723 | 0.73159 | 1.434162 |
| 35.36111 | -1.8241 | 0.971318 | -2.76594 | -2.07763 | 1.784265 | 1.408284 | 2.130645 | -0.1636 | 1.172912 | 1.484464 | -0.54277 | -0.19359 | -0.11088 | 0.882491 | 0.017017 | 0.652159 | 0.410362 | 0.859752 | 1.332359 | 0.089913 | 0.846373 | 0.023395 | 0.616543 | 1.51852 |
| 35.375 | -2.78169 | 0.904754 | -7.29593 | -2.51894 | 2.054477 | 0.330044 | 2.210892 | -0.28168 | 1.227098 | 1.60833 | -0.64492 | -0.30528 | -2.47145 | 0.651723 | -0.31149 | 0.410029 | 0.285054 | 1.138728 | 1.582979 | 0.123609 | 0.818816 | 0.186919 | 0.524411 | 1.30654 |
| 35.38889 | -0.96716 | 0.901416 | -11.331 | -0.37236 | 2.370011 | -0.50999 | 2.326169 | -0.28456 | 1.432023 | 1.686268 | -0.72825 | -0.10931 | -4.26053 | 0.516379 | -0.35983 | 0.240281 | 0.197274 | 1.12232 | 1.711184 | 0.082313 | 0.717372 | 0.3114 | 0.505253 | 0.997043 |
| 35.40278 | -1.04034 | 1.016269 | -14.9101 | 0.772173 | 2.579798 | -0.74984 | 2.43461 | -0.17199 | 1.728556 | 1.712069 | -1.05325 | 0.175338 | -4.74014 | 0.577306 | -0.17365 | 0.362589 | 0.476243 | 0.930122 | 1.703749 | 0.025891 | 0.636014 | 0.398869 | 0.550273 | 0.782979 |
| 35.41667 | -1.54361 | 1.176096 | -17.852 | 1.186939 | 2.766619 | -0.62459 | 2.517206 | -0.18677 | 2.016971 | 1.689929 | -1.57568 | 0.454346 | -3.85836 | 0.601772 | -0.06209 | 0.586468 | 0.877293 | 0.723781 | 1.604836 | -0.01939 | 0.569009 | 0.44799 | 0.474671 | 0.704946 |
| 35.43056 | -1.31952 | 1.180688 | -17.7038 | 1.210558 | 2.798303 | -0.7861 | 2.588982 | -0.31331 | 1.950723 | 1.571328 | -2.23915 | 0.84198 | -1.96633 | 0.712908 | -0.06891 | 0.60399 | 0.993943 | 0.494407 | 1.461109 | -0.02915 | 0.366545 | 0.330683 | 0.409122 | 0.781815 |
| 35.44444 | -0.41481 | 0.842387 | -13.6858 | 1.157675 | 2.898425 | -1.17934 | 2.807747 | -0.23295 | 1.675188 | 1.71687 | -2.32984 | 1.238916 | 0.108746 | 0.917608 | 0.313829 | 0.833382 | 0.918051 | 0.245732 | 1.359554 | 0.305043 | 0.214288 | 0.299584 | 0.588973 | 1.184407 |
| 35.45833 | 1.1733 | 0.667128 | -8.16539 | 1.219326 | 3.315835 | 0.334578 | 3.444602 | 0.214971 | 1.53586 | 2.469682 | -1.87459 | 1.759621 | 2.201627 | 1.202117 | 1.482196 | 1.928894 | 1.350946 | -0.239 | 1.362968 | 1.184395 | 0.306545 | 0.565552 | 1.264338 | 2.11385 |
| 35.47222 | 2.739364 | 0.818947 | 0.462859 | 1.23652 | 3.617404 | 1.32909 | 3.607911 | 1.003698 | 1.476955 | 3.060287 | -0.96872 | 1.970353 | 4.002065 | 1.157488 | 2.537277 | 3.36386 | 2.201725 | -0.9629 | 1.009047 | 2.259855 | 0.614898 | 0.75735 | 2.117509 | 2.834691 |
| 35.48611 | 3.674904 | 0.903049 | 9.300217 | 1.370946 | 3.124711 | 2.351219 | 3.067032 | 1.75947 | 1.451359 | 3.083913 | 0.185607 | 1.735916 | 6.534098 | 0.634002 | 2.978743 | 4.211423 | 2.859731 | -0.58309 | 0.996923 | 2.646085 | 0.97023 | 0.79153 | 3.357545 | 3.025525 |
| 35.5 | 4.66637 | 0.822699 | 13.46438 | 1.8827 | 3.131375 | 2.208551 | 2.96841 | 2.317369 | 1.55143 | 2.828632 | 0.931394 | 1.768214 | 8.773089 | 0.942073 | 3.065198 | 4.099428 | 2.889467 | 2.314603 | 2.274455 | 2.984423 | 1.519057 | 1.044161 | 4.755803 | 3.204707 |
| 35.51389 | 3.991223 | 0.545324 | 11.82799 | 3.344399 | 2.913498 | 1.556917 | 2.662479 | 2.03302 | 1.599589 | 2.661292 | 0.926204 | 2.008186 | 7.730429 | 0.218748 | 2.697334 | 3.577492 | 2.630972 | 4.471805 | 3.6283 | 3.330392 | 2.035622 | 1.422295 | 5.947628 | 2.92192 |
| 35.52778 | 2.803143 | 0.332148 | 9.300615 | 3.163517 | 2.067294 | 2.767572 | 2.181209 | 1.118428 | 1.507776 | 2.787848 | 0.71163 | 2.399535 | 6.812398 | 0.625943 | 1.660513 | 2.890384 | 2.133148 | 4.633254 | 3.771477 | 3.119659 | 2.255903 | 1.710895 | 6.52787 | 2.815326 |
| 35.54167 | 2.754581 | 0.918947 | 7.510878 | 3.136628 | 1.386768 | 3.082527 | 1.874642 | 0.273436 | 1.427889 | 3.159618 | 0.801581 | 2.64336 | 4.604674 | 0.940677 | 0.577641 | 2.033905 | 1.324548 | 4.204501 | 3.781379 | 2.484105 | 1.983118 | 1.826025 | 6.609543 | 2.455113 |
| 35.55556 | 3.404104 | 1.302868 | 5.850547 | 1.899674 | 0.889212 | 2.073308 | 1.630831 | -0.74465 | 1.543565 | 3.482817 | 0.864867 | 2.726327 | 1.290171 | 0.432678 | -0.3594 | 1.059716 | 0.234387 | 3.709555 | 3.501874 | 2.342536 | 1.616666 | 1.851961 | 5.987109 | 1.256916 |
| 35.56944 | 3.324656 | 2.042123 | 5.045934 | 2.807644 | 0.47659 | 1.646914 | 1.33426 | -1.44102 | 1.458368 | 3.375984 | 0.687189 | 2.773476 | -2.11964 | 0.066501 | -1.19015 | 0.136619 | -0.64109 | 2.871298 | 2.512717 | 1.919367 | 0.742668 | 1.771691 | 6.246423 | -0.2306 |
| 35.58333 | 2.688113 | 2.277002 | 3.479831 | 3.279885 | 0.312463 | 1.214243 | 1.160178 | -1.2615 | 1.191373 | 2.776324 | 0.216518 | 2.472334 | -4.09449 | 0.455131 | -1.89645 | -0.48756 | -1.14024 | 1.242184 | 1.279522 | 1.578518 | -0.5296 | 1.580299 | 5.747413 | -0.76575 |
| 35.59722 | 1.796699 | 1.546801 | 1.175149 | 1.318549 | 0.471281 | 0.904621 | 1.237195 | -0.4077 | 0.982132 | 2.13685 | 0.055066 | 1.811623 | -7.32123 | 0.833183 | -1.71431 | -0.74416 | -0.99045 | -0.14456 | 0.327439 | 1.781358 | -0.80468 | 1.324203 | 4.852274 | 0.929456 |
| 35.61111 | 0.765924 | 0.32872 | -1.38955 | -1.32492 | 0.817956 | 0.600075 | 1.401552 | 0.654057 | 0.808598 | 1.53961 | 0.342489 | 1.08479 | -7.09353 | 0.895823 | -1.20023 | -0.70478 | -0.48468 | 0.201055 | -0.15911 | 2.069975 | 0.528648 | 0.945199 | 3.342503 | 2.542772 |
| 35.625 | -0.16334 | -0.75062 | -4.25583 | -1.78166 | 1.167393 | 0.217902 | 1.499856 | 1.335168 | 0.615666 | 0.839828 | 0.548323 | 0.434159 | -5.06649 | 0.758572 | -0.49671 | -0.473 | 0.164523 | -0.09541 | -0.31304 | 2.104353 | 1.835623 | 0.509246 | 4.177251 | 3.86959 |

| | | | | | | | | | | | | | | | | | | | | | | | | |
|---|---|---|---|---|---|---|---|---|---|---|---|---|---|---|---|---|---|---|---|---|---|---|---|---|
| 35.63889 | -0.37499 | -0.44108 | -7.15813 | -1.1042 | 1.474104 | -0.84678 | 1.535435 | 1.480137 | 0.49339 | 0.437094 | 0.67153 | 0.083823 | -1.3432 | 0.479781 | -0.1504 | -0.20603 | 0.91369 | -0.73728 | -0.52834 | 1.804256 | 2.259338 | 0.055801 | 4.179989 | 4.889955 |
| 35.65278 | -0.72811 | 0.302674 | -10.1432 | -0.45758 | 1.745675 | -1.732 | 1.650503 | 1.217557 | 0.434146 | 0.407478 | 0.599259 | 0.000313 | 1.394764 | 0.263752 | -0.07361 | 0.105083 | 1.636216 | -0.72308 | 0.041523 | 1.486133 | 1.479999 | -0.19922 | 4.443186 | 4.542497 |
| 35.66667 | -0.87507 | 0.429504 | -13.2195 | -0.67297 | 1.974362 | -1.60192 | 1.8965 | 0.685604 | 0.484476 | 0.696739 | 0.386063 | 0.177626 | 2.61096 | 0.360552 | 0.196995 | 0.607371 | 2.043798 | 0.135582 | 1.132869 | 1.019124 | 1.029863 | -0.15264 | 3.908394 | 3.144583 |
| 35.68056 | -1.68983 | 0.20658 | -16.0635 | -0.33281 | 2.13413 | -1.43378 | 2.160018 | 0.220001 | 0.623961 | 0.967349 | 0.308598 | 0.370781 | 3.896153 | 0.68778 | 0.638748 | 1.239717 | 2.059133 | 1.027921 | 2.376161 | 0.449444 | 0.671522 | 0.134807 | 2.938306 | 1.252519 |
| 35.69444 | -1.19548 | 0.224395 | -12.9002 | 0.653865 | 2.138067 | -0.72505 | 2.256814 | 0.239188 | 0.770954 | 1.088443 | 0.367456 | 0.534856 | 4.319761 | 0.988148 | 1.072747 | 1.783188 | 1.948648 | 1.369406 | 2.997214 | -0.00501 | 0.654912 | 0.47959 | 1.262841 | -0.50067 |
| 35.70833 | -0.35859 | 0.331174 | -6.61792 | 1.038682 | 2.183662 | -0.14012 | 2.29998 | 0.393066 | 0.84871 | 1.229071 | 0.395938 | 0.634148 | 4.518007 | 1.256642 | 1.030956 | 2.090057 | 1.964392 | 1.221384 | 2.857575 | -0.31092 | 0.874365 | 0.743591 | -1.7753 | -1.81554 |
| 35.72222 | 0.244998 | 0.440812 | -1.77537 | 1.314663 | 2.355442 | 0.344332 | 2.398174 | 0.230726 | 0.897025 | 1.449151 | 0.353018 | 0.634891 | 4.354921 | 1.22678 | 1.077709 | 2.105155 | 2.121749 | 0.718994 | 2.138478 | -0.58254 | 0.889145 | 0.899374 | -3.5139 | -2.33607 |
| 35.73611 | 0.511251 | 0.536404 | 3.064586 | 1.666148 | 2.541466 | 0.956635 | 2.556831 | -0.2 | 0.941329 | 1.720142 | 0.228963 | 0.55058 | 3.447521 | 0.891301 | 0.982617 | 1.916747 | 2.114615 | -0.12665 | 1.35202 | -0.84902 | 0.649171 | 0.722969 | -5.47864 | -2.39208 |
| 35.75 | 0.546934 | 0.552839 | 7.441887 | 1.920767 | 2.72449 | 1.527651 | 2.710831 | -0.59062 | 0.915588 | 1.839016 | 0.011276 | 0.463653 | 1.886444 | 0.476188 | 0.613737 | 1.602366 | 1.890173 | -0.97274 | 0.645793 | -1.16846 | 0.55784 | 0.455621 | -7.38043 | -1.68188 |
| 35.76389 | 0.404144 | 0.492071 | 10.34293 | 1.886074 | 2.636491 | 1.925346 | 2.581225 | -0.95216 | 0.842866 | 1.758109 | -0.15361 | 0.377836 | -0.60724 | 0.220111 | -0.02261 | 1.13597 | 1.485122 | -1.4602 | 0.257615 | -1.6997 | 0.69243 | 0.272043 | -8.16564 | -0.36309 |
| 35.77778 | 0.423628 | 0.345561 | 11.92507 | 1.922158 | 2.512153 | 2.131955 | 2.176275 | -1.18773 | 0.796155 | 1.615811 | -0.29489 | 0.332068 | -3.94878 | -0.07215 | -0.96289 | 0.475898 | 1.171374 | -1.56778 | 0.096097 | -1.81977 | 0.588859 | 0.060109 | -8.01484 | 1.37681 |
| 35.79167 | 0.437102 | 0.072493 | 12.08 | 2.655545 | 2.445836 | 2.307561 | 1.82116 | -1.04631 | 0.726817 | 1.431249 | -0.41277 | 0.211024 | -5.01354 | -0.35456 | -1.94025 | -0.18973 | 0.930544 | -1.44797 | -0.08434 | -1.66124 | 0.136031 | -0.16917 | -6.86681 | 1.726477 |
| 35.80556 | 0.11501 | -0.31398 | 12.88786 | 3.989426 | 2.137333 | 3.401755 | 1.238714 | -0.71783 | 0.626591 | 1.006662 | -0.54152 | 0.037054 | -6.33314 | -0.64886 | -2.61993 | -0.57146 | 0.582113 | -1.30534 | -0.29706 | -1.5569 | -0.39862 | -0.32536 | -5.35494 | 1.036732 |
| 35.81944 | -0.46119 | -0.65034 | 10.72757 | 3.983041 | 1.538297 | 2.472849 | 0.402902 | -0.89801 | 0.425334 | 0.38249 | -0.72885 | -0.20982 | -6.4726 | -0.61827 | -2.69457 | -0.89372 | 0.249742 | -0.93754 | -0.30959 | -1.52785 | -0.68766 | -0.38808 | -4.76877 | -0.23455 |
| 35.83333 | 0.189712 | -0.47625 | 6.614042 | 1.622598 | 0.717444 | 1.231937 | -0.4354 | -1.71302 | 0.118005 | -0.36695 | -1.16956 | -0.5916 | -6.13706 | -0.40212 | -2.73597 | -1.13724 | -0.01123 | -0.48689 | -0.00592 | -1.36422 | -0.67473 | -0.39712 | -3.43293 | -1.02317 |
| 35.84722 | -0.66628 | -0.60377 | 2.115962 | -0.69036 | 0.144831 | -1.24318 | -1.22292 | -2.59163 | -0.25731 | -1.07226 | -1.80912 | -1.02778 | -5.59893 | -0.33045 | -2.49713 | -1.11717 | -0.16954 | -0.336 | 0.238489 | -1.02421 | -0.70546 | -0.39724 | -1.21339 | -1.16966 |
| 35.86111 | -2.13687 | -0.89389 | -1.12498 | -2.47013 | -0.31017 | -2.35752 | -2.02486 | -2.78412 | -0.64057 | -1.6864 | -2.37821 | -1.31325 | -5.79094 | -0.33082 | -1.42341 | -1.02643 | -0.17309 | -0.48418 | 0.438912 | -1.45986 | -0.72542 | -0.47686 | 1.206487 | -0.79826 |
| 35.875 | -2.61816 | -1.02745 | -4.02436 | -3.60313 | -0.84705 | -2.9838 | -2.3021 | -2.61232 | -0.96001 | -2.12812 | -2.66743 | -1.37589 | -5.32872 | -0.23734 | -1.09646 | -0.68239 | 0.055826 | -0.87949 | 0.622323 | 0.062739 | -0.60584 | -0.44334 | 3.042039 | -0.01181 |
| 35.88889 | -3.03159 | -0.56596 | -8.19099 | -3.45802 | -0.82171 | -2.96006 | -1.98681 | -2.23147 | -1.14436 | -2.18037 | -2.59893 | -1.30159 | -4.03952 | -0.13541 | -0.69014 | -0.09187 | 0.451227 | -1.46711 | 0.625511 | 0.293281 | -0.39291 | -0.18759 | 3.391641 | 0.461183 |
| 35.90278 | -3.27227 | -0.00814 | -15.65 | -1.52793 | -0.55606 | -2.95855 | -1.46608 | -1.80917 | -1.1586 | -2.09314 | -2.18291 | -1.20805 | -1.86946 | -0.14738 | -0.22547 | 0.657655 | 0.856349 | -1.82702 | 0.662773 | 0.175086 | -0.01566 | 0.143675 | 2.089528 | 0.598991 |
| 35.91667 | -2.62941 | 0.310651 | -18.8954 | -0.2151 | -0.45401 | -2.83474 | -1.13967 | -1.22413 | -1.1922 | -1.90056 | -1.43051 | -1.00328 | 0.20222 | -0.22989 | 0.263785 | 1.328862 | 1.174749 | -1.37056 | 1.042187 | -0.03411 | 0.460499 | 0.433789 | -0.08715 | 0.470968 |
| 35.93056 | -1.73419 | 0.358949 | -18.8527 | -0.00837 | -0.54125 | -3.15723 | -1.18095 | -0.28918 | -1.19349 | -1.73243 | -0.74969 | -0.68582 | 1.690438 | -0.28325 | 0.685145 | 1.869041 | 1.504546 | -0.51314 | 1.728483 | -0.21014 | 0.875768 | 0.665975 | -2.12245 | 0.228074 |
| 35.94444 | -0.88746 | 0.200045 | -14.8408 | -0.11246 | -0.9442 | -3.80873 | -1.4472 | 0.203144 | -1.40588 | -2.06674 | -0.98443 | -0.8393 | 1.394531 | -0.46982 | 0.535796 | 1.553166 | 0.850916 | -0.79723 | 1.337734 | -0.80437 | 0.869426 | 0.557575 | -4.44732 | -0.13358 |
| 35.95833 | -2.25613 | -0.1515 | -11.5435 | -0.56511 | -1.92179 | -4.74868 | -2.25789 | -1.07584 | -2.18927 | -3.11575 | -2.84104 | -2.44954 | -1.73177 | -1.10414 | -1.21471 | -0.64737 | -1.70526 | -3.10363 | -0.93663 | -2.48151 | -0.21352 | -0.23076 | -7.70707 | -1.62832 |
| 35.97222 | -4.15217 | -0.58987 | -10.8053 | -1.11848 | -2.86392 | -5.6417 | -2.9675 | -3.84411 | -3.35815 | -4.43043 | -6.49041 | -5.35154 | -5.7809 | -1.69933 | -4.28463 | -2.91889 | -3.2938 | -7.04545 | -4.34783 | -4.7639 | -1.90246 | -1.68968 | -10.414 | -4.31967 |
| 35.98611 | -4.71964 | -0.9322 | -9.89547 | -1.40363 | -2.92026 | -5.61219 | -3.0253 | -5.4851 | -4.13645 | -5.06111 | -7.88828 | -7.0141 | -9.31984 | -1.50269 | -5.85145 | -2.41645 | -1.66888 | -9.10311 | -6.15145 | -5.25455 | -3.08094 | -2.85845 | -11.0106 | -6.11285 |
| 36 | -3.39844 | -0.90114 | -10.2087 | -1.04817 | -2.17867 | -4.92052 | -2.4593 | -4.00396 | -4.54773 | -4.88999 | -5.75489 | -6.78451 | -6.18668 | -0.84783 | -4.48568 | -1.49583 | -0.50786 | -7.2684 | -4.85927 | -3.4411 | -2.57147 | -2.59679 | -7.04941 | -2.91971 |
| 36.01389 | -2.30466 | -0.47541 | -9.53542 | -0.24702 | -1.06691 | -3.65299 | -1.63416 | -1.60237 | -4.3491 | -4.07422 | -3.84336 | -5.60768 | -2.40514 | -0.18061 | -3.07956 | -0.39698 | 0.368626 | -5.11022 | -3.2329 | -1.99767 | -1.55653 | -1.97724 | -5.07341 | -0.31736 |
| 36.02778 | -1.68223 | 0.023393 | -7.38921 | 0.511484 | -0.34708 | -2.37985 | -1.09175 | -0.02837 | -3.65408 | -3.05392 | -2.77018 | -4.19257 | -0.76782 | 0.443966 | -2.26363 | 0.423505 | 1.332855 | -4.964 | -3.30462 | -1.21417 | -1.36606 | -1.58999 | -3.72768 | 0.737006 |
| 36.04167 | -1.25117 | 0.387125 | -5.11421 | 0.91597 | 0.06624 | -1.2955 | -0.66169 | 0.691908 | -2.72885 | -1.90594 | -2.43656 | -3.14079 | 0.015369 | 0.850146 | -1.33431 | 1.127625 | 2.392372 | -8.29971 | -6.61425 | -0.76522 | -2.03301 | -1.55157 | -2.99483 | 0.975251 |
| 36.05556 | -0.90647 | 0.655969 | -3.75528 | 1.042096 | 0.638125 | -0.41082 | 0.075311 | 1.162199 | -1.67691 | -0.91907 | -2.13965 | -2.31657 | 0.182918 | 0.895141 | -0.37847 | 1.36942 | 2.804991 | -9.54458 | -5.91838 | -0.62301 | -2.43665 | -1.40818 | -2.52566 | 1.075988 |
| 36.06944 | -0.68726 | 0.782215 | -2.96286 | 1.005278 | 1.115604 | 0.189915 | 0.736329 | 1.202569 | -0.7887 | -0.17433 | -2.03476 | -1.59371 | 0.24343 | 0.683126 | 0.159958 | 0.588175 | 2.4737 | -5.60834 | -3.30652 | -0.51318 | -1.8159 | -0.99042 | -1.92983 | 1.223353 |
| 36.08333 | -0.56071 | 0.794705 | -2.22451 | 0.880246 | 1.436664 | 0.427246 | 1.255546 | 0.884604 | -0.24717 | 0.219669 | -2.07098 | -1.04957 | 0.729586 | 0.502236 | 0.402423 | -0.11277 | 2.265404 | -2.98944 | -2.00836 | -0.39934 | -0.89885 | -0.61322 | -0.94008 | 1.75254 |
| 36.09722 | -0.4502 | 0.73687 | -1.77824 | 0.710474 | 1.571243 | 0.467239 | 1.459277 | 0.516233 | 0.005294 | 0.357068 | -2.17487 | -0.74091 | 1.740102 | 0.415269 | 0.517206 | -0.21644 | 2.228556 | -1.67295 | -1.08817 | -0.36825 | -0.29148 | -0.4549 | -0.19895 | 2.77826 |
| 36.11111 | -0.31351 | 0.650006 | -1.39999 | 0.561047 | 1.618558 | 0.456951 | 1.515279 | 0.156829 | 0.118143 | 0.350638 | -2.11959 | -0.70877 | 3.309041 | 0.308837 | 0.568577 | -0.13435 | 2.169489 | -1.01688 | -0.55439 | -0.40338 | -0.18759 | -0.52517 | 0.035997 | 3.409553 |
| 36.125 | -0.133 | 0.564697 | -1.0092 | 0.507857 | 1.742046 | 0.456023 | 1.63601 | -0.16346 | 0.258198 | 0.398698 | -2.38337 | -0.6203 | 6.217447 | 0.177599 | 0.213368 | -0.28863 | 1.527096 | -0.82798 | -0.25898 | -0.2678 | -0.19345 | -0.66412 | -0.01526 | 2.674647 |
| 36.13889 | 0.038126 | 0.525024 | -0.67877 | 0.540879 | 2.066383 | 0.518022 | 1.900042 | -0.547 | 0.516559 | 0.6615 | -2.64116 | -0.30772 | 5.297772 | 0.101881 | -0.13506 | -0.39238 | 0.858418 | -0.95109 | -0.067 | -0.16252 | -0.05764 | -0.6312 | 0.098825 | 1.888352 |
| 36.15278 | 0.125799 | 0.513952 | -0.36648 | 0.596531 | 2.485083 | 0.664637 | 2.231719 | -0.95084 | 0.757611 | 1.025951 | -2.67495 | 0.19765 | 4.5115 | 0.190582 | -0.35807 | -0.38346 | 0.168068 | -1.2084 | 0.099653 | -0.05677 | 0.254821 | -0.38759 | 0.540754 | 1.782709 |
| 36.16667 | 0.155502 | 0.543377 | -0.13246 | 0.65214 | 2.794009 | 0.866814 | 2.534299 | -1.04922 | 0.972058 | 1.348729 | -2.34081 | 0.704641 | 4.468841 | 0.394765 | -0.55669 | -0.39537 | -0.43516 | -1.40964 | 0.325878 | 0.018631 | 0.642583 | -0.07111 | 1.160542 | 2.033816 |
| 36.18056 | 0.150669 | 0.664868 | 0.019917 | 0.731088 | 2.906027 | 1.038486 | 2.709104 | -0.70282 | 1.170428 | 1.538906 | -1.70877 | 1.162043 | 4.199517 | 0.522816 | -0.8277 | -0.61884 | -1.17534 | -1.17559 | 0.752076 | -0.01536 | 0.965266 | 0.106714 | 1.532151 | 1.955381 |
| 36.19444 | 0.222067 | 0.867367 | 0.273781 | 0.799676 | 2.862346 | 1.06529 | 2.696607 | -0.05061 | 1.247859 | 1.642019 | -0.97221 | 1.486038 | 3.238517 | 0.560615 | -0.99291 | -0.70208 | -1.41716 | -0.58184 | 1.290438 | -0.08045 | 0.932413 | 0.193863 | 1.570549 | 1.731976 |
| 36.20833 | 0.485351 | 1.034701 | 0.678957 | 0.863312 | 2.819526 | 0.936602 | 2.555086 | 0.711608 | 1.242727 | 1.709655 | -0.33512 | 1.759569 | 2.625321 | 0.653698 | -1.01266 | -0.41219 | -0.83538 | 0.034922 | 1.699318 | 0.170647 | 0.990686 | 0.30549 | 1.659292 | 1.734576 |
| 36.22222 | 0.849317 | 1.158983 | 1.300685 | 0.928712 | 2.856319 | 0.70075 | 2.456323 | 1.333558 | 1.124316 | 1.755352 | 0.180571 | 1.917324 | 1.869563 | 0.723414 | -0.69234 | 0.201306 | 0.310715 | 0.392375 | 1.915538 | 0.493838 | 1.253882 | 0.359403 | 1.730182 | 2.165637 |
| 36.23611 | 1.112395 | 1.220631 | 2.007925 | 0.950489 | 2.92421 | 0.38228 | 2.337392 | 1.624866 | 0.966294 | 1.817843 | 0.568464 | 2.139879 | 0.952468 | 0.679199 | -0.09069 | 0.916324 | 1.581362 | 0.490852 | 1.901661 | 0.725469 | 1.474289 | 0.316412 | 1.758048 | 2.706207 |
| 36.25 | 1.118528 | 1.171303 | 2.612722 | 0.922597 | 2.76514 | 0.117682 | 2.172392 | 1.504893 | 0.74677 | 1.729261 | 0.803427 | 2.041878 | -0.11541 | 0.589812 | 0.433174 | 1.480694 | 2.690725 | 0.433884 | 1.620515 | 0.765942 | 1.522167 | 0.195813 | 2.079486 | 3.627407 |
| 36.26389 | 0.95919 | 1.055318 | 2.754429 | 0.885682 | 2.31692 | 0.052117 | 1.884266 | 1.07472 | 0.567002 | 1.615952 | 0.999512 | 1.652337 | -0.92501 | 0.519764 | 0.526304 | 1.783367 | 3.083276 | 0.253561 | 1.27619 | 0.736661 | 1.403541 | 0.049363 | 2.584475 | 4.80991 |
| 36.27778 | 0.787876 | 0.914029 | 3.065197 | 0.833682 | 1.929752 | 0.104365 | 1.65827 | 0.349626 | 0.48275 | 1.514789 | 1.153421 | 1.146786 | -1.27319 | 0.504458 | 0.338693 | 1.74006 | 2.730436 | -0.0073 | 0.824968 | 0.6435 | 1.274565 | 0.006242 | 2.771318 | 5.114851 |
| 36.29167 | 0.724293 | 0.730346 | 4.478627 | 0.734986 | 1.657682 | 0.054347 | 1.460734 | -0.37827 | 0.324432 | 1.371319 | 1.00501 | 0.584274 | -0.75595 | 0.567888 | 0.338096 | 1.442697 | 1.953776 | -0.29096 | 0.426453 | 0.470814 | 1.101249 | 0.154154 | 2.498001 | 4.417048 |
| 36.30556 | 0.694972 | 0.643052 | 6.421866 | 0.674018 | 1.532301 | -0.12036 | 1.347574 | -0.84801 | 0.092881 | 1.126261 | 0.441392 | 0.064412 | 0.196029 | 0.613691 | 0.679808 | 1.041687 | 1.086048 | -0.58854 | 0.114222 | 0.082294 | 0.770782 | 0.204611 | 2.043433 | 3.234283 |
| 36.31944 | 0.639294 | 0.65011 | 8.032358 | 0.692402 | 1.592395 | -0.35792 | 1.392719 | -1.13626 | -0.11407 | 0.954987 | -0.11539 | -0.30269 | 0.787019 | 0.638429 | 1.114776 | 0.770427 | 0.600722 | -0.85711 | -0.07111 | -0.28339 | 0.530073 | 0.138928 | 1.543102 | 1.912362 |
| 36.33333 | 0.536901 | 0.540955 | 8.166165 | 0.715489 | 1.807846 | -0.63735 | 1.545577 | -1.28201 | -0.1606 | 0.874673 | -0.49593 | -0.38508 | 1.026666 | 0.70897 | 1.45613 | 0.711314 | 0.692728 | -0.89571 | -0.03595 | -0.46988 | 0.348227 | 0.119147 | 0.966129 | 0.539878 |
| 36.34722 | 0.370547 | 0.307233 | 7.751773 | 0.653724 | 2.010958 | 0.05145 | 1.79013 | -1.30938 | -0.03471 | 0.843228 | -0.78393 | -0.37865 | 1.580106 | 0.780551 | 1.35114 | 0.731492 | 0.781202 | -0.76703 | 0.086357 | -0.36444 | 0.206901 | 0.130437 | 0.566331 | -0.31979 |
| 36.36111 | 0.193835 | 0.156709 | 8.198183 | 0.54783 | 2.219287 | 0.263534 | 2.016278 | -1.42636 | 0.189851 | 0.821711 | -1.06421 | -0.42055 | 2.277618 | 0.763235 | 0.802269 | 0.657025 | 0.840212 | -0.71344 | 0.039941 | -0.19617 | 0.202908 | 0.052161 | 0.355263 | -0.04518 |
| 36.375 | 0.152336 | 0.161797 | 8.388331 | 0.456552 | 2.329069 | 0.372725 | 2.183548 | -1.60424 | 0.410767 | 0.813272 | -1.30635 | -0.49209 | 2.461042 | 0.71297 | 0.131893 | 0.463545 | 0.922786 | -0.82385 | -0.08479 | -0.08699 | 0.437483 | -0.0507 | 0.290933 | 1.106355 |
| 36.38889 | 0.15929 | 0.268153 | 7.644072 | 0.470554 | 2.428002 | 0.521758 | 2.291666 | -1.63392 | 0.636787 | 0.790308 | -1.31516 | -0.49327 | 2.047444 | 0.704589 | 0.059258 | 0.306719 | 0.971521 | -0.75567 | 0.006679 | 0.026233 | 0.702208 | -0.23126 | 0.352692 | 2.478806 |
| 36.40278 | 0.198251 | 0.550989 | 7.260144 | 0.596742 | 2.503378 | 0.68683 | 2.33883 | -1.53941 | 0.939603 | 0.899659 | -1.19673 | -0.34958 | 1.595704 | 0.698449 | 0.531827 | 0.058111 | 0.919608 | -0.3777 | 0.270735 | 0.148043 | 0.854439 | -0.59513 | 0.313515 | 3.605415 |
| 36.41667 | 0.280316 | 0.991543 | 6.993892 | 0.786981 | 2.656605 | 0.792807 | 2.437511 | -1.28199 | 1.285081 | 1.180851 | -1.23419 | -0.11364 | 1.289718 | 0.708407 | 0.9809 | -0.0619 | 0.74123 | 0.005059 | 0.587547 | 0.256364 | 0.903342 | -0.47552 | 0.159574 | 3.197742 |
| 36.43056 | 0.395138 | 1.062282 | 6.590389 | 0.73237 | 2.600173 | 0.805607 | 2.492824 | -0.86054 | 1.319486 | 1.450021 | -1.45379 | 0.110351 | 0.658466 | 0.716862 | 0.829804 | 0.089058 | 0.751978 | 0.437082 | 1.019064 | 0.387925 | 1.141515 | 0.1259 | 0.083566 | 1.679893 |
| 36.44444 | 0.489488 | 0.611967 | 6.80148 | 0.393104 | 2.492736 | 0.959956 | 2.690657 | 0.092323 | 1.255409 | 1.793194 | -1.38926 | 0.451377 | 0.21771 | 0.823808 | 0.498111 | 0.929432 | 1.479253 | 1.359801 | 2.009382 | 0.702972 | 1.624576 | 0.720909 | 0.677979 | 0.407132 |
| 36.45833 | 0.653876 | 0.385515 | 9.918636 | 0.198653 | 2.656494 | 1.455227 | 3.468915 | 1.818724 | 1.721744 | 2.427215 | -0.42311 | 1.343702 | 1.304962 | 1.130988 | 1.240513 | 2.300726 | 2.477603 | 3.221712 | 3.93336 | 1.786628 | 2.279529 | 1.413076 | 2.460074 | 0.184868 |
| 36.47222 | 1.647526 | 0.883496 | 11.47427 | 1.114001 | 2.765295 | 1.974106 | 4.06715 | 4.250673 | 2.497038 | 3.077905 | 1.50953 | 2.429275 | 3.728416 | 1.383134 | 2.849683 | 3.797312 | 3.061675 | 5.142738 | 5.530588 | 3.181138 | 2.560613 | 1.814866 | 4.494164 | 1.18487 |
| 36.48611 | 2.581365 | 1.3605 | 7.481067 | 1.367253 | 2.721856 | 2.194965 | 3.769227 | 5.320553 | 2.903694 | 3.361073 | 3.238972 | 3.531705 | 4.908114 | 0.997839 | 4.142282 | 4.181438 | 3.330567 | 6.699261 | 6.046508 | 3.661966 | 2.627959 | 1.858686 | 4.026464 | 2.152863 |
| 36.5 | 2.250758 | 1.171667 | 4.105943 | 1.031221 | 2.883387 | 2.289706 | 3.415046 | 5.143245 | 2.996822 | 3.417795 | 3.731188 | 3.883546 | 5.465436 | 0.787993 | 4.996651 | 3.654765 | 3.944963 | 8.434001 | 6.76781 | 3.608052 | 3.331412 | 1.956476 | 3.73686 | 3.330109 |
| 36.51389 | 2.505178 | 0.793717 | 2.568639 | 1.359499 | 2.520504 | 2.280682 | 3.033053 | 3.887303 | 2.999931 | 3.151569 | 3.325738 | 4.221345 | 6.691433 | 1.033078 | 4.947384 | 3.001959 | 3.045498 | 9.004652 | 7.426969 | 3.400959 | 3.628829 | 1.841803 | 4.425615 | 3.938665 |
| 36.52778 | 3.146758 | 0.675782 | 2.271977 | 1.12108 | 2.062981 | 2.162371 | 2.645878 | 2.213585 | 2.823127 | 3.033685 | 2.785881 | 4.162793 | 7.387657 | 1.737499 | 3.601615 | 2.294485 | 2.50502 | 8.784758 | 7.600924 | 3.263753 | 3.254689 | 1.867557 | 5.212163 | 3.723084 |
| 36.54167 | 1.935313 | 0.768217 | 1.188041 | 0.720976 | 1.739985 | 2.302893 | 2.469546 | 0.436193 | 2.485903 | 3.209492 | 2.406182 | 4.235512 | 7.80085 | 1.099932 | 1.783323 | 2.185063 | 2.437527 | 8.82033 | 8.162746 | 2.515696 | 2.563213 | 2.152929 | 5.777994 | 3.725065 |
| 36.55556 | 0.232396 | 0.874677 | 0.05095 | 0.307276 | 1.518686 | 2.621108 | 2.660646 | -1.32314 | 2.346833 | 3.500255 | 1.929881 | 4.507931 | 6.384811 | -0.32133 | -0.16222 | 2.169599 | 2.215049 | 8.442341 | 7.842797 | 1.686533 | 1.456425 | 2.595406 | 7.641118 | 2.580436 |
| 36.56944 | -0.73687 | 0.87696 | -0.49133 | 0.177154 | 1.471453 | 2.758883 | 2.855913 | -2.78438 | 2.185231 | 3.605219 | 1.208029 | 3.889416 | 4.305862 | -1.04903 | -1.92228 | 2.051327 | 1.994823 | 7.329314 | 6.634489 | 1.011404 | 0.798494 | 2.651329 | 6.919107 | 0.546545 |
| 36.58333 | -1.20038 | 0.818877 | -0.5961 | 0.290141 | 1.350857 | 2.575021 | 2.674346 | -2.90907 | 1.79711 | 3.347205 | 0.278638 | 2.925038 | 0.526181 | -0.78068 | -2.475 | 1.747217 | 1.575699 | 5.45023 | 4.712247 | 0.839204 | 0.835685 | 2.285606 | 5.750206 | -1.40788 |
| 36.59722 | -1.35805 | 0.799006 | -0.67694 | 0.378875 | 1.159817 | 2.24756 | 2.402293 | -2.53828 | 1.447739 | 3.094985 | -0.20114 | 2.108229 | -3.33206 | 0.322395 | -1.64066 | 1.201277 | 0.934424 | 3.415862 | 2.874001 | 0.851972 | 0.740311 | 1.581904 | 5.306738 | -3.45336 |
| 36.61111 | -1.12881 | 0.852625 | -0.69176 | 0.537753 | 1.030062 | 1.905385 | 2.192584 | -1.30634 | 1.202953 | 2.889638 | -0.32234 | 1.69834 | -8.35097 | 0.734901 | 0.023015 | 0.370242 | 0.332789 | 1.856319 | 1.495857 | 1.531903 | 0.455628 | 1.206977 | 3.997501 | -3.46359 |
| 36.625 | -0.90042 | 0.873929 | -0.6851 | 0.757786 | 0.959507 | 1.409815 | 1.815122 | -0.19454 | 0.978105 | 2.459909 | -0.27626 | 1.228754 | -11.546 | 0.980531 | 1.402735 | -0.399 | 0.382236 | 1.180333 | 0.781897 | 2.266977 | 0.292284 | 0.736682 | 4.105758 | -2.27703 |
| 36.63889 | -0.82734 | 0.840193 | -0.8683 | 0.879322 | 0.910325 | 0.89933 | 1.339186 | 0.760478 | 0.724418 | 1.790115 | -0.16682 | 0.729856 | -6.84865 | 1.132365 | 1.385018 | -0.75489 | 0.72436 | 0.933064 | 0.46694 | 2.617031 | 0.430318 | 0.341436 | 4.876021 | 2.866941 |
| 36.65278 | -1.12999 | 0.787217 | -0.99352 | 0.96001 | 0.916331 | 0.557994 | 1.055437 | 1.783667 | 0.613663 | 1.255438 | 0.038352 | 0.396386 | -1.17683 | 1.090419 | 0.759755 | -0.65488 | 1.010439 | 0.690846 | 0.347518 | 2.47008 | 0.315961 | 0.055312 | 5.376985 | 6.063495 |
| 36.66667 | -1.47889 | 0.745031 | -1.17918 | 1.04904 | 1.060928 | 0.479562 | 1.012813 | 2.716844 | 0.556388 | 0.967467 | 0.025739 | 0.242531 | 0.403737 | 0.967749 | -0.36629 | -0.18763 | 1.193415 | 0.463034 | 0.360387 | 2.060959 | 0.324249 | -0.10905 | 5.18434 | 6.787654 |
| 36.68056 | -1.20149 | 0.710681 | -1.22651 | 1.126748 | 1.231426 | 0.609668 | 1.223926 | 2.994588 | 0.546651 | 0.871995 | -0.25971 | 0.163671 | 1.228019 | 0.914966 | -1.1422 | 0.316925 | 1.453798 | 0.208703 | 0.358758 | 1.251528 | 0.465518 | -0.15329 | 3.121022 | 5.163818 |
| 36.69444 | -1.03476 | 0.721314 | -1.43688 | 1.192441 | 1.423157 | 0.813642 | 1.508479 | 2.207666 | 0.573085 | 0.880391 | -0.65858 | 0.198616 | 2.797404 | 1.198141 | -1.47459 | 0.622456 | 1.94147 | -0.24108 | 0.303885 | 0.037319 | 0.414467 | -0.18807 | 0.312259 | 2.622439 |
| 36.70833 | -0.8005 | 0.75275 | -1.73761 | 1.276315 | 1.636854 | 1.026729 | 1.809234 | 1.099059 | 0.618197 | 0.975381 | -0.86357 | 0.33066 | 4.302405 | 1.925648 | -1.17518 | 0.93624 | 2.127746 | -0.6895 | 0.229199 | -0.70666 | 0.190304 | 0.018161 | -0.52444 | -0.57493 |
| 36.72222 | -0.4911 | 0.756174 | -1.45855 | 1.304918 | 1.877164 | 1.1592 | 2.075761 | -0.05025 | 0.612075 | 1.059969 | -0.98247 | 0.518066 | 4.346519 | 1.448577 | -0.54761 | 1.28805 | 2.236489 | -0.82356 | 0.41035 | -0.72558 | 0.30094 | 0.353216 | 1.028207 | -2.45597 |
| 36.73611 | -0.36018 | 0.709856 | -1.68045 | 1.23771 | 2.049213 | 1.183219 | 2.271128 | -1.28124 | 0.55736 | 0.959735 | -1.02654 | 0.623025 | 2.161985 | 0.207543 | -0.03199 | 1.550887 | 2.507946 | -0.43503 | 0.953871 | -0.42055 | 0.604794 | 0.75669 | 2.159359 | -3.60934 |
| 36.75 | -0.2722 | 0.655302 | -2.67366 | 1.186223 | 2.231331 | 1.163384 | 2.477185 | -2.39994 | 0.530869 | 0.75765 | -1.0985 | 0.682113 | 1.597203 | -0.50361 | 0.246055 | 1.758635 | 3.004899 | 0.178167 | 1.663331 | -0.26297 | 1.146661 | 0.980902 | 0.586366 | -5.26692 |
| 36.76389 | 0.039369 | 0.598331 | -3.17468 | 1.206875 | 2.316955 | 1.137676 | 2.639611 | -3.2267 | 0.503108 | 0.570386 | -1.28267 | 0.733585 | 2.622522 | 0.073584 | 0.240568 | 1.887968 | 3.706101 | 0.930453 | 2.152913 | -0.36943 | 1.266759 | 0.987417 | -4.45536 | -6.86403 |
| 36.77778 | 0.143988 | 0.568603 | -3.76321 | 1.139823 | 2.27477 | 1.084974 | 2.483528 | -2.97971 | 0.542442 | 0.261772 | -1.40246 | 0.666489 | -0.85081 | -0.02509 | -0.04835 | 1.922293 | 3.747917 | 1.422236 | 2.243524 | -0.65539 | 0.969214 | 0.756266 | -9.33952 | -4.9026 |
| 36.79167 | 0.070715 | 0.594657 | -4.66634 | 0.952157 | 2.123489 | 1.005609 | 2.241426 | -2.15838 | 0.568627 | 0.247367 | -1.23997 | 0.602733 | -0.54356 | -0.39523 | -0.33941 | 1.727264 | 3.173219 | 1.340504 | 1.961795 | -1.01044 | 0.328792 | 0.533967 | -10.4604 | -0.7723 |
| 36.80556 | -0.27755 | 0.640388 | -5.40659 | 0.752432 | 1.886453 | 0.940158 | 1.933032 | -1.186 | 0.637358 | 0.324481 | -0.82801 | 0.441606 | -3.81894 | -0.78733 | -0.07209 | 1.269371 | 2.345805 | 0.8062 | 1.614417 | -1.39652 | -0.10302 | 0.363454 | -10.3489 | 1.597354 |
| 36.81944 | -0.6757 | 0.586998 | -5.89418 | 0.583096 | 1.556727 | 0.764623 | 1.462052 | -0.50253 | 0.601127 | 0.422967 | -0.45303 | 0.155733 | -5.9261 | -1.12857 | 0.080031 | 0.576954 | 1.559388 | 0.159352 | 1.237784 | -1.79801 | -0.54357 | 0.165078 | -9.50643 | 2.102144 |
| 36.83333 | -0.98465 | 0.433356 | -6.23362 | 0.370325 | 1.076274 | 0.301812 | 0.701143 | -0.31064 | 0.385337 | 0.303316 | -0.46992 | -0.28773 | -5.69218 | -1.29642 | -0.61611 | -0.23104 | 0.347681 | -0.5578 | 0.892698 | -1.8754 | -0.8509 | -0.0815 | -7.85877 | 1.503846 |
| 36.84722 | -1.02831 | 0.29781 | -6.96961 | 0.206369 | 0.471629 | -0.45468 | -0.32591 | -0.44084 | -0.04026 | -0.18699 | -1.01708 | -0.84066 | -4.72406 | -1.41241 | -1.98206 | -1.12827 | -1.25265 | -1.35476 | 0.490925 | -1.54797 | -1.04787 | -0.28387 | -6.5909 | 1.13824 |
| 36.86111 | -0.83689 | 0.258536 | -8.2999 | 0.193724 | -0.191 | -1.51111 | -1.67538 | -0.92428 | -0.65493 | -0.8282 | -1.75361 | -1.45503 | -4.80506 | -1.36364 | -3.0749 | -1.70273 | -1.67184 | -2.12103 | -0.28941 | -1.01795 | -1.0457 | -0.41275 | -6.08809 | 0.883248 |
| 36.875 | -0.77511 | 0.260476 | -7.45511 | 0.189553 | -0.83056 | -2.67734 | -3.11468 | -1.6224 | -1.31444 | -1.4236 | -2.19334 | -2.0335 | -5.07236 | -1.06561 | -2.61251 | -1.58887 | -1.35418 | -2.79471 | -1.95687 | -1.21844 | -1.10218 | -0.40979 | -5.08813 | 0.713507 |
| 36.88889 | -0.81591 | 0.264696 | -5.93984 | 0.161656 | -0.81704 | -3.6644 | -3.93506 | -2.07573 | -1.8838 | -1.94463 | -2.15199 | -2.35767 | -4.27232 | -0.58669 | -1.02009 | -0.96623 | -0.514 | -1.37876 | -4.75723 | 0.640601 | -1.00962 | -0.4927 | -3.94197 | 0.986383 |
| 36.90278 | -0.71267 | 0.299432 | -4.88811 | 0.242471 | -0.50271 | -4.10616 | -4.14065 | -2.11536 | -2.28504 | -2.23058 | -1.6614 | -2.3279 | -2.80406 | -0.25205 | 0.399888 | -0.15884 | 0.025564 | -1.02486 | -4.0164 | 1.179765 | -0.99763 | -0.83232 | -2.69664 | 1.477042 |

**Supplementary Table S2.** Raw motion data from: (a) Hypoxia stress

| Time | Control1 | Control2 | Control3 | Control4 | Control5 | Control6 | Control7 | Control8 | Control9 | Control10 | Control11 | Control12 | Stress1 | Stress2 | Stress3 | Stress4 | Stress5 | Stress6 | Stress7 | Stress8 | Stress9 | Stress10 | Stress11 | Stress12 |
|---|---|---|---|---|---|---|---|---|---|---|---|---|---|---|---|---|---|---|---|---|---|---|---|---|
| 18 | 1.704239 | 1.08016 | 0.925252 | -0.19446 | 1.028643 | -0.06545 | 1.057191 | 0.111242 | 1.508922 | 0.34051 | -0.33673 | 1.194786 | NaN | NaN | NaN | NaN | NaN | NaN | NaN | NaN | NaN | NaN | NaN | NaN |
| 18.01389 | 0.930757 | 0.747874 | 0.549492 | -0.12995 | 0.453548 | -0.05781 | 0.604172 | 0.066766 | 2.083702 | 0.291365 | -0.2096 | 0.590316 | NaN | NaN | NaN | NaN | NaN | NaN | NaN | NaN | NaN | NaN | NaN | NaN |
| 18.02778 | 0.39559 | 0.361766 | 0.311262 | -0.09334 | 0.198664 | -0.05414 | 0.121145 | 0.027602 | 1.109271 | 0.172795 | -0.15927 | 0.271464 | NaN | NaN | NaN | NaN | NaN | NaN | NaN | NaN | NaN | NaN | NaN | NaN |
| 18.04167 | 0.16207 | 0.147618 | 0.268798 | -0.08175 | 0.064079 | -0.05901 | -0.01687 | -0.01657 | 0.65747 | 0.091338 | -0.14838 | 0.131999 | NaN | NaN | NaN | NaN | NaN | NaN | NaN | NaN | NaN | NaN | NaN | NaN |
| 18.05556 | 0.074911 | 0.050375 | 0.325899 | -0.09691 | -0.00537 | -0.09041 | -0.05505 | -0.04635 | 0.376281 | 0.023256 | -0.1755 | 0.062955 | NaN | NaN | NaN | NaN | NaN | NaN | NaN | NaN | NaN | NaN | NaN | NaN |
| 18.06944 | 0.055192 | 0.02494 | 0.203995 | -0.11126 | -0.02917 | -0.13727 | -0.04631 | -0.04882 | 0.246685 | -0.01027 | -0.21547 | 0.047379 | NaN | NaN | NaN | NaN | NaN | NaN | NaN | NaN | NaN | NaN | NaN | NaN |
| 18.08333 | 0.056574 | 0.026593 | 0.089797 | -0.10228 | -0.01348 | -0.15745 | -0.02734 | -0.01986 | 0.222816 | -0.00541 | -0.23382 | 0.068756 | NaN | NaN | NaN | NaN | NaN | NaN | NaN | NaN | NaN | NaN | NaN | NaN |
| 18.09722 | 0.059824 | 0.045358 | 0.081557 | -0.07961 | 0.01527 | -0.12834 | -0.00423 | 0.0199 | 0.228278 | 0.018049 | -0.22577 | 0.080956 | NaN | NaN | NaN | NaN | NaN | NaN | NaN | NaN | NaN | NaN | NaN | NaN |
| 18.11111 | 0.045375 | 0.065659 | 0.073654 | -0.0652 | 0.023402 | -0.09071 | 0.013326 | 0.033386 | 0.192289 | 0.037053 | -0.19554 | 0.057783 | NaN | NaN | NaN | NaN | NaN | NaN | NaN | NaN | NaN | NaN | NaN | NaN |
| 18.125 | 0.02139 | 0.079442 | 0.057402 | -0.06893 | 0.011471 | -0.07087 | 0.014878 | 0.019975 | 0.13742 | 0.037261 | -0.17174 | 0.018855 | NaN | NaN | NaN | NaN | NaN | NaN | NaN | NaN | NaN | NaN | NaN | NaN |
| 18.13889 | 0.008963 | 0.080796 | 0.04673 | -0.06972 | 0.009089 | -0.05966 | 0.009149 | 0.008193 | 0.103853 | 0.027927 | -0.17946 | 0.002514 | NaN | NaN | NaN | NaN | NaN | NaN | NaN | NaN | NaN | NaN | NaN | NaN |
| 18.15278 | 0.003819 | 0.050688 | 0.036568 | -0.06972 | 0.015249 | -0.05984 | 0.008771 | -0.00022 | 0.092099 | 0.012812 | -0.21316 | 0.005837 | NaN | NaN | NaN | NaN | NaN | NaN | NaN | NaN | NaN | NaN | NaN | NaN |
| 18.16667 | -0.01321 | 0.018738 | 0.016211 | -0.07995 | -0.00025 | -0.07153 | -0.00928 | -0.00829 | 0.084718 | -0.01078 | -0.25411 | 0.0087 | NaN | NaN | NaN | NaN | NaN | NaN | NaN | NaN | NaN | NaN | NaN | NaN |
| 18.18056 | -0.01945 | 0.02721 | 0.009778 | -0.08048 | -0.01731 | -0.07479 | -0.02469 | -0.00469 | 0.083888 | -0.02815 | -0.3019 | 0.011218 | NaN | NaN | NaN | NaN | NaN | NaN | NaN | NaN | NaN | NaN | NaN | NaN |
| 18.19444 | -0.0026 | 0.048985 | 0.033163 | -0.05297 | -0.00273 | -0.06531 | -0.00646 | 0.008404 | 0.106537 | -0.01736 | -0.37425 | 0.020139 | NaN | NaN | NaN | NaN | NaN | NaN | NaN | NaN | NaN | NaN | NaN | NaN |
| 18.20833 | 0.009384 | 0.0657 | 0.056563 | -0.05153 | 0.018799 | -0.06772 | 0.014703 | 0.012447 | 0.125973 | 0.012725 | -0.43515 | 0.014467 | NaN | NaN | NaN | NaN | NaN | NaN | NaN | NaN | NaN | NaN | NaN | NaN |
| 18.22222 | 0.010003 | 0.087489 | 0.05072 | -0.07467 | 0.022139 | -0.08054 | 0.010396 | 0.008033 | 0.115391 | 0.043162 | -0.53806 | -0.00292 | NaN | NaN | NaN | NaN | NaN | NaN | NaN | NaN | NaN | NaN | NaN | NaN |
| 18.23611 | 0.015943 | 0.107935 | 0.052361 | -0.09622 | 0.028348 | -0.07553 | -0.00377 | 0.007596 | 0.103025 | 0.074895 | -0.72658 | -0.00053 | NaN | NaN | NaN | NaN | NaN | NaN | NaN | NaN | NaN | NaN | NaN | NaN |
| 18.25 | 0.018439 | 0.082526 | 0.061451 | -0.08443 | 0.029605 | -0.06031 | -0.01678 | 0.003057 | 0.097998 | 0.077653 | -0.81523 | 0.013323 | NaN | NaN | NaN | NaN | NaN | NaN | NaN | NaN | NaN | NaN | NaN | NaN |
| 18.26389 | 0.008066 | 0.032642 | 0.075447 | -0.08553 | 0.017273 | -0.0584 | -0.01791 | -0.00575 | 0.100812 | 0.047143 | -0.77907 | 0.022876 | NaN | NaN | NaN | NaN | NaN | NaN | NaN | NaN | NaN | NaN | NaN | NaN |
| 18.27778 | 0.011657 | 0.037668 | 0.108907 | -0.07355 | 0.032789 | -0.04907 | 0.014092 | 0.005989 | 0.137052 | 0.039083 | -0.61168 | 0.033643 | NaN | NaN | NaN | NaN | NaN | NaN | NaN | NaN | NaN | NaN | NaN | NaN |
| 18.29167 | 0.028091 | 0.084608 | 0.123246 | -0.0531 | 0.068287 | -0.03067 | 0.035364 | 0.027744 | 0.170515 | 0.052787 | -0.41025 | 0.035659 | NaN | NaN | NaN | NaN | NaN | NaN | NaN | NaN | NaN | NaN | NaN | NaN |
| 18.30556 | 0.031134 | 0.110849 | 0.115185 | -0.05139 | 0.081225 | -0.02738 | 0.013488 | 0.033861 | 0.164698 | 0.0584 | -0.30044 | 0.033294 | NaN | NaN | NaN | NaN | NaN | NaN | NaN | NaN | NaN | NaN | NaN | NaN |
| 18.31944 | 0.027134 | 0.11108 | 0.099806 | -0.0654 | 0.078546 | -0.03749 | -0.0064 | 0.025439 | 0.153645 | 0.065891 | -0.24994 | 0.033051 | NaN | NaN | NaN | NaN | NaN | NaN | NaN | NaN | NaN | NaN | NaN | NaN |
| 18.33333 | 0.021826 | 0.10635 | 0.09607 | -0.07811 | 0.065741 | -0.04778 | 0.003967 | 0.017745 | 0.166954 | 0.091546 | -0.23466 | 0.016475 | NaN | NaN | NaN | NaN | NaN | NaN | NaN | NaN | NaN | NaN | NaN | NaN |
| 18.34722 | 0.024823 | 0.121411 | 0.107827 | -0.07071 | 0.064261 | -0.04893 | 0.029658 | 0.020742 | 0.199853 | 0.122711 | -0.23156 | -0.01254 | NaN | NaN | NaN | NaN | NaN | NaN | NaN | NaN | NaN | NaN | NaN | NaN |
| 18.36111 | 0.027848 | 0.093635 | 0.102695 | -0.0634 | 0.03704 | -0.06406 | 0.050683 | 0.020906 | 0.218442 | 0.120188 | -0.24793 | -0.05814 | NaN | NaN | NaN | NaN | NaN | NaN | NaN | NaN | NaN | NaN | NaN | NaN |
| 18.375 | 0.009145 | -0.019 | 0.050213 | -0.08972 | -0.04066 | -0.111 | 0.051974 | 0.004351 | 0.195087 | 0.065229 | -0.29653 | -0.12552 | NaN | NaN | NaN | NaN | NaN | NaN | NaN | NaN | NaN | NaN | NaN | NaN |
| 18.38889 | 0.001238 | -0.02401 | 0.014411 | -0.11804 | -0.03103 | -0.13048 | 0.08749 | -0.00246 | 0.217075 | 0.057546 | -0.31866 | -0.13185 | NaN | NaN | NaN | NaN | NaN | NaN | NaN | NaN | NaN | NaN | NaN | NaN |
| 18.40278 | 0.120459 | 0.363693 | 0.108934 | -0.08995 | 0.286925 | -0.06405 | 0.49662 | 0.153459 | 0.574782 | 0.263373 | -0.22936 | -0.03143 | NaN | NaN | NaN | NaN | NaN | NaN | NaN | NaN | NaN | NaN | NaN | NaN |
| 18.41667 | 0.351209 | 1.098077 | 0.394108 | -0.01156 | 0.927119 | 0.064998 | 1.462515 | 0.865333 | 1.297152 | 0.703257 | -0.06302 | 0.129413 | NaN | NaN | NaN | NaN | NaN | NaN | NaN | NaN | NaN | NaN | NaN | NaN |
| 18.43056 | 0.415777 | 1.439728 | 0.413162 | 0.000497 | 1.29237 | 0.202305 | 1.742592 | 0.874584 | 1.700223 | 1.236928 | 0.077115 | 0.203991 | NaN | NaN | NaN | NaN | NaN | NaN | NaN | NaN | NaN | NaN | NaN | NaN |
| 18.44444 | 0.255432 | 0.961569 | 0.01294 | -0.06386 | 0.937187 | 0.263495 | 1.020748 | 0.243244 | 1.337319 | 1.235061 | 0.160468 | 0.159501 | NaN | NaN | NaN | NaN | NaN | NaN | NaN | NaN | NaN | NaN | NaN | NaN |
| 18.45833 | 0.164228 | 0.425619 | -0.04542 | -0.03347 | 0.629301 | 0.204488 | 0.532155 | 0.185583 | 0.920984 | 1.022121 | 0.19765 | 0.14869 | NaN | NaN | NaN | NaN | NaN | NaN | NaN | NaN | NaN | NaN | NaN | NaN |
| 18.47222 | 0.225682 | 0.217633 | -0.00254 | 0.079623 | 0.592518 | 0.197243 | 0.33099 | 0.197956 | 0.719661 | 0.881827 | 0.209381 | 0.109176 | NaN | NaN | NaN | NaN | NaN | NaN | NaN | NaN | NaN | NaN | NaN | NaN |
| 18.48611 | 0.297751 | 0.142827 | 0.017153 | 0.133536 | 0.634243 | 0.179981 | 0.278175 | 0.185198 | 0.688481 | 0.859321 | 0.215956 | 0.037671 | NaN | NaN | NaN | NaN | NaN | NaN | NaN | NaN | NaN | NaN | NaN | NaN |
| 18.5 | 0.376033 | 0.136101 | 0.038503 | 0.136965 | 0.661974 | 0.165093 | 0.292784 | 0.140422 | 0.799074 | 0.797108 | 0.210865 | -0.01081 | NaN | NaN | NaN | NaN | NaN | NaN | NaN | NaN | NaN | NaN | NaN | NaN |
| 18.51389 | 0.443632 | 0.182972 | 0.081847 | 0.131852 | 0.686129 | 0.1601 | 0.264576 | 0.105498 | 0.955881 | 0.735573 | 0.200528 | -0.02315 | NaN | NaN | NaN | NaN | NaN | NaN | NaN | NaN | NaN | NaN | NaN | NaN |
| 18.52778 | 0.474295 | 0.258897 | 0.133516 | 0.118312 | 0.627135 | 0.145488 | 0.182809 | 0.084239 | 1.097203 | 0.703533 | 0.224568 | 0.00391 | NaN | NaN | NaN | NaN | NaN | NaN | NaN | NaN | NaN | NaN | NaN | NaN |
| 18.54167 | 0.482366 | 0.34679 | 0.188311 | 0.113196 | 0.572142 | 0.121836 | 0.076998 | 0.128943 | 1.233704 | 0.618925 | 0.301194 | 0.053348 | NaN | NaN | NaN | NaN | NaN | NaN | NaN | NaN | NaN | NaN | NaN | NaN |
| 18.55556 | 0.4084 | 0.441963 | 0.214806 | 0.119872 | 0.471913 | 0.075726 | -0.01389 | 0.237108 | 1.325311 | 0.535766 | 0.349553 | 0.075222 | NaN | NaN | NaN | NaN | NaN | NaN | NaN | NaN | NaN | NaN | NaN | NaN |
| 18.56944 | 0.326109 | 0.476866 | 0.20698 | 0.12526 | 0.354829 | 0.038906 | -0.07083 | 0.289394 | 1.326709 | 0.482486 | 0.32883 | 0.070882 | NaN | NaN | NaN | NaN | NaN | NaN | NaN | NaN | NaN | NaN | NaN | NaN |
| 18.58333 | 0.230973 | 0.449906 | 0.149113 | 0.107812 | 0.215178 | 0.040432 | -0.08496 | 0.227968 | 1.197509 | 0.434837 | 0.302638 | 0.06279 | NaN | NaN | NaN | NaN | NaN | NaN | NaN | NaN | NaN | NaN | NaN | NaN |
| 18.59722 | 0.154345 | 0.385849 | 0.094942 | 0.07087 | 0.097247 | 0.051501 | -0.05843 | 0.130695 | 0.98657 | 0.358391 | 0.283471 | 0.060241 | NaN | NaN | NaN | NaN | NaN | NaN | NaN | NaN | NaN | NaN | NaN | NaN |
| 18.61111 | 0.087946 | 0.339553 | 0.055265 | 0.095685 | 0.050179 | 0.060334 | -0.03231 | 0.103064 | 0.807501 | 0.235847 | 0.270377 | 0.043008 | NaN | NaN | NaN | NaN | NaN | NaN | NaN | NaN | NaN | NaN | NaN | NaN |
| 18.625 | 0.012852 | 0.31593 | 0.02827 | 0.104762 | -0.00469 | 0.058827 | -0.02945 | 0.09285 | 0.641135 | 0.117196 | 0.262236 | 0.009764 | NaN | NaN | NaN | NaN | NaN | NaN | NaN | NaN | NaN | NaN | NaN | NaN |
| 18.63889 | -0.04299 | 0.293938 | 0.015366 | 0.120803 | -0.1121 | 0.063135 | -0.02295 | 0.078835 | 0.485438 | 0.028446 | 0.259537 | 0.002697 | NaN | NaN | NaN | NaN | NaN | NaN | NaN | NaN | NaN | NaN | NaN | NaN |
| 18.65278 | -0.0679 | 0.320742 | 0.014606 | 0.109993 | -0.2231 | 0.068005 | 0.001436 | 0.07064 | 0.353323 | -0.03992 | 0.263513 | 0.018529 | NaN | NaN | NaN | NaN | NaN | NaN | NaN | NaN | NaN | NaN | NaN | NaN |
| 18.66667 | -0.07833 | 0.387995 | 0.014908 | 0.08728 | -0.28059 | 0.069117 | 0.017446 | 0.07239 | 0.252746 | -0.10076 | 0.261789 | 0.025974 | NaN | NaN | NaN | NaN | NaN | NaN | NaN | NaN | NaN | NaN | NaN | NaN |
| 18.68056 | -0.07513 | 0.400967 | 0.017972 | 0.080586 | -0.22059 | 0.103588 | 0.017116 | 0.08791 | 0.224325 | -0.09077 | 0.24306 | 0.025924 | NaN | NaN | NaN | NaN | NaN | NaN | NaN | NaN | NaN | NaN | NaN | NaN |
| 18.69444 | -0.04494 | 0.3271 | 0.020191 | 0.082739 | -0.12171 | 0.255278 | 0.005182 | 0.10956 | 0.239561 | 0.042385 | 0.217565 | 0.030766 | NaN | NaN | NaN | NaN | NaN | NaN | NaN | NaN | NaN | NaN | NaN | NaN |
| 18.70833 | -0.01504 | 0.237097 | 0.020973 | 0.11243 | -0.0618 | 0.352355 | -0.00827 | 0.106997 | 0.231158 | 0.181501 | 0.219326 | 0.031665 | NaN | NaN | NaN | NaN | NaN | NaN | NaN | NaN | NaN | NaN | NaN | NaN |
| 18.72222 | -0.00319 | 0.196929 | 0.029002 | 0.174707 | -0.05267 | 0.319168 | -0.035 | 0.099107 | 0.200847 | 0.268508 | 0.274586 | 0.008618 | NaN | NaN | NaN | NaN | NaN | NaN | NaN | NaN | NaN | NaN | NaN | NaN |
| 18.73611 | -0.03042 | 0.182281 | 0.013552 | 0.184723 | -0.07553 | 0.288953 | -0.08025 | 0.0762 | 0.167409 | 0.249483 | 0.426193 | -0.00915 | -0.40395 | -0.15196 | 0.048381 | 0.121193 | -0.75181 | 0.034001 | 0.021107 | -0.09174 | 0.021624 | 0.540095 | 0.030469 | 0.007208 |
| 18.75 | -0.0856 | 0.170477 | 0.004201 | 0.143438 | -0.15074 | 0.239653 | -0.13125 | 0.046087 | 0.11089 | 0.155082 | 0.627126 | -0.00271 | -0.02031 | -0.1607 | 0.023624 | 0.134477 | -0.75791 | -0.02176 | -0.07994 | -0.01041 | 0.055279 | 0.456081 | -0.05341 | -0.06793 |
| 18.76389 | -0.12848 | 0.204911 | 0.005339 | 0.115467 | -0.26518 | 0.208357 | -0.19078 | 0.012726 | 0.022004 | 0.000158 | 0.588528 | 0.018145 | 0.405094 | -0.13752 | 0.009676 | 0.140822 | -0.40215 | -0.07333 | -0.10451 | -0.00027 | 0.037103 | 0.159841 | -0.1233 | -0.14204 |
| 18.77778 | -0.1759 | 0.245067 | 0.002054 | 0.10242 | -0.40705 | 0.190889 | -0.26024 | 0.008288 | -0.08086 | -0.19513 | 0.410174 | 0.02652 | 0.541068 | -0.18034 | -0.02173 | 0.123623 | -0.05425 | -0.11011 | -0.0462 | -0.09782 | 0.00179 | -0.30959 | -0.14999 | -0.19305 |
| 18.79167 | -0.21403 | 0.20966 | -0.00339 | 0.071741 | -0.47538 | 0.17274 | -0.3828 | 0.026786 | -0.14751 | -0.2911 | 0.283999 | 0.000519 | 0.297171 | -0.31577 | -0.08397 | 0.080822 | 0.042855 | -0.09972 | -0.00419 | -0.25683 | -0.06022 | -0.79442 | -0.1413 | -0.22566 |
| 18.80556 | -0.28869 | 0.151289 | -0.00785 | 0.05298 | -0.47102 | 0.132246 | -0.48382 | 0.043241 | -0.20537 | -0.18031 | 0.235331 | -0.03195 | -0.18665 | -0.50109 | -0.12862 | 0.028484 | 0.049856 | -0.07555 | -0.03111 | -0.3753 | -0.16632 | -1.03988 | -0.11046 | -0.27208 |
| 18.81944 | -0.39467 | 0.125886 | -0.03215 | 0.04817 | -0.42819 | 0.103804 | -0.56156 | 0.040806 | -0.288 | -0.01807 | 0.233902 | -0.04862 | -0.69382 | -0.60615 | -0.11785 | 0.003539 | 0.055538 | -0.07652 | -0.12933 | -0.40103 | -0.24226 | -0.84508 | -0.11526 | -0.3766 |
| 18.83333 | -0.50646 | 0.177268 | -0.05041 | 0.042781 | -0.36999 | 0.089841 | -0.64441 | 0.035213 | -0.35329 | 0.040056 | 0.246682 | -0.0519 | -0.87411 | -0.59121 | -0.1015 | -0.01242 | 0.073848 | -0.10559 | -0.32987 | -0.36944 | -0.27156 | -0.4438 | -0.19203 | -0.45523 |
| 18.84722 | -0.52786 | 0.253431 | -0.04259 | 0.033388 | -0.31113 | 0.084161 | -0.70172 | 0.027629 | -0.35942 | -0.02112 | 0.259618 | -0.06627 | -0.53083 | -0.53868 | -0.08968 | -0.02204 | 0.118786 | -0.0799 | -0.5805 | -0.32418 | -0.23761 | -0.08577 | -0.27633 | -0.47014 |
| 18.86111 | -0.54267 | 0.192233 | -0.04189 | 0.030185 | -0.25963 | 0.086589 | -0.75774 | 0.00392 | -0.32648 | -0.17611 | 0.26519 | -0.08632 | -0.00615 | -0.50925 | -0.08252 | -0.01431 | 0.151756 | 0.001407 | -0.80592 | -0.31094 | -0.18103 | 0.111715 | -0.34822 | -0.43418 |
| 18.875 | -0.44727 | 0.053366 | -0.05828 | 0.034867 | -0.19478 | 0.081178 | -0.69593 | -0.02418 | -0.31038 | -0.36281 | 0.287744 | -0.09438 | 0.385352 | -0.42757 | -0.09013 | -0.00104 | 0.150044 | 0.042041 | -0.93467 | -0.29361 | -0.1068 | 0.125655 | -0.3908 | -0.44714 |
| 18.88889 | -0.32968 | 0.002927 | -0.06905 | 0.035678 | -0.16409 | 0.07996 | -0.5893 | -0.01856 | -0.32385 | -0.47423 | 0.322148 | -0.09313 | 0.390019 | -0.34341 | -0.12498 | -0.01965 | 0.122785 | -0.02112 | -0.80966 | -0.25608 | -0.08643 | 0.010687 | -0.37107 | -0.58443 |
| 18.90278 | -0.25472 | -0.10622 | -0.13273 | 0.034926 | -0.20206 | 0.021574 | -0.55227 | 0.040161 | -0.47445 | -0.57316 | 0.330404 | -0.17731 | -0.16577 | -0.3057 | -0.3231 | -0.15878 | -0.04261 | -0.20413 | -0.8002 | -0.22678 | -0.16089 | -0.40235 | -0.41467 | -0.76887 |
| 18.91667 | -0.30498 | -0.61512 | -0.28182 | -0.06734 | -0.36369 | -0.13218 | -0.83013 | 0.107118 | -0.9897 | -1.0425 | 0.186828 | -0.53102 | -1.61535 | -0.39609 | -0.68811 | -0.56537 | -0.45123 | -0.49226 | -1.62432 | -0.1712 | -0.48094 | -1.17493 | -0.75245 | -0.84655 |
| 18.93056 | -0.36074 | -1.33183 | -0.53246 | -0.22942 | -0.66352 | -0.48929 | -1.37278 | -0.15054 | -1.95015 | -1.93024 | -0.22623 | -1.15338 | -4.65512 | -0.47158 | -1.01069 | -1.16621 | -0.82865 | -0.77493 | -4.06943 | -0.21385 | -1.38758 | -2.18223 | -1.68642 | -1.01153 |
| 18.94444 | -0.32752 | -1.25265 | -0.72979 | -0.28737 | -0.56158 | -0.78604 | -1.89993 | -1.01335 | -3.12412 | -2.24336 | -0.79047 | -1.7309 | -5.75898 | -0.4163 | -1.00763 | -1.34113 | -0.47932 | -0.79722 | -3.68098 | -0.0257 | -2.57435 | -3.12018 | -2.88708 | -1.56707 |
| 18.95833 | -0.266 | -0.21864 | -0.1967 | -0.28926 | 0.081553 | -0.51033 | -1.15101 | -0.51076 | -1.05821 | -1.51319 | -1.21428 | -0.8252 | -2.44876 | -0.29056 | -0.54347 | -0.31601 | 0.426638 | -0.1963 | -2.72145 | 0.769522 | -0.80416 | -2.08632 | -1.91187 | -1.00798 |
| 18.97222 | -0.20872 | 0.559562 | 0.395414 | -0.28332 | 0.339484 | -0.28888 | 0.158057 | 0.446896 | 1.598898 | -0.16104 | -1.34083 | 0.480384 | 1.552392 | -0.20959 | 0.058965 | 0.401867 | 0.863849 | 0.196144 | -1.35822 | -0.27294 | 0.759325 | 0.285647 | 0.200974 | 0.328267 |
| 18.98611 | -0.15415 | 0.48369 | 0.203698 | -0.2475 | 0.151059 | -0.22339 | 0.67152 | 0.498934 | 1.832796 | 0.602676 | -1.35437 | 0.64028 | 5.144379 | -0.15737 | 0.282453 | 0.483598 | 0.626131 | 0.328875 | -0.52353 | -0.58105 | 0.93051 | 1.153603 | 2.367947 | 0.520692 |
| 19 | -0.11354 | 0.224621 | 0.080935 | -0.21974 | -0.03286 | -0.17114 | 0.445542 | 0.096213 | 0.750257 | 0.606167 | -1.18031 | 0.333425 | 4.358513 | -0.11227 | 0.236305 | 0.224841 | 0.296907 | 0.27364 | -0.29885 | -0.61336 | 0.512428 | 0.838472 | 1.325959 | 0.178099 |
| 19.01389 | -0.06921 | 0.1291 | 0.036103 | -0.16908 | -0.05668 | -0.14034 | 0.228866 | -0.02108 | 0.310711 | 0.369031 | -0.88476 | 0.128891 | 2.332259 | -0.04097 | 0.154271 | 0.10951 | 0.187813 | 0.180232 | -0.22227 | -0.52343 | 0.22608 | 0.4965 | 0.605483 | 0.01566 |
| 19.02778 | -0.03169 | 0.090408 | 0.00532 | -0.10804 | -0.03074 | -0.11821 | 0.100098 | -0.06746 | 0.126418 | 0.189024 | -0.65478 | 0.02887 | 0.925109 | 0.012427 | 0.083018 | 0.042192 | 0.176227 | 0.051492 | -0.19497 | -0.36261 | 0.03479 | 0.223237 | 0.248473 | -0.04297 |
| 19.04167 | -0.01 | 0.070177 | -0.00508 | -0.04503 | -0.01542 | -0.09707 | 0.025356 | -0.04506 | 0.052326 | 0.08942 | -0.54027 | 0.001914 | 0.220496 | 0.03181 | 0.057034 | 0.046313 | 0.109623 | -0.05319 | -0.16721 | -0.22861 | -0.07209 | 0.11028 | 0.06968 | -0.04012 |
| 19.05556 | 0.003074 | 0.092617 | -0.00195 | -0.02565 | 0.000699 | -0.07573 | 0.011876 | -0.02194 | 0.043722 | 0.054203 | -0.46492 | 0.014483 | 0.084632 | 0.028883 | 0.054547 | 0.065121 | 0.08068 | -0.07973 | -0.13596 | -0.16181 | -0.13442 | 0.124685 | 0.020289 | -0.00711 |
| 19.06944 | 0.012038 | 0.103275 | 0.017252 | -0.03191 | 0.010185 | -0.05449 | -0.00372 | -0.00206 | 0.011706 | 0.064809 | -0.35497 | 0.051563 | 0.10672 | 0.023335 | 0.0096 | 0.052452 | 0.150336 | -0.03523 | -0.10691 | -0.1756 | -0.18641 | 0.161952 | -0.00013 | -0.04776 |
| 19.08333 | -0.00382 | 0.03029 | 0.062528 | -0.02413 | -0.03167 | -0.05511 | -0.18622 | 0.023062 | -0.26717 | 0.00273 | -0.26283 | 0.090636 | -0.27416 | 0.081576 | -0.12783 | 0.014977 | 0.325873 | 0.021578 | -0.05486 | -0.23942 | -0.30592 | -0.01067 | -0.126 | -0.24226 |
| 19.09722 | -0.04078 | -0.0028 | 0.092838 | -0.02674 | -0.1794 | -0.09427 | -0.3959 | 0.016115 | -0.55545 | -0.10637 | -0.24375 | 0.026216 | -1.08927 | 0.219808 | -0.17108 | 0.011854 | 0.509425 | 0.073362 | 0.033665 | -0.21912 | -0.36879 | -0.18763 | -0.23694 | -0.2995 |
| 19.11111 | 0.022804 | 0.152977 | 0.036032 | -0.11205 | -0.02314 | -0.12129 | -0.12734 | -0.04098 | -0.07607 | 0.023907 | -0.19729 | -0.05993 | -0.30835 | 0.21305 | 0.125944 | 0.092697 | 0.361622 | 0.101391 | 0.071814 | -0.03669 | 0.005654 | 0.441535 | -0.00408 | 0.067807 |
| 19.125 | 0.153222 | 0.396231 | -0.0502 | -0.16802 | 0.222 | -0.0555 | 0.363518 | -0.03214 | 0.636754 | 0.329453 | -0.07301 | 0.007594 | 1.477041 | 0.069218 | 0.372735 | 0.146875 | 0.081824 | 0.095902 | 0.015267 | 0.010149 | 0.338614 | 1.208596 | 0.295647 | 0.375529 |
| 19.13889 | 0.201054 | 0.434003 | -0.06721 | -0.0308 | 0.174648 | 0.023837 | 0.429892 | 0.04023 | 0.601729 | 0.409463 | 0.010027 | 0.104179 | 1.425358 | 0.018761 | 0.21676 | 0.07418 | 0.079218 | 0.1061 | -0.02089 | -0.13439 | 0.169124 | 1.118012 | 0.248833 | 0.197619 |
| 19.15278 | 0.149823 | 0.329083 | -0.04695 | 0.010254 | 0.110433 | 0.018643 | 0.193336 | 0.0492 | 0.235493 | 0.15188 | 0.020868 | 0.104494 | 0.226806 | 0.062419 | 0.070495 | 0.037063 | 0.178574 | 0.115797 | 0.023613 | -0.18145 | -0.02671 | 0.7043 | 0.037652 | 0.004001 |
| 19.16667 | 0.081072 | 0.1965 | -0.0467 | -0.00881 | 0.034671 | -0.02404 | 0.007097 | 0.004999 | 0.030521 | -0.10857 | -0.00162 | 0.063059 | -0.5177 | 0.077584 | 0.006093 | 0.011072 | 0.18467 | 0.062051 | 0.024038 | -0.15375 | -0.05855 | 0.450401 | -0.12035 | -0.01621 |
| 19.18056 | 0.045443 | 0.044725 | -0.0487 | -0.02661 | 0.008478 | -0.04245 | -0.04852 | -0.01353 | -0.00202 | -0.15799 | -0.02197 | 0.029975 | -0.71623 | 0.069113 | -0.06189 | 0.004241 | 0.187625 | -0.02735 | -0.00882 | -0.18537 | -0.04877 | 0.351887 | -0.18449 | 0.008777 |
| 19.19444 | 0.059921 | 0.003323 | -0.01999 | -0.01136 | 0.038191 | -0.02308 | 0.001593 | 0.018973 | 0.052121 | -0.03005 | -0.02339 | 0.028103 | -0.39405 | 0.115346 | -0.05742 | 0.037192 | 0.2234 | -0.00059 | -0.0109 | -0.23577 | 0.005254 | 0.393708 | -0.11959 | 0.057185 |
| 19.20833 | 0.106914 | 0.103948 | 0.030523 | 0.027964 | 0.083422 | 0.013583 | 0.098393 | 0.074513 | 0.134505 | 0.135729 | 0.006035 | 0.070213 | 0.142746 | 0.206862 | 0.006194 | 0.088176 | 0.247651 | 0.167781 | 0.026095 | -0.24315 | 0.094963 | 0.535569 | 0.022628 | 0.134173 |
| 19.22222 | 0.144492 | 0.244255 | 0.050084 | 0.048433 | 0.099436 | 0.027931 | 0.165418 | 0.094588 | 0.187494 | 0.254779 | 0.027513 | 0.099488 | 0.585469 | 0.270001 | 0.057292 | 0.111288 | 0.208974 | 0.300137 | 0.059887 | -0.24659 | 0.154471 | 0.659658 | 0.151025 | 0.209916 |
| 19.23611 | 0.141209 | 0.277477 | 0.027105 | 0.033146 | 0.085292 | 0.014288 | 0.170434 | 0.077622 | 0.194984 | 0.270651 | 0.019122 | 0.0917 | 0.712481 | 0.272635 | 0.063466 | 0.106491 | 0.129473 | 0.287259 | 0.066017 | -0.26717 | 0.120659 | 0.71114 | 0.193338 | 0.261116 |
| 19.25 | 0.127346 | 0.253696 | 0.003123 | 0.023015 | 0.081108 | 0.002884 | 0.158404 | 0.064858 | 0.191078 | 0.219604 | 0.018839 | 0.07221 | 0.591211 | 0.249284 | 0.038897 | 0.097204 | 0.10813 | 0.213748 | 0.071268 | -0.26938 | 0.054135 | 0.724116 | 0.170455 | 0.262931 |
| 19.26389 | 0.118757 | 0.216298 | -0.00264 | 0.027339 | 0.089199 | 0.002833 | 0.158556 | 0.063061 | 0.186585 | 0.158695 | 0.031067 | 0.049123 | 0.418216 | 0.234181 | 0.022261 | 0.101973 | 0.135285 | 0.128671 | 0.094842 | -0.2165 | 0.030303 | 0.700832 | 0.142215 | 0.215015 |
| 19.27778 | 0.12258 | 0.184714 | -0.00407 | 0.02599 | 0.093269 | 0.005077 | 0.174587 | 0.058437 | 0.189872 | 0.127574 | 0.041633 | 0.066344 | 0.317011 | 0.240805 | 0.018246 | 0.113931 | 0.168657 | 0.084767 | 0.130709 | -0.13428 | 0.092524 | 0.71895 | 0.141263 | 0.194324 |
| 19.29167 | 0.144731 | 0.187909 | 0.003575 | 0.031153 | 0.107706 | 0.014744 | 0.214571 | 0.054314 | 0.235221 | 0.142921 | 0.0555 | 0.089806 | 0.346566 | 0.300563 | 0.025286 | 0.115064 | 0.157772 | 0.11435 | 0.164021 | -0.03514 | 0.215288 | 0.890959 | 0.170006 | 0.231851 |
| 19.30556 | 0.172787 | 0.202541 | 0.0119 | 0.040325 | 0.125829 | 0.024405 | 0.243588 | 0.048229 | 0.295961 | 0.182559 | 0.069598 | 0.085661 | 0.410595 | 0.364908 | 0.024011 | 0.100746 | 0.103831 | 0.196848 | 0.168031 | 0.09466 | 0.29359 | 1.167285 | 0.180015 | 0.272845 |
| 19.31944 | 0.203196 | 0.172013 | 0.0092 | 0.033075 | 0.124333 | 0.022628 | 0.233992 | 0.040559 | 0.315082 | 0.199803 | 0.080244 | 0.05813 | 0.383566 | 0.382333 | 0.023349 | 0.077276 | 0.026587 | 0.25291 | 0.16315 | 0.22744 | 0.274884 | 1.384729 | 0.13976 | 0.269913 |
| 19.33333 | 0.252105 | 0.141393 | 0.009067 | 0.022802 | 0.123323 | 0.024237 | 0.233689 | 0.047079 | 0.312637 | 0.209531 | 0.095351 | 0.050507 | 0.353278 | 0.365484 | 0.019626 | 0.065208 | -0.00298 | 0.230719 | 0.169923 | 0.300353 | 0.178236 | 1.488476 | 0.092889 | 0.306114 |
| 19.34722 | 0.316651 | 0.127687 | 0.016731 | 0.016528 | 0.119503 | 0.028277 | 0.258835 | 0.053756 | 0.335258 | 0.203503 | 0.101223 | 0.046714 | 0.375846 | 0.357129 | 0.009749 | 0.085646 | 0.000922 | 0.170256 | 0.207259 | 0.304125 | 0.092398 | 1.5395 | 0.090796 | 0.396248 |
| 19.36111 | 0.342293 | 0.053487 | 0.015428 | 0.003308 | 0.06298 | 0.002235 | 0.278513 | 0.038839 | 0.396951 | 0.163941 | 0.085146 | -0.02316 | 0.418354 | 0.399369 | 0.061022 | 0.159168 | 0.036016 | 0.192946 | 0.310199 | 0.343685 | 0.071258 | 1.776233 | 0.173986 | 0.495116 |
| 19.375 | 0.321209 | -0.06243 | 0.004059 | -0.01227 | -0.01137 | -0.0438 | 0.286562 | 0.025665 | 0.471494 | 0.080632 | 0.055702 | -0.14976 | 0.429118 | 0.458487 | 0.197802 | 0.261127 | 0.073521 | 0.291565 | 0.463338 | 0.460561 | 0.140406 | 1.943646 | 0.297067 | 0.53025 |
| 19.38889 | 0.300354 | -0.02897 | 0.004833 | -0.01698 | 0.010188 | -0.04262 | 0.333191 | 0.010005 | 0.559708 | 0.053662 | 0.065733 | -0.10837 | 0.394548 | 0.453871 | 0.307582 | 0.264663 | 0.168091 | 0.326317 | 0.572401 | 0.526308 | 0.171582 | 1.725256 | 0.32081 | 0.43937 |
| 19.40278 | 0.444997 | 0.308027 | 0.069956 | -0.0067 | 0.306457 | 0.057399 | 0.633991 | 0.023324 | 1.040211 | 0.267144 | 0.18581 | 0.444627 | 0.703546 | 0.557003 | 0.49256 | 0.273822 | 0.799825 | 0.274784 | 0.81316 | 0.502972 | 0.12436 | 1.623971 | 0.271707 | 0.408261 |
| 19.41667 | 0.924878 | 0.872826 | 0.218653 | 0.004616 | 0.78865 | 0.253474 | 1.559941 | 0.090838 | 2.299098 | 0.694685 | 0.531069 | 2.353106 | 1.66115 | 1.038783 | 0.853647 | 0.373888 | 2.123946 | 0.298131 | 1.485058 | 0.521302 | 0.07673 | 1.953966 | 0.270041 | 0.634889 |
| 19.43056 | 1.206838 | 1.168807 | 0.34443 | -0.05466 | 1.036047 | 0.452585 | 1.293581 | 0.106147 | 3.432819 | 1.050869 | 1.226247 | 6.328777 | 2.424725 | 1.732451 | 1.26259 | 0.385613 | 2.739889 | 0.326097 | 2.905796 | 0.697281 | 0.005873 | 1.871691 | 0.163766 | 0.698942 |
| 19.44444 | 0.992885 | 0.923481 | 0.371931 | 0.111105 | 0.904315 | 0.568662 | 1.117371 | 0.114173 | 3.013266 | 1.222256 | 2.628079 | 7.904811 | 1.653832 | 1.545404 | 0.908698 | 0.047218 | 2.397808 | 0.142592 | 3.222605 | 0.679745 | -0.19029 | 0.900637 | -0.01163 | 0.517006 |
| 19.45833 | 0.828623 | 0.662423 | 0.418892 | 0.589909 | 0.76953 | 0.529019 | 0.642353 | 0.138567 | 2.176547 | 1.216293 | 2.840682 | 5.737824 | 0.745925 | 1.029226 | 0.666221 | 0.023848 | 2.019266 | 0.072884 | 3.124855 | 0.576592 | -0.15473 | 0.146814 | 0.021213 | 0.37001 |
| 19.47222 | 0.856467 | 0.593959 | 0.642453 | 1.137731 | 0.81937 | 0.753464 | 0.443386 | 0.158042 | 1.799822 | 1.157879 | 3.041516 | 3.922197 | 0.320503 | 0.710976 | 0.700238 | 0.139265 | 1.624299 | 0.161169 | 2.45968 | 0.561338 | 0.174576 | -0.11585 | 0.20822 | 0.298926 |
| 19.48611 | 0.695713 | 0.563343 | 0.867403 | 1.415374 | 0.833708 | 0.801143 | 0.352665 | 0.125661 | 1.440169 | 0.934811 | 6.051561 | 2.387977 | 0.112178 | 0.436377 | 0.644024 | 0.189529 | 0.969208 | 0.215456 | 1.792246 | 0.584419 | 0.534204 | -0.03157 | 0.388072 | 0.196287 |
| 19.5 | 0.410099 | 0.56267 | 0.944978 | 1.254666 | 0.770179 | 0.797634 | 0.226809 | 0.070837 | 1.258631 | 0.553938 | 4.330519 | 1.57552 | -0.10745 | 0.177182 | 0.526173 | 0.153518 | 0.419106 | 0.146827 | 1.191561 | 0.593848 | 0.775549 | 0.3459 | 0.554053 | 0.044631 |

| | | | | | | | | | | | | | | | | | | | | | | | | |
|---|---|---|---|---|---|---|---|---|---|---|---|---|---|---|---|---|---|---|---|---|---|---|---|---|
| 19.51389 | 0.128522 | 0.681324 | 1.023406 | 0.882364 | 0.586281 | 0.756432 | 0.143201 | 0.037302 | 1.111163 | 0.339993 | 2.121178 | 1.063103 | -0.08826 | 0.027143 | 0.432564 | 0.097238 | 0.141788 | 0.104075 | 0.705126 | 0.503745 | 0.795309 | 0.539284 | 0.65414 | -0.09679 |
| 19.52778 | 0.070243 | 0.856072 | 0.853389 | 0.698121 | 0.322409 | 0.612552 | 0.137599 | 0.037908 | 0.876345 | 0.314056 | 2.411204 | 0.600816 | 0.19181 | 0.011765 | 0.391292 | 0.054255 | 0.1079 | 0.086158 | 0.288197 | 0.408982 | 0.672087 | 0.38285 | 0.657674 | -0.04768 |
| 19.54167 | 0.38522 | 0.978109 | 0.682478 | 0.583077 | -0.01942 | 0.620123 | 0.149737 | 0.074347 | 0.537687 | 0.37344 | 3.175691 | 0.325952 | 0.364559 | 0.046345 | 0.411905 | 0.042554 | 0.364374 | 0.108796 | 0.199162 | 0.428212 | 0.666838 | 0.063503 | 0.682291 | 0.329075 |
| 19.55556 | 0.901945 | 1.004342 | 0.544952 | 0.422835 | -0.24027 | 0.690974 | 0.125195 | 0.100215 | 0.271195 | 0.501177 | 3.721768 | 0.154 | 0.368656 | 0.008205 | 0.468813 | 0.056637 | 0.912994 | 0.10972 | 0.468373 | 0.486763 | 0.769612 | -0.26551 | 0.756604 | 0.848228 |
| 19.56944 | 1.120045 | 0.957603 | 0.467113 | 0.35536 | -0.20403 | 0.687536 | 0.113256 | 0.099624 | 0.179384 | 0.612471 | 3.499789 | 0.003835 | 0.269215 | -0.06891 | 0.500014 | 0.077402 | 1.47952 | 0.081565 | 0.746293 | 0.474925 | 0.725891 | -0.39726 | 0.772831 | 1.116592 |
| 19.58333 | 0.715034 | 0.901023 | 0.398605 | 0.322606 | 0.034645 | 0.541212 | 0.137186 | 0.096857 | 0.184006 | 0.627754 | 2.714291 | -0.09754 | 0.155759 | -0.02419 | 0.417544 | 0.103053 | 1.959091 | 0.059942 | 0.794482 | 0.397682 | 0.555856 | -0.24714 | 0.680326 | 1.041069 |
| 19.59722 | 0.187395 | 0.915305 | 0.320325 | 0.281113 | 0.211253 | 0.46327 | 0.157966 | 0.089098 | 0.150026 | 0.409423 | 1.874594 | 0.003354 | 0.012957 | 0.156717 | 0.2949 | 0.119381 | 2.179811 | 0.061347 | 0.55182 | 0.245924 | 0.274034 | 0.077425 | 0.502445 | 0.742813 |
| 19.61111 | -0.23213 | 0.926856 | 0.219543 | 0.238529 | 0.221316 | 0.478517 | 0.139454 | 0.056338 | 0.064753 | 0.102038 | 1.318327 | 0.092854 | -0.17637 | 0.281066 | 0.20555 | 0.105555 | 1.987283 | 0.059808 | 0.194481 | 0.063449 | 0.024629 | 0.251884 | 0.33036 | 0.429872 |
| 19.625 | -0.54542 | 0.865913 | 0.135117 | 0.197298 | 0.191526 | 0.496953 | 0.101865 | 0.022855 | -0.00828 | -0.18357 | 0.990363 | 0.111716 | -0.47436 | 0.291393 | 0.16351 | 0.075298 | 1.688403 | 0.040766 | -0.02025 | -0.10542 | -0.0297 | 0.258699 | 0.198472 | 0.182458 |
| 19.63889 | -0.75807 | 0.739263 | 0.092912 | 0.135512 | 0.208849 | 0.465702 | 0.066702 | 0.019423 | -0.03819 | -0.28874 | 0.799195 | 0.183521 | -0.48626 | 0.236538 | 0.198135 | 0.074533 | 1.469377 | 0.058335 | -0.02771 | -0.1956 | 0.132968 | 0.514016 | 0.082696 | 0.049327 |
| 19.65278 | -0.55087 | 0.624429 | 0.089326 | 0.074054 | 0.190586 | 0.389714 | 0.040155 | 0.025177 | -0.02201 | -0.28556 | 0.631572 | 0.300381 | -0.13538 | 0.15096 | 0.27219 | 0.081661 | 1.339434 | 0.093325 | 0.046003 | -0.23316 | 0.320998 | 0.725315 | -0.03923 | -0.01125 |
| 19.66667 | -0.01964 | 0.551508 | 0.115903 | 0.033025 | 0.115274 | 0.298302 | 0.043316 | 0.033866 | -0.01272 | -0.05404 | 0.469377 | 0.311793 | 0.296234 | 0.063275 | 0.267024 | 0.054842 | 1.058462 | 0.087278 | -0.04683 | -0.1692 | 0.44527 | 0.515911 | -0.15439 | -0.10644 |
| 19.68056 | 0.523429 | 0.497625 | 0.150172 | 0.041102 | 0.035301 | 0.209459 | 0.088035 | 0.038522 | 0.010556 | 0.314131 | 0.369376 | 0.270312 | 0.382289 | 0.030025 | 0.219249 | 0.009851 | 0.600173 | 0.026497 | -0.24229 | -0.03056 | 0.468241 | 0.101681 | -0.19419 | -0.22844 |
| 19.69444 | 0.658103 | 0.429482 | 0.16529 | 0.0711 | -0.02227 | 0.146331 | 0.130017 | 0.037178 | 0.023771 | 0.525615 | 0.350494 | 0.237211 | 0.161541 | 0.055921 | 0.221707 | -0.00279 | 0.170441 | -0.04317 | -0.2268 | 0.113297 | 0.416363 | -0.1854 | -0.18339 | -0.32018 |
| 19.70833 | 0.410594 | 0.279112 | 0.17272 | 0.063811 | -0.05353 | 0.135863 | 0.131611 | 0.031534 | -0.04023 | 0.514819 | 0.314429 | 0.187716 | -0.05199 | 0.059382 | 0.23891 | -0.00497 | -0.05147 | -0.12306 | -0.06044 | 0.234936 | 0.342979 | -0.15215 | -0.15741 | -0.27415 |
| 19.72222 | 0.116885 | 0.178737 | 0.148113 | 0.059249 | -0.0525 | 0.143958 | 0.118308 | 0.016696 | -0.16234 | 0.349763 | 0.263734 | 0.068934 | -0.24236 | 0.002429 | 0.20161 | -0.02413 | -0.31793 | -0.163 | -0.04789 | 0.335893 | 0.266692 | 0.030453 | -0.16384 | -0.30377 |
| 19.73611 | -0.18485 | 0.144902 | 0.103023 | 0.078682 | -0.03716 | 0.115891 | 0.103387 | 0.001437 | -0.30793 | 0.091489 | 0.223132 | -0.10247 | -0.39492 | -0.06177 | 0.148131 | -0.07004 | -0.74659 | -0.11496 | -0.11262 | 0.345826 | 0.134422 | -0.01325 | -0.20031 | -0.33895 |
| 19.75 | -0.45785 | 0.073761 | 0.067493 | 0.071197 | -0.00784 | 0.103332 | 0.055907 | 0.0111 | -0.42289 | -0.25737 | 0.227932 | -0.30761 | -0.57214 | -0.13322 | 0.078142 | -0.12394 | -1.32077 | -0.07107 | -0.00034 | 0.216079 | -0.05961 | -0.18019 | -0.23095 | -0.35226 |
| 19.76389 | -0.47203 | -0.02666 | 0.044965 | 0.018576 | 0.051477 | -0.1313 | -0.01538 | 0.02666 | -0.49982 | -0.57458 | 0.180902 | -0.60902 | -0.32898 | -0.257 | -0.00894 | -0.14149 | -1.74364 | -0.0648 | 0.223203 | 0.020864 | -0.07056 | -0.20157 | -0.19603 | -0.3231 |
| 19.77778 | -0.22057 | -0.14096 | 0.048406 | -0.0336 | 0.088902 | -0.24591 | -0.09659 | 0.025142 | -0.5515 | -0.75438 | 0.110715 | -1.04238 | 0.23744 | -0.39475 | -0.08462 | -0.12092 | -1.84727 | -0.12441 | 0.314407 | -0.21082 | 0.040376 | -0.16133 | -0.14994 | -0.30011 |
| 19.79167 | 0.0213 | -0.21926 | 0.058848 | -0.04607 | 0.045583 | -0.30161 | -0.22167 | 0.013069 | -0.6087 | -0.68184 | 0.051676 | -1.349 | 0.427883 | -0.4879 | -0.18092 | -0.09406 | -1.69924 | -0.18883 | 0.122048 | -0.4204 | 0.140155 | -0.3654 | -0.16039 | -0.36515 |
| 19.80556 | -0.01035 | -0.37791 | -0.01131 | -0.03961 | -0.11311 | -0.26402 | -0.26623 | -0.00567 | -0.70218 | -0.3849 | -0.00041 | -1.3189 | 0.091765 | -0.43577 | -0.33912 | -0.08536 | -1.53922 | -0.25746 | -0.18539 | -0.52392 | 0.175252 | -0.85951 | -0.22899 | -0.55722 |
| 19.81944 | -0.33136 | -0.61103 | -0.1743 | -0.02767 | -0.34437 | -0.18514 | -0.25772 | -0.0224 | -0.87501 | -0.20059 | -0.05706 | -1.22455 | -0.30125 | -0.27976 | -0.52963 | -0.08177 | -1.80044 | -0.33931 | -0.26157 | -0.63655 | 0.098249 | -1.48539 | -0.36536 | -0.78384 |
| 19.83333 | -0.82456 | -0.83542 | -0.37451 | 0.006235 | -0.55253 | -0.13348 | -0.28861 | -0.0251 | -1.01726 | -0.20481 | -0.08465 | -1.2784 | -0.71685 | -0.23631 | -0.71744 | -0.10547 | -2.37013 | -0.39379 | -0.17178 | -0.70619 | -0.0268 | -2.03689 | -0.54854 | -1.01757 |
| 19.84722 | -1.28658 | -0.97754 | -0.56477 | 0.03625 | -0.75192 | -0.12899 | -0.46338 | -0.04485 | -0.93016 | -0.41186 | -0.07661 | -1.4661 | -1.09164 | -0.43308 | -0.81516 | -0.12671 | -2.9234 | -0.33808 | -0.19644 | -0.78685 | -0.07239 | -2.33124 | -0.71823 | -1.39626 |
| 19.86111 | -1.37562 | -1.08827 | -0.60021 | 0.027738 | -0.76293 | -0.12595 | -0.69997 | -0.07118 | -0.74861 | -0.58491 | -0.07707 | -1.822 | -1.21332 | -0.74612 | -0.78859 | -0.13387 | -2.82442 | -0.24969 | -0.34539 | -0.78203 | -0.03002 | -2.26306 | -0.74986 | -1.63898 |
| 19.875 | -1.16214 | -1.03643 | -0.71429 | -0.01688 | -0.46117 | -0.12358 | -0.83418 | -0.07388 | -0.63498 | -0.78558 | -0.1074 | -2.06409 | -0.99431 | -1.0176 | -0.74261 | -0.1444 | -2.13487 | -0.22422 | -0.38695 | -0.72772 | -0.00168 | -1.87959 | -0.69593 | -1.7518 |
| 19.88889 | -0.95927 | -0.90046 | -0.8058 | -0.06536 | -0.35796 | -0.1092 | -0.77736 | -0.06484 | -0.60503 | -0.89595 | -0.17565 | -1.83126 | -0.8395 | -0.83419 | -0.67518 | -0.16308 | -1.41102 | -0.20874 | -0.21988 | -0.40401 | 0.022485 | -1.27789 | -0.6436 | -1.50961 |
| 19.90278 | -0.79598 | -0.71852 | -0.75831 | -0.08497 | -0.30663 | -0.15471 | -0.69181 | -0.04412 | -0.57444 | -0.96762 | -0.32905 | -1.38047 | -0.90039 | -0.51653 | -0.70524 | -0.22265 | -0.77658 | -0.20616 | -0.31876 | -0.01747 | 0.007169 | -0.81352 | -0.56646 | -1.14741 |
| 19.91667 | -0.67382 | -0.66615 | -0.72671 | -0.06812 | -0.23455 | -0.36194 | -0.69109 | -0.02537 | -0.52117 | -1.37707 | -0.61701 | -1.46824 | -1.31564 | -0.3941 | -0.85108 | -0.34441 | -0.51872 | -0.28116 | -1.29676 | 0.123616 | -0.05492 | -0.85877 | -0.53328 | -0.88809 |
| 19.93056 | -0.50966 | -0.61418 | -0.86283 | -0.01278 | -0.11856 | -1.11784 | -0.74762 | 0.053787 | -0.49604 | -2.3936 | -1.26457 | -2.10323 | -2.07822 | -0.37959 | -0.9985 | -0.40066 | -0.2152 | -0.31773 | -3.63898 | 0.028029 | -0.04886 | -1.37453 | -0.73007 | -0.71902 |
| 19.94444 | -0.45522 | -0.53496 | -1.04357 | -0.02823 | 0.008821 | -1.4641 | -0.81184 | 0.005618 | -0.51913 | -2.26743 | -3.05861 | -3.34587 | -3.56899 | -0.37016 | -1.00744 | -0.29323 | 0.233795 | -0.28955 | -5.12249 | -0.08873 | 0.008079 | -1.95032 | -1.62724 | -0.65738 |
| 19.95833 | -0.48899 | -0.43046 | -0.80182 | -0.36409 | -0.03654 | -0.72481 | -0.64194 | -0.33953 | -0.53668 | -0.99526 | -3.32166 | -2.39412 | -4.29811 | -0.5138 | -0.69456 | -0.13858 | 0.325901 | -0.26281 | -3.6185 | -0.26545 | -0.14932 | -1.8657 | -1.91062 | -0.7864 |
| 19.97222 | -0.46753 | -0.26081 | -0.51942 | -0.39293 | -0.15046 | -0.40823 | -0.15747 | -0.2558 | -0.45632 | 0.248308 | -1.47577 | 0.14164 | -1.16466 | -0.68676 | -0.27446 | -0.05728 | 0.153658 | -0.14662 | -1.32405 | -0.55624 | -0.38125 | -1.32644 | 0.520168 | -0.84747 |
| 19.98611 | -0.32274 | -0.12867 | -0.32313 | -0.24067 | -0.1564 | -0.21742 | 0.114711 | -0.17141 | -0.2909 | 0.83356 | -0.16577 | 1.074485 | 1.832231 | -0.50789 | 0.002475 | -0.01111 | 0.009171 | 0.114744 | -0.38217 | -0.73066 | -0.36919 | -0.88058 | 1.279634 | -0.60286 |
| 20 | -0.07046 | -0.09948 | -0.2259 | -0.16012 | -0.1212 | -0.13876 | 0.181946 | -0.10275 | -0.0788 | 0.524651 | 0.185598 | 0.943597 | 2.491923 | -0.13846 | 0.145049 | 0.014731 | -0.05295 | 0.456564 | -0.13718 | -0.69329 | -0.15497 | -0.52338 | 0.54555 | -0.02948 |
| 20.01389 | 0.20603 | -0.08021 | -0.15861 | -0.111 | -0.10942 | -0.10282 | 0.143143 | -0.06637 | 0.218016 | 0.16634 | 0.141606 | 0.689477 | 1.65145 | 0.233508 | 0.138102 | -0.00388 | 0.04984 | 0.551994 | -0.12702 | -0.46319 | -0.07178 | -0.26044 | 0.0974 | 0.423506 |
| 20.02778 | 0.422268 | -0.03194 | -0.1155 | -0.08944 | -0.12894 | -0.07746 | 0.014638 | -0.06151 | 0.497492 | -0.0119 | 0.028673 | 0.493264 | 0.894997 | 0.431569 | 0.115255 | -0.00087 | 0.102628 | 0.286025 | -0.1459 | -0.25166 | -0.11375 | -0.0786 | -0.05052 | 0.552105 |
| 20.04167 | 0.557633 | 0.007534 | -0.09089 | -0.07971 | -0.12531 | -0.05305 | -0.07604 | -0.07413 | 0.591681 | -0.10879 | -0.04369 | 0.318286 | 0.426244 | 0.43734 | 0.151624 | 0.048882 | -0.00788 | 0.101605 | -0.17302 | -0.09494 | -0.10489 | 0.042602 | -0.03149 | 0.480257 |
| 20.05556 | 0.563036 | 0.028137 | -0.06773 | -0.06499 | -0.07966 | -0.04131 | -0.05991 | -0.05907 | 0.456262 | -0.15291 | -0.05715 | 0.228203 | 0.139028 | 0.437958 | 0.256764 | 0.063623 | -0.07852 | -0.05333 | -0.17374 | -0.01438 | -0.12692 | 0.065369 | -0.00741 | 0.317044 |
| 20.06944 | 0.529216 | 0.02336 | -0.03921 | -0.04411 | -0.01554 | -0.03701 | 0.016114 | -0.01427 | 0.332575 | -0.18521 | -0.01816 | 0.23703 | -0.10306 | 0.428358 | 0.473449 | -0.00668 | 0.029062 | -0.21053 | -0.14251 | -0.02605 | -0.08777 | 0.010042 | -0.03687 | 0.187903 |
| 20.08333 | 0.479289 | 0.007553 | 0.013793 | -0.01184 | 0.030022 | -0.02904 | 0.028892 | 0.014554 | 0.372517 | -0.22425 | 0.034236 | 0.254956 | -0.23556 | 0.427539 | 0.462293 | -0.06807 | 0.259204 | -0.24358 | -0.0744 | -0.11491 | -0.00071 | -0.03978 | -0.10084 | 0.092246 |
| 20.09722 | 0.427086 | 0.006426 | 0.098637 | 0.010022 | 0.029843 | -0.01435 | -0.20114 | 0.015503 | 0.322149 | -0.28994 | -0.04973 | 0.012175 | -0.32458 | 0.59653 | 0.285195 | -0.06098 | 0.626855 | -0.20401 | 0.077117 | -0.24116 | 0.064096 | -0.12076 | -0.1907 | -0.03671 |
| 20.11111 | 0.40232 | 0.040322 | 0.157923 | 0.031002 | 0.010412 | -0.00815 | -0.37325 | 0.00482 | 0.222925 | -0.27672 | -0.10684 | -0.17465 | -0.28264 | 0.74581 | 0.171067 | 0.038632 | 0.635648 | -0.16569 | 0.181767 | -0.34497 | 0.138282 | -0.10261 | -0.15559 | -0.12863 |
| 20.125 | 0.377496 | 0.113368 | 0.131951 | 0.092054 | 0.04895 | -0.02262 | 0.098152 | 0.010716 | 0.670415 | 0.020373 | 0.190318 | 0.372247 | 0.190399 | 0.465479 | 0.211333 | 0.148979 | 0.040237 | -0.02072 | 0.063852 | -0.31264 | 0.216835 | 0.343958 | 0.140218 | -0.00571 |
| 20.13889 | 0.341437 | 0.214732 | 0.082458 | 0.114449 | 0.094489 | -0.0326 | 0.51842 | 0.008972 | 1.071842 | 0.27737 | 0.54366 | 0.950066 | 0.578447 | 0.213801 | 0.256501 | 0.144862 | -0.24881 | 0.14308 | -0.0775 | -0.22173 | 0.220341 | 0.696433 | 0.266409 | 0.165943 |
| 20.15278 | 0.232581 | 0.224525 | 0.092996 | 0.052699 | 0.098108 | -0.01523 | 0.266166 | -0.00869 | 0.8305 | 0.160026 | 0.524356 | 0.874378 | 0.471465 | 0.417801 | 0.211388 | 0.086002 | 0.122177 | 0.16463 | 0.019169 | -0.13256 | 0.139057 | 0.577891 | 0.11522 | 0.275927 |
| 20.16667 | 0.09098 | 0.17763 | 0.163522 | 0.053554 | 0.101921 | 0.007747 | 0.031163 | 0.045056 | 0.557382 | 0.004487 | 0.361026 | 0.666843 | 0.275419 | 0.570353 | 0.219436 | 0.076295 | 0.458939 | 0.041665 | 0.112815 | -0.11178 | 0.170553 | 0.447578 | 0.040966 | 0.391172 |
| 20.18056 | 0.019125 | 0.146561 | 0.220719 | 0.065765 | 0.131504 | 0.020264 | 0.049016 | 0.106542 | 0.525033 | 0.025435 | 0.231939 | 0.55248 | 0.201436 | 0.551289 | 0.241771 | 0.079853 | 0.638533 | -0.08607 | 0.125367 | -0.09533 | 0.271532 | 0.471055 | 0.055797 | 0.511923 |
| 20.19444 | 0.019084 | 0.163148 | 0.230682 | 0.055849 | 0.129071 | 0.018413 | 0.12311 | 0.102656 | 0.58115 | 0.073107 | 0.177918 | 0.413932 | 0.203525 | 0.435443 | 0.263332 | 0.086009 | 0.75888 | -0.20348 | 0.092069 | -0.09966 | 0.343401 | 0.501732 | 0.065837 | 0.585263 |
| 20.20833 | 0.11596 | 0.200546 | 0.209611 | 0.038113 | 0.133357 | 0.01233 | 0.215649 | 0.07166 | 0.767861 | 0.165687 | 0.220861 | 0.434357 | 0.250599 | 0.344652 | 0.291982 | 0.082236 | 0.831838 | -0.20922 | 0.0614 | -0.11062 | 0.315379 | 0.544098 | 0.077758 | 0.605246 |
| 20.22222 | 0.310746 | 0.224778 | 0.205036 | 0.028897 | 0.188816 | 0.012514 | 0.296676 | 0.066173 | 0.983976 | 0.273218 | 0.31408 | 0.572001 | 0.313811 | 0.333641 | 0.339321 | 0.085083 | 0.884311 | -0.05752 | 0.042407 | -0.03679 | 0.331908 | 0.640293 | 0.088386 | 0.558132 |
| 20.23611 | 0.524236 | 0.222613 | 0.223167 | 0.034532 | 0.252648 | 0.022062 | 0.363044 | 0.08727 | 1.00675 | 0.341126 | 0.621941 | 0.689375 | 0.321253 | 0.376642 | 0.328639 | 0.090052 | 0.898751 | 0.112928 | 0.05252 | 0.11959 | 0.511894 | 0.732335 | 0.098751 | 0.486932 |
| 20.25 | 0.629139 | 0.20884 | 0.253831 | 0.04821 | 0.257029 | 0.032637 | 0.440002 | 0.106005 | 0.924381 | 0.371358 | 1.189307 | 0.731977 | 0.292305 | 0.444195 | 0.289518 | 0.100509 | 0.896756 | 0.208831 | 0.074796 | 0.307631 | 0.663832 | 0.980174 | 0.103676 | 0.469548 |
| 20.26389 | 0.606762 | 0.22747 | 0.280742 | 0.053865 | 0.216649 | 0.047502 | 0.577725 | 0.101255 | 0.769295 | 0.36419 | 1.450118 | 0.772145 | 0.273583 | 0.604983 | 0.248885 | 0.11053 | 1.035962 | 0.213481 | 0.078196 | 0.515196 | 0.641419 | 1.173613 | 0.12352 | 0.544506 |
| 20.27778 | 0.577181 | 0.279999 | 0.305233 | 0.035012 | 0.178741 | 0.062278 | 0.685026 | 0.098601 | 0.674528 | 0.362032 | 1.317661 | 0.818953 | 0.28496 | 0.774467 | 0.236804 | 0.108443 | 1.386804 | 0.156535 | 0.066327 | 0.717767 | 0.540465 | 1.356296 | 0.158392 | 0.711851 |
| 20.29167 | 0.548993 | 0.35702 | 0.311917 | 0.033502 | 0.183985 | 0.089287 | 0.670904 | 0.095805 | 0.775157 | 0.386993 | 1.155173 | 0.907766 | 0.312595 | 0.873763 | 0.225354 | 0.070461 | 1.834647 | 0.108055 | 0.062324 | 1.004738 | 0.50163 | 1.469415 | 0.144769 | 0.893619 |
| 20.30556 | 0.531679 | 0.396342 | 0.322851 | 0.068475 | 0.236753 | 0.119104 | 0.646759 | 0.105998 | 1.028001 | 0.412542 | 1.074054 | 0.999651 | 0.391506 | 0.78347 | 0.241949 | 0.057961 | 2.074645 | 0.066428 | 0.06018 | 0.988167 | 0.555579 | 1.455551 | 0.136983 | 1.118182 |
| 20.31944 | 0.471385 | 0.37092 | 0.322152 | 0.136075 | 0.284783 | 0.121558 | 0.566308 | 0.092386 | 1.208227 | 0.463703 | 0.940184 | 1.071749 | 0.495135 | 0.527866 | 0.269117 | 0.123618 | 1.996281 | 0.049784 | 0.063851 | 0.652053 | 0.572927 | 1.362929 | 0.201902 | 1.106636 |
| 20.33333 | 0.382089 | 0.338805 | 0.397241 | 0.203905 | 0.29262 | 0.118695 | 0.580905 | 0.084788 | 1.247617 | 0.593501 | 0.926754 | 1.167621 | 0.491549 | 0.230024 | 0.263673 | 0.18073 | 1.819325 | 0.082282 | 0.068852 | 0.377599 | 0.512476 | 1.385432 | 0.252884 | 0.903922 |
| 20.34722 | 0.225303 | 0.37068 | 0.464051 | 0.248945 | 0.326035 | 0.158361 | 0.630349 | 0.091992 | 1.224699 | 0.743943 | 0.992536 | 1.275038 | 0.458624 | -0.02289 | 0.238325 | 0.184359 | 1.672908 | 0.174012 | 0.085032 | 0.112175 | 0.397134 | 1.483911 | 0.26848 | 0.632228 |
| 20.36111 | 0.091076 | 0.420583 | 0.459679 | 0.280554 | 0.312567 | 0.240367 | 0.685533 | 0.117588 | 1.115996 | 0.779168 | 0.9488 | 1.310263 | 0.478319 | 0.021339 | 0.294626 | 0.236361 | 1.521359 | 0.258935 | 0.168512 | -0.00028 | 0.375731 | 1.458555 | 0.358523 | 0.507348 |
| 20.375 | -0.00155 | 0.390088 | 0.361359 | 0.209591 | 0.240249 | 0.334556 | 0.580142 | 0.112647 | 0.925135 | 0.661615 | 0.826742 | 1.252102 | 0.557759 | 0.46607 | 0.416254 | 0.320404 | 1.364852 | 0.293221 | 0.297691 | 0.210914 | 0.4478 | 1.29279 | 0.466853 | 0.491862 |
| 20.38889 | 0.030834 | 0.45316 | 0.311762 | 0.120933 | 0.268745 | 0.353384 | 0.335155 | 0.07895 | 0.736333 | 0.402777 | 0.863172 | 1.261079 | 0.559725 | 0.987523 | 0.413115 | 0.326389 | 1.400911 | 0.238218 | 0.3742 | 0.566186 | 0.552144 | 1.081433 | 0.531455 | 0.485041 |
| 20.40278 | 0.291817 | 0.972345 | 0.486412 | 0.171823 | 0.705851 | 0.558854 | 0.138754 | 0.088875 | 0.749574 | 0.431131 | 1.136616 | 1.918885 | 0.41705 | 1.119452 | 0.222115 | 0.344544 | 2.658271 | 0.207719 | 1.004136 | 0.972272 | 0.561594 | 1.090125 | 0.669782 | 0.392971 |
| 20.41667 | 0.798111 | 1.973677 | 1.011831 | 0.321494 | 1.593638 | 1.034154 | 0.119227 | 0.157985 | 1.037074 | 1.428865 | 1.698576 | 3.341955 | 0.33135 | 0.965844 | 0.100531 | 0.481853 | 5.829791 | 0.349071 | 3.385387 | 1.216659 | 0.518157 | 1.389348 | 1.017694 | 0.410074 |
| 20.43056 | 1.692166 | 2.62391 | 1.437638 | 0.425133 | 1.605276 | 1.584704 | 0.2599 | 0.191298 | 1.145583 | 2.589799 | 2.110723 | 3.779788 | 0.193753 | 0.855786 | 0.003935 | 0.553898 | 8.443083 | 0.610229 | 5.480127 | 1.046473 | 0.385352 | 1.280756 | 1.395766 | 0.55111 |
| 20.44444 | 1.751072 | 2.446343 | 1.40754 | 0.405122 | 1.048661 | 1.583404 | 0.289885 | 0.168137 | 0.985261 | 2.901271 | 1.991536 | 2.564935 | -0.24237 | 0.874124 | -0.03407 | 0.367048 | 7.150548 | 0.703211 | 3.336445 | 0.527571 | 0.140658 | 0.192994 | 1.837193 | 0.749927 |
| 20.45833 | 0.982936 | 2.393358 | 1.267787 | 0.316774 | 0.880779 | 0.948767 | 0.08403 | 0.179775 | 0.895285 | 2.269593 | 1.708142 | 1.512333 | -0.15997 | 0.500193 | -0.03759 | 0.22502 | 4.347383 | 0.420455 | 1.722458 | 0.201286 | 0.246933 | -0.6535 | 2.150546 | 0.849061 |
| 20.47222 | 0.484232 | 2.71741 | 1.159931 | 0.250434 | 1.317171 | 0.616263 | -0.0883 | 0.168944 | 1.081729 | 1.43569 | 1.389185 | 0.860821 | -0.06143 | 0.249888 | 0.208517 | 0.306242 | 2.511891 | 0.251948 | 1.008954 | 0.187079 | 0.668041 | -0.87462 | 1.668284 | 0.654841 |
| 20.48611 | 0.000378 | 2.825467 | 0.879398 | 0.200664 | 2.805549 | 0.396035 | -0.27646 | 0.105995 | 0.895693 | 0.251179 | 0.966861 | 0.262326 | -0.11711 | 0.06032 | 0.346094 | 0.355236 | 0.971252 | 0.043709 | 0.694677 | 0.070837 | 0.804906 | -0.44267 | 1.167939 | 0.233916 |
| 20.5 | -0.51288 | 2.246307 | 0.493527 | 0.286118 | 2.065981 | 0.219676 | -0.36331 | 0.053753 | 0.341131 | -0.43033 | 0.65371 | -0.65091 | -0.36782 | -0.22361 | 0.383177 | 0.292353 | -1.06025 | -0.1219 | 0.399288 | -0.2146 | 0.61375 | -0.03665 | 0.90715 | -0.24536 |
| 20.51389 | -0.97507 | 1.446074 | 0.239217 | 0.44501 | 0.785822 | 0.094968 | -0.28493 | 0.049424 | -0.07461 | -0.10879 | 0.46094 | -1.45343 | -0.24401 | -0.47278 | 0.377166 | 0.261812 | -3.91038 | -0.03761 | 0.022066 | -0.45483 | 0.401426 | 0.044288 | 0.655503 | -0.69143 |
| 20.52778 | -0.88146 | 0.839947 | 0.08476 | 0.563204 | 0.207152 | 0.215847 | -0.00705 | 0.028979 | -0.36027 | 0.572735 | 0.294088 | -1.60671 | 0.003753 | -0.462 | 0.309195 | 0.202939 | -5.92911 | 0.08161 | -0.16126 | -0.35476 | 0.354297 | -0.36736 | 0.289083 | -1.04895 |
| 20.54167 | -0.08909 | 0.505963 | 0.131101 | 0.562253 | -0.13552 | 0.480984 | 0.285899 | 0.039548 | -0.55096 | 1.226991 | 0.136555 | -1.03829 | 0.275366 | -0.18684 | 0.169954 | 0.076727 | -3.45846 | 0.145414 | 0.181397 | 0.038935 | 0.396582 | -0.1213 | -0.26081 | -1.1018 |
| 20.55556 | 0.891652 | 0.328093 | 0.23567 | 0.539433 | -0.40247 | 0.820139 | 0.420195 | 0.058003 | -0.60682 | 1.571946 | -0.06976 | -0.27518 | 0.390785 | 0.485896 | -0.02005 | -0.08805 | 0.637146 | 0.15565 | 0.639577 | 0.44918 | 0.507561 | -1.6712 | -0.88403 | -0.74033 |
| 20.56944 | 1.739997 | 0.198042 | 0.50319 | 0.415024 | -0.34544 | 0.845671 | 0.261191 | 0.078437 | -0.54579 | 1.416352 | -0.21397 | -0.04605 | 0.291658 | 0.798838 | -0.2192 | -0.16478 | 3.73488 | 0.154038 | 0.649653 | 0.550523 | 0.552255 | -1.70791 | -1.18743 | -0.11383 |
| 20.58333 | 1.593293 | 0.143932 | 0.492827 | 0.265786 | -0.07644 | 0.412687 | -0.0094 | 0.060673 | -0.29415 | 0.565229 | -0.23594 | -0.19311 | 0.073433 | 0.644573 | -0.312 | -0.16851 | 4.604932 | 0.112211 | 0.305162 | 0.468026 | 0.700287 | -1.2981 | -0.7834 | 0.359417 |
| 20.59722 | 1.160418 | 0.102483 | 0.404499 | 0.174779 | 0.492899 | 0.007526 | -0.22023 | 0.037139 | 0.118516 | -0.39025 | -0.12996 | -0.24279 | -0.09232 | 0.162216 | -0.06751 | -0.09017 | 4.715829 | -0.04274 | 0.203125 | 0.393476 | 0.825465 | -0.60585 | -0.09245 | 0.477139 |
| 20.61111 | 0.482892 | 0.03505 | 0.317352 | 0.163264 | 1.1056 | -0.22681 | -0.40336 | 0.019089 | 0.514684 | -1.2042 | 0.000996 | -0.04448 | -0.29233 | -0.41052 | 0.124065 | -0.06025 | 3.170703 | -0.22496 | 0.389898 | 0.223189 | 0.740024 | 0.216219 | 0.505428 | 0.495917 |
| 20.625 | -0.2188 | -0.13889 | 0.225503 | 0.23145 | 1.217806 | -0.28113 | -0.43147 | -0.00203 | 0.652381 | -1.10368 | 0.177071 | 0.023041 | -0.30189 | -0.79773 | 0.256549 | -0.04402 | -0.62997 | -0.30822 | 0.413406 | -0.07247 | 0.386172 | 1.061901 | 0.991693 | 0.43161 |
| 20.63889 | -0.77334 | -0.334 | 0.216299 | 0.315168 | 0.845188 | -0.29411 | -0.287 | -0.00994 | 0.510067 | -0.42223 | 0.375626 | 0.004921 | -0.06691 | -0.81805 | 0.328851 | -0.01001 | -3.5302 | -0.18693 | 0.15206 | -0.34192 | 0.183874 | 1.621923 | 1.199904 | 0.232298 |
| 20.65278 | -1.18702 | -0.48246 | 0.196954 | 0.333106 | 0.269328 | -0.32798 | -0.19648 | 0.009697 | 0.148939 | 0.216694 | 0.735504 | 0.060921 | 0.19115 | -0.4227 | 0.273474 | 0.046325 | -5.17727 | 0.026072 | -0.16804 | -0.50711 | -0.08779 | 1.753499 | 0.986182 | -0.26145 |
| 20.66667 | -1.63626 | -0.54636 | 0.089444 | 0.288906 | -0.29656 | -0.15242 | -0.03041 | -0.00249 | -0.23083 | 0.530685 | 0.864224 | 0.290639 | 0.286872 | 0.128161 | 0.153484 | 0.002715 | -5.10488 | 0.120281 | -0.28273 | -0.48121 | 0.104506 | 1.339325 | 1.014071 | -0.88152 |
| 20.68056 | -1.20977 | -0.44913 | -0.05301 | 0.224201 | -0.50108 | 0.177938 | 0.208537 | -0.00861 | -0.50273 | 0.525298 | 0.658524 | 0.413184 | 0.284457 | 0.621747 | 0.029534 | -0.08968 | -4.14914 | 0.111552 | -0.22034 | -0.28754 | 0.385957 | 0.559043 | 1.44107 | -1.36462 |
| 20.69444 | -0.09319 | -0.27966 | -0.10534 | 0.200476 | -0.44093 | 0.682464 | 0.485399 | 0.028028 | -0.5183 | 0.246783 | 0.348746 | 0.162568 | 0.20443 | 0.817207 | -0.05038 | -0.11221 | -1.33643 | 0.032498 | -0.06182 | -0.00116 | 0.485466 | -0.24583 | 0.895436 | -1.41991 |
| 20.70833 | 0.815305 | -0.15 | -0.12393 | 0.193357 | -0.29934 | 0.882417 | 0.375197 | 0.039606 | -0.28179 | -0.43275 | 0.078986 | -0.26284 | 0.051838 | 0.802595 | -0.16625 | -0.04446 | 1.699249 | -0.05744 | 0.196653 | 0.317557 | 0.461444 | -0.86959 | 0.128978 | -1.09309 |
| 20.72222 | 1.670718 | -0.07675 | -0.14159 | 0.158262 | -0.35189 | 0.573125 | 0.070214 | 0.015797 | 0.067725 | -1.31697 | -0.0312 | -0.68659 | -0.10118 | 0.516379 | -0.28329 | 0.113998 | 3.846635 | -0.17845 | 0.420271 | 0.336125 | 0.341562 | -1.15919 | -0.49384 | -0.59736 |
| 20.73611 | 1.56841 | -0.03056 | -0.11676 | 0.100753 | -0.35792 | 0.251052 | -0.15485 | -0.00892 | 0.264491 | -1.48795 | -0.17928 | -0.98028 | -0.28242 | -0.00332 | -0.32178 | -0.04148 | 3.533468 | -0.2894 | 0.520318 | 0.099921 | 0.160142 | -1.23587 | -0.98793 | -0.13838 |
| 20.75 | 0.866921 | -0.03694 | -0.09837 | 0.064226 | -0.31873 | -0.01764 | -0.26264 | -0.04465 | 0.188203 | -0.81526 | -0.42025 | -0.89215 | -0.36276 | -0.38212 | -0.2126 | -0.3988 | 1.854496 | -0.37532 | 0.329778 | -0.10128 | 0.029806 | -1.07728 | -1.47542 | 0.046115 |
| 20.76389 | 0.211162 | -0.08563 | -0.07592 | 0.058124 | -0.19018 | -0.21767 | -0.22492 | -0.07437 | -0.04377 | -0.09949 | -0.44011 | -0.46201 | -0.37601 | -0.51817 | -0.26156 | -0.16608 | 0.073642 | -0.45625 | -0.08961 | -0.28193 | -0.18109 | -0.79116 | -1.58758 | -0.02582 |
| 20.77778 | -0.42233 | -0.1707 | -0.10666 | 0.025674 | -0.08376 | -0.32566 | -0.18112 | -0.01692 | -0.1972 | 0.267422 | -0.34539 | -0.13898 | -0.30147 | -0.44358 | -0.26341 | -0.10992 | -1.45386 | -0.46509 | -0.52063 | -0.44276 | -0.28088 | -0.44686 | -1.32628 | -0.31774 |
| 20.79167 | -1.06872 | -0.32005 | -0.18816 | -0.07794 | -0.10332 | -0.41126 | -0.15315 | 0.020899 | -0.18182 | 0.099217 | -0.21765 | -0.25116 | -0.2135 | -0.42067 | -0.19292 | -0.07854 | -2.64212 | -0.47239 | -0.68827 | -0.54132 | -0.03335 | -0.2977 | -0.78736 | -0.72764 |
| 20.80556 | -1.72574 | -0.45045 | -0.3237 | -0.20179 | -0.21048 | -0.47442 | -0.2042 | 0.03301 | -0.02292 | -0.35157 | -0.17246 | -0.6391 | -0.21289 | -0.47761 | -0.19438 | -0.07855 | -3.02379 | -0.42383 | -0.43764 | -0.61051 | 0.168441 | -0.3212 | -0.44913 | -1.08056 |
| 20.81944 | -2.1388 | -0.52231 | -0.48371 | -0.22383 | -0.303 | -0.38242 | -0.27052 | 0.02128 | 0.091257 | -0.86755 | -0.2316 | -0.91844 | -0.27097 | -0.51196 | -0.19189 | -0.11196 | -2.72674 | -0.31979 | -0.02036 | -0.81942 | 0.047558 | -0.39834 | -0.42153 | -1.24822 |
| 20.83333 | -1.85623 | -0.53259 | -0.52326 | -0.13515 | -0.30469 | -0.28107 | -0.3018 | -0.03082 | -0.05477 | -1.16515 | -0.33757 | -0.95071 | -0.34573 | -0.56802 | -0.17937 | -0.14983 | -2.20875 | -0.15805 | 0.26979 | -0.72074 | -0.20045 | -0.41613 | -0.49117 | -1.28688 |
| 20.84722 | -1.56329 | -0.71891 | -0.50644 | -0.05419 | -0.32647 | -0.09813 | -0.32556 | -0.12936 | -0.44161 | -1.223 | -0.42003 | -0.8228 | -0.38908 | -0.7919 | -0.20957 | -0.16343 | -2.08091 | -0.08563 | 0.358501 | -0.4605 | -0.4011 | -0.39959 | -0.52261 | -1.33795 |
| 20.86111 | -0.9185 | -1.01548 | -0.62251 | -0.0568 | -0.33332 | -0.05315 | -0.35303 | -0.25607 | -0.88538 | -0.65106 | -0.56396 | -0.68176 | -0.43398 | -1.09723 | -0.3349 | -0.11599 | -2.03902 | -0.23509 | 0.286727 | -0.19593 | -0.61053 | -0.42796 | -0.82346 | -1.47369 |
| 20.875 | -0.20408 | -1.23464 | -0.81223 | -0.09024 | -0.38497 | -0.05425 | -0.35864 | -0.32444 | -1.25796 | 0.134122 | -0.78335 | -0.61278 | -0.42587 | -1.05896 | -0.49814 | -0.06274 | -1.75129 | -0.59194 | 0.124926 | -0.07867 | -0.67142 | -0.55588 | -1.32739 | -1.56706 |
| 20.88889 | 0.397468 | -1.20938 | -0.99151 | -0.06454 | -0.34884 | -0.07263 | -0.30177 | -0.20921 | -1.36601 | 0.752935 | -0.90263 | -0.53667 | -0.32742 | -0.52665 | -0.64292 | 0.020478 | -0.90877 | -0.92706 | -0.08251 | -0.05935 | -0.40251 | -0.68902 | -1.76711 | -1.46594 |
| 20.90278 | 0.585997 | -0.99177 | -1.02502 | -0.04549 | -0.30892 | -0.11429 | -0.2691 | -0.03944 | -1.10343 | 0.79834 | -0.94116 | -0.56914 | -0.26476 | -0.00613 | -0.71579 | 0.052292 | -0.2471 | -1.00423 | -0.49037 | -0.15848 | -0.0637 | -0.76266 | -1.7825 | -1.12291 |
| 20.91667 | 0.121745 | -0.87736 | -1.10793 | -0.11268 | -0.28597 | -0.19332 | -0.4924 | 0.028268 | -1.07933 | -0.08457 | -0.95772 | -1.02679 | -0.51381 | -0.03033 | -0.84508 | 0.025138 | -0.17543 | -1.01181 | -1.32577 | -0.394 | -0.09189 | -0.83317 | -1.34495 | -0.9744 |
| 20.93056 | -0.61831 | -0.90974 | -1.11401 | -0.13692 | -0.21376 | -0.34911 | -0.88656 | -0.06215 | -1.2611 | -1.4494 | -1.01839 | -1.82422 | -0.97281 | -0.4889 | -0.86931 | 0.103791 | -0.19453 | -1.58067 | -2.64465 | -0.71764 | -0.06878 | -0.95488 | -0.89399 | -0.96416 |
| 20.94444 | -1.36567 | -1.13395 | -1.10198 | -0.01907 | -0.10028 | -0.57164 | -1.16511 | -0.07811 | -1.39198 | -2.43568 | -1.1224 | -2.7591 | -1.32962 | -0.88272 | -0.98576 | 0.271285 | -0.13087 | -1.66889 | -4.7933 | -1.12154 | 0.25285 | -1.22997 | -0.54698 | -1.00479 |
| 20.95833 | -1.98995 | -1.09385 | -1.1564 | 0.048702 | -0.17475 | -0.47478 | -1.27594 | -0.04249 | -1.08143 | -1.8591 | -1.06397 | -2.67882 | -1.35408 | -1.02408 | -1.14608 | 0.276708 | -0.10716 | -2.06095 | -6.1202 | -1.52145 | 0.359042 | -1.68237 | -0.52918 | -1.21159 |
| 20.97222 | -1.5307 | -0.80354 | -0.95514 | 0.149971 | -0.32282 | -0.34663 | -0.94221 | -0.05666 | -0.76618 | -0.5368 | -0.66293 | -1.67552 | -1.07994 | -0.8143 | -0.78768 | 0.082768 | 0.051382 | -1.482 | -2.37018 | -1.6692 | 0.037585 | -1.69848 | -0.57294 | -1.39589 |
| 20.98611 | -0.5967 | -0.41637 | -0.55395 | 0.159469 | -0.3402 | -0.15004 | -0.44205 | -0.10586 | -0.37127 | 0.441818 | -0.32614 | -0.64406 | -0.54566 | -0.25443 | -0.3117 | -0.06418 | 0.275651 | -0.3295 | 0.178717 | -1.17493 | -0.18365 | -1.21228 | -0.55164 | -1.0392 |
| 21 | 0.083644 | -0.06751 | -0.23875 | -0.02228 | -0.22613 | -0.07515 | -0.08803 | -0.15663 | 0.04195 | 0.797255 | -0.22191 | -0.04327 | 0.030421 | 0.211926 | 0.159694 | -0.05575 | 0.239786 | 0.569698 | 1.084332 | -0.68412 | -0.00022 | -0.61665 | -0.41028 | -0.34152 |
| 21.01389 | 0.487767 | 0.061914 | -0.09318 | -0.21739 | -0.16276 | -0.04789 | -0.01767 | -0.17597 | 0.19914 | 0.606692 | -0.20387 | 0.040371 | 0.277602 | 0.319168 | 0.610362 | -0.03615 | 0.144102 | 1.447617 | 0.952421 | -0.16544 | 0.093462 | -0.18641 | -0.32871 | 0.20163 |
| 21.02778 | 0.567391 | 0.067334 | 0.009397 | -0.40001 | -0.18879 | -0.02755 | -0.16127 | -0.18807 | 0.091078 | 0.139113 | -0.16148 | -0.06061 | 0.062277 | 0.140073 | 0.662644 | -0.07549 | 0.157666 | 1.818159 | 0.740836 | 0.213747 | 0.014202 | -0.04352 | -0.27482 | 0.331657 |
| 21.04167 | 0.470554 | -0.01517 | 0.022004 | -0.47702 | -0.23708 | -0.03632 | -0.23834 | -0.10076 | -0.04921 | -0.31263 | -0.17134 | -0.15448 | -0.17119 | 0.121088 | 0.401709 | -0.08825 | 0.122211 | 1.749774 | 0.459636 | 0.418635 | -0.05715 | -0.00043 | -0.21573 | 0.314638 |

| | | | | | | | | | | | | | | | | | | | | | | | | |
|---|---|---|---|---|---|---|---|---|---|---|---|---|---|---|---|---|---|---|---|---|---|---|---|---|
| 21.05556 | 0.358446 | -0.09134 | -0.04709 | -0.36467 | -0.18878 | -0.04786 | -0.14274 | 0.007815 | -0.12933 | -0.57637 | -0.217 | -0.20579 | -0.26239 | 0.26998 | 0.228439 | -0.08519 | 0.041378 | 1.570341 | 0.162717 | 0.543789 | -0.10862 | -0.00925 | -0.16168 | 0.297847 |
| 21.06944 | 0.313557 | -0.11714 | -0.09331 | -0.08867 | -0.03384 | -0.04135 | -0.06018 | 0.081607 | -0.13965 | -0.71339 | -0.25258 | -0.03543 | -0.29573 | 0.459734 | 0.143772 | -0.08441 | -0.01482 | 0.966181 | 0.051765 | 0.603283 | -0.08657 | -0.0393 | -0.08396 | 0.239313 |
| 21.08333 | 0.288073 | -0.14399 | -0.08907 | 0.117509 | 0.06409 | -0.03581 | -0.06714 | 0.021557 | -0.04288 | -0.69965 | -0.25151 | 0.207035 | -0.26069 | 0.548649 | 0.114561 | -0.03297 | -0.00152 | 0.445734 | 0.031057 | 0.474285 | 0.04189 | -0.01204 | -0.05202 | 0.066384 |
| 21.09722 | 0.228917 | -0.20373 | -0.16492 | -0.00211 | 0.008523 | -0.0614 | -0.08267 | -0.08658 | 0.107637 | -0.47897 | -0.30439 | 0.16265 | -0.24997 | 0.538365 | 0.019467 | 0.003479 | 0.052864 | -0.00765 | -0.21274 | 0.301868 | 0.178979 | -0.06631 | -0.06014 | -0.01304 |
| 21.11111 | 0.221186 | -0.19563 | -0.18839 | -0.19852 | -0.05573 | -0.07069 | -0.00889 | -0.1208 | 0.407726 | -0.06435 | -0.32214 | 0.027046 | -0.25213 | 0.453396 | -0.08327 | -0.05071 | 0.047101 | -0.3362 | -0.37308 | 0.270903 | 0.130141 | -0.17933 | -0.12622 | 0.196498 |
| 21.125 | 0.394397 | 0.000076 | 0.138458 | -0.02283 | -0.0229 | 0.029168 | 0.163042 | -0.06421 | 0.868983 | 0.517672 | -0.05906 | 0.22023 | -0.00893 | 0.524597 | -0.03031 | -0.09653 | 0.079337 | -0.42685 | 0.141861 | 0.448661 | -0.02507 | -0.02154 | -0.07782 | 0.616111 |
| 21.13889 | 0.648353 | 0.185529 | 0.430045 | 0.216599 | 0.060906 | 0.11755 | 0.250871 | 0.044235 | 1.00063 | 0.718305 | 0.133609 | 0.420295 | 0.26518 | 0.754468 | 0.164094 | -0.04957 | 0.116347 | -0.31966 | 0.628364 | 0.58109 | 0.031472 | 0.274405 | 0.030855 | 0.831772 |
| 21.15278 | 0.653531 | 0.180856 | 0.317822 | 0.188898 | 0.111671 | 0.094391 | 0.134005 | 0.140973 | 0.657358 | 0.257259 | -8.1E-05 | 0.326343 | 0.194195 | 0.775628 | 0.295034 | 0.00107 | 0.068775 | -0.25161 | 0.35652 | 0.520154 | 0.219746 | 0.345901 | 0.083822 | 0.698895 |
| 21.16667 | 0.434662 | 0.115695 | 0.194115 | 0.177301 | 0.179912 | 0.048045 | 0.024703 | 0.094106 | 0.361933 | -0.32178 | -0.14244 | 0.053208 | -0.04133 | 0.628152 | 0.320942 | 0.024315 | 0.047435 | -0.33684 | 0.025942 | 0.387892 | 0.320774 | 0.306771 | 0.127297 | 0.512949 |
| 21.18056 | 0.055874 | 0.014176 | 0.176755 | 0.205031 | 0.195444 | 0.043489 | 0.019067 | -0.09859 | 0.436227 | -0.68464 | -0.15284 | -0.07364 | -0.25523 | 0.420401 | 0.271555 | -0.00647 | 0.081927 | -0.45389 | -0.0587 | 0.278923 | 0.229962 | 0.26673 | 0.14802 | 0.530373 |
| 21.19444 | -0.28682 | -0.07938 | 0.199987 | 0.183742 | 0.144394 | 0.063287 | 0.161015 | -0.09767 | 0.862252 | -0.3957 | -0.05801 | 0.298782 | -0.26219 | 0.311557 | 0.192308 | -0.07504 | 0.150167 | -0.45113 | -0.0567 | 0.186 | 0.133643 | 0.234325 | 0.159248 | 0.643588 |
| 21.20833 | -0.31274 | -0.01372 | 0.258428 | 0.193853 | 0.127836 | 0.089407 | 0.317569 | 0.177149 | 1.367742 | 0.446113 | 0.062885 | 0.862594 | 0.004763 | 0.385444 | 0.256307 | -0.02961 | 0.295777 | -0.28674 | -0.01183 | 0.031977 | 0.223492 | 0.334198 | 0.232484 | 0.719601 |
| 21.22222 | -0.19366 | 0.089803 | 0.318377 | 0.211045 | 0.120565 | 0.097659 | 0.401316 | 0.167259 | 1.434571 | 1.063277 | 0.109162 | 1.004317 | 0.325827 | 0.453717 | 0.287596 | 0.040114 | 0.369113 | -0.11703 | 0.046393 | -0.02662 | 0.31795 | 0.448413 | 0.33953 | 0.855684 |
| 21.23611 | -0.04294 | 0.212858 | 0.40021 | 0.208388 | 0.140189 | 0.108636 | 0.465052 | 0.316985 | 1.141865 | 0.877079 | 0.115298 | 0.667312 | 0.44225 | 0.441696 | 0.293683 | 0.034111 | 0.401815 | 0.015211 | 0.101881 | 0.037563 | 0.408171 | 0.491137 | 0.43441 | 1.035014 |
| 21.25 | 0.092555 | 0.289496 | 0.538733 | 0.185248 | 0.170312 | 0.138049 | 0.540739 | 0.22894 | 0.942988 | 0.316248 | 0.123696 | 0.422513 | 0.335493 | 0.411295 | 0.36914 | 0.0195 | 0.677432 | 0.169666 | 0.202113 | 0.096609 | 0.523454 | 0.583061 | 0.565004 | 1.273514 |
| 21.26389 | 0.335963 | 0.296965 | 0.695712 | 0.190052 | 0.176788 | 0.194346 | 0.653164 | 0.085529 | 0.876013 | -0.16363 | 0.165745 | 0.686513 | 0.218669 | 0.365406 | 0.481451 | 0.00985 | 1.170286 | 0.381408 | 0.332533 | 0.123658 | 0.57701 | 0.701238 | 0.724106 | 1.560264 |
| 21.27778 | 0.571216 | 0.275192 | 0.791541 | 0.266595 | 0.187209 | 0.22547 | 0.794289 | -0.00394 | 0.859484 | 0.078073 | 0.212645 | 1.312616 | 0.24008 | 0.361159 | 0.567295 | -0.00368 | 1.569778 | 0.580765 | 0.40491 | 0.217939 | 0.675103 | 0.726275 | 0.871436 | 1.868688 |
| 21.29167 | 0.738229 | 0.279877 | 0.739157 | 0.335575 | 0.196362 | 0.297764 | 0.868878 | 0.077872 | 0.831895 | 1.127016 | 0.243474 | 1.791458 | 0.269041 | 0.346852 | 0.473673 | -0.03416 | 1.78159 | 0.562886 | 0.422816 | 0.434991 | 0.852557 | 0.686863 | 0.999552 | 2.022572 |
| 21.30556 | 0.783668 | 0.319438 | 0.661101 | 0.376276 | 0.185907 | 0.345491 | 0.841157 | 0.235769 | 0.766607 | 2.087024 | 0.275041 | 1.591314 | 0.283447 | 0.293652 | 0.354457 | -0.06292 | 2.093905 | 0.363219 | 0.452878 | 0.656632 | 0.986896 | 0.630249 | 1.178431 | 1.758073 |
| 21.31944 | 0.811411 | 0.396408 | 0.584503 | 0.351152 | 0.17415 | 0.335394 | 0.587925 | 0.245097 | 0.599272 | 1.786323 | 0.319662 | 1.100039 | 0.413093 | 0.298005 | 0.370349 | 0.060091 | 2.513594 | 0.155035 | 0.598706 | 0.760827 | 1.063028 | 0.694807 | 1.448396 | 1.083474 |
| 21.33333 | 0.840941 | 0.504223 | 0.478458 | 0.326211 | 0.20323 | 0.342135 | 0.406951 | 0.203208 | 0.498932 | 1.068525 | 0.390624 | 0.884451 | 0.595983 | 0.433118 | 0.426658 | 0.235973 | 2.765794 | 0.052832 | 0.84805 | 0.702094 | 1.075068 | 0.917536 | 1.698742 | 0.4813 |
| 21.34722 | 0.980538 | 0.58661 | 0.475288 | 0.262684 | 0.240842 | 0.333858 | 0.279744 | 0.177149 | 0.497829 | 0.264032 | 0.469006 | 1.030539 | 0.538265 | 0.665141 | 0.393367 | 0.245452 | 2.791302 | 0.146202 | 0.913269 | 0.599255 | 1.044892 | 0.966122 | 1.722735 | 0.335375 |
| 21.36111 | 1.071996 | 0.581315 | 0.544106 | 0.145737 | 0.245658 | 0.285589 | 0.268886 | 0.164768 | 0.553924 | -0.10586 | 0.49601 | 1.406671 | 0.417803 | 0.86502 | 0.434761 | 0.283077 | 2.709853 | 0.324188 | 0.822632 | 0.478703 | 1.022318 | 1.147463 | 1.570575 | 0.627029 |
| 21.375 | 1.047193 | 0.476248 | 0.487834 | 0.080612 | 0.199593 | 0.230767 | 0.164814 | 0.117549 | 0.518542 | 0.395524 | 0.44833 | 1.576275 | 0.501298 | 1.002961 | 0.482228 | 0.354942 | 2.658337 | 0.342647 | 0.775044 | 0.378123 | 0.967161 | 1.453445 | 1.456594 | 1.029746 |
| 21.38889 | 0.662073 | 0.535732 | 0.418392 | 0.102503 | 0.229754 | 0.270772 | -0.05146 | 0.098549 | 0.412523 | 1.172581 | 0.433458 | 1.186237 | 0.514569 | 0.903725 | 0.469861 | 0.365766 | 2.672072 | 0.238255 | 0.700864 | 0.331896 | 0.806852 | 1.859423 | 1.30394 | 1.312661 |
| 21.40278 | 0.268844 | 0.998794 | 0.628591 | 0.342338 | 0.55027 | 0.48259 | -0.24109 | 0.165508 | 0.263832 | 1.646353 | 0.789196 | 1.272249 | 0.260489 | 0.675126 | 0.51228 | 0.335063 | 4.192915 | 0.297484 | 1.053346 | 0.400046 | 0.553754 | 2.062447 | 1.292207 | 1.405197 |
| 21.41667 | 0.116758 | 2.056285 | 1.403594 | 0.833462 | 1.419989 | 0.962816 | -0.33763 | 0.266933 | 0.533679 | 2.2683 | 1.842907 | 2.672009 | -0.05286 | 0.806882 | 0.739379 | 0.426829 | 8.776912 | 0.739472 | 2.242104 | 0.614014 | 0.477536 | 2.826528 | 1.60229 | 1.790972 |
| 21.43056 | 0.382047 | 2.939217 | 1.951033 | 0.964765 | 1.512081 | 1.051427 | -0.09577 | 0.279152 | 1.030087 | 3.28115 | 2.528413 | 3.870128 | -0.26084 | 1.255092 | 1.020925 | 0.506822 | 12.33945 | 1.235218 | 3.521117 | 0.640689 | 0.432059 | 2.48357 | 1.720039 | 1.813913 |
| 21.44444 | 1.040952 | 2.360632 | 1.910113 | 0.556398 | 1.112596 | 0.548409 | 0.413689 | 0.23839 | 1.605592 | 3.338106 | 1.632274 | 2.950477 | -0.23367 | 1.587504 | 0.925961 | 0.415378 | 11.37258 | 0.835414 | 3.102632 | 0.399907 | 0.298959 | 0.921832 | 0.979852 | 0.839246 |
| 21.45833 | 1.771283 | 1.968446 | 1.516152 | 0.238237 | 1.084536 | 0.203484 | 0.112181 | 0.198942 | 1.953301 | 2.240251 | 1.124001 | 1.914048 | -0.14643 | 0.728811 | 0.570521 | 0.270256 | 8.454301 | 0.520157 | 2.028816 | 0.345779 | 0.731987 | 0.377808 | 0.121834 | -0.27148 |
| 21.47222 | 1.666148 | 1.737681 | 1.115424 | 0.143667 | 1.070418 | -0.03004 | -0.22538 | 0.21827 | 1.532994 | 1.006406 | 0.78651 | 1.074729 | -0.13523 | 0.091242 | 0.456923 | 0.31504 | 6.011777 | 0.339997 | 1.261454 | 0.473593 | 1.118215 | 0.240131 | -0.4849 | -0.66248 |
| 21.48611 | 0.9227 | 1.379225 | 0.658752 | 0.062422 | 0.885601 | -0.25712 | -0.65491 | 0.188945 | 0.872542 | -0.59629 | 0.404767 | 0.290223 | -0.2011 | -0.39061 | 0.184305 | 0.300703 | 3.586633 | 0.101689 | 0.196647 | 0.404573 | 1.264407 | 0.052738 | -0.50447 | -0.66794 |
| 21.5 | 0.368577 | 1.06062 | 0.346722 | -0.01008 | 0.634557 | -0.51461 | -0.89662 | 0.136565 | 0.460554 | -2.54222 | 0.009967 | -0.8456 | -0.27076 | -0.86043 | -0.0276 | 0.169307 | 0.690111 | -0.23397 | -0.32446 | 0.176671 | 0.722312 | -0.11396 | -0.16674 | -0.89515 |
| 21.51389 | 0.090178 | 0.552447 | 0.091552 | -0.0019 | 0.410076 | -0.68261 | -0.76689 | 0.066089 | -0.04313 | -2.99258 | -0.3239 | -1.50585 | -0.28135 | -1.08559 | -0.34416 | -0.03119 | -2.84364 | -0.65935 | -0.40562 | -0.17173 | 0.12932 | -0.18022 | 0.220283 | -1.1535 |
| 21.52778 | 1.082254 | 0.077842 | -0.13547 | 0.10395 | 0.183702 | -0.5308 | -0.17598 | -0.05117 | -0.57143 | -0.97033 | -0.45403 | -0.96716 | -0.18102 | -0.69117 | -0.60756 | -0.00819 | -5.95335 | -0.71516 | -0.03848 | -0.59737 | -0.22002 | -0.41317 | 0.375033 | -1.33887 |
| 21.54167 | 0.743395 | -0.49183 | -0.28248 | 0.255355 | -0.05868 | -0.13733 | 0.595401 | -0.12356 | -0.6221 | 1.139582 | -0.25806 | 0.172261 | -0.10152 | 0.029602 | -0.3239 | 0.06919 | -8.40867 | -0.29787 | 0.634721 | -0.5932 | -0.00506 | -0.71065 | 0.23222 | -0.9421 |
| 21.55556 | -0.68412 | -0.82205 | -0.28227 | 0.302736 | -0.25782 | 0.295479 | 0.881868 | -0.11804 | -0.3206 | 2.361255 | 0.161972 | 0.474679 | 0.032949 | 0.677215 | 0.034699 | 0.266829 | -8.16198 | 0.105981 | 1.095422 | -0.10728 | 0.156614 | -0.81091 | -0.08157 | -0.17447 |
| 21.56944 | -0.54458 | -0.98719 | -0.08906 | 0.241666 | -0.3781 | 0.665213 | 0.508744 | -0.06122 | 0.03065 | 1.773173 | 0.554357 | -0.03252 | 0.044812 | 0.735164 | 0.390851 | 0.188366 | -5.7161 | 0.295945 | 0.819333 | 0.495703 | 0.519079 | -0.78478 | -0.37035 | 0.661312 |
| 21.58333 | 0.481811 | -0.91693 | 0.157599 | 0.133617 | -0.39367 | 0.860938 | 0.006705 | -0.02495 | 0.110594 | 0.658244 | 0.742098 | -0.51235 | -0.06224 | 0.365075 | 0.598134 | 0.039181 | -2.82824 | 0.387782 | 0.142848 | 1.007548 | 0.9657 | -0.86679 | -0.6236 | 1.257216 |
| 21.59722 | 1.773771 | -0.71377 | 0.272955 | 0.048996 | -0.22861 | 0.711976 | -0.48242 | -0.0303 | -0.10106 | -0.26434 | 0.581935 | -0.96276 | -0.3365 | 0.09108 | 0.439172 | -0.09049 | -0.8053 | 0.272316 | -0.3024 | 1.104723 | 0.959523 | -1.08774 | -0.66923 | 1.021393 |
| 21.61111 | 2.266485 | -0.49871 | 0.219654 | -0.00985 | -0.03691 | 0.352543 | -0.94948 | -0.03177 | -0.21713 | -1.2748 | 0.377333 | -1.21476 | -0.70106 | -0.19288 | 0.172214 | -0.19067 | 0.023533 | -0.03426 | -0.2306 | 0.568995 | 0.346216 | -1.16703 | -0.86769 | 0.460068 |
| 21.625 | 1.354878 | -0.25307 | 0.201396 | -0.06515 | 0.137519 | 0.089463 | -0.96667 | -0.05384 | 0.118742 | -2.12705 | 0.158478 | -0.91105 | -0.63704 | -0.41233 | -0.13126 | -0.24304 | 0.086996 | -0.18161 | 0.143866 | 0.345028 | 0.078639 | -0.84233 | -0.83971 | 0.244752 |
| 21.63889 | 0.355248 | -0.1337 | 0.156739 | -0.05353 | 0.225425 | -0.18161 | -0.40033 | -0.04087 | 0.365029 | -2.0063 | -0.06508 | -0.07278 | -0.08706 | -0.30989 | -0.49805 | -0.11144 | 0.177843 | -0.34796 | 0.492502 | 0.094015 | -0.15197 | -0.18067 | -0.40749 | 0.167332 |
| 21.65278 | -0.09968 | -0.17057 | 0.035996 | 0.017471 | 0.257074 | -0.4803 | 0.162058 | -0.00858 | 0.38325 | -0.61074 | -0.20944 | 0.719952 | 0.367674 | 0.04158 | -0.71887 | 0.117139 | -0.03488 | -0.48106 | 0.506164 | -0.09811 | -0.32109 | 0.462796 | 0.438069 | -0.16468 |
| 21.66667 | -0.55656 | -0.23837 | -0.12758 | 0.120858 | 0.252599 | -0.70019 | 0.195846 | -0.00453 | 0.218263 | 0.749598 | -0.33615 | 1.225038 | 0.44617 | 0.606452 | -0.78718 | 0.221009 | -0.6523 | -0.44235 | 0.242399 | -0.27593 | -0.45673 | 0.70307 | 1.154922 | -0.64979 |
| 21.68056 | -0.84772 | -0.47762 | -0.31702 | 0.172302 | 0.158306 | -0.62401 | -0.03424 | 0.031071 | 0.105487 | 1.571492 | -0.40241 | 1.124618 | 0.159566 | 0.774541 | -0.46633 | 0.318761 | -1.73971 | -0.189 | -0.13371 | -0.57002 | -0.48776 | 0.637968 | 1.332717 | -1.25408 |
| 21.69444 | -1.174 | -0.709 | -0.50048 | 0.155568 | 0.011898 | -0.33166 | -0.24491 | 0.022176 | 0.038199 | 1.219266 | -0.31966 | 0.371882 | -0.16725 | 0.537383 | 0.015471 | 0.17133 | -3.22845 | 0.088275 | -0.52805 | -1.0609 | -0.36964 | 0.308633 | 0.86428 | -1.93522 |
| 21.70833 | -1.56047 | -0.97015 | -0.60156 | 0.120048 | -0.13466 | -0.15813 | -0.45925 | -0.03029 | -0.19115 | 0.263675 | -0.06479 | -0.53461 | -0.5012 | 0.120735 | 0.210749 | -0.09731 | -4.12942 | 0.207804 | -0.83644 | -1.35646 | -0.03145 | 0.024394 | 0.010021 | -2.39353 |
| 21.72222 | -1.67074 | -1.25464 | -0.55785 | 0.062276 | -0.25455 | -0.12833 | -0.81399 | -0.10659 | -0.71492 | -0.71256 | 0.250398 | -1.28301 | -0.80666 | -0.37271 | 0.262217 | -0.29145 | -3.51178 | 0.114088 | -0.60772 | -1.40998 | 0.178452 | -0.1736 | -1.0311 | -2.39316 |
| 21.73611 | -1.5189 | -1.44679 | -0.50521 | -0.04313 | -0.3522 | 0.02244 | -0.84163 | -0.21474 | -1.28108 | -1.8016 | 0.335726 | -1.29477 | -0.95803 | -0.91096 | 0.203392 | -0.45567 | -2.56698 | -0.02412 | -0.1396 | -1.05215 | 0.33991 | -0.3053 | -1.95373 | -1.76804 |
| 21.75 | -0.98884 | -1.17378 | -0.49844 | -0.09229 | -0.45105 | 0.206841 | -0.51371 | -0.37135 | -1.58859 | -2.57209 | 0.158009 | -0.86508 | -0.75896 | -1.58045 | 0.007638 | -0.4769 | -1.82831 | -0.17256 | 0.326472 | -0.48843 | 0.646212 | -0.51244 | -2.2769 | -0.84908 |
| 21.76389 | -0.5367 | -1.06213 | -0.56774 | -0.08887 | -0.53048 | 0.103726 | -0.42301 | -0.49921 | -1.35388 | -2.14154 | -0.1505 | -0.6284 | -0.52556 | -1.80865 | -0.29019 | -0.41166 | -1.41931 | -0.38351 | 0.454418 | -0.04098 | 0.778125 | -0.69626 | -2.29816 | -0.33315 |
| 21.77778 | -0.40029 | -1.26938 | -0.71735 | -0.0405 | -0.65074 | -0.25546 | -0.69802 | -0.46627 | -1.066 | -1.35352 | -0.51163 | -0.79136 | -0.53538 | -1.41928 | -0.68317 | -0.35414 | -1.01132 | -0.70655 | 0.124191 | 0.067453 | 0.126054 | -0.88244 | -2.13326 | -0.4819 |
| 21.79167 | -0.78284 | -1.62558 | -0.87574 | -0.01028 | -0.72234 | -0.58439 | -1.08914 | -0.35572 | -0.94489 | -0.44058 | -0.82374 | -1.08257 | -0.63856 | -1.16048 | -1.08167 | -0.25134 | -0.53307 | -1.00066 | -0.3616 | -0.07154 | -0.83221 | -0.99444 | -1.82545 | -1.03628 |
| 21.80556 | -1.21415 | -1.73757 | -1.04535 | -0.02833 | -0.78265 | -0.64345 | -1.40141 | -0.20111 | -1.13079 | 0.36993 | -1.02068 | -1.35763 | -0.54471 | -1.11728 | -1.26167 | -0.15348 | -0.64447 | -0.72759 | -0.67756 | -0.35303 | -0.50142 | -0.9794 | -1.41109 | -1.22118 |
| 21.81944 | -1.49332 | -1.71602 | -1.09991 | -0.08465 | -0.75051 | -0.47705 | -1.44372 | -0.11907 | -1.41788 | 0.357383 | -1.14846 | -1.58491 | -0.47579 | -1.0921 | -1.10496 | -0.17058 | -1.70808 | -0.41873 | -0.80471 | -0.60495 | -1.37935 | -0.86614 | -0.98614 | -1.22636 |
| 21.83333 | -1.32926 | -1.63925 | -1.0045 | -0.14088 | -0.52146 | -0.29531 | -1.1047 | -0.20132 | -1.53525 | -0.44208 | -1.21069 | -1.77379 | -0.49055 | -0.9044 | -0.76829 | -0.28 | -2.93791 | -0.20394 | -0.8708 | -0.7408 | -0.71476 | -0.63956 | -0.67367 | -1.22842 |
| 21.84722 | -1.073 | -1.27917 | -0.78548 | -0.1805 | -0.30417 | -0.21216 | -0.46606 | -0.40085 | -1.36038 | -1.53318 | -1.05442 | -1.66713 | -0.44078 | -0.64894 | -0.42542 | -0.31815 | -3.19277 | -0.07748 | -0.68164 | -0.74861 | -0.07983 | -0.30837 | -0.30685 | -1.22191 |
| 21.86111 | -0.92076 | -0.88981 | -0.61182 | -0.21547 | -0.13787 | -0.13831 | 0.111759 | -0.50578 | -0.89753 | -2.60367 | -0.78112 | -1.09741 | -0.32911 | -0.40654 | -0.24662 | -0.24505 | -2.72874 | -0.0202 | -0.31781 | -0.6728 | 0.431184 | -0.10867 | -0.08368 | -1.19711 |
| 21.875 | -1.00029 | -0.66717 | -0.50038 | -0.18873 | -0.12105 | -0.11132 | 0.239973 | -0.39472 | -0.53603 | -3.03471 | -0.55438 | -0.74031 | -0.45062 | -0.47722 | -0.26554 | -0.18739 | -1.66697 | -0.12368 | -0.13579 | -0.62741 | 0.374496 | -0.06664 | -0.21077 | -1.06195 |
| 21.88889 | -0.94681 | -0.57276 | -0.41152 | -0.07565 | -0.13665 | -0.15574 | 0.134214 | -0.22926 | -0.30394 | -2.25963 | -0.42298 | -0.45091 | -0.63186 | -0.59472 | -0.46421 | -0.19608 | -0.66081 | -0.29267 | -0.26798 | -0.62813 | 0.319298 | -0.00079 | -0.54766 | -0.89945 |
| 21.90278 | -0.73052 | -0.55792 | -0.40394 | -0.00357 | -0.12653 | -0.19513 | -0.03717 | -0.08214 | -0.30097 | -1.17827 | -0.39829 | -0.24145 | -0.62374 | -0.64764 | -0.73084 | -0.19011 | -0.16759 | -0.49857 | -0.72149 | -0.52811 | 0.129482 | 0.113028 | -0.63427 | -0.8208 |
| 21.91667 | -0.72496 | -0.5927 | -0.66742 | -0.15113 | -0.12434 | -0.26802 | -0.22851 | -0.04704 | -0.5724 | -0.66608 | -0.77105 | -0.29338 | -0.69275 | -0.81612 | -1.08179 | -0.1746 | -0.24028 | -0.74605 | -1.66943 | -0.33319 | -0.34848 | -0.00436 | -0.57273 | -0.97382 |
| 21.93056 | -1.05165 | -0.66403 | -1.15486 | -0.67131 | -0.08074 | -0.45426 | -0.36575 | -0.04899 | -0.89223 | -1.17941 | -1.82351 | -0.812 | -0.95156 | -1.39938 | -1.59883 | -0.11901 | -0.42518 | -1.14376 | -2.7785 | -0.24591 | -1.04349 | -0.44777 | -0.68076 | -1.24225 |
| 21.94444 | -1.34243 | -0.75323 | -1.52968 | -0.92758 | 0.010087 | -0.63594 | -0.47033 | -0.1462 | -0.85548 | -2.22766 | -2.82651 | -1.58374 | -1.25467 | -2.41209 | -2.15173 | 0.061343 | -0.38621 | -1.69094 | -3.72443 | -0.33082 | -1.88919 | -0.96732 | -0.98902 | -1.39509 |
| 21.95833 | -1.49029 | -1.14562 | -1.47537 | -0.6022 | -0.00466 | -0.54111 | -0.62692 | -0.38371 | -0.58528 | -1.93318 | -1.73771 | -1.91509 | -1.31928 | -3.05798 | -2.11426 | 0.122779 | -0.19406 | -1.98778 | -1.63366 | -0.71684 | -3.2292 | -1.26483 | -0.81591 | -1.39597 |
| 21.97222 | -1.13805 | -1.21934 | -0.87573 | 0.006224 | -0.07178 | -0.36966 | -0.75931 | -0.25019 | -0.39932 | -0.80841 | -0.08453 | -1.71686 | -1.13294 | -1.99699 | -1.51509 | 0.020009 | 0.147436 | -1.41684 | 0.330998 | -1.15576 | -2.27511 | -1.34445 | -0.33636 | -0.98567 |
| 21.98611 | -0.49858 | -0.87594 | -0.11511 | 0.514885 | -0.0874 | -0.21749 | -0.65812 | 0.167319 | -0.33006 | 0.21707 | 0.468362 | -0.89483 | -0.74477 | -0.00156 | -0.67728 | -0.02749 | 0.6142 | -0.49996 | 0.93925 | -1.22026 | -1.43291 | -1.04088 | 0.132541 | -0.56161 |
| 22 | -0.07249 | -0.39167 | 0.307499 | 0.639428 | -0.0515 | -0.11193 | -0.39588 | 0.650811 | -0.39172 | 1.448546 | 0.367802 | 0.226435 | -0.1995 | 1.638616 | 0.171907 | 0.069811 | 0.934303 | 0.105643 | 0.61118 | -0.93297 | -0.50238 | -0.54658 | 0.654308 | -0.33389 |
| 22.01389 | 0.16022 | 0.050814 | 0.295825 | 0.502558 | -0.01744 | -0.0532 | -0.2165 | 0.743569 | -0.28288 | 2.479216 | 0.188916 | 0.827712 | 0.269419 | 2.065476 | 0.751327 | 0.153946 | 0.964888 | 0.432514 | 0.287058 | -0.583 | 0.678267 | -0.16793 | 0.738499 | -0.27538 |
| 22.02778 | 0.21432 | 0.224033 | 0.180126 | 0.298655 | 0.039536 | -0.03844 | -0.29015 | 0.675402 | -0.15352 | 3.013766 | 0.07039 | 0.770939 | 0.466525 | 1.428645 | 0.916426 | 0.098591 | 0.69219 | 0.558765 | 0.088987 | -0.28243 | 1.581968 | -0.09988 | 0.455496 | -0.38143 |
| 22.04167 | 0.057522 | 0.178715 | 0.07697 | 0.144293 | 0.062929 | -0.04661 | -0.48957 | 0.374245 | -0.0931 | 2.715307 | -0.02254 | 0.398984 | 0.456546 | 0.831608 | 0.955051 | 0.02631 | 0.403435 | 0.601012 | -0.05657 | -0.08887 | 1.292135 | -0.12984 | 0.21267 | -0.40502 |
| 22.05556 | -0.07728 | 0.087714 | 0.04679 | 0.059293 | 0.036378 | -0.0441 | -0.32681 | 0.022296 | -0.08048 | 1.975205 | -0.04421 | -0.01821 | 0.44408 | 0.498346 | 0.892202 | 0.022451 | 0.224241 | 0.560698 | -0.0473 | 0.013742 | 1.066962 | -0.06317 | 0.056523 | -0.17017 |
| 22.06944 | -0.12441 | 0.141977 | 0.038623 | 0.028201 | -0.01595 | -0.02505 | 0.174686 | -0.28615 | -0.09989 | 1.245206 | -0.03421 | -0.27463 | 0.509704 | 0.396314 | 0.902839 | 0.072164 | 0.142455 | 0.511481 | 0.011384 | 0.093914 | 1.207367 | 0.046327 | -0.02938 | 0.243339 |
| 22.08333 | 0.02192 | 0.210593 | 0.021394 | 0.006863 | -0.08734 | -0.02979 | 0.384493 | -0.41678 | -0.12158 | 0.377492 | -0.07378 | -0.19534 | 0.48995 | 0.321572 | 0.916261 | 0.085685 | 0.10411 | 0.498957 | 0.021459 | 0.107859 | 1.227744 | 0.077976 | -0.14126 | 0.464324 |
| 22.09722 | 0.315334 | 0.248157 | -0.00637 | 0.00348 | -0.1201 | -0.05974 | 0.256527 | -0.28886 | -0.13854 | -0.46841 | -0.14988 | 0.20912 | 0.301795 | 0.182397 | 0.706431 | -0.01725 | 0.080922 | 0.549404 | -0.0637 | 0.085494 | 0.999592 | 0.014592 | -0.40752 | 0.236123 |
| 22.11111 | 0.461314 | 0.22276 | -0.00608 | 0.036854 | -0.07093 | -0.06785 | 0.132327 | -0.04427 | -0.12804 | -1.10604 | -0.1802 | 0.519694 | 0.121239 | 0.111798 | 0.64748 | -0.08453 | 0.058267 | 0.612045 | -0.13384 | 0.062762 | 0.688773 | -0.05779 | -0.50728 | -0.10209 |
| 22.125 | 0.456434 | 0.11955 | 0.002978 | 0.063699 | 0.014615 | -0.04514 | 0.000975 | 0.017981 | -0.12179 | -1.27978 | -0.15769 | 0.417329 | 0.105556 | -0.01473 | 0.775077 | -0.04249 | -0.03864 | 0.709682 | -0.1249 | 0.026678 | 0.407713 | -0.02015 | -0.25699 | -0.16517 |
| 22.13889 | 0.357377 | 0.010147 | 0.015629 | 0.044283 | 0.048357 | -0.02327 | -0.15273 | -0.10944 | -0.15011 | -0.63976 | -0.11574 | -0.05009 | 0.047802 | -0.35101 | 0.75345 | -0.01162 | -0.1541 | 0.710487 | -0.10249 | 0.015947 | 0.088181 | 0.032147 | -0.00783 | 0.107305 |
| 22.15278 | 0.127411 | -0.06839 | 0.00807 | -0.00904 | 0.03394 | -0.01862 | -0.18381 | -0.26291 | -0.17162 | 0.250925 | -0.09085 | -0.4821 | -0.10734 | -0.71107 | 0.567792 | -0.02591 | -0.19973 | 0.619844 | -0.15262 | 0.054044 | -0.22104 | 0.018691 | 0.04995 | 0.332702 |
| 22.16667 | -0.08875 | -0.09087 | -0.00158 | -0.01176 | -0.00662 | -0.02379 | -0.12571 | -0.2952 | -0.15875 | 0.437779 | -0.09376 | -0.60272 | -0.24643 | -0.71598 | 0.336544 | -0.04459 | -0.08343 | 0.475567 | -0.2334 | 0.105837 | -0.33806 | -0.01183 | -0.06221 | 0.185259 |
| 22.18056 | -0.18007 | -0.07301 | 0.018564 | 0.048165 | -0.03663 | -0.01752 | -0.10172 | -0.08361 | -0.10355 | -0.05676 | -0.09298 | -0.34219 | -0.18702 | -0.22528 | 0.160515 | 0.013371 | 0.120631 | 0.29925 | -0.22125 | 0.131703 | -0.26813 | 0.0143 | -0.1625 | -0.12124 |
| 22.19444 | -0.18389 | -0.08297 | 0.062655 | 0.109475 | -0.05572 | -0.00989 | -0.01966 | 0.249264 | -0.04364 | -0.52301 | -0.07682 | 0.059426 | -0.06761 | 0.118403 | 0.024199 | 0.05445 | 0.219908 | 0.15005 | -0.17604 | 0.114045 | -0.14792 | 0.009516 | -0.20673 | -0.17434 |
| 22.20833 | -0.09887 | -0.07124 | 0.153288 | 0.137643 | -0.03547 | -0.00729 | 0.178764 | 0.34553 | 0.008395 | -0.60409 | -0.00637 | 0.282314 | 0.002599 | -0.0638 | -0.11371 | 0.075101 | 0.139406 | 0.01857 | -0.12829 | 0.068692 | -0.01774 | -0.02826 | -0.04498 | 0.147891 |
| 22.22222 | 0.139507 | -0.03585 | 0.237534 | 0.171303 | 0.012372 | 0.000625 | 0.366962 | 0.165114 | 0.079572 | -0.1623 | 0.142463 | 0.247504 | 0.145912 | -0.27747 | -0.03016 | 0.132867 | 0.035197 | -0.0354 | 0.016799 | 0.030759 | 0.118387 | -0.01603 | 0.4866 | 0.548049 |
| 22.23611 | 0.507505 | 0.061474 | 0.307397 | 0.216703 | 0.04877 | 0.019455 | 0.479959 | -0.06244 | 0.195008 | 0.716541 | 0.280446 | 0.221203 | 0.179751 | -0.28592 | 0.132431 | 0.121459 | 0.02407 | 0.035995 | 0.226187 | 0.075046 | 0.145494 | 0.019927 | 0.975377 | 0.630961 |
| 22.25 | 0.888269 | 0.201523 | 0.384106 | 0.264952 | 0.093657 | 0.05802 | 0.561973 | -0.08007 | 0.358634 | 1.655479 | 0.293354 | 0.470348 | 0.287976 | 0.026035 | 0.291531 | 0.13983 | 0.16918 | 0.139889 | 0.482258 | 0.113057 | 0.386605 | 0.074739 | 1.404069 | 0.501372 |
| 22.26389 | 1.010153 | 0.283709 | 0.430671 | 0.302217 | 0.10431 | 0.089189 | 0.558724 | 0.265762 | 0.510228 | 1.672719 | 0.20682 | 0.773011 | 0.567224 | 0.519684 | 0.389833 | 0.259436 | 0.300041 | 0.135886 | 0.708321 | 0.075595 | 0.854823 | 0.108259 | 1.46703 | 0.593489 |
| 22.27778 | 0.685516 | 0.219231 | 0.424164 | 0.300717 | 0.06525 | 0.084282 | 0.480718 | 0.649978 | 0.582609 | 0.780784 | 0.113071 | 1.050956 | 0.55811 | 0.653852 | 0.302245 | 0.27808 | 0.24127 | 0.113538 | 0.687413 | 0.123418 | 0.945045 | 0.082879 | 1.149425 | 0.737233 |
| 22.29167 | 0.402266 | 0.167352 | 0.490389 | 0.33948 | 0.095348 | 0.117705 | 0.49768 | 0.715701 | 0.632361 | -0.27162 | 0.15977 | 1.04555 | 0.435619 | 0.594271 | 0.207736 | 0.194824 | 0.144271 | 0.183413 | 0.579724 | 0.287867 | 0.797972 | 0.110437 | 0.864284 | 0.811821 |
| 22.30556 | 0.329847 | 0.154195 | 0.572035 | 0.382183 | 0.172046 | 0.191652 | 0.603404 | 0.456326 | 0.655325 | -0.19682 | 0.311631 | 0.830061 | 0.4189 | 0.557859 | 0.30273 | 0.179114 | 0.100996 | 0.324296 | 0.570195 | 0.461351 | 0.645137 | 0.149936 | 0.919281 | 0.766127 |
| 22.31944 | 0.474572 | 0.13618 | 0.575195 | 0.344901 | 0.162438 | 0.232259 | 0.654794 | 0.007648 | 0.673062 | 1.118382 | 0.463606 | 0.617319 | 0.358115 | 0.596045 | 0.556247 | 0.182637 | 0.103338 | 0.470942 | 0.620091 | 0.546416 | 0.493065 | 0.192085 | 1.109776 | 0.746395 |
| 22.33333 | 0.781566 | 0.16948 | 0.53539 | 0.221073 | 0.093244 | 0.24719 | 0.708415 | -0.11883 | 0.734401 | 2.287613 | 0.468361 | 0.620793 | 0.2663 | 0.766919 | 0.83504 | 0.194839 | 0.12765 | 0.607403 | 0.587485 | 0.524759 | 0.442061 | 0.250428 | 1.210155 | 0.766818 |
| 22.34722 | 1.11664 | 0.229435 | 0.471388 | 0.104721 | 0.023513 | 0.258972 | 0.74255 | 0.22978 | 0.900353 | 2.032843 | 0.383544 | 0.738065 | 0.133774 | 1.044845 | 0.847799 | 0.227381 | 0.176429 | 0.63882 | 0.440227 | 0.526065 | 0.481952 | 0.305284 | 1.085111 | 0.765727 |
| 22.36111 | 1.262941 | 0.264436 | 0.507744 | 0.042315 | 0.013313 | 0.265839 | 0.646696 | 0.707463 | 1.011851 | 1.05415 | 0.354876 | 0.810079 | 0.136293 | 1.144978 | 0.801167 | 0.334129 | 0.350402 | 0.565347 | 0.319968 | 0.568342 | 0.716882 | 0.43951 | 1.015765 | 0.746922 |
| 22.375 | 1.084424 | 0.227098 | 0.605384 | 0.102484 | 0.046025 | 0.287279 | 0.509476 | 0.761451 | 1.049979 | 0.014776 | 0.402864 | 0.724848 | 0.607927 | 1.169459 | 0.958444 | 0.485674 | 0.592651 | 0.571621 | 0.362604 | 0.719263 | 1.091291 | 0.606251 | 1.109151 | 0.815284 |
| 22.38889 | 0.710566 | 0.221249 | 0.670643 | 0.109376 | 0.067729 | 0.279234 | 0.461287 | 0.454287 | 1.16199 | -0.82565 | 0.54602 | 0.615685 | 0.973071 | 0.896355 | 1.090545 | 0.469693 | 0.949888 | 0.53887 | 0.545296 | 0.839513 | 1.184603 | 0.637659 | 1.267582 | 0.947687 |
| 22.40278 | 0.48314 | 0.418231 | 0.833998 | 0.292013 | 0.111472 | 0.405157 | 0.454863 | 0.166287 | 2.021674 | 0.138396 | 0.961969 | 0.744063 | 0.955325 | 0.591722 | 1.115597 | 0.333347 | 2.841474 | 0.546059 | 1.133906 | 0.843014 | 0.902271 | 0.793826 | 1.641487 | 1.367609 |
| 22.41667 | 0.817734 | 0.939903 | 1.480043 | 0.87285 | 0.236813 | 0.738548 | 0.630638 | 0.225816 | 3.719094 | 2.577468 | 1.686513 | 1.442505 | 0.932369 | 0.736678 | 1.429455 | 0.339038 | 8.996785 | 0.645099 | 2.47973 | 0.908098 | 0.960308 | 1.535242 | 2.561349 | 2.910953 |
| 22.43056 | 1.45178 | 1.737731 | 2.154031 | 1.356069 | 0.248635 | 0.884539 | 1.040906 | 0.691352 | 3.174451 | 5.020569 | 1.618751 | 2.205899 | 0.970105 | 1.419414 | 1.927339 | 0.377106 | 15.32763 | 0.848202 | 3.449143 | 0.80773 | 1.134637 | 2.326089 | 3.151321 | 4.513806 |
| 22.44444 | 1.696098 | 1.613905 | 1.789835 | 1.072692 | 0.05148 | 0.66369 | 1.733618 | 0.903972 | 1.629741 | 4.525907 | 0.978919 | 2.507425 | 0.581138 | 2.824001 | 1.785143 | 0.490874 | 12.14288 | 0.606868 | 2.375428 | 0.68278 | 1.521531 | 2.569865 | 2.709093 | 3.309115 |
| 22.45833 | 1.695277 | 1.223246 | 1.619082 | 0.757119 | 0.007556 | 0.701703 | 0.948431 | 0.410325 | 1.181199 | 2.643174 | 0.619827 | 1.9143 | 0.351421 | 3.316128 | 1.314624 | 0.468786 | 8.238346 | 0.653883 | 0.650062 | 0.546483 | 1.112239 | 2.181751 | 1.706223 | 1.963983 |
| 22.47222 | 0.963981 | 0.945096 | 1.62666 | 0.494469 | 0.199753 | 0.658591 | -0.08166 | 0.230413 | 0.79382 | 1.19103 | 0.373852 | 0.859828 | 0.183714 | 2.375277 | 0.891409 | 0.422118 | 5.581586 | 0.649861 | -0.3139 | 0.548691 | 0.629016 | 1.971422 | 0.825892 | 1.296575 |
| 22.48611 | 0.161928 | 1.007558 | 1.275933 | 0.189718 | 0.268006 | 0.374447 | -0.98534 | 0.147167 | 0.231129 | 0.135177 | 0.205008 | 0.115233 | -0.2825 | 1.135286 | 0.382628 | 0.163075 | 3.728964 | 0.418588 | -1.5935 | 0.363416 | 0.230107 | 1.092446 | 0.042243 | 0.549277 |
| 22.5 | -0.74836 | 0.95972 | 0.880715 | -0.12682 | 0.196384 | 0.150509 | -1.4952 | -0.02922 | 0.09211 | -1.32796 | 0.125044 | -0.73494 | -0.90194 | -0.55292 | -0.40657 | -0.19449 | 1.482842 | 0.059838 | -2.59587 | 0.038701 | -0.45124 | 0.535256 | -0.72349 | -0.071 |
| 22.51389 | -1.46372 | 0.976319 | 0.468942 | -0.26045 | 0.193309 | -0.09031 | -1.56759 | -0.40934 | 0.327243 | -2.59354 | 0.159144 | -0.73305 | -0.95276 | -1.72849 | -1.20312 | -0.33606 | -1.21156 | -0.36443 | -0.96335 | -0.16094 | -1.22763 | 0.266464 | -0.87033 | -0.29906 |
| 22.52778 | -1.55307 | 1.159334 | 0.150232 | -0.15588 | 0.286789 | -0.19196 | -0.97577 | -0.82932 | 0.621743 | -2.73842 | 0.089909 | 0.021923 | -0.64487 | -1.4822 | -1.49453 | -0.14332 | -2.96603 | -0.52651 | 1.001828 | -0.40605 | -2.1591 | 0.347348 | -0.68745 | -0.22491 |
| 22.54167 | -1.07492 | 1.350536 | -0.07228 | 0.068459 | 0.328992 | -0.15922 | -0.02741 | -0.24428 | 0.46034 | -0.06956 | 0.072047 | 0.876498 | -0.27326 | -0.37046 | -0.93758 | 0.171647 | -3.82536 | -0.22275 | 2.874384 | -0.19735 | -1.75273 | 0.621141 | -0.42185 | 0.070712 |
| 22.55556 | -0.03477 | 1.593274 | -0.28186 | 0.293562 | 0.3119 | -0.06474 | 1.18114 | 0.4064 | 0.096397 | 2.503619 | 0.078301 | 1.490961 | 0.080363 | 0.827761 | 0.202536 | 0.47686 | -4.99959 | 0.301574 | 3.257813 | 0.399965 | -0.98434 | 0.756861 | -0.29825 | 0.299694 |
| 22.56944 | 1.081795 | 1.640993 | -0.46009 | 0.384195 | 0.240238 | 0.004599 | 2.158325 | 0.967363 | 0.065199 | 3.182892 | 0.108001 | 1.67386 | 0.57247 | 1.531885 | 1.087372 | 0.661621 | -6.41414 | 0.718074 | 2.368783 | 0.713521 | 0.265086 | 0.795795 | -0.00204 | 0.17805 |
| 22.58333 | 1.83049 | 1.563426 | -0.60646 | 0.414755 | 0.114337 | -0.03909 | 0.929891 | 0.561323 | 0.172188 | 2.249587 | 0.239096 | 1.114332 | 0.769453 | 1.538142 | 1.723197 | 0.52444 | -7.31121 | 0.903158 | 0.766047 | 0.436363 | 1.768486 | 0.532614 | 0.277206 | -0.11967 |

| | | | | | | | | | | | | | | | | | | | | | | | | |
|---|---|---|---|---|---|---|---|---|---|---|---|---|---|---|---|---|---|---|---|---|---|---|---|---|
| 22.59722 | 1.751551 | 1.312823 | -0.43995 | 0.42335 | 0.009907 | -0.04878 | 0.050542 | 0.205579 | 0.226561 | 0.791061 | 0.384446 | 0.11101 | 0.141518 | 1.020417 | 1.413453 | 0.215771 | -6.21658 | 0.968562 | -1.04462 | 0.163634 | 2.376947 | -0.11679 | 0.295279 | -0.20069 |
| 22.61111 | 1.499386 | 0.99862 | -0.06333 | 0.301727 | -0.03061 | -0.05771 | -0.83634 | 0.006819 | 0.096241 | -0.18036 | 0.479764 | -0.55771 | -0.58075 | 0.052246 | 0.779709 | -0.03003 | -3.79761 | 0.616201 | -2.07259 | 0.152893 | 1.571364 | -0.65321 | 0.194715 | -0.24731 |
| 22.625 | 1.090209 | 0.508792 | 0.045345 | 0.089952 | 0.006859 | 0.03578 | -1.89394 | -0.10486 | -0.4001 | -0.67403 | 0.443514 | -0.81654 | -1.06688 | -0.52413 | 0.18459 | -0.26056 | -1.86242 | 0.185527 | -1.3241 | 0.188601 | 0.758685 | -0.48314 | 0.171726 | -0.36901 |
| 22.63889 | 0.307297 | 0.094613 | 0.132645 | -0.14039 | 0.078832 | 0.180901 | -2.31414 | -0.2424 | -0.67532 | -1.50597 | 0.31397 | -1.01805 | -1.03529 | -0.91297 | -0.41069 | -0.44532 | -0.81338 | -0.23415 | -0.13139 | 0.121304 | 0.303949 | -0.09491 | 0.396553 | -0.03441 |
| 22.65278 | -0.66471 | -0.21449 | 0.170133 | -0.29595 | 0.10871 | 0.239077 | -1.34422 | -0.53255 | -0.65926 | -2.49653 | 0.106246 | -1.44587 | -0.66252 | -1.3529 | -1.12485 | -0.4489 | -0.64232 | -0.5773 | 0.544469 | 0.110861 | -0.14367 | 0.076044 | 0.586108 | 0.413187 |
| 22.66667 | -1.50186 | -0.42628 | 0.08611 | -0.28794 | 0.089264 | 0.223466 | 0.000736 | -0.86369 | -0.50539 | -3.08048 | -0.03509 | -1.11294 | -0.27038 | -1.50172 | -1.5981 | -0.19691 | -1.3893 | -0.713 | 0.326896 | 0.163296 | -1.17439 | 0.019399 | 0.696943 | 0.773747 |
| 22.68056 | -1.72528 | -0.65346 | -0.03963 | -0.06656 | -0.01452 | 0.142338 | 1.121635 | -0.4615 | -0.30094 | -1.97522 | -0.15031 | -0.22023 | 0.04453 | -1.11598 | -1.73417 | -0.0234 | -2.50994 | -0.73411 | -0.25428 | 0.240037 | -2.17055 | -0.03946 | 0.262348 | 0.818564 |
| 22.69444 | -1.55322 | -0.73205 | -0.15268 | 0.132447 | -0.09966 | 0.029379 | 1.082333 | -0.09844 | -0.20156 | -0.2671 | -0.26706 | 0.804239 | 0.280978 | -0.3279 | -1.4913 | 0.090696 | -3.10382 | -0.68607 | -1.00173 | 0.131391 | -2.2195 | -0.23542 | -0.60689 | 0.434887 |
| 22.70833 | -1.15833 | -0.7044 | -0.22427 | 0.276662 | -0.09296 | -0.18631 | 0.30767 | 0.324943 | -0.11631 | 1.354944 | -0.30748 | 1.448197 | 0.315103 | 0.300078 | -1.12944 | 0.06092 | -2.72066 | -0.58161 | -1.35068 | -0.28293 | -1.52772 | -0.42465 | -1.42548 | -0.41255 |
| 22.72222 | -0.23466 | -0.63953 | -0.32189 | 0.287263 | 0.01098 | -0.25188 | -0.54791 | 0.805933 | 0.118264 | 2.58754 | -0.24177 | 0.699112 | 0.253806 | 0.589701 | -0.62058 | -0.07535 | -2.38927 | -0.53167 | -1.15465 | -0.59233 | -0.3701 | -0.31894 | -1.85227 | -1.25127 |
| 22.73611 | 0.236989 | -0.59719 | -0.50082 | 0.135503 | 0.07592 | -0.32086 | -1.58766 | 0.504806 | 0.142686 | 1.692397 | -0.16406 | -0.05176 | 0.078678 | 0.410259 | -0.09258 | -0.27453 | -2.57224 | -0.4039 | -0.49818 | -1.06578 | 0.844486 | -0.08986 | -2.16097 | -1.42894 |
| 22.75 | -0.01719 | -0.71196 | -0.66112 | -0.13108 | 0.040376 | -0.37875 | -2.35941 | -0.1364 | -0.18786 | 0.171351 | -0.16739 | -0.63136 | -0.29636 | -0.06625 | 0.346608 | -0.53292 | -3.3633 | -0.16111 | 0.145083 | -0.85266 | 1.5161 | -0.00058 | -2.5549 | -1.41826 |
| 22.76389 | -0.40443 | -0.95031 | -0.76576 | -0.36207 | -0.02551 | -0.36288 | -1.3989 | -0.72316 | -0.62264 | -1.01868 | -0.29696 | -1.22747 | -0.62798 | -0.52421 | 0.42994 | -0.7455 | -4.29083 | 0.02466 | 0.443592 | -0.23731 | 1.428863 | -0.01833 | -2.05833 | -1.03923 |
| 22.77778 | -0.72461 | -1.23678 | -0.94665 | -0.38642 | -0.15752 | -0.29635 | -0.56304 | -1.09876 | -1.0462 | -2.378 | -0.55689 | -1.46745 | -0.81026 | -0.94006 | 0.388877 | -0.93982 | -5.55264 | 0.127095 | 0.310583 | 0.248868 | 0.786155 | -0.1246 | -0.99382 | -0.8501 |
| 22.79167 | -1.34047 | -1.40737 | -1.09942 | -0.24795 | -0.26671 | -0.28295 | 0.045205 | -0.77222 | -1.4991 | -4.22545 | -0.64474 | -1.2794 | -1.04868 | -1.46571 | 0.203253 | -1.03087 | -6.08472 | 0.147365 | -0.24855 | 0.216383 | -0.47106 | -0.50267 | -0.05635 | -1.32427 |
| 22.80556 | -2.27395 | -1.33133 | -1.18828 | -0.04848 | -0.22858 | -0.27499 | 0.145355 | -0.32451 | -1.45787 | -4.00501 | -0.4738 | -0.97017 | -1.07992 | -1.71954 | -0.23867 | -0.88217 | -4.54464 | 0.038888 | -0.91601 | -0.12906 | -1.82615 | -0.83836 | 0.357912 | -1.93255 |
| 22.81944 | -1.60166 | -1.10276 | -1.24692 | 0.092758 | -0.15202 | -0.27391 | -0.20238 | -0.3706 | -1.1176 | -2.72034 | -0.30687 | -0.7074 | -0.6993 | -1.60986 | -0.69548 | -0.54817 | -2.40713 | -0.20085 | -1.35082 | -0.43203 | -2.37486 | -0.84738 | 0.286355 | -2.14237 |
| 22.83333 | -0.87501 | -0.7263 | -1.2918 | 0.112953 | -0.09127 | -0.29055 | -0.58836 | 0.011983 | -0.93625 | -1.26442 | -0.26305 | -0.48242 | -0.25101 | -1.29095 | -1.22858 | -0.35285 | -0.82418 | -0.36954 | -1.83329 | -0.6364 | -2.5705 | -0.67769 | -0.22388 | -2.29954 |
| 22.84722 | -0.29724 | -0.09339 | -1.14917 | 0.060554 | -0.06462 | -0.29207 | -0.9775 | 0.362317 | -0.8993 | -0.00638 | -0.28257 | -0.38419 | 0.054484 | -1.02083 | -1.52646 | -0.30646 | 0.18585 | -0.40333 | -1.75714 | -0.75652 | -2.11171 | -0.51048 | -0.88883 | -2.34755 |
| 22.86111 | 0.114629 | 0.268713 | -0.89225 | -0.01479 | -0.0678 | -0.26609 | -1.04818 | 0.28101 | -0.89726 | 0.569722 | -0.3544 | -0.49871 | 0.356257 | -0.70927 | -1.26836 | -0.21915 | 0.412957 | -0.26352 | -1.12575 | -0.82768 | -1.17362 | -0.22054 | -1.34342 | -1.94062 |
| 22.875 | 0.119237 | 0.322652 | -0.62899 | -0.0691 | -0.08341 | -0.25817 | -0.64644 | 0.262575 | -0.77762 | 0.389279 | -0.48515 | -0.61658 | 0.724971 | -0.46293 | -0.68848 | -0.09064 | 0.252393 | -0.11256 | -0.24098 | -0.84696 | -0.36553 | 0.071582 | -1.42537 | -1.06141 |
| 22.88889 | -0.09923 | 0.224737 | -0.39143 | -0.08947 | -0.09521 | -0.25122 | 0.053673 | 0.143348 | -0.61706 | -0.54218 | -0.50975 | -0.61188 | 0.746105 | -0.231 | -0.36215 | 0.062449 | 0.302974 | 0.039334 | 0.215899 | -0.71056 | 0.199443 | -0.01532 | -1.09 | -0.17113 |
| 22.90278 | -0.42875 | 0.03809 | -0.35856 | -0.18255 | -0.11853 | -0.23563 | 0.488459 | -0.12574 | -0.63837 | -1.7144 | -0.53114 | -0.57323 | 0.300509 | -0.24436 | -0.37976 | -0.0177 | 0.195518 | -0.02454 | -0.22361 | -0.52632 | 0.099082 | -0.56451 | -0.69261 | 0.030569 |
| 22.91667 | -0.81268 | -0.35807 | -0.58514 | -0.39915 | -0.16803 | -0.23341 | 0.340766 | -0.38427 | -0.78556 | -2.67742 | -0.75006 | -0.58411 | -0.22878 | -0.62238 | -0.47257 | -0.21992 | -0.23103 | -0.31973 | -1.23548 | -0.53223 | -0.50814 | -1.08916 | -0.54651 | -0.55599 |
| 22.93056 | -1.30044 | -0.64063 | -0.89499 | -0.37063 | -0.18412 | -0.32192 | -0.18472 | -0.43149 | -1.05245 | -3.48389 | -1.49287 | -1.17517 | -0.73314 | -1.21019 | -0.48182 | -0.33142 | -0.56226 | -0.65749 | -2.4387 | -0.55627 | -1.7446 | -1.47023 | -0.8692 | -1.34331 |
| 22.94444 | -1.85023 | -0.63727 | -1.41253 | -0.2492 | -0.16071 | -0.50476 | -0.81261 | -0.69419 | -0.9995 | -4.67437 | -2.2442 | -2.1012 | -1.11811 | -1.94107 | -0.84505 | -0.21149 | -0.74498 | -1.03895 | -3.1194 | -0.63689 | -3.15868 | -1.62561 | -1.40421 | -2.17384 |
| 22.95833 | -2.26775 | -0.74947 | -1.76267 | -0.27309 | -0.16082 | -0.49047 | -1.54558 | -1.59016 | -0.69217 | -5.22213 | -2.21941 | -2.85766 | -1.2792 | -2.59749 | -1.5387 | -0.03015 | -0.70882 | -1.45411 | -2.55615 | -1.10879 | -3.66356 | -2.01443 | -1.58799 | -3.04789 |
| 22.97222 | -1.80713 | -0.9522 | -1.22519 | -0.24479 | -0.13165 | -0.3006 | -1.56688 | -2.89466 | -0.44437 | -3.81172 | -1.48993 | -2.47369 | -1.23195 | -1.88864 | -1.87268 | 0.061559 | -0.38476 | -1.46138 | -1.64324 | -1.73477 | -3.99347 | -2.51449 | -1.37926 | -2.2279 |
| 22.98611 | -0.89627 | -0.84112 | -0.35286 | -0.05001 | -0.03664 | -0.19554 | -0.89829 | -2.60467 | -0.38479 | -1.50098 | -0.72251 | -1.46743 | -0.9456 | -0.28419 | -1.20036 | 0.079336 | 0.303178 | -0.78425 | -1.1371 | -1.7439 | -2.61892 | -2.12946 | -0.98299 | 0.075926 |
| 23 | 0.01433 | -0.51888 | 0.399845 | 0.084471 | -0.05583 | -0.17903 | -0.03325 | -0.90597 | -0.37772 | 0.195898 | 0.145851 | -0.38972 | -0.56675 | 0.914983 | -0.3277 | 0.063585 | 0.801851 | -0.08583 | -0.32699 | -1.3082 | -0.9737 | -1.23799 | -0.20868 | 2.039278 |
| 23.01389 | 0.526551 | -0.24939 | 0.761142 | 0.087646 | -0.19307 | -0.13271 | 0.588936 | 0.497882 | -0.42901 | 1.431156 | 0.730648 | 0.421238 | -0.2237 | 1.333078 | 0.392601 | 0.047821 | 0.827568 | 0.351624 | 0.572717 | -0.8421 | 0.339126 | -0.64319 | 0.60001 | 2.290447 |
| 23.02778 | 0.539859 | -0.08493 | 0.722391 | 0.069034 | -0.24893 | -0.09278 | 0.655495 | 1.304651 | -0.52078 | 2.214044 | 0.808481 | 0.781428 | 0.111665 | 1.370576 | 0.912342 | 0.150554 | 0.513094 | 0.560506 | 1.584968 | -0.58602 | 1.265487 | -0.73757 | 1.329739 | 1.182976 |
| 23.04167 | 0.409612 | -0.05974 | 0.533676 | 0.065492 | -0.12915 | -0.07124 | 0.473389 | 1.549647 | -0.49087 | 2.461587 | 0.750452 | 0.89928 | 0.206169 | 1.312641 | 0.910895 | 0.138024 | 0.309851 | 0.509438 | 1.895042 | -0.42387 | 1.521357 | -0.98603 | 1.476438 | 0.407685 |
| 23.05556 | 0.360291 | -0.05614 | 0.347477 | 0.037628 | 0.070312 | -0.05407 | 0.356386 | 1.586944 | -0.29262 | 2.617162 | 0.548447 | 0.707173 | 0.046073 | 1.103904 | 0.643629 | -0.00031 | 0.272604 | 0.448648 | 1.595825 | -0.38678 | 1.23165 | -1.05483 | 1.039107 | 0.106513 |
| 23.06944 | 0.346531 | 0.037974 | 0.144372 | 0.00897 | 0.199734 | -0.05049 | 0.463651 | 1.433843 | -0.14219 | 2.467291 | 0.526112 | 0.3857 | 0.037943 | 0.990726 | 0.472541 | -0.01334 | 0.269128 | 0.389146 | 1.332871 | -0.33727 | 0.983522 | -0.84944 | 0.647948 | 0.12132 |
| 23.08333 | 0.346929 | 0.153412 | 0.020322 | -0.0074 | 0.110575 | -0.057 | 0.688263 | 1.218651 | -0.09543 | 2.217962 | 0.472869 | 0.166215 | 0.166726 | 0.940909 | 0.478878 | 0.056278 | 0.264064 | 0.32067 | 1.087672 | -0.22627 | 0.839883 | -0.54733 | 0.168213 | 0.294088 |
| 23.09722 | 0.377097 | 0.230536 | -0.00464 | 0.003711 | -0.0934 | -0.04325 | 0.739951 | 1.134188 | -0.107 | 2.024919 | 0.444601 | 0.11751 | 0.254487 | 0.901833 | 0.597638 | 0.10995 | 0.290174 | 0.307716 | 0.947606 | -0.13253 | 1.001996 | -0.32083 | -0.29436 | 0.418056 |
| 23.11111 | 0.502768 | 0.210172 | 0.014853 | 0.01376 | -0.30947 | -0.01711 | 0.454132 | 0.991097 | -0.06274 | 1.773553 | 0.329765 | 0.064634 | 0.219051 | 0.797319 | 0.593943 | 0.082878 | 0.33757 | 0.384006 | 0.800183 | -0.09874 | 1.034045 | -0.23469 | -0.68907 | 0.25816 |
| 23.125 | 0.612241 | 0.1543 | 0.01237 | 0.021348 | -0.26292 | -0.0126 | 0.022984 | 0.932404 | 0.021291 | 1.281597 | 0.222766 | -0.00217 | 0.164089 | 0.588087 | 0.562185 | 0.017714 | 0.35539 | 0.539543 | 0.659481 | -0.09378 | 0.9658 | -0.10831 | -0.65839 | -0.04788 |
| 23.13889 | 0.583723 | 0.096812 | 0.001834 | 0.024414 | -0.07586 | -0.01851 | -0.479 | 0.771967 | -0.01065 | 0.778242 | 0.149821 | -0.00378 | 0.415821 | 0.389521 | 0.774398 | 0.096152 | 0.267637 | 0.57809 | 0.769154 | -0.02929 | 0.826621 | 0.103352 | -0.16831 | -0.13675 |
| 23.15278 | 0.46897 | 0.066033 | -0.01309 | 0.004735 | 0.054011 | -0.03298 | -0.92286 | 0.641303 | -0.12062 | 0.211808 | 0.059959 | 0.048115 | 0.6474 | 0.197613 | 1.075758 | 0.134655 | 0.135859 | 0.582741 | 0.83778 | 0.047481 | 0.711616 | 0.166734 | 0.248762 | 0.050072 |
| 23.16667 | 0.3115 | 0.054023 | -0.07809 | -0.05766 | 0.057861 | -0.06102 | -0.82776 | 0.31998 | -0.119 | -0.20623 | 0.012549 | 0.128632 | 0.479152 | -0.01974 | 1.03456 | 0.012869 | 0.066982 | 0.589167 | 0.540762 | 0.048496 | 0.203138 | -0.02137 | 0.138687 | 0.157019 |
| 23.18056 | 0.157588 | 0.045167 | -0.14085 | -0.11739 | -0.01018 | -0.05242 | -0.41736 | 0.047826 | -0.00951 | -0.43828 | -0.03576 | 0.277884 | 0.274338 | -0.07741 | 0.856689 | -0.00722 | -0.00866 | 0.568737 | 0.083671 | 0.029502 | -0.34568 | -0.11015 | -0.21075 | -0.05243 |
| 23.19444 | 0.053264 | 0.05138 | -0.11525 | -0.12242 | -0.03034 | 0.015275 | 0.007385 | -0.09672 | 0.012989 | -0.59026 | -0.07436 | 0.541492 | 0.245915 | 0.055725 | 0.652085 | 0.066609 | -0.01795 | 0.441686 | -0.32457 | 0.026276 | -0.61536 | -0.10067 | -0.45862 | -0.38282 |
| 23.20833 | 0.037532 | 0.037536 | -0.02739 | -0.0706 | -0.00298 | 0.062856 | 0.189143 | -0.18271 | 0.012968 | -0.59185 | -0.07784 | 0.75605 | 0.239624 | 0.162613 | 0.445805 | 0.061205 | 0.032047 | 0.300583 | -0.54332 | 0.022761 | -0.72142 | -0.0462 | -0.37812 | -0.4519 |
| 23.22222 | 0.071893 | 0.039345 | 0.038566 | -0.00541 | 0.026069 | 0.058678 | 0.106623 | -0.23814 | 0.077049 | -0.25675 | -0.06082 | 0.662685 | 0.256789 | 0.088879 | 0.340058 | 0.04711 | 0.079617 | 0.176842 | -0.51578 | 0.056752 | -0.63345 | 0.044238 | 0.010093 | -0.11263 |
| 23.23611 | 0.142529 | 0.037593 | 0.057728 | 0.037333 | 0.012753 | 0.024404 | 0.02459 | -0.15605 | 0.160635 | 0.257554 | -0.03804 | 0.542507 | 0.179717 | -0.09591 | 0.248957 | 0.049076 | 0.103353 | 0.107953 | -0.19962 | 0.08292 | -0.37383 | 0.159284 | 0.541578 | 0.370037 |
| 23.25 | 0.249343 | 0.031468 | 0.080846 | 0.030945 | -0.05676 | -0.00253 | 0.135223 | 0.177991 | 0.237805 | 0.581357 | -1.5E-05 | 0.463196 | -0.09063 | -0.1254 | 0.032379 | -0.05597 | 0.161085 | 0.097953 | 0.10619 | 0.058001 | -0.15134 | 0.243355 | 0.858628 | 0.638486 |
| 23.26389 | 0.402495 | 0.025263 | 0.145362 | 0.021324 | -0.08071 | 0.007126 | 0.338291 | 0.52673 | 0.256224 | 0.694862 | 0.042351 | 0.506554 | -0.07176 | 0.177743 | 0.161446 | -0.04224 | 0.099713 | 0.09842 | 0.415138 | 0.098967 | 0.246971 | 0.250039 | 1.099921 | 0.61182 |
| 23.27778 | 0.537207 | 0.051941 | 0.204118 | 0.061896 | 0.012012 | 0.025585 | 0.532076 | 0.607887 | 0.244991 | 0.706265 | 0.073342 | 0.65715 | 0.329838 | 0.479845 | 0.749476 | 0.141517 | -0.04092 | 0.089291 | 0.832522 | 0.250909 | 0.683123 | 0.132322 | 1.12068 | 0.360244 |
| 23.29167 | 0.660338 | 0.095671 | 0.252456 | 0.132286 | 0.144275 | -0.05058 | 0.666082 | 0.455248 | 0.259155 | 0.864349 | 0.073777 | 0.859315 | 0.508276 | 0.502942 | 1.062721 | 0.219772 | -0.0788 | 0.13544 | 1.094276 | 0.33773 | 0.759298 | 0.055519 | 0.789187 | 0.163143 |
| 23.30556 | 0.779941 | 0.119506 | 0.307602 | 0.182592 | 0.264464 | -0.2429 | 0.675664 | 0.286066 | 0.2893 | 1.185124 | 0.083757 | 0.964674 | 0.311681 | 0.258996 | 0.811507 | 0.188982 | 0.030471 | 0.258426 | 0.970088 | 0.265885 | 0.459818 | 0.089319 | 0.320668 | 0.067249 |
| 23.31944 | 0.837671 | 0.11834 | 0.335382 | 0.211482 | 0.212001 | -0.26433 | 0.497131 | 0.162089 | 0.281013 | 1.294556 | 0.143887 | 0.890739 | 0.057738 | 0.09934 | 0.412107 | 0.158126 | 0.156342 | 0.372824 | 0.460788 | 0.15494 | 0.206 | 0.187705 | 0.063852 | 0.13027 |
| 23.33333 | 0.763777 | 0.084732 | 0.331136 | 0.189029 | 0.049366 | -0.01605 | 0.342099 | 0.12848 | 0.264107 | 1.120558 | 0.223049 | 0.646342 | -0.04608 | 0.171041 | 0.326731 | 0.152281 | 0.24112 | 0.42024 | 0.155983 | 0.108283 | 0.236332 | 0.256496 | 0.206715 | 0.273323 |
| 23.34722 | 0.768342 | 0.030934 | 0.404369 | 0.134213 | -0.12036 | 0.13803 | 0.347435 | 0.104494 | 0.322186 | 1.115296 | 0.321521 | 0.453151 | 0.203095 | 0.421915 | 0.54338 | 0.247406 | 0.165738 | 0.370544 | 0.37002 | 0.218024 | 0.574258 | 0.173565 | 0.69025 | 0.322405 |
| 23.36111 | 0.853465 | -0.0086 | 0.525562 | 0.125703 | -0.1692 | 0.215247 | 0.498416 | 0.097435 | 0.502659 | 1.335646 | 0.33097 | 0.400718 | 0.508604 | 0.752605 | 0.780555 | 0.343601 | 0.096336 | 0.366862 | 0.686775 | 0.384104 | 0.819355 | 0.034534 | 1.171611 | 0.279712 |
| 23.375 | 0.966349 | 0.006756 | 0.545986 | 0.171221 | -0.04589 | 0.254317 | 0.545089 | 0.304528 | 0.660198 | 1.631315 | 0.277357 | 0.37824 | 0.504329 | 0.992028 | 0.649708 | 0.359721 | 0.310617 | 0.436913 | 0.65821 | 0.470968 | 0.628842 | 0.107972 | 1.238857 | 0.200015 |
| 23.38889 | 0.979709 | 0.132863 | 0.396968 | 0.193705 | 0.199684 | 0.243155 | 0.421157 | 0.399942 | 0.760905 | 1.65809 | 0.287279 | 0.450205 | 0.443788 | 1.061593 | 0.570603 | 0.361044 | 0.551458 | 0.532405 | 0.644413 | 0.611057 | 0.370684 | 0.303089 | 1.027411 | 0.237397 |
| 23.40278 | 1.194756 | 0.466453 | 0.423661 | 0.274461 | 0.381756 | 0.314968 | 0.481425 | 0.412799 | 1.483954 | 1.768225 | 0.627537 | 1.005152 | 0.726709 | 1.139852 | 0.990439 | 0.355224 | 1.121016 | 0.640507 | 1.243799 | 1.106052 | 0.495543 | 0.485139 | 0.967928 | 0.787922 |
| 23.41667 | 1.918686 | 1.123048 | 0.942058 | 0.617571 | 0.361942 | 0.669998 | 0.952645 | 0.752701 | 2.922916 | 2.847547 | 1.328575 | 2.084174 | 1.45128 | 1.653813 | 1.626669 | 0.362 | 3.190733 | 1.122381 | 2.586909 | 2.487328 | 0.919695 | 0.905171 | 1.868074 | 2.396185 |
| 23.43056 | 2.089753 | 1.691448 | 1.34891 | 0.770456 | 0.273211 | 0.817851 | 1.315396 | 1.149103 | 3.204423 | 5.135376 | 1.548011 | 2.609192 | 1.685047 | 2.467914 | 1.682264 | 0.256104 | 7.764646 | 1.145393 | 3.881895 | 2.657645 | 1.401334 | 1.44315 | 3.230802 | 3.746791 |
| 23.44444 | 2.146461 | 1.331873 | 1.045245 | 0.649709 | 0.103342 | 0.716171 | 1.757284 | 1.249734 | 2.525679 | 6.602774 | 1.219772 | 2.098051 | 1.194838 | 3.257257 | 1.351777 | -0.12486 | 9.832285 | 0.922464 | 4.141604 | 1.696165 | 1.617851 | 1.676233 | 3.735759 | 3.300798 |
| 23.45833 | 2.057218 | 0.747484 | 0.75327 | 0.588864 | -0.01103 | 0.636799 | 0.874979 | 1.096812 | 2.30085 | 4.662307 | 0.939648 | 1.517677 | 0.806696 | 3.164753 | 1.431496 | -0.28586 | 9.902933 | 0.80515 | 3.494875 | 1.149906 | 1.984098 | 1.534442 | 2.74527 | 1.871084 |
| 23.47222 | 1.143233 | 0.872518 | 0.801903 | 0.501443 | -0.01732 | 0.375178 | -0.27177 | 0.669452 | 2.019442 | 2.109671 | 0.531851 | 1.005779 | 0.489644 | 2.181177 | 0.938266 | -0.49945 | 9.298517 | 0.819447 | 1.337777 | 0.860156 | 1.45762 | 0.950208 | 0.964942 | 0.978916 |
| 23.48611 | 0.262712 | 0.908913 | 1.135229 | 0.382233 | 0.001577 | -0.2493 | -1.42452 | 0.14695 | 1.562913 | 0.60784 | 0.236963 | 0.380822 | 0.051161 | 1.290903 | 0.445997 | -0.72975 | 6.743006 | 0.595976 | -0.76193 | 0.618471 | 0.585882 | 0.152226 | -0.63326 | 0.156633 |
| 23.5 | -0.36206 | 0.606576 | 1.361333 | 0.14854 | 0.114561 | -0.82674 | -1.61835 | -0.18219 | 1.090438 | -1.2029 | 0.105131 | 0.053263 | -0.31408 | 0.292265 | 0.041971 | -0.80853 | 5.352572 | 0.344616 | -2.85685 | 0.544243 | -0.15701 | -0.52765 | -1.80508 | -0.55416 |
| 23.51389 | -0.85853 | 0.651529 | 1.117553 | -0.04259 | 0.250586 | -0.57963 | -1.10305 | -0.357 | 1.436147 | -3.57738 | 0.169595 | 0.325808 | -0.39962 | -1.05758 | -0.55361 | -1.03424 | 4.029866 | 0.103923 | -3.68073 | 0.478395 | -0.82537 | -0.32153 | -2.34655 | -0.49051 |
| 23.52778 | -1.21 | 0.759485 | 0.594575 | -0.0831 | 0.244392 | -0.21261 | -0.23745 | -0.29435 | 1.546268 | -2.71541 | 0.25531 | 0.864901 | -0.27586 | -2.13032 | -0.96612 | -0.56241 | 3.062992 | -0.08035 | -1.86781 | 0.23813 | -1.053 | 0.677364 | -2.57097 | 0.304318 |
| 23.54167 | -1.28642 | 0.795227 | 0.336766 | -0.11358 | 0.142827 | 0.12979 | 0.912611 | -0.20038 | 1.256926 | -1.40666 | 0.337728 | 1.432622 | -0.05827 | -2.01932 | -1.06523 | 0.215309 | 1.930817 | -0.30247 | -0.87753 | -0.04163 | -1.24332 | 1.822834 | -1.63875 | 0.983406 |
| 23.55556 | -1.0669 | 0.699253 | 0.153347 | -0.14063 | -0.01414 | 0.278194 | 1.466377 | -0.07569 | 1.1679 | 0.721475 | 0.396995 | 1.708002 | 0.239417 | -0.70612 | -1.20683 | 0.754197 | -0.69056 | -0.37262 | -0.96638 | -0.14519 | -1.39495 | 2.578858 | -0.06066 | 1.319698 |
| 23.56944 | -0.34388 | 0.690243 | -0.00961 | -0.05547 | -0.11966 | 0.20484 | 1.514956 | 0.246843 | 1.085779 | 2.832031 | 0.450465 | 0.714956 | 0.546506 | 1.0332 | -0.66159 | 1.037226 | -4.37962 | -0.22348 | -0.69335 | -0.1477 | -1.05149 | 2.405666 | 1.094219 | 0.775616 |
| 23.58333 | 0.207989 | 0.833802 | 0.042902 | 0.143175 | -0.11419 | 0.087845 | 0.892811 | 0.581909 | 0.757827 | 3.443838 | 0.44941 | -0.0189 | 0.857165 | 2.94958 | -0.08435 | 1.150766 | -7.74002 | 0.028296 | 0.791943 | -0.16565 | 0.005707 | 1.892408 | 1.777916 | 0.122841 |
| 23.59722 | 0.671776 | 0.863882 | -0.00598 | 0.338945 | -0.06035 | -0.00321 | 0.21213 | 0.661274 | 0.600488 | 2.66809 | 0.418323 | -0.53931 | 0.954936 | 3.882409 | 0.509833 | 0.919022 | -9.59526 | 0.322371 | 2.26237 | -0.01374 | 0.631154 | 1.084248 | 2.400738 | -0.17035 |
| 23.61111 | 1.139985 | 0.570464 | -0.23122 | 0.401808 | 0.138455 | -0.07484 | -0.53008 | 0.593272 | -0.04377 | 0.393412 | 0.297013 | -0.66656 | 0.575439 | 2.733397 | 1.268529 | 0.311094 | -8.62354 | 0.697145 | 2.416241 | 0.241551 | 0.606925 | 0.159519 | 1.903714 | -0.5824 |
| 23.625 | 0.881772 | 0.19124 | -0.33999 | 0.327535 | 0.324214 | -0.07694 | -1.41094 | 0.37233 | -0.65935 | -1.37068 | 0.113978 | -0.7309 | 0.058362 | 1.212736 | 2.44956 | -0.39412 | -6.61635 | 1.044975 | 1.682387 | 0.60828 | 0.582416 | 0.002365 | 1.121601 | -1.03674 |
| 23.63889 | 0.200492 | 0.023864 | -0.4884 | 0.221105 | 0.331465 | 0.064403 | -1.60453 | 0.08863 | -1.11939 | -3.06163 | -0.07823 | -0.92029 | -0.42066 | -0.32781 | 1.637883 | -1.17813 | -3.56535 | 0.805205 | 0.625815 | 0.665302 | 0.464311 | 0.103123 | 0.456555 | -0.61738 |
| 23.65278 | -0.4678 | -0.13083 | -0.48123 | 0.141888 | 0.17441 | 0.30558 | -0.81453 | -0.27143 | -1.22031 | -4.32948 | -0.18326 | -0.59332 | -0.64655 | -2.05379 | 0.622708 | -1.40032 | 0.150207 | 0.357311 | -0.82452 | 0.738507 | 0.2477 | 0.868917 | -0.32792 | 0.302324 |
| 23.66667 | -0.96536 | -0.46943 | -0.43577 | 0.048508 | -0.11339 | 0.519911 | -0.05557 | -0.68319 | -0.74057 | -4.34708 | -0.27011 | 0.031175 | -0.64847 | -2.98802 | 0.058296 | -0.93877 | 2.839797 | 0.054836 | -1.92306 | 0.637278 | -0.12548 | 1.639455 | -0.58158 | 0.863045 |
| 23.68056 | -1.32872 | -0.94819 | -0.32633 | 0.030118 | -0.32485 | 0.570135 | 0.449575 | -0.91501 | -0.70506 | -3.74286 | -0.28744 | 0.681968 | -0.4278 | -3.26098 | -0.55487 | -0.46556 | 4.721655 | -0.21333 | -2.39808 | 0.381878 | -0.53165 | 1.778734 | -0.48539 | 0.918162 |
| 23.69444 | -1.64739 | -1.19743 | -0.28648 | 0.001627 | -0.37353 | 0.367183 | 0.35953 | -0.78732 | -0.40379 | -1.55688 | -0.14515 | 0.192097 | -0.30251 | -2.62253 | -1.26679 | -0.24267 | 4.995623 | -0.43511 | -2.91291 | 0.172318 | -0.85996 | 1.123571 | -0.70257 | 0.465373 |
| 23.70833 | -1.24652 | -1.03819 | -0.42173 | 0.033432 | -0.19708 | 0.123514 | 0.048761 | -0.49934 | 0.037928 | 0.28935 | 0.054944 | -0.35748 | -0.18089 | -1.321 | -1.34756 | -0.04804 | 2.967826 | -0.5837 | -3.13923 | -0.0542 | -1.11941 | 0.54132 | -1.22234 | -0.20318 |
| 23.72222 | -0.57257 | -0.7047 | -0.56245 | 0.124904 | 0.109134 | -0.10989 | -0.37324 | -0.22592 | 0.21301 | 0.759968 | 0.139792 | -0.51857 | -0.13196 | 0.078466 | -0.94703 | 0.165003 | -0.14026 | -0.63948 | -2.72688 | -0.2795 | -1.03966 | -0.06106 | -1.73533 | -0.94704 |
| 23.73611 | -0.08002 | -0.59015 | -0.61766 | 0.183222 | 0.385357 | -0.27079 | -1.00846 | -0.04602 | 0.034337 | 0.115769 | 0.064449 | -0.90948 | -0.25774 | 1.069687 | -0.77668 | 0.135823 | -3.02247 | -0.69554 | -2.21163 | -0.50826 | -1.04664 | -0.97397 | -2.02139 | -1.57608 |
| 23.75 | -0.06135 | -0.65522 | -0.7337 | 0.17391 | 0.309223 | -0.43775 | -1.41553 | 0.072246 | -0.10378 | -0.32933 | 0.017519 | -1.7001 | -0.25385 | 0.880433 | -0.84263 | -0.27334 | -5.12747 | -0.92238 | -1.05366 | -0.59222 | -0.70061 | -0.65581 | -1.631 | -1.35731 |
| 23.76389 | -0.29197 | -0.73647 | -0.84883 | 0.110615 | 0.034403 | -0.5047 | -1.12984 | 0.025477 | -0.31688 | -1.02117 | -0.11065 | -1.9339 | -0.14374 | -0.03193 | -0.99481 | -0.6451 | -6.98772 | -1.12793 | -0.1065 | -0.7217 | -0.23178 | -0.12257 | -0.93355 | -0.89239 |
| 23.77778 | -0.73077 | -0.80422 | -0.89239 | 0.006108 | -0.35066 | -0.44696 | -0.62143 | -0.13172 | -0.7488 | -2.48554 | -0.31393 | -1.58487 | -0.29928 | -1.05025 | -0.79556 | -0.96519 | -9.17801 | -0.9597 | -0.1802 | -1.17781 | -0.11669 | 0.107112 | -0.4329 | -0.77673 |
| 23.79167 | -1.2615 | -0.9821 | -0.81719 | -0.1239 | -0.36936 | -0.42451 | -0.24757 | -0.1931 | -1.17027 | -3.70324 | -0.53965 | -0.8606 | -0.46087 | -1.81986 | -0.51214 | -1.07242 | -8.81277 | -0.59228 | -0.38132 | -1.26326 | -0.09764 | -0.01253 | -0.34425 | -1.09925 |
| 23.80556 | -1.52991 | -1.1721 | -0.69747 | -0.20387 | -0.33819 | -0.5776 | -0.00282 | -0.29438 | -1.18925 | -5.12884 | -0.6826 | -0.04215 | -0.54716 | -2.14824 | -0.24982 | -1.02426 | -7.09516 | -0.20412 | -0.67214 | -1.07953 | 0.101075 | -0.28577 | -0.58236 | -1.46449 |
| 23.81944 | -1.97329 | -1.09928 | -0.60115 | -0.12912 | -0.02199 | -0.65727 | 0.053907 | -0.77297 | -1.14364 | -5.87877 | -0.70186 | 0.11722 | -0.50508 | -2.18939 | 0.02068 | -0.7842 | -5.36738 | 0.175003 | -1.24006 | -1.08772 | 0.021187 | -0.74432 | -0.72092 | -1.82333 |
| 23.83333 | -2.18746 | -0.77535 | -0.55124 | 0.038804 | 0.423789 | -0.67493 | -0.10275 | -0.56759 | -1.22868 | -3.5714 | -0.6598 | -0.05367 | -0.46328 | -1.90916 | 0.194132 | -0.18073 | -3.6467 | 0.493017 | -1.75717 | -0.84474 | -0.27552 | -1.38485 | -0.76409 | -2.23652 |
| 23.84722 | -1.42082 | -0.48989 | -0.54879 | 0.169547 | 0.566631 | -0.56946 | -0.3024 | -0.08714 | -1.06627 | -1.28533 | -0.44565 | -0.49484 | -0.4571 | -1.61224 | 0.038449 | 0.116714 | -1.7765 | 0.661751 | -1.65275 | -0.49186 | -0.5191 | -1.79101 | -0.79266 | -1.86957 |
| 23.86111 | -0.45614 | -0.22725 | -0.59816 | 0.159568 | 0.351804 | -0.34085 | -0.45769 | 0.187099 | -0.79263 | 0.545926 | -0.13769 | -0.8023 | -0.36901 | -1.17433 | 0.10264 | 0.167922 | -0.78218 | 0.617229 | -0.91013 | -0.19261 | -0.56154 | -1.63086 | -0.83075 | -1.21161 |
| 23.875 | 0.216854 | -0.05126 | -0.49374 | 0.042007 | 0.059368 | -0.16323 | -0.36824 | 0.285672 | -0.60776 | 1.649761 | 0.13702 | -1.00625 | -0.2107 | -0.92515 | 0.280146 | 0.183441 | -0.3137 | 0.594215 | -0.33015 | -0.07758 | -0.4847 | -1.20217 | -0.79217 | -0.7632 |
| 23.88889 | 0.463208 | 0.099559 | -0.29265 | -0.08247 | -0.24539 | -0.06502 | -0.20814 | 0.115595 | -0.64375 | 1.998542 | 0.268355 | -1.41408 | 0.012483 | -0.79329 | 0.3617 | 0.213113 | -0.03722 | 0.557437 | -0.2152 | 0.00833 | -0.40624 | -0.721 | -0.96663 | -0.51596 |
| 23.90278 | 0.226406 | 0.207524 | -0.27771 | -0.17847 | -0.39623 | -0.07972 | -0.15215 | -0.269 | -0.89856 | 1.218216 | 0.165923 | -1.83702 | 0.057709 | -0.65711 | 0.103207 | 0.085292 | 0.023919 | 0.382554 | -0.58117 | -0.04719 | -0.48369 | -0.65828 | -1.22694 | -0.55015 |
| 23.91667 | -0.37517 | 0.141529 | -0.60647 | -0.24867 | -0.38931 | -0.29444 | -0.41127 | -0.64459 | -1.16739 | -0.67874 | -0.2382 | -1.93884 | -0.25976 | -0.70706 | -0.45933 | -0.3593 | -0.09945 | -0.03658 | -1.20171 | -0.13338 | -0.7297 | -1.21043 | -1.54498 | -1.08347 |
| 23.93056 | -1.07402 | -0.06428 | -0.96556 | -0.24519 | -0.15366 | -0.71145 | -0.74623 | -0.66616 | -1.14031 | -2.53599 | -0.85473 | -1.88305 | -0.67615 | -1.17577 | -0.91448 | -0.93017 | -0.22109 | -0.52491 | -1.64611 | -0.3627 | -0.86877 | -1.91937 | -2.02877 | -1.93987 |
| 23.94444 | -1.86096 | -0.36612 | -1.18245 | -0.13862 | 0.49739 | -1.15087 | -1.07389 | -0.45709 | -0.75523 | -4.12303 | -1.52475 | -2.59199 | -0.94251 | -2.2066 | -1.19425 | -1.0102 | -0.65915 | -0.91441 | -2.25087 | -0.5721 | -1.38297 | -2.50669 | -2.76949 | -2.99378 |
| 23.95833 | -2.6598 | -0.6194 | -1.3509 | -0.26101 | 0.823839 | -1.02871 | -1.38854 | -1.002 | -0.43419 | -3.24494 | -1.6693 | -3.45806 | -0.91222 | -3.41756 | -2.06968 | -0.61796 | -1.99356 | -1.61679 | -2.17849 | -0.628 | -1.85535 | -2.8403 | -2.62103 | -3.2087 |
| 23.97222 | -2.23868 | -0.75377 | -1.23577 | -0.484 | 0.105025 | -0.68695 | -1.15454 | -1.94678 | -0.44998 | -1.19866 | -1.77512 | -2.77535 | -0.93117 | -3.11039 | -2.23134 | 0.270092 | -1.44847 | -1.79314 | -1.67246 | -0.66739 | -1.9482 | -2.959 | -1.91814 | -2.60951 |
| 23.98611 | -1.35252 | -0.59065 | -0.90504 | -0.36582 | -0.34564 | -0.46797 | -0.36346 | -1.52096 | -0.56451 | -0.20891 | -1.44362 | -2.27419 | -0.80106 | -1.02668 | -2.09008 | 0.372151 | -0.42506 | -1.15424 | -1.43437 | -0.94254 | -1.73524 | -2.06188 | -1.17367 | -0.97468 |
| 24 | -0.4373 | -0.25315 | -0.35646 | -0.07327 | -0.45364 | -0.3168 | 0.38535 | -0.78669 | -0.63675 | 0.046526 | -0.6689 | -1.06257 | -0.31414 | 0.758674 | -1.17667 | 0.17576 | 0.263018 | -0.34851 | -0.75128 | -1.01644 | -0.68312 | -0.68959 | -0.40673 | 1.022686 |
| 24.01389 | 0.149787 | 0.013207 | 0.163416 | 0.186796 | -0.14327 | -0.1759 | 0.858992 | 0.045782 | -0.57471 | 0.102542 | 0.112394 | -0.14038 | 0.189145 | 1.836373 | -0.19959 | 0.459793 | 0.728157 | 0.223493 | 0.178849 | -0.8765 | 0.048852 | 0.468161 | 0.36114 | 2.42423 |
| 24.02778 | 0.295818 | 0.133451 | 0.334072 | 0.322945 | 0.276517 | -0.09252 | 0.81893 | 0.7125 | -0.36815 | 0.104969 | 0.409019 | 0.360774 | 0.367191 | 2.246095 | 0.546708 | 1.120541 | 1.106693 | 0.549016 | 1.081206 | -0.5197 | 0.502995 | 0.913594 | 0.862791 | 2.740779 |
| 24.04167 | 0.284965 | 0.10472 | 0.265303 | 0.258093 | 0.385341 | -0.07356 | 0.650531 | 1.015458 | -0.17723 | 0.063549 | 0.423909 | 0.580472 | 0.300671 | 1.791814 | 0.754528 | 0.793152 | 0.764051 | 0.682345 | 1.314781 | -0.39225 | 0.49178 | 0.446699 | 0.802016 | 2.370999 |
| 24.05556 | 0.311822 | 0.029204 | -0.20047 | 0.109653 | 0.050206 | -0.10035 | 0.43276 | 0.835137 | -0.19573 | 0.016183 | 0.351224 | 0.736027 | 0.278023 | 1.445775 | 0.944011 | -0.08731 | -1.06245 | 0.796318 | 0.852318 | -0.4457 | -0.00757 | -0.18923 | 0.431076 | 1.377385 |
| 24.06944 | 0.219731 | -0.05513 | -0.73097 | -0.0514 | -0.08749 | -0.07905 | 0.300743 | 0.747827 | -0.19917 | 0.184549 | 0.605129 | 0.811148 | 0.383714 | 2.126995 | 1.839225 | -0.37905 | -2.31492 | 0.861215 | 0.368745 | -0.49201 | -0.62838 | -0.60122 | -0.163 | -0.01495 |
| 24.08333 | -0.29205 | -0.18433 | -0.04518 | -0.15003 | 0.214411 | 0.239421 | 1.34888 | 1.573724 | 0.363491 | 1.630979 | 1.234347 | 1.031523 | 0.435713 | 1.958096 | 1.696811 | 0.275584 | -0.51539 | 0.795551 | 0.595377 | -0.42582 | -0.60394 | -1.40153 | -0.39523 | -0.58006 |
| 24.09722 | -1.19442 | -0.30441 | 1.237064 | -0.75571 | 0.528278 | 0.392548 | 2.052542 | 1.856739 | 0.95377 | 3.294262 | 1.864247 | 2.152788 | 0.373281 | 0.979416 | 1.074835 | 0.065124 | 0.779512 | 0.500207 | 1.554756 | -0.12572 | -0.04562 | -2.42063 | 0.098748 | -0.14508 |
| 24.11111 | 0.092305 | 0.160409 | 0.173347 | -0.18564 | -0.36579 | 0.229621 | 0.816864 | 0.873143 | -0.43605 | 2.802308 | 0.568812 | 2.410812 | 0.390325 | 0.458889 | 0.807396 | 0.172399 | 0.585697 | 0.38276 | 1.798747 | 0.112853 | 0.068986 | -2.40829 | 0.483515 | -0.09852 |
| 24.125 | 1.319169 | 0.192571 | -1.10603 | 0.394706 | -0.06587 | -0.22026 | -1.53894 | -1.07106 | -1.72847 | -0.11097 | -0.3934 | 1.145172 | 0.330272 | 0.3508 | 0.590715 | 0.287189 | 0.253933 | 0.372986 | 0.959739 | 0.02325 | -0.04672 | -0.208 | 0.158124 | -0.27129 |

| | | | | | | | | | | | | | | | | | | | | | | | | |
|---|---|---|---|---|---|---|---|---|---|---|---|---|---|---|---|---|---|---|---|---|---|---|---|---|
| 24.13889 | 1.165424 | 0.206509 | -0.5253 | 0.107756 | -0.2153 | -0.0595 | -1.52933 | -1.26499 | -1.73475 | -0.19241 | -0.26109 | 1.05863 | 0.24603 | 0.321265 | 0.325311 | 0.242612 | -0.047 | 0.218803 | 0.274855 | -0.14887 | -0.15475 | 0.159686 | -0.45372 | -0.44626 |
| 24.15278 | 1.1587 | 0.231914 | 0.038551 | 0.030308 | 0.087384 | 0.128432 | -0.52516 | 0.109222 | -0.37099 | 0.947527 | 0.299376 | 1.671863 | 0.237817 | 0.253069 | 0.167724 | 0.040388 | -0.12709 | 0.068157 | -0.1515 | -0.20955 | -0.10341 | -0.50275 | -0.88298 | -0.70384 |
| 24.16667 | 1.359277 | 0.199264 | 0.147152 | -0.05414 | 0.065802 | 0.140351 | -0.58338 | 0.249443 | -0.11699 | 0.798261 | 0.25649 | 1.536457 | 0.285849 | 0.130293 | 0.09563 | -0.12542 | -0.09931 | -0.00051 | -0.46443 | -0.21327 | -0.12428 | -0.73627 | -0.96096 | -0.8886 |
| 24.18056 | 1.413376 | 0.142402 | 0.12003 | -0.15665 | 0.043751 | 0.102415 | -0.63448 | 0.092915 | -0.14961 | 0.376924 | 0.152535 | 1.133815 | 0.362365 | 0.034366 | 0.128368 | -0.16581 | -0.09237 | 0.021737 | -0.57601 | -0.15611 | -0.16146 | -0.36338 | -0.61485 | -0.68642 |
| 24.19444 | 1.266076 | 0.104762 | 0.171194 | -0.20103 | 0.069713 | 0.141127 | -0.39792 | -0.00367 | -0.14763 | 0.140063 | 0.178995 | 0.805555 | 0.460804 | 0.048534 | 0.308789 | -0.07374 | -0.02613 | 0.177869 | -0.3772 | 0.011162 | -0.09869 | 0.318149 | -0.07229 | -0.09256 |
| 24.20833 | 1.304809 | 0.161605 | 0.296944 | -0.14653 | 0.141578 | 0.29919 | -0.02364 | 0.09573 | -0.0701 | 0.294708 | 0.351298 | 0.593631 | 0.49796 | 0.132851 | 0.426466 | 0.061962 | 0.085112 | 0.334407 | -0.06667 | 0.152185 | -0.00789 | 1.010571 | 0.399972 | 0.55808 |
| 24.22222 | 1.331785 | 0.213219 | 0.332195 | -0.01049 | 0.250689 | 0.365887 | 0.310981 | 0.257227 | 0.058041 | 0.511227 | 0.478682 | 0.465311 | 0.414816 | 0.181577 | 0.347255 | 0.131485 | 0.108097 | 0.328823 | -0.05078 | 0.105266 | 0.012754 | 1.475901 | 0.620677 | 0.838805 |
| 24.23611 | 1.020599 | 0.185151 | 0.251914 | 0.129352 | 0.297623 | 0.259286 | 0.417546 | 0.247663 | 0.04116 | 0.506921 | 0.416998 | 0.172949 | 0.262666 | 0.212931 | 0.226545 | 0.114936 | 0.076928 | 0.215721 | -0.24648 | 0.041395 | 0.056417 | 0.944319 | 0.539946 | 0.630012 |
| 24.25 | 0.655116 | 0.107139 | 0.147536 | 0.158384 | 0.225956 | 0.145014 | 0.292236 | 0.084159 | -0.02932 | 0.339368 | 0.253164 | -0.14474 | 0.151836 | 0.289926 | 0.201269 | 0.057628 | 0.022347 | 0.101452 | -0.37099 | 0.030922 | 0.084028 | 0.097502 | 0.311349 | 0.182098 |
| 24.26389 | 0.382781 | 0.06371 | 0.079303 | 0.048479 | 0.06757 | 0.088512 | 0.045728 | -0.02008 | -0.05254 | -0.01691 | 0.111476 | -0.35179 | 0.127505 | 0.328376 | 0.24736 | -0.01711 | 0.02911 | 0.028536 | -0.26895 | 0.073496 | 0.064862 | -0.46387 | 0.232409 | 0.005015 |
| 24.27778 | 0.205004 | 0.028801 | 0.07023 | -0.06101 | -0.05317 | 0.114423 | -0.01488 | 0.148154 | 0.02349 | -0.36199 | 0.150552 | -0.32417 | 0.226502 | 0.258145 | 0.314455 | 0.001992 | 0.083194 | 0.036335 | 0.047908 | 0.160926 | 0.048912 | -0.27968 | 0.455425 | 0.356756 |
| 24.29167 | 0.150198 | -0.01504 | 0.148089 | -0.06642 | -0.07465 | 0.234742 | 0.19287 | 0.47891 | 0.276555 | -0.36214 | 0.375208 | 0.008545 | 0.361089 | 0.187792 | 0.371448 | 0.03863 | 0.110262 | 0.129203 | 0.405279 | 0.220863 | 0.120599 | 0.386861 | 0.893287 | 0.843303 |
| 24.30556 | 0.286105 | -0.03244 | 0.26861 | 0.100725 | 0.042154 | 0.421472 | 0.553064 | 0.925251 | 0.406993 | 0.087818 | 0.615837 | 0.376244 | 0.442693 | 0.242673 | 0.463316 | 0.077173 | 0.071453 | 0.243569 | 0.635627 | 0.263735 | 0.235046 | 0.931654 | 1.453601 | 1.035466 |
| 24.31944 | 0.581694 | 0.034971 | 0.381507 | 0.30242 | 0.201588 | 0.591095 | 0.907673 | 1.140836 | 0.328428 | 0.825847 | 0.685818 | 0.571379 | 0.436329 | 0.454945 | 0.5729 | 0.107002 | 0.050207 | 0.349824 | 0.757824 | 0.263891 | 0.295528 | 1.015616 | 1.617518 | 0.808517 |
| 24.33333 | 0.910576 | 0.115695 | 0.461494 | 0.366284 | 0.295968 | 0.684926 | 0.947336 | 0.952872 | 0.203835 | 1.540299 | 0.644673 | 0.580966 | 0.367852 | 0.625019 | 0.644711 | 0.159956 | 0.061845 | 0.399675 | 0.811887 | 0.22348 | 0.381349 | 0.595863 | 1.422019 | 0.400311 |
| 24.34722 | 1.11944 | 0.162312 | 0.466822 | 0.318323 | 0.209992 | 0.615639 | 0.730788 | 0.645118 | 0.157403 | 1.960063 | 0.547758 | 0.595417 | 0.331553 | 0.768183 | 0.732941 | 0.208531 | 0.185436 | 0.435012 | 1.060456 | 0.249361 | 0.537022 | 0.115207 | 1.192656 | 0.11152 |
| 24.36111 | 1.175742 | 0.181536 | 0.443803 | 0.228614 | 0.050476 | 0.490124 | 0.468735 | 0.289693 | 0.241926 | 2.144347 | 0.424595 | 0.664367 | 0.39379 | 0.814454 | 0.865875 | 0.300055 | 0.335445 | 0.424297 | 1.293022 | 0.330921 | 0.657121 | -0.26985 | 1.107366 | 0.038102 |
| 24.375 | 1.046648 | 0.157094 | 0.363607 | 0.132276 | -0.11843 | 0.441705 | 0.251609 | 0.021923 | 0.434189 | 2.078679 | 0.325221 | 0.703199 | 0.474897 | 0.678601 | 0.823892 | 0.398137 | 0.160029 | 0.357992 | 1.187602 | 0.370348 | 0.589479 | -0.35849 | 1.145065 | 0.243384 |
| 24.38889 | 0.80278 | 0.130718 | 0.332376 | 0.114989 | -0.14826 | 0.478697 | 0.214588 | -0.06024 | 0.655251 | 1.583245 | 0.311437 | 0.727736 | 0.633459 | 0.67076 | 0.777507 | 0.566958 | -0.45881 | 0.392237 | 1.191398 | 0.501701 | 0.527698 | 0.064586 | 1.634047 | 0.741821 |
| 24.40278 | 0.829656 | 0.272046 | 0.613608 | 0.328788 | 0.006955 | 0.811642 | 0.548276 | 0.075514 | 1.03527 | 1.191698 | 0.536885 | 0.795932 | 1.092531 | 1.135063 | 1.201284 | 1.094016 | -0.85933 | 0.734208 | 2.077327 | 1.054341 | 0.843852 | 0.564896 | 2.963161 | 1.551742 |
| 24.41667 | 1.432688 | 0.628938 | 1.575907 | 0.613628 | 0.250351 | 1.535203 | 1.224081 | 0.338795 | 1.697574 | 1.71851 | 1.067748 | 1.324256 | 1.592703 | 2.123894 | 1.521338 | 1.445703 | -0.64241 | 1.153279 | 3.123286 | 1.706104 | 1.151751 | 0.974205 | 4.28409 | 2.070135 |
| 24.43056 | 1.908404 | 0.862623 | 2.115819 | 0.738864 | 0.379969 | 2.020338 | 1.708021 | 0.742866 | 1.833807 | 3.196292 | 1.621826 | 1.828036 | 1.425836 | 2.789792 | 1.04286 | 1.125195 | -0.42359 | 0.996318 | 2.75966 | 1.488376 | 1.231418 | 0.909909 | 3.722149 | 1.88483 |
| 24.44444 | 2.447802 | 0.592291 | 1.53395 | 0.639403 | 0.443893 | 1.529465 | 1.685207 | 0.846346 | 1.409967 | 4.837334 | 1.788548 | 1.6739 | 1.468645 | 2.910037 | 0.824746 | 0.845461 | 0.06622 | 0.822901 | 1.946832 | 1.330432 | 1.09148 | 0.710792 | 2.443095 | 1.464279 |
| 24.45833 | 3.476647 | 0.224421 | 1.135528 | 0.608373 | 0.353663 | 1.130986 | 1.060419 | 0.875565 | 2.601795 | 4.896615 | 1.496559 | 1.160962 | 1.511639 | 2.629784 | 0.837769 | 0.406629 | 0.885284 | 0.655364 | 1.309916 | 1.557722 | 0.933623 | 0.523971 | 1.395068 | 1.049202 |
| 24.47222 | 2.626234 | 0.335156 | 1.236786 | 0.578887 | 0.21582 | 0.845992 | 0.288682 | 0.969099 | 3.275782 | 4.194284 | 0.908786 | 0.815821 | 1.289647 | 1.910958 | 0.603237 | 0.031192 | 1.610381 | 0.500388 | 0.70232 | 1.393322 | 0.632246 | -0.1076 | 0.516095 | 0.538777 |
| 24.48611 | 1.129779 | 0.489323 | 1.392978 | 0.497652 | 0.000737 | 0.629134 | -0.48443 | 0.691912 | 2.263445 | 2.311646 | 0.227031 | 0.476822 | 0.971636 | 1.463387 | 0.388291 | -0.20559 | 1.929344 | 0.437125 | 0.04227 | 1.146767 | 0.382682 | -0.69917 | 0.112484 | 0.399109 |
| 24.5 | 0.1721 | 0.494769 | 1.420584 | 0.425593 | -0.13019 | 0.41693 | -1.14066 | 0.257917 | 1.172731 | 0.225637 | -0.2837 | 0.165281 | 0.753 | 0.935578 | 0.259357 | -0.22518 | 2.322287 | 0.497872 | -0.40088 | 1.057875 | 0.292249 | -0.51414 | -0.05356 | 0.547454 |
| 24.51389 | -0.57238 | 0.415783 | 1.036634 | 0.504726 | -0.05468 | 0.309938 | -1.24679 | 0.092954 | 0.321434 | -1.66067 | -0.09865 | -0.15412 | 0.595762 | 0.298404 | 0.166867 | -0.13379 | 2.389401 | 0.518292 | -0.41229 | 0.966534 | 0.181688 | -0.0104 | -0.09694 | 0.794852 |
| 24.52778 | -1.18159 | 0.384572 | 0.901898 | 0.870106 | 0.019259 | 0.220087 | -1.08384 | -0.0378 | 1.221304 | -2.5405 | 0.462426 | -0.3109 | 0.366661 | -0.53105 | 0.159964 | 0.030352 | 1.641697 | 0.499788 | -0.23196 | 0.781676 | 0.027455 | 0.547677 | -0.26986 | 1.139716 |
| 24.54167 | -1.44971 | 0.341427 | 0.904389 | 0.848817 | 0.360034 | 0.070674 | -0.36761 | 0.047711 | 1.634529 | -3.51677 | 0.949781 | -0.34225 | 0.06712 | -1.44362 | 0.188083 | 0.270082 | 0.917641 | 0.47154 | -0.12929 | 0.514527 | -0.13746 | 1.025082 | -0.51528 | 1.375824 |
| 24.55556 | -1.31568 | 0.328403 | 0.757464 | 0.638584 | 0.364945 | -0.05237 | 0.469872 | -0.05534 | 1.4649 | -3.79609 | 0.876803 | 0.081065 | -0.15328 | -1.75019 | 0.171296 | 0.484666 | 0.755426 | 0.385893 | 0.062348 | 0.35324 | -0.20444 | 1.256379 | -0.45296 | 1.143342 |
| 24.56944 | -1.21582 | 0.34468 | 0.611796 | 0.610112 | 0.251869 | -0.01256 | 1.425617 | -0.34869 | 1.190615 | -2.79969 | 0.732547 | 0.703497 | -0.09885 | -1.24052 | 0.006273 | 0.637691 | 0.555237 | 0.169141 | 0.424401 | 0.268089 | 0.029997 | 1.14921 | -0.40749 | 0.728077 |
| 24.58333 | -0.78745 | 0.448049 | 0.541623 | 0.288933 | 0.110317 | 0.013963 | 1.603128 | -0.28939 | 0.901651 | 0.334633 | 0.582578 | 1.089992 | -0.0659 | -0.33843 | -0.19375 | 0.640609 | -0.16438 | -0.11762 | 0.820348 | 0.2197 | 0.299525 | 0.460866 | -0.3892 | 0.234935 |
| 24.59722 | 0.357096 | 0.617382 | 0.238805 | -0.12437 | -0.05913 | -0.09188 | 0.968163 | 0.140863 | 0.385158 | 3.844193 | 0.158393 | 1.019086 | 0.040378 | 0.554446 | -0.37023 | 0.667373 | -1.08443 | -0.26281 | 1.128102 | 0.139452 | 0.604271 | -0.33234 | -0.45082 | -0.08415 |
| 24.61111 | 1.122127 | 0.860986 | -0.11461 | -0.36435 | -0.25274 | 0.005113 | -0.0174 | 0.54777 | -0.03403 | 5.234736 | -0.41091 | 0.556872 | 0.076376 | 1.235827 | -0.28368 | 0.557389 | -2.0139 | -0.24891 | 1.309704 | -0.05863 | 0.788934 | -0.92754 | -0.53026 | -0.20529 |
| 24.625 | 1.673456 | 1.224517 | -0.37744 | -0.47428 | -0.19551 | 0.063404 | -0.9696 | 0.630098 | 0.212724 | 4.108068 | -0.27165 | -0.02135 | 0.22635 | 1.67232 | -0.03729 | 0.287899 | -2.35244 | -0.08782 | 1.200816 | -0.27326 | 0.834327 | -1.04099 | -0.46726 | -0.12103 |
| 24.63889 | 1.954033 | 1.375502 | -0.56412 | -0.40892 | -0.04331 | 0.084264 | -1.69229 | 0.543262 | 1.483529 | 3.206125 | 0.129942 | -0.1314 | 0.552268 | 2.30514 | 0.121021 | -0.02109 | -2.4255 | 0.107942 | 0.709142 | -0.27854 | 0.740788 | -1.17264 | -0.13042 | 0.039777 |
| 24.65278 | 1.94572 | 1.089924 | -0.64102 | -0.3147 | 0.158306 | 0.120892 | -1.69405 | 0.222462 | 1.004972 | 1.489718 | 0.381941 | -0.03294 | 0.736623 | 1.561022 | 0.083653 | -0.15147 | -2.29114 | 0.217877 | 0.150606 | -0.15163 | 0.406629 | -0.43266 | 0.281873 | 0.168115 |
| 24.66667 | 0.944422 | 0.643675 | -0.6481 | -0.3526 | 0.236753 | 0.075965 | -1.5445 | -0.01074 | -0.0003 | -0.48234 | 0.441229 | -0.29345 | 0.567643 | 0.491367 | 0.11947 | -0.1472 | -1.91647 | 0.177888 | -0.25552 | -0.13172 | 0.14359 | 0.856507 | 0.257446 | 0.210274 |
| 24.68056 | -0.14452 | 0.127044 | -0.66398 | -0.33052 | 0.075035 | 0.207737 | -0.84038 | -0.13976 | -0.93922 | -2.12416 | 0.301896 | -0.76775 | 0.283894 | -0.13921 | 0.235842 | -0.18363 | -1.31875 | 0.069028 | -0.68182 | -0.18132 | -0.08919 | 1.777193 | 0.055065 | 0.323865 |
| 24.69444 | -1.07017 | -0.19103 | -0.58441 | -0.44782 | -0.15726 | 0.222691 | -0.0302 | -0.26649 | -1.80303 | -3.81804 | -0.14419 | -1.18957 | -0.01159 | -0.64583 | 0.307107 | -0.2786 | -0.66229 | 0.021163 | -1.07284 | -0.17291 | -0.31422 | 1.400953 | -0.57617 | 0.431283 |
| 24.70833 | -1.6383 | -0.34231 | -0.52086 | -0.65813 | -0.42875 | 0.113415 | 0.496134 | -0.3771 | -1.54873 | -4.56839 | -0.54676 | -1.24517 | -0.33043 | -1.09532 | 0.249434 | -0.48904 | -0.42582 | 0.072435 | -1.25136 | -0.06006 | -0.42241 | 0.854885 | -1.04096 | 0.211726 |
| 24.72222 | -1.48516 | -0.45328 | -0.41856 | -0.88053 | -0.29482 | -0.01683 | 0.39317 | -0.4311 | -0.92324 | -4.60776 | -0.6818 | -0.78641 | -0.54198 | -1.40172 | 0.149256 | -0.5175 | -0.73342 | 0.118969 | -1.22933 | 0.106872 | -0.46433 | 0.129042 | -1.06879 | -0.23623 |
| 24.73611 | -1.12482 | -0.53453 | -0.32103 | -0.51926 | -0.03445 | -0.13087 | -0.03238 | -0.47514 | -0.69061 | -4.97824 | -0.26486 | -0.4044 | -0.65331 | -1.65735 | -0.10159 | -0.31834 | -1.06108 | 0.038263 | -1.03301 | 0.19336 | -0.52587 | -0.68593 | -0.90057 | -0.48952 |
| 24.75 | -1.04756 | -0.54211 | -0.31496 | -0.24339 | 0.260879 | -0.22117 | -0.51931 | -0.40375 | -0.70978 | -4.50008 | 0.017338 | 0.171555 | -0.4436 | -1.81012 | -0.51519 | -0.2675 | -1.01094 | -0.08728 | -0.82921 | 0.164535 | -0.57478 | -1.81193 | -0.43384 | -0.59858 |
| 24.76389 | -1.29526 | -0.51248 | -0.32411 | -0.50993 | 0.304434 | -0.29874 | -1.00342 | -0.21673 | -0.58329 | -3.4025 | 0.042704 | 0.779335 | -0.06653 | -2.12274 | -0.83254 | -0.27939 | -1.08886 | -0.25291 | -0.66216 | 0.025426 | -0.51285 | -2.113 | -0.08316 | -0.76842 |
| 24.77778 | -1.9349 | -0.53362 | -0.48821 | -1.03567 | 0.134166 | -0.34211 | -1.35788 | -0.0429 | -0.44098 | -2.09339 | -0.0693 | 1.020318 | 0.008346 | -1.969 | -0.83886 | -0.33884 | -1.45629 | -0.39012 | -0.52758 | -0.22097 | -0.36528 | -0.6377 | -0.0228 | -1.1155 |
| 24.79167 | -2.26147 | -0.64037 | -0.7858 | -1.4004 | -0.00724 | -0.33321 | -1.18948 | -0.13555 | -0.41693 | -0.97653 | -0.3153 | 0.288205 | -0.18141 | -1.16868 | -0.77963 | -0.26558 | -1.94083 | -0.48013 | -0.54243 | -0.45392 | -0.12651 | 0.982097 | -0.26592 | -1.12753 |
| 24.80556 | -2.34125 | -0.74188 | -1.17305 | -1.5399 | -0.09101 | -0.38646 | -0.96114 | -0.42327 | -0.61241 | -0.26553 | -0.58921 | -0.69515 | -0.57767 | -0.20536 | -0.65648 | -0.19853 | -2.46429 | -0.45042 | -0.60168 | -0.53715 | 0.129704 | 2.314613 | -0.5254 | -0.41372 |
| 24.81944 | -2.41665 | -0.76414 | -1.17349 | -1.30244 | -0.0592 | -0.42084 | -0.493 | -0.5387 | -0.90338 | -0.01694 | -0.84601 | -1.46272 | -0.96937 | 0.449738 | -0.56265 | -0.27724 | -2.77667 | -0.30018 | -0.66388 | -0.55171 | 0.148515 | 1.964926 | -0.71219 | 0.321246 |
| 24.83333 | -2.07111 | -0.73252 | -1.04327 | -1.1034 | 0.006474 | -0.51782 | -0.0855 | -0.65342 | -1.03267 | 0.040103 | -1.13458 | -1.6549 | -1.14906 | 0.663655 | -0.40235 | -0.51365 | -2.41032 | -0.18367 | -0.535 | -0.65833 | 0.061406 | 0.828792 | -0.85068 | -0.01452 |
| 24.84722 | -1.33899 | -0.5628 | -0.89321 | -0.4171 | 0.005954 | -0.58558 | 0.172743 | -1.00669 | -0.96531 | -0.01784 | -1.19008 | -1.75537 | -1.01796 | 0.663321 | -0.25906 | -0.56809 | -1.23135 | -0.18038 | -0.24318 | -0.72487 | -0.07765 | -0.42654 | -0.93744 | -0.6493 |
| 24.86111 | -0.35434 | -0.24857 | -0.70923 | -0.11004 | -0.03126 | -0.64699 | 0.227577 | -1.49499 | -0.79259 | -0.30717 | -0.71989 | -1.74397 | -0.47156 | 0.37503 | -0.41371 | -0.54637 | 0.130395 | -0.21038 | -0.07104 | -0.51444 | -0.19884 | -1.63415 | -0.75285 | -0.90905 |
| 24.875 | 0.330334 | -0.08788 | -0.53758 | -0.16543 | -0.05034 | -0.63608 | 0.072159 | -1.54674 | -0.54819 | -0.87069 | -0.10879 | -1.44263 | 0.288509 | -0.33857 | -0.5861 | -0.18421 | 1.118002 | -0.15507 | 0.006338 | -0.31279 | -0.34626 | -2.19607 | -0.45114 | -1.20419 |
| 24.88889 | 0.600913 | 0.069605 | -0.54083 | -0.21275 | -0.01284 | -0.44049 | -0.16254 | -1.15261 | -0.29776 | -1.4637 | 0.424094 | -0.73962 | 0.696632 | -1.1516 | -0.65766 | 0.005727 | 1.306717 | -0.17619 | -0.08394 | -0.12056 | -0.41923 | -1.70383 | -0.14151 | -1.33235 |
| 24.90278 | 0.375999 | 0.138864 | -0.60295 | -0.30925 | -0.00564 | -0.23664 | -0.44568 | -0.67572 | -0.13428 | -1.79843 | 0.498789 | 0.128108 | 0.214726 | -1.67716 | -0.66763 | -0.62045 | 0.427242 | -0.39671 | -0.48349 | -0.11369 | -0.46323 | -1.08013 | -0.24712 | -1.77652 |
| 24.91667 | -0.35369 | -0.01412 | -0.81471 | -0.54035 | -0.12501 | -0.32648 | -0.80542 | -0.4743 | -0.26674 | -1.92232 | -0.15861 | 0.386074 | -0.62603 | -1.80187 | -0.54896 | -1.45624 | -0.41066 | -0.71931 | -1.05884 | -0.35993 | -0.5829 | -0.85178 | -0.74096 | -2.08725 |
| 24.93056 | -1.24676 | -0.20547 | -1.04493 | -0.70314 | -0.25792 | -0.64394 | -1.21936 | -0.50525 | -0.66045 | -1.97517 | -1.07991 | -0.1674 | -1.16547 | -2.00396 | -0.68191 | -2.11921 | -1.56772 | -1.07948 | -1.7635 | -0.6142 | -0.79411 | -0.55231 | -1.66356 | -2.79472 |
| 24.94444 | -2.08346 | -0.42026 | -1.13601 | -0.78951 | -0.14403 | -1.05558 | -1.57646 | -0.695 | -1.30521 | -2.04951 | -1.87224 | -1.00411 | -1.10834 | -2.90363 | -1.7724 | -1.65237 | -2.25541 | -1.59435 | -2.08846 | -0.67901 | -1.03783 | -0.07745 | -2.32271 | -2.82027 |
| 24.95833 | -2.96992 | -0.70648 | -1.08391 | -1.04715 | -0.02579 | -1.30278 | -1.80131 | -1.03762 | -0.6013 | -1.79675 | -2.23945 | -1.7626 | -0.56282 | -3.03964 | -2.79683 | -0.65219 | -1.32858 | -1.79624 | -1.82431 | -0.49819 | -0.82577 | 0.595359 | -1.4841 | -1.7003 |
| 24.97222 | -2.75665 | -0.95684 | -1.04529 | -0.90321 | -0.00587 | -1.18946 | -1.3154 | -0.97435 | 0.391141 | -1.29863 | -1.98408 | -1.81672 | 0.028853 | -2.25298 | -2.12395 | 0.519321 | 0.653533 | -1.28449 | -1.57187 | -0.34151 | -0.72984 | 1.639334 | -0.60095 | 0.351409 |
| 24.98611 | -1.97804 | -0.67824 | -0.76487 | -0.41425 | 0.042428 | -0.84124 | -0.38877 | -0.28642 | 1.214484 | -0.50611 | -1.35914 | -1.49089 | 0.427304 | -0.86684 | -0.87414 | 1.329475 | 2.373678 | -0.61902 | -0.79977 | -0.05205 | -0.3187 | 2.489289 | 0.179366 | 1.323489 |
| 25 | -0.75289 | -0.21603 | -0.38503 | 0.0427 | 0.100098 | -0.37993 | 0.291377 | -0.02437 | 1.470995 | 0.283464 | -0.62732 | -0.85585 | 0.305305 | 0.245178 | 0.019924 | 1.597562 | 2.01777 | -0.13741 | 0.228945 | 0.108035 | 0.278818 | 2.493505 | 0.78473 | 1.396623 |
| 25.01389 | 0.191818 | 0.119038 | -0.06266 | 0.330739 | 0.208925 | -0.04911 | 0.640376 | -0.01974 | 1.061463 | 0.829093 | 0.376908 | -0.25249 | 0.115371 | 0.77277 | 0.418555 | 1.603329 | 0.837365 | 0.114552 | 0.855545 | -0.0367 | 0.638327 | 1.69496 | 1.458253 | 1.614595 |
| 25.02778 | 0.713628 | 0.226093 | 0.12641 | 0.478193 | 0.337856 | 0.120783 | 0.635145 | -0.03584 | 0.425162 | 0.954557 | 1.004614 | 0.136682 | -0.00491 | 0.814932 | 0.472274 | 1.067322 | -0.04118 | 0.156726 | 0.903507 | -0.17222 | 0.647729 | 0.761447 | 1.449422 | 1.015791 |
| 25.04167 | 0.785638 | 0.157789 | 0.189211 | 0.392731 | 0.275459 | 0.177269 | 0.485138 | -0.06392 | -0.02157 | 0.652892 | 0.993842 | 0.305042 | -0.11664 | 0.652753 | 0.256204 | 0.455485 | -0.66177 | 0.050895 | 0.684059 | -0.30779 | 0.55314 | 0.122012 | 0.953789 | 0.38256 |
| 25.05556 | 0.589656 | 0.085378 | 0.133145 | 0.211607 | 0.128072 | 0.124611 | 0.344144 | -0.0608 | -0.26703 | 0.214394 | 0.644944 | 0.31538 | -0.14942 | 0.492794 | 0.028098 | 0.085399 | -0.75282 | -0.09849 | 0.551681 | -0.33232 | 0.473279 | -0.3909 | 0.478907 | 0.043931 |
| 25.06944 | 0.306388 | 0.049697 | 0.11069 | 0.018562 | -0.06029 | 0.036065 | 0.285198 | -0.02499 | -0.38124 | 0.002557 | 0.425582 | 0.287483 | -0.07115 | 0.487133 | -0.06966 | 0.047211 | -0.27401 | -0.10439 | 0.484327 | -0.29256 | 0.434728 | -0.78761 | 0.23019 | -0.11944 |
| 25.08333 | 0.048555 | 0.048122 | 0.094891 | -0.14463 | -0.23942 | -0.04612 | 0.253256 | 0.025616 | -0.29265 | 0.006018 | 0.416888 | 0.323346 | 0.066222 | 0.697739 | 0.070044 | 0.146229 | 0.254558 | 0.108334 | 0.426642 | -0.18623 | 0.396442 | -0.69776 | 0.16959 | -0.3317 |
| 25.09722 | -0.04622 | 0.064165 | 0.095227 | -0.184 | -0.27619 | -0.0683 | 0.212629 | 0.098861 | 0.033885 | 0.08528 | 0.550781 | 0.481198 | 0.142178 | 0.824214 | 0.330264 | 0.234688 | 0.338804 | 0.398567 | 0.350569 | -0.11054 | 0.359917 | -0.03147 | 0.233489 | -0.57346 |
| 25.11111 | 0.14555 | 0.066187 | 0.107924 | -0.08528 | -0.04622 | -0.02018 | 0.182896 | 0.236259 | 0.264619 | 0.190357 | 0.66702 | 0.670544 | 0.158428 | 0.82796 | 0.756823 | 0.346272 | 0.050345 | 0.59831 | 0.262613 | -0.09534 | 0.313683 | 0.632094 | 0.276917 | -0.45666 |
| 25.125 | 0.463545 | 0.05967 | 0.118768 | 0.014952 | 0.201215 | 0.101505 | 0.112232 | 0.343786 | 0.260123 | 0.258867 | 0.658504 | 0.901106 | 0.119366 | 0.659745 | 1.004966 | 0.375555 | -0.32251 | 0.628357 | 0.217565 | -0.11887 | 0.354465 | 0.720091 | 0.184156 | -0.094 |
| 25.13889 | 0.677666 | 0.048414 | 0.099456 | -0.00599 | 0.199734 | 0.233756 | 0.012797 | 0.421793 | 0.171649 | 0.255916 | 0.615497 | 1.139229 | 0.106798 | 0.36026 | 1.028214 | 0.244747 | -0.51155 | 0.598262 | 0.158867 | -0.20467 | 0.397818 | -0.02659 | 0.010726 | 0.057948 |
| 25.15278 | 0.717813 | 0.047007 | 0.063925 | -0.14098 | 0.023723 | 0.284047 | -0.09894 | 0.547396 | 0.115564 | 0.254751 | 0.523887 | 1.303506 | 0.139347 | 0.066085 | 1.079851 | 0.073986 | -0.23193 | 0.606936 | 0.079098 | -0.29069 | 0.408893 | -1.09935 | -0.13375 | -0.09177 |
| 25.16667 | 0.553508 | 0.024557 | 0.024575 | -0.27249 | -0.16949 | 0.16422 | -0.22721 | 0.689188 | 0.10308 | 0.251753 | 0.451612 | 1.366587 | 0.208242 | -0.11562 | 1.052727 | -0.01523 | 0.28105 | 0.632153 | 0.04418 | -0.27548 | 0.405361 | -2.36324 | -0.14621 | -0.30059 |
| 25.18056 | 0.334508 | -0.00624 | 0.021469 | -0.27257 | -0.19754 | 0.038134 | -0.30147 | 0.681752 | 0.184254 | 0.197434 | 0.380418 | 1.236062 | 0.397095 | -0.14449 | 0.932063 | 0.120312 | 0.598365 | 0.65588 | 0.004687 | -0.19334 | 0.375948 | -2.17699 | 0.018586 | -0.46497 |
| 25.19444 | 0.138654 | -0.03186 | 0.03574 | -0.15718 | 0.054037 | -0.05404 | -0.23139 | 0.596865 | 0.2887 | 0.155114 | 0.330488 | 0.937655 | 0.543548 | -0.07659 | 0.864977 | 0.181925 | 0.476032 | 0.581571 | -0.0556 | -0.1475 | 0.323876 | -0.24865 | 0.287799 | -0.3957 |
| 25.20833 | -0.0355 | -0.06171 | 0.035833 | 0.055233 | 0.257982 | -0.02161 | -0.09709 | 0.459466 | 0.223343 | 0.144403 | 0.316399 | 0.711315 | 0.445071 | 0.033107 | 0.823031 | 0.154249 | 0.123105 | 0.53561 | -0.08494 | -0.05911 | 0.36079 | 1.717613 | 0.419556 | -0.09806 |
| 25.22222 | -0.19351 | -0.0698 | 0.037038 | 0.211746 | 0.241472 | 0.08567 | 0.014572 | 0.479371 | 0.209237 | 0.182195 | 0.278088 | 0.493556 | 0.292968 | 0.102168 | 0.85273 | 0.192062 | -0.15608 | 0.479616 | 0.020402 | 0.019096 | 0.467137 | 2.181155 | 0.525325 | 0.249249 |
| 25.23611 | -0.30318 | -0.06641 | 0.042864 | 0.309714 | 0.13919 | 0.242831 | 0.134538 | 0.543234 | 0.401141 | 0.228486 | 0.229582 | 0.293924 | 0.17017 | -0.02851 | 0.769984 | 0.24572 | -0.29288 | 0.362622 | 0.15767 | -0.01223 | 0.558035 | 0.590095 | 0.679654 | 0.374322 |
| 25.25 | -0.33853 | -0.07332 | 0.063833 | 0.362752 | 0.093171 | 0.366802 | 0.209566 | 0.622552 | 0.56061 | 0.218638 | 0.222823 | 0.071604 | 0.105827 | -0.35845 | 0.587071 | 0.247194 | -0.3071 | 0.225281 | 0.208525 | -0.08121 | 0.529988 | -1.38183 | 0.802026 | 0.252608 |
| 25.26389 | -0.30513 | -0.05676 | 0.093536 | 0.3504 | 0.120109 | 0.439542 | 0.241384 | 0.346134 | 0.447739 | 0.183054 | 0.219107 | -0.19711 | 0.151688 | -0.647 | 0.410226 | 0.065624 | -0.18247 | 0.167719 | 0.245696 | 0.003806 | 0.455348 | -2.7081 | 0.925575 | 0.005348 |
| 25.27778 | -0.24896 | -0.04889 | 0.150014 | 0.325203 | 0.175608 | 0.413385 | 0.277726 | 0.074446 | 0.202299 | 0.158832 | 0.243858 | -0.35905 | 0.238056 | -0.61788 | 0.259262 | -0.05362 | -0.05727 | 0.203735 | 0.29827 | 0.165408 | 0.364574 | -1.91648 | 0.97255 | -0.29352 |
| 25.29167 | -0.29627 | -0.07609 | 0.214924 | 0.300828 | 0.198498 | 0.281993 | 0.369024 | -0.03944 | 0.050407 | 0.145552 | 0.293578 | -0.31581 | 0.239091 | -0.29725 | 0.255271 | 0.11357 | -0.0894 | 0.285037 | 0.348168 | 0.346979 | 0.326547 | -0.10357 | 1.038213 | -0.41848 |
| 25.30556 | -0.28985 | -0.08525 | 0.254249 | 0.284907 | 0.188396 | 0.174561 | 0.453085 | -0.00877 | 0.144152 | 0.233799 | 0.398659 | -0.11839 | 0.228949 | 0.128186 | 0.349578 | 0.332389 | -0.2456 | 0.364265 | 0.44209 | 0.524669 | 0.349371 | 1.808335 | 1.108329 | -0.0808 |
| 25.31944 | -0.17151 | -0.0421 | 0.275466 | 0.356855 | 0.19059 | 0.267549 | 0.557647 | 0.106569 | 0.34779 | 0.462761 | 0.569551 | 0.144152 | 0.124399 | 0.469657 | 0.473135 | 0.555508 | -0.36668 | 0.377647 | 0.555911 | 0.520405 | 0.460278 | 2.805101 | 1.159603 | 0.644345 |
| 25.33333 | 0.080509 | 0.043176 | 0.284633 | 0.485702 | 0.184508 | 0.435223 | 0.540415 | 0.371534 | 0.62005 | 0.570154 | 0.586653 | 0.342816 | 0.063996 | 0.684179 | 0.521931 | 0.686592 | -0.33762 | 0.327911 | 0.69497 | 0.364741 | 0.557584 | 1.220658 | 1.128553 | 0.969123 |
| 25.34722 | 0.446744 | 0.09724 | 0.296077 | 0.607557 | 0.146063 | 0.601096 | 0.43714 | 0.515871 | 0.639638 | 0.47992 | 0.409161 | 0.449644 | 0.154408 | 0.837272 | 0.643675 | 0.81033 | -0.07209 | 0.341479 | 0.899381 | 0.264316 | 0.682036 | -0.41116 | 1.138943 | 0.907789 |
| 25.36111 | 0.748191 | 0.096338 | 0.313559 | 0.64893 | 0.133854 | 0.536174 | 0.370992 | 0.561413 | 0.464982 | 0.467171 | 0.225433 | 0.473049 | 0.363846 | 0.995639 | 0.744305 | 0.949119 | 0.177864 | 0.416233 | 1.002001 | 0.310476 | 0.808075 | -2.00179 | 1.110019 | 0.394984 |
| 25.375 | 0.897376 | 0.072734 | 0.286293 | 0.59084 | 0.135925 | 0.644577 | 0.355678 | 0.542738 | 0.143544 | 0.517036 | 0.215981 | 0.578932 | 0.459318 | 1.054959 | 0.656633 | 0.835151 | 0.133255 | 0.405056 | 0.843388 | 0.449376 | 0.893379 | -1.95989 | 1.002054 | -0.01399 |
| 25.38889 | 0.880958 | 0.077973 | 0.263633 | 0.559754 | 0.171292 | 0.649395 | 0.442576 | 0.371694 | -0.57739 | 0.663018 | 0.410006 | 0.766326 | 0.647412 | 1.012508 | 0.584724 | 0.665918 | -0.03741 | 0.41595 | 0.777059 | 0.679802 | 0.779163 | -0.27968 | 1.093507 | -0.21598 |
| 25.40278 | 1.100002 | 0.169285 | 0.331267 | 0.756253 | 0.278997 | 0.613779 | 0.727837 | 0.256723 | -1.85095 | 1.246838 | 0.743676 | 1.034611 | 1.174802 | 1.442854 | 0.904523 | 0.998611 | -0.03239 | 0.725541 | 1.058099 | 1.395323 | 0.660784 | 1.513641 | 1.720649 | 0.612408 |
| 25.41667 | 1.769752 | 0.364935 | 0.567594 | 1.299043 | 0.448711 | 1.216781 | 1.385648 | 0.062509 | -0.54056 | 2.500315 | 1.387962 | 1.762072 | 1.672256 | 2.070053 | 1.037299 | 1.370034 | 0.087364 | 1.090168 | 1.194579 | 1.885462 | 0.620918 | 2.197476 | 2.347982 | 2.220948 |
| 25.43056 | 2.69387 | 0.476934 | 0.751652 | 1.599321 | 0.493259 | 2.011729 | 1.760657 | -0.2294 | 1.389712 | 3.210859 | 2.145499 | 2.331864 | 2.18479 | 2.277793 | 0.864046 | 1.450927 | 0.105865 | 0.896121 | 0.634923 | 1.611991 | -0.12773 | 1.400106 | 2.180197 | 2.687971 |
| 25.44444 | 3.163647 | 0.374466 | 0.789483 | 1.182015 | 0.589617 | 1.560661 | 1.607753 | -0.03056 | 1.502404 | 3.012042 | 2.380005 | 2.150476 | 2.798317 | 2.652394 | 0.960425 | 1.289954 | 0.211426 | 0.823517 | 0.247571 | 1.334595 | -0.07512 | 1.255247 | 1.738793 | 2.039559 |
| 25.45833 | 3.743152 | 0.288304 | 0.564324 | 0.882489 | 0.495456 | 1.170844 | 1.345945 | 0.217767 | 1.696757 | 3.342059 | 1.909686 | 1.47914 | 2.58489 | 1.513493 | 1.098071 | 1.032936 | 0.178331 | 0.704739 | 0.464474 | 1.287899 | 0.126075 | 0.551633 | 1.197135 | 1.260599 |
| 25.47222 | 4.560006 | 0.219396 | 0.63888 | 0.775586 | 0.324971 | 1.541396 | 0.795675 | 0.409015 | 3.087447 | 3.585485 | 1.16603 | 1.085045 | 1.82255 | 0.833047 | 0.675302 | 0.71536 | -0.13567 | 0.347415 | 0.485959 | 1.133358 | 0.059642 | -0.67259 | 0.542554 | 0.679735 |
| 25.48611 | 3.53513 | 0.149225 | 0.471395 | 0.574113 | 0.161793 | 0.907005 | 0.275248 | 0.388974 | 3.496019 | 3.49411 | 0.438113 | 0.81811 | 1.234396 | 0.473673 | 0.293666 | 0.181489 | -0.64508 | 0.058401 | 0.24369 | 0.841639 | -0.02226 | -1.21613 | 0.173565 | 0.264375 |
| 25.5 | 2.218831 | 0.127148 | 0.250671 | 0.413287 | 0.016814 | 0.51654 | -0.13034 | 0.154213 | 2.26696 | 2.33968 | -0.47179 | 0.613844 | 0.477408 | 0.161776 | 0.048454 | -0.42839 | -1.03135 | -0.01616 | -0.02361 | 0.730991 | -0.01036 | -1.17693 | 0.174015 | -0.04256 |
| 25.51389 | 1.026638 | 0.09802 | 0.261407 | 0.18355 | -0.07263 | 0.320764 | -0.54206 | -0.17331 | 0.792653 | 0.697651 | -1.28521 | 0.339451 | -0.38321 | -0.45324 | -0.05784 | -1.26263 | -1.76131 | 0.193144 | -0.45658 | 0.676548 | -0.22921 | -1.82538 | 0.067551 | -0.17448 |
| 25.52778 | -0.17455 | 0.057892 | 0.586089 | -0.44162 | -0.01792 | 0.336822 | -0.8873 | -0.33831 | -0.78346 | -0.86904 | -1.14699 | -0.16805 | -1.2347 | -1.67807 | -0.26239 | -1.21159 | -1.39993 | 0.43089 | -0.76593 | 0.586122 | -0.32823 | -3.07526 | 0.040605 | -0.52077 |
| 25.54167 | -0.62846 | -0.00297 | 0.70309 | -0.69026 | 0.072746 | 0.240118 | -0.89457 | -0.33572 | -0.99573 | -0.99017 | -0.16997 | -0.76572 | -1.7573 | -2.84801 | -0.62038 | -0.36968 | -0.55698 | 0.469752 | -0.73842 | 0.494094 | -0.25831 | -1.28315 | -0.10078 | -0.50137 |
| 25.55556 | -1.1518 | 0.018407 | 0.839555 | -0.56447 | 0.288589 | -0.01621 | -0.38874 | -0.35522 | 0.50602 | -1.4714 | 1.084007 | -1.08306 | -1.63924 | -2.09574 | -1.03965 | 0.652301 | 0.045506 | 0.34333 | -0.56682 | 0.39278 | 0.118497 | 0.950076 | -0.20757 | 0.481717 |
| 25.56944 | -1.4505 | 0.114917 | 0.749678 | -0.08117 | 0.522568 | -0.17657 | 0.399441 | -0.5898 | 0.752795 | -2.37851 | 1.87061 | -0.94945 | -0.81768 | -1.16005 | -1.36242 | 1.00263 | -0.05197 | 0.129242 | -0.54286 | 0.014886 | 0.186656 | 2.34858 | -0.68684 | 1.291068 |
| 25.58333 | -2.01066 | 0.117629 | 0.680889 | 0.182877 | 0.558484 | -0.25115 | 1.253113 | -0.88272 | 0.446267 | -1.81931 | 1.331404 | -0.25862 | 0.010038 | 0.127957 | -1.14507 | 1.311593 | 0.22664 | -0.08045 | 0.10594 | -0.39993 | 0.251049 | 1.812307 | -0.69338 | 0.074156 |
| 25.59722 | -2.48177 | 0.049516 | 0.76931 | 0.292522 | 0.433501 | -0.47105 | 1.19084 | -0.1906 | 0.20305 | -0.5776 | 0.422728 | 0.447369 | 1.037759 | 1.378081 | -1.46747 | 1.383427 | -0.1509 | -0.34937 | 0.735184 | -0.63989 | 0.158338 | 1.135714 | -0.7214 | -0.59417 |
| 25.61111 | -1.6911 | -0.05305 | 0.591354 | 0.391493 | 0.194465 | -0.47105 | 0.74761 | 0.419495 | 0.44639 | 0.466209 | -0.14463 | 0.834039 | 2.427359 | 2.698882 | -2.08085 | 1.283655 | -0.85368 | -0.6082 | 1.122778 | -0.69183 | -0.02926 | 0.417041 | -1.42004 | -1.27758 |
| 25.625 | -0.3981 | -0.10263 | 0.365808 | 0.475192 | -0.04289 | -0.17197 | 0.150994 | 0.51542 | 0.85675 | 1.783605 | -0.52662 | 0.798394 | 2.402992 | 2.638327 | -1.57585 | 0.239471 | -1.45134 | -0.41751 | 1.119255 | -0.49086 | -0.28743 | -0.54661 | -1.47083 | -1.48883 |
| 25.63889 | -0.01963 | -0.03974 | 0.294315 | 0.520099 | -0.07003 | 0.189182 | -0.38313 | 0.464123 | 1.097209 | 2.99092 | -1.50731 | 0.600377 | 1.650076 | 1.635632 | -0.21868 | -0.48308 | -2.3466 | -0.02428 | 0.801497 | -0.29263 | -0.25657 | -1.17781 | -0.91587 | -1.11924 |
| 25.65278 | 0.412682 | 0.022061 | 0.117601 | 0.400523 | 0.106487 | 0.401068 | -0.71525 | 0.868402 | 0.772991 | 3.181629 | -1.46832 | 0.251161 | 0.736002 | 0.76588 | 0.613902 | -0.56411 | -2.81801 | 0.377361 | 0.629722 | 0.016355 | -0.10608 | -2.60653 | -0.69269 | -0.75145 |
| 25.66667 | 0.678321 | 0.062602 | -0.14794 | 0.092418 | 0.236081 | 0.356477 | -0.92319 | 0.463573 | 0.365581 | 2.569468 | -0.63976 | -0.17782 | -0.19254 | 0.136062 | 1.439835 | -0.58114 | -1.19121 | 0.671273 | 0.540258 | 0.230978 | 0.028319 | -4.55154 | -0.21805 | -0.49654 |

| | | | | | | | | | | | | | | | | | | | | | | | | |
|---|---|---|---|---|---|---|---|---|---|---|---|---|---|---|---|---|---|---|---|---|---|---|---|---|
| 25.68056 | 0.9759 | 0.011998 | -0.30744 | -0.327 | 0.240189 | 0.195363 | -1.01214 | 0.015109 | 0.153878 | 1.582295 | 0.282143 | -0.32754 | -0.67095 | -0.43926 | 0.70031 | -0.40592 | 0.490517 | 0.728026 | 0.209648 | 0.026618 | 0.185093 | -4.16249 | 0.276795 | -0.46005 |
| 25.69444 | 1.181142 | -0.05458 | -0.46292 | -0.77653 | 0.008174 | -0.0274 | -1.04865 | -0.22817 | 0.138812 | -0.15885 | 0.964738 | -0.40844 | -0.54879 | -1.09557 | 0.954397 | 0.031556 | 1.809186 | 0.559013 | -0.04887 | -0.16929 | 0.227185 | -1.45489 | 0.687946 | -0.68864 |
| 25.70833 | 0.927865 | -0.15213 | -0.56095 | -0.98442 | -0.09629 | -0.14243 | -0.81282 | -0.22845 | 0.116864 | -1.81946 | 0.896456 | -0.52575 | -0.42109 | -1.79469 | 1.228713 | -0.34716 | 1.882844 | 0.260387 | -0.23145 | -0.31859 | 0.142148 | 1.548595 | 0.88159 | -1.06336 |
| 25.72222 | 0.224706 | -0.19515 | -0.58044 | -0.97088 | -0.18947 | 0.048271 | -0.58637 | -0.09468 | 0.019216 | -2.82426 | 0.274924 | -0.83776 | -1.18231 | -2.22511 | 1.599854 | -1.32993 | 1.108061 | 0.030692 | -0.51306 | -0.37685 | -0.21625 | 4.741626 | 0.840245 | -0.2217 |
| 25.73611 | -0.80676 | -0.18336 | -0.59242 | -0.8762 | -0.34276 | 0.238531 | -0.32467 | 0.01529 | 0.100781 | -3.38975 | -0.44521 | -1.19914 | -1.60072 | -2.02971 | 0.890348 | -1.84099 | 0.174854 | -0.05904 | -0.88096 | -0.35249 | -0.57392 | 4.909135 | 0.410626 | 1.202537 |
| 25.75 | -1.82161 | -0.18714 | -0.4846 | -0.83364 | -0.22632 | 0.155765 | -0.13686 | 0.056714 | -0.08122 | -3.51743 | -0.909 | -1.55597 | -1.41028 | -2.51253 | 0.191683 | -1.33751 | -0.85017 | -0.24002 | -0.95255 | -0.21775 | -0.65423 | 3.190893 | -0.13088 | 1.488252 |
| 25.76389 | -2.18764 | -0.27568 | -0.28707 | -0.95869 | -0.02212 | -0.29606 | -0.09372 | -0.18026 | -0.29284 | -2.58726 | -1.11989 | -1.70702 | -0.74974 | -2.47297 | -0.3402 | -0.90738 | -1.26527 | -0.44209 | -0.8109 | -0.21106 | -0.35173 | 0.870336 | -0.66379 | 1.027303 |
| 25.77778 | -1.99956 | -0.39535 | -0.27429 | -1.18213 | 0.240492 | -0.77558 | -0.09404 | -0.55744 | -0.80666 | -1.33248 | -1.00076 | -1.48723 | -0.73289 | -1.61011 | -0.65585 | -0.57336 | -1.08841 | -0.59105 | -0.65792 | -0.27165 | -0.15714 | -1.3788 | -0.89624 | 0.232167 |
| 25.79167 | -1.64613 | -0.40628 | -0.39171 | -1.15385 | 0.350982 | -1.18568 | -0.02137 | -1.01975 | -1.33812 | -0.64767 | -0.61082 | -1.01767 | -0.87953 | -0.94675 | -0.75391 | -0.78337 | -0.50152 | -0.74869 | -0.56679 | -0.53898 | -0.08557 | -3.5972 | -0.90234 | -0.75213 |
| 25.80556 | -1.4144 | -0.28376 | -0.57812 | -0.88506 | 0.495203 | -1.05788 | 0.065367 | -1.39049 | -1.81735 | -0.5676 | -0.2841 | -0.75886 | -0.77297 | -0.5539 | -0.89351 | -0.96869 | 0.591658 | -0.8577 | -0.5652 | -0.95197 | -0.14213 | -3.24157 | -0.97066 | -1.49382 |
| 25.81944 | -1.25575 | -0.13116 | -0.93458 | -0.87024 | 0.428529 | -1.06956 | -0.02911 | -0.95628 | -2.06152 | -0.77543 | -0.28192 | -0.66273 | -0.37705 | -0.20398 | -1.43102 | -0.74289 | 1.474885 | -0.71613 | -0.54716 | -1.06215 | -0.18477 | -3.68208 | -1.09727 | -1.66151 |
| 25.83333 | -1.4092 | -0.01261 | -1.10662 | -0.87893 | 0.239277 | -1.06011 | -0.29266 | -0.53526 | -1.76698 | -0.88467 | -0.66156 | -0.55281 | 0.149206 | 0.216118 | -1.16396 | 0.081151 | 2.137292 | -0.35928 | -0.37302 | -1.00344 | -0.15922 | -1.89383 | -1.03471 | -1.40226 |
| 25.84722 | -0.83665 | 0.070356 | -0.7314 | -0.81678 | 0.044693 | -0.22906 | -0.58394 | -0.33577 | -1.00636 | -0.99976 | -0.92605 | -0.31078 | 0.410458 | 0.601984 | -0.73666 | 0.899855 | 1.968561 | -0.05184 | -0.00867 | -0.91586 | -0.15823 | 0.245721 | -0.85838 | -1.21351 |
| 25.86111 | -0.36208 | 0.075881 | -0.38589 | -0.71017 | -0.0144 | 0.033476 | -0.7626 | -0.08153 | -0.33251 | -1.29678 | -0.95655 | 0.008769 | 0.071861 | 0.838685 | -0.30312 | 0.853705 | 1.199358 | 0.148416 | 0.19698 | -0.79428 | -0.1543 | 2.75917 | -0.76593 | -0.84871 |
| 25.875 | -0.16936 | 0.054452 | -0.18195 | -0.54396 | 0.048302 | -0.4471 | -0.66153 | 0.057389 | -0.34508 | -1.21974 | -0.74383 | 0.327842 | -0.2471 | 0.854674 | 0.168117 | 0.837296 | 0.327553 | 0.314247 | 0.209732 | -0.67463 | -0.01276 | 4.902382 | -0.52467 | -0.38721 |
| 25.88889 | -0.10522 | 0.130348 | -0.0319 | -0.38227 | 0.085284 | -0.4914 | -0.27562 | -0.10509 | -0.55455 | -0.69354 | -0.34397 | 0.582114 | -0.30378 | 0.721688 | 0.379093 | 0.859624 | -0.51804 | 0.215331 | 0.089236 | -0.56789 | 0.03025 | 3.809063 | -0.64765 | -0.2168 |
| 25.90278 | -0.28306 | 0.05897 | -0.16818 | -0.33701 | -0.11992 | -0.57257 | -0.00766 | -0.30329 | -1.10679 | -0.52732 | -0.18026 | 0.619383 | -0.48998 | 0.155468 | 0.152378 | 0.389425 | -1.36091 | -0.22704 | -0.09009 | -0.55882 | -0.05035 | 1.405637 | -0.85611 | -0.46955 |
| 25.91667 | -0.80657 | -0.11207 | -0.4798 | -0.53146 | -0.46315 | -0.66618 | -0.06089 | -0.35995 | -1.52923 | -0.68314 | -0.42336 | 0.314297 | -0.37765 | -0.58619 | -0.36203 | -0.57207 | -1.60223 | -0.72372 | -0.07455 | -0.36201 | 0.101324 | -1.06495 | -1.21006 | -0.77576 |
| 25.93056 | -1.60018 | -0.19719 | -0.70716 | -0.70629 | -0.69714 | -0.71266 | -0.16442 | -0.463 | -1.98231 | -0.69656 | -0.89239 | -0.11843 | -0.02699 | -1.32438 | -0.43891 | -1.28688 | -0.84299 | -1.26627 | -0.09914 | -0.11273 | 0.389082 | -2.63307 | -1.53154 | -1.09605 |
| 25.94444 | -2.54174 | -0.24264 | -0.58185 | -0.87053 | -0.73616 | -0.90636 | -0.34251 | -0.39189 | -2.32284 | -0.68843 | -0.92937 | -0.70703 | -0.30268 | -1.71754 | -1.57893 | -1.10364 | 0.301223 | -1.17823 | -0.87393 | -0.2776 | 0.295858 | -3.56573 | -2.35085 | -1.63835 |
| 25.95833 | -3.34794 | -0.52886 | -0.3201 | -1.23016 | -0.60236 | -1.37497 | -0.84934 | -0.17582 | -2.28016 | -0.81272 | -0.99793 | -1.50573 | -0.26027 | -2.31636 | -2.65158 | -0.92532 | 0.963067 | -1.13185 | -1.46712 | -0.8289 | 0.034028 | -4.09133 | -2.83963 | -2.076 |
| 25.97222 | -3.09449 | -0.84347 | -0.30452 | -1.35625 | -0.2554 | -1.47313 | -1.10778 | -0.11981 | -1.64592 | -0.84139 | -1.50721 | -1.92805 | 0.253773 | -1.92409 | -2.16471 | -0.5108 | 1.201918 | -0.90057 | -1.34104 | -0.98955 | 0.004469 | -2.03416 | -2.21809 | -1.94694 |
| 25.98611 | -1.85346 | -0.83304 | -0.38885 | -0.9043 | 0.252883 | -1.19617 | -0.71716 | -0.04698 | -1.31559 | -0.76244 | -1.31967 | -1.66792 | 0.491144 | -1.21928 | -1.2666 | -0.22188 | 0.980045 | -0.50607 | -0.83673 | -0.87272 | -0.00132 | 0.394609 | -1.27138 | -1.2997 |
| 26 | -0.68153 | -0.58718 | -0.32472 | -0.25422 | 0.599483 | -0.65977 | -0.3033 | 0.074313 | -0.81874 | -0.927 | -0.06714 | -0.8343 | 0.601797 | -0.51451 | -0.59162 | 0.202609 | 0.921333 | -0.1812 | -0.36838 | -0.45469 | 0.009279 | 2.670703 | -0.42268 | -0.34813 |
| 26.01389 | 0.089505 | -0.31116 | -0.14723 | 0.199393 | 0.564847 | -0.0749 | -0.06383 | 0.259988 | -0.19032 | -1.52227 | 1.26444 | -0.30896 | 0.387173 | -0.09526 | -0.09038 | 0.597019 | 0.926944 | -0.18447 | -0.0852 | -0.02321 | 0.031288 | 4.365193 | 0.008233 | 0.166727 |
| 26.02778 | 0.441443 | -0.05536 | 0.053385 | 0.395932 | 0.407928 | 0.259876 | 0.025043 | 0.250909 | 0.502658 | -1.11748 | 2.340541 | -0.02855 | 0.126501 | 0.048136 | 0.322369 | 0.671312 | 0.954184 | -0.37168 | 0.024987 | 0.235269 | 0.067922 | 5.442283 | 0.166924 | 0.441563 |
| 26.04167 | 0.526906 | 0.125541 | 0.204513 | 0.318539 | 0.262151 | 0.144795 | 0.119237 | 0.243442 | 0.905118 | -0.1103 | 2.230622 | 0.069215 | 0.002343 | -0.03972 | 0.274333 | 0.577956 | 0.779056 | -0.56025 | 0.06824 | 0.210218 | 0.090684 | 5.217496 | 0.140263 | 0.698574 |
| 26.05556 | 0.428397 | 0.199325 | 0.189687 | 0.231375 | 0.076131 | -0.16267 | 0.225518 | 0.107924 | 1.188124 | 0.338026 | 1.565494 | 0.106555 | -0.0747 | -0.16604 | 0.24087 | 0.442068 | 0.545848 | -0.56975 | 0.057782 | 0.147514 | 0.11881 | 4.439579 | 0.085414 | 0.823083 |
| 26.06944 | 0.211663 | 0.166736 | 0.043208 | 0.120149 | -0.0952 | -0.51787 | 0.271512 | 0.123356 | 1.087062 | 0.304247 | 0.687132 | 0.085056 | -0.11438 | -0.18852 | -0.02893 | 0.586322 | 0.254359 | -0.44296 | -0.02267 | -0.03252 | 0.124342 | 2.74876 | 0.002787 | 1.185097 |
| 26.08333 | 0.026806 | 0.121188 | -0.12037 | 0.017031 | -0.21852 | -0.74694 | 0.231459 | 0.038656 | 0.472681 | 0.239633 | -0.02865 | 0.038805 | -0.08644 | -0.02765 | -0.00962 | 0.568409 | 0.113336 | -0.26497 | -0.09888 | -0.08755 | 0.164906 | 1.318921 | 0.096661 | 1.705923 |
| 26.09722 | -0.07137 | 0.099431 | -0.1625 | -0.01794 | -0.21426 | -0.50496 | 0.185084 | -0.14859 | 0.065176 | 0.200586 | -0.30097 | 0.038188 | -0.01787 | 0.259606 | 0.33745 | 0.587718 | 0.130789 | -0.10618 | -0.11616 | -0.02191 | 0.191259 | 0.26015 | 0.301779 | 1.684572 |
| 26.11111 | -0.03403 | 0.087316 | -0.0572 | 0.085666 | -0.14355 | -0.18994 | 0.181233 | -0.21017 | -0.19799 | 0.135753 | -0.07806 | 0.093439 | 0.08159 | 0.509035 | 0.50731 | 0.543971 | 0.244184 | 0.031475 | -0.03479 | 0.11077 | 0.15917 | -0.67411 | 0.602164 | 1.371745 |
| 26.125 | 0.078412 | 0.107296 | 0.112174 | 0.196206 | -0.02239 | 0.079907 | 0.196451 | -0.15953 | -0.31324 | 0.032928 | 0.431125 | 0.133688 | 0.15289 | 0.615945 | 0.556506 | 0.484461 | 0.382287 | 0.180882 | 0.089664 | 0.271569 | 0.078095 | -1.358 | 0.577578 | 0.85748 |
| 26.13889 | 0.241173 | 0.115674 | 0.216697 | 0.249823 | 0.064055 | 0.1947 | 0.160393 | -0.01019 | -0.23264 | -0.02574 | 0.858561 | 0.159501 | 0.120464 | 0.572886 | 0.600132 | 0.429137 | 0.359053 | 0.281977 | 0.206991 | 0.335579 | 0.039748 | -1.29261 | 0.648422 | 0.739645 |
| 26.15278 | 0.183716 | 0.101198 | 0.301305 | 0.233885 | 0.093361 | 0.181264 | 0.088808 | 0.176407 | -0.02595 | -0.01319 | 0.76777 | 0.20099 | 0.057969 | 0.456741 | 0.641847 | 0.372977 | 0.111266 | 0.30459 | 0.272663 | 0.232679 | 0.078324 | -0.31405 | 0.609877 | 0.788002 |
| 26.16667 | -0.08239 | 0.088523 | 0.3039 | 0.17487 | 0.086464 | 0.129949 | -0.00789 | 0.065785 | 0.075247 | -0.01515 | 0.069233 | 0.218261 | 0.066104 | 0.353071 | 0.684629 | 0.326207 | -0.18998 | 0.25529 | 0.278603 | 0.122652 | 0.120479 | 0.619564 | 0.493202 | 0.498179 |
| 26.18056 | -0.32174 | 0.070023 | 0.287992 | 0.11397 | 0.082007 | 0.15618 | -0.08713 | -0.05633 | 0.009983 | -0.00996 | -0.53679 | 0.224718 | 0.113168 | 0.269586 | 0.699893 | 0.173008 | -0.45519 | 0.201262 | 0.206396 | 0.153021 | 0.187641 | 0.336218 | 0.366248 | 0.083997 |
| 26.19444 | -0.38544 | 0.059031 | 0.25724 | 0.077558 | 0.062161 | 0.243932 | -0.12321 | -0.30326 | -0.20919 | 0.020666 | -0.56127 | 0.232089 | 0.129679 | 0.202309 | 0.673091 | -0.19992 | -0.62037 | 0.199299 | 0.119418 | -0.06026 | 0.183496 | -1.19489 | 0.202913 | -0.34674 |
| 26.20833 | -0.2605 | 0.051046 | 0.268353 | 0.021775 | 0.039544 | 0.320055 | -0.12148 | -0.35169 | -0.40137 | 0.050355 | -0.11631 | 0.253811 | 0.110768 | 0.151859 | 0.600515 | -0.52049 | -0.59387 | 0.272581 | 0.061009 | -0.1106 | 0.170519 | -3.24073 | -0.00569 | -0.75538 |
| 26.22222 | -0.08348 | 0.032828 | 0.242391 | -0.05062 | 0.003597 | 0.356439 | -0.11403 | -0.22455 | -0.36021 | 0.045876 | 0.352637 | 0.264491 | 0.044754 | 0.174544 | 0.567934 | -0.52783 | -0.39634 | 0.39257 | 0.03939 | 0.077187 | 0.097256 | -4.99124 | -0.22138 | -0.93878 |
| 26.23611 | 0.022279 | 0.014474 | 0.201775 | -0.07512 | -0.07244 | 0.352256 | -0.10141 | 0.000254 | 0.060295 | 0.060843 | 0.524935 | 0.281402 | -0.04485 | 0.254295 | 0.604977 | -0.43157 | -0.16726 | 0.525717 | 0.064966 | 0.306907 | 0.10209 | -4.51465 | -0.34961 | -0.7531 |
| 26.25 | 0.147361 | 0.038949 | 0.144884 | 0.001589 | -0.16236 | 0.398299 | -0.07217 | 0.267036 | 0.464415 | 0.128176 | 0.420708 | 0.322857 | -0.0429 | 0.3155 | 0.607408 | -0.21456 | 0.004388 | 0.541845 | 0.128485 | 0.344164 | 0.124167 | -1.10212 | -0.34359 | -0.52425 |
| 26.26389 | 0.233091 | 0.091337 | 0.102993 | 0.140427 | -0.16169 | 0.518465 | -0.07928 | 0.357058 | 0.552284 | 0.293073 | 0.279293 | 0.395061 | 0.098621 | 0.277704 | 0.590516 | 0.077517 | -0.02839 | 0.407323 | 0.176319 | 0.273569 | 0.179541 | 2.397075 | -0.25788 | -0.29678 |
| 26.27778 | 0.226679 | 0.115261 | 0.087058 | 0.276689 | -0.01081 | 0.590426 | -0.08356 | 0.447872 | 0.436277 | 0.50019 | 0.145552 | 0.44366 | 0.213681 | 0.161536 | 0.476349 | 0.284278 | -0.1742 | 0.292625 | 0.145759 | -0.07109 | 0.200076 | 3.834688 | -0.27178 | -0.3216 |
| 26.29167 | 0.242046 | 0.125883 | 0.12637 | 0.36567 | 0.125686 | 0.546806 | -0.01442 | 0.27309 | 0.237267 | 0.505872 | 0.095178 | 0.360638 | 0.22226 | -1.1E-05 | 0.430058 | 0.385101 | -0.2875 | 0.277126 | 0.107476 | -0.44898 | 0.185098 | 2.752564 | -0.18421 | -0.33931 |
| 26.30556 | 0.387496 | 0.136737 | 0.209999 | 0.499438 | 0.168159 | 0.440662 | 0.132696 | 0.02924 | -0.01462 | 0.373909 | 0.219188 | 0.317446 | 0.108156 | -0.10674 | 0.567091 | 0.30437 | -0.32829 | 0.288665 | 0.092218 | -0.55197 | 0.18412 | 0.833243 | 0.076569 | -0.23974 |
| 26.31944 | 0.695136 | 0.135684 | 0.21258 | 0.677865 | 0.173063 | 0.391743 | 0.296132 | -0.16728 | -0.25826 | 0.324894 | 0.483347 | 0.35438 | -0.01934 | -0.1024 | 0.73307 | 0.176618 | -0.21975 | 0.312995 | 0.122825 | -0.23239 | 0.152018 | -1.34798 | 0.31329 | -0.0123 |
| 26.33333 | 0.963557 | 0.118777 | 0.202052 | 0.814901 | 0.213077 | 0.465172 | 0.347034 | -0.14193 | -0.40034 | 0.412099 | 0.58806 | 0.371412 | -0.05337 | -0.00824 | 0.774372 | 0.169417 | -0.03317 | 0.360288 | 0.212032 | 0.171984 | 0.154226 | -2.777 | 0.554157 | 0.35387 |
| 26.34722 | 1.06588 | 0.094459 | 0.188669 | 0.785419 | 0.252942 | 0.626912 | 0.374127 | 0.076303 | -0.48221 | 0.609019 | 0.706669 | 0.350473 | 0.063201 | 0.215665 | 0.81926 | 0.261253 | 0.069251 | 0.405657 | 0.351446 | 0.476482 | 0.22543 | -2.24002 | 0.848006 | 0.866949 |
| 26.36111 | 0.920668 | 0.05557 | 0.182569 | 0.786968 | 0.203279 | 0.720883 | 0.346891 | 0.145511 | -0.85952 | 0.807496 | 0.753629 | 0.370261 | 0.244998 | 0.496943 | 0.903958 | 0.392107 | 0.043124 | 0.451059 | 0.429507 | 0.610584 | 0.327794 | -0.51268 | 1.230037 | 1.235845 |
| 26.375 | 0.711815 | 0.020347 | 0.181225 | 0.830323 | 0.084337 | 0.663476 | 0.350724 | 0.187228 | -0.90129 | 0.724911 | 0.4632 | 0.406987 | 0.328205 | 0.600997 | 0.96503 | 0.4634 | -0.08781 | 0.392134 | 0.31637 | 0.332182 | 0.330887 | 1.580588 | 1.437529 | 1.188007 |
| 26.38889 | 0.632496 | 0.042042 | 0.218795 | 0.915956 | 0.038516 | 0.567253 | 0.351753 | 0.384167 | -0.16216 | 0.628138 | 0.030821 | 0.442984 | 0.333251 | 0.780646 | 1.357268 | 0.55649 | -0.07981 | 0.413675 | 0.312737 | 0.014872 | 0.331647 | 3.595663 | 1.544581 | 1.130804 |
| 26.40278 | 0.787897 | 0.117547 | 0.345139 | 1.183033 | 0.151603 | 0.573098 | 0.464651 | 0.445578 | 0.7489 | 0.87139 | 0.238673 | 0.681245 | 0.517337 | 1.400517 | 2.766697 | 1.190122 | 0.302477 | 1.102847 | 0.705423 | 0.20712 | 0.565573 | 4.581096 | 1.873934 | 1.221207 |
| 26.41667 | 1.510437 | 0.262214 | 0.543556 | 2.026772 | 0.312906 | 0.80458 | 0.80092 | 0.637318 | 1.290256 | 1.724971 | 1.347709 | 1.250484 | 0.930286 | 2.001274 | 3.721314 | 1.498636 | 0.886235 | 2.31758 | 1.218548 | 0.735711 | 0.69342 | 4.235843 | 1.93403 | 1.151864 |
| 26.43056 | 2.675583 | 0.491775 | 0.674926 | 2.463683 | 0.399617 | 1.011378 | 1.062132 | 1.21637 | 0.911948 | 3.01234 | 2.829419 | 1.573864 | 1.136921 | 2.12758 | 2.988975 | 1.453422 | 1.575264 | 2.631045 | 1.138713 | 1.002165 | 0.502741 | 3.488243 | 1.119729 | 0.783193 |
| 26.44444 | 3.18704 | 0.327213 | 0.574365 | 1.833673 | 0.415643 | 0.813405 | 1.003812 | 2.162159 | 0.220167 | 4.067387 | 3.64699 | 1.520148 | 1.952994 | 2.725976 | 1.717602 | 1.295427 | 1.496739 | 2.073936 | 0.589072 | 1.220876 | 0.555997 | 3.140661 | 0.834648 | 0.533538 |
| 26.45833 | 3.257361 | 0.337048 | 0.406587 | 1.256519 | 0.486068 | 0.641009 | 0.818095 | 1.828455 | -0.53342 | 5.254834 | 2.846583 | 1.062281 | 3.079978 | 1.986161 | 1.5201 | 1.386473 | 0.67014 | 1.510339 | 0.656641 | 1.514143 | 1.066289 | 1.942885 | 0.852226 | 0.498993 |
| 26.47222 | 3.287322 | 0.431794 | 0.409867 | 1.132731 | 0.36823 | 0.37999 | 0.702197 | 1.256945 | -0.66352 | 5.07466 | 1.634323 | 0.657501 | 2.411702 | 1.458728 | 1.293762 | 0.728174 | -0.11686 | 0.632139 | 0.899619 | 1.53038 | 1.156626 | 0.457756 | 0.516831 | 0.045785 |
| 26.48611 | 2.869089 | 0.423237 | 0.471213 | 0.810615 | 0.169314 | -0.01969 | 0.507767 | 0.760287 | 1.20131 | 4.571307 | 0.411999 | 0.367284 | 1.793478 | 0.958849 | 1.085581 | 0.409831 | -0.92327 | -0.2043 | 0.858992 | 1.272506 | 0.823416 | -1.07085 | 0.264184 | -0.4333 |
| 26.5 | 2.498638 | 0.344591 | 0.539955 | 0.314958 | 0.046664 | -0.31512 | 0.307383 | 0.174595 | 1.128422 | 3.449971 | -0.68398 | 0.172313 | 1.248401 | 0.540903 | 1.107082 | 0.277429 | -1.4809 | -0.51297 | 0.569639 | 0.859477 | 0.595731 | -2.76833 | 0.3249 | -1.04334 |
| 26.51389 | 1.891807 | 0.22632 | 0.431053 | -0.11894 | -0.00853 | -0.39828 | 0.059577 | -0.40137 | 0.69696 | 1.552009 | -1.57706 | 0.076689 | 0.444106 | 0.150171 | 1.251637 | -0.2583 | -1.47454 | -0.55857 | -0.02745 | 0.438264 | 0.306261 | -3.68206 | 0.616968 | -1.39942 |
| 26.52778 | 1.234003 | 0.111326 | 0.380067 | -0.2368 | 0.000372 | -0.31967 | -0.17403 | -0.5367 | 0.200424 | -0.30411 | -2.11853 | -0.1928 | -0.31383 | -0.23128 | 1.201304 | -0.60062 | -1.27891 | -0.08045 | -0.32121 | 0.191951 | 0.361446 | -5.6889 | 0.819162 | -0.74935 |
| 26.54167 | 0.558009 | 0.027035 | 0.045398 | -0.20273 | 0.022781 | -0.10774 | -0.27795 | -0.81168 | -0.41192 | -1.90189 | -1.40314 | -0.55657 | -0.58616 | -0.71269 | 0.798193 | -0.98885 | -0.85327 | 0.567972 | -0.38662 | 0.096989 | 0.338787 | -4.40604 | 1.144032 | 0.424386 |
| 26.55556 | -0.26507 | -0.0558 | -0.00667 | -0.28236 | 0.094969 | 0.067553 | -0.02626 | -1.31636 | -0.73192 | -2.72216 | 0.120886 | -0.99545 | -0.65256 | -1.06548 | 0.563386 | -1.04466 | -0.44444 | 0.927928 | -0.47901 | 0.033733 | 0.390981 | -0.14068 | 1.364917 | 1.4244 |
| 26.56944 | -0.98077 | -0.14729 | -0.00554 | -0.30111 | 0.220377 | 0.074677 | 0.361304 | -0.65015 | 0.193789 | -3.46343 | 1.644941 | -1.28019 | -0.86803 | -1.35033 | 0.197385 | -0.69368 | -0.07402 | 1.096836 | -0.43876 | -0.0466 | 0.249479 | 3.408088 | 1.035166 | 1.632383 |
| 26.58333 | -1.73082 | -0.18606 | -0.09297 | 0.134492 | 0.430259 | 0.099922 | 0.688656 | -0.01393 | 0.554742 | -4.1139 | 2.452215 | -1.33769 | -1.7115 | -1.61141 | 0.085428 | -0.2357 | 0.205788 | 1.390957 | -0.52057 | -0.05239 | 0.301217 | 4.126234 | 0.305141 | 0.645172 |
| 26.59722 | -1.98507 | -0.21451 | 0.09926 | 0.732671 | 0.602199 | 0.037 | 0.723441 | -0.12693 | 0.046829 | -2.8555 | 1.777481 | -0.93474 | -1.65105 | -1.93805 | -0.54308 | 0.693669 | 0.045403 | 0.991774 | -0.504 | 0.066855 | 0.634593 | 3.661596 | 0.041673 | -0.02998 |
| 26.61111 | -1.77244 | -0.23663 | 0.34771 | 1.089715 | 0.527666 | -0.58101 | 0.396041 | -0.23566 | -0.60013 | -1.22281 | 0.330543 | -0.32635 | -0.8233 | -1.34612 | -0.89814 | 1.710112 | -0.53891 | 0.210159 | -0.01431 | -0.12292 | 0.677154 | 2.105456 | -0.11836 | -0.40495 |
| 26.625 | -1.35111 | -0.29353 | 0.525035 | 1.202873 | 0.282948 | -1.0442 | 0.011771 | -0.2045 | -0.42133 | -0.34542 | -0.8523 | 0.23252 | -0.15274 | -0.3565 | -0.66788 | 1.45142 | -1.06866 | -0.5174 | 0.554759 | -0.55557 | 0.584837 | 0.320792 | -0.11071 | -0.77052 |
| 26.63889 | -0.75024 | -0.30389 | 0.225609 | 0.952516 | -0.03592 | -0.91804 | -0.41772 | 0.087232 | 0.014509 | -0.08812 | -2.09073 | 0.650604 | 0.112873 | 0.62242 | -0.99824 | 0.588841 | -1.38959 | -1.50123 | 0.839639 | -0.75316 | 0.536806 | -1.37656 | -0.03968 | -0.79913 |
| 26.65278 | -1.05877 | -0.29148 | 0.292423 | 0.494514 | -0.4364 | -0.77757 | -0.70478 | 0.121192 | 0.36056 | 0.014897 | -3.18472 | 1.055178 | 0.681059 | 1.601085 | -1.21097 | -0.50316 | -1.6511 | -1.64893 | 0.647573 | -0.64807 | 0.605654 | -3.27513 | 0.134128 | -0.70779 |
| 26.66667 | -1.06508 | -0.24182 | 0.081242 | 0.088565 | -0.51255 | -0.46522 | -0.8888 | 0.152142 | 0.383533 | 0.021893 | -3.35949 | 1.01346 | 1.59375 | 2.094018 | -0.72007 | -1.65995 | -1.58431 | -1.22554 | -0.02546 | -0.78699 | 0.685314 | -3.92093 | 0.235489 | -0.86577 |
| 26.68056 | -0.46247 | -0.20396 | -0.0332 | -0.53979 | -0.40968 | -0.10899 | -0.89539 | 0.053508 | -0.09433 | -0.03937 | -2.70522 | 0.764278 | 1.005077 | 1.465705 | -0.04273 | -1.87668 | -1.26812 | -0.96931 | -0.20936 | -0.51442 | 0.670902 | -5.39158 | -0.04995 | -1.54249 |
| 26.69444 | 0.048502 | -0.18357 | 0.012245 | -0.96638 | -0.22499 | 0.117024 | -0.70259 | -0.23555 | -0.51463 | -0.11073 | -1.1613 | 0.365959 | 0.6357 | 0.855759 | 0.185718 | -1.65861 | -0.95493 | -0.42142 | -0.30461 | -0.06753 | 0.454863 | -3.39745 | -0.64541 | -1.45524 |
| 26.70833 | 0.236306 | -0.14387 | -0.09249 | -1.02772 | -0.0241 | 0.057408 | -0.3572 | -0.34234 | -0.56008 | -0.12228 | 0.310685 | -0.05105 | 0.317566 | 0.371663 | -0.17337 | -1.15457 | -0.6886 | 0.005891 | -0.28809 | 0.285053 | 0.15011 | -1.23887 | -1.27432 | -1.52539 |
| 26.72222 | 0.073229 | -0.07685 | -0.32726 | -0.71415 | 0.097901 | -0.09809 | -0.15528 | 0.235417 | -0.33463 | -0.1415 | 1.02906 | -0.38513 | -0.02735 | -0.09553 | -0.53889 | -0.58098 | -0.53711 | -0.12331 | -0.22337 | 0.442616 | 0.064851 | 0.831287 | -1.52289 | -1.07375 |
| 26.73611 | -0.50539 | -0.03056 | -0.48757 | -0.33241 | 0.198234 | -0.51091 | -0.05821 | 0.409069 | 0.162537 | -0.20372 | 1.001756 | -0.63996 | -0.66648 | -0.51171 | -0.59159 | -0.31702 | -0.65625 | -0.57619 | -0.1598 | 0.340262 | 0.118221 | 2.428753 | -1.60487 | -0.52322 |
| 26.75 | -1.12026 | 0.027479 | -0.63475 | -0.26309 | 0.196773 | -0.94992 | -0.10582 | -0.09286 | 0.168591 | -0.28262 | 0.664072 | -0.96709 | -1.34743 | -0.80506 | -0.6064 | -0.31066 | -0.9863 | -0.97378 | -0.42369 | -0.03817 | -0.11802 | 4.10648 | -1.52198 | -0.31498 |
| 26.76389 | -1.38179 | 0.072739 | -0.79804 | -0.69147 | -0.02649 | -1.15855 | -0.27695 | -0.90466 | -0.57914 | -0.34243 | -0.00412 | -0.95707 | -2.06219 | -1.00778 | -0.79305 | -0.62265 | -1.26619 | -1.23403 | -1.01081 | -0.53167 | -0.39452 | 5.430299 | -1.10779 | -0.27303 |
| 26.77778 | -2.09695 | 0.088721 | -0.91473 | -1.34647 | -0.34315 | -0.90377 | -0.48776 | -1.33861 | -1.09741 | -0.47754 | -1.04679 | -0.99432 | -2.63406 | -1.14259 | -1.25362 | -0.8201 | -1.01962 | -1.23966 | -1.42884 | -0.99176 | -0.76109 | 1.689297 | -0.62989 | -0.28673 |
| 26.79167 | -2.1643 | 0.045496 | -0.87533 | -1.8272 | -0.56254 | -0.54058 | -0.58292 | -1.38016 | -1.44575 | -0.63218 | -2.07646 | -0.95959 | -2.47916 | -1.62556 | -2.51764 | -1.02022 | -0.65798 | -1.16375 | -1.1974 | -1.26275 | -0.82083 | -2.21234 | -0.35335 | -0.40105 |
| 26.80556 | -1.7809 | -0.01442 | -0.65548 | -1.90226 | -0.51551 | -0.54954 | -0.51472 | -1.12547 | -1.56974 | -0.84389 | -2.5108 | -1.07617 | -1.80465 | -2.62975 | -2.98934 | -1.08678 | -0.25397 | -0.99874 | -0.83594 | -1.26086 | -0.55179 | -3.12361 | -0.4178 | -0.13752 |
| 26.81944 | -1.13162 | -0.06789 | -0.4892 | -1.76578 | -0.23389 | -0.80166 | -0.37918 | -0.64611 | -1.47818 | -0.77399 | -1.68561 | -0.83683 | -0.96553 | -2.39264 | -3.27057 | -1.46822 | 0.206692 | -0.81652 | -0.49731 | -1.03551 | -0.22136 | -2.49594 | -0.47375 | 0.276813 |
| 26.83333 | -0.75101 | -0.0654 | -0.3026 | -1.39382 | 0.144444 | -0.91534 | -0.15584 | -0.26307 | -1.17254 | -0.4428 | -0.25354 | -0.41599 | -0.29256 | -1.77353 | -2.81484 | -2.07024 | 0.617449 | -0.66161 | -0.25463 | -0.34733 | -0.18175 | -3.17718 | -0.40281 | 0.566077 |
| 26.84722 | -0.61205 | -0.07676 | -0.20517 | -0.83056 | 0.522492 | -0.79891 | 0.001135 | -0.06275 | -0.49706 | -0.1568 | 0.754194 | -0.05297 | 0.033147 | -1.13781 | -2.2982 | -1.51071 | 0.570482 | -0.712 | -0.06187 | 0.411663 | -0.23609 | -1.61246 | -0.22834 | 0.734397 |
| 26.86111 | -0.57159 | -0.15409 | -0.07477 | -0.26774 | 0.587428 | -0.48204 | 0.020852 | -0.06845 | 0.147742 | -0.17904 | 0.880909 | 0.085254 | 0.217398 | -0.43352 | -1.66378 | -0.2216 | 0.715206 | -0.70328 | 0.107399 | 0.885877 | -0.06025 | 0.138229 | 0.229731 | 0.747228 |
| 26.875 | -0.43313 | -0.17631 | 0.166563 | 0.038181 | 0.169195 | -0.15992 | 0.080496 | 0.045013 | 0.659061 | -0.33582 | 0.430033 | 0.137975 | 0.366425 | 0.153229 | -0.82595 | 0.704294 | 0.744249 | -0.59331 | 0.102336 | 1.002104 | 0.228321 | 1.970101 | 0.589033 | 0.774256 |
| 26.88889 | -0.3413 | -0.06003 | 0.393211 | 0.155518 | -0.25253 | 0.063646 | 0.108962 | 0.333948 | 0.869517 | -0.46583 | -0.08119 | 0.208225 | 0.279928 | 0.246419 | -0.40596 | 0.861621 | 0.403891 | -0.50783 | -0.16505 | 0.78399 | 0.436876 | 3.483272 | 0.34372 | 0.474216 |
| 26.90278 | -0.30029 | 0.006274 | 0.340154 | -0.03951 | -0.50501 | -0.05353 | -0.01059 | 0.469312 | 0.510507 | -0.52024 | -0.53106 | 0.197359 | -0.02903 | -0.14112 | -0.60453 | -0.06494 | -0.33231 | -0.56611 | -0.53692 | 0.19811 | 0.310941 | 3.239416 | -0.69798 | -0.26943 |
| 26.91667 | -0.53789 | 0.006072 | -0.03278 | -0.47701 | -0.661 | -0.58332 | -0.21016 | 0.026917 | -0.32251 | -0.48532 | -1.08364 | -0.00774 | -0.35143 | -0.55075 | -0.82892 | -1.24289 | -0.90904 | -0.56124 | -0.71319 | -0.30794 | 0.026402 | 0.21744 | -1.99433 | -1.09982 |
| 26.93056 | -1.16339 | 0.025363 | -0.45056 | -0.77785 | -0.72479 | -1.13108 | -0.29005 | -0.50337 | -1.49792 | -0.37979 | -1.76387 | -0.27017 | -0.54093 | -0.79407 | -0.8191 | -1.70934 | -1.12701 | -0.23527 | -0.64428 | -0.86703 | -0.18312 | -2.02879 | -2.50263 | -1.68089 |
| 26.94444 | -1.50555 | 0.085636 | -0.45316 | -0.93158 | -0.6103 | -1.17548 | -0.28569 | -0.34589 | -2.74853 | -0.35019 | -2.13756 | -0.41017 | -0.80904 | -1.27173 | -1.10087 | -1.53139 | -1.46284 | 0.105713 | -0.72747 | -0.83991 | -0.42074 | -2.98982 | -2.59514 | -1.93655 |
| 26.95833 | -1.89983 | -0.19567 | -0.19678 | -1.28926 | -0.58901 | -1.09209 | -0.47394 | -0.47429 | -4.1189 | -0.55164 | -1.80794 | -0.41554 | -0.88972 | -1.77642 | -1.70963 | -1.72178 | -1.3928 | 0.163471 | -1.07069 | -0.7634 | -0.40355 | -4.29409 | -2.17435 | -1.90937 |
| 26.97222 | -2.35923 | -0.71011 | -0.10703 | -2.09729 | -0.63583 | -0.97853 | -0.54282 | -0.33904 | -5.35997 | -0.69255 | -1.3021 | -0.41587 | -0.692 | -1.78008 | -1.59808 | -1.71809 | -0.8866 | -0.00212 | -0.96049 | -1.0127 | 0.110749 | -4.52895 | -1.29238 | -1.52578 |
| 26.98611 | -2.25554 | -0.8061 | -0.03521 | -2.1408 | -0.31801 | -0.60743 | -0.38478 | -0.20694 | -3.35934 | -0.57937 | -0.79752 | -0.32916 | -0.35348 | -1.35922 | -1.22557 | -1.28934 | -0.19057 | -0.21384 | -0.5854 | -0.96636 | 0.742068 | -2.75066 | -0.44409 | -0.56005 |
| 27 | -1.31499 | -0.62737 | 0.099136 | -1.46239 | 0.117601 | -0.12713 | -0.16807 | 0.198583 | 0.161622 | -0.3117 | -0.25007 | -0.12918 | 0.027726 | -0.90818 | -0.8086 | -0.87693 | 0.230418 | -0.22541 | -0.3453 | -0.56227 | 0.629245 | -0.62897 | -0.14113 | 0.102745 |
| 27.01389 | -0.45751 | -0.3921 | 0.212113 | -0.75496 | 0.358982 | 0.275503 | 0.001558 | 0.650308 | 1.809328 | -0.12894 | 0.316096 | 0.052944 | 0.242804 | -0.64696 | -0.52964 | -0.60239 | 0.538618 | -0.02986 | -0.26688 | -0.08062 | 0.280737 | 1.253003 | -0.36954 | 0.321732 |
| 27.02778 | 0.159168 | -0.1854 | 0.318415 | -0.37102 | 0.358498 | 0.50717 | 0.095323 | 0.724402 | 3.075585 | -0.05446 | 0.901444 | 0.054911 | 0.336818 | -0.58733 | -0.28979 | -0.41195 | 0.892167 | 0.110823 | -0.25133 | 0.303426 | -0.14951 | 2.559402 | -0.6414 | 0.329095 |
| 27.04167 | 0.469732 | -0.02286 | 0.455172 | -0.23311 | 0.307609 | 0.5386 | 0.083275 | 0.387425 | 2.973966 | 0.019892 | 1.54241 | -0.01428 | 0.312691 | -0.62863 | -0.15263 | -0.26554 | 1.017303 | 0.157068 | -0.219 | 0.549778 | -0.60375 | 3.322399 | -0.71682 | 0.198238 |
| 27.05556 | 0.459282 | 0.101863 | 0.654577 | -0.23382 | 0.287542 | 0.39166 | 0.036048 | 0.211218 | 2.588103 | 0.045604 | 1.700305 | -0.09273 | 0.247047 | -0.66217 | -0.0637 | -0.13526 | 0.938977 | 0.078962 | -0.02945 | 0.746328 | -1.02279 | 3.837477 | -0.71497 | -0.01328 |
| 27.06944 | 0.219345 | 0.190915 | 0.534143 | -0.25483 | 0.262914 | 0.228252 | 0.00268 | 0.177116 | 2.010005 | -0.01381 | 1.619773 | -0.11323 | 0.214247 | -0.5921 | 0.021898 | -0.02457 | 0.836329 | -0.0451 | 0.1405 | 0.807799 | -0.9043 | 4.005742 | -0.64613 | -0.16639 |
| 27.08333 | -0.12155 | 0.244518 | 0.191017 | -0.21732 | 0.236584 | 0.017353 | 0.002473 | 0.489319 | 1.262672 | -0.07466 | 1.38105 | -0.04376 | 0.181246 | -0.43878 | 0.030148 | -0.00037 | 0.716881 | -0.16284 | 0.128286 | 0.706401 | 0.127303 | 3.619483 | -0.55381 | -0.26485 |
| 27.09722 | -0.26203 | 0.312041 | 0.126707 | -0.23155 | 0.177114 | -0.07108 | 0.149656 | 0.190366 | 1.125244 | 0.138193 | 1.018193 | 0.150717 | 0.086122 | -0.00248 | -0.091 | -0.21808 | 0.217871 | -0.1169 | 0.092163 | 0.435723 | 1.275814 | 2.286508 | -0.57085 | -0.1518 |
| 27.11111 | 0.15497 | 0.50969 | 0.368359 | -0.47888 | 0.271579 | 0.237853 | 0.827064 | -1.22422 | 4.107879 | 1.385965 | 0.430392 | 0.829235 | 0.640534 | 1.427264 | -0.57421 | -1.18479 | -0.41998 | 0.399888 | 0.17698 | -0.01817 | 2.140754 | -0.53682 | -0.75752 | -0.00578 |
| 27.125 | -0.5941 | 0.201245 | 0.375747 | -0.77815 | -0.15655 | 0.125727 | 0.94316 | -1.71171 | 2.989558 | 1.172731 | 0.114027 | 0.810339 | 1.241285 | 0.50983 | -1.58112 | -1.7576 | -0.01069 | 1.252945 | -0.45947 | 0.37917 | 2.981451 | -2.40728 | -1.07594 | 0.747932 |
| 27.13889 | 0.893534 | -0.29321 | 0.260853 | 0.794875 | -0.06859 | -0.43476 | -1.5512 | 1.00078 | 0.139297 | -1.06638 | -0.37072 | -1.21639 | 0.995219 | -2.3993 | 0.478988 | 0.723702 | -1.7683 | 0.653135 | -0.23145 | 1.061604 | 0.869919 | 4.49931 | 0.81308 | 1.29025 |
| 27.15278 | 2.560261 | -0.91696 | 0.058887 | 2.641187 | 0.078218 | -0.55222 | -3.10159 | 3.227167 | -0.81445 | -3.09839 | -0.89013 | -3.07313 | -0.83007 | -3.16214 | 0.836183 | 3.094359 | -2.39639 | -0.85787 | 0.309898 | 0.418064 | -1.69134 | 3.75152 | 1.968421 | 0.29789 |
| 27.16667 | 0.672471 | -0.20137 | -0.38147 | 1.331002 | -0.32515 | -0.49565 | -1.60313 | 2.113082 | -3.17997 | -2.671 | -0.30497 | -1.55657 | -0.78129 | -1.73452 | 1.141064 | 1.510033 | 0.523194 | -0.33248 | -0.02549 | 0.540804 | -0.93849 | 1.359877 | 1.288736 | 0.213809 |
| 27.18056 | 0.521845 | 0.243108 | 0.435577 | 0.514073 | -0.19864 | 0.132843 | 0.374679 | 0.630721 | 0.760993 | 0.316703 | 0.461969 | 0.5292 | -0.04582 | -0.46324 | 0.435 | 0.560697 | 0.596475 | -0.0356 | 0.187758 | 0.273344 | 0.012932 | 0.4636 | 0.152193 | 0.04336 |
| 27.19444 | 0.238553 | 0.29857 | 0.492491 | 0.066676 | -0.01209 | 0.326999 | 0.872329 | 0.490189 | 1.811485 | 1.032535 | 0.501814 | 1.01723 | 0.044736 | -0.34578 | 0.127738 | 0.324967 | 0.253477 | -0.23572 | 0.058439 | -0.02411 | 0.395593 | -0.82483 | -0.75442 | -0.174 |
| 27.20833 | -0.11567 | 0.194919 | 0.150717 | -0.06976 | 0.014547 | 0.247235 | 0.332043 | 0.64819 | 1.072961 | 0.030401 | 0.19333 | 0.453266 | 0.047279 | -0.31896 | 0.04066 | 0.401093 | -0.02679 | -0.40991 | -0.12158 | -0.1596 | 0.562111 | -1.6689 | -0.81663 | -0.07088 |

| | | | | | | | | | | | | | | | | | | | | | | | | |
|---|---|---|---|---|---|---|---|---|---|---|---|---|---|---|---|---|---|---|---|---|---|---|---|---|
| 27.22222 | -0.19319 | 0.108693 | -0.10988 | -0.03126 | -0.07069 | 0.08459 | -0.39509 | 0.445363 | -0.27412 | -1.05067 | -0.34484 | -0.37404 | 0.124006 | -0.13732 | 0.028683 | 0.734496 | -0.13352 | -0.30857 | -0.22291 | -0.0517 | 0.454196 | -1.98038 | -0.16867 | 0.247071 |
| 27.23611 | -0.06696 | 0.016812 | -0.19168 | 0.17722 | -0.1162 | -0.01198 | -0.93386 | 0.080259 | -1.46483 | -1.57563 | -0.79039 | -0.98749 | 0.162445 | 0.093508 | 0.114065 | 0.95241 | -0.22061 | 0.017586 | -0.29812 | 0.156897 | 0.147403 | -1.91417 | 0.478568 | 0.436263 |
| 27.25 | 0.174446 | -0.07186 | 0.061556 | 0.276616 | -0.16211 | -0.02646 | -0.68774 | -0.02979 | -1.46083 | -1.01647 | -0.88043 | -0.7207 | 0.138102 | 0.345831 | 0.266621 | 1.062009 | -0.28005 | 0.300753 | -0.2987 | 0.499894 | -0.10777 | -1.37785 | 0.907384 | 0.481403 |
| 27.26389 | 0.296342 | 0.053751 | 0.548144 | 0.192282 | -0.14543 | 0.107098 | 0.14186 | 0.011019 | -0.31461 | 0.114296 | -0.50832 | 0.28927 | 0.092655 | 0.531298 | 0.44398 | 1.130696 | -0.16395 | 0.404919 | -0.12092 | 0.791432 | -0.21001 | -0.87121 | 1.144762 | 0.60036 |
| 27.27778 | 0.269017 | 0.157432 | 1.021046 | 0.120566 | -0.09353 | 0.194382 | 0.597242 | 0.115474 | 0.602895 | 0.727857 | 0.133679 | 0.695716 | 0.032225 | 0.403086 | 0.6092 | 1.141169 | 0.046619 | 0.325118 | -0.01758 | 0.779109 | -0.12263 | -0.41397 | 0.665743 | 0.613589 |
| 27.29167 | 0.338435 | 0.08152 | 0.77898 | 0.104106 | -0.1602 | 0.225662 | 0.105161 | 0.488664 | 0.627984 | 0.074578 | 0.608484 | 0.070535 | -0.12658 | -0.05549 | 0.618247 | 1.02963 | 0.148931 | 0.194689 | -0.09643 | 0.460607 | 0.173471 | -0.15973 | -0.26473 | 0.413147 |
| 27.30556 | 0.458712 | -0.03025 | 0.306063 | 0.160253 | -0.19075 | 0.310382 | -0.53621 | 0.614636 | -0.05063 | -0.78981 | 0.687719 | -0.92333 | -0.15166 | -0.40881 | 0.619614 | 0.931976 | 0.115071 | 0.004607 | -0.258 | 0.128601 | 0.583708 | 0.231236 | -0.73386 | 0.166029 |
| 27.31944 | 0.431319 | 0.034518 | 0.198868 | 0.116767 | -0.03795 | 0.42487 | -0.06597 | 0.293424 | 0.013491 | -0.35962 | 0.419973 | -0.61926 | 0.332681 | -0.00025 | 0.553318 | 1.062849 | -0.05558 | -0.1373 | -0.24031 | 0.155509 | 0.634241 | 0.973401 | -0.26603 | 0.199248 |
| 27.33333 | 0.026862 | 0.265973 | 0.221693 | 0.200745 | 0.125809 | 0.565558 | 0.929922 | -0.21574 | 0.784473 | 0.80958 | -0.18299 | 0.695027 | 0.757237 | 0.595835 | 0.507131 | 1.090174 | -0.19811 | 0.098032 | -0.02823 | 0.451651 | 0.428357 | 1.460746 | 0.607787 | 0.511408 |
| 27.34722 | 0.11132 | 0.180156 | 0.352607 | 0.595233 | 0.293575 | 0.615226 | 1.156335 | -0.13895 | 0.692131 | 1.104132 | -0.53427 | 1.053837 | 0.671958 | 0.701648 | 0.58294 | 0.813214 | -0.24197 | 0.347225 | 0.170981 | 0.713449 | 0.30299 | 1.26839 | 1.347974 | 0.732849 |
| 27.36111 | 0.561501 | 0.080836 | 0.304203 | 0.924566 | 0.31121 | 0.49456 | 0.471198 | 0.356421 | -0.47475 | 0.2939 | -0.53547 | 0.291601 | 0.413513 | 0.609957 | 0.559748 | 0.739038 | -0.25925 | 0.390541 | 0.222335 | 0.918671 | 0.166166 | 0.742241 | 1.577112 | 0.719495 |
| 27.375 | 0.821391 | -0.00951 | 0.301014 | 1.025104 | 0.203924 | 0.250699 | -0.00547 | 0.617382 | -1.1044 | -0.2181 | -0.20336 | -0.35469 | 0.270934 | 0.557032 | 0.475073 | 0.7209 | -0.19211 | 0.211225 | 0.141871 | 0.798557 | 0.046546 | 0.515145 | 0.671331 | 0.395061 |
| 27.38889 | 0.416368 | 0.078241 | 0.376148 | 0.566144 | 0.17138 | 0.381987 | 0.668675 | 0.005597 | 0.578143 | 1.342109 | -0.07944 | 0.704141 | 0.201789 | 0.509441 | 0.531493 | 0.933849 | 0.136572 | 0.052961 | 0.227532 | 0.64674 | 0.236491 | 1.28229 | -0.02741 | 0.255092 |
| 27.40278 | 0.010081 | 0.266458 | 0.448187 | 0.524761 | 0.206236 | 0.798694 | 1.801108 | -0.75934 | 2.807936 | 3.531173 | 0.168893 | 2.287891 | 0.214442 | 0.852144 | 1.047407 | 1.735346 | 0.801388 | 0.323497 | 0.83822 | 0.883308 | 0.737655 | 3.448745 | 0.614372 | 0.604873 |
| 27.41667 | 0.776092 | 0.266032 | 0.495361 | 2.039187 | 0.310493 | 1.236148 | 1.970363 | -0.17047 | 3.215963 | 4.22356 | 0.867952 | 2.254511 | 0.099377 | 1.38316 | 2.314163 | 2.274079 | 1.118112 | 1.253706 | 1.457486 | 1.106599 | 1.061689 | 5.657607 | 1.526139 | 1.119358 |
| 27.43056 | 2.3459 | 0.151526 | 0.661263 | 2.953964 | 0.420654 | 1.681478 | 1.066223 | 1.165103 | 1.621002 | 3.685701 | 1.759285 | 1.566733 | -0.12943 | 1.377859 | 3.274548 | 2.216118 | 0.881516 | 1.73975 | 1.18545 | 0.939305 | 0.969648 | 6.887108 | 1.795565 | 1.40406 |
| 27.44444 | 2.818525 | 0.11599 | 0.778809 | 2.300898 | 0.219579 | 1.36882 | 0.771171 | 1.848731 | 0.07631 | 3.033775 | 2.167439 | 1.184661 | -0.16235 | 1.140984 | 3.237231 | 2.115337 | 0.814693 | 1.757633 | 0.872913 | 0.902627 | 1.354173 | 7.075128 | 1.628651 | 1.541612 |
| 27.45833 | 3.436883 | 0.172076 | 0.817965 | 1.973254 | 0.137475 | 1.141687 | 0.721148 | 1.908776 | -0.16356 | 2.877616 | 1.845732 | 0.709658 | 0.017106 | 0.976887 | 2.862937 | 2.493801 | 0.947983 | 1.531637 | 1.199733 | 0.989103 | 1.670461 | 6.015181 | 1.853561 | 1.039404 |
| 27.47222 | 3.228105 | 0.23857 | 0.671972 | 1.704706 | 0.144053 | 0.821035 | 0.683742 | 1.368723 | 0.329304 | 3.011626 | 0.887052 | 0.395509 | 0.089848 | 0.799732 | 1.582487 | 1.856321 | 0.809485 | 0.686943 | 1.143619 | 1.043405 | 1.626455 | 4.774184 | 1.133024 | 0.195881 |
| 27.48611 | 2.115895 | 0.213686 | 0.432451 | 1.185815 | 0.095966 | 0.371275 | 0.588056 | 0.50537 | 0.734215 | 2.326038 | -0.20094 | 0.499165 | 0.101832 | 0.708603 | 0.833409 | 0.997596 | 0.727373 | 0.315383 | 0.781254 | 1.016668 | 1.272356 | 3.548496 | 0.251848 | -0.52727 |
| 27.5 | 1.385106 | 0.226883 | 0.236232 | 1.03404 | 0.119344 | 0.532406 | 0.679831 | -0.348 | 0.825423 | 2.040158 | -0.95715 | 0.71074 | 0.137882 | 0.720832 | 0.081879 | 0.485393 | 0.456744 | -0.04373 | 0.346379 | 0.699979 | 1.111963 | 1.697198 | 0.279794 | -0.81552 |
| 27.51389 | 0.44591 | 0.169298 | -0.02113 | 0.853377 | 0.253361 | 0.602254 | 0.700268 | -1.06822 | 0.917417 | 1.684462 | -1.48033 | 0.749074 | 0.174705 | 0.821524 | -0.29177 | 0.498215 | 0.069177 | -0.31872 | -0.04593 | 0.448281 | 0.749518 | -0.54888 | 0.071643 | -0.88659 |
| 27.52778 | -0.20234 | 0.062178 | -0.03122 | 0.33658 | 0.100136 | 0.344994 | 0.519816 | -1.26529 | 1.032583 | 1.412325 | -1.55945 | 0.553475 | 0.191937 | 0.906839 | 0.008249 | 1.051238 | -0.29157 | -0.17983 | -0.182 | 0.379841 | 0.296124 | -2.7675 | -0.2912 | -0.96504 |
| 27.54167 | -0.06959 | 0.016378 | -0.30313 | -0.094 | -0.09514 | 0.239436 | 0.248809 | -0.81281 | 1.22119 | 1.645152 | -0.91398 | 0.594861 | 0.277522 | 1.004115 | 1.11584 | 0.856409 | -0.51541 | -0.00083 | 0.209289 | 0.412973 | -0.18721 | -4.56472 | 0.216462 | -0.82422 |
| 27.55556 | -0.15576 | 0.014619 | -0.34369 | -0.43841 | -0.23546 | 0.229575 | -0.06197 | 0.256649 | 1.02447 | 1.434208 | 0.01647 | 0.334951 | 0.300308 | 0.98561 | 1.377763 | 0.588788 | -0.70371 | 0.268296 | 0.566331 | 0.350802 | -0.76758 | -6.3471 | 0.235588 | -0.49259 |
| 27.56944 | 0.178036 | 0.040105 | 0.088431 | -0.58077 | -0.18615 | 0.011401 | -0.2874 | 1.357866 | 0.434758 | 0.627574 | 0.998743 | -0.18307 | 0.269257 | 0.791405 | 1.130366 | 0.610036 | -0.68199 | 0.539439 | 0.607295 | 0.208389 | -1.26003 | -8.50491 | 0.123544 | 0.018728 |
| 27.58333 | 0.246867 | 0.029188 | 0.386811 | -0.35808 | 0.02844 | -0.0406 | -0.36046 | 1.514764 | 0.144428 | -0.4363 | 1.915086 | -0.62604 | 0.216043 | 0.442024 | 0.739007 | 0.504432 | -0.46983 | 0.808893 | 0.358347 | -0.01288 | -1.27619 | -6.90886 | 0.600319 | -0.24514 |
| 27.59722 | -0.55673 | -0.05195 | 0.287602 | -0.10484 | 0.336442 | 0.123164 | -0.20794 | 1.235658 | -0.11172 | -1.72447 | 1.605096 | -1.13284 | 0.229158 | 0.10089 | 0.431155 | -0.0826 | -0.35701 | 0.929624 | 0.146509 | 0.039422 | -0.68211 | -2.69339 | 0.333747 | -0.42066 |
| 27.61111 | -0.95623 | -0.11165 | 0.068215 | 0.262385 | 0.513467 | 0.179536 | 0.140116 | 0.442261 | -0.7379 | -3.01646 | 0.537059 | -1.24562 | 0.251956 | -0.20594 | 0.440361 | -0.58356 | -0.49757 | 0.462185 | -0.07924 | 0.289759 | -0.07832 | 1.528406 | -0.06143 | -0.04576 |
| 27.625 | -0.1239 | -0.09271 | -0.19303 | 0.899897 | 0.645383 | -0.19774 | 0.528735 | -0.57887 | -1.42901 | -2.62398 | -0.41711 | -0.91722 | 0.145381 | -0.33494 | 1.460854 | -0.37356 | -0.67494 | 0.751678 | -0.26145 | 0.41892 | 0.399904 | 4.475944 | -0.42742 | -0.13776 |
| 27.63889 | 0.423945 | -0.09398 | -0.34481 | 0.853077 | 0.547005 | -0.61407 | 0.712612 | -1.48025 | -1.39625 | -1.81458 | -1.24506 | -0.68331 | 0.107967 | -0.10845 | 1.438676 | 0.112774 | -0.72664 | 0.422841 | -0.24269 | 0.100896 | 0.513928 | 5.747603 | -0.00349 | -0.17497 |
| 27.65278 | 0.691102 | -0.11984 | -0.37508 | 0.983631 | 0.205353 | -0.68869 | 0.677058 | -1.45401 | -1.25406 | -1.10057 | -1.70321 | -0.46056 | 0.160431 | 0.2889 | 0.461682 | -0.28077 | -0.66388 | -0.01428 | -0.21463 | 0.132019 | -0.11563 | 6.140603 | -0.18855 | 0.125348 |
| 27.66667 | 0.784558 | -0.1358 | -0.48117 | 1.063258 | -0.09485 | -0.29505 | 0.476686 | 0.069371 | -0.20849 | -0.39493 | -2.02126 | -0.10397 | 0.156399 | 0.53836 | -1.167 | -0.61929 | -0.66623 | -0.44271 | -0.38366 | 0.02461 | -0.2805 | 4.908321 | -0.01816 | 0.311725 |
| 27.68056 | 0.646049 | -0.14545 | -0.71629 | 0.656868 | -0.20754 | -0.05898 | 0.270969 | 1.037424 | 0.582311 | 0.098917 | -1.89676 | 0.185177 | 0.114244 | 0.533687 | -2.01237 | -1.01583 | -0.77135 | -0.70557 | -0.57427 | -0.33181 | 0.198984 | 3.13557 | 0.99882 | 0.563016 |
| 27.69444 | 0.195465 | -0.18291 | -0.65678 | 0.159782 | -0.24442 | 0.046869 | -0.07335 | 0.959921 | 0.857002 | 0.150859 | -1.24514 | 0.005223 | 0.235117 | 0.434399 | -1.68506 | -1.04917 | -0.80116 | -1.00795 | -0.577 | -0.28342 | 0.198749 | 0.977943 | 1.446141 | 0.613122 |
| 27.70833 | -0.15627 | -0.16587 | -0.49341 | -0.37553 | -0.16907 | 0.013833 | -0.48452 | -0.2599 | 0.486467 | -0.14147 | -0.82148 | -0.4344 | 0.202947 | 0.272823 | -0.91003 | -0.76724 | -0.73178 | -0.77525 | -0.58071 | 0.509776 | -0.37557 | -1.45081 | 1.477545 | 0.38552 |
| 27.72222 | -0.56329 | -0.12647 | -0.57344 | -0.86082 | -0.21059 | 0.077295 | -0.93729 | -0.77205 | 0.256138 | -0.32196 | -0.5294 | -0.6853 | 0.037733 | -0.02024 | -0.59771 | -0.47705 | -0.53038 | -0.77669 | -0.5624 | 0.724199 | -0.89045 | -4.33023 | 0.536946 | -0.22729 |
| 27.73611 | -0.88717 | -0.11896 | -0.72557 | -1.21012 | -0.45822 | -0.07548 | -1.08146 | -0.84118 | -0.0755 | -0.20976 | -0.28519 | -0.41373 | -0.17679 | -0.34087 | -0.32674 | -0.46118 | -0.29347 | -1.17679 | -0.71174 | 0.502668 | -0.99428 | -6.33453 | -0.68608 | -1.10318 |
| 27.75 | -1.13446 | -0.1234 | -0.84092 | -1.2987 | -0.59265 | -0.508 | -1.1573 | -1.53355 | -0.53407 | -0.10119 | -0.12984 | -0.09164 | -0.35697 | -0.65152 | -0.00492 | -0.46432 | -0.16786 | -1.00386 | -0.63154 | 0.587443 | -0.77069 | -6.20061 | -1.4535 | -1.7218 |
| 27.76389 | -1.38702 | -0.21252 | -0.87432 | -1.13093 | -0.43073 | -0.78397 | -1.09388 | -1.58696 | -0.89015 | -0.39788 | 0.025748 | -0.16148 | -0.56642 | -0.91936 | 0.371491 | -0.51191 | -0.24838 | -0.91512 | -0.42031 | 0.342566 | -0.31736 | -6.21896 | -1.55646 | -2.14331 |
| 27.77778 | -1.71525 | -0.27806 | -0.76997 | -0.89624 | -0.25522 | -0.87581 | -0.78247 | -0.81743 | -0.9083 | -0.72023 | 0.146087 | -0.54376 | -0.77557 | -1.14794 | 0.424228 | -0.68413 | -0.42123 | -1.11719 | -0.82801 | -0.13916 | 0.135358 | -5.95371 | -2.45899 | -1.6721 |
| 27.79167 | -2.12299 | -0.28952 | -0.62317 | -0.92547 | -0.28501 | -0.85448 | -0.70351 | -0.23851 | -1.01124 | -0.7438 | 0.192302 | -0.7513 | -0.73842 | -1.03595 | -0.16467 | -0.98169 | -0.48811 | -1.35566 | -1.11991 | -0.17199 | 0.4542 | -4.47336 | -1.78971 | -0.95822 |
| 27.80556 | -2.25526 | -0.24413 | -0.45136 | -1.18924 | -0.39621 | -0.65927 | -0.62632 | 0.116849 | -0.7486 | -0.68599 | 0.094964 | -0.90271 | -0.46039 | -0.83316 | -1.14327 | -1.02051 | -0.55386 | -1.26667 | -1.4011 | -0.36375 | 0.223102 | -2.45113 | -0.87605 | -0.64523 |
| 27.81944 | -2.21831 | -0.23368 | -0.37165 | -1.24382 | -0.46904 | -0.63062 | -0.49352 | -0.29812 | -0.5315 | -0.70496 | -0.0632 | -0.70789 | -0.14953 | -0.60727 | -1.5377 | -1.09684 | -0.73488 | -0.76017 | -1.64486 | -0.27248 | 0.157497 | -0.37202 | 0.016866 | -0.16879 |
| 27.83333 | -2.04488 | -0.14862 | -0.23502 | -0.95342 | -0.4679 | -0.55808 | -0.25583 | -0.58034 | -0.42223 | -0.63008 | -0.06654 | -0.24668 | -0.09588 | -0.34389 | -0.97597 | -0.89857 | -0.79149 | -0.10699 | -1.727 | -0.00454 | 0.08026 | 1.695693 | 0.575442 | 0.568129 |
| 27.84722 | -1.56346 | 0.041013 | -0.02967 | -0.56022 | -0.27516 | -0.30214 | 0.126438 | -0.6745 | -0.50586 | -0.27402 | 0.00003 | 0.424101 | -0.05798 | -0.01033 | -0.37527 | -0.46896 | -0.57359 | 0.386402 | -1.47555 | 0.062854 | 0.130019 | 3.967569 | 0.486725 | 0.800522 |
| 27.86111 | -0.94911 | 0.167231 | 0.113767 | -0.25708 | -0.03812 | 0.00312 | 0.312289 | -0.68238 | -0.41359 | 0.061302 | 0.284044 | 0.855696 | 0.014381 | 0.134056 | 0.065689 | 0.0364 | -0.2326 | 0.565563 | -0.75289 | -0.33373 | 0.016021 | 4.002453 | 0.018475 | 0.453588 |
| 27.875 | -0.3169 | 0.219095 | 0.015154 | -0.16472 | 0.276478 | -0.01491 | 0.070545 | -0.5148 | 0.047518 | -0.45284 | 0.497205 | 0.452489 | 0.12 | 0.102587 | 0.190419 | 0.578095 | -0.02328 | 0.575604 | -0.1182 | -0.21577 | 0.157068 | 3.278802 | -0.63436 | -0.22549 |
| 27.88889 | 0.617905 | -0.02476 | -0.26688 | -0.461 | 0.493069 | -0.3516 | -0.87502 | 0.629292 | 0.800444 | -2.47726 | 0.733964 | -1.35434 | 0.082807 | -0.22683 | 0.05155 | 0.457035 | -0.19725 | 0.233423 | 0.125103 | 0.169128 | -0.12185 | 1.241027 | -1.3359 | -0.78141 |
| 27.90278 | 1.172375 | -0.34997 | -0.62217 | -0.47715 | 0.307276 | -0.74081 | -2.1586 | 1.643012 | 1.566483 | -5.17574 | 0.721166 | -3.35939 | -0.2279 | -0.83404 | -0.70693 | -0.41734 | -0.98068 | -0.70079 | -0.26552 | -0.03611 | -0.6957 | -1.66109 | -2.17002 | -1.16329 |
| 27.91667 | 0.633051 | -0.54659 | -0.72953 | -0.63935 | -0.01546 | -0.97452 | -2.10024 | 1.392519 | 0.479641 | -4.96095 | 0.118434 | -3.3335 | -0.60505 | -1.39338 | -1.26356 | -1.33131 | -2.03451 | -1.93596 | -0.82149 | -0.67029 | -1.06697 | -3.66279 | -3.44657 | -1.92538 |
| 27.93056 | -0.80294 | -0.09589 | -0.69363 | -0.86861 | -0.27288 | -0.91088 | -0.65553 | -0.01159 | -0.66654 | -2.42744 | -0.83498 | -1.0719 | -0.81936 | -1.7824 | -1.2665 | -1.65852 | -2.62135 | -2.9453 | -1.32889 | -0.94273 | -1.81534 | -3.61382 | -3.65716 | -3.08092 |
| 27.94444 | -1.43198 | -0.02878 | -0.72336 | -0.92981 | -0.23744 | -0.77912 | 0.025545 | -0.80455 | -0.58844 | -1.6565 | -1.07842 | -0.35046 | -0.73336 | -2.26749 | -0.8307 | -1.93751 | -2.69578 | -2.08504 | -1.01245 | -0.9493 | -2.75395 | -3.87091 | -1.7818 | -3.26603 |
| 27.95833 | -1.28235 | -0.6523 | -0.70388 | -1.49812 | -0.46906 | -0.62586 | -0.46612 | -0.78525 | 0.062189 | -2.32757 | -0.68144 | -0.96138 | -0.63886 | -2.17411 | -0.52913 | -2.46966 | -2.28622 | -1.29255 | -0.44709 | -1.01903 | -2.17881 | -3.98946 | 0.271953 | -1.85916 |
| 27.97222 | -1.10546 | -0.95451 | -0.46148 | -1.07208 | -0.60619 | -0.51561 | -0.73478 | -0.79514 | 0.158304 | -1.6469 | -0.3028 | -0.98741 | -0.4443 | -1.58652 | -0.31208 | -2.62008 | -1.47916 | 0.17557 | 0.05095 | -0.86237 | 0.322752 | -5.28943 | 1.695387 | 0.9065 |
| 27.98611 | -0.67718 | -0.83957 | -0.22207 | -0.40235 | -0.41486 | -0.32787 | -0.44176 | -0.70142 | -0.04402 | 0.119252 | -0.51467 | -0.24855 | -0.15522 | -0.77772 | -0.13547 | -1.5488 | -0.71712 | 1.989007 | 0.441389 | -0.35325 | 2.117722 | -4.42329 | 1.915008 | 2.934995 |
| 28 | 0.00823 | -0.43644 | -0.00593 | 0.395943 | -0.16599 | -0.05948 | -0.14154 | -0.18278 | -0.18454 | 1.276761 | -0.57992 | 0.270212 | 0.089809 | -0.34702 | 0.068583 | -0.76109 | -0.11232 | 3.972966 | 0.521486 | 0.367841 | 1.300247 | -2.38568 | 1.441369 | 3.841379 |
| 28.01389 | 0.266254 | -0.03501 | 0.149078 | 0.51978 | 0.113023 | 0.103527 | 0.280269 | 0.326747 | 0.354471 | 2.248222 | -0.37907 | 0.664017 | 0.369949 | -0.36479 | 0.085591 | -0.30042 | 0.335463 | 4.028751 | 0.500852 | 0.966752 | 0.207526 | -0.47662 | 1.29198 | 2.500768 |
| 28.02778 | 0.241601 | 0.164235 | 0.499013 | 0.388105 | 0.172507 | 0.215735 | 0.456635 | 0.633431 | 0.753713 | 1.932914 | 0.044489 | 0.479161 | 0.498283 | -0.61089 | 0.078222 | -0.16223 | 0.542477 | 2.289874 | 0.374421 | 1.085134 | 0.017463 | 1.217115 | 0.84068 | 1.55761 |
| 28.04167 | 0.482611 | 0.121696 | 0.860369 | 0.336176 | -0.04915 | 0.35579 | -0.00225 | 0.852389 | 0.620134 | 0.110272 | 0.472642 | -0.21016 | 0.466786 | -0.9288 | 0.152177 | -0.22382 | 0.52319 | 1.100656 | 0.130066 | 0.736012 | -0.0417 | 2.362115 | 0.223145 | 0.909116 |
| 28.05556 | 0.46532 | 0.182167 | 1.083626 | 0.153203 | -0.11227 | 0.440463 | 0.13472 | 0.555238 | 0.680442 | -0.60757 | 0.514422 | -0.3113 | 0.338097 | -1.16587 | 0.083654 | -0.27648 | 0.488782 | 0.120978 | 0.088206 | 0.247268 | -0.04473 | 3.044179 | -0.267 | 0.270356 |
| 28.06944 | 0.410134 | 0.243352 | 1.053268 | -0.06399 | -0.12719 | 0.415468 | 0.309498 | 0.060037 | 0.583669 | -0.54491 | 0.407264 | -0.02384 | 0.252738 | -1.22198 | 0.004331 | -0.24687 | 0.591535 | -0.44554 | 0.220692 | -0.07918 | -0.0061 | 3.482615 | -0.41231 | -0.23442 |
| 28.08333 | 0.248369 | 0.177281 | 0.679122 | -0.09997 | -0.14217 | 0.399452 | 0.235235 | -0.16391 | 0.504151 | -0.58742 | 0.358334 | 0.040806 | 0.221382 | -0.99774 | 0.076179 | -0.14562 | 0.691843 | -0.78811 | 0.314286 | -0.23916 | 0.088519 | 3.850224 | -0.25844 | -0.28244 |
| 28.09722 | 0.004641 | 0.12353 | 0.408388 | -0.15242 | -0.07483 | 0.278339 | 0.235867 | -0.31388 | 0.314924 | -0.26015 | 0.309529 | 0.263499 | 0.187075 | -0.61784 | 0.164645 | -0.01243 | 0.759928 | -0.90742 | 0.291682 | -0.36243 | 0.161974 | 4.01164 | -0.1431 | 0.015496 |
| 28.11111 | -0.00362 | -0.04369 | 0.257282 | -0.01984 | -0.08526 | 0.183402 | -0.05084 | -0.0254 | 0.258602 | -0.56234 | 0.285474 | 0.071938 | 0.146352 | -0.36074 | 0.234968 | 0.114586 | 0.87919 | -0.42444 | 0.208534 | -0.40403 | 0.223761 | 3.959686 | -0.09488 | 0.37161 |
| 28.125 | 0.168702 | -0.14966 | 0.326646 | 0.157843 | -0.06302 | 0.16563 | -0.31291 | 0.223916 | 0.226174 | -0.94076 | 0.362337 | -0.16871 | 0.168431 | -0.25668 | 0.299758 | 0.213095 | 1.077915 | 0.416873 | 0.268452 | -0.35131 | 0.247452 | 3.21209 | 0.092663 | 0.452471 |
| 28.13889 | 0.230751 | -0.06477 | 0.530105 | 0.256343 | 0.028194 | 0.158363 | -0.28575 | 0.43676 | 0.203254 | -0.68922 | 0.402714 | -0.16601 | 0.176522 | -0.26026 | 0.357863 | 0.291843 | 1.179776 | 0.851439 | 0.291385 | -0.21552 | 0.249064 | 2.608672 | 0.434449 | 0.199846 |
| 28.15278 | 0.171304 | 0.11243 | 0.708339 | 0.193131 | 0.140228 | 0.13925 | -0.06626 | 0.442687 | 0.238704 | 0.081396 | 0.365428 | 0.045579 | 0.241168 | -0.26197 | 0.344708 | 0.361692 | 1.001558 | 0.564564 | 0.24467 | -0.04094 | 0.200123 | 1.892385 | 0.661434 | -0.13932 |
| 28.16667 | -0.22662 | 0.493439 | 0.629139 | -0.18043 | 0.309657 | 0.083808 | 0.64901 | 0.049784 | 0.548324 | 2.069702 | 0.003839 | 0.880314 | 0.254479 | -0.18357 | 0.32567 | 0.432917 | 0.727615 | 0.00494 | 0.244527 | -0.01211 | 0.093899 | 1.373421 | 0.495847 | -0.29934 |
| 28.18056 | -0.44835 | 0.616861 | 0.233007 | -0.43826 | 0.365601 | 0.074966 | 0.988231 | -0.32321 | 0.874427 | 2.96787 | -0.25954 | 1.231471 | 0.269935 | -0.14081 | 0.366872 | 0.489148 | 0.518332 | -0.44657 | 0.15926 | -0.06838 | 0.042838 | 0.995592 | 0.098122 | -0.27808 |
| 28.19444 | -0.09541 | 0.362125 | 0.267529 | -0.06084 | 0.073275 | 0.082779 | 0.198433 | -0.17131 | 0.327025 | 0.823874 | -0.00246 | 0.272159 | 0.269764 | -0.21385 | 0.414988 | 0.607004 | 0.400415 | -0.76993 | 0.114472 | -0.05582 | 0.069523 | 0.613663 | -0.15934 | -0.01698 |
| 28.20833 | 0.409687 | -0.11372 | 0.535985 | 0.593627 | -0.23042 | 0.117618 | -0.90499 | 0.325341 | -0.53715 | -2.17521 | 0.372023 | -1.13622 | 0.262687 | -0.30779 | 0.433313 | 0.636864 | 0.287045 | -0.64769 | 0.143987 | 0.105887 | 0.054536 | 0.238044 | -0.13704 | 0.196794 |
| 28.22222 | 0.487479 | -0.16527 | 0.456237 | 0.958594 | -0.16671 | 0.059037 | -1.09698 | 0.577538 | -0.20285 | -2.2861 | 0.294887 | -1.2637 | 0.234042 | -0.33554 | 0.39712 | 0.585962 | 0.17844 | -0.08832 | 0.156708 | 0.319296 | -0.0483 | -0.27696 | 0.099915 | -0.06207 |
| 28.23611 | -0.14071 | 0.201897 | 0.117969 | 0.388566 | 0.2934 | -0.09841 | -0.03868 | 0.399195 | -0.16017 | 0.498828 | -0.15391 | 0.181982 | 0.206866 | -0.32183 | 0.383704 | 0.547552 | 0.142269 | 0.453512 | 0.126039 | 0.506185 | 0.07093 | -0.82584 | 0.378185 | -0.61388 |
| 28.25 | -0.56525 | 0.485455 | -0.00476 | -0.21764 | 0.55342 | -0.20447 | 0.754305 | 0.037637 | -0.15549 | 2.460685 | -0.50524 | 1.26132 | 0.202654 | -0.31902 | 0.422404 | 0.611395 | 0.181785 | 0.409812 | 0.161781 | 0.684851 | 0.31964 | -1.08661 | 0.556743 | -0.86638 |
| 28.26389 | -0.26224 | 0.353102 | 0.189742 | -0.1763 | 0.382998 | -0.05278 | 0.511819 | -0.03004 | -0.20305 | 1.625258 | -0.2404 | 0.700474 | 0.196283 | -0.2865 | 0.456278 | 0.798133 | 0.210683 | -0.04036 | 0.270331 | 0.483416 | 0.370027 | -0.98392 | 0.556215 | -0.46233 |
| 28.27778 | 0.269596 | 0.045783 | 0.334444 | 0.326879 | 0.027221 | 0.129463 | -0.26293 | 0.347804 | -0.04347 | -0.41351 | 0.194875 | -0.38477 | 0.175539 | -0.12361 | 0.425742 | 0.995935 | 0.184462 | -0.57841 | 0.269031 | 0.21596 | 0.209351 | -0.87797 | 0.513574 | 0.400582 |
| 28.29167 | 0.458657 | 0.024836 | 0.379467 | 0.533839 | -0.07253 | 0.227734 | -0.38583 | 0.588013 | 0.116597 | -0.75682 | 0.406607 | -0.50766 | 0.139826 | 0.103563 | 0.342529 | 0.99991 | 0.126718 | -1.04021 | 0.108189 | 0.018303 | 0.058642 | -0.90069 | 0.4634 | 0.784357 |
| 28.30556 | 0.163046 | 0.207215 | 0.357241 | 0.389552 | 0.152529 | 0.249973 | 0.213033 | 0.465651 | -0.08044 | 0.768005 | 0.110933 | 0.171931 | 0.104788 | 0.247439 | 0.289096 | 0.877139 | 0.083963 | -1.18163 | -0.08113 | -0.17197 | -0.1881 | -0.86015 | 0.400872 | 0.507495 |
| 28.31944 | 0.186479 | 0.268439 | 0.269712 | 0.605872 | 0.186306 | 0.266548 | 0.288544 | 0.398079 | -0.22667 | 0.884626 | -0.02699 | 0.261706 | 0.114301 | 0.237561 | 0.288559 | 0.765677 | 0.084479 | -0.82866 | -0.15964 | -0.25466 | -0.24653 | -0.50451 | 0.416262 | -0.23929 |
| 28.33333 | 0.441646 | 0.221065 | 0.285027 | 1.02679 | 0.124783 | 0.370903 | 0.074091 | 0.455887 | -0.26796 | 0.103407 | 0.127436 | -0.05163 | 0.146597 | 0.164992 | 0.351464 | 0.697444 | 0.126015 | -0.3553 | -0.00996 | 0.227894 | 0.121043 | -0.02623 | 0.54674 | -0.85135 |
| 28.34722 | 0.349964 | 0.308214 | 0.195293 | 0.643547 | 0.243195 | 0.473623 | 0.488797 | 0.358967 | -0.46389 | 0.838567 | -0.0885 | 0.376751 | 0.20846 | -0.02881 | 0.389599 | 0.840553 | 0.149225 | 0.000627 | 0.282653 | 0.580429 | 0.554269 | 0.308471 | 0.76316 | -0.73768 |
| 28.36111 | 0.238914 | 0.297253 | 0.203267 | 0.165788 | 0.253167 | 0.52078 | 0.717278 | 0.342644 | -0.52686 | 1.393542 | -0.31788 | 0.69867 | 0.233638 | -0.20676 | 0.47458 | 0.987071 | 0.101763 | 0.232257 | 0.185533 | 0.377743 | 0.75718 | 0.507111 | 0.830044 | 0.13563 |
| 28.375 | 0.258065 | 0.153568 | 0.317989 | 0.033396 | 0.045787 | 0.460784 | 0.41158 | 0.364674 | -0.23908 | 0.628614 | -0.08826 | 0.265651 | 0.221618 | -0.24871 | 0.569696 | 1.147755 | 0.096571 | 0.105607 | -0.12645 | 0.27103 | 0.354175 | 0.63997 | 0.631375 | 0.898149 |
| 28.38889 | 0.239426 | 0.107961 | 0.260171 | 0.120689 | -0.01093 | 0.392154 | 0.405396 | 0.367179 | -0.30979 | 0.831338 | -0.03318 | 0.245088 | 0.296889 | 0.02733 | 0.679629 | 1.395746 | 0.615899 | -0.02724 | -0.36756 | 0.412616 | 0.380261 | 1.006076 | 0.935627 | 1.464351 |
| 28.40278 | 0.228927 | 0.326348 | 0.119708 | 0.478042 | 0.421887 | 0.3688 | 1.16288 | 0.20371 | -0.964 | 2.416353 | -0.13975 | 0.982539 | 0.591428 | 0.8242 | 0.833282 | 2.059415 | 2.120012 | 0.77473 | 0.048548 | 0.672063 | 0.890641 | 1.503648 | 2.078051 | 2.23122 |
| 28.41667 | 0.688069 | 0.615031 | 0.382528 | 1.475146 | 0.688926 | 0.687752 | 1.63748 | 0.510203 | -1.39824 | 3.369761 | 0.244982 | 1.27996 | 0.860715 | 2.203713 | 1.571903 | 1.766731 | 2.6765 | 2.275547 | 0.854075 | 0.84561 | 1.902585 | 2.126485 | 2.282467 | 2.808994 |
| 28.43056 | 1.879638 | 0.468545 | 1.111752 | 3.161492 | 0.751759 | 1.363168 | 1.294515 | 1.215258 | 0.061273 | 2.681335 | 1.132576 | 0.8222 | 1.355217 | 3.342489 | 4.574775 | 0.652605 | 2.865942 | 2.722674 | 1.392757 | 0.108996 | 3.527346 | 3.153119 | 0.663279 | 2.425835 |
| 28.44444 | 2.864091 | 0.094937 | 1.796799 | 2.805484 | 0.558999 | 2.01767 | 0.650906 | 2.371751 | 2.139019 | 1.86813 | 1.084323 | 0.309947 | 0.998572 | 4.217212 | 11.96419 | 1.100701 | 3.146236 | 1.931054 | 0.697627 | -1.61347 | 16.08609 | 8.155815 | -1.91752 | 5.205358 |
| 28.45833 | 2.639962 | 0.033074 | 1.40772 | 0.863192 | 0.491988 | 1.578059 | 0.729599 | 2.500864 | 2.42598 | 1.190584 | 0.380256 | 0.173189 | -0.54493 | 2.510898 | NaN | -3.94272 | -0.69053 | -0.16787 | -7.5558 | -8.1599 | NaN | -14.9449 | -0.23437 | 8.937506 |
| 28.47222 | 2.053564 | 0.165794 | 0.439506 | 0.537019 | 0.541478 | 1.025444 | 0.934491 | 1.745869 | 1.065706 | 0.697056 | -0.0637 | 0.214237 | 0.380053 | 2.889321 | NaN | 4.200467 | 4.160999 | -0.06972 | 8.278029 | 6.615692 | NaN | 14.61022 | 5.212203 | -6.59119 |
| 28.48611 | 1.553881 | 0.180233 | -0.28333 | 0.887197 | 0.526916 | 0.719962 | 0.96999 | 0.906966 | -0.36004 | -0.08098 | -0.16007 | 0.356358 | -1.03979 | -4.67534 | 5.094023 | -0.62217 | 0.496112 | 0.503186 | -1.03961 | 2.08583 | -6.4189 | -5.65629 | 1.618816 | -5.58681 |
| 28.5 | 0.902235 | 0.13499 | -0.58471 | 0.775802 | 0.442696 | 0.509421 | 0.743599 | 0.144788 | -1.12821 | -0.98621 | -0.12862 | 0.41898 | -0.25205 | -1.84301 | 2.13889 | 0.646537 | 0.950312 | 0.883496 | -1.45642 | -0.72168 | -0.49476 | -0.70215 | 0.275591 | -1.83248 |
| 28.51389 | 0.481312 | 0.108929 | -0.45673 | 0.274059 | 0.327291 | 0.473792 | 0.473021 | -0.67395 | -0.86099 | -1.43644 | -0.16719 | 0.344331 | 0.205139 | 0.501139 | 0.291757 | 0.633221 | 0.048977 | 0.111626 | -0.93306 | -0.63528 | -0.34723 | 0.072982 | -0.13136 | -1.07292 |
| 28.52778 | 0.247622 | 0.109036 | -0.33219 | 0.378735 | 0.082409 | 0.286804 | 0.243855 | -1.05501 | -0.08802 | -0.99816 | -0.32584 | 0.306126 | 0.442419 | 0.816787 | -1.18847 | 0.407362 | -0.38906 | -0.63568 | -0.08822 | -0.59791 | -0.57709 | -2.34947 | 0.184834 | -0.73345 |
| 28.54167 | 0.163395 | 0.101468 | -0.11161 | 0.530743 | -0.0748 | 0.166491 | -0.07064 | -0.75719 | 0.64506 | -0.13257 | -0.5516 | 0.340205 | 0.692499 | 0.813473 | -1.11013 | 0.264152 | -0.6015 | -1.31025 | 0.920828 | -0.51822 | -0.62903 | -5.32941 | 0.531185 | -0.48153 |
| 28.55556 | 0.165094 | 0.069488 | 0.053487 | 0.752405 | -0.11607 | 0.160708 | -0.42691 | -0.13898 | 0.765365 | 0.577297 | -0.52745 | 0.290101 | 0.845534 | 0.905041 | -0.01142 | -0.18877 | -1.0382 | -1.60965 | 1.509152 | -0.34236 | -0.1413 | -7.84596 | 0.667496 | -0.25058 |
| 28.56944 | 0.336084 | 0.046138 | 0.018038 | 0.841615 | -0.08226 | 0.277536 | -0.79794 | 0.4477 | 0.566161 | 0.876399 | 0.014184 | 0.199361 | 0.558931 | 0.854888 | 0.711942 | -0.63492 | -1.67866 | -1.8731 | 1.424417 | 0.036394 | 0.575259 | -6.79939 | 0.763818 | -0.21954 |
| 28.58333 | 0.451938 | 0.042247 | 0.110181 | 0.762159 | -0.14053 | 0.358315 | -0.75298 | 1.37356 | 0.075822 | 0.699492 | 0.535372 | 0.038085 | 0.097576 | 0.646459 | 0.836311 | -0.79901 | -1.76966 | -0.97179 | 0.672264 | 0.775246 | 0.609019 | -7.84384 | 0.015695 | 0.236708 |
| 28.59722 | 0.459778 | 0.062434 | 0.256857 | 0.515215 | -0.14287 | 0.433398 | -0.62861 | 2.42952 | -0.43597 | 0.185729 | 0.640857 | -0.18055 | -0.35485 | 0.423013 | 0.517347 | -0.03749 | -1.30199 | 0.073673 | 0.063529 | 1.514976 | 0.373209 | -7.54462 | -0.42317 | 0.704992 |
| 28.61111 | 0.104702 | 0.047058 | 0.525421 | 0.434061 | -0.10319 | 0.482089 | -0.42813 | 2.787816 | -0.57641 | -0.3556 | 0.36941 | -0.37271 | -0.63821 | 0.142717 | 0.319585 | 0.238014 | -1.00948 | 1.093174 | -0.56588 | 2.020892 | -0.06707 | -4.93746 | -0.29621 | 0.774911 |
| 28.625 | -0.28819 | -0.01835 | 0.071894 | 0.135752 | 0.006138 | 0.48328 | 0.053691 | 2.692435 | -0.4801 | -0.76222 | 0.058031 | -0.51285 | -0.6739 | -0.13611 | 0.36235 | 0.48491 | -0.81169 | 1.62955 | -1.2651 | 2.094075 | -0.45704 | -1.90549 | -0.20319 | 0.40656 |
| 28.63889 | -0.62242 | -0.0677 | -0.54585 | -0.29513 | 0.142996 | 0.385381 | 0.608202 | 1.652564 | -0.19389 | -1.04783 | -0.00547 | -0.48544 | -0.2329 | -0.36282 | 0.639896 | 0.772755 | -0.52398 | 1.47814 | -1.91838 | 1.214381 | -0.58549 | -0.35462 | -0.25086 | 0.019553 |
| 28.65278 | -0.40474 | -0.0517 | -0.58522 | -0.37348 | 0.185706 | 0.266108 | 0.818951 | 0.024744 | -0.01141 | -1.34641 | 0.057858 | -0.26812 | 0.264949 | -0.53751 | 0.835985 | 1.160431 | 0.016555 | 0.691327 | -1.56345 | -0.30322 | 0.063531 | 0.277657 | -0.20332 | 0.209027 |
| 28.66667 | -0.05871 | 0.007148 | -0.6121 | 0.170029 | 0.258061 | 0.181609 | 0.854054 | -1.41838 | -0.03632 | -1.47705 | 0.057418 | -0.06672 | 0.612343 | -0.60397 | 1.601972 | 0.781594 | 0.430193 | 0.063695 | -0.91719 | -1.68598 | 0.162985 | 2.400816 | -0.00956 | -0.33635 |
| 28.68056 | 0.554624 | 0.01729 | -0.75397 | 0.761467 | 0.436744 | -0.02696 | 0.794759 | -2.31568 | -0.40333 | -1.37932 | 0.205886 | 0.042212 | 0.579894 | -0.39608 | 1.615688 | 0.28851 | 0.522734 | -0.57457 | -0.3123 | -2.40625 | 0.151778 | 6.63085 | 0.512337 | -0.68178 |
| 28.69444 | 0.735833 | -0.02253 | -0.52804 | 0.955354 | 0.553445 | -0.24637 | 0.649336 | -2.32343 | -0.70433 | -0.96037 | 0.329521 | 0.098875 | 0.619442 | -0.04899 | 0.762579 | -0.20809 | 0.32542 | -1.27351 | 0.310305 | -2.53837 | 0.211718 | 9.924224 | 0.929055 | -0.81705 |
| 28.70833 | 0.596018 | -0.05541 | -0.3299 | 0.644652 | 0.451506 | -0.45258 | 0.328429 | -1.72632 | -0.62767 | -0.72179 | -0.03677 | 0.089648 | 0.704919 | 0.327953 | -0.39844 | -0.705 | 0.073528 | -1.62511 | 0.801069 | -2.04597 | 0.268965 | 8.7225 | 1.16105 | -0.64965 |
| 28.72222 | 0.080333 | -0.09321 | -0.34471 | -0.04301 | 0.164089 | -0.46892 | 0.056831 | -1.42025 | -0.21568 | -0.75666 | -0.35725 | 0.041725 | 0.805932 | 0.586809 | -1.14637 | -1.14705 | -0.01844 | -1.29528 | 0.695603 | -1.40672 | -0.13694 | 7.286909 | 0.827511 | -0.42368 |
| 28.73611 | -0.63349 | -0.13728 | -0.33028 | -0.55016 | -0.1777 | -0.39128 | -0.06219 | -1.15876 | 0.178578 | -1.12767 | -0.56892 | -0.0655 | 0.668362 | 0.694206 | -1.20474 | -1.64952 | -0.06966 | -0.54694 | 0.424169 | -0.27834 | -0.2595 | 4.383067 | 0.533012 | -0.35573 |
| 28.75 | -1.22991 | -0.17566 | -0.20594 | -0.65443 | -0.39522 | -0.37528 | -0.17073 | -1.05559 | 0.40789 | -1.77803 | -0.567 | -0.22485 | 0.431425 | 0.671659 | -0.4801 | -2.26486 | -0.10157 | 0.174064 | -0.06887 | 0.818407 | 0.552979 | 1.335989 | 0.440097 | -0.52829 |

| | | | | | | | | | | | | | | | | | | | | | | | | |
|---|---|---|---|---|---|---|---|---|---|---|---|---|---|---|---|---|---|---|---|---|---|---|---|---|
| 28.76389 | -1.53213 | -0.24939 | -0.21576 | -0.57296 | -0.75133 | -0.39859 | -0.37199 | -1.12351 | 0.547494 | -2.41869 | -0.3766 | -0.34097 | 0.110564 | 0.527683 | -0.34847 | -2.64798 | -0.13181 | 0.213059 | -0.88166 | 1.232454 | 1.081492 | -2.67545 | 0.070674 | -0.75172 |
| 28.77778 | -1.53354 | -0.37231 | -0.35844 | -0.22367 | -0.74485 | -0.39732 | -0.51617 | -1.27638 | 0.174838 | -2.36186 | -0.21965 | -0.30423 | -0.30605 | 0.263863 | -0.22217 | -2.2573 | -0.03497 | 1.310116 | -0.98443 | 1.166909 | 0.345278 | -5.73254 | -0.73467 | -0.80612 |
| 28.79167 | -1.17831 | -0.41892 | -0.32549 | -0.45608 | -0.63937 | -0.34745 | -0.57811 | -1.25934 | -0.39216 | -1.92054 | -0.22074 | -0.15753 | -0.69322 | -0.04903 | -0.12467 | -2.41235 | 0.125937 | 0.623608 | -0.50807 | 0.848345 | -0.31671 | -8.86387 | -1.16216 | -0.70892 |
| 28.80556 | -0.82383 | -0.57297 | -0.17744 | -0.70233 | -0.49155 | -0.3371 | -0.53276 | -1.10434 | -0.51134 | -1.08649 | -0.08449 | -0.00273 | -0.96159 | -0.33218 | 0.0821 | -1.79723 | 0.359131 | 0.268759 | 0.106215 | 0.097099 | -0.83701 | -8.6649 | -1.47492 | -0.20906 |
| 28.81944 | -0.87776 | -0.40155 | 0.036367 | -0.93129 | -0.32097 | -0.28921 | -0.46428 | -0.83478 | -0.07811 | -0.28258 | 0.134896 | 0.108487 | -1.09728 | -0.50873 | 0.04863 | -1.43661 | 0.393177 | -0.01493 | 0.549951 | -0.42777 | -0.75208 | -6.76674 | -1.36971 | 0.261074 |
| 28.83333 | -0.93711 | -0.06353 | 0.0116 | -0.97154 | -0.11245 | -0.16662 | -0.38057 | -0.53299 | 0.59063 | 0.360227 | 0.377233 | 0.179716 | -0.93898 | -0.62364 | 0.088454 | -0.96145 | 0.304896 | -0.52752 | 0.775807 | -0.78139 | -0.14763 | -3.62544 | -0.89352 | 0.57902 |
| 28.84722 | -0.84082 | 0.182595 | 0.045299 | -0.81296 | -0.06176 | -0.05802 | -0.2973 | -0.20497 | 1.203646 | 0.830011 | 0.610171 | 0.268047 | -0.76365 | -0.68577 | 0.269972 | -0.48945 | 0.299351 | -0.75566 | 0.612016 | -0.89784 | 0.341092 | -2.03474 | -0.07574 | 0.649106 |
| 28.86111 | -0.58358 | 0.274201 | 0.046005 | -0.57575 | -0.15399 | 0.000771 | -0.2423 | -0.04581 | 1.479565 | 0.995486 | 0.732861 | 0.341384 | -0.74091 | -0.7642 | 0.390224 | -0.11688 | 0.406488 | -0.47169 | 0.132767 | -0.85977 | 0.693191 | -0.82436 | 0.573395 | 0.368649 |
| 28.875 | -0.2356 | 0.316361 | 0.027945 | -0.48143 | -0.26295 | 0.034703 | -0.1946 | 0.060293 | 1.364492 | 0.653271 | 0.569094 | 0.408575 | -0.73376 | -0.9737 | 0.165584 | 0.115134 | 0.383564 | -0.31875 | -0.14349 | -1.62856 | 1.052823 | 0.509675 | 0.748898 | -0.33591 |
| 28.88889 | -0.28636 | 0.254626 | -0.00099 | -0.77496 | -0.28434 | -0.02843 | -0.40289 | -0.0047 | 0.884979 | -0.32008 | 0.3471 | 0.318759 | -0.68185 | -1.25183 | -0.21392 | -0.0611 | 0.080248 | -0.43495 | -0.44958 | -1.80109 | 0.384781 | 1.115128 | 0.315871 | -1.19702 |
| 28.90278 | -0.9613 | -0.07996 | -0.06956 | -1.17472 | -0.16212 | -0.28311 | -0.61714 | -0.23458 | 0.022638 | -1.9181 | -0.01065 | -0.10366 | -0.90156 | -1.5031 | -0.92646 | -0.87704 | -0.56552 | -1.40126 | -1.07315 | -1.60216 | -0.34084 | 0.285848 | -0.63974 | -1.68315 |
| 28.91667 | -1.43175 | -0.28273 | -0.29794 | -1.24362 | 0.002243 | -0.51417 | -0.90534 | -0.61 | -1.03566 | -3.9605 | -0.40317 | -0.76125 | -1.06069 | -1.63293 | -1.46971 | -1.75101 | -1.20035 | -3.04364 | -1.4178 | -1.68386 | -0.98111 | -1.45093 | -2.08032 | -2.21023 |
| 28.93056 | -1.45217 | -0.18608 | -0.55151 | -1.75947 | 0.215086 | -0.54375 | -0.90584 | -0.67612 | -1.77693 | -5.30073 | -0.85369 | -1.45331 | -1.0972 | -1.90387 | -1.55013 | -2.05308 | -1.66791 | -4.01569 | -1.21888 | -1.45254 | -1.74856 | -2.91001 | -3.31937 | -3.52897 |
| 28.94444 | -1.58411 | -0.19287 | -0.48561 | -1.4623 | 0.189991 | -0.41274 | -0.71132 | -1.06874 | -1.71403 | -4.47663 | -0.9961 | -1.96043 | -0.99915 | -2.22795 | -1.22472 | -2.10159 | -2.1601 | -3.116 | -0.48589 | -1.24767 | -2.63951 | -4.44261 | -3.13467 | -3.94386 |
| 28.95833 | -1.65642 | -0.42998 | -0.30009 | -0.79662 | 0.054362 | -0.65966 | -0.49373 | -1.31859 | -1.4872 | -1.54731 | -0.70975 | -1.23533 | -0.72888 | -2.02674 | -0.74815 | -2.22286 | -2.51715 | -1.52377 | 0.513808 | -0.67615 | -2.92792 | -5.67138 | -1.81572 | -4.15767 |
| 28.97222 | -1.27996 | -0.32237 | -0.01988 | -0.15865 | 0.08752 | -0.59114 | -0.17303 | -1.1607 | -0.72183 | 1.378207 | -0.55522 | -0.05006 | -0.59993 | -0.88374 | -0.54858 | -2.24941 | -2.11608 | -0.05862 | 1.752786 | -0.07113 | -1.81286 | -4.93329 | 0.577916 | -1.81849 |
| 28.98611 | -0.83488 | -0.07449 | 0.335334 | 0.439066 | 0.238408 | -0.3061 | 0.056824 | -0.25079 | 0.288264 | 3.201602 | -0.50959 | 0.961671 | -0.29251 | 0.44319 | -0.76837 | -1.43227 | -1.25484 | 1.374658 | 1.079665 | 0.401983 | -0.04915 | -2.52647 | 1.96983 | 1.284242 |
| 29 | -0.29659 | 0.014986 | 0.497176 | 0.709576 | 0.327068 | -0.02401 | 0.186277 | 0.650438 | 1.33802 | 3.726008 | -0.46175 | 1.536023 | -0.02906 | 1.389161 | -0.82958 | -0.75586 | -0.5261 | 2.231573 | 0.355998 | 0.740393 | 1.118034 | -0.73381 | 2.881584 | 3.253681 |
| 29.01389 | 0.162499 | 0.155955 | 0.432052 | 0.604216 | 0.289881 | 0.177254 | 0.207344 | 0.739886 | 1.676208 | 3.356419 | -0.32595 | 1.501565 | 0.117978 | 1.299703 | -0.79616 | -0.38839 | 0.02871 | 3.001949 | -0.01926 | 0.45648 | 1.778316 | -0.09479 | 2.448002 | 3.358332 |
| 29.02778 | 0.394031 | 0.330922 | 0.372835 | 0.542035 | 0.188874 | 0.295807 | 0.227918 | 1.258169 | 1.183882 | 2.987166 | -0.17272 | 1.223024 | 0.129116 | 0.668784 | -0.28837 | -0.35185 | 0.315405 | 3.342958 | -0.11095 | 0.185251 | 1.670139 | -0.34667 | 1.943966 | 4.010339 |
| 29.04167 | 0.677139 | 0.424098 | 0.530843 | 0.572305 | 0.078243 | 0.352047 | 0.246094 | 1.116368 | 0.175535 | 2.519779 | -0.07884 | 0.825693 | 0.08171 | 0.082341 | 0.019467 | -0.38725 | 0.475064 | 3.741938 | 0.068535 | 0.259433 | 1.120733 | -0.29475 | 0.910026 | 2.45973 |
| 29.05556 | 0.807477 | 0.43888 | 0.670923 | 0.557696 | 0.036674 | 0.339858 | 0.249023 | 0.873261 | -0.24877 | 1.740524 | 0.013875 | 0.440414 | 0.009105 | -0.31685 | -0.13062 | -0.38156 | 0.561797 | 3.012735 | 0.230258 | 0.361367 | 0.398946 | -0.14039 | 0.243915 | 1.171553 |
| 29.06944 | 0.738076 | 0.497449 | 0.76353 | 0.580328 | 0.04758 | 0.290464 | 0.231608 | 0.430426 | -0.02148 | 0.959094 | 0.096794 | 0.137124 | -0.03966 | -0.60782 | -0.01993 | -0.3137 | 0.652774 | 1.642228 | 0.207831 | 0.384933 | -0.16986 | 0.082357 | -0.03468 | 0.815122 |
| 29.08333 | 0.811221 | 0.414057 | 0.607306 | 0.606094 | 0.061054 | 0.300895 | 0.247728 | 0.232274 | 0.240866 | 0.124372 | 0.168221 | -0.05922 | -0.00111 | -0.72909 | 0.176799 | -0.21842 | 0.865828 | 0.704922 | 0.151301 | 0.481891 | -0.43092 | 0.416053 | -0.16169 | 0.366071 |
| 29.09722 | 0.591648 | 0.293054 | 0.466345 | 0.678745 | 0.088484 | 0.339346 | 0.280256 | 0.113941 | 0.47056 | -0.40859 | 0.251701 | -0.14293 | 0.100723 | -0.6636 | 0.5081 | -0.07397 | 1.135379 | 0.214786 | 0.057814 | 0.5177 | -0.31905 | 0.953913 | -0.11608 | -0.04452 |
| 29.11111 | 0.381815 | 0.158017 | 0.456902 | 0.760129 | 0.09104 | 0.382366 | 0.312146 | -0.01075 | 0.625978 | -0.68856 | 0.358334 | -0.11834 | 0.220089 | -0.49488 | 0.678967 | 0.087065 | 1.421939 | -0.13581 | 0.049345 | 0.467257 | -0.1541 | 1.549862 | -0.05086 | -0.29779 |
| 29.125 | 0.212715 | 0.048673 | 0.423832 | 0.763484 | 0.048568 | 0.386814 | 0.287624 | 0.05646 | 0.748528 | -0.77665 | 0.437133 | -0.01309 | 0.307327 | -0.2129 | 0.735031 | 0.2006 | 1.532756 | -0.50498 | 0.052071 | 0.378451 | 0.162965 | 2.072712 | 0.094082 | -0.30413 |
| 29.13889 | -0.01177 | -0.03758 | 0.345139 | 0.666369 | 0.013564 | 0.358507 | 0.21403 | 0.246101 | 0.987574 | -0.4737 | 0.458776 | 0.096915 | 0.329749 | 0.015705 | 0.610811 | 0.227401 | 1.431016 | -0.61698 | 0.33313 | 0.274127 | 0.331567 | 2.454552 | 0.175996 | -0.14667 |
| 29.15278 | -0.16108 | -0.09058 | 0.307707 | 0.461145 | 0.011834 | 0.34015 | 0.104655 | 0.400904 | 0.9992 | -0.17978 | 0.375475 | 0.154903 | 0.281153 | -0.05041 | 0.481502 | 0.204667 | 1.121196 | -0.46025 | 0.417211 | 0.265464 | 0.462265 | 2.439008 | 0.241717 | 0.075466 |
| 29.16667 | -0.31521 | -0.08066 | 0.251978 | 0.258292 | -0.00711 | 0.309251 | 0.008877 | 0.336748 | 0.892411 | -0.0773 | 0.29996 | 0.18832 | 0.190476 | -0.50896 | 0.379456 | 0.127912 | 0.790862 | -0.07267 | 0.403329 | 0.331053 | 0.4626 | 2.260278 | 0.29158 | 0.231737 |
| 29.18056 | -0.41964 | -0.03389 | 0.17912 | 0.038131 | -0.08557 | 0.215177 | -0.09177 | 0.140065 | 0.629911 | -0.27677 | 0.232446 | 0.163658 | 0.059359 | -1.09714 | 0.308828 | 0.035569 | 0.53153 | 0.229119 | 0.218848 | 0.420857 | 0.442667 | 1.800692 | 0.348087 | 0.164198 |
| 29.19444 | -0.38689 | -0.11579 | 0.114824 | -0.0139 | -0.17332 | 0.106086 | -0.11792 | -0.22423 | 0.258538 | -0.58349 | 0.18102 | 0.142139 | -0.01012 | -1.48084 | 0.318841 | 0.02946 | 0.349622 | 0.293573 | 0.092882 | 0.428011 | 0.422568 | 1.518113 | 0.445337 | -0.1016 |
| 29.20833 | -0.35938 | -0.08228 | 0.12887 | 0.130666 | -0.16283 | 0.046613 | -0.0084 | -0.28343 | 0.065044 | -0.69906 | 0.175855 | 0.22216 | 0.058721 | -1.16022 | 0.386531 | 0.185038 | 0.322099 | 0.039363 | -0.02816 | 0.241104 | 0.449171 | 1.587323 | 0.495327 | -0.27164 |
| 29.22222 | -0.3259 | -0.09088 | 0.098839 | 0.419251 | -0.06949 | 0.018129 | 0.144858 | 0.081628 | 0.21425 | -0.47241 | 0.227501 | 0.386436 | 0.172463 | -0.57441 | 0.41652 | 0.455697 | 0.404422 | -0.48049 | -0.10344 | 0.03953 | 0.453022 | 1.591282 | 0.62592 | -0.28276 |
| 29.23611 | -0.31797 | -0.12689 | 0.091706 | 0.698868 | 0.006133 | 0.002404 | 0.239036 | 0.543618 | 0.520767 | 0.063365 | 0.298275 | 0.503283 | 0.280819 | -0.07241 | 0.438056 | 0.691467 | 0.467102 | -1.21375 | 0.070311 | -0.07541 | 0.415536 | 1.472329 | 0.79923 | -0.1153 |
| 29.25 | -0.14995 | -0.19499 | 0.134454 | 0.749381 | 0.007741 | 0.058928 | 0.261469 | 0.873571 | 0.814174 | 0.528598 | 0.32412 | 0.482421 | 0.335289 | 0.108863 | 0.434398 | 0.841939 | 0.425262 | -1.79226 | 0.295207 | -0.03231 | 0.365731 | 1.458715 | 0.864353 | 0.121167 |
| 29.26389 | 0.043455 | -0.13437 | 0.213948 | 0.540137 | -0.00694 | 0.180209 | 0.240003 | 0.823215 | 0.909659 | 0.722602 | 0.305708 | 0.391541 | 0.315196 | -0.28079 | 0.373988 | 0.909896 | 0.329278 | -1.9508 | 0.340744 | 0.170269 | 0.356211 | 1.559829 | 0.788052 | 0.205202 |
| 29.27778 | 0.294726 | 0.01848 | 0.265125 | 0.392596 | -0.0588 | 0.230121 | 0.188096 | 0.523167 | 0.73558 | 0.581083 | 0.281294 | 0.223924 | 0.236246 | -0.95932 | 0.355423 | 0.894655 | 0.20762 | -1.70069 | 0.144745 | 0.439108 | 0.320004 | 1.624855 | 0.593905 | 0.010144 |
| 29.29167 | 0.441429 | 0.120579 | 0.258936 | 0.269449 | -0.06426 | 0.185931 | 0.134248 | 0.294372 | 0.508543 | 0.225482 | 0.267742 | 0.044798 | 0.207615 | -1.33507 | 0.37549 | 0.802021 | 0.09171 | -0.98474 | 0.011813 | 0.671542 | 0.339419 | 1.543199 | 0.262937 | -0.30462 |
| 29.30556 | 0.464955 | 0.142731 | 0.235152 | 0.333233 | -0.0364 | 0.145973 | 0.143911 | 0.188905 | 0.342808 | -0.08266 | 0.253439 | -0.06784 | 0.279384 | -1.01415 | 0.404427 | 0.770742 | -0.00623 | -0.10395 | -0.12635 | 0.896399 | 0.360122 | 1.366783 | -0.01045 | -0.55706 |
| 29.31944 | 0.482644 | 0.161367 | 0.209182 | 0.521086 | -0.03004 | 0.140463 | 0.243572 | 0.302617 | 0.264057 | -0.16005 | 0.266216 | -0.00963 | 0.36801 | -0.54046 | 0.370848 | 0.789715 | -0.10069 | 0.121485 | -0.28789 | 0.702836 | 0.339941 | 1.216272 | 0.054772 | -0.64004 |
| 29.33333 | 0.580046 | 0.141557 | 0.108568 | 0.680715 | -0.08357 | 0.194614 | 0.418447 | 0.557764 | 0.185283 | -0.13605 | 0.298989 | 0.133929 | 0.460932 | -0.16524 | 0.340358 | 0.986496 | -0.17448 | -0.2847 | -0.36022 | 0.65251 | 0.43454 | 1.262142 | 0.479226 | -0.39078 |
| 29.34722 | 0.692435 | 0.11179 | -0.08088 | 0.780254 | -0.17313 | 0.307331 | 0.61317 | 0.790495 | 0.195701 | -0.03341 | 0.285587 | 0.256039 | 0.559667 | 0.079737 | 0.374784 | 1.313475 | -0.1721 | -0.91489 | 0.119741 | 0.70827 | 0.620307 | 1.287465 | 0.956591 | 0.150067 |
| 29.36111 | 0.730005 | 0.04066 | -0.01869 | 0.656943 | -0.2703 | 0.385863 | 0.701889 | 0.964316 | 0.309587 | 0.119198 | 0.232638 | 0.226218 | 0.589619 | 0.043858 | 0.424925 | 1.61833 | -0.13571 | -1.56141 | 0.142216 | 0.809627 | 0.744482 | 1.070844 | 1.145589 | 0.764411 |
| 29.375 | 0.746774 | 0.057064 | 0.073338 | 0.482696 | -0.30541 | 0.536456 | 0.744713 | 1.05313 | 0.525255 | 0.262168 | 0.220449 | 0.022128 | 0.549287 | -0.24503 | 0.382016 | 1.748058 | -0.1008 | -1.73651 | 0.013369 | 0.78401 | 0.734456 | 0.823602 | 1.009873 | 1.001954 |
| 29.38889 | 0.907214 | 0.22245 | 0.151142 | 0.756885 | -0.11152 | 0.866413 | 0.864465 | 1.445024 | 0.628061 | 0.579811 | 0.340632 | -0.14061 | 0.757674 | -0.24044 | 0.439911 | 2.06937 | 0.071245 | -0.58327 | -0.02449 | 0.834865 | 0.707504 | 0.774422 | 1.172537 | 0.86803 |
| 29.40278 | 1.266326 | 0.491294 | 0.309898 | 1.585021 | 0.359913 | 1.57627 | 0.952671 | 1.971444 | 1.262756 | 1.468696 | 0.574582 | -0.04717 | 1.624907 | 0.569882 | 0.800232 | 2.721583 | 0.572825 | 1.87737 | 0.223777 | 1.131819 | 0.881868 | 1.055545 | 1.979105 | 1.364429 |
| 29.41667 | 1.857868 | 0.400029 | 0.468068 | 1.926303 | 0.70739 | 1.436638 | 0.641617 | 2.676482 | 2.322134 | 2.737201 | 0.68604 | 0.209645 | 2.23364 | 1.936218 | 1.316288 | 2.807055 | 0.88436 | 4.387778 | 0.32728 | 1.313601 | 1.249285 | 1.562016 | 2.277174 | 2.665816 |
| 29.43056 | 2.004985 | 0.216881 | 0.606665 | 1.362478 | 0.929379 | 0.781931 | -0.11988 | 3.038035 | 3.035846 | 3.123491 | 0.962652 | 0.397106 | 1.531739 | 2.010297 | 1.416795 | 1.909507 | 0.443868 | 5.556637 | -0.02024 | 1.308363 | 1.483736 | 2.338786 | 2.486748 | 3.262797 |
| 29.44444 | 2.231319 | 0.215095 | 0.784462 | 0.528301 | 0.923324 | 0.448989 | -0.63522 | 3.484162 | 4.619332 | 3.190135 | 1.134231 | 0.398439 | 1.61432 | 1.040264 | 1.275074 | 2.149303 | 0.248297 | 4.477184 | -0.14219 | 1.380546 | 1.758658 | 4.525561 | 1.879387 | 2.050094 |
| 29.45833 | 2.68872 | 0.308101 | 1.292142 | 0.608608 | 0.771375 | 0.582084 | -0.43986 | 3.262922 | 3.777184 | 2.188365 | 0.967004 | 0.269309 | 2.047545 | 1.138788 | 1.401036 | 3.120529 | 0.434264 | 2.999063 | 0.059881 | 1.546312 | 2.13641 | 6.393504 | 1.275361 | 1.979675 |
| 29.47222 | 2.335526 | 0.275047 | 0.953493 | 0.974865 | 0.644095 | 0.74767 | 0.112032 | 2.216105 | 1.316455 | 1.185629 | 0.831683 | 0.233567 | 1.907279 | 1.297094 | 2.382717 | 2.531971 | 0.664332 | 1.802416 | 0.294714 | 1.35143 | 2.112143 | 10.03306 | 0.595153 | 1.214352 |
| 29.48611 | 1.857673 | 0.235669 | 0.622774 | 1.001594 | 0.438768 | 0.790625 | 0.518721 | 1.41797 | -0.99143 | 0.80456 | 0.692715 | 0.186311 | 1.161672 | 1.21704 | 2.763029 | 2.022216 | 0.565822 | 1.342085 | 0.26648 | 0.842104 | 0.623441 | 10.88591 | -0.04345 | 0.43225 |
| 29.5 | 1.098784 | 0.189925 | 0.346563 | 1.039374 | 0.139813 | 0.667401 | 0.662171 | 0.654289 | -3.12853 | 0.304736 | 0.472594 | 0.022306 | 0.397502 | 0.902544 | 3.435474 | 1.493676 | 0.346926 | 1.158329 | 0.12001 | -0.01297 | 0.790737 | 5.551563 | -0.70601 | -0.05894 |
| 29.51389 | 0.434775 | 0.103921 | 0.026637 | 1.238084 | -0.25878 | 0.402599 | 0.655204 | -0.23385 | -2.76127 | 0.488896 | 0.364893 | -0.46107 | 0.023608 | 0.505409 | 2.466204 | 1.120588 | 0.163057 | 0.180198 | 0.074658 | -0.92734 | -0.26836 | 3.376006 | -1.42428 | -0.60268 |
| 29.52778 | -0.01554 | 0.015682 | -0.22307 | 1.180331 | -0.61439 | 0.172765 | 0.580245 | -1.21633 | -1.16667 | 0.30778 | 0.430082 | -0.77298 | -0.37562 | 0.157656 | 1.551519 | 0.855076 | 0.00427 | -0.50921 | 0.167532 | -1.53735 | -3.46278 | 1.845711 | -1.16016 | -1.19457 |
| 29.54167 | 0.454127 | -0.01597 | -0.15751 | 1.07454 | -0.61565 | 0.077716 | 0.384705 | -1.95098 | 0.144624 | 0.558494 | 0.467867 | -0.72179 | -0.12131 | -0.39742 | 0.700512 | 0.54615 | -0.22666 | -1.53603 | 0.191819 | -1.41192 | -1.27975 | 0.578483 | 0.136675 | -1.55373 |
| 29.55556 | 0.598653 | -0.03563 | 0.002773 | 0.912818 | -0.38855 | 0.024395 | 0.107246 | -2.42513 | 0.953079 | 1.414943 | 0.370824 | -0.51105 | 0.190833 | -0.1664 | -0.17819 | 0.197072 | -0.51546 | -2.7583 | 0.041294 | -0.71164 | -1.15143 | -0.62463 | 1.331655 | -1.67253 |
| 29.56944 | 0.58172 | -0.08636 | 0.131667 | 0.736336 | -0.23845 | -0.08871 | -0.18282 | -2.66158 | 1.395161 | 1.572878 | 0.165588 | -0.16908 | 0.438605 | 0.819765 | -0.6264 | -0.55513 | -0.74125 | -2.84282 | -0.10724 | 0.149291 | 0.149007 | -1.93755 | 1.734858 | -0.37626 |
| 29.58333 | 0.471796 | -0.15979 | 0.187131 | 0.618522 | -0.13527 | -0.14959 | -0.23963 | -0.87734 | 1.254133 | 1.092971 | -0.05708 | 0.241285 | 0.805065 | 1.628709 | -0.18856 | -1.16504 | -0.97907 | -1.48505 | -0.41215 | 0.585597 | 1.63197 | -3.53477 | 1.784379 | 0.815334 |
| 29.59722 | 0.364422 | -0.15378 | 0.24192 | 0.510063 | -0.08883 | -0.06036 | -0.13 | 0.954224 | 0.469019 | 0.327934 | -0.19073 | 0.512829 | 1.18013 | 1.775133 | 0.502411 | -1.84877 | -1.05446 | -0.02035 | -0.39516 | 0.724881 | 0.481095 | -5.05064 | 1.437917 | 0.534733 |
| 29.61111 | 0.002562 | -0.15956 | 0.303138 | 0.310846 | -0.14025 | 0.106005 | -0.02402 | 3.383515 | -0.58891 | -0.59597 | -0.2276 | 0.52078 | 0.912423 | 1.197122 | 0.474681 | -2.60489 | -1.0997 | 1.31204 | -0.17945 | 0.843092 | 0.652761 | -8.28059 | 0.820163 | 0.254261 |
| 29.625 | -0.2508 | -0.32888 | 0.321116 | 0.181315 | -0.24271 | 0.12309 | 0.083778 | 4.657341 | -1.32763 | -1.12887 | -0.10764 | 0.183049 | 0.369475 | 0.487826 | 0.348839 | -1.15884 | -0.91021 | 1.233475 | -0.09257 | 1.1769 | -0.71922 | -9.39086 | 0.151925 | 0.157001 |
| 29.63889 | -0.2979 | -0.38582 | 0.354059 | 0.171409 | -0.16076 | 0.301757 | 0.19132 | 4.806477 | -1.23256 | -1.06534 | 0.119927 | -0.30494 | -0.32908 | 0.034746 | 0.475631 | 0.471169 | -0.61167 | 1.471102 | 0.017524 | 2.042084 | -1.16298 | -7.29048 | -0.64938 | -0.45834 |
| 29.65278 | -0.18923 | -0.20337 | 0.291012 | 0.252238 | -0.11928 | 0.53234 | 0.203263 | 4.448106 | -1.20875 | -0.87511 | 0.45038 | -0.82235 | -0.629 | -0.18866 | 0.619975 | 2.127115 | -0.43748 | 1.669155 | 0.141209 | 2.048016 | -0.74042 | -5.84089 | -0.86985 | -0.93246 |
| 29.66667 | -0.00958 | 0.034667 | 0.19866 | 0.318726 | -0.11377 | 0.591348 | 0.087662 | 3.144885 | -1.09537 | -0.78859 | 0.73686 | -1.23982 | -0.72922 | -0.35796 | 0.238868 | 3.098669 | -0.32902 | 1.810381 | 0.144724 | 1.745087 | 0.152094 | -3.48413 | -1.24492 | -0.74663 |
| 29.68056 | 0.331882 | 0.231962 | -0.18916 | 0.381349 | -0.09745 | 0.580845 | -0.09334 | 1.444195 | -0.94859 | -0.61093 | 0.784003 | -1.28838 | -0.70918 | -0.69981 | 0.956073 | 3.10038 | -0.19219 | 0.977109 | 0.111208 | 1.110233 | 1.185302 | -0.90468 | -1.066 | -1.24374 |
| 29.69444 | 0.574198 | 0.296577 | -0.60101 | 0.423578 | -0.05119 | 0.29247 | -0.30519 | 0.18774 | -0.47529 | -0.59825 | 0.687274 | -1.01363 | -0.49303 | -1.5736 | 1.618625 | 1.796889 | -0.05312 | 0.347535 | 0.109641 | 0.046117 | 0.393721 | 1.400274 | -0.22866 | -1.92986 |
| 29.70833 | 0.551441 | 0.219039 | -0.62605 | 0.322228 | -0.00715 | 0.064032 | -0.42356 | -0.89006 | -0.03592 | -0.73968 | 0.574727 | -0.57518 | -0.19715 | -2.59125 | 2.065896 | 0.88596 | -0.003 | -0.05456 | -0.08666 | -0.99791 | 0.517603 | 3.66329 | 0.790722 | -1.50867 |
| 29.72222 | 0.414415 | 0.149845 | -0.44086 | 0.120844 | 0.014002 | -0.12474 | -0.27422 | -2.50418 | -0.09232 | -1.14143 | 0.400613 | -0.45873 | -0.0125 | -2.4554 | 1.203883 | 0.220919 | -0.10977 | -1.41281 | -0.29695 | -1.27186 | -0.10807 | 4.255978 | 1.366441 | -1.06102 |
| 29.73611 | -0.21495 | -0.14425 | -0.39094 | -0.09922 | 0.016796 | -0.25067 | -0.28753 | -3.18548 | -0.73139 | -1.81077 | 0.231971 | -0.67863 | 0.025371 | -1.29376 | -0.23876 | -0.35971 | -0.23739 | -2.53179 | -0.49808 | -1.55216 | -0.65726 | 3.864314 | 1.76816 | 0.301297 |
| 29.75 | -0.82346 | -0.26905 | -0.33665 | -0.27919 | 0.006641 | -0.50162 | -0.34396 | -3.0765 | -0.78382 | -2.76062 | 0.094412 | -0.9414 | 0.192268 | -0.02259 | -1.41735 | -0.94618 | -0.33099 | -3.12848 | -0.69575 | -1.15547 | 0.050402 | 2.469882 | 1.674666 | 1.471066 |
| 29.76389 | -1.18385 | -0.22714 | -0.34672 | -0.39836 | -0.16379 | -0.70653 | -0.48831 | -2.72786 | -0.34564 | -2.8701 | 0.023544 | -0.97574 | 0.354219 | 1.06175 | -1.99024 | -1.564 | -0.44964 | -3.80981 | -0.98791 | -1.17609 | 1.104259 | 1.286455 | 0.843693 | 2.489158 |
| 29.77778 | -1.14476 | -0.12139 | -0.48631 | -0.49662 | -0.34917 | -0.95009 | -0.64145 | -2.42225 | 0.183267 | -2.71467 | -0.12367 | -0.81373 | 0.210189 | 1.753699 | -1.56651 | -2.41077 | -0.58589 | -4.0958 | -1.18252 | -0.49212 | 0.93291 | -0.26191 | -0.98563 | 2.698954 |
| 29.79167 | -1.16428 | -0.15115 | -0.70527 | -0.71774 | -0.57959 | -1.01507 | -0.32321 | -2.67377 | 0.522983 | -2.22298 | -0.43087 | -0.62354 | -0.25439 | 1.464839 | -1.56618 | -3.23523 | -0.77031 | -2.80447 | -1.03677 | -0.64529 | 0.366402 | -2.48371 | -2.83428 | 1.618634 |
| 29.80556 | -0.8745 | -0.19396 | -0.91911 | -1.07944 | -0.5612 | -0.97875 | -0.40969 | -2.17574 | 0.360108 | -1.44732 | -0.86122 | -0.11132 | -0.76691 | 0.926503 | -1.56177 | -3.49098 | -1.03082 | -1.02649 | -0.68401 | -0.62758 | -0.16237 | -4.58551 | -3.26188 | 0.117094 |
| 29.81944 | -0.60239 | -0.13669 | -0.52568 | -1.37642 | -0.41411 | -0.71146 | -0.81673 | -1.92726 | 0.342432 | -0.56265 | -1.20133 | 0.733468 | -1.04604 | 0.610509 | -0.99244 | -3.21857 | -1.0174 | 1.36707 | -0.11335 | -0.45252 | -0.03429 | -4.55327 | -2.50131 | -1.03639 |
| 29.83333 | -0.42699 | -0.15168 | 0.026995 | -1.52939 | -0.1941 | -0.34805 | -1.24787 | -2.05987 | 0.313274 | 0.106971 | -1.22064 | 1.183875 | -1.49091 | 0.378713 | -0.68356 | -2.89212 | -0.94201 | 1.602778 | 0.319343 | -0.61719 | 0.237092 | -3.74326 | -2.15878 | -1.3445 |
| 29.84722 | -0.26203 | -0.22554 | 0.56786 | -0.74941 | 0.112404 | -0.01395 | -1.1866 | -1.35939 | 0.076627 | 0.558517 | -0.35995 | 0.880466 | -1.47482 | -0.01416 | -0.58104 | -2.81063 | -0.86655 | 1.901974 | 0.294051 | -0.00332 | 0.400503 | -2.96909 | -1.49627 | -1.38699 |
| 29.86111 | -0.12638 | -0.13672 | 0.897274 | -0.83654 | 0.307972 | 0.353119 | -0.39358 | -0.66435 | 0.195709 | 0.880725 | -0.05562 | 0.4883 | -1.26538 | -0.49821 | -0.20619 | -2.24676 | -0.79116 | 2.833254 | -0.02883 | 0.474721 | 0.293747 | -0.54334 | -0.30864 | -2.50605 |
| 29.875 | 0.091503 | 0.050616 | 1.01284 | -0.7461 | 0.419706 | 0.592171 | 0.484766 | 0.153815 | 0.461425 | 0.787815 | 0.151144 | 0.334473 | -1.05578 | -1.01174 | -0.02534 | -1.37818 | -0.64169 | 4.08097 | -0.37797 | 0.781382 | 0.207804 | 0.072951 | 0.611767 | -2.3417 |
| 29.88889 | 0.171989 | 0.098342 | 1.16128 | -0.77712 | 0.299985 | 0.604327 | 0.494116 | 0.395149 | 0.617603 | 0.074532 | 0.264469 | 0.123284 | -1.26221 | -1.50479 | -0.58109 | -0.94662 | -0.63021 | 3.602456 | -0.61655 | 1.141558 | -0.13479 | 0.206226 | 0.895809 | -2.41317 |
| 29.90278 | -0.41423 | -0.11419 | 0.804145 | -0.5198 | -0.00106 | 0.198089 | -0.20362 | -0.1989 | 0.274453 | -1.34659 | -0.13089 | -0.45565 | -1.69305 | -2.15752 | -2.02413 | -0.86779 | -0.87831 | 0.930088 | -0.81013 | 0.590253 | -0.57042 | -0.2631 | -0.41672 | -2.22872 |
| 29.91667 | -1.1224 | -0.29123 | -0.04044 | -0.53357 | -0.27515 | -0.35402 | -0.44954 | -1.33569 | -0.56663 | -3.20028 | -0.7439 | -1.29524 | -1.99428 | -2.03082 | -2.2421 | -0.93573 | -1.33906 | -2.4081 | -1.55059 | -0.22222 | -0.86928 | -1.6663 | -2.34281 | -3.28841 |
| 29.93056 | -1.5067 | -0.55 | -0.65466 | -1.04508 | -0.38307 | -0.82941 | -0.21895 | -2.16066 | -1.28575 | -4.34627 | -1.08005 | -1.74344 | -1.85209 | -2.40451 | -1.67413 | -0.62144 | -1.83403 | -5.43662 | -2.12659 | -0.5469 | -1.46162 | -2.93344 | -3.8179 | -4.00151 |
| 29.94444 | -1.70026 | -0.79902 | -0.71101 | -1.37345 | -0.55007 | -1.1285 | -0.53025 | -1.87862 | -1.03525 | -3.07297 | -0.99134 | -1.63086 | -1.5923 | -2.87717 | -1.72677 | -0.41251 | -2.22797 | -6.61204 | -1.9519 | -2.54491 | -1.56479 | -4.40702 | -3.28387 | -3.09651 |
| 29.95833 | -1.96498 | -1.1657 | -0.47645 | -1.63542 | -0.64418 | -1.18175 | -0.6567 | -1.58702 | -0.73512 | -1.52247 | -0.74922 | -1.21471 | -1.59598 | -3.17986 | -1.08864 | -0.60242 | -1.79306 | -6.0814 | -1.07672 | -2.75201 | -1.19551 | -5.46514 | -2.10898 | -3.83952 |
| 29.97222 | -1.8678 | -1.07076 | -0.2005 | -1.55105 | -0.51591 | -0.88356 | -0.29199 | -1.67325 | -0.46339 | -0.625 | -0.6178 | -0.34484 | -0.84966 | -1.97419 | -0.29544 | -0.57464 | -1.02106 | -3.54223 | -0.5994 | -1.73548 | -1.16183 | -4.97432 | -0.88543 | -4.38768 |
| 29.98611 | -1.35301 | -0.46468 | 0.084831 | -1.18921 | -0.3306 | -0.70577 | -0.03935 | -1.56738 | -0.21234 | -0.24382 | -0.26381 | 0.492856 | -0.15245 | 0.013446 | -0.39376 | -0.62751 | -0.29721 | -2.02467 | 0.05978 | -0.57315 | -1.01801 | -2.03944 | -0.17729 | -1.70448 |
| 30 | -0.50312 | 0.218825 | 0.314273 | -0.78029 | -0.20185 | -0.53017 | 0.052745 | -1.03777 | 0.089394 | 0.074445 | 0.01974 | 1.131385 | 0.419744 | 1.522564 | 0.086082 | -0.52217 | -0.21576 | -1.12132 | 1.004589 | 0.438194 | -0.26567 | -0.79491 | 0.922097 | 1.365861 |
| 30.01389 | 0.157743 | 0.841846 | 0.413725 | -0.452 | -0.04172 | -0.32256 | 0.031003 | -0.4504 | 0.376172 | 0.661328 | 0.161192 | 1.572758 | 0.530203 | 1.999928 | 0.827465 | -0.35974 | -0.44852 | -0.27593 | 1.869215 | 0.679577 | 0.472501 | -0.47407 | 1.805464 | 3.379934 |
| 30.02778 | 0.631879 | 0.476394 | 0.48347 | -0.10973 | 0.161513 | -0.10027 | -0.0046 | -0.17521 | 0.52325 | 1.322986 | 0.239163 | 1.499088 | 0.550357 | 1.880833 | 1.153354 | -0.13372 | -0.36867 | 0.271599 | 2.338319 | 0.971691 | 0.929433 | -0.34126 | 2.650561 | 3.454696 |
| 30.04167 | 0.925787 | 0.615461 | 0.576654 | 0.374849 | 0.300537 | 0.069122 | 0.206927 | -0.18507 | 0.526344 | 2.947699 | 0.238847 | 1.648805 | 0.566241 | 1.57016 | 1.257364 | 0.065999 | 0.049361 | 0.724472 | 2.002093 | 0.878665 | 1.138743 | -0.18528 | 3.138815 | 2.622195 |
| 30.05556 | 0.851035 | 0.537673 | 0.591385 | 0.806355 | 0.359953 | 0.141226 | 0.539557 | -0.18239 | 0.592308 | 2.456326 | 0.186096 | 1.447768 | 0.847345 | 1.224245 | 1.1562 | 0.169864 | 0.269621 | 0.919989 | 1.236859 | 2.084166 | 0.856372 | -0.00528 | 2.752589 | 2.475058 |
| 30.06944 | 0.591615 | 0.378503 | 0.577902 | 0.990732 | 0.356157 | 0.155767 | 0.657487 | -0.04253 | 0.683573 | 1.852918 | 0.2735 | 1.199112 | 0.724086 | 0.925225 | 0.835352 | 0.239064 | 0.191865 | 1.287099 | 0.819346 | 2.333111 | 0.82978 | 0.16128 | 1.921484 | 1.906578 |
| 30.08333 | 0.317523 | 0.523408 | 0.563927 | 0.995383 | 0.340973 | 0.137801 | 0.448184 | 0.163986 | 0.831568 | 1.932213 | 0.520508 | 0.490335 | 0.589842 | 0.660601 | 0.224392 | 0.283543 | 0.105573 | 2.053653 | 0.195786 | 1.177443 | 0.845219 | 0.068116 | 1.338345 | 1.385164 |
| 30.09722 | 0.154078 | 0.55241 | 0.433322 | 1.036196 | 0.282239 | 0.153058 | 0.321114 | 0.35846 | 1.071448 | 1.394701 | 0.784637 | -0.05438 | 0.535424 | 0.468169 | 0.292441 | 0.379382 | 0.084643 | 2.379324 | -0.0869 | -0.15341 | 0.479781 | -0.18833 | 0.257186 | 0.935412 |
| 30.11111 | 0.576741 | 0.330071 | 0.416821 | 0.602963 | 0.231895 | 0.151641 | 0.66671 | 0.678106 | 1.079762 | 0.62518 | 0.656472 | -0.40085 | 0.498676 | 0.295144 | 0.835868 | 0.42621 | 0.102532 | 2.474595 | -0.20121 | -0.12147 | 0.405825 | -0.46804 | -0.44071 | 0.860567 |
| 30.125 | 1.064791 | 0.304644 | 0.654264 | -0.04321 | 0.187548 | 0.171718 | 0.471848 | 1.08476 | 1.030472 | 0.030722 | 0.701322 | -0.42763 | 0.53185 | 0.171069 | 0.915482 | 0.481852 | 0.12772 | 1.91559 | -0.25295 | -0.09536 | 0.168128 | -0.51379 | -0.75674 | 0.790572 |
| 30.13889 | 1.300399 | 0.24397 | 0.88446 | -0.22625 | 0.153024 | 0.298318 | 0.248516 | 1.573877 | 1.090713 | -0.06896 | 0.826646 | -0.28955 | 0.696378 | 0.193746 | 0.571596 | 0.475275 | 0.199547 | 1.491444 | -0.29404 | 0.232909 | -0.13314 | -0.36464 | -0.69255 | 0.653223 |
| 30.15278 | 1.086489 | 0.174956 | 1.098141 | -0.25425 | 0.132093 | 0.49701 | 0.149672 | 1.939932 | 1.150512 | 0.208071 | 0.63412 | -0.07083 | 0.826734 | 0.233778 | 0.428376 | 0.406954 | 0.362189 | 1.02376 | -0.3395 | 0.389424 | 0.063668 | -0.2344 | -0.20533 | 0.490749 |
| 30.16667 | 0.628007 | 0.05899 | 1.065807 | -0.12494 | 0.096197 | 0.589338 | 0.138312 | 1.716899 | 0.960161 | 0.53767 | 0.589673 | 0.121245 | 0.781797 | 0.219195 | 0.420914 | 0.336886 | 0.453472 | 0.562619 | -0.3497 | 0.364777 | 0.269653 | -0.22029 | 0.326573 | 0.388752 |
| 30.18056 | 0.278987 | -0.11371 | 0.842375 | -0.01035 | 0.043605 | 0.505868 | 0.108169 | 1.227419 | 0.856314 | 0.640332 | 0.499511 | 0.140633 | 0.608171 | 0.102527 | 0.39378 | 0.270039 | 0.426822 | 0.218909 | -0.31562 | 0.204068 | 0.305963 | -0.23706 | 0.53224 | 0.238906 |
| 30.19444 | -0.00362 | -0.35026 | 0.505619 | 0.113832 | 0.017515 | 0.315934 | 0.05868 | 0.806404 | 0.664925 | 0.336991 | 0.407279 | -0.01694 | 0.363536 | -0.13075 | 0.42174 | 0.257662 | 0.408923 | 0.014681 | -0.14182 | 0.01224 | 0.232854 | -0.22583 | 0.396606 | -0.03778 |
| 30.20833 | -0.21807 | -0.55212 | 0.212122 | 0.21927 | 0.007027 | 0.028751 | -0.03875 | 0.415826 | 0.392968 | -0.33824 | 0.247855 | -0.22246 | 0.086795 | -0.36406 | 0.505388 | 0.314182 | 0.512624 | -0.16062 | -0.01019 | -0.15043 | 0.144801 | -0.21764 | 0.12269 | -0.62013 |
| 30.22222 | -0.13038 | -1.03742 | -0.04035 | 0.255346 | -0.02342 | -0.27754 | -0.16884 | -0.00253 | 0.142356 | -0.90633 | 0.139146 | -0.42892 | 0.003006 | -0.22969 | 0.449299 | 0.33403 | 0.721744 | -0.39296 | -0.03391 | -0.15046 | 0.100768 | -0.12415 | -0.21554 | -1.52279 |
| 30.23611 | 0.010213 | -1.42749 | -0.14182 | 0.245322 | -0.02985 | -0.35184 | -0.26972 | -0.23322 | 0.068828 | -0.96077 | 0.173004 | -0.66907 | 0.156009 | 0.106471 | 0.152804 | 0.3031 | 0.928229 | -0.577 | -0.22773 | -0.10088 | -0.0537 | 0.066894 | -0.34522 | -1.90539 |
| 30.25 | 0.125967 | -1.17624 | -0.10272 | 0.226892 | 0.005213 | -0.14241 | -0.14333 | -0.07985 | 0.250886 | -0.56261 | 0.365195 | -0.52787 | 0.459272 | -0.64443 | -0.0254 | 0.13617 | 0.751778 | -0.45973 | -0.31701 | 0.243097 | -0.20534 | 0.261587 | -0.26975 | -1.28925 |
| 30.26389 | 0.39405 | -1.15521 | -0.01093 | 0.154251 | 0.082557 | 0.188261 | 0.160006 | 0.372263 | 0.579131 | 0.121752 | 0.521699 | -0.2035 | -0.003 | -0.85499 | 0.029346 | 0.193348 | 0.569373 | -0.19305 | -0.12375 | 0.645329 | -0.08183 | 0.390553 | 0.498193 | -0.03285 |
| 30.27778 | 0.574639 | -1.12935 | 0.126581 | 0.111546 | 0.121781 | 0.387041 | 0.361857 | 0.818854 | 0.514811 | 0.618696 | 0.607285 | 0.080654 | 0.132838 | -0.37368 | 0.162487 | 0.260334 | 0.615401 | -0.26039 | 0.22179 | 0.863736 | 0.009447 | 0.475077 | 1.108694 | 1.52594 |
| 30.29167 | 0.371846 | -1.1403 | 0.157421 | 0.076732 | 0.149642 | 0.446585 | 0.487661 | 1.103715 | 0.753915 | 0.823577 | 0.500929 | 0.146016 | 0.346723 | -0.11656 | 0.172368 | 0.309385 | 0.567202 | -0.45401 | 0.542015 | 0.8671 | 0.101832 | 0.385755 | 1.538414 | 2.449405 |

| | | | | | | | | | | | | | | | | | | | | | | | | |
|---|---|---|---|---|---|---|---|---|---|---|---|---|---|---|---|---|---|---|---|---|---|---|---|---|
| 30.30556 | 0.259399 | -0.75455 | 0.168247 | 0.079812 | 0.170464 | 0.421582 | 0.534564 | 1.18031 | 1.020594 | 0.896815 | 0.527133 | -0.03377 | 0.365593 | -0.24243 | 0.233997 | 0.375394 | 0.428902 | -0.16884 | 0.493153 | 0.727676 | 0.17756 | 0.21242 | 1.575566 | 2.217627 |
| 30.31944 | 0.17559 | -0.29725 | 0.132699 | 0.118224 | 0.173034 | 0.402507 | 0.377956 | 1.265314 | 1.168395 | 0.750217 | 0.556571 | -0.31423 | 0.207462 | -0.52314 | 0.304671 | 0.434829 | 0.342643 | 0.249732 | 0.370727 | 0.582202 | 0.219011 | 0.073597 | 1.09705 | 1.346651 |
| 30.33333 | 0.161372 | 0.062617 | 0.166037 | 0.224747 | 0.124381 | 0.398938 | -0.10516 | 1.371165 | 1.141077 | 0.180258 | 0.572914 | -0.4431 | 0.013248 | -0.81127 | 0.478439 | 0.423586 | 0.3287 | 0.574841 | 0.225775 | 0.46118 | 0.283589 | 0.060365 | 0.451469 | 0.585231 |
| 30.34722 | 0.37499 | 0.258553 | 0.1806 | 0.31336 | 0.055112 | 0.377873 | -0.67812 | 1.291003 | 0.998712 | -0.56376 | 0.480845 | -0.36723 | -0.13144 | -0.98327 | 0.61001 | 0.427069 | 0.338955 | 0.657344 | 0.040347 | 0.360031 | 0.315578 | 0.191349 | -0.34127 | 0.09053 |
| 30.36111 | 0.559226 | 0.314627 | 0.173353 | 0.346199 | -0.04817 | 0.290674 | -0.90927 | 1.236674 | 0.847145 | -0.7297 | 0.475951 | -0.10446 | -0.10678 | -0.95781 | 0.594801 | 0.391721 | 0.343659 | 0.450387 | -0.21637 | 0.421306 | 0.358741 | 0.329311 | -0.4262 | -0.16367 |
| 30.375 | 0.751641 | 0.446816 | 0.234846 | 0.23942 | -0.10325 | 0.296847 | -0.51443 | 1.315955 | 0.87752 | -0.18256 | 0.53114 | 0.143231 | 0.11286 | -0.84932 | 0.497737 | 0.359952 | 0.372273 | 0.131505 | -0.2642 | 0.671287 | 0.279478 | 0.382258 | -0.00452 | -0.1536 |
| 30.38889 | 0.947884 | 0.800628 | 0.39184 | 0.297341 | -0.00132 | 0.583827 | 0.479306 | 1.738543 | 1.231257 | 0.883961 | 0.694981 | 0.255793 | 0.594181 | -0.17664 | 0.400771 | 0.448026 | 0.60596 | 0.48832 | 0.22204 | 0.977793 | 0.24117 | 0.679642 | 0.890033 | 0.414716 |
| 30.40278 | 1.54634 | 1.496135 | 0.819973 | 0.752871 | 0.214355 | 1.444509 | 1.791353 | 2.537932 | 1.988701 | 2.103579 | 1.027766 | 0.524486 | 1.324161 | 1.331849 | 0.453421 | 0.748411 | 1.166137 | 0.977399 | 0.386715 | 1.47933 | 0.336665 | 1.819445 | 2.469466 | 1.463411 |
| 30.41667 | 1.932936 | 2.57282 | 1.243731 | 1.69141 | 0.279567 | 2.450588 | 2.938969 | 2.439629 | 2.497802 | 2.392081 | 1.010823 | 1.27264 | 2.140263 | 2.242802 | 0.468885 | 1.123045 | 1.666174 | 1.150441 | -0.08725 | 1.254303 | 0.395085 | 3.929435 | 3.558732 | 2.044037 |
| 30.43056 | 0.763588 | 3.472127 | 1.58864 | 1.22428 | 0.171877 | 1.750602 | 2.382901 | 2.395327 | 4.09507 | 2.464847 | 0.809902 | 2.125829 | 2.476114 | 1.590093 | 0.283914 | 1.37913 | 1.513534 | 1.330252 | -0.24524 | 0.720383 | 0.408391 | 4.171413 | 3.465371 | 1.769445 |
| 30.44444 | 2.154908 | 3.13085 | 1.339907 | 0.451694 | 0.240316 | 1.8127 | 2.037445 | 2.492028 | 5.255082 | 1.662699 | 0.455259 | 1.93519 | 2.895462 | 1.622669 | 0.174393 | 2.582335 | 1.571963 | 2.203158 | 0.284213 | 0.274501 | 0.473586 | 3.82126 | 2.744625 | 1.811324 |
| 30.45833 | 1.920394 | 2.762545 | 1.119098 | 0.765192 | 0.284137 | 1.573782 | 1.485304 | 3.204777 | 5.155907 | 1.331071 | 0.653645 | 1.107675 | 3.366405 | 1.541419 | 0.449087 | 4.370419 | 2.253558 | 2.949195 | 0.856514 | 0.398155 | 1.312102 | 4.06773 | 2.578913 | 2.082096 |
| 30.47222 | 1.630856 | 2.031318 | 1.031299 | 1.128659 | 0.26878 | 1.046803 | 0.92509 | 3.126364 | 2.668378 | 1.031734 | 0.9054 | 0.433829 | 2.709015 | 1.143412 | 1.001894 | 4.622238 | 2.430104 | 2.431626 | 0.958234 | 0.41813 | 1.081259 | 4.431518 | 2.01677 | 1.783652 |
| 30.48611 | 1.435743 | 1.412449 | 0.588147 | 1.257765 | 0.191053 | 0.300534 | 0.227804 | 2.860044 | -0.14498 | 0.622197 | 0.946891 | -0.03294 | 2.011437 | 0.348455 | 1.36211 | 3.748402 | 2.088471 | 1.973664 | 0.856627 | 0.445837 | 0.568924 | 4.364535 | 1.56174 | 1.284619 |
| 30.5 | 0.79654 | 0.936661 | 0.063691 | 1.27251 | 0.201346 | -0.14918 | -0.23748 | 2.751298 | -1.87529 | 0.410383 | 0.809648 | -0.40015 | 1.771442 | 0.064115 | 1.664495 | 0.7776 | 1.896898 | 1.080089 | 0.659792 | 0.573437 | 0.123002 | 4.660263 | 0.974121 | 0.377194 |
| 30.51389 | -0.25987 | 0.493341 | -0.33626 | 1.180996 | 0.231443 | -0.17262 | -0.37935 | 2.328284 | -2.58956 | 0.179773 | 0.450572 | -0.76915 | 1.321094 | -0.05364 | 1.720207 | 1.13083 | 1.673648 | 1.08444 | 0.598013 | 0.389085 | 0.45028 | 4.133673 | 0.380919 | -0.08321 |
| 30.52778 | -1.00135 | 0.067563 | -0.61954 | 1.085017 | 0.221247 | 0.14206 | -0.59503 | 2.018558 | -2.90043 | 0.085586 | 0.182845 | -0.87753 | 0.409242 | -0.61388 | 1.809661 | 1.627675 | 1.772138 | 0.884864 | 0.870259 | 0.136203 | 0.526422 | 3.584728 | -0.20489 | -0.50247 |
| 30.54167 | -0.82512 | -0.231 | -0.6064 | 0.846862 | 0.202106 | 0.386295 | -0.68125 | 1.373837 | -2.22527 | 0.113744 | 0.579362 | -0.77307 | 0.121529 | -1.60534 | 2.008163 | 1.72336 | 1.451581 | 0.969099 | 0.565444 | -0.04147 | -0.03176 | 3.023459 | -0.31048 | -0.83829 |
| 30.55556 | 0.151876 | -0.5068 | -0.93732 | 0.728568 | 0.206743 | 0.131224 | -1.27891 | 0.40828 | -1.12373 | 0.083979 | 0.867295 | -0.73027 | 0.412816 | -1.50798 | 2.17389 | 2.12836 | 0.945441 | 0.568113 | 0.166444 | -0.00706 | 0.400451 | 2.279933 | -0.62381 | -1.40815 |
| 30.56944 | 0.399728 | -0.6632 | -1.12702 | 0.618254 | 0.185885 | -0.0632 | -1.40981 | -0.48899 | -0.48816 | 0.017326 | 0.850335 | -0.37873 | 1.032947 | -0.94788 | 2.112995 | 2.228234 | 0.479114 | 0.018968 | -0.19127 | 0.368257 | -0.62384 | 0.936992 | -1.28771 | -2.01209 |
| 30.58333 | -0.44113 | -0.55133 | -0.57253 | 0.511215 | 0.211825 | -0.07932 | -0.53922 | -0.98547 | -0.32922 | -0.0779 | 0.392555 | -0.00898 | 1.316007 | 0.272559 | 1.802912 | 1.95585 | 0.076534 | -0.81166 | -0.34355 | 0.791845 | -0.38658 | -0.13652 | -1.0672 | -1.72861 |
| 30.59722 | -0.89267 | -0.66941 | 0.17864 | 0.435981 | 0.286175 | -0.24192 | 1.189687 | -0.83598 | -0.12219 | -0.20459 | -0.10756 | 0.420319 | 1.511199 | 1.523873 | 1.440794 | 1.62108 | -0.12758 | -1.11525 | -0.57862 | 0.882743 | -0.00862 | -0.95105 | 0.296532 | -0.87948 |
| 30.61111 | -1.04412 | -0.58952 | 0.860031 | 0.43767 | 0.346364 | -0.39336 | 1.672121 | -1.03331 | 0.301205 | -0.29531 | -0.19118 | 0.711411 | 1.47501 | 2.317148 | 1.003191 | 1.037304 | -0.39691 | 0.400885 | -0.68811 | 0.699501 | 0.367091 | -1.67405 | 1.396964 | -0.21949 |
| 30.625 | 0.259172 | -0.52624 | 1.21228 | 0.531321 | 0.302801 | -0.37858 | 0.821552 | -1.38604 | 0.039091 | -0.43573 | -0.22039 | 0.859123 | 1.668105 | 2.745629 | 0.638642 | 0.195448 | -0.67932 | 0.597216 | -0.83567 | 0.635433 | 0.488376 | -2.38546 | 1.679165 | -0.21918 |
| 30.63889 | 0.767438 | -0.30176 | 0.870385 | 0.487637 | 0.213515 | 0.055859 | -0.06122 | -0.79932 | -0.7229 | -0.63084 | -0.43182 | 0.771162 | 0.960026 | 2.021743 | 0.606229 | -0.56516 | -0.85968 | 0.054736 | -0.86785 | 0.762622 | 1.06586 | -2.99764 | 1.167557 | -0.90453 |
| 30.65278 | 0.815959 | -0.19728 | 0.576136 | 0.49172 | 0.119723 | 0.72434 | -0.50081 | 0.534533 | -1.02479 | -0.71825 | -0.29359 | 0.446412 | -0.26436 | 1.180854 | 0.538798 | -0.92658 | -1.03615 | -0.26156 | -1.17146 | 0.754071 | 0.75447 | -2.59373 | 0.14622 | -1.24012 |
| 30.66667 | 0.789945 | -0.18558 | 0.281986 | 0.426389 | 0.078595 | 0.567602 | -0.4402 | 1.650685 | -0.76481 | -0.64565 | 0.097725 | -0.157 | -1.50915 | 0.493686 | 0.254068 | -0.5033 | -1.05286 | -0.03795 | -1.41484 | 0.38607 | -0.44539 | -2.73981 | -0.59272 | -1.07684 |
| 30.68056 | 1.058301 | -0.0707 | -0.22859 | 0.311009 | 0.039729 | -0.11605 | -0.29231 | 2.129599 | -0.42538 | -0.54563 | 0.374983 | -0.91596 | -2.8192 | -0.10991 | -0.71587 | -0.19116 | -0.97296 | 0.745345 | -1.37058 | 0.042889 | -0.68644 | -2.59706 | -0.92452 | -0.87826 |
| 30.69444 | 1.661654 | 0.121218 | -0.89336 | 0.263716 | 0.018064 | -0.39829 | -0.30267 | 1.763951 | -0.2177 | -0.66281 | 0.631213 | -1.5219 | -2.94447 | -1.01064 | -1.17009 | -0.22672 | -0.87193 | 1.439114 | -1.1365 | -0.40558 | -0.82925 | -2.48086 | -1.69152 | -1.05106 |
| 30.70833 | 1.862623 | 0.182413 | -0.89938 | 0.270872 | -0.00758 | -0.38029 | -0.96124 | 0.893637 | -0.13427 | -0.58828 | 0.661891 | -1.84733 | -3.26772 | -1.92285 | -1.24122 | -0.47768 | -0.69764 | 0.842063 | -0.8582 | -0.45092 | -1.24178 | -2.21157 | -1.95041 | -1.3155 |
| 30.72222 | 1.054897 | 0.062646 | -0.76938 | 0.293771 | -0.114 | -0.38313 | -2.19338 | 0.224473 | -0.33358 | -0.75363 | 0.49114 | -1.97099 | -2.94631 | -3.78569 | -0.47305 | -0.97223 | -0.57771 | 1.171563 | -0.72352 | -0.03034 | -0.85389 | -1.27484 | -2.31891 | -1.56381 |
| 30.73611 | 0.29326 | -0.13899 | -0.12948 | 0.196426 | -0.15241 | -0.36836 | -2.17036 | -0.20849 | -0.52218 | -1.07943 | 0.132866 | -1.75513 | -2.55819 | -5.40428 | -0.62575 | -1.04025 | -0.59551 | 0.646763 | -0.95783 | 0.198976 | 0.159476 | 0.281035 | -2.76794 | -1.39089 |
| 30.75 | -0.63324 | -0.26852 | -0.04509 | 0.093987 | -0.22741 | -0.28318 | -1.98488 | -0.76999 | -0.79943 | -1.52305 | -0.26926 | -1.48877 | -2.28058 | -5.17645 | -0.76327 | -0.74766 | -0.57777 | -0.28532 | -0.32969 | 0.164738 | 0.857184 | 2.285996 | -2.47282 | -1.32858 |
| 30.76389 | -1.46546 | -0.33349 | -0.0493 | 0.009677 | -0.29671 | -0.33619 | -1.80115 | -1.61452 | -1.07458 | -2.02895 | -0.51864 | -1.26648 | -1.45825 | -3.74413 | -0.69089 | -0.26132 | -0.36408 | -1.17747 | 0.525945 | -0.02425 | 0.91026 | 2.172703 | -2.10784 | -1.70045 |
| 30.77778 | -2.24093 | -0.47928 | -0.1074 | -0.15128 | -0.37705 | -0.51782 | -1.59217 | -2.46088 | -1.20238 | -2.35791 | -0.55857 | -1.12417 | -0.80594 | -1.62829 | -0.7403 | -0.23846 | -0.01172 | -2.54644 | 1.215928 | -0.65714 | 0.474687 | 1.179892 | -1.70104 | -2.31166 |
| 30.79167 | -3.1081 | -0.90968 | -0.10533 | -0.24519 | -0.31429 | -0.65384 | -0.8337 | -2.43457 | -0.96477 | -2.44781 | -0.50298 | -0.86145 | -0.22734 | -0.1628 | -1.25743 | -0.44279 | 0.233633 | -3.65914 | 1.233555 | -0.57243 | -0.60902 | 0.664967 | -0.32824 | -3.52796 |
| 30.80556 | -2.3212 | -1.21481 | -0.02572 | -0.21558 | -0.14564 | -0.6476 | -0.46813 | -2.1053 | -0.86993 | -2.17938 | -0.51198 | -0.64995 | 0.247301 | 1.095692 | -1.12915 | -0.76756 | 0.344085 | -3.25392 | 0.581637 | -0.65815 | -0.90952 | 0.045374 | 0.10782 | -4.56246 |
| 30.81944 | -1.7236 | -1.62627 | 0.070985 | -0.21878 | 0.002784 | -0.55774 | -0.4052 | -2.41556 | -0.65496 | -1.78953 | -0.38757 | -0.30166 | 0.711219 | 2.410474 | -1.02912 | -1.30724 | 0.423165 | -1.26221 | 0.527244 | -0.74835 | -0.54884 | -1.04911 | 1.063291 | -4.92703 |
| 30.83333 | -0.86795 | -1.74285 | 0.137071 | -0.16509 | 0.094326 | -0.34956 | -0.43069 | -2.75225 | -0.291 | -0.95336 | -0.53519 | 0.158657 | 0.489944 | 3.866817 | -0.77842 | -1.46556 | 0.519092 | -0.69845 | 0.520563 | -0.75971 | 0.044383 | -1.83505 | 2.076706 | -2.87919 |
| 30.84722 | -0.45396 | -1.66758 | 0.225764 | -0.16763 | 0.199484 | -0.2137 | -0.31862 | -2.58812 | 0.30427 | -0.07134 | -0.52155 | 0.677079 | 0.392892 | 4.145679 | -0.44747 | -1.20238 | 0.491633 | -0.23665 | 0.183655 | -0.65856 | 0.647538 | -2.54339 | 2.677683 | 0.239422 |
| 30.86111 | -0.14735 | -1.55029 | 0.330284 | -0.16481 | 0.284401 | -0.15789 | -0.09695 | -1.96943 | 1.021147 | 0.715908 | 0.11721 | 1.081929 | 0.460864 | 3.568314 | 0.020864 | -0.68684 | 0.342621 | 0.101418 | -0.1675 | -0.79299 | 0.90713 | -2.42886 | 1.894729 | 2.902134 |
| 30.875 | 0.285706 | -1.11461 | 0.39933 | -0.09323 | 0.319417 | -0.09416 | 0.119508 | -1.24833 | 1.588237 | 1.322623 | -0.01775 | 1.269515 | 0.272297 | 2.323109 | 0.448518 | -0.2946 | 0.076349 | 0.249476 | -0.36556 | -0.34405 | 0.129066 | -0.91835 | 1.052746 | 4.251959 |
| 30.88889 | 0.508256 | -0.56667 | 0.359634 | -0.06965 | 0.248844 | -0.02509 | 0.17829 | -0.7758 | 1.396082 | 1.463446 | -0.09057 | 1.030704 | -0.41737 | 0.738367 | 0.522348 | 0.055156 | -0.40895 | 0.013458 | -0.60081 | -0.17358 | -0.19413 | -1.60463 | -0.06464 | 4.834281 |
| 30.90278 | 0.107175 | -0.26874 | 0.181901 | -0.23091 | 0.081025 | -0.17164 | -0.11243 | -0.97029 | 0.315795 | 0.689535 | -0.39477 | 0.267925 | -1.55866 | -1.2387 | 0.460255 | -0.24796 | -1.12885 | -0.70677 | -0.91235 | -0.30407 | -0.18534 | -2.55217 | -1.99267 | 1.878131 |
| 30.91667 | -0.73521 | -0.34898 | -0.05456 | -0.40517 | -0.05206 | -0.49699 | -0.72318 | -1.78919 | -1.54777 | -1.26583 | -0.9151 | -0.80816 | -4.08501 | -3.28401 | 0.568802 | -0.48083 | -1.61041 | -1.67909 | -1.45034 | -0.50663 | -0.26984 | -3.31813 | -4.48362 | -2.16739 |
| 30.93056 | -1.29901 | -0.17922 | -0.16868 | -0.40417 | -0.15401 | -0.7611 | -1.30862 | -2.70794 | -3.17341 | -2.93288 | -1.62586 | -2.10634 | -2.98127 | -5.27615 | 0.803418 | -0.52997 | -1.87229 | -2.17538 | -2.00148 | -0.5217 | -0.17286 | -4.70295 | -7.48605 | -5.78452 |
| 30.94444 | -1.54904 | -0.30884 | -0.26772 | -0.40005 | -0.29713 | -1.05843 | -1.34098 | -2.99326 | -3.60399 | -3.05889 | -2.49745 | -2.26937 | -2.10193 | -5.92268 | 0.822825 | -0.402 | -2.33454 | -1.42391 | -2.6095 | -0.40841 | -0.09682 | -5.75026 | -6.5948 | -4.58651 |
| 30.95833 | -1.70709 | -0.7576 | -0.22622 | -0.4902 | -0.48397 | -1.33073 | -1.07844 | -2.91678 | -2.13972 | -1.93857 | -2.25923 | -1.56174 | -1.52179 | -5.26069 | 0.237803 | -0.73548 | -3.10596 | -0.46873 | -3.39717 | -0.40747 | 0.320035 | -6.19055 | -5.26174 | -4.73248 |
| 30.97222 | -1.38819 | -1.56925 | 0.04249 | -0.54797 | -0.56468 | -0.80717 | -1.50946 | -2.39594 | -1.03011 | -1.31231 | -2.31587 | -1.28686 | -1.45815 | -3.6195 | -0.12253 | -0.85573 | -3.291 | -0.44158 | -3.73048 | -0.51691 | -0.53386 | -5.15273 | -3.67202 | -3.85304 |
| 30.98611 | -1.03438 | -2.2457 | 0.152975 | -0.60268 | -0.58351 | -0.55385 | -1.15819 | -1.90938 | -0.54472 | -1.09799 | -0.63038 | -1.07282 | -0.78511 | -2.32815 | -0.07039 | -0.53461 | -2.76607 | -0.0565 | -2.83421 | -0.70338 | -0.46154 | -3.56454 | -2.11357 | -2.81061 |
| 31 | -0.63328 | -2.28894 | 0.161324 | -0.55381 | -0.55481 | -0.44998 | -0.57198 | -1.10478 | 0.04714 | -0.83662 | 0.616878 | -0.61613 | -0.14481 | -1.23526 | 0.005901 | -0.02174 | -1.8796 | -0.30547 | -0.65761 | -0.40033 | -1.16427 | -2.43893 | 0.236613 | -2.01702 |
| 31.01389 | -0.25148 | -1.90123 | 0.13349 | -0.53202 | -0.47718 | -0.33409 | -0.33418 | -0.56401 | 0.564882 | -0.52376 | 1.526657 | -0.06861 | 0.272333 | -0.48843 | 0.082761 | 0.167491 | -1.0909 | -0.45925 | 0.828211 | -0.16804 | -0.91174 | -1.65474 | 1.001847 | -1.19204 |
| 31.02778 | -0.1143 | -1.30476 | 0.188314 | -0.49559 | -0.34144 | -0.28775 | -0.47836 | -0.23774 | 1.200134 | -0.25146 | 2.158658 | 0.38393 | 0.390727 | 0.081132 | 0.1407 | 0.094451 | -0.61314 | -0.41606 | 1.352486 | -0.06049 | -0.72785 | -1.31535 | 0.74576 | -0.5605 |
| 31.04167 | -0.01164 | -0.95174 | 0.246776 | -0.46173 | -0.23187 | -0.32128 | -0.72666 | -0.06002 | 1.147165 | -0.0469 | 1.319652 | 0.704692 | 0.361359 | 0.353423 | 0.234221 | -0.03313 | -0.44516 | -0.16763 | 1.009102 | 0.156121 | -0.45244 | -0.92242 | 0.325173 | -0.3699 |
| 31.05556 | 0.047219 | -0.79884 | 0.301979 | -0.41033 | -0.12718 | -0.32401 | -0.69672 | -0.11809 | 0.741544 | 0.009912 | 0.439244 | 0.897648 | 0.183987 | 0.412712 | 0.281536 | -0.16504 | -0.49563 | 0.071893 | 0.518543 | 0.086476 | -0.21126 | -0.55625 | 0.234369 | -0.46378 |
| 31.06944 | 0.301009 | -0.50199 | 0.349614 | -0.20641 | -0.01883 | -0.29336 | -0.29677 | -0.28742 | 0.214012 | 0.036449 | -0.05654 | 1.106106 | -0.0553 | 0.759923 | 0.31346 | -0.28802 | -0.38728 | 0.200614 | 0.36364 | 0.13515 | 0.080476 | -0.17506 | -0.1639 | -0.56061 |
| 31.08333 | 0.623538 | 0.148744 | 0.345428 | 0.133897 | 0.107631 | -0.13566 | 0.258961 | -0.39808 | -0.29295 | 0.344648 | -0.22339 | 1.163759 | -0.25407 | 1.319191 | 0.386529 | -0.31899 | -0.16128 | 0.357766 | 0.568804 | 0.077978 | 0.285812 | -0.07185 | -0.24374 | -0.15329 |
| 31.09722 | 0.988774 | 0.925397 | 0.346415 | 0.268213 | 0.236927 | 0.019737 | 0.905609 | -0.39342 | -0.45718 | 1.013158 | -1.23014 | 1.217455 | -0.23475 | 1.820591 | 0.538812 | -0.09187 | 0.049424 | 0.301706 | 0.760411 | 0.486237 | 0.434011 | -0.09292 | 0.088148 | 0.525803 |
| 31.11111 | 1.160535 | 1.8459 | 0.410983 | 0.203041 | 0.294764 | 0.150118 | 1.296634 | -0.17789 | -0.08864 | 1.714647 | -1.50988 | 1.747696 | 0.178052 | 2.375721 | 0.676246 | 0.374981 | 0.154515 | 0.264323 | 0.85595 | 1.027429 | 0.547195 | -0.26843 | 0.601247 | 1.076369 |
| 31.125 | 1.333418 | 2.621965 | 0.564222 | 0.175778 | 0.288026 | 0.229257 | 1.287223 | 0.029787 | 0.429717 | 2.382778 | -0.40491 | 1.81395 | 0.846044 | 2.679819 | 0.772558 | 1.00703 | 0.169108 | 0.303968 | 0.689085 | 1.393751 | 0.679453 | -0.36089 | 1.068137 | 1.475901 |
| 31.13889 | 1.307988 | 1.599959 | 0.60604 | 0.249006 | 0.301384 | 0.294709 | 1.238796 | 0.23295 | 1.395188 | 2.880528 | 0.938497 | 2.048599 | 1.603023 | 2.667472 | 0.847431 | 1.369814 | 0.17235 | 0.464535 | 0.30894 | 1.581625 | 0.93148 | -0.32353 | 1.481371 | 1.972443 |
| 31.15278 | 1.434784 | 1.770644 | 0.608396 | 0.242414 | 0.305196 | 0.327023 | 1.241848 | 0.46308 | 3.09498 | 2.401952 | 2.456195 | 1.919481 | 2.114143 | 2.799875 | 0.805848 | 1.525401 | 0.207934 | 0.656152 | 0.402752 | 1.696074 | 1.08471 | -0.21701 | 2.081243 | 2.131787 |
| 31.16667 | 1.455647 | 1.386649 | 0.572184 | 0.250452 | 0.258788 | 0.284742 | 1.355957 | 0.625607 | 2.862078 | 1.624608 | 2.245497 | 1.347222 | 1.570767 | 2.58601 | 0.761977 | 1.410021 | 0.251068 | 0.720453 | 0.567285 | 1.84012 | 1.194369 | -0.04964 | 3.190773 | 1.904582 |
| 31.18056 | 1.308013 | 1.338855 | 0.529019 | 0.296551 | 0.134167 | 0.208161 | 1.159542 | 0.583872 | 1.407005 | 0.549797 | 1.261605 | 0.783907 | 1.342814 | 2.115173 | 0.661195 | 1.158058 | 0.199423 | 0.656933 | 0.473773 | 1.902153 | 1.062405 | 0.103413 | 3.123047 | 1.505422 |
| 31.19444 | 1.17695 | 0.915478 | 0.469707 | 0.303119 | -0.02917 | 0.077902 | 0.588764 | 0.207292 | -0.24743 | -1.18155 | 0.402097 | 0.098378 | 1.096848 | 1.422532 | 0.445995 | 0.908123 | 0.046888 | 0.560466 | 0.070285 | 1.361971 | 0.788981 | 0.11767 | 2.32173 | 0.881431 |
| 31.20833 | 1.01299 | 0.614589 | 0.434928 | 0.146127 | -0.06724 | 0.074323 | 0.054138 | -0.16528 | -1.69481 | -1.5791 | -0.10797 | -0.35418 | 0.788192 | 0.919838 | 0.246777 | 0.704254 | -0.10794 | 0.459824 | -0.47495 | 0.820537 | 0.470414 | -0.01323 | 1.817377 | 0.364943 |
| 31.22222 | 0.763856 | 0.600437 | 0.383218 | 0.002653 | 0.082553 | 0.302684 | 0.052101 | -0.14227 | -1.69337 | -0.40474 | 0.090391 | -0.3387 | 0.595913 | 0.560341 | 0.151958 | 0.592973 | -0.10642 | 0.273396 | -0.67345 | 0.825 | 0.430699 | -0.0893 | 1.389411 | 0.794671 |
| 31.23611 | 0.646458 | 0.404862 | 0.401266 | -0.05844 | 0.185644 | 0.451116 | 0.005824 | 0.011605 | -1.18292 | 0.372381 | 0.276542 | -0.16185 | 0.525512 | 0.372793 | 0.12846 | 0.651396 | -0.0832 | 0.125007 | -0.57873 | 0.838175 | 0.39245 | -0.14506 | 1.267883 | 1.638727 |
| 31.25 | 0.670814 | -0.3766 | 0.414158 | -0.10622 | 0.143029 | 0.44931 | -0.26973 | 0.143017 | -0.05895 | 0.031131 | 0.629025 | -0.10049 | 0.511661 | 0.09156 | 0.077515 | 0.68758 | -0.16601 | 0.037122 | -0.50633 | 0.803218 | 0.264521 | -0.24147 | 0.850666 | 1.693487 |
| 31.26389 | 0.877632 | -1.01725 | 0.52645 | -0.13435 | 0.091971 | 0.328855 | -0.44629 | 0.232033 | 0.771826 | -0.08697 | 0.61081 | 0.080104 | 0.471112 | -0.13211 | 0.017037 | 0.573493 | -0.27337 | 0.070259 | -0.5057 | 0.52735 | 0.107092 | -0.19914 | 0.432397 | 0.894289 |
| 31.27778 | 0.871816 | -1.06847 | 0.524051 | -0.12721 | 0.064796 | 0.202315 | -0.31481 | 0.196692 | 1.074099 | -0.02291 | 0.466637 | 0.313522 | 0.473516 | -0.21319 | 0.021243 | 0.432304 | -0.32803 | 0.097051 | -0.6156 | 0.233655 | 0.045225 | -0.23549 | 0.206442 | 0.279059 |
| 31.29167 | 0.784478 | -1.19389 | 0.465538 | -0.11099 | 0.011718 | 0.0637 | -0.18703 | 0.112504 | 0.609291 | -0.43183 | 0.169591 | 0.241447 | 0.363164 | -0.22807 | 0.03143 | 0.2695 | -0.35936 | 0.002476 | -0.78916 | -0.17042 | -0.09802 | -0.37915 | 0.013844 | -0.2255 |
| 31.30556 | 0.509566 | -1.4941 | 0.351075 | -0.11362 | -0.02094 | 0.03285 | -0.18408 | 0.00632 | -0.01571 | -1.07391 | -0.05241 | -0.08709 | 0.180922 | -0.29735 | 0.037501 | 0.279798 | -0.33204 | -0.10046 | -0.9993 | -0.33628 | -0.16625 | -0.39529 | -0.16159 | -0.50849 |
| 31.31944 | 0.3172 | -1.58157 | 0.253516 | -0.07016 | -0.04157 | 0.132503 | -0.26801 | -0.08016 | -0.52085 | -1.59036 | -0.30581 | -0.3286 | 0.05545 | -0.35038 | 0.038664 | 0.382014 | -0.22167 | -0.22716 | -1.04408 | 0.365631 | -0.12806 | -0.35628 | -0.13368 | -0.54583 |
| 31.33333 | 0.157193 | -1.72646 | 0.204584 | -0.04909 | -0.04453 | 0.253181 | -0.41373 | -0.1222 | -0.74111 | -1.76281 | -0.4588 | -0.35209 | 0.052579 | -0.29393 | 0.033818 | 0.449292 | -0.11224 | -0.38972 | -0.95505 | 0.472227 | -0.11879 | -0.33883 | -0.07594 | -0.32352 |
| 31.34722 | 0.161804 | -1.86645 | 0.28852 | -0.06341 | -0.03285 | 0.270465 | -0.36164 | -0.17465 | -0.65214 | -1.22937 | -0.61075 | -0.14907 | 0.169517 | -0.10543 | -0.04396 | 0.464886 | -0.03033 | -0.4107 | -0.58595 | 0.313545 | -0.08546 | -0.32761 | 0.022076 | 0.003522 |
| 31.36111 | 0.215923 | -2.16381 | 0.362 | -0.0824 | -0.03467 | 0.231449 | -0.23268 | -0.20522 | -0.32207 | -0.5802 | -0.24328 | 0.07439 | 0.374421 | 0.2205 | -0.24246 | 0.481072 | 0.072828 | -0.19467 | -0.12397 | 0.210482 | 0.048983 | -0.28163 | 0.200281 | 0.387749 |
| 31.375 | 0.31383 | -2.06319 | 0.322747 | -0.07312 | -0.01839 | 0.221603 | -0.07543 | -0.08784 | 0.170238 | -0.17634 | 0.005369 | 0.226944 | 0.664934 | 0.634036 | -0.25924 | 0.485738 | 0.20723 | 0.089431 | 0.349138 | 0.157834 | 0.243137 | -0.08521 | 0.430956 | 0.741432 |
| 31.38889 | 0.740222 | -1.03358 | 0.436527 | 0.158286 | 0.113732 | 0.500484 | 0.541237 | 0.446873 | 0.646097 | 0.623502 | 0.09838 | 0.342875 | 1.344457 | 1.401577 | -0.184 | 0.555438 | 0.599947 | 0.403773 | 0.877505 | 0.220479 | 0.446345 | 0.236599 | 0.906713 | 1.936683 |
| 31.40278 | 1.50391 | 1.282507 | 0.684122 | 0.796287 | 0.414861 | 1.202926 | 1.935935 | 1.542772 | 1.06231 | 2.100396 | 0.414122 | 0.709839 | 2.571342 | 2.843514 | -0.14009 | 0.896527 | 1.581007 | 0.674722 | 0.760547 | 0.468029 | 0.728686 | 0.814706 | 1.937415 | 4.428061 |
| 31.41667 | 2.243783 | 3.090779 | 1.113415 | 1.544356 | 0.659845 | 1.581962 | 3.550787 | 2.800185 | 1.225083 | 3.256242 | 1.209283 | 1.231979 | 2.728167 | 4.687154 | -0.09845 | 1.281003 | 2.398692 | 0.651485 | 0.805469 | 1.098855 | 0.950135 | 1.434737 | 3.466882 | 4.624005 |
| 31.43056 | 2.471671 | 2.929799 | 1.470061 | 1.348663 | 0.614697 | 1.237294 | 3.207731 | 2.675593 | 0.862959 | 2.905206 | 2.063814 | 1.83745 | 2.028914 | 4.603854 | 0.137129 | 1.192822 | 2.72103 | 0.394169 | 0.610622 | 1.08876 | 0.822805 | 2.509662 | 3.082772 | 3.230514 |
| 31.44444 | 1.844458 | 2.831003 | 1.373615 | 0.607781 | 0.58033 | 0.594773 | 2.272138 | 2.174293 | 1.046487 | 2.71004 | 1.646178 | 2.344992 | 1.26973 | 4.251201 | 0.388567 | 0.810322 | 2.716753 | 0.87723 | 1.277042 | 0.648144 | 0.571977 | 3.734448 | 2.255328 | 2.631305 |
| 31.45833 | 0.848438 | 2.524474 | 1.014218 | 0.35939 | 0.647008 | 0.849167 | 2.143663 | 2.386084 | 2.807348 | 2.708975 | 1.024261 | 2.094461 | 1.952014 | 3.058647 | 0.286873 | 0.73013 | 2.84041 | 1.101021 | 1.742867 | 0.59081 | 0.688862 | 3.662192 | 1.900768 | 2.106232 |
| 31.47222 | 0.360169 | 2.784349 | 0.692535 | 0.296519 | 0.575601 | 1.446715 | 3.582966 | 2.633604 | 2.876276 | 1.764274 | 0.331003 | 1.13199 | 1.46343 | 1.475554 | 0.278865 | 0.714047 | 2.574238 | 1.05601 | 1.720398 | 0.460991 | 0.834946 | 4.625578 | 1.384894 | 1.352456 |
| 31.48611 | 1.540043 | 3.308035 | 0.338927 | 0.282734 | 0.440329 | 1.28385 | 3.491104 | 2.688444 | 2.562787 | 1.189398 | -0.50411 | 0.431021 | 1.367236 | -0.64868 | 0.505364 | 0.72179 | 1.848433 | 1.079653 | 1.785589 | 0.521568 | 0.959882 | 2.469837 | 1.017902 | 0.820852 |
| 31.5 | 1.638678 | 2.990862 | 0.016606 | 0.328033 | 0.280179 | 1.43867 | 1.826174 | 2.591413 | 1.838791 | 0.628405 | -0.52594 | -0.03338 | 0.830562 | -1.67646 | 0.973535 | 0.613264 | 1.309077 | 1.53373 | 1.672718 | 0.56052 | 0.484082 | 1.596176 | 1.368524 | 0.816845 |
| 31.51389 | 1.430182 | 2.174972 | -0.29677 | 0.409178 | 0.178511 | 0.739647 | 0.342033 | 1.952767 | 0.902984 | 0.28756 | 0.207824 | -0.52051 | 0.282121 | -1.48193 | 1.372194 | 0.504588 | 0.734248 | 1.388993 | 1.197119 | 0.802618 | 0.38176 | 1.275336 | 1.345683 | 0.825923 |
| 31.52778 | 1.242442 | 1.523212 | -0.54903 | 0.50415 | 0.060772 | 0.002322 | -1.02939 | 1.399914 | -0.31946 | -0.09355 | 0.02 | -0.91197 | -0.48305 | -1.55074 | 1.408895 | 0.464587 | 0.348319 | 0.935124 | 0.671463 | 1.090448 | 0.355994 | 0.856279 | 0.772174 | 0.602373 |
| 31.54167 | 0.532938 | 0.881271 | -0.90484 | 0.52632 | -0.06427 | -0.64785 | -2.2785 | 1.01187 | -1.18109 | -0.52957 | -0.34298 | -0.74302 | -0.97245 | -1.97489 | 1.202404 | 0.454135 | 0.111972 | 0.599858 | 0.387598 | 1.2039 | 0.124605 | 0.540333 | 0.279767 | 0.247542 |
| 31.55556 | -0.34689 | 0.226532 | -1.10538 | 0.407106 | -0.14538 | -0.47131 | -3.18916 | 0.817882 | -1.35227 | -0.75092 | 0.27089 | -0.9468 | -1.22703 | -1.96514 | 0.982935 | 0.412939 | -0.04771 | 0.766748 | 0.320423 | 1.208689 | -0.01127 | 0.191214 | -0.25888 | -0.01037 |
| 31.56944 | -0.00765 | -0.45812 | -1.28256 | 0.234854 | -0.13361 | -0.35053 | -2.62896 | 0.610015 | -1.15526 | -1.08158 | 0.80384 | -0.91009 | -1.27381 | -2.03527 | 0.770343 | 0.333091 | -0.12406 | 0.678842 | 0.07561 | 0.89156 | -0.04041 | -0.00701 | -0.18305 | -0.22974 |
| 31.58333 | -0.56853 | -1.1064 | -0.85315 | 0.031331 | -0.11288 | -0.12083 | -2.12844 | 0.488764 | -0.69608 | -1.32812 | 1.126724 | -0.89878 | -1.50374 | -2.55662 | 0.66941 | 0.267305 | -0.13549 | 0.408572 | -0.47866 | 0.583486 | -0.09522 | -0.29087 | -0.21896 | -0.53071 |
| 31.59722 | -1.29877 | -1.58826 | -0.20454 | -0.16663 | -0.09026 | 0.027116 | -2.56988 | 0.450173 | -0.46803 | -1.55629 | 1.000754 | -0.49629 | -1.51508 | -3.19719 | 0.554819 | 0.256007 | -0.0924 | 0.158637 | -0.74702 | 0.325021 | -0.09059 | -0.50996 | -0.46649 | -0.78573 |
| 31.61111 | -2.30043 | -1.65518 | 0.134636 | -0.14625 | -0.00775 | 0.081068 | -2.82173 | 0.577269 | -0.38487 | -1.68141 | 1.11404 | -0.05105 | -1.30983 | -3.05874 | 0.510363 | 0.239252 | -0.05267 | -0.11196 | -0.72575 | -0.00802 | -0.06153 | -0.47002 | -0.87388 | -0.95492 |
| 31.625 | -0.21182 | -1.68253 | 0.112427 | 0.141563 | 0.085832 | 0.173046 | -2.5622 | 0.859827 | -0.1694 | -1.90983 | 1.322855 | 0.161042 | -0.89987 | -2.43334 | 0.258354 | 0.301672 | -0.03565 | -0.46399 | -0.44178 | -0.27839 | 0.105618 | -0.27489 | -1.32763 | -1.15042 |
| 31.63889 | -0.50843 | -2.49068 | 0.00714 | 0.418379 | 0.119463 | 0.22628 | -1.98419 | 0.830362 | -0.38951 | -2.13057 | 1.018229 | 0.140551 | -0.60466 | -1.72353 | 0.286461 | 0.409886 | -0.03908 | -0.91517 | -0.10887 | -0.50115 | 0.144639 | 0.007958 | -2.01428 | -1.22118 |
| 31.65278 | -0.52138 | -2.853 | 0.000883 | 0.478477 | 0.032955 | 0.207418 | -1.45048 | 0.697496 | -0.68603 | -2.26047 | -0.28923 | 0.090533 | -0.0925 | -1.02066 | 0.217548 | 0.653863 | -0.04599 | -1.17126 | 0.097628 | -0.65764 | 0.209789 | 0.186474 | -1.91975 | -1.1044 |
| 31.66667 | -0.92199 | -3.06437 | -0.00507 | 0.304431 | -0.03416 | 0.213642 | -1.04104 | 0.512716 | -1.1396 | -2.5002 | -0.88288 | 0.118402 | 0.444471 | -0.52587 | 0.193215 | 0.864846 | -0.00952 | -0.77108 | 0.236736 | -0.50505 | 0.29318 | 0.36261 | -1.54554 | -0.82576 |
| 31.68056 | -0.21608 | -2.81606 | -0.09069 | 0.061527 | -0.03745 | 0.235253 | -0.75419 | 0.281011 | -1.11387 | -2.25982 | 0.357211 | -0.09984 | 0.779729 | -0.30699 | 0.281434 | 0.970751 | 0.033227 | -0.3702 | 0.33066 | -0.23269 | 0.3202 | 0.347018 | -1.17939 | -0.55171 |
| 31.69444 | 0.374451 | -2.92848 | -0.14228 | -0.16158 | -0.00383 | 0.226723 | -0.6826 | 0.152573 | -1.0845 | -1.91681 | -0.06593 | -0.53412 | 0.944449 | -0.16134 | 0.473927 | 0.934204 | -0.00855 | -0.01846 | 0.241927 | 0.1427 | 0.458065 | 0.23891 | -0.97793 | -0.41078 |
| 31.70833 | 0.314442 | -2.66058 | -0.15501 | -0.30071 | -0.02105 | 0.170447 | -0.82281 | -0.03555 | -0.75906 | -2.0501 | -0.02213 | -1.03027 | 0.941076 | -0.07193 | 0.473439 | 0.779108 | -0.12283 | 0.247911 | 0.227888 | 0.278514 | 0.499594 | 0.193663 | -0.63134 | -0.48595 |
| 31.72222 | 0.131392 | -1.97691 | -0.17557 | -0.19711 | -0.02149 | 0.01262 | -0.90079 | -0.1044 | -0.46936 | -2.43801 | -0.24707 | -1.50516 | 0.861221 | -0.13054 | 0.406739 | 0.571127 | -0.2833 | 0.634811 | 0.234889 | 0.052496 | 0.443275 | -0.01278 | -0.04898 | -0.64355 |
| 31.73611 | 0.171841 | -1.29613 | -0.47808 | 0.022046 | 0.053608 | -0.07475 | -1.10562 | -0.09055 | -0.10509 | -2.84905 | 0.079253 | -2.02197 | 0.487194 | -0.52229 | 0.45055 | 0.302393 | -0.44759 | 0.786831 | -0.0396 | -0.19429 | 0.310579 | -0.22625 | 0.186409 | -0.68023 |
| 31.75 | 0.276957 | -0.86377 | -0.81984 | 0.114832 | 0.074811 | -0.15886 | -1.43655 | -0.13387 | -0.16454 | -2.57248 | 0.522457 | -2.69261 | 0.038811 | -1.19029 | 0.310169 | -0.137 | -0.6717 | 0.602283 | -0.20155 | -0.0561 | 0.231203 | -0.54607 | -0.17923 | -0.71205 |
| 31.76389 | 0.267474 | -0.78349 | -0.76743 | 0.033356 | 0.023954 | -0.30363 | -1.95423 | -0.26708 | -0.59565 | -2.22416 | 0.492999 | -2.46559 | -0.20167 | -1.82442 | 0.085346 | -0.42009 | -0.87765 | -0.01739 | -0.71597 | -0.02618 | 0.160338 | -0.8997 | -0.94854 | -0.98065 |
| 31.77778 | 0.280729 | -0.79347 | -0.62572 | -0.12764 | 0.008062 | -0.45231 | -2.1893 | -0.52278 | -0.46859 | -1.70496 | 0.074997 | -2.0526 | 0.017181 | -2.27613 | -0.25342 | -0.50377 | -0.97162 | -0.55025 | -1.3455 | 0.021737 | -0.10375 | -1.12252 | -1.88244 | -1.67861 |
| 31.79167 | 0.275845 | -1.01121 | -0.62561 | -0.25335 | -0.00273 | -0.49053 | -1.96179 | -0.82947 | -0.08441 | -0.93879 | -0.01001 | -1.72959 | -0.08604 | -2.25922 | -0.43462 | -0.33359 | -0.98808 | 0.003415 | -2.06066 | -0.05338 | -0.03274 | -1.07326 | -2.31848 | -2.38657 |
| 31.80556 | 0.307851 | -1.00392 | -0.56716 | -0.29933 | -0.06669 | -0.43843 | -1.72557 | -1.05134 | 0.132226 | -0.41186 | 0.237741 | -1.26962 | -0.05234 | -1.79492 | -0.28087 | -0.05096 | -0.89334 | 0.016358 | -2.53536 | -0.32912 | -0.02717 | -0.90615 | -2.35153 | -2.59959 |
| 31.81944 | 0.261786 | -0.72205 | -0.47766 | -0.28673 | -0.11035 | -0.35271 | -1.33222 | -1.01624 | 0.214929 | -0.17436 | 0.26532 | -0.80603 | 0.193932 | -1.46407 | 0.001241 | -0.03227 | -0.77602 | 0.235172 | -2.53318 | -0.59706 | -0.12929 | -0.71473 | -1.86754 | -2.55361 |
| 31.83333 | 0.426995 | -0.5092 | -0.23643 | -0.19583 | -0.13569 | -0.15676 | -1.04665 | -0.791 | 0.250811 | -0.01865 | 0.436244 | -0.46948 | 0.48307 | -1.43117 | 0.135905 | -0.26871 | -0.8198 | 0.363927 | -2.22797 | -0.67395 | -0.1977 | -0.51585 | -2.14726 | -2.4525 |

| | | | | | | | | | | | | | | | | | | | | | | | | |
|---|---|---|---|---|---|---|---|---|---|---|---|---|---|---|---|---|---|---|---|---|---|---|---|---|
| 31.84722 | 0.698969 | -0.16097 | 0.094283 | -0.07742 | -0.14614 | 0.083065 | -0.75015 | -0.50029 | 0.277912 | 0.119642 | 0.47584 | -0.12525 | 0.646913 | -1.26729 | 0.211912 | -0.35321 | -0.82897 | 0.567341 | -1.28656 | -0.45441 | -0.16239 | -0.36908 | -2.35875 | -2.58896 |
| 31.86111 | 0.936451 | 0.521743 | 0.367346 | 0.023666 | -0.16479 | 0.287839 | -0.54929 | -0.20504 | 0.356978 | 0.257128 | 0.633176 | 0.195828 | 0.601772 | -0.54216 | 0.224159 | -0.19862 | -0.75735 | 0.445298 | -0.09034 | -0.22322 | -0.06585 | -0.12053 | -2.22694 | -2.9055 |
| 31.875 | 1.113076 | 1.118336 | 0.605391 | 0.035568 | -0.167 | 0.431553 | -0.39648 | -0.01442 | 0.451025 | 0.186256 | 0.590164 | 0.469743 | 0.639241 | 0.084811 | 0.158963 | -0.00644 | -0.66984 | 0.446126 | 0.628335 | -0.17281 | 0.029044 | 0.022324 | -1.91746 | -3.0591 |
| 31.88889 | 1.170725 | 1.117122 | 0.580904 | -0.23166 | -0.15784 | 0.391852 | -0.49924 | -0.15776 | 0.315805 | -0.33092 | 0.102106 | 0.460978 | 0.404018 | -0.20143 | -0.04636 | 0.032229 | -0.97794 | 0.258301 | 0.752926 | -0.15534 | -0.11072 | -0.66606 | -2.33871 | -3.73854 |
| 31.90278 | 0.624911 | 0.540967 | 0.230262 | -1.07279 | -0.22148 | 0.073738 | -0.99219 | -0.6323 | -0.19853 | -1.49935 | -0.58892 | 0.086695 | -0.64524 | -1.93388 | -0.58752 | -0.44615 | -1.84334 | -0.26514 | -0.24569 | -0.41924 | -0.62054 | -2.85692 | -3.8612 | -5.88395 |
| 31.91667 | -0.15405 | -0.5371 | -0.27098 | -1.58624 | -0.24237 | -0.29356 | -1.24483 | -0.98148 | -0.91941 | -2.96741 | -1.43878 | -0.65018 | -1.58664 | -3.82461 | -1.00253 | -0.89845 | -2.17292 | -0.50712 | -2.49743 | -0.45184 | -0.73139 | -2.69757 | -4.33008 | -6.64317 |
| 31.93056 | -0.50811 | -1.11611 | -0.58216 | -1.35473 | -0.14501 | -0.42118 | -1.32383 | -1.04466 | -1.32473 | -4.17767 | -1.89922 | -1.58837 | -1.68952 | -3.97514 | -0.64274 | -0.6381 | -1.43685 | -1.45514 | -3.34682 | -0.54809 | -0.48162 | -3.28045 | -4.0829 | -6.13405 |
| 31.94444 | -0.76356 | -1.34899 | -0.66304 | -1.10793 | -0.15517 | -0.47358 | -1.08767 | -1.0206 | -1.69776 | -2.64268 | -1.4626 | -1.75742 | -1.39788 | -3.20113 | -1.07345 | -0.20523 | -0.58821 | -1.81759 | -2.9387 | -0.74923 | -0.11622 | -1.83441 | -2.58154 | -3.65846 |
| 31.95833 | -1.01309 | -1.49276 | -0.58793 | -1.03685 | -0.35325 | -0.43128 | -0.96853 | -0.88018 | -1.41964 | 0.337057 | -1.15818 | -1.48871 | -0.90129 | -2.92646 | -0.87677 | -0.17054 | -0.63482 | -2.00554 | -2.30888 | -1.00815 | 0.097258 | -1.17271 | -0.89456 | -3.10708 |
| 31.97222 | -0.46847 | -1.58713 | -0.42255 | -0.57637 | -0.44817 | -0.2559 | -0.92716 | -0.7351 | -0.94871 | 1.243998 | -1.15016 | -0.77893 | 0.550742 | -2.18261 | -0.35798 | 0.456555 | -1.22421 | 0.332672 | -2.22313 | -0.96108 | 0.252537 | -0.78402 | 0.010772 | -2.38635 |
| 31.98611 | 0.308374 | -1.59922 | -0.15898 | 0.058183 | -0.33479 | -0.03158 | -0.49628 | -0.43739 | -0.48075 | 0.453046 | -1.6487 | -0.18444 | 1.634875 | -1.94987 | 0.245832 | 0.893341 | -1.21281 | 1.243917 | -2.06961 | -0.93859 | 0.38927 | -0.56159 | 0.077077 | -1.27481 |
| 32 | 0.877437 | -1.62019 | 0.13719 | 0.521168 | -0.18462 | 0.183944 | -0.15944 | -0.05921 | -0.17316 | -0.1778 | -1.60406 | 0.215215 | 1.578825 | -0.33417 | 0.358868 | 0.434239 | -1.05877 | 2.202778 | -0.53114 | -0.50751 | 0.38709 | -0.32819 | 0.252105 | -0.50817 |
| 32.01389 | 1.056469 | -1.33967 | 0.325441 | 0.627737 | -0.13537 | 0.347961 | 0.158865 | 0.297558 | 0.226023 | -0.35363 | -0.57328 | 0.367303 | 1.169329 | 1.094575 | 0.373597 | 0.114682 | -0.81033 | 2.209374 | 0.598219 | -0.02001 | 0.280209 | 0.221643 | 0.226186 | -0.17855 |
| 32.02778 | 1.102958 | -1.01068 | 0.324492 | 0.58278 | -0.13872 | 0.444251 | 0.243667 | 0.472737 | 0.393963 | -0.29816 | -0.16569 | 0.302006 | 1.139872 | 1.724376 | 0.382555 | 0.328454 | -0.58088 | 0.716746 | 0.992489 | 0.275672 | 0.198925 | 0.736696 | 0.316271 | 0.011761 |
| 32.04167 | 1.175863 | -0.82117 | 0.369633 | 0.462479 | -0.1445 | 0.423684 | 0.128412 | 0.497662 | 0.372713 | -0.22297 | 0.305693 | 0.274901 | 1.039743 | 1.947989 | 0.439079 | 0.441124 | -0.41351 | 0.824991 | 1.256648 | 0.367381 | 0.167514 | 0.941922 | 0.39532 | 0.21391 |
| 32.05556 | 1.427666 | -0.63482 | 0.604398 | 0.387887 | -0.11095 | 0.240544 | 0.015118 | 0.445839 | 0.31444 | -0.08304 | 0.305921 | 0.299251 | 0.964099 | 1.7604 | 0.33517 | 0.566834 | -0.205 | 0.535128 | 1.34118 | 0.367058 | 0.196942 | 0.775668 | 0.295138 | 0.321325 |
| 32.06944 | 1.334359 | -0.29123 | 0.746997 | 0.352742 | -0.06939 | 0.017796 | 0.029872 | 0.322739 | 0.168046 | 0.152822 | 0.09584 | 0.386432 | 0.893385 | 1.224878 | 0.165895 | 0.573627 | -0.02903 | 0.277245 | 1.651274 | 0.201962 | 0.255127 | 0.516548 | 0.279824 | 0.43562 |
| 32.08333 | 1.423355 | 0.138582 | 0.709507 | 0.37111 | -0.01052 | -0.09175 | 0.206881 | 0.178628 | 0.075766 | 0.389084 | 0.230637 | 0.460858 | 0.9206 | 0.905027 | 0.091068 | 0.533715 | 0.128192 | -0.09563 | 2.001555 | 0.099399 | 0.354795 | 0.363514 | 0.305746 | 0.550841 |
| 32.09722 | 1.527539 | 0.606396 | 0.646538 | 0.431485 | 0.090256 | -0.13264 | 0.470598 | 0.070711 | 0.307375 | 0.499168 | 0.528333 | 0.702853 | 1.251474 | 0.500829 | 0.117865 | 0.602066 | 0.229497 | -0.08192 | 2.434289 | 0.110884 | 0.421996 | 0.23196 | 0.443614 | 0.614524 |
| 32.11111 | 1.707747 | 0.939583 | 0.638765 | 0.505662 | 0.193854 | -0.10431 | 0.683654 | 0.06252 | 0.38312 | 0.538348 | 0.728674 | 0.975118 | 1.608111 | 0.295033 | 0.181036 | 0.649461 | 0.263467 | 0.319378 | 2.469633 | 0.178457 | 0.47943 | 0.18403 | 0.599874 | 0.632223 |
| 32.125 | 1.249418 | 1.117297 | 0.569029 | 0.543998 | 0.255576 | -0.00317 | 0.755598 | 0.140208 | 0.511702 | 0.445881 | 0.731651 | 1.13948 | 1.653549 | 0.240129 | 0.192643 | 0.707218 | 0.228208 | 0.380985 | 2.10699 | 0.340627 | 0.527713 | 0.113921 | 0.582301 | 0.580991 |
| 32.13889 | 0.886478 | 1.184928 | 0.529911 | 0.509779 | 0.286307 | 0.186438 | 0.698634 | 0.276245 | 0.597582 | 0.386161 | 1.182275 | 1.028713 | 1.395446 | 0.195806 | 0.217736 | 0.654472 | 0.133672 | 0.349955 | 1.569727 | 0.666948 | 0.613196 | 0.187362 | 0.629059 | 0.450348 |
| 32.15278 | 0.695336 | 1.083211 | 0.676269 | 0.458066 | 0.258395 | 0.363483 | 0.741989 | 0.457885 | 0.620145 | 0.43338 | 1.365032 | 1.039737 | 1.138153 | 0.126923 | 0.269416 | 0.627054 | 0.025738 | 0.44259 | 0.995495 | 1.071775 | 0.619048 | 0.218733 | 0.594789 | 0.329587 |
| 32.16667 | 0.247755 | 1.001874 | 0.776853 | 0.393051 | 0.195868 | 0.396532 | 0.705587 | 0.534904 | 0.45445 | 0.516209 | 1.119843 | 0.986252 | 0.983422 | -0.09018 | 0.323285 | 0.588028 | -0.03078 | 0.561128 | 0.591735 | 1.196739 | 0.623109 | 0.309295 | 0.603662 | 0.349299 |
| 32.18056 | -0.13789 | 1.031428 | 0.722369 | 0.333589 | 0.142269 | 0.321518 | 0.689905 | 0.449633 | 0.249937 | 0.647341 | 0.634574 | 0.877257 | 0.859073 | -0.40201 | 0.451298 | 0.598315 | -0.06772 | 0.576839 | 0.212765 | 1.20965 | 0.549945 | 0.287181 | 0.604473 | 0.588161 |
| 32.19444 | -0.01644 | 0.89672 | 0.513868 | 0.298756 | 0.096915 | 0.271111 | 0.628486 | 0.400216 | 0.300195 | 0.747055 | 0.02509 | 0.788506 | 0.866194 | -0.44791 | 0.490988 | 0.59301 | -0.0892 | 0.542535 | -0.10038 | 1.247887 | 0.483374 | 0.222048 | 0.450748 | 1.022516 |
| 32.20833 | 0.300637 | 0.826429 | 0.290575 | 0.266055 | 0.074261 | 0.343833 | 0.571942 | 0.221676 | 0.301675 | 0.7849 | -0.42506 | 0.640194 | 0.834404 | -0.12456 | 0.559587 | 0.602655 | -0.1077 | 0.592297 | -0.34848 | 1.18969 | 0.436507 | 0.107158 | 0.346239 | 1.503107 |
| 32.22222 | 0.442853 | 0.702083 | 0.159668 | 0.285833 | 0.064724 | 0.506261 | 0.646996 | -0.09991 | 0.307839 | 0.772865 | -0.47327 | 0.468221 | 0.828011 | 0.265306 | 0.668195 | 0.625977 | -0.22341 | 0.626375 | -0.14134 | 1.1493 | 0.427215 | -0.02453 | 0.425992 | 1.837525 |
| 32.23611 | 0.625718 | 0.515074 | 0.195227 | 0.295145 | 0.028472 | 0.728861 | 0.778641 | -0.44992 | 0.270651 | 0.705506 | 0.103801 | 0.253149 | 1.000894 | 0.530905 | 0.713645 | 0.657868 | -0.46975 | 0.625579 | 0.482895 | 1.058147 | 0.409839 | -0.21869 | 0.567068 | 2.054837 |
| 32.25 | 0.656664 | 0.345489 | 0.310777 | 0.314263 | 0.035014 | 0.938957 | 0.760551 | -0.50122 | 0.144297 | 0.588882 | 0.414379 | 0.074228 | 1.043779 | 0.686562 | 0.729256 | 0.581573 | -0.58809 | 0.557589 | 0.623066 | 0.847981 | 0.366867 | -0.50981 | 0.656653 | 1.913491 |
| 32.26389 | 0.609568 | 0.228016 | 0.364534 | 0.285437 | 0.048053 | 0.996509 | 0.536325 | -0.34956 | 0.108694 | 0.484268 | 0.509051 | 0.078077 | 0.866079 | 0.596646 | 0.716394 | 0.589039 | -0.52928 | 0.345071 | 0.068584 | 0.654491 | 0.256027 | -0.70119 | 0.752101 | 1.492286 |
| 32.27778 | 0.545739 | 0.109341 | 0.294177 | 0.170544 | 0.041735 | 0.909127 | 0.29771 | -0.27698 | 0.079698 | 0.35682 | 0.565113 | 0.155505 | 0.463387 | 0.486518 | 0.529872 | 0.581152 | -0.43674 | 0.16524 | -0.53006 | 0.556787 | 0.120382 | -0.69976 | 0.787286 | 1.335944 |
| 32.29167 | 0.331513 | -0.12607 | 0.210373 | 0.002604 | 0.002791 | 0.670897 | 0.066193 | -0.39072 | 0.118716 | 0.193866 | 0.475553 | 0.159007 | 0.317705 | 0.374765 | 0.433696 | 0.4923 | -0.37387 | -0.10564 | -1.01675 | 0.476721 | 0.058229 | -0.64522 | 0.900791 | 1.419141 |
| 32.30556 | 0.242214 | -0.18223 | 0.138151 | -0.06535 | -0.01442 | 0.475742 | -0.14319 | -0.50458 | 0.180089 | 0.03335 | 0.422376 | 0.082875 | 0.16344 | 0.228877 | 0.218954 | 0.476964 | -0.30897 | 0.091779 | -1.30008 | 0.306282 | 0.004478 | -0.73665 | 0.911958 | 1.335606 |
| 32.31944 | 0.315022 | -0.53231 | 0.294259 | -0.07343 | -0.0133 | 0.510763 | -0.12623 | -0.17907 | 0.363598 | -0.04064 | 0.778306 | 0.018621 | 0.183938 | 0.227924 | -0.00259 | 0.477788 | -0.25208 | 0.264771 | -1.23293 | 0.232685 | -0.03041 | -0.87846 | 0.886733 | 1.39949 |
| 32.33333 | 0.606681 | -0.79915 | 0.494743 | 0.019744 | 0.005376 | 0.557665 | 0.03892 | 0.415275 | 0.576896 | -0.00028 | 1.251292 | 0.048066 | 0.459009 | 0.30483 | 0.114959 | 0.566987 | -0.20518 | 0.681799 | -0.73061 | 0.175977 | -0.01343 | -0.79826 | 0.911082 | 1.691675 |
| 32.34722 | 0.857061 | -0.89761 | 0.644913 | 0.062363 | -0.05098 | 0.556014 | 0.235195 | 0.64731 | 0.845777 | 0.057718 | 1.63312 | 0.074828 | 0.759893 | 0.474053 | 0.435957 | 0.629022 | -0.17982 | 0.908748 | 0.053243 | 0.322573 | 0.061056 | -0.63624 | 1.023245 | 1.916787 |
| 32.36111 | 1.159222 | -0.8571 | 0.79671 | 0.012503 | -0.16387 | 0.42051 | 0.34894 | 0.603716 | 0.621005 | 0.071064 | 0.570554 | -0.00916 | 1.111826 | 0.545862 | 0.658998 | 0.695034 | -0.16935 | 1.119968 | 0.741306 | 0.533177 | 0.056408 | -0.5189 | 1.085043 | 1.724658 |
| 32.375 | 1.261654 | -0.71149 | 0.89289 | 0.002726 | -0.20014 | 0.322045 | 0.443937 | 0.360509 | 0.30277 | 0.11189 | 0.447082 | -0.10602 | 1.144528 | 0.53362 | 0.75807 | 0.720934 | -0.15978 | 1.257654 | 1.123937 | 0.672152 | 0.059718 | -0.3774 | 1.202177 | 1.755247 |
| 32.38889 | 1.484447 | -0.36437 | 0.919267 | 0.188957 | -0.13243 | 0.409345 | 0.856885 | 0.258863 | 0.438422 | 0.473191 | 0.667697 | -0.07517 | 1.208254 | 0.858808 | 0.821138 | 0.857807 | 0.020187 | 1.286776 | 1.583379 | 0.835243 | 0.313381 | 0.219332 | 1.95209 | 2.7498 |
| 32.40278 | 1.607807 | 0.399307 | 1.015395 | 0.686224 | 0.06865 | 0.682104 | 1.545396 | 0.64436 | 1.160474 | 1.083029 | 0.747451 | 0.237365 | 2.04389 | 1.745676 | 1.000394 | 1.27269 | 0.681411 | 1.539532 | 1.930195 | 1.19318 | 0.792601 | 1.579596 | 4.282154 | 5.366076 |
| 32.41667 | 1.881773 | 1.205985 | 0.954691 | 1.231904 | 0.239923 | 0.784769 | 2.893702 | 1.504705 | 1.605916 | 1.138353 | 1.337004 | 0.930883 | 3.220558 | 2.317134 | 1.129585 | 1.583018 | 1.565386 | 2.278831 | 3.210178 | 1.65348 | 0.989338 | 3.135076 | 5.09644 | 6.361741 |
| 32.43056 | 2.26236 | 1.841709 | 0.918419 | 1.262786 | 0.093026 | 0.499214 | 3.514357 | 2.210706 | 1.113118 | 1.109856 | 2.969458 | 1.841208 | 3.021052 | 1.981618 | 0.726269 | 1.69571 | 1.899783 | 3.088718 | 3.840282 | 1.710624 | 0.772008 | 3.650184 | 3.770718 | 6.657647 |
| 32.44444 | 2.457976 | 2.204717 | 1.239252 | 1.123049 | -0.04769 | -0.1419 | 3.515461 | 2.820729 | 0.592963 | 1.421433 | 3.400468 | 2.805136 | 2.920637 | 2.245038 | 0.818606 | 1.722228 | 2.038678 | 3.471434 | 5.527068 | 1.170697 | 1.035317 | 3.593052 | 2.868108 | 7.594922 |
| 32.45833 | 2.208668 | 3.590088 | 1.26601 | 1.129403 | 0.107141 | 0.052951 | 3.940451 | 2.856214 | 0.49175 | 2.190078 | 2.310354 | 2.351241 | 3.091316 | 2.717336 | 0.714282 | 1.512289 | 2.418917 | 2.584403 | 3.877449 | 0.56183 | 1.410892 | 3.673223 | 2.500921 | 7.412487 |
| 32.47222 | 1.67682 | 4.966351 | 0.859779 | 0.908514 | 0.160007 | 0.203313 | 4.114769 | 2.419366 | 1.327146 | 3.38479 | 0.496007 | 1.967862 | 2.653812 | 3.019381 | 1.023669 | 1.115519 | 2.485672 | 2.087953 | 2.559405 | 0.02172 | 1.294741 | 3.669399 | 2.146027 | 5.409989 |
| 32.48611 | 1.569293 | 3.340864 | 0.475178 | 0.52785 | 0.089179 | 0.169407 | 4.177933 | 2.087917 | 1.756066 | 3.156547 | 0.105984 | 1.68298 | 2.276008 | 2.568027 | 0.548336 | 1.030325 | 2.498888 | 1.791965 | 2.455287 | -0.10359 | 1.191281 | 3.52454 | 1.526073 | 3.213376 |
| 32.5 | 0.556613 | 2.628826 | -0.12552 | 0.544252 | 0.174269 | 0.26879 | 3.480008 | 1.226536 | 1.906148 | 2.874205 | -1.26903 | 1.143075 | 2.179302 | 2.171786 | -0.01621 | 1.142495 | 2.604547 | 1.528915 | 3.066869 | -0.04295 | 1.260234 | 2.300059 | 1.369119 | 2.190075 |
| 32.51389 | 0.69095 | 2.717835 | -0.6324 | 0.757102 | 0.087191 | 0.313768 | 5.500778 | 0.901013 | 1.916796 | 2.776227 | -1.59331 | 0.826209 | -0.00343 | 1.784105 | -0.27807 | 1.227516 | 2.63892 | 1.448091 | 3.011227 | 0.040403 | 0.950362 | 1.233381 | 1.560791 | 1.742563 |
| 32.52778 | 2.567266 | 2.362728 | 0.126664 | 0.704308 | 0.027185 | 0.74297 | 5.335098 | 1.637398 | 1.375383 | 2.025644 | -0.47178 | 0.48984 | 0.640171 | 1.306886 | 0.497828 | 1.407671 | 2.452525 | 2.362063 | 1.810952 | 0.418037 | 0.037689 | 0.518337 | 1.758671 | 1.051731 |
| 32.54167 | 2.752712 | 2.040451 | 0.126438 | 0.802494 | 0.10546 | 0.742073 | 2.783523 | 2.157082 | 0.849658 | 0.853343 | -0.5172 | 0.18707 | 0.425042 | 0.469365 | 0.837397 | 1.180216 | 1.955684 | 1.788623 | 0.925686 | 0.96737 | -0.51262 | 0.068686 | 1.108645 | -0.12331 |
| 32.55556 | 2.381825 | 1.721678 | -0.00583 | 0.758547 | 0.13497 | 0.620278 | 1.298347 | 1.430594 | 0.362486 | -0.59748 | -0.47225 | -0.04195 | -0.41752 | -0.66547 | 0.768147 | 0.788743 | 1.299488 | 1.871127 | 0.667203 | 1.162466 | -0.87405 | -0.31676 | 0.876543 | -1.05545 |
| 32.56944 | 1.824133 | 1.400127 | -0.1214 | 0.472059 | 0.089159 | 0.499784 | -0.70152 | 0.703416 | 0.048221 | -1.88339 | -0.85248 | -0.26697 | -1.57022 | -1.71837 | 0.650624 | 0.311094 | 0.539795 | 1.826023 | 0.466475 | 1.112817 | -0.5314 | -0.58137 | 0.990822 | -1.32812 |
| 32.58333 | 1.156841 | 1.153821 | -0.11245 | 0.111955 | -0.0453 | 0.281645 | -3.26851 | 0.288218 | -0.10408 | -2.65447 | 1.296418 | -0.62908 | -1.58643 | -2.5845 | 0.765716 | 0.119933 | -0.09016 | 1.689631 | -0.60062 | 1.217182 | -0.85355 | -0.77622 | 0.466523 | -1.61922 |
| 32.59722 | 0.582749 | 1.037562 | -0.17448 | -0.1946 | -0.1634 | 0.098043 | -3.52789 | 0.218291 | -0.3497 | -2.43177 | 2.234107 | -1.00216 | -0.73569 | -2.63067 | 0.929518 | 0.282993 | -0.54215 | 0.568809 | -1.53558 | 0.967768 | -0.94055 | -0.87291 | -0.23504 | -2.13914 |
| 32.61111 | -0.56139 | 0.652384 | -0.18729 | -0.44397 | -0.25505 | 0.070793 | -3.60064 | 0.33161 | -0.60531 | -1.99495 | 1.98706 | -1.18263 | -0.23818 | -1.95987 | 0.897296 | 0.357089 | -0.65817 | -0.2539 | -1.81295 | 0.799103 | -0.98714 | -0.84485 | -0.71887 | -2.85608 |
| 32.625 | -1.41906 | 0.100323 | -0.20958 | -0.47425 | -0.27103 | 0.127288 | -2.81984 | 0.590953 | -0.61271 | -2.17169 | 1.885568 | -0.95777 | -0.21248 | -1.66365 | 0.903811 | 0.386052 | -0.49814 | -0.59078 | -1.7522 | 0.171805 | -1.42973 | -0.63807 | -0.64083 | -4.45008 |
| 32.63889 | -1.25256 | -0.19892 | -0.17451 | -0.25599 | -0.23485 | 0.141918 | -2.04306 | 0.894998 | -0.45158 | -1.77192 | 0.687548 | -0.64715 | -0.05737 | -1.03369 | 1.014142 | 0.238914 | -0.2667 | -0.35926 | -0.96057 | -0.1765 | -0.77947 | -0.02607 | -1.13407 | -5.8033 |
| 32.65278 | -1.27942 | -0.52745 | -0.16504 | -0.16373 | -0.23062 | 0.017535 | -1.94769 | 1.015595 | -0.10168 | -1.72949 | 0.484409 | -0.50526 | 0.042315 | -0.41199 | 0.438848 | 0.187102 | -0.00235 | -0.07089 | 0.264426 | -0.22011 | 0.159611 | 0.750645 | -1.66324 | -5.15367 |
| 32.66667 | -1.17395 | -1.0463 | -0.1665 | -0.3177 | -0.32415 | -0.11035 | -2.35868 | 0.944269 | 0.090492 | -1.68775 | 0.808777 | -0.63545 | -0.28687 | -0.12522 | 0.261239 | 0.037652 | 0.220395 | -0.00442 | 1.385429 | -0.10085 | 1.142649 | 1.248821 | -0.82735 | -4.09129 |
| 32.68056 | -1.12678 | -1.88523 | -0.25731 | -0.4643 | -0.4069 | -0.04797 | -2.77137 | 0.917879 | 0.137763 | -1.84225 | 0.349671 | -0.85597 | 0.217423 | 0.202862 | -0.0207 | 0.306156 | 0.227515 | -0.12951 | 1.699145 | -0.60545 | 1.58106 | 0.620337 | 0.702148 | -3.07723 |
| 32.69444 | -1.1686 | -2.12248 | -0.20878 | -0.57657 | -0.48731 | 0.12286 | -3.33304 | 0.782419 | 0.183919 | -1.9787 | -1.00424 | -0.98664 | 0.506117 | 0.658489 | -0.07221 | 0.478268 | -0.06679 | -0.04498 | 1.056593 | -1.30656 | 1.412374 | 0.369034 | 1.45816 | -1.49835 |
| 32.70833 | -1.04202 | -2.12121 | -0.23587 | -0.64271 | -0.65979 | 0.229382 | -3.50972 | 0.480528 | -0.06246 | -2.00337 | -1.34346 | -0.87522 | 0.649151 | 0.757855 | -0.24619 | 0.664139 | -0.47244 | 0.149919 | 0.156973 | -1.34874 | 1.249958 | -0.12187 | 0.82602 | 0.269086 |
| 32.72222 | -0.84096 | -1.99478 | -0.34296 | -0.76012 | -0.9382 | 0.138353 | -4.34305 | 0.352614 | 0.096166 | -1.66429 | -1.27078 | -0.64854 | 0.023456 | 0.577224 | -0.21369 | 0.838804 | -0.66045 | -0.19981 | -0.22101 | -1.07365 | 1.382643 | -0.29154 | 0.482537 | 1.892472 |
| 32.73611 | -1.06038 | -1.72727 | -0.53654 | -0.81742 | -1.02217 | 0.00734 | -5.25805 | 0.582362 | 0.151561 | -1.40141 | -1.26864 | -0.37999 | -0.28029 | 0.396281 | -0.23238 | 1.127037 | -0.49776 | -0.10661 | -0.14258 | -1.06795 | 1.34669 | -0.49837 | 0.63219 | 1.834077 |
| 32.75 | -1.03829 | -1.45381 | -0.98894 | -0.80032 | -0.70736 | -0.14601 | -5.07662 | 0.766275 | 0.053491 | -1.28724 | -1.29823 | -0.4055 | 0.055193 | 0.385991 | 0.040867 | 1.274339 | -0.27263 | -0.112 | -0.41442 | -0.78814 | 0.963891 | -0.96268 | 0.895262 | 0.60229 |
| 32.76389 | -1.19678 | -1.92687 | -1.35235 | -0.79274 | -0.47312 | -0.28113 | -5.31071 | 0.663437 | -0.22704 | -1.37896 | -1.61742 | -0.67955 | 0.607201 | 0.27342 | 0.186195 | 1.174048 | -0.54802 | -0.71384 | -0.91455 | -0.22634 | 0.220367 | -1.47411 | 0.530624 | -0.44998 |
| 32.77778 | -0.88162 | -2.75914 | -1.61047 | -0.73791 | -0.334 | -0.40727 | -6.03712 | 0.411713 | -0.64488 | -1.6368 | -1.6289 | -0.99166 | 0.295061 | -0.20982 | -0.01853 | 1.013388 | -1.1169 | -0.35391 | -1.3175 | -0.1379 | -0.99678 | -1.84749 | -0.04026 | -1.67275 |
| 32.79167 | -0.36816 | -3.16036 | -1.7235 | -0.81154 | -0.30705 | -0.52962 | -6.33191 | 0.091512 | -0.71971 | -1.80108 | -1.37663 | -1.15124 | -0.71271 | -0.59319 | -0.46462 | 0.845422 | -1.66862 | 0.327318 | -1.47307 | -0.25297 | -1.55159 | -1.8434 | -0.60347 | -2.29282 |
| 32.80556 | -0.18081 | -2.60206 | -1.362 | -0.74176 | -0.27114 | -0.51652 | -6.81849 | -0.04924 | -0.31687 | -1.32354 | -1.10167 | -1.15421 | -0.58868 | -0.97638 | -1.70555 | 0.499942 | -1.99375 | -0.14863 | -1.76719 | -0.57292 | -1.67732 | -2.12097 | -0.77584 | -2.82149 |
| 32.81944 | 0.134303 | -1.90572 | -1.04345 | -0.56831 | -0.1777 | -0.45985 | -6.10298 | -0.16226 | 0.081557 | -0.81489 | -0.27318 | -1.06525 | -0.59547 | -1.42999 | -1.41567 | 0.294451 | -1.78276 | -1.47823 | -1.8998 | -0.71834 | -1.30548 | -2.07403 | -0.96607 | -3.53292 |
| 32.83333 | 0.214047 | -1.481 | -0.37569 | -0.4476 | -0.03379 | -0.36542 | -3.54655 | -0.27566 | 0.132156 | -0.34359 | 0.375687 | -0.8294 | -0.28835 | -1.77457 | -0.85104 | 0.385872 | -1.46341 | -2.01238 | -2.27462 | -0.72222 | -1.40711 | -2.18537 | -1.02802 | -3.34922 |
| 32.84722 | 0.481768 | -1.09734 | 0.096364 | -0.39816 | 0.112788 | -0.22356 | -2.10463 | -0.22909 | 0.021379 | -0.1318 | 1.361179 | -0.55984 | 0.185446 | -2.02346 | 0.03823 | 0.903884 | -1.4781 | -2.7845 | -3.25018 | -0.63455 | -1.04022 | -1.76036 | -1.18702 | -3.15492 |
| 32.86111 | 0.656329 | -0.49912 | 0.205172 | -0.25259 | 0.144241 | -0.09633 | -1.4852 | 0.007984 | 0.174601 | -0.00569 | 1.56606 | -0.24049 | 0.561271 | -2.12323 | 1.026888 | 0.604069 | -1.69665 | -3.20244 | -4.44227 | -0.26089 | -0.66404 | -1.12586 | -1.39552 | -3.73924 |
| 32.875 | 0.788544 | 0.098795 | 0.063354 | 0.057884 | 0.038223 | -0.13519 | -0.53134 | 0.273569 | 0.495204 | 0.088208 | 1.900913 | -0.03849 | 0.137251 | -1.45105 | 1.929525 | 0.484646 | -2.01693 | -2.56251 | -5.53882 | 0.337583 | -0.0895 | -0.54822 | -2.24904 | -3.83368 |
| 32.88889 | 0.894111 | 0.180454 | -0.38579 | 0.070242 | -0.22982 | -0.20514 | -0.07778 | 0.218956 | 0.476877 | -0.09049 | 1.37774 | -0.08375 | -0.55898 | -0.81872 | 2.545871 | -0.37143 | -2.37205 | -1.14367 | -5.02718 | 0.262668 | -0.17012 | -0.6103 | -1.72241 | -2.52435 |
| 32.90278 | 0.169699 | -0.24296 | -0.94304 | -0.54275 | -0.59938 | -0.40636 | 0.015209 | -0.58154 | 0.016033 | -0.95946 | -0.07115 | -0.34213 | -1.21879 | -1.19655 | 1.202203 | -1.57902 | -2.56883 | -0.61117 | -3.96624 | -0.45936 | -0.79895 | -1.76683 | -2.63671 | -4.48034 |
| 32.91667 | -0.45205 | -0.70717 | -1.55716 | -1.02685 | -0.6309 | -0.24252 | -0.45323 | -1.60262 | -0.80461 | -1.97194 | -1.62461 | -0.60179 | -1.64692 | -1.22892 | -0.94116 | -1.79934 | -2.41769 | -0.05181 | -1.78845 | -1.79568 | -1.37178 | -3.51404 | -3.02321 | -5.62105 |
| 32.93056 | -0.31816 | -0.95435 | -1.81943 | -0.79726 | -0.71564 | -0.00045 | 1.142604 | -2.4797 | -1.8204 | -2.16893 | -1.07806 | -0.96084 | -0.91746 | -0.40565 | -1.7997 | -1.33508 | -2.12816 | 0.376221 | -1.41606 | -2.40213 | -1.08824 | -4.19225 | -2.93378 | -4.52207 |
| 32.94444 | -0.54277 | -1.13364 | -2.16772 | -0.39573 | -0.77294 | -0.11472 | 0.630631 | -2.67811 | -2.10128 | -1.26508 | -10.4985 | -1.35607 | -0.269 | -0.88296 | -1.53729 | -0.73286 | -1.96481 | 0.506621 | -2.69869 | -2.71608 | -0.31048 | -3.52966 | -1.38378 | -1.75336 |
| 32.95833 | -0.87304 | -1.5588 | -2.33325 | -0.05726 | -0.67675 | -0.05245 | 0.106556 | -1.97147 | -1.94745 | -0.23296 | -5.90276 | -1.09635 | -0.00242 | -1.37112 | -0.86277 | -0.10722 | -2.20913 | 0.02004 | -4.7557 | -1.77817 | 0.129144 | -2.76936 | -0.90484 | -1.94505 |
| 32.97222 | -0.96922 | -1.92132 | -1.57253 | 0.278104 | -0.2092 | 0.154799 | 0.152304 | -0.80814 | -1.10068 | 0.198642 | -2.70151 | -0.46818 | -0.00212 | -1.06846 | 0.33278 | 0.239438 | -2.16619 | 0.329666 | -4.39646 | -0.25263 | 0.675522 | -2.00587 | -1.39525 | -2.27521 |
| 32.98611 | -0.8727 | -2.13661 | -0.59621 | 0.405548 | 0.442026 | 0.459882 | 0.301054 | 0.246156 | -0.30484 | 0.140969 | -1.13657 | 0.057758 | -0.07539 | -0.48508 | 1.159127 | 0.30274 | -1.64783 | 0.758075 | -4.96756 | 0.533516 | 0.706459 | -1.02368 | -1.70146 | -1.6244 |
| 33 | -0.55051 | -1.88855 | 0.405302 | 0.369634 | 0.688583 | 0.543037 | 0.270051 | 0.762679 | 0.12637 | -0.01617 | -0.2843 | 0.48301 | -0.16573 | -0.06289 | 1.359425 | 0.359509 | -1.20434 | 0.870488 | -4.30926 | 1.474285 | 0.676212 | 0.231693 | -1.11458 | -0.82368 |
| 33.01389 | -0.19423 | -1.66626 | 0.941203 | 0.285135 | 0.378249 | 0.423254 | 0.244479 | 0.8903 | 0.287224 | -0.20133 | 0.370554 | 0.780333 | -0.1798 | 0.184544 | 1.10824 | 0.392511 | -0.80567 | 0.729472 | -3.22412 | 1.06514 | 0.710346 | 1.246034 | -0.39636 | -0.27065 |
| 33.02778 | 0.003954 | -1.42992 | 0.758923 | 0.244378 | -0.10263 | 0.352766 | 0.157185 | 0.822208 | 0.1754 | -0.35552 | 0.498162 | 1.064095 | -0.07552 | 0.274456 | 0.941946 | 0.380669 | -0.48883 | 0.745708 | -2.37642 | 0.810129 | 0.82151 | 1.589795 | -0.00074 | -0.06899 |
| 33.04167 | -0.15341 | -1.37498 | 0.153803 | 0.257778 | -0.35514 | 0.318703 | 0.088441 | 0.541742 | 0.045294 | -0.43036 | 0.422604 | 1.225452 | 0.04509 | 0.281128 | 0.810847 | 0.290356 | -0.35604 | 0.251071 | -1.87655 | 0.558281 | 0.831065 | 1.414713 | 0.243923 | 0.077118 |
| 33.05556 | -0.15297 | -1.28108 | 0.015632 | 0.329824 | -0.32679 | 0.36219 | 0.03973 | 0.252183 | 0.036664 | -0.38719 | 0.588461 | 0.754408 | 0.071204 | 0.207487 | 0.749684 | 0.20119 | -0.35065 | 0.101792 | -1.52057 | 0.214164 | 0.59883 | 0.709361 | 0.355446 | 0.376018 |
| 33.06944 | -0.48489 | -1.1258 | 0.379951 | 0.399604 | -0.19555 | 0.510581 | 0.021393 | -0.05528 | 0.050367 | -0.26779 | 0.694816 | 0.443704 | 0.089059 | 0.245872 | 0.634051 | 0.154744 | -0.35395 | 0.265826 | -1.31902 | -0.1567 | 0.361088 | 0.372374 | 0.227117 | 0.169273 |
| 33.08333 | -0.40513 | -0.88867 | 0.774597 | 0.447843 | -0.08839 | 0.665326 | 0.044678 | -0.37072 | 0.088392 | -0.16911 | 1.31689 | 0.335371 | 0.165732 | 0.40936 | 0.481191 | 0.175742 | -0.30761 | 0.339318 | -1.01262 | -0.40338 | 0.157518 | 0.035318 | 0.242961 | -0.03389 |
| 33.09722 | -0.15506 | -0.7562 | 0.906374 | 0.465367 | 0.09627 | 0.732545 | 0.126986 | -0.44238 | 0.205856 | -0.07937 | 1.33117 | 0.297709 | 0.24719 | 0.624601 | 0.177127 | 0.210559 | -0.24534 | 0.23269 | -0.54662 | 0.093866 | 0.157414 | -0.03888 | 0.412407 | -0.22384 |
| 33.11111 | -0.00103 | -0.56428 | 0.96857 | 0.446207 | 0.338273 | 0.638496 | 0.212828 | -0.23518 | 0.275483 | 0.021118 | 0.486754 | 0.269013 | 0.306446 | 0.823271 | 0.028108 | 0.231766 | -0.17307 | 0.229888 | 0.022014 | 0.633216 | 0.338233 | -0.00288 | 0.52776 | -0.30129 |
| 33.125 | 0.317351 | -0.41512 | 0.982119 | 0.40794 | 0.51231 | 0.361552 | 0.275992 | 0.177176 | 0.321792 | 0.070032 | 0.421734 | 0.255741 | 0.397526 | 0.888922 | 0.165913 | 0.305881 | -0.08204 | 0.217451 | 0.505892 | 1.101122 | 0.606033 | 0.054847 | 0.533501 | -0.13704 |
| 33.13889 | 0.675948 | -0.28861 | 0.859493 | 0.330032 | 0.44922 | -0.06052 | 0.169492 | 0.507066 | 0.224184 | 0.029374 | 0.293689 | 0.232935 | 0.491264 | 0.735836 | 0.459317 | 0.405194 | -0.04575 | 0.091865 | 0.887656 | 1.201651 | 0.820292 | 0.179372 | 0.561626 | -0.05683 |
| 33.15278 | 0.699934 | -0.25132 | 0.621281 | 0.256801 | 0.166854 | -0.36679 | -0.08378 | 0.750091 | 0.138439 | -0.00608 | 0.178618 | 0.234698 | 0.43398 | 0.559555 | 0.727622 | 0.42596 | -0.0415 | 0.033644 | 1.110215 | 0.851664 | 1.016386 | 0.298527 | 0.659663 | 0.01117 |
| 33.16667 | 0.618219 | -0.24384 | 0.113417 | 0.21973 | -0.20544 | -0.17825 | -0.24323 | 0.854114 | 0.136897 | 0.013523 | 0.206805 | 0.263295 | 0.474499 | 0.606167 | 0.869423 | 0.414759 | -0.04823 | -0.01134 | 1.167931 | 0.530336 | 1.132958 | 0.382474 | 0.71519 | 0.122988 |
| 33.18056 | 0.541099 | -0.31251 | -0.57499 | 0.195148 | -0.4193 | 0.091025 | -0.03361 | 0.769094 | 0.253148 | -0.00942 | 0.134722 | 0.295962 | 0.893993 | 0.750154 | 0.80649 | 0.493432 | -0.01628 | 0.083614 | 1.179773 | 0.341539 | 1.134601 | 0.385282 | 0.700129 | 0.233179 |
| 33.19444 | 0.56736 | -0.36915 | -1.04081 | 0.170756 | -0.3437 | 0.253918 | 0.233207 | 0.680706 | 0.382106 | -0.15152 | 0.028818 | 0.321865 | 0.934771 | 0.833466 | 0.625646 | 0.565504 | 0.024684 | 0.151059 | 1.087341 | 0.377707 | 1.067548 | 0.355251 | 0.68063 | 0.260999 |
| 33.20833 | 0.49034 | -0.37581 | -0.75408 | 0.161988 | -0.10797 | 0.27093 | 0.359809 | 0.638568 | 0.336136 | -0.24513 | -0.04022 | 0.269401 | 0.989746 | 0.807645 | 0.495425 | 0.590234 | 0.043801 | 0.267227 | 0.871947 | 0.687978 | 0.9407 | 0.289956 | 0.632092 | 0.207865 |
| 33.22222 | 0.4142 | -0.32855 | -0.00151 | 0.141359 | 0.074424 | 0.246736 | 0.303356 | 0.537212 | 0.335118 | -0.20862 | -0.0468 | 0.178527 | 1.40042 | 0.798548 | 0.426228 | 0.628021 | 0.05925 | 0.34566 | 0.598271 | 0.931903 | 0.834815 | 0.29698 | 0.594185 | 0.1045 |
| 33.23611 | 0.376484 | -0.32273 | 0.591136 | 0.117097 | 0.07373 | 0.267104 | 0.207817 | 0.505716 | 0.324091 | -0.11393 | -0.0088 | 0.040059 | 1.28019 | 0.663763 | 0.462707 | 0.654674 | 0.064731 | 0.414391 | 0.417715 | 1.177951 | 0.814401 | 0.302962 | 0.524427 | 0.094927 |
| 33.25 | 0.277893 | -0.40265 | 0.66449 | 0.07209 | -0.08705 | 0.290786 | 0.222228 | 0.515591 | 0.16344 | -0.11566 | 0.033457 | -0.07728 | 1.188525 | 0.417781 | 0.572632 | 0.654905 | 0.072357 | 0.585409 | 0.419818 | 1.454688 | 0.790074 | 0.300494 | 0.49613 | 0.155036 |
| 33.26389 | 0.130236 | -0.46211 | 0.10991 | -0.02218 | -0.30964 | 0.272257 | 0.216142 | 0.486513 | -0.03235 | -0.16388 | -0.00214 | -0.16685 | 0.965963 | 0.340629 | 0.692632 | 0.63574 | 0.062339 | 0.461456 | 0.313035 | 1.224896 | 0.799834 | 0.297313 | 0.603542 | 0.122692 |
| 33.27778 | 0.099606 | -0.40301 | -0.31222 | -0.05039 | -0.40651 | 0.28176 | 0.241044 | 0.471103 | -0.09579 | -0.14259 | 0.041517 | -0.16237 | 0.724383 | 0.559785 | 0.825677 | 0.711955 | 0.029833 | 0.386353 | 0.21401 | 0.795469 | 1.141912 | 0.47603 | 0.709682 | 0.110648 |
| 33.29167 | 0.221349 | -0.35163 | -0.0704 | 0.043909 | -0.33829 | 0.41964 | 0.500276 | 0.556672 | -0.00902 | 0.051501 | 0.1088 | -0.0605 | 0.777379 | 0.943277 | 1.045981 | 0.787835 | 0.038191 | 0.49258 | 0.350071 | 0.566123 | 1.656131 | 0.574268 | 0.917918 | 0.223004 |
| 33.30556 | 0.305127 | -0.368 | 0.41318 | 0.164072 | -0.15502 | 0.575424 | 0.690967 | 0.658294 | 0.100563 | 0.238382 | 0.160115 | 0.06797 | 0.853103 | 1.192973 | 1.140057 | 0.785222 | 0.025158 | 0.674976 | 0.739755 | 0.629685 | 1.848848 | 0.495892 | 1.262446 | 0.32685 |
| 33.31944 | 0.282903 | -0.39899 | 0.782257 | 0.254089 | 0.195567 | 0.639115 | 0.767576 | 0.828005 | 0.260456 | 0.313489 | 0.26455 | 0.194774 | 0.980832 | 1.153969 | 1.258516 | 0.760782 | 0.006469 | 0.886043 | 1.481506 | 0.960619 | 1.566394 | 0.663951 | 1.374893 | 0.557223 |
| 33.33333 | 0.189869 | -0.38084 | 0.914593 | 0.296654 | 0.527097 | 0.63584 | 0.596501 | 1.0044 | 0.353703 | 0.287686 | 0.476821 | 0.266914 | 0.991131 | 1.053959 | 1.248788 | 0.726192 | 0.031545 | 1.048453 | 2.359137 | 1.413778 | 1.118724 | 0.642393 | 1.46928 | 0.82972 |
| 33.34722 | 0.16143 | -0.48345 | 0.852603 | 0.301721 | 0.548529 | 0.539746 | 0.360295 | 1.065457 | 0.303307 | 0.178589 | 0.657746 | 0.21497 | 0.80994 | 0.827666 | 1.252013 | 0.682993 | 0.03666 | 1.122755 | 2.392273 | 1.830426 | 0.690277 | 0.501252 | 1.295068 | 0.986167 |
| 33.36111 | 0.059962 | -0.69396 | 0.68612 | 0.240686 | 0.204582 | 0.424395 | 0.212507 | 1.002037 | 0.138348 | -0.01055 | 0.770594 | 0.049736 | 0.61817 | 0.632371 | 1.139874 | 0.64849 | 0.012596 | 1.056194 | 1.540301 | 1.812584 | 0.385897 | 0.334116 | 1.087624 | 0.831473 |
| 33.375 | 0.047204 | -0.83727 | 0.595058 | 0.184412 | -0.2193 | 0.378917 | 0.186828 | 0.900203 | 0.065959 | -0.07199 | 0.913116 | -0.10981 | 0.676225 | 0.621323 | 1.203491 | 0.635798 | -0.00956 | 0.994264 | 0.751946 | 1.55146 | 0.404591 | 0.212772 | 1.006432 | 0.730526 |

| | | | | | | | | | | | | | | | | | | | | | | | | |
|---|---|---|---|---|---|---|---|---|---|---|---|---|---|---|---|---|---|---|---|---|---|---|---|---|
| 33.38889 | 0.234069 | -0.62028 | 0.523782 | 0.221602 | -0.41805 | 0.475869 | 0.384888 | 0.795319 | 0.202319 | 0.24018 | 1.422687 | -0.01823 | 1.080849 | 1.178956 | 1.307148 | 0.819512 | 0.050945 | 1.008678 | 1.434381 | 1.316945 | 0.955123 | 0.485742 | 1.333562 | 1.60698 |
| 33.40278 | 0.905066 | 0.242812 | 0.531758 | 0.445201 | -0.11845 | 0.646254 | 0.592721 | 0.600428 | 0.482692 | 0.717034 | 2.125716 | 0.383991 | 1.777015 | 1.851915 | 1.473382 | 1.10603 | 0.366766 | 1.035179 | 4.025714 | 1.921013 | 2.084946 | 1.416851 | 2.384778 | 3.580402 |
| 33.41667 | 1.512987 | 1.263452 | 0.782124 | 0.56191 | 0.381009 | 0.684191 | 0.690724 | 0.518278 | 0.589162 | 0.742387 | 2.355566 | 0.824319 | 3.102345 | 2.498087 | 1.493112 | 1.339767 | 0.872825 | 1.108815 | 2.040845 | 2.403388 | 2.506834 | 2.342689 | 3.525804 | 4.569419 |
| 33.43056 | 1.835534 | 1.526294 | 1.383981 | 0.628279 | 0.261398 | 0.484563 | 0.546439 | 0.91205 | 0.600859 | 0.649148 | 2.775666 | 1.307358 | 3.209641 | 2.239166 | 1.571153 | 1.04973 | 1.071202 | 1.232925 | 1.032722 | 1.077498 | 2.793471 | 2.64214 | 4.213588 | 6.022782 |
| 33.44444 | 1.800832 | 1.076664 | 2.252635 | 0.821124 | 0.12423 | 0.523608 | 0.379992 | 1.470293 | 0.828265 | 0.991874 | 3.00204 | 1.917607 | 2.260556 | 3.221904 | 2.186029 | 1.641889 | 1.353999 | 1.505037 | 2.264929 | 0.673033 | 3.806398 | 3.344496 | 4.896666 | 8.823319 |
| 33.45833 | 1.89801 | 0.938491 | 2.748839 | 1.23824 | 0.342326 | 0.616858 | 0.17674 | 1.78184 | 0.722303 | 1.272206 | 4.224037 | 2.275812 | 2.6255 | 3.353929 | -0.37763 | 3.074711 | 1.759492 | 2.628492 | 0.214871 | 1.130898 | 5.052038 | 3.968154 | 5.424269 | 9.054826 |
| 33.47222 | 1.473471 | 1.400333 | 2.463714 | 1.277141 | 0.175884 | 0.656691 | -0.1341 | 1.844711 | 0.391529 | 0.812658 | 7.492945 | 2.243568 | 1.809194 | 2.141929 | 0.945414 | 2.473683 | 1.699904 | 3.48199 | 2.271375 | 1.729064 | 4.350509 | 3.929113 | 4.854494 | 8.702063 |
| 33.48611 | 0.837838 | 2.2237 | 2.310175 | 0.843501 | -0.34612 | 0.624017 | 0.291215 | 1.478854 | 0.287314 | 0.399507 | 8.492417 | 2.119178 | 1.278576 | 3.171109 | 0.744198 | 1.558116 | 1.181527 | 1.88877 | 1.997003 | 1.680874 | 3.577211 | 2.994142 | 3.217838 | 6.301872 |
| 33.5 | 0.471633 | 2.965882 | 2.128931 | 0.498096 | -0.39151 | 0.575377 | 1.265179 | 0.717342 | 0.382103 | 0.360177 | 6.03851 | 2.001596 | 1.372128 | 2.905568 | 0.058612 | 1.130737 | 0.713425 | 0.798045 | 1.494102 | 1.466861 | 3.800446 | 2.308969 | 0.580567 | 4.395415 |
| 33.51389 | 0.406787 | 3.121435 | 1.814756 | 0.194439 | -0.24102 | 0.637999 | 1.770123 | 0.172314 | 0.38449 | 0.304242 | 2.66192 | 1.594645 | 0.725533 | 2.380671 | -0.39035 | 1.053313 | 0.421864 | 0.165128 | 1.604599 | 1.157399 | 4.266886 | 1.482711 | -0.60905 | 3.229907 |
| 33.52778 | 0.567482 | 3.569086 | 1.418744 | 0.259138 | -0.37909 | 0.749773 | 1.744256 | 0.087661 | 0.54475 | 0.182139 | 0.223245 | 0.988208 | 0.521328 | 2.089577 | -1.1806 | 0.738294 | 0.272683 | -0.14519 | 1.862034 | 0.904214 | 3.040847 | 0.859209 | -0.7542 | 2.382307 |
| 33.54167 | 0.678815 | 3.844658 | 1.082276 | 0.338566 | -0.01617 | 0.712151 | 1.380507 | 0.169394 | 0.723895 | 0.173818 | -1.60617 | 0.25882 | 1.047647 | 0.733902 | -2.05487 | 0.394178 | 0.257892 | -0.06715 | 0.124867 | 0.464547 | 0.506447 | 0.205582 | -0.71645 | 1.36998 |
| 33.55556 | 0.790675 | 3.686841 | 0.866225 | 0.431372 | 0.160147 | 0.636645 | 1.429888 | -0.06945 | 0.783078 | 0.258893 | -7.38107 | -0.29481 | 0.246158 | -0.72095 | -2.29737 | -0.01653 | 0.263403 | 0.500675 | -1.6932 | 0.044065 | -1.80025 | -0.5413 | -0.07186 | 0.458002 |
| 33.56944 | 0.928488 | 3.420133 | 0.70462 | 0.297807 | -0.07054 | 0.562077 | 1.598287 | -0.32135 | 0.708753 | 0.389437 | -6.8045 | -0.7908 | -0.07119 | -1.85041 | -2.43172 | -0.2665 | 0.248144 | 0.173143 | -1.7774 | -0.12877 | -3.23385 | -0.96089 | 0.23505 | -1.14474 |
| 33.58333 | 1.021964 | 3.186421 | 0.417911 | 0.045314 | -0.37513 | 0.459651 | 1.821938 | -0.46628 | 0.272089 | 0.230189 | -3.56546 | -1.30219 | -0.51609 | -2.32273 | -2.43278 | -0.32356 | 0.168951 | -0.97107 | -1.76091 | -0.01288 | -3.37181 | -0.82506 | 0.117424 | -2.24718 |
| 33.59722 | 0.978471 | 3.054436 | 0.241317 | -0.11105 | -0.61613 | 0.431745 | 2.029358 | -0.27158 | -0.34996 | -0.10635 | -1.81495 | -1.50136 | -1.06522 | -2.44164 | -2.27695 | -0.26473 | 0.023942 | -1.1428 | -0.99311 | 0.309722 | -2.19149 | -0.7179 | 0.077604 | -3.37733 |
| 33.61111 | 0.751592 | 2.381209 | 0.307057 | -0.15516 | -0.87165 | 0.45832 | 1.944587 | 0.043317 | -1.09528 | -0.26134 | -1.47736 | -1.40113 | -0.73658 | -2.2702 | -1.55565 | -0.1289 | -0.11875 | -0.51604 | -1.23374 | 0.550347 | -1.43612 | -0.39492 | 0.158044 | -4.58227 |
| 33.625 | 0.258707 | 1.038438 | 0.224858 | -0.15564 | -0.96464 | 0.405713 | 1.505635 | 0.169322 | -0.74984 | -0.32626 | -0.01082 | -1.34036 | -0.48899 | -2.08447 | -0.23717 | -0.08173 | -0.16746 | -0.28494 | -1.83553 | 0.511794 | -1.50629 | -0.20923 | 0.299371 | -5.57508 |
| 33.63889 | -0.05808 | -0.07667 | 0.017149 | -0.08089 | -1.00941 | 0.376115 | 0.955566 | 0.204413 | -0.80401 | -0.38749 | 1.283199 | -1.17079 | -0.05379 | -1.90971 | 1.071542 | -0.10969 | -0.19641 | 0.133799 | -2.74309 | 0.192843 | -1.89713 | -0.27363 | 0.368053 | -5.14991 |
| 33.65278 | -0.40567 | -0.82804 | -0.2343 | -0.03549 | -1.0034 | 0.338253 | 0.759154 | 0.266726 | -0.54178 | -0.47079 | 2.872747 | -0.94582 | 0.059912 | -2.36369 | 2.140482 | -0.33613 | -0.28506 | 0.450878 | -3.50671 | -0.25022 | -2.33024 | -0.39122 | 0.138109 | -4.25624 |
| 33.66667 | -0.79854 | -1.2932 | -0.51858 | -0.09536 | -1.20048 | 0.3045 | 0.62026 | 0.326067 | -0.55834 | -0.49514 | 3.800821 | -1.03039 | 0.089556 | -2.21998 | 2.946118 | -0.40151 | -0.38013 | 1.275211 | -2.57624 | -0.131 | -2.37658 | -0.44772 | -0.1525 | -2.51916 |
| 33.68056 | -0.8882 | -1.03649 | -0.8275 | -0.1754 | -1.37613 | 0.229033 | 0.39976 | 0.432375 | -0.6687 | -0.56364 | 5.201402 | -1.39075 | -0.40037 | -1.87756 | 2.822267 | -0.37842 | -0.39845 | 2.028007 | -1.66588 | 0.01203 | -1.71967 | -0.47725 | -0.48704 | 0.040112 |
| 33.69444 | -0.90614 | -0.70928 | -1.47878 | -0.32648 | -1.5799 | 0.196864 | 0.120744 | 0.375698 | -0.46451 | -0.60349 | 4.548028 | -1.69842 | -0.38381 | -1.01495 | 2.805195 | -0.00214 | -0.44652 | 1.322616 | -0.9765 | 0.051901 | -0.68164 | -0.21639 | -0.85021 | 2.0655 |
| 33.70833 | -1.3211 | -0.8938 | -2.16283 | -0.39768 | -1.98632 | 0.125451 | -0.21202 | 0.26478 | -0.495 | -0.66999 | 10.95313 | -1.82762 | 0.065711 | -0.06555 | 2.738708 | 0.475605 | -0.58641 | 0.284535 | -0.77476 | -0.09348 | 0.052161 | -0.15232 | -0.95854 | 2.560021 |
| 33.72222 | -2.44718 | -0.98384 | -2.80832 | -0.34019 | -2.40762 | -0.00856 | -0.47231 | 0.11835 | -0.50023 | -0.71003 | 7.923407 | -1.77535 | 0.377245 | 0.61602 | 1.578683 | 0.694498 | -0.65107 | -0.28011 | -0.76887 | -0.47813 | 0.489092 | -0.14228 | -1.16229 | 3.784367 |
| 33.73611 | -3.30184 | -1.13622 | -3.04823 | -0.29105 | -2.63059 | -0.24629 | -0.73051 | 0.176794 | -0.3113 | -0.66602 | 4.394439 | -1.5948 | 0.149638 | 0.629003 | 0.373341 | 0.715626 | -0.65683 | -0.65822 | -0.8338 | -0.90154 | 0.664364 | 0.088979 | -1.6421 | 4.926957 |
| 33.75 | -3.17826 | -1.23489 | -3.06158 | -0.29193 | -2.14763 | -0.32873 | -1.02194 | 0.332412 | -0.02097 | -0.59691 | 1.325046 | -1.23733 | -0.59898 | 0.113761 | -1.05636 | 0.36187 | -0.75299 | -0.89326 | -0.9633 | -1.07978 | 0.387431 | 0.081225 | -1.79509 | 4.277389 |
| 33.76389 | -2.8375 | -1.61821 | -3.04415 | -0.20603 | -1.5042 | -0.33209 | -1.21707 | 0.502403 | 0.370172 | -0.54421 | -2.23672 | -0.93705 | -1.4385 | -0.49767 | -2.91482 | -0.12017 | -0.92159 | -0.89414 | -2.35546 | -0.94825 | -0.58044 | -0.32502 | -1.68147 | 1.410671 |
| 33.77778 | -1.94398 | -1.91932 | -2.79778 | -0.14372 | -1.57352 | -0.33914 | -1.24798 | 0.581978 | 0.660604 | -0.50768 | -7.11458 | -0.58762 | -1.79556 | -0.97728 | -4.24654 | -0.64101 | -1.15752 | -0.67951 | -6.45025 | -0.58729 | -2.13899 | -0.63618 | -1.34417 | -1.27197 |
| 33.79167 | -2.12932 | -2.28334 | -2.4114 | -0.16068 | -1.75487 | -0.47336 | -1.22593 | 0.638909 | 0.646057 | -0.46708 | -8.30276 | -0.1766 | -1.88228 | -1.29807 | -4.10082 | -1.01317 | -1.1088 | -0.50083 | -5.02749 | -0.22657 | -2.36011 | -0.87019 | -1.21295 | -3.78845 |
| 33.80556 | -1.86063 | -2.22155 | -2.31704 | -0.1762 | -1.23565 | -0.56881 | -1.79335 | 0.643971 | 0.719256 | -0.45508 | -5.15253 | 0.241415 | -1.85827 | -1.17952 | -2.89164 | -1.41134 | -0.8021 | -0.44874 | -4.48464 | -0.01757 | -1.87924 | -0.88741 | -1.37101 | -5.57635 |
| 33.81944 | -1.6378 | -1.92521 | -2.4528 | -0.14749 | -0.33082 | -0.72641 | -2.51578 | 0.62408 | 0.890989 | -0.45493 | -2.97978 | 0.532357 | -1.84168 | -0.85611 | -1.29083 | -1.3928 | -0.62739 | -0.02406 | -5.72683 | 0.128797 | -1.00022 | -0.86036 | -1.44982 | -5.42542 |
| 33.83333 | -1.57457 | -1.31136 | -2.87778 | -0.13618 | 0.296712 | -0.83354 | -2.93657 | 0.550171 | 1.346308 | -0.31311 | -2.84318 | 0.652725 | -1.91337 | -0.64525 | -0.44348 | -1.22013 | -0.53677 | 0.282781 | -5.56247 | 0.269313 | -0.52424 | -0.90982 | -1.63863 | -5.10556 |
| 33.84722 | -1.01694 | -0.56698 | -2.84374 | -0.08539 | 0.288168 | -0.93633 | -2.71818 | 0.502611 | 1.705582 | -0.13454 | -1.98525 | 0.910438 | -1.90607 | -0.76621 | -0.21837 | -0.68547 | -0.45079 | 0.593125 | -4.56974 | 0.362251 | -0.40977 | -0.91955 | -1.74897 | -4.9259 |
| 33.86111 | -0.73212 | 0.273159 | -2.25091 | 0.040061 | 0.04137 | -1.00713 | -2.56371 | 0.465099 | 1.851659 | 0.03532 | -0.81798 | 0.996882 | -1.99335 | -0.28908 | -0.16016 | -0.13334 | -0.30462 | 0.569827 | -3.76429 | 0.185709 | -0.04342 | -0.66343 | -1.49465 | -3.91342 |
| 33.875 | -0.55168 | 0.575565 | -1.23214 | 0.411653 | 0.1164 | -0.84221 | -1.52684 | 0.415828 | 1.95994 | 0.347413 | -0.39333 | 0.938516 | -2.21604 | 0.396722 | 0.804253 | 0.231121 | -0.25052 | -0.0783 | -3.49352 | -0.13277 | 0.631243 | -0.35152 | -0.91685 | -3.65061 |
| 33.88889 | -0.52899 | 0.014083 | -0.87714 | 0.856686 | 0.206277 | -0.95841 | -0.85331 | 0.24113 | 1.373636 | 0.193585 | -0.91292 | 0.64547 | -3.00794 | 0.62618 | 1.511165 | 0.171187 | -0.61312 | -0.98036 | -4.81372 | -0.6556 | 0.762496 | -0.63725 | -1.13402 | -3.99875 |
| 33.90278 | -0.95027 | -0.57127 | -1.29245 | 0.769834 | -0.0618 | -1.03292 | -1.26542 | -0.04049 | 0.045417 | -0.67441 | -1.73444 | 0.073899 | -3.51936 | -0.86862 | 0.236264 | -0.46569 | -1.27886 | -1.29303 | -5.59117 | -1.09395 | -0.85508 | -1.82864 | -2.24058 | -7.19211 |
| 33.91667 | -1.61108 | -1.22556 | -1.68206 | 0.186944 | -1.0633 | -0.66701 | -2.28023 | -0.09651 | -1.12873 | -1.64343 | -2.17076 | -0.47424 | -1.58899 | -4.30571 | -2.34443 | -1.2652 | -1.82851 | -1.25916 | -4.83352 | -1.62861 | -3.01486 | -3.07642 | -4.27811 | -8.98743 |
| 33.93056 | -1.70187 | -1.69401 | -1.68618 | -0.07663 | -1.72585 | -0.36831 | -2.66462 | -0.05588 | -1.92875 | -2.13046 | -1.9247 | -0.90735 | -1.05776 | -5.0322 | -4.23014 | -1.71313 | -1.29425 | -1.51048 | -5.60534 | -1.95342 | -3.62287 | -3.00564 | -4.99285 | -6.03895 |
| 33.94444 | -1.47562 | -2.26084 | -1.84823 | -0.0929 | -1.62857 | -0.20362 | -2.26153 | 0.188326 | -2.13555 | -1.63348 | -1.10566 | -0.91679 | -0.46103 | -3.17908 | -2.75356 | -1.19306 | -1.13596 | -0.81765 | -3.34266 | -1.10474 | -1.99418 | -1.90728 | -3.56607 | -2.83702 |
| 33.95833 | -1.54462 | -2.88591 | -2.24674 | -0.20442 | -0.28142 | -0.51095 | -1.64669 | 0.3955 | -2.11016 | -0.97609 | -0.34416 | -0.61854 | 0.355163 | -0.96063 | -0.07351 | -0.82592 | -0.52172 | 0.285241 | -0.08084 | -0.14968 | 0.070051 | -0.59217 | -0.86327 | -1.33554 |
| 33.97222 | -1.74114 | -3.25922 | -2.44768 | -0.19794 | 1.296937 | -0.43317 | -1.21318 | 0.120373 | -1.89007 | -0.40879 | 0.307435 | -0.34245 | 0.871233 | 0.801633 | 1.09001 | -0.30022 | -0.00709 | 1.671558 | 1.53544 | 0.487548 | 1.303836 | -0.10066 | -0.11311 | -1.14961 |
| 33.98611 | -1.52719 | -3.02414 | -1.98729 | -0.15165 | 2.182928 | -0.36464 | -0.9441 | -0.53281 | -1.15015 | -0.17521 | 0.397881 | -0.16511 | 1.039322 | 1.546607 | 1.107191 | 0.074931 | 0.123401 | 1.758777 | 1.432251 | 0.744242 | 1.453866 | -0.07696 | -0.31339 | -1.16873 |
| 34 | -1.64882 | -2.67919 | -1.3383 | -0.05183 | 1.833795 | -0.07976 | -0.63228 | -0.80837 | -0.68463 | -0.05036 | -0.05396 | -0.06167 | 1.0294 | 1.507581 | 1.460522 | 0.234971 | 0.286979 | 1.645344 | 1.086102 | 0.931956 | 1.266836 | 0.561904 | -0.21755 | -1.01605 |
| 34.01389 | -1.33264 | -2.36514 | -0.91751 | -0.01224 | 1.12699 | -0.01831 | -0.45387 | -0.55807 | -0.31153 | -0.0361 | -0.28721 | 0.072094 | 0.847512 | 1.166247 | 1.716147 | 0.462585 | 0.370771 | 1.257231 | 0.711417 | 0.910926 | 1.018446 | 1.61256 | -0.02259 | -1.04903 |
| 34.02778 | -1.04871 | -2.09151 | -0.70571 | 0.041619 | 0.714336 | 0.050705 | -0.31554 | -0.16586 | 0.006762 | -0.14059 | -0.05383 | 0.214895 | 0.651301 | 0.675666 | 1.4573 | 0.52276 | 0.371898 | 1.050273 | 0.389166 | 0.893578 | 0.698517 | 2.012803 | -0.02764 | -1.2208 |
| 34.04167 | -0.90025 | -1.69712 | -0.8089 | 0.101901 | 0.57049 | 0.014672 | -0.26445 | 0.066325 | 0.21222 | -0.21197 | 0.254588 | 0.262798 | 0.531357 | 0.330969 | 1.09358 | 0.534628 | 0.358271 | 0.944142 | 0.131626 | 0.882157 | 0.424111 | 2.107147 | -0.12331 | -0.66896 |
| 34.05556 | -0.63701 | -1.36418 | -1.16336 | 0.136052 | 0.325463 | 0.030864 | -0.34248 | 0.180645 | 0.251942 | -0.19428 | 0.637076 | 0.256134 | 0.472331 | 0.187349 | 0.773017 | 0.446286 | 0.273523 | 0.653552 | 0.009266 | 0.686358 | 0.26041 | 1.700119 | -0.20004 | 0.206292 |
| 34.06944 | -0.19523 | -1.18589 | -1.00412 | 0.212497 | -0.25991 | 0.08853 | -0.27726 | 0.243291 | 0.127213 | -0.07795 | 1.247478 | 0.262639 | 0.434919 | 0.122781 | 0.476733 | 0.31005 | 0.170951 | 0.474407 | -0.00557 | 0.434339 | 0.15264 | 1.124281 | -0.15918 | 0.258803 |
| 34.08333 | 0.00996 | -1.14572 | -0.36992 | 0.376088 | -0.68551 | 0.24036 | -0.04747 | 0.287019 | 0.108759 | 0.031053 | 1.778474 | 0.287505 | 0.404456 | 0.095822 | 0.247993 | 0.175424 | 0.042973 | 0.315539 | 0.006485 | 0.174993 | 0.171366 | 0.738121 | -0.05827 | 0.291314 |
| 34.09722 | 0.266444 | -1.16714 | 0.066491 | 0.474413 | -0.42954 | 0.300274 | 0.09845 | 0.277944 | 0.255325 | 0.017007 | 2.032505 | 0.281047 | 0.353274 | 0.041526 | 0.101567 | 0.078986 | -0.09601 | 0.110912 | -0.02244 | -0.0273 | 0.240945 | 0.497398 | -0.02497 | 0.230467 |
| 34.11111 | 0.279869 | -1.19374 | 0.254305 | 0.509951 | 0.237208 | 0.385565 | 0.045744 | 0.247216 | 0.488994 | -0.03902 | 2.014163 | 0.274293 | 0.322721 | -0.00509 | 0.060379 | -0.01755 | -0.17067 | -0.02817 | -0.00087 | -0.31349 | 0.336209 | 0.562217 | 0.039521 | -0.03827 |
| 34.125 | 0.293012 | -1.20218 | 0.35135 | 0.499012 | 0.776051 | 0.465452 | -0.14588 | 0.285043 | 0.512071 | -0.01138 | 2.017458 | 0.304651 | 0.358415 | 0.029085 | 0.15054 | -0.13608 | -0.18226 | 0.021675 | 0.131451 | -0.36319 | 0.518227 | 0.854372 | 0.158225 | -0.10394 |
| 34.13889 | 0.376671 | -1.06966 | 0.338546 | 0.491225 | 1.09323 | 0.523096 | -0.20208 | 0.383407 | 0.308503 | 0.057116 | 2.000583 | 0.334842 | 0.395272 | 0.064704 | 0.226208 | -0.15241 | -0.19761 | 0.080876 | 0.182606 | -0.04063 | 0.591628 | 1.191953 | 0.18296 | -0.19365 |
| 34.15278 | 0.375536 | -0.88686 | 0.207396 | 0.528951 | 1.188733 | 0.504828 | -0.11388 | 0.406392 | -0.01625 | 0.070725 | 1.914228 | 0.289997 | 0.354934 | 0.000238 | 0.208798 | -0.10498 | -0.41524 | 0.013368 | 0.039194 | 0.02788 | 0.498589 | 1.313513 | 0.053451 | -0.42479 |
| 34.16667 | 0.394615 | -0.6949 | 0.065564 | 0.453934 | 1.066823 | 0.433929 | -0.12004 | 0.317426 | -0.20405 | 0.065929 | 1.885432 | 0.237538 | 0.310007 | -0.07294 | 0.170779 | -0.09639 | -0.4976 | -0.0277 | -0.10965 | -0.00733 | 0.351831 | 1.193907 | -0.03174 | -0.50551 |
| 34.18056 | 0.381056 | -0.56394 | 0.0414 | 0.282605 | 0.363455 | 0.385523 | -0.13623 | 0.219368 | -0.07909 | 0.080089 | 1.842569 | 0.207913 | 0.34456 | -0.01725 | 0.199774 | -0.025 | -0.39416 | 0.010347 | -0.10265 | 0.120205 | 0.241497 | 0.922752 | 0.053594 | -0.38806 |
| 34.19444 | 0.505724 | -0.58968 | 0.123939 | 0.114435 | -0.62868 | 0.361359 | -0.09554 | 0.214478 | 0.04998 | 0.058362 | 1.754857 | 0.162296 | 0.387896 | 0.067656 | 0.218448 | 0.0381 | -0.2092 | 0.105236 | -0.04003 | 0.257981 | 0.191028 | 0.554892 | 0.156786 | -0.31642 |
| 34.20833 | 0.727606 | -0.75731 | 0.15129 | -0.02672 | -1.43357 | 0.348783 | -0.06892 | 0.289013 | 0.170769 | 0.007315 | 1.599225 | 0.0334 | 0.40085 | 0.062153 | 0.174179 | 0.0908 | -0.00058 | 0.218278 | -0.01985 | 0.281061 | 0.13057 | 0.323346 | 0.178844 | -0.42434 |
| 34.22222 | 0.811056 | -0.99525 | 0.036318 | -0.1304 | -1.76214 | 0.36815 | -0.03077 | 0.39467 | 0.266092 | -0.06515 | 1.367032 | -0.10312 | 0.369146 | -0.01206 | 0.13207 | 0.175261 | 0.173132 | 0.253686 | -0.04349 | 0.257543 | 0.125039 | 0.268021 | 0.153812 | -0.61225 |
| 34.23611 | 0.803893 | -1.22873 | -0.04543 | -0.18828 | -1.65351 | 0.366236 | 0.005316 | 0.448079 | 0.225661 | -0.13103 | 1.110341 | -0.21133 | 0.300754 | -0.12359 | 0.119613 | 0.200273 | 0.180379 | 0.175527 | -0.14919 | 0.206137 | 0.074179 | 0.068284 | 0.030801 | -0.78559 |
| 34.25 | 0.76736 | -1.40796 | -0.03274 | -0.20946 | -1.18774 | 0.415096 | -0.01677 | 0.464594 | 0.078498 | -0.16697 | 0.925491 | -0.30797 | 0.232391 | -0.21076 | 0.112407 | 0.15542 | 0.036213 | 0.142646 | -0.29019 | 0.113379 | -0.01086 | -0.24934 | -0.10098 | -0.89927 |
| 34.26389 | 0.519119 | -1.45625 | 0.041411 | -0.09423 | -0.4755 | 0.437967 | -0.0292 | 0.513127 | -0.06084 | -0.19108 | 0.804097 | -0.37915 | 0.251834 | -0.17342 | 0.14626 | 0.103148 | -0.13953 | 0.196122 | -0.2374 | 0.12632 | 0.060306 | -0.05172 | -0.1079 | -0.80636 |
| 34.27778 | 0.302801 | -1.43864 | 0.163993 | 0.009182 | 0.030599 | 0.43693 | -0.03691 | 0.624708 | -0.21641 | -0.15587 | 0.87371 | -0.44307 | 0.392989 | 0.049588 | 0.296721 | 0.158494 | -0.13875 | 0.341526 | 0.123258 | 0.241515 | 0.294364 | 0.603223 | 0.12364 | -0.41238 |
| 34.29167 | 0.157489 | -1.4766 | 0.272107 | 0.003042 | -0.00964 | 0.428884 | -0.06639 | 0.743599 | -0.37081 | -0.15215 | 0.526742 | -0.50293 | 0.580234 | 0.354654 | 0.64348 | 0.31186 | 0.061943 | 0.513804 | 0.643862 | 0.438671 | 0.701239 | 1.285511 | 0.463676 | 0.298074 |
| 34.30556 | -0.10034 | -1.46186 | 0.304349 | -0.08373 | -0.49848 | 0.42059 | -0.13852 | 0.815407 | -0.41134 | -0.14472 | 0.585317 | -0.46062 | 0.736319 | 0.52094 | 1.111485 | 0.444098 | 0.379455 | 0.639488 | 0.98799 | 0.582845 | 0.933586 | 1.72738 | 0.648536 | 0.885214 |
| 34.31944 | -0.14466 | -1.4545 | 0.269256 | -0.20953 | -0.80828 | 0.482888 | -0.2828 | 0.746096 | -0.27421 | -0.11788 | 0.434173 | -0.39626 | 0.831187 | 0.51542 | 1.453811 | 0.51209 | 0.580874 | 0.642302 | 0.980242 | 0.588652 | 0.897851 | 1.942052 | 0.714302 | 0.999526 |
| 34.33333 | -0.09197 | -1.39437 | 0.246139 | -0.15305 | -0.39849 | 0.557926 | -0.43465 | 0.717569 | -0.29496 | -0.05714 | 0.645022 | -0.38057 | 0.827584 | 0.49327 | 1.585447 | 0.520756 | 0.601668 | 0.645451 | 0.819305 | 0.532897 | 0.765104 | 2.097259 | 0.653813 | 1.182428 |
| 34.34722 | -0.13814 | -1.34494 | 0.238354 | -0.05 | 0.282195 | 0.501442 | -0.56899 | 0.710309 | -0.5403 | -0.04222 | 0.659789 | -0.47219 | 0.904238 | 0.514902 | 1.615681 | 0.508165 | 0.468984 | 0.55599 | 0.792908 | 0.526316 | 0.7627 | 2.243589 | 0.484067 | 1.196892 |
| 34.36111 | -0.19489 | -1.34106 | 0.349702 | -0.02388 | 0.758144 | 0.410923 | -0.69605 | 0.72022 | -0.78593 | -0.08482 | 0.900476 | -0.59025 | 0.991219 | 0.507016 | 1.509552 | 0.451714 | 0.371588 | 0.549251 | 0.887415 | 0.528705 | 0.87665 | 2.304842 | 0.282809 | 1.4921 |
| 34.375 | -0.02332 | -1.3151 | 0.494692 | 0.021245 | 0.974093 | 0.356766 | -0.77525 | 0.673988 | -0.94009 | -0.04215 | 1.392425 | -0.58099 | 1.129934 | 0.455593 | 1.481521 | 0.433045 | 0.289594 | 0.551584 | 0.901966 | 0.453177 | 1.049252 | 2.441947 | 0.178371 | 1.982327 |
| 34.38889 | 0.205395 | -1.09954 | 0.348697 | 0.032119 | 0.998327 | 0.318474 | -0.80706 | 0.657783 | -0.63808 | 0.194298 | 1.956162 | -0.50214 | 1.35508 | 0.670732 | 1.505429 | 0.515052 | 0.333951 | 0.625412 | 1.264263 | 0.470313 | 1.369988 | 2.900827 | 0.522803 | 2.140419 |
| 34.40278 | 0.824437 | -0.57238 | 0.122452 | 0.105108 | 1.217348 | 0.403776 | -0.66111 | 0.737441 | 0.079633 | 0.651064 | 2.773957 | -0.21315 | 1.694392 | 1.376088 | 1.766249 | 0.720715 | 0.663397 | 0.820067 | 2.341186 | 0.771033 | 1.883584 | 4.164859 | 1.540072 | 2.666279 |
| 34.41667 | 1.324324 | 0.036943 | 0.075131 | 0.375658 | 1.979602 | 0.368666 | -0.27844 | 0.66942 | 1.525257 | 1.070821 | 2.812068 | 0.279941 | 2.174835 | 1.875564 | 2.150812 | 1.180019 | 0.898177 | 0.63187 | 2.922109 | 0.857641 | 1.872045 | 5.800058 | 2.184725 | 2.742675 |
| 34.43056 | 1.21138 | 0.494467 | 0.115064 | 0.670974 | 3.249323 | 0.394081 | 0.359845 | 0.693749 | 2.983718 | 1.080157 | 2.950896 | 0.776436 | 1.942866 | 1.680487 | 1.794669 | 1.645012 | 0.86387 | 0.136669 | 2.23977 | 0.572923 | 1.325805 | 7.410114 | 2.035573 | 1.768738 |
| 34.44444 | 1.015417 | 0.916917 | 0.528494 | 0.976624 | 3.782494 | 0.499202 | 0.959096 | 1.277139 | 2.394653 | 0.79104 | 3.584525 | 1.227971 | 1.704501 | 1.38056 | 1.179167 | 1.454165 | 0.685874 | -0.19521 | 1.593032 | 0.272159 | 1.054111 | 6.858771 | 1.816114 | 2.56541 |
| 34.45833 | 0.97267 | 1.451359 | 1.17972 | 0.95701 | 2.031739 | 0.750281 | 1.058923 | 1.85535 | 1.805416 | 0.504679 | 3.293586 | 1.351213 | 1.452759 | 1.325032 | 0.765748 | 1.40968 | 0.666087 | 0.259321 | 2.118014 | 0.634366 | 1.197205 | 3.403124 | 1.770549 | 3.576628 |
| 34.47222 | 1.046102 | 1.700906 | 1.597098 | 0.638954 | -0.02632 | 0.746566 | 0.666028 | 1.964869 | 0.288362 | 0.366642 | 2.733958 | 1.485718 | 1.310219 | 1.192176 | 0.927185 | 1.44853 | 0.605171 | 0.738175 | 2.812792 | 0.592709 | 1.223351 | 1.389039 | 2.097144 | 3.544853 |
| 34.48611 | 1.189387 | 1.703049 | 1.657161 | 0.304753 | -1.66682 | 0.692867 | 0.141257 | 2.00707 | -1.11306 | 0.413591 | 2.158373 | 1.373505 | 1.148953 | 1.044846 | 1.259466 | 1.532678 | 0.531271 | 0.982468 | 3.074439 | 0.116873 | 1.266255 | 0.005558 | 2.018876 | 3.598489 |
| 34.5 | 1.273769 | 1.654445 | 1.677043 | 0.156098 | -2.45082 | 0.645227 | -0.12547 | 1.921028 | -1.43436 | 0.719045 | 1.440319 | 1.390443 | 1.11156 | 0.997203 | 1.807656 | 1.645892 | 0.445282 | 0.71124 | 2.957187 | -0.32123 | 1.685526 | -0.74778 | 1.790837 | 4.389279 |
| 34.51389 | 1.226501 | 1.750296 | 1.875932 | 0.199681 | -1.91369 | 0.568683 | -0.10219 | 1.968462 | -0.89565 | 0.827235 | 0.82491 | 1.308992 | 1.007173 | 1.051882 | 2.078839 | 1.528283 | 0.404344 | 0.427033 | 2.961805 | 0.005517 | 2.515795 | -1.05615 | 1.956158 | 4.155669 |
| 34.52778 | 1.251004 | 1.850882 | 2.260019 | 0.4103 | -0.68323 | 0.534849 | 0.126041 | 2.052037 | -0.27422 | 0.732935 | 0.415266 | 1.249744 | 0.947512 | 1.063319 | 2.28259 | 1.579709 | 0.387103 | -0.02888 | 3.132246 | 0.472651 | 3.102195 | -0.00822 | 1.703444 | 3.266049 |
| 34.54167 | 1.073843 | 1.807847 | 2.276486 | 0.55925 | 0.234357 | 0.506671 | 0.359686 | 1.892639 | 0.016935 | 0.488219 | 0.103194 | 0.94333 | 0.967447 | 1.032256 | 2.453459 | 1.7289 | 0.346427 | 0.171267 | 3.520764 | 0.815919 | 2.996487 | 1.726041 | 1.660449 | 2.513343 |
| 34.55556 | 1.063555 | 1.585814 | 1.811726 | 0.432913 | 0.471522 | 0.539908 | 0.481802 | 1.442488 | -0.28492 | 0.342916 | -0.08791 | 0.634735 | 0.938085 | 1.155152 | 2.52749 | 1.93399 | 0.278442 | 0.525438 | 3.366843 | 0.837596 | 2.427381 | 2.774929 | 1.693637 | 2.119846 |
| 34.56944 | 1.051266 | 1.667951 | 1.455455 | 0.274247 | 0.253648 | 0.496404 | 0.464419 | 0.97958 | -1.01574 | 0.272123 | -0.23635 | 0.255782 | 0.942221 | 1.238061 | 2.286349 | 1.74133 | 0.237149 | 1.060191 | 2.559852 | 0.879211 | 1.781561 | 2.876403 | 0.879759 | 1.97445 |
| 34.58333 | 1.154544 | 1.807738 | 1.212126 | 0.146058 | -0.30118 | 0.481616 | 0.407066 | 0.479041 | -1.61474 | 0.174017 | -0.36769 | -0.07783 | 0.862769 | 1.32973 | 1.916157 | 1.163755 | 0.165616 | 1.262355 | 1.654348 | 0.831657 | 1.174003 | 2.129271 | -0.42433 | 1.62609 |
| 34.59722 | 1.180674 | 1.721794 | 1.068152 | -0.02958 | -1.46092 | 0.527732 | 0.447796 | 0.009097 | -2.16034 | -0.10017 | -0.36505 | -0.35688 | 0.595714 | 0.983848 | 1.229938 | 0.603218 | 0.054307 | 0.821951 | 1.09921 | 0.779582 | 0.675689 | 0.856309 | -1.84731 | 0.873576 |
| 34.61111 | 1.183567 | 1.37267 | 0.961969 | -0.35617 | -3.01314 | 0.510394 | 0.491347 | -0.19071 | -2.04751 | -0.25267 | -0.31278 | -0.54806 | 0.467801 | 0.575055 | 0.619804 | 0.226012 | -0.04167 | 0.294752 | 0.602099 | 0.76233 | 0.289172 | -0.40207 | -1.96308 | 0.330843 |
| 34.625 | 1.231034 | 0.958526 | 0.834494 | -0.65268 | -3.62482 | 0.5107 | 0.386834 | -0.1724 | -1.66339 | -0.4482 | -0.08414 | -0.6938 | 0.514292 | 0.381771 | 0.337519 | 0.105275 | -0.07415 | 0.405801 | 0.159489 | 0.931959 | 0.082478 | -1.47072 | -2.88377 | 0.061827 |
| 34.63889 | 1.303339 | 0.618358 | 0.723456 | -0.67765 | -3.09069 | 0.462092 | 0.033939 | -0.06001 | -1.30753 | -0.53126 | 0.530011 | -0.95814 | 0.688553 | 0.205821 | 0.332305 | 0.199984 | -0.13685 | 0.409145 | -0.09139 | 1.087315 | -0.08164 | -2.20688 | -0.82433 | -0.14155 |
| 34.65278 | 1.488438 | 0.482001 | 0.447107 | -0.60738 | -2.06121 | 0.389909 | -0.4937 | 0.075049 | -1.36333 | -0.55866 | 1.614986 | -1.32969 | 0.865522 | -0.00039 | 0.493106 | 0.41449 | -0.18211 | 0.021873 | -0.256 | 0.814237 | -0.35282 | -3.22603 | -0.39386 | -0.13422 |
| 34.66667 | 1.652817 | 0.457417 | 0.217327 | -0.53055 | -1.05328 | 0.358839 | -0.85805 | 0.189193 | -1.42154 | -0.57067 | 1.788956 | -1.40237 | 0.945466 | -0.04091 | 0.625561 | 0.551691 | -0.25554 | -0.4521 | -0.42819 | 0.597736 | -0.6058 | -3.04652 | 0.049867 | -0.13353 |
| 34.68056 | 1.72437 | 0.308343 | 0.146517 | -0.55548 | -0.7014 | 0.398739 | -0.8999 | 0.225666 | -1.22955 | -0.77725 | 1.223945 | -1.51314 | 0.897367 | -0.01306 | 0.950111 | 0.380276 | -0.35958 | -0.61743 | -0.45255 | 0.326702 | -0.76192 | -2.57523 | 1.125451 | -0.35001 |
| 34.69444 | 1.639688 | -0.02809 | 0.239056 | -0.66737 | -0.96551 | 0.501477 | -0.90949 | 0.14905 | -0.76808 | -0.90866 | 0.895406 | -1.50723 | 0.911432 | -0.08218 | 1.049267 | -0.07202 | -0.47461 | -0.38981 | -0.61795 | 0.262472 | -0.73066 | -2.32057 | 2.072686 | -0.49023 |
| 34.70833 | 1.393124 | -0.52913 | 0.180237 | -0.72812 | -1.69177 | 0.472246 | -1.02798 | 0.048135 | -0.55128 | -0.92497 | 1.31019 | -1.4032 | 0.976007 | -0.22663 | 0.847373 | -0.68086 | -0.50114 | 0.384834 | -0.6537 | 0.665598 | -0.64274 | -2.29482 | 2.648355 | -0.84607 |
| 34.72222 | 0.98382 | -1.00035 | 0.173959 | -0.7967 | -2.09452 | 0.496059 | -1.10104 | -0.13645 | -0.54995 | -0.81747 | 1.406693 | -1.34105 | 1.078638 | -0.20142 | 0.367652 | -0.83055 | -0.48376 | 0.856355 | -0.85455 | 0.446129 | -0.46154 | -2.41811 | 2.168496 | -1.22778 |
| 34.73611 | 0.705945 | -1.35364 | 0.218625 | -0.8224 | -1.61991 | 0.373738 | -0.99831 | -0.36715 | -0.21899 | -0.76299 | 1.404058 | -1.27295 | 1.101042 | -0.14043 | -0.07744 | -0.66047 | -0.36352 | 1.015927 | -1.28013 | 0.604024 | -0.45818 | -3.38 | 1.372103 | -1.49198 |
| 34.75 | 0.427569 | -1.5699 | 0.290813 | -0.84232 | -1.68432 | 0.178884 | -0.86849 | -0.7546 | 0.143625 | -0.79324 | 1.195195 | -1.16159 | 0.843117 | -0.19851 | -0.59441 | -0.685 | -0.2882 | 0.793495 | -1.53595 | 0.55523 | -0.31045 | -4.26484 | -0.03564 | -1.91072 |
| 34.76389 | 0.097376 | -1.5525 | 0.232057 | -0.89059 | -2.20686 | 0.142075 | -0.7759 | -1.18141 | 0.344727 | -0.80725 | 0.749858 | -0.85666 | 0.535678 | -0.3211 | -0.82301 | -0.83728 | -0.33579 | 0.56771 | -1.44999 | 0.549558 | -0.4122 | -4.30087 | -1.53077 | -2.44353 |
| 34.77778 | -0.3071 | -1.59853 | 0.184781 | -0.88618 | -2.93188 | 0.112645 | -0.80567 | -0.89183 | 0.617646 | -0.75416 | 0.364561 | -0.50174 | 0.564676 | -0.31488 | -1.23703 | -0.8315 | -0.37746 | 0.268978 | -1.34294 | 0.439069 | -0.67664 | -4.06029 | -2.99427 | -2.70518 |
| 34.79167 | -0.69611 | -1.78134 | 0.200025 | -0.80813 | -3.81808 | 0.114178 | -0.74333 | -0.33699 | 0.883557 | -0.6634 | 0.042648 | -0.06271 | 0.740187 | -0.54321 | -1.51644 | -0.86083 | -0.30521 | -0.05997 | -1.01691 | 0.451918 | -0.61962 | -3.42814 | -4.37246 | -2.35986 |
| 34.80556 | -0.91159 | -1.6319 | 0.34116 | -0.60711 | -3.83092 | 0.126356 | -0.57079 | -0.04804 | 1.22077 | -0.67549 | -0.44574 | 0.274095 | 0.562026 | -0.70087 | -1.61047 | -0.89801 | -0.24845 | -0.26235 | -0.92137 | 0.396539 | -0.27187 | -2.73374 | -4.59283 | -2.92045 |
| 34.81944 | -0.92413 | -1.28022 | 0.439577 | -0.3366 | -3.41813 | 0.200052 | -0.25364 | 0.062037 | 1.382128 | -0.76304 | -0.77812 | 0.464669 | 0.325535 | -0.82796 | -1.36903 | -0.88592 | -0.33148 | -0.16546 | -0.90048 | 0.394845 | -0.31644 | -1.54321 | -2.97437 | -3.2848 |
| 34.83333 | -0.79611 | -1.05521 | 0.514551 | -0.08046 | -2.64787 | 0.348868 | 0.03843 | 0.102398 | 1.508302 | -0.56685 | -0.49221 | 0.527773 | 0.160555 | -0.77262 | -1.14598 | -0.81837 | -0.41647 | -0.13546 | -0.89723 | 0.490797 | -0.08544 | -0.83717 | -0.8233 | -4.01834 |
| 34.84722 | -0.54544 | -0.81528 | 0.33757 | 0.038183 | -2.16563 | 0.529001 | 0.248759 | 0.244411 | 1.671399 | -0.12195 | -0.41025 | 0.572641 | -0.31354 | -0.74031 | -1.09958 | -0.6586 | -0.44216 | -0.16336 | -0.88777 | 0.842416 | -0.1723 | -0.31287 | -0.89686 | -3.56649 |
| 34.86111 | -0.44318 | -0.50575 | 0.220012 | 0.165838 | -1.50701 | 0.448819 | 0.401788 | 0.415817 | 1.620958 | 0.25426 | -0.52026 | 0.504656 | -0.49154 | -0.58026 | -0.90191 | -0.60187 | -0.36551 | 0.004195 | -0.78863 | 1.047199 | -0.06353 | -0.13224 | -1.15589 | -2.95711 |
| 34.875 | -0.35921 | 0.279046 | 0.233626 | 0.339439 | -1.03587 | 0.207054 | 0.452511 | 0.407214 | 1.64724 | 0.241193 | -1.17205 | 0.279685 | -0.85839 | -0.02413 | -0.19958 | -0.185 | -0.28064 | 0.515193 | -1.13325 | 1.038239 | 0.27428 | 0.091123 | -0.8509 | -1.90172 |
| 34.88889 | -0.73594 | 1.305088 | 0.338286 | 0.341969 | -0.94101 | -0.1311 | 0.471966 | -0.16075 | 1.390783 | -0.4338 | -2.50147 | -0.32608 | -2.17279 | 0.877631 | 1.030455 | 0.502907 | 0.080968 | 0.701149 | -2.82969 | 0.684434 | 0.321109 | 0.279187 | -1.31625 | -0.76703 |
| 34.90278 | -0.91133 | 1.405917 | 0.177742 | 0.035573 | -1.62969 | -0.38538 | 0.134103 | -0.7926 | 0.476885 | -1.56144 | -3.44494 | -0.82552 | -2.23358 | 0.495532 | 1.438784 | 0.367032 | -0.18376 | 0.310041 | -4.90812 | 0.3426 | -0.7976 | -0.33989 | -1.23146 | -0.57484 |
| 34.91667 | -0.78334 | 0.386968 | 0.242234 | -0.15119 | -2.65505 | -0.36922 | -0.19294 | -0.89207 | -0.41975 | -2.90109 | -2.98277 | -0.81658 | -1.55381 | -0.67436 | 0.348614 | -0.57873 | -0.8165 | 0.016433 | -4.88201 | 0.068077 | -1.25375 | -1.7858 | -2.16897 | -2.39379 |

| | | | | | | | | | | | | | | | | | | | | | | | | |
|---|---|---|---|---|---|---|---|---|---|---|---|---|---|---|---|---|---|---|---|---|---|---|---|---|
| 34.93056 | -0.34795 | -0.5213 | 0.344449 | 0.115365 | -2.76366 | 0.014304 | -0.45462 | -0.57175 | -0.56918 | -3.26065 | -1.91415 | -0.4744 | -0.28756 | -1.04807 | -0.85124 | -0.58798 | -1.0894 | -0.08267 | -4.05637 | -0.2392 | -1.90908 | -1.99908 | -2.27388 | -2.41159 |
| 34.94444 | -0.03888 | -0.95856 | 0.250143 | 0.45567 | -2.12479 | 0.356491 | -0.62923 | -0.25057 | -0.49283 | -3.39047 | -0.82048 | -0.11744 | 0.627501 | -0.95144 | -1.28825 | 0.010949 | -1.29627 | 0.006513 | -2.72752 | -0.14218 | -1.55496 | -3.43041 | -2.41051 | -1.73107 |
| 34.95833 | 0.011494 | -1.20204 | 0.041824 | 0.52933 | -1.3093 | 0.459971 | -0.79618 | -0.10376 | -0.90838 | -3.54053 | 0.150428 | 0.046765 | 0.972749 | -0.77559 | -0.8625 | 0.111771 | -1.14067 | 0.228565 | -1.68298 | 0.397718 | -0.7665 | -3.36662 | -2.06956 | -1.91637 |
| 34.97222 | -0.04627 | -1.05663 | 0.011088 | 0.394192 | -0.38497 | 0.488646 | -0.72397 | -0.08907 | -0.92177 | -2.92234 | 0.764099 | -0.02618 | 0.866821 | -0.6591 | -0.12925 | 0.121695 | -1.26815 | 0.528421 | -1.00069 | -0.71475 | 0.012782 | -1.68053 | -2.24479 | -1.93532 |
| 34.98611 | 0.060778 | -0.95136 | 0.027415 | 0.334275 | 0.068557 | 0.461379 | -0.58338 | -0.11694 | -0.69019 | -1.90756 | 0.868424 | -0.18107 | 0.755703 | -0.50427 | 0.449522 | 0.34946 | -0.8588 | 0.64076 | -0.42357 | 0.240704 | 0.4912 | -0.64888 | -1.79606 | -1.03515 |
| 35 | 0.31632 | -0.94587 | 0.077697 | 0.39571 | 0.029057 | 0.465926 | -0.49488 | -0.08358 | -0.4481 | -0.92734 | 0.808744 | -0.18995 | 0.616875 | -0.35344 | 0.814527 | 0.668104 | 0.073893 | 0.762104 | -0.02732 | 1.376285 | 0.553654 | 0.4714 | -0.48307 | -0.40872 |
| 35.01389 | 0.502853 | -1.15227 | 0.23466 | 0.344947 | 0.033552 | 0.495946 | -0.3917 | 0.065626 | -0.32597 | -0.26248 | 0.652526 | -0.07379 | 0.449456 | -0.23372 | 0.967443 | 0.897964 | 0.796592 | 0.89393 | 0.17603 | 2.300343 | 0.435232 | 1.698296 | 0.28905 | 0.016796 |
| 35.02778 | 0.580055 | -1.29092 | 0.322851 | 0.314379 | 0.315665 | 0.470468 | -0.20863 | 0.252666 | -0.29797 | -0.05984 | 0.517303 | 0.088704 | 0.25362 | -0.18746 | 0.971273 | 1.019438 | 1.274575 | 0.857548 | 0.195826 | 1.842016 | 0.310223 | 2.516258 | 0.347068 | 0.563881 |
| 35.04167 | 0.565168 | -1.17784 | 0.314046 | 0.237705 | 0.734324 | 0.401199 | -0.00071 | 0.428424 | -0.29164 | -0.32461 | 0.422984 | 0.154419 | 0.107632 | -0.15656 | 0.883101 | 1.003301 | 1.238483 | 0.938888 | 0.169782 | -0.21301 | 0.105594 | 3.099868 | 0.3809 | 0.985996 |
| 35.05556 | 0.478679 | -0.93558 | 0.25417 | 0.226565 | 1.17996 | 0.330345 | 0.118758 | 0.550277 | -0.22963 | -0.83519 | 0.506162 | 0.215769 | 0.048635 | -0.09949 | 0.976268 | 1.059045 | 0.762956 | 0.803169 | 0.137711 | 1.233309 | -0.08263 | 3.688537 | 0.338153 | 0.774023 |
| 35.06944 | 0.451104 | -0.8842 | 0.179468 | 0.311145 | 1.520734 | 0.348331 | 0.158776 | 0.568823 | -0.15139 | -1.41515 | 0.689881 | 0.247877 | 0.040878 | -0.09014 | 1.082229 | 1.236945 | 0.592077 | 0.474426 | 0.126115 | 2.730188 | -0.09134 | 4.005489 | 0.232108 | 0.576394 |
| 35.08333 | 0.431033 | -0.93665 | 0.174979 | 0.351574 | 1.992329 | 0.343813 | 0.107975 | 0.53892 | 0.16111 | -1.66076 | 0.887326 | 0.236602 | 0.11778 | -0.08097 | 1.127379 | 1.254786 | 0.426475 | 0.611519 | 0.079302 | 1.281399 | 0.00207 | 4.065428 | 0.17564 | 0.31408 |
| 35.09722 | 0.495401 | -0.98253 | 0.255928 | 0.483909 | 2.227696 | 0.318899 | 0.015867 | 0.474382 | 0.262592 | -1.57745 | 1.025833 | 0.214135 | 0.253092 | -0.09729 | 1.124844 | 1.195307 | 0.309067 | 0.475031 | -0.02512 | 0.681108 | 0.055798 | 3.464842 | 0.128178 | 0.132313 |
| 35.11111 | 0.539955 | -0.95351 | 0.38647 | 0.608294 | 2.498339 | 0.351957 | -0.10295 | 0.325731 | 0.437487 | -1.32157 | 0.991384 | 0.197894 | 0.372808 | -0.12603 | 1.012885 | 0.908748 | 0.151783 | 0.325117 | -0.14258 | 0.175568 | 0.078933 | 2.708142 | 0.032941 | 0.472318 |
| 35.125 | 0.531646 | -0.94566 | 0.496569 | 0.623201 | 2.863101 | 0.453378 | -0.13333 | 0.233626 | 0.368337 | -1.05024 | 0.961541 | 0.214162 | 0.428719 | -0.11983 | 0.929189 | 0.640942 | -0.01217 | 0.110001 | -0.27125 | -0.25747 | 0.007126 | 2.620893 | 0.083528 | 0.784184 |
| 35.13889 | 0.417493 | -0.72265 | 0.569697 | 0.748693 | 2.814825 | 0.519775 | -0.07391 | 0.297438 | 0.292481 | -0.84896 | 0.796232 | 0.161999 | 0.468588 | -0.1323 | 0.92946 | 0.366534 | -0.11927 | 0.152366 | -0.3467 | -1.09835 | -0.02006 | 3.041962 | 0.200658 | 1.308244 |
| 35.15278 | 0.375415 | -0.41516 | 0.607124 | 0.792921 | 2.738465 | 0.550628 | -0.02861 | 0.379555 | 0.308593 | -0.78215 | 0.739803 | 0.148752 | 0.571437 | -0.14913 | 0.93306 | 0.127666 | -0.20336 | 0.440592 | -0.30149 | -0.44085 | 0.074972 | 3.754098 | 0.368748 | 1.444343 |
| 35.16667 | 0.356458 | -0.14375 | 0.563879 | 0.777803 | 1.912547 | 0.498361 | -0.04955 | 0.354759 | 0.382214 | -0.8352 | 0.670338 | 0.087918 | 0.628823 | -0.17345 | 0.954943 | 0.028749 | -0.236 | 0.795376 | -0.28296 | 0.097215 | 0.136052 | 3.225623 | 0.615074 | 1.516166 |
| 35.18056 | 0.292936 | -0.07617 | 0.387726 | 0.756549 | 1.096376 | 0.445167 | -0.12498 | 0.228466 | 0.395651 | -1.00779 | 0.632738 | -0.02621 | 0.62725 | -0.21843 | 0.858803 | 0.076267 | -0.29339 | 0.932638 | -0.36038 | 0.519201 | 0.001719 | 2.433347 | 0.704928 | 1.564917 |
| 35.19444 | 0.221548 | -0.02958 | 0.202919 | 0.759897 | 0.796699 | 0.358646 | -0.2225 | 0.118539 | 0.313509 | -1.38462 | 0.753129 | -0.07533 | 0.702721 | -0.21157 | 0.816004 | 0.179743 | -0.2125 | 1.036045 | -0.32822 | 1.127468 | -0.05161 | 1.620781 | 0.750157 | 1.449998 |
| 35.20833 | 0.285138 | -0.04159 | 0.11184 | 0.735927 | 0.649626 | 0.317225 | -0.27134 | 0.098129 | 0.364067 | -1.72996 | 0.981517 | -0.03845 | 0.805292 | -0.14641 | 0.986197 | 0.263626 | -0.10306 | 1.104176 | -0.13811 | 1.481271 | 0.123562 | 0.895266 | 1.054501 | 1.865407 |
| 35.22222 | 0.372378 | -0.07488 | 0.052473 | 0.421261 | 0.01902 | 0.25697 | -0.24748 | 0.105578 | 0.44266 | -1.82535 | 1.14568 | 0.004191 | 0.8443 | -0.09385 | 1.166244 | 0.42809 | 0.016765 | 1.120236 | 0.030488 | 1.478169 | 0.299139 | 0.157845 | 1.303197 | 2.101601 |
| 35.23611 | 0.413898 | -0.18374 | -0.04698 | 0.196262 | -0.5143 | 0.275768 | -0.21453 | 0.132566 | 0.508275 | -1.57928 | 1.251629 | -0.03567 | 0.848931 | -0.09797 | 1.173521 | 0.570319 | 0.084339 | 1.049234 | 0.11349 | 1.618139 | 0.345921 | -0.60116 | 1.39151 | 1.568243 |
| 35.25 | 0.449183 | -0.30636 | -0.1298 | 0.195143 | -0.36139 | 0.28526 | -0.17214 | 0.178208 | 0.54105 | -1.25567 | 1.225756 | -0.09321 | 0.707362 | -0.12702 | 1.192079 | 0.694173 | 0.103212 | 0.724355 | 0.087478 | 2.269548 | 0.362039 | -1.15053 | 1.238194 | 1.208356 |
| 35.26389 | 0.467124 | -0.37794 | -0.21154 | 0.135757 | -0.24036 | 0.289421 | -0.17326 | 0.163849 | 0.578773 | -1.01518 | 1.094636 | -0.12342 | 0.583788 | -0.3102 | 1.217719 | 0.643496 | 0.04859 | 0.072482 | -0.00359 | 3.444562 | 0.30016 | -1.39835 | 1.082233 | 0.746959 |
| 35.27778 | 0.459465 | -0.46567 | -0.2501 | -0.00465 | -0.56526 | 0.294225 | -0.17068 | 0.133969 | 0.74911 | -0.85204 | 1.052954 | -0.13224 | 0.469132 | -0.62254 | 1.449392 | 0.535964 | -0.04342 | 0.069989 | -0.06271 | 4.221035 | 0.249629 | -1.26168 | 0.922143 | -0.05966 |
| 35.29167 | 0.483035 | -0.5418 | -0.19487 | -0.0537 | -0.62893 | 0.383982 | -0.09841 | 0.212904 | 0.914097 | -0.87259 | 1.046366 | -0.13733 | 0.443973 | -0.84832 | 1.652056 | 0.421937 | -0.14613 | 0.22394 | -0.01268 | 4.505866 | 0.264887 | -0.90162 | 0.90198 | -0.89741 |
| 35.30556 | 0.534965 | -0.64052 | -0.14532 | 0.025745 | -0.28728 | 0.431338 | -0.02137 | 0.429914 | 0.843043 | -0.90775 | 1.156251 | -0.1501 | 0.431812 | -0.81258 | 1.823948 | 0.266013 | -0.16906 | 0.53539 | 0.135594 | 3.286287 | 0.343839 | -0.40317 | 0.993243 | -1.10483 |
| 35.31944 | 0.660922 | -0.74284 | -0.1603 | 0.039758 | 0.039322 | 0.437393 | 0.019261 | 0.637513 | 0.852152 | -0.90475 | 1.285766 | -0.14216 | 0.462703 | -0.59838 | 1.714321 | 0.273174 | -0.15737 | 0.734509 | 0.200144 | 1.952127 | 0.47304 | -0.02095 | 1.050181 | -0.94225 |
| 35.33333 | 0.751514 | -0.8408 | -0.14271 | 0.021609 | 0.214108 | 0.391404 | 0.070467 | 0.664121 | 0.556775 | -0.79089 | 1.510095 | -0.07216 | 0.45468 | -0.50872 | 1.646383 | 0.297307 | -0.16471 | 0.8106 | 0.100752 | 0.714275 | 0.489667 | 0.47919 | 0.99894 | -0.8185 |
| 35.34722 | 0.803278 | -0.93414 | -0.04012 | -0.04873 | 0.677846 | 0.238946 | 0.066872 | 0.681818 | 0.088417 | -0.69302 | 1.918177 | 0.258924 | 0.450273 | -0.48844 | 0.921165 | 0.367533 | -0.22316 | 1.026411 | -0.04694 | -0.37597 | 0.670644 | 1.287614 | 0.887811 | -0.42683 |
| 35.36111 | 0.896773 | -0.91746 | 0.09384 | 0.091106 | 1.490777 | 0.189973 | -0.02209 | 0.663242 | -0.16609 | -0.69721 | 2.112981 | 0.241497 | 0.51035 | -0.34681 | -0.43012 | 0.772297 | -0.21047 | 1.215954 | -0.28031 | -1.50801 | 0.972417 | 2.270208 | 0.695839 | 0.581152 |
| 35.375 | 1.070725 | -0.37689 | 0.173465 | 0.543839 | 2.035504 | 0.414587 | -0.1019 | 0.379851 | 0.877057 | -0.79688 | 1.261961 | -0.74466 | 0.579255 | -0.02422 | -1.66857 | 0.950998 | -0.20701 | 1.114466 | -0.12499 | -1.10817 | 0.935189 | 2.901262 | 0.725992 | 0.937737 |
| 35.38889 | 1.488426 | -0.02514 | 0.525713 | 0.905818 | 2.709357 | 0.456742 | -0.12464 | 0.567486 | 1.035673 | -0.54864 | 1.326901 | -0.98137 | 0.903528 | 0.385517 | -1.57041 | 0.583768 | 0.026838 | 0.738418 | 1.260131 | 0.200168 | 0.732004 | 2.40951 | 0.999088 | 0.741195 |
| 35.40278 | 1.732163 | -0.6976 | 1.356705 | 0.488669 | 4.099556 | -0.07939 | 0.014389 | 1.745144 | -1.4635 | 0.50785 | 3.357545 | 0.587265 | 2.048882 | 1.170572 | 0.671694 | 0.731117 | 1.091607 | -0.18608 | 3.851956 | -1.36223 | 0.104965 | 2.345546 | 1.740889 | 1.612458 |
| 35.41667 | 1.437403 | -3.68711 | 2.645642 | -1.58089 | 6.399976 | -2.73188 | -0.03841 | 3.260598 | -7.16245 | 0.448233 | 8.192095 | 4.060876 | 5.100341 | 2.647747 | -0.55886 | 1.94234 | 1.409419 | -2.90213 | 8.663653 | NaN | -3.33236 | 2.736648 | 3.3864 | 2.196523 |
| 35.43056 | 1.045684 | -13.6441 | 4.926499 | -4.59951 | 7.701188 | -2.83442 | 1.176752 | 1.061395 | -6.39806 | 0.967454 | 11.39029 | 5.169963 | 8.28764 | 0.551543 | 3.482768 | 4.169587 | -5.47345 | -8.11842 | NaN | NaN | -12.6726 | 11.37368 | 4.483293 | 8.569454 |
| 35.44444 | 1.630467 | NaN | 0.837862 | 0.230724 | 1.841161 | 0.041726 | 2.338915 | 3.368873 | 0.997261 | 1.154025 | 3.943658 | -1.67412 | -0.76524 | 2.119914 | NaN | 0.990568 | 2.789311 | 1.345589 | NaN | NaN | NaN | NaN | 3.218221 | NaN |
| 35.45833 | 0.99057 | 11.47655 | 0.179505 | 2.48764 | -4.46391 | 5.349113 | 1.738646 | -0.23299 | 7.804894 | 5.21948 | -5.7902 | -5.72255 | -8.39747 | -0.70815 | NaN | 4.665773 | 2.729613 | 11.40824 | -20.3714 | NaN | 10.46849 | 15.46273 | 5.799239 | -0.03733 |
| 35.47222 | 0.962738 | 3.431101 | -0.78831 | 4.135095 | -1.16477 | 2.156323 | 1.386404 | -1.35783 | 7.516228 | 2.806571 | -3.01479 | -3.00421 | -3.58885 | -1.07122 | -2.59606 | 7.006274 | -0.4251 | 4.409068 | -1.25025 | -2.77741 | 2.830068 | 5.086718 | 4.257649 | 2.317162 |
| 35.48611 | 1.167242 | 1.354448 | 1.198343 | 2.554108 | 2.369986 | 0.816487 | 1.232464 | 0.697119 | 1.502416 | 1.040994 | 1.803437 | -0.51423 | -0.6114 | 1.08808 | 1.730241 | 4.811106 | -0.05301 | 1.467398 | 2.101543 | -0.28886 | 0.979872 | 2.728651 | 2.52275 | 2.58717 |
| 35.5 | 1.31354 | 0.802598 | 1.687031 | 1.281019 | 1.938523 | 0.937521 | 0.992012 | 1.487133 | -0.14917 | 0.341395 | 2.861892 | 0.249818 | -0.02197 | 1.477392 | 1.200293 | 3.976067 | 0.010611 | 0.956602 | 2.993873 | -0.41786 | 0.86739 | 1.660614 | 2.930002 | 2.194395 |
| 35.51389 | 1.382085 | 1.070974 | 1.592906 | 0.322624 | -0.55283 | 1.075978 | 0.851046 | 1.530529 | -0.19997 | 0.238777 | 2.377264 | 0.362788 | 0.204247 | 1.546814 | 2.372232 | 3.087404 | 0.00868 | 0.621512 | 3.453439 | -0.81495 | 1.151778 | 1.052884 | 2.82374 | 2.428624 |
| 35.52778 | 1.376215 | 1.432321 | 1.125602 | -0.28369 | -2.72378 | 0.931419 | 0.685963 | 1.298687 | 0.20769 | 0.337327 | 1.375909 | 0.074119 | 0.57662 | 1.939012 | 1.577729 | 2.669084 | 0.04163 | 0.162776 | 3.756519 | -1.20811 | 2.059621 | 0.944147 | 0.599751 | 2.323377 |
| 35.54167 | 1.326782 | 1.686527 | 0.786995 | -0.73213 | -3.83058 | 0.674735 | 0.619552 | 1.208255 | 0.653628 | 0.201136 | 0.749969 | -0.18441 | 0.790635 | 2.861088 | 0.937115 | 2.87456 | 0.027138 | 0.423132 | 3.572632 | -2.29184 | 2.975549 | 0.785305 | -0.3857 | 2.114726 |
| 35.55556 | 1.23152 | 1.554323 | 0.67621 | -1.06989 | -4.68969 | 0.663432 | 0.378788 | 0.895583 | 0.436613 | -0.05499 | 0.601225 | -0.15451 | 0.87208 | 3.127257 | 0.470261 | 2.347374 | -0.01972 | 2.043547 | 2.571006 | -0.07164 | 2.94509 | 0.172043 | 1.559496 | 2.100319 |
| 35.56944 | 1.19879 | 1.426388 | 0.606352 | -1.30408 | -3.72295 | 0.758929 | 0.197176 | 1.008558 | 0.31368 | -0.43639 | 0.987466 | 0.110369 | 0.828272 | 2.53768 | -0.20587 | 0.90307 | -0.03022 | 2.123475 | 1.443064 | 1.161457 | 2.483619 | -0.75303 | 1.731723 | 2.798797 |
| 35.58333 | 1.14176 | 2.202012 | 0.496353 | -1.34408 | -3.44329 | 0.901346 | 0.261704 | 0.988547 | 0.690832 | -0.83465 | 1.209659 | -0.10299 | 0.792414 | 2.058084 | -0.81967 | -0.64507 | -0.00937 | 1.783508 | 0.513484 | 0.52117 | 1.922963 | -2.01985 | 1.61856 | 5.207903 |
| 35.59722 | 0.956179 | 3.828664 | 0.375797 | -1.27798 | -2.88621 | 0.893788 | 0.271322 | 0.715476 | 0.487008 | -1.17851 | 1.352824 | -0.30918 | 0.792003 | 1.712031 | -1.55526 | -1.53768 | -0.03595 | 1.395665 | 0.342202 | 0.861748 | 1.381852 | -3.31447 | -2.37549 | -2.76257 |
| 35.61111 | 0.829079 | 4.885248 | 0.357872 | -0.88721 | -1.15982 | 0.924768 | 0.1418 | 0.705989 | -0.32063 | -1.2996 | 1.793362 | -0.09626 | 0.810325 | 1.462485 | -1.36425 | -1.77146 | -0.06177 | 1.525251 | 0.726319 | 1.540642 | 0.766024 | -3.788 | -3.54238 | 1.696015 |
| 35.625 | 0.79903 | 4.179708 | 0.342693 | -0.11016 | 0.230074 | 0.922049 | 0.148176 | 0.742966 | -0.45828 | -1.12604 | 1.765806 | -0.19515 | 0.861439 | 1.286424 | -0.18768 | -1.31778 | -0.05109 | 1.997964 | 1.315505 | 1.990838 | 0.684515 | -4.09535 | -2.8134 | 1.17187 |
| 35.63889 | 1.070432 | 3.524758 | 0.405698 | 0.543723 | 1.125734 | 0.922087 | 0.154714 | 0.767643 | -0.40353 | -0.80665 | 1.606486 | -0.45233 | 0.898311 | 0.921098 | 0.635022 | -0.69585 | -0.03672 | 2.310772 | 1.738387 | 2.227799 | 0.987941 | -4.51448 | -2.21113 | -0.49407 |
| 35.65278 | 1.185795 | 2.885529 | 0.497651 | 0.938029 | 1.682038 | 1.035675 | 0.181629 | 0.751275 | -0.17286 | -0.53816 | 1.522201 | -0.73206 | 0.878733 | 0.581026 | 0.910217 | -0.0769 | -0.06121 | 2.117304 | 1.82181 | 2.092489 | 1.458259 | -3.43235 | -0.99441 | -0.63678 |
| 35.66667 | 1.344128 | 2.777301 | 0.484958 | 1.224289 | 1.176344 | 0.889034 | 0.340665 | 0.317029 | 0.367377 | -0.23544 | 0.846601 | -1.1213 | 0.767199 | 0.387786 | 0.902949 | 0.802491 | -0.06861 | 1.85636 | 1.818307 | 2.614454 | 1.657981 | -1.97107 | 0.399265 | 0.054155 |
| 35.68056 | 1.309296 | 2.091105 | 0.472542 | 0.808736 | 0.191694 | 0.906791 | 0.329783 | 0.11673 | 0.484726 | -0.25577 | 0.672685 | -1.00599 | 0.657193 | 0.179494 | 0.820722 | 1.503309 | -0.0306 | 1.377142 | 1.604327 | 3.223648 | 1.591945 | -1.75666 | 1.712886 | 0.287786 |
| 35.69444 | 1.136159 | 1.149703 | 0.524583 | -0.02012 | -0.95413 | 1.085642 | 0.186317 | 0.173939 | 0.355031 | -0.54329 | 0.936798 | -0.76726 | 0.550832 | -0.09906 | 0.715728 | 1.712235 | 0.002849 | 0.827392 | 0.918624 | 3.693489 | 1.063661 | -2.13314 | 2.686497 | 0.122519 |
| 35.70833 | 1.235076 | 0.92844 | 0.515857 | -0.61645 | -2.09842 | 1.089517 | 0.127587 | -0.17203 | 0.519848 | -0.84749 | 0.896984 | -0.70413 | 0.382473 | -0.41679 | 0.560294 | 1.443932 | -0.01827 | -0.00611 | 0.104642 | 1.539641 | 0.316134 | -2.77654 | 3.626815 | -0.34444 |
| 35.72222 | 1.55428 | 1.08988 | 0.488633 | -0.76891 | -2.21699 | 0.652988 | 0.031029 | -0.76726 | 0.589945 | -1.05458 | 0.618297 | -0.59706 | 0.289034 | -0.75539 | 0.384137 | 1.108388 | -0.11396 | -0.86817 | -0.2985 | -0.88084 | -0.37382 | -2.94293 | 3.559616 | -1.87594 |
| 35.73611 | 1.792134 | 1.458745 | 0.491946 | -0.67393 | -2.53596 | 0.238041 | -0.1772 | -0.81358 | 0.551069 | -1.01612 | 0.534453 | -0.53807 | 0.323829 | -1.16499 | 0.130381 | 0.987587 | -0.17656 | -1.25863 | -0.23104 | -1.21564 | -0.66278 | -1.95036 | 3.570696 | -1.15758 |
| 35.75 | 1.684789 | 1.473142 | 0.643566 | -0.70547 | -3.08211 | -0.0657 | -0.38679 | -0.79332 | 0.646041 | -0.9218 | 0.544791 | -0.5288 | 0.551095 | -1.62619 | -0.05325 | 0.887325 | -0.13606 | -0.8117 | 0.275123 | -1.32862 | -0.38179 | -1.57185 | 3.459955 | -2.00177 |
| 35.76389 | 1.340531 | 0.926808 | 0.737316 | -1.02192 | -3.68087 | -0.11722 | -0.61045 | -0.76157 | 0.731755 | -0.84156 | 0.552903 | -0.2912 | 0.484058 | -2.15183 | -0.56311 | 0.703072 | -0.09877 | -0.51318 | 0.599975 | 0.316077 | 0.573194 | -2.15602 | 2.91883 | -2.4718 |
| 35.77778 | 1.130909 | 0.245645 | 0.78049 | -1.11013 | -3.87354 | -0.07811 | -0.69311 | -0.70026 | 0.455849 | -0.80953 | 0.518057 | 0.047125 | 0.179019 | -2.66233 | -1.70189 | 0.479488 | -0.11026 | -0.36495 | 0.850269 | -0.02604 | 0.886982 | -1.89313 | 1.845252 | -2.52707 |
| 35.79167 | 1.035571 | 0.03878 | 0.722215 | -0.68426 | -3.88806 | -0.07605 | -0.52081 | -0.82382 | 0.442878 | -0.85267 | 0.273514 | 0.146721 | -0.25127 | -2.86308 | -2.17952 | 0.482071 | -0.13949 | -0.25412 | 0.820168 | -0.21651 | 0.692906 | -1.95942 | 0.199508 | -3.81058 |
| 35.80556 | 0.991267 | 0.04722 | 0.578072 | -0.56142 | -3.53967 | -0.09695 | -0.3147 | -0.72575 | 0.60399 | -0.76786 | 0.008228 | 0.129668 | -0.60801 | -2.77369 | -1.62908 | 0.040829 | -0.20454 | -0.49283 | 0.292322 | -2.36073 | 0.557338 | -1.65579 | -1.05862 | -3.60009 |
| 35.81944 | 0.829214 | -0.01117 | 0.479703 | -0.67007 | -3.3308 | 0.049382 | -0.20336 | -0.62936 | 0.278682 | -0.6617 | -0.05252 | 0.167865 | -0.87448 | -2.90961 | -1.51698 | -0.60524 | -0.32613 | -0.49367 | -0.315 | -2.91976 | 0.128459 | -1.60947 | -1.71739 | -3.80359 |
| 35.83333 | 0.694256 | -0.04755 | 0.428357 | -0.53528 | -2.70598 | 0.225167 | -0.01097 | -0.64211 | -0.19557 | -0.55834 | -0.07473 | 0.240511 | -1.02776 | -2.82888 | -1.26342 | -1.28704 | -0.37048 | -0.56647 | -0.34892 | -0.93219 | -0.26963 | -1.71284 | -1.84663 | -4.40829 |
| 35.84722 | 0.661599 | -0.00047 | 0.561294 | -0.19725 | -1.61515 | 0.39776 | 0.295101 | -0.66737 | -0.19187 | -0.42076 | -0.03156 | 0.272472 | -1.0377 | -2.26169 | -0.7363 | -1.53622 | -0.2388 | -0.62736 | 0.367046 | 1.040752 | -0.32949 | -1.55393 | -2.14633 | -3.57333 |
| 35.86111 | 0.7266 | 0.176107 | 0.703526 | 0.126613 | -0.95435 | 0.700902 | 0.534373 | -0.4898 | 0.056133 | -0.4848 | 0.018041 | 0.306734 | -0.82397 | -1.64668 | -0.71192 | -1.48777 | -0.11289 | -0.1003 | 1.033048 | 1.678259 | -0.15747 | -1.49767 | -2.12121 | -3.1157 |
| 35.875 | 0.700178 | 0.376343 | 0.71196 | 0.227545 | -0.94648 | 0.759108 | 0.665057 | -0.37385 | 0.537517 | -0.68844 | -0.15507 | -0.06794 | -0.68008 | -1.25332 | -0.51933 | -1.44568 | -0.06692 | 0.324506 | 0.987762 | 1.345917 | -0.03114 | -1.15411 | -2.229 | -2.93725 |
| 35.88889 | 0.47719 | 0.455309 | 0.156611 | 0.34396 | -1.70035 | 0.398865 | 0.673141 | -0.72454 | 1.245086 | -1.33827 | -0.73192 | -1.55053 | -0.60401 | -1.49883 | -0.54135 | -1.24321 | -0.15378 | 0.263705 | 0.008297 | 0.890617 | -0.16439 | -1.08699 | -2.34429 | -2.33692 |
| 35.90278 | -0.40565 | -0.01151 | -0.44965 | -0.01695 | -2.49039 | -0.18624 | 0.421239 | -1.26914 | 1.368053 | -2.44416 | -1.21476 | -2.43786 | -0.64195 | -2.00888 | -0.39097 | -1.72434 | -0.3523 | -0.2838 | -1.84839 | 0.495409 | -0.9834 | -1.81342 | -2.56932 | -2.08643 |
| 35.91667 | -1.25151 | -1.53541 | -0.10017 | -1.11471 | -2.80856 | 0.464286 | -0.16322 | -1.00968 | 0.097023 | -2.68292 | -0.0529 | 0.11471 | -0.54578 | -2.07123 | -0.2621 | -3.14568 | -0.62462 | -0.92103 | -4.31665 | 0.212784 | -2.1398 | -3.43148 | -2.31837 | -3.13702 |
| 35.93056 | -1.59409 | -2.77602 | 0.462215 | -0.44447 | -2.9687 | 0.258409 | -0.57151 | -0.7741 | -0.57739 | -2.47305 | 1.10197 | 2.203588 | 0.005981 | -1.58419 | -0.13313 | -3.94322 | -0.97108 | -0.88475 | -7.74207 | -0.41218 | -2.87579 | -4.84462 | -1.70147 | -4.67472 |
| 35.94444 | -1.46507 | -1.91238 | 0.016197 | 0.602561 | -4.40409 | 0.468182 | -0.4355 | -1.26956 | 0.356641 | -4.17018 | 0.425814 | -0.75525 | 0.520526 | -1.08599 | -0.10093 | -2.92208 | -1.23019 | -0.31231 | -5.49407 | -1.90219 | -2.2803 | -3.88728 | -1.63432 | -2.23407 |
| 35.95833 | -1.13896 | -0.65407 | -0.3021 | 0.792017 | -5.08012 | -1.04138 | -0.31692 | -1.55586 | 1.405218 | -5.47204 | -0.2749 | -3.10015 | 0.95264 | -0.80807 | -0.20327 | -2.0686 | -1.57282 | -0.26152 | -6.26097 | -4.03734 | -0.70282 | -3.96602 | -1.02525 | -1.06029 |
| 35.97222 | -1.26814 | -1.50035 | 0.031317 | 0.724117 | -3.76041 | -0.22942 | -0.32103 | -0.62897 | 0.333527 | -4.77034 | 0.976943 | -0.50077 | 1.403154 | -0.62774 | 0.111111 | -1.32345 | -1.33754 | 0.182717 | -4.42355 | -0.20858 | 0.413407 | -3.06228 | -0.70953 | -0.99772 |
| 35.98611 | -1.71875 | -2.79934 | 0.464388 | 0.638436 | -4.04987 | -0.08681 | -0.4088 | 0.417408 | -1.5207 | -2.92005 | 2.323389 | 2.700784 | 1.403977 | -0.51181 | 0.52798 | -0.95171 | -1.22111 | 0.541732 | -2.49986 | -0.02176 | 0.585168 | -2.81615 | -0.30902 | -1.79153 |
| 36 | -1.35795 | -1.89264 | 0.032361 | 0.700277 | -2.20601 | 0.451974 | -0.21298 | 0.313496 | -0.31224 | -2.27774 | 1.256906 | 1.187132 | 0.924525 | -0.50217 | 0.649657 | -0.31476 | -0.7084 | 0.637934 | -1.6781 | 0.312325 | 0.390413 | -2.01353 | -0.18432 | -1.00627 |
| 36.01389 | -0.42046 | -0.13674 | -0.40009 | 0.732925 | -0.81025 | 1.850163 | 0.059256 | -0.30317 | 0.871592 | -2.35957 | -0.16088 | -2.39599 | 0.442831 | -0.44638 | 0.633316 | 0.436477 | 0.010702 | 0.477345 | -1.10027 | 0.946603 | -0.01417 | -1.37564 | -0.11232 | -0.4223 |
| 36.02778 | -0.27659 | 0.39941 | -0.1946 | 0.84841 | -0.48874 | 1.72198 | 0.057445 | -0.00278 | 1.315195 | -2.11629 | -0.09245 | -2.11046 | 0.255998 | -0.31318 | 0.648208 | 1.02441 | 0.464622 | 0.482619 | -0.6528 | 1.257266 | -0.2757 | -0.82356 | 0.038237 | -0.70074 |
| 36.04167 | -0.3983 | -0.17859 | -0.11552 | 0.691059 | -0.14625 | 0.861502 | 0.087568 | 0.345381 | 0.80556 | -1.77361 | 0.08779 | -1.11726 | 0.19615 | -0.16799 | 0.672337 | 1.204292 | 0.381033 | 0.566434 | -0.45749 | 0.675637 | -0.53413 | -0.53942 | 0.061323 | -0.67976 |
| 36.05556 | -0.59257 | -0.98472 | -0.12197 | 0.542447 | -0.05428 | 0.41463 | 0.066452 | 0.687895 | 0.283576 | -1.8897 | 0.582884 | 0.080881 | 0.109121 | -0.27382 | 0.53585 | 1.271057 | 0.317798 | 1.082444 | -0.50863 | -0.3114 | -0.74145 | -0.20011 | -0.16795 | -0.70295 |
| 36.06944 | -0.65409 | -3.97834 | 0.050507 | 0.603806 | -0.8553 | -0.71948 | 0.057962 | 1.6531 | -1.36274 | -0.97433 | 2.05202 | 3.714728 | 0.027127 | -0.27914 | 0.395608 | 1.332349 | 0.167891 | 0.673572 | -0.48505 | 0.281151 | -0.68642 | 0.350168 | -0.40823 | -0.50731 |
| 36.08333 | -0.33214 | -4.44428 | 0.38357 | 0.674894 | -1.47702 | -0.00891 | 0.479955 | 2.278427 | -1.13049 | -0.55951 | 2.005138 | 4.647708 | 0.000345 | -0.18943 | 0.250887 | 1.416306 | 0.164366 | 0.335018 | -0.27073 | 0.485177 | -0.40783 | 0.777008 | -0.4061 | -0.06327 |
| 36.09722 | 0.078913 | -2.40038 | 0.478233 | 0.819871 | -0.074 | 0.880888 | 0.906307 | 1.914063 | 0.428643 | -1.04442 | 1.331929 | 1.866597 | 0.069392 | -0.1519 | 0.163576 | 1.579194 | 0.176049 | 0.412888 | -0.05402 | 0.643276 | -0.00229 | 1.786893 | -0.39313 | 0.541012 |
| 36.11111 | 0.326907 | -0.83835 | 0.447816 | 0.824447 | 1.258692 | 1.548923 | 0.88128 | 1.380241 | 1.197874 | -1.38565 | 0.681187 | -0.27646 | 0.390479 | -0.14578 | 0.04119 | 1.7459 | 0.303003 | 0.547395 | 0.130809 | 0.936071 | 0.371999 | 2.709169 | -0.25065 | 0.991047 |
| 36.125 | 0.578633 | 0.312035 | 0.497764 | 0.770995 | 1.896898 | 1.271447 | 0.696921 | 0.911402 | 1.515482 | -1.01668 | 0.235787 | -1.60217 | 0.32871 | -0.11576 | 0.033609 | 1.881827 | 0.268992 | 0.536793 | 0.218706 | 0.94073 | 0.535028 | 3.749687 | 0.017365 | 1.439207 |
| 36.13889 | 1.130857 | 2.165003 | 0.557375 | 0.433946 | 2.497675 | 1.240951 | 0.749942 | 0.226307 | 1.71355 | -0.06076 | -0.32993 | -3.63223 | 0.316298 | -0.10223 | 0.204222 | 2.053109 | 0.278275 | 0.629958 | 0.213351 | 1.033013 | 0.559685 | 4.342703 | 0.258292 | 1.021694 |
| 36.15278 | 1.036533 | 1.957655 | 0.69504 | 0.4668 | 2.368619 | 1.433795 | 0.630487 | 0.447921 | 1.15912 | 0.324797 | -0.38726 | -3.01695 | 0.366512 | -0.03972 | 0.460721 | 2.292856 | 0.402847 | 0.751828 | 0.247409 | 1.398905 | 0.518293 | 5.089528 | 0.49032 | 0.476962 |
| 36.16667 | 0.533124 | -0.91009 | 0.943298 | 0.902665 | 1.385865 | 0.716461 | 0.277393 | 1.660889 | 1.46666 | -0.00896 | 1.185514 | 1.182994 | 0.449136 | 0.041922 | 0.620928 | 2.479523 | 0.496186 | 0.986052 | 0.326937 | 1.666759 | 0.318514 | 4.944108 | 0.796536 | 1.952453 |
| 36.18056 | 0.329009 | -1.99601 | 0.924618 | 0.87843 | 0.555956 | 0.033269 | -0.0068 | 2.028703 | 1.641581 | -0.31791 | 1.738624 | 2.916334 | 0.491523 | 0.050926 | 0.625406 | 2.271746 | 0.555062 | 1.003661 | 0.330655 | 1.594822 | 0.067839 | 4.294252 | 1.044578 | 2.523459 |
| 36.19444 | 0.546128 | 0.39696 | 0.502161 | 0.713963 | 1.648512 | 0.802227 | -0.04763 | 1.070165 | 1.179101 | -0.6145 | 0.527403 | -0.40567 | 0.514174 | 0.005882 | 0.509726 | 2.049554 | 0.388669 | 0.93772 | 0.288876 | 1.456645 | -0.0089 | 3.279697 | 1.162026 | 3.207136 |
| 36.20833 | 0.856915 | 2.697713 | 0.11224 | 0.664465 | 3.008434 | 0.568985 | -0.0599 | 0.152225 | 0.447878 | -0.95048 | -0.27411 | -3.36255 | 0.595844 | -0.00383 | 0.371487 | 1.577952 | 0.257701 | 0.893728 | 0.295959 | 1.421738 | 0.060871 | 2.241182 | 1.210247 | 3.69451 |
| 36.22222 | 0.5215 | 1.773694 | 0.115837 | 0.811528 | 3.402003 | 1.275682 | -0.14518 | 0.395532 | 0.282589 | -1.52629 | -0.19573 | -2.56322 | 0.627742 | -0.0287 | -0.10597 | 1.028695 | 0.159337 | 0.90639 | 0.308967 | 1.547749 | 0.209832 | 1.029705 | 1.231907 | 2.274266 |
| 36.23611 | -0.03155 | 0.268051 | 0.089339 | 0.694005 | 2.402089 | 0.716299 | -0.19918 | 0.919504 | 0.51074 | -2.46759 | 0.560297 | -0.59965 | 0.659255 | -0.07804 | -0.83461 | 0.374846 | 0.019906 | 0.992845 | 0.251859 | 1.4674 | 0.173023 | -0.42235 | 1.173957 | 2.613208 |
| 36.25 | -0.25605 | -0.03521 | -0.15075 | 0.576484 | 1.349951 | 0.406571 | -0.1688 | 1.000607 | 0.435847 | -2.80914 | 0.596493 | -0.21996 | 0.661614 | -0.11222 | -0.99096 | -0.35285 | -0.17984 | 0.564478 | 0.050776 | 1.289554 | -0.14446 | -1.68424 | 1.043475 | 3.277567 |
| 36.26389 | -0.65707 | -1.08411 | -0.47776 | 0.286665 | 0.310036 | -0.02278 | -0.16753 | 1.399826 | 0.665564 | -3.37096 | 1.010924 | 1.114981 | 0.602824 | -0.14533 | -0.84185 | -0.99537 | -0.36568 | 0.133344 | -0.21396 | 1.371477 | -0.58761 | -2.3625 | 1.203154 | 3.478191 |
| 36.27778 | -1.11704 | -3.10286 | -0.90359 | -0.62323 | -1.69617 | -0.45215 | -0.18308 | 2.160065 | 1.622604 | -4.57389 | 1.698241 | 4.064064 | 0.515866 | -0.12727 | -0.31125 | -1.23372 | -0.43853 | -0.07798 | -0.33007 | 1.59999 | -0.49402 | -2.42287 | 1.44783 | 4.043933 |
| 36.29167 | -0.65134 | -2.24781 | -0.64262 | -0.6829 | -2.33425 | -0.64474 | -0.05361 | 2.094914 | 1.489125 | -3.75267 | 1.31925 | 3.039317 | 0.492847 | -0.04437 | 0.158543 | -1.00092 | -0.36171 | -0.05368 | -0.33811 | 1.756949 | -0.27072 | -2.42029 | 1.409488 | 3.643658 |
| 36.30556 | -0.04479 | 0.525326 | -0.06377 | 0.516598 | -0.41124 | 1.05713 | 0.094581 | 0.704355 | -0.24254 | -1.61895 | -0.1563 | -1.50351 | 0.450948 | 0.031144 | 0.308557 | -0.65954 | -0.36391 | -0.01475 | -0.37559 | 1.555868 | -0.15422 | -2.15635 | 1.191744 | 2.643508 |
| 36.31944 | 0.208668 | 1.391184 | 0.061803 | 1.084956 | 0.567866 | 1.870677 | 0.182667 | 0.267081 | -0.89576 | -0.76497 | -0.60454 | -3.31783 | 0.548914 | 0.039999 | 0.39294 | -0.25346 | -0.3121 | 0.148131 | -0.3896 | 1.215872 | -0.15488 | -1.61618 | 0.869046 | 1.552165 |
| 36.33333 | -0.11828 | 0.012029 | -0.19492 | 0.616953 | 0.486582 | 1.175738 | 0.13661 | 0.662423 | -0.23255 | -0.88532 | -0.37656 | -1.87318 | 0.954855 | 0.000688 | 0.341878 | 0.150111 | -0.20537 | 0.175542 | -0.36783 | 0.929743 | -0.1033 | -1.10036 | 0.531369 | 0.287564 |
| 36.34722 | -0.57346 | -1.36247 | -0.50795 | 0.175704 | 0.176112 | 0.730869 | 0.101936 | 1.028796 | 0.333136 | -1.30474 | 0.304192 | -0.2437 | 1.489663 | -0.05426 | 0.171507 | 0.313443 | -0.16739 | 0.124347 | -0.36361 | 0.740896 | -0.16734 | -0.87117 | 0.284898 | -0.73474 |
| 36.36111 | -0.60213 | -2.06568 | -0.67312 | -0.05671 | 0.141581 | 0.400234 | 0.084977 | 1.022524 | 0.531326 | -0.99709 | 0.691629 | 0.446128 | 1.597776 | -0.05977 | 0.01345 | 0.226533 | -0.16812 | 0.014512 | -0.22815 | 0.843769 | -0.2808 | -0.65045 | 0.305753 | -1.10084 |
| 36.375 | -0.54561 | -2.43588 | -0.43804 | 0.033505 | 0.284718 | 0.534029 | 0.242578 | 0.77851 | 0.827275 | -0.60549 | 1.077523 | 1.025305 | 1.581999 | 0.025206 | -0.25275 | 0.300505 | -0.11046 | 0.075376 | 0.057363 | 1.165238 | -0.41505 | -0.38742 | 0.468593 | -1.01612 |
| 36.38889 | -0.71296 | -3.49024 | 0.089319 | -0.20656 | 0.131249 | -0.10493 | 0.521484 | 0.615048 | 1.503968 | -0.69679 | 2.0487 | 2.447256 | 1.669363 | 0.221813 | -0.23184 | 0.83499 | 0.002911 | 0.555412 | 0.637539 | 1.599028 | -0.15348 | 0.275441 | 0.848665 | 0.150844 |
| 36.40278 | -0.58083 | -3.4672 | 0.520341 | -0.22284 | 0.563552 | 0.241615 | 0.537923 | 0.551463 | 1.702924 | 0.159144 | 2.802364 | 2.60818 | 2.115965 | 0.7027 | 0.352024 | 1.955866 | 0.237064 | 1.485143 | 1.795317 | 2.204431 | 0.429022 | 1.340678 | 1.668273 | 2.205267 |
| 36.41667 | -0.56559 | -2.88011 | 0.708187 | 0.024308 | 1.417742 | -0.15847 | 0.550423 | 0.817988 | 1.358826 | 2.104274 | 2.751607 | 1.1041 | 2.27391 | 0.954642 | 1.301344 | 3.157172 | 0.714831 | 2.344548 | 2.699471 | 2.06921 | 0.681626 | 2.268291 | 2.346755 | 2.67757 |
| 36.43056 | -0.61213 | -2.60157 | 0.514085 | 0.126399 | 1.250334 | 0.533152 | 0.798482 | 1.524681 | 0.604237 | 6.889932 | 2.561696 | -0.02093 | 1.719271 | 0.963207 | 2.237932 | 4.491415 | 0.52177 | 2.304405 | 2.531024 | 1.624932 | 0.508014 | 2.652839 | 2.033055 | 2.41982 |
| 36.44444 | -0.1727 | -1.16167 | 0.254501 | 0.649822 | 2.446749 | 1.023705 | 1.496175 | 2.354582 | 0.026383 | 12.15802 | 2.373704 | -0.60805 | 1.159237 | 1.273429 | 3.553127 | 5.449668 | 0.339115 | 1.973767 | 2.382707 | NaN | 0.751731 | 2.856901 | 1.014309 | 1.578876 |
| 36.45833 | 0.542508 | 0.037452 | 0.421401 | 1.798174 | 4.366897 | 1.673172 | 1.574761 | 2.322945 | -0.44801 | 7.985423 | 2.285608 | -0.90882 | 1.132287 | 1.587833 | 3.70799 | 6.320043 | 0.445942 | 1.804955 | 2.7925 | 1.650792 | 1.212755 | 2.932931 | 0.713419 | -0.27424 |

| | | | | | | | | | | | | | | | | | | | | | | | | |
|---|---|---|---|---|---|---|---|---|---|---|---|---|---|---|---|---|---|---|---|---|---|---|---|---|
| 36.47222 | 0.575953 | -0.29086 | 0.584633 | 1.698364 | 6.156508 | 1.404627 | 0.810215 | 1.779416 | -0.62164 | 3.599544 | 1.71108 | -0.62371 | 1.378819 | 1.542548 | 3.628771 | 6.217131 | 0.410347 | 1.599483 | 3.019968 | 1.360306 | 1.547058 | 2.730567 | 1.458531 | 1.710729 |
| 36.48611 | 0.417816 | -0.8676 | 0.724269 | 1.290887 | 5.109328 | 0.813361 | 0.272438 | 1.535471 | -0.51563 | 0.565418 | 1.999868 | 0.017757 | 1.540008 | 1.371524 | 4.439681 | 5.824044 | 0.107952 | 1.658268 | 2.93336 | 0.763503 | 1.330093 | 2.42424 | 1.867856 | 6.781678 |
| 36.5 | 0.75234 | -0.41261 | 0.960561 | 0.991382 | 3.396128 | 0.760315 | 0.298818 | 1.563673 | -0.38629 | -1.59103 | 1.628923 | 0.149333 | 1.396185 | 1.312197 | 4.253855 | 4.495517 | -0.05581 | 2.012198 | 2.710221 | -0.04594 | 1.067083 | 2.076937 | 1.930374 | 8.131537 |
| 36.51389 | 1.415123 | 0.831853 | 1.142289 | 0.956941 | 1.420875 | 1.053734 | 0.266657 | 1.441821 | -0.16562 | -2.89492 | 0.897927 | -0.32233 | 1.416035 | 1.29441 | 2.993798 | 3.784525 | -0.11435 | 2.372364 | 2.418171 | -1.2031 | 1.13855 | 1.826175 | 2.080266 | 5.623314 |
| 36.52778 | 1.570212 | 1.879108 | 1.02713 | 0.721378 | 0.035921 | 1.127131 | 0.202957 | 1.276991 | 0.044134 | -3.02339 | 0.681869 | -0.57515 | 1.30768 | 1.054512 | 2.547456 | 2.48228 | -0.09769 | 2.29073 | 2.073223 | -0.305 | 1.371229 | 1.488714 | 1.861895 | 4.585718 |
| 36.54167 | 1.343475 | 2.620406 | 0.73661 | 0.332448 | -0.76184 | 1.187438 | 0.13562 | 0.984932 | 0.206344 | -2.12044 | 0.54223 | -0.58594 | 1.356131 | 0.888996 | 1.632146 | 0.983455 | -0.13178 | 1.865198 | 1.486334 | 0.346632 | 1.381727 | 1.022128 | 0.079372 | 5.145768 |
| 36.55556 | 0.964659 | 2.970081 | 0.537169 | -0.04225 | -0.83224 | 0.952246 | 0.112377 | 0.814424 | 0.322156 | -1.69588 | 0.71444 | -0.38922 | 1.518173 | 1.066892 | 0.507924 | -0.21716 | -0.15964 | 1.687139 | 1.092688 | 0.672979 | 1.006484 | 0.449338 | -1.22955 | 4.467063 |
| 36.56944 | 0.484188 | 2.53563 | 0.403086 | -0.67528 | -1.36228 | 0.666638 | 0.16861 | 0.768736 | 0.404206 | -2.22834 | 0.900473 | -0.22343 | 1.557809 | 1.344809 | -0.08421 | -1.17328 | -0.23904 | 1.590965 | 0.781444 | 1.795879 | 0.483369 | 0.005455 | -1.1217 | 5.941777 |
| 36.58333 | 0.193502 | 2.804268 | 0.310743 | -1.2491 | -2.63496 | 0.516429 | 0.057452 | 0.542426 | 0.466271 | -3.75921 | 0.741684 | -0.38119 | 1.366139 | 1.657839 | -0.43098 | -2.16584 | -0.32659 | 1.594106 | 0.35674 | 2.201726 | -0.04714 | -0.1509 | -0.02904 | 5.359637 |
| 36.59722 | 0.353999 | 3.257666 | 0.288252 | -1.33725 | -3.99755 | 0.4468 | -0.10717 | 0.181392 | 0.495283 | -5.65833 | 0.424714 | -0.72912 | 1.001235 | 1.622578 | -0.65187 | -2.52246 | -0.2959 | 1.774506 | -0.10696 | 2.277451 | -0.40944 | -0.32909 | 0.266954 | 4.468548 |
| 36.61111 | 0.642156 | 4.330093 | 0.25701 | -0.82328 | -4.20987 | 0.450777 | -0.34128 | -0.16158 | 0.467372 | -6.91284 | 0.186158 | -1.15314 | 0.5708 | 1.201494 | -1.19417 | -2.61009 | -0.22406 | 1.654514 | -0.37369 | 2.34706 | -0.64287 | -0.53297 | 0.421967 | 3.588786 |
| 36.625 | 0.941045 | 4.687203 | 0.182063 | -0.10989 | -3.5617 | 0.358018 | -0.50515 | -0.30923 | 0.381854 | -7.1689 | 0.159712 | -1.22023 | 0.278897 | 0.816789 | -0.9812 | -2.69688 | -0.23109 | 1.606503 | -0.74799 | 2.357391 | -0.44201 | -0.69738 | 0.831785 | 2.613424 |
| 36.63889 | 0.878403 | 4.144474 | 0.384993 | -0.07571 | -2.75467 | 0.509013 | -0.46787 | -0.18808 | 0.319854 | -5.46454 | 0.539673 | -0.82342 | 0.14898 | 0.505472 | -1.06732 | -3.02303 | -0.23606 | 1.280482 | -0.8567 | 2.266983 | -0.28182 | -0.47919 | 0.617594 | 1.13511 |
| 36.65278 | 0.982851 | 3.102971 | 0.475891 | 0.195784 | -1.08304 | 0.647598 | -0.22784 | 0.007378 | 0.340104 | -4.98888 | 0.73298 | -0.61587 | 0.009357 | 0.320061 | -0.79601 | -2.79255 | -0.22142 | 1.060527 | -0.57685 | 2.343646 | 0.056356 | -0.05125 | 0.283349 | 0.39117 |
| 36.66667 | 1.188243 | 2.299185 | 0.416044 | 0.3793 | -1.22089 | 0.804011 | -0.05666 | 0.183988 | 0.412512 | -3.10656 | 0.610632 | -0.64936 | -0.01241 | 0.28869 | -0.64811 | -2.10589 | -0.14085 | 0.994503 | -0.25993 | 2.347429 | 0.149595 | 0.054008 | -0.6304 | -0.60504 |
| 36.68056 | 1.104802 | 1.449292 | 0.508649 | 0.358624 | -1.09195 | 0.943322 | 0.132371 | 0.425485 | 0.37922 | -3.8871 | 0.775571 | -0.39716 | 0.040421 | 0.260218 | -0.82815 | -1.22626 | -0.10827 | 0.989259 | 0.042287 | 2.610993 | 0.077093 | -0.07377 | -1.21897 | -1.5317 |
| 36.69444 | 0.839655 | 0.839223 | 0.62548 | 0.258733 | -1.21715 | 0.63246 | 0.182737 | 0.705982 | 0.409915 | -4.58308 | 1.178717 | -0.10967 | 0.030122 | 0.12755 | -0.45124 | -0.26827 | -0.14338 | 1.039434 | 0.270533 | 0.800843 | 0.050273 | -0.24326 | -1.78803 | -1.76925 |
| 36.70833 | 0.745454 | 0.807052 | 0.591762 | 0.271606 | -1.13739 | 0.558424 | 0.081775 | 0.953227 | 0.557843 | -3.46674 | 1.182458 | -0.13232 | -0.05702 | -0.11589 | -0.22859 | 0.016449 | -0.15483 | 0.85329 | 0.376681 | 1.5045 | -0.06242 | -0.5099 | -1.8879 | -1.369 |
| 36.72222 | 0.709399 | 0.649147 | 0.510132 | 0.22985 | -0.8331 | 0.504819 | -0.10794 | 1.092094 | 0.690328 | -1.27314 | 0.739054 | -0.31691 | -0.18891 | -0.45197 | -0.28379 | -0.15756 | -0.15915 | 0.532993 | 0.086187 | 1.166906 | -0.12204 | -0.56374 | -1.49905 | -0.58091 |
| 36.73611 | 0.612503 | 0.449679 | 0.567325 | 0.208424 | -0.561 | 0.38788 | -0.31414 | 1.091809 | 0.687684 | -0.41063 | 0.241443 | -0.42028 | -0.28009 | -0.84313 | -0.697 | -0.35234 | -0.14554 | 0.277153 | -0.33936 | 0.067022 | -0.22273 | -0.63391 | -1.1895 | -1.3077 |
| 36.75 | 0.550482 | 0.37938 | 0.639062 | 0.337278 | -0.47579 | 0.23239 | -0.52997 | 0.969583 | 0.819684 | -0.00988 | -0.0526 | -0.48325 | -0.29349 | -1.196 | -1.53842 | -0.21426 | -0.14966 | -0.12563 | -0.88138 | -1.83083 | -0.0766 | -0.35275 | -1.13 | -2.43456 |
| 36.76389 | 0.327488 | 0.293634 | 0.502548 | 0.371681 | -0.37084 | 0.255012 | -0.46349 | 0.773744 | 0.81968 | 0.639111 | -0.3355 | -0.52347 | -0.50307 | -1.48838 | -2.03575 | -0.35527 | -0.12007 | -0.41293 | -1.14031 | -7.00871 | -0.13769 | 0.070774 | -1.40128 | -4.2847 |
| 36.77778 | -0.23885 | 0.05593 | 0.212037 | 0.343209 | -0.53311 | 0.305621 | -0.28799 | 0.725846 | 0.812149 | 0.509336 | -0.48629 | -0.44612 | -0.68973 | -1.59769 | -2.2472 | -0.91783 | 0.052297 | -0.46107 | -1.13947 | NaN | 0.058803 | 0.152985 | -1.44916 | -6.04078 |
| 36.79167 | -0.67305 | -0.35915 | 0.433834 | 0.195631 | -1.01754 | 0.31394 | -0.1719 | 0.776699 | 0.867244 | 0.187714 | -0.47381 | -0.25419 | -0.91699 | -1.45367 | -1.99657 | -1.50446 | 0.334266 | -0.56165 | -0.91483 | -2.10956 | 0.142347 | 0.320366 | -1.20897 | -8.09132 |
| 36.80556 | -0.91018 | -0.74523 | 0.676498 | -0.04492 | -1.36612 | 0.163964 | -0.22225 | 0.824748 | 0.967269 | -0.12697 | -0.73161 | -0.11134 | -0.97491 | -1.45275 | -1.14711 | -2.18279 | 0.460552 | -0.55861 | -0.94644 | -0.52195 | 0.314153 | 0.454427 | -0.7063 | -9.60279 |
| 36.81944 | -0.95171 | -1.1696 | 0.825235 | -0.17766 | -1.32544 | 0.049492 | -0.31988 | 0.989189 | 0.977863 | -0.39103 | -1.1601 | -0.03005 | -1.10365 | -1.47783 | -0.3033 | -2.88028 | 0.388709 | -0.45885 | -1.07604 | 0.380132 | 0.41983 | 0.466821 | -0.15688 | -9.10335 |
| 36.83333 | -0.76176 | -1.34814 | 0.903289 | -0.15455 | -1.30951 | 0.1391 | -0.2653 | 1.074987 | 1.089484 | -0.98534 | -1.3816 | 0.0382 | -1.08772 | -1.55009 | 0.049896 | -3.17661 | 0.297839 | -0.72493 | -1.05468 | 1.044855 | 0.639295 | 0.544905 | 0.167063 | -7.99666 |
| 36.84722 | -0.62 | -1.318 | 0.938209 | 0.030701 | -1.34868 | 0.199159 | -0.1203 | 1.038121 | 1.290246 | -1.13523 | -1.22138 | 0.226879 | -1.00516 | -1.49723 | 0.024456 | -3.23249 | 0.243611 | -1.0398 | -0.74318 | 1.26826 | 0.832949 | 0.547196 | 0.378503 | -6.90221 |
| 36.86111 | -0.53398 | -1.14875 | 0.889015 | 0.312201 | -1.00799 | 0.425624 | 0.073313 | 0.85966 | 1.301651 | -0.77753 | -1.06021 | 0.200895 | -0.57212 | -1.46984 | -0.32913 | -2.58506 | 0.203102 | -1.21388 | -0.49597 | 1.377116 | 0.900754 | 0.422787 | 0.385789 | -6.26476 |
| 36.875 | -0.56066 | -0.52908 | 0.653912 | 0.604176 | -0.19495 | 0.874681 | 0.150882 | 0.496286 | 1.085875 | 0.382684 | -1.17638 | -0.63659 | -0.32573 | -1.48753 | -0.63592 | -1.69985 | -0.00542 | -1.25897 | -0.41562 | 1.259864 | 0.808086 | 0.749951 | 0.314473 | -5.02341 |
| 36.88889 | -0.7481 | -0.01422 | 0.315452 | 0.681534 | 0.128068 | 1.103116 | 0.116419 | 0.309436 | 0.819653 | 0.747216 | -1.20714 | -1.48211 | -0.57523 | -1.54614 | -0.6128 | -0.67481 | -0.26827 | -0.8877 | -0.91356 | 0.831639 | 0.303941 | 0.778519 | -0.16916 | -3.29728 |
| 36.90278 | -1.02216 | -0.43731 | 0.258396 | 0.310518 | -0.85584 | 0.237535 | 0.143057 | 0.68187 | 0.920687 | -0.37776 | -0.50558 | -0.79458 | -1.2363 | -1.9167 | -0.6868 | -1.15775 | -0.3837 | -1.07487 | -2.09226 | 0.531604 | -0.5176 | 0.147981 | -0.77159 | -2.85481 |
| 36.91667 | -1.22193 | -1.29172 | 0.361162 | -0.09329 | -2.36125 | -0.4858 | -0.02676 | 1.040135 | 1.203943 | -2.51801 | 0.021953 | -0.12359 | -1.68551 | -2.15811 | -0.29502 | -2.46909 | -1.44276 | -1.16491 | -3.24292 | 0.151356 | -1.17017 | -1.27222 | -1.16351 | -2.97057 |
| 36.93056 | -1.46913 | -1.18388 | 0.382753 | 0.162853 | -2.6376 | -0.70476 | -0.47972 | 0.685833 | 0.957496 | -2.96159 | -0.39536 | -0.87156 | -1.23394 | -1.91752 | -0.18683 | -3.28402 | -5.34623 | -0.55991 | -3.2404 | -0.04173 | -1.00086 | -0.7098 | -1.51012 | -2.22542 |
| 36.94444 | -1.61718 | -0.03413 | 0.261695 | 0.848728 | -2.87567 | -1.29941 | -0.9375 | -0.16156 | 0.3421 | -2.41448 | -0.92292 | -2.3563 | -0.30411 | -1.62646 | -0.27993 | -3.5912 | -13.7921 | 0.09742 | -2.42871 | -0.28864 | -0.46177 | -0.5707 | -1.60373 | -0.68423 |
| 36.95833 | -1.37262 | -0.66758 | 0.758422 | 0.688375 | -5.81317 | -1.18323 | -0.63058 | 0.374076 | 0.396611 | -2.45047 | -0.04805 | -1.1745 | 0.319334 | -1.32817 | -0.43541 | -3.37463 | -13.0464 | 0.508837 | -1.40515 | -0.41362 | -0.17706 | -0.4182 | -1.03662 | -0.32023 |
| 36.97222 | -0.543 | -3.73877 | 1.536562 | -0.57117 | -3.91194 | -1.79134 | 0.095426 | 2.207147 | 1.19335 | -5.27305 | 3.190052 | 2.706303 | 0.660755 | -1.0189 | -0.34487 | -2.23392 | -10.7567 | 0.521764 | -0.63242 | -0.14745 | -0.30918 | -0.39744 | -0.44747 | -1.1056 |
| 36.98611 | -0.31561 | -3.05603 | 1.327384 | -0.41843 | -2.69934 | -2.58555 | 0.13329 | 1.611977 | 0.866601 | -4.00188 | 3.004455 | 2.047423 | 0.849412 | -0.71808 | -0.27965 | -1.40335 | -7.47309 | 0.59667 | -0.43406 | 0.146774 | -0.52994 | -0.41431 | -0.14277 | -2.81713 |
| 37 | -0.5356 | 0.374639 | 0.800352 | 1.147852 | -0.67673 | -0.18826 | -0.4744 | -0.91753 | -0.53622 | 0.526611 | 0.066089 | -1.87893 | 0.823987 | -0.51129 | -0.27835 | -0.86287 | -4.64503 | 0.507476 | -0.40757 | 0.417823 | -0.52825 | -0.74273 | -0.03033 | -3.41447 |
| 37.01389 | -0.51599 | 0.674184 | 0.574904 | 1.272214 | -0.05227 | 0.239023 | -0.88237 | -1.27568 | -0.41432 | 1.265158 | -0.11609 | -2.20151 | 0.753888 | -0.35119 | -0.22933 | -0.39147 | -3.41091 | 0.562077 | -0.35492 | 0.696142 | -0.48506 | -1.18082 | -0.10567 | -3.0894 |
| 37.02778 | -0.24456 | -0.95415 | 0.605326 | 0.548148 | -0.55279 | -0.79235 | -0.75675 | -0.34764 | -0.15492 | -0.33511 | 0.864336 | -0.45146 | 0.756457 | -0.28209 | -0.1775 | -0.15793 | -2.57326 | 0.783268 | -0.37088 | 0.761996 | -0.5132 | -1.19434 | -0.35921 | -2.55722 |
| 37.04167 | -0.42768 | -0.86228 | 0.282567 | 0.535605 | -0.27483 | -0.50827 | -0.83823 | -0.24889 | -0.32606 | 0.082149 | -0.09669 | -0.69907 | 0.694708 | -0.32746 | -0.25724 | -0.1141 | -2.35505 | 0.908675 | -0.48875 | 0.415695 | -0.4837 | -1.13677 | -0.67918 | -2.1469 |
| 37.05556 | -0.44753 | -1.37388 | 0.245585 | 0.33449 | -0.29461 | -1.0813 | -0.46511 | 0.243119 | 0.1536 | -0.24383 | 0.005532 | -0.21545 | 0.463079 | -0.39446 | -0.25475 | -0.33255 | -2.41183 | 0.897489 | -0.56336 | -0.34662 | -0.41008 | -0.91275 | -0.9021 | -1.65374 |
| 37.06944 | -0.12164 | -3.06022 | 0.879801 | -0.41037 | -1.04072 | -1.29366 | 0.214461 | 1.334145 | 0.740881 | -2.3005 | 1.936078 | 1.862897 | 0.166617 | -0.39973 | -0.17364 | -0.60679 | -2.84842 | 0.781972 | -0.4885 | -0.75339 | -0.24339 | -0.66603 | -0.88098 | -1.35186 |
| 37.08333 | 0.12017 | -3.25554 | 1.112068 | -0.52771 | -0.96883 | -0.94908 | 0.399853 | 1.65745 | 1.196426 | -2.5522 | 2.65945 | 2.50348 | -0.10158 | -0.40598 | -0.08038 | -0.77123 | -3.22006 | 0.87609 | -0.33551 | -0.42487 | -0.20424 | -0.32818 | -0.72824 | -1.06498 |
| 37.09722 | 0.019651 | -2.9323 | 1.066883 | -0.43763 | -0.71994 | -0.64425 | 0.687508 | 1.625101 | 1.218214 | -1.93961 | 2.683388 | 2.102989 | -0.37323 | -0.44814 | -0.09297 | -0.69491 | -3.24818 | 0.939612 | -0.24595 | -0.14423 | -0.53542 | 0.00054 | -0.61277 | -0.79374 |
| 37.11111 | -0.25559 | -1.90545 | 0.73964 | -0.02418 | 0.204722 | -0.11372 | 0.680671 | 1.120629 | 0.8956 | -0.72793 | 1.868687 | 0.916348 | -0.62507 | -0.45576 | -0.40541 | -0.60637 | -2.81831 | 0.965073 | -0.20404 | 0.100508 | -1.08352 | 0.141052 | -0.58629 | -0.5798 |
| 37.125 | -0.72019 | -0.4887 | 0.251141 | 0.836205 | 1.924669 | -0.00295 | 0.326161 | 0.310957 | 0.388829 | 1.412648 | 0.072405 | -1.23721 | -0.73805 | -0.4269 | -1.02758 | -0.46771 | -2.17068 | 0.939214 | -0.09951 | 0.339857 | -1.5438 | 0.236233 | -0.60016 | -0.36681 |
| 37.13889 | -0.47785 | -0.67879 | 0.485218 | 0.601782 | 1.607901 | 0.16617 | 0.31633 | 0.520628 | 0.36737 | 0.819979 | 0.499213 | -0.67224 | -0.65789 | -0.30769 | -1.4886 | -0.26404 | -1.55982 | 0.942054 | 0.164622 | 0.58363 | -1.28362 | 0.370936 | -0.54 | -0.08088 |
| 37.15278 | 0.222155 | -1.97602 | 1.207895 | -0.23333 | 0.074436 | -0.29995 | 0.710802 | 1.301418 | 1.1559 | -1.52916 | 2.592621 | 1.646257 | -0.45692 | -0.18994 | -1.33063 | -0.09291 | -0.95363 | 0.906415 | 0.223491 | 0.730218 | -0.78234 | 0.587853 | -0.42119 | 0.246434 |
| 37.16667 | 0.168917 | -1.3346 | 1.051092 | -0.11362 | 0.388201 | -0.4119 | 0.472853 | 1.125717 | 0.894629 | -0.8612 | 2.155691 | 1.025827 | -0.35986 | -0.0791 | -0.64955 | 0.128692 | -0.07815 | 0.914255 | 0.369647 | 0.724687 | -0.46013 | 0.805875 | -0.32034 | 0.553225 |
| 37.18056 | -0.04377 | -0.0653 | 0.648606 | 0.3436 | 1.67712 | 0.020003 | -0.04129 | 0.731365 | 0.182893 | 0.5808 | 1.110504 | -0.46199 | -0.21437 | 0.045242 | -0.02647 | 0.417235 | 0.891524 | 1.011893 | 0.345706 | 0.672173 | -0.3453 | 1.036863 | -0.20002 | 0.994643 |
| 37.19444 | 0.272745 | -1.02943 | 1.011666 | 0.010973 | 1.177917 | -1.2752 | 0.206413 | 0.940674 | 0.668439 | -0.95591 | 2.43326 | 0.824525 | 0.035858 | 0.17061 | 0.242107 | 0.710133 | 1.813541 | 1.067008 | 0.4861 | 0.618766 | -0.2116 | 1.262783 | -0.04146 | 1.59098 |
| 37.20833 | 0.223767 | -1.23146 | 0.962126 | -0.0978 | 0.786413 | -1.0945 | 0.070691 | 0.825441 | 0.644204 | -1.20227 | 2.616018 | 0.990747 | 0.391615 | 0.238508 | 0.322165 | 0.888036 | 2.379179 | 1.147167 | 0.511368 | 0.711408 | -0.07236 | 1.381802 | 0.061241 | 1.940451 |
| 37.22222 | -0.38911 | 0.109055 | 0.260945 | 0.502328 | 1.642226 | -1.63868 | -0.52507 | 0.135416 | -0.25421 | 0.735596 | 1.042667 | -0.96344 | 0.871609 | 0.209841 | 0.308612 | 0.934649 | 2.426778 | 1.19816 | 0.467561 | 0.667017 | -0.03027 | 1.119375 | 0.10712 | 1.808355 |
| 37.23611 | -0.45331 | -0.17025 | 0.2351 | 0.518844 | 0.7686 | -1.78067 | -0.89633 | 0.067085 | -0.40503 | 0.484156 | 1.335929 | -0.83773 | 1.036634 | 0.158725 | 0.301856 | 0.985982 | 2.539089 | 1.098127 | 0.39469 | 0.486541 | -0.14524 | 0.768656 | 0.14918 | 1.640155 |
| 37.25 | -0.22281 | -1.10902 | 0.478809 | 0.164918 | -0.56235 | -1.79517 | -2.31224 | 0.343575 | 0.028092 | -0.72903 | 2.163393 | 0.252493 | 0.749175 | 0.132446 | 0.356394 | 1.053957 | 2.920854 | 0.901198 | 0.434081 | 0.388063 | -0.29368 | 0.703835 | 0.017248 | 1.809548 |
| 37.26389 | -0.30267 | -1.16545 | 0.423411 | 0.172039 | -0.77611 | -1.85247 | -3.59271 | 0.499027 | -0.09863 | -0.54418 | 1.917117 | 0.133309 | 0.439591 | 0.134942 | 0.446223 | 1.050589 | 3.003867 | 0.734254 | 0.418517 | 0.427143 | -0.381 | 0.697134 | -0.38999 | 2.288718 |
| 37.27778 | -0.4107 | -1.06186 | 0.449917 | 0.242273 | -0.74997 | -1.63739 | -2.23951 | 0.839807 | -0.08694 | -0.22949 | 1.861776 | -0.00736 | 0.462126 | 0.195282 | 0.578698 | 1.009742 | 2.879656 | 0.60232 | 0.478535 | 0.550139 | -0.19571 | 0.662508 | -1.04678 | 3.501479 |
| 37.29167 | -0.10526 | -1.6515 | 0.854028 | -0.12306 | -1.68837 | 0.498379 | -0.76951 | 1.243306 | 0.551042 | -1.33435 | 2.979979 | 1.322388 | 0.760025 | 0.263108 | 0.722646 | 0.98542 | 2.982175 | 0.513288 | 0.571059 | 0.661044 | 0.142676 | 0.588007 | -1.9063 | 3.938298 |
| 37.30556 | 0.348304 | -2.12781 | 1.315128 | -0.4403 | -2.53132 | 0.342755 | -0.49018 | 1.253283 | 1.046679 | -2.054 | 3.757837 | 2.402293 | 0.940198 | 0.278816 | 0.877854 | 1.033257 | 2.82148 | 0.459285 | 0.562879 | 0.650552 | 0.378179 | 0.460277 | -1.86329 | 3.078506 |
| 37.31944 | 0.114002 | -1.15384 | 0.862788 | -0.21464 | -1.86557 | -0.19894 | 0.034124 | 1.006745 | 0.573652 | -1.00081 | 2.68957 | 1.213195 | 0.85291 | 0.228614 | 1.011872 | 0.915281 | 2.693917 | 0.438031 | 0.326616 | 0.570373 | 0.298765 | 0.391708 | -1.03279 | 2.412504 |
| 37.33333 | -0.35964 | 0.167311 | 0.086898 | 0.518342 | -0.02007 | -0.07629 | 0.192578 | 0.407901 | -0.07445 | 0.721677 | 0.63766 | -1.17759 | 0.74036 | 0.196803 | 1.091726 | 1.099635 | 2.264001 | 0.434395 | 0.123507 | 0.475223 | 0.000245 | 0.302302 | -0.56828 | 2.135601 |
| 37.34722 | -0.69327 | 1.172942 | -0.45558 | 1.023002 | 1.580452 | -1.57073 | 0.47901 | -0.07115 | -0.96829 | 1.888511 | -0.99194 | -3.00643 | 0.60719 | 0.275939 | 1.246904 | 1.314641 | 2.25155 | 0.439265 | 0.134139 | 0.591504 | -0.23153 | 0.337688 | -0.34547 | 2.251632 |
| 37.36111 | -0.36535 | 0.90362 | -0.27332 | 0.937976 | 1.438233 | -1.77458 | 0.201029 | 0.117388 | -0.87013 | 1.31503 | -0.42115 | -2.58381 | 0.626758 | 0.361679 | 1.493708 | 1.489218 | 2.263181 | 0.472936 | 0.301402 | 0.940446 | -0.23099 | 0.548382 | -0.07116 | 2.461123 |
| 37.375 | 0.525362 | -0.81756 | 0.779711 | -0.1464 | -0.58208 | 1.296623 | -0.04789 | 0.953975 | 0.372926 | -0.70085 | 2.41686 | 0.357041 | 0.652375 | 0.364214 | 1.591012 | 1.500685 | 2.109179 | 0.487998 | 0.276886 | 1.131267 | -0.00719 | 0.551721 | 0.191672 | 2.251749 |
| 37.38889 | 1.651413 | -2.12052 | 1.60511 | -0.93585 | -1.68082 | 4.648595 | -0.28211 | 1.92802 | 1.452186 | -2.10039 | 4.782524 | 2.850316 | 1.012802 | 0.594826 | 1.698549 | 1.670099 | 1.92768 | 0.675876 | 0.47984 | 1.315593 | 0.277772 | 0.81078 | 0.739491 | 2.277471 |
| 37.40278 | 2.276862 | -1.35086 | 1.308967 | -0.61766 | -0.07355 | 3.733809 | 0.103693 | 2.225151 | 1.922383 | -1.69201 | 4.212451 | 2.456162 | 1.693737 | 1.189752 | 2.13902 | 2.177413 | 2.948601 | 1.097515 | 1.433896 | 1.765593 | 1.068778 | 1.135455 | 1.5176 | 3.199124 |
| 37.41667 | 2.35067 | -0.68322 | 0.816974 | 0.387615 | 3.014169 | 0.775656 | 0.167112 | 1.993321 | 1.010069 | -0.44735 | 1.783441 | 0.898072 | 1.999288 | 1.445223 | 2.348207 | 2.565446 | 3.969977 | 1.53936 | 2.446465 | 2.210633 | 2.25843 | 1.409227 | 1.819195 | 5.205397 |
| 37.43056 | 2.139578 | -0.56097 | 0.637988 | 1.039316 | 3.796007 | 0.35023 | 0.362093 | 1.505292 | 0.69573 | -0.12684 | 0.573816 | 0.134216 | 2.093481 | 1.109321 | 1.818143 | 2.667177 | 4.777846 | 1.338742 | 2.626292 | 1.18005 | 3.520046 | 1.585044 | 1.579629 | 3.791927 |
| 37.44444 | 1.57232 | -0.93697 | 0.662761 | 1.216001 | 3.921316 | 0.477116 | 0.7431 | 1.451571 | 0.862297 | -0.15126 | 0.309353 | 0.094345 | 2.506065 | 0.992184 | 1.472781 | 3.454386 | 5.456797 | 0.962378 | 2.564815 | 2.043623 | 3.986438 | 1.845207 | 1.177541 | 3.112774 |
| 37.45833 | 0.976409 | -0.84076 | 0.703469 | 1.177571 | 4.17722 | 1.309823 | 1.268651 | 1.738861 | 1.271802 | -0.1689 | 0.344925 | 0.048267 | 2.22251 | 1.14722 | 1.50289 | 5.063482 | 5.367696 | 0.930798 | 2.339636 | 3.861209 | 2.956319 | 1.962945 | 0.515126 | 2.146629 |
| 37.47222 | 0.72824 | -0.54644 | 0.656892 | 0.882922 | 3.323217 | 0.982001 | 1.691359 | 2.119553 | 1.545697 | -0.19545 | 0.715027 | 0.042564 | 1.237562 | 1.184888 | 1.498715 | 5.286033 | 4.87574 | 1.005118 | 1.93368 | -0.07389 | 2.731632 | 2.104392 | 0.83834 | 1.511469 |
| 37.48611 | 0.565412 | 0.26083 | 0.445572 | 0.904415 | 3.076016 | 0.846534 | 1.700336 | 2.128944 | 1.233141 | 0.083368 | 0.860078 | -0.28022 | 0.800477 | 1.140907 | 1.272898 | 5.722349 | 3.304621 | 1.071534 | 1.448909 | -0.36444 | 1.579208 | 1.605784 | 0.411823 | 2.225295 |
| 37.5 | 0.40268 | 0.945187 | 0.423626 | 1.04599 | 2.639802 | 0.640285 | 1.480448 | 1.912021 | 0.827593 | 0.274478 | 1.15992 | -0.43335 | 0.963112 | 1.006322 | 1.171567 | 5.437332 | 1.777943 | 1.238733 | 1.18119 | -0.5273 | 0.400101 | 1.437058 | -0.38512 | 3.606849 |
| 37.51389 | 0.465705 | 0.892048 | 0.621597 | 0.692966 | 1.533752 | 0.90337 | 1.482268 | 1.750696 | 0.779161 | 0.099366 | 1.594116 | -0.15467 | 1.075764 | 0.902016 | 1.357757 | 5.24834 | 1.243366 | 1.309035 | 1.100688 | 0.726922 | -0.21668 | 1.309001 | 0.07184 | 4.257489 |
| 37.52778 | 0.37996 | 0.506307 | 0.766992 | 0.557016 | 0.903938 | 1.14188 | 1.324569 | 1.71263 | 0.933987 | -0.02544 | 1.656319 | 0.0318 | 0.975214 | 0.784441 | 1.840046 | 5.190456 | 1.218663 | 1.19431 | 1.020549 | 0.996575 | -0.8519 | 1.200335 | 0.168906 | 3.836784 |
| 37.54167 | -0.03733 | 0.447604 | 0.634308 | 0.805557 | 1.09582 | 0.876884 | 1.091832 | 1.45669 | 0.820988 | 0.137024 | 1.35196 | -0.09399 | 0.569641 | 0.452067 | 1.855468 | 4.981241 | 1.267823 | 1.03078 | 0.674755 | 2.025608 | -1.63408 | 0.850427 | 0.653038 | 3.57635 |
| 37.55556 | -0.18337 | 0.173483 | 0.700365 | 0.787853 | 0.806278 | 0.895436 | 0.903889 | 1.381428 | 0.853904 | 0.109463 | 1.495099 | -0.04244 | 0.287369 | 0.210592 | 1.733914 | 3.280336 | 1.311523 | 0.947456 | 0.377675 | 0.957353 | -2.57454 | 0.814171 | 0.898917 | 3.258679 |
| 37.56944 | 0.161535 | -0.1416 | 0.971199 | 0.225385 | -0.46661 | 1.596316 | 0.944851 | 1.716716 | 1.208657 | -0.25744 | 2.028198 | 0.293025 | 0.370133 | 0.16597 | 1.506282 | 1.772397 | 1.427312 | 0.906864 | 0.218387 | -0.73216 | -1.89552 | 1.142318 | 0.941035 | 3.689168 |
| 37.58333 | 0.294079 | 0.061289 | 0.979195 | -0.14511 | -1.33782 | 1.384955 | 1.001251 | 1.742974 | 1.185984 | -0.17541 | 2.354085 | 0.260834 | 0.547607 | 0.156686 | 0.805187 | 0.255885 | 1.355491 | 0.812039 | 0.003556 | -2.34142 | -1.22347 | 1.509529 | 0.899732 | 3.672285 |
| 37.59722 | 0.132604 | 0.498097 | 0.705345 | -0.03526 | -1.43711 | 0.693723 | 0.873948 | 1.346449 | 0.724959 | 0.180007 | 2.179765 | -0.07246 | 0.564659 | 0.117652 | -0.10954 | -1.28498 | 1.159786 | 0.591377 | -0.08585 | -4.37229 | -0.94143 | 1.724658 | 0.772548 | 2.499867 |
| 37.61111 | 0.089943 | 0.686248 | 0.688124 | -0.12198 | -1.91727 | 0.744689 | 0.804181 | 1.220129 | 0.647499 | 0.086388 | 2.131027 | -0.10687 | 0.516537 | 0.073969 | -0.71186 | -2.25891 | 1.024524 | 0.320105 | -0.21008 | -0.21619 | -0.90357 | 1.703224 | 0.763814 | 0.934413 |
| 37.625 | 0.003886 | 0.781739 | 0.708674 | -0.03641 | -2.27564 | 0.668676 | 0.804904 | 1.209588 | 0.628386 | -0.02046 | 2.083349 | -0.19751 | 0.577718 | 0.062289 | -0.85303 | -2.48838 | 0.696161 | 0.222948 | -0.41518 | 0.756858 | -0.9711 | 1.678866 | 0.612117 | -0.12479 |
| 37.63889 | -0.15471 | 0.940839 | 0.619052 | 0.267491 | -2.35937 | 0.372214 | 0.806318 | 1.009596 | 0.453508 | 0.043361 | 1.767313 | -0.34682 | 0.776695 | 0.039453 | -1.23383 | -2.51037 | 0.136739 | 0.148155 | -0.89412 | 1.330317 | -1.25744 | 1.486241 | 0.563113 | -0.91637 |
| 37.65278 | -0.17551 | 0.455545 | 0.747867 | 0.347046 | -2.59908 | 0.567023 | 0.69977 | 1.223325 | 0.692755 | -0.25472 | 1.802102 | -0.27129 | 0.733635 | -0.07322 | -1.13881 | -2.42892 | -0.35714 | 0.062502 | -1.14545 | 1.857461 | -1.52205 | 1.031019 | 0.629138 | -1.45442 |
| 37.66667 | -0.11521 | -0.12084 | 0.878654 | 0.354349 | -2.72945 | 0.919411 | 0.669233 | 1.574337 | 0.946666 | -0.47958 | 1.807374 | -0.12439 | 0.610576 | -0.21121 | -1.02689 | -2.47696 | -0.82597 | 0.025165 | -1.07133 | 2.291312 | -1.62976 | 0.593733 | 0.568658 | -3.45413 |
| 37.68056 | -0.03831 | -0.2505 | 0.919059 | 0.355059 | -1.95075 | 0.692044 | 0.598863 | 1.643933 | 0.837271 | -0.35157 | 1.499543 | -0.16222 | 0.400935 | -0.28102 | -1.34736 | -2.74219 | -1.32756 | 0.099819 | -1.10373 | 2.565142 | -1.67894 | 0.033642 | 0.231398 | -5.79727 |
| 37.69444 | -0.08028 | -0.34795 | 0.935987 | 0.361521 | -0.94512 | 0.493643 | 0.56891 | 1.554214 | 0.607351 | -0.12571 | 1.150101 | -0.22098 | 0.095857 | -0.25646 | -1.1952 | -2.96517 | -1.66668 | 0.337733 | -0.89666 | 2.956623 | -1.62452 | -0.36831 | 0.210272 | -5.02 |
| 37.70833 | -0.14023 | -0.37801 | 0.896884 | 0.386896 | -0.17845 | 0.530329 | 0.603904 | 1.563171 | 0.560445 | 0.003204 | 0.983518 | -0.24987 | -0.12166 | -0.2624 | -1.20904 | -3.1634 | -1.73546 | 0.701067 | -0.5781 | 3.804154 | -1.68759 | -0.44338 | -0.23937 | -3.83623 |
| 37.72222 | -0.17527 | -0.33949 | 0.840835 | 0.438162 | 0.248736 | 0.604085 | 0.584976 | 1.527439 | 0.478688 | 0.106689 | 0.867441 | -0.23438 | -0.28259 | -0.30238 | -0.80129 | -3.41417 | -1.52688 | 1.030321 | -0.57475 | 2.814197 | -1.7354 | -0.35661 | -0.40822 | -3.98166 |
| 37.73611 | -0.1597 | -0.52114 | 0.814692 | 0.476692 | 0.283611 | 0.724584 | 0.63852 | 1.536659 | 0.514693 | 0.107214 | 0.636249 | -0.13089 | -0.42401 | -0.37491 | -1.21094 | -3.87008 | -1.44589 | 1.221541 | -0.63108 | 1.872357 | -1.85122 | -0.80178 | -0.74043 | -3.52776 |
| 37.75 | -0.10054 | -0.74234 | 0.833227 | 0.536728 | 0.045661 | 0.745725 | 0.74287 | 1.629349 | 0.643868 | 0.103012 | 0.510777 | -0.02993 | -0.6225 | -0.45519 | -0.51308 | -3.82157 | -1.36814 | 1.235383 | -0.613 | 1.228709 | -1.74398 | -1.01733 | -1.16029 | -4.49273 |
| 37.76389 | -0.02473 | -0.85045 | 0.875901 | 0.547629 | -0.3564 | 0.732687 | 0.800447 | 1.628006 | 0.76379 | 0.137014 | 0.136553 | 0.014331 | -0.824 | -0.65471 | -0.36729 | -3.96113 | -1.45576 | 1.096455 | -0.5135 | 0.776439 | -1.33543 | -1.01817 | -1.12069 | -3.37906 |
| 37.77778 | 0.112876 | -0.79792 | 0.859174 | 0.482444 | -0.61042 | 0.494849 | 0.679193 | 1.498413 | 0.749282 | 0.148076 | -0.15002 | 0.013053 | -0.94438 | -0.751 | -0.55992 | -4.08618 | -1.75741 | 0.883103 | -0.52766 | 0.456434 | -1.18623 | -0.68593 | -0.85892 | -1.97344 |
| 37.79167 | 0.128856 | -0.76622 | 0.802599 | 0.451945 | -0.72705 | 0.337422 | 0.484367 | 1.330172 | 0.574613 | 0.124364 | -0.30356 | 0.01472 | -0.98208 | -0.67294 | -0.92885 | -3.86569 | -2.44684 | 0.697534 | -0.4065 | 0.125566 | -1.27898 | -0.35104 | -0.55306 | -1.51761 |
| 37.80556 | 0.27295 | -0.70495 | 0.781324 | 0.437028 | -0.78043 | 0.388842 | 0.30078 | 1.359603 | 0.483305 | 0.092048 | -0.27267 | -0.04722 | -0.81964 | -0.62276 | -1.06923 | -3.94022 | -3.29554 | 0.499634 | -0.1579 | -0.13585 | -1.19243 | -0.05789 | -0.17613 | -1.70671 |
| 37.81944 | 0.332651 | -0.71108 | 0.78594 | 0.428857 | -0.90594 | 0.23446 | 0.140406 | 1.188905 | 0.503022 | 0.066135 | -0.35119 | -0.09908 | -0.86376 | -0.68431 | -1.15496 | -3.58726 | -3.18166 | 0.181088 | -0.14389 | -0.08883 | -0.93935 | 0.120231 | -0.00854 | -1.50313 |
| 37.83333 | 0.390128 | -0.64372 | 0.765232 | 0.455663 | -1.15233 | 0.096576 | 0.093212 | 1.046241 | 0.441324 | 0.038697 | -0.47335 | -0.10429 | -0.90965 | -0.90975 | -1.00572 | -2.65089 | -2.26137 | 0.097279 | -0.35312 | -0.15472 | -0.66395 | 0.405679 | 0.347377 | -0.99172 |
| 37.84722 | 0.37655 | -0.49833 | 0.784184 | 0.512573 | -1.18252 | 0.011238 | 0.217394 | 0.922503 | 0.53763 | 0.066168 | -0.51294 | -0.03841 | -0.68374 | -0.94485 | -0.82415 | -2.35945 | -1.54733 | -0.00205 | -0.4006 | 0.042204 | -0.31231 | 0.567396 | 0.311822 | -0.4729 |
| 37.86111 | 0.387477 | -0.34198 | 0.800693 | 0.698066 | -0.64832 | 0.130395 | 0.435192 | 0.798737 | 0.852771 | 0.122071 | -0.5624 | 0.012713 | -0.5163 | -0.85771 | -0.57081 | -1.86333 | -1.16649 | -0.06285 | -0.23844 | 0.221499 | 0.108689 | 0.680088 | 0.568509 | -0.02789 |
| 37.875 | 0.451881 | -0.10201 | 0.818272 | 0.859468 | 0.08852 | 0.418855 | 0.656676 | 0.693663 | 1.014567 | 0.163094 | -0.59266 | -0.00654 | -0.57506 | -1.02192 | -0.64322 | -1.09197 | -0.80972 | 0.009186 | -0.52688 | -0.08812 | 0.719231 | 1.295759 | 1.100913 | 0.693719 |
| 37.88889 | 0.49183 | 0.148501 | 0.787297 | 0.880335 | 0.339122 | 0.690836 | 0.729611 | 0.558117 | 0.985511 | 0.169715 | -0.72852 | -0.0181 | -0.8657 | -1.41388 | -0.78601 | -0.21802 | -0.62809 | -0.05732 | -1.34778 | -0.43496 | 0.996701 | 1.813457 | 1.272023 | 0.750002 |
| 37.90278 | 0.401632 | 0.132592 | 0.649722 | 0.583638 | -0.09583 | 0.515356 | 0.544952 | 0.630397 | 0.67595 | 0.251677 | -0.94584 | -0.17503 | -1.25349 | -1.98086 | -0.60135 | -0.35439 | -1.2023 | -0.13252 | -1.96058 | -0.5482 | -0.33059 | 0.963099 | 0.773524 | -1.84384 |
| 37.91667 | -0.36085 | 0.155433 | 0.403661 | 0.308738 | -0.47703 | -0.22751 | 0.217718 | 0.52734 | 0.040208 | 0.668644 | -1.53627 | -0.82153 | -1.24105 | -2.01548 | -0.26483 | -0.8707 | -1.73833 | -0.35974 | -2.20394 | -0.2253 | -2.05428 | -0.1645 | -0.05281 | -4.82034 |
| 37.93056 | -1.24059 | 0.304496 | 0.354573 | 0.232505 | -0.50157 | -1.1019 | -0.06766 | 0.581232 | -0.66094 | 1.059267 | -2.85847 | -1.5527 | -0.41632 | -1.54355 | 0.099911 | -0.59154 | -1.60258 | -0.49598 | -2.05006 | -0.02869 | -3.04381 | -0.62279 | -0.16995 | -4.12043 |
| 37.94444 | -1.58707 | -0.35053 | 0.735889 | -0.15643 | -1.12581 | -0.81378 | -0.25008 | 0.375593 | -0.62728 | 0.456527 | -3.25169 | -1.04468 | 0.447023 | -1.00488 | 0.279812 | 0.156809 | -1.1066 | -0.4615 | -1.64417 | 0.030621 | -2.57929 | -0.59764 | 0.323371 | -4.01216 |
| 37.95833 | -1.29186 | -0.01054 | 1.054504 | -0.4631 | -2.21298 | 0.534004 | -0.53895 | 0.453101 | -0.30862 | -0.54654 | -2.30453 | -0.05303 | 0.686323 | -0.67784 | 0.346326 | 0.360245 | -1.13683 | -0.43688 | -1.02384 | 0.162486 | -1.83683 | -0.86659 | 0.612362 | -2.92619 |
| 37.97222 | -0.6 | -0.80659 | 1.218928 | -0.2148 | -2.91496 | 1.173486 | -0.96085 | 0.552524 | -0.78693 | -0.82149 | -1.10444 | 0.285023 | 0.776501 | -0.45485 | 0.361508 | 0.375123 | -1.18722 | -0.26249 | -0.54084 | 0.328309 | -0.76552 | -0.91239 | 0.675885 | -1.86971 |
| 37.98611 | 0.053418 | 0.182055 | 1.235968 | -0.13699 | -2.87191 | 1.207287 | -0.90691 | 0.473269 | -1.0407 | -0.71225 | -0.12148 | 0.381865 | 0.712128 | -0.31048 | 0.36893 | 0.248597 | -1.06192 | -0.05942 | -0.32851 | 0.464848 | 0.20662 | -0.80883 | 0.566099 | -2.08931 |
| 38 | 0.348931 | -0.5905 | 1.01755 | 0.08828 | -2.52967 | 0.402677 | -0.52206 | 0.210943 | -1.14029 | -0.0357 | 0.111744 | -0.15513 | 0.679104 | -0.21863 | 0.37074 | 0.196009 | -0.71286 | -0.04864 | -0.29393 | 0.513073 | 0.723905 | -0.67612 | 0.468634 | -2.28559 |

| | | | | | | | | | | | | | | | | | | | | | | | | |
|---|---|---|---|---|---|---|---|---|---|---|---|---|---|---|---|---|---|---|---|---|---|---|---|---|
| 38.01389 | 0.29051 | -0.25181 | 0.668994 | 0.348328 | -2.124 | -0.5795 | -0.30415 | -0.06481 | -1.37316 | 0.82383 | -0.17305 | -0.90953 | 0.704825 | -0.14999 | 0.364019 | 0.162626 | -0.49425 | -0.03783 | -0.19902 | 0.533219 | 0.906137 | -0.46675 | 0.390931 | -2.16436 |
| 38.02778 | 0.296379 | -0.38158 | 0.505383 | 0.374019 | -1.9237 | -0.47459 | -0.17526 | 0.020221 | -0.88479 | 0.662434 | 0.045852 | -0.70481 | 0.750169 | -0.13359 | 0.379116 | 0.217395 | -0.37538 | 0.04937 | -0.09097 | 0.578575 | 0.928447 | -0.31811 | 0.334754 | -2.05451 |
| 38.04167 | 0.090809 | -0.73967 | 0.40324 | 0.312015 | -1.72262 | 0.105505 | -0.22346 | 0.343598 | -0.38711 | 0.142337 | 0.418474 | -0.2263 | 0.678368 | -0.16964 | 0.418731 | 0.256104 | -0.39452 | 0.127773 | -0.08832 | 0.495672 | 0.915632 | -0.28744 | 0.284 | -1.70661 |
| 38.05556 | -0.21605 | -0.66504 | 0.340717 | 0.235607 | -1.49108 | 0.301938 | -0.31003 | 0.440542 | -0.20543 | -0.20134 | 0.694147 | 0.031176 | 0.601552 | -0.25722 | 0.413081 | 0.282884 | -0.35717 | 0.183311 | -0.16099 | 0.348655 | 0.853719 | -0.23688 | 0.286805 | -1.28182 |
| 38.06944 | -0.06063 | -1.53673 | 0.532001 | 0.132923 | -1.45223 | 1.548817 | -0.26315 | 0.800526 | 0.682174 | -1.00721 | 1.382912 | 1.180678 | 0.595014 | -0.26438 | 0.427154 | 0.360445 | -0.23917 | 0.275841 | -0.196 | 0.297964 | 0.7031 | -0.03486 | 0.363309 | -0.8145 |
| 38.08333 | -0.0561 | -1.55494 | 0.703207 | 0.096316 | -1.06746 | 2.448601 | 0.003408 | 1.018574 | 1.235653 | -1.30375 | 1.498262 | 1.602395 | 0.456491 | -0.74775 | 0.294203 | 0.345661 | -0.17215 | 0.130758 | -0.00437 | 0.811939 | 0.516632 | -0.12396 | 0.490009 | -0.53743 |
| 38.09722 | -0.38562 | NaN | 1.144668 | -0.0799 | -1.20644 | 3.67685 | 0.047911 | 1.654127 | 1.368419 | -1.81329 | 1.441547 | 0.991746 | -0.33744 | -6.261 | -0.48549 | -0.04812 | -1.07035 | -1.24589 | 0.969333 | 4.595085 | 0.282593 | -2.22005 | 0.943278 | -2.39665 |
| 38.11111 | -0.1763 | NaN | 2.815146 | -0.52053 | -3.40019 | NaN | 0.156445 | 3.307054 | 2.397966 | -4.24396 | 2.865701 | 2.299794 | 3.096719 | -17.5466 | -5.32403 | 0.707685 | -3.23509 | -2.35825 | 17.25741 | 13.74256 | -0.21509 | -6.01253 | 2.252786 | -5.11668 |
| 38.125 | NaN | NaN | 0.77873 | -0.26572 | -3.70646 | NaN | 1.320402 | 2.061074 | -4.91185 | -1.7764 | -0.6774 | 2.648894 | 2.959673 | -5.74269 | NaN | 8.749509 | 3.40177 | 3.572297 | NaN | -6.47581 | -2.13921 | -4.28309 | NaN | NaN |
| 38.13889 | NaN | NaN | NaN | 1.740524 | 3.305294 | NaN | NaN | NaN | NaN | 6.227948 | -3.61116 | -5.7342 | 1.242776 | -1.81534 | NaN | NaN | -12.5744 | 0.667329 | NaN | 5.830151 | 1.04343 | NaN | NaN | 7.707698 |
