## Supplemental Table S2b for "Leaf movements as a quantitative metric for early stress detection"

**Supplementary Table S2.** Raw motion data from **(b)** salt stress applied at different locations on lettuce with their respective controls.

**Supplementary Table S2.** Raw motion data from (b) salt stress applied at UoC

| Time | control1 | control2 | control3 | control4 | control5 | control6 | control7 | control8 | control9 | Stress1 | Stress2 | Stress3 | Stress4 | Stress5 | Stress6 | Stress7 | Stress8 | Stress9 | Stress10 | Stress11 |
|---|---|---|---|---|---|---|---|---|---|---|---|---|---|---|---|---|---|---|---|---|
| 17.0125 | -0.48687 | 0.078617 | -0.42981 | -0.46974 | 2.447937 | 0.222383 | 0.143915 | -0.04743 | -0.23948 | 1.244356 | -0.06867 | -0.22985 | -0.07823 | 0.028838 | 0.589063 | -0.0934 | 0.26855 | 0.077907 | 0.398281 | -1.21983 |
| 17.02639 | -0.6039 | -0.11112 | -0.15331 | -0.18533 | 1.758853 | 0.190258 | 0.009515 | 0.011356 | -0.12157 | -1.0083 | 0.246199 | -0.13875 | -0.02623 | 0.127234 | 0.507332 | -0.05202 | 0.1003 | 0.148059 | 0.149912 | -0.67374 |
| 17.04028 | 0.159598 | 0.01656 | 0.04205 | 0.131415 | 0.222639 | 0.21073 | 0.160996 | 0.03524 | -0.00264 | -0.1686 | 0.540113 | 0.082068 | -0.00864 | 0.134778 | 0.284575 | -0.051 | 0.046815 | 0.09491 | -0.09653 | -0.15029 |
| 17.05417 | 0.204236 | -0.01466 | 0.028332 | 0.14769 | -0.03087 | 0.107505 | 0.121616 | 0.012355 | -0.03723 | -0.13237 | 0.219181 | 0.075314 | -0.03133 | 0.05448 | 0.030212 | -0.06712 | -0.01313 | 0.036863 | -0.22874 | -0.31643 |
| 17.06806 | 0.060567 | -0.05441 | 0.001689 | 0.12901 | -0.07589 | 0.038541 | 0.076528 | -0.01773 | -0.06354 | -0.23779 | -0.04138 | 0.040544 | -0.04031 | 0.007287 | -0.07035 | -0.07343 | -0.03852 | 0.012504 | -0.20353 | -0.48847 |
| 17.08194 | 0.028284 | -0.06217 | -0.03414 | 0.134165 | -0.07226 | 0.005757 | 0.09241 | -0.03186 | -0.05726 | -0.29694 | -0.12847 | 0.040564 | -0.03235 | 0.015087 | -0.01741 | -0.04865 | -0.02862 | 0.029822 | -0.00155 | -0.47151 |
| 17.09583 | 0.02279 | -0.06964 | -0.03946 | 0.110124 | -0.08106 | -0.04408 | 0.075978 | -0.03789 | -0.04945 | -0.2571 | -0.13567 | 0.04242 | -0.03757 | 0.042409 | 0.029958 | -0.05876 | -0.05321 | 0.019899 | 0.062824 | -0.36844 |
| 17.10972 | 0.008047 | -0.03306 | 0.01365 | 0.121775 | -0.0622 | -0.07422 | 0.058864 | -0.00988 | -0.01327 | -0.10273 | -0.08967 | 0.076775 | -0.03434 | 0.089113 | 0.043664 | -0.03902 | -0.06623 | 0.00307 | 0.079518 | -0.21071 |
| 17.12361 | 0.005155 | 0.014615 | 0.033154 | 0.180007 | 0.028663 | -0.07112 | 0.078143 | 0.015023 | 0.035247 | 0.049731 | 0.024478 | 0.119911 | -0.02208 | 0.118906 | 0.051605 | 0.038997 | -0.01576 | 0.015542 | 0.304329 | -0.19047 |
| 17.1375 | 0.008664 | 0.028227 | -0.07368 | 0.195737 | 0.150753 | -0.06542 | 0.096803 | -0.01577 | 0.060728 | 0.239852 | 0.157104 | 0.100272 | -0.04794 | 0.094456 | 0.060817 | 0.147428 | 0.069047 | 0.032448 | 0.693195 | -0.3707 |
| 17.15139 | -0.00348 | 0.017987 | -0.08974 | 0.165896 | 0.228696 | -0.03528 | 0.069474 | -0.01314 | 0.058845 | 0.289425 | 0.115057 | 0.060872 | -0.07261 | 0.045475 | 0.028343 | 0.169561 | 0.066042 | 0.025188 | 0.653913 | -0.65855 |
| 17.16528 | -0.01666 | 0.013459 | 0.141975 | 0.100149 | 0.156785 | 0.007878 | -0.00507 | 0.080626 | 0.028053 | -0.02969 | -0.10151 | 0.056045 | -0.00489 | -0.01167 | -0.05642 | 0.002684 | -0.06221 | -0.00816 | -0.07667 | -0.32852 |
| 17.17917 | -0.00082 | 0.026244 | 0.264254 | 0.030397 | 0.088752 | 0.046412 | -0.01206 | 0.131215 | 0.033814 | -0.22016 | -0.23358 | 0.070649 | 0.056798 | -0.04109 | -0.06763 | -0.09998 | -0.15589 | -0.02864 | -0.44611 | -0.0815 |
| 17.19306 | 0.010061 | 0.023087 | 0.142853 | 0.020987 | 0.099004 | 0.064843 | 0.035636 | 0.098994 | 0.079862 | 0.007556 | -0.20608 | 0.100285 | 0.042226 | -0.00206 | -0.03642 | 0.002201 | -0.10401 | 0.007518 | -0.03998 | 0.050818 |
| 17.20694 | 0.020074 | 0.044166 | -0.03613 | 0.101878 | 0.187516 | 0.089256 | 0.105357 | 0.043857 | 0.141382 | 0.343889 | -0.12444 | 0.127083 | -0.00657 | 0.054142 | 0.018501 | 0.232877 | 0.042751 | 0.049008 | 0.592276 | 0.068466 |
| 17.22083 | 0.011721 | 0.08798 | -0.19599 | 0.122863 | 0.315827 | 0.092372 | 0.15791 | -0.0291 | 0.174669 | 0.512526 | -0.06227 | 0.135811 | -0.05293 | 0.074832 | 0.104031 | 0.469207 | 0.155832 | 0.065184 | 1.079956 | 0.005046 |
| 17.23472 | -0.02915 | 0.072233 | -0.15195 | 0.058906 | 0.363493 | 0.076043 | 0.147001 | -0.0459 | 0.126372 | 0.510319 | -0.05683 | 0.087555 | -0.06807 | 0.045057 | 0.121987 | 0.407563 | 0.102098 | 0.029406 | 0.958953 | 0.056234 |
| 17.24861 | -0.04546 | 0.076558 | 0.044209 | 0.103105 | 0.349771 | 0.103064 | 0.151131 | 0.020948 | 0.087837 | 0.511423 | 0.023313 | 0.10699 | -0.03126 | 0.071206 | 0.087816 | 0.202463 | 0.030164 | -0.00585 | 0.500857 | 0.184574 |
| 17.2625 | -0.04231 | 0.086933 | 0.138659 | 0.207313 | 0.368352 | 0.141529 | 0.188457 | 0.079426 | 0.109466 | 0.585255 | 0.149554 | 0.125229 | 0.007099 | 0.120189 | 0.084467 | 0.135665 | 0.005089 | -0.01148 | 0.263607 | 0.174651 |
| 17.27639 | -0.0526 | 0.067857 | 0.131376 | 0.35238 | 0.353111 | 0.174395 | 0.247551 | 0.105367 | 0.133513 | 0.601693 | 0.162499 | 0.131753 | -0.02066 | 0.102648 | 0.062074 | 0.129205 | -0.01507 | -0.03101 | 0.213353 | 0.095684 |
| 17.29028 | -0.02861 | 0.080994 | 0.140971 | 0.547738 | 0.353004 | 0.266031 | 0.356905 | 0.150708 | 0.184205 | 0.620641 | 0.149406 | 0.166982 | -0.02323 | 0.085534 | 0.073139 | 0.161331 | 0.0097 | -0.02097 | 0.290196 | 0.169336 |
| 17.30417 | 0.008649 | 0.106385 | 0.168108 | 0.595242 | 0.36965 | 0.324274 | 0.41103 | 0.191375 | 0.212799 | 0.645527 | 0.240209 | 0.217977 | 0.012053 | 0.099684 | 0.101618 | 0.222594 | 0.05955 | 0.02051 | 0.401637 | 0.302605 |
| 17.31806 | -0.00503 | 0.078877 | 0.151868 | 0.487868 | 0.293784 | 0.244859 | 0.341217 | 0.153941 | 0.157718 | 0.6994 | 0.30239 | 0.187343 | 0.013651 | 0.066119 | 0.075437 | 0.228605 | 0.038616 | 0.018206 | 0.389201 | 0.396137 |
| 17.33194 | -0.03457 | 0.03918 | 0.11536 | 0.351657 | 0.208304 | 0.128121 | 0.256216 | 0.075402 | 0.089283 | 0.740221 | 0.2073 | 0.107803 | -0.01279 | 0.017502 | 0.027381 | 0.181801 | -0.00864 | -0.02361 | 0.304054 | 0.46372 |
| 17.34583 | -0.02014 | 0.049705 | 0.115172 | 0.244254 | 0.255103 | 0.069618 | 0.241957 | 0.068171 | 0.097918 | 0.693732 | 0.112788 | 0.061111 | 0.001494 | 0.036185 | 0.034031 | 0.160438 | 0.001381 | -0.027 | 0.293711 | 0.516997 |
| 17.35972 | 0.011528 | 0.084867 | 0.12986 | 0.221459 | 0.382062 | 0.080453 | 0.348536 | 0.126479 | 0.155384 | 0.630358 | 0.044472 | 0.059358 | 0.040949 | 0.109046 | 0.07799 | 0.167109 | 0.038725 | 0.005501 | 0.361025 | 0.559505 |
| 17.37361 | 0.012802 | 0.09542 | 0.145062 | 0.264573 | 0.410152 | 0.118467 | 0.559663 | 0.16683 | 0.17174 | 0.546465 | -0.00913 | 0.058077 | 0.047961 | 0.13935 | 0.071085 | 0.130488 | 0.048162 | 0.011427 | 0.36028 | 0.693828 |
| 17.3875 | -0.00694 | 0.089827 | 0.230111 | 0.37535 | 0.290809 | 0.169255 | 0.573136 | 0.118365 | 0.152519 | 0.465476 | -0.00374 | 0.033979 | 0.046218 | 0.079112 | -0.01623 | 0.009954 | 0.008442 | -0.02041 | 0.067023 | 0.99895 |
| 17.40139 | -0.0058 | 0.100497 | 0.291794 | 0.530748 | 0.148898 | 0.240424 | 0.409692 | 0.033488 | 0.170163 | 0.380868 | 0.129296 | 0.031919 | 0.052074 | 0.013142 | -0.07108 | -0.04753 | -0.02746 | -0.04297 | -0.19336 | 1.122558 |
| 17.41528 | 0.048025 | 0.159218 | 0.238355 | 0.649405 | 0.204433 | 0.319656 | 0.416394 | 0.036199 | 0.263883 | 0.542537 | 0.407647 | 0.125546 | 0.085107 | 0.059424 | 0.045414 | 0.084607 | 0.042389 | 0.020459 | 0.243731 | 0.655773 |
| 17.42917 | 0.107558 | 0.239874 | 0.180384 | 0.712794 | 0.375173 | 0.385638 | 0.473569 | 0.115757 | 0.328887 | 0.869092 | 0.457495 | 0.241784 | 0.139879 | 0.157134 | 0.18827 | 0.237149 | 0.154335 | 0.101513 | 0.88457 | 0.324652 |
| 17.44306 | 0.178705 | 0.368321 | 0.198763 | 0.915257 | 0.631799 | 0.5072 | 0.648531 | 0.186405 | 0.497371 | 1.627413 | 0.330544 | 0.406935 | 0.207471 | 0.33436 | 0.342827 | 0.389461 | 0.341424 | 0.166251 | 1.926213 | 0.309351 |
| 17.45694 | 0.299708 | 0.58507 | 0.462864 | 1.751094 | 0.8224 | 0.762385 | 1.061282 | 0.181288 | 0.865371 | 2.69395 | 0.224609 | 0.659443 | 0.282463 | 0.605427 | 0.476364 | 0.48721 | 0.589181 | 0.208215 | 4.13125 | 1.084426 |
| 17.47083 | 0.480573 | 0.629879 | 0.694009 | 2.558776 | 0.609811 | 0.921371 | 1.34276 | 0.163145 | 1.298217 | 3.04199 | 0.211982 | 0.693974 | 0.281915 | 0.694261 | 0.393155 | 0.407275 | 0.567291 | 0.184211 | 5.240313 | 1.600398 |
| 17.48472 | 0.517011 | 0.416922 | 0.587418 | 1.975934 | 0.407757 | 0.760667 | 1.314402 | 0.179334 | 1.162805 | 2.284328 | 0.235185 | 0.474485 | 0.15504 | 0.557179 | 0.203255 | 0.21363 | 0.356888 | 0.166769 | 3.224626 | 1.504222 |
| 17.49861 | 0.381514 | 0.270922 | 0.369339 | 1.124938 | 0.52508 | 0.565192 | 1.056064 | 0.156366 | 0.922653 | 1.424507 | 0.337879 | 0.325276 | 0.149138 | 0.374478 | 0.09688 | 0.030566 | 0.155532 | 0.13891 | 1.634614 | 1.079538 |
| 17.5125 | 0.233737 | 0.199355 | 0.16456 | 0.345862 | 0.587829 | 0.355433 | 0.658079 | 0.054943 | 0.738323 | 0.648628 | 0.473904 | 0.193359 | 0.179926 | 0.151775 | 0.057088 | -0.0837 | 0.03422 | 0.128243 | 1.029094 | 0.550129 |
| 17.52639 | 0.178773 | 0.173134 | -0.03013 | -0.07469 | 0.476781 | 0.184813 | 0.360385 | -0.04371 | 0.58857 | 0.342082 | 0.649531 | 0.088095 | 0.184761 | -0.00919 | 0.074401 | -0.11044 | 0.02025 | 0.132469 | 0.920529 | 0.149439 |
| 17.54028 | 0.158155 | 0.166901 | -0.11437 | 0.003862 | 0.39939 | 0.132437 | 0.280169 | -0.00794 | 0.496097 | 0.697713 | 0.652088 | 0.048306 | 0.151073 | -0.06694 | 0.185779 | -0.02063 | 0.086234 | 0.158423 | 0.982126 | -0.04859 |
| 17.55417 | 0.155957 | 0.212557 | -0.01943 | 0.446332 | 0.60107 | 0.238966 | 0.389598 | 0.163474 | 0.509874 | 1.297721 | 0.533669 | 0.079955 | 0.128573 | 0.010611 | 0.370159 | 0.15456 | 0.188885 | 0.167733 | 0.984336 | 0.084691 |
| 17.56806 | 0.216026 | 0.277765 | 0.13607 | 0.809704 | 0.931899 | 0.417992 | 0.582898 | 0.379727 | 0.55562 | 1.357759 | 0.511332 | 0.117443 | 0.133256 | 0.16316 | 0.391581 | 0.232636 | 0.225527 | 0.111584 | 0.634695 | 0.49168 |
| 17.58194 | 0.348982 | 0.252131 | 0.146273 | 0.799248 | 1.042531 | 0.501611 | 0.726185 | 0.399161 | 0.515533 | 0.776141 | 0.56762 | 0.128333 | 0.120559 | 0.253838 | 0.250779 | 0.189223 | 0.222763 | 0.059686 | 0.222565 | 0.724625 |
| 17.59583 | 0.443759 | 0.133373 | 0.056735 | 0.49439 | 0.887233 | 0.424928 | 0.823276 | 0.223112 | 0.421539 | 0.320623 | 0.517973 | 0.107283 | 0.118949 | 0.255058 | 0.142487 | 0.129276 | 0.263977 | 0.048839 | 0.032333 | 0.654354 |
| 17.60972 | 0.349209 | -0.01688 | -0.0053 | 0.152133 | 0.638566 | 0.314196 | 0.774977 | 0.073446 | 0.381013 | 0.03101 | 0.414084 | 0.055706 | 0.117732 | 0.198027 | 0.053503 | 0.043858 | 0.237191 | 0.083512 | -0.31207 | 0.628648 |
| 17.62361 | 0.162731 | -0.11572 | -0.05326 | -0.25405 | 0.351197 | 0.176282 | 0.566685 | -0.08743 | 0.356921 | -0.15434 | 0.310123 | -0.05372 | 0.098707 | 0.05745 | -0.04169 | -0.08082 | 0.090004 | 0.113704 | -0.8377 | 0.563661 |
| 17.6375 | 0.016976 | -0.14371 | -0.10636 | -0.60131 | 0.196592 | -0.00853 | 0.347482 | -0.1999 | 0.218081 | -0.16878 | 0.266886 | -0.10645 | 0.08866 | -0.08431 | -0.07347 | -0.16674 | -0.02651 | 0.111822 | -0.97238 | 0.338428 |
| 17.65139 | -0.06385 | -0.08434 | -0.15429 | -0.80709 | 0.103078 | -0.17829 | 0.126418 | -0.19582 | -0.06409 | 0.035393 | 0.369524 | -0.09092 | 0.095809 | -0.206 | -0.02132 | -0.14757 | -0.03194 | 0.099708 | -0.7712 | 0.077225 |
| 17.66528 | -0.11337 | 0.001197 | -0.12527 | -0.70322 | -0.00555 | -0.23555 | -0.12499 | -0.06889 | -0.33204 | 0.301425 | 0.555688 | -0.02422 | 0.096814 | -0.27348 | 0.088314 | -0.0135 | 0.015061 | 0.067612 | -0.36025 | -0.13565 |
| 17.67917 | -0.14002 | 0.047393 | 0.00307 | -0.32025 | -0.1247 | -0.20193 | -0.35368 | 0.069316 | -0.42847 | 0.32814 | 0.718924 | 0.051801 | 0.05771 | -0.18954 | 0.161359 | 0.08571 | 0.035629 | 0.009852 | 0.091896 | -0.16947 |
| 17.69306 | -0.12954 | 0.082561 | 0.152512 | 0.187479 | -0.23802 | -0.1183 | -0.50666 | 0.110284 | -0.30558 | 0.194136 | 0.713901 | 0.093824 | -0.01001 | -0.00904 | 0.126246 | 0.07144 | 0.029924 | -0.05746 | 0.390621 | -0.0238 |
| 17.70694 | -0.0781 | 0.128399 | 0.195204 | 0.596339 | -0.33584 | -0.02548 | -0.55404 | 0.073656 | -0.08448 | 0.098034 | 0.383489 | 0.081371 | -0.03402 | 0.102078 | 0.047848 | 0.038085 | 0.03841 | -0.07921 | 0.564198 | 0.065661 |
| 17.72083 | 0.022959 | 0.146805 | 0.096334 | 0.730342 | -0.47266 | 0.032071 | -0.54441 | -0.00265 | 0.118239 | -0.00921 | -0.13944 | 0.042307 | -0.0029 | 0.058022 | -0.03936 | 0.002388 | 0.037068 | -0.06341 | 0.425392 | -0.04107 |
| 17.73472 | 0.15071 | 0.077726 | -0.02366 | 0.538922 | -0.52652 | 0.029736 | -0.42276 | -0.06165 | 0.212806 | -0.22509 | -0.58518 | 0.006107 | 0.036427 | -0.01099 | -0.10266 | -0.04372 | 0.021852 | -0.04967 | -0.0796 | -0.21485 |
| 17.74861 | 0.232169 | -0.02896 | -0.09971 | 0.256906 | -0.4538 | -0.06419 | -0.22444 | -0.1062 | 0.157286 | -0.48831 | -0.72556 | -0.01023 | 0.015906 | -0.04543 | -0.10199 | -0.08889 | -0.00763 | -0.04117 | -0.29637 | -0.38246 |
| 17.7625 | 0.213797 | -0.10204 | -0.14535 | 0.013224 | -0.33541 | -0.23679 | -0.1165 | -0.10172 | -0.02439 | -0.76185 | -0.61795 | -0.03241 | -0.00029 | -0.09213 | -0.10037 | -0.13004 | -0.04413 | -0.02032 | -0.22276 | -0.51631 |
| 17.77639 | 0.122358 | -0.1188 | -0.20728 | -0.2654 | -0.24634 | -0.33459 | -0.15425 | -0.08468 | -0.21407 | -0.88567 | -0.40612 | -0.05987 | -0.03748 | -0.15807 | -0.11098 | -0.16426 | -0.05462 | 0.025737 | -0.12864 | -0.61009 |
| 17.79028 | 0.000452 | -0.08257 | -0.30175 | -0.64395 | -0.32106 | -0.28673 | -0.273 | -0.08439 | -0.2943 | -1.0236 | -0.25753 | -0.10079 | -0.09866 | -0.17181 | -0.13681 | -0.1633 | -0.05149 | 0.053672 | -0.29721 | -0.74188 |
| 17.80417 | -0.12457 | -0.08438 | -0.36999 | -1.0673 | -0.55737 | -0.21702 | -0.4473 | -0.10448 | -0.32209 | -1.29298 | -0.35302 | -0.12423 | -0.16473 | -0.11938 | -0.14883 | -0.1133 | -0.05571 | 0.048561 | -0.69604 | -1.02679 |
| 17.81806 | -0.25629 | -0.16151 | -0.33978 | -1.27327 | -0.92863 | -0.33622 | -0.67609 | -0.18578 | -0.42389 | -1.55499 | -0.82661 | -0.16066 | -0.25867 | -0.13563 | -0.1956 | -0.14983 | -0.10154 | 0.003264 | -1.21002 | -1.48166 |
| 17.83194 | -0.34733 | -0.2987 | -0.23329 | -1.36051 | -1.15229 | -0.58249 | -0.93161 | -0.2907 | -0.61863 | -1.72883 | -1.30675 | -0.17798 | -0.30107 | -0.22343 | -0.27849 | -0.27672 | -0.1858 | -0.07043 | -1.56446 | -1.80127 |
| 17.84583 | -0.31115 | -0.44312 | -0.20138 | -1.36505 | -1.14151 | -0.6888 | -1.12891 | -0.38474 | -0.67075 | -1.45529 | -1.45828 | -0.19346 | -0.27588 | -0.25865 | -0.33356 | -0.34247 | -0.24583 | -0.14401 | -1.71301 | -1.88512 |
| 17.85972 | -0.22487 | -0.54173 | -0.34957 | -1.39994 | -1.11014 | -0.6826 | -1.19821 | -0.46334 | -0.64808 | -1.32464 | -1.17901 | -0.21637 | -0.27011 | -0.22667 | -0.35665 | -0.33443 | -0.28044 | -0.20243 | -1.48002 | -1.60656 |
| 17.87361 | -0.26479 | -0.60774 | -0.54226 | -1.33018 | -1.02184 | -0.74358 | -1.2065 | -0.48428 | -0.65381 | -1.42629 | -0.92568 | -0.31357 | -0.34506 | -0.19005 | -0.40326 | -0.3437 | -0.29882 | -0.20303 | -1.41006 | -1.52439 |
| 17.8875 | -0.36565 | -0.53832 | -0.56411 | -1.15969 | -0.8507 | -0.80552 | -1.02274 | -0.42263 | -0.67879 | -1.50927 | -1.01221 | -0.46947 | -0.47679 | -0.18868 | -0.44718 | -0.36998 | -0.27973 | -0.19474 | -1.35514 | -1.5701 |
| 17.90139 | -0.40546 | -0.43129 | -0.4617 | -0.94516 | -0.56951 | -0.65256 | -0.76935 | -0.30632 | -0.60748 | -1.48801 | -1.23402 | -0.47302 | -0.51881 | -0.18944 | -0.44571 | -0.35372 | -0.22942 | -0.1739 | -1.19337 | -1.4214 |
| 17.91528 | -0.42627 | -0.36871 | -0.37952 | -0.73921 | -0.39494 | -0.53157 | -0.61703 | -0.27032 | -0.52822 | -1.59559 | -1.32194 | -0.41125 | -0.49097 | -0.16396 | -0.41108 | -0.32018 | -0.19949 | -0.16096 | -1.02301 | -1.00293 |
| 17.92917 | -0.38901 | -0.37582 | -0.32182 | -0.55294 | -0.39663 | -0.49455 | -0.66222 | -0.26254 | -0.58739 | -1.73461 | -1.37538 | -0.43983 | -0.39964 | -0.19955 | -0.44596 | -0.40087 | -0.29186 | -0.15175 | -1.31422 | -0.76263 |
| 17.94306 | -0.33991 | -0.47996 | -0.18017 | -0.50191 | -0.50991 | -0.43281 | -0.73736 | -0.16087 | -0.62161 | -1.78677 | -1.39343 | -0.52426 | -0.30825 | -0.25172 | -0.52308 | -0.53962 | -0.46296 | -0.1076 | -2.05795 | -0.59158 |
| 17.95694 | -0.16884 | -0.67414 | -0.1434 | -0.66155 | -0.64067 | -0.32899 | -0.761 | -0.05895 | -0.57884 | -1.79782 | -1.32908 | -0.54727 | -0.28271 | -0.26996 | -0.60582 | -0.4657 | -0.58869 | -0.08385 | -2.44803 | -0.92512 |
| 17.97083 | -0.32409 | -1.09694 | -0.20235 | -0.89889 | -0.69803 | -0.27091 | -0.73076 | -0.14214 | -0.42033 | -1.56559 | -1.16582 | -0.56029 | -0.37201 | -0.30505 | -0.80082 | -0.25986 | -0.62085 | -0.1142 | -2.3531 | -1.06071 |
| 17.98472 | -0.1463 | -0.69749 | -0.22179 | -0.897 | -0.61961 | -0.2355 | -0.50337 | -0.27711 | -0.22323 | -1.27708 | -0.7879 | -0.3856 | -0.32044 | -0.2039 | -0.52425 | -0.09887 | -0.40199 | -0.11571 | -0.94966 | -1.01969 |
| 17.99861 | 0.339236 | 0.440638 | -0.15923 | -0.61829 | -0.43399 | -0.16731 | -0.22458 | -0.27184 | -0.0383 | -0.80813 | -0.32121 | -0.05722 | -0.14177 | -0.04777 | 0.010079 | 0.062528 | 0.066858 | -0.07293 | 0.414741 | -0.78167 |
| 18.0125 | 1.358882 | 0.935654 | -0.113 | -0.30281 | -0.27264 | -0.04156 | -0.13616 | -0.16282 | 0.08907 | -0.21754 | -0.10655 | 0.053945 | 0.012955 | -0.00056 | 0.266499 | 0.128775 | 0.28713 | -0.00471 | 0.916211 | -0.34134 |
| 18.02639 | 1.057636 | 0.645364 | -0.01134 | -0.09903 | -0.09833 | 0.099483 | -0.04067 | 0.025018 | 0.11893 | 0.111987 | -0.13196 | 0.028585 | 0.081335 | -0.0745 | 0.186857 | -0.00778 | 0.128098 | -0.02627 | 0.596263 | 0.122537 |
| 18.04028 | 0.421711 | 0.32085 | 0.045355 | 0.084674 | 0.122912 | 0.287564 | 0.216206 | 0.171128 | 0.149301 | 0.331593 | -0.08062 | 0.048204 | 0.065054 | -0.11536 | 0.100039 | -0.04714 | -0.02393 | -0.0229 | 0.305391 | 0.348727 |
| 18.05417 | 0.343324 | 0.268388 | -0.10363 | 0.240733 | 0.306534 | 0.417322 | 0.45542 | 0.158397 | 0.159445 | 0.58335 | 0.147802 | 0.091936 | -0.049 | -0.03186 | 0.116322 | 0.211073 | 0.052891 | 0.042122 | 0.569874 | 0.056211 |
| 18.06806 | 0.180837 | 0.270929 | -0.3129 | 0.221538 | 0.358479 | 0.364435 | 0.444522 | 0.034403 | 0.084121 | 0.611249 | 0.245638 | 0.06713 | -0.19235 | 0.009716 | 0.064482 | 0.389368 | 0.139171 | 0.02696 | 0.876144 | -0.41072 |
| 18.08194 | -0.00873 | 0.069019 | -0.1893 | 0.132358 | 0.287844 | 0.143586 | 0.219837 | -0.01402 | -0.04393 | 0.196767 | 0.037738 | -0.04224 | -0.18272 | -0.07961 | -0.07059 | 0.090317 | -0.01072 | -0.04538 | 0.385686 | -0.40502 |
| 18.09583 | -0.12801 | -0.12443 | 0.136015 | 0.177372 | 0.222497 | -0.01361 | 0.07766 | 0.101853 | -0.09295 | -0.13031 | -0.1173 | -0.10149 | -0.02253 | -0.11695 | -0.12296 | -0.32689 | -0.19189 | -0.07303 | -0.45422 | 0.00419 |
| 18.10972 | -0.05263 | -0.05957 | 0.176821 | 0.197407 | 0.264856 | -0.01819 | 0.106299 | 0.145248 | -0.02059 | -0.06354 | -0.00765 | -0.02575 | 0.032425 | -0.0374 | -0.00703 | -0.23085 | -0.10719 | 0.023402 | -0.3452 | -0.05495 |
| 18.12361 | 0.072381 | 0.153981 | -0.12325 | 0.165406 | 0.189009 | -2.6E-05 | 0.147427 | 0.028745 | 0.062246 | 0.336397 | 0.275726 | 0.045764 | -0.14392 | 0.040558 | 0.017125 | 0.221969 | 0.100295 | 0.077333 | 0.480305 | -0.49192 |
| 18.1375 | 0.011134 | 0.118415 | -0.17177 | 0.022395 | 0.069049 | -0.08993 | 0.044057 | -0.0544 | 0.052133 | 0.384187 | 0.261194 | -0.00441 | -0.21386 | -0.03547 | -0.01665 | 0.247184 | 0.062006 | 0.013155 | 0.513909 | -0.58517 |
| 18.15139 | -0.10929 | -0.01503 | 0.022734 | 0.012224 | 0.029386 | -0.16491 | -0.02399 | 0.024553 | 0.010778 | 0.32483 | 0.127012 | -0.09204 | -0.03952 | -0.09244 | -0.03492 | -0.09104 | -0.09122 | -0.03821 | -0.08702 | -0.19301 |
| 18.16528 | -0.06317 | 0.024784 | 0.027382 | 0.087954 | 0.125024 | -0.07989 | 0.013806 | 0.114746 | 0.051626 | 0.348277 | 0.225064 | -0.05179 | 0.045657 | -0.0235 | 0.055227 | -0.06934 | -0.04012 | 0.006627 | 0.025503 | -0.02884 |
| 18.17917 | 0.027022 | 0.12569 | -0.13615 | 0.107686 | 0.203395 | 0.023183 | 0.017371 | 0.099768 | 0.108107 | 0.341497 | 0.322874 | 0.024283 | -0.03775 | 0.049654 | 0.130734 | 0.215903 | 0.117276 | 0.056963 | 0.572493 | -0.2652 |
| 18.19306 | -0.00167 | 0.124096 | -0.14486 | 0.075969 | 0.183701 | -0.004 | -0.05538 | 0.074193 | 0.084009 | 0.211439 | 0.232599 | 0.00685 | -0.08169 | 0.037225 | 0.116995 | 0.279238 | 0.11039 | 0.043703 | 0.621695 | -0.30311 |
| 18.20694 | -0.05141 | 0.107495 | -0.00504 | 0.082361 | 0.105467 | -0.03755 | -0.07748 | 0.080767 | 0.095154 | 0.231388 | 0.124002 | -0.02129 | -0.01417 | 0.047855 | 0.096337 | 0.242558 | 0.04632 | 0.025732 | 0.45372 | -0.09729 |
| 18.22083 | -0.06846 | 0.139594 | 0.143413 | 0.171495 | 0.108237 | 0.005032 | -0.00201 | 0.112638 | 0.19509 | 0.284852 | 0.196246 | 0.00592 | 0.092749 | 0.110296 | 0.119347 | 0.254202 | 0.034954 | 0.029855 | 0.498034 | 0.131785 |
| 18.23472 | -0.07559 | 0.175392 | 0.324306 | 0.329217 | 0.21016 | 0.081368 | 0.095867 | 0.160682 | 0.261327 | 0.28377 | 0.408597 | 0.019431 | 0.204642 | 0.146065 | 0.164687 | 0.183951 | -0.00805 | 0.023197 | 0.496916 | 0.461028 |

| | | | | | | | | | | | | | | | | | | | | |
|---|---|---|---|---|---|---|---|---|---|---|---|---|---|---|---|---|---|---|---|---|
| 18.24861 | -0.07387 | 0.16913 | 0.407201 | 0.413786 | 0.364134 | 0.184624 | 0.197934 | 0.193679 | 0.291013 | 0.343204 | 0.534547 | 0.042195 | 0.220467 | 0.155516 | 0.194137 | 0.115914 | -0.04148 | 0.008912 | 0.348088 | 0.741341 |
| 18.2625 | -0.02765 | 0.18627 | 0.176398 | 0.355268 | 0.419637 | 0.270002 | 0.31815 | 0.13242 | 0.25478 | 0.488764 | 0.378695 | 0.094424 | 0.075113 | 0.115387 | 0.178091 | 0.260212 | 0.035195 | -0.01025 | 0.519434 | 0.474823 |
| 18.27639 | -0.00845 | 0.200506 | -0.02741 | 0.322351 | 0.353967 | 0.279478 | 0.39064 | 0.055472 | 0.193472 | 0.597676 | 0.111685 | 0.09326 | -0.0506 | 0.031177 | 0.130549 | 0.381466 | 0.100929 | -0.04781 | 0.72137 | 0.139604 |
| 18.29028 | -0.05449 | 0.172402 | 0.05188 | 0.375263 | 0.251455 | 0.26665 | 0.450423 | 0.077286 | 0.23912 | 0.714306 | 0.032028 | 0.054799 | 0.038546 | 0.016156 | 0.098189 | 0.258144 | 0.053937 | -0.04007 | 0.537042 | 0.23957 |
| 18.30417 | -0.06831 | 0.176316 | 0.188987 | 0.491707 | 0.306639 | 0.302338 | 0.543519 | 0.197406 | 0.313079 | 0.737238 | 0.215213 | 0.083089 | 0.223633 | 0.088994 | 0.131076 | 0.1815 | 0.035258 | -0.00973 | 0.318249 | 0.58765 |
| 18.31806 | -0.02583 | 0.217849 | 0.27259 | 0.669071 | 0.504517 | 0.416305 | 0.660999 | 0.386546 | 0.362186 | 0.680783 | 0.494276 | 0.163694 | 0.295 | 0.172702 | 0.1663 | 0.234613 | 0.055121 | 0.008952 | 0.273319 | 0.8603 |
| 18.33194 | 0.010722 | 0.260601 | 0.347445 | 0.823963 | 0.678912 | 0.479431 | 0.768552 | 0.529826 | 0.3667 | 0.632978 | 0.825942 | 0.254173 | 0.260292 | 0.245705 | 0.165134 | 0.329598 | 0.105913 | 0.039567 | 0.404998 | 0.905796 |
| 18.34583 | 0.014981 | 0.285928 | 0.347083 | 0.859276 | 0.694056 | 0.417321 | 0.756474 | 0.507617 | 0.277368 | 0.699336 | 1.11155 | 0.292702 | 0.191375 | 0.248024 | 0.115413 | 0.364194 | 0.129852 | 0.043553 | 0.630375 | 0.785708 |
| 18.35972 | 0.01087 | 0.341298 | 0.314564 | 0.858788 | 0.676215 | 0.331888 | 0.609185 | 0.35471 | 0.220987 | 0.808028 | 0.937507 | 0.309736 | 0.197322 | 0.209075 | 0.069061 | 0.345851 | 0.129459 | 0.01646 | 0.730338 | 0.678967 |
| 18.37361 | 0.018622 | 0.401948 | 0.358419 | 0.956058 | 0.628449 | 0.283295 | 0.520729 | 0.254177 | 0.262332 | 0.981558 | 0.564697 | 0.251839 | 0.325636 | 0.215542 | 0.062844 | 0.358507 | 0.140485 | 0.018892 | 0.744363 | 0.793314 |
| 18.3875 | -0.02851 | 0.323335 | 0.508599 | 1.01879 | 0.432502 | 0.255518 | 0.412732 | 0.21097 | 0.358351 | 1.070437 | 0.251158 | 0.196993 | 0.41017 | 0.2455 | 0.034029 | 0.230126 | 0.065338 | 0.025429 | 0.554847 | 1.13807 |
| 18.40139 | -0.08448 | 0.149955 | 0.718545 | 1.003306 | 0.128831 | 0.280895 | 0.313184 | 0.227137 | 0.444608 | 0.945409 | 0.069758 | 0.113898 | 0.399907 | 0.234242 | -0.0214 | -0.04219 | -0.07883 | 0.038317 | 0.220704 | 1.67645 |
| 18.41528 | 0.006541 | 0.147034 | 0.759448 | 0.903805 | 0.083424 | 0.395211 | 0.310115 | 0.322432 | 0.474547 | 0.979019 | 0.225615 | 0.169988 | 0.349302 | 0.30074 | 0.059369 | 0.037668 | -0.05582 | 0.072664 | 0.293935 | 1.580021 |
| 18.42917 | 0.200036 | 0.319826 | 0.449205 | 0.761553 | 0.403074 | 0.49139 | 0.453083 | 0.371047 | 0.426107 | 1.158822 | 0.76795 | 0.373554 | 0.345964 | 0.421282 | 0.182362 | 0.419912 | 0.150004 | 0.12409 | 1.05282 | 1.096888 |
| 18.44306 | 0.506608 | 0.637304 | 0.122286 | 1.021288 | 0.874841 | 0.603467 | 0.771945 | 0.286994 | 0.52725 | 1.731832 | 1.707585 | 0.653656 | 0.464481 | 0.592311 | 0.26413 | 0.718319 | 0.40626 | 0.110737 | 2.086294 | 0.727829 |
| 18.45694 | 0.913902 | 1.048713 | 0.349303 | 2.012176 | 1.383386 | 0.814046 | 1.379197 | 0.267209 | 1.088676 | 2.870559 | 3.376086 | 0.908199 | 0.759302 | 0.821378 | 0.324874 | 0.756017 | 0.652393 | 0.101323 | 3.567157 | 1.26049 |
| 18.47083 | 1.246565 | 1.262599 | 0.764572 | 2.515862 | 1.704707 | 1.129105 | 1.709172 | 0.306709 | 1.777561 | 3.112852 | 4.623445 | 0.839958 | 0.690266 | 0.760399 | 0.3591 | 0.587238 | 0.746865 | 0.02215 | 3.750283 | 2.013677 |
| 18.48472 | 1.41382 | 0.967993 | 0.79667 | 1.772582 | 1.459386 | 0.869206 | 1.59823 | 0.290744 | 1.313247 | 1.552219 | 3.269493 | 0.543851 | 0.377581 | 0.436527 | 0.400988 | 0.433815 | 0.715826 | -0.00261 | 2.315715 | 1.919755 |
| 18.49861 | 1.112248 | 0.563704 | 0.710559 | 1.004984 | 1.009667 | 0.717348 | 1.600032 | 0.287438 | 0.962478 | 0.542708 | 1.606671 | 0.371898 | 0.300037 | 0.278372 | 0.589038 | 0.301426 | 0.580447 | 0.069141 | 0.964194 | 1.127726 |
| 18.5125 | 0.672199 | 0.099108 | 0.59253 | 0.302883 | 0.683498 | 0.476101 | 1.335434 | 0.24302 | 0.772487 | -0.00394 | 0.803183 | 0.174296 | 0.207383 | 0.139091 | 0.590073 | 0.113073 | 0.285327 | 0.059473 | -0.01489 | 0.326758 |
| 18.52639 | 0.371322 | -0.26155 | 0.459467 | -0.01832 | 0.471211 | 0.227287 | 0.817968 | 0.156101 | 0.552193 | 0.08621 | 0.345818 | 0.033853 | 0.070417 | 0.028586 | 0.458721 | -0.06846 | -0.00089 | -0.01728 | -0.43068 | -0.00964 |
| 18.54028 | 0.214935 | -0.39256 | 0.36036 | -0.0339 | 0.380729 | 0.141656 | 0.464353 | 0.086951 | 0.406147 | 0.899634 | 0.212229 | -0.08493 | 0.070463 | 0.007903 | 0.263214 | -0.09102 | -0.13567 | -0.04186 | -0.20636 | 0.258442 |
| 18.55417 | 0.069549 | -0.2828 | 0.218598 | -0.02902 | 0.42227 | 0.118874 | 0.271492 | 0.089523 | 0.321566 | 1.816656 | 0.372791 | -0.06476 | 0.102858 | 0.050961 | 0.079337 | 0.043688 | -0.04846 | 0.015386 | 0.429187 | 0.69122 |
| 18.56806 | 0.029713 | -0.02073 | 0.088489 | 0.084319 | 0.615591 | 0.129065 | 0.198261 | 0.17444 | 0.26721 | 1.86926 | 0.717551 | 0.076532 | 0.179803 | 0.079004 | 0.055722 | 0.197283 | 0.099712 | 0.022475 | 1.279699 | 0.897771 |
| 18.58194 | 0.18806 | 0.295872 | 0.100823 | 0.232704 | 0.769865 | 0.168931 | 0.146944 | 0.238599 | 0.242241 | 0.962932 | 0.967159 | 0.271294 | 0.205321 | 0.058775 | 0.162362 | 0.222509 | 0.115721 | 0.002945 | 1.785898 | 0.792866 |
| 18.59583 | 0.422664 | 0.550014 | 0.168633 | 0.370154 | 0.72577 | 0.203668 | 0.170334 | 0.24706 | 0.219419 | 0.105297 | 0.893134 | 0.237462 | 0.174317 | 0.04583 | 0.283165 | 0.179669 | 0.078712 | -0.02794 | 1.536304 | 0.29271 |
| 18.60972 | 0.44709 | 0.585755 | 0.177167 | 0.472577 | 0.501209 | 0.235272 | 0.300985 | 0.171431 | 0.167291 | -0.28925 | 0.531047 | 0.036793 | 0.169396 | 0.013005 | 0.289445 | 0.124956 | 0.07547 | -0.0076 | 0.686343 | -0.13385 |
| 18.62361 | 0.295651 | 0.27931 | 0.155991 | 0.339378 | 0.09074 | 0.204067 | 0.44456 | 0.034392 | 0.064447 | -0.2279 | 0.079776 | -0.05845 | 0.093011 | -0.03572 | 0.267304 | 0.046712 | 0.027148 | 0.0327 | -0.49552 | -0.17396 |
| 18.6375 | 0.140637 | -0.15966 | 0.142831 | 0.083841 | -0.37191 | 0.053416 | 0.605279 | -0.1292 | -0.05837 | 0.016894 | -0.31329 | -0.15171 | -0.00313 | -0.08895 | 0.191637 | -0.05969 | -0.08589 | 0.004676 | -1.48077 | 0.075163 |
| 18.65139 | 0.035105 | -0.45985 | 0.107241 | -0.06765 | -0.69068 | -0.11793 | 0.380664 | -0.27763 | -0.09577 | 0.267096 | -0.61281 | -0.26893 | -0.08939 | -0.15778 | -0.00017 | -0.13784 | -0.22827 | -0.0438 | -1.64365 | 0.437019 |
| 18.66528 | -0.10705 | -0.48944 | 0.005841 | -0.1066 | -0.57178 | -0.22462 | -0.07933 | -0.26626 | -0.07272 | 0.435641 | -0.66375 | -0.2235 | -0.06795 | -0.09228 | -0.16323 | -0.0518 | -0.16359 | -0.02345 | -0.76517 | 0.709926 |
| 18.67917 | -0.28845 | -0.34606 | -0.09269 | -0.11043 | -0.15415 | -0.30081 | -0.27696 | -0.15941 | -0.00278 | 0.571763 | -0.40305 | 0.033568 | -0.07695 | 0.032207 | -0.22788 | 0.115302 | 0.02827 | 0.053832 | 0.406394 | 0.79345 |
| 18.69306 | -0.30784 | -0.19937 | -0.09799 | -0.14571 | 0.237428 | -0.31779 | -0.34782 | -0.00397 | 0.08977 | 0.531377 | -0.05722 | 0.292391 | -0.0508 | 0.080873 | -0.22965 | 0.128887 | 0.117712 | 0.099543 | 0.963545 | 0.708597 |
| 18.70694 | -0.12755 | -0.09747 | -0.015 | -0.01334 | 0.413357 | -0.18204 | -0.41628 | 0.108891 | 0.154698 | 0.213951 | 0.260258 | 0.420089 | -0.00451 | 0.056552 | -0.10403 | 0.018098 | 0.065286 | 0.074829 | 0.900357 | 0.55391 |
| 18.72083 | 0.08205 | -0.02858 | 0.061352 | 0.168446 | 0.395475 | 0.023584 | -0.4676 | 0.117032 | 0.10883 | -0.29152 | 0.411034 | 0.351895 | 0.029258 | 0.025123 | 0.02268 | -0.07627 | -0.01358 | 0.04454 | 0.608511 | 0.278136 |
| 18.73472 | 0.104654 | -0.04726 | 0.123479 | 0.192794 | 0.212946 | 0.153345 | -0.48849 | 0.024025 | -0.02654 | -0.8932 | 0.204679 | 0.175804 | 0.023631 | -0.06693 | -0.00772 | -0.15206 | -0.04738 | 0.005326 | 0.076914 | -0.35771 |
| 18.74861 | -0.01121 | -0.13772 | 0.138792 | -0.03674 | -0.00169 | 0.196677 | -0.26422 | -0.08849 | -0.12873 | -1.28673 | -0.08303 | 0.009266 | -0.01588 | -0.12069 | -0.10451 | -0.16964 | -0.02932 | -0.0083 | -0.55734 | -1.06059 |
| 18.7625 | -0.08365 | -0.16317 | 0.055887 | -0.43319 | -0.17554 | 0.177849 | 0.088825 | -0.16884 | -0.18076 | -1.22775 | -0.27322 | -0.14216 | -0.04719 | -0.14404 | -0.19946 | -0.13109 | 0.000347 | -0.02591 | -1.1089 | -1.37528 |
| 18.77639 | -0.07902 | -0.16054 | -0.12454 | -0.91454 | -0.36442 | 0.015742 | 0.271348 | -0.27023 | -0.29476 | -0.62689 | -0.43911 | -0.36425 | -0.09527 | -0.19583 | -0.31403 | -0.13065 | -0.06662 | -0.09336 | -1.36254 | -1.11358 |
| 18.79028 | -0.10558 | -0.23481 | -0.32675 | -1.25633 | -0.48989 | -0.20589 | 0.134958 | -0.32452 | -0.42637 | -0.06313 | -0.57927 | -0.45799 | -0.16747 | -0.2456 | -0.31453 | -0.12279 | -0.15892 | -0.15099 | -1.27403 | -0.65038 |
| 18.80417 | -0.26192 | -0.38992 | -0.5106 | -1.37429 | -0.50351 | -0.38309 | -0.17027 | -0.32181 | -0.45945 | 0.209398 | -0.81176 | -0.41297 | -0.18408 | -0.30815 | -0.16726 | -0.11377 | -0.2058 | -0.11568 | -0.96985 | -0.5499 |
| 18.81806 | -0.53218 | -0.58383 | -0.70864 | -1.37129 | -0.47615 | -0.59311 | -0.69668 | -0.35616 | -0.5381 | -0.02315 | -1.1763 | -0.44017 | -0.13659 | -0.3879 | -0.14164 | -0.26077 | -0.25009 | -0.06549 | -0.92885 | -0.86477 |
| 18.83194 | -0.6421 | -0.76168 | -0.89398 | -1.22179 | -0.53474 | -0.7942 | -1.27271 | -0.36841 | -0.6988 | -0.69523 | -1.20284 | -0.43852 | -0.24389 | -0.44444 | -0.32204 | -0.48242 | -0.27783 | -0.08245 | -1.15753 | -1.44036 |
| 18.84583 | -0.46771 | -0.75089 | -0.90573 | -1.06012 | -0.68885 | -0.88587 | -1.67206 | -0.36283 | -0.84874 | -1.40414 | -1.10134 | -0.37339 | -0.38955 | -0.41413 | -0.54109 | -0.56419 | -0.28275 | -0.11409 | -1.39567 | -2.02837 |
| 18.85972 | -0.31189 | -0.74548 | -0.8045 | -1.01826 | -0.9385 | -0.73986 | -1.91034 | -0.46407 | -0.96791 | -2.15262 | -1.30019 | -0.37344 | -0.59186 | -0.4523 | -0.73287 | -0.59319 | -0.33165 | -0.17988 | -1.89618 | -2.31386 |
| 18.87361 | -0.37096 | -0.87892 | -0.69269 | -0.96918 | -1.09428 | -0.57617 | -1.83575 | -0.63212 | -1.04099 | -2.57477 | -1.77542 | -0.55407 | -0.69713 | -0.46284 | -0.8162 | -0.69673 | -0.43836 | -0.23987 | -2.28774 | -2.12324 |
| 18.8875 | -0.49252 | -1.03955 | -0.67629 | -0.8735 | -0.99602 | -0.46431 | -1.5161 | -0.71494 | -1.06797 | -2.36706 | -2.03749 | -0.66818 | -0.65049 | -0.39888 | -0.76528 | -0.6744 | -0.41451 | -0.2205 | -2.23343 | -1.43821 |
| 18.90139 | -0.44838 | -0.94487 | -0.66997 | -0.80267 | -0.84479 | -0.37754 | -0.92598 | -0.67173 | -1.0006 | -1.64207 | -1.91617 | -0.5945 | -0.58732 | -0.32539 | -0.57957 | -0.433 | -0.32725 | -0.2017 | -1.8022 | -0.84066 |
| 18.91528 | -0.32698 | -0.75337 | -0.65344 | -0.76634 | -0.65656 | -0.33223 | -0.42154 | -0.55869 | -0.76486 | -1.04767 | -1.53194 | -0.53503 | -0.59083 | -0.29362 | -0.45634 | -0.21014 | -0.28133 | -0.14528 | -1.25514 | -0.35088 |
| 18.92917 | -0.21165 | -0.57005 | -0.55574 | -0.64983 | -0.4786 | -0.27195 | -0.20732 | -0.44623 | -0.33822 | -0.67875 | -1.25143 | -0.50024 | -0.56022 | -0.26238 | -0.40119 | -0.17276 | -0.2861 | -0.01224 | -0.64379 | -0.18512 |
| 18.94306 | -0.18159 | -0.37813 | -0.43326 | -0.56192 | -0.42495 | -0.24515 | -0.32704 | -0.29442 | -0.04476 | -0.59585 | -1.23959 | -0.46747 | -0.41394 | -0.18795 | -0.38024 | -0.40066 | -0.35942 | 0.06206 | -0.45194 | -0.30216 |
| 18.95694 | -0.39473 | -0.31526 | -0.39572 | -0.73205 | -0.58743 | -0.3106 | -0.58118 | -0.18766 | -0.09427 | -0.82995 | -1.29448 | -0.43992 | -0.25413 | -0.20586 | -0.4119 | -0.64449 | -0.48048 | 0.00526 | -0.82787 | -0.76228 |
| 18.97083 | -0.91392 | -0.33775 | -0.55551 | -1.13508 | -0.87332 | -0.54234 | -0.94498 | -0.21241 | -0.43305 | -1.3294 | -1.23144 | -0.43447 | -0.29477 | -0.32221 | -0.52863 | -0.58979 | -0.43918 | -0.2129 | -1.18277 | -1.14313 |
| 18.98472 | -0.91673 | -0.33668 | -0.80254 | -1.32773 | -1.09824 | -0.85144 | -1.41851 | -0.33017 | -0.73786 | -1.78348 | -1.04339 | -0.50345 | -0.37029 | -0.39926 | -0.5357 | -0.43955 | -0.26551 | -0.36231 | -1.3297 | -1.53265 |
| 18.99861 | -0.27166 | -0.23321 | -0.7888 | -1.02184 | -1.03061 | -0.87334 | -1.45893 | -0.36484 | -0.57021 | -1.73209 | -0.84408 | -0.45356 | -0.239 | -0.26018 | -0.23597 | -0.37387 | -0.11732 | -0.11218 | -1.1385 | -1.25947 |
| 19.0125 | 0.518586 | -0.08303 | -0.62228 | -0.57977 | -0.70361 | -0.46488 | -1.07712 | -0.31834 | -0.26796 | -1.0743 | -0.67709 | -0.34949 | -0.00479 | -0.02725 | 0.19013 | -0.15812 | -0.00973 | 0.138159 | -0.58526 | -0.40584 |
| 19.02639 | 0.859792 | 0.035135 | -0.34958 | -0.32482 | -0.40722 | -0.1352 | -0.62256 | -0.24598 | -0.04882 | -0.33453 | -0.54751 | -0.24106 | 0.000142 | 0.101568 | 0.34539 | 0.025722 | 0.019757 | 0.160259 | -0.15155 | 0.27453 |
| 19.04028 | 0.492691 | 0.162327 | -0.11748 | -0.23571 | -0.08584 | 0.104705 | -0.14642 | -0.12879 | 0.034312 | 0.422673 | -0.2141 | -0.08957 | -0.07666 | 0.115287 | 0.326069 | 0.287473 | 0.072558 | 0.074924 | 0.274568 | 0.528279 |
| 19.05417 | 0.265223 | 0.250007 | -0.03241 | -0.15911 | 0.227543 | 0.268012 | 0.226864 | -0.0021 | -0.00886 | 0.912673 | 0.204492 | 0.064925 | -0.21194 | 0.041536 | 0.288695 | 0.643196 | 0.145229 | 0.003408 | 0.735672 | 0.432635 |
| 19.06806 | 0.081601 | 0.167828 | 0.024273 | -0.14854 | 0.345938 | 0.224114 | 0.24516 | 0.064147 | -0.12727 | 0.794039 | 0.273827 | 0.133764 | -0.30765 | -0.04531 | 0.109708 | 0.596971 | 0.095107 | -0.04852 | 0.629128 | 0.087509 |
| 19.08194 | -0.04032 | -0.00079 | 0.17749 | -0.1318 | 0.290633 | 0.042615 | 0.038682 | 0.097476 | -0.15129 | 0.449336 | 0.029346 | 0.024357 | -0.22559 | -0.04619 | -0.12891 | 0.14931 | -0.01274 | -0.07611 | -0.05688 | -0.12702 |
| 19.09583 | -0.07182 | -0.08301 | 0.234569 | 0.035855 | 0.233999 | -0.06024 | 0.052389 | 0.153273 | -0.13055 | 0.287216 | -0.08873 | -0.02739 | -0.06902 | -0.00524 | -0.19785 | -0.12548 | -0.04721 | -0.05672 | -0.36794 | -0.11747 |
| 19.10972 | -0.02053 | -0.06717 | 0.09247 | 0.2231 | 0.254454 | -0.04699 | 0.152327 | 0.167819 | -0.15902 | 0.266087 | -0.01456 | 0.00026 | -0.08925 | -0.01039 | -0.06619 | 0.047633 | -0.00954 | -0.0458 | -0.00229 | -0.19813 |
| 19.12361 | -0.01496 | -0.09833 | 0.045565 | 0.267614 | 0.243159 | -0.06336 | 0.148725 | 0.131238 | -0.15101 | 0.178827 | -0.08669 | -0.03768 | -0.1162 | -0.01183 | -0.01211 | 0.029928 | -0.02064 | -0.04874 | 0.00552 | -0.30319 |
| 19.1375 | -0.03997 | -0.16193 | 0.225748 | 0.290284 | 0.163889 | -0.07201 | 0.114909 | 0.126374 | 0.01446 | 0.123087 | -0.27828 | -0.11871 | 0.019658 | 0.0548 | -0.08176 | -0.33486 | -0.07517 | -0.02726 | -0.44819 | -0.17505 |
| 19.15139 | -0.03908 | -0.13662 | 0.273497 | 0.324779 | 0.144819 | 0.055984 | 0.249781 | 0.142689 | 0.20331 | 0.261041 | -0.24881 | -0.1068 | 0.075756 | 0.093634 | -0.07781 | -0.34739 | -0.06032 | 0.012965 | -0.3525 | -0.04976 |
| 19.16528 | -0.0218 | -0.05225 | 0.163332 | 0.324102 | 0.184709 | 0.170062 | 0.382061 | 0.141208 | 0.23366 | 0.501776 | -0.09731 | -0.03863 | 0.004439 | 0.06582 | 0.018793 | -0.03207 | 0.01425 | 0.002982 | 0.151495 | -0.1853 |
| 19.17917 | -0.00661 | 0.047246 | 0.056924 | 0.304322 | 0.266553 | 0.18133 | 0.371944 | 0.138719 | 0.097971 | 0.644159 | 0.047237 | 0.032566 | -0.08542 | 0.056759 | 0.132903 | 0.24248 | 0.088753 | 0.000799 | 0.558503 | -0.30324 |
| 19.19306 | -0.01874 | 0.12108 | -0.06153 | 0.254238 | 0.284824 | 0.126851 | 0.229884 | 0.096058 | -0.07745 | 0.501936 | 0.170811 | 0.072071 | -0.2107 | 0.023762 | 0.169974 | 0.439697 | 0.108792 | -0.02012 | 0.778449 | -0.39584 |
| 19.20694 | -0.07363 | 0.129298 | -0.09492 | 0.187643 | 0.151168 | 0.040474 | 0.076951 | 0.034186 | -0.12822 | 0.218228 | 0.163426 | 0.017291 | -0.2313 | -0.04086 | 0.124319 | 0.323476 | 0.053954 | -0.07354 | 0.561938 | -0.32045 |
| 19.22083 | -0.05751 | 0.160325 | 0.033332 | 0.200875 | 0.086389 | 0.03831 | 0.042273 | 0.07183 | 0.056269 | 0.002802 | 0.140219 | -0.02582 | -0.0356 | 0.032002 | 0.098055 | 0.063988 | 0.05013 | -0.03175 | 0.150614 | 0.031148 |
| 19.23472 | -0.01678 | 0.222845 | 0.187421 | 0.257033 | 0.118966 | 0.098058 | 0.103577 | 0.149123 | 0.304784 | 0.01498 | 0.170699 | -0.02048 | 0.173465 | 0.136651 | 0.083884 | -0.03828 | 0.068759 | 0.037394 | 0.038526 | 0.405552 |
| 19.24861 | -0.02463 | 0.210662 | 0.216936 | 0.265187 | 0.159034 | 0.110805 | 0.179403 | 0.159884 | 0.399408 | 0.246881 | 0.204235 | -0.00977 | 0.214244 | 0.157983 | 0.077186 | 0.066256 | 0.057258 | 0.025188 | 0.167035 | 0.633748 |
| 19.2625 | -0.041 | 0.195849 | 0.142877 | 0.270137 | 0.246334 | 0.122681 | 0.257727 | 0.140987 | 0.32408 | 0.48854 | 0.313727 | 0.043596 | 0.133726 | 0.14774 | 0.176737 | 0.370343 | 0.086517 | -0.02264 | 0.444083 | 0.639217 |
| 19.27639 | -0.02447 | 0.189262 | 0.170916 | 0.314327 | 0.307631 | 0.140057 | 0.23432 | 0.180762 | 0.235692 | 0.606203 | 0.319551 | 0.092545 | 0.11249 | 0.203188 | 0.338767 | 0.386799 | 0.102739 | -0.00697 | 0.409533 | 0.816208 |
| 19.29028 | -0.01977 | 0.1638 | 0.269387 | 0.342448 | 0.233438 | 0.162747 | 0.161536 | 0.217764 | 0.248285 | 0.598739 | 0.231799 | 0.087093 | 0.19266 | 0.2606 | 0.390425 | 0.010062 | 0.049232 | 0.068735 | 0.060074 | 1.139758 |
| 19.30417 | -0.01368 | 0.17643 | 0.229669 | 0.333562 | 0.192848 | 0.192879 | 0.223278 | 0.201755 | 0.265856 | 0.550521 | 0.322586 | 0.047748 | 0.177768 | 0.203744 | 0.335109 | 0.026594 | 0.019921 | 0.089959 | 0.20122 | 1.064458 |
| 19.31806 | 0.006694 | 0.238323 | 0.19487 | 0.369883 | 0.270245 | 0.258805 | 0.393264 | 0.222938 | 0.405596 | 0.612322 | 0.417423 | 0.019921 | 0.162834 | 0.150371 | 0.122358 | 0.205774 | 0.035733 | 0.045966 | 0.555978 | 0.849059 |
| 19.33194 | 0.051948 | 0.299435 | 0.349297 | 0.578177 | 0.387063 | 0.322527 | 0.56754 | 0.275547 | 0.631556 | 0.759069 | 0.339538 | -0.00418 | 0.265907 | 0.218154 | 0.113419 | 0.183231 | 0.075845 | 0.053488 | 0.468568 | 0.795095 |
| 19.34583 | 0.115629 | 0.386677 | 0.460994 | 0.790825 | 0.502224 | 0.39182 | 0.720578 | 0.293372 | 0.68931 | 0.910437 | 0.43961 | 0.06671 | 0.295715 | 0.293663 | 0.153427 | 0.414808 | 0.152954 | 0.077746 | 0.501352 | 0.611823 |
| 19.35972 | 0.165547 | 0.441468 | 0.41034 | 0.883301 | 0.550497 | 0.365054 | 0.835716 | 0.261448 | 0.495388 | 0.838614 | 0.63232 | 0.114917 | 0.214573 | 0.249694 | 0.25704 | 0.615003 | 0.182292 | 0.044488 | 0.58844 | 0.387589 |
| 19.37361 | 0.201862 | 0.429194 | 0.471391 | 0.857875 | 0.476719 | 0.277311 | 0.815787 | 0.272207 | 0.367245 | 0.618504 | 0.584565 | 0.054742 | 0.171182 | 0.222152 | 0.200648 | 0.325765 | 0.07549 | 0.017025 | 0.291511 | 0.457734 |
| 19.3875 | 0.26359 | 0.422274 | 0.533215 | 0.771628 | 0.391606 | 0.214503 | 0.666544 | 0.299281 | 0.314618 | 0.377415 | 0.573112 | 0.040299 | 0.164436 | 0.276394 | 0.076945 | 0.080957 | 0.011932 | 0.025922 | 0.140158 | 0.603683 |
| 19.40139 | 0.365186 | 0.399369 | 0.447083 | 0.731984 | 0.436109 | 0.174843 | 0.557403 | 0.34488 | 0.243376 | 0.329272 | 0.813536 | 0.143546 | 0.142076 | 0.298054 | 0.132884 | 0.255827 | 0.08549 | 0.0175 | 0.588707 | 0.500257 |
| 19.41528 | 0.490409 | 0.399277 | 0.356825 | 0.72208 | 0.545763 | 0.189382 | 0.571964 | 0.40658 | 0.26139 | 0.438 | 0.97163 | 0.258203 | 0.160767 | 0.303243 | 0.25751 | 0.502261 | 0.200909 | 0.01068 | 1.207729 | 0.464711 |
| 19.42917 | 0.621555 | 0.446486 | 0.297811 | 0.734425 | 0.660452 | 0.285024 | 0.688525 | 0.507128 | 0.365214 | 0.685307 | 0.999205 | 0.384383 | 0.212497 | 0.345328 | 0.352368 | 0.688555 | 0.326234 | 0.024847 | 1.546342 | 0.559303 |
| 19.44306 | 0.852816 | 0.734389 | 0.326029 | 1.001673 | 0.832844 | 0.503779 | 0.925155 | 0.568585 | 0.564339 | 1.398852 | 1.23838 | 0.630563 | 0.34805 | 0.459284 | 0.386335 | 0.865127 | 0.527759 | 0.023922 | 2.009859 | 0.994416 |
| 19.45694 | 1.346318 | 1.502618 | 0.808515 | 2.077373 | 1.09748 | 0.883991 | 1.503697 | 0.6732 | 1.025967 | 2.67996 | 2.090321 | 1.02426 | 0.814544 | 0.800214 | 0.312498 | 0.937061 | 0.866032 | 0.034393 | 3.14029 | 2.042696 |
| 19.47083 | 1.27483 | 1.918025 | 1.551187 | 2.95512 | 1.356271 | 1.268669 | 2.016355 | 0.721915 | 1.67648 | 3.291061 | 3.027389 | 1.300899 | 1.395084 | 1.0967 | 0.289506 | 0.805484 | 1.133868 | 0.103386 | 4.114257 | 3.145458 |
| 19.48472 | 0.785103 | 1.44752 | 1.259667 | 2.044538 | 1.255055 | 1.282773 | 1.716019 | 0.411055 | 1.837614 | 2.104905 | 2.996859 | 1.076359 | 1.234182 | 0.902216 | 0.352674 | 0.64643 | 0.953211 | 0.219395 | 3.167598 | 3.119394 |
| 19.49861 | 0.368499 | 0.897902 | 0.740122 | 1.020828 | 0.914383 | 0.899805 | 1.157652 | 0.076709 | 1.306105 | 0.969218 | 2.192158 | 0.739471 | 0.713846 | 0.553444 | 0.529193 | 0.47048 | 0.476849 | 0.178358 | 1.896709 | 2.19143 |

| | | | | | | | | | | | | | | | | | | | | |
|---|---|---|---|---|---|---|---|---|---|---|---|---|---|---|---|---|---|---|---|---|
| 19.5125 | -0.01284 | 0.259858 | 0.32556 | 0.238531 | 0.362762 | 0.50262 | 0.5423 | -0.18829 | 0.738186 | 0.307352 | 1.282514 | 0.327367 | 0.299305 | 0.284035 | 0.462115 | 0.151968 | 0.014655 | 0.100054 | 0.686449 | 1.303851 |
| 19.52639 | -0.08626 | -0.17797 | 0.081868 | -0.08836 | -0.05722 | 0.189112 | 0.001951 | -0.22176 | 0.338182 | 0.225904 | 0.602524 | 0.074247 | 0.093993 | 0.051285 | 0.338107 | -0.11435 | -0.15857 | 0.026895 | -0.19627 | 0.567764 |
| 19.54028 | 0.20188 | -0.29897 | 0.047113 | 0.181606 | -0.10518 | 0.070454 | -0.00459 | -0.08802 | 0.179659 | 0.55374 | 0.47587 | 0.045666 | 0.156009 | -0.03837 | 0.236623 | -0.14934 | 0.017788 | 0.018922 | -0.36342 | 0.376705 |
| 19.55417 | 0.464775 | -0.17639 | 0.13536 | 0.690962 | 0.266455 | 0.120565 | 0.326879 | 0.097815 | 0.314132 | 1.005314 | 0.685242 | 0.096079 | 0.361819 | 0.021112 | 0.240439 | -0.0418 | 0.225406 | 0.0298 | -0.11493 | 0.602487 |
| 19.56806 | 0.42056 | 0.107581 | 0.178984 | 0.895039 | 0.765085 | 0.179428 | 0.752456 | 0.308644 | 0.550458 | 1.182002 | 0.926119 | 0.150594 | 0.631547 | 0.137305 | 0.220789 | 0.137067 | 0.263801 | 0.013731 | 0.203536 | 0.849341 |
| 19.58194 | 0.347538 | 0.390396 | 0.147417 | 0.531068 | 1.067217 | 0.127539 | 0.883471 | 0.444746 | 0.709527 | 1.068577 | 0.96204 | 0.162997 | 0.799313 | 0.175129 | 0.102724 | 0.315091 | 0.177296 | -0.01648 | 0.450756 | 1.043327 |
| 19.59583 | 0.346245 | 0.535473 | 0.259402 | 0.008376 | 1.031385 | 0.069619 | 0.788087 | 0.380475 | 0.763978 | 0.762167 | 0.886117 | 0.140327 | 0.614947 | 0.139531 | 0.076806 | 0.424999 | 0.075431 | -0.04007 | 0.371723 | 1.072175 |
| 19.60972 | 0.13968 | 0.54658 | 0.483062 | -0.36947 | 0.796788 | 0.131448 | 0.675215 | 0.238866 | 0.637018 | 0.254455 | 0.962084 | 0.151368 | 0.282167 | 0.127642 | 0.17775 | 0.473664 | -0.04417 | -0.05541 | 0.144135 | 0.485613 |
| 19.62361 | -0.22894 | 0.450422 | 0.622177 | -0.46586 | 0.484412 | 0.233918 | 0.614272 | 0.166357 | 0.411835 | 0.10195 | 1.056214 | 0.254385 | -0.03514 | 0.15075 | 0.336177 | 0.395396 | -0.12501 | -0.05293 | -0.25555 | -0.47107 |
| 19.6375 | -0.46652 | 0.269767 | 0.580433 | -0.27988 | 0.102962 | 0.306201 | 0.534315 | 0.078464 | 0.148522 | 0.263124 | 1.009627 | 0.290829 | -0.21374 | 0.121737 | 0.455136 | 0.174455 | -0.16521 | -0.02458 | -0.56009 | -1.52823 |
| 19.65139 | -0.35037 | 0.046766 | 0.28438 | -0.00398 | -0.27698 | 0.250194 | 0.232503 | -0.05779 | -0.06307 | 0.411074 | 0.650279 | 0.174402 | -0.22799 | 0.051815 | 0.386597 | -0.02555 | -0.1133 | 0.021108 | -0.28232 | -1.64192 |
| 19.66528 | -0.04401 | -0.15721 | 0.103626 | 0.169265 | -0.46795 | 0.073542 | -0.19444 | -0.15797 | -0.20106 | 0.403527 | -0.0013 | 0.067934 | -0.22384 | -0.01194 | 0.205738 | -0.15732 | -0.03984 | 0.052666 | 0.327684 | -0.36958 |
| 19.67917 | 0.111074 | -0.30617 | 0.0481 | 0.242316 | -0.45331 | -0.08505 | -0.61099 | -0.18746 | -0.2337 | 0.233278 | -0.74941 | -0.01927 | -0.20979 | -0.03945 | -0.01891 | -0.25156 | -0.04204 | 0.039793 | 1.035045 | 1.341817 |
| 19.69306 | -0.04635 | -0.38699 | -0.01653 | 0.368294 | -0.28001 | -0.13784 | -0.82799 | -0.08741 | -0.20911 | 0.186685 | -1.00438 | -0.08831 | -0.09634 | -0.05254 | -0.16371 | -0.21941 | -0.11635 | 0.025497 | 1.453185 | 2.275821 |
| 19.70694 | -0.19086 | -0.40452 | -0.17589 | 0.406127 | -0.01954 | -0.14696 | -0.81658 | 0.045756 | -0.15572 | 0.279714 | -0.73 | -0.09933 | 0.080085 | -0.03651 | -0.17034 | -0.112 | -0.17967 | 0.015761 | 1.29885 | 1.677512 |
| 19.72083 | -0.11481 | -0.36504 | -0.27249 | 0.158294 | 0.213664 | -0.26543 | -0.59618 | 0.031122 | -0.1052 | 0.258251 | -0.47842 | -0.06316 | 0.053835 | -0.09185 | -0.16486 | -0.10343 | -0.16393 | -0.04532 | 0.759424 | 0.501027 |
| 19.73472 | -0.03258 | -0.21168 | -0.2247 | -0.27411 | 0.32078 | -0.4403 | -0.25749 | -0.09663 | -0.15597 | 0.054669 | -0.38216 | -0.06429 | -0.06265 | -0.17293 | -0.15674 | -0.09944 | -0.12481 | -0.08891 | 0.150722 | -0.54088 |
| 19.74861 | -0.18282 | -0.00025 | -0.18085 | -0.63247 | 0.261874 | -0.48448 | 0.0774 | -0.17164 | -0.22482 | -0.22021 | -0.28412 | -0.04779 | -0.16972 | -0.10075 | -0.15521 | 0.00949 | -0.08754 | -0.12366 | -0.35139 | -1.35133 |
| 19.7625 | -0.42998 | 0.030344 | -0.16548 | -0.82878 | 0.108311 | -0.31006 | 0.200971 | -0.23875 | -0.24739 | -0.57522 | -0.35658 | -0.06372 | -0.32718 | 0.01128 | -0.18863 | 0.00332 | -0.10456 | -0.08783 | -0.90226 | -1.72284 |
| 19.77639 | -0.39378 | -0.22625 | -0.21446 | -0.92738 | -0.10516 | -0.06575 | -0.04945 | -0.34156 | -0.21878 | -1.14671 | -0.72788 | -0.20353 | -0.43677 | -0.05401 | -0.24028 | -0.24681 | -0.2044 | -0.03648 | -1.55408 | -1.48027 |
| 19.79028 | -0.09459 | -0.51524 | -0.32143 | -0.90175 | -0.30511 | 0.079258 | -0.47916 | -0.39913 | -0.20811 | -1.46497 | -1.10751 | -0.38286 | -0.42848 | -0.21493 | -0.21846 | -0.50619 | -0.33676 | -0.01142 | -2.21625 | -0.87491 |
| 19.80417 | 0.159306 | -0.7094 | -0.50225 | -0.86468 | -0.52213 | -0.01055 | -0.87081 | -0.45216 | -0.32842 | -1.41268 | -1.52837 | -0.52748 | -0.40824 | -0.40429 | -0.09832 | -0.59387 | -0.42517 | -0.02208 | -2.70835 | -0.55651 |
| 19.81806 | 0.102993 | -0.9426 | -0.77812 | -1.02577 | -0.86222 | -0.27092 | -1.3661 | -0.51154 | -0.6545 | -1.45643 | -1.99177 | -0.58469 | -0.42799 | -0.66219 | -0.10339 | -0.55598 | -0.48836 | -0.08787 | -2.79598 | -0.78912 |
| 19.83194 | -0.21401 | -1.23405 | -1.13134 | -1.33425 | -1.32615 | -0.61705 | -2.00351 | -0.58863 | -1.11305 | -1.52314 | -2.34623 | -0.73369 | -0.45844 | -0.94206 | -0.31586 | -0.46175 | -0.52671 | -0.18083 | -2.53932 | -1.41421 |
| 19.84583 | -0.55629 | -1.25951 | -1.51285 | -1.53278 | -1.63547 | -0.89847 | -2.45457 | -0.58435 | -1.44536 | -1.42887 | -2.02055 | -0.85989 | -0.51772 | -1.201 | -0.5796 | -0.36137 | -0.56442 | -0.25456 | -2.22512 | -1.81229 |
| 19.85972 | -0.76816 | -1.02703 | -1.55662 | -1.42504 | -1.60911 | -1.09397 | -2.35308 | -0.53052 | -1.60069 | -1.07782 | -1.51185 | -0.85868 | -0.55892 | -1.08364 | -0.78518 | -0.32251 | -0.49428 | -0.1714 | -1.90692 | -1.59621 |
| 19.87361 | -0.89132 | -0.97483 | -1.33681 | -1.23175 | -1.3074 | -1.16102 | -1.90309 | -0.55594 | -1.48487 | -0.69759 | -1.42482 | -0.68332 | -0.65384 | -0.66028 | -0.93303 | -0.35675 | -0.40898 | -0.1013 | -1.49505 | -1.07775 |
| 19.8875 | -0.97432 | -1.03621 | -1.04407 | -1.10644 | -0.87384 | -1.05833 | -1.39071 | -0.60763 | -1.21812 | -0.5597 | -1.60031 | -0.51051 | -0.77384 | -0.2488 | -0.90852 | -0.38318 | -0.36862 | -0.07024 | -1.09083 | -0.47028 |
| 19.90139 | -0.68857 | -1.02145 | -0.8045 | -1.00878 | -0.5014 | -0.87695 | -1.04574 | -0.68437 | -0.95424 | -0.60805 | -1.99069 | -0.38452 | -0.83213 | -0.11515 | -0.84241 | -0.43871 | -0.36097 | -0.15077 | -0.76681 | -0.05232 |
| 19.91528 | -0.38653 | -0.90391 | -0.61225 | -0.95995 | -0.31909 | -0.65981 | -0.86603 | -0.66192 | -0.703 | -0.74708 | -2.39226 | -0.36071 | -0.79741 | -0.17461 | -0.67902 | -0.57009 | -0.37161 | -0.28786 | -0.59105 | -0.06658 |
| 19.92917 | -0.20977 | -0.68039 | -0.44899 | -0.85692 | -0.24959 | -0.4781 | -0.71205 | -0.50423 | -0.54451 | -0.7832 | -2.03856 | -0.34881 | -0.64196 | -0.24862 | -0.42841 | -0.55181 | -0.37531 | -0.38087 | -0.41287 | -0.29882 |
| 19.94306 | -0.15436 | -0.47961 | -0.38752 | -0.71224 | -0.2828 | -0.34604 | -0.57122 | -0.34859 | -0.52261 | -0.83912 | -1.40576 | -0.29285 | -0.49785 | -0.31534 | -0.27143 | -0.50784 | -0.36931 | -0.40495 | -0.38711 | -0.51047 |
| 19.95694 | -0.19563 | -0.37035 | -0.44003 | -0.78442 | -0.54011 | -0.35635 | -0.61618 | -0.33707 | -0.60387 | -1.05401 | -1.08923 | -0.29743 | -0.40986 | -0.46247 | -0.52246 | -0.65193 | -0.44995 | -0.4467 | -0.80977 | -0.83823 |
| 19.97083 | -0.28874 | -0.36598 | -0.58618 | -0.8732 | -0.84866 | -0.49297 | -0.84205 | -0.40071 | -0.65833 | -1.33499 | -1.0878 | -0.41935 | -0.32033 | -0.57007 | -1.09255 | -0.87787 | -0.43168 | -0.53051 | -1.35002 | -1.15324 |
| 19.98472 | -0.27057 | -0.33352 | -0.70425 | -0.68372 | -0.92276 | -0.66706 | -1.13006 | -0.42923 | -0.59285 | -1.63525 | -0.78017 | -0.57949 | -0.26372 | -0.55943 | -1.41876 | -0.80195 | -0.24884 | -0.52084 | -1.46455 | -1.44042 |
| 19.99861 | -0.17105 | -0.30707 | -0.66667 | -0.41632 | -0.88783 | -0.78132 | -1.2575 | -0.47516 | -0.43323 | -1.82037 | -0.32503 | -0.66545 | -0.29312 | -0.43047 | -0.95715 | -0.67207 | -0.13767 | -0.19361 | -1.40503 | -1.38311 |
| 20.0125 | -0.11572 | -0.26607 | -0.56806 | -0.38135 | -0.82549 | -0.63725 | -1.07081 | -0.47867 | -0.37843 | -1.55629 | -0.33689 | -0.69215 | -0.30861 | -0.35079 | -0.13301 | -0.63031 | -0.07773 | 0.219933 | -1.30356 | -1.23999 |
| 20.02639 | -0.02396 | -0.17414 | -0.51788 | -0.36346 | -0.77463 | -0.4311 | -0.85351 | -0.3944 | -0.34357 | -1.07522 | -0.8244 | -0.61703 | -0.21853 | -0.23607 | 0.509648 | -0.41017 | 0.019134 | 0.371348 | -0.91836 | -0.6221 |
| 20.04028 | 0.075993 | -0.02265 | -0.36732 | -0.23322 | -0.52306 | -0.16479 | -0.4649 | -0.2042 | -0.19874 | -0.50759 | -1.00787 | -0.36569 | -0.00712 | -0.03992 | 0.743974 | -0.11208 | 0.16664 | 0.371761 | -0.35959 | -0.06866 |
| 20.05417 | 0.065527 | 0.045 | -0.17814 | -0.07083 | -0.22785 | 0.040485 | -0.15059 | -0.05518 | -0.01724 | -0.02592 | -0.84481 | -0.14968 | 0.117862 | 0.129544 | 0.513424 | 0.143392 | 0.189843 | 0.265764 | 0.12521 | 0.254801 |
| 20.06806 | 0.011493 | 0.036868 | -0.07539 | 0.080778 | -0.00127 | 0.133753 | 0.061229 | -0.02033 | 0.048796 | 0.24675 | -0.68869 | -0.03422 | 0.034991 | 0.09926 | 0.202053 | 0.338207 | 0.109305 | 0.065064 | 0.403402 | 0.179 |
| 20.08194 | 0.005751 | 0.041165 | -0.07892 | 0.225765 | 0.112494 | 0.156838 | 0.225931 | -0.03903 | -0.01105 | 0.480795 | -0.4897 | 0.0518 | -0.16836 | -0.00349 | -0.00227 | 0.499568 | 0.061271 | -0.08615 | 0.597767 | 0.094275 |
| 20.09583 | -0.02584 | 0.00946 | 0.005355 | 0.183307 | 0.07381 | 0.113344 | 0.218379 | -0.01629 | 0.006428 | 0.496886 | -0.32926 | 0.044271 | -0.20453 | -0.03271 | -0.09638 | 0.335864 | 0.011587 | -0.08448 | 0.332305 | 0.056451 |
| 20.10972 | -0.0676 | -0.03505 | 0.098458 | 0.048134 | 0.023317 | 0.059778 | 0.099914 | 0.040705 | 0.08237 | 0.271509 | -0.26075 | -0.04431 | -0.06968 | -0.0068 | -0.09955 | 0.114834 | -0.02936 | -0.02923 | -0.08323 | -0.02363 |
| 20.12361 | -0.01787 | -0.03504 | 0.044855 | 0.017926 | 0.082284 | 0.044099 | 0.095009 | 0.065921 | 0.059807 | 0.258725 | -0.16031 | -0.05537 | -0.0358 | 0.015826 | 0.074173 | 0.172665 | 0.008846 | -0.03629 | -0.01044 | -0.04253 |
| 20.1375 | -0.02825 | -0.04485 | 0.074611 | -0.00327 | 0.182376 | 0.007077 | 0.106214 | 0.075127 | 0.023747 | 0.31518 | -0.17688 | -0.03184 | -0.04109 | 0.050303 | 0.283277 | 0.132892 | 0.020987 | -0.01696 | 0.093219 | -0.04526 |
| 20.15139 | -0.03592 | -0.00791 | 0.065449 | 0.117032 | 0.292972 | 0.014014 | 0.181055 | 0.098403 | -0.0017 | 0.592463 | -0.08181 | 0.030784 | -0.10413 | 0.085225 | 0.370713 | 0.159014 | 0.035767 | -0.02785 | 0.38926 | 0.013416 |
| 20.16528 | -0.00116 | 0.02737 | -0.07637 | 0.303615 | 0.403086 | 0.047814 | 0.286821 | 0.097809 | -0.01195 | 0.928702 | 0.20627 | 0.097257 | -0.20884 | 0.069577 | 0.278628 | 0.325294 | 0.028084 | -0.12917 | 0.893914 | 0.100668 |
| 20.17917 | -0.10434 | -0.06479 | 0.052199 | 0.210554 | 0.372567 | 0.020556 | 0.160541 | 0.074658 | 0.099957 | 0.702836 | 0.29088 | 0.02007 | -0.10001 | 0.038999 | 0.013765 | 0.081908 | -0.06262 | -0.12 | 0.604714 | 0.027111 |
| 20.19306 | -0.20949 | -0.14173 | 0.348071 | -0.00255 | 0.259598 | 0.003675 | -0.02155 | 0.117963 | 0.277085 | 0.292408 | 0.023495 | -0.07405 | 0.15208 | 0.094136 | -0.18376 | -0.34898 | -0.11828 | 0.061292 | -0.21006 | 0.072407 |
| 20.20694 | -0.16973 | -0.08144 | 0.495916 | 0.03061 | 0.205512 | 0.091347 | 0.058721 | 0.219632 | 0.389588 | 0.201384 | -0.31378 | -0.01376 | 0.275767 | 0.206143 | -0.02521 | -0.39769 | -0.06146 | 0.18192 | -0.46025 | 0.135701 |
| 20.22083 | -0.04685 | 0.034097 | 0.362782 | 0.204567 | 0.283976 | 0.181143 | 0.268589 | 0.265753 | 0.335477 | 0.369401 | -0.28979 | 0.113423 | 0.19072 | 0.254087 | 0.356355 | -0.01897 | 0.064519 | 0.146411 | -0.12687 | 0.133292 |
| 20.23472 | 0.041294 | 0.159931 | 0.106578 | 0.33505 | 0.324825 | 0.196627 | 0.398872 | 0.232258 | 0.185564 | 0.480324 | 0.193732 | 0.16199 | 0.049502 | 0.243776 | 0.665829 | 0.408021 | 0.135558 | 0.042799 | 0.280388 | 0.188899 |
| 20.24861 | 0.083028 | 0.249085 | 0.011887 | 0.362263 | 0.367228 | 0.198846 | 0.382896 | 0.183556 | 0.127522 | 0.465014 | 0.783276 | 0.123109 | -0.01287 | 0.182255 | 0.699205 | 0.611252 | 0.09999 | -0.03984 | 0.444319 | 0.381762 |
| 20.2625 | 0.039717 | 0.329071 | 0.127163 | 0.318562 | 0.379089 | 0.195311 | 0.292914 | 0.163152 | 0.206042 | 0.416411 | 1.008961 | 0.063998 | 0.040586 | 0.143884 | 0.378933 | 0.537315 | 0.038705 | -0.03275 | 0.426054 | 0.657565 |
| 20.27639 | 0.002209 | 0.409144 | 0.239887 | 0.35858 | 0.375231 | 0.205108 | 0.275276 | 0.17076 | 0.281751 | 0.578219 | 0.498122 | 0.100218 | 0.091073 | 0.178413 | -0.00692 | 0.510788 | 0.078313 | -0.01909 | 0.655657 | 0.876662 |
| 20.29028 | 0.063533 | 0.48006 | 0.273155 | 0.477174 | 0.38369 | 0.226469 | 0.32547 | 0.185082 | 0.262573 | 0.79315 | 0.04796 | 0.202875 | 0.080803 | 0.234804 | -0.09645 | 0.665856 | 0.169215 | -0.07282 | 1.15676 | 1.001289 |
| 20.30417 | 0.092163 | 0.495997 | 0.372725 | 0.443411 | 0.369523 | 0.259679 | 0.341969 | 0.265595 | 0.327116 | 0.786345 | -0.23205 | 0.198198 | 0.136259 | 0.272373 | 0.193248 | 0.620096 | 0.132637 | -0.08091 | 1.138298 | 0.97411 |
| 20.31806 | 0.08703 | 0.545314 | 0.485097 | 0.417455 | 0.365606 | 0.334449 | 0.365309 | 0.456472 | 0.451687 | 0.767617 | -0.01827 | 0.151957 | 0.272776 | 0.326998 | 0.587138 | 0.428214 | 0.074536 | 0.003249 | 0.728236 | 0.909733 |
| 20.33194 | 0.200797 | 0.74243 | 0.456681 | 0.606421 | 0.42343 | 0.411785 | 0.523462 | 0.537879 | 0.493496 | 0.820558 | 0.654696 | 0.19727 | 0.267596 | 0.386166 | 0.852587 | 0.648268 | 0.125003 | 0.011434 | 0.767829 | 0.735353 |
| 20.34583 | 0.236452 | 0.936832 | 0.387458 | 0.651631 | 0.415223 | 0.394009 | 0.590996 | 0.478225 | 0.494416 | 0.688377 | 1.301954 | 0.172274 | 0.238202 | 0.38835 | 0.663387 | 0.759325 | 0.145709 | -0.02836 | 0.843448 | 0.768184 |
| 20.35972 | 0.102442 | 1.020054 | 0.472751 | 0.500814 | 0.374251 | 0.3334 | 0.505944 | 0.429791 | 0.542474 | 0.548961 | 1.161126 | 0.082363 | 0.274899 | 0.340902 | 0.346333 | 0.540942 | 0.09105 | 0.007345 | 0.590265 | 1.04034 |
| 20.37361 | 0.149508 | 1.184549 | 0.525234 | 0.565205 | 0.456823 | 0.333267 | 0.526904 | 0.466977 | 0.550497 | 0.588709 | 0.618735 | 0.131121 | 0.226553 | 0.350869 | 0.174975 | 0.648781 | 0.149901 | 0.01266 | 0.703493 | 1.156824 |
| 20.3875 | 0.276068 | 1.320152 | 0.634213 | 0.764878 | 0.639578 | 0.371094 | 0.682399 | 0.544087 | 0.553476 | 0.746562 | 0.431948 | 0.214403 | 0.152631 | 0.438041 | 0.201838 | 0.819425 | 0.252465 | 0.007882 | 1.034097 | 1.087248 |
| 20.40139 | 0.321634 | 1.290392 | 0.895607 | 0.885 | 0.818767 | 0.406563 | 0.822109 | 0.568774 | 0.578742 | 0.870917 | 0.615507 | 0.211788 | 0.208011 | 0.538584 | 0.346005 | 0.715118 | 0.249786 | 0.072775 | 1.2304 | 1.059432 |
| 20.41528 | 0.310398 | 1.062538 | 1.116868 | 0.880882 | 0.913192 | 0.410612 | 0.899832 | 0.545734 | 0.593504 | 0.831279 | 1.016194 | 0.203114 | 0.360105 | 0.575985 | 0.500182 | 0.449896 | 0.193195 | 0.193689 | 1.157846 | 1.254829 |
| 20.42917 | 0.384208 | 0.745133 | 1.033124 | 0.881359 | 0.949801 | 0.459744 | 0.932699 | 0.537887 | 0.638979 | 0.748488 | 1.26452 | 0.280878 | 0.498964 | 0.596 | 0.579755 | 0.238093 | 0.226536 | 0.285382 | 1.087095 | 1.319735 |
| 20.44306 | 0.742606 | 0.752821 | 0.848352 | 1.275025 | 1.094904 | 0.649478 | 1.167138 | 0.705623 | 0.741275 | 1.023208 | 1.638358 | 0.588079 | 0.513412 | 0.743149 | 0.498515 | 0.547795 | 0.543035 | 0.262053 | 1.544091 | 1.508241 |
| 20.45694 | 1.295975 | 1.412019 | 0.957244 | 2.479406 | 1.532445 | 0.939895 | 1.872608 | 1.290568 | 1.055313 | 2.01647 | 3.000651 | 1.156673 | 0.621367 | 1.131982 | 0.438855 | 0.945775 | 1.078138 | 0.171284 | 2.793203 | 2.338458 |
| 20.47083 | 1.279163 | 1.456785 | 1.390595 | 3.092563 | 1.52553 | 1.168687 | 2.62329 | 1.293664 | 1.548222 | 2.855997 | 3.83108 | 1.611174 | 0.891088 | 1.296403 | 0.550568 | 0.979821 | 1.269302 | 0.147217 | 4.640775 | 3.668815 |
| 20.48472 | 0.816411 | 0.915708 | 1.291748 | 2.234927 | 1.165109 | 1.054812 | 1.791326 | 0.59662 | 1.666006 | 2.269562 | 2.789356 | 1.451253 | 0.832789 | 0.941382 | 0.639591 | 0.744707 | 0.946037 | 0.2188 | 4.508627 | 3.72755 |
| 20.49861 | 0.57238 | 0.706192 | 0.76956 | 1.217879 | 0.9261 | 0.697042 | 1.064182 | 0.290496 | 1.239226 | 1.276956 | 2.143844 | 1.030355 | 0.568335 | 0.53261 | 0.525059 | 0.53142 | 0.523784 | 0.161553 | 3.518886 | 3.056716 |
| 20.5125 | 0.456945 | 0.703085 | 0.483925 | 0.49138 | 0.467037 | 0.358343 | 0.426516 | 0.148498 | 0.587706 | 0.688535 | 1.744387 | 0.651538 | 0.28481 | 0.197649 | 0.182269 | 0.250227 | 0.353085 | 0.111015 | 2.605816 | 1.859009 |
| 20.52639 | 0.541805 | 0.625545 | 0.413027 | 0.068577 | -0.01858 | 0.121675 | -0.04357 | 0.044468 | 0.121004 | 0.534696 | 1.11643 | 0.373285 | 0.115868 | -0.04237 | -0.15285 | 0.013553 | 0.316048 | 0.007179 | 1.417977 | 0.897952 |
| 20.54028 | 0.74931 | 0.411097 | 0.357231 | 0.043751 | -0.3928 | 0.067183 | -0.20485 | -0.01367 | -0.01686 | 0.657364 | 0.320337 | 0.259138 | 0.160387 | -0.08776 | -0.29845 | -0.06227 | 0.265744 | -0.03477 | 0.405481 | 0.648232 |
| 20.55417 | 0.744717 | 0.154132 | 0.276459 | 0.244345 | -0.44623 | 0.145859 | -0.08397 | -0.05374 | 0.180027 | 0.847708 | -0.32906 | 0.187469 | 0.372842 | 0.018528 | 0.004564 | 0.005534 | 0.151632 | -0.01772 | -0.14309 | 0.432462 |
| 20.56806 | 0.417301 | 0.116172 | 0.181997 | 0.470353 | -0.0829 | 0.300341 | 0.253207 | -0.08303 | 0.463184 | 0.95042 | -0.31898 | 0.181332 | 0.548709 | 0.19054 | 0.273224 | 0.156029 | 0.060621 | 0.012974 | -0.02648 | 0.002726 |
| 20.58194 | 0.043592 | 0.372881 | 0.185852 | 0.604769 | 0.534667 | 0.469308 | 0.671564 | -0.07687 | 0.808566 | 1.115095 | 0.044646 | 0.237836 | 0.6722 | 0.271931 | 0.496927 | 0.318056 | 0.07485 | -0.01802 | 0.512338 | -0.37385 |
| 20.59583 | -0.23551 | 0.713986 | 0.233867 | 0.638325 | 1.027664 | 0.52564 | 1.096931 | 0.019352 | 1.084206 | 1.299378 | 0.601012 | 0.378755 | 0.58851 | 0.307234 | 0.547454 | 0.46731 | 0.19009 | -0.13886 | 1.015785 | -0.17954 |
| 20.60972 | -0.30056 | 0.797078 | 0.303348 | 0.628517 | 1.119409 | 0.469641 | 1.338304 | 0.220402 | 1.096052 | 1.37923 | 1.081093 | 0.486987 | 0.404362 | 0.274022 | 0.330433 | 0.556931 | 0.323259 | -0.20439 | 1.03711 | 0.332196 |
| 20.62361 | -0.24265 | 0.639288 | 0.391397 | 0.590529 | 0.931561 | 0.248364 | 1.393941 | 0.400265 | 0.780375 | 1.117095 | 1.327613 | 0.460544 | 0.198423 | 0.141305 | 0.170982 | 0.490419 | 0.236033 | -0.15603 | 0.517626 | 0.527138 |
| 20.6375 | -0.1784 | 0.323251 | 0.399832 | 0.472568 | 0.534707 | -0.03057 | 1.201431 | 0.520658 | 0.278079 | 0.401953 | 1.110089 | 0.331129 | 0.083866 | 0.005 | -0.09529 | 0.266644 | 0.091451 | -0.20465 | -0.24562 | 0.364171 |
| 20.65139 | 0.018311 | 0.004886 | 0.316475 | 0.326721 | 0.010161 | -0.17376 | 0.750186 | 0.5099 | -0.17344 | -0.11126 | 0.459251 | 0.137198 | 0.219008 | -0.04024 | -0.07426 | -0.09197 | 0.051948 | -0.16779 | -1.03241 | 0.137418 |
| 20.66528 | 0.244492 | -0.20733 | 0.179964 | 0.244753 | -0.30088 | -0.12794 | 0.158908 | 0.378388 | -0.33524 | -0.03679 | 0.043275 | 0.048295 | 0.402057 | -0.00604 | 0.260206 | -0.31243 | 0.123736 | -0.03845 | -1.36459 | 0.303718 |
| 20.67917 | 0.259104 | -0.31702 | -0.04584 | 0.042713 | -0.30685 | -0.02042 | -0.35631 | 0.19185 | -0.24062 | 0.291192 | 0.496266 | 0.035954 | 0.408186 | 0.065719 | 0.503668 | -0.31548 | 0.090832 | 0.106841 | -0.87718 | 0.328047 |
| 20.69306 | 0.071193 | -0.37047 | -0.17763 | -0.15933 | -0.147 | 0.108038 | -0.59367 | 0.012618 | 0.080808 | 0.437866 | 0.873634 | 0.033436 | 0.260892 | 0.086229 | 0.660057 | -0.28663 | -0.0446 | 0.152892 | 0.133587 | 0.036869 |
| 20.70694 | -0.07516 | -0.3119 | -0.1303 | -0.23554 | 0.10092 | 0.290999 | -0.641 | -0.06509 | 0.416129 | 0.434185 | 0.433838 | 0.031339 | 0.07397 | 0.006349 | 0.497023 | -0.37588 | -0.15632 | 0.194983 | 1.116648 | -0.4122 |
| 20.72083 | -0.13905 | -0.18332 | -0.11777 | -0.26114 | 0.239632 | 0.408078 | -0.61157 | -0.17487 | 0.612326 | 0.485427 | -0.47961 | -0.07079 | -0.23047 | -0.1608 | 0.030294 | -0.40411 | -0.22353 | 0.201465 | 1.520282 | -0.50755 |
| 20.73472 | -0.17396 | -0.17387 | -0.16086 | -0.32075 | -0.00949 | 0.272506 | -0.64314 | -0.35951 | 0.392298 | 0.413755 | -1.39411 | -0.27626 | -0.42294 | -0.42829 | -0.38505 | -0.37533 | -0.29695 | 0.138083 | 1.001986 | -0.14501 |
| 20.74861 | -0.15807 | -0.26816 | -0.16274 | -0.3321 | -0.51971 | -0.10776 | -0.66048 | -0.57116 | -0.10548 | 0.09922 | -2.14576 | -0.39644 | -0.22527 | -0.66936 | -1.05343 | -0.25259 | -0.30891 | 0.024779 | 0.229363 | 0.639545 |
| 20.7625 | -0.11863 | -0.40113 | -0.2433 | -0.26881 | -0.88053 | -0.49206 | -0.58148 | -0.67415 | -0.55542 | -0.15352 | -2.47325 | -0.45003 | -0.07942 | -0.6655 | -1.11695 | -0.08013 | -0.2003 | -0.08593 | -0.43203 | 1.262717 |

| | | | | | | | | | | | | | | | | | | | | |
|---|---|---|---|---|---|---|---|---|---|---|---|---|---|---|---|---|---|---|---|---|
| 20.77639 | -0.14856 | -0.48544 | -0.28325 | -0.28836 | -0.97478 | -0.72408 | -0.50324 | -0.6361 | -0.73159 | -0.11496 | -2.30004 | -0.25905 | -0.1044 | -0.58511 | -0.56145 | 0.055271 | -0.06225 | -0.16615 | -0.78362 | 0.771888 |
| 20.79028 | -0.35158 | -0.62 | -0.41596 | -0.4666 | -0.93238 | -0.67089 | -0.53111 | -0.50801 | -0.66205 | -0.00562 | -1.79409 | -0.04608 | -0.2988 | -0.4982 | -0.04542 | 0.015048 | 0.019806 | -0.27268 | -0.96459 | -0.21353 |
| 20.80417 | -0.76173 | -0.95134 | -0.55156 | -0.686 | -0.89849 | -0.44342 | -0.57595 | -0.35824 | -0.57335 | -0.23564 | -1.37394 | -0.05976 | -0.55215 | -0.44316 | 0.425294 | -0.25293 | 0.02442 | -0.43651 | -1.35517 | -1.45287 |
| 20.81806 | -1.08123 | -1.37786 | -0.50893 | -0.79399 | -0.85159 | -0.22106 | -0.60425 | -0.26037 | -0.62841 | -0.83619 | -1.45392 | -0.31219 | -0.74573 | -0.39282 | 0.639262 | -0.58237 | -0.12535 | -0.5288 | -1.84753 | -2.60655 |
| 20.83194 | -1.08494 | -1.76044 | -0.61843 | -0.8292 | -0.79124 | -0.14752 | -0.6899 | -0.29422 | -0.76847 | -1.35615 | -2.31016 | -0.71761 | -0.89777 | -0.41545 | 0.097332 | -0.90866 | -0.35448 | -0.42613 | -2.27065 | -2.99252 |
| 20.84583 | -0.82514 | -1.82771 | -0.841 | -0.94602 | -0.93235 | -0.25847 | -0.94943 | -0.43121 | -0.98889 | -1.64413 | -3.07186 | -1.05223 | -0.84047 | -0.76697 | -0.75105 | -1.24143 | -0.53532 | -0.32682 | -2.48072 | -2.4127 |
| 20.85972 | -0.55866 | -1.46653 | -1.20565 | -1.17131 | -1.11759 | -0.49179 | -1.31488 | -0.55797 | -1.21939 | -1.82922 | -3.31078 | -1.16874 | -0.67908 | -1.16936 | -1.63454 | -1.61239 | -0.60241 | -0.35652 | -2.67526 | -1.6918 |
| 20.87361 | -0.46334 | -1.14222 | -1.41013 | -1.2447 | -1.30719 | -0.65943 | -1.45778 | -0.67641 | -1.36466 | -1.95303 | -3.2197 | -1.15744 | -0.62099 | -1.34151 | -2.28735 | -1.75345 | -0.68614 | -0.55666 | -2.5729 | -0.96273 |
| 20.8875 | -0.54534 | -0.97172 | -1.36084 | -1.17919 | -1.35889 | -0.74438 | -1.39594 | -0.72844 | -1.38933 | -1.79825 | -2.52466 | -1.05205 | -0.52907 | -1.26797 | -2.07793 | -1.72928 | -0.75622 | -0.75889 | -2.12591 | -0.51167 |
| 20.90139 | -0.68174 | -0.93157 | -1.13919 | -1.013 | -1.24344 | -0.71411 | -1.2498 | -0.65053 | -1.19853 | -1.46268 | -1.2897 | -0.89374 | -0.58902 | -0.93123 | -1.24507 | -1.32711 | -0.78508 | -0.75005 | -1.45186 | -0.35201 |
| 20.91528 | -0.68963 | -0.85627 | -0.88801 | -0.77632 | -1.01687 | -0.6021 | -1.07934 | -0.50356 | -0.85404 | -1.03315 | -0.17922 | -0.77538 | -0.61634 | -0.62122 | -0.40203 | -0.78423 | -0.65543 | -0.55354 | -0.74376 | -0.43774 |
| 20.92917 | -0.56355 | -0.69638 | -0.61021 | -0.57136 | -0.79373 | -0.42548 | -0.9491 | -0.36637 | -0.46515 | -0.65785 | 0.083636 | -0.57364 | -0.58478 | -0.50351 | 0.270198 | -0.32005 | -0.49373 | -0.33054 | -0.32962 | -0.45186 |
| 20.94306 | -0.4068 | -0.50539 | -0.45241 | -0.53195 | -0.6252 | -0.28523 | -0.84666 | -0.31351 | -0.21734 | -0.51316 | -0.47298 | -0.42162 | -0.5611 | -0.52942 | 0.302241 | -0.24391 | -0.45319 | -0.23919 | -0.40369 | -0.46606 |
| 20.95694 | -0.35679 | -0.52804 | -0.64081 | -0.80067 | -0.6675 | -0.29146 | -0.83332 | -0.40821 | -0.29535 | -0.7138 | -1.38779 | -0.43774 | -0.67338 | -0.78078 | -0.29863 | -0.45925 | -0.51765 | -0.32025 | -0.91048 | -0.63606 |
| 20.97083 | -0.26219 | -0.57981 | -1.04256 | -1.31022 | -0.94745 | -0.44811 | -0.96905 | -0.63617 | -0.62291 | -0.99704 | -2.54998 | -0.54498 | -0.83095 | -1.12191 | -1.1519 | -0.55396 | -0.42905 | -0.66515 | -1.38654 | -0.91151 |
| 20.98472 | 0.378154 | -0.20667 | -0.86862 | -1.25527 | -0.9939 | -0.34855 | -0.9049 | -0.65437 | -0.45959 | -0.95788 | -3.72362 | -0.54469 | -0.71029 | -1.20535 | -2.14629 | -0.35953 | -0.30779 | -0.83909 | -1.28218 | -1.01318 |
| 20.99861 | 1.395135 | 0.572016 | 0.339912 | 0.262864 | -0.00957 | 0.692566 | 0.314411 | 0.286168 | 0.929383 | -0.54695 | -3.44696 | -0.17297 | -0.43658 | -1.06131 | -2.7868 | -0.13128 | -0.28394 | -0.46477 | -1.15491 | -0.8488 |
| 21.0125 | 1.263969 | 0.35641 | 0.911414 | 2.497541 | 1.414818 | 2.209719 | 1.397585 | 1.59092 | 3.243914 | -0.01003 | -1.59871 | 0.232929 | -0.26366 | -0.91154 | -1.96259 | -0.12107 | -0.20281 | 0.015185 | -0.28419 | -0.52312 |
| 21.02639 | 0.175368 | -0.93678 | -0.62974 | 0.833185 | 0.742056 | 0.484623 | -0.42583 | 0.283885 | 1.150351 | 0.465714 | 0.193631 | 0.089129 | -0.25261 | -0.88643 | -0.67232 | -0.33975 | -0.14321 | 0.328115 | 0.093729 | -0.50245 |
| 21.04028 | -0.44451 | -1.35342 | -1.70801 | -2.34989 | -1.51529 | -1.79676 | -1.96634 | -1.3824 | -1.804 | 0.391994 | 1.241567 | -0.14427 | -0.2838 | -0.90282 | 0.443623 | -0.42161 | -0.18915 | 0.453082 | -0.03204 | -0.62587 |
| 21.05417 | -0.4397 | -0.67904 | -0.88621 | -1.25377 | -1.13402 | -0.82074 | -1.02486 | -0.73642 | -0.82642 | -0.10083 | 1.09805 | -0.10666 | -0.38087 | -0.52011 | 1.147772 | -0.35019 | -0.17639 | 0.259458 | -0.43978 | -0.54827 |
| 21.06806 | -0.33499 | -0.22843 | -0.03416 | -0.17678 | -0.08375 | 0.018708 | -0.12973 | -0.14698 | -0.0151 | -0.32897 | 0.373243 | 0.032921 | -0.25707 | -0.10119 | 1.242478 | -0.14251 | -0.08206 | 0.114826 | -0.11997 | -0.20502 |
| 21.08194 | -0.32111 | -0.14367 | 0.231731 | 0.062204 | 0.339581 | 0.165292 | 0.02591 | -0.03371 | 0.023862 | -0.28434 | -0.35385 | 0.018042 | -0.06207 | 0.06395 | 0.723594 | 0.002976 | -0.11491 | -0.00659 | -0.07499 | 0.083518 |
| 21.09583 | -0.3809 | -0.14198 | 0.257955 | 0.111726 | 0.444973 | 0.111048 | -0.02062 | -0.04966 | -0.01745 | -0.08061 | -0.89781 | -0.03599 | 0.113213 | 0.0937 | 0.231036 | -0.01455 | -0.18517 | -0.11379 | -0.26351 | 0.094548 |
| 21.10972 | -0.32226 | -0.12842 | 0.203405 | 0.156217 | 0.306683 | 0.089195 | -0.03902 | -0.03924 | 0.080356 | 0.091937 | -1.03998 | -0.04786 | 0.217463 | 0.073548 | -0.02936 | -0.13277 | -0.16362 | -0.13315 | -0.31548 | -0.07024 |
| 21.12361 | -0.22811 | -0.132 | 0.133925 | 0.065129 | 0.099615 | 0.050185 | -0.12382 | -0.04231 | 0.177901 | 0.033714 | -0.69657 | -0.13852 | 0.213764 | 0.03209 | -0.12251 | -0.3167 | -0.17507 | -0.11957 | -0.28035 | -0.21599 |
| 21.1375 | -0.16 | -0.1207 | 0.107478 | -0.0793 | -0.04472 | -0.01259 | -0.26327 | -0.04905 | 0.187408 | -0.04004 | -0.34411 | -0.30707 | 0.232449 | -0.01956 | -0.14499 | -0.44325 | -0.28872 | -0.11286 | -0.26641 | -0.14612 |
| 21.15139 | -0.14611 | -0.06797 | 0.188945 | -0.10563 | -0.03065 | -0.00313 | -0.22473 | 0.018536 | 0.144817 | 0.225905 | -0.20936 | -0.31667 | 0.234726 | -0.06646 | -0.06394 | -0.29728 | -0.25674 | -0.09237 | -0.05692 | 0.222033 |
| 21.16528 | -0.165 | 0.00335 | 0.325928 | -0.06195 | 0.071384 | 0.036221 | -0.06152 | 0.089755 | 0.039456 | 0.410182 | -0.24385 | -0.17636 | 0.122081 | -0.12955 | 0.0598 | -0.09437 | -0.0369 | -0.08209 | 0.145768 | 0.424502 |
| 21.17917 | -0.13743 | 0.024328 | 0.382257 | -0.00469 | 0.201136 | 0.06173 | 0.015304 | 0.120778 | -0.06367 | 0.452825 | -0.40803 | -0.04355 | 0.058363 | -0.11196 | 0.124333 | 0.01015 | 0.182618 | -0.06047 | 0.075067 | 0.299661 |
| 21.19306 | 0.022304 | 0.068682 | 0.336426 | 0.146019 | 0.373034 | 0.125258 | 0.093491 | 0.184908 | -0.07288 | 0.571744 | -0.36521 | 0.147075 | 0.095079 | 0.039648 | 0.194689 | 0.250031 | 0.335258 | 0.002715 | 0.103902 | 0.208693 |
| 21.20694 | 0.147768 | 0.094761 | 0.254727 | 0.236834 | 0.442486 | 0.174742 | 0.146451 | 0.210465 | -0.01152 | 0.706648 | 0.007059 | 0.28133 | 0.177831 | 0.219358 | 0.24846 | 0.49005 | 0.343057 | 0.007302 | 0.333226 | 0.285236 |
| 21.22083 | 0.102883 | 0.043326 | 0.266175 | 0.158167 | 0.33761 | 0.162622 | 0.130321 | 0.165262 | 0.055729 | 0.753991 | 0.480019 | 0.226521 | 0.275348 | 0.277916 | 0.189578 | 0.503341 | 0.230102 | -0.07159 | 0.518728 | 0.448089 |
| 21.23472 | 0.037202 | -0.0313 | 0.308419 | 0.035204 | 0.175211 | 0.107813 | 0.076895 | 0.09846 | 0.086365 | 0.736381 | 0.796093 | 0.079773 | 0.26512 | 0.215869 | 0.073715 | 0.341124 | 0.131446 | -0.15051 | 0.533752 | 0.557025 |
| 21.24861 | 0.109545 | -0.01372 | 0.263012 | -0.03159 | 0.100325 | 0.067184 | 0.067557 | 0.063976 | 0.080615 | 0.691921 | 0.703928 | -0.02421 | 0.087978 | 0.165659 | 0.130213 | 0.206351 | 0.081429 | -0.14725 | 0.407649 | 0.578867 |
| 21.2625 | 0.231283 | 0.188338 | 0.161084 | -0.01277 | 0.189676 | 0.081418 | 0.178038 | 0.064918 | 0.150118 | 0.659682 | 0.286812 | 0.018596 | -0.10957 | 0.251794 | 0.385901 | 0.283229 | 0.137726 | -0.09786 | 0.245023 | 0.630554 |
| 21.27639 | 0.29954 | 0.412753 | 0.135556 | 0.148796 | 0.431623 | 0.165765 | 0.385274 | 0.123274 | 0.305596 | 0.719459 | 0.239406 | 0.240564 | -0.14512 | 0.46819 | 0.647378 | 0.637323 | 0.321167 | -0.047 | 0.267801 | 0.751643 |
| 21.29028 | 0.30198 | 0.577696 | 0.259273 | 0.355115 | 0.647158 | 0.285329 | 0.552207 | 0.208302 | 0.45331 | 0.86473 | 0.764653 | 0.461408 | 0.020408 | 0.580292 | 0.75456 | 0.977449 | 0.550386 | -0.00528 | 0.597486 | 0.868426 |
| 21.30417 | 0.297233 | 0.549614 | 0.428361 | 0.393576 | 0.67064 | 0.341932 | 0.551152 | 0.224277 | 0.572034 | 0.804348 | 1.414721 | 0.461229 | 0.259465 | 0.554455 | 0.684134 | 0.907386 | 0.596916 | 0.001983 | 0.866641 | 0.905369 |
| 21.31806 | 0.380077 | 0.667835 | 0.544333 | 0.365166 | 0.615455 | 0.376685 | 0.543499 | 0.231995 | 0.645497 | 0.691375 | 2.033552 | 0.404802 | 0.283081 | 0.540477 | 0.584017 | 0.791872 | 0.576526 | -0.02263 | 1.062026 | 0.864899 |
| 21.33194 | 0.447869 | 0.79585 | 0.574045 | 0.257401 | 0.55032 | 0.381828 | 0.421124 | 0.202689 | 0.648739 | 0.566285 | 1.937115 | 0.26154 | 0.238767 | 0.567952 | 0.510762 | 0.702703 | 0.519696 | 0.016846 | 0.846468 | 0.858317 |
| 21.34583 | 0.374224 | 0.877563 | 0.582932 | 0.016144 | 0.439752 | 0.330342 | 0.160745 | 0.119201 | 0.55542 | 0.366505 | 0.993845 | 0.000943 | 0.466997 | 0.646255 | 0.449109 | 0.48029 | 0.309208 | 0.110941 | 0.204637 | 1.13857 |
| 21.35972 | 0.419643 | 1.064553 | 0.516598 | 0.049361 | 0.475911 | 0.315409 | 0.150811 | 0.171886 | 0.461327 | 0.513604 | 0.288389 | 0.076862 | 0.532181 | 0.711257 | 0.656454 | 0.507857 | 0.306006 | 0.093526 | 0.22953 | 1.488374 |
| 21.37361 | 0.598928 | 1.434867 | 0.378084 | 0.289499 | 0.585147 | 0.318596 | 0.378004 | 0.399511 | 0.419763 | 0.90534 | 0.280666 | 0.392746 | 0.431446 | 0.736907 | 1.070671 | 0.889165 | 0.506157 | 0.066602 | 0.911608 | 1.470348 |
| 21.3875 | 0.60734 | 1.668381 | 0.368523 | 0.365984 | 0.599756 | 0.355456 | 0.48986 | 0.581116 | 0.544715 | 1.085229 | 0.741459 | 0.440231 | 0.495646 | 0.779062 | 1.290039 | 0.846307 | 0.484424 | 0.059512 | 1.311609 | 1.384112 |
| 21.40139 | 0.553416 | 1.465888 | 0.422353 | 0.400366 | 0.630878 | 0.431857 | 0.524933 | 0.626756 | 0.750631 | 0.981431 | 1.442531 | 0.297683 | 0.532691 | 0.734103 | 0.932692 | 0.68749 | 0.370317 | 0.058833 | 1.316906 | 1.156418 |
| 21.41528 | 0.758129 | 1.112607 | 0.476413 | 0.613233 | 0.854252 | 0.559375 | 0.665928 | 0.640316 | 0.905376 | 0.948895 | 2.448731 | 0.324727 | 0.32134 | 0.767371 | 0.601748 | 0.926247 | 0.417666 | 0.065455 | 1.462458 | 0.848109 |
| 21.42917 | 1.022598 | 0.926068 | 0.588326 | 0.87653 | 1.1862 | 0.654175 | 0.906465 | 0.734119 | 0.939236 | 0.985086 | 2.853943 | 0.516723 | 0.118947 | 0.924588 | 0.554685 | 1.224872 | 0.617899 | 0.131773 | 1.66213 | 0.697729 |
| 21.44306 | 1.280451 | 1.010291 | 0.86275 | 1.223256 | 1.488586 | 0.680205 | 1.095233 | 0.98872 | 0.916072 | 1.150832 | 2.038827 | 0.641466 | 0.235856 | 1.100918 | 0.516439 | 1.293774 | 0.759543 | 0.313264 | 1.919968 | 0.946495 |
| 21.45694 | 1.864683 | 1.484133 | 1.402701 | 2.291903 | 2.001411 | 0.75192 | 1.49398 | 1.68141 | 1.111904 | 1.886742 | 2.350632 | 0.885436 | 0.822052 | 1.376397 | 0.594947 | 1.352634 | 0.825164 | 0.660454 | 2.779798 | 2.057453 |
| 21.47083 | 1.66481 | 1.678847 | 2.079824 | 3.124033 | 2.319789 | 0.757431 | 2.734928 | 2.178402 | 1.423493 | 2.778305 | 3.323547 | 1.358391 | 1.446197 | 1.463343 | 1.055736 | 1.581537 | 0.874898 | 1.014715 | 4.227124 | 3.522092 |
| 21.48472 | 1.124536 | 1.174495 | 1.907232 | 2.571165 | 1.858353 | 0.678858 | 2.07966 | 1.2447 | 1.408869 | 2.623079 | 2.919054 | 1.489885 | 1.54917 | 1.213976 | 1.310852 | 1.628857 | 0.606891 | 0.760006 | 5.088726 | 3.608437 |
| 21.49861 | 0.962308 | 0.711628 | 1.558536 | 1.91967 | 1.396371 | 0.645173 | 1.539251 | 0.587313 | 1.156678 | 1.975412 | 2.917616 | 1.209378 | 1.230991 | 0.952912 | 1.191117 | 1.126431 | 0.267829 | 0.320965 | 4.143176 | 2.79032 |
| 21.5125 | 0.934174 | 0.611742 | 1.036672 | 1.440403 | 0.722107 | 0.546508 | 1.174252 | 0.349046 | 0.747448 | 1.120177 | 2.202505 | 0.962128 | 0.800307 | 0.718142 | 0.759366 | 0.246707 | 0.131385 | 0.196669 | 2.551299 | 1.698109 |
| 21.52639 | 0.979368 | 0.657867 | 0.587033 | 0.951104 | -0.02325 | 0.29938 | 0.585168 | 0.179059 | 0.194781 | 0.313423 | 0.881627 | 0.648653 | 0.324048 | 0.177451 | 0.18483 | -0.35001 | 0.0878 | 0.102926 | 0.707357 | 0.675488 |
| 21.54028 | 0.904133 | 0.767327 | 0.331277 | 0.542576 | -0.7205 | 0.066132 | 0.106832 | 0.086796 | -0.15737 | -0.4382 | -0.47626 | 0.29813 | -0.21041 | -0.48137 | -0.36887 | -0.93473 | 0.093013 | 0.001833 | -0.81885 | 0.075059 |
| 21.55417 | 0.593558 | 0.826286 | 0.100341 | 0.365652 | -1.06288 | 0.045166 | -0.17191 | 0.077307 | -0.03764 | -0.83552 | -1.16778 | 0.019577 | -0.43696 | -0.66728 | -0.84921 | -1.11923 | 0.102133 | 0.036747 | -1.38389 | 0.320765 |
| 21.56806 | 0.342616 | 0.673312 | -0.00551 | 0.308408 | -0.37641 | 0.114707 | -0.13196 | 0.146049 | 0.297774 | -0.41266 | -0.52559 | -0.16767 | 0.05656 | -0.34711 | -0.62845 | -0.46236 | 0.061458 | 0.1575 | -0.11123 | 0.90816 |
| 21.58194 | 0.214859 | 0.385137 | 0.20911 | 0.357478 | 0.850948 | 0.29488 | 0.155635 | 0.251393 | 0.720279 | 0.474191 | 0.60047 | 0.052425 | 0.613716 | 0.230088 | 0.127918 | 0.6159 | 0.089484 | 0.260718 | 1.710657 | 1.166714 |
| 21.59583 | 0.159886 | 0.205072 | 0.49917 | 0.331492 | 1.746357 | 0.391198 | 0.515257 | 0.300505 | 0.925283 | 1.294926 | 1.75294 | 0.330525 | 0.968648 | 0.703458 | 0.717372 | 1.681379 | 0.162661 | 0.252224 | 3.126438 | 0.812021 |
| 21.60972 | 0.045023 | 0.311078 | 0.690126 | 0.182742 | 1.720294 | 0.318604 | 0.773266 | 0.26244 | 0.700328 | 1.522425 | 2.472118 | 0.444698 | 1.124811 | 0.767381 | 1.041423 | 1.695341 | 0.223381 | 0.186312 | 3.016785 | 0.432676 |
| 21.62361 | -0.20216 | 0.579633 | 0.683685 | 0.099539 | 0.989892 | 0.210955 | 0.914571 | 0.254928 | 0.377876 | 1.210965 | 1.799697 | 0.421446 | 0.883807 | 0.62534 | 1.157321 | 1.133928 | 0.285728 | 0.197739 | 1.657656 | 0.355722 |
| 21.6375 | -0.50453 | 0.635918 | 0.378495 | 0.161388 | 0.313038 | 0.111147 | 0.844665 | 0.262901 | 0.111292 | 0.558612 | -0.10466 | 0.351712 | 0.580606 | 0.407315 | 0.886745 | 0.313758 | 0.25673 | 0.187251 | -0.24903 | 0.238001 |
| 21.65139 | -0.63587 | 0.26478 | 0.061766 | 0.357943 | -0.34172 | -0.02975 | 0.635596 | 0.248045 | -0.08847 | -0.33452 | -1.95217 | 0.246788 | 0.062607 | 0.056383 | 0.229584 | -0.57753 | 0.098933 | 0.033004 | -2.03207 | -0.12748 |
| 21.66528 | -0.35761 | -0.18641 | -0.09913 | 0.54238 | -0.95005 | -0.17753 | 0.448976 | 0.266217 | -0.1984 | -0.76364 | -2.41515 | 0.037912 | -0.74044 | -0.36577 | -0.37141 | -0.98612 | -0.12301 | -0.14103 | -2.6903 | 0.027382 |
| 21.67917 | 0.060662 | -0.56625 | -0.17639 | 0.602817 | -0.96549 | -0.19875 | 0.275052 | 0.300466 | -0.19107 | -0.47805 | -1.48717 | -0.30829 | -0.57069 | -0.84488 | -0.77501 | -0.62038 | -0.27035 | -0.17367 | -2.07947 | 0.412857 |
| 21.69306 | 0.389603 | -0.70496 | -0.2654 | 0.563219 | -0.17304 | -0.02599 | 0.09521 | 0.285412 | -0.01436 | 0.097192 | 0.377533 | -0.28705 | -0.35101 | -0.38773 | -0.65065 | 0.063522 | -0.28053 | -0.04261 | -0.8899 | 0.592691 |
| 21.70694 | 0.475127 | -0.56578 | -0.36991 | 0.530942 | 0.544094 | 0.223123 | 0.033863 | 0.123642 | 0.173992 | 0.469156 | 2.106241 | 0.163265 | 0.023171 | 0.317816 | -0.09324 | 0.655116 | -0.19711 | 0.162241 | 0.258701 | 0.43783 |
| 21.72083 | 0.266353 | -0.44966 | -0.20891 | 0.300607 | 0.744648 | 0.324128 | 0.057154 | -0.14712 | 0.144015 | 0.292827 | 2.746001 | 0.576233 | 0.718537 | 0.733842 | 0.432112 | 0.768175 | -0.21061 | 0.295973 | 0.870003 | 0.12882 |
| 21.73472 | -0.15079 | -0.42373 | 0.073053 | -0.21687 | 0.37979 | 0.231583 | -0.00665 | -0.39203 | -0.04738 | -0.22627 | 0.84486 | 0.527432 | 0.547156 | 0.361509 | 0.492628 | 0.058187 | -0.28876 | 0.156036 | 0.70631 | -0.36853 |
| 21.74861 | -0.4494 | -0.39306 | 0.161337 | -0.77338 | -0.27082 | 0.025244 | -0.32589 | -0.51841 | -0.20418 | -0.81899 | -1.70799 | 0.022958 | -0.16893 | -0.35996 | 0.003807 | -0.98828 | -0.41648 | -0.12916 | -0.11335 | -0.60655 |
| 21.7625 | -0.47763 | -0.33649 | -0.11444 | -1.12761 | -0.9895 | -0.26008 | -0.88541 | -0.44937 | -0.35144 | -1.01441 | -3.39478 | -0.59969 | -0.93031 | -1.16786 | -0.43711 | -1.68823 | -0.32964 | -0.29288 | -1.14512 | -0.33173 |
| 21.77639 | -0.40079 | -0.40244 | -0.56701 | -1.46548 | -1.75404 | -0.59855 | -1.51785 | -0.29573 | -0.63479 | -0.52064 | -4.02334 | -1.26835 | -1.69017 | -2.00619 | -0.89599 | -1.87529 | -0.22042 | -0.22154 | -1.88143 | 0.252076 |
| 21.79028 | -0.31153 | -0.56593 | -0.94962 | -1.70372 | -2.17372 | -0.65706 | -1.95711 | -0.22327 | -0.69874 | 0.107329 | -4.15056 | -1.92533 | -1.53442 | -2.35865 | -1.37219 | -1.27314 | -0.17317 | -0.242 | -1.74943 | 0.339003 |
| 21.80417 | -0.42119 | -0.82934 | -1.33447 | -1.61666 | -1.7868 | -0.55974 | -2.02997 | -0.30202 | -0.49992 | 0.106517 | -2.77012 | -1.92089 | -1.11121 | -1.69772 | -1.39122 | -0.48921 | -0.23404 | -0.46113 | -1.27256 | -0.24356 |
| 21.81806 | -0.66436 | -1.19454 | -1.40673 | -1.41822 | -1.30906 | -0.4669 | -1.78776 | -0.58377 | -0.45795 | -0.61382 | -1.51617 | -1.15421 | -0.39298 | -0.66665 | -1.10498 | 0.019122 | -0.32627 | -0.66717 | -1.18466 | -1.04222 |
| 21.83194 | -0.96326 | -1.5715 | -1.33583 | -1.19308 | -1.21364 | -0.53456 | -1.43497 | -1.02266 | -0.67104 | -1.53747 | -0.70965 | -0.59861 | 0.398726 | -0.06717 | -0.79681 | -0.11967 | -0.37344 | -1.02757 | -1.42169 | -1.4407 |
| 21.84583 | -1.21025 | -1.81678 | -1.14625 | -1.005 | -1.30457 | -0.81174 | -1.01897 | -1.45333 | -1.17774 | -1.81508 | -0.69795 | -0.33284 | 0.666236 | -0.12611 | -0.54355 | -0.75413 | -0.30408 | -1.08898 | -1.798 | -1.71777 |
| 21.85972 | -1.34983 | -2.03543 | -0.99058 | -0.649 | -1.40405 | -1.04157 | -0.72506 | -1.42724 | -1.55247 | -1.80541 | -1.84019 | -0.18761 | 0.090349 | -0.54628 | -0.40135 | -1.53274 | -0.3491 | -0.91873 | -1.98681 | -2.01354 |
| 21.87361 | -1.3092 | -1.91249 | -0.87788 | -0.20358 | -1.20276 | -1.10982 | -0.50698 | -0.97369 | -1.56371 | -1.41575 | -3.06952 | -0.09632 | -0.81305 | -1.09208 | -0.40188 | -2.104 | -0.36332 | -0.83174 | -1.7454 | -1.57748 |
| 21.8875 | -0.51729 | -1.18836 | -0.37966 | 0.146694 | -0.75798 | -0.48943 | 0.303133 | 0.077305 | -1.13886 | 0.158119 | -2.78901 | 0.385599 | -1.62894 | -0.93823 | -0.4064 | -1.50029 | 0.347417 | -0.55363 | 0.005401 | -0.34771 |
| 21.90139 | 0.327438 | -0.20814 | -0.23626 | 0.131969 | -1.08308 | 0.41384 | 1.827959 | 1.531409 | -0.91902 | 3.077884 | -1.42834 | 1.513765 | -3.23903 | 1.563663 | -0.09923 | -0.21476 | 1.53972 | 0.865922 | 3.850785 | 0.310481 |
| 21.91528 | NaN | NaN | NaN | NaN | NaN | NaN | NaN | NaN | NaN | NaN | NaN | NaN | NaN | NaN | NaN | NaN | NaN | NaN | NaN | NaN |
| 21.92917 | NaN | NaN | NaN | NaN | NaN | NaN | NaN | NaN | NaN | NaN | NaN | NaN | NaN | NaN | NaN | NaN | NaN | NaN | NaN | NaN |
| 21.94306 | NaN | NaN | NaN | NaN | NaN | NaN | NaN | NaN | NaN | NaN | NaN | NaN | NaN | NaN | NaN | NaN | NaN | NaN | NaN | NaN |
| 21.95694 | NaN | NaN | NaN | NaN | NaN | NaN | NaN | NaN | NaN | NaN | NaN | NaN | NaN | NaN | NaN | NaN | NaN | NaN | NaN | NaN |
| 21.97083 | NaN | NaN | NaN | NaN | NaN | NaN | NaN | NaN | NaN | NaN | NaN | NaN | NaN | NaN | NaN | NaN | NaN | NaN | NaN | NaN |
| 21.98472 | NaN | NaN | NaN | NaN | NaN | NaN | NaN | NaN | NaN | NaN | NaN | NaN | NaN | NaN | NaN | NaN | NaN | NaN | NaN | NaN |
| 21.99861 | NaN | NaN | NaN | NaN | NaN | NaN | NaN | NaN | NaN | NaN | NaN | NaN | NaN | NaN | NaN | NaN | NaN | NaN | NaN | NaN |
| 22.0125 | NaN | NaN | NaN | NaN | NaN | NaN | NaN | NaN | NaN | NaN | NaN | NaN | NaN | NaN | NaN | NaN | NaN | NaN | NaN | NaN |
| 22.02639 | NaN | NaN | NaN | NaN | NaN | NaN | NaN | NaN | NaN | NaN | NaN | NaN | NaN | NaN | NaN | NaN | NaN | NaN | NaN | NaN |

| | | | | | | | | | | | | | | | | | | | | |
|---|---|---|---|---|---|---|---|---|---|---|---|---|---|---|---|---|---|---|---|---|
| 22.04028 | NaN | NaN | NaN | NaN | NaN | NaN | NaN | NaN | NaN | NaN | NaN | NaN | NaN | NaN | NaN | NaN | NaN | NaN | NaN | NaN |
| 22.05417 | NaN | NaN | NaN | NaN | NaN | NaN | NaN | NaN | NaN | NaN | NaN | NaN | NaN | NaN | NaN | NaN | NaN | NaN | NaN | NaN |
| 22.06806 | NaN | NaN | NaN | NaN | NaN | NaN | NaN | NaN | NaN | NaN | NaN | NaN | NaN | NaN | NaN | NaN | NaN | NaN | NaN | NaN |
| 22.08194 | NaN | NaN | NaN | NaN | NaN | NaN | NaN | NaN | NaN | NaN | NaN | NaN | NaN | NaN | NaN | NaN | NaN | NaN | NaN | NaN |
| 22.09583 | NaN | NaN | NaN | NaN | NaN | NaN | NaN | NaN | NaN | NaN | NaN | NaN | NaN | NaN | NaN | NaN | NaN | NaN | NaN | NaN |
| 22.10972 | NaN | NaN | NaN | NaN | NaN | NaN | NaN | NaN | NaN | NaN | NaN | NaN | NaN | NaN | NaN | NaN | NaN | NaN | NaN | NaN |
| 22.12361 | NaN | NaN | NaN | NaN | NaN | NaN | NaN | NaN | NaN | NaN | NaN | NaN | NaN | NaN | NaN | NaN | NaN | NaN | NaN | NaN |
| 22.1375 | NaN | NaN | NaN | NaN | NaN | NaN | NaN | NaN | NaN | NaN | NaN | NaN | NaN | NaN | NaN | NaN | NaN | NaN | NaN | NaN |
| 22.15139 | NaN | NaN | NaN | NaN | NaN | NaN | NaN | NaN | NaN | NaN | NaN | NaN | NaN | NaN | NaN | NaN | NaN | NaN | NaN | NaN |
| 22.16528 | NaN | NaN | NaN | NaN | NaN | NaN | NaN | NaN | NaN | NaN | NaN | NaN | NaN | NaN | NaN | NaN | NaN | NaN | NaN | NaN |
| 22.17917 | NaN | NaN | NaN | NaN | NaN | NaN | NaN | NaN | NaN | NaN | NaN | NaN | NaN | NaN | NaN | NaN | NaN | NaN | NaN | NaN |
| 22.19306 | NaN | NaN | NaN | NaN | NaN | NaN | NaN | NaN | NaN | NaN | NaN | NaN | NaN | NaN | NaN | NaN | NaN | NaN | NaN | NaN |
| 22.20694 | NaN | NaN | NaN | NaN | NaN | NaN | NaN | NaN | NaN | NaN | NaN | NaN | NaN | NaN | NaN | NaN | NaN | NaN | NaN | NaN |
| 22.22083 | NaN | NaN | NaN | NaN | NaN | NaN | NaN | NaN | NaN | NaN | NaN | NaN | NaN | NaN | NaN | NaN | NaN | NaN | NaN | NaN |
| 22.23472 | NaN | NaN | NaN | NaN | NaN | NaN | NaN | NaN | NaN | NaN | NaN | NaN | NaN | NaN | NaN | NaN | NaN | NaN | NaN | NaN |
| 22.24861 | NaN | NaN | NaN | NaN | NaN | NaN | NaN | NaN | NaN | NaN | NaN | NaN | NaN | NaN | NaN | NaN | NaN | NaN | NaN | NaN |
| 22.2625 | NaN | NaN | NaN | NaN | NaN | NaN | NaN | NaN | NaN | NaN | NaN | NaN | NaN | NaN | NaN | NaN | NaN | NaN | NaN | NaN |
| 22.27639 | NaN | NaN | NaN | NaN | NaN | NaN | NaN | NaN | NaN | NaN | NaN | NaN | NaN | NaN | NaN | NaN | NaN | NaN | NaN | NaN |
| 22.29028 | NaN | NaN | NaN | NaN | NaN | NaN | NaN | NaN | NaN | NaN | NaN | NaN | NaN | NaN | NaN | NaN | NaN | NaN | NaN | NaN |
| 22.30417 | NaN | NaN | NaN | NaN | NaN | NaN | NaN | NaN | NaN | NaN | NaN | NaN | NaN | NaN | NaN | NaN | NaN | NaN | NaN | NaN |
| 22.31806 | NaN | NaN | NaN | NaN | NaN | NaN | NaN | NaN | NaN | NaN | NaN | NaN | NaN | NaN | NaN | NaN | NaN | NaN | NaN | NaN |
| 22.33194 | NaN | NaN | NaN | NaN | NaN | NaN | NaN | NaN | NaN | NaN | NaN | NaN | NaN | NaN | NaN | NaN | NaN | NaN | NaN | NaN |
| 22.34583 | NaN | NaN | NaN | NaN | NaN | NaN | NaN | NaN | NaN | NaN | NaN | NaN | NaN | NaN | NaN | NaN | NaN | NaN | NaN | NaN |
| 22.35972 | NaN | NaN | NaN | NaN | NaN | NaN | NaN | NaN | NaN | NaN | NaN | NaN | NaN | NaN | NaN | NaN | NaN | NaN | NaN | NaN |
| 22.37361 | NaN | NaN | NaN | NaN | NaN | NaN | NaN | NaN | NaN | NaN | NaN | NaN | NaN | NaN | NaN | NaN | NaN | NaN | NaN | NaN |
| 22.3875 | NaN | NaN | NaN | NaN | NaN | NaN | NaN | NaN | NaN | NaN | NaN | NaN | NaN | NaN | NaN | NaN | NaN | NaN | NaN | NaN |
| 22.40139 | NaN | NaN | NaN | NaN | NaN | NaN | NaN | NaN | NaN | NaN | NaN | NaN | NaN | NaN | NaN | NaN | NaN | NaN | NaN | NaN |
| 22.41528 | NaN | NaN | NaN | NaN | NaN | NaN | NaN | NaN | NaN | NaN | NaN | NaN | NaN | NaN | NaN | NaN | NaN | NaN | NaN | NaN |
| 22.42917 | NaN | NaN | NaN | NaN | NaN | NaN | NaN | NaN | NaN | NaN | NaN | NaN | NaN | NaN | NaN | NaN | NaN | NaN | NaN | NaN |
| 22.44306 | NaN | NaN | NaN | NaN | NaN | NaN | NaN | NaN | NaN | NaN | NaN | NaN | NaN | NaN | NaN | NaN | NaN | NaN | NaN | NaN |
| 22.45694 | NaN | NaN | NaN | NaN | NaN | NaN | NaN | NaN | NaN | NaN | NaN | NaN | NaN | NaN | NaN | NaN | NaN | NaN | NaN | NaN |
| 22.47083 | NaN | NaN | NaN | NaN | NaN | NaN | NaN | NaN | NaN | NaN | NaN | NaN | NaN | NaN | NaN | NaN | NaN | NaN | NaN | NaN |
| 22.48472 | NaN | NaN | NaN | NaN | NaN | NaN | NaN | NaN | NaN | NaN | NaN | NaN | NaN | NaN | NaN | NaN | NaN | NaN | NaN | NaN |
| 22.49861 | NaN | NaN | NaN | NaN | NaN | NaN | NaN | NaN | NaN | NaN | NaN | NaN | NaN | NaN | NaN | NaN | NaN | NaN | NaN | NaN |
| 22.5125 | NaN | NaN | NaN | NaN | NaN | NaN | NaN | NaN | NaN | NaN | NaN | NaN | NaN | NaN | NaN | NaN | NaN | NaN | NaN | NaN |
| 22.52639 | NaN | NaN | NaN | NaN | NaN | NaN | NaN | NaN | NaN | NaN | NaN | NaN | NaN | NaN | NaN | NaN | NaN | NaN | NaN | NaN |
| 22.54028 | NaN | NaN | NaN | NaN | NaN | NaN | NaN | NaN | NaN | NaN | NaN | NaN | NaN | NaN | NaN | NaN | NaN | NaN | NaN | NaN |
| 22.55417 | NaN | NaN | NaN | NaN | NaN | NaN | NaN | NaN | NaN | NaN | NaN | NaN | NaN | NaN | NaN | NaN | NaN | NaN | NaN | NaN |
| 22.56806 | NaN | NaN | NaN | NaN | NaN | NaN | NaN | NaN | NaN | NaN | NaN | NaN | NaN | NaN | NaN | NaN | NaN | NaN | NaN | NaN |
| 22.58194 | NaN | NaN | NaN | NaN | NaN | NaN | NaN | NaN | NaN | NaN | NaN | NaN | NaN | NaN | NaN | NaN | NaN | NaN | NaN | NaN |
| 22.59583 | NaN | NaN | NaN | NaN | NaN | NaN | NaN | NaN | NaN | NaN | NaN | NaN | NaN | NaN | NaN | NaN | NaN | NaN | NaN | NaN |
| 22.60972 | NaN | NaN | NaN | NaN | NaN | NaN | NaN | NaN | NaN | NaN | NaN | NaN | NaN | NaN | NaN | NaN | NaN | NaN | NaN | NaN |
| 22.62361 | NaN | NaN | NaN | NaN | NaN | NaN | NaN | NaN | NaN | NaN | NaN | NaN | NaN | NaN | NaN | NaN | NaN | NaN | NaN | NaN |
| 22.6375 | NaN | NaN | NaN | NaN | NaN | NaN | NaN | NaN | NaN | NaN | NaN | NaN | NaN | NaN | NaN | NaN | NaN | NaN | NaN | NaN |
| 22.65139 | NaN | NaN | NaN | NaN | NaN | NaN | NaN | NaN | NaN | NaN | NaN | NaN | NaN | NaN | NaN | NaN | NaN | NaN | NaN | NaN |
| 22.66528 | NaN | NaN | NaN | NaN | NaN | NaN | NaN | NaN | NaN | NaN | NaN | NaN | NaN | NaN | NaN | NaN | NaN | NaN | NaN | NaN |
| 22.67917 | NaN | NaN | NaN | NaN | NaN | NaN | NaN | NaN | NaN | NaN | NaN | NaN | NaN | NaN | NaN | NaN | NaN | NaN | NaN | NaN |
| 22.69306 | NaN | NaN | NaN | NaN | NaN | NaN | NaN | NaN | NaN | NaN | NaN | NaN | NaN | NaN | NaN | NaN | NaN | NaN | NaN | NaN |
| 22.70694 | NaN | NaN | NaN | NaN | NaN | NaN | NaN | NaN | NaN | NaN | NaN | NaN | NaN | NaN | NaN | NaN | NaN | NaN | NaN | NaN |
| 22.72083 | NaN | NaN | NaN | NaN | NaN | NaN | NaN | NaN | NaN | NaN | NaN | NaN | NaN | NaN | NaN | NaN | NaN | NaN | NaN | NaN |
| 22.73472 | NaN | NaN | NaN | NaN | NaN | NaN | NaN | NaN | NaN | NaN | NaN | NaN | NaN | NaN | NaN | NaN | NaN | NaN | NaN | NaN |
| 22.74861 | NaN | NaN | NaN | NaN | NaN | NaN | NaN | NaN | NaN | NaN | NaN | NaN | NaN | NaN | NaN | NaN | NaN | NaN | NaN | NaN |
| 22.7625 | NaN | NaN | NaN | NaN | NaN | NaN | NaN | NaN | NaN | NaN | NaN | NaN | NaN | NaN | NaN | NaN | NaN | NaN | NaN | NaN |
| 22.77639 | NaN | NaN | NaN | NaN | NaN | NaN | NaN | NaN | NaN | NaN | NaN | NaN | NaN | NaN | NaN | NaN | NaN | NaN | NaN | NaN |
| 22.79028 | NaN | NaN | NaN | NaN | NaN | NaN | NaN | NaN | NaN | NaN | NaN | NaN | NaN | NaN | NaN | NaN | NaN | NaN | NaN | NaN |
| 22.80417 | NaN | NaN | NaN | NaN | NaN | NaN | NaN | NaN | NaN | NaN | NaN | NaN | NaN | NaN | NaN | NaN | NaN | NaN | NaN | NaN |
| 22.81806 | NaN | NaN | NaN | NaN | NaN | NaN | NaN | NaN | NaN | NaN | NaN | NaN | NaN | NaN | NaN | NaN | NaN | NaN | NaN | NaN |
| 22.83194 | NaN | NaN | NaN | NaN | NaN | NaN | NaN | NaN | NaN | NaN | NaN | NaN | NaN | NaN | NaN | NaN | NaN | NaN | NaN | NaN |
| 22.84583 | NaN | NaN | NaN | NaN | NaN | NaN | NaN | NaN | NaN | NaN | NaN | NaN | NaN | NaN | NaN | NaN | NaN | NaN | NaN | NaN |
| 22.85972 | NaN | NaN | NaN | NaN | NaN | NaN | NaN | NaN | NaN | NaN | NaN | NaN | NaN | NaN | NaN | NaN | NaN | NaN | NaN | NaN |
| 22.87361 | NaN | NaN | NaN | NaN | NaN | NaN | NaN | NaN | NaN | NaN | NaN | NaN | NaN | NaN | NaN | NaN | NaN | NaN | NaN | NaN |
| 22.8875 | NaN | NaN | NaN | NaN | NaN | NaN | NaN | NaN | NaN | NaN | NaN | NaN | NaN | NaN | NaN | NaN | NaN | NaN | NaN | NaN |
| 22.90139 | NaN | NaN | NaN | NaN | NaN | NaN | NaN | NaN | NaN | NaN | NaN | NaN | NaN | NaN | NaN | NaN | NaN | NaN | NaN | NaN |
| 22.91528 | NaN | NaN | NaN | NaN | NaN | NaN | NaN | NaN | NaN | NaN | NaN | NaN | NaN | NaN | NaN | NaN | NaN | NaN | NaN | NaN |
| 22.92917 | NaN | NaN | NaN | NaN | NaN | NaN | NaN | NaN | NaN | NaN | NaN | NaN | NaN | NaN | NaN | NaN | NaN | NaN | NaN | NaN |
| 22.94306 | NaN | NaN | NaN | NaN | NaN | NaN | NaN | NaN | NaN | NaN | NaN | NaN | NaN | NaN | NaN | NaN | NaN | NaN | NaN | NaN |
| 22.95694 | NaN | NaN | NaN | NaN | NaN | NaN | NaN | NaN | NaN | NaN | NaN | NaN | NaN | NaN | NaN | NaN | NaN | NaN | NaN | NaN |
| 22.97083 | NaN | NaN | NaN | NaN | NaN | NaN | NaN | NaN | NaN | NaN | NaN | NaN | NaN | NaN | NaN | NaN | NaN | NaN | NaN | NaN |
| 22.98472 | NaN | NaN | NaN | NaN | NaN | NaN | NaN | NaN | NaN | NaN | NaN | NaN | NaN | NaN | NaN | NaN | NaN | NaN | NaN | NaN |
| 22.99861 | NaN | NaN | NaN | NaN | NaN | NaN | NaN | NaN | NaN | NaN | NaN | NaN | NaN | NaN | NaN | NaN | NaN | NaN | NaN | NaN |
| 23.0125 | NaN | NaN | NaN | NaN | NaN | NaN | NaN | NaN | NaN | NaN | NaN | NaN | NaN | NaN | NaN | NaN | NaN | NaN | NaN | NaN |
| 23.02639 | 0.098769 | -0.07769 | 0.427162 | 0.08003 | 0.226806 | 0.024487 | -0.12697 | 0.089296 | 0.135192 | -0.18175 | 0.625084 | -0.09943 | 0.650173 | -0.13611 | 0.152156 | -0.27467 | -0.12829 | 0.059278 | -0.09586 | 0.7863 |
| 23.04028 | -0.70128 | 1.065308 | -2.95756 | -1.45375 | -0.58011 | 0.026722 | -0.36079 | -1.99644 | 0.005793 | -0.97026 | -2.22323 | -1.56395 | -2.39365 | 0.095775 | -0.79412 | -0.38975 | -0.31751 | 0.602428 | -0.33466 | 2.566262 |
| 23.05417 | 0.515579 | -0.06253 | 4.7978 | 1.966011 | 1.5541 | 0.152258 | 1.163451 | 1.405037 | 1.551028 | -0.94798 | 7.310418 | -0.43965 | 4.349969 | -2.06301 | 1.995273 | 0.673356 | -1.80404 | -2.1412 | -0.41342 | 10.26671 |
| 23.06806 | -2.6818 | -1.80255 | 2.219028 | -1.32872 | -0.69847 | -0.43944 | -0.74249 | 0.041374 | 1.224545 | -2.77371 | 0.761216 | -3.72169 | 2.405357 | -2.3158 | 0.317504 | -2.80226 | -2.16269 | -0.19322 | -4.13705 | 1.256103 |
| 23.08194 | -1.31648 | -0.53655 | 0.952767 | -0.03221 | 0.181107 | 0.169057 | 0.062449 | -0.03646 | 0.965402 | -0.36892 | 0.35552 | -0.58166 | 0.457216 | -0.37462 | 0.115961 | -1.0517 | -0.65652 | -0.00924 | -0.72754 | 0.625603 |
| 23.09583 | -0.31183 | 0.080438 | 0.592752 | 0.77282 | 0.586658 | 0.285325 | 0.687275 | 0.247319 | 0.972875 | 1.2757 | 1.267199 | 0.683515 | 0.302015 | 0.412868 | 0.331874 | 0.236645 | -0.06834 | -0.01265 | 1.027896 | 1.146443 |
| 23.10972 | -0.16623 | 0.242419 | 0.515921 | 0.888339 | 0.724096 | 0.289656 | 0.663011 | 0.256113 | 0.878131 | 1.914093 | 1.674159 | 0.886272 | 0.271187 | 0.732379 | 0.373106 | 0.305469 | 0.060691 | -0.02953 | 1.35037 | 1.107661 |
| 23.12361 | -0.11276 | 0.315371 | 0.599491 | 0.874544 | 0.90805 | 0.221887 | 0.598889 | 0.240825 | 0.665301 | 1.707845 | 2.181198 | 0.971269 | 0.190416 | 0.845808 | 0.466279 | 0.200778 | 0.100019 | -0.07576 | 1.113192 | 0.694753 |
| 23.1375 | -0.07155 | 0.282511 | 0.787773 | 0.753643 | 1.050016 | 0.14705 | 0.507438 | 0.250048 | 0.454319 | 1.054914 | 2.075963 | 0.928752 | 0.000028 | 0.603493 | 0.540955 | 0.068222 | 0.135329 | -0.08033 | 0.631655 | 0.159599 |
| 23.15139 | -0.07921 | 0.123874 | 0.744543 | 0.562144 | 0.962288 | 0.090496 | 0.433441 | 0.207698 | 0.303451 | 0.297715 | 1.308352 | 0.681227 | -0.25272 | 0.197107 | 0.432451 | -0.04705 | 0.111242 | -0.0885 | 0.19755 | -0.16693 |
| 23.16528 | -0.19312 | -0.13902 | 0.441215 | 0.325725 | 0.625833 | 0.02033 | 0.247414 | 0.117009 | 0.248956 | -0.2833 | 0.318895 | 0.314437 | -0.26146 | -0.22429 | 0.271398 | -0.32405 | -0.04169 | -0.01022 | -0.21748 | -0.17883 |
| 23.17917 | -0.26959 | -0.46596 | -0.32066 | -0.10824 | 0.219406 | -0.34557 | -0.09396 | -0.14145 | 0.009321 | -0.00754 | -0.0323 | 0.252229 | 0.18905 | -0.39789 | 0.658454 | -0.14413 | -0.10123 | 0.380377 | -0.03604 | -0.0085 |
| 23.19306 | -0.34724 | -0.84847 | -1.57815 | -0.69098 | -0.2149 | -1.10648 | -0.74114 | -0.73058 | -0.7185 | 0.936907 | 0.354225 | 0.543519 | 0.747584 | -0.24797 | 1.572482 | 0.554491 | 0.049044 | 0.932745 | 0.787766 | 0.167942 |
| 23.20694 | -0.55425 | -0.83232 | -1.82483 | -0.99462 | -0.59822 | -1.38906 | -1.26465 | -1.09607 | -1.1964 | 0.970813 | 0.695232 | 0.461337 | 0.960519 | -0.19326 | 1.519836 | 0.538456 | 0.075305 | 0.941919 | 0.715205 | 0.346669 |
| 23.22083 | -0.53299 | -0.21218 | -0.81235 | -0.73643 | -0.58169 | -0.77826 | -1.08356 | -0.75887 | -0.83974 | 0.222471 | 0.679843 | 0.053774 | 0.709516 | -0.25888 | 0.419226 | -0.07682 | 0.020441 | 0.391829 | 0.038681 | 0.547851 |
| 23.23472 | -0.34986 | 0.2799 | -0.11347 | -0.30006 | -0.29714 | -0.23559 | -0.55468 | -0.20576 | -0.3148 | -0.0429 | 0.494153 | -0.0377 | 0.457651 | -0.15827 | 0.042394 | -0.16649 | 0.044713 | 0.055165 | -0.04404 | 0.727656 |
| 23.24861 | -0.1516 | 0.383495 | 0.052504 | 0.028541 | -0.16109 | -0.07009 | -0.21487 | 0.025392 | -0.06039 | 0.164815 | 0.320887 | 0.078248 | 0.273182 | 0.076213 | 0.284046 | 0.327319 | 0.15213 | 0.039119 | 0.248525 | 0.84108 |
| 23.2625 | 0.085047 | 0.34185 | 0.087178 | 0.184976 | -0.24558 | -0.01723 | -0.06089 | 0.034019 | 0.085895 | 0.397781 | 0.246837 | 0.176226 | 0.196923 | 0.32224 | 0.424875 | 0.961606 | 0.261202 | 0.099138 | 0.409026 | 0.976109 |
| 23.27639 | 0.232024 | 0.293695 | 0.118778 | 0.130647 | -0.40069 | 0.058714 | -0.02995 | 0.012972 | 0.251218 | 0.515806 | 0.220993 | 0.145855 | 0.357879 | 0.461779 | 0.379234 | 1.180761 | 0.25679 | 0.144633 | 0.234187 | 1.031766 |
| 23.29028 | 0.24441 | 0.303912 | 0.157475 | -0.02592 | -0.49956 | 0.173142 | -0.03838 | 0.006281 | 0.463016 | 0.577931 | 0.328936 | 0.052098 | 0.609891 | 0.436939 | 0.278856 | 0.909064 | 0.21445 | 0.128887 | -0.02425 | 1.043459 |

| | | | | | | | | | | | | | | | | | | | | |
|---|---|---|---|---|---|---|---|---|---|---|---|---|---|---|---|---|---|---|---|---|
| 23.30417 | 0.404389 | 0.499362 | 0.279249 | -0.05511 | -0.38929 | 0.329044 | 0.077158 | 0.10705 | 0.757917 | 0.906516 | 0.920778 | 0.146083 | 0.808643 | 0.520025 | 0.289544 | 0.688571 | 0.150577 | 0.137315 | 0.122893 | 1.175647 |
| 23.31806 | 0.690026 | 0.763088 | 0.463896 | 0.0457 | -0.04085 | 0.43197 | 0.244941 | 0.252335 | 0.884267 | 1.420648 | 1.826535 | 0.376198 | 0.952197 | 0.825596 | 0.401781 | 0.678302 | 0.162175 | 0.123508 | 0.612569 | 1.141134 |
| 23.33194 | 0.837749 | 0.828957 | 0.584952 | 0.139061 | 0.292937 | 0.386889 | 0.353488 | 0.353268 | 0.683776 | 1.488473 | 2.51458 | 0.611804 | 0.922726 | 1.093482 | 0.497078 | 0.7834 | 0.190741 | 0.076986 | 1.002802 | 0.850505 |
| 23.34583 | 0.800096 | 0.74804 | 0.69688 | 0.252775 | 0.562229 | 0.246671 | 0.453594 | 0.403941 | 0.450768 | 1.213967 | 2.755003 | 0.833783 | 0.862114 | 1.380373 | 0.583554 | 1.105099 | 0.214612 | 0.089564 | 1.309508 | 0.593736 |
| 23.35972 | 0.683728 | 0.798139 | 0.773252 | 0.385125 | 0.74142 | 0.20005 | 0.601808 | 0.454796 | 0.443943 | 0.917806 | 2.258627 | 0.988883 | 0.566917 | 1.547169 | 0.642093 | 1.395711 | 0.244232 | 0.14762 | 1.495972 | 0.646996 |
| 23.37361 | 0.628551 | 0.964073 | 0.797262 | 0.481572 | 0.735382 | 0.21406 | 0.698184 | 0.535648 | 0.708377 | 0.693252 | 1.453302 | 0.990126 | 0.263232 | 1.209635 | 0.638125 | 1.208642 | 0.30596 | 0.214146 | 1.351813 | 0.95806 |
| 23.3875 | 0.684853 | 1.164764 | 0.739947 | 0.453097 | 0.65514 | 0.248426 | 0.71238 | 0.665928 | 0.96086 | 0.660461 | 1.100935 | 0.902644 | 0.278389 | 0.896791 | 0.675776 | 0.602392 | 0.328349 | 0.267973 | 1.134779 | 1.113901 |
| 23.40139 | 0.797006 | 1.304892 | 0.626779 | 0.427361 | 0.593121 | 0.260033 | 0.717797 | 0.803548 | 1.000003 | 0.691815 | 1.060255 | 0.906344 | 0.409059 | 0.701471 | 0.653998 | 0.373102 | 0.439194 | 0.274983 | 0.995597 | 0.871096 |
| 23.41528 | 0.850885 | 1.420016 | 0.448961 | 0.727723 | 0.732949 | 0.258399 | 0.824714 | 0.963238 | 0.717289 | 0.95308 | 1.298305 | 1.059769 | 0.347362 | 0.838041 | 0.520715 | 1.058086 | 0.714912 | 0.318335 | 1.406755 | 0.321963 |
| 23.42917 | 0.889367 | 1.723207 | 0.474784 | 1.266087 | 1.182836 | 0.394682 | 1.174815 | 1.339005 | 0.426629 | 1.644525 | 1.976566 | 1.432237 | 0.270023 | 1.448925 | 0.580803 | 2.303696 | 1.363456 | 0.774802 | 2.591111 | -0.41091 |
| 23.44306 | 0.569009 | 2.576924 | 1.498937 | 1.557699 | 2.098029 | 0.815766 | 1.812929 | 2.022679 | 0.904594 | 2.740895 | 3.578193 | 2.092367 | 1.247313 | 2.950463 | 1.614335 | 3.592613 | 1.817019 | 2.115662 | 3.998493 | 0.088112 |
| 23.45694 | 0.826799 | 2.783811 | 2.801937 | 1.674539 | 3.064437 | 1.317792 | 2.707633 | 2.612022 | 2.237869 | 3.723191 | 4.606537 | 2.417085 | 2.307256 | 4.004641 | 2.626257 | 3.500146 | 1.469101 | 2.662504 | 4.68873 | 2.162722 |
| 23.47083 | 1.428794 | 1.736238 | 3.016159 | 1.83752 | 2.648908 | 0.838262 | 2.904751 | 1.593425 | 2.510195 | 3.742331 | 3.291146 | 1.972387 | 2.132694 | 3.721775 | 2.145506 | 2.912627 | 0.84795 | 1.997722 | 4.581825 | 2.610823 |
| 23.48472 | 1.555076 | 0.600659 | 2.330071 | 1.734523 | 1.723504 | 0.486669 | 2.328503 | 0.468686 | 2.034641 | 2.493504 | 2.196275 | 1.488339 | 1.258139 | 2.616939 | 1.5537 | 2.114382 | 0.546999 | 1.285415 | 3.106493 | 1.793918 |
| 23.49861 | 1.158122 | -0.25287 | 1.150467 | 1.190989 | 0.615055 | 0.258055 | 1.599607 | 0.049934 | 1.186903 | 0.837484 | 1.432957 | 1.117957 | 0.291417 | 1.694286 | 1.071379 | 1.355069 | 0.396966 | 0.219387 | 1.717214 | 0.688264 |
| 23.5125 | 0.782501 | -0.78713 | 0.534954 | 0.559131 | -0.27918 | 0.110263 | 0.711813 | -0.1741 | 0.448549 | -0.52924 | 0.634364 | 0.882473 | -0.43287 | 0.901611 | 0.515441 | 0.706281 | 0.319579 | -0.47537 | 0.662033 | 0.029757 |
| 23.52639 | 0.644297 | -0.76883 | 0.261058 | 0.2737 | -0.44929 | 0.12556 | 0.007155 | 0.00188 | 0.079174 | -1.35145 | 0.161763 | 0.773252 | -0.32325 | 0.234862 | -0.03816 | 0.203405 | 0.158384 | -0.43638 | 0.036643 | -0.12013 |
| 23.54028 | 0.777579 | -0.04749 | 0.23737 | 0.561642 | -0.0257 | 0.361691 | -0.3653 | 0.374351 | -0.03169 | -1.16131 | 0.636423 | 1.078684 | 0.203818 | -0.04116 | -0.08511 | -0.04879 | 0.18746 | -0.11313 | -0.04482 | -0.01335 |
| 23.55417 | 1.083361 | 0.962479 | 0.270125 | 1.172143 | 0.544911 | 0.482468 | -0.36769 | 0.715683 | 0.060114 | -0.40181 | 1.408004 | 1.140984 | 0.711807 | -0.08579 | 0.047002 | -0.1411 | 0.446312 | 0.211318 | 0.101348 | 0.506312 |
| 23.56806 | 1.819635 | 2.024805 | 0.327646 | 1.640358 | 1.086318 | 0.44906 | -0.13 | 0.773864 | 0.500883 | 0.596776 | 1.551741 | 0.932903 | 1.384229 | 0.034933 | 0.331217 | 0.038852 | 0.675745 | 0.686496 | 0.52979 | 1.298284 |
| 23.58194 | 2.020967 | 2.420319 | 0.30038 | 1.609451 | 1.229475 | 0.346887 | 0.118354 | 1.040352 | 0.894121 | 1.723149 | 1.115686 | 0.706263 | 1.859091 | 0.416388 | 0.569748 | 0.639581 | 0.74846 | 0.863998 | 1.164792 | 1.633982 |
| 23.59583 | 1.367164 | 2.271927 | 0.310282 | 1.279242 | 1.099068 | 0.296969 | 0.353902 | 1.207188 | 1.072229 | 2.656414 | 0.753234 | 0.537334 | 1.893828 | 0.747756 | 0.75133 | 1.430489 | 0.805547 | 0.735314 | 1.690991 | 1.219559 |
| 23.60972 | 0.625586 | 1.675259 | 0.40497 | 0.807084 | 0.878722 | 0.1378 | 0.622182 | 1.035771 | 0.911632 | 2.540156 | 0.696454 | 0.223172 | 1.311455 | 0.975843 | 0.682913 | 1.63921 | 0.683848 | 0.604774 | 1.964826 | 0.602384 |
| 23.62361 | 0.041362 | 0.825273 | 0.456687 | 0.311821 | 0.709108 | -0.0043 | 0.824653 | 0.608995 | 0.514809 | 1.632795 | 1.061519 | 0.025916 | 0.5849 | 0.750883 | 0.510362 | 1.250446 | 0.487837 | 0.561289 | 1.916055 | 0.273233 |
| 23.6375 | -0.68411 | 0.014916 | 0.38738 | -0.23692 | 0.481035 | -0.02477 | 0.77982 | 0.017018 | 0.181784 | 0.631905 | 1.327042 | 0.150547 | -0.09251 | 0.282456 | 0.381445 | 0.849933 | 0.244148 | 0.620909 | 1.563468 | 0.197693 |
| 23.65139 | -1.31237 | -0.59549 | 0.24452 | -0.65427 | 0.201927 | -0.03854 | 0.495534 | -0.47468 | -0.21539 | -0.21924 | 0.708995 | 0.508866 | -0.68714 | 0.067573 | 0.400065 | 0.290534 | -0.02758 | 0.453639 | 1.182259 | 0.09601 |
| 23.66528 | -1.04628 | -0.48197 | 0.246863 | -0.56689 | -0.19167 | 0.004527 | 0.21562 | -0.50174 | -0.27504 | -0.76942 | 0.232265 | 1.026848 | -0.61113 | 0.038992 | 0.346503 | -0.1112 | -0.18862 | -0.0436 | 0.469293 | -0.39275 |
| 23.67917 | -0.37354 | -0.22153 | 0.38067 | -0.33945 | -0.38204 | 0.021012 | -0.01065 | -0.20017 | -0.02427 | -1.21402 | -0.15041 | 1.283448 | 0.097413 | 0.305062 | 0.219978 | -0.29584 | -0.15229 | -0.33738 | -0.10029 | -0.56148 |
| 23.69306 | 0.637498 | -0.18833 | 0.518226 | -0.06849 | -0.29693 | 0.076755 | -0.02055 | 0.329777 | 0.187507 | -1.77858 | -0.85083 | 1.005663 | 0.651876 | 0.363451 | 0.273556 | -0.50971 | 0.224136 | -0.22895 | -0.29753 | 0.415609 |
| 23.70694 | 1.209333 | -0.44582 | 0.380459 | 0.316327 | -0.16722 | 0.112749 | -0.05337 | 0.924626 | 0.341188 | -0.99744 | -1.33963 | 0.421833 | 0.927538 | 0.111975 | 0.267207 | -0.33942 | 0.53371 | -0.08883 | -0.44124 | 1.409326 |
| 23.72083 | 1.109228 | -0.70841 | 0.036728 | 0.474764 | -0.27316 | 0.054806 | -0.13532 | 1.148447 | 0.265976 | 0.050519 | -1.47472 | -0.22693 | 1.022035 | -0.72378 | 0.29473 | 0.515467 | 0.326549 | -0.46307 | -0.48976 | 1.25827 |
| 23.73472 | 0.575881 | -0.70663 | -0.40872 | 0.317127 | -0.48177 | -0.01668 | -0.23688 | 0.687159 | 0.021396 | 0.957229 | -0.72293 | -0.82066 | 0.799057 | -1.00249 | 0.13724 | 1.126351 | -0.02357 | -0.78585 | -0.22187 | 0.428057 |
| 23.74861 | -0.12302 | -0.49164 | -0.56336 | 0.013506 | -0.32243 | -0.07834 | -0.08568 | 0.127498 | -0.12086 | 1.545458 | 0.244835 | -1.20467 | -0.18113 | 0.042521 | -0.12882 | 1.096367 | -0.26039 | -0.72833 | 0.314234 | -0.11056 |
| 23.7625 | -0.9803 | -0.40227 | -0.26891 | -0.36474 | 0.367246 | -0.26432 | 0.253654 | -0.48016 | -0.43334 | 1.345859 | 1.108991 | -1.12289 | -0.79966 | 1.269071 | -0.48738 | 1.136162 | -0.49644 | -0.28391 | 0.600342 | -0.37396 |
| 23.77639 | -1.58935 | -0.44557 | -0.19327 | -0.7381 | 0.845555 | -0.60534 | 0.140552 | -1.12478 | -0.84534 | 0.286856 | 1.517162 | -0.71112 | -1.64635 | 1.345967 | -0.79123 | 0.426994 | -0.628 | -0.09132 | 0.283401 | -1.13464 |
| 23.79028 | -1.14917 | -0.05761 | -0.40921 | -1.03047 | 0.482989 | -0.65303 | -0.51074 | -1.58816 | -0.98954 | -1.25672 | 1.371893 | -0.31845 | -2.29744 | 0.573222 | -0.98919 | -0.67966 | -0.98372 | -0.45021 | -0.64729 | -2.47918 |
| 23.80417 | -0.87623 | 0.129751 | -0.9806 | -1.09304 | -0.27525 | -0.43121 | -1.50122 | -1.81531 | -0.73422 | -3.15799 | 0.143517 | -0.19262 | -2.43984 | 0.00169 | -1.08388 | -1.40485 | -1.18746 | -1.18708 | -1.53014 | -3.4017 |
| 23.81806 | -0.62611 | -0.03075 | -1.68871 | -0.88192 | -1.17567 | -0.16523 | -2.28533 | -1.76693 | -0.58719 | -4.18628 | -1.74885 | -0.35606 | -1.82927 | -1.1733 | -0.79268 | -1.76586 | -0.7894 | -1.78228 | -2.26741 | -2.94821 |
| 23.83194 | -0.38726 | -0.24936 | -2.09275 | -0.79802 | -1.80254 | -0.20461 | -2.29611 | -1.53 | -0.93491 | -3.64471 | -3.79922 | -0.87243 | -0.76571 | -2.11349 | -0.7572 | -2.96014 | -0.43001 | -2.78313 | -3.24219 | -1.27694 |
| 23.84583 | -0.45821 | -0.72032 | -1.85229 | -0.80039 | -2.65575 | -0.67481 | -2.15317 | -1.22715 | -1.86314 | -2.77595 | -5.50614 | -1.77341 | 0.056852 | -2.99298 | -0.76424 | -4.33427 | -0.27086 | -2.7916 | -4.71318 | 0.377189 |
| 23.85972 | -0.81459 | -1.3228 | -1.28031 | -0.85224 | -2.34287 | -1.01097 | -1.97592 | -0.92283 | -2.10004 | -1.48908 | -5.83185 | -2.16196 | 0.881553 | -4.31917 | -0.76464 | -2.39044 | -0.31541 | -2.66184 | -4.81987 | 1.424429 |
| 23.87361 | -1.29362 | -1.85518 | -0.69288 | -0.50207 | -1.76851 | -0.92607 | -1.46088 | -0.70498 | -1.7779 | -0.36446 | -4.89737 | -1.94352 | 0.399161 | -4.5468 | -0.89982 | 0.972817 | -0.47861 | -2.01705 | -3.22839 | 1.472591 |
| 23.8875 | -1.64477 | -2.16171 | -0.35493 | -0.2496 | -0.82934 | -0.76657 | -1.00476 | -0.56887 | -1.02608 | 0.433874 | -3.07292 | -1.5332 | -0.49312 | -3.44034 | -1.22286 | 2.793657 | -0.67314 | -1.09961 | -1.30727 | 0.853539 |
| 23.90139 | -1.51106 | -2.23918 | -0.22273 | -0.14282 | -0.0535 | -0.47293 | -0.52936 | -0.4067 | -0.13521 | 0.645542 | -0.42708 | -0.87169 | -1.12826 | -2.11358 | -1.35663 | 2.007775 | -0.74578 | -0.37043 | 0.386563 | 0.069742 |
| 23.91528 | -1.28977 | -2.21693 | -0.17398 | -0.25916 | 0.959957 | -0.28827 | -0.40173 | -0.46654 | 0.571032 | 0.363732 | 2.04977 | -0.36107 | -1.39209 | 0.16533 | -1.24791 | 0.896146 | -0.90779 | -0.02169 | 1.837767 | -0.64972 |
| 23.92917 | -1.12121 | -2.02424 | -0.31647 | -0.90934 | 1.363648 | -0.34546 | -0.91074 | -0.87183 | 0.775828 | -0.7004 | 2.051774 | -0.44244 | -1.30716 | 1.716921 | -1.30372 | -0.4188 | -1.2588 | 0.037803 | 1.754521 | -1.26079 |
| 23.94306 | -1.13481 | -1.85794 | -0.82582 | -1.42284 | 0.486201 | -0.56898 | -1.58249 | -1.20126 | 0.098378 | -1.94083 | -0.87544 | -1.15924 | -1.56332 | 0.354061 | -1.74556 | -1.46323 | -1.67263 | -0.33814 | -0.81641 | -3.1019 |
| 23.95694 | -1.2003 | -1.89704 | -1.60633 | -1.49171 | -1.06697 | -0.81463 | -2.24768 | -1.5137 | -1.05981 | -4.93079 | -4.89906 | -2.00239 | -2.37113 | -1.79993 | -2.66277 | -2.59708 | -2.50739 | -0.97232 | -4.98479 | -7.17023 |
| 23.97083 | -0.78323 | -1.88492 | -1.82879 | -1.14765 | -1.99988 | -0.92315 | -2.23307 | -1.52251 | -1.82357 | -9.6045 | -9.6639 | -2.83467 | -3.25247 | -2.89533 | -4.07914 | -3.66905 | -2.15189 | -1.35459 | -10.4152 | -12.8001 |
| 23.98472 | -0.20106 | -1.5881 | -1.34548 | -0.7311 | -2.11802 | -0.81611 | -1.63313 | -1.19851 | -1.84072 | -5.89263 | -6.16881 | -2.44253 | -1.73741 | -1.9969 | -2.49023 | -3.67443 | -0.93823 | -0.79799 | -5.96386 | -6.94949 |
| 23.99861 | 0.26432 | -1.11989 | -1.17357 | -0.38796 | -1.8169 | -0.66514 | -1.30848 | -1.0299 | -1.74135 | -1.56983 | -0.7101 | -1.41782 | -0.40669 | -0.22274 | -0.49938 | -1.10268 | 0.031722 | -0.32532 | -1.07702 | -1.7198 |
| 24.0125 | 0.56306 | -0.71155 | -0.97894 | -0.42716 | -1.39231 | -0.41627 | -0.98052 | -0.87604 | -1.06871 | 1.666301 | 1.973291 | -0.03093 | 0.49525 | 1.058429 | 0.644734 | 0.335426 | 0.433646 | -0.03534 | 1.326933 | 1.507129 |
| 24.02639 | 0.700508 | -0.1718 | -0.58746 | -0.48691 | -0.71219 | -0.07624 | -0.69521 | -0.45888 | -0.24001 | 2.251423 | 2.108199 | 0.418845 | 0.544317 | 1.184054 | 0.942456 | 0.760489 | 0.419298 | 0.110955 | 1.693358 | 2.929926 |
| 24.04028 | 0.569745 | 0.234675 | -0.11996 | -0.36687 | -0.06191 | 0.130304 | -0.2299 | -0.05021 | 0.327835 | 1.47705 | 1.362588 | 0.353919 | 0.503893 | 0.866922 | 0.668696 | 0.640122 | 0.290135 | 0.141028 | 1.309499 | 2.661752 |
| 24.05417 | 0.27208 | 0.363652 | 0.323717 | -0.09148 | 0.277353 | 0.250485 | 0.077418 | 0.162723 | 0.857491 | 0.733998 | 0.61572 | 0.127518 | 0.3634 | 0.379283 | 0.210093 | 0.368742 | 0.073816 | 0.013938 | 0.966836 | 1.727474 |
| 24.06806 | 0.007715 | 0.29156 | 0.542577 | 0.224679 | 0.406301 | 0.306804 | 0.150467 | 0.181923 | 1.196125 | 0.387122 | 0.08244 | -0.03358 | 0.184382 | 0.007807 | -0.06329 | 0.186072 | -0.09692 | -0.08559 | 0.893151 | 1.001489 |
| 24.08194 | -0.12986 | 0.227182 | 0.641333 | 0.534841 | 0.479596 | 0.395819 | 0.314831 | 0.19401 | 1.18155 | 0.38945 | -0.01836 | -0.00294 | 0.076424 | -0.07083 | -0.13594 | 0.223011 | -0.10511 | -0.1139 | 1.057095 | 0.862824 |
| 24.09583 | -0.23219 | 0.169361 | 0.817205 | 0.816877 | 0.61739 | 0.500431 | 0.525525 | 0.230606 | 1.303323 | 0.586824 | 0.115561 | 0.093102 | 0.058785 | 0.013388 | -0.11033 | 0.296762 | -0.04441 | -0.09006 | 1.526263 | 0.944142 |
| 24.10972 | -0.35929 | 0.099499 | 0.768132 | 0.835915 | 0.807751 | 0.411414 | 0.619106 | 0.256497 | 1.167473 | 0.687756 | 0.149309 | 0.131649 | 0.031724 | 0.044958 | -0.08734 | 0.276598 | -0.02597 | -0.08748 | 1.70034 | 0.87484 |
| 24.12361 | -0.47646 | 0.005189 | 0.581173 | 0.625303 | 0.857998 | 0.247824 | 0.618819 | 0.248767 | 0.629362 | 0.592117 | 0.046127 | 0.113341 | -0.07156 | -0.03753 | -0.10337 | 0.176713 | -0.05834 | -0.12159 | 1.419785 | 0.655455 |
| 24.1375 | -0.47264 | -0.11354 | 0.52599 | 0.352868 | 0.839629 | 0.0656 | 0.565732 | 0.19242 | 0.200112 | 0.380288 | -0.10004 | 0.010428 | -0.18555 | -0.20284 | -0.08768 | -0.04405 | -0.13196 | -0.10612 | 0.998838 | 0.28267 |
| 24.15139 | -0.35592 | -0.19012 | 0.478764 | 0.116083 | 0.696816 | -0.1026 | 0.531855 | 0.087644 | 0.07457 | 0.453895 | -0.05176 | -0.05452 | -0.15783 | -0.25592 | 0.010159 | -0.21669 | -0.17681 | -0.05328 | 0.93092 | 0.096117 |
| 24.16528 | -0.15824 | -0.05849 | 0.33604 | 0.099363 | 0.547575 | -0.07734 | 0.51943 | 0.164169 | 0.015561 | 0.854757 | 0.315513 | 0.108876 | 0.081206 | -0.03804 | 0.192042 | -0.06992 | -0.08214 | 0.055446 | 1.564476 | 0.541218 |
| 24.17917 | 0.040839 | 0.192798 | 0.349684 | 0.146589 | 0.696391 | 0.121409 | 0.586077 | 0.432206 | 0.238512 | 1.23269 | 0.660931 | 0.317827 | 0.278787 | 0.172739 | 0.393143 | 0.152962 | 0.06558 | 0.210672 | 2.188347 | 1.118477 |
| 24.19306 | 0.136701 | 0.319529 | 0.383999 | 0.163862 | 0.791854 | 0.304984 | 0.568336 | 0.649368 | 0.448933 | 1.055933 | 0.701591 | 0.386975 | 0.282772 | 0.172046 | 0.406678 | 0.212892 | 0.141332 | 0.335436 | 2.085269 | 1.206957 |
| 24.20694 | 0.164218 | 0.285367 | 0.248167 | 0.127249 | 0.731519 | 0.351185 | 0.494265 | 0.53368 | 0.370509 | 0.570392 | 0.521069 | 0.29444 | 0.173741 | 0.123323 | 0.242675 | 0.15743 | 0.1277 | 0.385807 | 1.432847 | 0.83129 |
| 24.22083 | 0.15853 | 0.138911 | 0.033222 | -0.15769 | 0.463692 | 0.202321 | 0.242199 | 0.221182 | -0.01455 | 0.062856 | 0.185203 | 0.069124 | -0.00015 | -0.02494 | 0.001213 | -0.04888 | -0.01894 | 0.299623 | 0.605601 | 0.262898 |
| 24.23472 | 0.168541 | 0.076266 | -0.14287 | -0.55578 | 0.087415 | -0.03422 | -0.00206 | -0.08498 | -0.45225 | -0.371 | -0.18997 | -0.20933 | -0.24634 | -0.21542 | -0.22103 | -0.32426 | -0.19902 | 0.164107 | -0.18737 | -0.32555 |
| 24.24861 | 0.249077 | 0.203004 | -0.1696 | -0.7306 | -0.27692 | -0.22075 | -0.08421 | -0.13441 | -0.75656 | -0.65369 | -0.44013 | -0.3347 | -0.37982 | -0.2968 | -0.33359 | -0.4627 | -0.25335 | 0.059106 | -0.68964 | -0.69805 |
| 24.2625 | 0.355904 | 0.400792 | 0.079853 | -0.50504 | -0.47832 | -0.22246 | -0.03329 | 0.001167 | -0.70377 | -0.78715 | -0.50487 | -0.31002 | -0.35507 | -0.28513 | -0.34351 | -0.40346 | -0.20601 | 0.028606 | -0.84961 | -0.77737 |
| 24.27639 | 0.421409 | 0.517317 | 0.411107 | -0.14772 | -0.5155 | -0.07472 | 0.043758 | 0.14612 | -0.31592 | -0.83094 | -0.47284 | -0.24887 | -0.29335 | -0.25628 | -0.35641 | -0.27425 | -0.15809 | -0.01894 | -0.78433 | -0.74035 |
| 24.29028 | 0.425154 | 0.550505 | 0.708722 | 0.214677 | -0.36885 | 0.099089 | 0.232846 | 0.177672 | 0.188871 | -0.89708 | -0.4755 | -0.24496 | -0.32254 | -0.32397 | -0.37734 | -0.20184 | -0.15978 | -0.10173 | -0.69127 | -0.70782 |
| 24.30417 | 0.422213 | 0.589758 | 0.836304 | 0.627006 | -0.21365 | 0.353335 | 0.529559 | 0.149253 | 0.609347 | -0.90175 | -0.44716 | -0.18131 | -0.27903 | -0.26706 | -0.36452 | -0.07481 | -0.08585 | -0.14888 | -0.43104 | -0.56468 |
| 24.31806 | 0.498735 | 0.629816 | 0.733391 | 0.858004 | -0.05253 | 0.51306 | 0.751212 | 0.148187 | 0.92903 | -0.79607 | -0.37706 | -0.09623 | -0.14077 | -0.1704 | -0.295 | 0.044759 | -0.00194 | -0.12825 | -0.19148 | -0.40454 |
| 24.33194 | 0.572362 | 0.619665 | 0.580771 | 0.571926 | 0.130781 | 0.539413 | 0.693197 | 0.061208 | 1.253099 | -0.59759 | -0.40293 | -0.18303 | -0.10745 | -0.30473 | -0.20876 | -0.12847 | -0.08366 | -0.16698 | -0.25115 | -0.40307 |
| 24.34583 | 0.53805 | 0.572583 | 0.53161 | 0.296595 | 0.209269 | 0.389413 | 0.573733 | 0.022141 | 1.009169 | -0.4678 | -0.42363 | -0.26398 | -0.15164 | -0.41541 | -0.15346 | -0.29018 | -0.15995 | -0.18829 | -0.27344 | -0.37066 |
| 24.35972 | 0.563524 | 0.658927 | 0.426292 | 0.283885 | 0.123215 | 0.120152 | 0.485973 | 0.123028 | 0.618191 | -0.37836 | -0.3352 | -0.18061 | -0.09226 | -0.31475 | -0.1133 | -0.19979 | -0.09458 | -0.13055 | -0.04845 | -0.26183 |
| 24.37361 | 0.614942 | 0.825487 | 0.415011 | 0.336467 | 0.111999 | -0.05639 | 0.407084 | 0.301549 | 0.387796 | -0.25603 | -0.17072 | -0.06506 | -0.00489 | -0.23253 | -0.05044 | -0.10982 | -0.03683 | -0.05009 | 0.112339 | -0.1885 |
| 24.3875 | 0.51736 | 0.917865 | 0.442258 | 0.343912 | 0.074298 | -0.08132 | 0.363057 | 0.503949 | 0.326389 | -0.14399 | -0.02073 | 0.030013 | 0.060742 | -0.15978 | -0.04873 | -0.0383 | 0.006539 | -0.02452 | 0.269616 | -0.13579 |
| 24.40139 | 0.402123 | 0.953409 | 0.491508 | 0.293832 | 0.277575 | 0.091109 | 0.495911 | 0.744676 | 0.649203 | -0.09053 | 0.102821 | 0.127083 | 0.148275 | -0.00437 | -0.06872 | 0.137428 | 0.110174 | 0.025832 | 0.568365 | -0.04531 |
| 24.41528 | 0.456614 | 1.059986 | 0.570543 | 0.441945 | 0.473812 | 0.411584 | 0.802389 | 1.062295 | 1.108402 | 0.035079 | 0.353184 | 0.36148 | 0.396416 | 0.318692 | 0.04472 | 0.526759 | 0.330234 | 0.213939 | 1.172376 | 0.229149 |
| 24.42917 | 0.808371 | 1.541081 | 0.669788 | 0.684832 | 0.598718 | 0.741792 | 1.223519 | 1.420358 | 1.451593 | 0.313004 | 0.985737 | 0.73513 | 0.747405 | 0.854006 | 0.313888 | 1.380658 | 0.692964 | 0.522107 | 2.373518 | 0.875128 |
| 24.44306 | 1.884584 | 2.966388 | 1.292614 | 0.726991 | 1.521328 | 1.354325 | 1.602821 | 2.151147 | 1.940889 | 1.032621 | 1.929586 | 1.245661 | 1.149065 | 1.165216 | 0.966502 | 2.230695 | 1.009464 | 0.85775 | 3.53242 | 1.849318 |
| 24.45694 | 2.985391 | 3.35914 | 2.608307 | 0.635182 | 2.412873 | 1.540262 | 2.471348 | 2.788562 | 2.305986 | 2.181683 | 3.097496 | 1.453331 | 1.577988 | 1.302112 | 1.71951 | 2.38408 | 0.765685 | 0.966503 | 3.958565 | 2.724591 |
| 24.47083 | 2.27298 | 2.157509 | 2.231791 | 1.113053 | 2.393092 | 1.360068 | 2.626051 | 2.6052 | 2.032889 | 3.130332 | 3.80946 | 1.755636 | 2.331443 | 1.576182 | 2.142494 | 2.660304 | 0.532226 | 0.851023 | 4.444037 | 3.080331 |
| 24.48472 | 1.583234 | 1.305387 | 1.912484 | 1.686723 | 1.862958 | 0.826335 | 2.417286 | 1.863949 | 1.442765 | 3.296463 | 3.919406 | 2.356278 | 2.585277 | 1.671531 | 2.291098 | 3.060677 | 0.42953 | 0.872163 | 4.700052 | 3.163586 |
| 24.49861 | 0.964471 | 0.676226 | 1.338668 | 1.472743 | 1.05552 | 0.402508 | 1.757256 | 0.920189 | 0.900122 | 2.530105 | 3.753095 | 2.750764 | 2.545568 | 1.534066 | 1.923767 | 3.591602 | 0.271097 | 0.866251 | 4.049192 | 3.438506 |
| 24.5125 | 0.520424 | 0.310712 | 0.886279 | 0.929873 | 0.242221 | 0.317714 | 1.047213 | 0.155171 | 0.473817 | 1.743767 | 3.920066 | 2.717285 | 2.365589 | 1.706871 | 1.60222 | 2.560556 | 0.397412 | 1.009488 | 3.246611 | 4.013187 |
| 24.52639 | 0.379381 | 0.125292 | 0.705228 | 0.483419 | -0.43178 | 0.394426 | 0.58943 | -0.24288 | 0.237635 | 1.324892 | 4.200754 | 2.392385 | 2.27729 | 2.045377 | 1.660415 | 1.795322 | 0.439567 | 0.89868 | 2.732344 | 4.134424 |
| 24.54028 | 0.508735 | -0.0071 | 0.5409 | 0.086156 | -0.18044 | 0.481964 | 0.196479 | -0.31352 | 0.247592 | 0.988031 | 3.691021 | 1.936375 | 1.8829 | 1.668121 | 1.557005 | 1.232865 | 0.279333 | 0.698582 | 1.826482 | 3.25565 |
| 24.55417 | 0.838175 | 0.053855 | 0.468663 | 0.045542 | 0.515638 | 0.479476 | -0.09767 | -0.06346 | 0.598475 | 0.771215 | 2.961807 | 1.534638 | 1.71785 | 1.199264 | 1.351474 | 0.8549 | 0.137384 | 0.563267 | 1.036224 | 2.22157 |

| | | | | | | | | | | | | | | | | | | | | |
|---|---|---|---|---|---|---|---|---|---|---|---|---|---|---|---|---|---|---|---|---|
| 24.56806 | 1.354806 | 0.338737 | 0.409828 | 0.223864 | 1.148653 | 0.548911 | 0.100727 | 0.203866 | 0.926491 | 1.09155 | 2.971124 | 1.220816 | 1.691662 | 0.891495 | 1.316835 | 0.692295 | 0.095658 | 0.662878 | 0.683204 | 1.64217 |
| 24.58194 | 1.502573 | 0.557296 | 0.271325 | 0.359871 | 1.309037 | 0.488343 | 0.260885 | 0.510443 | 1.263366 | 1.663681 | 2.889135 | 0.953546 | 1.466817 | 0.7981 | 1.182823 | 0.569668 | -0.02725 | 0.728696 | 0.613623 | 1.339058 |
| 24.59583 | 0.874367 | 0.574272 | 0.24575 | 0.372511 | 0.553168 | 0.320625 | 0.311994 | 0.720656 | 1.326896 | 1.962928 | 2.38652 | 0.704462 | 1.069682 | 0.69245 | 0.942609 | 0.445706 | 0.006781 | 0.669094 | 0.931229 | 1.108253 |
| 24.60972 | 0.314173 | 0.666864 | 0.523521 | 0.577238 | 0.164561 | 0.102958 | 0.377228 | 1.023174 | 0.670728 | 1.804116 | 2.010502 | 0.799502 | 0.704994 | 0.700491 | 0.915192 | 0.478265 | 0.240863 | 0.549215 | 1.369715 | 0.971361 |
| 24.62361 | -0.19409 | 0.85639 | 0.693785 | 0.601849 | -0.16689 | -0.04243 | 0.375372 | 1.09673 | 0.172758 | 1.680093 | 1.870416 | 0.857826 | 0.455095 | 0.705177 | 0.899041 | 0.501431 | 0.425369 | 0.540151 | 1.405219 | 0.831064 |
| 24.6375 | -0.40473 | 0.99348 | 0.433497 | 0.496493 | -0.23107 | -0.15091 | 0.058744 | 0.523726 | -0.17281 | 1.523074 | 1.624454 | 0.676152 | 0.430836 | 0.610671 | 0.7452 | 0.394948 | 0.328996 | 0.497017 | 1.042861 | 0.810131 |
| 24.65139 | -0.25498 | 0.896395 | 0.111833 | 0.164703 | -0.18762 | -0.33902 | -0.41653 | 0.051413 | -0.591 | 1.211043 | 1.342816 | 0.469528 | 0.443979 | 0.440583 | 0.519434 | 0.201834 | 0.164602 | 0.360805 | 0.669822 | 0.696962 |
| 24.66528 | -0.28884 | 0.732375 | -0.02673 | -0.27128 | 0.026567 | -0.54257 | -0.53303 | -0.20913 | -0.96002 | 0.947584 | 1.008095 | 0.31145 | 0.381272 | 0.283569 | 0.305479 | 0.054681 | 0.015765 | 0.207546 | 0.413629 | 0.565455 |
| 24.67917 | -0.21511 | 0.66892 | -0.10032 | -0.73284 | 0.707291 | -0.55124 | -0.25052 | -0.2774 | -0.76225 | 0.784907 | 0.72002 | 0.207593 | 0.356628 | 0.242648 | 0.169744 | 0.015316 | -0.08838 | 0.138775 | 0.295483 | 0.522504 |
| 24.69306 | -0.05086 | 0.432285 | -0.28821 | -1.00297 | 1.413096 | -0.29794 | 0.170899 | -0.35602 | 0.059512 | 0.743412 | 0.621951 | 0.161335 | 0.454919 | 0.335067 | 0.127838 | 0.043072 | -0.14657 | 0.157997 | 0.231531 | 0.516856 |
| 24.70694 | 0.08156 | 0.4036 | -0.41371 | -1.04848 | 1.443271 | 0.023771 | 0.354828 | -0.45761 | 1.021499 | 0.740581 | 0.728826 | 0.181515 | 0.517128 | 0.497584 | 0.149239 | 0.118179 | -0.08897 | 0.207346 | 0.215879 | 0.438564 |
| 24.72083 | 0.137165 | 0.328489 | -0.23865 | -0.72351 | 0.363491 | 0.207551 | 0.365083 | -0.73808 | 1.03457 | 0.692275 | 0.81821 | 0.247622 | 0.415684 | 0.566235 | 0.217843 | 0.167756 | 0.003791 | 0.197652 | 0.273974 | 0.37025 |
| 24.73472 | 0.148043 | 0.047077 | -0.06284 | -0.14585 | -0.16506 | 0.079824 | 0.30744 | -0.84043 | 0.591953 | 0.561701 | 0.775421 | 0.270697 | 0.207811 | 0.509734 | 0.245461 | 0.168336 | 0.062194 | 0.148101 | 0.294295 | 0.272972 |
| 24.74861 | 0.008926 | -0.48161 | -0.03201 | 0.494857 | -0.69473 | -0.17264 | 0.306719 | -0.78624 | -0.15818 | 0.474321 | 0.690169 | 0.236573 | 0.043173 | 0.435301 | 0.266414 | 0.214518 | 0.13176 | 0.125259 | 0.316395 | 0.17808 |
| 24.7625 | 0.021812 | -1.17208 | -0.49328 | 0.773035 | -0.93445 | -0.47718 | -0.09461 | -0.72834 | -0.96337 | 0.46374 | 0.657932 | 0.184009 | 0.044809 | 0.442941 | 0.272571 | 0.24052 | 0.174766 | 0.151368 | 0.323166 | 0.139895 |
| 24.77639 | -0.21763 | -1.93355 | -1.28188 | 0.276191 | -1.44153 | -0.74551 | -0.94874 | -0.69142 | -1.54958 | 0.38133 | 0.568769 | 0.011053 | 0.160205 | 0.355048 | 0.169767 | 0.118381 | 0.107328 | 0.168658 | 0.16486 | 0.158607 |
| 24.79028 | -0.78169 | -2.38436 | -1.66498 | -0.62754 | -1.99768 | -0.83856 | -1.65054 | -0.56607 | -1.86047 | 0.255878 | 0.319986 | -0.09694 | 0.140618 | 0.162783 | 0.074193 | -0.00449 | 0.001172 | 0.137375 | -0.01026 | 0.140828 |
| 24.80417 | -1.47913 | -2.75522 | -2.0988 | -1.60466 | -1.90936 | -0.90189 | -2.08927 | -0.56719 | -2.56344 | 0.147002 | 0.123314 | -0.18044 | 0.017258 | 0.064488 | 0.059609 | -0.08932 | -0.06408 | 0.09706 | -0.12813 | 0.105628 |
| 24.81806 | -2.09427 | -2.98669 | -2.26995 | -2.32386 | -1.51091 | -0.76526 | -2.17221 | -0.7791 | -1.60473 | 0.093649 | 0.09338 | -0.31206 | -0.02722 | 0.150143 | -0.01283 | -0.16832 | -0.10995 | 0.084026 | -0.20892 | 0.18206 |
| 24.83194 | -2.3522 | -2.83499 | -2.25374 | -2.62499 | -1.01058 | -0.63514 | -2.26421 | -0.79423 | -0.74857 | 0.054985 | 0.179814 | -0.42528 | -0.04238 | 0.282179 | -0.1057 | -0.17688 | -0.0946 | 0.043207 | -0.30766 | 0.175699 |
| 24.84583 | -1.95726 | -2.08984 | -1.91831 | -2.27382 | -0.74656 | -0.47195 | -1.86763 | -0.75979 | -0.14085 | 0.075823 | 0.225537 | -0.40507 | -0.1022 | 0.253642 | -0.13902 | -0.17011 | -0.0387 | -0.04999 | -0.34884 | 0.004546 |
| 24.85972 | -1.85268 | -1.32439 | -1.36197 | -1.51538 | -0.29504 | -0.30318 | -1.0827 | -0.73389 | 0.120784 | 0.206382 | 0.175262 | -0.23607 | -0.10632 | 0.102136 | -0.08034 | -0.16905 | 0.021715 | -0.09389 | -0.33678 | -0.08962 |
| 24.87361 | -1.63044 | -0.51339 | -0.7027 | -0.46134 | 0.070667 | -0.12634 | -0.52624 | -0.53116 | 0.117879 | 0.425899 | 0.080714 | 0.019214 | -0.03276 | -0.03354 | -0.00172 | -0.04683 | 0.090725 | -0.07846 | -0.21222 | -0.08762 |
| 24.8875 | -1.40719 | 0.213855 | -0.17938 | 0.231036 | 0.061962 | -0.06738 | -0.07428 | -0.31955 | -0.02567 | 0.583131 | 0.007186 | 0.188621 | 0.15479 | -0.11457 | -0.0353 | 0.09769 | 0.125485 | -0.04447 | -0.07554 | -0.13617 |
| 24.90139 | -0.79917 | 0.476349 | 0.065004 | 0.198405 | 0.022191 | -0.17376 | 0.211532 | -0.55972 | -0.27122 | 0.500604 | -0.06196 | 0.002457 | 0.498073 | -0.23907 | -0.29575 | -0.00334 | -0.03559 | -0.01326 | -0.11723 | -0.19628 |
| 24.91528 | -0.01832 | 0.09802 | 0.019982 | -0.42665 | 0.096311 | -0.32118 | 0.215787 | -1.14862 | -0.60918 | 0.17022 | -0.11886 | -0.63535 | 0.904423 | -0.43087 | -0.81046 | -0.47331 | -0.4122 | -0.02321 | -0.3925 | 0.016583 |
| 24.92917 | 0.124943 | -0.57586 | -0.52623 | -1.35296 | 0.211613 | -0.45956 | 0.007846 | -1.63389 | -0.92637 | -0.30634 | -0.32659 | -1.27214 | 0.927599 | -0.71982 | -1.42012 | -1.24199 | -0.93408 | -0.09491 | -0.89343 | 0.0858 |
| 24.94306 | -0.56939 | -1.05366 | -1.16108 | -2.02524 | -0.4311 | -0.62023 | -0.75564 | -1.71482 | -1.34341 | -1.25814 | -1.54976 | -1.55706 | -0.06003 | -1.19013 | -1.78198 | -1.55756 | -1.09399 | -0.23121 | -1.83278 | -0.58621 |
| 24.95694 | -1.42311 | -1.32913 | -2.14151 | -2.62579 | -1.58932 | -1.09056 | -1.5481 | -1.71217 | -2.05096 | -3.3572 | -4.95975 | -1.3675 | -2.34013 | -1.70984 | -2.54618 | -1.52149 | -1.32143 | -0.24886 | -4.04359 | -3.22973 |
| 24.97083 | -2.08684 | -1.59749 | -2.9959 | -3.2039 | -2.37518 | -1.41797 | -1.66012 | -1.5658 | -2.73064 | -4.1781 | -6.53134 | -1.36568 | -3.69895 | -1.18977 | -2.16574 | -1.51926 | -0.91714 | -0.6806 | -4.9607 | -5.19887 |
| 24.98472 | -2.27068 | -1.96299 | -3.01118 | -3.03657 | -2.36202 | -1.41272 | -1.33239 | -1.26318 | -2.81935 | -1.90704 | -3.08611 | -1.10687 | -1.04845 | -0.47919 | -0.6081 | -0.55174 | -0.38353 | -0.61871 | -1.51615 | -1.85627 |
| 24.99861 | -1.93091 | -2.13175 | -2.63988 | -1.86101 | -2.01769 | -1.13128 | -1.17074 | -1.16495 | -2.719 | 0.790195 | 0.523339 | -0.5847 | 0.632649 | -0.03786 | 0.480368 | 0.673435 | 0.099276 | -0.26423 | 0.860031 | 1.625534 |
| 25.0125 | -1.38786 | -1.96703 | -1.88329 | -0.74197 | -1.69119 | -0.84021 | -1.18066 | -1.31126 | -1.83829 | 1.98071 | 1.975256 | -0.07412 | 1.299967 | 0.285195 | 0.839458 | 1.145986 | 0.32835 | -0.0401 | 1.414003 | 1.743528 |
| 25.02639 | -0.69971 | -1.33499 | -0.93 | 0.168222 | -1.02871 | -0.52207 | -0.79786 | -0.9318 | -1.07328 | 1.986645 | 1.993025 | 0.465457 | 1.493146 | 0.470553 | 0.828704 | 1.118014 | 0.313737 | 0.113205 | 1.54926 | 1.253781 |
| 25.04028 | -0.08384 | -0.6713 | -0.01011 | 0.692381 | -0.55128 | -0.26247 | -0.24443 | -0.51017 | -0.53154 | 1.416185 | 1.903116 | 0.803732 | 1.10041 | 0.432401 | 0.668297 | 0.726151 | 0.137307 | 0.073126 | 1.650333 | 0.908722 |
| 25.05417 | 0.314904 | -0.17808 | 0.684187 | 0.655033 | -0.15654 | 0.084747 | 0.310937 | -0.18636 | 0.104782 | 0.828591 | 1.584764 | 0.935185 | 0.609754 | 0.247981 | 0.496143 | 0.277808 | -0.04732 | 0.0085 | 1.643309 | 0.663312 |
| 25.06806 | 0.513574 | 0.193776 | 0.993356 | 0.412675 | 0.148831 | 0.440806 | 0.61759 | 0.179085 | 0.665098 | 0.497984 | 1.157115 | 0.902592 | 0.355266 | 0.090782 | 0.419766 | 0.079724 | -0.17232 | -0.0157 | 1.501122 | 0.531171 |
| 25.08194 | 0.630757 | 0.40115 | 1.032531 | 0.346373 | 0.470588 | 0.709584 | 0.541539 | 0.578389 | 1.194625 | 0.477499 | 0.875725 | 0.84761 | 0.334385 | 0.089221 | 0.527393 | 0.180625 | -0.19704 | 0.002748 | 1.260593 | 0.559102 |
| 25.09583 | 0.727369 | 0.516171 | 1.121976 | 0.571895 | 0.66283 | 0.758966 | 0.451362 | 0.732294 | 1.537804 | 0.617554 | 0.700651 | 0.761643 | 0.409637 | 0.251218 | 0.665913 | 0.360437 | -0.11588 | 0.06853 | 0.943982 | 0.518431 |
| 25.10972 | 0.716418 | 0.567734 | 1.134141 | 1.090993 | 0.746739 | 0.632576 | 0.345114 | 0.709278 | 1.609957 | 0.737107 | 0.588213 | 0.600856 | 0.498255 | 0.428077 | 0.637415 | 0.465179 | -0.01409 | 0.102954 | 0.65486 | 0.334315 |
| 25.12361 | 0.685861 | 0.56006 | 1.095438 | 1.2894 | 0.804185 | 0.533796 | 0.340649 | 0.620251 | 1.662052 | 0.781401 | 0.550141 | 0.541946 | 0.464008 | 0.502161 | 0.428354 | 0.449822 | 0.089901 | 0.092696 | 0.475163 | 0.166454 |
| 25.1375 | 0.597022 | 0.522059 | 0.947307 | 1.200438 | 0.605615 | 0.496221 | 0.595391 | 0.504728 | 1.486708 | 0.667539 | 0.555188 | 0.481018 | 0.39476 | 0.458892 | 0.25732 | 0.363781 | 0.159848 | 0.12731 | 0.353453 | 0.1605 |
| 25.15139 | 0.530862 | 0.415816 | 0.757258 | 1.07783 | 0.221229 | 0.580686 | 0.838709 | 0.382282 | 1.491075 | 0.460694 | 0.620243 | 0.454656 | 0.343313 | 0.398459 | 0.196516 | 0.290183 | 0.179544 | 0.230688 | 0.285346 | 0.366334 |
| 25.16528 | 0.487498 | 0.30277 | 0.538435 | 0.968763 | -0.1185 | 0.68287 | 0.83156 | 0.30153 | 1.530536 | 0.244957 | 0.768483 | 0.380531 | 0.330958 | 0.378233 | 0.207429 | 0.252056 | 0.16198 | 0.326411 | 0.260912 | 0.60783 |
| 25.17917 | 0.45477 | 0.180029 | 0.325576 | 0.751123 | -0.1877 | 0.526178 | 0.664595 | 0.286723 | 1.313599 | 0.035493 | 0.74846 | 0.260914 | 0.257898 | 0.324962 | 0.210508 | 0.180856 | 0.11554 | 0.359868 | 0.279296 | 0.542742 |
| 25.19306 | 0.419781 | 0.08662 | 0.113977 | 0.398328 | 0.082773 | 0.253327 | 0.439652 | 0.289024 | 0.829558 | -0.09924 | 0.466817 | 0.137913 | 0.165375 | 0.186764 | 0.125906 | 0.123013 | 0.079312 | 0.339448 | 0.232487 | 0.247228 |
| 25.20694 | 0.368873 | 0.070046 | -0.05237 | 0.115304 | 0.47079 | 0.023826 | 0.27815 | 0.233768 | 0.305664 | -0.01094 | 0.11608 | 0.030589 | 0.162081 | 0.087894 | 0.031377 | 0.114713 | 0.088855 | 0.302832 | 0.024251 | -0.02201 |
| 25.22083 | 0.284677 | 0.099553 | -0.17842 | -0.03902 | 0.681947 | -0.0544 | 0.14417 | 0.14362 | -0.00671 | 0.325068 | -0.06128 | -0.021 | 0.300481 | 0.063971 | 0.014991 | 0.168619 | 0.113659 | 0.26191 | -0.23796 | -0.14929 |
| 25.23472 | 0.169991 | 0.112406 | -0.25429 | 0.154152 | 0.657803 | -0.01499 | 0.009934 | 0.091316 | -0.06443 | 0.532754 | -0.09244 | 0.001244 | 0.353075 | 0.095547 | 0.037586 | 0.201062 | 0.139425 | 0.192325 | -0.32936 | -0.06138 |
| 25.24861 | 0.125697 | 0.174626 | -0.24128 | 0.463485 | 0.464479 | 0.155203 | -0.00817 | 0.140464 | 0.125351 | 0.414449 | -0.03007 | 0.107662 | 0.282375 | 0.180115 | 0.061673 | 0.190932 | 0.17839 | 0.115326 | -0.09127 | 0.21503 |
| 25.2625 | 0.198444 | 0.265745 | -0.15074 | 0.665301 | 0.323496 | 0.323881 | 0.112389 | 0.231934 | 0.419995 | 0.151453 | 0.056399 | 0.23308 | 0.201763 | 0.290144 | 0.079488 | 0.16196 | 0.240137 | 0.095848 | 0.303491 | 0.521951 |
| 25.27639 | 0.298372 | 0.302845 | 0.015188 | 0.516195 | 0.319139 | 0.342622 | 0.203573 | 0.273843 | 0.569544 | -0.01905 | 0.229586 | 0.280962 | 0.163402 | 0.366417 | 0.032911 | 0.087859 | 0.237356 | 0.135604 | 0.666186 | 0.615719 |
| 25.29028 | 0.441455 | 0.396166 | 0.241576 | 0.28626 | 0.388321 | 0.23725 | 0.314092 | 0.32242 | 0.527824 | -0.05845 | 0.44219 | 0.259003 | 0.080352 | 0.337057 | -0.0448 | 0.022829 | 0.155978 | 0.162056 | 0.759572 | 0.430948 |
| 25.30417 | 0.662958 | 0.632145 | 0.507919 | 0.195312 | 0.523982 | 0.085498 | 0.507972 | 0.434179 | 0.408074 | 0.006584 | 0.584063 | 0.260031 | -0.01867 | 0.257966 | -0.01782 | 0.087099 | 0.0957 | 0.170377 | 0.539709 | 0.122447 |
| 25.31806 | 0.715383 | 0.791587 | 0.706352 | 0.22353 | 0.608505 | -0.03433 | 0.608118 | 0.515559 | 0.265327 | 0.288841 | 0.609869 | 0.350777 | -0.03534 | 0.241714 | 0.16107 | 0.371421 | 0.139959 | 0.141813 | 0.281803 | -0.08028 |
| 25.33194 | 0.629169 | 0.851104 | 0.851784 | 0.211176 | 0.737883 | -0.07417 | 0.496366 | 0.514745 | 0.247616 | 0.59041 | 0.513199 | 0.408798 | 0.103986 | 0.226335 | 0.273088 | 0.608535 | 0.190577 | 0.114338 | -0.00149 | -0.11342 |
| 25.34583 | 0.718479 | 0.849185 | 0.968465 | 0.219887 | 0.955763 | 0.069113 | 0.401035 | 0.494295 | 0.516998 | 0.606422 | 0.312022 | 0.259295 | 0.35209 | 0.211577 | 0.147437 | 0.45897 | 0.136677 | 0.16569 | -0.38303 | 0.057427 |
| 25.35972 | 0.835459 | 0.887377 | 1.025714 | 0.426253 | 0.962264 | 0.261265 | 0.529252 | 0.458447 | 0.851353 | 0.459518 | 0.212866 | 0.191847 | 0.501823 | 0.272814 | 0.080167 | 0.251393 | 0.143194 | 0.246925 | -0.49823 | 0.329242 |
| 25.37361 | 0.974429 | 0.946893 | 0.916382 | 0.753232 | 0.709266 | 0.331983 | 0.768767 | 0.474346 | 1.028135 | 0.368244 | 0.284889 | 0.259967 | 0.429916 | 0.384888 | 0.194 | 0.296357 | 0.224805 | 0.286679 | -0.22734 | 0.561571 |
| 25.3875 | 0.940083 | 0.88577 | 0.700735 | 0.851201 | 0.233844 | 0.308837 | 0.917415 | 0.4361 | 0.969903 | 0.284436 | 0.277915 | 0.234638 | 0.265705 | 0.4027 | 0.24629 | 0.329359 | 0.199756 | 0.262205 | 0.183408 | 0.671628 |
| 25.40139 | 0.706997 | 0.7165 | 0.493246 | 0.634869 | -0.21515 | 0.250924 | 0.992839 | 0.444245 | 0.789479 | 0.210618 | 0.222359 | 0.192628 | 0.122547 | 0.29196 | 0.271403 | 0.357197 | 0.132304 | 0.218975 | 0.654454 | 0.576341 |
| 25.41528 | 0.307704 | 0.937314 | 0.35141 | 0.551743 | -0.58032 | 0.183098 | 1.010289 | 0.526202 | 0.602189 | 0.380865 | 0.436691 | 0.454636 | -0.00823 | 0.35685 | 0.59012 | 0.935393 | 0.167024 | 0.17341 | 1.190814 | 0.449673 |
| 25.42917 | 0.019685 | 1.728772 | 0.499205 | 0.738162 | -0.57909 | 0.369738 | 1.135114 | 0.796674 | 0.688825 | 1.232985 | 1.192154 | 1.112897 | 0.045661 | 1.047031 | 1.245368 | 2.014461 | 0.473357 | 0.277534 | 1.939943 | 0.09932 |
| 25.44306 | 0.247524 | 3.18636 | 1.165718 | 0.893007 | 0.818325 | 1.141875 | 1.653686 | 1.621685 | 1.559982 | 2.856157 | 2.429623 | 1.827646 | 0.927909 | 2.693896 | 1.872797 | 2.477148 | 0.700327 | 0.685308 | 2.248439 | 0.258097 |
| 25.45694 | 1.445714 | 3.318703 | 2.107786 | 0.742697 | 2.840204 | 1.672312 | 2.00572 | 2.616502 | 2.29748 | 4.21167 | 3.527853 | 1.783012 | 2.617221 | 2.849285 | 2.102746 | 2.196843 | 1.093469 | 1.327285 | 2.463317 | 1.830452 |
| 25.47083 | 2.097693 | 2.132704 | 1.874543 | 1.006929 | 2.513545 | 1.288119 | 1.980883 | 2.799189 | 2.007703 | 3.447509 | 4.004393 | 1.481387 | 2.69344 | 2.545613 | 2.016538 | 1.687766 | 0.501977 | 1.367598 | 2.588607 | 2.827491 |
| 25.48472 | 1.693926 | 1.593761 | 2.104704 | 1.590408 | 2.049954 | 0.957573 | 1.814154 | 2.220187 | 1.814644 | 2.616396 | 3.750186 | 1.548977 | 2.242521 | 2.383126 | 1.631608 | 1.369979 | 0.332012 | 1.344441 | 2.724265 | 2.680126 |
| 25.49861 | 1.157635 | 0.951428 | 2.148726 | 1.734709 | 1.29917 | 0.852476 | 1.564373 | 1.578725 | 1.529896 | 1.876681 | 2.878892 | 1.677769 | 1.827831 | 1.895864 | 1.357268 | 1.128196 | 0.271764 | 1.306295 | 2.43045 | 1.911082 |
| 25.5125 | 0.945835 | 0.480107 | 1.733433 | 1.300712 | 0.409684 | 0.768663 | 1.060422 | 0.76299 | 1.386259 | 1.338363 | 1.858409 | 1.570724 | 1.36589 | 1.30088 | 0.99422 | 0.776527 | 0.235296 | 1.110998 | 2.026219 | 1.273643 |
| 25.52639 | 1.089211 | -0.10388 | 1.502552 | 0.791157 | -0.27997 | 0.636461 | 0.738074 | -0.0119 | 0.911543 | 0.795483 | 1.072159 | 1.213452 | 0.71434 | 0.773745 | 0.502455 | 0.309703 | 0.173175 | 0.874899 | 1.448032 | 0.37401 |
| 25.54028 | 1.129262 | -0.30034 | 1.152412 | 0.339308 | -0.67523 | 0.584244 | 0.569629 | -0.32462 | 0.316726 | 0.38203 | 0.636781 | 0.888197 | 0.13816 | 0.413888 | 0.104735 | -0.01909 | 0.199468 | 0.735671 | 0.779582 | -0.19305 |
| 25.55417 | 0.924407 | -0.20228 | 0.796468 | 0.01729 | -0.98702 | 0.517511 | 0.354637 | -0.29218 | -0.06087 | 0.279631 | 0.563988 | 0.717596 | -0.29866 | 0.344817 | -0.16589 | -0.25263 | 0.238162 | 0.627376 | 0.25285 | -0.46662 |
| 25.56806 | 0.515924 | -0.10036 | 0.504755 | -0.15995 | -1.03713 | 0.304308 | -0.00121 | -0.15234 | -0.20692 | 0.246494 | 0.562092 | 0.435915 | -0.70356 | 0.521109 | -0.39981 | -0.61674 | 0.187375 | 0.607255 | -0.09406 | -0.67435 |
| 25.58194 | 0.069419 | 0.278382 | 0.100966 | -0.241 | -0.2771 | 0.141901 | 0.047586 | -0.03875 | -0.273 | 0.152781 | 0.372471 | 0.133118 | -0.93459 | 0.580049 | -0.37709 | -0.72256 | 0.124454 | 0.466129 | -0.39278 | -0.53724 |
| 25.59583 | -0.19099 | 0.780717 | -0.22394 | -0.18465 | 0.448395 | 0.209793 | 0.237826 | 0.104892 | -0.02028 | 0.019852 | 0.094847 | -0.03453 | -0.52703 | 0.256634 | -0.18529 | -0.45736 | 0.05313 | 0.189317 | -0.67893 | -0.50859 |
| 25.60972 | -0.16013 | 0.917758 | -0.33148 | -0.085 | 0.619581 | 0.218243 | 0.335308 | 0.042055 | 0.334011 | -0.3205 | -0.21946 | -0.23845 | 0.116722 | -0.27769 | -0.08433 | -0.14464 | -0.10906 | 0.009022 | -0.75336 | -0.53574 |
| 25.62361 | 0.29954 | 0.781074 | -0.27691 | -0.01791 | -0.49785 | 0.285398 | 0.246777 | -0.0591 | 0.64617 | -0.72511 | -0.41829 | -0.32736 | 0.625972 | -0.56193 | -0.05543 | 0.049237 | -0.19001 | -0.13814 | -0.5215 | -0.42515 |
| 25.6375 | 0.905113 | 0.576468 | -0.10973 | 0.065853 | -1.08317 | 0.360011 | 0.152607 | -0.09883 | 0.739299 | -0.82399 | -0.38122 | -0.29807 | 0.399603 | -0.57662 | -0.17815 | 0.03428 | -0.21266 | -0.30865 | 0.411463 | -0.17056 |
| 25.65139 | 1.025877 | 0.426644 | 0.106409 | -0.00511 | -0.63122 | 0.344317 | -0.01465 | -0.11528 | 0.618174 | -1.03348 | -0.23576 | -0.16954 | -0.03227 | -0.51708 | -0.29119 | -0.27552 | -0.25051 | -0.55801 | 0.450217 | -0.37632 |
| 25.66528 | 0.696855 | 0.638981 | 0.28903 | 0.102029 | -0.41866 | 0.356343 | -0.36324 | -0.11332 | 0.485356 | -1.37043 | -0.15559 | -0.01903 | -0.61374 | -0.61487 | -0.35468 | -0.54572 | -0.38154 | -0.84123 | 0.030519 | -0.79346 |
| 25.67917 | 0.194635 | 0.490742 | 0.460619 | 0.161688 | -0.0766 | 0.406705 | -0.71955 | 0.162747 | 0.279153 | -1.59332 | -0.09054 | 0.127285 | -1.12817 | -0.74603 | -0.07806 | -0.79969 | -0.41375 | -0.94921 | -0.43018 | -0.81922 |
| 25.69306 | -0.23583 | 0.311928 | 0.686287 | -0.08469 | 0.157333 | 0.286384 | -0.82571 | 0.330108 | 0.08968 | -1.25964 | -0.05135 | 0.158472 | -1.38981 | -1.08059 | 0.406185 | -0.90179 | -0.11036 | -0.61436 | -0.85503 | -0.46917 |
| 25.70694 | -0.54766 | 0.179311 | 0.704746 | -0.66524 | 0.627628 | -0.04978 | -0.45605 | 0.596498 | -0.12419 | -0.70675 | -0.10592 | -0.06261 | -1.16368 | -1.46235 | 0.486336 | -0.91899 | 0.328611 | -0.10324 | -1.33892 | 0.302221 |
| 25.72083 | -0.79066 | 0.245786 | 0.573601 | -1.13181 | 0.547716 | -0.3117 | -0.00323 | 0.230723 | -0.36524 | -0.24542 | -0.30646 | -0.27451 | -0.72836 | -1.57634 | 0.364926 | -0.55788 | 0.602087 | 0.163965 | -1.58693 | 0.804777 |
| 25.73472 | -0.74457 | 0.117105 | 0.333453 | -1.63025 | 0.96108 | -0.40637 | -0.12305 | -0.29956 | -0.64004 | 0.089288 | -0.54109 | -0.45346 | -0.45746 | -1.61414 | 0.191352 | -0.21345 | 0.577739 | 0.157212 | -1.11423 | 0.89509 |
| 25.74861 | -0.35607 | -0.34305 | -0.0511 | -2.26985 | 1.149765 | -0.4325 | -0.07239 | -0.78432 | -0.82336 | 0.447524 | -0.78255 | -0.64906 | -0.30616 | -1.36823 | -0.05429 | -0.01803 | 0.248805 | -0.05376 | -0.69311 | 0.625284 |
| 25.7625 | 0.067899 | -0.86597 | -0.64868 | -2.4004 | 0.836449 | -0.52356 | -0.2192 | -1.16177 | -0.61476 | 0.749396 | -1.15082 | -0.81144 | -0.27738 | -1.01042 | -0.24788 | 0.003199 | -0.20495 | -0.39674 | -0.25173 | -0.08302 |
| 25.77639 | -0.00039 | -1.2843 | -1.24763 | -2.04056 | -0.20023 | -0.82849 | -0.80931 | -1.13055 | -0.68302 | 0.743463 | -1.46394 | -0.91565 | -0.27657 | -0.84641 | -0.51317 | -0.04155 | -0.67456 | -0.62783 | 0.054356 | -0.69909 |
| 25.79028 | -0.53297 | -1.42855 | -1.8209 | -1.52373 | -1.3707 | -1.08205 | -1.43068 | -0.99263 | -0.47754 | 0.349054 | -1.43008 | -1.01348 | -0.24558 | -0.73083 | -0.80806 | 0.021516 | -0.95133 | -0.73136 | 0.273558 | -1.12562 |
| 25.80417 | -1.20469 | -1.57275 | -2.51115 | -1.11143 | -2.19323 | -1.24166 | -2.03377 | -0.91395 | -0.49106 | -0.20989 | -1.17385 | -1.15523 | -0.14677 | -0.73511 | -1.03032 | -0.01735 | -1.04088 | -0.91304 | 0.447851 | -1.40027 |
| 25.81806 | -1.75371 | -1.87566 | -3.16594 | -0.90384 | -2.47182 | -1.35365 | -2.50152 | -0.99096 | -0.86772 | -0.89081 | -0.99678 | -1.47606 | 0.004965 | -0.77343 | -1.22361 | -0.24005 | -0.88062 | -1.02603 | 0.309516 | -1.40976 |

| | | | | | | | | | | | | | | | | | | | | |
|---|---|---|---|---|---|---|---|---|---|---|---|---|---|---|---|---|---|---|---|---|
| 25.83194 | -1.7237 | -1.85399 | -3.39371 | -0.74264 | -2.50849 | -1.24355 | -2.20937 | -1.30174 | -1.24149 | -1.20317 | -0.92883 | -1.65903 | 0.102035 | -0.69222 | -1.4159 | -0.5936 | -0.58765 | -1.19651 | -0.05218 | -1.21978 |
| 25.84583 | -2.05839 | -1.77006 | -2.89095 | -0.59048 | -2.57224 | -0.98015 | -1.57451 | -1.38215 | -1.29855 | -1.10737 | -0.93498 | -1.61407 | 0.069618 | -0.59548 | -1.29732 | -0.7325 | -0.30038 | -1.18578 | -0.62486 | -1.13298 |
| 25.85972 | -2.07903 | -1.79012 | -2.06642 | -0.69436 | -2.43899 | -0.71327 | -1.16836 | -1.24843 | -1.28707 | -0.88627 | -0.95154 | -1.44175 | -0.08971 | -0.62003 | -1.03682 | -0.75114 | -0.14576 | -1.00959 | -1.32311 | -1.01438 |
| 25.87361 | -1.75778 | -1.67657 | -1.07959 | -0.75337 | -1.39233 | -0.23436 | -0.81215 | -0.74366 | -1.06516 | -0.61828 | -1.03265 | -1.08811 | -0.3587 | -0.81084 | -0.69945 | -0.68619 | -0.12194 | -0.86646 | -1.76216 | -0.87321 |
| 25.8875 | -1.17943 | -1.44367 | -0.16851 | -0.84525 | -0.49237 | 0.168986 | -0.3447 | -0.18208 | -0.4287 | -0.33196 | -1.03002 | -0.67575 | -0.50271 | -0.93455 | -0.34525 | -0.5132 | -0.12216 | -0.69235 | -1.90973 | -0.71692 |
| 25.90139 | -0.61785 | -1.10778 | 0.558222 | -0.96761 | 0.461394 | 0.169128 | -0.12528 | 0.241946 | 0.295468 | -0.21612 | -0.8317 | -0.54365 | -0.38102 | -0.85031 | -0.1014 | -0.39175 | -0.18324 | -0.5225 | -1.85351 | -0.48234 |
| 25.91528 | -0.27698 | -0.83334 | 0.923648 | -0.93677 | 1.039758 | -0.26793 | -0.35823 | 0.164514 | 0.597583 | -0.36526 | -0.53668 | -0.9123 | 0.045117 | -1.02292 | -0.20845 | -0.67621 | -0.47209 | -0.28007 | -1.76524 | -0.12598 |
| 25.92917 | -0.26559 | -0.84564 | 0.703985 | -0.97162 | 0.66809 | -0.82021 | -1.10449 | -0.43394 | 0.185978 | -0.901 | -0.79179 | -1.60912 | 0.208819 | -1.66026 | -0.69489 | -1.45299 | -0.77768 | -0.12718 | -2.02037 | -0.01434 |
| 25.94306 | -0.59256 | -1.00335 | -0.35927 | -1.16544 | -0.30119 | -1.30329 | -1.7722 | -1.24345 | -0.70184 | -1.41865 | -1.80201 | -1.8388 | -0.61582 | -2.54216 | -1.51185 | -2.00631 | -0.76625 | -0.21392 | -3.18426 | -1.39041 |
| 25.95694 | -0.98604 | -1.2863 | -1.39165 | -1.52068 | -1.31335 | -1.75116 | -2.45211 | -2.0192 | -1.59651 | -2.54041 | -4.05365 | -1.81875 | -2.1778 | -2.45411 | -2.10889 | -1.32942 | -0.79337 | -0.62902 | -4.73994 | -3.39973 |
| 25.97083 | -1.29704 | -1.4846 | -1.98666 | -1.63869 | -1.80686 | -1.79814 | -2.86828 | -2.25106 | -2.07257 | -3.5103 | -4.19936 | -1.32417 | -2.11492 | -1.71356 | -1.62601 | -1.40718 | -0.60735 | -1.05494 | -4.70506 | -3.32851 |
| 25.98472 | -1.5326 | -1.54727 | -1.87707 | -1.13598 | -1.55139 | -1.33025 | -2.32151 | -1.81558 | -1.76495 | -2.04208 | -1.46407 | -0.64992 | -0.70124 | -0.41251 | -0.82106 | -0.76591 | -0.58751 | -0.51511 | -2.49747 | -1.50208 |
| 25.99861 | -1.65986 | -1.6424 | -1.86408 | -0.73296 | -1.04981 | -0.97407 | -1.52494 | -1.46594 | -1.60719 | -0.61916 | -0.34733 | -0.36394 | 0.093166 | 0.338856 | -0.32426 | -0.30904 | -0.49909 | -0.30617 | -1.0637 | 0.139138 |
| 26.0125 | -1.48299 | -1.47127 | -1.64152 | -0.57155 | -0.45076 | -0.67662 | -0.85733 | -1.11702 | -2.04031 | 0.253164 | 0.088258 | -0.35705 | 0.394286 | 0.59475 | 0.019622 | -0.18201 | -0.23272 | -0.24173 | -0.02108 | 0.674178 |
| 26.02639 | -1.12374 | -1.04479 | -1.20065 | -0.32152 | 0.019253 | -0.41469 | -0.31524 | -0.65105 | -1.84809 | 0.713473 | 0.397287 | -0.41421 | 0.527505 | 0.27678 | 0.304184 | 0.058602 | 0.160021 | -0.09665 | 0.908926 | 0.909531 |
| 26.04028 | -0.71516 | -0.58939 | -0.58463 | -0.07437 | 0.227869 | -0.04953 | 0.252075 | -0.17374 | -0.64771 | 0.977149 | 0.685816 | -0.32838 | 0.475906 | 0.080311 | 0.273006 | 0.204971 | 0.411066 | -0.00507 | 1.556565 | 0.916647 |
| 26.05417 | -0.41768 | -0.14935 | -0.04444 | 0.035037 | 0.233845 | 0.427375 | 0.417711 | 0.133677 | 0.43449 | 0.960238 | 0.744576 | -0.03138 | 0.369727 | -0.04661 | 0.137315 | 0.209319 | 0.427798 | 0.001573 | 1.733314 | 0.710514 |
| 26.06806 | -0.23312 | 0.157195 | 0.385005 | 0.129759 | 0.232771 | 0.521401 | 0.412679 | 0.304709 | 1.036712 | 0.813973 | 0.48951 | 0.23331 | 0.26731 | -0.31457 | 0.071664 | 0.111806 | 0.26323 | -0.01291 | 1.718402 | 0.476745 |
| 26.08194 | -0.09895 | 0.410366 | 0.66406 | 0.416082 | 0.233543 | 0.298948 | 0.333664 | 0.31686 | 0.984004 | 0.65644 | 0.186305 | 0.352368 | 0.302117 | -0.60724 | 0.0274 | -0.02058 | 0.100873 | -0.01988 | 1.513428 | 0.266977 |
| 26.09583 | 0.017175 | 0.570846 | 0.77634 | 0.772831 | 0.261913 | 0.192601 | 0.374757 | 0.37455 | 0.752577 | 0.575085 | -0.05084 | 0.418061 | 0.333044 | -0.58567 | 0.024271 | 0.069487 | -0.01598 | -0.02928 | 1.310617 | 0.16142 |
| 26.10972 | 0.145716 | 0.675422 | 0.772268 | 1.114071 | 0.298905 | 0.19164 | 0.588601 | 0.490352 | 0.563695 | 0.420237 | -0.09247 | 0.584734 | 0.296862 | -0.22701 | 0.137223 | 0.459408 | -0.01392 | -0.08977 | 1.340103 | 0.101011 |
| 26.12361 | 0.230715 | 0.726184 | 0.904902 | 1.625914 | 0.30918 | 0.22143 | 0.849725 | 0.605002 | 0.546011 | 0.100104 | 0.037822 | 0.664947 | 0.293017 | -0.01287 | 0.260892 | 0.621307 | -0.00849 | -0.05098 | 1.175645 | 0.033233 |
| 26.1375 | 0.285422 | 0.620421 | 1.16181 | 1.919012 | 0.323895 | 0.19531 | 1.045058 | 0.69832 | 0.637396 | -0.31018 | 0.170294 | 0.447361 | 0.339783 | 0.032799 | 0.225734 | 0.248137 | -0.10975 | 0.052611 | 0.555613 | -0.00067 |
| 26.15139 | 0.370922 | 0.58806 | 1.251338 | 1.848688 | 0.331496 | 0.234718 | 1.208741 | 0.72187 | 0.722865 | -0.57621 | 0.189843 | 0.230842 | 0.293679 | 0.07269 | 0.052089 | -0.10433 | -0.29978 | 0.084089 | -0.10779 | -0.07376 |
| 26.16528 | 0.501776 | 0.671088 | 1.219492 | 1.5301 | 0.344249 | 0.392595 | 1.440317 | 0.688389 | 0.901884 | -0.52385 | 0.055059 | 0.088328 | 0.142615 | 0.132647 | -0.11429 | -0.09489 | -0.33674 | 0.085423 | -0.43386 | -0.05638 |
| 26.17917 | 0.612651 | 0.720913 | 0.977772 | 1.1607 | 0.315198 | 0.509099 | 1.409507 | 0.507616 | 1.076826 | -0.37813 | -0.20981 | 0.019901 | 0.085129 | 0.049884 | -0.21219 | -0.11964 | -0.31563 | 0.128399 | -0.71879 | 0.031543 |
| 26.19306 | 0.649023 | 0.758088 | 0.630502 | 0.65768 | 0.208709 | 0.495038 | 1.124476 | 0.188518 | 1.086039 | -0.18483 | -0.49849 | -0.02469 | 0.099448 | -0.12588 | -0.25444 | -0.25388 | -0.21085 | 0.189911 | -0.97301 | 0.157952 |
| 26.20694 | 0.68712 | 0.801242 | 0.399879 | 0.323899 | 0.107066 | 0.459003 | 0.708911 | -0.00967 | 0.945812 | 0.082212 | -0.58917 | 0.065227 | 0.025498 | -0.20254 | -0.12165 | -0.06925 | 0.06158 | 0.211775 | -0.86142 | 0.177565 |
| 26.22083 | 0.730627 | 0.834812 | 0.231786 | 0.193612 | 0.072906 | 0.458226 | 0.392004 | 0.050153 | 0.784701 | 0.306582 | -0.36593 | 0.238454 | 0.018656 | -0.09993 | 0.120724 | 0.306649 | 0.331223 | 0.218978 | -0.38136 | 0.075799 |
| 26.23472 | 0.696004 | 0.743466 | 0.049639 | 0.097718 | 0.043236 | 0.4366 | 0.202873 | 0.208366 | 0.72396 | 0.275152 | 0.093684 | 0.297684 | 0.175991 | 0.01542 | 0.2601 | 0.407054 | 0.397365 | 0.267861 | 0.296318 | 0.030672 |
| 26.24861 | 0.595909 | 0.632443 | -0.03612 | 0.000209 | 0.096102 | 0.368307 | 0.223139 | 0.278631 | 0.657122 | 0.027506 | 0.496735 | 0.255249 | 0.299265 | 0.083446 | 0.204959 | 0.311015 | 0.318408 | 0.28304 | 1.016491 | -0.00309 |
| 26.2625 | 0.60884 | 0.554418 | -0.03043 | -0.05545 | 0.14427 | 0.31714 | 0.349802 | 0.205583 | 0.64571 | -0.23652 | 0.630034 | 0.149528 | 0.374102 | 0.106081 | -0.04526 | 0.116812 | 0.175114 | 0.307865 | 1.385385 | 0.084594 |
| 26.27639 | 0.641853 | 0.538684 | -0.00418 | -0.02552 | 0.194517 | 0.282539 | 0.351489 | 0.105152 | 0.726162 | -0.30993 | 0.519135 | 0.028427 | 0.428537 | 0.033407 | -0.2392 | -0.02877 | 0.021035 | 0.319527 | 1.083173 | 0.354706 |
| 26.29028 | 0.615287 | 0.547881 | -0.00154 | 0.076823 | 0.169237 | 0.228436 | 0.294933 | 0.094789 | 0.773001 | -0.12804 | 0.309418 | -0.02521 | 0.406147 | 0.021234 | -0.17618 | 0.036094 | -0.01704 | 0.295609 | 0.564121 | 0.509948 |
| 26.30417 | 0.449797 | 0.604051 | -0.00745 | 0.236837 | 0.157401 | 0.170328 | 0.285629 | 0.218751 | 0.651111 | 0.202597 | 0.133352 | 0.00029 | 0.306944 | 0.074624 | 0.12636 | 0.200666 | 0.06162 | 0.249378 | -0.05567 | 0.443604 |
| 26.31806 | 0.334811 | 0.632886 | 0.008579 | 0.456624 | 0.155013 | 0.118566 | 0.321296 | 0.400744 | 0.45605 | 0.552463 | 0.121429 | 0.107189 | 0.221677 | 0.223955 | 0.475574 | 0.419487 | 0.249462 | 0.160423 | -0.71312 | 0.318961 |
| 26.33194 | 0.268464 | 0.58705 | 0.075751 | 0.696869 | 0.148242 | 0.074919 | 0.433146 | 0.591393 | 0.331003 | 0.663895 | 0.15472 | 0.206603 | 0.146946 | 0.451561 | 0.64191 | 0.600797 | 0.37431 | 0.14198 | -1.08377 | 0.236701 |
| 26.34583 | 0.338895 | 0.58298 | 0.199586 | 0.910761 | 0.22675 | 0.045873 | 0.53156 | 0.607972 | 0.341651 | 0.510554 | 0.116047 | 0.239803 | 0.121967 | 0.619597 | 0.570426 | 0.679737 | 0.416173 | 0.271747 | -1.01027 | 0.182318 |
| 26.35972 | 0.539364 | 0.675902 | 0.334348 | 1.121934 | 0.362952 | 0.058695 | 0.588963 | 0.630878 | 0.429725 | 0.363388 | 0.151118 | 0.2664 | 0.23064 | 0.599089 | 0.468339 | 0.660431 | 0.382687 | 0.583344 | -0.15173 | 0.361019 |
| 26.37361 | 0.658974 | 0.828231 | 0.498075 | 1.171432 | 0.377701 | 0.052317 | 0.64825 | 0.636766 | 0.473739 | 0.338372 | 0.287599 | 0.321898 | 0.382916 | 0.562993 | 0.355819 | 0.647167 | 0.29455 | 0.682045 | 1.127638 | 0.67053 |
| 26.3875 | 0.758513 | 0.962886 | 0.701455 | 1.058578 | 0.321882 | -0.00031 | 0.763574 | 0.561809 | 0.432446 | 0.377171 | 0.485084 | 0.362282 | 0.42022 | 0.672381 | 0.341461 | 0.78804 | 0.245995 | 0.524202 | 1.99653 | 0.823927 |
| 26.40139 | 0.850694 | 0.900541 | 0.893109 | 0.834485 | 0.389195 | 0.033002 | 0.875652 | 0.603249 | 0.468629 | 0.450971 | 0.804638 | 0.439964 | 0.447771 | 0.832853 | 0.470199 | 0.929769 | 0.280627 | 0.400806 | 2.147417 | 0.810542 |
| 26.41528 | 0.826989 | 0.912791 | 1.186388 | 0.786072 | 0.409957 | 0.231188 | 0.929459 | 0.756425 | 0.547417 | 0.739594 | 1.261942 | 0.739936 | 0.409498 | 1.195045 | 0.82434 | 1.381193 | 0.584809 | 0.484387 | 2.226286 | 0.711812 |
| 26.42917 | 0.297813 | 1.463249 | 1.724163 | 0.829758 | 0.642743 | 0.557846 | 1.141032 | 1.105231 | 0.587586 | 1.381665 | 1.933843 | 1.32034 | 0.345026 | 2.176145 | 1.299613 | 2.336149 | 1.387663 | 0.892137 | 2.381035 | 0.730411 |
| 26.44306 | 0.416182 | 2.701518 | 2.088685 | 1.090932 | 1.689256 | 1.111289 | 1.672298 | 1.733237 | 0.807921 | 2.43356 | 3.193217 | 2.188237 | 1.349198 | 2.801344 | 1.498916 | 2.830694 | 2.143854 | 1.571146 | 2.051658 | 1.336158 |
| 26.45694 | 1.671832 | 3.21951 | 2.333823 | 1.572052 | 2.978572 | 1.266373 | 2.473544 | 1.838314 | 1.040078 | 3.004803 | 5.901337 | 2.619642 | 2.248779 | 2.730742 | 1.971776 | 2.394744 | 1.2699 | 2.336702 | 2.36242 | 2.739463 |
| 26.47083 | 1.7547 | 2.458431 | 1.732196 | 1.548769 | 2.697986 | 0.967111 | 2.618913 | 2.20807 | 1.300564 | 2.741508 | 5.857196 | 2.14173 | 2.242633 | 3.080331 | 2.321209 | 1.995577 | 0.627005 | 2.341158 | 2.98609 | 2.948786 |
| 26.48472 | 1.239249 | 1.811654 | 1.174547 | 1.687414 | 2.063508 | 0.720773 | 2.75582 | 2.747934 | 1.72067 | 2.293411 | 4.116107 | 2.004922 | 1.821742 | 3.058451 | 2.499131 | 1.915276 | 0.412965 | 2.281424 | 3.162416 | 2.643245 |
| 26.49861 | 0.817687 | 0.779358 | 1.050715 | 1.312385 | 1.669208 | 0.779764 | 2.399233 | 2.19205 | 2.117846 | 1.846597 | 3.03025 | 1.931119 | 1.506224 | 2.546576 | 2.305739 | 1.631264 | 0.257375 | 1.92175 | 2.796191 | 2.230304 |
| 26.5125 | 0.475508 | -0.01655 | 1.09141 | 0.659907 | 1.147311 | 0.850933 | 1.613681 | 0.879316 | 2.097737 | 1.472169 | 2.004153 | 1.937042 | 1.238051 | 1.836043 | 2.048619 | 1.21212 | 0.259466 | 1.523522 | 2.499642 | 1.336341 |
| 26.52639 | 0.478945 | -0.49856 | 1.122247 | 0.138503 | 0.875886 | 0.863063 | 0.989758 | -0.11507 | 1.726061 | 1.19563 | 1.114259 | 1.735384 | 0.76991 | 1.141745 | 1.64762 | 0.791901 | 0.372321 | 1.042774 | 2.427047 | 0.563815 |
| 26.54028 | 0.804227 | -0.55254 | 1.094805 | -0.23938 | 0.736179 | 0.789283 | 0.52719 | -0.66665 | 1.101545 | 0.870344 | 0.286757 | 1.248113 | 0.081896 | 0.587742 | 0.933524 | 0.492501 | 0.535675 | 0.562386 | 1.717087 | 0.041008 |
| 26.55417 | 1.012502 | -0.41188 | 1.060333 | -0.29015 | 0.751439 | 0.669419 | 0.423031 | -0.69924 | 0.468965 | 0.521574 | -0.28931 | 0.791922 | -0.59378 | 0.070755 | 0.160428 | 0.122458 | 0.682476 | 0.320275 | 0.666754 | -0.20254 |
| 26.56806 | 1.225487 | 0.126214 | 0.969385 | -0.03152 | 0.805172 | 0.538942 | 0.274125 | -0.47605 | 0.055416 | 0.202678 | -0.49769 | 0.315761 | -0.98796 | -0.25569 | -0.43674 | 0.002507 | 0.728899 | 0.682829 | -0.168 | -0.38633 |
| 26.58194 | 1.251701 | 0.723672 | 0.719195 | 0.26472 | 0.69653 | 0.39785 | 0.081695 | -0.02535 | -0.28904 | -0.51359 | -0.48972 | -0.05307 | -0.92899 | -0.38259 | -0.83882 | -0.07116 | 0.568848 | 1.057281 | -0.58214 | -0.56022 |
| 26.59583 | 0.800872 | 1.272319 | 0.377458 | 0.375378 | 0.379225 | 0.157108 | -0.24085 | 0.538579 | -0.49456 | -1.16693 | -0.49331 | -0.31476 | -0.8304 | -0.59141 | -0.86321 | -0.1842 | 0.252242 | 1.075575 | -1.06648 | -0.55079 |
| 26.60972 | 0.142676 | 1.247853 | 0.04054 | 0.348502 | -0.07243 | -0.1588 | -0.26311 | 1.002584 | -0.77444 | -1.43155 | -0.59323 | -0.60611 | -0.83454 | -0.89567 | -0.75825 | -0.33927 | 0.07342 | 0.835111 | -1.5529 | -0.60702 |
| 26.62361 | -0.34075 | 0.734626 | -0.12854 | 0.258591 | -0.70089 | -0.29117 | 0.082259 | 1.019508 | -0.87041 | -1.05483 | -0.76669 | -0.82919 | -0.72397 | -1.37613 | -0.61231 | -0.5391 | -0.0677 | 0.472303 | -1.41927 | -0.39696 |
| 26.6375 | -0.30706 | 0.041784 | -0.08014 | 0.217498 | -1.48198 | -0.18129 | 0.560224 | 0.458677 | -0.1215 | -0.34124 | -0.97239 | -0.93382 | -0.5424 | -1.69543 | -0.50315 | -0.96072 | -0.2555 | -0.04707 | -0.44065 | -0.08223 |
| 26.65139 | 0.06076 | -0.43162 | 0.063467 | 0.070523 | -1.79115 | -0.02713 | 0.464913 | -0.02394 | 0.561204 | -0.52285 | -1.24298 | -0.86735 | -0.46809 | -1.92139 | -0.41453 | -1.31949 | -0.48085 | -0.57835 | 0.447979 | 0.038728 |
| 26.66528 | 0.087468 | -0.75093 | 0.012111 | -0.21147 | -1.70661 | 0.284764 | 0.34058 | -0.62112 | 0.950092 | -0.74229 | -1.29596 | -0.82448 | -0.63875 | -1.46876 | -0.31243 | -1.04293 | -0.58916 | -0.94737 | 0.334735 | -0.44937 |
| 26.67917 | 0.296616 | -0.60663 | -0.10726 | -0.52969 | -1.34558 | 0.645339 | 0.100528 | -0.97776 | 1.336281 | -1.04782 | -1.36559 | -0.91561 | -0.80754 | -0.86952 | -0.21891 | -0.79555 | -0.62791 | -0.96177 | 0.053775 | -1.11209 |
| 26.69306 | 0.462921 | -0.78994 | -0.1708 | -0.89765 | -0.47994 | 0.95378 | -0.50503 | -0.80894 | 1.647805 | -1.21796 | -1.29562 | -0.95793 | -0.78189 | -0.26516 | -0.19732 | -0.62162 | -0.57392 | -0.50368 | -0.57342 | -1.00208 |
| 26.70694 | 0.391959 | -0.08309 | -0.12555 | -1.05979 | 0.267794 | 0.967052 | -0.68974 | -0.39692 | 1.163728 | -1.19766 | -1.29354 | -0.85671 | -0.7314 | 0.29843 | -0.18304 | -0.37971 | -0.51082 | -0.0535 | -1.45113 | -0.75631 |
| 26.72083 | -0.06819 | 0.529569 | -0.14266 | -0.99147 | 0.657311 | 0.695554 | -0.8546 | -0.11642 | 0.382282 | -1.01127 | -1.45158 | -0.5649 | -0.66749 | 0.693446 | -0.36122 | -0.20164 | -0.43416 | -0.01987 | -2.15031 | -0.57125 |
| 26.73472 | -0.78443 | 0.90823 | -0.31014 | -1.15595 | 0.721667 | 0.460897 | -0.82229 | 0.316654 | -0.1215 | -0.62243 | -1.70357 | -0.30981 | -0.47563 | 0.673598 | -0.63896 | -0.12539 | -0.4043 | -0.15 | -2.34909 | -0.31582 |
| 26.74861 | -1.52861 | 0.463444 | -0.44195 | -1.52087 | 0.56304 | 0.198203 | -0.8548 | 0.749801 | -0.57123 | -0.03133 | -1.82799 | -0.17568 | -0.26293 | 0.447043 | -0.92062 | -0.34344 | -0.39072 | -0.47412 | -2.07229 | -0.30068 |
| 26.7625 | -1.89155 | -0.29694 | -0.82685 | -1.81985 | 0.352307 | -0.14252 | -0.91623 | 0.030477 | -1.05713 | 0.173764 | -1.84263 | -0.31587 | -0.07648 | -0.08243 | -1.1623 | -0.68056 | -0.52694 | -1.11354 | -1.71577 | -0.68965 |
| 26.77639 | -1.54757 | -1.03316 | -1.11535 | -2.02782 | 0.013148 | -0.57301 | -0.86921 | -1.06954 | -1.59298 | -0.03689 | -1.84987 | -0.69868 | -0.05397 | -0.87041 | -1.3553 | -1.07911 | -0.80977 | -1.8869 | -1.56422 | -1.17165 |
| 26.79028 | -1.56632 | -1.84283 | -1.18383 | -2.20822 | -0.46885 | -0.99856 | -0.56053 | -2.12931 | -1.63712 | -0.39558 | -1.84426 | -1.18014 | -0.24416 | -1.49707 | -1.54487 | -1.32898 | -1.0854 | -2.39997 | -1.8871 | -1.24109 |
| 26.80417 | -1.23075 | -2.00282 | -1.29408 | -2.16419 | -0.92427 | -1.30579 | -0.63466 | -2.83899 | -1.53389 | -0.81015 | -1.75388 | -1.65909 | -0.51131 | -1.98289 | -1.69608 | -1.52097 | -1.18947 | -2.44891 | -1.89465 | -1.23564 |
| 26.81806 | -1.16055 | -1.68326 | -1.47999 | -2.1171 | -1.17283 | -1.40304 | -1.02407 | -2.46516 | -1.4461 | -1.13789 | -1.56392 | -1.87057 | -0.79712 | -2.14687 | -1.72389 | -1.48381 | -1.03236 | -2.2478 | -1.63957 | -1.39126 |
| 26.83194 | -1.28994 | -1.21042 | -1.57082 | -2.26401 | -1.41724 | -1.36372 | -1.58639 | -1.80775 | -1.30699 | -1.27331 | -1.4543 | -1.80968 | -0.92467 | -2.45408 | -1.62552 | -1.29007 | -0.88555 | -2.05931 | -1.22968 | -1.54984 |
| 26.84583 | -1.34665 | -0.72316 | -1.63951 | -2.04467 | -1.58517 | -1.32814 | -1.91121 | -1.20099 | -1.0388 | -1.27908 | -1.37389 | -1.67666 | -0.91488 | -2.30934 | -1.49542 | -1.32247 | -0.75225 | -1.80445 | -0.89997 | -1.69372 |
| 26.85972 | -1.25686 | -0.57369 | -1.61654 | -1.51376 | -1.40724 | -1.14701 | -1.8029 | -0.61247 | -0.65434 | -1.29227 | -1.22436 | -1.57552 | -0.78932 | -1.90675 | -1.15751 | -1.37682 | -0.81345 | -1.64705 | -0.60791 | -1.59904 |
| 26.87361 | -0.93417 | -0.70289 | -1.29508 | -0.87068 | -0.97301 | -0.84694 | -1.35243 | -0.20727 | -0.35433 | -1.19756 | -1.10133 | -1.3686 | -0.70513 | -1.61771 | -0.90151 | -1.3058 | -0.76531 | -1.31907 | -0.42311 | -1.30744 |
| 26.8875 | -0.65466 | -0.82433 | -0.78654 | -0.04784 | -0.43564 | -0.44646 | -0.59858 | -0.01226 | -0.08704 | -0.95403 | -0.92133 | -0.94083 | -0.59413 | -1.36769 | -0.51865 | -0.98085 | -0.5203 | -0.92176 | -0.23578 | -0.85555 |
| 26.90139 | -0.53957 | -0.9197 | -0.32575 | 0.619658 | -0.09621 | 0.005266 | -0.02655 | -0.02533 | 0.054521 | -0.57656 | -0.72993 | -0.58647 | -0.4702 | -1.185 | -0.25025 | -0.68873 | -0.36519 | -0.63024 | -0.0921 | -0.44347 |
| 26.91528 | -0.48861 | -1.05249 | -0.3425 | 0.711672 | -0.04117 | 0.213886 | 0.159114 | -0.33517 | 0.039078 | -0.44018 | -0.61424 | -0.85675 | -0.27086 | -1.18355 | -0.34448 | -0.85047 | -0.6777 | -0.55551 | -0.18011 | -0.17016 |
| 26.92917 | -0.33079 | -1.3479 | -0.8612 | -0.11283 | -0.42843 | -0.07614 | -0.21429 | -0.71149 | -0.14409 | -0.89397 | -1.03124 | -1.55033 | -0.53674 | -1.61886 | -0.93083 | -1.67678 | -1.55737 | -0.56363 | -1.00011 | -0.32999 |
| 26.94306 | -0.55171 | -1.52973 | -1.36705 | -0.9095 | -1.20986 | -0.79318 | -1.14176 | -1.03876 | -0.69826 | -1.47522 | -2.50219 | -2.01673 | -1.70432 | -1.88974 | -1.68428 | -2.27995 | -1.61087 | -0.80944 | -2.44358 | -1.62965 |
| 26.95694 | -1.10754 | -1.63477 | -1.54432 | -1.54622 | -1.99808 | -1.16628 | -2.06628 | -1.52978 | -0.97196 | -2.46088 | -4.60914 | -2.11222 | -2.54792 | -2.61453 | -2.04589 | -2.62838 | -1.38643 | -1.35901 | -4.07092 | -3.07028 |
| 26.97083 | -1.4406 | -1.4064 | -1.52002 | -1.94198 | -2.09228 | -0.88072 | -2.76528 | -1.514 | -1.09091 | -2.75213 | -3.89765 | -1.65006 | -1.9232 | -2.3204 | -1.88005 | -2.57201 | -1.04022 | -1.44372 | -3.73602 | -2.7706 |
| 26.98472 | -1.44923 | -1.02516 | -1.39016 | -1.98344 | -1.55959 | -0.59639 | -2.80868 | -1.11882 | -1.1016 | -0.88695 | -1.82721 | -0.94124 | -0.64579 | -1.08828 | -1.21622 | -1.42992 | -0.91125 | -0.76507 | -1.93048 | -1.13618 |
| 26.99861 | -1.56039 | -1.20021 | -1.46877 | -2.26503 | -1.19158 | -0.53957 | -2.46354 | -0.94505 | -1.22445 | 0.464927 | -0.07044 | -0.33587 | 0.043771 | -0.00476 | -0.4437 | -0.37528 | -1.02936 | -0.27558 | -0.51915 | 0.065938 |
| 27.0125 | -1.51163 | -1.70052 | -1.46841 | -2.07708 | -0.81443 | -0.58651 | -1.80621 | -0.6767 | -1.43562 | 0.637061 | 0.843931 | -0.14715 | 0.264996 | 0.518055 | 0.191017 | 0.087432 | -0.73942 | -0.15788 | 0.007004 | 0.460367 |
| 27.02639 | -1.08255 | -1.44458 | -1.13295 | -1.6124 | -0.50318 | -0.48743 | -1.0561 | -0.22664 | -1.37403 | 0.485064 | 1.184693 | -0.19925 | 0.323492 | 0.621136 | 0.568271 | 0.21226 | -0.09164 | -0.19378 | 0.064898 | 0.412876 |
| 27.04028 | -0.45667 | -0.71437 | -0.42962 | -1.06693 | -0.16306 | -0.13327 | -0.30436 | 0.137428 | -0.73307 | 0.551348 | 1.192677 | 0.044176 | 0.380506 | 0.674991 | 0.718078 | 0.331708 | 0.340885 | -0.10305 | 0.282345 | 0.185368 |
| 27.05417 | -0.00912 | -0.02296 | 0.303903 | -0.58482 | -0.0202 | 0.213723 | 0.206922 | 0.357785 | 0.108833 | 0.642612 | 0.917019 | 0.27266 | 0.362989 | 0.789423 | 0.68933 | 0.35083 | 0.422059 | 0.01573 | 0.514656 | 0.275658 |
| 27.06806 | 0.232823 | 0.581372 | 0.71688 | -0.18627 | 0.130721 | 0.481763 | 0.642372 | 0.625483 | 0.864246 | 0.482809 | 0.449178 | 0.353003 | 0.168658 | 0.643145 | 0.544958 | 0.221545 | 0.330554 | -0.0053 | 0.648628 | 0.281513 |
| 27.08194 | 0.304136 | 0.996742 | 0.906725 | 0.140189 | 0.278399 | 0.400669 | 0.990408 | 0.896567 | 0.970896 | 0.201104 | 0.009859 | 0.377857 | -0.03466 | 0.372644 | 0.504402 | 0.03158 | 0.159312 | -0.01232 | 0.64002 | 0.106319 |

| | | | | | | | | | | | | | | | | | | | | |
|---|---|---|---|---|---|---|---|---|---|---|---|---|---|---|---|---|---|---|---|---|
| 27.09583 | 0.249494 | 0.958736 | 1.050311 | 0.39937 | 0.391624 | 0.301582 | 1.265764 | 1.050197 | 0.904774 | 0.054747 | -0.29134 | 0.506425 | -0.02724 | 0.119844 | 0.634473 | -0.0919 | 0.033621 | -0.00403 | 0.666615 | -0.0865 |
| 27.10972 | 0.132648 | 0.805239 | 1.099075 | 0.569511 | 0.494164 | 0.277068 | 1.356544 | 1.054048 | 0.719393 | -0.01459 | -0.46245 | 0.619299 | 0.016165 | -0.08238 | 0.626851 | -0.29206 | -0.09251 | -0.04962 | 0.650224 | -0.16514 |
| 27.12361 | 0.077254 | 0.555719 | 0.987549 | 0.758785 | 0.501253 | 0.210782 | 1.325041 | 0.832331 | 0.50358 | -0.06398 | -0.48683 | 0.588892 | 0.000832 | -0.19847 | 0.548743 | -0.45326 | -0.1756 | -0.13511 | 0.591828 | -0.1101 |
| 27.1375 | 0.202875 | 0.397656 | 0.843517 | 1.056414 | 0.454484 | 0.130037 | 1.129184 | 0.611436 | 0.384944 | 0.079213 | -0.26309 | 0.573079 | 0.061082 | -0.08627 | 0.509583 | -0.33003 | -0.11517 | -0.15116 | 0.653539 | 0.065106 |
| 27.15139 | 0.405252 | 0.250787 | 0.815367 | 1.349769 | 0.433019 | 0.03272 | 1.046133 | 0.458337 | 0.434901 | 0.374686 | 0.176428 | 0.648069 | 0.239716 | 0.214411 | 0.509143 | -0.10685 | -0.06252 | -0.10034 | 0.781626 | 0.254667 |
| 27.16528 | 0.669951 | 0.269327 | 0.896668 | 1.550245 | 0.454352 | 0.026717 | 1.071163 | 0.476008 | 0.604973 | 0.693998 | 0.584495 | 0.778286 | 0.47851 | 0.53071 | 0.358976 | -0.04102 | -0.09466 | -0.05865 | 0.938601 | 0.260544 |
| 27.17917 | 1.019473 | 0.380898 | 0.979291 | 1.732097 | 0.513304 | 0.062147 | 1.11347 | 0.429286 | 0.710757 | 0.790584 | 0.533686 | 0.821524 | 0.53999 | 0.613481 | 0.09522 | -0.09434 | -0.08664 | -0.0318 | 1.059131 | 0.11115 |
| 27.19306 | 1.156381 | 0.57223 | 0.870197 | 1.989887 | 0.548494 | -0.018 | 1.054454 | 0.353899 | 0.742967 | 0.542647 | 0.10751 | 0.608316 | 0.338425 | 0.396471 | -0.10164 | -0.13361 | -0.02076 | 0.068076 | 1.054545 | 0.121661 |
| 27.20694 | 1.053336 | 0.503872 | 0.727217 | 1.877475 | 0.650263 | -0.06784 | 0.901037 | 0.425655 | 0.716676 | 0.174217 | -0.19898 | 0.217571 | 0.149832 | -0.01067 | -0.12514 | -0.12798 | 0.012687 | 0.219693 | 0.68413 | 0.227344 |
| 27.22083 | 0.752954 | 0.385692 | 0.457658 | 1.334415 | 0.674742 | -0.04746 | 0.558168 | 0.510388 | 0.561339 | -0.07382 | -0.26889 | 0.058107 | 0.083705 | -0.29062 | 0.049559 | -0.00098 | 0.011781 | 0.257095 | 0.157754 | 0.245319 |
| 27.23472 | 0.45448 | 0.244838 | 0.222034 | 0.812764 | 0.608533 | -0.03003 | 0.220551 | 0.45189 | 0.450306 | -0.1923 | -0.30065 | 0.059108 | 0.003977 | -0.34324 | 0.206111 | 0.122792 | -0.04295 | 0.151725 | -0.33502 | 0.173747 |
| 27.24861 | 0.247744 | 0.291019 | 0.070834 | 0.389643 | 0.438173 | -0.03927 | 0.088894 | 0.269121 | 0.327615 | -0.19871 | -0.24016 | 0.035285 | -0.03928 | -0.13196 | 0.22592 | 0.17249 | -0.0514 | 0.052421 | -0.54277 | 0.076407 |
| 27.2625 | 0.280056 | 0.580498 | 0.040687 | 0.1878 | 0.302074 | -0.05555 | 0.139469 | 0.058435 | 0.217624 | 0.014871 | 0.105922 | 0.06861 | 0.068989 | 0.30591 | 0.248451 | 0.237591 | 0.094407 | 0.055859 | -0.41143 | 0.209109 |
| 27.27639 | 0.515015 | 0.943518 | 0.038743 | 0.057764 | 0.239083 | -0.09116 | 0.361042 | 0.036041 | 0.191011 | 0.202165 | 0.541167 | 0.085342 | 0.179303 | 0.504867 | 0.236986 | 0.178464 | 0.178188 | 0.12407 | -0.08409 | 0.424 |
| 27.29028 | 0.779818 | 1.220886 | 0.057635 | -0.0056 | 0.302746 | -0.05871 | 0.633157 | 0.230988 | 0.300091 | 0.247485 | 0.80978 | 0.106946 | 0.23441 | 0.3839 | 0.208329 | 0.106951 | 0.309353 | 0.197786 | 0.266796 | 0.532407 |
| 27.30417 | 0.95628 | 1.44702 | 0.137724 | 0.080782 | 0.484904 | 0.02444 | 0.841867 | 0.526657 | 0.492212 | 0.2837 | 0.818415 | 0.173571 | 0.336402 | 0.287455 | 0.308498 | 0.221316 | 0.434674 | 0.148202 | 0.634857 | 0.602998 |
| 27.31806 | 0.962491 | 1.397412 | 0.237463 | 0.197853 | 0.598914 | 0.068568 | 1.015316 | 0.716726 | 0.578654 | 0.253376 | 0.582409 | 0.170329 | 0.394507 | 0.252202 | 0.488172 | 0.445201 | 0.491247 | 0.134875 | 0.830375 | 0.518817 |
| 27.33194 | 0.838115 | 1.284402 | 0.310343 | 0.321954 | 0.491406 | 0.077732 | 1.197144 | 0.676255 | 0.564751 | 0.143016 | 0.13247 | 0.131134 | 0.333293 | 0.264164 | 0.638529 | 0.665344 | 0.49198 | 0.083948 | 0.81239 | 0.352475 |
| 27.34583 | 0.754082 | 1.16515 | 0.261695 | 0.493187 | 0.303403 | 0.135882 | 1.156818 | 0.587624 | 0.582182 | 0.03996 | -0.26295 | 0.060703 | 0.233854 | 0.242031 | 0.635915 | 0.683757 | 0.393169 | 0.083642 | 0.786926 | 0.307052 |
| 27.35972 | 0.763679 | 0.861106 | 0.128224 | 0.696057 | 0.204475 | 0.212814 | 0.929111 | 0.517093 | 0.47422 | 0.041094 | -0.21346 | -0.01054 | 0.219562 | 0.157217 | 0.406137 | 0.437207 | 0.159815 | 0.149639 | 0.819701 | 0.467577 |
| 27.37361 | 0.790734 | 0.467482 | 0.066975 | 0.750079 | 0.102058 | 0.205667 | 0.66385 | 0.38595 | 0.346799 | 0.084897 | 0.119084 | 0.001445 | 0.223963 | 0.149643 | 0.229889 | 0.265092 | 0.00131 | 0.200683 | 0.752257 | 0.542446 |
| 27.3875 | 0.75975 | 0.154593 | 0.141491 | 0.726876 | 0.048511 | 0.108619 | 0.379296 | 0.255442 | 0.212703 | 0.18109 | 0.411335 | 0.120201 | 0.251127 | 0.220398 | 0.208221 | 0.337663 | 0.060683 | 0.245766 | 0.474084 | 0.439696 |
| 27.40139 | 0.642748 | 0.043747 | 0.287409 | 0.706241 | 0.015584 | 0.08095 | 0.189938 | 0.291533 | 0.219544 | 0.37312 | 0.604993 | 0.40614 | 0.338313 | 0.475389 | 0.422879 | 0.698145 | 0.405066 | 0.282448 | 0.294029 | 0.349853 |
| 27.41528 | 0.28451 | 0.351323 | 0.551825 | 0.822194 | -0.15345 | 0.221234 | 0.192243 | 0.422943 | 0.479792 | 0.760013 | 0.852927 | 1.025872 | 0.582963 | 1.187826 | 1.003564 | 1.497989 | 1.108444 | 0.386344 | 0.382247 | 0.40863 |
| 27.42917 | 0.061047 | 1.199578 | 1.140802 | 1.197652 | -0.27242 | 0.548972 | 0.498684 | 0.753269 | 1.033188 | 1.152636 | 1.053451 | 1.996285 | 0.921221 | 2.175335 | 1.597327 | 2.538151 | 1.648269 | 0.599819 | 1.023246 | 0.630561 |
| 27.44306 | 1.076968 | 2.522639 | 1.782436 | 1.341109 | 0.572484 | 0.608655 | 1.266531 | 2.068106 | 1.881752 | 1.649936 | 2.021165 | 2.837283 | 1.814188 | 2.615132 | 2.066893 | 3.534305 | 1.461993 | 1.162131 | 2.310139 | 1.816287 |
| 27.45694 | 2.806752 | 3.232872 | 2.107419 | 1.969617 | 1.895377 | 0.305097 | 2.080636 | 3.503819 | 2.043922 | 2.317327 | 4.402382 | 2.641515 | 2.464172 | 3.560343 | 2.374796 | 3.93512 | 1.124606 | 1.776383 | 3.203618 | 3.574648 |
| 27.47083 | 2.828424 | 2.620161 | 1.707661 | 1.889769 | 2.304109 | 0.226763 | 2.07766 | 3.116131 | 1.359977 | 2.381205 | 5.4558 | 2.375267 | 1.969286 | 3.459755 | 2.578748 | 3.328783 | 0.906106 | 2.142536 | 3.652791 | 3.549659 |
| 27.48472 | 2.129734 | 2.13599 | 1.519849 | 2.129844 | 2.236854 | 0.455662 | 2.079104 | 2.57387 | 1.366142 | 2.141134 | 4.808513 | 2.135159 | 1.25087 | 4.054202 | 2.741033 | 2.61713 | 0.889426 | 2.321629 | 3.910217 | 2.817149 |
| 27.49861 | 1.402652 | 1.478419 | 1.550858 | 2.029248 | 1.901584 | 0.713261 | 2.250722 | 1.897623 | 1.603637 | 1.777733 | 3.884669 | 2.128219 | 1.051868 | 4.627678 | 2.454066 | 2.232694 | 0.789104 | 2.279459 | 4.115585 | 2.595192 |
| 27.5125 | 0.565148 | 0.703594 | 1.275385 | 1.156688 | 1.468262 | 0.685341 | 1.690907 | 0.785262 | 1.086938 | 1.488599 | 2.951477 | 2.119048 | 1.002539 | 4.623187 | 2.08526 | 2.059666 | 0.479104 | 1.993143 | 4.081889 | 2.129139 |
| 27.52639 | 0.021572 | 0.342043 | 0.81164 | 0.222962 | 1.086831 | 0.529507 | 0.904032 | -0.26098 | 0.678589 | 1.185103 | 1.896158 | 1.829125 | 0.760361 | 3.556157 | 1.862327 | 1.658422 | 0.106261 | 1.41162 | 3.621107 | 1.648541 |
| 27.54028 | -0.03228 | 0.187619 | 0.609958 | -0.38172 | 0.721994 | 0.453467 | 0.480063 | -0.6509 | 0.360132 | 0.92407 | 0.989665 | 1.348687 | 0.572011 | 2.649163 | 1.598846 | 1.098854 | -0.02288 | 0.985745 | 2.645318 | 1.286663 |
| 27.55417 | 0.080848 | 0.02323 | 0.605165 | -0.63073 | 0.781091 | 0.420715 | 0.279963 | -0.44239 | 0.203977 | 0.6621 | 0.403394 | 0.797717 | 0.396976 | 1.607462 | 1.093703 | 0.560292 | -0.01097 | 0.808605 | 1.377772 | 1.009524 |
| 27.56806 | 0.195099 | -0.31938 | 0.695233 | -0.6264 | 0.987842 | 0.444155 | 0.011303 | -0.00071 | 0.157356 | 0.526468 | 0.113099 | 0.351474 | 0.240394 | 0.984147 | 0.596471 | 0.55534 | 0.191361 | 0.889926 | 0.28267 | 0.963712 |
| 27.58194 | -0.11658 | -0.32985 | 0.542768 | -0.45469 | 1.030198 | 0.524102 | -0.23811 | 0.254092 | 0.339322 | 0.406558 | 0.359758 | 0.12234 | 0.365271 | 0.351157 | 0.075303 | 0.651486 | 0.506823 | 1.2239 | -0.30052 | 1.103672 |
| 27.59583 | -0.69076 | 0.053134 | 0.207817 | -0.24191 | 0.778913 | 0.486546 | -0.32425 | 0.234684 | 0.373324 | 0.171896 | 0.69496 | 0.022796 | 0.524524 | -0.06783 | -0.53696 | 0.5583 | 0.787865 | 1.354819 | -0.89572 | 0.900559 |
| 27.60972 | -0.70272 | 0.41022 | -0.24445 | -0.19761 | 0.241768 | 0.27482 | -0.18363 | 0.060977 | 0.189626 | -0.1105 | 0.745473 | -0.09495 | 0.403408 | -0.2866 | -0.95233 | 0.228234 | 0.601256 | 1.088271 | -0.90699 | 0.16542 |
| 27.62361 | -0.09062 | 0.765185 | -0.73564 | -0.32822 | -0.47977 | 0.057463 | -0.18805 | -0.16505 | 0.044323 | -0.45738 | 0.525196 | -0.41161 | 0.036813 | -0.47827 | -0.79325 | -0.24074 | 0.251645 | 0.598976 | -1.35827 | -0.28379 |
| 27.6375 | 0.564204 | 0.962459 | -1.1183 | -0.58576 | -1.25465 | -0.02646 | 0.013483 | -0.31133 | 0.181052 | -0.62962 | -0.05251 | -0.84989 | -0.39203 | -1.16234 | -0.32303 | -0.73994 | -0.01023 | 0.03571 | -1.16173 | -0.32507 |
| 27.65139 | 0.935265 | 0.677179 | -1.31241 | -0.76314 | -2.08919 | -0.00222 | 0.0559 | -0.50153 | 0.172456 | -0.59506 | -0.46952 | -1.26331 | -0.75814 | -2.1562 | 0.042973 | -1.15197 | -0.14477 | -0.61376 | -0.53801 | -0.15798 |
| 27.66528 | 1.015323 | 0.182867 | -1.17816 | -0.28932 | -2.02276 | -0.01971 | 0.029307 | -0.33084 | 0.081356 | -0.40614 | -0.67795 | -1.55349 | -1.00919 | -2.74762 | -0.03635 | -1.5271 | -0.20676 | -1.17815 | -0.00082 | -0.04679 |
| 27.67917 | 1.01918 | -0.2497 | -0.66275 | 0.477275 | -1.20911 | 0.015292 | 0.011874 | 0.109202 | 0.006366 | -0.11395 | -0.85412 | -1.57666 | -1.01096 | -3.16698 | -0.37336 | -1.73416 | -0.3744 | -1.38951 | 0.09417 | -0.17873 |
| 27.69306 | 0.846382 | -0.74958 | 0.035321 | 0.991782 | 0.130209 | 0.117696 | 0.161856 | 0.477006 | 0.01468 | 0.165982 | -1.02934 | -1.46752 | -0.79443 | -2.87359 | -0.52659 | -1.59483 | -0.58867 | -1.34511 | -0.27615 | -0.2859 |
| 27.70694 | 0.746109 | -0.89811 | 0.476229 | 0.846748 | 0.949721 | 0.233472 | 0.366409 | 0.67284 | 0.256442 | 0.224484 | -1.33406 | -1.22572 | -0.61427 | -2.06559 | -0.50515 | -1.38803 | -0.70965 | -1.276 | -1.01419 | -0.58065 |
| 27.72083 | 0.235146 | -0.71284 | 0.626035 | 0.668433 | 1.139504 | 0.384666 | 0.242992 | 0.551659 | 0.515606 | 0.09031 | -1.61311 | -0.95951 | -0.52932 | -1.56264 | -0.47869 | -1.36371 | -0.87749 | -1.03406 | -1.61403 | -0.97839 |
| 27.73472 | -0.54737 | -0.61649 | 0.56284 | -0.07008 | 0.687183 | 0.425795 | -0.42277 | 0.349199 | 0.525908 | -0.11741 | -1.78431 | -0.72259 | -0.5813 | -0.83823 | -0.69609 | -1.4392 | -1.1682 | -0.80697 | -2.19947 | -1.26134 |
| 27.74861 | -1.43987 | -0.23939 | 0.277858 | -1.39995 | -0.281 | 0.370847 | -1.05922 | -0.2425 | 0.333157 | -0.40488 | -1.86951 | -0.51134 | -0.64275 | -0.03097 | -1.0305 | -1.35357 | -1.27535 | -0.65017 | -2.37089 | -1.34767 |
| 27.7625 | -2.12082 | -0.06776 | -0.20766 | -2.00158 | -1.50826 | 0.214158 | -1.40536 | -1.54319 | 0.170143 | -0.72836 | -1.72548 | -0.32037 | -0.85723 | 0.306636 | -1.27712 | -1.39401 | -1.20492 | -0.64793 | -2.17605 | -1.48704 |
| 27.77639 | -2.56372 | -0.34017 | -0.84315 | -2.52931 | -2.18207 | -0.04283 | -1.64994 | -2.25092 | -0.13419 | -1.13659 | -1.65879 | -0.26695 | -1.10468 | 0.002305 | -1.24657 | -1.52341 | -1.12216 | -0.91975 | -1.93078 | -1.62448 |
| 27.79028 | -2.89906 | -0.8769 | -1.50106 | -2.51424 | -1.65482 | -0.25407 | -1.4378 | -1.91761 | -0.87856 | -1.61998 | -1.66203 | -0.53996 | -1.46831 | -0.7856 | -1.13057 | -1.67976 | -1.15394 | -1.22637 | -1.54965 | -1.65112 |
| 27.80417 | -2.91867 | -1.4379 | -2.03143 | -2.74101 | -1.20695 | -0.54727 | -1.15437 | -1.34577 | -1.79915 | -2.06452 | -1.81964 | -1.05013 | -1.71136 | -1.66498 | -1.13812 | -1.80044 | -1.02108 | -1.35394 | -1.12849 | -1.68632 |
| 27.81806 | -2.81293 | -2.0008 | -2.60198 | -2.98128 | -0.96944 | -0.77347 | -1.17038 | -0.97587 | -2.34057 | -2.0546 | -2.09999 | -1.65603 | -1.7826 | -2.49158 | -1.43205 | -2.01821 | -0.84034 | -1.38942 | -0.95154 | -1.81593 |
| 27.83194 | -2.21634 | -1.91572 | -2.84233 | -2.95062 | -0.91832 | -0.82299 | -1.17536 | -0.69868 | -2.12071 | -1.76322 | -2.01191 | -2.01241 | -1.49065 | -3.09323 | -1.67179 | -2.06735 | -0.88786 | -1.28578 | -0.90857 | -1.63159 |
| 27.84583 | -1.50801 | -1.62687 | -2.45364 | -2.46844 | -1.14267 | -0.87604 | -1.50175 | -0.39531 | -1.89193 | -1.45977 | -1.61145 | -1.98318 | -1.11836 | -3.19523 | -1.67535 | -1.70025 | -0.8882 | -1.02186 | -0.81604 | -1.36346 |
| 27.85972 | -0.89465 | -1.32987 | -1.87642 | -1.36986 | -1.12166 | -0.88537 | -1.5414 | -0.2947 | -1.64129 | -1.16001 | -1.02716 | -1.68856 | -0.81746 | -2.52504 | -1.53413 | -1.36234 | -0.67872 | -0.90017 | -0.67488 | -1.27902 |
| 27.87361 | -0.41637 | -0.26571 | -1.24977 | -0.32445 | -0.82883 | -0.81257 | -1.38027 | -0.29066 | -0.83379 | -0.73431 | -0.46327 | -1.27606 | -0.46747 | -1.44089 | -1.30745 | -1.1769 | -0.42477 | -0.67177 | -0.50486 | -1.22518 |
| 27.8875 | -0.03101 | 0.703235 | -0.80332 | 0.535512 | -0.35605 | -0.75997 | -0.86896 | -0.47473 | -0.28529 | -0.4913 | -0.09551 | -0.97428 | -0.28669 | -0.86503 | -1.15674 | -1.16028 | -0.31475 | -0.57804 | -0.40397 | -1.09601 |
| 27.90139 | 0.078501 | 1.020856 | -0.53917 | 1.288174 | 0.148093 | -0.68412 | -0.32628 | -0.55829 | 0.091589 | -0.30926 | -0.06656 | -0.73486 | -0.22302 | -0.71947 | -0.93349 | -1.12186 | -0.3321 | -0.5577 | -0.42031 | -0.75837 |
| 27.91528 | 0.394244 | 0.717229 | -0.48869 | 1.411263 | 0.594463 | -0.70802 | -0.07697 | -0.55274 | 0.11361 | -0.18327 | -0.25137 | -0.78065 | -0.311 | -0.86889 | -0.70614 | -1.39124 | -0.55598 | -0.61803 | -0.83917 | -0.51351 |
| 27.92917 | 0.80501 | -0.26191 | -1.06591 | 0.928114 | 0.547696 | -1.01847 | -0.57783 | -0.77319 | -0.40462 | -0.55147 | -0.81659 | -1.38087 | -0.94382 | -1.66742 | -0.80264 | -2.41184 | -1.01608 | -0.81485 | -1.91781 | -1.00002 |
| 27.94306 | 0.137857 | -1.23433 | -1.92402 | -0.47961 | -0.2812 | -1.33723 | -1.48601 | -0.98324 | -1.04225 | -1.44683 | -2.40975 | -2.39008 | -2.20909 | -1.97272 | -1.63435 | -3.01095 | -1.42613 | -1.10971 | -3.11632 | -2.09289 |
| 27.95694 | -1.1288 | -1.75018 | -2.10677 | -1.25847 | -1.2476 | -1.27497 | -1.82828 | -1.18925 | -1.40079 | -2.65571 | -3.76582 | -3.24118 | -3.04963 | -2.61352 | -2.43613 | -3.27619 | -1.22865 | -1.7144 | -4.3264 | -3.53687 |
| 27.97083 | -1.87874 | -2.34554 | -1.86354 | -1.64221 | -1.61755 | -0.9054 | -2.27073 | -1.28356 | -1.60962 | -2.70191 | -3.47364 | -2.66466 | -2.76981 | -2.21461 | -2.57746 | -3.38992 | -0.81107 | -1.6493 | -3.65792 | -3.73022 |
| 27.98472 | -2.02412 | -2.63633 | -1.45462 | -1.60289 | -1.4424 | -0.6008 | -2.25896 | -0.90965 | -1.19371 | -0.99365 | -1.45831 | -1.4115 | -0.81492 | -1.05127 | -1.81088 | -1.84533 | -0.57814 | -0.53819 | -1.4815 | -1.68172 |
| 27.99861 | -2.31966 | -2.82492 | -1.2672 | -1.64986 | -1.21552 | -0.38034 | -1.96117 | -0.50031 | -0.83264 | 0.312516 | -0.03312 | -0.61998 | 0.287716 | -0.19347 | -1.01795 | -0.65306 | -0.47909 | 0.047291 | 0.065043 | -0.01562 |
| 28.0125 | -2.37144 | -2.65426 | -1.22845 | -1.56445 | -1.08509 | -0.22917 | -1.71016 | -0.24515 | -0.31532 | 0.644141 | 0.394099 | -0.08725 | 0.68646 | 0.29498 | -0.16851 | 0.252058 | -0.39698 | -0.01471 | 0.695243 | 0.610663 |
| 28.02639 | -1.75626 | -1.97405 | -0.87134 | -1.39772 | -0.85904 | -0.06879 | -1.42537 | -0.05319 | 0.288643 | 0.702026 | 0.467537 | 0.374519 | 0.799614 | 0.521322 | 0.570695 | 0.938011 | -0.19788 | -0.14261 | 1.053486 | 0.789929 |
| 28.04028 | -0.83395 | -0.90717 | -0.06436 | -1.09299 | -0.57928 | 0.138288 | -0.72964 | 0.165812 | 0.46109 | 0.780656 | 0.559789 | 0.657848 | 0.708914 | 0.618594 | 1.132072 | 1.234136 | -0.01456 | -0.14727 | 1.084974 | 0.910774 |
| 28.05417 | -0.10074 | 0.195034 | 0.813133 | -0.89713 | -0.36928 | 0.196448 | 0.125837 | 0.301957 | 0.185021 | 0.783296 | 0.660557 | 0.732798 | 0.535046 | 0.679392 | 1.395232 | 1.259183 | 0.094075 | -0.0596 | 0.907225 | 0.919086 |
| 28.06806 | 0.288384 | 0.930961 | 1.249488 | -0.76545 | -0.19502 | 0.091153 | 0.7445 | 0.482675 | 0.284149 | 0.600149 | 0.65697 | 0.439328 | 0.326045 | 0.595341 | 1.312989 | 0.892591 | 0.086821 | 0.002955 | 0.599109 | 0.699504 |
| 28.08194 | 0.492003 | 1.149868 | 1.054897 | -0.5298 | 0.137486 | -0.0005 | 1.160354 | 0.729186 | 0.311245 | 0.348884 | 0.52184 | 0.101163 | 0.157604 | 0.432183 | 0.993706 | 0.320062 | 0.05447 | -0.03644 | 0.228412 | 0.39356 |
| 28.09583 | 0.42831 | 0.99781 | 0.97437 | -0.1949 | 0.424993 | -0.07059 | 1.439814 | 0.868564 | 0.261545 | 0.126095 | 0.45716 | -0.12393 | 0.047217 | 0.28818 | 0.676021 | -0.07588 | 0.010479 | -0.05054 | 0.022822 | 0.112502 |
| 28.10972 | 0.200467 | 0.955548 | 0.919536 | 0.121176 | 0.643174 | -0.17071 | 1.514467 | 0.870385 | 0.232982 | -0.07648 | 0.4467 | -0.30608 | -0.10505 | 0.141724 | 0.363046 | -0.34322 | -0.10392 | -0.08707 | -0.01292 | -0.20324 |
| 28.12361 | 0.302153 | 1.059608 | 0.92139 | 0.293952 | 0.622684 | -0.1354 | 1.650592 | 0.840912 | 0.322388 | -0.15998 | 0.406392 | -0.24844 | -0.16483 | 0.032197 | 0.250471 | -0.34236 | -0.135 | -0.21948 | 0.054374 | -0.44142 |
| 28.1375 | 0.344059 | 1.138258 | 0.809018 | 0.385848 | 0.462259 | 0.094456 | 1.806852 | 0.686881 | 0.460667 | -0.09272 | 0.330844 | 0.008202 | -0.04877 | -0.00593 | 0.3213 | -0.11629 | -0.08821 | -0.2594 | 0.136834 | -0.48899 |
| 28.15139 | 0.543605 | 1.127565 | 0.552088 | 0.460443 | 0.342663 | 0.271519 | 1.510892 | 0.559658 | 0.483788 | 0.069786 | 0.21722 | 0.217282 | 0.094197 | -0.11821 | 0.345586 | 0.068311 | -0.0154 | -0.21477 | 0.211499 | -0.36904 |
| 28.16528 | 0.728723 | 1.055851 | 0.332868 | 0.44181 | 0.270686 | 0.294131 | 1.327701 | 0.471065 | 0.49521 | 0.193775 | 0.115663 | 0.320421 | 0.112714 | -0.23586 | 0.303273 | 0.130263 | 0.038788 | -0.101 | 0.350896 | 0.001008 |
| 28.17917 | 0.7667 | 1.055164 | 0.150929 | 0.338427 | 0.233305 | 0.223214 | 1.172165 | 0.436072 | 0.467899 | 0.143987 | 0.062223 | 0.229498 | -0.05671 | -0.28293 | 0.212303 | -0.01618 | 0.092555 | -0.03705 | 0.373031 | 0.322152 |
| 28.19306 | 0.664784 | 0.808692 | -0.04124 | 0.192443 | 0.369301 | 0.085396 | 0.797422 | 0.394379 | 0.431455 | -0.11351 | -0.02404 | -0.01357 | -0.29642 | -0.3054 | 0.063964 | -0.40157 | 0.085624 | -0.09338 | 0.190248 | 0.364892 |
| 28.20694 | 0.508613 | 0.512711 | -0.0051 | 0.072446 | 0.420049 | 0.014597 | 0.603625 | 0.293799 | 0.486069 | -0.32391 | -0.18385 | -0.09382 | -0.38553 | -0.30203 | -0.0072 | -0.63512 | 0.019556 | -0.17971 | -0.02876 | 0.209445 |
| 28.22083 | 0.350636 | 0.361283 | 0.126838 | 0.040087 | 0.320052 | 0.009833 | 0.512145 | 0.175662 | 0.475077 | -0.27865 | -0.26076 | 0.064461 | -0.16051 | -0.1566 | 0.066676 | -0.40062 | -0.03546 | -0.18364 | -0.06111 | 0.159146 |
| 28.23472 | 0.230701 | 0.341373 | 0.176988 | 0.125929 | 0.192625 | 0.07296 | 0.47558 | 0.100164 | 0.479922 | -0.03081 | -0.10894 | 0.35707 | 0.25148 | 0.192843 | 0.331871 | 0.236913 | 0.031047 | -0.09439 | 0.157144 | 0.377617 |
| 28.24861 | 0.182322 | 0.619547 | 0.301892 | 0.230078 | 0.034557 | 0.242333 | 0.704399 | 0.053653 | 0.476727 | 0.262803 | 0.091094 | 0.902656 | 0.670585 | 0.595389 | 0.764138 | 1.011637 | 0.238043 | 0.07176 | 0.538975 | 0.7156 |
| 28.2625 | 0.153832 | 1.058923 | 0.597735 | 0.361805 | -0.07054 | 0.424778 | 0.830513 | 0.019484 | 0.644332 | 0.363283 | 0.212027 | 1.409013 | 0.932426 | 0.694124 | 1.16997 | 1.552329 | 0.418854 | 0.158078 | 0.876038 | 1.069379 |
| 28.27639 | 0.198721 | 1.181178 | 0.782959 | 0.607849 | 0.217565 | 0.338695 | 0.742009 | 0.165572 | 0.724574 | 0.315649 | 0.262388 | 1.381426 | 0.768992 | 0.495799 | 1.108799 | 1.304359 | 0.225968 | 0.093491 | 0.93282 | 1.10508 |
| 28.29028 | 0.556679 | 0.961404 | 0.615836 | 0.940171 | 0.652066 | 0.0093 | 0.35463 | 0.498224 | 0.604786 | 0.18877 | 0.473373 | 0.505251 | 0.223743 | 0.19466 | 0.807479 | 0.328598 | 0.025604 | 0.052643 | 0.738706 | 0.755396 |
| 28.30417 | 0.97753 | 0.825106 | 0.575882 | 0.923833 | 0.929986 | -0.24904 | 0.353474 | 0.668641 | 0.430402 | 0.040056 | 0.510944 | -0.11673 | -0.21394 | -0.07374 | 0.568481 | -0.33174 | -0.01744 | -0.0026 | 0.463463 | 0.361173 |
| 28.31806 | 1.139359 | 0.931204 | 0.666712 | 0.735369 | 0.799936 | -0.18496 | 0.60072 | 0.655276 | 0.300471 | -0.02232 | 0.314606 | 0.083461 | -0.13523 | -0.11379 | 0.581092 | -0.21241 | 0.00711 | -0.05246 | 0.33702 | 0.248931 |
| 28.33194 | 1.13176 | 1.082231 | 0.637793 | 0.588417 | 0.556687 | 0.021852 | 0.748377 | 0.52102 | 0.305093 | 0.128963 | 0.189659 | 0.477241 | 0.19015 | -0.1221 | 0.715603 | 0.113938 | 0.139161 | -0.04516 | 0.368326 | 0.364193 |
| 28.34583 | 1.030206 | 1.009791 | 0.476623 | 0.582369 | 0.430283 | 0.136866 | 0.882415 | 0.471871 | 0.324506 | 0.366836 | 0.275184 | 0.713085 | 0.56407 | -0.02507 | 0.746901 | 0.3807 | 0.223628 | 0.066315 | 0.461463 | 0.68781 |

| | | | | | | | | | | | | | | | | | | | | |
|---|---|---|---|---|---|---|---|---|---|---|---|---|---|---|---|---|---|---|---|---|
| 28.35972 | 0.971786 | 1.005898 | 0.281942 | 0.585721 | 0.211916 | 0.245501 | 1.00057 | 0.355005 | 0.42654 | 0.458004 | 0.331858 | 0.898494 | 0.742577 | 0.195227 | 0.741465 | 0.78578 | 0.270248 | 0.158327 | 0.544746 | 1.036536 |
| 28.37361 | 0.908408 | 1.275829 | 0.075444 | 0.608026 | -0.01902 | 0.331168 | 1.014737 | 0.215783 | 0.509663 | 0.405311 | 0.336239 | 0.961195 | 0.682196 | 0.368929 | 0.700599 | 1.212223 | 0.294395 | 0.160462 | 0.538368 | 1.123189 |
| 28.3875 | 0.936287 | 1.14838 | -0.07458 | 0.446265 | 0.106666 | 0.174023 | 0.710905 | 0.247635 | 0.448415 | 0.277554 | 0.298329 | 0.533223 | 0.341342 | 0.461696 | 0.448752 | 0.902409 | 0.252067 | 0.113353 | 0.401766 | 0.843145 |
| 28.40139 | 0.909428 | 0.832056 | -0.06442 | 0.187976 | 0.294436 | -0.00799 | 0.557854 | 0.292797 | 0.338541 | 0.25117 | 0.282954 | 0.090208 | 0.088373 | 0.450094 | 0.354449 | 0.387263 | 0.16694 | 0.123121 | 0.227455 | 0.578498 |
| 28.41528 | 0.629851 | 0.964554 | 0.326013 | 0.208884 | -0.04666 | 0.242287 | 0.560488 | 0.204692 | 0.440537 | 0.201179 | 0.308289 | 0.686375 | 0.399934 | 0.414807 | 0.806337 | 0.802066 | 0.362434 | 0.165044 | 0.247975 | 0.347426 |
| 28.42917 | 0.352079 | 1.322422 | 0.874116 | 0.65639 | -0.2766 | 0.594602 | 0.717645 | 0.189306 | 0.68324 | 0.393262 | 0.629499 | 1.830502 | 1.247582 | 0.553759 | 1.462229 | 1.731196 | 0.724149 | 0.354543 | 0.591976 | 0.67515 |
| 28.44306 | 1.066196 | 1.488981 | 1.681159 | 2.001444 | 0.776717 | 0.561333 | 1.491131 | 0.567008 | 1.000144 | 0.989651 | 1.868553 | 2.799107 | 2.672187 | 1.250183 | 2.172009 | 2.513691 | 0.932261 | 0.847982 | 1.271874 | 1.875837 |
| 28.45694 | 2.635263 | 1.68204 | 2.199696 | 3.074677 | 2.170434 | 0.375811 | 2.166171 | 1.4801 | 1.349978 | 1.651324 | 2.943564 | 2.870001 | 3.394612 | 2.310692 | 2.595461 | 2.88321 | 1.061285 | 1.18386 | 1.856005 | 3.077904 |
| 28.47083 | 2.94682 | 1.438096 | 1.900738 | 2.351318 | 3.045344 | 0.461844 | 2.390246 | 2.457663 | 1.318088 | 1.992784 | 3.485431 | 2.716966 | 3.031644 | 3.489378 | 2.961452 | 3.614753 | 1.13051 | 1.370016 | 2.118302 | 3.340941 |
| 28.48472 | 2.648329 | 1.108088 | 1.378227 | 2.322745 | 3.826604 | 0.183971 | 2.35116 | 3.272369 | 0.973667 | 1.963898 | 3.51645 | 2.556256 | 2.224764 | 3.826899 | 3.229972 | 4.11157 | 1.342112 | 1.273817 | 2.376799 | 3.153091 |
| 28.49861 | 2.169285 | 0.767864 | 1.25244 | 2.381527 | 3.53417 | 0.206009 | 2.194578 | 3.32025 | 0.825977 | 1.448753 | 3.105229 | 2.449881 | 1.588528 | 4.022885 | 2.89044 | 3.845413 | 1.055521 | 1.346694 | 2.738569 | 2.696605 |
| 28.5125 | 1.350969 | 0.294916 | 0.873376 | 2.54791 | 2.456693 | 0.287203 | 1.634928 | 1.872471 | 0.832169 | 1.069053 | 2.674737 | 2.350194 | 1.411945 | 4.091264 | 2.410174 | 3.43756 | 0.587748 | 1.356181 | 3.295989 | 2.21268 |
| 28.52639 | 0.605221 | 0.113234 | 0.493034 | 2.238204 | 1.60508 | 0.343383 | 1.049004 | 1.008431 | 0.817486 | 0.816664 | 2.187391 | 2.025157 | 1.230129 | 3.545295 | 2.053802 | 2.606318 | 0.298279 | 1.333929 | 3.297633 | 1.630149 |
| 28.54028 | 0.283765 | 0.546652 | 0.216626 | 2.121154 | 1.017609 | 0.467723 | 0.710265 | 0.515645 | 0.796532 | 0.790532 | 1.722387 | 1.525164 | 0.926477 | 3.117226 | 1.788471 | 1.683134 | 0.229462 | 1.333525 | 3.05631 | 0.920976 |
| 28.55417 | 0.430814 | 1.045143 | 0.204789 | 1.687733 | 0.388453 | 0.484012 | 0.354669 | 0.032982 | 0.577683 | 0.904424 | 1.147821 | 0.967949 | 0.676956 | 2.432247 | 1.507679 | 0.665652 | 0.036028 | 1.269535 | 2.56319 | 0.526485 |
| 28.56806 | 0.583308 | 1.170922 | 0.209028 | 0.776819 | -0.11862 | 0.477751 | 0.118369 | -0.63135 | 0.193685 | 0.882866 | 0.637576 | 0.397043 | 0.49591 | 1.016408 | 0.842497 | -0.21403 | -0.25506 | 1.00058 | 1.513153 | 0.599659 |
| 28.58194 | 0.541155 | 1.227178 | 0.079739 | 0.199285 | -0.52929 | 0.571947 | -0.00041 | -1.20658 | 0.24046 | 0.62647 | 0.35887 | 0.077944 | 0.334844 | 0.308659 | -0.04399 | -0.409 | -0.23 | 0.842652 | 0.466663 | 0.655302 |
| 28.59583 | 0.360044 | 1.099797 | -0.19078 | -0.30551 | -0.4696 | 0.802057 | -0.095 | -1.28295 | 0.510516 | 0.316401 | 0.415649 | -0.04598 | 0.228237 | 0.094072 | -0.8499 | -0.42623 | 0.120573 | 0.992671 | -0.3204 | 0.67271 |
| 28.60972 | 0.099855 | 0.824567 | -0.3577 | -1.19059 | -0.23674 | 0.803612 | -0.08972 | -0.77906 | 0.746895 | 0.010035 | 0.232702 | -0.09022 | 0.359039 | -0.18619 | -1.39741 | -0.5652 | 0.361777 | 1.142427 | -0.777 | 0.425689 |
| 28.62361 | -0.29647 | 0.373185 | -0.5903 | -1.80414 | 0.02347 | 0.530356 | -0.00677 | 0.046827 | 0.620505 | -0.34195 | 0.174622 | -0.23237 | 0.287096 | -0.48252 | -1.51314 | -0.62486 | 0.344132 | 1.063871 | -1.04086 | 0.015031 |
| 28.6375 | -0.6774 | -0.25788 | -0.86184 | -2.21157 | 0.059274 | 0.243288 | 0.029996 | 0.999932 | 0.523781 | -0.70181 | 0.082576 | -0.47818 | -0.05737 | -0.98036 | -1.22145 | -0.61061 | 0.13824 | 0.838444 | -1.26309 | -0.5772 |
| 28.65139 | -0.77837 | -1.07226 | -1.11445 | -2.51331 | -0.06329 | -0.12008 | -0.0927 | 0.943863 | 0.170181 | -1.11583 | -0.35499 | -0.84246 | -0.44708 | -1.70139 | -0.63693 | -0.64907 | -0.04599 | 0.542637 | -1.36648 | -1.06881 |
| 28.66528 | -0.61059 | -1.57952 | -1.13281 | -2.04248 | 0.024744 | -0.37726 | -0.21442 | 0.639818 | -0.12792 | -1.29041 | -0.84654 | -1.15464 | -0.69918 | -2.22622 | -0.07078 | -0.97273 | -0.01834 | 0.167707 | -1.78645 | -1.23398 |
| 28.67917 | -0.3106 | -1.45934 | -0.79611 | -1.21928 | -0.37251 | -0.40372 | -0.14355 | 0.195443 | -0.15052 | -1.15947 | -1.19233 | -1.07485 | -0.72442 | -2.58942 | 0.44193 | -1.47156 | 0.217108 | -0.24357 | -2.26375 | -1.47226 |
| 28.69306 | -0.0505 | -1.03444 | -0.25009 | 0.217549 | -0.77604 | -0.21747 | -0.02865 | -0.52881 | -0.08474 | -0.98394 | -1.35203 | -0.8719 | -0.57508 | -2.97833 | 0.90407 | -1.86407 | 0.474472 | -0.74855 | -2.35437 | -1.38893 |
| 28.70694 | 0.157084 | -0.39399 | 0.254921 | 1.279561 | -0.81883 | -0.09023 | 0.000515 | -1.2384 | 0.090936 | -0.7707 | -1.46369 | -0.79068 | -0.51695 | -3.17096 | 1.097202 | -2.34463 | 0.484017 | -1.28274 | -1.87907 | -1.15258 |
| 28.72083 | -0.00546 | 0.360976 | 0.36647 | 2.283573 | -0.71579 | -0.01897 | -0.14069 | -1.56069 | 0.07239 | -0.48186 | -1.51504 | -1.03185 | -0.64677 | -2.88133 | 0.620089 | -2.63354 | 0.093219 | -1.78036 | -1.00805 | -0.65223 |
| 28.73472 | -0.3808 | 1.057182 | 0.297648 | 1.928099 | -0.79449 | -0.05194 | -0.42233 | -1.18588 | 0.042514 | -0.26599 | -1.39896 | -1.44387 | -0.79927 | -2.16685 | -0.29646 | -2.18079 | -0.52805 | -2.15187 | 0.06993 | -0.01547 |
| 28.74861 | -0.89093 | 1.338413 | 0.250885 | 1.14993 | -1.08207 | -0.08037 | -0.80546 | -1.01056 | -0.02722 | -0.15015 | -1.09062 | -1.69023 | -0.83834 | -1.35611 | -0.90431 | -1.14913 | -1.1651 | -2.19778 | 0.547738 | 0.248514 |
| 28.7625 | -1.43627 | 0.736377 | 0.128271 | -0.01145 | -1.21384 | -0.18548 | -1.56805 | -0.77428 | -0.11344 | -0.18354 | -0.75533 | -1.73832 | -0.97353 | -0.84111 | -1.40208 | -0.49086 | -1.76446 | -1.99119 | 0.810563 | -0.00239 |
| 28.77639 | -1.6229 | -0.03744 | -0.23794 | -1.27128 | -0.62431 | -0.32824 | -1.59469 | -0.36817 | -0.31054 | -0.40908 | -0.66655 | -1.55446 | -1.31262 | -0.52419 | -1.75919 | -0.38071 | -2.04527 | -1.63259 | 0.439112 | -0.61848 |
| 28.79028 | -1.76431 | -0.79355 | -0.9033 | -1.88834 | -0.09642 | -0.46345 | -1.87378 | -0.28808 | -0.47859 | -0.72709 | -0.94933 | -1.3178 | -1.76457 | -0.41735 | -1.94611 | -0.60933 | -1.9951 | -1.20605 | -0.13906 | -1.26301 |
| 28.80417 | -2.00756 | -1.55536 | -1.52028 | -1.92736 | -0.66212 | -0.51839 | -1.77004 | -0.50411 | -0.57824 | -1.0473 | -1.38965 | -1.09834 | -2.1488 | -0.4964 | -2.34228 | -1.04085 | -1.91881 | -0.95581 | -0.71056 | -1.36384 |
| 28.81806 | -2.05919 | -1.63365 | -1.82163 | -2.24677 | -1.23518 | -0.58029 | -1.66771 | -0.83877 | -0.78455 | -1.11086 | -1.76622 | -1.1315 | -2.30838 | -0.83398 | -2.75823 | -1.59785 | -1.7512 | -0.97683 | -1.13227 | -1.29717 |
| 28.83194 | -2.06356 | -1.63242 | -1.729 | -2.33589 | -1.39601 | -0.66935 | -1.37004 | -1.13719 | -1.02144 | -0.99892 | -1.72625 | -1.40064 | -1.8658 | -1.0338 | -2.7234 | -1.85497 | -1.24783 | -0.97296 | -1.29117 | -0.90614 |
| 28.84583 | -1.8664 | -1.54553 | -1.45283 | -1.88348 | -1.589 | -0.81068 | -1.11154 | -1.17015 | -1.37056 | -0.88747 | -1.55057 | -1.7096 | -1.27368 | -1.02306 | -2.24092 | -1.78738 | -0.68997 | -0.95892 | -1.29339 | -0.73236 |
| 28.85972 | -1.55601 | -1.81845 | -1.33508 | -1.60591 | -1.11322 | -0.95189 | -0.84703 | -0.92726 | -2.10996 | -0.63468 | -1.53314 | -1.98901 | -0.79944 | -0.84593 | -1.7251 | -1.56102 | -0.2874 | -0.9054 | -1.08262 | -0.92983 |
| 28.87361 | -1.31227 | -1.31711 | -1.24327 | -1.29734 | -0.29956 | -0.97764 | -0.51332 | -0.598 | -1.86562 | -0.37003 | -1.2636 | -1.96862 | -0.42786 | -0.62627 | -1.21757 | -1.24541 | -0.04549 | -0.6475 | -0.76886 | -1.15383 |
| 28.8875 | -0.81769 | -0.73928 | -1.19109 | -0.79053 | 0.193923 | -0.78122 | -0.26121 | -0.33991 | -1.08413 | -0.19025 | -0.70172 | -1.76061 | -0.22636 | -0.35232 | -0.79256 | -0.97746 | 0.065748 | -0.36799 | -0.45369 | -1.08196 |
| 28.90139 | -0.27156 | -0.31424 | -0.8358 | -0.19825 | 0.504556 | -0.36341 | 0.021695 | 0.172038 | -0.30769 | -0.08305 | -0.05649 | -1.18321 | -0.13932 | -0.07652 | -0.41363 | -0.70209 | 0.176749 | -0.18061 | -0.09014 | -0.67807 |
| 28.91528 | 0.090742 | -0.29149 | -0.47802 | 0.309736 | 0.742095 | 0.06473 | 0.247822 | 0.587541 | 0.323409 | 0.121171 | 0.488266 | -0.87416 | -0.05506 | 0.169971 | -0.26351 | -0.83905 | 0.210163 | -0.0671 | 0.022973 | -0.25626 |
| 28.92917 | 0.290146 | -0.82675 | -0.90822 | 0.267169 | 0.598651 | -0.02148 | -0.06846 | 0.605387 | 0.519132 | 0.169623 | 0.543542 | -1.50941 | -0.41606 | 0.029609 | -0.71821 | -1.59183 | -0.18495 | -0.30683 | -0.53157 | -0.57695 |
| 28.94306 | -0.34793 | -1.54568 | -2.00038 | -0.49578 | -0.45187 | -0.66356 | -0.9858 | 0.112825 | 0.070052 | -0.58195 | -0.51441 | -2.59889 | -1.4883 | -1.1328 | -1.86787 | -2.64417 | -0.86428 | -0.92982 | -1.82034 | -1.85951 |
| 28.95694 | -1.38056 | -1.99023 | -2.71787 | -1.27698 | -1.6559 | -1.06519 | -1.79884 | -0.5004 | -0.42877 | -2.21291 | -2.09879 | -3.17644 | -3.0217 | -2.81531 | -3.11755 | -3.47273 | -1.24726 | -1.89181 | -3.2881 | -3.221 |
| 28.97083 | -1.75836 | -2.15046 | -2.51769 | -1.33574 | -1.92604 | -0.72589 | -1.81123 | -0.8686 | -0.5837 | -2.76646 | -2.41489 | -2.55931 | -3.21735 | -2.94999 | -3.00852 | -3.3926 | -0.94534 | -1.9661 | -3.10896 | -3.50422 |
| 28.98472 | -1.9733 | -2.32558 | -2.36661 | -1.15956 | -1.58122 | -0.41651 | -1.38748 | -0.93431 | -0.43607 | -1.23694 | -1.38395 | -1.73796 | -1.26734 | -1.95485 | -1.84774 | -2.17542 | -0.43395 | -1.20008 | -1.63604 | -2.08421 |
| 28.99861 | -1.67271 | -2.90015 | -2.12923 | -0.84753 | -1.22451 | -0.3282 | -1.29469 | -0.76056 | -0.29205 | 0.127341 | -0.23777 | -1.06196 | 0.223644 | -0.71363 | -1.14469 | -1.24921 | -0.26181 | -0.43625 | -0.45336 | -0.43559 |
| 29.0125 | -1.48528 | -3.27052 | -1.80961 | -0.17569 | -0.9803 | -0.11618 | -1.07357 | -0.42407 | -0.05144 | 0.620178 | 0.410252 | -0.32818 | 0.855544 | 0.266752 | -0.33034 | -0.3824 | -0.23991 | 0.08078 | 0.248857 | 0.630181 |
| 29.02639 | -1.15287 | -2.6946 | -1.55364 | -0.12454 | -0.63621 | 0.264195 | -0.91231 | -0.14503 | 0.416907 | 0.715314 | 0.528605 | 0.338554 | 0.96439 | 0.911546 | 0.396988 | 0.3472 | -0.21169 | 0.38169 | 0.717998 | 1.082442 |
| 29.04028 | -0.79645 | -1.74359 | -1.05927 | -0.36323 | -0.16447 | 0.466448 | -0.80169 | 0.106688 | 0.835491 | 0.805065 | 0.505736 | 0.788487 | 0.833655 | 1.147319 | 0.733471 | 0.849549 | -0.05656 | 0.637271 | 1.001084 | 1.296214 |
| 29.05417 | -0.25933 | -0.8397 | -0.24705 | -0.14705 | 0.259581 | 0.259374 | -0.62474 | 0.360839 | 0.963172 | 0.798367 | 0.41193 | 0.835422 | 0.5686 | 1.026559 | 0.806216 | 0.815183 | -0.00406 | 0.77422 | 1.13184 | 1.290807 |
| 29.06806 | 0.210561 | 0.05312 | 0.708003 | 0.127731 | 0.514766 | 0.196284 | -0.26051 | 0.570743 | 0.943444 | 0.706977 | 0.265827 | 0.913738 | 0.463324 | 0.936354 | 0.801568 | 0.636781 | 0.038503 | 0.64158 | 0.967531 | 1.107512 |
| 29.08194 | 0.400872 | 0.763827 | 1.276491 | 0.364169 | 0.582205 | 0.172038 | 0.177009 | 0.64877 | 0.791525 | 0.562425 | 0.019861 | 1.092058 | 0.612972 | 0.897663 | 0.911581 | 0.687617 | 0.001054 | 0.304351 | 0.62619 | 0.989294 |
| 29.09583 | 0.60196 | 0.972293 | 1.432642 | 0.482317 | 0.701189 | -0.08567 | 0.454206 | 0.733855 | 0.495381 | 0.434913 | -0.14723 | 0.773129 | 0.653074 | 0.631819 | 0.839526 | 0.360311 | -0.03206 | -0.21239 | 0.162813 | 0.500042 |
| 29.10972 | 0.734097 | 1.029507 | 1.322185 | 0.569851 | 0.771658 | -0.22327 | 0.50782 | 0.8533 | 0.295566 | 0.288586 | -0.05108 | 0.330583 | 0.365502 | 0.33214 | 0.573745 | -0.01904 | 0.065145 | -0.58287 | -0.12472 | -0.09031 |
| 29.12361 | 0.518529 | 1.27441 | 0.836125 | 0.49148 | 0.641104 | -0.03708 | 0.936185 | 0.895926 | 0.401204 | 0.164001 | 0.210558 | 0.307054 | 0.231624 | 0.216922 | 0.53411 | -0.00062 | 0.098568 | -0.50471 | -0.1505 | -0.30868 |
| 29.1375 | 0.268578 | 1.446685 | 0.850133 | 0.348261 | 0.54232 | 0.203668 | 1.249745 | 0.918 | 0.635406 | 0.069298 | 0.387415 | 0.29383 | 0.184425 | 0.121924 | 0.436858 | 0.018868 | 0.058141 | -0.37004 | -0.17441 | -0.40607 |
| 29.15139 | 0.379872 | 1.407083 | 0.723423 | 0.39959 | 0.366874 | 0.28582 | 1.342342 | 0.756061 | 0.65448 | 0.031087 | 0.347106 | 0.104914 | 0.06335 | 0.039633 | 0.25126 | -0.14878 | 0.025881 | -0.27104 | -0.12229 | -0.46874 |
| 29.16528 | 0.544206 | 1.41777 | 0.26801 | 0.679118 | 0.11876 | 0.237483 | 1.207818 | 0.787829 | 0.525324 | 0.056043 | 0.228821 | -0.07637 | -0.02562 | 0.104586 | 0.142446 | -0.18225 | -0.0047 | -0.25953 | 0.139857 | -0.43107 |
| 29.17917 | 0.724767 | 1.580085 | -0.12805 | 0.743766 | -0.08633 | 0.17401 | 1.286083 | 0.671372 | 0.434839 | 0.061545 | 0.186216 | -0.10539 | -0.03216 | 0.2276 | 0.207597 | 0.121783 | -0.02613 | -0.27978 | 0.453734 | -0.11315 |
| 29.19306 | 0.615715 | 1.676777 | -0.36757 | 0.637442 | -0.16922 | 0.135948 | 1.613498 | 0.462938 | 0.361064 | -0.03775 | 0.366488 | -0.1032 | -0.03001 | 0.186199 | 0.335982 | 0.430442 | -0.05746 | -0.1869 | 0.500458 | 0.26494 |
| 29.20694 | 0.924505 | 1.291758 | -0.63001 | 0.523664 | -0.09091 | -0.02769 | 1.60662 | 0.358403 | 0.149279 | -0.17092 | 0.627396 | -0.22061 | -0.15915 | -0.12539 | 0.343515 | 0.148068 | -0.05608 | -0.15131 | 0.081268 | 0.27721 |
| 29.22083 | 1.22268 | 0.746292 | -0.98885 | 0.448845 | 0.19035 | -0.17798 | 1.073741 | 0.279897 | -0.11075 | -0.15092 | 0.754371 | -0.37325 | -0.24889 | -0.4143 | 0.266777 | -0.38189 | 0.041102 | -0.13535 | -0.35434 | -0.07076 |
| 29.23472 | 1.007058 | 0.61647 | -0.78642 | 0.388704 | 0.268571 | -0.07017 | 0.808098 | 0.245909 | -0.00563 | -0.06573 | 0.641077 | -0.05469 | -0.01767 | -0.3913 | 0.324382 | -0.32392 | 0.024312 | 0.031443 | -0.2435 | -0.27034 |
| 29.24861 | 0.811945 | 0.796152 | -0.40633 | 0.34735 | 0.202956 | 0.170095 | 0.745526 | 0.152505 | 0.350341 | 0.034869 | 0.386957 | 0.381882 | 0.293272 | -0.17289 | 0.519092 | 0.153515 | -0.06786 | 0.178036 | 0.284898 | -0.21832 |
| 29.2625 | 0.746507 | 0.910392 | -0.02528 | 0.237525 | 0.118639 | 0.320227 | 0.670896 | 0.139962 | 0.614984 | 0.178126 | 0.2029 | 0.574899 | 0.323347 | 0.010543 | 0.666601 | 0.520008 | -0.07283 | 0.201135 | 0.774116 | -0.0034 |
| 29.27639 | 0.762235 | 0.90378 | 0.22046 | 0.099806 | 0.075959 | 0.280058 | 0.560842 | 0.182071 | 0.583682 | 0.262802 | 0.108663 | 0.555455 | 0.101401 | 0.131987 | 0.771658 | 0.683087 | -0.00945 | 0.124608 | 0.899683 | 0.305066 |
| 29.29028 | 0.608173 | 0.634824 | 0.335027 | 0.051335 | 0.095411 | 0.161141 | 0.318897 | 0.326971 | 0.419237 | 0.231397 | 0.216644 | 0.448226 | -0.09455 | 0.328971 | 0.851552 | 0.704066 | 0.086805 | 0.062441 | 0.713326 | 0.590592 |
| 29.30417 | 0.359741 | 0.456453 | 0.521471 | 0.079853 | 0.138316 | 0.122497 | 0.040482 | 0.484824 | 0.360592 | 0.150342 | 0.493952 | 0.383794 | -0.17517 | 0.512752 | 0.900409 | 0.620995 | 0.105219 | 0.006547 | 0.500298 | 0.69723 |
| 29.31806 | 0.269955 | 0.51561 | 0.740961 | 0.110302 | 0.171098 | 0.112589 | -0.00536 | 0.557403 | 0.358158 | 0.085594 | 0.792847 | 0.40176 | -0.10491 | 0.512447 | 0.882339 | 0.44711 | 0.050931 | 0.041504 | 0.41575 | 0.494703 |
| 29.33194 | 0.09128 | 0.634538 | 0.860014 | 0.170991 | 0.233908 | 0.049358 | -0.04946 | 0.623683 | 0.272226 | 0.085841 | 0.944118 | 0.460821 | 0.159502 | 0.393801 | 0.832606 | 0.387401 | -0.02625 | 0.160715 | 0.451591 | 0.269543 |
| 29.34583 | 0.322486 | 0.754808 | 0.935426 | 0.280496 | 0.332001 | -0.0166 | 0.216255 | 0.667721 | 0.146875 | 0.126294 | 0.765562 | 0.407293 | 0.409874 | 0.203904 | 0.727599 | 0.335208 | -0.0563 | 0.249558 | 0.457263 | 0.089209 |
| 29.35972 | 0.880169 | 0.855259 | 0.936144 | 0.402193 | 0.410235 | -0.05025 | 0.474234 | 0.622217 | 0.129175 | 0.223554 | 0.363756 | 0.245422 | 0.451722 | 0.04239 | 0.518835 | 0.118624 | 0.007383 | 0.237386 | 0.379422 | -0.05382 |
| 29.37361 | 1.187101 | 1.058529 | 0.83352 | 0.504206 | 0.414634 | -0.00991 | 0.744441 | 0.538679 | 0.229766 | 0.369698 | 0.088935 | 0.277281 | 0.393071 | 0.073437 | 0.440408 | 0.040176 | 0.051396 | 0.26416 | 0.450943 | 0.133483 |
| 29.3875 | 1.388071 | 1.266727 | 0.519012 | 0.632255 | 0.34519 | 0.093249 | 0.887985 | 0.450452 | 0.434803 | 0.392746 | 0.101406 | 0.489103 | 0.358454 | 0.28224 | 0.587659 | 0.255888 | 0.035711 | 0.286576 | 0.654862 | 0.605306 |
| 29.40139 | 1.350065 | 1.236023 | 0.186529 | 0.577837 | 0.057929 | 0.305375 | 0.831177 | 0.392157 | 0.779617 | 0.237983 | 0.221467 | 0.857935 | 0.407531 | 0.616187 | 0.872395 | 0.705829 | -0.03452 | 0.279203 | 0.811226 | 0.974209 |
| 29.41528 | 0.767974 | 1.267139 | 0.127088 | 0.520884 | -0.38068 | 0.612776 | 0.81479 | 0.383492 | 1.422277 | -0.00758 | 0.303116 | 1.522814 | 0.657818 | 0.942677 | 1.279843 | 1.46465 | -0.11271 | 0.362774 | 0.952871 | 1.223791 |
| 29.42917 | 0.755376 | 1.643056 | 0.549121 | 0.805492 | -0.48594 | 0.899137 | 0.905135 | 0.486334 | 1.713465 | 0.024558 | 0.534906 | 2.297703 | 1.254249 | 1.196171 | 1.838434 | 2.182113 | 0.138918 | 0.577968 | 1.217264 | 1.525083 |
| 29.44306 | 1.800337 | 2.355528 | 1.305048 | 2.005519 | 0.179204 | 0.771856 | 1.330393 | 0.755317 | 1.384006 | 0.919592 | 1.824492 | 2.373752 | 2.358063 | 1.386742 | 2.720454 | 2.644211 | 0.484748 | 0.84369 | 1.684621 | 1.880714 |
| 29.45694 | 3.380274 | 2.744782 | 1.848637 | 3.650402 | 1.588792 | 0.297409 | 2.041442 | 1.085739 | 0.681062 | 1.727327 | 2.570913 | 2.050743 | 2.916229 | 1.627667 | 3.048116 | 2.834175 | 1.010695 | 1.191144 | 2.155408 | 2.272941 |
| 29.47083 | 3.186587 | 3.149048 | 2.119534 | 3.312932 | 2.195152 | 0.481526 | 2.245781 | 1.409106 | 0.947964 | 1.756926 | 2.543897 | 2.019408 | 2.841953 | 2.071676 | 3.282886 | 3.377666 | 1.379464 | 1.366915 | 2.332042 | 2.468561 |
| 29.48472 | 2.610137 | 2.806373 | 1.681613 | 2.989305 | 2.272393 | 0.754613 | 2.113292 | 1.827057 | 1.158635 | 1.742219 | 2.451418 | 2.46896 | 2.532541 | 2.966633 | 3.797169 | 3.831348 | 1.603791 | 1.203978 | 2.08071 | 2.532062 |
| 29.49861 | 2.103616 | 2.252714 | 2.084611 | 2.94824 | 2.698441 | 0.854762 | 2.251535 | 1.904722 | 1.073536 | 1.826422 | 2.502522 | 2.835027 | 2.252304 | 3.645523 | 3.998861 | 3.790778 | 1.784794 | 1.140664 | 1.944949 | 2.462442 |
| 29.5125 | 1.359188 | 1.293564 | 2.162114 | 2.919906 | 2.245374 | 0.834176 | 2.029496 | 1.492808 | 0.812194 | 1.514066 | 2.140249 | 2.864086 | 1.907006 | 4.275895 | 3.674077 | 3.451592 | 1.615403 | 1.072395 | 1.811389 | 2.333614 |
| 29.52639 | 0.922438 | 0.574119 | 1.99285 | 2.457395 | 1.392287 | 0.796719 | 1.356953 | 0.960686 | 0.531954 | 1.048567 | 1.636138 | 2.696656 | 1.635014 | 3.489054 | 3.235855 | 2.862198 | 1.404731 | 1.089014 | 1.792862 | 2.183457 |
| 29.54028 | 0.821644 | 0.355499 | 1.568282 | 2.555841 | 0.914517 | 0.627794 | 0.5807 | 0.602428 | 0.354104 | 0.53442 | 1.179988 | 2.208275 | 1.433822 | 2.380159 | 2.324168 | 1.275401 | 1.156059 | 1.087759 | 1.464927 | 1.632014 |
| 29.55417 | 0.484428 | 0.279613 | 1.364141 | 2.766744 | 1.034623 | 0.548827 | 0.105231 | 0.425075 | 0.299482 | 0.136183 | 0.834555 | 1.559293 | 1.037509 | 1.353518 | 1.162294 | 0.050441 | 1.055087 | 1.169986 | 0.947997 | 1.147115 |
| 29.56806 | 0.331072 | 0.141646 | 0.92485 | 2.373802 | 1.340216 | 0.618237 | -0.09544 | 0.333919 | 0.084544 | -0.05188 | 0.570149 | 0.860183 | 0.551382 | 0.294928 | 0.13756 | -0.7802 | 0.785324 | 1.260761 | 1.021854 | 0.671328 |
| 29.58194 | 0.232549 | 0.131617 | 0.287774 | 1.278473 | 1.178998 | 0.576656 | -0.26063 | 0.063391 | -0.08407 | -0.30029 | 0.509607 | 0.327796 | 0.049145 | 0.07386 | -0.92546 | -0.90358 | 0.390493 | 1.30548 | 0.872188 | 0.351943 |
| 29.59583 | 0.273773 | 0.157415 | -0.18564 | 0.831615 | 0.818651 | 0.532059 | -0.16707 | 0.08877 | 0.241794 | -0.68775 | 0.222267 | 0.261221 | -0.11115 | 0.225477 | -1.99183 | -0.67292 | 0.156151 | 1.339478 | 0.472794 | 0.184284 |
| 29.60972 | 0.103824 | 0.196279 | -0.57807 | 0.537418 | 0.385358 | 0.61383 | -0.01966 | 0.136628 | 0.712904 | -0.84906 | 0.009343 | 0.148814 | -0.06053 | 0.333112 | -2.53212 | -0.81838 | 0.125959 | 1.408267 | -0.16019 | 0.224894 |

| | | | | | | | | | | | | | | | | | | | | |
|---|---|---|---|---|---|---|---|---|---|---|---|---|---|---|---|---|---|---|---|---|
| 29.62361 | -0.25082 | 0.425471 | -1.09514 | 0.210941 | -0.00846 | 0.546817 | 0.074675 | 0.072835 | 1.04643 | -0.8061 | -0.21555 | 0.076987 | 0.141341 | 0.344089 | -2.40874 | -0.8128 | 0.100813 | 1.051032 | -0.9765 | 0.274951 |
| 29.6375 | -0.74023 | 0.949084 | -1.16692 | -0.42333 | -0.10997 | 0.478579 | 0.17765 | 0.080199 | 1.136976 | -0.59704 | -0.45106 | -0.12851 | 0.203541 | 0.240561 | -2.11203 | -0.86392 | 0.18988 | 0.560853 | -1.39868 | 0.103628 |
| 29.65139 | -0.94071 | 1.583019 | -0.96694 | -1.09025 | 0.203878 | 0.300787 | 0.133305 | 0.01946 | 0.782695 | -0.28814 | -0.77324 | -0.49696 | -0.01932 | -0.18186 | -1.18307 | -1.18003 | 0.40493 | 0.2637 | -1.69763 | -0.26814 |
| 29.66528 | -1.09005 | 0.784367 | -0.8413 | -1.53164 | 0.391772 | -0.08683 | -0.00588 | -0.07017 | 0.151046 | -0.00723 | -0.93707 | -0.82615 | -0.53002 | -0.84516 | -0.25471 | -1.66919 | 0.653853 | 0.133416 | -2.05283 | -0.93734 |
| 29.67917 | -1.12141 | 0.439219 | -0.76408 | -1.68099 | 0.244545 | -0.48882 | -0.09809 | -0.09592 | -0.21401 | -0.01347 | -0.79989 | -1.02062 | -1.02195 | -1.70635 | 0.528702 | -1.93008 | 0.663155 | 0.188112 | -2.20224 | -1.36227 |
| 29.69306 | -1.14432 | -0.15238 | -0.77459 | -1.55187 | -0.05677 | -0.79587 | -0.05206 | -0.1539 | -0.48666 | -0.20289 | -0.75012 | -1.04205 | -1.50986 | -2.01829 | 1.038945 | -2.13535 | 0.476257 | 0.114525 | -2.08958 | -1.60088 |
| 29.70694 | -1.10964 | -1.11704 | -0.76422 | -1.90669 | -0.23941 | -1.09728 | -0.09703 | -0.33875 | -0.74565 | -0.15742 | -0.99444 | -1.08885 | -1.87497 | -2.34454 | 1.068304 | -2.48144 | 0.161451 | -0.10056 | -1.92115 | -1.90857 |
| 29.72083 | -1.14149 | -1.90555 | -0.63857 | -1.4735 | -0.3844 | -1.21409 | -0.14241 | -0.69517 | -0.67086 | 0.007775 | -1.21604 | -1.11095 | -1.9218 | -2.24777 | 0.723716 | -2.60707 | -0.36608 | -0.40482 | -1.54899 | -1.90083 |
| 29.73472 | -1.29675 | -1.75163 | -0.48576 | -0.31406 | -0.65108 | -1.05228 | -0.10901 | -0.71066 | -0.16381 | -0.09587 | -1.30594 | -1.17096 | -1.65794 | -1.85761 | 0.417407 | -2.32231 | -0.87002 | -0.6645 | -0.99811 | -1.11746 |
| 29.74861 | -1.37409 | -1.31381 | -0.50103 | 0.994079 | -1.025 | -0.81712 | -0.04319 | -0.51205 | 0.343177 | -0.33714 | -1.46436 | -1.39555 | -1.39176 | -1.65896 | 0.086045 | -1.94562 | -1.46932 | -1.04008 | -0.3993 | -0.21262 |
| 29.7625 | -1.39442 | -1.03356 | -0.60141 | 1.628713 | -1.11321 | -0.70315 | -0.2115 | -0.46326 | 0.469812 | -0.33916 | -1.50739 | -1.60901 | -1.11509 | -1.57389 | -0.47091 | -1.71292 | -2.00217 | -1.3107 | -0.04663 | 0.284209 |
| 29.77639 | -1.38575 | -0.87376 | -0.85973 | 1.358539 | -1.19577 | -0.81372 | -0.54812 | -0.51144 | 0.280628 | -0.29182 | -1.3406 | -1.40346 | -0.8698 | -1.54782 | -0.94675 | -1.29422 | -2.29511 | -1.38983 | 0.121017 | 0.046972 |
| 29.79028 | -1.35348 | -1.2139 | -1.22281 | 0.852357 | -1.26289 | -1.04426 | -0.82631 | -0.58625 | -0.18333 | -0.45525 | -1.23802 | -1.46958 | -0.68309 | -1.46031 | -1.46775 | -1.07361 | -2.01294 | -1.72906 | 0.227135 | -0.54273 |
| 29.80417 | -1.19565 | -1.45723 | -1.38438 | -0.01474 | -1.19674 | -1.42451 | -0.99083 | -0.83879 | -0.88251 | -0.66114 | -1.26398 | -1.93955 | -0.66629 | -1.39134 | -2.28021 | -1.15031 | -1.72125 | -2.22278 | 0.347887 | -1.22051 |
| 29.81806 | -1.44936 | -1.39556 | -1.42058 | -1.17539 | -1.10012 | -1.94814 | -1.53039 | -1.01054 | -1.21127 | -0.88138 | -1.24476 | -2.20974 | -0.69221 | -1.30989 | -2.99979 | -1.42929 | -1.33472 | -2.35274 | 0.305271 | -1.61348 |
| 29.83194 | -1.55869 | -1.12725 | -1.6624 | -1.66142 | -0.93475 | -2.2408 | -2.21494 | -1.03093 | -1.47507 | -1.13958 | -1.1429 | -2.09304 | -0.69032 | -1.10111 | -2.88892 | -1.75461 | -0.89804 | -2.23382 | 0.136815 | -1.40295 |
| 29.84583 | -1.96914 | -0.93518 | -1.63322 | -1.59907 | -0.78462 | -2.29607 | -2.32606 | -0.83283 | -1.09792 | -1.21251 | -1.01574 | -1.91674 | -0.67262 | -1.04393 | -2.38127 | -1.99324 | -0.79131 | -2.14974 | -0.14755 | -1.15082 |
| 29.85972 | -1.78122 | -0.97245 | -1.1401 | -1.0835 | -0.57052 | -2.02456 | -1.86364 | -0.54741 | -0.82926 | -1.13067 | -1.0831 | -1.92264 | -0.80531 | -1.06113 | -1.69722 | -2.09856 | -0.78164 | -1.85367 | -0.60936 | -0.6736 |
| 29.87361 | -0.9693 | -0.85016 | -0.70129 | -0.36239 | -0.13849 | -1.26482 | -1.37513 | -0.14425 | -0.37427 | -0.70868 | -1.04561 | -1.86877 | -0.84739 | -1.05055 | -1.32591 | -2.15479 | -0.78551 | -1.49625 | -0.87643 | -0.22885 |
| 29.8875 | -0.06406 | -0.35137 | -0.19664 | 0.121647 | 0.139706 | -0.42121 | -0.52614 | 0.143821 | 0.12521 | -0.21849 | -0.702 | -1.57398 | -0.66326 | -0.70168 | -1.14336 | -1.65842 | -0.57627 | -0.96166 | -0.69592 | 0.048043 |
| 29.90139 | 0.632219 | 0.12106 | 0.125826 | 0.195481 | 0.225111 | 0.057216 | 0.321175 | 0.316753 | 0.289084 | 0.055632 | -0.26012 | -1.19194 | -0.3747 | -0.30731 | -0.99034 | -0.86694 | -0.331 | -0.5017 | -0.38385 | 0.007827 |
| 29.91528 | 0.950643 | -0.02647 | 0.194074 | 0.015266 | 0.49006 | -0.17251 | 0.654198 | 0.388744 | 0.008381 | 0.32064 | 0.157407 | -1.10331 | -0.34182 | -0.16677 | -0.85219 | -0.54339 | -0.20726 | -0.36871 | 0.031217 | -0.23667 |
| 29.92917 | 0.714098 | -0.83783 | -0.24599 | -0.57243 | 0.462443 | -0.69945 | 0.057372 | 0.252921 | -0.46863 | 0.229681 | 0.212365 | -1.30446 | -0.66105 | -0.71378 | -1.11046 | -0.90971 | -0.39597 | -0.56982 | -0.09687 | -0.92589 |
| 29.94306 | -0.6555 | -1.74775 | -0.99776 | -2.09481 | -0.54891 | -1.32934 | -0.76503 | -0.20938 | -0.74689 | -0.83287 | -0.73333 | -1.75604 | -1.42466 | -2.0389 | -2.23407 | -1.98646 | -0.96325 | -1.27922 | -1.22224 | -1.97636 |
| 29.95694 | -1.87283 | -2.64635 | -1.66924 | -3.60767 | -1.59322 | -1.92263 | -1.39403 | -0.78022 | -0.86062 | -2.37154 | -2.1486 | -2.5807 | -2.82689 | -3.58215 | -4.11743 | -4.00316 | -1.32193 | -2.6002 | -3.21719 | -3.78451 |
| 29.97083 | -1.99429 | -3.11089 | -1.70533 | -3.93046 | -1.87416 | -2.15938 | -1.156 | -0.74597 | -0.70165 | -2.44557 | -2.24179 | -2.63767 | -2.71131 | -3.15666 | -3.22787 | -4.42558 | -0.69267 | -2.41661 | -3.68233 | -3.75726 |
| 29.98472 | -2.02203 | -2.61599 | -1.707 | -2.28339 | -2.01095 | -1.34829 | -0.37534 | -0.33763 | -0.37095 | -1.13363 | -1.39649 | -1.57694 | -0.84236 | -1.67106 | -1.19347 | -1.94935 | -0.54206 | -1.42057 | -2.58742 | -1.69001 |
| 29.99861 | -1.71777 | -1.95645 | -1.67692 | -1.22656 | -2.11287 | -0.66007 | -0.05247 | 0.095849 | -0.1279 | -0.11786 | -0.3419 | -0.72919 | 0.32969 | -0.3565 | -0.06397 | 0.139829 | -0.3697 | -0.52995 | -1.2548 | 0.139429 |
| 30.0125 | -1.20786 | -2.32581 | -1.57172 | -1.33247 | -1.48483 | -0.10402 | -0.26165 | 0.218267 | 0.448328 | 0.775881 | 0.625743 | -0.48871 | 0.607351 | 0.376586 | 0.468714 | 0.823844 | 0.034522 | 0.074618 | 0.388541 | 0.568595 |
| 30.02639 | -0.77004 | -2.56969 | -1.3403 | -1.09173 | -0.86119 | 0.496158 | -0.31228 | 0.281293 | 0.993754 | 1.249852 | 1.315891 | -0.77397 | 0.583526 | 0.518626 | 0.63963 | 0.829408 | 0.41761 | 0.420885 | 1.418795 | 0.464089 |
| 30.04028 | -0.53318 | -2.05858 | -1.23221 | -0.50013 | -0.63607 | 1.150163 | -0.09615 | 0.165224 | 1.214064 | 1.087415 | 1.459943 | -0.44297 | 0.82295 | 0.558692 | 0.727442 | 1.29683 | 0.295238 | 0.68897 | 1.713234 | 0.743892 |
| 30.05417 | -0.40947 | -1.17107 | -0.63948 | 0.05676 | -0.38485 | 1.520428 | 0.140005 | 0.001258 | 1.253935 | 0.795961 | 1.30957 | 0.215872 | 1.032468 | 0.739901 | 0.801597 | 1.798723 | 0.050744 | 0.962673 | 1.86648 | 0.898618 |
| 30.06806 | -0.35839 | -0.66192 | 0.330394 | 0.405198 | -0.1977 | 1.520334 | 0.269927 | -0.08893 | 1.290789 | 0.592987 | 0.791156 | 0.67294 | 0.892092 | 0.843559 | 0.637228 | 1.638771 | -0.15448 | 0.918681 | 1.74008 | 0.799572 |
| 30.08194 | -0.14508 | -0.32057 | 1.166929 | 0.719204 | 0.112575 | 1.795183 | 0.269905 | 0.015198 | 1.252454 | 0.513698 | 0.322345 | 0.820492 | 0.601435 | 0.698561 | 0.471872 | 1.080863 | -0.2284 | 0.743422 | 1.316242 | 0.638355 |
| 30.09583 | 0.476055 | 0.329558 | 1.592785 | 1.323688 | 0.84395 | 1.93424 | 0.915056 | 0.647736 | 1.314964 | 0.847662 | 0.238517 | 1.116843 | 0.636898 | 0.720668 | 0.646748 | 0.896905 | -0.02484 | 0.785744 | 1.061008 | 0.710443 |
| 30.10972 | 1.51924 | 1.474129 | 2.055896 | 2.225978 | 1.972356 | 2.026047 | 1.981828 | 1.658618 | 1.727772 | 1.514008 | 0.571615 | 1.84828 | 1.284616 | 1.212518 | 1.363063 | 1.3049 | 0.475776 | 1.063201 | 1.309932 | 1.254508 |
| 30.12361 | 2.158561 | 2.091781 | 2.251936 | 2.412944 | 2.382099 | 1.707706 | 2.306722 | 1.948883 | 1.556835 | 1.688146 | 0.8027 | 2.139313 | 1.688182 | 1.668287 | 1.783512 | 1.711682 | 0.696818 | 1.061674 | 1.456974 | 1.508503 |
| 30.1375 | 1.754111 | 1.426348 | 2.126437 | 1.445885 | 1.830634 | 1.037896 | 1.611511 | 1.212025 | 0.498281 | 1.104334 | 0.522356 | 1.748005 | 1.361033 | 1.37793 | 1.34173 | 1.377185 | 0.436515 | 0.538221 | 0.872747 | 0.962799 |
| 30.15139 | 1.269031 | 0.982742 | 2.044669 | 0.856199 | 1.32737 | 0.502801 | 0.928475 | 0.624086 | 0.051364 | 0.38473 | 0.065325 | 1.329725 | 0.733996 | 0.756238 | 0.686059 | 0.963607 | 0.062523 | 0.069906 | 0.426441 | 0.355266 |
| 30.16528 | 0.973775 | 0.983966 | 2.204749 | 0.829153 | 1.053972 | 0.8253 | 0.831935 | 0.355255 | 0.409793 | -0.11381 | -0.2052 | 1.130977 | 0.31464 | 0.343289 | 0.356016 | 0.946655 | -0.26615 | -0.11689 | 0.224078 | 0.20382 |
| 30.17917 | 0.794348 | 0.882918 | 1.61715 | 0.940273 | 1.04401 | 0.529743 | 0.733798 | 0.265744 | 0.533066 | -0.27128 | -0.32091 | 0.507692 | -0.09938 | -0.03905 | 0.052679 | 0.301605 | -0.30566 | -0.16081 | -0.14717 | -0.01128 |
| 30.19306 | 0.758081 | 0.616517 | 0.523791 | 0.898092 | 1.184253 | -0.42474 | 0.443482 | 0.419086 | 0.065351 | -0.12768 | -0.36638 | -0.38923 | -0.45954 | -0.27556 | -0.25618 | -0.92547 | 0.052912 | -0.21488 | -0.66763 | -0.31364 |
| 30.20694 | 0.462187 | 0.493886 | -0.53975 | 0.639301 | 1.082344 | -0.74289 | 0.24793 | 0.393755 | -0.18271 | -0.01784 | -0.47642 | -0.8849 | -0.4155 | -0.45774 | -0.43819 | -1.47681 | 0.478453 | -0.30331 | -0.94712 | -0.4027 |
| 30.22083 | 0.165915 | 0.577157 | -1.29402 | 0.382748 | 0.709766 | -0.59146 | 0.317964 | 0.267957 | -0.04532 | -0.0339 | -0.41018 | -0.88316 | -0.24392 | -0.38381 | -0.36634 | -1.07438 | 0.590858 | -0.36875 | -0.74199 | -0.23141 |
| 30.23472 | 0.228831 | 0.721096 | -1.54894 | 0.258252 | 0.293314 | -0.71403 | 0.544304 | 0.210762 | 0.159561 | 0.043078 | 0.033039 | -0.69322 | -0.17271 | -0.23293 | -0.11557 | -0.30914 | 0.431104 | -0.16228 | -0.13869 | 0.018534 |
| 30.24861 | 0.386945 | 0.72834 | -1.49821 | 0.195908 | 0.040556 | -0.5051 | 0.782408 | 0.24715 | 0.341992 | 0.145436 | 0.531723 | -0.42544 | -0.08218 | -0.00528 | 0.086579 | 0.26042 | 0.259353 | 0.093051 | 0.456028 | 0.173576 |
| 30.2625 | 0.505077 | 0.591811 | -1.39687 | 0.123089 | -0.11449 | -0.15788 | 0.925394 | 0.378038 | 0.394997 | 0.154836 | 0.731787 | -0.17291 | -0.01054 | 0.12655 | 0.079898 | 0.515974 | 0.169336 | 0.151296 | 0.750736 | 0.199848 |
| 30.27639 | 0.579047 | 0.555515 | -1.1324 | 0.18367 | -0.12211 | 0.211131 | 0.954884 | 0.497882 | 0.409383 | 0.153628 | 0.678824 | 0.035045 | 0.106103 | 0.197038 | 0.083359 | 0.692638 | 0.254304 | 0.037829 | 0.727341 | 0.194066 |
| 30.29028 | 0.611447 | 0.674576 | -0.61075 | 0.377776 | 0.043434 | 0.55735 | 1.025306 | 0.566266 | 0.549293 | 0.188964 | 0.549783 | 0.316183 | 0.312144 | 0.237405 | 0.322131 | 0.93328 | 0.451491 | -0.10359 | 0.570129 | 0.255919 |
| 30.30417 | 0.547959 | 0.79148 | -0.01648 | 0.479613 | 0.116493 | 0.723211 | 1.029774 | 0.552626 | 0.705351 | 0.229511 | 0.344126 | 0.487264 | 0.501692 | 0.207516 | 0.578929 | 1.003369 | 0.523824 | -0.18605 | 0.399373 | 0.254704 |
| 30.31806 | 0.357233 | 0.776866 | 0.35319 | 0.499667 | 0.137875 | 0.743225 | 1.015839 | 0.476024 | 0.827333 | 0.246276 | 0.198551 | 0.471354 | 0.535388 | 0.109394 | 0.588445 | 0.922837 | 0.36041 | -0.24555 | 0.272177 | 0.228 |
| 30.33194 | 0.213884 | 0.940282 | 0.675724 | 0.578197 | -0.01933 | 0.868775 | 0.997253 | 0.466999 | 1.043787 | 0.135097 | 0.097405 | 0.6979 | 0.506662 | 0.164152 | 0.525117 | 1.13016 | -0.09903 | -0.09445 | 0.424274 | 0.423757 |
| 30.34583 | 0.264415 | 1.105856 | 1.070505 | 0.641607 | -0.14996 | 0.942211 | 0.98985 | 0.529422 | 1.083016 | 0.023294 | 0.120749 | 0.984248 | 0.399612 | 0.254491 | 0.436252 | 1.331817 | -0.394 | 0.16943 | 0.734042 | 0.683772 |
| 30.35972 | 0.622338 | 0.90119 | 1.315357 | 0.474826 | 0.241613 | 0.46314 | 0.680468 | 0.537291 | 0.600189 | 0.197292 | 0.50255 | 0.593768 | -0.07158 | 0.038421 | 0.23219 | 0.666634 | -0.0453 | 0.37119 | 1.007913 | 0.444979 |
| 30.37361 | 0.985722 | 0.650684 | 1.355098 | 0.336335 | 0.704003 | -0.04552 | 0.309721 | 0.448428 | 0.144443 | 0.383979 | 0.789663 | 0.017937 | -0.2978 | -0.08923 | 0.295886 | -0.09963 | 0.563997 | 0.390012 | 0.734888 | 0.081967 |
| 30.3875 | 1.041047 | 0.636933 | 0.97249 | 0.223409 | 0.609996 | -0.15084 | 0.137762 | 0.426279 | 0.113597 | 0.25782 | 0.557542 | 0.05606 | 0.098638 | -0.01698 | 0.573371 | 0.053649 | 0.65199 | 0.248865 | 0.362611 | 0.139462 |
| 30.40139 | 0.659037 | 0.763971 | 0.498867 | 0.332308 | 0.242629 | 0.082521 | -0.00843 | 0.497226 | 0.323904 | 0.043927 | 0.16168 | 0.528015 | 0.634901 | 0.183989 | 0.824312 | 0.911375 | 0.199055 | 0.103451 | 0.110874 | 0.416976 |
| 30.41528 | 0.244091 | 0.945349 | 0.193271 | 0.580222 | -0.07077 | 0.45268 | 0.00068 | 0.697622 | 0.638708 | -0.13425 | -0.16487 | 1.214204 | 1.129688 | 0.503662 | 0.972955 | 2.059253 | -0.47282 | 0.088437 | 0.006148 | 0.824039 |
| 30.42917 | 0.308102 | 1.324306 | 0.244699 | 0.794772 | 0.263322 | 0.714637 | 0.421446 | 1.073363 | 0.713429 | 0.162772 | -0.00082 | 1.764149 | 1.305224 | 0.885582 | 1.101468 | 3.093608 | -0.72074 | 0.139869 | 0.374481 | 1.353365 |
| 30.44306 | 1.515713 | 1.722266 | 1.001458 | 0.993403 | 1.558436 | 0.455649 | 1.120464 | 1.544925 | 0.29775 | 1.780662 | 1.328747 | 1.783018 | 1.893957 | 1.691827 | 1.899185 | 3.684185 | 0.117373 | 0.639188 | 1.276765 | 2.152113 |
| 30.45694 | 3.221168 | 2.440498 | 2.009491 | 1.662569 | 2.517998 | 0.221006 | 2.049469 | 1.853285 | 0.146931 | 1.837805 | 2.465772 | 1.583763 | 2.726314 | 2.100977 | 3.021165 | 4.022363 | 1.132631 | 1.419324 | 2.283758 | 2.935401 |
| 30.47083 | 3.229488 | 2.623172 | 2.614847 | 3.705145 | 2.212048 | 0.325978 | 2.441701 | 1.600979 | 0.717225 | 1.807398 | 2.755717 | 1.702024 | 2.837346 | 2.470132 | 3.173233 | 4.115488 | 1.53012 | 1.778012 | 3.01258 | 3.086607 |
| 30.48472 | 2.60244 | 2.959397 | 3.057867 | 3.441013 | 1.733793 | 0.677321 | 2.533496 | 1.131621 | 2.249021 | 1.715417 | 2.673503 | 1.939921 | 2.553292 | 2.854651 | 3.583684 | 4.222378 | 1.63913 | 1.696239 | 3.475318 | 2.957526 |
| 30.49861 | 1.525825 | 2.800938 | 3.042539 | 3.730729 | 1.588478 | 0.698427 | 2.745933 | 0.88223 | 2.347542 | 1.526255 | 2.331676 | 2.082128 | 2.141749 | 3.517781 | 3.858019 | 4.332318 | 2.095222 | 1.349661 | 3.341248 | 2.557436 |
| 30.5125 | 0.453123 | 2.168055 | 2.798995 | 3.92163 | 1.457925 | 0.672802 | 2.543954 | 0.689704 | 1.703704 | 1.356385 | 1.848467 | 2.312133 | 2.003193 | 4.014669 | 3.58828 | 3.804735 | 2.084349 | 1.254963 | 2.770238 | 1.89886 |
| 30.52639 | -0.03466 | 1.8626 | 2.483576 | 3.674653 | 1.068052 | 0.681759 | 1.727318 | 0.537683 | 0.913835 | 1.065193 | 1.26249 | 2.548863 | 1.874823 | 4.062718 | 3.232356 | 3.25022 | 1.921757 | 1.151355 | 1.835984 | 1.343221 |
| 30.54028 | 0.034344 | 1.971971 | 2.139313 | 3.669259 | 1.209519 | 0.498021 | 1.137077 | 0.442044 | 0.062439 | 0.894098 | 0.797788 | 2.42165 | 1.543027 | 3.330775 | 2.651871 | 2.281273 | 1.457452 | 1.387596 | 0.922133 | 0.965691 |
| 30.55417 | -0.05607 | 2.036096 | 1.485279 | 3.420896 | 1.669812 | 0.337404 | 0.732113 | 0.51351 | -0.66997 | 0.703791 | 0.325854 | 1.891838 | 0.956455 | 2.247578 | 1.554827 | 0.915871 | 1.044299 | 1.41922 | 0.218034 | 0.435665 |
| 30.56806 | 0.173732 | 1.84937 | 0.608791 | 2.748969 | 1.586121 | 0.331098 | 0.184331 | 0.600584 | -0.86547 | 0.397546 | 0.035476 | 1.183964 | 0.277081 | 1.13897 | 0.659634 | -0.11366 | 0.653128 | 1.320857 | -0.47798 | 0.148417 |
| 30.58194 | 0.272181 | 1.625706 | -0.36321 | 1.953474 | 0.634226 | 0.360301 | -0.16198 | 0.769802 | -0.9779 | 0.085885 | -0.17614 | 0.752204 | -0.16117 | 0.422044 | 0.307268 | -0.59117 | 0.501701 | 1.186546 | -0.64411 | -0.00411 |
| 30.59583 | 0.29812 | 1.291625 | -1.08576 | 0.545358 | -0.44063 | 0.183052 | -0.23815 | 0.837892 | -1.03484 | -0.05834 | -0.10885 | 0.257051 | -0.33654 | 0.027234 | -0.12799 | -1.58242 | 0.557554 | 1.269899 | -0.54533 | -0.21017 |
| 30.60972 | 0.671448 | 0.865699 | -1.77565 | -0.56641 | -0.78215 | 0.148271 | -0.33829 | 0.834815 | -0.90159 | -0.21858 | 0.022769 | 0.155582 | -0.23353 | -0.34287 | -0.61024 | -2.3527 | 0.594134 | 1.283505 | -0.32137 | -0.52902 |
| 30.62361 | 1.133505 | 0.494259 | -1.94106 | -1.34506 | -0.64661 | 0.132441 | -0.14572 | 0.74895 | -0.13941 | -0.23675 | 0.000867 | 0.026648 | -0.19294 | -0.54184 | -0.76923 | -1.7282 | 0.280385 | 1.399127 | -0.08257 | -0.69544 |
| 30.6375 | 1.123351 | 0.217849 | -1.79817 | -1.45176 | -0.62632 | 0.043408 | -0.03784 | 0.576639 | 0.240092 | -0.29841 | -0.33831 | -0.05463 | -0.39005 | -0.44416 | -0.95882 | -0.55693 | -0.19868 | 1.262752 | 0.1027 | -0.52294 |
| 30.65139 | 0.551606 | 0.205813 | -1.07655 | -0.9365 | -0.50274 | 0.235356 | 0.20751 | 0.343271 | 0.527597 | -0.35053 | -0.69085 | -0.03468 | -0.58027 | -0.57024 | -0.98348 | 0.103012 | -0.38197 | 0.839693 | -0.24983 | -0.34435 |
| 30.66528 | 0.003231 | 0.295743 | -0.06923 | -0.38197 | 0.25812 | 0.431192 | 0.278533 | 0.267937 | 1.03085 | -0.29142 | -0.98726 | -0.20617 | -0.92295 | -0.9647 | -1.17138 | 0.089335 | -0.18947 | 0.404285 | -0.63593 | -0.70685 |
| 30.67917 | -0.72869 | 0.395817 | 0.419673 | 0.044835 | 0.920015 | 0.55632 | 0.303519 | 0.035601 | 1.487224 | -0.26802 | -1.03293 | -0.36783 | -1.04702 | -1.11898 | -1.29213 | -0.58709 | -0.04087 | -0.07702 | -1.19229 | -0.89414 |
| 30.69306 | -1.27068 | 0.333395 | 0.519194 | 0.228812 | 0.864759 | 0.584046 | 0.659818 | -0.15462 | 1.345673 | -0.59388 | -0.9836 | -0.56349 | -1.15675 | -1.04031 | -1.16587 | -1.11495 | -0.16944 | -0.42529 | -1.39742 | -0.85079 |
| 30.70694 | -1.86811 | -0.12346 | 0.616572 | 0.257911 | 0.643657 | 0.326015 | 0.960107 | -0.28477 | 0.771447 | -0.98561 | -0.90468 | -0.89748 | -1.6294 | -1.01081 | -0.60603 | -1.53044 | -0.29718 | -0.51328 | -1.37907 | -0.67939 |
| 30.72083 | -2.10244 | -0.69172 | 0.543909 | -0.61251 | 0.608652 | -0.45177 | 0.850797 | -0.45847 | 0.06151 | -0.93003 | -0.70791 | -1.34414 | -2.07041 | -0.95728 | -0.03093 | -1.88016 | -0.63839 | -0.48534 | -1.1739 | -0.65102 |
| 30.73472 | -1.58859 | -1.24022 | 0.275786 | -1.15184 | 0.270471 | -1.5784 | 0.48231 | -0.36961 | -0.73905 | -0.86328 | -0.60465 | -1.56347 | -2.0884 | -0.74543 | 0.411465 | -1.86213 | -0.99838 | -0.35798 | -0.80392 | -0.61412 |
| 30.74861 | -0.96937 | -1.80247 | -0.17203 | -2.0317 | -0.32618 | -2.63456 | 0.141407 | -0.41437 | -1.35666 | -0.53724 | -0.47674 | -1.62112 | -1.79693 | -0.65246 | 0.607325 | -1.87909 | -1.35081 | -0.44807 | -0.37686 | -0.60571 |
| 30.7625 | -0.69533 | -2.0474 | -0.6422 | -2.32775 | -0.55938 | -2.86729 | -0.19417 | -0.56377 | -1.44032 | -0.18494 | -0.29005 | -1.3308 | -1.42975 | -0.6859 | 0.363744 | -2.05744 | -1.50958 | -0.72976 | 0.048094 | -0.37812 |
| 30.77639 | -0.81821 | -1.94244 | -1.15582 | -2.50435 | -0.83964 | -2.81625 | -0.49281 | -0.76189 | -1.36711 | -0.03711 | -0.40414 | -0.89715 | -1.22866 | -0.94367 | 0.055272 | -2.10835 | -1.39144 | -1.14509 | -0.07284 | -0.04052 |
| 30.79028 | -1.16507 | -1.75225 | -1.56204 | -2.52665 | -1.23463 | -2.73188 | -0.86269 | -1.21611 | -1.29507 | -0.09629 | -0.82512 | -0.62637 | -1.17922 | -1.27212 | -0.20858 | -2.02506 | -1.2341 | -1.68129 | -0.48202 | 0.034015 |
| 30.80417 | -1.45473 | -1.2538 | -1.79243 | -1.9233 | -1.18646 | -2.30969 | -1.07335 | -1.6531 | -0.80139 | -0.03017 | -1.09733 | -0.59978 | -1.00767 | -1.62756 | -0.41709 | -1.79054 | -1.23563 | -2.10605 | -0.69226 | -0.16234 |
| 30.81806 | -1.6631 | -1.00381 | -2.04641 | -1.64801 | -1.31281 | -2.00647 | -1.33444 | -1.96314 | -0.33461 | 0.068968 | -0.98112 | -0.81693 | -0.78318 | -2.12806 | -0.57637 | -1.54055 | -1.30753 | -2.50691 | -0.76286 | -0.46913 |
| 30.83194 | -1.45692 | -1.08038 | -2.33648 | -1.68283 | -1.306 | -1.78351 | -1.15866 | -1.94013 | -0.12981 | 0.110187 | -0.86079 | -1.00378 | -0.56477 | -2.33297 | -0.66964 | -1.32265 | -1.27804 | -2.66175 | -0.63544 | -0.83595 |
| 30.84583 | -1.66402 | -1.46904 | -2.27289 | -1.07636 | -1.24587 | -1.51675 | -0.79468 | -1.69624 | -0.13134 | -0.05696 | -1.01973 | -1.04019 | -0.42458 | -2.2178 | -0.69629 | -1.00809 | -1.18989 | -2.79473 | -0.46936 | -1.14083 |
| 30.85972 | -1.31281 | -1.91232 | -1.59321 | -1.06972 | -0.53647 | -1.76331 | -0.59674 | -1.63324 | -0.25852 | -0.1746 | -1.00709 | -1.18493 | -0.47514 | -2.02865 | -0.97503 | -0.94736 | -1.29052 | -2.67497 | -0.17865 | -1.26324 |
| 30.87361 | -0.66055 | -1.80598 | -1.11623 | -0.7595 | 0.046461 | -2.01499 | -0.37771 | -1.28565 | -0.56218 | -0.099 | -0.74875 | -1.12702 | -0.57009 | -1.41199 | -1.12444 | -1.1312 | -1.32026 | -2.04019 | 0.242122 | -1.19177 |

| | | | | | | | | | | | | | | | | | | | | |
|---|---|---|---|---|---|---|---|---|---|---|---|---|---|---|---|---|---|---|---|---|
| 30.8875 | -0.1787 | -1.06911 | -0.85135 | 0.003603 | 0.362565 | -1.87403 | -0.16363 | -0.81691 | -0.77625 | -0.0549 | -0.45957 | -1.00806 | -0.50464 | -0.62881 | -0.94875 | -1.09656 | -0.97299 | -1.22141 | 0.44258 | -0.90411 |
| 30.90139 | 0.048803 | -0.37893 | -0.39343 | 0.825386 | 0.490563 | -1.23904 | 0.029101 | -0.50121 | -0.75244 | -0.00347 | -0.24148 | -0.84076 | -0.38189 | -0.14385 | -0.61642 | -0.99407 | -0.58539 | -0.55214 | 0.506402 | -0.46203 |
| 30.91528 | 0.217056 | -0.06686 | 0.173934 | 1.087025 | 0.802735 | -0.67348 | 0.225563 | -0.21191 | -0.6042 | 0.20644 | 0.02006 | -0.80028 | -0.43922 | 0.004826 | -0.69955 | -1.34182 | -0.47139 | -0.00713 | 0.607717 | -0.09263 |
| 30.92917 | 0.188293 | -0.53537 | 0.316235 | 0.318618 | 1.055297 | -0.53956 | 0.188109 | -0.24536 | -0.57806 | 0.208306 | 0.015983 | -1.02405 | -0.86858 | -0.39009 | -1.62408 | -1.2106 | -0.71458 | -0.30995 | 0.32486 | -0.34425 |
| 30.94306 | -0.47664 | -1.79233 | -0.49752 | -1.74799 | 0.393225 | -1.06103 | -0.34784 | -0.61975 | -0.85397 | -0.72124 | -0.84127 | -1.60442 | -1.73787 | -2.13124 | -2.86722 | -3.06073 | -1.36488 | -1.24233 | -1.34187 | -1.21221 |
| 30.95694 | -1.32715 | -2.76736 | -1.5176 | -4.0287 | -0.70745 | -1.72446 | -0.95142 | -1.16373 | -1.34955 | -2.69094 | -2.91112 | -2.49707 | -3.50767 | -4.36022 | -4.328 | -4.5179 | -2.28637 | -2.88767 | -3.62729 | -2.89363 |
| 30.97083 | -1.3823 | -2.77043 | -1.65899 | -3.58989 | -1.1907 | -2.38877 | -1.23967 | -1.05494 | -1.46929 | -3.23006 | -4.07957 | -2.06914 | -3.42374 | -4.24687 | -3.85109 | -3.81024 | -2.58126 | -2.70417 | -3.73098 | -3.37513 |
| 30.98472 | -0.93873 | -2.15686 | -1.61381 | -2.11453 | -1.3461 | -2.01291 | -0.71302 | -0.07879 | -0.71307 | -2.01917 | -2.53389 | -0.93071 | -1.43295 | -1.85292 | -1.70534 | -1.19457 | -1.717 | -1.70711 | -1.59755 | -2.12474 |
| 30.99861 | -0.32756 | -2.27832 | -1.8424 | -1.60315 | -1.13352 | -1.64888 | -0.2298 | 0.979873 | -0.29954 | -0.75769 | -0.56968 | -0.55516 | 0.093163 | -0.27123 | -0.55024 | 0.255496 | -0.39661 | -0.81362 | -0.28541 | -0.35685 |
| 31.0125 | -0.09057 | -2.75336 | -2.10849 | -1.88451 | -0.45301 | -0.98212 | 0.207377 | 1.654236 | -0.07906 | 0.203051 | 0.678869 | -0.45523 | 0.876334 | 0.681503 | -0.16056 | 0.932622 | 0.601435 | 0.059785 | 0.161051 | 0.857247 |
| 31.02639 | 0.124421 | -2.6573 | -1.93446 | -1.8289 | 0.334026 | -0.61475 | 0.452743 | 1.951135 | 0.19729 | 0.983616 | 1.424799 | -0.49242 | 1.100208 | 1.134645 | 0.235288 | 1.327597 | 1.357537 | 0.876371 | 0.351039 | 1.425378 |
| 31.04028 | 0.342111 | -2.3406 | -1.42882 | -1.7109 | 0.985913 | -0.1669 | 0.336656 | 1.705772 | 0.361076 | 1.319781 | 1.930614 | -0.46675 | 1.094906 | 1.122009 | 0.291231 | 1.307557 | 1.68392 | 1.467191 | 0.996651 | 1.138137 |
| 31.05417 | 0.40862 | -1.63027 | -1.03214 | -1.23596 | 0.89112 | 0.414506 | 0.429492 | 1.489047 | 0.61323 | 1.216233 | 1.66106 | -0.06282 | 1.050011 | 1.021559 | 0.491318 | 1.09714 | 1.472364 | 1.300092 | 1.444683 | 0.686236 |
| 31.06806 | 0.241203 | -1.09612 | -0.36079 | -0.43683 | 0.78913 | 1.061473 | 0.769464 | 0.953792 | 1.036963 | 0.963971 | 1.343104 | 0.26115 | 0.997116 | 0.866908 | 0.745036 | 1.141247 | 0.835985 | 0.735724 | 1.525418 | 0.340385 |
| 31.08194 | 0.049722 | -0.85232 | 0.33 | 0.275163 | 0.810066 | 1.714968 | 1.065846 | 0.583952 | 1.485822 | 0.729948 | 0.988293 | 0.317313 | 0.958276 | 0.901502 | 0.855153 | 1.21126 | 0.26147 | 0.474134 | 1.245361 | 0.371855 |
| 31.09583 | -0.39468 | -1.03199 | 0.540659 | 0.309422 | 0.581524 | 2.164982 | 0.993037 | -0.00445 | 1.159214 | 0.462374 | 0.636242 | 0.163343 | 0.936785 | 0.973038 | 0.811478 | 1.178669 | -0.13875 | 0.488409 | 1.067135 | 0.541081 |
| 31.10972 | -0.98694 | -1.59401 | 0.408362 | 0.110641 | -0.04407 | 1.93475 | 0.529929 | -0.79764 | 0.875821 | -0.15663 | -0.01451 | -0.40893 | 0.575257 | 0.782149 | 0.240468 | 0.537302 | -0.72827 | 0.046203 | 0.638806 | 0.304587 |
| 31.12361 | -0.94011 | -1.57653 | 0.640533 | 0.600875 | -0.35617 | 1.674618 | 0.512771 | -0.82537 | 0.429884 | -0.4835 | -0.44749 | -0.61075 | 0.304515 | 0.416495 | -0.30434 | -0.25654 | -1.17825 | -0.33214 | 0.464835 | -0.24634 |
| 31.1375 | -0.21848 | -0.80085 | 1.574011 | 1.522111 | 0.101005 | 1.852531 | 1.049233 | -0.13293 | 0.246487 | -0.10288 | -0.32177 | -0.18524 | 0.486376 | 0.501987 | -0.13276 | -0.32879 | -0.90736 | -0.2617 | 1.023518 | -0.09274 |
| 31.15139 | 0.294164 | -0.12434 | 1.859561 | 1.606845 | 0.524704 | 1.809545 | 1.217865 | 0.426029 | -0.00844 | 0.267469 | -0.05307 | 0.331735 | 0.7613 | 0.756502 | 0.262569 | -0.06301 | -0.42892 | -0.07917 | 1.232071 | 0.122539 |
| 31.16528 | 0.35024 | 0.394909 | 1.646013 | 1.726575 | 0.60612 | 1.850083 | 1.127759 | 0.742564 | -0.3583 | 0.229144 | -0.05128 | 0.573717 | 0.91307 | 0.691724 | 0.569883 | 0.293724 | -0.39896 | -0.07314 | 0.532575 | -0.01639 |
| 31.17917 | 0.230288 | 0.780889 | 1.551588 | 1.608208 | 0.137199 | 1.96941 | 0.933181 | 0.811951 | -0.59316 | 0.114612 | -0.36404 | 0.656102 | 0.763698 | 0.283465 | 0.633668 | 0.407199 | -0.73432 | -0.18016 | -0.4646 | -0.31596 |
| 31.19306 | 0.230147 | 0.742437 | 1.612639 | 1.073671 | -0.02593 | 1.400325 | 0.485563 | 0.537262 | -0.53526 | -0.07799 | -0.5386 | 0.280673 | 0.259192 | 0.05931 | 0.223216 | -0.1386 | -0.59688 | -0.25986 | -1.13648 | -0.57527 |
| 31.20694 | 0.251321 | 0.422204 | 1.000454 | 0.297449 | -0.04104 | 0.675168 | -0.00488 | 0.264252 | -0.40586 | -0.23525 | -0.53059 | -0.17037 | -0.22706 | -0.10848 | -0.27369 | -0.73084 | -0.07421 | -0.37094 | -1.22132 | -0.47254 |
| 31.22083 | 0.157318 | 0.064633 | 0.009045 | -0.65854 | -0.0192 | -0.24295 | -0.22544 | -0.05543 | -0.45789 | -0.41093 | -0.38273 | -0.62656 | -0.70155 | -0.34937 | -0.66798 | -1.13513 | 0.393322 | -0.39774 | -1.04058 | -0.34478 |
| 31.23472 | 0.194283 | -0.24826 | -1.05758 | -0.97353 | 0.369852 | -1.18268 | -0.40215 | -0.19765 | -0.4816 | -0.48859 | -0.03769 | -0.96383 | -1.11142 | -0.52856 | -0.89 | -1.17453 | 0.810164 | -0.23804 | -0.54865 | -0.19672 |
| 31.24861 | 0.285418 | 0.040498 | -1.2381 | -0.69369 | 0.511297 | -1.27832 | -0.13185 | -0.04359 | -0.24026 | -0.21895 | 0.224192 | -0.71665 | -0.84072 | -0.17574 | -0.48758 | -0.48003 | 0.92162 | -0.0451 | -0.21876 | 0.066656 |
| 31.2625 | 0.432721 | 0.758082 | -1.05598 | -0.04007 | 0.12767 | -0.75678 | 0.6176 | 0.375745 | 0.116985 | -0.13496 | 0.300883 | 0.021351 | 0.029701 | 0.445813 | 0.279901 | 0.599655 | 0.764467 | 0.083255 | -0.10476 | 0.571671 |
| 31.27639 | 0.71684 | 0.731454 | -1.27151 | -0.01356 | 0.366304 | -0.7734 | 0.801862 | 0.532095 | 0.250656 | 0.058261 | 0.56711 | 0.200389 | 0.237229 | 0.791704 | 0.389581 | 0.923936 | 0.713772 | 0.325295 | 0.223667 | 0.864807 |
| 31.29028 | 0.979128 | -0.08575 | -1.70653 | -0.85142 | 1.041975 | -1.22891 | 0.443383 | 0.286358 | 0.031441 | 0.447541 | 1.002376 | -0.4326 | -0.13913 | 0.70809 | -0.25048 | 0.365476 | 0.957196 | 0.42646 | 0.686046 | 0.812078 |
| 31.30417 | 0.789518 | -0.33103 | -1.46808 | -0.89533 | 0.997889 | -1.28902 | 0.529577 | 0.075069 | 0.084822 | 0.636173 | 0.918088 | -0.56955 | -0.10985 | 0.514964 | -0.44124 | 0.099235 | 1.052685 | 0.381773 | 0.672692 | 0.761036 |
| 31.31806 | 0.850097 | 0.174408 | -0.76335 | -0.15939 | 0.623258 | -0.3559 | 1.022912 | 0.190882 | 0.672219 | 0.614497 | 0.505157 | 0.012481 | 0.270042 | 0.672353 | 0.02275 | 0.359789 | 0.921991 | 0.208188 | 0.546047 | 0.875543 |
| 31.33194 | 0.742773 | 0.570458 | -0.05714 | 0.513452 | 0.65222 | 0.381447 | 1.367836 | 0.329714 | 1.029536 | 0.597227 | 0.23716 | 0.356619 | 0.394559 | 0.648926 | 0.361364 | 0.47216 | 0.895937 | 0.121107 | 0.412604 | 0.888995 |
| 31.34583 | 0.627299 | 0.768197 | 0.473412 | 1.052151 | 0.731338 | 0.788473 | 1.370397 | 0.324379 | 0.93988 | 0.491466 | 0.047255 | 0.501697 | 0.33498 | 0.576851 | 0.395232 | 0.438845 | 0.648322 | 0.063141 | 0.287307 | 0.68316 |
| 31.35972 | 0.494655 | 0.836757 | 0.707826 | 1.288286 | 0.641491 | 0.977502 | 1.330432 | 0.375515 | 0.612007 | 0.330222 | -0.00531 | 0.499008 | 0.225734 | 0.5001 | 0.302878 | 0.468342 | 0.183562 | 0.004744 | 0.239122 | 0.538163 |
| 31.37361 | 0.252553 | 0.715854 | 0.748216 | 0.952648 | 0.806087 | 0.917296 | 1.121303 | 0.289192 | 0.189963 | 0.249565 | 0.144994 | 0.239147 | -0.0465 | 0.252784 | 0.108592 | 0.325954 | -0.08293 | 0.167523 | 0.297556 | 0.460775 |
| 31.3875 | 0.106066 | 0.534685 | 0.476232 | 0.508489 | 0.696322 | 0.613431 | 0.746607 | 0.245562 | -0.16846 | 0.243362 | 0.304511 | -0.0126 | -0.25044 | 0.087955 | 0.019764 | 0.280288 | 0.115556 | 0.418741 | 0.390893 | 0.485246 |
| 31.40139 | 0.080712 | 0.61264 | 0.160592 | 0.383372 | 0.313767 | 0.455757 | 0.549688 | 0.241131 | -0.3512 | 0.271614 | 0.410555 | 0.057451 | -0.06465 | 0.213723 | 0.146808 | 0.477594 | 0.604144 | 0.545601 | 0.463755 | 0.595672 |
| 31.41528 | 0.248227 | 0.967419 | 0.046201 | 0.509703 | -0.14947 | 0.548545 | 0.569312 | 0.27436 | -0.32899 | 0.315187 | 0.467249 | 0.548396 | 0.550729 | 0.72912 | 0.564042 | 0.839371 | 1.054508 | 0.48007 | 0.513802 | 0.796477 |
| 31.42917 | 0.695839 | 1.632692 | 0.236075 | 1.11033 | -0.49639 | 0.822459 | 0.783075 | 0.537443 | -0.04625 | 0.472674 | 0.62273 | 1.389168 | 1.248782 | 1.614407 | 1.408867 | 1.673793 | 0.916329 | 0.328662 | 0.795919 | 1.22886 |
| 31.44306 | 1.891216 | 2.741255 | 0.984994 | 1.209016 | 0.165661 | 1.231085 | 1.064725 | 0.967127 | 0.050928 | 1.112192 | 1.245848 | 1.938283 | 2.042972 | 2.415019 | 2.488383 | 2.722506 | 1.041798 | 0.63174 | 1.572739 | 1.875581 |
| 31.45694 | 2.354418 | 3.475412 | 1.993206 | 1.997375 | 1.695781 | 0.887653 | 1.10555 | 1.526354 | 0.354474 | 1.660615 | 1.901562 | 2.01646 | 2.777989 | 2.455976 | 2.849816 | 3.748626 | 1.353026 | 1.375661 | 2.560967 | 3.103777 |
| 31.47083 | 1.799245 | 3.132151 | 2.594667 | 4.584227 | 2.192779 | 1.415824 | 1.42999 | 1.862616 | 1.114651 | 1.623796 | 1.857974 | 2.379663 | 3.156845 | 2.674207 | 2.664617 | 4.344712 | 1.427467 | 1.897645 | 3.035115 | 3.561546 |
| 31.48472 | 1.343156 | 3.354056 | 3.039759 | 7.308522 | 1.980768 | 1.214822 | 1.90574 | 2.375192 | 1.692738 | 1.432286 | 1.805805 | 2.448341 | 3.343465 | 3.354224 | 3.047526 | 5.195548 | 1.632828 | 2.07296 | 3.308027 | 3.428801 |
| 31.49861 | 0.810417 | 3.773311 | 3.266095 | 5.975768 | 2.027721 | 1.196511 | 1.555539 | 3.020689 | 2.514818 | 1.320317 | 1.895477 | 2.212623 | 2.883409 | 4.043306 | 3.52966 | 5.58321 | 1.931794 | 2.079161 | 3.735936 | 3.144192 |
| 31.5125 | 0.352718 | 3.151826 | 4.09695 | 4.672381 | 1.973273 | 1.166852 | 1.592922 | 2.787681 | 2.411908 | 1.203709 | 1.926884 | 1.970423 | 2.328506 | 3.870564 | 3.482154 | 4.769766 | 2.060863 | 1.899412 | 3.650156 | 2.410139 |
| 31.52639 | 0.417636 | 2.755613 | 3.453758 | 4.747882 | 1.023997 | 1.289053 | 1.915993 | 2.412826 | 1.673268 | 0.889786 | 1.726371 | 1.731911 | 2.199153 | 3.782492 | 3.268967 | 3.573198 | 1.965379 | 1.529237 | 2.480081 | 1.7126 |
| 31.54028 | 0.705346 | 3.134932 | 2.789485 | 4.681236 | 0.226912 | 1.128623 | 1.504536 | 2.055682 | 1.325561 | 0.577678 | 1.196924 | 1.881356 | 2.0426 | 3.513912 | 2.676178 | 2.268355 | 1.835452 | 1.060867 | 1.340559 | 1.073841 |
| 31.55417 | 1.000516 | 3.459378 | 1.656394 | 4.248294 | 0.361136 | 0.751368 | 0.83195 | 1.67033 | 0.284011 | 0.365276 | 0.747134 | 1.932441 | 1.334087 | 2.292831 | 1.650122 | 0.732268 | 1.449339 | 0.787437 | 0.193149 | 0.412934 |
| 31.56806 | 1.226784 | 3.54207 | 0.448137 | 3.998222 | 0.939485 | 0.242904 | 0.203113 | 1.093707 | -0.47521 | 0.230457 | 0.213971 | 1.634508 | 0.854742 | 1.044723 | 0.575782 | -0.65587 | 1.215691 | 0.552094 | -0.70535 | -0.14506 |
| 31.58194 | 1.035122 | 2.853848 | -0.89312 | 3.618105 | 1.273613 | -0.11697 | -0.58999 | 0.587806 | -1.27244 | -0.01746 | -0.27722 | 0.98051 | 0.309277 | 0.082617 | -0.12569 | -1.21327 | 0.795511 | 0.353277 | -1.00471 | -0.48182 |
| 31.59583 | 0.822167 | 1.955864 | -1.92708 | 2.570887 | 0.70384 | -0.23898 | -0.92395 | 0.629307 | -2.86889 | -0.27702 | -0.52991 | 0.387516 | -0.23844 | -0.27232 | -0.71997 | -1.86038 | 0.629491 | -0.08337 | -0.97837 | -0.55559 |
| 31.60972 | 0.962934 | 1.45646 | -2.75773 | 1.690088 | -0.09983 | -0.22047 | -0.7022 | 0.998825 | -2.55331 | -0.48528 | -0.65144 | -0.00011 | -0.66342 | -0.29139 | -1.85618 | -2.23116 | 0.750891 | -0.01029 | -0.90941 | -0.80585 |
| 31.62361 | 1.055179 | 0.954392 | -3.3367 | 1.021286 | 0.127656 | -1.008 | -0.69131 | 0.624476 | -1.38727 | -0.48219 | -0.3937 | -0.32361 | -0.82639 | -0.46937 | -3.65769 | -2.55348 | 0.699217 | 0.322 | -0.85901 | -1.21393 |
| 31.6375 | 0.650101 | 0.501386 | -2.79813 | 0.276841 | 0.792723 | -1.44822 | -0.50437 | -0.03411 | -0.64603 | -0.41569 | 0.065593 | -0.53042 | -0.93657 | -0.51263 | -2.83665 | -2.25437 | 0.351345 | 0.67222 | -0.40034 | -1.4048 |
| 31.65139 | 0.39372 | 0.336751 | -2.4655 | -0.58732 | 0.821112 | -0.7983 | 0.204296 | -0.35015 | 0.3569 | -0.50823 | 0.262871 | -0.34206 | -0.93317 | -0.39275 | -2.34838 | -0.87235 | 0.07418 | 0.861131 | 0.150323 | -1.10898 |
| 31.66528 | 0.30606 | 0.30831 | -1.79012 | -1.90716 | 0.843311 | -0.1515 | 0.793233 | -0.80546 | 0.883407 | -0.44592 | 0.247847 | -0.01624 | -1.09471 | -0.47386 | -1.74138 | 0.531107 | 0.022811 | 0.915526 | 0.240174 | -0.76183 |
| 31.67917 | 0.080394 | 0.294545 | -0.44038 | -3.14816 | 0.693895 | 0.262077 | 0.479459 | -1.34663 | 1.321097 | 0.021397 | 0.024676 | 0.135159 | -1.30869 | -0.77251 | -1.76401 | 1.270701 | -0.27506 | 0.957992 | -0.07992 | -0.44494 |
| 31.69306 | -0.52085 | 0.399564 | 0.759859 | -3.45753 | 0.142993 | 0.539952 | 0.405501 | -1.46977 | 1.527801 | 0.162932 | -0.40506 | 0.294024 | -1.24243 | -1.03728 | -1.37798 | 1.318384 | -0.48692 | 0.99725 | -0.82137 | -0.31347 |
| 31.70694 | -0.92567 | 0.290281 | 1.439499 | -3.13821 | -0.24845 | 0.540972 | 0.619979 | -1.3553 | 1.220937 | 0.018735 | -0.9838 | 0.351024 | -1.16772 | -0.97089 | -0.43387 | 0.937491 | -0.47889 | 0.697865 | -1.54917 | -0.24114 |
| 31.72083 | -1.58691 | -0.04512 | 1.37847 | -2.3975 | -0.59743 | 0.13008 | 0.829075 | -1.26582 | 0.880443 | -0.36608 | -1.4917 | 0.144381 | -1.12017 | -1.01073 | 0.401511 | 0.07356 | -0.32625 | -0.08719 | -1.56169 | -0.11114 |
| 31.73472 | -2.31102 | -0.46641 | 0.869111 | -2.11419 | -1.00885 | -0.56233 | 0.876104 | -1.38068 | 0.069311 | -0.71093 | -1.69383 | -0.30721 | -1.23876 | -1.36622 | 0.744555 | -0.9984 | -0.29846 | -0.93536 | -1.28451 | 0.043575 |
| 31.74861 | -2.24334 | -1.23784 | 0.229783 | -2.27026 | -1.30595 | -1.36016 | 0.470753 | -1.75568 | -1.00735 | -0.86319 | -1.52086 | -1.05627 | -1.50198 | -1.86313 | 0.506329 | -1.99149 | -0.65983 | -1.41217 | -1.00593 | -0.01819 |
| 31.7625 | -1.59421 | -2.06105 | -0.51866 | -2.20674 | -0.99581 | -1.88898 | -0.28654 | -2.03864 | -0.83508 | -0.75459 | -1.00179 | -1.65286 | -1.81588 | -2.33026 | -0.18469 | -2.5604 | -1.41705 | -1.37666 | -0.48505 | -0.37694 |
| 31.77639 | -1.11301 | -2.45226 | -1.15205 | -2.0322 | -0.48606 | -2.51021 | -1.18841 | -1.93875 | -0.96666 | -0.47705 | -0.50552 | -1.87495 | -1.97933 | -2.2049 | -0.8179 | -2.84845 | -1.99102 | -1.31386 | 0.064011 | -0.5747 |
| 31.79028 | -0.72463 | -2.64595 | -1.78128 | -2.05976 | -0.51634 | -2.76588 | -1.79163 | -1.8054 | -1.26246 | -0.18726 | -0.30869 | -1.82884 | -1.90091 | -1.9372 | -1.01175 | -2.98462 | -2.15532 | -1.524 | 0.13491 | -0.59655 |
| 31.80417 | -0.58628 | -2.88807 | -2.03973 | -2.4268 | -1.30548 | -2.91116 | -2.04681 | -1.93645 | -1.26609 | -0.33424 | -0.46675 | -1.7108 | -1.58262 | -2.01234 | -0.85519 | -2.27129 | -2.23048 | -2.00339 | -0.24002 | -0.79708 |
| 31.81806 | -0.73623 | -3.60041 | -2.29259 | -3.13902 | -2.23829 | -2.69986 | -2.08096 | -2.55902 | -1.61392 | -0.58399 | -0.70241 | -1.70393 | -1.60876 | -2.20489 | -0.75469 | -1.62154 | -2.40187 | -2.71446 | -0.71516 | -0.96656 |
| 31.83194 | -0.86201 | -4.03176 | -2.4568 | -3.69957 | -2.25754 | -2.84065 | -2.29081 | -2.66983 | -1.62824 | -0.51607 | -0.71991 | -1.68922 | -1.6447 | -1.9423 | -0.76434 | -1.57157 | -2.15752 | -3.12601 | -0.81216 | -1.12933 |
| 31.84583 | -0.85192 | -3.57927 | -2.38674 | -3.15049 | -1.5437 | -2.06367 | -2.3055 | -1.97135 | -1.50709 | -0.38656 | -0.65839 | -1.44824 | -1.50365 | -1.47971 | -0.70819 | -1.49382 | -1.57187 | -2.61421 | -0.84831 | -1.08661 |
| 31.85972 | -0.87228 | -3.2549 | -2.0856 | -2.61656 | -1.07979 | -1.78376 | -1.51063 | -1.22084 | -0.72373 | -0.47219 | -0.62792 | -1.05261 | -1.29128 | -1.17508 | -0.49151 | -1.10814 | -1.30666 | -1.98446 | -0.79462 | -0.74807 |
| 31.87361 | -0.79911 | -3.05642 | -1.63987 | -1.87579 | -0.71995 | -1.68498 | -0.83621 | -0.85928 | -0.26727 | -0.47927 | -0.59461 | -0.74033 | -1.13056 | -1.06328 | -0.30255 | -0.68777 | -1.1377 | -1.52348 | -0.73648 | -0.53845 |
| 31.8875 | -0.56129 | -2.80973 | -0.98899 | -0.82423 | -0.29968 | -1.24758 | -0.23709 | -0.52782 | 0.106506 | -0.35106 | -0.49062 | -0.52762 | -0.93882 | -0.89615 | -0.17346 | -0.2931 | -1.10581 | -0.99385 | -0.65651 | -0.18532 |
| 31.90139 | -0.24391 | -2.2088 | -0.49405 | 0.211545 | -0.0164 | -0.81483 | 0.275144 | -0.23048 | 0.433365 | -0.1498 | -0.30386 | -0.42121 | -0.64493 | -0.68778 | -0.10645 | 0.092472 | -0.98644 | -0.46646 | -0.28558 | -0.11822 |
| 31.91528 | 0.030354 | -1.64768 | -0.13227 | 0.65173 | 0.197424 | -0.13657 | 0.325961 | 0.167118 | 0.459921 | 0.286594 | -0.00604 | -0.55998 | -0.28261 | -0.37883 | -0.22001 | 0.118469 | -0.81217 | -0.0897 | 0.202318 | -0.11732 |
| 31.92917 | -0.02976 | -1.52534 | -0.12455 | -0.12727 | 0.304869 | -0.27852 | -0.34047 | 0.472909 | -0.09032 | 0.633662 | 0.027448 | -1.18475 | -0.46272 | -0.56243 | -0.90729 | -0.71767 | -0.71399 | -0.30088 | 0.376435 | -0.55985 |
| 31.94306 | -0.71606 | -2.24283 | -1.05009 | -2.85081 | -0.19919 | -1.52503 | -1.53953 | 0.317681 | -0.82239 | -0.39703 | -0.88669 | -2.2884 | -1.72264 | -1.88617 | -2.25035 | -2.17103 | -0.92144 | -1.49114 | -1.09851 | -1.67395 |
| 31.95694 | -1.49374 | -3.24599 | -1.9316 | -5.45308 | -1.14298 | -2.47537 | -2.49337 | -0.7878 | -1.1997 | -2.2964 | -2.53007 | -2.69541 | -3.615 | -3.06834 | -4.22953 | -3.98653 | -1.51291 | -2.43105 | -3.27425 | -3.38937 |
| 31.97083 | -1.735 | -2.72745 | -2.1301 | -5.06332 | -1.71269 | -2.63045 | -2.6332 | -1.45204 | -0.99928 | -2.71645 | -3.2294 | -2.12947 | -4.46185 | -3.17106 | -4.23544 | -4.55569 | -1.67273 | -2.23721 | -3.80683 | -3.4252 |
| 31.98472 | -1.43429 | -1.90259 | -1.79852 | -3.67999 | -1.59367 | -2.78024 | -1.97556 | -1.05701 | -0.95925 | -1.55078 | -2.13703 | -1.22229 | -2.63853 | -1.74869 | -2.03444 | -2.45952 | -0.99272 | -1.22803 | -1.98477 | -1.94181 |
| 31.99861 | -0.92848 | -1.39431 | -1.19351 | -3.44816 | -1.31427 | -2.83525 | -1.29481 | -0.17765 | -1.00498 | -0.51213 | -0.83243 | -0.6926 | -0.73224 | -0.51392 | -0.5074 | -0.62095 | -0.31455 | -0.26702 | -0.2327 | -0.4266 |
| 32.0125 | -0.21713 | -1.16099 | -0.71332 | -3.16103 | -0.80485 | -2.32142 | -0.76416 | 0.519325 | -0.83147 | -0.10209 | 0.010813 | -0.56954 | 0.754202 | 0.065426 | 0.161318 | 0.549382 | 0.370706 | 0.538913 | 0.637876 | 0.349158 |
| 32.02639 | 0.350785 | -0.87537 | -0.48099 | -2.87083 | -0.08618 | -1.38174 | -0.39099 | 0.766853 | -0.3641 | -0.00081 | 0.549765 | -0.33511 | 1.435782 | 0.260979 | 0.492981 | 1.086615 | 0.710899 | 0.70723 | 0.808368 | 0.412687 |
| 32.04028 | 0.707137 | -0.2522 | -0.28442 | -1.85378 | 0.11779 | -0.98621 | -0.11609 | 0.519153 | 0.143964 | -0.03994 | 0.916089 | 0.287901 | 1.207878 | 0.527522 | 0.955052 | 1.568376 | 0.666999 | 0.846809 | 0.965907 | 0.605365 |
| 32.05417 | 0.932033 | 0.40311 | -0.09642 | -0.17879 | 0.122828 | -0.4442 | 0.597141 | 0.100925 | 0.66243 | -0.22161 | 1.039751 | 1.043872 | 1.323777 | 0.970997 | 1.263475 | 1.983662 | 0.637408 | 0.545116 | 0.926559 | 0.728935 |
| 32.06806 | 1.400199 | 0.919618 | 0.48052 | 0.74927 | 0.626624 | 0.1293 | 0.636688 | 0.210486 | 0.99262 | 0.038507 | 0.892985 | 0.944393 | 1.203224 | 1.134472 | 1.025841 | 1.737453 | 0.534898 | 0.372473 | 0.54357 | 0.623938 |
| 32.08194 | 1.601569 | 0.995594 | 0.504349 | 0.927348 | 1.523936 | 0.500427 | 1.014359 | 0.973082 | 0.851323 | 0.722155 | 0.950705 | 0.426913 | 0.653847 | 1.013777 | 0.241673 | 1.132894 | 0.556627 | 0.526128 | 0.113302 | 0.434737 |
| 32.09583 | 1.411099 | 0.795954 | 0.521225 | 0.777937 | 2.474564 | 0.678611 | 1.271715 | 1.727578 | 0.648114 | 1.206721 | 1.149328 | -0.0291 | 0.207274 | 0.71684 | -0.35174 | 0.615045 | 0.382315 | 0.635625 | -0.17721 | 0.430371 |
| 32.10972 | 1.169439 | 0.556387 | 0.4568 | 1.234201 | 2.769528 | 0.809892 | 1.756317 | 1.635785 | 0.661524 | 1.058956 | 1.134853 | -0.04276 | 0.157188 | 0.468091 | -0.23195 | 0.620466 | 0.202528 | 0.386747 | -0.2836 | 0.345326 |
| 32.12361 | 0.879546 | 0.558269 | 0.432008 | 2.288318 | 2.466664 | 0.879373 | 2.729445 | 1.148588 | 0.696886 | 0.762529 | 0.873426 | 0.24162 | 0.504646 | 0.526883 | 0.296918 | 0.89424 | -0.0183 | 0.14559 | -0.11748 | 0.308307 |
| 32.1375 | 0.592096 | 0.446241 | 0.43454 | 2.871772 | 1.84423 | 1.403998 | 2.400224 | 1.096987 | 0.761574 | 0.592591 | 0.692964 | 0.442433 | 0.527917 | 0.526055 | 0.563633 | 0.724476 | -0.22958 | 0.022701 | 0.212293 | 0.466119 |

| | | | | | | | | | | | | | | | | | | | | |
|---|---|---|---|---|---|---|---|---|---|---|---|---|---|---|---|---|---|---|---|---|
| 32.15139 | 0.510617 | 0.221042 | 0.426335 | 2.371235 | 1.268823 | 2.440881 | 1.793172 | 1.13059 | 0.746848 | 0.657434 | 0.575608 | 0.381236 | 0.229972 | 0.386392 | 0.533331 | 0.357904 | -0.35081 | 0.064569 | 0.587824 | 0.382931 |
| 32.16528 | 0.450889 | 0.152546 | 0.524568 | 1.944628 | 0.777738 | 2.811745 | 1.640888 | 0.979884 | 0.664087 | 0.800539 | 0.373398 | 0.30414 | 0.10314 | 0.328944 | 0.338672 | 0.08975 | -0.4154 | 0.185432 | 0.77604 | 0.403092 |
| 32.17917 | 0.397204 | 0.200045 | 0.715578 | 1.869589 | 0.48238 | 2.951362 | 1.259846 | 0.760947 | 0.691228 | 0.949278 | 0.047406 | 0.391711 | 0.129807 | 0.32311 | 0.208211 | -0.09637 | -0.44373 | 0.299894 | 0.738054 | 0.427106 |
| 32.19306 | 0.251361 | 0.219102 | 0.804331 | 1.835982 | 0.421223 | 3.096525 | 0.84649 | 0.604539 | 0.646469 | 1.035994 | -0.19813 | 0.504686 | 0.2181 | 0.282648 | 0.19943 | -0.05363 | -0.26529 | 0.30979 | 0.411942 | 0.442737 |
| 32.20694 | 0.014568 | 0.099726 | 0.782063 | 1.456189 | 0.517981 | 2.709227 | 0.548461 | 0.455955 | 0.515625 | 0.893586 | -0.39613 | 0.569949 | 0.316264 | 0.163123 | 0.402374 | 0.167111 | -0.17846 | 0.183017 | 0.062286 | 0.436632 |
| 32.22083 | -0.26962 | -0.01304 | 0.558135 | 0.847949 | 0.475733 | 2.049079 | 0.201807 | 0.400295 | 0.34707 | 0.629527 | -0.4087 | 0.51258 | 0.449939 | 0.013127 | 0.661063 | 0.245599 | -0.22336 | 0.085775 | -0.1533 | 0.537601 |
| 32.23472 | -0.11051 | -0.00387 | 0.290826 | 0.587676 | 0.401722 | 1.463571 | -0.56366 | 0.410109 | -0.00933 | 0.516894 | -0.15202 | 0.099729 | 0.284082 | -0.27543 | 0.567108 | -0.06768 | -0.14349 | -0.12002 | -0.02116 | 0.401485 |
| 32.24861 | 0.004374 | 0.149587 | 0.055819 | 0.511124 | 0.231929 | 1.081831 | -0.88491 | 0.456011 | -0.04092 | 0.319707 | -0.03254 | -0.00896 | 0.128506 | -0.25333 | 0.45625 | -0.1598 | 0.012088 | -0.10928 | 0.195468 | 0.31215 |
| 32.2625 | -0.17265 | 0.29685 | -0.13914 | 0.176461 | 0.023569 | 0.852258 | -0.7212 | 0.41361 | 0.344443 | 0.097863 | -0.24435 | 0.30411 | 0.19755 | -0.1846 | 0.604488 | 0.06235 | 0.000584 | 0.023309 | 0.215279 | 0.393848 |
| 32.27639 | -0.02547 | 0.16923 | -0.34447 | 0.300014 | 0.218315 | 0.280739 | -0.68858 | 0.409657 | 0.060749 | 0.152393 | 0.131992 | 0.114176 | 0.024324 | -0.01655 | 0.382165 | 0.009354 | 0.199088 | 0.00946 | 0.159377 | 0.107418 |
| 32.29028 | 0.371925 | 0.026078 | -0.32862 | 0.675669 | 0.742668 | -0.19993 | -0.59687 | 0.462515 | -0.28691 | 0.430304 | 0.868894 | -0.30178 | 0.008524 | 0.155456 | 0.17842 | 0.00509 | 0.504559 | 0.021062 | 0.197241 | -0.06395 |
| 32.30417 | 0.368764 | 0.102846 | -0.18567 | 0.538747 | 0.872713 | -0.1337 | -0.04474 | 0.497189 | 0.016831 | 0.474857 | 0.937496 | -0.18431 | 0.357224 | 0.237183 | 0.360399 | 0.357088 | 0.499074 | 0.101578 | 0.188974 | -0.06012 |
| 32.31806 | 0.139046 | 0.241022 | -0.01198 | 0.195118 | 0.671821 | 0.185956 | 0.65458 | 0.472553 | 0.698811 | 0.311916 | 0.559753 | 0.303051 | 0.795257 | 0.424328 | 0.793984 | 0.772913 | 0.205454 | 0.232906 | 0.409161 | 0.106592 |
| 32.33194 | -0.1128 | 0.439192 | 0.119005 | 0.018188 | 0.620513 | 0.508116 | 1.164198 | 0.645552 | 1.220477 | 0.028909 | 0.191663 | 0.703599 | 1.006071 | 0.570569 | 1.223185 | 1.136103 | -0.10345 | 0.314197 | 0.567749 | 0.201569 |
| 32.34583 | 0.288879 | 0.453803 | 0.161781 | 0.331294 | 1.003161 | 0.570248 | 1.141771 | 0.738479 | 0.882135 | 0.041844 | 0.060021 | 0.513107 | 0.728461 | 0.567748 | 1.107809 | 1.051543 | -0.05715 | 0.240281 | 0.766566 | 0.213331 |
| 32.35972 | 1.080633 | 0.2048 | 0.283233 | 0.443242 | 1.309327 | 0.350169 | 0.754904 | 0.732263 | -0.16068 | 0.4038 | 0.236732 | -0.11221 | 0.07404 | 0.456928 | 0.391243 | 0.407507 | 0.16694 | 0.125744 | 0.663506 | 0.215911 |
| 32.37361 | 1.377299 | 0.040781 | 0.336248 | 0.48448 | 1.238934 | 0.422551 | 0.649398 | 0.615156 | -0.33722 | 0.539835 | 0.294869 | -0.32992 | -0.20256 | 0.340791 | -0.08626 | -0.00086 | 0.145184 | 0.029942 | 0.31607 | 0.319424 |
| 32.3875 | 0.9873 | 0.181818 | 0.248599 | 0.459447 | 0.959033 | 0.760501 | 0.891264 | 0.700087 | 0.195663 | 0.291976 | 0.173727 | -0.16451 | -0.06287 | 0.492481 | 0.101933 | 0.123406 | -0.01065 | 0.122582 | 0.264422 | 0.372684 |
| 32.40139 | 0.578344 | 0.247658 | 0.063068 | 0.338876 | 0.4633 | 0.882254 | 0.926191 | 0.450667 | 0.599396 | 0.14392 | 0.132217 | -0.01777 | 0.216113 | 0.607665 | 0.360699 | 0.368272 | -0.01021 | 0.121708 | 0.348665 | 0.388497 |
| 32.41528 | 0.171469 | 0.460155 | -0.09525 | 0.168698 | -0.11259 | 0.993135 | 1.095006 | 0.335008 | 1.15703 | -0.03444 | 0.209878 | 0.320182 | 0.70325 | 0.738212 | 0.81627 | 0.865365 | 0.128832 | 0.134946 | 0.699209 | 0.545213 |
| 32.42917 | -0.25724 | 1.076833 | -0.00742 | 0.161855 | -0.40555 | 1.228374 | 1.363561 | 0.438743 | 1.632796 | -0.03275 | 0.320118 | 0.798154 | 1.416278 | 1.010393 | 1.517934 | 1.671324 | 0.315389 | 0.222358 | 1.6717 | 1.045734 |
| 32.44306 | 0.155966 | 2.520104 | 0.785491 | 0.503671 | 0.377228 | 2.018068 | 1.418637 | 1.109658 | 1.447523 | 0.696553 | 0.778691 | 1.126231 | 2.114053 | 1.678514 | 2.307991 | 2.362063 | 0.525839 | 0.637758 | 2.780496 | 1.853591 |
| 32.45694 | 1.150219 | 2.801101 | 1.967807 | 1.189285 | 1.564291 | 2.976158 | 1.512638 | 1.879543 | 1.182842 | 1.413461 | 1.331776 | 1.551719 | 2.697918 | 1.808145 | 2.582481 | 2.76836 | 0.896179 | 1.205575 | 3.227132 | 2.548161 |
| 32.47083 | 1.46272 | 4.752599 | 2.841715 | 2.061266 | 2.150422 | 2.945692 | 1.57734 | 2.663603 | 1.353974 | 1.588989 | 1.364471 | 2.076511 | 3.015209 | 1.579124 | 2.289261 | 3.178397 | 1.110852 | 1.575491 | 2.878427 | 3.038754 |
| 32.48472 | 1.677199 | 4.621108 | 3.628101 | 3.130447 | 2.236905 | 2.899705 | 1.742201 | 2.924346 | 1.44356 | 1.580946 | 1.370465 | 1.9436 | 2.95405 | 1.655323 | 1.78564 | 3.219309 | 1.004019 | 1.659592 | 2.496604 | 2.998437 |
| 32.49861 | 1.604491 | 4.542646 | 3.857281 | 3.333492 | 2.285372 | 2.986037 | 1.999473 | 3.083706 | 1.124484 | 1.446649 | 1.533737 | 1.781243 | 2.746724 | 2.293409 | 1.665226 | 2.952749 | 0.997139 | 1.73676 | 2.356315 | 2.640371 |
| 32.5125 | 1.00197 | 4.391823 | 3.771724 | 2.971944 | 2.009991 | 3.085274 | 1.935037 | 2.632707 | 1.145627 | 1.3309 | 1.48333 | 1.405035 | 1.980467 | 2.890499 | 1.570024 | 2.99581 | 0.987959 | 1.726414 | 2.220238 | 2.119654 |
| 32.52639 | 0.284357 | 3.664082 | 3.724692 | 2.434925 | 1.544606 | 1.991776 | 2.609254 | 1.829753 | 1.469901 | 1.026862 | 1.176947 | 1.167033 | 1.105251 | 2.779344 | 1.277294 | 2.767225 | 1.051535 | 1.261992 | 1.344632 | 1.331284 |
| 32.54028 | 0.107306 | 3.823883 | 3.76073 | 2.694965 | 1.589289 | 1.665537 | 2.548411 | 1.124525 | 1.246694 | 0.581352 | 0.799772 | 1.051046 | 0.376121 | 2.398285 | 1.000206 | 2.226697 | 1.290666 | 0.624699 | 0.815105 | 0.861301 |
| 32.55417 | -0.15722 | 3.340723 | 2.992816 | 3.338193 | 2.127205 | 1.463664 | 2.148508 | 0.462933 | 0.886274 | 0.276319 | 0.482431 | 0.957034 | -0.08264 | 1.914324 | 0.41413 | 1.366602 | 1.556467 | 0.205009 | 0.547695 | 0.349158 |
| 32.56806 | -0.06844 | 3.032049 | 1.207268 | 2.958996 | 2.090599 | 1.12595 | 1.368541 | 0.160827 | 0.430306 | 0.124554 | 0.3375 | 0.872106 | -0.4072 | 1.197476 | -0.3079 | -0.1786 | 1.294982 | 0.324729 | 0.182592 | -0.36519 |
| 32.58194 | 0.1132 | 2.885221 | -0.1747 | 1.77026 | 1.438351 | 0.617897 | 0.425407 | 0.320592 | -0.08025 | -0.04396 | 0.079737 | 0.497927 | -0.47709 | 0.407746 | -0.89721 | -1.87799 | 1.094306 | 0.120757 | -0.27596 | -0.55191 |
| 32.59583 | 0.394933 | 2.805094 | -1.24764 | 1.452192 | 1.073248 | 0.025843 | -0.84201 | 0.340806 | -0.55763 | -0.31347 | -0.17885 | 0.106977 | -0.47192 | -0.00705 | -1.09925 | -2.76355 | 1.0995 | -0.33828 | -0.41623 | -0.62151 |
| 32.60972 | 0.586422 | 3.09175 | -2.56622 | 1.630331 | 1.04481 | -0.40155 | -2.04294 | 0.121561 | -0.62954 | -0.49821 | -0.40575 | -0.15011 | -0.47387 | -0.30271 | -1.0767 | -2.93424 | 1.14657 | -0.54972 | -0.01027 | -0.70897 |
| 32.62361 | 0.430245 | 2.802817 | -4.00655 | 1.676828 | 1.109059 | -0.74339 | -2.68691 | -0.11295 | -0.66849 | -0.53712 | -0.35656 | -0.33408 | -0.42195 | -0.70748 | -1.06988 | -2.43912 | 0.858424 | -0.38744 | 0.283656 | -0.98813 |
| 32.6375 | 0.084927 | 2.163816 | -4.16957 | 1.552806 | 1.074797 | -1.37721 | -1.90623 | -0.22952 | -0.67434 | -0.68407 | -0.19813 | -0.43885 | -0.21002 | -1.00498 | -0.85763 | -1.41093 | 0.612606 | -0.01728 | 0.265577 | -1.03866 |
| 32.65139 | 0.067637 | 1.664722 | -3.657 | 1.421101 | 1.004108 | -2.13876 | -1.2078 | -0.12817 | -0.66178 | -0.66639 | -0.00057 | -0.42119 | -0.20974 | -0.98999 | -0.64114 | -0.83937 | 0.291785 | 0.088765 | -0.08099 | -0.9231 |
| 32.66528 | 0.160011 | 1.106559 | -3.18043 | 0.939947 | 0.891326 | -1.9949 | -0.2238 | 0.235535 | -0.34508 | -0.45095 | 0.328391 | -0.14646 | -0.40834 | -0.85935 | -0.56994 | -0.68049 | -0.02361 | 0.141584 | -0.38077 | -0.60632 |
| 32.67917 | 0.352666 | 0.462876 | -2.71335 | 0.538382 | 0.584955 | -1.84136 | 0.199438 | 0.354694 | 0.51266 | -0.24929 | 0.48934 | 0.011741 | -0.71544 | -0.72708 | -0.42288 | -0.5423 | -0.27487 | 0.128514 | -0.62295 | -0.59057 |
| 32.69306 | 0.522543 | -0.03156 | -2.06562 | 0.361106 | 0.438209 | -1.31661 | 0.633681 | 0.140504 | 0.914083 | -0.11402 | 0.423179 | 0.133525 | -0.91637 | -0.5852 | -0.35927 | -0.19669 | -0.39878 | 0.143686 | -0.53041 | -0.54122 |
| 32.70694 | 0.374276 | -0.64662 | -0.8331 | 0.04443 | 0.486959 | -0.9064 | 0.671946 | -0.06894 | 0.950211 | 0.027225 | 0.210158 | 0.117334 | -0.87973 | -0.38819 | -0.72219 | -0.00626 | -0.50718 | 0.260514 | -0.17072 | -0.39637 |
| 32.72083 | 0.300377 | -1.28322 | -0.11213 | -0.44296 | 0.553388 | -0.55048 | 0.481556 | -0.40087 | 0.998649 | 0.021721 | -0.16762 | -0.02967 | -0.88513 | -0.27196 | -1.54806 | 0.096819 | -0.70351 | 0.291475 | 0.369118 | -0.43724 |
| 32.73472 | 0.061272 | -1.49717 | 0.05964 | -0.71062 | 0.479022 | -0.03342 | 0.407371 | -0.74409 | 0.978458 | -0.14294 | -0.59986 | -0.17571 | -0.7624 | -0.11872 | -2.11445 | -0.07724 | -0.9334 | 0.037683 | -0.10661 | -0.42405 |
| 32.74861 | -0.18121 | -1.68016 | 0.091564 | -0.69954 | 0.322542 | 0.181573 | 0.23158 | -1.16206 | 0.810421 | -0.43132 | -1.04668 | -0.30439 | -0.99958 | -0.1957 | -1.70146 | -0.1356 | -0.87755 | -0.31588 | -0.91822 | -0.59406 |
| 32.7625 | -0.54711 | -2.09441 | -0.22232 | -1.17863 | 0.149942 | -0.45057 | 0.017106 | -1.43393 | 0.31526 | -0.55407 | -1.4051 | -0.51717 | -1.34541 | -0.42291 | -1.06003 | -0.49879 | -0.92868 | -0.54076 | -1.82295 | -0.9789 |
| 32.77639 | -1.16189 | -2.96785 | -0.8072 | -2.08296 | 0.017146 | -1.75745 | -0.43795 | -1.53686 | -0.37304 | -0.45621 | -1.53734 | -0.97966 | -1.52178 | -0.80785 | -0.76009 | -1.13045 | -1.03099 | -0.64044 | -1.59419 | -1.32475 |
| 32.79028 | -1.5917 | -3.5211 | -1.29899 | -2.48068 | 0.021574 | -2.8802 | -0.69046 | -1.70397 | -0.97687 | -0.33272 | -1.49332 | -1.37501 | -1.6221 | -1.09623 | -0.65848 | -1.71513 | -1.07226 | -0.73259 | -1.12378 | -1.62362 |
| 32.80417 | -1.71368 | -3.53088 | -1.59213 | -1.97034 | 0.116815 | -3.41014 | -0.55276 | -1.83635 | -1.17697 | -0.41814 | -1.24783 | -1.47048 | -1.73303 | -1.37881 | -0.49577 | -1.85053 | -1.12707 | -1.06554 | -0.87214 | -1.73817 |
| 32.81806 | -1.88133 | -3.64149 | -1.73524 | -1.5369 | -0.12123 | -3.56483 | -0.50805 | -2.01121 | -1.39106 | -0.59923 | -0.9129 | -1.48471 | -1.86975 | -1.8662 | -0.47646 | -1.74952 | -1.18886 | -1.43115 | -0.92245 | -1.93886 |
| 32.83194 | -2.08859 | -3.91102 | -1.83707 | -1.80318 | -0.56103 | -3.80595 | -0.6164 | -2.06499 | -1.52212 | -0.69981 | -0.49042 | -1.63855 | -1.94747 | -2.07247 | -0.69374 | -1.75065 | -1.29691 | -1.46314 | -0.93557 | -1.84705 |
| 32.84583 | -2.31372 | -3.90539 | -1.57511 | -1.6718 | -0.95925 | -3.55834 | -0.83798 | -1.71577 | -1.25503 | -0.80291 | -0.12528 | -1.54123 | -1.97848 | -1.95422 | -0.89577 | -1.62516 | -1.29141 | -1.40334 | -0.81079 | -1.57318 |
| 32.85972 | -2.27077 | -3.85598 | -1.29629 | -1.23939 | -0.93056 | -2.65181 | -0.85237 | -1.24683 | -0.66608 | -0.75577 | 0.02471 | -1.26981 | -1.81349 | -1.72972 | -0.91643 | -1.36737 | -1.14617 | -1.33724 | -0.85026 | -1.4537 |
| 32.87361 | -1.59472 | -3.45949 | -1.35522 | -0.68436 | -0.61863 | -1.99459 | -0.91619 | -0.95858 | -0.05417 | -0.77792 | -0.00035 | -0.84161 | -1.49094 | -1.49931 | -0.88662 | -0.9433 | -0.96707 | -1.09382 | -0.78796 | -1.37804 |
| 32.8875 | -0.73691 | -2.7234 | -1.24741 | -0.19008 | -0.19285 | -1.80443 | -0.61629 | -0.65357 | 0.195567 | -0.82856 | -0.07597 | -0.50585 | -1.01077 | -1.13853 | -0.66016 | -0.40297 | -0.7047 | -0.74774 | -0.6608 | -1.12198 |
| 32.90139 | 0.003148 | -1.73935 | -0.82074 | 0.392369 | 0.294542 | -1.31974 | 0.011649 | -0.30775 | 0.334671 | -0.79513 | 0.003575 | -0.21809 | -0.45143 | -0.70563 | -0.25101 | 0.233357 | -0.42944 | -0.52324 | -0.5153 | -0.61415 |
| 32.91528 | 0.414066 | -0.99301 | -0.23424 | 0.477952 | 0.521871 | -1.0888 | 0.529935 | -0.035 | 0.210761 | -0.59854 | 0.065552 | -0.04668 | -0.07451 | -0.33646 | 0.009687 | 0.483111 | -0.2244 | -0.25753 | -0.36512 | -0.2564 |
| 32.92917 | 0.210448 | -1.14273 | -0.17643 | -0.19856 | 0.108884 | -1.41205 | 0.314292 | 0.000754 | -0.35625 | -0.49618 | -0.35939 | -0.30951 | -0.43615 | -0.5269 | -0.3995 | -0.34905 | -0.3394 | -0.3598 | -0.67858 | -0.6083 |
| 32.94306 | -0.89355 | -2.55402 | -1.00748 | -1.03468 | -0.69601 | -2.60743 | -0.8326 | -0.55375 | -1.22276 | -1.06685 | -1.24634 | -1.26229 | -1.81208 | -1.77877 | -1.63902 | -2.26156 | -0.99169 | -1.27691 | -2.10634 | -1.8698 |
| 32.95694 | -2.6179 | -3.38907 | -1.92701 | -1.45291 | -1.29118 | -3.76553 | -2.41275 | -1.27953 | -1.78539 | -2.05713 | -2.4804 | -2.2908 | -3.31224 | -3.31285 | -3.62205 | -4.12249 | -1.7912 | -2.08793 | -3.78829 | -2.96248 |
| 32.97083 | -3.7204 | -2.64344 | -1.79957 | -1.4334 | -1.16478 | -3.66203 | -3.28935 | -1.25401 | -1.51832 | -1.95966 | -2.61498 | -2.05781 | -3.61914 | -3.09799 | -3.79633 | -3.26518 | -2.37574 | -1.67135 | -3.52447 | -2.76766 |
| 32.98472 | -3.71824 | -2.06279 | -1.23794 | -0.6881 | -0.59013 | -2.4565 | -2.75022 | -0.85037 | -1.17817 | -0.78142 | -1.493 | -0.96019 | -2.09589 | -1.46463 | -1.84656 | -1.28529 | -1.6568 | -1.25312 | -1.85599 | -1.54317 |
| 32.99861 | -3.10346 | -1.95369 | -0.82065 | -0.33346 | -0.15486 | -1.80761 | -2.03155 | -0.36502 | -1.17532 | 0.164639 | -0.57469 | -0.00136 | -0.87013 | -0.08028 | -0.13863 | 0.267002 | -0.84276 | -0.57849 | -0.43034 | -0.33038 |
| 33.0125 | -2.32403 | -1.61108 | -0.42317 | -0.63437 | 0.110102 | -1.68173 | -1.35624 | 0.147657 | -1.09587 | 0.537529 | -0.03826 | 0.413673 | 0.313036 | 0.64534 | 0.760592 | 1.023547 | 0.065662 | 0.111302 | 0.312183 | 0.415763 |
| 33.02639 | -1.48683 | -1.11051 | -0.02426 | -0.92288 | 0.230727 | -1.29308 | -0.68089 | 0.544725 | -0.81638 | 0.551449 | 0.286278 | 0.586733 | 1.221908 | 0.920289 | 1.123393 | 1.402848 | 0.972541 | 0.715132 | 0.703206 | 0.657728 |
| 33.04028 | -0.79564 | -0.63018 | 0.261085 | -0.92219 | 0.179765 | -0.98063 | -0.12387 | 0.633244 | -0.57576 | 0.474541 | 0.580144 | 0.588693 | 1.602974 | 1.004012 | 1.18167 | 1.410828 | 1.387384 | 1.17057 | 1.156995 | 0.645893 |
| 33.05417 | -0.28202 | -0.18505 | 0.389779 | -0.63767 | 0.173168 | -0.79631 | 0.23107 | 0.534979 | -0.39495 | 0.324349 | 0.757435 | 0.44262 | 1.745256 | 0.998354 | 1.056851 | 1.253127 | 1.136654 | 1.532272 | 1.243958 | 0.710219 |
| 33.06806 | 0.172029 | 0.302987 | 0.346919 | -0.52052 | 0.188979 | -0.63163 | 0.56178 | 0.390724 | -0.25905 | 0.286567 | 0.754402 | 0.334186 | 1.609298 | 0.782165 | 0.596011 | 0.916903 | 0.835318 | 0.885825 | 0.838503 | 0.57776 |
| 33.08194 | 0.676097 | 0.769544 | 0.347336 | -0.43394 | 0.234878 | -0.42791 | 0.90623 | 0.31897 | -0.0086 | 0.2614 | 0.683009 | 0.240464 | 1.512956 | 0.570328 | 0.127293 | 0.6207 | 0.642977 | 0.530523 | 0.627822 | 0.469365 |
| 33.09583 | 1.234522 | 1.249618 | 0.461529 | -0.21773 | 0.372154 | -0.21277 | 1.289677 | 0.38068 | 0.270545 | 0.350183 | 0.621302 | 0.134344 | 1.464618 | 0.478467 | -0.28929 | 0.450064 | 0.588875 | 0.447306 | 0.438953 | 0.370101 |
| 33.10972 | 1.805329 | 1.624959 | 0.487592 | 0.327084 | 0.441466 | -0.03651 | 1.680056 | 0.431682 | 0.534762 | 0.35709 | 0.603962 | 0.074767 | 1.293658 | 0.426611 | -0.46544 | 0.390882 | 0.432354 | 0.489483 | 0.283796 | 0.198118 |
| 33.12361 | 2.472793 | 2.008889 | 0.555077 | 0.797847 | 0.49431 | 0.182162 | 1.909108 | 0.495799 | 0.810363 | 0.366572 | 0.720381 | 0.08457 | 1.140556 | 0.476737 | -0.33433 | 0.461112 | 0.21341 | 0.507211 | 0.219256 | 0.162795 |
| 33.1375 | 2.859678 | 2.198791 | 0.562802 | 0.843909 | 0.549721 | 0.437867 | 1.656259 | 0.59079 | 0.987773 | 0.452114 | 0.902835 | 0.071826 | 1.013081 | 0.548555 | 0.02374 | 0.548717 | 0.116566 | 0.566086 | 0.290936 | 0.357733 |
| 33.15139 | 2.96756 | 1.984472 | 0.496634 | 0.899925 | 0.591731 | 0.605347 | 1.296686 | 0.608901 | 1.04102 | 0.463022 | 1.052665 | 0.084242 | 0.849113 | 0.593591 | 0.387889 | 0.710186 | -0.01773 | 0.450466 | 0.452073 | 0.370407 |
| 33.16528 | 2.990604 | 1.62632 | 0.476013 | 1.041842 | 0.602192 | 0.640122 | 1.206929 | 0.580896 | 1.048684 | 0.386913 | 1.069115 | 0.300836 | 0.461127 | 0.62705 | 0.564053 | 0.820653 | -0.12697 | 0.219402 | 0.554259 | 0.148664 |
| 33.17917 | 2.58919 | 1.315786 | 0.464156 | 1.419628 | 0.611682 | 0.597558 | 1.351146 | 0.430917 | 1.008302 | 0.180017 | 0.719414 | 0.413183 | 0.006976 | 0.574381 | 0.48635 | 1.005721 | -0.20417 | 0.128203 | 0.534947 | -0.20683 |
| 33.19306 | 2.192002 | 1.173428 | 0.451605 | 1.847285 | 0.664192 | 0.62095 | 1.501571 | 0.312969 | 0.962961 | -0.01551 | 0.27044 | 0.386972 | -0.17 | 0.567111 | 0.488109 | 1.274383 | -0.18063 | 0.217374 | 0.564098 | -0.443 |
| 33.20694 | 1.767356 | 0.732406 | 0.247433 | 0.994351 | 0.580209 | 0.849595 | 1.289517 | 0.386511 | 0.718758 | 0.223782 | 0.051555 | 0.299892 | -0.34934 | 0.501367 | 0.447798 | 0.819375 | -0.13318 | 0.31918 | 0.388471 | 0.15545 |
| 33.22083 | 1.469153 | -0.05635 | -0.03425 | -0.6341 | 0.3975 | 1.034709 | 0.613806 | 0.448957 | 0.312186 | 0.778475 | -0.13737 | 0.139163 | -0.39907 | 0.389285 | 0.163443 | -0.35189 | -0.07245 | 0.332071 | 0.182138 | 1.029293 |
| 33.23472 | 1.053705 | -0.34852 | 0.071476 | -0.76734 | 0.34604 | 1.076077 | 0.346122 | 0.371512 | 0.159161 | 0.924978 | -0.25724 | 0.010585 | -0.37887 | 0.266013 | 0.064398 | -0.66435 | -0.01869 | 0.097209 | -0.18964 | 1.121421 |
| 33.24861 | 0.584716 | -0.31121 | 0.113952 | 0.116455 | 0.373524 | 0.816012 | 0.24566 | 0.09162 | 0.27437 | 0.638715 | -0.19623 | 0.06448 | -0.25553 | 0.192339 | 0.244863 | -0.17447 | 0.164284 | -0.0038 | 0.09084 | 0.52686 |
| 33.2625 | 0.043309 | -0.47395 | 0.114252 | 0.532099 | 0.351702 | 0.426378 | 0.198405 | -0.0178 | 0.266148 | 0.333754 | -0.04 | 0.168887 | -0.16281 | 0.002842 | 0.354204 | 0.077045 | 0.25118 | -0.12519 | 0.504655 | 0.091865 |
| 33.27639 | -0.3051 | -0.74613 | 0.101459 | 0.369005 | 0.293368 | 0.112724 | 0.211744 | 0.116145 | 0.157682 | 0.290005 | -0.00861 | 0.266207 | 0.0882 | -0.09973 | 0.361849 | 0.063006 | 0.181943 | -0.03257 | 0.517076 | 0.074223 |
| 33.29028 | -0.40997 | -0.6903 | 0.10505 | 0.372857 | 0.297827 | -0.02345 | 0.438013 | 0.178869 | 0.147617 | 0.358121 | 0.00858 | 0.378476 | 0.358393 | -0.05513 | 0.448478 | 0.235363 | 0.141979 | 0.220671 | 0.517339 | 0.090731 |
| 33.30417 | -0.33306 | -0.46235 | 0.094922 | 0.4371 | 0.29558 | -0.06638 | 0.493703 | 0.145049 | 0.173179 | 0.451394 | -0.0657 | 0.417457 | 0.492953 | 0.030432 | 0.588363 | 0.471831 | 0.096781 | 0.496972 | 0.509905 | 0.158615 |
| 33.31806 | -0.1122 | -0.19781 | 0.084762 | 0.101782 | 0.311921 | 0.017528 | 0.495736 | 0.203424 | 0.212985 | 0.676034 | -0.03272 | 0.472156 | 0.681629 | 0.199907 | 0.734838 | 0.508995 | 0.075508 | 0.717527 | 0.497944 | 0.491244 |
| 33.33194 | 0.267007 | 0.115517 | 0.156591 | -0.05575 | 0.447012 | 0.203396 | 0.696606 | 0.378718 | 0.298636 | 0.96597 | 0.128684 | 0.526034 | 0.949857 | 0.419126 | 0.830084 | 0.59701 | 0.127549 | 0.779447 | 0.722026 | 0.823561 |
| 33.34583 | 0.608432 | 0.534926 | 0.182866 | 0.35942 | 0.620408 | 0.395892 | 0.653355 | 0.449602 | 0.354508 | 0.921598 | 0.365735 | 0.446201 | 1.124212 | 0.55182 | 0.733619 | 0.841941 | 0.27695 | 0.682027 | 0.996564 | 0.797706 |
| 33.35972 | 1.059514 | 0.979537 | 0.132772 | 0.922007 | 0.732931 | 0.609403 | 0.395177 | 0.43528 | 0.47462 | 0.741263 | 0.600556 | 0.301412 | 1.174419 | 0.601383 | 0.649265 | 1.170385 | 0.433682 | 0.4981 | 1.234102 | 0.62266 |
| 33.37361 | 1.577247 | 1.41271 | 0.143082 | 1.164195 | 0.747074 | 0.785634 | 0.192745 | 0.402405 | 0.530673 | 0.630513 | 0.653057 | 0.273917 | 1.090455 | 0.613166 | 0.569741 | 1.175408 | 0.461986 | 0.488389 | 1.359359 | 0.519116 |
| 33.3875 | 1.778548 | 1.431961 | 0.134666 | 1.079485 | 0.642064 | 0.778548 | 0.137453 | 0.245133 | 0.484194 | 0.504906 | 0.508898 | 0.330698 | 0.81947 | 0.469464 | 0.48248 | 0.813126 | 0.31186 | 0.316579 | 1.156004 | 0.382181 |
| 33.40139 | 1.634804 | 1.158813 | 0.188139 | 0.94622 | 0.577009 | 0.692942 | 0.357171 | -0.07988 | 0.365723 | 0.389642 | 0.290172 | 0.473792 | 0.58937 | 0.307535 | 0.471426 | 0.48625 | 0.136876 | 0.128366 | 0.883137 | 0.346339 |

| | | | | | | | | | | | | | | | | | | | | |
|---|---|---|---|---|---|---|---|---|---|---|---|---|---|---|---|---|---|---|---|---|
| 33.41528 | 1.487274 | 0.933517 | 0.395808 | 1.557259 | 0.599929 | 0.671871 | 0.686922 | -0.28074 | 0.428204 | 0.306769 | 0.180367 | 0.736863 | 0.489425 | 0.299455 | 0.680782 | 0.680099 | 0.171145 | 0.094838 | 0.926 | 0.318143 |
| 33.42917 | 1.752393 | 1.068967 | 0.72109 | 2.781703 | 0.73903 | 0.848427 | 1.188053 | -0.07822 | 0.739157 | 0.404122 | 0.427677 | 1.063701 | 0.865743 | 0.557953 | 1.185664 | 1.547567 | 0.397446 | 0.127754 | 1.422736 | 0.566542 |
| 33.44306 | 2.33209 | 1.864213 | 1.23264 | 2.922088 | 0.994148 | 1.561266 | 1.491789 | 0.539057 | 1.024118 | 0.866239 | 0.852713 | 1.341697 | 2.044729 | 1.472427 | 1.835937 | 2.66349 | 0.67927 | 0.542189 | 2.264695 | 1.722518 |
| 33.45694 | 2.424187 | 1.804632 | 1.555533 | 2.204347 | 1.084093 | 3.286611 | 1.660406 | 1.174787 | 1.136067 | 1.554619 | 1.146826 | 1.441471 | 2.907642 | 2.146717 | 2.212795 | 2.918548 | 0.576644 | 1.111074 | 3.07516 | 2.755201 |
| 33.47083 | 2.088716 | 3.08286 | 1.703455 | 1.596971 | 0.97461 | 3.308814 | 1.859585 | 1.540876 | 1.117149 | 1.573007 | 1.288479 | 1.422216 | 3.13309 | 1.854389 | 1.726431 | 3.141745 | -0.04886 | 1.209517 | 2.788244 | 2.545803 |
| 33.48472 | 2.131453 | 3.30244 | 1.781908 | 1.062976 | 0.716919 | 2.344926 | 2.30469 | 1.715089 | 1.045218 | 1.062182 | 1.278957 | 1.258095 | 3.153602 | 1.380691 | 1.202833 | 2.989795 | -0.17868 | 0.990705 | 2.237873 | 2.288703 |
| 33.49861 | 2.110468 | 2.794497 | 1.637704 | 0.954086 | 0.681933 | 2.390742 | 1.57476 | 1.724749 | 1.068966 | 0.742995 | 1.22646 | 0.988034 | 3.039341 | 1.111712 | 0.78127 | 2.823599 | 0.300618 | 0.984777 | 1.588569 | 2.120923 |
| 33.5125 | 1.628829 | 2.254278 | 1.684561 | 1.268302 | 0.769531 | 3.185456 | 0.739123 | 1.410408 | 1.345123 | 0.454695 | 1.10595 | 0.760467 | 2.868321 | 1.191102 | 0.679982 | 3.017472 | 0.978439 | 0.990123 | 1.312237 | 1.978869 |
| 33.52639 | 0.7105 | 1.881 | 1.71604 | 1.286636 | 0.879665 | 3.583905 | 0.283647 | 1.156983 | 1.505186 | 0.312827 | 0.989606 | 0.560979 | 2.421805 | 1.540418 | 0.6441 | 3.136893 | 0.959612 | 0.839187 | 1.031115 | 1.503101 |
| 33.54028 | -0.1976 | 1.409863 | 1.615002 | 0.932054 | 0.973392 | 3.146238 | -0.31021 | 0.927114 | 1.521706 | 0.262344 | 0.864627 | 0.431843 | 1.592725 | 1.655754 | 0.363336 | 2.297672 | 0.944751 | 0.717747 | 0.764882 | 0.815059 |
| 33.55417 | -0.70385 | 1.41719 | 1.281219 | 0.478825 | 0.966078 | 2.568975 | -0.50381 | 0.572315 | 1.396054 | 0.259143 | 0.66934 | 0.490043 | 0.75666 | 1.359678 | 0.155063 | 1.049775 | 1.091454 | 0.581429 | 0.651626 | -0.04561 |
| 33.56806 | -0.71372 | 1.344414 | 0.879677 | 0.586234 | 0.839538 | 2.375837 | -0.08808 | 0.322621 | 1.295348 | 0.104983 | 0.523458 | 0.362486 | -0.06532 | 0.779349 | -0.11523 | -0.16582 | 1.072274 | 0.349794 | 0.67197 | -0.81441 |
| 33.58194 | -0.35018 | 1.369583 | 0.536148 | 0.690387 | 0.779167 | 2.188904 | 0.31294 | 0.29396 | 1.160533 | -0.11749 | 0.283206 | 0.035344 | -0.67184 | 0.342341 | -0.47014 | -1.25607 | 0.878861 | 0.073487 | 0.160979 | -1.06651 |
| 33.59583 | -0.04124 | 1.460557 | 0.207405 | 0.974132 | 0.860889 | 1.605854 | 0.353852 | 0.399392 | 0.974403 | -0.32627 | 0.033147 | -0.35095 | -1.30321 | -0.34188 | -0.83566 | -2.16483 | 0.960935 | 0.021781 | -0.49308 | -1.2154 |
| 33.60972 | 0.259844 | 2.298746 | -0.28894 | 1.250986 | 0.955029 | 0.627825 | 0.073722 | 0.577966 | 0.748731 | -0.40683 | -0.06784 | -0.62339 | -1.48477 | -0.60219 | -1.17947 | -1.98638 | 1.052153 | 0.099315 | -0.39909 | -1.27288 |
| 33.62361 | 0.538523 | 2.934262 | -0.82834 | 1.616686 | 0.922451 | -0.38749 | -0.37907 | 0.7261 | 0.34155 | -0.43607 | -0.17056 | -0.8036 | -1.18251 | -0.71242 | -1.40877 | -1.29477 | 0.884515 | -0.12251 | 0.0615 | -1.24383 |
| 33.6375 | 0.771726 | 2.984399 | -1.02505 | 1.672443 | 0.554015 | -1.12143 | -0.65815 | 0.72758 | -0.03515 | -0.42813 | -0.26923 | -0.69692 | -0.51511 | -0.72476 | -1.34948 | -0.8194 | 0.383151 | -0.11414 | 0.533027 | -0.88965 |
| 33.65139 | 0.830537 | 2.963415 | -0.90095 | 1.918782 | -0.01028 | -1.5028 | -0.7721 | 0.604706 | -0.25825 | -0.39319 | -0.33418 | -0.53681 | 0.039636 | -0.73138 | -0.99338 | -0.70808 | 0.122641 | -0.05057 | 0.64 | -0.63215 |
| 33.66528 | 0.684325 | 2.762537 | -0.67332 | 1.843467 | -0.3269 | -1.67765 | -0.70398 | 0.428088 | -0.34226 | -0.38862 | -0.31195 | -0.38249 | 0.623788 | -0.68038 | -0.61589 | -0.43223 | -0.06984 | 0.006368 | 0.46588 | -0.60525 |
| 33.67917 | 0.334798 | 2.179804 | -0.19938 | 1.802779 | -0.21072 | -1.62926 | -0.38041 | 0.235406 | -0.21075 | -0.35838 | -0.1754 | -0.12796 | 1.013188 | -0.46494 | -0.56153 | -0.03855 | -0.12222 | 0.150592 | 0.262465 | -0.5923 |
| 33.69306 | 0.01855 | 1.499668 | 0.20375 | 1.869388 | 0.051638 | -1.47326 | -0.07211 | 0.227042 | -0.09472 | -0.25448 | 0.035199 | 0.044076 | 1.134447 | -0.30731 | -0.41024 | 0.318859 | -0.20553 | 0.304648 | 0.042636 | -0.44676 |
| 33.70694 | -0.25664 | 0.989302 | 0.286924 | 1.548992 | 0.173749 | -1.44088 | 0.22361 | 0.3414 | -0.11436 | -0.15276 | 0.253088 | 0.099255 | 0.824367 | -0.31367 | -0.30371 | 0.525357 | -0.36096 | 0.282965 | -0.12568 | -0.42143 |
| 33.72083 | -0.42608 | 0.544451 | -0.15902 | 1.175311 | 0.126668 | -1.45412 | 0.214272 | 0.314396 | -0.20821 | -0.00407 | 0.327318 | -0.00259 | 0.262008 | -0.44314 | -0.34409 | 0.398689 | -0.51952 | -0.00818 | -0.21876 | -0.48046 |
| 33.73472 | -0.50146 | 0.03267 | -1.0029 | 0.654354 | 0.019094 | -1.56132 | -0.28582 | 0.147464 | -0.25812 | 0.114474 | 0.181874 | -0.17549 | -0.70599 | -0.57394 | -0.45915 | -0.03636 | -0.74832 | -0.3604 | -0.23549 | -0.22142 |
| 33.74861 | -0.53429 | -0.49081 | -1.79447 | 0.084163 | -0.06043 | -1.78236 | -1.06681 | -0.41872 | -0.35943 | 0.172922 | -0.09142 | -0.39815 | -1.90444 | -0.74391 | -0.48186 | -0.59527 | -0.44331 | -0.64301 | -0.2511 | 0.130172 |
| 33.7625 | -0.58962 | -0.9761 | -2.1284 | -0.18681 | -0.16238 | -2.06588 | -1.51813 | -1.06187 | -0.5133 | 0.162954 | -0.3363 | -0.596 | -2.56513 | -0.75892 | -0.35424 | -1.35321 | -0.43005 | -0.66731 | -0.23811 | 0.239102 |
| 33.77639 | -0.92697 | -1.43634 | -1.95779 | -0.1464 | -0.35675 | -2.3575 | -1.48073 | -1.35915 | -0.81735 | -0.19206 | -0.54494 | -0.72045 | -2.09879 | -0.6239 | -0.11975 | -1.75445 | -0.55274 | -0.84698 | -0.06132 | -0.33682 |
| 33.79028 | -1.43689 | -2.05208 | -1.675 | -0.26121 | -0.64096 | -2.75888 | -1.22778 | -1.6999 | -1.07067 | -0.67495 | -0.73509 | -0.72571 | -1.96134 | -0.51598 | -0.04998 | -2.026 | -0.43315 | -1.31396 | 0.034288 | -0.52061 |
| 33.80417 | -1.5158 | -2.72666 | -1.28112 | -0.88991 | -0.67929 | -2.3379 | -0.98406 | -2.05217 | -1.02843 | -0.73542 | -0.87034 | -0.69179 | -1.52949 | -0.50436 | -0.12314 | -1.96372 | -0.26555 | -1.34973 | 0.001044 | -0.28868 |
| 33.81806 | -1.6526 | -3.00803 | -0.96401 | -1.57281 | -0.48062 | -1.84948 | -0.52717 | -1.97665 | -0.65754 | -0.41052 | -0.97574 | -0.69367 | -1.18595 | -0.59831 | -0.2155 | -1.96511 | -0.18336 | -0.7076 | -0.08113 | 0.026923 |
| 33.83194 | -1.61608 | -2.66133 | -0.64679 | -1.67017 | -0.5169 | -1.46011 | -0.23442 | -1.78216 | -0.4426 | -0.157 | -1.04329 | -0.64901 | -0.86945 | -0.81253 | -0.23729 | -1.76011 | -0.27479 | -0.23979 | -0.40927 | -0.09858 |
| 33.84583 | -1.3685 | -2.5841 | -0.40078 | -1.53994 | -0.8907 | -1.46477 | -0.0516 | -1.82922 | -0.58398 | -0.19683 | -0.91722 | -0.58385 | -0.63309 | -0.99338 | -0.21388 | -1.0785 | -0.48585 | -0.2054 | -0.80553 | -0.81669 |
| 33.85972 | -1.15266 | -2.66236 | -0.13025 | -1.26942 | -0.80062 | -1.48324 | -0.06641 | -1.45619 | -0.63001 | -0.45561 | -0.74009 | -0.47795 | -0.532 | -1.02861 | -0.17898 | -0.61068 | -0.57955 | -0.29771 | -1.03758 | -1.2031 |
| 33.87361 | -0.77076 | -2.60227 | -0.09137 | -0.84338 | -0.24218 | -1.39625 | -0.2755 | -0.89588 | -0.41963 | -0.72424 | -0.49074 | -0.40446 | -0.48394 | -0.95247 | -0.12957 | -0.24496 | -0.59099 | -0.29071 | -1.1647 | -1.59536 |
| 33.8875 | -0.3609 | -2.41026 | -0.18372 | -0.47789 | 0.043571 | -0.92242 | -0.44149 | -0.4916 | -0.17016 | -0.72992 | -0.28634 | -0.34527 | -0.31121 | -0.89274 | -0.07927 | -0.25451 | -0.64089 | -0.42163 | -1.18603 | -1.49022 |
| 33.90139 | -0.00893 | -1.94742 | -0.1476 | -0.30866 | 0.034725 | -0.50518 | -0.33643 | -0.24838 | 0.149479 | -0.51028 | -0.1794 | -0.34601 | -0.14233 | -0.71058 | -0.13859 | -0.35703 | -0.63546 | -0.48904 | -0.90287 | -1.29077 |
| 33.91528 | 0.075572 | -1.30463 | 0.061009 | -0.18208 | 0.048267 | -0.43106 | 0.106777 | -0.11672 | 0.251575 | -0.26112 | -0.1706 | -0.29965 | -0.10615 | -0.40986 | -0.25244 | -0.45126 | -0.60201 | -0.48218 | -0.68103 | -0.91323 |
| 33.92917 | -0.20448 | -1.25252 | -0.20076 | -0.68435 | -0.18927 | -0.96973 | 0.141407 | -0.17785 | 0.156712 | -0.28324 | -0.61265 | -0.38585 | -0.71666 | -0.87376 | -0.64265 | -1.19286 | -0.88945 | -0.67245 | -0.84622 | -0.44207 |
| 33.94306 | -1.38598 | -2.01547 | -1.43221 | -2.38421 | -0.87394 | -2.25508 | -0.72938 | -0.61425 | -0.21026 | -0.75883 | -1.62604 | -1.05586 | -2.22972 | -1.94853 | -1.87798 | -2.67381 | -1.90405 | -1.32912 | -1.96126 | -0.50385 |
| 33.95694 | -3.15825 | -3.48835 | -3.01771 | -3.84417 | -1.55337 | -2.70807 | -1.92504 | -1.2338 | -0.3961 | -1.83625 | -2.74866 | -2.06751 | -3.91856 | -3.03383 | -3.99134 | -3.80775 | -2.92195 | -1.98492 | -3.31137 | -1.68745 |
| 33.97083 | -3.54516 | -4.23622 | -3.11812 | -2.94442 | -1.73706 | -2.26018 | -2.2332 | -1.18932 | -0.17058 | -2.02905 | -2.68772 | -2.2672 | -4.3395 | -2.67567 | -3.85356 | -3.56147 | -3.12727 | -1.6922 | -3.44296 | -2.24056 |
| 33.98472 | -2.94176 | -3.9981 | -2.6832 | -1.60823 | -1.44509 | -2.81964 | -2.11566 | -1.04514 | 0.103873 | -0.84134 | -1.30076 | -1.18357 | -2.03141 | -1.00364 | -1.98541 | -1.82325 | -2.07896 | -0.51825 | -2.0474 | -0.93703 |
| 33.99861 | -2.83734 | -3.30686 | -2.21099 | -1.43798 | -1.39438 | -3.39392 | -1.90289 | -1.52206 | -0.14671 | -0.08222 | -0.43993 | -0.00295 | -0.85006 | 0.40556 | -0.20184 | -0.27718 | -1.15871 | 0.096238 | -0.58188 | -0.26167 |
| 34.0125 | -2.54423 | -2.40847 | -1.07052 | -1.07247 | -0.79068 | -2.09699 | -1.11141 | -1.59637 | -0.47535 | 0.151652 | 0.027593 | 0.76689 | -0.14065 | 0.520054 | 0.88313 | 0.861094 | -0.11012 | 0.367067 | 0.28814 | 0.33485 |
| 34.02639 | -1.88836 | -1.25398 | 0.114549 | -0.64995 | -0.4095 | -1.17556 | -0.05971 | -0.9262 | -0.59325 | 0.301559 | 0.363808 | 1.158795 | 0.675034 | 0.760328 | 1.601347 | 1.488443 | 0.935624 | 0.72434 | 0.811665 | 1.019824 |
| 34.04028 | -1.17981 | -0.26007 | 0.66136 | -0.1856 | -0.13274 | -0.80784 | 0.67581 | -0.22163 | -0.60449 | 0.447825 | 0.555731 | 1.075384 | 1.083305 | 1.124966 | 1.772061 | 1.488378 | 1.555346 | 0.931307 | 0.719739 | 1.840619 |
| 34.05417 | -0.3293 | 0.397808 | 0.757895 | -0.16152 | -0.16893 | -0.56423 | 1.258724 | 0.168021 | -0.67765 | 0.548875 | 0.425813 | 0.677273 | 1.268097 | 0.939534 | 1.43693 | 0.976943 | 1.389314 | 0.811782 | 0.528261 | 1.855256 |
| 34.06806 | 0.595894 | 0.877421 | 0.77025 | 0.109171 | -0.07637 | -0.39985 | 1.701356 | 0.344629 | -0.61091 | 0.587824 | 0.389571 | 0.643981 | 1.518974 | 0.65774 | 1.538917 | 0.434032 | 1.052081 | 0.700077 | 0.35191 | 1.07486 |
| 34.08194 | 1.388585 | 1.110484 | 0.790012 | 0.326699 | 0.123874 | -0.43829 | 2.005528 | 0.199398 | -0.44703 | 0.336543 | 0.483174 | 0.56281 | 1.619611 | 0.544715 | 1.446413 | 0.594441 | 0.700358 | 0.479803 | 0.036957 | -0.18655 |
| 34.09583 | 1.850808 | 1.213942 | 0.744723 | 0.712531 | 0.234274 | -0.47449 | 2.040448 | -0.18814 | -0.23422 | -0.04282 | 0.57144 | 0.018331 | 1.528013 | 0.441864 | 0.794305 | 0.763037 | 0.635446 | 0.316996 | -0.24918 | -1.20301 |
| 34.10972 | 1.853836 | 1.21979 | 0.715564 | 0.513502 | 0.232349 | -0.21791 | 1.76119 | -0.12705 | -0.11405 | 0.103395 | 0.637634 | -0.47175 | 1.661493 | 0.253194 | -0.45722 | -0.21766 | 0.490484 | 0.672037 | -0.12806 | -1.17447 |
| 34.12361 | 1.882206 | 1.258164 | 0.834484 | 0.245068 | 0.244602 | 0.12258 | 1.580538 | 0.280151 | 0.154751 | 0.376489 | 0.752071 | -0.45918 | 1.613941 | 0.219906 | -1.15104 | -0.42371 | 0.595671 | 1.201772 | 0.15026 | 0.30356 |
| 34.1375 | 2.139424 | 1.470099 | 0.864608 | 0.842612 | 0.351588 | 0.120653 | 1.627809 | 0.472126 | 0.504739 | 0.345347 | 0.83067 | -0.40324 | 1.083909 | 0.229818 | -1.19068 | 0.373575 | 0.560217 | 0.841459 | 0.206367 | 0.586036 |
| 34.15139 | 2.386985 | 1.56523 | 0.744711 | 1.2439 | 0.353567 | 0.040479 | 1.236078 | 0.451609 | 0.650148 | 0.409982 | 0.780535 | -0.37412 | 0.699285 | 0.255586 | -0.95683 | 0.651291 | 0.254635 | 0.525125 | 0.260381 | 0.582541 |
| 34.16528 | 2.540779 | 1.605634 | 0.532291 | 1.18537 | 0.368637 | -0.29843 | 0.597924 | 0.384029 | 0.676411 | 0.568305 | 0.763392 | -0.12573 | 0.402932 | 0.337237 | -0.53162 | 0.602004 | -0.04352 | 0.489244 | 0.423978 | 0.424152 |
| 34.17917 | 2.85855 | 1.692775 | 0.356477 | 0.94153 | 0.429993 | -0.83268 | 0.332732 | 0.347596 | 0.60205 | 0.725137 | 0.774309 | 0.26275 | 0.053632 | 0.335545 | 0.128451 | 0.501978 | -0.03643 | 0.866121 | 0.5975 | 0.119561 |
| 34.19306 | 2.609451 | 1.22677 | 0.330485 | 0.388641 | 0.474326 | -0.9091 | 0.340422 | 0.39741 | 0.577598 | 0.981448 | 0.758276 | 0.468075 | -0.29718 | 0.153336 | 0.569237 | 0.002507 | 0.054327 | 1.322514 | 0.569511 | 0.26657 |
| 34.20694 | 1.802073 | 0.359701 | 0.074178 | 0.0683 | 0.351671 | -0.63431 | 0.340311 | 0.272745 | 0.512982 | 0.931346 | 0.61167 | 0.273897 | -0.60186 | 0.067574 | 0.521216 | -0.27648 | -0.13264 | 1.146535 | 0.245659 | 0.580038 |
| 34.22083 | 1.072193 | -0.2872 | -0.36629 | 0.268391 | 0.175561 | -0.43437 | 0.293918 | 0.010992 | 0.342515 | 0.446048 | 0.49225 | -0.17212 | -0.67381 | 0.343246 | 0.275903 | -0.07543 | -0.22274 | 0.275642 | -0.18107 | 0.469916 |
| 34.23472 | 0.880478 | -0.55834 | -0.51505 | 0.74877 | 0.21484 | -0.17955 | 0.332154 | -0.10087 | 0.219353 | -0.106 | 0.345847 | -0.50826 | -0.43328 | 0.721574 | -0.0085 | 0.362955 | -0.04419 | -0.54256 | -0.24761 | 0.398284 |
| 34.24861 | 0.588353 | -0.7135 | -0.41554 | 0.852085 | 0.390374 | 0.198452 | 0.440625 | -0.04305 | 0.280874 | -0.26241 | 0.312278 | -0.57239 | -0.16779 | 0.923157 | -0.23064 | 0.532913 | 0.372854 | -0.48416 | 0.048088 | 0.669116 |
| 34.2625 | 0.143099 | -0.86344 | -0.10226 | 0.183651 | 0.398561 | 0.435851 | 0.516948 | 0.086952 | 0.472728 | 0.27412 | 0.311722 | -0.30069 | 0.12316 | 0.825915 | -0.34026 | 0.020929 | 0.394229 | -0.2737 | 0.326935 | 1.109927 |
| 34.27639 | -0.11135 | -0.82866 | 0.07483 | 0.000325 | 0.309226 | 0.272111 | 0.641897 | -0.00542 | 0.411408 | 0.507222 | 0.221614 | 0.157135 | 0.218306 | 0.521944 | -0.1007 | 0.041953 | 0.326953 | 0.125411 | 0.361602 | 0.996584 |
| 34.29028 | -0.04023 | -0.94544 | 0.129855 | 0.600228 | 0.291288 | 0.135934 | 0.879955 | -0.26961 | 0.366864 | 0.361614 | 0.012174 | 0.539692 | 0.153923 | 0.213456 | 0.405986 | 0.852009 | 0.428247 | -0.19105 | 0.182513 | 0.434421 |
| 34.30417 | -0.26257 | -0.8439 | 0.475395 | 0.415452 | 0.26146 | 0.219695 | 1.005502 | -0.1684 | 0.410682 | 0.736251 | -0.14631 | 1.048974 | 0.193352 | 0.246454 | 1.009445 | 0.821554 | 0.557222 | 0.317921 | 0.245232 | 1.106212 |
| 34.31806 | -0.5933 | -0.43016 | 0.963615 | -0.30108 | 0.278622 | 0.35681 | 0.939069 | 0.139503 | 0.612332 | 1.496919 | -0.07524 | 1.421995 | 0.571528 | 0.544745 | 1.412553 | 0.265492 | 0.543938 | 1.497396 | 0.688757 | 1.889272 |
| 34.33194 | -0.48103 | -0.1223 | 0.983843 | -0.29731 | 0.459204 | 0.16692 | 0.955626 | 0.256699 | 0.510444 | 1.329499 | -0.06089 | 1.373356 | 0.785028 | 0.716573 | 1.283071 | 0.275822 | 0.46128 | 1.482141 | 0.777124 | 1.578526 |
| 34.34583 | -0.18027 | 0.247623 | 0.705877 | 0.071065 | 0.479744 | 0.071595 | 0.946231 | 0.169607 | 0.338205 | 0.81759 | 0.02499 | 1.025975 | 0.92801 | 0.786112 | 0.874082 | 0.426995 | 0.272724 | 0.835858 | 0.608746 | 0.898566 |
| 34.35972 | 0.074843 | 0.49016 | 0.363801 | 0.323512 | 0.588685 | 0.096246 | 0.824902 | 0.078389 | 0.306267 | 0.554734 | 0.147385 | 0.52054 | 0.991709 | 0.687844 | 0.14825 | 0.409895 | -0.00211 | 0.392134 | 0.532022 | 0.465719 |
| 34.37361 | 0.419683 | 0.70619 | 0.162239 | 0.47397 | 0.713763 | 0.300986 | 0.748725 | -0.00463 | 0.352355 | 0.688281 | 0.264207 | 0.105129 | 0.974066 | 0.501053 | -0.41459 | 0.339745 | -0.17531 | 0.304552 | 0.54443 | 0.390264 |
| 34.3875 | 0.76991 | 0.881622 | 0.132087 | 0.588436 | 0.842197 | 0.326622 | 0.81625 | -0.04876 | 0.521943 | 0.941536 | 0.321579 | 0.04033 | 0.886411 | 0.294314 | -0.51546 | 0.340146 | -0.12324 | 0.449863 | 0.511133 | 0.621659 |
| 34.40139 | 0.868244 | 0.894192 | 0.076407 | 0.623367 | 0.833351 | 0.173289 | 0.885854 | -0.11081 | 0.650885 | 0.832215 | 0.344959 | 0.283177 | 0.502785 | 0.161036 | -0.16213 | 0.368145 | 0.155493 | 0.442646 | 0.382691 | 0.85203 |
| 34.41528 | 0.891818 | 0.758757 | -0.20704 | 1.125566 | 0.706949 | 0.103428 | 1.049764 | -0.20842 | 0.809039 | 0.038758 | 0.393261 | 0.558419 | -0.11396 | 0.334931 | 0.368763 | 0.937253 | 0.625103 | -0.04171 | 0.060998 | 0.48562 |
| 34.42917 | 1.274649 | 0.898145 | -0.37212 | 2.161522 | 0.717563 | -0.06662 | 1.609873 | -0.20383 | 0.972243 | -0.999 | 0.49491 | 0.862872 | -0.73017 | 0.964774 | 1.056139 | 2.207852 | 0.938644 | -0.86277 | -0.36734 | 0.162208 |
| 34.44306 | 2.221119 | 1.367589 | 0.295795 | 2.676103 | 0.967579 | 0.800168 | 2.435409 | 0.632233 | 1.470438 | -0.80718 | 0.869755 | 1.084402 | 0.271945 | 1.826397 | 1.771509 | 3.507972 | 1.436428 | -1.21106 | -0.36986 | 0.180924 |
| 34.45694 | 2.938934 | 1.918484 | 1.379559 | 2.815996 | 1.648859 | 3.109526 | 2.659951 | 2.079255 | 1.606382 | 1.115219 | 1.451994 | 1.559126 | 2.054122 | 2.132697 | 2.498871 | 3.604899 | 1.28751 | 0.08601 | 1.258089 | 1.723644 |
| 34.47083 | 2.929946 | 2.637218 | 1.522399 | 2.547778 | 1.585052 | 2.91851 | 2.362159 | 2.423222 | 1.705575 | 1.732831 | 1.337267 | 1.854415 | 3.309185 | 1.804039 | 2.901484 | 3.310998 | 0.724356 | 0.932818 | 1.687016 | 2.559235 |
| 34.48472 | 2.758695 | 2.970501 | 1.429448 | 2.053317 | 1.183532 | 2.615682 | 1.377107 | 1.6988 | 1.723707 | 1.561401 | 0.985355 | 1.763711 | 4.157106 | 1.226888 | 2.498185 | 3.253854 | 0.374428 | 1.158893 | 1.438315 | 2.599746 |
| 34.49861 | 2.533745 | 2.3087 | 1.166699 | 1.572212 | 0.862049 | 2.307357 | 1.379623 | 0.99372 | 1.64371 | 1.2192 | 0.600748 | 1.441437 | 4.490135 | 0.991005 | 1.56831 | 2.902843 | 0.280875 | 1.03944 | 1.423187 | 2.065762 |
| 34.5125 | 2.008417 | 1.917049 | 0.910751 | 1.292974 | 0.72022 | 2.521179 | 1.376881 | 0.584317 | 1.567167 | 0.956727 | 0.526155 | 0.925495 | 4.66053 | 0.854953 | 0.866371 | 2.997847 | 0.317751 | 0.909506 | 1.433797 | 1.482976 |
| 34.52639 | 1.218354 | 1.419935 | 0.924481 | 0.968469 | 0.755968 | 2.886539 | 0.142682 | 0.221055 | 1.69904 | 0.795405 | 0.568946 | 0.539844 | 4.303679 | 0.652231 | 0.168901 | 3.179908 | 0.332272 | 0.752405 | 1.590578 | 1.123713 |
| 34.54028 | 0.539869 | 1.015219 | 0.919837 | 0.413154 | 0.57211 | 3.034372 | -0.69217 | 0.135534 | 1.879432 | 0.535067 | 0.471103 | 0.061257 | 2.68852 | 0.519615 | -0.47656 | 2.751699 | 0.459399 | 0.340449 | 1.658537 | 0.60878 |
| 34.55417 | -0.00895 | 0.410149 | 0.843623 | -0.17237 | 0.293315 | 3.11061 | -0.91192 | 0.312445 | 1.705528 | 0.14192 | 0.345713 | -0.28041 | 0.752658 | 0.185123 | -1.01253 | 1.738058 | 0.445684 | -0.09413 | 1.704261 | 1.140056 |
| 34.56806 | -0.67765 | 0.015165 | 0.733796 | -0.99038 | 0.260062 | 2.838337 | -1.23661 | 0.425736 | 1.614922 | -0.15958 | -0.11465 | -0.46991 | -1.36471 | -0.22619 | -0.97078 | 0.062967 | 0.43968 | -0.21802 | 1.375722 | 0.789799 |
| 34.58194 | -0.95349 | -0.29875 | 0.336958 | -1.25741 | 0.273681 | 2.040632 | -1.26234 | 0.514923 | 1.291887 | -0.60415 | -0.21253 | -0.76338 | -2.99605 | -0.48423 | -0.24753 | -1.4148 | 0.046967 | -0.3631 | 0.819122 | -0.40385 |
| 34.59583 | -0.9769 | -0.29669 | 0.037473 | -1.6461 | 0.204545 | 0.741365 | -1.0645 | 0.72683 | 0.991018 | -1.13108 | -0.06692 | -0.70932 | -3.98038 | -0.83977 | -0.19079 | -2.52582 | -0.00515 | -0.79205 | 0.480987 | -0.45096 |
| 34.60972 | -0.92274 | 0.039984 | -0.16919 | -1.89311 | 0.108934 | -0.38197 | -1.05167 | 0.829969 | 0.81838 | -1.13087 | -0.15544 | -0.65164 | -4.02708 | -1.14263 | -0.27971 | -2.29743 | 0.208485 | -0.86868 | 0.330977 | -0.59879 |
| 34.62361 | -0.73193 | 0.544103 | -0.41701 | -1.66774 | 0.025793 | -1.08806 | -1.03785 | 0.672784 | 0.644258 | -1.14829 | -0.26244 | -0.59499 | -3.04328 | -1.36014 | -0.56671 | -1.65756 | -0.24507 | -0.85819 | 0.52163 | -0.99557 |
| 34.6375 | -0.49484 | 1.023764 | -0.58825 | -1.1641 | 0.087767 | -1.71541 | -1.08656 | 0.392868 | 0.543658 | -1.24337 | -0.35704 | -0.74652 | -2.05491 | -1.57103 | -0.88878 | -1.21001 | -0.08954 | -0.40808 | 0.8515 | -1.75946 |
| 34.65139 | -0.27768 | 0.903781 | -0.43097 | -1.18293 | 0.299079 | -1.93719 | -0.9944 | 0.088817 | 0.420878 | -1.03305 | -0.29772 | -0.90026 | -0.59631 | -1.62988 | -0.87875 | -1.05039 | 0.193483 | 0.116113 | 0.946379 | -0.87543 |
| 34.66528 | 0.060522 | 1.655363 | -0.24726 | -0.39186 | 0.481008 | -1.6695 | -0.39045 | -0.09908 | 0.14219 | -0.92592 | -0.08948 | -0.95447 | 0.379017 | -1.52732 | -0.63181 | -0.47996 | 0.42512 | 0.218182 | 0.718468 | 0.263316 |

| | | | | | | | | | | | | | | | | | | | | |
|---|---|---|---|---|---|---|---|---|---|---|---|---|---|---|---|---|---|---|---|---|
| 34.67917 | 0.358836 | 2.524962 | -0.13781 | 0.384236 | 0.599139 | -1.39236 | 0.331005 | -0.0657 | -0.04675 | -0.72586 | 0.077762 | -1.0237 | 1.015814 | -1.17646 | -0.50887 | 0.011542 | 0.52592 | 0.157903 | 0.474862 | 0.28292 |
| 34.69306 | 0.267347 | 2.289167 | 0.006455 | 1.19444 | 0.700879 | -1.03109 | 0.990213 | 0.101893 | -0.00566 | -0.48171 | 0.192829 | -0.8387 | 1.420061 | -0.6646 | -0.36133 | 0.21449 | 0.560688 | 0.308297 | 0.386526 | 0.023225 |
| 34.70694 | 0.066706 | 1.760044 | 0.18813 | 1.644406 | 0.726107 | -0.37279 | 1.612895 | 0.248335 | 0.106612 | -0.3266 | 0.220725 | -0.41498 | 1.404222 | -0.22897 | -0.07577 | 0.418682 | 0.419331 | 0.293552 | 0.401863 | 0.362377 |
| 34.72083 | 0.08976 | 1.334623 | 0.28927 | 1.279907 | 0.663932 | -0.07784 | 2.062744 | 0.300753 | 0.147641 | -0.09894 | 0.087224 | -0.05487 | 1.045717 | -0.12154 | 0.053935 | 0.595362 | 0.29246 | 0.435015 | 0.404094 | 0.678088 |
| 34.73472 | 0.130901 | 0.643889 | 0.228928 | 0.784232 | 0.558719 | -0.80136 | 1.745683 | 0.245809 | 0.081166 | 0.145846 | -0.19092 | -0.01337 | 0.712601 | -0.29287 | -0.02391 | 0.676947 | 0.106116 | 0.315494 | 0.293565 | 0.615711 |
| 34.74861 | -0.07942 | 0.2222 | -0.04107 | 0.248552 | 0.486561 | -1.62486 | 0.662136 | 0.07598 | -0.17707 | 0.119883 | -0.40358 | -0.21309 | 0.288353 | -0.51344 | -0.35648 | 0.427626 | -0.25056 | -0.02468 | 0.03985 | 0.290898 |
| 34.7625 | -0.28616 | -0.16251 | -0.34923 | -0.29789 | 0.442953 | -2.03376 | -0.50319 | -0.27521 | -0.45558 | 0.077532 | -0.47701 | -0.51298 | -0.34071 | -0.67277 | -0.71354 | 0.028246 | -0.50925 | -0.50034 | -0.06101 | -0.2837 |
| 34.77639 | -0.76935 | -0.45505 | -0.57823 | -1.10267 | 0.478168 | -2.51698 | -1.16115 | -0.64699 | -0.63059 | 0.015193 | -0.39852 | -0.77469 | -1.01445 | -0.87215 | -0.79879 | -0.27592 | -0.75605 | -0.82441 | -0.3196 | -0.57276 |
| 34.79028 | -1.24893 | -0.75169 | -0.68219 | -2.08577 | 0.434392 | -2.77996 | -1.43628 | -1.00324 | -0.72194 | 0.065274 | -0.23738 | -0.88339 | -1.50788 | -0.78994 | -0.65606 | -0.48856 | -0.89354 | -0.91262 | -0.50528 | -0.47392 |
| 34.80417 | -1.47893 | -1.03064 | -0.86276 | -2.25438 | 0.260359 | -2.63035 | -1.29397 | -1.29676 | -0.72204 | 0.056846 | -0.08805 | -0.93542 | -1.67607 | -0.45904 | -0.48738 | -0.72605 | -1.12933 | -0.95237 | -0.63357 | -0.5682 |
| 34.81806 | -1.44763 | -1.37772 | -0.8778 | -2.5283 | 0.084773 | -2.43376 | -1.10328 | -1.29103 | -0.52078 | -0.04027 | -0.18396 | -0.87635 | -1.42605 | -0.19424 | -0.31498 | -1.02505 | -1.44643 | -0.79198 | -0.7725 | -0.66356 |
| 34.83194 | -1.453 | -1.73018 | -0.72465 | -2.56213 | 0.038044 | -2.446 | -0.9064 | -1.02686 | -0.12193 | -0.2395 | -0.53805 | -0.61381 | -1.34104 | -0.24123 | -0.17968 | -1.53312 | -1.61535 | -0.61474 | -1.40296 | -0.90981 |
| 34.84583 | -1.40381 | -1.91092 | -0.41633 | -2.26831 | 0.104295 | -2.32608 | -0.74246 | -0.69765 | 0.125587 | -0.47606 | -0.98583 | -0.25034 | -1.38488 | -0.34378 | -0.13075 | -1.8406 | -1.85515 | -0.29893 | -1.68308 | -1.53047 |
| 34.85972 | -1.2716 | -1.87151 | -0.2901 | -1.65875 | 0.21713 | -2.27743 | -0.62511 | -0.45206 | 0.281827 | -0.57668 | -1.17421 | 0.043915 | -1.4928 | -0.39879 | -0.07606 | -1.5888 | -1.54557 | -0.18477 | -1.19042 | -1.83967 |
| 34.87361 | -1.13601 | -1.78713 | -0.32941 | -1.02974 | 0.427296 | -2.01334 | -0.43874 | -0.34465 | 0.400494 | -0.54318 | -1.02879 | 0.212928 | -1.34059 | -0.41448 | -0.03048 | -1.07662 | -0.92057 | -0.30376 | -0.7274 | -1.26211 |
| 34.8875 | -1.08223 | -1.70766 | -0.2474 | -0.59629 | 0.581361 | -1.5662 | -0.03354 | -0.25439 | 0.487184 | -0.41381 | -0.70717 | 0.331261 | -1.36928 | -0.35857 | -0.06928 | -0.64905 | -0.09048 | -0.31976 | -0.46096 | -1.01293 |
| 34.90139 | -1.27272 | -1.78028 | 0.100251 | -0.40399 | 0.538284 | -1.13219 | 0.199376 | -0.09505 | 0.532136 | -0.29294 | -0.46932 | 0.315988 | -1.09439 | -0.22288 | -0.04013 | -0.63999 | 0.452577 | -0.05135 | -0.33045 | -0.30429 |
| 34.91528 | -1.65783 | -1.86511 | 0.405763 | -0.46808 | 0.3112 | -0.74748 | 0.137571 | 0.178707 | 0.354734 | -0.21712 | -0.31454 | 0.233321 | -0.81135 | -0.35346 | 0.022778 | -1.15302 | 0.763387 | 0.678488 | -0.2388 | 0.996041 |
| 34.92917 | -2.34952 | -2.21394 | 0.270082 | -1.3168 | -0.15119 | -0.97287 | -0.41652 | 0.370042 | -0.1354 | -0.35855 | -0.61943 | 0.191994 | -0.89179 | -0.85227 | -0.17504 | -2.15238 | 0.200386 | 1.411791 | -0.54305 | 1.658471 |
| 34.94306 | -2.75044 | -2.71309 | -1.12579 | -2.43166 | -0.7437 | -2.47726 | -1.11277 | 0.041735 | -0.925 | -1.23011 | -1.83298 | -0.34403 | -2.00545 | -1.74975 | -1.43793 | -3.58483 | -1.14412 | 0.4381 | -2.0582 | 0.175813 |
| 34.95694 | -2.68721 | -3.05548 | -2.29466 | -3.05244 | -0.96071 | -3.21732 | -1.73345 | -0.76941 | -1.50332 | -2.15413 | -3.46054 | -1.66961 | -3.89914 | -2.78312 | -3.14102 | -4.43198 | -2.94735 | -1.08789 | -3.8618 | -2.4435 |
| 34.97083 | -2.24263 | -2.6404 | -2.21261 | -2.18673 | -0.60266 | -3.18887 | -1.69744 | -1.18353 | -1.39746 | -2.36376 | -3.30772 | -2.84219 | -4.08558 | -3.19359 | -3.08825 | -3.1259 | -3.38901 | -1.17878 | -4.07543 | -3.6214 |
| 34.98472 | -1.76808 | -2.22383 | -1.71259 | -1.00177 | -0.08441 | -2.54158 | -1.49474 | -0.84202 | -0.88397 | -1.38059 | -1.18794 | -2.41886 | -1.63174 | -1.65148 | -1.53624 | -1.01642 | -1.93335 | -0.28102 | -2.31811 | -2.18067 |
| 34.99861 | -1.85839 | -1.67904 | -1.28458 | -0.45607 | 0.106595 | -2.17419 | -1.14919 | -0.41715 | -0.46966 | -0.25971 | 0.458706 | -1.1303 | -0.00801 | 0.280067 | -0.04934 | 0.397596 | -0.33592 | 0.487985 | -0.72569 | -0.94194 |
| 35.0125 | -2.39427 | -1.45476 | -0.9891 | -0.37054 | -0.12271 | -1.93315 | -0.76682 | -0.21938 | -0.32517 | 0.488556 | 1.15156 | 0.484669 | 0.828909 | 1.375625 | 0.70148 | 1.020545 | 0.607964 | 0.826314 | -0.02121 | -0.229 |
| 35.02639 | -2.29761 | -0.47574 | -0.58987 | -0.42566 | -0.356 | -1.6904 | -0.39641 | -0.09339 | -0.23696 | 0.844008 | 1.11799 | 1.976735 | 1.251768 | 1.532118 | 1.118102 | 1.136779 | 0.931565 | 0.960075 | 0.295405 | 0.495862 |
| 35.04028 | -1.75301 | 0.125835 | -0.27719 | -0.47288 | -0.43057 | -1.30207 | -0.02573 | 0.047181 | -0.11614 | 0.424354 | 0.9419 | 3.037044 | 1.336265 | 1.475065 | 1.47829 | 1.106206 | 0.793442 | 1.061379 | 0.480775 | 1.238795 |
| 35.05417 | -1.3337 | 0.140803 | -0.02856 | -0.51518 | -0.37522 | -0.91547 | 0.200794 | 0.151938 | 0.09673 | 0.415979 | 0.655388 | 3.047209 | 1.179816 | 1.5429 | 1.318685 | 1.054891 | 0.932912 | 0.857065 | 0.358952 | 1.92187 |
| 35.06806 | -1.24402 | -0.05717 | 0.087377 | -0.51321 | -0.27063 | -0.76276 | 0.400089 | 0.166375 | 0.32912 | 0.462821 | 0.418691 | 2.634311 | 0.921333 | 1.066035 | 1.23013 | 0.958036 | 1.095423 | 0.645323 | 0.265011 | 1.800926 |
| 35.08194 | -0.70477 | -0.00056 | 0.254016 | -0.31583 | -0.04855 | -0.49084 | 0.721115 | 0.173198 | 0.408849 | 0.395667 | 0.325486 | 1.954628 | 1.02465 | 0.712972 | 1.238305 | 0.77769 | 0.733924 | 0.377004 | 0.582043 | 1.044488 |
| 35.09583 | 0.162638 | 0.347742 | 0.527486 | -0.03921 | 0.128974 | -0.2138 | 1.202765 | 0.187531 | 0.3679 | 0.257189 | 0.303942 | 1.390009 | 1.359519 | 0.57879 | 1.404992 | 0.619712 | 0.357311 | -0.0863 | 0.556867 | 0.271071 |
| 35.10972 | 0.742356 | 0.598293 | 0.908076 | 0.137186 | 0.183824 | -0.07584 | 1.888048 | 0.129061 | 0.28007 | 0.345547 | 0.257285 | 0.913502 | 1.488558 | 0.230929 | 1.066197 | 0.509404 | 0.695605 | -0.27836 | 0.117429 | -0.0302 |
| 35.12361 | 0.958653 | 0.72512 | 0.413897 | 0.397174 | 0.212786 | -0.06955 | 1.67333 | 0.034871 | 0.205016 | 0.262731 | 0.195564 | 0.007619 | 1.483263 | 0.000794 | 0.341186 | 0.520278 | 0.798758 | -0.33422 | -0.32633 | 0.292428 |
| 35.1375 | 1.128773 | 0.741229 | 0.360297 | 0.665274 | 0.2417 | -0.06724 | 1.662552 | 0.013851 | 0.228925 | 0.176802 | 0.15215 | -0.84186 | 1.60263 | -0.2331 | -0.46878 | 0.401966 | 0.742241 | -0.09397 | -0.79009 | 0.513401 |
| 35.15139 | 1.143487 | 0.590866 | 0.547225 | 0.602706 | 0.218068 | 0.106309 | 2.115543 | 0.213934 | 0.251108 | 0.451794 | 0.236133 | -1.10504 | 1.7005 | -0.49502 | -0.9082 | 0.054447 | 0.831617 | 0.745164 | -0.82493 | 1.054283 |
| 35.16528 | 1.173731 | 0.669466 | 0.528056 | 0.506811 | 0.226565 | 0.174229 | 1.815046 | 0.370015 | 0.292815 | 0.551768 | 0.359253 | -1.24202 | 1.573047 | -0.58575 | -0.98583 | -0.17687 | 0.907929 | 1.303482 | -0.52205 | 1.529912 |
| 35.17917 | 1.350099 | 0.961526 | 0.33593 | 0.850828 | 0.327667 | 0.055229 | 1.000147 | 0.29199 | 0.294942 | 0.460579 | 0.37129 | -1.22346 | 1.252108 | -0.50776 | -0.99024 | -0.02616 | 0.831471 | 1.222104 | -0.49104 | 1.756443 |
| 35.19306 | 1.502511 | 1.172427 | 0.035262 | 0.973304 | 0.429585 | 0.246172 | -0.12611 | 0.238624 | 0.256378 | 0.28399 | 0.26789 | -0.75962 | 0.87312 | -0.38782 | -0.71755 | 0.164123 | 0.719027 | 0.793341 | -0.55009 | 1.458897 |
| 35.20694 | 1.539629 | 1.126876 | 0.087983 | 0.814313 | 0.481193 | 0.426017 | -0.89368 | 0.225766 | 0.229132 | 0.078084 | 0.293602 | -0.20364 | 0.337602 | -0.33656 | -0.47937 | 0.211508 | 0.29144 | 0.317841 | -0.5192 | 0.790773 |
| 35.22083 | 1.556502 | 1.018225 | -0.05915 | 0.370632 | 0.37665 | 0.677319 | -0.9025 | 0.182985 | 0.241685 | -0.06421 | 0.543431 | 0.206712 | -0.23508 | -0.46265 | -0.14793 | 0.097149 | -0.12229 | -0.16121 | -0.52168 | 0.11435 |
| 35.23472 | 1.681841 | 1.063588 | -0.09703 | 0.202829 | 0.30688 | 0.476932 | -0.78147 | 0.199098 | 0.325822 | -0.13673 | 0.871097 | 0.371942 | -0.59951 | -0.39201 | 0.222154 | 0.057569 | -0.36304 | -0.44906 | -0.29236 | -0.07494 |
| 35.24861 | 1.889243 | 1.080422 | -0.02496 | 0.298797 | 0.373775 | 0.358836 | -0.50135 | 0.178396 | 0.332513 | -0.05609 | 0.906568 | 0.209367 | -0.80365 | -0.06419 | 0.447953 | 0.14757 | -0.27404 | -0.4417 | 0.061072 | -0.19986 |
| 35.2625 | 1.639007 | 0.808313 | -0.05566 | 0.118086 | 0.423776 | 0.207227 | -0.36023 | 0.170566 | 0.199728 | 0.128021 | 0.664171 | -0.15581 | -0.73858 | -0.00143 | 0.392366 | 0.105074 | -0.13698 | -0.046 | 0.258776 | -0.06665 |
| 35.27639 | 1.195197 | 0.273854 | -0.07436 | -0.27477 | 0.368555 | 0.151856 | -0.56037 | 0.20739 | 0.011897 | 0.430517 | 0.279425 | -0.41995 | -0.36322 | -0.03458 | 0.227969 | 0.00727 | -0.18594 | 0.52114 | 0.362624 | 0.370538 |
| 35.29028 | 0.791502 | -0.10133 | -0.09052 | -0.43652 | 0.223239 | 0.173154 | -0.69763 | 0.202623 | -0.03143 | 0.584219 | -0.01714 | -0.48407 | 0.135743 | 0.138278 | 0.075929 | 0.137666 | -0.29686 | 0.649097 | 0.323182 | 1.222086 |
| 35.30417 | 0.505867 | -0.36859 | -0.04333 | -0.343 | 0.10379 | 0.241432 | -0.78786 | 0.119104 | 0.057114 | 0.455305 | -0.09906 | -0.30044 | 0.467151 | 0.403006 | -0.01423 | 0.372811 | -0.16825 | 0.163978 | 0.224169 | 0.969682 |
| 35.31806 | 0.308688 | -0.57623 | 0.139002 | -0.13437 | 0.070306 | 0.401885 | -0.64715 | 0.114804 | 0.145902 | 0.422862 | 0.152271 | 0.245266 | 0.750617 | 0.613794 | 0.073133 | 0.477801 | -0.11835 | 0.049961 | 0.181094 | 0.383825 |
| 35.33194 | 0.208316 | -0.68842 | 0.342097 | 0.006039 | 0.067927 | 0.494901 | -0.29643 | 0.155842 | 0.218391 | 0.290717 | 0.653701 | 1.089213 | 0.94132 | 0.744884 | 0.286913 | 0.458426 | -0.13739 | 0.174776 | 0.106683 | -0.11459 |
| 35.34583 | 0.204789 | -0.67927 | 0.598214 | 0.18222 | 0.13228 | 0.561358 | 0.277202 | 0.191826 | 0.318363 | 0.213975 | 0.854442 | 1.519701 | 0.993133 | 0.924624 | 0.563152 | 0.447629 | 0.033697 | 0.300525 | 0.041917 | 0.009289 |
| 35.35972 | 0.272352 | -0.53966 | 0.939295 | 0.253187 | 0.354091 | 0.678668 | 1.03314 | 0.250096 | 0.38897 | 0.376453 | 0.848456 | 1.6785 | 1.075869 | 1.215094 | 0.847999 | 0.462582 | 0.371783 | 0.525604 | 0.077745 | 1.000867 |
| 35.37361 | 0.332211 | -0.32646 | 1.095494 | 0.167229 | 0.564436 | 0.639482 | 1.722327 | 0.27615 | 0.398003 | 0.545796 | 0.509953 | 1.593714 | 0.95349 | 1.250107 | 0.916149 | 0.406608 | 0.691913 | 0.915424 | 0.067245 | 1.979143 |
| 35.3875 | 0.290565 | -0.09005 | 0.95742 | -0.09981 | 0.67676 | 0.290884 | 2.103939 | 0.239575 | 0.39637 | 0.475859 | 0.147914 | 0.951934 | 0.501999 | 0.814285 | 0.708903 | 0.378636 | 0.768532 | 0.765093 | -0.04471 | 1.952995 |
| 35.40139 | 0.248009 | 0.104916 | 0.635031 | -0.14089 | 0.774869 | -0.0154 | 2.418681 | 0.221401 | 0.397633 | 0.267479 | -0.07284 | 0.132337 | 0.135217 | 0.333088 | 0.272894 | 0.379855 | 0.661386 | 0.163475 | -0.05125 | 0.988407 |
| 35.41528 | 0.443926 | 0.427061 | 0.019477 | 0.29044 | 0.907346 | -0.16897 | 2.523453 | 0.087554 | 0.519917 | -0.12052 | 0.069289 | -0.48382 | -0.15286 | 0.005348 | -0.17967 | 0.678041 | 0.156978 | -0.47702 | -0.08035 | 0.132504 |
| 35.42917 | 1.029812 | 0.998235 | 0.076902 | 1.439038 | 1.199664 | -0.01217 | 2.778526 | -0.18711 | 0.941022 | -0.28571 | 0.687005 | -0.90633 | -0.10552 | 0.133695 | -0.1509 | 1.57627 | 0.108069 | -1.36232 | -0.17581 | -0.69834 |
| 35.44306 | 1.946935 | 1.741353 | 0.451843 | 2.60282 | 1.542033 | 1.139012 | 2.670721 | 0.305446 | 1.430084 | 0.590565 | 1.633048 | -0.24616 | 1.181419 | 0.794757 | 0.355004 | 2.413348 | 0.977107 | -1.71909 | 0.074219 | -0.26264 |
| 35.45694 | 2.651092 | 2.11299 | 1.054532 | 2.568185 | 1.894748 | 2.884249 | 2.88768 | 1.04086 | 1.701204 | 1.700469 | 2.152106 | 1.065149 | 2.934667 | 1.766597 | 1.187028 | 2.802216 | 1.577619 | -0.15014 | 1.106638 | 1.87275 |
| 35.47083 | 2.624644 | 1.812879 | 1.356553 | 2.185354 | 1.76168 | 3.244934 | 2.880066 | 1.216919 | 1.40739 | 2.452761 | 1.750321 | 2.39724 | 4.114483 | 2.483575 | 2.197346 | 2.982991 | 1.651202 | 1.416756 | 1.517465 | 3.210872 |
| 35.48472 | 2.360428 | 1.352983 | 1.379119 | 1.560112 | 1.330948 | 3.009485 | 2.64179 | 1.119285 | 1.201918 | 2.616785 | 1.301273 | 2.675657 | 4.979959 | 2.315872 | 2.582225 | 2.990326 | 1.579832 | 1.824806 | 1.606476 | 2.677499 |
| 35.49861 | 2.266927 | 1.069293 | 1.332157 | 0.851095 | 1.075634 | 2.893625 | 3.674026 | 1.090959 | 1.118186 | 2.253361 | 1.130546 | 2.157591 | 5.152268 | 1.791903 | 2.562566 | 2.784663 | 1.790007 | 2.02324 | 1.453345 | 2.451825 |
| 35.5125 | 2.042562 | 0.851744 | 1.081566 | 0.386386 | 0.897573 | 2.829775 | 2.346704 | 1.05562 | 1.16366 | 1.588063 | 0.991839 | 1.647014 | 5.05577 | 1.238067 | 2.183574 | 2.61949 | 0.957845 | 2.220773 | 1.247544 | 1.968306 |
| 35.52639 | 1.519523 | 0.673866 | 0.705281 | 0.157612 | 0.885912 | 2.872333 | 0.50698 | 0.923482 | 1.193878 | 0.921247 | 0.690225 | 1.053207 | 4.720921 | 0.483327 | 1.484228 | 2.441602 | 0.130702 | 1.861198 | 1.097592 | 1.618944 |
| 35.54028 | 1.067684 | 0.616981 | 0.49261 | 0.158811 | 0.653085 | 2.695529 | -0.88498 | 0.73634 | 1.111148 | 0.600817 | 0.193439 | 0.406183 | 3.716507 | -0.24213 | 0.322538 | 2.184551 | -0.49617 | 0.770516 | 0.878583 | 0.95924 |
| 35.55417 | 0.801716 | 0.827484 | 0.299236 | 0.035181 | 0.30059 | 2.358085 | -1.95901 | 0.616303 | 0.926026 | 0.345728 | -0.14685 | -0.1415 | 2.473928 | -0.36327 | -0.67627 | 1.718579 | -0.48833 | -0.53899 | 0.533147 | 0.392883 |
| 35.56806 | 0.620816 | 0.947127 | -0.00963 | -0.00378 | 0.194552 | 2.263823 | -2.41305 | 0.541665 | 0.707714 | -0.14991 | -0.43225 | -0.2512 | 0.872792 | -0.8074 | -1.14009 | 1.00458 | -0.04251 | -1.27486 | 0.311136 | -0.03797 |
| 35.58194 | 0.441923 | 0.390899 | -0.26222 | -0.0294 | 0.244039 | 1.954103 | -2.77514 | 0.494385 | 0.571128 | -0.31448 | -0.23445 | -0.37779 | -0.98288 | -0.99937 | -1.02679 | 0.326737 | -0.31658 | -1.58207 | 0.2166 | -0.33219 |
| 35.59583 | 0.128833 | -0.29817 | -0.41962 | -0.129 | 0.287086 | 1.405056 | -2.93839 | 0.431885 | 0.506336 | -0.26799 | 0.181934 | -0.76126 | -2.88269 | -0.67211 | -0.97233 | -0.36281 | -0.30903 | -1.56335 | 0.128061 | -0.7499 |
| 35.60972 | -0.0398 | -0.59503 | -0.33831 | -0.20295 | 0.381533 | 0.732849 | -3.13377 | 0.381152 | 0.439323 | -0.34112 | 0.10832 | -1.25715 | -4.14087 | -0.33281 | -0.86198 | -0.98903 | -0.35562 | -1.07265 | 0.032261 | -0.32789 |
| 35.62361 | -0.13023 | -0.56637 | -0.42858 | -0.22227 | 0.487952 | 0.060826 | -2.89136 | 0.420304 | 0.350885 | -0.52474 | -0.23549 | -1.13577 | -4.60022 | -0.44739 | -0.4097 | -1.53088 | -0.45838 | -0.75727 | -0.0538 | -0.05686 |
| 35.6375 | -0.4327 | -0.54568 | -0.36445 | -0.25997 | 0.486025 | -0.22564 | -2.70377 | 0.419295 | 0.294672 | -0.54671 | -0.4727 | -0.06102 | -4.03786 | -0.78134 | 0.452544 | -1.25739 | -0.71859 | -0.35231 | 0.090937 | 0.024337 |
| 35.65139 | -0.70701 | -0.21441 | -0.41618 | -0.31583 | 0.467422 | -0.22956 | -2.58623 | 0.327908 | 0.24042 | -0.55847 | -0.75343 | 0.763112 | -2.98613 | -1.01545 | 1.038768 | -1.06124 | -0.82187 | -0.03519 | 0.300744 | -0.8524 |
| 35.66528 | -0.88001 | 0.070036 | -0.27713 | -0.32128 | 0.495936 | -0.13089 | -1.93916 | 0.203599 | 0.254947 | -0.56113 | -1.1175 | 1.201832 | -1.36186 | -1.39985 | 1.162749 | -1.05999 | -0.47917 | -0.07657 | 0.409472 | -1.06388 |
| 35.67917 | -1.06991 | 0.16678 | -0.12761 | -0.21499 | 0.634386 | 0.183026 | -0.62064 | 0.117734 | 0.38109 | -0.56731 | -0.94186 | 1.250903 | 0.000396 | -1.64822 | 0.977205 | -1.15765 | 0.024987 | -0.05379 | 0.633735 | -0.15521 |
| 35.69306 | -1.68336 | 0.091951 | 0.172015 | -0.15265 | 0.76989 | 0.4265 | 0.972513 | 0.154038 | 0.56339 | -0.55384 | -0.38927 | 0.989437 | 1.0685 | -1.82546 | 0.616278 | -0.76474 | 0.567626 | -0.179 | 0.687284 | 0.260701 |
| 35.70694 | -2.47987 | -0.11024 | 0.281982 | -0.1783 | 0.83971 | 0.491492 | 2.363097 | 0.247914 | 0.715343 | -0.52662 | 0.120346 | 0.508291 | 1.6019 | -1.46874 | 0.20218 | -0.23943 | 0.659425 | -0.31773 | 0.829639 | 0.461596 |
| 35.72083 | -2.42868 | -0.5002 | 0.316738 | -0.1257 | 0.817717 | 0.565549 | 2.80986 | 0.335959 | 0.844607 | -0.5116 | 0.327049 | -0.04988 | 1.519898 | -0.82239 | -0.17059 | 0.16134 | 0.549378 | -0.63677 | 0.82092 | 0.640775 |
| 35.73472 | -1.97686 | -0.78048 | 0.355214 | -0.00221 | 0.757167 | 0.622135 | 2.353188 | 0.336816 | 0.789004 | -0.60874 | 0.143086 | -0.45685 | 0.945935 | -0.35849 | -0.40912 | 0.291754 | 0.277943 | -1.03643 | 0.473265 | 0.568647 |
| 35.74861 | -1.11034 | -0.58721 | 0.356755 | 0.080838 | 0.657314 | 0.578866 | 1.36038 | 0.295014 | 0.617771 | -0.77061 | -0.42081 | -0.74376 | 0.023706 | -0.08581 | -0.45225 | 0.141584 | 0.080037 | -1.24529 | 0.081954 | -0.0795 |
| 35.7625 | -0.52121 | -0.3584 | 0.296632 | 0.156906 | 0.561944 | 0.348396 | 0.394244 | 0.243431 | 0.341316 | -0.87638 | -1.13657 | -0.99738 | -1.02423 | 0.243202 | -0.45905 | -0.24205 | -0.22758 | -1.219 | -0.02401 | -0.37537 |
| 35.77639 | -0.4341 | -0.34259 | 0.202718 | 0.193376 | 0.56437 | -0.00971 | -0.40954 | 0.130835 | 0.0013 | -0.95814 | -1.46368 | -1.23087 | -2.23181 | 0.411227 | -0.51706 | -0.74913 | -0.5558 | -1.02182 | -0.11844 | -0.9381 |
| 35.79028 | -0.55071 | -0.43063 | 0.047536 | 0.088201 | 0.564185 | -0.51638 | -1.00872 | -0.02674 | -0.40206 | -0.8246 | -1.35401 | -1.27374 | -3.17457 | 0.03851 | -0.56246 | -1.24893 | -1.03253 | -0.71291 | -0.28197 | -1.57175 |
| 35.80417 | -0.62339 | -0.49641 | 0.004208 | -0.18964 | 0.469752 | -1.11958 | -1.22977 | -0.19473 | -0.82917 | -0.88253 | -1.10945 | -0.95125 | -3.35845 | -0.58639 | -0.7095 | -1.59139 | -1.4679 | -0.43318 | -0.30608 | -2.01516 |
| 35.81806 | -0.60441 | -0.59075 | -0.0831 | -0.59041 | 0.331185 | -1.42393 | -1.14988 | -0.32365 | -1.08012 | -0.97254 | -0.63649 | -0.91934 | -3.04672 | -1.15692 | -0.85568 | -1.72818 | -1.49313 | -0.3983 | -0.56619 | -1.57578 |
| 35.83194 | -0.65042 | -0.71128 | -0.2522 | -1.0177 | 0.218943 | -1.96203 | -0.9757 | -0.35148 | -1.09806 | -0.79732 | -0.15717 | -1.01712 | -2.71524 | -1.35457 | -0.74355 | -1.78021 | -1.82555 | -0.40582 | -0.5937 | -0.53656 |
| 35.84583 | -0.73318 | -0.70315 | -0.33257 | -1.43907 | -0.07729 | -2.34063 | -0.83582 | -0.33636 | -1.09523 | -0.59824 | 0.055694 | -1.21418 | -2.08702 | -1.12767 | -0.40975 | -1.43991 | -2.00001 | -0.51727 | -0.45528 | -1.18374 |
| 35.85972 | -0.81844 | -0.66983 | -0.63556 | -1.61944 | -0.75605 | -2.60169 | -0.78474 | -0.41932 | -1.21741 | -0.4407 | 0.033224 | -1.2063 | -1.56261 | -0.65998 | -0.15696 | -1.36063 | -2.04952 | -0.80426 | -0.4075 | -1.83289 |
| 35.87361 | -0.89299 | -0.66044 | -1.06128 | -1.76247 | -1.04551 | -2.87742 | -0.63221 | -0.56519 | -1.24489 | -0.29669 | -0.14195 | -1.04988 | -1.27938 | 0.129246 | -0.13157 | -1.16256 | -1.92398 | -0.73168 | -0.26118 | -2.16017 |
| 35.8875 | -0.97038 | -0.64373 | -0.96232 | -2.14269 | -0.73501 | -2.66092 | -0.48906 | -0.62974 | -1.23337 | -0.07601 | -0.43124 | -0.74066 | -0.98873 | 0.891191 | -0.16356 | -0.7521 | -1.50906 | -0.6418 | -0.12274 | -1.56909 |
| 35.90139 | -1.01837 | -0.71032 | -0.69865 | -1.95926 | -0.20639 | -2.33802 | -0.18055 | -0.52188 | -0.91637 | 0.200857 | -0.71146 | -0.33554 | -0.66856 | 1.245699 | -0.13475 | -0.6227 | -1.04403 | -0.1946 | -0.1963 | -0.79543 |
| 35.91528 | -1.21158 | -1.22524 | -0.35867 | -2.34 | -0.298 | -1.8229 | 0.085749 | -0.40594 | -0.44164 | 0.70115 | -0.53682 | 0.284008 | -0.41215 | 1.244811 | -0.18081 | -0.85221 | -0.38487 | 0.890978 | 0.034349 | 0.428037 |
| 35.92917 | -1.75287 | -2.04314 | -0.21043 | -3.20849 | -0.74205 | -1.76239 | -0.07859 | -0.40346 | -0.36472 | 0.937471 | -0.22787 | 0.768757 | -0.64629 | 0.799312 | -0.62045 | -1.82713 | -0.43225 | 1.72275 | 0.359343 | 1.290504 |

| | | | | | | | | | | | | | | | | | | | | |
|---|---|---|---|---|---|---|---|---|---|---|---|---|---|---|---|---|---|---|---|---|
| 35.94306 | -2.60201 | -2.64423 | -0.68237 | -3.64946 | -0.93915 | -2.84611 | -0.82726 | -0.88884 | -0.96717 | -0.32893 | -1.11314 | 0.188802 | -1.99553 | -0.66774 | -1.63616 | -3.18093 | -1.27617 | 0.800229 | -0.5023 | 0.38157 |
| 35.95694 | -3.18313 | -2.72231 | -1.60219 | -2.93554 | -0.68525 | -4.37067 | -1.75881 | -1.43204 | -2.22467 | -2.15685 | -2.69917 | -1.56911 | -3.87941 | -2.70863 | -2.99763 | -4.16156 | -1.93972 | -1.39444 | -2.42443 | -1.57518 |
| 35.97083 | -3.19794 | -2.42904 | -2.07122 | -1.50484 | -0.07138 | -4.03014 | -2.10052 | -1.66011 | -2.47361 | -2.99402 | -2.61606 | -3.07735 | -4.03605 | -3.55003 | -3.26331 | -3.06643 | -1.79816 | -1.94005 | -2.83482 | -1.86877 |
| 35.98472 | -2.44623 | -1.85546 | -1.60214 | -0.93294 | 0.587372 | -2.57099 | -1.66268 | -1.31764 | -1.61358 | -1.55956 | -0.85678 | -2.86793 | -2.01634 | -2.10961 | -1.98038 | -1.31976 | -1.0086 | -0.55485 | -1.52023 | -0.63962 |
| 35.99861 | -2.14351 | -1.5712 | -1.66895 | -0.66147 | 0.868481 | -2.02432 | -1.34447 | -1.0165 | -1.23869 | -0.15198 | 0.551645 | -1.527 | -0.47807 | -0.27769 | -0.78599 | -0.32053 | 0.060877 | 0.800085 | -0.65153 | 0.331294 |
| 36.0125 | -1.90481 | -1.26236 | -1.01599 | -0.51107 | 0.647001 | -1.73256 | -0.91207 | -0.74851 | -1.03488 | 0.467386 | 1.127176 | -0.12349 | 0.226518 | 1.020682 | -0.05601 | 0.338749 | 0.602912 | 1.458979 | -0.41229 | 0.71483 |
| 36.02639 | -1.58501 | -0.79664 | 0.020329 | -0.35257 | 0.355779 | -1.193 | -0.24312 | -0.41269 | -0.77057 | 0.822523 | 1.19143 | 1.142996 | 0.799821 | 1.664757 | 0.586175 | 0.982085 | 0.821765 | 1.715946 | -0.15123 | 0.567946 |
| 36.04028 | -1.23655 | -0.31162 | 0.617439 | -0.25915 | 0.105122 | -0.76066 | 0.323377 | -0.13647 | -0.53734 | 1.18743 | 1.092236 | 2.296592 | 1.07195 | 2.17068 | 1.409149 | 1.506991 | 0.66998 | 1.750078 | 0.090643 | 0.450398 |
| 36.05417 | -0.89732 | -0.0635 | 0.84764 | -0.20055 | 0.087624 | -0.48371 | 0.684559 | 0.036934 | -0.35437 | 1.16933 | 1.117722 | 3.049774 | 1.184995 | 2.249027 | 1.911284 | 1.456785 | 0.478687 | 1.859063 | 0.359453 | 0.681435 |
| 36.06806 | -0.56946 | 0.063869 | 0.944953 | -0.15686 | 0.280485 | -0.29492 | 1.10859 | 0.124667 | -0.12927 | 1.186869 | 1.216101 | 3.19794 | 1.251214 | 2.205003 | 1.939661 | 0.945036 | 0.518468 | 1.978393 | 0.5214 | 0.907589 |
| 36.08194 | -0.23815 | 0.248641 | 0.772258 | -0.19057 | 0.472795 | -0.12328 | 1.257206 | 0.208182 | 0.137321 | 0.77792 | 0.722051 | 3.182888 | 1.236369 | 2.224839 | 1.387701 | 0.465245 | 0.464985 | 1.671093 | 0.421302 | 1.198804 |
| 36.09583 | 0.12374 | 0.666155 | 0.629154 | -0.06585 | 0.571048 | 0.073741 | 1.493831 | 0.317229 | 0.418453 | 0.054364 | 0.233412 | 3.202668 | 1.181797 | 2.53596 | 1.143362 | 0.255211 | 0.428398 | 0.730337 | 0.099031 | 0.832905 |
| 36.10972 | 0.485578 | 0.940202 | 0.437586 | 0.119319 | 0.472911 | 0.18022 | 1.578235 | 0.45808 | 0.616466 | -0.14816 | -0.25648 | 2.935823 | 1.262907 | 2.464877 | 1.002929 | 0.066951 | 0.45333 | -0.47321 | -0.07071 | 0.388951 |
| 36.12361 | 0.74273 | 0.799932 | 0.469304 | 0.242465 | 0.295975 | 0.145828 | 1.594381 | 0.589713 | 0.646819 | -0.03621 | -0.66385 | 2.327463 | 1.421929 | 1.867159 | 1.214508 | 0.004088 | 0.534739 | -1.16009 | -0.07583 | 0.320355 |
| 36.1375 | 0.892995 | 0.674691 | 0.437597 | 0.221378 | 0.139587 | -0.00212 | 1.400136 | 0.679304 | 0.52866 | 0.016812 | -0.70868 | 1.412345 | 1.517486 | 0.845532 | 1.075321 | 0.341167 | 0.454923 | -1.4577 | -0.09105 | 0.620293 |
| 36.15139 | 1.009962 | 0.654269 | 0.403543 | 0.133648 | 0.129011 | -0.0687 | 0.959084 | 0.630676 | 0.320939 | 0.037633 | -0.53323 | 0.652784 | 1.591596 | -0.14547 | 0.525432 | 0.71705 | 0.524726 | -1.46557 | -0.13251 | 0.300268 |
| 36.16528 | 1.150981 | 0.612941 | 0.312876 | 0.07906 | 0.147022 | 0.00404 | 0.576507 | 0.589917 | 0.206028 | 0.175462 | -0.06798 | 0.13271 | 1.57308 | -1.02215 | -0.07141 | 0.789721 | 0.329698 | -0.9707 | -0.034 | 0.295666 |
| 36.17917 | 1.23422 | 0.456459 | 0.207271 | -0.00022 | 0.18468 | 0.139006 | 0.215272 | 0.601901 | 0.180022 | 0.528102 | 0.279828 | -0.2032 | 1.697185 | -1.55334 | -0.58246 | 0.458221 | 0.275554 | -0.40916 | 0.152262 | 0.724306 |
| 36.19306 | 1.204624 | 0.343335 | 0.096996 | -0.04548 | 0.236902 | 0.189336 | 0.144335 | 0.59051 | 0.070013 | 0.695875 | 0.182439 | -0.51976 | 1.654503 | -1.67317 | -0.90158 | 0.029531 | 0.124228 | 0.109403 | 0.099673 | 1.179515 |
| 36.20694 | 1.250783 | 0.470427 | 0.117833 | 0.105998 | 0.35486 | 0.161034 | 0.330804 | 0.505197 | -0.04447 | 0.655987 | -0.27285 | -0.84162 | 1.241822 | -1.56395 | -0.95689 | -0.23902 | 0.048134 | 0.567559 | -0.07499 | 1.32941 |
| 36.22083 | 1.567662 | 0.806216 | 0.275449 | 0.41169 | 0.517746 | 0.145516 | 0.597452 | 0.365484 | -0.07553 | 0.440445 | -0.55181 | -0.9741 | 0.737401 | -1.77423 | -0.85013 | -0.132 | 0.216418 | 0.74634 | -0.24813 | 0.986415 |
| 36.23472 | 1.50832 | 0.921385 | 0.327825 | 0.519113 | 0.496061 | 0.056693 | 0.806924 | 0.291016 | -0.14227 | 0.15441 | -0.30247 | -0.66653 | 0.103036 | -1.22925 | -0.60193 | 0.144935 | 0.343646 | 0.580071 | -0.27864 | 0.583536 |
| 36.24861 | 1.105974 | 0.79348 | 0.246436 | 0.402519 | 0.319398 | -0.11164 | 0.8372 | 0.285037 | -0.26819 | 0.205864 | 0.211755 | -0.08993 | -0.60487 | -0.13851 | -0.32238 | 0.45817 | 0.218488 | 0.353226 | 0.050488 | 0.21488 |
| 36.2625 | 0.906637 | 0.663645 | 0.197227 | 0.394492 | 0.276073 | -0.20409 | 0.670905 | 0.206097 | -0.32377 | 0.291396 | 0.504141 | 0.494085 | -1.16868 | 0.594333 | 0.036156 | 0.733494 | 0.093106 | 0.185828 | 0.298463 | -0.15355 |
| 36.27639 | 0.765579 | 0.469214 | 0.052525 | 0.408552 | 0.299355 | -0.23428 | 0.205908 | 0.117636 | -0.37561 | 0.326875 | 0.48837 | 0.910916 | -1.25128 | 0.593664 | 0.298448 | 0.739763 | 0.027374 | -0.06499 | 0.316879 | -0.15293 |
| 36.29028 | 0.489042 | 0.112659 | -0.15908 | 0.302302 | 0.255747 | -0.23407 | -0.38765 | 0.104992 | -0.37602 | 0.480215 | 0.55218 | 1.033222 | -0.88884 | 0.221296 | 0.25586 | 0.545906 | 0.111766 | -0.33761 | 0.282145 | 0.005642 |
| 36.30417 | 0.268275 | -0.18476 | -0.2528 | 0.191591 | 0.231702 | -0.10903 | -0.82497 | 0.153153 | -0.23755 | 0.717239 | 0.716475 | 0.92434 | -0.22752 | -0.1189 | 0.14587 | 0.456846 | 0.170914 | -0.39788 | 0.241965 | 0.526188 |
| 36.31806 | 0.039822 | -0.3666 | -0.31437 | 0.159781 | 0.241478 | 0.034249 | -1.05861 | 0.173245 | -0.15778 | 0.943448 | 0.733498 | 0.755288 | 0.627771 | -0.38036 | 0.112632 | 0.470714 | 0.21524 | -0.00147 | 0.250822 | 1.009813 |
| 36.33194 | -0.11367 | -0.43631 | -0.27352 | 0.068066 | 0.230993 | 0.136409 | -1.08337 | 0.194879 | -0.25897 | 1.055272 | 0.533094 | 0.705185 | 1.605093 | -0.44542 | 0.173999 | 0.436089 | 0.384042 | 0.763684 | 0.376887 | 1.486662 |
| 36.34583 | -0.15107 | -0.2855 | -0.20955 | 0.12661 | 0.292247 | 0.247361 | -1.00455 | 0.197772 | -0.38416 | 0.961322 | 0.379751 | 0.697739 | 2.159115 | 0.012844 | 0.352766 | 0.466679 | 0.836081 | 1.481985 | 0.374671 | 1.849123 |
| 36.35972 | -0.05563 | 0.016854 | 0.05082 | 0.388963 | 0.429221 | 0.342031 | -0.85968 | 0.148313 | -0.40828 | 0.617123 | 0.3836 | 0.710228 | 1.982206 | 0.866437 | 0.6188 | 0.71432 | 1.390545 | 1.646523 | 0.37986 | 1.365266 |
| 36.37361 | 0.112727 | 0.263204 | 0.294931 | 0.614722 | 0.522737 | 0.2749 | -0.47668 | 0.147731 | -0.33539 | 0.249555 | 0.540482 | 0.843481 | 1.568438 | 1.638762 | 0.859022 | 0.863769 | 1.741157 | 1.200274 | 0.509442 | 0.811678 |
| 36.3875 | 0.169189 | 0.259974 | 0.243027 | 0.57982 | 0.496459 | 0.133662 | -0.17516 | 0.222591 | -0.22051 | 0.092278 | 0.655692 | 1.053147 | 1.104368 | 1.860819 | 0.794174 | 0.707285 | 1.513256 | 0.457519 | 0.685765 | 0.657355 |
| 36.40139 | 0.142722 | 0.141365 | 0.035195 | 0.405065 | 0.445561 | 0.005734 | -0.08998 | 0.259424 | -0.1423 | -0.01819 | 0.549017 | 1.056039 | 0.611869 | 1.526642 | 0.5552 | 0.414569 | 0.883819 | -0.43084 | 0.577894 | 0.783915 |
| 36.41528 | 0.358426 | 0.510664 | 0.188315 | 0.467504 | 0.626354 | -0.00677 | -0.18729 | 0.178371 | 0.072501 | -0.05744 | 0.290392 | 0.780404 | 0.055739 | 0.811727 | 0.526064 | 0.550941 | 0.47444 | -1.0976 | 0.059654 | 0.542561 |
| 36.42917 | 1.116619 | 1.484532 | 0.862669 | 1.122051 | 1.041301 | 0.323348 | -0.19466 | -0.0328 | 0.555022 | 0.185178 | 0.13779 | 0.437106 | -0.18737 | 0.19697 | 0.829562 | 1.560644 | 0.741593 | -1.05936 | -0.54671 | 0.414515 |
| 36.44306 | 2.255692 | 2.293212 | 1.650147 | 1.799007 | 1.411634 | 1.04614 | 0.699941 | 0.10586 | 1.364969 | 0.993055 | 0.418314 | 0.667159 | 0.959081 | 0.260221 | 1.264684 | 2.590473 | 1.291787 | 0.144486 | -0.15981 | 1.728231 |
| 36.45694 | 2.833689 | 2.531436 | 2.294928 | 2.194343 | 1.358641 | 1.639394 | 2.011457 | 0.524615 | 1.973215 | 2.239175 | 1.273239 | 1.41999 | 2.651266 | 1.146004 | 1.620534 | 2.544869 | 2.164974 | 2.045353 | 0.997382 | 2.733899 |
| 36.47083 | 2.545901 | 2.379442 | 2.172295 | 1.766286 | 0.940885 | 1.304144 | 2.62997 | 0.665772 | 1.762382 | 2.49941 | 1.808548 | 2.202725 | 3.447403 | 2.488833 | 1.986326 | 3.151782 | 2.623891 | 3.139592 | 1.514081 | 3.338956 |
| 36.48472 | 1.889587 | 1.908227 | 2.052755 | 1.135466 | 0.465238 | 0.46434 | 2.944386 | 0.617968 | 0.997214 | 2.070612 | 1.630528 | 2.843772 | 3.943684 | 3.188358 | 2.324885 | 3.133034 | 2.361112 | 3.465212 | 1.383708 | 2.425287 |
| 36.49861 | 1.637503 | 1.505926 | 2.208227 | 0.700295 | 0.201248 | -0.03057 | 3.196515 | 0.529369 | 0.523633 | 1.536253 | 1.212148 | 3.233069 | 4.211238 | 2.991673 | 2.393163 | 2.87882 | 2.654527 | 4.305948 | 1.276829 | 1.142205 |
| 36.5125 | 1.659594 | 1.226301 | 2.028373 | 0.56683 | 0.130268 | 0.033408 | 3.083749 | 0.479126 | 0.265633 | 1.252957 | 0.74809 | 3.607859 | 4.323404 | 2.821637 | 2.338163 | 2.624751 | 1.552309 | 3.081852 | 1.196546 | 0.810303 |
| 36.52639 | 1.549624 | 1.051235 | 1.607838 | 0.532312 | 0.105091 | 0.314571 | 2.680776 | 0.470135 | 0.209884 | 1.101016 | 0.360685 | 3.071587 | 4.044662 | 2.628346 | 1.800572 | 2.288998 | 0.356491 | 1.013152 | 1.123661 | 0.583139 |
| 36.54028 | 1.270367 | 0.820245 | 1.175436 | 0.486334 | 0.04985 | 0.634548 | 2.103125 | 0.421921 | 0.215937 | 0.782227 | 0.035771 | 1.887857 | 3.425167 | 2.274023 | 1.000082 | 1.646078 | -0.71361 | -0.76245 | 0.948636 | 0.343809 |
| 36.55417 | 1.048776 | 0.380502 | 0.874055 | 0.426997 | 0.063421 | 0.847206 | 1.462797 | 0.356493 | 0.149378 | 0.595601 | -0.17248 | 0.299405 | 2.264787 | 1.254619 | 0.064591 | 1.240198 | -1.40943 | -1.64127 | 0.69961 | 0.491535 |
| 36.56806 | 0.898572 | -0.17856 | 0.719162 | 0.406075 | 0.163901 | 0.91183 | 0.742895 | 0.31576 | 0.237808 | 0.267653 | -0.25671 | -1.19035 | 1.062574 | 0.142513 | -0.6035 | 0.670204 | -1.04429 | -1.69472 | 0.612628 | 0.935451 |
| 36.58194 | 0.882785 | -0.32125 | 0.568743 | 0.370526 | 0.218788 | 0.803281 | 0.033104 | 0.261287 | 0.342309 | 0.19665 | -0.21253 | -2.06356 | -0.14296 | -0.92766 | -0.58833 | 0.643812 | -0.60856 | -1.52333 | 0.466193 | 0.984186 |
| 36.59583 | 0.774422 | -0.25325 | 0.085153 | 0.338451 | 0.181672 | 0.545469 | -0.6027 | 0.193667 | 0.329314 | 0.196162 | -0.01301 | -2.2406 | -1.3284 | -1.69221 | -0.18686 | 0.540384 | -0.16372 | -1.66644 | 0.214725 | 0.261913 |
| 36.60972 | 0.41089 | -0.40777 | -0.27338 | 0.332609 | 0.163158 | 0.239366 | -1.00305 | 0.140135 | 0.244164 | -0.10064 | -0.08784 | -2.17628 | -2.16759 | -2.24813 | -0.31417 | 0.130606 | 0.2593 | -1.51043 | 0.105138 | -0.77738 |
| 36.62361 | 0.002096 | -0.54197 | -0.35732 | 0.361476 | 0.246245 | 0.069424 | -1.19724 | 0.098726 | 0.223734 | -0.38532 | -0.18347 | -2.06184 | -2.63733 | -2.38583 | -0.60899 | -0.33208 | 0.374365 | -0.96543 | 0.319042 | -1.02098 |
| 36.6375 | -0.29185 | -0.55434 | -0.1886 | 0.416948 | 0.350988 | 0.014372 | -1.1192 | 0.08132 | 0.243496 | -0.64669 | -0.19723 | -1.34265 | -2.42256 | -2.3242 | -0.67389 | -0.73797 | -0.57631 | -0.65799 | 0.531306 | -0.60778 |
| 36.65139 | -0.61852 | -0.56734 | -0.25234 | 0.41793 | 0.38533 | -0.10024 | -1.30666 | 0.037801 | 0.24143 | -0.89339 | -0.31687 | -0.69839 | -2.37014 | -2.00071 | -0.39533 | -1.12259 | -1.56978 | -0.96997 | 0.553051 | -0.56083 |
| 36.66528 | -1.13322 | -0.66697 | -0.51799 | 0.378797 | 0.386094 | -0.31509 | -1.63696 | 0.000045 | 0.078547 | -0.89726 | -0.53508 | 0.0401 | -2.23495 | -1.18562 | -0.07225 | -1.1756 | -2.0502 | -1.48435 | 0.51525 | -1.28537 |
| 36.67917 | -1.65325 | -0.7845 | -0.76497 | 0.324795 | 0.472369 | -0.54783 | -1.77658 | -0.01972 | -0.15319 | -0.57714 | -0.67073 | 0.710791 | -1.97512 | -0.42632 | 0.178394 | -1.26017 | -1.78986 | -1.53507 | 0.445825 | -0.59386 |
| 36.69306 | -1.96884 | -0.82228 | -0.98025 | 0.293336 | 0.610281 | -0.61453 | -1.50081 | 0.022207 | -0.30452 | -0.15389 | -0.57458 | 1.191277 | -1.54954 | 0.111227 | 0.428389 | -1.21215 | -1.04556 | -1.29578 | 0.423334 | -0.06914 |
| 36.70694 | -1.95659 | -0.72784 | -0.79208 | 0.301215 | 0.673272 | -0.64453 | -1.0883 | 0.094032 | -0.28424 | 0.182037 | -0.25486 | 1.254039 | -0.98662 | 0.306297 | 0.517126 | -1.09815 | -0.05274 | -1.08964 | 0.371091 | 0.289875 |
| 36.72083 | -1.76313 | -0.55453 | -0.70669 | 0.219919 | 0.668954 | -0.7115 | -0.58759 | 0.127412 | -0.29144 | 0.196391 | 0.000881 | 1.009432 | -0.65587 | 0.053033 | 0.457551 | -1.03811 | 0.642397 | -0.86238 | 0.322254 | 0.332443 |
| 36.73472 | -1.42342 | -0.38751 | -0.54473 | 0.095511 | 0.640782 | -0.75207 | -0.21922 | 0.045282 | -0.28532 | -0.13314 | 0.055328 | 0.565941 | -0.56315 | -0.60781 | 0.111827 | -1.06818 | 0.809067 | -0.41503 | 0.222282 | -0.07578 |
| 36.74861 | -1.0004 | -0.25153 | -0.48728 | 0.080278 | 0.656326 | -0.76072 | -0.07003 | 0.019835 | -0.28345 | -0.3349 | -0.02521 | 0.07816 | -0.56791 | -1.13087 | -0.27505 | -1.09274 | 0.494024 | -0.32197 | 0.102456 | -0.74468 |
| 36.7625 | -0.68482 | -0.23149 | -0.65043 | 0.128142 | 0.662969 | -0.7268 | -0.15559 | 0.086458 | -0.2726 | -0.67339 | -0.19106 | -0.36686 | -0.69731 | -1.57087 | -0.67723 | -1.3502 | 0.186879 | -0.63826 | -0.01922 | -1.17466 |
| 36.77639 | -0.64832 | -0.42562 | -0.9983 | 0.140369 | 0.601256 | -0.77886 | -0.53312 | 0.090892 | -0.31839 | -0.82052 | -0.58802 | -0.82671 | -0.96345 | -1.84866 | -1.0204 | -1.66629 | 0.03765 | -1.07256 | -0.17464 | -1.34846 |
| 36.79028 | -0.7769 | -0.77821 | -1.34478 | 0.026611 | 0.507617 | -0.96842 | -1.01794 | 0.019806 | -0.56936 | -0.89571 | -0.96616 | -1.27806 | -1.30946 | -1.84188 | -1.25698 | -2.079 | -0.12711 | -1.21255 | -0.38947 | -1.59415 |
| 36.80417 | -1.05335 | -1.06099 | -1.73405 | -0.09757 | 0.438826 | -0.98978 | -1.20366 | -0.03744 | -0.76901 | -1.01812 | -1.05267 | -1.67433 | -1.65727 | -1.67372 | -1.2375 | -1.93001 | -0.34549 | -1.18122 | -0.60126 | -1.6375 |
| 36.81806 | -1.36419 | -1.23433 | -1.74238 | -0.11558 | 0.404638 | -0.92826 | -0.99786 | -0.02313 | -0.88971 | -1.18682 | -0.88258 | -1.94564 | -1.80797 | -1.31018 | -1.12163 | -1.71713 | -0.56882 | -1.05364 | -0.66057 | -1.60144 |
| 36.83194 | -1.54684 | -1.35774 | -1.50642 | -0.14952 | 0.38809 | -0.88994 | -0.62217 | 0.138479 | -0.99843 | -1.26117 | -0.67862 | -2.0561 | -1.78904 | -0.77348 | -0.91631 | -1.39206 | -0.80805 | -0.82284 | -0.6474 | -1.71876 |
| 36.84583 | -1.75763 | -1.42782 | -1.56692 | -0.22879 | 0.329613 | -0.92731 | -0.52914 | 0.18332 | -1.12738 | -1.23003 | -0.4977 | -2.02676 | -1.98543 | -0.43851 | -0.72664 | -1.04284 | -1.10606 | -0.71104 | -0.63022 | -1.87648 |
| 36.85972 | -1.84595 | -1.44019 | -1.38538 | -0.33056 | 0.292218 | -1.02093 | -0.26599 | 0.133982 | -1.20893 | -1.06791 | -0.32863 | -1.96341 | -2.1001 | -0.3244 | -0.47273 | -0.72445 | -1.26465 | -0.60444 | -0.56003 | -2.0127 |
| 36.87361 | -1.90011 | -1.41559 | -1.20747 | -0.3557 | 0.298851 | -0.99035 | 0.066643 | 0.096041 | -1.13231 | -0.7652 | -0.18161 | -1.92816 | -1.9134 | -0.31608 | -0.2082 | -0.57841 | -1.37376 | -0.63587 | -0.45455 | -1.64782 |
| 36.8875 | -1.73139 | -1.23181 | -0.89212 | -0.36369 | 0.329603 | -0.81825 | 0.300452 | 0.057636 | -0.86442 | -0.51216 | -0.13305 | -1.63059 | -1.58901 | -0.34671 | -0.02938 | -0.44713 | -1.34775 | -0.50957 | -0.36017 | -1.06275 |
| 36.90139 | -1.59764 | -0.95891 | -0.62975 | -0.65465 | 0.248078 | -0.56647 | 0.583665 | 0.163385 | -0.5112 | -0.22419 | -0.08279 | -1.03997 | -1.26237 | -0.36542 | 0.094623 | -0.50398 | -1.30517 | -0.73412 | -0.24651 | -0.62765 |
| 36.91528 | -1.68335 | -0.86847 | -0.41014 | -1.53286 | 0.043461 | -0.3561 | 0.96664 | 0.405593 | -0.28526 | 0.148172 | 0.180412 | -0.75084 | -0.65027 | -0.25148 | 0.102184 | -0.87424 | -1.08056 | -0.26968 | -0.47517 | 0.537684 |
| 36.92917 | -1.82993 | -1.25825 | -0.61317 | -2.62752 | -0.15962 | -0.50281 | 0.934876 | 0.465669 | -0.43675 | 0.264249 | 0.162519 | -0.89863 | -0.4057 | -0.27006 | -0.31716 | -1.75582 | -0.88368 | 0.419907 | -1.15092 | 1.448733 |
| 36.94306 | -2.51742 | -1.95451 | -1.46033 | -2.74567 | -0.17587 | -1.24075 | -0.22754 | 0.230195 | -1.13382 | -0.67574 | -0.87066 | -1.51722 | -1.72577 | -1.25553 | -1.63789 | -2.96543 | -1.828 | 0.078155 | -1.89637 | 0.15529 |
| 36.95694 | -2.74721 | -2.42908 | -2.85294 | -2.11825 | -0.0849 | -1.44291 | -1.44886 | 0.245022 | -1.37401 | -2.40686 | -2.59836 | -2.77006 | -4.02407 | -2.8165 | -3.53494 | -4.00143 | -3.20567 | -0.90638 | -2.90137 | -1.61941 |
| 36.97083 | -2.13109 | -1.92013 | -3.50141 | -0.82719 | 0.738201 | 0.241377 | -0.76634 | 1.102683 | 0.279294 | -3.13691 | -3.12567 | -3.79715 | -4.41505 | -3.48247 | -4.51453 | -3.55694 | -3.31109 | -1.27354 | -2.71533 | -2.52808 |
| 36.98472 | -1.76877 | -1.53329 | -2.5162 | -0.31953 | 1.020347 | 1.562188 | -0.37817 | 1.25281 | 1.143492 | -2.31763 | -1.84924 | -2.94597 | -2.97006 | -2.28271 | -3.23436 | -1.27811 | -2.26963 | -0.87525 | -1.49158 | -1.72579 |
| 36.99861 | -2.31746 | -1.55763 | -0.92638 | -0.0855 | 0.402716 | 0.170204 | -1.34187 | 0.417477 | 0.120791 | -0.99389 | -0.25643 | -0.9066 | -1.26534 | -0.87669 | -1.50195 | 0.881618 | -1.47188 | -0.30196 | -0.1056 | -0.47387 |
| 37.0125 | -2.74105 | -1.56696 | -0.66441 | -0.15556 | 0.160004 | -0.7299 | -1.60084 | -0.14044 | -0.89208 | 0.269681 | 0.811798 | 0.155915 | -0.25487 | 0.002668 | -0.34434 | 1.311121 | -0.56738 | 0.334809 | 0.461626 | 0.838071 |
| 37.02639 | -2.39857 | -1.05501 | -0.44425 | -0.45096 | 0.133416 | -0.99548 | -1.0416 | -0.25359 | -1.35752 | 1.366124 | 1.010983 | 0.652127 | 0.49864 | 0.604095 | 0.235831 | 0.943261 | 0.146952 | 1.174737 | 0.529847 | 1.620024 |
| 37.04028 | -1.94081 | -0.26452 | -0.22822 | -0.53895 | 0.097615 | -1.02103 | -0.4464 | -0.20522 | -1.45082 | 1.85462 | 0.8731 | 1.025761 | 1.063736 | 1.404982 | 0.7566 | 0.973487 | 0.588086 | 1.868351 | 0.593615 | 2.10189 |
| 37.05417 | -1.49933 | 0.628823 | 0.035656 | -0.33475 | 0.099923 | -0.91008 | -0.07809 | -0.11254 | -1.22932 | 1.504796 | 0.916431 | 1.498386 | 1.004911 | 1.902338 | 1.11435 | 1.62848 | 0.969428 | 2.105137 | 0.633817 | 1.790835 |
| 37.06806 | -1.1438 | 0.925632 | 0.323173 | -0.15863 | 0.101208 | -0.7263 | 0.30358 | 0.081847 | -1.00487 | 0.991388 | 0.327977 | 1.804907 | 0.902425 | 1.893569 | 1.27062 | 1.928254 | 1.035287 | 2.083402 | 0.253593 | 1.287764 |
| 37.08194 | -0.818 | 0.989592 | 0.503692 | -0.16817 | -0.00227 | -0.64058 | 0.720206 | 0.085865 | -0.85717 | 0.706145 | 0.317774 | 1.856774 | 0.73014 | 2.214009 | 1.327805 | 1.220303 | 1.132972 | 2.267008 | 0.01158 | 1.546532 |
| 37.09583 | -0.62646 | 0.824153 | 0.594207 | -0.17682 | -0.09429 | -0.59851 | 0.973882 | 0.041749 | -0.80092 | 0.612905 | 0.187001 | 1.689344 | 0.232686 | 2.355823 | 1.238475 | 0.14732 | 1.131731 | 2.198498 | -0.29119 | 1.863968 |
| 37.10972 | -0.54381 | 0.539816 | 0.632614 | -0.08543 | -0.02953 | -0.52686 | 1.066921 | 0.091398 | -0.57882 | 0.401529 | -0.23699 | 1.808162 | 0.048587 | 2.365964 | 1.120292 | -0.58765 | 0.993496 | 1.759697 | -0.52986 | 1.139229 |
| 37.12361 | -0.38502 | 0.354098 | 0.666131 | 0.031848 | 0.103732 | -0.36351 | 1.110601 | 0.179631 | -0.21982 | 0.277823 | -0.33876 | 1.718246 | 0.216402 | 2.073063 | 1.016971 | -0.69528 | 0.797696 | 1.402143 | -0.53563 | 0.399776 |
| 37.1375 | -0.20238 | 0.429735 | 0.448355 | 0.031682 | 0.071935 | -0.23071 | 1.097754 | 0.161364 | 0.028083 | 0.34143 | -0.81463 | 1.851938 | 0.501547 | 1.621901 | 1.115146 | -0.58128 | 0.660375 | 1.040224 | -0.47737 | 0.11273 |
| 37.15139 | -0.14248 | 0.289584 | 0.160944 | -0.07576 | -0.11016 | -0.2305 | 0.915695 | 0.044183 | 0.117756 | 0.453764 | -0.78536 | 1.775785 | 0.702827 | 0.910723 | 1.139628 | -0.17828 | 0.357655 | 0.56364 | -0.41664 | 0.415423 |
| 37.16528 | -0.07597 | -0.05511 | 0.13501 | -0.12656 | -0.18256 | -0.24909 | 0.672321 | 0.112056 | 0.119146 | 0.429299 | -0.5708 | 1.631273 | 0.715494 | 0.515762 | 1.051772 | 0.491848 | -0.07717 | -0.00512 | -0.16713 | 0.834665 |
| 37.17917 | 0.056736 | -0.1609 | 0.217902 | -0.05261 | -0.14684 | -0.21299 | 0.537874 | 0.197195 | 0.167151 | 0.28226 | -0.07374 | 1.305982 | 0.82818 | 0.214806 | 0.758755 | 0.9765 | -0.32652 | -0.49032 | 0.135451 | 1.215947 |
| 37.19306 | 0.136415 | -0.4053 | 0.144301 | 0.048039 | -0.11532 | -0.1385 | 0.545632 | 0.208574 | 0.200039 | 0.169162 | 0.378689 | 0.882363 | 1.030604 | -0.22161 | 0.242738 | 0.798792 | -0.43632 | -0.96474 | 0.263767 | 1.146414 |

| | | | | | | | | | | | | | | | | | | | | |
|---|---|---|---|---|---|---|---|---|---|---|---|---|---|---|---|---|---|---|---|---|
| 37.20694 | 0.130002 | -0.54813 | 0.191115 | 0.049519 | -0.1201 | -0.1138 | 0.529767 | 0.196971 | 0.150673 | 0.017308 | 0.44474 | 0.497086 | 1.086693 | -0.8684 | -0.21748 | 0.185264 | -0.59475 | -1.50397 | 0.115912 | 0.636658 |
| 37.22083 | -0.01281 | -0.48294 | 0.205058 | 0.003453 | -0.1461 | -0.20248 | 0.440173 | 0.166738 | -0.02765 | -0.30239 | 0.017145 | 0.088793 | 0.83936 | -1.36012 | -0.55656 | -0.43136 | -0.78587 | -1.97849 | -0.13334 | 0.261713 |
| 37.23472 | -0.10921 | -0.33573 | 0.136196 | -0.06652 | -0.12953 | -0.29623 | 0.334574 | 0.132098 | -0.22438 | -0.40367 | -0.47444 | -0.27523 | 0.431045 | -1.66814 | -0.5908 | -0.83624 | -0.89489 | -2.01322 | -0.2894 | 0.119679 |
| 37.24861 | -0.05836 | -0.08212 | 0.103531 | -0.03769 | -0.06422 | -0.28491 | 0.307865 | 0.131016 | -0.27435 | -0.29688 | -0.69826 | -0.43357 | 0.133079 | -1.96926 | -0.52987 | -0.74944 | -0.78137 | -1.63268 | -0.23583 | 0.50626 |
| 37.2625 | -0.02375 | 0.057698 | 0.077496 | -0.05232 | -0.03467 | -0.31648 | 0.282665 | 0.078987 | -0.27465 | -0.12001 | -0.65137 | -0.56474 | -0.022 | -1.78059 | -0.55316 | -0.30636 | -0.34056 | -0.7983 | -0.07061 | 1.18488 |
| 37.27639 | -0.133 | -0.11457 | -0.00956 | -0.07913 | -0.0825 | -0.41503 | 0.170727 | 0.01736 | -0.35857 | 0.014934 | -0.41883 | -0.59981 | -0.0759 | -1.27425 | -0.63633 | 0.244636 | -0.01998 | 0.261223 | 0.092265 | 1.744575 |
| 37.29028 | -0.24534 | -0.40938 | -0.13059 | -0.06145 | -0.09988 | -0.43417 | 0.026919 | -0.00969 | -0.439 | 0.151474 | -0.02873 | -0.49412 | -0.09179 | -0.27279 | -0.60117 | 0.775509 | -0.00143 | 1.093737 | 0.233441 | 1.548145 |
| 37.30417 | -0.33658 | -0.60887 | -0.20214 | -0.01072 | -0.09004 | -0.37675 | -0.09222 | -0.00827 | -0.42349 | 0.24525 | 0.383745 | -0.24609 | 0.065057 | 0.520378 | -0.4233 | 0.93154 | 0.036192 | 1.411611 | 0.331578 | 1.008125 |
| 37.31806 | -0.48044 | -0.70589 | -0.25532 | -0.01029 | -0.10333 | -0.36593 | -0.23364 | -0.0237 | -0.38767 | 0.257725 | 0.627165 | 0.020648 | 0.158847 | 0.594487 | -0.22041 | 0.772852 | 0.159735 | 1.332278 | 0.315682 | 0.375257 |
| 37.33194 | -0.86229 | -0.74952 | -0.22472 | 0.0009 | -0.11042 | -0.34057 | -0.25892 | -0.02977 | -0.3338 | 0.289487 | 0.770456 | 0.297471 | 0.31095 | 0.311381 | 0.091106 | 0.399722 | 0.40246 | 1.162346 | 0.290155 | 0.044277 |
| 37.34583 | -1.41254 | -0.76518 | -0.1632 | 0.086768 | -0.03117 | -0.21565 | -0.20369 | 0.045691 | -0.22634 | 0.45115 | 0.733041 | 0.619014 | 0.629337 | 0.220242 | 0.428476 | 0.106277 | 0.780734 | 1.05427 | 0.261874 | 0.076115 |
| 37.35972 | -1.54996 | -0.56993 | -0.12382 | 0.209234 | 0.105667 | -0.12867 | -0.11471 | 0.125197 | -0.12792 | 0.588195 | 0.42377 | 0.814654 | 0.961692 | 0.228883 | 0.670825 | -0.07349 | 0.973557 | 0.945537 | 0.247736 | 0.359345 |
| 37.37361 | -1.11763 | -0.10748 | -0.01389 | 0.260921 | 0.200727 | -0.21714 | -0.09862 | 0.233702 | -0.14269 | 0.446455 | 0.079236 | 0.872688 | 1.005914 | 0.083129 | 0.675266 | 0.042798 | 0.776601 | 0.82846 | 0.266001 | 0.822416 |
| 37.3875 | -0.63296 | 0.212854 | 0.01535 | 0.240212 | 0.180842 | -0.3719 | -0.11563 | 0.276088 | -0.23888 | 0.250286 | -0.09997 | 0.780598 | 0.866941 | -0.03476 | 0.547554 | 0.238847 | 0.439963 | 0.708705 | 0.177044 | 1.180367 |
| 37.40139 | -0.42547 | 0.202372 | -0.12682 | 0.150338 | 0.113419 | -0.42727 | -0.13685 | 0.239473 | -0.28189 | 0.123833 | -0.02054 | 0.559708 | 0.649706 | 0.159204 | 0.249248 | 0.308577 | 0.257608 | 0.721144 | -0.09906 | 1.191903 |
| 37.41528 | -0.16884 | 0.07534 | -0.22994 | 0.083884 | 0.109503 | -0.40106 | -0.15389 | 0.239851 | -0.23229 | 0.084833 | 0.200236 | 0.342437 | 0.440591 | 0.491172 | 0.053129 | 0.258424 | 0.276128 | 0.88394 | -0.22762 | 1.089685 |
| 37.42917 | 0.592227 | 0.358592 | 0.016269 | 0.268257 | 0.186618 | -0.19018 | 0.126786 | 0.322109 | 0.024973 | 0.332693 | 0.478386 | 0.54526 | 1.010897 | 1.155561 | 0.191307 | 0.274424 | 0.88617 | 1.234895 | 0.148897 | 1.382284 |
| 37.44306 | 2.015755 | 1.339191 | 0.700193 | 0.720459 | 0.332867 | 0.399946 | 0.844802 | 0.430258 | 0.667525 | 1.072777 | 0.885022 | 1.337126 | 3.01561 | 2.102845 | 0.798368 | 0.350101 | 2.545203 | 1.892491 | 0.954684 | 1.856704 |
| 37.45694 | 3.732003 | 2.67777 | 1.450184 | 0.993791 | 0.488292 | 0.975058 | 1.550845 | 0.58696 | 1.211559 | 1.893021 | 1.540222 | 1.97891 | 4.474958 | 2.694546 | 1.376902 | 0.783415 | 3.93035 | 2.417467 | 1.756998 | 2.48358 |
| 37.47083 | 3.804492 | 3.421823 | 1.717551 | 0.858199 | 0.483319 | 0.993312 | 1.80278 | 0.568404 | 1.118553 | 2.165161 | 1.788794 | 1.936437 | 4.319193 | 2.445512 | 1.422776 | 2.220161 | 3.986344 | 2.476827 | 2.015859 | 3.722544 |
| 37.48472 | 3.090582 | 3.576411 | 1.578724 | 0.603196 | 0.362947 | 0.694403 | 2.025932 | 0.448067 | 0.759193 | 1.750044 | 1.753431 | 1.902001 | 3.390899 | 2.399455 | 1.341848 | 2.611609 | 3.724463 | 2.54943 | 1.67691 | 3.82029 |
| 37.49861 | 2.828238 | 3.553926 | 1.527996 | 0.522352 | 0.277685 | 0.528157 | 2.178928 | 0.489558 | 0.491531 | 1.456916 | 1.381353 | 2.318227 | 3.276574 | 2.418481 | 1.424787 | 2.509149 | 3.732928 | 2.581092 | 1.233724 | 3.128008 |
| 37.5125 | 2.798726 | 3.221188 | 1.645284 | 0.445827 | 0.235446 | 0.745847 | 2.031833 | 0.5004 | 0.420986 | 1.491339 | 1.020241 | 2.839799 | 3.639957 | 2.196234 | 1.435034 | 2.167916 | 2.936507 | 2.688625 | 0.91639 | 3.32039 |
| 37.52639 | 2.642932 | 2.602212 | 1.533099 | 0.389798 | 0.214723 | 0.757102 | 1.691388 | 0.376302 | 0.303912 | 1.519119 | 0.568034 | 2.703871 | 3.731787 | 1.634491 | 1.286631 | 1.433413 | 2.001241 | 2.590262 | 0.69134 | 2.777402 |
| 37.54028 | 2.385438 | 1.79481 | 1.223787 | 0.285355 | 0.184694 | 0.609604 | 1.191176 | 0.218345 | 0.146298 | 1.414873 | -0.03684 | 1.960376 | 3.281285 | 1.111988 | 0.942872 | 0.687517 | 1.033063 | 1.793017 | 0.559877 | 1.277822 |
| 37.55417 | 2.021507 | 0.996102 | 0.825509 | 0.221694 | 0.128938 | 0.550162 | 0.651226 | 0.157621 | 0.042039 | 0.892413 | -0.50263 | 1.169057 | 2.467089 | 0.606721 | 0.44984 | 0.00081 | 0.141739 | 0.657395 | 0.348811 | 0.584461 |
| 37.56806 | 1.64849 | 0.35956 | 0.423107 | 0.222087 | 0.112632 | 0.651366 | 0.108647 | 0.110912 | 0.020698 | 0.689298 | -0.6888 | 0.445486 | 1.607266 | -0.02473 | -0.00277 | -0.63499 | -0.45229 | -0.26382 | 0.201015 | -0.09202 |
| 37.58194 | 1.295752 | -0.13868 | 0.156815 | 0.244267 | 0.100661 | 0.815474 | -0.19 | 0.088695 | 0.056626 | 0.552432 | -0.5324 | -0.25898 | 0.884355 | -0.61998 | -0.35276 | -0.70242 | -0.67085 | -0.82729 | 0.164693 | -0.33274 |
| 37.59583 | 0.957329 | -0.38509 | 0.081451 | 0.263108 | 0.068218 | 0.861954 | -0.3125 | 0.081969 | 0.182828 | 0.458745 | -0.35567 | -0.8873 | 0.259561 | -0.97767 | -0.47809 | -0.30426 | -1.0194 | -1.16786 | 0.21335 | -0.72363 |
| 37.60972 | 0.70418 | -0.52393 | 0.001122 | 0.25945 | 0.044113 | 1.104004 | -0.44525 | 0.082864 | 0.453112 | 0.272751 | -0.48614 | -1.28772 | -0.18924 | -1.20554 | -0.41987 | 0.00622 | -1.30931 | -1.38958 | 0.233031 | -1.06055 |
| 37.62361 | 0.337023 | -0.78129 | -0.31458 | 0.256294 | 0.036752 | 1.013835 | -0.73682 | 0.074469 | 0.302238 | 0.067647 | -1.01608 | -1.44282 | -0.4363 | -1.4983 | -0.43809 | -0.06971 | -1.32895 | -1.65563 | 0.142359 | -1.15492 |
| 37.6375 | -0.08693 | -0.93593 | -0.66887 | 0.250564 | 0.064185 | 0.387863 | -1.00691 | 0.002132 | 0.188838 | -0.00703 | -1.17262 | -1.50149 | -0.58929 | -1.63181 | -0.53877 | -0.50474 | -1.07941 | -2.11134 | 0.050912 | -1.27827 |
| 37.65139 | -0.41887 | -0.92621 | -0.78828 | 0.240806 | 0.145145 | 0.007816 | -1.07607 | -0.00193 | 0.113252 | 0.011276 | -1.04219 | -1.37214 | -0.75222 | -1.63193 | -0.58923 | -0.99514 | -0.90216 | -1.94601 | 0.145958 | -1.09906 |
| 37.66528 | -0.64432 | -0.87576 | -0.70851 | 0.230348 | 0.193142 | -0.05224 | -0.95932 | 0.051893 | 0.123624 | -0.04061 | -0.56515 | -1.28996 | -0.94814 | -1.50746 | -0.6347 | -1.33734 | -0.97552 | -1.29847 | 0.376501 | -0.95259 |
| 37.67917 | -0.75242 | -0.76757 | -0.50827 | 0.213573 | 0.169289 | 0.02116 | -0.72978 | 0.06049 | 0.18677 | -0.19854 | -0.14832 | -1.24045 | -1.15028 | -1.33938 | -0.72048 | -1.30127 | -0.98319 | -0.41861 | 0.483351 | -0.69489 |
| 37.69306 | -0.7562 | -0.64126 | -0.22857 | 0.200709 | 0.156591 | 0.075024 | -0.41867 | 0.042641 | 0.224977 | -0.35312 | 0.069872 | -1.0893 | -1.33043 | -0.87723 | -0.70954 | -1.03903 | -0.78335 | 0.601602 | 0.422729 | -0.59428 |
| 37.70694 | -0.59851 | -0.41833 | -0.03693 | 0.186291 | 0.170714 | 0.137338 | -0.187 | 0.013668 | 0.20888 | -0.37969 | 0.238769 | -0.86893 | -1.3674 | -0.21288 | -0.59218 | -0.43374 | -0.45599 | 1.329521 | 0.313683 | -0.41899 |
| 37.72083 | -0.34011 | -0.23349 | 0.040695 | 0.162172 | 0.20359 | 0.235692 | 0.001706 | 0.024364 | 0.21431 | -0.364 | 0.312153 | -0.58756 | -1.27064 | 0.381408 | -0.29948 | 0.285654 | -0.23188 | 1.713097 | 0.275956 | -0.27678 |
| 37.73472 | -0.09676 | -0.16054 | 0.011392 | 0.136288 | 0.21164 | 0.17706 | 0.117371 | 0.075586 | 0.15558 | -0.31912 | 0.0874 | -0.31125 | -0.96899 | 0.606217 | 0.012007 | 0.885414 | -0.14912 | 1.330323 | 0.26667 | -0.35252 |
| 37.74861 | 0.124888 | -0.15239 | -0.11008 | 0.076727 | 0.18456 | -0.03325 | 0.15432 | 0.083302 | -0.01291 | -0.26369 | -0.39655 | -0.19921 | -0.80879 | 0.52551 | 0.179155 | 0.913467 | -0.21056 | 0.465426 | 0.192402 | -0.67662 |
| 37.7625 | 0.249588 | -0.17136 | -0.22913 | 0.038377 | 0.114146 | -0.37509 | 0.116046 | 0.112593 | -0.27058 | -0.30497 | -0.89863 | -0.19938 | -0.87815 | 0.257227 | 0.146508 | 0.208709 | -0.50681 | -0.54588 | 0.030213 | -1.07178 |
| 37.77639 | 0.144514 | -0.3097 | -0.34598 | -0.03574 | 0.039663 | -0.67766 | 0.015549 | 0.100914 | -0.58426 | -0.4581 | -1.27059 | -0.33353 | -1.00073 | -0.06988 | -0.07285 | -1.04224 | -0.9293 | -1.46834 | -0.18941 | -1.3067 |
| 37.79028 | -0.11118 | -0.57579 | -0.47624 | -0.06612 | 0.053676 | -0.72152 | -0.03017 | 0.152295 | -0.87218 | -0.67731 | -1.34715 | -0.59081 | -1.15457 | -0.46308 | -0.41923 | -2.30545 | -1.21171 | -2.05984 | -0.46527 | -1.36332 |
| 37.80417 | -0.46786 | -0.72386 | -0.55171 | -0.07269 | 0.144366 | -0.56539 | 0.023391 | 0.157925 | -0.94635 | -0.90944 | -1.33374 | -0.88874 | -1.32048 | -0.84775 | -0.71851 | -2.95212 | -1.26203 | -2.15482 | -0.42058 | -1.20945 |
| 37.81806 | -0.7031 | -0.76073 | -0.49999 | -0.03904 | 0.156664 | -0.42942 | 0.061969 | 0.066468 | -0.90391 | -0.7019 | -1.13577 | -1.07473 | -1.79756 | -1.17135 | -0.83326 | -2.43164 | -1.24715 | -2.12226 | -0.58761 | -0.98099 |
| 37.83194 | -0.94592 | -0.88925 | -0.46793 | -0.07488 | 0.108272 | -0.31922 | 0.122743 | 0.002784 | -0.80683 | -0.88807 | -0.91909 | -1.18048 | -2.06397 | -1.29577 | -0.75968 | -1.7222 | -1.28518 | -1.93055 | -0.64506 | -0.87767 |
| 37.84583 | -1.00879 | -1.11705 | -0.44371 | -0.1338 | 0.059308 | -0.31284 | 0.187461 | -0.09716 | -0.83758 | -0.77591 | -0.85263 | -1.247 | -1.92151 | -1.37333 | -0.64608 | -0.95664 | -1.33079 | -1.22378 | -0.69418 | -1.19104 |
| 37.85972 | -0.77004 | -1.16872 | -0.41343 | -0.16897 | 0.024727 | -0.30314 | 0.238762 | -0.17891 | -0.79857 | -0.72807 | -0.60351 | -1.09688 | -1.62305 | -1.19242 | -0.43162 | -0.28963 | -1.11817 | -0.54154 | -0.70838 | -1.23264 |
| 37.87361 | -0.35098 | -1.02776 | -0.38876 | -0.1742 | 0.117296 | -0.25169 | 0.277648 | -0.18583 | -0.73587 | -0.69456 | -0.21959 | -0.8215 | -1.25883 | -0.86731 | -0.22613 | 0.303986 | -0.79264 | -0.06105 | -0.69154 | -1.14028 |
| 37.8875 | 0.07072 | -0.74344 | -0.26781 | -0.10731 | 0.283242 | -0.11594 | 0.363647 | -0.14605 | -0.64086 | -0.6054 | 0.128715 | -0.66958 | -0.93743 | -0.42812 | -0.12875 | 0.26366 | -0.33445 | 0.276853 | -0.46612 | -0.96189 |
| 37.90139 | 0.435569 | -0.48063 | -0.07462 | -0.06492 | 0.438084 | 0.019776 | 0.506364 | -0.01477 | -0.53233 | -0.3447 | 0.312338 | -0.60506 | -0.66292 | -0.06659 | -0.10278 | 0.190561 | 0.007486 | 0.4751 | -0.19418 | -0.59036 |
| 37.91528 | 0.500688 | -0.27611 | -0.01129 | -0.10962 | 0.487865 | 0.024258 | 0.564421 | 0.002123 | -0.46858 | -0.01933 | 0.265664 | -0.66309 | -0.43846 | 0.17736 | -0.18066 | -0.35767 | 0.115882 | 0.431307 | 0.01231 | -0.50863 |
| 37.92917 | -0.02647 | -0.43685 | -0.26201 | -0.33465 | 0.252189 | -0.33615 | 0.208644 | -0.22588 | -0.7675 | -0.06721 | -0.10447 | -0.96628 | -0.86038 | 0.024968 | -0.46948 | -1.21935 | -0.44386 | -0.10233 | -0.19062 | -0.81307 |
