## Supplemental Table S2c for "Leaf movements as a quantitative metric for early stress detection"

**Supplementary Table S2.** Raw motion data from: **(c)** 100 mM NaCl treatment on different CEA crops and their respective controls.

**Supplementary Table S2.** Raw motion data from: **(c)** 100 mM NaCl treatment in Mizuna

| Time | MizunaSalt1 | MizunaSalt10 | MizunaSalt11 | MizunaSalt12 | MizunaSalt2 | MizunaSalt3 | MizunaSalt4 | MizunaSalt5 | MizunaSalt6 | MizunaSalt7 | MizunaSalt8 | MizunaSalt9 | MizunaControl1 | MizunaControl10 | MizunaControl11 | MizunaControl12 | MizunaControl2 | MizunaControl3 | MizunaControl4 | MizunaControl5 | MizunaControl6 | MizunaControl7 | MizunaControl8 | MizunaControl9 |
|---|---|---|---|---|---|---|---|---|---|---|---|---|---|---|---|---|---|---|---|---|---|---|---|---|
| 18 | -0.889234 | -1.067249 | -0.291703 | -0.524097 | -1.331776 | -0.129562 | -0.273517 | -0.551585 | -0.139619 | -0.121616 | -0.2613 | -0.128362 | -0.721349 | -1.750728 | -0.214733 | -1.147701 | -2.828991 | -0.050361 | -0.379762 | -1.329151 | -0.237413 | -0.384475 | -0.216794 | -0.496667 |
| 18.01388889 | 0.020965 | 0.777096 | 0.063592 | 0.294019 | -0.477579 | -0.059443 | -0.124058 | -0.25013 | 0.09043 | 0.119537 | 0.190446 | 0.012438 | -0.021889 | 0.097695 | 0.034256 | 1.494603 | -1.631383 | 0.109609 | 0.20181 | -0.656747 | 0.257735 | 0.428433 | 0.395798 | 0.824471 |
| 18.02777778 | 0.658602 | 1.479638 | 0.076987 | 0.38416 | -0.044138 | -0.050984 | -0.097643 | -0.03864 | 0.199885 | 0.144646 | 0.464913 | -0.019601 | 0.265603 | 1.712224 | 0.21701 | 2.597248 | -0.124542 | 0.140183 | 0.467293 | -0.043107 | 0.457975 | 0.773087 | 0.731825 | 1.271464 |
| 18.04166667 | 0.844593 | 1.566676 | 0.028037 | 0.166832 | 0.102219 | -0.03713 | -0.107204 | 0.069345 | 0.164491 | 0.105575 | 0.43791 | -0.026631 | 0.367072 | 2.360561 | 0.163285 | 1.932748 | 0.766223 | 0.144968 | 0.469253 | 0.254373 | 0.375281 | 0.414887 | 0.728459 | 1.042163 |
| 18.05555556 | 0.617763 | 1.272229 | 0.039735 | 0.059503 | 0.086573 | -0.013929 | -0.103871 | 0.096041 | 0.166636 | 0.097298 | 0.326975 | -0.005945 | 0.309485 | 2.491374 | 0.00177 | 1.085224 | 0.825798 | 0.130045 | 0.348123 | 0.380948 | 0.259736 | 0.230576 | 0.5288 | 0.661846 |
| 18.06944444 | 0.356532 | 1.032059 | 0.072079 | 0.052824 | 0.007562 | -0.010797 | -0.102395 | 0.081323 | 0.121821 | 0.082506 | 0.266827 | 0.005864 | 0.248711 | 2.311534 | -0.087638 | 0.594122 | 0.462381 | 0.104608 | 0.256758 | 0.351153 | 0.166005 | 0.132482 | 0.321906 | 0.326801 |
| 18.08333333 | 0.196004 | 0.841001 | 0.108363 | 0.102225 | 0.025283 | 0.01012 | -0.072292 | 0.091863 | 0.131429 | 0.062757 | 0.249988 | 0.026794 | 0.217772 | 2.337221 | -0.145826 | 0.333569 | 0.358782 | 0.116113 | 0.246571 | 0.223688 | 0.097243 | 0.273893 | 0.217623 | 0.132979 |
| 18.09722222 | 0.147863 | 0.656997 | 0.099424 | 0.101447 | 0.090471 | 0.046731 | -0.026766 | 0.108239 | 0.100917 | 0.035912 | 0.216654 | 0.035866 | 0.169639 | 2.073883 | -0.174601 | 0.200455 | 0.314451 | 0.116164 | 0.210497 | 0.112096 | 0.045661 | 0.426545 | 0.138005 | 0.063503 |
| 18.11111111 | 0.103945 | 0.327068 | -0.002065 | -0.047975 | 0.045323 | 0.061899 | -0.02194 | 0.090701 | -0.000461 | -0.038536 | 0.109225 | -0.009439 | 0.034276 | 1.235003 | -0.219554 | 0.037977 | 0.046165 | 0.044797 | 0.078124 | -0.040624 | -0.011657 | 0.23188 | 0.041124 | -0.026113 |
| 18.125 | 0.03206 | 0.038829 | -0.101313 | -0.205725 | -0.062134 | 0.050541 | -0.05219 | 0.036387 | -0.101955 | -0.100804 | -0.02251 | -0.060454 | -0.094755 | 0.293492 | -0.258565 | -0.132409 | -0.285451 | -0.04136 | -0.075606 | -0.230029 | -0.059199 | -0.08387 | -0.036424 | -0.160964 |
| 18.13888889 | -0.055321 | -0.0801 | -0.091813 | -0.207373 | -0.168365 | 0.015789 | -0.090027 | -0.030594 | -0.138807 | -0.112657 | -0.062485 | -0.065047 | -0.135816 | -0.194236 | -0.250995 | -0.215059 | -0.436489 | -0.061685 | -0.163895 | -0.326734 | -0.069243 | -0.194523 | -0.056481 | -0.199279 |
| 18.15277778 | -0.096212 | -0.049648 | -0.03943 | -0.146209 | -0.208816 | -0.017276 | -0.106242 | -0.080689 | -0.115059 | -0.102708 | -0.033833 | -0.046944 | -0.115614 | -0.119797 | -0.23529 | -0.21229 | -0.307335 | -0.044223 | -0.15706 | -0.317083 | -0.053418 | -0.053509 | -0.049956 | -0.153875 |
| 18.16666667 | -0.070856 | 0.062055 | -0.014029 | -0.099459 | -0.210782 | -0.028144 | -0.093774 | -0.096786 | -0.067396 | -0.098191 | 0.021891 | -0.039911 | -0.041007 | 0.377261 | -0.221626 | -0.152971 | -0.077976 | -0.014663 | -0.109753 | -0.245098 | -0.032769 | 0.121714 | -0.040212 | -0.071975 |
| 18.18055556 | -0.029815 | 0.20438 | -0.006713 | -0.018989 | -0.195041 | -0.024415 | -0.062966 | -0.087846 | -0.011336 | -0.078222 | 0.092305 | -0.026816 | 0.05645 | 1.092397 | -0.195101 | -0.059896 | 0.119144 | 0.014223 | -0.045601 | -0.140724 | -0.013161 | 0.255229 | -0.019192 | 0.010854 |
| 18.19444444 | -0.007003 | 0.153089 | -0.014914 | -0.01657 | -0.175642 | 0.003872 | -0.026914 | -0.052498 | -0.007976 | -0.049841 | 0.086517 | -0.000915 | 0.083028 | 1.061315 | -0.178931 | -0.06432 | 0.195041 | 0.012452 | -0.026284 | -0.055382 | -0.011928 | 0.241451 | -0.018536 | 0.033451 |
| 18.20833333 | -0.036837 | -0.105695 | -0.050494 | -0.150656 | -0.233059 | 0.018215 | -0.020812 | -0.042392 | -0.055735 | -0.059492 | -0.015506 | 0.007382 | -0.005135 | 0.010523 | -0.190425 | -0.210107 | 0.100685 | -0.029004 | -0.091514 | -0.047306 | -0.038639 | -0.018765 | -0.063244 | -0.051069 |
| 18.22222222 | -0.104448 | -0.354129 | -0.091776 | -0.273002 | -0.339965 | -0.016104 | -0.054735 | -0.087398 | -0.142288 | -0.097405 | -0.099768 | -0.013701 | -0.114449 | -0.903643 | -0.217521 | -0.373248 | -0.053586 | -0.074773 | -0.185601 | -0.095145 | -0.076135 | -0.2059 | -0.121581 | -0.184583 |
| 18.23611111 | -0.11096 | -0.356514 | -0.101079 | -0.262495 | -0.392397 | -0.033368 | -0.063857 | -0.110486 | -0.163718 | -0.106278 | -0.101855 | -0.023923 | -0.135512 | -0.896436 | -0.229169 | -0.457307 | -0.246996 | -0.085011 | -0.225707 | -0.143309 | -0.089221 | -0.240366 | -0.135396 | -0.256667 |
| 18.25 | -0.053816 | -0.197253 | -0.044831 | -0.134722 | -0.367308 | -0.007734 | -0.020434 | -0.081047 | -0.082292 | -0.049325 | -0.023856 | 0.001664 | -0.064453 | -0.447071 | -0.186381 | -0.373199 | -0.22143 | -0.051211 | -0.16188 | -0.119808 | -0.068505 | -0.182286 | -0.083737 | -0.201433 |
| 18.26388889 | 0.005154 | 0.00611 | 0.023254 | 0.023832 | -0.222298 | 0.016909 | 0.029492 | -0.050656 | 0.027695 | 0.010743 | 0.06856 | 0.036178 | 0.021819 | 0.024792 | -0.102949 | -0.191137 | -0.015657 | 0.007844 | -0.062614 | -0.01776 | -0.039138 | -0.081415 | -0.004781 | -0.076372 |
| 18.27777778 | 0.03382 | 0.095041 | 0.032824 | 0.079303 | -0.114668 | 0.022031 | 0.043845 | -0.040843 | 0.063522 | 0.024502 | 0.096756 | 0.043127 | 0.060162 | 0.323072 | -0.053969 | -0.084807 | 0.157874 | 0.032779 | -0.002198 | 0.061104 | -0.026068 | 0.005891 | 0.048653 | -0.000683 |
| 18.29166667 | 0.038933 | 0.07126 | -0.007533 | 0.04246 | -0.158808 | 0.016048 | 0.026831 | -0.046357 | 0.036361 | 0.009087 | 0.07055 | 0.025267 | 0.058235 | 0.358475 | -0.06944 | -0.141833 | 0.137794 | 0.016166 | -0.015654 | 0.035754 | -0.034019 | -0.016572 | 0.050385 | -0.037522 |
| 18.30555556 | 0.033839 | 0.004849 | -0.050371 | -0.018795 | -0.317197 | 0.009338 | 0.007395 | -0.062318 | -0.008052 | -0.012561 | 0.028361 | 0.009439 | 0.026398 | 0.153828 | -0.118782 | -0.298273 | -0.028275 | -0.023441 | -0.084659 | -0.061178 | -0.063006 | -0.105165 | 0.015934 | -0.150896 |
| 18.31944444 | 0.008157 | -0.079524 | -0.079626 | -0.082567 | -0.542531 | -0.004344 | -0.01597 | -0.084602 | -0.049192 | -0.042427 | -0.016151 | -0.007222 | -0.034498 | -0.151005 | -0.175025 | -0.52514 | -0.247478 | -0.076905 | -0.168964 | -0.164031 | -0.109053 | -0.206096 | -0.017769 | -0.303764 |
| 18.33333333 | -0.034543 | -0.153376 | -0.088044 | -0.10343 | -0.722298 | -0.024142 | -0.038083 | -0.115883 | -0.079104 | -0.05791 | -0.050131 | -0.024314 | -0.07912 | -0.340666 | -0.204284 | -0.743064 | -0.404231 | -0.121177 | -0.221757 | -0.179084 | -0.156022 | -0.181189 | -0.018008 | -0.426875 |
| 18.34722222 | -0.053964 | -0.113133 | -0.06587 | -0.039579 | -0.715212 | -0.02979 | -0.029893 | -0.117459 | -0.056399 | -0.027404 | -0.047431 | -0.02171 | -0.06972 | -0.308564 | -0.188037 | -0.779972 | -0.324755 | -0.10956 | -0.216321 | -0.110064 | -0.166062 | -0.080333 | 0.023865 | -0.418008 |
| 18.36111111 | -0.036116 | 0.000106 | -0.021192 | 0.074395 | -0.469027 | -0.020123 | 0.001198 | -0.104895 | 0.008833 | 0.03454 | -0.014015 | -0.006334 | -0.027543 | -0.14782 | -0.130955 | -0.56283 | -0.049195 | -0.032754 | -0.127324 | -0.054072 | -0.146339 | -0.001826 | 0.080575 | -0.246432 |
| 18.375 | 0.004481 | 0.108632 | 0.01428 | 0.159784 | -0.165643 | -0.018431 | 0.020332 | -0.09644 | 0.065476 | 0.089637 | 0.021986 | 0.005272 | 0.01783 | 0.025059 | -0.057705 | -0.247184 | 0.252069 | 0.045286 | -0.007943 | -0.036617 | -0.093935 | 0.034452 | 0.124957 | 0.028971 |
| 18.38888889 | 0.060784 | 0.16234 | 0.029415 | 0.181433 | 0.006519 | -0.015204 | 0.034027 | -0.086684 | 0.113488 | 0.125452 | 0.05942 | 0.022653 | 0.045576 | 0.128368 | -0.006533 | -0.043061 | 0.394999 | 0.074356 | 0.081117 | -0.016679 | -0.043822 | 0.051342 | 0.142942 | 0.204562 |
| 18.40277778 | 0.106517 | 0.207187 | 0.009794 | 0.148053 | 0.035534 | 0.021337 | 0.06224 | -0.051261 | 0.139225 | 0.148333 | 0.109942 | 0.040917 | 0.005582 | 0.088525 | -0.037438 | -0.031452 | 0.338234 | 0.02753 | 0.065543 | -0.034267 | -0.050057 | 0.009028 | 0.095755 | 0.187463 |
| 18.41666667 | 0.142102 | 0.25975 | -0.034468 | 0.102601 | 0.021575 | 0.048258 | 0.070966 | -0.024621 | 0.160841 | 0.176923 | 0.132665 | 0.033195 | -0.078647 | 0.042665 | -0.125688 | -0.049788 | 0.220016 | -0.064033 | -0.008492 | -0.088009 | -0.109385 | -0.086988 | -0.003381 | 0.109236 |
| 18.43055556 | 0.136703 | 0.540905 | -0.007851 | 0.150187 | 0.296378 | -0.048639 | 0.005872 | -0.092956 | 0.24612 | 0.192334 | 0.147379 | -0.015798 | -0.006348 | 0.498455 | -0.139487 | 0.465582 | 0.511971 | -0.115144 | 0.025118 | 0.053044 | -0.096848 | -0.130571 | -0.046938 | 0.294351 |
| 18.44444444 | 0.157947 | 1.231448 | 0.196819 | 0.298424 | 1.072197 | -0.226329 | -0.060696 | -0.134875 | 0.394472 | 0.266673 | 0.201077 | -0.07618 | 0.405764 | 2.133363 | -0.013145 | 1.668658 | 1.526744 | -0.111588 | 0.243194 | 0.494335 | 0.074905 | 0.100804 | 0.103897 | 0.889343 |
| 18.45833333 | 0.197229 | 2.111929 | 0.373731 | 0.375091 | 1.941912 | -0.233997 | 0.041243 | 0.09939 | 0.464021 | 0.398661 | 0.377254 | -0.00604 | 1.128458 | 4.73479 | 0.089131 | 2.160714 | 1.744119 | -0.058731 | 0.542925 | 0.807827 | 0.320937 | 0.643991 | 0.416239 | 1.585494 |
| 18.47222222 | 0.2233 | 2.942098 | 0.349628 | 0.349892 | 1.746449 | -0.062293 | 0.222937 | 0.360878 | 0.561554 | 0.503706 | 0.969927 | 0.121408 | 1.494159 | 4.944733 | 0.113784 | 1.514075 | 1.319995 | 0.018089 | 0.788842 | 0.921952 | 0.543374 | 0.239748 | 0.655837 | 2.248028 |
| 18.48611111 | 0.168981 | 2.856291 | 0.302276 | 0.291265 | 1.029161 | 0.061661 | 0.256094 | 0.383412 | 0.504592 | 0.473338 | 1.147231 | 0.153321 | 1.228106 | 4.853411 | 0.201819 | 0.938313 | 0.784631 | 0.121241 | 0.789842 | 0.694236 | 0.574315 | 0.225441 | 0.725589 | 2.725016 |
| 18.5 | 0.136949 | 2.447983 | 0.218905 | 0.408153 | 0.450424 | 0.085137 | 0.253055 | 0.268737 | 0.469441 | 0.470829 | 1.137133 | 0.199295 | 0.950522 | 2.561889 | 0.190739 | 0.5061 | 0.465482 | 0.15771 | 0.575803 | 0.53237 | 0.451449 | 0.233018 | 0.709086 | 2.192329 |
| 18.51388889 | 0.040058 | 1.906345 | 0.159172 | 0.362853 | 0.234119 | 0.054837 | 0.189589 | 0.114068 | 0.27979 | 0.373338 | 0.86932 | 0.185989 | 0.94994 | 1.60647 | 0.16703 | 0.254376 | 0.252705 | 0.131852 | 0.368494 | 0.505442 | 0.32863 | 0.360863 | 0.584156 | 1.425251 |
| 18.52777778 | -0.029569 | 1.492716 | 0.092972 | 0.238725 | 0.21093 | 0.024746 | 0.153587 | 0.003668 | 0.238282 | 0.292933 | 0.673046 | 0.154658 | 1.036313 | 1.133123 | 0.155844 | 0.131031 | 0.164089 | 0.111304 | 0.217649 | 0.507742 | 0.238038 | 0.370865 | 0.437587 | 0.946938 |
| 18.54166667 | -0.029547 | 1.306749 | 0.073701 | 0.160861 | 0.15406 | 0.038648 | 0.142655 | 0.000347 | 0.239309 | 0.282134 | 0.630031 | 0.139791 | 1.286641 | 1.074473 | 0.221451 | 0.101139 | 0.227417 | 0.103587 | 0.18103 | 0.382113 | 0.256215 | 0.240976 | 0.423672 | 0.838972 |
| 18.55555556 | -0.00426 | 1.135575 | 0.090567 | 0.19121 | 0.153984 | 0.049212 | 0.150102 | 0.017086 | 0.291989 | 0.295565 | 0.548846 | 0.149025 | 1.163365 | 1.123883 | 0.304799 | 0.139491 | 0.4045 | 0.08569 | 0.176083 | 0.318538 | 0.338848 | 0.157801 | 0.480782 | 0.996529 |
| 18.56944444 | -0.000545 | 0.74665 | 0.099505 | 0.251917 | 0.064759 | 0.044717 | 0.153909 | 0.019749 | 0.230491 | 0.216137 | 0.409348 | 0.143114 | 0.876268 | 1.364771 | 0.337864 | 0.239985 | 0.456843 | 0.059745 | 0.171392 | 0.303247 | 0.385281 | 0.197605 | 0.475317 | 1.092947 |
| 18.58333333 | -0.011936 | 0.49836 | 0.106743 | 0.26973 | -0.064052 | 0.053975 | 0.173233 | 0.047961 | 0.148583 | 0.178029 | 0.235981 | 0.112852 | 0.67409 | 1.395855 | 0.306713 | 0.302789 | 0.418025 | 0.048776 | 0.160581 | 0.235181 | 0.308395 | 0.347566 | 0.402141 | 0.989796 |
| 18.59722222 | -0.016003 | 0.380156 | 0.122552 | 0.214244 | -0.11241 | 0.066143 | 0.152479 | 0.084815 | 0.071088 | 0.151716 | 0.150437 | 0.073421 | 0.571384 | 1.236482 | 0.253379 | 0.281889 | 0.306596 | 0.043291 | 0.14208 | 0.169422 | 0.244536 | 0.423711 | 0.331435 | 0.716143 |
| 18.61111111 | -0.041022 | 0.279245 | 0.117815 | 0.114511 | -0.081145 | 0.03622 | 0.097587 | 0.061181 | -0.016135 | 0.110238 | 0.089522 | 0.039705 | 0.578545 | 0.880282 | 0.15467 | 0.212832 | 0.231837 | 0.020491 | 0.101499 | 0.118588 | 0.144287 | 0.32059 | 0.257701 | 0.539042 |
| 18.625 | -0.052482 | 0.218143 | 0.083807 | 0.057762 | -0.005927 | -0.005471 | 0.090049 | 0.015134 | -0.073434 | 0.06993 | 0.071786 | 0.008087 | 0.624245 | 0.559267 | 0.09684 | 0.158434 | 0.287682 | -0.001821 | 0.059156 | 0.111247 | 0.054256 | 0.182954 | 0.232844 | 0.442786 |
| 18.63888889 | -0.009158 | 0.24497 | 0.045898 | 0.067701 | 0.049513 | -0.004326 | 0.128763 | 0.016173 | -0.074993 | 0.070129 | 0.088688 | 0.012955 | 0.534702 | 0.38378 | 0.061733 | 0.134073 | 0.279381 | 0.001543 | 0.055258 | 0.148159 | 0.039445 | 0.105844 | 0.227889 | 0.455105 |
| 18.65277778 | 0.058048 | 0.250828 | 0.028857 | 0.110177 | 0.049618 | 0.020462 | 0.175596 | 0.040464 | -0.036383 | 0.096691 | 0.090949 | 0.038475 | 0.388784 | 0.21983 | 0.048632 | 0.124125 | 0.273766 | 0.016425 | 0.08137 | 0.209189 | 0.052297 | 0.100981 | 0.243883 | 0.543332 |
| 18.66666667 | 0.107371 | 0.274325 | 0.028193 | 0.13156 | 0.051071 | 0.026122 | 0.151552 | 0.054349 | -0.036482 | 0.092233 | 0.100959 | 0.046521 | 0.266677 | 0.094901 | 0.068336 | 0.108482 | 0.253237 | 0.02262 | 0.10479 | 0.282801 | 0.048982 | 0.137933 | 0.251851 | 0.600439 |
| 18.68055556 | 0.110734 | 0.354889 | 0.032875 | 0.134039 | 0.096409 | 0.007855 | 0.098507 | 0.056391 | -0.051757 | 0.069909 | 0.147465 | 0.028186 | 0.148694 | 0.061825 | 0.061513 | 0.097724 | 0.186352 | 0.015666 | 0.096449 | 0.316862 | 0.023534 | 0.183754 | 0.273326 | 0.489364 |
| 18.69444444 | 0.088697 | 0.422956 | 0.040795 | 0.124817 | 0.168145 | 0.000346 | 0.069753 | 0.070858 | -0.024774 | 0.06847 | 0.186847 | 0.010909 | 0.124358 | 0.055784 | 0.03565 | 0.107297 | 0.173405 | 0.006279 | 0.082863 | 0.314434 | 0.011523 | 0.18534 | 0.335199 | 0.401329 |
| 18.70833333 | 0.08732 | 0.347752 | 0.024786 | 0.084324 | 0.201716 | 0.015425 | 0.058245 | 0.104512 | -0.013902 | 0.075011 | 0.148674 | 0.003991 | 0.145215 | 0.024106 | 0.000046 | 0.126512 | 0.207686 | 0.005977 | 0.086071 | 0.24032 | 0.010737 | 0.080351 | 0.30298 | 0.335932 |
| 18.72222222 | 0.093347 | 0.219694 | -0.018142 | 0.034345 | 0.181929 | 0.023169 | 0.003392 | 0.136283 | -0.04338 | 0.06429 | 0.094138 | 0.005653 | 0.080786 | -0.064694 | -0.034105 | 0.145125 | 0.191726 | 0.015278 | 0.070019 | 0.175627 | 0.023033 | -0.007618 | 0.216669 | 0.29051 |
| 18.73611111 | 0.077462 | 0.206655 | -0.052276 | -0.000615 | 0.165552 | 0.007915 | -0.055794 | 0.126162 | -0.076169 | 0.029569 | 0.095571 | 0.012978 | -0.091063 | 0.048643 | -0.054732 | 0.143357 | 0.140163 | 0.022759 | 0.041019 | 0.152964 | 0.044933 | -0.007392 | 0.13674 | 0.205783 |
| 18.75 | 0.066211 | 0.240511 | -0.065255 | -0.048574 | 0.174922 | -0.010029 | -0.025469 | 0.071603 | -0.074311 | -0.020518 | 0.099714 | -0.007562 | -0.249498 | 0.322709 | -0.057586 | 0.123891 | 0.23845 | 0.024276 | 0.001585 | 0.21412 | 0.05017 | 0.027793 | 0.084165 | 0.0924 |
| 18.76388889 | 0.048117 | 0.187991 | -0.076353 | -0.107372 | 0.158999 | -0.012539 | 0.015018 | 0.004306 | -0.027236 | -0.019608 | 0.061299 | -0.028895 | -0.249733 | 0.309358 | -0.091397 | 0.100546 | 0.170847 | 0.016443 | -0.059365 | 0.178714 | 0.049654 | 0.010953 | 0.039921 | -0.006115 |
| 18.77777778 | -0.10111 | 0.009141 | -0.102675 | -0.139682 | 0.131946 | -0.021406 | -0.022992 | -0.048669 | -0.00455 | 0.021368 | -0.012704 | -0.042145 | -0.09795 | 0.09416 | -0.151815 | 0.080235 | 0.119972 | -0.003763 | -0.13099 | 0.112107 | 0.047503 | -0.09228 | -0.01576 | -0.041311 |
| 18.79166667 | -0.704892 | -0.052097 | -0.133861 | -0.16161 | 0.116204 | -0.05474 | -0.073261 | -0.086453 | -0.068317 | -0.001748 | -0.091551 | -0.04727 | -0.072254 | -0.07909 | -0.162681 | 0.075367 | 0.055182 | -0.026499 | -0.104194 | 0.050925 | 0.043735 | -0.175392 | -0.046868 | -0.03191 |
| 18.80555556 | -1.083614 | -0.01642 | -0.13089 | -0.199815 | 0.183379 | -0.074268 | -0.05179 | -0.060894 | -0.139754 | -0.070258 | -0.049648 | -0.025604 | -0.295706 | 0.069441 | -0.102876 | 0.093509 | 0.059938 | -0.03107 | -0.033987 | 0.088245 | 0.063368 | -0.13879 | -0.010817 | -0.034 |
| 18.81944444 | -0.49468 | 0.087415 | -0.095981 | -0.28482 | 0.17326 | -0.033715 | 0.018124 | 0.035614 | -0.104738 | -0.068764 | 0.075321 | 0.009222 | -0.573986 | 0.238114 | -0.067868 | 0.112107 | 0.117585 | -0.012996 | 0.056999 | 0.164111 | 0.096466 | -0.090987 | 0.035872 | -0.064704 |
| 18.83333333 | -0.489 | 0.041633 | -0.10187 | -0.414991 | 0.196799 | 0.001527 | 0.033864 | 0.097876 | -0.102863 | -0.044221 | 0.146168 | 0.016505 | -0.585949 | 0.124916 | -0.07254 | 0.102365 | 0.206196 | -0.000519 | 0.073165 | 0.13608 | 0.097549 | -0.118759 | -0.015421 | -0.155749 |
| 18.84722222 | -0.515308 | -0.143849 | -0.19711 | -0.555132 | 0.572254 | -0.024751 | -0.048476 | 0.077195 | -0.2332 | -0.110075 | 0.042558 | -0.0201 | -0.368562 | -0.154016 | -0.109892 | 0.02575 | 0.175458 | -0.018697 | -0.00337 | 0.049544 | 0.054596 | -0.15585 | -0.16148 | -0.312793 |
| 18.86111111 | -0.533025 | -0.25934 | -0.267944 | -0.601971 | 0.44018 | -0.043243 | -0.122028 | 0.046237 | -0.36095 | -0.16204 | -0.114012 | -0.080122 | -0.302424 | -0.396376 | -0.159667 | -0.058717 | -0.104638 | -0.047426 | -0.051982 | -0.001077 | 0.00683 | -0.171549 | -0.337052 | -0.490714 |
| 18.875 | -0.523712 | -0.263378 | -0.301543 | -0.595452 | 0.210345 | -0.020183 | -0.139821 | 0.037496 | -0.339556 | -0.202354 | -0.144248 | -0.130386 | -0.516969 | -0.478542 | -0.261597 | -0.133069 | -0.318948 | -0.064679 | -0.080007 | -0.060578 | -0.032732 | -0.205761 | -0.410953 | -0.515061 |
| 18.88888889 | -0.476569 | -0.347751 | -0.332787 | -0.658324 | 0.048907 | -0.008816 | -0.161519 | 0.018239 | -0.296372 | -0.230723 | -0.09746 | -0.130263 | -0.859869 | -0.354737 | -0.316568 | -0.199831 | -0.445982 | -0.064517 | -0.094836 | -0.14392 | -0.060595 | -0.246808 | -0.425147 | -0.480007 |
| 18.90277778 | -0.53347 | -0.415508 | -0.358106 | -0.714648 | -0.11186 | -0.007895 | -0.175606 | -0.014159 | -0.260392 | -0.261629 | -0.113044 | -0.156112 | -0.877552 | -0.078711 | -0.367028 | -0.249983 | -0.655185 | -0.052445 | -0.10765 | -0.223336 | -0.039657 | -0.195953 | -0.465167 | -0.497596 |
| 18.91666667 | -0.610199 | -0.491467 | -0.38069 | -0.806962 | -0.321798 | 0.003973 | -0.169664 | -0.023638 | -0.295367 | -0.284088 | -0.182685 | -0.254479 | -0.659459 | 0.098814 | -0.34255 | -0.260439 | -0.990178 | -0.040765 | -0.108605 | -0.201733 | 0.002896 | -0.17053 | -0.500024 | -0.567779 |
| 18.93055556 | -0.54553 | -0.475122 | -0.44312 | -0.80563 | -0.467715 | 0.080628 | -0.08627 | 0.070996 | -0.300055 | -0.271826 | -0.105715 | -0.235607 | -0.607241 | 0.226492 | -0.313515 | -0.265152 | -1.123477 | 0.020348 | -0.022891 | -0.179225 | 0.076286 | -0.158049 | -0.387747 | -0.595785 |
| 18.94444444 | -0.360087 | -0.50716 | -0.476661 | -0.692707 | -0.557997 | 0.22034 | 0.069344 | 0.182274 | -0.22904 | -0.23657 | 0.052654 | 0.034721 | -0.810386 | 0.182072 | -0.226176 | -0.259731 | -0.733379 | 0.149966 | 0.110693 | -0.244963 | 0.142084 | -0.093703 | -0.270862 | -0.744291 |
| 18.95833333 | -0.397452 | -0.788028 | -0.494 | -0.556461 | -1.003536 | 0.233809 | 0.032415 | -0.039188 | -0.268847 | -0.340449 | 0.045052 | 0.259984 | -1.137501 | -0.579002 | -0.231784 | -0.420778 | -0.663084 | 0.145302 | -0.022178 | -0.30889 | 0.072289 | -0.172635 | -0.341274 | -1.50091 |
| 18.97222222 | -0.675549 | -1.102343 | -0.61114 | -0.624617 | -1.514788 | 0.063627 | -0.199256 | -0.438187 | -0.289985 | -0.511159 | -0.118009 | 0.154568 | -1.462576 | -3.467248 | -0.392989 | -0.697156 | -1.239867 | -0.141563 | -0.670489 | -0.690444 | -0.205049 | -0.404605 | -0.595452 | -2.722185 |
| 18.98611111 | -0.726902 | -1.467581 | -0.51678 | -0.593495 | -1.200529 | -0.066471 | -0.293304 | -0.386212 | -0.249066 | -0.436478 | -0.368269 | -0.056167 | -1.347711 | -6.376339 | -0.462176 | -0.767039 | -1.602987 | -0.229241 | -0.665848 | -1.079322 | -0.420396 | -0.564861 | -0.413393 | -2.504231 |
| 19 | -0.442321 | -0.414708 | -0.229451 | -0.243933 | -0.412086 | -0.075515 | -0.227773 | -0.208279 | -0.145117 | -0.231088 | -0.208063 | -0.097094 | -0.624177 | -3.38806 | -0.243724 | -0.390122 | -1.193407 | -0.065631 | -0.229428 | -0.880768 | -0.235994 | -0.516862 | -0.217056 | -0.866492 |
| 19.01388889 | -0.095686 | 0.166002 | -0.153268 | -0.065671 | -0.020204 | -0.068286 | -0.170604 | -0.144414 | -0.077333 | -0.144715 | -0.041907 | -0.084061 | -0.008495 | -0.583483 | -0.044609 | -0.069129 | -0.633727 | 0.038392 | 0.064981 | -0.325306 | 0.069322 | -0.030524 | -0.125693 | 0.388674 |
| 19.02777778 | 0.026513 | -0.06313 | -0.263765 | -0.182056 | -0.213615 | -0.084073 | -0.158616 | -0.158638 | -0.070747 | -0.186001 | -0.072464 | -0.155853 | 0.216715 | 0.6354 | -0.031874 | -0.066937 | -0.412067 | 0.055593 | 0.194839 | -0.022015 | 0.150429 | 0.238114 | -0.082856 | 0.877432 |
| 19.04166667 | -0.073887 | -0.213436 | -0.288805 | -0.254269 | -0.350703 | -0.068499 | -0.132795 | -0.13545 | -0.039897 | -0.166983 | -0.057584 | -0.192223 | 0.211063 | 1.336886 | -0.070908 | -0.141869 | -0.279462 | 0.062466 | 0.279697 | 0.201305 | 0.120829 | 0.203678 | -0.045256 | 0.864247 |
| 19.05555556 | -0.138736 | -0.165734 | -0.184727 | -0.184381 | -0.156217 | -0.029529 | -0.081209 | -0.062088 | 0.039682 | -0.052987 | 0.073897 | -0.121199 | 0.192437 | 2.380321 | -0.104855 | -0.142315 | -0.18778 | 0.096691 | 0.441802 | 0.447274 | 0.078427 | 0.161752 | -0.009744 | 0.721848 |
| 19.06944444 | -0.097623 | 0.010223 | -0.104532 | -0.081016 | -0.064681 | -0.013194 | -0.048733 | -0.009988 | 0.106611 | 0.034002 | 0.19666 | -0.054506 | 0.203218 | 3.524611 | -0.163545 | -0.152099 | -0.161638 | 0.131044 | 0.571326 | 0.528011 | 0.0362 | 0.192874 | -0.000626 | 0.584359 |
| 19.08333333 | -0.014133 | 0.200466 | -0.092185 | -0.023525 | -0.197567 | -0.008506 | -0.040988 | 0.002083 | 0.097569 | 0.057339 | 0.30651 | -0.036448 | 0.226078 | 4.345357 | -0.232765 | -0.206578 | 0.030003 | 0.165997 | 0.617466 | 0.494569 | 0.0097 | 0.335201 | 0.006456 | 0.652896 |
| 19.09722222 | 0.057501 | 0.108701 | -0.121698 | -0.026764 | -0.16961 | -0.009976 | -0.055561 | 0.027697 | 0.061055 | 0.026527 | 0.243928 | -0.038642 | 0.255836 | 3.864101 | -0.315682 | -0.329771 | 0.308934 | 0.162387 | 0.552494 | 0.451922 | -0.007507 | 0.340687 | 0.015953 | 0.680469 |
| 19.11111111 | -0.006969 | -0.350766 | -0.238244 | -0.165255 | 0.024964 | -0.069861 | -0.129173 | -0.062457 | -0.019465 | -0.066942 | -0.067901 | -0.091243 | 0.208717 | 3.849712 | -0.370711 | -0.386257 | 0.420931 | 0.078735 | 0.364794 | 0.144627 | -0.031475 | 0.10875 | 0.00223 | 0.476901 |
| 19.125 | -0.140224 | -0.845032 | -0.369392 | -0.320751 | -0.020531 | -0.160058 | -0.224772 | -0.181272 | -0.106643 | -0.206637 | -0.46199 | -0.166171 | 0.102759 | 1.907102 | -0.319319 | -0.380063 | 0.241493 | -0.04522 | 0.13032 | -0.245227 | -0.048899 | -0.192318 | -0.051544 | 0.054994 |
| 19.13888889 | -0.194118 | -0.936601 | -0.32068 | -0.307903 | -0.14715 | -0.108807 | -0.172391 | -0.13531 | -0.099261 | -0.176751 | -0.606923 | -0.115163 | 0.005412 | 0.852711 | -0.26353 | -0.331465 | 0.084731 | -0.108369 | -0.009706 | -0.385253 | -0.059127 | -0.413685 | -0.110939 | -0.254301 |
| 19.15277778 | -0.136055 | -0.252153 | -0.138534 | 0.001019 | 0.033422 | 0.03019 | -0.002512 | 0.021947 | 0.063078 | 0.034594 | -0.121065 | 0.014055 | 0.006228 | 1.148384 | -0.30345 | -0.24779 | 0.187678 | -0.049626 | 0.078015 | -0.095971 | -0.107298 | -0.227 | -0.124104 | 0.162761 |
| 19.16666667 | -0.022892 | 0.211733 | -0.019847 | 0.218373 | 0.399729 | 0.062775 | 0.101643 | 0.053348 | 0.237434 | 0.21996 | 0.319891 | 0.052826 | 0.092669 | 1.607653 | -0.357411 | -0.1622 | 0.416871 | 0.037374 | 0.212722 | 0.286421 | -0.153569 | 0.069449 | -0.114688 | 0.801821 |
| 19.18055556 | 0.152286 | 0.070415 | 0.09078 | 0.214704 | 0.516974 | 0.018824 | 0.112398 | -0.012682 | 0.291048 | 0.270281 | 0.224591 | 0.038469 | 0.145028 | 1.124898 | -0.346094 | 0.020814 | 0.351142 | 0.053845 | 0.196841 | 0.39542 | -0.154451 | 0.258598 | -0.118212 | 0.863833 |
| 19.19444444 | 0.131988 | -0.181283 | 0.145458 | 0.163078 | 0.360088 | -0.002709 | 0.088216 | -0.040633 | 0.205728 | 0.216319 | 0.247682 | 0.021342 | 0.087624 | 0.639906 | -0.32607 | 0.170492 | 0.086886 | 0.019108 | 0.107549 | 0.106994 | -0.112192 | 0.221389 | -0.144652 | 0.354641 |
| 19.20833333 | -0.053106 | -1.331403 | 0.033788 | -0.052281 | 0.296014 | 0.007915 | 0.06004 | 0.018642 | 0.157661 | 0.162367 | 0.312338 | 0.011987 | 0.047284 | 0.433666 | -0.304571 | 0.137444 | -0.035563 | -0.006417 | 0.05502 | 0.063792 | -0.121476 | 0.169885 | -0.15986 | 0.002742 |
| 19.22222222 | -0.066341 | -0.317445 | -0.028419 | 0.032787 | 0.264725 | 0.012877 | 0.03169 | 0.089593 | 0.157594 | 0.125199 | 0.224172 | 0.023611 | 0.147438 | 0.411886 | -0.225682 | 0.072635 | -0.055156 | -0.019137 | 0.056998 | 0.350089 | -0.137648 | 0.227838 | -0.152544 | 0.046538 |
| 19.23611111 | 0.128925 | 0.443111 | -0.073108 | 0.114618 | 0.218936 | -0.012691 | -0.003345 | 0.101047 | 0.105488 | 0.081467 | 0.25772 | 0.052656 | 0.343608 | 0.46207 | -0.056463 | 0.074068 | 0.068581 | -0.023901 | 0.0706 | 0.525643 | -0.155594 | 0.319931 | -0.150023 | 0.313236 |
| 19.25 | 0.203908 | 0.968 | -0.120664 | 0.132856 | 0.202113 | -0.03963 | -0.016475 | 0.116431 | 0.002455 | 0.051341 | 0.454011 | 0.088097 | 0.479103 | 0.565448 | 0.086078 | 0.052317 | 0.310001 | -0.030914 | 0.077649 | 0.642631 | -0.127552 | 0.337216 | -0.142821 | 0.607079 |
| 19.26388889 | 0.039875 | 1.327887 | -0.13552 | 0.159716 | 0.210933 | -0.026325 | 0.014464 | 0.190555 | -0.064144 | 0.066894 | 0.812456 | 0.111041 | 0.432532 | 0.739271 | 0.141177 | 0.017785 | 0.209319 | -0.066338 | 0.08568 | 1.004866 | -0.061373 | 0.37705 | -0.069156 | 1.005597 |
| 19.27777778 | -0.041088 | 1.201563 | -0.133921 | 0.150948 | 0.256323 | 0.014092 | 0.042483 | 0.188278 | -0.085879 | 0.055582 | 0.886338 | 0.095318 | 0.306499 | 0.790725 | 0.158019 | -0.005211 | -0.11828 | -0.06993 | 0.106177 | 0.472298 | 0.028459 | 0.392605 | 0.089174 | 1.089029 |
| 19.29166667 | 0.035532 | 0.759645 | -0.104996 | 0.064324 | 0.3036 | 0.030583 | 0.042628 | 0.088666 | -0.036237 | 0.016177 | 0.531654 | 0.06418 | 0.279553 | 0.651956 | 0.200519 | 0.009479 | -0.149345 | -0.03952 | 0.096923 | 0.523911 | 0.090771 | 0.406651 | 0.124807 | 0.662234 |
| 19.30555556 | 0.17518 | 0.385772 | -0.081055 | 0.038685 | 0.362322 | 0.030588 | 0.035388 | 0.026507 | 0.045721 | -0.001458 | 0.277052 | 0.077856 | 0.384311 | 0.446813 | 0.207027 | 0.039325 | 0.074044 | -0.016361 | 0.06074 | -0.029888 | 0.095779 | 0.351302 | 0.050542 | 0.276618 |
| 19.31944444 | 0.377078 | 0.296873 | -0.035141 | 0.061501 | 0.46496 | 0.040924 | 0.028394 | 0.048388 | 0.141252 | 0.05908 | 0.29539 | 0.143724 | 0.516902 | 0.28912 | 0.195644 | 0.023515 | 0.059841 | -0.012537 | 0.025188 | -0.510968 | 0.097695 | 0.239021 | -0.033393 | 0.168052 |
| 19.33333333 | 0.50702 | 0.338612 | 0.020051 | 0.07872 | 0.535879 | 0.062771 | 0.007011 | 0.078774 | 0.254456 | 0.20134 | 0.353122 | 0.197658 | 0.611534 | 0.294943 | 0.232977 | -0.016404 | -0.013962 | -0.019623 | -0.00028 | -0.765649 | 0.143215 | 0.116649 | -0.058994 | 0.192211 |
| 19.34722222 | 0.447892 | 0.495089 | 0.072016 | 0.126916 | 0.525912 | 0.064059 | -0.006835 | 0.103597 | 0.327179 | 0.330081 | 0.372109 | 0.214447 | 0.634671 | 0.354129 | 0.242232 | -0.04127 | 0.211228 | -0.023255 | -0.002599 | -0.210837 | 0.23755 | 0.175645 | -0.005564 | 0.292125 |
| 19.36111111 | 0.162947 | 0.518191 | 0.159314 | 0.17953 | 0.488112 | 0.042462 | 0.026431 | 0.177261 | 0.367354 | 0.371752 | 0.376025 | 0.176378 | 0.52097 | 0.486016 | 0.232025 | -0.072492 | 0.500344 | -0.014424 | 0.020845 | 0.617885 | 0.342193 | 0.378357 | 0.096545 | 0.430116 |
| 19.375 | 0.15465 | 0.473943 | 0.245217 | 0.169125 | 0.624026 | 0.051707 | 0.090658 | 0.268242 | 0.369636 | 0.378898 | 0.417022 | 0.202981 | 0.44638 | 0.685909 | 0.188393 | -0.143983 | 0.372234 | -0.00346 | 0.055854 | 1.040625 | 0.356489 | 0.48532 | 0.211188 | 0.515554 |
| 19.38888889 | 0.469791 | 0.369966 | 0.293292 | 0.140511 | 0.793336 | 0.06228 | 0.164062 | 0.283812 | 0.366435 | 0.365143 | 0.419937 | 0.220042 | 0.493107 | 0.978165 | 0.240108 | -0.136512 | 0.157649 | -0.0057 | 0.066097 | 1.00491 | 0.336288 | 0.406811 | 0.311213 | 0.532836 |
| 19.40277778 | 0.58421 | 0.276048 | 0.342914 | 0.123292 | 0.661694 | 0.081676 | 0.215078 | 0.260493 | 0.310551 | 0.335558 | 0.415999 | 0.218166 | 0.550375 | 1.054035 | 0.247334 | -0.094839 | 0.306278 | -0.027896 | 0.037641 | 0.795603 | 0.324602 | 0.305677 | 0.368732 | 0.561395 |
| 19.41666667 | 0.508494 | 0.212428 | 0.372563 | 0.104214 | 0.424881 | 0.09772 | 0.207033 | 0.30504 | 0.217699 | 0.303709 | 0.366693 | 0.257018 | 0.507869 | 0.784441 | 0.197394 | -0.087156 | 0.572001 | -0.078481 | -0.011764 | 0.663539 | 0.335151 | 0.326359 | 0.420754 | 0.645546 |
| 19.43055556 | 0.23185 | 0.201239 | 0.480699 | -0.005692 | 0.367585 | -0.029056 | 0.074724 | 0.246386 | 0.045353 | 0.17499 | 0.248799 | 0.191782 | 0.465603 | 0.993589 | 0.23579 | -0.019821 | 0.58421 | -0.16709 | -0.038592 | 0.728409 | 0.235447 | 0.28491 | 0.489864 | 0.893538 |
| 19.44444444 | 0.179072 | 0.099795 | 0.490889 | -0.157825 | 0.443831 | -0.289847 | -0.068799 | 0.135545 | -0.140884 | 0.019423 | 0.124455 | 0.002276 | 0.444148 | 1.220246 | 0.431742 | 0.157168 | 0.460641 | -0.24525 | -0.016117 | 0.815741 | 0.081655 | 0.131121 | 0.517866 | 1.289179 |
| 19.45833333 | 0.275286 | 0.175746 | 0.431303 | -0.189316 | 0.425745 | -0.357318 | 0.008106 | 0.315193 | -0.086438 | 0.092966 | 0.187395 | -0.106354 | 0.415466 | 0.903282 | 0.504927 | 0.339382 | 0.532162 | -0.204663 | 0.106739 | 0.327225 | 0.042347 | 0.201442 | 0.572488 | 1.471856 |
| 19.47222222 | 0.431832 | 0.580859 | 0.597534 | 0.020393 | 0.376152 | -0.151804 | 0.237216 | 0.369768 | 0.133101 | 0.434332 | 0.379526 | 0.007235 | 0.405288 | 1.049932 | 0.472391 | 0.465726 | 0.668216 | -0.043052 | 0.310569 | 0.26655 | 0.197293 | 0.283708 | 0.865885 | 1.453962 |
| 19.48611111 | 0.629025 | 0.930599 | 0.822881 | 0.279986 | 0.509608 | 0.051157 | 0.402877 | 0.365709 | 0.28702 | 0.622547 | 0.544225 | 0.196764 | 0.513472 | 1.872416 | 0.538068 | 0.62432 | 0.703806 | 0.113532 | 0.460919 | 0.231027 | 0.383843 | 0.275885 | 1.101958 | 1.747298 |
| 19.5 | 0.718723 | 0.992103 | 0.872853 | 0.460653 | 0.570886 | 0.12014 | 0.546555 | 0.413696 | 0.304449 | 0.487907 | 0.630734 | 0.323855 | 0.874575 | 2.574486 | 0.675701 | 0.806931 | 0.898625 | 0.201645 | 0.553386 | 0.474948 | 0.451137 | 0.27088 | 0.902311 | 1.91935 |
| 19.51388889 | 0.670536 | 0.933788 | 0.638742 | 0.591876 | 0.490327 | 0.103962 | 0.619759 | 0.423095 | 0.233581 | 0.364389 | 0.555959 | 0.29814 | 0.899611 | 1.501259 | 0.817993 | 0.822234 | 1.098129 | 0.319218 | 0.628092 | 0.668313 | 0.330585 | 0.31716 | 0.593142 | 1.737491 |
| 19.52777778 | 0.52447 | 0.863927 | 0.40808 | 0.629642 | 0.357903 | 0.055829 | 0.536177 | 0.328295 | 0.203311 | 0.323156 | 0.455247 | 0.215755 | 0.693545 | 0.683458 | 0.787774 | 0.718016 | 1.102874 | 0.481699 | 0.65926 | 0.648929 | 0.205283 | 0.369518 | 0.392276 | 1.223465 |
| 19.54166667 | 0.534328 | 0.789072 | 0.310044 | 0.559754 | 0.261945 | 0.046426 | 0.488124 | 0.311138 | 0.189611 | 0.306488 | 0.401577 | 0.171545 | 0.586264 | 0.583719 | 0.677253 | 0.517736 | 0.892634 | 0.486518 | 0.577559 | 0.570559 | 0.210049 | 0.394638 | 0.336978 | 0.710121 |
| 19.55555556 | 0.480864 | 0.591843 | 0.281828 | 0.417377 | 0.206204 | 0.062081 | 0.508136 | 0.352508 | 0.16491 | 0.294546 | 0.363328 | 0.141424 | 0.436692 | 1.156945 | 0.541241 | 0.365484 | 0.627048 | 0.406918 | 0.463707 | 0.52637 | 0.254244 | 0.285026 | 0.333142 | 0.453568 |
| 19.56944444 | 0.427492 | 0.459091 | 0.325024 | 0.29553 | 0.237936 | 0.070255 | 0.460073 | 0.308181 | 0.159297 | 0.282862 | 0.420203 | 0.114561 | 0.238798 | 1.56817 | 0.406078 | 0.269471 | 0.74754 | 0.304917 | 0.363966 | 0.544602 | 0.296961 | 0.253488 | 0.43235 | 0.302801 |
| 19.58333333 | 0.425043 | 0.504435 | 0.395509 | 0.268892 | 0.294098 | 0.062871 | 0.308785 | 0.232356 | 0.155231 | 0.304698 | 0.473329 | 0.150826 | 0.125711 | 1.558745 | 0.387908 | 0.19797 | 0.943899 | 0.253241 | 0.320174 | 0.523148 | 0.33045 | 0.392201 | 0.541589 | 0.387902 |
| 19.59722222 | 0.401315 | 0.503167 | 0.361711 | 0.366413 | 0.375426 | 0.056668 | 0.249 | 0.194556 | 0.157676 | 0.324585 | 0.41472 | 0.151818 | 0.093985 | 1.622289 | 0.385816 | 0.187541 | 0.643871 | 0.24106 | 0.256402 | 0.446434 | 0.187318 | 0.468359 | 0.555784 | 0.580983 |
| 19.61111111 | 0.371854 | 0.395059 | 0.264251 | 0.480339 | 0.483524 | 0.053243 | 0.230863 | 0.134989 | 0.176218 | 0.331073 | 0.313776 | 0.140277 | 0.088436 | 1.836712 | 0.35573 | 0.209101 | 0.527821 | 0.255814 | 0.234342 | 0.474573 | 0.154832 | 0.146801 | 0.533135 | 0.621576 |
| 19.625 | 0.371502 | 0.242665 | 0.183214 | 0.514483 | 0.531532 | 0.046896 | 0.178289 | 0.047365 | 0.196117 | 0.326742 | 0.244177 | 0.117366 | 0.110379 | 1.558365 | 0.25578 | 0.207445 | 0.700006 | 0.279471 | 0.248507 | 0.575863 | 0.23001 | 0.062358 | 0.588571 | 0.670188 |
| 19.63888889 | 0.366687 | 0.04869 | 0.213997 | 0.535476 | 0.533806 | 0.025737 | 0.129021 | -0.001899 | 0.189514 | 0.28983 | 0.188232 | 0.126079 | 0.128146 | 1.39276 | 0.139267 | 0.148231 | 0.833584 | 0.329248 | 0.30483 | 0.622841 | 0.298818 | 0.346896 | 0.597676 | 0.641967 |
| 19.65277778 | 0.244373 | -0.072052 | 0.213452 | 0.540321 | 0.561522 | 0.007835 | 0.118519 | -0.013132 | 0.128319 | 0.245973 | 0.161378 | 0.129319 | 0.067916 | 1.149145 | 0.022162 | 0.078138 | 0.593471 | 0.397806 | 0.37327 | 0.593437 | 0.250332 | 0.642195 | 0.595335 | 0.650947 |
| 19.66666667 | 0.074292 | 0.150928 | 0.142361 | 0.536191 | 0.593039 | 0.000672 | 0.088079 | -0.017491 | 0.065994 | 0.204561 | 0.23052 | 0.069716 | -0.032925 | 1.000593 | -0.077713 | 0.0058 | 0.416907 | 0.402601 | 0.394453 | 0.507421 | 0.161115 | 0.305662 | 0.601199 | 0.651897 |
| 19.68055556 | -0.019307 | 0.544775 | 0.124965 | 0.504274 | 0.505312 | 0.008002 | 0.003695 | -0.051506 | 0.025043 | 0.202232 | 0.317643 | 0.071984 | -0.104734 | 1.009862 | -0.159619 | -0.071035 | 0.417982 | 0.387354 | 0.338421 | 0.429062 | 0.157961 | -0.164421 | 0.535002 | 0.68585 |
| 19.69444444 | 0.08705 | 0.524252 | 0.134285 | 0.360546 | 0.478202 | 0.019577 | -0.112613 | -0.131576 | 0.003005 | 0.222903 | 0.289076 | 0.151032 | -0.167167 | 0.868393 | -0.215909 | -0.16302 | 0.427661 | 0.332797 | 0.306837 | 0.421028 | 0.184377 | -0.205416 | 0.458709 | 0.654057 |
| 19.70833333 | 0.146541 | 0.210953 | 0.117094 | 0.230622 | 0.527521 | 0.016921 | -0.20962 | -0.217745 | -0.031459 | 0.221923 | 0.122625 | 0.194117 | -0.230728 | 0.666538 | -0.239962 | -0.227913 | 0.255883 | 0.265745 | 0.233361 | 0.488817 | 0.162673 | 0.066951 | 0.428316 | 0.585383 |
| 19.72222222 | -0.007717 | 0.155121 | 0.089177 | 0.152745 | 0.561286 | -0.003245 | -0.241909 | -0.262573 | -0.060565 | 0.168497 | 0.105172 | 0.135394 | -0.262277 | 0.791534 | -0.264376 | -0.244113 | 0.060825 | 0.163841 | 0.143433 | 0.505325 | 0.129869 | 0.06231 | 0.400502 | 0.339724 |
| 19.73611111 | -0.226788 | 0.296399 | 0.115646 | 0.112657 | 0.503864 | -0.031176 | -0.227201 | -0.250125 | -0.055221 | 0.113293 | 0.206248 | 0.077307 | -0.307236 | 1.120818 | -0.262509 | -0.2495 | 0.063551 | 0.112829 | 0.119474 | 0.443114 | 0.169254 | -0.224132 | 0.284685 | 0.165929 |
| 19.75 | -0.281408 | 0.395071 | 0.191448 | -0.05079 | 0.397154 | -0.04588 | -0.179884 | -0.286855 | -0.047108 | 0.077705 | 0.271717 | 0.090182 | -0.391318 | 1.057442 | -0.331597 | -0.26279 | 0.158094 | 0.083474 | 0.114168 | 0.320544 | 0.198293 | -0.249837 | 0.163376 | 0.107455 |
| 19.76388889 | -0.366823 | 0.366518 | 0.189235 | -0.224658 | 0.21429 | -0.050462 | -0.156807 | -0.31176 | -0.064096 | 0.02752 | 0.290238 | 0.117199 | -0.499342 | 0.591222 | -0.349547 | -0.307586 | 0.076327 | 0.048766 | 0.095848 | 0.152619 | 0.213471 | -0.063842 | 0.09984 | -0.10986 |
| 19.77777778 | -0.538012 | 0.028267 | 0.071403 | -0.363287 | 0.040988 | -0.07612 | -0.163834 | -0.28774 | -0.099109 | -0.043726 | 0.067651 | 0.129471 | -0.60738 | -0.111586 | -0.350856 | -0.350178 | -0.335285 | -0.003414 | 0.066177 | 0.026889 | 0.214693 | 0.071471 | 0.079347 | -0.487672 |
| 19.79166667 | -0.475429 | -0.02406 | -0.01944 | -0.406616 | -0.050707 | -0.124006 | -0.15586 | -0.24202 | -0.141093 | -0.099081 | 0.123778 | 0.148662 | -0.67143 | -0.115524 | -0.336078 | -0.369138 | -0.369437 | -0.052123 | 0.020815 | -0.040329 | 0.142855 | -0.226159 | 0.106294 | -0.717621 |
| 19.80555556 | -0.391662 | -0.045013 | -0.025355 | -0.475814 | -0.13246 | -0.141767 | -0.160767 | -0.302862 | -0.19536 | -0.150584 | 0.20428 | 0.174823 | -0.745033 | -0.024309 | -0.427341 | -0.286726 | -0.112035 | -0.084254 | -0.006921 | -0.04627 | 0.161002 | -0.332348 | 0.120239 | -0.916655 |
| 19.81944444 | -0.487479 | 0.21451 | -0.042917 | -0.602193 | -0.271707 | -0.126513 | -0.134306 | -0.310355 | -0.237809 | -0.21558 | 0.312049 | 0.172731 | -0.708118 | -0.406857 | -0.504411 | -0.203538 | -0.026054 | -0.102598 | -0.032674 | -0.086423 | 0.15502 | -0.222008 | 0.110739 | -1.187129 |
| 19.83333333 | -0.785166 | 0.326583 | -0.159079 | -0.714147 | -0.409351 | -0.143965 | -0.119625 | -0.323789 | -0.22434 | -0.298974 | 0.354458 | 0.138269 | -0.755608 | -0.938597 | -0.522497 | -0.19136 | -0.609306 | -0.169512 | -0.043248 | -0.143653 | 0.101261 | -0.347567 | 0.078588 | -1.278932 |
| 19.84722222 | -0.960811 | 0.116925 | -0.252845 | -0.810048 | -0.393476 | -0.1167 | -0.139749 | -0.313836 | -0.198976 | -0.367687 | 0.278793 | 0.050129 | -0.908095 | -1.335063 | -0.391536 | -0.264876 | -0.905975 | -0.242973 | -0.079733 | -0.16822 | 0.0882 | -0.524301 | -0.062524 | -1.266789 |
| 19.86111111 | -0.940442 | 0.055459 | -0.279699 | -0.856937 | -0.434827 | -0.049166 | -0.178389 | -0.262693 | -0.193227 | -0.449614 | 0.275308 | -0.0392 | -1.147028 | -1.233551 | -0.333813 | -0.367643 | -0.766979 | -0.269783 | -0.108205 | -0.165196 | 0.04638 | -0.445504 | -0.175088 | -1.479859 |
| 19.875 | -1.017414 | 0.055455 | -0.306711 | -0.905867 | -0.64342 | -0.008338 | -0.220058 | -0.331317 | -0.217946 | -0.498861 | 0.220061 | -0.161919 | -1.115164 | -1.030956 | -0.404313 | -0.430402 | -0.650365 | -0.302298 | -0.164276 | -0.223186 | -0.032524 | -0.313911 | -0.36882 | -1.376051 |
| 19.88888889 | -1.170823 | -0.166691 | -0.437507 | -0.94302 | -0.845286 | -0.049126 | -0.283627 | -0.425159 | -0.258414 | -0.550307 | -0.020806 | -0.17369 | -1.143909 | -1.165578 | -0.49116 | -0.487351 | -0.971989 | -0.328605 | -0.212928 | -0.479899 | -0.141198 | -0.32913 | -0.538187 | -1.209684 |
| 19.90277778 | -1.448421 | -0.608519 | -0.55816 | -0.979424 | -0.911984 | -0.083568 | -0.34835 | -0.591236 | -0.280354 | -0.573074 | -0.340918 | -0.094919 | -1.428914 | -1.722139 | -0.57123 | -0.532611 | -1.345834 | -0.413397 | -0.299902 | -0.68286 | -0.182046 | -0.359046 | -0.788203 | -1.218366 |
| 19.91666667 | -1.74172 | -0.992851 | -0.673374 | -0.935689 | -1.002251 | -0.066343 | -0.357324 | -0.448546 | -0.303869 | -0.579394 | -0.547233 | -0.086409 | -1.624734 | -1.92288 | -0.637717 | -0.622838 | -1.421025 | -0.46032 | -0.497391 | -0.792934 | -0.261221 | -0.432656 | -1.05427 | -1.17451 |
| 19.93055556 | -1.750584 | -1.174429 | -0.754753 | -0.854736 | -0.991572 | 0.053844 | -0.23812 | -0.219487 | -0.206782 | -0.54609 | -0.504634 | 0.008971 | -1.605106 | -1.729398 | -0.740324 | -0.678801 | -1.423726 | -0.491612 | -0.733136 | -0.732175 | -0.433758 | -0.432508 | -1.295444 | -1.143469 |
| 19.94444444 | -1.591834 | -1.304949 | -0.680834 | -0.766439 | -0.94131 | 0.246415 | -0.019691 | 0.01524 | -0.036384 | -0.476255 | -0.440252 | 0.129988 | -1.483533 | -1.174558 | -0.840842 | -0.660014 | -1.916806 | -0.43699 | -0.778262 | -0.497282 | -0.514656 | -0.284089 | -1.349017 | -1.021499 |
| 19.95833333 | -1.631872 | -1.238839 | -0.642626 | -0.726415 | -1.308112 | 0.187397 | -0.077677 | -0.126876 | -0.047443 | -0.56811 | -0.52566 | 0.03798 | -1.566114 | -1.077369 | -0.696443 | -0.655022 | -2.520901 | -0.449196 | -0.919397 | -0.649938 | -0.532636 | -0.33848 | -1.290484 | -1.030296 |
| 19.97222222 | -2.115481 | -1.526484 | -1.172789 | -0.811993 | -2.252487 | -0.079764 | -0.426416 | -0.364394 | -0.274117 | -1.025788 | -1.23656 | -0.51449 | -1.755574 | -1.516316 | -0.587533 | -0.662313 | -3.035679 | -0.69507 | -1.222891 | -1.582304 | -0.825802 | -0.281466 | -1.465556 | -1.313577 |
| 19.98611111 | -2.129129 | -1.837869 | -0.782739 | -0.838108 | -2.025534 | -0.169685 | -0.569144 | -0.204372 | -0.522365 | -0.99624 | -1.470921 | -0.618568 | -1.459184 | -1.826515 | -0.555066 | -0.864492 | -1.726213 | -0.666698 | -1.054137 | -1.73203 | -0.820279 | -0.18274 | -1.393152 | -1.332091 |
| 20 | -0.710071 | -0.46983 | 0.385363 | -0.560866 | -0.448417 | -0.075021 | -0.309044 | 0.083637 | -0.215527 | 0.134925 | -0.151903 | 0.107361 | -0.239245 | -1.310613 | -0.198409 | -0.841591 | 0.631109 | -0.293193 | -0.478769 | -0.569892 | -0.184474 | -0.026314 | -0.739701 | -0.383529 |
| 20.01388889 | 0.753543 | 0.933058 | 0.387769 | -0.014799 | 0.211455 | 0.029901 | 0.000384 | 0.016675 | 0.126086 | 0.838341 | 0.938255 | 0.365437 | 0.817062 | -0.103408 | -0.026335 | -0.775383 | 1.346327 | 0.052552 | -0.039504 | 0.016787 | 0.637698 | 0.122358 | 0.212925 | 0.215301 |
| 20.02777778 | 0.619589 | 1.350969 | -0.289533 | -0.047684 | -0.107563 | -0.018088 | 0.289441 | -0.06675 | 0.016406 | 0.308323 | 0.773233 | -0.155688 | 0.966068 | 0.298354 | -0.60609 | -0.802602 | 0.967607 | 0.172969 | 0.132575 | -0.1601 | 1.049211 | 0.116242 | 0.447415 | 0.025544 |
| 20.04166667 | 0.208029 | -0.329901 | -0.644021 | -0.686184 | -0.22368 | -0.030595 | 0.333629 | -0.035461 | -0.114096 | -0.000575 | -0.386891 | -0.320202 | 0.625766 | -0.117825 | -0.531078 | -0.726346 | 0.96278 | 0.113219 | 0.194452 | -0.725354 | 0.559389 | -0.047997 | 0.183436 | -0.1203 |
| 20.05555556 | 0.123373 | -1.530566 | -0.56886 | -0.829938 | -0.009068 | 0.009258 | 0.194505 | 0.138821 | -0.292859 | 0.062811 | -0.779391 | -0.092162 | 0.427228 | -0.902491 | 0.043077 | -0.503772 | 0.95833 | 0.088966 | 0.207765 | -0.783697 | 0.118816 | -0.19802 | 0.191834 | -0.194301 |
| 20.06944444 | 0.174551 | -1.476555 | -0.399042 | -0.364588 | 0.05068 | 0.013181 | 0.026329 | 0.194588 | -0.145464 | 0.065137 | -0.752062 | -0.06868 | 0.434353 | -1.341787 | 0.429688 | -0.264616 | 0.43106 | 0.484868 | 0.528728 | -0.012536 | -0.009061 | -0.208273 | 0.619266 | -0.209875 |
| 20.08333333 | 0.372022 | -0.341473 | -0.163687 | 0.082799 | 0.048287 | 0.007512 | -0.090224 | 0.065171 | 0.106882 | -0.039118 | -0.47125 | -0.038326 | 0.576859 | -1.65854 | 0.155447 | 0.051505 | -0.067299 | 0.60695 | 0.643579 | 0.4818 | 0.053783 | -0.050861 | 0.856132 | -0.134288 |
| 20.09722222 | 0.551416 | 0.863639 | 0.060073 | 0.116299 | 0.156123 | 0.013366 | -0.065616 | -0.021478 | 0.100952 | -0.01349 | 0.115682 | 0.145814 | 0.634543 | -1.226078 | 0.118944 | 0.275659 | 0.078526 | 0.479281 | 0.53174 | -0.243239 | 0.036275 | 0.02502 | 0.193595 | -0.03424 |
| 20.11111111 | 0.494734 | 1.221093 | 0.149427 | 0.082534 | 0.266198 | 0.012579 | 0.067897 | 0.059573 | -0.201517 | -0.029029 | 0.610345 | 0.24388 | 0.585339 | 0.490752 | 0.232027 | 0.250628 | 0.586251 | 0.388655 | 0.439939 | -0.342009 | -0.141281 | -0.072974 | -0.464984 | -0.133506 |
| 20.125 | 0.39584 | 0.851101 | 0.188199 | 0.282358 | 0.307022 | 0.003813 | 0.155612 | 0.263664 | -0.297252 | -0.02971 | 0.568762 | 0.183101 | 0.654346 | 0.908293 | 0.242685 | 0.213644 | 0.656527 | 0.304977 | 0.321932 | 0.431523 | -0.208072 | -0.19012 | -0.485261 | 0.056858 |
| 20.13888889 | 0.468325 | 0.484804 | 0.223933 | 0.503541 | 0.399436 | -0.002885 | 0.10365 | 0.428783 | -0.092209 | 0.055632 | 0.390467 | 0.154117 | 0.724797 | -0.616583 | 0.246521 | 0.184927 | 0.669262 | 0.202578 | 0.208615 | 0.393961 | -0.099331 | -0.20251 | -0.317683 | 0.152613 |
| 20.15277778 | 0.473163 | 0.24444 | 0.22613 | 0.339848 | 0.456993 | -0.020882 | 0.038974 | 0.382266 | -0.015122 | 0.066216 | 0.32814 | 0.186011 | 0.654238 | -1.382817 | 0.316019 | 0.217707 | 1.006542 | 0.171597 | 0.162124 | -0.094121 | -0.070814 | -0.186502 | -0.493745 | 0.077886 |
| 20.16666667 | 0.219771 | 0.176307 | 0.284324 | 0.20802 | 0.401524 | -0.036215 | 0.118336 | 0.330397 | -0.025186 | 0.154647 | 0.31738 | 0.205285 | 0.531831 | -0.596594 | 0.323423 | 0.231481 | 1.369423 | 0.164532 | 0.165053 | 0.133989 | -0.069431 | -0.100677 | -0.613205 | 0.044185 |
| 20.18055556 | 0.097631 | 0.413088 | 0.331032 | 0.219123 | 0.356843 | -0.039646 | 0.286557 | 0.369243 | 0.037825 | 0.307732 | 0.523447 | 0.261853 | 0.504982 | 0.127073 | 0.277541 | 0.194988 | 0.993192 | 0.107246 | 0.1303 | 0.485644 | 0.017407 | -0.125703 | -0.216386 | 0.037931 |
| 20.19444444 | 0.348392 | 0.84222 | 0.362234 | 0.240078 | 0.459633 | -0.022851 | 0.360378 | 0.458428 | 0.146682 | 0.467615 | 0.87141 | 0.354097 | 0.594655 | -0.119227 | 0.262867 | 0.178964 | 0.569469 | 0.092196 | 0.131932 | 0.319904 | 0.117494 | 0.017019 | 0.302532 | 0.165187 |
| 20.20833333 | 0.692394 | 0.908972 | 0.423612 | 0.077342 | 0.540313 | 0.000268 | 0.208307 | 0.491019 | 0.224877 | 0.453914 | 0.942452 | 0.408771 | 0.688877 | -0.990766 | 0.290302 | 0.211524 | 0.618655 | 0.146465 | 0.192647 | 0.363888 | 0.152701 | 0.213036 | 0.55158 | 0.423778 |
| 20.22222222 | 0.558928 | 0.631796 | 0.520229 | -0.002595 | 0.534429 | 0.000732 | 0.059637 | 0.532245 | 0.275889 | 0.433174 | 0.813201 | 0.447204 | 0.680796 | -0.700091 | 0.298033 | 0.275097 | 0.752927 | 0.149204 | 0.215313 | 0.869163 | 0.161554 | 0.172968 | 0.583679 | 0.541139 |
| 20.23611111 | 0.234233 | 0.49127 | 0.62181 | 0.195547 | 0.548192 | -0.007202 | -0.020628 | 0.557021 | 0.280172 | 0.575313 | 0.818377 | 0.507499 | 0.598593 | 0.054175 | 0.273126 | 0.30568 | 0.457192 | 0.110928 | 0.176802 | 1.047341 | 0.242384 | 0.227395 | 0.555062 | 0.387042 |
| 20.25 | 0.203888 | 0.555926 | 0.671513 | 0.455279 | 0.500805 | 0.007376 | 0.010212 | 0.546733 | 0.250369 | 0.530785 | 0.874817 | 0.517383 | 0.597545 | 0.198209 | 0.280462 | 0.342143 | 0.48618 | 0.079475 | 0.143262 | 0.754926 | 0.411011 | 0.37352 | 0.507422 | 0.281322 |
| 20.26388889 | 0.399678 | 0.599636 | 0.584657 | 0.374811 | 0.421584 | 0.013533 | 0.071432 | 0.482518 | 0.219488 | 0.331693 | 0.882011 | 0.501158 | 0.64652 | -0.073917 | 0.310198 | 0.33157 | 0.761451 | -0.002632 | 0.083216 | 0.707299 | 0.485508 | 0.459603 | 0.498408 | 0.324319 |
| 20.27777778 | 0.617619 | 0.447954 | 0.460055 | 0.084436 | 0.386961 | -0.005182 | 0.070316 | 0.564597 | 0.184478 | 0.252906 | 0.907693 | 0.473945 | 0.623306 | 0.130669 | 0.340101 | 0.309312 | 0.761639 | -0.039791 | 0.048861 | 0.990885 | 0.422697 | 0.156349 | 0.427844 | 0.27302 |
| 20.29166667 | 0.611168 | 0.128422 | 0.435675 | 0.016247 | 0.449905 | 0.020906 | 0.121822 | 0.765349 | 0.155638 | 0.367352 | 0.811255 | 0.450102 | 0.674419 | 0.819843 | 0.403372 | 0.357412 | 0.701343 | 0.073272 | 0.126536 | 0.771883 | 0.396455 | -0.242204 | 0.392025 | 0.249306 |
| 20.30555556 | 0.559993 | 0.000424 | 0.466065 | 0.213614 | 0.638099 | 0.105142 | 0.245999 | 0.662307 | 0.156064 | 0.465237 | 0.701909 | 0.419389 | 1.0195 | 1.285903 | 0.42635 | 0.441607 | 0.921215 | 0.240132 | 0.218232 | 0.296635 | 0.400434 | -0.457364 | 0.501928 | 0.386903 |
| 20.31944444 | 0.669442 | 0.048807 | 0.468264 | 0.468538 | 0.838925 | 0.157929 | 0.506389 | 0.682273 | 0.18454 | 0.562374 | 0.689444 | 0.436533 | 1.340174 | 1.214243 | 0.43232 | 0.520019 | 1.29235 | 0.296566 | 0.204173 | 0.330614 | 0.404634 | -0.130011 | 0.69265 | 0.544517 |
| 20.33333333 | 0.945188 | 0.040836 | 0.552698 | 0.496345 | 0.856637 | 0.198773 | 0.664403 | 0.693747 | 0.243971 | 0.735319 | 0.686604 | 0.43881 | 1.291471 | 1.366943 | 0.542549 | 0.522383 | 1.13444 | 0.330993 | 0.22479 | 0.726165 | 0.458621 | 0.407955 | 0.926658 | 0.539581 |
| 20.34722222 | 1.196913 | 0.049002 | 0.729869 | 0.35835 | 0.732064 | 0.329393 | 0.731525 | 0.655881 | 0.324473 | 0.8876 | 0.637943 | 0.425086 | 1.134969 | 2.142303 | 0.672324 | 0.517875 | 0.835974 | 0.379941 | 0.232144 | 0.932736 | 0.494391 | 0.809396 | 1.15231 | 0.507036 |
| 20.36111111 | 1.043363 | 0.190238 | 0.945855 | 0.440921 | 0.667457 | 0.157398 | 0.805134 | 0.638987 | 0.402351 | 0.937296 | 0.746238 | 0.406156 | 1.105327 | 2.694035 | 0.710996 | 0.518537 | 1.049061 | 0.377884 | 0.248089 | 1.147933 | 0.584613 | 0.842606 | 1.230229 | 0.659858 |
| 20.375 | 0.841497 | 0.365886 | 0.975482 | 0.829731 | 0.722252 | 0.123046 | 0.984458 | 0.605164 | 0.444703 | 1.048505 | 0.953412 | 0.467151 | 1.024467 | 2.34319 | 0.752824 | 0.691272 | 1.517366 | 0.343681 | 0.258433 | 1.546314 | 0.785403 | 0.856748 | 1.420849 | 1.048941 |
| 20.38888889 | 1.032245 | 0.576525 | 0.935151 | 0.986712 | 0.787765 | 0.125587 | 0.954061 | 0.46851 | 0.486909 | 1.045593 | 0.924919 | 0.512223 | 1.050895 | 2.366764 | 0.872026 | 0.803067 | 1.693587 | 0.150882 | 0.141201 | 1.656316 | 0.892163 | 0.930602 | 1.410966 | 1.32433 |

| | | | | | | | | | | | | | | | | | | | | | | | | |
|---|---|---|---|---|---|---|---|---|---|---|---|---|---|---|---|---|---|---|---|---|---|---|---|---|
| 20.40277778 | 1.486523 | 0.951267 | 0.920916 | 0.775447 | 0.755601 | 0.153793 | 0.699797 | 0.301987 | 0.51357 | 1.017648 | 0.775385 | 0.490995 | 1.170555 | 2.958315 | 0.991771 | 0.853074 | 1.57374 | -0.032598 | -0.286095 | 1.277813 | 0.803892 | 0.643822 | 1.370913 | 1.41262 |
| 20.41666667 | 1.526119 | 1.433157 | 0.8872 | 0.681989 | 0.789412 | 0.229799 | 0.573349 | 0.229137 | 0.530906 | 0.966888 | 0.788383 | 0.530768 | 1.220353 | 3.413891 | 0.93129 | 0.85747 | 1.815421 | 0.109908 | -0.943153 | 1.195385 | 0.598306 | 0.474048 | 1.273113 | 1.576768 |
| 20.43055556 | 1.116054 | 1.521279 | 0.657159 | 0.750747 | 0.864494 | 0.222656 | 0.462944 | 0.122543 | 0.336151 | 0.648369 | 0.778884 | 0.371917 | 1.155042 | 2.71341 | 0.860124 | 0.862437 | 2.004615 | 0.263574 | -0.823999 | 1.525442 | 0.423625 | 0.235592 | 1.21683 | 1.762377 |
| 20.44444444 | 0.737415 | 1.092184 | 0.382257 | 0.625539 | 0.730942 | -0.041333 | 0.304328 | -0.065171 | -0.050875 | 0.236482 | 0.356201 | 0.022624 | 0.863092 | 2.043279 | 0.941764 | 0.863414 | 1.676806 | 0.243851 | -0.237494 | 1.28715 | 0.365361 | -0.127259 | 0.999339 | 1.234225 |
| 20.45833333 | 0.68238 | 0.654955 | 0.293025 | 0.528294 | 0.591043 | -0.158088 | 0.214235 | -0.072456 | -0.128508 | 0.195766 | 0.187728 | -0.095566 | 0.42011 | 1.665518 | 0.880101 | 0.839407 | 1.244593 | 0.246405 | 0.080644 | 0.38579 | 0.235478 | -0.149808 | 0.801142 | 0.779358 |
| 20.47222222 | 0.827517 | 0.72861 | 0.465813 | 0.672478 | 0.589858 | 0.000898 | 0.267153 | 0.090818 | 0.10435 | 0.530998 | 0.278184 | 0.097274 | 0.082902 | 1.873804 | 0.787932 | 0.722775 | 1.012055 | 0.34904 | 0.190098 | 0.073702 | 0.274254 | 0.144437 | 0.820376 | 1.068623 |
| 20.48611111 | 0.981257 | 1.166958 | 0.73288 | 0.897115 | 0.639568 | 0.179343 | 0.394143 | 0.215592 | 0.320338 | 0.736064 | 0.26073 | 0.188178 | 0.256756 | 3.009095 | 0.982417 | 0.959178 | 0.893708 | 0.496807 | 0.341093 | 0.910313 | 0.419654 | 0.412737 | 1.19976 | 1.969517 |
| 20.5 | 1.317019 | 1.574309 | 0.849874 | 0.938715 | 0.951111 | 0.254597 | 0.541165 | 0.295995 | 0.345425 | 0.690423 | 0.196005 | 0.086655 | 0.692816 | 3.331147 | 1.484027 | 1.370845 | 0.750344 | 0.684364 | 0.618163 | 2.182702 | 0.605608 | 0.49991 | 1.193883 | 2.411284 |
| 20.51388889 | 1.550348 | 1.135388 | 0.731703 | 0.783839 | 0.997128 | 0.238969 | 0.513145 | 0.269359 | 0.277953 | 0.543036 | 0.158746 | 0.037349 | 0.79002 | 2.591592 | 1.714286 | 1.515415 | 0.732032 | 0.773837 | 0.655358 | 2.113863 | 0.69499 | 0.591918 | 1.01129 | 2.20764 |
| 20.52777778 | 1.034579 | 0.646532 | 0.642542 | 0.613418 | 0.674246 | 0.16171 | 0.31217 | 0.149313 | 0.197822 | 0.382093 | 0.392821 | -0.001761 | 0.516672 | 1.664198 | 1.513963 | 1.362216 | 0.696431 | 0.732399 | 0.607808 | 1.546055 | 0.591626 | 0.56663 | 0.720992 | 1.346036 |
| 20.54166667 | 0.79585 | 0.708111 | 0.391973 | 0.557735 | 0.299003 | 0.098855 | 0.163764 | 0.055047 | 0.117181 | 0.192743 | 0.58809 | 0.019542 | 0.528028 | 0.869359 | 1.225147 | 1.107666 | 0.516262 | 0.546715 | 0.694278 | 1.163805 | 0.486423 | 0.430853 | 0.465811 | 0.522662 |
| 20.55555556 | 0.64839 | 0.659123 | 0.086135 | 0.480108 | 0.144767 | 0.064569 | 0.064453 | 0.046431 | 0.079528 | 0.016055 | 0.562196 | 0.003157 | 0.551463 | 0.668058 | 1.008083 | 0.790868 | 0.466536 | 0.378856 | 0.793232 | 0.976166 | 0.400549 | 0.414948 | 0.296191 | 0.620983 |
| 20.56944444 | 0.475405 | 0.563897 | -0.035277 | 0.382994 | 0.141562 | 0.12008 | -0.026734 | 0.063374 | 0.064791 | -0.071771 | 0.642535 | -0.019952 | 0.536911 | 0.868177 | 0.917565 | 0.593528 | 0.500962 | 0.293488 | 0.816578 | 0.833993 | 0.304636 | 0.45664 | 0.160116 | 0.982159 |
| 20.58333333 | 0.43146 | 0.631463 | -0.04324 | 0.313766 | 0.178189 | 0.192801 | -0.14911 | 0.003905 | 0.075132 | -0.045446 | 0.829702 | 0.001446 | 0.527093 | 1.052001 | 0.737249 | 0.572161 | 0.689664 | 0.300987 | 0.761046 | 0.519276 | 0.19624 | 0.334786 | -0.038799 | 1.33813 |
| 20.59722222 | 0.454519 | 0.53355 | -0.040894 | 0.243578 | 0.206498 | 0.189284 | -0.279409 | -0.075104 | 0.039292 | -0.079621 | 0.666328 | 0.043578 | 0.446224 | 1.16465 | 0.394075 | 0.530495 | 0.579265 | 0.394331 | 0.644019 | 0.089291 | 0.141154 | 0.172231 | -0.141357 | 1.471598 |
| 20.61111111 | 0.409843 | 0.44157 | 0.006536 | 0.184134 | 0.177012 | 0.11779 | -0.265473 | -0.110084 | 0.033489 | -0.069849 | 0.533531 | 0.067661 | 0.274424 | 1.350768 | 0.037967 | 0.397694 | 0.371333 | 0.489517 | 0.639308 | -0.043293 | 0.154951 | -0.017547 | -0.041346 | 1.663306 |
| 20.625 | 0.412281 | 0.3489 | 0.167768 | 0.191856 | 0.170397 | 0.037635 | -0.221902 | -0.156527 | 0.061547 | -0.010198 | 0.550633 | 0.064789 | 0.231005 | 1.486759 | -0.085526 | 0.259646 | 0.260096 | 0.525437 | 0.642587 | 0.194306 | 0.16366 | -0.164181 | 0.114496 | 1.678147 |
| 20.63888889 | 0.490049 | 0.281301 | 0.331176 | 0.169957 | 0.183711 | -0.019849 | -0.180508 | -0.159528 | 0.041145 | 0.03572 | 0.535206 | 0.06184 | 0.163597 | 1.494666 | -0.071557 | 0.133174 | 0.071158 | 0.518835 | 0.737917 | 0.591463 | 0.162986 | -0.165775 | 0.19413 | 1.551903 |
| 20.65277778 | 0.497579 | 0.099357 | 0.410453 | 0.168611 | 0.217217 | -0.069264 | -0.077457 | -0.133511 | 0.036647 | 0.014642 | 0.404793 | 0.07137 | -0.007577 | 1.403851 | -0.005686 | 0.042345 | -0.199941 | 0.439864 | 0.608251 | 0.879942 | 0.190097 | -0.089556 | 0.149252 | 1.373134 |
| 20.66666667 | 0.400115 | -0.111774 | 0.413018 | 0.170031 | 0.244568 | -0.041339 | -0.002137 | -0.092926 | 0.039627 | 0.062652 | 0.318323 | 0.076426 | -0.11517 | 0.985421 | 0.07123 | -0.007556 | -0.563654 | 0.397734 | 0.531706 | 1.000053 | 0.169753 | -0.145921 | 0.029487 | 0.993574 |
| 20.68055556 | 0.406241 | -0.215032 | 0.373837 | 0.13879 | 0.248162 | 0.007976 | 0.005067 | -0.071633 | 0.01647 | 0.105116 | 0.227745 | 0.063389 | -0.138317 | 0.602873 | 0.075472 | -0.076759 | -0.671281 | 0.385433 | 0.436081 | 0.930675 | 0.127297 | -0.289549 | -0.039882 | 0.720813 |
| 20.69444444 | 0.344414 | -0.028787 | 0.333602 | 0.098019 | 0.247386 | 0.025028 | -0.08495 | -0.083782 | 0.004122 | 0.085607 | 0.287276 | 0.04561 | -0.307314 | 0.493535 | -0.062801 | -0.170667 | -0.526342 | 0.321452 | 0.313406 | 0.608622 | 0.138583 | -0.353668 | -0.009832 | 0.651979 |
| 20.70833333 | 0.196802 | 0.220756 | 0.26037 | 0.051074 | 0.243109 | 0.033653 | -0.135041 | -0.087984 | 0.020401 | 0.072819 | 0.385871 | 0.069304 | -0.461719 | 0.586961 | -0.27451 | -0.240588 | -0.422953 | 0.176504 | 0.193586 | 0.314226 | 0.199881 | -0.191576 | -0.036174 | 0.624047 |
| 20.72222222 | 0.28656 | 0.427457 | 0.140786 | 0.043839 | 0.238791 | 0.038426 | -0.090363 | -0.07407 | 0.023413 | 0.049439 | 0.430396 | 0.111311 | -0.390115 | 0.663364 | -0.348649 | -0.239833 | -0.589753 | -0.037887 | 0.025379 | 0.265505 | 0.217011 | -0.130696 | -0.132602 | 0.550461 |
| 20.73611111 | 0.365618 | 0.432226 | 0.077186 | 0.064754 | 0.209026 | 0.031944 | -0.103977 | -0.120351 | -0.005109 | 0.070019 | 0.389418 | 0.115356 | -0.286467 | 0.568291 | -0.334542 | -0.245477 | -0.624041 | -0.229313 | -0.225032 | 0.287194 | 0.230355 | -0.176415 | -0.163803 | 0.411208 |
| 20.75 | 0.249404 | 0.526252 | 0.081619 | 0.06879 | 0.118304 | 0.009034 | -0.202873 | -0.184806 | 0.011505 | 0.0909 | 0.379268 | 0.069371 | -0.350296 | 0.276775 | -0.298814 | -0.29477 | -0.348535 | -0.339257 | -0.38099 | 0.085167 | 0.209522 | -0.261506 | -0.093829 | 0.084434 |
| 20.76388889 | 0.172689 | 0.337955 | 0.040264 | 0.057757 | -0.008781 | -0.005447 | -0.358236 | -0.243467 | 0.032562 | -0.000666 | 0.292122 | 0.042179 | -0.517268 | -0.160849 | -0.320132 | -0.343575 | -0.150681 | -0.419212 | -0.576271 | -0.147916 | 0.109635 | -0.179913 | -0.076352 | -0.450236 |
| 20.77777778 | 0.158736 | -0.01833 | -0.072525 | -0.017688 | -0.116329 | -0.003353 | -0.570614 | -0.359122 | 0.003127 | -0.209251 | 0.219421 | 0.049318 | -0.59535 | -0.560352 | -0.314317 | -0.400252 | -0.277577 | -0.420189 | -0.74335 | -0.176179 | 0.012163 | -0.037521 | -0.12232 | -0.816712 |
| 20.79166667 | 0.10088 | -0.287177 | -0.166574 | -0.152903 | -0.133853 | -0.004768 | -0.677271 | -0.481459 | -0.096139 | -0.414628 | 0.144773 | 0.037113 | -0.653844 | -0.683349 | -0.335691 | -0.496677 | -0.620079 | -0.478798 | -0.863785 | -0.159146 | -0.061453 | 0.025819 | -0.256694 | -0.996253 |
| 20.80555556 | 0.071601 | -0.442135 | -0.178124 | -0.270539 | -0.090857 | -0.025233 | -0.764877 | -0.563846 | -0.202514 | -0.504603 | -0.030781 | 0.043304 | -0.728524 | -0.616011 | -0.38469 | -0.565877 | -0.623464 | -0.570345 | -1.095326 | -0.298711 | -0.038184 | 0.027093 | -0.497043 | -0.935333 |
| 20.81944444 | 0.033153 | -0.647319 | -0.220359 | -0.33609 | -0.043232 | -0.038863 | -0.818476 | -0.669984 | -0.210872 | -0.612411 | -0.198443 | 0.103558 | -0.857321 | -0.558599 | -0.311806 | -0.500401 | -0.273096 | -0.61373 | -1.183307 | -0.492491 | 0.008091 | 0.074562 | -0.682192 | -0.963089 |
| 20.83333333 | -0.004854 | -0.979693 | -0.235459 | -0.427898 | -0.064378 | -0.033293 | -0.682485 | -0.707652 | -0.15513 | -0.827221 | -0.247923 | 0.159117 | -0.940616 | -0.701312 | -0.322713 | -0.40604 | -0.051828 | -0.597612 | -1.129661 | -0.505319 | 0.02224 | 0.143939 | -0.792239 | -1.106329 |
| 20.84722222 | -0.135595 | -1.004546 | -0.267086 | -0.545341 | -0.229268 | -0.050726 | -0.458862 | -0.614956 | -0.103396 | -1.067665 | -0.266177 | 0.127124 | -0.981535 | -1.006891 | -0.344095 | -0.386854 | -0.403309 | -0.575427 | -1.146488 | -0.334616 | -0.031358 | 0.086841 | -0.794056 | -1.433452 |
| 20.86111111 | -0.358476 | -0.820948 | -0.294231 | -0.673736 | -0.432246 | -0.095322 | -0.346878 | -0.579228 | -0.093982 | -1.026775 | -0.374881 | 0.054844 | -0.97737 | -1.199133 | -0.442137 | -0.449997 | -0.866089 | -0.658579 | -1.274467 | -0.44557 | -0.20296 | -0.047892 | -0.829941 | -1.470017 |
| 20.875 | -0.465257 | -0.712618 | -0.355708 | -0.770838 | -0.452598 | -0.103076 | -0.287138 | -0.580913 | -0.110838 | -0.926382 | -0.447518 | 0.032161 | -0.913482 | -1.173329 | -0.501812 | -0.466887 | -1.153649 | -0.711781 | -1.293277 | -0.70019 | -0.320923 | -0.07633 | -1.026436 | -1.41777 |
| 20.88888889 | -0.486836 | -0.873236 | -0.470795 | -0.75601 | -0.371241 | -0.117254 | -0.307674 | -0.549624 | -0.18359 | -0.938109 | -0.553487 | 0.017509 | -1.013962 | -1.356431 | -0.535242 | -0.411037 | -1.61525 | -0.756463 | -1.186967 | -0.99554 | -0.549074 | -0.126029 | -1.281666 | -1.48357 |
| 20.90277778 | -0.776676 | -1.034451 | -0.539446 | -0.710413 | -0.316579 | -0.187454 | -0.422029 | -0.618711 | -0.339111 | -1.000596 | -0.451109 | 0.021328 | -1.213307 | -1.805799 | -0.628013 | -0.336326 | -2.125108 | -0.754754 | -1.132684 | -0.911539 | -0.499186 | -0.209428 | -1.295383 | -1.444689 |
| 20.91666667 | -1.111489 | -1.066342 | -0.518128 | -0.696626 | -0.341369 | -0.223449 | -0.436254 | -0.610412 | -0.394602 | -1.0302 | -0.335413 | 0.068007 | -1.44857 | -1.987506 | -0.761184 | -0.358373 | -1.999048 | -0.814435 | -1.11716 | -1.011621 | -0.604607 | -0.220782 | -1.229317 | -1.417445 |
| 20.93055556 | -1.297421 | -1.070996 | -0.678621 | -0.752967 | -0.296853 | -0.265197 | -0.322326 | -0.493291 | -0.395373 | -0.919688 | -0.322186 | 0.177489 | -1.490832 | -1.593277 | -0.909774 | -0.449116 | -1.538641 | -0.827716 | -1.159092 | -1.130876 | -0.639234 | -0.124133 | -1.075519 | -1.310224 |
| 20.94444444 | -1.358848 | -1.115462 | -0.783303 | -0.876755 | -0.338781 | -0.072019 | -0.197675 | -0.259591 | -0.366005 | -0.886932 | -0.507876 | 0.278944 | -1.566591 | -1.42978 | -1.002571 | -0.500974 | -1.288058 | -0.752007 | -1.370942 | -1.766932 | -0.631233 | -0.02991 | -0.825476 | -1.304463 |
| 20.95833333 | -1.970715 | -1.160918 | -1.064874 | -1.206923 | -1.065654 | -0.036921 | -0.529722 | -0.387523 | -0.439802 | -1.178808 | -0.751506 | 0.113881 | -1.609634 | -2.492354 | -1.159937 | -0.570916 | -2.269121 | -0.770303 | -1.669615 | -2.964182 | -0.745867 | -0.037595 | -1.034258 | -1.589137 |
| 20.97222222 | -3.292099 | -1.574136 | -1.798232 | -1.764391 | -2.068233 | -0.254023 | -0.957541 | -0.634912 | -0.686603 | -1.908541 | -2.088462 | -0.422334 | -2.016013 | -3.93429 | -1.665539 | -0.865134 | -2.953171 | -0.876535 | -1.93816 | -3.966302 | -1.084366 | 0.090266 | -1.667049 | -2.608824 |
| 20.98611111 | -2.248771 | -2.177563 | -1.68974 | -1.93702 | -2.234858 | -0.378861 | -0.742989 | -0.527045 | -0.87203 | -1.85318 | -2.987273 | -0.849838 | -1.282783 | -4.214267 | -1.446978 | -1.1047 | -2.000898 | -0.933673 | -1.741486 | -3.930804 | -1.408979 | -0.117116 | -1.989595 | -2.404382 |
| 21 | -0.453557 | -0.510916 | -0.646187 | -1.196118 | -1.164023 | -0.275521 | -0.099887 | -0.067986 | -0.926874 | -0.381822 | -1.495801 | -0.671742 | -0.274685 | -2.669646 | -0.274172 | -0.876024 | -0.358849 | -0.462128 | -0.795126 | -1.649736 | -0.513965 | -0.136538 | -1.25976 | -1.314319 |
| 21.01388889 | 1.140367 | 1.791326 | 0.1213 | -0.086037 | -0.333165 | -0.081953 | 0.491742 | 0.269099 | 0.05722 | 1.2238 | 0.489801 | -0.280913 | 0.336967 | -1.579219 | 0.519102 | -0.585341 | 0.79538 | 0.52628 | 0.494455 | 0.777354 | 0.920345 | 0.524627 | 0.005 | 0.05507 |
| 21.02777778 | 1.564336 | 2.1832 | 0.046416 | 0.595504 | 0.097488 | 0.061144 | 0.744895 | 0.352428 | 0.515765 | 1.808357 | 1.847996 | -0.185819 | 0.675135 | -1.389842 | 0.480596 | -0.670877 | 1.647491 | 0.722538 | 0.915195 | 1.146387 | 1.080105 | 0.809571 | 1.034049 | 0.434569 |
| 21.04166667 | 1.35773 | 0.068528 | -0.3006 | 0.57145 | 0.229173 | 0.178191 | 0.702162 | 0.354528 | 0.261106 | 1.685891 | 1.238886 | -0.253732 | 1.013136 | -1.680986 | 0.247577 | -0.945493 | 1.689106 | 0.298972 | 0.528391 | -0.180251 | 0.009523 | 0.858697 | 0.859878 | 0.530064 |
| 21.05555556 | 1.040965 | -1.593328 | -0.210508 | -0.170785 | 0.20311 | 0.2383 | 0.51775 | 0.199483 | 0.028011 | 1.544118 | 1.052683 | -0.109953 | 1.008346 | -1.500347 | -0.075116 | -1.556903 | 1.31666 | 0.140459 | 0.312104 | -0.318267 | -0.381841 | 1.025399 | 0.466614 | 0.887462 |
| 21.06944444 | 0.763373 | -1.44039 | 0.037346 | -0.489374 | 0.298857 | 0.269797 | 0.315157 | 0.133924 | -0.10067 | 1.52154 | 0.730651 | 0.074074 | 0.777855 | -0.788715 | -0.082302 | -1.74603 | 1.445603 | -0.029584 | 0.467838 | 0.253189 | -0.308718 | 1.232635 | 0.091496 | 1.261402 |
| 21.08333333 | 0.549411 | -1.232326 | 0.078458 | -0.392513 | 0.494484 | 0.217662 | 0.160533 | 0.324858 | -0.149029 | 1.299659 | 0.388872 | 0.002405 | 0.75212 | -0.444109 | 0.011076 | -1.195945 | 1.425248 | -0.156994 | 0.71604 | 0.422553 | -0.554935 | 1.067915 | -0.018567 | 1.513048 |
| 21.09722222 | 0.426822 | -1.176842 | -0.070471 | -0.257895 | 0.594836 | 0.14543 | 0.02898 | 0.397837 | 0.003122 | 0.543987 | 0.286013 | 0.029772 | 0.863709 | -0.514558 | -0.359312 | -0.772059 | 0.92613 | -0.324756 | 0.493624 | 0.481147 | -0.87521 | 0.696734 | 0.18626 | 1.287688 |
| 21.11111111 | 0.248302 | -1.238719 | -0.250608 | -0.360522 | 0.573213 | 0.146718 | -0.038692 | 0.429865 | 0.061192 | -0.077457 | 0.054933 | 0.237116 | 0.682303 | -0.009947 | -0.23904 | -0.252829 | 0.845439 | -0.242118 | 0.510883 | 0.486226 | -0.721657 | 0.523476 | 0.206244 | 0.845312 |
| 21.125 | 0.101427 | -0.481503 | -0.387145 | -0.538049 | 0.362656 | 0.156648 | -0.189842 | 0.261601 | -0.018471 | -0.650576 | -0.282871 | 0.186876 | 0.24004 | 0.515624 | -0.326345 | 0.877511 | 1.452009 | 0.431244 | 1.76493 | -0.723804 | -0.458829 | 0.954957 | 0.121085 | 0.221594 |
| 21.13888889 | -0.032743 | 0.499212 | -0.390713 | -0.55985 | 0.046222 | 0.103657 | -0.38953 | 0.077301 | -0.227965 | -0.942462 | -0.578318 | -0.072477 | -0.802926 | 0.241122 | -0.785558 | 2.134408 | 2.538153 | 1.042663 | 3.26395 | -2.82318 | -0.899758 | 1.576256 | -0.177337 | -1.087468 |
| 21.15277778 | 0.077783 | 1.151391 | -0.352852 | -0.064645 | -0.083314 | 0.16213 | -0.233959 | 0.218298 | -0.056343 | -0.457542 | -0.056288 | -0.051039 | -1.716575 | -1.41785 | -0.139518 | 0.138625 | 0.296957 | -0.397442 | 1.317465 | -1.721342 | -1.383216 | -0.083656 | -1.538268 | -0.893799 |
| 21.16666667 | 0.294023 | 1.182838 | -0.274982 | 0.486203 | 0.234906 | 0.218083 | 0.135725 | 0.383987 | 0.283747 | 0.04607 | 0.323457 | 0.090148 | -0.135063 | -1.031419 | 0.692957 | -1.409408 | -0.750973 | -1.139441 | -1.471415 | 0.750542 | -0.416043 | -1.371032 | -0.878062 | -0.130171 |
| 21.18055556 | 0.658089 | 1.169852 | 0.081121 | 0.648575 | 0.802703 | 0.129168 | 0.308629 | 0.31161 | 0.306619 | 0.550011 | 0.496453 | 0.115758 | 0.570533 | -0.030206 | 0.887384 | -0.629864 | 0.613394 | -0.345113 | -0.392464 | 1.328936 | 0.368919 | -0.830938 | 0.22371 | 0.062617 |
| 21.19444444 | 1.084608 | 1.085144 | 0.608262 | 0.742257 | 0.936008 | 0.141967 | 0.368547 | 0.51867 | 0.289285 | 1.169817 | 1.451548 | 0.26225 | 0.4867 | 0.90625 | 0.984155 | 0.205306 | 1.834487 | 0.332721 | 0.71367 | 2.318858 | 0.425337 | -0.279086 | 0.738281 | 0.705127 |
| 21.20833333 | 0.711327 | 1.43416 | 0.767859 | 0.795528 | 0.615054 | 0.223711 | 0.466849 | 0.763684 | 0.328786 | 1.244156 | 2.072619 | 0.416642 | 0.227881 | 1.131253 | 0.651744 | 0.394176 | 1.639491 | 0.406891 | 0.486369 | 2.273906 | 0.329077 | 0.094581 | 0.671134 | 0.847294 |
| 21.22222222 | 0.241235 | 1.359903 | 0.63456 | 0.893426 | 0.437803 | 0.190934 | 0.573518 | 0.518872 | 0.492503 | 1.22701 | 1.391261 | 0.430604 | 0.287573 | 0.6809 | 0.257198 | 0.368595 | 1.09623 | 0.15459 | 0.114364 | 1.393113 | 0.493914 | 0.290106 | 0.759314 | 0.562332 |
| 21.23611111 | 0.282528 | 0.750303 | 0.647934 | 0.633036 | 0.704388 | 0.11586 | 0.503456 | 0.459061 | 0.535312 | 1.317483 | 0.944305 | 0.474451 | 0.622565 | 0.520044 | 0.172358 | 0.393515 | 1.18624 | 0.006177 | 0.258165 | 1.597547 | 0.728542 | 0.261652 | 0.726998 | 0.523981 |
| 21.25 | 0.467565 | 0.641663 | 0.772036 | 0.294279 | 0.972875 | 0.087837 | 0.483077 | 0.642907 | 0.249715 | 1.262431 | 1.163764 | 0.592739 | 0.915798 | 0.877183 | 0.339933 | 0.281998 | 1.118234 | 0.034914 | 0.663545 | 2.317431 | 0.718339 | 0.270122 | 0.462565 | 0.876745 |
| 21.26388889 | 0.565683 | 0.84381 | 0.763482 | 0.128389 | 0.811318 | 0.092311 | 0.71862 | 0.85722 | 0.098631 | 1.03918 | 1.503857 | 0.647004 | 1.083196 | 1.045775 | 0.486639 | 0.250812 | 0.565903 | 0.095378 | 0.780107 | 2.326618 | 0.588691 | 0.258087 | 0.327929 | 1.149687 |
| 21.27777778 | 0.673369 | 0.586748 | 0.790553 | 0.22067 | 0.656886 | 0.102353 | 0.775101 | 0.647667 | 0.128128 | 0.878079 | 1.118059 | 0.576743 | 1.024612 | 0.928274 | 0.433724 | 0.366845 | 0.433083 | 0.203708 | 0.740246 | 1.940794 | 0.688737 | 0.230343 | 0.390114 | 1.051099 |
| 21.29166667 | 0.793429 | 0.369515 | 0.728516 | 0.345362 | 0.835841 | 0.118529 | 0.595134 | 0.477299 | 0.151447 | 0.705105 | 0.947757 | 0.564591 | 0.97101 | 1.087525 | 0.314215 | 0.420978 | 0.707366 | 0.312158 | 0.794815 | 1.842879 | 0.841363 | 0.289718 | 0.702449 | 0.917071 |
| 21.30555556 | 0.913542 | 0.338396 | 0.639541 | 0.238426 | 0.867364 | 0.137935 | 0.559621 | 0.384897 | 0.286063 | 0.444869 | 1.426885 | 0.543129 | 0.968345 | 1.525395 | 0.237606 | 0.382397 | 0.558932 | 0.368383 | 0.909498 | 1.497609 | 0.796621 | 0.398437 | 0.947552 | 1.066517 |
| 21.31944444 | 0.73268 | 0.469984 | 0.518693 | 0.094254 | 0.486074 | 0.14643 | 0.567287 | 0.214337 | 0.527552 | 0.264465 | 1.547067 | 0.507194 | 1.06907 | 1.9842 | 0.343029 | 0.334123 | 0.23269 | 0.419009 | 1.010843 | 1.219241 | 0.879021 | 0.439855 | 0.939914 | 1.284547 |
| 21.33333333 | 0.383186 | 0.176471 | 0.440531 | 0.180021 | 0.241881 | 0.115929 | 0.5266 | 0.21888 | 0.559728 | 0.21579 | 1.197705 | 0.446313 | 1.127345 | 2.28514 | 0.560279 | 0.416017 | 0.301883 | 0.587186 | 1.038494 | 1.474189 | 1.133024 | 0.415625 | 0.833463 | 1.296585 |
| 21.34722222 | 0.471533 | 0.266786 | 0.442367 | 0.470551 | 0.468111 | 0.089564 | 0.598006 | 0.392382 | 0.265194 | 0.266568 | 0.797886 | 0.41153 | 1.016164 | 2.085272 | 0.701944 | 0.613086 | 0.586517 | 0.788017 | 1.030055 | 1.369857 | 1.182502 | 0.322586 | 0.84219 | 1.130507 |
| 21.36111111 | 0.94641 | 0.374117 | 0.537001 | 0.708324 | 0.726661 | 0.120171 | 0.721136 | 0.498745 | 0.172166 | 0.457073 | 0.49928 | 0.378015 | 1.021134 | 2.069986 | 0.708447 | 0.801138 | 0.756343 | 0.838842 | 1.108915 | 0.762242 | 1.143391 | 0.269733 | 1.043145 | 0.98083 |
| 21.375 | 1.260896 | 0.46472 | 0.710069 | 0.753946 | 0.768067 | 0.178589 | 0.859187 | 0.434087 | 0.385375 | 0.811339 | 0.60319 | 0.374031 | 1.028343 | 2.491309 | 0.686474 | 0.741042 | 1.246412 | 0.896007 | 1.194125 | 0.462857 | 1.056879 | 0.2623 | 1.524419 | 0.872266 |
| 21.38888889 | 1.430258 | 0.707602 | 0.818179 | 0.625378 | 0.980323 | 0.216696 | 0.900501 | 0.299651 | 0.747353 | 1.293371 | 1.05712 | 0.455547 | 0.994438 | 3.228805 | 0.698135 | 0.717369 | 1.99359 | 0.95431 | 1.382101 | 0.801028 | 1.295001 | 0.203822 | 1.703436 | 1.518621 |
| 21.40277778 | 1.649394 | 0.817418 | 0.887195 | 0.680474 | 1.381167 | 0.294056 | 0.869872 | 0.35714 | 0.920506 | 1.362198 | 1.531207 | 0.673866 | 1.208644 | 3.402097 | 0.882402 | 0.751638 | 2.149652 | 1.14841 | 1.478353 | 1.493323 | 1.570206 | 0.098377 | 1.452973 | 2.5759 |
| 21.41666667 | 1.693358 | 0.958548 | 1.043223 | 0.886998 | 1.527774 | 0.367549 | 0.924697 | 0.552058 | 0.820899 | 1.295814 | 1.74099 | 0.870229 | 1.451366 | 2.799041 | 1.125986 | 0.918068 | 1.61187 | 1.323922 | 1.527312 | 1.76744 | 1.212675 | 0.085051 | 1.488249 | 3.240107 |
| 21.43055556 | 1.688729 | 0.769542 | 0.979067 | 0.837696 | 1.363843 | 0.220445 | 0.820755 | 0.494985 | 0.484685 | 0.8175 | 1.257889 | 0.684814 | 1.065656 | 1.966937 | 1.144874 | 0.936549 | 1.091156 | 1.309331 | 1.490419 | 0.983718 | 0.607132 | 0.002792 | 1.214302 | 2.47832 |
| 21.44444444 | 1.444893 | 0.375954 | 0.723102 | 0.48221 | 1.158825 | -0.047484 | 0.548842 | 0.183837 | 0.213063 | 0.566708 | 0.653659 | 0.305564 | 0.50237 | 1.56526 | 1.068999 | 0.741507 | 0.79067 | 1.222595 | 1.31777 | 0.533795 | 0.347425 | -0.142534 | 0.858749 | 1.937747 |
| 21.45833333 | 1.116337 | 0.142977 | 0.543247 | 0.321036 | 1.303392 | -0.097241 | 0.350768 | 0.084976 | 0.245872 | 0.699018 | 0.478275 | 0.160989 | 0.214743 | 1.763653 | 0.988368 | 0.608904 | 0.798744 | 1.055071 | 1.194339 | 1.125867 | 0.437522 | -0.111377 | 0.777187 | 1.967734 |
| 21.47222222 | 1.25964 | 0.335193 | 0.594559 | 0.404244 | 1.446116 | 0.060127 | 0.398325 | 0.195922 | 0.555928 | 0.983526 | 0.580042 | 0.318014 | 0.367998 | 1.907158 | 1.014323 | 0.551629 | 1.381234 | 0.783461 | 0.780826 | 2.021532 | 0.545618 | 0.128858 | 0.827143 | 1.641603 |
| 21.48611111 | 1.714916 | 0.576915 | 0.715599 | 0.659259 | 1.486796 | 0.179595 | 0.488719 | 0.288517 | 0.749184 | 1.01214 | 0.762344 | 0.501889 | 0.542102 | 2.373476 | 1.091373 | 0.569105 | 1.903472 | 0.818786 | 0.55024 | 2.120378 | 0.581009 | 0.441644 | 0.858637 | 1.311405 |
| 21.5 | 2.091273 | 0.781177 | 0.738145 | 0.906438 | 1.457011 | 0.195134 | 0.672895 | 0.398083 | 0.917656 | 1.08719 | 0.885412 | 0.628264 | 0.715959 | 3.321363 | 1.370834 | 0.846041 | 1.743098 | 1.109873 | 0.684934 | 1.705782 | 0.662601 | 0.662138 | 0.79789 | 1.921938 |
| 21.51388889 | 1.706933 | 0.842618 | 0.729669 | 1.037379 | 1.378129 | 0.158775 | 0.758504 | 0.454019 | 0.82536 | 0.952275 | 1.125434 | 0.566254 | 0.753776 | 3.299123 | 1.527112 | 1.151749 | 1.38438 | 0.950785 | 0.655675 | 1.128787 | 0.766235 | 0.615189 | 0.829095 | 2.509896 |
| 21.52777778 | 1.225951 | 0.781498 | 0.606729 | 0.970837 | 1.167921 | 0.103773 | 0.608462 | 0.320888 | 0.53896 | 0.556885 | 0.956931 | 0.404227 | 0.539992 | 2.353498 | 1.167751 | 0.978307 | 0.97129 | 0.648839 | 0.494105 | 0.984194 | 0.625179 | 0.492115 | 1.00667 | 1.773186 |
| 21.54166667 | 1.284486 | 0.575347 | 0.439457 | 0.737899 | 0.960535 | 0.058514 | 0.490262 | 0.199397 | 0.264125 | 0.260943 | 0.668098 | 0.273759 | 0.329136 | 1.335015 | 0.687816 | 0.597226 | 0.369117 | 0.391724 | 0.386194 | 0.943734 | 0.592184 | 0.375176 | 0.986536 | 1.232282 |
| 21.55555556 | 0.95948 | 0.312973 | 0.305804 | 0.487513 | 0.596646 | 0.032365 | 0.382432 | 0.091592 | -0.026405 | -0.040433 | 0.354551 | 0.168512 | 0.551925 | 0.299733 | 0.380983 | 0.428594 | -0.169928 | 0.24784 | 0.333209 | 0.478475 | 0.441963 | 0.295163 | 0.735406 | 0.862386 |
| 21.56944444 | -0.055557 | -0.17796 | 0.082706 | 0.28161 | 0.188686 | 0.018077 | 0.186413 | -0.0499 | -0.396445 | -0.500432 | 0.017288 | 0.132464 | 0.776934 | 0.269523 | 0.242526 | 0.437131 | -0.56618 | 0.136183 | 0.316181 | -0.107374 | 0.147024 | 0.262193 | 0.234484 | 0.947733 |
| 21.58333333 | -0.888628 | -0.489135 | -0.095032 | 0.179706 | -0.123685 | 0.003092 | -0.029941 | -0.256282 | -0.542518 | -0.603214 | -0.175399 | 0.142629 | 0.814908 | 0.739023 | 0.219687 | 0.444268 | -0.519878 | 0.348408 | 0.446563 | -0.136794 | -0.114572 | 0.14652 | 0.009214 | 1.223012 |
| 21.59722222 | -0.875607 | -0.58983 | -0.141308 | 0.112399 | -0.213213 | 0.029946 | -0.090015 | -0.133659 | -0.456365 | -0.539396 | -0.281583 | 0.130098 | 0.641511 | 1.08939 | 0.169872 | 0.314122 | -0.165327 | 0.728598 | 0.733453 | 0.130943 | -0.173534 | 0.019207 | -0.089575 | 1.51087 |
| 21.61111111 | -0.477006 | -0.035325 | -0.139209 | 0.056171 | -0.284106 | 0.045076 | -0.022148 | -0.130709 | -0.386981 | -0.429291 | -0.025667 | 0.115029 | 0.451088 | 1.137266 | 0.066868 | 0.148711 | 0.145054 | 0.937581 | 0.936928 | 0.717371 | 0.083489 | -0.12191 | -0.160861 | 1.551284 |
| 21.625 | -0.130807 | 0.406567 | -0.014989 | 0.048778 | -0.286886 | 0.033917 | 0.10459 | -0.085399 | -0.175445 | -0.244771 | 0.383084 | 0.095839 | 0.412392 | 1.105964 | 0.011507 | 0.101288 | 0.123031 | 0.799671 | 0.872827 | 0.963306 | 0.269459 | -0.375025 | -0.17259 | 1.50794 |
| 21.63888889 | 0.169124 | 0.564785 | 0.186395 | 0.116743 | -0.197329 | 0.049577 | 0.112937 | -0.027946 | 0.047352 | 0.022288 | 0.622478 | 0.120476 | 0.398651 | 1.114227 | 0.046596 | 0.171143 | 0.068081 | 0.658349 | 0.73482 | 0.723046 | 0.368942 | -0.270119 | -0.142627 | 1.418055 |
| 21.65277778 | 0.53995 | 0.697555 | 0.333896 | 0.190791 | -0.036256 | 0.057994 | -0.021442 | 0.048583 | 0.201225 | 0.228195 | 0.663619 | 0.154681 | 0.215653 | 1.168387 | 0.056871 | 0.154421 | -0.107005 | 0.565051 | 0.540575 | 0.408912 | 0.415444 | -0.005675 | -0.102868 | 1.203101 |
| 21.66666667 | 0.898254 | 0.467861 | 0.380065 | 0.179883 | 0.10067 | 0.086821 | -0.095308 | 0.131213 | 0.306705 | 0.358463 | 0.579643 | 0.142394 | -0.022814 | 0.643944 | 0.126742 | 0.007008 | -0.179217 | 0.463451 | 0.446827 | 0.074678 | 0.473514 | 0.252288 | 0.078298 | 0.86482 |
| 21.68055556 | 1.024568 | 0.127215 | 0.392448 | 0.145865 | 0.210449 | 0.140051 | 0.029137 | 0.135575 | 0.417006 | 0.51161 | 0.540571 | 0.136825 | 0.068445 | -0.249853 | 0.184612 | -0.173712 | -0.057659 | 0.39389 | 0.397334 | -0.134139 | 0.471676 | 0.386518 | 0.278538 | 0.57532 |
| 21.69444444 | 0.705413 | -0.108499 | 0.350025 | 0.088943 | 0.169981 | 0.183942 | 0.089143 | 0.039167 | 0.464852 | 0.580842 | 0.40236 | 0.177133 | 0.193761 | -0.598336 | 0.149556 | -0.230081 | 0.07029 | 0.287615 | 0.277813 | -0.08387 | 0.474861 | 0.373487 | 0.289489 | 0.395517 |
| 21.70833333 | 0.251663 | -0.49186 | 0.214483 | -0.002921 | -0.028128 | 0.172589 | -0.071864 | -0.043283 | 0.428631 | 0.56773 | 0.117029 | 0.191191 | -0.113503 | -0.724347 | 0.056865 | -0.200216 | -0.000385 | 0.101657 | 0.030489 | 0.16648 | 0.490682 | 0.256757 | 0.319928 | 0.26271 |
| 21.72222222 | 0.031776 | -1.044624 | 0.02989 | -0.113185 | -0.116387 | 0.122057 | -0.281117 | -0.023804 | 0.438992 | 0.48505 | -0.099923 | 0.140446 | -0.464392 | -0.603625 | -0.04683 | -0.18497 | -0.1786 | -0.071063 | -0.128257 | 0.430967 | 0.533378 | 0.133446 | 0.416734 | 0.211693 |
| 21.73611111 | -0.101813 | -1.354463 | -0.083893 | -0.163531 | 0.092068 | 0.065791 | -0.366257 | -0.016327 | 0.400537 | 0.387952 | -0.208613 | 0.090337 | -0.466184 | -0.513849 | -0.119505 | -0.239463 | -0.139302 | -0.346651 | -0.332554 | 0.538487 | 0.598942 | 0.047419 | 0.510487 | 0.020759 |
| 21.75 | -0.223957 | -1.322115 | -0.175201 | -0.213689 | 0.282779 | 0.020999 | -0.308336 | -0.120194 | 0.282895 | 0.25561 | -0.194379 | 0.069726 | -0.251045 | -0.374058 | -0.160464 | -0.406444 | -0.037567 | -0.765676 | -0.721131 | 0.370921 | 0.46977 | -0.011281 | 0.230426 | -0.40764 |
| 21.76388889 | -0.477398 | -1.282373 | -0.266544 | -0.300805 | 0.19486 | -0.015957 | -0.225588 | -0.192233 | 0.055371 | 0.059582 | -0.197751 | 0.030104 | -0.158333 | -0.275219 | -0.165282 | -0.583411 | -0.258814 | -1.170057 | -1.152348 | 0.001139 | 0.226508 | -0.030059 | -0.060239 | -0.830614 |
| 21.77777778 | -0.676392 | -1.086389 | -0.275181 | -0.340015 | -0.057425 | -0.040796 | -0.25821 | -0.143877 | -0.256347 | -0.271585 | -0.233812 | -0.030847 | -0.453924 | -0.607248 | -0.206267 | -0.70127 | -0.859689 | -1.26529 | -1.299526 | -0.346904 | 0.033542 | -0.025862 | -0.173816 | -1.15836 |
| 21.79166667 | -0.688951 | -0.81673 | -0.303783 | -0.327013 | -0.214322 | -0.061077 | -0.405336 | -0.149894 | -0.577136 | -0.65054 | -0.281314 | -0.040302 | -0.707566 | -1.242723 | -0.242753 | -0.749667 | -1.218317 | -1.198705 | -1.280821 | -0.570087 | -0.127444 | -0.031189 | -0.317818 | -1.51333 |
| 21.80555556 | -0.708995 | -0.434347 | -0.320562 | -0.327684 | -0.271141 | -0.053144 | -0.579231 | -0.285074 | -0.579745 | -0.736196 | -0.246586 | 0.035256 | -0.680047 | -1.700205 | -0.234448 | -0.732948 | -1.104346 | -1.111045 | -1.243957 | -0.63501 | -0.248688 | -0.001955 | -0.403748 | -1.7864 |
| 21.81944444 | -0.888851 | -0.156641 | -0.328135 | -0.405107 | -0.308164 | -0.070148 | -0.745501 | -0.441547 | -0.599692 | -0.74675 | -0.391524 | 0.028668 | -0.459185 | -1.662198 | -0.260656 | -0.769983 | -0.941749 | -1.084793 | -1.271506 | -0.691109 | -0.431817 | 0.081865 | -0.483665 | -1.89027 |
| 21.83333333 | -1.302045 | -0.241275 | -0.336089 | -0.474468 | -0.369112 | -0.158192 | -0.90698 | -0.604418 | -0.805822 | -1.101563 | -0.743208 | -0.126741 | -0.790485 | -1.813421 | -0.408914 | -0.734211 | -0.948868 | -1.139051 | -1.332592 | -0.737818 | -0.826862 | 0.099311 | -0.945997 | -2.004314 |
| 21.84722222 | -1.632391 | -0.471297 | -0.356974 | -0.523789 | -0.514071 | -0.186806 | -0.990666 | -0.704366 | -0.930821 | -1.382762 | -1.026658 | -0.293468 | -1.074029 | -2.080883 | -0.554267 | -0.684287 | -1.190762 | -1.287302 | -1.340576 | -0.794937 | -0.992455 | 0.049044 | -1.060132 | -1.993346 |
| 21.86111111 | -1.754736 | -0.600523 | -0.389213 | -0.65362 | -0.632282 | -0.155513 | -1.066994 | -0.834684 | -0.934752 | -1.716418 | -1.20253 | -0.309556 | -1.071379 | -1.935633 | -0.661399 | -0.744241 | -1.617782 | -1.286791 | -1.381911 | -1.073312 | -0.873912 | -0.00132 | -1.354769 | -1.816837 |
| 21.875 | -1.82973 | -0.603741 | -0.424889 | -0.78839 | -0.696892 | -0.160109 | -1.066802 | -0.73918 | -0.828865 | -1.949609 | -1.255848 | -0.268382 | -1.129848 | -1.77767 | -0.715664 | -0.9079 | -1.868125 | -1.190173 | -1.33469 | -1.514851 | -0.89758 | 0.093098 | -1.309969 | -1.745309 |
| 21.88888889 | -1.879882 | -0.715609 | -0.403037 | -0.808589 | -0.786184 | -0.175137 | -0.987115 | -0.724406 | -0.799399 | -1.765014 | -1.021305 | -0.301093 | -1.403945 | -1.495148 | -0.727909 | -0.88453 | -1.702497 | -1.149688 | -1.413028 | -1.729706 | -0.950766 | 0.173235 | -1.302757 | -1.861093 |
| 21.90277778 | -1.963176 | -1.179023 | -0.421827 | -0.907392 | -0.910186 | -0.212056 | -1.001335 | -0.898125 | -0.728845 | -1.636443 | -0.968197 | -0.387252 | -1.422346 | -1.481602 | -0.742353 | -0.836793 | -1.579528 | -1.120345 | -1.528191 | -1.982608 | -1.153932 | 0.050042 | -1.346291 | -1.805658 |
| 21.91666667 | -2.153585 | -1.479743 | -0.504426 | -1.158311 | -1.130173 | -0.237725 | -1.077672 | -0.94406 | -0.597887 | -1.613168 | -1.249936 | -0.38924 | -1.297867 | -1.541698 | -0.849728 | -0.802414 | -1.453523 | -1.105855 | -1.461361 | -2.194078 | -1.346544 | -0.108698 | -1.39623 | -1.844828 |
| 21.93055556 | -2.352779 | -1.327856 | -0.601945 | -1.29345 | -1.294223 | -0.102858 | -1.010396 | -1.055831 | -0.36538 | -1.54376 | -1.469568 | -0.24823 | -1.326256 | -1.490925 | -0.971579 | -0.940683 | -1.685501 | -1.123828 | -1.490008 | -2.492661 | -1.692249 | -0.062124 | -1.336432 | -2.137087 |
| 21.94444444 | -2.677661 | -1.332315 | -0.729366 | -1.298987 | -1.532369 | 0.12281 | -0.892647 | -0.982529 | -0.044301 | -1.138713 | -1.425391 | -0.108501 | -1.529426 | -1.430081 | -1.015107 | -1.01789 | -1.772674 | -1.330829 | -1.954945 | -2.091262 | -1.347462 | -0.033585 | -1.227634 | -2.297231 |
| 21.95833333 | -3.25155 | -1.325813 | -0.771594 | -1.366642 | -2.30371 | 0.104956 | -0.805186 | -0.818857 | -0.117392 | -0.790873 | -1.626773 | -0.206865 | -1.946724 | -1.637727 | -1.04473 | -1.166933 | -2.265055 | -1.417537 | -1.906762 | -2.690311 | -1.231408 | 0.13648 | -1.166625 | -1.992035 |
| 21.97222222 | -4.603548 | -1.9337 | -1.251596 | -1.52393 | -3.38089 | -0.287773 | -0.959896 | -0.942766 | -0.554569 | -1.328268 | -2.962959 | -0.664778 | -2.87586 | -1.830779 | -1.303382 | -1.269446 | -3.103925 | -1.69773 | -2.23929 | -3.068844 | -1.434737 | -0.009916 | -1.447511 | -1.958737 |
| 21.98611111 | -3.878401 | -2.490169 | -1.51819 | -2.098391 | -3.26689 | -0.557026 | -1.03564 | -0.762104 | -0.878656 | -1.967373 | -4.810467 | -1.518915 | -2.022652 | -1.160752 | -1.282193 | -1.186328 | -2.537972 | -1.699395 | -2.664739 | -3.173925 | -1.261567 | -0.6464 | -1.943091 | -2.349732 |
| 22 | -1.232678 | -1.755495 | -1.048655 | -1.127502 | -1.400615 | -0.58564 | -0.536357 | -0.268534 | -0.600494 | -0.852041 | -3.191647 | -0.939841 | -0.940172 | -1.077748 | -0.560532 | -0.923071 | -1.191866 | -1.209128 | -1.494183 | -2.569819 | -0.393852 | -0.935141 | -1.706546 | -1.285477 |
| 22.01388889 | 0.538351 | -0.312758 | -0.541601 | 0.066848 | 0.067222 | -0.344396 | -0.073172 | 0.202231 | 0.291204 | 0.64508 | -0.941502 | -0.544056 | 0.208904 | -1.907435 | 0.207922 | -0.777849 | 0.33109 | -0.339335 | -0.30067 | -1.568758 | 0.468478 | -0.158298 | -0.432121 | -0.040365 |
| 22.02777778 | 1.637857 | 0.485316 | -0.276145 | 0.760651 | 0.925213 | -0.030657 | 0.140401 | 0.405744 | 0.348468 | 1.610635 | 0.973882 | -0.412045 | 1.015711 | -2.118994 | 0.241677 | -0.802406 | 1.241921 | 0.03575 | 0.197233 | 0.127494 | 0.93155 | 0.569804 | 0.60394 | 0.695294 |
| 22.04166667 | 2.349195 | 0.573933 | -0.341909 | 0.482919 | 1.487267 | 0.189616 | 0.253948 | 0.386422 | 0.237596 | 1.339062 | 1.31947 | -0.188308 | 1.199142 | -1.908772 | -0.022025 | -0.906431 | 0.965403 | -0.101567 | -0.125757 | 1.036964 | 0.741969 | 0.418539 | 0.754555 | 0.782876 |
| 22.05555556 | 2.738968 | 0.713925 | -0.345115 | 0.001216 | 1.526644 | 0.263921 | 0.265386 | 0.406127 | 0.086825 | 1.062303 | 1.011476 | 0.10745 | 0.981093 | -1.813952 | 0.228052 | -0.830617 | 1.116264 | -0.050192 | -0.142911 | 0.469539 | 0.979619 | -0.002825 | 0.4146 | 0.772374 |
| 22.06944444 | 3.13878 | 0.889142 | -0.186451 | -0.253437 | 1.269565 | 0.268193 | 0.211157 | 0.416658 | 0.001995 | 1.203854 | 1.476343 | 0.262653 | 0.952092 | -1.763988 | 0.453849 | -0.471316 | 1.582258 | 0.160646 | 0.343127 | 0.039077 | 1.598404 | 0.245107 | 0.287609 | 0.900564 |
| 22.08333333 | 3.221768 | 0.864252 | -0.031287 | -0.165464 | 1.293019 | 0.231128 | 0.113782 | 0.641138 | -0.04622 | 1.377385 | 2.221459 | 0.361103 | 0.914874 | -1.567013 | 0.264016 | -0.10194 | 1.438403 | 0.140516 | 0.712668 | 0.489056 | 1.388725 | 0.54453 | 0.373943 | 0.780438 |
| 22.09722222 | 2.315552 | 0.243036 | -0.04688 | -0.334973 | 1.407177 | 0.169725 | 0.039294 | 0.524747 | -0.055011 | 1.008821 | 1.577 | 0.372455 | 0.684019 | -1.206271 | 0.140858 | 0.015025 | 1.151411 | -0.132097 | 0.717371 | 0.650745 | 0.911997 | 0.247862 | 0.483674 | 0.475703 |
| 22.11111111 | 1.50468 | -0.380685 | -0.056994 | -0.366182 | 1.106086 | 0.134952 | 0.011821 | 0.4082 | -0.068631 | 0.793852 | 0.897347 | 0.270796 | 0.474771 | -0.472551 | 0.092982 | -0.034688 | 0.722278 | -0.120093 | 0.889299 | 0.186965 | 0.404458 | 0.0753 | 0.205362 | 0.707457 |
| 22.125 | 0.908255 | -0.559314 | 0.007848 | -0.215486 | 0.823643 | 0.07406 | -0.054454 | 0.627566 | -0.072331 | 0.840767 | 0.690303 | 0.09724 | 0.568916 | 0.043366 | 0.162679 | -0.204461 | 0.354742 | -0.294818 | 0.873453 | -0.103453 | 0.017037 | 0.37478 | 0.04385 | 0.794999 |
| 22.13888889 | 0.564694 | -0.139475 | -0.01451 | 0.108612 | 0.615439 | 0.034176 | -0.123852 | 0.683766 | -0.028981 | 0.754646 | 0.424787 | -0.006216 | 0.364351 | 0.379199 | 0.210392 | -0.147563 | 0.196695 | -0.228926 | 0.880591 | -0.168659 | -0.358498 | 0.431922 | 0.213652 | 0.068706 |
| 22.15277778 | 0.49635 | 0.123885 | -0.100731 | 0.283 | 0.377481 | -0.003736 | -0.106942 | 0.599552 | -0.001559 | 0.259754 | 0.025321 | 0.075441 | -0.087025 | 0.628144 | 0.178312 | -0.049566 | -0.029091 | 0.248441 | 1.210603 | -0.399163 | -0.622277 | 0.236821 | 0.303784 | -0.586002 |
| 22.16666667 | 0.29235 | 0.432262 | -0.16581 | 0.445069 | 0.12658 | -0.026803 | -0.049984 | 0.334828 | -0.048849 | -0.100679 | -0.20498 | 0.115603 | -0.13424 | 0.813855 | 0.193172 | 0.079162 | -0.12026 | 0.431244 | 1.121801 | -0.597228 | -0.723655 | 0.140704 | 0.337638 | -0.678657 |
| 22.18055556 | -0.02174 | 1.008895 | -0.162458 | 0.666722 | 0.148092 | -0.069052 | 0.112512 | 0.268564 | 0.012024 | -0.041716 | 0.034663 | 0.199758 | 0.143849 | 1.04228 | 0.284887 | 0.257451 | 0.141568 | 0.382407 | 0.818071 | -0.254529 | -0.642667 | 0.109816 | 0.557272 | -0.507789 |
| 22.19444444 | -0.138519 | 1.380839 | -0.176026 | 0.700037 | 0.412245 | -0.096933 | 0.272806 | 0.390598 | 0.00438 | 0.05422 | 0.565896 | 0.354754 | 0.350413 | 1.189877 | 0.30612 | 0.378052 | 0.646593 | 0.459912 | 0.783607 | 0.244776 | -0.34929 | 0.056521 | 0.587103 | -0.25362 |
| 22.20833333 | -0.001149 | 1.790664 | -0.163534 | 0.696245 | 0.600266 | -0.096051 | 0.410666 | 0.478724 | -0.009224 | -0.027387 | 0.952053 | 0.458171 | 0.344415 | 1.534379 | 0.332041 | 0.420758 | 1.095152 | 0.607398 | 0.847522 | 0.626576 | 0.0488 | 0.080811 | 0.548053 | 0.128641 |
| 22.22222222 | 0.190944 | 2.040197 | -0.080165 | 0.475042 | 0.448082 | 0.035385 | 0.565481 | 0.683711 | 0.071318 | 0.021402 | 1.122033 | 0.359009 | 0.28 | 1.930845 | 0.258321 | 0.428021 | 1.284462 | 0.380114 | 0.464811 | 0.714321 | 0.381968 | 0.276824 | 0.567003 | 0.523297 |
| 22.23611111 | 0.308026 | 1.552883 | -0.000324 | 0.305858 | 0.451606 | 0.17885 | 0.589758 | 0.527929 | 0.03227 | 0.118975 | 0.811161 | 0.39395 | 0.262048 | 1.964912 | 0.09741 | 0.386571 | 1.123932 | 0.184613 | 0.172442 | 0.657911 | 0.523898 | 0.32853 | 0.55227 | 0.815198 |
| 22.25 | 0.688175 | 0.708301 | 0.071105 | 0.050736 | 0.987013 | 0.201095 | 0.248961 | -0.040177 | -0.023351 | 0.430484 | 0.450339 | 0.549802 | 0.490175 | 1.811397 | -0.023493 | 0.328979 | 1.005156 | 0.250735 | 0.347525 | 0.661414 | 0.518172 | 0.062451 | 0.295788 | 1.273095 |
| 22.26388889 | 1.160193 | 0.355754 | 0.245424 | -0.038674 | 1.235859 | 0.227186 | 0.18575 | -0.266706 | 0.200419 | 1.036295 | 0.74919 | 0.726149 | 0.845735 | 1.665417 | 0.145429 | 0.291938 | 1.041071 | 0.511091 | 0.701808 | 0.726341 | 0.57689 | -0.087897 | 0.010919 | 1.614443 |
| 22.27777778 | 1.21043 | 0.780788 | 0.467399 | 0.311466 | 1.039797 | 0.295091 | 0.503657 | 0.260001 | 0.527706 | 1.344096 | 1.69175 | 0.870582 | 0.902831 | 1.199917 | 0.417558 | 0.40254 | 0.938638 | 0.58393 | 0.718609 | 0.998781 | 0.870783 | 0.429551 | 0.287007 | 1.536952 |
| 22.29166667 | 1.035623 | 0.786723 | 0.420139 | 0.58841 | 0.95255 | 0.274342 | 0.724651 | 0.600889 | 0.388386 | 0.578734 | 1.485031 | 0.737686 | 0.589944 | 0.575702 | 0.39946 | 0.473243 | 0.899064 | 0.426569 | 0.506535 | 1.069701 | 0.937738 | 0.647592 | 0.49757 | 1.153816 |
| 22.30555556 | 0.863214 | 0.460126 | 0.380057 | 0.517508 | 1.088811 | 0.274971 | 0.353188 | 0.247248 | 0.205157 | 0.109369 | 0.742189 | 0.674109 | 0.612097 | 0.026535 | 0.11545 | 0.311906 | 1.212586 | 0.659942 | 0.722529 | 1.227836 | 0.855302 | 0.475029 | 0.449764 | 0.624613 |
| 22.31944444 | 1.075923 | 0.431468 | 0.552496 | 0.439944 | 1.102948 | 0.346499 | 0.238368 | 0.052058 | 0.294619 | 0.220799 | 0.953888 | 0.801717 | 0.864834 | 0.514715 | 0.094139 | 0.08016 | 1.659729 | 1.040916 | 1.343625 | 1.292175 | 0.950876 | 0.804868 | 0.37867 | 0.767482 |
| 22.33333333 | 1.271627 | 0.186429 | 0.616679 | 0.586061 | 0.895982 | 0.348467 | 0.313661 | 0.290082 | 0.500029 | 0.293899 | 1.19839 | 0.951187 | 0.905975 | 1.060214 | 0.236158 | -0.017099 | 1.48714 | 1.135401 | 1.622502 | 0.857389 | 0.936464 | 0.919205 | 0.296816 | 1.060871 |
| 22.34722222 | 1.237597 | 0.288733 | 0.56295 | 0.599736 | 0.859071 | 0.263518 | 0.455789 | 0.554046 | 0.495316 | 0.312767 | 1.289063 | 1.043647 | 0.759792 | 1.022496 | 0.353778 | -0.001594 | 0.929668 | 1.07657 | 1.550956 | 0.652368 | 0.856255 | 0.444858 | 0.293989 | 1.258935 |
| 22.36111111 | 1.315725 | 0.879229 | 0.806345 | 0.570389 | 1.067809 | 0.22327 | 0.572139 | 0.66967 | 0.496452 | 0.450276 | 1.678317 | 1.243891 | 0.744454 | 0.96905 | 0.506852 | 0.045466 | 0.796054 | 1.139464 | 1.620092 | 0.723593 | 0.909227 | 0.323498 | 0.606951 | 1.566962 |
| 22.375 | 1.557739 | 1.287324 | 1.115666 | 0.476197 | 1.204866 | 0.233144 | 0.770396 | 0.661701 | 0.515217 | 0.718079 | 2.254986 | 1.475656 | 0.873433 | 1.066592 | 0.572593 | 0.070167 | 1.147176 | 1.14001 | 1.585592 | 0.665375 | 1.060698 | 0.579174 | 0.848816 | 1.795706 |
| 22.38888889 | 1.90748 | 1.17646 | 1.155553 | 0.449728 | 1.016833 | 0.218062 | 0.887887 | 0.624721 | 0.703873 | 1.043174 | 2.396986 | 1.640126 | 1.075409 | 1.206598 | 0.553404 | 0.088205 | 1.461705 | 1.25839 | 1.559833 | 0.36873 | 1.15391 | 0.716713 | 0.9453 | 1.795557 |
| 22.40277778 | 2.254172 | 1.151547 | 0.956253 | 0.483857 | 0.931294 | 0.21126 | 0.830473 | 0.571007 | 1.168217 | 1.488375 | 2.275286 | 1.720872 | 1.2037 | 1.365221 | 0.53892 | 0.106967 | 1.674268 | 1.384139 | 1.770561 | 0.347951 | 1.527623 | 0.452074 | 1.094232 | 1.919365 |
| 22.41666667 | 2.410438 | 1.242177 | 1.052925 | 0.579029 | 1.242815 | 0.202833 | 0.875075 | 0.573419 | 1.449564 | 1.587256 | 2.448927 | 1.864756 | 1.358789 | 1.485957 | 0.64863 | 0.080453 | 1.778586 | 1.369192 | 1.786593 | 0.549393 | 2.042599 | 0.297867 | 1.363883 | 2.222481 |
| 22.43055556 | 2.222823 | 1.147853 | 1.091993 | 0.619165 | 1.368466 | 0.04623 | 0.789345 | 0.511645 | 1.217348 | 1.229099 | 2.363903 | 1.78924 | 1.591033 | 1.49018 | 0.89561 | 0.118728 | 1.63193 | 1.354148 | 1.513154 | 1.011353 | 1.62692 | 0.334529 | 1.490259 | 2.185949 |
| 22.44444444 | 2.052846 | 1.075009 | 0.913976 | 0.486063 | 1.29285 | -0.20242 | 0.647669 | 0.306209 | 0.794158 | 0.741877 | 1.642938 | 1.352839 | 1.336019 | 1.217282 | 0.981785 | 0.252939 | 1.446271 | 1.33515 | 1.276046 | 1.23259 | 0.910149 | 0.461195 | 1.281474 | 1.793824 |
| 22.45833333 | 2.085885 | 0.895278 | 0.658702 | 0.417113 | 1.112357 | -0.185612 | 0.65875 | 0.337277 | 0.668687 | 0.676921 | 1.103497 | 0.906524 | 0.919384 | 0.782484 | 0.795481 | 0.274234 | 1.277117 | 1.054886 | 1.068779 | 0.851271 | 0.584474 | 0.1532 | 1.341415 | 1.27572 |
| 22.47222222 | 1.879906 | 0.753847 | 0.595128 | 0.543215 | 1.035642 | -0.022948 | 0.633843 | 0.466957 | 0.936154 | 1.139035 | 0.953893 | 0.764562 | 0.721763 | 0.433021 | 0.736609 | 0.055743 | 1.221099 | 0.761374 | 0.845697 | 0.303585 | 0.839431 | 0.042987 | 1.3406 | 0.963584 |
| 22.48611111 | 1.656588 | 0.907982 | 0.591107 | 0.850705 | 0.882539 | 0.101574 | 0.750758 | 0.460572 | 1.5148 | 1.512941 | 1.474287 | 1.150674 | 0.806672 | 0.383533 | 0.926932 | -0.112731 | 1.310397 | 0.832659 | 0.951651 | 0.157194 | 1.325317 | 0.177852 | 1.445638 | 0.965617 |
| 22.5 | 1.939227 | 0.681412 | 0.758607 | 1.054559 | 0.648287 | 0.189395 | 1.205435 | 0.357268 | 1.66148 | 1.455034 | 1.757595 | 1.310832 | 1.016131 | 0.638976 | 1.036564 | 0.121513 | 1.554933 | 1.461888 | 1.428583 | 0.577553 | 1.486513 | 0.463404 | 1.582428 | 1.288037 |
| 22.51388889 | 1.864381 | 0.718112 | 0.661654 | 0.967949 | 0.568432 | 0.240407 | 1.458493 | 0.340606 | 1.372514 | 1.01006 | 1.483849 | 1.131492 | 1.172329 | 0.89924 | 1.103493 | 0.475599 | 1.328304 | 1.96102 | 1.997859 | 1.338928 | 1.464622 | 0.37897 | 1.556871 | 1.49286 |
| 22.52777778 | 1.704944 | 0.834469 | 0.463695 | 0.811787 | 0.703252 | 0.1771 | 1.091438 | 0.327861 | 0.801934 | 0.553777 | 0.971807 | 0.785555 | 1.160022 | 1.032688 | 1.121981 | 0.629732 | 1.103494 | 1.362435 | 1.426585 | 1.553115 | 1.193841 | 0.137942 | 1.550262 | 1.288395 |
| 22.54166667 | 1.509515 | 0.732206 | 0.317066 | 0.730217 | 0.764052 | 0.099682 | 0.442208 | 0.276428 | 0.394425 | 0.299215 | 0.736728 | 0.505903 | 1.156441 | 1.016488 | 1.117676 | 0.544384 | 1.049099 | 0.734821 | 0.831841 | 0.898772 | 0.655247 | 0.112211 | 1.492526 | 1.10499 |
| 22.55555556 | 1.286122 | 0.542983 | 0.301823 | 0.583357 | 0.514652 | 0.049801 | 0.225512 | 0.21961 | 0.153835 | 0.098612 | 0.618375 | 0.456924 | 1.066489 | 0.955403 | 1.011801 | 0.411817 | 1.011471 | 0.311981 | 0.264539 | 0.223266 | 0.418725 | 0.250526 | 1.30877 | 1.018197 |
| 22.56944444 | 0.895669 | 0.396813 | 0.310894 | 0.405069 | 0.168437 | -0.016694 | 0.110092 | 0.067746 | -0.082396 | -0.119147 | 0.446402 | 0.351304 | 0.852005 | 0.790261 | 0.795269 | 0.106596 | 0.712462 | 0.102829 | -0.081497 | -0.21053 | 0.155747 | 0.374332 | 0.708351 | 0.948741 |
| 22.58333333 | 0.462726 | -0.064838 | 0.312586 | 0.262637 | -0.03004 | -0.08591 | -0.089624 | -0.110831 | -0.272171 | -0.227175 | 0.179647 | 0.134168 | 0.759384 | 0.588813 | 0.517936 | -0.17043 | 0.159388 | 0.078335 | -0.216595 | -0.436397 | -0.235597 | 0.433836 | 0.048498 | 0.846411 |
| 22.59722222 | -0.018274 | -0.355182 | 0.098116 | 0.135028 | -0.136509 | -0.086507 | -0.199723 | -0.203065 | -0.390259 | -0.296894 | -0.113267 | -0.180819 | 0.561762 | 0.428574 | 0.345294 | -0.223151 | -0.445201 | 0.187853 | -0.184285 | -0.531391 | -0.496796 | 0.369001 | -0.645518 | 0.820339 |
| 22.61111111 | -0.371783 | -0.296611 | 0.051474 | 0.107642 | -0.16372 | -0.088655 | -0.253336 | -0.238924 | -0.432991 | -0.495592 | -0.244601 | -0.395483 | 0.442244 | 0.406703 | 0.317882 | -0.161398 | -0.846693 | 0.34873 | -0.156845 | -0.611266 | -0.505595 | 0.252714 | -1.1217 | 0.879741 |
| 22.625 | -0.454595 | -0.14108 | -0.067685 | 0.154422 | -0.018284 | -0.004621 | -0.280099 | -0.115519 | -0.429954 | -0.690561 | -0.234832 | -0.491216 | 0.259753 | 0.398814 | 0.312838 | -0.194198 | -0.686896 | 0.402102 | -0.095466 | -0.618379 | -0.170988 | 0.070253 | -1.110652 | 0.803156 |
| 22.63888889 | -0.283501 | -0.190227 | 0.056207 | 0.150036 | 0.135348 | 0.0064 | -0.314212 | -0.209882 | -0.281735 | -0.491553 | -0.13774 | -0.444228 | 0.150231 | 0.309215 | 0.197632 | -0.313633 | -0.256521 | 0.348847 | 0.025208 | -0.504054 | 0.164325 | -0.096471 | -0.742433 | 0.52654 |
| 22.65277778 | -0.065301 | -0.0962 | 0.225103 | 0.121726 | 0.126806 | 0.008402 | -0.301254 | -0.180262 | -0.142134 | -0.311032 | 0.150075 | -0.094532 | 0.221109 | 0.175623 | 0.041209 | -0.386951 | 0.249868 | 0.257184 | 0.19751 | -0.273828 | 0.276514 | -0.273993 | -0.365657 | 0.17344 |
| 22.66666667 | 0.064707 | -0.003625 | 0.31687 | 0.042991 | -0.05333 | -0.000811 | -0.252856 | -0.031245 | -0.114331 | -0.227346 | 0.351819 | 0.154928 | 0.320197 | 0.093429 | 0.014301 | -0.156005 | 0.349241 | 0.328368 | 0.288529 | -0.010828 | 0.212461 | -0.234369 | -0.193339 | -0.074421 |
| 22.68055556 | 0.089892 | -0.213373 | 0.176172 | -0.041254 | -0.183736 | -0.022729 | -0.321345 | 0.078367 | 0.002375 | -0.12266 | 0.351976 | 0.211211 | 0.220134 | 0.11084 | 0.12346 | 0.14527 | -0.066894 | 0.324534 | 0.271565 | 0.14979 | 0.202317 | -0.131672 | -0.009152 | -0.138536 |
| 22.69444444 | 0.092653 | -0.512082 | 0.160563 | -0.134081 | -0.230033 | -0.029344 | -0.373425 | 0.03297 | 0.134827 | -0.046055 | 0.292308 | 0.172768 | -0.099022 | 0.192875 | 0.167131 | 0.178072 | -0.299028 | 0.267566 | 0.256288 | 0.108323 | 0.242416 | -0.080629 | -0.040273 | -0.116801 |
| 22.70833333 | 0.060942 | -0.613565 | 0.15152 | -0.224797 | -0.263063 | -0.050817 | -0.408284 | -0.072585 | 0.125078 | -0.062907 | 0.261883 | 0.053117 | -0.408598 | 0.245937 | 0.117129 | -0.04001 | -0.148003 | 0.183731 | 0.243882 | -0.113717 | 0.154287 | -0.130605 | -0.131701 | -0.111618 |
| 22.72222222 | -0.040234 | -0.677777 | 0.056858 | -0.222 | -0.315665 | -0.030536 | -0.22859 | -0.034195 | 0.065657 | -0.033406 | 0.191566 | -0.023817 | -0.527406 | 0.20795 | 0.08169 | -0.156614 | 0.010023 | 0.077074 | 0.153045 | -0.350201 | 0.129566 | -0.147498 | -0.2734 | -0.064862 |
| 22.73611111 | -0.359977 | -0.820419 | -0.067171 | -0.086704 | -0.339999 | -0.021853 | -0.085096 | 0.058501 | 0.026172 | 0.03475 | 0.030437 | -0.077936 | -0.391882 | 0.093899 | 0.057518 | -0.120516 | -0.174718 | -0.042168 | 0.01779 | -0.552408 | 0.138022 | -0.184831 | -0.388547 | -0.071213 |
| 22.75 | -0.766439 | -0.816083 | -0.039877 | -0.07596 | -0.388137 | -0.042984 | -0.124756 | 0.028231 | -0.051002 | 0.064381 | -0.132094 | -0.184011 | -0.222274 | -0.050518 | -0.041584 | -0.195336 | -0.668383 | -0.182076 | -0.17467 | -0.76219 | 0.171056 | -0.163418 | -0.53388 | -0.197248 |
| 22.76388889 | -0.986575 | -0.697198 | 0.066828 | -0.205803 | -0.363791 | -0.054115 | -0.156499 | -0.035364 | -0.025891 | 0.076564 | -0.199734 | -0.14942 | -0.329509 | -0.101791 | -0.162874 | -0.357287 | -0.757734 | -0.296135 | -0.327551 | -1.026784 | 0.350753 | -0.012333 | -0.625092 | -0.397791 |
| 22.77777778 | -1.036343 | -0.641032 | -0.074465 | -0.222996 | -0.414287 | -0.034749 | -0.1623 | -0.093434 | 0.015219 | 0.009688 | -0.224558 | -0.091218 | -0.630977 | -0.13408 | -0.169136 | -0.461773 | -0.905632 | -0.372418 | -0.331291 | -1.10546 | 0.491206 | 0.08993 | -0.536959 | -0.524364 |
| 22.79166667 | -1.022353 | -0.75378 | -0.195317 | -0.265673 | -0.508529 | -0.068493 | -0.251372 | -0.239099 | -0.173138 | -0.251021 | -0.571967 | -0.284666 | -0.938756 | -0.25597 | -0.165343 | -0.477493 | -1.050406 | -0.471002 | -0.329326 | -1.189793 | 0.32535 | 0.085189 | -0.447006 | -0.544202 |
| 22.80555556 | -1.099898 | -0.864652 | -0.400165 | -0.467295 | -0.618768 | -0.158709 | -0.394045 | -0.333223 | -0.54845 | -0.587033 | -1.131461 | -0.516105 | -1.551359 | -0.456299 | -0.27816 | -0.453928 | -1.365859 | -0.651577 | -0.460594 | -1.478236 | -0.131532 | 0.108693 | -0.527329 | -0.71236 |
| 22.81944444 | -1.272467 | -0.773064 | -0.561704 | -0.664063 | -0.549969 | -0.177347 | -0.44922 | -0.373308 | -0.743584 | -0.83438 | -1.328963 | -0.459607 | -1.538395 | -0.557084 | -0.471876 | -0.464837 | -1.676759 | -0.852156 | -0.705028 | -1.565997 | -0.714507 | 0.122477 | -0.474186 | -0.818001 |

| | | | | | | | | | | | | | | | | | | | | | | | |
|---|---|---|---|---|---|---|---|---|---|---|---|---|---|---|---|---|---|---|---|---|---|---|---|
| 22.83333333 | -1.315629 | -0.682966 | -0.67183 | -0.749684 | -0.365515 | -0.112969 | -0.464631 | -0.360064 | -0.771 | -1.070677 | -1.215644 | -0.386562 | -1.188432 | -0.615619 | -0.542977 | -0.473543 | -1.686961 | -0.900904 | -1.010143 | -1.649887 | -1.259315 | 0.117211 | -0.360832 | -0.837971 |
| 22.84722222 | -1.557729 | -0.725284 | -0.816977 | -0.881368 | -0.42454 | -0.06964 | -0.526328 | -0.49442 | -0.858868 | -1.224156 | -1.125831 | -0.635833 | -1.064947 | -0.54077 | -0.567308 | -0.464012 | -1.498285 | -0.994431 | -1.394805 | -1.469864 | -1.93741 | -0.049316 | -0.446277 | -0.892255 |
| 22.86111111 | -2.321022 | -0.736058 | -0.876536 | -1.026416 | -0.909198 | -0.047665 | -0.603806 | -0.650136 | -1.188057 | -1.344993 | -1.513015 | -1.130689 | -1.097567 | -0.474212 | -0.642251 | -0.330297 | -1.301761 | -1.246077 | -1.631936 | -1.470904 | -2.47152 | -0.145395 | -0.566884 | -0.83991 |
| 22.875 | -2.638295 | -0.890398 | -0.677774 | -1.033716 | -1.463227 | -0.056933 | -0.737062 | -0.556816 | -1.562221 | -1.722689 | -2.078167 | -1.218914 | -1.561857 | -0.435774 | -0.910683 | -0.088033 | -1.367057 | -1.665568 | -1.714326 | -1.439374 | -2.525048 | -0.225528 | -0.638996 | -0.807563 |
| 22.88888889 | -2.605594 | -1.335219 | -0.4417 | -1.198554 | -1.728031 | -0.11073 | -0.846026 | -0.543274 | -1.741215 | -2.180001 | -2.147481 | -1.135862 | -1.658004 | -0.422711 | -0.917631 | 0.179679 | -1.523273 | -1.977505 | -1.889805 | -1.619705 | -2.419825 | -0.256285 | -0.658175 | -0.989811 |
| 22.90277778 | -2.879289 | -1.501993 | -0.371332 | -1.400661 | -1.628121 | -0.100749 | -0.947774 | -0.580867 | -1.507924 | -2.187059 | -2.129872 | -1.082903 | -1.703393 | -0.398055 | -0.704689 | 0.056346 | -1.646265 | -1.952545 | -2.053164 | -1.777587 | -2.43275 | -0.172253 | -0.702025 | -1.551513 |
| 22.91666667 | -3.51942 | -1.312259 | -0.573564 | -1.523489 | -1.755016 | -0.052555 | -0.943848 | -0.589046 | -1.116411 | -1.901375 | -2.258918 | -1.187428 | -1.690555 | -0.541141 | -0.81348 | -0.171891 | -1.587582 | -1.870819 | -2.189317 | -2.331082 | -2.141117 | -0.178439 | -0.744583 | -2.280807 |
| 22.93055556 | -3.927187 | -1.139326 | -0.82609 | -1.696302 | -2.358514 | -0.001603 | -0.761932 | -0.532286 | -0.962714 | -1.850509 | -2.418225 | -1.198067 | -1.560522 | -0.655614 | -1.039591 | -0.367174 | -1.253373 | -1.860076 | -2.164125 | -2.262264 | -1.700915 | -0.140497 | -0.90841 | -2.330635 |
| 22.94444444 | -3.929021 | -1.183607 | -0.981922 | -1.833968 | -3.004955 | 0.002623 | -0.718756 | -0.72193 | -0.84478 | -1.895714 | -3.088688 | -1.175953 | -1.587203 | -0.788908 | -1.41017 | -0.240006 | -1.295506 | -1.838327 | -2.091606 | -1.816404 | -1.426771 | -0.317557 | -1.059542 | -2.392017 |
| 22.95833333 | -3.695942 | -1.10266 | -1.012053 | -1.930479 | -3.911274 | -0.139858 | -0.936267 | -1.101306 | -0.869741 | -1.856415 | -3.178878 | -1.351516 | -2.129462 | -1.010016 | -1.605709 | -0.101527 | -2.028228 | -1.695542 | -2.011714 | -2.176337 | -1.348451 | -0.411531 | -1.332728 | -1.932322 |
| 22.97222222 | -4.097884 | -1.123241 | -1.188354 | -2.020209 | -4.897634 | -0.461113 | -1.361245 | -1.280926 | -0.999941 | -2.181005 | -3.719606 | -1.667726 | -3.025489 | -1.167045 | -1.834577 | -0.079811 | -2.426049 | -2.035667 | -2.441338 | -1.096914 | -1.345779 | -0.678607 | -1.863961 | -1.579224 |
| 22.98611111 | -3.955967 | -2.205759 | -1.570755 | -2.422249 | -4.163807 | -0.617407 | -1.412327 | -1.236781 | -1.015826 | -2.531471 | -4.438851 | -1.772662 | -2.732862 | -1.10928 | -1.78926 | -0.143548 | -2.033672 | -2.584466 | -2.720071 | -0.98066 | -1.253048 | -1.284351 | -2.021303 | -1.945826 |
| 23 | -1.445021 | -2.236533 | -1.215634 | -2.14023 | -2.029907 | -0.571349 | -0.531925 | -0.702661 | -0.688723 | -1.187577 | -2.781156 | -1.331075 | -0.636392 | -0.769497 | -0.578927 | -0.212637 | -1.210636 | -1.46683 | -1.563121 | -1.131641 | -0.372426 | -0.980479 | -2.049079 | -1.828395 |
| 23.01388889 | 0.21647 | -0.702209 | -0.41614 | -0.634792 | -0.165325 | -0.199181 | 0.370772 | -0.064429 | -0.056567 | 0.300777 | -0.732568 | -0.664122 | 0.60829 | -0.352967 | -0.020757 | -0.486321 | -0.164015 | -0.059862 | -0.081196 | -0.903087 | 0.6802 | -0.256124 | -0.959991 | -0.640556 |
| 23.02777778 | 1.078085 | 0.778274 | -0.180195 | 0.793379 | 0.993201 | 0.114046 | 0.646106 | 0.21856 | 0.330734 | 1.154131 | 0.738237 | -0.275645 | 0.852508 | -0.107925 | -0.421968 | -0.827148 | 0.693133 | 0.669726 | 0.746253 | -0.678378 | 0.769622 | 0.08465 | 0.606007 | -0.149975 |
| 23.04166667 | 1.388258 | 0.93808 | -0.402864 | 0.848141 | 1.682069 | 0.05205 | 0.44644 | 0.353027 | 0.296706 | 1.114949 | 1.319388 | 0.00349 | 0.931988 | -0.244258 | -0.597365 | -1.087835 | 1.116756 | 0.317953 | 0.340897 | -0.160866 | 0.55774 | -0.130129 | 0.732804 | 0.009853 |
| 23.05555556 | 1.498255 | 1.089766 | -0.132589 | 0.654117 | 1.762489 | 0.054443 | 0.358318 | 0.534949 | 0.188363 | 1.145799 | 1.183361 | 0.33047 | 1.104665 | -0.405253 | 0.032685 | -1.201012 | 1.253469 | 0.226218 | 0.263457 | 0.096205 | 0.458124 | -0.361691 | 0.55 | 0.308283 |
| 23.06944444 | 1.702987 | 1.227929 | 0.114209 | 0.512128 | 0.983713 | 0.105714 | 0.379796 | 0.639163 | 0.38386 | 1.351179 | 1.21494 | 0.283214 | 1.104063 | -0.568863 | 0.17062 | -1.277505 | 1.276507 | 0.244957 | 0.374673 | -0.01645 | 1.012138 | -0.220098 | 0.49362 | 0.622474 |
| 23.08333333 | 1.549721 | 0.537793 | 0.088244 | 0.355439 | 0.49537 | 0.168821 | 0.290263 | 0.708979 | 0.598092 | 1.544523 | 1.076722 | 0.113465 | 1.006529 | -0.696044 | -0.170491 | -1.240552 | 1.054744 | 0.159037 | 0.24933 | -0.140204 | 1.201843 | 0.374919 | 0.399567 | 0.786467 |
| 23.09722222 | 1.746646 | 0.711883 | 0.037975 | 0.089205 | 0.579844 | 0.124488 | 0.111562 | 0.921594 | 0.995197 | 1.994549 | 0.97731 | 0.107439 | 0.781068 | -0.858239 | 0.024135 | -1.108306 | 1.007686 | 0.128586 | 0.19698 | -0.108215 | 1.068193 | 0.542346 | 0.451988 | 0.971589 |
| 23.11111111 | 1.949249 | 0.686665 | 0.150784 | -0.257984 | 0.60431 | 0.102172 | 0.184609 | 1.007082 | 0.859489 | 1.711558 | 0.815629 | 0.091899 | 0.726563 | -0.745129 | 0.617736 | -0.896903 | 1.148174 | 0.247419 | 0.522605 | 0.094961 | 1.103172 | 0.195742 | 0.567429 | 0.770912 |
| 23.125 | 1.860742 | 0.461593 | 0.322039 | -0.178157 | 0.363505 | 0.083623 | 0.328783 | 0.769817 | 0.400912 | 0.763761 | 0.263943 | -0.036141 | 0.570505 | -0.221013 | 0.408027 | -0.671154 | 0.745391 | 0.137852 | 0.595101 | 0.274656 | 1.111419 | 0.005022 | 0.435412 | 0.684458 |
| 23.13888889 | 1.664779 | 0.267932 | 0.525762 | 0.023712 | 0.289521 | 0.064667 | 0.35195 | 0.69848 | 0.188073 | 0.442303 | -0.003057 | 0.041666 | 0.432151 | 0.3085 | -0.0016 | -0.609844 | 0.22329 | -0.107188 | 0.276732 | 0.205839 | 0.754497 | 0.122026 | -0.209288 | 0.749781 |
| 23.15277778 | 1.481553 | 0.277625 | 0.576607 | 0.103785 | 0.416854 | 0.044163 | 0.313459 | 0.797561 | 0.116178 | 0.442535 | 0.292635 | 0.233542 | 0.419958 | 0.520438 | -0.033323 | -0.676371 | -0.002874 | -0.201178 | 0.147156 | 0.079679 | 0.330819 | -0.185702 | -0.427955 | 0.804297 |
| 23.16666667 | 0.993963 | 0.187505 | 0.265902 | 0.104745 | 0.45975 | 0.162985 | 0.277543 | 0.794794 | -0.001132 | 0.37712 | 0.354572 | 0.182873 | 0.285333 | 0.431054 | 0.243013 | -0.68531 | -0.048578 | -0.10466 | 0.227026 | -0.176058 | 0.0169 | -0.456747 | -0.539449 | 0.6724 |
| 23.18055556 | 0.591982 | -0.141323 | -0.238943 | 0.111811 | 0.651479 | 0.234026 | 0.315785 | 0.615276 | -0.068063 | 0.220549 | 0.216772 | 0.072611 | -0.090479 | 0.217249 | 0.666214 | -0.486255 | 0.193568 | -0.074285 | 0.245573 | -0.433915 | -0.352817 | -0.275083 | -0.102148 | 0.444061 |
| 23.19444444 | 0.560401 | -0.317252 | -0.453242 | 0.080133 | 0.916908 | 0.096051 | 0.411696 | 0.46891 | 0.063838 | 0.31282 | 0.417842 | 0.366801 | -0.113787 | 0.178403 | 0.7174 | -0.286334 | 0.718932 | 0.017685 | 0.120519 | -0.654887 | -0.420357 | -0.066273 | 0.478754 | 0.139416 |
| 23.20833333 | 0.518289 | -0.27943 | -0.388301 | -0.035876 | 0.773671 | 0.032831 | 0.483936 | 0.63414 | 0.187968 | 0.397949 | 0.806226 | 0.597358 | 0.218742 | 0.302242 | 0.185972 | -0.285341 | 1.082137 | 0.17728 | 0.245573 | -0.638753 | -0.197228 | 0.243752 | 0.874676 | -0.084738 |
| 23.22222222 | 0.019867 | -0.360557 | -0.380066 | -0.120596 | 0.392934 | 0.067654 | 0.348391 | 0.656404 | 0.225657 | 0.44614 | 1.039975 | 0.605605 | 0.287559 | 0.377873 | -0.30674 | -0.387039 | 1.119428 | 0.544216 | 0.90486 | -0.211837 | 0.191426 | 0.323699 | 0.878584 | -0.041838 |
| 23.23611111 | -0.159471 | 0.106049 | 0.103867 | 0.355915 | 0.372104 | 0.076102 | 0.25098 | 0.483953 | 0.267653 | 0.490195 | 1.179841 | 0.551686 | 0.146106 | 0.419878 | -0.263665 | -0.379768 | 1.028224 | 0.782878 | 1.158019 | 0.070301 | 0.334855 | 0.35869 | 0.692238 | 0.097735 |
| 23.25 | 0.128608 | 0.907797 | 0.585746 | 1.012574 | 0.422818 | 0.060796 | 0.388869 | 0.517984 | 0.25072 | 0.457839 | 1.362666 | 0.570609 | 0.242773 | 0.32589 | -0.100195 | -0.244806 | 0.892391 | 0.759317 | 0.823389 | -0.034135 | 0.490114 | 0.691389 | 0.565028 | 0.215358 |
| 23.26388889 | 0.46189 | 1.255018 | 0.890487 | 1.0664 | 0.261208 | 0.040006 | 0.508272 | 0.37378 | 0.263853 | 0.424632 | 1.44747 | 0.597437 | 0.613961 | 0.058553 | -0.108398 | -0.073264 | 0.727136 | 0.76847 | 0.832403 | -0.030782 | 0.773022 | 0.818908 | 0.477955 | 0.420521 |
| 23.27777778 | 0.724105 | 1.240859 | 0.839958 | 0.60717 | 0.489892 | 0.102272 | 0.48699 | 0.143127 | 0.414393 | 0.484781 | 1.563922 | 0.632671 | 0.847536 | -0.171597 | -0.020378 | 0.212092 | 0.657716 | 0.776852 | 0.783732 | 0.320587 | 0.853253 | 0.781043 | 0.359501 | 0.524018 |
| 23.29166667 | 0.926309 | 0.860877 | 0.823092 | 0.379366 | 1.352654 | 0.239885 | 0.480233 | 0.189074 | 0.566488 | 0.632206 | 1.707642 | 0.711388 | 0.549132 | -0.107203 | 0.099334 | 0.611448 | 0.537875 | 0.754838 | 0.818579 | 0.717529 | 0.761283 | 0.779018 | 0.396084 | 0.113276 |
| 23.30555556 | 1.064743 | 0.633409 | 0.671343 | 0.394285 | 2.073847 | 0.270982 | 0.455349 | 0.394122 | 0.748748 | 0.784056 | 1.837259 | 0.86194 | 0.682826 | 0.08791 | 0.139421 | 0.667157 | 0.573292 | 0.786627 | 0.99903 | 0.860898 | 0.868216 | 0.683683 | 0.563263 | 0.300485 |
| 23.31944444 | 1.255325 | 0.638457 | 0.676027 | 0.556088 | 1.988328 | 0.22169 | 0.569919 | 0.510725 | 0.980942 | 1.001245 | 1.940525 | 1.159139 | 1.045932 | 0.546816 | 0.314487 | 0.474713 | 0.920075 | 0.745046 | 1.047441 | 0.928002 | 1.303268 | 0.545084 | 0.667372 | 0.829554 |
| 23.33333333 | 1.512919 | 0.583026 | 0.84521 | 0.583096 | 1.699403 | 0.255709 | 0.814192 | 0.64631 | 0.978829 | 0.96721 | 1.970057 | 1.298125 | 1.254814 | 1.527369 | 0.800095 | 0.446959 | 1.236893 | 0.723249 | 0.930187 | 0.920829 | 1.630089 | 0.40375 | 0.649429 | 1.32733 |
| 23.34722222 | 1.69009 | 0.568029 | 0.699827 | 0.520258 | 1.615147 | 0.366759 | 0.844889 | 0.623992 | 0.84439 | 0.7662 | 1.583175 | 1.183017 | 1.247513 | 2.90097 | 1.041853 | 0.411626 | 1.276067 | 0.707031 | 0.889006 | 1.06965 | 1.464797 | 0.392085 | 0.616448 | 1.766598 |
| 23.36111111 | 1.792421 | 0.5803 | 0.60386 | 0.532396 | 1.564299 | 0.456543 | 0.630257 | 0.501967 | 0.757441 | 0.671367 | 1.312564 | 1.128442 | 1.140663 | 2.188506 | 0.927152 | 0.511349 | 1.140498 | 0.670297 | 1.003337 | 1.303685 | 1.193394 | 0.511961 | 0.588939 | 1.816293 |
| 23.375 | 1.83623 | 0.638642 | 0.681976 | 0.587839 | 1.640482 | 0.494904 | 0.468781 | 0.648979 | 0.82821 | 0.809 | 1.403351 | 1.141755 | 1.115898 | 1.001163 | 0.825112 | 0.722705 | 1.166822 | 0.828772 | 1.348846 | 1.298702 | 1.18248 | 0.571611 | 0.705517 | 1.733302 |
| 23.38888889 | 2.035855 | 0.591163 | 0.745819 | 0.571381 | 1.725454 | 0.56437 | 0.513797 | 0.933174 | 1.211452 | 1.127792 | 1.905755 | 1.149645 | 1.080198 | 0.759123 | 0.858016 | 0.819228 | 1.135736 | 1.089118 | 1.646242 | 1.107163 | 1.49219 | 0.527885 | 0.876045 | 1.760994 |
| 23.40277778 | 2.152908 | 0.302647 | 0.654285 | 0.559267 | 1.661875 | 0.610441 | 0.697206 | 0.747115 | 1.665646 | 1.416262 | 2.037937 | 1.31266 | 1.061627 | 0.670793 | 0.878296 | 0.806836 | 1.128917 | 1.120636 | 1.753115 | 0.909842 | 1.659333 | 0.490056 | 1.044645 | 1.767937 |
| 23.41666667 | 2.077 | -0.049778 | 0.582671 | 0.6235 | 1.565149 | 0.604708 | 0.819635 | 0.552078 | 1.679179 | 1.621667 | 2.071217 | 1.282828 | 1.1382 | 0.604929 | 0.882567 | 0.709404 | 1.381418 | 1.143951 | 1.713365 | 0.80957 | 1.526378 | 0.567028 | 1.210402 | 1.665546 |
| 23.43055556 | 1.737871 | -0.378403 | 0.538551 | 0.528853 | 1.498946 | 0.449579 | 0.707321 | 0.346762 | 1.461105 | 1.334773 | 1.952727 | 1.222637 | 1.395937 | 0.61469 | 0.893127 | 0.787795 | 1.467914 | 1.47168 | 1.818786 | 0.570068 | 1.2684 | 0.574456 | 1.235469 | 1.560529 |
| 23.44444444 | 1.742715 | -0.229578 | 0.485997 | 0.351041 | 1.102825 | 0.238719 | 0.503192 | 0.217073 | 0.963941 | 0.74713 | 1.484838 | 1.243434 | 1.468767 | 0.518042 | 0.925326 | 0.860515 | 1.550164 | 1.854423 | 1.867704 | 0.297209 | 0.806259 | 0.454789 | 1.436109 | 1.256556 |
| 23.45833333 | 1.57743 | -0.020052 | 0.348806 | 0.388354 | 1.052956 | 0.137651 | 0.468418 | 0.431644 | 0.562194 | 0.530022 | 1.142796 | 1.118868 | 1.23129 | 0.314265 | 0.738433 | 0.775445 | 1.727326 | 1.294554 | 1.239021 | 0.086636 | 0.813073 | 0.332475 | 1.486231 | 0.967105 |
| 23.47222222 | 1.352794 | 0.349772 | 0.529863 | 0.782365 | 1.251868 | 0.312775 | 0.553998 | 0.595115 | 0.726692 | 0.746684 | 1.173378 | 1.179037 | 1.163789 | 0.109941 | 0.604832 | 0.48202 | 1.74121 | 0.784283 | 0.972154 | 0.040905 | 1.144844 | 0.267561 | 1.435043 | 0.842263 |
| 23.48611111 | 1.327122 | 0.733021 | 1.049099 | 1.077995 | 1.207041 | 0.448691 | 0.701633 | 0.528768 | 1.295296 | 1.149866 | 1.466546 | 1.373197 | 1.388579 | 0.229857 | 0.62684 | 0.263635 | 1.938323 | 0.773899 | 1.145441 | 0.022614 | 1.543158 | 0.255131 | 1.718321 | 1.089358 |
| 23.5 | 1.724376 | 0.7849 | 1.174275 | 1.00035 | 0.930138 | 0.288804 | 1.047563 | 0.585927 | 1.763649 | 1.464192 | 1.861687 | 1.832814 | 2.009605 | 0.687097 | 0.915168 | 0.471372 | 1.993983 | 1.166396 | 1.44408 | -0.001102 | 1.810542 | 0.496481 | 1.866548 | 1.240215 |
| 23.51388889 | 1.991451 | 0.638326 | 1.088797 | 0.939406 | 0.747986 | 0.330552 | 1.161089 | 0.39286 | 1.556008 | 1.116583 | 1.700071 | 1.79762 | 2.271644 | 1.113661 | 1.063623 | 0.720288 | 1.20671 | 1.60519 | 1.380604 | 0.313361 | 1.883849 | 0.683521 | 1.893145 | 1.333472 |
| 23.52777778 | 1.791401 | 0.52341 | 0.766252 | 0.717589 | 0.751398 | 0.332381 | 1.00619 | 0.17258 | 0.776606 | 0.534546 | 1.235819 | 1.400297 | 1.681631 | 1.277241 | 1.007807 | 0.720248 | 0.545669 | 1.452549 | 0.923466 | 0.648197 | 1.080083 | 0.735389 | 1.638394 | 1.336141 |
| 23.54166667 | 1.498845 | 0.256574 | 0.472715 | 0.484808 | 0.794312 | 0.326914 | 0.738521 | 0.11059 | 0.556818 | 0.47224 | 0.809882 | 0.982122 | 0.893507 | 1.128507 | 0.849929 | 0.662433 | 0.450622 | 0.802544 | 0.411539 | 0.646464 | 0.981778 | 0.562063 | 1.494633 | 1.213271 |
| 23.55555556 | 1.470598 | 0.125582 | 0.290007 | 0.425391 | 0.738156 | 0.316887 | 0.634439 | 0.241923 | 0.531567 | 0.379567 | 0.571307 | 0.719431 | 0.526962 | 0.910075 | 0.726309 | 0.535233 | 0.652212 | 0.162263 | 0.294375 | 0.414617 | 0.809504 | 0.541867 | 1.440342 | 1.045339 |
| 23.56944444 | 1.243344 | -0.064768 | 0.249453 | 0.337711 | 0.560033 | 0.151673 | 0.344088 | 0.199656 | 0.242547 | 0.210828 | 0.464509 | 0.427444 | 0.58496 | 0.643616 | 0.679001 | 0.233041 | 0.763123 | -0.173797 | 0.421791 | 0.088071 | 0.311798 | 0.354832 | 1.042764 | 0.904454 |
| 23.58333333 | 0.645265 | -0.290386 | 0.059894 | 0.173685 | 0.347439 | -0.127465 | -0.02736 | 0.012685 | -0.262884 | -0.013629 | 0.16909 | -0.001194 | 0.59078 | 0.33373 | 0.575439 | -0.228626 | 0.447835 | -0.178487 | 0.604459 | -0.147751 | 0.057557 | 0.173353 | 0.475017 | 0.701579 |
| 23.59722222 | 0.174608 | -0.422454 | -0.099378 | 0.042217 | 0.084619 | -0.368186 | -0.295184 | 0.003182 | -0.59826 | -0.226787 | -0.249962 | -0.384516 | 0.494541 | 0.110209 | 0.32946 | -0.628382 | 0.108486 | -0.131679 | 0.674525 | -0.281043 | 0.030976 | 0.252183 | -0.315916 | 0.55137 |
| 23.61111111 | -0.145105 | -0.365121 | -0.12318 | 0.058441 | -0.142675 | -0.424721 | -0.565778 | -0.037588 | -0.911186 | -0.404294 | -0.523597 | -0.454024 | 0.592067 | 0.067006 | 0.18928 | -0.946056 | -0.202901 | -0.104773 | 0.538229 | -0.345828 | -0.082784 | 0.295993 | -0.718111 | 0.472164 |
| 23.625 | -0.397729 | -0.392057 | -0.143468 | 0.108912 | -0.279056 | -0.408566 | -0.792952 | -0.220809 | -1.157828 | -0.560622 | -0.499288 | -0.492806 | 0.830858 | 0.006265 | 0.050337 | -0.706646 | -0.324897 | 0.012568 | 0.289206 | -0.396858 | -0.317029 | 0.12413 | -0.7471 | 0.383456 |
| 23.63888889 | -0.571785 | -0.506446 | -0.132411 | 0.112846 | -0.319411 | -0.457432 | -0.730169 | -0.452192 | -1.227389 | -0.725252 | -0.372819 | -0.424885 | 0.835158 | -0.131819 | -0.043764 | -0.452274 | -0.616678 | 0.135025 | -0.058905 | -0.414337 | -0.76994 | -0.070848 | -0.665339 | 0.311122 |
| 23.65277778 | -0.640995 | -0.533009 | -0.318218 | 0.077816 | -0.3264 | -0.6578 | -0.509053 | -0.342083 | -1.068484 | -0.859344 | -0.282025 | -0.410413 | 0.279927 | -0.178256 | -0.202093 | -0.356188 | -0.685381 | 0.255094 | -0.281735 | -0.259473 | -0.58497 | -0.088352 | -0.439532 | 0.212629 |
| 23.66666667 | -0.537327 | -0.354649 | -0.152979 | 0.076425 | -0.194381 | -0.624118 | -0.555473 | -0.194384 | -0.635297 | -0.694401 | -0.173643 | -0.342975 | -0.177527 | -0.173592 | -0.199143 | -0.293157 | -0.598662 | 0.31194 | -0.204787 | 0.10469 | 0.093466 | 0.061676 | -0.445981 | -0.068436 |
| 23.68055556 | -0.074079 | -0.149089 | 0.355203 | 0.03641 | 0.020411 | -0.278055 | -0.536162 | -0.008291 | -0.250876 | -0.343248 | 0.042658 | -0.221984 | -0.37836 | -0.132951 | -0.070125 | -0.223983 | -0.426539 | 0.436472 | 0.132259 | 0.36393 | 0.474094 | 0.150285 | -0.538427 | -0.314375 |
| 23.69444444 | 0.163336 | -0.118743 | 0.55274 | -0.053499 | -0.049833 | -0.182039 | -0.462095 | -0.053009 | -0.42455 | -0.19156 | 0.040068 | -0.220281 | -0.376074 | -0.111136 | 0.033733 | -0.191464 | -0.053159 | 0.281501 | 0.120808 | 0.417819 | 0.130337 | 0.109933 | -0.412801 | -0.37157 |
| 23.70833333 | 0.003324 | -0.126712 | 0.347204 | -0.120808 | -0.387877 | -0.264036 | -0.445213 | -0.168349 | -0.52912 | -0.266342 | -0.085246 | -0.298665 | -0.253793 | -0.184151 | 0.097336 | -0.178629 | 0.149923 | 0.006103 | -0.342276 | 0.150929 | -0.306703 | 0.000954 | -0.522434 | -0.496265 |
| 23.72222222 | -0.295098 | -0.071332 | 0.180697 | -0.122498 | -0.592563 | -0.20548 | -0.335622 | -0.114855 | -0.4212 | -0.237849 | -0.097923 | -0.335569 | -0.280651 | -0.121024 | 0.105427 | -0.210926 | -0.142943 | -0.069838 | -0.452611 | -0.188533 | -0.34819 | 0.050912 | -0.581734 | -0.38131 |
| 23.73611111 | -0.510592 | -0.000151 | 0.206527 | -0.110076 | -0.645893 | -0.001385 | -0.153313 | -0.047285 | -0.458231 | -0.180796 | 0.020683 | -0.259877 | -0.402244 | 0.006371 | 0.098835 | -0.353386 | -0.532153 | 0.001583 | -0.308635 | -0.544159 | -0.177706 | 0.151863 | -0.497451 | -0.04746 |
| 23.75 | -0.64403 | -0.022946 | 0.287585 | -0.174118 | -0.628261 | 0.131749 | -0.039724 | -0.090251 | -0.657052 | -0.259334 | 0.082097 | -0.213057 | -0.498872 | -0.001148 | 0.074381 | -0.557724 | -0.571289 | 0.025466 | -0.26108 | -1.083857 | -0.047585 | 0.02146 | -0.557275 | 0.022128 |
| 23.76388889 | -0.731573 | -0.065555 | 0.271179 | -0.30139 | -0.851408 | 0.091203 | -0.224866 | -0.204155 | -0.596511 | -0.213535 | -0.02533 | -0.264495 | -0.599019 | -0.051037 | 0.065342 | -0.59332 | -0.563918 | -0.067469 | -0.306942 | -1.859673 | -0.192096 | -0.126702 | -0.681808 | 0.012645 |
| 23.77777778 | -0.790322 | -0.057514 | 0.196177 | -0.394749 | -1.168072 | -0.033077 | -0.495339 | -0.247495 | -0.368019 | -0.104229 | -0.177423 | -0.350439 | -0.849911 | -0.116276 | 0.022111 | -0.462177 | -0.73542 | -0.266403 | -0.46328 | -1.884648 | -0.56498 | -0.128236 | -0.67788 | 0.041189 |
| 23.79166667 | -0.963526 | 0.006113 | 0.063156 | -0.369322 | -1.297219 | -0.026901 | -0.681832 | -0.256523 | -0.270497 | -0.004873 | -0.377368 | -0.466324 | -1.046892 | -0.273964 | -0.039061 | -0.341949 | -1.005997 | -0.509818 | -0.678461 | -1.80184 | -0.729011 | -0.124556 | -0.65231 | 0.053607 |
| 23.80555556 | -1.264565 | 0.029242 | -0.234152 | -0.289645 | -1.366318 | 0.033846 | -0.726027 | -0.409664 | -0.319708 | -0.024187 | -0.787854 | -0.671656 | -1.407409 | -0.513779 | -0.135393 | -0.459434 | -1.110934 | -0.640101 | -0.724006 | -2.130015 | -0.944227 | -0.197223 | -0.752691 | -0.065507 |
| 23.81944444 | -1.581701 | -0.041441 | -0.651089 | -0.295176 | -1.411153 | -0.038437 | -0.663219 | -0.495894 | -0.338541 | -0.00977 | -1.121073 | -0.765769 | -1.749899 | -0.708021 | -0.251397 | -0.703939 | -0.986696 | -0.726902 | -0.788991 | -2.125856 | -1.209787 | -0.256766 | -0.96316 | -0.292799 |
| 23.83333333 | -1.763999 | -0.020724 | -0.984699 | -0.500679 | -1.326671 | -0.197409 | -0.600894 | -0.661911 | -0.255639 | 0.120887 | -1.127786 | -0.712581 | -2.080526 | -0.829301 | -0.359914 | -0.76706 | -0.89542 | -0.814922 | -0.822424 | -1.936146 | -1.415507 | -0.227883 | -0.91836 | -0.515902 |
| 23.84722222 | -1.846024 | -0.061005 | -1.008564 | -0.679678 | -1.438594 | -0.416248 | -0.716769 | -0.92437 | -0.334299 | -0.035078 | -1.491733 | -0.877926 | -2.241001 | -1.00781 | -0.418637 | -0.711068 | -1.083646 | -0.975093 | -1.05945 | -1.860207 | -1.444523 | -0.257045 | -0.674637 | -0.847998 |
| 23.86111111 | -1.912902 | -0.384833 | -1.329574 | -0.815209 | -1.803124 | -0.709153 | -1.00673 | -0.881802 | -0.68765 | -0.56543 | -1.953943 | -1.184974 | -1.775927 | -1.225406 | -0.524805 | -0.698778 | -1.125441 | -1.342449 | -1.357272 | -1.773527 | -1.587518 | -0.325362 | -0.659884 | -1.105966 |
| 23.875 | -2.085064 | -0.515357 | -1.833715 | -0.971349 | -2.025204 | -0.869507 | -1.141895 | -0.859019 | -0.961273 | -0.976379 | -2.15237 | -1.173131 | -1.527767 | -1.316843 | -0.594607 | -0.765323 | -1.132874 | -1.396718 | -1.71174 | -1.669145 | -1.548331 | -0.285167 | -0.903633 | -1.033324 |
| 23.88888889 | -2.46418 | -0.495819 | -2.144192 | -1.122559 | -2.053144 | -0.693633 | -1.116174 | -0.720381 | -1.273555 | -1.20576 | -2.272174 | -1.067239 | -1.741451 | -1.374249 | -0.711025 | -0.884444 | -1.236868 | -1.445258 | -1.956249 | -1.896223 | -1.629773 | -0.200962 | -1.313398 | -1.023854 |
| 23.90277778 | -2.751024 | -0.599602 | -1.976587 | -1.293736 | -2.29757 | -0.634367 | -1.201314 | -0.84207 | -1.546196 | -1.865083 | -2.554414 | -1.036676 | -2.25221 | -1.444572 | -0.780841 | -0.973258 | -1.461797 | -1.642946 | -2.239745 | -1.791278 | -1.705445 | -0.272963 | -1.475347 | -1.291218 |
| 23.91666667 | -2.935369 | -0.699781 | -2.193165 | -1.204953 | -2.438479 | -0.615617 | -1.223931 | -1.220775 | -1.717803 | -2.405426 | -2.493613 | -1.092341 | -2.567119 | -1.447268 | -0.997417 | -0.961989 | -1.588887 | -1.794495 | -2.410772 | -1.31702 | -1.708788 | -0.373359 | -1.073015 | -1.473688 |
| 23.93055556 | -2.852262 | -0.627802 | -1.850568 | -1.115938 | -2.451112 | -0.323264 | -1.022801 | -1.033201 | -1.665055 | -2.540768 | -2.306806 | -0.993402 | -2.673814 | -1.29039 | -1.081279 | -0.898759 | -1.784823 | -1.824848 | -2.524299 | -1.177835 | -1.73806 | -0.357963 | -0.920288 | -1.541281 |
| 23.94444444 | -3.197577 | -0.445938 | -1.762706 | -1.512779 | -3.259941 | 0.067788 | -0.906055 | -0.621863 | -1.719056 | -2.77754 | -2.79762 | -0.845904 | -2.918793 | -1.148774 | -0.938479 | -1.018212 | -2.03109 | -1.717539 | -2.494474 | -1.846654 | -1.824291 | -0.37121 | -0.945421 | -1.598886 |
| 23.95833333 | -3.80248 | -0.283326 | -1.63408 | -2.194487 | -4.901775 | 0.112065 | -1.101099 | -0.941744 | -1.383944 | -3.099512 | -3.294197 | -1.182204 | -3.041249 | -1.14581 | -1.099365 | -1.3016 | -2.435302 | -1.578657 | -2.227041 | -1.920594 | -1.900605 | -0.764576 | -1.049595 | -1.436685 |
| 23.97222222 | -4.153983 | -0.645421 | -1.855365 | -1.909454 | -5.88845 | -0.181223 | -1.614222 | -1.184498 | -1.533754 | -3.481106 | -4.095423 | -1.967484 | -3.212072 | -1.244938 | -1.591038 | -1.558121 | -3.197263 | -2.082739 | -2.279214 | -1.360134 | -1.629778 | -1.273315 | -1.510215 | -1.673777 |
| 23.98611111 | -3.328153 | -1.373083 | -2.27518 | -1.685122 | -4.203805 | -0.32572 | -1.634204 | -1.085859 | -1.819573 | -2.928304 | -4.162286 | -2.37155 | -2.786078 | -1.432087 | -1.63772 | -1.615498 | -2.432806 | -2.494389 | -2.180079 | -1.28475 | -1.379413 | -1.155538 | -1.661867 | -2.213956 |
| 24 | -1.520304 | -0.491156 | -1.227942 | -1.005486 | -2.292228 | -0.331869 | -0.955711 | -0.482195 | -1.168234 | -1.46923 | -2.069397 | -1.680925 | -1.239927 | -1.61845 | -0.375111 | -1.408087 | -1.660201 | -1.319481 | -1.295331 | -1.021733 | -0.56193 | -0.771644 | -1.179322 | -1.760186 |
| 24.01388889 | -0.132949 | 0.132012 | -0.18403 | 0.124677 | -0.777245 | -0.235822 | -0.489644 | 0.039976 | -0.443548 | -0.115646 | -0.287317 | -0.897006 | 0.258332 | -1.733529 | 0.774724 | -1.176301 | -0.817384 | -0.067487 | -0.218154 | -0.383051 | 1.109462 | -0.472284 | -0.048325 | -0.585386 |
| 24.02777778 | 0.71029 | -0.113507 | -0.020708 | 0.546482 | 0.456786 | -0.072802 | -0.085321 | 0.200908 | 0.125025 | 0.514481 | 0.411431 | -0.445095 | 1.153523 | -1.476071 | 0.838932 | -1.176267 | 0.329442 | 0.168231 | 0.075524 | 0.203424 | 1.402033 | 0.294974 | 0.488691 | 0.219716 |
| 24.04166667 | 0.945845 | 0.188476 | 0.061597 | 0.481703 | 1.23352 | 0.007151 | 0.039762 | 0.289047 | 0.400519 | 0.976458 | 0.974731 | -0.075363 | 1.38232 | -0.984823 | 0.943675 | -1.196695 | 0.79042 | 0.008187 | 0.268066 | 0.544438 | 0.704321 | 0.511239 | 0.689525 | 0.67853 |
| 24.05555556 | 0.904039 | 0.745408 | 0.214713 | 0.624613 | 1.557917 | 0.000892 | 0.082848 | 0.414702 | 0.502998 | 1.247377 | 1.41067 | 0.558786 | 1.552761 | -0.754933 | 1.426984 | -1.206651 | 0.297971 | 0.373293 | 0.686477 | 0.304981 | 0.686704 | 0.176304 | 1.094901 | 0.846951 |
| 24.06944444 | 0.879547 | 1.344626 | 0.349982 | 0.97598 | 1.858683 | -0.004885 | 0.179342 | 0.475129 | 0.796487 | 1.464298 | 1.271507 | 0.743017 | 1.98702 | -0.707209 | 1.016741 | -1.284167 | 0.170393 | 0.825666 | 0.823963 | -0.18697 | 1.422164 | 0.026176 | 1.037185 | 0.808296 |
| 24.08333333 | 1.10025 | 0.981604 | 0.882964 | 1.179553 | 2.019952 | 0.000767 | 0.208597 | 0.516626 | 1.329055 | 1.914557 | 0.928656 | 0.712708 | 1.682925 | -0.69427 | 0.338041 | -1.324048 | 0.8043 | 0.803023 | 0.485398 | -0.304504 | 1.648971 | 0.193198 | 0.69352 | 1.051548 |
| 24.09722222 | 1.289868 | 0.529925 | 1.048144 | 0.99674 | 2.160609 | 0.01844 | 0.347834 | 1.012915 | 1.766743 | 2.334931 | 1.051716 | 0.822774 | 1.676439 | -0.581828 | 0.275772 | -0.588713 | 0.64641 | 0.624427 | 0.483745 | -0.112682 | 1.053341 | 0.39322 | 0.405913 | 1.301118 |
| 24.11111111 | 2.578233 | 0.643186 | 0.338875 | 0.831476 | 3.300351 | -0.11981 | 0.543283 | 1.524876 | 1.234111 | 2.361786 | 2.128978 | 0.975912 | 1.58304 | -0.516136 | 0.159508 | 0.230468 | 0.092765 | 0.478531 | 0.650891 | -0.064318 | 0.467929 | 0.159248 | 0.518081 | 1.228915 |
| 24.125 | 3.841224 | 1.413285 | 1.230035 | 1.188547 | 3.779576 | -1.206262 | 0.60574 | 1.437215 | 0.063888 | 1.429289 | 2.515222 | 0.478397 | 1.224069 | -0.533087 | -0.749264 | 0.322909 | 0.035974 | 0.172406 | 0.505716 | -0.229483 | 0.556299 | -0.126177 | 0.636608 | 0.819252 |
| 24.13888889 | 0.119562 | -0.509452 | 0.625146 | -0.23947 | -0.845723 | -1.207035 | -1.655032 | -1.446099 | -1.236838 | -3.606612 | -0.369599 | -1.674301 | 1.890225 | -0.354179 | -0.936558 | 0.048776 | 0.291092 | -0.033695 | 0.345895 | -0.196241 | 0.930628 | 0.176285 | 0.425407 | 0.812807 |
| 24.15277778 | 1.623695 | 0.636266 | 0.77182 | -2.278998 | -2.187569 | -3.535278 | -1.727628 | 0.125819 | 1.117124 | -1.404753 | -0.949556 | -1.836029 | 1.27113 | -0.034455 | -0.35748 | 0.017299 | -0.169423 | -0.054491 | 0.344138 | -0.047272 | 1.048483 | 0.737507 | 0.22609 | 0.800339 |
| 24.16666667 | 3.008914 | 1.716171 | 1.578988 | -0.081414 | -0.774439 | -1.696395 | -1.193416 | 0.786736 | 0.183943 | -0.060603 | -0.361785 | -0.701507 | 0.930273 | 0.153882 | 0.202398 | 0.082638 | -0.25418 | -0.018512 | 0.408031 | 0.012953 | 0.651059 | 0.493909 | 0.098476 | 0.448747 |
| 24.18055556 | 3.26446 | 2.065003 | 1.911764 | 1.791178 | -0.267428 | 0.190287 | -1.136907 | 1.208594 | 0.024124 | 1.111205 | 1.85957 | 0.591856 | 0.496995 | 0.354414 | 0.478921 | 0.158064 | 0.300277 | -0.125562 | 0.349723 | 0.080308 | 0.284052 | -0.336446 | -0.140882 | 0.324445 |
| 24.19444444 | 2.407329 | 2.223141 | 2.226183 | 2.179093 | 0.984234 | 0.938977 | -1.037608 | 1.117702 | 0.414383 | 1.866375 | 2.944624 | 1.483782 | 0.25567 | 0.479522 | 0.488748 | 0.430907 | 0.238903 | -0.03128 | 0.251237 | -0.039156 | 0.263296 | -0.432641 | -0.213344 | 0.246494 |
| 24.20833333 | 1.87196 | 2.531598 | 2.182157 | 2.155852 | 1.525271 | 1.172429 | -0.457469 | 1.052839 | 0.902541 | 2.189131 | 2.984356 | 1.905711 | 0.347381 | 0.330495 | 0.377851 | 0.534815 | -0.193902 | 0.258978 | 0.146294 | -0.295196 | 0.185016 | -0.009269 | -0.06142 | -0.140095 |
| 24.22222222 | 1.855336 | 2.501663 | 1.844055 | 2.069738 | 1.723415 | 1.361773 | 0.004211 | 1.022117 | 1.07601 | 2.078795 | 3.069862 | 2.232277 | 0.463965 | 0.003879 | 0.335142 | 0.284564 | -0.382254 | 0.336174 | 0.087507 | -0.36535 | 0.00795 | 0.167366 | 0.033666 | -0.38642 |
| 24.23611111 | 2.31023 | 2.669229 | 1.68272 | 2.355146 | 1.756504 | 1.126323 | 0.256536 | 1.040882 | 0.986582 | 1.996557 | 2.829773 | 2.178 | 0.271297 | 0.009721 | 0.335464 | 0.399085 | -0.379031 | 0.181758 | -0.002757 | -0.228313 | -0.27144 | 0.089908 | 0.111174 | -0.562032 |
| 24.25 | 2.753494 | 2.572395 | 1.254649 | 2.164488 | 1.865838 | 0.669551 | 0.342247 | 0.960989 | 0.952586 | 1.883894 | 2.432888 | 1.863011 | -0.132444 | 0.463885 | 0.358927 | 0.816504 | -0.033978 | 0.028349 | 0.062947 | -0.023546 | -0.328052 | 0.296042 | 0.070302 | -0.603147 |
| 24.26388889 | 3.442608 | 2.983325 | 1.048823 | 1.885887 | 2.424662 | 0.434179 | 0.57593 | 1.033005 | 1.188388 | 2.20052 | 2.541481 | 1.514936 | -0.254622 | 0.759147 | 0.423535 | 0.871343 | 0.226285 | 0.207849 | 0.299769 | 0.450659 | -0.016798 | 0.798776 | 0.156123 | -0.204858 |
| 24.27777778 | 2.962047 | 3.251462 | 0.792293 | 1.477662 | 2.530714 | 0.44461 | 0.629684 | 0.850718 | 0.939411 | 2.261655 | 2.706523 | 1.178675 | 0.00702 | 0.692076 | 0.207066 | 0.598768 | 0.567089 | 0.494526 | 0.58779 | 0.845345 | 0.413568 | 0.757744 | 0.26357 | 0.264221 |
| 24.29166667 | 2.12589 | 3.206925 | 0.384269 | 1.077188 | 1.971253 | 0.548117 | 0.441094 | 0.337929 | 0.579246 | 1.577467 | 2.451355 | 1.089177 | 0.339374 | 0.558851 | -0.297351 | 0.612734 | 0.387936 | 0.553385 | 0.760855 | 0.656267 | 0.737031 | 0.469361 | 0.248533 | 0.29379 |
| 24.30555556 | 1.929009 | 2.227691 | 0.176673 | 0.812592 | 1.458291 | 0.567543 | 0.294778 | 0.118948 | 0.515139 | 1.007248 | 2.256268 | 0.974592 | 0.69255 | 0.661265 | -0.245901 | 0.821248 | 0.407469 | 0.601584 | 1.010802 | 0.665397 | 1.22062 | 0.74208 | 0.268488 | 0.135814 |
| 24.31944444 | 1.965937 | 2.062219 | 0.141105 | 0.896272 | 1.194495 | 0.499686 | 0.307665 | 0.248732 | 0.447584 | 0.819719 | 2.129064 | 0.830696 | 0.959917 | 0.671079 | 0.450865 | 0.936012 | 0.325113 | 0.946475 | 1.632262 | 0.744997 | 1.795571 | 1.024347 | 0.3511 | 0.206639 |
| 24.33333333 | 1.742567 | 1.932835 | 0.133414 | 0.969706 | 1.013179 | 0.434573 | 0.460924 | 0.446761 | 0.394733 | 0.836435 | 2.151407 | 0.703449 | 1.074957 | 0.508678 | 1.008382 | 1.044684 | 0.333514 | 1.160802 | 1.910472 | 0.586852 | 1.807992 | 0.831727 | 0.462445 | 0.48705 |
| 24.34722222 | 1.547512 | 1.789247 | 0.144161 | 0.666594 | 0.891754 | 0.413758 | 0.455512 | 0.498373 | 0.345003 | 0.869966 | 2.26751 | 0.334868 | 1.196392 | 0.345292 | 0.845682 | 1.084219 | 0.664925 | 1.054567 | 1.765546 | 0.562046 | 1.393678 | 0.700835 | 0.40501 | 0.783668 |
| 24.36111111 | 1.37518 | 1.686736 | 0.150815 | 0.479412 | 0.834012 | 0.427204 | 0.408666 | 0.538128 | 0.253421 | 0.897803 | 2.070051 | 0.393307 | 1.354713 | 0.409775 | 0.54267 | 1.06079 | 0.82269 | 1.084378 | 1.775132 | 0.515849 | 1.500279 | 0.805947 | 0.405731 | 1.006268 |
| 24.375 | 1.123362 | 1.590748 | 0.150033 | 0.371225 | 0.770315 | 0.378571 | 0.385841 | 0.534946 | 0.211657 | 1.067969 | 1.976844 | 0.420335 | 1.404263 | 0.678981 | 0.379711 | 1.068083 | 0.63984 | 1.294551 | 1.992244 | 0.414938 | 1.992218 | 0.89684 | 0.32131 | 1.046762 |
| 24.38888889 | 0.930375 | 1.628088 | 0.186088 | 0.41117 | 0.727315 | 0.309727 | 0.380321 | 0.491895 | 0.355398 | 1.341852 | 1.94485 | 0.37325 | 1.515286 | 0.848273 | 0.427707 | 1.142404 | 0.874712 | 1.451476 | 2.13445 | 0.28566 | 2.35414 | 0.928142 | 0.370699 | 0.891068 |
| 24.40277778 | 0.93591 | 1.668732 | 0.261368 | 0.51533 | 0.792582 | 0.326815 | 0.392977 | 0.447209 | 0.494165 | 1.317331 | 1.977142 | 0.416922 | 1.704526 | 0.778502 | 0.873439 | 1.136759 | 1.367278 | 1.269535 | 1.781336 | 0.233909 | 2.241467 | 1.047223 | 0.53317 | 0.839586 |
| 24.41666667 | 1.021588 | 1.880609 | 0.313569 | 0.755468 | 0.894358 | 0.372084 | 0.378244 | 0.40592 | 0.435549 | 1.069977 | 2.12847 | 0.442666 | 1.946364 | 0.606229 | 1.165868 | 1.130705 | 1.419678 | 1.173639 | 1.812543 | 0.238657 | 2.224503 | 1.389327 | 0.700025 | 1.009387 |
| 24.43055556 | 1.167286 | 1.92899 | 0.355674 | 0.896801 | 1.120665 | 0.307353 | 0.35083 | 0.32275 | 0.49257 | 0.988818 | 2.292612 | 0.465198 | 1.959423 | 0.520682 | 1.10527 | 1.234003 | 1.526128 | 1.305557 | 1.841613 | 0.216638 | 2.279365 | 1.757412 | 1.019041 | 1.411784 |
| 24.44444444 | 1.58623 | 1.994324 | 0.528906 | 0.787438 | 1.771607 | 0.246535 | 0.3475 | 0.437858 | 0.762759 | 1.150381 | 2.843078 | 0.533343 | 1.873988 | 0.568979 | 0.857038 | 1.266546 | 1.861215 | 1.338738 | 1.676444 | 0.238909 | 1.890086 | 1.84567 | 1.13116 | 2.131649 |
| 24.45833333 | 2.324785 | 2.093797 | 0.96015 | 1.021203 | 2.848574 | 0.419887 | 0.619205 | 0.889318 | 1.663562 | 1.882475 | 4.1783 | 0.805658 | 1.28721 | 0.430468 | 0.685713 | 0.846202 | 1.684066 | 1.020164 | 1.450438 | 0.175999 | 0.951127 | 1.471381 | 0.921899 | 2.232479 |
| 24.47222222 | 2.003013 | 1.928653 | 1.239308 | 1.139746 | 2.834469 | 0.606365 | 0.859343 | 1.175233 | 2.605602 | 2.534902 | 4.48515 | 1.23088 | 0.849131 | 0.217859 | 0.588245 | 0.333795 | 1.550177 | 0.732556 | 1.235177 | 0.172091 | 0.800701 | 0.889647 | 0.674925 | 1.512427 |
| 24.48611111 | 1.293704 | 1.448714 | 1.504909 | 1.289429 | 2.177856 | 0.531394 | 0.797671 | 1.139829 | 2.28659 | 2.123785 | 3.015212 | 1.327802 | 0.997035 | 0.652398 | 0.541959 | 0.316431 | 2.202995 | 0.910608 | 1.366112 | 0.275307 | 1.181772 | 0.812042 | 0.840326 | 1.284258 |
| 24.5 | 0.78646 | 0.810561 | 1.374626 | 1.23014 | 1.550016 | 0.274936 | 0.645053 | 0.868029 | 1.655963 | 1.475814 | 2.073161 | 1.08102 | 1.634615 | 1.792922 | 0.693892 | 0.963275 | 2.212233 | 1.155187 | 1.61182 | 0.343593 | 0.921102 | 1.156262 | 1.339776 | 1.811933 |
| 24.51388889 | 0.513613 | 0.488104 | 1.26682 | 1.060387 | 1.201178 | 0.124357 | 0.538942 | 0.773836 | 1.288434 | 1.058094 | 1.574228 | 0.812571 | 2.180415 | 2.181243 | 0.851119 | 1.595846 | 1.686271 | 1.423406 | 1.742929 | 0.463029 | 0.639387 | 1.087797 | 1.532323 | 2.358872 |
| 24.52777778 | 0.35831 | 0.253312 | 1.075291 | 0.766081 | 1.032073 | 0.125832 | 0.545419 | 0.717526 | 1.084153 | 0.803412 | 1.144491 | 0.553044 | 1.793341 | 2.133598 | 0.897362 | 1.426845 | 1.076831 | 1.262145 | 1.624084 | 0.377612 | 0.669757 | 0.870061 | 1.315805 | 2.750844 |
| 24.54166667 | 0.270863 | 0.123074 | 0.859468 | 0.519958 | 0.880033 | 0.135225 | 0.508467 | 0.649634 | 0.924597 | 0.59016 | 0.780118 | 0.347865 | 1.093578 | 2.30098 | 0.685734 | 0.823212 | 0.755454 | 0.724189 | 1.107112 | 0.3707 | 0.546104 | 0.472103 | 0.911997 | 2.265635 |
| 24.55555556 | 0.222642 | 0.10087 | 0.702901 | 0.406666 | 0.762463 | 0.123821 | 0.377752 | 0.450308 | 0.847341 | 0.444048 | 0.593464 | 0.288522 | 0.972638 | 1.939514 | 0.378473 | 0.449786 | 0.611302 | 0.230241 | 1.052805 | 0.379846 | 0.391456 | 0.087047 | 0.775925 | 1.691814 |
| 24.56944444 | 0.241346 | 0.134954 | 0.640349 | 0.374707 | 0.657783 | 0.075605 | 0.2605 | 0.289432 | 0.895029 | 0.433475 | 0.455756 | 0.334243 | 1.323742 | 0.765408 | 0.20327 | 0.502779 | 0.633119 | 0.047717 | 1.078173 | 0.247789 | 0.183338 | -0.202789 | 0.813766 | 1.066995 |
| 24.58333333 | 0.208898 | 0.223569 | 0.660624 | 0.344395 | 0.547709 | 0.018203 | 0.246639 | 0.279592 | 0.865331 | 0.466293 | 0.310469 | 0.285854 | 1.839104 | -0.373413 | 0.144283 | 0.583835 | 0.336489 | 0.204336 | 1.14114 | 0.025339 | -0.112353 | 0.07677 | 0.750956 | 0.430038 |
| 24.59722222 | 0.166763 | 0.380071 | 0.640361 | 0.269427 | 0.434966 | -0.006063 | 0.210493 | 0.245012 | 0.721618 | 0.480207 | 0.241711 | 0.169119 | 1.720901 | -0.473742 | 0.126004 | 0.366099 | 0.148561 | 0.391937 | 1.121579 | -0.195111 | -0.076915 | 0.386565 | 0.581763 | 0.458111 |
| 24.61111111 | 0.157641 | 0.437807 | 0.563436 | 0.176976 | 0.365976 | -0.015063 | 0.15403 | 0.216055 | 0.583065 | 0.459138 | 0.187761 | 0.103397 | 1.395361 | 0.402428 | 0.117404 | 0.062029 | 0.292656 | 0.340486 | 0.558579 | -0.401478 | -0.122602 | 0.264664 | 0.316857 | 0.808117 |
| 24.625 | 0.164144 | 0.278164 | 0.468336 | 0.187714 | 0.348967 | -0.045096 | 0.103886 | 0.221739 | 0.485382 | 0.387307 | 0.077765 | 0.060085 | 0.997947 | 0.924456 | 0.110629 | -0.063998 | 0.360759 | 0.149671 | 0.130746 | -0.542004 | -0.42879 | 0.162311 | -0.028253 | 0.927911 |
| 24.63888889 | 0.167211 | 0.223435 | 0.382209 | 0.287057 | 0.362795 | -0.077191 | 0.114359 | 0.227289 | 0.378887 | 0.232427 | 0.119608 | 0.050045 | 0.602769 | 0.689533 | 0.082936 | 0.004721 | 0.189358 | 0.04322 | -0.259673 | -0.588619 | -0.601608 | 0.132264 | -0.310211 | 0.834551 |
| 24.65277778 | 0.23771 | 0.242933 | 0.319751 | 0.272203 | 0.357833 | -0.088447 | 0.142235 | 0.230355 | 0.313728 | 0.13472 | 0.214802 | 0.082209 | 0.153321 | 0.550887 | 0.067017 | 0.075629 | 0.037264 | -0.026633 | -0.494811 | -0.502574 | -0.266803 | 0.216037 | -0.164059 | 0.803647 |
| 24.66666667 | 0.35205 | 0.176112 | 0.39386 | 0.067833 | 0.301013 | -0.062514 | 0.088263 | 0.196253 | 0.31395 | 0.12692 | 0.193799 | 0.090434 | -0.195502 | 0.968051 | 0.020367 | 0.029406 | 0.262574 | -0.111866 | -0.353458 | -0.300603 | -0.168944 | 0.170659 | -0.092505 | 0.825552 |
| 24.68055556 | 0.440592 | 0.133676 | 0.594242 | -0.05032 | 0.23717 | -0.027706 | 0.019228 | 0.195602 | 0.375668 | 0.190266 | 0.125961 | 0.063448 | -0.314985 | 1.292613 | -0.049284 | -0.010598 | 0.41641 | -0.05847 | -0.039181 | -0.109294 | -0.259705 | -0.147065 | -0.018824 | 0.717715 |
| 24.69444444 | 0.408883 | 0.188511 | 0.729575 | 0.059075 | 0.200098 | -0.00462 | -0.020089 | 0.152182 | 0.549536 | 0.351717 | 0.090405 | 0.008433 | -0.152071 | 1.046525 | -0.010627 | 0.031057 | 0.19477 | 0.058143 | 0.448237 | 0.066399 | -0.326449 | -0.195964 | 0.062998 | 0.549875 |
| 24.70833333 | 0.340975 | 0.243689 | 0.683075 | 0.236739 | 0.175459 | -0.023087 | -0.032495 | 0.087153 | 0.718114 | 0.478596 | 0.117488 | -0.03375 | 0.025167 | 0.767402 | 0.03776 | 0.011005 | 0.208649 | 0.180693 | 0.732695 | 0.136407 | -0.371323 | -0.026159 | 0.100186 | 0.51727 |
| 24.72222222 | 0.329504 | 0.203701 | 0.525139 | 0.265382 | 0.168678 | -0.03021 | -0.002544 | 0.064136 | 0.75364 | 0.428392 | 0.164419 | 0.012439 | 0.010262 | 0.832917 | -0.027818 | -0.091394 | 0.265202 | 0.154698 | 0.673661 | 0.157862 | -0.405484 | 0.268821 | 0.096832 | 0.565249 |
| 24.73611111 | 0.333481 | 0.153168 | 0.438036 | 0.092443 | 0.179479 | -0.011634 | 0.007906 | 0.083333 | 0.748056 | 0.310872 | 0.216548 | 0.110989 | -0.280199 | 0.915986 | -0.140418 | -0.228139 | 0.208342 | -0.029324 | 0.398928 | 0.016233 | -0.392683 | 0.47759 | 0.189481 | 0.569066 |
| 24.75 | 0.348563 | 0.155131 | 0.451692 | -0.054596 | 0.21277 | 0.010316 | 0.039205 | 0.133185 | 0.832602 | 0.304115 | 0.288405 | 0.216024 | -0.641132 | 0.785252 | -0.186219 | -0.284912 | -0.220805 | -0.245676 | 0.082926 | -0.586768 | -0.477017 | 0.248958 | 0.222109 | 0.476294 |
| 24.76388889 | 0.369785 | 0.232963 | 0.534124 | 0.03299 | 0.263371 | 0.003475 | 0.052124 | 0.168515 | 0.878335 | 0.278303 | 0.389246 | 0.295117 | -0.931341 | 0.654359 | -0.204107 | -0.191944 | -0.536206 | -0.362651 | -0.190337 | -1.481877 | -0.576462 | -0.018562 | 0.156429 | 0.41861 |
| 24.77777778 | 0.388804 | 0.328986 | 0.643971 | 0.176431 | 0.315876 | -0.043737 | 0.031875 | 0.167371 | 0.862543 | 0.195661 | 0.445901 | 0.336774 | -1.309352 | 0.333914 | -0.155078 | -0.164176 | -0.302839 | -0.407498 | -0.318356 | -1.603418 | -0.479386 | -0.061896 | 0.043509 | 0.398573 |
| 24.79166667 | 0.417392 | 0.33133 | 0.76135 | 0.098306 | 0.377854 | -0.061154 | 0.019132 | 0.192114 | 0.789832 | 0.126255 | 0.490358 | 0.372449 | -1.888376 | -0.041221 | -0.099425 | -0.343427 | -0.233447 | -0.466813 | -0.300551 | -1.458809 | -0.359574 | 0.09653 | -0.050776 | 0.268797 |
| 24.80555556 | 0.432581 | 0.371908 | 0.902649 | -0.093555 | 0.439679 | 0.012751 | 0.061527 | 0.268468 | 0.75225 | 0.126265 | 0.554508 | 0.406556 | -2.458603 | -0.083174 | -0.170584 | -0.540219 | -0.747627 | -0.513437 | -0.252168 | -1.518867 | -0.32457 | 0.067362 | -0.036806 | 0.134432 |
| 24.81944444 | 0.420881 | 0.424053 | 0.924391 | -0.016244 | 0.459115 | 0.082486 | 0.131441 | 0.330469 | 0.652227 | 0.129702 | 0.567769 | 0.393832 | -2.591992 | -0.023397 | -0.243311 | -0.552197 | -1.193503 | -0.587102 | -0.357747 | -1.653247 | -0.018568 | -0.029584 | 0.008688 | -0.167961 |
| 24.83333333 | 0.374559 | 0.296838 | 0.877878 | 0.241083 | 0.45666 | 0.081004 | 0.170646 | 0.349455 | 0.457951 | 0.095486 | 0.487429 | 0.364123 | -2.4594 | -0.260831 | -0.232641 | -0.534532 | -0.984095 | -0.419628 | -0.455982 | -0.727222 | 0.40748 | -0.151354 | 0.010209 | -0.559028 |
| 24.84722222 | 0.300678 | 0.200338 | 0.704525 | 0.338877 | 0.456432 | 0.009645 | 0.155475 | 0.33243 | 0.256198 | 0.071887 | 0.418797 | 0.361301 | -2.348802 | -0.650877 | -0.298093 | -0.621973 | -0.790587 | -0.261856 | -0.435053 | 0.928029 | 0.492155 | -0.177055 | -0.006721 | -0.996572 |
| 24.86111111 | 0.2359 | -0.051421 | 0.576226 | 0.090933 | 0.504698 | -0.076013 | 0.142314 | 0.298109 | 0.091909 | -0.029604 | 0.356638 | 0.37509 | -2.491415 | -0.983068 | -0.470669 | -0.830562 | -1.136452 | -0.707399 | -0.672393 | -2.186494 | -0.005993 | -0.475631 | 0.157411 | -1.227112 |
| 24.875 | 0.268764 | -0.183298 | 0.511254 | -0.088319 | 0.63165 | -0.138962 | 0.079077 | 0.230138 | -0.066734 | -0.190855 | 0.303348 | 0.365726 | -2.80474 | -1.445013 | -0.635675 | -0.896371 | -1.623795 | -1.17686 | -0.79908 | -2.546613 | -0.561562 | -0.756223 | 0.044575 | -1.066038 |
| 24.88888889 | 0.308672 | -0.216834 | 0.523079 | -0.14049 | 0.773934 | -0.192375 | -0.126311 | -0.061439 | -0.19161 | -0.336119 | 0.329573 | 0.307083 | -2.893493 | -2.34759 | -0.729253 | -0.939811 | -1.58636 | -1.079034 | -0.41107 | -2.38891 | -0.402425 | -0.569057 | -0.185186 | -1.015843 |
| 24.90277778 | 0.275272 | -0.238761 | 0.646837 | -0.148302 | 0.867621 | -0.223586 | -0.230821 | -0.336391 | -0.33754 | -0.622103 | 0.305784 | 0.214115 | -2.725622 | -3.336668 | -0.750766 | -1.028946 | -1.129686 | -1.046125 | -0.469628 | -2.887243 | -0.433609 | -0.521639 | -0.386183 | -1.034057 |
| 24.91666667 | 0.244928 | -0.193563 | 0.844399 | -0.290806 | 0.951022 | -0.178569 | -0.086595 | -0.288938 | -0.343664 | -0.671961 | 0.372634 | 0.19176 | -2.688825 | -3.442806 | -0.855258 | -1.084496 | -1.699967 | -1.143507 | -0.904478 | -3.24437 | -0.678225 | -0.514784 | -0.361905 | -0.853408 |
| 24.93055556 | 0.203214 | 0.064384 | 1.045658 | -0.598695 | 1.021388 | -0.023457 | 0.06058 | 0.021844 | -0.308999 | -0.581521 | 0.520083 | 0.238612 | -2.779436 | -2.195503 | -1.023324 | -1.035993 | -2.465283 | -1.31325 | -1.667076 | -3.359929 | -0.659084 | -0.316361 | -0.383392 | -0.486915 |
| 24.94444444 | 0.009168 | 0.206787 | 0.862956 | -0.696892 | 0.521222 | 0.070955 | 0.004497 | -0.018313 | -0.181655 | -0.557278 | 0.382129 | 0.157016 | -2.865876 | -1.248916 | -1.064039 | -0.976995 | -2.516306 | -1.52313 | -2.080724 | -3.817865 | -0.848479 | 0.23164 | -0.79506 | -0.197538 |
| 24.95833333 | -0.630907 | 0.219038 | 0.324356 | -0.578203 | -0.651094 | -0.037408 | -0.339666 | -0.497583 | -0.256697 | -0.777705 | -0.399959 | -0.221007 | -2.687129 | -1.878954 | -0.978472 | -1.262548 | -2.823489 | -1.8188 | -2.303383 | -3.040819 | -0.961379 | 0.169917 | -1.282143 | -0.496844 |
| 24.97222222 | -1.540145 | 0.15316 | -0.28335 | -0.624659 | -1.57865 | -0.14876 | -0.449723 | -0.894569 | -0.578405 | -1.156787 | -1.441771 | -0.562167 | -2.632224 | -3.297827 | -1.187404 | -1.68949 | -3.525867 | -2.391642 | -2.729429 | -1.618776 | -1.570097 | -0.134572 | -2.042882 | -1.398949 |
| 24.98611111 | -1.585753 | -0.045624 | -0.374854 | -0.738266 | -1.164825 | -0.167898 | -0.36444 | -0.70821 | -0.700733 | -1.034065 | -0.855979 | -0.476063 | -2.928244 | -3.182753 | -1.405969 | -1.796545 | -2.806817 | -2.636006 | -2.807694 | -0.992445 | -2.47946 | -0.707021 | -2.466838 | -1.558218 |
| 25 | -0.70498 | -0.148111 | 0.0016 | -0.650864 | -0.252009 | -0.12723 | -0.380565 | -0.215481 | -0.651253 | -0.549715 | 0.024465 | -0.385244 | -2.380922 | -0.175366 | -1.017714 | -1.210086 | -1.014097 | -1.139736 | -1.576683 | -1.010442 | -1.138478 | -0.478383 | -1.449429 | -0.691202 |
| 25.01388889 | -0.208772 | -0.088361 | 0.047779 | -0.760835 | -0.056897 | -0.095209 | -0.400841 | -0.078449 | -0.603565 | -0.266458 | -0.033863 | -0.503382 | -1.368764 | 0.687697 | -0.073885 | -0.985231 | 0.035291 | 0.718794 | -0.399362 | -0.458723 | 1.28144 | 0.302961 | -0.671548 | -0.067345 |
| 25.02777778 | -0.112558 | -0.095607 | -0.001952 | -0.732329 | -0.158019 | -0.025064 | -0.254758 | -0.145242 | -0.437227 | -0.057689 | -0.16323 | -0.41788 | -0.766223 | -0.387341 | 0.302499 | -1.10878 | 0.672184 | 0.898947 | -0.393261 | 0.086013 | 1.37649 | 0.235634 | -0.156803 | -0.03869 |
| 25.04166667 | -0.084978 | 0.051959 | -0.001821 | -0.635987 | 0.025831 | 0.062752 | -0.03606 | -0.095843 | -0.131084 | 0.133079 | -0.008554 | -0.225421 | -0.494438 | -1.366624 | 0.253376 | -1.119251 | 1.346751 | 0.286151 | -0.897458 | 0.313601 | 0.83345 | -0.009736 | 0.211314 | -0.0629 |
| 25.05555556 | 0.021189 | 0.564614 | -0.042732 | -0.493605 | 0.348452 | 0.097165 | 0.030208 | 0.016909 | 0.073903 | 0.258546 | 0.138093 | -0.203304 | 0.059833 | -0.872816 | 0.444761 | -1.079041 | 1.841795 | 0.577839 | -0.306966 | 0.374467 | 1.226001 | 0.101405 | 0.383735 | 0.060699 |
| 25.06944444 | 0.054617 | 1.182073 | -0.017375 | -0.411061 | 0.62229 | 0.07459 | -0.067184 | 0.156433 | 0.116202 | 0.385367 | 0.191601 | -0.219561 | 0.877025 | 0.11355 | 0.963516 | -1.172695 | 1.571365 | 1.383892 | 0.280226 | 0.421458 | 1.223265 | 0.331236 | 0.225315 | 0.201632 |
| 25.08333333 | -0.013804 | 0.976288 | -0.019749 | -0.226713 | 0.811151 | 0.103496 | -0.135379 | 0.340607 | 0.193155 | 0.540782 | 0.27117 | -0.173673 | 1.66686 | 0.165208 | 1.249502 | -1.164223 | 1.221626 | 1.04567 | -0.098569 | 0.46777 | 0.994073 | 0.372615 | 0.372506 | 0.302465 |
| 25.09722222 | 0.094712 | 0.420201 | -0.032198 | -0.245143 | 0.957592 | 0.061073 | -0.098981 | 0.541548 | 0.352843 | 0.633696 | 0.387811 | -0.055172 | 1.790994 | -0.599402 | 1.348495 | -0.833433 | 0.423074 | 0.447498 | -0.52276 | 0.474814 | 1.267118 | 0.34195 | 0.57071 | 0.287284 |
| 25.11111111 | 0.406823 | -0.054086 | 0.016957 | -0.39139 | 1.088684 | -0.031054 | -0.051021 | 0.614458 | 0.437196 | 0.557789 | 0.561018 | 0.031755 | 1.27768 | -1.196818 | 1.131459 | -0.513228 | -0.090296 | 0.470194 | -0.36345 | 0.473008 | 1.391279 | 0.442923 | 0.747866 | 0.286466 |
| 25.125 | 0.522723 | 0.176131 | 0.149333 | -0.410487 | 1.28677 | -0.075021 | -0.034688 | 0.574243 | 0.26202 | 0.319252 | 0.651378 | 0.028635 | 1.08486 | -0.976339 | 1.155906 | -0.454497 | 0.208252 | 0.52375 | -0.224472 | 0.522498 | 0.763762 | 0.448401 | 0.364575 | 0.213726 |
| 25.13888889 | 0.379787 | 0.423672 | 0.254971 | -0.349491 | 1.47681 | -0.032849 | -0.093172 | 0.553975 | -0.162066 | -0.018889 | 0.596823 | -0.021705 | 1.468391 | -0.351949 | 0.82862 | -0.386942 | 0.462631 | 0.29632 | -0.332238 | 0.209062 | 0.467538 | 0.28708 | -0.185387 | 0.245371 |
| 25.15277778 | 0.134969 | 0.28593 | 0.278559 | -0.315811 | 1.46464 | -0.069999 | -0.09567 | 0.521776 | -0.508609 | -0.257914 | 0.445897 | -0.062437 | 1.535995 | 0.170389 | 0.642042 | -0.205927 | 0.79116 | -0.03265 | -0.09991 | -0.264867 | 0.850188 | 0.227619 | -0.349089 | 0.450741 |
| 25.16666667 | -0.034245 | 0.201687 | 0.344096 | -0.206732 | 1.274051 | 0.018222 | 0.021637 | 0.497625 | -0.661528 | -0.302056 | 0.24715 | -0.095008 | 2.256427 | 0.590853 | 0.320405 | -0.032268 | 0.743197 | -0.017385 | 0.224407 | -0.737293 | 0.929989 | 0.27245 | -0.395221 | 0.672128 |
| 25.18055556 | -0.087031 | 0.229404 | 0.379173 | -0.132668 | 1.03346 | 0.148783 | 0.122214 | 0.458054 | -0.69314 | -0.155688 | 0.075589 | -0.163052 | 2.273688 | 0.926294 | 0.238192 | 0.102126 | 0.636859 | 0.017124 | 0.233238 | -0.55408 | 0.806744 | 0.192201 | -0.227193 | 0.723074 |
| 25.19444444 | -0.118935 | 0.338 | 0.403413 | -0.101631 | 0.806862 | 0.091101 | 0.196727 | 0.335471 | -0.671404 | -0.050999 | -0.108652 | -0.276702 | 1.663254 | 0.930407 | 0.319856 | 0.195424 | 1.028375 | 0.07123 | 0.355905 | -0.056728 | 1.085841 | 0.015958 | 0.03628 | 0.711563 |
| 25.20833333 | -0.186858 | 0.250436 | 0.319907 | -0.106685 | 0.632409 | 0.146065 | 0.245923 | 0.259196 | -0.756361 | -0.133766 | -0.294829 | -0.320901 | 1.415585 | 0.669737 | 0.102115 | 0.305835 | 1.024943 | 0.223635 | 0.68954 | 0.171084 | 1.010744 | -0.074111 | 0.288095 | 0.770181 |
| 25.22222222 | -0.337891 | 0.027978 | 0.161068 | -0.266069 | 0.500015 | 0.205798 | 0.217718 | 0.243626 | -0.876261 | -0.272338 | -0.402062 | -0.246382 | 1.434668 | 0.665885 | -0.24627 | 0.394917 | 0.287732 | 0.622259 | 0.815056 | 0.066975 | 0.403502 | -0.09364 | 0.422904 | 0.754856 |
| 25.23611111 | -0.466754 | -0.010419 | 0.032927 | -0.237826 | 0.366974 | 0.143241 | 0.134706 | 0.183516 | -0.904102 | -0.343513 | -0.392257 | -0.154882 | 1.282391 | 0.801419 | -0.229253 | 0.421346 | -0.22251 | 0.67003 | 0.684335 | 0.029698 | -0.017943 | -0.030251 | 0.435317 | 0.53775 |
| 25.25 | -0.423709 | 0.145131 | -0.048222 | 0.124291 | 0.300691 | 0.0667 | 0.092484 | 0.103835 | -0.846407 | -0.298664 | -0.187477 | -0.030638 | 1.038271 | 0.796155 | 0.133945 | 0.393387 | -0.149388 | 0.511689 | 0.68884 | 0.118566 | 0.078416 | -0.006528 | 0.372655 | 0.41481 |

| | | | | | | | | | | | | | | | | | | | | | | | | |
|---|---|---|---|---|---|---|---|---|---|---|---|---|---|---|---|---|---|---|---|---|---|---|---|---|
| 25.26388889 | -0.17204 | 0.153347 | -0.064493 | 0.262103 | 0.392569 | 0.110004 | 0.166673 | 0.13079 | -0.78139 | -0.280149 | 0.072256 | 0.09309 | 0.866607 | 1.024523 | 0.307985 | 0.395026 | 0.28013 | 0.472485 | 0.701612 | 0.333911 | 0.310664 | 0.042632 | 0.26908 | 0.494008 |
| 25.27777778 | 0.060588 | -0.029539 | -0.08152 | -0.124238 | 0.498989 | 0.203625 | 0.293953 | 0.333555 | -0.745484 | -0.276478 | 0.243184 | 0.239585 | 0.857998 | 1.5495 | 0.301945 | 0.334974 | 0.741076 | 0.573388 | 0.773795 | 0.63426 | 0.248683 | 0.278215 | 0.157693 | 0.581278 |
| 25.29166667 | 0.14188 | -0.381726 | -0.171094 | -0.630212 | 0.508373 | 0.249227 | 0.383582 | 0.50819 | -0.695871 | -0.25594 | 0.332358 | 0.333331 | 1.083467 | 1.722219 | 0.371114 | 0.175144 | 0.67477 | 0.629522 | 0.891437 | 0.808717 | 0.132087 | 0.451526 | 0.142633 | 0.584753 |
| 25.30555556 | 0.191925 | -0.60249 | -0.149074 | -0.523303 | 0.486139 | 0.229402 | 0.393753 | 0.581792 | -0.629046 | -0.205092 | 0.392876 | 0.368123 | 1.325975 | 1.321336 | 0.546653 | 0.143817 | 0.124511 | 0.541519 | 0.978841 | 0.924962 | 0.319263 | 0.546267 | 0.327213 | 0.597501 |
| 25.31944444 | 0.31312 | -0.278182 | -0.000488 | 0.06155 | 0.514203 | 0.214198 | 0.511558 | 0.725184 | -0.501292 | -0.079516 | 0.637296 | 0.452851 | 1.312386 | 1.154549 | 0.60055 | 0.270126 | -0.179206 | 0.45706 | 0.975463 | 1.000854 | 0.700993 | 0.581834 | 0.564173 | 0.614185 |
| 25.33333333 | 0.433706 | 0.354321 | 0.13813 | 0.304693 | 0.597238 | 0.181829 | 0.569443 | 0.886258 | -0.398751 | 0.058125 | 0.923494 | 0.64938 | 1.39966 | 1.406968 | 0.322701 | 0.383776 | 0.14959 | 0.519374 | 0.86877 | 1.25984 | 0.874566 | 0.563247 | 0.67795 | 0.682568 |
| 25.34722222 | 0.517566 | 0.629813 | 0.220743 | 0.074067 | 0.695392 | 0.102533 | 0.583731 | 0.829591 | -0.347803 | 0.163824 | 1.150477 | 0.669918 | 1.666865 | 1.721 | 0.089327 | 0.365277 | 0.734325 | 0.620301 | 0.934051 | 1.472589 | 1.035203 | 0.542184 | 0.753404 | 0.904801 |
| 25.36111111 | 0.639306 | 0.643276 | 0.272883 | 0.01148 | 0.802227 | 0.070292 | 0.589705 | 0.710381 | -0.169691 | 0.334086 | 1.28625 | 0.730322 | 1.848406 | 2.051192 | 0.186174 | 0.362022 | 1.37561 | 0.654062 | 1.046456 | 1.578231 | 1.275586 | 0.494007 | 0.817367 | 1.104777 |
| 25.375 | 0.864586 | 0.752601 | 0.335012 | 0.510588 | 0.89816 | 0.116565 | 0.623431 | 0.72619 | 0.027184 | 0.518988 | 1.399166 | 0.799281 | 1.768724 | 2.283905 | 0.511263 | 0.410649 | 1.483658 | 0.715662 | 1.009408 | 1.808423 | 1.453644 | 0.496377 | 0.728062 | 1.289633 |
| 25.38888889 | 0.923871 | 0.819354 | 0.468589 | 1.105653 | 0.979636 | 0.152445 | 0.505642 | 0.717367 | 0.115175 | 0.656937 | 1.372104 | 0.808325 | 1.660716 | 2.579439 | 0.75049 | 0.459005 | 1.32587 | 0.868215 | 0.98046 | 1.605027 | 1.573054 | 0.555544 | 0.749588 | 1.299165 |
| 25.40277778 | 0.897972 | 0.741524 | 0.643639 | 1.10923 | 1.067132 | 0.157078 | 0.357229 | 0.667593 | 0.134893 | 0.63154 | 1.302388 | 0.698987 | 1.644288 | 2.941579 | 0.849603 | 0.445562 | 1.529802 | 1.129585 | 1.069324 | 1.815191 | 1.848982 | 0.787167 | 0.957991 | 1.726752 |
| 25.41666667 | 0.787251 | 0.536269 | 0.670224 | 0.743684 | 1.240151 | 0.195454 | 0.401679 | 0.695572 | 0.236416 | 0.623649 | 1.386552 | 0.789521 | 1.739357 | 2.901258 | 0.870395 | 0.437958 | 2.066293 | 1.240462 | 1.258921 | 2.285684 | 1.92936 | 1.095388 | 1.149491 | 2.245284 |
| 25.43055556 | 0.789617 | 0.478035 | 0.652539 | 0.728056 | 1.214003 | 0.15389 | 0.411428 | 0.53827 | 0.46829 | 0.574914 | 1.562684 | 0.973454 | 1.778365 | 2.02572 | 0.679829 | 0.467172 | 2.285475 | 1.188881 | 1.247145 | 2.455885 | 2.064779 | 1.240725 | 1.245474 | 1.949715 |
| 25.44444444 | 1.089085 | 0.301845 | 0.673429 | 0.905942 | 1.118588 | -0.005457 | 0.38589 | 0.330417 | 0.736629 | 0.477331 | 1.659121 | 1.009925 | 1.538872 | 1.533929 | 0.578793 | 0.521611 | 1.990654 | 1.004598 | 1.035687 | 2.789561 | 2.298519 | 1.154931 | 1.093487 | 1.660375 |
| 25.45833333 | 1.609046 | 0.368567 | 0.704375 | 0.791466 | 1.233993 | 0.040596 | 0.5723 | 0.352191 | 0.900999 | 0.487964 | 1.865459 | 0.964402 | 1.098543 | 1.158842 | 0.408511 | 0.487309 | 1.801587 | 0.734779 | 1.104569 | 2.28515 | 1.808353 | 0.847371 | 0.874661 | 1.482929 |
| 25.47222222 | 1.674849 | 0.693361 | 0.764481 | 0.559508 | 1.275812 | 0.195719 | 0.48874 | 0.40512 | 0.733484 | 0.478548 | 1.809162 | 0.953333 | 0.760293 | 0.816685 | 0.343489 | 0.303185 | 1.969647 | 0.57324 | 0.981638 | 2.056657 | 1.46582 | 0.685417 | 0.833918 | 1.424724 |
| 25.48611111 | 1.270315 | 1.173266 | 0.900417 | 0.662932 | 0.867407 | 0.278482 | 0.149588 | 0.308418 | 0.427606 | 0.389859 | 1.817553 | 1.036903 | 1.07334 | 0.700241 | 0.458353 | 0.23398 | 2.452358 | 0.669689 | 0.848658 | 2.225491 | 1.59673 | 0.768834 | 1.24716 | 1.324347 |
| 25.5 | 1.067945 | 1.545763 | 1.115288 | 0.874121 | 0.598534 | 0.416403 | -0.011741 | 0.387252 | 0.567893 | 0.765603 | 2.212715 | 1.208049 | 1.961618 | 0.98549 | 0.568775 | 0.577033 | 2.254688 | 1.087106 | 0.918242 | 2.877805 | 1.825532 | 0.856124 | 1.657441 | 1.525473 |
| 25.51388889 | 0.946865 | 1.463054 | 1.230039 | 0.464444 | 0.52925 | 0.579934 | 0.293486 | 0.52412 | 0.803384 | 1.072175 | 2.254296 | 1.246541 | 2.505149 | 1.620377 | 0.601798 | 1.06498 | 1.562583 | 1.084368 | 0.990987 | 1.272701 | 1.821193 | 0.564934 | 1.989596 | 1.814982 |
| 25.52777778 | 0.816086 | 1.09734 | 0.970524 | 0.443464 | 0.487589 | 0.649298 | 0.788273 | 0.481538 | 0.883757 | 1.019817 | 1.983974 | 1.20805 | 2.088794 | 2.436901 | 0.538842 | 1.16022 | 0.975914 | 0.910013 | 0.907539 | 0.543905 | 1.50941 | 0.492179 | 1.955881 | 2.051058 |
| 25.54166667 | 0.644786 | 0.781199 | 0.752565 | 0.431166 | 0.447678 | 0.619413 | 0.721639 | 0.255775 | 0.971108 | 0.950764 | 1.969867 | 1.512386 | 1.657522 | 2.787523 | 0.462098 | 0.880095 | 1.145652 | 0.515106 | 0.489827 | 0.536706 | 1.108026 | 0.483226 | 1.297198 | 2.12975 |
| 25.55555556 | 0.502678 | 0.592432 | 0.537543 | 0.417265 | 0.518405 | 0.677989 | 0.454468 | -0.044132 | 0.964577 | 0.969405 | 2.125392 | 1.941152 | 1.252161 | 2.620371 | 0.330263 | 0.510409 | 1.017383 | 0.111287 | 0.133653 | 0.267501 | 0.701105 | 0.509718 | 0.698578 | 2.095038 |
| 25.56944444 | 0.566866 | 0.400843 | 0.340485 | 0.381776 | 0.540512 | 0.771401 | 0.285355 | -0.20458 | 0.471904 | 0.754439 | 2.084031 | 2.192337 | 1.169751 | 1.80083 | 0.135431 | 0.339323 | 0.585371 | 0.194995 | 0.278304 | 0.189804 | 0.667931 | 0.576635 | 0.329629 | 1.706275 |
| 25.58333333 | 0.645399 | 0.171024 | 0.174884 | 0.224432 | 0.435995 | 0.639951 | 0.160659 | -0.223128 | -0.220539 | 0.330207 | 1.681772 | 1.961279 | 1.194862 | 1.043785 | 0.102223 | 0.285354 | 0.120841 | 0.271365 | 0.640986 | 0.046591 | 0.283302 | 0.637104 | 0.127027 | 1.222379 |
| 25.59722222 | 0.498939 | 0.032763 | 0.087852 | -0.400363 | 0.239951 | 0.294206 | -0.070923 | -0.40888 | -0.579368 | -0.039851 | 1.129657 | 1.262848 | 1.053807 | 0.559435 | 0.073788 | 0.22247 | -0.232169 | 0.184776 | 1.057467 | -0.083772 | -0.016756 | 0.619983 | 0.08754 | 0.855196 |
| 25.61111111 | 0.365414 | -0.126818 | 0.117895 | -0.171112 | 0.093583 | 0.132304 | -0.261666 | -0.717062 | -0.419107 | 0.082575 | 0.556702 | 0.773429 | 0.753247 | 0.083652 | 0.082561 | 0.108109 | 0.089083 | 0.208158 | 1.282526 | -0.113221 | -0.437687 | 0.424505 | 0.243961 | 0.487568 |
| 25.625 | 0.279203 | -0.165003 | 0.237899 | -0.019189 | 0.044433 | 0.217351 | -0.173681 | -0.596154 | 0.107418 | 0.540999 | 0.445834 | 0.704486 | 0.508279 | -0.506899 | 0.099544 | 0.065782 | 0.772443 | 0.259365 | 1.078529 | -0.040569 | -0.503717 | 0.219814 | 0.373274 | 0.113594 |
| 25.63888889 | 0.236088 | -0.109683 | 0.330524 | 0.158974 | -0.041073 | 0.313973 | -0.177225 | -0.404942 | 0.520273 | 0.811861 | 0.497629 | 0.803237 | 0.469476 | -0.923493 | 0.014666 | 0.160147 | 0.793692 | 0.13041 | 0.940239 | 0.057919 | -0.722856 | 0.157075 | 0.141293 | -0.053948 |
| 25.65277778 | 0.203638 | -0.116685 | 0.334882 | 0.305071 | -0.143998 | 0.303855 | -0.211152 | -0.440366 | 0.590734 | 0.667912 | 0.491307 | 0.687126 | 0.598298 | -0.767345 | -0.040398 | 0.29604 | 0.323432 | -0.020426 | 0.557486 | 0.113843 | -0.849503 | 0.043073 | -0.098152 | 0.209635 |
| 25.66666667 | 0.242818 | -0.108587 | 0.293008 | 0.163699 | -0.240268 | 0.201956 | -0.278038 | -0.60469 | 0.516612 | 0.423798 | 0.287912 | 0.366226 | 0.623351 | -0.475108 | -0.027195 | 0.254642 | 0.145971 | 0.069587 | -0.087717 | 0.131095 | -0.552439 | 0.025148 | -0.527709 | 0.468405 |
| 25.68055556 | 0.273859 | 0.06293 | 0.11499 | -0.174139 | -0.344177 | 0.181517 | -0.28514 | -0.754033 | 0.557111 | 0.378985 | -0.035623 | -0.020602 | 0.451389 | -0.326246 | 0.029118 | 0.109014 | 0.375132 | 0.150968 | -0.453977 | -0.709212 | -0.620255 | 0.060991 | -0.713633 | 0.512508 |
| 25.69444444 | 0.286946 | 0.31392 | -0.068565 | -0.334694 | -0.446896 | 0.207026 | -0.199572 | -0.584317 | 0.866049 | 0.552096 | -0.362761 | -0.403612 | 0.324537 | -0.071262 | -0.099259 | 0.071092 | 0.097992 | 0.182722 | -0.240373 | -1.225788 | -0.625554 | 0.111846 | -0.641695 | 0.601645 |
| 25.70833333 | 0.241679 | 0.359801 | -0.075596 | -0.154718 | -0.425999 | 0.177051 | -0.047162 | -0.295984 | 1.204925 | 0.62895 | -0.355969 | -0.411524 | 0.346511 | 0.174336 | -0.161341 | 0.096293 | -0.192275 | 0.249935 | -0.049904 | -1.787648 | -0.678024 | 0.157063 | -0.683924 | 0.651108 |
| 25.72222222 | 0.190224 | 0.289415 | 0.02063 | 0.175936 | -0.304 | -0.076982 | -0.082282 | -0.088781 | 1.199287 | 0.517763 | -0.158069 | -0.214566 | 0.423814 | 0.275023 | -0.075507 | 0.06521 | 0.133041 | 0.284223 | -0.037187 | -1.722932 | -0.504394 | 0.196492 | -0.777147 | 0.609376 |
| 25.73611111 | 0.221405 | 0.360643 | 0.126019 | 0.177037 | -0.11911 | -0.237248 | -0.189106 | 0.063997 | 0.954305 | 0.300453 | 0.033074 | -0.129608 | 0.046597 | 0.295537 | -0.067507 | 0.008671 | 0.633879 | 0.252591 | -0.122639 | -1.354929 | -0.3112 | 0.249725 | -0.73758 | 0.545495 |
| 25.75 | 0.228754 | 0.368828 | 0.055455 | -0.10719 | -0.022325 | -0.259911 | -0.181507 | 0.107293 | 0.686073 | 0.158264 | 0.022238 | -0.210051 | -0.4294 | 0.325991 | -0.193847 | -0.058095 | 0.256992 | 0.141363 | -0.094439 | -1.174811 | -0.250907 | 0.267547 | -0.493269 | 0.496671 |
| 25.76388889 | 0.168608 | 0.128154 | -0.224846 | -0.463285 | -0.146269 | -0.369105 | -0.108961 | 0.037885 | 0.477516 | -0.027685 | -0.312561 | -0.329854 | -0.682654 | 0.499671 | -0.250613 | -0.156056 | -0.78688 | -0.054051 | -0.149114 | -1.294906 | -0.26276 | 0.172298 | -0.246077 | 0.470609 |
| 25.77777778 | 0.093732 | -0.204571 | -0.431223 | -0.520407 | -0.357962 | -0.613135 | -0.086625 | -0.121704 | 0.439497 | -0.156618 | -0.410977 | -0.210178 | -1.027888 | 0.794347 | -0.111921 | -0.195608 | -0.547775 | -0.197498 | -0.291896 | -1.604661 | -0.330459 | 0.112233 | -0.289946 | 0.510684 |
| 25.79166667 | 0.115483 | -0.207341 | -0.448957 | -0.491418 | -0.507203 | -0.924777 | -0.219577 | -0.150118 | 0.205185 | -0.290519 | -0.441177 | -0.14114 | -1.615859 | 0.874103 | 0.094988 | -0.227752 | -0.062291 | -0.267007 | -0.247383 | -1.726075 | -0.433141 | 0.129868 | -0.484628 | 0.492653 |
| 25.80555556 | 0.153415 | -0.210162 | -0.475275 | -0.049764 | -0.604548 | -1.160303 | -0.343169 | -0.135829 | -0.154756 | -0.440221 | -0.651738 | -0.307196 | -2.261215 | 0.550472 | 0.082185 | -0.375482 | -0.391671 | -0.323579 | -0.111099 | -1.751405 | -0.384029 | -0.012195 | -0.87682 | 0.243619 |
| 25.81944444 | 0.144395 | -0.468731 | -0.646544 | -0.148276 | -0.670864 | -1.187373 | -0.417734 | -0.279797 | -0.377643 | -0.450251 | -1.030619 | -0.576447 | -2.597799 | 0.115142 | -0.052019 | -0.527637 | -1.585418 | -0.358549 | -0.356217 | -1.815484 | -0.393052 | -0.185743 | -0.925553 | -0.059798 |
| 25.83333333 | 0.053887 | -0.852015 | -0.845255 | -0.770582 | -0.771504 | -0.894586 | -0.468534 | -0.534751 | -0.51663 | -0.385643 | -1.303438 | -0.554918 | -2.621072 | -0.184084 | -0.136452 | -0.508422 | -2.059278 | -0.584933 | -0.298833 | -1.882068 | -0.463716 | -0.338749 | -0.917786 | -0.310368 |
| 25.84722222 | -0.128378 | -0.978385 | -0.969374 | -1.200233 | -0.609224 | -0.743438 | -0.642979 | -0.768355 | -0.800021 | -0.51371 | -1.374275 | -0.624165 | -2.813137 | -0.395906 | -0.258461 | -0.479262 | -1.573553 | -0.820841 | -0.171408 | -1.569383 | -0.733179 | -0.343166 | -1.045534 | -0.4073 |
| 25.86111111 | -0.128351 | -1.053119 | -0.713831 | -0.942259 | -0.346768 | -0.7244 | -0.834839 | -0.783895 | -1.207209 | -0.775236 | -1.628042 | -1.0487 | -3.240018 | -0.677463 | -0.347051 | -0.539858 | -1.850827 | -0.823366 | -0.011419 | -1.051659 | -0.739914 | -0.384479 | -1.355299 | -0.449332 |
| 25.875 | -0.343251 | -1.248994 | -0.455175 | -0.321215 | -0.516756 | -0.862378 | -0.844881 | -0.876982 | -1.563323 | -1.108699 | -1.938477 | -1.185363 | -3.596388 | -0.903637 | -0.671245 | -0.687371 | -2.728456 | -0.820894 | 0.134112 | -1.418597 | -1.034894 | -0.593392 | -1.225964 | -0.725331 |
| 25.88888889 | -1.250788 | -1.550381 | -0.627437 | 0.238916 | -0.889481 | -0.974056 | -0.985906 | -1.142486 | -1.70136 | -1.424578 | -2.250823 | -1.098214 | -3.43355 | -0.99909 | -0.967957 | -0.933816 | -2.552253 | -0.793288 | 0.134889 | -2.440972 | -0.776092 | -0.826416 | -1.398032 | -0.959726 |
| 25.90277778 | -1.837737 | -1.356672 | -1.086752 | 0.061899 | -1.076027 | -0.843782 | -1.13469 | -1.291704 | -1.833437 | -1.410325 | -2.253289 | -1.283447 | -3.565261 | -0.907344 | -1.156273 | -1.158485 | -2.232643 | -0.935258 | -0.201798 | -3.297648 | -0.561347 | -0.964822 | -1.535746 | -1.206332 |
| 25.91666667 | -1.893683 | -1.261106 | -1.424503 | -0.555571 | -1.11395 | -0.630699 | -1.110919 | -1.473753 | -1.792835 | -1.224176 | -2.000235 | -1.164986 | -3.826159 | -1.001063 | -1.263997 | -1.351465 | -2.285849 | -1.169639 | -0.58705 | -2.957072 | -0.514668 | -1.175314 | -1.421962 | -1.24622 |
| 25.93055556 | -1.887183 | -1.097373 | -1.592767 | -1.078153 | -1.005954 | -0.365858 | -1.031391 | -1.461776 | -1.476584 | -1.033871 | -1.743304 | -1.021697 | -4.202664 | -1.045105 | -1.304559 | -1.417126 | -3.19729 | -1.311605 | -1.044752 | -2.21436 | -0.695026 | -1.726079 | -1.362024 | -1.260651 |
| 25.94444444 | -1.550641 | -1.144727 | -1.761006 | -1.206772 | -1.008836 | -0.238375 | -0.809763 | -1.225523 | -1.20441 | -1.0586 | -1.794655 | -0.917556 | -4.554129 | -0.92267 | -1.053939 | -1.380934 | -3.385763 | -1.497172 | -1.234868 | -1.412203 | -0.970286 | -2.034986 | -1.359267 | -1.131591 |
| 25.95833333 | -2.214923 | -1.502698 | -1.94504 | -1.305982 | -1.605693 | -0.289735 | -0.940653 | -1.298532 | -1.182619 | -1.265323 | -2.377205 | -1.091331 | -4.043126 | -1.190904 | -0.481108 | -1.285694 | -3.552151 | -1.751277 | -1.476113 | -0.381824 | -1.264313 | -2.466754 | -1.724304 | -1.600763 |
| 25.97222222 | -3.868475 | -2.153349 | -2.227419 | -1.467036 | -2.740817 | -0.684397 | -1.327442 | -1.661672 | -1.545471 | -1.52478 | -2.966824 | -1.360127 | -3.94897 | -2.633233 | -0.593932 | -1.33863 | -3.137047 | -2.291255 | -1.625429 | -0.47534 | -1.978072 | -2.931177 | -1.851936 | -2.601451 |
| 25.98611111 | -2.538621 | -2.593712 | -2.353114 | -1.563632 | -2.449314 | -1.159578 | -1.303744 | -1.802096 | -1.361712 | -1.272491 | -2.856924 | -1.289954 | -2.841545 | -3.123495 | -1.355043 | -1.304643 | -2.521795 | -2.551319 | -1.699035 | -0.752294 | -2.181816 | -2.23444 | -2.199248 | -3.001023 |
| 26 | -0.684211 | -1.853386 | -1.596471 | -1.715008 | -1.223037 | -1.15674 | -0.870264 | -1.359313 | -1.073189 | -0.812118 | -1.557965 | -0.925412 | -2.520459 | -2.470867 | -1.240005 | -1.342069 | -1.395 | -1.789605 | -1.915326 | -0.501389 | -1.322683 | -1.737291 | -1.846839 | -2.464676 |
| 26.01388889 | -0.092828 | -0.517202 | -0.919302 | -1.412551 | -0.136199 | -0.876913 | -0.294179 | -0.83887 | -0.705749 | -0.408048 | -0.507115 | -0.721714 | -1.626392 | -1.832706 | -0.73485 | -1.318464 | -0.540831 | -0.083308 | -1.381715 | -0.19483 | -0.38246 | -1.341973 | -1.10911 | -1.689664 |
| 26.02777778 | -0.022728 | 0.28936 | -0.555459 | -0.79658 | 0.615484 | -0.559361 | -0.078753 | -0.386575 | -0.211427 | -0.140106 | -0.156094 | -0.791355 | -0.579343 | -0.600149 | -0.315154 | -1.347789 | 0.085877 | 0.676271 | -1.08851 | -0.059397 | -0.226266 | -0.840578 | -0.494763 | -0.469223 |
| 26.04166667 | -0.094941 | 0.770642 | -0.430577 | 0.095322 | 0.593029 | -0.356325 | -0.030855 | -0.019036 | 0.18082 | 0.160364 | -0.024126 | -0.909598 | -0.088261 | -0.047294 | -0.128782 | -1.2264 | 1.030313 | 0.404463 | -1.065704 | -0.086885 | -0.054259 | -0.366409 | 0.315467 | 0.144688 |
| 26.05555556 | -0.033062 | 1.116134 | -0.063211 | 0.866638 | 0.130542 | -0.242401 | 0.056198 | 0.143926 | 0.468916 | 0.483417 | 0.323085 | -0.946794 | 0.065637 | -0.025954 | 0.626883 | -1.138148 | 1.598981 | 0.306942 | -0.857417 | -0.161703 | 0.483961 | -0.385028 | 0.901477 | 0.186783 |
| 26.06944444 | 0.226202 | 1.288513 | 0.226593 | 1.098132 | 0.155978 | -0.166997 | -0.049382 | 0.203416 | 0.544935 | 0.713062 | 0.647937 | -0.641543 | 0.662635 | 0.437505 | 1.414015 | -1.144945 | 1.574287 | 0.673175 | -0.495798 | -0.21073 | 0.807571 | -0.041013 | 1.018526 | 0.64027 |
| 26.08333333 | 0.379272 | 1.183578 | 0.316599 | 0.672705 | 0.488984 | -0.071661 | -0.06076 | 0.544724 | 0.602454 | 0.788012 | 0.7932 | -0.145694 | 1.707604 | 1.298063 | 1.346146 | -0.953639 | 1.106088 | 0.817929 | -0.01823 | -0.20612 | 1.126327 | 0.316369 | 0.945838 | 1.366757 |
| 26.09722222 | 0.301215 | 1.038705 | 0.283057 | 0.267211 | 0.530394 | 0.070749 | -0.084979 | 0.823043 | 0.633144 | 0.810901 | 0.853145 | 0.137254 | 1.96226 | 2.238575 | 0.835626 | -0.504962 | 1.674747 | 1.632751 | 1.036415 | -0.669015 | 1.566308 | 1.165767 | 0.728426 | 1.962749 |
| 26.11111111 | 0.295868 | 0.889906 | 0.254002 | 0.048294 | 0.273925 | 0.147148 | -0.063284 | 1.034744 | 0.650139 | 0.839065 | 0.808966 | 0.088286 | 1.908931 | 2.719382 | 0.099654 | -0.000422 | 2.279955 | 2.603504 | 1.857914 | -2.067207 | 2.126263 | 2.50535 | 0.27327 | 2.381098 |
| 26.125 | 0.422624 | 0.671355 | 0.361521 | -0.064744 | 0.051805 | 0.166205 | 0.173512 | 1.055669 | 0.871836 | 0.939618 | 0.703156 | 0.069291 | 3.771137 | 0.053308 | -0.018078 | -1.194906 | 1.08894 | 0.274123 | -0.780641 | -0.787239 | 0.58995 | 0.965412 | 0.299099 | 0.958657 |
| 26.13888889 | 0.406452 | 0.553819 | 0.476142 | -0.339065 | -0.124155 | 0.02313 | 0.033532 | 0.802076 | 0.952509 | 0.900085 | 0.481806 | 0.116189 | 2.188971 | -2.649841 | 0.036784 | -1.572004 | -1.165888 | -2.417249 | -2.795257 | 0.647271 | -1.601056 | -1.212767 | 0.256655 | -1.67939 |
| 26.15277778 | 0.300878 | 0.669493 | 0.214451 | -0.817664 | -0.006136 | -0.196955 | -0.249803 | 0.557091 | 0.87265 | 0.74793 | 0.360071 | 0.050994 | 1.665634 | -3.283676 | 1.134862 | -0.782099 | -1.113617 | -1.544994 | -1.849274 | -0.132251 | -0.158125 | -0.356588 | 0.318719 | -1.936103 |
| 26.16666667 | 0.326709 | 1.05199 | -0.287152 | -1.077998 | 0.482691 | -0.223606 | -0.276219 | 0.426236 | 0.873142 | 0.820439 | 0.43221 | -0.040779 | 2.135724 | -1.897883 | 1.321209 | -0.425777 | 0.161169 | -0.127512 | -0.720614 | -0.366197 | 0.961827 | -0.19092 | 0.27729 | -0.84771 |
| 26.18055556 | 0.342184 | 1.212162 | -0.699443 | -1.24322 | 0.82349 | -0.128514 | -0.007475 | 0.340204 | 0.809979 | 0.859024 | 0.473874 | -0.173277 | 2.213043 | -0.573759 | 1.282287 | -0.295449 | 0.574073 | 0.219434 | -0.385416 | -0.411232 | 1.071868 | -0.221279 | 0.363926 | 0.104116 |
| 26.19444444 | 0.312917 | 1.169689 | -0.63527 | -0.865752 | 1.032751 | -0.179648 | 0.076141 | 0.206123 | 0.534433 | 0.682997 | 0.348714 | -0.337843 | 1.750215 | 0.144827 | 0.70704 | -0.289462 | -0.241938 | 0.220469 | -0.204496 | -0.36608 | 0.99101 | -0.357642 | 0.451531 | 0.362275 |
| 26.20833333 | 0.236549 | 1.42213 | -0.332152 | -0.259435 | 1.228889 | -0.233306 | -0.09412 | -0.040851 | 0.208014 | 0.454138 | 0.470487 | -0.355726 | 1.079005 | 0.614561 | 0.317043 | -0.128718 | -1.198457 | 0.259812 | -0.024646 | -0.331892 | 0.791442 | -0.730068 | 0.492702 | 0.429143 |
| 26.22222222 | 0.055033 | 1.744614 | -0.30164 | -0.101918 | 1.364021 | -0.281733 | -0.218931 | -0.133141 | 0.062563 | 0.303454 | 0.779532 | -0.205641 | 1.296353 | 0.976699 | 0.367645 | 0.067281 | -0.55307 | 0.43264 | 0.039338 | -0.345662 | 0.723548 | -0.719156 | 0.414472 | 0.504699 |
| 26.23611111 | 0.008718 | 1.602026 | -0.360201 | -0.024946 | 1.520568 | -0.298111 | -0.028077 | 0.209903 | 0.185063 | 0.336971 | 1.166044 | 0.063914 | 2.20457 | 0.816598 | 0.392081 | 0.250449 | 0.420448 | 0.425896 | 0.056164 | -0.370458 | 1.072456 | -0.200807 | 0.304193 | 0.41897 |
| 26.25 | 0.260676 | 1.339339 | -0.249147 | 0.150068 | 1.299591 | -0.291475 | 0.30819 | 0.547708 | 0.298455 | 0.382396 | 1.322188 | 0.375501 | 2.388256 | 0.453153 | 0.121432 | 0.372229 | 0.316732 | 0.454275 | 0.245159 | -0.250667 | 1.45372 | -0.190042 | 0.772895 | 0.37638 |
| 26.26388889 | 0.342685 | 1.015189 | -0.070219 | 0.30113 | 0.92986 | -0.292883 | 0.492782 | 0.497602 | 0.220113 | 0.229703 | 1.238049 | 0.481726 | 1.855264 | 0.32031 | -0.208802 | 0.279375 | -0.344497 | 0.584907 | 0.601496 | -0.100029 | 0.77164 | -0.767388 | 1.202209 | 0.432412 |
| 26.27777778 | 0.257198 | 0.81937 | 0.154162 | 0.566731 | 0.675245 | -0.250906 | 0.496642 | 0.158956 | 0.007238 | 0.029818 | 0.945739 | 0.437321 | 1.406663 | 0.153706 | -0.257592 | 0.16946 | -0.951449 | 0.548563 | 0.657275 | -0.010858 | -0.100737 | -0.93981 | 0.759799 | 0.346854 |
| 26.29166667 | 0.157715 | 0.731458 | 0.327492 | 0.90242 | 0.642033 | -0.173129 | 0.543416 | -0.045792 | 0.031222 | 0.059397 | 0.79069 | 0.55024 | 1.372019 | -0.232407 | 0.054747 | 0.118075 | -0.867109 | 0.35284 | 0.534103 | 0.019276 | -0.655732 | -0.492176 | 0.189304 | 0.047083 |
| 26.30555556 | 0.102154 | 0.665287 | 0.508531 | 0.937649 | 0.634634 | -0.138319 | 0.714162 | 0.049175 | 0.475911 | 0.301804 | 1.028893 | 0.716117 | 1.566231 | -0.332884 | 0.488516 | 0.153682 | 0.038746 | 0.422051 | 0.594956 | 0.094074 | -0.137948 | -0.116101 | 0.211341 | -0.026108 |
| 26.31944444 | 0.16933 | 0.649032 | 0.71415 | 0.814992 | 0.606485 | -0.099948 | 0.957873 | 0.305874 | 0.900857 | 0.532753 | 1.298604 | 0.720865 | 1.852968 | 0.003139 | 0.764173 | 0.245605 | 1.036722 | 0.705013 | 0.839149 | 0.203402 | 0.970853 | -0.084265 | 0.589726 | 0.216564 |
| 26.33333333 | 0.280726 | 0.507673 | 0.981351 | 0.795483 | 0.484504 | -0.012814 | 1.131444 | 0.54103 | 0.963786 | 0.592304 | 1.452958 | 0.729308 | 1.769319 | 0.329946 | 0.770536 | 0.248934 | 2.005999 | 0.855431 | 1.053232 | 0.208395 | 1.444929 | 0.058085 | 0.614007 | 0.44153 |
| 26.34722222 | 0.388896 | 0.274892 | 1.210303 | 0.980671 | 0.408711 | 0.029955 | 1.158649 | 0.566242 | 1.003018 | 0.601755 | 1.600111 | 0.922774 | 1.765023 | 0.421395 | 0.808509 | 0.252121 | 2.652447 | 0.817229 | 1.049575 | 0.280362 | 1.690358 | 0.301151 | 0.611754 | 0.505935 |
| 26.36111111 | 0.459096 | 0.222677 | 1.500001 | 1.18745 | 0.565747 | 0.028565 | 1.156828 | 0.573399 | 1.078537 | 0.651064 | 1.827965 | 1.131575 | 1.852264 | 0.736042 | 0.838121 | 0.253694 | 2.618944 | 0.788047 | 1.080711 | 0.361742 | 1.921492 | 0.4849 | 0.697071 | 0.690462 |
| 26.375 | 0.585409 | 0.3985 | 1.434787 | 1.010194 | 0.703511 | 0.158287 | 1.105093 | 0.745536 | 1.278139 | 0.729692 | 2.12377 | 1.304275 | 1.941107 | 1.177738 | 0.722304 | 0.264072 | 2.154177 | 0.802247 | 1.215578 | 0.322302 | 1.8545 | 0.528645 | 1.059421 | 0.957522 |
| 26.38888889 | 0.692119 | 0.667605 | 1.317682 | 1.050372 | 0.808197 | 0.335164 | 0.903505 | 0.910641 | 1.412716 | 0.788288 | 2.242373 | 1.348017 | 1.677874 | 1.342797 | 0.540701 | 0.278899 | 1.460746 | 0.803478 | 1.262246 | 0.214388 | 1.971726 | 0.636946 | 1.559016 | 1.179131 |
| 26.40277778 | 0.810246 | 0.801211 | 1.374882 | 0.839648 | 1.009635 | 0.486104 | 0.844497 | 1.242573 | 1.485747 | 0.83868 | 2.533888 | 1.285625 | 1.713835 | 1.632155 | 0.640453 | 0.349118 | 0.926696 | 0.864988 | 1.370276 | 0.166321 | 2.023778 | 0.665594 | 1.806548 | 1.426075 |
| 26.41666667 | 0.977723 | 0.703363 | 1.368019 | 0.747834 | 1.191013 | 0.564274 | 0.856056 | 1.249882 | 1.531524 | 0.888882 | 2.632735 | 1.218402 | 2.165131 | 2.107839 | 1.101177 | 0.376245 | 0.703874 | 0.943765 | 1.508569 | 0.165755 | 2.015643 | 0.765148 | 1.842924 | 1.62853 |
| 26.43055556 | 0.961858 | 0.27659 | 1.276579 | 0.823363 | 1.091227 | 0.470929 | 0.806869 | 0.900772 | 1.507572 | 0.766904 | 2.550915 | 1.164479 | 2.280792 | 2.046014 | 1.244387 | 0.316205 | 0.839769 | 0.939162 | 1.24373 | 0.252488 | 1.686458 | 1.01028 | 1.560117 | 1.510673 |
| 26.44444444 | 0.839764 | -0.196641 | 1.218545 | 0.615722 | 0.77846 | 0.304913 | 0.602498 | 0.681633 | 1.437 | 0.532109 | 2.174379 | 1.137638 | 2.26552 | 2.006911 | 1.118171 | 0.208458 | 1.210083 | 0.694982 | 0.254753 | 0.560672 | 1.438778 | 1.306484 | 1.306343 | 1.427811 |
| 26.45833333 | 0.831056 | -0.133988 | 0.84122 | 0.330929 | 0.715946 | 0.602464 | 0.658913 | 0.749077 | 1.539682 | 0.762458 | 2.008342 | 1.096146 | 1.744824 | 2.149977 | 0.776412 | 0.156856 | 1.602608 | 0.495431 | -0.346906 | 1.110788 | 1.350071 | 1.281575 | 1.126433 | 1.460738 |
| 26.47222222 | 0.652581 | 0.426918 | 0.504754 | 0.311359 | 0.739946 | 1.193695 | 0.770972 | 0.821577 | 1.485753 | 1.228078 | 1.605649 | 1.20718 | 1.621955 | 1.849484 | 0.785352 | 0.182576 | 1.759746 | 0.520586 | -0.248646 | 1.026337 | 1.222886 | 0.971635 | 1.098267 | 1.199901 |
| 26.48611111 | 0.434361 | 0.868057 | 0.429477 | 0.523926 | 0.711992 | 1.402911 | 0.652982 | 0.596244 | 1.411874 | 1.37087 | 1.536521 | 1.258498 | 1.566086 | 1.380064 | 0.882742 | 0.20684 | 2.308243 | 0.67343 | 0.193332 | 0.989359 | 1.112101 | 0.648606 | 1.14416 | 0.861037 |
| 26.5 | 0.480474 | 1.239732 | 0.613019 | 0.744299 | 0.955612 | 1.188697 | 0.461182 | 0.395332 | 1.371129 | 1.183179 | 2.061304 | 1.567952 | 1.794508 | 1.50739 | 0.658685 | 0.425218 | 2.993543 | 1.010151 | 0.501539 | 1.305847 | 1.018811 | 0.980543 | 1.429401 | 0.999883 |
| 26.51388889 | 0.652061 | 1.297153 | 0.886348 | 0.513981 | 1.023403 | 1.009921 | 0.585885 | 0.534693 | 1.44009 | 0.862226 | 2.584219 | 1.941944 | 1.94395 | 1.897844 | 0.550681 | 0.629392 | 2.540134 | 1.304448 | 0.716048 | 0.686147 | 0.731822 | 1.627253 | 1.221574 | 1.377259 |
| 26.52777778 | 0.622416 | 0.88975 | 0.917912 | 0.093087 | 0.863007 | 0.775446 | 0.644796 | 0.751104 | 1.365218 | 0.391886 | 2.204133 | 1.818327 | 1.417871 | 1.721641 | 0.337709 | 0.743865 | 1.734683 | 1.27911 | 0.72022 | 0.452935 | 1.165909 | 2.04142 | 1.240968 | 1.438675 |
| 26.54166667 | 0.420884 | 0.448232 | 0.648486 | -0.222181 | 0.660485 | 0.482271 | 0.776124 | 0.627482 | 1.307533 | 0.250824 | 1.365937 | 1.28492 | 1.109116 | 1.441099 | 0.281738 | 0.683681 | 1.318061 | 0.908864 | 0.812493 | 1.453139 | 1.030238 | 1.547724 | 1.15262 | 1.135272 |
| 26.55555556 | 0.280803 | 0.013163 | 0.389142 | -0.179947 | 0.419716 | 0.384304 | 0.964059 | 0.737659 | 1.193001 | 0.618863 | 1.161628 | 1.102001 | 0.900975 | 1.305995 | 0.400162 | 0.595304 | 1.012475 | 0.47829 | 0.790146 | 2.302734 | 1.224155 | 1.09624 | 1.202944 | 1.145988 |
| 26.56944444 | 0.24797 | 0.039482 | 0.162104 | 0.11513 | 0.233783 | 0.555348 | 0.890391 | 0.604777 | 0.779538 | 1.005042 | 1.460761 | 1.142186 | 0.997522 | 1.62173 | 0.326956 | 0.395736 | 0.615725 | 0.342595 | 1.023899 | 1.41255 | 1.731024 | 0.963088 | 1.136855 | 1.078646 |
| 26.58333333 | 0.279606 | 0.134112 | 0.250562 | 0.352184 | 0.102371 | 0.578625 | 0.595538 | 0.454241 | 0.382032 | 0.956921 | 1.096133 | 1.021124 | 0.969993 | 1.290453 | 0.215157 | 0.194009 | 0.295504 | 0.210929 | 1.050202 | 1.44325 | 1.7945 | 0.862789 | 0.633075 | 0.663173 |
| 26.59722222 | 0.251499 | 0.21083 | 0.471508 | 0.43251 | 0.080889 | 0.309631 | 0.213598 | 0.317493 | 0.393377 | 0.803468 | 0.506442 | 0.646918 | 0.646891 | 0.444235 | 0.00574 | 0.026423 | 0.118189 | 0.054354 | 0.972058 | 0.919746 | 1.54319 | 0.881951 | 0.485317 | 0.454619 |
| 26.61111111 | 0.206489 | 0.286902 | 0.70639 | 0.21596 | 0.050101 | 0.012632 | -0.098969 | 0.191972 | 0.439471 | 0.685809 | -0.101906 | 0.003331 | 0.22925 | -0.217201 | -0.015515 | 0.046345 | -0.222087 | -0.090913 | 0.701909 | 0.334716 | 1.118389 | 0.679787 | 0.320052 | 0.290079 |
| 26.625 | 0.194387 | 0.17619 | 0.650969 | 0.010674 | -0.028744 | -0.068451 | -0.209485 | 0.030615 | 0.242342 | 0.539692 | -0.513181 | -0.224349 | 0.081015 | -0.350011 | 0.053257 | 0.109078 | 0.078761 | -0.20301 | 0.49435 | 0.252183 | 0.597425 | 0.371323 | 0.474317 | 0.227543 |
| 26.63888889 | 0.075422 | -0.113228 | 0.20498 | 0.273454 | -0.219369 | 0.017882 | -0.086755 | -0.167355 | -0.049873 | 0.330274 | -0.370758 | -0.342483 | 0.01431 | -0.35736 | 0.077961 | 0.158605 | 0.428659 | -0.004518 | 0.418056 | -0.222437 | -0.024857 | 0.144569 | 0.219636 | 0.169859 |
| 26.65277778 | -0.057859 | -0.193596 | -0.3346 | 0.58094 | -0.47086 | 0.090884 | -0.053034 | -0.466551 | -0.229319 | 0.215999 | -0.153963 | -0.360073 | -0.100678 | -0.536092 | 0.022449 | 0.125779 | 0.500835 | -0.102998 | 0.178909 | -0.831329 | -0.471521 | 0.009218 | -0.315714 | 0.107763 |
| 26.66666667 | -0.00165 | 0.167114 | -0.676525 | 0.421342 | -0.470768 | 0.199294 | -0.119634 | -0.662454 | -0.180724 | 0.18688 | 0.04143 | -0.305557 | -0.167544 | -0.16399 | 0.005487 | 0.140515 | 0.404619 | -0.217387 | -0.176114 | -1.11875 | -0.373682 | 0.008842 | -0.438484 | 0.134731 |
| 26.68055556 | 0.17458 | 0.600261 | -0.513709 | 0.037437 | -0.230584 | 0.346938 | -0.019464 | -0.419629 | 0.019551 | 0.259394 | 0.416102 | 0.000708 | 0.001345 | 0.334071 | 0.086972 | 0.210553 | 0.585342 | -0.077817 | -0.194505 | -1.385404 | 0.026495 | 0.090329 | -0.624181 | 0.264746 |
| 26.69444444 | 0.279177 | 0.660536 | -0.248578 | -0.204901 | -0.030614 | 0.395448 | 0.134922 | -0.002281 | 0.246288 | 0.286378 | 0.56915 | 0.315215 | 0.070316 | 0.512737 | 0.270285 | 0.179899 | 0.948627 | 0.00791 | -0.093046 | -1.437966 | -0.107499 | -0.020946 | -0.601232 | 0.360487 |
| 26.70833333 | 0.214556 | 0.374286 | -0.246619 | -0.223996 | -0.058481 | 0.479039 | 0.120666 | 0.070246 | 0.396814 | 0.383391 | 0.498012 | 0.410568 | -0.066942 | 0.520502 | 0.32127 | 0.111258 | 0.853427 | -0.221676 | -0.28757 | -1.658544 | -0.208224 | -0.048538 | -0.622123 | 0.380504 |
| 26.72222222 | 0.085504 | 0.028557 | -0.527204 | 0.347699 | -0.241507 | 0.502078 | -0.112252 | -0.204413 | 0.434829 | 0.395119 | 0.479548 | 0.32847 | -0.310138 | 0.474303 | 0.143327 | -0.044546 | 0.662989 | -0.733117 | -0.697144 | -1.750797 | -0.330591 | -0.126554 | -0.898648 | 0.371376 |
| 26.73611111 | -0.009365 | 0.008889 | -0.856559 | 0.267941 | -0.43494 | 0.355978 | -0.143731 | -0.271145 | 0.449234 | 0.329988 | 0.604125 | 0.25019 | -0.451967 | 0.273744 | 0.030748 | -0.136759 | 0.660854 | -0.912249 | -0.988198 | -1.69815 | -0.688108 | -0.225561 | -1.468478 | 0.292145 |
| 26.75 | -0.102949 | 0.179413 | -0.375524 | 0.049699 | -0.488936 | 0.202904 | 0.072103 | 0.063666 | 0.427715 | 0.301787 | 0.513341 | 0.153766 | -0.496706 | 0.156976 | 0.043199 | -0.039396 | 0.814128 | -0.394427 | -0.646562 | -1.763927 | -0.844011 | -0.077527 | -1.907067 | 0.301057 |
| 26.76388889 | -0.119359 | 0.30934 | -0.033767 | -0.239138 | -0.301643 | 0.188213 | 0.23675 | 0.345404 | 0.357178 | 0.290115 | 0.144632 | -0.051478 | -0.609441 | 0.172082 | 0.018424 | 0.117343 | 0.441807 | 0.344261 | 0.105185 | -2.043148 | -0.939868 | -0.092347 | -1.810789 | 0.366324 |
| 26.77777778 | -0.18755 | 0.145384 | -0.267683 | -0.359238 | 0.002414 | 0.208298 | 0.100037 | 0.170244 | 0.119626 | 0.185642 | -0.318283 | -0.313057 | -0.91277 | -0.066009 | -0.148697 | 0.084933 | -0.139193 | 1.028711 | 0.916264 | -1.914158 | -0.376826 | -0.350293 | -1.583715 | 0.185862 |
| 26.79166667 | -0.214567 | -0.197985 | -0.791183 | -0.640196 | 0.251533 | 0.163563 | -0.210828 | -0.143358 | -0.1044 | -0.002266 | -0.499582 | -0.371443 | -1.0928 | -0.373964 | -0.14596 | -0.110392 | -0.419555 | 1.364551 | 1.062755 | -1.630451 | -0.473453 | -0.444483 | -1.477932 | -0.002713 |
| 26.80555556 | -0.237797 | -0.34111 | -1.206306 | -0.591092 | 0.086423 | -0.052722 | -0.352582 | -0.205221 | -0.187738 | -0.052084 | -0.435895 | -0.246204 | -0.809846 | -0.335159 | 0.034708 | -0.321443 | -0.288494 | 1.087572 | 0.759158 | -1.169777 | -0.436336 | -0.3183 | -0.902248 | 0.07046 |
| 26.81944444 | -0.356335 | -0.410075 | -1.182639 | -0.350003 | -0.284283 | -0.403741 | -0.450676 | -0.186663 | -0.267225 | 0.013849 | -0.621463 | -0.295465 | -0.591836 | -0.198223 | 0.061629 | -0.458545 | -0.42323 | 0.823885 | 0.410638 | -1.278993 | 0.124431 | -0.374652 | -0.492736 | 0.192555 |
| 26.83333333 | -0.495351 | -0.323231 | -0.982504 | -0.318885 | -0.567364 | -0.601675 | -0.642034 | -0.385331 | -0.39517 | 0.045465 | -1.024244 | -0.519934 | -0.932373 | -0.38795 | -0.012525 | -0.536464 | -1.141067 | 0.338505 | -0.095574 | -1.264949 | 0.377125 | -0.713694 | -0.675161 | 0.046059 |
| 26.84722222 | -0.579026 | -0.156127 | -0.837664 | -0.63101 | -0.719651 | -0.56812 | -0.817536 | -0.652157 | -0.603061 | -0.126098 | -1.218324 | -0.648468 | -1.643565 | -0.765201 | -0.108382 | -0.590378 | -2.442768 | -0.115669 | -0.447865 | -0.913722 | 0.398951 | -1.003043 | -0.785514 | -0.215132 |
| 26.86111111 | -0.663997 | -0.218381 | -0.928039 | -0.85462 | -0.799682 | -0.545201 | -0.826253 | -0.655439 | -0.804556 | 0.018854 | -1.089985 | -0.595684 | -1.906137 | -0.999751 | -0.195644 | -0.656529 | -2.604665 | -0.301729 | -0.524546 | -0.388828 | 0.1027 | -1.069944 | -0.943504 | -0.360244 |
| 26.875 | -0.726829 | -0.581115 | -1.278559 | -0.605401 | -0.845074 | -0.644772 | -0.839735 | -0.640806 | -0.997345 | -0.003941 | -1.22277 | -0.580686 | -2.157928 | -0.9649 | -0.251973 | -0.631148 | -2.260668 | -0.488671 | -0.521889 | -0.151748 | -0.634249 | -0.818512 | -0.903861 | -0.294374 |
| 26.88888889 | -0.859635 | -0.950869 | -1.835434 | -0.283439 | -0.79336 | -0.793526 | -0.839625 | -0.771834 | -1.168655 | -0.154387 | -1.41753 | -0.645569 | -2.629898 | -0.465273 | -0.242057 | -0.638607 | -2.533645 | -0.738468 | -0.622618 | -0.208363 | -1.206751 | -0.672843 | -1.08365 | -0.185108 |
| 26.90277778 | -0.972655 | -1.234977 | -1.764768 | -0.337702 | -0.875559 | -1.042844 | -0.960248 | -1.052776 | -1.373036 | -0.495155 | -1.638086 | -0.694176 | -3.046117 | 0.109733 | -0.36726 | -0.776006 | -2.836293 | -1.1524 | -0.854835 | -0.311301 | -1.356174 | -0.984089 | -1.307404 | 0.081031 |
| 26.91666667 | -1.014278 | -1.076347 | -1.749059 | -0.664065 | -1.111494 | -1.152161 | -0.971778 | -1.006938 | -1.447261 | -0.70663 | -1.544559 | -0.82264 | -3.129538 | 0.206623 | -0.482077 | -0.945766 | -2.596181 | -1.324483 | -0.870466 | -0.273467 | -1.102439 | -1.382744 | -1.254447 | 0.114318 |
| 26.93055556 | -1.072401 | -0.892234 | -1.941794 | -0.897118 | -1.615836 | -1.080744 | -0.829184 | -0.923736 | -1.368602 | -0.809189 | -1.400827 | -0.904512 | -3.262933 | -0.093558 | -0.627708 | -0.915204 | -1.988167 | -1.169555 | -0.624602 | -0.058767 | -0.951245 | -1.461768 | -0.829112 | 0.056971 |
| 26.94444444 | -1.182974 | -0.927509 | -2.201746 | -0.933862 | -2.185291 | -0.888886 | -0.790041 | -0.910138 | -1.426556 | -0.966565 | -1.535371 | -0.914986 | -3.500924 | -0.554825 | -1.067292 | -0.811514 | -2.612337 | -1.753852 | -0.650695 | 0.034981 | -1.202285 | -1.65145 | -0.706042 | -0.201545 |
| 26.95833333 | -1.814303 | -1.387884 | -2.341103 | -1.087377 | -3.572236 | -1.084212 | -1.411003 | -1.352413 | -1.910329 | -1.574388 | -2.191159 | -1.097258 | -4.067311 | -1.590138 | -1.116109 | -0.813816 | -4.105284 | -1.548206 | -1.142054 | -0.183971 | -1.895653 | -1.519148 | -1.79707 | -0.84535 |
| 26.97222222 | -3.186081 | -2.428002 | -2.570891 | -1.288635 | -5.150322 | -1.502889 | -2.46942 | -2.380649 | -2.676127 | -2.402901 | -3.055231 | -0.972489 | -3.811906 | -2.32026 | -1.210177 | -1.017376 | -4.692721 | -1.216298 | -1.292458 | -0.146968 | -2.868836 | -1.393187 | -2.51698 | -0.839775 |
| 26.98611111 | -2.768781 | -3.074062 | -2.956036 | -1.577761 | -5.371937 | -1.859754 | -2.848687 | -1.919411 | -2.297191 | -2.209957 | -3.269344 | -0.905764 | -4.682287 | -2.728083 | -1.273781 | -1.110796 | -3.405395 | -1.414401 | -1.183934 | -0.034541 | -3.030496 | -0.5749 | -3.234099 | -1.332151 |
| 27 | -1.336145 | -2.527373 | -2.045213 | -1.985219 | -3.698365 | -1.77113 | -1.828328 | -1.011346 | -1.106258 | -1.515631 | -2.428536 | -0.511575 | -3.726568 | -2.889032 | -1.225351 | -0.805513 | -1.437375 | -1.122854 | -0.79762 | 0.143001 | -1.921328 | 0.100481 | -3.002397 | -1.595105 |
| 27.01388889 | -0.506361 | -0.938719 | -0.864259 | -1.629271 | -1.933558 | -1.091789 | -0.996874 | -0.379787 | -0.530129 | -0.99558 | -1.246122 | -0.24979 | -2.458585 | -2.226981 | -0.786882 | -0.501833 | -0.454733 | -0.487969 | -0.540318 | 0.204013 | -0.6126 | -0.172601 | -1.729328 | -1.238393 |
| 27.02777778 | -0.392866 | 0.224659 | -0.08308 | -0.739528 | -0.280473 | -0.51459 | -0.499885 | -0.300961 | -0.137432 | -0.678393 | -0.398511 | -0.311281 | -0.84847 | -1.287894 | -0.890767 | -0.564318 | 0.033774 | -0.185217 | -0.423501 | 0.040492 | -0.104588 | -0.586969 | -1.110078 | -0.859134 |
| 27.04166667 | -0.705217 | 0.25692 | 0.205971 | 0.142327 | 0.242297 | -0.288132 | -0.164214 | -0.074952 | 0.196627 | -0.257253 | 0.191661 | -0.101546 | -0.057191 | -0.733657 | -0.805925 | -0.784749 | 0.605279 | -0.019222 | -0.072423 | -0.057565 | -0.236895 | -0.608333 | -0.896027 | -0.508227 |
| 27.05555556 | -0.572732 | 0.145038 | 0.467909 | 0.331674 | 0.607751 | -0.053527 | 0.150033 | 0.390355 | 0.516503 | 0.335702 | 0.642176 | 0.125693 | -0.156222 | -0.269817 | 0.188035 | -0.815157 | 0.68641 | 0.232512 | -0.005095 | 0.134596 | -0.026863 | -0.112455 | -1.184228 | -0.10506 |
| 27.06944444 | -0.042197 | 0.303909 | 0.624104 | 0.286421 | 1.540299 | 0.304649 | 0.470245 | 0.453832 | 0.704108 | 0.759365 | 0.932342 | 0.273695 | -0.117579 | -0.038777 | 0.683282 | -0.411146 | 1.149747 | 0.661288 | -0.220212 | 0.242278 | 0.099648 | 0.52257 | -1.01068 | 0.200604 |
| 27.08333333 | 0.472272 | 0.675645 | 0.706739 | 0.685441 | 1.968086 | 0.605337 | 0.587441 | 0.407889 | 0.78057 | 0.963122 | 1.374766 | 0.525641 | 0.921097 | 0.111242 | 0.686342 | -0.238267 | 1.570478 | 0.713308 | -0.212652 | 0.039054 | 0.05402 | 0.098877 | -0.592869 | 0.333449 |
| 27.09722222 | 0.721397 | 0.964988 | 0.843888 | 0.989024 | 1.717568 | 0.571055 | 0.497927 | 0.878646 | 1.007976 | 0.830306 | 1.875995 | 0.667918 | 1.581336 | 0.613311 | 1.059768 | -0.594933 | 1.919341 | 0.712548 | 0.30425 | -0.310689 | 0.357823 | -0.591532 | 0.033385 | 0.730467 |
| 27.11111111 | 0.744416 | 0.885671 | 0.780842 | 0.26256 | 1.493702 | 0.370418 | 0.35613 | 1.160906 | 1.352501 | 0.631536 | 1.773051 | 0.612116 | 1.310267 | 0.878156 | 1.388405 | -0.868474 | 1.519379 | 1.00035 | 1.326609 | -0.467548 | 0.878798 | -0.559687 | 0.742444 | 1.076387 |
| 27.125 | 0.622218 | 0.690337 | 0.639855 | 0.005014 | 1.398687 | 0.203667 | 0.287591 | 1.065556 | 1.444187 | 0.678086 | 1.132993 | 0.44439 | 1.305094 | 0.207443 | 1.111542 | -0.6473 | 0.808944 | 1.219303 | 2.112402 | -0.408958 | 0.895314 | -0.559922 | 1.294722 | 0.73982 |
| 27.13888889 | 0.462369 | 0.462959 | 0.509018 | -0.208073 | 1.362661 | 0.041485 | 0.14436 | 0.931054 | 1.274431 | 0.68537 | 0.597464 | 0.324995 | 1.697594 | -0.678607 | 1.161576 | -0.309264 | 0.199393 | 0.994112 | 0.774357 | -0.347893 | 0.875507 | -0.673925 | 1.252031 | 0.118051 |
| 27.15277778 | 0.471423 | 0.341141 | 0.515525 | -0.450994 | 1.208195 | -0.059708 | -0.019983 | 0.963053 | 1.077218 | 0.516622 | 0.357948 | 0.27243 | 1.294605 | -1.086956 | 1.209078 | -0.396446 | -0.070638 | 0.354463 | 0.25935 | -0.405132 | 1.276982 | -0.612626 | 0.755067 | -0.158928 |
| 27.16666667 | 0.445596 | 0.226349 | 0.698541 | -0.518342 | 1.475149 | -0.020382 | 0.01508 | 1.076313 | 0.883313 | 0.293344 | 0.324417 | 0.232682 | 0.907599 | -1.154764 | 0.685948 | -0.524541 | 0.515448 | 0.381469 | 0.861361 | -0.403868 | 1.361421 | -0.149611 | 0.599843 | -0.021123 |
| 27.18055556 | 0.281675 | 0.321883 | 0.92881 | -0.259018 | 1.56803 | 0.142997 | 0.113706 | 1.17053 | 0.570659 | 0.181775 | 0.41808 | 0.254083 | 1.506727 | -0.812368 | 0.136515 | -0.568724 | 1.157096 | 0.624425 | 1.388204 | -0.399083 | 0.740303 | -0.018581 | 1.436792 | 0.217261 |
| 27.19444444 | 0.128485 | 0.484255 | 1.052832 | -0.222773 | 1.599351 | 0.164917 | 0.205421 | 1.296664 | 0.279432 | 0.117581 | 0.52559 | 0.274373 | 1.724427 | -0.214299 | -0.020006 | -0.288174 | 0.780915 | 0.378231 | 0.497555 | -0.320909 | 0.581958 | 0.22534 | 1.681055 | 0.43659 |
| 27.20833333 | 0.080718 | 0.697108 | 0.968568 | -0.502466 | 1.693178 | 0.118945 | 0.185723 | 1.069579 | 0.128929 | 0.0384 | 0.572945 | 0.30389 | 1.68226 | 0.259414 | -0.00115 | -0.234161 | -0.02677 | 0.519917 | 0.576388 | -0.312958 | 0.672752 | 0.326032 | 0.836864 | 0.51374 |
| 27.22222222 | 0.243823 | 1.020293 | 1.05478 | -0.547805 | 1.791423 | 0.203077 | 0.178541 | 0.713477 | 0.054328 | 0.01365 | 0.650382 | 0.332089 | 1.362058 | 0.55595 | 0.173215 | -0.388775 | -0.803064 | 0.592752 | 0.97071 | -0.437231 | 0.624362 | -0.077671 | 1.140941 | 0.583281 |
| 27.23611111 | 0.442372 | 0.824948 | 1.247634 | -0.224497 | 1.67937 | 0.379158 | 0.283939 | 0.556951 | 0.096595 | 0.089123 | 0.677109 | 0.443306 | 1.581169 | 0.781207 | 0.006844 | -0.460154 | -0.642021 | 0.005786 | 0.115346 | -0.47034 | 0.286635 | -0.088595 | 2.072163 | 0.668102 |
| 27.25 | 0.531587 | 0.399522 | 1.338173 | 0.085694 | 1.53342 | 0.523287 | 0.50048 | 0.4393 | 0.201335 | 0.198698 | 0.816339 | 0.601777 | 1.421157 | 1.047961 | -0.55263 | -0.222449 | 0.385685 | 0.036642 | -0.163316 | -0.36964 | 0.235087 | 0.283881 | 2.301446 | 0.814227 |
| 27.26388889 | 0.635103 | 0.35768 | 1.437834 | 0.276415 | 1.523171 | 0.636042 | 0.68588 | 0.350688 | 0.352501 | 0.261623 | 1.038652 | 0.727469 | 1.031216 | 1.121156 | -0.79658 | -0.14546 | 0.909677 | 0.692368 | 0.612266 | -0.334352 | 0.525297 | 0.012845 | 1.972306 | 0.994046 |
| 27.27777778 | 0.764008 | 0.492694 | 1.444196 | 0.45546 | 1.568136 | 0.721496 | 0.687991 | 0.266665 | 0.426375 | 0.20753 | 1.166872 | 0.753723 | 1.084553 | 0.70321 | -0.254712 | -0.242017 | 0.871131 | 0.945995 | 1.153608 | -0.478795 | 0.916502 | -0.522544 | 1.40826 | 0.636049 |
| 27.29166667 | 0.853232 | 0.50873 | 1.682959 | 0.542069 | 1.49848 | 0.695857 | 0.637494 | 0.244298 | 0.453316 | 0.135147 | 1.206204 | 0.637238 | 1.555024 | 0.40514 | 0.231561 | -0.159262 | 0.351763 | 0.758866 | 1.08834 | -0.550638 | 0.918058 | -0.465221 | 1.313972 | 0.359499 |
| 27.30555556 | 0.932998 | 0.553028 | 1.85034 | 0.618814 | 1.28305 | 0.550451 | 0.59255 | 0.291364 | 0.664313 | 0.231996 | 1.336751 | 0.568708 | 1.90845 | 0.359362 | 0.428146 | 0.168476 | -0.031839 | 0.59808 | 0.892521 | -0.335772 | 0.531326 | -0.125257 | 2.007423 | 0.428189 |
| 27.31944444 | 1.090829 | 0.573074 | 1.794335 | 0.805084 | 1.066817 | 0.386122 | 0.715079 | 0.523496 | 0.801335 | 0.347802 | 1.331462 | 0.634642 | 2.302665 | 0.294573 | 0.654468 | 0.29629 | -0.577456 | 0.775622 | 0.910302 | -0.169955 | 0.576332 | 0.162835 | 1.863986 | 0.369653 |
| 27.33333333 | 1.194907 | 0.62607 | 1.574644 | 0.802682 | 0.954157 | 0.293345 | 0.811606 | 0.688801 | 1.004026 | 0.500443 | 1.336362 | 0.750232 | 2.792048 | 0.087658 | 0.993805 | 0.256474 | -0.112141 | 1.211329 | 1.299964 | -0.074326 | 0.878355 | -0.010953 | 1.742519 | 0.167576 |
| 27.34722222 | 1.012564 | 0.61098 | 1.523819 | 0.589734 | 0.702182 | 0.370585 | 0.833161 | 0.573004 | 1.229635 | 0.707799 | 1.441871 | 0.809563 | 2.464831 | 0.10068 | 1.151834 | 0.184385 | 0.72985 | 1.234226 | 1.02646 | 0.132539 | 1.092698 | -0.039329 | 1.935642 | 0.143231 |
| 27.36111111 | 0.971175 | 0.640555 | 1.560412 | 0.798693 | 0.427357 | 0.71116 | 0.894364 | 0.40951 | 1.336673 | 0.824885 | 1.549344 | 0.790814 | 1.557324 | 0.642956 | 1.397898 | 0.436403 | 1.453989 | 1.012357 | 0.420807 | 0.261579 | 1.256422 | 0.265777 | 2.001983 | 0.432835 |
| 27.375 | 0.888501 | 0.758122 | 1.429339 | 1.304512 | 0.328523 | 1.095418 | 1.02869 | 0.579348 | 1.32042 | 0.856833 | 1.643315 | 0.848386 | 1.274457 | 1.37028 | 1.703358 | 0.598529 | 1.591217 | 1.184432 | 0.552367 | 0.238728 | 1.557605 | 0.362119 | 2.029123 | 0.765121 |
| 27.38888889 | 0.908289 | 0.768539 | 1.243189 | 1.313561 | 0.404312 | 1.222934 | 1.238483 | 1.142592 | 1.297409 | 0.907601 | 1.622513 | 0.910116 | 1.557176 | 1.7662 | 1.840801 | 0.420084 | 1.609647 | 1.411412 | 0.761467 | 0.332734 | 1.935162 | 0.43603 | 1.941374 | 0.919742 |
| 27.40277778 | 1.11688 | 0.682263 | 1.207478 | 0.783986 | 0.591142 | 0.91205 | 1.211134 | 1.412788 | 1.376796 | 1.111707 | 1.636661 | 1.016932 | 1.719267 | 2.080129 | 1.824703 | 0.260137 | 1.376643 | 1.435357 | 0.68294 | 0.540241 | 1.859755 | 0.546171 | 1.820166 | 1.131205 |
| 27.41666667 | 1.130481 | 0.43268 | 1.267013 | 0.32194 | 0.756154 | 0.727266 | 1.030773 | 1.47914 | 1.388109 | 1.208433 | 1.585287 | 1.157528 | 2.0339 | 2.197801 | 1.795648 | 0.168092 | 0.764139 | 1.588543 | 0.972665 | 0.683875 | 1.553153 | 0.448213 | 2.113003 | 1.398325 |
| 27.43055556 | 1.016155 | 0.383392 | 1.322335 | 0.291421 | 0.974404 | 0.669579 | 0.82556 | 1.196055 | 1.267188 | 0.982222 | 1.694419 | 1.131251 | 2.313416 | 1.843139 | 1.872259 | 0.202131 | 0.379735 | 1.05265 | 0.734001 | 0.782713 | 1.410342 | 0.52973 | 2.871851 | 1.258105 |
| 27.44444444 | 0.991666 | 0.509329 | 1.085475 | 0.522512 | 1.033793 | 0.766799 | 0.75932 | 0.987877 | 1.099468 | 0.674475 | 1.846611 | 1.053344 | 1.755157 | 1.346838 | 1.702625 | 0.224937 | -0.253693 | 0.462935 | 0.519157 | 0.976349 | 1.532606 | 0.704308 | 2.741541 | 0.960727 |
| 27.45833333 | 1.289146 | 0.656772 | 0.754636 | 0.52773 | 1.075316 | 1.241876 | 0.895619 | 0.981212 | 1.130071 | 0.721721 | 2.264983 | 1.280001 | 1.37709 | 1.057166 | 1.052071 | 0.174836 | 0.07191 | 0.518577 | 0.371673 | 0.847252 | 1.380133 | 0.508479 | 1.770909 | 0.751272 |
| 27.47222222 | 1.46722 | 0.503883 | 0.359203 | 0.352666 | 0.931903 | 1.861383 | 0.795923 | 0.869481 | 1.393702 | 1.013122 | 2.027272 | 1.616344 | 1.133193 | 0.771669 | 0.703848 | 0.161004 | 0.374515 | 1.072638 | 0.407056 | 0.635694 | 1.520585 | 0.407515 | 1.873141 | 0.594568 |
| 27.48611111 | 1.401969 | 0.220865 | 0.383813 | 0.409157 | 1.009055 | 2.251455 | 0.605079 | 0.715081 | 1.4505 | 1.318418 | 1.675872 | 1.620723 | 1.714814 | 0.526129 | 0.62894 | 0.251043 | 0.611889 | 1.469759 | 0.284135 | 0.697426 | 1.529753 | 0.451995 | 1.51807 | 0.271019 |
| 27.5 | 1.617277 | -0.017441 | 0.849295 | 0.56761 | 1.309483 | 1.967418 | 0.647704 | 0.68547 | 1.436572 | 1.402223 | 1.798679 | 1.700736 | 2.520246 | 0.347481 | 0.494938 | 0.545003 | 1.419885 | 1.313947 | 0.246693 | 1.300224 | 1.318564 | 0.80568 | 1.50681 | -0.100569 |
| 27.51388889 | 1.522556 | -0.301372 | 1.282136 | 0.421318 | 1.359336 | 1.673964 | 0.867447 | 0.810852 | 1.397884 | 1.271996 | 1.702548 | 1.578383 | 1.97393 | 0.077023 | 0.605666 | 0.820047 | 2.543817 | 0.839822 | 0.395463 | 1.854683 | 0.947013 | 1.227314 | 1.6544 | -0.438986 |
| 27.52777778 | 1.100929 | -0.37113 | 1.095527 | -0.124451 | 0.876539 | 1.435958 | 0.575416 | 0.939975 | 1.302926 | 1.186477 | 1.421246 | 1.587416 | 0.742259 | -0.464841 | 0.544846 | 0.773779 | 2.227461 | 0.548452 | 0.590192 | 2.674103 | 1.280211 | 1.052682 | 1.039661 | -0.524732 |
| 27.54166667 | 0.776797 | -0.235038 | 0.641215 | -0.436013 | 0.387345 | 1.19022 | 0.250991 | 0.64073 | 1.342912 | 1.205578 | 0.913005 | 1.278876 | 1.148607 | 0.130052 | 0.425004 | 0.629027 | 1.391358 | 0.601831 | 0.562804 | 3.028851 | 1.713101 | 0.714787 | 0.746667 | 0.094363 |
| 27.55555556 | 0.80579 | -0.162225 | 0.980302 | -0.310074 | 0.385458 | 1.069198 | 0.288885 | 0.473774 | 1.122446 | 0.950709 | 0.902513 | 1.1215 | 1.155351 | 1.086626 | 0.611655 | 0.951856 | 0.76232 | 0.887001 | 0.617171 | 2.282409 | 1.865987 | 0.594761 | 1.047195 | 0.744918 |
| 27.56944444 | 0.960985 | -0.154047 | 1.343414 | -0.041446 | 0.567176 | 0.882836 | 0.207129 | 0.373071 | 0.606735 | 0.626194 | 1.150878 | 1.171009 | 0.798043 | 1.62792 | 0.526344 | 1.337718 | 0.776921 | 0.827885 | 0.809218 | 1.519294 | 1.540287 | 0.441184 | 1.086857 | 1.175287 |
| 27.58333333 | 0.909526 | -0.21793 | 0.829815 | -0.082065 | 0.5863 | 0.572591 | 0.104338 | 0.41064 | 0.192284 | 0.461736 | 0.964798 | 0.880036 | 0.385745 | 1.565265 | 0.206006 | 1.027458 | 1.39186 | 0.833989 | 1.119159 | 2.084584 | 1.168624 | 0.291328 | 0.963906 | 1.231311 |
| 27.59722222 | 0.738256 | -0.277787 | -0.060126 | -0.296017 | 0.376042 | 0.258862 | 0.059122 | 0.327486 | -0.086282 | 0.35642 | 0.63963 | 0.674045 | 0.235299 | 1.171198 | 0.023403 | 0.573032 | 1.243033 | 0.665824 | 1.14128 | 2.531213 | 0.901951 | 0.335263 | 1.029344 | 1.118054 |
| 27.61111111 | 0.514627 | -0.333238 | -0.331495 | -0.408772 | 0.182764 | 0.049923 | -0.130207 | 0.089151 | -0.038241 | 0.259318 | 0.460705 | 0.511133 | 0.139689 | 0.169559 | -0.169266 | 0.252184 | 0.861373 | 0.098527 | 0.92771 | 2.247585 | 0.557008 | 0.379628 | 1.094825 | 0.852029 |
| 27.625 | 0.233898 | -0.360313 | -0.025926 | -0.21334 | 0.189809 | -0.011974 | -0.255514 | -0.145781 | 0.001023 | 0.188848 | 0.114743 | 0.042679 | -0.118112 | -0.88763 | -0.166018 | -0.103633 | 0.629883 | -0.481026 | 0.820604 | 1.306868 | 0.445872 | 0.273862 | 0.7732 | 0.396958 |
| 27.63888889 | 0.090635 | -0.121303 | 0.21194 | -0.034699 | 0.240602 | 0.029233 | -0.078519 | -0.238736 | -0.161714 | 0.108116 | -0.008231 | -0.497558 | -0.23582 | -1.528499 | 0.019428 | -0.104478 | 1.078498 | -0.543295 | 0.542721 | 0.791214 | 0.353754 | 0.017337 | 0.814631 | -0.002419 |
| 27.65277778 | 0.057219 | -0.00154 | 0.001871 | -0.061316 | 0.221274 | -0.008106 | 0.127048 | -0.148934 | -0.344183 | 0.018935 | -0.251238 | -0.816701 | 0.072005 | -1.399462 | 0.164841 | 0.048276 | 1.329701 | -0.326725 | 0.290515 | -0.043083 | 0.240624 | -0.270368 | 0.738375 | -0.129111 |
| 27.66666667 | -0.011729 | 0.059247 | -0.280252 | -0.162951 | 0.135286 | -0.027043 | 0.066883 | -0.167654 | -0.314732 | 0.000321 | -0.239452 | -0.777036 | 0.584747 | -1.215007 | 0.201172 | 0.380158 | 1.34554 | 0.058339 | 0.227719 | -0.766783 | 0.714522 | -0.234772 | 0.218897 | -0.10572 |
| 27.68055556 | 0.010806 | -0.015136 | -0.352565 | -0.125366 | 0.061283 | 0.041362 | -0.079377 | -0.288165 | -0.03409 | -0.03685 | 0.008612 | -0.526883 | 0.047024 | -1.050082 | 0.127047 | 0.211809 | 1.080414 | 0.349052 | 0.204397 | -1.56815 | 0.6003 | -0.212463 | -0.186195 | -0.221157 |

| | | | | | | | | | | | | | | | | | | | | | | | | |
|---|---|---|---|---|---|---|---|---|---|---|---|---|---|---|---|---|---|---|---|---|---|---|---|---|
| 27.69444444 | -0.066902 | -0.028103 | -0.227187 | 0.16967 | -0.008833 | 0.120045 | -0.105819 | -0.257054 | 0.230041 | -0.027987 | 0.307077 | -0.273415 | -0.173135 | -1.110658 | 0.035202 | 0.047683 | 0.554778 | 0.101489 | 0.234581 | -1.914618 | 0.573656 | -0.1924 | 0.233672 | -0.311883 |
| 27.70833333 | -0.144047 | -0.020417 | -0.160317 | 0.521121 | -0.045908 | 0.175687 | 0.058686 | -0.102426 | 0.368964 | 0.100643 | 0.440254 | -0.010522 | -0.153628 | -0.944921 | 0.067341 | 0.03993 | 0.457191 | -0.299238 | 0.268372 | -1.738172 | 0.644406 | -0.174139 | 0.165766 | -0.298278 |
| 27.72222222 | 0.007569 | -0.09183 | -0.21514 | 0.52527 | -0.008748 | 0.210702 | 0.12931 | 0.057852 | 0.395921 | 0.189347 | 0.363734 | -0.008233 | -0.083592 | -0.653209 | 0.052457 | 0.002749 | 0.908992 | -0.294651 | 0.329881 | -1.32658 | 0.660014 | -0.234721 | 0.047257 | -0.086848 |
| 27.73611111 | 0.303291 | -0.163982 | -0.478358 | 0.154308 | 0.061622 | 0.204361 | 0.009854 | -0.034258 | 0.318154 | 0.066308 | 0.275017 | -0.032775 | -0.047083 | -0.357739 | 0.013238 | -0.036382 | 1.102853 | -0.193822 | 0.384265 | -1.0023 | 0.773977 | -0.283474 | -0.023521 | 0.000349 |
| 27.75 | 0.320387 | -0.309833 | -0.849795 | -0.192983 | 0.034165 | 0.145522 | -0.157768 | -0.285384 | 0.188457 | -0.055351 | 0.221112 | -0.116988 | -0.250575 | -0.730086 | 0.046019 | -0.039687 | 0.640387 | -0.34306 | 0.378756 | -0.788123 | 0.820803 | -0.237169 | 0.168227 | -0.045366 |
| 27.76388889 | 0.023394 | -0.578669 | -0.895411 | -0.393587 | -0.005592 | 0.006906 | -0.248379 | -0.410267 | 0.051541 | -0.057697 | 0.006707 | -0.111253 | -0.173913 | -1.622254 | 0.149635 | -0.005569 | 0.32409 | -0.726394 | 0.054604 | -0.88582 | 0.620971 | -0.271449 | 0.016437 | -0.354188 |
| 27.77777778 | -0.36498 | -0.658372 | -0.832091 | -0.216555 | 0.085048 | -0.153994 | -0.314707 | -0.319247 | -0.002989 | 0.017828 | -0.091531 | -0.057531 | -0.092222 | -1.517544 | 0.081608 | -0.111456 | 0.259411 | -0.717686 | -0.100032 | -1.268797 | 0.438077 | -0.42181 | -0.425196 | -0.200271 |
| 27.79166667 | -0.607641 | -0.553769 | -0.590636 | -0.058553 | 0.224827 | -0.300155 | -0.271145 | -0.150405 | 0.054277 | 0.043098 | -0.141376 | -0.114779 | -0.292437 | -1.453138 | -0.03392 | -0.110298 | 0.563307 | -0.878702 | -0.106695 | -1.607615 | 0.113778 | -0.479601 | -0.712533 | -0.077522 |
| 27.80555556 | -0.187135 | -0.434951 | -0.414462 | -0.240239 | 0.190817 | -0.338901 | -0.261437 | -0.120268 | 0.020957 | -0.200797 | -0.212894 | -0.346207 | -1.129503 | -1.316094 | -0.216138 | -0.249873 | 0.529286 | -1.410481 | -0.332702 | -1.725816 | -0.218832 | -0.423734 | -0.836368 | 0.145301 |
| 27.81944444 | 0.09531 | -0.498589 | -0.504726 | -0.610249 | 0.017895 | -0.278822 | -0.238527 | -0.259268 | -0.115948 | -0.508218 | -0.281484 | -0.543259 | -1.600689 | -1.237262 | -0.300076 | -0.446599 | 0.344045 | -1.96317 | -0.751179 | -1.885988 | -0.45736 | -0.534231 | -0.845661 | 0.239211 |
| 27.83333333 | -0.202357 | -0.790007 | -0.874151 | -0.849931 | -0.107532 | -0.282345 | -0.252922 | -0.432871 | -0.254093 | -0.640765 | -0.387809 | -0.476016 | -1.62194 | -1.144584 | -0.404216 | -0.593906 | 0.361895 | -2.236332 | -1.268991 | -2.690679 | -0.999346 | -0.848627 | -1.011597 | 0.153307 |
| 27.84722222 | -0.89389 | -0.980732 | -1.373231 | -0.652637 | -0.265689 | -0.466367 | -0.314559 | -0.55546 | -0.388467 | -0.600172 | -0.64955 | -0.523363 | -1.576354 | -0.920438 | -0.588862 | -0.659601 | 0.312974 | -2.237891 | -1.35344 | -3.055109 | -1.125964 | -1.06562 | -0.87647 | -0.011359 |
| 27.86111111 | -1.240233 | -0.91372 | -1.927083 | -0.428105 | -0.222791 | -0.754234 | -0.43047 | -0.733211 | -0.594136 | -0.601033 | -0.921394 | -0.679684 | -1.654267 | -0.830675 | -0.555247 | -0.72233 | 0.069342 | -1.971343 | -1.080974 | -2.744499 | -1.041497 | -1.108447 | -0.885577 | -0.006215 |
| 27.875 | -1.177406 | -0.942947 | -2.100682 | -0.64893 | -0.67449 | -0.962944 | -0.429138 | -0.852387 | -0.723577 | -0.837592 | -1.392693 | -0.986248 | -2.142553 | -1.009877 | -0.612216 | -0.837688 | -0.774318 | -2.025889 | -1.225798 | -2.351847 | -1.126201 | -1.137559 | -0.966564 | 0.037048 |
| 27.88888889 | -1.175908 | -0.788249 | -1.985653 | -1.1829 | -1.368372 | -1.036255 | -0.533081 | -1.140533 | -0.776698 | -1.118134 | -1.779063 | -1.309534 | -2.769775 | -1.241168 | -0.759393 | -0.829068 | -1.563975 | -2.230347 | -1.6616 | -1.847294 | -1.460757 | -1.230602 | -0.999823 | 0.053622 |
| 27.90277778 | -1.084204 | -0.694416 | -2.040878 | -1.440836 | -1.682044 | -1.176746 | -0.723745 | -1.545096 | -0.942699 | -1.283233 | -1.700697 | -1.345429 | -3.487121 | -1.130625 | -0.643673 | -0.818551 | -1.792994 | -2.317702 | -1.79222 | -1.712755 | -1.512662 | -1.353925 | -1.565832 | 0.028958 |
| 27.91666667 | -1.171185 | -0.903976 | -2.59198 | -0.944852 | -1.954147 | -1.304618 | -0.683295 | -1.465066 | -1.132748 | -1.316714 | -1.638585 | -1.146561 | -3.257398 | -1.432246 | -0.632029 | -0.902507 | -1.86759 | -2.752412 | -2.09381 | -1.415105 | -1.377529 | -1.294009 | -1.488897 | -0.225765 |
| 27.93055556 | -1.210656 | -1.215232 | -2.98042 | -0.462145 | -2.855421 | -1.178559 | -0.594819 | -1.482904 | -0.960831 | -0.947158 | -1.644761 | -0.954944 | -2.299651 | -2.13983 | -0.907696 | -0.935766 | -1.955064 | -3.077378 | -2.172197 | -1.205882 | -1.077876 | -0.922799 | -0.922363 | -0.642202 |
| 27.94444444 | -1.270179 | -1.050843 | -2.670745 | -0.542722 | -2.829093 | -1.012315 | -0.26325 | -1.372824 | -0.66231 | -0.594563 | -1.721893 | -0.813603 | -2.492134 | -2.747459 | -1.021136 | -0.710415 | -2.437225 | -2.821559 | -2.317127 | -1.041067 | -0.526504 | -0.742615 | -0.648807 | -0.729502 |
| 27.95833333 | -1.734775 | -1.24948 | -2.70622 | -1.125877 | -2.614638 | -1.492458 | -0.453433 | -1.703475 | -0.957759 | -0.92103 | -2.221929 | -1.1588 | -4.070157 | -2.905047 | -1.419903 | -0.669831 | -3.352129 | -2.452906 | -2.835449 | -0.982485 | -0.055006 | -1.310701 | -1.25111 | -0.716491 |
| 27.97222222 | -2.475737 | -1.738889 | -3.5645 | -1.706078 | -3.14883 | -2.659556 | -1.10969 | -2.345791 | -1.630785 | -1.656883 | -2.756577 | -1.569946 | -5.98218 | -3.666544 | -1.617265 | -0.56614 | -3.903032 | -2.636276 | -3.132223 | -0.976703 | -0.081171 | -2.214925 | -1.573641 | -0.757367 |
| 27.98611111 | -2.129385 | -2.295876 | -3.825121 | -1.755676 | -3.207431 | -2.901204 | -1.096756 | -2.082727 | -1.555908 | -1.528534 | -2.98252 | -1.589138 | -6.735591 | -5.298009 | -1.275595 | -0.825235 | -3.406145 | -2.20119 | -2.704116 | -1.763407 | -0.250496 | -2.733833 | -1.397034 | -1.11415 |
| 28 | -1.592089 | -2.326007 | -2.880011 | -1.479079 | -3.071071 | -2.557425 | -0.810183 | -0.993052 | -1.2119 | -1.184432 | -2.463727 | -0.982456 | -3.72129 | -5.054823 | -0.682853 | -0.865982 | -2.177062 | -1.15095 | -1.304137 | -2.024179 | -0.613585 | -2.298753 | -1.103286 | -1.956858 |
| 28.01388889 | -1.197054 | -1.53515 | -1.593363 | -1.141765 | -1.848985 | -2.247052 | -0.401399 | -0.271572 | -1.006384 | -1.129332 | -2.539779 | -1.213603 | -1.242612 | -3.528955 | -0.365626 | -0.137918 | -1.297622 | 0.269768 | 0.320062 | -2.016069 | -0.461873 | -1.730482 | -0.522119 | -1.599952 |
| 28.02777778 | -0.828383 | -0.517725 | -1.183176 | -0.653285 | -0.848933 | -1.8159 | -0.393954 | -0.311338 | -0.796873 | -1.078134 | -2.210674 | -1.290329 | -0.214783 | -2.198682 | -0.242585 | 0.460512 | -0.40637 | 1.229692 | 1.428344 | -1.254218 | -0.32204 | -1.03397 | -0.31512 | -0.75122 |
| 28.04166667 | -0.667205 | -0.026026 | -0.97122 | -0.08365 | -0.486955 | -1.131696 | -0.135004 | -0.216422 | -0.473279 | -0.74486 | -1.78966 | -1.083579 | -0.755334 | -1.51663 | -0.262312 | 0.490289 | -0.14366 | 0.963049 | 0.868967 | -0.451188 | -0.450936 | -0.837892 | -0.363843 | -0.379716 |
| 28.05555556 | -0.598124 | 0.446419 | -0.560958 | 0.541374 | -0.210114 | -0.446128 | 0.362623 | 0.31714 | -0.046234 | -0.131648 | -1.228087 | -0.59984 | -1.185133 | -0.841559 | -0.100208 | 0.731781 | -0.175861 | 0.484128 | 0.343381 | 0.010887 | -0.473267 | -0.553886 | -0.757965 | -0.080693 |
| 28.06944444 | -0.415108 | 0.882457 | -0.024173 | 0.897985 | 0.536709 | 0.182562 | 0.755744 | 0.545981 | 0.281658 | 0.312414 | -0.445452 | -0.215773 | -1.266563 | -0.05694 | 0.339373 | 0.632964 | -0.53071 | 0.938442 | 0.806348 | 0.167113 | -0.42961 | -0.097241 | -0.725254 | 0.289528 |
| 28.08333333 | 0.04498 | 1.254219 | 0.321442 | 0.867972 | 1.321987 | 0.556376 | 0.818155 | 0.521984 | 0.484348 | 0.442222 | 0.418377 | 0.111958 | -1.056454 | 0.787373 | 0.704662 | 0.460042 | -1.23 | 1.568655 | 1.650978 | 0.07432 | -0.175595 | 0.625892 | -0.11283 | 1.203926 |
| 28.09722222 | 0.609882 | 1.320743 | 0.523127 | 0.698365 | 1.867971 | 0.625151 | 0.999069 | 0.678858 | 0.702863 | 0.513775 | 0.888136 | 0.203206 | -0.317832 | 1.016111 | 1.019358 | 0.303689 | -1.131227 | 1.698951 | 1.99174 | 0.011896 | -0.000601 | 0.29169 | 0.122683 | 1.620771 |
| 28.11111111 | 1.055764 | 1.340022 | 0.719737 | 0.519427 | 2.003452 | 0.645696 | 0.930327 | 0.716058 | 0.857869 | 0.539549 | 1.102787 | 0.355132 | 0.050421 | 1.640458 | 1.276241 | 0.123553 | -0.796573 | 1.162432 | 1.559561 | 0.030461 | -0.250272 | -0.815592 | 0.316216 | 1.643979 |
| 28.125 | 0.990396 | 1.067112 | 1.011578 | 0.266319 | 1.710406 | 0.651118 | 0.424651 | 0.579756 | 0.654281 | 0.324697 | 1.24079 | 0.549326 | -0.116871 | 2.42521 | 0.734665 | -0.417504 | -0.737476 | 1.06445 | 1.208548 | -0.024064 | -0.496842 | -1.657506 | 0.707104 | 2.065169 |
| 28.13888889 | 0.617969 | 0.838731 | 1.116864 | 0.209392 | 1.549848 | 0.502915 | 0.114528 | 0.83876 | 0.343552 | -0.011266 | 1.310397 | 0.611176 | -0.293165 | 2.3248 | 0.657307 | -1.022234 | -0.849554 | 1.253048 | 1.672103 | -0.501892 | -0.147337 | -0.819426 | 0.906889 | 2.050927 |
| 28.15277778 | 0.29449 | 0.668449 | 0.9173 | 0.321915 | 1.138739 | 0.236874 | 0.192032 | 1.258945 | 0.285876 | -0.08823 | 1.295298 | 0.528262 | -0.27238 | 2.038915 | 0.655607 | -1.006617 | -1.045334 | 1.520906 | 2.272544 | -0.349806 | 0.213046 | -0.213208 | 1.015014 | 1.510976 |
| 28.16666667 | 0.159927 | 0.587719 | 0.535069 | 0.42799 | 1.041901 | -0.097708 | 0.272424 | 1.461622 | 0.21093 | 0.019161 | 0.915413 | 0.218393 | -0.605657 | 1.940419 | 0.243338 | -0.860559 | -1.31249 | 1.68683 | 2.377294 | -0.347567 | 0.342508 | 0.100193 | 0.894701 | 1.336589 |
| 28.18055556 | 0.05479 | 0.398043 | 0.193774 | 0.323297 | 1.741666 | -0.540236 | 0.038923 | 1.212874 | -0.147943 | -0.054229 | 0.308616 | -0.233866 | -1.163757 | 1.196389 | -0.165588 | -0.756869 | -1.440275 | 1.12997 | 1.570189 | -0.303981 | -0.043013 | -0.199563 | 0.774241 | 0.747784 |
| 28.19444444 | 0.042527 | 0.476865 | 0.106033 | 0.048844 | 2.954553 | -0.700964 | -0.093949 | 1.111276 | -0.349882 | -0.082461 | 0.080407 | -0.25536 | -1.210767 | 0.419786 | 0.168718 | -0.761972 | -1.15792 | 0.443613 | 0.777759 | -0.324272 | -0.17384 | -0.555614 | 0.693882 | 0.154008 |
| 28.20833333 | 0.314335 | 0.569337 | 0.373311 | -0.060204 | 4.074221 | -0.368509 | 0.343707 | 1.359489 | 0.01013 | 0.132846 | 0.582749 | 0.373166 | -0.621857 | 0.350547 | 0.45219 | -0.350589 | 0.001985 | 0.419382 | 0.592347 | -0.353405 | 0.392723 | -0.659847 | -0.139461 | 0.107176 |
| 28.22222222 | 0.534564 | 0.426929 | 0.716101 | 0.109077 | 3.418385 | -0.101632 | 0.693099 | 1.362303 | 0.345762 | 0.309433 | 0.9577 | 0.725017 | -0.512506 | 0.172637 | -0.083088 | 0.120977 | 0.295691 | 0.337648 | 0.246761 | -0.389204 | 0.602476 | -0.631953 | -0.524569 | -0.004726 |
| 28.23611111 | 0.502037 | 0.287644 | 0.582784 | 0.026578 | 3.191442 | -0.113079 | 0.640659 | 1.271581 | 0.088327 | 0.243229 | 0.75169 | 0.50748 | -0.641805 | -0.106115 | -0.464901 | 0.225438 | 0.002606 | -0.225783 | -0.420358 | -0.417043 | 0.207108 | -0.59776 | -0.22216 | -0.183264 |
| 28.25 | 0.676858 | 0.547023 | 0.654848 | -0.229373 | 3.853103 | -0.132833 | 0.556014 | 1.219865 | -0.300309 | 0.131048 | 0.631069 | 0.390361 | 0.2543 | 0.179646 | -0.160164 | 0.441954 | 0.261681 | -0.274716 | -0.766552 | -0.434176 | 0.034121 | -0.549467 | 0.271489 | -0.12348 |
| 28.26388889 | 0.98708 | 0.901449 | 1.58253 | -0.294631 | 3.773109 | 0.029838 | 0.652918 | 1.233521 | -0.239859 | 0.115628 | 0.853891 | 0.465868 | 1.256887 | 0.20068 | 0.118463 | 0.458017 | 0.337334 | 0.060469 | -0.720237 | -0.52569 | 0.456105 | -0.579607 | 0.69879 | 0.147253 |
| 28.27777778 | 0.83796 | 0.771137 | 2.048904 | -0.033156 | 3.385919 | 0.26618 | 0.819598 | 1.268104 | 0.133439 | 0.184887 | 0.960056 | 0.46702 | 1.058962 | -0.215064 | 0.091109 | 0.384636 | 0.212971 | -0.295233 | -0.807126 | -0.521574 | 0.770979 | -0.829505 | 0.074514 | 0.088754 |
| 28.29166667 | 0.619512 | 0.52319 | 1.817772 | 0.379172 | 2.900461 | 0.540915 | 0.887361 | 1.00478 | 0.350272 | 0.234258 | 0.876081 | 0.502294 | 0.433014 | -0.194781 | 0.034359 | 0.414197 | -0.293193 | 0.125667 | 0.021551 | -0.471402 | 0.643852 | -0.872613 | -0.515097 | -0.252888 |
| 28.30555556 | 0.593776 | 0.488502 | 1.905493 | 0.578104 | 2.022176 | 0.839525 | 0.829471 | 0.461838 | 0.300174 | 0.251998 | 0.807964 | 0.48943 | 0.59196 | -0.000215 | 0.52589 | 0.72372 | -0.567195 | 0.707497 | 0.576235 | -0.408815 | 0.588551 | -0.66604 | 0.04893 | 0.067345 |
| 28.31944444 | 0.632184 | 0.570479 | 2.204464 | 0.46468 | 1.17462 | 0.977462 | 0.617955 | 0.090479 | 0.161844 | 0.381006 | 0.928368 | 0.487664 | 1.433699 | -0.112462 | 0.653095 | 0.464436 | -0.060217 | 0.464774 | -0.004645 | -0.345413 | 0.551921 | -0.424011 | 1.033556 | 0.298835 |
| 28.33333333 | 0.605131 | 0.664124 | 2.63135 | 0.325635 | 0.858746 | 0.957364 | 0.595478 | 0.266506 | 0.174784 | 0.660568 | 1.038739 | 0.79621 | 2.311093 | -0.608379 | 0.693606 | -0.063463 | 0.54952 | 0.270591 | -0.274969 | -0.231145 | 0.738181 | -0.0997 | 1.451536 | 0.741449 |
| 28.34722222 | 0.699261 | 0.716125 | 2.748098 | 0.455778 | 1.231833 | 0.931861 | 0.757221 | 0.728646 | 0.505202 | 0.94315 | 1.139285 | 1.005717 | 2.741511 | -0.377621 | 1.209177 | 0.254779 | 0.851311 | 0.969121 | 0.68665 | -0.067997 | 1.174678 | 0.044971 | 1.419262 | 1.284813 |
| 28.36111111 | 0.855857 | 0.691628 | 2.00826 | 0.622331 | 1.642001 | 1.120471 | 0.945054 | 0.97206 | 0.855521 | 1.133673 | 1.237049 | 0.933293 | 2.768693 | 0.52218 | 1.926033 | 0.659546 | 1.179965 | 1.665524 | 1.24936 | 0.014698 | 1.445252 | -0.040644 | 1.569022 | 1.375727 |
| 28.375 | 0.934257 | 0.77756 | 1.392127 | 0.603346 | 1.047874 | 1.360363 | 1.117184 | 0.856954 | 0.986222 | 1.220543 | 1.321311 | 0.864779 | 2.945058 | 0.461993 | 2.120104 | 0.4973 | 1.497666 | 1.749496 | 0.895799 | -0.010606 | 1.243402 | -0.08953 | 1.686481 | 0.628012 |
| 28.38888889 | 0.97339 | 0.835816 | 1.249171 | 0.534776 | 0.576494 | 1.417517 | 1.180003 | 0.756659 | 0.937796 | 1.219795 | 1.494294 | 0.938482 | 3.146545 | -0.063558 | 2.388365 | 0.338661 | 1.505163 | 1.595785 | 0.663747 | -0.009058 | 1.016391 | 0.038411 | 1.705062 | -0.072312 |
| 28.40277778 | 1.05906 | 0.728131 | 1.42479 | 0.572916 | 0.602675 | 1.204582 | 1.124097 | 0.91553 | 0.865657 | 1.163974 | 1.390307 | 1.015706 | 2.991939 | -0.279674 | 2.766274 | 0.209626 | 1.233442 | 1.533626 | 0.781161 | 0.106588 | 0.965437 | 0.215598 | 1.864815 | -0.378646 |
| 28.41666667 | 1.069253 | 0.608631 | 1.721938 | 0.610687 | 0.79111 | 1.022348 | 1.121311 | 0.939862 | 0.939896 | 1.081878 | 1.460901 | 1.15446 | 3.231185 | -0.184938 | 2.946223 | 0.277381 | 0.999292 | 1.454304 | 0.830727 | 0.244774 | 0.943503 | 0.26833 | 1.810379 | -0.153838 |
| 28.43055556 | 1.001684 | 0.56046 | 1.72123 | 0.483484 | 0.769169 | 0.885864 | 1.033754 | 0.754826 | 1.021916 | 0.940867 | 1.602958 | 1.249637 | 3.893863 | 0.009485 | 2.88324 | 0.178701 | 0.725349 | 1.461425 | 0.648817 | 0.267378 | 0.985398 | 0.254591 | 1.503172 | -0.330586 |
| 28.44444444 | 0.98735 | 0.491025 | 1.391565 | 0.287819 | 0.31029 | 0.788627 | 0.68879 | 0.461848 | 0.948659 | 0.787952 | 1.711175 | 1.210026 | 4.435911 | 0.548301 | 2.410398 | 0.18411 | 0.438121 | 1.417003 | 0.45175 | 0.218855 | 1.414265 | 0.08146 | 1.295897 | -0.216706 |
| 28.45833333 | 1.089644 | 0.496739 | 1.418622 | 0.388372 | 0.088162 | 0.975602 | 0.456604 | 0.597619 | 0.785003 | 0.819789 | 1.808599 | 1.196492 | 4.209118 | 1.607398 | 1.933721 | 0.278374 | -0.084796 | 1.265865 | 0.227207 | 0.096284 | 1.533911 | -0.013472 | 1.838091 | 0.243643 |
| 28.47222222 | 0.937922 | 0.447459 | 1.37027 | 0.601298 | 0.422096 | 1.151488 | 0.4828 | 0.547383 | 0.654551 | 0.928352 | 1.869978 | 1.397314 | 3.070915 | 1.769189 | 1.795237 | 0.313125 | -0.34785 | 1.460069 | 0.024905 | 0.126315 | 1.404963 | 0.168731 | 2.633067 | 0.863282 |
| 28.48611111 | 0.786499 | 0.436906 | 1.04999 | 0.753771 | 1.001723 | 1.225771 | 0.66757 | 0.50883 | 0.675216 | 1.005003 | 1.775213 | 1.419149 | 2.608664 | 0.99603 | 1.95882 | 0.309617 | -0.433678 | 1.604072 | 0.123888 | 0.245227 | 1.844914 | 0.271079 | 1.832769 | 1.213877 |
| 28.5 | 0.944863 | 0.285907 | 1.08193 | 0.742397 | 1.247634 | 1.07635 | 0.854968 | 0.797989 | 0.881163 | 1.193871 | 1.830567 | 1.640966 | 3.453313 | 0.677884 | 2.005584 | 0.430901 | -0.405112 | 1.638295 | 0.321492 | 0.727107 | 2.145479 | 0.322484 | 1.555776 | 1.214127 |
| 28.51388889 | 1.005252 | -0.337398 | 1.139043 | 0.568991 | 1.127522 | 0.989627 | 0.819566 | 1.088058 | 1.120145 | 1.6754 | 1.585759 | 1.917389 | 3.570264 | 1.23713 | 1.743165 | 0.488591 | 0.248637 | 1.930703 | 0.415647 | 1.174258 | 2.287057 | 0.360733 | 2.588015 | 1.277623 |
| 28.52777778 | 0.836602 | -0.512622 | 0.52275 | 0.151815 | 0.601315 | 0.883561 | 0.666561 | 0.914539 | 1.322233 | 2.05997 | 1.745158 | 1.971059 | 1.990212 | 1.562201 | 0.919364 | 0.372642 | 0.960282 | 1.626242 | 0.313788 | 0.824396 | 2.391068 | 0.415863 | 1.908926 | 1.243422 |
| 28.54166667 | 0.63245 | -1.061187 | -0.183799 | -0.207799 | 0.463999 | 0.623409 | 0.562179 | 0.493941 | 1.210866 | 1.785875 | 1.148474 | 1.68267 | 0.567691 | 1.360918 | 0.937705 | 0.269487 | 1.428668 | 1.221016 | 0.215176 | 0.785753 | 2.310583 | 0.50588 | 1.41854 | 1.353232 |
| 28.55555556 | 0.679575 | -0.584397 | -0.1733 | -0.160698 | 0.553297 | 0.525359 | 0.636595 | 0.594264 | 1.031368 | 1.698188 | 0.572283 | 0.418874 | 0.434252 | 0.677908 | 1.099941 | 0.240382 | 1.434794 | 0.947082 | 0.351314 | 0.889815 | 1.921633 | 0.552901 | 0.921902 | 1.64256 |
| 28.56944444 | 0.8005 | -0.109965 | 0.482424 | 0.044214 | 0.518014 | 0.624726 | 0.721298 | 0.707419 | 0.643647 | 1.193509 | 0.032789 | -0.042792 | 0.325279 | -0.513474 | 0.995801 | 0.228697 | 1.421001 | 0.810302 | 0.611909 | 0.800926 | 1.656176 | 0.449682 | 0.682548 | 1.946197 |
| 28.58333333 | 0.693394 | 0.098881 | 0.368392 | 0.130911 | 0.427125 | 0.432658 | 0.410124 | 0.298483 | 0.176451 | 0.586707 | 0.031864 | 0.098789 | 0.524296 | 0.767826 | 0.351508 | 0.201435 | 1.29331 | 0.463672 | 0.556184 | 0.505774 | 0.906572 | 0.319037 | 0.530902 | 2.136299 |
| 28.59722222 | 0.351858 | 0.217 | -0.387456 | 0.10844 | 0.236798 | 0.218346 | 0.050768 | 0.064115 | -0.020295 | 0.02035 | 0.205008 | 0.382059 | 0.809346 | 1.243679 | -0.008247 | 0.160453 | 0.58707 | -0.016157 | 0.407689 | 0.268914 | 0.342241 | 0.250535 | 0.488477 | 1.857245 |
| 28.61111111 | 0.002044 | 0.026362 | -1.015031 | -0.091105 | -0.157163 | -0.053621 | -0.122828 | -0.023681 | -0.048388 | -0.291681 | 0.132597 | 0.267635 | 0.875312 | 0.983912 | -0.008355 | 0.06311 | -0.495666 | -0.383367 | 0.37849 | 0.013796 | -0.035724 | 0.138002 | 0.413324 | 1.172221 |
| 28.625 | -0.314804 | -0.444572 | -0.87838 | -0.374737 | -0.636408 | -0.234113 | -0.148931 | -0.271751 | -0.002559 | -0.071539 | -0.443843 | -0.148173 | 0.953715 | 0.450481 | -0.248389 | -0.046325 | -0.513097 | -0.552682 | 0.363218 | -0.077801 | -0.525364 | -0.115239 | 0.182975 | 0.534315 |
| 28.63888889 | -0.365892 | -0.157483 | -0.447743 | -0.449896 | -0.65466 | -0.175054 | 0.017415 | -0.485448 | 0.01092 | 0.265935 | -0.812888 | -0.669168 | 0.887346 | 0.026964 | -0.111414 | -0.016152 | -0.173912 | -0.857958 | 0.384274 | 0.01207 | -0.350225 | -0.327025 | 0.208457 | -0.015199 |
| 28.65277778 | -0.173994 | 0.188425 | -0.091427 | -0.208597 | -0.344579 | -0.028269 | 0.124021 | -0.390626 | -0.099946 | 0.191218 | -0.954945 | -0.936632 | 0.652799 | -0.367677 | 0.332023 | 0.034598 | -0.525133 | -0.169137 | 0.281423 | 0.121546 | 0.28042 | -0.241236 | 0.218099 | -0.578123 |
| 28.66666667 | -0.061315 | 0.416019 | 0.11698 | 0.014982 | -0.343962 | 0.036217 | 0.096438 | -0.264619 | -0.17374 | -0.114233 | -0.708603 | -0.833741 | 0.204895 | -0.243231 | 0.450662 | 0.006573 | -0.723287 | 0.355862 | 0.193084 | 0.294147 | 0.955658 | -0.039677 | -0.103982 | -1.117341 |
| 28.68055556 | 0.005643 | 0.545036 | 0.14114 | -0.007685 | -0.57913 | 0.092147 | 0.014315 | -0.124601 | -0.081266 | -0.183153 | -0.230164 | -0.520313 | 0.146041 | 0.074426 | 0.002305 | -0.032602 | 0.658129 | 0.689981 | 0.155149 | 0.110405 | 0.980138 | 0.060493 | -0.148682 | -1.010412 |
| 28.69444444 | 0.059486 | 0.527918 | 0.353613 | -0.072127 | -0.712912 | 0.169765 | 0.05622 | -0.031269 | 0.147736 | -0.027128 | 0.13997 | -0.257095 | 0.039133 | -0.734971 | -0.331609 | -0.073816 | 1.255257 | 0.515298 | 0.109869 | -0.184259 | -0.067271 | -0.027004 | 0.067443 | -0.945948 |
| 28.70833333 | 0.164637 | 0.309879 | 0.656592 | -0.029279 | -0.459105 | 0.22643 | 0.320999 | 0.003092 | 0.36348 | 0.09343 | -0.006518 | -0.249882 | -0.007219 | -1.56103 | -0.041063 | -0.068671 | 1.866352 | 0.009588 | 0.066785 | -0.545386 | -0.844003 | -0.176678 | 0.345769 | -0.521222 |
| 28.72222222 | 0.330019 | 0.174564 | 0.653293 | 0.08631 | -0.146738 | 0.282952 | 0.450184 | 0.121489 | 0.456142 | 0.13228 | -0.280432 | -0.428625 | -0.087136 | -2.210801 | 0.133799 | -0.002939 | 1.713232 | -0.435594 | 0.008378 | -0.5811 | -0.159708 | -0.248704 | 0.402687 | -0.107798 |
| 28.73611111 | 0.446005 | 0.311235 | 0.468402 | 0.131247 | -0.284771 | 0.335936 | 0.281507 | 0.108214 | 0.496813 | 0.107339 | -0.018405 | -0.290899 | -0.170189 | -1.58581 | 0.104727 | 0.049501 | 1.175543 | -0.757277 | 0.03165 | -0.536281 | 0.622433 | -0.136812 | 0.382551 | -0.315656 |
| 28.75 | 0.394561 | 0.438784 | 0.182445 | -0.028146 | -0.823705 | 0.301282 | -0.050915 | -0.029581 | 0.381534 | -0.005215 | 0.33044 | -0.044599 | 0.033603 | -0.908274 | 0.509874 | 0.065662 | 1.056961 | -0.987086 | 0.033658 | -0.67971 | 0.767316 | -0.154849 | 0.403542 | -0.660152 |
| 28.76388889 | 0.281043 | 0.482784 | 0.060218 | -0.250345 | -0.571587 | 0.117606 | -0.096739 | 0.025223 | 0.197499 | -0.133515 | 0.397691 | 0.019711 | -0.207192 | -1.022039 | 1.003885 | 0.098765 | 1.252625 | -1.522317 | 0.000251 | -0.753891 | 0.51614 | -0.200312 | 0.183135 | -0.795912 |
| 28.77777778 | 0.152168 | 0.499416 | -0.025145 | -0.357608 | -0.2308 | -0.075917 | -0.016279 | 0.256075 | 0.112632 | -0.220267 | 0.284583 | -0.104383 | -1.550227 | -1.966234 | 0.943314 | 0.147293 | 0.882049 | -2.274696 | -0.100094 | -0.793378 | 0.487279 | -0.243117 | -0.382698 | -0.821369 |
| 28.79166667 | -0.003516 | 0.354716 | -0.319706 | -0.385373 | -0.50507 | -0.159331 | -0.254189 | 0.049886 | 0.090422 | -0.156664 | -0.053753 | -0.429424 | -1.562513 | -1.863011 | 0.722113 | 0.078539 | 0.407702 | -2.171959 | -0.089564 | -0.637064 | 0.540215 | -0.15817 | -0.992096 | -0.637088 |
| 28.80555556 | -0.108673 | 0.174858 | -0.56372 | -0.41668 | -1.262721 | -0.176569 | -0.658311 | -0.394414 | 0.000116 | -0.199918 | -0.4094 | -0.824764 | -1.207238 | -0.284779 | 0.378028 | -0.043413 | 0.385596 | -2.259268 | -0.248357 | -0.554247 | 0.138443 | -0.19695 | -1.156198 | -0.241301 |
| 28.81944444 | -0.128831 | 0.125695 | -0.739411 | -0.319456 | -2.079877 | -0.134996 | -0.782098 | -0.760142 | -0.213301 | -0.530643 | -0.494163 | -0.951434 | -0.666497 | 1.149374 | -0.09001 | -0.13784 | 0.512742 | -2.825858 | -0.662855 | -0.477776 | -0.505205 | -0.268132 | -1.211025 | 0.095103 |
| 28.83333333 | -0.164413 | -0.013563 | -0.925061 | -0.323276 | -2.508616 | -0.145868 | -0.765646 | -0.985334 | -0.48462 | -1.094972 | -0.464243 | -0.741457 | -0.247521 | 2.270196 | -0.349309 | -0.197008 | 0.246928 | -3.463666 | -0.973238 | -0.538424 | -0.511106 | -0.289051 | -1.620046 | 0.195127 |
| 28.84722222 | -0.235123 | -0.415934 | -1.281845 | -0.501977 | -3.008325 | -0.221373 | -0.77391 | -1.222737 | -0.644291 | -1.528274 | -0.645362 | -0.674717 | -1.078003 | 2.213459 | -0.530354 | -0.319005 | 0.186331 | -3.610681 | -1.30877 | -0.492528 | -0.365636 | -0.265751 | -2.120784 | 0.32364 |
| 28.86111111 | -0.25293 | -0.969759 | -1.653131 | -0.601146 | -3.525131 | -0.361599 | -0.719325 | -1.614347 | -0.687178 | -1.639345 | -1.200942 | -0.755384 | -2.461303 | 2.311627 | -0.582028 | -0.515515 | -0.007274 | -3.796054 | -1.588507 | -0.417946 | -0.838723 | -0.221217 | -1.958044 | 0.342348 |
| 28.875 | -0.383744 | -1.175254 | -2.246374 | -0.712709 | -3.305675 | -0.545874 | -0.990108 | -1.921905 | -0.890886 | -1.632932 | -2.004674 | -0.904098 | -2.96092 | 1.965176 | -0.778954 | -0.697919 | 0.043039 | -3.894795 | -2.042049 | -0.290133 | -1.337593 | -0.303665 | -1.859311 | 0.079993 |
| 28.88888889 | -0.605009 | -0.835273 | -2.482549 | -0.692426 | -4.17868 | -0.894003 | -1.242761 | -2.095236 | -1.159578 | -1.801705 | -2.507512 | -1.078969 | -2.952118 | 1.912863 | -0.864064 | -0.90636 | 0.088309 | -3.381816 | -1.89007 | -0.266293 | -1.041294 | -0.415581 | -2.619541 | -0.003353 |
| 28.90277778 | -0.864228 | -0.34907 | -2.269167 | -0.657895 | -5.152774 | -1.184519 | -1.24921 | -1.888542 | -1.320594 | -1.817581 | -2.347874 | -1.112916 | -2.875826 | 1.484622 | -0.870469 | -0.946783 | -0.098786 | -2.039798 | -1.558999 | -0.655006 | -0.902291 | -0.391597 | -1.928266 | -0.004716 |
| 28.91666667 | -1.471828 | -0.578415 | -2.089751 | -0.789087 | -4.992446 | -1.419893 | -1.094368 | -1.426404 | -1.319531 | -2.119259 | -2.292064 | -1.326749 | -2.544619 | 0.920878 | -0.989016 | -1.070639 | -0.492606 | -1.533754 | -2.015941 | -0.776579 | -0.530617 | -0.355521 | -1.886916 | -0.050968 |
| 28.93055556 | -2.0367 | -1.15609 | -2.070916 | -0.938025 | -4.631579 | -1.360063 | -0.797267 | -1.081697 | -1.355201 | -2.480166 | -2.368343 | -1.512226 | -2.860877 | -0.385883 | -0.984109 | -1.097413 | -0.587339 | -1.256051 | -3.045982 | -0.765153 | 0.025951 | -0.346631 | -1.875485 | -0.534294 |
| 28.94444444 | -2.342527 | -1.171381 | -2.120523 | -1.00912 | -4.026218 | -1.326971 | -0.805966 | -0.763489 | -1.306332 | -3.039139 | -2.452132 | -1.335356 | -3.932978 | -1.302134 | -0.913422 | -1.409819 | -0.78833 | -1.099359 | -2.70856 | -0.448157 | -0.029507 | -0.32886 | -1.818927 | -1.228227 |
| 28.95833333 | -2.325964 | -1.187198 | -2.147 | -0.975685 | -4.026221 | -1.59683 | -1.174681 | -0.772114 | -1.631524 | -3.60464 | -2.972147 | -1.541408 | -4.7338 | -1.816009 | -1.15165 | -1.683106 | -1.500809 | -1.36314 | -3.247072 | 0.231934 | -0.482802 | -0.460241 | -2.583364 | -1.619941 |
| 28.97222222 | -2.835771 | -1.888526 | -2.523654 | -1.28057 | -4.446283 | -2.145198 | -1.536753 | -1.151291 | -2.162657 | -3.140812 | -3.424883 | -1.743026 | -6.028763 | -1.849241 | -1.881433 | -1.520338 | -2.150075 | -2.580937 | -4.169151 | 0.199149 | -1.027856 | -0.792944 | -3.437528 | -1.486773 |
| 28.98611111 | -3.40985 | -2.504812 | -2.694872 | -1.691022 | -3.37072 | -2.187929 | -1.468585 | -1.299754 | -1.992572 | -2.623757 | -2.954796 | -1.455764 | -6.188086 | -1.932415 | -2.33201 | -1.439288 | -2.1828 | -3.135319 | -4.954043 | -0.214161 | -1.072735 | -1.307096 | -2.796879 | -1.682961 |
| 29 | -2.142604 | -2.1254 | -2.050179 | -1.551438 | -2.070732 | -1.830542 | -0.938334 | -1.01284 | -1.510076 | -2.157053 | -2.032667 | -0.73985 | -5.372333 | -2.186112 | -1.194762 | -1.185466 | -1.426861 | -1.674398 | -4.20216 | -0.756811 | -0.574499 | -1.159556 | -1.659552 | -1.334388 |
| 29.01388889 | -0.868706 | -1.324297 | -1.554191 | -1.171747 | -0.884053 | -1.327941 | -0.408702 | -0.819337 | -1.06008 | -1.5421 | -1.400005 | -0.427308 | -3.49326 | -1.645551 | -0.61364 | -1.056776 | -1.263249 | -0.953495 | -2.446456 | -1.098929 | 0.286658 | -0.4997 | -0.818558 | -0.706661 |
| 29.02777778 | -0.280614 | -0.413805 | -1.37876 | -0.447816 | -0.081961 | -1.135512 | -0.33239 | -0.776199 | -0.779161 | -1.240989 | -1.017553 | -0.356133 | -2.30761 | -0.797092 | -0.694771 | -1.400275 | -1.474913 | -0.717662 | -0.830844 | -0.120787 | 0.280665 | -0.753043 | -0.801602 | 0.119538 |
| 29.04166667 | -0.126996 | -0.367751 | -1.20551 | 0.009636 | 0.278409 | -1.032554 | -0.522606 | -0.778819 | -0.709598 | -0.93601 | -0.844645 | -0.350058 | -1.672138 | 0.44672 | -0.532367 | -1.015875 | -1.618509 | -0.468145 | -1.059318 | 0.129082 | 0.018511 | -0.814362 | -0.897691 | 0.683573 |
| 29.05555556 | 0.326323 | -0.273588 | -0.909915 | 0.175077 | 0.59127 | -0.55645 | -0.37906 | -0.553609 | -0.520196 | -0.230704 | -0.699092 | -0.223403 | -0.904126 | 1.121649 | 0.037401 | -0.129543 | -1.019588 | -0.555858 | -2.273411 | -0.521814 | 0.041102 | 0.061376 | -0.627078 | 0.951774 |
| 29.06944444 | 1.10939 | 0.590381 | -0.45384 | 0.308961 | 1.08121 | 0.071192 | 0.110042 | -0.211241 | -0.296715 | 0.47716 | -0.183951 | 0.138309 | 0.014445 | 0.984187 | 0.480149 | 0.564046 | -0.588768 | 0.238919 | -0.993174 | -0.312643 | 0.404509 | 1.025831 | 0.077339 | 0.90893 |
| 29.08333333 | 1.413477 | 1.282098 | -0.041249 | 0.65812 | 1.582882 | 0.46702 | 0.41618 | -0.00752 | -0.095714 | 0.672452 | 0.514669 | 0.422802 | 0.531982 | 0.414666 | 0.447763 | 0.962574 | 0.004605 | 1.215333 | 1.694789 | 0.616583 | 0.449416 | 1.181094 | 0.248301 | 0.159725 |
| 29.09722222 | 1.675146 | 1.371532 | 0.243839 | 0.824447 | 1.880581 | 0.476923 | 0.404542 | 0.119626 | -0.024763 | 0.92648 | 0.93761 | 0.457631 | 0.722048 | -0.015804 | 1.036822 | 1.150271 | 0.42122 | 1.041149 | 2.73875 | 0.838742 | 0.357328 | 0.569614 | 0.426342 | 0.415388 |
| 29.11111111 | 1.999788 | 1.302013 | 0.433279 | 0.782524 | 2.094944 | 0.42012 | 0.262733 | 0.229451 | -0.015859 | 1.142642 | 0.857593 | 0.379512 | 0.538097 | 0.287552 | 1.741394 | 1.306356 | 0.456475 | 0.646565 | 1.679717 | 0.364045 | 0.151383 | -0.266506 | 0.711064 | 1.077519 |
| 29.125 | 2.073615 | 1.475735 | 0.676023 | 0.742202 | 2.413439 | 0.354207 | 0.221743 | 0.255885 | -0.111685 | 1.036132 | 0.64565 | 0.441504 | 0.240993 | -0.038498 | 1.627153 | 0.568756 | 0.477401 | 1.293727 | 2.089851 | 0.384386 | -0.178846 | -0.535803 | 0.693234 | 0.79078 |
| 29.13888889 | 2.047718 | 1.819589 | 0.838518 | 0.564269 | 2.736277 | 0.298849 | 0.27486 | 0.199977 | -0.077039 | 0.770117 | 0.720687 | 0.658689 | -0.150973 | -0.221895 | 1.065275 | -0.055233 | 0.503323 | 2.256074 | 3.642904 | -0.176302 | 0.197189 | 0.035698 | 0.863462 | 0.083933 |
| 29.15277778 | 1.489209 | 2.101443 | 0.704989 | 0.284854 | 2.870065 | 0.267407 | 0.171564 | 0.132319 | 0.108538 | 0.423767 | 0.945425 | 0.715054 | -0.334447 | 0.50267 | 0.756281 | -0.032681 | 1.526684 | 2.171925 | 3.371202 | -0.176986 | 0.550729 | -0.09347 | 1.623555 | 0.287305 |
| 29.16666667 | 1.184553 | 2.182433 | 0.647506 | 0.208505 | 2.893383 | 0.437865 | 0.24209 | 0.18924 | 0.350727 | 0.347694 | 1.103035 | 0.682951 | -0.299262 | 0.174684 | 0.567376 | 0.360256 | 1.954439 | 2.38448 | 3.324286 | 0.106368 | 0.497702 | -0.436724 | 2.479483 | 0.773427 |
| 29.18055556 | 1.227168 | 1.592395 | 0.891156 | 0.288357 | 3.068515 | 0.669864 | 0.427561 | 0.47103 | 0.523834 | 0.507763 | 0.854513 | 0.520335 | -0.416776 | -0.528793 | 0.220325 | 0.25547 | 1.75539 | 2.504403 | 4.014094 | 0.114238 | 0.530763 | 0.139794 | 1.937084 | 0.900289 |
| 29.19444444 | 1.070074 | 0.485933 | 0.926342 | 0.464861 | 3.002586 | 0.524921 | 0.414535 | 0.609616 | 0.460079 | 0.421566 | 0.112672 | 0.109693 | -0.621667 | 0.177671 | -0.171069 | -0.051656 | 1.304551 | 2.085304 | 4.059575 | 0.061971 | 0.735964 | 0.474279 | 1.106325 | 1.045236 |
| 29.20833333 | 0.737994 | -0.036881 | 0.721199 | 0.464032 | 3.103513 | 0.250975 | 0.290651 | 0.607076 | 0.324413 | 0.093265 | -0.318536 | -0.101073 | -0.265317 | 1.085262 | 0.025111 | 0.010742 | 0.954223 | 1.843053 | 3.487598 | 0.291764 | 0.692655 | 0.489623 | 1.150747 | 0.823973 |
| 29.22222222 | 0.516728 | -0.030598 | 0.671098 | 0.429148 | 3.534413 | 0.116375 | 0.243834 | 0.632397 | 0.278732 | -0.058536 | -0.196547 | -0.031463 | -0.197 | 0.550459 | 0.331857 | 0.505422 | 0.617264 | 2.340877 | 2.666526 | 0.285084 | 0.162206 | 0.202296 | 1.57684 | 0.729822 |
| 29.23611111 | 0.426643 | 0.384298 | 0.609287 | 0.253523 | 4.041449 | 0.09022 | 0.271734 | 0.761351 | 0.328324 | 0.109962 | 0.399359 | 0.065454 | -0.284239 | 0.420873 | 0.312482 | 0.580554 | 0.337959 | 1.93448 | 2.273491 | 0.026161 | -0.109336 | -0.065792 | 1.752737 | 0.995001 |
| 29.25 | 0.51869 | -0.03613 | 0.528146 | 0.14937 | 4.428179 | 0.130708 | 0.392652 | 0.834733 | 0.381385 | 0.324654 | 0.139897 | 0.147738 | -0.004289 | 0.768837 | 0.466925 | 0.383347 | 0.440736 | 1.846079 | 2.531437 | -0.111919 | 0.424101 | -0.083189 | 1.720442 | 1.043695 |
| 29.26388889 | 0.680106 | 0.073246 | 0.531772 | 0.16116 | 4.38888 | 0.20956 | 0.756418 | 0.910847 | 0.466112 | 0.531644 | 0.311178 | 0.286492 | 0.259342 | 0.484368 | 0.977626 | 0.587014 | 0.650143 | 2.055271 | 3.124443 | -0.096585 | 0.934079 | 0.008342 | 1.62021 | 0.54422 |
| 29.27777778 | 0.725752 | 0.310396 | 0.570502 | 0.26333 | 3.840726 | 0.294831 | 1.199844 | 0.981488 | 0.553312 | 0.753403 | 0.637339 | 0.485842 | 0.210412 | 0.353139 | 1.441828 | 0.787857 | 0.910958 | 2.621702 | 3.036103 | -0.055034 | 0.844805 | 0.081076 | 1.483327 | 0.620072 |
| 29.29166667 | 0.658108 | -0.026574 | 0.727267 | 0.337168 | 3.939591 | 0.355354 | 1.250199 | 1.11256 | 0.590222 | 1.016648 | 0.840485 | 0.677863 | 0.111943 | 1.124672 | 1.724389 | 1.144125 | 0.963903 | 2.421847 | 1.959076 | -0.024556 | 0.942833 | 0.126887 | 1.536517 | 1.344592 |
| 29.30555556 | 0.624633 | -0.146822 | 1.049664 | 0.464413 | 4.084431 | 0.287448 | 1.307613 | 1.509409 | 0.618251 | 1.209964 | 0.930179 | 0.806999 | -0.144017 | 1.162921 | 1.972203 | 1.373565 | 0.741084 | 3.137899 | 2.562846 | 0.15549 | 1.595725 | 0.440538 | 1.741967 | 2.088191 |
| 29.31944444 | 0.743076 | -0.230335 | 1.579972 | 1.394525 | 5.534443 | -0.040169 | 1.675208 | 2.195478 | 0.876184 | 1.462548 | 1.723003 | 1.219612 | -0.345177 | -0.468393 | 2.444642 | 1.074825 | 0.789494 | 4.906237 | 5.406895 | 0.618048 | 2.050076 | 2.040798 | 1.128116 | 2.131427 |
| 29.33333333 | 1.192605 | 1.465827 | 1.286044 | 3.377181 | 8.882266 | 0.247158 | 2.509267 | 2.775662 | 1.20144 | 2.157555 | 3.453196 | 2.113873 | 0.51228 | -0.747732 | 3.448495 | 1.98636 | 0.909122 | 7.187935 | 8.388581 | 1.916989 | 1.724129 | 2.93779 | 0.993513 | 3.098339 |
| 29.34722222 | 1.357601 | 2.182246 | 3.620074 | 0.724771 | 3.544011 | 1.56196 | 1.759494 | 0.454766 | 1.440571 | 2.290798 | 2.395668 | 1.466572 | 2.389275 | 2.602207 | 3.569591 | -0.27001 | -0.652297 | 3.924373 | 3.187681 | -0.531777 | 1.478364 | 0.175553 | 1.249417 | 3.691302 |
| 29.36111111 | 0.238623 | 2.010738 | 4.163764 | -2.226191 | -0.965983 | 1.268181 | 0.147035 | -0.836979 | 0.033391 | 0.659334 | -0.452044 | -0.437136 | 1.278031 | 2.824796 | 0.409816 | -1.517494 | 0.111535 | -4.171124 | -4.295074 | -1.700347 | 0.492343 | -1.752206 | 0.554744 | -1.635306 |
| 29.375 | 0.091983 | 2.274 | 2.834159 | -0.740713 | 0.126741 | 0.922003 | 0.142955 | -0.437971 | 0.098078 | 0.753441 | 0.463726 | 0.163425 | 0.950636 | 0.292253 | 0.084126 | 0.555897 | 0.575543 | -3.255647 | -3.028035 | -0.603036 | 1.023713 | -1.22457 | -0.227028 | -1.470979 |
| 29.38888889 | 0.624831 | 2.625305 | 2.442557 | 0.345919 | 1.112081 | 0.875921 | 0.721534 | 0.41155 | 0.729025 | 1.637684 | 1.674391 | 1.108159 | 1.118322 | 0.432802 | 1.624714 | 1.298583 | 0.827641 | -1.090855 | -0.864936 | -0.111036 | 1.176678 | -0.101398 | 0.480118 | -0.000039 |
| 29.40277778 | 1.087848 | 1.954709 | 2.271602 | 0.549277 | 1.412432 | 1.061286 | 1.156516 | 0.504488 | 0.942861 | 2.10621 | 1.807418 | 1.442482 | 1.334582 | 1.05784 | 2.681965 | 1.044763 | 0.921793 | 1.328454 | 0.712887 | 0.061726 | 1.36937 | -0.018361 | 0.991634 | 1.152525 |
| 29.41666667 | 1.599562 | 1.531868 | 2.347855 | 0.592792 | 1.366321 | 1.26775 | 1.445778 | 0.531167 | 0.961275 | 2.10826 | 1.646381 | 1.657865 | 1.423602 | 1.064941 | 2.946154 | 0.876755 | 1.035792 | 3.07128 | 1.683134 | 0.236164 | 1.812157 | -0.129088 | 0.908001 | 1.389593 |
| 29.43055556 | 1.793133 | 0.837421 | 2.296043 | 0.50619 | 1.124435 | 1.234862 | 1.190106 | 0.505184 | 0.973106 | 1.616602 | 1.310328 | 1.516692 | 1.506384 | 0.997239 | 2.598678 | 0.622976 | 0.925021 | 3.673448 | 1.951483 | 0.692655 | 1.972251 | -0.065678 | 1.729755 | 1.162888 |
| 29.44444444 | 1.568993 | 0.423935 | 2.159653 | 0.417792 | 0.480938 | 1.096823 | 0.796631 | 0.290415 | 0.984264 | 1.033908 | 1.019899 | 1.187499 | 1.503816 | 0.886836 | 1.938345 | 0.531699 | 0.54418 | 3.078282 | 1.753275 | 1.423609 | 2.186908 | 0.059105 | 2.541738 | 1.477294 |
| 29.45833333 | 1.397524 | 0.695977 | 2.27298 | 0.387198 | 0.582424 | 1.058045 | 0.508965 | 0.189236 | 1.083411 | 1.112395 | 0.952645 | 0.856083 | 1.221282 | 0.894786 | 1.525704 | 0.694498 | 0.180059 | 1.70315 | 0.769845 | 1.975734 | 1.647794 | 0.157674 | 1.882637 | 1.883906 |
| 29.47222222 | 0.960018 | 0.875665 | 1.819817 | 0.549134 | 0.425199 | 1.15119 | 0.341061 | 0.145075 | 1.021111 | 1.429855 | 0.757079 | 0.569073 | 0.935676 | 0.422745 | 1.179864 | 0.456905 | 0.189195 | 1.258122 | 0.285864 | 1.097241 | 1.01734 | 0.15855 | 1.79968 | 1.310766 |
| 29.48611111 | 0.660757 | 1.051662 | 1.1389 | 0.682339 | 0.375685 | 0.870947 | 0.427152 | 0.207766 | 0.837065 | 1.472498 | 0.807882 | 0.622826 | 1.028017 | 0.717155 | 1.535393 | 0.757916 | 0.847203 | 1.535594 | 0.25309 | 0.067545 | 1.223068 | 0.078408 | 1.659478 | 0.80514 |
| 29.5 | 1.032728 | 1.278778 | 1.046203 | 0.420478 | 0.630501 | 1.022802 | 0.742571 | 0.26652 | 1.043722 | 1.641127 | 1.302245 | 1.209531 | 1.676214 | 0.964109 | 1.764383 | 0.758882 | 1.774061 | 2.434546 | 0.346652 | -0.404841 | 1.861013 | 0.083384 | 2.362739 | 0.795918 |
| 29.51388889 | 1.639245 | 1.114858 | 1.34625 | 0.174378 | 0.924415 | 1.051209 | 1.007215 | 0.497678 | 1.458083 | 2.094625 | 1.711266 | 1.889846 | 1.664936 | 1.010084 | 1.2963 | 0.586224 | 2.018642 | 3.070068 | 0.292325 | 0.65285 | 2.782132 | 0.237616 | 2.820286 | 1.161029 |
| 29.52777778 | 1.333644 | 0.528507 | 1.40746 | 0.24452 | 0.646126 | 0.938776 | 0.806741 | 0.617286 | 1.615496 | 2.164175 | 1.513939 | 2.149742 | 1.114162 | 1.024212 | 0.80121 | 0.294938 | 1.626856 | 2.893822 | 0.46644 | 1.946762 | 1.980299 | 0.232413 | 2.11497 | 1.308398 |
| 29.54166667 | 0.538831 | 0.14077 | 1.072093 | 0.131908 | 0.393213 | 0.722093 | 0.58622 | 0.690099 | 1.504806 | 1.717165 | 1.048078 | 2.048451 | 0.933298 | 0.944681 | 0.408019 | 0.067284 | 1.359544 | 1.889064 | 0.453285 | 2.015879 | 1.183134 | 0.171476 | 2.542239 | 0.678688 |
| 29.55555556 | 0.239361 | 0.092986 | 0.934115 | 0.177732 | 0.237691 | 0.762558 | 0.658337 | 0.740981 | 1.343436 | 1.273304 | 0.548272 | 1.435614 | 1.379301 | 0.632377 | 0.739439 | -0.007281 | 0.97351 | 1.586172 | 0.277085 | 1.100434 | 1.359762 | 0.117349 | 1.548947 | 0.189242 |
| 29.56944444 | 0.56805 | 0.283614 | 1.145778 | 0.300306 | 0.321104 | 0.938662 | 0.753434 | 0.726007 | 1.28485 | 1.089019 | 0.214069 | 0.688477 | 1.072499 | 0.473384 | 0.851451 | -0.145295 | -0.248388 | 1.147145 | 0.187941 | 0.350019 | 1.154064 | 0.105308 | 1.130944 | 0.12058 |
| 29.58333333 | 0.660484 | 0.328428 | 0.999141 | 0.299855 | 0.300974 | 0.705608 | 0.471921 | 0.389245 | 1.032756 | 1.047747 | 0.363615 | 0.405784 | 0.256008 | 0.511306 | 0.036279 | -0.090636 | -1.967394 | 0.793704 | 0.214334 | 0.65699 | 0.591413 | 0.030756 | 0.438938 | 0.208757 |
| 29.59722222 | 0.482753 | -0.011215 | 0.072667 | 0.058638 | 0.328931 | 0.27114 | -0.070193 | 0.071386 | 0.551196 | 0.719226 | 0.406897 | 0.290965 | -0.183912 | 0.544109 | -0.802317 | -0.177174 | -1.965215 | 0.55499 | 0.314634 | 1.355726 | 0.38047 | 0.009944 | 0.225948 | 0.186303 |
| 29.61111111 | 0.066932 | -0.433188 | -0.501434 | -0.218367 | 0.041099 | -0.050839 | -0.267195 | -0.155903 | 0.2821 | 0.293335 | 0.245674 | 0.167918 | -0.116015 | 0.590262 | -1.064057 | -0.302585 | -1.543358 | 0.594001 | 0.369765 | 0.609269 | -0.038493 | 0.017971 | 0.049028 | -0.039169 |
| 29.625 | 0.100763 | -0.186509 | -0.554431 | -0.318645 | -0.223938 | -0.168529 | -0.274073 | -0.203725 | 0.112258 | -0.027912 | 0.03644 | -0.121492 | 0.097661 | 0.621967 | -0.874405 | -0.685673 | -1.439879 | 0.377033 | 0.331515 | 0.063691 | -0.553384 | 0.009731 | -0.053245 | -0.255686 |
| 29.63888889 | -0.15905 | -0.140996 | -0.257194 | -0.23419 | -0.557711 | -0.19003 | -0.01279 | -0.192981 | 0.030167 | -0.237734 | -0.279966 | -0.311565 | -0.125778 | 0.609141 | -0.64933 | -0.966919 | -1.577824 | 0.079445 | 0.258969 | -0.308643 | -0.06979 | 0.031182 | -0.221778 | -0.21341 |
| 29.65277778 | -0.338495 | -0.118593 | -0.168786 | -0.179485 | -0.913756 | -0.246383 | -0.009786 | -0.446254 | 0.013367 | -0.381532 | -0.297444 | -0.342365 | -0.203685 | 0.677122 | -0.325618 | -0.811585 | -0.74115 | -0.162807 | 0.161622 | -0.088746 | 0.628385 | 0.103344 | -0.423411 | -0.066056 |
| 29.66666667 | -0.230804 | 0.022379 | -0.170478 | -0.126621 | -0.647593 | -0.335406 | -0.287514 | -0.661125 | -0.088073 | -0.345077 | -0.087061 | -0.295613 | -0.183585 | 0.62899 | 0.019607 | -0.95191 | 0.644445 | -0.188531 | 0.107679 | -0.422242 | 0.685817 | 0.117518 | -0.271848 | 0.027318 |
| 29.68055556 | -0.10876 | 0.407897 | -0.096253 | -0.105249 | -1.000891 | -0.363297 | -0.31553 | -0.394329 | -0.056707 | -0.122133 | -0.033133 | -0.336124 | -0.045805 | 0.591838 | 0.218112 | -0.905651 | 1.06275 | -0.281738 | 0.109407 | -0.625519 | 0.564324 | 0.05943 | -0.256029 | -0.007275 |
| 29.69444444 | -0.014312 | 0.62728 | -0.11208 | -0.064851 | -1.166729 | -0.18328 | 0.10699 | -0.077622 | -0.07364 | -0.188533 | -0.109375 | -0.385647 | -0.03675 | 0.618506 | 0.029698 | -0.796759 | 0.702425 | -0.553591 | 0.084943 | -1.14772 | 0.007351 | 0.022349 | -0.373212 | 0.026721 |
| 29.70833333 | 0.154876 | 0.573907 | 0.211059 | -0.135197 | -1.149693 | 0.050616 | 0.545516 | -0.052066 | -0.137531 | -0.504564 | -0.202288 | -0.435648 | -0.139822 | 0.730308 | -0.231155 | -0.57263 | 0.451941 | -0.821318 | 0.047709 | -1.957164 | -0.55365 | 0.027843 | -0.344172 | -0.171001 |
| 29.72222222 | 0.273143 | 0.201271 | 0.528871 | -0.13219 | -1.004162 | 0.125101 | 0.492358 | -0.007633 | -0.19876 | -0.640476 | -0.147194 | -0.342551 | -0.170212 | 0.648778 | -0.220478 | -0.257244 | 0.154388 | -0.569416 | 0.057136 | -3.120312 | -0.843797 | 0.051511 | -0.287854 | -0.731277 |
| 29.73611111 | 0.251203 | 0.164639 | 0.573497 | 0.050241 | -0.935045 | 0.162266 | 0.382524 | 0.095411 | -0.18178 | -0.547233 | 0.128254 | 0.011334 | 0.052894 | 0.519841 | -0.035842 | -0.600348 | -0.405569 | -0.420569 | 0.097853 | -3.163055 | -0.684051 | 0.002336 | -0.574275 | -0.669946 |
| 29.75 | 0.095586 | 0.312984 | 0.112683 | 0.140821 | -1.009693 | 0.046539 | 0.27773 | 0.125308 | -0.146127 | -0.491346 | 0.162711 | 0.127926 | 0.347894 | 0.357234 | -0.132023 | -0.874805 | -0.842549 | -0.556051 | 0.114175 | -1.231104 | -0.73283 | -0.079596 | -0.711846 | -0.382147 |
| 29.76388889 | -0.103962 | 0.310856 | -0.482657 | -0.115374 | -1.202998 | -0.19156 | -0.018898 | -0.150503 | -0.208081 | -0.538979 | -0.095204 | -0.236364 | 0.731094 | 0.236969 | -0.084038 | -0.947559 | -0.688789 | -0.902412 | 0.097253 | 0.40042 | -0.870265 | 0.004259 | -0.740581 | -0.355887 |
| 29.77777778 | -0.200039 | -0.054459 | -0.937965 | -0.585369 | -1.684224 | -0.426686 | -0.272304 | -0.294065 | -0.397166 | -0.78587 | -0.526559 | -0.93739 | 1.54418 | 0.074272 | 0.378819 | -0.543896 | -0.327014 | -1.199243 | 0.115264 | 0.816782 | -1.080567 | 0.031401 | -0.588232 | -0.698693 |
| 29.79166667 | -0.203971 | -0.132695 | -1.058451 | -0.787051 | -1.818885 | -0.399391 | -0.395435 | -0.250989 | -0.401279 | -1.029237 | -0.635158 | -1.328469 | 1.169926 | 0.12906 | 0.928934 | -0.246691 | -0.542753 | -1.469026 | 0.093731 | -0.417198 | -1.12904 | -0.009564 | -0.705689 | -0.831147 |
| 29.80555556 | -0.222187 | 0.335173 | -0.860025 | -0.696172 | -1.55244 | -0.29501 | -0.396067 | -0.169839 | -0.189654 | -1.009437 | -0.432894 | -0.981202 | 0.070541 | 0.232325 | 0.87783 | -0.32006 | -1.111269 | -1.727615 | 0.024283 | -1.967341 | -0.50288 | -0.056945 | -1.124983 | -0.465913 |
| 29.81944444 | -0.321251 | -0.081229 | -0.81136 | -0.591104 | -1.784052 | -0.339539 | -0.579315 | -0.370192 | -0.151325 | -1.041932 | -0.493236 | -0.7932 | 0.214467 | 0.250228 | 0.388342 | -0.36929 | -1.193115 | -1.619047 | -0.206002 | -1.97884 | -0.282289 | -0.002241 | -1.437658 | -0.109586 |
| 29.83333333 | -0.402999 | -0.362463 | -1.190186 | -0.494225 | -3.207582 | -0.534432 | -1.180192 | -0.684855 | -0.338016 | -1.103236 | -0.69437 | -0.809843 | 0.534254 | 0.308479 | 0.038578 | -0.469053 | -0.797692 | -1.929442 | -0.361311 | -1.171338 | -0.393634 | 0.010874 | -1.474652 | -0.076159 |
| 29.84722222 | -0.377766 | -0.608028 | -1.639114 | -0.580359 | -4.613034 | -0.774795 | -1.68444 | -0.768054 | -0.464636 | -1.050527 | -0.856833 | -0.845878 | 0.816373 | 0.410837 | 0.106351 | -0.478274 | -0.43446 | -2.755498 | -0.700739 | -0.706014 | -0.784983 | 0.062744 | -1.123167 | -0.216675 |
| 29.86111111 | -0.439423 | -0.905876 | -1.598941 | -0.707366 | -5.409128 | -0.869756 | -1.71842 | -0.875222 | -0.418431 | -0.910691 | -0.939978 | -1.014713 | -0.133423 | 0.451629 | 0.144977 | -0.494754 | -1.206751 | -3.228927 | -0.729559 | -1.43928 | -1.333912 | 0.031857 | -0.657003 | -0.238128 |
| 29.875 | -0.633907 | -1.397479 | -1.620636 | -0.905187 | -5.474959 | -0.986044 | -1.46258 | -1.033291 | -0.648716 | -0.809964 | -1.081475 | -1.309302 | -1.240676 | 0.280692 | -0.137543 | -0.569951 | -2.228911 | -2.797265 | -0.474652 | -2.6616 | -1.89606 | 0.01842 | -0.389734 | -0.336348 |
| 29.88888889 | -0.789768 | -2.08327 | -1.58263 | -1.034045 | -5.734815 | -0.954753 | -1.52323 | -1.178476 | -1.027192 | -1.110056 | -1.443631 | -1.803537 | -1.905169 | -0.041187 | -0.77945 | -0.759657 | -2.365851 | -2.583505 | -0.613192 | -2.774287 | -2.243289 | 0.141657 | -0.497345 | -1.104141 |
| 29.90277778 | -0.967664 | -2.542496 | -1.572712 | -1.289987 | -6.357951 | -0.972458 | -1.49308 | -1.234013 | -1.260494 | -1.727273 | -1.79789 | -2.133174 | -2.559077 | -0.228631 | -1.318493 | -1.093798 | -1.41166 | -2.343495 | -0.90415 | -1.847877 | -1.775825 | 0.162923 | -1.120368 | -1.793429 |
| 29.91666667 | -1.17194 | -2.066113 | -1.847302 | -1.306078 | -6.237759 | -0.988114 | -1.512877 | -1.158033 | -1.264701 | -2.439478 | -1.941463 | -2.209667 | -3.055307 | -0.339882 | -1.533351 | -1.009974 | -0.341411 | -2.774019 | -1.73454 | -0.412888 | -1.606676 | 0.081245 | -1.941029 | -1.810155 |
| 29.93055556 | -1.069363 | -1.280858 | -1.999105 | -1.259796 | -5.578413 | -0.967667 | -1.644815 | -1.064715 | -1.06336 | -2.011512 | -2.034573 | -2.129001 | -3.433501 | -0.511928 | -1.708608 | -1.062466 | 0.960363 | -3.094127 | -1.855572 | 0.251092 | -1.424408 | 0.135085 | -2.386208 | -1.692378 |
| 29.94444444 | -0.874741 | -0.938367 | -1.999981 | -0.931876 | -5.149893 | -0.953545 | -1.171596 | -0.957029 | -0.947079 | -1.501931 | -1.933214 | -2.136525 | -3.957784 | -0.045504 | -1.472394 | -0.679489 | 1.747809 | -4.02757 | -2.038626 | 0.569299 | -1.329276 | 0.039597 | -2.26665 | -1.818245 |
| 29.95833333 | -1.443846 | -1.596732 | -2.382762 | -0.943826 | -5.251666 | -1.280789 | -1.132403 | -1.105203 | -1.583693 | -1.74722 | -2.379315 | -2.431662 | -5.094473 | -0.13311 | -1.478482 | -0.795219 | 1.182314 | -3.310833 | -3.759623 | 0.416542 | -1.771294 | -0.080243 | -2.090815 | -1.735883 |
| 29.97222222 | -2.51839 | -2.641578 | -3.060519 | -1.623184 | -6.03864 | -2.285694 | -1.614317 | -1.470255 | -2.513679 | -3.007581 | -3.869483 | -3.067695 | -5.984887 | -1.492792 | -1.273648 | -0.763514 | -0.830128 | -2.387177 | -3.324388 | 0.126595 | -1.998249 | -0.378115 | -2.02627 | -1.702823 |
| 29.98611111 | -2.122994 | -2.796361 | -3.414537 | -2.167336 | -5.530715 | -2.173895 | -1.570035 | -1.640679 | -2.357226 | -3.417028 | -4.290181 | -2.770944 | -5.947542 | -1.660809 | -1.339169 | -0.652993 | -1.98309 | -1.871028 | -2.580545 | -0.386793 | -1.271828 | -0.00531 | -1.399257 | -1.009963 |
| 30 | -1.304219 | -1.812709 | -2.589291 | -2.72148 | -3.360793 | -1.394962 | -1.009619 | -1.33324 | -1.794271 | -2.845891 | -3.224899 | -2.006249 | -4.784425 | -2.37467 | -0.733342 | -1.860511 | -2.350373 | -1.622616 | -3.638275 | -0.609488 | -0.337549 | -0.057939 | -1.267271 | -1.194469 |
| 30.01388889 | -0.823843 | -0.364226 | -1.722265 | -1.494043 | -2.289773 | -1.024479 | -0.384727 | -1.087151 | -1.479935 | -2.463685 | -2.315114 | -1.674594 | -2.975899 | -0.793868 | -0.967721 | -1.041367 | -3.232346 | -1.200107 | -1.665446 | -0.583587 | 0.235527 | -0.154425 | -1.386025 | -0.148527 |
| 30.02777778 | -0.650247 | 0.728137 | -1.026968 | -0.236001 | -1.455794 | -0.820826 | -0.009456 | -1.03635 | -1.228875 | -2.285042 | -1.611697 | -1.324822 | -1.583192 | -0.038085 | -1.21623 | -1.094629 | -2.882625 | -0.156164 | 0.914761 | -0.028288 | 0.037726 | -0.139302 | -1.371245 | 0.350127 |
| 30.04166667 | -0.415194 | 1.136017 | -0.689621 | 0.278318 | -0.747475 | -0.467785 | 0.234669 | -0.872423 | -0.96112 | -1.678919 | -1.08412 | -0.979177 | -1.103348 | 0.844196 | -1.147444 | -1.004558 | -2.770483 | -0.111562 | 1.263684 | 0.520829 | -0.029458 | -0.442716 | -0.764447 | 0.329712 |
| 30.05555556 | -0.236989 | 0.989269 | -0.569101 | 0.060934 | -0.181806 | -0.000794 | 0.540592 | -0.471104 | -0.625268 | -0.401094 | -0.596743 | -0.622414 | -0.277855 | 1.471137 | -0.143951 | 0.045701 | -2.407446 | -0.34606 | -1.202286 | 0.949904 | 0.008124 | 0.677792 | -0.417551 | 0.677719 |
| 30.06944444 | -0.023965 | 1.158458 | -0.390667 | 0.060273 | 0.216751 | 0.372873 | 0.709453 | -0.328025 | -0.226861 | 0.777831 | -0.220729 | -0.338969 | 0.411 | 1.652668 | 0.566042 | 0.51327 | -1.585928 | -0.12863 | -1.655168 | 0.663153 | 0.313357 | 0.83386 | -0.267196 | 0.81772 |
| 30.08333333 | 0.277732 | 1.520847 | -0.109604 | 0.534417 | 0.680786 | 0.472795 | 0.792776 | 0.097099 | 0.211335 | 1.437515 | 0.046721 | 0.006506 | 0.570255 | 0.818569 | 0.485855 | 0.677326 | -0.777146 | 0.763062 | -0.241345 | 0.151642 | 0.204873 | 0.55704 | -0.448984 | 0.584361 |
| 30.09722222 | 0.623508 | 1.422019 | 0.29584 | 0.92784 | 1.180676 | 0.433771 | 0.764792 | 0.587499 | 0.544191 | 1.608732 | 0.299799 | 0.362926 | 0.986496 | 0.014963 | 0.360595 | 0.781592 | -0.37858 | 1.630624 | 1.551286 | 0.267182 | 0.453574 | 0.280152 | -0.590173 | 0.616764 |
| 30.11111111 | 0.85343 | 0.908201 | 0.580813 | 0.919521 | 1.387566 | 0.492532 | 0.642733 | 0.647973 | 0.657164 | 1.573143 | 0.538536 | 0.562668 | 1.256812 | -0.313686 | 0.806774 | 1.227297 | 0.199731 | 1.871114 | 2.499006 | 0.255326 | 0.792739 | 0.230538 | -0.394681 | 0.557266 |

| | | | | | | | | | | | | | | | | | | | | | | | | |
|---|---|---|---|---|---|---|---|---|---|---|---|---|---|---|---|---|---|---|---|---|---|---|---|---|
| 30.125 | 0.824873 | 0.780554 | 0.73734 | 0.992649 | 1.518131 | 0.465452 | 0.644222 | 0.508615 | 0.514002 | 1.474707 | 0.625305 | 0.674861 | 1.3155 | -0.50832 | 1.444634 | 1.722536 | 1.195111 | 1.732015 | 2.354675 | 0.377448 | 1.245154 | -0.013159 | 0.076542 | 0.53402 |
| 30.13888889 | 0.635338 | 1.439296 | 0.987469 | 1.584753 | 1.625559 | 0.46446 | 0.589638 | 0.441169 | 0.353519 | 1.417765 | 0.735728 | 0.827306 | 1.127043 | 0.221782 | 1.563529 | 1.878435 | 1.584866 | 1.644254 | 2.409933 | -0.087791 | 0.629308 | 0.069195 | 0.655323 | 1.060031 |
| 30.15277778 | 0.564444 | 1.75932 | 1.198843 | 1.993513 | 1.665988 | 0.456873 | 0.384155 | 0.449154 | 0.276111 | 1.306854 | 0.747408 | 0.763759 | 1.116765 | 1.180846 | 1.212629 | 1.172846 | 1.163193 | 1.589854 | 2.711941 | -0.682026 | 0.109084 | 0.127045 | 1.238502 | 1.629898 |
| 30.16666667 | 0.603038 | 1.54876 | 1.189229 | 1.652747 | 1.642356 | 0.489409 | 0.326381 | 0.347777 | 0.214075 | 1.111471 | 0.624986 | 0.677906 | 1.321962 | 1.374274 | 1.18989 | 0.377303 | 1.049977 | 1.59133 | 2.823805 | -0.479885 | 0.444528 | 0.348888 | 1.609225 | 1.669884 |
| 30.18055556 | 0.575261 | 1.503967 | 1.00323 | 1.318989 | 1.513366 | 0.494095 | 0.29035 | 0.316698 | 0.154941 | 0.895182 | 0.379178 | 0.639942 | 1.346196 | 0.310119 | 0.880368 | 0.231359 | 1.14585 | 1.594001 | 2.910036 | 0.008696 | 0.363554 | 1.100093 | 1.704986 | 1.232036 |
| 30.19444444 | 0.532181 | 1.998524 | 0.831585 | 1.398704 | 1.56934 | 0.403156 | 0.273176 | 0.385873 | 0.119151 | 0.67089 | 0.161571 | 0.443811 | 0.426871 | -0.024999 | 0.195664 | 0.043013 | 0.770876 | 1.550308 | 2.481429 | 0.405273 | -0.523633 | 0.939901 | 1.512721 | 0.729441 |
| 30.20833333 | 0.54864 | 2.640585 | 0.881944 | 1.599013 | 1.484709 | 0.203389 | 0.244689 | 0.375492 | -0.005857 | 0.429114 | -0.114205 | 0.05016 | 0.444148 | 0.379674 | -0.546814 | -0.092244 | -0.189955 | 1.503275 | 2.156029 | 0.077076 | -0.40646 | 0.298194 | 1.381394 | 0.29121 |
| 30.22222222 | 0.602557 | 1.904356 | 0.810998 | 1.242126 | 1.355483 | 0.0863 | 0.236657 | 0.323348 | -0.12669 | 0.301172 | -0.329525 | -0.415682 | 0.80399 | 0.826341 | -0.536451 | -0.042082 | -0.995054 | 1.570008 | 2.079919 | -0.325408 | 0.679676 | 0.095909 | 1.694201 | 0.182326 |
| 30.23611111 | 0.65959 | 1.095926 | 0.664037 | 0.707329 | 1.4659 | 0.11627 | 0.303666 | 0.427307 | -0.14903 | 0.308777 | -0.118514 | -0.069931 | 0.670802 | 1.188926 | -0.012749 | 0.148487 | -1.301673 | 1.624177 | 2.098354 | -0.231267 | 1.219799 | 0.559874 | 1.794288 | 0.572891 |
| 30.25 | 0.660228 | 1.369339 | 0.649588 | 0.49547 | 1.880707 | 0.177561 | 0.414795 | 0.641804 | -0.113055 | 0.25575 | 0.063054 | 0.252442 | 0.596131 | 1.082973 | 0.19071 | 0.428845 | -0.923257 | 1.730717 | 2.394595 | 0.067 | 0.533137 | 1.02625 | 1.424385 | 0.79704 |
| 30.26388889 | 0.702764 | 1.457607 | 0.817384 | 0.719979 | 2.094111 | 0.234484 | 0.445436 | 0.691673 | -0.007385 | 0.123742 | 0.105049 | 0.435443 | 0.62992 | 0.996296 | 0.078795 | 1.01438 | -0.12343 | 1.548424 | 2.184317 | -0.029329 | 0.300599 | 1.087006 | 1.391135 | 0.553019 |
| 30.27777778 | 0.845508 | 0.721022 | 1.134069 | 0.762235 | 2.073468 | 0.374489 | 0.479359 | 0.689831 | 0.241337 | 0.348111 | 0.189854 | 0.736663 | 0.650734 | 1.320752 | 0.238143 | 1.304032 | 0.087079 | 1.263002 | 1.922817 | -0.160716 | 0.600767 | 0.395637 | 1.662307 | 0.369034 |
| 30.29166667 | 0.913475 | 0.241639 | 1.321539 | 0.71786 | 1.933579 | 0.517351 | 0.567249 | 0.714633 | 0.46861 | 0.735961 | 0.29294 | 1.003115 | 0.940804 | 1.481607 | 0.824838 | 1.431998 | -0.827064 | 1.61088 | 2.036674 | -0.010936 | 0.513762 | 0.253921 | 1.540846 | 0.772276 |
| 30.30555556 | 0.802122 | 0.006539 | 1.520018 | 0.718774 | 1.93621 | 0.567713 | 0.627482 | 0.660055 | 0.530607 | 0.921785 | 0.36445 | 0.986446 | 1.088901 | 1.762056 | 1.379285 | 1.419821 | -1.042465 | 2.140322 | 2.130258 | 0.193987 | 0.720746 | 0.303595 | 1.162274 | 0.813569 |
| 30.31944444 | 0.640764 | 0.108693 | 1.927353 | 0.86174 | 2.388108 | 0.541926 | 0.572483 | 0.520656 | 0.517522 | 0.989155 | 0.514196 | 0.93447 | 0.856927 | 1.880241 | 1.774918 | 1.507894 | -0.839606 | 2.458843 | 2.289309 | 0.300266 | 1.669705 | 0.309882 | 1.192073 | 0.690258 |
| 30.33333333 | 0.511793 | 0.091293 | 2.323367 | 0.964103 | 2.879311 | 0.622584 | 0.550646 | 0.476233 | 0.628946 | 1.38986 | 0.743265 | 1.03499 | 0.558875 | 1.738004 | 2.138239 | 1.570674 | 0.413261 | 2.23157 | 2.01496 | 0.336283 | 2.583715 | 0.179526 | 0.931798 | 0.245873 |
| 30.34722222 | 0.468269 | 0.077608 | 2.426097 | 0.729173 | 2.784829 | 0.813517 | 0.558538 | 0.533858 | 0.936231 | 1.760455 | 0.994434 | 1.308793 | 0.827072 | 1.614559 | 2.329081 | 1.497452 | 1.24546 | 2.313054 | 2.242607 | 0.589419 | 2.037518 | 0.147518 | 1.550982 | 0.197339 |
| 30.36111111 | 0.481323 | 0.021859 | 2.595771 | 0.504666 | 3.016072 | 1.034133 | 0.671393 | 0.729 | 1.276458 | 1.937323 | 1.251936 | 1.641841 | 1.147087 | 1.646313 | 2.732228 | 1.627835 | 0.828273 | 2.732745 | 2.181934 | 1.268241 | 1.999368 | 0.268311 | 1.839553 | 0.488262 |
| 30.375 | 0.530212 | 0.816537 | 2.862855 | 0.505256 | 3.554338 | 1.111958 | 0.885965 | 1.032421 | 1.535505 | 2.002656 | 1.553376 | 1.887772 | 0.891598 | 2.090142 | 3.457034 | 1.56967 | -0.166259 | 2.515502 | 1.506477 | 2.243192 | 2.559411 | 0.230189 | 2.90547 | 1.023787 |
| 30.38888889 | 0.550558 | 1.699949 | 3.258807 | 0.661539 | 4.728886 | 1.184436 | 0.994348 | 1.014597 | 1.640624 | 2.120448 | 1.732793 | 1.942591 | 0.639145 | 2.057581 | 3.513935 | 1.609073 | -0.754089 | 2.449993 | 1.864637 | 3.619347 | 2.301246 | 0.069782 | 3.593915 | 1.803224 |
| 30.40277778 | 0.525323 | 2.264262 | 3.093395 | 0.757604 | 5.106317 | 0.992503 | 0.858134 | 0.959554 | 1.529145 | 2.092786 | 1.806292 | 1.923088 | 1.23493 | 2.191026 | 3.310963 | 1.253702 | -0.194408 | 2.454422 | 1.870005 | 4.599338 | 2.442122 | -0.115151 | 3.007309 | 2.248436 |
| 30.41666667 | 0.520498 | 2.187048 | 2.799175 | 0.735424 | 5.006768 | 0.851383 | 0.704382 | 0.963633 | 1.399265 | 1.904312 | 1.795434 | 1.982114 | 2.014405 | 2.173265 | 3.405861 | 0.978589 | 0.920453 | 2.858671 | 2.008557 | 5.385124 | 2.362809 | -0.25001 | 2.462321 | 2.54063 |
| 30.43055556 | 0.46765 | 1.729727 | 2.264429 | 0.712691 | 5.227981 | 0.744589 | 0.588331 | 0.902625 | 1.158166 | 1.560386 | 1.800911 | 2.158691 | 1.77889 | 2.388277 | 3.539675 | 0.716089 | 1.671157 | 2.310237 | 1.438754 | 4.662187 | 2.111217 | -0.42197 | 2.496257 | 3.142293 |
| 30.44444444 | 0.505242 | 1.111432 | 2.06418 | 0.590677 | 5.437476 | 0.748358 | 0.495363 | 0.782568 | 1.036614 | 1.180921 | 2.012591 | 2.484581 | 1.484107 | 3.206878 | 3.720009 | 0.551373 | 1.571275 | 1.363071 | 0.409182 | 3.180154 | 1.906722 | -0.586614 | 2.712871 | 3.117232 |
| 30.45833333 | 0.892179 | 0.753547 | 2.2584 | 0.443102 | 5.203283 | 1.062631 | 0.599429 | 0.875268 | 1.201429 | 1.145032 | 2.287749 | 2.654016 | 1.637383 | 3.23923 | 3.556712 | 0.475032 | 2.427593 | 1.641359 | -0.102796 | 0.623088 | 1.979498 | -0.606389 | 2.912537 | 3.140617 |
| 30.47222222 | 1.335005 | 0.779806 | 2.069412 | 0.362056 | 3.32411 | 0.889015 | 0.607263 | 0.954068 | 1.353164 | 1.188851 | 1.949931 | 2.026078 | 2.018943 | 3.05549 | 2.640457 | 0.535646 | 2.127218 | 2.063607 | 0.062503 | -1.384982 | 1.720021 | -0.770492 | 2.927995 | 2.302716 |
| 30.48611111 | 1.958774 | 1.093246 | 1.433467 | 0.351372 | 2.752567 | 0.601243 | 0.494765 | 0.894691 | 1.306731 | 1.089478 | 1.50422 | 1.52208 | 2.300729 | 2.664051 | 2.427756 | 0.66883 | 2.083542 | 2.501486 | 0.467844 | -0.674578 | 1.645708 | -0.20582 | 2.48576 | 1.286319 |
| 30.5 | 2.637483 | 1.629525 | 1.301024 | 0.321157 | 3.377587 | 0.587926 | 0.934494 | 0.766749 | 1.254877 | 1.165455 | 1.557556 | 1.688045 | 2.212112 | 2.398498 | 2.307866 | 0.806627 | 1.652726 | 3.256885 | 0.858749 | 1.582427 | 2.03375 | 0.159492 | 2.385857 | 0.535069 |
| 30.51388889 | 2.240809 | 1.501423 | 1.523888 | 0.246785 | 3.309519 | 0.563352 | 1.278298 | 0.49854 | 1.298617 | 1.402881 | 1.82661 | 2.096281 | 1.505705 | 2.335383 | 1.566502 | 1.015103 | 1.257311 | 3.20868 | 1.154616 | 2.115055 | 2.225247 | 0.036775 | 2.362958 | 0.337601 |
| 30.52777778 | 2.292913 | 1.033238 | 1.155644 | 0.30926 | 2.64432 | 0.459778 | 0.56255 | 0.311087 | 1.331096 | 1.636985 | 2.054453 | 2.307487 | 0.084217 | 2.095873 | 0.664496 | 0.835969 | 0.140694 | 2.390038 | 0.736658 | 1.204372 | 1.73117 | 0.063704 | 1.894175 | 0.192958 |
| 30.54166667 | 1.715915 | 0.713 | 0.575825 | 0.064251 | 1.932863 | 0.369995 | 0.591152 | 0.439465 | 1.247695 | 1.683248 | 1.968643 | 2.106736 | -0.152862 | 2.331515 | 0.810131 | 0.563414 | 0.018233 | 2.665701 | 0.315897 | 1.233853 | 0.764635 | -0.134066 | 1.960654 | -0.172539 |
| 30.55555556 | 0.94973 | 0.345089 | 0.313093 | -0.080434 | 0.946186 | 0.522712 | 0.792161 | 0.414582 | 1.06122 | 1.505329 | 1.500268 | 1.685611 | 0.235972 | 1.916501 | 1.261112 | 0.293591 | 1.379515 | 1.962076 | 0.241983 | 1.178182 | 0.898979 | -0.022354 | 1.547372 | -0.253226 |
| 30.56944444 | 0.618589 | 0.212632 | 0.367198 | 0.193653 | 0.818287 | 0.50577 | 0.686343 | 0.640765 | 0.935863 | 1.069285 | 0.724404 | 1.213623 | 0.887904 | 0.7395 | 0.995909 | 0.120752 | 3.111442 | 1.264905 | 0.083594 | 0.7863 | 1.226389 | -0.121384 | 1.191204 | 0.230832 |
| 30.58333333 | 0.673585 | 0.340377 | 0.331346 | 0.144719 | 0.903622 | 0.44879 | 0.463838 | 0.764301 | 0.756669 | 0.684995 | 0.338645 | 0.940996 | 1.158111 | 0.293288 | 0.21679 | 0.072852 | 0.910352 | 0.810437 | -0.002527 | 0.816589 | 0.623449 | -0.392167 | 0.601368 | 0.716693 |
| 30.59722222 | 0.359793 | 0.209724 | -0.27984 | -0.385247 | 0.592791 | 0.35476 | -0.008102 | 0.152505 | 0.431737 | 0.243059 | 0.355901 | 0.619026 | 0.731106 | 0.634051 | -0.857004 | -0.036956 | -1.68271 | 0.452717 | -0.066703 | 0.396036 | -0.104071 | -0.394535 | 0.197161 | 0.940974 |
| 30.61111111 | -0.038173 | -0.057527 | -0.791875 | -0.755647 | -0.196334 | 0.211601 | -0.272372 | -0.670981 | 0.028564 | -0.185692 | 0.262756 | 0.337058 | 0.196172 | 0.677515 | -1.277765 | -0.18082 | -2.874376 | 0.290351 | -0.072893 | -0.098525 | -0.844524 | -0.525845 | -0.28739 | 0.812637 |
| 30.625 | -0.040543 | -0.278773 | -1.015831 | -0.695301 | -0.653284 | 0.105189 | -0.397264 | -0.969246 | -0.070097 | -0.371422 | 0.24355 | 0.355074 | -0.040488 | 0.201638 | -1.362574 | -0.286836 | -3.697219 | 0.154399 | 0.017102 | -0.244247 | -0.98105 | -0.362439 | -0.575728 | 0.67262 |
| 30.63888889 | -0.058051 | -0.058541 | -1.067734 | -0.621422 | -0.888161 | 0.348712 | -0.171947 | -0.456093 | 0.073928 | -0.336081 | 0.381024 | 0.620702 | 0.126997 | -0.240819 | -0.850829 | -0.267692 | -4.211118 | 0.087333 | 0.058916 | -0.705009 | -0.5206 | 0.010726 | -0.863965 | 0.245058 |
| 30.65277778 | 0.076275 | -0.194381 | -1.268836 | -0.723286 | -1.045035 | 0.488142 | -0.172924 | -0.046595 | 0.133894 | -0.420712 | 0.501585 | 0.894256 | 0.353001 | -0.138473 | -0.519095 | -0.203927 | -3.510774 | -0.201843 | 0.044963 | -1.646383 | -0.056708 | 0.490603 | -0.873622 | -0.244838 |
| 30.66666667 | 0.245954 | -0.055214 | -1.317518 | -0.537497 | -1.370596 | 0.329435 | -0.591919 | -0.432196 | 0.073379 | -0.556127 | 0.359994 | 0.682269 | 0.850714 | 0.041622 | -0.090736 | -0.159331 | -2.397237 | -0.352545 | 0.064069 | -2.515175 | 0.86155 | 0.481192 | -0.673566 | -0.48274 |
| 30.68055556 | 0.119192 | 0.335897 | -0.930805 | -0.214034 | -1.929751 | -0.044074 | -0.537092 | -0.777184 | 0.010645 | -0.281552 | 0.192553 | 0.31137 | 0.39845 | -0.1819 | 0.330814 | -0.127807 | -2.915693 | -0.263964 | 0.108738 | -2.400885 | 1.48604 | -0.141675 | -0.292316 | -0.419378 |
| 30.69444444 | -0.110251 | 0.518663 | -0.233788 | -0.051173 | -2.584404 | -0.26541 | -0.193406 | -0.270473 | 0.031545 | 0.323583 | 0.017256 | 0.030448 | -0.096705 | -0.51948 | 0.507329 | -0.09108 | -3.057862 | -0.017515 | 0.105527 | -1.885903 | 1.118231 | -0.365784 | -0.120881 | -0.472197 |
| 30.70833333 | -0.269553 | 0.421006 | 0.322837 | -0.051606 | -2.745962 | -0.203088 | 0.160507 | 0.294409 | 0.077561 | 0.66765 | 0.003279 | -0.022226 | -0.425483 | -0.358155 | 0.297555 | -0.072292 | -2.921057 | -0.147602 | -0.009845 | -1.518849 | 0.73412 | -0.379009 | -0.016908 | -0.422161 |
| 30.72222222 | -0.257128 | 0.582518 | 0.354496 | -0.060537 | -2.87033 | 0.079834 | 0.312973 | 0.392285 | 0.105972 | 0.570875 | 0.267346 | 0.284107 | -0.713484 | 0.168826 | -0.083595 | -0.143112 | -2.081142 | -0.342728 | -0.024914 | -1.34515 | 0.751867 | -0.433661 | 0.160091 | -0.268172 |
| 30.73611111 | -0.13551 | 0.516452 | 0.008277 | -0.069188 | -3.030396 | 0.201136 | 0.166005 | 0.044557 | 0.347802 | 0.358182 | 0.599995 | 0.722371 | -1.055193 | 0.412759 | 0.249669 | -0.180048 | -1.311399 | -0.428093 | -0.036522 | -0.663418 | 0.45127 | -0.236252 | 0.356966 | -0.211647 |
| 30.75 | -0.073748 | -0.028515 | -0.352906 | -0.24411 | -2.821339 | 0.225344 | 0.01032 | -0.232235 | 0.435518 | 0.271525 | 0.819874 | 0.996499 | -1.05412 | 0.212801 | 0.532172 | -0.108874 | -0.729558 | -0.452164 | -0.106548 | -0.412224 | 0.064628 | -0.244208 | 0.51455 | -0.355536 |
| 30.76388889 | -0.021134 | -0.32723 | -0.651579 | -0.453196 | -2.460024 | 0.20359 | 0.07683 | -0.143589 | 0.284315 | 0.239529 | 0.801688 | 0.996262 | -1.092938 | -0.087028 | 0.884948 | -0.107817 | -0.649713 | -0.55248 | -0.202118 | -0.04112 | 0.032694 | -0.612929 | 0.155427 | -0.570089 |
| 30.77777778 | 0.003742 | -0.33552 | -0.762124 | -0.572761 | -2.648527 | 0.25674 | 0.211299 | 0.085631 | 0.156336 | 0.068785 | 0.653888 | 0.569782 | -1.095569 | -0.176047 | 1.082594 | -0.155994 | -0.141994 | -0.899948 | -0.206347 | 0.075205 | 0.054477 | -0.331553 | -0.637517 | -0.806114 |
| 30.79166667 | -0.068143 | -0.35205 | -0.787603 | -0.760942 | -3.300551 | 0.133586 | 0.231509 | 0.120802 | 0.309629 | 0.12429 | 0.324081 | -0.096627 | -1.232067 | -0.056098 | 0.912385 | -0.16776 | -0.013986 | -0.924733 | -0.049378 | 0.055246 | -0.203025 | -0.145753 | -1.008764 | -0.921952 |
| 30.80555556 | -0.078132 | -1.024362 | -1.027333 | -1.044914 | -3.47606 | 0.005721 | 0.019102 | -0.078626 | 0.460773 | 0.121516 | -0.26319 | -0.769486 | -1.519661 | 0.289663 | 0.793343 | -0.134585 | -0.965533 | -1.009907 | -0.120609 | 0.025114 | -0.715226 | -0.274993 | -1.407827 | -0.962561 |
| 30.81944444 | 0.10293 | -1.458714 | -1.203832 | -1.060953 | -2.72583 | 0.014049 | -0.237625 | -0.316753 | 0.181883 | -0.147469 | -0.827947 | -0.185824 | -1.808132 | 0.613036 | 0.590659 | -0.099089 | -1.343851 | -1.405258 | -0.291996 | 0.113115 | -1.094425 | -0.328416 | -1.603307 | -0.971465 |
| 30.83333333 | 0.15566 | -0.824905 | -1.079667 | -1.029933 | -2.133496 | -0.026843 | -0.521378 | -0.549288 | -0.198823 | -0.567158 | -0.691704 | 0.112406 | -1.189568 | 0.416866 | 0.047409 | -0.226228 | -0.897838 | -1.369242 | -0.45316 | 0.077473 | -1.365137 | -0.346171 | -1.913309 | -0.675574 |
| 30.84722222 | -0.021302 | -0.206924 | -1.276166 | -0.985172 | -2.617711 | -0.342001 | -0.829474 | -0.809467 | -0.582782 | -1.015048 | -0.248062 | 0.037709 | -1.338385 | -0.248839 | -0.59815 | -0.3922 | -1.026483 | -1.316291 | -0.42533 | 0.02025 | -1.223217 | -0.375318 | -2.02928 | -1.132999 |
| 30.86111111 | -0.225039 | -1.031007 | -1.472373 | -1.078582 | -3.715755 | -0.662759 | -0.762634 | -1.020811 | -0.808266 | -1.374075 | -0.156077 | -0.132731 | -1.444635 | -0.768709 | -0.857098 | -0.310888 | -1.346899 | -1.444655 | -0.281821 | -0.074125 | -1.380093 | -0.243118 | -2.002068 | -1.209314 |
| 30.875 | -0.35969 | -1.277438 | -1.832643 | -1.513164 | -4.556031 | -0.955999 | -0.588058 | -0.717119 | -0.729346 | -1.357125 | -0.293887 | -0.3268 | -1.768179 | -0.862115 | -0.84358 | -0.224238 | -1.741575 | -1.463662 | -0.257054 | -0.212072 | -1.44395 | -0.070303 | -2.314575 | -0.782403 |
| 30.88888889 | -0.643837 | -1.943293 | -2.03203 | -2.483567 | -4.793883 | -1.106297 | -0.388398 | -0.564845 | -0.743624 | -1.50147 | -0.569861 | -0.678966 | -1.995668 | -0.971749 | -1.234337 | -0.277863 | -2.40517 | -1.240446 | -0.180154 | -0.317781 | -1.382733 | -0.05268 | -1.887576 | -0.948131 |
| 30.90277778 | -1.198308 | -2.654997 | -1.795638 | -3.868595 | -5.113033 | -1.401655 | -0.596321 | -0.79872 | -1.070521 | -2.35775 | -0.88065 | -1.111361 | -2.37755 | -1.395934 | -1.149701 | -0.648698 | -3.046713 | -1.259643 | -0.347588 | -0.427631 | -1.244181 | -0.04514 | -1.326759 | -1.449546 |
| 30.91666667 | -1.490751 | -2.762048 | -1.751419 | -4.223667 | -5.003695 | -1.572498 | -0.827385 | -1.202266 | -1.370148 | -3.191301 | -1.153705 | -1.562481 | -2.317245 | -1.726771 | -1.823715 | -1.053422 | -1.986405 | -1.828981 | -0.66823 | -0.674869 | -1.043021 | -0.086612 | -1.159535 | -1.116805 |
| 30.93055556 | -1.325741 | -2.859851 | -1.535302 | -4.255165 | -4.550383 | -1.397867 | -0.73016 | -1.066374 | -1.176954 | -2.760201 | -1.142576 | -1.868368 | -2.261785 | -1.76698 | -2.688376 | -1.711125 | -0.435396 | -2.527352 | -1.026587 | -0.62374 | -1.048731 | -0.085229 | -0.918072 | -0.938256 |
| 30.94444444 | -1.287954 | -2.454606 | -1.708186 | -4.271181 | -4.330877 | -1.217373 | -0.508672 | -0.740167 | -0.818742 | -2.505398 | -1.319235 | -2.016534 | -2.082427 | -2.415364 | -3.164487 | -2.671217 | -0.764073 | -3.415195 | -1.285783 | -0.910887 | -1.600566 | -0.781193 | -1.568651 | -1.270949 |
| 30.95833333 | -1.83681 | -2.792067 | -2.303212 | -4.742993 | -4.679862 | -1.255568 | -0.702706 | -0.47552 | -0.905625 | -2.972173 | -1.408996 | -2.070924 | -3.274112 | -1.878983 | -3.724215 | -2.573928 | -1.636517 | -3.906615 | -1.695585 | -1.841632 | -1.747137 | -1.254851 | -2.588112 | -1.113338 |
| 30.97222222 | -2.403744 | -3.433489 | -2.561559 | -4.919093 | -5.234407 | -1.847118 | -1.409841 | -0.625451 | -1.396763 | -3.658595 | -2.289277 | -2.982159 | -3.625716 | -1.002828 | -3.718819 | -2.010377 | -1.799151 | -4.168181 | -1.417439 | -1.508495 | -1.215742 | -0.411816 | -1.979014 | -0.647788 |
| 30.98611111 | -1.755517 | -3.298333 | -2.463553 | -5.021915 | -6.118798 | -2.229834 | -1.358171 | -0.869078 | -1.424046 | -3.625896 | -2.7711 | -2.221924 | -3.849336 | -1.67407 | -2.314965 | -1.191662 | -1.113515 | -3.43085 | -1.726528 | -1.24748 | -0.878043 | -0.716278 | -1.139207 | -0.290822 |
| 31 | -1.413336 | -2.646267 | -1.566378 | -3.890208 | -4.949354 | -1.565928 | -1.00854 | -0.744148 | -0.99357 | -2.410999 | -3.221817 | -1.260358 | -2.448307 | -2.711993 | -3.123101 | -1.355158 | -1.07291 | -2.959319 | -2.02453 | -0.668403 | -0.68382 | 0.057594 | -1.603916 | -1.016667 |
| 31.01388889 | -1.670381 | -1.700041 | -0.716881 | -2.553346 | -3.662975 | -0.849611 | -0.866376 | -0.236705 | -0.500018 | -1.305853 | -2.890596 | -0.665796 | 0.222949 | -3.40923 | -2.452015 | -1.771094 | -1.134205 | -1.911632 | -1.651276 | 0.316309 | 0.022908 | 1.271223 | -1.246912 | -1.402351 |
| 31.02777778 | -1.001818 | -0.89543 | -0.45521 | -1.272013 | -2.888615 | -0.465901 | -0.693665 | -0.266633 | -0.341378 | -0.778875 | -2.242835 | -0.936608 | 0.211115 | -1.777968 | -1.524723 | -1.935539 | -1.250264 | -0.722011 | -1.015906 | 0.709222 | 0.548574 | 0.389648 | -1.43546 | -0.757762 |
| 31.04166667 | -0.723464 | -0.73767 | -0.504172 | -0.50195 | -2.241312 | -0.316931 | -0.647146 | -0.50259 | -0.370713 | -0.608819 | -1.713871 | -0.741782 | 0.028361 | -0.012449 | -0.654809 | -1.058572 | -1.359905 | 0.101431 | -0.284226 | 0.558724 | 0.457799 | -0.675193 | -1.360437 | 0.119801 |
| 31.05555556 | -0.376607 | -0.655051 | -0.452479 | -0.527369 | -1.213147 | -0.115824 | -0.359453 | -0.457228 | -0.201332 | -0.30141 | -1.566289 | -0.545823 | -0.12761 | 0.88087 | 0.473816 | 0.84824 | -0.952041 | -0.197366 | -0.43438 | 1.495499 | 0.460535 | -0.287452 | -1.324394 | 0.613331 |
| 31.06944444 | 0.161125 | -0.344645 | -0.166302 | -0.748322 | -0.416221 | 0.133695 | -0.052235 | -0.066761 | 0.152406 | 0.155314 | -1.396578 | -0.46941 | -0.239094 | 1.295743 | 1.311003 | 2.626275 | -1.077523 | -0.445528 | -0.534065 | 1.222848 | 0.503163 | 0.341754 | -0.780866 | 0.705736 |
| 31.08333333 | 0.431188 | 0.463065 | 0.108664 | -0.287406 | -0.24744 | 0.38734 | 0.155587 | 0.181353 | 0.34552 | 0.408635 | -0.632038 | -0.225699 | 0.395413 | 1.656525 | 1.234915 | 3.348491 | -0.746628 | 0.002546 | -0.486501 | 0.59861 | 0.325417 | 0.634204 | -0.197905 | 0.687114 |
| 31.09722222 | 0.53898 | 1.263181 | 0.202265 | 0.441034 | -0.437006 | 0.658904 | 0.308774 | 0.286773 | 0.427684 | 0.603927 | 0.429126 | 0.191441 | 0.620645 | 1.639849 | 1.321651 | 3.0227 | -1.018027 | 0.534811 | 0.181247 | 0.36697 | 0.488419 | 0.68983 | -0.425275 | 0.632133 |
| 31.11111111 | 0.494866 | 1.640622 | 0.173361 | 0.984038 | -0.438304 | 0.748282 | 0.355457 | 0.366337 | 0.522034 | 0.962284 | 1.066597 | 0.519408 | 1.381837 | 1.283602 | 1.542636 | 2.308519 | -1.2515 | 0.995874 | 0.838764 | 0.135934 | 0.779109 | -0.182818 | -1.403318 | 0.359851 |
| 31.125 | 0.369583 | 1.6371 | 0.275973 | 1.408555 | -0.078376 | 0.855315 | 0.3689 | 0.583831 | 0.58486 | 1.376864 | 1.169228 | 0.683104 | 2.748005 | 0.721857 | 1.610117 | 1.194208 | -0.681786 | 1.02963 | 0.887315 | -0.245819 | 0.893647 | -0.20359 | -2.085579 | -0.045772 |
| 31.13888889 | 0.320767 | 1.535195 | 0.464028 | 1.591616 | 0.30104 | 1.010512 | 0.391447 | 0.717102 | 0.601131 | 1.618887 | 1.094889 | 0.658259 | 2.369667 | 0.240025 | 1.613762 | 0.687557 | 0.051371 | 0.968329 | 0.868769 | -0.124132 | 1.219498 | 0.253193 | -0.893853 | -0.411475 |
| 31.15277778 | 0.41629 | 1.293551 | 0.465043 | 1.122431 | 0.811328 | 1.027998 | 0.364606 | 0.641943 | 0.580053 | 1.688273 | 1.259356 | 0.677751 | 1.272412 | -0.242732 | 1.855781 | 0.453776 | 0.815579 | 1.05235 | 1.205061 | -0.084792 | 1.212506 | 0.445826 | 0.491366 | -0.480936 |
| 31.16666667 | 0.470684 | 0.963348 | 0.353714 | 0.385462 | 1.199022 | 0.87326 | 0.333345 | 0.668725 | 0.516548 | 1.547832 | 1.582744 | 0.77494 | 0.893108 | -0.308835 | 2.080561 | 0.209532 | 1.343713 | 1.1146 | 1.297087 | 0.210309 | 1.196638 | -0.158949 | 0.595604 | -0.388216 |
| 31.18055556 | 0.386611 | 0.791842 | 0.324828 | 0.00494 | 1.338441 | 0.673914 | 0.21651 | 0.705333 | 0.453098 | 1.277841 | 1.711245 | 1.103593 | 0.908554 | 0.595452 | 1.913563 | 0.122873 | 1.245099 | 1.014301 | 1.047715 | 0.831543 | 1.459658 | -0.558808 | -0.576774 | 0.169858 |
| 31.19444444 | 0.159982 | 0.916256 | 0.347179 | 0.168449 | 1.215318 | 0.527544 | 0.029059 | 0.538812 | 0.384848 | 0.971138 | 1.540787 | 1.016328 | 2.116839 | 1.091068 | 1.464582 | 0.382422 | 1.20842 | 0.935401 | 0.819275 | 0.839597 | 1.75115 | -0.451056 | -0.579316 | 0.574581 |
| 31.20833333 | 0.019685 | 1.014782 | 0.393162 | 0.543035 | 0.990617 | 0.432285 | -0.082904 | 0.429541 | 0.394003 | 0.822696 | 1.590946 | 1.012413 | 2.746369 | 1.355439 | 1.193291 | 0.784045 | 2.73823 | 1.16123 | 1.128625 | 0.251644 | 0.486535 | -0.365638 | -0.140034 | 0.573354 |
| 31.22222222 | 0.097291 | 0.916906 | 0.435256 | 0.572837 | 1.76996 | 0.37618 | -0.084645 | 0.422097 | 0.43868 | 0.721401 | 1.776427 | 1.164356 | 2.569472 | 1.262102 | 0.977447 | 0.903963 | 1.968439 | 1.668135 | 1.706065 | 0.145172 | 0.4093 | 0.074299 | -0.098303 | 0.426474 |
| 31.23611111 | 0.064034 | 1.003557 | 0.400859 | 0.731293 | 2.46984 | 0.232322 | 0.023274 | 0.339897 | 0.416811 | 0.356668 | 1.623946 | 1.108559 | 2.101139 | 0.905594 | 0.624209 | 0.776524 | 0.145634 | 2.190722 | 2.271378 | 0.473548 | 1.86357 | 0.289116 | 0.292245 | 0.381955 |
| 31.25 | -0.052443 | 1.142836 | 0.314252 | 0.73526 | 2.636946 | 0.131636 | 0.086028 | 0.291224 | 0.280005 | -0.45981 | 1.217676 | 0.856839 | 1.952559 | 0.436556 | 0.295625 | 0.687596 | -0.890012 | 2.381532 | 2.242049 | 0.569854 | 1.475225 | 0.455192 | 0.605122 | 0.320483 |
| 31.26388889 | -0.075222 | 1.091783 | 0.220385 | 0.222687 | 2.580324 | 0.1708 | -0.002807 | 0.430378 | 0.157187 | -0.897146 | 0.810321 | 0.688311 | 2.139079 | 0.421472 | 0.28071 | 0.524336 | 0.007301 | 2.543303 | 2.177196 | 0.394611 | 1.437135 | 0.430041 | 1.004407 | 0.50843 |
| 31.27777778 | 0.044895 | 1.293123 | 0.245093 | 0.056412 | 2.948045 | 0.402312 | 0.017914 | 0.835015 | 0.237903 | -0.832575 | 0.739141 | 0.660961 | 2.425499 | 0.782375 | 0.478638 | 0.764755 | 1.282846 | 2.41081 | 1.876203 | 0.362498 | 0.735356 | 0.142501 | 1.399869 | 0.649892 |
| 31.29166667 | 0.179434 | 1.186593 | 0.281612 | -0.062908 | 3.404102 | 0.583625 | 0.191015 | 0.921019 | 0.34977 | -0.15255 | 0.571964 | 0.574827 | 2.363226 | 1.380702 | 0.518418 | 0.974264 | 0.914323 | 1.629835 | 1.140623 | 0.622472 | 0.106121 | -0.165901 | 1.591424 | 0.916689 |
| 31.30555556 | 0.120177 | 0.864753 | 0.269573 | -0.373668 | 3.423701 | 0.561891 | 0.363302 | 0.652744 | 0.364768 | 0.339893 | 0.132127 | 0.16883 | 1.898388 | 1.208765 | 0.297077 | 1.020002 | 1.002249 | 1.14749 | 0.888136 | 0.807061 | 0.241036 | -0.205523 | 1.899759 | 1.029856 |
| 31.31944444 | 0.026367 | 0.081467 | 0.332801 | -0.952309 | 3.597848 | 0.535172 | 0.51135 | 0.384845 | 0.353901 | 0.922706 | -0.243313 | -0.217476 | 1.560477 | 0.994432 | 0.562365 | 1.324616 | 0.529005 | 1.274697 | 1.076855 | 0.700555 | 1.152744 | -0.003482 | 1.844145 | 1.132191 |
| 31.33333333 | 0.067653 | 0.023169 | 0.48391 | -1.012695 | 3.761366 | 0.702131 | 0.610172 | 0.400177 | 0.387432 | 1.53232 | -0.246307 | -0.229103 | 0.441802 | 1.186676 | 1.187093 | 1.743335 | -0.896035 | 1.195363 | 1.025419 | 0.791203 | 1.883352 | 0.478911 | 1.268918 | 0.791501 |
| 31.34722222 | 0.219586 | -0.049231 | 0.646794 | -0.815638 | 4.075718 | 0.920364 | 0.657096 | 0.654842 | 0.461848 | 1.823086 | -0.125234 | -0.150516 | -0.124385 | 1.885989 | 1.995621 | 2.034682 | -1.903318 | 1.241754 | 1.045255 | 0.645389 | 2.437358 | 0.562091 | 1.748251 | 1.579534 |
| 31.36111111 | 0.426174 | 0.086501 | 0.871278 | -0.242827 | 4.238809 | 1.102417 | 0.743548 | 0.715144 | 0.562027 | 2.008228 | 0.165902 | 0.138023 | 0.758769 | 2.567055 | 3.21025 | 2.557863 | -0.961998 | 1.761168 | 1.652197 | 0.569513 | 2.46729 | 0.302376 | 2.678964 | 1.847818 |
| 31.375 | 0.555463 | 0.503659 | 0.968216 | 0.339902 | 4.560115 | 1.201556 | 1.0016 | 0.847981 | 0.650488 | 1.943903 | 0.538066 | 0.515704 | 1.736048 | 2.561899 | 3.828544 | 2.52072 | 1.017005 | 1.968979 | 1.862367 | 0.962588 | 2.454918 | 0.0765 | 2.953361 | 1.715544 |
| 31.38888889 | 0.455026 | 1.08397 | 0.872912 | 0.955918 | 4.698622 | 1.392147 | 1.12872 | 0.773211 | 0.747629 | 2.035408 | 0.775041 | 0.716901 | 2.483524 | 2.490362 | 4.439798 | 2.624971 | 3.056731 | 1.700301 | 1.534181 | 1.413102 | 2.498247 | 0.417295 | 2.595955 | 1.428557 |
| 31.40277778 | 0.376007 | 1.499962 | 1.168232 | 1.683037 | 5.042747 | 1.715727 | 1.232508 | 0.702756 | 0.78502 | 1.963096 | 0.972135 | 0.816052 | 2.740162 | 2.421442 | 4.59682 | 2.028966 | 3.866743 | 1.862487 | 1.54797 | 1.726223 | 2.117216 | 0.369744 | 2.824664 | 1.480453 |
| 31.41666667 | 0.479042 | 1.793051 | 1.485981 | 2.329385 | 5.20758 | 1.893954 | 1.311453 | 0.685198 | 0.837649 | 1.886075 | 1.196831 | 1.106704 | 2.738216 | 2.514226 | 3.690183 | 1.555141 | 3.208464 | 2.355263 | 1.688277 | 2.25209 | 2.112676 | 0.106446 | 3.57966 | 1.608699 |
| 31.43055556 | 0.613738 | 1.72188 | 1.510202 | 2.505264 | 5.370005 | 1.580044 | 1.22713 | 0.520191 | 0.905031 | 1.770662 | 1.417499 | 1.43031 | 2.788358 | 2.808058 | 3.194626 | 1.595787 | 3.42216 | 2.531846 | 1.623181 | 3.136138 | 1.671081 | -0.12694 | 3.431569 | 1.975807 |
| 31.44444444 | 0.64297 | 1.189918 | 1.246111 | 2.130684 | 4.997307 | 1.535561 | 1.280346 | 0.402943 | 0.972838 | 1.783291 | 1.53887 | 1.951791 | 2.909871 | 3.327152 | 3.627354 | 2.111687 | 3.289726 | 2.609021 | 1.545381 | 4.377298 | 1.610484 | -0.189121 | 2.911629 | 2.361206 |
| 31.45833333 | 0.606042 | 0.77186 | 0.777603 | 1.682326 | 3.580846 | 1.67402 | 1.306221 | 0.512038 | 1.219469 | 2.035218 | 1.485716 | 2.218013 | 2.907421 | 3.753824 | 3.49044 | 2.106395 | 1.540518 | 2.621021 | 1.327132 | 4.331927 | 1.622141 | -0.116706 | 3.303934 | 2.467663 |
| 31.47222222 | 0.335687 | 0.33542 | 0.490734 | 0.731652 | 2.715204 | 1.457018 | 1.060886 | 0.495294 | 1.249965 | 1.905787 | 1.275852 | 1.903633 | 2.706078 | 3.284485 | 3.416659 | 1.76767 | 0.634926 | 2.52962 | 1.08979 | 2.51407 | 1.630172 | -0.340564 | 3.132085 | 2.350107 |
| 31.48611111 | 0.109237 | -0.145312 | 0.377459 | -0.275421 | 2.462144 | 0.879303 | 0.647159 | 0.434022 | 1.136411 | 1.563614 | 0.713354 | 0.962849 | 2.457116 | 3.180544 | 4.294821 | 1.878805 | 1.794099 | 2.905359 | 1.592449 | 1.058265 | 2.239575 | -0.83836 | 2.779068 | 2.547018 |
| 31.5 | 0.278447 | -1.139425 | 0.714927 | -0.483813 | 3.351246 | 0.428133 | 0.291349 | 0.403676 | 1.195812 | 1.646714 | 0.668568 | 0.087076 | 2.927747 | 2.851818 | 4.033233 | 1.36672 | 3.834809 | 3.009817 | 1.749264 | 0.867829 | 1.803184 | 0.175102 | 2.17843 | 2.574426 |
| 31.51388889 | 0.251256 | -1.986039 | 1.010843 | -1.147211 | 3.022037 | 0.401755 | -0.370496 | 0.295475 | 1.341551 | 1.988316 | 0.679432 | -1.015268 | 2.803075 | 2.464413 | 3.763707 | 0.838188 | 4.351967 | 2.475485 | 1.611516 | 1.722594 | 1.686797 | 1.093022 | 1.584607 | 2.161909 |
| 31.52777778 | -0.76073 | -2.294875 | 1.112133 | -1.58939 | 1.5935 | 0.388697 | -0.447833 | 0.263247 | 1.324022 | 1.756111 | 0.56228 | -0.395017 | 2.066876 | 2.154046 | 3.223514 | 0.307214 | 1.74306 | 2.344028 | 1.414348 | 2.407198 | 1.545313 | 0.620393 | 1.29813 | 1.824388 |
| 31.54166667 | -1.021601 | -4.204502 | 0.457674 | -1.302888 | 1.22562 | 0.27005 | -0.743983 | 0.272281 | 1.408829 | 1.937062 | 0.094525 | 0.326462 | 1.914329 | 1.820583 | 2.938608 | 0.349855 | 3.624378 | 2.383893 | 0.869933 | 1.529448 | 1.514361 | 0.208884 | 1.450967 | 1.418483 |
| 31.55555556 | -0.694884 | -4.086155 | 0.323662 | -1.115841 | 1.272456 | 0.285246 | -0.787545 | 0.018695 | 1.580015 | 2.285538 | -0.105649 | 0.37015 | 2.038107 | 1.48403 | 2.975413 | 0.20254 | 4.178777 | 1.942915 | 0.580694 | 1.816698 | 1.09104 | 0.351701 | 1.912027 | 1.174334 |
| 31.56944444 | -0.5442 | -3.614665 | 0.328621 | -1.611025 | 1.130245 | 0.526777 | -0.433627 | 0.104173 | 1.340698 | 1.961727 | -0.306735 | 0.004478 | 2.276749 | 1.3981 | 2.791903 | -0.024234 | 3.261511 | 1.304552 | 0.344997 | 1.107886 | 0.744075 | 0.317261 | 1.97513 | 1.08115 |
| 31.58333333 | -0.172371 | -2.852373 | 0.05524 | -2.323142 | 0.70262 | 0.459387 | -0.01501 | 0.176901 | 0.898773 | 1.312167 | -0.227458 | -0.038619 | 1.953625 | 0.832716 | 2.123483 | -0.069618 | 2.875667 | 1.023559 | 0.180414 | 1.278388 | 0.616242 | 0.311347 | 1.270504 | 0.967008 |
| 31.59722222 | -0.379612 | -1.310558 | -0.091214 | -2.315643 | 0.121573 | -0.09241 | -0.132974 | -0.007662 | 0.287767 | 0.220593 | -0.020212 | -0.098428 | 1.757889 | 0.0427 | 1.276094 | -0.154772 | 0.10928 | 0.787478 | 0.172814 | 0.60906 | 0.104194 | 0.229281 | 0.589052 | 0.835427 |
| 31.61111111 | -0.672633 | -0.852101 | -0.177797 | -2.246182 | -0.509148 | -0.338155 | -0.445873 | -0.366699 | -0.199978 | -0.649845 | -0.156501 | -0.25378 | 1.018787 | -0.502834 | 0.859656 | -0.290191 | -2.676039 | 0.47122 | 0.084699 | 0.661042 | -0.927173 | 0.05467 | 0.101315 | 0.603379 |
| 31.625 | -0.860095 | -1.04896 | -0.200263 | -2.179077 | -1.069494 | -0.526937 | -0.760614 | -0.684773 | -0.330277 | -0.829175 | -0.208528 | -0.081324 | 0.50045 | -0.756068 | -0.139219 | -0.252991 | -4.457325 | 0.478976 | -0.011926 | 0.049112 | -2.358005 | 0.158836 | -0.558944 | 0.390373 |
| 31.63888889 | -1.057292 | -0.792684 | -0.235269 | -2.391845 | -1.568106 | -0.560991 | -0.874944 | -0.634206 | -0.238412 | -0.62931 | 0.037281 | 0.316336 | -0.204849 | -0.51251 | -0.300491 | -0.17456 | -5.329123 | 0.437419 | 0.018027 | 0.061754 | -2.262299 | 0.165384 | -0.048721 | 0.385462 |
| 31.65277778 | -1.082222 | -0.542818 | -0.248476 | -2.838548 | -1.746622 | -0.461928 | -0.799565 | -0.475955 | -0.062089 | -0.602198 | 0.332199 | 0.630616 | -0.645959 | 0.075697 | -0.130505 | -0.149826 | -4.830362 | 0.332732 | -0.034794 | -0.248212 | -1.298643 | 0.149103 | 0.299061 | 0.506863 |
| 31.66666667 | -1.124765 | -0.737851 | 0.206071 | -2.931859 | -2.294884 | -0.390693 | -0.577557 | -0.419479 | -0.041656 | -1.043832 | 0.589172 | 0.633045 | -0.513012 | 0.474056 | -0.697815 | -0.13535 | -3.41225 | 0.082236 | -0.255344 | -0.866559 | -0.318973 | -0.036443 | 0.160354 | 0.646211 |
| 31.68055556 | -1.087298 | -0.996485 | 0.569986 | -2.514516 | -3.357454 | -0.433096 | -0.451749 | -0.411307 | 0.007186 | -1.098622 | 0.732204 | 0.659092 | 0.01924 | 0.464224 | -1.571754 | -0.185667 | -1.797173 | -0.007263 | -0.368142 | -1.579078 | -0.086673 | -0.19836 | -0.403574 | 0.385835 |
| 31.69444444 | -0.795413 | -0.693924 | 0.39807 | -1.470807 | -4.236129 | -0.576865 | -0.410531 | -0.291166 | 0.140079 | -0.610638 | 0.847884 | 0.719306 | 0.327709 | 0.450882 | -0.875592 | -0.162152 | -1.048869 | 0.033765 | -0.158323 | -2.163225 | -0.13304 | -0.118837 | -0.682406 | 0.204447 |
| 31.70833333 | -0.446751 | -0.378176 | 0.08767 | -0.693189 | -4.548874 | -0.730241 | -0.297654 | 0.015084 | 0.152621 | 0.060025 | 0.752504 | 0.74262 | 0.269757 | 0.833504 | 0.084323 | -0.081234 | -0.9546 | 0.326208 | -0.146026 | -1.592403 | 0.244006 | 0.149563 | -0.717354 | 0.306127 |
| 31.72222222 | -0.352028 | -0.125694 | -0.044784 | -0.753838 | -4.28321 | -0.815503 | -0.053696 | 0.432151 | 0.173864 | 0.601539 | 0.60724 | 0.904371 | 0.176606 | 1.389108 | 0.436349 | -0.059071 | -0.567399 | 0.266843 | -0.130682 | -0.471372 | 0.522763 | -0.676792 | -0.48266 | 0.492014 |
| 31.73611111 | -0.31665 | -1.022154 | 0.110503 | -0.921325 | -4.310076 | -0.761601 | 0.211393 | 0.502001 | 0.255945 | 0.977159 | 0.498559 | 1.366756 | 0.153448 | 1.973448 | 0.40149 | -0.040448 | -1.264299 | -0.231464 | -0.251021 | 0.063121 | 0.728497 | -1.464552 | -0.886982 | 0.82725 |
| 31.75 | -0.232942 | -2.14015 | 0.258387 | -1.140264 | -4.2572 | -0.587758 | 0.426579 | 0.342554 | 0.328549 | 0.961054 | -0.477338 | 0.969577 | -0.047606 | 2.054289 | 0.182996 | 0.096681 | -0.896503 | -0.605632 | -0.289158 | -0.242737 | 0.749959 | -1.404011 | -1.454804 | 1.035821 |
| 31.76388889 | 0.130838 | -1.81577 | 0.036482 | -1.941851 | -4.404278 | -0.452447 | 0.518936 | 0.209371 | 0.3814 | 0.617398 | -0.931912 | 0.128511 | -1.097814 | 1.54543 | 0.122524 | 0.15391 | -0.548088 | -0.650906 | -0.182737 | -0.941243 | 0.141017 | -0.794371 | -0.785665 | 1.200895 |
| 31.77777778 | 0.350792 | -0.8623 | -0.255844 | -1.320429 | -4.390876 | -0.365592 | 0.601965 | 0.296821 | 0.460128 | 0.614496 | -0.466458 | 0.068087 | -1.84738 | 0.87215 | 0.100265 | 0.000104 | -0.757669 | -0.765335 | -0.039594 | -1.097895 | -0.533111 | -0.240301 | -0.969 | 1.486311 |
| 31.79166667 | 0.321778 | 0.089541 | -0.261764 | -0.704421 | -3.932252 | -0.346217 | 0.688584 | 0.437255 | 0.495742 | 1.059788 | 0.690503 | 0.37228 | -1.760195 | 0.389905 | 0.019286 | -0.185795 | -1.354238 | -1.102716 | -0.119154 | -0.690993 | -0.380559 | 0.041499 | -1.649744 | 1.115941 |
| 31.80555556 | 0.0262 | 1.275302 | -0.169069 | -0.02924 | -3.501071 | -0.375593 | 0.452044 | 0.349247 | 0.435506 | 1.053097 | 1.840239 | 0.981706 | -1.19533 | 0.642606 | -0.212289 | -0.466803 | -1.355158 | -2.081145 | -0.576631 | -0.250873 | 0.217321 | 0.154328 | -1.939982 | 0.791253 |
| 31.81944444 | -0.186357 | 1.478024 | -0.395453 | 0.378696 | -3.406575 | -0.46897 | 0.216994 | 0.159539 | 0.365313 | 0.526276 | 1.926224 | 0.676213 | -1.061631 | 0.615934 | -0.949497 | -0.798887 | -1.102664 | -2.462383 | -0.939473 | -0.084357 | -0.023188 | 0.128097 | -1.445729 | 0.469699 |
| 31.83333333 | -0.123004 | 1.461037 | -0.631276 | 0.516513 | -3.821819 | -0.495205 | 0.100927 | -0.035457 | 0.389651 | -0.044452 | 0.329877 | -0.463044 | -1.619831 | 0.234669 | -1.97251 | -1.122632 | -0.804734 | -2.231252 | -0.841239 | -0.320259 | -0.845206 | -0.165503 | -2.006702 | 0.105553 |
| 31.84722222 | -0.017267 | 1.57627 | -0.645081 | 0.846307 | -4.615387 | -0.397361 | -0.05046 | -0.186915 | 0.280238 | -0.743793 | -0.386491 | -1.204067 | -1.638758 | 0.023353 | -2.64128 | -1.202116 | -1.537979 | -1.896351 | -0.536084 | -0.536083 | -1.077231 | -0.400013 | -2.051385 | -0.205959 |
| 31.86111111 | -0.134086 | 1.752183 | -0.730184 | 0.851636 | -5.043016 | -0.378073 | -0.568941 | -0.490827 | -0.048988 | -1.463638 | -0.247903 | -1.172394 | -1.161756 | -0.261718 | -2.619365 | -1.052924 | -2.632273 | -1.299775 | -0.506278 | -0.56866 | -0.959313 | -0.402915 | -2.371311 | -0.348777 |
| 31.875 | -0.468252 | 1.948877 | -0.975764 | 1.12246 | -5.127028 | -0.588967 | -0.919019 | -0.854252 | -0.446064 | -1.676326 | -0.475588 | -1.600315 | -1.469514 | -0.294385 | -2.751157 | -1.351209 | -3.339225 | -1.505997 | -0.65446 | -0.251891 | -0.998448 | -0.335196 | -3.431803 | -0.199768 |
| 31.88888889 | -0.764098 | 1.535346 | -1.267371 | 0.948074 | -4.961749 | -0.86027 | -0.87891 | -1.288647 | -0.705666 | -1.597208 | -1.388966 | -1.814746 | -2.087071 | -0.53747 | -2.415142 | -1.520702 | -3.866361 | -1.64636 | -0.819169 | -0.257386 | -0.668832 | -0.083079 | -3.615124 | -0.023369 |
| 31.90277778 | -0.778024 | 1.033075 | -1.110789 | 0.444518 | -5.157525 | -0.993463 | -0.597328 | -1.379497 | -0.716141 | -1.779939 | -2.829696 | -2.615512 | -2.350431 | -0.664113 | -2.128077 | -1.535274 | -3.664082 | -1.544349 | -0.668462 | -0.812324 | -0.477207 | -0.221234 | -3.554048 | 0.088502 |
| 31.91666667 | -0.667338 | 0.601714 | -1.011161 | 0.066592 | -5.913644 | -1.14435 | -0.300335 | -1.099781 | -0.634958 | -3.04816 | -3.958187 | -2.800672 | -2.451201 | -0.977972 | -1.807025 | -1.390508 | -4.007549 | -1.728835 | -0.602707 | -1.534262 | -1.472003 | -1.443631 | -4.090534 | 0.001916 |
| 31.93055556 | -0.613827 | 0.481404 | -1.071732 | -0.047259 | -6.472806 | -1.261579 | -0.324822 | -1.111872 | -0.60775 | -3.530199 | -3.669014 | -2.182104 | -2.993609 | -1.107967 | -1.792853 | -1.43672 | -4.474645 | -2.185072 | -0.495215 | -1.941034 | -2.284915 | -0.263991 | -4.412941 | -0.214379 |
| 31.94444444 | -0.412704 | 0.255332 | -0.95838 | 0.039074 | -6.263942 | -1.41426 | -0.412794 | -1.331302 | -0.540107 | -3.515658 | -3.448636 | -2.057103 | -3.457146 | -1.05093 | -2.105723 | -1.611017 | -5.386405 | -2.303255 | -0.442351 | -2.437508 | -1.31943 | -0.026115 | -5.157262 | -0.693829 |
| 31.95833333 | -0.352599 | -0.499304 | -0.795986 | -0.187967 | -5.196558 | -2.186611 | -0.988217 | -1.816347 | -0.922108 | -4.139917 | -4.283654 | -2.274712 | -3.993862 | -1.420458 | -2.113684 | -1.590269 | -7.009707 | -2.219221 | -0.02218 | -3.243288 | -0.115853 | -0.195642 | -6.046228 | -1.436986 |
| 31.97222222 | -0.567068 | -1.34583 | -0.898694 | -0.631767 | -5.196225 | -3.041066 | -1.940894 | -2.275749 | -1.401211 | -4.969558 | -5.422364 | -2.884573 | -3.897356 | -1.906862 | -2.465152 | -1.53444 | -5.062577 | -1.865708 | -0.278478 | -3.518485 | -1.208513 | 1.248945 | -6.191643 | -1.448467 |
| 31.98611111 | -0.751441 | -2.000887 | -0.670952 | -0.486251 | -6.737413 | -3.022021 | -2.227338 | -2.027063 | -1.306519 | -5.721044 | -6.099733 | -2.674929 | -3.900768 | -1.889249 | -2.702338 | -1.848774 | -3.066288 | -2.339627 | -1.152748 | -1.956042 | -1.24191 | 0.886293 | -5.424787 | -0.539991 |
| 32 | -0.349024 | -0.822289 | 0.088409 | -0.453833 | -6.536021 | -1.859623 | -1.440154 | -1.598167 | -0.746625 | -4.059529 | -3.969369 | -1.643142 | -3.855706 | -1.290292 | -2.032264 | -1.927279 | -1.827393 | -2.104467 | -1.115677 | -1.347006 | -0.632107 | -0.824138 | -2.247817 | -0.188515 |
| 32.01388889 | 0.076868 | 0.004646 | 0.574719 | 0.336772 | -5.405374 | -0.979349 | -0.577192 | -1.018529 | -0.427848 | -3.047546 | -2.220429 | -0.995181 | -3.046431 | -0.648784 | -0.799664 | -1.649944 | -1.247604 | -1.206075 | -0.334074 | -1.118264 | 0.215267 | -1.627267 | 2.121401 | -0.121638 |
| 32.02777778 | 0.274894 | 0.219638 | 0.329399 | 1.032297 | -4.788887 | -0.529695 | -0.220288 | -0.568478 | -0.435679 | -2.296686 | -1.397448 | -0.811324 | -2.014192 | -0.713961 | -0.181467 | -1.253367 | -1.202722 | -1.048988 | -0.333883 | -0.229799 | 0.446965 | -2.47862 | 3.290734 | -0.471613 |
| 32.04166667 | 0.449714 | 0.129552 | 0.56108 | 1.046363 | -4.150772 | -0.376808 | -0.136612 | -0.368485 | -0.520507 | -1.963353 | -1.233749 | -0.823336 | -0.945395 | -0.037554 | -0.096668 | -0.496412 | -0.294336 | -1.275243 | -0.41755 | 0.571522 | -0.001831 | -1.839741 | 1.047253 | -0.230593 |
| 32.05555556 | 0.679875 | 0.043655 | 0.547504 | 0.778757 | -3.335675 | -0.428588 | -0.008553 | -0.239392 | -0.399333 | -1.805974 | -1.215586 | -0.895384 | -0.063261 | 0.481354 | 0.388792 | 0.622943 | 0.999457 | -1.264756 | -0.500833 | 0.63761 | 0.474623 | -1.56995 | 0.002266 | 0.369988 |
| 32.06944444 | 0.452404 | 0.189549 | 0.915511 | 0.941639 | -2.032035 | -0.433856 | 0.210602 | -0.18374 | -0.27835 | -1.438035 | -0.967802 | -0.961209 | 0.732056 | 0.329721 | 0.985779 | 1.896536 | 2.032607 | -0.529899 | 0.024972 | 0.632763 | 1.171264 | -0.934516 | -0.602392 | 0.808759 |
| 32.08333333 | 0.45615 | 0.411239 | 0.984551 | 1.244986 | -0.603259 | -0.215063 | 0.242421 | 0.120312 | -0.080608 | -0.902685 | -0.488888 | -0.879984 | 1.219925 | -0.216506 | 0.94606 | 2.117812 | 2.874916 | -0.103264 | 0.206198 | 0.154135 | 1.196028 | 0.548602 | -0.973 | 0.626692 |
| 32.09722222 | 0.690592 | 0.766159 | 0.650666 | 1.209386 | 0.482902 | 0.111826 | 0.253161 | 0.462876 | 0.096709 | -0.387951 | 0.208019 | -0.492631 | 1.233866 | -0.07056 | 1.012369 | 2.596266 | 3.136034 | -0.003621 | 0.090189 | -0.527866 | 0.90579 | 0.894229 | -1.457644 | 0.412054 |
| 32.11111111 | 0.583821 | 0.895116 | 0.34449 | 0.80234 | 1.151897 | 0.369679 | 0.427566 | 0.775535 | 0.308389 | 0.098228 | 1.240744 | -0.024901 | 1.22582 | 0.280694 | 1.008412 | 1.996861 | 3.536746 | 0.11482 | 0.149404 | -0.411482 | 1.213523 | 0.310719 | -2.131779 | -0.378005 |
| 32.125 | 0.47198 | 1.033707 | 0.267128 | 0.774747 | 1.603676 | 0.453911 | 0.647926 | 0.870817 | 0.364739 | 0.522148 | 1.254401 | 0.262345 | 1.432029 | 0.735687 | 1.213521 | 2.183191 | 4.391394 | 0.427258 | 0.583387 | -0.174888 | 1.426222 | 1.088163 | -2.468045 | -0.318707 |
| 32.13888889 | 0.483874 | 0.907393 | 0.840534 | 0.947364 | 1.947574 | 0.477579 | 0.598841 | 0.970942 | 0.330065 | 0.801125 | 0.928104 | 0.383874 | 1.488641 | 0.946226 | 1.10824 | 1.387383 | 4.657649 | 0.630029 | 0.718013 | 0.323828 | 0.96025 | 0.9918 | -1.434044 | -0.632868 |
| 32.15277778 | 0.497572 | 0.830766 | 0.866041 | 1.063485 | 2.092272 | 0.552122 | 0.422206 | 1.121137 | 0.305412 | 0.965517 | 0.972945 | 0.37613 | 1.267251 | 0.755808 | 0.496548 | 0.421241 | 3.963057 | 0.479743 | 0.324074 | 0.444549 | 0.42695 | 0.14082 | -0.092399 | 0.240783 |
| 32.16666667 | 0.496239 | 0.936191 | 0.656911 | 1.059995 | 2.16434 | 0.57585 | 0.323946 | 1.251039 | 0.272964 | 0.989291 | 1.049218 | 0.390483 | 0.463038 | 0.977792 | -0.187964 | -0.631774 | 3.669207 | 0.206078 | -0.113878 | 0.442005 | 1.345654 | -1.00972 | 0.073573 | 1.016675 |
| 32.18055556 | 0.479261 | 1.012297 | 0.370994 | 1.08309 | 2.230829 | 0.504452 | 0.380667 | 1.530562 | 0.216323 | 0.918529 | 0.803242 | 0.412334 | -0.281601 | 1.101604 | -0.289911 | -0.630953 | 3.39993 | 0.055321 | 0.010479 | 1.052122 | 1.799145 | -0.544038 | -0.32062 | 0.932059 |
| 32.19444444 | 0.40212 | 0.961713 | 0.542201 | 1.200647 | 2.044102 | 0.439815 | 0.458192 | 1.529985 | 0.170551 | 0.872952 | 0.699692 | 0.35969 | -0.654072 | 0.987219 | 0.073659 | -0.527004 | 2.491084 | -0.280534 | -0.090018 | 1.092648 | 2.495091 | 0.544321 | -0.0882 | 1.063342 |
| 32.20833333 | 0.275341 | 0.954371 | 0.81792 | 1.260544 | 1.668087 | 0.343106 | 0.335221 | 1.249204 | 0.199728 | 0.730683 | 0.835322 | 0.285303 | -0.536014 | 0.646952 | 0.165943 | -0.724489 | 1.54771 | -0.274346 | -0.242442 | 0.538572 | 2.641845 | 1.383183 | 0.750644 | 1.194761 |
| 32.22222222 | 0.206433 | 0.926217 | 0.730412 | 1.146613 | 1.335744 | 0.078828 | 0.048886 | 0.757126 | 0.175878 | 0.36931 | 0.608875 | 0.162456 | -0.303479 | 0.409638 | -0.284525 | -1.100965 | 0.347074 | 0.029731 | -0.362818 | 0.101046 | 1.49743 | 0.710808 | 0.566979 | 0.478186 |
| 32.23611111 | 0.17259 | 0.727379 | 0.347446 | 1.018966 | 1.30292 | 0.010111 | 0.079266 | 0.567362 | 0.076864 | 0.266176 | 0.339463 | 0.011681 | 0.237222 | -0.128726 | -0.809159 | -1.22487 | 0.349695 | 0.421791 | 0.032867 | 0.361493 | 1.23737 | 0.950454 | -0.204618 | 0.490672 |
| 32.25 | 0.307269 | 1.021816 | 0.663413 | 1.583694 | 1.932482 | 0.442387 | 0.457388 | 1.010449 | 0.241272 | 0.827735 | 0.869379 | -0.003684 | 2.227641 | 0.217706 | -0.58013 | -0.694666 | 2.789369 | 1.551725 | 1.304824 | 0.632109 | 1.231725 | 2.678458 | 0.293822 | 0.906032 |
| 32.26388889 | 0.471021 | 1.612141 | 1.007404 | 2.631583 | 2.247085 | 0.806218 | 0.623432 | 1.107969 | 0.542032 | 1.169768 | 1.485359 | -0.021752 | 3.096109 | 0.425067 | -0.074045 | 0.156559 | 3.375122 | 1.756775 | 1.552085 | 0.758423 | 0.831507 | 2.38502 | 1.082975 | 0.831261 |
| 32.27777778 | 0.446335 | 1.54891 | 1.107293 | 2.486126 | 1.869807 | 0.510027 | 0.205835 | 0.628703 | 0.520354 | 0.72423 | 1.071863 | 0.074132 | 1.784447 | 0.539185 | 0.000803 | 0.160994 | 1.283147 | 0.872627 | 0.782041 | 0.43943 | -0.312558 | 1.082147 | 1.116481 | 0.860444 |
| 32.29166667 | 0.365355 | 1.225609 | 0.86798 | 1.707703 | 2.030118 | 0.145363 | -0.038883 | 0.476318 | 0.356804 | 0.436226 | 0.619724 | -0.050535 | 1.542657 | 0.507034 | 0.162398 | 0.636333 | 1.477095 | 0.677484 | 0.748949 | 0.373359 | 0.660474 | 0.764905 | 1.342746 | 0.46154 |
| 32.30555556 | 0.267451 | 1.031243 | 0.675848 | 1.420122 | 2.856662 | 0.13031 | 0.067673 | 0.702478 | 0.275978 | 0.681384 | 0.360055 | -0.130804 | 1.759228 | 0.630189 | 0.970093 | 1.837769 | 2.750106 | 0.878142 | 1.184248 | 0.756778 | 2.19334 | 0.953161 | 1.672525 | 0.544006 |
| 32.31944444 | 0.309671 | 0.903725 | 0.69982 | 1.400553 | 3.348515 | 0.200625 | 0.235809 | 0.822706 | 0.296197 | 0.837911 | 0.399432 | -0.088682 | 1.425004 | 0.761873 | 1.711488 | 2.448137 | 2.628027 | 0.725653 | 1.069333 | 1.058938 | 1.689508 | 1.466428 | 1.76133 | 0.762026 |
| 32.33333333 | 0.476171 | 0.898454 | 0.847181 | 1.294895 | 2.944878 | 0.27232 | 0.31587 | 0.732655 | 0.440636 | 0.875705 | 0.765676 | 0.001747 | 0.437758 | 0.872155 | 1.927076 | 1.525355 | 1.067606 | 0.223059 | 0.627468 | 1.134907 | 1.63159 | 0.417255 | 1.851275 | 1.132341 |
| 32.34722222 | 0.528834 | 1.034321 | 0.976654 | 1.253733 | 2.760148 | 0.35736 | 0.416396 | 0.798473 | 0.595431 | 1.175061 | 1.454211 | 0.140449 | -0.188713 | 0.908786 | 2.031738 | 1.719414 | 0.108119 | -0.213709 | 0.258279 | 1.054716 | 1.928234 | -0.159906 | 2.166151 | 1.433147 |
| 32.36111111 | 0.55464 | 1.416308 | 1.177061 | 1.621687 | 3.397389 | 0.537945 | 0.594242 | 1.140989 | 0.762455 | 1.895793 | 2.375194 | 0.242576 | 0.13886 | 0.922947 | 2.12133 | 1.620636 | 0.348835 | -0.019421 | 0.523518 | 0.911243 | 0.843413 | 0.105061 | 1.93447 | 1.048601 |
| 32.375 | 0.533699 | 1.883434 | 1.366259 | 2.5849 | 4.315523 | 0.69348 | 0.969031 | 1.542387 | 0.908885 | 2.634698 | 3.213051 | 0.284173 | 0.791176 | 0.929523 | 1.874613 | 0.993079 | 1.118449 | 0.548225 | 0.910067 | 0.827373 | -0.822752 | 1.406607 | 1.146662 | 0.727918 |
| 32.38888889 | 0.538008 | 2.286788 | 1.424832 | 3.446112 | 4.769585 | 0.829755 | 1.044489 | 1.578852 | 1.04904 | 2.832254 | 3.647217 | 0.470565 | 1.582049 | 1.225929 | 1.828202 | 0.996403 | 2.257901 | 1.208213 | 1.268662 | 1.33197 | -0.947537 | 2.87921 | 1.386135 | 0.835367 |
| 32.40277778 | 0.634501 | 2.481358 | 1.487792 | 3.374367 | 5.092454 | 1.231065 | 1.175313 | 1.838176 | 1.281252 | 2.852545 | 3.807364 | 0.941653 | 2.524728 | 1.531497 | 2.169097 | 1.346672 | 3.710767 | 2.153524 | 1.867152 | 2.698732 | 0.667548 | 3.891968 | 2.662133 | 1.624073 |
| 32.41666667 | 0.779631 | 2.902532 | 1.597666 | 3.772427 | 5.626598 | 2.046746 | 1.62567 | 1.992249 | 1.450679 | 2.924894 | 4.352447 | 1.365169 | 3.600119 | 2.102703 | 2.765297 | 1.593262 | 5.270955 | 2.735863 | 2.337943 | 2.050735 | 1.322566 | 3.37451 | 3.749089 | 2.33772 |
| 32.43055556 | 0.848695 | 3.102799 | 1.622786 | 5.14353 | 6.232636 | 2.303507 | 1.885069 | 1.645181 | 1.423871 | 3.12865 | 4.19402 | 1.206536 | 4.469897 | 3.049053 | 3.513776 | 1.733157 | 6.63259 | 2.768866 | 2.37313 | 2.308483 | 2.088579 | 3.205494 | 3.734156 | 2.594743 |
| 32.44444444 | 0.683098 | 2.756718 | 1.362716 | 3.679008 | 7.359423 | 2.014176 | 1.682695 | 1.370355 | 1.170609 | 2.882488 | 3.824388 | 1.246481 | 4.980195 | 3.859712 | 3.81288 | 2.412107 | 8.749221 | 3.126508 | 2.515722 | 1.97263 | 3.239192 | 2.248835 | 3.721597 | 2.838595 |
| 32.45833333 | 0.616862 | 2.460543 | 1.288079 | 2.678762 | 6.705032 | 1.767053 | 1.512173 | 1.48341 | 0.997074 | 2.77259 | 3.361159 | 1.503078 | 4.626744 | 4.294152 | 3.857742 | 2.586434 | 9.387057 | 3.148331 | 2.153329 | 2.337902 | 2.955205 | 2.635824 | 4.095831 | 3.705975 |
| 32.47222222 | 0.678528 | 1.769127 | 1.258796 | 2.389639 | 5.561457 | 1.498865 | 1.177921 | 1.936048 | 0.782118 | 2.535814 | 3.150757 | 2.058706 | 3.712404 | 4.305056 | 3.90361 | 2.072501 | 6.754379 | 2.567166 | 1.574062 | 1.32436 | 3.33261 | 2.94721 | 3.149013 | 4.030019 |
| 32.48611111 | 0.783396 | 1.047282 | 1.436431 | 1.856043 | 5.844752 | 1.284326 | 0.881121 | 2.120983 | 0.618883 | 2.304224 | 2.910837 | 2.178071 | 4.164089 | 3.304342 | 3.877914 | 1.67361 | 5.496596 | 2.437298 | 1.650782 | 0.766829 | 3.276837 | 0.900807 | 2.33642 | 3.15 |
| 32.5 | 1.015869 | 0.670045 | 2.00907 | 1.341785 | 6.539355 | 1.502907 | 0.946244 | 1.890953 | 0.484663 | 2.202638 | 2.506097 | 1.961845 | 4.740169 | 3.166215 | 3.631924 | 1.318402 | 5.811932 | 2.856534 | 1.903844 | 1.353892 | 2.871725 | 0.385312 | 1.893611 | 2.284419 |
| 32.51388889 | 1.298802 | 0.494365 | 2.135587 | 0.700155 | 6.203696 | 1.780533 | 1.047758 | 1.429511 | 0.683126 | 2.403526 | 2.029419 | 1.654802 | 4.118862 | 3.294531 | 3.352961 | 1.055022 | 5.741447 | 3.023963 | 1.811134 | 1.59514 | 2.461295 | 0.913493 | 2.413536 | 1.937107 |
| 32.52777778 | 1.387509 | 0.787078 | 1.4418 | 0.899869 | 5.395705 | 1.609223 | 0.699828 | 1.045502 | 0.962016 | 2.376664 | 2.089918 | 1.994143 | 3.103364 | 3.433433 | 3.233424 | 0.658458 | 5.867417 | 2.119928 | 1.092221 | 0.947271 | 2.042042 | 1.319947 | 3.779148 | 2.745953 |
| 32.54166667 | 1.426377 | 0.694885 | 0.919577 | 1.213609 | 4.257305 | 1.359488 | 0.498729 | 0.474756 | 1.104432 | 2.263149 | 2.352279 | 2.279918 | 1.687347 | 3.193099 | 3.486964 | 0.227569 | 6.173208 | 1.691413 | 0.809168 | 0.802897 | 1.627193 | 2.665956 | 4.573004 | 3.446147 |

| | | | | | | | | | | | | | | | | | | | | | | | | |
|---|---|---|---|---|---|---|---|---|---|---|---|---|---|---|---|---|---|---|---|---|---|---|---|---|
| 32.55555556 | 1.331709 | 0.601049 | 0.602708 | 1.174276 | 2.895808 | 1.973908 | -0.179465 | 0.467273 | 1.250797 | 2.477998 | 1.791655 | 2.216029 | 1.101815 | 2.291679 | 3.689667 | -0.117182 | 7.957255 | 1.920462 | 0.264759 | 0.808762 | 1.332049 | 2.783396 | 4.408436 | 3.445366 |
| 32.56944444 | 1.184894 | 0.035338 | 0.441448 | 0.669247 | 1.692279 | 2.118808 | -0.039764 | 0.464376 | 1.065958 | 2.396675 | 1.517567 | 2.136355 | 0.590148 | 1.772326 | 4.149328 | -0.219038 | 7.378179 | 2.304565 | 0.14881 | 0.800814 | 1.024423 | 1.910291 | 4.828753 | 2.853125 |
| 32.58333333 | 1.128411 | -0.1438 | 0.382026 | -0.412677 | 0.712939 | 1.073339 | -0.163185 | 0.154056 | 0.739337 | 1.801317 | 1.602994 | 1.733987 | 0.372189 | 0.958525 | 2.918813 | -0.308615 | 4.54414 | 2.066163 | 0.227258 | 0.743675 | 0.680746 | 1.61132 | 3.347905 | 1.099938 |
| 32.59722222 | 0.703433 | 0.264913 | 0.133513 | -0.883227 | 0.637246 | -0.517118 | -0.642382 | -0.226312 | 0.408641 | 1.133581 | 0.780019 | 0.968627 | -0.461353 | 0.322579 | 0.823888 | -0.41672 | 0.874106 | 1.430842 | 0.093673 | 0.443001 | 0.557482 | 0.564717 | 2.104382 | 0.085765 |
| 32.61111111 | 0.362949 | 1.009366 | -0.308717 | -0.686098 | 0.491627 | -1.153839 | -1.313251 | -0.489133 | 0.12076 | 0.568501 | -0.092037 | 0.121164 | -1.013317 | -0.020645 | -0.732756 | -0.446525 | -1.902076 | 1.260233 | -0.111848 | -0.389938 | 0.020318 | 0.445746 | 1.59553 | -0.123019 |
| 32.625 | 0.287607 | 0.655752 | -0.584067 | -0.319245 | 0.330959 | -0.480174 | -1.636282 | -0.693417 | -0.231542 | 0.46765 | -0.948143 | -0.716623 | -0.586809 | -0.501135 | -1.358824 | -0.486941 | -2.846421 | 0.274235 | -0.308758 | -0.762529 | -0.790953 | 0.533952 | 0.517368 | -0.427506 |
| 32.63888889 | 0.163016 | -0.010438 | -0.615076 | -0.457175 | -0.063006 | -0.15055 | -1.62182 | -0.982152 | -0.540697 | 0.220893 | -1.813699 | -1.384246 | 0.183021 | -1.290739 | -1.313649 | -0.287142 | -2.302263 | -0.483744 | -0.35678 | 0.248674 | -1.224756 | -0.019907 | -1.398866 | -0.916534 |
| 32.65277778 | 0.148757 | -0.231588 | -0.390909 | -0.779422 | -0.096762 | -0.419698 | -1.816748 | -1.493162 | -0.567982 | -0.04237 | -2.023714 | -1.29193 | 0.510435 | -1.112692 | -0.773593 | -0.295476 | -0.990776 | -0.890375 | -0.379544 | -0.173527 | -0.94135 | -0.331029 | -4.016276 | -1.240811 |
| 32.66666667 | 0.120748 | 0.219356 | -0.127094 | -0.680987 | -0.001506 | -0.329373 | -1.2699 | -2.066458 | -0.473825 | -0.884876 | -1.495495 | -1.258084 | 0.400622 | -0.803833 | -0.351259 | -0.252852 | -1.014266 | -0.984876 | -0.396094 | -0.59849 | -1.18481 | -0.248318 | -5.98078 | -1.162893 |
| 32.68055556 | 0.046023 | 0.498943 | 0.018419 | -1.024579 | -0.049182 | -0.397659 | -0.590299 | -2.047929 | -0.234059 | -1.566069 | -0.939891 | -1.354251 | 0.067313 | -0.370866 | 0.036676 | -0.225768 | -1.576108 | -0.434483 | -0.47454 | -1.105516 | -1.635545 | -0.844298 | -5.487442 | -0.732223 |
| 32.69444444 | 0.188647 | 0.358819 | -0.023 | -1.075937 | -0.429174 | -0.367382 | -0.320269 | -1.908583 | 0.099863 | -1.139337 | -0.537346 | -1.441411 | -0.023139 | -0.153691 | -0.030856 | -0.034724 | -2.672416 | 0.133388 | -0.524328 | -1.275467 | -1.858409 | -1.130454 | -3.430419 | -0.391721 |
| 32.70833333 | 0.222802 | 0.020698 | -0.224922 | -0.017095 | -1.208197 | -0.373868 | -0.438435 | -1.763676 | 0.340916 | -0.677088 | -0.787579 | -1.227699 | 0.083176 | 0.30231 | -0.221759 | 0.000071 | -2.733983 | 0.175 | -0.450277 | -1.095859 | -1.201844 | -1.0325 | -2.505679 | -0.352867 |
| 32.72222222 | 0.058963 | -0.062874 | -0.193673 | 0.992689 | -1.159396 | -0.500281 | -0.474711 | -1.279921 | 0.321825 | -0.78234 | -0.310986 | -0.34389 | 0.360992 | 0.767269 | -0.102215 | -0.053429 | -2.58041 | -0.340912 | -0.345118 | -0.87293 | -0.791869 | -0.811514 | -2.441849 | -0.587114 |
| 32.73611111 | -0.010788 | 0.506913 | 0.25904 | 1.171643 | -0.965507 | -0.584704 | -0.356341 | -0.813832 | 0.226028 | -0.876358 | 0.773781 | 0.434394 | 0.039599 | 1.145171 | 0.169681 | -0.197823 | -2.78123 | -0.639662 | -0.307157 | -0.681546 | -0.006433 | -0.802654 | -2.180863 | -0.680788 |
| 32.75 | -0.049785 | 0.774062 | 0.532609 | 0.835269 | -1.236637 | -0.459474 | -0.179162 | -0.503993 | 0.172167 | -0.924657 | 0.680499 | 0.473278 | -0.692806 | 1.386287 | 0.292011 | -0.494994 | -2.996434 | -0.431388 | -0.3788 | -0.768703 | -0.258466 | -0.627926 | -2.047776 | -0.203409 |
| 32.76388889 | 0.121407 | 0.559639 | 0.22255 | 0.390842 | -1.857952 | -0.270208 | 0.031622 | -0.365208 | 0.169409 | -1.011528 | 0.535116 | 0.231605 | -1.079347 | 1.061849 | 0.137326 | -0.59205 | -3.374959 | 0.133402 | -0.453485 | -1.296656 | -0.803932 | -0.703991 | -2.321938 | 0.69289 |
| 32.77777778 | 0.142825 | 0.349632 | -0.298839 | 0.429463 | -2.016527 | -0.213052 | 0.178181 | -0.216839 | 0.100248 | -1.15443 | 0.693001 | 0.484335 | -1.143398 | 0.3968 | 0.188601 | -0.648266 | -2.809259 | 0.52977 | -0.399788 | -1.896153 | -1.074622 | -0.640363 | -3.124272 | -0.260044 |
| 32.79166667 | 0.235114 | 0.178343 | -0.449004 | 0.577808 | -2.946958 | -0.306537 | 0.35545 | -0.157613 | -0.061938 | -1.035866 | 0.889667 | 0.623721 | -1.377586 | 0.509638 | -0.017085 | -0.833828 | -2.219367 | -0.191227 | -0.455884 | -1.965583 | -1.35684 | -0.51455 | -3.179061 | -0.229096 |
| 32.80555556 | 0.215373 | 0.239618 | -0.237935 | 0.173688 | -4.153303 | -0.442442 | 0.394145 | -0.259477 | -0.271817 | -0.728117 | 1.259443 | 0.85532 | -1.621377 | 0.366026 | -0.247492 | -0.890121 | -2.04097 | -1.318501 | -1.048641 | -1.217091 | -0.989728 | -0.591439 | -2.978758 | -0.493622 |
| 32.81944444 | 0.151224 | 0.340336 | -0.116674 | -0.576097 | -4.633763 | -0.522784 | 0.318409 | -0.236318 | -0.318915 | -0.36825 | 1.501843 | 1.14977 | -1.445417 | -0.178622 | -0.504522 | -1.029944 | -1.751651 | -1.858707 | -1.261357 | -0.834939 | -0.453614 | -0.852894 | -2.6563 | -1.140504 |
| 32.83333333 | 0.101764 | 0.09151 | -0.2695 | -1.009363 | -4.75314 | -0.499549 | 0.077765 | -0.130824 | -0.279623 | -0.276131 | 0.634709 | 0.608149 | -1.70847 | -0.221066 | -0.55861 | -1.249882 | -1.632887 | -2.081049 | -1.412485 | -1.316615 | -0.735962 | -0.801637 | -1.882625 | -1.551536 |
| 32.84722222 | 0.004542 | -0.299104 | -0.562347 | -1.218599 | -5.115357 | -0.487826 | -0.212608 | -0.223097 | -0.218539 | -0.459303 | -0.901104 | -0.430853 | -2.669564 | -0.298961 | -0.48342 | -1.241988 | -2.409151 | -2.343762 | -1.929554 | -0.945632 | -1.53052 | -0.737421 | -2.475809 | -1.550021 |
| 32.86111111 | -0.257217 | -0.650844 | -0.71162 | -1.17853 | -5.556689 | -0.635779 | -0.416799 | -0.647781 | -0.28272 | -0.747861 | -1.314332 | -0.528108 | -3.693888 | -0.057401 | -0.519734 | -1.250163 | -3.161144 | -2.574094 | -2.468484 | -1.70744 | -2.198768 | -1.031641 | -2.417828 | -1.329507 |
| 32.875 | -0.555303 | -0.970546 | -0.970217 | -1.406094 | -6.103538 | -0.877333 | -0.6888 | -0.955426 | -0.375443 | -1.299036 | -1.733244 | -0.861836 | -3.945506 | 0.005989 | -0.781261 | -1.734487 | -3.720491 | -3.163016 | -2.819138 | -2.498419 | -3.404492 | -1.022822 | -2.256769 | -1.319504 |
| 32.88888889 | -0.623684 | -1.349188 | -1.066661 | -1.84608 | -6.200769 | -1.211446 | -0.876283 | -1.179355 | -0.520639 | -2.197814 | -2.318249 | -1.16542 | -4.218475 | -1.005762 | -0.816912 | -2.601198 | -4.824594 | -3.820453 | -2.794327 | -3.064531 | -3.735354 | -0.761029 | -2.927871 | -1.537819 |
| 32.90277778 | -0.545884 | -1.430075 | -1.112187 | -1.769734 | -5.599409 | -1.609608 | -0.848523 | -1.380766 | -0.575796 | -2.805096 | -2.37637 | -1.116547 | -4.533968 | -1.578941 | -1.259136 | -2.750683 | -5.810361 | -3.986512 | -2.546832 | -3.238795 | -4.040183 | -0.57281 | -3.59428 | -1.668756 |
| 32.91666667 | -0.525002 | -1.962401 | -1.200107 | -2.21107 | -5.582356 | -1.682551 | -0.856582 | -1.464149 | -0.594479 | -2.606935 | -2.575532 | -0.995823 | -5.022247 | -1.841417 | -1.79633 | -3.13418 | -5.540502 | -3.472538 | -2.514863 | -3.672138 | -3.530068 | -0.886216 | -3.386127 | -1.314769 |
| 32.93055556 | -0.534254 | -1.973747 | -1.339352 | -2.351892 | -6.218089 | -1.790674 | -0.982824 | -2.193117 | -0.537772 | -2.378 | -3.697643 | -1.539979 | -6.413436 | -1.9102 | -1.732779 | -3.729608 | -5.833331 | -3.124522 | -2.318348 | -3.179435 | -1.908539 | -1.022904 | -3.264982 | -2.193085 |
| 32.94444444 | -0.597275 | -1.620681 | -1.177052 | -2.578224 | -6.304664 | -1.88598 | -1.218065 | -2.874884 | -0.548439 | -2.215669 | -3.893536 | -1.446117 | -6.771705 | -2.00659 | -1.273474 | -4.014825 | -6.437688 | -3.020411 | -2.423201 | -2.205491 | -1.64567 | -0.690896 | -3.709881 | -2.589338 |
| 32.95833333 | -0.823228 | -2.456251 | -1.075738 | -3.107124 | -6.272348 | -2.513054 | -1.45409 | -3.149238 | -1.062692 | -2.774587 | -4.28444 | -1.827 | -6.713686 | -1.498669 | -1.285514 | -4.457067 | -5.463831 | -3.346553 | -2.648021 | -2.617569 | -1.377836 | -0.355257 | -1.614228 | -2.10183 |
| 32.97222222 | -1.221023 | -3.210391 | -1.751572 | -3.149212 | -7.465629 | -3.318218 | -1.544883 | -2.857302 | -1.587495 | -3.656854 | -5.879651 | -2.883568 | -5.677916 | -1.886779 | -1.433543 | -1.783607 | -3.212998 | -3.921195 | -2.800662 | -3.333951 | -1.026018 | -0.780242 | -1.849265 | -4.156836 |
| 32.98611111 | -1.415116 | -3.149078 | -2.160166 | -1.349537 | -9.307826 | -3.098884 | -1.03902 | -3.418862 | -1.673332 | -3.412806 | -6.071809 | -2.624967 | -5.213592 | -1.520363 | -1.542462 | -1.06191 | -1.299708 | -4.528798 | -2.888271 | -3.32872 | -0.481324 | -1.457694 | -3.322228 | -4.459094 |
| 33 | -0.729107 | -1.464558 | -1.571283 | 0.300321 | -7.216622 | -2.16463 | -0.313326 | -2.781038 | -1.012226 | -2.223803 | -3.975971 | -2.022577 | -5.947326 | -1.588593 | -2.265629 | -2.620775 | 0.03442 | -3.248873 | -1.876492 | -3.180164 | -0.988599 | -1.986652 | -2.788427 | -1.437181 |
| 33.01388889 | -0.084 | 0.1474 | -0.85922 | 1.134841 | -5.218813 | -1.285408 | -0.131365 | -2.563288 | -0.503683 | -1.425667 | -1.563417 | -0.990017 | -3.674227 | -0.568601 | -2.364978 | -3.148054 | 1.474055 | -0.946353 | -0.728979 | -1.795056 | -0.369587 | -2.187335 | -2.207752 | -2.29908 |
| 33.02777778 | -0.028771 | 1.004255 | -0.359777 | 1.79993 | -4.017282 | -0.839306 | -0.211269 | -2.373432 | -0.316859 | -1.111045 | 0.020817 | -0.576144 | -1.739659 | -0.941197 | -2.524522 | -3.448371 | 2.45018 | 0.031339 | -0.221541 | -1.135858 | -0.743301 | -3.520672 | -1.690001 | -2.754186 |
| 33.04166667 | -0.155897 | 1.573798 | -0.113748 | 2.89907 | -3.466761 | -0.76377 | -0.404254 | -1.893451 | -0.254753 | -0.875248 | 0.927936 | -0.549716 | -1.664682 | -0.900799 | -1.491034 | -2.515893 | 2.088211 | -0.604666 | -0.614511 | -0.300571 | -1.077476 | -3.004003 | -1.284461 | -2.467365 |
| 33.05555556 | -0.054893 | 1.65837 | -0.002049 | 3.18903 | -3.027063 | -0.544756 | -0.361084 | -1.017514 | -0.230944 | -0.580792 | 1.389283 | -0.492348 | -2.209929 | -0.072525 | 0.339003 | -0.170729 | 0.748487 | -1.559673 | -0.620337 | 0.844791 | -0.155241 | -1.457259 | -0.641531 | -1.729756 |
| 33.06944444 | 0.328768 | 1.723322 | 0.193674 | 2.799319 | -2.054499 | -0.224847 | -0.19041 | -0.149132 | 0.016808 | -0.340358 | 2.018796 | -0.232087 | -0.65502 | 0.52767 | 1.279037 | 2.291776 | 0.301578 | -0.340324 | 0.605897 | 1.528405 | 0.686742 | 1.661659 | -0.057767 | -0.891991 |
| 33.08333333 | 0.702128 | 1.489232 | 0.479312 | 3.094217 | -0.584703 | 0.079779 | 0.053751 | 0.202338 | 0.195679 | -0.169322 | 2.184274 | -0.031795 | 2.595931 | 0.871862 | 1.166558 | 3.251968 | 2.139359 | 1.154529 | 1.375334 | 1.261385 | 0.731093 | 3.766954 | 0.097569 | -0.59748 |
| 33.09722222 | 0.854933 | 1.41731 | 0.601552 | 3.927638 | 0.722659 | 0.29989 | 0.25662 | 0.51485 | 0.278055 | 0.031546 | 2.494154 | 0.061852 | 3.833907 | 0.930134 | 1.392374 | 3.217077 | 3.775759 | 1.689737 | 1.269794 | 1.481717 | 0.46975 | 3.634534 | -0.077561 | -0.058928 |
| 33.11111111 | 0.681995 | 1.431224 | 0.522535 | 4.036869 | 1.100606 | 0.449321 | 0.346795 | 1.223995 | 0.344926 | 0.177593 | 2.607264 | 0.069385 | 3.091411 | 0.552387 | 1.664557 | 2.720328 | 3.949944 | 1.005219 | 0.786738 | 3.138014 | 0.170733 | 3.251043 | -0.450883 | -0.160145 |
| 33.125 | 0.713682 | 1.383262 | 0.378107 | 3.713118 | 1.156855 | 0.594799 | 0.385695 | 2.102898 | 0.466358 | 0.292159 | 2.137384 | -0.040063 | 2.288157 | 0.576716 | 1.79933 | 2.582469 | 3.704581 | 0.724104 | 0.597012 | 4.034879 | 0.53023 | 2.836955 | -0.758069 | -0.130919 |
| 33.13888889 | 0.727945 | 1.438857 | 0.340989 | 3.436497 | 1.601049 | 0.710087 | 0.419857 | 2.313361 | 0.585108 | 0.371466 | 1.817148 | -0.18527 | 1.997646 | 0.670494 | 1.504863 | 2.544252 | 3.056821 | 0.596862 | 0.564404 | 2.971586 | 1.172871 | 2.720778 | -0.91121 | 0.208776 |
| 33.15277778 | 0.654774 | 1.632183 | 0.346724 | 3.104853 | 2.305343 | 0.711297 | 0.311702 | 2.091005 | 0.655975 | 0.414472 | 1.819576 | -0.182285 | 1.654981 | 0.671829 | 1.048275 | 1.836902 | 1.579582 | -0.005845 | -0.209383 | 1.758286 | 1.601631 | 2.550033 | -0.672774 | 0.427257 |
| 33.16666667 | 0.532071 | 1.749156 | 0.215316 | 1.887826 | 2.439487 | 0.491373 | 0.120461 | 1.895734 | 0.5968 | 0.322283 | 1.798684 | -0.197635 | 0.709878 | 0.674717 | 0.806104 | 0.962734 | 0.114113 | -0.739686 | -0.734986 | 1.432079 | 1.531249 | 1.491755 | -0.029576 | 0.76096 |
| 33.18055556 | 0.441535 | 1.538019 | 0.088943 | 0.673585 | 2.046972 | 0.244057 | 0.08578 | 1.905072 | 0.472119 | 0.219276 | 1.356069 | -0.276188 | -0.213361 | 0.588611 | 0.446961 | 0.271349 | -0.581444 | -0.779615 | -0.575948 | 1.999481 | 1.142048 | 1.492482 | 0.413089 | 0.72252 |
| 33.19444444 | 0.450898 | 1.194159 | 0.102506 | 0.342709 | 1.820897 | 0.124098 | 0.080578 | 1.97122 | 0.411001 | 0.186297 | 0.830949 | -0.26962 | -0.787601 | 0.280842 | 0.092311 | -0.027491 | 0.031805 | -0.123169 | 0.027893 | 2.674622 | 1.182699 | 1.761967 | 0.170559 | 0.126521 |
| 33.20833333 | 0.431424 | 1.098655 | 0.185565 | 0.717874 | 2.083872 | 0.073053 | 0.037915 | 1.520247 | 0.358916 | 0.153865 | 0.826873 | -0.195339 | -1.067852 | -0.105212 | -0.145535 | -0.303183 | 1.059284 | 0.275893 | 0.183171 | 2.350554 | 1.555835 | 1.471828 | -0.113902 | -0.231812 |
| 33.22222222 | 0.381224 | 1.063476 | 0.189604 | 0.890334 | 2.385255 | 0.023537 | 0.005193 | 1.197567 | 0.272145 | 0.068594 | 1.007164 | -0.231406 | -1.20737 | -0.283906 | -0.327007 | -0.798481 | 1.23733 | -0.1525 | -0.449651 | 1.574351 | 1.713922 | 0.691971 | -0.38496 | -0.328108 |
| 33.23611111 | 0.354313 | 0.969531 | 0.142172 | 0.670572 | 2.299418 | -0.020272 | 0.014958 | 1.286271 | 0.178135 | -0.001204 | 0.974938 | -0.275419 | -1.40732 | -0.292888 | -0.299178 | -0.506757 | -0.094963 | -0.250076 | -0.223783 | 1.399771 | 1.697115 | 1.133727 | -0.524556 | -0.376681 |
| 33.25 | 0.343216 | 0.976576 | 0.156006 | 0.926361 | 1.982737 | -0.075497 | 0.008767 | 1.139406 | 0.190144 | -0.058637 | 0.965122 | -0.221537 | -1.525598 | -0.046731 | 0.28596 | -0.119848 | -1.393098 | 0.74196 | 0.648836 | 2.095699 | 1.856606 | 1.959558 | -0.506621 | 0.04253 |
| 33.26388889 | 0.335145 | 1.2018 | 0.238659 | 1.956593 | 1.885803 | -0.073877 | 0.048557 | 0.932689 | 0.288644 | -0.010813 | 1.350593 | -0.04488 | -1.003352 | 0.469502 | 1.187402 | 0.515733 | -0.842694 | 1.630208 | 1.325083 | 2.994614 | 1.936119 | 1.77548 | 0.001624 | 1.104387 |
| 33.27777778 | 0.386724 | 1.547135 | 0.336192 | 3.144455 | 2.221778 | 0.093037 | 0.179055 | 1.129345 | 0.380088 | 0.139158 | 2.108864 | 0.145654 | -0.184651 | 0.999283 | 1.856499 | 0.887152 | 0.538407 | 0.746429 | 0.834353 | 2.314913 | 1.877755 | 1.72748 | 0.562149 | 1.870539 |
| 33.29166667 | 0.555999 | 1.773728 | 0.387758 | 3.188555 | 2.783476 | 0.306075 | 0.302408 | 1.517291 | 0.464258 | 0.247632 | 2.820855 | 0.307727 | 0.182737 | 1.197961 | 1.770286 | 1.336585 | 1.479404 | -0.015969 | 0.397692 | 1.628344 | 1.780702 | 1.444721 | 0.754448 | 1.677525 |
| 33.30555556 | 0.788435 | 1.709542 | 0.39821 | 2.720217 | 3.192561 | 0.422708 | 0.350962 | 1.651318 | 0.590994 | 0.254619 | 2.63824 | 0.487521 | 0.346518 | 0.996494 | 1.606073 | 1.355229 | 1.988943 | 0.35436 | 1.079557 | 2.113606 | 2.360845 | 1.640278 | 0.595123 | 1.070918 |
| 33.31944444 | 0.879231 | 1.813693 | 0.38035 | 2.677186 | 3.529452 | 0.516719 | 0.425296 | 1.844927 | 0.725656 | 0.387797 | 2.468674 | 0.780398 | 1.230936 | 0.96503 | 1.891367 | 1.723711 | 2.64141 | 1.150543 | 2.312064 | 3.152151 | 2.576755 | 1.538987 | 0.751652 | 0.993416 |
| 33.33333333 | 0.818987 | 2.292825 | 0.407447 | 2.66982 | 3.827865 | 0.661001 | 0.570984 | 2.310211 | 0.756897 | 0.414146 | 3.147206 | 1.290542 | 2.389418 | 1.342924 | 2.4232 | 2.117701 | 2.450605 | 1.401962 | 3.048652 | 3.906723 | 2.433034 | -0.232403 | 1.141546 | 1.641781 |
| 33.34722222 | 0.703127 | 2.828035 | 0.459044 | 2.968026 | 3.763386 | 0.793968 | 0.658856 | 2.73799 | 0.741343 | 0.423303 | 4.040215 | 1.701777 | 2.806841 | 1.56732 | 2.599005 | 2.273512 | 1.51687 | 1.007947 | 2.929134 | 3.293918 | 3.676065 | -0.494575 | 1.338712 | 2.08549 |
| 33.36111111 | 0.698438 | 3.079834 | 0.59767 | 3.571292 | 3.549007 | 0.754026 | 0.708222 | 2.440971 | 0.809802 | 0.37388 | 4.290539 | 1.860365 | 2.244554 | 1.479311 | 2.427119 | 2.521185 | 0.283829 | 1.099682 | 2.785043 | 3.282658 | 4.127094 | 0.580487 | 1.302687 | 1.849419 |
| 33.375 | 0.753517 | 2.866709 | 0.861835 | 3.345638 | 3.509853 | 0.679933 | 0.863305 | 2.609763 | 0.880457 | 0.380418 | 3.838005 | 1.744733 | 2.075736 | 1.588336 | 2.543657 | 2.983586 | 0.230524 | 1.679944 | 2.663825 | 4.001548 | 4.75324 | 1.451913 | 1.615269 | 1.951087 |
| 33.38888889 | 0.767626 | 2.651392 | 1.075732 | 3.247412 | 3.936138 | 0.684 | 1.094079 | 2.686488 | 0.919695 | 0.658597 | 3.571306 | 2.372986 | 2.650385 | 2.047833 | 3.024525 | 3.60014 | 1.224343 | 2.169772 | 2.468672 | 4.189663 | 4.22927 | 1.169423 | 1.941165 | 2.403054 |
| 33.40277778 | 0.805302 | 2.474043 | 1.018634 | 3.507116 | 4.477877 | 0.884662 | 1.406921 | 2.479366 | 0.99953 | 1.25966 | 3.659748 | 2.684958 | 2.982001 | 2.401966 | 3.25824 | 3.770568 | 2.604767 | 2.030098 | 2.708983 | 3.139707 | 3.900427 | 2.068964 | 1.931109 | 2.540161 |
| 33.41666667 | 0.930348 | 2.316876 | 0.995772 | 3.608595 | 4.72459 | 0.992601 | 1.25493 | 2.229706 | 0.892678 | 1.62191 | 4.039467 | 2.845053 | 3.348372 | 2.575738 | 3.269213 | 3.42568 | 4.114392 | 1.420442 | 2.507349 | 3.429648 | 3.456541 | 2.332208 | 1.999219 | 2.73524 |
| 33.43055556 | 0.984877 | 2.294888 | 1.118329 | 3.373661 | 4.909913 | 1.133256 | 0.87791 | 1.772963 | 0.618883 | 1.503611 | 3.747403 | 2.640153 | 3.828907 | 3.070305 | 3.20808 | 2.876363 | 5.596688 | 1.928545 | 2.670517 | 4.548891 | 3.010437 | 1.26913 | 2.250282 | 3.35056 |
| 33.44444444 | 0.931681 | 2.609953 | 1.288638 | 2.880987 | 5.139456 | 1.431082 | 0.539154 | 1.551047 | 0.572033 | 1.617804 | 3.418484 | 2.710726 | 4.848981 | 4.031476 | 3.315983 | 1.748414 | 6.117762 | 2.996869 | 3.035719 | 4.94812 | 1.809872 | 0.788478 | 2.579907 | 4.037883 |
| 33.45833333 | 0.956396 | 2.932807 | 1.394254 | 2.386297 | 5.898456 | 1.824495 | 0.848261 | 2.369436 | 0.652775 | 1.908515 | 3.640161 | 3.281251 | 4.761344 | 4.199231 | 2.858784 | 1.557554 | 5.493914 | 2.970871 | 3.215113 | 4.436949 | 2.225467 | 0.721172 | 2.584734 | 3.965195 |
| 33.47222222 | 1.051186 | 2.374522 | 1.409088 | 1.835564 | 5.118188 | 2.082879 | 1.272605 | 1.78462 | 0.78172 | 2.012391 | 2.518927 | 2.719936 | 3.574956 | 2.391857 | 2.801224 | 2.426402 | 3.674673 | 2.515259 | 3.067791 | 3.504838 | 3.026373 | 1.657371 | 2.401799 | 3.116361 |
| 33.48611111 | 1.220561 | 1.113341 | 1.38214 | 1.179558 | 4.89257 | 2.184346 | 1.507437 | 1.279246 | 0.895068 | 2.390542 | 1.455626 | 1.855153 | 3.051379 | 1.221818 | 3.42562 | 1.985155 | 3.29686 | 2.506302 | 3.34173 | 2.890436 | 3.520307 | 0.994307 | 2.710181 | 2.550233 |
| 33.5 | 1.310065 | 0.602908 | 1.358235 | 0.918488 | 6.63035 | 2.230034 | 1.504263 | 1.261079 | 0.996028 | 2.976912 | 1.429992 | 1.992112 | 3.231884 | 1.637026 | 4.017574 | 2.015067 | 3.673199 | 2.869877 | 3.769813 | 3.178023 | 1.980877 | 1.517745 | 2.775925 | 2.746245 |
| 33.51388889 | 0.847007 | 0.826736 | 1.32986 | 0.837033 | 8.723776 | 2.300495 | 1.116143 | 1.066296 | 0.974524 | 3.077829 | 2.332294 | 2.740113 | 2.809597 | 2.201315 | 4.276983 | 2.366082 | 2.805227 | 3.421299 | 3.405453 | 2.858759 | 1.051582 | 0.618486 | 2.301368 | 3.033919 |
| 33.52777778 | 0.68472 | 1.127579 | 1.113815 | 1.020772 | 7.727664 | 2.594177 | 0.964434 | 1.08323 | 0.876197 | 2.839214 | 2.774692 | 2.781557 | 2.769793 | 3.164132 | 3.998585 | 1.605653 | 1.216579 | 3.941998 | 1.885095 | 1.611284 | 2.265521 | -0.50954 | 2.252115 | 2.214154 |
| 33.54166667 | 0.403136 | 1.258693 | 0.956764 | 1.313432 | 5.826561 | 3.068496 | 0.852858 | 1.504374 | 0.924011 | 2.941773 | 2.41455 | 2.442322 | 3.773301 | 3.849551 | 4.152344 | 1.54053 | 1.034884 | 3.338988 | 2.395931 | -1.294165 | 3.165432 | -0.798815 | 2.313299 | 2.184582 |
| 33.55555556 | 0.226982 | 2.57322 | 0.872462 | 1.851921 | 4.19557 | 2.969916 | 1.007357 | 1.369916 | 1.027852 | 3.117197 | 1.913259 | 2.178567 | 4.346046 | 3.588004 | 4.067574 | 1.136134 | 1.876894 | 3.191755 | 2.558031 | -1.806268 | 3.195216 | -0.492827 | 2.061545 | 2.967194 |
| 33.56944444 | 0.290152 | 3.46569 | 0.738019 | 2.132972 | 3.779184 | 2.078035 | 0.219942 | 0.823496 | 1.005311 | 2.680006 | 1.620877 | 2.054907 | 3.925901 | 1.049321 | 4.993722 | 0.75428 | 2.151745 | 2.623417 | 1.817603 | -0.965931 | 2.56402 | -0.096689 | 1.777121 | 1.850388 |
| 33.58333333 | 0.386859 | 2.101689 | 0.534887 | 1.643548 | 3.7238 | 1.555077 | -0.510672 | 0.408368 | 0.873781 | 1.982461 | 1.604298 | 2.144828 | 3.847131 | -1.280414 | 5.017691 | 0.900873 | 1.315656 | 1.354464 | 0.829258 | -0.194753 | 2.656331 | 0.105597 | 1.363331 | 1.100027 |
| 33.59722222 | 0.144092 | 0.385717 | 0.3625 | 0.632341 | 3.561507 | 0.832498 | -0.252735 | 0.058965 | 0.611082 | 1.307805 | 1.367757 | 2.30136 | 2.993584 | -0.926608 | 4.587979 | 0.926908 | 0.176557 | -0.495417 | -0.850545 | 0.051571 | 1.937251 | -0.485823 | 0.767014 | 1.775658 |
| 33.61111111 | -0.055886 | -0.507169 | 0.120389 | -0.526299 | 3.463388 | -0.074672 | 0.014547 | -0.099854 | 0.325933 | 0.515093 | 0.73638 | 1.759675 | 1.193175 | 0.140428 | 2.90129 | 0.254034 | -0.646963 | -2.38079 | -1.900085 | -0.377268 | -0.065906 | -0.672299 | 0.114681 | 0.598277 |
| 33.625 | -0.191547 | -0.37516 | -0.118053 | -1.223577 | 2.254779 | -0.606108 | 0.016045 | -0.155586 | 0.24757 | -0.115552 | 0.098268 | 0.7402 | -0.85507 | 0.36613 | 0.41874 | -0.074158 | -1.325326 | -4.285061 | -2.071981 | -0.783635 | -0.669901 | -0.855357 | -0.533881 | 0.134215 |
| 33.63888889 | -0.222371 | 0.287708 | -0.351071 | -0.827767 | -0.164709 | -0.604799 | -0.563021 | -0.253299 | 0.210353 | -0.490451 | -0.775819 | -0.482445 | -1.997776 | 0.760356 | -0.674391 | -0.179103 | -1.643826 | -4.997813 | -3.654215 | -0.936635 | -0.698057 | -1.145582 | -0.90051 | -0.168779 |
| 33.65277778 | -0.039784 | 0.685378 | -0.413202 | 0.044884 | -2.189072 | -0.366188 | -0.948361 | -0.407496 | 0.091972 | -0.540923 | -1.217287 | -1.572116 | -1.683622 | 1.316176 | 0.25959 | -0.150015 | -1.007883 | -3.451685 | -3.424698 | -0.848004 | -1.620952 | -0.895431 | -1.103142 | 0.056229 |
| 33.66666667 | 0.062098 | 0.565066 | -0.303514 | 0.284333 | -2.852085 | 0.13886 | -0.001872 | -0.44262 | 0.120983 | -0.497856 | -1.2512 | -2.222591 | -0.886409 | 1.076277 | 1.133346 | -0.216266 | -0.759263 | -2.268327 | -3.461986 | -0.707917 | -3.03601 | -0.674966 | -1.356221 | 0.502049 |
| 33.68055556 | -0.038703 | -0.059185 | -0.377162 | -0.653887 | -3.212333 | 0.259583 | 1.074467 | -0.555616 | 0.283359 | -0.796185 | -1.251217 | -2.101223 | -0.410002 | 0.011734 | 1.345024 | -0.048174 | -0.956679 | -1.224501 | -2.620761 | -0.062966 | -1.753053 | -0.656601 | -1.547875 | 0.613773 |
| 33.69444444 | -0.059574 | -0.605736 | -0.205191 | -1.736394 | -3.297456 | 0.293395 | 0.627646 | -0.468006 | 0.352771 | -1.156219 | -0.723525 | -0.876893 | 0.043728 | -1.930255 | 0.946905 | -0.228102 | -1.004468 | -0.491094 | -1.72865 | 0.295267 | -1.405353 | -1.029046 | -1.17148 | 0.619225 |
| 33.70833333 | 0.032692 | -0.272465 | 0.076811 | -1.181808 | -3.556038 | 0.460562 | -0.173167 | -0.393181 | 0.443889 | -1.07658 | 0.245759 | 0.542122 | 0.005915 | -1.663073 | 0.26423 | -0.600346 | 0.353799 | -0.59214 | -1.155736 | 0.53041 | -1.398982 | -1.205222 | -0.998444 | 0.369639 |
| 33.72222222 | 0.200479 | 0.276258 | 0.254692 | -0.428316 | -3.397022 | 0.123878 | -1.022599 | -0.310238 | 0.434859 | -0.586472 | 0.541433 | 0.815152 | 0.106096 | -1.64947 | -1.013092 | -1.042145 | 1.077175 | -1.028596 | -1.585882 | 0.445076 | -0.841846 | -1.48069 | -1.35302 | -0.386943 |
| 33.73611111 | 0.159714 | 0.225016 | 0.260324 | 0.13332 | -3.474116 | -0.182236 | -1.215997 | -0.448649 | 0.464985 | -0.794306 | -0.075536 | -0.04603 | -0.197785 | -2.079834 | -2.231152 | -1.857401 | 0.315842 | -1.190296 | -2.131799 | -0.195552 | -0.728886 | -1.921055 | -1.986122 | -0.973536 |
| 33.75 | 0.28226 | -0.110082 | 0.138604 | -0.609821 | -3.891894 | -0.335997 | -1.047204 | -0.373877 | 0.423226 | -0.817409 | -0.416138 | -0.527389 | -0.935199 | -2.434592 | -2.777608 | -2.351192 | -0.849314 | -1.212936 | -2.067596 | -0.364436 | -0.213296 | -1.781381 | -2.051122 | -1.767409 |
| 33.76388889 | 0.521144 | -0.12135 | 0.053072 | -2.040209 | -3.530642 | -0.485029 | -0.605047 | -0.203203 | 0.313203 | -0.830099 | -0.037465 | -0.132979 | -2.235994 | -3.328019 | -3.323151 | -2.395317 | -1.476687 | -1.47436 | -1.471469 | -0.134349 | 0.6376 | -0.918625 | -2.108625 | -1.895862 |
| 33.77777778 | 0.633943 | 0.013718 | 0.020454 | -2.211303 | -3.538523 | -0.436027 | -0.026871 | -0.038416 | 0.269762 | -1.122045 | 0.654961 | 0.85663 | -3.599643 | -3.767041 | -3.124285 | -1.883068 | -0.707335 | -1.673462 | -1.241957 | 0.121412 | 0.842637 | -0.175504 | -2.085056 | -2.46357 |
| 33.79166667 | 0.40313 | -0.161933 | -0.015998 | -1.414012 | -3.771228 | -0.122578 | 0.183778 | -0.156683 | 0.226514 | -0.813833 | 1.086255 | 1.450403 | -3.887929 | -4.635602 | -3.346644 | -1.401594 | -0.451257 | -1.317223 | -1.270101 | -0.29519 | 0.237984 | -0.202219 | -2.001281 | -3.328785 |
| 33.80555556 | 0.23102 | -0.394702 | -0.111311 | -1.073048 | -4.271055 | 0.092151 | 0.655632 | -0.147854 | 0.216036 | -0.162419 | 0.604587 | 0.553773 | -3.203239 | -4.68247 | -2.564646 | -0.890886 | -0.687187 | -0.81371 | -1.178764 | -1.306954 | -0.45634 | -0.37024 | -1.917641 | -3.520973 |
| 33.81944444 | 0.353611 | -0.300569 | -0.073091 | -1.478457 | -4.334428 | 0.225765 | 1.10263 | 0.101712 | 0.258409 | 0.29015 | -0.086562 | -0.678599 | -2.837046 | -3.268188 | -2.355017 | -0.771985 | -1.098795 | -0.56078 | -1.459271 | -1.13251 | -0.825655 | -0.553595 | -1.76087 | -2.586498 |
| 33.83333333 | 0.471826 | -0.791963 | -0.029446 | -2.288091 | -4.46469 | 0.047639 | 1.321442 | 0.042807 | 0.20144 | 0.157962 | -0.708014 | -1.709602 | -3.157081 | -2.513764 | -2.994365 | -1.133158 | -1.09896 | -0.858567 | -1.898427 | -0.906589 | -0.367214 | -0.972669 | -1.85537 | -2.103445 |
| 33.84722222 | 0.282591 | -1.286902 | -0.157215 | -2.530469 | -4.545593 | -0.413907 | 1.019398 | -0.314592 | -0.030906 | -0.290919 | -0.927826 | -2.295046 | -3.912083 | -2.386242 | -2.563537 | -0.540003 | -1.425093 | -1.371642 | -2.041428 | -1.97733 | 0.583782 | -1.017444 | -2.408787 | -2.073297 |
| 33.86111111 | -0.035967 | -1.456444 | -0.363668 | -2.534424 | -4.667681 | -0.649918 | 0.320819 | -0.422945 | -0.215661 | -0.429503 | -1.090539 | -2.637474 | -4.068194 | -2.579318 | -2.946608 | -0.6632 | -1.589708 | -2.073498 | -1.971498 | -2.809344 | -0.195357 | -0.609592 | -2.54738 | -2.719453 |
| 33.875 | -0.177565 | -1.393431 | -0.540905 | -2.343428 | -4.215015 | -0.701161 | -0.305676 | -0.639229 | -0.156847 | -0.318073 | -1.368815 | -2.583542 | -3.812551 | -3.231959 | -3.567894 | -1.039882 | -1.952275 | -1.976145 | -1.730889 | -2.922436 | -0.501188 | -0.168694 | -2.38342 | -3.508351 |
| 33.88888889 | -0.361313 | -1.402741 | -0.783724 | -2.472304 | -4.164443 | -0.708146 | -0.317087 | -0.447953 | -0.218737 | -0.923956 | -1.424584 | -2.571625 | -3.333681 | -3.764558 | -3.649072 | -1.724306 | -2.647911 | -0.9499 | -1.849087 | -2.956532 | -0.267594 | -0.247541 | -2.627914 | -4.200662 |
| 33.90277778 | -0.652575 | -1.733208 | -1.045953 | -3.114161 | -4.518275 | -0.908502 | -0.381874 | -0.626609 | -0.416556 | -1.236577 | -1.802667 | -2.553365 | -2.998424 | -4.902787 | -4.039798 | -2.007366 | -3.51238 | -1.335353 | -2.438659 | -3.476978 | -1.276215 | -0.184924 | -2.913284 | -5.059835 |
| 33.91666667 | -0.948296 | -2.39989 | -1.293211 | -3.822122 | -5.061187 | -1.169056 | -0.784137 | -1.24117 | -0.623221 | -1.869224 | -2.152525 | -2.637721 | -3.222318 | -4.638695 | -3.87657 | -2.03253 | -3.584784 | -2.179013 | -3.101751 | -4.920809 | -1.712832 | -0.678437 | -2.469904 | -5.340066 |
| 33.93055556 | -0.881712 | -2.688831 | -1.487216 | -4.020388 | -6.527588 | -1.299954 | -1.356164 | -1.914225 | -0.732492 | -2.600872 | -2.863622 | -3.219882 | -2.544568 | -5.698715 | -3.728437 | -2.049966 | -3.4638 | -3.362616 | -3.306539 | -7.036102 | -2.364717 | -1.468136 | -2.171349 | -5.546118 |
| 33.94444444 | -0.583928 | -2.987204 | -1.67823 | -4.977165 | -7.555459 | -1.541551 | -1.631971 | -2.6458 | -0.800921 | -2.876635 | -3.111928 | -3.884375 | -3.048405 | -5.325199 | -3.660101 | -2.118824 | -4.684408 | -5.266831 | -4.442608 | -6.462476 | -4.181393 | -1.726282 | -2.403318 | -5.449184 |
| 33.95833333 | -0.579833 | -3.305004 | -1.505817 | -4.84477 | -8.039201 | -2.006097 | -1.67284 | -2.638637 | -1.012402 | -3.73601 | -3.755159 | -4.617886 | -3.19072 | -4.222157 | -3.52124 | -2.046482 | -3.488373 | -5.38293 | -5.163118 | -4.611131 | -8.836453 | -1.683137 | -3.31125 | -8.369276 |
| 33.97222222 | -1.061914 | -3.24256 | -1.665701 | -4.84022 | -9.884341 | -2.711221 | -1.878226 | -3.898653 | -1.619659 | -3.75706 | -4.428525 | -4.986917 | -4.696411 | -6.676393 | -3.153001 | -1.432706 | -3.562973 | -5.243052 | -6.135086 | -1.198463 | -2.772163 | -2.260797 | -4.76112 | 0.278319 |
| 33.98611111 | -0.923396 | -2.410441 | -1.704623 | -4.672844 | -9.099357 | -2.95374 | -1.66092 | -3.773327 | -1.529801 | -3.606219 | -4.182835 | -4.838432 | -4.66452 | -5.265819 | -2.483781 | -1.477314 | -3.869156 | -2.378666 | -4.116998 | -2.440142 | -2.788991 | -0.809108 | -4.638873 | -5.508522 |
| 34 | -0.490924 | -1.304904 | -1.275924 | -2.912769 | -8.159258 | -2.386201 | -0.646434 | -3.783478 | -1.006358 | -2.76659 | -3.103227 | -3.681552 | -2.893043 | -5.658307 | -2.472182 | -1.611093 | -2.477042 | -2.084128 | -3.25747 | -3.099627 | -3.566119 | -0.961571 | -4.226936 | -8.585076 |
| 34.01388889 | -0.234635 | -0.408072 | -0.838409 | -0.88925 | -6.208624 | -1.680385 | -0.234652 | -3.552302 | -0.474803 | -2.280293 | -2.092611 | -2.520023 | -2.564351 | -3.384943 | -1.88238 | -0.769015 | -1.276868 | -0.532047 | -2.302319 | -3.822537 | -3.302522 | -2.20816 | -3.478773 | -6.086795 |
| 34.02777778 | -0.1486 | 0.15479 | -0.550761 | 0.124657 | -4.648731 | -1.337762 | -0.196409 | -2.651467 | -0.292911 | -1.884562 | -1.496597 | -1.800103 | -1.882515 | -3.77739 | -2.426883 | -1.746507 | -0.245442 | 0.425563 | -1.151169 | -4.299028 | -2.223437 | -1.469442 | -2.853563 | -5.546947 |
| 34.04166667 | -0.121545 | 0.222325 | -0.372973 | 0.532543 | -3.576374 | -1.108573 | -0.164194 | -1.791549 | -0.120828 | -1.290474 | -0.88826 | -1.196502 | -1.268089 | -3.618799 | -2.976646 | -1.767987 | -0.448181 | -0.817884 | -1.956542 | -4.00719 | -1.975886 | -1.882053 | -2.294272 | -6.422787 |
| 34.05555556 | -0.028565 | 0.551453 | -0.308609 | 0.401094 | -1.961201 | -0.937667 | 0.068041 | -0.931126 | 0.069063 | -0.730284 | -0.411008 | -0.732219 | -1.197511 | -4.879949 | -2.125753 | -1.559028 | -0.61692 | -2.237861 | -2.395881 | -1.47387 | -2.271656 | -2.169264 | -1.554353 | -6.127671 |
| 34.06944444 | 0.088426 | 0.963239 | -0.267929 | 0.306311 | -0.058481 | -0.420692 | 0.244027 | -0.08576 | 0.165189 | -0.405495 | -0.028922 | -0.250059 | -0.366651 | -4.427041 | -1.768705 | -1.328814 | 0.627245 | -0.818162 | -0.492321 | 1.831278 | -2.562064 | -0.816559 | -1.99066 | -4.78003 |
| 34.08333333 | 0.123292 | 0.915396 | -0.088753 | 0.770142 | 1.432704 | -0.196008 | 0.357142 | 0.64816 | 0.249217 | -0.237797 | 0.386247 | 0.404248 | 1.073633 | -4.631594 | -1.855912 | -1.120704 | 1.940131 | 1.395886 | 2.398322 | 3.827188 | -0.51754 | 0.317414 | -2.218803 | -4.958551 |
| 34.09722222 | 0.210962 | 0.791856 | 0.211741 | 1.783922 | 2.643566 | 0.066057 | 0.523577 | 1.500446 | 0.413465 | 0.092252 | 0.991027 | 1.13933 | 2.798 | -5.015507 | -1.508358 | -0.745014 | 1.567187 | 2.611344 | 3.870772 | 3.427136 | 0.380567 | 0.750985 | -2.14734 | -4.974469 |
| 34.11111111 | 0.34837 | 0.764161 | 0.411908 | 3.045251 | 3.298697 | 0.440906 | 0.751244 | 2.256072 | 0.579548 | 0.406318 | 1.330661 | 1.653389 | 3.707471 | -4.327972 | -0.580856 | -0.144293 | 1.440124 | 2.949072 | 3.346238 | 3.816915 | 0.662671 | 0.794696 | -1.683589 | -3.866289 |
| 34.125 | 0.374827 | 0.70798 | 0.404208 | 3.590695 | 3.455093 | 0.72182 | 0.934174 | 2.405917 | 0.610545 | 0.439711 | 1.419019 | 1.710528 | 3.308794 | -3.440101 | -0.118269 | 0.270534 | 1.392838 | 3.206704 | 3.354231 | 5.908677 | 1.648765 | 1.496783 | -1.282097 | -2.597365 |
| 34.13888889 | 0.407234 | 0.491817 | 0.344317 | 2.932748 | 3.133296 | 0.692452 | 0.854174 | 2.104982 | 0.617697 | 0.372111 | 1.127049 | 1.697584 | 2.0622 | -2.868597 | -0.057211 | 0.37702 | 0.344567 | 3.392028 | 2.850479 | 6.352448 | 3.145803 | 1.86762 | -1.116963 | -2.078612 |
| 34.15277778 | 0.5868 | 0.622929 | 0.38753 | 2.605376 | 3.1804 | 0.562488 | 0.867588 | 2.129966 | 0.639878 | 0.540663 | 0.947799 | 1.338661 | 1.393693 | -2.275163 | -0.168636 | 0.335179 | 0.274471 | 2.168893 | 1.323393 | 3.445701 | 3.434394 | 1.308206 | -1.087526 | -1.276538 |
| 34.16666667 | 1.070667 | 1.079627 | 0.174517 | 3.348712 | 3.059285 | 0.36025 | 1.737812 | 3.643494 | 0.715415 | 2.485787 | 1.109847 | 0.729019 | 0.486953 | -2.578889 | -0.677976 | -0.27896 | 0.567204 | -0.19642 | -0.586939 | 0.271338 | 2.183379 | 0.04971 | -1.33422 | -1.623822 |
| 34.18055556 | 1.365465 | 2.182404 | -0.19216 | 4.248498 | 2.126676 | -0.585857 | 4.33403 | 8.206315 | 0.826801 | 3.733012 | 1.255651 | -0.034189 | -0.962323 | -3.961559 | -1.386422 | -1.047709 | -0.895207 | -1.495345 | -2.082318 | -1.285651 | 0.828887 | -2.345068 | -1.717334 | -2.685191 |
| 34.19444444 | 0.571007 | 0.834167 | 0.152653 | 2.981111 | 6.326823 | -0.690135 | 0.842819 | 3.067142 | 0.421385 | 0.860731 | -0.014817 | -0.228692 | -0.574659 | -3.811465 | -1.031777 | -0.820125 | -1.2524 | 0.916 | -0.24176 | 2.159924 | 1.539558 | -1.790351 | -1.590883 | -2.93308 |
| 34.20833333 | -0.195479 | -1.347469 | 0.805835 | 1.377792 | 3.32611 | 0.407643 | -1.615377 | -3.496357 | -0.463339 | -2.794281 | 0.210774 | 1.056965 | 0.807 | -2.383274 | -0.166692 | 0.019542 | 0.685299 | 2.149011 | 1.723885 | 6.424564 | 2.301538 | 0.408297 | -1.171701 | -2.249127 |
| 34.22222222 | 0.03372 | 0.898617 | 1.109175 | 2.645612 | 0.523233 | 0.462669 | -0.437589 | -1.003957 | 0.03116 | -1.872239 | -0.115103 | 0.47519 | 0.674918 | -1.951182 | -0.105108 | -0.117486 | 1.36546 | 0.834464 | 0.516301 | 2.477138 | 1.882242 | 0.679765 | -1.099983 | -1.692799 |
| 34.23611111 | 0.206511 | 1.922585 | 0.88043 | 3.77338 | 1.775587 | 0.12967 | 0.30578 | 0.904999 | 0.441602 | -0.456711 | 0.099648 | 0.300408 | 0.174892 | -1.012778 | -0.18305 | -0.425886 | 0.817296 | -2.136871 | -1.924856 | -1.206203 | 0.998852 | 0.130196 | -0.851217 | -0.95496 |
| 34.25 | 0.353468 | 1.914144 | 0.677993 | 4.048937 | 2.809171 | -0.075858 | 0.560856 | 1.53993 | 0.423988 | -0.193184 | -0.060778 | 0.075336 | -0.017333 | -0.493124 | -0.01171 | -0.12031 | 0.918702 | -1.491942 | -1.39361 | -0.713639 | 1.369506 | 0.106393 | -0.301994 | -0.490166 |
| 34.26388889 | 0.455046 | 1.885283 | 0.632732 | 3.928475 | 3.213954 | 0.037703 | 0.832866 | 1.852847 | 0.478228 | 0.087342 | 0.266711 | 0.43889 | 0.193249 | -0.401612 | 0.127173 | 0.102662 | 1.136893 | 1.099642 | 0.988467 | 3.24826 | 3.271554 | 1.516945 | -0.038224 | -0.428083 |
| 34.27777778 | 0.479991 | 1.689157 | 0.590751 | 3.779612 | 2.610526 | 0.280475 | 0.89567 | 1.508068 | 0.567985 | 0.1344 | 0.6055 | 0.867056 | 0.491188 | -0.017394 | 0.134637 | -0.03996 | 0.824048 | 2.047706 | 2.173713 | 4.582565 | 4.240769 | 2.40242 | 0.057665 | -0.243551 |
| 34.29166667 | 0.520515 | 1.285339 | 0.390486 | 3.248236 | 2.563964 | 0.359869 | 0.68854 | 1.21393 | 0.540965 | 0.093621 | 0.498895 | 0.67526 | 0.317442 | 0.105813 | -0.088998 | -0.187418 | 0.28937 | 0.136331 | 0.934935 | 2.483389 | 4.077099 | 2.304426 | 0.035222 | -0.077544 |
| 34.30555556 | 0.736859 | 0.991546 | 0.250637 | 3.011973 | 3.207059 | 0.201815 | 0.606295 | 1.454315 | 0.473128 | 0.376227 | 0.188835 | 0.16451 | 0.01227 | -0.248964 | -0.135986 | -0.281386 | -0.802769 | -0.937383 | -0.030845 | 1.313788 | 2.903095 | 1.722045 | 0.026849 | -0.095756 |
| 34.31944444 | 0.912392 | 0.979167 | 0.202656 | 2.736048 | 3.482 | 0.067501 | 0.740891 | 2.00865 | 0.496758 | 0.808117 | 0.138006 | -0.005445 | -0.251522 | 0.050943 | 0.221236 | 0.118967 | -1.532986 | 0.516436 | 0.993734 | 2.119794 | 2.410323 | 2.09693 | 0.136849 | 0.242983 |
| 34.33333333 | 0.835207 | 1.010055 | 0.231148 | 2.675655 | 3.064667 | 0.168592 | 0.97461 | 2.283175 | 0.57459 | 0.782812 | 0.6094 | 0.640956 | 0.341463 | 1.5739 | 1.024487 | 1.063699 | -0.682554 | 2.517358 | 2.44483 | 4.245067 | 2.120974 | 2.11561 | 0.328496 | 1.182838 |
| 34.34722222 | 0.769031 | 1.174464 | 0.353407 | 2.697011 | 2.783121 | 0.543051 | 0.992684 | 2.234101 | 0.685471 | 0.688819 | 1.637367 | 2.282062 | 1.457783 | 3.228734 | 1.674744 | 1.852645 | 1.000826 | 2.353232 | 2.483387 | 6.461753 | 1.791308 | 2.206981 | 0.542468 | 2.179469 |
| 34.36111111 | 0.973708 | 1.500058 | 0.60983 | 3.159942 | 3.185571 | 1.015485 | 0.908159 | 2.403748 | 0.720038 | 0.655268 | 2.452764 | 3.1752 | 2.025132 | 3.932527 | 1.863171 | 2.170203 | 2.725028 | 2.092497 | 2.539958 | 7.544494 | 2.407138 | 3.210013 | 0.645213 | 2.537419 |
| 34.375 | 1.246312 | 1.879709 | 0.776383 | 3.616919 | 3.922629 | 1.27389 | 0.744861 | 2.071339 | 0.701418 | 0.784402 | 2.474969 | 3.458809 | 2.025561 | 3.65763 | 0.862917 | 1.375474 | 3.21458 | 3.003901 | 3.060805 | 7.443786 | 3.172486 | 3.896575 | 0.472015 | 2.169675 |
| 34.38888889 | 1.393255 | 2.204894 | 0.971584 | 3.703522 | 4.725108 | 1.414348 | 0.953899 | 2.152158 | 0.86065 | 1.182304 | 2.723589 | 3.798544 | 2.157812 | 2.88144 | -0.106063 | 0.629078 | 2.925711 | 4.170611 | 3.608284 | 6.949009 | 3.856152 | 3.468946 | 0.331117 | 1.949458 |
| 34.40277778 | 1.302449 | 2.214425 | 1.208078 | 3.654975 | 5.29336 | 1.593918 | 1.230052 | 2.508191 | 1.046256 | 1.310414 | 3.25635 | 3.950124 | 2.691668 | 3.105002 | 0.659617 | 0.531488 | 3.060357 | 3.401005 | 3.316328 | 8.242389 | 5.106128 | 4.038204 | 0.555796 | 2.526374 |
| 34.41666667 | 1.464512 | 1.978264 | 1.452551 | 3.758393 | 5.409185 | 1.638184 | 1.380809 | 2.658904 | 1.173094 | 1.497682 | 3.308593 | 3.950389 | 2.648624 | 3.501007 | 3.033188 | 1.800291 | 2.945364 | 2.784403 | 3.59089 | 8.795615 | 3.133699 | 3.347464 | 1.040636 | 3.079811 |
| 34.43055556 | 2.018348 | 1.84806 | 1.403824 | 3.065979 | 5.762306 | 1.662639 | 1.508555 | 2.826099 | 1.16468 | 1.47825 | 3.066022 | 3.71482 | 2.5342 | 4.537757 | 3.875353 | 2.407243 | 2.89516 | 2.162982 | 3.281883 | 7.11414 | 1.603737 | 2.869475 | 1.509174 | 3.51366 |
| 34.44444444 | 2.325086 | 2.252877 | 1.122398 | 3.232412 | 6.015979 | 1.948538 | 1.529354 | 2.335446 | 1.000337 | 1.38705 | 2.714447 | 3.941604 | 3.164635 | 6.752308 | 5.103179 | 3.099527 | 3.440526 | 2.49811 | 4.447185 | 3.809298 | 1.434465 | 1.71477 | 2.046609 | 5.16954 |
| 34.45833333 | 2.178289 | 2.521048 | 1.542446 | 3.331387 | 6.127433 | 2.543643 | 1.529502 | 2.133022 | 0.889164 | 1.756835 | 2.577285 | 3.126912 | 3.341814 | 8.112977 | 5.442115 | 2.881098 | 4.256333 | 3.156241 | 4.134513 | 4.155327 | 7.73252 | 0.754967 | 2.120691 | 6.464304 |
| 34.47222222 | 1.846898 | 2.374533 | 2.044259 | 2.66482 | 5.106422 | 2.483621 | 1.463758 | 2.007287 | 0.905078 | 2.230479 | 2.110829 | 2.541669 | 2.506563 | 7.153579 | 4.769063 | 2.798781 | 4.448375 | 2.777466 | 3.990529 | 4.827905 | 5.592858 | 0.907083 | 1.994457 | 6.074947 |
| 34.48611111 | 1.465088 | 1.67324 | 1.57387 | 1.959605 | 3.972748 | 1.867031 | 1.547788 | 1.998525 | 0.82781 | 2.073999 | 1.524046 | 1.930924 | 2.044396 | 6.921449 | 5.119782 | 3.140736 | 4.218961 | 2.946164 | 3.565617 | 3.964081 | 4.60765 | 2.1074 | 2.237561 | 5.591016 |
| 34.5 | 1.510787 | 1.197091 | 1.132507 | 1.398747 | 3.742146 | 1.410211 | 1.367409 | 2.139775 | 0.870946 | 2.113854 | 1.382598 | 1.646288 | 2.136837 | 7.087128 | 6.28319 | 3.754698 | 4.222534 | 2.809093 | 2.761629 | 3.232347 | 4.676267 | 2.224736 | 2.495331 | 5.680412 |
| 34.51388889 | 1.846695 | 1.634027 | 0.970863 | 1.741491 | 4.333906 | 1.627433 | 1.014738 | 2.416599 | 0.892569 | 1.997485 | 1.952473 | 1.631777 | 2.916553 | 7.479991 | 6.647491 | 3.84443 | 3.506349 | 2.855484 | 2.549768 | 3.318603 | 4.888646 | 2.531648 | 2.714202 | 6.406491 |
| 34.52777778 | 2.236533 | 1.699119 | 1.053577 | 1.546453 | 4.658192 | 1.873594 | 0.637365 | 2.256879 | 0.679913 | 1.694557 | 2.195723 | 1.794781 | 3.248228 | 7.556499 | 5.299975 | 4.144021 | 3.007299 | 3.05067 | 2.479392 | 3.765253 | 2.730216 | 2.863846 | 2.433561 | 6.283638 |
| 34.54166667 | 2.243861 | 1.110821 | 1.161947 | 1.225275 | 4.412428 | 1.781919 | 0.811919 | 1.690642 | 0.699463 | 1.893023 | 1.479546 | 0.701072 | 2.814411 | 6.964574 | 3.944915 | 3.294793 | 2.66389 | 2.549556 | 1.291353 | 3.826199 | 0.752648 | 3.65173 | 2.04541 | 5.761474 |
| 34.55555556 | 1.702995 | 0.530394 | 1.069685 | 0.361004 | 3.718168 | 1.44484 | 0.512545 | 1.382442 | 0.494641 | 1.956203 | 1.375028 | 0.502585 | 2.401875 | 5.456964 | 3.236841 | 2.830187 | 2.640301 | 1.737157 | 0.071732 | 3.319579 | 0.367137 | 2.590322 | 2.041221 | 5.302202 |
| 34.56944444 | 1.217893 | 0.022303 | 0.857138 | -0.44524 | 2.836822 | 1.10518 | -0.015299 | 1.189079 | 0.451356 | 2.146614 | 1.0572 | 0.213061 | 2.122547 | 3.027396 | 2.943887 | 2.59115 | 2.021454 | 1.410434 | 0.416531 | 2.942 | -0.245237 | 1.53036 | 1.826966 | 3.021915 |
| 34.58333333 | 0.590017 | -0.21354 | 0.536617 | -0.178353 | 1.821499 | 0.675721 | -0.290061 | 0.99806 | 0.709136 | 1.886209 | 0.786439 | 0.190984 | 1.942791 | 1.934444 | 2.737204 | 2.127393 | 0.583237 | -1.240097 | -0.055725 | 2.335378 | 0.135814 | 1.778707 | 1.418571 | 2.146158 |
| 34.59722222 | 0.027458 | -0.585793 | 0.231589 | 0.682767 | 0.732841 | 0.289261 | -0.147245 | 0.406858 | 0.80657 | 1.183724 | 0.632612 | 0.437742 | 1.610347 | 1.573935 | 2.179312 | 1.701461 | 0.099389 | -2.718464 | -0.821953 | 1.27875 | -0.128432 | 2.034683 | 0.912343 | 1.792082 |
| 34.61111111 | -0.202121 | -1.089972 | -0.021582 | 0.943593 | 0.056881 | -0.030177 | 0.128753 | -0.081782 | 0.69565 | 0.638947 | 0.390948 | 0.159238 | 1.019284 | 1.611928 | 1.181278 | 0.520165 | -0.002179 | -2.526812 | -1.423443 | 0.086471 | -0.368938 | 0.894988 | 0.342196 | 1.642158 |
| 34.625 | -0.223805 | -1.309517 | -0.075601 | -0.350596 | -0.410591 | -0.425338 | 0.110288 | -0.442547 | 0.396915 | 0.085521 | -0.070562 | -0.218065 | -0.284991 | 0.822472 | -0.408185 | -1.299323 | -0.121609 | -3.129908 | -1.649809 | -0.994591 | -0.600987 | -0.670398 | -0.413679 | 0.702392 |
| 34.63888889 | -0.254664 | -0.999112 | -0.003221 | -1.280968 | -1.551071 | -0.628736 | 0.000811 | -0.47008 | 0.171336 | -0.318478 | -0.2738 | -0.198397 | -1.642727 | 0.15197 | -1.99275 | -2.270649 | -1.662447 | -2.399824 | -2.039962 | -2.550434 | -0.591953 | -1.940031 | -0.905325 | -0.218338 |
| 34.65277778 | -0.435392 | -0.178736 | -0.09779 | -1.306073 | -2.871026 | -0.43236 | -0.100646 | -0.56769 | 0.039431 | -0.659021 | -0.456845 | -0.031423 | -1.820468 | 0.466253 | -1.084121 | -1.163821 | -3.285083 | -0.878607 | -1.781931 | -3.433437 | 0.165029 | -1.348446 | -1.058085 | -0.223242 |
| 34.66666667 | -0.493367 | 0.813229 | -0.317565 | -0.817907 | -3.321616 | -0.083981 | -0.388486 | -0.700581 | 0.01924 | -0.919637 | -0.489783 | 0.035592 | -1.618652 | 0.827382 | 0.491205 | -0.342357 | -2.993068 | 1.09395 | -1.568894 | -2.837125 | -0.313385 | -1.520492 | -0.949369 | 0.275093 |
| 34.68055556 | -0.32078 | 1.746245 | -0.523566 | 0.143508 | -3.292489 | 0.066021 | -0.559699 | -0.870994 | 0.037307 | -0.663127 | -0.504962 | -0.169157 | -0.821401 | 1.110439 | 1.129301 | 0.038086 | -4.073312 | 1.757687 | -0.080744 | -2.4312 | 1.343815 | -1.306204 | -1.057674 | 0.479276 |
| 34.69444444 | -0.227054 | 1.550376 | -0.52832 | 0.482344 | -3.109693 | -0.135745 | -0.620479 | -0.930513 | 0.161341 | -0.926057 | -0.42985 | -0.301617 | -0.364742 | 0.930157 | 0.797374 | -0.19531 | -6.515439 | 1.201291 | 0.454575 | -2.171723 | 0.12531 | -1.867659 | -1.142298 | 0.400833 |
| 34.70833333 | -0.421053 | 0.247412 | -0.467454 | -0.906433 | -2.858241 | -0.441744 | -0.38745 | -0.873026 | 0.053293 | -1.665661 | -0.52205 | -0.257894 | -0.412055 | 0.713227 | 0.602171 | -0.210097 | -6.674286 | 0.386032 | -0.008887 | -2.096384 | 0.301884 | -1.931771 | -1.044891 | 0.231103 |
| 34.72222222 | -0.816107 | -0.560255 | -0.36611 | -2.005922 | -2.827568 | -0.533579 | -0.090813 | -0.653151 | -0.02464 | -1.686241 | -0.569935 | -0.322399 | -0.824659 | 0.540644 | 1.279472 | 0.609539 | -4.681794 | 0.045691 | -0.783308 | -1.756499 | 0.794481 | -1.824055 | -0.644643 | 0.020668 |
| 34.73611111 | -1.070651 | -0.411961 | -0.242579 | -1.919518 | -3.041564 | -0.322748 | 0.035044 | -0.582482 | 0.030529 | -1.477309 | -0.416961 | -0.439193 | -0.822026 | -0.000977 | 2.61443 | 1.685249 | -1.455005 | -0.318906 | -1.422482 | -1.751775 | 1.505321 | -1.459999 | -0.428291 | -0.452377 |
| 34.75 | -0.885437 | 0.323494 | -0.173247 | -0.717964 | -3.443652 | -0.267255 | -0.120694 | -0.730877 | -0.015402 | -1.265693 | -0.736602 | -0.888293 | -0.454856 | -1.288772 | 2.333402 | 1.454736 | -0.355321 | -0.635813 | -1.724914 | -1.856962 | 0.021154 | -1.59185 | -0.895154 | -1.471019 |
| 34.76388889 | -0.415291 | 0.824665 | -0.233386 | 0.342904 | -3.063016 | -0.310746 | -0.405044 | -0.826192 | -0.060838 | -1.13395 | -1.16006 | -1.316888 | -0.046624 | -1.248045 | 0.727734 | 0.692772 | -0.788202 | -0.777026 | -1.906061 | -2.158241 | -0.50651 | -1.436594 | -1.101137 | -1.620742 |
| 34.77777778 | -0.052592 | 0.013868 | -0.29644 | -0.118346 | -2.224893 | -0.250881 | -0.468266 | -0.657069 | -0.034941 | -1.084012 | -0.987092 | -1.176547 | 0.070001 | -0.083691 | 0.344044 | 0.375072 | -0.457215 | -1.184842 | -1.880778 | -1.70957 | 0.065816 | -1.578893 | -0.574259 | -0.684941 |
| 34.79166667 | 0.036509 | -1.320237 | -0.301361 | -1.538006 | -1.293653 | -0.159217 | -0.246412 | -0.555424 | 0.103599 | -1.189773 | -0.303629 | -0.348125 | 0.131994 | 0.226738 | 0.661299 | 0.462921 | -0.196213 | -1.05141 | -1.542538 | -1.108566 | 0.989346 | -1.101514 | -0.193862 | -0.239607 |
| 34.80555556 | 0.188515 | -1.292673 | -0.225405 | -2.179026 | -1.241676 | -0.094061 | -0.259656 | -0.97073 | 0.230702 | -1.290584 | 0.402603 | 0.578244 | -0.03468 | -0.468 | 0.857129 | 0.689133 | -0.283901 | -1.01074 | -1.254324 | -0.931702 | 1.05466 | -0.634476 | -0.047983 | -0.711601 |
| 34.81944444 | 0.524101 | -0.612691 | -0.190112 | -2.043416 | -1.190186 | -0.082755 | -0.40798 | -1.343418 | 0.294167 | -0.591592 | 0.413644 | 0.433315 | -0.62599 | -1.134039 | 0.212764 | 0.291432 | -1.50919 | -1.207601 | -1.474045 | -1.542615 | 0.655439 | -0.966843 | -0.444685 | -1.352753 |
| 34.83333333 | 0.808411 | -0.496114 | -0.306303 | -2.021887 | -1.189903 | -0.150559 | -0.531233 | -1.466514 | 0.233277 | -0.697348 | -0.284779 | -0.289064 | -1.902588 | -2.013386 | -1.282128 | -1.19549 | -2.846617 | -1.610633 | -2.087496 | -2.513745 | -0.058317 | -1.704214 | -0.948361 | -2.233948 |
| 34.84722222 | 0.551648 | -1.15302 | -0.476408 | -2.305428 | -1.483813 | -0.265062 | -0.470583 | -1.587719 | 0.109908 | -1.088401 | -0.723201 | -0.786694 | -2.079521 | -3.229084 | -2.210723 | -1.978896 | -3.681586 | -1.512296 | -2.527033 | -2.997444 | -1.508185 | -1.177868 | -1.148976 | -3.168822 |
| 34.86111111 | -0.101684 | -1.601764 | -0.525746 | -2.781211 | -1.536169 | -0.364678 | -0.521373 | -1.528364 | 0.010151 | -0.888933 | -0.676094 | -0.466351 | -2.291633 | -3.79817 | -1.80584 | -1.255218 | -4.211991 | -1.705108 | -2.212484 | -2.71946 | -1.502032 | -0.954183 | -1.115292 | -3.720827 |
| 34.875 | -0.720321 | -1.601562 | -0.523872 | -3.301825 | -1.690315 | -0.443363 | -0.534554 | -1.780183 | -0.018504 | -0.73651 | -0.369265 | 0.123596 | -2.268944 | -4.854553 | -1.873768 | -1.200799 | -3.650468 | -1.717914 | -1.790578 | -2.734113 | -1.021238 | -1.316943 | -1.146002 | -4.172987 |
| 34.88888889 | -1.046359 | -1.53651 | -0.608314 | -3.416177 | -1.984586 | -0.49203 | -0.723515 | -2.045338 | -0.062803 | -0.781134 | -0.53746 | -0.022542 | -1.790677 | -7.742834 | -3.450601 | -1.569597 | -2.787032 | -1.93991 | -1.686141 | -3.863307 | -1.350129 | -1.482704 | -1.167837 | -5.831458 |
| 34.90277778 | -1.074651 | -1.970337 | -0.814045 | -3.455579 | -2.349979 | -0.417429 | -0.972495 | -2.357082 | -0.350067 | -1.227819 | -0.957327 | -0.878932 | -1.500105 | -5.963172 | -4.496054 | -1.597343 | -2.003041 | -2.472079 | -1.969343 | -5.673024 | -2.096475 | -1.476355 | -1.364575 | -5.467393 |
| 34.91666667 | -1.340702 | -1.999603 | -0.930785 | -3.566128 | -3.082019 | -0.265749 | -1.101856 | -2.68389 | -0.691259 | -1.865022 | -1.424007 | -1.766871 | -1.637658 | -5.565428 | -4.446113 | -1.962323 | -2.11681 | -2.997978 | -2.40926 | -6.496801 | -2.438165 | -1.894 | -0.931845 | -4.885063 |
| 34.93055556 | -1.66591 | -2.000176 | -0.969308 | -3.701255 | -3.519405 | -0.524933 | -1.051753 | -2.637386 | -0.73938 | -2.069745 | -1.53198 | -1.817078 | -2.489524 | -6.58574 | -4.989637 | -2.205657 | -2.228381 | -3.832755 | -2.680401 | -6.677438 | -3.740979 | -2.714455 | -0.903709 | -5.017183 |
| 34.94444444 | -1.610687 | -2.081842 | -1.0781 | -4.01439 | -3.585186 | -1.177192 | -0.984591 | -2.693981 | -0.592076 | -1.835996 | -1.255418 | -1.855158 | -4.182745 | -7.410007 | -4.89573 | -2.038659 | -3.062288 | -4.053493 | -3.296289 | -3.729801 | -3.380833 | -4.032846 | -1.206778 | -5.660868 |
| 34.95833333 | -1.499384 | -2.418032 | -1.396351 | -4.668061 | -4.379806 | -2.293697 | -1.35493 | -3.498768 | -0.736295 | -2.07457 | -1.575343 | -2.94553 | -6.933522 | -10.133508 | -4.854019 | -2.322507 | -4.96788 | -2.698999 | -4.257327 | 1.147478 | -3.267244 | -3.520116 | -2.880215 | -5.984064 |
| 34.97222222 | -1.303953 | -2.812599 | -1.281253 | -5.285099 | -4.754551 | -3.395141 | -1.537323 | -4.372604 | -1.363582 | -3.314065 | -2.688323 | -4.098479 | -8.125654 | -10.814257 | -4.562657 | -2.105222 | -9.631442 | -2.21662 | -5.715018 | -1.578175 | -3.583455 | -2.218245 | -5.092134 | -2.740003 |

| | | | | | | | | | | | | | | | | | | | | | | | | |
|---|---|---|---|---|---|---|---|---|---|---|---|---|---|---|---|---|---|---|---|---|---|---|---|---|
| 34.98611111 | -0.852577 | -2.322183 | -0.891037 | -4.905697 | -5.537662 | -2.508559 | -1.109496 | -4.301286 | -1.670825 | -4.20867 | -2.394453 | -3.678376 | -6.186029 | -5.393536 | -3.633964 | -2.408835 | -6.512489 | -2.807595 | -4.676551 | -4.908941 | -4.197471 | 0.07314 | -3.817532 | -4.024909 |
| 35 | -0.21342 | -1.425702 | -0.676086 | -2.906264 | -3.751973 | -1.109277 | -0.350983 | -3.750786 | -1.284551 | -4.420873 | -1.943992 | -1.887381 | -4.581815 | -4.459539 | -3.468726 | -3.331242 | -3.569306 | -1.695093 | -3.021188 | -3.048501 | -4.173209 | -0.35893 | -2.641326 | -4.228048 |
| 35.01388889 | 0.200261 | -0.64773 | -0.445547 | -1.662714 | -2.609898 | -0.416614 | -0.141322 | -4.165492 | -1.085349 | -4.184015 | -1.703032 | -1.516239 | -2.297145 | -4.551862 | -2.10884 | -2.704348 | -1.930404 | -0.46978 | -1.040654 | -1.75513 | -4.620944 | -1.070113 | -1.990294 | -3.878599 |
| 35.02777778 | 0.22163 | -0.024269 | -0.252969 | -0.713802 | -2.101298 | -0.001167 | -0.083024 | -4.273745 | -0.702648 | -3.316498 | -0.988484 | -1.065553 | -0.859203 | -4.289524 | -2.334404 | -2.150964 | -0.558952 | 0.54556 | 0.59947 | -2.254506 | -6.115336 | -1.266771 | -1.908895 | -3.778941 |
| 35.04166667 | 0.279041 | 0.260989 | -0.132151 | 0.041165 | -1.84065 | 0.578253 | -0.161004 | -4.157236 | -0.14606 | -2.199111 | -0.570248 | -0.84517 | -0.236053 | -4.416166 | -1.992074 | -1.847454 | -0.115757 | 1.828837 | 1.914044 | -2.388727 | -4.86689 | -0.836512 | -1.937786 | -3.646189 |
| 35.05555556 | 0.302274 | 0.197311 | -0.034565 | 0.353562 | -1.052125 | 0.779248 | -0.119613 | -3.401919 | 0.127245 | -1.225516 | -0.359767 | -0.734601 | 0.141654 | -4.434178 | -1.460105 | -0.871407 | 0.375641 | 2.093562 | 2.293922 | -1.099244 | -1.956709 | -0.738658 | -1.760995 | -3.636849 |
| 35.06944444 | 0.283371 | 0.025433 | 0.054722 | 0.392085 | 0.192028 | 0.431464 | -0.084675 | -2.127896 | 0.285255 | -0.631431 | -0.130127 | -0.455102 | 0.709815 | -4.345767 | -1.042498 | 0.198924 | 0.709996 | 1.681518 | 1.962683 | 0.3644 | -0.457167 | -0.419501 | -1.555826 | -3.390081 |
| 35.08333333 | 0.403454 | 0.054964 | 0.12791 | 0.552834 | 1.140306 | 0.303169 | 0.0981 | -1.410699 | 0.401927 | -0.259732 | 0.35398 | 0.237195 | 1.754729 | -3.192588 | -0.734846 | 0.531318 | 1.164229 | 2.243864 | 2.069951 | 1.334144 | 0.294892 | 0.201001 | -1.360847 | -2.308151 |
| 35.09722222 | 0.651634 | 0.171345 | 0.199661 | 1.10485 | 1.582588 | 0.469905 | 0.401785 | -0.777369 | 0.560459 | 0.130827 | 0.805359 | 1.007746 | 2.724654 | -1.562016 | -0.842299 | 0.354713 | 1.865239 | 4.160942 | 3.08201 | 1.942389 | 0.809088 | 0.759948 | -1.131557 | -1.118852 |
| 35.11111111 | 0.775863 | 0.462695 | 0.333287 | 1.721103 | 1.801523 | 0.631674 | 0.67907 | 0.16938 | 0.757116 | 0.534836 | 1.059967 | 1.549129 | 3.249538 | -1.312583 | -0.662311 | 0.311821 | 2.200733 | 5.874895 | 3.196898 | 1.913813 | 1.134148 | 1.176972 | -1.144219 | -0.986342 |
| 35.125 | 0.638079 | 0.486256 | 0.397013 | 2.402796 | 2.173819 | 0.67286 | 0.805663 | 1.231528 | 0.845916 | 0.703859 | 1.036268 | 1.816565 | 3.022617 | -2.366498 | 0.25621 | 0.937017 | 1.576243 | 5.412586 | 2.535326 | 1.714925 | 0.898167 | 1.150446 | -1.435155 | -1.665742 |
| 35.13888889 | 0.525007 | 0.443272 | 0.339221 | 2.769118 | 2.438705 | 0.596197 | 0.881734 | 1.884135 | 0.797357 | 0.632257 | 0.933905 | 1.851821 | 2.943722 | -3.43959 | 1.486336 | 1.821442 | 0.663261 | 3.766581 | 1.360781 | 1.584588 | -0.036602 | 0.610524 | -1.668127 | -2.429415 |
| 35.15277778 | 0.616717 | 0.373575 | 0.337966 | 2.331899 | 2.50052 | 0.504265 | 0.934431 | 2.143149 | 0.687932 | 0.552655 | 0.959456 | 1.663279 | 2.050675 | -3.91588 | 2.118881 | 2.474737 | 0.481648 | 1.647032 | -0.192543 | 1.149997 | -0.420091 | -0.009353 | -1.458148 | -2.568649 |
| 35.16666667 | 0.694784 | 0.221264 | 0.377404 | 1.778455 | 2.25161 | 0.426151 | 0.86635 | 2.465962 | 0.553893 | 0.417463 | 0.868419 | 1.336158 | 1.327056 | -3.550323 | 2.049244 | 2.381948 | 0.494711 | 0.381484 | -1.103139 | -0.03272 | 0.042092 | -0.199652 | -1.067541 | -2.192095 |
| 35.18055556 | 0.622777 | 0.105664 | 0.390073 | 1.654265 | 1.953455 | 0.322748 | 0.673858 | 2.621466 | 0.451087 | 0.080934 | 0.707633 | 0.942795 | 0.504851 | -2.898598 | 1.751376 | 1.747442 | 0.340835 | 0.531654 | -0.821753 | -0.835912 | 0.739014 | -0.14092 | -0.823257 | -1.821257 |
| 35.19444444 | 0.534499 | 0.234651 | 0.434313 | 2.011718 | 1.691027 | 0.203292 | 0.45116 | 2.723762 | 0.338425 | -0.283156 | 0.45083 | 0.486531 | -0.179239 | -2.32276 | 1.469518 | 1.216464 | 0.353592 | 1.813307 | 0.143116 | -0.221242 | 0.87371 | -0.20595 | -0.853527 | -1.585932 |
| 35.20833333 | 0.526358 | 0.490679 | 0.490898 | 2.062664 | 1.560257 | 0.105694 | 0.24578 | 2.726641 | 0.18583 | -0.483824 | 0.206386 | 0.04645 | -0.116843 | -1.679427 | 1.174646 | 0.600593 | 0.454106 | 2.170078 | 0.380881 | 0.725748 | 0.348539 | -0.399247 | -0.755735 | -1.303308 |
| 35.22222222 | 0.539515 | 0.60839 | 0.47431 | 1.511103 | 1.505762 | 0.150812 | 0.218597 | 2.483542 | 0.117977 | -0.608995 | 0.065048 | -0.310288 | 0.018969 | -0.934734 | 0.861897 | 0.210892 | 0.28683 | 0.69594 | -0.253847 | 1.110393 | -0.133708 | -0.089111 | -0.459496 | -0.841806 |
| 35.23611111 | 0.560178 | 0.443673 | 0.388622 | 0.792594 | 1.401395 | 0.216387 | 0.374968 | 2.185636 | 0.109024 | -0.593849 | -0.028029 | -0.428597 | -0.027105 | -0.39476 | 0.780118 | 0.163922 | -0.130608 | -0.868941 | -1.272054 | 0.758228 | -0.071534 | 0.123023 | -0.16187 | -0.399036 |
| 35.25 | 0.504863 | 0.298265 | 0.293369 | 0.166174 | 1.217569 | 0.175931 | 0.502042 | 2.158735 | 0.098676 | -0.40521 | -0.067375 | -0.494903 | -0.123672 | -0.425343 | 0.757779 | 0.254691 | -0.38458 | -0.710182 | -1.248208 | 0.22698 | -0.017832 | 0.098067 | -0.065393 | -0.268323 |
| 35.26388889 | 0.433461 | 0.620166 | 0.320243 | -0.089522 | 1.131074 | 0.133259 | 0.480055 | 2.532115 | 0.091723 | -0.224614 | -0.154502 | -0.53343 | 0.219059 | -0.472652 | 0.778498 | 0.43484 | -0.121111 | 0.820504 | -0.312122 | 0.200839 | 0.192487 | 0.279151 | -0.11073 | -0.239943 |
| 35.27777778 | 0.416431 | 1.163691 | 0.438857 | 0.531775 | 1.30103 | 0.297059 | 0.426806 | 2.942378 | 0.145347 | 0.008601 | -0.114151 | -0.321699 | 1.15196 | 0.105887 | 1.040234 | 0.842688 | 0.523762 | 2.750437 | 0.677666 | 1.185226 | 0.988544 | 1.257499 | -0.034191 | 0.016431 |
| 35.29166667 | 0.539964 | 1.521892 | 0.557826 | 1.642746 | 1.70637 | 0.545162 | 0.428497 | 2.738316 | 0.341277 | 0.197831 | 0.134177 | 0.144216 | 1.962215 | 1.035922 | 1.329966 | 1.288186 | 0.991218 | 3.178472 | 1.467076 | 2.59385 | 1.854639 | 2.303068 | 0.072883 | 0.367663 |
| 35.30555556 | 0.673621 | 1.58956 | 0.636754 | 2.442607 | 2.157981 | 0.843433 | 0.488224 | 2.33786 | 0.541237 | 0.372259 | 0.338853 | 0.576635 | 2.026663 | 1.595571 | 1.383765 | 1.254991 | 0.977253 | 2.295734 | 1.674529 | 3.989918 | 2.432982 | 2.563526 | 0.086125 | 0.492922 |
| 35.31944444 | 0.744063 | 1.674792 | 0.585879 | 2.51559 | 2.500549 | 1.188958 | 0.542311 | 2.390965 | 0.641766 | 0.54069 | 0.381912 | 0.731019 | 1.728036 | 1.327935 | 1.14857 | 0.891046 | 0.893249 | 1.951067 | 2.029612 | 4.048604 | 3.131334 | 2.481033 | 0.105811 | 0.466912 |
| 35.33333333 | 0.753542 | 1.753754 | 0.553733 | 2.623729 | 2.669648 | 1.348398 | 0.591744 | 2.740985 | 0.685698 | 0.752848 | 0.392925 | 0.746853 | 1.294215 | 1.041493 | 1.135007 | 0.774181 | 0.427882 | 2.443018 | 2.666395 | 3.524359 | 3.482582 | 2.182254 | 0.231636 | 0.595182 |
| 35.34722222 | 0.851967 | 1.954425 | 0.561077 | 3.276774 | 2.849148 | 1.124423 | 0.621352 | 2.973357 | 0.760626 | 0.971642 | 0.599215 | 1.004088 | 1.199868 | 1.231158 | 1.601247 | 1.170108 | 0.232982 | 3.450831 | 3.251831 | 4.717926 | 3.46501 | 2.433428 | 0.438022 | 0.818884 |
| 35.36111111 | 1.047573 | 2.33722 | 0.636567 | 4.262937 | 2.99629 | 1.090174 | 0.693403 | 3.186426 | 0.86776 | 1.138343 | 0.948088 | 1.616442 | 1.543528 | 1.597443 | 2.201095 | 1.769296 | 0.500024 | 4.420244 | 4.046312 | 6.224463 | 3.509404 | 2.977309 | 0.634374 | 1.059339 |
| 35.375 | 1.326075 | 2.924063 | 0.766445 | 4.856129 | 3.353813 | 1.266214 | 0.696678 | 3.447537 | 0.993498 | 1.134793 | 1.24256 | 2.153081 | 1.898464 | 1.751769 | 2.469432 | 2.099029 | 0.998838 | 5.045778 | 5.356422 | 6.176992 | 4.285438 | 3.985627 | 0.698711 | 1.098325 |
| 35.38888889 | 1.551531 | 3.075185 | 0.867574 | 4.756081 | 3.74017 | 1.40778 | 0.72572 | 3.700532 | 0.935392 | 1.284022 | 1.460178 | 2.542507 | 1.269843 | 1.656044 | 2.455554 | 2.246019 | 1.317495 | 4.563811 | 5.374015 | 5.15037 | 5.280451 | 4.905706 | 0.662802 | 0.980641 |
| 35.40277778 | 1.695586 | 3.325914 | 0.884428 | 4.657949 | 3.728368 | 1.409218 | 0.886527 | 3.407053 | 0.77974 | 1.384808 | 1.710202 | 2.750643 | 0.881537 | 1.554394 | 2.472519 | 2.262796 | 1.265467 | 4.21335 | 4.832439 | 3.828318 | 5.775646 | 4.843791 | 0.75453 | 0.97571 |
| 35.41666667 | 1.690149 | 3.228724 | 0.941137 | 4.952826 | 3.738025 | 1.371272 | 0.792033 | 3.541885 | 0.886383 | 1.243641 | 1.841073 | 2.810598 | 1.089222 | 1.718922 | 2.779828 | 2.572876 | 1.424797 | 4.65508 | 4.51678 | 5.173258 | 5.176493 | 4.627911 | 1.183423 | 1.284541 |
| 35.43055556 | 1.826384 | 3.309297 | 1.122703 | 5.029054 | 4.160117 | 1.303823 | 0.627666 | 3.084145 | 1.109719 | 1.140737 | 2.11455 | 3.067243 | 2.153407 | 2.401 | 3.208663 | 2.985117 | 1.984773 | 5.161386 | 4.146575 | 5.031831 | 4.422805 | 4.474192 | 1.752014 | 1.898132 |
| 35.44444444 | 2.001865 | 2.908724 | 1.408595 | 5.570353 | 4.399853 | 1.591647 | 0.428125 | 2.569247 | 1.172785 | 1.197776 | 2.287994 | 3.197826 | 3.305804 | 3.596515 | 3.723037 | 3.353415 | 3.172231 | 6.197753 | 3.754605 | 6.532762 | 5.286712 | 4.133574 | 2.269047 | 2.911442 |
| 35.45833333 | 2.164409 | 3.442335 | 1.79557 | 5.385767 | 4.607771 | 2.191477 | 0.593702 | 1.488466 | 1.30054 | 1.209447 | 1.945035 | 2.637226 | 3.066883 | 4.207902 | 3.798185 | 3.551059 | 3.526023 | 6.494886 | 3.769601 | 9.867211 | 6.129124 | 4.535786 | 2.331983 | 3.511958 |
| 35.47222222 | 2.087959 | 3.621107 | 1.814922 | 4.596238 | 4.082572 | 2.303476 | 0.453215 | 1.568818 | 1.402917 | 1.25447 | 1.524264 | 2.468353 | 2.364137 | 3.949662 | 4.49662 | 3.256017 | 3.536724 | 5.86637 | 3.9436 | 8.640341 | 6.288089 | 5.74964 | 2.209062 | 3.601301 |
| 35.48611111 | 1.898036 | 2.849238 | 1.553474 | 3.188521 | 3.33485 | 1.925296 | 0.803098 | 2.154876 | 1.475222 | 1.115881 | 1.092578 | 1.956051 | 1.996598 | 3.863997 | 6.219751 | 2.742292 | 2.951395 | 5.393395 | 3.620022 | 6.217617 | 4.827949 | 6.096031 | 2.373209 | 3.834133 |
| 35.5 | 2.034993 | 2.319327 | 1.112289 | 2.558544 | 3.305667 | 1.447521 | 1.05389 | 1.80094 | 1.384367 | 0.93543 | 1.52996 | 1.792232 | 1.492318 | 4.106958 | 7.148762 | 2.232413 | 2.955334 | 4.543683 | 2.749244 | 5.309193 | 4.756764 | 6.076311 | 2.829855 | 4.380829 |
| 35.51388889 | 2.310106 | 2.244932 | 1.19609 | 2.226366 | 3.51061 | 1.257927 | 1.089982 | 1.463066 | 1.301566 | 0.898183 | 2.428623 | 2.004529 | 1.172094 | 4.515353 | 7.143282 | 2.379122 | 2.748378 | 4.246795 | 3.432332 | 5.899116 | 3.442604 | 6.817399 | 3.195538 | 4.400848 |
| 35.52777778 | 2.731452 | 2.387456 | 1.660231 | 2.866463 | 3.455774 | 1.461118 | 0.924801 | 1.39068 | 1.097497 | 0.972104 | 2.285171 | 1.056391 | 0.956364 | 4.626417 | 6.605845 | 3.116733 | 2.294371 | 3.657711 | 3.492451 | 6.077426 | 1.267189 | 6.474599 | 3.42797 | 3.879763 |
| 35.54166667 | 2.587713 | 1.944879 | 2.033549 | 2.720988 | 2.921202 | 1.790533 | 1.175692 | 1.448997 | 0.895536 | 1.170861 | 1.847629 | 0.56336 | 1.146034 | 3.813551 | 4.618093 | 2.891509 | 1.839191 | 3.705676 | 2.426352 | 4.758563 | 1.618521 | 5.455372 | 2.969197 | 3.24018 |
| 35.55555556 | 1.984201 | 0.997163 | 1.793881 | 1.493187 | 2.891818 | 2.558565 | 1.62454 | 1.200141 | 0.818158 | 1.217484 | 1.80544 | 1.167716 | 1.105102 | 2.959242 | 3.890689 | 2.110105 | 2.14121 | 2.808365 | 2.24353 | 3.500032 | 1.277305 | 2.994909 | 2.456243 | 2.555234 |
| 35.56944444 | 0.41406 | 0.672179 | 1.247532 | 0.551233 | 2.122371 | 3.038033 | 1.618467 | 1.414593 | 0.82048 | 1.422552 | 1.558046 | 1.454855 | 1.238768 | 2.639076 | 3.752588 | 2.114752 | 2.617729 | 1.983912 | 2.18478 | 3.320108 | 2.392075 | 1.704438 | 2.043228 | 2.140104 |
| 35.58333333 | -1.062353 | 0.741566 | 0.784831 | 0.177078 | 1.071048 | 2.792149 | 1.579026 | 1.334381 | 0.836976 | 1.613251 | 1.02296 | 1.125017 | 1.259253 | 2.77258 | 3.218212 | 1.562407 | 2.372838 | 1.281592 | 2.175483 | 2.780307 | 2.629963 | 1.450549 | 1.313208 | 2.328108 |
| 35.59722222 | -1.22377 | 0.123769 | 0.578262 | 0.070004 | 0.371764 | 2.249175 | 1.174412 | 0.928703 | 0.727864 | 1.481183 | 0.255137 | 0.48988 | 1.021552 | 3.442779 | 2.381527 | -0.169944 | 2.108438 | 0.891726 | 1.751904 | 2.008514 | 2.994456 | 1.221171 | 1.676783 | 2.753247 |
| 35.61111111 | -0.888943 | -0.631704 | 0.372456 | -0.302118 | 0.297415 | 2.385457 | 1.050069 | 0.509957 | 0.385957 | 0.85442 | -0.039465 | 0.175826 | 0.716962 | 4.177813 | 1.172769 | -0.374962 | 1.4351 | 0.14213 | 0.844228 | 1.327348 | 1.332095 | 1.305977 | 1.920488 | 3.166709 |
| 35.625 | -1.044665 | -1.263268 | 0.080586 | -0.938772 | 0.288777 | 1.7362 | 0.683645 | 0.179086 | 0.170413 | 0.330643 | -0.003165 | 0.226204 | 0.405752 | 3.908467 | 0.395348 | -0.292887 | 0.705386 | -0.922423 | -0.663569 | 0.810639 | 0.804595 | 1.098171 | 1.553954 | 2.99113 |
| 35.63888889 | -1.655841 | -1.313153 | -0.208762 | -1.504274 | -0.491348 | 0.173564 | 0.213042 | 0.026816 | 0.199619 | 0.630325 | -0.09391 | 0.151202 | 0.045377 | 2.824272 | -0.323825 | 0.282576 | 0.329544 | -1.598091 | -2.10958 | -0.044701 | -0.389843 | 0.739389 | 1.123953 | 2.139516 |
| 35.65277778 | -1.871949 | -0.476841 | -0.378809 | -1.800349 | -1.280239 | -1.153285 | -0.017171 | -0.156122 | 0.241661 | 0.579926 | -0.368538 | -0.235139 | -0.157522 | 2.2875 | -0.586042 | 0.402298 | 0.323142 | -1.692003 | -2.964527 | -0.946165 | -0.116318 | 0.418349 | 1.016221 | 1.625377 |
| 35.66666667 | -1.338149 | -0.449883 | -0.41983 | -2.106294 | -1.838847 | -1.23643 | -0.183305 | -0.054411 | 0.251429 | 0.135505 | -0.562741 | -0.547164 | -0.22298 | 1.909508 | -0.806706 | -0.094928 | 0.207472 | -0.842062 | -2.458814 | -2.134518 | 0.746951 | 0.525029 | 1.062279 | 1.273951 |
| 35.68055556 | -0.689207 | -0.324838 | -0.455066 | -2.484582 | -1.983595 | -1.212909 | -0.239323 | -0.010163 | 0.238626 | -0.154429 | -0.614507 | -0.752833 | 0.300271 | 0.939637 | -0.779917 | -0.339653 | -0.319027 | -0.421653 | -2.955033 | -2.624057 | 0.189545 | 0.676833 | 0.688297 | 0.351938 |
| 35.69444444 | -0.355936 | -0.738024 | -0.589819 | -2.435942 | -2.146024 | -1.358809 | -0.513458 | -0.088819 | 0.187546 | -0.322757 | -0.268351 | -0.781436 | 0.613516 | -0.563562 | -0.586647 | 0.464774 | -1.24147 | -1.547359 | -4.072114 | -2.803296 | -0.564731 | 0.446474 | 0.015345 | -0.817093 |
| 35.70833333 | -0.439819 | -0.12618 | -0.745171 | -1.599294 | -2.356406 | -1.114921 | -0.770259 | -0.377657 | 0.074422 | -0.678602 | 0.114487 | -0.455396 | -0.186931 | -1.699481 | 0.059295 | 0.135716 | -1.488361 | -2.111533 | -4.333389 | -2.735592 | -1.111054 | -0.188405 | -0.449387 | -1.647294 |
| 35.72222222 | -0.601059 | 0.481315 | -0.77119 | -1.041523 | -2.337521 | -0.386185 | -0.932646 | -0.520296 | 0.111776 | -0.888567 | 0.213496 | 0.032806 | -0.11591 | -2.067857 | 0.859235 | 0.15574 | -0.242781 | -2.4322 | -2.773045 | -1.764601 | -1.559913 | -0.704898 | -0.640308 | -1.844864 |
| 35.73611111 | -0.655048 | 0.303937 | -0.697416 | -1.030071 | -2.165126 | -0.144081 | -0.956458 | -0.589715 | 0.350845 | -0.791147 | -0.055719 | 0.157773 | -0.084072 | -1.726023 | 0.976051 | 0.344636 | 0.800664 | -1.977419 | -1.813204 | -1.324469 | -2.164228 | -0.505159 | -0.524553 | -1.476419 |
| 35.75 | -0.676627 | -0.578634 | -0.542778 | -1.333781 | -2.380611 | 0.030677 | -0.854237 | -0.825143 | 0.427291 | -0.32513 | -0.359704 | -0.208368 | -0.267562 | -1.101186 | 0.42088 | 0.274703 | 1.30945 | -1.45946 | -1.485978 | -2.771968 | -2.485289 | -0.581228 | -0.440194 | -0.877398 |
| 35.76388889 | -0.719571 | -1.22554 | -0.425828 | -1.102384 | -2.305721 | 0.005253 | -1.312659 | -0.898358 | 0.271808 | -0.171834 | -0.521993 | -0.510412 | -1.085643 | -0.974217 | -0.015069 | -0.128293 | 0.891346 | -2.372137 | -2.023668 | -4.115947 | -1.849895 | -1.047895 | -0.704764 | -0.722474 |
| 35.77777778 | -0.56161 | -1.140713 | -0.339734 | -0.57788 | -1.96384 | 0.064412 | -1.126437 | -0.543379 | 0.189826 | -0.283586 | -0.559374 | -0.455711 | -0.768569 | -1.072296 | -0.05106 | -0.210265 | 0.284961 | -2.844592 | -2.674996 | -5.018061 | -1.33048 | -1.626606 | -0.865311 | -0.840924 |
| 35.79166667 | -0.254736 | -0.861151 | -0.377904 | -0.59518 | -1.528794 | 0.055561 | -0.821884 | -0.209846 | -0.018311 | -0.756041 | -0.440547 | -0.176248 | -0.169278 | -1.216878 | -0.169418 | -0.235505 | -0.470371 | -1.214875 | -1.677494 | -2.521398 | -0.558497 | -2.108423 | -0.895561 | -1.003008 |
| 35.80555556 | -0.290166 | -1.029435 | -0.529498 | -0.884419 | -1.420494 | 0.07595 | -0.524956 | -0.550768 | -0.253323 | -1.530961 | -0.298332 | -0.092817 | -0.50443 | -1.516624 | -0.491328 | -0.527816 | -0.821212 | -0.065546 | -1.087464 | 1.350863 | 0.086256 | -2.903055 | -0.859594 | -1.290913 |
| 35.81944444 | -0.512173 | -0.935802 | -0.692991 | -1.220835 | -1.67727 | 0.039245 | -0.32328 | -0.728154 | -0.350293 | -1.365069 | -0.176758 | -0.198216 | -1.142823 | -1.40477 | -0.718386 | -0.68392 | -0.969285 | 0.896106 | -0.544936 | 2.865437 | -3.573446 | -2.884383 | -0.989348 | -1.331988 |
| 35.83333333 | -0.661557 | -0.741405 | -0.685534 | -1.302085 | -1.750812 | -0.040099 | -0.03744 | -0.887891 | -0.248691 | -0.480781 | -0.052132 | -0.212232 | -1.188274 | -0.617785 | -0.601942 | -0.452413 | -1.003825 | -0.056462 | -0.855711 | 0.170646 | -4.634579 | -2.313884 | -0.961433 | -1.000362 |
| 35.84722222 | -0.692202 | -1.111323 | -0.712318 | -1.649693 | -1.786111 | -0.129331 | -0.074325 | -0.897988 | -0.064347 | -0.56507 | -0.212966 | -0.137697 | -0.952508 | 0.143249 | -0.643681 | -0.010422 | -1.6721 | -0.545915 | -1.094637 | -2.959983 | -11.617198 | -2.059619 | -0.840967 | -0.565339 |
| 35.86111111 | -0.566886 | -1.651554 | -0.785249 | -2.168069 | -1.715643 | -0.107994 | -0.059576 | -1.343108 | -0.147673 | -0.891676 | -0.359108 | -0.140554 | -0.863904 | 0.368246 | -1.176715 | -0.606706 | -2.598029 | -0.083253 | -1.253411 | -3.884399 | -3.346677 | -2.026672 | -1.003359 | -0.32118 |
| 35.875 | -0.567609 | -2.090208 | -0.943066 | -2.666795 | -1.804097 | -0.0245 | -0.213725 | -1.700951 | -0.33314 | -1.061295 | -0.369334 | 0.079939 | -0.846978 | 0.263616 | -1.501749 | -0.812157 | -3.126248 | 0.53563 | -1.624099 | -4.976601 | -1.395526 | -3.548964 | -1.14078 | -0.213245 |
| 35.88888889 | -0.941536 | -2.270657 | -1.139644 | -3.082169 | -2.199067 | -0.084876 | -0.39028 | -1.733703 | -0.488321 | -1.455144 | -0.546723 | 0.484342 | -1.04157 | 0.358512 | -0.995369 | -0.668139 | -2.676614 | -0.424812 | -1.759427 | -6.053041 | -1.186447 | -4.106867 | -1.390294 | 0.095745 |
| 35.90277778 | -1.313768 | -2.101473 | -1.234148 | -3.541317 | -2.785136 | -0.161167 | -0.587581 | -2.195485 | -0.539726 | -1.862918 | -0.539761 | 0.297244 | -1.617332 | 1.080989 | -0.096556 | -0.146932 | -1.808408 | -1.783782 | -2.605095 | -6.558515 | -1.403115 | -4.437785 | -1.730669 | 0.771794 |
| 35.91666667 | -1.496012 | -2.578558 | -1.279795 | -4.564392 | -3.002042 | -0.202429 | -0.346253 | -2.548534 | -0.561303 | -1.969084 | -0.639513 | -0.403999 | -1.853909 | 1.465186 | 0.535398 | 0.036248 | -1.919608 | -2.699207 | -3.567541 | -5.809645 | -1.623374 | -3.300881 | -1.910378 | 0.923475 |
| 35.93055556 | -1.505174 | -3.464735 | -1.283493 | -5.627337 | -2.984834 | -0.149108 | -0.41136 | -3.073661 | -0.367157 | -1.597372 | -0.879543 | -0.825789 | -2.102895 | 0.609456 | 0.270429 | -0.129193 | -3.187 | -3.636415 | -3.640715 | -6.674522 | -2.122793 | -3.396092 | -2.075805 | 0.434492 |
| 35.94444444 | -1.317384 | -4.575324 | -1.623409 | -6.143914 | -3.07368 | -0.397177 | -0.230346 | -3.26981 | -0.370454 | -1.381802 | -1.013548 | -1.200859 | -2.60797 | -1.32185 | -0.755751 | -1.348005 | -5.402717 | -4.676947 | -2.940491 | -6.648474 | -1.713928 | -3.90129 | -2.849459 | -1.031931 |
| 35.95833333 | -1.457139 | -4.69423 | -2.778404 | -5.861635 | -3.177373 | -1.329336 | 0.034097 | -3.008344 | -0.811594 | -2.470672 | -1.465782 | -2.590439 | -3.719698 | -3.303014 | -2.448807 | -2.690772 | -6.475716 | -5.049775 | -4.452132 | -5.579776 | -1.819977 | -4.133664 | -3.120563 | -2.498941 |
| 35.97222222 | -1.266236 | -3.808322 | -3.319222 | -6.376944 | -4.600256 | -2.458765 | -0.250617 | -2.788448 | -1.407518 | -3.865402 | -2.453701 | -3.888464 | -4.259691 | -4.420793 | -4.455695 | -4.068834 | -5.112491 | -5.76999 | -5.566302 | -4.707906 | -3.292521 | -6.0061 | -3.631493 | -3.238083 |
| 35.98611111 | -0.842706 | -3.751413 | -2.357688 | -4.847063 | -4.235979 | -2.150434 | -0.39148 | -1.958367 | -1.16581 | -4.000255 | -2.399047 | -3.324523 | -4.663471 | -3.597422 | -3.988696 | -3.245975 | -3.31872 | -3.286991 | -4.00431 | -5.9573 | -5.75085 | -9.378177 | -3.315467 | -2.830212 |
| 36 | -0.28149 | -2.862006 | -1.442692 | -3.117504 | -3.149616 | -1.232033 | -0.244928 | -2.255901 | -0.686767 | -3.906349 | -1.71385 | -1.873498 | -4.523297 | -2.342095 | -2.617275 | -2.303787 | -2.490713 | -3.487678 | -2.955242 | -7.4422 | -4.140011 | -4.671899 | -3.203223 | -1.976398 |
| 36.01388889 | 0.035068 | -1.723323 | -0.852185 | -1.908797 | -2.11332 | -0.510835 | -0.080204 | -3.315408 | -0.414667 | -3.290139 | -1.269477 | -1.204198 | -4.263305 | -1.321875 | -2.046095 | -1.891833 | -2.500792 | -5.073417 | -3.185439 | -6.661854 | -3.64868 | -4.072039 | -2.307789 | -1.303895 |
| 36.02777778 | 0.216608 | -0.800884 | -0.605641 | -1.356538 | -1.436495 | -0.290745 | -0.185777 | -3.241655 | -0.340474 | -3.290016 | -0.949376 | -0.966171 | -4.397379 | -1.106693 | -1.858245 | -1.66745 | -1.439408 | -3.926269 | -3.190301 | -4.737891 | -3.783206 | -3.502962 | -1.261631 | -0.833125 |
| 36.04166667 | 0.157652 | -0.453889 | -0.421316 | -0.969449 | -0.925968 | -0.37431 | -0.458068 | -3.246175 | -0.304143 | -2.88723 | -0.916551 | -0.904379 | -4.027383 | -2.028807 | -1.878171 | -2.079996 | -0.216823 | -0.437982 | -1.759327 | -2.82064 | -3.133292 | -2.903886 | -0.062376 | -1.00581 |
| 36.05555556 | 0.146979 | -0.35047 | -0.179651 | -0.350451 | -0.454232 | -0.443153 | -0.596521 | -2.664177 | -0.158137 | -1.903714 | -0.84937 | -1.043691 | -3.109836 | -3.657382 | -1.88819 | -1.736286 | 0.408102 | 2.448828 | 0.64163 | -0.628049 | -2.279236 | -1.440056 | 0.587272 | -1.950872 |
| 36.06944444 | 0.25203 | 0.003667 | -0.01829 | 0.320567 | 0.027362 | -0.427903 | -0.500389 | -1.415965 | -0.024008 | -0.844669 | -0.467822 | -0.924163 | -2.660902 | -4.058899 | -1.718153 | -1.123868 | 0.444925 | 2.500304 | 1.31601 | 0.605006 | -1.548887 | -1.233003 | 0.142047 | -2.238844 |
| 36.08333333 | 0.330089 | 0.776264 | 0.075787 | 0.857878 | 0.851286 | -0.415017 | -0.401821 | -0.240315 | 0.086985 | -0.050033 | -0.053753 | -0.47286 | -2.351109 | -3.150469 | -1.431725 | -0.718056 | 0.314424 | 1.377823 | 0.980146 | 1.384185 | -1.31835 | -2.669496 | -0.42523 | -1.465171 |
| 36.09722222 | 0.43709 | 1.333405 | 0.010743 | 0.981172 | 1.751957 | -0.317826 | -0.227691 | 0.190934 | 0.267212 | 0.394362 | 0.193946 | -0.046215 | -2.128668 | -2.74333 | -1.200084 | -0.34273 | 0.304518 | 1.103758 | 0.986261 | 2.607715 | -1.037547 | -3.007639 | -0.816317 | -1.092314 |
| 36.11111111 | 0.470249 | 1.319738 | -0.043529 | 0.920642 | 1.99122 | -0.120385 | 0.022324 | 0.557796 | 0.435914 | 0.684655 | 0.29196 | 0.115878 | -1.859946 | -2.688477 | -0.940769 | -0.010153 | 0.904986 | 2.132115 | 1.710749 | 2.952309 | -1.214822 | -2.995989 | -1.1221 | -1.262133 |
| 36.125 | 0.448886 | 1.270667 | -0.00695 | 1.166106 | 1.908954 | 0.077875 | 0.274163 | 1.387428 | 0.471129 | 0.891121 | 0.384961 | 0.314713 | -1.412693 | -2.604467 | -0.951331 | 0.151605 | 1.530491 | 3.451532 | 1.87254 | 2.072797 | -0.766302 | -3.117944 | -1.376763 | -0.932172 |
| 36.13888889 | 0.378328 | 1.515382 | 0.161344 | 1.52469 | 2.18701 | 0.213213 | 0.486299 | 2.336975 | 0.482707 | 1.083909 | 0.595809 | 0.716415 | -0.851086 | -2.240614 | -0.872478 | 0.119426 | 1.831622 | 3.450143 | 2.095225 | 1.337679 | -0.485191 | -2.91921 | -1.241575 | -0.415021 |
| 36.15277778 | 0.247857 | 1.667251 | 0.328532 | 1.520973 | 2.442877 | 0.329761 | 0.646665 | 2.517677 | 0.580294 | 1.111904 | 0.831266 | 1.147478 | -0.727836 | -1.825148 | -0.969355 | -0.115627 | 1.696557 | 2.692449 | 0.66117 | -0.029004 | -0.572607 | -3.401599 | -0.660987 | -0.365278 |
| 36.16666667 | 0.28546 | 1.792991 | 0.40692 | 1.382912 | 2.45074 | 0.395183 | 0.709756 | 2.43871 | 0.576889 | 0.966282 | 0.993232 | 1.383184 | -1.24805 | -1.95365 | -1.037427 | -0.204997 | 0.489856 | 0.967962 | -1.345316 | -1.668181 | -0.742203 | -3.10886 | -0.78001 | -0.790387 |
| 36.18055556 | 0.492471 | 2.722359 | 0.434255 | 1.604191 | 2.404434 | 0.422257 | 0.705546 | 2.593385 | 0.495874 | 0.824877 | 1.06658 | 1.352905 | -1.543955 | -2.172018 | -1.088684 | -0.214711 | 0.152271 | 0.252295 | -2.492823 | -2.493082 | -0.149124 | -2.455631 | -0.623151 | -1.308904 |
| 36.19444444 | 0.608974 | 3.365583 | 0.393985 | 1.775036 | 2.057868 | 0.365545 | 0.689899 | 2.748398 | 0.426508 | 0.742638 | 1.03273 | 1.224998 | -1.311064 | -1.51378 | -0.928153 | -0.489787 | 0.003999 | 0.835095 | -2.063914 | -2.282436 | 0.061278 | -2.084449 | -0.038529 | -1.171048 |
| 36.20833333 | 0.561757 | 2.621273 | 0.39955 | 1.202771 | 1.523578 | 0.276437 | 0.678572 | 2.69464 | 0.344287 | 0.643724 | 0.968921 | 1.09378 | -1.017775 | -0.741369 | -0.735946 | -0.636021 | 0.063503 | 1.244285 | -1.171934 | -1.306036 | -0.3919 | -2.619143 | -0.313454 | -0.973673 |
| 36.22222222 | 0.478537 | 1.76989 | 0.448774 | 0.247654 | 1.073202 | 0.156848 | 0.626534 | 2.423084 | 0.290485 | 0.367007 | 0.821544 | 0.910082 | -0.886341 | -0.648894 | -0.617048 | -0.912418 | -0.313763 | 0.473256 | -0.739886 | -0.700155 | -1.151099 | -3.258573 | -0.942815 | -0.994314 |
| 36.23611111 | 0.489786 | 1.581745 | 0.537193 | -0.417094 | 0.843389 | 0.036871 | 0.480165 | 2.318011 | 0.187341 | 0.219524 | 0.639592 | 0.736553 | -0.225348 | -0.690857 | -0.561821 | -1.173473 | -0.242808 | -0.255516 | -0.476157 | -0.237282 | -1.245749 | -3.473142 | -1.107825 | -1.074769 |
| 36.25 | 0.488032 | 2.235273 | 0.687183 | 0.306248 | 0.847026 | -0.028892 | 0.343793 | 2.542404 | 0.172066 | 0.231551 | 0.450068 | 0.667175 | 1.069591 | -0.314759 | -0.331994 | -0.652331 | 1.005308 | 0.55599 | 0.047684 | 0.469527 | 0.285992 | -1.545572 | -0.404874 | -0.680101 |
| 36.26388889 | 0.439129 | 2.583482 | 0.802235 | 1.561193 | 0.8537 | 0.015551 | 0.375982 | 2.900033 | 0.307244 | 0.422026 | 0.477264 | 0.783734 | 1.650692 | -0.01036 | 0.042715 | 0.244017 | 1.97214 | 1.82712 | 0.159777 | 0.780718 | 1.558043 | -0.525727 | 0.149595 | -0.469698 |
| 36.27777778 | 0.404901 | 2.03007 | 0.647711 | 1.975534 | 0.916816 | 0.098427 | 0.608877 | 3.211303 | 0.405453 | 0.49584 | 0.533826 | 0.842565 | 0.846688 | -0.319997 | 0.126663 | 0.496097 | 1.875812 | 1.738438 | -0.698698 | 0.344529 | 1.248977 | -0.758671 | 0.063931 | -0.715495 |
| 36.29166667 | 0.460314 | 2.508745 | 0.497605 | 2.084933 | 0.982988 | 0.099104 | 0.717817 | 3.178075 | 0.350693 | 0.447899 | 0.514429 | 0.6895 | 0.072507 | -0.819487 | 0.09291 | 0.443362 | 1.684965 | 0.871523 | -0.999118 | -0.08688 | 1.22986 | -0.763792 | -0.053147 | -1.098014 |
| 36.30555556 | 0.551195 | 4.176202 | 0.552971 | 2.680743 | 1.114739 | 0.061493 | 0.761789 | 3.434494 | 0.287531 | 0.403033 | 0.432352 | 0.483683 | 0.345401 | -0.687123 | 0.207182 | 0.561435 | 2.101592 | 0.608591 | 0.060328 | 0.262021 | 2.311213 | -1.539832 | -0.055809 | -1.115916 |
| 36.31944444 | 0.66841 | 4.257545 | 0.727869 | 3.423649 | 1.655529 | 0.203387 | 0.907994 | 3.813363 | 0.342248 | 0.538761 | 0.46933 | 0.534939 | 1.428828 | 0.178087 | 0.282312 | 0.996928 | 2.455375 | 1.412581 | 1.155651 | 1.855744 | 2.134987 | -2.380756 | 0.17326 | -1.031893 |
| 36.33333333 | 0.787129 | 3.500195 | 1.20581 | 4.539137 | 4.464629 | 1.314702 | 1.737689 | 5.920482 | 0.845617 | 0.733793 | 0.451866 | 1.432646 | 3.858361 | 1.326137 | -0.427235 | 2.312862 | 1.612442 | 2.731722 | 0.085674 | 7.268775 | 0.318741 | 4.148872 | 2.873294 | -0.279505 |
| 36.34722222 | 2.15205 | 5.673165 | 2.558226 | 8.356116 | 13.037891 | 5.106059 | 5.017049 | 12.583909 | 1.455082 | 1.230333 | 2.451646 | 4.796451 | 10.540318 | 6.707429 | -0.487239 | 4.901254 | -5.205848 | 1.535734 | -2.229668 | 19.526404 | -1.964323 | 15.419679 | 14.295918 | 3.126744 |
| 36.36111111 | 6.784194 | NaN | 7.396838 | 6.020635 | NaN | 7.046543 | 3.492749 | NaN | 25.855455 | -17.036397 | NaN | NaN | 10.412773 | NaN | NaN | NaN | NaN | 16.726476 | NaN | NaN | NaN | NaN | NaN | 32.906947 |
| 36.375 | NaN | NaN | NaN | NaN | NaN | NaN | NaN | NaN | NaN | -5.730273 | NaN | NaN | NaN | NaN | NaN | NaN | NaN | NaN | NaN | NaN | NaN | NaN | NaN | NaN |
| 36.38888889 | NaN | NaN | NaN | NaN | NaN | NaN | NaN | NaN | NaN | -0.71305 | NaN | NaN | NaN | NaN | NaN | NaN | NaN | NaN | NaN | NaN | NaN | NaN | NaN | NaN |

**Supplementary Table S2.** Raw motion data from: **(c)** 100 mM NaCl treatment in Radish

| Time | RadishSalt1 | RadishSalt10 | RadishSalt11 | RadishSalt12 | RadishSalt2 | RadishSalt3 | RadishSalt4 | RadishSalt5 | RadishSalt6 | RadishSalt7 | RadishSalt8 | RadishSalt9 | RadishControl1 | RadishControl10 | RadishControl11 | RadishControl12 | RadishControl2 | RadishControl3 | RadishControl4 | RadishControl5 | RadishControl6 | RadishControl7 | RadishControl8 | RadishControl9 |
|---|---|---|---|---|---|---|---|---|---|---|---|---|---|---|---|---|---|---|---|---|---|---|---|---|
| 18 | 0.557695 | -0.494876 | 0.07075 | 0.942106 | 1.00062 | 0.01119 | 1.216815 | 0.012765 | -0.043603 | 0.630301 | -0.038872 | 0.121593 | -0.565465 | -0.486906 | -0.381353 | -1.59045 | -0.607432 | -1.318567 | -0.613557 | -1.205897 | -0.447911 | -0.332326 | -0.401487 | -0.844672 |
| 18.01389 | 0.862077 | 0.388146 | 0.125194 | 1.188092 | 1.177016 | 0.014952 | 1.638644 | 0.188223 | -0.048015 | 1.208809 | 0.004443 | 0.186202 | -0.366809 | -0.366382 | -0.072231 | 0.233124 | -0.425736 | -0.335646 | -0.686252 | -0.690514 | -0.288939 | -0.315197 | -0.265248 | -0.501504 |
| 18.02778 | 0.930782 | 0.974449 | 0.336783 | 1.492864 | 1.016206 | 0.006428 | 1.937747 | 0.230921 | -0.036757 | 1.343647 | 0.002379 | 0.130735 | -0.2735 | -0.317277 | -0.05846 | 0.801079 | -0.15606 | 0.27707 | -0.802798 | -0.371659 | -0.206991 | -0.392276 | -0.251468 | -0.147128 |
| 18.04167 | 0.815285 | 1.079083 | 0.640843 | 1.772128 | 1.079296 | 0.004594 | 2.148567 | 0.178781 | 0.022898 | 0.868595 | -0.006329 | 0.074758 | -0.231794 | -0.293087 | 0.035663 | 0.591073 | -0.023974 | 0.646529 | -0.355745 | 0.059344 | -0.115354 | -0.124256 | -0.214797 | -0.219257 |
| 18.05556 | 0.673104 | 1.186255 | 0.85584 | 1.938205 | 1.831349 | 0.056943 | 2.141763 | 0.192415 | 0.120441 | 0.510746 | 0.009997 | 0.11032 | -0.24018 | -0.321533 | 0.0728 | 0.35002 | -0.039931 | 0.662811 | -0.017689 | 0.375799 | -0.068365 | 0.115274 | -0.20463 | -0.419223 |
| 18.06944 | 0.60079 | 1.379566 | 0.841055 | 2.003488 | 2.513 | 0.131901 | 2.123032 | 0.171961 | 0.269585 | 0.474178 | 0.036168 | 0.214629 | -0.296388 | -0.389432 | 0.009468 | 0.1698 | -0.11338 | 0.458023 | 0.10831 | 0.467984 | -0.073561 | 0.197989 | -0.223163 | -0.548758 |
| 18.08333 | 0.507381 | 1.788187 | 0.77034 | 2.174787 | 1.811759 | 0.177226 | 1.871108 | 0.116073 | 0.455005 | 0.586878 | 0.06337 | 0.286097 | -0.341765 | -0.331527 | -0.197368 | 0.074352 | -0.156151 | 0.258936 | 0.143025 | 0.516706 | -0.067116 | 0.203844 | -0.222583 | -0.596209 |
| 18.09722 | 0.388999 | 2.180898 | 0.688272 | 2.25992 | 0.944233 | 0.176452 | 1.22882 | 0.173642 | 0.512746 | 0.487025 | 0.117116 | 0.257485 | -0.399901 | -0.24368 | -0.615607 | -0.237023 | -0.132301 | 0.129046 | 0.010375 | 0.44267 | -0.082348 | 0.008843 | -0.240947 | -0.486603 |
| 18.11111 | 0.249733 | 2.031263 | 0.588847 | 2.412452 | 0.460097 | 0.109485 | 0.645381 | 0.329192 | 0.41885 | 0.57542 | 0.187666 | 0.161684 | -0.475812 | -0.170324 | -0.939475 | -0.495095 | -0.101172 | 0.01774 | -0.036111 | 0.304171 | -0.111283 | -0.162856 | -0.158658 | -0.388006 |
| 18.125 | 0.241323 | 1.354216 | 0.497266 | 2.77569 | 0.510662 | 0.046546 | 0.210052 | 0.543743 | 0.171201 | 0.788307 | 0.210731 | 0.09912 | -0.560047 | -0.088813 | -0.817119 | -0.441096 | -0.081685 | -0.047679 | 0.092808 | 0.199097 | -0.109833 | -0.035359 | 0.006508 | -0.409708 |
| 18.13889 | 0.45967 | 1.298939 | 0.589132 | 3.0979 | 1.141262 | 0.06251 | 0.137493 | 1.16699 | 0.089691 | 0.865718 | 0.193096 | 0.206673 | -0.528248 | -0.039522 | -0.76245 | -0.491257 | -0.038946 | -0.118691 | -0.031076 | -0.023049 | -0.108798 | -0.122441 | 0.113686 | -0.255212 |
| 18.15278 | 0.694526 | 1.624741 | 0.792818 | 2.64449 | 1.672087 | 0.140552 | 0.420196 | 1.848706 | 0.252227 | 0.719313 | 0.182148 | 0.415793 | -0.395234 | -0.089724 | -0.908678 | -0.664859 | 0.001881 | -0.170967 | -0.313509 | -0.353167 | -0.127753 | -0.429224 | 0.134626 | -0.037723 |
| 18.16667 | 0.780593 | 1.60844 | 0.777127 | 1.810212 | 2.104515 | 0.204344 | 1.009717 | 2.100406 | 0.525881 | 0.447111 | 0.267431 | 0.616702 | -0.405448 | -0.182879 | -0.90578 | -0.594766 | 0.011406 | -0.159435 | -0.366053 | -0.408555 | -0.150945 | -0.465872 | 0.103342 | -0.079069 |
| 18.18056 | 0.884266 | 1.574753 | 0.646737 | 1.802707 | 2.378102 | 0.328769 | 1.497239 | 1.613636 | 0.77713 | 0.179654 | 0.39247 | 0.506175 | -0.513538 | -0.204104 | -0.637629 | -0.294269 | -0.020333 | -0.083061 | -0.169114 | -0.217136 | -0.184517 | -0.291035 | 0.07397 | -0.241555 |
| 18.19444 | 1.012563 | 2.071624 | 0.570396 | 1.787498 | 2.786999 | 0.539181 | 1.643388 | 0.747583 | 0.936386 | 0.141111 | 0.467666 | 0.634926 | -0.519854 | -0.081314 | -0.41747 | -0.012879 | -0.104813 | 0.016238 | 0.058981 | 0.056732 | -0.202254 | -0.039934 | 0.03616 | -0.366042 |
| 18.20833 | 0.955502 | 3.427466 | 0.410102 | 2.334833 | 2.488402 | 0.681148 | 1.435233 | 0.125453 | 1.085352 | 0.373023 | 0.473725 | 1.112747 | -0.407245 | 0.011129 | -0.367812 | 0.102516 | -0.173791 | 0.01675 | 0.131449 | 0.176107 | -0.207328 | 0.049361 | -0.039276 | -0.389852 |
| 18.22222 | 0.619584 | 3.983933 | 0.383084 | 3.518173 | 1.610205 | 0.644528 | 0.961309 | -0.10548 | 1.090775 | 0.608305 | 0.494048 | 0.857418 | -0.360178 | -0.023373 | -0.462199 | -0.012932 | -0.163194 | -0.100435 | -0.053335 | -0.019492 | -0.258528 | -0.164523 | -0.135695 | -0.292414 |
| 18.23611 | 0.227856 | 1.996952 | 0.275856 | 2.780252 | 0.234479 | 0.445023 | 0.538511 | -0.042885 | 1.050402 | 0.586222 | 0.333377 | 0.522419 | -0.360555 | -0.107826 | -0.441721 | -0.131155 | -0.143074 | -0.167744 | -0.181743 | -0.186262 | -0.263665 | -0.299015 | -0.166274 | -0.184963 |
| 18.25 | -0.108601 | -0.476414 | 0.06843 | 0.739717 | -1.548551 | 0.286622 | 0.228663 | -0.195655 | 1.237118 | 0.405926 | 0.278946 | 0.218989 | -0.355094 | -0.150307 | -0.299322 | -0.109509 | -0.134433 | -0.135135 | -0.133442 | -0.170216 | -0.186029 | -0.196121 | -0.116894 | -0.098226 |
| 18.26389 | -0.050007 | -2.429786 | 0.253754 | -0.125767 | -1.498071 | 0.153195 | 0.191356 | -0.366498 | 1.24874 | 0.160479 | 0.151549 | -0.020775 | -0.339996 | -0.144623 | -0.102643 | 0.029692 | -0.193873 | -0.039352 | 0.021771 | -0.009375 | -0.043701 | 0.049096 | -0.040573 | -0.028448 |
| 18.27778 | 0.324214 | -1.142575 | 0.344114 | -0.010456 | 0.398745 | 0.120852 | 0.307846 | -0.607702 | 0.975759 | -0.002384 | 0.151478 | 0.05056 | -0.315657 | -0.189892 | -0.02876 | 0.042665 | -0.208586 | 0.011046 | 0.079219 | 0.065685 | 0.035276 | 0.157101 | -0.019084 | 0.090484 |
| 18.29167 | 0.514146 | 2.077483 | 0.243406 | 0.657704 | 2.022827 | 0.143656 | 0.488581 | -0.193893 | 0.747337 | -0.002079 | 0.195224 | 0.393423 | -0.293065 | -0.29996 | -0.114969 | -0.132233 | -0.120528 | -0.05693 | -0.105501 | -0.087684 | 0.008285 | -0.039537 | -0.051629 | 0.28436 |
| 18.30556 | 0.330486 | 3.739393 | 0.045994 | 1.00146 | 1.726019 | 0.156475 | 0.549349 | 0.493866 | 0.46949 | 0.114389 | 0.163326 | 0.753784 | -0.266529 | -0.344817 | -0.141257 | -0.227196 | -0.089678 | -0.112197 | -0.213225 | -0.186786 | -0.02948 | -0.160359 | -0.058154 | 0.30532 |
| 18.31944 | 0.203366 | 1.10261 | -0.045236 | 0.303306 | 1.295622 | 0.168622 | 0.583196 | 0.653347 | 0.379146 | 0.187477 | 0.255887 | 0.516371 | -0.244938 | -0.337275 | -0.14675 | -0.287738 | -0.087025 | -0.150926 | -0.282968 | -0.250678 | -0.059212 | -0.239273 | -0.066781 | 0.266248 |
| 18.33333 | 0.327087 | -2.153311 | 0.06332 | -1.108971 | 0.847218 | 0.225019 | 0.610094 | 0.193649 | 0.314921 | 0.1904 | 0.355256 | -0.310004 | -0.219403 | -0.261336 | -0.081813 | -0.26634 | -0.147715 | -0.147791 | -0.259449 | -0.230338 | -0.063756 | -0.227602 | -0.054039 | 0.286102 |
| 18.34722 | 0.622359 | -2.921131 | 0.25325 | -0.798328 | 0.555981 | 0.246368 | 0.437533 | -0.354647 | 0.122504 | 0.268488 | 0.412156 | -0.779625 | -0.164772 | -0.105099 | 0.171778 | -0.053311 | -0.242747 | -0.056946 | -0.036115 | -0.043807 | -0.018728 | 0.024367 | -0.013354 | 0.282294 |
| 18.36111 | 0.410965 | -0.419648 | 0.320893 | 0.575972 | 0.706122 | 0.188205 | 0.190231 | -0.25272 | -0.102032 | 0.334401 | 0.199882 | -0.265328 | -0.155903 | -0.125674 | 0.365689 | 0.025426 | -0.242163 | -0.01427 | 0.014546 | -0.039627 | -0.028653 | 0.051939 | 0.003082 | 0.359888 |
| 18.375 | 0.335079 | 2.398837 | 0.24298 | 2.052292 | 0.83462 | 0.053547 | 0.165408 | -0.337584 | -0.24415 | 0.404069 | 0.136583 | 0.066313 | -0.202538 | -0.324191 | 0.422433 | -0.084841 | -0.125954 | -0.051463 | -0.179327 | -0.259555 | -0.096119 | -0.189149 | 0.022849 | 0.533367 |
| 18.38889 | 0.241169 | 1.552765 | 0.065141 | 1.744346 | 0.5123 | -0.009995 | 0.452125 | 0.085137 | -0.192523 | 0.467448 | 0.225738 | 0.435429 | -0.115861 | -0.298903 | 0.59623 | 0.004993 | -0.026932 | -0.015107 | -0.097138 | -0.183123 | -0.078841 | -0.095172 | 0.063334 | 0.635438 |
| 18.40278 | 0.157175 | -0.625376 | -0.020312 | 0.421664 | -0.092528 | 0.11152 | 0.62155 | 0.887612 | -0.054895 | 0.495508 | 0.213792 | 0.116681 | 0.081694 | -0.023432 | 0.892246 | 0.311294 | -0.054693 | 0.09026 | 0.226448 | 0.160657 | 0.022456 | 0.298888 | 0.092831 | 0.589102 |
| 18.41667 | 0.173726 | -1.321608 | 0.012683 | -0.608174 | -0.449983 | 0.223058 | 0.485452 | 1.171378 | 0.131873 | 0.413769 | 0.313937 | -0.240112 | 0.16264 | 0.074076 | 0.976035 | 0.418935 | -0.110338 | 0.144365 | 0.270112 | 0.253569 | 0.059873 | 0.405195 | 0.088516 | 0.639006 |
| 18.43056 | 0.396589 | 0.258244 | -0.004149 | -0.788271 | -0.191507 | 0.234973 | 0.30048 | 0.836165 | 0.300865 | 0.348127 | 0.254688 | 0.050767 | 0.091147 | -0.128023 | 0.993298 | 0.468789 | -0.113923 | 0.190508 | 0.072081 | 0.057816 | 0.012018 | 0.212918 | 0.066216 | 0.576576 |
| 18.44444 | 0.648223 | 2.05605 | -0.405645 | -0.015144 | 0.114572 | 0.147798 | 0.148474 | -0.190001 | 0.260379 | 0.269303 | 0.291539 | 0.46563 | -0.053532 | -0.620741 | 1.307296 | 0.610371 | 0.007121 | 0.3444 | 0.03964 | -0.047205 | -0.08774 | 0.056462 | -0.056984 | 0.577303 |
| 18.45833 | 0.493438 | 1.957056 | -0.358678 | 0.48086 | 0.101606 | 0.076155 | 0.131671 | 0.046996 | 0.09116 | 0.194606 | 0.309769 | 0.589594 | -0.094704 | -1.187704 | 1.354081 | 0.58294 | 0.30737 | 0.526325 | 0.176862 | 0.034213 | -0.107692 | -0.07664 | -0.247591 | 0.639363 |
| 18.47222 | 0.164435 | 1.092861 | -0.169582 | 0.33262 | -0.023684 | 0.088369 | 0.126045 | 0.393496 | 0.032143 | 0.162996 | 0.402915 | 0.135937 | 0.096314 | -1.206869 | 1.067441 | 0.471827 | 0.524283 | 0.622475 | 0.327912 | 0.019329 | -0.016694 | -0.041658 | -0.109637 | 0.996092 |
| 18.48611 | 0.123625 | 0.7989 | -0.026895 | -0.011224 | 0.019992 | 0.068262 | 0.06713 | 0.588735 | 0.107909 | 0.157184 | 0.168199 | 0.082995 | 0.34104 | -0.827586 | 0.880992 | 0.467462 | 0.579156 | 0.549151 | 0.389359 | 0.073032 | 0.161339 | 0.167305 | 0.202453 | 1.241278 |
| 18.5 | 0.191316 | 0.758941 | 0.233845 | 0.064444 | 0.255618 | -0.013349 | -0.053895 | 0.253372 | 0.15286 | 0.201392 | 0.089456 | 0.348932 | 0.547196 | -0.266751 | 0.861441 | 0.493555 | 0.542227 | 0.45063 | 0.447127 | 0.239644 | 0.425437 | 0.425791 | 0.380453 | 1.309235 |
| 18.51389 | 0.140132 | 0.616395 | 0.563734 | 0.555906 | 0.320198 | -0.05036 | -0.080693 | -0.10353 | 0.057421 | 0.246456 | 0.063947 | 0.62187 | 0.709009 | 0.238422 | 0.904764 | 0.552054 | 0.52284 | 0.324443 | 0.430916 | 0.263212 | 0.595744 | 0.664906 | 0.473299 | 1.290669 |
| 18.52778 | 0.066064 | 0.076209 | 0.12334 | 0.56398 | 0.385279 | -0.025127 | -0.057955 | 0.08239 | -0.174574 | 0.225414 | 0.024607 | 0.434126 | 0.937278 | 0.264766 | 0.6752 | 0.769039 | 0.751371 | 0.228068 | 0.276585 | 0.14667 | 0.574475 | 0.586411 | 0.658402 | 1.232554 |
| 18.54167 | 0.051191 | -0.777211 | -0.090816 | -0.054831 | 0.370964 | -0.069052 | -0.086572 | 0.410374 | -0.408653 | 0.188939 | 0.048993 | -0.168197 | 1.282412 | 0.328716 | 0.540893 | 1.110131 | 0.963552 | 0.327188 | 0.194596 | 0.225038 | 0.437378 | 0.413254 | 0.994765 | 1.350338 |
| 18.55556 | 0.063404 | -1.171615 | -0.229491 | -0.36741 | 0.014151 | -0.11415 | -0.264342 | 0.190873 | -0.456487 | 0.232326 | 0.081276 | -0.1784 | 1.615437 | 0.55354 | 0.557156 | 1.459702 | 0.911713 | 0.570489 | 0.222158 | 0.421397 | 0.352611 | 0.40481 | 1.170856 | 1.586621 |
| 18.56944 | 0.026648 | -0.567459 | -0.135683 | -0.198485 | -0.42467 | -0.124487 | -0.457953 | -0.555005 | -0.42833 | 0.28323 | 0.024747 | -0.214359 | 1.433057 | 0.532739 | 0.202553 | 1.153573 | 0.709259 | 0.590906 | 0.23105 | 0.403652 | 0.251195 | 0.187015 | 0.790461 | 1.228179 |
| 18.58333 | -0.027575 | -0.085242 | 0.047529 | -0.471257 | -0.572769 | -0.098407 | -0.41151 | -0.851309 | -0.588534 | 0.279089 | -0.007676 | -0.703408 | 0.96205 | 0.373439 | -0.330161 | 0.556884 | 0.49452 | 0.435097 | 0.235304 | 0.243734 | 0.096785 | -0.035683 | 0.112201 | 0.456668 |
| 18.59722 | -0.029571 | 0.11671 | -0.001798 | -1.371315 | -0.541221 | -0.074705 | -0.245191 | -0.377215 | -0.946198 | 0.242972 | 0.049169 | -1.266506 | 0.622201 | 0.096612 | -0.950628 | 0.079587 | 0.404245 | 0.300656 | 0.180804 | 0.082442 | -0.011646 | -0.175717 | -0.193118 | -0.286657 |
| 18.61111 | -0.228972 | -0.443191 | -0.791051 | -2.415286 | -0.703958 | -0.147476 | -0.197792 | -0.09242 | -0.890329 | 0.14894 | -0.087267 | -1.337708 | 0.361936 | -0.039066 | -0.869885 | -0.198227 | 0.39251 | 0.189742 | 0.07876 | 0.028843 | -0.007449 | -0.107918 | -0.22081 | -0.857449 |
| 18.625 | -1.021538 | -2.064667 | -1.41809 | -4.118719 | -1.461761 | -0.3903 | -0.6242 | -0.26487 | -0.768886 | -0.170633 | -0.612647 | -1.479841 | 0.28281 | 0.168106 | -0.260823 | -0.377609 | 0.315024 | 0.15984 | 0.041879 | 0.127766 | 0.167962 | 0.205846 | -0.055438 | -0.762031 |
| 18.63889 | -1.046793 | -1.552567 | -0.602241 | -4.145418 | -1.980347 | -0.618818 | -0.713982 | -0.187249 | -0.790676 | -0.569196 | -0.952199 | -1.22806 | 0.367174 | 0.426895 | 0.303964 | -0.544072 | 0.272121 | 0.053063 | 0.019027 | 0.091617 | 0.208999 | 0.208091 | 0.228331 | 0.022469 |
| 18.65278 | -0.131082 | 0.573619 | 0.784011 | -0.846121 | -0.662169 | -0.505259 | -0.36272 | 1.026439 | -0.65614 | -0.31074 | -0.566529 | -0.537047 | 0.385389 | 0.442625 | 0.454273 | -0.513273 | 0.246478 | -0.057612 | 0.032317 | -0.017082 | -0.055547 | -0.186308 | 0.314291 | 0.931659 |
| 18.66667 | 0.382193 | 0.870566 | 1.720059 | 1.164239 | 0.122588 | -0.210675 | -0.713414 | 1.282577 | -0.364314 | 0.056874 | -0.287421 | 0.001003 | 0.280492 | 0.144303 | 0.313502 | -0.054377 | 0.15239 | -0.030669 | 0.088472 | 0.02446 | -0.367011 | -0.503075 | 0.343467 | 1.311924 |
| 18.68056 | 0.201496 | -0.648848 | 1.46962 | 1.602345 | -0.838593 | -0.031385 | -1.07455 | -0.475077 | -0.063444 | 0.131452 | -0.044749 | 0.225187 | 0.115976 | -0.068201 | -0.096049 | 0.564454 | 0.0943 | 0.026355 | 0.071914 | 0.106343 | -0.265445 | -0.34479 | 0.282744 | 0.617239 |
| 18.69444 | 0.086824 | -3.100685 | 0.217126 | 0.391171 | -1.60023 | -0.060152 | -1.143899 | -3.269821 | 0.040852 | 0.047979 | -0.290447 | -0.024876 | -0.00813 | 0.004247 | -0.59974 | 0.717746 | 0.091788 | 0.068472 | 0.039753 | 0.178884 | 0.157123 | 0.311802 | 0.213671 | -0.302973 |
| 18.70833 | 0.060501 | -4.651356 | -0.599585 | -1.400622 | -1.27139 | -0.326266 | -1.065841 | -4.093133 | -0.109348 | -0.082596 | -0.228027 | -0.042546 | 0.025332 | 0.561763 | -0.499969 | 0.149314 | 0.073548 | 0.098745 | 0.063324 | 0.20187 | 0.762254 | 1.050757 | 0.289224 | -0.381117 |
| 18.72222 | -0.342761 | -2.180838 | -1.231228 | -3.092417 | -0.962438 | -0.482988 | -0.958944 | -1.128476 | -0.437459 | -0.180214 | -0.184888 | 0.12111 | 0.114794 | 0.865387 | -0.303388 | -0.393662 | 0.051188 | 0.090275 | 0.119763 | 0.124142 | 0.609673 | 0.597847 | 0.372211 | 0.324212 |
| 18.73611 | -0.295288 | -1.431866 | -1.438938 | -4.822167 | -1.434196 | -0.382303 | -0.903268 | 1.687797 | -0.729833 | -0.186486 | -0.173247 | 0.08417 | 0.034618 | 0.264299 | -0.321441 | -0.3769 | 0.073192 | 0.025684 | 0.110068 | -0.043609 | -0.189819 | -0.284176 | 0.263286 | 0.804686 |
| 18.75 | 0.285348 | -2.98053 | -0.576061 | -5.383522 | -2.124632 | -0.250572 | -1.095359 | 0.63337 | -0.729855 | -0.34097 | -0.289999 | -0.36703 | -0.182504 | -0.332816 | -0.351932 | 0.05265 | 0.120253 | -0.078906 | -0.018132 | -0.232415 | -0.678057 | -0.872996 | 0.187374 | 0.811381 |
| 18.76389 | 0.098826 | -4.721672 | -0.015673 | -4.288573 | -1.615094 | -0.327896 | -1.196905 | -2.313838 | -0.743661 | -0.800826 | -0.226811 | -0.682201 | -0.210138 | -0.582267 | -0.030444 | 0.404965 | 0.174296 | -0.130757 | -0.116463 | -0.286423 | -0.332023 | -0.462009 | 0.1316 | 0.535646 |
| 18.77778 | -0.743145 | -4.142716 | 0.196423 | -3.525478 | -0.632883 | -0.459843 | -0.888337 | -2.924735 | -0.688137 | -0.729567 | -0.346676 | -0.121147 | 0.086408 | -0.007928 | 0.513439 | 0.32657 | 0.134973 | -0.11367 | -0.128788 | -0.206174 | 0.293436 | 0.550233 | 0.047879 | 0.28955 |
| 18.79167 | -1.213805 | -1.416448 | 0.057118 | -2.903281 | 0.310967 | -0.347939 | -0.513369 | -0.445345 | -0.845049 | -0.681269 | -0.332073 | 0.287851 | 0.263452 | 0.507788 | 0.903675 | -0.007687 | 0.043821 | -0.102416 | -0.091522 | -0.120651 | 0.465204 | 0.586015 | -0.179821 | 0.189855 |
| 18.80556 | -0.251752 | 0.490673 | 0.134777 | -1.477904 | 0.678695 | -0.226279 | -0.160419 | 2.201989 | -0.689619 | -0.39714 | -0.306831 | -0.338538 | 0.018091 | 0.058701 | 0.319482 | -0.112449 | 0.017241 | -0.054907 | -0.016932 | -0.067167 | -0.032949 | -0.043972 | -0.361151 | 0.076342 |
| 18.81944 | 0.345679 | 0.689816 | 0.924849 | 1.191925 | 0.273127 | -0.220984 | 0.381768 | 2.359357 | 0.023832 | 0.112619 | -0.191565 | -0.881213 | -0.306214 | -0.430896 | -0.327228 | 0.157305 | 0.039629 | 0.001248 | 0.023856 | -0.088075 | -0.434145 | -0.593368 | -0.157767 | 0.14685 |
| 18.83333 | 0.138599 | 0.791432 | 1.714387 | 3.185138 | -0.027341 | -0.205894 | 0.938106 | 0.633313 | 0.66962 | 0.268014 | 0.008957 | -0.108063 | -0.330512 | -0.678113 | -0.502864 | 0.506578 | 0.172403 | -0.007497 | -0.025258 | -0.12592 | -0.278414 | -0.508516 | 0.30379 | 0.428152 |
| 18.84722 | -0.320542 | 0.958899 | 0.901127 | 2.795114 | 0.652598 | -0.207705 | 1.227231 | -0.274072 | 0.860187 | 0.308367 | 0.150801 | 0.783212 | -0.067464 | -0.329689 | -0.234554 | 0.389134 | 0.29546 | -0.026551 | -0.099642 | -0.038019 | 0.144396 | 0.091451 | 0.592121 | 0.589186 |
| 18.86111 | -0.245286 | 1.061975 | -0.374918 | 2.086106 | 1.901484 | -0.182428 | 1.388031 | 0.290198 | 0.5186 | 0.698394 | 0.240671 | 0.718786 | 0.106118 | 0.244895 | -0.061145 | -0.087072 | 0.254457 | -0.038227 | -0.122028 | 0.091995 | 0.18522 | 0.207723 | 0.416432 | 0.483079 |
| 18.875 | 0.195008 | 0.913894 | -1.148099 | 2.758643 | 2.440801 | 0.000706 | 1.30564 | 1.591068 | 0.387331 | 0.750209 | 0.350943 | 0.115807 | -0.061166 | 0.103295 | -0.264254 | -0.386293 | 0.136391 | 0.021619 | -0.054992 | 0.212898 | -0.067226 | -0.043208 | 0.064721 | 0.103042 |
| 18.88889 | 0.359841 | 0.657304 | -1.02755 | 4.104404 | 1.96825 | 0.119597 | 0.834238 | 1.023279 | 0.661654 | 0.091955 | 0.423856 | 0.011307 | -0.288238 | -0.135725 | -0.547513 | -0.342691 | -0.014871 | 0.11094 | 0.017209 | 0.231038 | -0.273286 | -0.293622 | -0.062196 | -0.195225 |
| 18.90278 | 0.096215 | 0.418781 | -0.186663 | 3.90556 | 0.708803 | -0.080818 | 0.494564 | -0.574814 | 1.129139 | -0.16737 | 0.444903 | 0.288346 | -0.262707 | -0.323574 | -0.35595 | -0.25709 | -0.044485 | 0.088 | 0.000843 | 0.053334 | -0.104413 | -0.17041 | 0.048253 | -0.019799 |
| 18.91667 | -0.221922 | 0.319418 | 0.520155 | 2.760475 | -0.247207 | -0.39718 | 0.532181 | -2.071452 | 1.647653 | -0.132228 | 0.483871 | 0.436039 | -0.106364 | -0.135552 | -0.002213 | -0.342688 | 0.080902 | -0.003827 | -0.07349 | -0.133769 | 0.065769 | -0.034069 | 0.110756 | 0.280462 |
| 18.93056 | -0.196644 | 0.541786 | 1.007933 | 1.87435 | 0.046333 | -0.441355 | 0.623071 | -1.056576 | 1.452231 | 0.038519 | 0.487773 | 0.218949 | 0.000999 | -0.118518 | -0.08924 | -0.65362 | 0.077787 | -0.132413 | -0.15707 | -0.343035 | -0.012295 | -0.201802 | 0.009634 | 0.348578 |
| 18.94444 | 0.097236 | 1.222884 | 0.734302 | 1.701261 | 1.362431 | -0.199643 | 0.562602 | 0.440738 | 0.405052 | 0.157011 | 0.381618 | 0.097363 | -0.022759 | -0.597086 | -0.589759 | -1.049659 | -0.124375 | -0.279343 | -0.194824 | -0.644566 | -0.216852 | -0.456394 | -0.219711 | 0.098752 |
| 18.95833 | 0.284857 | 1.845604 | 0.021402 | 1.064373 | 2.40037 | 0.102732 | 0.674623 | 0.725477 | -0.531567 | 0.104258 | 0.284914 | 0.228717 | -0.337583 | -0.92688 | -1.096324 | -1.486417 | -0.500027 | -0.455656 | -0.280119 | -0.96457 | -0.690821 | -0.670928 | -0.613439 | -0.585476 |
| 18.97222 | 0.192068 | 1.79061 | -0.363597 | -0.417244 | 2.13679 | 0.201224 | 0.922664 | 0.157158 | -1.026135 | 0.00569 | 0.102765 | 0.258757 | -0.820564 | -1.288349 | -1.679332 | -2.102827 | -0.953177 | -0.70567 | -0.675117 | -1.432442 | -0.908071 | -0.419871 | -1.069398 | -1.352093 |
| 18.98611 | 0.034933 | 1.284809 | -0.103906 | -2.021204 | 0.849256 | 0.135664 | 1.064136 | -0.176697 | -0.672698 | -0.04332 | -0.014493 | 0.10104 | -1.028061 | -1.184369 | -1.612654 | -1.993306 | -0.924456 | -0.616475 | -0.921895 | -1.335519 | -0.578403 | 0.007856 | -1.109278 | -1.577451 |
| 19 | 0.071157 | 0.684809 | 0.411913 | -2.982988 | -0.033097 | 0.058063 | 0.991718 | 0.023981 | -0.066776 | -0.025593 | -0.05615 | 0.140304 | -0.769707 | -0.698558 | -0.863715 | -0.76848 | -0.429031 | -0.086003 | -0.70828 | -0.397223 | -0.758517 | -0.070081 | -0.787563 | -1.222867 |
| 19.01389 | 0.231016 | 0.026594 | 0.684869 | -2.358807 | -0.050524 | 0.138027 | 0.751213 | 0.230502 | 0.397013 | 0.089399 | 0.003775 | 0.316089 | -0.400012 | -0.45481 | -0.455778 | 0.069994 | -0.136348 | 0.109656 | -0.445375 | -0.00783 | -0.633561 | -0.198291 | -0.488148 | -0.924914 |
| 19.02778 | 0.30041 | -0.826605 | 0.710242 | -0.583596 | 0.599974 | 0.268525 | 0.605129 | 0.212984 | 0.46281 | 0.182927 | 0.064101 | 0.323429 | -0.121898 | -0.580122 | -0.421203 | 0.048244 | -0.20285 | -0.017723 | -0.420179 | -0.245249 | -0.178968 | -0.451351 | -0.218169 | -0.575282 |
| 19.04167 | 0.285602 | -1.560674 | 0.817807 | 1.18262 | 1.300296 | 0.330925 | 0.806583 | 0.288729 | 0.319879 | 0.203017 | 0.069278 | 0.15971 | -0.070079 | -0.627868 | -0.336987 | -0.012789 | -0.189503 | -0.039051 | -0.279709 | -0.220539 | -0.109353 | -0.611722 | 0.009311 | -0.276141 |
| 19.05556 | 0.347293 | -1.265531 | 0.800118 | 2.356701 | 1.757236 | 0.288069 | 1.201223 | 0.708947 | 0.240981 | 0.218423 | 0.049446 | -0.04404 | -0.206533 | -0.55116 | -0.280385 | 0.0401 | -0.070008 | 0.015823 | -0.043329 | 0.031506 | -0.073436 | -0.534992 | 0.133997 | -0.182854 |
| 19.06944 | 0.601892 | -0.083974 | 0.404157 | 2.361665 | 2.048464 | 0.249349 | 1.62893 | 1.425946 | 0.273664 | 0.180012 | 0.063627 | 0.022887 | -0.352721 | -0.503006 | -0.382902 | 0.055269 | -0.039121 | 0.006326 | 0.086641 | 0.128647 | 0.002234 | -0.398802 | 0.084704 | -0.263019 |
| 19.08333 | 1.048765 | 1.544422 | -0.158191 | 1.913579 | 1.638382 | 0.309764 | 1.854914 | 1.857181 | 0.223126 | 0.164829 | 0.118548 | 0.4122 | -0.391915 | -0.413032 | -0.338961 | 0.084609 | 0.025865 | 0.002334 | 0.187285 | 0.242741 | 0.104689 | -0.133366 | -0.023667 | -0.419189 |
| 19.09722 | 1.032451 | 2.589321 | -0.128894 | 1.387535 | 0.75432 | 0.356541 | 1.403966 | 1.549211 | -0.015437 | 0.136185 | 0.21091 | 0.683004 | -0.35543 | -0.324021 | -0.149015 | 0.077637 | 0.118157 | -0.002317 | 0.213902 | 0.280357 | 0.116528 | 0.040733 | -0.078468 | -0.461197 |
| 19.11111 | 0.451029 | 2.327981 | 0.285334 | 0.865747 | -0.076965 | 0.204309 | 0.426907 | 0.926145 | -0.418933 | 0.128076 | 0.169852 | 0.459359 | -0.327416 | -0.352434 | 0.005847 | -0.024497 | 0.030644 | -0.097448 | 0.037103 | 0.004946 | -0.01445 | -0.081758 | -0.086512 | -0.25743 |
| 19.125 | 0.032154 | 1.779943 | 0.706008 | 0.592945 | -0.376015 | -0.0396 | -0.290978 | 0.310568 | -0.869463 | 0.216144 | 0.045917 | 0.122254 | -0.323019 | -0.310827 | 0.015417 | -0.14002 | -0.163636 | -0.216867 | -0.199912 | -0.31111 | -0.212664 | -0.304013 | -0.094397 | -0.016218 |
| 19.13889 | 0.22016 | 1.797401 | 1.126161 | 1.25451 | -0.26967 | -0.052795 | -0.205332 | 0.02439 | -0.781609 | 0.30715 | 0.027939 | 0.231219 | -0.330924 | -0.141775 | -0.078456 | -0.137313 | -0.252469 | -0.252998 | -0.306363 | -0.372971 | -0.294443 | -0.318165 | -0.108244 | 0.109101 |
| 19.15278 | 0.905426 | 1.841152 | 1.669783 | 2.675435 | -0.07065 | 0.319971 | 0.606621 | 0.220892 | 0.121415 | 0.251555 | 0.385005 | 0.724264 | -0.418596 | 0.065311 | -0.112529 | 0.154987 | -0.168905 | -0.151961 | -0.167843 | -0.063833 | -0.157365 | -0.214666 | -0.130276 | 0.093066 |
| 19.16667 | 1.835643 | 2.031597 | 1.883901 | 4.756099 | 0.78542 | 0.85476 | 1.592323 | 0.840447 | 1.229937 | 0.201378 | 1.083948 | 1.261252 | -0.452281 | 0.233426 | -0.011261 | 0.61517 | 0.113288 | -0.031556 | 0.143093 | 0.439344 | -0.031748 | -0.239921 | -0.107424 | 0.008783 |
| 19.18056 | 2.21288 | 1.851806 | 1.905196 | 5.093659 | 2.220428 | 1.20244 | 2.028477 | 1.544241 | 2.041508 | 0.293695 | 1.582678 | 1.433248 | -0.31715 | 0.216553 | -0.048048 | 0.715949 | 0.3121 | -0.00404 | 0.242161 | 0.555136 | -0.136628 | -0.395952 | -0.035868 | 0.024255 |
| 19.19444 | 1.851382 | 0.951446 | 1.690141 | 3.341399 | 2.953065 | 1.27253 | 2.188632 | 1.753993 | 1.872823 | 0.454698 | 1.937612 | 1.93856 | -0.152211 | 0.036399 | -0.351538 | 0.476401 | 0.244594 | -0.059938 | 0.073544 | 0.210248 | -0.236664 | -0.401663 | -0.002755 | 0.017691 |
| 19.20833 | 1.740188 | -0.328653 | 1.181038 | 1.881579 | 1.878338 | 1.086706 | 2.554838 | 1.543515 | 1.68944 | 0.553733 | 2.222057 | 1.801018 | -0.092058 | -0.059959 | -0.606092 | 0.270323 | 0.075429 | -0.092572 | -0.067179 | -0.063878 | -0.215673 | -0.295682 | -0.016616 | -0.109339 |
| 19.22222 | 1.318896 | -0.535923 | 0.626016 | 1.310378 | 0.564916 | 0.689999 | 3.144941 | 1.192998 | 1.526972 | 0.527128 | 1.639864 | 1.507062 | -0.11808 | -0.141249 | -0.456664 | 0.145938 | -0.103042 | -0.131055 | -0.198028 | -0.219608 | -0.147305 | -0.270303 | 0.014579 | -0.132595 |
| 19.23611 | 0.785347 | 0.487503 | 0.185623 | 1.275982 | -0.404582 | 0.279741 | 3.284744 | 0.90996 | 1.102477 | 0.39554 | 1.499039 | 1.49135 | -0.162593 | -0.210218 | 0.074222 | 0.048863 | -0.086128 | -0.174707 | -0.309011 | -0.31237 | -0.097725 | -0.324026 | 0.122989 | -0.001747 |
| 19.25 | 0.500965 | 1.339735 | -0.375085 | 1.831659 | -0.952862 | 0.041251 | 2.929555 | 0.635614 | 0.928004 | 0.253611 | 1.077007 | 1.144085 | -0.140419 | -0.157566 | 0.66181 | 0.121173 | 0.131603 | -0.097701 | -0.175625 | -0.115789 | -0.009417 | -0.187369 | 0.231269 | 0.019747 |
| 19.26389 | 0.352315 | 1.204624 | -1.202031 | 2.465173 | -0.905825 | 0.034866 | 1.676577 | 0.318777 | 0.833288 | 0.116948 | 0.78541 | 0.640001 | -0.054371 | -0.08626 | 0.572133 | 0.143424 | 0.236196 | 0.013658 | 0.057434 | 0.129119 | 0.165515 | -0.006328 | 0.244 | -0.017884 |
| 19.27778 | 0.254161 | 0.686411 | -1.484009 | 2.495958 | -0.674974 | 0.118038 | 0.409937 | 0.018525 | 0.650256 | -0.132598 | 0.274005 | 0.283052 | 0.051608 | -0.049215 | 0.151832 | 0.012413 | 0.146589 | 0.07883 | 0.143769 | 0.212827 | 0.248346 | 0.094546 | 0.222431 | 0.152884 |
| 19.29167 | 0.130321 | 0.604233 | -0.439574 | 1.842904 | -0.419067 | 0.215245 | 0.142859 | 0.084179 | 0.88339 | -0.201332 | 0.305709 | 0.277672 | 0.163107 | -0.04567 | -0.239395 | -0.156646 | 0.053244 | 0.019314 | 0.010669 | 0.028724 | 0.09955 | 0.18428 | 0.267368 | 0.666767 |
| 19.30556 | 0.138471 | 0.891668 | 0.899991 | 1.686243 | -0.152854 | 0.20638 | 0.341815 | 0.492396 | 1.445525 | -0.182785 | 0.395503 | 0.655853 | 0.279897 | -0.022032 | -0.292592 | -0.173226 | 0.012914 | -0.093142 | -0.150565 | -0.169368 | -0.105975 | 0.303569 | 0.280941 | 0.993838 |
| 19.31944 | 0.070226 | 1.199973 | 1.932756 | 0.939978 | 0.689514 | 0.040073 | 0.297498 | 0.278818 | 2.064438 | -0.097319 | 0.409931 | 1.210198 | 0.332594 | 0.081532 | 0.151823 | 0.078027 | 0.115427 | -0.052501 | -0.034805 | -0.004794 | -0.157137 | 0.548225 | 0.202655 | 0.729793 |
| 19.33333 | 0.156578 | 0.202489 | 1.415588 | 0.060879 | 1.614654 | -0.061654 | 0.199285 | -0.016245 | 2.081632 | -0.002081 | 1.002344 | 1.044238 | 0.392033 | 0.11539 | 0.72896 | 0.285307 | 0.184709 | 0.022642 | 0.046746 | 0.090531 | -0.061409 | 0.765822 | 0.121245 | 0.402731 |
| 19.34722 | 0.280261 | -0.215215 | 1.10093 | -0.443498 | 1.732925 | 0.184306 | 0.61611 | -0.271599 | 1.433763 | 0.191178 | 2.037561 | 0.666795 | 0.480999 | 0.047365 | 0.977697 | 0.264875 | 0.075579 | 0.002401 | -0.064867 | -0.181434 | 0.025429 | 0.704702 | 0.148127 | 0.284632 |
| 19.36111 | 0.229026 | -0.270501 | 0.110324 | -0.710909 | 1.443952 | 0.538761 | 1.734677 | -0.992853 | 0.481683 | 0.386136 | 2.514757 | 0.315691 | 0.620697 | 0.094575 | 0.76519 | 0.241753 | -0.044547 | 0.014036 | -0.212509 | -0.463963 | 0.081295 | 0.454128 | 0.296395 | 0.412322 |
| 19.375 | -0.197104 | 0.256979 | -0.9613 | -1.188908 | 0.75173 | 0.5333 | 2.367007 | -0.563421 | 0.035082 | 0.351173 | 1.710861 | -0.011335 | 0.655235 | 0.245043 | 0.501952 | 0.349109 | -0.001851 | 0.084698 | -0.185202 | -0.364215 | 0.159983 | 0.314067 | 0.513129 | 0.633361 |
| 19.38889 | -0.712327 | 0.509529 | -2.501496 | -1.609315 | 0.250006 | 0.312988 | 0.920761 | 0.338003 | -0.119638 | 0.291418 | 0.934047 | -0.172336 | 0.566384 | 0.355069 | 0.428935 | 0.57528 | 0.242879 | 0.180694 | 0.067719 | 0.085211 | 0.274168 | 0.390104 | 0.574664 | 0.587773 |
| 19.40278 | -0.607336 | 0.107841 | -2.258278 | -2.173147 | -0.86785 | 0.277144 | -0.220758 | 0.053257 | -0.422989 | 0.27668 | 0.428177 | 0.170653 | 0.521079 | 0.358737 | 0.364498 | 0.82533 | 0.479355 | 0.279777 | 0.304639 | 0.522179 | 0.453686 | 0.6724 | 0.490763 | 0.369329 |
| 19.41667 | -0.27938 | -0.569495 | -1.291517 | -0.969323 | -1.682692 | 0.416903 | -1.161549 | -0.690245 | -1.265156 | 0.176497 | -0.1056 | 0.639369 | 0.546154 | 0.358207 | 0.241277 | 0.807987 | 0.488247 | 0.310145 | 0.25834 | 0.509794 | 0.627762 | 0.807815 | 0.406469 | 0.444251 |
| 19.43056 | 0.102 | -0.554361 | -0.177671 | -0.007137 | -2.280915 | 0.34964 | -1.465996 | -1.260709 | -1.307362 | 0.099046 | -0.747189 | 0.776695 | 0.569208 | 0.484028 | 0.135914 | 0.621443 | 0.279198 | 0.264522 | 0.091572 | 0.372577 | 0.588424 | 0.658108 | 0.287889 | 0.759036 |
| 19.44444 | -0.11176 | -0.312509 | 0.739818 | -0.010204 | -2.323601 | 0.167869 | -1.136807 | -1.208869 | -0.658045 | -0.116175 | -1.72248 | 1.027109 | 0.473287 | 0.808597 | 0.391858 | 0.571076 | 0.318386 | 0.329148 | 0.07865 | 0.672999 | 0.395728 | 0.26829 | 0.216581 | 0.925306 |
| 19.45833 | -0.343632 | -0.233374 | 0.998855 | -0.087288 | -1.817254 | 0.069061 | -0.516042 | -0.522589 | -0.15748 | -0.20449 | -1.584249 | 0.780478 | 0.429017 | 0.746331 | 0.910265 | 0.50805 | 0.661867 | 0.499435 | 0.242873 | 1.153991 | 0.222686 | -0.077439 | 0.197091 | 0.745436 |
| 19.47222 | -0.108083 | 0.137774 | 1.10378 | -0.351843 | -0.599283 | 0.207841 | -0.291574 | 0.275045 | 0.19119 | -0.16859 | -0.400263 | 1.349011 | 0.663014 | 0.61374 | 1.592744 | 0.485503 | 0.9263 | 0.755652 | 0.40196 | 1.17276 | 0.178649 | -0.074605 | 0.37361 | 0.431639 |
| 19.48611 | 0.508348 | 0.37306 | 1.161206 | -1.172296 | 0.082387 | 0.347076 | -0.218784 | 0.636093 | 0.580855 | -0.101119 | 0.873386 | 0.452951 | 0.978844 | 0.8215 | 2.217018 | 0.70929 | 0.988155 | 0.865579 | 0.455763 | 0.9904 | 0.270868 | 0.125466 | 0.559489 | 0.328521 |
| 19.5 | 0.589165 | 0.449867 | 0.71178 | -1.983477 | 0.862843 | 0.295279 | -0.044859 | 0.217897 | 0.534222 | -0.021688 | 1.000337 | 0.129987 | 1.246186 | 1.610873 | 2.054246 | 0.981342 | 0.947194 | 0.763237 | 0.418088 | 1.032556 | 0.378321 | 0.414704 | 0.609756 | 0.616027 |
| 19.51389 | 0.274565 | 0.689232 | -0.048511 | -2.093431 | 0.791786 | 0.148769 | 0.311514 | -0.294908 | 0.052372 | -0.143584 | 0.450469 | 0.097786 | 1.450643 | 2.570728 | 0.99471 | 1.172391 | 0.894617 | 0.691834 | 0.410859 | 1.238308 | 0.657226 | 0.889043 | 0.766161 | 1.402051 |
| 19.52778 | 0.01059 | 0.569013 | -1.60752 | -3.048571 | 0.46406 | 0.038841 | 0.714856 | -0.310186 | -0.393999 | -0.143369 | -0.149608 | -0.062891 | 1.460601 | 3.235749 | -0.228587 | 1.267605 | 0.889459 | 0.715269 | 0.423006 | 1.256128 | 0.715207 | 1.135468 | 1.107047 | 2.309771 |
| 19.54167 | -0.405524 | 0.484244 | -2.801686 | -2.771897 | 0.129382 | -0.022693 | 0.742643 | -0.252811 | -0.7179 | -0.082262 | -0.417673 | -0.457237 | 1.277683 | 3.663652 | -0.514588 | 1.190546 | 0.896864 | 0.674975 | 0.418357 | 0.950152 | 0.556034 | 0.648863 | 1.682508 | 2.487375 |
| 19.55556 | -0.615653 | 0.364993 | -3.04277 | -1.733581 | -0.172678 | -0.083618 | -0.270658 | -0.878116 | -0.893717 | -0.067916 | -0.454387 | -0.903431 | 1.105996 | 3.416374 | 0.333461 | 1.060003 | 0.803516 | 0.554131 | 0.369025 | 0.93506 | 0.351915 | 0.236427 | 0.909472 | 1.482492 |
| 19.56944 | -0.910215 | -0.475625 | -2.713958 | -1.003845 | -0.538052 | -0.152693 | -1.297016 | -1.644264 | -1.214304 | -0.089944 | -0.496204 | -0.353125 | 0.900844 | 2.25137 | 0.870748 | 0.959259 | 0.569526 | 0.422434 | 0.277597 | 1.219442 | -0.032879 | -0.238371 | -0.051952 | 0.615016 |
| 19.58333 | -0.944841 | -1.366384 | -1.851294 | -1.004273 | -1.401926 | -0.225758 | -1.478611 | -1.411916 | -1.914385 | 0.024829 | -0.591144 | -0.072227 | 0.533776 | 0.845343 | -0.214858 | 0.736263 | 0.452197 | 0.227008 | 0.17504 | 1.106376 | -0.285962 | -0.454458 | -0.714014 | 0.109842 |
| 19.59722 | -0.859667 | -1.233393 | -1.570885 | -1.277744 | -2.084947 | -0.348812 | -0.903922 | -0.912746 | -1.98519 | 0.118087 | -0.484804 | -0.767469 | 0.18045 | -0.21107 | -0.259979 | 0.38867 | 0.399703 | 0.104757 | 0.160545 | 0.742117 | -0.054463 | -0.182366 | -0.643357 | 0.041324 |
| 19.61111 | -0.955244 | -0.945156 | -1.754832 | -2.156665 | -3.067321 | -0.647118 | -0.756709 | -1.525005 | -1.708798 | 0.133616 | -0.589886 | -0.766201 | 0.039387 | -1.222962 | 0.040715 | 0.090292 | 0.476145 | 0.083058 | 0.267537 | 0.490471 | 0.347564 | 0.212877 | 0.247169 | 0.262673 |
| 19.625 | -2.15699 | -1.765142 | -2.139524 | -5.125566 | -5.164903 | -1.04136 | -1.06055 | -4.493546 | -1.874411 | -0.172009 | -2.052137 | -1.087323 | 0.299114 | -1.795064 | 0.332477 | -0.166046 | 0.590455 | 0.038048 | 0.334727 | 0.413946 | 0.555942 | 0.324775 | 1.267751 | 0.775268 |
| 19.63889 | -2.104168 | -3.790076 | -1.779697 | -7.059905 | -5.398776 | -1.125242 | -1.187495 | -4.285586 | -1.318583 | -0.799322 | -2.520159 | -2.429612 | 0.696505 | -1.33111 | 0.822702 | -0.019838 | 0.545221 | -0.032179 | 0.15891 | 0.54775 | 0.663539 | 0.218078 | 1.659039 | 1.095745 |
| 19.65278 | -0.864508 | -3.03757 | -0.041366 | -4.899486 | -4.396438 | -0.531424 | -1.097302 | -0.305177 | -0.458162 | -0.673048 | -1.525981 | -1.874199 | 0.812566 | 0.158395 | 0.907173 | 0.324379 | 0.4538 | 0.037544 | -0.00087 | 0.616805 | 0.300052 | -0.095515 | 0.963478 | 0.842922 |
| 19.66667 | 0.012705 | -1.37032 | 1.338506 | -3.151769 | -1.544191 | -0.247464 | -1.086017 | 2.760564 | -0.176455 | 0.371025 | -0.94362 | -1.322643 | 0.684003 | 1.590014 | 0.273163 | 0.698674 | 0.426193 | 0.136156 | 0.048143 | 0.36838 | 0.022466 | -0.140814 | 0.134667 | 0.32388 |
| 19.68056 | 0.266855 | -0.212945 | 1.875443 | -2.203215 | 1.506485 | -0.461207 | -1.069008 | 1.990093 | -0.149217 | 0.819525 | -0.699705 | -0.926614 | 0.522848 | 1.775254 | 0.122192 | 0.674782 | 0.413005 | 0.120711 | 0.174766 | 0.044717 | -0.229339 | -0.060461 | -0.266087 | 0.340723 |
| 19.69444 | 0.171813 | 0.179681 | 1.920101 | -0.142247 | 2.760205 | -0.724593 | -0.856016 | 0.718446 | 0.039522 | 0.546386 | -0.573461 | -0.534815 | 0.476077 | 0.530903 | 0.161719 | 0.570726 | 0.391821 | 0.084117 | 0.197092 | -0.199761 | -0.201915 | -0.241178 | 0.090986 | 0.782536 |
| 19.70833 | 0.091071 | 0.224956 | 1.281229 | 2.437178 | 2.558145 | -0.652756 | -0.747003 | -0.316691 | -0.137477 | 0.209077 | -0.374401 | 0.063023 | 0.490819 | -0.429919 | -0.015139 | 0.604644 | 0.381922 | 0.052211 | 0.045542 | -0.203679 | 0.443885 | 0.284584 | 1.091474 | 1.234174 |
| 19.72222 | -0.184015 | 0.335601 | -0.413123 | 4.235741 | 2.925426 | -0.367096 | -0.729589 | -1.053034 | -0.945862 | 0.039576 | -0.180312 | 0.103928 | 0.4512 | -0.564739 | -0.289794 | 0.698285 | 0.403057 | 0.050086 | -0.04656 | -0.060559 | 0.819478 | 0.519017 | 1.067768 | 1.202347 |
| 19.73611 | -0.387173 | 0.088913 | -2.43076 | 3.20161 | 3.571009 | -0.271432 | -0.795743 | -1.241403 | -2.006449 | -0.115189 | -0.105721 | -0.422355 | 0.268801 | -0.340729 | -0.440138 | 0.59366 | 0.460645 | -0.012326 | -0.021538 | 0.141797 | 0.53476 | 0.288962 | 0.71237 | 0.650851 |
| 19.75 | 0.040514 | -1.459972 | -2.074387 | 0.321577 | 2.519198 | -0.375618 | -0.943727 | -1.690715 | -1.569116 | -0.441571 | -0.186586 | -0.74475 | 0.099641 | 0.262977 | -0.404555 | 0.274358 | 0.521283 | -0.18616 | 0.044075 | 0.224175 | 0.306968 | 0.197823 | 0.12757 | 0.598854 |
| 19.76389 | 0.590097 | -3.44239 | -0.634193 | -2.579367 | 0.386889 | -0.164032 | -1.292844 | -2.325979 | -1.026312 | -0.823804 | -0.384819 | -1.056806 | 0.008113 | 0.473403 | -0.148984 | -0.18106 | 0.518514 | -0.409032 | 0.023052 | 0.098567 | 0.304642 | 0.281661 | -0.549163 | 0.058799 |
| 19.77778 | 0.491982 | -5.009173 | 0.289688 | -4.980297 | -1.544631 | 0.039573 | -1.275813 | -2.26287 | -0.698942 | -1.575647 | -0.549613 | -1.167201 | -0.019155 | 0.471466 | 0.22535 | -0.38792 | 0.442824 | -0.537781 | -0.012916 | 0.073922 | 0.367499 | 0.315501 | 0.066486 | 0.157323 |
| 19.79167 | -0.163633 | -3.70881 | 0.419289 | -5.945329 | -2.936542 | -0.176821 | -0.844112 | -1.119056 | -0.620954 | -1.755524 | -0.387569 | -0.737934 | 0.018553 | 0.12044 | 0.524373 | -0.533758 | 0.377696 | -0.424393 | 0.021782 | 0.273104 | 0.480159 | 0.276942 | 1.307169 | 0.076763 |
| 19.80556 | -0.652959 | -0.618271 | -0.186891 | -4.567342 | -3.023805 | -0.218296 | -0.354737 | -0.100742 | -0.798609 | -1.455572 | -0.253065 | -0.123285 | 0.008242 | 0.072047 | 0.655326 | -0.567517 | 0.403394 | 0.036982 | 0.083825 | 0.447693 | 0.380566 | 0.194491 | 1.64084 | 0.551634 |
| 19.81944 | -0.609382 | 1.300093 | -0.851987 | -2.234326 | -2.145859 | -0.057651 | 0.206261 | 0.261148 | -1.204658 | -1.153428 | -0.16842 | 0.152073 | -0.048895 | 0.337481 | 0.761285 | -0.287319 | 0.448471 | 0.428093 | 0.135681 | 0.588177 | 0.13613 | 0.013237 | 1.035965 | 0.84778 |
| 19.83333 | -0.141806 | 1.29364 | -0.465941 | -0.638881 | -0.643033 | -0.20856 | 0.530684 | 0.252405 | -1.339329 | -0.543654 | -0.109228 | 0.105369 | -0.109592 | 0.410027 | -0.364984 | 0.101915 | 0.466479 | 0.6367 | 0.148133 | 0.621119 | -0.129422 | -0.220077 | 0.59971 | 0.488541 |
| 19.84722 | 0.447991 | 0.824722 | 0.600187 | 0.202081 | 0.716823 | -0.34422 | 0.636173 | 0.659029 | -0.616357 | 0.59711 | -0.051775 | 0.172117 | -0.132713 | 0.463624 | 0.284551 | 0.202464 | 0.449643 | 0.560904 | 0.084779 | 0.419953 | -0.379256 | -0.400214 | 0.391492 | 0.172129 |
| 19.86111 | 0.605667 | 1.002834 | 1.352065 | 0.767131 | 1.548851 | 0.011405 | 0.539814 | 1.384179 | 0.309792 | 1.201437 | 0.074579 | 0.32429 | -0.128794 | -0.204488 | 0.491643 | -0.006205 | 0.471101 | 0.287605 | -0.000632 | 0.135428 | -0.581726 | -0.627551 | 0.052781 | 0.05197 |
| 19.875 | 0.04367 | 1.116528 | 0.895116 | 0.938632 | 1.613584 | 0.162943 | 0.474294 | 1.402277 | 0.944504 | 0.987237 | 0.227886 | 0.303417 | -0.023596 | -0.895672 | 0.450893 | 0.050752 | 0.4716 | 0.052283 | -0.01781 | 0.097247 | -0.609576 | -0.667242 | -0.370754 | 0.340062 |
| 19.88889 | -0.567317 | 0.604385 | -0.086125 | 0.353078 | 1.029604 | -0.166621 | 0.296221 | 0.836149 | 0.938169 | 0.45979 | 0.295943 | 0.153102 | -0.084426 | -0.771545 | 0.281317 | 0.43635 | 0.405844 | -0.018431 | 0.048256 | 0.201717 | -0.443315 | -0.484972 | -0.483722 | 0.716146 |
| 19.90278 | -0.293353 | -0.037557 | -0.972824 | -0.099664 | 0.477819 | -0.326912 | 0.112675 | 0.637022 | 0.905904 | 0.276074 | 0.339733 | 0.034973 | -0.348291 | -0.395819 | -0.014524 | 0.751456 | 0.316365 | -0.150788 | 0.070458 | -0.016759 | -0.422882 | -0.371666 | -0.413861 | 0.402309 |
| 19.91667 | 0.400095 | -0.461555 | -1.169814 | 0.452252 | 0.318053 | -0.276006 | -0.092958 | 0.588422 | 1.110687 | 0.488429 | 0.430942 | 0.017619 | -0.512796 | -0.178278 | -0.466893 | 0.85364 | 0.266814 | -0.515963 | 0.058827 | -0.442777 | -0.589607 | -0.397159 | -0.632292 | 0.116402 |
| 19.93056 | 0.816679 | -0.41998 | -0.257751 | 1.11562 | 0.220818 | -0.250738 | -0.400318 | 0.413754 | 1.259836 | 0.813504 | 0.475239 | 0.085748 | -0.384889 | -0.522814 | -0.87032 | 0.686826 | 0.365415 | -1.057395 | 0.073523 | -0.588319 | -0.6017 | -0.467815 | -0.754253 | -0.042349 |
| 19.94444 | 0.663648 | -0.063184 | 0.802166 | 1.357944 | -0.02068 | -0.11168 | -0.497148 | 0.222294 | 1.23866 | 0.955501 | 0.416278 | 0.161069 | -0.341143 | -1.205912 | -0.672896 | -0.110238 | 0.497846 | -1.296259 | 0.053069 | -0.658591 | -0.640627 | -0.652244 | -0.52514 | -0.04463 |
| 19.95833 | 0.263291 | 0.222917 | 1.440976 | 1.7313 | -0.162823 | 0.160349 | -0.287486 | 0.230275 | 1.232283 | 1.127507 | 0.32081 | 0.248723 | -0.767558 | -2.475411 | -0.591897 | -0.991435 | 0.367961 | -1.852254 | -0.239683 | -1.059265 | -1.078022 | -1.170597 | -0.439459 | -0.616849 |
| 19.97222 | 0.019351 | 0.380868 | 1.429016 | 2.160185 | 0.008608 | 0.251427 | 0.11421 | 0.098129 | 0.898576 | 0.694056 | 0.268301 | 0.405489 | -1.126584 | -3.631491 | -1.170531 | -1.376848 | 0.112172 | -2.619576 | -0.584959 | -1.700972 | -1.467615 | -2.120858 | 0.258401 | -0.566414 |
| 19.98611 | 0.04306 | 0.441117 | 1.050785 | 2.598165 | 0.290515 | 0.096655 | 0.358038 | -0.089938 | 0.52871 | 0.1049 | 0.20177 | 0.493386 | -0.812964 | -3.251218 | -1.229559 | -1.929851 | -0.051171 | -1.954775 | -0.601457 | -1.690571 | -0.961294 | -1.769407 | 1.561183 | 0.045015 |
| 20 | 0.271402 | 0.164196 | 0.665072 | 3.290758 | 0.41698 | 0.083744 | 0.444877 | -0.087628 | 0.266796 | -0.455235 | 0.145621 | 0.35557 | -0.263568 | -1.602758 | -0.420981 | -1.149849 | -0.130504 | -0.926342 | -0.448897 | -1.086088 | -0.090526 | -0.438649 | 1.533952 | 0.507305 |
| 20.01389 | 0.579273 | 0.060892 | 0.474067 | 3.52654 | 0.430694 | 0.256452 | 0.537148 | 0.157241 | 0.095781 | -0.652553 | 0.172925 | 0.254734 | 0.260617 | -0.21999 | 0.45053 | 0.415375 | -0.159087 | 0.00467 | -0.241314 | -0.3672 | 0.806993 | 0.830842 | -0.297592 | 0.501879 |
| 20.02778 | 0.768115 | 0.084805 | 0.419343 | 2.483318 | 0.417978 | 0.324238 | 0.586693 | 0.432914 | 0.029274 | -0.541652 | 0.235947 | 0.229429 | 0.217387 | 0.914686 | 0.674445 | 1.287508 | -0.06337 | 0.791157 | 0.059533 | 0.355082 | 1.044522 | 1.043804 | -1.366794 | 0.555765 |
| 20.04167 | 0.818638 | 0.035542 | 0.531948 | 1.274475 | 0.471405 | 0.2763 | 0.670446 | 0.566214 | 0.046794 | -0.225836 | 0.250763 | 0.13833 | -0.334823 | 1.674757 | -0.563685 | -0.05585 | 0.002651 | 1.238306 | 0.161957 | 0.677237 | 0.550108 | 0.214523 | -2.867828 | -0.873064 |
| 20.05556 | 0.86156 | 0.147343 | 0.92479 | 0.671807 | 0.940824 | 0.324388 | 0.760509 | 0.935715 | 0.11749 | 0.20731 | 0.280853 | 0.137856 | -0.987942 | 1.487539 | -1.046208 | -1.006467 | 0.023459 | 1.363505 | -0.004741 | 0.457726 | 0.169497 | -0.313703 | -2.91407 | -2.0149 |
| 20.06944 | 0.992489 | 0.522471 | 1.479196 | 0.41162 | 1.612926 | 0.411505 | 0.845385 | 1.464854 | 0.194712 | 0.626982 | 0.319146 | 0.268793 | -1.155386 | 0.822955 | -0.400721 | -1.196546 | 0.018947 | 0.854158 | -0.28179 | -0.004313 | 0.169454 | -0.252444 | -0.915024 | -1.783504 |
| 20.08333 | 1.095036 | 1.129949 | 1.830836 | 0.533463 | 1.987917 | 0.464336 | 0.922974 | 1.479952 | 0.343594 | 0.593612 | 0.330573 | 0.337805 | -0.909721 | 0.274619 | 0.095484 | -0.713164 | 0.03773 | 0.357012 | -0.354509 | -0.406232 | 0.309922 | 0.00619 | 1.069118 | -0.352584 |
| 20.09722 | 0.892888 | 1.959667 | 2.039176 | 1.272014 | 1.927026 | 0.457932 | 1.021221 | 1.252674 | 0.426968 | 0.420103 | 0.375519 | 0.362076 | -0.62923 | -0.115524 | 0.157947 | -0.19876 | 0.04791 | -0.024148 | -0.194165 | -0.64051 | 0.072233 | -0.044587 | 2.373126 | 1.00649 |
| 20.11111 | 0.499213 | 2.20761 | 1.614334 | 1.751255 | 1.630959 | 0.286507 | 0.902973 | 1.03006 | 0.247859 | 0.284818 | 0.418087 | 0.447316 | -0.625132 | -0.35128 | 0.290373 | 0.233969 | 0.071384 | -0.391877 | -0.009979 | -0.745605 | -0.337084 | -0.427815 | 1.646358 | 0.997506 |
| 20.125 | 0.169141 | 1.395142 | 0.883407 | 1.067435 | 1.178042 | -0.029589 | 0.494903 | 0.322695 | -0.073786 | 0.218382 | 0.413656 | 0.377639 | -0.655505 | -0.559902 | 0.495014 | 0.227016 | 0.098956 | -0.710276 | -0.132192 | -0.773251 | -0.50461 | -0.744994 | -0.25341 | -0.014878 |
| 20.13889 | 0.086458 | 0.295016 | 0.352036 | 0.076206 | 0.233254 | -0.18227 | 0.190301 | -0.870398 | -0.018246 | 0.08726 | 0.28359 | 0.119106 | -0.467343 | -0.653766 | 0.449086 | 0.02299 | 0.11184 | -0.571303 | -0.20375 | -0.333854 | -0.458659 | -0.48095 | -1.361247 | -0.393152 |
| 20.15278 | 0.146764 | -0.08184 | 0.213778 | -0.1668 | -0.777873 | 0.068008 | 0.364347 | -1.705737 | 0.558121 | -0.233191 | 0.325221 | 0.142883 | -0.163439 | -0.471627 | -0.215292 | 0.12307 | 0.091892 | -0.083225 | 0.011494 | 0.415414 | -0.378646 | 0.102684 | -1.680077 | -0.47469 |
| 20.16667 | 0.324213 | 0.360527 | 0.316629 | 0.065256 | -1.098093 | 0.580967 | 1.136889 | -1.283376 | 0.69748 | -0.416492 | 0.48086 | 0.479223 | -0.009288 | -0.107761 | -0.765187 | 0.242535 | 0.116631 | 0.242503 | 0.04773 | 0.653113 | -0.245258 | 0.39689 | -1.171777 | -0.133215 |
| 20.18056 | 0.666687 | 1.301875 | 0.537384 | 1.014097 | -0.324964 | 0.947757 | 2.268078 | -0.119001 | 0.970295 | -0.505421 | 0.707173 | 0.906966 | 0.038099 | 0.251997 | -0.620092 | 0.472739 | 0.165084 | 0.35391 | -0.080372 | 0.557667 | 0.003055 | 0.325176 | 0.511437 | 0.596364 |
| 20.19444 | 1.142958 | 2.233098 | 0.661832 | 2.695228 | 1.41551 | 1.164756 | 3.083748 | 1.224262 | 1.207486 | -0.469072 | 0.679254 | 1.290065 | 0.132485 | 0.491638 | 0.187376 | 0.934596 | 0.148599 | 0.415729 | -0.078543 | 0.633965 | 0.287359 | 0.296684 | 2.325696 | 1.315509 |
| 20.20833 | 1.981299 | 3.160265 | 1.325704 | 3.85897 | 2.213331 | 1.188003 | 3.31702 | 2.749979 | 1.588922 | -0.399394 | 0.927032 | 1.610447 | 0.333456 | 0.56052 | 0.545086 | 1.06307 | 0.144816 | 0.359371 | -0.079122 | 0.55765 | 0.554121 | 0.47975 | 2.740803 | 1.299051 |
| 20.22222 | 2.909972 | 3.477475 | 2.41071 | 3.547346 | 2.299636 | 0.837799 | 2.999544 | 4.534036 | 1.879496 | -0.33716 | 0.785035 | 1.626173 | 0.517659 | 0.523802 | 0.513402 | 0.821926 | 0.193412 | 0.169253 | -0.128913 | 0.241509 | 0.616478 | 0.667918 | 0.997832 | -0.212343 |
| 20.23611 | 2.654695 | 3.216924 | 2.241212 | 2.736786 | 1.630908 | 0.117778 | 2.2579 | 3.560437 | 2.050969 | 0.107632 | 1.771337 | 1.594114 | 0.496068 | 0.438584 | 0.459945 | 0.441476 | 0.243219 | -0.067356 | -0.262741 | -0.084446 | 0.401192 | 0.502152 | -1.432922 | -0.868089 |
| 20.25 | 1.683864 | 1.907976 | 1.493391 | 0.909171 | 0.813764 | -0.226406 | 2.017725 | 0.68786 | 2.628594 | 0.188344 | 2.204029 | 1.44494 | 0.370224 | 0.330242 | 0.44326 | 0.354053 | 0.24238 | -0.184876 | -0.271806 | -0.098631 | 0.332174 | 0.353054 | -2.000045 | -1.065744 |
| 20.26389 | 0.939358 | 0.999225 | 1.325018 | -0.588312 | 0.12731 | -0.347636 | 1.453303 | -0.21499 | 3.183899 | 0.005386 | 0.982034 | 1.231987 | 0.344701 | 0.167907 | 0.437508 | 0.587074 | 0.168588 | -0.058217 | -0.025723 | 0.092897 | 0.539481 | 0.551946 | -1.674889 | -1.300484 |
| 20.27778 | 0.366784 | 0.228992 | 0.322841 | 0.103861 | -0.018053 | -0.166872 | 0.642305 | -0.662367 | 2.756504 | 0.154745 | 0.049742 | 1.049118 | 0.391122 | 0.160933 | 0.425917 | 0.698145 | 0.184233 | 0.147636 | 0.126693 | 0.085965 | 0.614018 | 0.452675 | -0.577938 | -0.521984 |
| 20.29167 | -0.101358 | -0.425102 | -0.651669 | 1.584543 | 0.087024 | 0.229714 | 0.225012 | -0.729561 | 1.676883 | 0.280527 | -1.00713 | 0.831754 | 0.449213 | 0.326941 | 0.368523 | 0.704714 | 0.197147 | 0.418008 | 0.238242 | 0.220221 | 0.48053 | 0.264086 | 0.650216 | 0.978459 |
| 20.30556 | -0.104102 | -0.800304 | -1.045422 | 3.870864 | 0.308772 | -0.413149 | 0.392818 | -0.740933 | 0.281167 | 0.414515 | -1.991707 | 0.750475 | 0.469793 | 0.468433 | 0.113004 | 0.629276 | 0.130391 | 0.699257 | 0.382395 | 0.595522 | 0.403204 | 0.23805 | 1.411532 | 2.178473 |
| 20.31944 | -0.251103 | -0.496195 | -0.545754 | 4.842991 | 0.359249 | -0.818768 | 0.850048 | -1.043222 | -0.604988 | 0.34908 | -1.988319 | 1.082256 | 0.506018 | 0.417937 | -0.188511 | 0.331483 | 0.222177 | 0.673712 | 0.323736 | 0.632402 | 0.35413 | 0.355307 | 1.432766 | 1.981277 |
| 20.33333 | -0.034416 | 0.289806 | -0.199355 | 1.681942 | 0.101814 | -0.892306 | 0.60186 | -1.036404 | -1.29926 | 0.095887 | -1.467205 | 1.565166 | 0.695151 | 0.332464 | -0.103692 | 0.071115 | 0.330189 | 0.597858 | 0.268613 | 0.559762 | 0.387499 | 0.357911 | 1.110806 | 1.065167 |
| 20.34722 | 0.536128 | 1.02863 | -0.277231 | -0.307296 | -0.278065 | -1.033575 | -0.024913 | -0.525101 | -1.672264 | -0.006475 | -0.647961 | 1.934319 | 1.022505 | 0.509707 | 0.400652 | 0.192347 | 0.30105 | 0.677011 | 0.338417 | 0.729465 | 0.652903 | 0.464343 | 0.818414 | 0.202581 |
| 20.36111 | 0.69675 | 0.781592 | 0.403084 | -1.189345 | -0.98817 | -0.907975 | -0.294062 | 0.090516 | -1.231485 | -0.159302 | 0.229566 | 1.395647 | 1.206655 | 0.844304 | 0.893193 | 0.534772 | 0.222422 | 0.695793 | 0.285257 | 0.790912 | 1.035326 | 0.787033 | 0.904499 | -0.457252 |
| 20.375 | 0.394589 | 0.300809 | 0.865539 | -1.960128 | -0.389649 | -0.018643 | 0.263985 | 1.335157 | -0.479943 | -0.471773 | 0.957855 | 0.950446 | 1.076661 | 1.001978 | 1.180393 | 0.890455 | 0.228927 | 0.642896 | 0.186729 | 0.651151 | 1.232242 | 0.866352 | 0.017309 | -0.665224 |
| 20.38889 | 0.151025 | -0.067549 | 1.055335 | -1.709902 | 0.422245 | 0.489322 | 0.895801 | 1.98881 | -0.103357 | -0.5243 | 1.223291 | 0.403089 | 0.917885 | 0.923347 | 1.318773 | 1.08728 | 0.245874 | 0.63861 | 0.301684 | 0.784519 | 1.167464 | 0.710786 | 0.125238 | 0.055104 |
| 20.40278 | 0.166903 | -0.290372 | 0.445808 | -0.707317 | 0.709038 | 0.774316 | 0.853249 | 0.516203 | -0.029525 | -0.189774 | 1.447714 | 0.080481 | 0.87987 | 0.835768 | 1.341475 | 0.850472 | 0.222649 | 0.557923 | 0.492635 | 0.888379 | 0.867925 | 0.411798 | 1.152457 | 1.353951 |
| 20.41667 | 0.403568 | -0.509732 | -0.841672 | 0.316321 | 0.940533 | 0.917811 | 0.183355 | -0.861588 | -0.499286 | 0.001064 | 1.636299 | -0.093573 | 0.808483 | 0.981959 | 1.331906 | 0.477288 | 0.202976 | 0.444622 | 0.454918 | 0.72916 | 0.459052 | 0.028041 | 2.359048 | 2.206689 |
| 20.43056 | 0.402992 | -0.640371 | -1.242675 | 0.743662 | 0.979351 | 0.918457 | -0.970471 | -3.366945 | -0.628425 | 0.088441 | 0.60394 | -0.104438 | 0.668936 | 1.438336 | 1.290057 | 0.56284 | 0.127241 | 0.505732 | 0.333264 | 0.816413 | 0.068651 | -0.248148 | 1.76887 | 1.771543 |
| 20.44444 | 0.09698 | -0.484691 | -1.739949 | -0.500904 | 0.501047 | 0.596578 | -2.701101 | -2.174311 | -0.126977 | 0.194777 | -0.491725 | 0.164975 | 0.497407 | 1.708979 | 1.00484 | 0.863326 | 0.000175 | 0.638771 | 0.242955 | 0.921471 | -0.339081 | -0.424433 | 0.902119 | 1.066007 |
| 20.45833 | -0.30855 | -0.281242 | -1.842028 | -2.092917 | 0.123999 | 0.216907 | -0.913099 | -0.471047 | 0.116827 | 0.304164 | -0.846673 | 0.330041 | 0.421645 | 1.504528 | 0.799486 | 0.731852 | 0.198667 | 0.659864 | 0.101891 | 0.672023 | -0.45103 | -0.385557 | 0.589966 | 0.626841 |

| | | | | | | | | | | | | | | | | | | | | | | | | |
|---|---|---|---|---|---|---|---|---|---|---|---|---|---|---|---|---|---|---|---|---|---|---|---|---|
| 20.47222 | -0.493575 | -0.166514 | -1.999693 | -2.47336 | 0.035856 | -0.155663 | 0.313592 | 0.185852 | -0.189289 | 0.405653 | -0.979225 | -0.06679 | 0.608972 | 1.29 | 0.859611 | 0.588014 | 0.433524 | 0.810987 | 0.193601 | 0.53189 | -0.115102 | 0.036501 | 0.299171 | 0.366041 |
| 20.48611 | -0.19243 | -0.021478 | -1.508702 | -2.459524 | -0.331031 | -0.649789 | 0.563551 | -0.357282 | -0.586969 | 0.569017 | -1.80177 | -0.534869 | 1.022682 | 1.558394 | 1.292951 | 1.033063 | 0.523329 | 1.108941 | 0.459747 | 0.819488 | 0.410881 | 0.48243 | 0.037183 | 0.163719 |
| 20.5 | -0.057255 | 0.37058 | -0.946717 | -4.158637 | -0.590252 | -1.253653 | 0.346834 | -0.84283 | -0.278975 | 0.809262 | -2.05862 | -0.572522 | 1.520898 | 2.351919 | 1.933308 | 1.878697 | 0.527498 | 1.018201 | 0.872883 | 1.347322 | 0.894689 | 0.812608 | 0.146062 | 0.179284 |
| 20.51389 | -0.178084 | 0.437869 | -0.483943 | -10.066437 | -0.448502 | -1.517754 | 0.518366 | -0.55149 | 0.271061 | 0.779175 | -1.857429 | -0.33791 | 1.750433 | 3.290734 | 1.812074 | 2.478264 | 0.470297 | 0.673971 | 1.403151 | 1.531615 | 1.110434 | 0.890337 | 0.067847 | 0.955377 |
| 20.52778 | -0.248217 | -0.114041 | 0.904058 | -8.356978 | -0.572113 | -1.343093 | 1.459261 | 0.188865 | 0.441276 | 0.264658 | -1.904042 | -0.077912 | 1.613251 | 3.367496 | 1.110081 | 1.396799 | 0.433384 | 0.307817 | 1.686976 | 1.448953 | 0.821976 | 0.683812 | 0.644634 | 2.256335 |
| 20.54167 | 0.219244 | -0.18289 | 1.621332 | -5.018474 | -0.877174 | -1.505077 | 1.740985 | 0.710155 | 0.561431 | -0.409897 | -1.184597 | -0.143559 | 1.462053 | 2.258682 | 0.602616 | 0.842831 | 0.384441 | 0.137665 | 1.970234 | 0.972051 | 0.777951 | 0.326073 | 1.859323 | 2.466796 |
| 20.55556 | 0.112431 | -0.046155 | 0.297856 | -2.893228 | -0.517575 | -0.199192 | 0.69584 | 0.425295 | 0.914081 | -0.951802 | -0.191482 | -0.360996 | 1.401168 | 0.702179 | 0.13617 | 0.535222 | 0.305488 | 0.249517 | 2.02843 | 0.614401 | 1.058801 | 0.1615 | 2.624135 | 1.471232 |
| 20.56944 | -0.065832 | -0.116513 | -1.152733 | -1.224578 | -0.382243 | 0.423508 | -0.19861 | -0.059947 | 1.548794 | -1.611129 | 0.107937 | -0.384365 | 1.113603 | -0.713638 | -0.020225 | 0.032358 | 0.281331 | 0.284474 | 0.811475 | 0.353347 | 1.305084 | 0.199642 | 1.556519 | 0.466404 |
| 20.58333 | -0.397125 | -0.527165 | -1.99998 | 0.131421 | -0.448161 | 0.862632 | -0.814812 | -0.521988 | 0.274884 | -3.584305 | -0.462712 | -0.501833 | 0.564658 | -1.914828 | -0.404096 | -0.396326 | 0.313454 | -0.127445 | -0.105992 | 0.072317 | 1.123468 | 0.250625 | 0.690415 | -0.294364 |
| 20.59722 | -0.615886 | -0.991185 | -2.936523 | 0.457627 | -0.396302 | 1.432503 | -1.455177 | -1.124995 | -0.555201 | -2.87077 | -0.468225 | -0.808324 | 0.105824 | -2.663814 | -0.252311 | -0.509525 | 0.290968 | -0.589577 | -0.680909 | 0.010322 | 0.444313 | 0.184751 | 0.126951 | -0.813979 |
| 20.61111 | -1.435934 | -1.46712 | -3.671999 | -1.355994 | -0.593071 | 1.256357 | -2.094823 | -1.860143 | -2.04558 | -2.469194 | -0.604477 | -1.287094 | -0.013674 | -2.982647 | -0.399978 | -0.321353 | 0.210633 | -0.4515 | -1.034655 | 0.043942 | -0.282416 | 0.092639 | 0.02401 | -0.732688 |
| 20.625 | -2.452332 | -2.364054 | -3.87161 | -4.577494 | -1.168776 | 0.315939 | -2.579609 | -3.366724 | -3.813951 | -1.541776 | -1.146838 | -2.393261 | 0.28276 | -2.430452 | 0.157747 | -0.034643 | 0.210814 | 0.421868 | -1.204763 | -0.416115 | -0.798261 | -0.059744 | 0.374723 | -0.217203 |
| 20.63889 | -3.560924 | -3.249192 | -3.847661 | -7.724084 | -1.6576 | -0.010565 | -0.985033 | -3.754717 | -4.489567 | -3.468519 | -0.684054 | -3.668184 | 0.527225 | -0.566735 | 0.649681 | 0.373253 | 0.343569 | 1.076631 | -0.815954 | -0.480544 | -0.832255 | -0.319273 | 0.798462 | 0.38924 |
| 20.65278 | -2.636077 | -2.853234 | -1.922488 | -3.867912 | -0.471562 | -0.019277 | -0.88671 | -1.488195 | -4.098891 | -1.972438 | -0.011383 | -1.732628 | 0.558426 | 1.732272 | 0.415924 | 0.930352 | 0.415762 | 0.797819 | -0.363444 | -0.569261 | -0.167169 | -0.46147 | 0.763539 | 1.095395 |
| 20.66667 | -1.717109 | -3.230731 | -1.466715 | -0.571997 | 0.320915 | -0.130145 | -1.667541 | 0.4586 | -2.852969 | 0.364443 | -0.033548 | -0.76677 | 0.537919 | 2.774275 | 0.865138 | 1.374854 | 0.359334 | 0.361023 | 0.07354 | -0.168329 | 0.460909 | -0.355267 | 0.810693 | 1.472689 |
| 20.68056 | -0.382303 | -2.823279 | -0.991851 | 0.527097 | -0.015448 | 0.126095 | -1.792083 | 0.868657 | -1.967338 | 2.340247 | -0.27502 | -0.55386 | 0.593871 | 1.850237 | 0.823944 | 1.749069 | 0.255109 | 0.300497 | 0.483671 | 0.393248 | 0.590178 | -0.058765 | 0.726338 | 1.199381 |
| 20.69444 | 0.364105 | -1.545644 | 0.012195 | 0.429727 | -0.417395 | 0.856094 | -1.210666 | 0.41849 | -1.458285 | 3.774387 | -0.446057 | -0.343194 | 0.600908 | 1.321582 | 0.536056 | 1.399794 | 0.183493 | 0.363457 | 0.882105 | 0.844779 | 0.714297 | 0.194746 | 0.538471 | 0.802196 |
| 20.70833 | 0.223218 | 0.17324 | 0.65494 | 0.791437 | -0.055079 | 1.18305 | -0.649343 | 0.585012 | -0.856868 | 3.710947 | -0.402212 | 0.284205 | 0.48086 | 1.4511 | 0.227786 | 0.818201 | 0.206936 | 0.302587 | 0.992358 | 0.758771 | 0.775781 | 0.173679 | 0.789293 | 0.654578 |
| 20.72222 | -0.352105 | 1.802307 | 0.295753 | 1.7064 | 0.315028 | 1.021619 | -0.500286 | 0.78047 | -0.162388 | 3.601247 | -0.200384 | 0.48513 | 0.277457 | 1.677831 | -0.030232 | 0.433964 | 0.242492 | 0.037429 | 0.754805 | 0.545588 | 0.716837 | 0.064514 | 0.673464 | 1.009728 |
| 20.73611 | -0.817085 | 2.384726 | -0.465047 | 1.695623 | 0.687575 | 0.37282 | -0.562585 | 0.71815 | 0.382484 | 3.56597 | -0.001656 | 0.44852 | 0.154372 | 1.0813 | -0.31864 | 0.117172 | 0.178927 | -0.214135 | 0.451839 | 0.424725 | 0.464461 | 0.072084 | 0.535035 | 0.67895 |
| 20.75 | -0.837302 | 1.254144 | -0.693072 | 0.616749 | 1.046654 | -1.133731 | -0.574626 | 0.399962 | 1.101548 | 0.239599 | 0.102331 | 0.359676 | 0.032322 | 0.06365 | -1.043113 | -0.035733 | 0.096144 | -0.492585 | 0.323569 | 0.518532 | 0.17457 | 0.091309 | 0.126951 | 0.250253 |
| 20.76389 | -0.544038 | -0.779458 | -0.489724 | -0.608352 | 0.518311 | -2.897972 | -0.761464 | -0.20234 | 0.418813 | -1.216708 | -0.014908 | 0.177857 | -0.142158 | -0.889498 | -1.387894 | 0.182198 | 0.084641 | -1.712361 | 0.261349 | 0.436319 | -0.323776 | -0.052065 | -0.449104 | -0.038542 |
| 20.77778 | -0.27235 | -2.506692 | -0.123093 | -1.769083 | 0.007416 | -2.80997 | -0.897734 | -0.846947 | -0.752428 | -1.469416 | -0.209453 | -0.083555 | -0.329684 | -1.113938 | -0.714501 | 0.358812 | 0.12036 | -2.354272 | 0.092696 | 0.160085 | -0.963536 | -0.268787 | -0.862444 | -0.006891 |
| 20.79167 | -0.278801 | -1.926269 | 0.279381 | -2.692256 | -0.217545 | -1.263672 | -0.984857 | -1.27083 | -1.313775 | -1.81406 | -0.340554 | -0.315739 | -0.393979 | -0.753485 | 0.343529 | 0.426988 | 0.154953 | -1.849596 | -0.303914 | -0.1429 | -1.465278 | -0.438886 | -0.969987 | 0.215985 |
| 20.80556 | -0.681037 | -1.446505 | 0.421002 | -2.578856 | -0.48657 | -1.302667 | -1.268222 | -1.495458 | -1.917657 | -4.77238 | -0.332015 | -0.627916 | -0.487852 | -0.59867 | 1.146925 | 0.304141 | 0.188169 | -0.934813 | -0.90047 | -0.430074 | -1.63662 | -0.759932 | -0.863271 | -0.047517 |
| 20.81944 | -0.954172 | -0.984933 | 0.298395 | -1.412992 | -0.750131 | -1.036292 | -1.88675 | -1.152207 | -1.639306 | -3.902828 | -0.16804 | -0.709263 | -0.558194 | -0.611632 | 1.054507 | 0.249358 | 0.209941 | -0.472994 | -1.310249 | -0.896667 | -1.292034 | -0.86926 | -0.393215 | 0.233546 |
| 20.83333 | -0.654666 | -0.715993 | 0.258804 | -0.367215 | -0.616313 | -0.658322 | -0.964258 | -0.326685 | -0.909698 | -2.293787 | -0.070533 | -0.274006 | -0.459131 | -0.632818 | 1.004185 | 0.076173 | 0.154525 | 0.107317 | -1.305328 | -1.23857 | -0.796646 | -0.631964 | 0.03595 | 0.446496 |
| 20.84722 | -0.182314 | -0.177775 | 0.638696 | -0.074351 | -0.252163 | 0.224063 | 0.410777 | 0.660327 | -0.253264 | -1.159006 | 0.060361 | 0.198075 | -0.220812 | -0.864071 | 0.897268 | -0.476658 | 0.083286 | 1.238354 | -0.922463 | -0.885549 | -0.913459 | -0.71258 | 0.26816 | -0.067755 |
| 20.86111 | 0.177668 | 0.156141 | 1.262586 | -0.464843 | -0.082746 | 0.669484 | 1.753488 | 1.232561 | 0.655595 | 0.412457 | 0.214337 | 0.39199 | -0.105435 | -0.877336 | 0.634349 | -1.068079 | 0.001981 | 2.320417 | -0.725433 | -0.476937 | -0.947723 | -0.679715 | -0.054398 | -0.072337 |
| 20.875 | 0.332 | 0.3895 | 1.477797 | -0.759887 | -0.146869 | 1.246033 | 0.926815 | 1.284635 | 1.391934 | 2.898667 | 0.286642 | 0.399612 | -0.154621 | -0.782384 | 0.080362 | -0.949533 | -0.057724 | 1.976618 | -0.632844 | -0.187514 | -0.191691 | -0.26315 | -0.24374 | -0.09192 |
| 20.88889 | 0.153105 | 0.68553 | 1.169092 | -0.404562 | -0.190575 | 1.094587 | -0.167321 | 1.465075 | 0.940218 | 2.255187 | 0.304234 | 0.373327 | -0.28398 | -0.590284 | -1.16648 | -0.727983 | -0.022863 | 1.339055 | -0.679502 | -0.252782 | 0.519161 | 0.185952 | -0.820905 | -0.233187 |
| 20.90278 | -0.155053 | 0.97237 | 0.773269 | 0.222564 | 0.085903 | -0.299131 | -0.398455 | 1.602118 | 0.702537 | 2.153666 | 0.321008 | 0.330902 | -0.222677 | -0.540075 | -1.933734 | -0.770457 | 0.017145 | 0.630893 | -0.692216 | -0.51739 | 0.760473 | 0.190013 | -1.02044 | -0.217739 |
| 20.91667 | -0.106231 | 1.02684 | 0.723166 | 0.627884 | 0.482406 | -0.243401 | -0.344117 | 1.297003 | 0.565328 | 1.786138 | 0.372128 | 0.255623 | 0.01199 | -1.098837 | -2.841238 | -0.944611 | -0.004854 | 0.431837 | -0.700165 | -0.571732 | 0.449299 | 0.029379 | -1.2574 | -0.75405 |
| 20.93056 | 0.277075 | 1.041249 | 0.956678 | 0.753942 | 0.638975 | 0.44662 | -0.210629 | 0.796792 | -0.246133 | 1.147633 | 0.4274 | 0.201996 | 0.12282 | -1.38117 | -2.796125 | -1.053235 | 0.071087 | 0.101561 | -0.558273 | -0.407937 | -0.074092 | -0.025638 | -1.84381 | -0.934637 |
| 20.94444 | 0.597892 | 0.934059 | 0.894187 | 0.794047 | 0.519059 | 0.318537 | -0.142197 | 0.349649 | -0.143634 | 0.497928 | 0.333371 | 0.127513 | -0.19846 | -1.409322 | -3.213376 | -1.102767 | 0.169397 | -0.353208 | -0.523051 | -0.480095 | -0.833786 | -0.311644 | -2.411683 | -1.116004 |
| 20.95833 | 0.562818 | 0.537372 | 0.543427 | 0.950221 | 0.338959 | 0.092697 | -0.085621 | -0.01863 | 0.098416 | -0.496481 | 0.257597 | 0.075301 | -0.927478 | -1.414119 | -3.128245 | -1.471951 | 0.001663 | -1.013147 | -1.031536 | -1.410149 | -2.027796 | -1.12678 | -3.321698 | -1.447186 |
| 20.97222 | 0.351251 | -0.186763 | 0.320536 | 1.067632 | 0.261689 | 0.072675 | 0.07413 | -0.330996 | 0.482413 | -0.989551 | 0.241402 | 0.070627 | -1.622892 | -1.416026 | -2.762906 | -2.086189 | -0.322426 | -1.142969 | -1.502958 | -3.959874 | -1.980237 | -1.48784 | -3.452719 | -2.597492 |
| 20.98611 | 0.129098 | -0.792008 | 0.226301 | 0.830023 | 0.292156 | 0.041428 | 0.278276 | -0.304822 | 1.024371 | -0.079941 | 0.226218 | 0.088005 | -1.4232 | -1.224532 | -2.095562 | -2.154043 | -0.390871 | -0.507869 | -0.98046 | -2.70877 | -0.665664 | -1.27139 | -1.793226 | -2.603359 |
| 21 | 0.032631 | -0.895545 | 0.217332 | 0.267547 | 0.381852 | 0.071351 | 0.380296 | 0.214919 | 1.701294 | 0.524841 | 0.192563 | 0.089247 | -0.850683 | -1.191366 | -0.781707 | -1.833336 | -0.308227 | 0.074816 | -0.591412 | -1.23052 | 0.270561 | -0.785724 | -0.955462 | -1.696818 |
| 21.01389 | 0.077855 | -0.349893 | 0.390694 | -0.205532 | 0.39863 | 0.067802 | 0.417836 | 0.742913 | 2.40435 | 0.030738 | 0.146093 | 0.074259 | -0.376 | -0.86596 | -0.402994 | -1.755985 | -0.348518 | 0.019436 | -0.533903 | -0.595288 | 0.363931 | -0.263391 | -0.489696 | -0.595449 |
| 21.02778 | 0.197561 | 0.599802 | 0.846496 | -0.352093 | 0.367845 | 0.094664 | 0.548927 | 0.946637 | 2.554612 | 0.845096 | 0.135551 | 0.065422 | -0.01729 | -0.21019 | -0.317202 | -1.667479 | -0.363634 | -0.31651 | -0.396392 | -0.172773 | 0.386472 | 0.052138 | 0.069951 | 0.219073 |
| 21.04167 | 0.332358 | 1.534578 | 1.34752 | 0.121532 | 0.481437 | 0.339915 | 0.973842 | 0.877911 | 2.105972 | 2.655725 | 0.16969 | 0.065418 | 0.000611 | 0.166929 | 0.209564 | -1.293912 | -0.226884 | -0.559037 | -0.25 | 0.027843 | 0.380906 | 0.184235 | 0.560266 | 0.429444 |
| 21.05556 | 0.458698 | 2.244296 | 1.438461 | 1.038974 | 0.681657 | 0.640411 | 1.109215 | 0.958858 | 1.504683 | 2.992752 | 0.234145 | 0.06829 | -0.266545 | -0.0726 | 1.035912 | -0.737626 | -0.051406 | -0.72447 | -0.204276 | 0.157729 | 0.588111 | 0.285587 | 0.722887 | 0.308575 |
| 21.06944 | 0.579396 | 2.563154 | 1.364939 | 1.817283 | 0.818847 | 0.646907 | 1.198869 | 0.954962 | 0.481365 | 1.806685 | 0.274252 | 0.061229 | -0.436458 | -0.58918 | 1.210309 | -0.242142 | -0.004288 | -0.761059 | -0.171688 | 0.286225 | 1.093326 | 0.439778 | 0.529303 | -0.078573 |
| 21.08333 | 0.775221 | 2.573604 | 1.425498 | 2.618385 | 0.845343 | 0.92894 | 1.469303 | 0.878146 | 0.684577 | 1.874903 | 0.369536 | 0.083581 | -0.416885 | -1.098981 | 1.197361 | -0.003512 | -0.019473 | -0.531892 | -0.10957 | 0.304228 | 1.076206 | 0.501166 | -0.159146 | -0.954692 |
| 21.09722 | 0.577505 | 2.565068 | 1.351259 | 3.198204 | 0.767554 | 1.105124 | 1.389917 | 0.96695 | 0.415692 | -0.31586 | 0.377311 | 0.130845 | -0.395229 | -1.194982 | 1.360756 | -0.234259 | 0.019341 | -0.197862 | -0.157029 | 0.179473 | 0.703184 | 0.440842 | -0.96217 | -2.037113 |
| 21.11111 | 0.298161 | 2.595775 | 0.855053 | 2.860695 | 0.436543 | -0.206178 | 0.871302 | 0.947632 | 0.274812 | -0.668361 | 0.21928 | 0.122344 | -0.420084 | -0.689557 | 1.480573 | -0.554406 | 0.000783 | 0.09305 | -0.232179 | -0.110618 | 0.119417 | 0.241259 | -1.396071 | -2.523559 |
| 21.125 | NaN | NaN | NaN | NaN | NaN | NaN | NaN | NaN | NaN | NaN | NaN | NaN | -0.467501 | -0.14035 | 0.646652 | -0.728705 | -0.006423 | 0.257522 | -0.123476 | -0.462131 | -0.470632 | -0.072006 | -1.131429 | -1.916507 |
| 21.13889 | 0.051115 | -0.31048 | 0.09277 | -0.030128 | -0.285569 | 0.007475 | -0.257313 | 0.34057 | 0.421379 | 0.888851 | 0.483652 | 0.622811 | -0.638036 | 0.139192 | 0.203112 | -0.870967 | 0.095199 | 0.354558 | 0.058219 | -0.609638 | -0.962816 | -0.475989 | -0.852862 | -1.013139 |
| 21.15278 | 0.099603 | 0.567485 | -0.012467 | 0.090926 | 0.163864 | -0.093326 | -0.481474 | 0.447798 | 0.180159 | 0.423983 | 0.364326 | 0.269954 | -0.844348 | 0.487188 | 0.389977 | -0.777372 | 0.100355 | 0.445848 | 0.12578 | -0.348473 | -1.200775 | -0.521014 | -0.582799 | -0.169592 |
| 21.16667 | 0.29598 | 1.042303 | 0.1333 | 0.303192 | 0.582006 | 0.084054 | -0.206871 | 0.614433 | 0.193955 | 0.238192 | 0.273539 | 0.414119 | -0.732419 | 0.835621 | 0.334381 | -0.449276 | -0.13072 | 0.457893 | 0.20513 | 0.110112 | -1.003507 | -0.221061 | -0.202309 | 0.467967 |
| 21.18056 | 0.439699 | 0.516877 | 0.263615 | 0.382615 | 0.720704 | 0.358726 | 0.204912 | 0.459327 | 0.40943 | 0.365884 | 0.187956 | 0.586341 | -0.290897 | 0.867244 | 0.396885 | -0.186017 | -0.136597 | 0.012884 | -0.103087 | 0.484109 | -0.351833 | 0.138494 | 0.397468 | 1.056329 |
| 21.19444 | 0.406452 | -0.16982 | 0.346256 | 0.389418 | 0.609074 | 0.51323 | 0.549232 | 0.052129 | 0.56828 | 0.553977 | 0.014903 | 0.643265 | -0.082697 | 0.654803 | 0.476156 | 0.127957 | -0.032077 | -0.131855 | 0.092972 | 0.775409 | 0.304751 | 0.301551 | 0.808324 | 1.376627 |
| 21.20833 | 0.209023 | -0.187389 | 0.447413 | 0.408433 | 0.395445 | 0.458469 | 0.64903 | -0.103818 | 0.518545 | 0.691052 | -0.266781 | 0.756467 | -0.043585 | 0.215587 | 0.454013 | 0.468365 | -0.076536 | 0.247007 | 0.326683 | 0.788173 | 0.609623 | 0.36868 | 0.722224 | 1.125281 |
| 21.22222 | 0.096361 | 0.393523 | 0.591634 | 0.380771 | 0.260684 | 0.333646 | 0.613345 | 0.185458 | 0.392232 | 0.845502 | -0.384162 | 1.125762 | 0.021483 | -0.313232 | 0.358693 | 0.680283 | -0.139621 | 0.221623 | 0.258767 | 0.51948 | 0.497515 | 0.316483 | 0.429901 | 0.498085 |
| 21.23611 | 0.167738 | 0.841099 | 0.641643 | 0.407435 | 0.278214 | 0.270044 | 0.624045 | 0.72095 | 0.406708 | 0.997739 | -0.411347 | 1.441207 | 0.103573 | -0.488863 | 0.122664 | 0.64932 | -0.137207 | -0.196536 | -0.055964 | 0.249325 | 0.424063 | 0.142354 | 0.259325 | -0.077053 |
| 21.25 | 0.248888 | 0.899962 | 0.562224 | 0.614415 | 0.520386 | 0.301975 | 0.782728 | 1.206277 | 0.540903 | 1.190203 | -0.269952 | 1.285148 | 0.05703 | -0.236367 | 0.211383 | 0.513477 | -0.088641 | -0.266449 | -0.123802 | 0.200571 | 0.376824 | 0.000905 | 0.102566 | -0.533733 |
| 21.26389 | 0.276054 | 0.613163 | 0.510159 | 0.921332 | 0.866178 | 0.415275 | 0.859694 | 1.507236 | 0.756281 | 1.358062 | -0.037654 | 1.169986 | -0.071048 | 0.067609 | 0.730905 | 0.510302 | -0.089492 | -0.102512 | 0.017188 | 0.353797 | 0.37577 | 0.010911 | -0.052529 | -0.650708 |
| 21.27778 | 0.321796 | 0.176771 | 0.637978 | 1.135232 | 1.117413 | 0.519699 | 0.811885 | 1.56607 | 0.883942 | 1.411956 | 0.196287 | 1.296904 | -0.08402 | 0.176369 | 1.115976 | 0.677596 | -0.21735 | -0.072015 | -0.002998 | 0.477483 | 0.380193 | 0.174845 | -0.150252 | -0.356855 |
| 21.29167 | 0.439788 | -0.114801 | 0.867967 | 1.389617 | 1.446833 | 0.572361 | 0.711295 | 1.362307 | 0.856536 | 1.444282 | 0.358837 | 1.352394 | 0.138522 | 0.135261 | 0.806188 | 0.747366 | -0.264517 | -0.094169 | -0.116964 | 0.388985 | 0.29969 | 0.221331 | -0.09083 | 0.395713 |
| 21.30556 | 0.689392 | 0.279764 | 1.158494 | 1.977971 | 2.0465 | 0.55609 | 0.77639 | 0.666826 | 0.82145 | 1.865784 | 0.476784 | 1.26735 | 0.341309 | 0.134651 | 0.433871 | 0.642891 | -0.074728 | -0.068151 | -0.097021 | 0.221562 | 0.233892 | 0.044804 | 0.21975 | 1.079726 |
| 21.31944 | 0.979608 | 1.145033 | 1.475586 | 2.483618 | 2.426575 | 0.626005 | 0.94072 | 0.088546 | 0.96459 | 2.897282 | 0.579153 | 1.094997 | 0.494125 | 0.399011 | 0.553069 | 0.84894 | 0.017954 | 0.149584 | 0.08786 | 0.231826 | 0.254939 | -0.049902 | 0.495739 | 1.336356 |
| 21.33333 | 1.106186 | 1.550961 | 1.670823 | 2.651805 | 2.272229 | 0.762825 | 0.843798 | -0.127209 | 1.11415 | 3.536191 | 0.700466 | 1.047795 | 0.555287 | 0.676585 | 0.823516 | 1.144287 | -0.026188 | 0.259444 | 0.148043 | 0.387837 | 0.292239 | -0.024684 | 0.668547 | 1.582517 |
| 21.34722 | 0.881895 | 1.032506 | 1.670436 | 2.313428 | 1.627759 | 0.701418 | 0.501085 | 0.038695 | 0.84997 | 2.615226 | 0.508848 | 1.050402 | 0.488143 | 0.777898 | 0.937017 | 1.126941 | -0.041679 | -0.030071 | 0.03281 | 0.487267 | 0.34106 | -0.068087 | 0.830329 | 1.67218 |
| 21.36111 | 0.529874 | 0.311912 | 1.528923 | 1.940845 | 0.988142 | 0.339683 | 0.218143 | 0.539316 | 0.439413 | 1.116771 | 0.13035 | 0.892043 | 0.418242 | 0.698486 | 0.883911 | 0.899993 | -0.05747 | -0.288545 | 0.029247 | 0.529069 | 0.426131 | -0.135981 | 0.913772 | 1.48504 |
| 21.375 | 0.335716 | 0.119751 | 1.419599 | 1.760784 | 0.835939 | 0.074164 | 0.185058 | 1.424923 | 0.225978 | 0.211547 | -0.017562 | 0.875257 | 0.36755 | 0.515792 | 0.782925 | 0.638968 | -0.045771 | -0.335054 | 0.164617 | 0.614768 | 0.471153 | -0.224144 | 0.872541 | 1.046294 |
| 21.38889 | 0.357663 | 0.266502 | 1.357608 | 1.707985 | 0.927332 | -0.009949 | 0.371641 | 2.302887 | 0.262889 | -0.01307 | 0.021429 | 0.947248 | 0.381745 | 0.401216 | 1.006795 | 0.643642 | -0.027083 | -0.41578 | 0.415363 | 0.914484 | 0.505828 | -0.198543 | 0.715817 | 0.664355 |
| 21.40278 | 0.588828 | 0.43741 | 1.354154 | 1.986494 | 1.151437 | 0.0177 | 0.61551 | 2.753927 | 0.576426 | 0.55271 | 0.037646 | 0.899977 | 0.492064 | 0.643956 | 1.594578 | 0.998993 | -0.06155 | 0.450446 | 0.936096 | 1.307999 | 0.574718 | 0.026599 | 0.568714 | 0.408278 |
| 21.41667 | 1.038438 | 0.849005 | 1.761352 | 2.908949 | 1.321677 | 0.351118 | 1.089495 | 2.910188 | 1.189966 | 1.426578 | 0.040327 | 1.046588 | 0.574235 | 1.215854 | 2.283958 | 1.208077 | -0.105703 | 0.976142 | 1.181456 | 1.237999 | 0.625832 | 0.272272 | 0.316494 | 0.201889 |
| 21.43056 | 1.478511 | 1.463935 | 2.593558 | 3.94923 | 1.470069 | 0.75398 | 1.029626 | 2.192721 | 1.896761 | 2.242793 | 0.132659 | 1.371326 | 0.481643 | 1.410547 | 2.141318 | 1.056525 | -0.160178 | 0.884643 | 0.817924 | 0.586027 | 0.527635 | 0.369297 | -0.034128 | -0.40017 |
| 21.44444 | 1.854984 | 2.00394 | 3.165354 | 4.113962 | 1.509385 | 0.665222 | 0.543721 | 1.335048 | 2.088561 | 3.59662 | 0.369813 | 1.958636 | 0.22816 | 1.149677 | 1.838705 | 0.695413 | -0.229366 | 0.405835 | 0.188011 | 0.028155 | 0.33022 | 0.2979 | -0.259699 | -0.640867 |
| 21.45833 | 1.923038 | 2.379096 | 3.537215 | 3.874451 | 1.391716 | 0.353436 | 0.012693 | 0.707419 | 1.875194 | 4.502089 | 0.672249 | 2.804921 | 0.069885 | 0.873019 | 1.638336 | 0.683137 | -0.288715 | 0.235544 | -0.080546 | -0.069527 | 0.239525 | 0.278835 | -0.003172 | -0.158887 |
| 21.47222 | 1.691881 | 2.550967 | 3.448643 | 2.825828 | 1.264918 | 0.074344 | -0.362825 | 0.075576 | 1.238108 | 3.10894 | 0.612603 | 2.200497 | 0.152447 | 0.86213 | 1.751787 | 1.189935 | -0.565007 | 0.458945 | 0.073944 | 0.233346 | 0.233611 | 0.351156 | 0.535589 | 0.433958 |
| 21.48611 | 1.320022 | 2.287376 | 2.765538 | 1.497517 | 1.124956 | -0.099997 | -0.531903 | -0.530995 | 0.391393 | 1.154831 | -0.041368 | 0.991037 | 0.480073 | 1.130003 | 2.543752 | 2.074167 | -0.489446 | 0.748922 | 0.490023 | 0.723466 | 0.047046 | 0.447869 | 1.17952 | 1.216001 |
| 21.5 | 0.833092 | 1.782614 | 1.800202 | 0.34433 | 0.78584 | -0.447347 | -0.714095 | -0.833547 | -0.349187 | 0.007486 | -0.048603 | 0.199603 | 0.814499 | 1.889859 | 3.182055 | 2.574597 | -0.041985 | 0.732365 | 1.080173 | 1.110234 | -0.34928 | 0.50498 | 1.610141 | 2.145074 |
| 21.51389 | 0.603118 | 1.419994 | 0.978677 | -0.407844 | 0.578238 | -0.309895 | -0.678253 | -0.782486 | -0.295111 | -0.629573 | 0.236354 | -0.051187 | 1.042771 | 3.301376 | 3.825012 | 2.490138 | 0.352472 | 0.568882 | 1.881237 | 1.350159 | -0.45624 | 0.639709 | 1.598928 | 3.304833 |
| 21.52778 | 0.507607 | 1.258978 | 0.525473 | -0.671482 | 0.630173 | 0.209002 | -0.135528 | -0.318536 | 0.237124 | -0.736406 | 0.447009 | -0.028548 | 1.254699 | 3.17041 | 3.384094 | 2.337643 | 0.621361 | 0.515488 | 2.534264 | 1.390406 | 0.227047 | 0.766233 | 1.444762 | 3.816105 |
| 21.54167 | 0.591889 | 1.202873 | 0.296081 | -0.604797 | 0.938168 | 0.930987 | 0.633475 | 0.359318 | 1.077881 | -0.24352 | 0.51687 | 0.39888 | 1.505211 | 2.054413 | 1.884092 | 1.895321 | 0.889218 | 0.497413 | 2.750715 | 1.258052 | 1.042763 | 0.831507 | 1.281313 | 3.154769 |
| 21.55556 | 0.777835 | 1.178105 | -0.021173 | -0.438316 | 1.420254 | 1.167732 | 1.007146 | 0.976598 | 1.395858 | 0.406702 | 0.400781 | 0.603005 | 1.676796 | 1.206839 | 0.695238 | 1.245593 | 0.71679 | 0.446233 | 2.233012 | 0.869125 | 1.865328 | 0.758544 | 0.972533 | 2.051802 |
| 21.56944 | 0.923807 | 1.454021 | -0.404927 | -0.61885 | 1.751812 | 0.915403 | 1.346748 | 1.391215 | 0.979802 | 0.863068 | 0.296636 | 0.511685 | 1.520363 | 0.153956 | -0.242826 | 0.594825 | 0.34183 | 0.152288 | 0.945239 | 0.335098 | 2.052766 | 0.741658 | 0.482998 | 0.800252 |
| 21.58333 | 0.981404 | 1.744047 | -0.671086 | -0.669108 | 1.650398 | 0.939843 | 1.089848 | 0.229925 | 0.863317 | 0.868635 | 0.231929 | 0.372503 | 1.302407 | -1.060865 | -1.047706 | 0.114169 | 0.194367 | -0.109909 | 0.130853 | 0.110439 | 1.202233 | 0.534921 | -0.094422 | -0.384917 |
| 21.59722 | 0.835691 | 1.796264 | -0.770797 | -0.359276 | 1.198425 | 1.035659 | 0.516909 | -0.351002 | 0.748607 | 0.346012 | 0.183172 | 0.471027 | 1.338077 | -2.170295 | -1.155678 | -0.130479 | 0.106567 | -0.135434 | -0.464299 | -0.185724 | 0.227533 | 0.203645 | -0.578216 | -1.336199 |
| 21.61111 | 0.488427 | 1.626643 | -0.827495 | -0.102987 | 0.685439 | 1.31939 | 0.247487 | 0.213855 | 0.601369 | -0.174586 | 0.228114 | 0.896466 | 1.25439 | -2.797944 | -1.234527 | -0.235557 | 0.056696 | 0.125492 | -0.654042 | -0.248179 | -0.446928 | -0.074222 | -0.852257 | -1.768044 |
| 21.625 | -0.020223 | 1.252058 | -0.950431 | -0.31719 | 0.159241 | 1.279966 | 0.0522 | 0.633314 | 0.417486 | -0.271765 | 0.328527 | 1.165066 | 1.208504 | -2.359967 | -0.78509 | -0.254491 | 0.029992 | 0.426272 | -0.210904 | 0.007204 | -0.413169 | -0.249604 | -0.892955 | -1.477384 |
| 21.63889 | -0.387521 | 0.629431 | -0.875837 | -0.862864 | -0.273841 | 0.802998 | -0.170304 | 0.218963 | 0.161634 | 0.051294 | 0.386642 | 0.98423 | 1.208684 | -0.965749 | 0.169235 | -0.166529 | 0.040217 | 0.492384 | 0.145072 | 0.197812 | -0.51898 | -0.311262 | -0.747443 | -0.589875 |
| 21.65278 | -0.473683 | -0.254837 | -0.602588 | -0.992174 | -0.458895 | 0.190204 | -0.399718 | -0.443398 | -0.0786 | 0.484064 | 0.230157 | 0.488546 | 1.132355 | 0.350074 | 0.944277 | 0.263215 | 0.142713 | 0.336229 | 0.568543 | 0.430592 | -0.367849 | -0.292964 | -0.147699 | 0.397332 |
| 21.66667 | -0.169432 | -0.724907 | -0.427954 | -0.853842 | -0.627936 | -0.512102 | -0.668284 | -0.541973 | -0.249797 | 0.69725 | -0.031739 | -0.087267 | 1.045235 | 1.25389 | 1.434087 | 1.077463 | 0.250947 | 0.27414 | 0.93715 | 0.41268 | -0.248586 | -0.166968 | 1.078292 | 1.43871 |
| 21.68056 | 0.589352 | -0.691189 | -0.554657 | -0.948154 | -0.749876 | -1.008124 | -0.99825 | -0.471879 | -0.375305 | 0.507549 | -0.206643 | -0.57798 | 1.087064 | 1.299978 | 0.991775 | 1.897096 | 0.275998 | 0.482754 | 1.077218 | 0.278308 | 0.008339 | 0.093951 | 1.758168 | 2.175914 |
| 21.69444 | 1.226212 | -0.818818 | -0.755649 | -1.210034 | -0.789114 | -1.12444 | -1.138503 | 0.075682 | -0.399236 | -0.000747 | -0.185513 | -0.80471 | 1.056076 | 0.881052 | -0.30122 | 2.209695 | 0.222133 | 0.74021 | 1.036736 | 0.35345 | 0.471333 | 0.432473 | 1.929631 | 2.370337 |
| 21.70833 | 0.94454 | -1.09835 | -1.039658 | -1.233983 | -0.684697 | -0.943437 | -0.725581 | 0.090136 | -0.378179 | -0.382545 | -0.112742 | -0.758526 | 0.725761 | 0.875633 | -1.591498 | 1.993423 | 0.159966 | 0.608112 | 0.958197 | 0.440419 | 0.7794 | 0.606212 | 1.977153 | 2.319751 |
| 21.72222 | 0.03169 | -0.966504 | -0.944887 | -1.032287 | -0.491022 | -0.56572 | -0.100636 | -0.508592 | -0.227605 | -0.264264 | 0.105569 | -0.549011 | 0.09294 | 1.242163 | -3.056663 | 1.7594 | 0.204715 | 0.014681 | 0.882791 | 0.542569 | 0.780053 | 0.511708 | 2.498136 | 2.451152 |
| 21.73611 | -0.542703 | -0.862337 | -0.743244 | -0.864042 | -0.474389 | -0.344887 | 0.158453 | -0.807053 | -0.398557 | -0.085427 | 0.216735 | -0.472609 | -0.232037 | 1.445174 | -5.063832 | 1.733976 | 0.199589 | -0.375929 | 1.903036 | 0.737377 | 0.79468 | 0.384777 | 3.573172 | 2.442784 |
| 21.75 | -0.719231 | -1.189482 | -0.59965 | -0.8367 | -0.549067 | -0.294926 | 0.210051 | 0.157474 | -0.652892 | -0.334737 | 0.180907 | -0.544503 | -0.107354 | 1.006072 | -5.009905 | 1.507248 | 0.08523 | -0.133163 | 1.01834 | 0.614367 | 0.576152 | 0.269613 | 2.947677 | 1.109455 |
| 21.76389 | -0.878132 | -1.189662 | -0.407688 | -0.823901 | -0.69846 | -0.343135 | 0.320992 | 0.878716 | -0.51996 | -1.216602 | 0.019066 | -0.667522 | 0.028605 | 0.046367 | -3.309149 | 0.839513 | -0.027509 | 0.086079 | -1.306169 | 0.198999 | -0.139874 | 0.04853 | 1.183281 | -0.05722 |
| 21.77778 | -1.116715 | -0.258073 | -0.353398 | -0.953126 | -0.915429 | -0.307314 | 0.468597 | 0.874475 | -0.339404 | -2.337906 | -0.101935 | -0.614078 | -0.180333 | -1.01705 | -1.805461 | 0.193129 | -0.049466 | -0.062904 | -3.045392 | -0.138198 | -0.775845 | -0.211066 | -0.642858 | -0.919744 |
| 21.79167 | -1.195027 | 0.63201 | -0.492067 | -1.170471 | -1.159646 | -0.390069 | 0.29514 | -0.014939 | -0.485234 | -2.79775 | -0.046065 | -0.200148 | -0.573713 | -1.406365 | -0.480381 | -0.57915 | -0.00584 | -0.223834 | -3.345718 | -0.707806 | -1.506834 | -0.464411 | -2.119743 | -1.806312 |
| 21.80556 | -1.040871 | 0.336892 | -0.67365 | -1.122957 | -1.327161 | -0.664946 | -0.015009 | -0.6905 | -0.896363 | -2.401604 | -0.027361 | 0.0382 | -0.985775 | -0.954849 | 0.600007 | -1.127524 | 0.009692 | -0.744904 | -1.906337 | -0.599457 | -1.839301 | -0.620255 | -3.045649 | -1.789677 |
| 21.81944 | -0.939635 | -1.111051 | -0.924505 | -1.001798 | -1.249551 | -0.971528 | -0.354776 | -0.999781 | -1.30102 | -2.4317 | -0.156708 | -0.420509 | -0.776803 | -0.497483 | 1.601447 | -1.599912 | -0.014955 | -0.859781 | -1.001367 | -0.330784 | -1.53932 | -0.868054 | -2.67789 | -1.421259 |
| 21.83333 | -0.163408 | -2.272102 | -1.175239 | -1.058973 | -1.020106 | -1.146195 | -0.542438 | -1.020142 | -1.622663 | -2.704439 | -0.26705 | -1.259923 | -0.334882 | -0.269747 | 2.390714 | -2.231894 | -0.028089 | -0.924082 | -0.391947 | -0.136532 | -1.191347 | -1.318092 | -1.367578 | -0.464148 |
| 21.84722 | 0.199394 | -2.118229 | -1.240368 | -1.224035 | -0.902517 | -1.15212 | -0.459383 | -1.364158 | -1.766295 | -2.474516 | -0.293314 | -1.515136 | 0.015144 | 0.009204 | 2.776387 | -3.067733 | -0.015984 | -0.310212 | 0.092103 | -0.078536 | -0.621397 | -1.437986 | -0.686904 | 0.396829 |
| 21.86111 | 0.011915 | -1.957313 | -1.571177 | -1.899724 | -1.377786 | -1.218538 | -0.367101 | -2.189114 | -1.69548 | -2.083319 | -0.323285 | -1.359081 | 0.150764 | 0.014206 | 1.72757 | -3.293815 | 0.008614 | -0.595718 | 0.153231 | -0.194013 | -0.336484 | -1.059875 | 0.212292 | 0.480122 |
| 21.875 | -0.525631 | -2.522958 | -2.6402 | -3.861688 | -2.715525 | -2.192204 | -0.962987 | -3.690232 | -2.355069 | -2.993213 | -0.520618 | -2.343155 | 0.085627 | -0.407943 | 0.070571 | -2.61492 | 0.018836 | -0.206854 | -0.129364 | -0.231677 | -0.224675 | -0.556698 | 0.948618 | 0.823953 |
| 21.88889 | -1.181118 | -3.131253 | -2.794534 | -4.508898 | -4.224171 | -2.939187 | -2.305269 | -4.516757 | -3.236409 | -4.19067 | -0.652375 | -3.409898 | -0.071091 | -0.846753 | 0.311301 | -1.80247 | 0.016006 | -0.266569 | -0.224876 | -0.153819 | -0.102636 | -0.022177 | 1.520063 | 1.413976 |
| 21.90278 | -1.353441 | -2.329635 | -2.03997 | -2.746509 | -4.528668 | -2.37746 | -2.310636 | -2.765316 | -2.610225 | -3.482129 | -0.552094 | -3.003607 | -0.250545 | -1.001657 | 0.893703 | -0.583881 | -0.020368 | -0.380689 | -0.149499 | -0.026487 | -0.113137 | 0.242875 | 1.903226 | 2.012207 |
| 21.91667 | -1.248049 | -1.264973 | -1.349228 | -1.32021 | -3.513591 | -1.554682 | -2.031779 | -1.299519 | -1.634456 | -2.761869 | -0.32451 | -1.618685 | -0.448149 | -0.772253 | 1.262584 | 0.388283 | -0.03884 | -0.179145 | -0.133461 | -0.030188 | -0.27244 | 0.185667 | 0.944802 | 1.637026 |
| 21.93056 | -1.115957 | -0.721444 | -0.559375 | -0.784467 | -2.338062 | -1.167656 | -1.56836 | -0.147655 | -0.900498 | -2.091037 | -0.091803 | -0.483431 | -0.386587 | -0.567081 | 1.272731 | 0.936105 | 0.032833 | 0.088557 | -0.301424 | -0.169746 | -0.551982 | -0.098186 | -0.206156 | 1.451195 |
| 21.94444 | -1.135076 | -1.451829 | -0.12268 | -0.911112 | -2.142704 | -0.730507 | -1.267261 | 0.366943 | -0.394883 | -1.690011 | 0.190298 | 0.38308 | -0.534809 | -0.629572 | -0.45506 | 2.139912 | 0.107427 | -0.124314 | -0.63119 | -0.479201 | -0.994858 | -0.515897 | -0.520421 | 0.948507 |
| 21.95833 | -0.96657 | -2.849626 | -0.043062 | -1.094456 | -1.294867 | -0.30885 | -0.488299 | 0.167135 | -0.078844 | -1.305043 | 0.40926 | 0.980482 | -1.489911 | -1.15045 | -1.099195 | 2.53013 | 0.001473 | -0.981379 | -1.313659 | -1.379239 | -1.928832 | -1.172961 | -1.555652 | -1.29758 |
| 21.97222 | -0.682866 | -2.91879 | -0.226385 | -0.969567 | -0.01034 | -0.010833 | 0.359664 | -0.471225 | 0.142246 | -0.754294 | 0.461273 | 1.693901 | -2.452854 | -1.470723 | -1.892001 | 0.556288 | -0.222536 | -1.137057 | -1.332914 | -2.053338 | -1.966757 | -1.736183 | -2.519839 | -2.179459 |
| 21.98611 | -0.364316 | -2.189383 | -0.420816 | -0.877179 | 0.323758 | 0.226326 | 0.943783 | 0.213428 | 0.165846 | -0.319594 | 0.411956 | 1.980125 | -2.625791 | -1.410124 | -3.234774 | -1.539382 | -0.397605 | -0.156822 | -0.645589 | -1.688367 | -1.100733 | -1.962031 | -2.925343 | -3.335932 |
| 22 | -0.018574 | -0.309686 | -0.473434 | -0.876752 | -0.106908 | 0.430515 | 0.740777 | 1.892418 | 0.015846 | -0.180657 | 0.334758 | 1.287135 | -2.683755 | -1.641884 | -3.03247 | -2.821132 | -0.733156 | 0.717112 | 0.043597 | -1.142187 | -0.506979 | -1.963206 | -2.672431 | -3.657465 |
| 22.01389 | 0.325736 | 0.468579 | -0.548056 | -0.899358 | -0.491795 | 0.395185 | 0.091234 | 1.898496 | -0.158089 | -0.071844 | 0.242713 | 0.639595 | -1.880546 | -1.633436 | -2.106108 | -2.801706 | -0.799377 | 0.819428 | 0.197476 | -0.902086 | -0.228422 | -1.155046 | -1.957678 | -2.564472 |
| 22.02778 | 0.717423 | 0.111687 | -0.616364 | -0.912059 | -0.465447 | 0.247281 | -0.52013 | 0.070318 | -0.337473 | 0.111268 | 0.060178 | -0.321131 | -0.88368 | -1.42262 | -1.021105 | -2.20395 | -0.424158 | 0.212563 | -0.096543 | -0.690992 | -0.069551 | -0.251634 | -0.930291 | -0.849944 |
| 22.04167 | 0.724279 | -0.44471 | -0.753236 | -0.824254 | -0.204708 | 0.066179 | -0.611925 | -1.541716 | -0.432639 | 0.067696 | -0.213101 | -1.068037 | -0.29212 | -1.190926 | 0.014338 | -2.116255 | -0.003149 | -0.30145 | -0.385075 | -0.536838 | 0.083442 | 0.509101 | 0.456658 | 0.171351 |
| 22.05556 | 0.527004 | -0.539652 | -0.858086 | -0.592684 | -0.103439 | -0.169391 | -0.528218 | -1.773129 | -0.400357 | -0.109028 | -0.347944 | -1.455493 | -0.014002 | -1.031289 | 0.714619 | -2.314781 | 0.297319 | -0.599837 | -0.591254 | -0.405788 | 0.363556 | 0.916806 | 1.555092 | 1.295404 |
| 22.06944 | 0.16491 | -0.312555 | -0.785335 | -0.406126 | 0.182358 | -0.335041 | -0.4453 | -0.583677 | -0.348425 | -0.176851 | -0.262124 | -1.437915 | 0.024784 | -0.731193 | 0.842653 | -2.108443 | 0.360066 | -0.758437 | -0.509653 | -0.313409 | 0.424503 | 1.210016 | 1.653238 | 2.024543 |
| 22.08333 | -0.165748 | -0.035452 | -0.572268 | -0.345786 | 0.600772 | -0.471091 | -0.344716 | -0.36831 | -0.306431 | -0.092504 | -0.065945 | -1.08784 | -0.026646 | -0.257109 | 0.953354 | -1.064551 | 0.269571 | -0.767678 | -0.42611 | -0.297335 | 0.263238 | 1.323893 | 1.038872 | 1.564295 |
| 22.09722 | -0.39245 | 0.262952 | -0.338239 | -0.30391 | 0.766318 | -0.512828 | -0.035656 | -1.19609 | -0.290064 | -0.018836 | 0.098 | -0.690223 | -0.070694 | 0.164928 | 0.886192 | -0.345831 | 0.111124 | -0.619442 | -0.247526 | -0.329499 | 0.058103 | 0.95726 | 0.430408 | 0.546116 |
| 22.11111 | -0.352484 | 0.627339 | -0.253104 | -0.241748 | 0.584874 | -0.473057 | 0.276435 | -0.909047 | -0.286234 | -0.052543 | 0.17583 | -0.400169 | -0.135194 | 0.34813 | 0.689462 | -0.277681 | -0.072344 | -0.408099 | -0.037776 | -0.360094 | -0.145476 | 0.448935 | -0.218668 | -0.214118 |
| 22.125 | -0.356627 | 0.771977 | -0.294649 | -0.249396 | 0.334615 | -0.388141 | 0.188672 | 0.721783 | -0.271742 | -0.096344 | 0.211902 | -0.242026 | -0.202229 | 0.164653 | 0.462676 | -0.691152 | -0.141859 | -0.25853 | 0.07854 | -0.490132 | -0.315493 | 0.015002 | -0.851862 | -0.883321 |
| 22.13889 | -0.392322 | -0.194652 | -0.267523 | -0.354202 | 0.268059 | -0.340681 | 0.059078 | 1.077684 | -0.280556 | -0.181134 | 0.237817 | -0.189234 | -0.290276 | -0.212785 | 0.373092 | -1.203033 | -0.159222 | -0.209609 | -0.015735 | -0.589695 | -0.445216 | -0.320866 | -1.392426 | -1.422569 |
| 22.15278 | -0.289087 | -0.953249 | -0.108836 | -0.392673 | 0.466941 | -0.303792 | 0.071364 | -0.038919 | -0.191979 | -0.204517 | 0.221452 | -0.089126 | -0.34138 | -0.481768 | 0.427056 | -1.237439 | -0.237259 | -0.050423 | -0.043868 | -0.44349 | -0.521641 | -0.493967 | -1.580572 | -1.58504 |
| 22.16667 | 0.07561 | -0.327791 | 0.056285 | -0.351284 | 0.811531 | -0.268016 | 0.034353 | -0.884761 | 0.000066 | -0.106446 | 0.137306 | 0.053207 | -0.296528 | -0.508191 | 0.520632 | -0.79451 | -0.281602 | 0.235245 | 0.100668 | -0.10988 | -0.601315 | -0.508954 | -1.35809 | -1.094198 |
| 22.18056 | 0.389633 | 0.716982 | 0.15417 | -0.283399 | 0.808517 | -0.171614 | -0.104491 | -0.32421 | 0.128124 | 0.011771 | 0.072551 | 0.095015 | -0.176584 | -0.302153 | 0.604702 | -0.250933 | -0.255671 | 0.38231 | 0.156462 | 0.160089 | -0.645678 | -0.377396 | -1.034927 | -0.106609 |
| 22.19444 | 0.229656 | 0.870342 | 0.131769 | -0.258384 | 0.246357 | -0.060403 | -0.207211 | 1.085926 | 0.069725 | 0.034824 | 0.045788 | 0.058553 | -0.079017 | 0.14143 | 0.675638 | 0.141749 | -0.216667 | 0.21263 | -0.029256 | 0.209342 | -0.60417 | -0.210936 | -0.566477 | 0.14965 |
| 22.20833 | -0.166878 | 0.609364 | 0.04475 | -0.184025 | -0.18975 | 0.019559 | -0.080154 | 0.935733 | -0.048009 | 0.003388 | -0.005541 | 0.048208 | -0.046604 | 0.665224 | 0.604304 | 0.284186 | -0.22431 | 0.032649 | -0.289587 | 0.128485 | -0.454835 | -0.009842 | -0.006964 | 0.259162 |
| 22.22222 | -0.288859 | 0.891367 | 0.128469 | 0.066457 | 0.058647 | 0.113196 | 0.199542 | 0.66579 | 0.02032 | 0.148891 | -0.052596 | 0.186319 | -0.016474 | 0.838532 | 0.34128 | 0.170767 | -0.319567 | 0.173267 | -0.306837 | 0.114666 | -0.331183 | 0.179498 | 0.516103 | 0.563268 |
| 22.23611 | 0.047729 | 1.145445 | 0.344596 | 0.282168 | 0.427511 | 0.19331 | 0.364404 | 0.804702 | 0.240722 | 0.469394 | -0.011928 | 0.399311 | 0.101013 | 0.569658 | 0.185454 | 0.150352 | -0.255332 | 0.44371 | 0.01814 | 0.172021 | -0.303078 | 0.228881 | 0.886054 | 1.304739 |
| 22.25 | 0.428183 | 0.901912 | 0.438746 | 0.354894 | 0.474332 | 0.207994 | 0.322653 | 0.996166 | 0.371344 | 0.702168 | 0.110828 | 0.575868 | 0.194772 | 0.218487 | 0.253672 | 0.4792 | 0.234795 | 0.425554 | 0.261249 | 0.201193 | -0.304418 | 0.109213 | 1.28403 | 2.166477 |
| 22.26389 | 0.530129 | 0.487515 | 0.463661 | 0.542409 | 0.547362 | 0.31046 | 0.325861 | 0.902589 | 0.419408 | 0.83186 | 0.257109 | 0.799895 | 0.173843 | 0.078259 | 0.438195 | 0.885772 | 0.508788 | 0.210903 | 0.226273 | 0.223551 | -0.295867 | 0.010084 | 1.527287 | 2.488733 |
| 22.27778 | 0.39081 | 0.410508 | 0.583503 | 0.904334 | 0.892785 | 0.543927 | 0.467615 | 0.573665 | 0.472563 | 0.941539 | 0.355344 | 1.131921 | 0.055322 | 0.123466 | 0.455444 | 0.734233 | 0.208283 | 0.028527 | 0.105867 | 0.226273 | -0.229749 | 0.040667 | 1.419496 | 1.533265 |
| 22.29167 | 0.26145 | 0.503888 | 0.696986 | 1.112097 | 1.201307 | 0.767031 | 0.606223 | 0.438585 | 0.545715 | 0.985767 | 0.424367 | 1.532259 | 0.007951 | 0.15088 | 0.374406 | 0.277232 | -0.056597 | -0.009744 | 0.175394 | 0.18778 | -0.141355 | 0.148984 | 0.893375 | 0.527653 |
| 22.30556 | 0.304609 | 0.575934 | 0.734469 | 1.290516 | 1.208974 | 0.854032 | 0.722669 | 0.935385 | 0.594811 | 0.976423 | 0.489617 | 1.928673 | 0.120004 | 0.224643 | 0.379668 | 0.055452 | 0.043164 | 0.144367 | 0.379793 | 0.186623 | 0.082043 | 0.267196 | 0.486442 | 0.065974 |
| 22.31944 | 0.544271 | 0.744714 | 0.792574 | 1.489046 | 1.229026 | 0.842975 | 0.837599 | 1.628456 | 0.679555 | 1.022593 | 0.535403 | 2.071339 | 0.307545 | 0.263771 | 0.47318 | 0.169796 | 0.223567 | 0.264413 | 0.413452 | 0.271291 | 0.241611 | 0.310148 | 0.193128 | -0.006655 |
| 22.33333 | 0.837416 | 0.945479 | 0.998877 | 1.77361 | 1.392464 | 0.972613 | 0.923496 | 2.043826 | 0.818017 | 1.218524 | 0.526098 | 2.031465 | 0.468681 | -0.002269 | 0.615842 | 0.52865 | 0.260381 | 0.309958 | 0.300431 | 0.35684 | 0.196616 | 0.214721 | -0.173419 | 0.094205 |
| 22.34722 | 0.856605 | 0.795074 | 1.105184 | 1.843261 | 1.324251 | 0.966871 | 0.740135 | 1.817836 | 0.799576 | 1.466663 | 0.402962 | 1.672926 | 0.570856 | 0.041314 | 0.960766 | 1.238494 | 0.217747 | 0.394711 | 0.331692 | 0.460496 | 0.146139 | 0.129018 | -0.310958 | 0.095466 |
| 22.36111 | 0.331542 | 0.255508 | 0.907024 | 1.453769 | 0.905852 | 0.773177 | 0.45789 | 0.859574 | 0.511256 | 1.310392 | 0.1835 | 1.075948 | 0.586185 | 0.633497 | 1.383771 | 1.96529 | 0.123457 | 0.659669 | 0.52658 | 0.545685 | 0.189753 | 0.17353 | -0.269855 | 0.112609 |
| 22.375 | -0.12236 | -0.026171 | 0.616272 | 0.937865 | 0.491182 | 0.515905 | 0.32673 | -0.310583 | 0.123431 | 0.78643 | 0.079079 | 0.649973 | 0.529243 | 0.342283 | 1.44627 | 1.871612 | 0.021438 | 0.965309 | 0.666626 | 0.549173 | 0.212249 | 0.184368 | -0.118998 | 0.187878 |
| 22.38889 | -0.206874 | 0.219251 | 0.55754 | 0.95147 | 0.365883 | 0.349138 | 0.33551 | -1.158264 | -0.058125 | 0.310689 | 0.121033 | 0.706242 | 0.543152 | -0.046842 | 1.264701 | 1.432403 | -0.036931 | 1.129237 | 0.66852 | 0.475599 | 0.155099 | 0.140324 | 0.242646 | 0.80544 |
| 22.40278 | 0.199601 | 0.662831 | 0.991602 | 1.918847 | 0.747695 | 0.532331 | 0.464303 | -0.856624 | 0.215999 | 0.566134 | 0.205564 | 1.275898 | 0.667274 | 0.349039 | 1.047207 | 1.200047 | -0.0116 | 1.081763 | 0.64196 | 0.466172 | 0.098493 | 0.138165 | 1.008529 | 1.980411 |
| 22.41667 | 0.688393 | 1.221505 | 1.76901 | 3.088097 | 1.435977 | 1.033158 | 0.970822 | 0.441941 | 0.754836 | 1.609338 | 0.288862 | 2.078108 | 0.709178 | 1.202639 | 1.019739 | 1.153516 | -0.016283 | 1.050829 | 0.622624 | 0.480868 | 0.102984 | 0.158881 | 2.365177 | 3.046321 |
| 22.43056 | 1.096484 | 1.587896 | 2.557685 | 3.579194 | 1.881341 | 1.585018 | 1.600883 | 2.224466 | 1.321327 | 2.811176 | 0.396654 | 1.692346 | 0.492697 | 1.117186 | 0.960766 | 1.035459 | -0.257527 | 1.093938 | 0.57074 | 0.433654 | 0.028725 | 0.178238 | 2.672692 | 2.661127 |
| 22.44444 | 1.511735 | 1.594434 | 2.845844 | 3.403668 | 2.004909 | 1.734845 | 1.755252 | 3.098092 | 1.720189 | 3.341902 | 0.424956 | 1.021456 | 0.088154 | 0.468749 | 0.690389 | 0.491341 | -0.879066 | 0.882209 | 0.504533 | 0.247581 | -0.299084 | 0.133859 | 2.010702 | 1.617209 |
| 22.45833 | 1.866956 | 1.680222 | 2.935667 | 3.406901 | 2.266685 | 1.431887 | 1.563351 | 3.328426 | 1.921076 | 3.538738 | 0.538828 | 0.937453 | -0.134326 | 0.073469 | 0.231313 | 0.035612 | -2.111654 | 0.539413 | 0.38887 | 0.051845 | -0.50402 | 0.100549 | 1.416536 | 0.539421 |
| 22.47222 | 1.532875 | 1.744097 | 2.590173 | 3.479867 | 2.163136 | 1.101461 | 1.362397 | 2.406056 | 1.76023 | 3.471919 | 0.820523 | 1.53534 | 0.152296 | 0.555836 | -0.048499 | 0.269924 | -2.556255 | 0.321379 | 0.413796 | 0.08744 | -0.176336 | 0.21233 | 0.835306 | -0.169114 |
| 22.48611 | 0.98054 | 1.791008 | 2.098697 | 2.970098 | 1.606057 | 1.089542 | 1.108792 | 1.359361 | 1.580199 | 2.59686 | 0.965612 | 2.010809 | 0.612248 | 1.559184 | 0.58705 | 1.128955 | -1.479466 | 0.281064 | 0.569336 | 0.41667 | 0.129233 | 0.528655 | 0.183889 | -0.663664 |
| 22.5 | 0.499433 | 2.0133 | 1.651215 | 1.866748 | 1.257419 | 1.080849 | 0.845518 | -0.040153 | 1.240651 | 1.421994 | 0.875651 | 1.758133 | 0.871889 | 2.568833 | 1.823328 | 1.995547 | -0.294992 | 0.331206 | 0.875669 | 0.847712 | 0.432223 | 1.034107 | -0.425502 | -0.757285 |
| 22.51389 | 0.430957 | 2.116121 | 1.196773 | 1.077506 | 1.253376 | 0.986128 | 0.524616 | -1.811946 | 0.960945 | 0.193503 | 0.809965 | 1.447356 | 0.937254 | 3.625201 | 2.188092 | 2.565907 | 0.739598 | 0.429067 | 1.279884 | 1.035498 | 0.712364 | 1.327656 | -0.83484 | -0.184715 |
| 22.52778 | 0.674603 | 2.180599 | 0.895182 | 0.881973 | 1.598176 | 0.785389 | 0.17443 | -1.185411 | 0.687999 | -0.644748 | 0.777936 | 1.14672 | 1.121041 | 4.449963 | 1.80914 | 2.790724 | 1.835322 | 0.622709 | 1.446444 | 1.142137 | 0.839329 | 1.319813 | -0.960134 | 0.968592 |
| 22.54167 | 1.021326 | 2.107795 | 0.659077 | 0.941867 | 2.078154 | 0.612083 | -0.016795 | 0.178335 | 0.606914 | -0.765723 | 0.853921 | 1.069752 | 1.528515 | 3.895895 | 1.755559 | 2.941256 | 2.751495 | 0.889655 | 1.527569 | 1.382731 | 1.104247 | 1.253607 | -0.471599 | 2.028382 |
| 22.55556 | 1.068722 | 1.990251 | 0.608122 | 0.654276 | 2.572647 | 0.612635 | 0.039501 | 0.869107 | 0.794752 | -0.231882 | 1.102455 | 1.241625 | 1.672591 | 2.215627 | 1.95104 | 2.202879 | 2.572507 | 1.081858 | 1.424745 | 1.634073 | 1.297547 | 1.269227 | 0.091943 | 2.187897 |
| 22.56944 | 0.901883 | 2.004742 | 0.743085 | 0.103938 | 2.74208 | 0.71864 | 0.291006 | 0.227575 | 0.912614 | 0.197277 | 1.127013 | 1.179768 | 1.400111 | 0.767249 | 1.974369 | 1.34466 | 1.69429 | 0.946694 | 1.099253 | 1.168755 | 1.298739 | 1.114383 | 0.697947 | 1.824711 |
| 22.58333 | 0.677903 | 1.973222 | 0.854321 | -0.136848 | 2.414977 | 0.859531 | 0.501632 | -0.123609 | 0.889299 | 0.46114 | 0.796687 | 0.968775 | 0.983334 | -0.345773 | 1.975789 | 0.755599 | 0.916024 | 0.569764 | 0.650685 | 0.784774 | 1.122243 | 0.853176 | 1.055796 | 1.173795 |
| 22.59722 | 0.720374 | 1.649381 | 0.564162 | 0.036712 | 1.87981 | 0.952533 | 0.753741 | 0.112637 | 0.788689 | 0.578787 | 0.459196 | 0.751907 | 0.745912 | -1.055574 | 2.484963 | 0.398593 | 0.387549 | 0.33108 | 0.384269 | 0.599559 | 0.902364 | 0.642421 | 0.671971 | 0.175633 |
| 22.61111 | 0.919441 | 1.618835 | 0.209864 | 0.190086 | 1.178127 | 0.936007 | 1.029376 | 0.407114 | 0.68003 | 0.49157 | 0.103836 | 0.51372 | 0.720361 | -1.215152 | 2.668992 | 0.210159 | 0.162586 | 0.333506 | 0.313679 | 0.470081 | 0.740414 | 0.406111 | 1.092306 | -0.390557 |
| 22.625 | 1.318441 | 0.824264 | 0.124221 | 0.028777 | 0.798062 | 0.961915 | 1.054978 | 0.403699 | 0.635147 | 0.145466 | -0.130653 | 0.389668 | 0.848219 | -1.205043 | 2.59604 | 0.588702 | 0.144846 | 0.680635 | 0.28738 | 0.443533 | 0.640383 | 0.165432 | 1.978583 | 0.718501 |
| 22.63889 | 1.050624 | -0.087658 | 0.386453 | -0.201613 | 0.708735 | 1.174461 | 0.823973 | -0.005191 | 0.735671 | -0.104354 | -0.193548 | 0.245769 | 1.006991 | -1.58901 | 1.460227 | 1.582372 | 0.156253 | 1.162279 | 0.517207 | 0.451759 | 0.669443 | 0.116854 | 2.502967 | 0.798806 |
| 22.65278 | 0.395107 | -0.503142 | 0.460278 | -0.300937 | 0.597911 | 1.374999 | 0.531989 | -1.234487 | 0.926588 | 0.033375 | -0.35697 | 0.000826 | 1.178954 | -1.652512 | -0.06576 | 2.492753 | 0.154715 | 1.158535 | 1.097093 | 0.590159 | 0.622097 | 0.244272 | 2.539085 | 1.30337 |
| 22.66667 | 0.209753 | -0.80172 | -0.042792 | -0.296722 | 0.300686 | 1.278419 | 0.227304 | -2.857965 | 0.786942 | 0.54242 | -0.437703 | -0.319092 | 1.180561 | -0.745381 | -1.189935 | 2.888185 | 0.108709 | 0.832395 | 1.522316 | 0.807013 | 0.654044 | 0.317888 | 1.569076 | 1.180184 |
| 22.68056 | 0.703702 | -0.786416 | -0.241782 | -0.280648 | -0.023897 | 0.68111 | 0.008423 | -1.13085 | 0.469753 | 1.129834 | -0.421981 | -0.432319 | 0.827066 | 0.527754 | -2.59548 | 1.652779 | 0.073915 | 0.366544 | 1.198841 | 0.49386 | 0.63247 | 0.220466 | 0.283337 | 0.622015 |
| 22.69444 | 1.277894 | -0.663667 | -0.37718 | -0.382723 | -0.020073 | 0.437861 | 0.032835 | 0.714265 | 0.265906 | 0.951119 | -0.376906 | -0.505541 | 0.579212 | 1.549373 | -2.294266 | 1.038111 | 0.096416 | -0.39541 | 0.624716 | 0.315357 | 0.613682 | 0.038112 | -1.073354 | -0.045365 |
| 22.70833 | 1.19096 | -0.619763 | -0.375514 | -0.567121 | 0.271693 | 0.463836 | 0.183791 | 0.806093 | 0.308765 | 0.603499 | -0.368078 | -0.727946 | 0.603184 | 1.427649 | -1.556069 | 0.935105 | 0.141333 | 0.438621 | 0.388922 | 0.135265 | 0.462372 | -0.130217 | -1.194058 | -0.445472 |
| 22.72222 | 0.315102 | -0.847946 | -0.4049 | -0.70999 | 0.299607 | 0.581104 | 0.144924 | -0.422785 | 0.46145 | 0.495992 | -0.281049 | -1.02276 | 0.610426 | 0.551301 | -1.586342 | 0.641252 | 0.150104 | 0.480735 | 0.216198 | -0.092742 | 0.124392 | -0.204662 | -0.751402 | -0.563963 |
| 22.73611 | 0.002605 | -1.110014 | -0.55555 | -0.84401 | 0.067753 | 0.48518 | -0.072471 | -1.073411 | 0.403949 | 0.566547 | -0.005062 | -1.016086 | 0.423255 | 0.07563 | -1.69365 | 0.464079 | 0.102029 | 0.000781 | -0.130641 | -0.223173 | -0.178704 | -0.278795 | -0.482597 | -0.48858 |
| 22.75 | 0.485313 | -0.759319 | -0.773154 | -1.002277 | -0.158185 | 0.139907 | -0.152304 | -0.38744 | 0.174601 | 0.601202 | 0.183191 | -0.909347 | 0.109172 | 0.312982 | -0.821552 | 0.171007 | 0.047854 | -0.403103 | -0.400034 | -0.429543 | -0.596054 | -0.295349 | -0.14975 | -0.805861 |
| 22.76389 | 1.025671 | -0.207238 | -0.826569 | -1.048164 | -0.537313 | -0.120893 | 0.126584 | 1.199119 | -0.052819 | 0.367785 | 0.120321 | -0.845053 | -0.078497 | 0.89738 | -0.222452 | -0.177014 | 0.009943 | -0.620905 | -0.738663 | -0.393254 | -1.00944 | -0.314368 | -0.044548 | -1.105406 |
| 22.77778 | 0.189701 | -0.057502 | -0.748663 | -0.938782 | -0.831168 | -0.076575 | 0.495665 | 2.612056 | -0.072653 | 0.016174 | 0.025675 | -0.829329 | -0.051867 | 1.022706 | -0.040289 | -0.423465 | 0.026954 | -0.466876 | -1.059851 | -0.405935 | -1.026931 | -0.31726 | -0.356178 | -0.664832 |
| 22.79167 | -0.591254 | -0.61484 | -0.666452 | -1.009239 | -0.751521 | -0.018618 | 0.604358 | 1.359939 | -0.041398 | -0.266527 | 0.011728 | -0.961388 | -0.020323 | 0.292415 | 0.739001 | -0.845507 | 0.054517 | -0.191795 | -1.163311 | -0.336164 | -0.650743 | -0.237899 | -0.209169 | -0.169499 |
| 22.80556 | -0.759692 | -1.25983 | -1.019521 | -1.3285 | -0.58405 | -0.229377 | 0.343867 | -1.275692 | -0.276537 | -0.605349 | -0.091959 | -1.050921 | -0.161566 | -0.794134 | 1.990875 | -0.696403 | 0.049516 | 0.122974 | -1.12803 | -0.093787 | -0.231228 | -0.117654 | 0.890779 | -0.031353 |
| 22.81944 | -0.592472 | -1.623712 | -0.998689 | -1.603029 | -0.46729 | -0.664079 | 0.006451 | -2.845225 | -0.695665 | -1.124096 | -0.386656 | -1.202165 | -0.349357 | -1.547916 | 2.100297 | -1.151428 | 0.050103 | 0.288363 | -1.029238 | 0.112601 | 0.030042 | -0.003718 | 2.707032 | 0.313379 |
| 22.83333 | -1.003772 | -2.092967 | -1.04335 | -1.495865 | -0.725506 | -1.023328 | -0.281352 | -1.635007 | -0.85114 | -1.716049 | -0.495019 | -0.788353 | -0.355594 | -1.523486 | 2.629641 | -1.788547 | 0.058172 | 0.291079 | -0.5406 | 0.237496 | 0.136709 | 0.046532 | 4.631017 | 0.319776 |
| 22.84722 | -1.502101 | -2.549566 | -1.217316 | -1.420843 | -1.182097 | -1.044042 | -0.251987 | -0.976279 | -0.882024 | -1.425854 | -0.348284 | -0.079291 | -0.099072 | -1.078987 | 3.300457 | -1.724781 | 0.041535 | 0.225398 | 0.026183 | 0.380102 | 0.177494 | 0.11402 | 3.658824 | 0.319834 |
| 22.86111 | -1.876574 | -2.585232 | -1.49673 | -2.062098 | -1.683901 | -0.966978 | -0.341427 | -1.423035 | -1.278658 | -1.440883 | -0.348598 | -0.099998 | 0.047902 | -0.868778 | 1.668929 | -1.829356 | 0.01784 | 0.131199 | 0.240401 | 0.540971 | 0.32466 | 0.216216 | 1.526114 | 0.845502 |
| 22.875 | -2.681082 | -2.709904 | -1.958602 | -3.462586 | -2.818538 | -1.424191 | -0.954106 | -3.616505 | -2.33909 | -2.46528 | -0.783687 | -1.224356 | -0.121926 | -1.032127 | -1.64702 | -1.513229 | -0.003145 | 0.102968 | 0.145669 | 0.359576 | 0.495936 | 0.367574 | 0.169424 | 1.440525 |
| 22.88889 | -3.080547 | -3.388821 | -2.572264 | -4.396907 | -3.816853 | -2.352739 | -1.722995 | -4.75863 | -2.889365 | -3.137092 | -1.292794 | -3.287793 | -0.309533 | -1.250748 | 0.245284 | -0.181006 | 0.023294 | -0.016675 | -0.190252 | 0.101429 | 0.584788 | 0.353959 | -1.217619 | 1.606778 |
| 22.90278 | -2.298969 | -3.491526 | -2.354663 | -4.147207 | -2.957931 | -2.520235 | -2.202784 | -2.527392 | -2.35767 | -2.574611 | -1.163571 | -3.776046 | -0.334118 | -1.131924 | 0.493574 | 1.314653 | 0.044234 | -0.473141 | -0.507695 | 0.13952 | 0.527315 | 0.369813 | -1.456182 | 1.120795 |
| 22.91667 | -1.546291 | -2.321404 | -1.406229 | -3.172637 | -2.533352 | -1.690488 | -2.263008 | -0.661174 | -1.712904 | -2.308798 | -0.916423 | -3.07102 | -0.347539 | -0.674255 | 0.767348 | 1.651344 | 0.019527 | -1.351733 | -0.863944 | 0.213788 | 0.353591 | 0.235823 | -1.961239 | -0.461455 |
| 22.93056 | -1.282069 | -0.380707 | -0.923847 | -2.029725 | -2.65572 | -1.275487 | -2.352899 | -0.077481 | -1.407869 | -2.40633 | -0.468655 | -2.645778 | -0.317417 | 0.049926 | 0.359713 | 1.811016 | 0.036367 | -2.256731 | -1.472879 | 0.089139 | 0.236174 | 0.144371 | -1.975545 | -1.460549 |
| 22.94444 | -1.258215 | 0.710526 | -0.722398 | -1.483622 | -3.299893 | -1.562219 | -2.481706 | 0.687689 | -1.408958 | -2.305597 | -0.188193 | -1.60903 | -0.371382 | 0.624013 | -0.359708 | 1.666286 | 0.103482 | -1.850112 | -1.895569 | -0.232887 | -0.315812 | -0.012932 | -1.738859 | -2.421506 |
| 22.95833 | -1.127583 | 0.029603 | -0.649341 | -1.201655 | -3.146464 | -1.39346 | -1.897348 | 1.412437 | -1.130607 | -1.835732 | -0.200435 | -0.605002 | -0.652079 | 0.640045 | -1.012066 | 0.630952 | -0.019884 | -1.235956 | -2.382132 | -1.147741 | -1.003529 | -0.282145 | -1.934968 | -3.305733 |

| | | | | | | | | | | | | | | | | | | | | | | | | |
|---|---|---|---|---|---|---|---|---|---|---|---|---|---|---|---|---|---|---|---|---|---|---|---|---|
| 22.97222 | -0.629012 | -1.636712 | -0.459559 | -0.98861 | -1.240716 | -0.786861 | -0.972992 | 1.492468 | -0.536456 | -1.054756 | -0.296129 | 0.402775 | -1.06171 | 0.142542 | -1.566733 | -2.021664 | -0.437987 | -1.306988 | -2.490181 | -2.905097 | -1.236004 | -0.481791 | -2.295807 | -2.389786 |
| 22.98611 | -0.400087 | -2.154717 | -0.257383 | -0.88152 | 0.348158 | -0.316141 | -0.034238 | 0.721892 | -0.417496 | -0.675455 | -0.102967 | 1.389111 | -1.406576 | -0.452168 | -2.086249 | -3.974805 | -0.526096 | -1.530529 | -2.541752 | -2.881515 | -1.167497 | -0.444933 | -1.692304 | -2.12474 |
| 23 | -0.541754 | -1.114661 | 0.092742 | -0.926828 | -0.203545 | -0.276984 | 0.4426 | -0.371972 | -0.78491 | -0.470938 | 0.204973 | 1.345901 | -1.517718 | -1.135184 | -2.035864 | -3.860803 | -0.374568 | -0.796876 | -1.537598 | -2.125824 | -0.801929 | -0.401375 | -0.969433 | -2.045244 |
| 23.01389 | -0.722294 | -0.08346 | 0.150025 | -0.82572 | -2.338188 | -0.451107 | 0.831205 | -1.000593 | -0.999492 | -0.131668 | 0.108049 | 0.928856 | -1.358358 | -1.542418 | -1.717941 | -3.507098 | -0.398175 | -0.039863 | -0.973466 | -1.393959 | -0.663423 | -0.458534 | -0.39285 | -1.537667 |
| 23.02778 | -0.848784 | 0.362146 | 0.122376 | -0.567598 | -4.265612 | -0.702995 | 0.855729 | -0.963065 | -0.850039 | -0.018536 | -0.071418 | 0.310455 | -0.977101 | -1.463893 | -1.229353 | -3.2466 | -0.258044 | 0.130292 | -0.608879 | -0.814904 | -0.59616 | -0.268473 | 1.039356 | -0.017865 |
| 23.04167 | -0.787274 | -0.117669 | -0.024458 | -0.36913 | -3.166036 | -0.748513 | 0.711003 | -0.66974 | -0.536642 | 0.041847 | -0.038951 | -0.231622 | -0.568159 | -1.542161 | -0.663271 | -1.177754 | -0.089572 | -0.008569 | -0.496265 | -0.2473 | -0.353257 | 0.075752 | 2.492047 | 1.816454 |
| 23.05556 | -0.446776 | -1.201543 | -0.162325 | -0.233355 | -0.703224 | -0.452276 | -0.079222 | -0.129044 | -0.291878 | -0.053003 | -0.074632 | -0.781098 | -0.252121 | -1.94512 | -0.298073 | 0.537387 | -0.019283 | -0.268283 | -0.504807 | 0.334504 | -0.162972 | 0.255922 | 2.794365 | 3.755076 |
| 23.06944 | -0.143172 | -2.108114 | -0.293942 | -0.188813 | -0.09618 | -0.350826 | -0.952145 | 0.257713 | -0.220982 | -0.067038 | -0.151414 | -1.237929 | -0.119358 | -1.937199 | -0.302189 | 0.203306 | -0.056401 | -0.495706 | -0.554439 | 0.668056 | -0.20123 | 0.10053 | 2.002494 | 3.396666 |
| 23.08333 | -0.187134 | -1.601061 | -0.362788 | -0.121847 | -1.244291 | -0.339566 | -1.448893 | 0.38425 | -0.101371 | 0.072994 | -0.054414 | -1.239645 | -0.079666 | -1.392853 | -0.85768 | -0.629431 | -0.035572 | -0.494852 | -0.441166 | 0.607556 | -0.328115 | -0.013571 | 1.009021 | 1.561613 |
| 23.09722 | -0.435973 | -0.653987 | -0.366348 | -0.016745 | -1.683594 | -0.162405 | -0.991355 | 0.077061 | -0.054455 | 0.193535 | 0.06017 | -0.87739 | 0.022982 | -0.647673 | -1.588902 | -1.247656 | 0.114692 | -0.407411 | -0.265635 | 0.309698 | -0.165236 | 0.073046 | 0.136388 | -0.351216 |
| 23.11111 | -0.43372 | -0.129426 | -0.182433 | 0.095422 | -0.521575 | 0.128587 | -0.342376 | 0.060301 | -0.168413 | 0.140086 | 0.065512 | -0.558683 | 0.11427 | 0.02928 | -1.339986 | -0.980854 | 0.180376 | -0.275177 | -0.092263 | 0.115649 | -0.088662 | 0.031898 | -0.137122 | -0.995772 |
| 23.125 | -0.065665 | -0.084095 | -0.005883 | 0.152334 | 1.235903 | 0.384373 | 0.045007 | 0.384503 | -0.356384 | -0.089875 | 0.020753 | -0.403626 | 0.065745 | 0.48035 | 1.900897 | -0.690658 | 0.120117 | -0.224451 | -0.183034 | -0.056463 | -0.274874 | -0.013816 | -0.303264 | -1.304073 |
| 23.13889 | 0.144441 | -0.150449 | 0.088116 | 0.16722 | 1.02075 | 0.357453 | 0.150422 | 0.366196 | -0.421001 | -0.246361 | -0.038376 | -0.275335 | -0.098837 | 0.431569 | 1.182802 | -0.524297 | 0.104071 | -0.295552 | -0.352971 | -0.189861 | -0.407627 | 0.01748 | -0.560223 | -2.250187 |
| 23.15278 | -0.024111 | -0.074827 | 0.194431 | 0.12134 | -0.676804 | 0.273564 | 0.301286 | -0.215561 | -0.336579 | -0.174582 | -0.08992 | -0.160212 | -0.172975 | -0.028903 | -1.199831 | -0.289468 | 0.083972 | -0.284287 | -0.37078 | -0.237137 | -0.481681 | -0.022716 | -0.792297 | -3.181769 |
| 23.16667 | -0.108132 | -0.000299 | 0.272525 | 0.169381 | -1.649713 | 0.294503 | 0.635197 | -0.882073 | -0.210138 | -0.025783 | -0.091765 | -0.055103 | -0.149782 | -0.37125 | -0.885903 | -0.066044 | 0.065523 | -0.110192 | -0.071834 | -0.188618 | -0.578779 | -0.139534 | -0.73457 | -2.588027 |
| 23.18056 | 0.185638 | 0.03094 | 0.321227 | 0.158279 | 0.020274 | 0.293741 | 0.802266 | -0.880805 | -0.131412 | -0.002038 | -0.031206 | -0.116408 | -0.211279 | -0.411663 | -0.027061 | 0.395291 | 0.048521 | 0.066979 | 0.21252 | -0.048967 | -0.674923 | -0.134362 | -0.509003 | -1.586847 |
| 23.19444 | 0.413144 | 0.015948 | 0.305096 | 0.132366 | 1.929456 | 0.220008 | 0.521472 | -0.387875 | -0.0739 | -0.092267 | 0.061989 | -0.178956 | -0.222148 | -0.353207 | 1.053546 | 1.064661 | 0.013276 | 0.042571 | 0.131103 | 0.091698 | -0.613128 | 0.035728 | -0.392318 | -0.596264 |
| 23.20833 | 0.409077 | -0.018947 | 0.279685 | 0.159802 | 1.674681 | 0.216125 | 0.23579 | 0.014425 | 0.036871 | -0.064415 | 0.169083 | -0.0271 | -0.096387 | -0.245744 | 1.378809 | 1.240027 | -0.018004 | -0.065042 | -0.151109 | 0.084023 | -0.51577 | 0.171479 | -0.196209 | 0.585377 |
| 23.22222 | 0.344037 | 0.196347 | 0.308268 | 0.235481 | 0.302646 | 0.340052 | 0.109332 | 0.180536 | 0.214231 | 0.116188 | 0.317793 | 0.292848 | 0.043688 | -0.045774 | 0.199463 | 0.971694 | -0.031439 | 0.064966 | -0.128609 | 0.025052 | -0.446671 | 0.281662 | 0.090379 | 1.421395 |
| 23.23611 | 0.23607 | 0.655762 | 0.429278 | 0.312539 | -0.77372 | 0.460584 | 0.190717 | 0.533952 | 0.434849 | 0.279023 | 0.384641 | 0.565928 | 0.074147 | 0.114107 | -0.132307 | 0.648045 | -0.008207 | 0.190182 | 0.140211 | 0.080734 | -0.40555 | 0.36704 | 0.462802 | 2.078294 |
| 23.25 | 0.184464 | 0.963704 | 0.460539 | 0.458376 | -0.837233 | 0.541018 | 0.441005 | 1.126194 | 0.529546 | 0.385061 | 0.292196 | 0.733574 | 0.090559 | 0.090872 | -0.359339 | 0.323535 | -0.0379 | 0.101211 | 0.20384 | 0.176289 | -0.30232 | 0.349599 | 0.764972 | 2.417692 |
| 23.26389 | 0.381268 | 0.799429 | 0.451851 | 0.709254 | 0.168898 | 0.592946 | 0.784239 | 1.686917 | 0.581222 | 0.520874 | 0.11133 | 0.85545 | 0.171922 | 0.010921 | -0.444038 | 0.130269 | -0.153841 | -0.035348 | 0.163049 | 0.16472 | -0.170111 | 0.197927 | 0.960199 | 1.950994 |
| 23.27778 | 0.621769 | 0.638224 | 0.455683 | 0.9337 | 1.315216 | 0.62297 | 1.140286 | 1.963315 | 0.675413 | 0.628498 | 0.008088 | 1.041622 | 0.262211 | -0.032565 | -0.510673 | 0.201924 | -0.240615 | -0.200151 | 0.158603 | 0.069912 | -0.104604 | 0.051028 | 1.025038 | 1.366409 |
| 23.29167 | 0.724422 | 0.641368 | 0.54153 | 1.031667 | 1.792829 | 0.738235 | 1.391359 | 2.18053 | 0.845818 | 0.712023 | 0.08341 | 1.288836 | 0.309042 | -0.093083 | -0.302407 | 0.439721 | -0.136597 | -0.164961 | 0.325861 | 0.16451 | -0.038087 | 0.032975 | 0.825166 | 0.983361 |
| 23.30556 | 0.692196 | 0.652121 | 0.689977 | 1.091356 | 1.434828 | 0.904384 | 1.666206 | 2.212187 | 1.007563 | 0.752829 | 0.278547 | 1.452343 | 0.354824 | -0.135279 | 0.093973 | 0.74307 | 0.055568 | 0.060641 | 0.367382 | 0.239663 | 0.20361 | 0.166482 | 0.599462 | 0.646521 |
| 23.31944 | 0.72523 | 0.515546 | 0.844682 | 1.206868 | 1.23671 | 1.100158 | 1.528267 | 2.044761 | 1.079735 | 0.930999 | 0.527482 | 1.53082 | 0.449832 | -0.020101 | 0.396858 | 0.918256 | 0.165875 | 0.29299 | 0.583287 | 0.30077 | 0.447352 | 0.241664 | 0.377748 | 0.393442 |
| 23.33333 | 0.90743 | 0.350112 | 0.961608 | 1.311548 | 1.432275 | 1.23703 | 1.256224 | 1.899959 | 1.030273 | 1.158952 | 0.654547 | 1.665861 | 0.505472 | 0.126165 | 0.802477 | 0.886216 | 0.204564 | 0.414676 | 0.65126 | 0.332621 | 0.583441 | 0.225764 | 0.166819 | 0.221521 |
| 23.34722 | 0.967557 | 0.159548 | 0.921087 | 1.428754 | 1.636849 | 1.290201 | 0.93909 | 1.344444 | 0.904678 | 1.251957 | 0.587356 | 1.642543 | 0.466562 | 0.162171 | 1.167051 | 0.799681 | 0.195148 | 0.423694 | 0.591258 | 0.327772 | 0.609469 | 0.186144 | 0.007886 | 0.188085 |
| 23.36111 | 0.829802 | -0.118271 | 0.611801 | 1.463747 | 1.474423 | 0.985438 | 0.428167 | 0.651436 | 0.513869 | 1.011648 | 0.373123 | 1.493837 | 0.420976 | 0.168953 | 1.324239 | 1.037805 | 0.198791 | 0.454356 | 0.553529 | 0.314844 | 0.6451 | 0.167557 | -0.064044 | 0.157362 |
| 23.375 | 0.910653 | -0.392219 | 0.364117 | 1.421271 | 1.363813 | 0.462576 | -0.048869 | 0.102187 | 0.095579 | 0.77776 | 0.275321 | 1.274934 | 0.430048 | 0.245885 | 1.429958 | 1.393735 | 0.211315 | 0.631302 | 0.649933 | 0.364304 | 0.693391 | 0.190002 | 0.066372 | 0.351992 |
| 23.38889 | 1.175406 | -0.400756 | 0.368378 | 1.5857 | 1.480976 | 0.289605 | -0.053408 | 0.059672 | -0.068163 | 0.66591 | 0.475262 | 1.232541 | 0.509333 | 0.240751 | 1.521948 | 1.387521 | 0.235513 | 0.840928 | 0.824041 | 0.415257 | 0.748396 | 0.219622 | 0.385092 | 0.873794 |
| 23.40278 | 1.249644 | 0.119217 | 0.981518 | 2.040166 | 1.958755 | 0.508782 | 0.361599 | 0.824799 | 0.109653 | 0.938763 | 0.718343 | 1.264364 | 0.519701 | 0.503117 | 1.515716 | 1.307891 | 0.196998 | 0.929371 | 1.01093 | 0.4157 | 0.843706 | 0.192156 | 0.715098 | 1.552135 |
| 23.41667 | 0.856367 | 0.321337 | 1.436897 | 2.84075 | 2.487254 | 1.098761 | 1.060513 | 1.736992 | 0.749914 | 1.727542 | 0.917651 | 1.39388 | 0.454086 | 1.077293 | 1.412939 | 1.377836 | 0.094089 | 0.896628 | 1.204109 | 0.450589 | 0.931295 | 0.173022 | 0.915502 | 1.918092 |
| 23.43056 | 0.696875 | 0.628133 | 1.762059 | 3.040577 | 2.297948 | 2.001879 | 1.979255 | 2.611158 | 1.694819 | 2.526821 | 1.119814 | 1.543596 | 0.283258 | 1.118647 | 1.273814 | 1.216794 | -0.019739 | 0.810591 | 1.179168 | 0.437236 | 0.830856 | 0.141101 | 1.013802 | 2.218755 |
| 23.44444 | 1.324658 | 1.257358 | 1.908307 | 2.790215 | 2.363954 | 2.492111 | 3.031548 | 2.564698 | 2.346503 | 2.987312 | 1.225423 | 1.454144 | -0.367888 | 0.997568 | 1.039796 | 0.919073 | -0.626312 | 0.591776 | 0.947206 | 0.285638 | 0.525425 | 0.069944 | 0.912913 | 1.844042 |
| 23.45833 | 1.696783 | 1.805643 | 1.816785 | 2.432066 | 2.051013 | 2.131987 | 3.13862 | 2.008361 | 2.252798 | 3.462204 | 0.785489 | 1.142836 | -1.377687 | 0.872529 | 0.600654 | 0.474905 | -3.128772 | 0.423564 | 0.694372 | 0.20266 | 0.109739 | 0.043814 | 0.719636 | 0.746328 |
| 23.47222 | 1.877696 | 1.581208 | 1.367382 | 1.694792 | 1.477871 | 1.740988 | 2.008845 | 0.586535 | 1.778873 | 3.279473 | 0.267983 | 0.797344 | -1.37808 | 0.487917 | 0.202519 | 0.505668 | -6.311158 | 0.468298 | 0.659289 | 0.362186 | -0.149971 | 0.195962 | 0.397671 | -0.165794 |
| 23.48611 | 0.290003 | 1.21611 | 0.91057 | 0.850266 | 0.531604 | 1.812541 | 1.318283 | 0.445506 | 1.287587 | 2.610867 | 0.027347 | 0.728165 | -0.471307 | 0.419812 | 0.380371 | 1.473928 | -5.197364 | 0.707237 | 0.897351 | 0.649117 | 0.014681 | 0.371716 | 0.346694 | -0.611256 |
| 23.5 | 0.885077 | 1.355198 | 0.543458 | 0.179706 | 1.171246 | 1.976088 | 1.377155 | 0.79676 | 0.942573 | 1.799332 | 0.199072 | 0.957484 | 0.31783 | 1.113279 | 1.035904 | 2.996062 | -1.66118 | 1.051928 | 1.369196 | 1.074537 | 0.234101 | 0.459578 | 0.576419 | -0.15003 |
| 23.51389 | 1.773242 | 1.608578 | 0.43622 | -0.200708 | 2.194404 | 1.592157 | 1.055047 | 0.803553 | 0.710531 | 0.968543 | 0.516216 | 1.352039 | 0.848563 | 2.292891 | 1.615293 | 3.511711 | 0.870044 | 1.520177 | 1.986903 | 1.462152 | 0.283527 | 0.549258 | 1.025529 | 0.99947 |
| 23.52778 | 1.120132 | 1.589956 | 0.43382 | 0.00926 | 1.910586 | 1.238822 | 0.377088 | 0.562161 | 0.570755 | 0.190424 | 0.792509 | 1.811241 | 1.447834 | 3.167365 | 1.907549 | 3.172186 | 3.327403 | 1.576479 | 2.344694 | 1.589003 | 0.245208 | 0.654941 | 1.446444 | 2.431663 |
| 23.54167 | 0.122431 | 1.690282 | 0.411739 | 0.583198 | 1.45556 | 0.7899 | 0.145543 | 0.48859 | 0.434901 | -0.354732 | 0.909717 | 1.601071 | 2.016081 | 2.843157 | 1.913384 | 3.079286 | 5.158561 | 1.448611 | 2.207778 | 1.502396 | 0.281159 | 0.680622 | 1.641372 | 3.31256 |
| 23.55556 | 0.381805 | 2.003294 | 0.518005 | 1.161686 | 1.260231 | 0.645249 | 0.57209 | 0.744493 | 0.281054 | -0.430535 | 1.035688 | 1.438266 | 1.921668 | 1.804545 | 1.662709 | 2.724162 | 4.455338 | 1.049407 | 1.792155 | 1.018307 | 0.223405 | 0.589865 | 1.715303 | 2.511603 |
| 23.56944 | 1.261671 | 2.183849 | 0.688288 | 1.502619 | 1.613344 | 0.882516 | 1.401411 | 1.282439 | 0.247063 | -0.02522 | 1.414697 | 1.73007 | 1.629851 | 0.861113 | 1.39105 | 1.610322 | 2.812031 | 0.572413 | 1.221395 | 0.542834 | 0.296238 | 0.440976 | 1.644147 | 1.189811 |
| 23.58333 | 1.033205 | 1.370114 | 1.06262 | 1.617382 | 2.169429 | 0.977131 | 1.736574 | 1.70901 | 0.381159 | 0.461959 | 1.762397 | 2.062393 | 1.140016 | 0.262483 | 1.127563 | 1.109754 | 1.346016 | 0.023448 | 0.759726 | 0.221819 | 0.617224 | 0.349246 | 1.382342 | 0.490584 |
| 23.59722 | -0.034801 | 0.542803 | 1.426998 | 1.938106 | 2.279143 | 0.928677 | 1.154403 | 0.746811 | 0.637559 | 0.807109 | 1.853293 | 2.056421 | 0.724794 | 0.012639 | 0.717305 | 0.840549 | 0.324966 | -0.481556 | 0.438374 | 0.222253 | 1.059538 | 0.364385 | 1.031459 | 0.1546 |
| 23.61111 | -0.75606 | -0.239532 | 1.474819 | 2.200287 | 1.829913 | 0.864207 | 0.29595 | -0.108113 | 0.79264 | 0.651819 | 1.25572 | 1.330888 | 0.599461 | -0.124506 | 0.557992 | 0.330951 | -0.009418 | -0.550356 | 0.18001 | 0.373989 | 1.290887 | 0.241087 | 0.656071 | 0.120067 |
| 23.625 | -0.295716 | -0.485237 | 1.088372 | 2.050022 | 1.000418 | 1.230182 | 0.082944 | -0.873169 | 0.794762 | 0.266188 | 0.533219 | 0.530998 | 0.631818 | -0.253493 | 0.728365 | 0.102592 | -0.107107 | 0.133923 | 0.220961 | 0.45105 | 1.152948 | 0.100042 | 0.35196 | 0.137533 |
| 23.63889 | 0.866933 | -0.105895 | 0.747245 | 1.543718 | 0.479885 | 1.720028 | 0.101242 | -1.064843 | 0.719566 | 0.068677 | 0.077948 | 0.077717 | 0.7127 | -0.167368 | 0.934651 | 0.415025 | -0.124383 | 1.197286 | 0.463863 | 0.519026 | 0.933205 | 0.039312 | 0.144716 | 0.67627 |
| 23.65278 | 1.535951 | 0.65477 | 0.215992 | 0.971437 | 0.89806 | 1.527197 | 0.21094 | -0.730869 | 0.615171 | 0.216668 | -0.129762 | -0.093775 | 0.893681 | 0.226785 | 0.980738 | 1.117514 | -0.177482 | 1.496625 | 0.456579 | 0.696083 | 0.809883 | 0.029277 | 0.051216 | 1.678459 |
| 23.66667 | 1.675872 | 1.210837 | -0.107624 | 0.358221 | 1.179411 | 0.94038 | 0.289468 | -0.597721 | 0.535289 | 0.70935 | -0.297385 | -0.263387 | 1.000827 | 0.75986 | 1.184163 | 1.501483 | -0.149122 | 1.185389 | 0.37345 | 0.939591 | 0.657127 | -0.031387 | 0.245221 | 2.680569 |
| 23.68056 | 1.306612 | 0.909957 | -0.29985 | -0.225333 | 0.847213 | 0.21631 | 0.269122 | -0.621698 | 0.601987 | 1.179175 | -0.263587 | -0.389439 | 1.061549 | 1.225584 | 1.545485 | 1.803452 | 0.022126 | 1.006517 | 0.419293 | 0.756775 | 0.507057 | -0.192798 | 0.700087 | 2.092139 |
| 23.69444 | 1.504625 | 0.052217 | -0.274349 | -0.803163 | 0.391883 | 0.057642 | 0.148645 | -0.998899 | 0.529271 | 1.021583 | -0.164093 | -0.323436 | 0.880042 | 1.485531 | 1.115854 | 1.425946 | 0.155217 | 0.889331 | 0.144872 | 0.374781 | 0.56211 | -0.229936 | 0.969785 | 1.60005 |
| 23.70833 | 1.590213 | -0.559952 | -0.165748 | -1.183656 | 0.249289 | -0.09559 | -0.028101 | -0.718802 | 0.250812 | 0.632367 | -0.116617 | -0.105154 | 0.628608 | 1.504226 | 0.358521 | 1.110806 | 0.083353 | 0.660174 | -0.381168 | 0.117224 | 0.720473 | -0.081377 | 0.862875 | 1.169936 |
| 23.72222 | 1.964125 | -0.41422 | -0.199041 | -1.357481 | 0.425597 | -0.105752 | -0.168206 | 0.051537 | 0.027912 | 0.474764 | -0.02453 | -0.109978 | 0.137844 | 1.593519 | -0.337202 | 0.906054 | -0.058183 | 0.0481 | -0.717051 | -0.108423 | 0.70055 | 0.10159 | 0.572109 | 0.901803 |
| 23.73611 | 1.444788 | 0.645201 | -0.371288 | -1.482812 | 0.211634 | -0.142978 | -0.071458 | 0.237732 | 0.161099 | 0.660702 | 0.019015 | -0.268551 | -0.047602 | 1.318248 | -0.797404 | 0.738177 | -0.126696 | -0.728537 | -0.695235 | -0.377811 | 0.218338 | 0.002596 | 0.613058 | 0.623776 |
| 23.75 | 0.475705 | 1.700877 | -0.694549 | -1.632018 | 0.019425 | -0.340379 | -0.071347 | 0.006058 | 0.425936 | 0.950157 | 0.091005 | -0.197017 | -0.062425 | 0.619028 | -0.914591 | 0.526166 | -0.109503 | -1.186836 | -0.741254 | -0.392518 | -0.651465 | -0.210159 | 0.882622 | 0.428266 |
| 23.76389 | -0.168172 | 1.414527 | -0.755753 | -1.939888 | -0.079721 | -0.52049 | -0.382647 | 0.089385 | 0.517845 | 0.846205 | 0.230531 | -0.024871 | -0.10768 | 0.032267 | -0.674473 | 0.223842 | -0.101985 | -0.781896 | -0.889611 | -0.224597 | -1.585158 | -0.307822 | 0.764408 | 0.338283 |
| 23.77778 | -0.582607 | 0.412407 | -0.486083 | -1.864286 | -0.051237 | -0.406633 | -0.732522 | 0.546714 | 0.396413 | 0.230757 | 0.425211 | 0.219257 | -0.261511 | -0.409728 | -0.654957 | -0.023458 | -0.210595 | -0.402601 | -1.02644 | -0.149206 | -1.363983 | -0.066136 | 0.145272 | 0.230211 |
| 23.79167 | -0.792743 | -0.003939 | -0.343906 | -1.720225 | 0.034458 | -0.185808 | -0.674313 | 1.088324 | 0.203957 | -0.283915 | 0.559815 | 0.230478 | -0.500918 | -0.6844 | -0.162784 | -0.290702 | -0.305845 | -0.079511 | -0.963706 | -0.195018 | -0.670217 | 0.066856 | -0.435762 | 0.218282 |
| 23.80556 | -0.921831 | -0.024041 | -0.432752 | -1.77696 | 0.144914 | 0.071259 | -0.134201 | 1.055333 | 0.033463 | -0.33479 | 0.536435 | 0.057971 | -0.572192 | -0.880982 | 0.497886 | -0.479345 | -0.295187 | -0.034027 | -0.51305 | -0.298671 | -0.207948 | -0.086937 | -0.860455 | 0.203048 |
| 23.81944 | -1.341636 | 0.138093 | -0.526469 | -1.868486 | -0.185134 | 0.387047 | 0.325184 | 0.350994 | -0.045302 | -0.096537 | 0.326214 | -0.167639 | -0.47555 | -1.079387 | 0.732902 | -0.560139 | -0.220418 | 0.008936 | 0.011121 | -0.175964 | -0.073615 | -0.189939 | -1.183375 | 0.065295 |
| 23.83333 | -1.716744 | -0.082943 | -0.505488 | -1.507814 | -1.11564 | 0.614268 | 0.195106 | -0.551873 | -0.148388 | -0.105524 | 0.121265 | -0.50874 | -0.306722 | -1.271033 | 0.818089 | -0.469771 | -0.124417 | 0.246397 | 0.11933 | 0.287705 | 0.115454 | -0.071545 | -1.201612 | -0.191232 |
| 23.84722 | -1.751026 | -0.985077 | -0.265814 | -1.264238 | -1.979479 | 0.595566 | -0.171281 | -1.373566 | -0.404668 | -0.376692 | -0.10882 | -0.99985 | -0.243303 | -1.121894 | 0.850691 | -0.653787 | -0.079963 | 0.372598 | -0.067459 | 0.52308 | 0.51416 | 0.066426 | -1.030191 | -0.404898 |
| 23.86111 | -1.090602 | -1.948524 | -0.116537 | -1.71058 | -1.345649 | 0.191523 | -0.448656 | -2.277764 | -0.814431 | -1.131782 | -0.478445 | -1.549543 | -0.376801 | -0.908608 | 0.41593 | -2.129619 | -0.135486 | 0.203256 | -0.097745 | 0.270221 | 0.907865 | -0.007966 | -1.283141 | -0.5423 |
| 23.875 | -1.328699 | -2.505946 | -0.286423 | -2.910952 | -1.311578 | -1.024882 | -1.79577 | -2.79248 | -1.741188 | -3.007963 | -1.267011 | -2.284942 | -0.581817 | -0.897147 | -0.372611 | -2.591238 | -0.243781 | -0.182544 | -0.055051 | 0.029967 | 1.135376 | -0.039269 | -1.523546 | -1.279736 |
| 23.88889 | -2.838102 | -2.687969 | -0.744775 | -4.569777 | -2.794463 | -1.88262 | -2.512671 | -3.853629 | -2.381583 | -4.788821 | -2.132272 | -2.890645 | -0.72865 | -0.743385 | -1.29583 | -2.576963 | -0.330749 | -0.581889 | 0.037763 | 0.025696 | 1.084354 | 0.111718 | -0.838056 | -0.817618 |
| 23.90278 | -2.324123 | -2.805267 | -0.982909 | -3.531699 | -3.593948 | -1.338587 | -1.957026 | -3.193383 | -2.343853 | -3.603396 | -1.176237 | -3.290147 | -0.828529 | -0.141572 | -1.966195 | -1.904468 | -0.444056 | -0.385628 | -0.064176 | -0.01134 | 0.885672 | 0.258064 | -0.148933 | -0.640307 |
| 23.91667 | -1.173808 | -2.631361 | -1.0122 | -2.351758 | -4.251996 | -0.707542 | -1.164935 | -2.181821 | -1.644836 | -2.49125 | -0.885753 | -2.529385 | -1.044395 | 0.460078 | -1.839065 | -1.786512 | -0.520539 | -0.124562 | -0.308498 | -0.164398 | 0.530297 | 0.281315 | 0.121027 | -0.927628 |
| 23.93056 | -0.487256 | -1.570768 | -1.156384 | -1.693828 | -3.938128 | -0.319368 | -1.526823 | -1.030093 | -0.94098 | -1.810197 | -0.781689 | -1.633038 | -1.160988 | 0.562216 | -1.497209 | -1.815694 | -0.66509 | -0.499014 | -0.603304 | -0.333957 | 0.075204 | 0.143675 | 0.038545 | -0.794781 |
| 23.94444 | -0.804608 | 0.538302 | -1.498553 | -1.406966 | -2.448842 | -0.295839 | -1.972384 | -0.056295 | -0.510743 | -1.236202 | -0.810402 | -0.941054 | -1.488153 | 0.210048 | -1.613271 | -1.646157 | -1.38553 | -1.246573 | -1.011337 | -0.565765 | -0.390267 | 0.117475 | -0.172819 | -0.823564 |
| 23.95833 | -0.81086 | 1.372395 | -1.543183 | -0.873394 | -1.621885 | -0.477228 | -1.939597 | 0.609888 | -0.365515 | -0.88691 | -0.808742 | -0.21346 | -2.221802 | -0.586698 | -1.593308 | -1.896176 | -3.778153 | -3.247335 | -2.264492 | -1.022479 | -0.939766 | 0.10227 | -0.635681 | -1.965633 |
| 23.97222 | -0.232922 | 0.926788 | -1.035505 | -0.662459 | -1.517719 | -0.677394 | -2.056042 | 1.361977 | -0.396687 | -0.614174 | -0.367055 | 0.594566 | -3.355585 | -1.11489 | -1.372225 | -2.696935 | -6.576026 | -3.210087 | -2.617636 | -1.98935 | -1.446786 | -0.053291 | -1.407614 | -2.905888 |
| 23.98611 | 0.662857 | 0.447164 | -0.521494 | -0.811841 | -0.964393 | -0.668773 | -2.156763 | 1.805112 | -0.646162 | -0.447202 | 0.133471 | 1.479248 | -3.315301 | -1.291102 | -1.148339 | -2.039658 | -7.401746 | -1.42732 | -1.678995 | -2.375955 | -1.290305 | -0.195124 | -1.462861 | -2.235555 |
| 24 | 0.951248 | -0.12499 | -0.215289 | -0.774855 | 0.090452 | -0.389589 | -2.069938 | 1.172219 | -1.039126 | -0.407106 | 0.274957 | 1.442337 | -2.186679 | -1.045815 | -0.760036 | -0.818772 | -8.341218 | -0.541687 | -1.091358 | -1.471762 | -1.242291 | -0.290199 | -1.154603 | -1.731988 |
| 24.01389 | 0.248666 | -0.058184 | 0.033885 | -0.310842 | 0.217345 | -0.191178 | -1.285037 | 0.437226 | -1.142252 | -0.204297 | 0.029063 | 0.803422 | -1.119569 | -0.966635 | -0.392412 | -0.598674 | -5.559452 | -0.388126 | -0.796243 | -0.683613 | -1.119502 | -0.154411 | -1.205318 | -1.235019 |
| 24.02778 | -0.118748 | 1.628073 | 0.293802 | -0.244333 | -0.21779 | 0.079058 | -0.151919 | 1.207237 | -0.391797 | 0.597828 | 0.094207 | 0.782082 | -0.794037 | -0.991032 | -0.209932 | -0.825958 | -2.218397 | -0.364113 | -0.614527 | -0.621509 | -0.96739 | 0.07087 | -1.174383 | -0.725992 |
| 24.04167 | -1.657323 | 3.99614 | -1.556739 | -1.755602 | -2.203133 | 1.154606 | -0.015225 | 1.656252 | 0.747791 | 0.439872 | 1.31127 | 1.053506 | -0.731989 | -0.992091 | -0.024008 | -0.713998 | -0.377363 | -0.195031 | -0.355119 | -0.616801 | -0.601171 | 0.135517 | -0.754611 | -0.599575 |
| 24.05556 | -5.882148 | -3.126757 | -4.907569 | -5.716314 | -5.554543 | -0.159509 | -1.416613 | -0.197145 | -1.690361 | -4.75369 | -0.925038 | 1.412104 | -0.474145 | -1.213398 | 0.155199 | -0.402778 | 1.008603 | -0.055672 | -0.12132 | -0.334668 | -0.150578 | 0.0743 | -0.30003 | -0.398127 |
| 24.06944 | -1.675369 | -1.427785 | -2.519602 | -0.640491 | -5.536546 | -1.527636 | -1.356273 | -2.802106 | -0.860814 | -2.080529 | -2.440968 | 0.648405 | -0.129469 | -1.374387 | 0.267436 | -0.099672 | 1.729172 | -0.119252 | -0.002391 | -0.164502 | 0.096024 | 0.005212 | 0.051417 | -0.07405 |
| 24.08333 | 1.310507 | 1.184272 | 1.102399 | 2.582986 | -0.089807 | -0.101216 | 0.158987 | -0.172623 | 1.069564 | 0.896648 | -0.386489 | 1.572508 | -0.097772 | -1.060498 | 0.201615 | -0.194533 | 1.579487 | -0.321761 | -0.069911 | -0.256505 | 0.089016 | 0.036527 | 0.00053 | 0.304237 |
| 24.09722 | 3.709287 | 2.042385 | 1.070652 | 3.22786 | 3.720012 | 1.085883 | 0.654472 | 2.280804 | 1.827226 | 2.213196 | 1.163773 | 3.509651 | -0.284608 | -0.502359 | 0.08936 | -0.53176 | 0.837676 | -0.473633 | -0.220483 | -0.408779 | 0.031552 | 0.091731 | -0.157731 | 0.245166 |
| 24.11111 | 5.133272 | 3.015624 | 0.654498 | 2.274132 | 6.612252 | 0.963305 | 1.025673 | 2.890756 | 1.324848 | 2.285514 | 2.151567 | 3.77163 | -0.392187 | -0.200767 | 0.071083 | -0.838249 | -0.233859 | -0.561311 | -0.325497 | -0.390011 | 0.082565 | 0.05223 | 0.036195 | -0.07995 |
| 24.125 | 4.39135 | 2.774474 | 0.33191 | 0.873291 | 6.22382 | 0.643782 | 1.474621 | 2.060479 | 1.20364 | 2.023137 | 1.994811 | 2.643985 | -0.387935 | -0.239165 | 0.088271 | -0.927378 | -1.689421 | -0.627919 | -0.462799 | -0.475131 | 0.116772 | -0.008202 | 0.142745 | -0.517298 |
| 24.13889 | 2.465823 | 1.745836 | 0.155635 | 0.322184 | 4.097914 | 0.388349 | 1.639099 | 1.25002 | 1.15868 | 1.694221 | 1.209476 | 1.836354 | -0.299204 | -0.285295 | 0.170763 | -0.622499 | -2.325927 | -0.579418 | -0.53647 | -0.771802 | -0.007268 | -0.007732 | 0.073673 | -0.943576 |
| 24.15278 | 1.661448 | 1.102216 | 0.144005 | 0.361685 | 2.95923 | 0.288097 | 1.566515 | 0.809535 | 1.0906 | 1.40947 | 0.754956 | 1.089121 | -0.205116 | -0.310471 | 0.388225 | -0.184073 | -0.803629 | -0.349716 | -0.402631 | -0.723966 | -0.139157 | 0.059447 | -0.265084 | -1.35637 |
| 24.16667 | 1.109357 | 0.421382 | 0.299147 | 0.428374 | 2.088201 | 0.367065 | 1.329071 | 0.724842 | 1.043378 | 1.153367 | 0.543426 | 0.709582 | -0.102211 | -0.250589 | 0.491318 | 0.173589 | 0.056344 | -0.069187 | -0.247088 | -0.186149 | -0.133729 | 0.115339 | -0.333018 | -1.574435 |
| 24.18056 | 0.713252 | 0.382344 | 0.538459 | 0.464413 | 1.748091 | 0.570752 | 1.307766 | 0.82113 | 1.154267 | 1.034726 | 0.536581 | 0.883549 | 0.000839 | -0.132656 | 0.352768 | 0.540615 | 0.078023 | 0.06145 | -0.126703 | 0.134741 | -0.086367 | 0.120767 | -0.208709 | -1.382881 |
| 24.19444 | 0.529929 | 0.400351 | 0.554266 | 0.573985 | 1.412937 | 0.694199 | 1.166494 | 0.862133 | 1.113742 | 0.881427 | 0.539636 | 0.940726 | -0.054712 | -0.025786 | 0.201283 | 0.970559 | -0.125763 | 0.029024 | -0.105907 | -0.022372 | -0.124581 | 0.155569 | -0.08348 | -0.859483 |
| 24.20833 | 0.404706 | 0.322136 | 0.370886 | 0.603514 | 0.981949 | 0.663074 | 0.897333 | 0.724369 | 0.807205 | 0.600796 | 0.443219 | 0.746599 | -0.24874 | -0.114295 | 0.162611 | 1.398817 | -0.086362 | 0.015827 | -0.190028 | -0.276157 | -0.245083 | 0.168933 | -0.1265 | -0.39526 |
| 24.22222 | 0.311321 | 0.293511 | 0.216338 | 0.568821 | 0.711464 | 0.55495 | 0.741815 | 0.616697 | 0.547917 | 0.450464 | 0.359384 | 0.529412 | -0.336593 | -0.463975 | 0.272771 | 1.437094 | 0.178137 | 0.032879 | -0.270928 | -0.39332 | -0.27665 | 0.160957 | -0.00343 | 0.058798 |
| 24.23611 | 0.277865 | 0.258046 | 0.148207 | 0.393713 | 0.583457 | 0.489769 | 0.646347 | 0.511203 | 0.458774 | 0.390363 | 0.322281 | 0.417443 | -0.164091 | -0.689411 | 0.420159 | 0.839272 | 0.356025 | 0.080226 | -0.297768 | -0.477195 | -0.120786 | 0.19886 | 0.235414 | 0.490929 |
| 24.25 | 0.244781 | 0.19647 | 0.143103 | 0.222851 | 0.455921 | 0.429918 | 0.552636 | 0.386078 | 0.434192 | 0.348538 | 0.296211 | 0.336248 | -0.008803 | -0.528917 | 0.491397 | 0.38681 | 0.427356 | 0.151895 | -0.206075 | -0.475473 | 0.113339 | 0.280171 | 0.560504 | 0.910268 |
| 24.26389 | 0.204177 | 0.152894 | 0.174345 | 0.211815 | 0.373733 | 0.448824 | 0.535705 | 0.305262 | 0.428506 | 0.333547 | 0.270732 | 0.288756 | 0.116117 | -0.254691 | 0.583889 | 0.549934 | 0.632233 | 0.266321 | -0.014165 | -0.342011 | 0.226106 | 0.306981 | 0.861642 | 1.298987 |
| 24.27778 | 0.183985 | 0.145589 | 0.199179 | 0.224392 | 0.343496 | 0.501957 | 0.557559 | 0.263983 | 0.454429 | 0.342664 | 0.262363 | 0.282988 | 0.246854 | -0.193399 | 0.61627 | 0.879758 | 0.737121 | 0.307644 | 0.109618 | -0.104201 | 0.191378 | 0.286113 | 0.911902 | 1.073364 |
| 24.29167 | 0.171353 | 0.135568 | 0.189356 | 0.223571 | 0.310866 | 0.551552 | 0.527347 | 0.221662 | 0.427358 | 0.351019 | 0.236685 | 0.280205 | 0.317325 | -0.18517 | 0.463084 | 0.974203 | 0.531006 | 0.203362 | 0.099142 | 0.149909 | 0.151131 | 0.209287 | 0.841922 | 0.819094 |
| 24.30556 | 0.162568 | 0.123989 | 0.197967 | 0.259327 | 0.258352 | 0.764826 | 0.43741 | 0.19017 | 0.302978 | 0.37452 | 0.199664 | 0.28413 | 0.311883 | 0.004303 | 0.349204 | 1.021156 | 0.229387 | 0.067117 | 0.057927 | 0.356015 | 0.193816 | 0.150464 | 0.730495 | 0.791236 |
| 24.31944 | 0.19907 | 0.173018 | 0.327327 | 0.355619 | 0.254532 | 1.458735 | 0.345363 | 0.243348 | 0.175742 | 0.467299 | 0.223251 | 0.359479 | 0.311263 | 0.182998 | 0.302243 | 1.049983 | 0.04515 | 0.033277 | 0.065697 | 0.529983 | 0.289565 | 0.111062 | 0.532585 | 0.836432 |
| 24.33333 | 0.240248 | 0.27309 | 0.486871 | 0.42948 | 0.285605 | 1.453508 | 0.336297 | 0.330252 | 0.333255 | 0.57255 | 0.291458 | 0.456495 | 0.378583 | 0.189414 | 0.313494 | 1.187983 | 0.153448 | 0.155431 | 0.143009 | 0.700001 | 0.34533 | 0.065989 | 0.433195 | 0.759749 |
| 24.34722 | 0.152581 | 0.240986 | 0.545404 | 0.401573 | 0.231512 | 1.086166 | 0.337635 | 0.262366 | 0.589768 | 0.568544 | 0.261628 | 0.494669 | 0.542118 | 0.151666 | 0.236826 | 1.435989 | 0.535275 | 0.344818 | 0.27235 | 0.729352 | 0.362336 | 0.007557 | 0.362824 | 0.766858 |
| 24.36111 | 0.047008 | 0.070826 | 0.607335 | 0.436241 | 0.390647 | 0.877851 | 0.368012 | 0.143461 | 0.861673 | 0.611534 | 0.160302 | 0.557301 | 0.655883 | 0.154288 | 0.078218 | 1.532821 | 0.789378 | 0.443246 | 0.37182 | 0.674426 | 0.368415 | -0.056733 | 0.311437 | 0.885171 |
| 24.375 | 0.138926 | 0.059915 | 0.911638 | 0.629445 | 1.026363 | 0.914001 | 0.617273 | 0.263045 | 1.306817 | 0.915022 | 0.239193 | 0.835837 | 0.617871 | 0.23277 | -0.059258 | 1.375964 | 0.7795 | 0.45819 | 0.40428 | 0.641055 | 0.368262 | -0.11223 | 0.285897 | 0.900868 |
| 24.38889 | 0.308545 | 0.153611 | 1.159069 | 0.774433 | 1.427153 | 0.959352 | 0.785292 | 0.377029 | 1.554861 | 1.099597 | 0.485223 | 1.025786 | 0.53228 | 0.371115 | -0.084608 | 1.192322 | 0.603898 | 0.523427 | 0.426062 | 0.578874 | 0.367756 | -0.152775 | 0.365387 | 0.888391 |
| 24.40278 | 0.285385 | 0.161478 | 1.111551 | 0.773715 | 1.059036 | 0.788762 | 0.668601 | 0.263141 | 1.28904 | 0.918943 | 0.674958 | 0.971231 | 0.452037 | 0.434097 | -0.027125 | 1.056167 | 0.26169 | 0.669654 | 0.611375 | 0.591155 | 0.443667 | -0.109478 | 0.523657 | 0.795869 |
| 24.41667 | 0.248761 | 0.075144 | 0.769741 | 0.557151 | 0.51029 | 0.585407 | 0.525821 | 0.097301 | 0.98063 | 0.64114 | 0.648005 | 0.733209 | 0.370453 | 0.416939 | -0.046252 | 1.069871 | -0.036001 | 0.877843 | 1.023208 | 0.629388 | 0.600736 | -0.039271 | 0.71865 | 0.797737 |
| 24.43056 | 0.189385 | -0.007286 | 0.464675 | 0.381142 | 0.197054 | 0.367731 | 0.303073 | -0.001516 | 0.816263 | 0.473226 | 0.51292 | 0.463365 | 0.389064 | 0.346618 | -0.136989 | 1.321576 | 0.04689 | 0.871222 | 1.1454 | 0.540345 | 0.523221 | -0.042863 | 0.815448 | 0.831495 |
| 24.44444 | 0.133509 | -0.067985 | 0.324285 | 0.253383 | 0.065017 | 0.128105 | 0.006744 | -0.075374 | 0.7666 | 0.390039 | 0.305578 | 0.251056 | 0.634273 | 0.259194 | -0.254427 | 1.080507 | 0.527323 | 0.752331 | 1.07142 | 0.445362 | 0.242638 | -0.144233 | 0.460396 | 0.639352 |
| 24.45833 | 0.097237 | -0.11283 | 0.330752 | 0.204023 | 0.057401 | 0.036687 | -0.086008 | -0.087405 | 0.606524 | 0.368084 | 0.167308 | 0.164146 | 0.974105 | 0.264773 | -0.393564 | 0.342672 | 0.754483 | 0.504328 | 0.862986 | 0.557481 | -0.030105 | -0.194391 | 0.189642 | 0.387692 |
| 24.47222 | 0.054681 | -0.09568 | 0.432596 | 0.189411 | 0.230476 | 0.289456 | 0.234079 | -0.058147 | 0.703727 | 0.333835 | 0.098262 | 0.184483 | 1.215898 | 0.322209 | -0.450285 | -0.138288 | -0.108744 | 0.367704 | 0.776862 | 0.83252 | -0.08006 | -0.094347 | 0.163562 | 0.29058 |
| 24.48611 | 0.022053 | 0.017219 | 0.472561 | 0.14046 | 0.340342 | 0.426901 | 0.537704 | 0.028397 | 0.797308 | 0.337359 | 0.07551 | 0.311398 | 1.214224 | 0.64793 | 0.024872 | 1.2386 | -0.755512 | 0.608392 | 0.84543 | 1.075732 | 0.177053 | 0.036737 | 0.490881 | 0.824599 |
| 24.5 | -0.007999 | 0.323439 | 0.384424 | 0.079092 | 0.292824 | 0.427369 | 0.630208 | 0.087155 | 0.758678 | 0.332946 | 0.065683 | 0.676816 | 1.060521 | 1.726055 | 1.070201 | 3.284836 | -0.396471 | 1.17386 | 1.221037 | 1.392432 | 0.695845 | 0.129446 | 1.466336 | 2.221474 |
| 24.51389 | -0.043894 | -0.042567 | 0.26486 | 0.042187 | 0.244069 | 0.352945 | 0.551282 | 0.081141 | 0.653969 | 0.255324 | 0.058603 | 0.419674 | 0.894799 | 3.262674 | 2.081655 | 3.786834 | 0.302935 | 1.647343 | 1.431698 | 1.646827 | 1.18202 | 0.120424 | 2.150098 | 3.583357 |
| 24.52778 | -0.05056 | 0.072517 | 0.146038 | 0.009111 | 0.230431 | 0.251486 | 0.391241 | 0.058654 | 0.506621 | 0.172864 | 0.074848 | 0.254348 | 0.994959 | 3.485513 | 2.408188 | 2.749911 | 1.5268 | 1.777656 | 1.415814 | 1.71445 | 1.380958 | 0.14098 | 2.041952 | 3.685435 |
| 24.54167 | -0.003502 | 0.035581 | 0.059852 | -0.006061 | 0.090572 | 0.195476 | 0.240905 | 0.029861 | 0.374345 | 0.112592 | 0.101312 | 0.122887 | 1.424795 | 2.864639 | 2.417838 | 2.903192 | 3.310886 | 1.552001 | 1.259217 | 1.64189 | 1.303362 | 0.343684 | 2.501965 | 3.452909 |
| 24.55556 | 0.072181 | -0.033148 | 0.019315 | -0.004157 | -0.022687 | 0.169169 | 0.135845 | -0.006842 | 0.316177 | 0.091704 | 0.113766 | 0.07809 | 1.817506 | 2.300096 | 1.529736 | 2.884441 | 4.136319 | 1.120591 | 1.162422 | 1.360732 | 1.256516 | 0.354826 | 2.142402 | 2.555633 |
| 24.56944 | 0.14948 | -0.07851 | 0.018059 | -0.006335 | -0.119704 | 0.162509 | 0.059054 | -0.033257 | 0.294066 | 0.09781 | 0.117014 | 0.051378 | 1.749267 | 1.587208 | 0.311666 | 2.5472 | 3.919768 | 0.793057 | 0.981564 | 0.885813 | 1.329925 | 0.358155 | 1.300932 | 2.49113 |
| 24.58333 | 0.189436 | -0.115779 | 0.023141 | -0.025198 | -0.246366 | 0.16742 | -0.001085 | -0.049763 | 0.295294 | 0.110544 | 0.116025 | 0.017445 | 1.474509 | 0.871946 | 0.286083 | 2.095902 | 3.449738 | 0.744363 | 0.751694 | 0.414513 | 1.178077 | 0.325435 | 0.532007 | 1.800609 |
| 24.59722 | 0.168098 | -0.110096 | 0.028066 | 0.008435 | -0.258309 | 0.209391 | -0.008742 | -0.033748 | 0.34448 | 0.143726 | 0.110665 | 0.019475 | 1.214 | 0.277187 | 1.158559 | 1.878915 | 2.952499 | 0.624012 | 0.561496 | 0.096472 | 0.720653 | 0.280785 | -0.0575 | 0.968637 |
| 24.61111 | 0.071429 | -0.059775 | 0.038057 | 0.104581 | -0.334511 | 0.280178 | 0.032659 | 0.001802 | 0.411748 | 0.173762 | 0.116744 | 0.058942 | 1.025188 | -0.294757 | 2.278322 | 1.531908 | 2.58122 | 0.418651 | 0.390735 | -0.035979 | 0.262278 | 0.258571 | -0.59719 | 0.47219 |
| 24.625 | -0.052622 | -0.011984 | 0.046673 | 0.188103 | -0.379369 | 0.306027 | 0.088232 | 0.045874 | 0.420284 | 0.174314 | 0.126317 | 0.095114 | 0.954897 | -0.679351 | 2.643626 | 1.0326 | 2.191969 | 0.401315 | 0.290801 | 0.124665 | -0.093269 | 0.213836 | -1.090887 | 0.610474 |
| 24.63889 | -0.096502 | 0.015088 | 0.077969 | 0.21206 | -0.287178 | 0.306925 | 0.140546 | 0.114509 | 0.365911 | 0.178827 | 0.153483 | 0.125825 | 0.956563 | -0.691776 | 2.366308 | 1.06738 | 1.851233 | 0.665699 | 0.355516 | 0.466906 | 0.210671 | 0.135317 | -1.610577 | 0.499941 |
| 24.65278 | -0.00683 | 0.038211 | 0.146831 | 0.213846 | 0.016572 | 0.290484 | 0.178884 | 0.192367 | 0.340093 | 0.195752 | 0.181001 | 0.171521 | 0.957055 | -0.43315 | 2.228435 | 1.390606 | 1.307323 | 1.142683 | 0.504804 | 0.68899 | 0.55726 | 0.098436 | -1.774219 | -0.133464 |
| 24.66667 | 0.083199 | 0.053447 | 0.216855 | 0.222066 | 0.297171 | 0.274896 | 0.201937 | 0.252554 | 0.308569 | 0.210929 | 0.206663 | 0.193272 | 0.960581 | 0.025343 | 0.880508 | 1.557264 | 0.642926 | 1.323171 | 0.567476 | 0.817749 | 0.863942 | 0.122342 | -1.727699 | -0.872117 |
| 24.68056 | 0.138738 | 0.035621 | 0.235991 | 0.219204 | 0.427305 | 0.259607 | 0.202234 | 0.264573 | 0.28502 | 0.187647 | 0.212571 | 0.178503 | 0.73187 | 0.693806 | -0.484108 | 2.053419 | 0.102167 | 1.122989 | 0.508495 | 0.816211 | 0.871781 | 0.122436 | -0.568337 | -0.641934 |
| 24.69444 | 0.180376 | -0.017379 | 0.236383 | 0.189101 | 0.530561 | 0.239505 | 0.189805 | 0.220787 | 0.24003 | 0.126979 | 0.200758 | 0.145762 | 0.333919 | 1.408988 | -1.893966 | 2.331506 | -0.24011 | 0.61664 | 0.342965 | 0.865561 | 0.902506 | 0.153889 | 0.123759 | -0.08627 |
| 24.70833 | 0.271908 | -0.012178 | 0.211864 | 0.16278 | 0.472252 | 0.247486 | 0.203911 | 0.176572 | 0.215483 | 0.088708 | 0.190366 | 0.146306 | 0.092957 | 2.008835 | -2.922282 | 1.214236 | -0.247372 | 0.270388 | 0.144994 | 0.78445 | 0.966137 | 0.151957 | 0.733023 | 1.012275 |
| 24.72222 | 0.435909 | 0.081156 | 0.20886 | 0.184603 | 0.438765 | 0.32001 | 0.316754 | 0.180272 | 0.271437 | 0.129094 | 0.18959 | 0.188335 | 0.241438 | 2.241529 | -2.008847 | 0.849491 | -0.05213 | 0.039109 | 0.070167 | 0.615773 | 0.961358 | 0.16967 | 1.082897 | 1.174232 |
| 24.73611 | 0.451784 | 0.2074 | 0.23377 | 0.205982 | 0.411306 | 0.386687 | 0.528279 | 0.204285 | 0.339271 | 0.210513 | 0.156276 | 0.233402 | 0.855023 | 1.625373 | -1.502712 | 0.633855 | 0.250495 | -0.219327 | 0.091394 | 0.183378 | 0.667367 | 0.189931 | 0.891881 | 0.949976 |
| 24.75 | 0.338678 | 0.284168 | 0.258802 | 0.188636 | 0.425466 | 0.376462 | 0.724706 | 0.213825 | 0.346912 | 0.246062 | 0.0822 | 0.238081 | 2.0163 | 0.654252 | -0.14727 | 0.701023 | 0.532136 | -0.75452 | 0.023332 | -0.16349 | 0.330631 | 0.156175 | 1.159787 | 0.762351 |
| 24.76389 | 0.248493 | 0.294364 | 0.243055 | 0.189108 | 0.475582 | 0.308862 | 0.788666 | 0.214326 | 0.307914 | 0.20897 | 0.043303 | 0.217082 | 2.801481 | -0.057783 | 1.132203 | 0.694322 | 0.599217 | -1.118673 | -0.121245 | -0.048252 | 0.050335 | 0.08921 | 1.883175 | 1.216245 |
| 24.77778 | 0.247879 | 0.247807 | 0.203271 | 0.216615 | 0.721849 | 0.233977 | 0.769918 | 0.180809 | 0.279313 | 0.144197 | 0.07922 | 0.214259 | 2.554768 | -0.216559 | 1.591612 | 0.550083 | 0.436908 | -0.898414 | -0.325082 | 0.061684 | -0.105268 | 0.026642 | 1.773279 | 1.262802 |
| 24.79167 | 0.212386 | 0.213609 | 0.143853 | 0.232562 | 1.128358 | 0.156151 | 0.669038 | 0.111367 | 0.187193 | 0.092794 | 0.118971 | 0.224662 | 1.486812 | -0.07502 | 1.541761 | 0.491769 | 0.226245 | -0.841231 | -0.406865 | -0.136183 | -0.076796 | -0.035754 | 1.393725 | 1.132729 |
| 24.80556 | 0.140807 | 0.201995 | 0.076168 | 0.219784 | 0.866177 | 0.050256 | 0.53094 | 0.060579 | 0.063781 | 0.061885 | 0.088009 | 0.222445 | 0.695363 | -0.13658 | 1.969681 | 0.526564 | 0.152276 | -0.814023 | -0.349835 | -0.421701 | 0.023575 | -0.04138 | 0.782056 | 1.157268 |
| 24.81944 | 0.042515 | 0.223995 | 0.034082 | 0.243527 | 0.559077 | -0.081295 | 0.377855 | 0.080741 | -0.00864 | 0.071547 | 0.024906 | 0.205206 | 0.110738 | -0.676453 | 1.746903 | 0.418599 | 0.004525 | -0.719367 | -0.127893 | -0.438649 | 0.097448 | 0.016618 | 0.22259 | 0.616157 |
| 24.83333 | 0.040134 | 0.256438 | 0.077568 | 0.321305 | 0.44421 | -0.133828 | 0.285849 | 0.159343 | 0.117449 | 0.177484 | 0.014575 | 0.207737 | -0.433774 | -1.363536 | 1.30443 | 0.154653 | -0.193892 | -0.341502 | 0.187058 | -0.168713 | 0.143838 | 0.068772 | -0.052018 | 0.350792 |
| 24.84722 | 0.240681 | 0.308265 | 0.149854 | 0.389455 | 0.44194 | -0.05146 | 0.35111 | 0.320195 | 0.331914 | 0.387381 | 0.099563 | 0.285814 | -1.012648 | -1.759368 | 0.978857 | -0.270734 | -0.429333 | 0.267068 | 0.219895 | 0.169867 | 0.382265 | 0.136368 | -0.228037 | 0.501473 |
| 24.86111 | 0.4279 | 0.313069 | 0.125341 | 0.344629 | 0.48955 | 0.085367 | 0.382299 | 0.495591 | 0.406062 | 0.535403 | 0.214091 | 0.332293 | -1.619465 | -1.845464 | 0.893083 | -0.641628 | -0.694236 | 0.691915 | 0.138671 | 0.27138 | 0.664722 | 0.255837 | -0.293364 | 0.471898 |
| 24.875 | 0.467979 | 0.032701 | -0.094107 | 0.105646 | 0.171844 | 0.000926 | 0.038585 | 0.39408 | 0.087451 | 0.274942 | 0.183008 | 0.099749 | -2.349748 | -1.723321 | 0.499724 | -1.033165 | -0.892292 | 0.525017 | 0.116626 | 0.19193 | 0.562736 | 0.364423 | -0.583526 | 0.100929 |
| 24.88889 | 0.010651 | -0.486507 | -0.405389 | -0.142332 | -0.442305 | -0.196881 | -0.403529 | 0.085842 | -0.473905 | -0.28249 | -0.045339 | -0.21651 | -2.892248 | -1.268625 | -0.144412 | -1.005903 | -1.347117 | -0.137574 | 0.200776 | 0.129273 | 0.693647 | 0.393932 | -0.956365 | -0.103614 |
| 24.90278 | -0.163651 | -0.626813 | -0.474864 | -0.136478 | -0.401314 | -0.10257 | -0.318374 | 0.000691 | -0.832268 | -0.511511 | -0.328848 | -0.216502 | -3.096909 | -0.576963 | -0.761857 | -1.513364 | -2.625555 | -0.337467 | 0.144858 | 0.102334 | 0.805934 | 0.30491 | -0.89567 | 0.024961 |
| 24.91667 | 0.018265 | -0.271125 | -0.317221 | -0.040261 | 0.157009 | 0.13415 | -0.031599 | 0.183936 | -0.723504 | -0.303192 | -0.558612 | -0.0781 | -2.254787 | -0.107138 | -1.883517 | -2.225396 | -3.756719 | -0.254978 | -0.033067 | -0.007166 | 0.240025 | 0.20081 | -0.902656 | -0.308007 |
| 24.93056 | 0.1692 | 0.220865 | -0.059376 | 0.024983 | 0.207114 | 0.33413 | 0.010171 | 0.285533 | -0.443158 | -0.104673 | -0.605291 | -0.047229 | -1.217718 | 0.078573 | -2.160625 | -2.214857 | -5.018087 | -0.306271 | -0.208849 | -0.262661 | -0.258977 | 0.204474 | -0.570831 | -1.325423 |
| 24.94444 | 0.280324 | 0.540932 | 0.153875 | 0.142555 | 0.046558 | 0.664664 | 0.008872 | 0.249229 | -0.261143 | -0.035688 | -0.530212 | -0.021334 | -0.900294 | -0.058796 | -1.382151 | -2.292931 | -3.579504 | -0.684635 | -0.483588 | -0.539795 | -0.417636 | 0.340005 | -0.248664 | -2.034722 |
| 24.95833 | 0.368783 | 0.675695 | 0.316613 | 0.283513 | -0.0361 | 0.774235 | 0.155703 | 0.272436 | -0.122658 | 0.004603 | -0.413879 | 0.032058 | -1.051892 | -0.476222 | -0.638698 | -1.924582 | -2.525962 | -2.176862 | -1.554157 | -1.37575 | -1.015893 | 0.510523 | -0.491811 | -2.627459 |
| 24.97222 | 0.460999 | 0.755846 | 0.38593 | 0.405902 | 0.017534 | 0.55036 | 0.407112 | 0.317682 | 0.017615 | 0.024606 | -0.290103 | 0.09384 | -1.088937 | -0.849306 | -0.883912 | -2.633147 | -1.230611 | -3.69734 | -2.49627 | -2.246955 | -1.478185 | 0.555586 | -0.931015 | -2.360993 |
| 24.98611 | 0.559925 | 0.874572 | 0.464214 | 0.54977 | 0.107665 | 0.266463 | 0.522696 | 0.304461 | 0.148095 | 0.029141 | -0.113056 | 0.149856 | -0.28496 | -1.181018 | -1.42571 | -3.285678 | 0.804011 | -2.017766 | -1.72906 | -2.693981 | -0.8828 | 0.167674 | -0.739918 | -1.870825 |
| 25 | 0.586933 | 0.944032 | 0.570262 | 0.637212 | 0.205944 | -0.126941 | 0.21496 | 0.223847 | 0.238243 | 0.082692 | 0.065609 | 0.211974 | 1.078189 | -1.166698 | -1.800744 | -2.916352 | 3.707478 | -0.875605 | -1.222304 | -1.469713 | -0.418903 | 0.144071 | -0.063467 | -1.165711 |
| 25.01389 | 0.500249 | 0.806364 | 0.49525 | 0.552255 | 0.177677 | -0.480338 | -0.220159 | 0.102111 | 0.240212 | 0.095485 | 0.229372 | 0.225634 | 1.257584 | -0.950891 | -1.675845 | -1.849969 | 6.147089 | -0.583754 | -0.829843 | 0.096981 | -0.380016 | 0.066695 | 0.239288 | -0.123174 |
| 25.02778 | 0.343218 | 0.662995 | 0.150312 | 0.356139 | 0.026452 | -0.595075 | -0.374433 | 0.013658 | 0.138309 | 0.067982 | 0.320887 | 0.199167 | 0.960158 | -0.582693 | -0.727592 | -1.112082 | 8.03467 | -0.522416 | -0.764083 | -0.054145 | -0.233527 | 0.001425 | 0.17514 | 1.061062 |
| 25.04167 | 0.193826 | 0.535109 | -0.224202 | 0.117999 | -0.187495 | -0.61296 | -0.361469 | -0.024843 | -0.029352 | 0.016716 | 0.349574 | 0.122005 | 2.244904 | -0.580668 | 0.614183 | 0.225918 | 6.010112 | -0.390456 | -0.821716 | -1.336958 | 0.193857 | -0.056905 | 0.165446 | 1.272432 |
| 25.05556 | 0.092457 | 0.369061 | -0.196918 | 0.050863 | -0.309718 | -0.415096 | -0.317054 | -0.03691 | -0.201135 | -0.034766 | 0.316503 | 0.05407 | 2.326903 | -0.828753 | 1.170355 | 1.321329 | 2.90169 | -0.239384 | -0.588281 | -2.463576 | 0.624435 | -0.037484 | 0.618801 | 1.14829 |
| 25.06944 | 0.032373 | 0.282263 | 0.171985 | 0.149789 | -0.33413 | -0.007021 | -0.212835 | -0.017727 | -0.327178 | -0.037615 | 0.284433 | 0.072306 | -0.326479 | -1.263509 | 0.535967 | 1.24925 | 0.724798 | -0.069668 | -0.27548 | -2.258735 | 0.132234 | 0.122296 | 0.992467 | 0.868407 |
| 25.08333 | -0.029059 | 0.259067 | 0.398117 | 0.317554 | -0.360112 | 0.52994 | -0.113478 | 0.005333 | -0.408479 | -0.014326 | 0.246889 | 0.106547 | -2.287622 | -1.133297 | 0.109032 | 0.498647 | -1.267642 | -0.053622 | -0.212217 | -1.092116 | -0.297193 | 0.288113 | 0.811953 | 0.291592 |
| 25.09722 | -0.100934 | 0.226759 | 0.308995 | 0.395983 | -0.357987 | 1.070212 | -0.073938 | -0.014878 | -0.46562 | 0.022449 | 0.172946 | 0.107002 | -2.985745 | -0.387832 | 0.232903 | -0.122493 | -3.604297 | -0.311625 | -0.346209 | -0.238666 | -0.482966 | 0.11041 | 0.174382 | 0.322566 |
| 25.11111 | -0.134123 | 0.263676 | 0.17873 | 0.375434 | -0.237626 | 0.932797 | -0.058366 | -0.048353 | -0.479849 | 0.074329 | 0.102772 | 0.086809 | -2.470568 | 0.281515 | 0.460573 | -0.306415 | -5.756411 | -0.52138 | -0.34904 | 0.29054 | 0.122505 | 0.334993 | 0.028048 | 1.037873 |
| 25.125 | -0.141382 | 0.311509 | 0.297795 | 0.361132 | 0.043669 | 0.547691 | -0.002553 | -0.044079 | -0.417881 | 0.132471 | 0.053263 | 0.066079 | -2.529475 | 0.636996 | 0.404666 | -0.101147 | -7.066814 | -0.552806 | -0.254815 | 0.145286 | 0.420737 | 0.559958 | 0.240704 | 1.011127 |
| 25.13889 | -0.16626 | 0.243471 | 0.59666 | 0.391203 | 0.159478 | 0.37971 | 0.074728 | -0.045754 | -0.356719 | 0.126225 | 0.02258 | 0.058812 | -1.905965 | 0.772153 | 0.229232 | -0.07556 | -6.208888 | -0.481977 | -0.232425 | -0.207834 | 0.780993 | 0.221703 | 0.210133 | 0.066615 |
| 25.15278 | -0.19724 | 0.143268 | 0.654284 | 0.436635 | 0.001556 | 0.30699 | 0.136638 | -0.072094 | -0.25444 | 0.067488 | 0.008472 | 0.03992 | -0.71566 | 0.725827 | 0.223021 | -0.286194 | -3.527535 | -0.250627 | -0.163329 | -0.082226 | 0.621297 | -0.067365 | 0.174006 | -0.810863 |
| 25.16667 | -0.177898 | 0.172135 | 0.45118 | 0.431093 | -0.062158 | 0.248241 | 0.182599 | -0.088792 | -0.161671 | 0.020061 | 0.000373 | 0.009486 | 0.203609 | 0.320839 | 0.240175 | -0.476603 | -1.330777 | 0.098873 | 0.076314 | 0.527426 | 0.1193 | -0.115181 | -0.015869 | -1.476934 |
| 25.18056 | -0.096895 | 0.313792 | 0.227542 | 0.378405 | 0.036044 | 0.152808 | 0.174932 | -0.091962 | -0.100637 | 0.02942 | -0.021002 | -0.009134 | 0.722485 | -0.359863 | 0.098426 | -0.179853 | 0.583279 | 0.327447 | 0.20046 | 0.460795 | -0.041483 | -0.173802 | -0.052351 | -1.56565 |
| 25.19444 | -0.034396 | 0.397002 | 0.171661 | 0.400458 | 0.16703 | 0.097292 | 0.149476 | -0.093222 | -0.074182 | 0.071738 | -0.02621 | 0.011762 | 1.270099 | -0.920528 | -0.095993 | 0.510084 | 2.687227 | 0.266556 | 0.15064 | -0.639117 | -0.11172 | -0.387625 | -0.172905 | -1.781626 |
| 25.20833 | 0.032652 | 0.368701 | 0.233694 | 0.433441 | 0.141468 | 0.122618 | 0.121717 | -0.088396 | -0.036764 | 0.133507 | 0.016067 | 0.073838 | 1.824482 | -0.810196 | -0.133086 | 0.83653 | 4.458139 | 0.200559 | 0.133039 | -1.673819 | -0.242835 | -0.6266 | -0.31564 | -1.813543 |
| 25.22222 | 0.066234 | 0.316737 | 0.297954 | 0.427303 | 0.024005 | 0.235324 | 0.08725 | -0.126562 | -0.002105 | 0.174341 | 0.067351 | 0.137259 | 1.882112 | -0.331698 | 0.016439 | 0.782939 | 5.283352 | 0.277553 | 0.225146 | -1.414926 | -0.250813 | -0.560998 | -0.64305 | -1.263815 |
| 25.23611 | 0.130797 | 0.349036 | 0.371411 | 0.408144 | 0.07847 | 0.374602 | 0.055235 | -0.149809 | 0.0475 | 0.219318 | 0.093938 | 0.196615 | 1.780468 | -0.159046 | 0.220312 | 0.619617 | 3.875032 | 0.347233 | 0.248842 | -0.999728 | -0.172592 | -0.351461 | -0.585334 | -0.624416 |
| 25.25 | 0.22464 | 0.415571 | 0.510279 | 0.409716 | 0.348652 | 0.429966 | 0.036547 | -0.151671 | 0.109068 | 0.254575 | 0.163866 | 0.266896 | 1.295934 | -0.45725 | 0.319751 | 0.514107 | 2.187416 | 0.123822 | 0.192185 | -1.018648 | -0.091645 | -0.193407 | -0.191653 | -0.398902 |
| 25.26389 | 0.252899 | 0.407412 | 0.606674 | 0.407899 | 0.560732 | 0.402119 | 0.046231 | -0.147989 | 0.153515 | 0.264932 | 0.270767 | 0.331668 | 0.518363 | -0.795537 | 0.326267 | 0.49311 | 0.666161 | -0.032504 | 0.236996 | -0.921414 | -0.068446 | -0.201897 | 0.209276 | -0.310557 |
| 25.27778 | 0.245616 | 0.349989 | 0.570077 | 0.393434 | 0.482549 | 0.292789 | 0.085309 | -0.142899 | 0.210978 | 0.259513 | 0.359575 | 0.370314 | -0.06359 | -0.82094 | 0.36263 | 0.379361 | -0.29626 | -0.092624 | 0.370316 | -0.346666 | -0.087343 | -0.209648 | 0.433638 | 0.031496 |
| 25.29167 | 0.196484 | 0.299475 | 0.425181 | 0.337648 | 0.377966 | 0.190935 | 0.142308 | -0.138186 | 0.297214 | 0.261547 | 0.41318 | 0.369291 | -0.327529 | -0.456737 | 0.410922 | 0.313056 | -0.769506 | 0.018433 | 0.522558 | 0.238705 | -0.024306 | -0.003663 | 0.675675 | 0.683956 |
| 25.30556 | 0.162706 | 0.254964 | 0.376594 | 0.296617 | 0.386413 | 0.130254 | 0.188451 | -0.143469 | 0.358884 | 0.271258 | 0.404349 | 0.324339 | -0.480251 | -0.125242 | 0.44837 | 0.428174 | -1.066817 | 0.137785 | 0.630166 | 0.570213 | 0.0973 | 0.181102 | 0.881286 | 1.233322 |
| 25.31944 | 0.189183 | 0.268322 | 0.426867 | 0.330789 | 0.58214 | 0.11311 | 0.255109 | -0.108328 | 0.439832 | 0.318557 | 0.354591 | 0.314651 | -0.587857 | 0.016774 | 0.330605 | 0.493666 | -0.992567 | 0.205454 | 0.650183 | 0.731524 | 0.196304 | 0.118821 | 0.940105 | 1.18661 |
| 25.33333 | 0.265248 | 0.292582 | 0.474236 | 0.347463 | 0.793307 | 0.101195 | 0.346464 | -0.06063 | 0.534095 | 0.361113 | 0.31248 | 0.344427 | -0.506234 | 0.000023 | 0.160064 | 0.426551 | -0.703239 | 0.28437 | 0.64045 | 0.773492 | 0.282574 | -0.000703 | 0.738092 | 1.165905 |
| 25.34722 | 0.16801 | 0.164266 | 0.355796 | 0.241814 | 0.586915 | -0.045308 | 0.283428 | -0.180096 | 0.580119 | 0.269622 | 0.154684 | 0.291041 | -0.193968 | -0.015074 | 0.268341 | 0.574628 | -0.332379 | 0.351218 | 0.620109 | 0.619425 | 0.386161 | -0.087961 | 0.437808 | 1.125405 |
| 25.36111 | 0.024671 | -0.019134 | 0.158566 | 0.210302 | 0.191499 | -0.213061 | 0.118729 | -0.340873 | 0.527411 | 0.162725 | -0.169777 | 0.124548 | 0.21646 | 0.008186 | 0.54092 | 0.89527 | 0.016536 | 0.405607 | 0.699112 | 0.470118 | 0.453161 | -0.133156 | 0.563733 | 0.794556 |
| 25.375 | 0.087863 | -0.016187 | 0.181618 | 0.382276 | 0.148383 | -0.119375 | 0.134969 | -0.196562 | 0.414421 | 0.309631 | -0.330252 | 0.118959 | 0.44163 | 0.083027 | 0.759281 | 1.174896 | 0.409706 | 0.521415 | 0.803886 | 0.582347 | 0.459148 | -0.106552 | 0.697658 | 0.665449 |
| 25.38889 | 0.312571 | 0.052766 | 0.363503 | 0.5674 | 0.439091 | 0.061544 | 0.263404 | 0.140698 | 0.265879 | 0.514524 | -0.2419 | 0.206151 | 0.552168 | 0.126367 | 0.881939 | 1.451165 | 0.708988 | 0.668062 | 0.873174 | 0.987102 | 0.502292 | -0.032416 | 0.927863 | 0.674028 |
| 25.40278 | 0.6656 | 0.02314 | 0.51602 | 0.525229 | 1.056987 | 0.096992 | 0.367985 | 0.495864 | 0.278125 | 0.518501 | -0.078553 | 0.20871 | 0.581608 | 0.186525 | 0.944166 | 1.51546 | 0.852937 | 0.797011 | 0.94825 | 1.437594 | 0.551155 | 0.063218 | 0.778507 | 0.646229 |
| 25.41667 | 1.019098 | 0.067254 | 0.61807 | 0.316275 | 1.883237 | 0.120507 | 0.532529 | 0.792676 | 0.54678 | 0.582118 | 0.159247 | 0.219771 | 0.508723 | 0.262447 | 0.82153 | 1.285845 | 0.807605 | 0.833303 | 0.908763 | 1.718338 | 0.557618 | 0.173446 | 0.726128 | 0.689937 |
| 25.43056 | 1.34044 | 0.158871 | 0.707281 | 0.16236 | 2.564089 | 0.199391 | 0.755352 | 0.912052 | 0.8891 | 0.624748 | 0.344384 | 0.240208 | 0.320721 | 0.279037 | 0.568472 | 1.027266 | 0.513678 | 0.760917 | 0.722433 | 1.593996 | 0.46714 | 0.204792 | 0.684439 | 0.823411 |
| 25.44444 | 1.317791 | 0.166608 | 0.709316 | 0.162636 | 2.726971 | 0.331159 | 0.808714 | 0.823997 | 0.977457 | 0.508641 | 0.343206 | 0.2209 | 0.112973 | 0.214378 | 0.445411 | 0.580564 | 0.011572 | 0.524048 | 0.374896 | 1.259522 | 0.275683 | 0.036072 | 0.258463 | 0.874523 |
| 25.45833 | 1.139346 | 0.147152 | 0.632269 | 0.263339 | 2.560675 | 0.446621 | 0.806584 | 0.703845 | 1.027102 | 0.359991 | 0.193799 | 0.225313 | 0.061147 | 0.058478 | 0.467219 | 0.086432 | -0.490549 | 0.301643 | 0.270811 | 0.90492 | -0.029917 | -0.366965 | -0.051706 | 0.739596 |

| | | | | | | | | | | | | |
|---|---|---|---|---|---|---|---|---|---|---|---|---|
| 25.47222 | 0.87257 | 0.128957 | 0.52217 | 0.342034 | 1.844142 | 0.583478 | 0.835866 | 0.544381 | 1.109679 | 0.248703 | 0.183887 | 0.256857 |
| 25.48611 | 0.404391 | 0.137206 | 0.62619 | 0.335304 | 0.831788 | 0.652872 | 0.790742 | 0.426904 | 1.042589 | 0.131012 | 0.267415 | 0.310877 |
| 25.5 | -0.128892 | 0.213273 | 0.902734 | 0.268259 | -0.252185 | 0.590267 | 0.578066 | 0.345325 | 0.830231 | 0.04785 | 0.348606 | 0.348641 |
| 25.51389 | -0.524431 | 0.291419 | 1.038326 | 0.17005 | -1.083473 | 0.426324 | 0.396729 | 0.243713 | 0.560155 | 0.014691 | 0.546204 | 0.360211 |
| 25.52778 | -0.579264 | 0.271735 | 0.772928 | 0.109503 | -1.470035 | 0.31275 | 0.344818 | 0.147477 | 0.315313 | 0.026546 | 0.906414 | 0.357687 |
| 25.54167 | -0.541667 | 0.18229 | 0.347379 | 0.07727 | -1.345949 | 0.231903 | 0.482764 | 0.111297 | 0.101335 | 0.071474 | 1.165287 | 0.332455 |
| 25.55556 | -0.195125 | 0.056481 | -0.08484 | 0.026234 | -0.992084 | 0.169762 | 0.79739 | 0.081364 | -0.041987 | 0.177211 | 1.12262 | 0.323951 |
| 25.56944 | -0.077377 | -0.040966 | -0.314194 | -0.008104 | -0.384684 | 0.153803 | 1.298588 | 0.009093 | -0.046859 | 0.331229 | 0.893153 | 0.23376 |
| 25.58333 | 0.224107 | -0.127317 | -0.169266 | 0.04014 | 0.600525 | 0.160931 | 1.311707 | -0.080434 | 0.125025 | 0.451901 | 0.65795 | 0.108203 |
| 25.59722 | 0.770677 | -0.152919 | 0.238005 | 0.107507 | 2.2211 | 0.156859 | 1.045264 | -0.107674 | 0.367722 | 0.465045 | 0.317601 | 0.033853 |
| 25.61111 | 1.056414 | -0.117186 | 0.650241 | 0.126276 | 3.047689 | 0.181721 | 0.761667 | -0.027393 | 0.464846 | 0.449041 | -0.007098 | 0.070193 |
| 25.625 | 0.819035 | 0.069586 | 0.850651 | 0.092785 | 2.901532 | 0.208158 | 0.492767 | 0.09523 | 0.416416 | 0.408179 | -0.230744 | 0.138683 |
| 25.63889 | 0.881056 | 0.265099 | 0.728368 | 0.081242 | 2.975864 | 0.229413 | 0.208707 | 0.182719 | 0.349582 | 0.34775 | -0.31519 | 0.198549 |
| 25.65278 | 1.02272 | 0.353025 | 0.603972 | 0.072069 | 2.064456 | 0.239681 | -0.063668 | 0.218369 | 0.355405 | 0.293317 | -0.132706 | 0.296667 |
| 25.66667 | 0.854374 | 0.290687 | 0.507179 | 0.045183 | 0.947059 | 0.223677 | -0.292286 | 0.198861 | 0.327709 | 0.185196 | 0.05093 | 0.33933 |
| 25.68056 | 0.271689 | 0.183205 | 0.385036 | 0.015453 | 0.240381 | 0.214233 | -0.438064 | 0.212109 | 0.300203 | 0.040544 | 0.160544 | 0.313515 |
| 25.69444 | -0.324759 | 0.105415 | 0.267152 | 0.014376 | -0.405411 | 0.242714 | -0.477479 | 0.301647 | 0.329601 | -0.07496 | 0.264516 | 0.311891 |
| 25.70833 | -0.828682 | 0.055362 | 0.107363 | 0.065083 | -1.722998 | 0.296429 | -0.370325 | 0.338804 | 0.332329 | -0.102053 | 0.466436 | 0.304978 |
| 25.72222 | -0.914128 | 0.001061 | -0.086889 | 0.152587 | -2.810007 | 0.333372 | -0.074359 | 0.216918 | 0.179696 | -0.150475 | 0.821262 | 0.270573 |
| 25.73611 | -1.273538 | -0.025784 | -0.26225 | 0.23813 | -3.461729 | 0.369249 | 0.272731 | 0.087092 | 0.061852 | -0.073816 | 0.984817 | 0.236945 |
| 25.75 | -1.42164 | 0.004117 | -0.326817 | 0.305893 | -3.836311 | 0.397307 | 0.572907 | 0.076548 | -0.091793 | 0.109275 | 0.749361 | 0.233417 |
| 25.76389 | -1.475542 | 0.042716 | -0.198129 | 0.314191 | -3.850926 | 0.359815 | 0.685981 | 0.065659 | -0.458663 | 0.249098 | 0.465961 | 0.253071 |
| 25.77778 | -1.067521 | 0.145995 | 0.06084 | 0.321591 | -2.222521 | 0.323573 | 0.706254 | 0.02331 | -0.992457 | 0.236852 | 0.306685 | 0.278707 |
| 25.79167 | -0.745282 | 0.219643 | 0.176217 | 0.302854 | -1.379005 | 0.254339 | 0.709363 | -0.095728 | -1.280783 | 0.147431 | 0.365091 | 0.26299 |
| 25.80556 | -0.408322 | 0.109006 | 0.079829 | 0.292939 | -0.761314 | 0.105962 | 0.63975 | -0.295372 | -0.939292 | 0.121816 | 0.454659 | 0.150786 |
| 25.81944 | 0.07418 | -0.102277 | -0.058592 | 0.267392 | 0.216485 | -0.056668 | 0.406712 | -0.399669 | -0.575104 | 0.174248 | 0.358061 | 0.043269 |
| 25.83333 | 0.753006 | -0.226289 | -0.006573 | 0.24063 | 1.663755 | -0.040941 | 0.069032 | -0.299236 | -0.473053 | 0.310653 | 0.191756 | 0.068325 |
| 25.84722 | 1.311592 | -0.110938 | 0.149646 | 0.179894 | 2.998719 | 0.104934 | -0.200747 | 0.013247 | -0.308975 | 0.37174 | 0.078734 | 0.263178 |
| 25.86111 | 1.081278 | -0.027738 | -0.021265 | -0.031243 | 2.425446 | 0.275571 | -0.44678 | 0.304932 | -0.076428 | 0.253658 | 0.011556 | 0.496565 |
| 25.875 | 0.502577 | -0.31567 | -0.688398 | -0.536536 | 0.575907 | 0.17444 | -1.072819 | 0.172791 | -0.127214 | -0.267179 | -0.184941 | 0.425412 |
| 25.88889 | -0.467973 | -0.909599 | -1.576008 | -1.233345 | -1.913067 | -0.054209 | -1.906418 | -0.308418 | -0.364914 | -1.048751 | -0.380728 | 0.024313 |
| 25.90278 | -1.182692 | -1.188143 | -1.747451 | -1.291479 | -3.27794 | 0.030188 | -2.197268 | -0.740208 | -0.125326 | -1.461283 | -0.844642 | -0.600734 |
| 25.91667 | -1.367316 | -1.01255 | -1.850039 | -0.672129 | -3.762732 | 0.150983 | -1.628725 | -0.854632 | -0.029619 | -1.259863 | -0.792967 | -0.861696 |
| 25.93056 | -1.299558 | -0.798091 | -1.776175 | -0.080937 | -3.358572 | 0.085093 | -0.964413 | -0.695407 | 0.400332 | -1.015633 | -0.424324 | -0.911286 |
| 25.94444 | -1.211974 | -0.603305 | -1.20084 | 0.100298 | -3.452372 | 0.172767 | -0.447286 | -0.454028 | 0.517356 | -0.634594 | -0.057262 | -0.763133 |
| 25.95833 | -0.96439 | -0.235226 | -0.858429 | 0.067436 | -3.616101 | 0.375822 | -0.013289 | -0.209438 | 0.277762 | -0.403533 | 0.303319 | -0.463185 |
| 25.97222 | -0.638054 | 0.136922 | -0.541483 | 0.1629 | -2.887214 | 0.271328 | 0.413783 | 0.011422 | -0.033395 | -0.319244 | 0.635797 | -0.048344 |
| 25.98611 | -0.357025 | 0.409802 | -0.29878 | 0.433156 | -2.097527 | 0.125747 | 0.996411 | 0.144278 | -0.113709 | -0.111074 | 0.866924 | 0.41974 |
| 26 | -0.194443 | 0.488002 | -0.225451 | 0.722917 | -1.771322 | 0.130685 | 1.496792 | 0.276496 | 0.012423 | 0.180922 | 0.819292 | 0.686305 |
| 26.01389 | -0.14223 | 0.454438 | -0.177397 | 0.90484 | -1.588422 | 0.213791 | 1.316861 | 0.382331 | -0.220478 | 0.398106 | 0.264636 | 0.699065 |
| 26.02778 | -0.161846 | 0.50948 | -0.277327 | 0.780647 | -1.118823 | 0.195074 | 0.624698 | 0.367539 | -0.687311 | 0.458865 | -0.264756 | 0.625435 |
| 26.04167 | -0.073668 | 0.600381 | -0.547476 | 0.477438 | -0.236672 | 0.088707 | -0.115665 | 0.197828 | -1.204244 | 0.330078 | -0.682922 | 0.463964 |
| 26.05556 | 0.13043 | 0.596576 | -0.714554 | 0.194174 | 0.22275 | 0.101118 | -0.741971 | -0.074992 | -1.266274 | 0.191387 | -0.806421 | 0.150038 |
| 26.06944 | 0.256137 | 0.494809 | -0.513182 | 0.06095 | -0.118869 | 0.113933 | -0.86659 | -0.373165 | -1.000441 | 0.201041 | -0.701889 | -0.021365 |
| 26.08333 | 0.160057 | 0.361098 | -0.314761 | 0.006174 | -0.618029 | 0.082694 | -0.769425 | -0.832277 | -0.739852 | 0.254645 | -0.595067 | -0.089369 |
| 26.09722 | -0.02005 | 0.296514 | -0.291183 | 0.008198 | -0.662387 | -0.002649 | -0.744717 | -1.516044 | -0.391544 | 0.25922 | -0.607176 | -0.086865 |
| 26.11111 | -0.189874 | 0.244569 | -0.321953 | 0.114557 | -0.292183 | 0.116845 | -0.687834 | -1.742849 | 0.038962 | 0.238612 | -0.35476 | -0.023007 |
| 26.125 | -0.276706 | 0.230149 | -0.34241 | 0.286073 | 0.053292 | 0.355433 | -0.44924 | -1.306479 | 0.450393 | 0.264731 | 0.172529 | 0.048281 |
| 26.13889 | -0.249751 | 0.2969 | -0.199573 | 0.37399 | 0.342986 | 0.455282 | -0.153251 | -0.910509 | 0.798883 | 0.245964 | 0.745917 | 0.262528 |
| 26.15278 | -0.063232 | 0.417959 | 0.181477 | 0.367233 | 0.874653 | 0.484272 | 0.142936 | -0.619579 | 0.971653 | 0.220684 | 1.167692 | 0.515234 |
| 26.16667 | 0.206138 | 0.498284 | 0.554274 | 0.39798 | 1.624113 | 0.465875 | 0.386391 | -0.50846 | 0.877284 | 0.22761 | 1.166819 | 0.628462 |
| 26.18056 | 0.389154 | 0.407038 | 0.784921 | 0.523646 | 2.136786 | 0.343609 | 0.616078 | -0.395057 | 0.700716 | 0.211531 | 1.125829 | 0.575995 |
| 26.19444 | 0.448016 | 0.180875 | 1.005745 | 0.656454 | 1.961896 | 0.168336 | 0.799549 | -0.12488 | 0.739598 | 0.24567 | 1.16131 | 0.53126 |
| 26.20833 | 0.474793 | 0.052092 | 1.237341 | 0.731665 | 1.567637 | 0.107429 | 1.01305 | 0.233269 | 0.925372 | 0.343381 | 1.094429 | 0.596591 |
| 26.22222 | 0.487936 | 0.258971 | 1.239334 | 0.803643 | 1.32621 | 0.095385 | 1.124721 | 0.461789 | 0.904375 | 0.313497 | 0.815886 | 0.713767 |
| 26.23611 | 0.522954 | 0.626222 | 1.281824 | 0.828661 | 1.362603 | 0.128999 | 1.089801 | 0.584236 | 0.664939 | 0.231975 | 0.441734 | 0.786008 |
| 26.25 | 0.59986 | 0.676068 | 1.219787 | 0.813059 | 1.524104 | 0.175935 | 1.112644 | 0.824279 | 0.485853 | 0.272142 | 0.089996 | 0.737004 |
| 26.26389 | 0.663029 | 0.468726 | 1.143648 | 0.780752 | 1.530036 | 0.193832 | 1.0642 | 1.271862 | 0.491478 | 0.463743 | -0.029207 | 0.598389 |
| 26.27778 | 0.629237 | 0.247743 | 1.223633 | 0.718836 | 1.42547 | 0.183061 | 0.933641 | 1.304642 | 0.65461 | 0.539853 | 0.282906 | 0.4664 |
| 26.29167 | 0.595449 | 0.318995 | 1.351681 | 0.653333 | 1.297717 | 0.168588 | 0.898807 | 1.134834 | 0.851144 | 0.611167 | 0.682716 | 0.450383 |
| 26.30556 | 0.629589 | 0.590021 | 1.344472 | 0.60593 | 1.405462 | 0.174079 | 0.974025 | 1.028869 | 0.964673 | 0.698266 | 1.037003 | 0.496941 |
| 26.31944 | 0.752762 | 0.717164 | 1.115663 | 0.54127 | 1.651816 | 0.179605 | 1.097599 | 0.908397 | 0.883357 | 0.646037 | 1.291443 | 0.467411 |
| 26.33333 | 0.851472 | 0.629226 | 0.87369 | 0.389799 | 1.942612 | 0.170775 | 1.034907 | 0.783984 | 0.906334 | 0.491826 | 1.36273 | 0.403503 |
| 26.34722 | 0.783084 | 0.426266 | 0.648504 | 0.21988 | 1.716438 | 0.040042 | 0.759184 | 0.586675 | 0.844251 | 0.226068 | 1.108011 | 0.2746 |
| 26.36111 | 0.563215 | 0.369676 | 0.644054 | 0.156329 | 1.215133 | -0.124626 | 0.547602 | 0.337911 | 0.80757 | 0.065166 | 0.717588 | 0.24294 |
| 26.375 | 0.544103 | 0.630631 | 1.003924 | 0.269357 | 0.857239 | -0.165761 | 0.472199 | 0.253353 | 0.933143 | 0.082149 | 0.531379 | 0.482657 |
| 26.38889 | 0.79909 | 0.994073 | 1.3488 | 0.34845 | 0.949313 | -0.088255 | 0.604264 | 0.365025 | 0.740017 | 0.175559 | 0.432404 | 0.565975 |
| 26.40278 | 1.472822 | 1.018816 | 1.158608 | 0.361716 | 1.754605 | -0.065908 | 0.64394 | 0.677191 | 0.390848 | 0.048134 | 0.252534 | 0.585751 |
| 26.41667 | 2.377417 | 0.741949 | 0.717016 | 0.601207 | 3.057678 | 0.185012 | 0.832806 | 1.38934 | 0.328076 | -0.040928 | 0.076255 | 0.582702 |
| 26.43056 | 2.392908 | 0.694401 | 0.266742 | 0.707876 | 2.925805 | 0.671599 | 1.01213 | 1.755048 | 0.503396 | -0.006417 | -0.070072 | 0.443715 |
| 26.44444 | 1.520337 | 0.570223 | 0.218372 | 0.63097 | 1.451105 | 0.912975 | 0.954202 | 1.298531 | 0.761564 | 0.036659 | -0.183823 | 0.36272 |
| 26.45833 | 0.562877 | 0.251062 | 0.865749 | 0.489052 | -0.097958 | 1.104086 | 0.871434 | 0.922662 | 1.066158 | 0.040099 | -0.003464 | 0.463834 |
| 26.47222 | -0.07142 | 0.182138 | 1.221755 | 0.432342 | -1.001817 | 1.01073 | 0.723502 | 0.763303 | 1.199915 | 0.054058 | 0.374863 | 0.669056 |
| 26.48611 | -0.550528 | 0.391392 | 0.817712 | 0.408687 | -0.693661 | 0.687718 | 0.483896 | 0.537762 | 0.946757 | 0.051896 | 0.654017 | 0.729395 |
| 26.5 | -0.371757 | 0.683509 | 0.097789 | 0.445648 | 0.126866 | 0.394525 | 0.309158 | 0.097301 | 0.669007 | 0.063973 | 0.58472 | 0.614138 |
| 26.51389 | -0.222513 | 0.809358 | -0.342883 | 0.593143 | 0.325136 | 0.297676 | 0.413305 | -0.261621 | 0.465701 | 0.08086 | 0.563815 | 0.52071 |
| 26.52778 | -0.810171 | 0.637357 | -0.68897 | 0.777051 | -0.875726 | 0.339487 | 0.765501 | -0.565599 | 0.369515 | 0.041685 | 0.358452 | 0.532723 |
| 26.54167 | -0.942039 | 0.335181 | -0.056855 | 0.6481 | -1.780767 | 0.367831 | 1.288469 | -0.889863 | 0.579615 | -0.050393 | 0.271578 | 0.6572 |
| 26.55556 | -0.229144 | -0.019922 | 0.5062 | 0.410835 | -0.998489 | 0.412098 | 2.107848 | -1.169505 | 0.811893 | -0.160972 | 0.457659 | 0.848679 |
| 26.56944 | 0.54001 | -0.256772 | 0.783824 | 0.260633 | 1.291637 | 0.517682 | 1.926078 | -1.327253 | 0.762249 | -0.145008 | 0.505645 | 0.91109 |
| 26.58333 | 1.168852 | -0.242179 | 0.861098 | 0.284558 | 3.22102 | 0.697285 | 1.264757 | -1.146498 | 0.651739 | 0.039668 | 0.410684 | 0.971971 |
| 26.59722 | 1.535108 | 0.001138 | 1.038578 | 0.442736 | 3.677794 | 0.927379 | 0.837247 | -0.755718 | 0.548435 | 0.37631 | 0.446966 | 1.09046 |
| 26.61111 | 1.064649 | 0.26103 | 1.279613 | 0.538735 | 2.280949 | 1.20077 | 0.608544 | -0.515921 | 0.516287 | 0.653028 | 0.410937 | 1.083062 |
| 26.625 | 0.518625 | 0.354597 | 1.112634 | 0.457791 | -0.013393 | 1.051058 | 0.287657 | -0.298404 | 0.53864 | 0.694001 | 0.219168 | 0.852753 |
| 26.63889 | 0.120602 | 0.282249 | 1.085891 | 0.396248 | -1.027958 | 0.752985 | -0.133661 | -0.089666 | 0.562093 | 0.602872 | -0.220559 | 0.62358 |
| 26.65278 | 0.553803 | 0.163418 | 0.832758 | 0.343616 | 0.068614 | 0.349114 | -0.671262 | 0.016877 | 0.483814 | 0.452834 | -0.07058 | 0.471319 |
| 26.66667 | 1.226732 | 0.116266 | 0.594568 | 0.271939 | 2.318995 | 0.091581 | -1.08652 | 0.054108 | 0.268919 | 0.399443 | 0.263328 | 0.390956 |
| 26.68056 | 1.348129 | 0.109312 | 0.278309 | 0.215352 | 3.921581 | -0.068534 | -0.986589 | 0.187894 | 0.004261 | 0.255164 | 0.521324 | 0.421272 |
| 26.69444 | 1.261243 | 0.132464 | 0.063922 | 0.187655 | 2.852537 | -0.240935 | -0.64872 | 0.447572 | -0.265673 | -0.049112 | 0.726029 | 0.415395 |
| 26.70833 | 0.802403 | 0.196894 | -0.123505 | 0.169797 | 1.114426 | -0.412049 | -0.151021 | 0.815692 | -0.506642 | -0.263505 | 0.963537 | 0.409893 |
| 26.72222 | 0.093317 | 0.23984 | -0.231098 | 0.122894 | -0.085936 | -0.406725 | 0.301398 | 1.16516 | -0.569662 | -0.16322 | 0.842659 | 0.417577 |
| 26.73611 | -0.972411 | 0.14934 | -0.246003 | 0.022364 | -0.443386 | -0.173219 | 0.581321 | 1.318577 | -0.447823 | 0.037589 | 0.3647 | 0.414771 |
| 26.75 | -1.713106 | 0.335225 | -0.230897 | -0.09219 | -0.838965 | 0.151823 | 0.75841 | 1.321545 | -0.375851 | 0.097373 | 0.236149 | 0.474886 |
| 26.76389 | -2.330404 | 0.409078 | -0.20039 | -0.153732 | -2.178932 | 0.379636 | 0.952724 | 1.315653 | -0.427022 | 0.015392 | 0.54158 | 0.522008 |
| 26.77778 | -1.555928 | 0.358394 | -0.040561 | -0.216285 | -3.228808 | 0.573479 | 1.251344 | 1.046357 | -0.429086 | 0.011081 | 0.546347 | 0.533789 |
| 26.79167 | -1.212747 | 0.380512 | 0.124925 | -0.236802 | -4.544478 | 0.626778 | 1.378535 | 0.628192 | -0.247669 | 0.162339 | 0.425074 | 0.502402 |
| 26.80556 | -1.297159 | 0.55052 | 0.149571 | -0.220052 | -4.532776 | 0.517266 | 1.10801 | 0.153608 | 0.041646 | 0.272006 | 0.266875 | 0.466169 |
| 26.81944 | -1.090005 | 0.357093 | 0.106798 | -0.315975 | -3.635615 | 0.368347 | 0.641983 | -0.217762 | 0.271737 | 0.180448 | 0.228878 | 0.457549 |
| 26.83333 | -0.863543 | 0.709547 | 0.054317 | -0.398166 | -2.494077 | 0.293537 | 0.287544 | -0.864061 | 0.392004 | -0.064051 | 0.284323 | 0.413293 |
| 26.84722 | -0.908171 | 0.971648 | 0.133987 | -0.386394 | -2.644233 | 0.308637 | -0.089486 | -1.094356 | 0.41496 | -0.192674 | 0.328822 | 0.404846 |
| 26.86111 | -1.020852 | 0.885503 | 0.11575 | -0.545802 | -3.013378 | 0.102792 | -0.810149 | -0.954009 | -0.016968 | -0.47807 | -0.174545 | 0.059843 |
| 26.875 | -1.186821 | -0.072302 | -0.334811 | -1.128781 | -1.969881 | -0.595295 | -1.987537 | -1.268108 | -1.318619 | -1.309563 | -1.122633 | -1.157641 |
| 26.88889 | -1.213248 | -1.163 | -0.820307 | -1.526549 | -1.311467 | -1.25083 | -2.960701 | -2.99152 | -2.156588 | -2.158642 | -1.342591 | -2.104903 |
| 26.90278 | -0.917245 | -1.55208 | -0.896218 | -1.312447 | -1.228989 | -1.247114 | -3.125042 | -2.415451 | -1.155842 | -2.645165 | -0.78369 | -1.879076 |
| 26.91667 | -0.824762 | -0.765055 | -0.893575 | -0.71888 | -1.838315 | -0.541377 | -1.9415 | -1.354717 | -0.500684 | -2.752029 | -0.304583 | -1.334779 |
| 26.93056 | -0.782171 | -0.676946 | -0.957924 | -0.361948 | -2.282427 | -0.127442 | -0.995359 | -0.291831 | -0.394328 | -1.894933 | -0.04258 | -1.098352 |
| 26.94444 | -0.443838 | -0.896729 | -0.913578 | -0.202089 | -1.52513 | -0.012795 | -0.638158 | 0.41619 | -0.475479 | -1.02208 | 0.124271 | -0.804414 |
| 26.95833 | 0.004425 | -0.952566 | -1.00647 | -0.196059 | -0.307771 | -0.10065 | -0.583872 | 0.742647 | -0.489313 | -0.431887 | 0.339235 | -0.459065 |
| 26.97222 | 0.242784 | -1.081777 | -1.111084 | -0.21991 | 0.442581 | -0.207656 | -0.732104 | 0.997531 | -0.446546 | 0.069545 | 0.870304 | -0.215338 |
| 26.98611 | 0.188211 | -0.657094 | -1.08173 | 0.05515 | 0.128291 | -0.201097 | -0.689823 | 1.065084 | -0.455102 | 0.484805 | 1.308496 | 0.057617 |
| 27 | -0.163017 | -0.264909 | -0.682726 | 0.352464 | -1.374084 | -0.103824 | -0.238697 | 0.569217 | -0.364087 | 0.706549 | 1.969301 | 0.37468 |
| 27.01389 | -0.566426 | 0.227946 | -0.250225 | 0.540836 | -3.060095 | 0.104112 | 0.362184 | 0.05779 | -0.223759 | 0.632512 | 1.946868 | 0.427295 |
| 27.02778 | -0.41396 | 0.503235 | -0.051366 | 0.562019 | -2.535618 | 0.23865 | 0.722328 | -0.220587 | -0.191465 | 0.205205 | 1.255594 | 0.469248 |
| 27.04167 | -0.084641 | 0.470599 | -0.051196 | 0.324369 | -0.74938 | 0.262592 | 0.567402 | -0.731859 | -0.136377 | -0.580531 | 0.457928 | 0.361183 |
| 27.05556 | 0.127664 | 0.056247 | -0.133273 | 0.11715 | 0.326618 | 0.144199 | 0.168271 | -1.192913 | -0.124347 | -1.298389 | -0.34997 | 0.110334 |
| 27.06944 | 0.238692 | -0.251583 | -0.339391 | 0.006508 | 0.158241 | -0.017491 | -0.133098 | -1.277173 | -0.020593 | -1.606323 | -1.092882 | -0.179043 |
| 27.08333 | 0.264103 | -0.257221 | -0.618189 | 0.13611 | -0.579111 | -0.177025 | -0.244325 | -1.03188 | 0.015672 | -1.53041 | -1.530732 | -0.390803 |
| 27.09722 | 0.241189 | -0.014601 | -0.899891 | 0.271254 | -0.506612 | -0.264744 | -0.338135 | -0.755637 | 0.000719 | -1.220154 | -1.859155 | -0.512149 |
| 27.11111 | 0.187578 | 0.440291 | -0.783126 | 0.326466 | 0.595473 | -0.255943 | -0.484765 | -0.538973 | 0.007088 | -1.072753 | -1.832381 | -0.591184 |
| 27.125 | 0.187919 | 0.772491 | -0.60092 | 0.428065 | 1.958608 | -0.235739 | -0.621603 | -0.549523 | 0.082359 | -0.850042 | -1.562046 | -0.476043 |
| 27.13889 | 0.187903 | 0.91824 | -0.405664 | 0.612506 | 2.003315 | -0.227472 | -0.691743 | -0.477682 | 0.145302 | -0.418737 | -0.915655 | -0.167948 |
| 27.15278 | 0.123766 | 0.953944 | -0.252773 | 0.816799 | 0.576099 | -0.152351 | -0.577317 | 0.091147 | 0.087122 | 0.024016 | -0.342335 | 0.159448 |
| 27.16667 | 0.068088 | 0.877507 | -0.11415 | 1.068188 | -0.584582 | -0.055533 | -0.325474 | 0.705629 | 0.031898 | 0.16888 | 0.091161 | 0.392311 |
| 27.18056 | 0.13605 | 0.816102 | 0.139802 | 1.188806 | -0.849553 | 0.033978 | -0.050919 | 0.906017 | 0.158136 | 0.097662 | 0.544591 | 0.678107 |
| 27.19444 | 0.285881 | 0.824665 | 0.45644 | 1.267892 | 0.40152 | 0.113989 | 0.191436 | 0.861091 | 0.450015 | 0.164162 | 1.084723 | 1.078663 |
| 27.20833 | 0.319374 | 0.835833 | 0.735714 | 1.295613 | 1.934703 | 0.245863 | 0.555466 | 0.888052 | 0.785294 | 0.458628 | 1.256106 | 1.399524 |
| 27.22222 | 0.432092 | 0.713088 | 0.964023 | 1.122146 | 2.54236 | 0.376767 | 0.87212 | 1.177276 | 1.062327 | 0.803061 | 1.332699 | 1.49748 |
| 27.23611 | 0.473933 | 0.50376 | 1.124643 | 0.97252 | 1.960297 | 0.478977 | 1.044388 | 1.172387 | 1.178546 | 0.811966 | 1.430774 | 1.573264 |
| 27.25 | 0.382239 | 0.360312 | 1.237361 | 0.862544 | 1.295058 | 0.56518 | 1.018517 | 1.207324 | 1.210634 | 0.613732 | 1.336531 | 1.713233 |
| 27.26389 | 0.387118 | 0.421268 | 1.380249 | 0.874512 | 0.965516 | 0.600239 | 0.990895 | 1.128379 | 1.337681 | 0.462603 | 1.160538 | 1.539985 |
| 27.27778 | 0.397909 | 0.66274 | 1.469302 | 0.920164 | 1.188204 | 0.665255 | 1.017604 | 1.075723 | 1.421665 | 0.581756 | 1.032893 | 1.28711 |
| 27.29167 | 0.396683 | 0.861977 | 1.609295 | 0.901747 | 1.472616 | 0.775327 | 1.039011 | 0.960534 | 1.247068 | 0.856872 | 0.981564 | 1.041977 |
| 27.30556 | 0.454392 | 0.82988 | 1.656173 | 0.826102 | 1.787838 | 0.867782 | 1.020191 | 0.900093 | 1.016257 | 1.142663 | 1.077944 | 0.934738 |
| 27.31944 | 0.575798 | 0.695133 | 1.614338 | 0.786846 | 1.873655 | 0.964635 | 1.125374 | 1.20844 | 0.874464 | 1.458365 | 1.076792 | 0.929453 |
| 27.33333 | 0.685366 | 0.719183 | 1.408918 | 0.857554 | 1.752989 | 0.954489 | 1.220204 | 2.277322 | 0.954065 | 1.689071 | 1.024816 | 0.87745 |
| 27.34722 | 0.709177 | 0.797464 | 1.201254 | 0.861989 | 1.472186 | 0.776728 | 1.207246 | 2.979697 | 0.965956 | 1.651415 | 0.513878 | 0.593094 |
| 27.36111 | 0.607675 | 0.869381 | 0.992213 | 0.854927 | 1.380558 | 0.4625 | 1.176707 | 2.777278 | 0.809811 | 1.451401 | 0.214614 | 0.265142 |
| 27.375 | 0.615575 | 1.118652 | 1.017337 | 1.089888 | 1.499058 | 0.177451 | 1.124203 | 2.079467 | 0.398393 | 1.746172 | 0.207326 | 0.170173 |
| 27.38889 | 0.737311 | 1.167059 | 1.04436 | 1.26566 | 1.308935 | -0.014844 | 0.848172 | 1.505998 | -0.226844 | 1.665724 | 0.231223 | 0.079168 |
| 27.40278 | 0.970433 | 0.871417 | 1.063194 | 1.539031 | 1.461031 | -0.082069 | 0.575496 | 1.300878 | -0.463872 | 1.391033 | 0.27188 | -0.070288 |
| 27.41667 | 1.563058 | 0.354628 | 0.988088 | 2.031432 | 2.097078 | 0.064658 | 0.483197 | 1.158436 | -0.048261 | 1.24363 | 0.305327 | -0.348005 |
| 27.43056 | 1.796365 | -0.019543 | 0.779066 | 1.853283 | 2.054634 | 0.400702 | 0.473181 | 0.840768 | 0.41054 | 1.059537 | 0.290436 | -0.548317 |
| 27.44444 | 1.437014 | -0.088953 | 0.467061 | 1.220066 | 1.199768 | 0.93811 | 0.598266 | 0.710185 | 0.995194 | 0.947129 | 0.599336 | 0.114937 |
| 27.45833 | 1.126044 | 0.217352 | 0.456927 | 0.815373 | 0.652291 | 1.52178 | 0.929188 | 1.156984 | 1.5914 | 0.900766 | 0.661478 | 1.104673 |
| 27.47222 | 1.211387 | 0.839317 | 0.56487 | 0.645315 | 0.759521 | 2.004394 | 1.54006 | 1.91791 | 1.940612 | 0.911725 | 0.462255 | 1.333965 |
| 27.48611 | 1.508167 | 1.358462 | 0.207571 | 0.39751 | 0.92059 | 1.965956 | 1.676568 | 1.816933 | 1.700949 | 0.730126 | 1.289721 | 1.77795 |
| 27.5 | 1.550701 | 1.561143 | -0.365138 | 0.292185 | 1.528291 | 1.370082 | 1.58102 | 0.016491 | 1.436967 | 0.244328 | 0.843077 | 1.74366 |
| 27.51389 | 1.024977 | 1.310472 | -0.529352 | 0.265477 | 1.686654 | 0.694755 | 1.095285 | -0.870117 | 1.073386 | -0.215907 | 0.562155 | 0.917161 |
| 27.52778 | 0.298397 | 1.002769 | -0.335593 | 0.500968 | 1.477643 | 0.097371 | 0.739248 | 0.177047 | 0.828877 | -0.247506 | 0.338748 | 0.377465 |
| 27.54167 | -0.313722 | 0.854694 | 0.087107 | 0.660272 | 1.087353 | -0.199817 | 0.865204 | 1.064062 | 0.623385 | 0.164366 | -0.122019 | 0.283225 |
| 27.55556 | -0.321453 | 0.680208 | 0.520785 | 0.540383 | 0.442822 | -0.105572 | 0.956848 | 2.151685 | 0.403955 | 0.774892 | -0.299704 | 0.270222 |
| 27.56944 | 0.202712 | 0.525113 | 0.734871 | 0.339565 | 0.03888 | 0.316529 | 0.953062 | 2.252133 | 0.248214 | 1.142429 | -0.214642 | 0.33834 |
| 27.58333 | 1.210162 | 0.450466 | 0.505183 | 0.132937 | 0.680423 | 0.795961 | 0.911799 | 2.022473 | 0.081352 | 0.989832 | 0.026018 | 0.183097 |
| 27.59722 | 2.196653 | 0.478688 | 0.338865 | 0.219046 | 1.809322 | 1.047006 | 1.029088 | 2.381791 | 0.002432 | 0.828739 | 0.29439 | 0.167795 |
| 27.61111 | 2.192052 | 0.971313 | 0.64202 | 0.347964 | 2.368414 | 0.910826 | 1.332141 | 2.968988 | -0.078098 | 0.737773 | 0.438877 | 0.735578 |
| 27.625 | 1.562485 | 1.386388 | 1.02813 | 0.43398 | 2.539337 | 0.772727 | 1.590665 | 2.698879 | 0.020389 | 0.712522 | 0.640757 | 1.561087 |
| 27.63889 | 1.481809 | 0.967394 | 1.516026 | 0.261442 | 2.491197 | 0.915931 | 1.239055 | 0.547239 | 0.302994 | 0.806787 | 0.902207 | 2.222456 |
| 27.65278 | 1.818823 | 0.754964 | 1.473064 | 0.069217 | 2.271562 | 1.211341 | 0.568117 | -0.921331 | 0.557126 | 0.814009 | 1.231892 | 2.749269 |
| 27.66667 | 1.378411 | 0.503888 | 1.211385 | 0.054651 | 0.402578 | 1.199898 | 0.192599 | -1.14194 | 0.619502 | 0.610853 | 1.63092 | 2.270964 |
| 27.68056 | 1.3421 | -0.048976 | 0.592772 | 0.118435 | -0.552778 | 0.906768 | -0.05827 | 0.086338 | 0.52278 | 0.358118 | 1.71588 | 1.502476 |
| 27.69444 | 0.778879 | -0.57654 | 0.087649 | 0.200519 | 0.168878 | 0.629733 | -0.016026 | 2.210428 | 0.380281 | 0.225644 | 1.840219 | 1.1388 |
| 27.70833 | 0.60638 | -0.449388 | -0.218306 | 0.262941 | 1.417606 | 0.444385 | 0.038801 | 2.13143 | 0.57822 | 0.28634 | 1.980135 | 0.826883 |
| 27.72222 | 0.415924 | -0.643768 | -0.159733 | 0.237914 | 1.297652 | 0.257637 | -0.093078 | 1.525568 | 1.058411 | 0.491746 | 1.666829 | 0.522364 |
| 27.73611 | -0.15097 | -0.008477 | 0.187615 | 0.114753 | 0.293831 | -0.025733 | -0.581235 | 0.754796 | 1.47164 | 0.582855 | 0.836757 | 0.400252 |
| 27.75 | -0.660287 | 0.468285 | 0.207036 | -0.037301 | -0.796898 | -0.376224 | -0.958869 | 0.37562 | 1.469374 | 0.535019 | -0.320034 | -0.212345 |
| 27.76389 | -0.705268 | 0.667772 | -0.437303 | -0.234236 | -0.762798 | -0.812628 | -0.948072 | -0.205561 | 0.220679 | 0.387698 | -0.839068 | -0.680344 |
| 27.77778 | -0.515674 | 0.792926 | -1.11081 | -0.647021 | 0.453943 | -0.949537 | -0.5894 | -1.051225 | -0.873362 | 0.22975 | -0.999085 | -1.114807 |
| 27.79167 | -0.513899 | 0.774572 | -1.296089 | -0.767931 | 1.387784 | -0.543363 | -0.259537 | -1.450521 | -2.174913 | 0.051573 | -1.639629 | -1.634334 |
| 27.80556 | -0.866618 | 0.68499 | -0.713369 | -0.43185 | 0.561861 | -0.142066 | -0.080426 | -1.434273 | -3.795679 | -0.007799 | -1.68627 | -0.995074 |
| 27.81944 | -1.130775 | 0.450874 | -0.300972 | -0.437767 | -1.217289 | 0.074926 | 0.120424 | -1.567128 | -3.262561 | -0.028305 | -0.750477 | -0.302701 |
| 27.83333 | -1.726922 | 0.245221 | -0.136112 | -0.532071 | -2.953803 | 0.262069 | 0.32009 | -1.160985 | -2.759148 | -0.094497 | -0.439454 | -0.066616 |
| 27.84722 | -2.065306 | 0.341708 | -0.179339 | -0.601996 | -2.817778 | 0.628654 | 0.498493 | -0.890221 | -2.724625 | -0.301657 | 0.046147 | 0.057475 |
| 27.86111 | -1.786762 | 0.280355 | -0.480146 | -0.955555 | -0.932226 | 0.797063 | 0.463555 | -0.962791 | -2.793878 | -0.762681 | 0.299 | 0.118947 |
| 27.875 | -1.644589 | -0.47561 | -0.983792 | -2.564408 | -0.218886 | 0.108356 | -0.32804 | -2.74431 | -3.096763 | -1.72227 | 0.593493 | -0.119871 |
| 27.88889 | -1.857192 | -1.099961 | -0.693506 | -4.04298 | -1.753077 | -1.193218 | -1.385217 | -5.617366 | -3.194124 | -2.74078 | 0.610868 | -0.204471 |
| 27.90278 | -1.086348 | -1.279221 | 0.211429 | -2.550238 | -3.591832 | -1.818985 | -1.573373 | -3.141617 | -2.098058 | -3.678598 | 0.524837 | 0.219494 |
| 27.91667 | -0.746155 | -1.677432 | 0.613663 | -0.926764 | -4.356251 | -1.836004 | -1.331123 | -0.015798 | -1.168871 | -2.869699 | 0.250004 | 0.621175 |
| 27.93056 | -0.637947 | -1.503381 | 0.539969 | -0.654526 | -2.864785 | -1.608065 | -0.804656 | 0.446079 | -0.580599 | -1.030597 | 0.023677 | 0.595111 |
| 27.94444 | -0.459546 | -1.413842 | 0.171231 | -1.164421 | -0.956922 | -0.92658 | -0.67763 | 0.53944 | 0.357428 | -0.051139 | -0.614659 | 0.403543 |
| 27.95833 | -0.094041 | -1.323737 | -0.215528 | -1.47655 | 0.171577 | -0.182922 | -0.765765 | 0.525662 | 1.069107 | 0.20472 | -0.842604 | 0.14807 |

| | | | | | | | | | | | | |
|---|---|---|---|---|---|---|---|---|---|---|---|---|
| 25.47222 | 0.274125 | 0.062155 | 0.924413 | 0.462024 | -0.816197 | 0.297588 | 0.561116 | 1.148462 | -0.132427 | -0.255005 | -0.002399 | 0.543699 |
| 25.48611 | 0.541282 | 0.342216 | 1.384614 | 1.714262 | -0.452668 | 0.679002 | 1.231249 | 1.815923 | 0.064849 | 0.188578 | 0.099586 | 0.658636 |
| 25.5 | 0.746209 | 1.285143 | 1.174236 | 3.171035 | 0.269843 | 1.671473 | 2.066563 | 2.411493 | 0.455871 | 0.811032 | 0.65588 | 1.325879 |
| 25.51389 | 1.343519 | 3.03692 | 0.320618 | 3.480987 | 0.903879 | 2.955392 | 2.791052 | 2.691122 | 0.630985 | 1.181356 | 1.119423 | 2.838007 |
| 25.52778 | 1.994564 | 4.158167 | -0.049755 | 3.660741 | 1.409107 | 2.942205 | 2.106843 | 2.124926 | 0.427594 | 0.685053 | 1.810718 | 3.275538 |
| 25.54167 | 2.452699 | 3.783195 | 0.150507 | 3.804931 | 1.968149 | 2.053994 | 1.305978 | 1.650045 | 0.057276 | 0.308263 | 2.112051 | 1.971584 |
| 25.55556 | 2.355313 | 2.403032 | 0.515636 | 4.158883 | 2.175937 | 1.104306 | 0.400646 | 1.168639 | -0.028682 | 0.260418 | 1.713654 | 0.779537 |
| 25.56944 | 2.123484 | 1.389691 | 1.263383 | 3.377783 | 1.812092 | 0.304597 | 0.170017 | 0.886224 | 0.440163 | 0.360439 | 1.201858 | 0.616354 |
| 25.58333 | 1.536644 | 0.856249 | 1.85552 | 2.209643 | 1.518026 | -0.212262 | 0.424361 | 0.732549 | 0.578594 | 0.463614 | 0.531724 | 1.029317 |
| 25.59722 | 1.206252 | 0.55069 | 1.482819 | 1.592667 | 1.379012 | -0.484276 | 0.649455 | 0.558604 | 0.445238 | 0.393169 | 0.116571 | 1.257245 |
| 25.61111 | 1.329964 | 0.167047 | 0.049222 | 1.478361 | 1.409663 | -0.937781 | 0.440619 | 0.326628 | 0.290075 | 0.18189 | -0.190387 | 0.735941 |
| 25.625 | 1.572248 | -0.27169 | -1.789711 | 1.201508 | 1.520084 | -0.944599 | 0.142552 | 0.177647 | 0.005027 | 0.067241 | -0.270516 | 0.039961 |
| 25.63889 | 1.810007 | -0.400427 | -2.321143 | 1.159746 | 1.292134 | 0.151522 | 0.256186 | 0.273016 | 0.257225 | 0.002729 | -0.163011 | 0.071743 |
| 25.65278 | 1.815866 | 0.006427 | -1.504824 | 1.457991 | 0.979929 | 1.564774 | 0.527541 | 0.560361 | 0.586748 | 0.085923 | 0.208842 | 0.933326 |
| 25.66667 | 1.855582 | 0.825457 | -0.088013 | 1.890917 | 0.665759 | 2.392189 | 0.723898 | 0.777414 | 0.86496 | 0.208654 | 0.403537 | 1.88214 |
| 25.68056 | 1.659528 | 1.800465 | 1.741087 | 1.954701 | 0.467535 | 2.05663 | 0.546358 | 0.724757 | 1.288699 | 0.19775 | 0.678666 | 1.967522 |
| 25.69444 | 1.268349 | 2.276615 | 2.70631 | 1.582414 | 0.341438 | 1.40929 | 0.446772 | 0.407897 | 1.041826 | 0.147068 | 0.816003 | 1.297632 |
| 25.70833 | 0.827321 | 2.085661 | 1.508816 | 1.362129 | 0.190173 | 0.911935 | 0.525009 | -0.070235 | 0.570441 | 0.089636 | 1.099828 | 0.534877 |
| 25.72222 | 0.359361 | 2.225707 | 1.889713 | 1.546367 | 0.043527 | 0.554164 | 0.513959 | -0.466702 | 0.284012 | -0.02513 | 1.461544 | 0.486792 |
| 25.73611 | 0.000411 | 2.35699 | 1.54835 | 1.322055 | -0.051263 | -0.127479 | 0.649411 | -0.496269 | 0.144931 | 0.075266 | 1.715849 | 0.699449 |
| 25.75 | -0.293607 | 1.712883 | 1.580767 | 1.177001 | -0.07583 | -1.319607 | 0.711494 | -0.220052 | 0.313875 | 0.098666 | 1.345572 | 0.624285 |
| 25.76389 | -0.451049 | 0.954332 | 1.007325 | 0.833745 | -0.032733 | -2.13718 | 0.507923 | 0.077671 | 0.593846 | -0.131522 | 0.613773 | -0.062275 |
| 25.77778 | -0.290542 | 0.08154 | -0.604875 | 0.52646 | 0.068249 | -0.715254 | 0.65662 | 0.145297 | 0.655237 | -0.551232 | -0.089279 | -0.591616 |
| 25.79167 | -0.308739 | -0.565811 | -1.727059 | 0.1356 | 0.110903 | 0.85431 | 0.765861 | 0.135282 | 0.606824 | -0.107017 | -0.58756 | -0.374821 |
| 25.80556 | -0.70601 | -0.521229 | -1.947963 | -0.344594 | 0.012034 | 1.146083 | 0.534307 | 0.011931 | 0.608637 | 0.544705 | -0.259692 | 0.329911 |
| 25.81944 | -1.107307 | -0.670083 | -2.034797 | -0.535356 | -0.146228 | 0.01884 | 0.299998 | 0.203942 | 0.734652 | 1.177041 | 0.028599 | 0.652611 |
| 25.83333 | -1.072902 | -1.267491 | -0.483786 | -0.84738 | -0.264973 | -1.331357 | 0.356925 | 0.679756 | 0.822787 | 1.075399 | 0.073155 | 0.223941 |
| 25.84722 | -0.797764 | -2.212076 | -0.525414 | -0.777582 | -0.36494 | -0.876867 | 0.414233 | 0.733089 | 0.72328 | 0.597547 | 0.258855 | -0.587284 |
| 25.86111 | -0.557179 | -2.575109 | -0.094493 | -0.755588 | -0.468146 | 0.458656 | 0.294918 | 0.28006 | 0.535161 | 0.419826 | 0.733809 | -0.615866 |
| 25.875 | -0.530062 | -1.960339 | 0.276214 | -1.466634 | -0.57067 | 1.90368 | 0.049688 | -0.012404 | 0.477556 | 0.352166 | 1.309807 | -0.321399 |
| 25.88889 | -0.697301 | -1.802981 | 0.944358 | -2.001623 | -0.734291 | 1.291584 | -0.417018 | -0.09674 | 0.499901 | 0.184623 | 1.254952 | -0.276316 |
| 25.90278 | -0.853552 | -1.875038 | 0.931389 | -1.714439 | -0.893451 | -0.778399 | -0.718257 | -0.480278 | 0.528445 | -0.252769 | -0.160343 | -1.156919 |
| 25.91667 | -0.666828 | -1.707121 | 1.273597 | -1.844946 | -0.833081 | -3.011937 | -1.376302 | -1.284927 | 0.592219 | -0.375477 | -1.413127 | -1.805263 |
| 25.93056 | -0.233918 | -1.044811 | 1.261262 | -2.489669 | -0.687544 | -2.637614 | -1.715751 | -1.785213 | 0.683688 | -0.739203 | -1.489349 | -1.895961 |
| 25.94444 | -0.278186 | -0.548824 | 0.963291 | -2.26303 | -0.894597 | -1.89918 | -1.708412 | -1.778473 | 0.847338 | -0.815378 | -1.46551 | -1.442438 |
| 25.95833 | -1.285008 | -0.627532 | -0.153612 | -1.510529 | -1.711287 | -2.577284 | -2.651477 | -3.173065 | 0.270446 | -1.81619 | -2.037383 | -1.737583 |
| 25.97222 | -1.889805 | -0.638133 | -0.842167 | -1.987519 | -2.185338 | -3.298354 | -3.282075 | -4.488368 | -0.647497 | -1.258087 | -1.547935 | -1.819748 |
| 25.98611 | -1.746189 | -0.856482 | -0.671381 | -1.984025 | -1.217829 | -2.747285 | -2.841829 | -3.775371 | -0.734307 | -0.510681 | -1.029915 | -1.081253 |
| 26 | -1.02193 | -0.908954 | -0.755837 | -2.116265 | -0.149441 | -1.412095 | -2.022021 | -1.905975 | -0.769012 | 0.264717 | -0.600712 | -0.094609 |
| 26.01389 | -0.423216 | -0.560512 | -1.107427 | -1.831345 | 0.21799 | -0.654092 | -1.293556 | -0.271712 | -0.576996 | 0.998519 | 0.000906 | 1.097122 |
| 26.02778 | -0.315275 | -0.346242 | -1.486222 | -2.015463 | 0.228602 | -0.414667 | -0.925824 | 0.121542 | -0.42045 | 1.683439 | 0.200445 | 2.170543 |
| 26.04167 | -0.335306 | -0.624466 | -1.34324 | -1.422401 | 0.322047 | -0.309175 | -0.951164 | 0.076951 | -0.297919 | 0.948172 | 0.468522 | 1.377019 |
| 26.05556 | -0.222228 | -1.07453 | 0.073741 | 0.11673 | 0.41609 | -0.246744 | -0.757273 | 0.254187 | -0.231089 | 0.271945 | 0.385593 | 0.445572 |
| 26.06944 | 0.111754 | -1.485319 | 0.857843 | 0.940551 | 0.496477 | -0.183793 | -0.347004 | 0.18954 | -0.107557 | -0.774768 | 0.920022 | -0.403474 |
| 26.08333 | 0.215034 | -1.313524 | 0.246077 | 0.484465 | 0.588974 | -0.1319 | -0.169071 | -0.535217 | 0.27572 | -2.036095 | 1.688085 | -0.952249 |
| 26.09722 | -0.049108 | -0.833993 | -0.338495 | -0.326268 | 0.597832 | -0.013233 | -0.112762 | -1.628599 | 0.480268 | -1.416048 | 1.588696 | -0.062535 |
| 26.11111 | -0.476238 | 0.078203 | -0.843781 | -0.718061 | 0.618629 | 0.152379 | -0.149989 | -1.833128 | 0.11529 | 0.170356 | 0.474825 | 0.628019 |
| 26.125 | -0.594294 | 0.755155 | -1.125669 | -0.380078 | 0.492637 | 0.01989 | -0.300134 | -1.302995 | 0.11504 | 1.488714 | -0.131075 | 1.22224 |
| 26.13889 | -0.273717 | 0.66265 | -0.289867 | 0.189634 | 0.300659 | -0.462824 | -0.355408 | -0.481122 | 0.405118 | 0.722613 | -0.050507 | 1.229677 |
| 26.15278 | 0.05631 | 0.038443 | 0.721185 | 0.268966 | 0.181276 | -0.47367 | 0.007777 | 0.516157 | 0.20046 | -0.667326 | -0.103337 | -0.065469 |
| 26.16667 | 0.106926 | -0.44647 | 0.909703 | 0.072683 | 0.086578 | -0.035886 | 0.381493 | 1.414548 | 0.019125 | -1.39683 | 0.016575 | -0.409465 |
| 26.18056 | 0.186818 | -0.702054 | 0.53193 | 0.213326 | -0.106486 | 0.239243 | 0.331272 | 1.287573 | 0.009045 | -0.66467 | 0.474687 | -0.402843 |
| 26.19444 | 0.269715 | -0.985658 | 0.295302 | 0.35994 | -0.288436 | 0.12245 | 0.071991 | 0.182094 | -0.066755 | -0.118134 | 0.864503 | -0.4357 |
| 26.20833 | 0.2531 | -1.30529 | 0.197793 | 0.108173 | -0.287414 | -0.057269 | 0.100625 | -1.006682 | -0.290202 | -0.100954 | 0.781871 | -0.661512 |
| 26.22222 | 0.141161 | -1.314966 | 0.335188 | -0.075004 | -0.097769 | 0.032115 | 0.18952 | -1.453494 | -0.590047 | -0.395456 | 0.49193 | -0.763298 |
| 26.23611 | 0.060068 | -1.097694 | 0.606098 | 0.021884 | 0.134866 | -0.036082 | 0.018512 | -1.160522 | -0.739923 | -0.496687 | 0.692356 | -0.590763 |
| 26.25 | 0.004095 | -0.841748 | 0.54255 | 0.31845 | 0.283587 | -0.26267 | -0.188839 | -0.864986 | -0.766015 | -0.347377 | 0.651338 | -0.124202 |
| 26.26389 | -0.194533 | -0.732892 | 0.226367 | 0.466576 | 0.34659 | -0.475096 | -0.153764 | -0.711641 | -0.784651 | -0.160802 | 0.218944 | 0.180381 |
| 26.27778 | -0.219053 | -0.626707 | -0.066891 | 0.39082 | 0.332433 | -0.461262 | 0.014006 | -0.393074 | -0.8256 | -0.113933 | 0.052902 | 0.214633 |
| 26.29167 | -0.025974 | -0.534542 | -0.065069 | 0.435072 | 0.351796 | -0.249326 | 0.289397 | 0.286299 | -0.634622 | -0.239353 | 0.042873 | 0.035649 |
| 26.30556 | 0.195642 | -0.558288 | 0.094683 | 0.788443 | 0.401065 | 0.018276 | 0.586924 | 0.836644 | -0.41103 | -0.525714 | 0.294353 | -0.139294 |
| 26.31944 | 0.358979 | -0.56688 | 0.190838 | 1.167409 | 0.39738 | 0.250868 | 0.775688 | 0.879903 | -0.298841 | -0.686575 | 0.44512 | -0.346109 |
| 26.33333 | 0.513162 | -0.379192 | 0.328624 | 1.403549 | 0.370325 | 0.396573 | 0.772495 | 0.663982 | -0.224905 | -0.484151 | 0.593046 | -0.190526 |
| 26.34722 | 0.654661 | -0.031685 | 0.377345 | 1.593054 | 0.361922 | 0.445494 | 0.738852 | 0.406029 | -0.167608 | -0.156135 | 0.645776 | 0.244661 |
| 26.36111 | 0.781172 | 0.236449 | 0.276509 | 1.502478 | 0.369451 | 0.489092 | 0.64172 | 0.354862 | -0.105178 | 0.092107 | 0.813386 | 0.638568 |
| 26.375 | 0.842499 | 0.186925 | 0.103548 | 1.162503 | 0.328785 | 0.560376 | 0.57428 | 0.558008 | 0.000636 | 0.091234 | 0.957425 | 0.515705 |
| 26.38889 | 0.601028 | -0.027515 | 0.110556 | 0.832375 | 0.304281 | 0.687139 | 0.64803 | 0.861975 | 0.086628 | -0.015144 | 0.759041 | 0.179313 |
| 26.40278 | 0.449314 | -0.105053 | 0.38377 | 0.78557 | 0.317542 | 0.743165 | 0.909903 | 1.02426 | 0.141482 | 0.07977 | 0.671431 | 0.251846 |
| 26.41667 | 0.360573 | 0.062053 | 0.661259 | 0.996554 | 0.226729 | 0.791145 | 1.165027 | 1.091939 | 0.14723 | 0.513807 | 0.567059 | 0.937671 |
| 26.43056 | 0.21985 | 0.155333 | 0.629438 | 0.997873 | 0.049512 | 0.753947 | 0.991112 | 1.064483 | 0.101732 | 1.254996 | 0.343643 | 1.414657 |
| 26.44444 | 0.084467 | 0.104263 | 0.428477 | 0.439751 | -0.02271 | 0.630088 | 0.607586 | 0.359012 | -0.035291 | 0.429896 | -0.371098 | 1.046828 |
| 26.45833 | 0.175672 | -0.001483 | 0.517254 | -0.34264 | -0.011541 | 0.598614 | 0.372098 | -0.423612 | -0.331006 | -0.597941 | -0.41809 | 0.600249 |
| 26.47222 | 0.509373 | 0.028615 | 0.981379 | -0.084019 | -0.014043 | 0.774978 | 0.553623 | -0.136197 | -0.506571 | -1.126346 | 0.237545 | 0.614537 |
| 26.48611 | 1.017574 | 0.370357 | 0.960422 | 1.536499 | 0.203336 | 1.170454 | 1.125495 | 1.434966 | -0.196684 | -0.547158 | 0.907819 | 0.84927 |
| 26.5 | 1.479547 | 1.570403 | 0.489583 | 3.677454 | 0.429938 | 1.823474 | 2.180325 | 3.633266 | 0.548369 | 0.788055 | 1.330012 | 1.432424 |
| 26.51389 | 1.683314 | 3.343328 | -0.007519 | 5.320584 | 0.590385 | 2.369302 | 3.068614 | 4.727718 | 1.285432 | 2.723964 | 1.692478 | 1.917381 |
| 26.52778 | 2.009279 | 3.715379 | 0.017471 | 5.146938 | 0.8873 | 2.112526 | 3.092241 | 4.54294 | 1.539263 | 2.384442 | 2.161441 | 2.213211 |
| 26.54167 | 2.583489 | 3.069998 | 0.448274 | 5.085145 | 1.306877 | 1.17271 | 2.730322 | 4.933809 | 1.688197 | 1.856057 | 2.054795 | 1.407651 |
| 26.55556 | 2.888946 | 1.546957 | 0.706369 | 3.737025 | 1.615435 | 0.91375 | 2.143674 | 3.741101 | 1.783988 | 0.896442 | 1.437861 | 0.223228 |
| 26.56944 | 2.727218 | 0.468187 | 0.426114 | 2.884418 | 1.693683 | 0.483097 | 0.958996 | 2.065805 | 1.510992 | 0.409044 | 1.027463 | -0.366068 |
| 26.58333 | 2.312634 | 0.138281 | 0.347656 | 1.869618 | 1.535783 | 0.110062 | 0.08651 | 0.778891 | 1.070079 | 0.216927 | 0.980446 | 0.134279 |
| 26.59722 | 1.951709 | 0.15138 | 0.481524 | 1.257547 | 1.214146 | -0.150988 | -0.224521 | -0.260524 | 0.92734 | 0.319877 | 0.930636 | 0.875442 |
| 26.61111 | 1.687575 | 0.332945 | 0.767114 | 0.494649 | 0.999704 | -0.311845 | -0.123219 | -0.38974 | 0.778519 | 0.553182 | 1.060598 | 1.607037 |
| 26.625 | 1.653917 | 0.538829 | 1.23847 | 0.122314 | 0.780649 | -0.157738 | 0.407639 | 0.511424 | 0.535527 | 0.510822 | 0.924361 | 1.096801 |
| 26.63889 | 2.01813 | 0.725413 | 1.337235 | 0.411058 | 0.644948 | 0.518144 | 1.00386 | 1.641028 | 0.402693 | 0.158229 | 0.642688 | 0.913649 |
| 26.65278 | 2.491706 | 0.713996 | 1.662165 | 1.485612 | 0.561592 | 1.451367 | 1.06123 | 1.795414 | 0.380135 | 0.146897 | 0.307415 | 0.782027 |
| 26.66667 | 2.950115 | 0.873196 | 2.174253 | 2.329833 | 0.521713 | 1.857771 | 1.365389 | 1.179999 | 0.461205 | 0.200916 | 0.022212 | 1.058885 |
| 26.68056 | 3.225798 | 1.381312 | 1.762004 | 2.702616 | 0.428677 | 1.376455 | 1.148606 | 0.799416 | 0.633948 | 0.269397 | 0.474734 | 1.527579 |
| 26.69444 | 3.04362 | 2.152548 | 1.067256 | 3.263985 | 0.38279 | 0.773226 | 1.07614 | 0.355267 | 0.768264 | 0.257804 | 0.569149 | 1.377895 |
| 26.70833 | 2.997852 | 2.58914 | 0.726973 | 2.745121 | 0.376918 | 0.712006 | 0.939705 | -0.623563 | 0.923718 | 0.150841 | -0.087698 | 0.902993 |
| 26.72222 | 2.369897 | 2.469807 | 1.486645 | 2.089727 | 0.299386 | 1.571032 | 0.624565 | -1.764351 | 0.686239 | -0.197338 | -0.803781 | 0.627462 |
| 26.73611 | 1.449519 | 1.925851 | 1.955388 | 1.725148 | 0.243118 | 3.013109 | 0.438456 | -3.50913 | 0.576498 | -0.234828 | -0.693446 | 0.951965 |
| 26.75 | 0.630317 | 1.928724 | 1.223903 | 1.623367 | 0.234745 | 2.213382 | 0.181187 | -4.339469 | 0.688772 | -0.352807 | -0.355374 | 1.692901 |
| 26.76389 | 0.297593 | 2.02676 | -0.056912 | 1.547871 | 0.24613 | 0.700724 | 0.062377 | -3.499184 | 0.73063 | -0.277413 | 0.101145 | 1.442691 |
| 26.77778 | 0.3005 | 1.503612 | -1.929253 | 1.428212 | 0.219216 | 0.216346 | 0.329818 | -1.831702 | -0.079054 | -1.254613 | -0.286287 | 0.742177 |
| 26.79167 | 0.018869 | 0.680652 | -2.038376 | 0.620077 | 0.15436 | -0.397699 | 0.286753 | -1.271803 | -0.097293 | -1.173006 | -0.745703 | 0.33188 |
| 26.80556 | -0.680999 | 0.036215 | -1.12346 | -0.103795 | 0.158358 | -0.987436 | 0.18945 | -1.104057 | 0.311772 | -0.31881 | -0.298311 | 0.426718 |
| 26.81944 | -1.136355 | -0.376027 | 0.96365 | -0.439916 | 0.237002 | -0.79501 | 0.28679 | -0.16506 | 0.63846 | 0.662016 | 0.297194 | 0.667677 |
| 26.83333 | -1.342297 | -0.77489 | 1.645775 | -0.860226 | 0.27392 | -0.989326 | 0.525209 | 1.490979 | 1.025047 | 1.947301 | 0.678151 | 0.653526 |
| 26.84722 | -1.372944 | -1.215333 | 1.147791 | -0.939808 | 0.240642 | -0.601932 | 1.017676 | 4.224623 | 1.2117 | 1.925818 | 1.066376 | 0.143742 |
| 26.86111 | -1.089089 | -2.007543 | 0.755866 | -0.650254 | 0.181728 | 0.057157 | 1.347104 | 5.446134 | 1.292866 | 1.436298 | 1.469751 | -0.925282 |
| 26.875 | -0.788818 | -2.908533 | -0.036123 | -1.654631 | 0.135958 | 0.019839 | 1.459346 | 5.35332 | 1.087805 | 0.869738 | 2.384784 | -1.377486 |
| 26.88889 | -0.602942 | -2.747692 | -1.400234 | -1.93976 | 0.119058 | -0.339655 | 1.041606 | 2.707622 | 0.76651 | 0.784328 | 1.932815 | -0.337549 |
| 26.90278 | -0.744943 | -1.623259 | -2.555041 | -1.551239 | 0.102914 | -0.855487 | 0.03825 | 0.238265 | 0.685797 | 0.90354 | 0.282838 | 0.600288 |
| 26.91667 | -1.278749 | -1.361082 | -2.410302 | -2.431479 | 0.01997 | -1.25335 | -1.006598 | -1.276319 | 0.831344 | 0.814736 | 0.027208 | 0.108687 |
| 26.93056 | -1.77507 | -1.967745 | -1.136058 | -2.771489 | -0.179598 | -1.137266 | -1.500634 | -1.992039 | 0.347862 | 0.249484 | 0.242834 | -0.717998 |
| 26.94444 | -1.788864 | -2.374119 | -0.68518 | -2.607811 | -0.731599 | -1.415632 | -1.509916 | -3.640117 | -0.329658 | -0.208424 | -1.037416 | -0.944809 |
| 26.95833 | -2.22404 | -2.629317 | -1.786477 | -2.714462 | -2.400475 | -2.305485 | -2.392126 | -5.456343 | -0.97033 | -0.939366 | -1.542082 | -0.179837 |
| 26.97222 | -2.224407 | -2.74266 | -2.581944 | -3.701436 | -2.341072 | -2.246719 | -3.023648 | -6.251674 | -1.742766 | -2.180635 | -1.341111 | -1.718739 |
| 26.98611 | -1.195059 | -2.473549 | -2.888617 | -3.578519 | -0.803223 | -1.463777 | -1.666445 | -4.940986 | -0.835369 | -3.173017 | -1.092609 | -1.776546 |
| 27 | -0.541479 | -2.759728 | -2.149201 | -4.530461 | -0.178334 | -0.695866 | -0.49344 | -2.241697 | -0.805857 | -4.213224 | -0.465352 | -1.772087 |
| 27.01389 | -0.554815 | -2.20533 | -1.685902 | -3.80935 | -0.057813 | -0.504299 | -0.20938 | -0.141893 | -0.337922 | -1.060841 | 0.830017 | -0.79806 |
| 27.02778 | -0.533139 | -1.785789 | -2.178488 | -2.649511 | -0.072854 | -0.860722 | -0.639037 | 0.685622 | -0.568659 | 1.733712 | 0.512452 | 0.647872 |
| 27.04167 | -0.297768 | -2.003625 | -1.231445 | -1.425929 | 0.014124 | -0.906293 | -1.02685 | 0.922748 | -0.380007 | 2.93867 | 0.026931 | 1.165859 |
| 27.05556 | 0.350449 | -1.495112 | 0.331791 | -0.218621 | 0.225285 | -0.175948 | -0.995 | 0.619535 | -0.06998 | 1.358359 | -0.049113 | -0.098598 |
| 27.06944 | 1.028226 | -0.178857 | 1.43832 | 0.683748 | 0.374873 | -0.137163 | -0.462266 | -0.431242 | 0.372023 | -0.14993 | 0.149886 | -0.786938 |
| 27.08333 | 0.689837 | 0.354266 | 1.883048 | 0.431704 | 0.290068 | 0.105766 | -0.136297 | -1.701136 | 0.501304 | -0.663735 | 0.499174 | -0.781116 |
| 27.09722 | 0.0861 | 0.372803 | 1.210118 | -0.227841 | 0.083431 | 1.007291 | -0.014553 | -2.108569 | 0.424638 | -0.084545 | 0.87011 | -0.120274 |
| 27.11111 | -0.303958 | 0.679625 | 0.215527 | -0.200653 | -0.031296 | 0.607433 | -0.084358 | -2.026178 | 0.370342 | 0.818454 | 0.395335 | 0.435558 |
| 27.125 | -0.699169 | 1.060746 | -0.288356 | 0.217591 | -0.060159 | 0.27783 | -0.017906 | -1.508966 | 0.657539 | 1.761207 | 0.436968 | 0.82906 |
| 27.13889 | -0.918336 | 0.44159 | 0.136476 | 0.547008 | -0.195319 | 0.186735 | 0.086559 | -0.945677 | 0.371529 | 1.290589 | 0.117416 | 0.591867 |
| 27.15278 | -0.667416 | -0.371357 | 0.360699 | 0.544929 | -0.492051 | 0.157461 | -0.025328 | -0.740533 | -0.200097 | 0.117043 | -0.235468 | 0.084688 |
| 27.16667 | -0.37875 | -0.526784 | 0.66119 | 0.291887 | -0.821709 | 0.217257 | -0.222445 | -0.237093 | -0.464272 | -0.377766 | -0.26966 | 0.480245 |
| 27.18056 | -0.208089 | -0.30635 | 1.176731 | 0.053927 | -1.025542 | 0.123437 | -0.199296 | 0.581864 | -0.546194 | -0.351487 | -0.158176 | 1.164562 |
| 27.19444 | -0.147213 | -0.53622 | 1.389854 | -0.041832 | -1.145257 | -0.007637 | -0.048618 | 0.949796 | -0.634387 | -0.367522 | -0.093449 | 0.899212 |
| 27.20833 | -0.074323 | -0.852291 | 1.206534 | -0.002723 | -1.219619 | 0.078753 | 0.031484 | 0.219473 | -0.488196 | -0.446329 | -0.155898 | 0.096693 |
| 27.22222 | -0.099829 | -0.892883 | 0.834932 | 0.001279 | -1.16085 | 0.159903 | 0.071978 | -1.004366 | -0.388034 | -1.122201 | -0.118315 | -0.943481 |
| 27.23611 | -0.283107 | -0.881706 | 0.59009 | -0.12369 | -1.091087 | 0.22138 | 0.083979 | -1.760253 | -0.204582 | -3.02177 | -0.002618 | -1.491654 |
| 27.25 | -0.578392 | -0.789606 | 0.521809 | -0.229357 | -1.134909 | 0.361076 | 0.114136 | -1.86684 | -0.118241 | -3.156778 | -0.000294 | -1.208739 |
| 27.26389 | -0.854225 | -0.566585 | 0.510265 | -0.099182 | -1.350675 | 0.573429 | 0.162161 | -1.581937 | -0.198677 | -1.704031 | 0.095013 | -0.556556 |
| 27.27778 | -0.829544 | -0.308204 | 0.586036 | 0.178643 | -1.489372 | 0.690324 | 0.205145 | -0.96358 | -0.358939 | -0.423879 | 0.193482 | -0.061584 |
| 27.29167 | -0.558888 | -0.335738 | 0.805622 | 0.488749 | -1.320307 | 0.617511 | 0.242516 | -0.111154 | -0.355518 | 0.181137 | 0.258893 | 0.161113 |
| 27.30556 | -0.455382 | -0.459158 | 1.124562 | 0.731425 | -1.140945 | 0.389008 | 0.303396 | 0.778973 | 0.087995 | 0.10181 | 0.27413 | -0.087529 |
| 27.31944 | -0.511234 | -0.38601 | 1.109723 | 0.875739 | -1.209899 | 0.222097 | 0.340002 | 1.358846 | 0.223127 | -0.173278 | 0.445375 | -0.358888 |
| 27.33333 | -0.385428 | -0.219421 | 0.844653 | 1.08752 | -1.116176 | 0.275473 | 0.321786 | 1.350032 | 0.092867 | -0.212012 | 0.58653 | -0.354469 |
| 27.34722 | -0.060639 | -0.089792 | 0.64286 | 1.6662 | -0.798617 | 0.485915 | 0.318027 | 1.091363 | -0.166295 | 0.022248 | 0.673674 | -0.145194 |
| 27.36111 | 0.299096 | 0.134995 | 0.804379 | 1.823345 | -0.428646 | 0.625783 | 0.360568 | 1.051327 | -0.366387 | 0.303141 | 0.681959 | 0.149986 |
| 27.375 | 0.493524 | 0.303239 | 0.94522 | 1.496379 | -0.219121 | 0.601706 | 0.401777 | 1.172085 | -0.450822 | 0.587887 | 0.627679 | 0.450794 |
| 27.38889 | 0.509525 | 0.230498 | 0.876307 | 1.385941 | -0.151483 | 0.584536 | 0.381824 | 1.191475 | -0.525522 | 0.974445 | 0.604797 | 0.642131 |
| 27.40278 | 0.35651 | 0.234993 | 0.927802 | 1.51603 | -0.145154 | 0.842837 | 0.42455 | 1.117119 | -0.351972 | 0.915395 | 0.633994 | 0.695278 |
| 27.41667 | 0.21135 | 0.284893 | 1.116313 | 1.780855 | -0.194957 | 0.950616 | 0.489864 | 1.028102 | -0.28792 | 0.347927 | 0.748041 | 0.552692 |
| 27.43056 | 0.143747 | 0.292549 | 1.186719 | 1.931506 | -0.294701 | 0.883928 | 0.492493 | 0.946849 | -0.318256 | -0.08735 | 0.924128 | 0.336466 |
| 27.44444 | 0.064396 | 0.341778 | 1.106406 | 1.502086 | -0.555068 | 0.565724 | 0.399978 | 0.83878 | -0.777343 | -0.451662 | 0.867381 | 0.146571 |
| 27.45833 | 0.000055 | 0.447419 | 0.910135 | 1.01101 | -1.188024 | 0.207065 | 0.301475 | 0.528158 | -1.066001 | -0.924883 | 0.539291 | 0.090282 |
| 27.47222 | 0.04164 | 0.647831 | 0.676904 | 0.716939 | -2.412141 | 0.171651 | 0.357285 | 0.420127 | -1.297317 | -1.848534 | 0.287785 | 0.170021 |
| 27.48611 | 0.383881 | 1.023385 | 0.862987 | 1.039352 | -2.439207 | 0.379005 | 0.67027 | 1.469121 | -0.382522 | -0.950896 | 0.500519 | 0.576622 |
| 27.5 | 1.244804 | 2.143811 | 1.232552 | 3.118444 | -0.79363 | 0.569686 | 1.184986 | 3.301613 | 0.490378 | 0.670521 | 0.669508 | 1.246002 |
| 27.51389 | 1.483628 | 3.635143 | 1.261659 | 3.770876 | 0.361479 | 0.859685 | 1.757097 | 4.646817 | 1.070862 | 2.363943 | 1.456108 | 1.970542 |
| 27.52778 | 1.409628 | 4.318269 | 1.112772 | 4.336965 | 1.389107 | 1.166864 | 2.29788 | 4.699895 | 1.47031 | 3.48644 | 2.574463 | 1.933703 |
| 27.54167 | 1.913812 | 3.188689 | 0.883104 | 4.57684 | 2.607835 | 1.620571 | 3.000302 | 4.239198 | 0.815938 | 2.557861 | 3.104805 | 1.48018 |
| 27.55556 | 2.333833 | 1.569254 | 0.991625 | 4.1439 | 3.494954 | 2.442345 | 3.348472 | 3.94145 | 0.552939 | 2.910339 | 2.712932 | 1.218008 |
| 27.56944 | 2.171649 | 0.500338 | 1.522516 | 3.776987 | 3.40154 | 2.611245 | 1.950003 | 2.671634 | 1.277996 | 1.598727 | 1.308646 | 1.120008 |
| 27.58333 | 1.667952 | -0.010945 | 1.96481 | 2.810873 | 2.886796 | 1.817127 | 0.724806 | 1.179735 | 1.216827 | 0.391176 | 0.088753 | 0.740458 |
| 27.59722 | 1.049652 | -0.35552 | 1.964371 | 2.031191 | 2.312345 | 1.559533 | 0.3996 | 0.059681 | 0.858239 | -0.074964 | -1.109235 | 1.150393 |
| 27.61111 | 0.731943 | -0.754947 | 1.251098 | 1.181948 | 1.963412 | 0.705331 | 0.380833 | -0.123561 | 0.459361 | -0.138179 | -1.004964 | 1.160006 |
| 27.625 | 0.631352 | -1.368564 | -0.117382 | -0.127781 | 1.881601 | 0.204762 | 0.560467 | 0.798341 | 0.163572 | 0.09839 | -0.229911 | 1.095957 |
| 27.63889 | 0.771559 | -1.458061 | -0.950124 | 0.150065 | 1.858841 | 0.336492 | 0.846573 | 2.323812 | -0.028589 | 0.179412 | 0.568289 | 1.00225 |
| 27.65278 | 0.967761 | -1.060928 | 0.036563 | 0.90252 | 1.663117 | 0.872713 | 0.988636 | 2.948056 | 0.288091 | 1.363231 | 1.165372 | 0.943551 |
| 27.66667 | 1.412009 | -0.323542 | 1.708481 | 2.325776 | 1.659172 | 1.111533 | 0.809825 | 0.920532 | 0.265594 | 0.598927 | 0.889974 | 0.950323 |
| 27.68056 | 1.61346 | 0.180867 | 3.775754 | 2.765558 | 1.879412 | 0.961505 | 0.693319 | -0.818869 | 0.600505 | 0.443175 | 0.980591 | 0.619593 |
| 27.69444 | 1.673074 | 0.677704 | 5.356132 | 1.743725 | 2.132182 | 0.729385 | 0.828038 | -1.74259 | 0.613304 | 0.384608 | 0.591671 | 0.356251 |
| 27.70833 | 1.553282 | 1.670223 | 5.557759 | 2.316761 | 1.902872 | 1.129971 | 0.91723 | -1.371785 | 0.488978 | -0.102097 | 0.202527 | 1.211164 |
| 27.72222 | 1.309001 | 2.593051 | 3.279573 | 1.835785 | 1.515134 | 1.279724 | 0.641481 | -1.311301 | 0.631703 | -0.066063 | -0.525216 | 1.589536 |
| 27.73611 | 1.018838 | 2.784715 | 0.448231 | 1.882942 | 1.109903 | 0.771373 | 0.102397 | -2.615659 | 0.191569 | 0.101146 | -0.712054 | 0.836332 |
| 27.75 | 0.798303 | 2.353997 | -1.8126 | 1.961939 | 0.795344 | 0.217145 | -0.218059 | -3.761419 | 0.207263 | 0.141414 | -1.028563 | 0.768613 |
| 27.76389 | 0.801584 | 1.893008 | -2.22908 | 1.352446 | 0.599092 | 0.256632 | -0.419923 | -3.697529 | 0.073473 | -0.4164 | -1.430102 | 1.066919 |
| 27.77778 | 0.807312 | 1.177973 | -1.425913 | 0.789048 | 0.43334 | -0.025109 | -0.147888 | -3.142715 | -0.812136 | -1.197428 | -1.166118 | 0.752951 |
| 27.79167 | 0.685194 | 0.477609 | -0.625545 | 0.639611 | 0.237288 | -1.415653 | -0.376541 | -2.65359 | -0.374192 | -0.273525 | -0.700572 | 0.737708 |
| 27.80556 | 0.368355 | -0.092001 | 0.459032 | 0.147703 | -0.099855 | -0.969641 | 0.005837 | -1.785524 | -0.08223 | -0.273857 | -0.491288 | 0.511375 |
| 27.81944 | 0.034019 | -0.142431 | 0.959986 | -0.80213 | -0.492123 | -0.267469 | 0.223558 | -0.781702 | -0.265467 | -0.934552 | -0.721335 | -0.380256 |
| 27.83333 | -0.169724 | -0.89848 | 1.228563 | -1.562317 | -0.888894 | -0.105836 | 0.218311 | 0.679278 | -0.549016 | -1.53924 | -0.72167 | -1.307583 |
| 27.84722 | -0.200927 | -1.866931 | 1.916231 | -2.006472 | -1.158436 | 0.093786 | 0.534748 | 2.298327 | -0.410988 | -1.18148 | -0.145019 | -0.636374 |
| 27.86111 | -0.078088 | -2.147453 | 2.318456 | -2.263927 | -1.0624 | 0.200575 | 1.455586 | 4.35008 | 0.02063 | -0.258867 | 0.73932 | 0.241053 |
| 27.875 | 0.071112 | -2.583788 | 2.654635 | -2.597057 | -0.817891 | 0.907588 | 2.218288 | 4.612831 | 0.52132 | 0.645168 | 1.324925 | 0.827613 |
| 27.88889 | 0.006345 | -2.321111 | -0.199604 | -2.72549 | -0.657606 | 1.544571 | 1.83127 | 4.312637 | 0.869521 | 1.101392 | 1.85048 | 0.487259 |
| 27.90278 | 0.07171 | -3.08657 | -1.353843 | -2.407036 | -0.591694 | 1.22457 | 1.259856 | 3.112241 | 1.144288 | 1.011612 | 2.124923 | -0.368674 |
| 27.91667 | 0.140648 | -3.334287 | -2.337279 | -2.592936 | -0.609147 | 0.834932 | 0.669524 | 1.318541 | 1.29386 | 0.75907 | 2.026262 | -0.388562 |
| 27.93056 | 0.067797 | -1.77558 | -2.689431 | -1.816122 | -0.720993 | 0.378995 | 0.172251 | 0.060864 | 1.063277 | 0.957985 | 1.742048 | 0.700358 |
| 27.94444 | -0.243408 | -0.886398 | -3.1618 | -1.178813 | -0.895186 | -0.402929 | -0.68052 | -1.206519 | 1.187142 | 1.64979 | 1.190901 | 1.122315 |
| 27.95833 | -1.050275 | -1.03861 | -2.206528 | -1.622843 | -1.639437 | -1.91069 | -2.345561 | -3.652496 | -0.089987 | 0.085754 | 0.397918 | 0.631075 |

| | | | | | | | | | | | | | | | | | | | | | | | | |
|---|---|---|---|---|---|---|---|---|---|---|---|---|---|---|---|---|---|---|---|---|---|---|---|---|
| 27.97222 | 0.133974 | -1.616613 | -0.450028 | -1.260089 | 0.355479 | -0.204075 | -1.122388 | -0.143409 | 1.392529 | 0.306527 | -0.957591 | -0.468138 | -1.790325 | -1.482446 | -1.864911 | -2.720156 | -1.317358 | -2.62305 | -4.130755 | -7.37677 | -1.009556 | -1.055492 | -0.492439 | -0.375275 |
| 27.98611 | 0.193188 | -1.672858 | -0.477634 | -0.60536 | 0.097518 | -0.623481 | -1.557169 | -0.072274 | 1.615028 | 0.864988 | -0.74335 | -1.429845 | -0.853506 | -1.942839 | -1.596649 | -3.04358 | -0.243946 | -1.742322 | -3.923376 | -6.851986 | -1.217337 | -2.283233 | -0.548894 | -1.165017 |
| 28 | 0.14602 | -0.381076 | -0.415739 | -0.055649 | -0.140091 | -0.947874 | -1.979168 | 1.043484 | 1.438049 | 1.702752 | -0.425266 | -2.991441 | -0.358056 | -1.989411 | -0.843119 | -2.835511 | 0.272121 | -1.773148 | -3.115019 | -5.570939 | -1.134114 | -3.426845 | -1.729922 | -2.558932 |
| 28.01389 | 0.089793 | 0.822772 | -0.423336 | 0.146399 | -0.300668 | -0.728182 | -1.855386 | 2.939566 | 1.496948 | 2.22205 | -0.082692 | -2.136702 | -0.058863 | -1.245118 | -0.132046 | -2.722777 | 0.436241 | -1.352525 | -1.423364 | -2.457627 | -0.261409 | -1.760964 | -1.510158 | -1.588197 |
| 28.02778 | 0.048277 | 0.736345 | -0.503767 | 0.092583 | -0.487272 | -0.487705 | -1.105603 | 1.187668 | 0.920544 | 1.167059 | 0.18266 | -1.094601 | 0.205923 | -0.454051 | -0.479206 | -2.445486 | 0.404 | -0.517423 | -0.555119 | -0.635254 | 0.130452 | -0.397014 | -0.928349 | 0.268244 |
| 28.04167 | 0.014837 | 0.351402 | -0.532075 | -0.083349 | -0.805555 | -0.08889 | -0.258159 | -0.959034 | 0.381525 | -0.757547 | 0.397485 | -0.67345 | 0.489709 | -0.594302 | -1.380075 | -2.01877 | 0.401504 | -0.232907 | -0.380955 | 0.642782 | 0.415165 | 1.281733 | -0.533326 | 1.023036 |
| 28.05556 | 0.013247 | -0.084483 | -0.484616 | -0.144256 | -1.141281 | 0.1651 | 0.192596 | -0.840798 | -0.040267 | -2.470351 | 0.523573 | -0.684136 | 0.76369 | -1.556869 | -0.558844 | -1.927641 | 0.433493 | 0.087949 | -0.027511 | 0.574884 | 0.461979 | 0.412998 | -1.236976 | -0.129546 |
| 28.06944 | -0.031972 | -0.169903 | -0.438384 | 0.056219 | -1.106612 | 0.157237 | -0.010008 | -0.255939 | -0.254602 | -2.463667 | 0.468696 | -0.530203 | 0.865123 | -2.354108 | 0.186006 | -1.007253 | 0.521385 | 0.523115 | 0.19777 | -0.707504 | 0.361402 | -1.209974 | -2.252809 | -1.097852 |
| 28.08333 | -0.059463 | -0.053015 | -0.253112 | 1.004881 | 0.807023 | 0.404001 | -0.042496 | -1.371224 | 0.028527 | 0.486166 | 0.018909 | -0.050004 | 0.392567 | -1.823642 | 0.529221 | -0.156125 | 0.430658 | 0.484544 | -0.179535 | -2.21 | 0.362173 | -1.175184 | -1.273687 | -0.806381 |
| 28.09722 | -2.3347 | 0.778882 | -0.435698 | 3.449077 | 5.19277 | 1.353035 | 0.810439 | -5.71292 | 1.001016 | 4.540979 | 0.873819 | 0.068897 | -0.061351 | -0.71926 | 0.479523 | 0.243071 | 0.31233 | -0.356554 | -0.682525 | -2.799972 | 0.174902 | -0.36165 | 0.78023 | -0.010431 |
| 28.11111 | -4.688369 | 1.283214 | 2.36651 | 3.579229 | 3.688761 | 2.31777 | -0.022601 | -7.941232 | -0.265773 | -1.313004 | 3.980654 | -0.935475 | -0.27646 | 0.339114 | -0.279794 | 0.270655 | 0.217959 | -0.954736 | -0.997444 | -2.48502 | -0.250516 | 0.378359 | 2.905769 | 0.614694 |
| 28.125 | 3.774258 | -3.100946 | -5.203876 | -5.283333 | -6.771593 | -1.532213 | -0.232339 | 4.112667 | -0.489652 | -0.892522 | -4.837276 | -0.348236 | -0.339871 | 1.115043 | -0.487752 | -0.025429 | 0.122409 | -1.231485 | -1.320318 | -1.772122 | -0.465223 | 0.904544 | 3.933337 | 1.360155 |
| 28.13889 | 0.654206 | -0.105423 | -0.723162 | -2.991781 | -5.733572 | -1.885517 | -0.44886 | 4.65865 | -0.851275 | -6.377429 | -1.060833 | -0.024702 | -0.140571 | 1.012475 | -0.115606 | -0.464746 | -0.053152 | 0.005463 | -0.620754 | -1.35208 | -0.359987 | 0.133263 | 3.013879 | 1.321952 |
| 28.15278 | -0.000082 | 0.561893 | -0.444477 | -0.614383 | -1.648774 | -0.595347 | -0.128665 | 1.283104 | -0.213319 | -2.171734 | -0.350144 | 0.013047 | -0.050217 | 0.418675 | 0.182409 | -0.596212 | -0.39905 | 0.459469 | -0.006324 | -1.299323 | -0.238945 | -1.17181 | 1.828777 | -0.022047 |
| 28.16667 | 0.057428 | 0.191559 | -0.067069 | 0.354828 | -0.392762 | -0.195669 | -0.133356 | 0.289564 | 0.149969 | 0.494186 | -0.094172 | -0.053287 | 0.917943 | 0.07614 | 0.304543 | -0.171504 | -1.372517 | -0.072599 | 0.728048 | -0.992176 | -0.684575 | -1.684583 | 0.838627 | -1.319257 |
| 28.18056 | 0.157757 | -0.061529 | 0.134587 | 0.603033 | -0.174423 | -0.218476 | -0.05091 | 0.106266 | 0.165606 | 1.217604 | 0.046329 | -0.118678 | 1.158025 | -0.518714 | -0.255234 | -1.113031 | -3.054698 | -0.926328 | 0.341596 | -1.523784 | -0.738526 | -1.241494 | 0.868691 | -1.858295 |
| 28.19444 | 0.322825 | -0.182448 | 0.216154 | 1.025896 | 0.080193 | -0.148917 | 0.100271 | 0.087937 | 0.091041 | 1.565718 | 0.15503 | -0.069646 | -1.027058 | -0.993921 | 0.290472 | -1.295919 | -1.443804 | 1.886511 | 0.877534 | -3.030357 | -1.015897 | -0.358799 | -0.364071 | -0.596587 |
| 28.20833 | 0.454468 | 0.065734 | 0.263457 | 1.580569 | 0.376667 | 0.034973 | 0.212373 | 0.199643 | 0.143797 | 1.830093 | 0.261537 | 0.212068 | -1.471967 | -1.16285 | -1.334378 | -0.286873 | 1.834525 | 2.924637 | -0.102265 | -2.34459 | -0.291849 | -0.1869 | -1.684256 | 0.44631 |
| 28.22222 | 0.412209 | 0.527531 | 0.328253 | 1.894045 | 0.50995 | 0.132203 | 0.245931 | 0.301555 | 0.274942 | 1.785347 | 0.376292 | 0.580175 | -0.242507 | -0.679222 | -0.369527 | -0.223193 | 1.075719 | 0.258543 | -0.873352 | -0.843343 | -0.510448 | -0.311076 | -1.964626 | -0.337157 |
| 28.23611 | 0.23665 | 0.862304 | 0.425176 | 1.908292 | 0.354399 | 0.216296 | 0.288682 | 0.326555 | 0.367171 | 1.503802 | 0.568492 | 0.937075 | 0.483203 | -0.652037 | 0.305799 | -0.169148 | -0.027171 | -0.918722 | -0.372774 | -0.532825 | -0.562033 | -0.690586 | -1.500399 | -0.933803 |
| 28.25 | 0.057567 | 0.896336 | 0.569978 | 1.859617 | 0.001695 | 0.362358 | 0.387925 | 0.600826 | 0.41427 | 0.946057 | 0.726276 | 1.252215 | 0.573078 | -0.593729 | 0.932954 | -0.226226 | -0.161628 | -0.241014 | -0.158287 | -0.223514 | -0.708802 | -0.973744 | -1.388465 | -1.000931 |
| 28.26389 | 0.105185 | 0.83007 | 0.70297 | 2.032232 | -0.050766 | 0.582223 | 0.506442 | 0.458794 | 0.506098 | 0.351204 | 0.889079 | 1.495659 | 0.372699 | -0.255063 | 1.054826 | -0.082409 | -0.150002 | 0.327174 | -0.086684 | -0.068726 | -0.908592 | -0.976406 | -1.471096 | -1.10806 |
| 28.27778 | 0.369203 | 0.839491 | 0.854899 | 2.119477 | 0.330669 | 0.698816 | 0.58926 | 0.425585 | 0.687533 | 0.008818 | 1.062076 | 1.751054 | 0.018335 | -0.127343 | 0.254931 | 0.097055 | -0.167059 | 0.390974 | -0.059344 | -0.229189 | -0.714796 | -0.429618 | -1.473392 | -0.998408 |
| 28.29167 | 0.574274 | 0.961667 | 0.942646 | 1.809286 | 0.728133 | 0.759067 | 0.523691 | 0.740313 | 1.004315 | 0.24123 | 1.184907 | 1.849891 | -0.242446 | -0.167573 | 0.264786 | 0.379613 | -0.256755 | 0.433496 | 0.17692 | -0.239353 | -0.361286 | -0.109978 | -0.912625 | -0.564364 |
| 28.30556 | 0.652412 | 1.222107 | 1.035618 | 1.70187 | 0.988416 | 0.85425 | 0.525418 | 1.271396 | 1.1675 | 1.020281 | 1.268429 | 1.898246 | -0.407225 | -0.201214 | 1.214054 | 0.737426 | -0.260683 | 0.532204 | 0.576637 | 0.181995 | 0.162052 | 0.226282 | -0.068088 | -0.035743 |
| 28.31944 | 0.696851 | 1.379347 | 1.113822 | 1.424664 | 1.201885 | 1.02465 | 0.598648 | 1.412848 | 1.268896 | 1.989175 | 1.416451 | 1.90619 | -0.439274 | -0.262078 | 2.223633 | 1.086625 | -0.103459 | 0.451405 | 0.847846 | 0.60222 | 0.190432 | 0.451106 | 0.618796 | 0.293948 |
| 28.33333 | 0.699046 | 1.379169 | 1.219868 | 1.268397 | 1.375834 | 1.165753 | 0.708626 | 1.574679 | 1.165219 | 2.539666 | 1.445902 | 2.019564 | -0.374005 | -0.314102 | 1.15208 | 1.26254 | 0.083574 | 0.241456 | 0.692378 | 0.817825 | -0.049565 | 0.596172 | 1.073491 | 0.220834 |
| 28.34722 | 0.490938 | 1.156373 | 1.213638 | 1.1419 | 1.355061 | 1.065488 | 0.618869 | 1.495927 | 0.858668 | 2.380113 | 1.249587 | 1.98718 | -0.173109 | -0.240339 | -0.084725 | 1.1466 | 0.234872 | 0.15432 | 0.409211 | 1.006385 | -0.277009 | 0.755932 | 1.560406 | 0.191007 |
| 28.36111 | 0.439217 | 0.739405 | 1.074857 | 0.973214 | 1.198881 | 0.854406 | 0.534823 | 1.212245 | 0.456387 | 2.021945 | 0.935789 | 1.50622 | 0.098355 | -0.022508 | -0.469449 | 1.011227 | 0.301767 | 0.308539 | 0.248503 | 1.198042 | -0.105022 | 0.972272 | 2.140886 | 0.420505 |
| 28.375 | 0.554957 | 0.47713 | 1.0059 | 1.042269 | 1.140149 | 0.822257 | 0.707357 | 1.310673 | 0.163494 | 2.075854 | 0.69059 | 1.136381 | 0.384168 | 0.271708 | -0.300276 | 1.042658 | 0.317942 | 0.640184 | 0.299347 | 1.416339 | 0.226464 | 1.060331 | 2.464706 | 0.731568 |
| 28.38889 | 0.552967 | 0.407172 | 0.785572 | 1.12212 | 0.868498 | 0.680443 | 0.777855 | 1.08678 | -0.079223 | 1.859273 | 0.493649 | 0.784552 | 0.670895 | 0.653061 | 0.689943 | 1.070401 | 0.29679 | 0.920439 | 0.444228 | 1.672902 | 0.405275 | 0.785153 | 2.217933 | 0.789907 |
| 28.40278 | 0.487803 | 0.277356 | 0.602409 | 1.362854 | 0.623545 | 0.518137 | 0.546351 | 0.832553 | -0.304891 | 1.498 | 0.267734 | 0.291253 | 0.860514 | 0.696229 | 1.429878 | 1.148764 | 0.267275 | 1.024105 | 0.658936 | 1.980263 | 0.397785 | 0.384485 | 1.732895 | 0.540173 |
| 28.41667 | 0.626649 | 0.224802 | 1.048164 | 2.043219 | 0.772045 | 0.734312 | 0.532207 | 0.859972 | -0.303052 | 1.283915 | 0.067777 | -0.110191 | 0.939982 | 0.794296 | 2.26356 | 1.326822 | 0.287657 | 0.791194 | 0.845435 | 2.423586 | 0.329082 | 0.133556 | 1.39099 | 0.332564 |
| 28.43056 | 0.855576 | 0.344198 | 1.195434 | 2.2071 | 1.030412 | 1.095478 | 0.657475 | 1.10107 | 0.003961 | 1.21899 | 0.003322 | -0.463933 | 0.930056 | 0.869752 | 2.364051 | 1.329549 | 0.327075 | 0.496143 | 0.980614 | 2.15217 | -0.00669 | -0.110865 | 1.31243 | 0.229995 |
| 28.44444 | 0.897854 | 0.753504 | 1.144496 | 1.677927 | 1.129226 | 1.308607 | 0.597694 | 1.956459 | 0.573397 | 1.030366 | 0.103443 | -0.477896 | 0.884871 | 1.164462 | 2.157138 | 0.979872 | 0.05251 | 0.021106 | 1.035573 | 1.013002 | -0.824559 | -0.779352 | 1.204073 | 0.071553 |
| 28.45833 | 0.904103 | 1.154931 | 1.225392 | 1.376776 | 1.206724 | 1.489683 | 0.58141 | 2.173885 | 1.084107 | 0.75263 | 0.378369 | 0.016885 | 0.564188 | 0.864021 | 1.441068 | 0.595653 | -0.463466 | -0.379937 | 1.021943 | 0.267672 | -1.426541 | -2.187583 | 1.033523 | -0.100771 |
| 28.47222 | 1.000038 | 1.404725 | 1.397244 | 1.096922 | 1.411922 | 1.528927 | 0.676722 | 1.546645 | 1.454326 | 0.530034 | 0.72265 | 0.895285 | 0.260601 | 0.347642 | 0.861268 | 0.880507 | -0.448177 | -0.262996 | 1.00332 | -0.06863 | -1.522015 | -3.654787 | 0.704329 | -0.232202 |
| 28.48611 | 1.208834 | 2.134039 | 0.939648 | 0.713175 | 1.786762 | 1.448161 | 0.741653 | 1.166854 | 1.988923 | 0.384152 | 1.029943 | 1.523846 | 0.304471 | 0.354361 | 0.595494 | 1.619578 | -0.344101 | 0.07493 | 1.198386 | 0.468562 | -0.903609 | -2.962905 | 0.784776 | -0.198278 |
| 28.5 | 1.451101 | 2.333438 | 0.10883 | 0.546159 | 2.019444 | 1.375805 | 0.722992 | 1.793126 | 1.989736 | 0.200792 | 1.400099 | 1.455063 | 0.546095 | 0.964399 | 0.196545 | 2.216376 | 0.164891 | 0.535712 | 1.970573 | 1.753534 | 0.480514 | -0.580304 | 0.983602 | 0.384812 |
| 28.51389 | 1.495334 | 2.542428 | -0.065611 | 0.516663 | 1.795161 | 1.119053 | 0.652397 | 1.377131 | 1.837196 | 0.114506 | 1.203793 | 1.213019 | 0.59412 | 2.308129 | 0.086544 | 1.742611 | 0.762143 | 1.297271 | 2.897717 | 3.240463 | 1.615809 | 1.779545 | 1.211783 | 1.402452 |
| 28.52778 | 1.432383 | 1.913307 | 0.16911 | 0.635234 | 1.430065 | 0.729978 | 0.452807 | 1.52696 | 1.60242 | 0.129922 | 0.710718 | 1.038051 | 0.716427 | 2.942611 | 1.426388 | 2.318756 | 1.371484 | 2.235585 | 2.9731 | 4.387352 | 1.569037 | 4.177579 | 1.608573 | 2.556825 |
| 28.54167 | 1.51477 | 0.832305 | 0.649817 | 0.831134 | 1.186967 | 0.415366 | 0.309658 | 0.782311 | 1.389487 | 0.470225 | 0.40012 | 0.88601 | 1.194583 | 2.297417 | 2.554728 | 3.037432 | 2.017357 | 2.869177 | 2.26413 | 4.197245 | 0.612732 | 2.203195 | 2.140211 | 2.682013 |
| 28.55556 | 1.739323 | 0.199322 | 1.480212 | 0.848568 | 1.238148 | 0.158504 | 0.306301 | 0.691507 | 1.466853 | 1.080121 | 0.402834 | 1.336672 | 1.572348 | 1.531038 | 2.555131 | 2.678409 | 2.518139 | 2.350097 | 1.631731 | 3.616356 | 0.972972 | 3.657058 | 1.60992 | 2.385799 |
| 28.56944 | 2.007079 | 0.239196 | 1.606333 | 0.652101 | 1.336229 | -0.012662 | 0.402641 | 2.494876 | 1.371302 | 1.504275 | 0.627275 | 1.463842 | 1.74069 | 0.480679 | 1.850119 | 1.895098 | 2.359554 | 1.717377 | 1.382223 | 2.91328 | 1.908715 | 3.220002 | 0.652895 | 1.391742 |
| 28.58333 | 2.110647 | 0.499968 | 1.505482 | 0.382295 | 1.327676 | 0.030623 | 0.523637 | 3.03238 | 1.023123 | 1.132407 | 0.852213 | 1.229932 | 1.675228 | -0.232102 | 1.046256 | 2.94574 | 2.015649 | 0.682453 | 0.546338 | 1.604747 | 1.948783 | 1.550744 | 0.014341 | 0.494798 |
| 28.59722 | 1.650684 | 0.675086 | 1.255201 | 0.329523 | 1.212907 | 0.462415 | 0.79024 | 2.809735 | 0.798882 | 1.143033 | 1.054645 | 0.913505 | 1.559972 | -0.741943 | 0.331285 | 1.399327 | 1.781645 | 0.467678 | 0.066052 | 0.46769 | 0.723974 | 0.270971 | -0.183266 | 0.015818 |
| 28.61111 | 1.149139 | 1.124987 | 1.09671 | 0.485385 | 0.811041 | 0.932891 | 1.098118 | 2.991709 | 0.752389 | 1.272832 | 1.124758 | 0.656215 | 1.366277 | -0.699801 | -0.173573 | 0.352736 | 1.232978 | 0.266872 | 0.861588 | 0.659732 | 0.055403 | -0.324867 | -0.530211 | -0.093622 |
| 28.625 | 0.92977 | 0.512239 | 1.091182 | 0.456386 | 0.344345 | 1.086695 | 1.288505 | 3.294632 | 1.074808 | 1.394904 | 0.958878 | 0.642697 | 1.182184 | -0.499432 | -0.288565 | 0.044659 | 0.707669 | 0.470072 | 1.034166 | 1.451665 | 0.103346 | -0.123348 | -0.576184 | 0.234089 |
| 28.63889 | 0.802242 | 0.658224 | 0.841341 | 0.321216 | 0.110577 | 0.90177 | 1.393634 | 3.2335 | 0.798097 | 1.37061 | 1.000875 | 0.796915 | 1.140963 | -0.497051 | 0.133776 | 0.375582 | 0.479232 | 1.222459 | 1.616967 | 2.298885 | 0.299176 | 0.258457 | 0.426892 | 0.632526 |
| 28.65278 | 0.666797 | 0.540897 | 0.541558 | 0.095788 | 0.2408 | 0.870279 | 1.510591 | 2.411273 | 0.658662 | 1.254672 | 1.130072 | 0.834066 | 1.207677 | 0.073107 | 0.53972 | 0.576332 | 0.374746 | 1.727591 | 2.114656 | 3.93598 | 0.685141 | 0.910262 | 1.325963 | 1.046059 |
| 28.66667 | 0.493077 | 0.29555 | 0.68326 | -0.092353 | 0.572854 | 0.630917 | 1.489255 | 1.973374 | 0.42861 | 0.98791 | 1.255556 | 0.740262 | 1.317816 | 1.123569 | 0.747327 | 1.091999 | 0.231486 | 1.493822 | 2.23275 | 5.394261 | 1.260579 | 3.104611 | 1.84801 | 1.415943 |
| 28.68056 | 0.313841 | 1.510816 | 0.925188 | -0.068801 | 0.733131 | 0.700451 | 1.362305 | 1.526689 | 0.187877 | 0.68587 | 1.580906 | 0.966931 | 1.552701 | 2.000069 | 0.966716 | 2.505637 | 0.182249 | 0.54102 | 2.012856 | 3.807459 | 1.041306 | 2.718432 | 2.765645 | 1.956986 |
| 28.69444 | 0.43662 | 1.279488 | 0.862521 | -0.011775 | 0.579561 | 0.669241 | 1.420815 | 0.128958 | -0.310504 | 0.623801 | 1.636129 | 1.395961 | 1.788457 | 2.228177 | 0.791374 | 2.453396 | 0.251172 | 0.127609 | 1.211237 | 1.442611 | 0.793435 | 1.322039 | 2.301196 | 2.060319 |
| 28.70833 | 0.613543 | 0.864154 | 0.228545 | 0.022713 | 0.321376 | 0.604682 | 1.311159 | -0.974103 | -0.406665 | -0.135487 | 1.172475 | 1.588222 | 1.781824 | 1.747909 | 0.496717 | 2.735142 | 0.264682 | 0.047407 | 0.532627 | -0.329191 | 0.401558 | 0.332945 | 1.353343 | 1.446833 |
| 28.72222 | 0.690871 | 1.995532 | -0.318693 | 0.117729 | 0.1686 | 0.662827 | 0.663993 | -1.846101 | -0.13469 | -0.936363 | 0.501751 | 1.517096 | 1.646154 | 1.439631 | 0.182947 | 2.082934 | 0.193845 | 1.082909 | -0.252244 | -2.080618 | 0.053934 | -0.245612 | 0.379995 | 0.613412 |
| 28.73611 | 0.775689 | 0.682396 | -0.710575 | 0.071749 | 0.176961 | 0.700278 | 0.174913 | -1.172951 | 0.095489 | -0.856334 | 0.224981 | 1.40978 | 1.503971 | 1.356271 | 0.207962 | 1.277572 | 0.070488 | 1.914572 | -1.113587 | -3.563051 | -0.523379 | -0.692231 | -0.317902 | -0.101426 |
| 28.75 | 0.821515 | 1.320366 | -0.888953 | -0.138988 | -0.040732 | 0.71949 | 0.155821 | -0.487966 | 0.224126 | -0.440466 | 0.57309 | 1.475485 | 1.340003 | 1.039933 | 0.433843 | 1.155974 | -0.122468 | 0.733041 | -1.183124 | -4.388271 | -0.831235 | -1.087489 | -0.430557 | -0.5155 |
| 28.76389 | 0.750431 | 1.705007 | -1.097984 | -0.497999 | -0.477872 | 0.603168 | 0.151499 | 0.213378 | 0.166407 | 0.499166 | 0.705601 | 1.22512 | 1.22096 | 0.73773 | 0.905807 | 2.006512 | -0.270949 | -0.492198 | -1.398688 | -4.268462 | -1.238269 | -1.545643 | -0.330236 | -0.554459 |
| 28.77778 | 0.431587 | 1.5939 | -0.973276 | -0.902352 | -0.901176 | 0.445821 | -0.190554 | 0.97909 | -0.117893 | 1.164673 | 0.424243 | 0.908141 | 0.961241 | 0.080254 | 0.718757 | 2.141335 | 0.000871 | -0.932864 | -0.95371 | -4.830328 | -1.271872 | -3.471994 | -0.714724 | -0.715936 |
| 28.79167 | 0.255138 | 1.236046 | -1.114906 | -0.952389 | -1.120336 | 0.105307 | -0.272709 | 1.534981 | -1.410349 | 1.332068 | -0.197052 | 0.584151 | 0.780801 | -0.463866 | -0.214209 | 1.504311 | 0.321626 | -0.364968 | -0.970435 | -4.456763 | -1.096613 | -2.388052 | -1.082984 | -0.655728 |
| 28.80556 | 0.591837 | 0.676187 | -0.922512 | -1.223237 | -0.76287 | -0.352898 | -0.246947 | 2.043367 | -1.195299 | 0.618393 | -0.804369 | 0.239428 | 0.941056 | -0.292573 | -0.94582 | 0.962887 | 0.158616 | -0.423741 | -1.599993 | -4.310481 | -0.567079 | -2.801883 | -1.389225 | -0.383181 |
| 28.81944 | 0.890113 | 0.098321 | -0.711201 | -1.339781 | -0.385401 | -0.634759 | -0.21271 | 3.463881 | -0.857361 | 0.556484 | -1.295241 | -0.153605 | 1.076663 | -0.84268 | -0.781441 | 0.278426 | -0.040236 | -0.531329 | -1.532583 | -4.505628 | 0.44962 | -2.360953 | -1.500571 | -0.297722 |
| 28.83333 | 0.851119 | -0.183611 | -0.420772 | -1.324899 | 0.164333 | -1.127689 | -0.16355 | 4.63761 | -0.734281 | 0.59136 | -1.88486 | -0.905201 | 1.027305 | -1.448137 | -0.574614 | -0.239439 | -0.11457 | -0.959635 | -0.942857 | -3.389415 | 0.185236 | -2.406767 | -0.86643 | -0.260137 |
| 28.84722 | 0.52213 | -0.32472 | -0.009647 | -1.311637 | 0.698809 | -1.492655 | 0.350374 | 3.346579 | -0.676533 | 0.117395 | -2.076003 | -1.535656 | 0.916812 | -2.380781 | 0.423863 | -1.169636 | -0.105561 | -0.429293 | 0.359455 | -0.65262 | -0.198118 | -2.16789 | -0.247482 | -0.249481 |
| 28.86111 | 0.027838 | -0.526716 | 0.071646 | -1.861983 | 0.817115 | -1.433706 | 1.169515 | 1.512365 | -1.220366 | 0.00861 | -1.491567 | -1.418368 | 0.72757 | -3.03745 | 1.362521 | -2.679316 | -0.01583 | 0.47118 | 1.253599 | 1.70049 | -0.117725 | -1.123275 | 0.523359 | 0.007693 |
| 28.875 | -0.566574 | -0.518316 | -0.553116 | -3.455616 | -0.119732 | -1.417933 | -0.380795 | -1.86239 | -2.252948 | -0.527987 | -0.972169 | -1.042291 | 0.47696 | -2.515916 | 1.330039 | -3.053285 | 0.111569 | 1.383185 | 1.828874 | 2.380335 | 0.200946 | 0.012322 | 1.590575 | 0.654814 |
| 28.88889 | -0.649109 | -1.856321 | -1.053704 | -4.81132 | -1.582188 | -1.471009 | -1.979343 | -5.399633 | -2.586254 | -1.667276 | -0.927049 | -0.393126 | 0.298917 | -1.947836 | 1.084694 | -2.881231 | 0.083225 | 2.605351 | 2.06235 | 2.461842 | 0.237414 | 1.383516 | 2.228396 | 1.249751 |
| 28.90278 | -1.67194 | -2.507838 | -0.208433 | -2.857132 | -2.058164 | -1.307259 | -2.593451 | -4.485347 | -2.478729 | -2.978798 | -0.167067 | 0.236919 | 0.316591 | -1.79358 | -0.482465 | -2.811051 | -0.010102 | 2.734423 | 2.219319 | 3.598417 | -0.029199 | 1.588006 | 2.309208 | 1.114515 |
| 28.91667 | -2.691162 | -3.165314 | 0.677629 | -0.481672 | -1.923338 | -1.042299 | -2.447613 | 0.687766 | -2.190799 | -3.978664 | 0.289897 | 0.485867 | 0.204587 | -2.338067 | -2.06958 | -2.836964 | 0.030419 | 1.630657 | 1.51578 | 4.10132 | -0.291815 | 0.677832 | 1.554578 | 0.221814 |
| 28.93056 | -2.44638 | -1.002158 | 0.584091 | -0.233783 | -1.323168 | -0.787297 | -1.22985 | 3.777565 | -1.616353 | -3.224717 | 0.47039 | 0.464073 | -0.458875 | -2.65021 | -2.171916 | -2.501304 | 0.565981 | 1.44951 | 1.169255 | 3.42286 | 0.572132 | 0.084769 | 0.538883 | -0.069983 |
| 28.94444 | -1.808295 | -1.675129 | -0.147413 | -1.725609 | -0.635874 | -0.504009 | -0.608433 | 1.969135 | -1.288163 | -2.948765 | 0.465382 | 0.116935 | -1.54539 | -2.366785 | -1.567134 | -2.80856 | 1.365246 | 1.302222 | 0.373311 | 0.442764 | 0.621417 | 0.033581 | -0.617204 | 0.98158 |
| 28.95833 | -1.664569 | -2.35821 | -0.485543 | -2.629073 | -0.634361 | -0.45963 | -0.098485 | 0.303814 | -1.147374 | -3.093002 | 0.189202 | -0.507785 | -1.818013 | -0.805008 | 0.08531 | -2.803571 | 1.011968 | 0.117591 | -0.962432 | -3.78964 | 0.169124 | -0.468134 | -1.188111 | 1.286996 |
| 28.97222 | -1.345332 | -2.557354 | -0.4872 | -2.48458 | -0.718132 | -0.073428 | -0.113194 | 0.017979 | -1.01072 | -2.741597 | -0.26746 | -1.084785 | -1.354583 | -0.954687 | -0.811722 | -2.695967 | 0.328654 | -0.917399 | -2.957362 | -6.458934 | -0.143995 | -1.257051 | -2.173377 | 0.322213 |
| 28.98611 | -0.622534 | -2.440896 | -0.336445 | -2.179322 | -0.627532 | 0.474892 | -0.123882 | -0.09848 | -0.597263 | -1.020572 | -0.572981 | -1.059643 | -0.608461 | -1.880368 | -0.909168 | -1.552788 | -0.064602 | -1.16857 | -3.705448 | -7.130664 | 0.616528 | -2.202443 | -2.759021 | -1.322554 |
| 29 | -0.086772 | -1.420973 | -0.274362 | -1.707609 | -0.272547 | 0.403306 | -0.190243 | 2.00491 | -0.093604 | 1.558814 | -0.339245 | -0.923569 | -0.478655 | -2.090428 | -0.517765 | -2.874993 | -0.112856 | -1.015623 | -1.964402 | -6.862821 | 0.928074 | -3.598718 | -2.889221 | -2.215305 |
| 29.01389 | -0.180461 | 0.049446 | -0.217059 | -1.366446 | -0.431948 | -0.106753 | -0.316056 | 2.832696 | 0.395905 | 3.654195 | 0.031467 | -0.563678 | -0.493567 | -1.742638 | 0.995815 | -2.02168 | -0.006653 | -0.156001 | 0.169738 | -6.131098 | -0.228642 | -3.479447 | -4.56988 | -2.110219 |
| 29.02778 | -0.546899 | 0.642244 | -0.44374 | -1.121298 | -0.719023 | -0.733389 | -0.377888 | 1.905388 | 0.592594 | 2.728246 | 0.195399 | -0.240669 | -0.613377 | -0.90159 | 0.684628 | -2.000089 | 0.004011 | 0.074843 | 0.649963 | -2.954352 | -0.617586 | -0.581687 | -3.970315 | -0.493989 |
| 29.04167 | -0.71213 | 0.425791 | -0.687882 | -0.998172 | -0.749756 | -1.034965 | -0.440437 | 0.605719 | 0.551544 | 2.682267 | 0.120333 | -0.138832 | -0.520708 | -0.35192 | 0.781838 | -1.743349 | -0.156532 | -0.193761 | -0.544391 | -0.183098 | -0.203965 | 1.778442 | -1.429749 | 0.814714 |
| 29.05556 | -0.676528 | 0.048155 | -0.989739 | -0.892022 | -0.529518 | -0.766646 | -0.670199 | -0.485662 | 0.440613 | 2.170356 | -0.132042 | -0.324997 | -0.206134 | -0.25573 | -0.150122 | -1.793546 | -0.243054 | -0.29833 | -1.833977 | 0.976461 | 0.57007 | 0.903042 | 0.936501 | 0.908502 |
| 29.06944 | -0.698391 | -0.033229 | -1.223154 | -0.805466 | -0.35585 | -0.470543 | -0.743188 | -0.553696 | 0.259775 | 1.232933 | -0.48914 | -0.784122 | 0.191516 | -0.355033 | -2.013631 | -1.722425 | -0.136082 | 0.037012 | -2.576778 | -0.001693 | 1.159051 | -0.257238 | 2.538316 | 0.214611 |
| 29.08333 | -0.720576 | 0.011487 | -1.300602 | -0.752219 | -0.295095 | -0.678104 | -0.695367 | 0.503188 | 0.127992 | -0.258314 | -0.800947 | -1.177534 | 0.292245 | -0.380977 | -1.654048 | -1.530288 | -0.007475 | -0.07787 | -1.574702 | -1.569256 | 0.928649 | -0.626763 | 3.576548 | -0.183427 |
| 29.09722 | -0.588182 | 0.077037 | -1.272777 | -0.803689 | -0.253766 | -0.840208 | -0.975919 | 0.865421 | 0.073984 | -1.786445 | -0.787611 | -1.00424 | 0.174971 | -0.162793 | -1.038903 | 0.098852 | 0.115487 | -0.267161 | -0.940804 | -2.018219 | 0.969531 | -0.491578 | 3.291061 | -0.424231 |
| 29.11111 | -0.189122 | -0.285251 | -1.187909 | -0.835479 | -0.261142 | -0.57708 | -0.854352 | 0.110324 | -0.018207 | -2.964513 | -0.613445 | -0.5728 | 0.097436 | 0.11362 | 0.48039 | -0.070932 | 0.11246 | -0.503202 | -0.949902 | -1.0447 | 0.669232 | -0.447774 | 2.447604 | -0.480083 |
| 29.125 | 0.172695 | -0.76541 | -1.046255 | -0.857536 | -0.323533 | -0.745905 | -0.80231 | -0.479167 | -0.193317 | -3.379198 | -0.524542 | -0.386441 | 0.038564 | 0.604729 | 1.157056 | -0.178441 | 0.100581 | -0.570585 | -1.009899 | -0.147509 | 0.525688 | -0.748166 | 1.214358 | -0.312311 |
| 29.13889 | 0.223404 | -1.221396 | -0.893696 | -0.720189 | -0.484181 | -0.783904 | -0.687119 | -0.410347 | -0.377415 | -3.047451 | -0.490994 | -0.479783 | -0.253801 | 0.530785 | 0.431922 | -0.132968 | 0.118714 | -0.565673 | -0.791202 | 0.416874 | 0.234771 | -1.256797 | 0.212508 | -0.483449 |
| 29.15278 | -0.057764 | -1.073606 | -0.845915 | -0.448189 | -0.696833 | -0.618807 | -0.560775 | -0.545263 | -0.514504 | -1.697726 | -0.37334 | -0.565438 | -0.396511 | 0.087634 | -0.685056 | -0.421488 | 0.053673 | -0.751532 | -0.083886 | 0.60467 | -0.166066 | -2.035376 | -0.334781 | -1.034914 |
| 29.16667 | -0.381994 | -0.303135 | -0.491747 | 0.004484 | -0.788644 | -0.442595 | -0.393676 | -0.608094 | -0.639106 | -0.245891 | -0.301121 | -0.656811 | -0.474605 | 0.065779 | -0.15735 | -0.346579 | -0.027207 | -0.579431 | 0.123583 | 0.068592 | -0.594439 | -2.212886 | -0.724053 | -1.402995 |
| 29.18056 | -0.308078 | 0.56749 | -0.008061 | 0.427885 | -0.623759 | -0.199551 | -0.177632 | -0.409351 | -0.781225 | 1.070482 | -0.249518 | -0.721003 | -0.422392 | 0.496523 | 0.328828 | 0.084124 | -0.113 | -0.10518 | -0.399254 | -1.115722 | -0.854539 | -0.722761 | -0.933684 | -0.754511 |
| 29.19444 | 0.24505 | 0.837564 | 0.314846 | 0.732267 | -0.294657 | -0.041289 | 0.027895 | 0.231505 | -0.900978 | 2.179744 | -0.214882 | -0.547112 | -0.250142 | 0.782219 | -0.408908 | 0.064226 | -0.157546 | 0.144625 | -0.863289 | -1.770755 | -0.565767 | 0.394764 | -0.912294 | 0.153247 |
| 29.20833 | 0.784043 | 0.652125 | 0.481294 | 1.015494 | 0.01008 | 0.075212 | 0.160926 | 0.735318 | -0.641104 | 3.467031 | -0.183539 | -0.251826 | -0.379238 | 0.578869 | -2.317786 | -0.378883 | -0.155812 | 0.141404 | -0.68358 | -1.984736 | -0.164013 | 1.55069 | -0.891144 | 0.621248 |
| 29.22222 | 1.04558 | 1.209938 | 0.773791 | 1.625503 | 0.762062 | -0.083455 | -0.073908 | 0.291975 | 0.698009 | 3.950997 | -0.62943 | -0.174609 | -0.47692 | 0.07739 | -2.43862 | -0.312079 | -0.213049 | 0.138142 | -0.565322 | -1.49419 | 0.152826 | 0.931576 | -2.065223 | 0.35279 |
| 29.23611 | 0.514099 | 2.214506 | 1.341201 | 2.965563 | 2.164923 | 0.578343 | -0.300957 | -0.463051 | 1.984797 | 4.375825 | -0.392085 | 0.049247 | -0.335193 | -0.15491 | -1.248015 | -0.723097 | -0.299313 | 0.169892 | -0.668735 | -1.108156 | 0.382024 | -0.204575 | -2.885429 | 0.118809 |
| 29.25 | -0.463869 | 1.360638 | 0.815715 | 3.027056 | -0.246396 | 0.534479 | 0.699359 | 1.917901 | -0.112262 | 2.424847 | 0.513495 | 1.083225 | -0.088364 | -0.089365 | 1.167111 | -0.733646 | -0.236035 | 0.28248 | -0.696779 | -1.111816 | 0.465203 | -0.404171 | -2.717884 | 0.195492 |
| 29.26389 | -0.709724 | -0.827829 | -0.088325 | -0.14678 | -2.764402 | 0.113402 | 0.949488 | 1.351145 | -2.251802 | -1.584833 | 1.422301 | 0.7683 | 0.070619 | 0.189255 | 2.589313 | -0.421078 | -0.088255 | 0.328723 | -0.416741 | -1.097258 | 0.423013 | -0.472639 | -1.711303 | 0.293762 |
| 29.27778 | -0.488044 | -0.16236 | 0.202501 | 0.415319 | -0.927996 | -0.033237 | 0.619893 | -0.2195 | -1.227988 | -1.16378 | 0.607767 | 0.407716 | 0.252194 | 0.136136 | 2.203343 | 0.030283 | -0.061856 | 0.276851 | -0.076561 | -1.432044 | 0.130713 | -0.773116 | -0.686152 | -0.135787 |
| 29.29167 | 0.342126 | 0.995084 | 0.784423 | 1.877346 | 0.022271 | 0.383817 | 0.559694 | 0.773718 | 0.341344 | 0.668404 | 0.280815 | 0.562231 | 0.384907 | -0.308684 | 0.636086 | 0.112591 | -0.129282 | 0.180648 | 0.075503 | -1.780911 | -0.072214 | -1.498906 | -0.375706 | -0.729429 |
| 29.30556 | 0.650006 | 1.10827 | 1.251442 | 2.50065 | -0.051555 | 0.472686 | 0.497848 | 0.829526 | 0.524021 | 0.781226 | 0.523717 | 0.662843 | 0.653484 | -0.552103 | 0.09036 | 0.106221 | -0.350182 | 0.077977 | 0.180799 | -1.78755 | -0.119406 | -2.024638 | -0.622728 | -1.272134 |
| 29.31944 | 0.581853 | 1.036371 | 1.611346 | 2.500222 | -0.204831 | 0.443988 | 0.387486 | 0.96859 | 0.487301 | 0.565747 | 0.73366 | 0.52797 | 0.94699 | -0.619939 | -0.166445 | 0.454028 | -0.606988 | -0.115563 | 0.163308 | -2.539974 | -0.050486 | -1.855933 | -1.003926 | -1.563243 |
| 29.33333 | 0.587635 | 1.084118 | 1.766827 | 2.39529 | 0.098402 | 0.636931 | 0.377896 | 1.013267 | 0.762405 | 1.143678 | 0.585455 | 0.698703 | 0.779917 | 0.858108 | -0.929118 | 1.409742 | -0.557955 | -0.157647 | 0.236395 | -1.853877 | 0.023988 | -0.289255 | 0.231019 | -0.781859 |
| 29.34722 | 0.620031 | 1.315962 | 1.785903 | 2.574928 | 0.502578 | 0.78303 | 0.34954 | 0.937975 | 0.969156 | 2.096686 | 0.427659 | 0.684235 | 0.482794 | 1.611883 | -1.386997 | 1.670717 | -0.140803 | -0.013831 | -0.005831 | 0.336995 | -0.019533 | 0.599353 | 1.616227 | 0.210832 |
| 29.36111 | 0.725828 | 1.582063 | 1.44023 | 2.417552 | 0.73787 | 0.751822 | 0.194568 | 0.700998 | 0.895655 | 2.595725 | 0.313371 | 0.204215 | 0.327233 | 1.4162 | -0.287777 | 1.008304 | 0.074947 | 0.193717 | -0.219906 | -0.30723 | -0.130719 | 0.598509 | 1.648017 | 1.025999 |
| 29.375 | 1.026769 | 1.644476 | 0.83605 | 2.453548 | 0.930877 | 0.686337 | 0.249206 | 0.651877 | 0.734668 | 2.765191 | 0.289223 | 0.27159 | 0.343267 | 0.793425 | 1.452634 | 0.605095 | 0.078149 | 0.525773 | -0.061992 | 0.140354 | -0.120362 | 0.235653 | 1.375144 | 0.855605 |
| 29.38889 | 1.13717 | 1.282946 | 0.456946 | 3.06296 | 0.755662 | 0.318326 | 0.391099 | 0.900808 | 0.472827 | 2.115576 | 0.348903 | 0.722224 | 0.428525 | 0.878297 | 2.583341 | 0.576777 | 0.081539 | 0.776743 | 0.349564 | 1.157854 | 0.043428 | 0.294214 | 1.100211 | 0.990744 |
| 29.40278 | 1.093643 | 0.692362 | 0.281167 | 3.599601 | 0.602716 | 0.035707 | 0.367928 | 1.213634 | 0.17359 | 1.12833 | 0.276174 | 0.746502 | 0.465228 | 1.201342 | 2.905864 | 1.432991 | 0.071576 | 0.676433 | 0.68679 | 2.703599 | 0.309265 | 0.791925 | 0.885731 | 0.660548 |
| 29.41667 | 1.324728 | 0.210377 | 0.399818 | 4.127647 | 0.755679 | 0.07026 | 0.404687 | 1.166166 | 0.476363 | 1.333172 | 0.367517 | 0.702907 | 0.415394 | 1.663332 | 2.910362 | 2.182387 | 0.042342 | 0.487731 | 0.769911 | 4.055412 | 0.440028 | 1.286241 | 0.958609 | 0.586508 |
| 29.43056 | 1.488077 | 0.086819 | 1.161324 | 4.145755 | 1.080924 | 0.39748 | 0.421078 | 0.967166 | 1.090517 | 1.754576 | 0.599516 | 0.933563 | 0.35856 | 1.017933 | 3.061998 | 2.99258 | 0.029695 | 0.339891 | 0.594766 | 3.943329 | 0.163234 | 0.774268 | 1.174419 | -0.147444 |
| 29.44444 | 1.354927 | 0.617032 | 1.316842 | 3.182861 | 1.174209 | 0.802444 | 0.475975 | 0.856915 | 1.241785 | 1.816481 | 0.917231 | 1.360914 | 0.3128 | 0.420363 | 3.09275 | 2.767373 | 0.499071 | 0.232304 | 0.304653 | 2.466929 | -0.207232 | -0.395017 | 1.090301 | 0.178627 |
| 29.45833 | 1.16259 | 1.216526 | 1.524526 | 1.962649 | 1.117236 | 1.11552 | 0.514454 | 0.627042 | 1.163763 | 1.477643 | 1.348219 | 1.780285 | 0.325421 | 0.014792 | 2.119705 | 1.702196 | 0.81954 | 0.21379 | 0.2233 | 1.419934 | -0.334953 | -1.564603 | 0.569745 | -0.4458 |
| 29.47222 | 0.91513 | 1.510278 | 2.089965 | 1.00699 | 1.061515 | 1.498873 | 0.631195 | 0.225926 | 1.167296 | 1.217732 | 1.889625 | 2.035296 | 0.487326 | 0.01431 | 1.672359 | 2.008783 | 0.888757 | 0.361643 | 0.32871 | 1.185701 | -0.229552 | -2.416069 | 0.441406 | -0.611031 |
| 29.48611 | 0.899494 | 3.029076 | 2.124415 | 0.11999 | 0.788727 | 1.696332 | 0.70861 | 0.03164 | 1.337598 | 0.940243 | 2.400553 | 1.810039 | 0.79696 | 0.830684 | 1.383867 | 3.542243 | 0.885225 | 0.74261 | 0.732018 | 1.637763 | 0.202389 | -0.823833 | 0.747866 | 0.686055 |
| 29.5 | 0.813585 | 3.306465 | 1.148187 | -0.470197 | 0.002671 | 1.707879 | 0.663282 | 0.47365 | 1.438242 | 0.530246 | 2.107966 | 1.340277 | 0.960097 | 1.81536 | 2.103691 | 3.138352 | 1.333184 | 1.213748 | 1.548117 | 2.999598 | 0.432268 | 0.614542 | 1.332388 | 1.376154 |
| 29.51389 | 0.75484 | 2.741667 | 0.278877 | -0.753473 | 0.406208 | 1.583568 | 0.7343 | 0.889991 | 1.283104 | 0.222846 | 1.658798 | 1.308381 | 0.9707 | 1.574565 | 3.10912 | 2.115297 | 1.313223 | 1.080752 | 2.247802 | 3.82502 | 0.845426 | 2.634322 | 2.596341 | 1.269265 |
| 29.52778 | 0.811589 | 2.303259 | -0.85524 | -1.748977 | 0.354623 | 1.364606 | 0.785766 | 0.976035 | 0.982405 | -0.061471 | 1.385947 | 1.422509 | 1.020622 | 1.636762 | 3.771316 | 1.731934 | 0.157758 | 1.284846 | 3.228131 | 4.042835 | 0.606361 | 3.641087 | 3.918941 | 3.093683 |
| 29.54167 | 1.136095 | 1.350615 | 0.30841 | 0.12326 | 0.230626 | 1.03191 | 0.726487 | 1.360416 | 0.622101 | -0.212847 | 1.337403 | 1.406924 | 1.162473 | 1.359974 | 4.663688 | 1.471207 | 0.026149 | 1.204414 | 2.998295 | 3.170193 | 0.572764 | 1.91055 | 4.351559 | 3.675909 |
| 29.55556 | 1.359403 | 0.22718 | 1.45934 | 2.348113 | 0.913371 | 0.866662 | 0.491015 | 2.14889 | 0.56341 | 0.22672 | 1.058125 | 1.411155 | 1.168368 | 0.901613 | 4.918359 | 1.746069 | 0.229129 | 0.401393 | 1.981301 | 2.610627 | 0.56654 | 3.355415 | 3.408098 | 2.932103 |
| 29.56944 | 1.399367 | -0.732661 | 1.097982 | 1.910213 | 1.731972 | 0.889197 | 0.514122 | 2.507923 | 0.734011 | 1.138219 | 0.401739 | 1.053945 | 0.887329 | 0.216039 | 2.840776 | 1.907975 | -0.203038 | -0.201084 | 1.49523 | 2.771739 | 0.591741 | 2.016784 | 1.440015 | 1.384244 |
| 29.58333 | 1.22024 | -1.708568 | 0.805125 | 1.43207 | 2.017974 | 0.905481 | 0.749567 | 1.619906 | 0.911665 | 1.495526 | -0.375839 | 0.557057 | 0.581841 | -0.317187 | 1.222486 | 1.372631 | -0.914127 | -0.256781 | 1.123934 | 1.954347 | 0.506385 | 0.421509 | 0.10829 | 0.024336 |
| 29.59722 | 1.135335 | -0.987856 | 0.407404 | 1.569327 | 1.677326 | 0.81095 | 0.97687 | 0.989103 | 1.028559 | 1.185849 | -0.657802 | 0.064645 | 0.330482 | -1.81651 | 0.058625 | 0.847492 | -0.745763 | 0.253581 | 0.535389 | 0.718073 | 0.330262 | -0.731644 | -0.982613 | -1.172179 |
| 29.61111 | 1.307336 | 0.616451 | 0.248833 | 2.112831 | 0.640066 | 0.728566 | 0.768104 | -0.633568 | 1.143539 | 1.001012 | -0.684223 | -0.221806 | 0.155639 | -2.69325 | -1.385966 | 0.639177 | -0.685522 | 0.774937 | 0.248617 | -0.283887 | 0.17617 | -1.523286 | -1.692475 | -1.515552 |
| 29.625 | 1.456399 | 1.681999 | 0.356009 | 2.563551 | 0.881002 | 0.664799 | 0.737771 | 0.860411 | 1.318738 | 1.016073 | -0.680277 | -0.37151 | 0.086731 | -1.640046 | -3.279613 | 0.609582 | -0.800245 | 0.821408 | 0.651068 | -0.016037 | 0.07637 | -1.517654 | -1.651952 | -1.372651 |
| 29.63889 | 1.549591 | 1.825726 | 0.522375 | 1.981713 | 1.411234 | 0.574621 | 0.95976 | 0.931518 | 1.800617 | 0.842985 | -0.2435 | -0.268463 | 0.114408 | -0.681858 | -3.141766 | 0.998739 | -0.184481 | 0.899895 | 1.620416 | 0.917091 | -0.053789 | -0.587061 | -1.661543 | -1.000062 |
| 29.65278 | 1.487371 | 1.481907 | 0.823056 | 0.812056 | 2.077726 | 0.422497 | 1.730987 | 0.04244 | 1.927723 | 0.784234 | -0.041429 | -0.185305 | 0.210736 | 0.03323 | 0.248118 | 1.61104 | 0.684152 | 1.238425 | 2.44548 | 0.986388 | 0.101503 | 0.550602 | -1.185396 | -0.0953 |
| 29.66667 | 1.517082 | 1.524817 | 1.095297 | -0.350835 | 2.232026 | 0.46734 | 1.68801 | -0.047309 | 1.600269 | 1.076655 | 0.215602 | 0.125573 | 0.33496 | 1.174705 | 3.46058 | 2.288326 | 0.787058 | 1.285421 | 2.187442 | 1.332853 | 0.612748 | 2.278957 | -0.045473 | 1.463857 |
| 29.68056 | 1.100383 | 1.827852 | 1.229949 | -0.619571 | 3.123978 | 0.608785 | 1.555914 | -1.258845 | 1.116686 | 1.180733 | 0.913285 | 0.295198 | 0.540108 | 2.248492 | 2.399874 | 3.408757 | 0.557156 | 1.194217 | 1.582953 | 1.948109 | 1.781571 | 3.747689 | 0.882085 | 3.031726 |
| 29.69444 | 0.278788 | 3.465071 | 1.225092 | -0.696242 | 2.462642 | 0.716145 | 1.986996 | -1.400285 | 0.555652 | 1.149124 | 1.254748 | 0.327061 | 0.810188 | 2.046721 | -0.057689 | 3.569791 | 0.556211 | 1.156183 | 1.381712 | 2.120467 | 0.82236 | 1.699167 | 2.095859 | 2.352068 |
| 29.70833 | -0.266675 | 3.532221 | 1.301822 | -0.897529 | 0.349381 | 0.587934 | 2.832386 | 0.522416 | -0.185894 | 0.98411 | 1.184674 | 0.27687 | 0.997285 | 1.117113 | -0.550205 | 2.298486 | 0.926754 | 1.204905 | 1.547671 | 1.605492 | -1.355138 | -0.551018 | 2.049463 | 0.399724 |
| 29.72222 | -0.705918 | 1.281255 | 1.045275 | -1.256438 | -0.07556 | 0.376316 | 3.087661 | -0.08976 | -0.375794 | 0.860635 | 0.980042 | 0.462582 | 0.979442 | 0.755514 | 0.921846 | 1.54785 | 0.811717 | 1.142437 | 1.422615 | 1.433119 | -1.634118 | -1.377764 | 1.809935 | -0.525615 |
| 29.73611 | -1.169805 | -1.127428 | 0.473384 | -1.85611 | -1.151483 | 0.282023 | 1.107415 | 0.57134 | 0.034616 | 0.456868 | 1.189563 | 0.661468 | 0.792631 | 0.809055 | 2.305484 | 1.860089 | 0.01747 | 0.848853 | 0.981993 | 1.533049 | -0.591252 | -1.125322 | 1.948462 | -0.658062 |
| 29.75 | -1.321039 | -2.079674 | -0.188503 | -1.76799 | -2.728798 | 0.21988 | -0.478518 | -0.262147 | -0.02089 | 0.443853 | 1.504291 | 0.729336 | 0.577847 | 0.725939 | 3.035992 | 2.300954 | -0.684061 | 0.188053 | 0.544325 | 0.970679 | 0.472972 | -0.30743 | 1.596259 | -0.524594 |
| 29.76389 | -1.035429 | -2.399195 | -0.495002 | -2.289964 | -2.998838 | 0.269291 | -1.526857 | 0.022028 | 0.427978 | 0.243369 | 1.855188 | 0.731437 | 0.432342 | 1.036549 | -0.893597 | 2.638265 | -0.741956 | -0.334803 | 0.249799 | -0.256834 | 0.348737 | -0.248896 | 0.980049 | -0.601431 |
| 29.77778 | -0.746411 | -0.92242 | -0.598257 | -3.668748 | -3.228102 | 0.30897 | -1.139257 | 0.636291 | 0.752407 | -0.877518 | 1.419559 | 0.81418 | 0.354919 | 0.686357 | -2.923071 | 2.321164 | -0.473516 | -0.653997 | -0.477379 | -1.173495 | -0.478084 | -1.414385 | 0.545753 | -1.603494 |
| 29.79167 | -0.510728 | -1.543616 | -0.814944 | -3.788793 | -2.945458 | 0.142229 | -0.255598 | 2.030098 | 0.927125 | -1.775053 | 1.434178 | 1.000248 | 0.344412 | 0.258831 | -0.296244 | 1.502255 | -0.237273 | -1.089073 | -1.271787 | -2.022303 | -0.881929 | -3.742974 | 0.139915 | -2.260594 |
| 29.80556 | 0.344151 | -1.672394 | -1.242682 | -2.495565 | -3.0594 | 0.014706 | -0.842459 | 2.583297 | 0.767515 | -1.8451 | 1.272091 | 0.931739 | 0.326122 | -0.536652 | 0.440598 | 0.770894 | -0.344625 | -1.104329 | -0.948875 | -2.773656 | -0.835827 | -4.688361 | 0.038053 | -1.915811 |
| 29.81944 | 1.232855 | -1.356577 | -1.572029 | -0.757125 | -1.637206 | 0.041369 | -1.013657 | 1.776263 | 0.428609 | -0.661541 | 1.438093 | 1.073645 | 0.334727 | -1.052272 | 1.094868 | -0.030629 | -0.755707 | -0.392842 | -0.095214 | -2.804392 | -0.824167 | -4.002915 | -0.190088 | -1.0851 |
| 29.83333 | 1.336446 | -1.328735 | -1.429037 | 0.943072 | -1.325703 | -0.060925 | -0.367979 | 1.168528 | 0.102221 | 0.696562 | 1.636296 | 2.027646 | 0.384521 | -1.153115 | 0.472804 | -1.505588 | -1.004187 | 0.105331 | 0.503465 | -2.617847 | -0.854635 | -3.106653 | -0.4678 | -0.598764 |
| 29.84722 | 0.919881 | -1.020894 | -0.951264 | 2.309209 | -0.060564 | -0.063149 | -0.795333 | 1.65649 | -0.081362 | 1.317553 | 0.761615 | 2.673412 | 0.456907 | -0.994404 | -1.711671 | -3.142808 | -0.849384 | 0.161447 | 0.890601 | -2.183611 | -0.471032 | -2.419208 | -0.89876 | -0.35107 |
| 29.86111 | 0.175185 | -1.206562 | -0.879804 | 1.750223 | 0.347161 | -0.097149 | -0.690568 | 2.139474 | -0.688819 | 0.390659 | 0.046104 | 1.425671 | 0.475051 | -1.1102 | -2.010631 | -4.15741 | -0.232607 | 0.700402 | 1.286295 | -0.909416 | -0.010038 | -2.904118 | -1.378174 | -0.614316 |
| 29.875 | -1.270634 | -2.078561 | -1.730937 | -1.340998 | -1.45589 | -0.784808 | -1.032993 | 0.038377 | -2.03592 | -1.366892 | -0.98445 | -0.202212 | 0.378273 | -1.379067 | -0.20848 | -4.51834 | 0.076272 | 1.332626 | 1.557853 | 0.420627 | 0.289164 | -4.184571 | -2.003306 | -0.736622 |
| 29.88889 | -3.016065 | -2.438178 | -2.451252 | -3.115489 | -4.397398 | -1.522946 | -1.175233 | -1.805291 | -2.372708 | -2.887244 | -1.381765 | -1.186288 | 0.233383 | -1.575419 | 0.120432 | -4.706838 | 0.052909 | 1.94105 | 1.041366 | 0.803508 | 0.430608 | -3.360356 | -1.61553 | -0.335628 |
| 29.90278 | -3.227314 | -1.317538 | -1.650593 | -0.758563 | -3.287899 | -1.20331 | -0.392103 | -3.391359 | -1.942712 | -4.040569 | -0.66508 | -1.062346 | 0.178135 | -1.413844 | -0.410615 | -3.872362 | 0.042503 | 1.821397 | 0.3941 | 0.54278 | 0.888019 | -0.881926 | -1.641936 | 0.402295 |
| 29.91667 | -2.526904 | -0.715131 | -0.628916 | 0.977701 | -1.600991 | -0.792893 | 0.780418 | 1.632253 | -1.541799 | -3.202189 | -0.209865 | -0.806459 | 0.234605 | -1.252484 | -2.269883 | -3.980193 | 0.11305 | 1.699479 | 0.174792 | -0.015197 | 1.156629 | 1.159589 | -1.473677 | 0.869985 |
| 29.93056 | -2.039392 | 0.666677 | -0.311489 | 0.700687 | -1.003253 | -0.601859 | 2.043389 | 1.550843 | -1.271152 | -3.217393 | -0.391362 | -0.957998 | 0.19398 | -2.030695 | -3.914371 | -3.932596 | 0.20656 | 0.788963 | -1.066689 | -0.940051 | 0.771859 | 0.916389 | -1.404079 | 0.642613 |
| 29.94444 | -1.697301 | 1.243983 | -0.384068 | 0.714743 | -1.118565 | -0.323573 | 2.257511 | 0.215994 | -0.638692 | -3.114973 | -1.002422 | -1.684734 | -0.39137 | -2.188016 | -4.345213 | -3.324214 | 0.11759 | -0.529231 | -3.58777 | -2.639629 | 0.024542 | -0.534234 | -2.016808 | 0.062536 |
| 29.95833 | -1.49194 | 1.341354 | -0.565734 | -0.003475 | -1.2795 | -0.033433 | 2.052217 | -0.30314 | -0.455739 | -1.721339 | -1.41722 | -1.723868 | -1.11522 | -2.031532 | -4.667219 | -4.33827 | -0.066719 | -1.592737 | -4.34829 | -2.764147 | -0.765999 | -1.995516 | -1.715303 | -1.091201 |
| 29.97222 | -1.200831 | -0.340995 | -0.437751 | -0.76773 | -0.689858 | 0.323378 | 0.976815 | -2.115054 | -0.349966 | -1.581666 | -1.644759 | -1.593906 | -0.826515 | -2.310073 | -2.380805 | -1.021516 | -0.281411 | -1.784172 | -3.757491 | -2.629343 | 0.03103 | -2.589085 | -1.335473 | -1.874129 |
| 29.98611 | -0.692596 | -1.4434 | -0.037516 | -0.92197 | 1.038386 | 0.605531 | 0.595017 | -0.973886 | -0.014053 | -1.764637 | -1.215616 | -1.098187 | -0.231338 | -2.689593 | -1.267237 | 0.011662 | -0.549981 | -1.123348 | -2.198788 | -4.427788 | 1.513103 | -2.992366 | 0.134128 | -1.986799 |
| 30 | -0.278119 | -2.05388 | 0.018269 | -2.09266 | 0.896937 | 0.561493 | -0.158249 | 2.520796 | 0.359436 | -1.077446 | -0.666591 | -0.5707 | 0.170246 | -1.454316 | -0.939834 | -0.033226 | 0.668908 | -0.251408 | -0.318985 | -2.138277 | 1.687906 | -2.091302 | 1.544612 | -1.280389 |
| 30.01389 | -0.263498 | -2.819898 | -0.148125 | -2.837202 | -1.366018 | 0.252324 | -0.365441 | 2.106284 | 0.309781 | -0.186057 | -0.377951 | -0.547512 | 0.232342 | -0.27443 | 0.936568 | -0.401066 | 0.292478 | 0.265381 | -0.205482 | -0.468577 | 0.286748 | -0.007503 | 0.684655 | -0.216467 |
| 30.02778 | -0.493452 | -2.793986 | -0.265028 | -3.631335 | -1.209215 | -0.162901 | -0.326582 | 0.224946 | 0.120889 | 0.272502 | -0.07315 | -0.603178 | 0.096351 | 0.853044 | 1.837665 | -1.450554 | 0.032689 | 0.977676 | 0.235998 | -1.279895 | 0.167251 | 2.113593 | 0.412941 | 0.807788 |
| 30.04167 | -0.589062 | -1.708072 | -0.389039 | -3.675241 | -0.982885 | -0.249494 | -0.612683 | -1.052241 | 0.207093 | 0.902705 | 0.030664 | -0.475029 | 0.001562 | 0.242027 | 0.668406 | -1.631266 | 0.038308 | 0.510442 | 1.402406 | -0.093699 | 0.533381 | 2.792541 | 0.706235 | 0.883752 |
| 30.05556 | -0.653241 | -0.642338 | -0.715687 | -3.381052 | -1.454399 | -0.086662 | -0.351662 | -1.282868 | 0.561663 | 0.788672 | -0.050563 | -0.24549 | -0.026139 | 0.031353 | -1.934697 | -1.225591 | -0.006318 | -0.233936 | 1.619261 | 0.558932 | 0.604055 | 1.923461 | -0.703766 | 0.125146 |
| 30.06944 | -0.604248 | -0.80188 | -0.970944 | -3.628157 | -1.733045 | -0.076894 | -0.26476 | -0.533223 | 0.551668 | -0.403066 | -0.11258 | -0.253364 | -0.018301 | 0.356574 | -2.432206 | -0.989214 | -0.080201 | -0.168105 | 0.440329 | 0.213614 | -0.562849 | 1.252783 | -1.644166 | -0.140741 |
| 30.08333 | -0.660444 | -0.905524 | -1.18177 | -3.151566 | -1.172699 | -0.283302 | -0.936068 | 1.233866 | 0.516399 | -1.632985 | -0.205192 | -0.326683 | 0.038703 | 0.989931 | -0.298108 | -0.560123 | -1.136052 | 0.129427 | -0.962606 | 0.37675 | -0.694723 | 2.57095 | -1.381004 | 0.421255 |
| 30.09722 | -0.663087 | -0.622594 | -1.458044 | -2.438351 | -0.949991 | -0.360944 | -1.054887 | 3.506095 | 0.510398 | -2.437844 | -0.267403 | -0.266993 | -0.105491 | 0.784199 | 0.568888 | -0.349596 | -0.455928 | 0.258637 | -1.266877 | 1.031459 | -0.053835 | 2.890139 | -1.6844 | 0.996702 |
| 30.11111 | -0.612423 | -0.460779 | -1.790303 | -1.86187 | -0.726037 | -0.303409 | -0.406755 | 2.291203 | 0.369945 | -2.23925 | -0.203207 | -0.26979 | -0.132463 | 0.705029 | -1.135547 | -0.52409 | -0.036096 | 0.078407 | -0.298352 | 2.076699 | 0.307582 | 1.614007 | -1.67255 | 0.511539 |
| 30.125 | -0.627934 | -0.328984 | -1.813755 | -1.492856 | -0.448649 | -0.118769 | 0.363869 | 1.071423 | 0.295662 | -1.639559 | -0.051519 | -0.133171 | 0.044182 | 0.578468 | -2.17372 | -0.178848 | 0.125279 | -0.403381 | 1.039102 | 3.07314 | 0.486381 | 0.707574 | -1.402483 | 0.03488 |
| 30.13889 | -0.650814 | -0.179383 | -1.461654 | -1.10832 | -0.156623 | -0.010561 | 1.392644 | -0.075265 | 0.190903 | -1.144386 | 0.171655 | -0.038982 | 0.1408 | 0.892302 | -1.43447 | -0.495252 | 0.237068 | -0.691521 | 1.525089 | 3.124405 | 0.554579 | -0.347845 | -1.290039 | -0.079903 |
| 30.15278 | -0.718298 | -0.208491 | -1.140073 | -0.673875 | 0.012347 | -0.127894 | 1.553933 | -1.186899 | -0.049914 | -0.744717 | 0.314827 | -0.036237 | -0.087745 | 1.51851 | 0.887394 | -0.463266 | -0.050031 | -0.718359 | 0.878284 | 2.637138 | 0.764443 | -1.521984 | -0.695065 | 0.324574 |
| 30.16667 | -0.562628 | -0.140363 | -0.795346 | -0.126576 | 0.072612 | -0.202159 | 0.841046 | -0.571483 | -0.144188 | 0.173651 | 0.358967 | -0.002639 | -0.283345 | 1.204414 | 1.975608 | -0.422939 | -0.016534 | -0.454734 | 0.242021 | 2.737629 | 0.969517 | -1.709647 | 0.194149 | 0.179653 |
| 30.18056 | -0.203715 | 0.221229 | -0.367717 | 1.120478 | -0.170847 | -0.109805 | 0.334831 | 0.736546 | 0.030279 | 1.551971 | 0.360912 | 0.137564 | -0.418301 | 0.484616 | 1.396172 | -1.0103 | 0.276857 | -0.034404 | -0.032147 | 2.319896 | 0.833587 | -0.881032 | 1.201264 | 0.128539 |
| 30.19444 | 0.08598 | 0.471101 | 0.102898 | 1.432999 | 0.924561 | -0.078986 | 0.1288 | 1.28621 | -0.020344 | 2.170664 | 0.247913 | 0.242179 | -0.574039 | 0.45268 | 0.222046 | -0.853999 | 0.176146 | 0.203771 | -0.208991 | 0.439554 | 0.301645 | -0.500903 | 1.755709 | 0.101043 |
| 30.20833 | 0.239262 | 0.400471 | 0.437311 | 0.94367 | 1.180835 | -0.077878 | 0.206464 | 0.072559 | -0.229808 | 1.989488 | 0.045107 | 0.18447 | -0.687351 | -0.224108 | -0.852519 | -1.026517 | -0.225437 | -0.232619 | -0.702816 | -1.336644 | -0.228788 | -0.460368 | 1.844061 | -0.020366 |
| 30.22222 | 0.315042 | 0.512214 | 0.717898 | 0.479563 | 1.853637 | 0.024584 | 0.058066 | -1.647529 | -0.245569 | 2.185811 | -0.054927 | 0.245662 | -0.740839 | -0.284821 | -0.740134 | -1.276648 | -0.499904 | -0.701339 | -1.275262 | -2.630511 | -0.659196 | -0.40078 | 1.656118 | 0.049609 |
| 30.23611 | 0.448789 | 0.558066 | 0.937336 | -0.131991 | 2.171131 | 0.154651 | -0.181792 | -2.361682 | -0.037264 | 2.483845 | 0.085392 | 0.379782 | -0.605659 | 0.322172 | -0.148164 | -0.703349 | -0.568477 | -0.807719 | -1.153653 | -2.237046 | -0.561612 | -0.462581 | 0.993685 | 0.113614 |
| 30.25 | 0.211139 | 0.403562 | 1.12704 | -0.499515 | 1.239073 | 0.110112 | -0.765041 | -0.205241 | -0.071911 | 1.767284 | 0.000598 | 0.325985 | -0.216337 | 0.393039 | -0.67524 | -0.604109 | -0.416943 | -0.508651 | -0.000952 | -1.448583 | -0.53084 | -0.581486 | 0.203112 | -0.201952 |
| 30.26389 | -0.081516 | 0.617811 | 1.240597 | 0.016841 | 0.493693 | 0.146654 | -1.209492 | 2.700317 | -0.083112 | 0.948916 | 0.075752 | 0.34606 | 0.340713 | -0.019514 | -1.710165 | -1.069821 | -0.156303 | -0.016049 | 0.750416 | -0.842755 | -0.412045 | -0.490395 | -0.604018 | -0.36125 |
| 30.27778 | 0.029056 | 1.017248 | 1.355206 | 0.991999 | 0.379916 | 0.255186 | -1.116776 | 2.897351 | 0.027245 | 0.815913 | 0.253294 | 0.48462 | 0.577687 | 0.150179 | -2.084774 | -1.485473 | 0.086898 | 0.243728 | 0.909513 | -0.02117 | -0.208264 | -0.192506 | -1.350677 | -0.111732 |
| 30.29167 | 0.177818 | 1.190808 | 1.384961 | 1.503018 | 0.514862 | 0.338049 | -0.002368 | 1.425003 | 0.098796 | 0.866542 | 0.372983 | 0.538695 | 0.548704 | 0.201148 | -0.46148 | -0.249091 | 0.18483 | 0.270078 | 0.595841 | 0.704579 | -0.012518 | 0.011361 | -1.880281 | 0.146635 |
| 30.30556 | 0.280613 | 1.355523 | 1.643762 | 1.71288 | 0.505295 | 0.337199 | 0.576619 | 0.452431 | 0.040124 | 1.422193 | 0.322388 | 0.463423 | 0.466834 | 0.067766 | 1.858449 | 1.10099 | 0.223306 | 0.237421 | 0.455261 | 0.914171 | 0.054038 | -0.013452 | -1.736979 | 0.53604 |
| 30.31944 | 0.289408 | 1.575532 | 1.851743 | 2.204071 | 0.395795 | 0.344772 | 0.846548 | -0.09789 | 0.181256 | 2.03257 | 0.284886 | 0.391191 | 0.432526 | 0.430205 | 2.670714 | 1.729554 | 0.150024 | 0.221015 | 0.519829 | 0.986836 | 0.043187 | -0.157038 | -1.442225 | 0.771645 |
| 30.33333 | 0.314607 | 2.040461 | 2.229724 | 3.48676 | 0.008237 | 0.300458 | 1.184199 | -0.231066 | 0.391462 | 1.796918 | 0.253275 | 0.279845 | 0.391655 | 0.975484 | 1.369742 | 1.14137 | 0.018813 | 0.245621 | 0.645006 | 1.616493 | 0.131854 | -0.256165 | -1.079694 | 1.250159 |
| 30.34722 | 0.269739 | 2.088064 | 2.388468 | 4.921915 | 0.162715 | 0.31905 | 1.328457 | -0.191344 | 0.69368 | 1.997233 | 0.260176 | 0.256583 | 0.270756 | 1.422205 | 0.190078 | 0.968396 | -0.038315 | 0.086106 | 0.726637 | 2.006004 | 0.233476 | -0.206584 | -1.447712 | 1.641907 |
| 30.36111 | 0.27976 | 2.843622 | 1.954738 | 4.798824 | 0.972902 | 0.379856 | 1.496473 | 0.403647 | 0.737351 | 3.358594 | 0.403085 | 0.575581 | 0.214005 | 1.773205 | -0.1713 | 1.214836 | -0.06707 | -0.060382 | 0.704843 | 1.877491 | 0.19499 | -0.061383 | -1.130917 | 2.403416 |
| 30.375 | 0.429644 | 2.490622 | 1.370621 | 4.32649 | 1.375528 | 0.280738 | 1.566228 | 0.838972 | 0.605018 | 3.111252 | 0.316828 | 0.608289 | 0.176717 | 1.867843 | 0.191684 | 1.56894 | -0.088179 | -0.018835 | 0.696574 | 1.53394 | 0.033261 | 0.131748 | -0.398821 | 1.986927 |
| 30.38889 | 0.492129 | 1.640899 | 0.47751 | 3.923815 | 1.075651 | 0.038633 | 1.598335 | 0.829936 | 0.335296 | 1.718395 | -0.182569 | 0.237612 | 0.181152 | 1.965767 | 0.538022 | 1.910234 | -0.090966 | 0.21085 | 0.920374 | 2.151717 | -0.086257 | 0.324056 | 0.560401 | 1.666894 |
| 30.40278 | 0.55644 | 1.014227 | 0.086498 | 3.369487 | 0.768969 | -0.195352 | 0.857311 | 0.241605 | 0.196172 | 0.84911 | -0.476871 | -0.01844 | 0.14588 | 1.957951 | -0.053701 | 2.003009 | 0.089981 | 0.44712 | 1.078583 | 2.470061 | -0.143395 | 0.337672 | 1.523714 | 2.624078 |
| 30.41667 | 1.214747 | 0.033308 | 0.605804 | 3.334665 | 0.938537 | 0.05106 | 0.563087 | -0.257946 | 0.451992 | 0.675301 | -0.281218 | 0.051884 | 0.051362 | 1.534351 | -0.609377 | 1.263825 | 0.443099 | 0.583167 | 0.830684 | 2.314864 | -0.166013 | 0.146148 | 1.771986 | 2.444126 |
| 30.43056 | 1.38913 | -0.479257 | 0.841521 | 2.252323 | 1.345062 | 0.529297 | 0.299707 | -0.413012 | 0.688302 | 0.778344 | 0.104335 | 0.317469 | -0.04106 | 1.240884 | -1.093547 | 1.011506 | 0.950557 | 0.598506 | 0.528145 | 3.01728 | -0.145863 | -0.178563 | 2.600364 | 1.715921 |
| 30.44444 | 1.190269 | -0.010675 | 1.104536 | 1.398831 | 1.506839 | 1.06857 | 0.460842 | -0.036576 | 0.832881 | 1.205959 | 0.605829 | 0.628478 | -0.071119 | 0.784533 | -0.496859 | 2.008172 | 1.511888 | 0.565906 | 0.246931 | 3.38559 | -0.056934 | -0.482067 | 3.063101 | 1.950535 |
| 30.45833 | 0.833985 | 1.67589 | 2.467783 | 1.860741 | 1.286897 | 1.328815 | 0.809537 | 0.60751 | 0.687761 | 1.793985 | 0.894898 | 0.856256 | -0.019573 | 0.224148 | 0.399547 | 2.857525 | 1.887465 | 0.413888 | 0.132114 | 3.140356 | -0.107795 | -0.6615 | 2.570215 | 0.911967 |

| | | | | | | | | | | | | | | | | | | | | | | | | |
|---|---|---|---|---|---|---|---|---|---|---|---|---|---|---|---|---|---|---|---|---|---|---|---|---|
| 30.47222 | 0.326682 | 3.586972 | 2.661196 | 0.502173 | 0.397709 | 1.31079 | 1.347019 | 0.80536 | 0.501606 | 2.268473 | 0.962113 | 1.06506 | 0.180544 | -0.079413 | 1.430087 | 2.858096 | 1.637926 | 0.287396 | 0.655171 | 3.614945 | -0.110304 | -0.643727 | 2.437028 | 0.301679 |
| 30.48611 | 0.202303 | 2.839057 | 1.730699 | -0.84708 | -0.347951 | 1.408554 | 1.443055 | 1.862276 | 0.674118 | 3.688782 | 1.27238 | 1.464605 | 0.62097 | -0.004813 | 2.944473 | 3.336657 | 2.515901 | 0.37694 | 1.667714 | 4.110615 | 0.044283 | 0.06699 | 2.07924 | 0.852993 |
| 30.5 | 0.546479 | 2.910798 | 0.181963 | -1.580014 | -0.077843 | 1.557906 | 0.842133 | 2.059615 | 0.93391 | 4.517332 | 1.659758 | 2.135286 | 0.830921 | 0.47573 | 5.192829 | 2.06343 | 2.072828 | 0.736072 | 2.6826 | 3.835492 | 0.34275 | 2.446294 | 1.722791 | 1.833194 |
| 30.51389 | 0.176123 | 2.689192 | -1.247376 | -1.743209 | 0.318624 | 1.419994 | 0.316405 | 1.461673 | 0.875832 | 4.319721 | 1.201579 | 2.486152 | 0.929703 | 1.064153 | 5.831424 | 0.963988 | 1.254009 | 1.197514 | 3.366266 | 2.282469 | 0.642599 | 3.919698 | 1.358753 | 2.485798 |
| 30.52778 | 0.14891 | 2.045271 | -1.606798 | -1.531688 | 0.907137 | 1.179378 | -0.234098 | 0.715011 | 0.808673 | 2.881933 | 1.017001 | 1.697249 | 1.025256 | 1.320488 | 5.911335 | 1.059953 | 0.568206 | 1.614523 | 3.819116 | 1.035341 | 0.879115 | 4.111242 | 1.424172 | 1.93363 |
| 30.54167 | 0.113307 | 1.347403 | -1.539656 | -0.976698 | 1.605642 | 1.052746 | -0.637714 | -0.218085 | 0.428538 | -0.19242 | 1.23461 | 1.193316 | 1.023595 | 0.854026 | 4.95169 | 1.55705 | 0.629744 | 1.481087 | 3.584273 | 2.026932 | 0.683742 | 3.723783 | 1.600353 | 1.678434 |
| 30.55556 | -0.045082 | 0.672343 | -0.265905 | 0.814052 | 1.660967 | 0.926337 | -0.924672 | -1.043788 | 0.389769 | -0.562122 | 1.477872 | 0.90271 | 0.813672 | 0.436417 | 2.127419 | 0.479915 | 1.046828 | 0.396849 | 2.814733 | 3.036986 | 0.781018 | 1.838498 | 1.680514 | 1.784536 |
| 30.56944 | -0.103165 | 0.487494 | 0.681987 | 2.074663 | 0.814517 | 0.739067 | -0.322998 | -0.347892 | 0.600502 | 0.661095 | 1.3726 | 0.866456 | 0.598649 | 0.472289 | 0.388348 | 0.293254 | 0.443546 | -0.310894 | 2.079345 | 2.712187 | 1.160333 | -0.422177 | 0.446229 | 1.186906 |
| 30.58333 | 0.197548 | 1.336914 | 1.482441 | 2.987732 | 0.659082 | 0.715701 | 0.659344 | 0.600288 | 0.773499 | 1.415421 | 1.314393 | 0.877379 | 0.368922 | 0.810477 | -0.914344 | -0.194392 | -0.22383 | -0.519193 | 1.502704 | 0.152541 | 0.595847 | -1.851014 | -1.249689 | -0.085233 |
| 30.59722 | 0.460781 | 2.114249 | 1.899188 | 3.396349 | 1.063641 | 0.721149 | 1.912287 | 0.82931 | 1.044703 | 2.493581 | 1.165583 | 0.825165 | 0.080917 | 0.045749 | -2.167568 | -0.175043 | -1.285642 | -0.242718 | 0.55529 | -2.15213 | -0.052146 | -2.421101 | -2.2043 | -1.253624 |
| 30.61111 | 0.641235 | 1.670348 | 1.493074 | 2.63682 | 0.502357 | 0.650523 | 2.911435 | 0.671018 | 1.048633 | 3.94308 | 0.967946 | 0.811261 | 0.015233 | -0.351236 | -2.529013 | 0.980749 | -1.671656 | -0.029144 | -0.7925 | -1.902965 | -0.381789 | -2.09952 | -2.580074 | -1.833759 |
| 30.625 | 0.941353 | 1.392257 | 1.53911 | 1.796282 | -1.031507 | 0.491574 | 2.483417 | 2.517792 | 0.507771 | 2.908262 | 0.921613 | 0.907586 | 0.217044 | -0.395655 | -1.701133 | 2.577495 | -1.527401 | 0.372088 | -0.97791 | -0.754291 | -0.64039 | -2.15151 | -2.532029 | -2.198819 |
| 30.63889 | 1.453391 | 0.605104 | 2.208863 | 2.087141 | -0.931868 | 0.35861 | 0.890108 | 1.665375 | 0.155329 | 1.547812 | 0.872357 | 1.004952 | 0.605669 | -0.965079 | 0.62297 | 2.741439 | -0.716579 | 0.67692 | -0.053604 | 0.234339 | -0.402134 | -1.880441 | -1.913973 | -1.984792 |
| 30.65278 | 1.318762 | -0.774833 | 2.416821 | 1.157036 | 0.56728 | 0.289858 | 0.421973 | 0.878263 | -0.128638 | 1.040046 | 1.03896 | 1.050914 | 1.126319 | -0.874978 | 4.534736 | 1.083085 | 0.047146 | 1.122567 | 1.322664 | 0.507067 | -0.165176 | -0.574909 | -1.214664 | -0.847393 |
| 30.66667 | 1.238087 | -0.151521 | 2.055783 | 0.492409 | 0.815524 | 0.426753 | 0.407462 | 0.59675 | -0.380918 | 1.088191 | 1.483772 | 1.116069 | 1.480601 | 0.824758 | 1.726117 | 0.884965 | 0.780794 | 1.57917 | 1.454041 | 1.170739 | 1.013359 | 1.031501 | -0.354206 | 0.807959 |
| 30.68056 | 1.203814 | 2.185247 | 1.682191 | 0.233054 | 1.727703 | 0.706619 | 0.170337 | -0.241203 | -0.128435 | 1.895979 | 1.603697 | 1.17212 | 1.483016 | 1.714056 | 2.1126 | 0.18975 | 0.945672 | 1.437758 | 1.741238 | 1.040284 | 1.961733 | 2.57028 | 1.580777 | 2.577589 |
| 30.69444 | 0.370243 | 4.373349 | 0.988238 | -0.217029 | 3.231624 | 0.774539 | -0.062841 | -0.117211 | 0.10458 | 4.300797 | 1.579303 | 1.102787 | 1.46815 | 1.763176 | 2.065033 | -0.156186 | 0.614155 | 1.419468 | 0.810456 | 0.549027 | 2.483855 | 3.566442 | 4.186025 | 2.916354 |
| 30.70833 | -0.342295 | 5.618118 | 0.635508 | -0.61438 | 4.367736 | 0.685615 | 0.077162 | -1.734624 | 0.070072 | 3.73392 | 1.535814 | 1.280447 | 1.36432 | 1.569857 | 0.766213 | -0.367189 | 0.093867 | 1.52632 | -0.349713 | 0.230307 | 3.27514 | 3.36612 | 4.037141 | 1.820421 |
| 30.72222 | -0.397656 | 5.984111 | 0.576366 | -0.125794 | 2.81616 | 0.688935 | -1.178006 | -1.463803 | 0.211045 | 2.804293 | 1.348617 | 1.555699 | 1.214717 | 0.730775 | 1.19633 | -0.892789 | -0.08483 | 1.939786 | -0.736878 | -0.050344 | 2.519438 | 0.813053 | 2.775244 | 1.331534 |
| 30.73611 | -0.203747 | 3.939158 | 0.685197 | 0.437323 | 1.738236 | 0.773442 | -1.243111 | 1.907355 | 0.545694 | 0.797792 | 0.994333 | 1.457092 | 0.666772 | 0.06385 | 1.658337 | -0.961256 | -0.192483 | 1.542516 | -0.461656 | -0.286935 | 1.216344 | -0.770532 | 1.743441 | 0.481088 |
| 30.75 | -0.104371 | 1.489974 | 0.49966 | 0.857432 | 0.97273 | 0.796405 | -0.571762 | 3.26181 | -0.059085 | -3.023397 | 1.144327 | 0.482586 | 0.123794 | -0.350171 | -1.194116 | -1.283162 | -0.615903 | 0.232236 | -0.045218 | -0.09232 | 0.023704 | -1.096033 | 1.092797 | -0.856639 |
| 30.76389 | -0.14875 | 0.252207 | -0.18773 | 1.158564 | -0.030775 | 0.614273 | 0.281024 | 2.306787 | -0.307924 | -6.145682 | 1.163514 | -0.351137 | -0.716224 | -0.250691 | -1.660237 | -2.446276 | -1.200741 | -0.958453 | 0.551865 | 0.574163 | -1.28145 | -0.537367 | 0.846356 | -0.995821 |
| 30.77778 | -0.576824 | 0.195271 | -0.465097 | -1.506241 | -1.162606 | 0.597975 | 0.166805 | 1.116683 | 0.292624 | -6.650262 | 0.622339 | -0.818223 | -1.027595 | 0.782036 | -0.439904 | -2.91528 | -1.170136 | -1.452917 | 0.183546 | 0.613273 | -1.789354 | -0.318574 | 0.68607 | -0.632625 |
| 30.79167 | -0.767275 | -1.07593 | -0.41981 | -3.083349 | -2.542719 | 0.873252 | -0.187481 | 0.545782 | 0.750129 | -7.606895 | -0.163923 | -0.621609 | -0.819164 | 1.25442 | 0.119868 | -3.144869 | -0.619651 | -1.424622 | -0.631597 | -0.01563 | -2.097104 | -0.720369 | 1.312251 | -0.87158 |
| 30.80556 | -0.78786 | -2.400081 | 0.117041 | 0.8694 | -3.802999 | 0.757456 | -0.067943 | -0.31169 | 0.7657 | -5.831316 | -0.277365 | -0.242587 | -0.796174 | 0.605218 | -0.979819 | -3.093901 | -0.556536 | -0.914929 | -1.403711 | 0.145964 | -1.520855 | -2.021485 | 1.022667 | -1.176159 |
| 30.81944 | -0.722739 | -2.983804 | -0.678634 | -1.110559 | -4.444037 | 0.649922 | 0.555234 | 0.250229 | 0.705188 | -4.122726 | -0.429432 | 0.030231 | -0.463501 | -0.669748 | -1.14648 | -2.057783 | -0.88799 | -0.589582 | -1.24869 | -0.856789 | -1.425107 | -3.240004 | 0.167107 | -1.647613 |
| 30.83333 | -0.649012 | -4.433983 | -1.733734 | -6.25815 | -3.869762 | 0.484441 | 0.798988 | 1.138386 | 0.577843 | -4.941577 | -0.331055 | 0.100741 | 0.096369 | -2.013715 | -5.418915 | -1.805439 | -1.162609 | -0.134608 | -0.602964 | -1.188155 | -1.475212 | -3.753004 | -1.579228 | -2.469703 |
| 30.84722 | -0.422836 | -4.340109 | -1.874778 | -9.359011 | -3.372198 | 0.265882 | 0.909505 | 1.316274 | 0.310002 | -3.843766 | 0.010814 | 0.095453 | 0.555393 | -2.474117 | -4.844939 | -1.331552 | -0.673594 | 0.211661 | 0.023299 | -1.233177 | -1.281259 | -4.193835 | -3.051598 | -2.464048 |
| 30.86111 | -0.069128 | -3.584267 | -2.340131 | -4.436043 | -2.366081 | -0.08993 | 0.330303 | 1.306106 | 0.025029 | -1.494756 | -0.211578 | 0.083122 | 1.266004 | -2.270887 | -0.569202 | -1.256335 | -0.097311 | 0.191874 | 0.170258 | -1.77055 | -1.061792 | -4.497608 | -3.585716 | -2.163319 |
| 30.875 | 0.743023 | -3.582728 | -1.944514 | -5.255172 | 1.172373 | -0.860983 | -0.150382 | 0.323519 | -0.469708 | -0.084047 | -0.755982 | -0.013679 | 1.075079 | -2.311598 | 2.208387 | -1.433139 | -0.011142 | 0.546293 | 0.165965 | -2.819092 | -0.632523 | -3.204192 | -3.437544 | -1.371382 |
| 30.88889 | 0.060605 | -2.339083 | -2.032978 | -2.729002 | 0.322342 | -2.287835 | -0.869976 | -0.498607 | -0.791707 | 0.21221 | -1.27278 | 0.105675 | 1.407649 | -1.792283 | -0.443774 | -1.985151 | -0.020256 | 0.891298 | 0.203576 | -3.580899 | -0.216479 | -2.965744 | -2.727803 | -0.578601 |
| 30.90278 | -0.305751 | -0.617497 | -1.369958 | -1.935907 | 1.067659 | -3.207056 | -1.923965 | -1.153241 | -0.655639 | 2.8453 | -1.254695 | 0.367238 | 1.362681 | -1.8028 | -2.639027 | -2.705642 | 0.066363 | 0.8224 | 0.183303 | -3.390861 | 0.310522 | -3.368363 | -2.415614 | -1.086061 |
| 30.91667 | -0.455103 | 0.972358 | -0.52752 | -0.156592 | 0.753121 | -2.308757 | -1.768062 | -1.250175 | -0.751201 | 6.390409 | -0.792148 | 0.377939 | 1.224557 | -2.042984 | -3.986373 | -3.065468 | 0.200217 | 0.571352 | -0.452607 | -3.029623 | 0.576814 | -2.50372 | -2.906615 | -1.306063 |
| 30.93056 | 0.16886 | 0.19476 | 0.159191 | 0.642033 | 2.378071 | -0.745031 | -0.669606 | 2.095936 | -0.197107 | 6.865174 | -0.789824 | 0.195947 | 0.586512 | -1.626692 | -3.212294 | -3.650107 | 0.158077 | 0.058364 | -1.312569 | -3.911989 | 0.157913 | -1.689544 | -3.479455 | -1.037962 |
| 30.94444 | 0.088168 | -0.312234 | 0.216451 | 1.475075 | 0.510897 | -0.390343 | -1.287752 | 4.187172 | 0.549501 | 7.154987 | -0.632916 | 0.214189 | -0.660684 | -1.941014 | -2.907876 | -4.355379 | -0.232821 | -1.247717 | -2.820629 | -5.974543 | -0.854655 | -1.62181 | -4.079012 | -0.962807 |
| 30.95833 | -0.392636 | -0.494115 | 0.196365 | 2.236956 | -0.598464 | -0.399684 | -2.234858 | 1.2584 | 0.556757 | 3.286659 | -0.234292 | 0.063235 | -1.341334 | -2.017622 | -5.378843 | -4.006132 | -1.241167 | -2.850893 | -4.436768 | -7.680287 | -0.905259 | -1.990745 | -3.984932 | -1.549912 |
| 30.97222 | -0.794614 | -0.32107 | 0.030562 | 2.324336 | -0.6047 | -0.504207 | -2.454778 | -1.760317 | 1.004522 | 3.280637 | -0.005545 | 0.130212 | -1.413678 | -2.383015 | -4.06318 | -2.010052 | -2.55661 | -3.207073 | -6.176902 | -7.536998 | -0.954628 | -1.748597 | -2.702451 | -1.658698 |
| 30.98611 | -0.529197 | 0.528915 | -0.10861 | 4.231758 | -0.125735 | -0.65512 | -1.557877 | -2.365365 | 1.055462 | 2.455705 | 0.063875 | 0.512479 | -1.393379 | -2.207359 | -1.28842 | -2.885237 | -2.790988 | -1.498511 | -3.643036 | -4.929362 | -0.241949 | -1.164824 | -1.143978 | -0.776194 |
| 31 | 0.100917 | 2.137754 | -0.154429 | 3.096968 | 0.32448 | -0.732849 | -0.690089 | -1.017537 | 0.201516 | 0.931379 | 0.250825 | 0.52118 | -0.320688 | -1.637507 | -1.227926 | -2.289278 | -1.469983 | 0.851318 | -1.718802 | -2.844734 | 0.094799 | -1.263828 | 0.17834 | -0.14691 |
| 31.01389 | 0.707523 | 3.270258 | -0.218269 | 1.248992 | -0.121177 | -0.459486 | 0.404701 | -0.661404 | -1.061233 | -1.998225 | 0.542396 | 0.114405 | 0.649426 | -1.000558 | 1.34325 | 0.093092 | 0.833734 | 1.514816 | -0.214985 | -1.209461 | 0.0708 | -0.817869 | 1.502203 | 0.219367 |
| 31.02778 | 0.993486 | 1.100057 | -0.139898 | -1.012593 | 0.515116 | 0.03675 | 0.625455 | -0.159207 | -1.6175 | -4.15055 | 0.606706 | -0.231239 | 0.897949 | -0.246441 | 2.304081 | 0.087052 | 1.415759 | 1.328352 | 0.634871 | 0.522239 | 0.171944 | 0.423025 | 3.04911 | 0.630032 |
| 31.04167 | 0.515856 | -0.375293 | -0.042469 | -3.856778 | 1.091431 | 0.464742 | 0.459103 | -0.563604 | -1.12766 | -5.032319 | 0.576793 | -0.107685 | 0.28227 | 0.421677 | 5.378439 | 0.540339 | 1.445406 | 1.185285 | 1.223198 | 1.977579 | 0.097111 | 2.413648 | 3.749723 | 0.752138 |
| 31.05556 | 0.266463 | -1.214368 | 0.093844 | -4.847608 | 1.373717 | 0.814383 | 0.341581 | 0.130897 | -0.802026 | -4.516028 | 0.300989 | -0.161341 | 0.371482 | 0.880868 | 2.851473 | 0.519059 | 1.361483 | 1.067697 | 2.61659 | 2.403579 | -0.064777 | 2.241495 | 1.57741 | 0.413939 |
| 31.06944 | 0.338741 | -2.283234 | -0.08461 | -4.417875 | 1.494146 | 0.618409 | 0.065577 | -0.254944 | -0.597059 | -2.50509 | 0.000075 | -0.829886 | 0.733556 | 0.545393 | -2.53502 | -0.672264 | 0.354995 | 0.623896 | 1.279474 | 1.193997 | 0.086668 | 1.229586 | 0.615279 | -0.013126 |
| 31.08333 | -0.019109 | -1.739186 | -0.229829 | -2.658995 | 1.520563 | 0.199967 | 0.078312 | -0.126619 | -0.257002 | -3.767143 | -0.306188 | -0.734268 | 0.028432 | 0.884569 | -1.14003 | -0.602086 | -1.317336 | 0.21523 | -0.810481 | -0.062167 | 0.551387 | 1.051682 | -0.685442 | -0.146713 |
| 31.09722 | -0.299283 | -1.058486 | -0.446119 | -1.572371 | 2.841345 | 0.163177 | -0.205509 | -0.041182 | 0.268804 | -5.388638 | -0.525832 | -0.310907 | -0.506928 | 0.771786 | 2.167294 | -1.121383 | -2.175359 | 0.29622 | -2.106457 | -0.481107 | 0.787181 | 0.866636 | -1.6173 | 0.255575 |
| 31.11111 | -0.238114 | -0.732561 | -0.986172 | -1.280642 | 1.955741 | 0.338474 | 0.005575 | 0.01801 | 0.429888 | -5.863874 | -0.580761 | -0.490042 | -0.366346 | 1.567835 | 0.647929 | -0.603515 | -2.162073 | 0.623154 | -1.805824 | 0.115514 | 0.779333 | 1.134229 | -1.642543 | 0.302174 |
| 31.125 | -0.19948 | -1.202113 | -0.991841 | -1.025707 | 1.332243 | 0.388659 | -0.770993 | 1.551867 | 0.57748 | -4.20591 | -0.667479 | -0.496577 | -0.105674 | 1.292996 | -1.747715 | 0.039572 | -2.239152 | 0.395933 | -0.860394 | 0.908511 | 0.023114 | 0.718895 | -2.292666 | 0.291157 |
| 31.13889 | -0.097607 | -1.662769 | -0.78432 | -0.798009 | 1.072742 | 0.483967 | -0.705262 | 2.055221 | 0.680102 | -2.460724 | -0.646981 | -0.3016 | 0.037678 | -0.378611 | -3.44051 | 0.471572 | -1.318076 | -0.422199 | -0.20158 | 1.32181 | -0.45292 | -0.649839 | -2.754172 | 0.013453 |
| 31.15278 | -0.057751 | -2.034143 | -1.024708 | -0.591069 | 0.584391 | 0.348911 | -0.342389 | 1.319558 | 0.522322 | -0.783896 | -0.447952 | -0.24852 | 0.096935 | -1.01159 | -1.991402 | 0.181217 | -0.367244 | -0.620552 | 0.323174 | 1.618903 | -0.488634 | -1.688873 | -3.088391 | -0.015181 |
| 31.16667 | -0.050791 | -1.907435 | -1.190008 | -0.291132 | 0.396684 | 0.202261 | -0.0381 | 0.335354 | 0.401115 | 1.254288 | -0.188542 | -0.242347 | 0.057023 | 0.513595 | 0.585024 | -0.170072 | -0.075392 | -0.373636 | 0.891089 | 1.526724 | -0.018003 | -1.534 | -2.707572 | -0.020998 |
| 31.18056 | -0.044927 | -1.474915 | -0.866557 | -0.071186 | 0.434144 | 0.242614 | 0.481051 | -0.050977 | 0.431831 | 2.933226 | -0.018372 | -0.036113 | 0.163345 | 1.534792 | 3.033956 | -0.113442 | 0.477672 | -0.087272 | 1.74991 | 1.625789 | 0.20089 | -1.563871 | -2.111347 | 0.323665 |
| 31.19444 | -0.056647 | -1.075143 | -0.484595 | 0.302117 | 0.046049 | 0.329257 | 0.949424 | 0.333062 | 0.502601 | 3.386448 | 0.107924 | 0.091798 | 0.268633 | 0.580577 | 2.983854 | -0.28281 | 1.538025 | 0.118051 | 2.315561 | 1.468426 | 0.317652 | -1.431284 | -1.434454 | 0.651667 |
| 31.20833 | -0.125399 | -0.635449 | -0.386783 | 0.218166 | 0.573837 | 0.228056 | 0.390008 | 0.930309 | 0.802044 | 3.643845 | 0.172291 | 0.10891 | 0.239958 | 0.309617 | 0.596128 | 0.222484 | 1.331923 | 0.074171 | 2.004769 | 1.101225 | 0.375421 | 0.536198 | -0.733404 | 0.475319 |
| 31.22222 | -0.21865 | -0.035569 | -0.266253 | 0.819095 | 0.486267 | 0.124945 | 0.140974 | 0.546174 | 0.834164 | 0.453748 | 0.234525 | 0.110037 | -0.015894 | 2.552247 | -1.392962 | -0.982949 | 0.181574 | -0.074226 | 1.202851 | 1.026629 | 0.27876 | 1.737315 | -0.473133 | 0.372637 |
| 31.23611 | 0.160628 | 1.114582 | 0.050961 | 0.420877 | 1.287626 | 0.265269 | 0.420366 | -0.399428 | 0.846626 | 1.755121 | 0.362238 | 0.343943 | -0.2952 | 3.265129 | -2.049236 | -0.724977 | -0.905684 | -0.088935 | 0.40882 | 1.076383 | 0.33352 | 0.823858 | -0.367812 | 0.512604 |
| 31.25 | 0.305036 | 1.811427 | 0.376924 | 0.190801 | 1.078651 | 0.458485 | 0.729091 | -0.788322 | 0.694565 | 2.119866 | 0.350385 | 0.558612 | -0.469535 | 1.52786 | -1.656669 | -0.258195 | -1.479671 | -0.004355 | -0.24675 | 1.313633 | 0.59683 | -1.029649 | -0.251689 | -0.122789 |
| 31.26389 | 0.263162 | 1.974412 | 0.565416 | 0.245539 | 1.645038 | 0.432442 | 0.157085 | -0.814619 | 0.655457 | 2.402543 | 0.162311 | 0.470297 | -0.442146 | 0.794247 | -1.664783 | -0.205489 | -1.569094 | -0.038132 | -0.479985 | 1.485476 | 0.43552 | -0.635085 | -0.054098 | 0.037983 |
| 31.27778 | 0.083936 | 1.737149 | 0.632517 | 0.228145 | 2.359531 | 0.228111 | -0.643419 | -1.377882 | 0.676717 | 2.715946 | 0.255892 | 0.24981 | -0.327088 | 1.6403 | -2.31952 | 0.112756 | -1.322989 | -0.184977 | -0.229761 | 1.325249 | 0.485058 | 1.116913 | 0.114976 | 0.642104 |
| 31.29167 | -0.006312 | 1.504348 | 0.557389 | 0.391144 | 2.449284 | 0.017786 | -0.021288 | -1.42387 | 0.641697 | 2.415516 | 0.507326 | 0.27078 | -0.119249 | 2.255794 | -2.59587 | 0.122397 | -0.702724 | -0.37065 | 0.347835 | 1.592379 | 0.063672 | 1.799121 | 0.111681 | 1.172379 |
| 31.30556 | 0.13034 | 2.244572 | 0.796131 | 0.652295 | 2.484255 | 0.036453 | 0.044224 | -0.878772 | 0.56311 | 2.006207 | 0.794698 | 0.434625 | 0.074965 | 2.084166 | -1.755028 | -0.495085 | -0.021648 | -0.222481 | 0.776008 | 2.677189 | 0.353859 | 1.380672 | 0.194544 | 1.652892 |
| 31.31944 | 0.167625 | 3.73137 | 1.260972 | 1.520156 | 2.624703 | 0.418294 | -0.168896 | 0.008112 | 0.500636 | 2.715568 | 0.583632 | 0.633185 | 0.206548 | 1.324277 | -0.13936 | 0.132769 | 0.400894 | 0.088407 | 0.795666 | 1.416781 | 0.324014 | 1.844238 | 0.332024 | 1.410069 |
| 31.33333 | -0.021635 | 5.053558 | 1.383997 | 2.883938 | 1.301707 | 0.761703 | -0.114774 | 0.816514 | 0.432464 | 3.460603 | 0.513532 | 0.79272 | 0.286144 | 1.183943 | 1.402851 | -0.14239 | 0.684617 | 0.280459 | 0.766868 | 1.31356 | 0.123943 | 2.570583 | 0.549722 | 1.220704 |
| 31.34722 | -0.161393 | 5.602653 | 1.555526 | 3.309801 | 0.719075 | 0.71709 | 0.410801 | 1.002508 | 0.268447 | 2.621259 | 0.305523 | 0.684187 | 0.373679 | 1.538989 | 2.20915 | -0.122869 | 0.791045 | 0.260317 | 0.824483 | 1.634032 | 0.026233 | 3.0905 | 0.832416 | 1.092165 |
| 31.36111 | -0.187289 | 5.085179 | 1.886756 | 2.866089 | 0.363191 | 0.51913 | 0.863065 | 0.618726 | 0.083884 | 1.60496 | 0.093281 | 0.403986 | 0.460219 | 1.832994 | 3.1999 | 0.254419 | 0.771517 | 0.232166 | 0.921547 | 1.941356 | -0.077929 | 2.817489 | 1.116743 | 0.989198 |
| 31.375 | -0.087003 | 3.560295 | 2.136122 | 2.639297 | 0.280257 | 0.392154 | 1.497919 | 0.630802 | 0.055659 | 0.60324 | 0.039026 | 0.223413 | 0.54675 | 1.670454 | 3.716832 | 0.990655 | 0.89784 | 0.122794 | 1.035582 | 2.254686 | -0.394117 | 3.43907 | 1.081131 | 0.931876 |
| 31.38889 | 0.020214 | 1.63336 | 1.653654 | 2.145924 | 0.23739 | 0.437196 | 1.998542 | 0.798849 | -0.039093 | -0.999916 | 0.005423 | -0.179423 | 0.517741 | 1.349558 | 3.814546 | 0.76161 | 1.204463 | 0.075006 | 1.171795 | 2.600012 | -0.305783 | 4.868498 | 0.952087 | 1.201577 |
| 31.40278 | -0.123989 | 0.443484 | 1.115282 | 1.900387 | 0.296175 | 0.834336 | 2.228761 | 0.694038 | -0.268458 | -1.771383 | -0.048496 | -0.181404 | 0.477269 | 1.430216 | 3.617629 | 0.408459 | 1.220951 | 0.059896 | 1.240302 | 3.160428 | 0.183047 | 5.133928 | 0.790041 | 2.161894 |
| 31.41667 | -0.052876 | -0.874534 | 1.168957 | 2.569217 | 0.516531 | 0.985135 | 1.58875 | 0.616604 | -0.21952 | -1.53866 | 0.062207 | -0.042001 | 0.492127 | 1.997952 | 2.864122 | 1.001198 | 1.076777 | 0.094346 | 1.319532 | 3.90305 | 0.305119 | 4.774358 | 0.934873 | 3.382444 |
| 31.43056 | 0.020111 | -2.443037 | 1.667196 | 3.334997 | 0.838264 | 0.71502 | 0.826699 | 0.648246 | 0.036053 | -1.006121 | 0.256995 | 0.144139 | 0.593348 | 2.735261 | 2.713571 | 1.528791 | 1.084154 | 0.176797 | 1.316016 | 5.113844 | 0.037896 | 5.269162 | 0.863042 | 3.850829 |
| 31.44444 | 0.171564 | -2.479394 | 2.017677 | 3.600227 | 1.156492 | 0.749617 | 0.323722 | 0.785075 | 0.312374 | 0.115634 | 0.400901 | 0.346801 | 0.818295 | 1.378821 | 1.79558 | 1.456885 | 1.208933 | 0.249716 | 1.867639 | 3.653218 | -0.150664 | 3.164039 | 0.770508 | 1.788077 |
| 31.45833 | 0.674543 | 0.830111 | 1.693399 | 4.719074 | 1.713016 | 1.343715 | 0.772514 | 0.849262 | 0.541196 | 1.585901 | 0.636912 | 0.567256 | 0.713738 | -0.110259 | 0.889405 | 2.068451 | 0.545276 | 0.290325 | 2.205321 | 1.651389 | -0.197146 | 1.581328 | 0.597593 | 0.59886 |
| 31.47222 | 1.080588 | 3.930223 | 1.557859 | 5.278337 | 2.430739 | 2.287475 | 1.612481 | 0.671472 | 0.7053 | 4.317573 | 0.941214 | 0.754047 | 0.561433 | -0.288942 | 0.253855 | 2.883249 | -0.516582 | 0.330986 | 3.146036 | 0.428635 | 0.148986 | 1.540553 | 1.926776 | 1.063891 |
| 31.48611 | 1.462955 | 4.510685 | 1.387431 | 2.983271 | 3.569365 | 2.351479 | 1.703694 | -0.413598 | 1.078493 | 5.867654 | 1.308624 | 0.80684 | 0.477849 | -0.123674 | 0.270508 | 2.99815 | 0.471949 | 0.48724 | 4.163864 | 0.680056 | 0.458436 | 3.689095 | 1.779358 | 0.611667 |
| 31.5 | 1.143582 | 4.213451 | 0.531907 | 1.101593 | 3.703629 | 1.949098 | 1.53972 | -1.428752 | 1.638398 | 5.246752 | 1.754631 | 0.729007 | 0.607975 | -0.029103 | -0.603485 | 1.743388 | 2.356836 | 0.681805 | 3.584921 | 2.08826 | 0.299463 | 5.330928 | 0.884921 | -0.645755 |
| 31.51389 | 0.878681 | 3.300968 | -0.114393 | -1.482505 | 1.846109 | 1.502873 | 0.636252 | -1.300817 | 2.09336 | 4.530841 | 1.9319 | 0.672464 | 0.473143 | 0.153248 | -0.313448 | 1.901919 | 1.534985 | 1.08742 | 3.326954 | 2.0031 | 0.172279 | 4.263191 | -0.498457 | -0.325067 |
| 31.52778 | 0.32006 | 2.462891 | 0.016436 | -2.819224 | 0.652482 | 0.938515 | -0.055966 | -0.782326 | 1.807696 | 4.171863 | 1.645351 | 0.717974 | 0.491034 | 0.785233 | 2.434078 | 1.075643 | 1.492409 | 1.565932 | 3.649855 | 5.587614 | 0.300873 | 2.892298 | -0.358065 | 0.480355 |
| 31.54167 | 0.161225 | 2.210561 | 0.28215 | -2.4533 | 0.251763 | 0.644856 | 0.218765 | -0.349962 | 1.481503 | 4.472175 | 1.347037 | 0.752394 | 0.412653 | 1.71418 | 3.780896 | 3.501706 | 1.395531 | 1.560022 | 4.248349 | 5.354775 | 0.647949 | 2.035775 | -0.111758 | 1.014761 |
| 31.55556 | 0.052842 | 2.476786 | 0.356127 | -1.204212 | -0.040836 | 0.617759 | 1.171902 | -0.125521 | 1.210676 | 5.02208 | 1.382095 | 0.789022 | 0.613815 | 2.719718 | 4.425124 | 5.605561 | 1.05227 | 1.192779 | 3.498663 | 5.002513 | 0.933523 | 0.545094 | 0.185841 | 0.02104 |
| 31.56944 | -0.203654 | 4.041869 | 0.46395 | 0.245595 | -0.864562 | 0.774932 | 1.985646 | 0.173324 | 1.407318 | 4.50065 | 1.802359 | 0.863362 | 0.458716 | 1.736229 | 6.065317 | 5.35041 | 0.671154 | 0.449606 | 1.43948 | 4.333267 | 0.869631 | -0.366741 | 1.407476 | 0.959934 |
| 31.58333 | -0.846159 | 5.345129 | 0.627361 | 1.530484 | -3.20785 | 0.291999 | 0.599074 | 0.330322 | 1.441561 | 2.929569 | 1.360137 | 0.725175 | 0.060487 | 1.071324 | 3.324022 | 7.512828 | -0.389627 | -0.24926 | -0.738179 | 2.905184 | 0.546225 | -1.098639 | 0.890071 | -0.295756 |
| 31.59722 | -0.309987 | 5.731773 | 0.745309 | 2.675481 | -5.045022 | -0.200194 | -0.274889 | 0.094066 | 1.449751 | 2.09438 | 0.880606 | 0.64633 | -0.257446 | -0.103063 | 0.145008 | 4.242356 | -1.422729 | -0.548434 | -1.65209 | 2.106209 | 0.362669 | -2.84159 | -1.164784 | -2.687315 |
| 31.61111 | 0.415485 | 4.464188 | 0.815761 | 3.272568 | -2.183998 | 0.113586 | 0.151431 | 0.136565 | 1.613764 | 2.15308 | 0.653419 | 0.857958 | -0.354703 | -1.321337 | -1.520269 | -0.089194 | -2.380318 | -0.431151 | -1.771051 | 0.94362 | 0.279044 | -4.751406 | -0.7801 | -4.428084 |
| 31.625 | 0.745581 | 4.225183 | 1.000445 | 4.437983 | -2.165501 | 0.36195 | 0.837352 | 0.322021 | 1.572498 | 2.946161 | 0.399272 | 1.124897 | 0.273493 | -0.853339 | -1.84204 | -5.085288 | -3.052526 | -0.129993 | -0.695444 | 1.307932 | 0.372627 | -3.906697 | -0.599221 | -2.72678 |
| 31.63889 | 0.77797 | 2.528802 | 1.415338 | 4.234346 | 0.29225 | 0.265837 | 0.463049 | 0.757562 | 1.601772 | 3.244684 | 0.330038 | 1.267556 | 0.823786 | 0.115071 | -2.153432 | -8.149906 | -1.574618 | 0.568465 | 0.380386 | 1.990583 | 0.449019 | -1.812274 | -0.646607 | -1.04655 |
| 31.65278 | 0.718834 | -0.117292 | 1.511679 | 3.519559 | 3.323376 | 0.20666 | 0.008208 | 1.459231 | 2.317286 | 1.708652 | 0.309382 | 1.184227 | 1.000195 | 0.997644 | -1.468091 | -3.705274 | -0.291391 | 1.248114 | 1.153923 | 2.502305 | 0.405501 | 0.303867 | -0.319344 | 0.300106 |
| 31.66667 | 0.090042 | -0.973263 | 1.831789 | 3.924302 | 4.665079 | 0.214571 | 0.197589 | 2.363572 | 2.8055 | 0.180296 | 0.436158 | 0.903741 | 1.141342 | 1.714875 | 0.988403 | -0.584321 | 0.318905 | 1.592963 | 2.235777 | 2.242532 | 0.336673 | 0.580677 | 0.190672 | 1.034327 |
| 31.68056 | -0.092144 | -2.07956 | 1.959036 | 3.720736 | 4.57647 | 0.381674 | 0.665457 | 2.077701 | 2.386329 | -0.792035 | 0.568821 | 0.826837 | 1.408161 | 1.452049 | 3.04997 | -0.477017 | 0.326933 | 1.844501 | 3.312993 | 2.137194 | 0.263211 | 0.051398 | 0.633547 | 1.361132 |
| 31.69444 | -0.033975 | 1.14289 | 1.778321 | 2.901791 | 5.844123 | 0.183703 | -0.356462 | 1.477808 | 1.530323 | -0.763222 | 0.705863 | 1.060001 | 1.289509 | 1.165379 | 2.915714 | -1.241015 | -0.125114 | 1.4713 | 3.416226 | 2.090585 | 0.251179 | -0.286684 | 1.065146 | 1.147196 |
| 31.70833 | 0.714737 | 1.93298 | 2.23478 | 1.735974 | 3.983 | 0.235358 | -2.228061 | 2.105954 | 0.970822 | -0.309062 | 0.801461 | 1.469495 | 1.097966 | 1.056493 | 3.852716 | -3.194708 | 0.404267 | 0.954574 | 2.751475 | 1.851332 | 0.4927 | -1.33457 | 1.671868 | 0.944559 |
| 31.72222 | 1.077769 | 1.837165 | 1.376092 | 0.215392 | 2.414583 | 0.435323 | -1.59251 | 2.658022 | 1.072815 | -0.342301 | 0.835227 | 1.656478 | 1.059333 | 1.120847 | 2.312352 | -4.012556 | 1.43289 | 0.768225 | 1.850115 | 0.912281 | 0.645307 | -1.710015 | 1.787173 | 0.655178 |
| 31.73611 | 0.668494 | 1.765914 | 0.219674 | -2.873175 | 1.585966 | 0.265667 | 0.978657 | 2.118696 | 1.669979 | -1.522606 | 1.15421 | 1.506616 | 0.741166 | 1.225479 | 0.711775 | -3.136148 | 1.330616 | 0.55068 | 0.414326 | -0.355223 | 0.786332 | -1.047139 | 1.56651 | 0.670185 |
| 31.75 | 0.271116 | 1.94695 | -0.407961 | -6.207644 | 1.133823 | 0.087895 | 2.492617 | 1.102525 | 1.450016 | -3.338278 | 1.411039 | 1.16998 | 0.037979 | 0.763007 | 0.43534 | -2.844405 | 0.594309 | 0.33325 | -0.961534 | -0.39419 | 1.120795 | -0.855707 | 1.378693 | 0.631195 |
| 31.76389 | -0.003921 | 1.344854 | -0.04279 | -5.022229 | 0.770146 | 0.079449 | 2.067548 | -0.262502 | 0.817277 | -4.295755 | 1.397611 | 0.797195 | -0.577845 | -0.509221 | 0.026552 | -2.346345 | 0.011339 | -0.248817 | -2.16423 | -0.040478 | 1.651787 | -0.966761 | 1.196321 | -0.119003 |
| 31.77778 | 0.084825 | 2.19527 | 0.035186 | -4.990363 | 0.008 | 0.396069 | 0.960462 | -1.780053 | -0.37569 | -4.742011 | 1.141728 | 0.457176 | -1.117542 | -1.119723 | -0.613963 | -1.507405 | -0.609754 | -0.540697 | -3.398059 | -0.084974 | 2.943261 | -2.642409 | 0.826678 | -0.525489 |
| 31.79167 | 1.204066 | 3.296017 | -0.284222 | -3.812401 | -0.639028 | 0.006853 | 1.207549 | -1.639082 | -1.038141 | -4.694297 | 0.874896 | 0.333843 | -1.762969 | -0.712737 | -1.238113 | -1.412174 | -1.264994 | -0.296151 | -4.307747 | -0.569452 | 2.228839 | -3.79494 | 0.799719 | -0.792677 |
| 31.80556 | 1.475299 | 2.357807 | -0.773103 | -3.961023 | -1.128976 | -0.112425 | -0.150859 | -0.921106 | -1.101995 | -3.931521 | 0.573817 | 0.281223 | -0.994813 | -0.588326 | -1.266173 | -0.925193 | -1.488164 | 0.312245 | -3.966122 | -0.957525 | 0.560292 | -4.488414 | 0.508075 | -0.861483 |
| 31.81944 | 0.806062 | 0.557686 | -1.20564 | -3.882259 | -0.302248 | 0.308053 | 0.333344 | -1.290696 | -1.358377 | -3.556739 | 0.434041 | 0.164335 | -0.637492 | -0.76313 | -1.334912 | -0.802802 | -1.424784 | 0.698729 | -3.455376 | -1.419268 | -0.388985 | -5.085206 | -0.208789 | -1.009125 |
| 31.83333 | -0.588148 | 5.280612 | -1.555264 | -5.033299 | -1.414259 | 0.558428 | 0.198326 | -2.608011 | -2.127328 | -3.739495 | 0.204535 | -0.13106 | -0.524309 | -0.426609 | -0.764418 | -0.778779 | -1.985004 | 0.47956 | -2.840717 | -1.527508 | -1.61234 | -6.202866 | -1.388592 | -1.620693 |
| 31.84722 | -0.315674 | 3.883116 | -1.574451 | -4.468556 | -3.734588 | 0.458326 | -0.047136 | -3.196034 | -1.338171 | -4.699283 | 0.139787 | -0.380345 | -0.286337 | -0.620883 | -0.323064 | -0.716016 | -2.599913 | -0.166162 | -2.475065 | -1.591202 | -2.190247 | -8.413979 | -1.331725 | -1.884667 |
| 31.86111 | -1.035394 | -3.332289 | -1.679283 | -3.215764 | -5.741302 | 0.209633 | 0.102514 | -2.211566 | -0.481416 | -7.021795 | 0.108338 | -0.562233 | 0.206262 | -0.263812 | -2.021138 | -0.232705 | -2.061551 | -0.50919 | -2.363432 | -2.398204 | -1.451015 | -6.93195 | -0.510254 | -1.628353 |
| 31.875 | -2.540951 | -5.569891 | -3.182253 | -2.597692 | -6.488686 | -0.347076 | 0.149637 | -1.977679 | 0.104063 | -8.381788 | -0.429057 | -0.874226 | 0.881554 | -0.646985 | -0.874119 | -0.824298 | -0.903093 | 0.390322 | -1.64936 | -2.910564 | -1.071754 | -6.672198 | 0.328143 | -1.4921 |
| 31.88889 | -2.298258 | -6.410626 | -3.158672 | -2.631037 | -5.012642 | -0.942621 | 0.027684 | -1.939345 | -0.482626 | -6.201101 | -1.077806 | -0.822079 | 1.024929 | -0.293463 | 2.241081 | -0.844122 | -0.51835 | 1.410896 | -0.382596 | -2.949099 | -0.807093 | -6.642799 | 1.145388 | -0.96105 |
| 31.90278 | -1.857236 | -1.616298 | -2.060131 | -2.606337 | -6.338371 | -0.712724 | 0.326494 | -0.654431 | -0.735991 | -7.148814 | -1.426101 | -0.370336 | 1.172775 | 0.157232 | 1.609169 | -0.649774 | -0.090726 | 1.299139 | 0.120633 | -2.292542 | 0.013545 | -6.311205 | 1.556358 | -0.185396 |
| 31.91667 | -1.268145 | -0.384486 | -1.185121 | -2.292078 | -7.597191 | -0.262992 | 0.525572 | 0.820536 | 0.269348 | -1.615303 | -0.98066 | -0.04517 | 1.290613 | 0.010788 | 0.531553 | -0.698606 | 0.206561 | 0.670045 | -0.305555 | -1.684263 | 0.69455 | -5.479033 | 1.587796 | 0.24494 |
| 31.93056 | -0.278658 | -0.593509 | -0.897738 | -0.664636 | -2.441035 | -0.501912 | 0.249285 | 1.885044 | 0.553014 | 0.452408 | -0.36434 | -0.081536 | 1.331151 | -0.368644 | -1.308319 | -1.142332 | -0.076353 | 0.207115 | -1.730832 | -1.642081 | 1.110925 | -3.967955 | 1.467716 | 0.250027 |
| 31.94444 | -0.021644 | -1.224601 | -0.678825 | 0.032461 | 0.129599 | -0.120963 | -1.030626 | 2.406399 | 0.28191 | 2.44039 | -0.391152 | -0.188347 | 0.326238 | -0.641576 | -4.412417 | -1.941838 | -1.290136 | -0.631382 | -2.593647 | -2.151843 | 1.318514 | -3.320763 | 0.355073 | -0.262639 |
| 31.95833 | -0.331754 | -1.780507 | -0.282834 | 0.729292 | -0.247878 | 0.128772 | -1.019469 | 2.500985 | 0.250269 | 4.45613 | -0.648057 | -0.167074 | -0.957288 | -0.748719 | -4.514299 | -3.523573 | -2.394696 | -2.448892 | -4.661754 | -2.162273 | 1.08856 | -3.813253 | -1.432997 | -0.898636 |
| 31.97222 | -0.11405 | -1.127161 | 0.008037 | 0.528729 | -0.552704 | 0.184273 | -0.03022 | 1.929953 | 0.657409 | 4.896837 | -0.752953 | -0.048369 | -1.302402 | -1.081427 | -3.218881 | -4.391572 | -2.992167 | -2.072221 | -5.616782 | -1.986333 | 0.424495 | -3.439415 | -2.245854 | -0.891484 |
| 31.98611 | 0.029505 | 0.616869 | 0.07533 | -0.311225 | -0.996852 | 0.159982 | 0.615282 | 0.143136 | 0.627858 | 6.149865 | -0.486242 | 0.23646 | -0.987512 | -1.122234 | -1.111654 | -3.565442 | -3.076324 | -1.641937 | -3.782014 | -1.414027 | -0.005129 | -2.272689 | -1.110056 | -0.597862 |
| 32 | 0.073956 | 2.987937 | 0.068701 | -0.679483 | -1.10918 | 0.059679 | 1.470356 | 1.900595 | 0.74247 | 7.351998 | 0.19672 | 0.715733 | -0.523856 | -0.93088 | -2.365513 | -1.987332 | -2.29638 | -0.483821 | -1.840963 | -1.644242 | -0.221136 | -1.685595 | -0.957596 | -0.427974 |
| 32.01389 | -0.016954 | 3.388685 | 0.105051 | 0.369256 | -1.424059 | 0.029135 | 1.200566 | 2.168576 | 0.58456 | 7.082559 | 0.556876 | 0.773727 | 0.386991 | -0.599598 | -2.03167 | -0.373347 | -0.185051 | -0.483356 | -0.731782 | -1.820106 | -0.339438 | -1.306762 | -1.556351 | -0.27176 |
| 32.02778 | 0.261287 | 2.278001 | 0.177187 | 0.512301 | 0.428355 | 0.22097 | 0.158541 | 1.92685 | 0.945176 | 6.520716 | 0.571291 | 0.739509 | 1.281321 | -0.396963 | 0.594112 | -0.118769 | 1.727132 | -0.81954 | -0.789444 | -1.417981 | -0.177138 | -0.628661 | -2.022558 | -0.118975 |
| 32.04167 | 0.511098 | 1.643054 | 0.344179 | 0.721002 | 1.098319 | 0.402377 | 0.026917 | 0.632985 | 1.686992 | 5.259246 | 0.487455 | 0.722658 | 1.558524 | -0.08225 | 1.697894 | -0.428785 | 2.028013 | -0.029925 | -0.977228 | -0.912577 | -0.100695 | 0.750623 | -2.071794 | 0.223447 |
| 32.05556 | 1.191178 | 1.654567 | 0.597213 | 0.496161 | 2.490771 | 0.419528 | 0.244121 | -0.297741 | 1.422557 | 3.322136 | 0.211509 | 0.607284 | 1.523245 | 0.365427 | 1.077507 | -0.559101 | 1.916176 | 1.264983 | 0.294244 | -0.520928 | 0.052296 | 2.512201 | -2.99232 | 0.705496 |
| 32.06944 | 2.014585 | 2.314828 | 0.469467 | 0.12842 | 2.832136 | 0.36692 | 0.151521 | -1.315827 | 0.216442 | 1.111313 | 0.065551 | 0.529192 | 0.938963 | 0.282188 | -0.192802 | -0.462403 | 1.718753 | 1.438261 | 0.95738 | -0.564826 | 0.118966 | 3.690871 | -3.488925 | 0.706729 |
| 32.08333 | 1.919896 | 3.185024 | 0.405054 | 0.037622 | 2.969922 | 0.290806 | -0.071868 | -0.945859 | -1.05578 | -0.601421 | 0.015996 | 0.585266 | 0.329787 | -0.118916 | 0.280085 | -0.6714 | 1.501466 | 0.691833 | 0.560532 | -0.797436 | 0.233467 | 3.227322 | -3.008366 | 0.102182 |
| 32.09722 | 0.569988 | 2.422038 | 0.329932 | 0.156269 | 2.141012 | 0.299234 | -0.314901 | -0.193752 | -1.580696 | -1.3873 | -0.107836 | 0.553317 | -0.10134 | -0.182352 | 0.708908 | -0.315352 | 1.292676 | -0.291056 | 0.319262 | -0.411636 | 0.1115 | 3.897297 | -2.551297 | -0.29687 |
| 32.11111 | -0.41029 | 0.434621 | 0.291837 | 0.334644 | 3.025255 | 0.364324 | 0.43614 | -0.073496 | -0.807347 | -2.524777 | -0.192328 | 0.320594 | -0.1479 | -0.390882 | 0.868418 | 0.122852 | 1.448502 | -0.654058 | -0.898957 | 0.200549 | -0.296292 | 6.163747 | -1.850557 | -0.101363 |
| 32.125 | -0.72015 | -1.359878 | 0.093153 | 0.201185 | 3.555516 | 0.425109 | 0.458748 | 0.68983 | -0.823852 | -5.414045 | -0.402374 | -0.011668 | -0.103409 | 0.373419 | -0.347328 | -0.432214 | 1.576134 | -0.486134 | -1.380315 | 0.596917 | -0.64503 | 9.150385 | -0.954386 | 0.343863 |
| 32.13889 | -0.454438 | -2.914354 | -0.163369 | -0.115676 | 2.40495 | -0.057184 | -0.820352 | 0.566164 | -0.830284 | -7.868537 | -0.637887 | -0.48357 | 0.015343 | 0.856362 | -1.974435 | -0.74924 | 1.103573 | -0.477794 | -0.878587 | 0.776222 | -0.101817 | 6.819218 | -0.33629 | 0.429453 |
| 32.15278 | -0.83818 | -4.635943 | -0.538142 | -0.453492 | 1.18606 | -0.396465 | -1.273512 | 1.256725 | -0.422852 | -5.132566 | -0.900119 | -1.092509 | -0.094619 | 0.850998 | -2.780802 | -1.211774 | -0.091444 | -0.428244 | 0.378371 | 0.813107 | 0.169625 | 5.160403 | -0.169678 | 0.329542 |
| 32.16667 | -0.823957 | -6.14952 | -0.664883 | -0.518477 | 0.360905 | -0.33578 | -0.826484 | 0.497983 | -0.526467 | -8.619863 | -0.865469 | -1.374194 | -0.209621 | 1.864763 | -1.470508 | -2.226361 | -0.707561 | -0.382855 | 0.728793 | 0.943828 | 0.121065 | 4.784436 | -0.055141 | 0.473031 |
| 32.18056 | -0.402645 | -5.041298 | -0.59107 | -0.301904 | -0.044892 | -0.129321 | 0.049309 | 0.513719 | -0.68595 | -7.48605 | -0.43147 | -0.625753 | -0.045956 | 0.899756 | 1.34115 | -1.749774 | -0.912036 | -0.190549 | -0.039625 | 0.885517 | 0.083743 | 3.196063 | 0.351191 | 0.55521 |
| 32.19444 | -0.289568 | -3.442523 | -0.530138 | -0.102079 | 0.802371 | -0.340499 | -0.818323 | 1.77612 | -0.363887 | -5.56438 | -0.026628 | -0.507436 | 0.224029 | -0.247296 | 4.633028 | -1.532114 | -1.101701 | -0.006663 | -0.617705 | 0.376828 | 0.271313 | 0.502393 | 0.786881 | 0.346789 |
| 32.20833 | 0.446566 | -2.18968 | -0.491875 | -0.123909 | 1.029715 | -0.16141 | -1.337599 | 1.089953 | 0.052237 | -1.84725 | 0.187368 | -0.31189 | 0.191028 | -0.976977 | 3.546056 | -1.03627 | -1.631639 | 0.059605 | -0.334548 | -0.635756 | 0.435494 | -2.270527 | 0.941095 | 0.395532 |
| 32.22222 | 0.777016 | -1.239746 | -0.460497 | -0.19526 | 1.052602 | 0.057753 | -0.131092 | -0.13626 | 0.356145 | 1.635458 | 0.318837 | -0.17795 | 0.035625 | -0.733133 | 0.964717 | -1.287876 | -2.243964 | -0.024408 | 0.242625 | -0.901469 | -0.034788 | -3.563222 | 1.135643 | 0.176899 |
| 32.23611 | 1.299703 | -0.616192 | -0.420557 | -0.164885 | 2.084776 | 0.100329 | 0.303123 | -0.626734 | 0.707122 | 4.304602 | 0.527853 | -0.004279 | 0.344891 | -0.025252 | -0.103598 | -0.30482 | -1.289672 | -0.111751 | 0.821548 | 0.497746 | -0.349738 | 2.576552 | 1.169106 | 0.388474 |
| 32.25 | 1.387868 | 0.116083 | -0.252239 | 0.045077 | 2.975853 | 0.158267 | 0.133664 | -0.55524 | 1.074351 | 4.633956 | 0.825579 | 0.142425 | 0.585234 | -1.238345 | 0.301305 | 1.469276 | -0.470004 | -0.212166 | 1.085644 | 1.224117 | 0.073606 | 4.029425 | 1.259043 | 1.144588 |
| 32.26389 | 1.178649 | 0.503898 | -0.056563 | 0.35573 | 3.640175 | 0.104945 | -0.091797 | -0.507844 | 1.220998 | 4.702272 | 0.945467 | 0.262819 | 0.278619 | 0.099859 | 0.8851 | 1.616923 | -0.570691 | -0.176173 | 0.740851 | 0.496358 | -0.014413 | 1.585335 | 0.869049 | 0.898828 |
| 32.27778 | 0.762087 | 0.5687 | 0.118862 | 0.637316 | 4.080545 | 0.068208 | -0.154689 | -0.481914 | 1.543326 | 3.878723 | 1.18427 | 0.290455 | 0.012214 | 0.060064 | 0.101695 | 1.040639 | -0.660469 | 0.00105 | 0.842067 | 0.061916 | -0.094062 | 0.089543 | 0.308671 | 0.273969 |
| 32.29167 | 0.33303 | 0.777163 | 0.302882 | 0.889638 | 3.917992 | 0.16141 | -0.191259 | -0.411874 | 1.402497 | 3.146006 | 1.441409 | 0.342487 | -0.079764 | -0.228355 | -1.367492 | 1.053999 | 0.022531 | 0.170346 | 0.814128 | 0.386711 | 0.083798 | 0.385554 | -0.123313 | 0.04696 |
| 32.30556 | -0.164792 | 1.838916 | 0.551875 | 1.196945 | 3.809788 | 0.269076 | 0.841661 | 0.055248 | 1.507834 | 5.271222 | 1.428181 | 0.512134 | -0.175306 | -0.27361 | -2.616351 | 1.844683 | 0.660304 | 0.137626 | 0.588727 | 0.988006 | 0.159216 | 1.873451 | -0.666443 | 0.172584 |
| 32.31944 | 0.094189 | 3.224875 | 0.69281 | 1.443396 | 2.930518 | 0.325336 | 0.341922 | 0.419739 | 1.493567 | 5.668352 | 1.090392 | 0.684576 | -0.208054 | 0.152064 | -3.085314 | 1.723141 | 0.65655 | 0.146381 | 0.279418 | 1.32245 | 0.312718 | 3.479278 | -1.297777 | 0.402742 |
| 32.33333 | 0.078919 | 3.553323 | 0.654469 | 1.376127 | 1.76791 | 0.322574 | 0.467299 | 0.821109 | 1.487852 | 5.53117 | 0.541159 | 0.56736 | -0.126835 | 1.224881 | -1.659778 | 1.033543 | 0.655271 | 0.135001 | 0.244819 | 1.29576 | 0.02358 | 4.076585 | -1.414406 | 0.781244 |
| 32.34722 | -0.256276 | 1.986421 | 0.576916 | 1.26986 | 1.728898 | 0.156646 | 0.639178 | 0.687044 | 1.52817 | 4.79144 | 0.148076 | 0.113442 | -0.041444 | 2.095807 | 0.325703 | 1.103161 | 0.96563 | 0.068872 | 0.370279 | 1.236655 | -0.164247 | 3.878435 | -1.492424 | 0.916175 |
| 32.36111 | -0.242008 | 0.752196 | 0.524058 | 1.098555 | 1.991246 | 0.166045 | 1.249616 | -0.042001 | 1.679331 | 2.730651 | -0.194481 | -0.250343 | -0.010957 | 1.515381 | 1.477131 | 1.118602 | 0.965868 | 0.009318 | 0.435117 | 1.202992 | -0.210175 | 3.375859 | -1.370825 | 0.506649 |
| 32.375 | -0.031437 | 0.380667 | 0.417681 | 0.939502 | 1.305969 | 0.2075 | 0.995957 | -0.592888 | 1.602774 | 2.580043 | -0.381889 | -0.397571 | 0.075696 | -0.031764 | 1.610864 | 1.142192 | 0.998373 | 0.06957 | 0.514902 | 1.324043 | -0.201464 | 3.514107 | -1.179652 | 0.36336 |
| 32.38889 | 0.332341 | 1.056161 | 0.473166 | 1.101704 | 2.786869 | 0.706332 | 1.605111 | -0.491939 | 1.040475 | 4.669689 | -0.623201 | -0.329543 | 0.372283 | 0.329662 | 1.464842 | 1.654646 | 2.409225 | 0.273831 | 0.704591 | 1.48677 | -0.065755 | 5.018034 | -0.868001 | 0.773898 |
| 32.40278 | 0.598993 | 1.854963 | 1.083207 | 1.833227 | 2.632401 | 1.554207 | 1.871159 | 1.246303 | 0.601892 | 4.597329 | -0.695451 | -0.186115 | 0.760929 | 1.635161 | 1.850899 | 2.663536 | 3.439904 | 0.549105 | 0.823858 | 1.773641 | 0.205927 | 5.675887 | -0.53345 | 1.60963 |
| 32.41667 | 0.655784 | 1.859282 | 2.24161 | 3.351729 | 0.990361 | 1.218893 | 0.15475 | 0.927311 | 0.565789 | 2.183839 | -0.497991 | -0.013994 | 1.115816 | 1.248 | 2.675617 | 3.712765 | 3.185766 | 0.662007 | 0.808041 | 2.0575 | 0.2999 | 6.341399 | -0.144114 | 1.665913 |
| 32.43056 | 0.631119 | 2.625544 | 2.828396 | 4.475458 | -0.303086 | 0.299897 | 0.040311 | 1.175697 | 0.679157 | 0.859778 | -0.076902 | 0.27968 | 1.389739 | 0.786358 | 3.018562 | 4.186628 | 3.564213 | 0.653137 | 0.739882 | 2.213179 | -0.090621 | 6.348289 | -0.258651 | 0.972228 |
| 32.44444 | 1.32756 | 4.640372 | 2.578976 | 4.903418 | -1.027958 | -0.620505 | -0.361109 | 0.704515 | 0.712065 | 0.209195 | 0.468749 | 0.922163 | 1.531126 | 0.19694 | 3.199851 | 2.803022 | 3.22072 | 0.63363 | 0.852708 | 1.962454 | -0.660743 | 6.10894 | 0.790732 | 0.813007 |
| 32.45833 | 0.989238 | 5.140236 | 2.496484 | 5.348737 | -4.085235 | -0.966487 | -0.18871 | 0.437228 | 0.841328 | 0.291802 | 1.012183 | 1.413532 | 1.402514 | 0.059999 | 2.840729 | 1.11225 | 2.033762 | 0.814776 | 1.484346 | 1.179491 | -0.510611 | 4.255312 | 1.487464 | 1.042829 |
| 32.47222 | -0.096995 | 4.06967 | 3.125407 | 4.80003 | -6.882305 | -0.391865 | 0.572805 | 0.583255 | 1.076777 | -0.666909 | 1.365058 | 1.535546 | 1.515955 | 0.604076 | 2.060331 | 0.934947 | 1.813152 | 1.344405 | 3.008979 | 1.045452 | 0.187901 | 4.130305 | 1.375142 | 1.426024 |
| 32.48611 | 0.924274 | 2.11784 | 3.054056 | 5.033801 | -6.117054 | -0.013848 | 0.764157 | 0.743029 | 1.301599 | -0.954963 | 1.338881 | 1.181656 | 1.536925 | 2.319194 | 2.008803 | 2.615647 | 2.343763 | 2.122698 | 4.538372 | 1.846251 | 0.669421 | 6.393509 | 1.100681 | 2.159057 |
| 32.5 | -0.182201 | 0.750103 | 2.138325 | 4.33695 | -8.322204 | 0.172278 | 1.250203 | -0.431375 | 1.512413 | -0.189698 | 1.257171 | 0.916706 | 1.132889 | 1.336065 | 2.036222 | 4.912569 | 3.577759 | 2.500299 | 3.101151 | 3.38641 | 0.862087 | 4.566728 | 1.225706 | 1.431267 |
| 32.51389 | -1.81106 | 1.051073 | 1.382731 | 3.107629 | -9.222954 | 0.290657 | 1.318731 | -0.471158 | 1.297146 | 0.587543 | 1.109147 | 0.799701 | 0.65886 | 0.654244 | 1.957098 | 4.635217 | 2.638873 | 2.264583 | 2.874637 | 4.456753 | 0.88986 | 3.543911 | 1.650113 | 0.427412 |
| 32.52778 | -1.602222 | 1.429787 | 1.271493 | 2.235232 | -5.295785 | 0.340744 | 1.177031 | -0.327986 | 0.629378 | 1.554324 | 0.892735 | 0.965333 | 0.588205 | 1.760257 | 3.520916 | 4.78018 | 1.613759 | 2.31011 | 2.958643 | 4.748725 | 1.032516 | 0.593708 | 2.237447 | 0.615312 |
| 32.54167 | -1.42422 | 1.966685 | 1.095984 | 1.61172 | -0.787532 | 0.388319 | 0.934069 | -0.041837 | -0.275523 | 2.770181 | 0.834803 | 1.277441 | 0.62078 | 1.894742 | 3.543697 | 2.898702 | 1.374639 | 2.282216 | 3.690978 | 4.329696 | 1.44989 | -1.022604 | 2.193379 | 0.847229 |
| 32.55556 | -0.675315 | 1.621449 | 0.7491 | 1.101848 | 0.367767 | 0.447253 | 0.689949 | 0.655155 | -0.951003 | 2.728054 | 0.840741 | 1.421892 | 0.106149 | 1.639709 | 1.849837 | 2.31194 | 1.885142 | 2.044973 | 3.792266 | 4.137995 | 1.737293 | 1.658705 | 2.801162 | 0.580599 |
| 32.56944 | -0.20752 | -0.030583 | 0.187528 | 0.737492 | 0.106836 | 0.527409 | 0.435925 | 0.869929 | -1.29498 | 1.670649 | 1.104674 | 1.15777 | -0.557659 | 1.495123 | 0.307404 | 3.913073 | 2.733738 | 1.528737 | 3.057566 | 3.525534 | 1.130449 | 4.16271 | 4.797636 | 0.169124 |
| 32.58333 | 1.745437 | -1.057291 | -0.459011 | 0.663829 | 5.38992 | 0.578947 | 0.189727 | 1.150361 | -1.15633 | 1.733862 | 1.311971 | 0.982152 | -0.992093 | -0.402596 | -0.959036 | 2.298691 | 0.954162 | 0.520829 | 2.0991 | 2.901624 | 0.585442 | 2.495732 | 4.266192 | -0.874594 |
| 32.59722 | 2.255824 | -0.001216 | -0.78869 | 0.180932 | 6.827506 | 0.579964 | -0.012279 | 0.755906 | -1.127228 | 3.234952 | 1.47927 | 1.238169 | -1.288565 | -1.541794 | -1.140013 | 3.33225 | -1.066831 | -0.708116 | 1.156822 | 2.484334 | 0.072693 | -0.41772 | 3.397551 | -0.9077 |
| 32.61111 | 1.093759 | 2.096712 | -0.662986 | -0.255619 | 10.987666 | 0.729862 | -0.910545 | -0.111119 | -0.886023 | 4.813707 | 1.417731 | 1.938325 | -0.881258 | -0.800027 | -1.47898 | 2.665756 | -2.679979 | -1.407251 | 0.23576 | 2.286257 | -0.246247 | -0.490421 | 2.147683 | -0.449064 |
| 32.625 | 1.18191 | 3.494668 | -0.367066 | -1.304041 | 8.762417 | 0.740183 | -0.0693 | -3.156157 | 0.001276 | 4.851016 | 1.292017 | 2.599928 | -0.091983 | -0.042664 | -1.636254 | 2.32893 | -2.816286 | -1.276796 | -0.132862 | 2.092642 | -0.146987 | 0.570705 | 1.063684 | 0.47073 |
| 32.63889 | -0.628446 | 5.081603 | 0.125401 | -1.625738 | 3.135248 | 0.79721 | 0.355389 | -1.647205 | 1.576415 | 4.658699 | 1.115005 | 3.270444 | 0.733081 | 0.68367 | -2.040512 | 2.183024 | -1.14567 | 0.467337 | 1.034681 | 2.326385 | 0.271804 | 0.328004 | 0.51163 | 1.517614 |
| 32.65278 | -1.568225 | 6.557832 | 0.515673 | -1.171319 | 1.414291 | 0.267739 | -0.998458 | 1.622232 | 3.348381 | 4.505389 | 0.891721 | 2.319924 | 1.749106 | -0.119056 | -1.812532 | 1.454391 | 0.4852 | 1.866521 | 1.077662 | 2.509495 | 0.964674 | 2.032558 | -0.528931 | 1.101091 |
| 32.66667 | -0.771515 | 2.841612 | 1.031108 | -1.402294 | 1.517628 | 0.018956 | -1.390241 | 2.954968 | 5.127338 | 3.884236 | 0.994228 | 1.595329 | 1.717661 | -0.935911 | -1.647944 | 1.52704 | 1.222253 | 1.394811 | -0.122434 | 2.259822 | 1.827552 | 0.88721 | -1.325389 | 0.495507 |
| 32.68056 | 0.091079 | 2.594392 | 1.58772 | -2.190533 | 1.177135 | -0.575967 | -2.755204 | 5.129372 | 5.854116 | 2.876828 | 1.264128 | 1.382103 | 1.074561 | -1.058416 | -0.731703 | 0.054779 | 0.895655 | 0.541248 | -0.844344 | 1.79363 | 2.768118 | 1.224938 | -1.859184 | 0.33334 |
| 32.69444 | 0.190042 | 2.658989 | 1.563145 | -0.243912 | -1.415527 | -0.627937 | -3.619469 | 3.378805 | 5.023332 | 1.510274 | 0.865443 | 1.458564 | 0.607806 | -0.774934 | -0.75362 | -1.036201 | 0.666847 | 0.367823 | -1.298717 | 1.198947 | 3.600727 | 0.472234 | -1.853153 | 0.415352 |
| 32.70833 | 0.041241 | 2.926477 | 1.623931 | 1.066029 | -2.16429 | -0.015088 | -1.876525 | 1.742288 | 3.576993 | 1.394985 | 0.85463 | 1.721393 | 0.117302 | -0.359822 | -0.170741 | -2.160124 | 0.744054 | 0.832483 | -0.729519 | 1.063393 | 3.78234 | 0.02818 | -1.328907 | 0.109499 |
| 32.72222 | 0.06014 | -0.044742 | 1.928614 | 1.13946 | -1.079121 | -0.010944 | -0.340073 | 2.584984 | 2.051191 | 1.967671 | 0.818522 | 1.736419 | -0.146106 | 0.741874 | 0.451796 | -2.934418 | 0.674147 | 1.294567 | -0.165664 | 1.407118 | 2.812211 | 0.503878 | -1.038644 | 1.242057 |
| 32.73611 | 0.445424 | -6.613926 | 2.020592 | 1.093763 | 0.594181 | -0.009011 | -0.400639 | 1.181506 | 1.285953 | 2.677797 | 0.80694 | 1.177415 | -0.376762 | 2.090375 | 0.967811 | -2.052646 | 0.438295 | 1.312143 | -0.638877 | 1.638464 | 1.91257 | 0.791973 | -1.350549 | 2.314086 |
| 32.75 | 0.724702 | -8.086058 | 1.717504 | 1.588721 | 0.981977 | 0.009938 | 0.215002 | -0.191912 | 1.227079 | 1.298084 | 0.818981 | 0.252527 | -0.683814 | 2.859662 | 0.24512 | -1.013107 | 0.400352 | 0.920821 | -2.170342 | 0.820497 | 1.238454 | 0.51113 | -0.727381 | 2.699101 |
| 32.76389 | 0.582232 | -6.575684 | 1.439433 | 2.054896 | 0.866848 | 0.039417 | 0.23784 | -0.909236 | 1.35727 | 0.449549 | 0.40265 | -0.533301 | -0.611485 | 1.440924 | -1.603588 | -0.618593 | 0.123829 | 0.63748 | -2.721723 | 0.319073 | 0.70599 | -0.538704 | -0.371456 | 0.790587 |
| 32.77778 | 0.420348 | -5.425165 | 1.241011 | 1.69534 | 1.026726 | 0.072015 | -0.198366 | -0.44647 | 1.183989 | -0.669461 | 0.445885 | -1.497004 | -0.556396 | 0.141609 | -2.514689 | -0.322998 | -0.71765 | -0.05694 | -2.749322 | 0.620508 | 0.660696 | -2.303751 | 0.01942 | -1.225502 |
| 32.79167 | 0.286807 | -4.934142 | 0.931666 | 1.004613 | 1.880827 | 0.094658 | -0.814205 | -0.348277 | -0.181487 | -2.352897 | 0.491992 | -1.84506 | -1.21562 | -0.929957 | -2.250369 | -0.234005 | -1.160619 | 0.13768 | -2.859719 | 1.033476 | 0.237745 | -2.296498 | 0.402348 | -1.792365 |
| 32.80556 | 0.184886 | -3.72059 | 0.372129 | 0.721903 | 3.214824 | 0.092258 | -0.893808 | -0.487868 | -0.738966 | -3.279166 | 0.620467 | -0.904893 | -1.781989 | -1.929022 | -2.465473 | -0.615144 | -0.534011 | 1.131741 | -3.331367 | 0.970426 | -1.061614 | -3.030237 | -0.247523 | -1.958808 |
| 32.81944 | 0.088466 | -1.8329 | -0.245986 | 0.428227 | 4.179287 | 0.152083 | -0.580363 | -0.894507 | -0.858395 | -4.971566 | 0.64226 | -0.714269 | -2.203946 | -2.619274 | -1.460145 | -1.477806 | -1.48361 | 1.858981 | -3.263574 | 0.644139 | -1.991916 | -4.443963 | -0.311003 | -1.739575 |
| 32.83333 | 0.293115 | -6.295548 | -0.816754 | -0.478494 | -1.812699 | 0.335043 | -0.354396 | -0.616067 | -1.499376 | -4.929239 | 0.312992 | -0.764665 | -1.481144 | -2.425908 | -1.150711 | -1.820137 | -2.349764 | 2.059347 | -3.175798 | 0.181627 | -1.89148 | -5.979115 | -0.040951 | -2.155311 |
| 32.84722 | -0.279704 | -7.334994 | -1.497374 | -1.616369 | -5.018254 | 0.49421 | 0.147363 | -1.711836 | -0.49486 | -2.69547 | 0.346241 | -0.358621 | -0.367068 | -1.651524 | -1.418957 | -1.390114 | -2.297407 | 2.233258 | -2.965874 | -0.21134 | -1.801814 | -6.935082 | 0.68713 | -2.285188 |
| 32.86111 | -0.510229 | -5.159228 | -2.364026 | -2.916789 | -0.723577 | 0.363115 | 0.186575 | -0.909639 | 0.509146 | -2.724443 | -0.378819 | -1.310279 | 0.5295 | -1.014723 | -2.070992 | -1.006336 | -2.076656 | 1.855407 | -2.427713 | -0.588592 | -1.998564 | -6.496348 | 0.666613 | -1.681513 |
| 32.875 | -0.613477 | -5.973289 | -1.724866 | -3.522489 | -1.271229 | -0.172844 | 0.306844 | 0.033781 | -1.863246 | -5.687221 | -1.166165 | -1.619083 | 1.146828 | -0.975619 | -2.082338 | -0.829758 | -1.837982 | 1.963551 | -2.545761 | -1.541455 | -1.869854 | -7.165185 | -0.087024 | -1.402628 |
| 32.88889 | -0.636941 | -4.841261 | -0.376914 | -3.551133 | -4.417064 | -0.537816 | -0.308738 | 2.811227 | -0.909325 | -7.613243 | -1.045538 | -0.886144 | 1.5803 | -1.335877 | -0.876673 | -0.811115 | -1.949516 | 2.619095 | -2.262478 | -2.913273 | -1.897968 | -9.231299 | 1.787013 | -1.515193 |
| 32.90278 | -0.469701 | -1.546395 | -1.041703 | -1.903589 | -6.034721 | -0.544256 | -0.772263 | 3.605809 | -0.172649 | -4.971352 | -0.641711 | -0.311172 | 2.093111 | -1.219164 | 0.106921 | -1.126078 | -2.077776 | 2.734479 | -1.927801 | -3.640304 | -1.748547 | -9.456449 | 3.479714 | -1.229617 |
| 32.91667 | -0.418705 | -0.246405 | -2.054174 | -1.201511 | -6.179167 | -0.303735 | -1.075505 | 2.27122 | 0.294021 | -4.788886 | -0.17626 | -0.191014 | 2.523948 | -0.670232 | 0.65058 | -1.98592 | -2.086337 | 0.471598 | -0.407554 | -4.569666 | -1.455363 | -7.323845 | 2.889036 | -0.510369 |
| 32.93056 | -0.321298 | -0.61026 | -1.692791 | -1.113472 | -3.58873 | 0.171072 | -0.443893 | 1.223024 | 0.38885 | -1.282709 | -0.481266 | -0.298384 | 2.215617 | -0.664927 | 1.026387 | -3.639011 | -2.550943 | -0.358366 | 1.113901 | -4.744574 | -1.359667 | -6.050991 | 3.080819 | -0.040789 |
| 32.94444 | -0.336976 | -0.990932 | -1.094187 | -0.71803 | -3.108813 | 0.534619 | 0.413173 | -0.976341 | 0.271544 | 1.865246 | -0.570694 | -0.327634 | 0.363914 | 0.269571 | 0.290534 | -5.572648 | -3.946635 | -3.367787 | 0.171041 | -5.110945 | -1.496973 | -7.073313 | 2.279874 | 0.098837 |
| 32.95833 | -0.090118 | -0.2772 | -0.665783 | -0.357891 | -4.926975 | 0.498767 | 0.918792 | -0.857968 | 0.245289 | 5.002685 | -0.583874 | -0.070262 | -1.2615 | 0.717708 | -0.894017 | -5.32853 | -5.223986 | -5.280632 | -1.417483 | -4.697084 | -1.102643 | -7.477423 | 0.050729 | 0.078646 |

| | | | | | | | | | | | | | | | | | | | | | | | | |
|---|---|---|---|---|---|---|---|---|---|---|---|---|---|---|---|---|---|---|---|---|---|---|---|---|
| 32.97222 | 0.11398 | 1.090723 | -0.719347 | -0.566468 | -5.705616 | 0.427119 | 0.931301 | -0.428545 | 0.440394 | 6.610966 | -0.479857 | 0.35751 | -1.357446 | 0.012939 | -1.814973 | -4.904453 | -5.547627 | -4.416647 | -2.617132 | -4.139975 | -0.866508 | -5.967488 | -1.068177 | 0.370604 |
| 32.98611 | 0.281114 | 2.754948 | -0.777587 | -1.110766 | -0.376432 | 0.394196 | 1.036991 | -0.007428 | 1.103998 | 9.35336 | -0.376421 | 0.449643 | -1.217734 | -0.210242 | -1.748855 | -3.032743 | -3.765788 | -3.548916 | -1.672063 | -1.874648 | -0.517876 | -3.079637 | -1.242631 | 0.24279 |
| 33 | 0.316178 | 3.995293 | -0.628538 | -1.190872 | 2.56154 | 0.259285 | 1.086844 | -0.009807 | 2.084024 | 11.14536 | -0.296716 | 0.238651 | -1.189808 | 0.200681 | -1.777884 | -0.579356 | -2.438613 | -0.788307 | -0.313952 | -0.183175 | -0.787159 | -0.146272 | -1.048812 | -0.015045 |
| 33.01389 | 0.267742 | 4.085965 | -0.381309 | -0.392826 | 2.562221 | 0.152608 | 1.063788 | 0.025248 | 2.458025 | 15.807202 | -0.085313 | 0.146941 | -0.543703 | 0.898201 | -1.310007 | 0.783303 | -0.55573 | -0.006224 | 0.222497 | 0.612031 | -0.407175 | 1.597741 | -1.205866 | 0.035638 |
| 33.02778 | 0.097597 | 4.228959 | -0.351468 | -0.171514 | 1.416546 | 0.106005 | 1.105751 | 0.196918 | 2.139836 | 15.962761 | 0.222397 | 0.343598 | 0.602881 | 1.226363 | -0.977658 | 1.755627 | 1.125861 | 1.620213 | 0.49206 | 0.531779 | 0.080176 | 2.166041 | -1.038079 | 0.277942 |
| 33.04167 | -0.011801 | 4.464217 | -0.341546 | -0.286397 | 0.481277 | 0.14819 | 1.154218 | 0.478843 | 2.541604 | 7.509893 | 0.582887 | 0.444278 | 1.864571 | 2.035921 | 0.340597 | 1.551875 | 1.54556 | 1.522507 | 0.220677 | 0.222071 | 0.538273 | 3.51687 | -2.560355 | 0.598093 |
| 33.05556 | -0.015799 | 1.370961 | -0.13 | -0.535371 | -0.253243 | 0.345488 | 1.323758 | 1.117878 | 2.395374 | 5.181194 | 0.698492 | 0.549169 | 2.517809 | 3.066054 | -0.039382 | 1.337684 | 1.40791 | -0.595463 | 0.071404 | 0.129756 | 0.802187 | 7.169482 | -4.952269 | 1.104036 |
| 33.06944 | -0.082403 | 2.543968 | 0.165246 | -0.370684 | -0.845695 | 0.134678 | 1.374018 | 1.790827 | 2.219955 | 6.657521 | 0.611939 | 0.551607 | 2.301959 | 1.590934 | -1.685573 | 1.306671 | 1.389116 | -0.282084 | 0.441512 | 0.478747 | 1.623544 | 8.148295 | -4.374614 | 1.388093 |
| 33.08333 | -0.186218 | 3.435263 | 0.286822 | 0.03545 | -0.560526 | -0.042234 | 1.305873 | 1.516291 | 1.204825 | 4.198994 | 0.417077 | 0.435411 | 1.986513 | 0.877669 | -2.507405 | 1.921696 | 1.580274 | 0.078003 | 0.661003 | 0.889008 | 1.402563 | 8.037489 | -1.992673 | 1.350144 |
| 33.09722 | -0.228197 | 1.667779 | 0.118882 | 0.414657 | 0.608321 | -0.138208 | 1.032419 | 1.056795 | 0.031934 | 0.82224 | 0.243392 | 0.389442 | 1.569939 | 0.979815 | -1.93088 | 2.144974 | 1.492273 | -0.764579 | 0.325334 | 1.046012 | 0.954139 | 7.186996 | -0.092854 | 1.180726 |
| 33.11111 | -0.1293 | 0.450488 | -0.115824 | 0.531402 | -0.933893 | -0.128386 | 0.795642 | 0.556599 | -1.053958 | -2.37164 | 0.040194 | 0.137055 | 1.257307 | 1.280939 | -0.513135 | 1.435821 | 1.437332 | -0.307507 | -0.044494 | 1.011661 | 1.142274 | 8.177226 | -0.052386 | 1.649446 |
| 33.125 | 0.219233 | -0.755632 | -0.190112 | 0.51927 | 0.087902 | 0.045472 | 1.005835 | 0.254109 | -1.861125 | -5.077621 | -0.11764 | -0.035485 | 1.244362 | 1.501105 | -0.102648 | 1.034898 | 1.46095 | -0.458661 | -0.502523 | 0.953807 | 0.927187 | 9.112792 | 0.119029 | 1.616535 |
| 33.13889 | 0.254309 | -2.155642 | -0.110851 | 0.206627 | 1.3057 | 0.197976 | 1.693747 | 0.25459 | -2.195635 | -4.421598 | -0.219083 | -0.238828 | 0.98939 | 0.48581 | -0.455049 | 1.860811 | 1.520074 | -0.93643 | -0.711542 | 0.791782 | 0.039094 | 7.916699 | 0.347883 | 1.419475 |
| 33.15278 | -0.282516 | -3.291328 | -0.060644 | -0.136884 | 0.14251 | 0.5768 | 2.537951 | 0.56926 | -2.760728 | -5.04261 | -0.231788 | -0.28004 | 0.568597 | -0.851003 | -0.494327 | 2.618694 | 1.48644 | -0.733459 | 0.100803 | 0.738874 | -0.372757 | 4.653661 | -0.005258 | 1.325298 |
| 33.16667 | -0.810584 | -4.1143 | -0.047389 | -0.580619 | -0.472924 | 0.423782 | 2.927437 | 0.592748 | -2.801285 | -6.853379 | -0.115825 | -0.399782 | 0.328572 | -1.120577 | 0.372507 | 2.507576 | 1.255659 | -0.373773 | 0.982912 | 0.771736 | -0.648482 | 2.32028 | -0.449984 | 1.156973 |
| 33.18056 | -0.55581 | -5.776084 | -0.106935 | -0.703095 | 0.558264 | 0.616019 | 2.423597 | 0.311831 | -2.388207 | -9.053992 | 0.026082 | -0.555035 | 0.367826 | -1.284779 | 0.999792 | 1.633232 | 0.826672 | -0.001288 | 0.935975 | 0.592635 | -0.705141 | 1.594164 | -0.731411 | 0.853809 |
| 33.19444 | 0.189464 | -4.989521 | -0.164907 | -0.630157 | 0.426243 | 0.561788 | 2.055827 | -0.011419 | -1.803978 | -8.431756 | 0.159494 | -0.670276 | 0.22687 | -1.565633 | 0.84957 | 0.443134 | 0.392645 | -0.455189 | 0.332183 | 0.443438 | -0.887489 | 0.800312 | -0.816282 | 0.489858 |
| 33.20833 | 0.215179 | -2.687224 | -0.237053 | -0.477616 | -3.214361 | 0.646827 | 2.02204 | 0.003963 | -1.524293 | -6.851084 | 0.183089 | -0.603595 | -0.216338 | -0.797812 | 0.303818 | -0.999259 | 0.204623 | -1.101817 | -0.238571 | 0.567774 | -0.950482 | 0.077014 | -0.327891 | 0.329503 |
| 33.22222 | 0.156009 | -0.258591 | -0.277899 | -0.095508 | -4.301681 | 0.591515 | 1.777354 | -0.002348 | -0.954521 | -4.567476 | 0.169606 | -0.361471 | -0.6722 | 0.109137 | -1.070427 | -1.901131 | 0.228536 | -0.474905 | -0.086586 | 0.709059 | -0.651766 | -0.065829 | -0.02263 | 0.316924 |
| 33.23611 | 0.146993 | 2.302756 | -0.269697 | 0.214132 | -1.423232 | 0.477075 | 1.352445 | 0.00122 | -0.459854 | -3.362234 | 0.155146 | -0.240537 | -0.775187 | 1.048913 | -1.197409 | -1.472594 | 0.080076 | 0.257214 | 0.209961 | 0.596768 | -0.154585 | 0.339091 | 0.259736 | 0.498611 |
| 33.25 | 0.131196 | 3.927385 | -0.193273 | 0.122755 | 0.133162 | 0.469453 | 1.078082 | -0.006553 | -0.061904 | -1.194081 | 0.139733 | 0.242375 | -0.639721 | 2.030786 | -0.966725 | -0.914231 | 0.073573 | 0.83382 | 0.437974 | 0.481748 | 0.198722 | 1.509589 | 0.722813 | 1.003775 |
| 33.26389 | 0.105198 | 4.646641 | -0.118109 | 0.089269 | 1.050896 | 0.554423 | 1.209909 | -0.03516 | 0.282666 | 0.955201 | 0.134407 | 0.740992 | -0.185565 | 1.631042 | -0.866311 | -0.003512 | 0.230136 | 0.707381 | 0.429438 | 0.612624 | 0.303782 | 2.700951 | 0.976715 | 1.456329 |
| 33.27778 | 0.075859 | 2.875932 | -0.03592 | 0.405597 | 2.198218 | 0.743096 | 1.399437 | -0.102693 | 0.60436 | 2.877984 | 0.155358 | 0.809447 | 0.248117 | 1.975478 | -0.409784 | 0.63909 | 0.508193 | 0.302368 | 0.424186 | 0.962191 | 0.150399 | 2.707767 | 1.149555 | 1.048095 |
| 33.29167 | 0.125828 | 0.81424 | 0.078029 | 0.575683 | 4.145386 | 0.857235 | 1.32987 | -0.146056 | 0.922828 | 4.633454 | 0.178076 | 0.665824 | 0.495805 | 2.55798 | -0.049497 | 1.208433 | 0.57443 | 0.328776 | 0.634646 | 1.286997 | 0.097984 | 2.483187 | 0.710097 | 0.767432 |
| 33.30556 | 0.268292 | -1.828598 | 0.130829 | 0.693714 | 6.42178 | 0.769884 | 1.062006 | -0.013669 | 1.256005 | 5.767515 | 0.275491 | 0.384058 | 0.573784 | 2.41257 | -0.261362 | 1.82966 | 0.561516 | 0.81557 | 0.804713 | 1.205144 | 0.233448 | 3.068981 | 0.346307 | 0.903524 |
| 33.31944 | 0.413754 | -1.322698 | 0.145887 | 0.740706 | 9.60499 | 0.659618 | 0.701631 | 0.043641 | 1.487051 | 5.780395 | 0.413025 | 0.101903 | 0.593549 | 2.451785 | 0.240887 | 2.492781 | 0.587172 | 1.308792 | 0.874813 | 1.019613 | 0.376203 | 4.10543 | 0.414031 | 1.236882 |
| 33.33333 | 0.405997 | -1.865941 | 0.109446 | 0.428424 | 10.286838 | 0.569381 | 0.304189 | 0.005801 | 1.655597 | 4.705846 | 0.505505 | 0.319242 | 0.612357 | 2.00191 | 0.647684 | 2.588495 | 0.625954 | 1.383197 | 0.982531 | 1.204028 | 0.249895 | 4.842084 | 0.791419 | 1.399243 |
| 33.34722 | 0.238616 | -3.08416 | -0.10995 | 0.033106 | 11.189105 | 0.431229 | 0.013063 | -0.300349 | 1.969582 | 4.770445 | 0.394687 | 0.361459 | 0.709255 | 0.999969 | 0.49418 | 2.283061 | 0.685368 | 0.917268 | 0.895604 | 1.470976 | 0.153179 | 4.829612 | 1.165067 | 1.47391 |
| 33.36111 | 0.28112 | -2.529475 | -0.587523 | -0.04853 | 10.479614 | 0.40714 | 0.090603 | -0.772898 | 2.712958 | 4.569659 | 0.310981 | 0.069851 | 0.867123 | 0.766057 | 0.594806 | 2.104231 | 0.764077 | 0.50059 | 0.623172 | 1.638641 | 0.155948 | 5.116404 | 1.369439 | 1.837822 |
| 33.375 | 0.678864 | -0.420642 | -1.070845 | 0.061172 | 5.797514 | 0.412845 | 0.465674 | -1.265605 | 2.631673 | 4.513323 | 0.137071 | -0.210288 | 1.092615 | 0.463806 | 1.368749 | 2.002726 | 1.21567 | 0.413808 | 0.569929 | 2.12533 | 0.184029 | 5.438558 | -0.066954 | 2.082641 |
| 33.38889 | 1.467033 | 1.525241 | -1.358939 | 0.488774 | 4.781108 | 0.64475 | 0.691107 | -0.735132 | 1.997577 | 3.794888 | -0.0604 | -0.139102 | 1.22039 | 0.769604 | 1.983199 | 1.451926 | 1.911989 | 0.531313 | 0.775132 | 2.700762 | 0.196794 | 5.253481 | 0.072926 | 1.786142 |
| 33.40278 | 2.428364 | 2.976571 | -0.179184 | 2.188997 | 2.08529 | 0.989436 | -0.473456 | -0.044115 | 1.69544 | 3.890745 | -0.06231 | 0.111484 | 1.409468 | 0.550413 | 2.642381 | 1.05407 | 2.381122 | 0.665756 | 0.965281 | 2.81235 | 0.240162 | 5.512325 | 0.238755 | 1.413002 |
| 33.41667 | 1.372821 | 2.653656 | 1.774074 | 3.266844 | -0.223506 | 0.909311 | -0.256046 | 0.211298 | 1.611221 | 3.900012 | 0.086935 | 0.160617 | 1.4152 | 0.283665 | 2.90083 | 0.897095 | 2.845109 | 0.72052 | 0.93238 | 2.944133 | 0.194442 | 6.144976 | -0.018158 | 1.641943 |
| 33.43056 | 1.134033 | 2.603734 | 3.100239 | 3.538723 | -0.320202 | -0.367715 | -0.085405 | -0.033545 | 1.511557 | 3.752172 | 0.362039 | 0.376028 | 1.515847 | 1.887427 | 2.720062 | 0.360705 | 3.234575 | 0.77336 | 0.802365 | 3.646587 | 0.086598 | 5.475969 | 0.933089 | 2.186487 |
| 33.44444 | 1.137099 | 2.556801 | 4.040508 | 3.752474 | 0.337471 | 0.293103 | 0.391736 | -0.378812 | 1.264072 | 4.4978 | 0.721517 | 0.836477 | 1.197521 | 2.171212 | 1.713472 | 0.6102 | 2.846277 | 0.792188 | 0.586891 | 3.118639 | -0.206623 | 4.189352 | 1.52686 | 2.383066 |
| 33.45833 | -0.340509 | 6.936751 | 4.763041 | 2.587614 | -1.456258 | 0.514834 | 0.60091 | -0.097063 | 0.907225 | 5.142074 | 0.953443 | 1.177967 | 1.006959 | 2.830303 | 2.213489 | 0.280562 | 1.881424 | 1.010317 | 0.437141 | 1.896782 | -0.206528 | 3.898937 | 2.181329 | 2.197575 |
| 33.47222 | -1.008193 | 6.918775 | 4.792729 | 2.093909 | -4.993811 | 0.341938 | 0.900207 | 0.249864 | 0.494615 | 4.997811 | 1.104412 | 1.537355 | 0.677123 | 3.241302 | 1.721578 | 0.382864 | 1.345451 | 1.740193 | 1.19733 | 1.875437 | 0.174564 | 2.7256 | 1.996043 | 2.372804 |
| 33.48611 | 1.00483 | 3.999795 | 3.508127 | 1.802947 | -7.083288 | -0.069047 | 0.942853 | 0.499056 | 0.661994 | 5.262732 | 1.360595 | 2.297289 | -0.255286 | 3.450862 | 2.006889 | 0.441849 | 0.84832 | 3.098232 | 3.081186 | 2.725698 | 0.936242 | 4.025331 | 2.084138 | 2.990334 |
| 33.5 | 2.441621 | 1.140077 | 2.180419 | 1.188498 | -8.723598 | -0.38194 | 1.12445 | 0.625663 | 0.88355 | 5.205908 | 1.324795 | 2.125575 | 2.03164 | 2.943082 | 1.925616 | 1.400952 | 0.860641 | 4.02467 | 4.332918 | 3.581886 | 1.259953 | 5.99383 | 2.470992 | 3.011322 |
| 33.51389 | 0.709127 | -6.284039 | 1.275185 | 0.048372 | -5.285959 | -0.155421 | 0.94875 | 0.796426 | 0.904481 | 4.565156 | 1.189659 | 2.581337 | 5.452312 | 2.458611 | 1.603398 | 1.319653 | 0.870905 | 3.500605 | 3.162241 | 3.980348 | 1.720922 | 4.359322 | 2.799293 | 2.750099 |
| 33.52778 | -2.446275 | -8.527197 | 1.140911 | 0.064522 | 1.997217 | 0.272703 | 1.203033 | 0.975639 | 0.765622 | 3.397811 | 0.94057 | 1.863456 | 4.165438 | 2.864875 | 2.299636 | 1.780982 | 1.131188 | 2.67698 | 2.007348 | 4.26307 | 2.383039 | 3.690885 | 3.752361 | 3.177605 |
| 33.54167 | -3.536256 | 1.750549 | 2.308977 | 0.065375 | 8.184034 | 0.513452 | 1.011635 | 0.39973 | 0.293392 | 2.41531 | 0.833453 | 0.959694 | 2.908029 | 2.183881 | 3.583969 | 1.650894 | 1.311579 | 1.962618 | 4.000939 | 4.289006 | 2.508481 | 4.55794 | 2.224976 | 3.101543 |
| 33.55556 | -4.272277 | 10.508162 | 3.210978 | 0.351444 | 11.433445 | -0.62936 | 0.770354 | 0.446211 | -0.334069 | 1.623863 | 0.781955 | 0.177367 | 1.159793 | 1.221502 | 3.211348 | 1.530334 | 0.865061 | 1.387868 | 4.265837 | 4.070884 | 2.275031 | 4.124814 | 0.343178 | 2.256026 |
| 33.56944 | -3.042777 | 8.891811 | 2.891627 | 0.56857 | 10.769492 | -0.504822 | 0.599042 | 0.982529 | -0.676438 | 1.386754 | 0.77906 | 0.385075 | 0.40181 | 0.342337 | 1.591647 | 0.815932 | -0.262352 | 0.347307 | 3.775952 | 3.680064 | 0.620147 | 3.29589 | -0.325478 | 1.494057 |
| 33.58333 | -3.855997 | 0.516588 | 1.943385 | 0.596588 | 5.413697 | -0.064872 | 0.487164 | 1.350668 | -0.132923 | 1.719761 | 0.842122 | 0.65468 | 0.184541 | -0.262073 | 0.161423 | -0.389878 | -0.851132 | 0.234133 | 2.470834 | 3.284641 | 0.023189 | 3.061998 | -0.240431 | 0.794406 |
| 33.59722 | -2.813203 | -1.71821 | 1.36977 | 0.596941 | -0.338286 | 0.6401 | 0.814174 | 1.622075 | 0.643994 | 2.963789 | 0.845902 | 0.685916 | -1.365593 | -0.108582 | -0.698937 | 0.045176 | -0.280403 | -0.27274 | 0.521679 | 2.717057 | 0.363978 | 2.359557 | -1.398596 | 0.442194 |
| 33.61111 | -0.779847 | 0.271604 | 2.955678 | 0.585019 | -0.445214 | -0.162347 | 1.281352 | 1.53931 | 1.242336 | 5.152512 | 0.774517 | 1.539282 | -4.003141 | 0.669107 | -1.231426 | 0.063242 | 0.013908 | -0.637606 | -0.995908 | 2.404671 | 1.114527 | 0.812243 | -3.88806 | 0.433306 |
| 33.625 | 0.852811 | 1.102347 | 3.32464 | 0.503502 | 1.670072 | 0.25463 | 0.302113 | 1.49523 | 1.832065 | 6.619235 | 0.722525 | 2.305696 | -8.122573 | 1.54492 | -1.049237 | -0.470057 | -0.287371 | -0.895674 | -1.419481 | 2.384096 | 1.689621 | -0.706174 | -6.092581 | 0.919388 |
| 33.63889 | 2.697992 | 2.186095 | 1.676056 | 0.371713 | -1.509955 | 0.407109 | 0.005429 | 1.442391 | 2.906333 | 6.887792 | 0.689579 | 2.430023 | -8.16376 | 1.030323 | 0.000634 | 0.304358 | -0.48466 | -0.516905 | -1.456506 | 2.651979 | 1.890158 | -1.746582 | -3.584871 | 1.026874 |
| 33.65278 | 5.110539 | 2.766511 | 0.512092 | 2.102206 | -5.215397 | 0.473092 | 0.609563 | 1.121741 | 3.59413 | 6.487263 | 0.729646 | 2.105708 | -6.710006 | 0.24753 | 0.78308 | 1.823013 | -1.953713 | -0.164913 | -1.56833 | 3.118459 | 1.964225 | -1.836475 | -1.705033 | 0.162922 |
| 33.66667 | 7.361603 | 3.100359 | -0.793777 | 3.469775 | -3.675694 | 0.757756 | 0.81356 | 0.582272 | 3.824039 | 5.341606 | 0.976402 | 1.660904 | -2.342324 | -0.703423 | 1.241915 | 2.752626 | -2.770605 | 0.468957 | -2.1792 | 3.178652 | 2.539698 | -0.555787 | 1.145996 | 0.057403 |
| 33.68056 | 7.355639 | 4.836113 | -1.910371 | 3.002807 | -3.021496 | 0.894644 | 1.027337 | 0.273497 | 5.339962 | 4.451805 | 1.089373 | 1.440889 | 0.445846 | 0.174447 | 0.930229 | 2.291495 | 0.199471 | 1.777306 | -2.441156 | 2.76141 | 2.75758 | 0.573146 | 2.908698 | 0.719505 |
| 33.69444 | 7.869209 | 2.414597 | -2.304384 | 0.390717 | -0.758558 | 0.754491 | 0.682341 | 0.204949 | 5.822357 | 3.907336 | 1.144564 | 1.060758 | 1.473454 | 0.558461 | 0.340155 | 0.758066 | 2.116893 | 2.448278 | -1.915071 | 2.303385 | 3.299562 | 1.212127 | 2.376531 | 1.431351 |
| 33.70833 | 7.264124 | 6.934334 | -1.373114 | 0.341464 | 4.809156 | 0.576439 | 0.321619 | 0.339526 | 4.400704 | 3.303997 | 1.155575 | -0.04061 | 1.449784 | 0.964684 | 0.644324 | -0.321924 | 2.666189 | 1.9166 | -2.381941 | 1.492683 | 3.525058 | 0.977361 | 1.81776 | 2.337649 |
| 33.72222 | 5.921213 | NaN | -1.067591 | -0.111254 | 8.047285 | 0.198376 | -0.083676 | 0.749731 | 5.030496 | 2.605431 | 1.123596 | -0.053527 | 1.309028 | 2.283828 | 1.200908 | -1.161704 | 2.969725 | 1.366959 | -3.090021 | 0.811338 | 3.101473 | 1.211582 | 1.943504 | 2.984494 |
| 33.73611 | 7.060893 | -1.135071 | 0.107675 | -0.302926 | 9.520228 | -0.187655 | 0.198753 | 1.236885 | 4.418826 | 1.777561 | 1.065421 | 0.568058 | 2.207972 | 1.981345 | 0.378438 | -1.060165 | 3.306503 | 1.073886 | -3.145579 | 0.382752 | 2.476986 | 2.051579 | 1.590498 | 2.749047 |
| 33.75 | 3.345315 | 3.599516 | 0.34396 | -1.140158 | 11.471343 | -0.246165 | 0.805977 | 1.514259 | 3.790535 | 0.837261 | 1.047036 | -0.22933 | 2.715856 | 1.391029 | -0.079815 | -0.895228 | 4.149837 | 0.656385 | -2.907817 | 0.352676 | 2.706965 | 2.436639 | 0.053006 | 2.096896 |
| 33.76389 | 1.507734 | 12.926239 | 0.123511 | -1.666482 | 11.806263 | -1.107691 | 1.353859 | 2.47497 | 3.656478 | -0.107372 | 0.765626 | -1.443324 | 1.459828 | 1.266539 | -0.430865 | -0.69448 | 3.897612 | 0.793974 | -1.676344 | 0.391012 | 3.492504 | 2.164832 | -1.151146 | 2.126625 |
| 33.77778 | 0.720378 | 14.117179 | 0.355663 | -1.809484 | 2.853203 | -0.77611 | 0.763588 | 2.220243 | 1.789894 | -0.051136 | 0.51655 | -1.631518 | -0.459325 | 0.929004 | -0.991193 | -0.588455 | 2.499226 | 1.140044 | -1.601129 | 0.309267 | 4.66017 | 0.953198 | -1.109725 | 2.347093 |
| 33.79167 | 1.535326 | 9.144992 | 1.261717 | -0.889314 | 7.387912 | -1.313882 | 1.996409 | 3.037131 | -0.59313 | -0.061276 | 0.06333 | -1.164692 | -2.134689 | 0.451223 | -1.129566 | -0.374199 | 0.490891 | 0.819108 | -2.289448 | -0.093881 | 3.809058 | -0.263246 | -0.889583 | 1.4779 |
| 33.80556 | -0.668429 | 0.354102 | 0.563699 | 0.327104 | 7.213851 | -0.694799 | 1.063902 | 1.936411 | -2.603093 | -0.742843 | -0.427742 | -1.49991 | -2.706316 | -0.161591 | -2.06351 | 0.13186 | -1.530176 | 0.283439 | -2.069089 | -0.16853 | 1.045585 | -1.250831 | -1.293518 | 0.241962 |
| 33.81944 | 0.379325 | 0.102436 | -6.895534 | 0.603497 | 5.045417 | 0.779862 | -0.108494 | 0.682924 | -3.324998 | -1.300788 | -0.459244 | -1.899262 | -3.762867 | -0.367727 | -2.29034 | 0.000267 | -1.297985 | -0.110476 | -1.506087 | -0.28994 | -0.105241 | -1.747093 | -1.126704 | -0.654837 |
| 33.83333 | -1.684392 | -1.0756 | -6.68559 | -2.102511 | 0.392117 | 1.711076 | -0.572246 | 0.500903 | -3.459653 | -1.521884 | -0.145833 | -2.212253 | -2.859308 | -1.206045 | -1.95774 | -0.670701 | -1.185716 | 0.435561 | -1.692532 | -0.75097 | -0.558198 | -2.273199 | -1.201871 | -1.88038 |
| 33.84722 | 0.670977 | -3.977969 | -5.821701 | -1.946005 | -0.751453 | 1.052483 | -0.082186 | 1.260458 | 0.439718 | -2.484985 | -0.280571 | -2.08188 | -2.263841 | -1.244632 | -1.40923 | -0.760846 | -1.290716 | 1.096302 | -1.048656 | -0.90164 | 0.235658 | -2.97958 | -1.471513 | -1.65602 |
| 33.86111 | -0.828091 | -4.391187 | -6.571973 | -0.665651 | -2.569857 | -0.190067 | -0.715241 | 2.375361 | -1.157424 | -7.375742 | -1.235134 | -2.012273 | -2.304646 | 0.20819 | -1.264514 | 0.333347 | -1.841193 | 1.097646 | -0.548666 | -0.071523 | 0.491628 | -3.417637 | -2.864772 | -0.581027 |
| 33.875 | -2.950714 | -7.553674 | -7.452256 | 0.934703 | -2.339681 | -0.478507 | -0.57401 | 2.517714 | -1.418137 | -10.95672 | -2.163969 | -1.97147 | -2.758249 | 0.168116 | -0.941182 | 1.23981 | -2.185648 | 0.431461 | -0.124734 | 0.611227 | -0.761246 | -4.121984 | -3.20658 | 0.216104 |
| 33.88889 | -5.122018 | -5.995593 | -5.940094 | 3.827688 | -2.326336 | -0.16173 | -0.442556 | 1.339543 | -2.132445 | -11.104916 | -1.227741 | -1.419138 | -2.914912 | -0.854004 | -0.304985 | 1.73107 | -1.700586 | 0.243459 | -0.136017 | 0.060817 | -1.885378 | -6.007383 | -3.013493 | -1.156487 |
| 33.90278 | -3.089613 | -3.009012 | -3.588113 | 3.25572 | -2.977334 | 0.104363 | -0.303216 | 0.952845 | -1.92248 | -10.731526 | -0.301401 | -0.745217 | -2.644551 | -1.632884 | -0.908392 | 1.85895 | -1.591121 | -0.057909 | -0.410653 | -0.997463 | -2.219145 | -6.080528 | -1.507836 | -2.075049 |
| 33.91667 | -1.427351 | -1.011883 | -3.405571 | 2.605824 | -0.855994 | 0.738372 | 0.174314 | -0.003919 | -0.976699 | -8.197089 | 0.146647 | -0.358116 | -1.789364 | -2.154936 | 0.302933 | 1.482508 | -2.717211 | 0.135884 | -0.145092 | -1.696272 | -1.649215 | -8.660044 | 0.069387 | -2.395798 |
| 33.93056 | -0.722754 | 0.308434 | -3.467103 | 1.526897 | -1.104914 | 1.746234 | 0.443341 | -0.513152 | -0.525829 | -5.941545 | 0.33077 | -0.177518 | -0.884599 | -1.794311 | 2.552376 | 0.346409 | -4.701657 | 0.437706 | 1.209798 | -1.975283 | -2.003218 | -8.736871 | 0.496229 | -2.681795 |
| 33.94444 | -0.831329 | 1.001583 | -2.739626 | 1.283969 | -2.358369 | 1.191899 | -0.664011 | -0.656761 | -0.289961 | -1.250191 | 0.253739 | -0.17886 | -0.667959 | -1.364642 | 2.631604 | -0.721779 | -6.347757 | -0.163626 | 1.878898 | -2.522031 | -6.080021 | -9.171833 | 0.452341 | -2.928963 |
| 33.95833 | -0.640059 | 0.797184 | -1.009889 | 0.86108 | -2.765519 | 0.543596 | -0.610014 | -0.452817 | -0.095829 | 2.176525 | 0.218469 | -0.186385 | -0.983104 | -2.426185 | 1.875088 | -1.284196 | -3.515897 | -1.173748 | 1.099938 | -2.712525 | -9.066817 | -7.314307 | -1.747583 | -2.420069 |
| 33.97222 | -0.01375 | 0.436378 | 0.759233 | 1.06093 | -2.111495 | 0.319681 | 0.04162 | -0.345361 | 0.059073 | 3.378505 | 0.192791 | -0.064936 | -1.131659 | -2.113777 | 1.104003 | -1.167419 | -2.357766 | -1.734082 | -0.44736 | -1.9739 | -5.039165 | -7.658157 | -2.81746 | -2.165831 |
| 33.98611 | 0.159024 | 0.348881 | 2.485817 | 1.211066 | 0.855411 | 0.671769 | 0.104873 | -0.397719 | 0.32665 | 4.541179 | 0.182839 | 0.165132 | -0.18942 | -1.950485 | 1.437734 | -0.651569 | -2.359281 | -2.611878 | -0.404603 | -1.016323 | -2.899096 | -6.599301 | -3.520593 | -1.978387 |
| 34 | 0.448379 | 0.768609 | 3.935835 | 0.727754 | 1.304207 | 1.137807 | -0.062903 | 0.392914 | 0.909732 | 6.987937 | 0.205908 | 0.380292 | 0.370758 | -1.54087 | 2.377776 | 0.51274 | -1.113324 | -1.095917 | -1.240246 | -0.379318 | -1.29508 | -3.335004 | -2.672878 | -1.832602 |
| 34.01389 | 1.65856 | 1.148369 | 4.163509 | -0.342316 | -1.11622 | 1.371697 | -1.136264 | 0.693746 | 1.803168 | 8.985071 | 0.174512 | 0.460062 | 0.527992 | -1.376464 | 2.39786 | 1.903438 | 0.843681 | 0.305295 | -1.18551 | 0.620748 | 0.44481 | 0.482322 | -2.727095 | -1.441607 |
| 34.02778 | 4.249722 | 2.115075 | 4.243982 | -0.468688 | 1.965728 | 0.815422 | -0.246091 | -0.010024 | 2.595427 | 7.14848 | 0.083895 | 0.475571 | 0.595632 | -0.751199 | 3.351391 | 2.502877 | 2.797722 | 0.018366 | -1.414486 | 0.078158 | 1.166573 | 2.369808 | -1.959212 | -0.251504 |
| 34.04167 | 4.464258 | 2.686676 | 3.139884 | -0.668815 | 18.3866 | 0.191849 | -0.206352 | -0.783085 | 2.729245 | 8.865045 | 0.044736 | 0.554829 | 0.52072 | 0.504435 | 3.202906 | 2.278651 | 2.187053 | 0.311607 | -1.178465 | -0.077657 | 1.055272 | 2.339256 | 0.570467 | 1.172614 |
| 34.05556 | 3.346565 | 2.647462 | 1.001223 | -1.10765 | 9.325615 | -0.390597 | -0.309087 | -0.011742 | 2.715833 | 4.904106 | -0.056515 | 0.69995 | 0.745383 | 1.471829 | 1.196788 | 1.624274 | 1.692113 | 1.223141 | -0.794415 | -0.661775 | 0.749571 | 3.330829 | 2.55354 | 2.288673 |
| 34.06944 | 0.630145 | 2.134503 | -1.279207 | -1.688055 | 0.136281 | -0.430191 | -0.442385 | 0.07189 | 2.114642 | 0.887941 | -0.206641 | 0.687293 | 1.230455 | 2.120457 | -0.236145 | 1.516551 | 1.226827 | 0.507019 | -0.875462 | -0.851291 | 0.377823 | 4.033441 | 5.310263 | 3.029823 |
| 34.08333 | 1.468323 | 1.550895 | -3.001449 | -0.169523 | -4.453307 | -0.425332 | -0.194333 | -0.13629 | 1.427399 | 0.697925 | -0.386994 | 0.613421 | 1.015722 | 2.532362 | -1.617059 | 1.355782 | -0.744262 | 0.196793 | -0.60207 | -1.381182 | 0.809347 | 4.00942 | 5.123598 | 3.473031 |
| 34.09722 | -2.669272 | 1.127112 | -4.018854 | -1.817906 | -13.511171 | -0.216198 | -0.172423 | -0.423106 | 0.943482 | 2.222058 | -0.412484 | -0.251696 | 0.581685 | 2.112543 | -2.468895 | 0.963038 | 0.788497 | 0.141044 | -0.274281 | -2.002999 | 1.480818 | 4.296195 | 0.502763 | 3.166321 |
| 34.11111 | -0.046092 | 0.730464 | -3.744931 | -1.485721 | -5.974701 | -0.224619 | -0.401862 | -0.965672 | 0.528174 | 3.638193 | -0.248133 | -0.655467 | 0.28623 | 2.230716 | -2.73394 | 0.506074 | 0.492763 | 0.475858 | -0.55165 | -1.815368 | 1.315977 | 5.81736 | 0.492289 | 2.546007 |
| 34.125 | -0.465422 | -0.646119 | -2.535863 | -2.03012 | 2.822075 | -1.110751 | -0.821215 | -1.249308 | -0.406582 | 1.734109 | -0.194551 | -0.538866 | 0.544433 | 1.463607 | -2.337704 | 0.335402 | 0.240616 | 0.300089 | -0.407306 | -1.172219 | 0.235929 | 5.79718 | 0.522241 | 2.529934 |
| 34.13889 | -1.05946 | -1.745877 | -0.910744 | -1.081516 | -0.37975 | 0.525943 | -0.778954 | -0.65032 | -1.743223 | 0.278111 | -0.260922 | -0.205011 | 0.317515 | 1.292698 | -1.619279 | 0.536389 | 0.578449 | -0.017524 | -1.132803 | -0.126017 | -0.597399 | 5.850157 | 0.073199 | 3.641819 |
| 34.15278 | -1.405592 | -0.329206 | 0.092226 | 0.139741 | -2.813921 | -3.408355 | -0.886772 | -0.174439 | -1.306327 | -0.122376 | -0.279853 | -0.122217 | -1.713048 | -0.484422 | -0.843836 | 1.713225 | 1.737413 | 1.0713 | -4.046732 | 1.799742 | -0.120335 | 3.094953 | -1.438667 | 2.46748 |
| 34.16667 | -1.867735 | 0.877677 | 1.166782 | 1.202204 | -0.981531 | -3.061531 | -0.923655 | 0.124482 | -0.925034 | 0.773387 | -0.331378 | -0.074101 | -2.403828 | -1.451302 | -0.548724 | 1.206601 | -1.302957 | 3.118118 | -8.044335 | 2.633116 | 0.563903 | 2.121904 | -2.564664 | -0.645815 |
| 34.18056 | -1.726516 | 2.533187 | 2.965416 | 1.796178 | 1.250096 | -2.788157 | -0.865831 | 0.38859 | -1.166301 | 1.010301 | -0.493374 | -0.057683 | -0.820497 | -0.789456 | 1.988237 | -0.792115 | -2.52043 | 1.478624 | -1.202987 | 0.442023 | -1.406066 | -0.023279 | -0.227597 | -1.010252 |
| 34.19444 | -1.402815 | 2.170591 | 5.116036 | 1.240145 | 2.601227 | -2.174838 | -0.633258 | 0.606138 | -1.411365 | 0.979389 | -0.489523 | -0.050446 | 1.371573 | 1.123109 | 2.118126 | -1.044252 | 0.518767 | -2.746259 | 5.123824 | 2.158962 | -2.019443 | 3.791687 | -0.174518 | 2.832105 |
| 34.20833 | -0.801685 | 1.117441 | 4.729771 | 0.634791 | 4.059869 | -1.648515 | -0.442453 | 0.760553 | -1.501122 | 0.685427 | -0.257221 | 0.280394 | 0.984428 | 0.804841 | 1.91874 | 0.076084 | 0.962863 | -1.679641 | 1.737155 | 3.015266 | -1.105659 | 4.713653 | 2.172494 | 1.817568 |
| 34.22222 | -0.260201 | 1.28562 | 4.338575 | 0.648757 | 4.337912 | -1.281722 | -0.133023 | 0.804678 | -1.321518 | -0.124288 | -0.080247 | 0.902628 | 0.392412 | -0.675527 | 1.094536 | 0.701761 | -0.281055 | -0.309862 | -0.550596 | 1.17657 | -0.423875 | 3.192286 | -1.29594 | -1.314736 |
| 34.23611 | 0.305447 | 0.973281 | 2.449732 | 0.670064 | 0.825468 | -1.143143 | 0.303239 | 1.010212 | -1.301936 | -0.091753 | 0.0163 | 1.088374 | 0.245567 | -0.52318 | 0.699866 | 0.989724 | -0.444039 | 0.432329 | -0.459458 | 1.13698 | -0.097082 | 0.700708 | -2.072406 | -2.634516 |
| 34.25 | 1.388266 | 0.298152 | 1.289044 | 0.795766 | -0.070866 | -0.80802 | 0.396796 | 1.160824 | -1.186934 | -0.078708 | 0.161664 | 1.354957 | 0.265512 | -0.454453 | 0.207053 | 1.098726 | 0.424717 | 0.497871 | -0.452747 | 0.524236 | 0.416935 | 0.353098 | -2.088125 | -2.101484 |
| 34.26389 | 3.012827 | -0.641 | 1.071871 | 1.025871 | 3.444825 | -0.213262 | 0.592068 | 1.103148 | -0.984327 | -0.097428 | 0.2469 | 1.644117 | 0.290378 | 0.447847 | -0.483535 | 0.888454 | 1.301393 | 0.562216 | -0.330462 | 0.771829 | 0.611542 | 1.767569 | 0.196389 | -0.756232 |
| 34.27778 | 4.155004 | -0.635561 | 0.366751 | 1.255044 | 11.439482 | 0.211324 | 0.710353 | 0.789454 | -0.667678 | 0.336935 | 0.236507 | 1.359459 | 0.253279 | 0.721611 | -0.845669 | 0.710956 | 1.277679 | 0.508527 | -0.307184 | 0.996242 | 0.327916 | 2.495479 | 1.061223 | -0.204356 |
| 34.29167 | 7.173159 | 1.163185 | -0.181146 | 1.176658 | 9.075113 | -0.035028 | 0.547969 | 0.392744 | -0.309041 | 1.21186 | 0.094864 | 0.131574 | 0.120366 | 0.300358 | -1.038587 | 0.574138 | 0.752788 | 0.448143 | -0.132683 | 0.74426 | 0.010683 | 1.471979 | -0.092614 | -0.689453 |
| 34.30556 | 5.944418 | 2.245039 | -0.551022 | 0.835252 | 2.256779 | 0.321144 | 0.355913 | 0.379693 | 0.063709 | 2.131991 | 0.039295 | -0.59273 | 0.197348 | 0.229982 | -1.411015 | 0.355064 | 0.720736 | 0.356696 | -0.119561 | 0.534857 | -0.037014 | 0.655623 | -0.972765 | -0.879712 |
| 34.31944 | 2.219611 | 2.853266 | -0.907254 | 0.440832 | -1.971243 | 0.971242 | 0.077227 | 0.416384 | 0.292429 | 2.795813 | 0.148988 | -1.077668 | 0.683365 | 0.70592 | -1.51158 | 0.266195 | 1.35921 | 0.21541 | -0.160896 | 0.733411 | 0.209798 | 0.926673 | 0.191138 | -0.024784 |
| 34.33333 | -1.475335 | 2.424951 | -1.386049 | 0.340586 | -3.660341 | 0.715624 | -0.078387 | 0.156503 | 0.359564 | 2.721801 | 0.206972 | -1.510034 | 1.106947 | 1.491572 | -1.117649 | 0.452184 | 1.54911 | 0.127132 | -0.119407 | 1.474048 | 0.319255 | 2.224008 | 2.60624 | 1.513081 |
| 34.34722 | -2.423531 | 1.285473 | -1.669429 | 0.015116 | -5.1025 | 0.404379 | -0.257933 | -0.362881 | 0.532628 | 2.323789 | 0.164284 | -0.95445 | 1.010966 | 1.995628 | -0.200135 | 0.924399 | 1.056151 | -0.01858 | -0.129576 | 1.965172 | 0.16859 | 3.434288 | 3.788704 | 2.776098 |
| 34.36111 | -1.051157 | 1.287382 | -1.211091 | -0.109354 | -3.016995 | 0.196916 | -0.468047 | -0.720894 | 1.144741 | 2.296475 | 0.146892 | -0.574477 | 0.939768 | 2.30575 | 0.528975 | 1.019979 | 1.228676 | 0.089844 | 0.046776 | 2.132064 | 0.163703 | 4.186752 | 4.898748 | 2.798046 |
| 34.375 | -0.320425 | 1.593381 | -0.368174 | 0.422255 | -0.497854 | 0.007403 | -0.651622 | -0.806079 | 1.707174 | 1.579724 | 0.177172 | -0.337531 | 1.709065 | 2.943001 | 0.627821 | 0.935087 | 2.821345 | 0.481154 | 0.35624 | 2.84729 | 0.504419 | 5.229277 | 4.374025 | 2.593331 |
| 34.38889 | 0.271871 | 1.951993 | 0.662391 | 1.238416 | -2.651767 | -0.163761 | -0.443939 | -0.647748 | 1.456631 | 1.153822 | 0.161819 | -0.312985 | 2.584046 | 4.359707 | 0.492034 | 0.797901 | 4.675756 | 0.696845 | 0.57457 | 3.350722 | 0.734823 | 7.559437 | 2.375206 | 3.860018 |
| 34.40278 | 0.612175 | 2.777897 | 1.856526 | 1.675952 | -4.818942 | 0.295214 | -0.047964 | -0.330755 | 1.155001 | 1.090185 | 0.144441 | -0.249996 | 4.091772 | 4.716176 | 0.457009 | 0.790043 | 4.869077 | 0.818004 | 0.743253 | 2.993109 | 0.893625 | 11.61338 | 1.428843 | 3.984917 |
| 34.41667 | 0.781913 | 3.476508 | 2.211359 | 1.492756 | -3.489539 | 0.757326 | 0.334265 | 0.228531 | 0.908015 | 1.301317 | 0.250234 | -0.116661 | 5.506874 | 5.141385 | 0.718072 | 1.13345 | 4.190737 | 0.838769 | 1.087528 | 2.979574 | 0.926062 | 9.78712 | 2.961745 | 3.827215 |
| 34.43056 | 0.882447 | 3.494533 | 3.948511 | 1.550449 | 0.507397 | 1.103455 | 0.739884 | 1.306216 | 0.785964 | 1.461682 | 0.446164 | 0.158573 | 7.629666 | 5.606973 | 1.248137 | 1.637296 | 4.42008 | 0.779746 | 1.549709 | 3.147457 | 1.137589 | 9.254685 | 3.037693 | 4.290729 |
| 34.44444 | 1.604843 | 3.568765 | 7.198335 | 1.388462 | 3.853869 | 1.195315 | 0.902997 | 2.209047 | 0.812369 | 2.041289 | 0.500437 | 0.495417 | 7.476956 | 4.288057 | 1.970905 | 1.838091 | 3.809943 | 0.762335 | 2.14996 | 2.427573 | 1.410604 | 6.707606 | 2.191966 | 2.930406 |
| 34.45833 | 1.770825 | 4.75049 | 7.188528 | 1.488622 | 3.456068 | 1.099292 | 1.358633 | 1.979519 | 0.794869 | 2.540434 | 0.394206 | 0.629127 | 6.956682 | 2.695081 | 1.590721 | 1.627152 | 2.351089 | 0.998198 | 2.154789 | 1.59139 | 1.324576 | 3.887211 | 2.582217 | 1.510072 |
| 34.47222 | 0.655461 | 4.107886 | 5.195282 | 1.424448 | 5.371205 | 0.509245 | 0.864826 | 1.383529 | 1.080855 | 3.233836 | 0.332033 | 0.629552 | 5.263738 | 2.249251 | 0.97477 | 1.397423 | 1.16832 | 1.264603 | 2.464408 | 1.472144 | 0.959634 | 3.447373 | 2.047212 | 1.236623 |
| 34.48611 | 0.09783 | 3.542299 | 3.647301 | 2.710233 | 3.767847 | 1.103358 | 1.161538 | 0.73509 | 1.79585 | 3.586856 | 0.487025 | 1.016273 | 2.589253 | 3.06886 | 0.697297 | 1.519613 | 1.435597 | 1.137174 | 3.886085 | 2.253108 | 0.802732 | 4.90407 | 2.052714 | 1.689416 |
| 34.5 | -1.741908 | 2.342093 | 2.523193 | 2.735729 | 2.371329 | 1.814935 | 1.446276 | 0.741574 | 2.316626 | 2.902539 | 0.573486 | 1.144397 | 4.502174 | 3.837998 | 0.645027 | 1.547956 | 1.445192 | 1.303637 | 3.694881 | 3.281729 | 1.169129 | 6.286064 | 2.909259 | 2.58003 |
| 34.51389 | -2.159226 | 0.773705 | 1.814672 | 1.665481 | 3.782435 | 2.022962 | 1.586235 | 0.960539 | 2.287178 | 1.681573 | 0.655015 | 1.058045 | 4.255126 | 5.086595 | 0.421163 | 2.013464 | 2.077782 | 2.025485 | 4.097449 | 3.707866 | 2.236647 | 6.145401 | 2.003578 | 2.786424 |
| 34.52778 | -1.835437 | -0.281648 | 0.653031 | -0.405237 | 4.10759 | 1.461092 | 1.5431 | 1.094734 | 2.123925 | 1.195582 | 0.714297 | 1.000542 | 2.714361 | 5.305579 | 1.654599 | 2.385181 | 1.672666 | 2.521817 | 4.288581 | 3.556717 | 2.160504 | 4.23493 | 2.321052 | 2.859709 |
| 34.54167 | -1.253137 | -1.067028 | -0.906704 | -1.727222 | 3.708695 | 1.202677 | 1.254619 | 1.03916 | 2.418062 | 2.646269 | 0.682803 | 0.837411 | 1.578997 | 3.189876 | 2.7628 | 2.417851 | 1.27111 | 2.093159 | 3.945157 | 3.203242 | 1.795878 | 1.822681 | 2.338989 | 1.969341 |
| 34.55556 | 0.456037 | -3.023953 | -3.899811 | -1.065415 | 4.626991 | 0.847798 | 0.82043 | 0.83147 | 2.899025 | 3.313427 | 0.692478 | 1.682976 | 1.205138 | 2.24194 | 2.096126 | 1.872314 | 1.356761 | 1.162618 | 3.691222 | 2.697988 | 0.936235 | 0.815085 | 1.323805 | 1.680785 |
| 34.56944 | 2.982286 | -4.629037 | -5.422107 | -2.087108 | 3.600846 | 0.932801 | 0.504228 | 0.659721 | 3.390631 | 3.039379 | 0.662249 | 1.124909 | 0.891844 | 1.781058 | 1.316726 | 1.390008 | 1.333091 | 0.998126 | 3.768495 | 2.674037 | 0.167479 | 0.466366 | 0.184057 | 1.26761 |
| 34.58333 | 2.4553 | 0.011349 | -2.961569 | -1.848757 | 0.695386 | 0.96336 | 0.535865 | 0.615127 | 3.592699 | 3.044554 | 0.647738 | 1.506531 | 0.768102 | 0.479276 | 0.605753 | 0.893393 | 0.460952 | 0.737585 | 2.521282 | 2.657508 | -0.437147 | 0.141135 | -0.215558 | 0.173794 |
| 34.59722 | 2.491877 | 2.087383 | -0.540384 | 0.432837 | -4.028527 | 0.710655 | 0.624438 | 0.773976 | 3.138658 | 3.21129 | 0.466585 | -0.018923 | 0.174429 | -0.30716 | -0.023822 | 0.670658 | -0.576586 | 0.171288 | 1.347141 | 2.689015 | -0.643667 | -0.567847 | -0.149984 | -0.662037 |
| 34.61111 | 0.573747 | 4.25437 | 1.763289 | 1.788298 | -3.061827 | -0.121088 | 0.356383 | 0.903603 | 2.413548 | 3.579007 | 0.443041 | -1.203971 | -1.773007 | -0.845144 | -0.493248 | 0.630533 | -1.289144 | 0.109904 | 0.533701 | 3.14276 | -0.911326 | -0.98891 | -0.94626 | -0.854904 |
| 34.625 | -1.063362 | 5.487706 | 4.113271 | 1.355029 | -12.135761 | -0.52155 | 0.547334 | 1.247819 | 1.755976 | 4.699532 | 0.349241 | -1.73852 | -5.655995 | -0.526851 | -0.535537 | 0.767476 | -0.635534 | 0.341163 | 0.127576 | 3.350557 | -0.899208 | -1.920533 | -2.370206 | -0.574484 |
| 34.63889 | -4.866698 | 5.430657 | 7.321734 | 0.547256 | -17.266448 | -1.748536 | 0.041573 | 1.219655 | 0.952436 | 5.333888 | 0.381827 | -1.53767 | -4.261869 | -0.849242 | -0.247189 | 0.845941 | -0.897081 | 0.549252 | 0.037133 | 3.47818 | -1.151118 | -2.819171 | -4.365616 | -0.687425 |
| 34.65278 | -3.298814 | 6.423555 | 7.645088 | -0.0006 | -6.841199 | -1.876892 | -0.060225 | 1.761046 | 0.060133 | 5.089584 | 0.261713 | -0.338329 | -4.263639 | -1.480673 | 0.050228 | 0.651608 | -1.733708 | 0.460701 | -0.488894 | 4.084207 | -0.602243 | -2.667446 | -4.760093 | -0.955144 |
| 34.66667 | 0.843789 | 4.558971 | 10.452716 | -0.469394 | -5.229349 | -1.62951 | 0.694957 | 2.227228 | -0.764727 | 4.593069 | 0.078247 | 0.819819 | -1.798895 | -1.758417 | 0.412938 | 0.448272 | -0.963368 | 0.22425 | -1.519489 | 3.439796 | -0.300476 | -0.140272 | -4.448863 | -0.585592 |
| 34.68056 | 0.628383 | 3.326791 | 15.559579 | -0.385295 | -6.267024 | -0.393956 | 2.404313 | 3.168717 | -1.331759 | 3.907643 | -0.062873 | 1.984705 | 1.191179 | -1.283478 | 0.436317 | 0.176784 | 1.077916 | 1.075936 | -2.688944 | 2.074871 | 0.018009 | 0.787985 | -2.812621 | 0.278662 |
| 34.69444 | 0.148492 | 3.474411 | 9.386579 | -0.004959 | -0.13814 | -0.230713 | 2.462457 | 1.662865 | -1.832676 | 3.279064 | -0.032486 | 2.253893 | 3.0762 | -0.342147 | 0.580383 | 0.194274 | 2.188919 | 1.264549 | -3.182891 | 1.070621 | 0.310365 | 0.912572 | -2.35844 | 1.067118 |
| 34.70833 | -0.154202 | 3.378695 | 7.158227 | 1.098729 | 4.846956 | -0.16694 | 1.59358 | 0.433623 | -2.273829 | 3.343357 | 0.016024 | 1.415192 | 4.342208 | 0.659604 | 1.021473 | 0.325641 | 2.488513 | 0.919295 | -3.395593 | 0.652619 | 0.946479 | 1.506997 | -2.444439 | 0.949391 |
| 34.72222 | -0.306403 | 2.305716 | 4.644575 | 0.320008 | 6.9149 | 0.399472 | 1.437183 | 0.614281 | -2.310277 | 2.287684 | 0.180788 | 1.61615 | 4.983138 | 1.041983 | 1.606414 | 0.417509 | 2.430762 | 0.595148 | -3.795335 | 0.087451 | 1.708311 | 1.195387 | -2.599713 | 0.728918 |
| 34.73611 | -0.261925 | 0.899717 | 1.554462 | -3.11531 | 7.10564 | 0.966798 | 0.645077 | 0.94995 | -1.418631 | 1.278025 | 0.487576 | 2.42153 | 4.151227 | 0.835281 | 1.545026 | 0.327819 | 0.304341 | 0.536924 | -4.612991 | -0.350201 | 1.889799 | 0.700437 | -1.801612 | -0.68007 |
| 34.75 | 0.273144 | -1.73381 | -0.442887 | -5.661174 | 3.080647 | 1.350825 | 0.610253 | 0.971256 | -0.65866 | 1.602911 | 0.788155 | 3.795598 | 3.666661 | 0.51588 | 1.105478 | 0.109401 | -0.030332 | 0.793562 | -2.95553 | -0.648371 | 1.806459 | 0.694762 | -1.352997 | -0.108521 |
| 34.76389 | 0.899664 | -3.127187 | -1.425663 | -7.220331 | 3.400359 | 1.205154 | 0.869479 | 1.001262 | -0.031498 | 1.614877 | 0.033038 | 2.044525 | 2.453488 | 1.0845 | 0.345446 | 0.026357 | -0.133428 | 1.067136 | -1.467855 | -0.842641 | 2.342683 | 1.031777 | -1.437996 | 0.66375 |
| 34.77778 | -1.546465 | -2.470457 | -2.37946 | -6.84257 | 3.242479 | 0.77518 | 1.809176 | 1.038536 | -0.176285 | 1.131437 | 0.069035 | 1.68894 | 1.367774 | 1.578684 | 0.243795 | -0.027004 | 0.097833 | 1.159029 | -0.573743 | -0.00276 | 2.188144 | 0.891371 | -1.452283 | 0.915481 |
| 34.79167 | -3.146785 | -1.695597 | -3.663121 | -3.017623 | 4.003295 | 0.55464 | 2.858092 | 1.1427 | -0.714634 | 0.530408 | 0.198183 | 0.559136 | -0.055047 | 1.335637 | 0.204728 | -0.313784 | 0.274574 | 1.549685 | -0.022569 | 0.079524 | 1.577979 | 0.510279 | -1.259185 | 0.511518 |
| 34.80556 | 5.740757 | 0.389595 | -3.119695 | -5.487151 | 4.591811 | 1.561156 | 3.230411 | 1.033671 | -1.31532 | -0.232675 | -0.102925 | 0.322396 | -1.942507 | 0.546816 | 0.192492 | -0.704589 | -0.400285 | 1.932995 | 0.213917 | -0.123673 | 1.212141 | 0.087346 | -1.653363 | -0.053675 |
| 34.81944 | 3.448225 | 3.048028 | -2.298063 | -3.924735 | 2.931978 | 2.166912 | 2.274669 | 0.399985 | -1.321663 | -0.828961 | -0.162955 | -0.31621 | -3.749473 | -0.118213 | -0.363785 | -1.184128 | -0.685529 | 1.596095 | -0.15249 | -0.226925 | 1.160499 | 0.030261 | -1.902003 | -0.378688 |
| 34.83333 | 2.084388 | 3.923271 | -2.728103 | -2.710211 | 0.317341 | 2.770003 | 1.384312 | 1.111269 | -0.94982 | -0.674305 | 0.374625 | -1.665784 | -4.200542 | -0.213436 | -0.848558 | -0.805925 | -0.860347 | 0.717432 | -0.161051 | -0.978283 | 0.61202 | -0.738714 | -1.705735 | -0.187856 |
| 34.84722 | 2.089493 | 4.095158 | -1.369128 | -4.511722 | -2.543554 | 0.909162 | 1.01276 | 0.184727 | -0.521752 | -0.720804 | -0.119224 | -2.132688 | -4.803942 | -0.219345 | -0.972514 | -0.38622 | -0.729904 | 0.007446 | -0.143485 | -1.734176 | 0.215155 | -1.335806 | -1.706414 | -0.148891 |
| 34.86111 | 2.877296 | 3.227155 | 1.247562 | -8.904126 | -2.95743 | -1.376962 | -0.362937 | -1.649338 | -0.704439 | -1.884056 | -0.622602 | -1.033386 | -4.862099 | -0.498573 | -0.745357 | -0.193454 | -0.541196 | 0.223271 | -0.067499 | -1.236275 | -0.132572 | -2.082299 | -1.790685 | -0.049053 |
| 34.875 | -2.101339 | 2.295386 | 4.744375 | -7.334851 | -3.956027 | -3.144658 | -2.479359 | -3.145059 | -1.692787 | -5.951819 | -0.868481 | -1.135208 | -5.232697 | -0.834529 | -0.456954 | -0.311361 | -1.389013 | 1.141387 | 0.087453 | -1.087978 | -0.019578 | -3.015797 | -2.332796 | -0.489496 |
| 34.88889 | -4.094498 | 0.687774 | 7.946449 | 0.626408 | -3.5576 | -3.31362 | -3.299538 | -3.627767 | -1.086067 | -9.589545 | -0.574415 | 0.100735 | -5.604115 | -1.619414 | -0.413752 | -0.260169 | -1.359538 | 1.638608 | 0.39278 | -1.824059 | -0.192198 | -3.045842 | -2.139884 | -1.262835 |
| 34.90278 | -1.979498 | 0.11666 | 8.243632 | 1.495879 | -3.91972 | -2.131766 | -2.067669 | -2.794985 | -0.843387 | -3.982025 | -0.0142 | 0.380703 | -3.274483 | -1.575102 | -0.765424 | -0.490667 | -1.011466 | 0.974152 | 0.706988 | -2.265651 | -0.403522 | -4.251366 | -1.532223 | -1.213658 |
| 34.91667 | -0.304519 | -0.463964 | 8.892782 | 1.149295 | -3.570123 | -1.647945 | -1.08942 | -1.324487 | -0.630796 | -1.233995 | 0.172902 | 0.060881 | -3.27763 | -1.596652 | -1.608333 | -1.11177 | -0.548815 | 0.259464 | 0.736827 | -2.270772 | -0.382981 | -4.423925 | -1.01112 | -1.579085 |
| 34.93056 | 0.416725 | -0.730845 | 8.980932 | 0.122543 | -2.869812 | -1.833898 | -1.328088 | -0.111692 | 0.247981 | 0.549806 | 0.234699 | -0.59381 | -4.138524 | -1.542315 | -3.186089 | -2.245132 | -0.997079 | 0.158932 | 0.592928 | -0.370428 | -0.845289 | -3.546529 | -0.496295 | -1.784183 |
| 34.94444 | 0.482507 | -0.568754 | 10.037074 | -1.012368 | -1.708723 | -2.812653 | -1.895095 | 0.342694 | 0.301747 | 0.172784 | 0.356459 | -0.167185 | -3.499759 | -1.261151 | -3.989592 | -2.781272 | -2.0827 | -0.085805 | 0.144535 | 0.148205 | -1.730496 | -4.194593 | -1.100443 | -2.29564 |
| 34.95833 | 0.481842 | 1.221142 | 6.141082 | -2.210982 | -1.654147 | -3.370345 | -2.157652 | 0.115725 | 0.451682 | -0.488786 | 0.564377 | 0.43503 | -3.604881 | -0.811842 | -4.086241 | -1.735611 | -2.85139 | -1.420376 | -0.86839 | -0.282605 | -4.136192 | -6.793975 | -1.845741 | -1.739655 |
| 34.97222 | 0.764137 | 2.170532 | 3.849329 | -1.027779 | -0.002522 | -2.734345 | -1.699413 | 0.114772 | 0.649958 | -0.430539 | 0.682061 | 0.468571 | -2.314791 | -2.218466 | -4.71605 | -1.057757 | -2.901876 | -1.712993 | -1.635787 | -0.716832 | -3.912461 | -4.921521 | -9.506464 | -1.360119 |
| 34.98611 | 1.09692 | 1.73534 | 4.393181 | 0.651377 | 2.862204 | -1.437956 | -0.927444 | 0.325424 | 0.842017 | 0.263569 | 0.550682 | 1.257791 | -0.82685 | -2.970395 | -3.46056 | 0.023463 | -1.622336 | -1.612582 | -1.771442 | -1.068556 | -5.098788 | -5.616443 | -10.276579 | -3.352503 |
| 35 | 1.025163 | 1.155597 | 2.785898 | 0.720464 | 4.204909 | -0.341685 | -0.380994 | 0.216846 | 0.894114 | 1.19403 | 0.23541 | 2.410448 | -0.405547 | -1.241419 | -2.400916 | -0.093689 | 0.137746 | -0.880986 | -1.252851 | -1.486509 | -5.132183 | -3.402495 | -3.132811 | -2.042053 |
| 35.01389 | 0.618987 | 0.600297 | 1.226648 | 1.031066 | 1.898921 | 0.358885 | -0.170851 | -0.234404 | 0.722374 | 1.34877 | 0.000762 | 2.354325 | -0.174791 | -0.430638 | -2.101413 | 0.012619 | 1.237865 | -0.109314 | -0.81631 | -1.581676 | -2.149774 | -1.038402 | -3.569089 | -1.111739 |
| 35.02778 | 0.209867 | -0.19618 | -1.071301 | 0.411987 | -0.210669 | 0.68537 | -0.174412 | 0.461367 | 0.61912 | 2.003293 | 0.109981 | 1.641858 | 0.128743 | -0.611601 | -1.25915 | 0.519904 | 1.612512 | 0.531658 | -0.420501 | -1.717933 | 0.913881 | -0.043474 | -1.200789 | -1.406745 |
| 35.04167 | 0.055771 | -0.771847 | 0.108419 | -0.382898 | -2.0822 | 0.600102 | -0.184005 | 0.683521 | 0.564573 | 2.573752 | 0.135473 | 1.649208 | 0.428617 | -0.42544 | 0.55581 | 1.53137 | 1.522683 | 0.304815 | -0.356385 | -1.73464 | 2.227801 | 0.326357 | 1.534306 | -0.827701 |
| 35.05556 | 0.397581 | -1.155997 | -0.750015 | -0.708312 | -2.723296 | 0.544829 | 0.001384 | 0.688539 | 0.514003 | 2.697603 | 0.381556 | 2.35528 | 0.416743 | 0.767589 | 1.889901 | 2.16711 | 0.868293 | 0.048787 | -0.744264 | -1.512495 | 1.023253 | 1.33669 | 3.075557 | 0.610366 |
| 35.06944 | 0.889066 | -0.590555 | -0.676589 | 0.290422 | -0.866667 | 0.955836 | 0.781376 | 0.64887 | 0.247338 | 2.529714 | 0.496737 | 2.028919 | 0.087841 | 1.761229 | 2.171607 | 2.845635 | 0.329746 | -0.446615 | -1.317936 | -1.055428 | -0.225954 | 2.59487 | 1.511239 | 1.356496 |
| 35.08333 | 2.824548 | 0.299249 | -0.760554 | 1.299892 | 1.458878 | 0.597521 | 0.853986 | 0.651357 | 0.075446 | 2.528755 | 0.19867 | 0.959531 | -0.11814 | 2.331941 | 1.806302 | 2.640045 | 0.212256 | -1.072811 | -1.456134 | -0.595505 | -0.471388 | 4.33326 | 2.059453 | 2.016718 |
| 35.09722 | 0.975337 | 0.447571 | -0.734928 | 1.902513 | 0.156363 | 0.346954 | 0.821569 | 0.411083 | -0.046415 | 2.367036 | -0.250541 | -0.357961 | 0.037981 | 2.355603 | 1.756116 | 2.420634 | 0.206501 | -0.34879 | -0.863574 | -0.259287 | 0.032665 | 5.409755 | 1.349614 | 2.265468 |
| 35.11111 | -0.654927 | 1.66047 | -0.212433 | 1.796182 | -1.94498 | 0.135692 | 0.718666 | 0.260131 | 0.210186 | 1.471044 | -0.61936 | -1.457617 | 0.097721 | 2.136888 | 2.119993 | 2.356885 | 0.423662 | 0.599395 | -0.208326 | -0.1318 | 1.218304 | 4.639312 | 3.89084 | 2.060515 |
| 35.125 | -2.784312 | 1.842263 | 0.613775 | 1.823051 | -3.763508 | 0.279485 | 0.566806 | 0.058094 | 0.34908 | 0.326372 | -0.809471 | -2.138935 | 0.380214 | 1.910744 | 2.447718 | 2.04325 | 0.676438 | 0.99984 | 0.107578 | -0.097999 | 2.865124 | 4.223228 | 3.284097 | 1.945229 |
| 35.13889 | -1.895463 | 1.84044 | 1.563758 | 2.221876 | -1.931898 | 0.194575 | 0.329463 | -0.293426 | 0.555975 | 0.488079 | -0.635287 | -2.18055 | 0.570619 | 1.808747 | 2.836337 | 1.829378 | 0.577341 | 1.250323 | 0.00247 | -0.203798 | 3.213648 | 3.537821 | 1.552945 | 1.968113 |
| 35.15278 | -0.374305 | 1.972672 | 0.334812 | 1.615811 | 1.218091 | 0.0512 | 0.305704 | -0.6741 | 0.636412 | 0.06473 | -0.214771 | -1.744576 | 0.27825 | 1.694759 | 2.796986 | 1.553317 | -0.010992 | 1.438605 | -0.346417 | -0.074494 | 2.304122 | 3.423515 | 1.377019 | 2.260405 |
| 35.16667 | -0.101173 | 2.209071 | 2.179937 | 0.880651 | 1.282071 | -0.087657 | 0.294637 | -0.208784 | 0.68246 | -0.591932 | -0.236892 | -1.399852 | -0.305853 | 1.664178 | 2.263447 | 1.507983 | -0.671706 | 0.86305 | -0.608185 | 0.108519 | 1.165892 | 3.618337 | 1.121907 | 2.209789 |
| 35.18056 | 0.035252 | 2.279777 | 3.978265 | 0.224816 | 2.537834 | -0.153114 | 0.24927 | -0.121495 | 0.660935 | -0.268447 | -0.380793 | -1.189376 | -0.742932 | 1.711952 | 1.769424 | 1.69436 | -1.115628 | 0.249005 | -0.75907 | 0.17653 | -0.032976 | 2.51629 | 1.152669 | 1.790105 |
| 35.19444 | 0.218797 | 2.465586 | 3.23343 | 0.251322 | 1.112164 | -0.07402 | 0.188775 | -0.140345 | 0.58767 | 0.327232 | -0.335319 | -0.820762 | -0.829189 | 1.458082 | 1.358711 | 1.473843 | -1.370499 | -0.453798 | -0.822891 | 0.088815 | -0.366634 | 1.317785 | 0.929029 | 1.457091 |
| 35.20833 | 0.189406 | 2.282385 | 2.092752 | 1.23062 | -1.235645 | 0.016652 | 0.084367 | 0.005407 | 0.541068 | 0.748669 | -0.166553 | -0.305821 | -0.669149 | 1.347586 | 1.130768 | 1.218179 | -1.38037 | -0.825118 | -0.81996 | 0.336532 | -1.024749 | 0.510146 | 0.877265 | 1.252521 |
| 35.22222 | 0.151298 | 1.459474 | 1.250435 | 1.446269 | -0.956103 | 0.057025 | 0.004961 | 0.208382 | 0.512569 | 0.834867 | -0.074978 | 0.215836 | -0.508696 | 1.48187 | 1.05613 | 1.238219 | -1.163648 | -0.913557 | -0.631029 | 1.018079 | -1.251071 | 0.841817 | 0.713328 | 1.070744 |
| 35.23611 | 0.097679 | 0.789175 | 0.496543 | 0.911254 | -0.162887 | 0.148518 | 0.043412 | 0.463046 | 0.389357 | 0.681704 | -0.04822 | 0.678241 | -0.398211 | 1.818311 | 0.827275 | 1.243083 | -0.850417 | -0.806468 | -0.448031 | 1.367167 | -1.219756 | 1.291601 | 0.715101 | 1.157586 |
| 35.25 | 0.107789 | 0.461087 | 0.412522 | 0.628431 | 0.829805 | 0.396886 | 0.189727 | 0.831947 | 0.236093 | 0.461739 | -0.015101 | 1.115368 | -0.268504 | 1.618793 | 0.362692 | 1.241799 | -0.439346 | -0.677776 | -0.326425 | 0.949505 | -0.868324 | 1.326837 | 1.232662 | 1.02291 |
| 35.26389 | 0.228296 | 0.436149 | -0.195776 | 0.828019 | 1.671942 | 0.800976 | 0.428195 | 1.22234 | 0.134836 | 0.393377 | -0.002052 | 1.54816 | 0.05334 | 1.447627 | -0.153906 | 1.307258 | 0.123121 | -0.24366 | -0.187997 | 0.369533 | 0.061455 | 2.095972 | 1.969514 | 1.050516 |
| 35.27778 | 0.500321 | 0.495592 | 0.51262 | 1.124531 | 0.820169 | 1.293892 | 0.706265 | 1.408622 | 0.132016 | 0.501406 | -0.053098 | 1.341425 | 0.405792 | 1.573606 | -0.519855 | 1.457483 | 0.718501 | 0.313958 | 0.089732 | 0.28712 | 0.858609 | 3.524426 | 2.439848 | 1.462891 |
| 35.29167 | 0.712837 | 0.417782 | 1.958881 | 1.410827 | -0.16803 | 1.516855 | 0.798273 | 1.280336 | 0.223558 | 0.612605 | -0.023324 | 1.126793 | 0.443531 | 1.498114 | -0.619251 | 1.608369 | 0.911817 | 0.628888 | 0.316156 | 0.452828 | 0.864048 | 4.141601 | 4.794251 | 1.590381 |
| 35.30556 | 0.899966 | 0.112982 | 3.875215 | 1.589969 | -1.318066 | 1.409276 | 0.64812 | 0.920154 | 0.298616 | 0.722385 | 0.045433 | 1.067945 | 0.189009 | 1.437174 | -0.753325 | 1.537618 | 0.650861 | 0.572704 | 0.305231 | 0.457607 | 0.389213 | 3.486553 | 5.935226 | 1.613332 |
| 35.31944 | 1.140133 | -0.295372 | 5.445875 | 1.371509 | 0.767848 | 1.425703 | 0.428405 | 0.557832 | 0.354889 | 0.853435 | 0.01806 | 1.036544 | 0.10214 | 1.427698 | -0.965015 | 1.37201 | 0.369621 | 0.312894 | 0.165046 | 0.329704 | 0.171602 | 2.963518 | 4.544381 | 1.274885 |
| 35.33333 | 1.608678 | -0.811997 | 4.92144 | 1.173583 | 1.885021 | 1.80017 | 0.333898 | 0.584139 | 0.377801 | 1.220734 | -0.04987 | 1.096606 | 0.233056 | 1.392655 | -1.036665 | 1.350663 | 0.523364 | 0.133991 | 0.141166 | 0.294232 | -0.129043 | 3.282314 | 5.176457 | 1.3911 |
| 35.34722 | 2.092458 | -1.208485 | 4.714526 | 0.696448 | 2.147072 | 2.58947 | 0.366588 | 0.847086 | 0.417323 | 2.035207 | -0.139094 | 1.042656 | 0.463592 | 1.239436 | -0.764476 | 1.508454 | 1.264801 | 0.139146 | 0.318492 | 0.489579 | 0.296019 | 4.066632 | 6.560477 | 1.492907 |
| 35.36111 | 2.640347 | -1.204209 | 3.296969 | 0.067868 | 3.877392 | 4.016275 | 0.518461 | 1.102626 | 0.578571 | 2.328549 | -0.311585 | 0.422013 | 0.876754 | 1.367823 | -0.481089 | 1.49114 | 2.24254 | 0.200937 | 0.517839 | 0.791575 | 1.034429 | 4.86363 | 6.405353 | 1.732609 |
| 35.375 | 2.883787 | -1.104678 | 1.565959 | -0.455664 | 5.397152 | 4.441813 | 0.568263 | 1.103729 | 0.817393 | 1.189653 | -0.390096 | -0.359197 | 1.583034 | 1.815991 | -0.07229 | 1.201065 | 2.177159 | 0.182362 | 0.5987 | 1.00508 | 1.582013 | 5.209223 | 7.170478 | 1.958086 |
| 35.38889 | 2.229452 | -0.669174 | 1.95103 | -0.44552 | 2.245091 | 3.869868 | 0.018072 | 0.934282 | 0.603433 | -0.543132 | -0.261746 | -0.491437 | 1.776874 | 2.04269 | 0.187116 | 1.048214 | 1.537698 | 0.142432 | 0.636784 | 1.02346 | 1.54463 | 5.302908 | 6.415823 | 1.912873 |
| 35.40278 | 1.033626 | -0.04898 | 2.789413 | -0.036557 | -3.758009 | 4.040416 | 0.036361 | 0.788754 | 0.501996 | -1.902674 | -0.012463 | -0.059615 | 1.839223 | 2.2247 | 0.274944 | 1.186094 | 1.762327 | 0.239875 | 0.728285 | 1.09791 | 1.475075 | 5.45274 | 6.555136 | 1.992997 |
| 35.41667 | 0.526205 | 0.749218 | 3.204903 | 0.395885 | -9.520682 | 4.48554 | 0.303528 | 1.487361 | 0.548614 | -1.668448 | 0.12084 | 0.085925 | 2.242386 | 2.777439 | 0.169215 | 1.653347 | 2.123503 | 0.462937 | 0.846173 | 1.397763 | 1.530184 | 5.035646 | 7.299185 | 2.38524 |
| 35.43056 | 0.73317 | 1.13883 | 3.231823 | 0.894425 | -2.80557 | 4.431244 | 0.508539 | 1.561646 | 0.525679 | -1.960372 | 0.05702 | 0.291075 | 3.327409 | 3.035006 | 0.431711 | 2.039038 | 2.310538 | 0.639668 | 0.94165 | 2.060555 | 2.189265 | 5.248293 | 7.47726 | 2.97401 |
| 35.44444 | 0.859108 | 1.371006 | 3.82116 | 1.40179 | -0.966464 | 3.79421 | 0.536663 | 1.722725 | 0.465131 | -1.732086 | 0.006091 | 0.346788 | 3.5849 | 1.908826 | 0.898322 | 1.836192 | 2.608728 | 0.638518 | 1.011102 | 2.367423 | 3.134451 | 4.778051 | 7.085633 | 1.928036 |
| 35.45833 | 1.478091 | 1.758648 | 4.226527 | 1.800729 | 0.195235 | 3.374599 | 0.60285 | 1.637503 | 0.460372 | -0.535833 | -0.013549 | 0.023499 | 2.543728 | 2.035617 | 1.170912 | 0.133329 | 2.468221 | 0.619535 | 1.225123 | 2.30132 | 2.791318 | 3.152531 | 5.678226 | 2.025043 |

| | | | | | | | | | | | | | | | | | | | | | | | | |
|---|---|---|---|---|---|---|---|---|---|---|---|---|---|---|---|---|---|---|---|---|---|---|---|---|
| 35.47222 | 1.820016 | 2.480081 | 3.907365 | 1.75728 | -0.298421 | 3.216213 | 0.732371 | 1.448612 | 0.429517 | 0.727753 | 0.207253 | -0.470956 | 1.412114 | 3.15789 | 1.262665 | 0.083402 | 2.924242 | 0.35849 | 1.238386 | 2.0199 | 2.459995 | 3.422729 | 4.434968 | 2.824166 |
| 35.48611 | 2.186456 | 2.748759 | 2.61428 | 1.311608 | -2.915312 | 3.135375 | 0.765472 | 1.711709 | 0.39805 | 1.751827 | 0.155541 | -0.346126 | 2.762245 | 3.454536 | 0.685812 | 0.843794 | 3.654828 | 0.369771 | 1.500202 | 2.26417 | 2.61476 | 3.373746 | 3.894279 | 3.754817 |
| 35.5 | 2.097194 | 2.242952 | 0.241341 | 0.458422 | -4.778273 | 2.711503 | 0.692113 | 0.86925 | 0.557843 | 2.294098 | 0.50445 | 0.751451 | 3.625749 | 3.390496 | 0.039028 | 2.046144 | 3.801812 | 0.858656 | 2.369817 | 2.439227 | 3.051988 | 4.260201 | 4.728699 | 4.773145 |
| 35.51389 | 1.058428 | 1.130231 | -1.65275 | -0.46632 | -4.865422 | 2.406817 | 0.425723 | 0.060182 | 0.882421 | 2.943397 | 0.832532 | 1.504986 | 2.940013 | 3.179099 | -0.107617 | 2.132 | 4.239539 | 1.923635 | 2.722716 | 2.852253 | 4.804854 | 3.557633 | 5.381053 | 4.735335 |
| 35.52778 | 0.588455 | 0.449009 | -3.475486 | -0.282335 | -3.098302 | 1.541289 | 0.338337 | 0.664795 | 1.106536 | 2.633924 | 0.873185 | 1.322804 | 2.86558 | 3.255382 | 0.588676 | 2.545622 | 3.06515 | 2.918469 | 2.570614 | 2.521305 | 6.849151 | 2.597412 | 4.336645 | 4.07784 |
| 35.54167 | -0.335433 | 0.420312 | -3.048593 | 0.352291 | 0.348551 | 0.780948 | 0.723043 | 1.541573 | 1.121777 | 2.639546 | 0.971193 | 0.881652 | 4.593878 | 3.256583 | 1.075332 | 2.861722 | 2.088736 | 2.661287 | 2.296886 | 1.969413 | 5.285666 | 1.891356 | 3.903192 | 3.715603 |
| 35.55556 | -1.185606 | 0.7089 | -0.058255 | 1.236119 | 1.858843 | 0.460492 | 1.138389 | 2.153272 | 1.251428 | 2.013167 | 0.835459 | 0.725767 | 4.745592 | 2.82628 | 1.329483 | 2.21658 | 1.517007 | 1.204316 | 2.243915 | 1.749973 | 3.357693 | 1.454397 | 3.58835 | 3.184045 |
| 35.56944 | -1.092042 | 1.456191 | 1.687956 | 2.117801 | 0.062331 | 0.427425 | 1.102429 | 2.568079 | 1.389592 | 1.665131 | 0.767775 | 1.032006 | 3.686765 | 1.843014 | 1.570229 | 1.532353 | 1.393429 | -0.353271 | 2.136888 | 1.461978 | 1.499008 | 1.310422 | 2.667472 | 2.70365 |
| 35.58333 | -0.215786 | 2.043379 | 2.8437 | 3.178651 | 2.596504 | 0.216239 | 0.902068 | 2.86442 | 1.602671 | 1.409167 | 0.926864 | 1.385343 | 2.772265 | 1.220475 | 1.545095 | 1.79282 | 1.632895 | -1.316188 | 1.919683 | 1.578285 | -0.468424 | 1.113124 | 1.76635 | 2.438333 |
| 35.59722 | 0.422735 | 2.288652 | 2.266804 | 3.776528 | 2.617937 | 0.482736 | 0.60545 | 3.007892 | 1.869991 | 0.797568 | 0.740857 | 1.808183 | 2.216137 | 1.094161 | 1.02737 | 2.198115 | 2.406717 | -0.560102 | 1.743351 | 1.509192 | -0.846211 | 0.966013 | 1.185221 | 2.114577 |
| 35.61111 | -0.065333 | 2.420981 | 3.81897 | 3.329341 | 1.997138 | 0.503521 | 0.451279 | 3.585189 | 2.319224 | 0.340233 | 0.686822 | 2.099428 | 0.832188 | 1.215712 | 0.268155 | 2.801195 | 2.238504 | 0.696584 | 1.652967 | 1.495035 | -0.252492 | 0.726217 | 0.32845 | 1.939493 |
| 35.625 | 1.1019 | 3.031635 | 6.049999 | 1.63296 | 1.805381 | 0.545033 | 0.363907 | 4.293868 | 2.877389 | 1.025722 | 0.644476 | 2.21764 | -0.376231 | 0.99549 | -0.133105 | 3.19779 | -0.093676 | 1.529575 | 1.51217 | 1.454655 | 0.134102 | 0.413137 | -0.612752 | 1.505509 |
| 35.63889 | 0.339671 | 2.812626 | 6.767583 | -0.472153 | 3.709828 | 0.818545 | 0.397168 | 3.761464 | 3.115246 | 1.39277 | 0.56656 | 2.141852 | -1.900809 | 0.806519 | -0.231796 | 2.540909 | -2.1768 | 1.548733 | 1.330623 | 0.874308 | -0.380613 | -0.084346 | -1.362881 | 1.154304 |
| 35.65278 | 4.195315 | 2.220985 | 5.78017 | -1.447805 | 2.965836 | 1.070108 | 0.279173 | 3.392006 | 2.739771 | 2.071578 | 0.595465 | 2.222108 | -2.217887 | 0.734791 | -0.036095 | 2.068618 | -2.618706 | 1.354346 | 1.061739 | 0.670997 | -1.838831 | -0.754582 | -1.330787 | 0.701993 |
| 35.66667 | 4.421324 | 1.713653 | 5.66081 | -1.501503 | -1.698413 | 0.831795 | -0.00617 | 3.081235 | 1.869235 | 2.424164 | 0.828686 | 2.160289 | -2.206847 | 0.128312 | 0.2552 | 2.021369 | -3.176951 | 0.865514 | 0.849596 | 1.568452 | -2.7025 | -0.921028 | -2.399287 | 0.321464 |
| 35.68056 | -9.680549 | 1.418283 | 6.141155 | -1.458654 | -5.586932 | 0.669105 | 0.360502 | 2.616851 | 1.155909 | 3.007538 | 1.475452 | 1.745661 | -1.369574 | -0.611374 | 0.534932 | 2.774097 | -2.939576 | 1.199342 | 1.088569 | 1.698313 | -2.883676 | -0.632988 | -2.87029 | 0.123087 |
| 35.69444 | -9.250745 | 1.423203 | 5.956468 | -1.053987 | -13.111571 | 0.368988 | 0.675811 | 2.347421 | 0.759991 | 3.113754 | 1.737317 | 0.747898 | -0.571713 | -0.068541 | 0.648968 | 2.709124 | -1.507545 | 1.470812 | 1.363442 | 1.484658 | -2.590198 | -0.484925 | -2.468055 | 0.308013 |
| 35.70833 | -2.918645 | 1.570363 | 4.400032 | -0.267525 | -27.848294 | 0.868299 | 0.978133 | 1.950242 | 0.462617 | 2.531278 | 1.332958 | -0.447096 | 0.313778 | 0.244281 | 0.741239 | 2.385183 | -0.038812 | 1.124499 | 1.377998 | 1.502009 | -1.631316 | -0.777519 | -1.894116 | 0.723734 |
| 35.72222 | -1.621656 | 2.352369 | 2.381448 | 0.616488 | -4.411425 | 1.539278 | 1.380381 | 1.746324 | -0.262196 | 2.512508 | 1.20072 | -3.613895 | 1.028953 | -0.292329 | 0.9407 | 1.99426 | 0.393537 | 0.602394 | 1.235404 | 1.311645 | -0.719309 | -1.073494 | -1.698278 | 0.991172 |
| 35.73611 | -0.092208 | 1.461332 | 1.64999 | 1.559583 | -0.79615 | 2.095281 | 2.087204 | 2.037558 | -1.470066 | 2.819714 | -0.898243 | -4.430662 | 1.402403 | -0.442639 | 1.180144 | 1.662184 | 0.317615 | -0.00217 | 1.179219 | 1.076788 | -0.094871 | -1.010825 | -2.075699 | 1.050082 |
| 35.75 | 0.249795 | 5.296702 | -1.249493 | 3.627547 | -4.794025 | 2.63989 | 2.379408 | 2.454857 | -1.48853 | 3.936189 | -1.319468 | -7.117929 | 1.38324 | -0.170256 | 0.906217 | 1.33565 | 0.44131 | -0.275273 | 1.190619 | 0.920707 | 0.310473 | -1.284002 | -2.336185 | 1.001554 |
| 35.76389 | -0.05958 | 3.876649 | 0.34746 | 5.140151 | -5.226256 | 2.053487 | 2.281792 | 2.284807 | 0.711642 | 3.889953 | -0.528451 | -2.93884 | 0.985031 | 0.295536 | 1.363559 | 1.009199 | 0.482722 | -0.182073 | 1.504374 | 0.763023 | -0.1766 | -1.548937 | -3.008713 | 0.63356 |
| 35.77778 | -0.688988 | 2.206976 | 0.033543 | 3.951369 | -2.648127 | 1.469414 | 2.409931 | 1.42798 | 2.727725 | 3.01283 | -0.312556 | -0.865535 | 0.360389 | 0.81773 | 1.146081 | 0.959811 | 0.390862 | 0.286181 | 2.000732 | 0.720729 | 0.437287 | -1.050926 | -2.419715 | 0.446643 |
| 35.79167 | -1.151559 | -0.099174 | -3.676211 | 3.473082 | -2.370559 | 1.426343 | 2.221726 | 0.21481 | 3.587407 | 1.398106 | 1.318675 | 1.098561 | -0.199745 | 0.772227 | 0.808637 | 0.785162 | 0.478078 | 0.918334 | 2.328608 | 0.37128 | 0.830172 | -0.459035 | -2.893652 | 0.235805 |
| 35.80556 | -1.756303 | -1.057283 | -5.308136 | 1.752596 | -2.581955 | 2.226212 | 1.805435 | -1.126785 | 3.435268 | -0.527985 | 1.104828 | 3.475755 | -0.873206 | 0.874582 | 1.020587 | 0.495459 | 1.033824 | 1.652858 | 2.292792 | -0.776367 | 0.476349 | -0.236316 | -3.620088 | 0.098228 |
| 35.81944 | -1.738705 | -1.214955 | -0.366366 | 0.819592 | 0.015007 | 3.443049 | 1.713572 | -1.33624 | 2.251248 | -1.948108 | 2.423993 | 5.733364 | -2.717562 | 1.182225 | 1.082865 | 0.495304 | 1.395406 | 1.561723 | 2.123287 | -1.169205 | -0.399993 | -0.15687 | -4.157088 | 0.309803 |
| 35.83333 | -1.992305 | 1.169943 | 5.725793 | -0.431642 | 2.157923 | 2.236027 | 1.074248 | -1.380756 | 0.756292 | -2.220629 | 0.985791 | 2.956157 | -4.937335 | 1.034607 | 0.717771 | 0.667965 | 1.512101 | 1.536088 | 1.911695 | -0.732693 | -1.285943 | 0.372478 | -4.383183 | 0.661314 |
| 35.84722 | -2.428132 | 1.30702 | 1.531544 | -2.119002 | 2.03301 | 0.45074 | 0.995511 | -1.788318 | 2.708147 | -1.812696 | -0.798567 | 2.78226 | -6.937633 | 0.956996 | -0.268855 | 0.471757 | 1.280276 | 1.629353 | 1.569139 | -0.040108 | -1.799532 | 0.908447 | -4.979142 | 0.895582 |
| 35.86111 | -3.014964 | 0.462502 | -6.259811 | -3.781137 | -3.22755 | -2.06151 | 0.341751 | -3.031545 | 2.903681 | -1.341005 | -3.07317 | 2.468242 | -6.359182 | 0.149498 | -1.174872 | 0.175521 | 1.425887 | 1.704656 | 1.350865 | 0.416232 | -1.994252 | 1.021471 | -4.925928 | 1.136272 |
| 35.875 | -2.52051 | 2.134686 | -1.576326 | -2.899551 | -4.026625 | -3.931156 | -0.776793 | -3.60363 | 1.81342 | -4.673195 | -5.498832 | 1.094419 | -4.09748 | -0.253506 | -1.525267 | -0.269452 | 1.061383 | 1.411803 | 1.175626 | 0.039144 | -2.501652 | 1.071065 | -4.381308 | 0.882719 |
| 35.88889 | -2.856669 | 2.546955 | -1.44153 | 0.64318 | -4.655409 | -5.444311 | -0.882142 | -2.938465 | 1.044481 | -3.8274 | -2.541409 | -1.348492 | -3.324853 | -0.599311 | -1.698153 | -0.35172 | 0.870459 | 1.321565 | 1.143886 | -0.597778 | -2.985254 | 0.984642 | -3.21303 | 0.623726 |
| 35.90278 | -1.890295 | 1.596652 | -1.625625 | 3.476937 | -3.161899 | -4.491657 | -0.563222 | -1.664094 | 1.395736 | 0.774797 | -2.701173 | -0.461427 | -3.278362 | -0.79016 | -1.611835 | -0.928272 | 0.015211 | 1.562043 | 1.096355 | -1.477888 | -3.3652 | 0.663187 | -2.691835 | 0.641464 |
| 35.91667 | -1.26598 | 0.675755 | 0.87934 | 4.791111 | -0.948729 | -4.576774 | -0.81643 | -0.202871 | -0.219191 | 1.596362 | -1.35958 | 0.411041 | -2.330234 | -0.851188 | -1.620342 | -1.707788 | -1.047656 | 1.719796 | 1.243458 | -1.495705 | -3.068476 | -0.521495 | -2.299158 | 0.508183 |
| 35.93056 | -0.538118 | 0.041599 | 1.251773 | 4.751862 | 2.148651 | -5.084052 | -1.384098 | 0.497117 | -0.257345 | 0.13516 | -0.828254 | 0.803307 | -1.708805 | -1.211925 | -1.746833 | -2.993505 | -1.088861 | 0.882264 | 0.928763 | -1.258704 | -3.062119 | -2.414034 | -2.936598 | -0.291403 |
| 35.94444 | -0.073758 | -0.180708 | 1.265991 | 3.205831 | 4.037549 | -5.766167 | -1.984473 | 0.477206 | 0.138208 | -0.086583 | -0.589573 | 0.794298 | -2.604723 | -1.565127 | -2.307326 | -4.4619 | -1.23752 | -0.693483 | -0.183637 | -1.457232 | -3.881657 | -4.24211 | -3.900028 | -1.526403 |
| 35.95833 | 0.093436 | 0.516631 | 2.508395 | 1.584326 | 3.94512 | -4.032775 | -2.183568 | 0.340053 | -0.499564 | -0.070241 | -0.203064 | 0.595039 | -5.287117 | -1.449398 | -3.25578 | -5.711592 | -2.034019 | -1.376348 | -1.354322 | -1.85991 | -4.321192 | -4.854255 | -4.497734 | -1.509064 |
| 35.97222 | 0.510811 | 0.719356 | 3.819856 | 0.310139 | 2.963032 | -0.908909 | -2.09617 | 0.292622 | -0.554421 | 0.231982 | 0.438726 | 0.354408 | -6.600999 | -1.101956 | -4.28226 | -3.227646 | -2.109891 | -1.306179 | -1.241261 | -1.769373 | -3.684943 | -3.464945 | -3.98887 | -0.836489 |
| 35.98611 | 1.028015 | 0.540156 | 4.443722 | -0.092785 | 2.121122 | 0.12466 | -1.416919 | -0.129032 | 1.065971 | -0.442359 | 0.805831 | 0.498733 | -4.999472 | -0.71605 | -3.652982 | -2.101382 | -2.459951 | -0.67077 | -0.503637 | -0.552311 | -2.70312 | -3.01692 | -1.668677 | -0.595745 |
| 36 | 1.983623 | 0.550638 | 4.924798 | 0.886578 | 1.560751 | 0.937148 | -0.635721 | -0.857853 | 1.015686 | -1.030482 | 0.723584 | 0.629311 | -4.52994 | -0.67956 | -2.507132 | -0.97986 | -2.53555 | -0.144434 | -0.036344 | -0.227639 | -2.017002 | -1.12304 | -0.42336 | -0.462914 |
| 36.01389 | 1.920512 | 2.197805 | 3.719285 | 1.094427 | 0.900667 | 1.105164 | 0.355621 | -1.582733 | 0.14263 | -1.455071 | 0.575828 | 0.692033 | -4.132926 | -0.714558 | -1.527808 | -0.629136 | -2.688171 | 0.035972 | -0.062696 | 0.060953 | -0.7349 | 0.21931 | 0.427515 | -0.569428 |
| 36.02778 | 1.329951 | 3.063064 | 2.675807 | 1.465944 | -0.155302 | 0.215553 | 0.696324 | -1.93447 | -0.793249 | -1.016554 | 0.783379 | 0.917218 | -3.154109 | -0.484057 | -0.971637 | 0.272406 | -2.189967 | -0.126219 | -0.339807 | 0.335994 | 0.663693 | 0.741998 | 0.70009 | -0.552911 |
| 36.04167 | 0.829654 | 3.340006 | 3.260696 | 1.327151 | -0.90773 | -1.765689 | 0.644005 | -1.706468 | -0.243083 | -0.885795 | 1.594767 | 1.596915 | -2.0369 | 0.298195 | -0.230641 | 1.875463 | -1.401141 | 0.18347 | -0.08113 | 0.680831 | 1.353998 | 0.917621 | 0.762365 | -0.08595 |
| 36.05556 | 0.475789 | 2.818263 | 4.015039 | 1.585716 | -1.035052 | -2.643539 | 0.327927 | -1.168365 | -0.331719 | -0.582146 | 2.609275 | 1.969347 | -1.365931 | 1.202245 | 0.770942 | 3.73191 | -1.31302 | 0.308429 | 0.139909 | 0.60666 | 1.400729 | 1.012667 | 0.873377 | 0.503507 |
| 36.06944 | -0.082086 | 2.58797 | 4.21388 | 0.910944 | -2.564555 | -1.861757 | -0.687929 | -0.355953 | 0.156086 | 0.768215 | 2.95465 | 1.612908 | -0.974935 | 1.435806 | 1.798748 | 3.291609 | -0.633325 | 0.558428 | 0.406464 | 0.476063 | 1.151092 | 1.121515 | 0.9813 | 0.32472 |
| 36.08333 | -0.865409 | 2.143144 | 3.135576 | -1.638095 | -3.365249 | -1.024988 | -0.860037 | 0.20628 | 0.794442 | 0.553047 | 3.813049 | 1.485981 | -0.824061 | 1.088455 | 1.949781 | 2.168797 | 0.036312 | 0.563955 | 0.582065 | 0.275713 | 0.832393 | 0.847513 | 0.386063 | -0.445659 |
| 36.09722 | -1.677875 | 1.067901 | 2.838244 | -1.042414 | -2.364735 | -1.389286 | -0.551358 | 0.295092 | 1.041251 | -0.078054 | 2.838017 | 0.128187 | -0.690601 | 0.695663 | 1.917958 | 1.855085 | 0.415401 | 0.797843 | 0.958919 | 0.479568 | 0.803459 | 0.180876 | 0.141773 | -0.998098 |
| 36.11111 | -2.044809 | 0.531743 | 2.956806 | 6.390267 | -0.665717 | -1.930667 | -0.361187 | 0.345498 | 0.542758 | -0.423381 | 0.982202 | 0.114627 | -0.230183 | 0.945111 | 1.882433 | 1.93317 | 1.165661 | 0.492104 | 0.998953 | 0.54232 | 1.396984 | -0.444399 | 0.507472 | -0.524971 |
| 36.125 | -1.870224 | 0.105938 | 1.99252 | 1.346055 | 0.506056 | -2.120237 | 0.039393 | 0.380967 | -0.832382 | 0.004955 | -0.564434 | -0.005691 | 0.141724 | 1.864422 | 1.877453 | 2.160962 | 2.022177 | -0.121058 | 1.028216 | 0.500607 | 1.73279 | -0.285906 | 1.043678 | 0.79281 |
| 36.13889 | -1.39933 | 0.072123 | 1.046616 | 0.885647 | 0.605346 | -1.635002 | 0.504656 | 0.577736 | -2.060856 | 1.11581 | -1.77618 | 0.037999 | -0.310292 | 1.013079 | 1.929369 | 2.225352 | 2.417497 | -0.677657 | 0.797296 | 0.325608 | 1.849978 | 0.128798 | 1.265988 | 0.810725 |
| 36.15278 | -1.098063 | 0.521138 | 0.467752 | 3.841519 | 0.757318 | -1.503008 | 0.427388 | 0.749727 | -1.601694 | 1.711955 | -3.662165 | 0.053922 | -1.303248 | 0.672556 | 2.009064 | 1.931219 | 1.463046 | -0.192028 | 0.801279 | 0.08956 | 1.547971 | 0.689587 | 0.751598 | 0.787542 |
| 36.16667 | -0.746008 | 1.164691 | 0.265452 | 4.736966 | 0.915297 | -1.145848 | 0.261328 | 0.811938 | -0.217881 | 1.904519 | -4.319486 | -0.087283 | -2.092303 | 0.554842 | 1.937468 | 1.858886 | 0.299325 | 0.461548 | 1.113158 | -0.01239 | 0.931534 | 1.066314 | 0.370701 | 1.040456 |
| 36.18056 | -0.194015 | 0.552479 | 0.146713 | -1.520831 | 0.160444 | -0.243389 | 0.333398 | 1.000085 | 0.643247 | 1.008333 | -3.684867 | -0.853524 | -2.217364 | 0.851862 | 1.582852 | 1.565645 | -0.404176 | 0.758721 | 1.123254 | 0.009645 | 0.984442 | 0.93088 | -0.064217 | 1.012267 |
| 36.19444 | 0.234034 | -0.106268 | 0.479641 | -1.551157 | 0.392754 | -0.805578 | 0.22912 | 0.932547 | 1.439245 | 0.52422 | -2.8145 | -0.603631 | -1.351985 | 0.99982 | 0.997533 | 1.046057 | -0.825845 | 0.455286 | 0.963318 | -0.129899 | 1.124738 | 0.455487 | -0.609305 | 0.82607 |
| 36.20833 | 0.63601 | -0.320563 | 0.881294 | 0.757826 | 0.430022 | -0.616326 | 0.175706 | 0.537622 | 1.236918 | 1.348672 | -1.241215 | 0.389045 | -1.165199 | 1.245552 | 0.559259 | 0.610535 | -0.978405 | 0.249569 | 0.857376 | -0.201904 | 1.702839 | -0.213927 | -0.764682 | 0.807216 |
| 36.22222 | 0.271555 | -0.188872 | 1.117699 | 1.342055 | 0.176432 | 0.396342 | 0.284702 | 0.109924 | 0.555109 | 0.273725 | -0.668226 | 0.906415 | -1.125643 | 1.632493 | 0.560343 | 0.682435 | -1.007309 | 0.371467 | 0.951935 | -0.052273 | 2.681975 | -0.405592 | -0.457979 | 1.105715 |
| 36.23611 | 0.304852 | 0.358487 | 1.824788 | 0.418498 | 0.334693 | 1.07383 | 0.390292 | -0.769566 | -0.088365 | -1.448308 | -0.442197 | 1.190667 | -0.523507 | 1.509281 | 0.615822 | 1.221992 | -0.685802 | 0.669808 | 1.1355 | 0.380321 | 3.054607 | 0.263376 | -0.002176 | 1.50479 |
| 36.25 | 1.662037 | 0.205579 | 3.952205 | -0.021367 | -0.082338 | -1.028481 | 0.579672 | -5.539454 | -0.531468 | -2.209043 | -0.155655 | 2.091961 | 0.195935 | 0.908504 | 0.390013 | 1.42802 | -0.286245 | 0.910351 | 1.184322 | 0.632055 | 4.233164 | 0.988891 | 0.238138 | 1.142968 |
| 36.26389 | 2.325829 | -4.796644 | 5.76295 | -2.870418 | -2.913718 | -9.950403 | 3.466266 | -15.173122 | 0.010638 | -3.897558 | 0.339226 | 1.367194 | 0.472007 | 0.372818 | -0.04452 | 0.994022 | 0.287126 | 0.677822 | 0.961614 | 0.582063 | 3.700659 | 1.062145 | 0.461004 | 0.488029 |
| 36.27778 | -12.461326 | NaN | 32.449576 | NaN | NaN | NaN | NaN | -19.539414 | 0.249437 | 4.775064 | NaN | -16.640052 | 0.360795 | 0.137257 | -0.271892 | 0.614655 | 0.737856 | 0.618804 | 0.840005 | 0.420335 | 2.267233 | 0.730777 | 0.294492 | 0.060172 |
| 36.29167 | NaN | NaN | NaN | NaN | NaN | NaN | NaN | NaN | NaN | 2.679513 | NaN | NaN | 0.183682 | 0.023277 | -0.270785 | 0.621122 | 0.848872 | 0.470306 | 1.150943 | 0.399254 | 0.93557 | 0.601466 | 0.076085 | -0.248716 |
| 36.30556 | NaN | NaN | NaN | NaN | NaN | NaN | NaN | NaN | NaN | -0.098633 | NaN | NaN | 0.189301 | -0.225012 | -0.128877 | 0.731881 | 0.749794 | 0.467088 | 1.346044 | 0.460428 | 0.200721 | 0.690065 | -0.177957 | -0.581941 |
| 36.31944 | NaN | NaN | NaN | NaN | NaN | NaN | NaN | NaN | NaN | NaN | NaN | NaN | -0.27484 | -0.2835 | -0.134857 | 0.824131 | 0.661515 | 0.070305 | 1.563657 | 0.233056 | -0.230776 | 1.482788 | -0.828371 | -0.733493 |
| 36.33333 | NaN | NaN | NaN | NaN | NaN | NaN | NaN | NaN | NaN | NaN | NaN | NaN | -2.890337 | 0.796929 | -0.770309 | 0.84448 | -0.398279 | -2.308941 | 1.836273 | -1.512433 | -2.263471 | 6.915168 | -2.942012 | -1.258982 |
| 36.34722 | NaN | NaN | NaN | NaN | NaN | NaN | NaN | NaN | NaN | NaN | NaN | NaN | -8.49866 | 0.475349 | -5.494706 | -1.859243 | -2.532623 | -6.17924 | -1.494794 | -4.113131 | -3.81463 | 10.967559 | -8.135741 | -1.244831 |
| 36.36111 | NaN | NaN | NaN | NaN | NaN | NaN | NaN | NaN | NaN | NaN | NaN | NaN | -23.080546 | 5.199341 | 14.11583 | NaN | NaN | -17.341699 | -12.127084 | NaN | 6.782179 | 2.134585 | NaN | NaN |
| 36.375 | NaN | NaN | NaN | NaN | NaN | NaN | NaN | NaN | NaN | NaN | NaN | NaN | NaN | NaN | NaN | NaN | NaN | NaN | NaN | NaN | NaN | -0.828035 | NaN | NaN |
| 36.38889 | NaN | NaN | NaN | NaN | NaN | NaN | NaN | NaN | NaN | NaN | NaN | NaN | NaN | NaN | NaN | NaN | NaN | NaN | NaN | NaN | NaN | -0.103655 | NaN | NaN |

**Supplementary Table S2.** Raw motion data from: **(c)** 100 mM NaCl treatment in Amaranth

| Time | Amaranth 100mMNaCl1 | Amaranth 100mMNaCl10 | Amaranth 100mMNaCl11 | Amaranth 100mMNaCl12 | Amaranth 100mMNaCl2 | Amaranth 100mMNaCl3 | Amaranth 100mMNaCl4 | Amaranth 100mMNaCl5 | Amaranth 100mMNaCl6 | Amaranth 100mMNaCl7 | Amaranth 100mMNaCl8 | Amaranth 100mMNaCl9 | Amaranth Control1 | Amaranth Control10 | Amaranth Control11 | Amaranth Control12 | Amaranth Control2 | Amaranth Control3 | Amaranth Control4 | Amaranth Control5 | Amaranth Control6 | Amaranth Control7 | Amaranth Control8 | Amaranth Control9 |
|---|---|---|---|---|---|---|---|---|---|---|---|---|---|---|---|---|---|---|---|---|---|---|---|---|
| 18 | 0.350558 | 0.149029 | -0.206832 | 0.041471 | 0.423121 | 0.335366 | 0.364606 | 0.389818 | 0.089492 | 0.300196 | 0.215653 | 0.179756 | -0.067626 | 0.024806 | -0.297207 | 0.047323 | -0.197359 | 0.029746 | 0.121529 | 0.126831 | -0.056917 | 0.098258 | 0.054394 | -0.06796 |
| 18.01389 | 0.345844 | 0.104771 | 0.069686 | 0.539447 | 0.672112 | -0.015711 | 0.473994 | 0.216636 | 0.024917 | 0.518819 | 0.292189 | 0.203259 | -0.117417 | 0.279021 | 0.044147 | 0.295398 | -0.107355 | 0.09258 | 0.611138 | 0.543028 | 0.044223 | 0.34294 | 0.21918 | 0.026003 |
| 18.02778 | 0.218139 | -0.431929 | 0.102807 | 0.412528 | 0.608831 | -0.771426 | 0.337519 | -0.132201 | -0.403048 | 0.456815 | 0.026742 | -0.127599 | -0.252145 | 0.273779 | 0.144617 | 0.363195 | -0.689838 | 0.069225 | 0.890973 | 0.749213 | 0.063367 | 0.458512 | 0.274475 | -0.116593 |
| 18.04167 | -0.096277 | -0.279891 | -0.496913 | -1.001495 | -0.24047 | -0.335594 | -0.120711 | -0.072548 | -0.984344 | -0.350161 | -0.687133 | -0.277994 | -0.064456 | -0.173428 | -0.229509 | 0.046471 | -0.766732 | 0.105092 | 0.128907 | -0.12424 | 0.045395 | 0.125373 | 0.047894 | -0.089823 |
| 18.05556 | -0.148533 | 0.436369 | -0.435186 | -1.033651 | -0.419843 | 0.804974 | -0.378728 | 0.380555 | -0.525654 | -0.309721 | -0.74396 | 0.163334 | 0.221482 | -0.24954 | -0.46987 | -0.108756 | -0.098667 | 0.148266 | -0.634539 | -0.772134 | 0.122148 | -0.13772 | -0.049551 | 0.049434 |
| 18.06944 | 0.042254 | 0.678549 | 0.206707 | -0.142062 | 0.02173 | 0.728415 | -0.037005 | 0.441164 | -0.400125 | 0.107467 | -0.289401 | 0.448274 | 0.156643 | 0.01531 | -0.308558 | 0.03334 | 0.260974 | 0.107897 | -0.369176 | -0.286583 | 0.128891 | -0.004171 | 0.056229 | 0.227618 |
| 18.08333 | 0.305071 | 0.669165 | 0.562937 | 0.294337 | 0.49514 | 0.507755 | 0.355749 | 0.418852 | -0.303786 | 0.390636 | 0.073562 | 0.475932 | 0.034855 | 0.136162 | -0.225078 | 0.087946 | 0.221339 | 0.048044 | -0.015682 | 0.109307 | 0.085593 | 0.109461 | 0.089309 | 0.274054 |
| 18.09722 | 0.478553 | 0.711828 | 0.607173 | 0.409405 | 0.574804 | 0.507864 | 0.533917 | 0.50947 | 0.124437 | 0.461686 | 0.261529 | 0.487897 | -0.010734 | 0.061055 | -0.283959 | -0.018328 | 0.102359 | -0.00274 | 0.009749 | 0.073787 | 0.016272 | 0.029115 | 0.01557 | 0.149205 |
| 18.11111 | 0.478788 | 0.691971 | 0.575005 | 0.567907 | 0.513359 | 0.511724 | 0.535822 | 0.51725 | 0.387716 | 0.477208 | 0.337245 | 0.426316 | -0.012021 | 0.003502 | -0.266295 | -0.123018 | 0.123487 | -0.042552 | -0.034117 | 0.010304 | -0.034254 | -0.039404 | -0.035249 | 0.043221 |
| 18.125 | 0.375969 | 0.558663 | 0.433658 | 0.64811 | 0.416873 | 0.388615 | 0.438724 | 0.403678 | 0.473583 | 0.3584 | 0.313798 | 0.23979 | -0.026704 | -0.019517 | -0.18457 | -0.17267 | 0.131981 | -0.071568 | -0.078406 | -0.012122 | -0.060711 | -0.06681 | -0.054183 | -0.043076 |
| 18.13889 | 0.199463 | 0.360271 | 0.193061 | 0.552655 | 0.245257 | 0.198656 | 0.269741 | 0.215413 | 0.442308 | 0.134375 | 0.191613 | -0.045037 | -0.049303 | -0.050473 | -0.136876 | -0.213882 | 0.054556 | -0.092977 | -0.130985 | -0.037603 | -0.091947 | -0.080287 | -0.063954 | -0.130761 |
| 18.15278 | 0.02783 | 0.175104 | -0.061592 | 0.337571 | 0.023073 | 0.053514 | 0.075072 | 0.048553 | 0.258118 | -0.08007 | 0.026064 | -0.344298 | -0.036301 | -0.059657 | -0.108435 | -0.217646 | 0.054838 | -0.084573 | -0.180513 | -0.075973 | -0.114667 | -0.070489 | -0.061301 | -0.19163 |
| 18.16667 | -0.003038 | 0.134531 | -0.156789 | 0.202123 | -0.054184 | 0.06953 | 0.020306 | 0.034197 | 0.083887 | -0.139982 | -0.015955 | -0.481876 | -0.03151 | -0.059175 | -0.104603 | -0.228309 | 0.062131 | -0.088314 | -0.203323 | -0.08656 | -0.133282 | -0.061968 | -0.061958 | -0.217874 |
| 18.18056 | 0.121493 | 0.250932 | -0.024265 | 0.251969 | 0.085553 | 0.191086 | 0.137923 | 0.146368 | 0.056819 | 0.119209 | 0.115176 | -0.419369 | -0.066786 | -0.071023 | -0.1289 | -0.256532 | -0.016269 | -0.10841 | -0.196608 | -0.061727 | -0.157543 | -0.074976 | -0.081225 | -0.236941 |
| 18.19444 | 0.220314 | 0.333436 | 0.101391 | 0.291784 | 0.202034 | 0.259242 | 0.22664 | 0.221329 | 0.063141 | 0.213321 | 0.218223 | -0.399657 | -0.077355 | -0.080406 | -0.141539 | -0.235592 | -0.093316 | -0.107899 | -0.157291 | -0.045767 | -0.157881 | -0.090229 | -0.091854 | -0.24858 |
| 18.20833 | 0.157585 | 0.196404 | 0.002882 | 0.123056 | 0.117564 | 0.168441 | 0.132123 | 0.147347 | -0.018509 | 0.106083 | 0.126221 | -0.526897 | -0.043103 | -0.070044 | -0.113844 | -0.183665 | -0.115533 | -0.083697 | -0.115554 | -0.059138 | -0.139582 | -0.092114 | -0.073375 | -0.221892 |
| 18.22222 | -0.036949 | -0.096249 | -0.215178 | -0.200436 | -0.097782 | -0.058123 | -0.080833 | -0.052611 | -0.167662 | -0.11337 | -0.089677 | -0.752741 | -0.02113 | -0.061798 | -0.08336 | -0.151571 | -0.087998 | -0.077486 | -0.103677 | -0.083686 | -0.131721 | -0.090408 | -0.054247 | -0.17533 |
| 18.23611 | -0.157572 | -0.22942 | -0.258016 | -0.356348 | -0.202361 | -0.201802 | -0.173556 | -0.173139 | -0.222166 | -0.271541 | -0.161616 | -0.698584 | -0.040341 | -0.084166 | -0.100898 | -0.191359 | -0.059043 | -0.078931 | -0.100364 | -0.109471 | -0.122435 | -0.094536 | -0.06653 | -0.163136 |
| 18.25 | -0.102989 | -0.124067 | -0.107181 | -0.233312 | -0.110701 | -0.139019 | -0.083453 | -0.107026 | -0.130003 | -0.201695 | -0.039937 | -0.44577 | -0.090795 | -0.099578 | -0.137371 | -0.231265 | -0.070206 | -0.087582 | -0.08539 | -0.100297 | -0.116622 | -0.092249 | -0.098686 | -0.160965 |
| 18.26389 | -0.021924 | -0.041106 | -0.015517 | -0.10535 | -0.026492 | -0.044099 | -0.000774 | -0.02036 | -0.071594 | -0.158452 | 0.051258 | -0.33262 | -0.121988 | -0.084125 | -0.142207 | -0.202798 | -0.116139 | -0.110153 | -0.092306 | -0.088391 | -0.132491 | -0.094462 | -0.115381 | -0.153263 |
| 18.27778 | -0.046305 | -0.088645 | -0.050838 | -0.145239 | -0.072933 | -0.067144 | -0.042009 | -0.03824 | -0.113126 | 0.016521 | 0.001835 | -0.302062 | -0.091068 | -0.061154 | -0.116899 | -0.125731 | -0.123501 | -0.112384 | -0.114517 | -0.108897 | -0.12476 | -0.098619 | -0.101378 | -0.125568 |
| 18.29167 | -0.120789 | -0.177362 | -0.115973 | -0.22694 | -0.150928 | -0.136557 | -0.128811 | -0.103228 | -0.124291 | -0.037433 | -0.093059 | -0.282296 | -0.032332 | -0.015696 | -0.066312 | -0.046993 | -0.042875 | -0.079248 | -0.108671 | -0.113403 | -0.079884 | -0.074524 | -0.063352 | -0.074171 |
| 18.30556 | -0.179736 | -0.260707 | -0.170716 | -0.264322 | -0.195006 | -0.192818 | -0.191283 | -0.168181 | -0.127106 | -0.132618 | -0.162764 | -0.255009 | -0.007681 | 0.020611 | -0.025649 | -0.006166 | 0.008259 | -0.053135 | -0.054191 | -0.091301 | -0.046496 | -0.044513 | -0.039139 | -0.038041 |
| 18.31944 | -0.247206 | -0.314125 | -0.205611 | -0.282455 | -0.226586 | -0.251074 | -0.233798 | -0.231173 | -0.119272 | -0.207913 | -0.207492 | -0.220528 | -0.060197 | -0.014251 | -0.058476 | -0.060497 | -0.06012 | -0.064485 | -0.012986 | -0.069515 | -0.056545 | -0.044529 | -0.066742 | -0.057108 |
| 18.33333 | -0.208179 | -0.227792 | -0.118203 | -0.15776 | -0.153758 | -0.195959 | -0.167166 | -0.176142 | -0.017138 | -0.018455 | -0.138137 | -0.087096 | -0.103627 | -0.056331 | -0.097262 | -0.115557 | -0.106471 | -0.075295 | -0.020541 | -0.047 | -0.070503 | -0.055458 | -0.095372 | -0.078806 |
| 18.34722 | -0.033216 | -0.03677 | 0.075002 | 0.055354 | 0.016646 | -0.022994 | -0.005509 | -0.001935 | 0.178461 | 0.157682 | -0.01743 | 0.130264 | -0.05031 | -0.031326 | -0.054358 | -0.059436 | -0.046328 | -0.04257 | -0.046121 | -0.038856 | -0.039208 | -0.043861 | -0.060358 | -0.037867 |
| 18.36111 | 0.095097 | 0.085547 | 0.192908 | 0.176631 | 0.12699 | 0.100348 | 0.100145 | 0.121901 | 0.309879 | 0.063555 | -0.012121 | 0.275796 | -0.001917 | 0.00478 | -0.006409 | 0.000292 | 0.004677 | -0.007456 | -0.063115 | -0.032598 | -0.005171 | -0.024945 | -0.017447 | 0.007392 |
| 18.375 | 0.137887 | 0.125289 | 0.252233 | 0.21576 | 0.162707 | 0.141806 | 0.139256 | 0.156575 | 0.343309 | 0.065043 | -0.021932 | 0.329011 | -0.022504 | 0.001939 | -0.007486 | -0.00976 | -0.001375 | -0.013309 | -0.064925 | -0.029333 | -0.010944 | -0.024108 | -0.020722 | 0.002714 |
| 18.38889 | 0.148014 | 0.134283 | 0.302828 | 0.248127 | 0.165592 | 0.139581 | 0.150663 | 0.149106 | 0.322076 | 0.064686 | 0.041947 | 0.326334 | -0.052497 | -0.012434 | -0.023949 | -0.042352 | -0.032587 | -0.038124 | -0.057251 | -0.037985 | -0.034302 | -0.038109 | -0.044064 | -0.033562 |
| 18.40278 | 0.089774 | 0.082182 | 0.266268 | 0.19242 | 0.111582 | 0.084691 | 0.098944 | 0.087145 | 0.257234 | 0.013944 | 0.056508 | 0.240245 | -0.060052 | -0.023928 | -0.042232 | -0.059891 | -0.04982 | -0.056613 | -0.064922 | -0.048582 | -0.054129 | -0.051147 | -0.061075 | -0.058618 |
| 18.41667 | 0.014927 | 0.012544 | 0.212694 | 0.096539 | 0.054955 | 0.024806 | 0.035481 | 0.024689 | 0.212037 | -0.057322 | 0.033196 | 0.164139 | -0.054178 | -0.020493 | -0.046777 | -0.050125 | -0.034836 | -0.056487 | -0.078344 | -0.054475 | -0.055423 | -0.059039 | -0.064669 | -0.055429 |
| 18.43056 | -0.016112 | -0.036335 | 0.181349 | 0.080999 | 0.032984 | -0.022423 | -0.000865 | -0.007288 | 0.206138 | -0.103245 | 0.017998 | 0.117446 | -0.05357 | -0.036777 | -0.050939 | -0.053751 | 0.002864 | -0.061624 | -0.090301 | -0.063638 | -0.052229 | -0.076948 | -0.071636 | -0.049391 |
| 18.44444 | -0.118456 | -0.191698 | 0.039414 | 0.141596 | -0.035445 | -0.181501 | -0.124532 | -0.161277 | 0.065079 | -0.256795 | -0.105116 | -0.067026 | -0.130369 | -0.220562 | -0.177454 | -0.228101 | 0.027768 | -0.187777 | -0.242163 | -0.140691 | -0.137849 | -0.221087 | -0.174455 | -0.113797 |
| 18.45833 | -0.30523 | -0.383051 | -0.212086 | -0.03094 | -0.153845 | -0.40986 | -0.266116 | -0.449592 | -0.13021 | -0.484332 | -0.291631 | -0.3541 | -0.291751 | -0.58594 | -0.428964 | -0.590408 | 0.002184 | -0.427793 | -0.433303 | -0.17781 | -0.347654 | -0.479807 | -0.355846 | -0.212705 |
| 18.47222 | -0.382205 | -0.487631 | -0.239226 | -0.341812 | -0.053594 | -0.851022 | -0.251193 | -0.689644 | -0.226459 | -0.288931 | -0.34902 | -0.347576 | -0.458413 | -0.611534 | -0.558564 | -0.569665 | 0.075569 | -0.615748 | -0.055472 | 0.337524 | -0.527764 | -0.349455 | -0.344313 | -0.098576 |
| 18.48611 | -0.276249 | -1.212812 | 0.116017 | 0.675887 | 0.675285 | -1.875237 | -0.195775 | -1.093099 | -0.370222 | 0.371599 | -0.338356 | -0.277224 | -0.757424 | 0.138994 | -0.575236 | -0.257852 | 0.437282 | -0.756297 | 0.822591 | 1.231115 | -0.658433 | -0.074387 | -0.149234 | 0.305683 |
| 18.5 | -0.304351 | -2.35475 | 0.322956 | 0.971625 | 0.999293 | -2.382975 | -0.214221 | -1.235085 | -0.224192 | 0.893679 | -0.319367 | -0.515216 | -0.939458 | -0.164506 | 0.145615 | -0.001775 | 0.501402 | -0.627523 | 1.056356 | 1.353507 | -0.624266 | -0.10894 | -0.285615 | -0.236719 |
| 18.51389 | -0.423209 | -1.630864 | 0.094142 | 0.152409 | -0.123074 | -0.266163 | -0.161091 | -0.350941 | 0.392344 | 0.295733 | -0.059111 | -0.378444 | -0.306508 | -0.545124 | 0.738969 | -0.035272 | 0.092749 | -0.063998 | -0.027578 | 0.006592 | -0.065412 | -0.055383 | -0.261062 | -0.678176 |
| 18.52778 | -0.826136 | 0.242748 | -1.063102 | -1.164827 | -2.047373 | 1.311903 | -0.810988 | 0.190999 | 0.248521 | -1.416865 | -0.642618 | -0.192671 | 0.637845 | -0.213101 | 0.167715 | 0.109795 | -0.328724 | 0.488677 | -1.163278 | -1.637781 | 0.628444 | -0.134288 | -0.042101 | 0.032047 |
| 18.54167 | -2.419729 | -0.938981 | -3.0205 | -2.390589 | -3.410147 | -0.038504 | -2.783099 | -1.035785 | -1.276973 | -3.003558 | -2.025365 | -1.258569 | 0.954185 | -0.454801 | 0.135282 | 0.130005 | -0.470852 | 0.537219 | -1.14111 | -1.858641 | 0.728487 | -0.361241 | 0.032555 | 0.336282 |
| 18.55556 | -2.98886 | -2.059545 | -3.407912 | -3.159852 | -3.312778 | -2.041318 | -2.969844 | -2.191495 | -2.264563 | -3.353784 | -2.202064 | -1.826225 | 0.357588 | -0.247147 | 0.234342 | 0.067389 | -0.192122 | 0.184749 | -0.442785 | -0.938271 | 0.328998 | -0.282738 | -0.027651 | 0.238449 |
| 18.56944 | -1.715757 | -1.331393 | -1.823658 | -1.856319 | -1.901311 | -1.380831 | -1.630343 | -1.304217 | -1.288772 | -1.668716 | -1.060406 | -0.917325 | 0.020066 | 0.102451 | 0.353438 | 0.043588 | -0.017509 | -0.018465 | -0.02363 | -0.157232 | 0.124785 | -0.052411 | -0.065035 | 0.305107 |
| 18.58333 | -0.648244 | -0.246383 | -0.519414 | -0.603061 | -0.668045 | -0.344333 | -0.589106 | -0.412709 | 0.0375 | -0.554911 | -0.167516 | -0.094015 | 0.082907 | 0.228528 | 0.341118 | 0.154295 | 0.127722 | 0.01628 | 0.104213 | 0.049949 | 0.19054 | 0.071248 | 0.004781 | 0.315171 |
| 18.59722 | -0.129298 | 0.26364 | 0.095763 | 0.016879 | -0.123136 | 0.116151 | -0.113527 | 0.051511 | 0.866989 | -0.144903 | 0.271292 | 0.147374 | 0.171179 | 0.163514 | 0.234271 | 0.194295 | 0.171832 | 0.051897 | 0.106197 | -0.001691 | 0.256722 | 0.058663 | 0.039563 | 0.181201 |
| 18.61111 | 0.086869 | 0.49689 | 0.390072 | 0.290325 | 0.102472 | 0.254974 | 0.093346 | 0.234253 | 0.804469 | 0.051114 | 0.39833 | 0.260166 | 0.151471 | 0.108409 | 0.196076 | 0.159258 | 0.162149 | 0.048758 | 0.058895 | -0.058835 | 0.27506 | 0.020788 | 0.027617 | 0.08614 |
| 18.625 | 0.263423 | 0.724822 | 0.620065 | 0.515534 | 0.30451 | 0.37434 | 0.277545 | 0.36927 | 0.643164 | 0.277215 | 0.458477 | 0.468841 | 0.124329 | 0.132534 | 0.270401 | 0.162079 | 0.15957 | 0.061254 | 0.029259 | -0.056756 | 0.232692 | 0.019571 | 0.03173 | 0.113803 |
| 18.63889 | 0.438571 | 0.933776 | 0.7839 | 0.732567 | 0.504897 | 0.531528 | 0.451454 | 0.519967 | 0.540119 | 0.486077 | 0.557148 | 0.709747 | 0.089642 | 0.177634 | 0.326062 | 0.162669 | 0.133371 | 0.077816 | 0.022523 | -0.0224 | 0.177617 | 0.02933 | 0.038663 | 0.14266 |
| 18.65278 | 0.557168 | 1.000851 | 0.869682 | 0.896462 | 0.626059 | 0.618152 | 0.569624 | 0.609984 | 0.473632 | 0.564069 | 0.684104 | 0.882634 | 0.025505 | 0.14662 | 0.274553 | 0.090446 | 0.052184 | 0.05112 | -0.002494 | -0.013086 | 0.107885 | 0.000594 | 0.005848 | 0.129885 |
| 18.66667 | 0.598029 | 0.958476 | 0.880426 | 0.941104 | 0.644289 | 0.637494 | 0.611453 | 0.628854 | 0.416296 | 0.563077 | 0.764499 | 0.932473 | -0.030377 | 0.088904 | 0.155602 | 0.019113 | -0.018064 | 0.017022 | -0.028155 | -0.011679 | 0.033427 | -0.033594 | -0.028874 | 0.102913 |
| 18.68056 | 0.633578 | 0.831233 | 0.88671 | 0.933842 | 0.645494 | 0.661378 | 0.636019 | 0.649722 | 0.440598 | 0.584186 | 0.837487 | 0.946327 | -0.012579 | 0.064594 | 0.104929 | -0.022447 | -0.023163 | 0.008521 | -0.024202 | -0.00263 | 0.011707 | -0.036732 | -0.0167 | 0.093101 |
| 18.69444 | 0.734471 | 0.846735 | 1.011663 | 1.020976 | 0.742901 | 0.77455 | 0.731479 | 0.764167 | 0.624831 | 0.693162 | 0.929291 | 1.037145 | -0.005448 | 0.039853 | 0.064259 | -0.102809 | -0.01321 | 0.005747 | -0.003021 | 0.021828 | 0.010503 | -0.024059 | -0.009417 | 0.070738 |
| 18.70833 | 0.893673 | 0.994346 | 1.274914 | 1.206774 | 0.92312 | 0.972888 | 0.895665 | 0.950238 | 0.83 | 0.867706 | 1.041771 | 1.188558 | -0.003413 | 0.038155 | 0.035906 | -0.128599 | 0.002237 | 0.001449 | 0.022814 | 0.060745 | 0.007422 | -0.00652 | -0.004319 | 0.052596 |
| 18.72222 | 0.950387 | 1.039509 | 1.392213 | 1.22238 | 0.981134 | 1.042477 | 0.953334 | 0.994877 | 0.888534 | 0.915002 | 1.014866 | 1.222369 | 0.041942 | 0.057773 | 0.050104 | -0.063162 | 0.022677 | 0.020045 | 0.043659 | 0.05778 | 0.02319 | -0.002396 | 0.020551 | 0.052719 |
| 18.73611 | 0.742038 | 0.807556 | 1.126553 | 0.947277 | 0.774242 | 0.816434 | 0.741302 | 0.761979 | 0.684658 | 0.712736 | 0.710454 | 0.984336 | 0.015084 | 0.0525 | 0.017619 | -0.058613 | -0.001831 | 0.004993 | 0.032412 | 0.030875 | 0.014035 | -0.021248 | 0.000655 | 0.023706 |
| 18.75 | 0.378872 | 0.41793 | 0.639019 | 0.517954 | 0.42885 | 0.434271 | 0.376982 | 0.39038 | 0.330366 | 0.374528 | 0.303951 | 0.55551 | -0.090501 | 0.023532 | -0.04603 | -0.125414 | -0.055745 | -0.054291 | -0.014656 | 0.015238 | -0.042928 | -0.048494 | -0.061316 | -0.035206 |
| 18.76389 | 0.136676 | 0.147396 | 0.290185 | 0.231916 | 0.197253 | 0.173068 | 0.136658 | 0.141578 | 0.078504 | 0.160154 | 0.091552 | 0.242171 | -0.110704 | 0.023358 | -0.036607 | -0.148073 | -0.054207 | -0.068685 | -0.031994 | -0.003134 | -0.078918 | -0.053361 | -0.066382 | -0.052398 |
| 18.77778 | 0.02056 | 0.012678 | 0.076837 | 0.083644 | 0.068834 | 0.036048 | 0.020607 | 0.020727 | -0.057003 | -0.045537 | 0.045527 | 0.078782 | -0.059406 | 0.021044 | 0.002116 | -0.13709 | -0.031064 | -0.036374 | -0.013708 | -0.03734 | -0.085582 | -0.047626 | -0.032081 | -0.044836 |
| 18.79167 | -0.085785 | -0.093811 | -0.112646 | -0.056194 | -0.05941 | -0.081048 | -0.080021 | -0.074946 | -0.14147 | -0.384118 | 0.008363 | -0.021766 | -0.077405 | -0.083425 | -0.048427 | -0.175346 | -0.077637 | -0.047629 | -0.024682 | -0.075748 | -0.121218 | -0.075769 | -0.057691 | -0.084988 |
| 18.80556 | -0.101355 | -0.087695 | -0.15425 | -0.083317 | -0.079513 | -0.086034 | -0.086646 | -0.06913 | -0.116126 | -0.219297 | 0.027039 | -0.022521 | -0.121705 | -0.17727 | -0.112645 | -0.201953 | -0.114 | -0.078458 | -0.048821 | -0.065412 | -0.157976 | -0.092025 | -0.100936 | -0.113062 |
| 18.81944 | -0.028348 | 0.00368 | -0.066437 | 0.012771 | 0.001944 | 0.010234 | -0.021667 | 0.014111 | -0.03368 | -0.127558 | 0.084147 | 0.049697 | -0.10729 | -0.150211 | -0.090469 | -0.156217 | -0.074954 | -0.06544 | -0.045124 | -0.022927 | -0.141189 | -0.06102 | -0.084883 | -0.075317 |
| 18.83333 | -0.041217 | -0.019407 | -0.069201 | 0.003977 | -0.010933 | -0.006845 | -0.047931 | -0.009507 | -0.09705 | -0.16602 | 0.01832 | 0.03041 | -0.083036 | -0.093328 | -0.052017 | -0.087856 | -0.057159 | -0.044685 | -0.039391 | -0.037147 | -0.121275 | -0.04834 | -0.05481 | -0.053159 |
| 18.84722 | -0.157231 | -0.195869 | -0.231668 | -0.144633 | -0.160336 | -0.168473 | -0.186303 | -0.165548 | -0.321741 | -0.324481 | -0.164755 | -0.10506 | -0.066856 | -0.065672 | -0.0485 | -0.03711 | -0.074154 | -0.041346 | -0.039359 | -0.090211 | -0.109686 | -0.066724 | -0.044818 | -0.068384 |
| 18.86111 | -0.243172 | -0.324897 | -0.376819 | -0.219666 | -0.278292 | -0.269321 | -0.285369 | -0.270133 | -0.492142 | -0.413234 | -0.294573 | -0.221398 | -0.074122 | -0.022489 | -0.068873 | -0.030667 | -0.080305 | -0.041012 | -0.030981 | -0.07953 | -0.103221 | -0.055366 | -0.045642 | -0.079185 |
| 18.875 | -0.216246 | -0.30829 | -0.389834 | -0.120327 | -0.254296 | -0.221898 | -0.270341 | -0.239446 | -0.52039 | -0.380946 | -0.294585 | -0.228128 | -0.114515 | 0.023799 | -0.097505 | -0.042953 | -0.084288 | -0.0438 | -0.027952 | -0.026865 | -0.102783 | -0.02462 | -0.049239 | -0.084525 |
| 18.88889 | -0.132868 | -0.26054 | -0.313416 | 0.038167 | -0.165025 | -0.133962 | -0.192367 | -0.155844 | -0.388676 | -0.297468 | -0.22638 | -0.185606 | -0.132685 | 0.023397 | -0.117322 | -0.045967 | -0.099957 | -0.045263 | -0.035174 | -0.007545 | -0.114298 | -0.019833 | -0.047187 | -0.093271 |
| 18.90278 | -0.049281 | -0.218282 | -0.203121 | 0.040418 | -0.075445 | -0.066443 | -0.107406 | -0.086504 | -0.214203 | -0.187166 | -0.14198 | -0.138054 | -0.116925 | 0.003275 | -0.109286 | -0.030534 | -0.117869 | -0.030017 | -0.025423 | -0.009632 | -0.110234 | -0.019996 | -0.030015 | -0.090313 |
| 18.91667 | -0.001941 | -0.199785 | -0.13644 | -0.134068 | -0.036253 | -0.043435 | -0.063941 | -0.06848 | -0.110753 | -0.16058 | -0.081412 | -0.134151 | -0.103042 | -0.011017 | -0.081884 | -0.001369 | -0.142618 | -0.008077 | -0.006439 | -0.033868 | -0.078421 | -0.020493 | -0.008427 | -0.09353 |
| 18.93056 | -0.023709 | -0.260354 | -0.184743 | -0.490239 | -0.089985 | -0.103293 | -0.11188 | -0.132492 | -0.157909 | -0.257098 | -0.129883 | -0.204182 | -0.090495 | -0.079116 | -0.085641 | 0.003607 | -0.161227 | -0.00632 | -0.009733 | -0.076719 | -0.072658 | -0.042194 | -0.006817 | -0.105233 |
| 18.94444 | -0.079544 | -0.321408 | -0.272729 | -0.774548 | -0.151834 | -0.165085 | -0.164862 | -0.184644 | -0.248504 | -0.315444 | -0.222424 | -0.288066 | -0.095832 | -0.177222 | -0.131358 | -0.03005 | -0.180567 | -0.02289 | -0.018772 | -0.06477 | -0.084848 | -0.045802 | -0.023289 | -0.152214 |
| 18.95833 | -0.066311 | -0.262623 | -0.246884 | -0.775004 | -0.11117 | -0.108127 | -0.116297 | -0.125425 | -0.247255 | -0.221024 | -0.210601 | -0.274474 | -0.102515 | -0.186738 | -0.168644 | -0.057426 | -0.199122 | -0.033682 | -0.010777 | 0.000399 | -0.09411 | -0.018565 | -0.03189 | -0.146829 |
| 18.97222 | 0.02255 | -0.133284 | -0.1219 | -0.587703 | -0.006672 | 0.024199 | 0.002019 | 0.011575 | -0.178999 | -0.075069 | -0.119538 | -0.154468 | -0.086012 | -0.153841 | -0.168986 | -0.050968 | -0.210064 | -0.033096 | -0.003306 | 0.030354 | -0.08466 | -0.003379 | -0.025328 | -0.119839 |
| 18.98611 | 0.117154 | -0.008471 | 0.004041 | -0.387886 | 0.087647 | 0.11804 | 0.104632 | 0.133275 | -0.114896 | -0.009392 | -0.027118 | -0.013095 | -0.05964 | -0.133087 | -0.13345 | -0.017165 | -0.198151 | -0.019367 | 0.005216 | 0.022098 | -0.059902 | 0.005141 | -0.008444 | -0.095605 |
| 19 | 0.195609 | 0.092753 | 0.169551 | -0.13052 | 0.249592 | 0.095687 | 0.221005 | 0.183738 | 0.018784 | 0.07027 | 0.120807 | 0.091955 | -0.02646 | 0.005847 | -0.003145 | 0.123835 | -0.117442 | 0.045509 | 0.145194 | 0.113399 | 0.015418 | 0.097957 | 0.070369 | -0.024627 |
| 19.01389 | 0.137249 | -0.169294 | 0.420498 | 0.402638 | 0.544684 | -0.471305 | 0.395205 | -0.183323 | -0.354586 | 0.250655 | 0.350935 | -0.112911 | -0.13746 | 0.383289 | 0.278695 | 0.33703 | 0.062316 | 0.059067 | 0.648637 | 0.589503 | 0.051419 | 0.328562 | 0.20671 | 0.011073 |
| 19.02778 | -0.094432 | -1.011752 | 0.438322 | 0.746971 | 0.808597 | -1.96358 | 0.356554 | -1.139274 | -1.600737 | 0.190608 | 0.277415 | -0.80824 | -0.504132 | 0.690242 | 0.492539 | 0.411895 | 0.21021 | -0.1005 | 1.239584 | 1.300574 | -0.123211 | 0.564774 | 0.266452 | -0.204211 |
| 19.04167 | -0.344718 | -1.068772 | -0.048819 | -0.147084 | 0.315507 | -2.141232 | -0.159419 | -1.170828 | -2.074987 | -0.518389 | -0.335902 | -1.039311 | -0.301347 | 0.525021 | 0.475706 | 0.301664 | 0.119537 | -0.078165 | 0.877273 | 0.985064 | -0.061404 | 0.400042 | 0.194646 | 0.251163 |
| 19.05556 | -0.132711 | 0.800664 | -0.130475 | -0.976108 | -0.67742 | 0.989528 | -0.330761 | 0.51976 | 0.178478 | -0.51819 | -0.475424 | 0.499406 | 0.418077 | 0.071354 | 0.050207 | 0.288603 | -0.238679 | 0.248192 | -0.480068 | -0.528575 | 0.322039 | 0.118053 | 0.098453 | 0.833029 |
| 19.06944 | 0.288231 | 1.329643 | 0.35025 | -0.521483 | -0.583885 | 2.067776 | -0.012528 | 1.492334 | 1.104035 | -0.03639 | -0.114928 | 1.088884 | 0.687404 | -0.03343 | -0.143578 | 0.165819 | -0.257855 | 0.410518 | -0.977477 | -1.066347 | 0.436408 | -0.032244 | 0.102124 | 0.588402 |
| 19.08333 | 0.390382 | 0.369076 | 0.441033 | 0.174797 | 0.045548 | 0.978661 | 0.247656 | 0.767467 | 0.221754 | 0.41851 | 0.108306 | 0.56378 | 0.241308 | 0.03181 | 0.003719 | 0.039469 | -0.066357 | 0.204129 | -0.505478 | -0.461782 | 0.244051 | 0.004982 | 0.000485 | 0.118808 |
| 19.09722 | 0.307498 | -0.136036 | 0.412312 | 0.439065 | 0.313088 | 0.345531 | 0.324583 | 0.340564 | -0.111934 | 0.930799 | 0.123794 | 0.367826 | -0.004348 | 0.012366 | 0.000747 | -0.045893 | 0.00181 | 0.019348 | -0.143278 | -0.061451 | 0.059231 | 0.018165 | -0.02685 | -0.054243 |
| 19.11111 | 0.299087 | -0.096646 | 0.32999 | 0.354887 | 0.284384 | 0.270287 | 0.249484 | 0.332248 | 0.195936 | 1.170214 | -0.020351 | 0.403962 | -0.034766 | -0.037423 | -0.065286 | -0.120286 | 0.044524 | -0.037972 | -0.07976 | -0.023219 | 0.009602 | -0.021624 | -0.037415 | -0.059176 |
| 19.125 | 0.162644 | -0.24105 | 0.120997 | 0.134151 | 0.122125 | 0.186033 | 0.083247 | 0.266589 | 0.387291 | 0.834514 | -0.25025 | 0.304039 | -0.033646 | -0.070231 | -0.09354 | -0.208917 | 0.040011 | -0.050053 | -0.089169 | -0.051183 | -0.003964 | -0.063589 | -0.04388 | -0.17489 |
| 19.13889 | -0.013242 | -0.535605 | -0.063383 | -0.061447 | -0.007537 | -0.039284 | -0.030995 | -0.001584 | 0.103098 | 0.402474 | -0.398726 | 0.027 | -0.022459 | -0.083959 | -0.09928 | -0.268585 | 0.004491 | -0.043861 | -0.085152 | -0.085252 | -0.042268 | -0.081635 | -0.044993 | -0.369505 |
| 19.15278 | -0.073672 | -0.590051 | -0.106732 | -0.144065 | -0.019757 | -0.166673 | -0.036714 | -0.157912 | -0.13594 | 0.116445 | -0.415018 | -0.122551 | -0.015583 | -0.084105 | -0.103274 | -0.275562 | -0.020197 | -0.026247 | -0.058827 | -0.106659 | -0.065585 | -0.077308 | -0.042116 | -0.510599 |
| 19.16667 | -0.032731 | -0.223927 | -0.037269 | -0.192873 | -0.020208 | -0.081588 | -0.026456 | 0.009539 | 0.109656 | -0.140591 | -0.28979 | 0.020153 | -0.018444 | -0.062661 | -0.109907 | -0.221163 | -0.015117 | -0.014692 | -0.033284 | -0.089302 | -0.075824 | -0.059638 | -0.034844 | -0.487983 |
| 19.18056 | -0.017859 | 0.066656 | -0.076869 | -0.311321 | -0.064709 | -0.024952 | -0.069896 | 0.117236 | 0.315871 | -0.387014 | -0.155633 | 0.117117 | -0.025047 | -0.042442 | -0.117589 | -0.179789 | -0.001032 | -0.025192 | -0.037159 | -0.059212 | -0.107696 | -0.050638 | -0.033739 | -0.397424 |
| 19.19444 | -0.105486 | -0.037074 | -0.224759 | -0.413704 | -0.08479 | -0.185012 | -0.095709 | -0.062646 | 0.109681 | -0.510843 | -0.087906 | -0.05123 | -0.028765 | -0.035588 | -0.130022 | -0.140726 | 0.002782 | -0.039179 | -0.05126 | -0.032618 | -0.140264 | -0.051629 | -0.037102 | -0.312023 |
| 19.20833 | -0.177149 | -0.179791 | -0.328527 | -0.439817 | -0.067909 | -0.337458 | -0.089439 | -0.216183 | -0.024815 | -0.432609 | -0.066232 | -0.20582 | -0.036263 | -0.028682 | -0.146562 | -0.126942 | -0.009197 | -0.040272 | -0.055018 | -0.004002 | -0.156254 | -0.047445 | -0.037292 | -0.215054 |
| 19.22222 | -0.173703 | -0.118803 | -0.398095 | -0.424027 | -0.099584 | -0.342114 | -0.119485 | -0.170706 | 0.07727 | -0.316465 | -0.077087 | -0.165816 | -0.042052 | -0.023436 | -0.153062 | -0.11615 | -0.022745 | -0.032883 | -0.054227 | 0.002762 | -0.147561 | -0.043141 | -0.034211 | -0.116281 |
| 19.23611 | -0.189214 | -0.083222 | -0.474711 | -0.37949 | -0.147512 | -0.257723 | -0.165782 | -0.12175 | 0.012704 | -0.226943 | -0.110776 | -0.090979 | -0.034608 | -0.01295 | -0.149886 | -0.104497 | -0.012974 | -0.019511 | -0.042703 | -0.007492 | -0.088081 | -0.038433 | -0.02697 | -0.050644 |
| 19.25 | -0.211271 | -0.056806 | -0.467393 | -0.233703 | -0.109448 | -0.163483 | -0.135122 | -0.094145 | -0.121985 | -0.130967 | -0.080921 | -0.000786 | -0.01143 | -0.007114 | -0.142796 | -0.074633 | 0.000569 | -0.00814 | -0.031068 | -0.032405 | -0.024041 | -0.041514 | -0.015767 | -0.030349 |
| 19.26389 | -0.334084 | -0.000334 | -0.388147 | -0.081263 | -0.04323 | -0.087212 | -0.066265 | -0.037313 | -0.172517 | -0.072684 | -0.016558 | 0.100099 | -0.000487 | -0.028602 | -0.168276 | -0.054144 | -0.027653 | -0.014026 | -0.047754 | -0.069086 | -0.015084 | -0.061586 | -0.020765 | -0.045119 |
| 19.27778 | -0.031786 | -0.04549 | -0.405731 | -0.149444 | -0.081477 | -0.104452 | -0.097617 | -0.068312 | -0.354118 | -0.14727 | -0.054753 | 0.040074 | -0.031654 | -0.064102 | -0.227563 | -0.074793 | -0.069289 | -0.039333 | -0.072866 | -0.067021 | -0.045423 | -0.080277 | -0.051412 | -0.054863 |
| 19.29167 | -0.169182 | -0.196904 | -0.487738 | -0.377893 | -0.196857 | -0.211636 | -0.213866 | -0.199719 | -0.628357 | -0.301588 | -0.164617 | -0.150144 | -0.042386 | -0.075876 | -0.248347 | -0.088451 | -0.069331 | -0.050269 | -0.070375 | -0.030729 | -0.041794 | -0.074484 | -0.06024 | -0.027482 |
| 19.30556 | -0.302943 | -0.342026 | -0.569494 | -0.565858 | -0.329657 | -0.340064 | -0.334538 | -0.336674 | -0.783806 | -0.434731 | -0.247325 | -0.360189 | -0.019525 | -0.073073 | -0.217686 | -0.06261 | -0.061392 | -0.049688 | -0.060534 | -0.025796 | -0.011866 | -0.068134 | -0.045952 | 0.027997 |
| 19.31944 | -0.466185 | -0.413374 | -0.643259 | -0.595691 | -0.466284 | -0.439366 | -0.442551 | -0.440898 | -0.687035 | -0.512924 | -0.308471 | -0.566969 | -0.03033 | -0.059231 | -0.174163 | -0.041759 | -0.057219 | -0.063606 | -0.060218 | -0.032775 | -0.015387 | -0.063009 | -0.04607 | 0.091709 |
| 19.33333 | -0.51067 | -0.399687 | -0.59841 | -0.422596 | -0.497924 | -0.437239 | -0.441464 | -0.455578 | -0.391777 | -0.491059 | -0.298849 | -0.649405 | -0.021077 | -0.01211 | -0.106836 | -0.019707 | -0.024864 | -0.059638 | -0.034794 | -0.00968 | -0.037887 | -0.036233 | -0.03197 | 0.156989 |
| 19.34722 | -0.350661 | -0.254566 | -0.321369 | -0.129663 | -0.36364 | -0.293347 | -0.26125 | -0.319931 | -0.076684 | -0.343865 | -0.1138 | -0.477392 | 0.022577 | 0.013187 | -0.03842 | -0.00713 | -0.019865 | -0.030797 | 0.00757 | -0.005509 | -0.061112 | -0.015057 | -0.011186 | 0.192219 |
| 19.36111 | -0.106494 | -0.035443 | 0.053458 | 0.078387 | -0.151663 | -0.053014 | -0.002563 | -0.101929 | 0.20561 | -0.111094 | 0.125043 | -0.181829 | 0.049185 | 0.04759 | 0.022842 | 0.010333 | -0.033779 | -0.005293 | 0.018478 | -0.024473 | -0.076593 | -0.001705 | -0.000493 | 0.203504 |
| 19.375 | 0.071328 | 0.157852 | 0.330589 | 0.137075 | 0.00775 | 0.164812 | 0.20354 | 0.061073 | 0.374013 | 0.085131 | 0.275758 | 0.049131 | 0.077004 | 0.130722 | 0.098716 | 0.051313 | -0.01461 | 0.020195 | -0.000705 | -0.027863 | 0.036383 | 0.018999 | 0.016333 | 0.179172 |
| 19.38889 | 0.092015 | 0.313101 | 0.421213 | 0.089052 | 0.029626 | 0.211652 | 0.253791 | 0.073564 | 0.358014 | 0.13471 | 0.233548 | 0.132323 | 0.0529 | 0.15513 | 0.14046 | 0.037901 | -0.003348 | 0.02547 | -0.031936 | -0.017786 | 0.181217 | 0.017829 | 0.004996 | 0.137914 |
| 19.40278 | 0.002725 | 0.418938 | 0.390142 | 0.0087 | -0.027926 | 0.099861 | 0.188654 | -0.017911 | 0.252793 | 0.084984 | 0.137795 | 0.095445 | 0.001126 | 0.133877 | 0.160784 | 0.001825 | -0.00453 | 0.00847 | -0.058709 | -0.014706 | 0.206995 | -0.002705 | -0.022789 | 0.107349 |
| 19.41667 | -0.097754 | 0.441554 | 0.372628 | -0.043448 | -0.078557 | -0.034896 | 0.151207 | -0.105143 | 0.155214 | 0.033497 | 0.083766 | 0.032628 | 0.011008 | 0.103708 | 0.19709 | 0.02431 | -0.011492 | 0.009968 | -0.045041 | -0.011211 | 0.21313 | -0.009664 | -0.02116 | 0.101045 |
| 19.43056 | -0.155151 | 0.403752 | 0.431593 | 0.054711 | -0.059137 | -0.089662 | 0.183281 | -0.124734 | 0.141603 | 0.047186 | 0.099352 | -0.039727 | 0.044776 | 0.073045 | 0.22648 | 0.047027 | -0.011007 | 0.003585 | -0.022647 | 0.015504 | 0.215707 | -0.015392 | -0.024465 | 0.087412 |
| 19.44444 | -0.128079 | 0.290951 | 0.489978 | 0.426349 | 0.06943 | -0.081254 | 0.245199 | -0.127852 | 0.177473 | 0.126752 | 0.144596 | -0.135307 | 0.009813 | -0.071654 | 0.155387 | -0.06395 | 0.006887 | -0.115278 | -0.15486 | -0.015384 | 0.141821 | -0.133336 | -0.10983 | 0.020725 |
| 19.45833 | -0.138455 | 0.11116 | 0.409347 | 0.498921 | 0.116655 | -0.111836 | 0.254156 | -0.208488 | 0.143941 | 0.101351 | 0.194084 | -0.247071 | -0.124316 | -0.420443 | -0.052462 | -0.364798 | -0.044835 | -0.342257 | -0.415871 | -0.164557 | -0.061016 | -0.393041 | -0.296799 | -0.127524 |
| 19.47222 | -0.310098 | -0.112068 | 0.133186 | -0.073514 | -0.061029 | -0.232244 | 0.160881 | -0.332553 | 0.176665 | -0.142785 | 0.132753 | -0.463314 | -0.165295 | -0.56414 | -0.178627 | -0.412605 | -0.151586 | -0.390273 | -0.433159 | -0.233185 | -0.152872 | -0.40611 | -0.35097 | -0.206251 |
| 19.48611 | -0.653592 | -0.452961 | -0.052204 | -0.128076 | -0.339072 | -0.502837 | -0.135506 | -0.447282 | 0.105711 | -0.397413 | -0.107667 | -0.728021 | -0.085538 | -0.261284 | -0.031795 | -0.137486 | -0.117026 | -0.213341 | -0.224037 | -0.173262 | -0.055081 | -0.185132 | -0.210973 | -0.125456 |
| 19.5 | -0.901495 | -0.659734 | -0.151185 | -0.140901 | -0.583502 | -0.715635 | -0.375651 | -0.520447 | 0.137601 | -0.558217 | -0.27243 | -0.962428 | -0.009264 | 0.013116 | 0.130007 | 0.088823 | -0.011022 | -0.056309 | -0.063532 | -0.105244 | 0.073581 | -0.035361 | -0.068474 | 0.005363 |
| 19.51389 | -0.65905 | -0.496107 | -0.053002 | -0.042444 | -0.492904 | -0.581234 | -0.29693 | -0.328497 | 0.44084 | -0.433606 | -0.127619 | -0.795198 | 0.042428 | 0.116975 | 0.215238 | 0.191838 | 0.061809 | 0.019078 | 0.004683 | -0.077519 | 0.159045 | 0.029646 | 0.009272 | 0.084536 |
| 19.52778 | -0.310536 | -0.211669 | 0.047479 | -0.06147 | -0.362846 | -0.437992 | -0.195809 | -0.126715 | 0.756737 | -0.373067 | -0.006771 | -0.519309 | 0.121859 | 0.113206 | 0.274338 | 0.297982 | 0.071413 | 0.071661 | 0.074997 | -0.161485 | 0.251046 | 0.001343 | 0.067508 | 0.068922 |
| 19.54167 | -0.42615 | -0.321365 | -0.286655 | -0.525916 | -0.687134 | -0.656748 | -0.48778 | -0.400613 | 0.507038 | -0.758989 | -0.314105 | -0.646803 | 0.172665 | 0.02874 | 0.277208 | 0.284162 | 0.03663 | 0.068304 | 0.10793 | -0.326745 | 0.323386 | -0.115956 | 0.065304 | -0.042594 |
| 19.55556 | -0.667911 | -0.504286 | -0.659262 | -0.774035 | -1.00478 | -0.786993 | -0.755848 | -0.65141 | 0.151317 | -0.949779 | -0.633885 | -0.856789 | 0.074186 | 0.01291 | 0.181719 | 0.099015 | 0.058247 | -0.006898 | 0.086057 | -0.245651 | 0.237539 | -0.110336 | 0.005231 | -0.066862 |
| 19.56944 | -0.380245 | -0.211829 | -0.405685 | -0.268985 | -0.752462 | -0.330379 | -0.40078 | -0.329932 | 0.432857 | -0.419021 | -0.32581 | -0.529759 | -0.029348 | 0.148384 | 0.134559 | -0.000214 | 0.170259 | -0.034586 | 0.09473 | 0.037483 | 0.106725 | 0.051089 | -0.007788 | 0.083004 |
| 19.58333 | 0.19147 | 0.228003 | 0.179704 | 0.382725 | -0.145667 | 0.277264 | 0.191662 | 0.219151 | 0.963434 | 0.241779 | 0.266671 | 0.085037 | -0.011442 | 0.205837 | 0.160228 | 0.064958 | 0.225056 | 0.01151 | 0.122004 | 0.137793 | 0.069194 | 0.13028 | 0.043537 | 0.237415 |
| 19.59722 | 0.513077 | 0.33385 | 0.558014 | 0.68519 | 0.228835 | 0.555757 | 0.50316 | 0.501324 | 1.047764 | 0.523597 | 0.591843 | 0.430546 | 0.015899 | 0.182145 | 0.125981 | 0.108319 | 0.191499 | 0.0399 | 0.108457 | 0.097571 | 0.055156 | 0.095118 | 0.054935 | 0.286846 |
| 19.61111 | 0.517713 | 0.107488 | 0.636492 | 0.730179 | 0.327531 | 0.554479 | 0.513491 | 0.518928 | 0.796048 | 0.507428 | 0.598213 | 0.472033 | 0.005259 | 0.125905 | 0.040603 | 0.053309 | 0.146152 | 0.040015 | 0.092407 | 0.103865 | 0.011485 | 0.051358 | 0.025088 | 0.265662 |
| 19.625 | 0.417631 | -0.066083 | 0.542964 | 0.675006 | 0.352317 | 0.4651 | 0.418237 | 0.434309 | 0.446229 | 0.410674 | 0.465193 | 0.407831 | 0.013609 | 0.128932 | -0.015538 | 0.028187 | 0.107417 | 0.048281 | 0.104024 | 0.141113 | 0.009638 | 0.035643 | 0.015013 | 0.224874 |
| 19.63889 | 0.308439 | -0.100351 | 0.444701 | 0.615232 | 0.341696 | 0.356886 | 0.305764 | 0.329154 | 0.090233 | 0.314198 | 0.336486 | 0.336595 | 0.018424 | 0.177728 | -0.032759 | 0.037336 | 0.070315 | 0.055931 | 0.122121 | 0.122622 | 0.047601 | 0.028688 | 0.021812 | 0.182465 |
| 19.65278 | 0.241999 | -0.076357 | 0.371183 | 0.504858 | 0.307655 | 0.244028 | 0.215223 | 0.254502 | -0.146943 | 0.24471 | 0.224933 | 0.27773 | 0.012548 | 0.236777 | -0.039854 | 0.041198 | 0.060817 | 0.048881 | 0.114852 | 0.043615 | 0.047584 | 0.017067 | 0.022099 | 0.117708 |
| 19.66667 | 0.230455 | 0.036831 | 0.367419 | 0.44434 | 0.316865 | 0.192665 | 0.193987 | 0.250345 | -0.162404 | 0.23991 | 0.200408 | 0.286759 | 0.007226 | 0.249297 | -0.025521 | 0.032473 | 0.067606 | 0.035718 | 0.100205 | 0.003147 | 0.006509 | 0.014072 | 0.017423 | 0.064537 |
| 19.68056 | 0.199388 | 0.186664 | 0.328035 | 0.433089 | 0.31853 | 0.278776 | 0.19937 | 0.267015 | -0.004263 | 0.269482 | 0.226794 | 0.332574 | -0.014796 | 0.201853 | -0.015393 | 0.014782 | 0.044035 | 0.014598 | 0.073809 | 0.022601 | -0.035794 | 0.017531 | -0.001877 | 0.04742 |
| 19.69444 | 0.120613 | 0.23786 | 0.256813 | 0.402778 | 0.254714 | 0.307015 | 0.176162 | 0.244543 | 0.098925 | 0.25286 | 0.262498 | 0.329211 | -0.017927 | 0.117132 | -0.018769 | 0.005863 | 0.014575 | -0.007016 | 0.039476 | 0.056515 | -0.046746 | 0.02341 | -0.015317 | 0.078806 |
| 19.70833 | 0.065703 | 0.19781 | 0.187431 | 0.320491 | 0.169914 | 0.255288 | 0.136715 | 0.171069 | 0.110751 | 0.157447 | 0.252255 | 0.247018 | 0.004329 | 0.011291 | -0.025861 | -0.006323 | -0.004459 | -0.023982 | -0.008965 | 0.043442 | -0.035882 | 0.00443 | -0.016874 | 0.116639 |
| 19.72222 | 0.090757 | 0.19321 | 0.212344 | 0.298596 | 0.155906 | 0.182764 | 0.137021 | 0.147053 | 0.116825 | 0.11292 | 0.247157 | 0.17204 | 0.011879 | -0.014905 | -0.017581 | 0.003625 | -0.004473 | -0.031634 | -0.037872 | 0.026279 | -0.010597 | -0.001791 | -0.00314 | 0.127121 |
| 19.73611 | 0.145121 | 0.246956 | 0.274564 | 0.339258 | 0.210486 | 0.177221 | 0.170122 | 0.194607 | 0.183376 | 0.160644 | 0.28457 | 0.181774 | 0.014217 | 0.084822 | 0.001783 | 0.061557 | 0.019334 | -0.022625 | -0.016419 | 0.028259 | 0.019999 | 0.027652 | 0.026759 | 0.103889 |
| 19.75 | 0.109492 | 0.226591 | 0.241865 | 0.269477 | 0.18923 | 0.123121 | 0.132563 | 0.166775 | 0.154243 | 0.125313 | 0.269715 | 0.154078 | -0.025976 | 0.125957 | -0.065449 | 0.027695 | -0.008114 | -0.044135 | -0.023224 | -0.00942 | -0.001633 | 0.006141 | -0.0048 | -0.024381 |
| 19.76389 | 0.001174 | 0.109196 | 0.105328 | 0.109559 | 0.080399 | 0.022107 | 0.026352 | 0.049412 | 0.06593 | 0.001415 | 0.227089 | 0.039015 | -0.058006 | 0.093665 | -0.139729 | -0.045487 | -0.041623 | -0.057872 | -0.044595 | -0.04051 | -0.040551 | -0.031242 | -0.050887 | -0.169681 |
| 19.77778 | -0.058897 | 0.026842 | 0.022877 | 0.005255 | 0.021721 | -0.041284 | -0.044829 | -0.025407 | 0.002904 | -0.070942 | -0.009343 | -0.035994 | -0.013501 | 0.032472 | -0.118119 | -0.014408 | -0.023057 | -0.009703 | -0.021962 | -0.038556 | -0.035506 | -0.023772 | -0.02982 | -0.081847 |
| 19.79167 | -0.066082 | 0.049955 | 0.050167 | -0.010818 | 0.044454 | -0.04496 | -0.054818 | -0.017879 | 0.003132 | -0.064586 | -0.075337 | -0.006952 | 0.002463 | -0.091659 | -0.07891 | 0.003588 | -0.031788 | 0.027818 | -0.001395 | -0.034902 | -0.036008 | -0.015902 | -0.016915 | 0.126361 |
| 19.80556 | -0.014523 | 0.178765 | 0.111697 | 0.045829 | 0.117385 | 0.043428 | -0.000277 | 0.068387 | 0.096511 | 0.034813 | -0.005062 | 0.108032 | -0.016668 | -0.236587 | -0.069922 | -0.033992 | -0.050925 | 0.022812 | 0.001044 | -0.014078 | -0.06835 | -0.007041 | -0.02674 | 0.194942 |
| 19.81944 | 0.125649 | 0.36634 | 0.142569 | 0.164626 | 0.157999 | 0.18633 | 0.10426 | 0.195782 | 0.240893 | 0.183566 | 0.050145 | 0.255548 | -0.003079 | -0.26398 | -0.058055 | -0.027444 | -0.033526 | 0.01742 | -0.004649 | -0.009469 | -0.115155 | -0.002722 | -0.011583 | 0.095609 |
| 19.83333 | 0.237376 | 0.500608 | 0.0416 | 0.205192 | 0.056939 | 0.243687 | 0.157184 | 0.246242 | 0.300965 | 0.246173 | -0.008371 | 0.32433 | 0.012669 | -0.141348 | -0.07859 | 0.002097 | -0.029749 | 0.014764 | -0.034087 | -0.039129 | -0.250675 | -0.023738 | 0.005808 | 0.007762 |
| 19.84722 | 0.196737 | 0.508318 | -0.232216 | 0.096575 | -0.14473 | 0.19817 | 0.094997 | 0.186506 | 0.240227 | 0.186912 | -0.102395 | 0.239823 | -0.018139 | -0.025581 | -0.181916 | -0.011545 | -0.051694 | -0.008429 | -0.079058 | -0.046989 | -0.319725 | -0.042891 | -0.00669 | -0.141037 |
| 19.86111 | 0.07005 | 0.413127 | -0.566603 | -0.126614 | -0.243991 | 0.102422 | -0.024223 | 0.07821 | 0.120881 | 0.070709 | -0.137511 | 0.089903 | -0.088195 | -0.018211 | -0.344106 | -0.068171 | -0.073342 | -0.040037 | -0.108104 | -0.019895 | -0.141575 | -0.044928 | -0.030323 | -0.26166 |
| 19.875 | -0.011997 | 0.343163 | -0.79875 | -0.316314 | -0.281497 | 0.025672 | -0.112215 | -0.010822 | 0.014517 | -0.027468 | -0.159763 | -0.026525 | -0.111456 | -0.159985 | -0.555515 | -0.074007 | -0.089048 | -0.041288 | -0.084292 | -0.011415 | -0.297083 | -0.072135 | -0.026046 | -0.227602 |
| 19.88889 | 0.007353 | 0.309534 | -0.788788 | -0.274923 | -0.246153 | 0.023774 | -0.121446 | -0.018381 | 0.000775 | -0.047473 | -0.216496 | -0.04179 | -0.103487 | -0.315584 | -0.584791 | -0.043891 | -0.096586 | -0.004917 | -0.029017 | -0.01099 | -0.221685 | -0.07578 | 0.006205 | -0.0615 |
| 19.90278 | 0.034328 | 0.199478 | -0.673618 | -0.352652 | -0.254004 | 0.024385 | -0.169871 | -0.01604 | 0.022631 | -0.046962 | -0.109549 | -0.053945 | -0.102204 | -0.522518 | -0.520839 | -0.064838 | -0.12514 | 0.012252 | 0.007419 | 0.000234 | -0.193376 | -0.062261 | 0.013609 | -0.045471 |
| 19.91667 | -0.012564 | -0.155893 | -0.527636 | -0.580016 | -0.307317 | -0.038084 | -0.279795 | -0.092441 | 0.026726 | -0.103467 | -0.133727 | -0.153923 | -0.120994 | -0.516973 | -0.384442 | -0.118123 | -0.178964 | -0.012394 | -0.020004 | 0.025436 | -0.271565 | -0.054073 | -0.0153 | -0.09603 |
| 19.93056 | -0.063207 | -0.637501 | -0.383206 | -0.920208 | -0.347316 | -0.073196 | -0.346912 | -0.183026 | 0.057672 | -0.145422 | -0.143802 | -0.232989 | -0.102267 | -0.558612 | -0.322217 | -0.125951 | -0.207468 | -0.03005 | -0.069465 | 0.050578 | -0.362077 | -0.040813 | -0.031231 | -0.168238 |
| 19.94444 | -0.058203 | -0.91866 | -0.308618 | -1.100646 | -0.349882 | -0.097107 | -0.298055 | -0.183941 | 0.19314 | -0.159707 | -0.307242 | -0.211014 | -0.062231 | -0.513481 | -0.305044 | -0.085188 | -0.198833 | -0.004769 | -0.073555 | 0.054576 | -0.400578 | -0.029317 | -0.000528 | -0.241835 |
| 19.95833 | -0.04499 | -0.8786 | -0.271544 | -1.11281 | -0.362708 | -0.123161 | -0.252214 | -0.163749 | 0.324013 | -0.181521 | -0.741105 | -0.16223 | -0.040824 | -0.445806 | -0.367354 | -0.024617 | -0.184557 | 0.025358 | -0.040826 | 0.031809 | -0.34722 | -0.037076 | 0.033383 | -0.258793 |
| 19.97222 | -0.08126 | -0.702972 | -0.271793 | -1.068832 | -0.383281 | -0.140979 | -0.294721 | -0.175339 | 0.337823 | -0.200566 | -0.913292 | -0.142629 | -0.046539 | -0.449447 | -0.468324 | -0.027964 | -0.186671 | 0.00218 | -0.064185 | 0.014652 | -0.271029 | -0.068241 | 0.009828 | -0.256547 |
| 19.98611 | -0.100869 | -0.514175 | -0.308944 | -0.947183 | -0.36282 | -0.109347 | -0.338615 | -0.147074 | 0.195669 | -0.158967 | -0.771868 | -0.115687 | -0.07172 | -0.331627 | -0.419664 | -0.058815 | -0.179682 | -0.026716 | -0.086122 | 0.041731 | -0.194929 | -0.060593 | -0.020263 | -0.187913 |
| 20 | 0.005195 | -0.269185 | -0.260868 | -0.631203 | -0.248398 | 0.008278 | -0.259894 | -0.007771 | 0.128561 | -0.004438 | -0.417215 | 0.006389 | -0.013685 | -0.079706 | -0.151942 | 0.053081 | -0.11432 | 0.055 | -0.016762 | 0.102099 | -0.046087 | 0.034111 | 0.059073 | -0.057176 |
| 20.01389 | 0.167004 | -0.124437 | -0.120606 | -0.167932 | -0.105259 | 0.102689 | -0.102824 | 0.181631 | 0.094789 | 0.174361 | 0.004683 | 0.171636 | 0.144373 | 0.386919 | 0.251956 | 0.379353 | 0.00341 | 0.257526 | 0.204886 | 0.18024 | 0.218584 | 0.253554 | 0.252782 | 0.089282 |
| 20.02778 | -0.082345 | -0.416564 | -0.379573 | -0.335621 | -0.306462 | -0.220375 | -0.229888 | -0.041571 | -0.303012 | -0.087611 | -0.113991 | -0.083898 | 0.250914 | 0.715877 | 0.487552 | 0.564349 | 0.078848 | 0.361792 | 0.341813 | 0.228073 | 0.345744 | 0.42831 | 0.35964 | 0.192541 |
| 20.04167 | -0.78565 | -1.122474 | -1.082949 | -1.255822 | -0.813325 | -0.896552 | -0.673166 | -0.760473 | -0.973444 | -0.873247 | -0.909233 | -0.864484 | 0.127489 | 0.552062 | 0.325028 | 0.300671 | 0.058849 | 0.213655 | 0.200395 | 0.178252 | 0.224599 | 0.330723 | 0.215137 | 0.185799 |
| 20.05556 | -0.863422 | -1.007563 | -1.055075 | -1.16282 | -0.80442 | -0.876432 | -0.627766 | -0.879513 | -1.10746 | -1.022533 | -1.156969 | -0.980381 | -0.033038 | 0.203745 | 0.067723 | 0.000894 | 0.062416 | 0.022966 | 0.005332 | 0.139163 | -0.012177 | 0.16472 | 0.031385 | 0.116751 |
| 20.06944 | -0.233679 | -0.234963 | -0.303772 | -0.471043 | -0.186006 | -0.192087 | -0.200227 | -0.273575 | -0.628155 | -0.447534 | -0.59727 | -0.268169 | -0.075619 | -0.147949 | -0.093904 | -0.143439 | 0.068583 | -0.030212 | -0.043192 | 0.136981 | -0.222999 | 0.063277 | -0.021868 | 0.023961 |
| 20.08333 | 0.161181 | -0.124407 | -0.019855 | -0.409447 | 0.124865 | 0.137749 | -0.112079 | 0.095526 | -0.174844 | -0.014253 | -0.255906 | 0.17649 | -0.062182 | -0.274025 | -0.149832 | -0.157644 | 0.044998 | -0.013922 | -0.010789 | 0.116417 | -0.43719 | -0.023998 | -0.009341 | -0.040531 |
| 20.09722 | 0.176903 | -0.445206 | -0.134578 | -0.347458 | 0.075392 | 0.066699 | -0.012519 | 0.083922 | -0.668817 | 0.078162 | -0.20724 | 0.118262 | -0.029556 | -0.044148 | -0.084357 | -0.114474 | 0.037582 | 0.000264 | 0.010241 | 0.103234 | -0.263225 | -0.02789 | 0.01309 | -0.035089 |
| 20.11111 | 0.160046 | -0.340637 | -0.113362 | 0.108066 | 0.040784 | 0.032043 | 0.250957 | 0.089497 | -0.736861 | 0.174525 | -0.09659 | 0.036159 | 0.001158 | 0.224094 | 0.023226 | -0.073232 | 0.037365 | -0.00744 | 0.000263 | 0.102585 | 0.040445 | 0.023797 | 0.016437 | -0.027937 |
| 20.125 | 0.144835 | 0.056762 | -0.128612 | 0.247464 | 0.021698 | 0.054814 | 0.268389 | 0.112893 | -0.710084 | 0.223199 | -0.016028 | 0.014569 | 0.001023 | 0.177578 | 0.08211 | -0.065033 | 0.010574 | -0.031797 | -0.030848 | 0.067134 | 0.112999 | 0.023391 | -0.010562 | -0.05349 |
| 20.13889 | 0.093676 | 0.327649 | -0.328666 | -0.00323 | -0.026685 | 0.088206 | 0.062477 | 0.082133 | -1.198144 | 0.099802 | -0.081089 | -0.019524 | -0.012525 | -0.015552 | 0.067241 | -0.068216 | -0.009169 | -0.033183 | -0.039075 | 0.009934 | 0.018741 | -0.025683 | -0.026857 | -0.079632 |
| 20.15278 | 0.077758 | 0.300057 | -0.467224 | -0.044382 | -0.043055 | 0.132819 | 0 | 0.064742 | -0.908626 | -0.040775 | -0.162234 | -0.006321 | -0.022332 | -0.009208 | -0.008907 | -0.068013 | 0.006698 | -0.01167 | -0.020017 | -0.016221 | 0.017889 | -0.043563 | -0.023299 | -0.108534 |
| 20.16667 | 0.091313 | -0.023854 | -0.513233 | 0.073119 | -0.031595 | 0.12897 | 0.035573 | 0.066078 | -0.38668 | -0.091053 | -0.160676 | 0.06897 | -0.044715 | 0.098227 | -0.132568 | -0.082984 | 0.01811 | -0.014002 | -0.013987 | -0.020238 | 0.061812 | -0.037505 | -0.02857 | -0.128977 |
| 20.18056 | 0.083078 | -0.251541 | -0.521447 | 0.0172 | -0.052096 | 0.086421 | 0.033627 | 0.048411 | -0.16508 | -0.116687 | -0.110084 | 0.132252 | -0.072986 | 0.065517 | -0.271975 | -0.11126 | 0.000082 | -0.042743 | -0.036152 | -0.027695 | 0.078622 | -0.045459 | -0.051397 | -0.108473 |
| 20.19444 | 0.062586 | -0.173153 | -0.513698 | -0.027489 | -0.072473 | 0.061208 | 0.060638 | 0.026287 | -0.14519 | -0.149363 | -0.015949 | 0.207763 | -0.066921 | -0.000321 | -0.364008 | -0.103742 | -0.020462 | -0.050355 | -0.053282 | -0.018221 | 0.133946 | -0.0414 | -0.042901 | -0.063078 |
| 20.20833 | 0.015564 | -0.038005 | -0.471686 | 0.081452 | -0.087996 | -0.016848 | 0.047588 | -0.036646 | -0.231997 | -0.237843 | 0.064974 | 0.266436 | -0.021127 | 0.012887 | -0.320744 | -0.066959 | -0.042742 | -0.033025 | -0.052175 | -0.02317 | 0.121088 | -0.039008 | -0.016831 | -0.07834 |
| 20.22222 | -0.017675 | 0.027734 | -0.387348 | 0.184582 | -0.067799 | -0.078413 | 0.020874 | -0.072537 | -0.254022 | -0.30092 | 0.111618 | 0.282178 | -0.004811 | -0.033479 | -0.1852 | -0.071162 | -0.094031 | -0.025889 | -0.039371 | -0.046617 | 0.040984 | -0.059581 | -0.026034 | -0.190572 |
| 20.23611 | 0.098148 | 0.140457 | -0.07603 | 0.218844 | 0.108048 | 0.073907 | 0.178706 | 0.080959 | -0.069772 | -0.114114 | 0.222745 | 0.378134 | -0.0061 | -0.058978 | -0.017385 | -0.086073 | -0.10402 | -0.022353 | -0.007138 | -0.033517 | 0.020492 | -0.039665 | -0.034818 | -0.207981 |
| 20.25 | 0.230669 | 0.168745 | 0.242401 | 0.186957 | 0.282777 | 0.291719 | 0.296456 | 0.248911 | 0.128269 | 0.087516 | 0.29755 | 0.38135 | 0.0181 | -0.003186 | 0.068633 | -0.076741 | -0.058486 | -0.011686 | 0.018301 | -0.015083 | 0.01864 | -0.001317 | -0.023214 | -0.095111 |
| 20.26389 | 0.081401 | -0.08679 | 0.149648 | -0.080241 | 0.10855 | 0.182384 | 0.035362 | 0.09478 | 0.001566 | -0.075925 | 0.079443 | 0.104113 | 0.025934 | 0.014463 | 0.014699 | -0.095082 | -0.064381 | -0.018934 | 0.00793 | -0.05521 | -0.020104 | -0.025122 | -0.03419 | -0.013424 |
| 20.27778 | -0.213482 | -0.406189 | -0.225659 | -0.406659 | -0.237363 | -0.097084 | -0.355884 | -0.188128 | -0.227853 | -0.422342 | -0.247363 | -0.156128 | -0.017015 | -0.007902 | -0.078473 | -0.118331 | -0.113054 | -0.038641 | -0.01086 | -0.092021 | -0.033542 | -0.066824 | -0.052638 | -0.026894 |
| 20.29167 | -0.303217 | -0.502367 | -0.472286 | -0.467741 | -0.339476 | -0.182568 | -0.492698 | -0.252307 | -0.256474 | -0.517384 | -0.317718 | -0.226307 | -0.04025 | -0.001169 | -0.116846 | -0.092171 | -0.123293 | -0.038051 | -0.007509 | -0.07993 | 0.004553 | -0.057855 | -0.039226 | -0.081037 |
| 20.30556 | -0.170076 | -0.291777 | -0.303959 | -0.160988 | -0.145655 | -0.081834 | -0.324931 | -0.216794 | -0.189791 | -0.268781 | -0.057071 | -0.153569 | -0.001053 | 0.040111 | -0.105666 | -0.057606 | -0.093696 | -0.017348 | 0.007768 | -0.04291 | 0.046368 | -0.024747 | -0.01161 | -0.105232 |
| 20.31944 | 0.098543 | 0.199648 | 0.257919 | 0.299579 | 0.173977 | 0.17689 | 0.041868 | -0.01438 | -0.000973 | 0.120569 | 0.38353 | 0.025111 | 0.059354 | 0.116097 | -0.05563 | -0.012387 | -0.055128 | 0.00445 | 0.024318 | 0.005166 | 0.090492 | 0.005997 | 0.011754 | -0.056068 |
| 20.33333 | 0.309526 | 0.541941 | 0.68036 | 0.514036 | 0.365386 | 0.42123 | 0.30233 | 0.264348 | 0.24933 | 0.319575 | 0.617398 | 0.099195 | 0.087788 | 0.157464 | 0.00674 | 0.052011 | -0.025029 | 0.015255 | 0.034489 | 0.033126 | 0.148255 | 0.031085 | 0.023161 | 0.034608 |
| 20.34722 | 0.175495 | 0.404697 | 0.614408 | 0.272254 | 0.202192 | 0.321357 | 0.168615 | 0.173942 | 0.147964 | 0.149489 | 0.407473 | -0.080026 | 0.078416 | 0.125189 | 0.04759 | 0.081811 | -0.001016 | 0.006461 | 0.018085 | 0.015289 | 0.188102 | 0.033386 | 0.011433 | 0.091125 |
| 20.36111 | -0.114397 | 0.089449 | 0.449713 | -0.044874 | -0.091526 | 0.063158 | -0.096945 | -0.139176 | -0.14202 | -0.085795 | 0.137601 | -0.259982 | 0.020986 | 0.095407 | 0.05939 | 0.053907 | -0.006132 | -0.023326 | -0.016174 | -0.027774 | 0.184118 | 0.007077 | -0.019581 | 0.075129 |
| 20.375 | -0.202817 | 0.075917 | 0.556946 | -0.034899 | -0.124122 | 0.067271 | -0.121467 | -0.221357 | -0.212803 | -0.041636 | 0.172664 | -0.249754 | -0.04087 | 0.158084 | 0.080707 | 0.073842 | -0.014817 | -0.035728 | -0.022809 | -0.040962 | 0.17893 | -0.017307 | -0.029107 | 0.041656 |
| 20.38889 | -0.128263 | 0.339743 | 0.741812 | 0.142788 | 0.117766 | 0.230443 | 0.016696 | -0.121537 | -0.129332 | 0.176042 | 0.348587 | -0.185114 | -0.034086 | 0.275842 | 0.122023 | 0.20557 | 0.007486 | -0.008814 | 0.002274 | -0.012722 | 0.189064 | 0.004066 | -0.001181 | 0.05855 |
| 20.40278 | -0.144944 | 0.492238 | 0.69573 | 0.141398 | 0.330491 | 0.164641 | 0.035161 | -0.135324 | -0.179604 | 0.202616 | 0.417678 | -0.254896 | 0.031941 | 0.292895 | 0.13716 | 0.291222 | 0.02097 | -0.001927 | -0.003333 | -0.012513 | 0.215637 | 0.008681 | 0.012869 | 0.080767 |
| 20.41667 | -0.3604 | 0.388882 | 0.528347 | -0.071868 | 0.326287 | -0.167051 | -0.136427 | -0.343026 | -0.436207 | -0.006065 | 0.209309 | -0.46374 | 0.080077 | 0.22649 | 0.149562 | 0.2205 | 0.02856 | -0.034667 | -0.030681 | -0.028181 | 0.259907 | -0.024001 | -0.00612 | 0.06631 |
| 20.43056 | -0.567329 | 0.23043 | 0.408354 | -0.201579 | 0.258782 | -0.444709 | -0.284859 | -0.505226 | -0.664437 | -0.20982 | 0.053524 | -0.697759 | 0.070361 | 0.228269 | 0.162578 | 0.052787 | 0.05743 | -0.067327 | -0.048688 | -0.004228 | 0.392319 | -0.039839 | -0.050481 | 0.04167 |
| 20.44444 | -0.542254 | 0.188741 | 0.484969 | 0.091099 | 0.219711 | -0.438463 | -0.205503 | -0.458637 | -0.683083 | -0.219035 | 0.141288 | -0.779086 | 0.013008 | 0.111993 | 0.09173 | -0.142184 | 0.117578 | -0.148724 | -0.14844 | -0.017743 | 0.530388 | -0.124687 | -0.142367 | 0.018654 |
| 20.45833 | -0.423088 | 0.121144 | 0.538938 | 0.258628 | 0.149199 | -0.220607 | -0.031811 | -0.34973 | -0.57828 | -0.175323 | 0.290294 | -0.740905 | -0.121029 | -0.126189 | -0.077606 | -0.405937 | 0.104015 | -0.292399 | -0.30242 | -0.113562 | 0.49812 | -0.329148 | -0.31652 | -0.113914 |
| 20.47222 | -0.17104 | 0.24294 | 0.594487 | 0.414099 | 0.235217 | 0.101571 | 0.2211 | -0.022527 | -0.240347 | 0.013446 | 0.390081 | -0.255281 | -0.096019 | -0.126446 | -0.146788 | -0.354922 | 0.04436 | -0.278135 | -0.23806 | -0.164178 | 0.354211 | -0.313388 | -0.303222 | -0.075296 |
| 20.48611 | 0.331643 | 0.690288 | 1.027562 | 1.195541 | 0.571307 | 0.496424 | 0.552877 | 0.50709 | 0.251427 | 0.498792 | 0.601893 | 0.293622 | 0.029972 | 0.235896 | 0.012013 | -0.036463 | 0.10062 | -0.097278 | -0.010732 | -0.140635 | 0.453458 | -0.091327 | -0.110705 | 0.034466 |
| 20.5 | 0.609 | 0.942348 | 1.317851 | 1.458893 | 0.676023 | 0.648366 | 0.689436 | 0.807712 | 0.467352 | 0.752274 | 0.68788 | 0.467779 | 0.0676 | 0.482888 | 0.090639 | 0.091768 | 0.146501 | 0.009477 | 0.111933 | -0.084905 | 0.460122 | 0.008256 | -0.000857 | 0.106266 |
| 20.51389 | 0.403643 | 0.708607 | 1.099302 | 1.153271 | 0.311842 | 0.353074 | 0.423439 | 0.534374 | 0.229718 | 0.497574 | 0.367196 | 0.215182 | 0.041009 | 0.363045 | 0.054089 | 0.045101 | 0.115337 | 0.012264 | 0.120211 | -0.082046 | 0.400099 | 0.002506 | 0.000008 | 0.083053 |
| 20.52778 | -0.186364 | 0.14816 | 0.498653 | 0.506543 | -0.337946 | -0.348875 | -0.250635 | -0.178596 | -0.402201 | -0.261943 | -0.278589 | -0.348734 | 0.070352 | 0.223114 | 0.052933 | 0.039345 | 0.059053 | 0.028093 | 0.114711 | -0.196219 | 0.355015 | -0.08096 | 0.00007 | 0.027618 |
| 20.54167 | -0.879278 | -0.533446 | -0.365597 | -0.300966 | -1.029618 | -1.184016 | -1.057636 | -0.968807 | -1.101359 | -1.16645 | -0.977227 | -0.883316 | 0.135698 | 0.119356 | 0.132558 | 0.082612 | 0.030069 | 0.070589 | 0.131373 | -0.354109 | 0.340981 | -0.171247 | 0.021776 | -0.000318 |
| 20.55556 | -0.74489 | -0.36028 | -0.420317 | -0.244659 | -0.890379 | -0.940304 | -0.864511 | -0.746262 | -0.753858 | -0.913624 | -0.733706 | -0.563948 | 0.097294 | 0.191862 | 0.228916 | 0.073825 | 0.10811 | 0.061504 | 0.131393 | -0.260371 | 0.278306 | -0.105185 | 0.024398 | 0.065713 |
| 20.56944 | 0.106824 | 0.594347 | 0.436669 | 0.753245 | 0.150333 | 0.181315 | 0.128607 | 0.189529 | 0.341499 | 0.209483 | 0.299916 | 0.340204 | 0.013514 | 0.33796 | 0.288999 | 0.033939 | 0.271401 | 0.02911 | 0.11554 | 0.007681 | 0.202851 | 0.061396 | 0.014828 | 0.158966 |
| 20.58333 | 0.572069 | 1.116336 | 0.968424 | 1.300461 | 0.76464 | 0.670121 | 0.624363 | 0.606905 | 0.834996 | 0.707211 | 0.886008 | 0.746424 | 0.021263 | 0.351891 | 0.294754 | 0.039822 | 0.339576 | 0.013734 | 0.104509 | 0.054093 | 0.153314 | 0.093144 | 0.019949 | 0.168942 |
| 20.59722 | 0.351981 | 0.883228 | 0.808907 | 0.951699 | 0.559998 | 0.37164 | 0.364257 | 0.326434 | 0.524074 | 0.370457 | 0.631934 | 0.448929 | -0.003898 | 0.247127 | 0.240867 | 0.046103 | 0.290654 | -0.028672 | 0.072007 | -0.03313 | 0.012696 | 0.015188 | -0.012864 | 0.096686 |
| 20.61111 | -0.062335 | 0.463806 | 0.386402 | 0.417778 | 0.103279 | -0.066093 | -0.076311 | -0.078442 | 0.082935 | -0.117661 | 0.121823 | -0.01561 | -0.100636 | 0.098697 | 0.203851 | -0.001722 | 0.199981 | -0.080416 | 0.033132 | -0.046539 | -0.076887 | -0.030795 | -0.076708 | 0.057111 |
| 20.625 | -0.288658 | 0.111085 | 0.051181 | 0.055198 | -0.216693 | -0.304241 | -0.33909 | -0.259917 | -0.130038 | -0.365939 | -0.203775 | -0.25943 | -0.15826 | -0.009217 | 0.178003 | -0.023242 | 0.077628 | -0.107933 | -0.00458 | -0.034186 | 0.048714 | -0.045211 | -0.115536 | 0.026693 |
| 20.63889 | -0.278521 | -0.110634 | -0.15077 | -0.096532 | -0.316618 | -0.286343 | -0.380364 | -0.260072 | -0.114493 | -0.354049 | -0.265435 | -0.311414 | -0.14276 | -0.050393 | 0.181195 | -0.024851 | -0.009743 | -0.113476 | -0.042475 | -0.046772 | -0.01554 | -0.060881 | -0.114297 | 0.003568 |
| 20.65278 | -0.161697 | -0.281492 | -0.193066 | -0.171411 | -0.270849 | -0.176537 | -0.285432 | -0.180825 | -0.004873 | -0.23538 | -0.137345 | -0.32767 | -0.093932 | -0.00278 | 0.208027 | -0.008663 | -0.023654 | -0.09124 | -0.052934 | -0.055866 | -0.031697 | -0.056248 | -0.083767 | 0.012618 |
| 20.66667 | -0.106404 | -0.415291 | -0.186524 | -0.289441 | -0.213003 | -0.121429 | -0.220748 | -0.138291 | 0.039317 | -0.165735 | -0.120167 | -0.348404 | -0.069448 | 0.068102 | 0.193015 | -0.002477 | -0.008678 | -0.063301 | -0.038014 | -0.046851 | -0.018427 | -0.033712 | -0.055249 | 0.034903 |
| 20.68056 | -0.148842 | -0.510032 | -0.176509 | -0.400322 | -0.187209 | -0.159869 | -0.239312 | -0.177375 | -0.012383 | -0.198649 | -0.17859 | -0.267124 | -0.053706 | 0.086285 | 0.128369 | -0.000535 | -0.001545 | -0.037353 | -0.017724 | -0.031421 | -0.001884 | -0.015289 | -0.036747 | 0.061037 |
| 20.69444 | -0.189724 | -0.503399 | -0.228011 | -0.440028 | -0.172461 | -0.210892 | -0.253271 | -0.216152 | -0.070325 | -0.245708 | -0.192941 | -0.106854 | -0.018085 | 0.03605 | 0.074775 | 0.018022 | -0.003067 | -0.008195 | 0.007438 | -0.016311 | 0.018401 | -0.004741 | -0.017048 | 0.068518 |
| 20.70833 | -0.165968 | -0.376252 | -0.173999 | -0.318384 | -0.151179 | -0.195676 | -0.207761 | -0.186325 | -0.060039 | -0.229524 | -0.220815 | -0.003299 | -0.004279 | -0.057112 | 0.012245 | 0.007791 | -0.014427 | -0.00536 | 0.013776 | -0.015457 | 0.004557 | -0.01284 | -0.021652 | 0.038863 |
| 20.72222 | -0.091408 | -0.21711 | -0.064892 | -0.10793 | -0.16297 | -0.114752 | -0.117711 | -0.108651 | 0.042717 | -0.14144 | -0.090192 | -0.021774 | -0.025173 | -0.169779 | -0.03995 | -0.02466 | -0.036908 | -0.032017 | -0.002524 | -0.013588 | -0.03081 | -0.033159 | -0.045859 | -0.023148 |
| 20.73611 | -0.007969 | -0.125605 | -0.095395 | 0.071487 | -0.156812 | -0.025579 | -0.037331 | -0.031814 | 0.209662 | -0.051711 | 0.029701 | -0.092986 | -0.047283 | -0.19643 | -0.065861 | -0.048803 | -0.051633 | -0.052801 | -0.018559 | 0.006935 | -0.047226 | -0.042381 | -0.053286 | -0.071502 |
| 20.75 | 0.045646 | -0.084359 | -0.266917 | 0.140697 | -0.115195 | 0.015099 | -0.010461 | 0.001716 | 0.307224 | -0.01686 | 0.095714 | -0.114461 | -0.034498 | -0.098219 | -0.076216 | -0.047939 | -0.036125 | -0.038419 | -0.032891 | 0.016917 | -0.028092 | -0.023887 | -0.030692 | -0.071148 |
| 20.76389 | 0.099898 | -0.038718 | -0.325918 | 0.140521 | -0.117157 | 0.033976 | -0.003064 | 0.017997 | 0.324829 | 0.001376 | 0.026537 | -0.076325 | 0.003537 | 0.008795 | -0.084398 | -0.011256 | -0.000717 | -0.005496 | -0.030724 | -0.006888 | -0.005308 | 0.001579 | -0.003005 | -0.039003 |
| 20.77778 | 0.159994 | 0.040295 | -0.119864 | 0.154572 | -0.158446 | 0.092942 | 0.046257 | 0.084216 | 0.355818 | 0.086062 | -0.019327 | 0.016623 | 0.022826 | -0.003526 | -0.088028 | 0.03839 | 0.020237 | 0.019766 | -0.025854 | -0.049303 | -0.008565 | 0.015981 | 0.00809 | 0.000072 |
| 20.79167 | 0.179255 | 0.105701 | 0.118816 | 0.137195 | -0.211775 | 0.122994 | 0.106378 | 0.175947 | 0.407352 | 0.197801 | 0.093694 | 0.135842 | 0.017191 | -0.170446 | -0.101367 | 0.057487 | 0.01466 | 0.037089 | -0.016161 | -0.036061 | -0.042992 | 0.025511 | 0.01223 | 0.037104 |
| 20.80556 | 0.151472 | 0.073428 | 0.156157 | -0.03601 | -0.24925 | 0.020832 | 0.05813 | 0.206846 | 0.386215 | 0.228626 | 0.069848 | 0.169693 | -0.023319 | -0.436337 | -0.164154 | 0.024356 | -0.006417 | 0.019622 | -0.04414 | 0.036162 | -0.108829 | 0.019149 | 0.009797 | 0.04592 |
| 20.81944 | 0.121366 | -0.056321 | 0.029667 | -0.345875 | -0.177817 | -0.051237 | -0.061154 | 0.16007 | 0.297583 | 0.174621 | 0.072664 | 0.125571 | -0.090212 | -0.599883 | -0.278343 | -0.027594 | -0.04075 | -0.015719 | -0.071335 | 0.088354 | -0.170142 | 0.003516 | -0.002954 | 0.002595 |
| 20.83333 | 0.057858 | -0.311176 | -0.226911 | -0.663153 | -0.123047 | -0.069857 | -0.157666 | 0.064738 | 0.186349 | 0.062518 | 0.019845 | 0.021094 | -0.161606 | -0.600827 | -0.410322 | -0.088398 | -0.092839 | -0.040168 | -0.093062 | 0.111527 | -0.192545 | -0.006621 | -0.02267 | -0.06982 |
| 20.84722 | -0.066819 | -0.824136 | -0.530294 | -0.846434 | -0.165644 | -0.123943 | -0.218346 | -0.071121 | 0.080288 | -0.108724 | -0.061969 | -0.130075 | -0.194606 | -0.513573 | -0.522768 | -0.120342 | -0.160897 | -0.024738 | -0.08722 | 0.05192 | -0.202557 | -0.007467 | -0.031714 | -0.154178 |
| 20.86111 | -0.191399 | -1.468534 | -0.801524 | -1.123893 | -0.314559 | -0.226114 | -0.221682 | -0.189257 | 0.05754 | -0.278382 | -0.430839 | -0.251267 | -0.200102 | -0.427076 | -0.597148 | -0.119616 | -0.261412 | -0.02275 | -0.125324 | -0.096769 | -0.697112 | -0.027886 | -0.031064 | -0.207195 |
| 20.875 | -0.264117 | -1.400047 | -0.928892 | -1.092675 | -0.501465 | -0.371489 | -0.229923 | -0.245688 | 0.149087 | -0.381202 | -0.808983 | -0.318308 | -0.194148 | -0.434307 | -0.583865 | -0.148822 | -0.325115 | -0.034175 | -0.219255 | -0.212196 | -0.6407 | -0.055966 | -0.030612 | -0.198004 |
| 20.88889 | -0.334101 | -1.011202 | -0.935341 | -0.90307 | -0.676771 | -0.490919 | -0.376334 | -0.299067 | 0.316403 | -0.473271 | -0.856776 | -0.390386 | -0.146314 | -0.372189 | -0.502429 | -0.217099 | -0.33638 | -0.031856 | -0.234816 | -0.202715 | -0.093448 | -0.062305 | -0.022533 | -0.188261 |
| 20.90278 | -0.472742 | -1.061935 | -1.010766 | -0.774231 | -0.879382 | -0.589556 | -0.661335 | -0.398351 | 0.477884 | -0.609853 | -0.841134 | -0.501729 | -0.126571 | -0.366297 | -0.473416 | -0.306212 | -0.361411 | -0.048975 | -0.162624 | -0.091573 | 0.015684 | -0.072683 | -0.039415 | -0.244383 |
| 20.91667 | -0.596898 | -1.2772 | -1.075266 | -0.692357 | -1.016678 | -0.758529 | -0.877702 | -0.466821 | 0.25392 | -0.739223 | -0.86257 | -0.580401 | -0.108032 | -0.378387 | -0.427861 | -0.349712 | -0.369498 | -0.065303 | -0.097239 | 0.054283 | -0.154293 | -0.082521 | -0.050703 | -0.229526 |
| 20.93056 | -0.613876 | -1.386245 | -1.065241 | -0.668876 | -1.141393 | -1.106174 | -0.798188 | -0.439127 | -0.02682 | -0.739307 | -0.91544 | -0.559563 | -0.051227 | -0.327306 | -0.342227 | -0.253854 | -0.355278 | -0.04827 | -0.093644 | 0.133587 | -0.279049 | -0.070596 | -0.019912 | -0.181573 |
| 20.94444 | -0.548651 | -1.224596 | -0.893065 | -0.728813 | -1.236917 | -1.163898 | -0.60943 | -0.383657 | -0.093403 | -0.639107 | -1.098278 | -0.475591 | -0.049016 | -0.234233 | -0.222237 | -0.135321 | -0.320947 | -0.04748 | -0.145428 | 0.126365 | -0.413024 | -0.060916 | -0.010411 | -0.146142 |
| 20.95833 | -0.41459 | -0.979366 | -0.597083 | -0.776071 | -1.034982 | -0.736461 | -0.405915 | -0.305716 | -0.107682 | -0.449704 | -1.490327 | -0.347518 | -0.076344 | -0.145546 | -0.102803 | -0.091611 | -0.276637 | -0.039902 | -0.150632 | 0.116684 | -0.644457 | -0.034339 | -0.005124 | -0.153145 |
| 20.97222 | -0.268156 | -0.825475 | -0.306794 | -0.799202 | -0.593445 | -0.246286 | -0.230003 | -0.284155 | -0.011931 | -0.302135 | -1.072927 | -0.217254 | -0.075203 | -0.137687 | 0.006342 | -0.125685 | -0.229878 | -0.009027 | -0.107341 | 0.122136 | -0.672136 | -0.020132 | 0.024266 | -0.219329 |
| 20.98611 | -0.180639 | -0.690381 | -0.281463 | -0.842805 | -0.282142 | -0.066745 | -0.139537 | -0.230837 | 0.131905 | -0.243765 | -0.431928 | -0.185863 | -0.087929 | -0.220012 | 0.037555 | -0.147559 | -0.186864 | -0.001023 | -0.082869 | 0.113975 | -0.479182 | -0.02005 | 0.026424 | -0.321935 |
| 21 | -0.061088 | -0.430813 | -0.360033 | -0.77574 | -0.120664 | -0.020159 | -0.07775 | -0.201985 | 0.22406 | -0.156867 | 0.009712 | -0.148579 | -0.057407 | -0.178567 | 0.103283 | 0.014949 | -0.092296 | 0.063884 | 0.014361 | 0.132401 | -0.253952 | 0.069106 | 0.081818 | -0.277901 |
| 21.01389 | 0.142199 | -0.004925 | -0.283607 | -0.217089 | 0.062063 | 0.147026 | 0.027314 | -0.118372 | 0.318063 | 0.006186 | 0.074396 | 0.008312 | 0.087766 | 0.219136 | 0.317955 | 0.263802 | 0.04874 | 0.252098 | 0.243528 | 0.197699 | 0.013761 | 0.292745 | 0.266055 | -0.012768 |
| 21.02778 | 0.146919 | 0.34901 | -0.332403 | 0.209152 | 0.140823 | 0.22274 | 0.048275 | -0.091196 | 0.312294 | 0.042658 | -0.329493 | 0.111124 | 0.181978 | 0.467725 | 0.475925 | 0.159865 | 0.140686 | 0.36276 | 0.380986 | 0.286162 | 0.136318 | 0.482151 | 0.348679 | 0.275928 |
| 21.04167 | -0.098854 | 0.337332 | -0.559468 | 0.230414 | 0.005464 | 0.262665 | -0.070086 | -0.290113 | 0.118005 | -0.197237 | -0.729383 | -0.067714 | 0.09364 | 0.180942 | 0.339049 | -0.096236 | 0.112951 | 0.232959 | 0.244093 | 0.283759 | -0.012148 | 0.397993 | 0.218374 | 0.321378 |
| 21.05556 | -0.098666 | 0.2989 | -0.369777 | 0.548664 | -0.039985 | 0.682705 | -0.076537 | -0.372713 | 0.065847 | -0.32648 | -0.846603 | -0.235885 | -0.028464 | -0.338033 | 0.094803 | -0.211002 | 0.089023 | 0.076356 | 0.0635 | 0.22282 | -0.233052 | 0.206323 | 0.069628 | 0.122238 |
| 21.06944 | 0.088474 | 0.228743 | -0.140167 | 0.248981 | 0.099732 | 1.276896 | 0.076556 | -0.268297 | 0.249749 | -0.177813 | -0.452127 | -0.096208 | -0.080644 | -0.843982 | -0.121411 | -0.413834 | 0.086607 | 0.02122 | -0.002041 | 0.23198 | -0.487835 | 0.060148 | 0.023741 | -0.104483 |
| 21.08333 | 0.104226 | -0.070365 | -0.319127 | -0.770314 | 0.116934 | 0.895563 | 0.133474 | -0.199076 | 0.344196 | 0.082909 | -0.033881 | 0.151618 | -0.090641 | -0.584169 | -0.258371 | -0.483685 | 0.079863 | 0.011284 | -0.011337 | 0.122302 | -0.778096 | -0.035943 | 0.023647 | -0.195794 |
| 21.09722 | 0.090746 | -0.136908 | -0.275253 | -0.930206 | 0.08822 | 0.339925 | 0.062582 | -0.125251 | 0.280633 | 0.403959 | 0.209122 | 0.289467 | -0.047029 | 0.071656 | -0.266858 | 0.017426 | 0.066836 | 0.01122 | 0.002259 | 0.001674 | -0.610566 | -0.019628 | 0.022753 | -0.093947 |
| 21.11111 | 0.15756 | 0.219715 | 0.192884 | -0.402642 | 0.126031 | 0.089014 | 0.014995 | 0.006075 | 0.197011 | 0.727262 | 0.193861 | 0.349777 | 0.048056 | 0.441719 | -0.195594 | 0.643361 | 0.044277 | -0.001374 | -0.002035 | -0.048912 | -0.221537 | 0.001906 | -0.000978 | 0.016551 |
| 21.125 | 0.15525 | 0.378632 | 0.574359 | -0.363579 | 0.096086 | 0.108685 | -0.011607 | 0.139927 | 0.12608 | 0.817002 | 0.047913 | 0.209735 | 0.176494 | 0.352393 | -0.102208 | 0.903986 | 0.031644 | -0.00444 | -0.035117 | -0.054021 | 0.14197 | -0.019341 | -0.009557 | -0.029307 |
| 21.13889 | 0.034646 | 0.024446 | 0.583191 | -0.510239 | 0.023876 | 0.164815 | -0.064771 | 0.164646 | 0.04407 | 0.54005 | -0.085528 | 0.029572 | 0.221953 | 0.291922 | -0.011178 | 0.566428 | 0.010617 | 0.006441 | -0.057505 | 0.021303 | 0.373462 | -0.055872 | -0.004324 | -0.183138 |
| 21.15278 | -0.065742 | -0.3506 | 0.163221 | -0.319721 | 0.049403 | 0.164016 | -0.106255 | 0.035864 | -0.034481 | 0.137825 | -0.161617 | -0.07024 | 0.076004 | 0.331433 | 0.019643 | 0.062809 | -0.028951 | -0.006917 | -0.049275 | -0.039812 | 0.323244 | -0.083375 | -0.039159 | -0.222232 |
| 21.16667 | -0.042682 | -0.263682 | -0.593789 | 0.051701 | 0.187711 | 0.13839 | -0.139051 | -0.092349 | -0.067223 | -0.205418 | -0.132345 | -0.102132 | -0.074311 | 0.35277 | 0.021443 | -0.433991 | -0.04035 | -0.025268 | -0.013508 | -0.05156 | 0.165951 | -0.117568 | -0.049768 | -0.096631 |
| 21.18056 | 0.008569 | -0.117358 | -0.990553 | 0.150678 | 0.24426 | 0.011894 | -0.17422 | -0.099736 | -0.116253 | -0.459785 | 0.026617 | -0.122066 | -0.060032 | 0.35615 | 0.065694 | -0.496276 | 0.007891 | -0.020615 | 0.017039 | -0.02765 | 0.097926 | -0.113704 | -0.020108 | 0.012439 |
| 21.19444 | -0.052804 | -0.260416 | -0.739077 | 0.002484 | 0.154819 | -0.122516 | -0.202466 | -0.038235 | -0.184222 | -0.617685 | 0.195862 | -0.14491 | 0.07443 | 0.12458 | 0.154147 | -0.016283 | 0.082905 | -0.003471 | 0.040193 | -0.008267 | 0.199225 | -0.006051 | 0.015743 | 0.046923 |
| 21.20833 | -0.114481 | -0.284423 | -0.361168 | 0.082583 | 0.159204 | -0.140174 | -0.168737 | 0.041119 | -0.167799 | -0.676418 | 0.261421 | -0.093799 | 0.149154 | -0.018237 | 0.240469 | 0.319877 | 0.135297 | 0.009908 | 0.057437 | -0.007959 | 0.095552 | 0.108614 | 0.038184 | 0.049101 |
| 21.22222 | -0.056203 | 0.040648 | -0.027703 | 0.273092 | 0.189107 | -0.1687 | -0.065883 | 0.065896 | -0.097274 | -0.563911 | 0.274938 | -0.019207 | 0.106278 | 0.142805 | 0.239905 | 0.155714 | 0.144825 | 0.006844 | 0.068677 | -0.029249 | -0.155927 | 0.072191 | 0.035342 | 0.041993 |
| 21.23611 | 0.009422 | 0.368908 | 0.18709 | 0.15639 | 0.07434 | -0.226658 | 0.02499 | 0.010839 | -0.085281 | -0.371113 | 0.297226 | 0.0051 | 0.025999 | 0.213675 | 0.152583 | -0.114745 | 0.124093 | 0.005453 | 0.075032 | -0.049856 | -0.167547 | -0.013463 | 0.014171 | 0.023435 |
| 21.25 | 0.00968 | 0.33903 | 0.374165 | -0.05708 | -0.022397 | -0.099731 | 0.114477 | -0.064843 | -0.073265 | -0.184041 | 0.303549 | -0.001595 | 0.01649 | 0.235596 | 0.116615 | -0.09308 | 0.124518 | 0.023363 | 0.088439 | -0.035106 | 0.086701 | -0.036962 | 0.012863 | 0.02601 |

| | | | | | | | | | | | | | | | | | | | | | | | | |
|---|---|---|---|---|---|---|---|---|---|---|---|---|---|---|---|---|---|---|---|---|---|---|---|---|
| 21.26389 | -0.062283 | 0.087128 | 0.389145 | -0.092315 | -0.040204 | 0.188309 | 0.187011 | -0.107903 | -0.050987 | -0.085921 | 0.23359 | -0.036323 | 0.028361 | 0.415885 | 0.142713 | -0.01962 | 0.148037 | 0.037231 | 0.105738 | 0.026336 | 0.385057 | -0.018662 | 0.021822 | 0.056677 |
| 21.27778 | -0.144198 | 0.071647 | 0.20386 | -0.048394 | -0.029245 | 0.363752 | 0.220122 | -0.0996 | -0.058553 | -0.059753 | 0.251775 | -0.043856 | 0.012865 | 0.473413 | 0.147773 | -0.139368 | 0.14649 | 0.034497 | 0.108267 | 0.032919 | 0.412284 | -0.010596 | 0.030738 | 0.080215 |
| 21.29167 | -0.115973 | 0.199931 | -0.026045 | 0.031098 | 0.027245 | 0.202388 | 0.194278 | -0.051303 | -0.100413 | -0.04836 | 0.219257 | -0.022404 | -0.005336 | 0.28484 | 0.113262 | -0.246006 | 0.079333 | 0.018082 | 0.08238 | -0.03716 | 0.225039 | -0.026164 | 0.025839 | 0.064098 |
| 21.30556 | -0.009581 | 0.093599 | -0.167723 | 0.153952 | 0.06163 | -0.04011 | 0.118887 | -0.044659 | -0.16227 | -0.081718 | 0.033178 | -0.061396 | -0.023193 | 0.107479 | 0.05275 | -0.147765 | -0.02476 | -0.017221 | 0.053955 | -0.054381 | 0.031787 | -0.042254 | -0.01612 | 0.027031 |
| 21.31944 | 0.056704 | 0.009551 | -0.021211 | 0.266285 | 0.094056 | -0.120893 | 0.099825 | -0.002588 | -0.10665 | -0.060537 | 0.000166 | -0.050054 | 0.024334 | 0.093414 | 0.052584 | 0.059191 | -0.055581 | -0.024634 | 0.062229 | 0.007899 | 0.007883 | 0.000888 | -0.017537 | 0.063591 |
| 21.33333 | 0.038146 | 0.303435 | 0.333047 | 0.40653 | 0.180095 | 0.023374 | 0.171232 | 0.143388 | 0.050359 | 0.058484 | 0.335668 | 0.078373 | 0.095233 | 0.213023 | 0.088605 | 0.249452 | 0.003694 | 0.01582 | 0.067524 | 0.057511 | 0.117683 | 0.12533 | 0.035948 | 0.167576 |
| 21.34722 | 0.007931 | 0.586107 | 0.45763 | 0.554498 | 0.225166 | 0.186047 | 0.160528 | 0.17239 | 0.092282 | 0.06338 | 0.673173 | 0.097197 | 0.07329 | 0.252663 | 0.060481 | 0.232274 | 0.043096 | 0.033044 | 0.048294 | 0.060986 | 0.259909 | 0.13243 | 0.044092 | 0.229449 |
| 21.36111 | 0.035293 | 0.531498 | 0.312433 | 0.520701 | 0.264209 | 0.37918 | 0.105475 | 0.07919 | -0.004037 | -0.021935 | 0.574923 | 0.00847 | 0.080319 | 0.237144 | 0.078302 | 0.117676 | 0.062436 | 0.019494 | 0.044132 | 0.038933 | 0.394182 | 0.031457 | 0.030667 | 0.261096 |
| 21.375 | 0.059704 | 0.498317 | 0.178477 | 0.358323 | 0.378248 | 0.513526 | 0.157092 | 0.060704 | -0.150929 | -0.005603 | 0.532388 | 0.006624 | 0.15959 | 0.332599 | 0.183935 | 0.199513 | 0.089474 | 0.015599 | 0.047154 | 0.00773 | 0.443907 | 0.007971 | 0.040397 | 0.310146 |
| 21.38889 | 0.049627 | 0.713602 | 0.240687 | 0.301974 | 0.521075 | 0.446586 | 0.245731 | 0.112961 | -0.212776 | 0.05082 | 0.671555 | 0.062953 | 0.185455 | 0.495732 | 0.264717 | 0.438428 | 0.089229 | 0.018472 | 0.05385 | -0.0423 | 0.346974 | 0.044181 | 0.029619 | 0.323923 |
| 21.40278 | 0.126683 | 1.133419 | 0.512615 | 0.456228 | 0.666202 | 0.236478 | 0.340679 | 0.149282 | 0.280328 | 0.071848 | 0.934192 | 0.090103 | 0.196871 | 0.607213 | 0.367856 | 0.658721 | 0.127042 | 0.030671 | 0.076392 | -0.026252 | 0.24383 | 0.090489 | 0.035523 | 0.310981 |
| 21.41667 | 0.266031 | 1.511414 | 0.795269 | 0.546113 | 0.720677 | 0.048286 | 0.368815 | 0.125857 | 0.083003 | 0.060118 | 1.047147 | 0.024547 | 0.199842 | 0.670585 | 0.534892 | 0.758945 | 0.238247 | 0.061554 | 0.118984 | 0.032119 | 0.200327 | 0.142562 | 0.076168 | 0.405661 |
| 21.43056 | 0.280882 | 1.538131 | 0.955854 | 0.527103 | 0.600176 | -0.08498 | 0.331524 | 0.128389 | -0.070253 | 0.183472 | 0.939842 | -0.06539 | 0.12708 | 0.685149 | 0.643945 | 0.548061 | 0.314876 | 0.063846 | 0.139958 | 0.043952 | 0.210414 | 0.146622 | 0.039097 | 0.527986 |
| 21.44444 | 0.304151 | 1.112133 | 1.117605 | 0.737603 | 0.572189 | 0.047956 | 0.41167 | 0.272774 | 0.026793 | 0.626592 | 0.970151 | -0.039953 | 0.025467 | 0.533148 | 0.550403 | 0.108682 | 0.38708 | -0.039318 | -0.008791 | -0.041476 | 0.20335 | -0.006718 | -0.072677 | 0.395114 |
| 21.45833 | 0.39176 | 0.593378 | 1.253672 | 1.193498 | 0.723882 | 0.468299 | 0.5546 | 0.486567 | 0.319418 | 0.842911 | 1.112892 | 0.091167 | -0.057499 | 0.304903 | 0.336395 | -0.409195 | 0.399717 | -0.208683 | -0.260438 | -0.174331 | 0.290854 | -0.262421 | -0.215884 | 0.140326 |
| 21.47222 | 0.384654 | 0.305346 | 1.070238 | 1.548192 | 0.843199 | 0.89608 | 0.605923 | 0.577218 | 0.630703 | 0.700768 | 1.101148 | 0.33749 | -0.038381 | 0.154484 | 0.119091 | -0.22607 | 0.267558 | -0.221359 | -0.274361 | -0.249135 | 0.547144 | -0.287919 | -0.248185 | 0.129155 |
| 21.48611 | 0.343848 | 0.499307 | 1.059875 | 1.770151 | 0.839915 | 0.866737 | 0.519425 | 0.505375 | 0.753283 | 0.534685 | 0.761975 | 0.363476 | 0.027675 | 0.458193 | 0.226161 | -0.064574 | 0.176163 | -0.08479 | -0.09474 | -0.293524 | 1.032846 | -0.109582 | -0.128975 | 0.340727 |
| 21.5 | 0.34641 | 0.772611 | 0.996314 | 1.397118 | 0.651103 | 0.547327 | 0.412574 | 0.355271 | 0.730766 | 0.393447 | 0.45655 | 0.273389 | -0.123582 | 0.627794 | 0.459166 | 0.120135 | 0.215321 | -0.033992 | 0.071687 | -0.216494 | 1.059826 | 0.016358 | -0.042701 | 0.622568 |
| 21.51389 | 0.233302 | 0.812618 | 0.670364 | 0.987406 | 0.349141 | 0.310747 | 0.304947 | 0.170732 | 0.750964 | 0.281768 | 0.361843 | 0.215137 | -0.252798 | 0.553899 | 0.557937 | 0.138699 | 0.215664 | -0.052492 | 0.134178 | -0.07172 | 0.576122 | 0.062088 | -0.004376 | 0.578765 |
| 21.52778 | -0.08363 | 0.28424 | 0.065922 | 0.551309 | -0.218407 | -0.174219 | -0.134475 | -0.222725 | 0.402663 | -0.207346 | 0.005298 | -0.088462 | -0.034057 | 0.432677 | 0.56864 | 0.1864 | 0.131023 | 0.005545 | 0.138344 | -0.073852 | 0.322437 | 0.032156 | 0.06064 | 0.388971 |
| 21.54167 | -0.769988 | -0.61555 | -0.819555 | -0.046984 | -0.927615 | -0.995281 | -0.924007 | -0.892967 | -0.347118 | -1.071463 | -0.75139 | -0.668352 | 0.151467 | 0.204028 | 0.466936 | 0.213625 | 0.012337 | 0.045189 | 0.141102 | -0.215303 | 0.24408 | -0.098337 | 0.057104 | 0.41384 |
| 21.55556 | -1.204821 | -1.322201 | -1.251262 | -0.412546 | -1.17187 | -1.397289 | -1.279033 | -1.300636 | -0.683632 | -1.402162 | -1.09604 | -0.95446 | 0.060712 | 0.142686 | 0.30276 | 0.150659 | -0.053089 | -0.004564 | 0.10158 | -0.204913 | 0.178582 | -0.106134 | -0.001765 | 0.349294 |
| 21.56944 | -0.981052 | -1.216222 | -0.956623 | -0.154699 | -0.729996 | -1.012468 | -0.944587 | -0.97725 | -0.322314 | -0.974168 | -0.84312 | -0.360704 | -0.024706 | 0.375127 | 0.246882 | 0.058342 | 0.041714 | -0.032067 | 0.055308 | -0.040566 | 0.159359 | 0.02373 | -0.003877 | 0.360028 |
| 21.58333 | -0.518117 | -0.717373 | -0.493426 | 0.260655 | -0.215683 | -0.528896 | -0.499854 | -0.562652 | 0.17698 | -0.4936 | -0.5381 | 0.125005 | -0.007795 | 0.528413 | 0.341517 | 0.041102 | 0.153388 | -0.006495 | 0.043211 | -0.022667 | 0.303788 | 0.101842 | 0.039141 | 0.372744 |
| 21.59722 | -0.150247 | -0.248448 | -0.08429 | 0.696853 | 0.150359 | -0.192785 | -0.184065 | -0.233315 | 0.336363 | -0.16643 | -0.32098 | 0.209485 | 0.016844 | 0.425411 | 0.468725 | 0.057749 | 0.20699 | 0.012343 | 0.039434 | -0.081998 | 0.243141 | 0.104203 | 0.060914 | 0.288196 |
| 21.61111 | 0.002515 | 0.110141 | 0.145134 | 0.844884 | 0.287874 | -0.02325 | -0.046588 | -0.062135 | 0.278962 | -0.020042 | -0.14098 | 0.129068 | 0.017934 | 0.210564 | 0.461564 | 0.067314 | 0.192045 | 0.016403 | 0.037311 | -0.063296 | 0.233735 | 0.097034 | 0.061779 | 0.227789 |
| 21.625 | -0.037959 | 0.299628 | 0.147086 | 0.455943 | 0.147384 | -0.037481 | -0.066336 | -0.063485 | 0.027781 | -0.073802 | -0.098132 | -0.197096 | -0.00009 | 0.060182 | 0.283662 | 0.06297 | 0.089064 | 0.011239 | 0.049664 | -0.043207 | 0.388446 | 0.065566 | 0.040611 | 0.151835 |
| 21.63889 | -0.099343 | 0.445065 | 0.097732 | 0.122886 | -0.013854 | -0.094364 | -0.12722 | -0.121522 | -0.195943 | -0.14546 | 0.129795 | -0.591895 | -0.005281 | -0.063009 | 0.150913 | 0.072229 | -0.023752 | 0.019486 | 0.086215 | 0.006546 | 0.299851 | 0.03922 | 0.030665 | 0.138755 |
| 21.65278 | -0.131251 | 0.37374 | 0.050307 | 0.096561 | -0.032211 | -0.124349 | -0.171291 | -0.174999 | -0.214636 | -0.17179 | 0.069799 | -0.527403 | -0.008002 | -0.170177 | 0.06938 | 0.061402 | -0.07684 | 0.016387 | 0.122524 | 0.082154 | 0.174827 | 0.040524 | 0.024322 | 0.253887 |
| 21.66667 | -0.180134 | 0.095309 | -0.051349 | 0.044706 | 0.016633 | -0.205936 | -0.241857 | -0.247015 | -0.033977 | -0.233109 | -0.202011 | -0.600781 | -0.017527 | -0.202806 | -0.013567 | 0.022033 | -0.055274 | -0.016077 | 0.110504 | 0.096177 | 0.100077 | 0.037286 | 0.013065 | 0.388003 |
| 21.68056 | -0.271589 | -0.200944 | -0.213461 | -0.171744 | -0.050555 | -0.366613 | -0.354983 | -0.349326 | 0.162195 | -0.356258 | -0.467929 | -0.639309 | -0.030907 | -0.173785 | -0.052908 | -0.015336 | 0.01328 | -0.04757 | 0.049057 | 0.037446 | 0.025448 | 0.010438 | -0.010563 | 0.315662 |
| 21.69444 | -0.355435 | -0.47964 | -0.331999 | -0.433547 | -0.138673 | -0.506408 | -0.429172 | -0.432207 | 0.244384 | -0.459907 | -0.682585 | -0.682369 | -0.017804 | -0.108891 | 0.003414 | 0.006961 | 0.091202 | -0.026617 | 0.018035 | -0.018106 | -0.033744 | -0.001497 | -0.014288 | 0.145392 |
| 21.70833 | -0.380962 | -0.643011 | -0.309424 | -0.467372 | -0.161326 | -0.51144 | -0.373873 | -0.420137 | 0.025859 | -0.451581 | -0.666783 | -0.660678 | 0.01065 | -0.04134 | 0.121304 | 0.079443 | 0.146878 | 0.031079 | 0.024988 | -0.052838 | -0.016211 | 0.018895 | 0.004799 | -0.038435 |
| 21.72222 | -0.355824 | -0.673212 | -0.169634 | -0.383985 | -0.22203 | -0.414514 | -0.274207 | -0.339605 | -0.29631 | -0.356897 | -0.488791 | -0.531501 | -0.000087 | -0.106568 | 0.187178 | 0.122236 | 0.138926 | 0.046469 | -0.003104 | -0.063687 | -0.003391 | 0.030216 | 0.01335 | -0.263091 |
| 21.73611 | -0.248904 | -0.365338 | 0.050225 | -0.258074 | -0.225993 | -0.222688 | -0.141611 | -0.182593 | -0.400594 | -0.167774 | 0.057586 | -0.296594 | -0.006192 | -0.193156 | 0.173251 | 0.152321 | 0.142179 | 0.055979 | -0.009238 | 0.025091 | 0.01527 | 0.044333 | 0.052836 | -0.31194 |
| 21.75 | 0.059236 | 0.311216 | 0.401511 | 0.131297 | -0.157716 | 0.147181 | 0.134025 | 0.143798 | -0.268674 | 0.181321 | 0.540848 | 0.142112 | 0.026635 | -0.128403 | 0.158885 | 0.141626 | 0.197593 | 0.089189 | 0.035484 | 0.138272 | 0.067518 | 0.061713 | 0.1202 | 0.149054 |
| 21.76389 | 0.566174 | 1.177309 | 0.871602 | 0.815819 | 0.118896 | 0.655992 | 0.593978 | 0.645064 | 0.005384 | 0.660522 | 0.851499 | 0.765177 | 0.011448 | -0.027039 | 0.114406 | 0.091869 | 0.207507 | 0.09257 | 0.057604 | 0.186424 | 0.093914 | 0.050471 | 0.11557 | 0.42302 |
| 21.77778 | 0.997773 | 1.701558 | 1.216854 | 1.414389 | 0.576145 | 1.032833 | 1.000194 | 1.093978 | -0.016199 | 1.070389 | 0.734626 | 1.290034 | -0.041378 | 0.055276 | 0.035325 | 0.045346 | 0.167918 | 0.070368 | 0.058984 | 0.16417 | 0.085428 | 0.028503 | 0.078529 | 0.32704 |
| 21.79167 | 1.126327 | 1.470428 | 1.242496 | 1.493102 | 0.936102 | 1.064684 | 1.123749 | 1.266142 | -0.298376 | 1.208337 | 0.629659 | 1.451226 | -0.004261 | 0.011143 | -0.023427 | 0.07477 | 0.155365 | 0.069895 | 0.069489 | 0.102505 | 0.068835 | 0.032583 | 0.053907 | 0.130579 |
| 21.80556 | 1.142373 | 1.126633 | 1.13171 | 1.204242 | 1.082328 | 0.893945 | 1.060643 | 1.228548 | 0.034368 | 1.160427 | 0.540323 | 1.382358 | 0.049003 | -0.145255 | -0.066077 | 0.140248 | 0.123241 | 0.070593 | 0.071812 | 0.042803 | 0.00092 | 0.031054 | 0.022357 | -0.102003 |
| 21.81944 | 0.999237 | 0.638927 | 0.862332 | 0.715303 | 0.780309 | 0.755306 | 0.857396 | 1.031744 | 0.746009 | 0.96445 | 0.615933 | 1.120988 | 0.032417 | -0.417947 | -0.172203 | 0.273659 | 0.022947 | 0.04109 | 0.032902 | 0.013679 | -0.118423 | 0.002167 | -0.045585 | -0.454003 |
| 21.83333 | 0.639375 | -0.016363 | 0.430752 | 0.061485 | 0.168014 | 0.458008 | 0.421938 | 0.598221 | 1.113367 | 0.551618 | 0.836089 | 0.565465 | -0.01774 | -0.687651 | -0.259971 | 0.220666 | -0.12274 | -0.006724 | -0.034782 | 0.014943 | -0.207133 | -0.028392 | -0.072791 | -0.649569 |
| 21.84722 | 0.191894 | -0.688394 | -0.059742 | -0.54583 | -0.425323 | 0.032453 | -0.064123 | 0.107909 | 0.563933 | 0.052203 | 0.298647 | -0.163469 | -0.055508 | -0.482684 | -0.436458 | 0.03902 | -0.264385 | -0.042577 | -0.094598 | 0.036806 | -0.452102 | -0.056224 | -0.051024 | -0.423588 |
| 21.86111 | -0.214658 | -1.517343 | -0.252605 | -1.019047 | -0.866773 | -0.410897 | -0.469678 | -0.306602 | -0.491116 | -0.379955 | -0.466858 | -0.846057 | -0.063221 | -0.345504 | -0.664718 | -0.05767 | -0.367509 | -0.060382 | -0.145679 | 0.009706 | -1.223316 | -0.098766 | -0.053904 | -0.120215 |
| 21.875 | -0.597856 | -1.786532 | -0.326609 | -1.18166 | -0.86507 | -0.819219 | -0.77843 | -0.606682 | -1.30696 | -0.681617 | -1.046567 | -1.470909 | -0.049526 | -0.401258 | -0.504884 | -0.098537 | -0.378409 | -0.06794 | -0.182289 | -0.053726 | -0.023731 | -0.127111 | -0.068997 | -0.034373 |
| 21.88889 | -0.970469 | -1.61312 | -0.620154 | -1.199451 | -1.109385 | -1.21909 | -1.11368 | -0.891087 | -1.737613 | -0.924712 | -1.408244 | -2.091096 | -0.012148 | -0.52077 | -0.11104 | -0.143388 | -0.344587 | -0.075755 | -0.195474 | -0.100854 | 0.314009 | -0.102059 | -0.086525 | -0.202386 |
| 21.90278 | -1.359355 | -1.717316 | -1.131192 | -1.467517 | -1.310139 | -1.582606 | -1.481904 | -1.142909 | -1.994786 | -1.180924 | -1.472718 | -2.447316 | 0.019999 | -0.537291 | 0.023819 | -0.247736 | -0.377209 | -0.092206 | -0.184343 | -0.08836 | 0.18638 | -0.0877 | -0.124636 | -0.431135 |
| 21.91667 | -1.496425 | -1.754823 | -1.417506 | -1.655643 | -1.381147 | -1.514397 | -1.454817 | -1.196826 | -1.945423 | -1.200417 | -1.26857 | -2.340321 | 0.050991 | -0.363649 | -0.203813 | -0.358924 | -0.432794 | -0.094147 | -0.141071 | -0.015194 | 0.067116 | -0.060304 | -0.124026 | -0.391107 |
| 21.93056 | -1.170956 | -1.510006 | -1.339597 | -1.578503 | -1.238656 | -1.149963 | -1.083139 | -0.988135 | -1.853382 | -0.936189 | -1.144059 | -1.79453 | 0.04088 | -0.390043 | -0.538907 | -0.429994 | -0.438879 | -0.05376 | -0.106389 | 0.051851 | 0.006151 | -0.020734 | -0.026359 | -0.274382 |
| 21.94444 | -0.883025 | -1.26883 | -1.272965 | -1.580726 | -1.120423 | -0.818228 | -0.776868 | -0.694261 | -1.422414 | -0.655708 | -1.062045 | -1.23862 | -0.022883 | -0.582181 | -0.903734 | -0.257439 | -0.327841 | -0.006572 | -0.119812 | 0.125283 | -0.093278 | -0.003243 | 0.017656 | -0.311712 |
| 21.95833 | -0.654827 | -1.267976 | -1.318957 | -1.738173 | -1.002964 | -0.525468 | -0.572626 | -0.447911 | -1.192329 | -0.45266 | -1.022008 | -0.828939 | -0.070742 | -0.713074 | -0.816216 | -0.282123 | -0.149093 | 0.035134 | -0.123464 | 0.218052 | -0.228882 | -0.004995 | 0.031989 | -0.583611 |
| 21.97222 | -0.475868 | -1.370719 | -1.308858 | -1.492757 | -0.721185 | -0.367101 | -0.3655 | -0.210611 | -1.329916 | -0.303832 | -0.851638 | -0.549119 | -0.054429 | -0.64759 | -0.667615 | -0.73504 | -0.047342 | 0.076749 | -0.096781 | 0.172219 | -0.342475 | -0.03166 | 0.060237 | -0.521127 |
| 21.98611 | -0.448171 | -1.20073 | -1.251578 | -1.09726 | -0.466521 | -0.362274 | -0.198905 | -0.087665 | -1.632511 | -0.237484 | -0.626329 | -0.422157 | -0.014339 | -0.50844 | -0.458354 | -0.935295 | -0.042125 | 0.097931 | -0.061588 | 0.112745 | -0.33185 | 0.001805 | 0.056424 | -0.101314 |
| 22 | -0.460931 | -0.696686 | -1.248694 | -0.703571 | -0.263329 | -0.316474 | -0.105467 | -0.182071 | -1.449624 | -0.198228 | -0.423627 | -0.437695 | 0.098841 | -0.380402 | 0.259979 | -0.758888 | -0.002766 | 0.161742 | 0.026023 | 0.135874 | -0.22899 | 0.095499 | 0.127721 | 0.157325 |
| 22.01389 | -0.414353 | -0.327213 | -1.188399 | -0.162077 | -0.197662 | -0.141711 | -0.091445 | -0.299624 | -1.08303 | -0.155268 | -0.281468 | -0.595365 | 0.220908 | -0.116682 | 1.275528 | -0.11259 | 0.094037 | 0.309208 | 0.210188 | 0.233849 | -0.00365 | 0.291308 | 0.293876 | 0.15795 |
| 22.02778 | -0.481825 | -0.40954 | -1.148694 | -0.228605 | -0.346821 | -0.068305 | -0.154548 | -0.453499 | -0.708362 | -0.132884 | -0.27302 | -0.73446 | 0.136319 | 0.023494 | 1.432068 | 0.261166 | 0.143421 | 0.355426 | 0.327919 | 0.328886 | 0.068731 | 0.468254 | 0.35754 | 0.017957 |
| 22.04167 | -0.337078 | -0.237904 | -0.674086 | -0.992521 | 0.015355 | -0.063527 | 0.061477 | -0.35406 | -0.341397 | 0.065469 | -0.114353 | -0.307177 | -0.110448 | -0.517114 | 0.434122 | 0.022549 | 0.168646 | 0.206316 | 0.220371 | 0.349186 | -0.280597 | 0.385023 | 0.197995 | 0.132795 |
| 22.05556 | 0.223372 | 0.659476 | 0.717606 | -0.594513 | 0.960227 | -0.125773 | 0.583819 | 0.032987 | 0.028835 | 0.452809 | 0.318645 | 0.693437 | -0.290483 | -1.34267 | -0.272131 | -0.338478 | 0.309081 | 0.040459 | 0.019386 | 0.410254 | -0.798311 | 0.208601 | 0.052123 | 0.10228 |
| 22.06944 | 0.595711 | 1.177221 | 1.72275 | -0.723975 | 0.914399 | -0.414257 | 0.755859 | 0.277612 | 0.057904 | 0.639192 | 0.313473 | 1.31848 | -0.408931 | -1.720249 | -0.844375 | -0.978537 | 0.311266 | -0.021822 | -0.085089 | 0.420679 | -1.385933 | 0.05424 | 0.02351 | -0.278865 |
| 22.08333 | 0.56023 | 0.661076 | 0.511543 | -0.870667 | -0.029787 | 0.123072 | 0.597079 | 0.296118 | -0.171105 | 0.436687 | -0.06375 | 0.888455 | -0.348564 | -0.971965 | -0.629586 | -1.492038 | 0.04526 | 0.001443 | -0.077786 | 0.268073 | -1.487163 | -0.098591 | 0.079274 | -0.393786 |
| 22.09722 | 0.493282 | 0.308633 | -0.63402 | 0.149788 | -0.304503 | 0.466003 | 0.623808 | 0.502567 | 0.028024 | 0.23438 | 0.31525 | 0.441797 | -0.004328 | 0.12276 | 0.146919 | -0.837688 | -0.103933 | 0.05034 | -0.012452 | 0.11223 | -0.960308 | -0.110747 | 0.151291 | -0.050453 |
| 22.11111 | 0.519141 | 0.398303 | -0.782555 | 0.978802 | -0.097332 | 0.816698 | 0.728498 | 0.624561 | 0.477441 | 0.284332 | 0.693936 | 0.5824 | 0.30367 | 0.808125 | 0.597941 | 0.101988 | 0.137804 | 0.058848 | 0.03134 | 0.053886 | -0.324153 | -0.03506 | 0.157672 | 0.188514 |
| 22.125 | 0.61885 | 0.407689 | -0.321644 | 1.175613 | 0.013579 | 1.464339 | 0.648302 | 0.797106 | 0.646589 | 0.514351 | 0.956336 | 0.907656 | 0.274481 | 0.777769 | 0.229342 | 0.619856 | 0.353083 | 0.056067 | 0.044566 | 0.02672 | 0.212684 | 0.062471 | 0.04583 | 0.211076 |
| 22.13889 | 0.777209 | 0.188455 | 0.407968 | 0.72628 | 0.044165 | 1.528258 | 0.634262 | 1.035819 | 0.675843 | 0.688406 | 0.993051 | 0.883679 | -0.012246 | 0.494088 | -0.6553 | 0.437757 | 0.161281 | 0.086241 | 0.064635 | -0.008786 | 0.521326 | 0.112246 | -0.014763 | -0.063252 |
| 22.15278 | 0.896484 | 0.065725 | 1.136792 | 0.780786 | 0.373136 | 0.973422 | 0.853517 | 1.086903 | 1.118419 | 0.813343 | 0.969766 | 0.803268 | -0.189528 | 0.39041 | -0.85011 | 0.105394 | -0.137355 | 0.088026 | 0.068781 | -0.030945 | 0.445288 | 0.044762 | 0.038759 | -0.337274 |
| 22.16667 | 0.861682 | 0.261235 | 1.283185 | 1.270839 | 0.84657 | 0.38956 | 0.88795 | 0.916797 | 2.020763 | 0.876053 | 0.9629 | 0.855645 | -0.133504 | 0.3424 | -0.052119 | 0.02234 | -0.120894 | 0.036897 | 0.074261 | -0.03816 | 0.188974 | -0.00613 | 0.117075 | -0.045601 |
| 22.18056 | 0.725456 | 0.407736 | 0.725214 | 1.265838 | 1.058305 | -0.122057 | 0.614471 | 0.719396 | 1.979344 | 0.858277 | 0.803656 | 0.823825 | -0.060294 | 0.170845 | 0.621667 | 0.257986 | 0.049981 | 0.007195 | 0.12703 | -0.039526 | 0.01365 | 0.057534 | 0.131496 | 0.458796 |
| 22.19444 | 0.71661 | 0.315539 | 0.209135 | 0.845263 | 0.977816 | -0.083163 | 0.613539 | 0.64526 | 1.332078 | 0.79135 | 0.247079 | 0.739705 | 0.085731 | 0.063313 | 0.518497 | 0.506584 | 0.094341 | 0.015089 | 0.162692 | -0.034038 | -0.036354 | 0.097834 | 0.07808 | 0.418744 |
| 22.20833 | 0.752574 | 0.409567 | 0.119345 | 0.604638 | 0.535803 | 0.334003 | 0.682117 | 0.576626 | 1.24307 | 0.64013 | 0.325176 | 0.595062 | 0.30464 | 0.171972 | 0.254134 | 0.444625 | 0.08038 | 0.006083 | 0.132583 | -0.045961 | 0.055279 | 0.049385 | 0.032598 | -0.030672 |
| 22.22222 | 0.616094 | 0.952331 | 0.346911 | 0.52904 | 0.048919 | 0.686867 | 0.335172 | 0.365979 | 1.483841 | 0.380344 | 1.05298 | 0.341418 | 0.250136 | 0.381423 | 0.074719 | 0.390066 | 0.122113 | -0.014375 | 0.106985 | -0.089561 | 0.325102 | 0.049399 | 0.029004 | 0.022088 |
| 22.23611 | 0.468682 | 1.508626 | 0.747182 | 0.308939 | 0.112702 | 0.946241 | 0.087347 | 0.227804 | 1.481756 | 0.196095 | 1.695725 | 0.137221 | -0.02221 | 0.614659 | -0.095026 | 0.533257 | 0.134743 | -0.010305 | 0.117188 | -0.146488 | 0.400396 | 0.124036 | 0.045951 | 0.657658 |
| 22.25 | 0.524541 | 0.953787 | 0.89806 | 0.406532 | 0.475033 | 1.022628 | 0.310844 | 0.315457 | 1.24703 | 0.212382 | 0.848459 | 0.069203 | -0.083403 | 0.827898 | -0.14528 | 0.438915 | 0.086429 | 0.004286 | 0.102456 | -0.172179 | 0.227624 | 0.155849 | 0.026419 | 1.13443 |
| 22.26389 | 0.496424 | 0.064321 | 0.620841 | 0.67478 | 0.427035 | 0.723041 | 0.500384 | 0.364186 | 1.11246 | 0.240949 | 0.098367 | 0.003165 | 0.166477 | 0.847001 | 0.232205 | -0.133911 | 0.100552 | 0.003911 | 0.081921 | -0.121054 | 0.13752 | 0.127059 | 0.01347 | 0.890971 |
| 22.27778 | 0.328295 | 0.07104 | 0.405896 | 0.799653 | 0.316021 | 0.124699 | 0.423776 | 0.305691 | 1.105498 | 0.242792 | -0.91734 | -0.005051 | 0.29848 | 0.734959 | 0.452953 | -0.151664 | 0.183361 | 0.003621 | 0.085696 | -0.037022 | 0.199009 | 0.055512 | 0.038775 | 0.068104 |
| 22.29167 | 0.299912 | 1.008913 | 0.409553 | 0.801673 | 0.649936 | -0.390373 | 0.367052 | 0.31829 | 1.130577 | 0.322313 | 0.252023 | 0.140889 | 0.151729 | 0.525509 | 0.269881 | 0.269777 | 0.182385 | 0.001266 | 0.082022 | -0.04595 | 0.320941 | -0.002743 | 0.035612 | -0.305805 |
| 22.30556 | 0.424881 | 1.689348 | 0.324605 | 0.710785 | 0.81821 | -0.68917 | 0.310229 | 0.367815 | 0.735272 | 0.38493 | 1.086741 | 0.217769 | -0.093283 | 0.280022 | 0.032873 | 0.390683 | 0.071072 | -0.030615 | 0.035389 | -0.079562 | 0.344908 | -0.044985 | -0.013202 | -0.063895 |
| 22.31944 | 0.538834 | 1.244567 | 0.163259 | 0.51691 | 0.368036 | -0.691999 | 0.222933 | 0.343098 | 0.14942 | 0.470722 | 0.751603 | 0.209221 | -0.262213 | 0.190387 | -0.070957 | 0.025863 | 0.009887 | -0.065604 | 0.001342 | -0.041747 | 0.343794 | -0.0515 | -0.044679 | 0.405297 |
| 22.33333 | 0.453595 | 0.307086 | 0.074251 | 0.366294 | -0.060239 | -0.141513 | 0.183898 | 0.209829 | 0.018472 | 0.542433 | 0.347825 | 0.241261 | -0.183389 | 0.225119 | 0.208679 | -0.115438 | 0.023816 | -0.067828 | -0.007144 | -0.016522 | 0.339757 | -0.014316 | -0.033526 | 0.531797 |
| 22.34722 | 0.179017 | -0.202999 | 0.05353 | 0.327727 | 0.048399 | 0.394203 | 0.142973 | 0.003996 | 0.28412 | 0.350815 | 0.250881 | 0.245455 | 0.079759 | 0.189411 | 0.748126 | -0.050512 | 0.046941 | -0.054634 | -0.025756 | -0.096845 | 0.277916 | 0.020426 | -0.002819 | 0.203839 |
| 22.36111 | -0.079582 | -0.162113 | 0.103225 | 0.37242 | 0.308692 | 0.476621 | 0.060788 | -0.191954 | 0.522123 | -0.001977 | 0.247987 | 0.31706 | 0.272628 | 0.113285 | 0.771855 | -0.203354 | 0.020927 | -0.051858 | -0.052245 | -0.165037 | 0.20988 | -0.018725 | -0.010534 | -0.183668 |
| 22.375 | -0.184859 | 0.411668 | 0.215557 | 0.487817 | 0.084755 | 0.450707 | -0.009435 | -0.238541 | 0.605002 | -0.123798 | 0.455074 | 0.561704 | 0.21894 | 0.301903 | 0.205611 | -0.280742 | -0.013823 | -0.061089 | -0.061497 | -0.095252 | 0.245945 | -0.081297 | -0.045747 | -0.183116 |
| 22.38889 | -0.060407 | 1.074283 | 0.319688 | 0.597832 | -0.26974 | 0.414618 | -0.009325 | -0.103502 | 0.724743 | -0.104919 | 0.64871 | 0.766096 | 0.038132 | 0.718918 | -0.173461 | 0.022186 | 0.022004 | -0.064732 | -0.032154 | 0.043561 | 0.343018 | -0.059341 | -0.044696 | 0.187954 |
| 22.40278 | 0.238102 | 1.222252 | 0.360133 | 0.623983 | -0.033044 | 0.21889 | 0.144925 | 0.032446 | 1.038855 | -0.10314 | 0.677477 | 0.520535 | -0.039208 | 0.839368 | -0.09437 | 0.465322 | 0.103098 | -0.045043 | -0.002676 | 0.094429 | 0.36875 | 0.018712 | -0.007528 | 0.56887 |
| 22.41667 | 0.432357 | 0.660152 | 0.481154 | 0.677593 | 0.6691 | 0.006492 | 0.37788 | -0.006102 | 1.828109 | -0.114814 | 0.708419 | -0.030419 | 0.032975 | 0.581061 | 0.348857 | 0.858694 | 0.169618 | -0.013176 | -0.006008 | 0.0628 | 0.422757 | 0.061128 | 0.019703 | 0.669776 |
| 22.43056 | -0.01272 | 0.103088 | 0.771197 | 0.853642 | 1.027269 | -0.21199 | 0.369419 | -0.143094 | 2.594346 | -0.157556 | 0.859185 | -0.156536 | 0.112463 | 0.553421 | 1.130198 | 1.111641 | 0.210199 | -0.013765 | -0.036728 | -0.031724 | 0.558076 | 0.03901 | 0.010998 | 0.629784 |
| 22.44444 | -0.478387 | -0.006265 | 0.980371 | 1.213071 | 0.771358 | -0.430545 | -0.04058 | -0.250614 | 2.044957 | -0.296665 | 0.975791 | -0.10424 | 0.012011 | 0.66636 | 1.745157 | 0.905263 | 0.232377 | -0.133473 | -0.160634 | -0.165408 | 0.477188 | -0.140311 | -0.110989 | 0.570319 |
| 22.45833 | -0.392362 | 0.137214 | 0.896851 | 1.186455 | 0.314051 | -0.463082 | -0.343575 | -0.335511 | 1.860875 | -0.460642 | 1.021347 | -0.021058 | -0.25363 | 0.695699 | 0.77098 | 0.255313 | 0.20324 | -0.316721 | -0.328584 | -0.303197 | 0.078324 | -0.404503 | -0.29161 | 0.330597 |
| 22.47222 | -0.193346 | 0.306327 | 0.691317 | 0.746614 | 0.308424 | -0.285421 | -0.335907 | -0.34805 | 1.979681 | -0.471465 | 0.894144 | 0.012449 | -0.380029 | 0.467176 | 0.105316 | 0.044518 | 0.153929 | -0.325757 | -0.315053 | -0.394345 | -0.062771 | -0.389348 | -0.305104 | 0.280209 |
| 22.48611 | 0.626667 | 0.560881 | 0.471963 | 0.462143 | 0.9723 | -0.309524 | -0.310833 | -0.451291 | 1.638316 | -0.472424 | 0.26555 | -0.04272 | -0.211766 | 0.555772 | 0.293265 | 0.369728 | 0.225626 | -0.169723 | -0.12691 | -0.378464 | 0.141332 | -0.202393 | -0.127305 | 0.260476 |
| 22.5 | 0.096412 | 0.590591 | 0.239968 | 0.539184 | 1.336905 | -0.455529 | -0.506562 | -0.66544 | 0.984198 | -0.546375 | -0.031604 | -0.021617 | -0.042123 | 0.573942 | -0.102276 | 0.392987 | 0.265747 | -0.040412 | 0.049858 | -0.300323 | 0.431009 | -0.091076 | 0.008371 | 0.240683 |
| 22.51389 | -0.474642 | 0.370441 | 0.143164 | 0.890262 | 1.305813 | -0.495045 | -0.875901 | -0.753775 | 0.425068 | -0.641875 | -0.304278 | -0.002856 | 0.06134 | 0.57887 | -0.338851 | 0.173708 | 0.204725 | 0.019003 | 0.102282 | -0.196124 | 0.779169 | -0.057663 | 0.036367 | 0.283973 |
| 22.52778 | -0.858761 | -0.200471 | -0.12683 | 1.055921 | -0.28155 | -0.895169 | -1.423107 | -0.939776 | -0.299984 | -1.15918 | -0.420223 | -0.505665 | 0.1292 | 0.412744 | 0.615842 | 0.208567 | 0.073113 | 0.042724 | 0.114976 | -0.337239 | 1.00192 | -0.095196 | 0.053278 | 0.385645 |
| 22.54167 | -1.584328 | -1.10215 | -0.982452 | -0.056336 | -1.259132 | -1.854146 | -2.078199 | -1.740498 | -1.377592 | -2.141616 | -1.227993 | -1.505465 | 0.144653 | 0.263684 | 1.32919 | 0.164392 | -0.072588 | 0.009327 | 0.107584 | -0.459477 | 1.102416 | -0.150211 | 0.037688 | 0.685893 |
| 22.55556 | -1.687109 | -1.313147 | -1.202933 | -1.607175 | -2.049288 | -1.995992 | -1.927659 | -1.821349 | -1.694425 | -2.078939 | -1.587055 | -1.701345 | 0.014292 | 0.191435 | 0.501773 | 0.198517 | -0.121165 | -0.066856 | 0.047266 | -0.392044 | 0.754663 | -0.115717 | -0.004251 | 0.602404 |
| 22.56944 | -0.952709 | -0.6272 | -0.257195 | -0.89563 | -1.337785 | -1.060658 | -0.971933 | -0.910266 | -0.945997 | -1.02459 | -0.964903 | -0.962459 | -0.126407 | 0.197831 | 0.055561 | 0.053575 | -0.04022 | -0.086043 | -0.011772 | -0.13494 | 0.301085 | 0.037308 | -0.013849 | 0.534565 |
| 22.58333 | -0.44334 | -0.071148 | 0.556637 | -0.162296 | -0.58739 | -0.4185 | -0.311479 | -0.230187 | -0.295288 | -0.369018 | -0.444868 | -0.393976 | -0.133883 | 0.149965 | -0.075385 | -0.008333 | 0.029371 | -0.053072 | -0.034001 | -0.059444 | 0.168079 | 0.123672 | 0.014284 | 0.423211 |
| 22.59722 | -0.355854 | 0.06583 | 0.64429 | -0.077969 | -0.304671 | -0.300464 | -0.196215 | 0.107145 | -0.128161 | -0.275511 | -0.381663 | -0.261932 | -0.106157 | 0.101942 | -0.193565 | -0.009485 | -0.016222 | -0.062111 | -0.056791 | -0.16006 | 0.180791 | 0.065811 | -0.006508 | 0.176422 |
| 22.61111 | -0.423171 | 0.120384 | 0.539302 | -0.026493 | -0.088366 | -0.385273 | -0.298486 | 0.131847 | -0.110442 | -0.327635 | -0.332264 | -0.337965 | -0.089446 | 0.131762 | -0.148026 | 0.045863 | -0.081113 | -0.07735 | -0.057069 | -0.166616 | 0.172964 | 0.025539 | -0.027684 | -0.016522 |
| 22.625 | -0.285189 | 0.116062 | 0.472919 | 0.215028 | 0.108209 | -0.467825 | -0.338012 | -0.023244 | -0.004976 | -0.2895 | -0.265477 | -0.466005 | -0.046527 | 0.174423 | 0.022538 | 0.055773 | -0.113259 | -0.053003 | -0.023676 | -0.075235 | 0.16099 | 0.009087 | -0.013323 | -0.133154 |
| 22.63889 | -0.216451 | 0.091492 | 0.30241 | 0.421445 | -0.072408 | -0.439086 | -0.308099 | -0.097759 | 0.067041 | -0.232125 | -0.183651 | -0.512419 | -0.04435 | 0.134075 | 0.081295 | 0.025464 | -0.179122 | -0.041075 | -0.006641 | 0.010921 | 0.090241 | -0.021864 | -0.032303 | -0.196101 |
| 22.65278 | -0.149134 | 0.135531 | 0.164215 | 0.525063 | -0.205821 | -0.256713 | -0.207064 | -0.106077 | 0.129784 | -0.148054 | -0.148295 | -0.287224 | -0.075729 | 0.050945 | 0.068544 | -0.028735 | -0.259008 | -0.037232 | 0.00227 | 0.060793 | -0.024528 | -0.034139 | -0.054821 | -0.08345 |
| 22.66667 | 0.008913 | 0.21148 | 0.145176 | 0.392196 | 0.010533 | -0.018044 | -0.030632 | -0.016078 | 0.24539 | -0.015922 | -0.180263 | 0.016988 | -0.040394 | -0.03371 | 0.095926 | -0.004304 | -0.264769 | -0.007018 | 0.02367 | 0.055662 | -0.084955 | -0.022325 | -0.030875 | 0.156298 |
| 22.68056 | 0.154342 | 0.193621 | 0.117221 | 0.092093 | 0.127356 | 0.266326 | 0.086357 | 0.047485 | 0.479525 | 0.041359 | -0.201197 | 0.159008 | 0.01653 | -0.145667 | 0.106965 | 0.040371 | -0.174284 | 0.029684 | 0.024928 | -0.003103 | -0.15625 | -0.020921 | 0.019738 | 0.442106 |
| 22.69444 | 0.144 | 0.095859 | 0.00025 | -0.038295 | -0.021895 | 0.352335 | 0.090213 | 0.015148 | 0.555683 | -0.002953 | -0.237591 | 0.243054 | -0.007853 | -0.239877 | 0.06067 | 0.004443 | 0.001765 | 0.019703 | -0.017215 | -0.059757 | -0.316065 | -0.026926 | 0.028085 | 0.505482 |
| 22.70833 | 0.082722 | 0.05854 | -0.104446 | -0.078689 | -0.13322 | 0.281178 | 0.067466 | -0.017906 | 0.493422 | -0.023188 | -0.298115 | 0.347131 | -0.05523 | -0.208394 | 0.016287 | -0.051979 | 0.274293 | -0.01601 | -0.060965 | -0.063702 | -0.469686 | -0.004603 | -0.026487 | 0.233664 |
| 22.72222 | 0.094375 | 0.108043 | -0.174893 | -0.035294 | -0.172501 | 0.276781 | 0.077903 | -0.001784 | 0.417128 | 0.024996 | -0.255889 | 0.382927 | -0.052175 | -0.089628 | 0.040158 | -0.049346 | 0.504271 | -0.04324 | -0.081335 | -0.025935 | -0.247266 | 0.01335 | -0.04895 | -0.084257 |
| 22.73611 | 0.124084 | 0.160377 | -0.315198 | 0.08718 | -0.073255 | 0.232299 | 0.078559 | 0.000791 | 0.084459 | 0.06017 | -0.097053 | 0.203622 | -0.036382 | 0.004736 | 0.093332 | -0.004356 | 0.531859 | -0.032203 | -0.076481 | 0.060226 | -0.078709 | 0.008231 | -0.061913 | -0.337443 |
| 22.75 | 0.116231 | 0.184632 | -0.428883 | 0.210947 | 0.075922 | 0.128327 | 0.041001 | -0.037151 | -0.566852 | 0.050098 | -0.001118 | 0.036854 | -0.032019 | 0.046753 | 0.116553 | 0.103084 | 0.381311 | 0.027105 | -0.048089 | 0.157119 | 0.116114 | -0.017461 | -0.068446 | -0.486457 |
| 22.76389 | 0.08721 | 0.184174 | -0.331304 | 0.284218 | 0.202093 | 0.03677 | 0.004422 | -0.051399 | -1.052464 | 0.030242 | 0.011483 | 0.08234 | -0.020453 | 0.02174 | 0.086922 | 0.230621 | 0.262741 | 0.071816 | -0.031673 | 0.190073 | 0.303912 | -0.047322 | -0.050927 | -0.625251 |
| 22.77778 | 0.062053 | 0.119006 | 0.03557 | 0.392592 | 0.113838 | 0.02387 | 0.014538 | -0.073654 | -0.738608 | 0.031671 | -0.007924 | 0.221388 | 0.012716 | -0.02309 | 0.042259 | 0.344233 | 0.158392 | 0.047895 | -0.026566 | 0.148084 | 0.297154 | -0.074324 | -0.062716 | -0.547961 |
| 22.79167 | 0.04795 | -0.122613 | 0.494494 | 0.553023 | -0.079248 | 0.096102 | 0.057476 | -0.04519 | -0.276498 | 0.057172 | -0.069516 | 0.310774 | -0.007809 | -0.083021 | 0.025661 | 0.254794 | 0.030552 | -0.001245 | -0.02756 | 0.127289 | 0.043924 | -0.093985 | -0.040045 | -0.212263 |
| 22.80556 | 0.04011 | -0.495136 | 0.714041 | 0.506945 | -0.004055 | 0.062524 | 0.09287 | 0.003351 | 0.324718 | 0.075863 | -0.174184 | 0.41472 | -0.067386 | -0.137918 | 0.016624 | 0.147245 | -0.130801 | -0.049784 | -0.044724 | 0.036006 | -0.194294 | -0.070258 | -0.010636 | 0.214435 |
| 22.81944 | 0.000365 | -0.579992 | 0.457289 | -0.00891 | 0.084298 | 0.037874 | 0.12868 | 0.076203 | 1.082169 | 0.086581 | -0.231846 | 0.390208 | -0.077534 | -0.158476 | -0.015993 | 0.0624 | -0.35094 | -0.058178 | -0.062757 | -0.096551 | -0.407419 | -0.055401 | 0.001539 | 0.409229 |
| 22.83333 | -0.04487 | -0.375767 | -0.052972 | -0.145804 | -0.119123 | -0.040688 | 0.143993 | 0.073802 | 0.728721 | 0.07307 | -0.218731 | 0.121566 | -0.034204 | -0.177816 | -0.106635 | 0.087927 | -0.554579 | -0.031859 | -0.058213 | -0.153332 | -0.575147 | -0.02083 | -0.002103 | 0.314169 |
| 22.84722 | -0.13818 | -0.355942 | -0.575762 | 0.073801 | -0.255037 | -0.139758 | 0.07239 | -0.02408 | -0.139495 | -0.002555 | -0.308528 | -0.414248 | -0.008733 | -0.110623 | -0.288023 | -0.005951 | -0.544054 | -0.010884 | -0.066427 | -0.056082 | -0.655615 | -0.008387 | -0.02867 | 0.028269 |
| 22.86111 | -0.248982 | -0.50505 | -0.891439 | -1.309581 | -0.286668 | -0.263401 | -0.138819 | -0.155958 | -0.99976 | -0.131442 | -0.370758 | -0.808041 | -0.031955 | -0.036031 | -0.457696 | 0.082867 | -0.45975 | -0.04015 | -0.107857 | 0.142063 | -0.533714 | -0.026015 | -0.075098 | -0.335383 |
| 22.875 | -0.342609 | -0.672974 | -0.903686 | -1.743264 | -0.578161 | -0.358428 | -0.365327 | -0.246803 | -0.832563 | -0.247805 | -0.449118 | -0.725602 | -0.106835 | -0.099108 | -0.481517 | 0.198934 | -0.404889 | -0.078687 | -0.141424 | 0.26268 | -0.28009 | -0.062764 | -0.127781 | -0.630284 |
| 22.88889 | -0.37315 | -0.812728 | -0.823051 | -1.582445 | -0.871922 | -0.34871 | -0.415042 | -0.270944 | -0.569466 | -0.295478 | -0.493778 | -0.494818 | -0.168581 | -0.361158 | -0.22491 | 0.079911 | -0.27959 | -0.082797 | -0.136635 | 0.188845 | -0.112231 | -0.084459 | -0.189117 | -0.690346 |
| 22.90278 | -0.372298 | -0.928176 | -0.768878 | -1.051768 | -0.941707 | -0.285217 | -0.382611 | -0.216469 | -1.18675 | -0.276561 | -0.675779 | -0.459621 | -0.18096 | -0.686357 | -0.048841 | -0.109535 | -0.11773 | -0.083712 | -0.12128 | -0.002191 | -0.15244 | -0.107742 | -0.238232 | -0.783237 |
| 22.91667 | -0.35473 | -1.066047 | -0.689076 | -0.702594 | -0.869514 | -0.225334 | -0.36515 | -0.146031 | -0.975704 | -0.210448 | -0.737191 | -0.637031 | -0.110081 | -0.99894 | -0.064203 | -0.330971 | -0.102563 | -0.069199 | -0.119985 | -0.073658 | -0.358827 | -0.062941 | -0.178052 | -0.866905 |
| 22.93056 | -0.308626 | -1.202184 | -0.639351 | -0.524397 | -0.728561 | -0.190708 | -0.333201 | -0.041683 | -0.674609 | -0.119658 | -0.62219 | -0.799028 | 0.022654 | -1.518022 | -0.175211 | -0.701418 | -0.19447 | -0.057769 | -0.147271 | 0.05415 | -0.53354 | -0.005212 | -0.078739 | -0.544882 |
| 22.94444 | -0.296635 | -1.181848 | -0.590297 | -0.829744 | -0.741192 | -0.189064 | -0.309958 | -0.016771 | -0.563812 | -0.06359 | -0.527049 | -0.788609 | 0.120667 | -1.845474 | -0.422518 | -0.892468 | -0.237069 | -0.016883 | -0.18038 | 0.057941 | -0.537498 | 0.020433 | 0.002939 | -0.492012 |
| 22.95833 | -0.423093 | -0.970848 | -0.730337 | -1.117319 | -1.029996 | -0.215168 | -0.328371 | -0.155565 | -0.860466 | -0.096347 | -0.527971 | -0.746989 | 0.076596 | -1.856012 | -0.756449 | -1.164209 | -0.216475 | -0.023833 | -0.201256 | -0.050306 | -0.419795 | -0.01893 | -0.028315 | -0.487845 |
| 22.97222 | -0.606056 | -0.815745 | -0.884118 | -1.244746 | -1.528807 | -0.245539 | -0.424794 | -0.131953 | -1.287317 | -0.154143 | -0.492699 | -0.688081 | -0.042943 | -1.757648 | -0.831687 | -0.875239 | -0.224465 | -0.058124 | -0.181029 | -0.073444 | -0.278606 | -0.049533 | -0.096425 | -0.536818 |
| 22.98611 | -0.78215 | -0.789144 | -0.922802 | -1.182936 | -1.674881 | -0.125997 | -0.424305 | -0.145357 | -1.619024 | -0.124493 | -0.439364 | -0.497672 | -0.098782 | -1.217663 | -0.713013 | -0.600361 | -0.199119 | -0.045312 | -0.133335 | -0.060876 | -0.137748 | -0.054999 | -0.094049 | -0.508832 |
| 23 | -0.702692 | -0.634959 | -0.643683 | -0.447708 | -1.188871 | 0.100086 | -0.176948 | -0.143064 | -2.036689 | 0.051601 | -0.26912 | -0.215525 | -0.042007 | -0.52708 | -0.426622 | -0.181983 | -0.057181 | 0.074022 | 0.009303 | -0.01111 | -0.003646 | 0.043599 | 0.025113 | -0.393454 |
| 23.01389 | -0.411184 | -0.21422 | -0.150198 | 0.400821 | -0.269405 | 0.292868 | 0.115317 | -0.027175 | -1.845722 | 0.340552 | 0.081267 | 0.193699 | 0.078424 | 0.07699 | -0.084712 | 0.283955 | 0.159827 | 0.265935 | 0.221118 | 0.165011 | 0.155535 | 0.249336 | 0.221437 | -0.166614 |
| 23.02778 | -0.194503 | 0.089681 | -0.087955 | 0.313749 | 0.409195 | 0.318573 | 0.271248 | 0.046749 | -0.574449 | 0.567108 | 0.289402 | 0.608384 | 0.060783 | 0.214467 | 0.099258 | 0.354905 | 0.200602 | 0.322323 | 0.326666 | 0.29242 | 0.113649 | 0.40976 | 0.297687 | -0.016238 |
| 23.04167 | 0.00968 | 0.066499 | -0.561378 | -0.172439 | 0.657842 | 0.069193 | 0.274863 | 0.049365 | -0.310456 | 0.621127 | 0.061016 | 0.845088 | -0.195983 | -0.467872 | -0.107282 | -0.040317 | 0.023137 | 0.149834 | 0.178765 | 0.291115 | -0.24652 | 0.275418 | 0.140579 | -0.412279 |
| 23.05556 | 0.612957 | 0.278313 | -0.931007 | -0.801584 | 0.965278 | -0.359887 | 0.333878 | 0.120349 | -1.222057 | 0.743802 | -0.495647 | 1.230906 | -0.417852 | -1.618485 | -0.282315 | 0.045728 | 0.048308 | 0.002269 | -0.004642 | 0.244929 | -0.675412 | 0.065541 | -0.090732 | -1.241478 |
| 23.06944 | 1.109308 | 1.623691 | -0.023339 | -1.477012 | 1.606717 | -0.431475 | 0.384626 | 0.203152 | -2.449767 | 0.794625 | -0.331551 | 1.991387 | -0.222631 | -2.082858 | -0.23719 | 0.548237 | 0.420024 | 0.025628 | -0.013073 | 0.15591 | -0.979755 | 0.128762 | -0.108957 | -1.211791 |
| 23.08333 | 0.763166 | 5.432768 | 4.588965 | -1.677626 | 2.648837 | 0.171403 | 0.19747 | 0.591012 | -3.495019 | 0.634624 | 0.012002 | 3.481696 | 0.374316 | -0.906134 | -0.109115 | 1.10066 | 0.640744 | 0.076719 | 0.082915 | -0.02982 | -1.192986 | 0.661675 | 0.071808 | -0.582992 |
| 23.09722 | -0.458179 | 3.600446 | 3.806553 | -0.716724 | 1.881972 | 0.58401 | 0.010382 | 0.294078 | -3.536539 | 0.416582 | -0.064874 | 3.072488 | 0.33377 | 0.212042 | -0.165276 | -0.28654 | 0.18605 | -0.0653 | 0.091162 | 0.096559 | -0.842434 | 0.708449 | 0.16751 | -0.076939 |
| 23.11111 | 0.373459 | 0.13173 | -0.687562 | -0.03161 | -0.040526 | 0.924986 | 0.183241 | 0.286816 | 0.053266 | 0.242098 | 0.036739 | 0.507725 | 0.08945 | 0.848488 | -0.023886 | -0.381508 | -0.24978 | -0.03699 | 0.003043 | 0.043128 | -0.242286 | 0.227628 | 0.158355 | 0.305795 |
| 23.125 | 0.404665 | 0.283819 | -1.371821 | 0.083179 | -0.670635 | 1.128049 | 0.273575 | 0.856854 | 3.061291 | 0.130575 | 0.199937 | -0.926464 | 0.132348 | 0.815205 | 0.115643 | -0.28886 | -0.36522 | 0.012759 | -0.053869 | -0.032965 | 0.21721 | 0.072127 | 0.092161 | 0.462031 |
| 23.13889 | 0.862695 | 0.916345 | 0.132799 | -0.193477 | -0.679498 | 1.521688 | 0.148235 | 0.904514 | 4.906491 | 0.074758 | 0.233429 | -1.328386 | 0.048936 | 0.327452 | -0.059759 | -0.614843 | -0.368373 | -0.082324 | -0.070119 | -0.236914 | 0.4742 | 0.046081 | -0.013642 | 0.340788 |
| 23.15278 | 1.246735 | 0.796523 | 0.080932 | -0.547248 | -0.768635 | 1.17785 | -0.161534 | 0.317763 | 5.044277 | 0.021113 | 0.089483 | -0.660958 | -0.328531 | 0.005264 | -0.449485 | -1.131835 | -0.235472 | -0.240405 | -0.074397 | -0.297799 | 0.484382 | 0.007167 | -0.121309 | 0.196647 |
| 23.16667 | 1.609837 | 0.338133 | -0.940426 | -0.684585 | -0.704904 | 0.392562 | -0.256531 | -0.115844 | 3.502842 | -0.014461 | 0.039269 | 0.302265 | -0.451017 | 0.164296 | -0.60768 | -1.166364 | 0.041369 | -0.260718 | -0.009176 | -0.071934 | 0.335022 | 0.010968 | -0.092683 | 0.105562 |
| 23.18056 | 1.329794 | 0.094122 | -0.56145 | -0.085979 | 0.050208 | 0.015719 | 0.129459 | 0.166563 | 1.060839 | -0.003678 | 0.381036 | 0.907255 | -0.117954 | 0.788319 | -0.481259 | -0.522503 | 0.347233 | -0.173867 | 0.072992 | 0.22166 | 0.191318 | 0.071546 | 0.038341 | 0.116052 |
| 23.19444 | 0.316594 | 0.11728 | -0.682758 | 0.211991 | 1.115612 | 0.016931 | 0.438842 | 0.964307 | -0.771571 | -0.056547 | 0.657451 | 0.854227 | 0.227067 | 1.507658 | -0.12204 | 0.245352 | 0.479572 | -0.024217 | 0.099114 | 0.257665 | 0.165337 | 0.148838 | 0.095236 | 0.252578 |
| 23.20833 | -0.458174 | -0.107301 | -1.362792 | -0.104461 | 0.668235 | -0.222812 | 0.008623 | 0.819405 | -1.004906 | -0.438769 | 0.255835 | 0.142514 | 0.399068 | 1.541592 | 0.230522 | 1.144685 | 0.373114 | 0.071036 | 0.104346 | 0.163819 | 0.266356 | 0.209151 | 0.03496 | 0.588426 |
| 23.22222 | 0.018197 | -0.638144 | -1.601817 | -0.371571 | -0.753065 | -0.913133 | -0.737884 | -0.336001 | 0.155261 | -1.016677 | -0.626181 | -1.331163 | 0.282544 | 0.951221 | 0.314018 | 1.689402 | 0.158827 | -0.010797 | 0.06264 | 0.18546 | 0.469774 | 0.228963 | -0.00325 | 0.944772 |
| 23.23611 | 1.445823 | -0.909712 | -1.047781 | -0.054959 | -0.818621 | -1.46341 | -0.681734 | -1.233163 | 1.813141 | -1.013703 | -0.495416 | -3.975116 | -0.015001 | 0.352408 | 0.176818 | 1.398164 | -0.011338 | -0.096601 | -0.020359 | 0.265496 | 0.689489 | 0.155592 | 0.018072 | 1.013548 |
| 23.25 | 1.305927 | -1.05371 | -0.105919 | 0.263708 | 0.677055 | -0.539126 | 0.083131 | -2.094118 | 3.509438 | -0.460796 | 0.21962 | -4.152734 | -0.099738 | 0.162527 | 0.183448 | 0.850788 | 0.058783 | -0.101253 | -0.03803 | 0.192725 | 0.633388 | 0.091602 | 0.025008 | 0.530585 |
| 23.26389 | 0.283906 | -1.609002 | 0.390073 | 0.191573 | 1.413787 | 0.295921 | 0.439792 | -1.619484 | 2.455026 | -0.094025 | 0.319749 | -2.616987 | -0.015976 | 0.314375 | 0.236679 | 0.127505 | 0.245773 | -0.008588 | 0.043563 | 0.020849 | 0.360485 | 0.051865 | 0.011235 | -0.031056 |
| 23.27778 | -0.390062 | -2.256282 | 0.519069 | 0.243962 | 0.496801 | 0.536665 | 0.207703 | -0.841456 | 0.034194 | 0.03859 | -0.167126 | -1.482985 | -0.012658 | 0.375813 | 0.230668 | -0.519269 | 0.214297 | -0.008521 | 0.092673 | -0.007777 | 0.231332 | -0.009108 | 0.02583 | -0.178109 |
| 23.29167 | -0.346071 | -2.009973 | 0.452219 | 0.37346 | -0.117278 | 0.405731 | -0.170891 | -0.538177 | -1.324224 | 0.033359 | -0.141747 | 0.175094 | -0.09037 | 0.237083 | 0.068695 | -0.558354 | 0.035136 | -0.082729 | 0.038395 | 0.027348 | 0.082457 | -0.134981 | 0.014322 | 0.043122 |
| 23.30556 | -0.489219 | -1.850413 | -0.344727 | -0.300547 | -0.497655 | -0.219044 | -0.764182 | -0.869462 | -1.741161 | -0.643421 | -0.478248 | 0.356883 | -0.116579 | 0.248221 | -0.347294 | -0.413297 | -0.029576 | -0.130201 | -0.013627 | 0.049814 | -0.128384 | -0.263628 | -0.12561 | 0.304991 |
| 23.31944 | -1.051009 | -2.54298 | -1.43427 | -1.731651 | -1.199394 | -1.099976 | -1.533881 | -1.610217 | -1.436504 | -1.897178 | -1.582202 | -0.981965 | -0.046506 | 0.677004 | -0.428281 | -0.257389 | 0.031482 | -0.072804 | 0.057096 | 0.08463 | -0.080917 | -0.185399 | -0.218636 | 0.444248 |
| 23.33333 | -0.710969 | -2.403005 | -0.78426 | -1.547489 | -1.418898 | -0.799265 | -1.182125 | -1.37243 | 0.184423 | -1.647032 | -1.72956 | -1.102978 | 0.056897 | 1.244802 | 0.03714 | 0.225577 | 0.12448 | 0.063367 | 0.171842 | 0.120761 | 0.338485 | 0.18031 | -0.006728 | 0.461615 |
| 23.34722 | 0.211715 | -1.07536 | 0.908058 | -0.044974 | -0.715083 | 0.025931 | -0.135382 | -0.550877 | 1.794188 | -0.368906 | -0.027432 | 0.058955 | -0.001808 | 1.201215 | 0.438264 | 0.616148 | 0.158395 | 0.075928 | 0.144802 | 0.096693 | 0.792341 | 0.501927 | 0.206006 | 0.346238 |
| 23.36111 | 0.379988 | -0.222894 | 1.42311 | 0.56894 | 0.088854 | -0.186776 | 0.146784 | -0.076349 | 1.315784 | 0.122877 | 1.131988 | 0.182864 | -0.142785 | 0.871404 | 0.386186 | 0.474974 | 0.087692 | -0.027865 | 0.006552 | 0.07019 | 1.15543 | 0.376138 | 0.16019 | 0.191309 |
| 23.375 | 0.156915 | 0.194641 | 1.205524 | 0.674812 | 0.87495 | -0.556464 | 0.058881 | 0.222158 | 0.598452 | 0.086623 | 1.280781 | -0.465781 | -0.107774 | 0.824863 | 0.245305 | 0.317931 | 0.033942 | -0.073269 | -0.005677 | 0.16941 | 1.051815 | 0.176664 | 0.075539 | 0.418944 |
| 23.38889 | 0.118752 | 0.285292 | 0.859193 | 0.734563 | 1.186569 | -0.849971 | 0.140647 | 0.287773 | 0.367549 | -0.012538 | 0.870346 | -0.330453 | 0.046465 | 0.651719 | 0.283723 | 0.194827 | 0.080985 | -0.021851 | 0.057306 | 0.424681 | 0.5491 | 0.088003 | 0.053511 | 0.532117 |
| 23.40278 | 0.260167 | -0.008843 | 0.52618 | 0.977604 | 1.167199 | -0.570595 | 0.281869 | 0.091961 | 0.347248 | -0.057647 | 0.249068 | 0.545126 | 0.080358 | 0.458888 | 0.434491 | 0.054188 | 0.200597 | 0.01188 | 0.055118 | 0.332615 | 0.349985 | 0.094793 | 0.057773 | 0.322037 |
| 23.41667 | 0.422083 | -0.284276 | 0.345097 | 1.183179 | 1.00551 | -0.043921 | 0.307098 | 0.057343 | 0.741101 | -0.022061 | 0.150592 | 0.658635 | 0.038656 | 0.553115 | 0.569048 | -0.030585 | 0.289738 | 0.020444 | 0.007209 | 0.120762 | 0.244717 | 0.115808 | 0.056331 | 0.137411 |
| 23.43056 | 0.436606 | -0.155964 | 0.53052 | 1.036341 | 0.718634 | 0.599709 | 0.256549 | 0.162289 | 1.572198 | -0.032503 | 0.551567 | 0.468404 | 0.055588 | 0.750989 | 0.54563 | -0.083862 | 0.289361 | 0.046622 | -0.033466 | -0.098059 | 0.171871 | 0.07023 | 0.026212 | 0.163741 |
| 23.44444 | 0.432352 | 0.079001 | 0.852516 | 0.65814 | -0.020699 | 0.749041 | 0.209613 | 0.127741 | 2.739534 | -0.150892 | 1.029129 | 0.520782 | 0.044833 | 0.786746 | 0.374089 | -0.20822 | 0.297728 | 0.018933 | -0.119182 | -0.262057 | 0.091628 | -0.086548 | -0.074228 | 0.251995 |
| 23.45833 | 0.409358 | 0.105559 | 0.868361 | 0.300252 | -0.051224 | 0.598927 | 0.054735 | -0.14602 | 3.572128 | -0.346289 | 1.120126 | 0.690266 | -0.107043 | 0.648761 | 0.109568 | -0.471466 | 0.340947 | -0.102492 | -0.279014 | -0.359638 | -0.015643 | -0.309496 | -0.237809 | 0.278749 |
| 23.47222 | 0.447753 | -0.037647 | 0.616465 | -0.019581 | 1.298333 | 0.535767 | -0.11372 | -0.237019 | 2.98804 | -0.435339 | 0.693096 | 0.844877 | -0.241412 | 0.483165 | 0.157396 | -0.501631 | 0.362762 | -0.107272 | -0.202979 | -0.248316 | 0.002242 | -0.272494 | -0.241128 | 0.545299 |
| 23.48611 | 0.592682 | -0.040149 | 0.764633 | 0.629612 | 2.508174 | 0.298211 | -0.03576 | -0.225731 | 2.357976 | -0.312491 | 0.564323 | 0.828755 | -0.131948 | 0.330561 | 0.819983 | -0.17021 | 0.414676 | 0.114787 | 0.038853 | -0.106944 | 0.241646 | -0.029768 | -0.060075 | 0.751131 |
| 23.5 | 0.561937 | 0.094391 | 0.970576 | 1.148553 | 2.894694 | 0.204793 | 0.054074 | -0.326652 | 1.841788 | -0.239113 | 0.813528 | 0.569998 | 0.001279 | 0.462785 | 1.41853 | 0.079693 | 0.451244 | 0.235712 | 0.173995 | -0.15897 | 0.42811 | 0.097351 | 0.102418 | 0.738545 |
| 23.51389 | 0.419549 | 0.337577 | 0.887525 | 0.727568 | 1.189812 | -0.008845 | 0.003067 | -0.336712 | 1.300606 | -0.27748 | 1.184073 | 0.450381 | 0.095407 | 0.68504 | 1.36979 | 0.200834 | 0.426558 | 0.227988 | 0.143928 | -0.366759 | 0.699768 | 0.119617 | 0.204477 | 0.552622 |
| 23.52778 | 0.237579 | 0.527327 | 0.473816 | 0.064974 | 0.829541 | -0.410075 | -0.334008 | -0.531881 | 0.691558 | -0.580075 | 1.012698 | 0.612757 | 0.23167 | 0.657805 | 1.037639 | 0.362998 | 0.376095 | 0.190352 | 0.141579 | -0.50002 | 0.959698 | 0.050246 | 0.210989 | 0.618188 |
| 23.54167 | -0.169871 | -0.026785 | 0.06977 | -0.637235 | 0.204958 | -1.061718 | -0.891658 | -1.066486 | -0.590492 | -1.206415 | 0.077994 | 0.57545 | 0.38355 | 0.486802 | 0.71747 | 0.431508 | 0.249116 | 0.078029 | 0.128532 | -0.618337 | 1.049131 | -0.018431 | 0.197071 | 0.766976 |
| 23.55556 | -0.461778 | -0.804723 | -0.115909 | -1.153809 | -1.896296 | -1.411823 | -0.899185 | -1.238155 | -2.098299 | -1.506811 | -0.777725 | -0.091044 | 0.30233 | 0.230404 | 0.474667 | 0.674153 | 0.110798 | 0.010978 | 0.075116 | -0.388245 | 0.706068 | 0.097455 | 0.132069 | 0.714575 |
| 23.56944 | -0.543762 | -0.754689 | 0.584634 | -0.811029 | -1.532387 | -1.093711 | -0.787891 | -0.534194 | -1.05213 | -1.130078 | -0.918545 | -0.617462 | 0.090373 | 0.026035 | 0.214852 | 0.312458 | 0.066669 | 0.01797 | 0.020674 | 0.119016 | 0.476957 | 0.359297 | 0.065913 | 0.612053 |
| 23.58333 | -0.225177 | -0.259842 | 1.465851 | -0.348335 | -0.017759 | -0.582256 | -0.488453 | 0.551122 | -0.922186 | -0.599171 | -0.547519 | -0.485336 | 0.052389 | -0.024238 | 0.193563 | 0.010211 | 0.046675 | 0.083237 | 0.033771 | 0.43481 | 0.430066 | 0.419485 | 0.096337 | 0.27409 |
| 23.59722 | 0.011861 | 0.165138 | 1.295519 | 0.322265 | 0.930551 | -0.235144 | -0.19698 | 0.99811 | 0.440143 | -0.276565 | -0.34219 | -0.232915 | 0.123155 | 0.109187 | 0.278216 | 0.085764 | 0.004861 | 0.086464 | 0.041134 | 0.417949 | 0.360937 | 0.329982 | 0.112862 | 0.054182 |
| 23.61111 | -0.029539 | 0.258765 | 0.916454 | 0.628268 | 0.826045 | -0.12522 | -0.140243 | 0.498199 | 0.876146 | -0.231026 | -0.190173 | -0.223464 | 0.115304 | 0.099119 | 0.318608 | -0.225357 | -0.100503 | 0.014639 | 0.005311 | 0.189331 | 0.238846 | 0.227245 | 0.066362 | 0.056694 |
| 23.625 | -0.117662 | 0.174353 | 0.580516 | 0.394182 | 0.044392 | -0.10037 | -0.160988 | -0.034166 | 1.093809 | -0.233976 | -0.000575 | -0.279901 | -0.032224 | 0.132745 | 0.298068 | -0.350936 | -0.232878 | -0.051427 | -0.024342 | 0.010286 | 0.085364 | 0.106835 | 0.01821 | -0.064547 |
| 23.63889 | -0.014131 | 0.194111 | 0.490722 | 0.222351 | -0.486087 | 0.026053 | -0.057903 | -0.11128 | 0.601387 | -0.103824 | -0.056918 | -0.145631 | -0.178538 | 0.280476 | 0.27024 | -0.228816 | -0.259659 | -0.043354 | -0.027944 | -0.102697 | -0.060292 | 0.039949 | 0.014157 | 0.204485 |
| 23.65278 | 0.112731 | 0.204096 | 0.409231 | 0.048083 | -0.088787 | 0.15531 | -0.029592 | -0.065422 | 0.412031 | -0.026568 | -0.051019 | 0.066717 | -0.149276 | 0.35026 | 0.209738 | -0.129095 | -0.089662 | 0.038226 | -0.006214 | -0.165705 | -0.161282 | 0.034069 | 0.010206 | 0.339196 |
| 23.66667 | 0.05297 | 0.048882 | 0.202442 | -0.138386 | 0.309571 | 0.101137 | -0.130974 | -0.144459 | -0.685722 | -0.119588 | -0.073061 | 0.170006 | 0.006182 | 0.364712 | 0.10962 | -0.07959 | 0.102274 | 0.037395 | 0.023599 | -0.172688 | -0.161681 | 0.061461 | -0.012837 | 0.362638 |
| 23.68056 | -0.071894 | -0.099905 | -0.033842 | -0.00607 | -0.130657 | -0.017941 | -0.236554 | -0.268757 | -2.938225 | -0.213863 | -0.163079 | 0.209353 | 0.160673 | 0.215522 | 0.01047 | -0.020854 | 0.285389 | 0.001549 | 0.054656 | -0.108657 | -0.046771 | 0.080332 | 0.04949 | 0.225357 |
| 23.69444 | -0.101265 | -0.007819 | -0.138082 | 0.065184 | -0.704085 | -0.075118 | -0.218553 | -0.251689 | -2.557095 | -0.176942 | -0.26274 | 0.285997 | 0.27187 | 0.058074 | -0.009697 | 0.080264 | 0.395562 | 0.017166 | 0.093945 | -0.025089 | 0.037656 | 0.095467 | 0.053492 | 0.295526 |
| 23.70833 | -0.084927 | 0.255418 | -0.205804 | 0.067783 | -0.393089 | -0.037155 | -0.12434 | -0.145683 | -1.719992 | -0.083937 | -0.345696 | 0.343402 | 0.23163 | -0.135306 | 0.038517 | 0.245092 | 0.397673 | 0.077509 | 0.121657 | 0.017163 | 0.020853 | 0.096418 | 0.054704 | 0.116457 |
| 23.72222 | -0.108143 | 0.474325 | -0.379968 | -0.227004 | 0.242406 | -0.001854 | -0.030742 | -0.149627 | -0.438802 | -0.041076 | -0.378953 | 0.231967 | 0.135964 | -0.267399 | 0.102674 | 0.343139 | 0.27993 | 0.103721 | 0.10581 | 0.02122 | -0.057826 | 0.097006 | -0.016233 | -0.042493 |
| 23.73611 | -0.194981 | 0.497971 | -0.687616 | -0.227522 | 0.368024 | -0.02638 | -0.000651 | -0.299967 | 0.924344 | -0.053586 | -0.318945 | 0.061819 | 0.00367 | -0.306875 | 0.17556 | 0.227992 | 0.145993 | 0.092431 | 0.064434 | 0.020709 | -0.165729 | 0.092516 | -0.080937 | -0.184289 |
| 23.75 | -0.237181 | 0.410406 | -0.910719 | 0.211131 | 0.407211 | -0.032112 | 0.021508 | -0.443772 | 2.298119 | -0.033946 | -0.153925 | 0.14611 | -0.11835 | -0.268645 | 0.231968 | 0.10737 | 0.073381 | 0.057451 | 0.025028 | 0.022002 | -0.254546 | 0.073031 | -0.074427 | -0.21134 |
| 23.76389 | -0.141031 | 0.317578 | -0.851339 | 0.214075 | 0.417018 | 0.009485 | 0.026568 | -0.397251 | 2.534885 | 0.078734 | 0.117426 | 0.550343 | -0.227425 | -0.158654 | 0.251873 | 0.022725 | 0.025205 | 0.02493 | 0.010435 | 0.081299 | -0.306811 | 0.058757 | -0.046214 | -0.241356 |
| 23.77778 | -0.088567 | 0.264968 | -0.730215 | 0.446391 | 0.164292 | -0.059275 | 0.122693 | -0.279258 | 1.526487 | 0.154393 | 0.263385 | 0.866681 | -0.181564 | -0.048698 | 0.240304 | -0.005598 | -0.035158 | 0.002865 | 0.022255 | 0.136915 | -0.279835 | 0.027951 | -0.023353 | -0.264173 |
| 23.79167 | -0.08282 | 0.123683 | -0.551294 | 0.570013 | -0.136417 | -0.19316 | 0.211569 | -0.136051 | 0.49464 | 0.128999 | 0.241743 | 0.77755 | -0.212803 | 0.013007 | 0.172201 | 0.041258 | -0.177686 | -0.003857 | 0.039686 | 0.098382 | -0.220246 | -0.031853 | 0.023413 | -0.332749 |
| 23.80556 | 0.001139 | 0.203767 | -0.264154 | 0.292186 | -0.58932 | -0.164344 | 0.30505 | 0.052961 | 0.058007 | 0.15514 | 0.197633 | 0.629012 | -0.120367 | 0.016617 | 0.059699 | 0.150806 | -0.319177 | 0.001646 | 0.049905 | -0.028848 | -0.223097 | -0.095718 | 0.001703 | -0.315433 |
| 23.81944 | 0.161511 | 0.048969 | -0.058322 | 0.36236 | -1.171891 | 0.026223 | 0.411771 | 0.256394 | -0.235418 | 0.271999 | 0.18335 | 0.632961 | -0.097176 | 0.00551 | -0.059039 | 0.195598 | -0.401738 | 0.001129 | 0.048221 | -0.163769 | -0.322233 | -0.103712 | -0.013602 | -0.087694 |
| 23.83333 | 0.257336 | -0.223731 | -0.104056 | 0.522594 | -0.694036 | 0.114642 | 0.428542 | 0.299599 | -0.654991 | 0.36695 | 0.164425 | 0.622614 | 0.006042 | -0.081506 | -0.070226 | 0.150959 | -0.491872 | -0.013629 | 0.026249 | -0.230994 | -0.399171 | -0.078199 | -0.013105 | 0.137559 |
| 23.84722 | 0.175528 | -0.203656 | -0.281457 | 0.180964 | 0.391928 | 0.142807 | 0.377743 | 0.166143 | -1.010961 | 0.399075 | 0.130792 | 0.542728 | 0.116663 | -0.20487 | 0.036843 | 0.148764 | -0.221769 | -0.060257 | -0.015223 | -0.225261 | -0.377408 | -0.04218 | -0.025017 | 0.183904 |
| 23.86111 | -0.089617 | 0.004662 | -0.517284 | 0.001337 | 1.32021 | 0.219251 | 0.357938 | 0.045455 | -0.950778 | 0.394098 | 0.050204 | 0.438487 | 0.171186 | -0.320992 | 0.071992 | 0.209853 | 0.186316 | -0.09413 | -0.037411 | -0.198805 | -0.276871 | -0.012845 | -0.038041 | 0.115396 |
| 23.875 | -0.488107 | 0.154074 | -0.742597 | -0.270676 | 1.282966 | 0.335261 | 0.268246 | 0.090309 | -0.881347 | 0.355989 | -0.036943 | -0.966117 | 0.16051 | -0.41424 | -0.061764 | 0.180727 | 0.312581 | -0.088091 | -0.037244 | -0.184125 | -0.171532 | 0.004675 | -0.059028 | -0.080446 |
| 23.88889 | -0.832894 | -0.125502 | -0.855635 | -0.064781 | 0.309773 | 0.344241 | 0.1672 | 0.131519 | -0.793609 | 0.318896 | -0.022087 | -0.633508 | 0.063978 | -0.468966 | -0.305274 | -0.067977 | 0.063875 | -0.086553 | -0.065031 | -0.18532 | -0.137557 | -0.037697 | -0.108184 | -0.430799 |
| 23.90278 | -0.935874 | -0.466251 | -0.857213 | -0.198417 | -0.659952 | 0.252182 | 0.070275 | 0.143064 | -0.96195 | 0.304979 | -0.035615 | 0.707781 | -0.083454 | -0.62229 | -0.500252 | -0.195536 | -0.24214 | -0.086649 | -0.090967 | -0.201274 | -0.230372 | -0.137547 | -0.157543 | -0.814661 |
| 23.91667 | -0.810205 | -0.488298 | -0.936136 | -0.493709 | -1.224049 | 0.065194 | 0.200689 | 0.225605 | -1.243045 | 0.283055 | -0.128924 | 2.011634 | -0.174951 | -0.89166 | -0.500459 | -0.241609 | -0.668371 | -0.142924 | -0.119371 | -0.213984 | -0.442088 | -0.232089 | -0.173718 | -0.947088 |
| 23.93056 | -0.651822 | -0.475973 | -1.049106 | -0.580795 | -2.223601 | -0.08039 | 0.255053 | 0.275571 | -1.491024 | 0.222303 | -0.25981 | 2.090513 | -0.222019 | -1.136411 | -0.340959 | -0.391565 | -1.100515 | -0.230993 | -0.156408 | -0.268151 | -0.505162 | -0.357296 | -0.143331 | -0.955439 |
| 23.94444 | -0.627499 | -0.597219 | -1.148755 | -0.524446 | -3.327765 | -0.225719 | 0.178559 | 0.272583 | -1.899579 | 0.161159 | -0.418045 | 0.152963 | -0.16636 | -1.374654 | -0.230482 | -0.597218 | -1.040532 | -0.289275 | -0.204574 | -0.257046 | -0.378126 | -0.351487 | -0.079885 | -0.785819 |
| 23.95833 | -0.673029 | -0.618585 | -1.554301 | -0.291601 | -3.322229 | -0.378664 | 0.124013 | 0.17203 | -2.964255 | 0.137321 | -0.49752 | -1.055092 | -0.141793 | -1.457374 | -0.369567 | -0.659081 | -0.722313 | -0.299567 | -0.274512 | -0.087249 | -0.260273 | -0.229501 | -0.056416 | -0.610586 |
| 23.97222 | -0.689422 | -0.55568 | -1.844121 | -0.341851 | -2.342282 | -0.529029 | 0.123809 | -0.052238 | -3.068789 | 0.120707 | -0.422628 | -2.378223 | -0.11348 | -1.750189 | -0.673908 | -0.685583 | -0.427739 | -0.27172 | -0.283695 | 0.029405 | -0.359589 | -0.148739 | -0.084941 | -0.507888 |
| 23.98611 | -0.660997 | -0.604025 | -1.987686 | -0.200413 | -1.26824 | -0.521245 | 0.155078 | -0.27027 | -2.517272 | 0.1265 | -0.386544 | -2.02964 | -0.073125 | -1.857356 | -0.733308 | -0.593997 | -0.167127 | -0.207898 | -0.210777 | -0.158907 | -0.411466 | -0.105771 | -0.124599 | -0.476897 |
| 24 | -0.554325 | -0.609517 | -1.458983 | 0.263806 | -0.316249 | -0.254838 | 0.241953 | -0.283297 | -1.483337 | 0.258764 | -0.232237 | -1.107741 | 0.082302 | -1.492054 | -0.38449 | -0.394304 | 0.091425 | -0.064325 | 0.011797 | -0.265032 | -0.293161 | -0.010697 | -0.063245 | -0.300462 |
| 24.01389 | -0.309256 | -0.381442 | -0.638615 | 0.927651 | 0.390542 | 0.130811 | 0.368081 | -0.077682 | -0.26907 | 0.517968 | -0.021688 | -0.49003 | 0.274335 | -0.670468 | 0.013032 | 0.117427 | 0.266431 | 0.159171 | 0.215177 | -0.121026 | 0.014842 | 0.158327 | 0.111712 | 0.021469 |
| 24.02778 | -0.258159 | -0.376042 | -0.76817 | 0.771516 | 0.552775 | 0.173241 | 0.301699 | -0.057109 | 0.581747 | 0.568347 | -0.070182 | -0.153485 | 0.290802 | -0.630518 | -0.010826 | 0.416207 | 0.262996 | 0.266238 | 0.337481 | 0.075419 | 0.074268 | 0.286568 | 0.202887 | 0.013285 |
| 24.04167 | -0.630197 | -0.707569 | -1.418024 | -0.423597 | 0.021698 | -0.197848 | 0.008099 | -0.346589 | 0.606339 | 0.255786 | -0.754961 | -0.031881 | 0.065483 | -1.798025 | -0.446564 | 0.357355 | 0.027906 | 0.085171 | 0.146475 | 0.157045 | -0.163055 | 0.061156 | 0.021636 | -0.544129 |
| 24.05556 | -0.612069 | -0.690883 | -1.578132 | -1.672872 | -0.297604 | -0.555938 | -0.096364 | -0.385709 | 0.380963 | 0.163477 | -1.616102 | 0.52864 | -0.176362 | -2.773972 | -1.458176 | 0.526057 | -0.250386 | -0.114354 | -0.113149 | 0.178321 | -0.587648 | -0.577188 | -0.40833 | -1.573914 |
| 24.06944 | -0.384872 | -0.414355 | -1.800179 | -2.185606 | -0.470557 | -1.077324 | -0.088445 | -0.207706 | 0.534119 | 0.169173 | -1.785192 | 0.986414 | -0.297094 | -3.069294 | -2.175552 | 0.452865 | -0.381938 | -0.167774 | -0.236147 | 0.029403 | -1.437333 | -1.031816 | -0.48719 | -1.874456 |
| 24.08333 | -0.574474 | -0.403973 | -2.54584 | -2.424069 | -1.006701 | -1.685586 | -0.341728 | -0.298518 | 1.548998 | -0.022842 | -1.155561 | 0.453737 | -0.408599 | -2.004295 | -1.822181 | -1.080601 | -0.237918 | -0.175473 | -0.248607 | -0.166259 | -1.953318 | -0.567508 | -0.296251 | -1.22364 |
| 24.09722 | -1.05628 | -0.523922 | -2.447215 | -2.702975 | -1.299115 | -1.662973 | -0.331749 | -0.344021 | -0.023515 | -0.175806 | -0.828818 | 0.051248 | -0.329297 | -0.750203 | -0.925847 | -1.310551 | 0.052464 | -0.156709 | -0.155945 | -0.185935 | -1.317605 | -0.033157 | -0.086768 | -0.815671 |
| 24.11111 | -0.314717 | -0.53619 | -1.124176 | -1.121555 | -0.880514 | -1.320367 | -0.01885 | -0.111172 | -0.661337 | -0.279066 | -0.415699 | -0.247354 | -0.048892 | 0.421387 | -0.158122 | -0.49033 | 0.391287 | -0.069004 | -0.054401 | 0.049231 | -0.543919 | 0.391556 | 0.13677 | -0.115854 |
| 24.125 | -0.305421 | -0.532914 | -0.20483 | -0.573792 | -0.498821 | -1.251919 | 0.149925 | 0.065494 | -0.502906 | -0.304129 | -0.140021 | -0.223188 | 0.206522 | 1.479402 | 0.366677 | 0.10769 | 0.632879 | 0.014434 | -0.006225 | 0.351833 | -0.038516 | 0.55808 | 0.288537 | 0.560495 |
| 24.13889 | -0.287144 | -0.588225 | 0.236065 | -0.305255 | -0.384784 | -1.05253 | 0.143238 | 0.109364 | -0.316681 | -0.062592 | -0.004397 | -0.039643 | 0.296987 | 1.619899 | 0.481739 | 0.483257 | 0.535287 | 0.055804 | -0.003646 | 0.268505 | 0.297932 | 0.442562 | 0.241538 | 1.055262 |
| 24.15278 | -0.047746 | -0.791371 | 0.391193 | -0.153854 | -0.301742 | -0.645424 | 0.067501 | 0.094987 | -0.333344 | 0.060569 | 0.018101 | 0.104631 | 0.128025 | 1.106547 | 0.438327 | 0.755864 | 0.078359 | 0.043067 | 0.016017 | -0.071238 | 0.565751 | 0.267442 | 0.099371 | 1.150529 |
| 24.16667 | 0.18317 | -1.00234 | 0.490522 | -0.15762 | -0.261256 | -0.231119 | 0.001723 | 0.04921 | -0.641911 | 0.048472 | -0.045642 | 0.18357 | -0.133419 | 0.889955 | 0.481115 | 1.028382 | -0.326243 | 0.002344 | 0.064885 | -0.206079 | 0.599876 | 0.110773 | 0.011325 | 1.123025 |
| 24.18056 | 0.352913 | -0.972024 | 0.746133 | -0.146989 | -0.14204 | -0.087356 | 0.001158 | 0.044041 | -1.094601 | 0.019005 | -0.119878 | 0.099982 | -0.264681 | 1.051284 | 0.503295 | 0.921986 | -0.329557 | -0.054135 | 0.082862 | 0.062938 | 0.259687 | -0.003601 | -0.023352 | 0.330005 |
| 24.19444 | 0.574744 | -0.641963 | 1.170594 | -0.006962 | 0.166622 | -0.1015 | 0.029369 | 0.038887 | -1.129744 | -0.022165 | -0.019282 | -0.081324 | -0.093159 | 1.354297 | 0.319137 | -0.007831 | -0.022023 | -0.079078 | 0.018138 | 0.233836 | -0.131212 | -0.000789 | 0.016838 | 0.174616 |
| 24.20833 | 0.868675 | -0.097313 | 1.508574 | 0.121143 | 0.601491 | -0.15477 | 0.030545 | -0.00649 | -0.925975 | -0.082083 | 0.291303 | -0.09094 | 0.275776 | 1.473477 | 0.031454 | 0.031469 | 0.161308 | -0.052852 | -0.040754 | -0.006692 | -0.288469 | 0.096788 | 0.094769 | 0.560361 |
| 24.22222 | 1.182316 | 0.658969 | 1.253184 | 0.231927 | 1.187367 | -0.235625 | 0.030986 | -0.030664 | -0.57465 | -0.126976 | 0.724689 | 0.114834 | 0.440121 | 1.146323 | -0.20142 | 0.284102 | 0.161283 | -0.009485 | -0.012877 | -0.24987 | -0.069194 | 0.206885 | 0.098366 | 0.795667 |
| 24.23611 | 1.104308 | 1.55884 | 0.469582 | 0.445974 | 1.594857 | -0.309577 | 0.053347 | 0.052451 | -0.269153 | -0.144958 | 1.116568 | 0.333333 | 0.109415 | 0.878931 | -0.02311 | 0.1026 | 0.110322 | 0.07749 | 0.058678 | -0.222232 | 0.337632 | 0.260279 | 0.05349 | 0.112152 |
| 24.25 | 0.49366 | 2.281925 | 0.074863 | 0.801221 | 1.741467 | -0.325488 | 0.095218 | 0.236684 | 0.021636 | -0.130487 | 1.256182 | 0.604307 | -0.191543 | 0.717084 | 0.41804 | -0.315733 | 0.102511 | 0.182714 | 0.107585 | 0.016268 | 0.091793 | 0.230744 | 0.029748 | -0.545663 |
| 24.26389 | -0.256865 | 2.361402 | -0.301698 | 0.764506 | 1.600921 | -0.394538 | 0.079168 | 0.258857 | 0.238024 | -0.155715 | 1.085188 | 0.945999 | -0.072235 | 0.555133 | 0.780291 | -0.470861 | 0.156419 | 0.18172 | 0.148137 | 0.099648 | -0.100799 | 0.12675 | 0.044926 | -0.448855 |
| 24.27778 | -1.064545 | 1.90523 | -0.566213 | 0.19094 | 1.153031 | -0.526529 | -0.012676 | 0.104129 | 0.359987 | -0.23337 | 0.649109 | 1.110164 | 0.222679 | 0.678575 | 0.67902 | -0.339828 | 0.180333 | 0.124287 | 0.103305 | -0.008476 | 0.243877 | -0.023269 | 0.049009 | 0.117535 |
| 24.29167 | -1.132277 | 1.382188 | -0.168057 | -0.193084 | 0.643558 | -0.517921 | -0.056896 | 0.06793 | 0.482834 | -0.226501 | 0.380824 | 1.062916 | 0.177659 | 0.996529 | 0.283015 | -0.129019 | 0.103256 | 0.147565 | 0.091141 | -0.060271 | 0.710903 | -0.127386 | -0.012497 | 0.341261 |
| 24.30556 | -0.694739 | 0.858058 | 0.983621 | -0.327371 | 0.277027 | -0.382862 | -0.064681 | 0.135532 | 0.569643 | -0.163585 | 0.57162 | 0.766328 | -0.104489 | 1.156238 | 0.272625 | -0.041934 | 0.10302 | 0.2002 | 0.09752 | 0.072183 | 0.757505 | -0.032061 | -0.043237 | 0.709857 |
| 24.31944 | -0.155213 | 0.315182 | 1.77583 | -0.249341 | 0.153182 | -0.286926 | -0.057767 | 0.221192 | 0.665912 | -0.134199 | 0.705197 | 0.218915 | -0.145062 | 1.193162 | 0.350858 | -0.061397 | 0.20053 | 0.174829 | 0.109682 | 0.239337 | 0.562632 | 0.130846 | 0.008332 | 0.863007 |
| 24.33333 | 0.257877 | 0.09641 | 1.882563 | 0.015771 | 0.248846 | -0.185091 | -0.007098 | 0.339296 | 0.704193 | -0.094909 | 0.781566 | -0.183724 | 0.046668 | 0.87617 | 0.318025 | -0.022381 | 0.278867 | 0.08904 | 0.094256 | 0.27199 | 0.503726 | 0.212603 | 0.07254 | 0.334341 |
| 24.34722 | 0.495077 | 0.377786 | 1.431676 | 0.288181 | 0.560969 | -0.099873 | 0.071256 | 0.40385 | 0.738659 | -0.045594 | 0.819433 | -0.193814 | 0.161276 | 0.50172 | 0.095746 | 0.169423 | 0.241926 | 0.02582 | 0.000723 | 0.282607 | 0.752058 | 0.203432 | 0.068765 | 0.121639 |
| 24.36111 | 0.492494 | 0.713075 | 0.156366 | 0.192567 | 0.712053 | -0.100251 | 0.116974 | 0.35475 | 0.543945 | -0.036893 | 0.86408 | 0.221824 | 0.091008 | 0.595037 | -0.07594 | 0.309851 | 0.075426 | -0.027715 | -0.039875 | 0.250922 | 0.997159 | 0.165306 | 0.048178 | 0.228738 |
| 24.375 | 0.366019 | 0.896843 | -0.750982 | -0.292502 | 0.614227 | -0.135499 | 0.174442 | 0.240858 | 0.264219 | -0.036843 | 0.631094 | 0.714181 | 0.066763 | 0.92072 | 0.048415 | 0.202829 | -0.121866 | -0.046347 | -0.014418 | 0.29471 | 0.764843 | 0.153209 | 0.051905 | 0.290828 |
| 24.38889 | 0.184928 | 0.987787 | -0.586359 | -0.113718 | 0.597524 | -0.091162 | 0.225213 | 0.206201 | 0.35408 | 0.011074 | 0.364089 | 0.8638 | 0.192156 | 0.86373 | 0.22179 | 0.003485 | -0.119834 | 0.044723 | 0.008747 | 0.263444 | 0.635805 | 0.184345 | 0.079303 | 0.28292 |
| 24.40278 | -0.094301 | 1.00537 | 0.375966 | 1.040406 | 0.64233 | -0.026649 | 0.184525 | 0.247429 | 0.684301 | 0.033048 | 0.082519 | 0.67695 | 0.346622 | 0.649765 | 0.291074 | -0.092229 | 0.17844 | 0.172925 | 0.059534 | 0.21165 | 0.603501 | 0.233722 | 0.10266 | 0.488217 |
| 24.41667 | [illegible] | [illegible] | [illegible] | [illegible] | [illegible] | [illegible] | [illegible] | [illegible] | [illegible] | [illegible] | [illegible] | [illegible] | [illegible] | [illegible] | [illegible] | [illegible] | [illegible] | [illegible] | [illegible] | [illegible] | [illegible] | [illegible] | [illegible] | [illegible] |
| 24.43056 | [illegible] | [illegible] | [illegible] | [illegible] | [illegible] | [illegible] | [illegible] | [illegible] | [illegible] | [illegible] | [illegible] | [illegible] | [illegible] | [illegible] | [illegible] | [illegible] | [illegible] | [illegible] | [illegible] | [illegible] | [illegible] | [illegible] | [illegible] | [illegible] |
| 24.44444 | [illegible] | [illegible] | [illegible] | [illegible] | [illegible] | [illegible] | [illegible] | [illegible] | [illegible] | [illegible] | [illegible] | [illegible] | [illegible] | [illegible] | [illegible] | [illegible] | [illegible] | [illegible] | [illegible] | [illegible] | [illegible] | [illegible] | [illegible] | [illegible] |
| 24.45833 | [illegible] | [illegible] | [illegible] | [illegible] | [illegible] | [illegible] | [illegible] | [illegible] | [illegible] | [illegible] | [illegible] | [illegible] | [illegible] | [illegible] | [illegible] | [illegible] | [illegible] | [illegible] | [illegible] | [illegible] | [illegible] | [illegible] | [illegible] | [illegible] |
| 24.47222 | [illegible] | [illegible] | [illegible] | [illegible] | [illegible] | [illegible] | [illegible] | [illegible] | [illegible] | [illegible] | [illegible] | [illegible] | [illegible] | [illegible] | [illegible] | [illegible] | [illegible] | [illegible] | [illegible] | [illegible] | [illegible] | [illegible] | [illegible] | [illegible] |
| 24.48611 | [illegible] | [illegible] | [illegible] | [illegible] | [illegible] | [illegible] | [illegible] | [illegible] | [illegible] | [illegible] | [illegible] | [illegible] | [illegible] | [illegible] | [illegible] | [illegible] | [illegible] | [illegible] | [illegible] | [illegible] | [illegible] | [illegible] | [illegible] | [illegible] |
| 24.5 | [illegible] | [illegible] | [illegible] | [illegible] | [illegible] | [illegible] | [illegible] | [illegible] | [illegible] | [illegible] | [illegible] | [illegible] | [illegible] | [illegible] | [illegible] | [illegible] | [illegible] | [illegible] | [illegible] | [illegible] | [illegible] | [illegible] | [illegible] | [illegible] |
| 24.51389 | [illegible] | [illegible] | [illegible] | [illegible] | [illegible] | [illegible] | [illegible] | [illegible] | [illegible] | [illegible] | [illegible] | [illegible] | [illegible] | [illegible] | [illegible] | [illegible] | [illegible] | [illegible] | [illegible] | [illegible] | [illegible] | [illegible] | [illegible] | [illegible] |
| 24.52778 | [illegible] | [illegible] | [illegible] | [illegible] | [illegible] | [illegible] | [illegible] | [illegible] | [illegible] | [illegible] | [illegible] | [illegible] | [illegible] | [illegible] | [illegible] | [illegible] | [illegible] | [illegible] | [illegible] | [illegible] | [illegible] | [illegible] | [illegible] | [illegible] |
| 24.54167 | [illegible] | [illegible] | [illegible] | [illegible] | [illegible] | [illegible] | [illegible] | [illegible] | [illegible] | [illegible] | [illegible] | [illegible] | [illegible] | [illegible] | [illegible] | [illegible] | [illegible] | [illegible] | [illegible] | [illegible] | [illegible] | [illegible] | [illegible] | [illegible] |

| | | | | | | | | | | | | | | | | | | | | | | | | |
|---|---|---|---|---|---|---|---|---|---|---|---|---|---|---|---|---|---|---|---|---|---|---|---|---|
| 24.55556 | -0.579776 | -0.449814 | 0.20277 | 0.117189 | -0.10993 | -0.837783 | -0.783247 | -0.576521 | 0.8135 | -0.8851 | 0.105173 | -0.653032 | 0.300937 | 0.320014 | 0.835323 | 0.065407 | 0.140953 | 0.219269 | 0.107578 | -1.228921 | 0.717803 | 0.229613 | 0.072489 | 0.620118 |
| 24.56944 | -0.197399 | 0.185069 | 0.611077 | 0.091628 | 0.31165 | -0.310525 | -0.367216 | -0.022221 | 0.730819 | -0.289413 | 0.158368 | -0.182294 | 0.257658 | 0.336764 | 0.655583 | 0.384808 | -0.015995 | 0.177788 | 0.075625 | -0.561789 | 1.06607 | 0.327755 | 0.027393 | 0.597671 |
| 24.58333 | 0.223284 | 0.66735 | 1.201091 | 0.391784 | 0.61195 | 0.103612 | 0.056877 | 0.75215 | 0.642982 | 0.157842 | 0.308831 | 0.148512 | 0.273269 | 0.156664 | 0.650788 | 0.791627 | -0.026081 | 0.176344 | 0.081506 | -0.043498 | 0.883995 | 0.330385 | 0.083314 | 0.71498 |
| 24.59722 | 0.045172 | 0.565438 | 1.122681 | 0.579849 | 0.52651 | 0.023247 | 0.089512 | 0.813268 | 0.384006 | 0.056911 | 0.176456 | 0.082012 | 0.271975 | -0.117451 | 0.568753 | 0.460241 | -0.016282 | 0.076724 | 0.025709 | 0.232578 | 0.59206 | 0.270321 | 0.114092 | 0.543167 |
| 24.61111 | -0.59998 | 0.198396 | 0.762387 | 0.285138 | 0.500242 | -0.228527 | -0.101867 | 0.428619 | 0.077595 | -0.209457 | -0.10538 | -0.15384 | 0.221668 | -0.003093 | 0.393344 | 0.100944 | 0.019881 | -0.09576 | -0.04002 | 0.337546 | 0.405992 | 0.237736 | 0.051883 | 0.225247 |
| 24.625 | -0.833667 | 0.028855 | 0.774782 | -0.00494 | 0.575862 | -0.257589 | -0.192291 | 0.258571 | -0.020884 | -0.235368 | -0.205739 | -0.225025 | 0.090763 | 0.324882 | 0.263064 | -0.064966 | 0.124328 | -0.165527 | -0.053807 | 0.393233 | 0.195126 | 0.228064 | 0.00064 | 0.146262 |
| 24.63889 | -0.521326 | 0.066894 | 0.946989 | -0.041148 | -1.035433 | -0.135726 | -0.114249 | 0.302258 | 0.038078 | -0.11671 | -0.134228 | -0.159607 | -0.001917 | 0.546151 | 0.255958 | -0.138938 | 0.150577 | -0.096045 | -0.046068 | 0.360895 | -0.04192 | 0.195691 | -0.015154 | -0.002591 |
| 24.65278 | -0.239147 | 0.064174 | 0.788358 | -0.027831 | -1.205861 | -0.061989 | -0.032079 | 0.214041 | 0.064672 | -0.063633 | -0.053538 | -0.074671 | -0.060303 | 0.37555 | 0.263983 | -0.194498 | -0.002113 | -0.071497 | -0.020733 | 0.132428 | -0.26029 | 0.122218 | -0.005085 | 0.264806 |
| 24.66667 | -0.105222 | 0.021745 | 0.494777 | -0.042342 | -0.226274 | -0.022543 | 0.019524 | 0.044035 | 0.036805 | -0.037262 | -0.021857 | -0.014887 | -0.058541 | 0.034294 | 0.199885 | -0.279229 | -0.035448 | 0.034786 | 0.029596 | -0.092914 | -0.311488 | 0.075635 | 0.002852 | 0.160976 |
| 24.68056 | 0.013592 | 0.04937 | 0.396026 | 0.025721 | 0.374796 | 0.062571 | 0.097516 | -0.043158 | 0.050327 | 0.054866 | 0.042635 | 0.084765 | 0.019624 | -0.200809 | -0.000015 | -0.395431 | 0.04071 | 0.156035 | 0.089692 | -0.240436 | -0.285471 | 0.061659 | -0.047276 | -0.015165 |
| 24.69444 | 0.157595 | 0.111565 | 0.401659 | 0.122838 | 0.506898 | 0.158914 | 0.197466 | -0.025278 | 0.101994 | 0.164399 | 0.159912 | 0.145198 | 0.113531 | -0.290858 | -0.114995 | -0.293161 | 0.106471 | 0.157572 | 0.078769 | -0.274095 | -0.148284 | 0.049217 | -0.002896 | 0.200759 |
| 24.70833 | 0.22878 | 0.138165 | 0.43417 | 0.131575 | 0.365507 | 0.205069 | 0.210813 | 0.023874 | 0.152409 | 0.207899 | 0.245492 | 0.119585 | 0.139291 | -0.263443 | -0.069849 | -0.18836 | 0.087951 | 0.128827 | 0.048386 | -0.157551 | 0.053062 | 0.036557 | 0.024136 | -0.112165 |
| 24.72222 | 0.226191 | 0.114911 | 0.660001 | 0.113556 | 0.228516 | 0.202922 | 0.159443 | 0.071616 | 0.168141 | 0.199893 | 0.266759 | 0.117015 | 0.09848 | -0.054382 | 0.111775 | -0.196216 | 0.050638 | 0.072759 | 0.022294 | -0.025404 | 0.144317 | 0.043195 | 0.101255 | 0.009135 |
| 24.73611 | 0.192066 | 0.0926 | 0.82985 | 0.092321 | 0.15352 | 0.189493 | 0.094391 | 0.075273 | 0.140869 | 0.173142 | 0.243206 | 0.159158 | 0.03715 | 0.058154 | 0.301674 | -0.201439 | 0.018416 | 0.015278 | -0.005262 | 0.094743 | 0.087924 | 0.071293 | 0.062676 | -0.009023 |
| 24.75 | 0.146474 | 0.031221 | -1.429249 | 0.068295 | 0.08213 | 0.173203 | 0.039511 | 0.027484 | 0.095863 | 0.131049 | 0.187902 | 0.140364 | -0.037002 | 0.096908 | 0.394687 | -0.16648 | -0.017106 | -0.052069 | -0.040806 | 0.158065 | -0.044785 | 0.101975 | -0.003684 | 0.045614 |
| 24.76389 | 0.089477 | -0.055288 | -1.555776 | 0.050763 | 0.006146 | 0.154655 | 0.007463 | -0.013402 | 0.039615 | 0.113098 | 0.11998 | 0.055655 | -0.133838 | 0.200706 | 0.379914 | -0.08902 | -0.033188 | -0.12736 | -0.069318 | 0.13186 | -0.20241 | 0.074698 | -0.0022 | 0.179501 |
| 24.77778 | 0.047514 | -0.197055 | -1.07239 | 0.018766 | -0.038132 | 0.126688 | -0.024951 | -0.031987 | -0.013588 | 0.110643 | 0.072541 | -0.005129 | -0.219861 | 0.08746 | 0.260287 | -0.048141 | -0.097469 | -0.200553 | -0.076722 | 0.006154 | -0.399251 | -0.043281 | -0.046726 | -0.038248 |
| 24.79167 | 0.013575 | -0.331452 | -0.746471 | -0.031896 | -0.110311 | 0.078624 | -0.067154 | -0.054568 | -0.094868 | 0.084024 | 0.034578 | -0.095062 | -0.301291 | -0.160749 | 0.147976 | 0.024184 | -0.217875 | -0.20286 | -0.073018 | -0.156566 | -0.624925 | -0.285522 | -0.123676 | -0.377188 |
| 24.80556 | -0.041285 | -0.446231 | -0.64893 | -0.075931 | -0.243806 | 0.052335 | -0.029897 | -0.029571 | -0.181679 | 0.056294 | 0.008906 | -0.223694 | -0.45033 | -0.426763 | 0.016405 | 0.129908 | -0.278891 | -0.173974 | -0.053693 | -0.217168 | -0.730264 | -0.493685 | -0.091012 | -0.483174 |
| 24.81944 | -0.08052 | -0.489373 | -0.392768 | -0.099393 | -0.284772 | 0.084074 | 0.012255 | 0.029464 | -0.24563 | 0.08436 | 0.016343 | -0.29493 | -0.321537 | -0.481074 | -0.153647 | 0.096829 | -0.296156 | -0.083867 | -0.022101 | -0.079109 | -0.631054 | -0.287221 | -0.072009 | -0.511659 |
| 24.83333 | -0.092159 | -0.515291 | -0.09971 | -0.126039 | -0.272655 | 0.132178 | 0.07049 | 0.003758 | -0.303693 | 0.136937 | 0.045745 | -0.270811 | -0.154801 | -0.425323 | -0.579292 | -0.053873 | -0.858052 | 0.004282 | 0.001567 | -0.044514 | -0.439994 | -0.161035 | -0.055636 | -0.603599 |
| 24.84722 | -0.131208 | -0.545133 | 0.065633 | -0.267844 | -0.264489 | 0.154511 | 0.088774 | -0.150107 | -0.392609 | 0.16902 | 0.057865 | -0.251208 | 0.027918 | -0.254298 | -0.928221 | -0.316852 | -0.453098 | 0.005228 | -0.001964 | -0.294268 | -0.270918 | -0.009162 | -0.028635 | -0.51539 |
| 24.86111 | -0.191822 | -0.601989 | 0.118057 | -0.44767 | -0.276786 | 0.157978 | 0.060546 | -0.277617 | -0.578759 | 0.17645 | 0.053606 | -0.214771 | 0.140021 | -0.120907 | -0.83706 | -0.554203 | 0.054015 | -0.028965 | -0.013057 | -0.53865 | -0.190107 | 0.10031 | -0.019784 | -0.413133 |
| 24.875 | -0.28356 | -0.646756 | 0.095353 | -0.456832 | -0.311541 | 0.157749 | 0.004763 | -0.340766 | -0.860852 | 0.161674 | 0.032809 | -0.225007 | 0.083672 | -0.235587 | -0.228857 | -0.849782 | 0.349731 | -0.054222 | -0.025868 | -0.368686 | -0.233503 | 0.000064 | -0.008663 | -0.311877 |
| 24.88889 | -0.36718 | -0.573565 | -0.025922 | -0.468848 | -0.367441 | 0.15637 | -0.050418 | -0.100972 | -1.095796 | 0.130167 | -0.010966 | -0.367338 | -0.075869 | -0.399344 | 0.562645 | -0.844153 | 0.174016 | -0.092917 | -0.044102 | -0.097802 | -0.32254 | -0.215423 | -0.021947 | -0.291199 |
| 24.90278 | -0.456962 | -0.4471 | -0.243144 | -0.585792 | -0.491101 | 0.1346 | -0.090881 | 0.082501 | -1.527178 | 0.107065 | -0.075946 | -0.663505 | -0.264358 | -0.658504 | 1.11899 | -0.568796 | -0.331021 | -0.159995 | -0.081618 | 0.056279 | -0.532135 | -0.473751 | -0.061231 | -0.402434 |
| 24.91667 | -0.613188 | -0.403788 | -0.404885 | -0.715242 | -0.667696 | 0.08657 | -0.165056 | 0.129771 | -1.987505 | 0.082491 | -0.166893 | -1.065649 | -0.372256 | -0.93518 | 0.755448 | -0.364958 | -0.690282 | -0.187128 | -0.108988 | 0.026493 | -0.757655 | -0.708819 | -0.121299 | -0.519694 |
| 24.93056 | -0.83122 | -0.434061 | -0.427827 | -0.867223 | -0.813736 | 0.037866 | -0.269072 | 0.059973 | -2.353336 | 0.047302 | -0.27099 | -1.30851 | -0.431756 | -0.941258 | -0.165635 | -0.209694 | -0.774438 | -0.211154 | -0.12308 | -0.028467 | -0.869829 | -0.717108 | -0.175625 | -0.545911 |
| 24.94444 | -0.941441 | -0.442985 | -0.437305 | -0.957098 | -0.940446 | 0.012482 | -0.345472 | -0.29868 | -2.715589 | 0.023734 | -0.386122 | -1.087577 | -0.262851 | -1.061827 | -0.656945 | -0.171646 | -1.10432 | -0.260133 | -0.157682 | 0.007495 | -0.810086 | -0.498312 | -0.232741 | -0.337646 |
| 24.95833 | -0.889362 | -0.404038 | -0.490022 | -0.828376 | -0.961354 | 0.008491 | -0.382525 | -0.433462 | -2.50753 | 0.022056 | -0.385049 | -0.648151 | -0.084779 | -1.420348 | -1.539152 | -0.28984 | -1.605381 | -0.338036 | -0.223867 | -0.084364 | -0.75505 | -0.341711 | -0.277227 | -0.530326 |
| 24.97222 | -0.736754 | -0.313367 | -0.559533 | -0.619394 | -0.931037 | 0.011221 | -0.382218 | -0.383764 | -2.01185 | 0.018342 | -0.397382 | -0.217883 | 0.009738 | -1.894513 | -2.708383 | -0.395967 | -1.75683 | -0.326852 | -0.21444 | -0.217093 | -0.393662 | -0.265289 | -0.303538 | -0.887012 |
| 24.98611 | -0.443616 | -0.155663 | -0.651867 | -0.336232 | -0.837099 | 0.037659 | -0.350444 | -0.296011 | -1.527068 | 0.038654 | -0.362759 | 0.065095 | -0.011126 | -2.104449 | -2.737995 | -0.357662 | -1.131067 | -0.271177 | -0.132897 | -0.311301 | -0.101119 | -0.241178 | -0.230221 | -1.031345 |
| 25 | 0.004846 | 0.169964 | -0.631147 | 0.147793 | -0.554073 | 0.123604 | -0.098591 | -0.097695 | -0.978461 | 0.190334 | -0.225878 | 0.314492 | 0.006227 | -2.084976 | -1.870454 | -0.042166 | -0.455781 | -0.188649 | -0.024896 | -0.326424 | 0.086978 | -0.134305 | -0.070394 | -0.875023 |
| 25.01389 | 0.545427 | 0.547116 | -0.38358 | 0.826162 | -0.142912 | 0.234741 | 0.142296 | 0.188724 | -0.355207 | 0.452658 | -0.013343 | 0.489773 | 0.104442 | -1.29832 | -0.769841 | 0.437108 | 0.070544 | -0.001005 | 0.18232 | -0.213128 | 0.154764 | 0.094616 | 0.16184 | -0.499944 |
| 25.02778 | 0.569612 | 0.287379 | -0.436627 | 0.725228 | -0.105729 | 0.082561 | 0.018105 | 0.17276 | -0.235676 | 0.32785 | -0.162937 | 0.227076 | 0.129205 | -1.409268 | -0.307916 | 0.615371 | 0.150221 | 0.092594 | 0.307282 | -0.090243 | -0.015159 | 0.123354 | 0.213059 | -0.324517 |
| 25.04167 | -0.057556 | -0.558984 | -1.238543 | -0.325862 | -0.53396 | -0.487326 | -0.443609 | -0.294686 | -0.712775 | -0.356094 | -0.76502 | -0.546978 | -0.045483 | -2.634078 | -0.529088 | 0.352975 | -0.153714 | -0.108416 | 0.159414 | 0.010171 | -0.427197 | -0.239132 | -0.060418 | -0.776907 |
| 25.05556 | -0.133696 | -0.696497 | -1.751433 | -0.551545 | -0.721981 | -0.504562 | -0.491585 | -0.367443 | -0.857266 | -0.4632 | -0.850894 | -0.882744 | -0.233332 | -3.128222 | -1.457233 | 0.063037 | -0.441747 | -0.327157 | -0.116952 | 0.076424 | -0.724098 | -0.63647 | -0.449578 | -1.683144 |
| 25.06944 | 0.363245 | -0.18722 | -2.35761 | -0.253949 | -0.716366 | 0.055269 | -0.079996 | -0.047214 | -0.385558 | 0.049493 | -0.404184 | -0.652537 | -0.257174 | -3.318052 | -2.262494 | -0.232864 | -0.500852 | -0.317147 | -0.238083 | 0.029833 | -1.101871 | -0.614788 | -0.679326 | -2.392084 |
| 25.08333 | 0.589488 | -0.360285 | -2.692622 | -0.440196 | -0.837322 | 0.280247 | 0.05592 | -0.105886 | -0.203995 | 0.269284 | -0.22296 | -0.508336 | -0.268779 | -1.81805 | -1.718708 | -0.622881 | -0.472698 | -0.262351 | -0.279401 | 0.060319 | -1.775409 | -0.300562 | -0.415954 | -1.897441 |
| 25.09722 | 0.404215 | -0.635524 | -1.349237 | -0.945839 | -0.743761 | 0.087642 | -0.046467 | -0.415239 | -0.355137 | 0.077096 | -0.353446 | 0.127662 | -0.286975 | -0.506415 | -0.983567 | -0.805444 | -0.093089 | -0.211216 | -0.196383 | 0.120311 | -1.991744 | -0.034228 | -0.067969 | -0.819076 |
| 25.11111 | 0.128045 | -0.617004 | -0.463484 | -0.892022 | -0.516833 | -0.151974 | -0.081883 | -0.462854 | -0.298114 | -0.171347 | -0.548462 | 0.677459 | -0.119373 | 0.397131 | -0.220962 | -0.716629 | 0.445824 | -0.109938 | -0.035028 | 0.221482 | -1.653358 | 0.262014 | 0.146142 | 0.316955 |
| 25.125 | -0.078948 | -0.479093 | -0.144545 | -0.188821 | -0.322156 | -0.2253 | -0.015666 | -0.258142 | 0.05747 | -0.266843 | -0.626241 | 0.912479 | 0.168282 | 1.143603 | 0.442393 | -0.202962 | 0.806756 | 0.024709 | 0.079706 | 0.182203 | -1.074991 | 0.52148 | 0.272182 | 1.04379 |
| 25.13889 | -0.114463 | -0.344028 | 0.074139 | 0.439504 | -0.112481 | -0.126528 | 0.059035 | -0.053852 | 0.674067 | -0.181205 | -0.534961 | 0.784731 | 0.455046 | 1.396123 | 1.161148 | 0.39833 | 0.52608 | 0.14945 | 0.140239 | -0.036774 | -0.277689 | 0.529161 | 0.262713 | 1.291261 |
| 25.15278 | -0.107547 | -0.178358 | 0.206824 | 0.082144 | 0.00904 | -0.071014 | -0.004067 | -0.075176 | 0.953627 | -0.119595 | -0.455574 | 0.412863 | 0.558609 | 1.094791 | 1.607566 | 1.275771 | 0.178461 | 0.173524 | 0.12988 | -0.275351 | 0.260559 | 0.272099 | 0.178365 | 1.196694 |
| 25.16667 | -0.156174 | -0.022379 | 0.303533 | -0.587993 | -0.000498 | -0.113743 | -0.182845 | -0.163431 | 0.773389 | -0.147702 | -0.348714 | 0.073442 | 0.071044 | 0.313105 | 1.349563 | 1.730992 | 0.109028 | 0.051753 | 0.088801 | -0.273049 | 0.428441 | 0.038856 | 0.086093 | 0.883405 |
| 25.18056 | -0.210634 | -0.235586 | 0.402511 | -1.708968 | -0.131244 | -0.207802 | -0.338488 | -0.349713 | 0.556733 | -0.231308 | -0.15509 | -0.15565 | -0.522198 | 0.070807 | 0.563709 | 0.268525 | 0.076165 | -0.030612 | -0.040223 | -0.071181 | 0.389826 | 0.005374 | 0.099273 | 0.881772 |
| 25.19444 | -0.267346 | -0.731964 | 0.213167 | -1.909034 | -0.235786 | -0.318117 | -0.346216 | -0.434445 | 0.185863 | -0.330755 | -0.091284 | -0.319996 | -0.135066 | 0.898752 | -0.3048 | -0.355273 | -0.287525 | 0.111721 | -0.053888 | 0.165558 | 0.334 | 0.151687 | 0.211155 | 1.544779 |
| 25.20833 | -0.207273 | -0.683259 | 0.05209 | -1.178854 | -0.059977 | -0.326154 | -0.18341 | -0.306089 | -0.036858 | -0.330033 | -0.181518 | -0.287692 | 0.428839 | 1.458294 | -0.564453 | -0.154575 | -0.505795 | 0.324479 | 0.001152 | 0.338659 | 0.436672 | 0.248196 | 0.248082 | 1.263782 |
| 25.22222 | -0.007832 | 0.220002 | 0.24411 | -0.357627 | 0.398939 | -0.27699 | -0.044968 | -0.068921 | -0.124975 | -0.275162 | -0.267491 | 0.08756 | 0.363015 | 1.058328 | -0.536442 | 0.15774 | -0.248043 | 0.346266 | -0.022027 | 0.369192 | 0.709479 | 0.230021 | 0.199218 | 0.041501 |
| 25.23611 | 0.109205 | 1.4267 | 0.536703 | 0.647498 | 0.77209 | -0.301076 | -0.112229 | 0.046751 | -0.087499 | -0.298862 | -0.161053 | 0.700885 | 0.07268 | 1.030136 | 0.283199 | -0.301311 | 0.067242 | 0.171509 | -0.019462 | 0.089769 | 1.019579 | 0.255446 | 0.112222 | -0.640783 |
| 25.25 | 0.119475 | 1.232028 | 0.591133 | 1.752398 | 0.644945 | -0.347412 | -0.259991 | -0.06785 | -0.011179 | -0.345191 | 0.136539 | 1.571796 | 0.008906 | 1.304391 | 1.11471 | -0.828448 | 0.026666 | 0.185865 | 0.050947 | -0.244444 | 1.233219 | 0.3048 | 0.093367 | 0.109201 |
| 25.26389 | 0.173826 | 0.378517 | 0.510153 | 1.747188 | 0.372132 | -0.319757 | -0.264851 | -0.18869 | 0.018088 | -0.324943 | 0.43048 | 1.481428 | 0.133804 | 1.447426 | 2.057681 | -0.308784 | -0.175923 | 0.21365 | 0.076477 | -0.135859 | 1.148934 | 0.244693 | 0.106252 | 1.471383 |
| 25.27778 | 0.444098 | 0.353163 | 0.360862 | 0.387686 | 0.070928 | -0.249906 | -0.144123 | -0.111293 | -0.001711 | -0.254169 | 0.548683 | 0.463627 | 0.175748 | 1.239785 | 0.88773 | -0.204601 | -0.231852 | 0.030346 | 0.048998 | -0.017185 | 0.791263 | 0.055803 | 0.122253 | 1.679878 |
| 25.29167 | 0.610749 | 1.251791 | 0.197487 | -0.410597 | -0.126741 | -0.264005 | -0.108968 | 0.038702 | -0.068599 | -0.252641 | 0.372613 | -0.334547 | 0.056972 | 0.75542 | -0.466428 | -0.254789 | -0.040037 | -0.084225 | 0.027129 | -0.118416 | 0.627191 | -0.06255 | 0.109901 | 0.462249 |
| 25.30556 | 0.221863 | 1.422489 | 0.042137 | -0.275106 | 0.223718 | -0.309493 | -0.204309 | 0.129408 | -0.019347 | -0.293262 | 0.173828 | -0.662298 | -0.058553 | 0.461083 | -0.753589 | -0.081066 | 0.184028 | -0.012481 | 0.064106 | -0.285681 | 0.619524 | 0.033045 | 0.031024 | -0.327863 |
| 25.31944 | -0.272225 | 0.950826 | -0.073215 | 0.394422 | 0.843233 | -0.23473 | -0.160128 | 0.163963 | 0.346225 | -0.220333 | 0.172072 | -0.355182 | -0.091272 | 0.972246 | -0.913423 | 0.202611 | 0.281423 | 0.032457 | 0.106404 | 0.060247 | 0.692807 | 0.220303 | -0.020721 | -0.137201 |
| 25.33333 | -0.470545 | 0.629051 | -0.023647 | 0.644065 | 0.855262 | -0.13548 | -0.022578 | -0.055922 | 0.705282 | -0.120907 | 0.355808 | 0.159505 | 0.01547 | 1.358895 | 0.122941 | 0.311241 | 0.274229 | -0.013663 | 0.091368 | 0.44012 | 0.835275 | 0.220457 | 0.01741 | 0.532744 |
| 25.34722 | -0.307984 | 0.640686 | 0.238978 | 0.4651 | 0.439557 | -0.161585 | 0.000284 | -0.306776 | 0.745735 | -0.152805 | 0.480328 | 0.450357 | 0.17255 | 0.926428 | 0.593036 | 0.266444 | 0.124404 | 0.007353 | 0.101539 | 0.484802 | 0.762438 | 0.08873 | 0.055167 | 0.704872 |
| 25.36111 | -0.094925 | 0.58398 | 0.45719 | 0.372469 | 0.374685 | -0.258155 | -0.114446 | -0.407893 | 0.463917 | -0.251664 | 0.486496 | 0.378867 | 0.195325 | 1.013641 | 0.54781 | 0.250051 | -0.090522 | 0.125148 | 0.121431 | 0.206432 | 0.503477 | 0.081838 | 0.020336 | 0.285224 |
| 25.375 | 0.121581 | 0.289135 | 0.538089 | 0.588689 | 0.5481 | -0.25632 | -0.09742 | -0.283846 | 0.352148 | -0.248605 | 0.450781 | 0.049595 | 0.125667 | 0.984808 | 0.365271 | 0.329304 | -0.115351 | 0.197664 | 0.128442 | -0.120496 | 0.480124 | 0.182288 | 0.010612 | 0.024439 |
| 25.38889 | 0.262764 | 0.247973 | 0.596532 | 0.966238 | 0.669872 | -0.177375 | 0.100383 | 0.021791 | 0.396504 | -0.163745 | 0.457499 | -0.256833 | 0.138007 | 0.974512 | 0.223642 | 0.526829 | 0.182715 | 0.116107 | 0.133921 | 0.05566 | 0.694308 | 0.194216 | 0.0635 | 0.277585 |
| 25.40278 | 0.200798 | 0.500615 | 0.91316 | 1.379701 | 0.875328 | -0.094046 | 0.321367 | 0.281779 | 0.577191 | -0.081589 | 0.420022 | -0.080592 | 0.241283 | 0.937388 | 0.20313 | 0.808397 | 0.577422 | 0.041958 | 0.15878 | 0.379653 | 0.923925 | 0.222665 | 0.102422 | 0.813647 |
| 25.41667 | 0.222746 | 0.790311 | 1.126313 | 1.425295 | 1.238219 | -0.010602 | 0.47796 | 0.373872 | 1.024761 | -0.014507 | 0.333826 | 0.511203 | 0.350722 | 0.851393 | 0.309219 | 1.024341 | 0.733761 | 0.073044 | 0.24416 | 0.571969 | 1.037457 | 0.358314 | 0.135287 | 0.937888 |
| 25.43056 | 0.230231 | 0.558911 | 0.863781 | 1.130722 | 1.435778 | -0.082256 | 0.471776 | 0.33553 | 1.75833 | -0.11477 | 0.237226 | 0.839966 | 0.341648 | 1.163347 | 0.450718 | 1.064857 | 0.525301 | 0.164297 | 0.292699 | 0.383043 | 0.957275 | 0.465843 | 0.142986 | 0.654639 |
| 25.44444 | 0.054007 | 0.181881 | 0.333651 | 0.902556 | 1.250418 | -0.348722 | 0.310187 | 0.172997 | 2.169392 | -0.420137 | 0.064585 | 0.519393 | 0.140219 | 1.651387 | 0.413702 | 1.112939 | 0.27768 | 0.118615 | 0.085362 | 0.230312 | 0.756922 | 0.45042 | 0.033299 | 0.802362 |
| 25.45833 | -0.179629 | -0.207286 | -0.202251 | 0.473059 | 0.921958 | -0.552924 | 0.143211 | -0.249066 | 2.41643 | -0.724213 | -0.060558 | 0.116913 | -0.119005 | 1.168999 | 0.286639 | 1.502289 | 0.218483 | -0.09119 | -0.202935 | 0.247502 | 0.589906 | 0.2534 | -0.138121 | 1.009133 |
| 25.47222 | -0.289368 | -0.741235 | -0.17993 | 0.192614 | 0.234346 | -0.410879 | 0.043495 | -0.339271 | 2.394809 | -0.667465 | -0.130126 | -0.957842 | -0.141068 | 0.506805 | 0.283483 | 0.152267 | 0.50135 | -0.181342 | -0.296547 | 0.184485 | 0.676157 | 0.214502 | -0.13901 | 1.506558 |
| 25.48611 | -0.122153 | -0.432641 | -0.070758 | 0.455944 | 0.127259 | -0.270682 | -0.062976 | -0.002978 | 2.06379 | -0.417205 | -0.006273 | -0.691873 | 0.076081 | 0.570695 | 0.615563 | 0.798244 | 1.154272 | -0.054902 | -0.095625 | -0.410564 | 0.99805 | 0.331604 | 0.030585 | 1.811075 |
| 25.5 | -0.080342 | 0.023234 | -0.064108 | 0.523184 | 0.291674 | -0.285029 | -0.165262 | 0.273045 | 1.856821 | -0.31729 | 0.022063 | -0.746501 | 0.084472 | 0.870268 | 0.984859 | 0.95441 | 1.508989 | 0.228109 | 0.124544 | -1.03732 | 1.061837 | 0.277042 | 0.136089 | 1.554302 |
| 25.51389 | -0.093687 | 0.191349 | -0.106919 | 0.256827 | 0.205056 | -0.272077 | -0.20824 | 0.351699 | 1.7267 | -0.280678 | -0.049686 | -0.338411 | 0.059199 | 1.104847 | 1.245585 | 0.902721 | 1.35922 | 0.487762 | 0.205669 | -0.95223 | 0.978841 | 0.176786 | 0.101848 | 1.031971 |
| 25.52778 | -0.183975 | 0.106743 | -0.214551 | 0.016515 | 0.043206 | -0.424311 | -0.414394 | 0.17957 | 1.335315 | -0.449519 | -0.284101 | 0.476986 | 0.229923 | 1.297399 | 1.37017 | 0.850075 | 1.253834 | 0.626518 | 0.166299 | -0.418786 | 0.8961 | 0.143813 | 0.096589 | 0.551261 |
| 25.54167 | -0.547394 | -0.534565 | -0.506147 | -0.344397 | -0.243384 | -0.854827 | -0.843631 | -0.123976 | 0.531165 | -0.904061 | -0.703813 | 0.442191 | 0.360718 | 0.868491 | 1.146785 | 0.754504 | 0.986632 | 0.603322 | 0.140409 | -0.166205 | 1.027343 | 0.127671 | 0.069535 | -0.295222 |
| 25.55556 | -0.778916 | -0.732003 | -0.66076 | -0.439635 | -0.209815 | -0.896311 | -0.911476 | -0.092133 | -0.121773 | -0.958486 | -0.825847 | -0.413035 | 0.299656 | 0.379815 | 0.654556 | 0.575398 | 0.435601 | 0.437644 | 0.124639 | -0.171255 | 1.158107 | 0.137183 | 0.001606 | 0.2291 |
| 25.56944 | -0.639918 | 0.012756 | -0.374526 | -0.101767 | 0.106121 | -0.425594 | -0.481458 | 0.250201 | -0.09678 | -0.445324 | -0.442838 | -0.309653 | 0.132636 | 0.256752 | 0.312491 | 0.425683 | 0.230909 | 0.263663 | 0.136198 | 0.12982 | 1.00696 | 0.185201 | -0.031756 | 0.098159 |
| 25.58333 | -0.556423 | 0.526218 | -0.012922 | 0.135996 | 0.085847 | -0.051971 | -0.096748 | 0.518105 | -0.049967 | -0.05389 | -0.049629 | 0.023625 | 0.055147 | 0.077134 | 0.328387 | 0.210132 | 0.217729 | 0.193771 | 0.154509 | 0.424318 | 0.786554 | 0.148476 | -0.02066 | -0.011256 |
| 25.59722 | -0.389694 | 0.55289 | 0.231954 | 0.192402 | 0.071907 | 0.093109 | 0.069083 | 0.482954 | -0.083797 | 0.094158 | 0.052632 | 0.167065 | 0.067843 | -0.009707 | 0.544295 | 0.021981 | 0.225411 | 0.086144 | 0.105766 | 0.252547 | 0.531813 | 0.092351 | 0.042803 | 0.244794 |
| 25.61111 | -0.07085 | 0.420068 | 0.356247 | 0.152795 | 0.095844 | 0.138388 | 0.107533 | 0.325114 | -0.23174 | 0.137965 | 0.074742 | 0.154141 | 0.09859 | 0.229093 | 0.816292 | -0.141266 | 0.147118 | 0.013329 | 0.067678 | -0.074957 | 0.311587 | 0.072862 | 0.051451 | 0.207296 |
| 25.625 | 0.042822 | 0.162592 | 0.26486 | 0.03697 | 0.00225 | 0.089004 | 0.045418 | 0.131484 | -0.392028 | 0.083801 | 0.094761 | 0.00112 | 0.048626 | 0.569894 | 1.071553 | -0.347677 | -0.037023 | 0.028375 | 0.04334 | -0.433083 | 0.150061 | 0.112529 | 0.052593 | 0.215016 |
| 25.63889 | -0.06182 | -0.183942 | 0.095294 | -0.064452 | -0.168044 | 0.006885 | -0.036055 | -0.055725 | -0.358423 | -0.00636 | 0.045111 | -0.21729 | -0.056477 | 0.43611 | 1.36887 | -0.380868 | -0.272439 | -0.005029 | 0.009167 | -0.739636 | 0.026198 | 0.192998 | -0.011362 | 0.431139 |
| 25.65278 | -0.128708 | -0.466203 | -0.063435 | -0.132253 | -0.451451 | -0.075978 | -0.104307 | -0.221906 | -0.341268 | -0.101778 | -0.047495 | -0.397596 | -0.136923 | -0.10947 | 0.952799 | -0.378732 | -0.439257 | -0.01317 | 0.018162 | -0.387493 | -0.093076 | 0.088086 | -0.022531 | 0.272929 |
| 25.66667 | -0.110519 | -0.481908 | -0.139195 | -0.135507 | -0.544772 | -0.115579 | -0.154131 | -0.350587 | -0.269486 | -0.145028 | -0.121551 | -0.490418 | -0.092644 | -0.230338 | 0.259739 | -0.559686 | -0.394707 | -0.05009 | 0.068401 | -0.119186 | -0.24904 | 0.008959 | -0.058159 | -0.021052 |
| 25.68056 | -0.062339 | -0.278059 | -0.101652 | -0.035568 | -0.3085 | -0.086347 | -0.145411 | -0.431574 | -0.143516 | -0.101137 | -0.079314 | -0.312496 | 0.057078 | -0.342843 | -0.28867 | -0.333869 | -0.194075 | 0.091237 | 0.081404 | 0.043726 | -0.419049 | 0.005274 | -0.078143 | -0.608705 |
| 25.69444 | -0.047515 | -0.089244 | -0.042191 | 0.037019 | -0.063655 | -0.045773 | -0.096408 | -0.400755 | 0.065619 | -0.050913 | -0.105712 | 0.032233 | 0.176716 | -0.344611 | -0.781579 | 0.102603 | -0.051221 | 0.25281 | -0.004642 | -0.002879 | -0.460861 | 0.036909 | -0.010986 | -0.391644 |
| 25.70833 | -0.043787 | -0.051977 | -0.035448 | -0.000123 | -0.124519 | -0.027568 | -0.034804 | -0.303308 | 0.142046 | -0.036105 | -0.12928 | 0.331254 | 0.1998 | 0.007358 | -1.452682 | 0.44213 | -0.004433 | 0.217025 | -0.108387 | -0.149914 | -0.383744 | -0.008058 | 0.001917 | -0.184354 |
| 25.72222 | -0.010798 | -0.143331 | -0.067 | -0.099664 | -0.083994 | -0.037531 | -0.001964 | -0.222278 | -0.077759 | -0.04954 | 0.012606 | 0.477747 | 0.146097 | 0.180213 | -1.632088 | 0.555489 | -0.014689 | 0.137673 | -0.12394 | -0.211569 | -0.343118 | -0.013442 | 0.021183 | -0.04647 |
| 25.73611 | 0.051006 | -0.191696 | -0.115216 | -0.171501 | 0.284771 | -0.044979 | -0.006109 | -0.145956 | -0.34013 | -0.056932 | 0.052052 | 0.329341 | 0.067208 | 0.188676 | -0.7034 | 0.393712 | -0.025028 | 0.092673 | -0.045662 | -0.090077 | -0.318089 | 0.025601 | -0.048665 | 0.011395 |
| 25.75 | 0.089637 | -0.227018 | -0.187846 | -0.219145 | 0.098675 | -0.048021 | -0.041855 | -0.035085 | -0.421171 | -0.061898 | -0.183347 | 0.071738 | 0.011425 | 0.286869 | 0.434219 | 0.171289 | -0.006547 | 0.061887 | 0.036366 | 0.126965 | -0.32482 | 0.008285 | -0.01839 | 0.119126 |
| 25.76389 | 0.065083 | -0.36989 | -0.310654 | -0.257488 | -0.358958 | -0.085838 | -0.076476 | 0.041171 | -0.476489 | -0.098197 | -0.301303 | -0.185974 | -0.054822 | -0.191739 | 1.376088 | -0.020531 | -0.028196 | -0.004721 | 0.041352 | 0.188072 | -0.344636 | -0.091341 | -0.001841 | 0.218122 |
| 25.77778 | -0.027046 | -0.576171 | -0.485731 | -0.286579 | -0.256451 | -0.10925 | -0.108013 | 0.060235 | -0.575636 | -0.118494 | -0.179395 | -0.347788 | -0.176122 | -0.244166 | 1.584382 | -0.443725 | -0.097429 | -0.184783 | 0.005465 | 0.315524 | -0.402831 | -0.181257 | -0.060083 | 0.089458 |
| 25.79167 | -0.135158 | -0.613734 | -0.61393 | -0.315779 | -0.093721 | -0.079363 | -0.102893 | 0.008807 | -0.737684 | -0.092389 | 0.036231 | -0.394859 | -0.325241 | -0.34449 | 1.52718 | -0.597166 | -0.305382 | -0.245133 | -0.03301 | 0.438429 | -0.565338 | -0.225866 | -0.139939 | -0.102907 |
| 25.80556 | -0.280028 | -0.655421 | -0.715363 | -0.527761 | -0.295561 | -0.110468 | -0.139429 | -0.185547 | -1.085817 | -0.128493 | 0.030985 | -0.61972 | -0.400175 | -0.423568 | 0.870571 | -0.774525 | -0.538792 | -0.237629 | -0.061366 | 0.472758 | -0.737528 | -0.172703 | -0.114394 | -0.208555 |
| 25.81944 | -0.457352 | -0.718193 | -0.947583 | -0.92847 | -0.612358 | -0.240128 | -0.28109 | -0.496531 | -1.476856 | -0.262735 | -0.296685 | -1.087427 | -0.396293 | -0.411779 | 0.060003 | -0.452501 | -0.400272 | -0.166478 | -0.09857 | 0.341727 | -0.925344 | -0.08933 | -0.074918 | -0.209857 |
| 25.83333 | -0.57432 | -0.816773 | -1.070576 | -1.288572 | -0.895702 | -0.32539 | -0.396967 | -0.783436 | -1.705667 | -0.350769 | -0.563597 | -1.663751 | -0.206008 | -0.449512 | -0.536589 | 0.114649 | -0.150266 | -0.075103 | -0.140699 | 0.194404 | -0.842834 | -0.020558 | -0.066486 | -0.291406 |
| 25.84722 | -0.636768 | -0.89518 | -1.037725 | -1.3995 | -1.353022 | -0.320832 | -0.463242 | -0.935621 | -1.800091 | -0.337337 | -0.536685 | -1.987255 | 0.002736 | -0.498706 | -1.339179 | 0.346986 | 0.003978 | -0.003387 | -0.140616 | -0.05411 | -0.492935 | -0.012299 | -0.066222 | -0.488034 |
| 25.86111 | -0.736912 | -0.938614 | -0.977097 | -1.206039 | -1.825767 | -0.313051 | -0.555419 | -1.050903 | -1.882988 | -0.323224 | -0.444187 | -1.503294 | 0.136435 | -0.582393 | -2.053102 | 0.296394 | -0.079641 | -0.008034 | -0.095859 | -0.22191 | -0.195536 | -0.199361 | -0.036182 | -0.708279 |
| 25.875 | -0.925397 | -1.026743 | -0.960392 | -0.954124 | -2.104001 | -0.336995 | -0.680911 | -1.082795 | -1.735324 | -0.356642 | -0.46561 | -0.891467 | 0.072178 | -0.767422 | -2.111498 | -0.351783 | -0.257354 | -0.046894 | -0.050989 | -0.199414 | -0.2689 | -0.421132 | -0.019333 | -0.719168 |
| 25.88889 | -1.058498 | -1.054278 | -0.973141 | -0.795381 | -2.243616 | -0.362972 | -0.710953 | -0.805734 | -1.42244 | -0.409164 | -0.603667 | -0.377142 | -0.096501 | -0.83851 | -1.115732 | -0.665166 | -0.425785 | -0.068527 | -0.026629 | -0.150405 | -0.68629 | -0.426595 | -0.051377 | -0.613366 |
| 25.90278 | -1.100743 | -1.011011 | -1.067174 | -0.463641 | -2.137999 | -0.381162 | -0.838763 | -0.640405 | -1.225469 | -0.438909 | -0.626315 | 0.051236 | -0.180974 | -0.904485 | -0.304703 | -1.013597 | -0.71134 | -0.110703 | -0.028075 | -0.227198 | -1.145765 | -0.360915 | -0.122378 | -0.604448 |
| 25.91667 | -1.055762 | -0.865923 | -0.999332 | -0.021904 | -1.85434 | -0.377651 | -0.82826 | -0.580307 | -1.271404 | -0.408011 | -0.667135 | 0.370641 | -0.218605 | -0.783252 | 0.259501 | -0.395229 | -1.219987 | -0.29267 | -0.125593 | -0.329761 | -0.984751 | -0.253688 | -0.22951 | -0.762345 |
| 25.93056 | -0.927682 | -0.642019 | -0.684976 | 0.341611 | -1.441567 | -0.277277 | -0.635947 | -0.66258 | -1.687644 | -0.259805 | -0.611224 | 0.559136 | -0.154449 | -0.810345 | 0.22305 | -0.173052 | -1.299713 | -0.336753 | -0.133778 | -0.225784 | -0.73301 | -0.133437 | -0.287929 | -0.930483 |
| 25.94444 | -0.662357 | -0.410998 | -0.183103 | 0.484048 | -1.134431 | -0.045449 | -0.347106 | -0.449362 | -1.731554 | -0.00516 | -0.433517 | 0.685519 | -0.210742 | -1.015915 | 0.13798 | 0.026917 | -1.151606 | -0.257127 | -0.064767 | -0.15784 | -0.72824 | -0.076456 | -0.275802 | -1.217918 |
| 25.95833 | -0.348889 | -0.332565 | 0.275809 | 0.343367 | -1.124679 | 0.169373 | -0.111791 | -0.314062 | -1.129625 | 0.195177 | -0.274554 | 0.690404 | -0.166776 | -1.375653 | -0.201537 | -0.082855 | -0.94699 | -0.203199 | -0.083763 | -0.065922 | -0.898284 | -0.195893 | -0.246891 | -1.541737 |
| 25.97222 | -0.189335 | -0.577989 | 0.474207 | 0.057711 | -1.210375 | 0.212491 | -0.075785 | -0.372436 | -0.714597 | 0.214475 | -0.26332 | 0.644849 | -0.177261 | -1.375781 | -0.465865 | -0.269014 | -0.693872 | -0.189546 | -0.116892 | 0.009374 | -0.989122 | -0.388649 | -0.197917 | -1.452754 |
| 25.98611 | -0.077608 | -0.784113 | 0.450831 | -0.105776 | -1.098677 | 0.172474 | -0.111984 | -0.53988 | -0.399003 | 0.173322 | -0.267828 | 0.470119 | -0.224045 | -1.007346 | -0.511771 | -0.400413 | -0.407893 | -0.130355 | -0.116016 | 0.05562 | -0.917669 | -0.410746 | -0.150507 | -1.212527 |
| 26 | 0.118645 | -0.403897 | 0.513466 | 0.094605 | -0.497775 | 0.23277 | -0.00944 | -0.389172 | -0.102793 | 0.271054 | -0.151495 | 0.371356 | -0.26043 | -0.670255 | -0.659847 | -0.665252 | -0.200524 | -0.005158 | -0.038276 | 0.197353 | -0.649078 | -0.015292 | -0.029298 | -0.789553 |
| 26.01389 | 0.288472 | 0.043351 | 0.674708 | 0.423786 | -0.027461 | 0.311874 | 0.193273 | -0.093558 | 0.11348 | 0.421755 | 0.055501 | 0.358707 | -0.069614 | -0.267285 | -0.62923 | -1.17235 | -0.10223 | 0.14781 | 0.15186 | 0.345943 | -0.202365 | 0.445066 | 0.164011 | -0.282089 |
| 26.02778 | 0.412719 | 0.775592 | 0.669021 | 0.77474 | 0.278752 | 0.454322 | 0.419665 | 0.19485 | 0.226435 | 0.568983 | 0.243287 | 0.48712 | 0.130139 | 0.850653 | -0.992044 | -1.703812 | -0.199827 | 0.095627 | 0.423954 | 0.553974 | -0.012835 | -0.225855 | 0.32673 | -0.521279 |
| 26.04167 | 0.680685 | 1.385825 | 0.959521 | 1.196789 | 0.672846 | 0.816265 | 0.827862 | 0.526801 | 0.151911 | 0.877821 | 0.607342 | 0.663516 | 0.073589 | 1.994886 | -2.686362 | -1.371344 | -0.736365 | -0.270745 | 0.477855 | 0.588746 | -0.234445 | -0.581366 | 0.400095 | -0.116037 |
| 26.05556 | 0.697166 | 0.514769 | 0.354413 | 0.633192 | 0.913893 | 0.931943 | 0.872972 | 0.606275 | -0.526817 | 1.061561 | 0.607995 | 0.328169 | -0.165868 | -1.124913 | -2.348999 | -0.281755 | -1.470072 | -0.461786 | 0.049731 | -0.226071 | -0.528933 | -0.945481 | -0.469588 | -0.837996 |
| 26.06944 | 0.495806 | -0.662827 | -0.501885 | -0.555043 | 0.877038 | 0.604292 | 0.42008 | 0.142963 | -0.971223 | 0.787057 | 0.073469 | -0.635261 | -0.274694 | -3.819207 | -2.448886 | 0.644679 | -1.233019 | -0.206702 | -0.255306 | -0.212436 | -0.771513 | -0.385824 | -1.144226 | -0.723838 |
| 26.08333 | 0.429537 | -0.522068 | -1.094437 | -1.002771 | 1.193062 | 0.425702 | 0.270385 | -0.111557 | -0.414504 | 0.509098 | -0.318418 | -1.013603 | -0.210129 | -1.788112 | -1.730824 | 0.410731 | -0.432846 | -0.164322 | -0.336891 | -0.219777 | -1.262929 | 0.244089 | -0.405479 | -0.121559 |
| 26.09722 | 0.337134 | -0.422067 | -0.771362 | -0.721903 | 1.840075 | 0.295625 | 0.281079 | -0.035685 | 0.361841 | 0.323203 | -0.202671 | -0.603418 | -0.094846 | -0.216384 | -0.328021 | -0.457043 | 0.14718 | -0.239472 | -0.14894 | -0.029475 | -1.517477 | 0.400572 | 0.120504 | -0.525472 |
| 26.11111 | 0.338354 | -0.554827 | -0.323663 | -0.483235 | 2.077251 | 0.138949 | 0.301733 | 0.15557 | 0.921316 | 0.15444 | -0.10272 | -0.380662 | 0.22949 | 0.664912 | 0.8553 | -0.881274 | 0.61341 | -0.039129 | 0.123312 | 0.328308 | -0.62863 | 0.70788 | 0.351702 | -0.026988 |
| 26.125 | 0.608028 | -0.479601 | -0.097032 | -0.381405 | 1.608302 | 0.086472 | 0.356412 | 0.373748 | 1.04289 | 0.092449 | -0.13331 | -0.193036 | 0.723937 | 1.190367 | 1.843578 | -0.716096 | 0.979673 | 0.223761 | 0.340758 | 0.352943 | 0.010782 | 1.153749 | 0.458455 | 0.583718 |
| 26.13889 | 0.721932 | -0.32778 | -0.12763 | -0.340439 | 1.046982 | 0.067178 | 0.35395 | 0.494894 | 0.814319 | 0.062339 | -0.254434 | 0.006551 | 1.034573 | 1.536393 | 2.416988 | -0.375072 | 0.846181 | 0.393219 | 0.427749 | 0.060475 | 0.091677 | 1.029508 | 0.358275 | 1.438483 |
| 26.15278 | 0.453977 | -0.346369 | -0.240635 | -0.230343 | 0.461979 | 0.099943 | 0.419834 | 0.622263 | 0.579551 | 0.08852 | -0.232062 | 0.323261 | 0.503953 | 0.515408 | 2.080591 | 0.122109 | 0.729696 | 0.463154 | 0.329602 | -0.208655 | 0.012084 | 0.411307 | 0.225981 | 2.276078 |
| 26.16667 | 0.212113 | -0.139614 | -0.256729 | -0.114348 | 0.029636 | 0.206889 | 0.57832 | 0.726551 | 0.412848 | 0.185573 | 0.144688 | 0.932462 | 0.008913 | -0.785688 | 0.649708 | 1.001755 | 0.691088 | 0.278931 | 0.184441 | -0.235768 | 0.155133 | -0.263122 | 0.186265 | 1.627649 |
| 26.18056 | 0.174756 | -0.046849 | -0.187413 | -0.114924 | 0.507925 | 0.245158 | 0.653614 | 0.770188 | 0.252721 | 0.223178 | 0.605639 | 1.939479 | -0.01964 | -0.43594 | -0.5775 | 1.301775 | 0.396854 | 0.115186 | 0.17844 | 0.05052 | 0.685688 | -0.61689 | 0.279433 | 0.316435 |
| 26.19444 | 0.499975 | -0.058131 | -0.164615 | -0.182498 | 1.245668 | 0.190691 | 0.622272 | 0.703181 | 0.245634 | 0.180618 | 0.756267 | 2.070476 | 0.204785 | 1.186158 | -1.863805 | 0.146347 | -0.200651 | 0.072645 | 0.163387 | 0.084918 | 1.181637 | 0.204549 | 0.33669 | -0.105128 |
| 26.20833 | 0.873566 | 0.17681 | -0.134002 | -0.092264 | 1.423859 | 0.134476 | 0.615141 | 0.618897 | 0.444198 | 0.136633 | 0.383333 | 1.588741 | 0.12898 | 1.566699 | -1.13423 | -0.534447 | -0.584856 | 0.150213 | 0.131683 | -0.186944 | 1.221926 | 0.750453 | 0.257989 | 0.623516 |
| 26.22222 | 1.03946 | 0.30164 | -0.140865 | 0.200273 | 1.071262 | 0.109922 | 0.665648 | 0.64569 | 0.675538 | 0.109945 | 0.070279 | 0.545624 | 0.036379 | 0.72168 | 0.840455 | 0.072739 | -0.190724 | 0.215327 | -0.023989 | -0.020518 | 1.243887 | 0.997057 | 0.284877 | 2.332616 |
| 26.23611 | 0.771372 | 0.398409 | -0.384083 | 0.619202 | 0.78937 | 0.112471 | 0.787939 | 0.847742 | 0.790173 | 0.083589 | 0.336117 | -0.715036 | 0.268256 | 0.443037 | 2.650245 | 1.059751 | 0.329588 | 0.192156 | -0.154837 | 0.403795 | 1.396107 | 0.742121 | 0.350556 | 1.590004 |
| 26.25 | 0.381135 | 0.126279 | -0.919319 | 1.011873 | 0.494744 | 0.127696 | 0.917128 | 0.9633 | 0.719946 | 0.067049 | 0.931049 | -0.803436 | 0.563889 | 1.195965 | 2.285604 | 0.848932 | 0.64513 | 0.094096 | -0.092467 | 0.461052 | 0.923224 | 0.54175 | 0.348769 | -0.036085 |
| 26.26389 | 0.235164 | -0.141424 | -1.126713 | 0.977109 | 0.632843 | 0.135131 | 0.944818 | 0.69876 | 0.667879 | 0.078424 | 1.284104 | 0.391513 | 0.486466 | 1.591829 | 0.215004 | 0.212034 | 0.625567 | 0.13314 | 0.008718 | 0.055942 | 0.33758 | 0.194419 | 0.278302 | 0.171554 |
| 26.27778 | 0.288257 | -0.055654 | -0.715502 | 0.569198 | 0.939058 | 0.107781 | 0.825537 | 0.382101 | 0.762554 | 0.086257 | 0.989374 | 1.36828 | 0.206864 | 1.116301 | -1.427217 | 0.294832 | -0.019685 | 0.391797 | 0.06344 | -0.103688 | 0.47758 | 0.113501 | 0.29757 | 1.214004 |
| 26.29167 | 0.409246 | 0.762942 | -0.383612 | 0.077888 | 0.823259 | 0.068808 | 0.698575 | 0.356474 | 0.954178 | 0.051692 | 0.790948 | 1.122286 | -0.003458 | 1.03684 | -0.897254 | 0.844243 | -0.704369 | 0.463691 | 0.254597 | 0.206643 | 0.599197 | 0.52161 | 0.262346 | 1.895634 |
| 26.30556 | 0.465759 | 1.829938 | -0.288635 | 0.025465 | 0.816314 | 0.119326 | 0.58761 | 0.478128 | 1.054354 | 0.096857 | 0.928361 | 0.896804 | -0.108744 | 0.713771 | 0.612999 | 0.40731 | -0.682928 | 0.062226 | 0.547965 | 0.370586 | 0.529921 | 0.79413 | 0.064758 | 0.399885 |
| 26.31944 | 0.448054 | 1.764703 | -0.076076 | 0.623439 | 0.69194 | 0.317492 | 0.698499 | 0.588555 | 1.113108 | 0.297597 | 1.073443 | 1.074927 | -0.11459 | 0.745489 | 1.574117 | -0.67746 | -0.11716 | -0.252199 | 0.68577 | 0.153622 | 0.294429 | 0.726142 | 0.023347 | -0.753119 |
| 26.33333 | 0.500804 | 0.907935 | 0.14496 | 1.002456 | 0.926265 | 0.473855 | 0.830284 | 0.588441 | 1.32414 | 0.459263 | 1.095169 | 1.381762 | -0.035035 | 1.165762 | 1.483969 | -1.006232 | 0.251959 | 0.014123 | 0.364124 | -0.145429 | 0.200886 | 0.542332 | 0.182111 | -0.283251 |
| 26.34722 | 0.668236 | 0.447598 | 0.256752 | 0.92885 | 0.981668 | 0.396764 | 0.783723 | 0.492376 | 1.614116 | 0.391116 | 1.009969 | 1.149175 | 0.100443 | 1.501664 | 0.99086 | -0.401207 | 0.271677 | 0.402206 | 0.127109 | -0.11325 | 0.378867 | 0.411294 | 0.272371 | 0.421859 |
| 26.36111 | 0.745542 | 0.466793 | 0.109957 | 0.795445 | 1.14465 | 0.18854 | 0.462765 | 0.350381 | 1.674476 | 0.184143 | 0.801089 | 0.469557 | 0.212296 | 1.84535 | 0.944646 | -0.105556 | 0.230546 | 0.504275 | 0.113645 | 0.294378 | 0.555358 | 0.387616 | 0.180108 | 0.986468 |
| 26.375 | 0.614472 | 0.515982 | -0.082047 | 0.699143 | 1.429961 | 0.032101 | 0.201156 | 0.315042 | 1.128088 | 0.024428 | 0.548626 | -0.177676 | 0.272916 | 1.709291 | 0.965298 | 0.330099 | 0.307493 | 0.349775 | 0.271298 | 0.544432 | 0.679636 | 0.390059 | 0.120437 | 1.207397 |
| 26.38889 | 0.533147 | 0.38326 | -0.148575 | 0.634522 | 1.511237 | -0.008441 | 0.215526 | 0.443061 | 0.574164 | -0.015771 | 0.388966 | -0.204663 | 0.357322 | 1.310689 | 0.565343 | 0.94318 | 0.494033 | 0.277956 | 0.353155 | 0.347366 | 0.635157 | 0.30914 | 0.16872 | 0.759255 |
| 26.40278 | 0.679898 | 0.397237 | -0.100914 | 0.504653 | 1.758191 | 0.025066 | 0.368682 | 0.558919 | 0.898706 | 0.006406 | 0.379327 | 0.046111 | 0.450699 | 0.742996 | 0.048137 | 1.382837 | 0.759809 | 0.391243 | 0.303954 | -0.013695 | 0.41762 | 0.257226 | 0.227861 | 0.411454 |
| 26.41667 | 0.738865 | 0.750276 | -0.055464 | 0.411215 | 1.510331 | 0.02758 | 0.055414 | 0.575999 | 1.792753 | -0.005621 | 0.451689 | 0.058518 | 0.48728 | 1.170755 | 0.1808 | 1.556382 | 0.852381 | 0.605765 | 0.267587 | -0.154049 | 0.248282 | 0.403025 | 0.201827 | 0.640768 |
| 26.43056 | 0.724392 | 1.372623 | -0.101089 | 0.424264 | 2.009589 | -0.037281 | 0.117695 | 0.669889 | 2.586896 | -0.077537 | 0.538715 | -0.113087 | 0.478616 | 2.175606 | 0.469676 | 2.032489 | 0.59066 | 0.716682 | 0.296719 | 0.282415 | 0.206925 | 0.77975 | 0.160598 | 1.1089 |
| 26.44444 | 0.440729 | 1.930067 | -0.276416 | 0.468673 | 1.079677 | -0.136467 | 0.011524 | 0.647094 | 2.755643 | -0.196859 | 0.414293 | 0.301966 | 0.454174 | 2.750783 | 0.547288 | 2.399245 | 0.521837 | 0.511416 | 0.227647 | 0.760829 | 0.267611 | 1.012128 | 0.076833 | 1.177646 |
| 26.45833 | 0.12675 | 1.379041 | -0.428274 | 0.338107 | -0.168895 | -0.21444 | -0.195641 | 0.47393 | 2.823646 | -0.369722 | 0.115171 | 0.857311 | 0.177942 | 2.500755 | 0.637313 | 1.327653 | 0.726235 | 0.088582 | 0.010903 | 0.671096 | 0.190709 | 0.759233 | -0.120578 | 1.135324 |
| 26.47222 | -0.103917 | 0.83248 | -0.296242 | 0.235287 | 1.900425 | -0.118069 | -0.117556 | 0.178294 | 2.744193 | -0.390786 | -0.049635 | 0.65413 | 0.067314 | 1.800212 | 0.222611 | -0.140002 | 0.979226 | 0.225468 | -0.003235 | 0.306138 | 0.098841 | 0.411197 | -0.153068 | 1.454396 |
| 26.48611 | -0.088183 | 0.672087 | -0.181672 | 0.43932 | -1.643736 | -0.046454 | 0.17416 | 0.20155 | 1.617227 | -0.215087 | 0.030445 | 0.325726 | 0.302348 | 1.043401 | -0.251337 | 0.921585 | 1.128184 | 0.660929 | 0.11626 | 0.148156 | 0.264027 | 0.491195 | 0.001144 | 1.585955 |
| 26.5 | -0.09835 | 0.566812 | -0.126472 | 0.594274 | -1.330093 | -0.046707 | 0.27037 | 0.282273 | 1.531462 | -0.108142 | 0.166961 | -0.841061 | 0.360233 | 1.021832 | -0.150649 | 1.552009 | 1.688319 | 1.014493 | 0.187746 | 0.066326 | 0.463746 | 0.915003 | 0.097808 | 1.784509 |
| 26.51389 | -0.169055 | 0.085131 | -0.128701 | 0.555079 | 1.106361 | -0.089649 | 0.174334 | 0.275312 | 1.403937 | -0.125192 | 0.125358 | -2.514121 | 0.178229 | 1.222386 | 1.693437 | 1.989285 | 1.563435 | 0.935752 | 0.167756 | 0.193355 | 0.600008 | 1.325646 | 0.178625 | 1.003899 |
| 26.52778 | -0.384678 | -0.239068 | -0.351419 | 0.246243 | 0.7129 | -0.401092 | -0.338946 | 0.169909 | 1.246588 | -0.43171 | -0.228627 | -2.109691 | 0.335806 | 1.34327 | 2.568425 | 1.679942 | 1.011065 | 0.382569 | 0.075924 | 0.248974 | 0.689597 | 0.785221 | 0.220158 | 0.640545 |
| 26.54167 | -0.910785 | -0.5954 | -1.031276 | -0.556581 | -1.527992 | -1.140434 | -1.146024 | -0.119319 | 0.523979 | -1.170749 | -0.993823 | -0.48835 | 0.331565 | 0.845934 | 2.507078 | 1.41251 | 0.682749 | 0.234538 | 0.052799 | -0.184872 | 0.726443 | 0.200467 | 0.06965 | 0.776638 |
| 26.55556 | -1.509886 | 0.341889 | -1.836222 | -1.466982 | -2.111039 | -1.823306 | -1.643638 | -0.419747 | -0.49079 | -1.829812 | -1.623123 | -0.48643 | 0.184038 | 0.495206 | 1.662124 | 0.579059 | 0.656657 | 0.406566 | 0.0446 | 0.144445 | 0.764528 | -0.189706 | -0.011376 | 0.760776 |
| 26.56944 | -1.44916 | 0.891002 | -1.824594 | -1.376369 | -1.079802 | -1.530556 | -1.314825 | -0.229843 | -0.821306 | -1.559497 | -1.364263 | -0.404806 | 0.029894 | -0.075603 | 0.514223 | 0.206067 | 0.690286 | 0.428399 | 0.101438 | 0.4153 | 0.920427 | -0.103095 | 0.041601 | 0.633573 |
| 26.58333 | -0.686384 | 0.343054 | -1.028424 | -0.643517 | 0.300601 | -0.744908 | -0.5112 | 0.234448 | -0.485079 | -0.779678 | -0.593873 | 0.12068 | 0.006716 | -0.227463 | 0.100854 | 0.314731 | 0.53311 | 0.375498 | 0.176473 | 0.227548 | 0.846686 | -0.126635 | 0.167746 | 0.592631 |
| 26.59722 | 0.034009 | 0.105874 | -0.359173 | -0.194342 | 0.686767 | -0.190009 | -0.022445 | 0.18013 | -0.159986 | -0.206719 | -0.132862 | 0.614063 | 0.072148 | 0.201286 | -0.669531 | 0.220201 | 0.358523 | 0.390565 | 0.190747 | -0.21934 | 0.620379 | -0.078543 | 0.184831 | 0.418602 |
| 26.61111 | 0.333392 | 0.086953 | -0.162571 | 0.04185 | -1.135783 | 0.007492 | 0.132796 | -0.161227 | -0.044031 | 0.008877 | 0.01651 | 0.79377 | 0.158519 | 0.502631 | -1.167846 | -0.115873 | 0.136427 | 0.281445 | 0.141747 | -0.141865 | 0.391263 | 0.350893 | 0.188582 | 0.256339 |
| 26.625 | 0.527312 | 0.133133 | -0.046093 | 0.178833 | -1.119732 | 0.138427 | 0.065761 | -0.345544 | -0.018353 | 0.158764 | 0.146194 | 0.944487 | 0.212357 | 0.436738 | -1.195132 | -0.169707 | -0.130022 | 0.019148 | 0.104192 | -0.209506 | 0.191395 | 0.721731 | 0.106075 | 0.208484 |
| 26.63889 | 0.645933 | 0.142567 | 0.230942 | 0.403733 | 0.840909 | 0.363588 | -0.106974 | -0.212295 | -0.071148 | 0.406956 | 0.282337 | 1.251671 | 0.245529 | 0.325123 | -0.680203 | -0.377767 | -0.47976 | -0.342262 | 0.096798 | -0.220891 | 0.153026 | 0.53641 | 0.008427 | -0.096124 |
| 26.65278 | 0.585777 | 0.098991 | 0.388824 | 0.511992 | 2.285334 | 0.49316 | -0.344802 | -0.009378 | -0.305586 | 0.533937 | 0.465917 | 1.253014 | 0.226226 | 0.003176 | 0.364018 | -1.003043 | -0.877187 | -0.344388 | 0.077725 | -0.023692 | 0.33262 | 0.011271 | -0.000515 | -0.031717 |
| 26.66667 | 0.38649 | 0.121936 | 0.308191 | 0.420271 | 2.866163 | 0.430177 | -0.561636 | 0.112331 | -0.641835 | 0.470734 | 0.431444 | 0.935792 | 0.14687 | -0.30448 | 1.173049 | -1.528939 | -0.921984 | -0.17619 | 0.058272 | 0.119501 | 0.892701 | -0.214132 | -0.015306 | 0.114975 |
| 26.68056 | 0.263363 | 0.251356 | 0.252782 | 0.359951 | -0.541833 | 0.408687 | -0.157487 | 0.232432 | -0.504723 | 0.450039 | 0.39965 | 0.653213 | 0.033496 | -0.072117 | 1.323689 | -0.849125 | -0.446729 | 0.097203 | 0.03472 | 0.306363 | 0.874067 | -0.400004 | 0.084317 | 0.659089 |
| 26.69444 | 0.299933 | 0.395956 | 0.311005 | 0.389525 | 1.653092 | 0.491755 | 0.49752 | 0.490685 | -0.080632 | 0.517239 | 0.397837 | 0.565542 | -0.034525 | 0.137108 | 1.203532 | 0.728906 | 0.218592 | 0.455259 | -0.013828 | 0.550061 | 0.427637 | -0.420757 | 0.16724 | 0.219206 |
| 26.70833 | 0.342743 | 0.464011 | 0.347091 | 0.37954 | 3.111338 | 0.522748 | 0.773818 | 0.869645 | 0.460672 | 0.536626 | 0.463482 | 0.512522 | -0.054163 | 0.070231 | 0.716665 | 0.874116 | 0.592432 | 0.373892 | -0.047394 | 0.515616 | 0.302676 | -0.243883 | 0.019377 | 0.070346 |
| 26.72222 | 0.361309 | 0.469039 | 0.311897 | 0.240347 | -2.986545 | 0.47889 | 0.783622 | 1.292166 | 0.796487 | 0.491682 | 0.449334 | 0.481166 | -0.043403 | -0.218957 | 0.186061 | 0.393542 | 0.535366 | 0.160263 | -0.067544 | 0.214596 | 0.174195 | -0.012852 | -0.11014 | 0.025853 |
| 26.73611 | 0.418249 | 0.447614 | 0.285821 | -0.162021 | -2.57967 | 0.43794 | 0.805881 | 1.23148 | 1.201951 | 0.452436 | 0.316718 | 0.472159 | -0.016993 | -0.412911 | -0.375648 | 0.391085 | 0.387531 | -0.107603 | -0.05016 | -0.188071 | 0.039753 | 0.208735 | -0.11751 | -0.184982 |
| 26.75 | 0.425846 | 0.411448 | 0.290105 | -0.762196 | -0.956532 | 0.419922 | 0.818588 | 0.668521 | 0.881002 | 0.431888 | 0.28806 | 0.429647 | -0.02092 | -0.287177 | -0.915483 | 0.232531 | 0.211115 | -0.081941 | -0.016826 | -0.517022 | -0.033225 | 0.243519 | -0.313128 | -0.147848 |
| 26.76389 | 0.335703 | 0.352749 | 0.279014 | -1.156872 | 1.12975 | 0.39083 | 0.680803 | 0.189039 | 0.41837 | 0.400083 | 0.356188 | 0.315911 | -0.099446 | -0.230895 | -1.481825 | 0.192765 | 0.071036 | -0.07699 | 0.013615 | -0.614019 | -0.204115 | 0.032873 | -0.387452 | -0.212868 |
| 26.77778 | 0.08105 | 0.276991 | 0.229339 | -1.362244 | 1.914127 | 0.323095 | 0.393843 | -0.266933 | 0.146359 | 0.328565 | 0.307507 | 0.120584 | -0.245081 | -0.057178 | -1.894815 | -0.057792 | -0.154835 | -0.114044 | 0.031775 | -0.396121 | -0.479162 | -0.163491 | -0.363081 | 0.056489 |
| 26.79167 | -0.188368 | 0.191295 | 0.157698 | -1.325063 | 1.236912 | 0.20694 | 0.140006 | -0.715361 | -0.187852 | 0.215138 | 0.044506 | -0.069372 | -0.364685 | 0.095734 | -1.836195 | -0.092502 | -0.595922 | -0.186928 | 0.066224 | 0.034961 | -0.769476 | -0.361962 | -0.145497 | -0.119288 |
| 26.80556 | -0.479881 | 0.132672 | 0.091843 | -1.192309 | -0.539151 | 0.090679 | -0.051489 | -0.663918 | -0.409931 | 0.102952 | -0.10111 | -0.255027 | -0.361897 | 0.063948 | -1.630299 | -0.182141 | -1.012689 | -0.237 | -0.036778 | 0.407636 | -0.934097 | -0.463067 | -0.094378 | -0.156901 |
| 26.81944 | -0.798275 | 0.111604 | 0.048864 | -0.84329 | -1.312114 | 0.043345 | -0.150257 | -0.360053 | -0.58253 | 0.053713 | -0.056939 | -0.37717 | -0.232253 | -0.118632 | -1.101922 | -1.352671 | -1.267173 | -0.417308 | -0.116195 | 0.47247 | -0.989223 | -0.351241 | -0.080252 | -0.111841 |
| 26.83333 | -0.847777 | 0.121025 | 0.036914 | -0.457841 | -1.776133 | 0.075576 | -0.323505 | -0.286643 | -0.685947 | 0.081427 | -0.030604 | -0.440182 | -0.055857 | -0.366508 | 0.114252 | -0.348024 | -0.805372 | -0.444508 | -0.106668 | 0.288953 | -0.802639 | -0.217315 | -0.10178 | -0.233707 |
| 26.84722 | -0.593687 | 0.158129 | 0.076119 | -0.201776 | -1.568971 | 0.121288 | -0.557001 | -0.388385 | -0.840353 | 0.136142 | -0.01806 | -0.454247 | 0.003729 | -0.606351 | 1.099587 | 0.540527 | -0.32326 | -0.375884 | -0.081779 | 0.097725 | -0.440814 | -0.155909 | -0.07812 | -0.36749 |
| 26.86111 | -0.399189 | 0.188673 | 0.123082 | -0.021485 | -0.062583 | 0.132274 | -0.525695 | -0.470697 | -1.026138 | 0.166149 | 0.018809 | -0.43603 | -0.061901 | -0.663025 | 1.503693 | -0.089067 | -0.019088 | -0.228785 | -0.079596 | -0.087729 | -0.087432 | -0.263011 | -0.055135 | -0.389175 |
| 26.875 | -0.333701 | 0.210068 | 0.145273 | 0.099962 | 1.110326 | 0.128153 | -0.315702 | -0.555616 | -1.149796 | 0.173939 | 0.004441 | -0.437506 | -0.136668 | -0.579904 | 0.083899 | -0.21285 | -0.024486 | -0.229929 | -0.108727 | -0.265576 | -0.090875 | -0.479067 | -0.113139 | -0.385761 |
| 26.88889 | -0.393148 | 0.254507 | 0.164103 | 0.191688 | 1.1877 | 0.149368 | -0.261654 | -0.593921 | -1.112457 | 0.193181 | -0.13132 | -0.452906 | -0.146107 | -0.507242 | -1.257102 | -0.530532 | -0.279369 | -0.320597 | -0.136498 | -0.324787 | -0.359109 | -0.634379 | -0.162487 | -0.403047 |
| 26.90278 | -0.547412 | 0.314228 | 0.199975 | 0.302771 | -0.887593 | 0.202384 | -0.263789 | -0.615358 | -1.047014 | 0.230242 | -0.260937 | -0.447476 | -0.160694 | -0.523985 | -2.540116 | -0.94074 | -0.696861 | -0.413549 | -0.149246 | -0.314274 | -0.588082 | -0.630987 | -0.2123 | -0.496731 |
| 26.91667 | -0.71529 | 0.335994 | 0.221214 | 0.398443 | -2.159766 | 0.240346 | -0.359918 | -0.631663 | -1.007755 | 0.245039 | -0.295145 | -0.374511 | -0.139328 | -0.520739 | -2.854042 | -1.51582 | -1.050348 | -0.530383 | -0.162623 | -0.313951 | -0.778821 | -0.716808 | -0.282252 | -0.765267 |
| 26.93056 | -0.732443 | 0.279584 | 0.184622 | 0.392372 | -2.453617 | 0.214003 | -0.445026 | -0.723853 | -0.89915 | 0.206781 | -0.346606 | -0.240732 | -0.363104 | -0.325708 | -1.707316 | -1.051357 | -1.449638 | -0.585446 | -0.152498 | -0.37077 | -0.95892 | -0.589137 | -0.356609 | -1.112732 |
| 26.94444 | -0.635353 | 0.173959 | 0.114539 | 0.333401 | -1.961938 | 0.142549 | -0.453629 | -0.878587 | -0.628399 | 0.139418 | -0.287771 | -0.220873 | -0.57581 | -0.542389 | -1.096921 | -1.060097 | -1.667319 | -0.578412 | -0.147819 | -0.451685 | -1.107187 | -0.705375 | -0.362203 | -1.376703 |
| 26.95833 | -0.4597 | 0.074179 | 0.057007 | 0.246134 | -1.237093 | 0.086385 | -0.36035 | -0.970802 | -0.413744 | 0.080722 | -0.280847 | -0.231735 | -0.751185 | -1.184344 | -1.744757 | -1.195234 | -1.54925 | -0.54897 | -0.168658 | -0.35049 | -0.975693 | -0.616559 | -0.400745 | -1.51482 |
| 26.97222 | -0.25701 | 0.027028 | 0.032024 | 0.197659 | -0.471343 | 0.059862 | -0.266279 | -1.09626 | -0.218269 | 0.047747 | -0.242005 | -0.259486 | -0.682669 | -1.994895 | -0.79327 | -0.732987 | -1.388883 | -0.474999 | -0.173034 | -0.196336 | -0.928671 | -0.518424 | -0.45877 | -2.058451 |
| 26.98611 | -0.120833 | 0.017161 | 0.050292 | 0.178659 | 0.054752 | 0.063939 | -0.202251 | -1.026671 | -0.006813 | 0.060791 | -0.210498 | -0.337647 | -0.451512 | -2.420347 | 0.837594 | 0.12458 | -1.152827 | -0.326068 | -0.200735 | -0.268777 | -0.721228 | -0.497189 | -0.420516 | -1.978009 |
| 27 | -0.029547 | 0.118684 | 0.155238 | 0.300275 | 0.433784 | 0.10741 | -0.135665 | -0.688546 | 0.148078 | 0.162416 | -0.105516 | -0.31957 | -0.123258 | -1.54526 | 2.717885 | 0.656446 | -0.845918 | -0.105778 | -0.106682 | -0.120906 | -0.499303 | -0.402688 | -0.266817 | -1.518402 |
| 27.01389 | 0.125565 | 0.265496 | 0.306388 | 0.532754 | 0.568356 | 0.156101 | -0.046784 | -0.296499 | 0.236064 | 0.32542 | 0.029722 | -0.152205 | 0.269041 | -0.504632 | 2.375872 | 0.789243 | -0.422961 | 0.102502 | 0.126161 | 0.201451 | -0.237069 | -0.206584 | -0.076845 | -0.762006 |
| 27.02778 | 0.102396 | 0.043256 | 0.218181 | 0.38277 | 0.310611 | 0.090604 | -0.112142 | -0.383216 | 0.032347 | 0.272158 | -0.073487 | -0.228254 | 0.450523 | -0.426419 | 0.488055 | 0.717813 | -0.198366 | 0.012059 | 0.204627 | 0.37341 | 0.012351 | -0.143101 | -0.044069 | -0.345593 |
| 27.04167 | -0.184801 | -0.356799 | -0.161309 | -0.131932 | -0.142851 | -0.121711 | -0.34996 | -0.69039 | -0.602529 | -0.029767 | -0.412141 | -0.629642 | 0.207644 | -1.801624 | -0.08678 | 0.109821 | -0.392648 | -0.528347 | -0.095072 | 0.317205 | -0.125506 | -0.555919 | -0.614546 | -0.838069 |
| 27.05556 | -0.107668 | -0.252639 | -0.169706 | -0.18394 | -0.220085 | -0.018892 | -0.513588 | -1.048392 | -1.29414 | 0.028476 | -0.51135 | -0.589459 | 0.06945 | -3.821852 | -1.057722 | -0.504695 | -0.715953 | -1.031741 | -0.542408 | 0.160961 | -0.406322 | -1.393602 | -0.701999 | -1.758214 |
| 27.06944 | 0.45147 | 0.441597 | 0.24055 | 0.077893 | -0.283109 | 0.401053 | -0.506709 | -1.19635 | -1.894089 | 0.372779 | -0.590843 | -0.261442 | 0.326449 | -4.651318 | -2.229834 | -0.270585 | -0.806486 | -1.035266 | -0.889705 | -0.047204 | -0.792035 | -2.096391 | -0.620391 | -2.158732 |
| 27.08333 | 1.00962 | 0.782502 | 0.515083 | 0.180107 | -0.523722 | 0.579555 | -0.285075 | -0.500301 | -1.803933 | 0.543651 | -0.56127 | -0.056653 | 0.418626 | -3.507163 | -2.755555 | 0.093239 | -0.606371 | -0.546988 | -0.94723 | -0.066982 | -1.861751 | -1.106026 | -0.327547 | -1.471099 |
| 27.09722 | 0.88668 | 0.466655 | 0.529261 | 0.086871 | 0.184645 | 0.523197 | 0.137505 | 0.031033 | -1.208174 | 0.508185 | -0.097354 | -0.240111 | 0.016519 | -1.241134 | -2.433536 | -0.395589 | -0.239741 | -0.138313 | -0.410317 | -0.012077 | -2.105563 | -0.025585 | -0.10455 | -0.716869 |
| 27.11111 | 0.85476 | 0.278605 | 0.433087 | -0.052316 | 0.830687 | 0.399183 | 0.506199 | 0.314999 | -0.697897 | 0.386282 | 0.236265 | -0.471695 | -0.323149 | 0.550162 | -1.878313 | -0.795336 | 0.269395 | 0.148143 | -0.084404 | 0.062484 | -1.191454 | 0.617061 | 0.073131 | -0.303728 |
| 27.125 | 0.849772 | 0.218644 | 0.32919 | -0.189669 | 1.052663 | 0.281754 | 0.665952 | 0.538538 | -0.408413 | 0.264577 | 0.454862 | -0.297244 | -0.361237 | 1.568617 | -0.099694 | -1.069356 | 0.669815 | 0.321617 | 0.153613 | 0.06721 | -0.427151 | 0.895906 | 0.212051 | 0.075646 |
| 27.13889 | 0.819441 | 0.11704 | 0.276619 | -0.31454 | 1.204083 | 0.221098 | 0.641027 | 0.798633 | -0.18807 | 0.191448 | 0.438816 | 0.200481 | -0.206494 | 2.407535 | 1.36457 | -0.987254 | 0.630189 | 0.330125 | 0.391721 | -0.083399 | 0.20994 | 1.378269 | 0.244757 | 0.617126 |
| 27.15278 | 0.770173 | -0.01596 | 0.235078 | -0.488518 | 1.220274 | 0.189709 | 0.535269 | 0.984343 | 0.019322 | 0.141556 | 0.107983 | 0.587264 | -0.001985 | 2.462484 | 2.572992 | -0.638419 | 0.717305 | 0.28658 | 0.456324 | -0.379089 | 0.76906 | 1.102991 | 0.144876 | 1.500158 |
| 27.16667 | 0.846664 | -0.085438 | 0.186435 | -0.59052 | 0.927987 | 0.163338 | 0.468019 | 0.970001 | 0.051027 | 0.117144 | -0.271599 | 0.643924 | 0.183233 | 1.588557 | 2.687833 | -0.18735 | 1.324613 | 0.362663 | 0.195036 | -0.677346 | 0.989027 | 0.146246 | 0.100531 | 1.690932 |
| 27.18056 | 0.83261 | -0.060314 | 0.172422 | -0.597235 | 0.631908 | 0.146863 | 0.557393 | 0.834177 | -0.079858 | 0.117247 | -0.213753 | 0.528107 | 0.320911 | 0.80042 | 2.517222 | 0.232958 | 1.611505 | 0.798151 | -0.048003 | -0.751672 | 0.931513 | -0.632411 | 0.255117 | 0.424044 |
| 27.19444 | 0.434326 | 0.044033 | 0.195728 | -0.493686 | 0.670407 | 0.159435 | 0.744853 | 0.68802 | -0.22399 | 0.141274 | 0.593811 | 0.402806 | 0.352494 | 0.127428 | 1.515454 | 0.754616 | 0.693255 | 0.954925 | 0.002189 | -0.456579 | 0.526331 | -0.664151 | 0.51263 | -0.562195 |
| 27.20833 | -0.006307 | 0.177218 | 0.239811 | -0.385672 | 0.983724 | 0.213253 | 0.886508 | 0.785816 | -0.185734 | 0.192834 | 1.404712 | 0.155671 | 0.152867 | -0.642251 | 0.327322 | 1.956174 | -0.179429 | 0.336056 | 0.20949 | 0.009083 | 0.294319 | -0.220083 | 0.257698 | -0.297962 |
| 27.22222 | 0.051795 | 0.240684 | 0.228651 | -0.248157 | 1.052261 | 0.237926 | 0.971412 | 1.161774 | 0.011961 | 0.205885 | 1.570706 | -0.247036 | -0.202977 | -0.880272 | -0.404674 | 2.484515 | -0.403492 | -0.251363 | 0.216514 | 0.483742 | 0.183699 | -0.060227 | 0.07481 | 0.722626 |
| 27.23611 | 0.656884 | 0.030815 | 0.005377 | -0.247091 | 0.518284 | 0.106057 | 0.805702 | 1.391545 | 0.097644 | 0.058319 | 1.106019 | -0.37307 | -0.470789 | -0.246739 | -1.952498 | 0.768105 | 0.016613 | -0.19028 | 0.041609 | 0.581112 | 0.033527 | 0.043763 | 0.224182 | 1.797055 |
| 27.25 | 0.984134 | -0.27198 | -0.2559 | -0.302385 | 0.248696 | -0.075935 | 0.563943 | 1.121331 | 0.118518 | -0.131056 | 0.45812 | -0.442795 | -0.345961 | 0.257097 | -1.69529 | -1.475971 | -0.288701 | 0.067601 | -0.043339 | 0.154781 | -0.084489 | 0.369445 | 0.249825 | 0.883905 |
| 27.26389 | 0.631772 | -0.237436 | -0.119708 | -0.084727 | 0.354119 | -0.030348 | 0.735982 | 0.859453 | 0.277695 | -0.06652 | 0.344204 | -0.270768 | -0.098708 | 0.643864 | 0.247111 | -1.715328 | -0.530387 | 0.085472 | 0.077312 | -0.365194 | 0.253736 | 0.661599 | 0.181482 | -0.054168 |
| 27.27778 | 0.284685 | -0.052746 | 0.15099 | 0.250671 | 0.604339 | 0.109462 | 1.092035 | 0.900118 | 0.513209 | 0.101701 | 0.609021 | 0.71174 | -0.086601 | 1.334136 | 1.828711 | -0.407122 | -0.277619 | -0.097413 | 0.224914 | -0.23012 | 0.759418 | 0.305703 | 0.086966 | 0.124292 |
| 27.29167 | 0.096556 | -0.036459 | 0.158709 | 0.326615 | 0.932041 | 0.121601 | 0.877905 | 1.091245 | 0.577305 | 0.122274 | 0.764998 | 1.501859 | -0.252751 | 1.565266 | 2.082894 | 0.083386 | 0.08492 | -0.060264 | 0.209929 | 0.179761 | 0.863298 | -0.112811 | 0.110174 | 0.538226 |
| 27.30556 | 0.054286 | -0.005958 | 0.108513 | 0.240418 | 1.074756 | 0.085856 | 0.297246 | 1.254714 | 0.582286 | 0.100586 | 0.74976 | 1.853878 | -0.24685 | 1.686652 | 1.071719 | 0.117524 | 0.238263 | 0.307288 | 0.289713 | 0.76237 | 0.923592 | -0.129859 | 0.245951 | 0.947648 |
| 27.31944 | 0.15305 | 0.09837 | 0.108095 | 0.129963 | 0.685032 | 0.073373 | 0.101414 | 0.948315 | 0.582117 | 0.093196 | 0.862779 | 1.524152 | -0.071622 | 1.327277 | 0.467453 | -0.023371 | 0.136814 | 0.457428 | 0.523995 | 0.739656 | 1.004171 | 0.209513 | 0.270031 | 1.037918 |
| 27.33333 | 0.3525 | 0.164207 | 0.067173 | 0.013527 | 0.562768 | 0.062401 | 0.258918 | 0.626205 | 0.546359 | 0.073225 | 1.034585 | 1.243918 | 0.111182 | 0.408254 | 0.255686 | -0.251864 | -0.055215 | 0.399567 | 0.441589 | 0.582164 | 0.983448 | 0.498667 | 0.110325 | 0.073835 |
| 27.34722 | 0.488658 | 0.048943 | -0.074596 | -0.164978 | 0.888371 | -0.022364 | 0.568622 | 0.494006 | 0.521563 | -0.034512 | 0.814923 | 1.128134 | 0.268176 | 1.136776 | 0.316144 | -0.444456 | -0.159521 | 0.173817 | 0.214145 | -0.193921 | 0.699584 | 0.557503 | 0.095483 | 0.37524 |
| 27.36111 | 0.480952 | -0.203801 | -0.267032 | -0.394538 | 1.129934 | -0.170545 | 0.685182 | 0.456524 | 0.558268 | -0.213464 | 0.328276 | 1.027209 | 0.359601 | 1.983516 | 0.502662 | -0.183393 | 0.132363 | 0.172605 | 0.088991 | -0.577844 | 0.407616 | 0.447651 | 0.203295 | 0.910864 |
| 27.375 | 0.604311 | -0.333063 | -0.307351 | -0.468632 | 0.876798 | -0.219919 | 0.490126 | 0.494398 | 0.853856 | -0.273442 | 0.102813 | 0.791742 | 0.37399 | 2.703168 | 0.549512 | 0.371889 | 0.685733 | 0.397313 | 0.136433 | -0.685881 | 0.471077 | 0.327303 | 0.296977 | 0.568172 |
| 27.38889 | 0.8306 | -0.291453 | -0.203501 | -0.418515 | 0.650253 | -0.166795 | 0.231156 | 0.563478 | 1.375995 | -0.203218 | 0.113659 | 0.672877 | 0.386987 | 2.411985 | 0.420833 | 0.987596 | 1.117886 | 0.653293 | 0.277305 | -0.114414 | 0.808813 | 0.111745 | 0.28477 | 0.7581 |
| 27.40278 | 0.892528 | -0.189907 | -0.1734 | -0.538395 | 0.887976 | -0.157917 | 0.08445 | 0.580557 | 1.893812 | -0.166969 | 0.131532 | 0.62357 | 0.432932 | 1.433913 | 0.338321 | 1.528821 | 1.086258 | 0.689242 | 0.262201 | 0.414098 | 1.038939 | -0.033817 | 0.125189 | 0.845469 |
| 27.41667 | 0.807073 | -0.034276 | -0.202674 | -0.702169 | 1.442102 | -0.163308 | 0.127317 | 0.665447 | 2.276335 | -0.152968 | 0.21013 | 0.620085 | 0.558409 | 1.470086 | 0.513252 | 1.903874 | 0.802386 | 0.470026 | 0.122263 | 0.900365 | 1.144253 | 0.030516 | 0.093928 | 0.640611 |
| 27.43056 | 0.850492 | 0.131621 | -0.184007 | -0.801877 | 1.720041 | -0.123732 | 0.413355 | 0.719625 | 2.37524 | -0.107226 | 0.401769 | 0.732417 | 0.653311 | 2.206413 | 1.094472 | 1.999716 | 0.615537 | 0.296916 | 0.038752 | 1.041207 | 1.30673 | 0.297969 | 0.141797 | 0.895769 |
| 27.44444 | 0.84373 | 0.217821 | -0.224271 | -0.937856 | 1.248342 | -0.148352 | 0.793789 | 0.605548 | 2.391498 | -0.141824 | 0.451412 | 1.03069 | 0.632666 | 2.122743 | 1.446266 | 1.714584 | 0.535629 | 0.312237 | -0.082635 | 0.325916 | 1.287372 | 0.608324 | 0.275541 | 1.697462 |
| 27.45833 | 0.477322 | 0.08351 | -0.348504 | -1.279922 | 0.688874 | -0.212392 | 0.774644 | 0.508787 | 2.490006 | -0.298591 | 0.240616 | 1.034347 | 0.687767 | 2.336032 | 1.039353 | 1.657523 | 0.615791 | 0.439867 | -0.260129 | -0.683777 | 0.918554 | 0.556662 | 0.201693 | 1.516195 |
| 27.47222 | -0.139359 | -0.182392 | -0.277445 | -1.295865 | 0.232581 | -0.073173 | 0.288682 | 0.69729 | 2.393906 | -0.292489 | 0.075946 | 0.838449 | 0.766825 | 3.107126 | 0.66642 | 1.49084 | 0.784019 | 0.674773 | -0.214192 | -0.47779 | 0.530953 | 0.244042 | 0.267596 | 1.671017 |
| 27.48611 | -0.193089 | -0.272452 | -0.111897 | -0.859454 | 0.505621 | 0.072152 | 0.160912 | 0.836161 | 2.57351 | -0.085476 | 0.079125 | 0.407425 | 0.861388 | 1.683273 | 0.68999 | 1.674336 | 1.263785 | 0.880228 | -0.040258 | 0.092246 | 0.536067 | 0.233301 | 0.225629 | 1.632314 |
| 27.5 | 0.06898 | -0.360154 | -0.000599 | -0.464664 | 0.876273 | 0.112475 | 0.142493 | 0.780339 | 2.187808 | 0.056087 | 0.167108 | 0.059644 | 0.619893 | 1.363832 | 1.004461 | 1.360819 | 2.036547 | 0.728945 | 0.051494 | 0.412117 | 0.828101 | 0.070542 | 0.260305 | 0.783685 |
| 27.51389 | 0.15088 | -0.560332 | 0.01199 | -0.077943 | 1.117066 | 0.072294 | 0.212355 | 0.480779 | 1.617627 | 0.048995 | 0.179547 | -0.144201 | 0.405995 | 1.586616 | 1.04175 | 0.267502 | 2.128074 | 0.668837 | 0.136425 | 0.501605 | 1.171125 | -0.207682 | 0.369401 | 0.602328 |
| 27.52778 | -0.250957 | -1.025949 | -0.436998 | -0.326308 | 1.017798 | -0.507793 | -0.287725 | 0.01201 | 1.16397 | -0.539779 | -0.345428 | -0.149197 | 0.272612 | 1.373033 | 1.040687 | 0.462626 | 1.466996 | 0.769139 | 0.142201 | 0.258386 | 1.333398 | -0.255043 | 0.436928 | 0.694905 |
| 27.54167 | -1.361819 | -1.94616 | -1.530752 | -1.595631 | -0.490031 | -1.81012 | -1.471668 | -0.791439 | 0.039703 | -1.878795 | -1.578395 | -0.614123 | 0.25795 | 0.449357 | 1.13717 | 0.7544 | 0.778325 | 0.761066 | 0.047032 | -0.447558 | 1.455988 | -0.135821 | 0.519802 | 1.560754 |
| 27.55556 | -1.904764 | -2.080472 | -1.850785 | -2.26329 | -1.431779 | -2.067811 | -1.827822 | -0.966359 | -0.695743 | -2.115884 | -1.832177 | -0.887353 | 0.077579 | 0.402426 | 0.839064 | 0.747247 | 0.426117 | 0.649427 | -0.015478 | -0.492944 | 1.081643 | -0.019763 | 0.314175 | 1.673484 |
| 27.56944 | -1.136094 | -0.941694 | -0.900209 | -1.509148 | -0.426423 | -0.800436 | -0.870874 | -0.58063 | -0.102583 | -0.865424 | -0.68429 | -0.280627 | -0.041761 | 0.648147 | 0.742623 | 0.643554 | 0.243433 | 0.333531 | -0.009073 | 0.159487 | 0.915073 | 0.333015 | 0.102148 | 0.669773 |
| 27.58333 | -0.002508 | 0.217479 | -0.026243 | -1.089297 | 0.919297 | 0.125661 | 0.174267 | -0.650172 | 0.554688 | 0.099209 | 0.278245 | 0.3926 | -0.022276 | 0.644916 | 0.826714 | 0.538227 | 0.316946 | 0.159046 | 0.046261 | 0.392874 | 0.928362 | 0.237418 | 0.11313 | 0.439078 |
| 27.59722 | 0.711478 | 0.845823 | 0.343939 | -0.769494 | 1.198602 | 0.441656 | 0.447386 | -0.742238 | 0.784649 | 0.452644 | 0.540457 | 0.144959 | 0.003958 | 0.240091 | 0.856708 | 0.371204 | 0.23351 | 0.024471 | 0.109971 | -0.001513 | 0.304258 | 0.097557 | -0.105325 | 0.566697 |
| 27.61111 | 0.901796 | 0.936307 | 0.330654 | -0.187501 | 0.578837 | 0.384709 | 0.391206 | -0.400433 | 0.688407 | 0.420534 | 0.425191 | -1.013213 | -0.004463 | 0.13287 | 0.713976 | 0.005134 | 0.043303 | -0.076875 | 0.071612 | -0.158408 | 0.142917 | 0.252146 | 0.030781 | -0.273104 |
| 27.625 | 0.711828 | 0.602931 | 0.161796 | 0.31716 | 0.483124 | 0.191503 | 0.16344 | -0.027363 | 0.412554 | 0.223264 | 0.199257 | -0.849067 | -0.035127 | -0.143885 | 0.592299 | -0.098059 | -0.116813 | -0.153898 | -0.003792 | 0.11872 | 0.027718 | 0.197036 | 0.136064 | -0.788746 |
| 27.63889 | 0.340669 | 0.20187 | 0.036919 | 0.605572 | 0.745541 | 0.056425 | -0.039835 | 0.353967 | 0.156422 | 0.098694 | -0.06873 | -0.238777 | -0.086995 | -0.304278 | 0.382171 | -0.445967 | -0.198052 | -0.167421 | -0.07226 | 0.432275 | 0.017126 | -0.003508 | 0.123089 | -0.802414 |
| 27.65278 | 0.113763 | -0.104979 | 0.016008 | 0.681638 | 0.064736 | 0.049662 | -0.141913 | 0.772682 | 0.061304 | 0.097819 | -0.047202 | 0.400178 | -0.137256 | 0.206906 | 0.023523 | -1.122186 | -0.240153 | -0.079313 | -0.054432 | 0.5347 | 0.133062 | -0.240152 | 0.119987 | -0.297492 |
| 27.66667 | -0.00847 | -0.300679 | 0.049948 | 0.645781 | -0.291938 | 0.083443 | -0.104828 | 0.802291 | 0.046757 | 0.123005 | 0.183665 | 1.383143 | -0.106707 | 0.649684 | -0.056618 | -2.01543 | -0.20306 | 0.073152 | 0.010057 | 0.065755 | 0.322254 | -0.427391 | -0.029917 | 0.750038 |
| 27.68056 | -0.172986 | -0.397812 | 0.074064 | 0.564797 | 0.318457 | 0.094894 | -0.042793 | 0.524128 | -0.050721 | 0.129943 | 0.309205 | 1.949766 | -0.064935 | 0.582353 | -0.381739 | -1.338062 | -0.135818 | 0.184815 | 0.036467 | -0.196094 | 0.570598 | -0.505929 | -0.090712 | 1.944968 |
| 27.69444 | -0.270988 | -0.383556 | 0.063904 | 0.474714 | 0.59448 | 0.088407 | -0.012695 | 0.185909 | -0.135422 | 0.111671 | 0.18481 | 1.198849 | -0.010346 | 0.204092 | -0.690603 | 0.56154 | -0.031381 | 0.273006 | -0.027065 | -0.329087 | 0.628361 | -0.352691 | -0.248196 | 0.540697 |
| 27.70833 | -0.239649 | -0.262127 | 0.041592 | 0.382486 | 0.205917 | 0.076232 | 0.007816 | -0.168714 | -0.158121 | 0.075972 | 0.013548 | 0.5552 | 0.014338 | 0.148876 | -0.915648 | 2.134871 | 0.061821 | 0.235301 | -0.092802 | -0.326599 | 0.418444 | -0.218769 | -0.11007 | 0.384234 |
| 27.72222 | -0.187389 | -0.14329 | 0.002897 | 0.2595 | -0.764376 | 0.04021 | -0.002889 | -0.397051 | -0.169972 | 0.028783 | -0.11584 | 0.254845 | 0.041295 | -0.032938 | -0.905802 | 1.686758 | 0.087366 | -0.024127 | -0.028958 | -0.127125 | 0.101292 | -0.124191 | -0.007269 | 0.286974 |
| 27.73611 | -0.166905 | -0.113534 | -0.038026 | 0.098406 | -0.825378 | -0.01588 | -0.049576 | -0.499265 | -0.208942 | -0.024452 | -0.119422 | 0.036341 | 0.058453 | -0.224379 | -0.545593 | 0.542792 | 0.032194 | -0.217869 | 0.013923 | 0.332798 | -0.263434 | -0.045619 | 0.135054 | 0.158844 |
| 27.75 | -0.180718 | -0.096389 | -0.060949 | -0.118865 | -0.591021 | -0.070799 | -0.119375 | -0.455192 | -0.258749 | -0.068293 | -0.042027 | -0.213447 | 0.025924 | -0.585972 | -0.122878 | 0.24599 | -0.092278 | -0.103722 | 0.026997 | 0.614818 | -0.611619 | -0.008467 | 0.162093 | 0.547924 |
| 27.76389 | -0.170459 | -0.075959 | -0.081421 | -0.476665 | -0.668438 | -0.088407 | -0.219866 | -0.326569 | -0.280114 | -0.07667 | 0.03147 | -0.512485 | -0.027634 | -0.421551 | 0.402967 | -0.02933 | -0.307244 | 0.014917 | 0.03827 | 0.603387 | -0.895398 | 0.016694 | 0.198771 | -0.67588 |
| 27.77778 | -0.183933 | -0.042612 | -0.076123 | -0.758264 | -0.560995 | -0.056841 | -0.299398 | -0.227146 | -0.266183 | -0.04428 | 0.06297 | -0.749867 | -0.055333 | 0.016075 | 1.035376 | 0.060365 | -0.572386 | 0.069158 | 0.007661 | 0.385313 | -0.981699 | 0.050938 | 0.214507 | -0.714897 |
| 27.79167 | -0.290772 | 0.003498 | -0.054594 | -0.624996 | -0.467706 | -0.025005 | -0.371594 | -0.118543 | -0.271548 | -0.016976 | -0.001656 | -0.929572 | -0.065659 | 0.386751 | 1.843872 | 0.103601 | -0.929258 | -0.06437 | 0.041909 | 0.389306 | -0.83796 | 0.003413 | 0.092242 | -0.190439 |
| 27.80556 | -0.54764 | 0.041169 | -0.042387 | -0.355345 | -0.422104 | -0.016011 | -0.41085 | 0.009467 | -0.317059 | -0.004526 | -0.150399 | -0.876243 | -0.076457 | 0.195211 | 2.124119 | -0.035619 | -0.851147 | -0.155112 | 0.067774 | 0.253664 | -0.490293 | -0.141855 | -0.180102 | 0.324544 |
| 27.81944 | -0.702896 | 0.083707 | 0.005943 | -0.103071 | -0.616925 | 0.014287 | -0.370238 | 0.134226 | -0.351351 | 0.032432 | -0.24643 | -0.647901 | -0.050773 | -0.014982 | 0.561708 | -0.240358 | -0.531208 | -0.216926 | 0.091614 | -0.169614 | 0.042774 | -0.286037 | -0.396385 | 0.49468 |
| 27.83333 | -0.565273 | 0.126023 | 0.067828 | 0.171843 | -0.833 | 0.06501 | -0.339798 | 0.065608 | -0.408893 | 0.081333 | -0.201219 | -0.483812 | -0.068164 | -0.35791 | -0.857691 | -1.832764 | -0.061932 | -0.307107 | 0.12027 | -0.423056 | 0.422102 | -0.427791 | -0.173188 | 0.192898 |

| | | | | | | | | | | | | | | | | | | | | | | |
|---|---|---|---|---|---|---|---|---|---|---|---|---|---|---|---|---|---|---|---|---|---|---|
| 27.84722 | -0.376434 | 0.139219 | 0.089875 | 0.364263 | -1.073778 | 0.084067 | -0.413569 | -0.046983 | -0.562244 | 0.096226 | -0.058796 | -0.515887 | -0.090861 | -0.707931 | -2.123441 | -1.576807 | 0.201859 | -0.272039 | 0.113343 | -0.218075 | 0.296349 | -0.479723 | -0.219435 | -0.406833 |
| 27.86111 | -0.297194 | 0.100538 | 0.054846 | 0.38412 | -1.242853 | 0.061668 | -0.585259 | -0.284566 | -0.827088 | 0.067314 | 0.023145 | -0.54957 | -0.042644 | -0.710716 | -1.975617 | -0.255213 | 0.008448 | -0.321246 | 0.094345 | -0.112792 | -0.336663 | -0.371642 | -0.270134 | -0.579342 |
| 27.875 | -0.375552 | 0.032263 | 0.004404 | 0.321122 | -0.766865 | 0.026018 | -0.680965 | -0.503518 | -1.019006 | 0.022734 | -0.116068 | -0.522415 | 0.008054 | -0.798166 | -2.359711 | 1.10728 | -0.423578 | -0.41341 | 0.073294 | 0.00058 | -1.197465 | -0.386929 | -0.337665 | -0.645774 |
| 27.88889 | -0.560383 | -0.004195 | -0.011855 | 0.27538 | -0.376191 | 0.01006 | -0.651204 | -0.711022 | -1.126455 | 0.003465 | -0.405521 | -0.429899 | 0.006514 | -0.805066 | -1.693028 | 1.361467 | -0.971682 | -0.727747 | -0.008705 | -0.155851 | -1.548952 | -0.541125 | -0.2801 | -0.728384 |
| 27.90278 | -0.841415 | -0.00035 | 0.005371 | 0.23658 | -0.383928 | 0.016434 | -0.555026 | -0.892859 | -1.112176 | 0.011218 | -0.611526 | -0.465843 | -0.063482 | -0.752962 | -0.050029 | 0.405809 | -1.669857 | -0.819565 | -0.039737 | -0.702122 | -1.479054 | -0.569275 | -0.34004 | -0.848096 |
| 27.91667 | -1.174557 | -0.008518 | 0.00632 | 0.183516 | -0.829614 | 0.003275 | -0.653205 | -1.019376 | -1.049302 | -0.001076 | -0.762078 | -0.539452 | -0.240231 | -0.87551 | 0.951221 | 0.038195 | -2.143576 | -0.647886 | -0.107792 | -0.375553 | -0.983011 | -0.64812 | -0.438458 | -1.101678 |
| 27.93056 | -1.305021 | -0.027433 | -0.002145 | 0.121239 | -1.005499 | -0.022271 | -0.847421 | -1.092921 | -0.928041 | -0.019641 | -0.836964 | -0.640064 | -0.42495 | -1.60722 | 0.880811 | -0.562758 | -2.070321 | -0.474712 | -0.16779 | 0.311743 | -1.165379 | -1.072859 | -0.62113 | -1.296051 |
| 27.94444 | -1.023697 | -0.053893 | -0.000854 | 0.084788 | -0.940673 | -0.031601 | -0.823386 | -1.083096 | -0.801726 | -0.01236 | -0.792066 | -0.801261 | -0.459432 | -2.342773 | 0.350848 | -1.654369 | -1.75806 | -0.429411 | -0.22224 | 0.157556 | -0.343871 | -1.047224 | -0.61187 | -1.343459 |
| 27.95833 | -0.617552 | -0.100946 | -0.018519 | 0.049773 | -1.009561 | -0.040715 | -0.597552 | -1.081608 | -0.707232 | -0.017344 | -0.637514 | -0.971865 | -0.44154 | -2.180839 | 0.054332 | -2.87531 | -1.471942 | -0.369753 | -0.245341 | -0.1224 | 0.087657 | -0.982639 | -0.65207 | -1.589631 |
| 27.97222 | -0.220157 | -0.133594 | -0.046497 | 0.008038 | -1.141276 | -0.049544 | -0.414076 | -1.077001 | -0.65475 | -0.042748 | -0.415904 | -1.04876 | -0.522422 | -2.607945 | -0.200733 | -2.737958 | -0.922212 | -0.27847 | -0.276285 | -0.397465 | -0.301712 | -0.865108 | -0.600962 | -1.787839 |
| 27.98611 | 0.083041 | -0.106977 | -0.033113 | 0.001078 | -1.032952 | -0.037886 | -0.352284 | -0.966275 | -0.647041 | -0.040416 | -0.151941 | -0.894697 | -0.602462 | -2.674245 | -0.464187 | -1.837535 | -0.558737 | -0.13372 | -0.273386 | -0.562302 | -1.261672 | -0.760794 | -0.540013 | -1.909836 |
| 28 | 0.298442 | 0.065142 | 0.106064 | 0.160742 | -0.730956 | 0.04706 | -0.274588 | -0.686554 | -0.515024 | 0.082659 | 0.082384 | -0.570095 | -0.621032 | -1.788283 | -0.887271 | -0.998244 | -0.367481 | 0.0485 | -0.111156 | -0.704044 | -1.231351 | -0.409925 | -0.423204 | -1.190043 |
| 28.01389 | 0.440882 | 0.30214 | 0.302622 | 0.458796 | -0.470925 | 0.163132 | -0.128722 | -0.40588 | -0.282708 | 0.291214 | 0.286306 | -0.16153 | -0.390698 | -0.812322 | -1.46269 | -0.352622 | -0.148697 | 0.271342 | 0.137726 | -0.474887 | -0.947305 | 0.062898 | -0.15973 | -0.376267 |
| 28.02778 | 0.251409 | 0.112123 | 0.184844 | 0.447936 | -0.551835 | 0.079201 | -0.243081 | -0.581626 | -0.294332 | 0.241233 | 0.211249 | -0.075732 | -0.353347 | -0.62556 | -1.639464 | 0.146932 | -0.064732 | 0.226977 | 0.295266 | 0.195532 | -0.699486 | 0.284364 | -0.095791 | -0.191551 |
| 28.04167 | -0.281373 | -0.430712 | -0.300518 | -0.032927 | -0.905298 | -0.261276 | -0.642297 | -1.222256 | -0.710666 | -0.136181 | -0.254806 | -0.399064 | -0.651197 | -1.758176 | -0.982295 | -0.009445 | -0.506133 | -0.291983 | 0.139939 | 0.547321 | -0.169774 | -0.135212 | -0.501714 | -0.550697 |
| 28.05556 | -0.509175 | -0.500567 | -0.405542 | -0.110416 | -1.229801 | -0.228601 | -0.923527 | -2.208473 | -1.229516 | -0.176076 | -0.456929 | -0.422643 | -1.702058 | -3.14258 | -0.329065 | -0.608554 | -1.525459 | -1.172548 | -0.346007 | 0.476138 | 0.215348 | -1.165527 | -1.133377 | -1.776262 |
| 28.06944 | -0.538445 | -0.161342 | -0.020805 | 0.161147 | -1.204867 | 0.128829 | -0.974248 | -3.344126 | -1.799491 | 0.128341 | -0.458957 | -0.004152 | -1.487457 | -4.547753 | 0.767578 | -0.618028 | -2.220018 | -1.350288 | -0.948539 | 0.198945 | -0.15705 | -1.59422 | -0.974287 | -2.443681 |
| 28.08333 | -0.495549 | 0.032014 | 0.237905 | 0.297798 | -0.683486 | 0.299538 | -0.708082 | -3.423847 | -1.914286 | 0.300471 | -0.539204 | 0.578148 | -0.542169 | -3.965845 | 1.57399 | -0.011883 | -1.708437 | -0.912369 | -1.139884 | -0.206655 | -1.511638 | -0.720546 | -0.331936 | -1.709938 |
| 28.09722 | -0.332502 | -0.03138 | 0.205111 | 0.192725 | -0.115184 | 0.241969 | -0.115964 | -1.927826 | -1.463168 | 0.237454 | -0.445119 | 0.792985 | -0.040554 | -1.903685 | 0.496893 | 0.253813 | -0.736347 | -0.412431 | -0.799533 | -0.316285 | -3.097428 | 0.175776 | 0.017373 | -1.074985 |
| 28.11111 | -0.350457 | -0.17966 | 0.089461 | -0.037181 | 0.09763 | 0.131178 | 0.26466 | -0.697204 | -1.156918 | 0.114026 | -0.251057 | 0.35861 | 0.147643 | 0.144743 | -0.397351 | -0.046439 | 0.11404 | -0.045979 | -0.378889 | -0.303676 | -2.438517 | 0.817835 | 0.172075 | -0.493303 |
| 28.125 | -0.208089 | -0.30893 | 0.015023 | -0.288211 | -0.152971 | 0.057546 | 0.318294 | -0.057017 | -0.651133 | 0.03537 | -0.103795 | 0.014995 | 0.137833 | 1.132987 | -0.930599 | -0.107522 | 0.718186 | 0.293037 | -0.07155 | -0.189641 | -0.937458 | 0.935028 | 0.194718 | -0.04599 |
| 28.13889 | 0.105863 | -0.407951 | 0.002253 | -0.432893 | -0.26857 | 0.026876 | 0.216798 | 0.345109 | -0.163888 | 0.008892 | -0.105623 | -0.001092 | 0.131065 | 1.782907 | -1.589727 | 0.026055 | 1.178989 | 0.329433 | 0.171626 | 0.116932 | 0.018401 | 1.116757 | 0.252642 | 0.441355 |
| 28.15278 | 0.496302 | -0.424036 | 0.008033 | -0.381266 | 0.159667 | 0.005277 | 0.10392 | 0.675087 | 0.012508 | 0.001539 | -0.228839 | 0.190778 | 0.174355 | 2.46533 | -1.375454 | 0.168603 | 1.309354 | 0.337528 | 0.264589 | 0.12974 | 0.655572 | 1.218968 | 0.338155 | 1.005568 |
| 28.16667 | 0.95716 | -0.424017 | 0.00541 | -0.272922 | 0.751871 | -0.006309 | -0.032241 | 0.803155 | -0.127258 | -0.000404 | -0.308151 | 0.392926 | 0.16081 | 1.876258 | -1.268849 | 0.257359 | 1.190033 | 0.250955 | 0.196145 | -0.003624 | 0.823204 | 0.737626 | 0.53886 | 1.460751 |
| 28.18056 | 1.205955 | -0.385022 | 0.034222 | -0.137696 | 0.974142 | 0.03694 | -0.011624 | 0.777443 | -0.271722 | 0.033865 | -0.083463 | 0.39873 | 0.032935 | 0.626734 | -1.158532 | 0.295342 | 1.238051 | 0.380196 | 0.054361 | -0.127822 | 0.74508 | -0.071572 | 0.502343 | 1.048678 |
| 28.19444 | 0.905923 | -0.358361 | 0.085753 | -0.01512 | 0.964232 | 0.099763 | 0.326826 | 0.784663 | -0.273459 | 0.092568 | 0.496491 | 0.122204 | -0.063485 | -0.325669 | -1.026843 | 0.445355 | 1.582682 | 0.610133 | -0.041415 | -0.411026 | 0.786137 | -0.76647 | 0.345915 | 0.25583 |
| 28.20833 | 0.386018 | -0.333132 | 0.099176 | 0.033717 | 0.933602 | 0.111275 | 0.656066 | 0.716416 | -0.23966 | 0.110724 | 1.016059 | -0.151442 | 0.0644 | -0.408881 | -0.642926 | 0.838323 | 1.781896 | 0.489442 | -0.039931 | -0.481691 | 0.861496 | -1.079006 | 0.004959 | -0.295256 |
| 28.22222 | -0.007088 | -0.336795 | 0.083874 | 0.003737 | 0.862797 | 0.090717 | 0.631593 | 0.731899 | -0.161905 | 0.08956 | 1.006719 | -0.194345 | 0.295017 | -0.098595 | 0.117943 | 1.921251 | 0.996515 | 0.173025 | 0.076315 | -0.074902 | 0.73986 | -0.581024 | -0.1027 | -0.039143 |
| 28.23611 | 0.075235 | -0.363034 | 0.050288 | -0.051606 | 0.764142 | 0.050824 | 0.484808 | 0.779575 | 0.020613 | 0.046389 | 0.676625 | -0.300392 | 0.272668 | -0.032574 | 0.686196 | 2.250766 | -0.590846 | -0.17248 | 0.08852 | 0.32962 | 0.402119 | 0.038995 | 0.041547 | 0.3717 |
| 28.25 | 0.630343 | -0.435296 | -0.024267 | -0.114846 | 0.486026 | -0.028976 | 0.515207 | 0.860952 | 0.225208 | -0.025815 | 0.470735 | -0.374229 | 0.135668 | -0.121757 | 0.763291 | 0.769386 | -0.569453 | -0.368446 | 0.052888 | 0.307343 | -0.044822 | 0.444686 | 0.476254 | 0.598467 |
| 28.26389 | 1.102284 | -0.51549 | -0.113306 | -0.192858 | 0.591734 | -0.10763 | 0.699468 | 1.092007 | 0.32348 | -0.106486 | 0.599065 | -0.532118 | 0.29613 | 0.303399 | 0.643554 | -1.356605 | 0.738654 | -0.118745 | 0.083987 | 0.120212 | -0.383989 | 0.785106 | 0.479194 | 0.174498 |
| 28.27778 | 1.168724 | -0.53591 | -0.160752 | -0.264706 | 0.833067 | -0.142927 | 0.904088 | 1.511193 | 0.370501 | -0.153094 | 0.844771 | -0.643976 | 0.505638 | 1.07943 | 0.251315 | -2.390126 | 0.82579 | 0.329452 | 0.171528 | 0.098788 | -0.396056 | 0.787889 | 0.202084 | -0.137583 |
| 28.29167 | 0.962607 | -0.501858 | -0.158856 | -0.294324 | 0.870262 | -0.140953 | 1.097788 | 1.646222 | 0.511359 | -0.153122 | 0.893463 | -0.383358 | 0.236876 | 1.289467 | -0.334206 | -1.039411 | -0.103571 | 0.667567 | 0.147836 | 0.492356 | 0.050747 | 0.541097 | -0.153452 | 0.251742 |
| 28.30556 | 0.769101 | -0.457621 | -0.134988 | -0.26652 | 0.784231 | -0.120126 | 1.139898 | 1.232854 | 0.716552 | -0.12806 | 0.883582 | 0.072356 | -0.182452 | 1.259232 | -0.163704 | 0.044781 | -0.250494 | 0.517259 | 0.092344 | 0.889385 | 0.710893 | 0.216833 | -0.181868 | 1.031352 |
| 28.31944 | 0.671898 | -0.412986 | -0.092775 | -0.197374 | 1.022825 | -0.073804 | 0.863743 | 0.878955 | 0.87448 | -0.085359 | 0.793587 | 0.771261 | -0.338088 | 1.410208 | 0.642793 | 0.434894 | 0.658604 | 0.257855 | 0.166605 | 0.541768 | 1.184237 | -0.016423 | 0.419364 | 1.41259 |
| 28.33333 | 0.661076 | -0.360485 | -0.016546 | -0.094393 | 1.276018 | -0.003008 | 0.64679 | 0.906215 | 0.96425 | -0.008772 | 0.636247 | 1.247608 | -0.129335 | 1.563869 | 1.269704 | 0.318631 | 1.194584 | 0.309212 | 0.294816 | -0.54283 | 1.336864 | 0.048817 | 0.902152 | 1.319925 |
| 28.34722 | 0.700525 | -0.325719 | 0.052381 | -0.012363 | 1.300474 | 0.062103 | 0.702783 | 1.082927 | 0.882576 | 0.05949 | 0.629763 | 1.50465 | 0.220722 | 2.018236 | 1.106949 | 0.132567 | 0.418442 | 0.344054 | 0.330735 | -0.785899 | 1.190403 | 0.353246 | 0.60369 | 0.61876 |
| 28.36111 | 0.742025 | -0.342091 | 0.047572 | -0.0256 | 1.255892 | 0.063346 | 1.012584 | 1.109682 | 0.73665 | 0.050434 | 0.57576 | 1.679869 | 0.385484 | 1.68837 | 0.51771 | 0.014434 | -0.558346 | 0.381927 | 0.122053 | -0.713853 | 0.793248 | 0.603023 | 0.278897 | 0.494844 |
| 28.375 | 0.808236 | -0.328062 | 0.021709 | -0.082096 | 1.225111 | 0.03525 | 1.058944 | 1.029345 | 0.796577 | 0.009163 | 0.571927 | 1.680142 | 0.319176 | 1.077247 | 0.197799 | 0.008601 | -0.196453 | 0.438326 | -0.013102 | 0.090139 | 0.624229 | 0.644638 | 0.118613 | 1.074487 |
| 28.38889 | 0.990542 | -0.25394 | 0.075062 | -0.051877 | 1.308778 | 0.078339 | 0.692764 | 0.928618 | 1.057281 | 0.052416 | 0.632446 | 1.291588 | 0.268184 | 1.68945 | 0.348596 | 0.307837 | 1.045785 | 0.562254 | 0.087635 | 0.341017 | 0.939047 | 0.544576 | 0.238996 | 1.803427 |
| 28.40278 | 1.278954 | -0.151727 | 0.238354 | 0.121568 | 1.522231 | 0.21816 | 0.295672 | 1.022655 | 1.414562 | 0.211189 | 0.764171 | 0.752106 | 0.380374 | 2.496991 | 0.713103 | 0.960748 | 1.616327 | 0.697232 | 0.262105 | 0.24754 | 1.33085 | 0.482687 | 0.484821 | 2.167773 |
| 28.41667 | 1.561315 | 0.000233 | 0.44304 | 0.329623 | 1.765491 | 0.38099 | 0.163873 | 1.35084 | 1.644334 | 0.396051 | 0.81168 | 1.000666 | 0.529809 | 2.981623 | 0.988285 | 1.680845 | 0.980076 | 0.945247 | 0.260502 | 0.222963 | 1.629728 | 0.517637 | 0.599234 | 1.679882 |
| 28.43056 | 1.558397 | 0.211415 | 0.49882 | 0.359078 | 2.027596 | 0.457876 | 0.479917 | 1.611508 | 1.776352 | 0.45282 | 0.709515 | 2.243661 | 0.575002 | 3.424342 | 0.834218 | 1.789142 | 0.684238 | 0.937854 | 0.10967 | -0.182497 | 1.752659 | 0.482579 | 0.506467 | 1.439141 |
| 28.44444 | 1.297037 | 0.266157 | 0.246383 | 0.104542 | 2.294046 | 0.345768 | 0.854248 | 1.564973 | 1.693259 | 0.274972 | 0.477089 | 2.576009 | 0.48335 | 2.855389 | 0.607104 | 1.656172 | 0.870083 | 0.771035 | -0.122409 | -0.519961 | 1.633551 | 0.338651 | 0.306542 | 1.583549 |
| 28.45833 | 0.95829 | 0.003497 | -0.131231 | -0.431887 | 2.247046 | 0.057792 | 0.877141 | 1.422229 | 1.361491 | -0.044515 | 0.264424 | 1.890903 | 0.367892 | 1.999571 | 0.656188 | 1.816986 | 1.286691 | 0.589482 | -0.300812 | -0.754131 | 1.242629 | 0.268671 | 0.117153 | 1.639114 |
| 28.47222 | 0.922734 | -0.169534 | -0.140382 | -0.450551 | 1.995189 | 0.030873 | 0.691491 | 0.886203 | 0.944531 | -0.095605 | 0.26301 | 1.536387 | 0.423742 | 1.546909 | 1.197329 | 1.921469 | 1.804273 | 0.509545 | -0.243679 | -0.48602 | 0.818715 | 0.39929 | 0.008032 | 1.206181 |
| 28.48611 | 1.22215 | -0.134999 | 0.045357 | -0.128975 | 1.29188 | 0.130781 | 0.407192 | 0.873778 | 1.120657 | 0.059995 | 0.412904 | 1.399463 | 0.704241 | 1.943101 | 1.543598 | 1.86373 | 2.11707 | 0.634494 | -0.124005 | 0.707382 | 0.898642 | 0.924278 | 0.328718 | 1.118727 |
| 28.5 | 1.006367 | -0.240099 | 0.088456 | -0.028174 | 0.914629 | 0.093737 | 0.101976 | 0.532057 | 1.303658 | 0.082293 | 0.348074 | 0.865599 | 1.07259 | 1.981494 | 1.456536 | 1.420901 | 1.824783 | 0.67726 | -0.101364 | 0.716868 | 1.041291 | 1.106486 | 0.969519 | 1.63772 |
| 28.51389 | 0.582428 | -0.342794 | -0.046117 | -0.179061 | 0.670739 | -0.073858 | -0.164015 | 0.647695 | 1.195833 | -0.070238 | 0.08317 | -0.527987 | 1.169922 | 1.437707 | 1.396614 | 1.018495 | 1.721655 | 0.737357 | -0.164744 | 0.589913 | 1.088347 | 0.903766 | 1.034667 | 2.15936 |
| 28.52778 | 0.047615 | -0.631083 | -0.414192 | -0.642816 | 0.252986 | -0.518977 | -0.493997 | 0.687168 | 0.720531 | -0.553333 | -0.488243 | -2.324154 | 0.45832 | 1.244986 | 1.102348 | 0.677134 | 1.25992 | 0.76377 | -0.139584 | 0.211222 | 0.845969 | 0.632336 | 0.850632 | 1.858305 |
| 28.54167 | -0.578878 | -1.23197 | -0.995184 | -1.41234 | 0.029065 | -1.243506 | -0.957607 | 0.098064 | -0.086405 | -1.325473 | -1.151152 | -3.685873 | 0.266347 | 1.340653 | 0.935254 | 0.328088 | 0.965186 | 0.535923 | 0.007477 | 0.041757 | 0.849636 | 0.425479 | 0.761893 | 1.297023 |
| 28.55556 | -0.696019 | -1.395257 | -1.186045 | -1.591345 | -0.141676 | -1.407433 | -0.974852 | -0.417096 | -0.522038 | -1.457117 | -1.125829 | -1.151176 | 0.153715 | 1.142729 | 0.756742 | 0.269258 | 0.779518 | 0.378713 | -0.052493 | 0.250939 | 1.268134 | 0.259481 | 0.057268 | 1.30146 |
| 28.56944 | -0.309642 | -0.865171 | -0.819128 | -0.983097 | -0.06608 | -0.82317 | -0.399935 | -0.573607 | -0.377431 | -0.830979 | -0.513583 | 0.012663 | 0.008466 | 1.056555 | 1.07379 | 0.43216 | 0.661119 | 0.198897 | -0.029429 | 0.25465 | 0.811919 | 0.051918 | -0.32558 | 1.413438 |
| 28.58333 | 0.059069 | -0.341249 | -0.348683 | -0.411705 | 0.171508 | -0.278758 | 0.126122 | -0.615724 | -0.165801 | -0.272562 | -0.084342 | 0.69966 | 0.029536 | 0.574966 | 1.496272 | 0.487408 | 0.611472 | 0.212213 | 0.28826 | 0.221842 | 0.551631 | -0.071507 | -0.467942 | 0.291607 |
| 28.59722 | 0.186404 | -0.102064 | -0.066774 | -0.152832 | 0.118665 | -0.032415 | 0.245396 | -0.81415 | -0.006163 | -0.033254 | 0.033874 | 2.273457 | 0.074502 | 0.13653 | 1.724007 | 0.503186 | 0.320945 | 0.161639 | 0.222098 | 0.090539 | 0.429834 | -0.130913 | -0.314133 | -0.342504 |
| 28.61111 | 0.015941 | -0.10238 | -0.033099 | -0.142154 | 0.267153 | -0.034061 | 0.019763 | -0.216675 | 0.048129 | -0.033955 | 0.017577 | 1.545268 | 0.005205 | -0.280565 | 1.236163 | 0.449897 | 0.093892 | 0.035796 | 0.120786 | 0.501192 | 0.344696 | 0.023068 | 0.684794 | 0.438452 |
| 28.625 | -0.235416 | -0.170204 | -0.105354 | -0.200307 | 0.262298 | -0.091248 | -0.280791 | 0.078038 | -0.026574 | -0.096968 | -0.105011 | 0.261917 | -0.062761 | -0.561231 | 0.378243 | -0.0317 | -0.0764 | -0.104725 | 0.082756 | 0.685863 | 0.255102 | 0.256754 | 0.581273 | 0.377416 |
| 28.63889 | -0.182213 | -0.174286 | -0.151837 | -0.222087 | -0.035215 | -0.10437 | -0.338435 | 0.092457 | -0.156679 | -0.120882 | -0.179558 | -0.581877 | -0.05297 | -0.46236 | -0.324467 | -0.553005 | -0.196852 | -0.10903 | 0.039187 | 0.048973 | 0.151738 | 0.387539 | -0.016039 | 0.183275 |
| 28.65278 | -0.269864 | -0.173377 | -0.169768 | -0.231277 | 0.237394 | -0.111484 | -0.373985 | -0.04154 | -0.196176 | -0.136579 | -0.121019 | -1.310751 | -0.03542 | -0.371 | -0.467413 | -0.81917 | -0.250122 | -0.486606 | -0.141924 | -0.892056 | 0.026138 | 0.548542 | 0.122426 | -0.159325 |
| 28.66667 | -0.354092 | -0.134165 | -0.126517 | -0.186668 | 0.323637 | -0.090742 | -0.241909 | -0.144808 | 0.003768 | -0.109622 | -0.041862 | -1.81788 | -0.023637 | 0.13547 | -0.262642 | -0.574795 | -0.202078 | -0.549107 | 0.072542 | -1.11848 | -0.243454 | 0.496641 | -0.17158 | 0.840614 |
| 28.68056 | -0.123327 | -0.037751 | -0.012116 | -0.07353 | -0.367513 | -0.009937 | 0.060361 | -0.122032 | 0.233107 | -0.015539 | 0.043839 | -1.974931 | 0.005552 | 0.359216 | -0.251322 | -0.480739 | -0.055741 | 0.004762 | 0.232736 | -0.715128 | -0.291701 | 0.291056 | -0.313506 | 0.196296 |
| 28.69444 | 0.177013 | 0.068632 | 0.114636 | 0.056768 | -1.076764 | 0.095398 | 0.285516 | -0.098776 | 0.389422 | 0.099227 | 0.140701 | -1.107859 | 0.01519 | 0.006994 | -0.276735 | -0.218431 | -0.151065 | -0.19623 | 0.194039 | 0.026695 | 0.074992 | -0.014032 | 0.0679 | -0.4863 |
| 28.70833 | 0.297259 | 0.137365 | 0.189633 | 0.136361 | -0.343493 | 0.171259 | 0.379624 | -0.141435 | 0.424886 | 0.168924 | 0.168883 | 0.070554 | 0.030411 | 0.096384 | -0.244573 | 0.288457 | -0.256903 | -0.215353 | 0.076084 | 0.647527 | 0.419009 | -0.142107 | 0.451509 | 0.005908 |
| 28.72222 | 0.247619 | 0.110644 | 0.158371 | 0.108893 | 0.472856 | 0.154669 | 0.312795 | -0.235218 | 0.36444 | 0.134249 | 0.089221 | 1.094386 | 0.057743 | -0.103216 | -0.107571 | 0.488617 | -0.199973 | 0.126526 | 0.131472 | 0.919911 | 0.895192 | -0.113221 | 0.333295 | -0.159279 |
| 28.73611 | 0.132726 | 0.023593 | 0.060567 | 0.027764 | 0.884415 | 0.070594 | 0.058551 | -0.311389 | 0.261389 | 0.032746 | -0.01558 | 1.595225 | 0.013905 | -0.399804 | -0.116195 | 1.263355 | 0.203774 | -0.003125 | 0.096263 | 0.822714 | 0.655428 | -0.080897 | 0.210914 | 0.017588 |
| 28.75 | -0.012363 | -0.040373 | -0.023065 | -0.043713 | 0.414725 | -0.012916 | -0.303744 | -0.223852 | 0.170738 | -0.062406 | -0.042824 | 1.539154 | 0.007021 | -0.448558 | -0.023325 | 0.678401 | 0.268376 | 0.131949 | 0.066537 | 0.44848 | 0.274281 | -0.044637 | -0.294074 | 0.104989 |
| 28.76389 | -0.45877 | -0.076883 | -0.075181 | -0.094266 | 0.229338 | -0.076407 | -0.366649 | -0.059628 | 0.087229 | -0.134021 | -0.042385 | 1.603057 | 0.016979 | -0.266682 | -0.001882 | 0.318008 | 0.087319 | 0.005545 | 0.026373 | 0.378282 | -0.121244 | -0.038143 | -0.43954 | -0.718338 |
| 28.77778 | -0.806956 | -0.097922 | -0.091889 | -0.111382 | 0.217372 | -0.101988 | -0.201774 | 0.040532 | 0.004751 | -0.165877 | -0.066757 | 0.772132 | 0.022948 | -0.433284 | -0.043088 | 0.0676 | -0.173512 | 0.399749 | -0.066349 | 0.411481 | -0.543554 | -0.061009 | -0.280958 | -0.708931 |
| 28.79167 | -0.828938 | -0.106307 | -0.085291 | -0.10879 | -1.154863 | -0.101502 | -0.080518 | 0.035171 | -0.073163 | -0.150796 | -0.106734 | 0.030027 | 0.001634 | -0.208277 | -0.024182 | -0.158705 | -0.392579 | 0.12833 | -0.094209 | 0.033106 | -0.95104 | -0.140068 | -0.110953 | -0.692345 |
| 28.80556 | -0.737814 | -0.12223 | -0.090931 | -0.119427 | -2.455547 | -0.110533 | -0.029044 | -0.058329 | -0.183268 | -0.145712 | -0.155639 | -0.464876 | -0.064359 | -0.113006 | 0.201142 | -0.320346 | -0.457798 | -0.226005 | -0.0733 | -0.457407 | -0.973386 | -0.326379 | 0.062873 | -1.061492 |
| 28.81944 | -0.491376 | -0.138437 | -0.111957 | -0.128811 | -2.181942 | -0.124986 | -0.111547 | -0.234494 | -0.33445 | -0.176467 | -0.232087 | -0.822419 | -0.130059 | -0.37432 | 0.646047 | -0.290537 | -0.506239 | -0.036685 | -0.079647 | -0.996581 | -0.791241 | -0.565027 | -0.019649 | -0.490594 |
| 28.83333 | -0.462693 | -0.128921 | -0.119369 | -0.105047 | -1.418033 | -0.11799 | -0.319669 | -0.47121 | -0.515418 | -0.171601 | -0.232809 | -1.076112 | -0.146937 | -0.616945 | 0.716397 | -0.380435 | -0.749267 | -0.01859 | 0.013315 | -0.878488 | -0.345652 | -0.709688 | -0.206371 | 0.270841 |
| 28.84722 | -0.531278 | -0.095079 | -0.095861 | -0.054805 | -1.064728 | -0.085148 | -0.565535 | -0.752806 | -0.757021 | -0.120699 | -0.257275 | -1.078117 | -0.1269 | -0.821928 | 0.318383 | -1.475533 | -0.814201 | -0.137591 | 0.309474 | -0.578633 | 0.416542 | -0.583624 | -0.350636 | 0.462733 |
| 28.86111 | -0.674017 | -0.059338 | -0.049806 | -0.005839 | -0.978705 | -0.043208 | -0.789666 | -1.006627 | -0.877766 | -0.063106 | -0.307244 | -1.00264 | -0.168357 | -0.732528 | -0.411738 | -1.505946 | -0.889425 | -0.203162 | 0.397405 | -0.670729 | 0.519774 | -0.392031 | -0.386349 | -0.269672 |
| 28.875 | -0.839716 | -0.028352 | -0.004751 | 0.034989 | -0.967799 | -0.005623 | -1.180369 | -0.884093 | -0.893789 | -0.018482 | -0.219527 | -1.282148 | -0.360513 | -0.563931 | -0.833028 | -0.101653 | -0.819422 | -0.299133 | 0.156123 | -0.317411 | 0.330128 | -0.213683 | -0.339453 | -0.791113 |
| 28.88889 | -1.013783 | -0.00771 | 0.021655 | 0.054834 | -1.113262 | 0.024357 | -1.63411 | -0.836292 | -0.866184 | 0.015515 | -0.077977 | -1.513397 | -0.415925 | -0.618715 | -1.222555 | 1.481378 | -0.727095 | -0.07225 | -0.113639 | 0.29203 | -0.252847 | -0.177959 | -0.400173 | -1.451134 |
| 28.90278 | -1.211544 | 0.001076 | 0.045443 | 0.066431 | -1.188069 | 0.044327 | -1.721056 | -0.807477 | -0.797501 | 0.038674 | -0.104772 | -1.516728 | -0.512507 | -1.260182 | -1.881269 | 1.439333 | -0.840645 | -0.457116 | -0.153566 | 0.571427 | -0.983078 | -0.32806 | -0.4144 | -1.911619 |
| 28.91667 | -1.607691 | 0.020061 | 0.077377 | 0.089088 | -1.077574 | 0.060723 | -1.491562 | -0.663521 | -0.626629 | 0.067451 | -0.263354 | -1.267989 | -0.520184 | -2.296575 | -2.609332 | -0.137659 | -1.226065 | -0.879814 | -0.005387 | 0.386611 | -1.221394 | -0.520216 | -0.430771 | -1.522637 |
| 28.93056 | -1.98112 | 0.060847 | 0.113578 | 0.115017 | -1.103076 | 0.086501 | -0.975477 | -0.57729 | -0.515114 | 0.109186 | -0.366996 | -1.079907 | -0.386285 | -3.352731 | -2.027492 | -0.551784 | -1.687048 | -0.986561 | 0.054159 | -0.019131 | -0.940091 | -0.774426 | -0.461265 | -1.141219 |
| 28.94444 | -1.638974 | 0.111566 | 0.160932 | 0.150804 | -1.131267 | 0.138858 | -0.512965 | -0.416716 | -0.408144 | 0.161848 | -0.692576 | -0.842863 | -0.329547 | -3.25841 | -1.580148 | -1.048008 | -1.844863 | -0.856961 | 0.016203 | -0.436191 | -1.346585 | -1.049256 | -0.537134 | -1.050437 |
| 28.95833 | -1.052909 | 0.137686 | 0.197643 | 0.176765 | -1.084789 | 0.185753 | -0.191159 | -0.340618 | -0.287332 | 0.198592 | -0.770235 | -0.600214 | -0.333496 | -3.15385 | -1.016274 | -2.316332 | -1.842241 | -0.676636 | -0.061554 | -0.973117 | -1.257338 | -0.965277 | -0.589859 | -1.368642 |
| 28.97222 | -0.480186 | 0.100726 | 0.164253 | 0.13585 | -0.848821 | 0.164854 | -0.100693 | -0.492488 | -0.120135 | 0.169378 | -0.698264 | -0.474306 | -0.416229 | -3.323242 | -0.268442 | -3.274259 | -1.743128 | -0.472928 | -0.155359 | -1.256219 | -0.564461 | -0.657663 | -0.66088 | -1.867605 |
| 28.98611 | 0.206487 | 0.046702 | 0.100138 | 0.073042 | -0.627776 | 0.10894 | -0.099111 | -0.433448 | 0.011161 | 0.11486 | -0.494056 | -0.409757 | -0.557698 | -3.309575 | 0.3841 | -2.928799 | -1.377852 | -0.281496 | -0.235866 | -1.095821 | 0.463772 | -0.469268 | -0.575659 | -2.274896 |
| 29 | 0.861901 | 0.124267 | 0.175777 | 0.176711 | -0.339079 | 0.13935 | -0.021201 | 0.043817 | 0.159138 | 0.188953 | -0.170267 | -0.248778 | -0.625922 | -3.213595 | 0.630579 | -1.208191 | -0.795161 | -0.096764 | -0.188812 | -0.481367 | 0.83176 | -0.379375 | -0.491129 | -1.969138 |
| 29.01389 | 1.081403 | 0.325008 | 0.394235 | 0.47448 | 0.064553 | 0.266312 | 0.183918 | 0.392118 | 0.354148 | 0.40246 | 0.169448 | 0.062783 | -0.5431 | -1.915667 | 1.226987 | 0.870104 | -0.225902 | 0.166529 | 0.044545 | 0.210939 | 0.291735 | -0.175291 | -0.173217 | -1.265462 |
| 29.02778 | 0.571201 | 0.315115 | 0.407523 | 0.619392 | 0.013443 | 0.261965 | 0.1906 | 0.037825 | 0.220293 | 0.423417 | 0.168323 | 0.142633 | -0.611965 | -1.317705 | 1.47742 | 2.020173 | -0.110466 | -0.058092 | 0.096017 | 0.551525 | -0.747737 | -0.045033 | -0.246945 | -0.74808 |
| 29.04167 | -0.316392 | -0.005978 | 0.026335 | 0.195583 | -0.338496 | -0.007885 | -0.197311 | -1.083107 | -0.319379 | 0.096129 | -0.203766 | -0.15973 | -0.830914 | -2.340466 | 1.590226 | 0.863436 | -0.738885 | -0.772535 | -0.101492 | 0.323803 | -1.968574 | -0.428731 | -0.680242 | -1.200692 |
| 29.05556 | -1.07138 | -0.127442 | -0.166017 | -0.06779 | -0.625333 | -0.054494 | -0.626831 | -2.009614 | -0.700995 | -0.018366 | -0.472639 | -0.38339 | -1.831643 | -3.966488 | 0.642435 | -0.812415 | -2.360286 | -1.626332 | -0.42244 | -0.147969 | -1.995044 | -1.346201 | -1.328835 | -2.516714 |
| 29.06944 | -1.659723 | 0.030048 | 0.010721 | 0.034462 | -0.969932 | 0.125581 | -1.140245 | -2.201541 | -1.013526 | 0.132048 | -0.643552 | -0.393195 | -1.973393 | -5.582349 | -0.622087 | -1.832538 | -3.976048 | -1.832976 | -0.723164 | -0.873955 | -1.007185 | -1.901231 | -1.34009 | -2.930995 |
| 29.08333 | -1.521253 | 0.092168 | 0.106101 | 0.088258 | -1.183329 | 0.179406 | -1.32114 | -2.028642 | -1.57801 | 0.178114 | -0.887797 | -0.484966 | -1.526405 | -4.66527 | 0.139549 | -1.330262 | -2.455945 | -1.571892 | -0.886684 | -1.142948 | -0.98766 | -1.200542 | -0.600613 | -2.831381 |
| 29.09722 | -0.238648 | 0.065055 | 0.063546 | 0.047326 | -0.796558 | 0.139204 | -0.440975 | -1.986868 | -1.86238 | 0.116488 | -0.655162 | -0.604209 | -1.196498 | -2.855419 | -1.395568 | -0.448236 | -0.898117 | -0.967985 | -0.712011 | -0.694515 | -2.443192 | -0.322274 | -0.066101 | -2.384307 |
| 29.11111 | 0.523959 | 0.035832 | 0.038438 | 0.015682 | -0.197343 | 0.126476 | 0.079392 | -1.520852 | -1.903011 | 0.07319 | -0.114998 | -0.898142 | -0.855922 | -0.768057 | -1.557793 | 0.224128 | 0.084325 | -0.349257 | -0.397879 | -0.135934 | -2.385284 | 0.400357 | 0.284448 | -1.458373 |
| 29.125 | 0.881025 | -0.050358 | 0.017198 | -0.020864 | 0.14742 | 0.092117 | 0.244569 | -0.895644 | -1.680075 | 0.035984 | 0.194829 | -1.109025 | -0.537782 | 0.550282 | -0.732119 | 0.723724 | 0.668023 | -0.018266 | -0.280908 | 0.372576 | -1.64153 | 0.900658 | 0.404881 | -0.742242 |
| 29.13889 | 0.825828 | -0.121026 | -0.001799 | -0.050232 | 0.151356 | 0.045912 | 0.190592 | -0.539193 | -1.260497 | 0.009532 | 0.223264 | -0.809565 | -0.273916 | 0.90151 | 0.258325 | 1.201344 | 1.219831 | 0.106199 | -0.120408 | 0.731196 | -0.827931 | 1.23705 | 0.456488 | -0.056259 |
| 29.15278 | 0.571439 | -0.120303 | 0.004264 | -0.034315 | -0.018016 | 0.034715 | 0.152761 | -0.086754 | -0.972204 | 0.013401 | 0.084506 | -0.230985 | -0.172192 | 1.294377 | 0.807797 | 1.715105 | 2.304252 | 0.297976 | -0.036961 | 0.655857 | 0.143296 | 1.27867 | 0.493765 | 0.942907 |
| 29.16667 | 0.515701 | -0.064332 | 0.04654 | 0.034591 | -0.112352 | 0.055445 | 0.21636 | 0.326088 | -0.957158 | 0.060483 | -0.126238 | 0.426607 | -0.586055 | 1.615763 | 0.973561 | 2.116514 | 2.670479 | 0.548831 | 0.367661 | 0.323649 | 0.872571 | 1.029635 | 0.577999 | 2.135613 |
| 29.18056 | 0.843946 | 0.03598 | 0.129911 | 0.133321 | 0.154298 | 0.10521 | 0.492006 | 0.504353 | -1.143182 | 0.141801 | 0.006404 | 1.276774 | -1.984531 | 1.737102 | 0.766057 | 1.379261 | 1.654441 | 0.723318 | 0.687483 | -0.076823 | 1.582427 | 0.580289 | 0.629629 | 2.524207 |
| 29.19444 | 1.023427 | 0.105151 | 0.195159 | 0.197105 | 0.762038 | 0.168508 | 0.974807 | 0.233194 | -1.268968 | 0.198382 | 0.585984 | 1.373746 | -2.83518 | 1.878535 | 0.410521 | 0.174997 | 1.123645 | 1.294974 | 0.213165 | -0.481752 | 2.548868 | -0.020334 | 0.30389 | 1.329965 |
| 29.20833 | 0.638211 | 0.139126 | 0.219218 | 0.219994 | 1.480634 | 0.227411 | 1.21449 | 0.036928 | -1.200997 | 0.223074 | 1.088775 | 1.399227 | -0.560011 | 1.809662 | 0.100446 | -0.628536 | 0.456303 | 1.481221 | -0.129038 | -0.824841 | 2.356214 | -0.538855 | 0.268459 | -0.028495 |
| 29.22222 | 0.325609 | 0.195882 | 0.269724 | 0.273787 | 1.755772 | 0.290107 | 0.912407 | 0.165678 | -0.952928 | 0.282006 | 1.010576 | 1.455274 | 0.219604 | 1.65839 | -0.108142 | -1.023878 | 1.117342 | 0.980174 | -0.124399 | -0.702719 | 1.640044 | -0.737274 | 0.172901 | -0.231872 |
| 29.23611 | 0.464351 | 0.239309 | 0.310169 | 0.312969 | 1.517078 | 0.305862 | 0.667972 | 0.469479 | -0.527626 | 0.326561 | 0.60096 | 1.57392 | 1.065679 | 0.933202 | -0.027051 | -0.580162 | 1.755997 | 0.597272 | 0.330533 | -0.239017 | 0.046107 | -0.617979 | 0.081408 | 0.348821 |
| 29.25 | 0.763951 | 0.149361 | 0.208989 | 0.204979 | 1.1736 | 0.202071 | 0.685041 | 0.66589 | -0.260617 | 0.228504 | 0.425616 | 1.378025 | 1.156192 | -0.137599 | 0.502141 | -0.458047 | 0.388272 | 0.520711 | 0.748542 | 0.159978 | -0.864869 | -0.329658 | 0.434196 | 0.747141 |
| 29.26389 | 1.0507 | 0.004894 | 0.024206 | 0.009776 | 0.780312 | 0.039294 | 0.887986 | 0.417209 | -0.225566 | 0.036196 | 0.564544 | 0.976008 | 0.812942 | -0.107344 | 1.401034 | -0.446818 | -0.596776 | 0.205517 | 0.380363 | 0.502176 | -1.515361 | 0.139155 | 0.78644 | -0.085503 |
| 29.27778 | 1.122926 | -0.073024 | -0.100746 | -0.112572 | 0.157142 | -0.06764 | 1.064941 | 0.213449 | -0.271982 | -0.091821 | 1.140924 | 0.797433 | 0.264885 | 0.426147 | 1.680887 | 0.057415 | -0.758707 | -0.161277 | 0.018126 | 0.726159 | -0.959468 | 0.726367 | 0.82471 | -0.498811 |
| 29.29167 | 1.049904 | -0.077161 | -0.117924 | -0.105856 | -0.102656 | -0.094924 | 1.228673 | 0.306505 | -0.277173 | -0.099588 | 1.387538 | 0.019794 | -0.234636 | 1.115307 | 0.964005 | 0.806244 | 0.630604 | -0.004454 | -0.174711 | 0.693279 | 0.285917 | 0.958726 | 0.379395 | 0.307065 |
| 29.30556 | 0.804128 | 0.011494 | -0.022994 | 0.004909 | 0.385837 | -0.012739 | 1.369097 | 0.550296 | -0.103526 | -0.000447 | 0.964718 | -0.503688 | -0.596172 | 1.368594 | 0.435682 | 0.871132 | 1.743884 | 0.399285 | 0.09685 | 0.593003 | 1.541919 | 0.39983 | 0.052933 | 1.807909 |
| 29.31944 | 0.721749 | 0.111388 | 0.101646 | 0.136064 | 0.942491 | 0.113887 | 1.451286 | 0.66151 | 0.265423 | 0.123747 | 0.49505 | -0.757092 | -0.772058 | 0.128564 | 0.047855 | 0.13193 | 1.324884 | 0.517892 | 0.244487 | 0.516178 | 1.880109 | -0.045491 | 0.022133 | 2.805875 |
| 29.33333 | 0.638127 | 0.096908 | 0.122644 | 0.174296 | 1.087667 | 0.150316 | 1.304265 | 0.392461 | 0.580462 | 0.149659 | 0.372168 | -0.406692 | -0.736679 | 0.544967 | -0.084964 | -0.338612 | 1.042656 | 0.394391 | 0.145452 | 0.186959 | 1.101179 | 0.006345 | 0.522202 | 1.558011 |
| 29.34722 | 0.764547 | 0.033518 | 0.064782 | 0.130336 | 1.240104 | 0.102509 | 0.983441 | 0.21545 | 0.643593 | 0.094229 | 0.332242 | 0.238848 | -0.555288 | 1.950095 | 0.341147 | 0.075818 | 0.587415 | 0.278974 | 0.002543 | 0.034243 | 0.449356 | 0.461051 | 0.789004 | -0.070577 |
| 29.36111 | 1.290752 | 0.015435 | 0.029379 | 0.092687 | 1.544144 | 0.066794 | 0.70342 | 0.336476 | 0.586441 | 0.045357 | 0.357858 | 0.460763 | -0.304661 | 2.916086 | 0.931048 | 0.601481 | 0.776689 | 0.354619 | -0.09256 | 0.260262 | 0.07997 | 0.730127 | 0.632082 | -0.435264 |
| 29.375 | 1.911782 | -0.017767 | -0.018071 | 0.040163 | 1.602318 | -0.002283 | 0.494959 | 0.39893 | 0.55369 | -0.019002 | 0.432064 | 1.249086 | -0.050918 | 3.178398 | 1.29123 | 0.381028 | 0.516106 | 0.488242 | -0.063811 | 0.441721 | -0.027468 | 0.500682 | 0.316253 | 0.573566 |
| 29.38889 | 2.117577 | -0.04726 | -0.06067 | 0.002195 | 1.314242 | -0.045999 | 0.398239 | 0.324765 | 0.660259 | -0.048037 | 0.399487 | 2.247566 | 0.20893 | 1.65537 | 1.549969 | -0.12758 | -0.024788 | 0.553295 | -0.022348 | -0.114057 | 0.211431 | 0.276591 | 0.181581 | 2.095123 |
| 29.40278 | 1.947673 | 0.044322 | 0.004269 | 0.061697 | 0.914663 | 0.040918 | 0.460751 | 0.516874 | 0.89695 | 0.029641 | 0.304896 | 2.356771 | 0.40931 | 0.595084 | 1.707914 | -0.426405 | 0.101785 | 0.731634 | 0.038317 | -1.276463 | 0.646142 | 0.384643 | 0.37138 | 2.228736 |
| 29.41667 | 1.618154 | 0.128261 | 0.079782 | 0.098613 | 0.99497 | 0.10407 | 0.637191 | 1.005039 | 1.335476 | 0.075792 | 0.274075 | 2.284889 | 0.520283 | 2.223247 | 1.787081 | -0.167958 | 0.913845 | 0.873494 | 0.099662 | -1.812472 | 1.000795 | 0.681078 | 0.723316 | 1.262734 |
| 29.43056 | 1.204674 | -0.030587 | -0.032304 | -0.037327 | 1.443029 | -0.031802 | 0.809794 | 1.448501 | 1.627091 | -0.059471 | 0.256829 | 2.160832 | 0.54181 | 4.340132 | 1.585515 | 0.534334 | 1.891272 | 0.949938 | 0.155084 | -1.451359 | 1.322678 | 0.785704 | 0.878475 | 0.74241 |
| 29.44444 | 1.350942 | -0.3836 | -0.383027 | -0.345572 | 1.595996 | -0.314267 | 0.700332 | 1.696535 | 1.279393 | -0.356675 | -0.065031 | 2.380059 | 0.454738 | 4.523501 | 1.160315 | 1.406622 | 2.332039 | 0.933735 | 0.002289 | 0.904386 | 1.434389 | 0.608086 | 1.549779 | 1.361406 |
| 29.45833 | 1.296601 | -0.66859 | -0.734354 | -0.943382 | 0.895318 | -0.5664 | 0.384026 | 1.70793 | 0.893374 | -0.669015 | -0.360528 | 2.141866 | 0.19002 | 4.625417 | 0.406729 | 2.323743 | 1.452721 | 0.985413 | -0.339181 | 1.188753 | 1.537249 | 0.470604 | 0.290385 | 2.143405 |
| 29.47222 | 0.976645 | -0.528529 | -0.52209 | -0.676078 | 0.426267 | -0.407827 | -0.056296 | 1.322018 | 1.118422 | -0.559074 | -0.355603 | 0.974081 | 0.018362 | 4.310233 | 0.280136 | 2.2212 | 0.353786 | 1.085429 | -0.390177 | -0.319618 | 1.263027 | 0.694006 | 0.107964 | 2.629658 |
| 29.48611 | 1.024587 | -0.202516 | -0.131443 | -0.168517 | 0.952778 | -0.10966 | -0.067302 | 1.119619 | 1.540906 | -0.182194 | -0.077175 | 0.924492 | 0.053215 | 3.495427 | 0.700414 | 2.2299 | 0.345471 | 1.41005 | -0.036586 | -0.709171 | 1.287024 | 0.925305 | 0.485538 | 2.62289 |
| 29.5 | 1.18697 | 0.021814 | 0.097436 | 0.067372 | 1.70978 | 0.071035 | 0.149078 | 1.251976 | 1.709308 | 0.055379 | 0.185606 | 0.867155 | 0.10787 | 2.002477 | 1.127859 | 1.349852 | 1.269326 | 1.444505 | 0.16269 | 0.930753 | 1.287324 | 0.838105 | 0.905067 | 2.011581 |
| 29.51389 | 1.079525 | 0.127976 | 0.201397 | 0.181391 | 2.006006 | 0.182672 | 0.26633 | 1.310405 | 1.602916 | 0.171723 | 0.297169 | 1.82901 | 0.254144 | 2.314369 | 1.419875 | 0.877169 | 1.545254 | 1.373196 | 0.249804 | 1.189435 | 1.228405 | 0.735775 | 0.89673 | 1.147821 |
| 29.52778 | 0.587329 | -0.222099 | -0.023427 | -0.090634 | 1.200259 | -0.145205 | 0.031782 | 1.172316 | 1.315428 | -0.176811 | -0.004961 | 0.818157 | 0.577194 | 0.853552 | 1.640743 | 1.166743 | 1.896413 | 1.297016 | 0.216605 | 0.713282 | 1.104802 | 0.76708 | 0.999397 | 1.778262 |
| 29.54167 | -0.110251 | -1.092374 | -0.700438 | -0.859811 | 0.005451 | -1.01864 | -0.644062 | 0.530429 | 0.753618 | -1.072656 | -0.742286 | 0.032774 | 0.975229 | 0.742206 | 2.365508 | 1.360165 | 0.717012 | 0.94331 | 0.119507 | -0.006547 | 1.022175 | 0.694 | 0.540555 | 2.019319 |
| 29.55556 | -0.427814 | -1.470033 | -1.102932 | -1.264458 | -0.486558 | -1.369862 | -0.912489 | 0.232616 | 0.37388 | -1.385958 | -1.105961 | -1.179064 | 0.895364 | 0.788069 | 1.962563 | 0.86944 | 0.423895 | 0.816699 | 0.017969 | -0.243119 | 0.674076 | 0.347892 | 0.209141 | 1.377073 |
| 29.56944 | 0.005847 | -0.774098 | -0.6657 | -0.659806 | 0.222457 | -0.660113 | -0.326096 | 0.164983 | 0.13041 | -0.643991 | -0.592503 | -1.707319 | 0.489853 | 0.383486 | 1.626099 | 0.575649 | 0.174649 | 0.938888 | -0.017788 | -0.029369 | 0.487763 | 0.143741 | 0.614002 | 1.527431 |
| 29.58333 | 0.710184 | 0.028476 | 0.05443 | 0.110225 | 1.046893 | 0.127722 | 0.345957 | 0.097009 | 0.230105 | 0.148371 | 0.123447 | 0.261755 | 0.328668 | -0.354785 | 1.587768 | 0.196074 | 0.118574 | 0.775884 | 0.062016 | -0.573275 | 0.29436 | 0.077778 | 0.916145 | 1.152078 |
| 29.59722 | 0.985446 | 0.380085 | 0.466195 | 0.465937 | 1.171469 | 0.463684 | 0.549579 | 0.162421 | 0.452641 | 0.481944 | 0.493071 | 0.534907 | 0.0163 | -0.206419 | 1.147487 | -0.011925 | 0.327496 | 0.56008 | 0.115768 | -0.950961 | 0.10958 | -0.206691 | 0.606176 | 0.724004 |
| 29.61111 | 0.404668 | 0.318722 | 0.447145 | 0.402458 | 0.955203 | 0.399421 | 0.45511 | 0.353453 | 0.719517 | 0.406473 | 0.471718 | 1.102967 | -0.441855 | 0.712651 | 0.653104 | -0.090613 | 0.18764 | 0.019888 | -0.054675 | -0.230288 | 0.177176 | -0.262592 | 0.44853 | -0.327803 |
| 29.625 | -0.293111 | 0.072061 | 0.168776 | 0.137298 | 0.41784 | 0.153836 | 0.221833 | 0.510019 | 0.718835 | -0.137427 | 0.20799 | 1.013686 | -0.848666 | 0.464242 | -0.466354 | -0.300628 | -0.236375 | 0.107157 | -0.010407 | 0.625347 | 0.165231 | 0.167307 | 0.54867 | -1.372448 |
| 29.63889 | -0.518081 | -0.135263 | -0.102444 | -0.103845 | -0.049854 | -0.059053 | -0.038764 | 0.449137 | 0.427386 | -0.195767 | -0.069885 | 0.74723 | -1.047173 | 0.165023 | -0.779514 | -0.686706 | -0.381895 | 0.064666 | 0.009071 | 0.1503 | 0.581815 | 0.605148 | 0.536785 |
| 29.65278 | -0.432137 | -0.19945 | -0.216312 | -0.213455 | 0.206529 | -0.155531 | -0.22484 | 0.260947 | 0.164256 | -0.192742 | -0.196029 | 0.535499 | -0.917618 | 0.022795 | -0.211381 | -0.661597 | -0.493949 | 0.238171 | -0.032817 | 0.238169 | 0.377737 | 0.625685 | 0.457678 | 1.613263 |
| 29.66667 | -0.183325 | -0.14972 | -0.185063 | -0.185218 | 0.18778 | -0.148929 | -0.281572 | 0.02094 | 0.15665 | -0.165103 | -0.121235 | 0.744024 | -0.576136 | -0.07418 | 0.062103 | -0.036511 | -0.194954 | -0.285135 | 0.21547 | -0.236543 | 0.333541 | 0.335418 | 0.054041 | 0.902528 |
| 29.68056 | 0.076791 | -0.04935 | -0.091528 | -0.088322 | -0.38687 | -0.077275 | -0.154996 | -0.185387 | 0.231572 | -0.074469 | -0.053384 | 0.489002 | -0.188202 | -0.176933 | 0.325487 | 0.474971 | 0.19037 | -0.488085 | 0.129397 | -0.704156 | 0.048559 | 0.139401 | -0.328921 | -0.010159 |
| 29.69444 | -0.036386 | -0.03685 | -0.056293 | -0.063438 | -0.467582 | -0.065676 | -0.154025 | -0.43528 | 0.288424 | -0.058527 | -0.024253 | -0.403446 | 0.260934 | 0.020054 | 0.293266 | 0.663948 | 0.265017 | -0.388394 | -0.011518 | -0.107377 | -0.15038 | 0.077254 | 0.058909 | -0.471263 |
| 29.70833 | -0.184721 | -0.14455 | -0.135329 | -0.159302 | -0.555308 | -0.159774 | -0.324025 | -0.657708 | 0.233275 | -0.157309 | -0.084003 | -0.338595 | 0.716793 | 0.227709 | -0.288567 | 0.669981 | 0.175086 | 0.168744 | -0.017122 | -0.079463 | -0.140427 | 0.184238 | 0.041789 | -0.49816 |
| 29.72222 | -0.416043 | -0.286804 | -0.262456 | -0.280628 | -0.981739 | -0.254488 | -0.489397 | -0.524233 | 0.168111 | -0.26986 | -0.197725 | -0.367579 | 0.977722 | 0.240127 | -0.763842 | 0.581905 | 0.195543 | 0.501517 | -0.057017 | -0.018381 | -0.10711 | 0.19785 | -0.006826 | 0.086323 |
| 29.73611 | -0.831565 | -0.341923 | -0.345652 | -0.355654 | -0.727491 | -0.30068 | -0.486603 | -0.085693 | 0.183264 | -0.32441 | -0.288453 | -0.388026 | 1.003874 | 0.252129 | -0.848878 | 0.230102 | 0.360609 | 0.235441 | -0.127428 | 0.408441 | -0.040039 | 0.04082 | 0.046575 | -1.40422 |
| 29.75 | -0.838027 | -0.29808 | -0.330272 | -0.352694 | -0.035969 | -0.286635 | -0.267403 | 0.256507 | 0.307666 | -0.303695 | -0.296541 | -0.533579 | 1.169249 | 0.174872 | -0.609671 | -0.042721 | 0.460676 | 0.103445 | 0.02736 | 0.589289 | -0.159628 | -0.069184 | 0.303305 | -1.224156 |
| 29.76389 | -0.763912 | -0.226452 | -0.264762 | -0.29355 | 0.383382 | -0.236399 | -0.03224 | 0.227142 | 0.442077 | -0.244033 | -0.239929 | -0.774739 | 1.244058 | -0.05296 | 0.004269 | -0.203179 | 0.304456 | 0.139 | 0.19094 | 0.582297 | -0.428479 | -0.174413 | -0.002662 | 0.421424 |
| 29.77778 | -0.325378 | -0.206825 | -0.248501 | -0.265106 | -0.247792 | -0.226197 | 0.050414 | 0.060467 | 0.486283 | -0.233626 | -0.202063 | -0.933441 | 1.013011 | -0.296715 | 0.644865 | -0.356464 | -0.013093 | 0.231698 | 0.178152 | 0.267562 | 0.215953 | -0.238742 | 0.245699 | 0.882229 |
| 29.79167 | -0.230979 | -0.216752 | -0.251853 | -0.270851 | -1.392861 | -0.232472 | 0.067395 | -0.125072 | 0.331091 | -0.244623 | -0.242736 | -1.041229 | 0.781866 | -0.414534 | 1.308824 | -0.558208 | -0.234573 | 0.096968 | 0.037 | 0.157571 | 0.52289 | -0.228461 | 0.3291 | 1.370271 |
| 29.80556 | -0.478313 | -0.17806 | -0.212985 | -0.248577 | -0.810945 | -0.193429 | 0.030819 | -0.26647 | 0.252384 | -0.207554 | -0.226076 | -0.757434 | 0.565098 | -0.326099 | 0.825109 | -0.710487 | -0.503092 | -0.471079 | -0.034945 | 0.035815 | -0.017485 | -0.207474 | -0.210183 | 0.442243 |
| 29.81944 | -0.839726 | -0.188278 | -0.191618 | -0.234698 | -0.540112 | -0.165593 | -0.157325 | -0.445107 | 0.159533 | -0.179371 | -0.131359 | -0.411842 | 0.416428 | -0.006365 | 0.433213 | -0.597617 | -0.855226 | -0.606327 | 0.009754 | -0.425675 | -0.836615 | -0.188736 | 0.072081 | 0.841892 |
| 29.83333 | -1.120924 | -0.27714 | -0.228141 | -0.268059 | -0.824676 | -0.195916 | -0.48341 | -1.030909 | -0.003614 | -0.21747 | -0.123183 | -0.383336 | 0.325579 | 0.020791 | 0.315796 | -0.098596 | -1.118714 | -0.191647 | -0.196917 | -0.060229 | -1.159507 | -0.18498 | 0.137215 | -0.700729 |
| 29.84722 | -1.447616 | -0.336389 | -0.25289 | -0.291896 | -0.996476 | -0.228865 | -0.715717 | -1.585827 | -0.206703 | -0.257463 | -0.241409 | -0.719179 | 0.23966 | -0.187304 | 0.164996 | 0.450158 | -1.509017 | 0.004225 | -0.111494 | 0.186505 | -0.816985 | -0.186589 | 0.097136 | -1.767275 |
| 29.86111 | -1.419013 | -0.294983 | -0.215295 | -0.255633 | -0.894761 | -0.207187 | -0.896938 | -1.037285 | -0.434692 | -0.23558 | -0.285826 | -0.7399 | 0.12588 | -0.292863 | -0.604317 | 0.534474 | -1.007403 | -0.171389 | 0.013398 | -0.058305 | -0.786816 | -0.156758 | -0.076834 | -1.81904 |
| 29.875 | -1.653127 | -0.227899 | -0.161656 | -0.204277 | -1.541249 | -0.17179 | -1.096249 | -0.898987 | -0.596476 | -0.190036 | -0.250389 | -0.727209 | 0.064076 | -0.801974 | -1.948003 | 0.061015 | -0.384482 | -0.414579 | 0.020299 | -0.534365 | 0.071854 | -0.082877 | -0.815998 | -1.978028 |
| 29.88889 | -2.151656 | -0.184682 | -0.139464 | -0.176937 | -2.315966 | -0.159051 | -1.324053 | -0.880883 | -0.674011 | -0.164118 | -0.315891 | -0.827676 | 0.020758 | -2.17793 | -2.006908 | -0.329487 | -0.232665 | -0.738293 | 0.033652 | -1.03018 | 0.102526 | -0.173856 | -0.897737 | -2.045858 |
| 29.90278 | -2.527628 | -0.154645 | -0.137631 | -0.16582 | -2.398032 | -0.141295 | -1.284321 | -1.078632 | -0.805123 | -0.149104 | -0.283007 | -1.076683 | -0.085855 | -2.774039 | -2.548786 | -0.807932 | -0.474129 | -0.934676 | 0.127584 | -0.939213 | -0.367868 | -0.372883 | -0.635564 | -1.804426 |
| 29.91667 | -1.827669 | -0.146648 | -0.13342 | -0.148368 | -2.128653 | -0.111486 | -1.359802 | -1.284879 | -0.94095 | -0.134865 | -0.270955 | -1.628396 | -0.314815 | -2.92248 | -2.948746 | -1.914719 | -0.815061 | -0.634428 | 0.271986 | -0.859969 | -0.287558 | -0.673067 | -0.658165 | -1.597166 |
| 29.93056 | -0.880595 | -0.134617 | -0.098555 | -0.098959 | -2.164347 | -0.080632 | -1.341146 | -1.804553 | -1.055123 | -0.104708 | -0.168109 | -2.117389 | -0.553924 | -2.691852 | -2.376158 | -3.194933 | -1.136581 | -0.502563 | 0.186792 | -0.643947 | -0.158054 | -0.954984 | -0.688877 | -2.081051 |
| 29.94444 | -0.395581 | -0.060349 | -0.029927 | -0.016436 | -1.948947 | -0.036871 | -1.180318 | -2.835959 | -1.133198 | -0.04555 | -0.046643 | -2.239103 | -0.880949 | -3.365931 | -1.16397 | -2.349939 | -1.985309 | -0.889568 | -0.111694 | 0.013346 | -0.642594 | -1.074863 | -0.824442 | -1.823243 |
| 29.95833 | -0.434101 | 0.072599 | 0.034704 | 0.057988 | -1.557073 | 0.020403 | -0.956211 | -2.817573 | -1.120873 | 0.020963 | -0.010081 | -2.036161 | -1.362779 | -4.608875 | 0.189144 | -0.938474 | -2.388415 | -1.169543 | -0.232487 | 1.022321 | -1.772029 | -0.676904 | -0.877355 | -1.330428 |
| 29.97222 | -0.605677 | 0.178333 | 0.102532 | 0.133493 | -1.034903 | 0.089472 | -0.40787 | -2.038849 | -0.937572 | 0.094249 | -0.010488 | -1.588196 | -1.725061 | -5.241191 | 1.439259 | -0.063017 | -2.548346 | -1.21147 | -0.129579 | 1.594497 | -1.280962 | -0.469252 | -0.91912 | -1.195402 |
| 29.98611 | -0.589284 | 0.237886 | 0.206523 | 0.240416 | -0.488928 | 0.181733 | 0.105782 | -0.91292 | -0.67101 | 0.198828 | 0.029291 | -1.067482 | -1.452187 | -4.350954 | 2.572673 | 0.027108 | -2.20719 | -1.179466 | -0.048694 | 1.259269 | -1.259802 | -0.536658 | -0.921864 | -1.514179 |
| 30 | -0.390291 | 0.340608 | 0.355752 | 0.406823 | -0.082936 | 0.30343 | 0.435464 | 0.053221 | -0.405377 | 0.352555 | 0.131994 | -0.509592 | -1.059296 | -2.776225 | 2.854747 | 0.100502 | -1.257183 | -1.07058 | 0.153397 | 1.367161 | -0.384335 | -0.505054 | -0.624041 | -1.505388 |
| 30.01389 | -0.175654 | 0.539242 | 0.570234 | 0.671632 | 0.13102 | 0.46953 | 0.616139 | 0.577971 | -0.191938 | 0.583656 | 0.345647 | 0.105309 | -0.747002 | -1.961137 | 2.797936 | 0.702578 | -0.341465 | -0.746273 | 0.419311 | 1.078665 | 0.796382 | -0.075357 | 0.023573 | -1.003982 |
| 30.02778 | 0.059316 | 0.892026 | 0.908114 | 1.172335 | 0.2603 | 0.766528 | 0.849061 | 0.686315 | 0.010583 | 0.898639 | 0.60559 | 0.63431 | -0.513429 | -2.347114 | 1.624553 | 0.866356 | -0.02432 | -0.521983 | 0.359822 | -0.092557 | 1.799049 | 0.212567 | 0.1793 | -0.954019 |
| 30.04167 | 0.208091 | 1.187503 | 1.128028 | 1.34152 | 0.447546 | 1.034503 | 0.847028 | 0.65386 | 0.211494 | 1.127139 | 0.840073 | 0.933229 | -0.299132 | -3.893712 | -0.126852 | -0.049208 | -0.631984 | -0.728889 | -0.05938 | -0.11566 | 1.286502 | -0.303617 | -0.620728 | -2.167958 |
| 30.05556 | -0.144161 | 1.11652 | 1.008456 | 1.09325 | 0.296058 | 1.03094 | 0.363679 | 0.382696 | -0.127152 | 1.096916 | 0.756187 | 0.731715 | -0.064273 | -5.108644 | -1.289547 | -1.064926 | -1.987623 | -1.318841 | -0.376451 | 0.805225 | -0.247786 | -1.312329 | -1.669289 | -3.771219 |
| 30.06944 | -0.957023 | 0.8076 | 0.767668 | 0.812936 | -0.112318 | 0.819672 | -0.744498 | 0.090283 | -0.977214 | 0.858874 | 0.353771 | -0.135975 | 0.02605 | -5.133143 | -1.409027 | -1.781134 | -2.489701 | -1.6624 | -0.710641 | 1.495217 | -0.808708 | -1.640601 | -2.151269 | -4.305602 |
| 30.08333 | -1.371563 | 0.597151 | 0.64474 | 0.643349 | -0.513478 | 0.648222 | -1.229888 | 0.529692 | -1.397329 | 0.670176 | -0.03898 | -0.19699 | -0.141582 | -4.356549 | -1.44316 | -2.36287 | -1.364417 | -0.842609 | -0.632311 | 0.226023 | -1.108449 | -1.565339 | -2.245511 | -4.269547 |
| 30.09722 | -0.886413 | 0.457631 | 0.520153 | 0.489178 | -0.760601 | 0.509225 | -0.39017 | 0.974981 | -1.599499 | 0.515868 | -0.120465 | -0.072803 | -0.381525 | -3.566139 | -2.767025 | -0.613335 | -0.565025 | -0.156165 | -0.33434 | -0.901154 | -2.39609 | -0.764765 | -0.785593 | -3.256565 |
| 30.11111 | -0.166279 | 0.29728 | 0.327531 | 0.290239 | -0.719888 | 0.354634 | 0.253142 | 0.671132 | -1.737972 | 0.334528 | -0.08613 | -0.159256 | -0.512809 | -1.002703 | -0.937422 | -0.289454 | -0.043101 | -0.175887 | -0.27376 | -1.261541 | -3.45884 | 0.110974 | 0.010082 | -1.725434 |
| 30.125 | 0.386667 | 0.194549 | 0.204492 | 0.170804 | -0.099093 | 0.246385 | 0.638306 | -0.114969 | -1.442694 | 0.216715 | 0.037429 | 0.018783 | -0.470557 | 0.655093 | 0.335204 | 0.112914 | 0.469172 | -0.110993 | -0.242994 | -0.784027 | -2.177863 | 0.693614 | 0.477698 | -0.452471 |
| 30.13889 | 0.817924 | 0.150759 | 0.155933 | 0.122977 | 0.575474 | 0.173672 | 0.703136 | -0.704603 | -0.739945 | 0.156865 | 0.120546 | 0.168833 | -0.274178 | 1.315271 | 0.933054 | 0.608334 | 1.137798 | 0.286105 | -0.021089 | -0.052773 | -0.708267 | 0.961916 | 0.661818 | 0.694749 |
| 30.15278 | 1.208715 | 0.102654 | 0.117343 | 0.091917 | 0.738418 | 0.12742 | 0.507437 | -0.794954 | -0.083178 | 0.122215 | 0.167842 | 0.120751 | 0.01745 | 1.513753 | 1.04674 | 1.112524 | 2.058089 | 0.769676 | 0.335694 | 0.286998 | 1.016249 | 1.243374 | 0.600306 | 1.96785 |
| 30.16667 | 1.712894 | 0.077283 | 0.091479 | 0.080525 | 0.599184 | 0.114554 | 0.497571 | -0.596968 | 0.197959 | 0.113326 | 0.163832 | 0.166464 | 0.289652 | 1.382138 | 1.033196 | 1.550932 | 2.922176 | 0.917431 | 0.358732 | 0.273965 | 2.288683 | 0.759223 | 0.688086 | 2.758291 |
| 30.18056 | 2.281378 | 0.107553 | 0.096917 | 0.091893 | 0.432744 | 0.129142 | 0.891488 | -0.459009 | -0.01222 | 0.124387 | 0.143282 | 0.334276 | 0.503057 | 0.942081 | 0.836387 | 1.998762 | 1.862621 | 0.565654 | 0.104816 | -0.247007 | 2.558412 | 0.135706 | 0.680659 | 2.90909 |
| 30.19444 | 2.235784 | 0.166839 | 0.135588 | 0.126006 | 0.619087 | 0.16127 | 1.335657 | -0.361398 | -0.526186 | 0.155785 | 0.228168 | 0.613053 | 0.630938 | 0.60143 | 0.388932 | 1.938234 | 0.534006 | 0.092211 | -0.190557 | -0.942264 | 2.511228 | -0.053123 | 0.354773 | 1.777747 |
| 30.20833 | 1.495117 | 0.202253 | 0.175494 | 0.163278 | 0.879044 | 0.191835 | 1.25299 | -0.300802 | -0.889559 | 0.188273 | 0.411303 | 1.099475 | 0.670948 | 0.609483 | -0.142661 | 0.637055 | -0.402389 | 0.109855 | -0.158538 | -1.003135 | 0.926644 | -0.158865 | 0.220523 | 0.187775 |
| 30.22222 | 0.590974 | 0.107776 | 0.085127 | 0.090493 | 0.770282 | 0.119202 | 0.702755 | -0.074217 | -0.982188 | 0.116481 | 0.485058 | 1.464533 | 0.522581 | 0.544005 | -0.353123 | -0.338029 | 0.48048 | 0.427671 | 0.242622 | -0.230473 | -0.02746 | -0.107115 | 0.311749 | -0.675943 |
| 30.23611 | -0.173597 | -0.055319 | -0.098344 | -0.071604 | 0.641262 | -0.031341 | 0.372575 | 0.188179 | -1.137463 | -0.042033 | 0.354355 | 1.170007 | 0.209955 | 0.494083 | 0.021701 | -1.639443 | 1.155975 | 0.727009 | 0.65458 | 0.267349 | -0.629747 | -0.077426 | 0.444112 | -0.426825 |
| 30.25 | -0.176777 | -0.026767 | -0.068151 | -0.051301 | 0.576794 | -0.023516 | 0.754537 | 0.269429 | -1.140047 | -0.031935 | 0.277961 | 0.993265 | 0.03209 | 0.415268 | 0.648393 | -1.288025 | 0.714023 | 0.985371 | 0.597441 | 0.048162 | -1.071098 | -0.218746 | 0.873551 | 0.475221 |
| 30.26389 | 0.37576 | 0.168086 | 0.159412 | 0.140965 | 0.986399 | 0.159208 | 1.296512 | 0.367904 | -0.818585 | 0.162547 | 0.348515 | 1.250251 | 0.130341 | 0.565076 | 0.826406 | -0.195452 | -0.205654 | 0.613831 | 0.270774 | -0.21217 | -0.658416 | -0.164609 | 1.030733 | 0.888233 |
| 30.27778 | 1.06997 | 0.277905 | 0.287082 | 0.243908 | 1.598337 | 0.286791 | 1.1492 | 0.409788 | -0.407509 | 0.288277 | 0.47306 | 1.201956 | 0.134878 | 0.86777 | 0.439695 | 0.429794 | -0.499543 | -0.021164 | -0.08958 | 0.161173 | 0.03946 | -0.061467 | 0.629227 | 0.679122 |
| 30.29167 | 1.808687 | 0.218063 | 0.223827 | 0.172145 | 1.359787 | 0.245801 | 0.889862 | 0.354071 | -0.260779 | 0.231405 | 0.584731 | 0.594627 | -0.189217 | 0.869524 | 0.010907 | 1.067836 | -0.204468 | -0.522433 | -0.134024 | 1.121578 | 0.615554 | 0.112758 | 0.138976 | 1.024098 |
| 30.30556 | 1.811183 | 0.096823 | 0.081135 | 0.024581 | 0.274697 | 0.107526 | 0.785611 | 0.225021 | -0.18284 | 0.0862 | 0.444805 | -0.14195 | -0.37544 | 0.350013 | -0.085881 | 1.254677 | 0.67382 | -0.807659 | 0.21566 | 2.152376 | 1.078837 | 0.351061 | -0.182735 | 1.31712 |
| 30.31944 | 1.254506 | 0.025307 | 0.01278 | -0.041171 | -0.77688 | 0.029021 | 1.066811 | 0.206931 | 0.09729 | 0.029071 | 0.186844 | -0.929907 | -0.260764 | 0.22181 | 0.137977 | 0.599508 | 0.945926 | -0.240168 | 0.606781 | 1.157044 | 0.909012 | 0.536479 | 0.027369 | 1.688443 |
| 30.33333 | 0.997971 | 0.021816 | 0.031085 | -0.009835 | -1.057758 | 0.015182 | 1.27861 | 0.37981 | 0.414483 | 0.06126 | 0.139483 | -1.048843 | -0.10344 | 0.968119 | 0.548626 | 0.055268 | 0.60981 | 0.67162 | 0.233505 | -0.120886 | 0.37543 | 0.511672 | 0.84846 | 1.657344 |
| 30.34722 | 0.937587 | 0.021804 | 0.022686 | -0.00568 | 0.13428 | 0.006162 | 0.984184 | 0.564552 | 0.516576 | 0.049572 | 0.123596 | -0.56622 | -0.044218 | 2.04386 | 1.063261 | -0.088306 | 0.260871 | 1.218302 | 0.057752 | -1.145304 | 0.102312 | 0.458797 | 1.522073 | 1.797923 |
| 30.36111 | 0.912177 | -0.035788 | -0.06498 | -0.083754 | 1.877737 | -0.063236 | 0.572825 | 0.752774 | 0.525194 | -0.038304 | 0.011224 | 0.183226 | -0.034507 | 2.849285 | 1.347436 | 0.205175 | 0.393603 | 1.490755 | -0.026178 | -0.9584 | -0.162809 | 0.448858 | 1.131299 | 1.208149 |
| 30.375 | 0.891159 | -0.094662 | -0.156097 | -0.165449 | 2.653124 | -0.123691 | 0.317235 | 0.907381 | 0.612604 | -0.116399 | -0.045179 | 1.172282 | 0.091789 | 2.645094 | 1.449975 | 0.827561 | 0.839614 | 0.95386 | -0.009869 | -0.239077 | -0.138902 | 0.494951 | 0.385039 | 0.830066 |
| 30.38889 | 0.966652 | -0.09253 | -0.126745 | -0.143828 | 2.168245 | -0.099056 | 0.330982 | 1.142066 | 0.864965 | -0.094307 | 0.047773 | 1.932018 | 0.324239 | 1.458171 | 1.717721 | 1.279364 | 1.040015 | 0.201496 | -0.065124 | 0.465547 | 0.234924 | 0.53619 | 0.0312 | 0.957747 |
| 30.40278 | 0.972453 | -0.114998 | -0.075543 | -0.113146 | 1.77831 | -0.078096 | 0.462371 | 1.245306 | 1.247279 | -0.070744 | 0.091922 | 2.527369 | 0.501955 | 1.158483 | 1.912786 | 1.324619 | 0.911857 | 0.213484 | 0.000338 | 1.1036 | 0.622281 | 0.554948 | 0.353196 | 1.621003 |
| 30.41667 | 0.761743 | -0.195217 | -0.161821 | -0.193626 | 1.89528 | -0.159009 | 0.51953 | 1.116874 | 1.623036 | -0.154546 | 0.003307 | 2.586766 | 0.505179 | 1.79489 | 2.049263 | 1.08785 | 1.102652 | 0.551914 | 0.02429 | 1.256854 | 0.824115 | 0.479645 | 1.010761 | 2.282253 |
| 30.43056 | 1.105478 | -0.193128 | -0.254875 | -0.256266 | 2.174827 | -0.199059 | 0.437899 | 1.056671 | 1.917039 | -0.214899 | -0.055557 | 2.071694 | 0.495135 | 2.357774 | 1.934478 | 0.953645 | 1.138067 | 0.931216 | -0.04594 | 0.964349 | 0.70936 | 0.535406 | 1.457763 | 2.580773 |
| 30.44444 | 1.970324 | -0.199654 | -0.342054 | -0.312998 | 2.144215 | -0.201178 | 0.254757 | 1.036299 | 1.959158 | -0.261611 | -0.132208 | 1.873015 | 0.503515 | 3.504471 | 1.567489 | 1.013795 | 1.096094 | 1.269469 | -0.150863 | 0.447865 | 0.314316 | 0.54741 | 1.319395 | 2.036074 |
| 30.45833 | 2.378747 | -0.342042 | -0.47666 | -0.681364 | 1.736669 | -0.317957 | 0.04339 | 0.994859 | 1.83645 | -0.432844 | -0.166878 | 1.592099 | 0.502707 | 5.988632 | 1.552096 | 1.061152 | 0.821181 | 1.288347 | -0.358114 | -0.213449 | 0.214441 | 0.51342 | 0.863344 | 1.34668 |
| 30.47222 | 2.119387 | -0.382337 | -0.406486 | -0.578426 | 1.33616 | -0.327371 | -0.07207 | 1.136493 | 1.831176 | -0.469317 | -0.18172 | 1.344615 | 0.720854 | 6.580383 | 1.718445 | 1.287583 | 0.49917 | 1.203264 | -0.381183 | -0.235319 | 0.312819 | 0.60779 | 0.312445 | 0.830884 |
| 30.48611 | 1.258753 | -0.353037 | -0.317418 | -0.354457 | 1.19625 | -0.276437 | 0.003189 | 1.540755 | 1.853528 | -0.340107 | -0.143215 | 0.734813 | 0.933837 | 5.53449 | 1.6362 | 1.428391 | 0.469745 | 1.200467 | -0.259341 | 0.264313 | 0.599734 | 0.943704 | 0.218685 | 1.524158 |
| 30.5 | 1.005161 | -0.236883 | -0.24902 | -0.265856 | 1.12443 | -0.22027 | 0.099378 | 1.544488 | 1.762872 | -0.240012 | -0.109557 | -0.547359 | 1.068845 | 4.150116 | 1.682407 | 1.413217 | 1.020457 | 0.925447 | -0.064177 | 0.539107 | 0.654456 | 0.945904 | 0.725931 | 2.771237 |
| 30.51389 | 1.501888 | -0.162752 | -0.199388 | -0.239891 | 0.788955 | -0.190723 | 0.127921 | 1.599181 | 1.505385 | -0.22471 | -0.104798 | -0.620809 | 1.123814 | 2.154738 | 1.787713 | 0.771596 | 1.423863 | 0.882128 | 0.123948 | 0.101666 | 0.605289 | 0.895357 | 1.539132 | 3.44484 |
| 30.52778 | 1.507954 | -0.517964 | -0.394532 | -0.500182 | 0.718664 | -0.561792 | -0.180104 | 1.296893 | 1.04439 | -0.605307 | -0.367651 | 0.993073 | 0.935816 | 1.692753 | 1.999347 | 0.901257 | 1.162665 | 1.023242 | 0.129319 | -0.823414 | 0.729794 | 0.790041 | 1.75855 | 2.61721 |
| 30.54167 | 0.074983 | -1.340434 | -0.973624 | -1.174953 | 0.312455 | -1.424805 | -0.92328 | 0.199703 | 0.213044 | -1.454606 | -1.026043 | 1.876135 | 0.772981 | 2.07997 | 2.345953 | 1.208609 | 0.689482 | 0.138017 | -0.013354 | -0.622129 | 0.449424 | 0.614487 | 1.647097 | 1.727977 |
| 30.55556 | -0.269626 | -1.824515 | -1.486323 | -1.693645 | -0.224128 | -1.898777 | -1.481963 | -0.74657 | -0.502961 | -1.855359 | -1.470422 | 0.361882 | 0.631188 | 1.19504 | 1.863463 | 0.949675 | -0.13042 | 0.126906 | -0.089383 | -0.496719 | 0.370248 | 0.404969 | 1.379733 | 1.623875 |
| 30.56944 | -0.575512 | -1.385067 | -1.297047 | -1.326993 | 0.067394 | -1.327999 | -1.142999 | -0.793276 | -0.621461 | -1.285415 | -1.141828 | -0.421309 | 0.491891 | 0.180843 | 1.057236 | 0.699831 | 0.138131 | 0.964444 | -0.012573 | 0.064179 | 0.12444 | 0.358343 | 1.463002 | 2.301496 |
| 30.58333 | -1.24436 | -0.647185 | -0.677838 | -0.623811 | 0.145506 | -0.567054 | -0.56071 | -0.681054 | -0.456748 | -0.541996 | -0.505777 | -0.06288 | 0.65345 | 0.044023 | 1.220853 | 0.370247 | 0.239346 | 1.44942 | 0.093454 | 0.394905 | 0.002428 | 0.183369 | 1.717204 | 2.078572 |
| 30.59722 | -0.500058 | -0.167472 | -0.171493 | -0.146934 | 0.212449 | -0.109075 | -0.200235 | -0.792699 | -0.239968 | -0.106775 | -0.049006 | 0.374564 | 0.712131 | 0.00948 | 0.489622 | 0.18882 | -0.149486 | 0.566849 | 0.05651 | 0.475598 | 0.104941 | 0.065659 | 0.709568 | -0.050896 |
| 30.61111 | 0.265969 | 0.004242 | 0.051535 | 0.03829 | 0.812998 | 0.062399 | -0.005445 | -0.74515 | -0.004003 | 0.045977 | 0.114956 | 0.486768 | 0.578648 | 0.368623 | 0.133216 | 0.432814 | -0.059949 | -0.155004 | -0.272744 | 0.531997 | 0.182848 | 0.059192 | -0.202858 | -0.481804 |
| 30.625 | 1.033921 | -0.0538 | 0.003223 | -0.017616 | 1.209362 | 0.010538 | 0.113456 | -0.616771 | 0.082558 | -0.018164 | 0.023343 | 0.066163 | 0.451424 | 0.528845 | -0.625589 | 0.128384 | 0.625914 | 0.000895 | -0.334303 | 0.583602 | 0.14097 | 0.183594 | -0.647898 | -1.13422 |
| 30.63889 | 1.2072 | -0.144373 | -0.119077 | -0.1224 | 0.308601 | -0.077823 | 0.221436 | -0.536648 | -0.002546 | -0.117443 | -0.121109 | -0.237994 | 0.386459 | 0.265815 | -1.510339 | -0.317512 | 0.778379 | 0.1245 | -0.096311 | 0.068475 | 0.098989 | 0.354842 | -0.010095 | -0.196642 |
| 30.65278 | 0.188736 | -0.122698 | -0.11923 | -0.113728 | 0.582913 | -0.070422 | 0.183549 | -0.416572 | -0.062835 | -0.103057 | -0.12868 | -0.305177 | 0.306668 | 0.04715 | -1.09961 | -0.371059 | 0.313188 | -0.398673 | -0.050671 | 0.046131 | 0.051015 | 0.440679 | -0.293474 | 1.588358 |
| 30.66667 | -0.095038 | -0.037798 | -0.038402 | -0.037863 | 1.336616 | -0.019138 | -0.067832 | -0.317308 | -0.036502 | -0.027655 | -0.037017 | -0.250913 | 0.173206 | -0.184322 | -0.281919 | 0.005566 | -0.294881 | -0.457134 | -0.016533 | 0.036785 | 0.037042 | 0.405438 | -0.613377 | 2.525341 |
| 30.68056 | -0.147341 | 0.007344 | 0.015708 | 0.00805 | -0.921994 | 0.002306 | -0.334521 | -0.438795 | -0.004118 | 0.007378 | 0.055237 | -0.668319 | 0.056427 | -0.21724 | 0.39619 | 0.329473 | -0.56826 | 0.168934 | 0.076053 | -0.473471 | -0.037114 | 0.250854 | -0.022353 | 1.121658 |
| 30.69444 | -0.382597 | -0.006332 | 0.009862 | -0.004679 | -0.362993 | -0.029986 | -0.384563 | -1.142036 | -0.029452 | -0.021905 | 0.089791 | -1.880227 | -0.00671 | -0.19961 | 0.684117 | 0.281991 | -0.507427 | -0.050162 | 0.111248 | -0.228711 | -0.107191 | 0.025666 | 0.363667 | -0.422891 |
| 30.70833 | -0.557159 | -0.001144 | -0.002467 | -0.008232 | 1.329754 | -0.042545 | -0.228199 | -1.58419 | -0.038649 | -0.030048 | 0.096675 | -0.78489 | -0.037827 | 0.290166 | 0.249235 | 0.13355 | -0.223053 | 0.125991 | 0.19713 | 0.231265 | -0.074229 | -0.125774 | 0.472595 | -0.883742 |
| 30.72222 | 0.19901 | 0.04174 | 0.021662 | 0.032557 | 0.227889 | 0.010009 | -0.008877 | -1.322057 | 0.009746 | 0.01682 | 0.075235 | 0.00345 | -0.02603 | 0.472085 | 0.027822 | 0.048979 | 0.482186 | 0.446778 | 0.299942 | 0.442164 | 0.022376 | -0.091924 | 0.66532 | -2.831517 |
| 30.73611 | 0.207713 | 0.068824 | 0.057006 | 0.067334 | -2.518633 | 0.059458 | 0.22669 | -0.732372 | 0.068078 | 0.050249 | 0.045002 | 0.498301 | -0.013316 | 0.430632 | 0.065601 | 0.080693 | 0.897069 | 0.647887 | 0.387195 | 0.917732 | -0.107006 | -0.046565 | 0.049799 | -1.505791 |
| 30.75 | -0.058197 | 0.075447 | 0.080643 | 0.078907 | -1.855771 | 0.081053 | 0.403803 | -0.462107 | 0.116502 | 0.063044 | 0.045785 | 0.719466 | -0.036043 | 0.08218 | 0.16784 | 0.081911 | 0.736786 | 0.366794 | 0.292666 | 1.552091 | -0.237486 | 0.00751 | 0.39397 | 0.063396 |
| 30.76389 | -0.379381 | 0.09588 | 0.104114 | 0.092786 | 0.575982 | 0.105165 | 0.36403 | -0.414817 | 0.164638 | 0.091023 | 0.07313 | 0.575847 | -0.08313 | -0.291244 | -0.234193 | -0.063838 | 0.230162 | 0.430697 | 0.145414 | 2.002472 | -0.114133 | -0.00609 | -0.059622 | 0.901794 |
| 30.77778 | -0.635513 | 0.12649 | 0.128777 | 0.114191 | 1.867354 | 0.127272 | 0.152833 | -0.461661 | 0.202414 | 0.117457 | 0.098129 | 0.434849 | -0.164399 | -0.42584 | -0.904305 | -0.219255 | 0.064296 | 0.597778 | 0.014424 | 1.307141 | 0.038028 | -0.043118 | -0.973876 | 1.35741 |
| 30.79167 | -0.684309 | 0.117555 | 0.115529 | 0.099905 | 0.008916 | 0.119513 | -0.142909 | -0.531155 | 0.18946 | 0.104182 | 0.118771 | -0.135416 | -0.267425 | -0.656099 | -1.157118 | -0.380703 | 0.021099 | 0.661682 | -0.052922 | -0.24993 | 0.221365 | -0.027991 | 0.147852 | 1.630699 |
| 30.80556 | -0.801581 | 0.077883 | 0.073586 | 0.060066 | -0.629901 | 0.08871 | -0.411329 | -0.586684 | 0.150942 | 0.072963 | 0.122082 | -0.761383 | -0.33628 | -0.470608 | -1.568462 | -0.476167 | -0.38183 | 0.652647 | -0.01799 | -0.654036 | 0.274942 | 0.011002 | -0.009754 | -0.174552 |
| 30.81944 | -0.483894 | 0.069913 | 0.067431 | 0.05907 | -0.184482 | 0.078961 | -0.412619 | -0.574807 | 0.154101 | 0.069456 | 0.133777 | -1.148248 | -0.353993 | 0.000978 | -1.30566 | -0.399702 | -0.708191 | 0.568415 | -0.008448 | -1.185621 | 0.078476 | -0.06776 | -0.073822 | -0.959751 |
| 30.83333 | -0.234789 | 0.110971 | 0.117647 | 0.111518 | 0.633942 | 0.115855 | -0.303923 | -0.466323 | 0.213476 | 0.110477 | 0.175125 | -1.536263 | -0.358228 | 0.181575 | -0.736628 | -0.165 | -0.477604 | 0.119526 | 0.027473 | -2.383235 | -0.040837 | -0.229514 | 0.019576 | -0.847006 |
| 30.84722 | -0.048337 | 0.172473 | 0.193675 | 0.184733 | -0.096677 | 0.180958 | -0.076119 | -0.143558 | 0.27135 | 0.175935 | 0.198782 | -1.583806 | -0.407319 | 0.083194 | 0.162053 | 0.124801 | 0.056117 | -0.237807 | 0.005431 | -1.442068 | 0.079674 | -0.3773 | 0.027525 | -0.70544 |
| 30.86111 | -0.330457 | 0.229791 | 0.257886 | 0.244586 | -1.217837 | 0.242647 | -0.016302 | -0.015028 | 0.233016 | 0.234031 | 0.185429 | -1.493737 | -0.409287 | 0.03648 | 0.181587 | -0.077078 | 0.161357 | -0.167392 | 0.000817 | -0.486662 | -0.021454 | -0.650828 | -0.055882 | -1.632452 |
| 30.875 | -1.063138 | 0.286428 | 0.308896 | 0.291911 | -1.356196 | 0.300819 | -0.112646 | 0.155167 | 0.116633 | 0.286595 | 0.193363 | -1.400965 | -0.48443 | -0.242801 | -0.308061 | -0.291432 | -0.206734 | 0.02054 | 0.091045 | -0.385915 | -0.042437 | -0.617478 | -0.283525 | -1.326667 |
| 30.88889 | -1.78543 | 0.32684 | 0.342816 | 0.318075 | -1.038286 | 0.333561 | -0.250133 | 0.171759 | 0.009157 | 0.319319 | 0.180722 | -1.378087 | -0.587959 | -0.803662 | -0.589176 | 0.013121 | -0.256238 | -0.405254 | 0.102602 | 0.213502 | -0.088474 | -0.411734 | -0.574009 | -0.851852 |
| 30.90278 | -2.376298 | 0.330208 | 0.333822 | 0.302276 | -0.90133 | 0.323333 | -0.50015 | -0.144343 | -0.060567 | 0.291556 | 0.072353 | -1.249132 | -0.845617 | -1.051955 | -1.413983 | 0.010677 | -0.034593 | -0.248028 | 0.042002 | 0.265874 | -0.050064 | -0.262399 | -0.917931 | -0.724748 |
| 30.91667 | -2.438944 | 0.309972 | 0.300893 | 0.271116 | -0.718722 | 0.301009 | -0.999438 | -0.576729 | -0.134911 | 0.220153 | 0.195083 | -1.249779 | -0.831365 | -1.58682 | -1.148002 | -0.197852 | -0.367042 | -0.168719 | -0.061124 | 0.508434 | -0.297796 | -0.283688 | -1.068304 | -0.93055 |
| 30.93056 | -1.859088 | 0.281496 | 0.273289 | 0.247364 | -0.622157 | 0.278657 | -1.308961 | -0.792702 | -0.252301 | 0.165111 | 0.27335 | -1.143416 | -0.796305 | -2.295621 | -0.627067 | -0.379132 | -1.03518 | -0.25962 | -0.121933 | 0.996657 | -0.477913 | -0.425403 | -1.228821 | -1.016228 |
| 30.94444 | -1.233119 | 0.237916 | 0.239515 | 0.215055 | -0.814805 | 0.235055 | -1.164121 | -0.843841 | -0.379844 | 0.158752 | 0.130069 | -0.849214 | -0.875774 | -4.379988 | -0.391894 | -0.522754 | -1.886848 | -0.323319 | -0.124579 | 1.400024 | -0.288381 | -0.53832 | -1.754267 | -1.157121 |
| 30.95833 | -0.758503 | 0.176789 | 0.182182 | 0.163988 | -0.99793 | 0.166888 | -1.093487 | -0.908134 | -0.573513 | 0.143797 | 0.073402 | -0.636741 | -0.928219 | -5.266431 | -1.156098 | -1.652517 | -2.569245 | -0.380018 | -0.063869 | 1.558804 | 0.236966 | -0.617867 | -1.605076 | -1.542869 |
| 30.97222 | -0.390601 | 0.109476 | 0.111043 | 0.10058 | -1.133059 | 0.090307 | -0.840672 | -1.063706 | -0.787973 | 0.106174 | -0.00617 | -0.526478 | -0.95307 | -3.747881 | -1.90663 | -2.451878 | -2.370263 | -0.531774 | -0.065466 | 0.819007 | 0.328403 | -0.678968 | -1.430577 | -1.770095 |
| 30.98611 | -0.184088 | 0.058324 | 0.063312 | 0.057574 | -1.108038 | 0.039417 | -0.484319 | -1.013012 | -0.88975 | 0.076025 | -0.050276 | -0.501852 | -0.984187 | -1.86452 | -2.646932 | -3.002776 | -1.758783 | -1.014747 | -0.174804 | -0.858674 | 0.297744 | -0.720683 | -1.21938 | -1.964395 |
| 31 | -0.098664 | 0.118292 | 0.133044 | 0.144987 | -0.881623 | 0.079374 | -0.185438 | -0.658924 | -0.851918 | 0.167772 | 0.007662 | -0.472287 | -0.833774 | -1.910894 | -2.761834 | -2.224315 | -0.948553 | -1.02239 | -0.166138 | -1.446567 | 0.043188 | -0.506959 | -0.995928 | -1.750354 |
| 31.01389 | -0.018463 | 0.276258 | 0.306165 | 0.370209 | -0.683522 | 0.182348 | 0.050199 | -0.349599 | -0.675234 | 0.35929 | 0.133446 | -0.232122 | -0.610333 | -2.422057 | -2.519186 | -0.803485 | -0.222294 | -0.6477 | 0.171149 | -0.33795 | -0.230069 | -0.081999 | -0.441904 | -1.334373 |
| 31.02778 | -0.077805 | 0.360482 | 0.362962 | 0.555797 | -1.064724 | 0.230708 | 0.012364 | -0.487911 | -0.559017 | 0.408824 | 0.132265 | -0.01516 | -0.60808 | -2.2043 | -2.073823 | -0.084099 | -0.127087 | -0.575138 | 0.433164 | 1.141153 | -0.903336 | 0.104753 | -0.37157 | -1.791381 |
| 31.04167 | -0.454955 | 0.353614 | 0.210828 | 0.389021 | -1.694934 | 0.196825 | -0.339426 | -0.793091 | -0.767739 | 0.258103 | 0.006412 | -0.066174 | -0.620915 | -3.223852 | -0.748351 | -0.347855 | -0.574732 | -1.383071 | 0.200441 | 2.148289 | -1.416205 | -0.362802 | -1.280177 | -2.982102 |
| 31.05556 | -0.887004 | 0.357358 | 0.170483 | 0.259992 | -2.448376 | 0.272469 | -0.702815 | -1.029637 | -1.152115 | 0.276165 | -0.101837 | -0.472404 | -0.800732 | -5.367782 | -0.333573 | -1.090888 | -0.832263 | -1.954127 | -0.207855 | 1.534101 | -0.927852 | -1.272342 | -2.358989 | -4.095869 |
| 31.06944 | -1.020665 | 0.380969 | 0.266456 | 0.30589 | -4.63979 | 0.369806 | -1.129479 | -1.351079 | -2.147178 | 0.367177 | -0.573142 | -1.045353 | -1.161041 | -6.937373 | -0.338143 | -1.988886 | -0.834636 | -2.114295 | -0.681331 | 0.213554 | -0.500921 | -2.153845 | -3.416861 | -4.804041 |
| 31.08333 | -0.918775 | 0.251865 | 0.199712 | 0.213564 | -3.129405 | 0.264307 | -1.114038 | -1.164247 | -3.050782 | 0.254877 | -0.669517 | -1.312875 | -1.936876 | -6.021889 | -0.735545 | -1.198328 | -0.430841 | -1.577085 | -0.704928 | -0.674897 | -0.028406 | -2.058909 | -3.031199 | -4.935258 |
| 31.09722 | -0.434581 | 0.011492 | -0.01701 | 0.002518 | -0.405233 | 0.037481 | -0.674106 | -0.571561 | -2.576331 | 0.018119 | -0.343047 | -0.964561 | -2.013975 | -3.553015 | -1.336898 | -0.420151 | -0.226701 | -0.844782 | -0.686653 | -1.873836 | -0.328487 | -1.246831 | -1.706297 | -3.500347 |
| 31.11111 | -0.119143 | -0.158536 | -0.147761 | -0.128989 | 1.263188 | -0.117552 | -0.119365 | -0.127105 | -1.564353 | -0.129052 | -0.236174 | -0.39818 | -1.638524 | -1.559846 | -0.9029 | 0.199288 | -0.15461 | -0.507987 | -0.298246 | -2.149295 | -1.088467 | 0.143541 | -0.458287 | -1.634914 |
| 31.125 | -0.070452 | -0.132502 | -0.098063 | -0.095419 | 1.474835 | -0.105273 | 0.414538 | 0.10127 | -0.357125 | -0.10707 | -0.072224 | 0.019325 | -1.273675 | -0.647229 | -0.187921 | 0.76317 | -0.033884 | -0.187908 | -0.109279 | -1.21305 | -1.134394 | 0.912365 | 0.373922 | -0.389872 |

| | | | | | | | | | | | | | | | | | | | | | | | | |
|---|---|---|---|---|---|---|---|---|---|---|---|---|---|---|---|---|---|---|---|---|---|---|---|---|
| 31.13889 | -0.033484 | -0.015903 | 0.011453 | 0.002303 | 0.840373 | -0.00554 | 0.734711 | 0.391888 | 0.760758 | -0.0117 | 0.052456 | 0.146965 | -0.966606 | -0.10202 | 0.158445 | 0.848072 | 0.136064 | 0.316614 | -0.058269 | -0.278493 | -0.445516 | 1.223054 | 0.816301 | 0.268757 |
| 31.15278 | 0.17575 | 0.045403 | 0.066949 | 0.04596 | 0.260078 | 0.067567 | 0.706259 | 0.674589 | 1.414968 | 0.059999 | 0.112712 | 0.276897 | -0.72656 | 0.619828 | 0.324137 | 0.511331 | 0.400938 | 0.780754 | 0.299254 | 0.237448 | 0.395693 | 1.13402 | 0.967995 | 0.954264 |
| 31.16667 | 0.664718 | 0.151597 | 0.167779 | 0.123762 | 0.053831 | 0.167817 | 0.703172 | 0.832975 | 1.194399 | 0.169285 | 0.208656 | 0.884587 | -0.486117 | 1.466873 | 0.212142 | 0.344989 | 0.662182 | 0.876771 | 1.010699 | 0.423319 | 1.263389 | 0.761513 | 1.569986 | 1.883562 |
| 31.18056 | 1.320192 | 0.376383 | 0.376528 | 0.326165 | 0.061196 | 0.357238 | 0.964765 | 0.827392 | 0.620076 | 0.350714 | 0.406386 | 1.512319 | -0.289396 | 1.880965 | -0.000855 | 0.630432 | 0.85493 | 1.211563 | 0.811916 | 0.562429 | 2.030803 | 0.483502 | 1.746882 | 2.428491 |
| 31.19444 | 1.579426 | 0.524495 | 0.518885 | 0.49094 | -0.288678 | 0.497864 | 1.329626 | 0.594668 | -0.003523 | 0.479327 | 0.530605 | 1.77348 | -0.025228 | 2.088447 | -0.173928 | 1.626766 | 0.738115 | 1.794534 | 0.336533 | 0.237634 | 1.845057 | 0.313368 | 1.016312 | 2.171281 |
| 31.20833 | 1.178937 | 0.466663 | 0.473626 | 0.464921 | -0.891465 | 0.464595 | 1.374269 | 0.21194 | -0.515818 | 0.456657 | 0.472915 | 2.025143 | 0.47409 | 1.674689 | -0.089779 | 2.340978 | 0.584463 | 1.087055 | -0.070155 | -0.358779 | 0.515016 | 0.280284 | 0.340831 | 1.18445 |
| 31.22222 | 0.53157 | 0.301202 | 0.31012 | 0.313431 | -1.173388 | 0.317529 | 1.133098 | -0.186139 | -0.63421 | 0.317504 | 0.341496 | 2.299879 | 1.133255 | 1.554651 | 0.066121 | 1.782959 | 0.558379 | 0.282294 | -0.345039 | -0.685652 | -0.23653 | 0.244923 | -0.495543 | 0.058987 |
| 31.23611 | -0.217698 | 0.134279 | 0.107883 | 0.120367 | -0.676561 | 0.144641 | 0.877241 | -0.510889 | -0.497602 | 0.127381 | 0.164213 | 1.766972 | 1.717473 | 1.500655 | 0.088935 | 0.387984 | 0.694647 | -0.001293 | -0.105789 | -0.33717 | -0.379621 | 0.217923 | 0.007901 | -0.322644 |
| 31.25 | -0.503971 | 0.002448 | -0.07901 | -0.061184 | -0.068174 | 0.003671 | 0.760278 | -0.720541 | -0.47537 | -0.027722 | 0.002601 | 0.268107 | 2.238769 | 1.626011 | -0.030696 | -0.704651 | 0.664358 | 0.68747 | 0.677825 | 1.099138 | -0.530471 | 0.183679 | 0.922456 | 0.248928 |
| 31.26389 | 0.070949 | -0.122284 | -0.210677 | -0.16884 | 0.440186 | -0.096886 | 0.808112 | -0.68652 | -0.579619 | -0.106184 | -0.074998 | 0.205001 | 2.783776 | 1.597569 | -0.143939 | -1.961951 | 0.205508 | 1.172625 | 0.554646 | 1.611584 | -1.019771 | 0.132292 | 1.951976 | 0.98622 |
| 31.27778 | 0.901527 | -0.13528 | -0.193179 | -0.145644 | 0.874429 | -0.108974 | 0.907182 | -0.489031 | -0.575761 | -0.089046 | -0.033419 | 1.809497 | 2.223198 | 0.876519 | -0.122242 | -0.4794 | -0.381391 | 1.210378 | -0.02738 | 1.769796 | -0.301766 | 0.048694 | 1.734437 | 1.645219 |
| 31.29167 | 1.431444 | 0.068518 | 0.050649 | 0.05258 | 0.915869 | 0.065245 | 1.159216 | -0.079339 | -0.234142 | 0.085885 | 0.176702 | 3.65079 | 1.612506 | 0.101087 | 0.104997 | 1.316428 | -0.643834 | 1.465791 | -0.336375 | 0.641807 | 0.372756 | 0.003813 | 0.876582 | 1.888422 |
| 31.30556 | 1.764118 | 0.356762 | 0.37624 | 0.343811 | 0.491 | 0.352223 | 1.428057 | 0.387331 | 0.312806 | 0.371406 | 0.61704 | 3.114348 | 1.163883 | -0.962187 | 0.887179 | 2.426245 | -0.358484 | 0.831732 | -0.392503 | -0.834968 | 0.733677 | 0.091332 | 0.36721 | 1.673009 |
| 31.31944 | 1.978896 | 0.508051 | 0.541343 | 0.501137 | 0.096711 | 0.51981 | 1.468458 | 0.632583 | 0.727285 | 0.556804 | 0.861105 | 0.952892 | 0.834447 | -0.655342 | 1.709959 | 1.328574 | 0.080563 | 0.190456 | -0.130974 | -1.788963 | 0.465813 | 0.249484 | -0.205661 | 1.351994 |
| 31.33333 | 1.970348 | 0.453175 | 0.463511 | 0.411896 | 0.309141 | 0.469873 | 1.323131 | 0.602532 | 0.889852 | 0.519291 | 0.689885 | -1.30456 | 0.433015 | 1.128811 | 1.83786 | 0.144396 | 0.388575 | -0.380004 | 0.086351 | -1.282038 | 0.1754 | 0.370453 | 0.633498 | 0.65785 |
| 31.34722 | 1.944932 | 0.313173 | 0.299664 | 0.232643 | 1.245958 | 0.326473 | 1.099279 | 0.409198 | 0.910336 | 0.367989 | 0.377301 | -2.180802 | 0.211632 | 3.225596 | 1.996761 | -0.692987 | 0.638317 | -0.700147 | 0.112433 | -0.098902 | -0.089822 | 0.432773 | 2.271724 | 0.621773 |
| 31.36111 | 1.668347 | 0.145245 | 0.135463 | 0.055601 | 2.543981 | 0.142256 | 0.983399 | 0.249982 | 0.802868 | 0.162447 | 0.16323 | -1.206485 | 0.121804 | 3.053402 | 2.031887 | -0.857228 | 0.730384 | -0.055838 | -0.042721 | 0.407986 | -0.133565 | 0.478119 | 2.646883 | 0.977081 |
| 31.375 | 1.063277 | -0.101797 | -0.090663 | -0.16014 | 3.183409 | -0.115625 | 0.853197 | 0.167634 | 0.630767 | -0.093897 | -0.031888 | -0.034671 | 0.031698 | 2.174686 | 1.628971 | -0.132608 | 0.749145 | 0.145456 | -0.120626 | -0.07185 | 0.072923 | 0.47302 | 1.435317 | 1.301326 |
| 31.38889 | 0.614647 | -0.36338 | -0.369271 | -0.398699 | 2.499609 | -0.370269 | 0.575631 | 0.155204 | 0.485348 | -0.351602 | -0.230466 | 1.032117 | -0.061695 | 1.592169 | 1.110165 | 0.54324 | 0.805784 | -0.286386 | 0.037294 | 0.483918 | 0.221341 | 0.436031 | 0.299419 | 1.995223 |
| 31.40278 | 0.538283 | -0.527959 | -0.572117 | -0.561713 | 1.624833 | -0.51149 | 0.390628 | 0.277153 | 0.424332 | -0.517518 | -0.363267 | 1.588761 | -0.123099 | 1.337574 | 0.828065 | 0.85816 | 0.92882 | 0.337406 | -0.38501 | 0.092915 | 0.201447 | 0.467176 | -0.755903 | 2.123602 |
| 31.41667 | 1.090457 | -0.533675 | -0.581222 | -0.570803 | 0.693307 | -0.518382 | 0.307341 | 0.60981 | 0.535876 | -0.555648 | -0.380089 | 1.597894 | -0.117367 | 1.687562 | 0.751834 | 0.79154 | 1.151955 | 0.211615 | -0.077717 | -0.340913 | 0.190074 | 0.53349 | -0.511615 | 0.874678 |
| 31.43056 | 1.974136 | -0.415653 | -0.46756 | -0.478938 | -0.143413 | -0.453813 | 0.374498 | 1.203227 | 0.8198 | -0.495439 | -0.276795 | 1.456642 | -0.07956 | 3.690284 | 0.790038 | 0.7265 | 1.344481 | 0.120685 | -0.11561 | 0.023285 | 0.314135 | 0.566896 | 0.489536 | 1.614196 |
| 31.44444 | 2.349905 | -0.383301 | -0.443992 | -0.429089 | -0.222863 | -0.402723 | 0.404762 | 1.659151 | 1.25623 | -0.414793 | -0.191812 | 1.35943 | -0.090782 | 4.435542 | 1.164363 | 0.923289 | 1.770596 | -0.569417 | 0.004388 | 0.762923 | 0.292858 | 0.517666 | 1.030359 | 2.535833 |
| 31.45833 | 2.187376 | -0.376396 | -0.41821 | -0.599331 | 0.739049 | -0.300951 | 0.380823 | 1.793268 | 1.902138 | -0.346526 | -0.128497 | 1.501967 | -0.135777 | 2.615026 | 0.351035 | 1.237282 | 2.040107 | -0.32488 | -0.277198 | 1.197379 | -0.009532 | 0.555525 | 0.998598 | 2.284389 |
| 31.47222 | 1.865358 | -0.091928 | -0.093423 | -0.267144 | 2.972028 | 0.003622 | 0.416474 | 1.703318 | 2.492697 | -0.120637 | 0.011556 | 1.583405 | -0.083948 | 2.524587 | 0.430223 | 1.37306 | 1.669272 | 0.297816 | -0.38102 | 1.043759 | 0.147732 | 0.543214 | 0.387228 | 2.77475 |
| 31.48611 | 1.604583 | 0.135262 | 0.1128 | 0.053301 | 4.819403 | 0.154923 | 0.485985 | 1.382072 | 2.706161 | 0.084458 | 0.157015 | 0.906662 | 0.161178 | 2.872267 | 0.651815 | 1.5282 | 1.162257 | 1.276234 | -0.136094 | 0.568206 | 0.534456 | 0.730573 | 0.327937 | 2.999211 |
| 31.5 | 1.929458 | 0.168829 | 0.139911 | 0.109361 | 4.881386 | 0.135276 | 0.449711 | 1.341237 | 2.59027 | 0.14308 | 0.252105 | -0.484628 | 0.414486 | 3.502618 | 1.311742 | 1.419887 | 0.587305 | 2.412453 | 0.178096 | -0.212625 | 0.476192 | 0.894618 | 0.714232 | 1.737284 |
| 31.51389 | 2.330317 | 0.164593 | 0.166821 | 0.159767 | 2.76831 | 0.14456 | 0.350218 | 1.412916 | 2.339465 | 0.173031 | 0.319051 | -0.878196 | 0.579128 | 3.852197 | 1.135148 | 0.710688 | 0.351659 | 2.1596 | 0.231626 | -0.41999 | 0.391372 | 0.844163 | 1.015711 | 1.058627 |
| 31.52778 | 2.336318 | -0.131729 | 0.002478 | -0.083747 | 0.92549 | -0.162223 | 0.05585 | 1.199204 | 2.009651 | -0.140072 | 0.096677 | 0.345906 | 0.719075 | 4.032092 | 0.65835 | 0.602881 | 0.423086 | 1.256225 | 0.073177 | -0.569778 | 0.398525 | 0.746955 | 1.517949 | 3.118155 |
| 31.54167 | 1.419998 | -0.879267 | -0.536136 | -0.745212 | -0.053506 | -0.941296 | -0.551441 | 0.302529 | 1.30999 | -0.902542 | -0.541317 | 1.597529 | 0.636857 | 2.465673 | 0.968991 | 0.758373 | 0.594201 | 0.463048 | 0.025745 | -0.548246 | 0.217198 | 0.597923 | 2.310527 | 3.500686 |
| 31.55556 | 0.87813 | -1.205034 | -0.875785 | -1.026933 | -0.196644 | -1.230451 | -0.760128 | -0.418284 | 0.547452 | -1.165041 | -0.873221 | 1.285242 | 0.358552 | 0.994722 | 1.525499 | 0.754455 | 0.723854 | 0.596204 | 0.079067 | -0.579654 | 0.248999 | 0.314963 | 2.019265 | 1.72967 |
| 31.56944 | 0.573916 | -0.699217 | -0.631835 | -0.615674 | 0.777017 | -0.681916 | -0.341151 | -0.270232 | 0.044408 | -0.627796 | -0.545033 | 0.718028 | 0.181734 | -0.179302 | 2.657669 | 0.731799 | 0.711929 | 1.08638 | -0.342934 | 0.292814 | 0.116686 | 0.304934 | 2.318533 | 2.165412 |
| 31.58333 | 1.12877 | -0.228198 | -0.278035 | -0.215233 | 1.553193 | -0.208228 | 0.104656 | 0.06863 | -0.121104 | -0.176647 | -0.158389 | 0.750592 | 0.34447 | -0.699178 | 3.564831 | 0.651558 | 0.256059 | 1.373685 | -0.856975 | 0.193866 | -0.132752 | 0.247556 | 2.918343 | 1.827956 |
| 31.59722 | 0.941944 | -0.145112 | -0.178195 | -0.146336 | 1.216141 | -0.146446 | 0.16607 | 0.064267 | -0.201021 | -0.12706 | -0.072877 | 0.65844 | -0.361273 | 0.445821 | 2.510797 | 0.082495 | -0.825346 | 0.382469 | -0.295339 | -0.26142 | -0.404895 | -0.153718 | 0.720466 | 0.584123 |
| 31.61111 | 0.134351 | -0.190758 | -0.228379 | -0.207539 | 0.132971 | -0.220557 | 0.012443 | -0.088456 | -0.207261 | -0.209689 | -0.155129 | 0.495064 | -0.184895 | 1.055549 | 1.055495 | -0.25314 | -0.95965 | 0.202967 | 0.502968 | -0.513633 | -0.066309 | -0.170123 | -0.454871 | -1.801721 |
| 31.625 | -0.73671 | -0.194233 | -0.23012 | -0.209405 | -1.53562 | -0.221878 | 0.089092 | -0.215539 | -0.154032 | -0.218816 | -0.189178 | 0.368055 | 0.119715 | 0.391687 | -0.381161 | -0.134019 | 0.169608 | 0.081647 | 0.47068 | 0.174952 | -0.018426 | -0.119242 | -1.250835 | -0.78269 |
| 31.63889 | -0.813444 | -0.178455 | -0.194804 | -0.184241 | -2.093787 | -0.1892 | 0.218826 | -0.415242 | -0.136694 | -0.199324 | -0.170681 | 0.285118 | 0.071386 | 0.038883 | -1.130132 | -0.407431 | 0.739045 | 0.52118 | 0.2461 | 0.899447 | -0.014118 | 0.078471 | -0.479899 | 1.734547 |
| 31.65278 | 0.051453 | -0.18369 | -0.196566 | -0.182302 | -0.334075 | -0.171736 | 0.111408 | -0.707449 | -0.134478 | -0.189642 | -0.164263 | 0.129713 | 0.437749 | -0.3432 | -1.496797 | -0.773931 | 0.0456 | 1.157513 | -0.000443 | 1.032039 | 0.078435 | 0.296736 | 0.153492 | 4.026566 |
| 31.66667 | 0.45267 | -0.162047 | -0.197868 | -0.165053 | 1.376029 | -0.141732 | -0.063093 | -1.073797 | -0.085193 | -0.157106 | -0.142235 | 0.010622 | 0.362803 | -0.350643 | -1.128572 | -0.393317 | 0.504366 | 0.692713 | -0.345595 | 1.024415 | 0.196694 | 0.421047 | -0.734063 | 1.466642 |
| 31.68056 | 0.019021 | -0.121114 | -0.156421 | -0.123653 | 2.153293 | -0.11733 | -0.182361 | -1.26574 | -0.015429 | -0.11492 | -0.072139 | -0.051636 | 0.24858 | 0.105484 | -0.401266 | 0.115367 | 0.927269 | 0.778022 | 0.147544 | 0.689309 | 0.182765 | 0.322866 | -0.35929 | -0.377113 |
| 31.69444 | -0.641685 | -0.111596 | -0.127436 | -0.114317 | 1.152096 | -0.135625 | -0.382117 | -1.236772 | 0.035989 | -0.114972 | -0.018383 | -0.114852 | 0.487425 | 0.582082 | 0.113347 | 0.495956 | 0.7293 | 0.477049 | -0.091438 | 0.188069 | 0.194077 | 0.179303 | 0.247064 | -0.828978 |
| 31.70833 | -1.307238 | -0.138091 | -0.142374 | -0.143298 | 0.271239 | -0.161006 | -0.540443 | -1.202454 | 0.072151 | -0.140118 | -0.040682 | -0.215971 | 0.375237 | 0.502649 | 1.889579 | 0.66885 | 0.015011 | -0.93106 | -0.556215 | -0.333638 | 0.147016 | 0.074692 | 0.435511 | -2.882882 |
| 31.72222 | -1.402819 | -0.058493 | -0.07197 | -0.067235 | -0.21843 | -0.063883 | -0.64187 | -1.073982 | 0.171995 | -0.054562 | -0.039839 | -0.226848 | 0.134312 | 0.111114 | 2.325116 | 0.508571 | -0.103556 | -0.716004 | -1.445323 | -0.630629 | 0.098755 | 0.015254 | -0.01925 | -0.088674 |
| 31.73611 | -1.142822 | 0.130806 | 0.114435 | 0.127403 | -0.686069 | 0.136559 | -0.409462 | -0.688717 | 0.319746 | 0.136293 | 0.062711 | -0.130061 | -0.133425 | -0.166577 | 1.786245 | 0.190779 | -0.035899 | -0.264265 | -0.045316 | -0.832679 | 0.173036 | -0.03295 | -0.581979 | -0.602457 |
| 31.75 | -0.993784 | 0.233843 | 0.218789 | 0.229539 | -1.079125 | 0.242919 | 0.154197 | -0.45707 | 0.354288 | 0.231747 | 0.143937 | -0.089095 | -0.397889 | -0.183156 | 0.86947 | -0.120236 | -0.003851 | 0.183843 | 1.202653 | -0.449884 | 0.218263 | -0.033122 | -0.138211 | 0.331012 |
| 31.76389 | -0.536639 | 0.182698 | 0.175609 | 0.182911 | -0.595489 | 0.20018 | 0.671367 | -0.453441 | 0.246547 | 0.179954 | 0.12749 | -0.133599 | -0.605345 | -0.326018 | -0.01322 | -0.278721 | -0.048754 | 0.388832 | 0.707569 | 0.203155 | 0.142297 | -0.036477 | 0.029925 | 1.112502 |
| 31.77778 | -0.207427 | 0.128192 | 0.121861 | 0.124681 | 0.289127 | 0.138347 | 0.720829 | -0.543577 | 0.145131 | 0.120641 | 0.104977 | -0.136029 | -0.646991 | -0.461275 | -1.382402 | -0.471431 | 0.029746 | -0.074432 | -0.060814 | 0.430816 | 0.083407 | -0.118852 | -0.691352 | -0.452569 |
| 31.79167 | -0.116739 | 0.123666 | 0.115662 | 0.115832 | 1.116557 | 0.128013 | 0.449307 | -0.594808 | 0.123139 | 0.115974 | 0.139218 | -0.157012 | -0.421227 | -0.312968 | -2.534548 | -0.525012 | 0.106019 | 0.001093 | -0.079905 | 0.524388 | -0.151608 | -0.154737 | 0.035837 | -0.360306 |
| 31.80556 | -0.324965 | 0.107567 | 0.093473 | 0.094142 | 1.147638 | 0.096068 | -0.068312 | -0.57332 | 0.088312 | 0.094161 | 0.169329 | -0.3684 | -0.202273 | -0.035379 | -2.933095 | -0.360399 | 0.054462 | 0.141677 | 0.125363 | 0.399025 | -0.220266 | -0.083233 | 0.439825 | 0.536191 |
| 31.81944 | -0.609667 | 0.056379 | 0.043723 | 0.034718 | 0.492815 | 0.022772 | -0.621219 | -0.500692 | -0.009946 | 0.030963 | 0.138096 | -0.699808 | -0.334576 | 0.012468 | -3.168477 | -0.137534 | -0.079308 | 0.094869 | 0.187696 | -0.4599 | 0.118042 | -0.166505 | 0.13881 | 0.009965 |
| 31.83333 | -0.768465 | 0.03694 | 0.029664 | 0.008447 | -0.283105 | 0.004758 | -1.199819 | -0.433997 | -0.07542 | 0.011353 | 0.091761 | -0.877114 | -0.74364 | 0.059531 | -3.013909 | 0.078453 | -0.099019 | 0.270484 | 0.258022 | -0.906576 | -0.014265 | -0.295706 | 0.029807 | -2.423352 |
| 31.84722 | -0.956306 | 0.079836 | 0.07962 | 0.057812 | -1.136968 | 0.072609 | -1.591015 | -0.390109 | -0.068002 | 0.071142 | 0.062722 | -0.917835 | -1.496883 | 0.056732 | -2.162124 | 0.146916 | -0.156808 | 0.344882 | 0.305636 | -0.950218 | -0.126542 | -0.417282 | 0.004494 | -0.381992 |
| 31.86111 | -1.429985 | 0.144284 | 0.153472 | 0.137729 | -2.304036 | 0.154593 | -2.494163 | -0.368123 | -0.051797 | 0.145692 | 0.09748 | -1.091013 | -0.626699 | -0.16725 | -1.534461 | 0.105142 | -0.338825 | 0.114473 | 0.340636 | -0.763555 | -0.250693 | -0.512656 | -0.423677 | -0.360111 |
| 31.875 | -1.695124 | 0.202117 | 0.215594 | 0.203271 | -3.260236 | 0.207124 | -2.796803 | -0.374495 | -0.066811 | 0.206471 | 0.193996 | -1.374607 | -0.272312 | -0.507959 | -0.778948 | -0.09857 | -0.767181 | 0.133551 | 0.250323 | -0.20459 | -0.267948 | -0.505241 | -0.414874 | 0.92838 |
| 31.88889 | -2.22789 | 0.245978 | 0.248335 | 0.238766 | -2.42143 | 0.225864 | -1.527552 | -0.386513 | -0.136152 | 0.24551 | 0.225784 | -1.739515 | -0.218079 | -1.617404 | -0.41182 | -0.340911 | -1.151445 | 0.05463 | 0.162733 | -0.085884 | -0.373411 | -0.478069 | 0.120243 | -0.610515 |
| 31.90278 | -3.18116 | 0.244256 | 0.243741 | 0.233001 | -1.648895 | 0.211375 | 0.43817 | -0.366817 | -0.265035 | 0.238088 | 0.114834 | -2.519004 | -0.486007 | -2.313698 | -0.243629 | -0.489019 | -1.242468 | -0.409053 | 0.057178 | 0.451894 | -0.345627 | -0.523563 | 0.365794 | -1.412946 |
| 31.91667 | -2.863925 | 0.200365 | 0.194802 | 0.180981 | -1.272362 | 0.16557 | 0.602127 | -0.340552 | -0.454706 | 0.181288 | 0.111157 | -2.890002 | -0.967516 | -2.466186 | -0.54741 | -0.543428 | -1.884381 | -0.558398 | -0.015576 | 0.134866 | -0.31747 | -0.652813 | 0.014074 | -1.856706 |
| 31.93056 | -2.81349 | 0.103835 | 0.079308 | 0.068791 | -1.243282 | 0.075248 | -0.284 | -0.385114 | -0.686729 | 0.079486 | 0.039477 | -2.922605 | -1.126619 | -2.177203 | -0.950981 | -0.710602 | -2.709473 | -0.424386 | -0.013766 | -0.260039 | -0.475212 | -0.839338 | -0.591194 | -2.966595 |
| 31.94444 | -2.864814 | 0.031498 | -0.004926 | -0.004796 | -1.310431 | 0.010446 | -0.912935 | -0.292647 | -0.894623 | 0.013617 | -0.158671 | -2.58618 | -0.800226 | -2.062591 | -1.017873 | -0.91811 | -3.282843 | -0.512498 | -0.034785 | -0.432028 | -0.422497 | -0.838744 | -1.085676 | -3.239764 |
| 31.95833 | -2.481173 | 0.08188 | 0.069947 | 0.065352 | -1.549413 | 0.059044 | -1.270528 | -0.037227 | -0.965956 | 0.073067 | -0.028379 | -1.992979 | -0.477325 | -1.954666 | -0.936916 | -0.916246 | -2.927263 | -0.558439 | 0.0189 | -0.230006 | -0.327034 | -0.824313 | -1.874327 | -3.517624 |
| 31.97222 | -1.705168 | 0.132885 | 0.149033 | 0.12463 | -1.601053 | 0.103603 | -1.377907 | 0.088243 | -1.032512 | 0.123389 | 0.094979 | -1.794198 | -0.293589 | -1.103228 | -0.465288 | -0.644747 | -2.099094 | -0.496748 | -0.071757 | -0.385384 | -0.030672 | -0.820289 | -2.118971 | -2.913746 |
| 31.98611 | -1.107201 | 0.074511 | 0.074249 | 0.05089 | -1.164642 | 0.035799 | -1.361201 | -0.121311 | -1.187056 | 0.051636 | 0.022267 | -1.836357 | -0.332067 | 0.323227 | -0.09226 | -0.219905 | -1.116165 | -0.864512 | -0.28489 | -1.442339 | 0.103652 | -0.678123 | -1.602676 | -3.046999 |
| 32 | -0.398377 | 0.129759 | 0.108139 | 0.120661 | -0.556047 | 0.058091 | -1.15803 | -0.4447 | -1.063371 | 0.101536 | 0.020413 | -1.507255 | -0.421151 | -0.051277 | 0.04057 | 0.174102 | -0.536464 | -1.390014 | -0.27865 | -2.00053 | 0.169125 | -0.366525 | -0.759864 | -3.035083 |
| 32.01389 | 0.449363 | 0.380934 | 0.372255 | 0.465377 | -0.015922 | 0.244934 | -0.465957 | -0.442943 | -0.684516 | 0.367241 | 0.197122 | -0.804374 | -0.32313 | -2.009934 | 0.309367 | 0.539105 | -0.278021 | -1.160308 | 0.074115 | -1.55205 | -0.051862 | 0.110554 | 0.19946 | -2.198628 |
| 32.02778 | 0.981357 | 0.388374 | 0.370092 | 0.565261 | -0.147649 | 0.27194 | -0.054654 | -0.414054 | -0.47454 | 0.379646 | 0.1044 | -0.355241 | -0.06212 | -2.086954 | 0.624106 | 0.339833 | -0.298849 | -1.078241 | 0.507647 | -0.777655 | -0.187531 | 0.334051 | 0.227143 | -1.677339 |
| 32.04167 | 0.524595 | 0.006482 | -0.178882 | -0.097721 | -1.103883 | -0.097694 | -0.452805 | -0.787701 | -0.751863 | -0.095689 | -0.362164 | -0.361736 | 0.065641 | -2.893394 | 0.39263 | -0.871544 | -0.807124 | -1.584158 | 0.365116 | 0.096296 | -0.281581 | -0.126125 | -0.604056 | -2.152747 |
| 32.05556 | -0.090413 | -0.254572 | -0.490112 | -0.438675 | -2.201053 | -0.293201 | -0.982677 | -1.206889 | -1.007408 | -0.321092 | -0.563503 | -0.506295 | -0.008259 | -4.530752 | -0.138385 | -2.044262 | -1.620724 | -2.257427 | -0.128294 | 0.962838 | -0.544752 | -0.988678 | -1.982771 | -3.543877 |
| 32.06944 | -0.008588 | -0.162518 | -0.277286 | -0.228481 | -2.297143 | -0.125338 | -1.039486 | -1.295892 | -1.254462 | -0.135842 | -0.370729 | -0.15261 | -0.172115 | -5.438964 | -0.25869 | -2.731929 | -1.756411 | -2.922879 | -0.34012 | 1.57988 | -0.250604 | -2.035858 | -2.945447 | -4.424157 |
| 32.08333 | 0.319258 | -0.0263 | -0.035326 | -0.005703 | -1.074269 | 0.050356 | -0.588142 | -1.208468 | -1.62019 | 0.039668 | -0.313779 | 0.627684 | -0.315822 | -5.05013 | -0.815121 | -2.19947 | -1.513208 | -2.179322 | -0.164719 | 0.976565 | 0.18154 | -2.842721 | -2.976664 | -4.214245 |
| 32.09722 | 0.403475 | 0.029815 | 0.075741 | 0.089183 | 0.386498 | 0.112387 | -0.097382 | -0.783426 | -1.810428 | 0.10415 | -0.219906 | 0.620051 | -0.435978 | -3.553074 | -1.608393 | -0.969046 | -1.284619 | -1.557846 | 0.02583 | 0.184944 | 0.107253 | -1.70067 | -2.195628 | -2.782089 |
| 32.11111 | 0.563394 | 0.124747 | 0.197569 | 0.189833 | 0.977426 | 0.192796 | 0.469399 | -0.223627 | -1.3761 | 0.190971 | -0.014443 | 0.858783 | -0.550319 | -2.492474 | -1.027008 | 0.146124 | -0.586272 | -0.849614 | -0.020083 | -0.231828 | -0.220831 | -0.340729 | -1.078114 | -1.135451 |
| 32.125 | 0.729439 | 0.245622 | 0.286581 | 0.266778 | 1.135632 | 0.269786 | 0.940518 | 0.070671 | -0.67059 | 0.260723 | 0.147865 | 0.735262 | -0.573334 | -1.653121 | -0.481588 | 1.028701 | 0.011594 | -0.213089 | -0.07904 | -0.216038 | -0.738163 | 0.232486 | -0.210323 | -0.288189 |
| 32.13889 | 0.750925 | 0.266881 | 0.254684 | 0.244959 | 0.77468 | 0.256491 | 1.183747 | 0.01533 | -0.151374 | 0.24317 | 0.175446 | -0.058313 | -0.414756 | -0.437591 | -0.61442 | 1.666872 | 0.444396 | 0.263879 | -0.01624 | 0.099667 | -0.50731 | 0.492635 | 0.526304 | 0.484026 |
| 32.15278 | 0.610656 | 0.175957 | 0.162438 | 0.158423 | -0.157426 | 0.177189 | 1.136721 | -0.078923 | -0.082989 | 0.168734 | 0.161477 | -0.42666 | -0.133595 | 0.847927 | -1.258025 | 1.0707 | 0.718223 | 0.773224 | 0.206973 | 0.277217 | -0.095808 | 1.039284 | 1.107674 | 1.609811 |
| 32.16667 | 0.472017 | 0.042096 | 0.070436 | 0.06493 | -0.78531 | 0.088253 | 0.826786 | -0.025146 | -0.317376 | 0.080767 | 0.130632 | -0.212828 | 0.018073 | 2.084006 | -2.093312 | 0.01453 | 0.640526 | 1.490843 | 0.676771 | 0.031957 | 0.243369 | 1.38808 | 1.731353 | 2.463174 |
| 32.18056 | 0.335857 | -0.021645 | 0.010507 | 0.00249 | -0.764785 | 0.017265 | 0.632382 | -0.054087 | -0.591966 | 0.02022 | 0.017361 | 0.775205 | -0.082316 | 2.932014 | -2.788211 | -0.750684 | 0.313676 | 1.867569 | 0.714603 | -0.431885 | 0.702308 | 1.175275 | 2.074086 | 2.434773 |
| 32.19444 | 0.081301 | -0.001645 | 0.012048 | -0.003955 | -0.457307 | 0.006023 | 0.907899 | -0.226297 | -0.691164 | 0.012056 | -0.016452 | 1.916447 | -0.30104 | 3.165009 | -2.351974 | -0.695817 | 0.112548 | 1.650603 | 0.372472 | -0.63565 | 0.432246 | 0.799153 | 1.53018 | 1.883312 |
| 32.20833 | -0.17931 | -0.013049 | 0.014609 | -0.005841 | -0.267625 | 0.013836 | 1.11293 | -0.445857 | -0.606221 | 0.007006 | 0.006907 | 2.120158 | -0.447324 | 2.698258 | -1.300337 | 0.131471 | 0.34956 | 0.866815 | -0.300812 | -0.314074 | 0.18619 | 0.567915 | 1.126344 | 1.330035 |
| 32.22222 | -0.427477 | -0.063007 | -0.027087 | -0.052181 | -0.425133 | -0.009561 | 0.919571 | -0.616199 | -0.537986 | -0.029027 | -0.148391 | 1.396145 | -0.409846 | 1.576717 | -0.513997 | 0.625671 | 0.563927 | -0.130985 | -0.507228 | 0.401499 | -0.064459 | 0.562658 | 0.599887 | 1.01043 |
| 32.23611 | -0.691975 | -0.051592 | -0.029248 | -0.057147 | -0.464057 | -0.005912 | 0.651356 | -0.628127 | -0.507365 | -0.022845 | -0.173109 | 1.235041 | -0.170081 | 1.553978 | 0.004579 | 0.894577 | 0.279856 | -0.266524 | -0.274988 | 1.310601 | -0.28759 | 0.73402 | 0.154179 | 1.166466 |
| 32.25 | -0.55752 | 0.044588 | 0.057828 | 0.030606 | -0.423082 | 0.071066 | 0.723115 | -0.473656 | -0.508545 | 0.05574 | 0.005221 | 1.605275 | 0.118691 | 2.744966 | 0.140652 | 0.642274 | 0.072823 | 0.49991 | -0.012469 | 1.890067 | -0.953719 | 0.775645 | 0.722005 | 1.231591 |
| 32.26389 | 0.190497 | 0.15341 | 0.16855 | 0.132907 | -0.480303 | 0.158664 | 1.424085 | -0.313562 | -0.501614 | 0.144089 | 0.154763 | 1.766268 | 0.285204 | 3.6345 | 0.047928 | 0.23861 | 0.088073 | 1.271707 | 0.083309 | 1.173125 | -0.833707 | 0.530859 | 1.68742 | 1.036607 |
| 32.27778 | 1.163227 | 0.159886 | 0.17323 | 0.12968 | -0.375822 | 0.138773 | 2.063511 | -0.264829 | -0.608319 | 0.140584 | 0.155872 | 1.08307 | 0.429483 | 2.601507 | -0.050291 | 0.28421 | 0.150132 | 1.702725 | 0.073671 | 0.326661 | -0.3241 | 0.087226 | 1.935797 | 1.044827 |
| 32.29167 | 2.061924 | 0.00923 | 0.012073 | -0.018678 | 0.093387 | -0.023428 | 1.399879 | -0.351527 | -0.74511 | 0.001361 | 0.037646 | 0.521174 | 0.610976 | -0.06742 | -0.09223 | 0.635625 | 0.199931 | 1.705075 | 0.006096 | -0.087611 | 0.393132 | -0.177527 | 1.170339 | 1.693692 |
| 32.30556 | 2.418859 | -0.197298 | -0.179593 | -0.187429 | 0.742945 | -0.179797 | 0.661158 | -0.468961 | -0.813088 | -0.16171 | -0.13862 | 0.392246 | 0.767538 | -1.866083 | -0.057402 | 0.740433 | 0.191818 | 1.033478 | 0.05269 | 0.22255 | 1.062712 | -0.204483 | 0.533269 | 2.57913 |
| 32.31944 | 1.974438 | -0.323937 | -0.272126 | -0.285881 | 1.068454 | -0.242553 | 0.152799 | -0.594063 | -0.803716 | -0.256959 | -0.206696 | 0.344242 | 0.898812 | -1.689913 | 0.274544 | 0.603277 | 0.212538 | 0.267585 | 0.091279 | 1.033446 | 0.82174 | 0.032342 | -0.47195 | 2.07039 |
| 32.33333 | 1.1668 | -0.319004 | -0.273876 | -0.328699 | 0.946344 | -0.269119 | 0.057086 | -0.656025 | -0.735388 | -0.297283 | -0.220041 | 0.073726 | 1.022408 | -0.210547 | 0.920479 | 0.46679 | 0.293245 | -0.360293 | -0.017171 | 2.242849 | 0.369049 | 0.593828 | -0.361653 | 0.950878 |
| 32.34722 | 0.407123 | -0.192324 | -0.180813 | -0.24751 | 0.811048 | -0.204622 | 0.454363 | -0.581978 | -0.515198 | -0.223662 | -0.127358 | -0.084546 | 1.101698 | 2.070411 | 1.770067 | 0.480062 | 0.541431 | -0.187754 | -0.042973 | 2.220322 | 0.195768 | 1.132015 | 1.614429 | 1.523627 |
| 32.36111 | -0.038978 | -0.014959 | -0.00128 | -0.038869 | 0.985847 | -0.038705 | 1.014832 | -0.481998 | -0.155223 | -0.030603 | 0.047142 | -0.12261 | 1.236694 | 4.202798 | 2.000563 | 0.568341 | 0.896603 | 0.440636 | 0.036728 | 0.92284 | 0.057002 | 1.290015 | 2.810315 | 1.417898 |
| 32.375 | 0.15303 | 0.111979 | 0.150687 | 0.120921 | 1.49172 | 0.104823 | 1.2386 | -0.366489 | 0.106197 | 0.12035 | 0.198656 | -0.317098 | 1.354019 | 4.625048 | 1.927972 | 0.685393 | 1.318094 | 1.395222 | 0.081826 | -0.044387 | -0.024937 | 1.15663 | 2.223632 | 1.196007 |
| 32.38889 | 0.912327 | 0.179421 | 0.198677 | 0.167311 | 2.16123 | 0.166114 | 1.432828 | -0.219294 | 0.226608 | 0.166308 | 0.227887 | -0.015489 | 1.174277 | 3.991219 | 1.811472 | 0.705108 | 1.655166 | 2.2002 | 0.073516 | -0.210847 | 0.081609 | 1.114099 | 1.302814 | 2.454548 |
| 32.40278 | 1.768786 | 0.224018 | 0.224121 | 0.200223 | 2.634143 | 0.212362 | 1.828487 | -0.034757 | 0.371517 | 0.19954 | 0.232126 | 0.869758 | 0.874214 | 3.047956 | 1.588951 | 0.607205 | 1.666609 | 1.684496 | 0.115911 | 0.461734 | 0.299404 | 1.11115 | 0.356074 | 1.968378 |
| 32.41667 | 2.195647 | 0.284543 | 0.294649 | 0.246955 | 2.727124 | 0.270436 | 1.848912 | 0.110488 | 0.559316 | 0.253538 | 0.282121 | 2.051054 | 0.658482 | 4.098372 | 1.552351 | 0.635212 | 1.514441 | 0.877969 | 0.30909 | 1.495434 | 0.468653 | 1.022301 | -0.454991 | 0.959422 |
| 32.43056 | 2.320319 | 0.33095 | 0.336953 | 0.295904 | 2.272298 | 0.318744 | 1.824298 | 0.226767 | 0.757923 | 0.314412 | 0.392169 | 3.105419 | 0.463037 | 4.347792 | 1.470463 | 0.993948 | 1.358689 | 0.196327 | 0.355469 | 2.515732 | 0.530026 | 0.761915 | 0.332142 | 2.114727 |
| 32.44444 | 2.381241 | 0.268791 | 0.234404 | 0.262886 | 1.764914 | 0.294151 | 1.62659 | 0.278308 | 0.894002 | 0.293598 | 0.380904 | 2.808033 | 0.261061 | 3.454811 | 1.513332 | 1.528679 | 1.293831 | -0.593518 | 0.056513 | 1.778265 | 0.354849 | 0.593928 | 0.801301 | 2.897347 |
| 32.45833 | 2.580393 | 0.140546 | 0.077808 | -0.000675 | 1.397516 | 0.200544 | 1.201902 | 0.1755 | 0.844624 | 0.162789 | 0.237846 | 2.595269 | 0.18955 | 3.256664 | 1.843499 | 1.561463 | 1.269225 | -0.219089 | -0.486999 | -0.364859 | 0.424732 | 0.403349 | 1.198188 | 1.843646 |
| 32.47222 | 2.663388 | 0.227299 | 0.16775 | -0.068944 | 1.558311 | 0.270437 | 0.744586 | 0.040341 | 0.764649 | 0.162254 | 0.202499 | 1.553993 | 0.654254 | 4.003759 | 2.387475 | 0.981114 | 1.68491 | 0.44858 | -0.632694 | -2.248954 | 0.833904 | 0.337254 | 1.044539 | 1.813998 |
| 32.48611 | 2.818037 | 0.336705 | 0.276209 | 0.166397 | 2.764803 | 0.346705 | 0.701961 | 0.069797 | 0.821682 | 0.264091 | 0.251484 | 3.747086 | 1.588262 | 2.322888 | 2.916639 | 0.826013 | 2.176671 | 1.643053 | -0.20275 | -1.969268 | 1.081979 | 0.632468 | 1.433013 | 2.992364 |
| 32.5 | 2.7961 | 0.243145 | 0.194837 | 0.094587 | 3.23598 | 0.231166 | 0.55826 | -0.01168 | 0.886732 | 0.171524 | 0.177425 | 2.998519 | 1.305907 | 2.161172 | 3.181902 | 0.861693 | 2.260717 | 1.773025 | 0.08506 | -0.511855 | 1.189212 | 0.881042 | 1.723147 | 2.776672 |
| 32.51389 | 2.05293 | -0.107246 | -0.100568 | -0.177446 | 2.4797 | -0.073788 | 0.203206 | -0.308367 | 0.732679 | -0.117751 | -0.074536 | 0.395565 | 0.811392 | 2.214519 | 2.763205 | 0.849472 | 1.414294 | 0.702887 | 0.202467 | 0.684499 | 0.983779 | 0.699941 | 1.263138 | 1.218114 |
| 32.52778 | 1.318678 | -0.677456 | -0.545096 | -0.589627 | 1.470549 | -0.619099 | -0.353318 | -0.735824 | 0.612406 | -0.615552 | -0.499931 | -1.269486 | 0.772544 | 1.161674 | 1.883687 | 0.874498 | 0.606927 | 0.160503 | 0.375155 | 1.066958 | 0.728645 | 0.4069 | 0.948311 | 2.306192 |
| 32.54167 | 0.888136 | -1.333879 | -1.073509 | -1.157466 | 0.363388 | -1.324508 | -1.031358 | -1.39843 | 0.284796 | -1.274965 | -1.070824 | 1.148974 | 1.117119 | 0.319906 | 1.620744 | 1.060686 | 0.182436 | 0.453357 | 0.373197 | 0.355242 | 0.50855 | 0.18808 | 1.566649 | 1.028518 |
| 32.55556 | 0.538093 | -1.483495 | -1.272879 | -1.345384 | -0.373939 | -1.434989 | -1.204811 | -1.544239 | -0.148248 | -1.407471 | -1.268299 | 2.411875 | 0.812087 | -0.049252 | 0.995604 | 0.724654 | -0.175652 | 1.581895 | 0.255723 | -0.499926 | 0.582507 | 0.187083 | 2.2273 | 0.45333 |
| 32.56944 | 0.529564 | -1.098257 | -1.0755 | -1.042042 | -0.461686 | -1.021568 | -0.768023 | -1.170994 | -0.298213 | -1.02231 | -0.97334 | 1.770259 | 0.25595 | -0.02324 | 1.253542 | 0.524109 | -0.054687 | 2.663834 | 0.101331 | -1.369081 | 0.792707 | 0.241547 | 2.755343 | 0.918404 |
| 32.58333 | 0.692227 | -0.818476 | -0.873478 | -0.802204 | -0.453472 | -0.75211 | -0.430449 | -0.900796 | -0.221607 | -0.771612 | -0.704682 | 1.646521 | 0.340622 | 0.075908 | 1.538827 | 0.684568 | 0.399812 | 1.07966 | 0.05321 | -1.179745 | 0.623523 | 0.143015 | 2.067396 | 1.599229 |
| 32.59722 | 0.220038 | -0.712052 | -0.768107 | -0.689373 | -0.436188 | -0.643971 | -0.360239 | -0.797814 | -0.052177 | -0.666585 | -0.575048 | 1.022801 | 0.357173 | 0.041241 | 1.15177 | 0.726631 | 1.287808 | 0.053177 | 0.025071 | -1.095581 | 0.252413 | 0.052514 | 0.67906 | 1.282804 |
| 32.61111 | 0.155698 | -0.556075 | -0.591124 | -0.526171 | 0.428426 | -0.486137 | -0.294071 | -0.649232 | 0.116272 | -0.507359 | -0.4415 | 0.266916 | 0.408561 | 0.145016 | 1.009146 | 0.263732 | 1.450342 | -1.370333 | 0.133004 | -0.669001 | -0.028712 | 0.07881 | 0.782031 | 1.009721 |
| 32.625 | 0.012469 | -0.383662 | -0.382428 | -0.349767 | 1.495936 | -0.332999 | -0.22108 | -0.505947 | 0.179364 | -0.352605 | -0.318943 | -0.086677 | 0.415528 | -0.979902 | -0.050297 | 0.127601 | -0.47556 | -0.991799 | -0.480219 | -0.411027 | -0.021447 | 0.120599 | 1.625097 | 0.009302 |
| 32.63889 | 0.636374 | -0.275456 | -0.258732 | -0.237607 | 1.803765 | -0.239812 | -0.261523 | -0.470964 | 0.20496 | -0.254658 | -0.256692 | -0.970515 | 0.220612 | -1.300038 | -0.367832 | -0.159447 | -0.981737 | 0.204192 | -0.033025 | -0.69009 | -0.127754 | -0.015396 | 0.847894 | 0.010296 |
| 32.65278 | 0.897172 | -0.160281 | -0.13921 | -0.113342 | 0.028346 | -0.119911 | -0.040074 | -0.45848 | 0.291965 | -0.130638 | -0.152511 | -2.172632 | -0.845288 | -0.358771 | -0.229344 | -0.365072 | -0.25258 | 0.149503 | 0.903053 | -0.263827 | -0.078639 | -0.047176 | 1.061453 | 0.324493 |
| 32.66667 | 0.403991 | -0.098418 | -0.064398 | -0.053048 | -1.387634 | -0.065356 | 0.170577 | -0.523318 | 0.330243 | -0.075218 | -0.055665 | -4.166704 | -0.19756 | -0.054428 | -0.878094 | -0.356498 | 0.109313 | -0.664632 | 0.393175 | 0.211351 | -0.323573 | 0.045383 | 0.394771 | 0.953173 |
| 32.68056 | -1.318458 | -0.167476 | -0.136317 | -0.140554 | -2.883426 | -0.161137 | 0.032234 | -0.710872 | 0.251463 | -0.170351 | -0.075897 | -4.924375 | 0.174727 | 0.114391 | -0.864026 | -0.209523 | 0.164551 | -0.747809 | -0.23883 | 0.535304 | -0.26148 | 0.083595 | -0.343765 | 0.016372 |
| 32.69444 | -1.686228 | -0.235919 | -0.215057 | -0.212966 | -2.637176 | -0.237043 | -0.089837 | -0.917799 | 0.242515 | -0.240755 | -0.088775 | -2.845378 | -0.068482 | -0.05766 | -0.069762 | 0.048609 | 0.564719 | -0.278562 | -0.179873 | 0.716774 | -0.266921 | 0.076313 | -0.120349 | 0.58161 |
| 32.70833 | -0.995772 | -0.23078 | -0.207501 | -0.193992 | -0.357317 | -0.207156 | -0.22718 | -1.153496 | 0.303676 | -0.203266 | -0.042351 | -0.930164 | 0.172173 | -0.179397 | 0.692402 | 0.283324 | 0.47363 | -0.028926 | -0.035543 | 0.30973 | -0.401914 | 0.19476 | -0.327483 | 0.381119 |
| 32.72222 | -0.305534 | -0.168913 | -0.125716 | -0.119194 | 1.323491 | -0.129227 | -0.296492 | -1.059882 | 0.367058 | -0.115554 | 0.011206 | 0.998665 | 0.184033 | 0.303045 | 1.444385 | 0.393614 | 1.107827 | -0.577311 | 0.349095 | -0.150098 | 0.012278 | 0.027163 | 0.588302 | 0.932277 |
| 32.73611 | -0.629341 | -0.076379 | -0.010817 | -0.026328 | 2.630643 | -0.045709 | -0.208068 | -0.710765 | 0.433962 | -0.027152 | 0.041356 | 2.736441 | 0.144499 | 1.354415 | 1.567598 | 0.439544 | 1.111513 | -1.004496 | 0.804227 | -0.220989 | 0.662533 | -0.254267 | 0.655325 | -0.70456 |
| 32.75 | -1.271621 | -0.029712 | 0.025439 | -0.003494 | 2.461094 | -0.007744 | -0.07368 | -0.430463 | 0.41926 | 0.002439 | -0.00569 | 3.289464 | 0.381636 | 1.418215 | 1.448252 | 0.334259 | 1.728897 | 0.541356 | 0.417626 | -0.120065 | 0.57272 | -0.297633 | 0.506659 | 0.215164 |
| 32.76389 | -1.378958 | 0.005667 | 0.026828 | 0.006439 | 0.705333 | 0.035692 | 0.00671 | -0.230144 | 0.384063 | 0.030143 | -0.033515 | 2.582257 | -0.052274 | 0.535453 | 0.310276 | -0.006824 | 0.691742 | 0.497064 | -0.071123 | -0.048998 | 0.2848 | -0.169645 | 0.883516 | -2.332078 |
| 32.77778 | -1.071245 | 0.076018 | 0.088154 | 0.072842 | -0.651937 | 0.117131 | 0.116101 | -0.044053 | 0.413121 | 0.105264 | 0.028014 | 1.478835 | -0.378045 | 0.174351 | -0.748575 | -0.288755 | -1.210781 | 0.39313 | -0.202859 | 0.123109 | 0.154067 | -0.078518 | -0.188476 | -0.937557 |
| 32.79167 | -1.299063 | 0.115036 | 0.134495 | 0.117083 | -2.120086 | 0.151706 | 0.142325 | 0.046927 | 0.42302 | 0.147179 | 0.093797 | 0.390708 | -0.36649 | 0.189685 | -1.176921 | -0.557327 | -0.282376 | 0.673197 | -0.289808 | -0.93362 | 0.066964 | 0.016652 | -0.094395 | -1.617154 |
| 32.80556 | -2.017857 | 0.100302 | 0.111195 | 0.094448 | -3.943418 | 0.119598 | 0.010882 | 0.032919 | 0.347481 | 0.115712 | 0.115919 | -0.521868 | -0.178199 | 0.287579 | -1.314362 | -0.782845 | 0.348001 | 0.761047 | -0.111102 | -1.202673 | -0.03793 | 0.083492 | 1.007247 | 0.006815 |
| 32.81944 | -2.194733 | 0.069683 | 0.0634 | 0.053322 | -4.746076 | 0.072285 | -0.274013 | -0.003712 | 0.225059 | 0.066842 | 0.122036 | -1.105906 | -0.528219 | 0.062539 | -1.150327 | -0.535301 | 0.025461 | 1.044377 | -0.234264 | -0.354583 | -0.219787 | 0.097981 | 0.403135 | 1.008201 |
| 32.83333 | -2.080674 | 0.044785 | 0.029444 | 0.023717 | -1.750191 | 0.024857 | -0.616809 | -0.024894 | 0.11833 | 0.033066 | 0.124145 | -1.568773 | -0.981223 | -0.132095 | -0.217236 | -0.181193 | -0.032984 | 0.397592 | -0.255078 | 0.327167 | -0.636061 | -0.030913 | 0.290246 | 1.394345 |
| 32.84722 | -2.981154 | 0.044993 | 0.036273 | 0.01478 | -0.253466 | -0.017395 | -0.926304 | -0.027329 | 0.070209 | 0.018441 | 0.141058 | -1.667703 | -0.365895 | -0.516167 | 0.8709 | 0.260393 | -0.045062 | 0.305741 | -0.062691 | 0.558808 | -1.146213 | -0.233231 | 0.621926 | 1.645537 |
| 32.86111 | -2.008857 | 0.059811 | 0.070497 | 0.029687 | 0.878575 | -0.024332 | -1.398306 | -0.010149 | 0.034918 | 0.022154 | 0.139344 | -1.86154 | 0.832755 | -0.394516 | 0.312504 | -0.649888 | -0.050247 | -0.48713 | 0.361487 | 0.392654 | -1.365801 | -0.404823 | 0.620321 | -0.626041 |
| 32.875 | 0.285111 | 0.059056 | 0.074178 | 0.044937 | 0.425385 | 0.005423 | -1.964036 | -0.002966 | -0.046819 | 0.031966 | 0.049232 | -1.94158 | 0.902833 | -0.260779 | -0.13979 | -1.268018 | -0.789806 | -0.502643 | 0.525649 | -0.042702 | -1.008808 | -0.425574 | -0.126271 | -0.447219 |
| 32.88889 | 1.273432 | 0.074563 | 0.086038 | 0.078751 | -1.676325 | 0.07667 | -2.005818 | 0.037215 | -0.154576 | 0.075761 | -0.005557 | -1.860959 | -0.52524 | -0.028926 | -0.748835 | 0.197714 | -0.639347 | -0.645977 | 0.292814 | 0.325672 | -0.689208 | -0.396983 | 0.179528 | -0.056879 |
| 32.90278 | 0.285702 | 0.116621 | 0.136893 | 0.13531 | -3.375322 | 0.151933 | -1.833957 | 0.106615 | -0.263784 | 0.139148 | -0.13637 | -1.6752 | -1.454951 | 0.299781 | -0.752635 | 0.531216 | -0.402516 | -0.585982 | -0.057286 | -0.445859 | -0.813944 | -0.35248 | 0.651795 | -0.054522 |
| 32.91667 | -1.395187 | 0.117963 | 0.134539 | 0.137518 | -3.199634 | 0.146546 | -1.814285 | 0.115064 | -0.461296 | 0.137835 | 0.012143 | -1.618576 | -2.046504 | -1.174745 | -0.77435 | 0.599149 | -0.376509 | -0.495299 | -0.266854 | -1.189929 | -1.028608 | -0.468678 | 0.577073 | -0.182925 |
| 32.93056 | -1.545065 | 0.069239 | 0.066484 | 0.07255 | -1.975075 | 0.082025 | -2.145215 | 0.060337 | -0.575379 | 0.068393 | 0.109165 | -1.712843 | -2.341673 | -2.680536 | -0.723686 | -0.607495 | -0.496558 | 0.240204 | -0.204441 | -1.079198 | -1.154843 | -0.653753 | 0.286568 | -1.065509 |
| 32.94444 | -2.152363 | 0.019822 | 0.008872 | 0.003444 | -0.627732 | 0.035336 | -2.057921 | 0.015324 | -0.693595 | 0.005961 | 0.039382 | -1.953806 | -2.15102 | -3.029164 | -0.881591 | -1.238698 | -0.785674 | 0.67926 | -0.338714 | -0.262488 | -1.222184 | -0.827599 | 0.697963 | 0.148421 |
| 32.95833 | -2.842235 | -0.007548 | -0.017856 | -0.034979 | 0.332939 | 0.011267 | -1.453081 | -0.013482 | -0.830925 | -0.016623 | -0.078923 | -2.212921 | -1.346457 | -2.622952 | -1.162286 | -0.687151 | -0.959554 | 0.666828 | -0.69346 | -0.094723 | -0.980468 | -0.855952 | 0.452487 | 1.309962 |
| 32.97222 | -2.708972 | -0.004545 | -0.021865 | -0.03645 | 0.627659 | -0.002718 | -0.914365 | -0.037164 | -1.090911 | -0.01233 | -0.107925 | -2.309313 | -1.117302 | -4.568149 | -1.682968 | 0.512197 | -1.085911 | 0.791416 | -0.764448 | -0.892509 | -0.514293 | -0.750273 | -0.860975 | 0.079717 |
| 32.98611 | -2.284066 | 0.021727 | 0.003692 | 0.00509 | 0.168853 | 0.009583 | -0.541622 | -0.038415 | -1.005962 | 0.009097 | -0.042575 | -2.129039 | -0.660594 | -3.158779 | -2.764677 | 1.874966 | -1.254886 | 0.645858 | -0.978539 | -1.282181 | -0.28298 | -0.468778 | -2.284557 | -2.544773 |
| 33 | -1.527252 | 0.12198 | 0.119093 | 0.159362 | -0.578725 | 0.080663 | -0.222255 | 0.047991 | -0.683652 | 0.112908 | 0.06291 | -1.545774 | 1.00232 | -2.354768 | -0.857755 | 2.094707 | -0.77267 | 0.991971 | 1.093808 | -1.299547 | -0.018659 | -0.154219 | -2.299218 | -4.038075 |
| 33.01389 | -0.730039 | 0.362682 | 0.373283 | 0.496559 | -1.064847 | 0.245176 | 0.191629 | 0.260915 | -0.222831 | 0.351626 | 0.246646 | -0.778678 | 2.918229 | -1.832081 | 1.596285 | 1.043459 | -0.501598 | -0.11181 | 1.387598 | -0.487453 | 0.213856 | 0.049736 | -2.599905 | -3.886203 |
| 33.02778 | -0.215859 | 0.713746 | 0.719661 | 0.900106 | -1.39537 | 0.546108 | 0.462918 | 0.495624 | 0.044253 | 0.636167 | 0.458729 | -0.343602 | 2.137206 | -0.748321 | 2.275397 | -0.180212 | -0.392654 | 0.18292 | 0.953499 | 0.428553 | 0.384097 | 0.146327 | 0.858657 | -4.36731 |
| 33.04167 | 0.054865 | 0.98603 | 0.88081 | 0.874051 | -1.803005 | 0.75893 | 0.385307 | 0.554588 | 0.321405 | 0.779388 | 0.612778 | -0.19723 | -0.074199 | -1.337135 | 0.632248 | -0.774065 | -1.035841 | 0.572849 | 0.137341 | 0.292092 | 0.534958 | -0.210052 | -0.000742 | -4.575472 |
| 33.05556 | -0.111846 | 0.931966 | 0.733425 | 0.693963 | -2.485959 | 0.779815 | -0.152883 | 0.395579 | 0.3924 | 0.75112 | 0.468446 | -0.014822 | -1.4974 | -3.431742 | -1.164229 | -1.342477 | -1.949998 | -0.297956 | -0.454477 | -0.613262 | 0.088901 | -1.090758 | -1.824962 | -6.173033 |
| 33.06944 | -0.979738 | 0.679254 | 0.564461 | 0.572224 | -2.772922 | 0.672042 | -1.200651 | 0.273534 | 0.190972 | 0.638293 | 0.144771 | 0.400721 | 0.058458 | -6.032602 | -0.638089 | -1.992281 | -3.271592 | -0.883461 | -0.137797 | -0.823449 | -0.044837 | -2.41185 | -2.735101 | -5.663492 |
| 33.08333 | -2.025583 | 0.534231 | 0.520354 | 0.504006 | -1.378489 | 0.580153 | -0.54091 | 0.328397 | -0.17937 | 0.541717 | 0.180631 | 1.108377 | -0.250167 | -6.82948 | -0.336055 | -2.086277 | -3.569346 | -1.62084 | -0.189363 | -0.553691 | 0.144109 | -3.130805 | -3.190575 | -4.082463 |
| 33.09722 | -1.394595 | 0.442318 | 0.456922 | 0.419921 | 0.539888 | 0.474169 | 0.016974 | 0.370831 | -0.595595 | 0.439053 | 0.209286 | 0.675237 | -0.657109 | -6.281681 | -1.748827 | -0.733488 | -2.603253 | -1.237025 | -0.651167 | -0.487229 | 0.304439 | -1.568099 | -2.243584 | -2.204648 |
| 33.11111 | -0.217413 | 0.288305 | 0.293738 | 0.270089 | 1.840301 | 0.316268 | 0.399327 | 0.253124 | -0.950073 | 0.296884 | 0.096075 | -0.636847 | -0.952233 | -4.637952 | -2.061445 | 0.978934 | -1.225639 | -0.83401 | -0.686329 | -0.379061 | -0.00014 | -0.046132 | -0.664817 | -0.669009 |
| 33.125 | 0.815041 | 0.126725 | 0.122596 | 0.120399 | 2.183991 | 0.156718 | 0.702939 | 0.093398 | -1.026699 | 0.155896 | 0.010344 | -0.144416 | -1.086244 | -3.022651 | -1.76123 | 1.745565 | -0.45823 | -0.174594 | -0.410843 | -0.003963 | -0.373103 | 0.307267 | -0.016815 | 0.11768 |
| 33.13889 | 1.615044 | 0.009095 | 0.015236 | 0.025056 | 1.687178 | 0.047989 | 0.770227 | -0.026913 | -0.834278 | 0.054264 | -0.026186 | -0.111813 | -1.092783 | -1.609641 | -1.414455 | 2.033521 | 0.028407 | 0.355968 | 0.271473 | 0.409276 | -0.330288 | 0.356043 | 0.427885 | 0.908956 |
| 33.15278 | 2.070213 | 0.011764 | 0.028463 | 0.029837 | 0.913954 | 0.050049 | 0.573006 | -0.050968 | -0.545114 | 0.054385 | 0.009583 | -0.664519 | -1.06939 | -0.321153 | -1.727166 | 1.037728 | 0.624203 | 1.000036 | 0.714621 | 0.666018 | 0.04541 | 0.622279 | 0.947992 | 1.965689 |
| 33.16667 | 2.310924 | 0.140953 | 0.155931 | 0.135756 | 0.077499 | 0.167347 | 0.369372 | 0.043302 | -0.339659 | 0.155873 | 0.119547 | -0.19127 | -0.922539 | 0.71939 | -1.948967 | 0.206173 | 1.390668 | 1.740077 | 1.1642 | 1.064724 | 0.329664 | 1.36973 | 1.643562 | 2.979831 |
| 33.18056 | 2.059117 | 0.273285 | 0.297751 | 0.268686 | -0.548738 | 0.297375 | 0.428803 | 0.191515 | -0.387152 | 0.27476 | 0.233226 | -0.208887 | -0.623052 | 1.400856 | -1.403783 | 0.216393 | 1.523499 | 2.137167 | 1.084186 | 1.638152 | 0.425589 | 1.220541 | 2.070166 | 2.980021 |
| 33.19444 | 1.494253 | 0.381362 | 0.422796 | 0.400266 | -0.247003 | 0.400617 | 0.779439 | 0.334106 | -0.470267 | 0.397327 | 0.337533 | -0.443952 | -0.245025 | 1.788656 | -1.042227 | 0.807941 | 0.723205 | 1.833367 | 0.668983 | 1.201139 | 0.326619 | 0.868171 | 1.928233 | 1.683383 |
| 33.20833 | 1.085614 | 0.496306 | 0.538645 | 0.524925 | 0.589071 | 0.499446 | 1.366744 | 0.448431 | -0.399172 | 0.520542 | 0.456225 | 1.684049 | 0.030657 | 2.357535 | -0.63054 | 1.533769 | 0.267776 | 1.640964 | -0.279684 | 0.166462 | 0.127451 | 0.660475 | 1.585467 | 0.657038 |
| 33.22222 | 0.692991 | 0.511539 | 0.547372 | 0.530212 | 1.157785 | 0.50773 | 1.860271 | 0.466157 | -0.216186 | 0.537423 | 0.47302 | 3.373902 | 0.088507 | 3.596905 | -0.359893 | 2.319738 | 0.370686 | 0.888036 | -0.883495 | -0.989696 | 0.032495 | 0.787433 | 0.84817 | -0.139076 |
| 33.23611 | 0.337282 | 0.373369 | 0.387581 | 0.356958 | 0.819005 | 0.36577 | 1.838319 | 0.337835 | -0.217892 | 0.378269 | 0.301962 | 3.198509 | 0.005676 | 3.271938 | -0.055587 | 1.617285 | 0.671703 | -0.140465 | -0.566331 | -1.051687 | 0.058804 | 0.777454 | 0.321568 | -0.020871 |
| 33.25 | 0.120131 | 0.164354 | 0.139397 | 0.103331 | 0.051321 | 0.146166 | 1.100191 | 0.124922 | -0.4394 | 0.139483 | 0.073962 | 2.255775 | -0.120381 | 1.709931 | 0.172071 | 0.365813 | 0.83109 | -0.316521 | -0.046635 | -0.531112 | 0.072693 | 0.571456 | 0.822114 | 1.0394 |
| 33.26389 | 0.105183 | -0.058094 | -0.123083 | -0.154079 | -0.789596 | -0.074354 | 0.329617 | -0.096399 | -0.641627 | -0.092256 | -0.136854 | 0.917247 | -0.163744 | 0.762649 | 0.451195 | -0.198121 | 0.807292 | 0.243299 | 0.164279 | 0.310396 | -0.363468 | 0.501933 | 1.869805 | 2.068925 |
| 33.27778 | 0.292061 | -0.194493 | -0.286397 | -0.324492 | -1.149323 | -0.203975 | 0.153189 | -0.219747 | -0.754338 | -0.235663 | -0.225415 | -0.117178 | 0.18016 | 0.686766 | 0.590644 | 0.257208 | 0.405662 | 0.286175 | 0.739512 | 1.210222 | -0.664353 | 0.157542 | 3.005778 | 2.214629 |
| 33.29167 | 0.684496 | -0.114163 | -0.189714 | -0.260293 | -0.420312 | -0.133894 | 0.803391 | -0.14068 | -0.625781 | -0.17421 | -0.111824 | -0.635811 | 0.059329 | 1.222175 | 0.280704 | 1.302743 | -0.187736 | 0.521406 | 0.904484 | 1.803377 | -0.590656 | -0.102344 | 2.437259 | 1.797384 |
| 33.30556 | 0.968584 | -0.019049 | -0.039747 | -0.113594 | 1.173249 | -0.024531 | 1.742491 | -0.043002 | -0.384397 | -0.03962 | 0.013446 | -0.73532 | 0.143751 | 2.480913 | -0.054355 | 2.913724 | -0.369418 | 0.791888 | 1.216186 | 1.610188 | -0.40418 | -0.046867 | 1.051046 | 2.05298 |
| 33.31944 | 0.914872 | -0.147241 | -0.103449 | -0.119157 | 2.315148 | -0.091581 | 2.212069 | -0.13176 | -0.266357 | -0.062817 | -0.01013 | -0.379624 | 0.418073 | 1.239364 | -0.376153 | 3.902336 | -0.480018 | 0.941299 | 1.105049 | 1.027032 | 0.042791 | 0.262198 | 0.158821 | 2.374735 |
| 33.33333 | 0.646491 | -0.270684 | -0.230008 | -0.223979 | 2.327063 | -0.215606 | 1.957405 | -0.256724 | -0.298871 | -0.181052 | -0.082649 | 0.220998 | 0.850018 | -1.115215 | -0.054725 | 3.408521 | -0.451358 | 0.931425 | 0.701868 | 0.405435 | 0.045612 | 0.481177 | -0.274268 | 1.729974 |
| 33.34722 | 0.100544 | -0.248043 | -0.276723 | -0.300724 | 1.481019 | -0.263201 | 1.105294 | -0.270375 | -0.36175 | -0.253442 | -0.182201 | 0.452602 | 1.477115 | -2.040334 | 0.317028 | 4.279284 | 0.153498 | 1.241887 | 0.223847 | -0.017407 | -0.09637 | 0.362318 | 0.741679 | 1.848196 |
| 33.36111 | -0.680689 | -0.209586 | -0.258019 | -0.296029 | 0.858097 | -0.24504 | 0.417981 | -0.244115 | -0.359733 | -0.244814 | -0.242904 | 0.311135 | 1.029109 | -0.53611 | 1.597547 | 4.148108 | 1.022332 | 1.008322 | 0.094829 | 0.130122 | -0.20378 | 0.231095 | 2.067686 | 0.867622 |
| 33.375 | -1.284761 | -0.192764 | -0.197662 | -0.226397 | 0.698694 | -0.197196 | 0.024352 | -0.209878 | -0.300867 | -0.202756 | -0.228585 | 0.160511 | 0.551654 | 2.342811 | 3.068759 | 1.628481 | 1.743484 | 1.082988 | 0.12046 | 0.704329 | 0.123312 | 0.262034 | 2.53777 | 1.266696 |
| 33.38889 | -1.044868 | -0.20132 | -0.170986 | -0.175393 | 1.127311 | -0.17482 | -0.273975 | -0.191649 | -0.242121 | -0.179359 | -0.182485 | 0.139019 | 0.287839 | 5.038241 | 3.158678 | 0.48026 | 2.198887 | 1.207935 | 0.159129 | 1.585566 | 0.317423 | 0.479702 | 1.769457 | 1.966018 |
| 33.40278 | 0.636617 | -0.233107 | -0.19 | -0.176632 | 1.881207 | -0.177034 | -0.327733 | -0.192729 | -0.191301 | -0.170036 | -0.187808 | 0.224913 | 0.615672 | 5.252207 | 2.193909 | 0.127997 | 2.025152 | 1.218958 | 0.321944 | 2.025634 | 0.339845 | 0.823457 | 1.290363 | 2.011462 |
| 33.41667 | 0.792015 | -0.293853 | -0.263871 | -0.266665 | 2.482264 | -0.237269 | -0.001882 | -0.241811 | -0.204694 | -0.229125 | -0.247761 | 0.416281 | 1.351633 | 5.374594 | 1.274857 | 0.709151 | 1.597935 | 1.125325 | 0.516708 | 1.750953 | 0.384018 | 1.088818 | 1.015741 | 0.927562 |
| 33.43056 | 1.662881 | -0.451034 | -0.480027 | -0.492304 | 3.166614 | -0.428322 | 0.946003 | -0.430476 | -0.360489 | -0.439205 | -0.439675 | 0.875805 | 1.155685 | 6.173517 | 1.0639 | 1.46404 | 2.328349 | 1.124301 | 0.479603 | 1.175858 | 0.532594 | 1.083111 | 0.580422 | 0.762352 |
| 33.44444 | 2.410396 | -0.651855 | -0.752749 | -0.700282 | 3.742145 | -0.624834 | 0.843787 | -0.652446 | -0.471557 | -0.661011 | -0.52765 | 2.01225 | 0.172464 | 7.030899 | 1.455484 | 2.18922 | 2.67577 | 1.037228 | 0.122682 | 0.468087 | 0.530166 | 0.926443 | 0.27007 | 0.817697 |
| 33.45833 | 0.899898 | -0.788051 | -0.876121 | -0.917112 | 3.526933 | -0.69965 | -0.861863 | -0.808279 | -0.441141 | -0.770754 | -0.505127 | 3.470732 | -0.059779 | 7.463238 | 2.155624 | 2.638646 | 2.813118 | 0.672833 | -0.577029 | -0.517861 | 0.419376 | 0.664744 | 0.365674 | 1.270565 |
| 33.47222 | -0.517003 | -0.584942 | -0.549826 | -0.75996 | 3.258673 | -0.46666 | -1.073016 | -0.568555 | -0.306692 | -0.58421 | -0.337314 | 3.312723 | 0.068093 | 6.338637 | 2.859872 | 2.727167 | 3.332677 | 0.228066 | -0.704612 | -1.137029 | 0.628858 | 0.440485 | 0.986844 | 3.255519 |
| 33.48611 | -0.965033 | -0.138451 | -0.109639 | -0.217088 | 3.088562 | -0.072312 | 0.117944 | -0.137709 | 0.125123 | -0.181661 | -0.137176 | 2.637363 | 0.504519 | 3.528238 | 3.655966 | 3.419878 | 3.368025 | 0.420375 | -0.18724 | -0.900309 | 0.887321 | 0.535565 | 1.414166 | 3.548931 |
| 33.5 | -0.765142 | 0.248439 | 0.259179 | 0.177572 | 3.276103 | 0.228941 | 0.294512 | 0.204171 | 0.703196 | 0.180757 | 0.176567 | 1.913196 | 1.143785 | 2.845135 | 3.764883 | 3.539499 | 2.565954 | 0.998387 | 0.470236 | -0.288578 | 1.152376 | 0.752392 | 1.566314 | 3.042285 |
| 33.51389 | 0.16896 | 0.38923 | 0.424107 | 0.324767 | 3.082998 | 0.343519 | 0.72697 | 0.379649 | 1.614828 | 0.324458 | 0.33486 | 0.788218 | 0.858423 | 2.401381 | 3.626612 | 3.299766 | 1.626747 | 1.126602 | 0.776749 | 0.245817 | 1.227065 | 0.698717 | 1.349747 | 1.947359 |
| 33.52778 | 1.511255 | 0.080548 | 0.228833 | 0.108867 | 2.421092 | 0.064492 | 0.519863 | 0.148233 | 1.914847 | 0.0598 | 0.117142 | -1.135972 | 0.351622 | 1.40635 | 3.293528 | 2.524462 | 0.423076 | 0.981231 | 0.873266 | 0.568888 | 1.203704 | 0.45812 | 1.155451 | 0.198426 |
| 33.54167 | 2.719022 | -0.543792 | -0.191019 | -0.28853 | 1.810655 | -0.496923 | -0.65734 | -0.427392 | 1.164872 | -0.432571 | -0.321534 | -1.931663 | 1.185117 | 0.251434 | 3.034141 | 1.047509 | -0.130214 | 1.166385 | 0.728471 | 0.885171 | 1.337504 | 0.223632 | 1.612592 | 1.787444 |
| 33.55556 | 3.018315 | -0.850341 | -0.515835 | -0.544591 | 1.351475 | -0.783584 | -0.425852 | -0.734831 | 0.675076 | -0.67543 | -0.61408 | -0.213828 | 1.674564 | -0.045976 | 1.251978 | -0.735907 | -0.280869 | 0.959653 | 0.324205 | 0.97503 | 0.732311 | 0.092917 | 1.260795 | 3.106738 |
| 33.56944 | 2.847791 | -0.644294 | -0.526732 | -0.498769 | 0.800135 | -0.582506 | -0.432683 | -0.54438 | 0.327471 | -0.537415 | -0.548543 | 0.693312 | 2.859923 | -0.269555 | -0.578763 | 0.641957 | -0.088575 | 0.608888 | -0.386555 | 0.495996 | 0.695894 | 0.278458 | 0.612538 | 4.97938 |
| 33.58333 | 2.689265 | -0.323721 | -0.306502 | -0.291406 | 0.039377 | -0.287648 | 0.365798 | -0.25588 | 0.549706 | -0.293221 | -0.307622 | 1.314893 | 4.451506 | -0.294046 | -0.622286 | 1.959965 | 0.016235 | 0.416632 | -0.233084 | 0.385423 | 0.458709 | 0.222537 | 1.962446 | -0.367559 |
| 33.59722 | 1.385038 | -0.089009 | -0.067133 | -0.078183 | -1.088549 | -0.077803 | 1.396019 | -0.053948 | 1.192126 | -0.083503 | -0.082526 | -0.159995 | 3.047787 | 0.094961 | 0.851466 | 1.183818 | 0.05662 | 0.655522 | 0.498711 | 0.157461 | 0.234368 | 0.114006 | 4.014601 | -0.125105 |
| 33.61111 | 1.194267 | 0.089835 | 0.109694 | 0.092548 | -0.793808 | 0.083805 | 0.617638 | 0.114152 | 1.895594 | 0.090582 | 0.08997 | -1.200368 | 1.894522 | 0.492404 | -0.466597 | 0.005898 | 0.065516 | 0.698669 | 0.792974 | -0.616682 | 0.020642 | 0.112221 | 2.156518 | -1.230243 |
| 33.625 | 1.351283 | 0.191914 | 0.203306 | 0.20117 | 0.32605 | 0.178569 | -0.361171 | 0.197684 | 1.646926 | 0.183805 | 0.183791 | 0.065885 | 1.48084 | 0.501471 | -0.213218 | -0.581783 | -0.137863 | -0.216446 | 0.438386 | -1.126658 | 0.001429 | 0.221322 | 1.165751 | -2.611556 |
| 33.63889 | 1.354099 | 0.232708 | 0.245375 | 0.251661 | 1.344212 | 0.224563 | -0.926608 | 0.209729 | 0.803867 | 0.226593 | 0.223756 | 0.666762 | 0.255065 | 0.065701 | 0.22424 | 0.004676 | -0.819152 | -0.422504 | 0.199914 | -0.021778 | 0.233655 | 0.193243 | 0.762139 | -0.350719 |
| 33.65278 | 1.108015 | 0.215002 | 0.240387 | 0.224415 | 1.604294 | 0.218456 | -0.289787 | 0.187442 | 0.519671 | 0.21995 | 0.235389 | -0.135475 | -0.933806 | -0.367278 | 1.371631 | -0.17304 | -1.612876 | 0.050393 | -0.371987 | 0.583853 | 0.508032 | -0.075443 | -0.459911 | 0.086821 |
| 33.66667 | 0.082617 | 0.149379 | 0.168346 | 0.140949 | -0.096744 | 0.152288 | 0.015322 | 0.14466 | 0.446939 | 0.155642 | 0.189414 | -0.638634 | -0.027861 | -0.488291 | 0.469034 | -1.833522 | -3.082354 | 0.099651 | -0.4689 | -0.424036 | 0.272839 | -0.143312 | 0.66897 | 0.193058 |
| 33.68056 | -0.423022 | 0.116594 | 0.115779 | 0.108782 | -1.636917 | 0.109335 | 0.125362 | 0.133112 | 0.490647 | 0.110594 | 0.151853 | -1.916085 | 1.193654 | -0.3349 | -0.003434 | -4.124537 | -3.731782 | -0.16113 | -0.721149 | -0.948194 | -0.373707 | 0.022557 | 1.104475 | 0.005713 |
| 33.69444 | -1.40921 | 0.077054 | 0.084781 | 0.086546 | -2.456198 | 0.078049 | 0.106291 | 0.12509 | 0.595568 | 0.063377 | 0.135119 | -1.402627 | 2.58887 | -0.048853 | -0.321971 | -4.77521 | -4.282651 | 0.316414 | -0.770647 | -2.661738 | -1.041898 | 0.09051 | -0.339143 | 0.152123 |
| 33.70833 | -1.527621 | -0.05076 | 0.012518 | 0.00291 | -1.274484 | 0.009981 | -0.067253 | 0.083515 | 0.624931 | -0.009842 | 0.08943 | -0.539783 | 1.078689 | 0.29584 | -0.617146 | -2.780243 | -5.443102 | 0.081818 | -1.000558 | -1.488165 | -0.631977 | 0.067987 | 0.318799 | -1.392134 |
| 33.72222 | -0.881161 | -0.023467 | 0.023517 | 0.009597 | 0.745808 | 0.023702 | 0.066255 | 0.111809 | 0.687305 | 0.021372 | 0.115707 | 0.242011 | 0.719775 | 0.598686 | -0.140202 | -0.27952 | -4.966415 | 0.111778 | -0.962451 | 0.117435 | -0.44923 | 0.007225 | -0.132515 | -1.469259 |
| 33.73611 | 1.390488 | 0.09294 | 0.156362 | 0.139534 | 2.813907 | 0.149918 | 0.132309 | 0.186679 | 0.815507 | 0.159613 | 0.215266 | 0.946712 | 0.062769 | 0.450394 | -0.06592 | 2.402768 | -1.482405 | 0.063148 | -0.738538 | 0.977785 | -0.378956 | -0.040965 | 0.153412 | 1.150293 |
| 33.75 | 2.21937 | 0.155955 | 0.21844 | 0.178913 | 3.511071 | 0.206051 | 0.220063 | 0.199357 | 0.74529 | 0.201799 | 0.200293 | 1.326191 | -0.595935 | 0.052486 | -0.239341 | 3.265335 | 0.836289 | -0.645713 | -0.052928 | 0.572853 | -0.169373 | -0.075877 | -1.223567 | 0.323304 |
| 33.76389 | 1.493955 | 0.111618 | 0.149174 | 0.09492 | 1.969537 | 0.130442 | 0.056056 | 0.109667 | 0.464874 | 0.105236 | 0.066121 | 1.578919 | -0.807175 | -0.456306 | 0.432664 | 3.225687 | 3.45922 | -0.685214 | 0.498577 | -0.788021 | 0.378085 | -0.061016 | -1.458792 | 1.363894 |
| 33.77778 | 1.158155 | 0.06689 | 0.07773 | 0.043896 | 0.125035 | 0.063702 | 0.062046 | 0.055409 | 0.25921 | 0.045471 | -0.02238 | 1.715268 | -0.751724 | -0.732039 | 0.111994 | 1.977824 | 5.038842 | -1.480687 | 0.24164 | -0.607027 | 0.937921 | -0.024691 | -0.428942 | 0.735795 |
| 33.79167 | 1.502203 | 0.078108 | 0.073804 | 0.068999 | -1.88886 | 0.077681 | 0.037377 | 0.073153 | 0.23947 | 0.080498 | -0.010107 | 1.36007 | 0.871574 | -0.320756 | -0.078873 | 0.74116 | 4.414906 | -0.750973 | -0.006326 | -0.267376 | 0.764866 | 0.011937 | 0.613998 | 0.343977 |
| 33.80556 | 1.68866 | 0.1011 | 0.090808 | 0.092499 | -3.875839 | 0.101991 | -0.104027 | 0.081191 | 0.26348 | 0.108809 | 0.023593 | 1.161818 | 1.304276 | -0.10125 | 0.862294 | -0.446693 | 2.022914 | 0.77893 | 0.087886 | -0.829415 | 0.280114 | -0.01726 | 1.948311 | 0.34017 |
| 33.81944 | 1.841256 | 0.068299 | 0.076481 | 0.064117 | -4.919374 | 0.075854 | -0.290227 | 0.021389 | 0.222722 | 0.072737 | 0.017207 | 0.086212 | 1.048895 | 0.235448 | 0.3627 | -1.465196 | -0.103557 | 1.312615 | -0.340674 | -1.120599 | -0.134765 | -0.087153 | 1.084446 | -1.528782 |
| 33.83333 | 2.084086 | -0.004924 | 0.011316 | -0.009745 | -5.144813 | -0.004591 | -0.491998 | -0.053248 | 0.145264 | -0.020671 | -0.000841 | -1.283169 | 0.386066 | 0.662764 | -0.101999 | -2.266787 | -1.877037 | 1.090016 | -0.705707 | -1.082104 | -0.616598 | -0.202147 | 1.190355 | -2.420405 |
| 33.84722 | 1.952351 | -0.052314 | -0.054082 | -0.064674 | -2.47327 | -0.062233 | -0.692873 | -0.11322 | 0.0969 | -0.082863 | 0.021246 | -0.907804 | -0.2925 | 0.419105 | -0.083374 | -2.408671 | -3.074039 | 0.860657 | -0.328087 | -0.749651 | -0.825747 | -0.214473 | 0.789882 | -1.715527 |
| 33.86111 | 0.582254 | -0.071772 | -0.087321 | -0.080486 | 1.33916 | -0.070894 | -0.825795 | -0.132421 | 0.081431 | -0.082903 | 0.176384 | -1.46276 | 0.033282 | 0.062482 | -0.428461 | -3.072169 | -3.071332 | 0.776844 | -0.167059 | -0.426912 | -0.977241 | -0.192446 | 0.392484 | -2.028944 |
| 33.875 | -1.462967 | -0.140979 | -0.135171 | -0.141111 | 4.622421 | -0.124232 | -1.104814 | -0.166403 | -0.018144 | -0.129907 | 0.094683 | -1.077225 | -1.017011 | -0.21018 | -1.277133 | -3.923966 | -3.147513 | 0.562568 | -0.291734 | -0.649533 | -1.176384 | -0.210812 | -0.006358 | -0.136952 |
| 33.88889 | -2.700759 | -0.28312 | -0.295991 | -0.289141 | 4.696045 | -0.260932 | -1.229116 | -0.282052 | -0.314992 | -0.275358 | -0.507574 | -0.943835 | -2.101764 | -0.331329 | -1.183253 | -4.099285 | -3.253657 | 0.352626 | -0.293977 | -0.535111 | -1.401863 | -0.242915 | -0.078592 | -2.626403 |
| 33.90278 | -3.269626 | -0.424745 | -0.444807 | -0.452345 | 1.217497 | -0.411755 | -2.746252 | -0.441024 | -0.69164 | -0.438943 | -0.663352 | -0.784774 | -2.072902 | -1.324443 | -1.26754 | -3.66707 | -3.647545 | 0.151303 | -0.066355 | 0.136538 | -1.325253 | -0.337937 | -0.88793 | -3.950084 |
| 33.91667 | -3.885137 | -0.454734 | -0.48303 | -0.493392 | -3.374429 | -0.448858 | -3.165249 | -0.477035 | -0.935821 | -0.469955 | -0.629928 | -0.915311 | -1.051145 | -2.188209 | -0.596499 | -2.96289 | -3.023788 | -0.095079 | 0.244237 | 0.750547 | -1.31723 | -0.436815 | -1.976561 | -2.690366 |
| 33.93056 | -3.885143 | -0.358426 | -0.390266 | -0.390402 | -7.901615 | -0.352692 | -2.785614 | -0.365846 | -0.885695 | -0.354657 | -0.45427 | -1.093676 | -0.255209 | -2.135241 | -0.318635 | -2.32419 | -1.850816 | -0.030158 | 0.343365 | 1.034921 | -1.557468 | -0.471533 | -0.809903 | -2.020244 |
| 33.94444 | -3.354846 | -0.269943 | -0.288582 | -0.288237 | -8.486177 | -0.257547 | -2.723191 | -0.263636 | -1.012446 | -0.257946 | -0.233702 | -1.050088 | -0.452513 | -2.334362 | -1.343847 | -1.09438 | -0.752454 | 0.113481 | 0.07699 | -0.589816 | -1.59195 | -0.582319 | 1.05017 | -2.577715 |
| 33.95833 | -3.362123 | -0.268226 | -0.259452 | -0.27234 | -5.866318 | -0.245451 | -2.770832 | -0.248891 | -1.344895 | -0.247176 | -0.206146 | 0.037295 | -1.682727 | -3.331598 | -2.319034 | -2.235607 | -0.62961 | -0.924451 | -0.442833 | -1.582014 | -1.290403 | -0.560882 | 1.640205 | -3.416283 |
| 33.97222 | -3.880797 | -0.310763 | -0.271799 | -0.308921 | -3.416349 | -0.28752 | -2.256958 | -0.283912 | -1.598085 | -0.284194 | -0.281331 | -0.061539 | -2.971483 | -4.021689 | -2.903145 | -2.270984 | -0.472995 | -0.096677 | -0.579453 | -0.602801 | -1.03634 | -0.446972 | 0.577576 | -3.718557 |
| 33.98611 | -4.167476 | -0.331795 | -0.293412 | -0.326135 | -1.77466 | -0.320884 | -2.013011 | -0.31924 | -1.55603 | -0.308997 | -0.335078 | -0.725668 | -3.845239 | -2.915111 | -1.689125 | -1.84608 | -0.529118 | 0.9471 | -0.393188 | 0.356274 | -0.636757 | -0.34375 | 0.661718 | -3.543776 |
| 34 | -4.015137 | -0.222721 | -0.204524 | -0.168703 | -0.9531 | -0.231009 | -1.586859 | -0.22567 | -1.305056 | -0.185218 | -0.271046 | -1.207339 | -3.624906 | -2.621787 | -0.515123 | -0.992603 | -0.696629 | 0.221741 | -0.036419 | 0.39047 | -0.306179 | -0.205536 | 0.240388 | -2.171513 |
| 34.01389 | -2.4838 | 0.050965 | 0.070142 | 0.234884 | -0.579489 | 0.007311 | -0.995226 | 0.032866 | -0.917918 | 0.118932 | -0.058799 | -1.033968 | -1.340659 | -1.47776 | -0.309027 | -0.058471 | -0.634981 | -0.024069 | 0.157692 | -0.013175 | -0.048545 | 0.104023 | 0.018102 | -0.348594 |
| 34.02778 | -0.952254 | 0.408769 | 0.40472 | 0.626078 | -0.494389 | 0.322488 | -0.583622 | 0.36993 | -0.310808 | 0.405502 | 0.13338 | -0.342481 | 0.04585 | -0.240532 | -0.628143 | 0.186869 | -0.990262 | -0.107347 | 0.172389 | -0.535891 | 0.299676 | 0.291451 | -0.15936 | 0.114987 |
| 34.04167 | -0.893217 | 0.674437 | 0.54478 | 0.549732 | -0.558341 | 0.518969 | -0.329459 | 0.611367 | 0.093111 | 0.508144 | 0.238439 | 0.165596 | 0.006546 | -1.413572 | -1.136517 | -0.590933 | -3.643365 | 0.079134 | -0.215601 | -1.553541 | 0.209852 | -0.175373 | -0.605899 | -1.528336 |
| 34.05556 | -0.273651 | 0.666434 | 0.453233 | 0.4248 | -1.178432 | 0.530227 | -0.441736 | 0.594905 | 0.247341 | 0.494668 | 0.123103 | 0.450433 | -1.582275 | -3.449885 | -1.890209 | -1.347713 | -5.412938 | -0.43189 | -1.069302 | -2.166394 | -0.410479 | -1.07205 | -2.153634 | -4.478695 |
| 34.06944 | 0.095047 | 0.421599 | 0.305178 | 0.316965 | -2.578225 | 0.389078 | -0.665741 | 0.413525 | 0.40477 | 0.390443 | -0.277602 | -1.012714 | -1.893354 | -5.071732 | -2.082188 | -2.677201 | -6.09882 | -1.100199 | -0.866214 | -2.186252 | -0.324449 | -1.952374 | -3.918098 | -6.86404 |
| 34.08333 | 0.546254 | 0.190877 | 0.141122 | 0.164004 | -2.176973 | 0.221534 | -0.526923 | 0.231424 | 0.476559 | 0.223873 | -0.253907 | -1.055112 | -0.355673 | -6.874123 | -2.776947 | -3.054576 | -5.955919 | -1.363365 | -0.201569 | -1.392747 | 0.130938 | -2.26847 | -3.386066 | -6.555681 |
| 34.09722 | 0.763989 | 0.087902 | 0.047994 | 0.080738 | -0.269646 | 0.14641 | -0.273609 | 0.181898 | 0.177758 | 0.148727 | 0.148421 | -0.579493 | -0.057054 | -6.735716 | -3.271227 | -1.457258 | -3.405505 | -1.581715 | -0.1468 | -0.943708 | 0.257149 | -1.707072 | -1.598699 | -4.862772 |
| 34.11111 | 0.714057 | 0.228128 | 0.248763 | 0.285588 | 2.081484 | 0.293226 | 0.380524 | 0.316797 | 0.113276 | 0.343686 | 0.365864 | -0.604987 | -0.25067 | -5.498848 | -3.107965 | 0.081541 | -1.218785 | -1.787885 | 0.146202 | -0.49065 | 0.157011 | -0.850269 | 0.234644 | -2.330554 |
| 34.125 | 0.756675 | 0.373052 | 0.482875 | 0.498257 | 3.530247 | 0.382975 | 1.162107 | 0.36843 | 0.221192 | 0.491373 | 0.459504 | -1.261509 | -0.119364 | -4.400216 | -1.579707 | 1.336541 | 0.040854 | -1.54998 | 0.661539 | -0.215823 | -0.039989 | 0.233893 | 0.928196 | -0.127127 |
| 34.13889 | 1.296967 | 0.258369 | 0.384072 | 0.382929 | 3.956403 | 0.182374 | 1.677095 | 0.178802 | 0.211708 | 0.32156 | 0.275852 | -0.493459 | 0.214396 | -3.174644 | -0.813086 | 2.172679 | 1.260263 | -0.185974 | 0.880883 | -0.003864 | -0.085309 | 0.835029 | 0.918521 | 1.966117 |
| 34.15278 | -0.120176 | 0.074778 | 0.151803 | 0.123893 | 3.515479 | -0.014994 | 1.835912 | 0.010275 | 0.250351 | 0.103816 | 0.078679 | -0.137773 | 0.87637 | -0.647668 | 0.104092 | 2.491927 | 2.227678 | 0.779466 | 0.369373 | 0.174312 | 0.073569 | 0.811168 | 0.937216 | 3.767132 |
| 34.16667 | -0.705425 | -0.042551 | -0.015378 | -0.060685 | 1.672509 | -0.070288 | 1.441713 | -0.062987 | 0.319737 | -0.018506 | -0.004586 | -0.38324 | 1.691228 | 1.401345 | 0.386338 | 2.702956 | 1.91768 | 1.624038 | 0.157602 | 1.074928 | 0.215746 | 0.320396 | 1.3502 | 4.18549 |
| 34.18056 | 0.690967 | -0.098466 | -0.072582 | -0.096538 | -0.868908 | -0.0586 | 0.861831 | -0.079536 | 0.243037 | -0.044303 | 0.087148 | -0.441377 | 1.349932 | 2.532786 | 0.251677 | 2.86305 | 0.701254 | 1.545667 | 0.02752 | 2.245589 | 0.260559 | -0.340892 | 1.582958 | 3.252817 |
| 34.19444 | 2.011785 | -0.084157 | -0.042975 | -0.041501 | -2.542787 | -0.021134 | 0.206779 | -0.045751 | 0.037585 | -0.003013 | 0.08529 | 0.827993 | 0.572078 | 2.54941 | -0.255249 | 2.784221 | 0.30133 | 0.806314 | -0.037644 | 2.325453 | 0.504378 | 0.219977 | 1.26812 | 1.827661 |
| 34.20833 | 2.133501 | -0.072818 | -0.036339 | -0.031582 | -2.542023 | -0.015351 | -0.079833 | -0.02503 | -0.19134 | -0.004723 | -0.015999 | 1.403411 | -0.069101 | 2.226196 | -0.942161 | 2.438163 | 1.237251 | 0.383585 | -0.235651 | 0.876287 | 0.765931 | 1.084819 | 0.830742 | 1.172219 |
| 34.22222 | 1.505278 | -0.081133 | -0.047708 | -0.056874 | -1.536118 | -0.032168 | -0.187916 | -0.031527 | -0.254615 | -0.036086 | -0.051913 | 0.711875 | -0.645823 | 2.408612 | -1.64487 | 1.029719 | 1.80668 | 0.391232 | -0.764575 | -0.754213 | 0.728885 | 1.633022 | 0.494927 | 0.23794 |
| 34.23611 | 1.232716 | -0.049569 | -0.001936 | -0.035748 | -0.409594 | -0.020082 | 0.070201 | -0.002295 | -0.080526 | -0.012627 | -0.122426 | 0.176757 | 0.67764 | 2.815587 | -1.812194 | -0.078298 | 0.812584 | 0.397336 | -0.150725 | -1.806075 | 0.456989 | 1.193929 | 0.190549 | -0.642963 |
| 34.25 | 1.919014 | 0.024995 | 0.069828 | 0.0325 | -1.744848 | 0.023667 | 0.55076 | 0.052445 | -0.130117 | 0.05913 | -0.172997 | -0.060139 | 4.111034 | 2.579338 | -1.413426 | -0.254736 | -0.072952 | 0.38287 | 0.276067 | -1.207393 | 0.091543 | 0.521072 | -0.309591 | -0.143585 |
| 34.26389 | -0.944927 | 0.172907 | 0.18788 | 0.166622 | -0.756454 | 0.15857 | 0.868137 | 0.184446 | -1.175388 | 0.190777 | 0.079146 | 0.612792 | 3.823664 | 2.033407 | -1.101619 | 0.348619 | -0.160621 | 0.381132 | 0.215669 | 0.231537 | -0.416268 | -0.277352 | -0.105562 | 1.097823 |
| 34.27778 | -0.61105 | 0.353077 | 0.373572 | 0.346678 | 0.87184 | 0.347739 | 1.126083 | 0.364984 | -1.374758 | 0.364241 | 0.365401 | 1.598466 | 0.437436 | 1.797782 | -0.672157 | 1.185202 | 0.579632 | 0.844414 | 0.062816 | 1.41135 | -0.496595 | -0.450303 | 0.377492 | 1.858138 |
| 34.29167 | 1.731284 | 0.285667 | 0.367948 | 0.336317 | 2.444177 | 0.311025 | 1.119678 | 0.324954 | -0.422841 | 0.344797 | 0.494786 | 2.20847 | -2.43825 | 1.759485 | -0.077494 | 1.722162 | 1.193968 | 1.038551 | 0.070392 | 1.417665 | -0.373842 | 0.290895 | 0.917022 | 2.174955 |
| 34.30556 | 1.868205 | 0.050768 | 0.147583 | 0.129937 | 3.625654 | 0.10384 | 0.794859 | 0.101846 | 0.295407 | 0.147587 | 0.582841 | 2.436932 | -0.263578 | 0.799459 | 0.964417 | 1.11451 | 0.508603 | 1.209468 | 0.468142 | 0.666897 | 0.069968 | 1.050886 | 1.621409 | 1.215914 |
| 34.31944 | 1.038133 | 0.007242 | 0.062385 | 0.037325 | 2.98146 | 0.069603 | 0.663838 | 0.057588 | 0.801295 | 0.074463 | 0.378478 | 1.900769 | 0.881159 | 0.339392 | 2.097547 | 0.687526 | -0.418759 | 1.079518 | 0.581327 | 0.050206 | 0.387523 | 1.359908 | 1.669786 | -0.529151 |
| 34.33333 | -0.071906 | 0.090879 | 0.131709 | 0.080176 | 1.312631 | 0.154507 | 0.799855 | 0.144654 | 0.908106 | 0.123447 | 0.195533 | 1.190768 | 3.531491 | 0.816226 | 2.458708 | 1.38217 | -0.606233 | 1.00689 | 0.55962 | 0.046362 | 0.655641 | 0.805807 | 1.060349 | -1.314928 |
| 34.34722 | 0.935768 | 0.076801 | 0.115096 | 0.069369 | 0.21289 | 0.122428 | 1.083532 | 0.119033 | 0.769776 | 0.105625 | 0.104459 | 0.708911 | 3.860405 | 2.425376 | 2.719542 | 2.54454 | 0.230679 | 0.814069 | 0.135648 | 0.607177 | 0.564192 | 0.288384 | 0.551042 | -0.410034 |
| 34.36111 | 1.834648 | -0.030922 | 0.029465 | 0.01685 | -1.257011 | 0.027917 | 1.402537 | 0.012439 | 0.534271 | 0.041081 | -0.010393 | 0.255499 | 2.875581 | 4.072524 | 2.447671 | 3.100055 | 1.323086 | 0.728849 | -0.085464 | 1.165029 | 0.200929 | 0.225175 | 0.823939 | 1.687846 |
| 34.375 | -0.57058 | -0.121198 | -0.003457 | -0.008648 | 0.934071 | -0.014515 | 1.650028 | -0.046073 | 0.45426 | 0.000511 | -0.056252 | 0.188267 | 1.531496 | 3.523849 | 2.010353 | 3.017859 | 1.988298 | 0.880132 | -0.014685 | 1.279031 | 0.094101 | 0.549806 | 1.28214 | 3.642286 |
| 34.38889 | -0.480001 | -0.132833 | -0.023668 | -0.044904 | 2.40598 | -0.026593 | 1.883572 | -0.063097 | 0.570824 | -0.031084 | -0.04772 | -0.615819 | 0.095174 | 2.283779 | 1.538764 | 2.46813 | 1.53775 | 1.150214 | -0.059756 | 1.169483 | 0.121685 | 0.886731 | 1.923129 | 3.687502 |
| 34.40278 | 0.934694 | -0.083938 | -0.042175 | -0.06579 | 4.194574 | -0.027787 | 2.008183 | -0.05312 | 0.794378 | -0.038323 | -0.008087 | -1.35702 | -1.31785 | 1.589446 | 1.392153 | 1.871528 | 0.769865 | 1.186622 | -0.067858 | 1.138373 | 0.068299 | 0.92757 | 1.915274 | 2.966594 |
| 34.41667 | 1.951995 | -0.04471 | -0.01437 | -0.031267 | 5.448799 | -0.020812 | 2.113878 | -0.042329 | 1.095648 | -0.020859 | 0.047451 | 0.944449 | -2.841695 | 1.747713 | 1.732208 | 1.493312 | 0.743973 | 1.097753 | 0.171721 | 1.200464 | 0.007813 | 0.780345 | 1.572362 | 2.645509 |

| | | | | | | | | | | | | | | | | | | | | | | | | |
|---|---|---|---|---|---|---|---|---|---|---|---|---|---|---|---|---|---|---|---|---|---|---|---|---|
| 34.43056 | 2.490779 | -0.038271 | 0.003117 | -0.000188 | 6.06921 | -0.007703 | 2.094106 | -0.037669 | 1.449634 | -0.007569 | 0.047427 | 3.599247 | -1.331827 | 2.594907 | 2.361651 | 1.404252 | 1.487895 | 1.132013 | 0.518597 | 0.910702 | -0.041602 | 0.633629 | 1.336207 | 2.352707 |
| 34.44444 | 2.578256 | -0.1478 | -0.122696 | -0.078876 | 4.710482 | -0.05598 | 1.670992 | -0.101237 | 1.784273 | -0.081444 | -0.050805 | 4.38426 | 0.175489 | 3.851965 | 2.272521 | 2.021068 | 2.279685 | 1.094793 | 0.467201 | 0.181537 | -0.153359 | 0.660111 | 1.193222 | 1.89445 |
| 34.45833 | 2.128432 | -0.36545 | -0.375827 | -0.448338 | 2.306202 | -0.227457 | 1.48466 | -0.260439 | 1.937206 | -0.309725 | -0.199235 | 3.45095 | 1.24518 | 4.807416 | 2.141434 | 3.026666 | 2.884741 | 0.769554 | -0.053017 | -0.67651 | -0.374025 | 0.659181 | 1.568633 | 2.287331 |
| 34.47222 | 1.492061 | -0.453509 | -0.448802 | -0.660308 | 1.180378 | -0.324659 | 1.401153 | -0.362113 | 1.936597 | -0.448234 | -0.346505 | 2.683736 | 1.873074 | 4.124036 | 2.352935 | 4.338171 | 1.19901 | 0.579997 | -0.245972 | -0.855256 | -0.383685 | 0.720397 | 1.245206 | 1.517907 |
| 34.48611 | 1.160393 | -0.372702 | -0.351236 | -0.402341 | 2.569508 | -0.278932 | 1.490811 | -0.340699 | 1.832906 | -0.357414 | -0.316242 | 1.944502 | 1.792198 | 1.667187 | 3.39971 | 5.503777 | 1.742894 | 0.684674 | -0.172744 | -0.420538 | 0.150839 | 0.817211 | 2.44106 | 2.342998 |
| 34.5 | 1.18694 | -0.17943 | -0.123432 | -0.150349 | 5.168436 | -0.125401 | 1.368796 | -0.154815 | 1.629321 | -0.134449 | -0.113936 | 2.100752 | 1.658437 | 2.835261 | 4.163619 | 5.228989 | 0.952496 | 1.725756 | 0.426364 | -0.076945 | 1.015081 | 0.554634 | 2.85791 | 3.717753 |
| 34.51389 | 1.271223 | 0.007448 | 0.137166 | 0.095688 | 7.032144 | 0.076797 | 1.454955 | 0.024491 | 1.443085 | 0.096326 | 0.102898 | 2.003836 | 1.852295 | 3.736807 | 3.975516 | 3.269111 | 0.480122 | 2.575486 | 0.54041 | 0.781822 | 1.377253 | 0.457721 | 2.6769 | 3.600509 |
| 34.52778 | 1.715934 | -0.117423 | 0.200129 | 0.142677 | 4.91564 | -0.009355 | 0.898781 | -0.048501 | 1.321864 | 0.053249 | 0.10859 | 1.989092 | 2.284588 | 1.844914 | 2.993148 | 2.433575 | 0.239199 | 2.10179 | 0.391335 | 2.007489 | 1.003391 | 0.43275 | 2.188689 | 2.739666 |
| 34.54167 | 2.125656 | -0.615988 | -0.080573 | -0.139989 | 1.197462 | -0.510142 | 0.256566 | -0.454754 | 0.940691 | -0.381395 | -0.227881 | 2.087751 | 3.11539 | 0.725741 | 2.166375 | 2.612546 | -0.126933 | 1.86528 | 0.138112 | 3.407585 | 1.057998 | 0.453386 | 1.261252 | 2.293969 |
| 34.55556 | 1.222776 | -0.952644 | -0.498793 | -0.5343 | 0.464828 | -0.905042 | -0.328417 | -0.753938 | 0.357747 | -0.760647 | -0.634922 | 1.406768 | 3.331944 | 0.81148 | 1.687568 | 2.799403 | -0.219314 | 0.718773 | -0.141868 | 2.479907 | 0.440778 | 0.418649 | 1.616455 | 3.332179 |
| 34.56944 | 0.403397 | -0.708261 | -0.555667 | -0.555254 | 0.227436 | -0.710403 | -0.741933 | -0.533969 | -0.130641 | -0.619108 | -0.659708 | 0.44282 | 2.188591 | 0.556728 | 1.994341 | 2.557574 | -0.172174 | 0.160094 | -0.13406 | 2.038424 | 0.435395 | 0.404437 | 2.454043 | 4.894883 |
| 34.58333 | 0.927279 | -0.221633 | -0.224503 | -0.218738 | -0.544378 | -0.23936 | -0.764367 | -0.176974 | -0.263609 | -0.194235 | -0.303522 | 0.22893 | 0.731859 | 0.087825 | 3.363568 | 2.542737 | -0.019589 | 0.304947 | -0.016051 | 1.413069 | 0.565838 | 0.351573 | 2.682549 | 4.548277 |
| 34.59722 | 0.975352 | 0.090198 | 0.075757 | 0.071575 | -1.437415 | 0.065847 | -0.134669 | 0.071682 | -0.265825 | 0.103397 | 0.0324 | 0.463566 | 2.021253 | 0.131556 | 2.202998 | 1.465711 | 0.055654 | 0.510665 | -0.024598 | -0.212989 | 0.607924 | 0.386094 | 1.850351 | 3.243906 |
| 34.61111 | 0.286397 | 0.219987 | 0.20669 | 0.197438 | 0.430002 | 0.193693 | 0.538184 | 0.105747 | -0.063873 | 0.230841 | 0.182765 | 0.91241 | 2.446278 | -0.424785 | 1.032001 | -0.074769 | 0.119594 | 0.801981 | -0.180216 | -1.811979 | 0.186355 | 0.317388 | 0.512934 | -0.490102 |
| 34.625 | 0.122935 | 0.279286 | 0.274159 | 0.265606 | 2.018952 | 0.269953 | 0.667751 | 0.091669 | 0.224648 | 0.298489 | 0.220551 | 1.616873 | 0.857382 | -0.135675 | 0.461368 | -1.311634 | 0.090989 | 1.247928 | -0.23831 | -0.64911 | -0.266048 | 0.185866 | 0.699925 | -2.350592 |
| 34.63889 | -0.405995 | 0.328864 | 0.340623 | 0.329014 | 0.866497 | 0.348447 | -0.035281 | 0.1927 | 0.472524 | 0.35813 | 0.248472 | 1.451111 | -2.166494 | -0.365265 | 0.168499 | -1.54962 | 0.060938 | 0.454645 | -0.155196 | 1.71482 | -0.557052 | 0.038924 | -0.659985 | -0.447154 |
| 34.65278 | -0.362669 | 0.379202 | 0.396649 | 0.364021 | -0.297821 | 0.40462 | -0.073362 | 0.289444 | 0.626362 | 0.394488 | 0.290243 | -2.183752 | -2.588503 | 0.541993 | -0.542056 | -1.314878 | -1.160946 | -0.456884 | -0.023153 | 1.951765 | -0.702982 | -0.061115 | -0.632878 | 1.756555 |
| 34.66667 | 2.260655 | 0.431655 | 0.451727 | 0.399632 | 0.659216 | 0.436929 | 0.373981 | 0.367229 | 0.743332 | 0.425341 | 0.353898 | -0.837424 | -1.017657 | 0.016512 | -0.666963 | -0.481857 | -0.45161 | -0.303992 | 0.041367 | 0.360641 | -0.50423 | -0.085576 | -0.601233 | 0.547944 |
| 34.68056 | 2.601317 | 0.440854 | 0.48291 | 0.422818 | 0.863139 | 0.448317 | 0.758242 | 0.410304 | 0.810113 | 0.445838 | 0.404775 | -0.204395 | 1.015978 | 0.368934 | 0.318776 | -0.097754 | 0.66584 | 0.317041 | 0.005368 | -0.946545 | 0.402255 | 0.111823 | 0.356972 | -0.170869 |
| 34.69444 | 1.207178 | 0.373754 | 0.427513 | 0.382321 | -2.168358 | 0.412349 | 0.728653 | 0.390349 | 0.73626 | 0.412595 | 0.387534 | 0.568354 | 3.713319 | 0.450238 | 0.012104 | -0.121562 | 0.355298 | 0.050453 | -0.16182 | -3.818928 | 1.771054 | 0.310043 | 0.946641 | -1.28216 |
| 34.70833 | 1.247381 | 0.329256 | 0.356139 | 0.339305 | -4.793489 | 0.380666 | 0.605812 | 0.369077 | 0.613439 | 0.375959 | 0.36787 | 0.249263 | 3.367598 | -0.363093 | -0.561837 | -0.89982 | -0.941991 | -0.195941 | -0.229251 | -2.942733 | 1.357151 | 0.330195 | -0.134155 | -3.251402 |
| 34.72222 | 1.799072 | 0.316763 | 0.323721 | 0.31506 | -2.587251 | 0.351086 | 0.354464 | 0.345104 | 0.501675 | 0.338754 | 0.364284 | -0.21747 | 2.031609 | -2.091336 | -0.470984 | -2.284931 | -1.523423 | 0.471547 | -0.230384 | -0.672207 | 0.836664 | 0.084898 | 0.522233 | -1.441755 |
| 34.73611 | 1.834748 | 0.251952 | 0.261295 | 0.243228 | -0.229315 | 0.253801 | 0.170436 | 0.270518 | 0.333216 | 0.244034 | 0.28119 | -0.354378 | 1.233607 | -0.474664 | 0.63556 | -3.174594 | -0.145855 | 0.31146 | -0.26471 | 0.332074 | 0.181626 | -0.241099 | 0.390853 | 0.171906 |
| 34.75 | 0.835032 | 0.179735 | 0.186212 | 0.169903 | 1.198785 | 0.171608 | 0.074103 | 0.173152 | 0.190606 | 0.167574 | 0.142354 | -0.433314 | 2.507971 | 2.112793 | 1.005103 | -2.791583 | 0.303638 | 0.240083 | -0.348046 | 0.56302 | -0.410009 | -0.141471 | -1.258073 | 1.991574 |
| 34.76389 | 0.193989 | 0.158327 | 0.149839 | 0.142721 | 2.071078 | 0.164994 | -0.051894 | 0.117459 | 0.116737 | 0.154226 | 0.085867 | -0.679081 | 2.075105 | 0.971078 | 1.485781 | -1.421382 | 0.375543 | 0.045304 | -0.029664 | 0.91105 | -0.775517 | -0.214206 | -0.90269 | 3.524187 |
| 34.77778 | 0.346216 | 0.077802 | 0.07712 | 0.065262 | 2.958097 | 0.101276 | -0.190281 | 0.01992 | -0.003478 | 0.085369 | 0.034062 | -0.811953 | 0.887797 | -0.240595 | 2.05446 | -0.036436 | 0.415973 | -0.327283 | 0.204134 | 0.839935 | -0.411166 | -0.311666 | 1.094156 | 1.38439 |
| 34.79167 | -0.466501 | -0.048189 | -0.035879 | -0.06251 | 2.952702 | -0.026994 | -0.305607 | -0.083916 | -0.173661 | -0.038519 | -0.03455 | -0.755225 | 0.244586 | -0.675185 | 1.538793 | 1.180918 | 0.051171 | -0.692885 | 0.258794 | 0.753903 | -0.637213 | -0.339984 | 0.608732 | 0.752527 |
| 34.80556 | -1.996003 | -0.055018 | -0.060661 | -0.081005 | 0.803244 | -0.049015 | -0.29098 | -0.079173 | -0.29051 | -0.057343 | -0.027169 | -0.703476 | 0.49188 | -0.285482 | 0.613417 | 1.636103 | -0.014074 | -0.53924 | -0.181445 | 0.790293 | 1.017783 | -0.328694 | 0.528818 | -1.361509 |
| 34.81944 | -2.055058 | 0.039586 | 0.007036 | 0.009413 | 0.351149 | 0.03622 | -0.215034 | -0.015377 | -0.279662 | 0.025472 | 0.04841 | -0.837147 | 0.272736 | -0.429272 | -0.691666 | 0.587133 | -0.020609 | -0.840121 | -0.114704 | 0.240336 | 1.184703 | -0.166857 | 1.151762 | -1.370731 |
| 34.83333 | -1.531985 | 0.103151 | 0.066152 | 0.069855 | 0.248729 | 0.087509 | -0.200003 | 0.012734 | -0.393761 | 0.077952 | 0.082109 | -1.048295 | -0.663624 | -1.276702 | -1.308095 | -1.248818 | -0.126079 | -0.483761 | -0.354204 | -0.180288 | 0.20045 | 0.087898 | 1.707172 | -3.371238 |
| 34.84722 | -1.183673 | 0.088945 | 0.080799 | 0.063654 | -0.29438 | 0.065123 | -0.536199 | 0.004938 | -0.642415 | 0.064317 | 0.037446 | -1.436398 | -1.048962 | -1.495121 | -1.533632 | -2.532907 | -1.393077 | -0.287501 | -0.435665 | -0.100215 | -0.80643 | 0.188146 | 1.001028 | -0.568364 |
| 34.86111 | -0.936788 | 0.019435 | 0.051208 | 0.017059 | -3.971616 | 0.009888 | -0.768521 | 0.0048 | -0.922728 | 0.01416 | -0.003668 | -1.750839 | -0.515859 | -0.962418 | -1.892967 | -4.68962 | -2.315547 | -0.821688 | -0.293722 | 0.016358 | -2.08899 | 0.103864 | 0.286997 | 0.436694 |
| 34.875 | -1.091847 | -0.036103 | 0.004944 | -0.026742 | -4.382818 | -0.019373 | -1.500391 | 0.047675 | -1.415756 | -0.021565 | 0.108331 | -1.916538 | 0.790945 | -0.075716 | -2.255453 | -5.909525 | -3.55439 | 0.246739 | -0.016775 | 0.194825 | -1.417065 | -0.151278 | 0.246052 | 1.018466 |
| 34.88889 | -1.577503 | -0.030608 | -0.010774 | -0.029952 | -2.904972 | -0.002636 | -1.920519 | 0.086366 | -1.822459 | -0.011377 | 0.071692 | -1.968693 | 1.272319 | 0.874321 | -1.074685 | -3.175537 | -3.436234 | 0.283914 | 0.030113 | 0.443655 | -0.059805 | -0.246754 | 1.210528 | 2.281732 |
| 34.90278 | -2.395205 | -0.009044 | -0.017556 | -0.029965 | -1.830444 | 0.005141 | -2.109245 | 0.07292 | -2.181977 | -0.001545 | -1.24997 | -1.677784 | -1.058837 | 1.425296 | -0.589766 | -0.495057 | -3.92362 | -0.004518 | 0.195363 | 0.308111 | 0.648966 | -0.238757 | 0.886757 | 1.843831 |
| 34.91667 | -2.888659 | -0.033503 | -0.063703 | -0.069896 | -2.269593 | -0.03852 | -3.037263 | -0.00957 | -2.391179 | -0.041506 | -1.394709 | -1.184117 | -2.481316 | 0.603205 | 0.252782 | 1.100106 | -4.323033 | -0.500979 | 0.253178 | 0.062096 | 0.131857 | -0.234675 | 0.916734 | -0.078653 |
| 34.93056 | -2.836673 | -0.083242 | -0.110173 | -0.107476 | -4.986114 | -0.092478 | -3.736963 | -0.050049 | -2.50643 | -0.092812 | -0.475752 | -0.819329 | -3.771187 | -0.384476 | 1.014602 | 1.325508 | -2.468704 | -0.925762 | 0.264843 | -0.121046 | -0.590765 | -0.563387 | 0.464105 | -0.663617 |
| 34.94444 | -2.611544 | -0.12852 | -0.125358 | -0.117362 | -7.068425 | -0.135064 | -3.593506 | -0.123834 | -2.569817 | -0.12888 | 0.717213 | -0.75083 | -3.18182 | -1.869576 | -0.776015 | 0.84106 | -2.17704 | -1.588111 | -0.116192 | -0.347245 | -0.760179 | -0.691854 | -0.299366 | -3.009206 |
| 34.95833 | -2.610817 | -0.19652 | -0.171296 | -0.16299 | -5.603582 | -0.204564 | -2.96387 | -0.212296 | -2.474393 | -0.194935 | 1.104983 | -0.836348 | -1.872882 | -2.51891 | -2.498547 | -0.008913 | -2.502491 | -2.045863 | -0.268069 | -0.713497 | -0.529678 | -0.517464 | -1.882208 | -2.976395 |
| 34.97222 | -2.379543 | -0.266406 | -0.255535 | -0.250771 | -3.624225 | -0.277901 | -2.324188 | -0.261579 | -2.277495 | -0.272185 | 0.806833 | -0.947736 | -0.032184 | -3.36113 | -2.623524 | -0.755715 | -0.77342 | -1.566233 | -0.283112 | -1.249227 | -1.398973 | -0.288718 | -3.282663 | -2.742993 |
| 34.98611 | -1.901071 | -0.249108 | -0.259203 | -0.249323 | -1.876459 | -0.256985 | -2.078444 | -0.241782 | -1.866431 | -0.251986 | -0.039549 | -0.99058 | 1.378816 | -3.878645 | -2.725081 | -1.003692 | -0.931465 | -0.44926 | -0.066017 | -1.145653 | -2.006475 | -0.290861 | -1.314445 | -2.174972 |
| 35 | -1.430564 | -0.078592 | -0.079015 | -0.027838 | -0.50136 | -0.09762 | -1.740476 | -0.082635 | -1.397468 | -0.061582 | -0.120729 | -0.826947 | 2.048798 | -2.787811 | -4.240616 | -0.850819 | -2.4811 | 0.614486 | 0.053296 | 0.0744 | -2.196704 | -0.566319 | 0.314905 | 0.213025 |
| 35.01389 | -1.1758 | 0.174729 | 0.196431 | 0.330293 | 0.315488 | 0.109766 | -0.936637 | 0.140446 | -0.948302 | 0.210674 | 0.078635 | -0.542256 | 1.492974 | -1.210854 | -3.343811 | -0.847924 | -4.755672 | 1.417493 | 0.326933 | 1.065611 | -0.753637 | -0.356906 | 1.865092 | 1.872489 |
| 35.02778 | -1.010393 | 0.256326 | 0.198201 | 0.376076 | 0.346911 | 0.141597 | -0.463098 | 0.187469 | -0.764462 | 0.213675 | 0.001625 | -0.510313 | -0.669086 | -0.34847 | -3.103298 | -1.037422 | -4.157785 | 1.038247 | 0.353534 | 1.454593 | -0.384119 | 0.054686 | 2.621338 | 0.149775 |
| 35.04167 | -1.050909 | 0.107227 | -0.188429 | -0.167547 | -0.261734 | -0.056814 | -0.701943 | 0.014762 | -0.919086 | -0.141016 | -0.520722 | -0.896777 | -3.06963 | -1.789203 | -2.577086 | -1.776436 | -3.09117 | -0.5273 | 0.310485 | 0.467638 | 0.087438 | -0.114698 | -0.042837 | -1.173007 |
| 35.05556 | -0.754662 | 0.033639 | -0.361442 | -0.335139 | -0.610895 | -0.075129 | -1.070299 | -0.061052 | -0.735143 | -0.235015 | -0.668838 | -1.015165 | -5.167069 | -3.251197 | -2.378226 | -2.676533 | -2.38084 | -2.409708 | -0.04699 | -0.621587 | 0.243032 | -0.754511 | -3.151116 | -3.315937 |
| 35.06944 | 0.502755 | 0.063985 | -0.142058 | -0.105263 | -0.753709 | 0.09121 | -0.969851 | 0.061657 | 0.170862 | -0.018862 | -0.46788 | -0.823939 | -5.853323 | -3.193122 | -1.964831 | -4.279691 | -1.910734 | -2.584867 | -0.086506 | -1.80824 | 0.002525 | -1.4398 | -3.617963 | -5.692375 |
| 35.08333 | -0.230371 | 0.080157 | 0.020899 | 0.045423 | -0.708678 | 0.14726 | -0.554023 | 0.105231 | 1.22748 | 0.094803 | -0.579241 | -0.41018 | -5.82977 | -3.049955 | -1.48155 | -4.549112 | -3.446275 | -3.3825 | -0.138397 | -1.570186 | -0.81241 | -1.951305 | -3.838737 | -5.888304 |
| 35.09722 | -0.517167 | 0.081615 | 0.030447 | 0.029065 | 0.121657 | 0.106041 | 0.019392 | 0.07276 | 1.202832 | 0.056123 | -0.201059 | 0.361331 | -3.189861 | -3.899026 | -2.254663 | -2.161383 | -2.28827 | -3.205171 | -0.406328 | -0.687058 | -0.712358 | -1.929276 | -2.43178 | -4.086236 |
| 35.11111 | -0.884168 | 0.02099 | -0.018866 | -0.025299 | 1.238706 | 0.034329 | 0.530671 | 0.005079 | 0.612432 | -0.010143 | -0.071796 | 0.091667 | -1.372943 | -4.698242 | -1.613435 | -0.254935 | -1.21249 | -1.630509 | -0.410129 | -0.334638 | -0.289136 | -0.799862 | -0.764317 | -2.371243 |
| 35.125 | -0.186631 | -0.131233 | -0.107682 | -0.079427 | 1.967194 | -0.08137 | 0.873794 | -0.109889 | -0.144666 | -0.079743 | -0.017028 | -0.184166 | -0.347525 | -4.438442 | -1.046512 | 0.887308 | -0.269883 | -0.209463 | -0.159821 | -0.459187 | 0.231415 | 0.040166 | 0.300048 | -1.401424 |
| 35.13889 | 0.954323 | -0.200759 | -0.165034 | -0.113394 | 1.793718 | -0.155856 | 1.203755 | -0.169457 | -0.77825 | -0.121917 | -0.004726 | -0.72977 | 0.920115 | -2.525884 | -0.648502 | 1.451918 | 1.13911 | 0.528307 | -0.115827 | -0.437199 | 0.712577 | 0.436517 | 1.324164 | -0.770411 |
| 35.15278 | 0.36076 | -0.126078 | -0.134673 | -0.106072 | 1.499669 | -0.123128 | 1.200387 | -0.175686 | -0.893212 | -0.116445 | 0.003084 | -1.399817 | 3.483618 | -0.670683 | -0.209709 | 1.543685 | 3.5302 | 1.15611 | 0.284201 | 0.148122 | 1.36997 | 0.25054 | 2.053022 | 0.026904 |
| 35.16667 | -0.472221 | -0.036864 | -0.065067 | -0.070541 | 1.389222 | -0.049437 | 0.690269 | -0.3572 | -0.591115 | -0.076157 | 0.094156 | -1.582454 | 5.177196 | 1.441802 | 0.946237 | 1.688535 | 4.952409 | 1.58475 | 0.334705 | 1.166841 | 1.51296 | -0.326644 | 2.168972 | 0.936828 |
| 35.18056 | 0.863691 | -0.029067 | -0.012657 | -0.023793 | 1.008565 | -0.006942 | 0.101612 | -0.354941 | -0.341754 | -0.023827 | 0.524694 | -1.114238 | 3.497794 | 3.25419 | 1.555513 | 2.51638 | 3.283571 | 1.853797 | 0.426806 | 2.043022 | 1.090578 | -0.557742 | 2.397178 | 1.653514 |
| 35.19444 | 1.104182 | -0.054161 | 0.017616 | 0.027249 | 0.247733 | 0.011674 | 0.076098 | -0.008698 | -0.122897 | 0.032069 | 0.624139 | -0.725552 | 0.417792 | 2.666046 | 1.628585 | 3.838284 | 1.160122 | 1.798879 | 0.403541 | 1.720016 | 0.636113 | -0.60668 | 2.133913 | 1.594512 |
| 35.20833 | 2.056322 | -0.00145 | 0.05431 | 0.071906 | -0.213073 | 0.053898 | 0.497029 | 0.359401 | 0.159546 | 0.083074 | 0.489144 | -0.96246 | -1.724813 | 2.386049 | 0.691421 | 3.573663 | -0.129893 | 1.578306 | 0.277385 | 0.874376 | 0.482097 | -0.265016 | 1.574459 | 0.864855 |
| 35.22222 | 1.118212 | 0.118802 | 0.107217 | 0.095507 | -0.075076 | 0.122906 | 1.250697 | 0.35316 | 0.592954 | 0.11719 | 0.172605 | -1.031073 | -2.25461 | 2.404196 | -0.591781 | 1.955376 | 0.238515 | 0.575375 | -0.059546 | 0.123418 | 0.566347 | -0.15228 | 1.333129 | -0.113907 |
| 35.23611 | 1.198866 | 0.204727 | 0.149832 | 0.105911 | 0.019134 | 0.17926 | 2.150299 | 0.181322 | 1.038921 | 0.132953 | -0.044689 | -0.230704 | -3.369711 | 1.5462 | -1.790505 | 0.810048 | 1.179895 | 0.269302 | -0.238073 | -0.34569 | 0.526558 | -0.421016 | 1.575621 | -0.571367 |
| 35.25 | -0.191454 | 0.17245 | 0.158122 | 0.128282 | -0.566067 | 0.175322 | 1.974747 | 0.172623 | 1.228129 | 0.140511 | -0.239269 | 0.509617 | 0.505609 | 0.864728 | -0.567429 | -0.475802 | 1.902208 | 0.144568 | 0.251623 | -1.605313 | 0.556905 | -0.115863 | 1.211906 | -0.257396 |
| 35.26389 | 0.374024 | 0.045982 | 0.109598 | 0.13097 | -0.854245 | 0.111698 | 1.165596 | 0.088469 | 1.017003 | 0.125878 | -0.246763 | 0.702301 | 3.00612 | 0.214014 | 0.979792 | -1.057583 | 1.284139 | 0.258076 | 0.141196 | 1.911005 | 0.611634 | 0.825972 | 0.313181 | 1.07477 |
| 35.27778 | 1.244557 | 0.008319 | 0.056952 | 0.087633 | -0.547395 | 0.065819 | 0.59452 | 0.036536 | 0.589254 | 0.079779 | -0.037028 | 0.449637 | 5.442948 | 0.891739 | 2.480107 | -0.247228 | -0.170821 | 0.706282 | 0.514751 | 4.867707 | 0.494491 | 1.900095 | -0.345809 | 2.32557 |
| 35.29167 | 1.240758 | 0.043233 | 0.038038 | 0.027201 | 0.070384 | 0.03885 | 0.470882 | 0.019781 | 0.176869 | 0.022364 | 0.181135 | 0.045596 | 4.841009 | 1.899819 | 3.225928 | 1.580175 | -0.4079 | 1.0254 | 0.34879 | 3.692685 | 0.223214 | 1.480585 | -0.117698 | 3.382034 |
| 35.30556 | 1.780944 | -0.005784 | 0.016224 | -0.021246 | 0.861964 | -0.013855 | 0.655205 | -0.048521 | -0.063512 | -0.03282 | 0.443775 | -0.150115 | 3.0497 | 2.765499 | 2.233 | 2.15569 | 0.24023 | 1.388143 | -0.106155 | 0.353953 | -0.010214 | 0.097416 | 0.189846 | 3.266458 |
| 35.31944 | -0.072782 | -0.043622 | 0.001917 | -0.035475 | 1.628307 | -0.032367 | 0.886514 | -0.070507 | -0.149858 | -0.043436 | 0.519015 | -0.052027 | 1.794168 | 2.74397 | 0.509541 | 0.919787 | 0.579086 | 1.827034 | 0.156621 | -2.805277 | -0.129033 | -0.360752 | 0.678399 | 2.686746 |
| 35.33333 | -2.086271 | -0.01344 | 0.014887 | 0.005249 | 2.268196 | 0.01136 | 0.993377 | -0.021606 | -0.102702 | 0.010712 | 0.318795 | -1.128815 | 0.261035 | 2.852047 | -0.477283 | -0.499428 | 1.100503 | 1.510534 | 0.736323 | -2.596813 | 0.053123 | 0.396521 | 1.24283 | 2.342921 |
| 35.34722 | -0.250286 | -0.006834 | 0.032841 | 0.061948 | 2.372827 | 0.04459 | 1.016027 | 0.009932 | 0.065338 | 0.069623 | 0.143072 | -0.645719 | -1.455557 | 2.82382 | 0.25683 | -0.337677 | 2.964827 | 1.045496 | 0.321305 | 0.621327 | 0.530906 | 1.467318 | 2.227902 | 2.575237 |
| 35.36111 | 1.57377 | 0.00867 | 0.042115 | 0.063808 | 1.91097 | 0.054889 | 1.051942 | 0.017598 | 0.294182 | 0.063955 | 0.075767 | 0.802529 | 0.579251 | 2.994442 | 1.892613 | 1.313789 | 5.264587 | 0.648364 | -0.192781 | 2.622075 | 1.065758 | 2.519601 | 2.206527 | 2.806084 |
| 35.375 | 2.833399 | 0.03288 | 0.013815 | -0.004595 | 1.560656 | 0.037888 | 1.178237 | 0.020569 | 0.468661 | -0.008618 | 0.002697 | 2.010605 | 1.952752 | 2.412267 | 3.762067 | 2.862439 | 4.46915 | 0.444652 | 0.013441 | 2.531747 | 1.16105 | 2.569205 | 1.550764 | 2.619974 |
| 35.38889 | 3.17487 | 0.018287 | -0.013758 | -0.041016 | 1.28417 | 0.017713 | 1.258261 | 0.014203 | 0.641451 | -0.046465 | -0.072619 | 1.986021 | 2.571666 | 1.641171 | 3.719743 | 3.111453 | 1.459555 | 0.749228 | 0.537914 | 1.610737 | 0.864336 | 1.109043 | 0.921326 | 2.365133 |
| 35.40278 | 2.689297 | -0.025521 | -0.002081 | -0.009181 | 1.236523 | -0.00402 | 1.387203 | 0.025573 | 0.855539 | -0.021277 | -0.060418 | 2.593961 | 1.909778 | 0.950618 | 2.651797 | 2.512393 | -0.473645 | 0.881287 | 0.497697 | 0.936985 | 0.4994 | -0.051122 | 0.650782 | 2.149454 |
| 35.41667 | 1.627499 | -0.080832 | -0.018611 | -0.016584 | 1.478184 | -0.072522 | 1.68065 | 0.004472 | 1.00257 | -0.037597 | -0.026 | 2.759349 | 1.285908 | 0.605622 | 1.685076 | 1.896095 | -2.597975 | 1.355516 | 0.633424 | 0.119301 | 0.16925 | -0.156104 | 0.689727 | 2.077732 |
| 35.43056 | 2.514073 | -0.080987 | -0.045641 | -0.050914 | 2.424539 | -0.060568 | 1.997586 | -0.01999 | 0.999977 | -0.054886 | -0.039392 | 2.50531 | 0.394633 | 0.465818 | 1.41434 | 1.621703 | -1.565472 | 1.308534 | 0.696189 | 0.383658 | 0.078899 | 0.594025 | 0.909386 | 2.074953 |
| 35.44444 | 3.776662 | -0.067357 | -0.081524 | -0.082767 | 4.696869 | 0.033575 | 2.344449 | -0.059871 | 0.902105 | -0.053814 | -0.025689 | 1.923612 | -1.418823 | 1.14764 | 1.302151 | 2.183356 | 1.770275 | 0.894101 | 0.441704 | 1.198112 | 0.082927 | 1.569133 | 1.098005 | 2.455312 |
| 35.45833 | 4.439403 | -0.104114 | -0.120875 | -0.237244 | 7.59312 | 0.050929 | 2.611618 | -0.141465 | 0.664992 | -0.088648 | 0.008147 | 1.639736 | -2.969461 | 2.362574 | 1.432406 | 3.32135 | 4.192368 | 0.968817 | 0.413036 | 1.466609 | 0.153005 | 1.920669 | 1.294813 | 3.882916 |
| 35.47222 | 3.865077 | -0.0004 | 0.018153 | -0.20734 | 8.498225 | 0.093848 | 2.729209 | -0.070186 | 0.403508 | -0.031518 | 0.089533 | 1.859562 | -3.043786 | 4.315745 | 2.349462 | 4.506199 | 2.830741 | 1.177348 | 0.216771 | 1.66641 | 0.148245 | 1.29684 | 1.689942 | 4.520607 |
| 35.48611 | 2.52845 | 0.147436 | 0.149202 | 0.066487 | 6.4508 | 0.167855 | 2.720993 | 0.069415 | 0.381676 | 0.102841 | 0.17872 | 2.39766 | -0.94247 | 6.185639 | 3.681311 | 4.622517 | -0.272735 | 1.582162 | 0.001993 | 1.687304 | 0.061129 | 1.146707 | 1.677513 | 5.103044 |
| 35.5 | 1.391195 | 0.18599 | 0.176514 | 0.169695 | 3.102618 | 0.176603 | 2.137539 | 0.107646 | 0.43125 | 0.163301 | 0.220647 | 2.079192 | 2.096618 | 5.395385 | 4.472917 | 3.933598 | -0.538635 | 2.029205 | -0.394109 | 1.725378 | 0.562539 | 0.99764 | 1.594233 | 4.961232 |
| 35.51389 | 1.282444 | 0.091877 | 0.116443 | 0.10522 | 1.345566 | 0.087431 | 1.467722 | 0.038347 | 0.405851 | 0.086581 | 0.169278 | 1.943128 | 4.101533 | 3.456496 | 4.709484 | 3.129617 | -0.313151 | 2.046387 | -0.384166 | 1.20795 | 1.470897 | 0.811941 | 2.022747 | 3.04721 |
| 35.52778 | 0.452683 | -0.353268 | -0.137622 | -0.169981 | 1.114728 | -0.315461 | 0.767004 | -0.342505 | 0.100421 | -0.265618 | -0.172595 | 1.807102 | 4.088916 | 3.560086 | 4.789007 | 2.834138 | 0.245973 | 1.601506 | -0.311265 | 1.349498 | 1.010107 | 0.533178 | 2.208281 | 1.85168 |
| 35.54167 | -0.202031 | -1.160948 | -0.671443 | -0.709176 | 1.743859 | -1.069604 | -0.053673 | -1.041117 | -0.493938 | -0.905064 | -0.814773 | 1.399294 | 3.639125 | 3.163217 | 3.849828 | 3.569271 | -0.025954 | 1.293095 | -0.362401 | 1.203684 | 0.251873 | 0.245512 | 2.089073 | 1.756935 |
| 35.55556 | -0.455033 | -1.569346 | -1.102674 | -1.115217 | 1.655123 | -1.449835 | -0.64106 | -1.369764 | -0.937527 | -1.246973 | -1.201178 | 0.854719 | 2.862645 | 1.78825 | 2.222953 | 5.467546 | -0.21492 | 0.766862 | -0.398036 | 0.67598 | -0.477745 | 0.343115 | 2.589052 | 2.921205 |
| 35.56944 | -0.210496 | -1.134449 | -0.963106 | -0.943915 | 0.643827 | -1.062449 | -0.661429 | -1.010044 | -0.91179 | -0.931926 | -0.950182 | 0.652616 | 0.615164 | 1.012054 | 1.296946 | 4.314555 | -0.293905 | 0.657649 | -0.280887 | 1.505399 | -0.550521 | 0.597141 | 2.67317 | 3.206827 |
| 35.58333 | -1.077631 | -0.539272 | -0.526865 | -0.519744 | -0.609397 | -0.531194 | -0.274795 | -0.52649 | -0.7119 | -0.485484 | -0.501642 | 0.953077 | -0.788615 | 0.242213 | 1.641277 | 2.917671 | -0.129315 | 0.027265 | 0.07638 | 2.643381 | -0.005886 | 0.727317 | 1.757298 | 2.343226 |
| 35.59722 | -0.849953 | -0.257399 | -0.270956 | -0.272868 | -0.215817 | -0.281616 | 0.053541 | -0.285336 | -0.663704 | -0.254223 | -0.256066 | 1.816207 | 0.066696 | 0.080952 | 2.485745 | 0.537507 | 0.43085 | -0.228796 | 0.175646 | 2.370513 | 0.916465 | 0.511196 | 0.262926 | 0.906093 |
| 35.61111 | 1.004758 | -0.150148 | -0.162817 | -0.162404 | 0.143959 | -0.162821 | 0.221312 | -0.178246 | -0.773251 | -0.145661 | -0.155686 | 1.784833 | 0.401377 | 0.264566 | 1.120306 | -1.52306 | -0.079509 | 0.267237 | 0.131423 | 1.13786 | 0.656989 | 0.035815 | -0.899343 | -2.287881 |
| 35.625 | 2.009339 | -0.085056 | -0.074491 | -0.073722 | 0.373347 | -0.082125 | 0.350392 | -0.103263 | -0.958686 | -0.07837 | -0.096649 | 0.025863 | -0.007132 | -0.116029 | 0.399498 | -4.144733 | -0.105692 | 0.561853 | -0.256373 | 0.30242 | 0.283497 | -0.13846 | 0.259723 | -3.112107 |
| 35.63889 | 2.11698 | -0.070028 | -0.061304 | -0.050224 | -0.313322 | -0.071961 | 0.327046 | -0.083796 | -0.746307 | -0.067797 | -0.105471 | -0.687262 | 0.375864 | -0.183929 | -0.445407 | -2.470614 | -0.067136 | -0.412613 | -0.329712 | -0.463328 | 0.098855 | 0.412008 | 1.458862 | -2.388923 |
| 35.65278 | 0.951487 | -0.064188 | -0.081748 | -0.055283 | -0.858249 | -0.065979 | 0.096439 | -0.079607 | -0.487715 | -0.054324 | -0.118125 | 0.23478 | -0.064156 | 0.183352 | 0.453105 | -0.164964 | -0.015987 | 0.992183 | 0.277998 | -0.504338 | 0.192303 | 0.410202 | 0.37444 | 0.661933 |
| 35.66667 | 0.873665 | -0.007105 | -0.018468 | 0.000967 | -1.736675 | 0.00311 | -0.251196 | -0.021914 | -0.240166 | 0.011604 | -0.038737 | 0.357325 | -1.747101 | 0.278045 | 1.397899 | 2.572341 | 0.064139 | 0.939399 | 0.897679 | -0.334808 | 0.835309 | 0.442006 | -0.121886 | 0.962021 |
| 35.68056 | 0.870025 | 0.037994 | 0.05636 | 0.05451 | -3.374553 | 0.049385 | -0.332577 | 0.029112 | -0.104444 | 0.052182 | 0.02996 | -0.408874 | -2.561931 | 0.255021 | 0.785933 | 3.501415 | 0.092518 | 1.318868 | 0.537531 | -0.71736 | -0.528475 | 0.09248 | -1.637651 | 0.504606 |
| 35.69444 | -0.337746 | 0.024325 | 0.054985 | 0.049065 | -3.925734 | 0.037839 | -0.246205 | 0.020791 | 0.057176 | 0.040038 | 0.033272 | -0.084543 | -1.781764 | 0.300365 | 0.100913 | 2.550194 | -0.010774 | 0.358365 | 0.083827 | -0.378724 | -0.26326 | -0.267277 | -1.999703 | -0.387196 |
| 35.70833 | -0.757014 | 0.043892 | 0.075123 | 0.077365 | -0.91378 | 0.072436 | -0.18162 | 0.044797 | 0.402943 | 0.075376 | 0.085876 | -0.968108 | 0.31221 | -0.192264 | -0.56843 | 0.268195 | -0.291641 | 0.45221 | -0.015221 | 1.021723 | -0.228375 | -0.376051 | -2.219482 | -0.657316 |
| 35.72222 | 0.803832 | 0.118667 | 0.148203 | 0.152337 | 1.278704 | 0.150163 | -0.085482 | 0.125446 | 0.828893 | 0.156488 | 0.185969 | -1.712055 | -1.09775 | -0.992951 | -0.568676 | -1.529245 | 0.10015 | 0.336132 | 0.008312 | 0.907791 | -0.564804 | -0.605369 | -2.015468 | -1.454815 |
| 35.73611 | 0.25547 | 0.161457 | 0.177294 | 0.18024 | 1.397324 | 0.174156 | -0.055672 | 0.170622 | 1.245952 | 0.189862 | 0.209477 | -2.289581 | -0.319095 | -0.812635 | -0.543338 | -3.291438 | 0.19175 | -0.13167 | -0.218351 | 0.69101 | 0.077626 | -0.138636 | -0.718702 | -0.893057 |
| 35.75 | -0.53156 | 0.149794 | 0.144165 | 0.146081 | -1.65264 | 0.141276 | -0.09482 | 0.152359 | 1.436264 | 0.154922 | 0.130845 | -2.682841 | 0.706999 | 0.378291 | 0.260719 | -4.167701 | 0.048774 | -0.265578 | -0.342853 | 0.461641 | 0.536209 | -0.152217 | 2.012522 | -0.31331 |
| 35.76389 | 1.754003 | 0.11913 | 0.096126 | 0.095424 | -1.295096 | 0.097706 | -0.281505 | 0.116713 | 1.121069 | 0.101967 | 0.023764 | -2.667643 | 0.613297 | 0.189208 | 0.406089 | -3.473385 | -0.104265 | 0.005881 | 0.31001 | 0.006609 | 0.770753 | -0.174605 | 2.78594 | 0.263847 |
| 35.77778 | 0.676546 | 0.10884 | 0.07413 | 0.074535 | -0.636899 | 0.075982 | -0.417282 | 0.093084 | 0.674031 | 0.083433 | -0.01133 | -3.00199 | 1.695117 | 0.141632 | -0.719879 | -1.98987 | -0.092547 | 0.422325 | 0.933651 | 0.0581 | 0.777357 | -0.018781 | 0.313797 | 0.545479 |
| 35.79167 | 0.216648 | 0.111566 | 0.081857 | 0.083526 | -0.252501 | 0.079605 | -0.376655 | 0.082981 | 0.339815 | 0.08794 | 0.017232 | -1.228834 | 2.545953 | 0.677648 | -1.332808 | -0.127858 | -0.053074 | 0.828689 | 0.365277 | -0.38212 | 1.289811 | 0.269174 | 0.394397 | -0.294041 |
| 35.80556 | 1.421226 | 0.093006 | 0.083198 | 0.077294 | -1.11743 | 0.076498 | -0.555278 | 0.06419 | 0.135758 | 0.079722 | 0.040917 | 0.240755 | 1.222163 | 0.607801 | 0.063283 | 1.22152 | 0.019671 | 1.800623 | 0.108138 | -0.789061 | 2.782713 | 0.408429 | 0.145384 | -1.474784 |
| 35.81944 | 2.773309 | 0.072824 | 0.067084 | 0.055645 | -2.367847 | 0.060104 | -0.780133 | 0.040782 | 0.015724 | 0.065091 | 0.037786 | -1.666678 | -0.832 | 0.538802 | 0.250549 | 0.884813 | -0.528565 | 1.904391 | -0.566866 | -1.263338 | 2.853999 | -0.125105 | -0.388449 | -0.978513 |
| 35.83333 | 3.026045 | 0.087044 | 0.069435 | 0.061348 | -3.392542 | 0.064683 | -0.760665 | 0.053944 | -0.051746 | 0.074231 | 0.044947 | -1.600899 | -1.228621 | 0.3171 | 0.097888 | -0.660384 | -0.365475 | 1.145699 | -0.574662 | -0.360206 | 0.857631 | -0.763394 | -1.196589 | -1.709875 |
| 35.84722 | 2.044215 | 0.107675 | 0.101837 | 0.089643 | -3.772069 | 0.086885 | -0.609414 | 0.081019 | -0.124859 | 0.092688 | 0.08643 | 0.028079 | -0.21938 | -0.043192 | -0.069875 | -2.793678 | 1.35798 | 0.23807 | -0.44954 | 0.572067 | -0.268732 | -1.15637 | -0.852536 | -1.306335 |
| 35.86111 | 1.601347 | 0.080524 | 0.10004 | 0.079386 | -3.920609 | 0.075959 | -0.844508 | 0.084095 | -0.271191 | 0.071729 | 0.122712 | 0.77626 | -0.530619 | -0.378377 | 0.286818 | -5.613451 | 2.177634 | -1.051841 | -0.186354 | 0.641565 | -0.840894 | -1.17506 | 0.341488 | -0.299327 |
| 35.875 | 1.111793 | 0.048303 | 0.0647 | 0.061786 | -3.979411 | 0.060375 | -1.244863 | 0.073911 | -0.408856 | 0.055665 | 0.201407 | 1.31512 | -0.995004 | -0.411203 | 0.562625 | -5.856509 | 1.260109 | -1.799847 | 0.274353 | 0.52729 | -1.538223 | -0.858289 | -0.234636 | 0.194265 |
| 35.88889 | 0.846171 | 0.089488 | 0.093762 | 0.11306 | -4.221666 | 0.103053 | -1.518579 | 0.109214 | -0.490801 | 0.107115 | 0.144297 | 1.568115 | -1.429006 | -0.213513 | -0.519475 | -4.413878 | 0.010195 | -2.129001 | 0.13069 | 0.132962 | -0.79293 | -0.420125 | -0.202646 | 1.844858 |
| 35.90278 | 0.736784 | 0.168874 | 0.162815 | 0.19416 | -4.294747 | 0.17311 | -1.639032 | 0.166464 | -0.545418 | 0.175662 | -0.451699 | 1.529316 | -0.875813 | 0.134925 | -0.103563 | -2.601391 | -0.072841 | -2.645174 | -0.021467 | -0.685271 | -0.01306 | -0.037948 | -0.365102 | 0.615937 |
| 35.91667 | -0.860765 | 0.204681 | 0.210723 | 0.229116 | -4.196658 | 0.219348 | -1.741085 | 0.207117 | -0.610411 | 0.204533 | -1.272457 | 1.692591 | 0.689402 | 0.483201 | -1.014036 | -1.329306 | -0.023531 | -1.831791 | -0.009327 | -1.088072 | 0.579424 | -0.214751 | -0.814544 | -0.806974 |
| 35.93056 | -0.54391 | 0.192259 | 0.23289 | 0.229485 | -3.590469 | 0.239376 | -2.093404 | 0.219202 | -0.640741 | 0.207941 | -1.561465 | 2.093979 | 1.586548 | 0.438777 | -1.859316 | -1.199093 | 0.064198 | -0.489282 | 0.103805 | -1.398892 | 0.593887 | -0.305865 | -0.126505 | 0.50057 |
| 35.94444 | -0.361227 | 0.179907 | 0.246445 | 0.232839 | -2.307838 | 0.240422 | -2.254685 | 0.216561 | -0.638924 | 0.214176 | -0.822764 | 1.644359 | 1.605522 | 0.736136 | -1.288277 | -1.807179 | 0.106411 | 0.396512 | -0.171771 | -0.589504 | -0.156212 | -0.457875 | -1.11061 | -0.449663 |
| 35.95833 | -0.951681 | 0.179125 | 0.239796 | 0.230365 | -1.270772 | 0.229767 | -2.272087 | 0.201216 | -0.694684 | 0.209584 | -0.055777 | 0.262054 | -0.970656 | 0.271106 | -1.611277 | -2.459149 | 0.017148 | 1.342127 | -0.432892 | 0.166509 | -1.607256 | -0.453582 | -0.493014 | -1.51282 |
| 35.97222 | -1.228873 | 0.197489 | 0.22958 | 0.226709 | -0.699475 | 0.230432 | -2.282122 | 0.183915 | -0.75198 | 0.207071 | 0.33532 | -0.569962 | -3.984392 | -0.275477 | -3.302169 | -3.086777 | -0.551796 | 1.014832 | -0.932021 | 0.92992 | -0.780805 | -0.431366 | 0.673842 | -2.149749 |
| 35.98611 | -1.713707 | 0.215653 | 0.234657 | 0.232413 | -0.530327 | 0.228297 | -1.922467 | 0.158552 | -0.7418 | 0.217527 | 0.555985 | -0.574773 | -3.533424 | -0.239685 | -3.726248 | -3.368475 | -2.360418 | -0.134562 | -0.759978 | 1.152248 | 1.164752 | -0.122303 | 2.743355 | -2.623453 |
| 36 | -1.848691 | 0.258589 | 0.301821 | 0.322184 | -0.286174 | 0.228074 | -1.35225 | 0.196128 | -0.662078 | 0.279363 | 0.30785 | -0.818604 | -1.556976 | -1.268485 | -3.292283 | -1.929874 | -0.856353 | -1.404446 | -0.386222 | 0.928691 | 1.719478 | -0.628029 | 2.606471 | -1.888855 |
| 36.01389 | -1.398619 | 0.389358 | 0.455333 | 0.567902 | 0.45869 | 0.273717 | -0.835133 | 0.304163 | -0.453191 | 0.430575 | 0.348897 | -0.812076 | 0.053741 | -0.052334 | -2.941562 | -0.240087 | 2.266077 | -1.669649 | 0.195637 | 0.308314 | 2.005387 | -1.815694 | 0.652217 | -2.567567 |
| 36.02778 | -1.287334 | 0.463241 | 0.419394 | 0.562205 | 0.514758 | 0.278809 | -0.505139 | 0.323119 | -0.367849 | 0.408393 | 0.241085 | -1.078269 | -0.712853 | 1.438441 | -3.096849 | 0.173266 | 2.78648 | -1.662496 | 0.720889 | -0.494096 | 1.265041 | -2.965892 | -1.746798 | -0.142691 |
| 36.04167 | -1.32508 | 0.401067 | 0.070597 | 0.066143 | -0.20318 | 0.173062 | -0.812799 | 0.210509 | -0.367277 | 0.10715 | -0.262002 | -1.852874 | -4.471701 | -0.352008 | -3.357225 | -0.833347 | 1.2496 | -2.128121 | 0.892103 | -0.941514 | -1.062608 | -3.044244 | -3.721878 | -0.436165 |
| 36.05556 | -1.318157 | 0.450476 | 0.047969 | 0.048073 | -0.618573 | 0.311312 | -1.620889 | 0.13624 | -0.009376 | 0.156027 | -0.218374 | -2.87432 | -6.337817 | -2.111999 | -5.046003 | -2.102916 | 0.888495 | -3.478261 | 0.492734 | -0.939945 | -1.428935 | -2.548205 | -5.912127 | -3.847556 |
| 36.06944 | -1.017058 | 0.604819 | 0.414681 | 0.439776 | -0.773289 | 0.598818 | -1.645922 | 0.315027 | 0.620868 | 0.509371 | 0.462161 | -4.176251 | -4.705255 | -4.687397 | -4.840582 | -3.939836 | -0.316988 | -4.725105 | -0.225694 | -0.597478 | -0.897692 | -2.079296 | -5.872411 | -6.768071 |
| 36.08333 | -0.530929 | 0.608466 | 0.573138 | 0.584496 | -0.144497 | 0.648062 | -1.127359 | 0.590299 | 0.869173 | 0.612883 | 0.975036 | -2.874227 | -5.145726 | -5.684579 | -3.169247 | -4.818922 | -1.35778 | -3.973544 | -0.290885 | -1.474784 | -0.340966 | -1.023591 | -3.659337 | -7.085525 |
| 36.09722 | -1.066576 | 0.463043 | 0.450428 | 0.449464 | 1.193089 | 0.49001 | -0.274868 | 0.476236 | 0.698259 | 0.460225 | 0.454083 | -1.295118 | -3.37921 | -4.449422 | -2.982898 | -3.438316 | -3.381585 | -2.731584 | -0.526609 | -1.733139 | 0.009935 | 0.040867 | -1.857286 | -4.978831 |
| 36.11111 | -1.174067 | 0.273271 | 0.277928 | 0.281363 | 2.602855 | 0.300455 | 0.098205 | 0.306912 | 0.693072 | 0.279048 | 0.213718 | 0.09874 | -1.745407 | -2.981253 | -2.486253 | -1.483409 | -2.701192 | -1.383254 | -0.369892 | -1.157029 | 0.026509 | 0.892334 | -0.791938 | -3.174078 |
| 36.125 | -0.76155 | 0.11107 | 0.135337 | 0.143584 | 2.70697 | 0.148708 | 0.301061 | 0.047221 | 0.748394 | 0.132768 | 0.128639 | -0.081598 | -1.582032 | -2.867608 | -1.757927 | 0.073845 | -0.68204 | -1.022216 | -0.199982 | -0.762171 | 0.179492 | 1.26917 | 0.009499 | -2.263955 |
| 36.13889 | 0.053478 | 0.033762 | 0.05194 | 0.053033 | 1.797444 | 0.07282 | 0.563253 | -0.115597 | 0.517852 | 0.043293 | 0.070896 | -1.433631 | -0.783484 | -2.284063 | -0.806092 | 1.724898 | 1.389265 | -0.591541 | -0.086536 | -0.294032 | 0.515633 | 0.891315 | 0.618146 | -1.541297 |
| 36.15278 | 0.070787 | 0.000079 | 0.017947 | 0.012985 | 1.048625 | 0.033807 | 0.845498 | -0.126097 | 0.126613 | 0.013847 | 0.046414 | -0.745238 | 1.333874 | -1.560242 | 0.365101 | 3.055306 | 2.688488 | 0.093864 | -0.031677 | 0.28152 | 0.432422 | 0.420952 | 0.848104 | -0.633417 |
| 36.16667 | -0.774848 | -0.055347 | -0.022872 | -0.022429 | 0.45828 | -0.018625 | 0.855779 | -0.155139 | -0.173809 | -0.018569 | 0.042891 | -0.584692 | 4.721177 | -0.981021 | 0.818957 | 3.537837 | 2.157615 | 1.244898 | -0.099858 | 1.041692 | 0.205252 | -0.427656 | 1.178785 | 0.113088 |
| 36.18056 | -0.635642 | -0.103383 | -0.068458 | -0.058434 | -0.251426 | -0.067961 | 0.508631 | -0.115451 | -0.360286 | -0.074398 | 0.129512 | -0.696443 | 4.685422 | -0.423134 | 1.883491 | 3.535959 | 1.042606 | 3.136066 | -0.276147 | 1.576701 | -0.01629 | -1.53319 | 1.768974 | 0.852078 |
| 36.19444 | -1.158525 | -0.101645 | -0.078766 | -0.073574 | -1.091193 | -0.076548 | 0.235203 | -0.077435 | -0.48678 | -0.100147 | 0.389358 | -1.595982 | 2.955434 | -0.015468 | 2.491648 | 3.146268 | 0.380455 | 3.42301 | -0.224086 | 1.443127 | -0.174387 | -2.613574 | 2.394009 | 1.272173 |
| 36.20833 | -1.223169 | -0.060539 | -0.057065 | -0.07441 | -1.737163 | -0.047497 | 0.611261 | -0.080168 | -0.579831 | -0.075659 | 0.47456 | -1.232311 | 0.687554 | 0.279605 | 1.784196 | 2.156191 | -0.253359 | 2.71838 | 0.287525 | 1.067398 | 0.150201 | -1.728785 | 1.852575 | 1.167478 |
| 36.22222 | -1.027124 | -0.018669 | -0.028789 | -0.068139 | -1.435422 | -0.009651 | 1.210414 | -0.067611 | -0.633031 | -0.045397 | 0.325659 | -0.42316 | -1.224105 | 0.010047 | 0.646705 | 0.775414 | -0.930165 | 1.379851 | 0.791229 | 0.510212 | 0.769222 | -0.313156 | 0.923897 | 0.869184 |
| 36.23611 | -0.505537 | -0.000336 | -0.014557 | -0.053539 | -0.664177 | 0.009614 | 1.81418 | -0.025226 | -0.61615 | -0.027466 | 0.180804 | -0.319499 | -2.487835 | 0.21032 | -0.710258 | -0.683521 | 0.449527 | -1.259873 | 0.322858 | -0.043092 | 0.249584 | 0.616321 | 0.051978 | 0.371802 |
| 36.25 | -0.473372 | -0.037832 | -0.028541 | -0.03911 | -0.009324 | -0.015027 | 1.569014 | -0.064363 | -0.545547 | -0.013794 | 0.051804 | -0.481099 | -3.502969 | 0.20948 | -0.927309 | -1.475596 | 1.333315 | -1.566479 | -0.773434 | -0.251149 | 0.287046 | 1.119556 | -0.491664 | 0.133272 |
| 36.26389 | -0.323097 | -0.06442 | -0.048962 | -0.050113 | 0.046061 | -0.055356 | 0.762597 | -0.116992 | -0.439346 | -0.021801 | -0.034274 | -0.076124 | -0.835969 | 0.167792 | 0.16305 | -0.63524 | 0.633454 | -0.307809 | -1.040901 | -0.255216 | 0.423548 | 1.187155 | -0.493566 | 0.402463 |
| 36.27778 | -0.62931 | -0.021953 | -0.040674 | -0.06564 | -0.317272 | -0.052729 | 0.175283 | -0.102681 | -0.388656 | -0.04069 | -0.045346 | 0.343418 | 1.893121 | 0.050057 | 0.527889 | 1.140266 | 0.172017 | 0.554776 | -0.961193 | 0.036311 | -0.198282 | 1.904767 | 0.158149 | 0.942553 |
| 36.29167 | -3.305133 | -0.017628 | -0.040041 | -0.061536 | -0.116147 | -0.029176 | -0.039667 | -0.099247 | -0.542673 | -0.050746 | -0.018299 | 0.647803 | 4.512067 | 0.278497 | 1.024621 | 2.567505 | -0.042413 | 1.046899 | 0.030086 | 0.796958 | -0.316 | 2.221357 | 1.047028 | 1.825006 |
| 36.30556 | -3.25296 | -0.053366 | -0.07132 | -0.078668 | 0.6445 | -0.026726 | 0.268739 | -0.116064 | -0.752444 | -0.076398 | -0.0059 | 0.974235 | 4.524045 | 0.670719 | 1.578489 | 2.602274 | -0.118935 | 1.537521 | 0.762635 | 1.874741 | 0.285801 | 1.05573 | 1.638003 | 2.459154 |
| 36.31944 | 0.253228 | -0.043883 | -0.083669 | -0.104587 | 1.286968 | -0.028969 | 0.973049 | -0.109361 | -0.776754 | -0.094145 | 0.011067 | 1.215724 | 2.490101 | 1.524868 | 2.102246 | 1.931022 | -0.171236 | 1.930704 | 0.605586 | 1.784026 | 0.827905 | 0.609208 | 1.707955 | 3.37671 |
| 36.33333 | 2.216558 | -0.047623 | -0.073571 | -0.098527 | 1.744284 | -0.046098 | 1.381855 | -0.116106 | -0.638628 | -0.077926 | 0.043944 | 1.215345 | 1.846344 | 1.636385 | 2.893658 | 1.116618 | -0.232671 | 2.315536 | 0.773199 | 0.989419 | 0.473704 | 1.453265 | 1.335784 | 3.385839 |
| 36.34722 | 2.671571 | -0.079382 | -0.07767 | -0.088759 | 2.128685 | -0.08075 | 2.210078 | -0.178758 | -0.510539 | -0.064714 | 0.020374 | 1.016351 | 1.719315 | 1.083427 | 2.770436 | 0.452914 | -0.499774 | 2.567462 | 0.671426 | 1.018519 | 0.285839 | 1.077441 | 1.427311 | 2.57232 |
| 36.36111 | 2.259988 | -0.108076 | -0.086057 | -0.084313 | 2.457539 | -0.099827 | 2.380786 | -0.248759 | -0.360329 | -0.057613 | -0.031215 | 0.802337 | 1.323413 | 0.783682 | 1.809992 | 0.393525 | -0.732696 | 2.250277 | 0.098039 | 1.087314 | 0.290245 | 0.370241 | 1.763384 | 2.119115 |
| 36.375 | 1.167238 | -0.094827 | -0.057961 | -0.04949 | 2.606299 | -0.070532 | 1.315479 | -0.206657 | -0.055942 | -0.029139 | -0.028656 | 0.808319 | 0.739173 | 0.755488 | 0.986535 | 1.141313 | -0.372093 | 1.345027 | -0.166002 | 0.989166 | 0.315964 | 0.530503 | 2.000906 | 2.097315 |
| 36.38889 | 2.389709 | 0.002417 | 0.00911 | 0.008396 | 2.662945 | -0.005724 | 0.564481 | -0.065084 | 0.398333 | 0.015464 | 0.015014 | 0.939398 | 0.008634 | 0.756545 | 1.440443 | 2.341223 | 0.35515 | 0.834357 | -0.127402 | 0.892438 | 0.40504 | 0.873875 | 2.357577 | 2.475389 |
| 36.40278 | 2.604028 | 0.080226 | 0.060917 | 0.052021 | 2.702979 | 0.041928 | 0.285018 | 0.050115 | 0.89217 | 0.054536 | 0.055044 | 1.096294 | -0.678926 | 0.887984 | 2.299085 | 3.25574 | 0.345954 | 1.225703 | 0.261994 | 1.001389 | 0.488019 | 1.392745 | 2.638393 | 2.813124 |
| 36.41667 | 1.928281 | 0.06747 | 0.06969 | 0.065251 | 2.929975 | 0.082023 | 0.646991 | 0.089166 | 1.363275 | 0.064594 | 0.062059 | 1.206647 | -0.465174 | 0.927595 | 3.536116 | 3.387809 | -0.150927 | 0.900548 | 0.316016 | 1.32682 | 0.664475 | 2.002487 | 2.237251 | 3.04997 |
| 36.43056 | 1.394331 | 0.014434 | 0.022658 | 0.024632 | 3.174367 | 0.089318 | 0.603872 | 0.041074 | 1.722207 | 0.020322 | 0.031381 | 1.364451 | -0.054212 | 1.141322 | 4.90372 | 3.202491 | -0.480413 | 0.427599 | -0.04559 | 1.610961 | 0.699141 | 2.575299 | 2.114303 | 3.39032 |
| 36.44444 | -0.434254 | -0.1666 | -0.185364 | -0.145729 | 3.654395 | -0.047621 | 0.966146 | -0.122683 | 1.624329 | -0.155633 | -0.087196 | 1.558554 | -0.27969 | 1.23962 | 4.532996 | 3.165089 | -0.892696 | 0.203548 | -0.716729 | 1.088383 | 0.615202 | 2.575507 | 2.346604 | 3.603211 |
| 36.45833 | -0.93304 | -0.441344 | -0.480436 | -0.575847 | 4.576448 | -0.283877 | 1.996317 | -0.421302 | 1.193374 | -0.421075 | -0.258794 | 1.8931 | -0.484636 | 1.603826 | 3.479023 | 3.28708 | -0.4381 | 0.233654 | -1.27422 | 0.161783 | 0.616362 | 1.972682 | 2.433455 | 3.385297 |
| 36.47222 | 1.378157 | -0.415797 | -0.43323 | -0.653221 | 5.7059 | -0.291184 | 3.408484 | -0.48074 | 0.817354 | -0.418949 | -0.332151 | 2.100723 | -0.192939 | 3.048759 | 2.67205 | 3.556157 | -0.779042 | 1.031407 | -0.683849 | -0.115797 | 0.476823 | 0.762396 | 2.923122 | 2.832714 |
| 36.48611 | 3.733414 | -0.211467 | -0.229015 | -0.296788 | 6.083457 | -0.135053 | 3.113537 | -0.314639 | 0.732567 | -0.243084 | -0.216788 | 2.531763 | 0.784177 | 3.451671 | 2.685187 | 4.869675 | -1.133862 | 2.249659 | 0.001448 | 0.273615 | 0.471872 | -0.210174 | 3.567927 | 3.348366 |
| 36.5 | 5.087535 | -0.077451 | -0.123888 | -0.173819 | 4.702341 | -0.078294 | 1.818458 | -0.196885 | 0.777416 | -0.16386 | -0.117692 | 2.672672 | 1.837732 | 4.036105 | 3.377547 | 6.205585 | -0.250601 | 3.753409 | 0.477779 | 0.732053 | 0.553854 | -0.633891 | 3.633451 | 4.135522 |
| 36.51389 | 4.848907 | -0.178981 | -0.164247 | -0.202517 | 1.780952 | -0.183219 | 1.629879 | -0.270074 | 0.703481 | -0.222354 | -0.12252 | 2.305494 | 2.188067 | 3.460969 | 4.750332 | 5.563907 | 0.602095 | 3.428476 | 0.791101 | 1.017805 | 0.469066 | 0.176811 | 2.744509 | 5.32362 |
| 36.52778 | 3.056532 | -0.729265 | -0.459807 | -0.485966 | 0.673582 | -0.676739 | 0.911844 | -0.712773 | 0.222635 | -0.605033 | -0.501903 | 1.691083 | 1.971041 | 2.548805 | 4.245465 | 4.275738 | 1.225535 | 1.945422 | 0.788163 | 0.679623 | 0.523818 | 1.98899 | 2.548641 | 4.77085 |
| 36.54167 | 0.63188 | -1.670809 | -1.068037 | -1.093377 | 1.204851 | -1.545935 | -0.68108 | -1.550027 | -0.564498 | -1.352706 | -1.162148 | 0.617552 | 1.841093 | 0.00286 | 3.401883 | 2.444359 | 1.036146 | 0.094316 | 0.40105 | 0.18615 | 0.78335 | 2.566114 | 2.236394 | 3.98682 |
| 36.55556 | -0.424674 | -2.068865 | -1.545428 | -1.55029 | 0.429194 | -1.970783 | -1.651337 | -1.86029 | -1.028677 | -1.692971 | -1.572482 | -0.073904 | 1.351962 | 0.160016 | 1.195536 | 0.988157 | 0.359157 | -0.627319 | 0.181386 | 0.070876 | 0.585007 | 1.441667 | 1.685281 | 3.866096 |
| 36.56944 | -0.142414 | -1.481145 | -1.363031 | -1.309665 | -0.562672 | -1.43447 | -1.306281 | -1.360299 | -0.817973 | -1.24632 | -1.293287 | 0.193557 | 0.314747 | -0.006171 | 0.610608 | 0.573004 | -0.0544 | -0.398943 | 0.647298 | 0.32926 | 0.740658 | 0.608873 | 1.616562 | 2.70167 |
| 36.58333 | 0.64062 | -0.706116 | -0.785536 | -0.725125 | -0.996562 | -0.718547 | -0.535755 | -0.687356 | -0.396387 | -0.623627 | -0.715779 | 1.004893 | 0.118652 | 1.077021 | 1.838137 | 1.53761 | -0.195623 | -0.40155 | 1.193869 | 0.791376 | 1.748214 | 0.459122 | 1.018206 | 3.352996 |
| 36.59722 | 1.369932 | -0.275259 | -0.339043 | -0.308621 | -0.913641 | -0.287206 | 0.2413 | -0.291504 | -0.129183 | -0.252791 | -0.296604 | 1.514852 | 0.237831 | 2.448124 | 1.196055 | 2.936638 | -0.862772 | 0.924784 | 1.564067 | 0.99103 | 1.622635 | 0.455559 | 1.852026 | 5.30485 |
| 36.61111 | 0.976972 | -0.072334 | -0.082238 | -0.069396 | -0.502227 | -0.061051 | 0.744808 | -0.093128 | -0.024496 | -0.062712 | -0.059285 | 1.211995 | 0.348709 | 1.120564 | 1.180309 | 2.619623 | 0.86293 | 1.706608 | 1.655108 | 0.884595 | 1.027898 | 0.592473 | 0.418747 | 1.566574 |
| 36.625 | -1.726036 | 0.052963 | 0.086978 | 0.090011 | -0.032491 | 0.068396 | 0.774536 | 0.021165 | 0.028125 | 0.06182 | 0.082479 | 0.095773 | 1.108706 | 0.926921 | -0.14526 | 1.163187 | 0.22188 | 1.217254 | 1.656619 | 0.681492 | 1.809643 | 0.215751 | 0.19749 | -1.968833 |
| 36.63889 | -2.900432 | 0.134724 | 0.159828 | 0.163431 | 0.169351 | 0.133133 | 0.441077 | 0.053237 | 0.034335 | 0.140342 | 0.123097 | -0.351721 | 1.742876 | 0.286698 | -0.507694 | -0.88893 | 0.070037 | 0.305269 | 3.262066 | 0.388697 | 0.593054 | 0.32189 | -0.006718 | -3.779881 |
| 36.65278 | -2.053551 | 0.246773 | 0.209246 | 0.222029 | 0.369261 | 0.212117 | 0.305667 | 0.103272 | 0.079132 | 0.217496 | 0.150861 | 0.14757 | 2.053565 | -1.884254 | -0.424165 | -1.562433 | 0.162052 | 0.019757 | 2.834296 | 0.274373 | -0.641851 | 0.534635 | 0.910418 | -2.023649 |
| 36.66667 | -0.204626 | 0.395798 | 0.304191 | 0.310342 | 0.35622 | 0.327529 | -0.060624 | 0.259322 | 0.163755 | 0.316877 | 0.257049 | 1.306823 | 1.641573 | -0.21754 | 0.224679 | -1.438534 | 0.065583 | 0.464218 | 2.026066 | -0.056176 | -2.717397 | -0.052828 | -0.123281 | -0.179039 |
| 36.68056 | 1.318602 | 0.449255 | 0.350114 | 0.345162 | -0.165402 | 0.367632 | -0.377536 | 0.391294 | 0.173337 | 0.35519 | 0.329485 | 2.028886 | -0.059254 | 2.422613 | 1.437202 | -1.717693 | 0.053206 | -0.016916 | 0.956415 | -0.497326 | -1.650146 | 0.097782 | -0.89745 | 2.501806 |
| 36.69444 | 1.860544 | 0.364094 | 0.304118 | 0.30085 | -0.583892 | 0.309362 | -0.368993 | 0.35711 | 0.081868 | 0.304907 | 0.305458 | 1.719542 | -2.750159 | 1.068213 | 2.341188 | 0.012958 | 0.040624 | 0.548942 | 1.157219 | -0.248125 | 0.755708 | 0.436231 | -0.241912 | 2.846215 |
| 36.70833 | 2.258919 | 0.244822 | 0.245215 | 0.244784 | -0.414042 | 0.243093 | -0.617057 | 0.278726 | -0.047321 | 0.241706 | 0.258855 | 1.369594 | -1.570363 | 0.415916 | 1.101278 | 1.792257 | -0.211018 | 1.399613 | 0.96737 | -0.017471 | 1.44937 | 1.015529 | -0.542343 | 1.002447 |
| 36.72222 | 1.416335 | 0.150124 | 0.228491 | 0.226646 | 0.296358 | 0.23199 | -0.600363 | 0.237417 | -0.15342 | 0.218951 | 0.250475 | 0.369658 | 0.61398 | 0.508941 | 0.196917 | 2.574977 | -0.263103 | 1.636503 | -0.075098 | 0.356208 | 0.186569 | 0.896011 | -0.738229 | -0.204812 |
| 36.73611 | -0.102621 | 0.087948 | 0.257804 | 0.255637 | 0.771126 | 0.266877 | -0.103556 | 0.238576 | -0.200097 | 0.241002 | 0.29134 | -2.465812 | 2.764038 | 0.540927 | -1.636192 | 2.512627 | -0.151565 | 0.317513 | -0.393345 | -0.053079 | -0.351903 | 0.635115 | -0.290264 | 0.348277 |
| 36.75 | -1.096566 | 0.084116 | 0.28801 | 0.282747 | 0.36636 | 0.297297 | 0.35267 | 0.260736 | -0.149725 | 0.266918 | 0.318135 | -3.902576 | 2.244679 | 0.444938 | -0.85563 | 1.053031 | 0.180862 | -0.049299 | -0.898503 | -1.128412 | 0.830838 | 0.468459 | 0.419439 | 0.411774 |
| 36.76389 | -0.369323 | 0.1563 | 0.267899 | 0.262138 | -0.49424 | 0.276678 | 0.34056 | 0.266121 | -0.158376 | 0.251093 | 0.232634 | -3.60483 | 2.006575 | 0.160572 | -0.470082 | -0.543717 | 0.254702 | 0.076166 | -0.939895 | -0.918291 | 1.469753 | -0.192397 | 0.639604 | 0.010851 |
| 36.77778 | -0.067625 | 0.269769 | 0.235575 | 0.236525 | -2.629358 | 0.239354 | 0.06427 | 0.202714 | -0.185354 | 0.22659 | 0.159983 | -2.179129 | 1.797841 | -0.180258 | 0.660153 | -1.648236 | 0.113743 | 0.371997 | -0.12009 | 0.241299 | 0.207085 | -0.258393 | -0.218798 | 0.446873 |
| 36.79167 | -0.01469 | 0.352342 | 0.226241 | 0.229745 | -2.559065 | 0.22428 | 0.301011 | 0.170578 | -0.222654 | 0.220224 | 0.151988 | -0.703113 | 1.676413 | -0.41594 | 1.669802 | -2.181055 | 0.136478 | -0.128295 | -0.34902 | 0.546903 | -0.110898 | -0.320982 | -0.402313 | 0.147155 |
| 36.80556 | 0.438841 | 0.352989 | 0.223597 | 0.229444 | -1.974531 | 0.225177 | -0.011799 | 0.159383 | -0.249908 | 0.218992 | 0.162157 | 0.200903 | 1.542242 | -0.548713 | -0.229434 | -2.516617 | 0.146996 | -0.34213 | -0.340751 | 0.40792 | 1.514255 | 0.091834 | -0.158446 | 0.555248 |
| 36.81944 | 0.714416 | 0.319223 | 0.232607 | 0.247396 | -1.271961 | 0.237931 | -0.648259 | 0.175782 | -0.239586 | 0.2285 | 0.189929 | 0.11375 | 1.35272 | -0.853971 | -1.429824 | -2.39192 | -0.009343 | -1.107558 | 0.018272 | 0.008245 | 2.772023 | 0.101508 | -0.05913 | 0.672807 |
| 36.83333 | -0.210832 | 0.282824 | 0.239594 | 0.247866 | -0.319684 | 0.236596 | -0.938471 | 0.207946 | -0.245777 | 0.222126 | 0.216813 | -0.208051 | 0.746656 | -0.5195 | -1.261997 | -1.56718 | -0.056212 | -1.164334 | 0.445888 | -0.252213 | 3.308684 | 0.038975 | -0.797559 | -0.676749 |
| 36.84722 | -1.025263 | 0.221249 | 0.227465 | 0.20804 | 0.962069 | 0.195614 | -1.2162 | 0.234585 | -0.283779 | 0.177854 | 0.207695 | -0.439889 | 0.610201 | 0.076948 | -1.968082 | -1.101112 | -0.053263 | -0.438057 | 0.912628 | -0.651342 | 4.053688 | -0.069684 | -1.134984 | -1.643491 |
| 36.86111 | -1.879183 | 0.153015 | 0.199991 | 0.167807 | 1.482854 | 0.147152 | -1.420674 | 0.222186 | -0.323786 | 0.142057 | 0.182363 | -0.796335 | 0.854098 | 0.261228 | -1.186031 | -1.640961 | -0.210645 | -0.081753 | 1.283892 | -1.050917 | 4.834258 | -0.574259 | -0.061988 | -0.113653 |
| 36.875 | -2.289743 | 0.111462 | 0.177893 | 0.157422 | 0.400107 | 0.136373 | -1.951746 | 0.238221 | -0.35355 | 0.145872 | 0.165022 | -1.169105 | -1.360034 | 0.492413 | 0.161882 | -2.947278 | -0.501091 | -0.173515 | 0.597089 | -0.923381 | 2.769055 | -1.035319 | 0.784186 | 0.71032 |
| 36.88889 | -2.25531 | 0.119574 | 0.186493 | 0.178003 | -2.024584 | 0.163236 | -3.008427 | 0.24566 | -0.368701 | 0.170092 | 0.179679 | -1.227496 | -0.97219 | 0.444569 | -1.053676 | -4.0379 | 1.049674 | -0.549489 | -0.373034 | -0.814234 | 0.334948 | -0.785318 | -0.713754 | -0.540472 |
| 36.90278 | -1.731538 | 0.130409 | 0.188538 | 0.182763 | -3.480801 | 0.170381 | -3.691954 | 0.208912 | -0.388714 | 0.170133 | 0.287612 | -1.131991 | -1.585323 | -0.04205 | -1.591939 | -4.628935 | 2.740622 | -0.928174 | -0.815848 | -0.376458 | 0.234537 | 0.491958 | -0.536947 | -1.741667 |
| 36.91667 | -1.128214 | 0.097806 | 0.142227 | 0.131707 | -3.809391 | 0.126052 | -3.852008 | 0.120971 | -0.428342 | 0.125115 | 0.302083 | -1.056061 | 0.33373 | -0.794867 | -2.17336 | -6.324285 | 0.949263 | -0.761762 | -1.047201 | -0.072521 | -3.37888 | 0.690548 | 0.280536 | -1.388 |
| 36.93056 | -0.881131 | 0.038732 | 0.06634 | 0.052336 | -3.340019 | 0.056942 | -3.530398 | 0.02134 | -0.478743 | 0.061608 | -0.304733 | -0.998643 | 0.818857 | -1.59796 | -2.052679 | -5.381283 | 1.240893 | -0.198851 | -1.625403 | -0.113467 | -3.21119 | 0.418876 | -0.06102 | -0.280735 |
| 36.94444 | -0.65777 | -0.02663 | -0.014711 | -0.024658 | -2.969989 | -0.016467 | -3.310924 | -0.036362 | -0.524331 | -0.009483 | -0.362838 | -0.843682 | 0.511665 | -1.783337 | -1.603314 | -3.301881 | 0.275974 | 0.023949 | -1.437428 | -0.24559 | -1.676796 | 0.074655 | 0.475047 | 0.183467 |
| 36.95833 | -0.771547 | -0.080378 | -0.07688 | -0.082166 | -2.897326 | -0.074503 | -2.896356 | -0.085112 | -0.590381 | -0.075952 | -0.312952 | -0.709583 | -0.539838 | -2.105256 | -1.7111 | -1.74658 | 0.087987 | 0.883119 | -0.820124 | -0.141169 | -0.905624 | -0.883234 | 0.178003 | 0.086293 |
| [illegible] | [illegible] | [illegible] | [illegible] | [illegible] | [illegible] | [illegible] | [illegible] | [illegible] | [illegible] | [illegible] | [illegible] | [illegible] | [illegible] | [illegible] | [illegible] | [illegible] | [illegible] | [illegible] | [illegible] | [illegible] | [illegible] | [illegible] | [illegible] | [illegible] |
| [illegible] | [illegible] | [illegible] | [illegible] | [illegible] | [illegible] | [illegible] | [illegible] | [illegible] | [illegible] | [illegible] | [illegible] | [illegible] | [illegible] | [illegible] | [illegible] | [illegible] | [illegible] | [illegible] | [illegible] | [illegible] | [illegible] | [illegible] | [illegible] | [illegible] |
| 37 | -1.5357 | -0.029687 | -0.017113 | 0.015729 | -0.980412 | -0.056342 | -0.828189 | -0.026834 | -0.626557 | -0.016363 | -0.319717 | -0.358793 | -4.780784 | -2.488781 | -0.143738 | -0.999196 | -0.031659 | -1.020348 | 1.690774 | -1.123883 | 0.466599 | -2.42858 | -1.744509 | -2.245535 |
| 37.01389 | -1.335493 | 0.141645 | 0.189548 | 0.30744 | -0.482551 | 0.068866 | -0.283701 | 0.157393 | -0.339161 | 0.186323 | -0.230971 | -0.028711 | -6.075952 | -1.934716 | 0.005353 | -0.771477 | -0.220154 | -1.685701 | 1.803021 | -1.630978 | 1.005971 | -1.94769 | -2.422373 | -5.159107 |
| 37.02778 | -1.327265 | 0.258052 | 0.202497 | 0.340143 | -1.068806 | 0.115494 | -0.055934 | 0.218487 | -0.298719 | 0.1845 | -0.359176 | -0.092497 | -5.218797 | -1.780358 | -0.039838 | -1.949014 | -0.251415 | -1.336022 | 1.430276 | -1.609843 | 0.115605 | -2.146305 | -1.74903 | -3.108005 |
| 37.04167 | -1.539044 | 0.229627 | -0.1964 | -0.209889 | -2.086333 | -0.069824 | -0.433962 | 0.027737 | -0.627274 | -0.171976 | -0.780595 | -0.646894 | -4.199998 | -1.860573 | -1.939471 | -3.225458 | -0.702289 | -1.819175 | 0.84559 | -1.54502 | -0.351944 | -1.86532 | -1.77443 | -3.372443 |
| 37.05556 | -1.616294 | 0.121331 | -0.404266 | -0.45345 | -2.117301 | -0.116761 | -0.828604 | -0.121305 | -0.827972 | -0.339393 | -0.948643 | -0.862239 | -2.754672 | -2.741934 | -4.597948 | -4.798714 | -2.725852 | -2.781363 | 0.347158 | -1.677315 | -1.965262 | -1.592341 | -2.393729 | -5.349144 |
| 37.06944 | -1.172117 | 0.115592 | -0.188882 | -0.200268 | -1.101324 | 0.043836 | -0.725568 | -0.012178 | -0.795164 | -0.099079 | -0.89525 | -0.857681 | -2.413452 | -4.252614 | -5.414178 | -4.719797 | -1.904707 | -3.11502 | -0.77972 | -1.949656 | -3.60719 | -2.057566 | -2.628371 | -7.469214 |
| 37.08333 | -1.500649 | 0.156055 | 0.04728 | 0.052697 | -0.207001 | 0.152124 | -0.428246 | 0.120503 | -0.714649 | 0.113032 | -0.480543 | -0.43848 | -1.660694 | -5.013516 | -3.115615 | -4.278137 | -0.672603 | -3.444883 | -1.444358 | -1.705235 | -3.010772 | -2.340277 | -1.916382 | -7.263576 |
| 37.09722 | -2.586314 | 0.148841 | 0.090191 | 0.08927 | 0.338634 | 0.143773 | -0.362169 | 0.130924 | -0.611493 | 0.130903 | -0.013663 | -0.375469 | -0.381733 | -3.563268 | -1.052146 | -2.652084 | -1.717044 | -2.885737 | -0.957214 | -1.271518 | -1.88094 | -1.508921 | -3.229199 | -5.293331 |
| 37.11111 | -2.472942 | 0.102616 | 0.068583 | 0.065454 | 0.894815 | 0.101029 | -0.113481 | 0.090699 | -0.423431 | 0.097421 | -0.039178 | -0.61025 | -0.515427 | -2.697918 | -1.046382 | -0.819857 | -0.571984 | -2.171057 | -0.267966 | -0.809379 | -0.827716 | -0.349766 | -3.006144 | -3.519836 |
| 37.125 | -1.639106 | 0.086163 | 0.068149 | 0.06323 | 1.132066 | 0.089812 | 0.291197 | 0.011443 | -0.221554 | 0.080737 | -0.080593 | -0.503844 | -0.1118 | -2.313385 | -0.925171 | 0.393101 | 0.78317 | -2.297373 | 0.106609 | -0.555479 | 0.207065 | 0.171224 | -1.707151 | -2.02375 |
| 37.13889 | -0.822412 | 0.08046 | 0.058541 | 0.04404 | 1.141752 | 0.078831 | 0.678499 | -0.073867 | -0.104276 | 0.051197 | -0.117679 | -0.144044 | 0.652948 | -1.689305 | 0.020848 | 1.306969 | 2.168485 | -2.373898 | 0.004202 | -0.497522 | 1.326139 | 0.779588 | -0.803806 | -1.00018 |
| 37.15278 | -0.181582 | 0.036489 | 0.027521 | 0.011922 | 1.110206 | 0.047085 | 0.901942 | -0.119048 | -0.092862 | 0.023097 | -0.068979 | 0.155594 | 2.196765 | -1.101118 | 1.169223 | 2.045708 | 2.201831 | -0.896533 | -0.310271 | 0.096352 | 2.257862 | 1.264178 | -0.382482 | -0.275073 |
| 37.16667 | 0.379358 | -0.04145 | -0.009405 | -0.009755 | 0.701475 | 0.008147 | 0.953839 | -0.101761 | -0.123058 | 0.006282 | 0.077228 | 0.185496 | 3.125394 | -0.145854 | 1.957663 | 2.441171 | 1.249758 | 0.518207 | -0.46157 | 0.440355 | 2.152636 | 1.475912 | 0.293023 | 0.444126 |
| 37.18056 | 0.784528 | -0.111789 | -0.03922 | -0.017123 | 0.094662 | -0.026242 | 0.729429 | -0.102381 | -0.124912 | -0.015354 | 0.306495 | -0.13151 | 2.535205 | 0.825116 | 2.671604 | 1.510513 | 0.518112 | 2.337792 | -0.333791 | 0.63218 | 1.142669 | 1.693846 | 0.576783 | 0.87916 |
| 37.19444 | 1.20652 | -0.10885 | -0.032639 | -0.006138 | -0.350344 | -0.028097 | 0.596069 | -0.070879 | -0.092767 | -0.020567 | 0.712878 | -0.66497 | 1.657264 | 1.452719 | 2.224282 | 0.647683 | 0.102854 | 3.425761 | 0.031783 | 0.835983 | 0.786781 | 1.149074 | 1.249124 | 1.729915 |
| 37.20833 | 1.602861 | -0.020099 | 0.005267 | 0.008195 | -0.439613 | 0.013301 | 0.771352 | -0.024905 | -0.044743 | -0.001009 | 1.220919 | -0.843018 | 0.416834 | 1.445258 | 0.877065 | 1.03545 | -0.917334 | 3.09788 | 0.494839 | 1.333238 | 0.428267 | 0.317613 | 0.940938 | 2.836767 |
| 37.22222 | 1.789912 | 0.062751 | 0.048894 | 0.028725 | -0.050323 | 0.060278 | 1.270542 | 0.000928 | 0.005447 | 0.037937 | 0.945824 | -0.656874 | 0.601833 | 1.256761 | -0.193018 | 1.935371 | -1.468232 | 1.700296 | 0.467085 | 1.527777 | 0.189674 | 0.135591 | -0.855302 | 2.930546 |
| 37.23611 | 1.384762 | 0.097107 | 0.093518 | 0.068511 | 0.212919 | 0.081244 | 1.637316 | 0.012923 | 0.045784 | 0.08727 | 0.990051 | -0.52918 | -0.36994 | 0.576325 | -0.932235 | 3.306002 | -0.360588 | 0.153636 | -0.011023 | 1.210491 | -0.048872 | 0.707454 | -2.748875 | 2.214985 |
| 37.25 | 0.966175 | 0.121319 | 0.12952 | 0.103653 | 0.028591 | 0.096279 | 1.638219 | -0.002506 | 0.055019 | 0.119569 | 0.89716 | -0.521524 | -3.166966 | 0.094239 | -0.53757 | 2.244979 | 1.482708 | -1.174022 | -0.286129 | 0.944983 | -0.075159 | 1.619107 | -1.419541 | 1.273484 |
| 37.26389 | 0.79515 | 0.145512 | 0.14764 | 0.122684 | -0.46803 | 0.127271 | 1.315923 | 0.021112 | 0.052906 | 0.130852 | 0.597725 | -0.557438 | -4.159337 | 0.492513 | 0.711051 | -0.33759 | 2.006143 | -1.797095 | -0.18889 | 0.461687 | 0.163438 | 2.585925 | -0.643266 | 0.739005 |
| 37.27778 | 0.574501 | 0.148459 | 0.155442 | 0.139303 | -0.820374 | 0.154898 | 0.840555 | 0.037117 | 0.05387 | 0.138965 | 0.305697 | -0.289561 | -2.327758 | 0.281253 | 1.669256 | -2.253623 | 1.797417 | -0.739379 | 0.177441 | 0.30489 | 0.653987 | 2.997638 | 0.144846 | 1.081769 |
| 37.29167 | 0.308556 | 0.13189 | 0.158943 | 0.156979 | -0.834245 | 0.155111 | 0.519519 | 0.040166 | 0.055561 | 0.153646 | 0.160325 | -0.021263 | -0.087624 | 0.175186 | 1.659479 | -1.870536 | 1.65792 | 0.973282 | -0.089216 | 0.3721 | 1.157933 | 2.856646 | 0.656803 | 1.78182 |
| 37.30556 | 0.112328 | 0.116553 | 0.148052 | 0.151915 | -0.423837 | 0.13066 | 0.42491 | 0.034867 | 0.036651 | 0.153331 | 0.094348 | 0.318574 | 2.867785 | -0.045613 | 1.737604 | 1.215864 | -0.84296 | 2.00282 | -0.296761 | 0.647721 | 1.319488 | 1.874718 | 1.433643 | 2.226889 |
| 37.31944 | -0.062275 | 0.097372 | 0.103301 | 0.099855 | 0.261694 | 0.095773 | 0.569206 | 0.013912 | -0.046494 | 0.101636 | 0.070504 | 0.602755 | 2.78371 | -0.066696 | 2.266391 | 4.765608 | -1.197902 | 1.958629 | 0.366257 | 0.949292 | 0.833643 | 0.951956 | 1.846187 | 2.209221 |
| 37.33333 | -0.294593 | 0.08899 | 0.065751 | 0.054343 | 1.086164 | 0.076675 | 0.719635 | -0.00724 | -0.142833 | 0.049651 | 0.116769 | 0.749925 | 1.307327 | 0.10791 | 2.965015 | 5.321632 | 0.445356 | 1.661561 | 1.127923 | 1.103604 | 0.523431 | 0.229951 | 1.740215 | 2.646534 |
| 37.34722 | -0.334633 | 0.117916 | 0.074883 | 0.056984 | 1.749944 | 0.090473 | 0.821705 | 0.027425 | -0.186716 | 0.054492 | 0.189799 | 0.795522 | 1.196591 | -0.270192 | 3.198926 | 3.717511 | 2.888507 | 1.394106 | 1.639947 | 0.888279 | 0.299686 | -0.01028 | 1.508128 | 3.081603 |
| 37.36111 | -0.06762 | 0.128722 | 0.094283 | 0.073202 | 1.989965 | 0.100197 | 0.751112 | 0.05113 | -0.185947 | 0.088379 | 0.178585 | 0.83471 | 1.471432 | -0.469816 | 3.846466 | 1.016626 | 4.098207 | 1.023862 | 1.238218 | 0.728417 | 0.250242 | 0.143044 | 1.493264 | 3.167042 |
| 37.375 | 0.540906 | 0.111031 | 0.090042 | 0.070521 | 1.851673 | 0.095881 | 0.595128 | 0.059858 | -0.157079 | 0.100797 | 0.105722 | 0.893761 | 1.933839 | -0.15694 | 4.498874 | -0.617305 | 2.543735 | 1.047754 | 0.8143 | 0.590377 | 0.45975 | 0.881912 | 1.882269 | 3.015095 |
| 37.38889 | 1.119889 | 0.215766 | 0.076671 | 0.065754 | 1.77325 | 0.08562 | 0.64004 | 0.059396 | -0.137107 | 0.069634 | 0.050758 | 0.858581 | 2.330622 | 0.347815 | 3.790395 | -0.14246 | 0.784368 | 1.039614 | 0.61168 | 0.622869 | 0.843294 | 1.668643 | 2.408334 | 2.681432 |
| 37.40278 | 1.667747 | 0.824315 | 0.125661 | 0.211254 | 1.739783 | 0.058638 | 0.996282 | 0.0522 | -0.104601 | 0.039835 | 0.062507 | 0.7833 | 2.047823 | 1.174435 | 3.425014 | 1.602291 | -0.051751 | 1.280094 | 0.578317 | 0.868023 | 1.133048 | 1.67682 | 2.561644 | 2.415707 |
| 37.41667 | 2.117453 | 1.413111 | 0.195442 | 0.530339 | 1.68057 | 0.043237 | 1.403198 | 0.059647 | -0.072587 | 0.043819 | 0.096226 | 0.843967 | 1.479976 | 1.496504 | 2.96864 | 3.72802 | -2.067322 | 1.326433 | 0.424296 | 1.137065 | 1.276541 | 1.246469 | 2.393689 | 2.786135 |
| 37.43056 | 1.929414 | 0.020984 | 0.015116 | 0.00283 | 1.700208 | 0.019643 | 1.672167 | -0.003536 | -0.183377 | 0.017171 | -0.015828 | 1.147288 | -0.160927 | 1.044577 | 3.943805 | 5.847171 | -1.033801 | 0.792631 | 0.614501 | 1.178075 | 1.296292 | 0.193345 | 2.418901 | 2.132409 |
| 37.44444 | 1.337699 | -1.464356 | -0.262347 | -0.598846 | 2.305591 | -0.032561 | 1.636395 | -0.122049 | -0.301185 | -0.086541 | -0.14095 | 1.384205 | -1.68909 | 0.974947 | 3.454468 | 6.655868 | 2.96774 | 0.465086 | 0.925034 | 0.52562 | 0.786341 | -0.987422 | 2.317529 | 2.360611 |
| 37.45833 | 1.347999 | -0.923484 | -0.287655 | -0.573964 | 0.85626 | -0.099352 | 1.814611 | -0.250131 | -0.347464 | -0.189393 | -0.124395 | 1.815507 | -2.362364 | 1.639889 | 2.636072 | 5.368765 | 5.182955 | 0.349677 | 0.818455 | -0.055113 | 0.130854 | -1.410688 | 2.582089 | 3.638307 |
| 37.47222 | 1.736363 | -0.172654 | -0.066884 | -0.35779 | 2.959715 | -0.017564 | 2.7233 | -0.199162 | -0.285078 | -0.110626 | -0.043863 | 2.443305 | -1.716208 | 2.38432 | 3.22211 | 3.725881 | 5.662015 | 0.671969 | 0.810346 | -0.122333 | -0.092968 | -0.620399 | 3.607402 | 3.5176 |
| 37.48611 | 1.950977 | -0.002739 | 0.045454 | -0.052199 | 2.585313 | 0.072444 | 3.792332 | -0.080292 | -0.179116 | -0.012913 | 0.057799 | 3.370845 | 1.006422 | 1.986327 | 3.932832 | 3.080277 | 2.943423 | 1.003809 | 0.623658 | 0.219418 | 0.788821 | 1.138822 | 3.817836 | 3.738356 |
| 37.5 | 2.6592 | -0.009211 | 0.034062 | -0.023694 | 4.258028 | 0.045886 | 4.003245 | -0.036849 | -0.17781 | 0.002592 | 0.081277 | 3.763907 | 3.21561 | 2.293298 | 4.088002 | 3.119255 | 0.973524 | 1.018721 | 1.022176 | 0.466925 | 2.163336 | 2.883152 | 3.855323 | 4.483316 |
| 37.51389 | 2.449881 | -0.069592 | -0.002407 | -0.03943 | 2.525325 | -0.03232 | 4.124705 | -0.094099 | -0.219982 | -0.041473 | 0.063182 | 3.679673 | 3.976918 | 2.487778 | 3.789085 | 3.884889 | -0.196321 | 0.747026 | 1.002347 | 0.56078 | 3.392846 | 2.977251 | 3.517258 | 4.712651 |
| 37.52778 | 2.295667 | -0.618266 | -0.330781 | -0.348907 | -1.238824 | -0.546105 | 3.532676 | -0.645344 | -0.635412 | -0.481883 | -0.366187 | 3.235322 | 3.139659 | 1.903374 | 4.454459 | 4.468707 | 0.144677 | 0.469834 | 1.320707 | 0.257698 | 3.448955 | 3.026986 | 2.749275 | 4.972901 |
| 37.54167 | 1.020127 | -1.746547 | -1.123691 | -1.131547 | -0.432556 | -1.599824 | 2.244637 | -1.668982 | -1.536335 | -1.413895 | -1.201084 | 2.60816 | 2.303916 | 0.343207 | 4.893693 | 6.071334 | 2.064565 | 0.598659 | 2.262792 | 0.272513 | 1.989001 | 3.445509 | 2.357798 | 6.135737 |
| 37.55556 | -0.2195 | -2.125939 | -1.604796 | -1.576555 | 0.256003 | -1.977459 | 0.799613 | -1.997473 | -1.87642 | -1.749906 | -1.624832 | 1.426056 | 1.371765 | -0.391761 | 3.283077 | 6.323637 | 0.336625 | 0.679356 | 1.959789 | 0.26241 | 0.359039 | 2.395806 | 3.109451 | 5.692987 |
| 37.56944 | -0.562571 | -1.33209 | -1.206954 | -1.110798 | 1.375631 | -1.240121 | 0.502874 | -1.211562 | -1.260362 | -1.092315 | -1.150232 | 1.138955 | 1.33491 | -0.079778 | 1.17664 | 4.240471 | 0.044981 | 1.006744 | 1.964804 | -0.030012 | 0.059853 | 1.238844 | 3.955052 | 4.206649 |
| 37.58333 | -0.411899 | -0.552877 | -0.619612 | -0.542362 | 0.920228 | -0.515766 | 1.096912 | -0.517219 | -0.645797 | -0.473505 | -0.568066 | 1.818693 | 0.466359 | 0.896369 | -0.165549 | 2.700521 | -0.112157 | 2.010018 | 2.73868 | 0.239941 | 0.023235 | 0.834151 | 4.850969 | 2.596406 |
| 37.59722 | -0.222986 | -0.203837 | -0.264932 | -0.218238 | -0.081868 | -0.186439 | 1.066903 | -0.196121 | -0.337467 | -0.180586 | -0.226598 | 2.412575 | 0.041264 | 0.888021 | -0.966891 | 2.109804 | -0.641956 | 2.199255 | 2.147596 | 0.303837 | 0.520238 | 0.725969 | 3.588104 | 2.145216 |
| 37.61111 | -0.112543 | -0.060636 | -0.096673 | -0.07125 | -1.313445 | -0.051411 | 0.574112 | -0.081552 | -0.204141 | -0.062262 | -0.073381 | 2.297448 | 0.429757 | -0.075045 | -2.20242 | 0.204576 | 0.307508 | 2.201601 | 0.370452 | -0.087941 | 1.068294 | 0.515101 | -0.14959 | -0.475059 |
| 37.625 | 0.454719 | -0.004214 | -0.020899 | -0.013457 | 0.415618 | 0.002548 | 0.326912 | -0.059047 | -0.141318 | -0.013014 | -0.026783 | 1.30132 | 2.268201 | -0.383591 | -1.467376 | -0.958249 | 0.031418 | 0.58838 | -1.258824 | -0.572309 | 2.047337 | 0.28493 | -1.369151 | -1.341536 |
| 37.63889 | 0.383875 | 0.001285 | -0.012699 | -0.01248 | 0.890164 | -0.00173 | 0.107504 | -0.082606 | -0.131964 | -0.011529 | -0.040928 | 0.095748 | 3.550448 | 0.098648 | 3.474776 | -1.674394 | -0.215319 | -0.668042 | -1.582895 | -0.616949 | 1.726446 | 0.024517 | -1.28312 | 0.744414 |
| 37.65278 | 0.262472 | -0.012859 | -0.046721 | -0.045717 | -0.74324 | -0.037803 | -0.559062 | -0.106131 | -0.162485 | -0.04404 | -0.069107 | -0.936187 | 4.21233 | 0.507958 | 5.218041 | -1.493986 | -0.708096 | -0.229068 | -0.070068 | 0.144003 | 4.053337 | -0.846378 | 3.214874 | 2.191352 |
| 37.66667 | -0.218882 | -0.002114 | -0.061332 | -0.057445 | -2.600058 | -0.055282 | -1.090888 | -0.082063 | -0.175637 | -0.063974 | -0.068204 | -2.091276 | 1.696083 | 1.410657 | 2.860722 | -1.286255 | -0.62213 | 1.252296 | 1.133838 | 0.533836 | -0.776946 | -1.020025 | 3.685428 | 2.064156 |
| 37.68056 | -0.441501 | 0.024776 | -0.049214 | -0.040323 | -3.019628 | -0.046203 | -0.791435 | -0.034268 | -0.150596 | -0.057541 | -0.043399 | -2.484688 | -1.630258 | -0.025062 | -0.636758 | 0.86514 | -0.25982 | 1.593247 | 1.036994 | 1.091329 | -2.346741 | -0.35358 | 1.710193 | 1.28791 |
| 37.69444 | -0.501403 | 0.04681 | -0.011476 | -0.00616 | -2.695471 | -0.016735 | -0.152937 | 0.025382 | -0.101381 | -0.018912 | 0.004899 | -1.662501 | -4.737807 | 0.031057 | -2.895767 | 3.030777 | 0.476457 | 1.697948 | 0.802875 | 1.418767 | 1.916466 | 0.482432 | -0.401259 | 0.391009 |
| 37.70833 | -0.59221 | 0.098055 | 0.099548 | 0.09286 | -2.218802 | 0.078246 | 0.384819 | 0.140958 | -0.00042 | 0.089632 | 0.118471 | -0.66379 | -3.064209 | 0.28023 | -5.252733 | 4.034444 | 0.218451 | 0.681801 | 0.404361 | 1.501104 | 3.017421 | 0.799378 | -2.043963 | -1.194223 |

| | | | | | | | | | | | | | | | | | | | | | | | | |
|---|---|---|---|---|---|---|---|---|---|---|---|---|---|---|---|---|---|---|---|---|---|---|---|---|
| 37.72222 | -0.689175 | 0.186572 | 0.281574 | 0.265227 | -1.206246 | 0.249903 | 0.647948 | 0.320208 | 0.169515 | 0.259332 | 0.287671 | -0.197418 | 0.334423 | 0.310965 | -2.477557 | 3.211799 | 0.075041 | 0.627486 | 0.090462 | 0.721877 | -1.777659 | 0.440046 | -1.763575 | -1.289408 |
| 37.73611 | -0.733639 | 0.283032 | 0.462151 | 0.435912 | -0.316203 | 0.427114 | 0.88904 | 0.495171 | 0.341829 | 0.419749 | 0.448101 | -0.546556 | 2.794917 | -0.60192 | -0.652097 | 0.760627 | 0.172924 | 0.32361 | -0.507885 | 0.146235 | -2.969879 | 0.13218 | -1.992919 | 0.927186 |
| 37.75 | -0.777342 | 0.389347 | 0.582199 | 0.550963 | 0.342128 | 0.541156 | 0.904934 | 0.606261 | 0.454177 | 0.529744 | 0.56119 | -0.90439 | 2.825447 | 1.098101 | -0.364867 | -0.90071 | 0.273117 | -0.651179 | -0.760036 | -0.002434 | -3.117598 | 0.122888 | -1.590656 | -0.781499 |
| 37.76389 | -0.694802 | 0.516473 | 0.62291 | 0.592295 | 0.634337 | 0.569239 | 0.615364 | 0.625239 | 0.490186 | 0.57457 | 0.581259 | -0.577598 | 0.357727 | 0.441116 | -0.090181 | -1.953434 | 0.175837 | -2.028327 | -0.975203 | -0.00227 | 0.774174 | 0.1139 | -0.450602 | -1.153556 |
| 37.77778 | -0.336814 | 0.597274 | 0.588162 | 0.556403 | 0.546722 | 0.529913 | 0.043167 | 0.542133 | 0.458178 | 0.546817 | 0.508737 | -0.274652 | -1.17794 | 0.117399 | 0.77478 | -3.026083 | -0.001305 | -1.415411 | -1.270952 | 0.037281 | 2.626707 | 0.086939 | -0.496693 | 0.538696 |
| 37.79167 | 0.204584 | 0.585023 | 0.509305 | 0.474807 | 0.045918 | 0.452433 | -1.121743 | 0.427271 | 0.380933 | 0.467123 | 0.405967 | -0.301802 | -2.323803 | 0.070643 | 1.454086 | -3.640952 | -0.199574 | -0.484087 | -0.183417 | -0.008943 | 0.514265 | -0.328271 | 0.490654 | 1.1609 |
| 37.80556 | 0.456303 | 0.486888 | 0.410872 | 0.37671 | -0.550679 | 0.354024 | -2.760639 | 0.313047 | 0.2888 | 0.367978 | 0.312128 | -0.614019 | -2.58316 | 0.102382 | 1.667094 | -4.204603 | -0.22745 | 0.436321 | 0.57157 | -0.171689 | -1.366693 | -0.693257 | 1.20844 | 1.089968 |
| 37.81944 | -0.13173 | 0.358312 | 0.303003 | 0.275918 | -1.26318 | 0.260981 | -3.451806 | 0.221533 | 0.201128 | 0.268474 | 0.23959 | -1.040761 | -1.324448 | -0.058567 | 2.049828 | -3.879115 | -0.131587 | -0.041883 | 1.321407 | -0.259273 | -2.487479 | -0.654358 | 1.667627 | 1.356967 |
| 37.83333 | -0.688329 | 0.214097 | 0.194362 | 0.174995 | -2.799959 | 0.170963 | -3.124672 | 0.15414 | 0.111371 | 0.174187 | 0.157375 | -0.976887 | -0.755738 | -0.284061 | 2.04617 | -2.295608 | 0.136986 | -0.244138 | 1.078853 | -0.715273 | -2.389895 | -0.156907 | -0.536922 | 0.925304 |
| 37.84722 | -1.419291 | 0.074405 | 0.104166 | 0.078938 | -4.209784 | 0.086861 | -2.104801 | 0.093345 | 0.022643 | 0.092023 | 0.086236 | -0.807707 | -0.743263 | -0.476775 | 1.203176 | -0.844793 | 0.167376 | -0.217416 | 0.701496 | -0.487482 | -1.595425 | 0.194448 | -0.251101 | 0.692561 |
| 37.86111 | -1.922426 | -0.006267 | 0.053927 | 0.021299 | -4.051682 | 0.03637 | -1.813457 | 0.060393 | -0.033454 | 0.044178 | 0.055631 | -0.923172 | -1.546516 | -0.505855 | -1.94249 | -0.755227 | 0.030047 | -0.450899 | -0.211843 | -0.63665 | -0.804376 | 0.081638 | -0.447418 | -0.154677 |
| 37.875 | -1.695 | -0.044541 | 0.018597 | -0.008837 | -3.101471 | 0.006556 | -1.35457 | 0.018646 | -0.069212 | 0.011079 | 0.040116 | -1.276124 | -1.461699 | -0.390809 | -3.981354 | -1.47089 | -0.18811 | -0.6888 | -0.680758 | -0.694175 | -0.14302 | -0.193102 | -0.816908 | -0.977555 |
| 37.88889 | -1.180309 | -0.066734 | -0.012514 | -0.032056 | -2.138916 | -0.015752 | -1.828709 | -0.029224 | -0.101123 | -0.019755 | 0.011317 | -1.742819 | 0.333815 | -0.220973 | -2.27824 | -2.527686 | 0.176647 | -0.485843 | -0.111821 | -0.705157 | 0.164667 | -0.606836 | -0.333714 | -2.154837 |
| 37.90278 | -0.938234 | -0.048954 | -0.002678 | -0.022946 | -1.652356 | -0.008078 | -2.65685 | -0.045903 | -0.099087 | -0.022012 | 0.051226 | -2.173717 | -0.173843 | 0.159124 | -0.717069 | -3.659851 | -0.010018 | -0.530391 | 0.444431 | -0.712393 | 0.340249 | -0.903548 | 0.054669 | -0.487686 |
| 37.91667 | -0.944854 | -0.005078 | 0.026518 | 0.006155 | -1.260786 | 0.014933 | -3.038479 | -0.005577 | -0.071031 | 0.00079 | 0.111084 | -2.264452 | 0.911438 | 0.350546 | -1.519332 | -4.113605 | -0.138665 | -0.276199 | 0.301809 | -0.386185 | 0.934057 | -1.413244 | -0.955174 | 0.429005 |
| 37.93056 | -1.040419 | 0.017555 | 0.039459 | 0.024855 | -1.051682 | 0.026967 | -2.599247 | 0.029143 | -0.051138 | 0.019436 | -0.158832 | -1.930211 | 1.427658 | 0.251078 | 2.971835 | -3.624329 | -0.153222 | 0.034746 | -0.684503 | 0.440752 | 0.569306 | -0.822466 | -0.328768 | 0.631125 |
| 37.94444 | -1.070932 | 0.028152 | 0.046496 | 0.036558 | -0.811875 | 0.032221 | -2.219064 | 0.043392 | -0.043437 | 0.024718 | -0.305355 | -1.358671 | -0.108879 | 0.079256 | 6.551459 | -2.59515 | -0.111566 | 0.325433 | -0.710299 | 0.89986 | 0.109914 | -1.232565 | 0.141648 | 0.4567 |
| 37.95833 | -1.089015 | 0.031461 | 0.034433 | 0.026183 | -0.551467 | 0.02031 | -2.015547 | 0.037492 | -0.051538 | 0.01015 | -0.257604 | -1.108047 | -0.781126 | -0.091287 | 5.548499 | -2.777037 | -0.109282 | -0.74419 | 0.121833 | 0.75788 | -0.244155 | -2.028954 | 1.318372 | 1.083738 |
| 37.97222 | -1.148269 | 0.002226 | -0.01161 | -0.018333 | -0.455388 | -0.017085 | -2.190619 | 0.007288 | -0.07964 | -0.023189 | -0.325963 | -1.017685 | -1.390888 | -0.262533 | -0.792002 | -2.557203 | -0.096218 | -1.570489 | 0.986782 | -0.243898 | 0.447232 | -0.477474 | 2.936399 | 0.841957 |
| 37.98611 | -1.175846 | -0.009644 | -0.017201 | -0.02289 | -0.565026 | -0.020916 | -1.982563 | -0.000596 | -0.079549 | -0.019195 | -0.407258 | -0.850487 | -2.023103 | -0.4332 | -5.683453 | -2.922347 | -0.001712 | -2.160377 | 1.658652 | -1.400396 | 3.934122 | 1.87306 | 3.728967 | -0.28451 |

**Supplementary Table S2.** Raw motion data from: **(c)** 100 mM NaCl treatment in Tomato

| Time | Tomato Control1 | Tomato Control10 | Tomato Control11 | Tomato Control12 | Tomato Control2 | Tomato Control3 | Tomato Control4 | Tomato Control5 | Tomato Control6 | Tomato Control7 | Tomato Control8 | Tomato Control9 | Tomato 100mMNaCl1 | Tomato 100mMNaCl10 | Tomato 100mMNaCl11 | Tomato 100mMNaCl12 | Tomato 100mMNaCl2 | Tomato 100mMNaCl3 | Tomato 100mMNaCl4 | Tomato 100mMNaCl5 | Tomato 100mMNaCl6 | Tomato 100mMNaCl7 | Tomato 100mMNaCl8 | Tomato 100mMNaCl9 |
|---|---|---|---|---|---|---|---|---|---|---|---|---|---|---|---|---|---|---|---|---|---|---|---|---|
| 18 | -0.840506 | -0.016154 | 0.151624 | 0.080564 | -0.65267 | 0.054369 | -0.012195 | 0.022118 | -0.234927 | -0.174012 | 0.026872 | -0.133871 | -0.371128 | -0.187633 | -0.377296 | -0.47298 | -0.265356 | -0.176035 | -0.187729 | 0.012637 | -0.640103 | -0.044978 | -0.187611 | -0.174933 |
| 18.01389 | -1.212601 | 0.116613 | 0.368856 | -0.313434 | -0.760362 | 0.048798 | 0.128125 | -0.000482 | -0.328754 | -0.089545 | -0.034663 | -0.03531 | -0.406183 | -0.182898 | -0.346196 | -0.437838 | -0.234746 | -0.299318 | -0.266388 | -0.080192 | -0.994585 | -0.059761 | -0.291471 | -0.236824 |
| 18.02778 | -1.673898 | 0.147207 | 0.531025 | -1.084823 | -0.721797 | 0.035402 | 0.299591 | -0.136618 | -0.635912 | -0.074529 | -0.256356 | -0.055309 | -0.547823 | -0.222436 | -0.386521 | -0.437797 | -0.22127 | -0.536011 | -0.428173 | -0.255227 | -1.333041 | -0.113142 | -0.436743 | -0.366718 |
| 18.04167 | -1.409844 | -0.074271 | -0.024064 | -1.05389 | -1.130363 | -0.128796 | 0.046441 | -0.209758 | -0.87785 | -0.346642 | -0.439346 | -0.542359 | -0.910891 | -0.35161 | -0.673758 | -0.568137 | -0.371117 | -0.575324 | -0.5187 | -0.557033 | -1.28803 | -0.207192 | -0.449624 | -0.447034 |
| 18.05556 | 0.304108 | -0.430195 | -0.982101 | 0.593533 | -1.695912 | -0.213851 | -0.398427 | -0.010023 | -0.638088 | -0.522788 | -0.169911 | -0.726109 | -1.171532 | -0.416241 | -0.821492 | -0.60641 | -0.561118 | -0.182266 | -0.455189 | -0.856084 | -0.723332 | -0.187612 | -0.376138 | -0.263264 |
| 18.06944 | 0.764986 | -0.203125 | -1.525657 | 0.65865 | -1.411561 | -0.195254 | -0.403127 | 0.139698 | -0.386213 | -0.471603 | 0.104309 | -0.492878 | -1.034463 | -0.250108 | -0.642164 | -0.367331 | -0.463778 | 0.153151 | -0.282226 | -0.871878 | -0.358346 | -0.004835 | -0.187374 | 0.019776 |
| 18.08333 | 0.211129 | -0.005472 | -2.176081 | -0.031067 | -1.187571 | -0.039781 | -0.100425 | 0.083567 | -0.38873 | -0.302153 | 0.044003 | -0.543469 | -0.841706 | -0.085749 | -0.492889 | -0.225859 | -0.375337 | 0.049903 | -0.250704 | -1.014247 | -0.580835 | 0.037501 | -0.079629 | 0.047629 |
| 18.09722 | -0.294704 | -0.119215 | -2.34942 | -0.663382 | -1.187227 | -0.050414 | 0.015352 | -0.142406 | -0.453987 | -0.320765 | -0.028505 | -0.850695 | -0.906623 | -0.106586 | -0.553432 | -0.372102 | -0.442044 | -0.180603 | -0.371242 | -1.059068 | -1.03478 | -0.051502 | -0.127418 | -0.056729 |
| 18.11111 | -0.53843 | -0.408549 | -1.107237 | -0.990517 | -1.067949 | -0.135575 | -0.087001 | -0.391798 | -0.491348 | -0.420343 | -0.061866 | -1.282281 | -1.087472 | -0.172059 | -0.643061 | -0.673496 | -0.525092 | -0.249085 | -0.435688 | -0.818474 | -1.330593 | -0.11643 | -0.1933 | -0.084481 |
| 18.125 | -0.446752 | -0.674511 | -0.527836 | -0.979269 | -0.815123 | -0.218185 | -0.243202 | -0.438264 | -0.51813 | -0.513894 | -0.098633 | -1.309211 | -1.177721 | -0.2007 | -0.643026 | -0.92042 | -0.53306 | -0.210791 | -0.420805 | -0.483038 | -1.263851 | -0.114168 | -0.233573 | -0.055517 |
| 18.13889 | -0.1993 | -0.695146 | -0.202634 | -0.745456 | -0.538088 | -0.202201 | -0.329714 | -0.356654 | -0.453353 | -0.533818 | -0.12453 | -0.972542 | -1.164403 | -0.202373 | -0.56235 | -0.953557 | -0.497377 | -0.170954 | -0.375809 | -0.221275 | -1.082567 | -0.114586 | -0.295864 | -0.054523 |
| 18.15278 | -0.077123 | -0.591378 | -0.055738 | -0.486497 | -0.356215 | -0.162162 | -0.35132 | -0.342186 | -0.345146 | -0.477489 | -0.134604 | -0.672137 | -0.986163 | -0.218676 | -0.461252 | -0.817555 | -0.419645 | -0.16507 | -0.329485 | -0.119219 | -0.874031 | -0.125608 | -0.320547 | -0.096066 |
| 18.16667 | -0.072708 | -0.469888 | -0.015431 | -0.35921 | -0.259841 | -0.0931 | -0.27827 | -0.269023 | -0.269979 | -0.388461 | -0.1353 | -0.420701 | -0.709825 | -0.212322 | -0.340308 | -0.623044 | -0.337273 | -0.162402 | -0.269145 | -0.093831 | -0.757371 | -0.129047 | -0.291406 | -0.127298 |
| 18.18056 | -0.081151 | -0.320415 | 0.01316 | -0.401078 | -0.125663 | -0.013038 | -0.149531 | -0.161065 | -0.186686 | -0.30591 | -0.113112 | -0.278396 | -0.515533 | -0.19424 | -0.220418 | -0.446227 | -0.25919 | -0.143876 | -0.216695 | -0.099045 | -0.662505 | -0.12155 | -0.247559 | -0.131119 |
| 18.19444 | -0.087974 | -0.221373 | 0.030071 | -0.353493 | -0.073133 | 0.021771 | -0.071097 | -0.16375 | -0.150349 | -0.286063 | -0.113241 | -0.191729 | -0.416249 | -0.191842 | -0.130114 | -0.319791 | -0.203143 | -0.11425 | -0.176005 | -0.106064 | -0.594157 | -0.111195 | -0.211391 | -0.116246 |
| 18.20833 | -0.142887 | -0.206207 | 0.031893 | -0.267274 | -0.196121 | 0.022755 | -0.068166 | -0.202687 | -0.194125 | -0.301576 | -0.125172 | -0.152548 | -0.356116 | -0.177753 | -0.072864 | -0.2284 | -0.151131 | -0.100129 | -0.14222 | -0.10899 | -0.514662 | -0.104274 | -0.169922 | -0.093784 |
| 18.22222 | -0.254416 | -0.2789 | 0.007001 | -0.247704 | -0.414558 | -0.003112 | -0.116139 | -0.255343 | -0.276683 | -0.361424 | -0.146856 | -0.156986 | -0.312465 | -0.173961 | -0.053748 | -0.174384 | -0.126677 | -0.126507 | -0.141621 | -0.12916 | -0.476898 | -0.114411 | -0.147922 | -0.094323 |
| 18.23611 | -0.301585 | -0.312257 | -0.052773 | -0.187784 | -0.571779 | -0.019671 | -0.16797 | -0.319722 | -0.283839 | -0.389847 | -0.120546 | -0.151267 | -0.301304 | -0.175242 | -0.073541 | -0.165179 | -0.127729 | -0.153677 | -0.151123 | -0.169343 | -0.446599 | -0.12302 | -0.151529 | -0.104714 |
| 18.25 | -0.272935 | -0.21646 | -0.06807 | -0.040798 | -0.633247 | 0.024006 | -0.15428 | -0.283412 | -0.209563 | -0.322218 | -0.057705 | -0.091394 | -0.297321 | -0.14676 | -0.092705 | -0.163663 | -0.130117 | -0.150562 | -0.134845 | -0.173192 | -0.397757 | -0.107508 | -0.152979 | -0.09617 |
| 18.26389 | -0.306538 | -0.146532 | -0.035018 | 0.039637 | -0.74807 | 0.034579 | -0.159783 | -0.279564 | -0.221508 | -0.288735 | -0.049153 | -0.037706 | -0.269394 | -0.10017 | -0.084047 | -0.143673 | -0.109644 | -0.137105 | -0.124284 | -0.153428 | -0.345332 | -0.085578 | -0.130904 | -0.083095 |
| 18.27778 | -0.352941 | -0.195813 | 0.014728 | 0.047742 | -0.853559 | -0.025851 | -0.228867 | -0.362326 | -0.351899 | -0.359172 | -0.090605 | -0.045208 | -0.226139 | -0.068155 | -0.058039 | -0.115783 | -0.083155 | -0.130834 | -0.132127 | -0.165903 | -0.331123 | -0.080655 | -0.104254 | -0.083585 |
| 18.29167 | -0.334583 | -0.232625 | 0.079859 | 0.00412 | -0.843505 | -0.067379 | -0.241035 | -0.373579 | -0.476285 | -0.402529 | -0.112767 | -0.085666 | -0.18001 | -0.049058 | -0.026523 | -0.093465 | -0.052866 | -0.114285 | -0.11517 | -0.191924 | -0.30011 | -0.077947 | -0.079917 | -0.074133 |
| 18.30556 | -0.258459 | -0.22403 | 0.097334 | -0.075129 | -0.751748 | -0.061033 | -0.184463 | -0.314015 | -0.489645 | -0.401123 | -0.115489 | -0.133718 | -0.152573 | -0.049337 | -0.018037 | -0.092953 | -0.043603 | -0.09975 | -0.098543 | -0.213342 | -0.273109 | -0.084963 | -0.069361 | -0.068031 |
| 18.31944 | -0.182567 | -0.18739 | 0.043661 | -0.121579 | -0.70526 | -0.047704 | -0.122664 | -0.276372 | -0.382262 | -0.368463 | -0.122349 | -0.165436 | -0.166933 | -0.075046 | -0.063566 | -0.13522 | -0.085129 | -0.125176 | -0.109396 | -0.24247 | -0.273093 | -0.097578 | -0.089052 | -0.086001 |
| 18.33333 | -0.11724 | -0.120758 | 0.012269 | -0.094866 | -0.629478 | -0.019782 | -0.04787 | -0.177287 | -0.223461 | -0.291595 | -0.10194 | -0.138682 | -0.161522 | -0.070208 | -0.088202 | -0.148185 | -0.10747 | -0.131921 | -0.098956 | -0.236232 | -0.257582 | -0.078148 | -0.094443 | -0.085417 |
| 18.34722 | -0.064789 | -0.100349 | 0.007203 | -0.049778 | -0.519059 | 0.00022 | 0.01169 | -0.130353 | -0.109182 | -0.214475 | -0.066257 | -0.081428 | -0.115067 | -0.050768 | -0.049537 | -0.104641 | -0.067501 | -0.101679 | -0.066935 | -0.201357 | -0.195317 | -0.05083 | -0.060598 | -0.058326 |
| 18.36111 | -0.029632 | -0.142223 | -0.019093 | -0.042632 | -0.442957 | -0.007541 | 0.025078 | -0.190155 | -0.068949 | -0.182984 | -0.056128 | -0.050461 | -0.081978 | -0.046411 | -0.023525 | -0.07882 | -0.035887 | -0.085931 | -0.05342 | -0.176278 | -0.131354 | -0.045177 | -0.03806 | -0.042237 |
| 18.375 | -0.017367 | -0.169988 | -0.02344 | -0.064956 | -0.391101 | -0.026964 | -0.022555 | -0.288906 | -0.094713 | -0.168437 | -0.072744 | -0.058203 | -0.068515 | -0.044812 | -0.026495 | -0.07939 | -0.035229 | -0.076461 | -0.056138 | -0.167252 | -0.100263 | -0.048593 | -0.039555 | -0.039165 |
| 18.38889 | -0.032732 | -0.154341 | 0.008836 | -0.081139 | -0.344052 | -0.057679 | -0.097748 | -0.285103 | -0.153377 | -0.155281 | -0.090609 | -0.084252 | -0.064104 | -0.04987 | -0.034759 | -0.085616 | -0.050162 | -0.076319 | -0.065209 | -0.161472 | -0.108135 | -0.056438 | -0.047935 | -0.048218 |
| 18.40278 | -0.022359 | -0.092789 | 0.071854 | -0.058182 | -0.260788 | -0.036421 | -0.082588 | -0.132743 | -0.133696 | -0.090631 | -0.064564 | -0.070578 | -0.032932 | -0.047715 | -0.032342 | -0.079013 | -0.054586 | -0.076918 | -0.057561 | -0.144333 | -0.091081 | -0.046579 | -0.044992 | -0.050767 |
| 18.41667 | 0.016372 | -0.029437 | 0.12004 | -0.06916 | -0.186459 | 0.023498 | 0.019801 | -0.002006 | -0.052534 | -0.004893 | -0.008395 | -0.009856 | 0.031215 | -0.034073 | -0.017014 | -0.047847 | -0.045695 | -0.0615 | -0.035589 | -0.112634 | -0.032923 | -0.016968 | -0.029196 | -0.037434 |
| 18.43056 | 0.035226 | -0.020677 | 0.099909 | -0.11153 | -0.219251 | 0.049258 | 0.083188 | -0.017707 | 0.000778 | 0.024631 | 0.010029 | 0.029188 | 0.07027 | -0.02963 | -0.010113 | -0.027637 | -0.041803 | -0.047321 | -0.02794 | -0.096637 | 0.011319 | 0.000002 | -0.017787 | -0.024566 |
| 18.44444 | 0.042727 | -0.045375 | 0.022493 | -0.118447 | -0.374752 | 0.025184 | 0.076291 | -0.082433 | 0.010889 | -0.000952 | -0.017256 | 0.00748 | 0.073339 | -0.034015 | -0.02347 | -0.040217 | -0.043975 | -0.040913 | -0.026103 | -0.094939 | 0.024651 | 0.005229 | -0.016869 | -0.021567 |
| 18.45833 | -0.030509 | -0.099229 | -0.081614 | -0.229405 | -0.624113 | -0.040293 | 0.035743 | -0.139268 | -0.064365 | -0.062587 | -0.110933 | -0.088899 | 0.071362 | -0.034231 | -0.04268 | -0.073371 | -0.040893 | -0.060483 | -0.019661 | -0.083315 | -0.002325 | 0.019154 | -0.031571 | -0.048303 |
| 18.47222 | -0.354814 | -0.098209 | -0.005819 | -0.475263 | -0.84806 | -0.027561 | 0.127807 | -0.253792 | -0.196522 | 0.021067 | -0.272227 | -0.148203 | 0.139679 | 0.005401 | -0.04047 | -0.120018 | 0.079462 | -0.186789 | -0.001875 | -0.050311 | -0.076601 | 0.025683 | -0.070694 | -0.146863 |
| 18.48611 | -0.901294 | -0.022482 | 0.496072 | -0.83133 | -0.939852 | 0.209296 | 0.386978 | -0.3897 | -0.236478 | 0.355297 | -0.398167 | -0.108621 | 0.359618 | 0.09489 | -0.002632 | -0.183963 | 0.212598 | -0.414908 | 0.039811 | 0.012821 | -0.169618 | 0.012355 | -0.125639 | -0.271594 |
| 18.5 | -1.242391 | 0.205547 | 0.975812 | -0.866684 | -0.788826 | 0.223843 | 0.510324 | -0.251888 | -0.154833 | 0.532577 | -0.385798 | -0.029119 | 0.368246 | 0.132084 | 0.056906 | -0.152019 | -0.103369 | -0.500411 | 0.036505 | 0.034869 | -0.14938 | 0.046536 | -0.13151 | -0.230865 |
| 18.51389 | -0.281051 | -0.058319 | 0.399326 | -0.436418 | -0.338106 | -0.057056 | 0.090667 | -0.026534 | -0.092199 | -0.097592 | -0.048521 | -0.075492 | -0.075776 | -0.093341 | -0.006645 | -0.053157 | -0.41488 | -0.16241 | -0.074661 | -0.068328 | -0.070621 | 0.069164 | -0.10321 | -0.042037 |
| 18.52778 | 0.642869 | -1.116516 | -1.044702 | 0.025092 | -0.52578 | -0.835288 | -1.198735 | -0.7281 | -0.709941 | -1.642876 | -0.303966 | -0.635462 | -0.284593 | -0.456632 | -0.081432 | -0.034551 | -0.307951 | 0.221261 | -0.31122 | -0.474031 | -0.224781 | -0.189139 | -0.094451 | -0.0498 |
| 18.54167 | -0.382025 | -2.541827 | -1.464598 | -0.74628 | -1.490233 | -1.641969 | -2.559468 | -2.535842 | -1.977034 | -3.146378 | -1.34506 | -1.670115 | -0.443142 | -0.622788 | 0.052769 | -0.047532 | -0.249942 | 0.175342 | -0.480722 | -0.989412 | -0.657345 | -0.53076 | -0.170242 | -0.226391 |
| 18.55556 | -1.502324 | -3.149178 | -0.827057 | -1.072419 | -1.801401 | -1.778141 | -2.661577 | -3.042025 | -2.432813 | -3.337497 | -1.966659 | -1.95141 | -0.496695 | -0.382273 | 0.0863 | -0.196234 | -0.445541 | -0.101954 | -0.360608 | -1.230499 | -1.160281 | -0.600196 | -0.390361 | -0.312545 |
| 18.56944 | -1.245101 | -2.26155 | -0.190103 | -0.660403 | -1.03018 | -1.152708 | -1.713899 | -1.719708 | -2.032969 | -2.542158 | -1.59126 | -1.187275 | -0.48371 | -0.005911 | -0.025532 | -0.306024 | -0.583304 | -0.195146 | -0.061213 | -0.917974 | -1.294491 | -0.3232 | -0.562926 | -0.186032 |
| 18.58333 | -0.521402 | -1.269156 | 0.531663 | -0.241312 | -0.075129 | -0.522301 | -0.934608 | -0.686246 | -1.596141 | -1.89156 | -0.852898 | -0.061723 | -0.380603 | 0.149271 | -0.057637 | -0.245535 | -0.517971 | -0.137539 | 0.090239 | -0.440888 | -1.270987 | -0.018966 | -0.531493 | -0.034953 |
| 18.59722 | -0.138685 | -0.63319 | 0.882235 | -0.039002 | 0.723075 | -0.146943 | -0.479172 | -0.270915 | -1.042658 | -1.065731 | -0.345438 | 0.281214 | -0.19695 | 0.015841 | -0.045351 | -0.16105 | -0.432772 | -0.098942 | 0.055874 | -0.221666 | -0.996121 | -0.007765 | -0.381496 | -0.017255 |
| 18.61111 | -0.009894 | -0.241611 | 0.72981 | -0.006808 | 1.754855 | 0.014281 | -0.223529 | -0.105841 | -0.569906 | -0.462122 | -0.160565 | 0.000156 | -0.049898 | -0.135021 | -0.061627 | -0.116931 | -0.364366 | -0.084725 | -0.005148 | -0.11597 | -0.569591 | -0.045753 | -0.272645 | -0.043398 |
| 18.625 | 0.086586 | 0.004582 | 0.552194 | 0.029853 | 2.090286 | 0.087212 | -0.05276 | 0.068769 | -0.335795 | -0.158869 | -0.077731 | 0.009846 | 0.040865 | -0.202881 | -0.059104 | -0.059106 | -0.227548 | -0.058457 | -0.012282 | 0.014673 | -0.262862 | -0.068234 | -0.164126 | -0.031297 |
| 18.63889 | 0.182089 | 0.158205 | 0.516655 | 0.106533 | 1.579651 | 0.160103 | 0.030613 | 0.222152 | -0.348591 | -0.064541 | 0.028432 | 0.134793 | 0.108768 | -0.201484 | -0.045286 | -0.017633 | -0.102003 | -0.045356 | 0.001341 | 0.21792 | -0.116599 | -0.076519 | -0.08479 | -0.013439 |
| 18.65278 | 0.227784 | 0.185192 | 0.593711 | 0.190601 | 1.131485 | 0.214098 | 0.015533 | 0.251745 | -0.416585 | -0.107572 | 0.192669 | 0.293665 | 0.165666 | -0.143941 | -0.046907 | -0.032203 | -0.064321 | -0.059717 | 0.008762 | 0.501038 | -0.069726 | -0.068274 | -0.030086 | -0.020223 |
| [illegible] | [illegible] | [illegible] | [illegible] | [illegible] | [illegible] | [illegible] | [illegible] | [illegible] | [illegible] | [illegible] | [illegible] | [illegible] | [illegible] | [illegible] | [illegible] | [illegible] | [illegible] | [illegible] | [illegible] | [illegible] | [illegible] | [illegible] | [illegible] | [illegible] |

| | | | | | | | | | | | | | | | | | | | | | | | | |
|---|---|---|---|---|---|---|---|---|---|---|---|---|---|---|---|---|---|---|---|---|---|---|---|---|
| 21.48611 | 0.970288 | 1.182734 | 1.768318 | 2.119701 | 1.427814 | 0.932726 | 0.991453 | 0.801138 | 1.008983 | 1.077078 | 0.993318 | 1.789523 | 1.659727 | 0.503643 | 1.16514 | 0.939782 | 1.502407 | 1.118814 | 1.14258 | 1.198743 | 1.64714 | 0.401569 | 0.73544 | 0.523969 |
| 21.5 | 0.96794 | 1.761487 | 1.965251 | 1.720431 | 1.835935 | 0.981865 | 1.035523 | 0.981244 | 1.10739 | 1.173871 | 1.256086 | 2.331604 | 1.841082 | 0.646558 | 1.529505 | 1.288753 | 1.773233 | 1.0501 | 1.503789 | 1.848105 | 2.121082 | 0.44278 | 1.020976 | 0.677871 |
| 21.51389 | 1.041712 | 1.999235 | 2.482755 | 1.952412 | 2.196677 | 1.049987 | 1.070831 | 1.259966 | 1.422235 | 1.472704 | 1.333192 | 2.612047 | 1.791959 | 0.727798 | 1.906356 | 1.46136 | 1.86242 | 0.941448 | 1.714198 | 2.233675 | 2.182892 | 0.428287 | 1.168765 | 0.751053 |
| 21.52778 | 1.248259 | 1.510804 | 3.117495 | 2.651124 | 2.028627 | 1.143061 | 1.024711 | 0.981237 | 1.486334 | 1.536538 | 1.353339 | 2.536397 | 1.647704 | 0.588496 | 2.385145 | 1.575733 | 1.812096 | 0.989143 | 1.265937 | 1.400637 | 2.425809 | 0.261419 | 0.982887 | 0.68607 |
| 21.54167 | 1.134946 | 1.069821 | 2.498399 | 1.790679 | 1.376484 | 0.507982 | 0.42518 | 0.113168 | 0.782543 | 0.780633 | 1.049296 | 2.066397 | 1.606392 | 0.212094 | 2.488542 | 1.700179 | 1.629599 | 1.058329 | 0.718448 | 0.612627 | 2.120154 | 0.111089 | 0.864002 | 0.370097 |
| 21.55556 | 1.001457 | 1.403546 | 1.568985 | 1.16659 | 1.289649 | 0.007749 | 0.040074 | 0.098534 | 0.195116 | 0.395826 | 0.73506 | 1.751279 | 1.978775 | 0.032132 | 2.088278 | 2.115729 | 1.549868 | 0.94775 | 0.730349 | 0.72187 | 1.812722 | 0.182955 | 0.893124 | 0.081105 |
| 21.56944 | 1.193565 | 1.891978 | 1.658774 | 2.274053 | 2.104195 | 0.213714 | 0.731447 | 0.927439 | 0.698795 | 1.031711 | 0.76233 | 2.278494 | 2.790987 | 0.26355 | 1.60996 | 2.364147 | 1.364329 | 0.630225 | 1.159506 | 1.419065 | 1.935962 | 0.447454 | 1.159946 | 0.144368 |
| 21.58333 | 1.860375 | 1.257484 | 1.04286 | 1.786376 | 2.788212 | 0.411583 | 0.957194 | 0.964092 | 1.515185 | 1.823369 | 1.057595 | 2.561775 | 2.020234 | 0.392764 | 1.20165 | 1.942092 | 0.557673 | 0.500026 | 1.2089 | 1.449683 | 1.908366 | 0.427326 | 0.676727 | 0.248142 |
| 21.59722 | 1.56773 | 0.743658 | 1.318286 | 1.053351 | 1.98941 | 0.665802 | 0.4008 | 0.730173 | 1.524628 | 1.311128 | 0.911656 | 1.819643 | 1.330853 | 0.343863 | 0.728945 | 1.378265 | 0.45609 | 0.460392 | 0.831028 | 1.647772 | 1.500455 | 0.283442 | 0.327257 | 0.242393 |
| 21.61111 | 1.080873 | 0.660738 | 0.886655 | 0.478049 | 1.17077 | 0.755424 | 0.288113 | 0.435079 | 0.249877 | 0.344941 | 0.56638 | 0.892947 | 0.897592 | 0.331746 | 0.930612 | 0.786041 | 0.63985 | 0.600096 | 0.338753 | 2.178156 | 0.611178 | 0.198859 | 0.228695 | 0.235566 |
| 21.625 | 0.18776 | 0.462026 | 0.304963 | -0.046572 | 0.641741 | 0.448537 | 0.790269 | 0.065013 | 0.363171 | 0.336522 | 0.453377 | 0.428183 | 0.474251 | 0.337061 | 0.732806 | 0.476276 | 0.695331 | 0.795479 | 0.225882 | 2.091093 | 0.339415 | 0.196091 | 0.339629 | 0.246222 |
| 21.63889 | -0.009212 | 0.311054 | 0.262128 | -0.317237 | 0.524069 | 0.141586 | 0.584774 | 0.097398 | 0.535011 | 0.429761 | 0.442157 | 0.384295 | 0.301253 | 0.39626 | 0.380616 | 0.619624 | 0.626979 | 0.615229 | 0.067417 | 1.795053 | 0.604859 | 0.209844 | 0.451809 | 0.208455 |
| 21.65278 | 0.249532 | 0.476594 | 0.945514 | -0.086333 | 0.101327 | -0.05631 | 0.394771 | 0.239545 | 0.376971 | 0.288781 | 0.656183 | 0.777571 | 0.344168 | 0.406842 | 0.161287 | 0.842669 | 0.610039 | 0.125911 | -0.077076 | 1.018186 | 1.330534 | 0.238382 | 0.355576 | 0.197316 |
| 21.66667 | 0.884135 | 0.616287 | 0.879115 | 0.122595 | 0.528522 | 0.020441 | 0.309358 | 0.266424 | 0.005039 | 0.095188 | 0.41896 | 0.784799 | 0.314113 | 0.142231 | 0.18916 | 0.591556 | 0.575776 | -0.229807 | -0.115966 | 0.384994 | 1.164148 | 0.255609 | 0.186584 | 0.072748 |
| 21.68056 | 0.548431 | 0.33593 | 0.597861 | 0.067258 | 0.823079 | 0.224838 | 0.392425 | 0.258476 | 0.073925 | 0.174938 | 0.226178 | 0.228284 | 0.370251 | -0.120367 | 0.508609 | 0.082834 | 0.342355 | -0.148994 | 0.023 | -0.128508 | 0.493452 | 0.207345 | 0.117768 | -0.081283 |
| 21.69444 | 0.00768 | -0.028881 | 0.422926 | 0.262583 | 0.733263 | 0.220502 | 0.26603 | 0.247536 | 0.464447 | 0.341953 | -0.020789 | -0.106586 | 0.28487 | -0.191865 | 0.437588 | -0.208786 | -0.08157 | -0.091746 | -0.029385 | -0.248118 | -0.02478 | 0.146121 | 0.092248 | -0.146827 |
| 21.70833 | 0.047479 | -0.156271 | 0.252258 | 0.919848 | 0.580039 | 0.166385 | 0.261889 | 0.080148 | 0.587658 | 0.308487 | -0.00552 | -0.113634 | 0.227425 | -0.011666 | 0.159728 | -0.219108 | -0.429582 | -0.066181 | -0.142701 | 0.394226 | -0.20077 | 0.125228 | 0.076695 | -0.004047 |
| 21.72222 | 0.741811 | 0.004426 | 0.125154 | 0.998505 | 0.821665 | 0.140591 | 0.098899 | 0.008532 | 0.330526 | 0.110893 | -0.000398 | -0.110302 | 0.188579 | 0.175967 | -0.008134 | -0.00522 | -0.259183 | 0.019531 | -0.334298 | 0.900992 | -0.095497 | 0.121208 | 0.063291 | 0.12383 |
| 21.73611 | 0.598367 | 0.06051 | 0.135487 | 0.35203 | 0.517761 | 0.065441 | 0.069315 | -0.000995 | 0.060514 | -0.031717 | -0.102814 | -0.265258 | 0.194814 | 0.160971 | -0.239392 | 0.304627 | 0.121353 | 0.129499 | 0.160896 | 0.479637 | -0.172842 | 0.102275 | 0.044885 | 0.011371 |
| 21.75 | 0.090963 | 0.083524 | 0.159807 | -0.155798 | 0.110319 | 0.051698 | 0.162282 | 0.168015 | 0.117122 | 0.127411 | -0.037553 | -0.143465 | 0.189854 | 0.076714 | -0.287221 | 0.410698 | 0.15082 | 0.032842 | 0.831028 | 0.684117 | -0.223654 | 0.084851 | 0.042392 | 0.022273 |
| 21.76389 | 0.582736 | 0.304984 | 0.360118 | -0.029712 | 0.944804 | 0.172684 | 0.305861 | 0.353322 | 0.444 | 0.510277 | 0.296022 | 0.302959 | 0.024311 | -0.072413 | 0.045741 | 0.155034 | -0.16855 | -0.013324 | 0.581716 | -0.286201 | -0.085984 | 0.055467 | 0.093282 | 0.037382 |
| 21.77778 | 1.238488 | 0.658466 | 0.52859 | 0.312716 | 1.960158 | 0.53272 | 0.316399 | 0.580705 | 0.718737 | 0.629304 | 0.329124 | 0.355459 | -0.242446 | -0.117597 | 0.204696 | -0.091036 | -0.311179 | -0.107343 | -0.336038 | -0.688505 | 0.142633 | 0.022954 | 0.17982 | 0.09936 |
| 21.79167 | 0.50127 | 0.655799 | 0.575279 | 0.690414 | 0.620524 | 0.712294 | 0.742515 | 0.495149 | 0.718189 | 0.530318 | 0.216991 | -0.063646 | -0.255918 | -0.038691 | -0.289935 | -0.092035 | -0.111088 | 0.128151 | -0.374101 | -0.937527 | -0.033975 | -0.006216 | 0.161087 | 0.192864 |
| 21.80556 | -0.44645 | 0.500086 | 0.621981 | 0.960014 | -0.780515 | 0.689687 | 0.845191 | 0.491174 | 0.668217 | 0.544653 | 0.398341 | -0.064436 | 0.083092 | 0.027242 | -0.824171 | 0.027912 | 0.075405 | 0.216493 | 0.152187 | -0.66726 | 0.251988 | 0.015621 | 0.161716 | 0.066386 |
| 21.81944 | 0.408341 | 0.306634 | 0.591218 | 0.61564 | 0.401379 | 0.748611 | 0.964696 | 0.509137 | 0.687639 | 0.691761 | 0.691348 | 0.653648 | 0.346935 | -0.027674 | -0.653232 | -0.11296 | -0.016568 | 0.110311 | 0.313385 | 0.004022 | -0.179233 | 0.014969 | 0.030127 | -0.003557 |
| 21.83333 | 1.807809 | 0.138369 | 0.366302 | -0.190473 | 2.601748 | 0.114038 | 0.284354 | 0.24361 | 0.609448 | 0.722997 | 0.86608 | 0.933158 | 0.276889 | -0.292195 | 0.048108 | -0.12221 | -0.195307 | -0.373758 | -0.131838 | 0.234006 | -0.087843 | -0.037828 | -0.102937 | -0.169814 |
| 21.84722 | 1.516011 | 0.13859 | 0.223602 | -0.500637 | 1.749949 | 0.108323 | 0.198281 | 0.111191 | 0.33788 | 0.248971 | 0.233647 | 0.06801 | -0.094245 | -0.397688 | 0.50188 | 0.26011 | -0.127146 | -0.501964 | -0.432053 | 0.016556 | 0.057376 | -0.057005 | -0.229484 | -0.074044 |
| 21.86111 | -0.84087 | 0.005021 | 0.362601 | 0.223714 | -1.667248 | 0.005204 | 0.076003 | 0.0023 | -0.388322 | -0.358836 | -0.19188 | -0.926708 | -0.357834 | -0.12436 | 0.072174 | 0.399949 | 0.071263 | 0.002074 | 0.069005 | -0.480196 | -0.177238 | -0.030014 | -0.232739 | 0.024968 |
| 21.875 | -1.267113 | -0.208808 | 0.104634 | 0.527172 | -1.722433 | 0.161966 | 0.153575 | -0.060835 | -0.755741 | -0.451561 | -0.330852 | -0.578251 | -0.289528 | -0.027931 | -0.713067 | 0.003225 | 0.067074 | 0.036577 | 0.132601 | -1.248217 | -0.212791 | 0.013865 | -0.196232 | -0.139397 |
| 21.88889 | -0.266084 | -0.383173 | -0.842791 | -0.124831 | 0.667803 | -0.094349 | -0.240994 | -0.224315 | -0.441906 | 0.000823 | -0.10565 | 0.225297 | -0.153563 | -0.179984 | -0.82645 | -0.499085 | -0.119811 | -0.250291 | 0.200272 | -1.524162 | -0.975333 | 0.04206 | -0.259382 | -0.256325 |
| 21.90278 | 0.149104 | -0.361561 | -0.760433 | -0.756831 | 1.137739 | -0.186183 | -0.497272 | -0.324024 | 0.134143 | 0.090458 | -0.327245 | -0.129916 | -0.201847 | -0.182897 | -0.589052 | -1.269903 | -0.355702 | -0.437442 | -0.445775 | -1.213437 | -0.836182 | 0.002608 | -0.406612 | -0.14588 |
| 21.91667 | -0.231951 | -0.496548 | 0.018968 | -0.461225 | -1.090612 | -0.211034 | -0.217729 | -0.283435 | -0.23252 | -0.686688 | -0.780055 | -1.386735 | -0.53352 | 0.088057 | -0.419734 | -1.050054 | -0.502336 | -0.204625 | -0.962139 | -0.719144 | -0.630484 | -0.054217 | -0.640747 | -0.150191 |
| 21.93056 | -0.654132 | -0.792123 | 0.068083 | -0.018503 | -2.075396 | -0.404076 | -0.315859 | -0.111569 | -0.655978 | -0.986302 | -0.665152 | -1.34485 | -0.859621 | 0.128062 | -0.551865 | -0.38922 | -0.51786 | 0.377506 | -0.63439 | -1.090974 | -0.680037 | -0.107236 | -0.552091 | -0.223681 |
| 21.94444 | -0.967261 | -0.915954 | -0.665267 | -0.352277 | -0.650632 | -0.273785 | -0.401501 | -0.29643 | -1.25589 | -0.609118 | -0.315788 | -0.674169 | -1.02237 | -0.341644 | -0.890984 | -0.762108 | -0.157585 | -0.040533 | -0.671293 | -0.921032 | -1.469337 | -0.14021 | -0.476779 | -0.58992 |
| [illegible] | [illegible] | [illegible] | [illegible] | [illegible] | [illegible] | [illegible] | [illegible] | [illegible] | [illegible] | [illegible] | [illegible] | [illegible] | [illegible] | [illegible] | [illegible] | [illegible] | [illegible] | [illegible] | [illegible] | [illegible] | [illegible] | [illegible] | [illegible] | [illegible] |

| | | | | | | | | | | | | | | | | | | | | | | | | |
|---|---|---|---|---|---|---|---|---|---|---|---|---|---|---|---|---|---|---|---|---|---|---|---|---|
| 25 | -2.157105 | -2.473026 | -1.452987 | 0.864045 | -2.102563 | -0.403538 | -1.598826 | -0.216474 | -2.51713 | -1.301029 | -1.435164 | -2.950605 | -2.116464 | -0.142738 | -0.969643 | -0.567023 | -0.459394 | -0.993979 | -0.712786 | -0.106806 | -1.94153 | -0.501595 | -0.946054 | -0.898259 |
| 25.01389 | -2.078796 | -2.725785 | -0.372254 | -0.722883 | -1.244329 | -1.003208 | -1.748691 | -1.303588 | -1.341794 | -1.33275 | -1.920978 | -2.518236 | -1.900676 | 0.25373 | -1.121338 | -0.8156 | -0.50589 | -0.876097 | -0.77435 | -0.293915 | -1.962719 | -0.260358 | -0.986585 | -0.639054 |
| 25.02778 | -1.922387 | -2.330893 | -0.17624 | -3.955344 | -1.388476 | -1.735903 | -1.760189 | -1.97224 | -0.669807 | -0.930704 | -2.055447 | -2.10861 | -1.639997 | 0.384312 | -1.219612 | -0.872952 | -0.33917 | -0.538309 | -0.534668 | -0.19319 | -1.107875 | -0.02807 | -0.739318 | -0.371088 |
| 25.04167 | -2.02845 | -1.646393 | -0.388089 | -4.885381 | -2.518341 | -2.041837 | -1.348636 | -1.332401 | -1.114228 | -0.868606 | -2.19658 | -2.223985 | -1.174094 | 0.31397 | -0.914334 | -0.566927 | 0.008432 | -0.001669 | -0.299478 | 0.039 | 0.191407 | 0.133779 | -0.086424 | 0.20429 |
| 25.05556 | -2.235331 | -1.333235 | -1.085999 | -2.012775 | -3.978942 | -1.26042 | -1.212429 | -0.178105 | -1.966618 | -0.9238 | -1.699514 | -1.95925 | -0.916024 | 0.173518 | -0.831397 | -0.293919 | -0.050938 | 0.042252 | -0.029688 | 0.165408 | 0.330419 | -0.032281 | -0.00143 | 0.223593 |
| 25.06944 | -2.341737 | -1.078489 | -1.315756 | 0.221152 | -3.951055 | -0.588531 | -1.096846 | 0.213196 | -2.674152 | -0.955014 | -1.175133 | -1.421031 | -1.159778 | -0.070305 | -1.131332 | -0.316224 | -0.43249 | -0.392467 | -0.295498 | 0.242199 | -0.588859 | -0.364438 | -0.666788 | -0.521007 |
| 25.08333 | -1.15597 | -0.717361 | -1.531314 | 0.152844 | -1.647423 | -0.207451 | -1.080742 | -0.196204 | -2.672325 | -0.9549 | -0.720308 | -0.870167 | -1.332905 | -0.174669 | -1.013573 | -0.301626 | -0.527982 | -0.485797 | -0.401762 | 0.133273 | -0.842939 | -0.281122 | -0.86741 | -1.122156 |
| 25.09722 | -0.289218 | -0.683399 | -1.620528 | -1.513184 | -1.364644 | -0.209512 | -1.231356 | -0.529242 | -2.794368 | -0.95834 | -0.659384 | -0.777673 | -1.223231 | -0.214958 | -0.587831 | -0.295717 | -0.430093 | -0.282885 | -0.319257 | -0.067588 | -0.693968 | -0.033704 | -0.760386 | -1.139396 |
| 25.11111 | -1.267824 | -1.073192 | -1.838412 | -3.461928 | -2.775238 | -0.677709 | -1.934775 | -0.341842 | -3.413089 | -1.309239 | -1.206794 | -1.298662 | -1.328084 | -0.339743 | -0.388645 | -0.381734 | -0.372051 | -0.06156 | -0.232926 | -0.089572 | -0.573404 | -0.025322 | -0.603852 | -0.98741 |
| 25.125 | -2.239919 | -1.465124 | -1.81764 | -3.325992 | -4.058054 | -1.09644 | -2.051141 | -0.870977 | -3.200903 | -1.369582 | -1.770438 | -1.84561 | -1.628397 | -0.455772 | -0.515979 | -0.481135 | -0.375989 | -0.071736 | -0.188952 | -0.119407 | -0.606729 | -0.235745 | -0.594146 | -0.92728 |
| 25.13889 | -1.908196 | -1.405957 | -1.482768 | -2.21507 | -3.560987 | -0.997915 | -1.502985 | -2.003814 | -2.138085 | -1.043909 | -1.674643 | -1.66476 | -1.623167 | -0.496885 | -0.67777 | -0.535412 | -0.372757 | -0.271757 | -0.259532 | -0.172602 | -0.690391 | -0.438399 | -0.651236 | -0.931911 |
| 25.15278 | -1.293909 | -1.209253 | -1.212479 | -1.553386 | -1.827074 | -0.698297 | -0.949304 | -2.353407 | -1.303734 | -0.812196 | -1.182082 | -1.33342 | -1.402092 | -0.490219 | -0.715545 | -0.550164 | -0.40035 | -0.43044 | -0.322718 | -0.204761 | -0.703869 | -0.575673 | -0.691403 | -0.912822 |
| 25.16667 | -0.953363 | -1.01761 | -0.999273 | -1.409579 | -0.801805 | -0.605255 | -0.762985 | -1.789836 | -0.881606 | -0.801638 | -0.978892 | -1.169134 | -1.236442 | -0.470078 | -0.749478 | -0.545337 | -0.461127 | -0.483617 | -0.358127 | -0.187013 | -0.663492 | -0.540414 | -0.694108 | -0.833548 |
| 25.18056 | -0.624914 | -0.871245 | -0.831644 | -1.495021 | -0.689732 | -0.694454 | -0.747863 | -1.2564 | -0.823576 | -0.906464 | -0.967026 | -1.11038 | -1.214708 | -0.455435 | -0.808829 | -0.543128 | -0.494845 | -0.43162 | -0.363365 | -0.194751 | -0.63801 | -0.387987 | -0.65818 | -0.656262 |
| 25.19444 | -0.211548 | -0.80444 | -0.732101 | -1.574682 | -1.473099 | -0.769779 | -0.7548 | -1.182935 | -0.966433 | -1.025894 | -0.834377 | -0.978135 | -1.375208 | -0.437482 | -0.935844 | -0.554508 | -0.492965 | -0.392566 | -0.378958 | -0.287673 | -0.655374 | -0.243753 | -0.669294 | -0.532116 |
| 25.20833 | -0.365269 | -0.849014 | -0.685343 | -1.354541 | -1.844539 | -0.693807 | -0.678508 | -1.165132 | -1.039692 | -1.03165 | -0.651155 | -0.831028 | -1.517731 | -0.440301 | -1.015405 | -0.593634 | -0.547095 | -0.454066 | -0.504778 | -0.436256 | -0.679807 | -0.204182 | -0.714677 | -0.515868 |
| 25.22222 | -0.765686 | -0.872176 | -0.704926 | -1.010239 | -1.308739 | -0.575546 | -0.519244 | -0.890349 | -0.996457 | -0.983709 | -0.614304 | -0.832051 | -1.565147 | -0.455881 | -1.004479 | -0.686373 | -0.673081 | -0.586065 | -0.678526 | -0.526864 | -0.747298 | -0.231739 | -0.728292 | -0.557907 |
| 25.23611 | -0.708419 | -0.785496 | -0.708338 | -0.820442 | -0.712787 | -0.580287 | -0.392024 | -0.520273 | -0.907462 | -0.926062 | -0.619046 | -0.84332 | -1.391695 | -0.481876 | -0.921283 | -0.793017 | -0.799128 | -0.714443 | -0.754589 | -0.525712 | -0.789049 | -0.27293 | -0.723898 | -0.561529 |
| 25.25 | -0.332202 | -0.67253 | -0.625276 | -0.801093 | -0.381138 | -0.626765 | -0.450788 | -0.419724 | -0.762901 | -0.855258 | -0.484466 | -0.695768 | -1.097967 | -0.449007 | -0.755873 | -0.761678 | -0.817083 | -0.728003 | -0.691464 | -0.413511 | -0.713302 | -0.252783 | -0.612441 | -0.455986 |
| 25.26389 | -0.366061 | -0.615797 | -0.578696 | -0.863149 | -0.440959 | -0.565801 | -0.578997 | -0.742778 | -0.627937 | -0.777553 | -0.367945 | -0.549874 | -0.807374 | -0.335004 | -0.609356 | -0.610717 | -0.722611 | -0.585548 | -0.598789 | -0.278239 | -0.548119 | -0.204688 | -0.380785 | -0.299288 |
| 25.27778 | -0.643247 | -0.664155 | -0.5929 | -0.901284 | -0.510589 | -0.521625 | -0.621991 | -1.02185 | -0.634531 | -0.759646 | -0.423475 | -0.568564 | -0.551612 | -0.246362 | -0.473635 | -0.480364 | -0.699688 | -0.44751 | -0.524966 | -0.182492 | -0.399701 | -0.157955 | -0.179752 | -0.187476 |
| 25.29167 | -0.610943 | -0.645396 | -0.568883 | -0.737753 | -0.331809 | -0.576744 | -0.582313 | -0.808895 | -0.667179 | -0.740955 | -0.526357 | -0.641253 | -0.388221 | -0.194194 | -0.301576 | -0.439131 | -0.724792 | -0.347041 | -0.461156 | -0.134591 | -0.302252 | -0.134619 | -0.134152 | -0.136348 |
| 25.30556 | -0.375575 | -0.432925 | -0.443183 | -0.38705 | -0.109258 | -0.531901 | -0.508275 | -0.391951 | -0.509722 | -0.580545 | -0.491675 | -0.58086 | -0.274247 | -0.121925 | -0.136521 | -0.458756 | -0.7045 | -0.326811 | -0.412471 | -0.081782 | -0.24746 | -0.146933 | -0.187441 | -0.13315 |
| 25.31944 | -0.301702 | -0.197063 | -0.315494 | -0.145552 | -0.081041 | -0.323856 | -0.365098 | -0.195266 | -0.153799 | -0.338805 | -0.307093 | -0.357911 | -0.049015 | -0.019833 | -0.110429 | -0.397635 | -0.602649 | -0.308842 | -0.381385 | 0.025436 | -0.173268 | -0.146229 | -0.15987 | -0.057695 |
| 25.33333 | -0.212293 | -0.089552 | -0.225778 | -0.059475 | -0.138536 | -0.139011 | -0.191559 | -0.266534 | 0.10168 | -0.225733 | -0.136312 | -0.188077 | 0.26842 | 0.030633 | -0.175961 | -0.190881 | -0.526934 | -0.155161 | -0.273129 | 0.115576 | -0.035274 | -0.078135 | 0.02163 | 0.104594 |
| 25.34722 | -0.030622 | -0.036211 | -0.130938 | 0.116066 | -0.165491 | -0.114704 | -0.069831 | -0.187931 | 0.100286 | -0.246601 | -0.086188 | -0.161111 | 0.3231 | -0.008257 | -0.199322 | 0.010296 | -0.474184 | 0.043871 | -0.150943 | 0.116186 | 0.068273 | -0.035349 | 0.105994 | 0.148912 |
| 25.36111 | -0.03193 | -0.061101 | -0.121246 | 0.30601 | -0.30518 | -0.215164 | -0.06648 | 0.049007 | 0.028101 | -0.255808 | -0.095374 | -0.174268 | 0.157676 | -0.007414 | -0.053106 | 0.113583 | -0.300314 | 0.148576 | -0.036641 | 0.090901 | 0.127537 | -0.021155 | 0.03984 | 0.109861 |
| 25.375 | -0.090724 | -0.132659 | -0.142486 | 0.383451 | -0.269257 | -0.212821 | -0.083463 | -0.015638 | 0.115645 | -0.181977 | -0.068239 | -0.12558 | 0.265445 | 0.109226 | 0.176456 | 0.191783 | 0.019557 | 0.209747 | 0.031872 | 0.134514 | 0.227284 | 0.022616 | 0.094397 | 0.236621 |
| 25.38889 | 0.073104 | -0.113419 | -0.042383 | 0.427182 | 0.206243 | -0.083369 | -0.002552 | -0.262917 | 0.34643 | -0.060111 | -0.017144 | -0.04134 | 0.760063 | 0.196212 | 0.25324 | 0.354392 | 0.35907 | 0.28424 | 0.08542 | 0.223729 | 0.341877 | 0.124006 | 0.408926 | 0.574628 |
| 25.40278 | 0.276854 | -0.034488 | 0.077482 | 0.460372 | 0.713057 | -0.04105 | 0.104927 | -0.145431 | 0.510189 | 0.033904 | 0.033517 | 0.015059 | 1.18615 | 0.250716 | 0.190862 | 0.575629 | 0.568151 | 0.361307 | 0.178897 | 0.245545 | 0.459148 | 0.263741 | 0.66316 | 0.839006 |
| 25.41667 | 0.358123 | 0.060311 | 0.175868 | 0.560567 | 0.747164 | -0.027624 | 0.189633 | 0.428784 | 0.677439 | 0.155869 | 0.154593 | 0.097235 | 1.048913 | 0.322942 | 0.332058 | 0.710373 | 0.635082 | 0.404948 | 0.365268 | 0.199275 | 0.643539 | 0.390025 | 0.54076 | 0.890829 |
| 25.43056 | 0.434438 | 0.124405 | 0.348699 | 0.645959 | 0.576547 | 0.118145 | 0.27858 | 0.56639 | 0.936405 | 0.331693 | 0.310624 | 0.220163 | 1.048559 | 0.417024 | 0.896347 | 0.732497 | 0.681032 | 0.490697 | 0.567909 | 0.308488 | 0.917591 | 0.49404 | 0.556528 | 1.058596 |
| 25.44444 | 0.60369 | 0.183412 | 0.513035 | 0.711448 | 0.771272 | 0.270406 | 0.390993 | 0.310849 | 1.228974 | 0.440787 | 0.369354 | 0.328869 | 1.735747 | 0.460736 | 1.597458 | 0.762779 | 0.908863 | 0.672498 | 0.695503 | 0.575916 | 1.028134 | 0.585236 | 1.054143 | 1.545398 |
| 25.45833 | 0.857836 | 0.392659 | 0.697323 | 0.932325 | 1.44204 | 0.359632 | 0.621674 | 0.366219 | 1.541721 | 0.559978 | 0.444551 | 0.520569 | 2.653797 | 0.466512 | 1.922809 | 1.131619 | 1.376058 | 0.884312 | 0.845261 | 0.659744 | 1.185364 | 0.638583 | 1.698358 | 2.087942 |
| 25.47222 | 1.178975 | 0.767129 | 1.115934 | 1.323955 | 2.306961 | 0.468776 | 0.915261 | 1.004003 | 1.933366 | 0.839487 | 0.729406 | 0.861089 | 2.97894 | 0.717129 | 2.229639 | 1.724571 | 1.96097 | 1.213596 | 1.168678 | 0.465714 | 1.693491 | 0.7497 | 1.846363 | 2.209621 |
| 25.48611 | 1.54641 | 1.138622 | 1.877519 | 1.970042 | 3.101351 | 0.708031 | 1.217209 | 1.758977 | 2.352003 | 1.156396 | 1.221484 | 1.350095 | 2.899211 | 0.999783 | 2.889105 | 1.997187 | 2.196367 | 1.591596 | 1.669304 | 0.240602 | 2.547692 | 0.925978 | 1.667919 | 2.149272 |
| 25.5 | 2.176847 | 1.49384 | 2.727528 | 2.710976 | 3.910831 | 1.05079 | 1.54524 | 1.407967 | 2.649491 | 1.392794 | 1.709772 | 1.826192 | 3.020927 | 0.943869 | 3.114285 | 1.673813 | 1.943799 | 2.044334 | 1.797454 | 0.296417 | 2.888591 | 0.971591 | 1.672664 | 2.058551 |
| 25.51389 | 2.900273 | 2.031675 | 3.132465 | 3.40893 | 4.816887 | 1.447102 | 2.065365 | 1.225696 | 3.209278 | 1.581881 | 2.131007 | 2.338689 | 3.908006 | 0.81235 | 3.004759 | 1.110756 | 1.569605 | 2.322531 | 1.605813 | 0.317428 | 3.022375 | 0.885796 | 1.845648 | 2.027586 |
| 25.52778 | 3.459541 | 2.736845 | 3.092271 | 3.787577 | 5.256654 | 1.443432 | 2.363645 | 1.605652 | 3.690133 | 1.55405 | 2.002773 | 2.755609 | 3.734311 | 0.772044 | 2.434619 | 0.800882 | 1.486579 | 2.028103 | 1.155312 | -0.012435 | 2.475058 | 0.772159 | 1.731599 | 1.75342 |
| 25.54167 | 3.099531 | 2.488849 | 2.502005 | 3.590209 | 4.399193 | 1.039645 | 2.041792 | 1.502897 | 2.831636 | 1.03913 | 1.544842 | 2.707366 | 2.322105 | 0.498294 | 1.462365 | 0.371627 | 1.569449 | 1.241753 | 0.861287 | -0.379293 | 1.371146 | 0.577373 | 1.249707 | 0.118081 |
| 25.55556 | 2.300022 | 2.132933 | 2.22812 | 3.483481 | 3.097449 | 0.942758 | 1.34262 | 1.371559 | 1.396431 | 0.811907 | 1.474184 | 2.438945 | 0.813897 | 0.109215 | 0.97523 | 0.121872 | 1.604291 | 0.24248 | 0.437594 | -0.393899 | 0.856361 | 0.742627 | -0.05684 | -0.699432 |
| 25.56944 | 1.631784 | 2.352816 | 1.950366 | 4.37343 | 2.700734 | 1.176089 | 1.399133 | 1.522907 | 1.100439 | 1.391357 | 1.959137 | 2.634178 | 0.301289 | -0.137132 | 1.601397 | 0.518076 | 1.610618 | -0.288937 | 0.083036 | -0.072095 | 1.564837 | 0.981276 | -0.346237 | -1.003478 |
| 25.58333 | 1.674471 | 2.291409 | 0.621603 | 4.561785 | 2.376901 | 0.9505 | 1.963285 | 0.996129 | 1.870557 | 1.827185 | 2.228426 | 2.585319 | 0.833465 | -0.096532 | 2.319677 | 1.188283 | 1.344482 | -0.247445 | -0.103938 | 0.047032 | 2.307322 | 0.826194 | 0.231804 | -0.171911 |
| 25.59722 | 2.823412 | 1.911187 | 0.314995 | 3.48605 | 1.710856 | 0.446052 | 2.214165 | 0.23429 | 2.501243 | 1.979137 | 1.714757 | 2.392556 | 1.012625 | -0.039074 | 2.122446 | 1.706062 | 0.722352 | 0.332066 | -0.044137 | -0.071425 | 2.021965 | 0.449201 | 1.086263 | 0.216235 |
| 25.61111 | 4.712126 | 1.177929 | 0.632284 | 2.011375 | 2.148177 | 0.448027 | 2.101407 | 0.254461 | 2.96758 | 1.986044 | 1.14573 | 1.370145 | 0.958677 | 0.253359 | 1.599282 | 1.524265 | 0.405298 | 1.258177 | 0.413737 | -0.171344 | 1.54312 | 0.260803 | 1.719391 | -0.070095 |
| 25.625 | 4.52969 | 0.217332 | 1.459188 | 1.26221 | 2.031795 | 0.433567 | 1.964407 | 0.627175 | 2.29642 | 2.013317 | 0.299097 | 0.427066 | 0.184569 | 0.582388 | 1.324433 | 0.724982 | 0.245155 | 1.50627 | 1.301365 | -0.217451 | 1.570604 | 0.167097 | 1.026047 | -0.384359 |
| 25.63889 | 2.588856 | 0.26082 | -0.542901 | -0.153462 | 1.509611 | 0.295294 | 0.437548 | 0.924567 | 0.51414 | 0.666091 | -0.028163 | 0.611633 | -0.13719 | 0.229554 | 0.899558 | 0.107573 | 0.247488 | 0.828307 | 1.173242 | -0.094928 | 0.927147 | 0.012447 | 0.184746 | -0.187657 |
| 25.65278 | 0.236111 | 1.109629 | -1.235605 | -0.367702 | 1.178838 | 0.422357 | 0.110021 | 0.518823 | -0.811964 | -0.484551 | 0.285432 | 1.58002 | 0.257493 | -0.117626 | 0.97544 | 0.178702 | 0.482153 | -0.534114 | 0.405042 | 0.09321 | -0.103083 | 0.016286 | -1.007669 | 0.480522 |
| 25.66667 | -1.318598 | 0.962395 | -1.285672 | 0.612924 | 1.082234 | 0.511908 | 0.087084 | 0.247545 | -0.216937 | -0.223243 | 1.033716 | 2.286188 | 1.218192 | 0.085159 | 0.674985 | 0.320896 | 0.62707 | -0.946062 | -0.254701 | 0.318621 | 0.016536 | 0.184505 | -0.902329 | 0.529524 |
| 25.68056 | -0.135136 | 0.080181 | 0.566271 | 1.986183 | 0.510609 | 0.599289 | 1.489612 | 0.067343 | 1.233163 | 0.638058 | 1.134349 | 0.850673 | 1.36885 | 0.374238 | 0.415873 | 0.235889 | 0.329246 | -0.307872 | -0.15982 | 0.424043 | 0.143036 | 0.164279 | 0.020013 | 0.015181 |
| 25.69444 | 1.826799 | -0.273101 | 0.672367 | 0.726162 | 0.438924 | -0.169455 | 1.508799 | 0.031056 | 1.936519 | 0.982824 | 0.164174 | -0.332539 | 0.659402 | 0.365522 | 0.038448 | -0.48478 | 0.032967 | 0.176271 | -0.054872 | 0.230523 | 0.275208 | 0.11118 | 0.168164 | -0.373962 |
| 25.70833 | 1.847395 | 0.486715 | 0.079083 | -0.504341 | 0.287529 | -0.403132 | 0.737442 | 0.062797 | 0.394939 | 0.411612 | -0.848819 | -0.149588 | 0.13515 | 0.049601 | 0.182215 | -0.355306 | -0.769485 | 0.779917 | -0.370455 | -0.011484 | 0.430772 | 0.068419 | 0.134842 | -0.07569 |
| 25.72222 | -0.274631 | 0.712745 | -1.262766 | 0.140796 | -0.406104 | -0.000135 | -0.100687 | 0.131389 | -1.260418 | -0.78107 | -0.565765 | 0.441991 | 0.047851 | 0.043692 | 0.289623 | 0.2608 | -0.283883 | 0.894821 | -0.344012 | 0.057082 | 0.282375 | 0.05389 | 0.303451 | 0.511093 |
| 25.73611 | -1.645236 | -0.33126 | -0.343152 | -0.117402 | 0.222896 | 0.029163 | -0.174669 | 0.100818 | -1.387559 | -1.042613 | 0.171186 | -0.212416 | -0.27182 | 0.452063 | -0.194401 | 0.781118 | 0.52412 | -0.052875 | -0.38724 | 0.331868 | 0.322676 | 0.171275 | -0.18118 | 0.571682 |
| 25.75 | -0.84079 | -1.05942 | -0.505097 | 0.141366 | 0.133356 | 0.39306 | -0.281053 | 0.044668 | -0.186404 | 0.102257 | 0.819233 | -1.05477 | 0.481939 | 0.633506 | -0.019902 | 0.587213 | 1.307052 | -0.438606 | -0.107682 | 0.521574 | 0.049863 | -0.049161 | -0.00505 | 0.230395 |
| 25.76389 | 0.702175 | -0.176923 | -0.329991 | 0.539666 | -0.46053 | 0.115924 | 0.343479 | -0.134652 | 1.072032 | 0.995456 | 0.457771 | -0.951787 | 0.174981 | 0.057526 | 0.879351 | -0.402776 | 0.521187 | -0.722085 | 0.034742 | 0.436268 | 0.327159 | -0.063476 | 0.525984 | -0.041301 |
| 25.77778 | 0.912468 | 0.772311 | -0.13272 | 1.421332 | -0.490772 | -0.055395 | 0.572093 | -0.145333 | 0.757729 | 0.169297 | 0.138708 | 0.200022 | -0.192225 | -0.359965 | 0.727876 | -1.524181 | -0.762175 | -1.122002 | -0.236036 | 0.155988 | 0.169644 | 0.106136 | 0.483112 | -0.29461 |
| 25.79167 | 0.863304 | 1.184339 | 0.492168 | 0.983749 | 0.287344 | 0.194 | 0.155565 | 0.051537 | -0.974865 | -0.742316 | 0.263613 | 1.205829 | 0.502381 | -0.355168 | -0.078968 | -1.024102 | -1.336426 | -0.754103 | -0.435578 | -0.008216 | -0.478996 | -0.001014 | -0.089415 | 0.009933 |
| 25.80556 | 0.675017 | 0.306911 | 0.708205 | 0.254441 | 0.718402 | -0.290898 | -0.790202 | 0.1617 | -1.359575 | -0.646492 | 0.148601 | 0.57741 | 0.970353 | 0.003955 | -0.891559 | 0.187172 | -0.763952 | -0.021828 | -0.188511 | -0.042664 | -0.221858 | 0.155368 | -0.583277 | 0.395829 |
| 25.81944 | -1.352738 | -0.55241 | 0.822164 | 0.883806 | 0.083877 | -0.764098 | -0.997885 | 0.015791 | 0.396559 | 0.645132 | -0.272375 | -1.024641 | 1.073018 | 0.37125 | -1.075591 | 0.852706 | 0.359211 | 0.234607 | 0.034658 | 0.161293 | -0.769741 | -0.034077 | -0.354809 | 0.433801 |
| 25.83333 | -1.823612 | -0.216682 | 0.074517 | 1.669275 | -0.986253 | -0.674446 | 0.365972 | -0.236748 | 1.463111 | 0.959226 | -0.340161 | -1.335312 | 0.080172 | 0.312352 | -0.209041 | 0.404805 | 1.314493 | 0.036513 | 0.020759 | 0.22218 | -0.54257 | -0.177535 | 0.242469 | 0.136706 |
| 25.84722 | -0.031114 | 0.447739 | 0.012841 | 0.214351 | -0.408486 | 0.126481 | 0.278233 | -0.292967 | 0.880031 | -0.295976 | -0.333708 | 0.052603 | -0.733878 | 0.043752 | 0.796511 | -0.212148 | 1.590487 | -0.260961 | 0.199919 | 0.182615 | -0.243843 | -0.07791 | 0.855364 | -0.238206 |
| 25.86111 | 2.426573 | 0.694418 | 1.021276 | 0.632349 | 0.762742 | 0.50324 | 0.040972 | -0.029226 | -0.427489 | -0.622947 | 0.247838 | 0.928449 | -0.680577 | -0.178867 | 0.306298 | -0.568668 | 0.375524 | -0.250288 | 0.254448 | -0.02949 | 0.266321 | -0.094469 | 0.035831 | -0.607883 |
| 25.875 | 1.540802 | -0.276317 | 1.128364 | -1.377348 | 0.628575 | -0.139967 | -0.831254 | 0.069936 | -1.211372 | -0.403581 | 0.330006 | 0.417224 | -1.375034 | -0.425985 | -0.381607 | -0.198649 | -0.820139 | -0.193161 | 0.10707 | -0.205539 | 0.345144 | -0.081322 | -0.474983 | -0.753993 |
| 25.88889 | -1.677457 | -0.961766 | 1.004624 | -0.328819 | -0.341746 | -0.16465 | -0.790565 | -0.172329 | -1.055106 | -0.036348 | 0.275837 | -0.351182 | -0.071412 | -0.456002 | -1.57223 | 0.182607 | -1.320736 | -0.769099 | -0.497605 | -0.147226 | -0.467074 | -0.14076 | -0.779064 | -0.174625 |
| 25.90278 | -3.037501 | -0.066743 | 0.372031 | 3.155436 | -1.259545 | -0.34186 | 0.748758 | -0.285217 | -0.745496 | -0.41909 | -0.280902 | 0.05181 | 1.629187 | -0.396933 | -1.399611 | 0.131028 | -1.322095 | -1.044646 | -0.437032 | 0.078223 | -1.28715 | -0.075179 | -0.376279 | 0.37826 |
| 25.91667 | -1.225859 | 0.714923 | -0.065394 | 1.384395 | -1.015034 | 0.446788 | 0.501374 | -0.023653 | -0.393927 | -0.380977 | -0.514434 | 0.329343 | 0.677112 | -0.139168 | -0.287255 | 0.077655 | -0.91821 | -0.951883 | -0.146339 | 0.141888 | -1.526605 | 0.116339 | 0.292562 | 0.644918 |
| 25.93056 | 0.598411 | 0.3763 | -0.557679 | -0.113418 | -0.799615 | 0.613358 | 0.380056 | 0.304278 | 0.285733 | 0.097511 | -0.33472 | -0.186947 | -1.068895 | -0.04443 | 0.225029 | -0.135738 | -0.614853 | -1.299119 | 0.06031 | 0.062111 | -1.722561 | -0.049714 | 0.129735 | 0.202388 |
| 25.94444 | -0.531695 | -0.40211 | -0.517046 | -0.729584 | -0.843179 | -0.112002 | -0.46415 | 0.197174 | -0.437278 | 0.176857 | 0.127495 | -0.695936 | -3.139829 | -0.130228 | -0.937844 | -0.299047 | -0.30949 | -1.570782 | -0.349614 | -0.032398 | -1.176789 | -0.099966 | -0.557848 | -0.072846 |
| 25.95833 | -2.17758 | -1.016196 | 0.244293 | 0.852162 | -0.535238 | 0.205058 | -0.236306 | -0.259831 | -2.114168 | -0.977937 | -0.003406 | -0.961828 | -3.855513 | -0.393578 | -2.219252 | -0.577694 | -0.340626 | -1.404342 | -0.839274 | -0.090224 | -0.309529 | -0.011559 | -0.917674 | 0.005126 |
| 25.97222 | -1.701892 | -0.946788 | 0.331281 | 4.17808 | -0.912449 | 0.077327 | -0.068988 | -0.888268 | -2.882913 | -1.28778 | -1.112235 | -1.278812 | -3.830132 | -0.820304 | -2.486451 | -1.13077 | -1.014782 | -1.074791 | -1.363273 | -0.001325 | -0.661115 | -0.097535 | -0.945746 | -0.788399 |
| 25.98611 | -1.543242 | -0.787496 | -0.27487 | 1.032823 | -2.621958 | -0.27504 | -0.553442 | 0.0273 | -1.994678 | -0.574424 | -1.554586 | -2.179699 | -2.42098 | -0.583255 | -2.363751 | -1.434702 | -2.06735 | -1.133873 | -1.300685 | 0.099738 | -1.841524 | -0.409496 | -1.233756 | -1.261691 |
| 26 | -2.211043 | -0.858809 | -0.804129 | -3.68129 | -3.340337 | -0.760869 | -1.563168 | 0.598816 | -0.905064 | -0.239638 | -1.386948 | -2.552351 | -2.252382 | -0.811889 | -2.330842 | -1.071779 | -2.726476 | -1.290251 | -1.245476 | 0.149886 | -2.709597 | -0.74801 | -1.296359 | -1.285507 |
| 26.01389 | -2.343118 | -1.181016 | -1.18051 | -4.192626 | -1.649657 | -1.222429 | -1.985488 | -0.45019 | -0.595415 | -1.064199 | -0.671361 | -2.296935 | -2.376896 | -0.925184 | -2.828231 | -0.528345 | -2.306814 | -1.054055 | -0.917121 | 0.132297 | -2.997547 | -0.884433 | -1.397294 | -1.270297 |
| 26.02778 | -1.613825 | -1.151839 | -1.390385 | -2.317921 | -1.064792 | -0.786061 | -2.041299 | -1.969706 | -1.734053 | -1.121082 | -0.346602 | -1.639017 | -2.835406 | -0.810909 | -3.38775 | -0.127499 | -0.960841 | -1.011974 | -1.092737 | 0.369932 | -2.51584 | -0.700736 | -1.608911 | -1.250828 |
| 26.04167 | -0.644918 | -0.216411 | -0.642269 | -0.803017 | -1.097039 | -0.132825 | -0.683087 | -0.854087 | -2.009147 | 0.146322 | -0.170085 | -0.535321 | -2.646199 | -0.418197 | -1.908925 | -0.022905 | -0.746423 | -0.72865 | -1.0923 | 0.6853 | -1.349874 | -0.503884 | -1.286821 | -0.996485 |
| 26.05556 | -0.632743 | 0.673385 | 0.00894 | 0.00008 | -1.024164 | 0.675688 | 0.409766 | 1.331827 | -0.899753 | 1.35072 | -0.222918 | -0.10865 | -1.991933 | -0.109218 | -0.80984 | -0.145099 | -0.361854 | -0.493317 | -0.703474 | 0.890916 | -0.444287 | -0.676904 | -0.618325 | -0.549124 |
| 26.06944 | -0.709744 | 0.712074 | 0.386929 | -0.074421 | -0.961839 | 0.854143 | 0.678012 | 2.098668 | -0.249452 | 1.279213 | -0.149455 | 0.084365 | -1.408764 | -0.02553 | -0.295885 | -0.095727 | -0.355276 | -0.540012 | -0.415985 | 0.101451 | -0.213384 | -0.382781 | -0.451725 | -0.355166 |
| 26.08333 | -0.092901 | 0.545974 | 0.205749 | -0.340969 | -1.325005 | 0.667021 | 0.310597 | 1.470039 | -0.296864 | 0.543298 | 0.498775 | 0.618146 | -0.960002 | 0.16803 | -0.119595 | 0.064405 | -0.964843 | -0.559332 | -0.226404 | -0.638908 | -0.188457 | -0.135347 | -0.575531 | -0.459387 |
| 26.09722 | -0.140714 | 0.080952 | -0.528173 | -0.926001 | -2.613677 | 0.009521 | -0.595273 | 0.653114 | -0.994983 | -0.103846 | 0.430388 | 0.514499 | -0.781582 | 0.342928 | -0.08486 | 0.060479 | -0.425238 | -0.424631 | -0.048049 | -0.600572 | -0.164886 | -0.100587 | -0.532626 | -0.553598 |
| 26.11111 | -1.117542 | -0.516218 | -1.145725 | -1.429682 | -3.732494 | -0.559187 | -1.092592 | -0.12225 | -2.063527 | -0.412015 | -0.4396 | -0.31382 | -0.595144 | 0.256737 | -0.260209 | -0.109474 | -0.309241 | -0.360341 | -0.029842 | -0.424765 | -0.266729 | -0.081794 | -0.4117 | -0.55177 |
| 26.125 | -1.257141 | -0.780167 | -1.103026 | -1.693686 | -3.268694 | -0.799672 | -1.293655 | -1.38018 | -2.53304 | -0.6058 | -1.159317 | -1.020593 | -0.569095 | -0.042527 | -0.452331 | -0.211193 | -0.201757 | -0.309079 | -0.082914 | -0.328219 | -0.372673 | -0.044805 | -0.444674 | -0.562492 |
| 26.13889 | -0.787788 | -0.738036 | -1.013678 | -1.681917 | -2.102505 | -0.895074 | -1.306278 | -1.879694 | -2.157858 | -0.825601 | -1.462409 | -1.105633 | -0.637553 | -0.299561 | -0.529354 | -0.208931 | -0.030716 | -0.237204 | -0.09996 | -0.320968 | -0.39629 | -0.455582 | -0.440625 | -0.554555 |
| 26.15278 | -0.521471 | -0.530842 | -0.8845 | -1.436612 | -1.682823 | -0.738453 | -1.230263 | -1.850076 | -1.361608 | -0.693622 | -1.191968 | -0.941749 | -0.758296 | -0.333304 | -0.615167 | -0.276774 | -0.060687 | -0.240529 | -0.132822 | -0.362376 | -0.428318 | -0.627347 | -0.458851 | -0.481861 |
| 26.16667 | -0.559406 | -0.312333 | -0.650023 | -1.226724 | -1.406168 | -0.586571 | -1.022221 | -1.61355 | -0.629775 | -0.438913 | -0.831992 | -0.671567 | -0.756676 | -0.269434 | -0.725573 | -0.478381 | -0.259357 | -0.26257 | -0.188637 | -0.366087 | -0.44966 | -0.499608 | -0.475923 | -0.459523 |
| 26.18056 | -0.352441 | -0.181173 | -0.327622 | -1.149883 | -0.690023 | -0.400293 | -0.562262 | -1.359408 | -0.459175 | -0.439139 | -0.533935 | -0.451923 | -0.55236 | -0.263818 | -0.693155 | -0.638597 | -0.383819 | -0.24535 | -0.223464 | -0.274534 | -0.460031 | -0.347798 | -0.465963 | -0.442211 |
| 26.19444 | 0.378906 | -0.126563 | -0.191425 | -0.942761 | -0.320283 | -0.125488 | -0.13205 | -1.171914 | -0.409908 | -0.388288 | -0.475542 | -0.401856 | -0.314255 | -0.337828 | -0.514259 | -0.60676 | -0.361294 | -0.181633 | -0.248944 | -0.184478 | -0.495995 | -0.204318 | -0.465803 | -0.428207 |
| 26.20833 | 0.785099 | -0.008237 | 0.03417 | -0.479444 | -0.553764 | 0.114272 | 0.036285 | -0.452173 | -0.003273 | -0.034686 | -0.420986 | -0.34098 | -0.237365 | -0.394637 | -0.355549 | -0.34848 | -0.272205 | -0.125654 | -0.229181 | -0.168969 | -0.44697 | -0.125543 | -0.461008 | -0.362723 |
| 26.22222 | 0.51234 | 0.233756 | 0.340436 | 0.106664 | -0.452152 | 0.256273 | 0.061889 | 0.455367 | 0.429027 | 0.320168 | 0.042549 | 0.016061 | -0.190934 | -0.34023 | -0.251895 | -0.096484 | -0.211802 | -0.100533 | -0.248114 | -0.134765 | -0.343078 | -0.152683 | -0.36872 | -0.189535 |
| 26.23611 | 0.281058 | 0.339014 | 0.506841 | 0.391631 | 0.146143 | 0.288497 | 0.159458 | 0.597528 | 0.394666 | 0.276875 | 0.43669 | 0.250104 | -0.038827 | -0.133729 | -0.063347 | 0.014272 | -0.144404 | -0.004403 | -0.212758 | -0.010231 | -0.217392 | -0.185453 | -0.195117 | 0.025236 |
| 26.25 | 0.480556 | 0.183096 | 0.402319 | 0.253168 | 0.419743 | 0.223981 | 0.32004 | 0.136288 | 0.157996 | 0.042022 | 0.308626 | 0.088636 | 0.125702 | 0.047846 | 0.198144 | 0.043803 | -0.07484 | 0.219017 | -0.110048 | 0.129719 | -0.115319 | -0.120272 | -0.006367 | 0.180039 |
| 26.26389 | 0.494762 | 0.040027 | 0.301553 | 0.080481 | 0.227735 | 0.169239 | 0.437053 | 0.030604 | 0.180858 | -0.007508 | 0.069887 | -0.060235 | 0.122872 | 0.058475 | 0.333049 | 0.083127 | 0.012256 | 0.553582 | 0.00933 | 0.184044 | 0.001318 | 0.043343 | 0.122641 | 0.177577 |
| 26.27778 | -0.091019 | -0.018944 | 0.149352 | 0.057301 | 0.085443 | 0.101986 | 0.343348 | 0.311412 | 0.250641 | -0.031273 | 0.161445 | 0.0366 | 0.124517 | 0.00649 | 0.295201 | 0.123489 | 0.121215 | 0.738116 | 0.060851 | 0.119755 | 0.134992 | 0.213789 | 0.166792 | 0.112886 |
| 26.29167 | -0.196169 | 0.006356 | 0.041061 | 0.062103 | 0.28006 | 0.073809 | 0.178029 | 0.517177 | 0.346003 | -0.050473 | 0.246393 | 0.091926 | 0.167998 | 0.018805 | 0.221435 | 0.107427 | 0.137528 | 0.695581 | 0.040028 | 0.0893 | 0.162322 | 0.304775 | 0.152664 | 0.106094 |
| 26.30556 | 0.423711 | 0.135643 | 0.254469 | 0.164493 | 0.686661 | 0.207287 | 0.318532 | 0.514487 | 0.648223 | 0.235337 | 0.227332 | 0.089572 | 0.250607 | 0.037467 | 0.141445 | 0.130077 | 0.001701 | 0.596904 | 0.039837 | 0.130315 | 0.0912 | 0.492348 | 0.087854 | 0.115581 |
| 26.31944 | 0.521023 | 0.340909 | 0.538783 | 0.420674 | 1.016763 | 0.420818 | 0.637168 | 0.654079 | 0.825554 | 0.620176 | 0.40558 | 0.236048 | 0.286346 | 0.051446 | 0.108916 | 0.278182 | -0.221246 | 0.441292 | 0.116314 | 0.193117 | 0.086423 | 0.333281 | 0.035111 | 0.111601 |
| 26.33333 | 0.269535 | 0.597682 | 0.220018 | 0.653812 | 1.169966 | 0.544038 | 0.83859 | 1.060209 | 0.692065 | 0.668696 | 0.654776 | 0.541348 | 0.284394 | 0.100354 | 0.083398 | 0.395665 | -0.206644 | 0.291087 | 0.140442 | 0.158576 | 0.168856 | -0.120196 | 0.03753 | 0.182357 |
| 26.34722 | 0.579708 | 0.726773 | 0.160201 | 0.894349 | 1.182685 | 0.607052 | 0.863475 | 1.169033 | 0.622697 | 0.505006 | 0.611184 | 0.617219 | 0.259995 | 0.097274 | 0.034384 | 0.367144 | 0.08652 | 0.222612 | 0.086893 | 0.012827 | 0.146607 | -0.394092 | 0.047519 | 0.24465 |
| 26.36111 | 0.724459 | 0.55756 | 0.540194 | 0.943169 | 1.142504 | 0.552243 | 0.69741 | 0.658331 | 0.763914 | 0.530168 | 0.372309 | 0.373261 | 0.236307 | 0.074895 | 0.055382 | 0.345519 | 0.255156 | 0.220145 | 0.086498 | -0.085462 | 0.059529 | -0.350911 | 0.052784 | 0.256581 |
| 26.375 | 0.270399 | 0.304948 | 0.495257 | 0.840761 | 0.995314 | 0.332392 | 0.417107 | 0.263934 | 0.678149 | 0.538439 | 0.339038 | 0.262789 | 0.251123 | 0.151868 | 0.214275 | 0.413295 | 0.127236 | 0.241983 | 0.178156 | -0.086041 | 0.089483 | -0.198036 | 0.109178 | 0.302633 |
| 26.38889 | 0.272629 | 0.252651 | 0.540416 | 0.846119 | 0.763817 | 0.164318 | 0.248627 | 0.449462 | 0.487113 | 0.369323 | 0.391035 | 0.383232 | 0.398177 | 0.336937 | 0.487539 | 0.486019 | 0.095649 | 0.335661 | 0.237015 | -0.023486 | 0.277005 | -0.023815 | 0.283602 | 0.421623 |
| 26.40278 | 0.826638 | 0.380924 | 0.854825 | 0.939093 | 0.728021 | 0.316906 | 0.421982 | 0.899498 | 0.671174 | 0.366944 | 0.461323 | 0.515576 | 0.761964 | 0.506406 | 0.85809 | 0.567368 | 0.392122 | 0.547701 | 0.318946 | 0.082898 | 0.474676 | 0.14567 | 0.609729 | 0.678321 |
| 26.41667 | 0.924782 | 0.479719 | 1.031703 | 0.939283 | 0.958616 | 0.60286 | 0.675543 | 0.839117 | 1.064055 | 0.581902 | 0.7084 | 0.660949 | 1.274504 | 0.575199 | 1.159013 | 0.671905 | 0.631076 | 0.841912 | 0.556591 | 0.191742 | 0.661506 | 0.263484 | 0.915519 | 1.040018 |
| 26.43056 | 0.67768 | 0.448481 | 0.984702 | 0.897787 | 1.247094 | 0.668935 | 0.677902 | 0.469625 | 1.126023 | 0.646397 | 0.913096 | 0.790958 | 1.914416 | 0.63226 | 1.297182 | 0.769177 | 0.751771 | 1.234714 | 0.894216 | 0.183408 | 0.872399 | 0.384366 | 1.052903 | 1.278251 |
| 26.44444 | 0.862581 | 0.519504 | 1.087965 | 1.13876 | 1.577175 | 0.744079 | 0.699871 | 0.453242 | 1.188287 | 0.613169 | 1.038118 | 0.947996 | 2.573688 | 0.719451 | 1.607761 | 0.822631 | 1.07759 | 1.401575 | 1.227883 | 0.085374 | 1.234289 | 0.601637 | 1.157978 | 1.471774 |
| 26.45833 | 1.478613 | 0.7831 | 1.656771 | 1.746 | 1.951775 | 1.026298 | 1.038122 | 0.906511 | 1.936014 | 0.869924 | 1.199283 | 1.223543 | 3.302569 | 0.770995 | 2.458961 | 0.878139 | 1.614438 | 1.80866 | 1.54584 | 0.09471 | 1.665046 | 0.81784 | 1.493444 | 1.915834 |
| 26.47222 | 1.857125 | 1.06239 | 2.493721 | 2.545936 | 2.231374 | 1.373545 | 1.512976 | 1.160881 | 2.852625 | 1.241606 | 1.427049 | 1.62947 | 4.062882 | 0.783823 | 3.127444 | 0.989433 | 1.843692 | 2.309448 | 1.485698 | 0.005291 | 2.059115 | 0.847357 | 1.887824 | 2.294884 |
| 26.48611 | 2.29115 | 1.423128 | 3.215998 | 3.492009 | 2.848734 | 1.811687 | 2.116066 | 1.194159 | 3.250444 | 1.536199 | 1.932352 | 2.529206 | 4.322406 | 0.865806 | 2.838771 | 1.078219 | 1.687013 | 2.406987 | 1.503586 | 0.004037 | 2.476603 | 0.843755 | 2.08194 | 2.372173 |
| 26.5 | 3.329441 | 2.059641 | 3.823095 | 4.276025 | 4.177012 | 2.227356 | 2.995332 | 1.510191 | 3.363303 | 1.742657 | 2.628699 | 3.777069 | 4.597548 | 0.786378 | 2.701095 | 1.116431 | 1.617926 | 2.450723 | 1.616997 | 0.040028 | 2.755267 | 0.813994 | 2.237919 | 2.532568 |
| 26.51389 | 4.563116 | 2.448787 | 4.145147 | 5.165224 | 4.993332 | 2.522754 | 3.840494 | 1.956273 | 3.698045 | 1.933223 | 2.880308 | 4.414097 | 4.225836 | 0.687056 | 3.025625 | 1.013131 | 1.7723 | 2.056174 | 1.623435 | 0.047391 | 2.877563 | 0.703096 | 2.369502 | 2.371301 |
| 26.52778 | 4.375069 | 2.431955 | 3.953851 | 5.552237 | 3.795786 | 2.312335 | 3.96157 | 1.720407 | 3.892265 | 1.974404 | 2.789014 | 4.377882 | 3.774933 | 0.654792 | 3.105599 | 0.714703 | 1.693711 | 1.287444 | 1.056728 | -0.147044 | 2.154102 | 0.382654 | 2.411456 | 1.760892 |
| 26.54167 | 3.647296 | 2.022051 | 2.866237 | 3.970815 | 2.893597 | 1.888721 | 3.034585 | 0.882636 | 3.14224 | 1.761753 | 2.213476 | 3.59361 | 2.706906 | 0.163958 | 2.521078 | 0.307715 | 0.886575 | 0.579951 | 0.340604 | -0.567694 | 1.32906 | 0.10963 | 1.698474 | 1.081442 |
| 26.55556 | 2.056839 | 1.028135 | 1.475037 | 1.215822 | 1.580774 | 1.375779 | 1.578824 | 0.230156 | 1.324156 | 1.123413 | 1.36313 | 1.389663 | 0.808043 | -0.383849 | 1.292553 | 0.241954 | -0.634637 | -0.249531 | 0.279488 | -0.961444 | 1.08617 | 0.055855 | 0.715902 | 0.358812 |
| 26.56944 | 0.077696 | 0.253195 | 0.777588 | -0.43405 | 0.396293 | 0.646696 | 0.312536 | -0.08323 | -0.166226 | 0.066349 | 1.037787 | -0.217215 | -0.191992 | -0.593339 | 1.067262 | 0.707749 | -1.691745 | 0.2359 | 0.718552 | -1.208478 | 1.37064 | 0.302055 | 0.371448 | 0.46684 |
| 26.58333 | 0.230063 | 0.55815 | 0.911241 | 0.992191 | 1.46331 | 0.028348 | 1.335232 | -0.064338 | 0.727778 | 0.364114 | 1.801564 | 0.897562 | 0.187163 | -0.149724 | 1.812181 | 1.618123 | -0.974799 | 1.622615 | 0.917328 | -0.869167 | 1.512942 | 0.455707 | 0.516099 | 1.017956 |
| 26.59722 | 3.695614 | 1.294588 | 1.520643 | 1.387487 | 2.117894 | -0.541709 | 2.162073 | 0.047095 | 2.550198 | 1.552677 | 2.590238 | 2.04799 | 1.140344 | 0.118382 | 1.951761 | 1.723498 | 0.701667 | 1.669911 | 0.709191 | -0.114642 | 1.410766 | 0.253398 | 1.008502 | 0.848051 |
| 26.61111 | 5.197839 | 1.213203 | 1.912242 | 1.387486 | 1.405785 | 0.331253 | 2.088115 | 0.176411 | 2.232772 | 2.455687 | 1.680687 | 1.949673 | 2.089131 | 0.647595 | 0.694504 | 0.817314 | 1.985117 | 1.135095 | 0.571004 | -0.169729 | 1.265674 | 0.082641 | 0.836443 | -0.0139 |
| 26.625 | 4.460761 | 1.127637 | 1.135681 | 2.583229 | 0.868135 | 1.854239 | 1.664542 | 0.452825 | 1.184663 | 1.560653 | 0.674884 | 1.868152 | 1.933401 | 0.257689 | 0.525565 | 0.261155 | 1.808461 | 0.876353 | 0.49073 | -0.325688 | 1.040342 | 0.270712 | 0.251702 | -0.370141 |
| 26.63889 | 2.928279 | -0.125484 | -0.239795 | 0.606013 | 1.04448 | 0.657706 | 0.589696 | 0.746851 | 0.327011 | 0.220655 | -0.136188 | 0.509488 | 0.571886 | 0.02109 | 0.459102 | 0.218532 | 1.50769 | 0.445016 | 0.222885 | -0.161163 | 0.81654 | 0.258843 | -0.212099 | -0.046736 |
| 26.65278 | -0.081291 | -0.354206 | -0.497563 | -0.866647 | 0.876572 | 0.283175 | -0.223882 | 0.60799 | -0.784596 | -0.56079 | -0.215127 | 1.239603 | -0.180546 | -0.080042 | 0.285862 | 0.365572 | 0.622289 | -0.074269 | 0.0174 | 0.067438 | 0.143201 | 0.100141 | 0.053261 | 0.073508 |
| 26.66667 | -0.885908 | 0.950033 | 0.081631 | -0.220198 | 0.632205 | 0.394317 | 0.256979 | 0.446116 | 0.181195 | 0.125254 | 1.490577 | 2.081971 | -0.054158 | -0.035909 | 0.495727 | 0.212708 | -0.409578 | -0.528083 | -0.157874 | 0.147355 | -0.012623 | 0.12016 | 0.007697 | 0.21125 |
| 26.68056 | -0.254476 | 1.023018 | 0.747916 | 0.226548 | 0.522689 | 1.162265 | 2.166052 | 0.444697 | 1.025533 | 1.132522 | 2.53399 | 2.938655 | 0.174603 | 0.000412 | 1.135811 | -0.199169 | -1.040427 | -0.096661 | -0.284858 | 0.055965 | -0.540357 | 0.08133 | 0.39442 | 0.143534 |
| 26.69444 | 1.60364 | 0.298364 | 0.654795 | 0.930786 | 0.640159 | 1.491259 | 1.89801 | 0.459245 | 0.967324 | 0.989536 | 1.036098 | 1.472985 | 0.411991 | -0.068569 | 1.07131 | -0.334896 | -1.122275 | 0.339612 | -0.464547 | -0.020327 | -0.233642 | -0.00405 | 0.505866 | 0.005475 |
| 26.70833 | 1.440029 | 0.2734 | -0.486586 | 1.920911 | 0.644244 | 0.147344 | -0.331978 | 0.494499 | -0.181988 | -0.035671 | -0.812619 | -0.932603 | 1.250537 | 0.111914 | -0.75991 | -0.085969 | -0.040505 | 0.484596 | -0.55587 | -0.076526 | 0.496512 | -0.121913 | 0.049714 | 0.135984 |
| 26.72222 | 0.432272 | 0.096122 | -0.337588 | -1.107568 | 0.343639 | -0.543456 | -0.145169 | 0.470651 | -1.366175 | -0.213236 | -1.765807 | -1.284184 | 2.036186 | 0.336227 | -1.522595 | 0.541643 | 0.996915 | 0.827541 | -0.314237 | -0.118255 | 0.630042 | -0.134829 | -0.72228 | 0.4283 |
| 26.73611 | -1.153542 | 0.426611 | -0.099924 | -1.094385 | 0.192126 | -0.159781 | -0.830341 | 0.396379 | -0.684018 | 0.267476 | -0.408013 | -0.045316 | 1.507637 | 0.277166 | -1.040515 | 0.900916 | 1.417777 | 1.125157 | 0.352687 | -0.109434 | -0.147263 | -0.112499 | -0.581289 | -0.251932 |
| 26.75 | -1.014165 | 0.32538 | 0.821198 | 0.13192 | 0.483195 | 0.432994 | 0.070649 | 0.32962 | 0.030135 | 0.570986 | 1.017839 | 0.632463 | 0.306837 | 0.058828 | 0.545145 | 0.077498 | 0.056498 | 0.850177 | 0.717994 | 0.074546 | -0.981002 | 0.044936 | 0.057273 | -0.118113 |
| 26.76389 | 0.974318 | 0.599492 | 0.607237 | 1.507639 | 0.669734 | 0.784631 | 1.146833 | 0.343606 | 0.589281 | 0.192817 | 0.424841 | 0.652252 | 0.63026 | -0.412216 | 1.114566 | -0.840745 | -1.390027 | 0.037603 | 0.686721 | 0.17133 | -0.826339 | 0.094284 | -0.109338 | -0.133833 |
| 26.77778 | 1.293478 | 0.936204 | -0.120788 | 1.247974 | 0.489221 | 0.686224 | 1.153535 | 0.389004 | 1.439672 | 0.674591 | -0.231126 | -0.299192 | -0.603767 | -0.535718 | -0.168566 | -1.052515 | -1.61932 | -0.797536 | -0.039621 | 0.092487 | -0.799152 | -0.044617 | 0.083741 | -0.197086 |
| 26.79167 | -0.163338 | 0.079071 | -1.677481 | -0.121115 | 0.00605 | -0.010462 | 0.161156 | 0.332075 | 0.998696 | 0.6578 | -0.034433 | -0.064023 | -1.170915 | -0.236036 | -0.590327 | -0.43343 | -0.834996 | -0.418659 | -0.595533 | -0.036571 | 0.108482 | 0.005623 | 0.301417 | -0.169989 |
| 26.80556 | -0.602945 | -0.160703 | -1.110396 | -1.420082 | -0.021742 | -0.441102 | -0.530394 | -0.100727 | -1.166291 | 0.486757 | 0.281891 | 0.525658 | -1.987545 | -0.001123 | -1.402652 | 0.326685 | 0.058542 | 0.235186 | -0.994823 | -0.054533 | 0.736795 | 0.122498 | 0.514168 | -0.002657 |
| 26.81944 | 0.291782 | -0.352122 | 0.12678 | 0.035975 | 0.244787 | -0.412509 | -0.152025 | -0.252083 | -1.674722 | -0.430667 | -0.188463 | -0.567049 | -2.697855 | 0.265087 | -0.33356 | 0.957466 | 0.255807 | 0.307336 | -0.913907 | 0.158414 | 0.823724 | 0.092341 | 0.209874 | 0.001662 |
| 26.83333 | -0.047217 | 0.142077 | 0.285729 | 1.52986 | 0.364872 | 0.123358 | 0.508502 | -0.209532 | -0.138478 | -0.672822 | -0.040964 | -0.596634 | -1.451491 | 0.328946 | 0.78733 | 0.309657 | 0.569924 | -0.293054 | -0.846435 | 0.305524 | -0.183883 | 0.066882 | -0.103061 | -0.037179 |
| 26.84722 | -0.594038 | 1.29342 | 0.136893 | -0.647016 | -0.005822 | -0.172477 | 0.448333 | 0.152855 | 1.620284 | 0.374004 | 0.414996 | 0.568433 | -0.713676 | 0.159059 | 1.091143 | -0.211128 | -0.20434 | -0.567298 | -0.575218 | 0.207209 | -0.835132 | 0.097482 | -0.522655 | -0.165778 |
| 26.86111 | -0.243152 | 0.661413 | -0.242933 | -0.878515 | 0.045913 | -0.038994 | 0.464582 | 0.094825 | 1.926028 | 1.074567 | 0.553669 | 0.555507 | -0.228932 | -0.11376 | 0.111673 | -0.938209 | -0.831778 | -1.004879 | -0.782702 | -0.028756 | -0.885456 | 0.099346 | -0.40359 | -0.374901 |
| 26.875 | 0.986191 | -0.974642 | -0.436263 | -0.429916 | 0.156201 | -0.841186 | -0.020602 | -0.097097 | 0.048746 | 0.641779 | -0.379201 | -0.996971 | 0.824424 | -0.482416 | -1.143385 | -0.809202 | -0.332927 | -1.905919 | -0.42163 | -0.122892 | -0.45401 | 0.178327 | 0.189141 | 0.240181 |
| 26.88889 | 0.663607 | -0.840194 | -0.266837 | 1.387718 | 0.067992 | -0.240927 | -0.381744 | -0.051089 | -1.986029 | -0.598117 | -0.876206 | -1.144783 | 2.323486 | -0.616143 | -1.544984 | -0.114078 | -0.158283 | -1.448612 | -0.120888 | 0.007716 | 0.39894 | 0.234325 | 0.595902 | 0.47629 |
| 26.90278 | -0.461315 | 0.366896 | 0.333511 | 0.894235 | -0.095103 | 0.472298 | 0.050088 | 0.215043 | -0.940412 | 0.12288 | -0.183588 | -0.294006 | 3.491176 | -0.33672 | -0.880083 | 0.528633 | 0.228819 | -0.044017 | 0.440999 | 0.044167 | 0.671125 | 0.188482 | -0.103793 | 0.256111 |
| 26.91667 | -1.43492 | 0.822889 | -0.343828 | -2.034461 | -0.069413 | -0.142647 | 1.089437 | 0.341944 | -0.003731 | 0.20595 | 0.48826 | -0.288033 | 1.75743 | 0.207377 | 0.389011 | 0.588485 | 0.599536 | 1.212679 | 0.632477 | -0.102724 | -0.159096 | 0.122266 | -0.622167 | 0.037426 |
| 26.93056 | -1.740183 | -0.352921 | -1.989169 | -1.171258 | -0.254098 | -0.298402 | -0.065283 | -0.229532 | 0.326133 | 0.316146 | -0.034799 | -1.608015 | -0.455878 | 0.434793 | -0.056201 | -0.019065 | -0.220947 | 0.650355 | -0.080153 | -0.236644 | -0.84981 | 0.032093 | -0.719739 | -0.338175 |
| 26.94444 | -1.745185 | -1.724146 | -0.897882 | -0.435422 | -0.653072 | 0.046383 | -0.806502 | -0.379454 | -0.265043 | -0.300143 | -1.072693 | -2.494932 | -1.844401 | -0.276782 | -1.509449 | -0.783531 | -1.605049 | -1.11923 | -0.478449 | -0.233846 | -1.64034 | -0.18424 | -0.175967 | -0.999883 |
| 26.95833 | -1.266892 | -0.63989 | -0.110909 | 0.573685 | -0.732458 | -0.151022 | -1.67951 | -0.243588 | -1.264872 | -0.563026 | -1.684501 | -0.766144 | -3.106228 | -1.316771 | -1.895363 | -1.150307 | -2.023443 | -3.295081 | -1.006502 | -0.111546 | -1.828402 | -0.341773 | 0.299565 | -1.032789 |
| 26.97222 | -0.534651 | -0.005709 | -0.829734 | 1.466159 | -0.313579 | -0.96385 | -1.159296 | 0.363683 | -2.044956 | -0.350228 | -0.930104 | 0.208975 | -3.752169 | -1.912732 | -2.402374 | -1.13037 | -1.185867 | -3.998056 | -1.744559 | 0.015725 | -1.177705 | -0.425444 | -0.102676 | -0.901921 |
| 26.98611 | -1.028929 | -1.389909 | -2.582451 | -2.039109 | -0.125486 | -1.313285 | -0.400371 | 0.492159 | -1.83943 | -0.302313 | -0.844811 | -1.929039 | -4.805091 | -1.531044 | -1.742685 | -0.592327 | -1.384601 | -2.926062 | -2.045059 | 0.012976 | -0.814242 | -0.61253 | -1.008352 | -0.922872 |
| 27 | -1.451913 | -2.366014 | -3.493169 | -3.729869 | -0.915874 | -0.758918 | -1.376225 | -0.174108 | -1.147737 | -0.49505 | -1.592048 | -4.263086 | -4.907811 | -1.370275 | -1.538055 | -0.350863 | -1.880309 | -1.804382 | -1.329177 | -0.092704 | -1.330094 | -0.818297 | -1.6704 | -1.672456 |
| 27.01389 | -2.544533 | -1.479061 | -3.110711 | -3.541122 | -2.329575 | -0.720883 | -2.096828 | -2.041998 | -1.425531 | -0.912272 | -2.292767 | -3.443285 | -4.391278 | -0.87249 | -2.273681 | -0.519192 | -1.604433 | -1.244362 | -0.642135 | -0.154958 | -2.293334 | -0.742581 | -1.865015 | -2.001204 |
| 27.02778 | -2.658906 | -0.734192 | -1.35616 | -0.990495 | -2.410926 | -1.142721 | -2.510333 | -3.774345 | -2.350232 | -0.883153 | -2.759279 | -1.61546 | -3.992747 | -0.527465 | -2.902869 | -0.745127 | -0.955745 | -1.322683 | -0.436944 | -0.096647 | -2.728433 | -0.498699 | -1.659594 | -1.863582 |
| 27.04167 | -2.315689 | -0.780824 | -1.241995 | -1.292097 | -1.493043 | -1.803516 | -2.358695 | -2.351052 | -2.447088 | -0.782482 | -1.944941 | -1.136798 | -2.41488 | -0.108961 | -2.335324 | -0.875186 | -0.696852 | -1.175271 | -0.159667 | -0.111485 | -1.6798 | -0.276978 | -1.053366 | -1.187789 |
| 27.05556 | -2.475196 | -1.354455 | -1.795342 | -1.944887 | -0.991757 | -1.961976 | -1.458301 | 0.133266 | -1.705607 | -0.611048 | -1.105904 | -1.99311 | -1.28174 | -0.022736 | -1.152891 | -0.756686 | -0.75727 | -0.778801 | -0.234935 | -0.252967 | -0.654181 | -0.259502 | -0.54684 | -0.783017 |
| 27.06944 | -1.509411 | -1.216471 | -1.406028 | -2.493736 | -1.052479 | -1.272275 | -0.403615 | 1.393096 | -0.84159 | -0.324821 | -0.586646 | -2.219617 | -0.710041 | -0.033301 | -0.48647 | -0.258941 | -0.311402 | -0.707913 | -0.678171 | -0.294544 | -0.486146 | -0.44978 | -0.256557 | -0.520285 |
| 27.08333 | -0.621503 | -0.592396 | -0.437709 | -1.917969 | -1.673989 | -0.238591 | -0.217185 | 0.974362 | -0.540255 | -0.269357 | -0.199222 | -1.218575 | -0.066868 | 0.090932 | -0.019811 | 0.519498 | 0.045662 | -0.556532 | -0.700343 | -0.134813 | -0.431004 | -0.464159 | -0.08622 | -0.253019 |
| 27.09722 | -0.563326 | -0.127684 | -0.71584 | -1.508507 | -3.191191 | -0.019576 | -1.187538 | 0.176567 | -0.976926 | -0.408007 | -0.325408 | -0.301897 | -0.023669 | 0.128618 | 0.124562 | 0.378563 | 0.174362 | -0.15404 | -0.254403 | 0.173859 | -0.331611 | -0.330597 | -0.090045 | -0.108588 |
| 27.11111 | -1.535461 | -0.182062 | -1.796868 | -1.974999 | -3.696246 | -0.607697 | -2.271223 | -0.478458 | -2.062729 | -0.565531 | -0.929088 | -0.511812 | -0.679937 | 0.05917 | -0.125226 | -0.050129 | 0.103394 | 0.078742 | 0.158262 | 0.306553 | -0.380952 | -0.188767 | -0.430729 | -0.286718 |
| 27.125 | -1.682191 | -0.470447 | -1.872669 | -2.441821 | -3.624306 | -1.191028 | -2.284826 | -1.14172 | -2.722078 | -0.639149 | -1.409635 | -1.504594 | -1.142077 | 0.029853 | -0.54291 | -0.384333 | -0.063125 | -0.012531 | 0.064194 | 0.099408 | -0.538917 | -0.19137 | -0.784971 | -0.517778 |
| 27.13889 | -0.921387 | -0.671296 | -1.008952 | -2.134488 | -3.110523 | -0.940277 | -1.487375 | -1.528739 | -2.260864 | -0.366483 | -1.345898 | -1.91668 | -1.189861 | 0.019858 | -0.782694 | -0.361243 | -0.329907 | -0.328113 | -0.270439 | -0.027977 | -0.667529 | -0.305445 | -0.977635 | -0.738877 |
| 27.15278 | -0.593353 | -0.606447 | -1.083886 | -1.541445 | -2.366437 | -0.433132 | -0.761317 | -1.883266 | -1.354731 | -0.21825 | -1.306849 | -1.580704 | -1.121927 | -0.165881 | -0.813886 | -0.210831 | -0.494857 | -0.576804 | -0.372043 | -0.069754 | -0.652542 | -0.41903 | -1.026854 | -0.974683 |
| 27.16667 | -0.719544 | -0.493244 | -0.958454 | -1.222583 | -1.963298 | -0.323009 | -0.595355 | -1.485344 | -0.97075 | -0.555117 | -1.23261 | -1.021663 | -1.118934 | -0.362091 | -0.847426 | -0.174124 | -0.422299 | -0.600131 | -0.320499 | -0.068138 | -0.587466 | -0.472052 | -1.072251 | -1.108401 |
| 27.18056 | -0.384464 | -0.313668 | -1.08258 | -0.94562 | -1.540097 | -0.618833 | -0.692343 | -1.149164 | -0.770824 | -0.679467 | -1.124086 | -0.610998 | -1.207172 | -0.440284 | -0.939754 | -0.399004 | -0.369056 | -0.519157 | -0.177966 | -0.108077 | -0.587845 | -0.462567 | -1.030043 | -1.105023 |
| 27.19444 | 0.095029 | -0.14777 | -0.734883 | -0.798689 | -1.088242 | -0.822621 | -0.709105 | -1.294251 | -0.356595 | -0.278518 | -0.890619 | -0.520871 | -1.390658 | -0.4424 | -1.017707 | -0.699991 | -0.627806 | -0.454592 | -0.156571 | -0.240023 | -0.65833 | -0.371863 | -0.988739 | -1.213866 |
| 27.20833 | -0.001276 | -0.076547 | -0.311505 | -0.74637 | -0.667415 | -0.698552 | -0.668983 | -1.621039 | -0.151164 | -0.047097 | -0.659765 | -0.492911 | -1.522292 | -0.428438 | -1.064111 | -0.843935 | -1.184176 | -0.498733 | -0.21699 | -0.390501 | -0.759491 | -0.239632 | -0.995048 | -1.261982 |
| 27.22222 | -0.214449 | -0.129362 | -0.295134 | -0.585531 | -0.277399 | -0.401766 | -0.511676 | -1.345986 | -0.476787 | -0.412607 | -0.376688 | -0.381587 | -1.451511 | -0.445152 | -1.043801 | -0.714893 | -1.327696 | -0.619795 | -0.355201 | -0.471769 | -0.845328 | -0.201114 | -1.030381 | -1.163598 |
| 27.23611 | -0.066047 | -0.1899 | -0.479658 | -0.388437 | -0.056584 | -0.322891 | -0.445347 | -0.637777 | -0.566997 | -0.518336 | -0.194831 | -0.232174 | -1.251809 | -0.480744 | -0.883858 | -0.53503 | -0.92895 | -0.731575 | -0.540609 | -0.475032 | -0.864816 | -0.308201 | -0.889442 | -0.901973 |
| 27.25 | 0.092565 | -0.099353 | -0.399115 | -0.361125 | -0.044202 | -0.340166 | -0.295917 | 0.003502 | -0.262307 | -0.156088 | -0.059715 | -0.173818 | -0.878457 | -0.496758 | -0.563508 | -0.473654 | -0.496838 | -0.622116 | -0.529096 | -0.346828 | -0.791024 | -0.412018 | -0.586305 | -0.624051 |
| 27.26389 | 0.083062 | 0.023833 | -0.054745 | -0.384732 | 0.035537 | -0.22442 | -0.120802 | 0.438487 | -0.077589 | 0.011436 | 0.064811 | -0.105118 | -0.317085 | -0.476728 | -0.245323 | -0.454893 | -0.003314 | -0.188871 | -0.249027 | -0.041862 | -0.55635 | -0.295017 | -0.232582 | -0.432371 |
| 27.27778 | 0.06328 | 0.029389 | 0.075358 | -0.202489 | 0.121937 | -0.03396 | -0.019061 | 0.582089 | -0.251761 | -0.269593 | 0.128448 | -0.111746 | 0.122581 | -0.442371 | -0.13691 | -0.409227 | 0.518607 | 0.253986 | -0.064257 | 0.182474 | -0.301642 | -0.114518 | 0.0092 | -0.217123 |
| 27.29167 | 0.116561 | -0.003521 | 0.073104 | -0.030395 | -0.003231 | 0.143763 | 0.098211 | 0.505323 | -0.105298 | -0.1288 | 0.21456 | -0.08037 | 0.0753 | -0.378329 | -0.198597 | -0.359631 | 0.491951 | 0.27935 | -0.14107 | 0.110358 | -0.205385 | -0.097345 | 0.029133 | -0.067382 |
| 27.30556 | 0.074389 | 0.034621 | 0.244363 | -0.145589 | -0.207827 | 0.223468 | 0.303406 | 0.412477 | 0.261326 | 0.30972 | 0.216931 | -0.037747 | -0.165062 | -0.321704 | -0.288501 | -0.361388 | 0.23546 | 0.066888 | -0.307391 | -0.028146 | -0.185258 | -0.189648 | -0.121528 | -0.107764 |
| 27.31944 | -0.129037 | -0.006708 | 0.160855 | -0.240431 | -0.279607 | 0.120531 | 0.286106 | 0.306445 | 0.204236 | 0.16124 | 0.045101 | -0.100915 | -0.207396 | -0.324126 | -0.319777 | -0.377054 | 0.003228 | -0.048255 | -0.331383 | -0.082392 | -0.113186 | -0.25823 | -0.269351 | -0.234102 |
| 27.33333 | -0.129031 | -0.057799 | -0.207847 | -0.130036 | -0.114794 | 0.010296 | 0.101955 | 0.147755 | -0.179102 | -0.353424 | 0.013465 | -0.073714 | -0.024298 | -0.343699 | -0.315672 | -0.279514 | -0.058164 | 0.010486 | -0.254799 | -0.073602 | -0.043368 | -0.199161 | -0.220809 | -0.217945 |
| 27.34722 | 0.104264 | 0.021919 | -0.015677 | -0.001928 | 0.155485 | 0.110808 | 0.182345 | 0.044079 | -0.10685 | -0.172459 | 0.211939 | 0.066269 | 0.091504 | -0.283985 | -0.351853 | -0.061749 | -0.006215 | 0.095349 | -0.164405 | -0.055897 | -0.034658 | -0.069511 | -0.082384 | -0.063366 |
| 27.36111 | 0.267031 | 0.158626 | 0.260975 | 0.117392 | 0.41025 | 0.379906 | 0.474709 | 0.070769 | 0.345461 | 0.325573 | 0.33757 | 0.160004 | -0.077669 | -0.075198 | -0.387116 | 0.126977 | 0.000661 | 0.098068 | -0.07148 | -0.063568 | -0.027588 | -0.000514 | -0.080422 | 0.04939 |
| 27.375 | 0.322271 | 0.220839 | 0.338972 | 0.352298 | 0.669613 | 0.516298 | 0.533857 | 0.168012 | 0.561571 | 0.262121 | 0.380322 | 0.264504 | -0.261132 | 0.190352 | -0.266893 | 0.109507 | -0.032852 | 0.093152 | 0.094901 | -0.090395 | 0.045154 | -0.001725 | -0.081304 | 0.152524 |
| 27.38889 | 0.438341 | 0.256939 | 0.397554 | 0.629433 | 0.849123 | 0.515384 | 0.507947 | 0.323572 | 0.557032 | 0.060004 | 0.515501 | 0.478293 | -0.100529 | 0.294876 | -0.030553 | 0.039123 | 0.130763 | 0.153715 | 0.30442 | -0.137394 | 0.155292 | 0.017943 | 0.09584 | 0.339599 |
| 27.40278 | 0.619157 | 0.292977 | 0.818331 | 0.688521 | 1.012384 | 0.560419 | 0.643446 | 0.542689 | 0.725314 | 0.368983 | 0.622258 | 0.565434 | 0.33549 | 0.218228 | 0.150597 | 0.223175 | 0.370077 | 0.209475 | 0.480565 | -0.214174 | 0.28473 | 0.103964 | 0.401257 | 0.558309 |
| 27.41667 | 0.848613 | 0.334979 | 1.08231 | 0.701373 | 1.457693 | 0.744518 | 0.819944 | 0.9255 | 1.015193 | 0.741463 | 0.810438 | 0.674485 | 0.79143 | 0.195611 | 0.283652 | 0.608599 | 0.519155 | 0.350967 | 0.59449 | -0.27128 | 0.470532 | 0.235581 | 0.580247 | 0.716932 |
| 27.43056 | 1.078827 | 0.498315 | 1.053546 | 1.084012 | 1.886786 | 1.006965 | 0.895983 | 1.431227 | 1.179432 | 0.667827 | 1.103153 | 0.993725 | 1.231118 | 0.394667 | 0.521686 | 0.73809 | 0.691285 | 0.730348 | 0.686704 | -0.203864 | 0.641221 | 0.346602 | 0.743074 | 1.034758 |
| 27.44444 | 1.216454 | 0.689398 | 1.264154 | 1.704071 | 1.95318 | 1.323587 | 1.142936 | 1.559462 | 1.448471 | 0.619709 | 1.387715 | 1.374991 | 1.669163 | 0.68885 | 0.942154 | 0.652842 | 1.068878 | 1.272599 | 0.856864 | 0.088828 | 0.825651 | 0.446742 | 1.115435 | 1.650391 |
| 27.45833 | 1.43408 | 0.862611 | 1.930526 | 2.108064 | 2.090618 | 1.581246 | 1.399269 | 1.132276 | 1.979617 | 0.866147 | 1.702728 | 1.690296 | 2.133704 | 0.884317 | 1.464557 | 0.697145 | 1.73749 | 1.96445 | 1.05867 | 0.46015 | 1.091598 | 0.582528 | 1.706224 | 2.302669 |
| 27.47222 | 1.939111 | 1.033359 | 2.544111 | 2.32703 | 2.48114 | 1.763029 | 1.649824 | 0.944749 | 2.618972 | 1.17458 | 1.983306 | 1.984829 | 3.07333 | 1.077235 | 1.974631 | 0.795325 | 1.987185 | 2.606941 | 1.170768 | 0.609431 | 1.473363 | 0.745252 | 2.322181 | 2.735978 |
| 27.48611 | 2.67598 | 1.321846 | 2.872487 | 2.836185 | 3.279068 | 2.038608 | 2.030794 | 1.608455 | 2.904368 | 1.151125 | 2.339663 | 2.507141 | 4.508742 | 1.424221 | 2.474137 | 0.868938 | 1.668 | 2.580812 | 1.321869 | 0.528439 | 2.040002 | 0.992164 | 2.583571 | 3.129715 |
| 27.5 | 3.424287 | 1.733053 | 3.486529 | 4.021539 | 4.187831 | 2.558703 | 2.813233 | 3.247775 | 3.331293 | 1.042797 | 2.728872 | 3.351873 | 5.422386 | 1.739383 | 3.102821 | 0.829887 | 1.625989 | 3.281245 | 1.740651 | 0.373054 | 2.788982 | 1.109307 | 2.563334 | 3.206522 |
| 27.51389 | 4.303741 | 2.092249 | 4.447186 | 5.054495 | 4.521984 | 2.808741 | 3.802778 | 3.495635 | 4.152835 | 1.351385 | 2.724109 | 4.314704 | 5.73098 | 1.683492 | 3.620915 | 0.783894 | 1.590587 | 3.593711 | 2.012407 | 0.153601 | 3.028686 | 0.803354 | 2.791976 | 3.343583 |
| 27.52778 | 4.731244 | 1.94666 | 4.626684 | 5.211515 | 4.01827 | 2.180793 | 3.989781 | 1.509703 | 4.250252 | 1.372186 | 2.07569 | 4.59359 | 5.989142 | 1.36817 | 3.562686 | 0.936945 | 1.729474 | 3.130592 | 1.949041 | -0.036193 | 2.624548 | 0.451527 | 2.963054 | 3.127144 |
| 27.54167 | 3.719315 | 0.967041 | 3.068333 | 3.283945 | 2.381665 | 1.008661 | 2.172788 | -0.845009 | 2.688496 | 0.608754 | 1.31219 | 3.314761 | 3.967441 | 0.99069 | 3.092678 | 1.025756 | 1.089151 | 2.34904 | 1.372544 | -0.24206 | 1.572274 | 0.105228 | 1.878077 | 2.382279 |
| 27.55556 | 1.668976 | 0.375808 | 0.712454 | 1.082856 | 0.343694 | 0.487293 | 0.125922 | -1.674438 | 0.540224 | 0.001081 | 0.514694 | 1.097248 | -0.329328 | 0.28401 | 2.090036 | 0.866688 | -0.01981 | 1.054814 | 0.836977 | -0.239263 | 0.689607 | -0.124657 | 0.555609 | 0.908271 |
| 27.56944 | -0.154447 | 0.504084 | -0.163619 | -0.521475 | 0.094651 | 0.67182 | 0.172159 | -0.3202 | -0.623597 | 0.076265 | 0.541471 | -0.127806 | -1.658972 | -0.311486 | 1.514527 | 0.709973 | -1.094404 | 0.174613 | 0.502461 | 0.031135 | 0.972204 | 0.06197 | 0.175279 | 0.301084 |
| 27.58333 | 0.290217 | 1.011039 | 0.909406 | 0.520679 | 0.830777 | 1.428354 | 1.344558 | 1.128318 | 0.83186 | 0.889837 | 1.975246 | 1.712984 | -0.122043 | -0.739716 | 1.692412 | 0.746751 | -0.756145 | 0.285145 | -0.003976 | 0.261569 | 1.757626 | 0.637062 | 1.017439 | 1.086406 |
| 27.59722 | 1.822071 | 1.342582 | 1.553501 | 2.207069 | 1.488875 | 1.632565 | 3.296275 | 1.603056 | 2.321374 | 1.601635 | 3.719927 | 3.247018 | 1.688832 | 0.045583 | 2.702394 | 0.844279 | 0.169213 | 1.226905 | -0.245563 | 0.259371 | 2.143259 | 0.720948 | 1.544071 | 1.843278 |
| 27.61111 | 2.806355 | 1.62829 | 2.733427 | 3.151241 | 1.154582 | 1.772637 | 3.000279 | 0.726494 | 2.43021 | 1.494468 | 2.494691 | 3.849988 | 1.818509 | 0.815963 | 2.499574 | 0.718407 | 0.709618 | 1.817654 | -0.031421 | 0.10437 | 1.925478 | 0.249303 | 1.65488 | 0.981029 |
| 27.625 | 4.069466 | 1.103365 | 3.094885 | 3.04742 | 0.743176 | 1.438119 | 1.66709 | 0.381367 | 1.469592 | 0.902886 | 1.147997 | 2.627943 | 1.052128 | 1.26289 | 1.28606 | 0.319643 | 1.835616 | 1.037021 | 0.096068 | -0.128562 | 1.535327 | 0.054354 | 1.006411 | 0.049917 |
| 27.63889 | 3.259806 | 0.846072 | 1.248626 | 2.227356 | 0.456781 | 0.643628 | 0.271093 | 0.345845 | 1.196988 | 0.772743 | 0.19858 | 0.32135 | 1.864582 | 0.557129 | 1.464611 | -0.072727 | 1.861356 | 0.00696 | -0.21542 | -0.234254 | 1.069452 | 0.019684 | 0.630677 | -0.12045 |
| 27.65278 | 0.679344 | 0.440642 | -0.771225 | -0.944749 | 0.281042 | -0.460377 | 0.683745 | 0.573941 | 0.000688 | 0.664058 | -0.481051 | -1.234318 | 1.094783 | 0.319158 | 0.846684 | 0.012133 | 1.183526 | -0.629163 | -0.093719 | 0.003703 | 0.68916 | -0.029153 | 1.045778 | 0.277301 |
| 27.66667 | -1.064688 | 0.489607 | -1.302421 | -1.824823 | 0.270731 | -0.360456 | 1.211662 | 0.558458 | -0.587312 | 0.175341 | 0.828032 | 0.481651 | 0.672083 | 0.229137 | -0.169773 | 0.355041 | 0.267747 | -0.801417 | 0.070159 | 0.293502 | 0.231969 | 0.003302 | 0.289797 | 0.347852 |
| 27.68056 | -1.269471 | 0.489609 | -0.106553 | -0.220774 | 0.33686 | 0.908163 | 1.159989 | 0.457085 | 0.000291 | -0.070176 | 2.264869 | 2.933745 | -0.004082 | -0.086412 | 0.037664 | 0.275923 | -0.551458 | -0.784396 | -0.093172 | 0.195052 | -0.257251 | 0.092185 | -0.126732 | 0.117287 |
| 27.69444 | 0.504735 | -0.130063 | 0.960933 | 1.410117 | 0.518182 | 1.203281 | 0.444062 | 0.40621 | 0.433547 | 0.084435 | 1.611951 | 2.497354 | -1.280286 | -0.360086 | 0.701792 | -0.011944 | -1.117989 | 0.027507 | -0.267414 | -0.033236 | 0.004805 | 0.176339 | 0.224202 | 0.030713 |
| 27.70833 | 2.595506 | -0.085629 | 1.341219 | 2.821481 | 0.475355 | 0.057765 | -0.107696 | 0.35081 | -0.174461 | -0.193364 | 0.133216 | 1.117825 | 0.15256 | -0.2252 | -0.023186 | -0.088525 | -1.136148 | 0.969663 | -0.105867 | -0.119353 | 0.526775 | 0.192324 | 0.603965 | 0.104482 |
| 27.72222 | 1.686475 | 0.254012 | -0.953737 | -0.470072 | 0.044934 | -1.53018 | -1.179212 | 0.134791 | 0.127406 | 0.159328 | -0.737517 | -1.419011 | 1.475643 | 0.237639 | -0.644301 | 0.235684 | -0.147106 | 1.316106 | -0.131232 | -0.010771 | 0.504063 | 0.175584 | 0.401468 | 0.095639 |
| 27.73611 | -1.054088 | 0.343674 | -3.123274 | -3.468873 | -0.40346 | -1.341403 | -0.825966 | -0.037126 | -0.156103 | 0.301756 | -0.793225 | -2.164229 | 1.929826 | 0.462845 | -1.022071 | 0.567954 | 0.915536 | 0.046848 | -0.229995 | 0.138777 | 0.299174 | 0.144767 | -1.095468 | 0.298886 |
| 27.75 | -2.611111 | 0.117197 | -1.4383 | -1.35607 | -0.195182 | -0.138352 | 0.360802 | -0.182786 | -1.016616 | 0.266582 | -0.26131 | -1.406496 | 0.165659 | 0.242496 | -0.137766 | 0.348898 | 1.40564 | -0.918334 | -0.3487 | 0.068938 | 0.093823 | 0.011456 | -0.429885 | 0.406907 |
| 27.76389 | -1.940551 | -0.007864 | 0.630721 | 1.402585 | 0.319362 | 0.593329 | 1.318109 | -0.273113 | -0.375179 | 0.137679 | 0.290461 | 0.530847 | 0.121043 | -0.209099 | 0.368539 | -0.145704 | 0.783913 | -0.687577 | -0.031692 | -0.155555 | 0.055176 | -0.043549 | 0.44205 | 0.365445 |
| 27.77778 | 1.124242 | 0.093777 | 1.626759 | 3.620128 | 0.656604 | 0.750473 | 0.94222 | -0.264368 | 1.027057 | 0.210048 | 1.002938 | 2.911736 | 1.013096 | -0.288525 | 1.247034 | -0.584373 | -0.044144 | 0.270232 | 0.350577 | -0.22061 | 0.088324 | 0.00768 | 0.066349 | 0.190387 |
| 27.79167 | 2.617243 | 0.253454 | 0.564693 | 0.91153 | 0.192666 | 0.171003 | 0.026082 | -0.132067 | 1.367301 | 0.290242 | 0.512637 | 2.079646 | 1.222094 | 0.03233 | 1.165055 | -0.808827 | -0.697995 | 0.907883 | 0.495157 | -0.066999 | 0.032355 | 0.103204 | 0.373673 | -0.028965 |
| 27.80556 | 1.424372 | 0.243598 | -1.323173 | -0.983469 | -0.172483 | -0.799594 | -0.557678 | 0.099364 | 1.1302 | 0.383365 | -0.903841 | -1.353513 | 0.394804 | 0.174127 | -0.954842 | -0.30313 | -0.783409 | -0.082743 | 0.356701 | 0.124299 | 0.011198 | 0.115416 | 1.288316 | 0.33636 |
| 27.81944 | -0.92776 | 0.028805 | -2.1786 | 0.031798 | -0.097681 | -1.078206 | 0.282347 | 0.191299 | -0.121098 | -0.011816 | -1.245982 | -3.367318 | -0.507253 | -0.13656 | -3.112851 | 0.324922 | -0.638441 | -1.50923 | -0.057639 | 0.172307 | -0.076175 | 0.058046 | -0.069965 | -0.187693 |
| 27.83333 | -1.468584 | 0.166213 | -0.297756 | 0.661991 | 0.299807 | -0.425145 | 0.647215 | -0.139856 | -0.989328 | -0.061802 | 0.05048 | -0.882634 | -1.025175 | -0.368218 | -1.084152 | 0.91058 | -0.476038 | -1.448113 | -0.470153 | 0.002597 | 0.048964 | -0.013695 | -0.207237 | -0.132124 |
| 27.84722 | -0.202961 | 0.19639 | 0.747065 | 0.246793 | 0.335725 | -0.10249 | 0.438446 | -0.119834 | -0.130492 | 0.338143 | 1.955071 | 1.739096 | -2.309911 | -0.213598 | -0.009917 | 0.623819 | -0.392703 | -0.159998 | -0.68359 | -0.158 | 0.131428 | -0.077087 | -0.55998 | -0.030913 |
| 27.86111 | -0.264716 | -0.227817 | 0.925486 | 0.775134 | 0.055202 | -0.239968 | 0.126789 | 0.061421 | 0.29962 | 0.119152 | 0.858654 | 1.482158 | -0.562215 | 0.011577 | 0.441957 | 0.093061 | 0.127479 | 0.460512 | -0.934442 | -0.131746 | 0.022068 | -0.086708 | -0.284354 | -0.020243 |
| 27.875 | -0.28942 | 0.000151 | -0.472527 | 1.577766 | 0.030928 | 0.112656 | -0.347532 | 0.062338 | 0.600861 | 0.08854 | -0.989398 | -0.847611 | 1.750568 | 0.175776 | 0.902347 | -0.472069 | 0.672253 | -0.110998 | -0.677819 | 0.034542 | -0.206506 | -0.020618 | -0.39266 | 0.005656 |
| 27.88889 | 1.254927 | 0.044526 | -1.807224 | 1.839078 | 0.220028 | -0.776795 | -1.025271 | -0.188287 | 1.060728 | -0.224859 | -1.851647 | -2.429527 | 1.130132 | -0.176607 | 0.09539 | -0.851724 | 0.70561 | -0.929199 | -0.104893 | 0.100715 | -0.376288 | -0.023755 | -0.752849 | -0.362201 |
| 27.90278 | 1.221875 | -0.492586 | -1.078183 | 0.246712 | 0.131794 | -0.255794 | -0.807671 | -0.568255 | 0.432697 | 0.195663 | -0.716947 | -1.603296 | -1.602926 | -0.637522 | -1.58241 | -0.542536 | -0.352508 | -1.672562 | 0.667961 | -0.027558 | -0.429179 | -0.143642 | -0.432471 | -0.445575 |
| 27.91667 | -0.761247 | -0.8698 | -0.949955 | -1.014962 | -0.142881 | -0.214726 | 0.036361 | -0.321601 | -0.422122 | 0.199115 | 0.903759 | -0.039374 | -2.770429 | -1.362069 | -1.943322 | 0.010218 | -1.587602 | -2.189427 | 0.812581 | -0.172855 | -0.495041 | -0.228428 | 0.150326 | -0.351892 |
| 27.93056 | -3.012952 | -0.198736 | 0.389409 | 0.703496 | -0.339551 | -0.902323 | -0.02606 | 0.004669 | -1.548957 | -0.415314 | 0.169191 | -0.240565 | -1.44309 | -1.335867 | -1.70249 | 0.589522 | -1.979557 | -2.062139 | 0.021837 | -0.167587 | -0.951389 | -0.23321 | 0.21731 | -0.318028 |
| 27.94444 | -1.882519 | 0.414133 | 0.092709 | 2.395621 | -0.193516 | -0.195429 | -0.015438 | 0.091286 | -0.460132 | -0.430237 | -1.438491 | -1.995791 | -0.161541 | -0.593091 | -1.476311 | 0.511771 | -1.241091 | -1.831553 | -0.304056 | -0.064727 | -0.935536 | -0.217686 | -0.241431 | -0.279417 |
| 27.95833 | 0.609287 | -0.478357 | -0.721699 | 3.741519 | -0.074315 | 0.018725 | -0.993961 | -0.207517 | 0.296433 | -0.19734 | -2.045723 | -3.270032 | -0.417797 | -0.109455 | -1.029392 | -0.100961 | -0.525059 | -1.924137 | -0.758999 | 0.028559 | -0.755898 | -0.25634 | -0.933707 | -0.285166 |
| 27.97222 | 0.583847 | -1.844333 | -1.917509 | -1.102331 | -0.31132 | -0.727552 | -2.684872 | -0.447663 | -0.065786 | -0.005999 | -1.885976 | -3.450031 | -2.372146 | -0.219845 | -1.764146 | -0.990837 | -0.170257 | -2.140285 | -0.689734 | 0.000823 | -0.923942 | -0.418717 | -1.562343 | -0.850289 |
| 27.98611 | -1.852551 | -1.538667 | -2.459015 | -3.71591 | -0.79025 | -1.074121 | -2.942803 | -0.391564 | -1.256833 | -0.609827 | -1.436722 | -2.277274 | -3.886449 | -0.694736 | -3.062092 | -1.91425 | -0.886126 | -2.61749 | -0.701377 | -0.071761 | -1.256822 | -0.796983 | -1.839338 | -1.79451 |
| 28 | -3.741588 | -0.237416 | -1.491714 | -4.48106 | -1.059631 | -1.368396 | -1.295679 | -0.285946 | -2.513894 | -1.022765 | -0.835981 | -1.506505 | -4.701801 | -1.03438 | -3.429466 | -1.503869 | -2.915026 | -2.705642 | -0.515757 | -0.076948 | -1.736808 | -1.060329 | -2.151742 | -2.209917 |
| 28.01389 | -2.642045 | -0.3664 | -0.262943 | -2.853644 | -1.188214 | -0.892759 | 0.398553 | -0.364372 | -2.904235 | -0.435279 | -1.803918 | -3.240952 | -3.0201 | -1.440847 | -3.253047 | -0.68389 | -2.661028 | -2.681473 | -0.265699 | -0.03416 | -2.015419 | -0.959789 | -2.046201 | -2.678735 |
| 28.02778 | -0.518123 | -1.846881 | -0.760961 | -0.358307 | -1.463925 | -1.179939 | -0.240316 | -0.380915 | -1.126967 | 0.41928 | -4.322464 | -5.055065 | -2.705021 | -1.219771 | -3.075037 | -0.383778 | -2.09349 | -2.401915 | -0.607588 | 0.043965 | -2.301939 | -0.796005 | -1.46115 | -2.234173 |
| 28.04167 | -0.900087 | -2.427646 | -2.961918 | -0.376645 | -1.369043 | -1.485877 | -2.701942 | -0.259224 | -1.213458 | -0.357131 | -3.947394 | -4.544422 | -2.566101 | -0.665353 | -2.17293 | -0.364055 | -1.934728 | -1.62548 | -0.870712 | 0.19277 | -1.581624 | -0.478178 | -0.732312 | -1.427697 |
| 28.05556 | -3.002571 | -1.695556 | -4.303293 | -0.81484 | -1.690439 | -1.375854 | -3.350273 | -0.422091 | -1.171546 | -1.432556 | -1.687821 | -2.007858 | -2.021789 | -0.355109 | -1.183706 | -0.44668 | -1.900225 | -0.940119 | -0.842211 | 0.160848 | -0.718747 | -0.121883 | -0.497542 | -1.022319 |
| 28.06944 | -3.766655 | -0.598725 | -3.284073 | -0.819365 | -1.805756 | -1.304693 | -2.634624 | -1.552083 | -1.365082 | -1.567298 | 0.369691 | -0.048094 | -1.860332 | -0.315006 | -0.642205 | -0.528831 | -1.153452 | -0.635869 | -0.62592 | -0.04872 | -0.679656 | 0.046043 | -0.624096 | -0.785812 |
| 28.08333 | -1.887702 | 0.42929 | -0.947834 | -0.631082 | -1.817442 | -0.77315 | -1.402131 | -1.598556 | -1.156632 | -0.712208 | 0.871679 | 0.631609 | -1.065509 | -0.377631 | -0.352725 | -0.271246 | -0.230849 | -0.483482 | -0.362813 | -0.251018 | -0.800507 | 0.13112 | -0.820796 | -0.633043 |
| 28.09722 | 0.488303 | 0.522927 | 0.401966 | -1.222985 | -2.200767 | -0.264675 | -0.832236 | -0.981588 | -0.445654 | 0.250491 | 0.066605 | 0.70071 | -0.208792 | -0.416206 | -0.199055 | 0.035905 | 0.342086 | -0.000892 | -0.314377 | -0.335111 | -0.493911 | 0.168977 | -0.347532 | -0.355589 |
| 28.11111 | 0.088589 | -0.01259 | -1.347244 | -2.576494 | -2.757346 | -0.859343 | -1.657532 | -1.075969 | -0.531521 | 0.330009 | -1.361442 | -0.485515 | 0.204316 | -0.394305 | -0.227724 | 0.026327 | 0.502096 | 0.280831 | -0.069481 | -0.39441 | -0.055679 | 0.05014 | 0.10172 | -0.176434 |
| 28.125 | -1.624654 | -0.5313 | -3.227902 | -3.474579 | -2.961031 | -1.861055 | -2.510892 | -2.103922 | -1.276021 | -0.256694 | -2.279123 | -1.768351 | 0.166501 | -0.236819 | -0.437161 | 0.018687 | 0.379601 | 0.062009 | 0.019857 | -0.390444 | -0.114148 | -0.112906 | -0.328909 | -0.221578 |
| 28.13889 | -2.036122 | -0.510193 | -2.253773 | -2.898268 | -2.93649 | -1.911011 | -2.439287 | -2.693792 | -2.053206 | -0.760844 | -1.911516 | -1.716105 | -0.204797 | -0.053207 | -0.724745 | 0.075185 | 0.211825 | -0.308213 | -0.155736 | -0.389172 | -0.371091 | -0.3016 | -0.690191 | -0.300978 |
| 28.15278 | -0.867387 | -0.337669 | -1.140397 | -1.693873 | -2.269459 | -1.069389 | -1.631841 | -2.437855 | -2.248753 | -1.088448 | -1.077729 | -1.267387 | -0.583796 | -0.078333 | -0.956165 | 0.108374 | -0.126407 | -0.61223 | -0.326452 | -0.372985 | -0.496212 | -0.458921 | -0.957771 | -0.417825 |
| 28.16667 | -0.221002 | -0.5302 | -0.412103 | -0.777969 | -1.573803 | -0.628132 | -1.14685 | -2.133841 | -1.748243 | -0.915749 | -0.67184 | -1.167109 | -0.851866 | -0.25624 | -1.064441 | 0.019511 | -0.425434 | -0.708918 | -0.355211 | -0.358078 | -0.485378 | -0.567518 | -1.094311 | -0.655457 |
| 28.18056 | -0.33764 | -0.552721 | -0.617107 | -0.369827 | -1.006828 | -0.698567 | -1.015456 | -1.90303 | -0.85888 | -0.442861 | -0.916522 | -1.301782 | -1.103587 | -0.427922 | -1.07693 | -0.140361 | -0.754336 | -0.7099 | -0.30712 | -0.33133 | -0.312179 | -0.623099 | -0.736588 | -0.86951 |
| 28.19444 | -0.461144 | -0.151335 | -1.124191 | -0.303481 | -0.639605 | -0.784455 | -1.074978 | -1.783238 | -0.427641 | -0.229535 | -1.364608 | -1.221651 | -1.305163 | -0.513119 | -1.057693 | -0.272746 | -1.134222 | -0.85589 | -0.325114 | -0.382179 | -0.35652 | -0.626574 | -0.726644 | -1.035312 |
| 28.20833 | -0.226207 | -0.07857 | -1.12538 | -0.340535 | -0.568708 | -0.733916 | -0.983927 | -1.680566 | -0.650893 | -0.412174 | -1.370935 | -0.880696 | -1.645047 | -0.505266 | -1.085894 | -0.30902 | -1.606073 | -1.053736 | -0.512875 | -0.511741 | -0.648427 | -0.565353 | -1.135163 | -1.16504 |
| 28.22222 | -0.132066 | -0.463195 | -0.506975 | -0.391148 | -0.629827 | -0.657131 | -0.783013 | -1.335265 | -0.966407 | -0.588562 | -0.911536 | -0.640716 | -1.807262 | -0.496239 | -1.032893 | -0.28851 | -2.246365 | -1.063704 | -0.773876 | -0.605565 | -0.968115 | -0.477626 | -1.540304 | -1.169599 |
| 28.23611 | -0.320989 | -0.668751 | -0.248729 | -0.415211 | -0.491407 | -0.664497 | -0.696308 | -0.819991 | -0.76674 | -0.402216 | -0.505075 | -0.554894 | -1.517735 | -0.449103 | -0.839281 | -0.245304 | -2.259803 | -0.791157 | -0.915533 | -0.620156 | -1.033019 | -0.319613 | -1.322774 | -0.897181 |
| 28.25 | -0.412818 | -0.35562 | -0.418126 | -0.39553 | -0.210634 | -0.574195 | -0.626463 | -0.366781 | -0.457494 | -0.215321 | -0.468191 | -0.496457 | -1.000927 | -0.401335 | -0.549591 | -0.159333 | -1.512379 | -0.465461 | -0.520435 | -0.464593 | -0.755541 | -0.18556 | -0.686072 | -0.558176 |
| 28.26389 | -0.252678 | -0.125022 | -0.506562 | -0.479292 | -0.183909 | -0.45353 | -0.476551 | -0.172593 | -0.456616 | -0.280758 | -0.466601 | -0.357386 | -0.59215 | -0.331917 | -0.314289 | -0.038726 | -0.69044 | -0.263259 | -0.287771 | -0.297058 | -0.579562 | -0.121999 | -0.254324 | -0.276872 |
| 28.27778 | -0.150218 | -0.219008 | -0.372666 | -0.696093 | -0.48334 | -0.315451 | -0.260965 | -0.08993 | -0.511344 | -0.338041 | -0.319889 | -0.290734 | -0.412813 | -0.286693 | -0.222252 | -0.022955 | -0.292577 | -0.162704 | -0.344126 | -0.181163 | -0.490859 | -0.097574 | -0.201969 | -0.137817 |
| 28.29167 | -0.316857 | -0.209135 | -0.319096 | -0.783459 | -0.833067 | -0.225009 | -0.160512 | 0.023523 | -0.314054 | -0.224634 | -0.176098 | -0.290529 | -0.339999 | -0.33272 | -0.234587 | -0.198349 | -0.439786 | -0.087343 | -0.444466 | -0.152074 | -0.372627 | -0.063824 | -0.317002 | -0.118159 |
| 28.30556 | -0.444729 | 0.055252 | -0.325629 | -0.713479 | -0.941692 | -0.053845 | -0.078252 | 0.110327 | -0.181328 | -0.17329 | 0.02659 | -0.08923 | -0.173838 | -0.376269 | -0.238614 | -0.39968 | -0.703834 | -0.011625 | -0.362389 | -0.143087 | -0.263236 | 0.022296 | -0.261811 | -0.169889 |
| 28.31944 | -0.290193 | 0.110957 | -0.108387 | -0.619675 | -0.789458 | 0.061626 | 0.05021 | 0.090357 | -0.320975 | -0.259644 | 0.286908 | 0.201394 | 0.028259 | -0.42589 | -0.220601 | -0.438479 | -0.567332 | 0.028328 | -0.181872 | -0.093759 | -0.237994 | 0.075803 | -0.17174 | -0.242297 |
| 28.33333 | -0.163093 | -0.1171 | -0.050804 | -0.466811 | -0.53418 | 0.032217 | 0.149239 | 0.033694 | -0.335269 | -0.20707 | 0.530982 | 0.233973 | 0.026772 | -0.473426 | -0.233443 | -0.25732 | -0.049613 | 0.032787 | -0.178804 | -0.000316 | -0.184578 | 0.023987 | -0.206394 | -0.2499 |
| 28.34722 | -0.160732 | -0.106574 | -0.212681 | -0.13276 | -0.277752 | 0.057288 | 0.248781 | -0.001332 | -0.005345 | 0.007641 | 0.541884 | 0.207586 | -0.084988 | -0.429221 | -0.200235 | -0.152742 | 0.526018 | 0.065609 | -0.253113 | 0.111346 | 0.03633 | -0.046114 | -0.313843 | -0.235254 |
| 28.36111 | 0.00223 | 0.236881 | 0.083533 | 0.315642 | 0.042107 | 0.283364 | 0.384332 | 0.135651 | 0.23869 | 0.094207 | 0.644368 | 0.430263 | -0.053448 | -0.317588 | -0.173759 | -0.233382 | 0.782967 | 0.106001 | -0.2105 | 0.168299 | 0.163782 | -0.040047 | -0.261569 | -0.077776 |
| 28.375 | 0.280176 | 0.292281 | 0.592214 | 0.420896 | 0.455896 | 0.484301 | 0.574966 | 0.46904 | 0.28106 | 0.077697 | 1.051647 | 0.815924 | 0.15294 | -0.172547 | -0.141749 | -0.242926 | 0.752892 | 0.082905 | -0.002463 | 0.14691 | 0.111934 | 0.027098 | -0.088867 | 0.124627 |
| 28.38889 | 0.464275 | 0.202855 | 0.752139 | 0.350324 | 0.903373 | 0.64435 | 0.74318 | 0.715714 | 0.49146 | 0.213216 | 1.445831 | 1.015931 | 0.3491 | -0.105559 | 0.012812 | 0.022932 | 0.679466 | 0.030225 | 0.191689 | 0.135433 | 0.14611 | 0.103352 | 0.028714 | 0.299101 |
| 28.40278 | 0.701522 | 0.440002 | 0.882244 | 0.428731 | 1.321704 | 0.853856 | 0.825749 | 0.730665 | 0.850812 | 0.445802 | 1.422624 | 1.098861 | 0.427264 | -0.027859 | 0.342982 | 0.395091 | 0.806552 | 0.097699 | 0.315514 | 0.183892 | 0.471354 | 0.251696 | 0.201651 | 0.48069 |
| [illegible] | [illegible] | [illegible] | [illegible] | [illegible] | [illegible] | [illegible] | [illegible] | [illegible] | [illegible] | [illegible] | [illegible] | [illegible] | [illegible] | [illegible] | [illegible] | [illegible] | [illegible] | [illegible] | [illegible] | [illegible] | [illegible] | [illegible] | [illegible] | [illegible] |
| [illegible] | [illegible] | [illegible] | [illegible] | [illegible] | [illegible] | [illegible] | [illegible] | [illegible] | [illegible] | [illegible] | [illegible] | [illegible] | [illegible] | [illegible] | [illegible] | [illegible] | [illegible] | [illegible] | [illegible] | [illegible] | [illegible] | [illegible] | [illegible] | [illegible] |
| [illegible] | [illegible] | [illegible] | [illegible] | [illegible] | [illegible] | [illegible] | [illegible] | [illegible] | [illegible] | [illegible] | [illegible] | [illegible] | [illegible] | [illegible] | [illegible] | [illegible] | [illegible] | [illegible] | [illegible] | [illegible] | [illegible] | [illegible] | [illegible] | [illegible] |
| [illegible] | [illegible] | [illegible] | [illegible] | [illegible] | [illegible] | [illegible] | [illegible] | [illegible] | [illegible] | [illegible] | [illegible] | [illegible] | [illegible] | [illegible] | [illegible] | [illegible] | [illegible] | [illegible] | [illegible] | [illegible] | [illegible] | [illegible] | [illegible] | [illegible] |
| [illegible] | [illegible] | [illegible] | [illegible] | [illegible] | [illegible] | [illegible] | [illegible] | [illegible] | [illegible] | [illegible] | [illegible] | [illegible] | [illegible] | [illegible] | [illegible] | [illegible] | [illegible] | [illegible] | [illegible] | [illegible] | [illegible] | [illegible] | [illegible] | [illegible] |
| [illegible] | [illegible] | [illegible] | [illegible] | [illegible] | [illegible] | [illegible] | [illegible] | [illegible] | [illegible] | [illegible] | [illegible] | [illegible] | [illegible] | [illegible] | [illegible] | [illegible] | [illegible] | [illegible] | [illegible] | [illegible] | [illegible] | [illegible] | [illegible] | [illegible] |
| [illegible] | [illegible] | [illegible] | [illegible] | [illegible] | [illegible] | [illegible] | [illegible] | [illegible] | [illegible] | [illegible] | [illegible] | [illegible] | [illegible] | [illegible] | [illegible] | [illegible] | [illegible] | [illegible] | [illegible] | [illegible] | [illegible] | [illegible] | [illegible] | [illegible] |

| | | | | | | | | | | | | | | | | | | | | | | | | |
|---|---|---|---|---|---|---|---|---|---|---|---|---|---|---|---|---|---|---|---|---|---|---|---|---|
| 28.51389 | 4.014422 | 1.481413 | 4.769185 | 4.739028 | 4.063439 | 2.993687 | 3.298968 | 2.501056 | 3.96321 | 1.21793 | 2.657034 | 3.725496 | 6.299304 | 1.511993 | 3.784876 | 0.590518 | 1.941937 | 4.237835 | 1.379655 | 0.021096 | 2.656104 | 0.789366 | 3.381264 | 2.751752 |
| 28.52778 | 3.984441 | 1.410176 | 4.925106 | 5.201143 | 2.95403 | 2.859237 | 3.709116 | 1.751664 | 3.613627 | 1.038942 | 1.988237 | 4.124368 | 6.356309 | 1.373798 | 3.674169 | 0.528834 | 2.078104 | 3.817273 | 1.009361 | -0.226058 | 2.312594 | 0.26874 | 3.50462 | 2.583198 |
| 28.54167 | 3.933345 | 0.963131 | 4.761934 | 5.350018 | 1.632704 | 2.055889 | 3.860853 | -0.017362 | 2.194843 | 0.357898 | 1.305216 | 2.689398 | 6.123664 | 1.403663 | 4.041773 | 0.336644 | 1.738098 | 4.191229 | 0.807089 | -0.338658 | 1.826332 | 0.024882 | 2.910375 | 2.5512 |
| 28.55556 | 2.793528 | 0.651564 | 1.783452 | 3.199794 | 0.705692 | 1.131838 | 1.304031 | -0.461253 | 0.642055 | 0.130982 | 0.903193 | 0.676775 | 2.581337 | 1.748895 | 3.550088 | 0.299904 | -0.447042 | 3.535791 | 0.46172 | -0.262656 | 1.248025 | 0.033012 | 1.16574 | 2.597804 |
| 28.56944 | 1.173441 | 0.819349 | -0.241871 | 0.492924 | 0.515786 | 1.113177 | -0.682985 | 0.082611 | -0.256629 | 0.496572 | 1.818262 | 0.960433 | -1.652907 | 0.569324 | 1.950511 | 0.609641 | -1.950074 | 1.682103 | 0.372414 | -0.021845 | 0.89728 | 0.185567 | -0.390765 | 1.864443 |
| 28.58333 | 0.882839 | 1.059355 | -0.861118 | 0.828088 | 0.398991 | 1.279553 | 0.380984 | 0.82272 | 0.699894 | 0.609296 | 2.537063 | 3.607259 | -1.329653 | 0.189142 | 1.097784 | 1.138263 | -1.933765 | 0.39905 | 0.694473 | 0.10593 | 1.270126 | 0.244862 | -0.759612 | 0.71448 |
| 28.59722 | 2.636117 | 0.907627 | 0.379869 | 2.702885 | 0.534821 | 0.916095 | 2.320276 | 0.848871 | 1.805361 | 0.966429 | 3.187496 | 4.759222 | 0.838423 | 0.480961 | 1.480761 | 1.081151 | -0.409774 | 0.374376 | 0.696989 | 0.085206 | 1.003123 | 0.025592 | -0.418304 | 0.179532 |
| 28.61111 | 5.214214 | 0.559867 | 2.656459 | 4.713848 | 0.798293 | 1.303145 | 4.120698 | 0.440596 | 2.478665 | 1.04498 | 2.953052 | 3.877216 | 2.467064 | 0.688672 | 1.183218 | 0.499968 | 0.831888 | 1.087657 | 0.521693 | 0.030242 | 0.421086 | -0.217316 | 0.016752 | 0.579353 |
| 28.625 | 5.902116 | 0.395367 | 3.634521 | 4.075943 | 0.687017 | 1.909536 | 3.575082 | 0.248723 | 1.890696 | 0.609712 | 1.664113 | 2.909294 | 2.235469 | 0.609458 | 0.7655 | -0.085525 | 1.739959 | 1.568944 | 0.394003 | -0.015044 | 0.679203 | -0.188855 | 0.487892 | 0.243593 |
| 28.63889 | 4.420476 | 0.91474 | 3.221235 | 4.111703 | 0.56151 | 0.708643 | 1.097429 | 0.351681 | 1.380146 | 0.564019 | -0.125331 | 0.516584 | 2.342662 | 0.15769 | 0.652958 | -0.3958 | 1.60909 | 0.61252 | 0.012031 | -0.011758 | 1.003057 | 0.016362 | -0.63238 | 0.082145 |
| 28.65278 | 1.799406 | 0.464752 | 0.15341 | 0.27807 | 0.445785 | -0.878264 | -0.694128 | 0.48657 | 0.526388 | 0.380097 | -0.971871 | -1.903319 | 2.575283 | 0.186489 | -0.597965 | -0.147249 | 1.212749 | -0.204413 | 0.090416 | 0.046811 | 0.873393 | 0.146076 | -0.189054 | -0.823901 |
| 28.66667 | -0.185784 | 0.062585 | -2.53232 | -3.20203 | 0.323412 | -0.834522 | -1.26765 | 0.446697 | -0.258727 | 0.316955 | 0.513332 | -0.599272 | 1.04517 | 0.139958 | -1.075893 | 0.212186 | 0.658766 | 0.23788 | 0.266463 | 0.182183 | 0.155826 | 0.141033 | -0.279618 | -0.203084 |
| 28.68056 | -1.01531 | -0.310033 | -1.047746 | -2.390304 | 0.2074 | -0.117534 | 0.547542 | 0.368034 | 0.966996 | 0.362536 | 1.935681 | 2.548229 | -0.133569 | 0.081726 | 0.476924 | 0.420422 | 0.113435 | 0.106223 | 0.072512 | 0.200503 | -0.00225 | 0.156297 | -0.707999 | 0.383824 |
| 28.69444 | 0.137519 | -0.147882 | 0.741792 | -0.148301 | 0.241257 | 1.847577 | 2.17709 | 0.305902 | 0.467261 | 0.262539 | 2.236156 | 4.458951 | -1.579222 | -0.003888 | 0.789852 | 0.344151 | -0.480507 | -0.135719 | 0.138019 | 0.110954 | -0.088639 | 0.110885 | -0.165951 | -0.526808 |
| 28.70833 | 1.986528 | 0.350629 | 1.396192 | 2.26726 | 0.371334 | 0.633394 | 1.517006 | 0.285504 | 0.509561 | 0.170654 | -0.171246 | 1.868037 | 0.138189 | -0.197057 | 1.545451 | 0.088755 | -0.682108 | -0.02523 | -0.048217 | -0.045724 | 0.109191 | 0.025257 | 0.609959 | -0.374933 |
| 28.72222 | 2.018137 | 0.755491 | 0.31293 | 3.321917 | 0.318242 | -0.515439 | -0.493992 | 0.242813 | 0.203766 | 0.120348 | -1.960231 | -1.55701 | 2.391868 | -0.210225 | 0.998679 | 0.065438 | -0.704139 | 0.562608 | -0.611574 | -0.14672 | 0.397472 | 0.035444 | 0.539654 | -0.267135 |
| 28.73611 | -0.226336 | 0.350025 | -1.748171 | -0.664841 | 0.053955 | -0.796868 | -1.624003 | 0.051207 | -0.383086 | 0.212164 | -1.428054 | -2.959057 | 2.183198 | -0.154809 | -0.889579 | 0.209804 | -0.260963 | -0.061189 | -0.376269 | 0.10508 | 0.381992 | 0.117258 | 0.531813 | 0.136714 |
| 28.75 | -2.330711 | -0.251823 | -2.671842 | -4.326515 | -0.080466 | 0.059404 | -0.883918 | -0.051179 | -0.69509 | 0.264738 | 0.75799 | -0.540978 | 1.189901 | 0.122157 | -1.914857 | 0.263196 | 0.347381 | -0.851845 | -0.022869 | 0.239494 | 0.470831 | 0.151074 | 0.742169 | 0.348803 |
| 28.76389 | -2.393013 | -0.551808 | -1.196414 | -2.053329 | -0.039567 | 0.649063 | 0.173945 | -0.056288 | -0.11034 | 0.205825 | 1.399727 | 2.082973 | -1.025437 | 0.371218 | -1.041596 | 0.193419 | 0.51676 | 0.140434 | 0.061544 | 0.139677 | 0.604491 | 0.117214 | 0.387591 | -0.005899 |
| 28.77778 | 0.371707 | 0.039875 | -0.364887 | 0.432489 | 0.028557 | 0.55197 | 0.251009 | -0.051814 | 0.489055 | 0.12674 | 0.370775 | 1.7623 | -1.588755 | 0.433429 | 0.176119 | 0.098055 | 0.241989 | -0.095637 | 0.316089 | -0.00819 | 0.405048 | 0.100965 | 0.117025 | 0.014414 |
| 28.79167 | 2.65472 | 0.731405 | 1.819991 | 3.250475 | -0.025826 | 0.538527 | 1.377952 | -0.112144 | 0.775866 | 0.123335 | -0.834241 | -0.690094 | -1.343829 | 0.134092 | 0.785672 | -0.112081 | -0.370682 | 0.550477 | 0.768001 | -0.076773 | 0.177953 | 0.093257 | 0.032715 | -0.077984 |
| 28.80556 | 1.938803 | 0.288844 | 1.470952 | 2.028207 | -0.085082 | -0.145122 | 0.931205 | -0.165471 | 0.467495 | 0.14528 | -1.183316 | -3.112846 | 0.459546 | -0.128753 | 0.99062 | -0.243729 | -0.708409 | -0.061033 | 0.248088 | 0.035793 | -0.110909 | 0.054218 | 0.006645 | -0.058036 |
| 28.81944 | -0.337604 | -0.331524 | -0.396666 | -0.861342 | 0.051634 | -0.783943 | 0.208076 | -0.011426 | -0.408617 | 0.027313 | -0.227198 | -1.827317 | 1.095288 | -0.368364 | 0.132713 | -0.202997 | -0.622002 | -0.046095 | -0.191921 | 0.212434 | -0.35994 | -0.025776 | -0.059781 | 0.038529 |
| 28.83333 | -0.176129 | -0.104843 | -3.094949 | -2.660664 | 0.198141 | -0.610331 | -0.575974 | 0.09103 | -1.367606 | -0.190311 | 1.223655 | 1.440877 | 1.077492 | -0.455666 | -0.013246 | -0.015789 | -0.389038 | -0.958941 | -0.310608 | 0.216334 | -0.149353 | -0.058478 | -0.243638 | 0.170362 |
| 28.84722 | 0.090593 | 0.343276 | -2.513941 | -0.813147 | 0.094446 | -0.410887 | -0.262701 | 0.092607 | -0.552319 | -0.19334 | 1.169642 | 3.319548 | 0.536944 | -0.235119 | 0.083715 | 0.176318 | 0.044189 | -1.836797 | -0.25463 | 0.137616 | -0.010038 | 0.014464 | 0.527596 | -0.042925 |
| 28.86111 | 0.017249 | 0.451328 | 0.264748 | 1.350573 | -0.070771 | 0.191114 | -0.011423 | -0.160739 | 0.312537 | 0.136561 | -0.427928 | 0.437449 | -0.308575 | 0.109677 | -0.09064 | 0.113908 | 0.589614 | -0.720856 | -0.46503 | 0.069196 | 0.181585 | 0.162858 | 0.649097 | -0.047208 |
| 28.875 | -0.889648 | -0.323607 | 1.933968 | 2.090324 | -0.278631 | 0.937513 | 0.414744 | -0.321168 | 0.662513 | 0.248577 | -1.993696 | -2.62832 | -1.335513 | 0.335151 | -1.044122 | -0.123807 | 0.535435 | 0.463602 | -0.334335 | 0.048247 | 0.266819 | 0.142099 | -0.292221 | -0.266313 |
| 28.88889 | -1.529752 | -0.542742 | -0.313958 | -0.107238 | -0.12822 | -0.190711 | 0.498379 | -0.186891 | 0.124497 | -0.192642 | -1.128541 | -1.46273 | -2.470183 | 0.064396 | -1.731117 | -0.3069 | 0.032047 | 1.100625 | 0.423259 | 0.202118 | 0.065549 | 0.030217 | -0.931977 | 0.052147 |
| 28.90278 | -1.110459 | -0.011597 | -2.301668 | 0.081259 | 0.080184 | -1.432527 | -0.555658 | -0.026311 | -1.205134 | -0.60461 | 0.549267 | 0.779927 | -1.838083 | -0.493344 | -1.535705 | -0.289406 | -0.396422 | 0.200211 | 0.227586 | 0.315666 | -0.483466 | -0.061484 | -1.395429 | -0.224845 |
| 28.91667 | -0.124755 | 0.2217 | -1.95497 | -0.287521 | 0.208001 | -1.137982 | -1.805461 | 0.01529 | -1.484451 | -0.412197 | 0.661298 | 0.075613 | -1.09252 | -0.739603 | -0.460777 | -0.101205 | -0.524527 | -0.256827 | -0.244115 | 0.171912 | -0.976428 | -0.099376 | -0.597148 | -0.067015 |
| 28.93056 | -0.652198 | -0.01457 | 0.548239 | 0.656082 | 0.088119 | 0.178023 | -1.840414 | -0.064623 | -0.36289 | 0.034324 | -0.134323 | -0.936139 | -0.708667 | -0.782816 | 0.155332 | 0.10918 | -0.672617 | -0.723527 | -0.100745 | -0.164698 | -0.869044 | -0.03834 | -0.273967 | -0.16848 |
| 28.94444 | -1.52674 | -0.371127 | 1.011423 | 3.675803 | -0.183076 | 0.184129 | -0.337861 | -0.159769 | 0.062783 | 0.046668 | -2.038354 | -2.741294 | -1.468682 | -1.011686 | -0.493756 | 0.102856 | -1.099572 | -1.389579 | 0.164458 | -0.244677 | -0.652144 | -0.0813 | -0.373645 | -0.803512 |
| 28.95833 | -1.755006 | -0.464176 | 0.459622 | 5.501538 | -0.444255 | -0.424713 | 0.172506 | -0.243101 | 0.037577 | -0.662927 | -3.118693 | -2.662771 | -2.508155 | -0.759245 | -1.337888 | -0.16543 | -2.162094 | -2.206334 | -0.263383 | -0.074867 | -0.525649 | -0.166244 | -0.814149 | -0.890411 |
| 28.97222 | -1.520955 | -0.335201 | 0.531548 | 4.107024 | -0.541493 | -1.571579 | -1.349419 | -0.316081 | -0.781506 | -1.115021 | -2.993758 | -2.178011 | -4.067962 | -0.462995 | -2.312326 | -0.424113 | -3.261364 | -2.270232 | -0.056248 | 0.02132 | -0.563646 | -0.413188 | -1.205087 | -1.315247 |
| 28.98611 | -2.336391 | -1.009601 | -0.657712 | 2.036148 | -0.364523 | -1.485236 | -3.470641 | -0.38065 | -0.768653 | -0.867132 | -1.936521 | -3.011157 | -5.022043 | -0.339963 | -3.00935 | -0.895424 | -3.395968 | -2.760368 | 0.179373 | -0.01503 | -1.22752 | -0.87434 | -1.701182 | -1.66037 |
| 29 | -3.420352 | -1.747028 | -2.668482 | -0.462261 | -0.164373 | -1.621559 | -3.730987 | -0.359078 | -1.101917 | -0.25546 | -1.447112 | -2.97411 | -4.46951 | -0.246644 | -2.517319 | -1.433252 | -1.485327 | -3.244141 | -0.135126 | -0.133182 | -2.205425 | -1.3237 | -2.105286 | -1.772431 |
| 29.01389 | -2.092137 | -1.217129 | -3.24513 | -1.528883 | -0.440048 | -1.608109 | -1.952712 | -0.291337 | -1.24077 | -0.195677 | -1.773721 | -2.68917 | -2.334352 | -0.179685 | -1.951384 | -1.159075 | -0.535757 | -3.331248 | -0.576677 | -0.218005 | -2.796058 | -1.231598 | -2.499156 | -2.21469 |
| 29.02778 | -1.685413 | -0.648906 | -2.21871 | -0.577269 | -1.043076 | -1.685997 | -1.068681 | -0.198616 | -2.468605 | -1.211105 | -2.392614 | -3.543281 | -1.544084 | -0.284848 | -1.659816 | -0.575808 | -0.895459 | -2.497684 | -1.074 | -0.142758 | -2.840995 | -0.673935 | -2.32711 | -3.010767 |
| 29.04167 | -2.224111 | -0.912733 | -1.496575 | -0.117307 | -1.418708 | -1.759868 | -2.218243 | -0.015301 | -2.484982 | -1.791269 | -2.985333 | -3.512186 | -1.799466 | -0.399057 | -1.538425 | -0.282196 | -2.476514 | -1.874526 | -1.041981 | -0.015931 | -2.247608 | -0.239664 | -1.55881 | -2.721413 |
| 29.05556 | -2.949798 | -1.401904 | -1.398861 | -0.034375 | -1.60002 | -1.936442 | -3.128355 | 0.034312 | -1.229264 | -1.458728 | -2.846561 | -2.82949 | -2.445033 | -0.345642 | -1.421325 | -0.293626 | -2.916582 | -1.118136 | -0.791313 | -0.058243 | -1.395463 | -0.137893 | -1.303691 | -2.159162 |
| 29.06944 | -2.595442 | -1.851266 | 0.520796 | 0.890127 | -1.217678 | -1.948404 | -2.972745 | -0.179983 | -0.396431 | -0.894087 | -2.012478 | -1.773198 | -2.253747 | -0.283565 | -1.431661 | -0.402475 | -1.331254 | -0.774858 | -0.597044 | -0.116638 | -0.836913 | -0.190354 | -0.917693 | -1.458757 |
| 29.08333 | -0.502152 | -0.865764 | 1.351884 | 1.273117 | -0.9487 | -1.14926 | -1.537801 | -0.243453 | 0.093073 | -0.168132 | -0.811958 | -0.489552 | -1.426171 | -0.302284 | -1.258352 | -0.356438 | -0.495992 | -0.618805 | -0.661383 | -0.121925 | -0.500096 | -0.047906 | -0.175816 | -0.751768 |
| 29.09722 | 0.662107 | 0.795271 | 0.097824 | 0.403785 | -2.018496 | -0.184667 | -0.211278 | -0.187315 | 0.36967 | 0.412976 | -0.658031 | 0.230499 | -1.004285 | -0.292282 | -0.874011 | -0.134383 | -0.47106 | -0.304118 | -0.441472 | -0.103289 | 0.00208 | 0.184366 | 0.405067 | -0.321259 |
| 29.11111 | -0.23981 | 0.497958 | -1.429356 | -1.009392 | -2.739716 | -0.590026 | -0.953523 | -0.608209 | -0.215038 | 0.266581 | -1.262806 | -0.508946 | -0.705292 | -0.282999 | -0.559109 | 0.091871 | -0.80089 | 0.129434 | -0.145341 | -0.118527 | 0.393019 | 0.227267 | 0.396386 | -0.154377 |
| 29.125 | -1.538087 | -0.288219 | -1.97572 | -1.687591 | -2.802851 | -1.195955 | -1.760046 | -1.878018 | -1.127918 | -0.348956 | -1.646242 | -1.399949 | -0.577231 | -0.270336 | -0.503258 | 0.25723 | -1.186797 | 0.252521 | -0.062731 | -0.111668 | -0.013048 | 0.03471 | -0.03413 | -0.345931 |
| 29.13889 | -2.062968 | -1.198228 | -1.907036 | -1.341808 | -3.099164 | -1.365257 | -1.884519 | -2.986167 | -1.65406 | -0.712938 | -1.456885 | -1.618964 | -0.769935 | -0.278921 | -0.744578 | 0.163874 | -1.532204 | -0.106846 | -0.205688 | -0.140651 | -0.473646 | -0.283997 | -0.480987 | -0.5793 |
| 29.15278 | -1.631241 | -1.47504 | -1.587882 | -0.791347 | -2.743743 | -1.042996 | -1.432741 | -2.813931 | -1.547745 | -0.774497 | -1.220479 | -1.338777 | -1.06289 | -0.307272 | -0.890977 | 0.020543 | -1.695023 | -0.481546 | -0.375952 | -0.208916 | -0.727224 | -0.485822 | -0.766775 | -0.673031 |
| 29.16667 | -1.049708 | -0.661848 | -1.332442 | -0.51528 | -1.822478 | -0.367183 | -0.782572 | -1.543429 | -1.324898 | -0.747861 | -0.988884 | -0.980668 | -1.188309 | -0.319202 | -0.833039 | -0.107066 | -1.685749 | -0.611467 | -0.370523 | -0.219131 | -0.44765 | -0.491139 | -0.928705 | -0.697424 |
| 29.18056 | -0.718519 | -0.203433 | -1.444826 | -0.349615 | -1.414829 | -0.00518 | -0.43766 | -1.087674 | -1.20481 | -0.566994 | -0.962445 | -0.964226 | -1.238529 | -0.345528 | -0.825203 | -0.19658 | -1.744365 | -0.550181 | -0.254469 | -0.177737 | -0.221259 | -0.475715 | -1.112383 | -0.84712 |
| 29.19444 | -0.593346 | -0.157246 | -2.063569 | -0.306382 | -1.172904 | -0.187751 | -0.481528 | -1.35165 | -1.046814 | -0.364367 | -1.091358 | -1.150904 | -1.430851 | -0.397499 | -1.012079 | -0.198941 | -2.02733 | -0.494428 | -0.279945 | -0.117456 | -0.402104 | -0.557409 | -1.385363 | -1.150726 |
| 29.20833 | -0.51903 | -0.473481 | -2.189869 | -0.43995 | -0.964495 | -0.735973 | -0.537016 | -1.485326 | -0.99837 | -0.423 | -1.135827 | -1.112592 | -1.764659 | -0.428125 | -1.222012 | -0.197938 | -2.492368 | -0.611554 | -0.468175 | -0.181208 | -0.950428 | -0.729414 | -1.682505 | -1.477986 |
| 29.22222 | -0.243339 | -0.429443 | -1.205254 | -0.582421 | -0.76195 | -0.852597 | -0.514552 | -1.037362 | -1.107076 | -0.659003 | -1.019102 | -0.893315 | -2.262301 | -0.467992 | -1.217153 | -0.285621 | -3.208052 | -0.744522 | -0.522265 | -0.303299 | -1.083155 | -0.828139 | -1.715268 | -1.463019 |
| 29.23611 | 0.049528 | -0.103578 | -0.495331 | -0.616354 | -0.633565 | -0.539888 | -0.274377 | -0.527995 | -0.958683 | -0.629752 | -0.866437 | -0.724722 | -2.336765 | -0.499067 | -0.96243 | -0.418979 | -3.77147 | -0.657686 | -0.24342 | -0.363918 | -0.834688 | -0.700445 | -1.440361 | -1.20966 |
| 29.25 | -0.041994 | -0.290404 | -0.498315 | -0.549091 | -0.468444 | -0.442015 | -0.259535 | -0.22188 | -0.643819 | -0.324933 | -0.609081 | -0.674933 | -1.867387 | -0.512385 | -0.669868 | -0.451755 | -3.799933 | -0.409155 | 0.082076 | -0.327467 | -0.720869 | -0.479209 | -1.035511 | -0.963415 |
| 29.26389 | -0.25439 | -0.690244 | -0.511562 | -0.448332 | -0.199313 | -0.617428 | -0.235197 | -0.057491 | -0.51127 | -0.188169 | -0.327923 | -0.573022 | -1.290329 | -0.546131 | -0.456196 | -0.375094 | -2.875736 | -0.166759 | 0.183243 | -0.254444 | -0.580908 | -0.286861 | -0.496703 | -0.708995 |
| 29.27778 | -0.28715 | -0.501155 | -0.32965 | -0.439256 | -0.040283 | -0.387262 | -0.120787 | 0.111454 | -0.636497 | -0.292459 | -0.138985 | -0.335367 | -0.887774 | -0.482033 | -0.316119 | -0.199547 | -2.055898 | -0.084341 | 0.053909 | -0.158836 | -0.380115 | -0.171485 | -0.082872 | -0.458815 |
| 29.29167 | -0.109472 | -0.017589 | -0.287849 | -0.453162 | -0.261911 | -0.127989 | 0.013498 | 0.23901 | -0.56233 | -0.306342 | -0.175481 | -0.209179 | -0.60161 | -0.356554 | -0.219857 | -0.091425 | -1.409255 | -0.166179 | -0.140162 | -0.089124 | -0.190358 | -0.148593 | 0.068049 | -0.287443 |
| 29.30556 | -0.11604 | -0.131933 | 0.029767 | -0.365827 | -0.591713 | -0.272738 | 0.041718 | 0.314601 | -0.224029 | -0.092162 | -0.153394 | -0.24036 | -0.310016 | -0.297397 | -0.200719 | -0.157046 | -1.128033 | -0.240914 | -0.245708 | -0.097831 | -0.301658 | -0.183336 | 0.035116 | -0.237015 |
| 29.31944 | -0.161507 | -0.353311 | 0.461316 | -0.254446 | -0.581739 | -0.11426 | 0.036039 | 0.386961 | -0.120691 | -0.011442 | 0.207432 | -0.141851 | -0.182356 | -0.388255 | -0.197506 | -0.292626 | -1.121775 | -0.230752 | -0.261363 | -0.108654 | -0.327103 | -0.207702 | -0.092496 | -0.278945 |
| 29.33333 | 0.081343 | -0.154457 | 0.258822 | -0.192836 | -0.173809 | 0.271311 | 0.181301 | 0.376918 | -0.195939 | -0.090798 | 0.661081 | 0.206596 | -0.181969 | -0.491376 | -0.170123 | -0.319737 | -0.934824 | -0.209362 | -0.2737 | -0.091573 | -0.307419 | -0.170484 | -0.195906 | -0.33252 |
| 29.34722 | 0.366819 | 0.175069 | -0.271713 | 0.029172 | 0.457185 | 0.386378 | 0.444015 | 0.310201 | -0.121286 | -0.041774 | 0.89487 | 0.446068 | -0.001678 | -0.472509 | -0.087981 | -0.193447 | -0.309086 | -0.21302 | -0.284948 | -0.014015 | -0.050394 | -0.088316 | -0.117102 | -0.277571 |
| 29.36111 | 0.422903 | 0.05625 | -0.071036 | 0.210159 | 0.946415 | 0.309065 | 0.526851 | 0.254571 | 0.067242 | 0.044112 | 0.861748 | 0.404085 | 0.306594 | -0.323834 | -0.025132 | -0.055952 | 0.561706 | -0.147704 | -0.219794 | 0.082073 | -0.022916 | -0.040386 | 0.027783 | -0.079809 |
| 29.375 | 0.452496 | -0.11656 | 0.7516 | 0.153277 | 0.90906 | 0.493174 | 0.511167 | 0.254273 | 0.161096 | 0.017387 | 0.811058 | 0.371578 | 0.441611 | -0.221094 | -0.001218 | -0.040925 | 1.220077 | -0.027986 | -0.127604 | 0.135384 | 0.016527 | -0.032992 | 0.032948 | 0.091333 |
| 29.38889 | 0.6815 | 0.242896 | 1.423506 | 0.15493 | 0.692832 | 0.744635 | 0.70484 | 0.470696 | 0.421007 | 0.142982 | 1.035001 | 0.715771 | 0.294641 | -0.224397 | 0.064824 | -0.051802 | 1.139685 | 0.088331 | -0.000243 | 0.13187 | 0.044115 | 0.031718 | -0.079499 | 0.115116 |
| 29.40278 | 1.00652 | 0.757113 | 1.740633 | 0.427961 | 0.832282 | 0.862719 | 1.072495 | 0.85849 | 0.854475 | 0.452183 | 1.468871 | 1.221233 | 0.262096 | -0.170245 | 0.248965 | 0.041669 | 0.902522 | 0.179127 | 0.188066 | 0.185221 | 0.398189 | 0.221363 | -0.019447 | 0.16749 |
| 29.41667 | 1.226468 | 0.647639 | 2.107302 | 0.639824 | 1.211451 | 1.289971 | 1.239691 | 1.114467 | 1.167051 | 0.570276 | 1.696155 | 1.328983 | 0.55753 | 0.073139 | 0.448531 | 0.207303 | 1.171313 | 0.39157 | 0.425085 | 0.268313 | 0.751064 | 0.499441 | 0.361246 | 0.435081 |
| 29.43056 | 1.292106 | 0.422119 | 2.556965 | 0.702406 | 1.421044 | 1.655548 | 1.219837 | 1.125267 | 1.184028 | 0.433136 | 1.605749 | 1.20415 | 1.055704 | 0.413717 | 0.665475 | 0.372523 | 1.95304 | 0.751096 | 0.612423 | 0.322047 | 0.870169 | 0.771619 | 0.829164 | 0.870547 |
| 29.44444 | 1.664721 | 0.575663 | 3.262783 | 0.910231 | 1.61326 | 1.559604 | 1.359588 | 1.239728 | 1.381713 | 0.500471 | 1.758342 | 1.442676 | 1.494346 | 0.536824 | 0.992563 | 0.46479 | 2.596937 | 1.24123 | 0.697834 | 0.372824 | 1.220616 | 0.961988 | 1.284885 | 1.36479 |
| 29.45833 | 2.49767 | 1.062219 | 3.982219 | 1.450052 | 2.06861 | 1.677204 | 1.785474 | 1.796927 | 2.013984 | 0.929336 | 2.262586 | 2.115582 | 2.010094 | 0.500811 | 1.430815 | 0.535194 | 3.221674 | 1.789904 | 0.845094 | 0.384562 | 1.606917 | 1.196688 | 1.792807 | 1.875811 |
| 29.47222 | 3.157201 | 1.436851 | 4.765124 | 2.249841 | 2.87172 | 2.402739 | 2.34652 | 2.537622 | 2.708791 | 1.336378 | 2.268656 | 2.59086 | 2.841161 | 0.588568 | 2.058215 | 0.747293 | 4.007699 | 2.639251 | 1.223228 | 0.461907 | 2.24955 | 1.494702 | 2.447081 | 2.410133 |
| 29.48611 | 3.392219 | 1.370414 | 5.562805 | 3.010094 | 3.6208 | 2.729477 | 3.059193 | 2.660365 | 3.094561 | 1.415774 | 2.330436 | 2.729979 | 4.016481 | 0.828894 | 2.85062 | 0.890336 | 4.518521 | 3.868881 | 1.506233 | 0.489788 | 2.462165 | 1.414268 | 2.937081 | 3.231075 |
| 29.5 | 3.783933 | 1.325951 | 6.415537 | 3.839602 | 3.57253 | 2.885127 | 3.945888 | 2.095444 | 3.597017 | 1.405433 | 3.109178 | 3.15835 | 5.193854 | 0.930102 | 3.632083 | 0.869546 | 3.911253 | 4.879172 | 1.264182 | 0.287821 | 2.548715 | 1.029274 | 3.030493 | 3.536605 |
| 29.51389 | 4.715636 | 1.854738 | 6.260402 | 4.509689 | 2.808071 | 3.325844 | 5.113246 | 0.780383 | 4.116807 | 1.467905 | 3.936798 | 4.091626 | 6.910287 | 0.773781 | 4.163912 | 0.762293 | 3.617318 | 4.577633 | 0.727933 | 0.057669 | 2.582096 | 0.697264 | 3.180614 | 3.529159 |
| 29.52778 | 4.521589 | 2.145309 | 5.672375 | 4.034437 | 1.861993 | 3.171354 | 5.465207 | -0.074471 | 3.900402 | 1.324493 | 4.032663 | 4.995854 | 5.979982 | 0.258905 | 4.376637 | 0.710783 | 3.520867 | 3.862096 | 0.474127 | -0.139625 | 2.416661 | 0.264932 | 3.686658 | 2.983909 |
| 29.54167 | 3.752524 | 1.345073 | 4.433393 | 3.493098 | 0.797989 | 2.130601 | 4.224915 | -0.823325 | 2.73197 | 0.752285 | 2.2995 | 3.604336 | 5.148696 | 0.642657 | 3.578902 | 0.563 | 2.188469 | 2.986452 | 0.216587 | -0.372545 | 1.87571 | -0.130164 | 3.432918 | 2.444601 |
| 29.55556 | 1.873191 | 0.324365 | 2.013117 | 2.126337 | 0.168081 | 0.792494 | 0.782077 | -0.942532 | 1.048983 | 0.225504 | -0.335659 | 0.67153 | 2.302755 | 1.080059 | 1.827292 | 0.304026 | -0.311229 | 1.723039 | 0.248052 | -0.359606 | 1.387252 | -0.140398 | 1.941434 | 2.394471 |
| 29.56944 | 0.843938 | 0.82626 | 0.001499 | 0.654539 | -0.220204 | -0.160219 | -1.225825 | -0.23123 | 0.20202 | 0.149282 | -0.823933 | -0.018555 | -1.588864 | 0.886434 | 0.448743 | 0.118351 | -1.964861 | 0.7647 | 0.333129 | -0.035774 | 1.07225 | 0.122006 | 0.559033 | 2.386275 |
| 29.58333 | 0.774947 | 1.629345 | 0.350349 | 0.670308 | -0.168123 | 0.601328 | -0.876835 | 0.500511 | 0.891741 | 0.401327 | 1.818149 | 2.591291 | -1.853616 | 0.399998 | 0.33224 | 0.154275 | -2.891218 | 0.428952 | 0.332418 | 0.128674 | 0.99252 | 0.379347 | -0.212543 | 1.648555 |
| 29.59722 | 1.672876 | 1.555753 | 3.90725 | 2.572638 | 0.32129 | 1.170362 | 1.652381 | 0.710536 | 2.111277 | 0.620585 | 5.059562 | 5.539612 | 0.681309 | 0.160169 | 0.83471 | 0.344622 | -0.343417 | 0.516286 | 0.268727 | 0.073443 | 1.168462 | 0.290017 | -0.251149 | 1.283294 |
| 29.61111 | 3.090501 | 0.950887 | 6.729652 | 4.207753 | 0.762487 | 0.541709 | 3.00748 | 0.564011 | 2.142556 | 0.912644 | 4.235281 | 5.860713 | 2.085769 | 0.381414 | 2.203652 | 0.592843 | 1.866019 | 0.96062 | 0.203695 | -0.041119 | 1.146974 | -0.083939 | 0.50467 | 0.582967 |
| 29.625 | 4.020231 | 0.465441 | 5.088424 | 4.070329 | 0.901465 | 0.754358 | 2.596374 | 0.456118 | 2.20653 | 1.02533 | 2.512869 | 3.191239 | 1.244479 | 0.683886 | 2.524846 | 0.613413 | 3.968787 | 0.892035 | 0.216295 | -0.099793 | 1.178332 | -0.273381 | 1.06162 | -0.168678 |
| 29.63889 | 3.518824 | 0.208535 | 3.081317 | 2.987069 | 0.690367 | 0.892655 | 1.193094 | 0.438576 | 1.361476 | 0.707022 | 0.151637 | 0.090066 | 0.691828 | 0.841145 | 2.20371 | 0.365169 | 2.362843 | 0.118809 | 0.257778 | -0.095627 | 0.851035 | -0.135944 | -1.018719 | 0.002752 |
| 29.65278 | 2.088149 | 0.745039 | -0.250397 | 0.417623 | 0.444812 | -0.118587 | 0.2873 | 0.533161 | -0.20167 | 0.290297 | -1.919749 | -1.501927 | 1.975468 | 0.445916 | 0.238232 | 0.060256 | 0.705068 | -0.048295 | -0.069703 | 0.050632 | 0.990831 | 0.010279 | -1.998491 | 0.173498 |
| 29.66667 | 0.376884 | 0.883744 | -2.166062 | -1.413108 | 0.046718 | 0.067294 | 0.147084 | 0.488005 | -0.487834 | 0.067898 | -0.637078 | 0.64085 | 1.427264 | 0.113382 | -0.049266 | -0.019666 | 0.455443 | -0.085507 | -0.056559 | 0.229764 | 1.127346 | 0.046005 | -0.326046 | 0.241698 |
| 29.68056 | -2.11905 | 0.398806 | -1.13151 | -1.183237 | -0.138108 | 0.756645 | 0.303852 | 0.336986 | -0.525763 | 0.055143 | 1.328461 | 2.508077 | 0.550984 | -0.102117 | 0.04324 | 0.128356 | 0.179466 | -0.441244 | -0.120364 | 0.206628 | 0.583129 | 0.136054 | 0.087415 | 0.036083 |
| 29.69444 | -1.549119 | 0.137962 | 1.438516 | 0.669534 | -0.047191 | 1.203504 | 0.875294 | 0.135877 | -0.072671 | 0.23356 | 2.03686 | 3.370067 | -0.103223 | -0.162099 | 0.606096 | 0.143875 | -0.273876 | 0.151194 | -0.23259 | 0.137082 | -0.060787 | 0.208963 | 0.117542 | -0.079932 |
| 29.70833 | -0.026055 | 0.016073 | 1.518546 | 2.280053 | 0.034355 | -0.055594 | 0.759432 | -0.018095 | 0.747018 | 0.343472 | 1.492357 | 0.83367 | -0.554798 | -0.428205 | 1.384623 | -0.024718 | -0.465681 | 0.297637 | -0.125034 | 0.054136 | 0.238496 | 0.176244 | 0.612664 | -0.182406 |
| 29.72222 | 1.54575 | 0.012876 | -1.364021 | 1.785677 | 0.09383 | -1.08843 | -0.664276 | -0.050151 | 1.004019 | 0.182362 | -0.607322 | -1.859719 | -0.302593 | -0.44244 | 2.052148 | -0.064962 | -0.361061 | 0.340238 | -0.107994 | -0.007186 | 0.309934 | 0.065109 | 0.664365 | -0.041852 |
| 29.73611 | 1.238948 | 0.095918 | -4.458143 | -0.811902 | 0.08227 | -1.22789 | -0.751313 | -0.024996 | -0.215748 | -0.011838 | -1.508726 | -1.778688 | 1.222166 | -0.297817 | 0.978775 | 0.065652 | 0.040583 | 0.167898 | -0.012735 | -0.002153 | -0.002338 | -0.051863 | -0.184428 | -0.017018 |
| 29.75 | 0.995669 | 0.084687 | -5.005297 | -2.777129 | 0.071762 | 0.115171 | 0.237561 | -0.036393 | -1.532364 | -0.141603 | -0.804207 | -0.768402 | 1.670158 | 0.01145 | -1.456207 | 0.215501 | 0.474574 | -0.937758 | 0.357913 | 0.037641 | 0.191625 | -0.015237 | -0.622489 | 0.232013 |
| 29.76389 | -0.451002 | -0.027297 | -1.906633 | -1.335273 | 0.090953 | 0.897206 | 0.371449 | -0.092036 | -1.889583 | -0.241905 | 2.080542 | 1.277611 | 0.338325 | 0.329521 | -1.94195 | 0.206634 | 0.539589 | -0.934357 | 0.357356 | 0.086544 | 0.644757 | 0.137209 | -0.498468 | 0.437764 |
| 29.77778 | -1.74917 | 0.049475 | 1.310414 | 1.015892 | 0.039883 | 0.191671 | -0.802073 | -0.040095 | -0.583693 | -0.096206 | 3.33951 | 2.714318 | -1.1975 | 0.706775 | -0.799425 | 0.161775 | 0.19068 | 0.248656 | 0.17189 | 0.076706 | 0.871252 | 0.204666 | 0.745248 | 0.146935 |
| 29.79167 | -1.817019 | 0.123192 | 2.776237 | 1.723956 | -0.174403 | -0.549127 | -0.253411 | 0.111482 | 0.493129 | 0.135254 | 1.091427 | 0.658151 | -1.278279 | 0.409521 | 0.50628 | 0.002618 | -0.318705 | 0.952057 | -0.064838 | 0.064876 | 0.52685 | 0.162707 | 1.307634 | -0.203935 |
| 29.80556 | 0.131643 | -0.155246 | 1.376887 | 0.797253 | -0.13098 | -0.90506 | 1.118708 | 0.132277 | 1.504479 | 0.289805 | -2.562314 | -2.060065 | -0.017764 | 0.110126 | 2.031453 | -0.039157 | -0.573727 | 0.733985 | -0.274307 | 0.111769 | -0.156777 | 0.110942 | 0.991407 | -0.059848 |
| 29.81944 | 0.691244 | -0.329362 | -1.262473 | -0.475722 | 0.02505 | 0.568988 | 2.22828 | 0.043038 | 1.479593 | 0.282199 | -2.876335 | -2.678366 | 1.359467 | 0.002423 | 1.657257 | 0.015093 | -0.365373 | -0.364838 | -0.508902 | 0.123752 | -0.535182 | 0.054524 | 0.452588 | 0.031073 |
| 29.83333 | 1.170736 | 0.01287 | -3.047652 | -0.58943 | 0.150021 | 0.744104 | 0.263092 | 0.006896 | -0.873921 | -0.019985 | -0.730218 | -0.841321 | 2.170116 | -0.16212 | -0.287516 | 0.098304 | 0.016622 | -1.63587 | -0.511288 | 0.060697 | -0.53904 | -0.015761 | 0.226755 | 0.198089 |
| 29.84722 | 1.272193 | 0.38805 | -2.247562 | -1.421791 | 0.111692 | -0.03289 | -1.364167 | 0.05127 | -1.932797 | -0.358494 | 2.17084 | 1.008452 | 0.390429 | -0.209751 | -1.125655 | 0.14004 | 0.382926 | -2.175527 | 0.165115 | 0.025655 | -0.105609 | -0.025113 | -0.472023 | 0.029992 |
| 29.86111 | -0.337577 | 0.339067 | -1.809019 | -1.028191 | 0.05064 | -0.678221 | -1.30119 | 0.052661 | -1.357181 | -0.365463 | 2.814769 | 1.560746 | -0.777235 | -0.107057 | -0.61109 | 0.138674 | 0.421394 | -1.757648 | 0.283417 | -0.018755 | 0.287629 | 0.019223 | -0.530209 | 0.095976 |
| 29.875 | -2.071651 | -0.075719 | -1.182152 | -0.477652 | -0.073169 | 0.477847 | 0.583416 | -0.144371 | 0.510442 | -0.006355 | 0.701874 | 0.170197 | -1.412367 | -0.09866 | 0.421965 | 0.056534 | -0.104636 | -0.672883 | 0.122197 | 0.041716 | 0.513741 | 0.089267 | -0.394937 | -0.445174 |
| 29.88889 | -1.927392 | -0.421886 | 0.078709 | 0.41231 | -0.302199 | -0.32902 | 0.514313 | -0.232389 | 0.995353 | 0.258441 | -3.258024 | -2.353723 | -1.763777 | -0.149478 | 0.815647 | -0.054491 | -1.357565 | 0.519705 | 0.086776 | 0.185031 | 0.470588 | 0.16249 | -0.337096 | -0.432149 |
| 29.90278 | -1.361929 | -0.274367 | 0.70868 | 0.010368 | -0.447723 | -1.68209 | -0.838197 | -0.076489 | 0.642524 | 0.157183 | -2.951211 | -2.123588 | -1.036056 | -0.286395 | 0.533441 | -0.167137 | -1.180196 | 0.933701 | 0.036855 | 0.153722 | -0.226802 | 0.036732 | -0.974731 | -0.232624 |
| 29.91667 | -0.557948 | 0.076144 | -1.7239 | -1.317933 | -0.287503 | -1.913122 | -1.895154 | 0.072821 | -0.832298 | -0.119099 | -0.373681 | -0.752598 | -0.398146 | -0.590535 | -0.591086 | -0.229614 | 0.290937 | 0.592007 | -0.02278 | 0.012871 | -0.540093 | -0.148596 | -0.514077 | 0.08579 |
| 29.93056 | -1.511994 | -0.030163 | -6.140593 | -2.79008 | -0.011123 | -0.499291 | -1.345383 | 0.023766 | -1.371009 | -0.349374 | 1.823576 | 0.167453 | -1.655978 | -0.55996 | -1.536513 | -0.192526 | 0.130096 | 0.193144 | -0.026992 | -0.124369 | -1.076351 | 0.069337 | 0.037764 | 0.066613 |
| 29.94444 | -0.569433 | -0.599859 | -5.072072 | -2.023084 | 0.059646 | -0.073353 | 0.431694 | -0.197501 | -1.597311 | -0.630621 | 1.259616 | -0.51512 | -3.45331 | -0.536775 | -1.919882 | -0.120863 | -1.572215 | -0.658135 | 0.043959 | -0.159906 | -1.858623 | -0.001202 | -0.57526 | -0.508082 |
| 29.95833 | 0.463377 | -1.548046 | -2.054109 | -1.459186 | -0.076237 | -0.423746 | -0.350185 | -0.267374 | -1.758901 | -0.765642 | -2.788095 | -2.437931 | -2.201012 | -0.492032 | -1.591335 | -0.085225 | -4.195035 | -1.93872 | -0.054279 | -0.072541 | -1.610054 | -0.511407 | -1.770269 | -1.099081 |
| 29.97222 | 0.504527 | -1.188008 | -0.579384 | -1.582958 | -0.19012 | -1.951533 | -2.088091 | -0.211123 | -2.268815 | -0.912408 | -6.115325 | -4.297952 | -0.623684 | -0.442818 | -1.620773 | -0.075357 | -5.397502 | -2.750677 | -0.240899 | 0.012329 | -1.43814 | -0.560121 | -1.534636 | -1.634421 |
| 29.98611 | -0.964406 | -0.856978 | -1.596185 | -2.670541 | -0.1095 | -1.668088 | -3.109825 | -0.164081 | -1.857795 | -1.083912 | -4.343242 | -4.656854 | -1.6276 | -0.278027 | -2.38524 | -0.125796 | -3.842301 | -2.712334 | -0.15413 | -0.00522 | -1.629605 | -0.583045 | -1.550238 | -0.658701 |
| 30 | -2.279809 | -1.612227 | -4.296706 | -3.755738 | -0.045366 | -1.199933 | -3.399981 | -0.219454 | -0.959585 | -0.668185 | -1.144053 | -3.180876 | -2.906866 | -0.310356 | -3.644889 | -0.642719 | -3.159624 | -2.657862 | -0.093895 | -0.047442 | -2.106186 | -0.606273 | -1.633836 | -0.459995 |
| 30.01389 | -2.543124 | -2.868434 | -5.512913 | -5.038539 | -0.233709 | -1.328276 | -3.892594 | -0.30591 | -1.26358 | -0.456247 | 0.292548 | -2.617663 | -3.129722 | -0.565414 | -3.160388 | -0.968856 | -2.465675 | -3.348789 | -0.222877 | -0.045091 | -2.238035 | -0.74551 | -1.762009 | -1.4675 |
| 30.02778 | -1.424152 | -4.415815 | -4.317434 | -5.075983 | -0.46507 | -1.128412 | -3.54225 | -0.189495 | -2.522915 | -0.977541 | -1.095092 | -3.390734 | -2.486145 | -0.539692 | -2.449894 | -0.646295 | -1.713771 | -3.612756 | -0.435468 | -0.162276 | -2.423909 | -1.05809 | -2.257944 | -2.590328 |
| 30.04167 | -2.123103 | -2.5459 | -3.188276 | -4.290921 | -0.631134 | -1.352725 | -2.230135 | 0.133261 | -3.635868 | -1.823275 | -2.957907 | -3.810268 | -2.949086 | -0.452588 | -1.659409 | -0.267416 | -2.749981 | -2.065243 | -0.386152 | -0.210569 | -2.002811 | -1.056831 | -1.933477 | -2.427302 |
| 30.05556 | -2.212588 | 0.556519 | -1.976656 | -3.254364 | -1.045208 | -1.886407 | -1.601465 | 0.417328 | -2.16819 | -1.70385 | -3.599051 | -2.693271 | -3.213616 | -0.335875 | -1.272465 | -0.120358 | -1.831334 | -1.304141 | -0.335154 | -0.194161 | -1.288973 | -0.855016 | -2.116458 | -1.941782 |
| 30.06944 | -1.883202 | 1.89495 | -0.698702 | -1.56641 | -1.237709 | -2.216542 | -1.930566 | 0.419507 | -0.396978 | -0.817025 | -3.044374 | -1.598827 | -2.106423 | -0.259489 | -0.933324 | -0.133645 | -0.799031 | -1.00047 | -0.369549 | -0.177592 | -1.074751 | -0.695885 | -1.582136 | -1.343625 |
| 30.08333 | -0.378125 | 0.436816 | -0.297072 | 0.142013 | -1.467688 | -1.119823 | -1.51653 | 0.10963 | 0.317828 | 0.044187 | -1.496215 | -0.943547 | -0.799335 | -0.250396 | -0.569502 | -0.13824 | 0.146071 | -1.011689 | -0.276778 | -0.191536 | -0.819271 | -0.432608 | -0.81178 | -0.813852 |
| 30.09722 | 0.193434 | -0.816492 | -1.441346 | 0.448633 | -3.141462 | -0.346866 | -0.531534 | -0.274441 | 0.285412 | 0.137393 | -1.025668 | -0.996877 | -0.407592 | -0.229204 | -0.341868 | -0.08136 | -0.173685 | -0.790959 | -0.251811 | -0.231626 | -0.382021 | -0.067458 | -0.570997 | -0.497749 |
| 30.11111 | -0.682254 | -0.171276 | -2.818634 | -0.709313 | -4.729127 | -0.806684 | -0.761484 | -0.450459 | -0.747643 | -0.382238 | -1.27027 | -0.854541 | -0.469117 | -0.151658 | -0.124509 | -0.023334 | -1.072114 | -0.340177 | -0.227591 | -0.230133 | 0.15517 | 0.247214 | -0.464118 | -0.300047 |
| 30.125 | -0.91363 | 0.76004 | -3.109342 | -1.9012 | -4.3785 | -0.980388 | -1.026531 | -0.663467 | -1.844267 | -0.75684 | -1.498671 | -0.610674 | -0.705588 | -0.120291 | -0.284397 | 0.016083 | -1.566615 | -0.168331 | -0.222082 | -0.183112 | 0.149427 | 0.275326 | -0.404079 | -0.251233 |
| 30.13889 | -1.27695 | 0.359609 | -2.601837 | -2.289107 | -2.987315 | -0.732768 | -1.535994 | -1.120079 | -1.395249 | -0.491412 | -1.442008 | -0.827854 | -0.659233 | -0.120142 | -0.467242 | 0.002574 | -1.567452 | -0.113811 | -0.204821 | -0.166107 | -0.105242 | 0.157259 | -0.353877 | -0.236869 |
| 30.15278 | -1.450835 | -0.783158 | -1.746394 | -1.347052 | -1.674713 | -0.148408 | -1.416262 | -1.317141 | -0.425055 | -0.121891 | -1.149394 | -1.012531 | -0.698081 | -0.16102 | -0.457844 | -0.02363 | -1.325735 | -0.176252 | -0.2198 | -0.184704 | -0.258972 | 0.037727 | -0.253439 | -0.226312 |
| 30.16667 | -1.496558 | -1.510045 | -1.233644 | -0.29844 | -0.731906 | 0.196896 | -0.797659 | -0.889088 | -0.121706 | -0.144486 | -0.912907 | -1.103253 | -0.388507 | -0.187471 | -0.270083 | -0.063833 | -1.221615 | -0.280352 | -0.272447 | -0.20267 | -0.223622 | -0.025669 | -0.216897 | -0.232995 |
| 30.18056 | -1.064795 | -1.107881 | -1.012063 | -0.160061 | -0.18016 | 0.151691 | -0.350718 | -0.803245 | -0.287341 | -0.315885 | -0.882445 | -0.980143 | -0.454135 | -0.204227 | -0.247882 | -0.134253 | -1.308229 | -0.356557 | -0.366676 | -0.223004 | 0.003044 | -0.090121 | -0.329885 | -0.367814 |
| 30.19444 | -0.435505 | -0.695158 | -0.957857 | -0.071081 | -0.151255 | 0.136419 | -0.187128 | -1.014681 | -0.538081 | -0.326146 | -1.053674 | -0.999409 | -1.208428 | -0.22239 | -0.524222 | -0.240305 | -1.545126 | -0.416557 | -0.478272 | -0.209913 | -0.051661 | -0.273208 | -0.629766 | -0.646698 |
| 30.20833 | -0.163165 | -0.746177 | -0.848273 | -0.2827 | -0.263005 | 0.297969 | -0.209271 | -1.433143 | -0.364952 | -0.306367 | -1.310303 | -1.242852 | -1.70127 | -0.278514 | -0.832288 | -0.312403 | -1.981615 | -0.541018 | -0.525587 | -0.162805 | -0.541581 | -0.539225 | -1.007683 | -0.914471 |
| 30.22222 | -0.319034 | -0.64245 | -0.509499 | -0.29332 | -0.043918 | 0.086912 | -0.178092 | -1.72822 | -0.331253 | -0.525591 | -1.420712 | -1.308946 | -2.029825 | -0.391457 | -0.898607 | -0.349945 | -2.881327 | -0.750377 | -0.536884 | -0.131377 | -1.064144 | -0.711765 | -1.425495 | -1.132875 |
| 30.23611 | -0.184306 | 0.011944 | -0.10722 | -0.540858 | 0.394731 | -0.433937 | -0.419674 | -1.363665 | -0.645121 | -0.727684 | -0.990518 | -0.849691 | -2.164926 | -0.52227 | -0.82679 | -0.368242 | -3.843885 | -0.773206 | -0.476042 | -0.221303 | -1.038749 | -0.565253 | -1.661013 | -1.257161 |
| 30.25 | 0.233309 | 0.78007 | 0.039279 | 0.090231 | 0.756494 | -0.298503 | -0.291557 | -0.549227 | -0.520657 | -0.489307 | -0.553178 | -0.180165 | -1.997783 | -0.540187 | -0.651248 | -0.296075 | -4.050648 | -0.470258 | -0.340832 | -0.351994 | -0.49933 | -0.2677 | -1.298465 | -1.048053 |
| 30.26389 | 0.326513 | 0.630012 | -0.008244 | 0.290498 | 0.896566 | 0.199769 | 0.150084 | -0.098202 | 0.03052 | -0.105966 | -0.224175 | 0.103412 | -1.321403 | -0.471837 | -0.448034 | -0.206143 | -2.992858 | -0.184462 | -0.253822 | -0.272706 | -0.287545 | -0.116076 | -0.66822 | -0.518277 |
| 30.27778 | 0.039 | -0.026803 | 0.074405 | -0.135244 | 0.686092 | 0.053751 | 0.156634 | -0.058254 | 0.091447 | -0.064155 | 0.058841 | -0.020461 | -0.759991 | -0.397485 | -0.234386 | -0.173388 | -2.047058 | -0.139381 | -0.306126 | -0.172303 | -0.389628 | -0.053882 | -0.136701 | -0.031249 |
| 30.29167 | -0.101475 | -0.355009 | 0.308285 | -0.464242 | 0.096663 | -0.415785 | -0.321338 | -0.145724 | -0.399143 | -0.209019 | 0.122218 | -0.148267 | -0.6624 | -0.334135 | -0.166118 | -0.178941 | -1.282376 | -0.246159 | -0.372285 | -0.119929 | -0.43394 | -0.015116 | 0.214283 | 0.217249 |
| 30.30556 | -0.047176 | 0.115706 | 0.313639 | -0.291835 | -0.511167 | -0.492787 | -0.274075 | 0.023196 | -0.483273 | -0.127434 | 0.109909 | 0.114412 | -0.66516 | -0.28798 | -0.251588 | -0.174243 | -0.679776 | -0.260103 | -0.369317 | -0.093428 | -0.34993 | -0.008861 | 0.285516 | 0.185469 |
| 30.31944 | -0.114996 | 0.486588 | 0.028015 | -0.553071 | -0.730185 | -0.06626 | 0.268762 | 0.412067 | -0.135327 | -0.000313 | 0.291072 | 0.377874 | -0.307566 | -0.300321 | -0.279457 | -0.188483 | -0.161568 | -0.121622 | -0.315594 | -0.094496 | -0.270699 | -0.095438 | 0.181042 | 0.092792 |
| 30.33333 | -0.131206 | 0.117174 | -0.26232 | -0.837114 | -0.492983 | 0.163703 | 0.579957 | 0.69711 | -0.049889 | -0.074929 | 0.682473 | 0.412709 | 0.202017 | -0.337139 | -0.161642 | -0.229372 | 0.490535 | -0.083468 | -0.264616 | -0.08272 | -0.210029 | -0.16801 | 0.156202 | 0.105166 |
| 30.34722 | 0.150598 | -0.414957 | -0.140891 | -0.602275 | 0.049897 | 0.330515 | 0.373696 | 0.816479 | -0.196586 | -0.109374 | 0.985948 | 0.342621 | 0.397175 | -0.328792 | -0.069343 | -0.24183 | 1.309984 | -0.23117 | -0.20663 | -0.011825 | -0.06222 | -0.179462 | 0.233685 | 0.159738 |
| 30.36111 | 0.403247 | -0.101469 | 0.289642 | -0.073553 | 0.598993 | 0.668455 | 0.355103 | 0.710961 | 0.008346 | 0.057433 | 1.036164 | 0.525626 | 0.382669 | -0.299333 | -0.096411 | -0.196676 | 2.070314 | -0.377515 | -0.157423 | 0.089721 | -0.049614 | -0.293531 | 0.213604 | 0.100456 |
| 30.375 | 0.556141 | 0.609055 | 0.797612 | -0.017723 | 1.104617 | 0.732962 | 0.734599 | 0.666921 | 0.535789 | 0.323366 | 1.145314 | 1.001344 | 0.471798 | -0.233569 | -0.129284 | -0.131628 | 2.675412 | -0.316415 | -0.152001 | 0.189854 | -0.109971 | -0.434073 | 0.079234 | -0.020744 |
| 30.38889 | 0.878645 | 0.698459 | 1.362663 | 0.111595 | 1.46564 | 0.685731 | 0.921173 | 0.784538 | 0.745505 | 0.386772 | 1.486062 | 1.471956 | 0.711454 | -0.152719 | -0.022668 | -0.041734 | 2.923609 | -0.05554 | -0.056285 | 0.287019 | 0.196049 | -0.255537 | 0.060362 | -0.010631 |
| 30.40278 | 1.192827 | 0.348598 | 1.896634 | 0.673699 | 1.37808 | 0.971384 | 0.917379 | 1.110757 | 0.519875 | 0.221601 | 1.770591 | 1.551358 | 0.905587 | -0.060073 | 0.158823 | 0.028038 | 2.774944 | 0.312316 | 0.149011 | 0.26902 | 0.753646 | 0.180492 | 0.243368 | 0.178148 |
| 30.41667 | 1.295554 | 0.57332 | 2.014206 | 0.999422 | 0.937875 | 1.269341 | 1.036269 | 1.048703 | 0.729417 | 0.293961 | 1.792758 | 1.484742 | 0.81353 | -0.010952 | 0.260658 | -0.006555 | 2.011121 | 0.565627 | 0.281887 | 0.198331 | 0.840633 | 0.338709 | 0.432556 | 0.399144 |
| 30.43056 | 1.497356 | 1.199384 | 1.742126 | 1.133534 | 0.743095 | 1.398534 | 1.382487 | 0.881545 | 1.497743 | 0.700798 | 1.853149 | 1.769998 | 0.784998 | 0.058069 | 0.345164 | -0.029666 | 1.381137 | 0.730783 | 0.373918 | 0.137843 | 0.611406 | 0.247662 | 0.533471 | 0.586017 |
| 30.44444 | 2.052025 | 1.176696 | 2.069509 | 1.597168 | 1.230811 | 1.624845 | 1.709502 | 1.031705 | 2.122891 | 1.035361 | 2.045194 | 2.190644 | 1.326679 | 0.269197 | 0.715675 | 0.130188 | 1.783829 | 1.08128 | 0.580555 | 0.198248 | 0.876551 | 0.488247 | 0.775086 | 0.980451 |
| 30.45833 | 2.561142 | 0.852203 | 3.604392 | 2.264566 | 2.186354 | 2.051651 | 2.449077 | 1.248602 | 2.523533 | 0.99447 | 2.483088 | 2.583322 | 2.37558 | 0.568247 | 1.27016 | 0.387701 | 3.288794 | 1.74689 | 0.759261 | 0.36824 | 1.834162 | 1.076601 | 1.372363 | 1.676238 |
| 30.47222 | 3.11204 | 0.920733 | 4.97846 | 2.757088 | 3.289607 | 2.376767 | 3.315011 | 1.250211 | 2.808087 | 0.901611 | 3.386368 | 3.170778 | 3.592681 | 0.653596 | 1.850127 | 0.551215 | 5.729215 | 2.500806 | 0.831699 | 0.556757 | 2.227137 | 1.458143 | 2.295351 | 2.329347 |
| 30.48611 | 3.704523 | 1.230318 | 5.99773 | 3.485685 | 3.777976 | 2.623292 | 3.853614 | 0.694099 | 3.070575 | 1.117135 | 4.024062 | 3.50748 | 5.096822 | 0.636435 | 2.448572 | 0.59023 | 7.337326 | 3.432004 | 0.870164 | 0.620407 | 2.113732 | 1.399366 | 2.90958 | 2.481707 |
| 30.5 | 4.311607 | 1.809091 | 6.160676 | 4.514158 | 2.960644 | 2.854185 | 4.067053 | 0.320955 | 3.73359 | 1.538972 | 4.420668 | 3.66768 | 6.31343 | 0.685857 | 3.020004 | 0.60597 | 6.703913 | 4.17301 | 0.724362 | 0.530874 | 2.295301 | 1.153668 | 3.108833 | 2.762657 |
| 30.51389 | 4.18087 | 1.879087 | 6.34493 | 5.070673 | 1.352671 | 3.02221 | 4.407315 | 0.298503 | 4.383552 | 1.614588 | 4.995694 | 4.164042 | 6.186191 | 0.835228 | 3.766416 | 0.631084 | 5.97452 | 3.797873 | 0.608632 | 0.430273 | 2.882825 | 0.806506 | 3.272126 | 3.118767 |
| 30.52778 | 3.886883 | 1.29425 | 6.809882 | 5.212381 | 0.972408 | 3.064183 | 4.645474 | -0.103294 | 3.533273 | 1.091114 | 5.055072 | 3.906327 | 6.433411 | 0.836723 | 3.923717 | 0.655404 | 5.608016 | 3.55861 | 0.496097 | 0.089186 | 2.777758 | 0.455728 | 3.575585 | 3.462119 |
| 30.54167 | 2.665964 | 0.396413 | 5.393033 | 3.884056 | 0.572927 | 2.234051 | 3.908505 | -0.942512 | 1.91383 | 0.424951 | 3.655442 | 2.69896 | 4.434713 | 1.010966 | 3.686325 | 0.640187 | 2.826932 | 3.796544 | 0.23232 | -0.230059 | 2.135766 | 0.10512 | 3.664627 | 3.568056 |
| 30.55556 | 0.862433 | -0.178442 | 1.110442 | 1.148096 | -0.326982 | 0.552502 | 0.997068 | -1.156593 | 1.009967 | -0.001451 | -0.496631 | 0.583383 | 0.201657 | 0.8599 | 2.408533 | 0.560131 | -1.409725 | 2.46272 | 0.002258 | -0.364858 | 1.949078 | -0.040318 | 1.948522 | 2.597996 |
| 30.56944 | -0.037996 | 0.290686 | -1.204876 | 0.115859 | -0.943508 | 0.025468 | -0.952295 | -0.423834 | 0.810897 | 0.019453 | -3.009171 | -0.058504 | -3.710768 | 0.424343 | 0.779406 | 0.284599 | -5.039193 | 1.671576 | 0.047696 | -0.180774 | 1.358716 | 0.004312 | -0.161329 | 1.229329 |
| 30.58333 | 0.650455 | 1.431367 | -0.615777 | 1.734561 | -0.660516 | 0.732184 | 0.156699 | 0.21891 | 1.062787 | 0.185527 | 0.445228 | 2.92568 | -2.833186 | -0.188319 | -0.293555 | 0.092946 | -5.554287 | 1.31899 | 0.135124 | 0.010369 | 1.260024 | 0.215904 | -0.561878 | 1.08597 |
| 30.59722 | 2.249365 | 1.54068 | 3.798967 | 3.121678 | 0.109523 | 1.685505 | 1.719092 | 0.287334 | 2.574802 | 0.20176 | 4.977339 | 6.011406 | 0.509008 | -0.556736 | 0.514245 | 0.148511 | 0.684777 | 0.890868 | 0.227138 | 0.069326 | 1.260071 | 0.326883 | -0.165164 | 0.778798 |
| 30.61111 | 2.928485 | 0.946518 | 6.346357 | -0.17215 | 0.670396 | 3.159415 | 3.359594 | 0.059585 | 3.32825 | 0.543774 | 5.891159 | 5.811855 | 1.898298 | 0.287209 | 1.407877 | 0.30128 | 4.879921 | 0.553742 | 0.243353 | 0.109211 | 1.327074 | 0.173649 | -0.063344 | 0.413978 |
| 30.625 | 3.508504 | 0.642831 | 6.01516 | -1.636413 | 0.553396 | 1.497567 | 3.514448 | 0.033457 | 3.561347 | 0.982728 | 5.017296 | 3.706861 | 1.095853 | 0.879068 | 1.659836 | 0.318536 | 6.488231 | 0.322236 | 0.19738 | 0.029478 | 0.862925 | -0.000472 | 0.733564 | 0.303927 |
| 30.63889 | 3.390571 | 0.235903 | 4.517089 | -1.57265 | 0.394108 | 1.126347 | 2.571683 | 0.244429 | 2.563814 | 0.851526 | 2.491817 | 1.424416 | -0.786732 | 1.20451 | 2.192156 | 0.336042 | 2.027755 | 0.77328 | 0.095428 | -0.049347 | 0.663152 | -0.123661 | 0.695154 | -0.146613 |
| 30.65278 | 2.384333 | 0.070137 | 1.001531 | -1.053905 | 0.017607 | -0.706851 | -0.054627 | 0.48021 | 0.926226 | 0.644052 | -1.066887 | -0.356427 | 0.676885 | 0.576796 | 1.306095 | 0.384191 | 0.194183 | 0.272289 | 0.16634 | -0.026991 | 0.120709 | -0.049033 | 0.69029 | -0.474404 |
| 30.66667 | 0.157667 | 0.259844 | -0.098934 | 3.377558 | -0.082431 | -1.079905 | -0.866158 | 0.555134 | 0.606779 | 0.363039 | -1.960876 | -0.20075 | -0.421271 | 0.96966 | -0.703925 | 0.267265 | 0.234094 | 0.251 | 0.13873 | 0.023272 | -0.288481 | 0.013505 | 0.445804 | -0.160504 |
| 30.68056 | -1.302646 | 0.546616 | -1.130624 | 5.763724 | -0.041546 | -0.108898 | -0.729535 | 0.451195 | 0.044333 | 0.140329 | 0.885735 | 1.320724 | -1.42104 | 0.370474 | -1.562287 | 0.054043 | 0.135319 | -0.173522 | -0.016328 | 0.09561 | 0.196417 | 0.070412 | -0.376313 | 0.443601 |
| 30.69444 | -0.808602 | 0.647019 | 0.06244 | 3.192295 | 0.111473 | 1.686762 | 0.613235 | 0.205141 | -1.063268 | -0.021616 | 3.133151 | 2.133867 | -0.199077 | -0.230197 | -0.081096 | 0.06305 | -0.210922 | -0.103331 | 0.046326 | 0.117112 | 0.789741 | 0.120755 | -2.261272 | 0.247594 |
| 30.70833 | 0.381205 | 0.370424 | 1.882148 | 0.940303 | 0.227735 | 0.448763 | 1.827154 | 0.017542 | -0.385428 | -0.008908 | 2.00861 | 0.633492 | 0.497333 | -0.448219 | 0.774507 | -0.093057 | -0.648113 | -0.471405 | 0.015891 | 0.020122 | 0.773947 | 0.145742 | -1.767915 | -0.260241 |
| 30.72222 | 1.867252 | -0.086482 | 2.84927 | 2.153436 | 0.178371 | 0.837429 | 2.060413 | 0.064012 | 1.085872 | 0.242009 | -1.491152 | -1.85692 | 0.987731 | -0.436224 | 1.257308 | -0.198158 | -0.573691 | -0.76696 | -0.019278 | -0.095106 | 0.525861 | 0.132712 | 1.005756 | -0.102602 |
| 30.73611 | 1.256712 | -0.393729 | 2.420057 | 2.066837 | 0.06307 | -0.639102 | 0.293945 | 0.190391 | 1.463794 | 0.431305 | -3.132505 | -2.261733 | 1.039303 | -0.533341 | 1.028938 | -0.035725 | 0.048625 | -0.350815 | 0.039082 | -0.071748 | 0.328602 | 0.05411 | 2.112382 | 0.078581 |
| 30.75 | -0.388901 | -0.217627 | -0.845961 | 1.242967 | 0.024934 | -1.490195 | -0.456983 | 0.225705 | 0.567444 | 0.2501 | -2.064982 | -0.519576 | -0.095216 | -0.162385 | 0.276211 | 0.062297 | 0.541236 | 0.062431 | 0.175266 | 0.049337 | 0.109306 | 0.012726 | 1.149631 | 0.218391 |
| 30.76389 | -1.248322 | 0.207424 | -4.273846 | -1.202142 | 0.065796 | -1.178414 | -0.974516 | 0.215302 | -0.90288 | -0.116828 | -0.462294 | 0.712546 | -2.276832 | 0.293198 | -0.928863 | 0.132375 | 0.544865 | 0.453331 | 0.158732 | 0.092462 | 0.442604 | 0.059565 | 0.345428 | 0.213267 |
| 30.77778 | -1.252282 | 0.593035 | -4.023955 | -0.303842 | 0.094677 | 0.856953 | -0.320606 | 0.147158 | -2.501289 | -0.264469 | 1.552236 | 0.995222 | -2.684262 | 0.562021 | -0.881597 | 0.164588 | 0.229492 | 0.03944 | 0.014565 | 0.042802 | 0.190669 | 0.119619 | -0.119326 | -0.044651 |
| 30.79167 | -0.475538 | 0.463288 | 0.463384 | 2.414492 | 0.039564 | 1.060191 | 0.588767 | 0.017377 | -1.191162 | -0.143109 | 0.952932 | -0.283767 | -2.734586 | 0.57463 | 0.284309 | 0.1381 | -0.156674 | -0.234449 | -0.051955 | 0.000547 | 0.032938 | 0.113813 | 0.08255 | -0.051421 |
| 30.80556 | 0.201449 | -0.061618 | 1.931673 | 1.840647 | 0.048113 | 0.359557 | 1.051622 | 0.009535 | 0.976766 | 0.134813 | -2.059175 | -1.838883 | -0.093086 | 0.077462 | 1.395913 | 0.120252 | -0.545406 | -0.210555 | -0.003492 | -0.000334 | 0.061739 | 0.097639 | 0.739657 | -0.030784 |
| 30.81944 | -0.243256 | -0.206548 | 0.696433 | -0.394818 | 0.129905 | -0.630836 | 1.077453 | 0.087537 | 2.530252 | 0.394061 | -3.496966 | -1.717778 | 1.324687 | -0.458775 | 1.601072 | -0.005372 | -0.648943 | -0.587423 | 0.031847 | 0.036624 | -0.345434 | 0.036212 | 2.575167 | 0.341973 |
| 30.83333 | -0.663311 | -0.189554 | -2.570439 | -1.040482 | 0.217953 | -0.800685 | -0.141553 | 0.178546 | 1.448389 | 0.426342 | -1.14885 | -0.362192 | 1.621406 | -0.817337 | 0.353119 | -0.177444 | -0.240824 | -0.620914 | -0.056955 | 0.015626 | -0.531489 | -0.167025 | 2.155684 | 0.235514 |
| 30.84722 | -0.404194 | 0.183889 | -3.175653 | -0.943152 | 0.252813 | -0.71026 | -1.524759 | 0.139524 | -0.699377 | 0.175605 | 2.188074 | 1.051799 | 0.355167 | -0.980226 | -0.632047 | -0.155478 | 0.141465 | -0.886087 | -0.098687 | -0.02099 | 0.23059 | 0.029738 | 0.082486 | -0.144319 |
| 30.86111 | 0.386699 | 0.584458 | -0.651162 | 1.209413 | 0.230931 | -1.24394 | -1.770016 | 0.064157 | -1.402371 | -0.168477 | 3.940483 | 0.8648 | -1.11463 | -0.692308 | -0.311969 | 0.007058 | 0.38504 | -1.708692 | -0.073269 | -0.0438 | 1.27875 | 0.145144 | -1.664441 | -0.281622 |
| 30.875 | -0.018615 | 0.450199 | -0.605243 | 4.038439 | 0.148517 | 0.955577 | 1.283412 | -0.00861 | -0.785879 | -0.290866 | 0.080534 | 0.127832 | -2.060574 | -0.30848 | -0.10633 | 0.126557 | 0.349794 | -1.487875 | -0.172532 | -0.013907 | 1.043779 | 0.123867 | -1.340927 | -0.368807 |
| 30.88889 | -0.76858 | 0.070235 | -1.647268 | 3.973606 | -0.020717 | 0.562811 | 1.389089 | 0.015627 | 0.589352 | -0.049801 | -3.808082 | -1.36052 | -4.441697 | 0.061014 | 0.173219 | 0.150133 | -0.769033 | 0.123143 | -0.025974 | 0.046018 | 0.269532 | 0.077736 | -0.801997 | 0.04419 |
| 30.90278 | -1.490995 | -0.210179 | -0.475738 | 1.24751 | -0.092264 | -0.515199 | -0.504702 | 0.079595 | 0.85285 | 0.306574 | -2.437769 | -0.795495 | -6.284474 | 0.276543 | 0.117191 | 0.125621 | -1.701753 | 1.303364 | 0.075533 | 0.031437 | -0.775714 | 0.002571 | -0.606113 | 0.276369 |
| 30.91667 | -1.359737 | -0.327023 | 0.029441 | -1.729534 | 0.067944 | -1.493883 | -0.872284 | 0.155749 | 0.276749 | 0.371488 | 1.717605 | 0.304375 | -5.104094 | 0.000438 | -0.000761 | -0.002961 | -2.24233 | 0.140855 | 0.017895 | -0.021247 | -1.96859 | -0.102511 | 0.42098 | 0.319807 |
| 30.93056 | -0.963867 | -0.109269 | -2.682128 | -3.048477 | 0.231725 | -1.416098 | -0.709989 | 0.190849 | -0.50918 | 0.136857 | 1.763014 | 0.028535 | -4.584627 | -0.438783 | -1.324282 | -0.239135 | -2.150759 | -0.707417 | -0.076294 | -0.04393 | -1.152214 | -0.115704 | 2.72993 | 0.138202 |
| 30.94444 | -1.39741 | 0.015083 | -4.665703 | -2.024072 | 0.183604 | -0.217936 | 0.304612 | 0.071961 | -1.842073 | -0.158023 | -0.05455 | -0.544913 | -4.072123 | -0.656487 | -1.878009 | -0.409075 | -2.584601 | -1.498452 | -0.173375 | 0.021076 | -0.040845 | -0.036289 | 2.109881 | -0.240431 |
| 30.95833 | -1.556354 | -0.039469 | -4.354404 | 0.205029 | 0.001575 | 0.526238 | -0.181435 | -0.115723 | -2.671593 | -0.413115 | -3.388567 | -1.902143 | -3.427254 | -0.90074 | -1.801265 | -0.324907 | -3.256036 | -2.739304 | -0.227346 | 0.053222 | 0.273365 | -0.029832 | -1.274993 | -1.266133 |
| 30.97222 | -1.601914 | -0.972253 | -1.795281 | 1.516597 | -0.100151 | -0.732326 | -2.451726 | -0.196649 | -2.64035 | -0.643299 | -4.435301 | -2.835825 | -1.134441 | -0.873092 | -1.075603 | -0.127122 | -3.888528 | -2.530575 | -0.148531 | 0.018028 | -1.151589 | -0.356062 | -3.677256 | -1.871509 |
| 30.98611 | -2.538491 | -1.634796 | -1.345231 | 0.303972 | -0.061578 | -2.533645 | -4.054417 | -0.168898 | -2.288118 | -0.91246 | -3.568908 | -3.165843 | -1.748996 | -0.698609 | -0.703732 | -0.028572 | -2.166445 | -1.419861 | -0.080074 | -0.043791 | -2.190134 | -0.637191 | -3.678612 | -1.702422 |
| 31 | -2.792433 | -1.377187 | -2.652907 | -3.24119 | -0.051194 | -3.314337 | -4.019043 | -0.265797 | -1.65337 | -0.531163 | -1.989023 | -2.685099 | -2.621133 | -0.472574 | -1.452777 | -0.188819 | -0.858519 | -1.210212 | -0.086802 | -0.036103 | -2.231647 | -0.856988 | -2.335283 | -1.372071 |
| 31.01389 | -2.9372 | -1.74997 | -4.345233 | -6.059433 | -0.070023 | -2.936935 | -3.020201 | -0.439093 | -1.454508 | -0.330737 | -2.242763 | -2.012382 | -3.799022 | -0.167949 | -2.120237 | -0.594425 | 1.442142 | -1.574319 | -0.260649 | 0.00021 | -2.001267 | -0.886305 | -1.19616 | -1.305171 |
| 31.02778 | -2.737031 | -3.210577 | -3.539895 | -5.565492 | -0.195239 | -2.181757 | -2.666571 | -0.277987 | -2.683778 | -0.793976 | -3.887108 | -3.555377 | -4.41443 | 0.175832 | -2.118461 | -0.953484 | 1.12158 | -2.080258 | -0.265774 | -0.0213 | -2.132843 | -0.80491 | -1.282716 | -2.22932 |
| 31.04167 | -3.0113 | -3.356649 | -2.987533 | -4.932593 | -0.41531 | -1.642401 | -1.999014 | 0.009677 | -3.231353 | -1.744331 | -5.175153 | -4.454653 | -5.585111 | -0.005265 | -2.086473 | -0.854115 | -3.430673 | -2.190491 | -0.279231 | -0.257114 | -2.464095 | -0.680427 | -1.738242 | -2.650982 |
| 31.05556 | -4.33139 | -1.345456 | -2.466 | -4.239278 | -1.150767 | -2.610161 | -2.406443 | 0.194158 | -2.741741 | -1.769526 | -4.780745 | -3.534261 | -5.650899 | -0.192419 | -1.728221 | -0.318003 | -4.492177 | -1.256013 | -0.291235 | -0.34249 | -1.667614 | -0.582245 | -1.81121 | -1.997371 |
| 31.06944 | -3.538036 | 0.439682 | -1.629657 | -2.172721 | -1.832013 | -2.90437 | -3.020386 | 0.069935 | -1.615779 | -0.790793 | -4.340991 | -1.963748 | -2.869347 | -0.220123 | -1.236945 | 0.040232 | -2.135679 | -0.880122 | -0.312862 | -0.432478 | -1.373149 | -0.374491 | -1.150993 | -1.382427 |
| 31.08333 | -1.011005 | 0.617829 | -1.477017 | 0.161518 | -1.573056 | -1.897206 | -2.106553 | -0.064537 | -0.264461 | 0.278198 | -2.640383 | -0.917335 | 0.00953 | -0.270667 | -0.753875 | 0.298634 | -0.665074 | -0.595092 | -0.185468 | -0.322169 | -0.914523 | -0.160135 | -0.472763 | -0.879841 |
| 31.09722 | -0.101844 | -0.067906 | -1.634799 | 0.135507 | -1.045649 | -1.123023 | -0.776881 | -0.169825 | 0.376498 | 0.560871 | -1.282285 | -0.852489 | 0.510071 | -0.266384 | -0.270104 | 0.241091 | -0.391943 | -0.329213 | 0.021077 | -0.157979 | -0.17378 | 0.048265 | -0.159032 | -0.516171 |
| 31.11111 | -1.095313 | -0.485333 | -2.017215 | -0.853812 | -0.745906 | -0.54932 | -0.833193 | -0.13607 | -0.170202 | -0.044475 | -1.61521 | -1.347194 | 0.353236 | -0.201884 | -0.05909 | 0.031077 | -0.684969 | -0.080065 | 0.116763 | -0.103464 | 0.269533 | 0.133623 | -0.265869 | -0.391681 |
| 31.125 | -1.890991 | -0.495283 | -2.180452 | -2.02251 | -1.292948 | -0.738298 | -2.015738 | -0.147218 | -1.810734 | -1.063632 | -2.471645 | -1.742297 | -0.056759 | -0.176391 | -0.278487 | -0.152352 | -0.997748 | -0.194904 | 0.025443 | -0.067657 | 0.099478 | 0.039848 | -0.5194 | -0.449477 |
| 31.13889 | -1.869356 | -0.518584 | -2.367551 | -1.236643 | -1.704287 | -1.48381 | -2.802966 | -0.755419 | -2.625291 | -1.358297 | -3.201556 | -1.945218 | -0.533894 | -0.185575 | -0.539347 | -0.27241 | -1.324734 | -0.440624 | -0.069193 | -0.096939 | -0.166338 | -0.079506 | -0.726544 | -0.596685 |
| 31.15278 | -1.645238 | -0.842554 | -2.222094 | -1.982417 | -1.827816 | -1.405389 | -2.292711 | -1.084767 | -1.961666 | -0.992611 | -2.76062 | -1.98359 | -0.548786 | -0.180089 | -0.613084 | -0.293521 | -1.551318 | -0.58556 | -0.079867 | -0.118695 | -0.368704 | -0.154638 | -0.796935 | -0.62456 |
| 31.16667 | -1.525422 | -1.369709 | -1.949157 | -1.286375 | -2.21709 | -1.02267 | -1.580331 | -1.07616 | -1.081088 | -0.407257 | -1.78042 | -2.083703 | -0.317741 | -0.16558 | -0.574138 | -0.247563 | -1.685553 | -0.661039 | -0.104314 | -0.108164 | -0.48189 | -0.201074 | -0.819459 | -0.535353 |
| 31.18056 | -1.251465 | -1.73205 | -1.836682 | -0.860345 | -1.995894 | -0.969129 | -1.362491 | -1.112927 | -0.520914 | -0.098428 | -1.649535 | -2.256594 | -0.342425 | -0.180314 | -0.663784 | -0.212051 | -1.860826 | -0.730109 | -0.167823 | -0.103481 | -0.370532 | -0.24107 | -0.810147 | -0.490753 |
| 31.19444 | -1.120848 | -1.212637 | -1.845733 | -0.823659 | -1.122716 | -1.017804 | -1.057626 | -1.199101 | -0.280662 | -0.111862 | -1.883396 | -2.10107 | -0.867048 | -0.235543 | -0.943817 | -0.186222 | -2.42215 | -0.886897 | -0.242903 | -0.158315 | -0.233977 | -0.310313 | -0.852921 | -0.744078 |
| 31.20833 | -1.018809 | -0.687562 | -1.604648 | -0.921617 | -0.37063 | -0.446003 | -0.520681 | -1.2333 | -0.501303 | -0.179597 | -1.445522 | -1.565043 | -1.39953 | -0.343753 | -1.214103 | -0.226098 | -3.417196 | -1.026954 | -0.305472 | -0.259234 | -0.419919 | -0.421617 | -0.995798 | -1.19392 |
| 31.22222 | -0.403039 | -0.788172 | -1.206291 | -0.785793 | 0.0818 | 0.089652 | -0.333274 | -1.000362 | -0.669891 | -0.256463 | -1.180187 | -1.344806 | -1.534497 | -0.439984 | -1.267278 | -0.292219 | -4.617099 | -1.069575 | -0.322984 | -0.443058 | -0.742622 | -0.535888 | -1.284878 | -1.515922 |
| 31.23611 | 0.059294 | -0.904226 | -0.765048 | -0.409881 | 0.413185 | -0.37998 | -0.513866 | -0.628646 | -0.247028 | -0.260117 | -1.374407 | -1.377284 | -1.257881 | -0.502336 | -1.141293 | -0.31823 | -5.808663 | -0.97442 | -0.353087 | -0.61111 | -1.113113 | -0.559623 | -1.732698 | -1.349452 |
| 31.25 | -0.082552 | -0.259146 | -0.362494 | -0.205567 | 0.579549 | -0.537918 | -0.172301 | -0.185797 | 0.037467 | -0.10252 | -1.034456 | -0.852002 | -1.39666 | -0.404938 | -0.916797 | -0.294878 | -6.364359 | -0.831664 | -0.359603 | -0.571397 | -1.076548 | -0.452202 | -1.681365 | -0.99015 |
| 31.26389 | -0.359034 | 0.361028 | -0.187316 | -0.463929 | 0.35881 | -0.180237 | 0.156852 | -0.044322 | -0.233345 | -0.014178 | -0.148769 | -0.063702 | -1.611721 | -0.27734 | -0.737408 | -0.224332 | -5.05365 | -0.639235 | -0.300924 | -0.493131 | -0.613392 | -0.26799 | -1.210312 | -0.683976 |
| 31.27778 | -0.266869 | 0.330378 | -0.305586 | -0.561939 | -0.122451 | 0.304245 | 0.163038 | -0.202147 | -0.353615 | 0.012028 | 0.37871 | 0.322054 | -1.138179 | -0.248339 | -0.564969 | -0.168464 | -3.438263 | -0.293872 | -0.149281 | -0.306498 | -0.253024 | -0.132402 | -0.809829 | -0.484999 |
| 31.29167 | -0.073421 | -0.167554 | -0.596996 | -0.315011 | -0.530812 | -0.175153 | -0.161026 | -0.359939 | 0.101224 | 0.14471 | 0.113688 | -0.007063 | -0.630377 | -0.312301 | -0.430182 | -0.144601 | -2.739166 | -0.083979 | 0.011696 | -0.296518 | -0.148158 | -0.05547 | -0.607817 | -0.443857 |
| 31.30556 | -0.065372 | -0.162298 | -0.652642 | -0.498026 | -0.850652 | -0.163208 | -0.057804 | -0.176298 | 0.301375 | 0.301619 | -0.390129 | -0.27329 | -0.977423 | -0.397624 | -0.300113 | -0.171689 | -3.046252 | -0.196235 | -0.008815 | -0.361757 | -0.282504 | -0.032419 | -0.603946 | -0.571573 |
| 31.31944 | 0.014311 | 0.425236 | -0.397454 | -0.772233 | -1.127523 | 0.351335 | 0.346042 | -0.052937 | 0.20924 | 0.291042 | -0.274804 | 0.09717 | -1.553228 | -0.443846 | -0.26499 | -0.232766 | -3.370905 | -0.422793 | -0.149811 | -0.372882 | -0.376669 | -0.02957 | -0.644012 | -0.720723 |
| 31.33333 | 0.234565 | 0.830145 | -0.142892 | -0.380745 | -1.003112 | 0.585304 | 0.364217 | 0.029823 | 0.278356 | 0.14544 | 0.435508 | 0.833929 | -1.079574 | -0.411027 | -0.278566 | -0.237062 | -2.454873 | -0.544683 | -0.241191 | -0.335196 | -0.274322 | -0.009992 | -0.485036 | -0.627606 |
| 31.34722 | 0.343378 | 0.261141 | -0.041105 | 0.336325 | -0.167645 | 0.30031 | 0.442418 | 0.195241 | 0.13919 | 0.043578 | 1.130044 | 0.972215 | -0.118854 | -0.346708 | -0.209398 | -0.18382 | -0.789163 | -0.503713 | -0.216971 | -0.163754 | -0.152261 | 0.013826 | -0.146279 | -0.365944 |
| 31.36111 | 0.486001 | -0.154433 | -0.008579 | 0.6779 | 0.994898 | 0.439071 | 0.599122 | 0.560679 | 0.013525 | 0.028586 | 1.462214 | 0.764057 | 0.344401 | -0.318521 | -0.05623 | -0.142705 | 1.152097 | -0.335511 | -0.12597 | 0.016978 | -0.093392 | 0.044141 | 0.208885 | -0.08117 |
| 31.375 | 0.974356 | 0.283784 | 0.185936 | 0.710012 | 1.935094 | 0.852343 | 0.826569 | 0.913102 | 0.405152 | 0.13266 | 1.373306 | 0.949976 | 0.443088 | -0.236985 | 0.111253 | -0.102215 | 3.045173 | -0.142377 | -0.042882 | 0.236131 | 0.01868 | 0.041349 | 0.432155 | 0.15969 |
| 31.38889 | 1.428238 | 0.977827 | 0.59709 | 0.821519 | 2.152738 | 0.910334 | 0.866728 | 1.163993 | 1.057838 | 0.374573 | 1.478488 | 1.460328 | 0.865835 | -0.137978 | 0.199973 | -0.045417 | 4.017713 | 0.003505 | -0.062574 | 0.311241 | 0.260064 | 0.01478 | 0.491598 | 0.324507 |
| 31.40278 | 1.576049 | 0.766785 | 1.115132 | 0.90653 | 1.892181 | 0.852125 | 0.912708 | 1.250497 | 1.217022 | 0.539542 | 2.459711 | 1.936905 | 1.666921 | -0.064097 | 0.376387 | 0.048848 | 4.479132 | 0.227908 | -0.055267 | 0.418075 | 0.736519 | 0.082698 | 0.589173 | 0.426611 |
| 31.41667 | 1.692944 | 0.319014 | 1.764951 | 0.995475 | 1.493703 | 1.140221 | 1.104573 | 1.288065 | 1.211623 | 0.452269 | 3.566466 | 2.322034 | 2.452767 | 0.023434 | 0.780199 | 0.201818 | 5.601566 | 0.691832 | 0.139952 | 0.657962 | 1.194442 | 0.309996 | 0.834141 | 0.608474 |
| 31.43056 | 2.016233 | 0.495265 | 2.699559 | 1.265236 | 1.355136 | 1.555623 | 1.355986 | 1.153008 | 1.554293 | 0.484968 | 3.755819 | 2.48252 | 2.786162 | 0.154402 | 1.207907 | 0.314172 | 6.873491 | 1.264926 | 0.361956 | 0.92641 | 1.278467 | 0.480328 | 1.234963 | 0.892713 |
| 31.44444 | 2.552063 | 1.080167 | 3.392141 | 2.311353 | 1.720759 | 1.857542 | 1.80585 | 1.064488 | 2.171012 | 0.812017 | 4.297761 | 2.802856 | 2.810295 | 0.270658 | 1.463589 | 0.351874 | 7.151255 | 1.672437 | 0.490098 | 0.986984 | 1.23069 | 0.526888 | 1.674689 | 1.329933 |
| 31.45833 | 3.213595 | 1.356854 | 3.559374 | 3.014265 | 2.207457 | 2.292825 | 2.631111 | 0.771802 | 2.902467 | 1.251387 | 5.177595 | 3.27049 | 3.161285 | 0.390429 | 1.75444 | 0.470037 | 6.770973 | 2.052662 | 0.534487 | 0.86305 | 1.75222 | 0.601993 | 2.039263 | 1.789477 |
| 31.47222 | 3.867101 | 1.28628 | 4.069396 | 3.72668 | 2.768253 | 3.067416 | 3.905547 | 0.381562 | 3.678848 | 1.388093 | 5.836783 | 3.482072 | 3.640672 | 0.524962 | 2.162068 | 0.641079 | 6.49538 | 2.602775 | 0.567944 | 0.86295 | 2.790969 | 0.705967 | 2.518347 | 2.148877 |
| 31.48611 | 4.018177 | 1.190156 | 4.929993 | 4.829791 | 2.853272 | 3.666016 | 4.888859 | 0.165356 | 3.525797 | 1.141495 | 6.499667 | 3.395281 | 4.25495 | 0.548813 | 2.660594 | 0.67274 | 7.873295 | 2.937114 | 0.508124 | 1.4168 | 2.946369 | 0.760242 | 2.618518 | 2.470615 |
| 31.5 | 3.933922 | 1.319613 | 6.016462 | 6.075738 | 2.026722 | 3.919478 | 4.882304 | 0.188179 | 3.481946 | 1.116197 | 5.635159 | 3.662445 | 4.867966 | 0.52353 | 3.319085 | 0.598523 | 9.633119 | 2.884629 | 0.63282 | 0.711256 | 2.90271 | 0.61316 | 2.94255 | 2.789961 |
| 31.51389 | 4.002646 | 1.568682 | 6.284912 | 6.482369 | 1.012576 | 3.099794 | 4.97056 | 0.150793 | 4.421298 | 1.726588 | 4.564735 | 4.220286 | 4.86457 | 0.436662 | 4.105356 | 0.61291 | 8.958908 | 2.852643 | 0.417797 | 0.23988 | 2.54792 | 0.457098 | 3.348706 | 2.877951 |
| 31.52778 | 4.012501 | 1.520773 | 5.570119 | 6.774042 | 0.851115 | 2.486066 | 4.331601 | -0.252482 | 4.176574 | 1.489933 | 5.325569 | 4.772915 | 5.637759 | 0.178338 | 4.493884 | 0.675937 | 7.281647 | 1.924055 | 0.076977 | 0.059756 | 2.557537 | 0.160284 | 3.909351 | 2.249037 |
| 31.54167 | 3.891069 | 0.8698 | 3.610971 | 5.254513 | 0.24572 | 1.553324 | 3.959748 | -0.941626 | 2.56207 | 0.374766 | 4.809095 | 4.119889 | 5.324966 | 0.435064 | 3.410984 | 0.701417 | 3.252158 | 0.955315 | -0.123877 | -0.209598 | 2.566155 | -0.169799 | 4.509594 | 1.96557 |
| 31.55556 | 1.916568 | -0.059938 | 1.231485 | 2.040294 | -0.517658 | 0.204813 | 2.181556 | -0.901878 | 0.472751 | -0.424097 | 0.613686 | 2.356229 | 2.062877 | 0.861322 | 2.257557 | 0.430219 | -2.572914 | 0.375885 | -0.126069 | -0.358109 | 2.279245 | -0.082727 | 3.568833 | 1.741808 |
| 31.56944 | -0.000889 | -0.303516 | 0.187133 | -0.37639 | -0.531273 | -0.951479 | 0.471432 | -0.330058 | -0.209155 | -0.33614 | -1.578277 | 1.708481 | -2.946762 | 0.564228 | 1.072631 | 0.055288 | -7.294158 | 0.374679 | -0.053821 | -0.179673 | 2.046181 | 0.059164 | 1.159935 | 1.046965 |
| 31.58333 | 0.978127 | -0.04057 | 2.23482 | 1.766807 | -0.238625 | 0.545191 | 1.47131 | 0.102018 | 1.242758 | 0.289812 | 0.481962 | 2.780627 | -3.697225 | 0.157042 | -0.0563 | -0.19508 | -8.668017 | 1.347474 | -0.053023 | -0.065431 | 1.751848 | 0.075995 | 1.390431 | 0.636152 |
| 31.59722 | 2.939619 | 0.352488 | 2.325338 | 3.907782 | 0.062199 | 0.617446 | 2.671438 | -0.063947 | 3.517422 | 0.608961 | 4.478644 | 4.241712 | -1.232871 | -0.238989 | -0.530234 | -0.157813 | -1.512553 | 0.975228 | -0.018849 | -0.206029 | 1.361272 | 0.036034 | 1.382776 | 1.334867 |
| 31.61111 | 4.038136 | 0.687034 | 0.997909 | 4.290609 | 0.327089 | 1.748009 | 3.055574 | -0.220869 | 4.039776 | 0.641514 | 5.292877 | 4.941435 | 1.771006 | -0.226938 | -0.907065 | 0.000091 | 3.73824 | 0.388097 | 0.040795 | -0.03224 | 0.814708 | 0.105515 | 1.085742 | 1.208708 |
| 31.625 | 4.608427 | 0.626107 | 2.936211 | 2.402462 | 0.424793 | 4.110326 | 3.500117 | -0.121819 | 3.455127 | 0.899154 | 5.116082 | 4.850183 | 2.42102 | 0.271901 | 0.305679 | 0.145602 | 5.624552 | -0.063305 | 0.139251 | 0.095061 | 0.459151 | -0.026156 | 0.698954 | 0.657214 |
| 31.63889 | 5.997893 | 0.388898 | 1.178309 | 1.288104 | 0.360399 | 0.932887 | 2.07044 | 0.091126 | 2.34566 | 0.833277 | 2.638648 | 2.448537 | 2.641204 | 0.646308 | 1.502675 | 0.207092 | 2.225139 | 0.084417 | 0.223303 | 0.256201 | 0.49378 | -0.082972 | 1.693476 | 0.134003 |
| 31.65278 | 4.629998 | -0.029836 | -0.816499 | 1.789562 | 0.169072 | -0.902043 | -0.000395 | 0.399189 | 1.278531 | 0.542883 | -0.726209 | -0.544022 | 1.665682 | 0.962683 | 2.502951 | 0.267087 | 1.592404 | 0.299744 | 0.300871 | 0.055345 | 0.639526 | -0.066532 | 0.929329 | -0.101819 |
| 31.66667 | 1.679053 | -0.133244 | -0.884873 | 2.062102 | -0.011435 | -0.280977 | -0.583507 | 0.640934 | 0.520838 | 0.193964 | -2.42777 | -1.797899 | 1.677156 | 0.87476 | 2.238536 | 0.334625 | 0.505202 | 0.561916 | 0.316841 | -0.076597 | -0.387453 | -0.051087 | -0.755633 | -0.019675 |
| 31.68056 | -0.706411 | -0.048408 | 0.737722 | 0.858208 | 0.012875 | -0.963638 | -0.250679 | 0.551379 | -0.236607 | -0.17598 | -0.971872 | -0.303676 | 0.613816 | 0.312723 | 1.083595 | 0.312193 | 0.258324 | 0.508071 | 0.182899 | -0.053283 | 0.213301 | -0.014878 | 1.515355 | 0.039762 |
| 31.69444 | -2.686719 | 0.027805 | 2.136976 | -0.08755 | 0.095014 | -0.306304 | 0.696702 | 0.268959 | -0.436117 | -0.318682 | 1.975694 | 1.314923 | -1.272532 | 0.491425 | -0.525545 | 0.300543 | 0.04883 | 0.713315 | -0.000554 | 0.131914 | 0.984614 | 0.054556 | 1.115198 | 0.482272 |
| 31.70833 | -2.211834 | 0.339953 | 1.036827 | 0.34437 | 0.09147 | 0.854389 | 1.702568 | 0.108802 | -0.220748 | -0.141365 | 2.185825 | 0.497966 | -1.575961 | 0.044438 | -0.702957 | 0.129951 | -0.413712 | 0.46982 | -0.271458 | 0.174225 | 0.193964 | 0.065973 | 0.645747 | 0.533422 |
| 31.72222 | -0.88475 | 0.458572 | 0.88987 | 1.916246 | -0.038024 | 0.668941 | 0.455146 | 0.060474 | -0.173938 | 0.123511 | 0.95396 | 0.240966 | -0.626076 | -0.453663 | -1.099103 | -0.059885 | -0.752848 | 0.318726 | -0.248573 | 0.111798 | 0.006526 | 0.08353 | 0.126404 | -0.011976 |
| 31.73611 | -0.138457 | 0.214892 | 2.094031 | 1.590001 | -0.182187 | -2.161409 | -1.510632 | 0.125311 | -0.027379 | 0.279646 | -0.193634 | 0.999262 | 0.309169 | -0.532622 | -0.603432 | -0.191978 | -0.619293 | 0.149992 | 0.002792 | -0.000137 | -0.00241 | 0.105221 | 1.078244 | -0.094572 |
| 31.75 | 0.262395 | -0.040704 | 0.502207 | -0.765185 | -0.065386 | -2.127419 | -1.011669 | 0.180951 | -0.39019 | 0.215303 | -1.776621 | 0.21266 | 2.107165 | -0.815596 | 0.782498 | -0.251926 | -0.232377 | -0.304106 | 0.014583 | -0.047933 | -0.043411 | 0.074008 | 1.203415 | 0.000411 |
| 31.76389 | 0.845916 | -0.080231 | -0.037947 | -3.142654 | 0.14791 | -1.073322 | -0.413241 | 0.150287 | -0.317494 | 0.091961 | -1.933043 | -1.10052 | 2.55276 | -0.789765 | 1.690014 | -0.226157 | 0.051381 | -0.571071 | 0.020653 | 0.041748 | 0.160521 | 0.080933 | 0.812838 | 0.312644 |
| 31.77778 | 1.312791 | -0.074595 | -3.17782 | -2.748802 | 0.293547 | -0.474089 | 0.187385 | 0.079015 | -0.762594 | 0.084396 | 0.509472 | -0.065155 | -0.741497 | -0.183906 | 1.361472 | -0.068938 | 0.176591 | -0.532135 | 0.005163 | 0.102909 | 0.103688 | 0.099509 | -0.458865 | 0.318299 |
| 31.79167 | 0.680125 | 0.32905 | -2.181432 | 0.500528 | 0.312893 | -0.479849 | 0.310054 | 0.020103 | -0.721455 | 0.121433 | 3.432169 | 1.658459 | -2.747949 | 0.302336 | 0.851836 | 0.09745 | 0.271613 | -0.096787 | -0.021201 | 0.090954 | 0.002081 | 0.095326 | -1.138365 | 0.082535 |
| 31.80556 | -0.85697 | 0.53029 | 1.977668 | 2.221542 | 0.092836 | -0.587684 | 0.363936 | -0.022299 | 0.554926 | 0.140379 | 3.918031 | 2.035005 | -0.74682 | 0.868415 | 0.040429 | 0.208024 | -0.036294 | 0.201246 | -0.015755 | -0.045711 | 0.007677 | 0.068332 | -0.503123 | -0.070029 |
| 31.81944 | -1.105637 | 0.386486 | 1.462892 | 3.011217 | -0.275594 | -0.791265 | 0.341281 | -0.027719 | 0.492345 | 0.123711 | 0.320044 | 0.699467 | -0.282326 | 1.182907 | -0.636853 | 0.164442 | -0.03978 | 0.041832 | -0.006822 | -0.0525 | -0.190108 | -0.009445 | 0.909988 | -0.167203 |
| 31.83333 | -0.007588 | -0.216569 | 1.256628 | 1.017596 | -0.655334 | 0.219651 | 0.073048 | -0.027446 | -0.219012 | 0.099148 | -4.469145 | -2.586554 | 0.044407 | 0.958065 | -0.471113 | 0.003838 | -0.145349 | 0.152353 | 0.053825 | -0.032018 | -0.684044 | -0.035923 | 0.253894 | -0.090928 |
| 31.84722 | -0.82444 | -0.733698 | -0.741855 | -1.240409 | -0.870801 | -0.264503 | 0.28623 | 0.153085 | -1.066216 | 0.11023 | -2.041084 | -2.850176 | 0.900745 | 0.507545 | -0.358368 | -0.111381 | -0.19087 | -0.160009 | 0.141919 | -0.011022 | -0.724264 | 0.006729 | -0.843266 | 0.033518 |
| 31.86111 | -1.077001 | -0.636899 | 0.368494 | -2.418514 | 0.274592 | -0.515071 | -0.348778 | 0.320357 | -0.786689 | 0.054251 | 0.147089 | -1.025122 | 0.40247 | -0.049709 | -0.05495 | -0.063098 | -0.265179 | 0.199758 | 0.215132 | -0.000889 | -0.061702 | 0.055003 | -0.990497 | 0.086555 |
| 31.875 | 1.809648 | -0.260023 | 0.562583 | -1.782205 | 1.338075 | -0.076505 | -0.348232 | 0.299944 | 0.51726 | 0.011764 | 1.026558 | -0.416377 | -0.947893 | -0.647402 | 0.280117 | 0.00816 | -0.752233 | 0.154071 | 0.140669 | -0.055852 | 1.271166 | 0.06166 | 0.822531 | 0.086594 |
| 31.88889 | 2.596754 | 0.368494 | -0.1825 | 0.067275 | 0.812257 | -0.213205 | 0.124359 | 0.089056 | 1.060234 | 0.056075 | -0.457336 | 0.07583 | -0.500457 | -0.913772 | 0.695328 | 0.012538 | -2.263483 | 0.008138 | -0.011183 | 0.036499 | 1.050452 | 0.070842 | 0.338784 | 0.057408 |
| 31.90278 | 0.549342 | 0.687052 | 0.597852 | 1.591138 | 0.215253 | 0.700726 | -0.324287 | -0.149464 | 0.393528 | 0.223662 | -2.085822 | 0.050532 | -0.045163 | -0.893757 | -0.369067 | -0.109932 | -4.367191 | -0.719267 | -0.121211 | 0.128496 | 0.610838 | 0.055463 | -1.004668 | -0.251156 |
| 31.91667 | -2.547065 | 0.666432 | 1.749093 | 0.986136 | -0.168046 | 0.216872 | 0.15982 | -0.288432 | -1.043029 | 0.286968 | -0.025003 | -0.219202 | -0.412687 | -0.842546 | -1.694278 | -0.206684 | -5.710643 | -0.922109 | -0.118361 | 0.136079 | 0.544141 | 0.049544 | 0.229058 | -0.487115 |
| 31.93056 | -2.851745 | 0.023179 | 1.26819 | -0.209123 | -0.357298 | -1.691612 | 0.114647 | -0.276827 | -1.116623 | -0.127982 | 0.578173 | -0.316405 | -3.831025 | -0.81697 | -2.288582 | -0.156411 | -4.460027 | -0.419217 | -0.130088 | 0.043744 | -1.627435 | 0.036559 | 1.177193 | -0.044311 |
| 31.94444 | -1.733813 | -0.337527 | -0.167815 | -3.910596 | -0.134826 | -0.693492 | -1.676812 | -0.137327 | -1.158157 | 0.000472 | 1.134079 | 1.037056 | -4.572128 | -0.664814 | -1.033772 | -0.061816 | -5.357184 | -0.176317 | -0.210575 | -0.090195 | -2.237779 | -0.132035 | 0.147597 | 0.226597 |
| 31.95833 | -0.692778 | -0.271792 | -0.401592 | -5.53041 | 0.17319 | -2.140303 | -3.22285 | -0.039971 | -1.900099 | -0.01094 | 0.608379 | -0.299303 | -2.729965 | -0.236289 | -0.153381 | 0.029298 | -4.132911 | -0.402991 | -0.244126 | -0.019643 | -1.364362 | -0.311617 | 1.017481 | 0.258908 |
| 31.97222 | -0.964273 | -0.132975 | -1.81234 | -5.679843 | 0.275453 | -1.319746 | -2.174425 | -0.139559 | -3.239824 | -0.410367 | -2.691301 | -2.125372 | -3.169009 | -0.298797 | -0.602888 | -0.01823 | -4.329618 | -0.769147 | -0.247185 | -0.048796 | -0.988761 | -0.222127 | -1.013237 | -0.596886 |
| 31.98611 | -3.829154 | -0.421203 | -2.911015 | -4.819005 | -0.213947 | 0.226474 | 0.060167 | -0.271815 | -3.233973 | -0.607267 | -4.730093 | -2.712844 | -4.948281 | -0.262962 | -1.07625 | -0.22341 | -7.314275 | -0.695167 | -0.112152 | -0.009573 | -1.904433 | -0.276242 | -2.007046 | -0.975846 |
| 32 | -5.053699 | -1.827884 | -4.668256 | -4.910041 | -1.1392 | -0.592667 | 0.148179 | -0.413438 | -3.517864 | -1.509389 | -5.432366 | -4.669616 | -5.18913 | -0.194011 | -1.218008 | -0.546278 | -5.660444 | -1.190292 | -0.047038 | -0.158266 | -3.401484 | -0.543228 | -1.820703 | -1.766459 |
| 32.01389 | -2.154734 | -3.204695 | -3.207835 | -3.587185 | -1.002581 | -2.884583 | -2.003033 | -0.30023 | -2.819316 | -1.896543 | -5.189471 | -6.456137 | -2.587994 | -0.250931 | -1.203559 | -0.779651 | -1.771461 | -2.773849 | 0.113969 | -0.810038 | -3.702471 | -0.584865 | -1.76421 | -1.999014 |

| | | | | | | | | | | | | | | | | | | | | | | | | |
|---|---|---|---|---|---|---|---|---|---|---|---|---|---|---|---|---|---|---|---|---|---|---|---|---|
| 32.02778 | -1.600095 | -2.43441 | -0.916199 | -4.157075 | -0.124776 | -3.311166 | -5.051542 | -0.089247 | -3.29819 | -1.947527 | -4.07484 | -5.025401 | -0.083728 | -0.226242 | -1.581157 | -0.67716 | -0.140587 | -2.047964 | 0.023148 | -1.560157 | -2.992281 | -0.57002 | -1.542625 | -1.812501 |
| 32.04167 | -3.250929 | -2.271361 | -1.584445 | -6.129275 | -0.173505 | -2.806563 | -5.16424 | 0.062027 | -3.871015 | -2.223574 | -4.052899 | -3.468051 | -1.42949 | -0.164066 | -1.575856 | -0.424747 | -4.983362 | -1.137245 | 0.036522 | -0.605559 | -2.380828 | -0.843197 | -1.541504 | -1.703263 |
| 32.05556 | -6.095325 | -1.59778 | -3.708453 | -5.883909 | -1.532686 | -3.429367 | -4.335559 | 0.075844 | -2.270519 | -1.979512 | -4.617763 | -3.187116 | -3.480514 | -0.308657 | -1.654565 | -0.149595 | -9.771228 | -1.838561 | -0.066977 | 0.147181 | -1.832713 | -0.723173 | -1.923928 | -1.651471 |
| 32.06944 | -7.589864 | -0.448069 | -4.442532 | -3.258296 | -3.849049 | -4.051625 | -2.689845 | -0.091749 | -0.622772 | -0.727287 | -3.586917 | -2.412629 | -2.779716 | -0.30724 | -1.840463 | -0.059281 | -4.896397 | -1.739519 | -0.096104 | -0.014192 | -1.261819 | -0.433064 | -1.488613 | -1.299341 |
| 32.08333 | -4.425075 | 0.273514 | -1.857221 | -1.083725 | -3.963837 | -2.895823 | -2.126024 | -0.232149 | -0.022386 | -0.187175 | -2.978242 | -1.704243 | -1.363528 | -0.235606 | -1.600038 | -0.001856 | -0.265047 | -1.17039 | -0.100351 | -0.000663 | -0.512681 | -0.146573 | -0.90376 | -0.764038 |
| 32.09722 | -1.68661 | 0.589233 | -0.42141 | -0.891782 | -2.736761 | -0.981785 | -2.952506 | -0.325565 | -0.381305 | -0.46179 | -3.571615 | -1.691544 | -0.857594 | -0.180778 | -0.918585 | 0.003838 | 0.147724 | -0.67558 | -0.11962 | -0.073316 | -0.153691 | -0.028079 | -0.779676 | -0.737726 |
| 32.11111 | -1.345237 | 0.314518 | -0.756969 | -2.173487 | -1.995992 | -0.662704 | -3.506828 | -0.377744 | -1.277402 | -0.61674 | -3.595347 | -1.937695 | -0.671905 | -0.162597 | -0.663351 | -0.0482 | -0.210191 | -0.485512 | -0.136248 | -0.156116 | -0.109176 | -0.036346 | -0.557727 | -0.768219 |
| 32.125 | -2.223952 | -0.847278 | -2.113384 | -3.030077 | -1.561872 | -1.073434 | -2.931457 | -0.671132 | -1.661654 | -0.64828 | -2.713059 | -1.961269 | -0.637245 | -0.180847 | -0.324115 | -0.096012 | -0.607738 | -0.462977 | -0.125561 | -0.175148 | -0.174834 | -0.08944 | -0.284446 | -0.753568 |
| 32.13889 | -2.777555 | -1.342835 | -2.628726 | -2.677219 | -1.424853 | -1.432704 | -2.425924 | -0.870069 | -2.024557 | -0.693243 | -2.094054 | -2.002317 | -0.970714 | -0.176504 | -0.280984 | -0.125182 | -1.083613 | -0.62222 | -0.113461 | -0.186633 | -0.240412 | -0.124349 | -0.244265 | -0.721691 |
| 32.15278 | -2.382871 | -1.728482 | -2.007884 | -2.162292 | -1.374027 | -1.186493 | -1.785662 | -1.118316 | -1.680188 | -0.651302 | -1.662515 | -2.036933 | -1.320648 | -0.166895 | -0.456985 | -0.138557 | -1.346744 | -0.684697 | -0.141424 | -0.200347 | -0.285186 | -0.154588 | -0.376023 | -0.759537 |
| 32.16667 | -1.956781 | -2.411425 | -1.386122 | -2.006867 | -0.951195 | -0.937648 | -1.189562 | -1.106527 | -1.324252 | -0.501012 | -1.147746 | -1.974074 | -1.099892 | -0.198883 | -0.491665 | -0.138955 | -1.425931 | -0.614485 | -0.179808 | -0.197606 | -0.328523 | -0.180465 | -0.496848 | -0.72699 |
| 32.18056 | -1.91515 | -2.363655 | -1.691698 | -1.442285 | -0.559805 | -0.980728 | -1.068426 | -0.986779 | -1.817702 | -0.446027 | -0.983384 | -1.776824 | -0.787683 | -0.218981 | -0.422897 | -0.150612 | -1.481343 | -0.632163 | -0.189455 | -0.169521 | -0.410859 | -0.182546 | -0.543056 | -0.651396 |
| 32.19444 | -1.898917 | -1.671666 | -2.123107 | -0.960605 | -0.572267 | -0.677355 | -1.16919 | -1.121662 | -1.553325 | -0.509221 | -1.0914 | -1.494256 | -0.683671 | -0.207514 | -0.359765 | -0.203094 | -1.717763 | -0.885573 | -0.206749 | -0.181009 | -0.522256 | -0.218204 | -0.616533 | -0.663313 |
| 32.20833 | -1.693506 | -0.79176 | -1.521452 | -0.564102 | -0.654209 | -0.556401 | -0.981752 | -1.393093 | -0.655978 | -0.494057 | -0.926915 | -0.992925 | -0.923763 | -0.24788 | -0.528681 | -0.297856 | -2.122615 | -1.230184 | -0.246772 | -0.243356 | -0.595957 | -0.32854 | -0.856325 | -0.98047 |
| 32.22222 | -1.212226 | -0.695771 | -0.880275 | -0.291064 | -0.576137 | -0.936172 | -0.709802 | -1.378012 | -0.515932 | -0.501438 | -0.54273 | -0.60808 | -1.237164 | -0.36747 | -0.726474 | -0.387059 | -2.711785 | -1.371839 | -0.239889 | -0.378456 | -0.604462 | -0.441799 | -1.210026 | -1.412145 |
| 32.23611 | -0.755892 | -0.848521 | -0.651402 | -0.17847 | -0.437204 | -0.736392 | -0.612127 | -1.078384 | -0.679982 | -0.484479 | -0.04935 | -0.335032 | -1.36801 | -0.442982 | -0.777404 | -0.323 | -3.527691 | -1.325841 | -0.220797 | -0.559318 | -0.537335 | -0.440425 | -1.396492 | -1.494885 |
| 32.25 | -0.430885 | -0.352935 | -0.491217 | -0.194664 | -0.43636 | 0.044486 | -0.305946 | -0.718261 | -0.179433 | -0.290653 | 0.17047 | -0.038175 | -1.222626 | -0.44093 | -0.740764 | -0.247936 | -4.470981 | -1.102415 | -0.191523 | -0.698644 | -0.40672 | -0.358547 | -1.271695 | -1.263324 |
| 32.26389 | -0.295033 | 0.416881 | -0.033611 | 0.018917 | -0.454415 | 0.082434 | -0.000386 | -0.412341 | 0.228761 | -0.116496 | 0.175579 | 0.133579 | -0.987653 | -0.427426 | -0.610324 | -0.20764 | -4.652469 | -0.862812 | -0.168742 | -0.786266 | -0.309096 | -0.271114 | -1.008142 | -1.016648 |
| 32.27778 | -0.246889 | 0.641819 | 0.236304 | 0.322784 | -0.486371 | -0.440836 | -0.085698 | -0.153105 | -0.174186 | -0.269019 | 0.144187 | 0.111687 | -0.809693 | -0.410243 | -0.445956 | -0.218889 | -3.833048 | -0.604241 | -0.155756 | -0.695106 | -0.279813 | -0.231913 | -0.754553 | -0.783501 |
| 32.29167 | -0.172522 | 0.172428 | -0.005704 | 0.213319 | -0.61436 | -0.374965 | -0.34318 | -0.082625 | -0.523301 | -0.527432 | -0.085755 | -0.13818 | -0.75799 | -0.364179 | -0.357956 | -0.297314 | -3.117846 | -0.416974 | -0.176151 | -0.634767 | -0.306133 | -0.26172 | -0.58588 | -0.581523 |
| 32.30556 | -0.124024 | -0.284446 | -0.299972 | -0.135018 | -0.767242 | -0.13913 | -0.37131 | -0.177773 | -0.282679 | -0.515411 | -0.484547 | -0.391885 | -0.804669 | -0.341731 | -0.390956 | -0.352644 | -2.891646 | -0.409329 | -0.192873 | -0.683239 | -0.331424 | -0.201136 | -0.440634 | -0.377914 |
| 32.31944 | -0.019476 | -0.153605 | 0.047807 | -0.073225 | -0.519488 | -0.245658 | -0.068521 | -0.237118 | 0.009579 | -0.229685 | -0.253128 | -0.317171 | -0.621168 | -0.256064 | -0.359853 | -0.250352 | -1.956063 | -0.229689 | -0.163001 | -0.568373 | -0.275992 | -0.082817 | -0.220054 | -0.112274 |
| 32.33333 | 0.332015 | 0.38889 | 0.47224 | 0.40883 | 0.071025 | -0.001796 | 0.083943 | -0.160222 | 0.203687 | 0.011916 | 0.412794 | 0.287017 | -0.173903 | -0.179818 | -0.16712 | -0.11299 | -0.477127 | -0.017837 | -0.125248 | -0.370423 | -0.162009 | 0.016615 | 0.09364 | 0.176357 |
| 32.34722 | 0.895898 | 0.663658 | 0.513223 | 0.95612 | 0.623202 | 0.553922 | 0.063564 | 0.108114 | 0.429334 | 0.077491 | 0.662356 | 0.77394 | 0.057034 | -0.12058 | 0.002723 | -0.124391 | 0.548865 | 0.059454 | -0.104037 | -0.239942 | -0.109948 | 0.022184 | 0.30261 | 0.277858 |
| 32.36111 | 1.313096 | 0.42104 | 0.325802 | 1.140502 | 1.065633 | 0.968383 | 0.549376 | 0.692483 | 0.536961 | 0.128687 | 0.808406 | 0.762375 | 0.080483 | -0.092 | 0.03965 | -0.183857 | 1.280948 | 0.023867 | -0.091612 | -0.137083 | -0.070059 | -0.008306 | 0.364739 | 0.203008 |
| 32.375 | 1.466001 | 0.274801 | 0.643984 | 1.102453 | 1.605316 | 0.975226 | 1.138941 | 1.468295 | 0.437725 | 0.248116 | 1.416844 | 0.905234 | 0.173841 | -0.051764 | 0.084756 | -0.151416 | 2.280221 | 0.145378 | -0.04576 | 0.047602 | 0.069858 | 0.011637 | 0.455226 | 0.15869 |
| 32.38889 | 1.45701 | 0.676493 | 1.459773 | 1.201071 | 1.931272 | 1.295581 | 1.355449 | 1.788884 | 0.647842 | 0.567321 | 1.927482 | 1.576069 | 0.425094 | -0.010402 | 0.305698 | -0.098665 | 3.106311 | 0.396224 | 0.00557 | 0.225191 | 0.280187 | 0.064852 | 0.549678 | 0.207735 |
| 32.40278 | 1.673727 | 1.392761 | 2.01308 | 1.718469 | 2.155739 | 1.68384 | 1.624966 | 1.758733 | 1.086323 | 0.995688 | 2.365891 | 2.358625 | 0.575779 | 0.035682 | 0.764928 | -0.012325 | 3.498163 | 0.612689 | 0.024639 | 0.406015 | 0.536699 | 0.123095 | 0.69651 | 0.363043 |
| 32.41667 | 2.120523 | 1.775946 | 2.237422 | 2.493842 | 2.462329 | 1.735856 | 2.023433 | 1.731771 | 1.252744 | 1.299886 | 3.085264 | 2.733998 | 0.82854 | 0.153544 | 1.21061 | 0.108472 | 4.585945 | 0.841533 | 0.076869 | 0.692035 | 0.828948 | 0.198547 | 0.855724 | 0.6524 |
| 32.43056 | 2.432468 | 1.399119 | 2.664481 | 2.394392 | 2.314911 | 2.10332 | 2.052555 | 1.1826 | 1.382954 | 1.301422 | 3.469514 | 2.651824 | 1.172837 | 0.289774 | 1.384011 | 0.21121 | 5.525392 | 1.278974 | 0.139557 | 0.921785 | 1.164321 | 0.284757 | 1.165106 | 0.99612 |
| 32.44444 | 2.879243 | 1.159727 | 3.189216 | 2.37173 | 2.390406 | 2.975882 | 2.217349 | 0.684474 | 1.706967 | 1.441983 | 3.704666 | 2.803543 | 1.575976 | 0.405107 | 1.51466 | 0.331882 | 6.591136 | 1.917171 | 0.175384 | 1.062878 | 1.545983 | 0.418652 | 1.543239 | 1.352203 |
| 32.45833 | 3.767677 | 1.712056 | 3.175542 | 2.809494 | 3.032646 | 3.581806 | 2.872805 | 0.55312 | 2.274981 | 1.794677 | 4.297955 | 3.519798 | 1.938028 | 0.521168 | 2.052006 | 0.290735 | 8.281084 | 2.512128 | 0.182512 | 1.346623 | 2.023821 | 0.527213 | 1.901031 | 1.902362 |
| 32.47222 | 4.579127 | 2.479032 | 3.327375 | 3.360136 | 3.278215 | 3.103632 | 3.718649 | 0.236443 | 3.188764 | 2.54983 | 5.48687 | 4.697285 | 2.19709 | 0.602983 | 2.820886 | 0.431612 | 8.829022 | 2.787628 | 0.189639 | 1.321387 | 2.482994 | 0.599639 | 2.253228 | 2.487819 |
| 32.48611 | 5.080323 | 2.663011 | 4.607674 | 4.304205 | 2.716945 | 3.19137 | 4.986013 | 0.033607 | 4.151709 | 3.436093 | 6.401311 | 5.327854 | 2.501234 | 0.5286 | 3.169367 | 0.405766 | 9.208841 | 2.801715 | 0.15225 | 1.323202 | 2.82519 | 0.737876 | 2.64601 | 3.105353 |
| 32.5 | 5.617853 | 1.783104 | 6.241793 | 5.441886 | 1.797265 | 3.547348 | 5.727113 | -0.136109 | 4.383031 | 3.306902 | 5.796343 | 4.969426 | 3.091958 | 0.479911 | 2.981349 | 0.37661 | 10.868324 | 2.749231 | 0.223668 | 1.525358 | 3.075292 | 0.757384 | 3.189746 | 3.129648 |
| 32.51389 | 5.745359 | 1.239773 | 7.737911 | 5.523516 | 1.258079 | 3.495232 | 5.524517 | -0.339934 | 4.880707 | 2.513485 | 5.977392 | 4.919479 | 3.950297 | 0.345972 | 2.539066 | 0.421433 | 12.22906 | 2.939024 | 0.346409 | 0.953124 | 3.292884 | 0.614066 | 3.355621 | 2.328189 |
| 32.52778 | 4.489485 | 0.864203 | 7.73429 | 6.309411 | 0.804994 | 2.390562 | 5.3091 | -0.767615 | 5.061337 | 1.734712 | 6.753341 | 5.391972 | 4.304148 | 0.43774 | 2.216793 | 0.530203 | 13.08592 | 1.537212 | 0.294045 | 0.668146 | 2.221085 | 0.438667 | 2.905024 | 1.711777 |
| 32.54167 | 2.374646 | 0.806832 | 4.961768 | 6.881982 | 0.069757 | 1.951713 | 5.095334 | -1.329448 | 3.565464 | 0.595423 | 6.03291 | 5.881246 | 4.611092 | 0.678496 | 2.656001 | 0.654881 | 10.94033 | 0.717574 | 0.190692 | 0.604824 | 1.913399 | 0.153184 | 2.837683 | 1.738553 |
| 32.55556 | 0.996475 | 0.094998 | 2.106474 | 3.210798 | -0.911714 | 1.503971 | 2.724513 | -1.14856 | 2.454537 | 0.304926 | 0.446992 | 5.255364 | 3.254595 | 1.138713 | 1.989457 | 0.66444 | 4.699845 | 0.767378 | 0.099833 | -0.007192 | 2.455418 | -0.017591 | 2.15085 | 1.246139 |
| 32.56944 | -0.201005 | -0.365826 | 1.340104 | 0.013191 | -1.903991 | 1.198751 | 1.088326 | -0.400521 | 1.224635 | 0.144133 | -4.035443 | 2.421067 | 1.123756 | 1.042625 | 0.927985 | 0.355094 | -2.975205 | 0.852673 | -0.04824 | -0.164768 | 2.591386 | 0.103332 | 1.480936 | 0.836459 |
| 32.58333 | 0.00905 | -0.551054 | 1.960792 | 1.78362 | -1.752926 | 0.878766 | 0.822376 | -0.060517 | 0.611595 | -0.612873 | -2.530462 | 0.546512 | 0.54367 | 0.600918 | 0.46634 | 0.085565 | -4.137294 | 0.743184 | -0.001914 | -0.081494 | 1.702333 | 0.123106 | 1.273086 | 0.133277 |
| 32.59722 | 1.255029 | -0.281767 | 4.598271 | 3.367713 | 0.52678 | 0.5247 | 1.923514 | -0.174698 | 0.782022 | -0.307153 | 1.321243 | 0.595511 | 2.031569 | 0.082472 | 1.121164 | -0.104578 | -0.992577 | 0.413234 | -0.012328 | -0.041616 | 0.523022 | 0.003302 | 0.568317 | -0.402555 |
| 32.61111 | 2.044168 | 0.275477 | 1.782364 | 3.926143 | 1.724159 | 0.331122 | 3.046514 | -0.489257 | 1.694018 | 0.232347 | 5.192636 | 2.278628 | 3.103643 | -0.070231 | 1.757577 | -0.10519 | 4.356831 | 0.211871 | 0.063425 | -0.039924 | 0.73103 | -0.013092 | 1.405361 | -0.631562 |
| 32.625 | 2.313067 | 0.806809 | -4.367041 | 2.267955 | 0.591549 | 0.574473 | 3.328432 | -0.426593 | 3.702403 | 0.387511 | 4.563832 | 4.526749 | 2.210387 | -0.029201 | 1.341005 | 0.126725 | 6.670737 | 0.249238 | 0.132798 | -0.016554 | 0.6663 | -0.102593 | 1.344387 | -0.132823 |
| 32.63889 | 3.136722 | 0.732211 | -0.858585 | 1.436052 | 0.068133 | 1.134617 | 3.508752 | 0.008623 | 3.647297 | 0.69283 | 3.588828 | 4.166534 | 1.397807 | 0.121838 | -0.103827 | 0.314186 | 5.249359 | 0.190782 | 0.13418 | -0.035535 | 0.524986 | -0.151672 | -0.53492 | 0.256432 |
| 32.65278 | 4.916245 | 0.322599 | 1.703285 | 2.53816 | 0.104726 | 0.705026 | 2.244741 | 0.215177 | 1.874953 | 0.632533 | -1.152812 | 0.103595 | 1.474791 | 0.303746 | -1.341006 | 0.382686 | 1.68685 | 0.84378 | 0.128853 | -0.00037 | 0.77383 | -0.031243 | -1.833225 | 0.509107 |
| 32.66667 | 4.710146 | -0.185042 | 0.908702 | 1.489558 | 0.256847 | -0.013174 | 0.601783 | 0.361499 | 0.323614 | 0.33036 | -1.412491 | -2.050962 | 1.10941 | 0.549338 | -0.974884 | 0.37649 | 0.61124 | 1.000001 | 0.195312 | 0.029006 | 0.932004 | 0.027834 | -1.002979 | 0.267525 |
| 32.68056 | 1.611782 | -0.520556 | 0.727412 | -0.240769 | 0.283103 | 0.046837 | -0.705588 | 0.363705 | -0.871655 | -0.056898 | 1.722571 | -0.113222 | -0.634167 | 0.756014 | 0.540466 | 0.237065 | -0.307157 | 0.457595 | 0.259093 | -0.004431 | 0.612339 | -0.010407 | -0.388097 | 0.183822 |
| 32.69444 | -1.761643 | -0.35239 | -0.616982 | -1.455151 | 0.192284 | -1.322967 | -1.095205 | 0.315778 | -0.950046 | -0.397043 | 2.711707 | 0.143416 | -1.411508 | 0.513052 | 2.145142 | 0.127438 | -1.506491 | 0.324546 | 0.178437 | 0.005536 | -0.201214 | -0.033121 | -1.099728 | 0.071327 |
| 32.70833 | -2.285136 | -0.175564 | -0.482784 | 0.434221 | -0.035128 | -1.15709 | -0.111162 | 0.182412 | -0.298697 | -0.440591 | 0.656668 | -1.852542 | -1.604869 | -0.066193 | 1.704203 | 0.063807 | -1.398758 | 0.224386 | 0.096153 | 0.061493 | -0.72484 | 0.033579 | 0.169386 | 0.133101 |
| 32.72222 | -1.296497 | 0.163809 | -1.088225 | 2.010629 | -0.21129 | -0.114112 | 1.058636 | 0.015188 | -0.143587 | -0.17872 | 0.87241 | -2.170634 | 0.112068 | -0.456395 | 0.742925 | -0.009198 | 0.403122 | 0.061893 | 0.012987 | 0.112897 | -0.654108 | 0.106213 | 0.663691 | -0.242042 |
| 32.73611 | -0.297181 | 0.538339 | 3.059729 | 1.707787 | -0.212941 | 0.605342 | 1.615583 | -0.047343 | -0.204047 | 0.16691 | 0.097092 | -0.822854 | 0.063031 | -0.85969 | 0.103408 | -0.146136 | 1.962643 | -0.011874 | -0.059826 | 0.093493 | -1.074491 | 0.139562 | -0.205196 | -0.059553 |
| 32.75 | -0.277287 | 0.667821 | 1.311709 | -0.396777 | -0.078244 | 0.43149 | 0.177441 | 0.130677 | 0.267762 | 0.254593 | -1.828698 | 2.220802 | 0.356462 | -0.97937 | -0.34496 | -0.291796 | 1.500243 | -0.160143 | -0.038959 | -0.004908 | -0.581165 | 0.129411 | -0.110611 | 0.25904 |
| 32.76389 | -0.70881 | 0.099017 | 3.179609 | -6.082192 | -0.02569 | -0.795226 | -1.88935 | 0.320624 | -0.067421 | 0.342873 | -0.938968 | 3.470963 | 0.118229 | -0.69057 | -0.667471 | -0.276225 | 0.11083 | -0.420006 | -0.221751 | -0.067455 | 0.370289 | 0.105614 | 0.008356 | 0.17276 |
| 32.77778 | -1.289953 | -0.199206 | 0.054614 | -5.414233 | 0.078751 | -2.360701 | -3.448472 | 0.437158 | 0.076219 | 0.319147 | 1.028281 | 2.277465 | -1.520989 | -0.159842 | -0.074426 | -0.107182 | -0.32907 | -0.455436 | -0.171325 | -0.098659 | 0.848735 | 0.060542 | -0.47126 | 0.110247 |
| 32.79167 | -1.615369 | -0.238826 | 2.790238 | -2.724643 | 0.263644 | -1.893582 | -1.298779 | 0.269062 | -0.466751 | 0.147021 | 2.588488 | 0.998582 | -3.224329 | 0.200412 | 0.152126 | 0.140399 | -0.417806 | -0.018688 | -0.061755 | -0.136999 | 0.558753 | 0.010677 | -2.358613 | -0.18312 |
| 32.80556 | -0.705893 | -0.064552 | 2.705463 | 2.318952 | 0.288155 | 1.400095 | 1.604598 | 0.140147 | -0.215762 | 0.036501 | 2.300753 | 0.450568 | -2.331624 | 0.499461 | 0.130678 | 0.284075 | -0.722019 | 0.10705 | 0.050794 | -0.06196 | 0.116996 | 0.003768 | -5.617705 | -0.895875 |
| 32.81944 | 0.349188 | 0.11319 | 3.052786 | 4.545868 | 0.11989 | 1.149556 | 2.873379 | -0.006543 | 0.39106 | -0.023808 | -2.74737 | -0.122796 | -1.312518 | 0.737746 | -0.14773 | 0.30676 | -1.992751 | 0.190065 | 0.100757 | 0.067589 | -0.298502 | 0.023344 | -2.037081 | -0.844999 |
| 32.83333 | 1.142172 | 0.332495 | 3.312845 | 3.682005 | -0.033597 | 2.52142 | 1.769531 | -0.085794 | 0.173507 | -0.035931 | -4.84216 | -1.178314 | -1.654086 | 0.553495 | -0.221759 | 0.202915 | -2.917294 | 0.153304 | 0.077799 | 0.140844 | -1.071273 | 0.050404 | 0.06461 | -0.224345 |
| 32.84722 | 0.649961 | 0.217982 | 2.677362 | 0.879781 | -0.024581 | 3.508532 | 0.864625 | 0.091788 | 0.12564 | 0.146089 | -2.685769 | -1.566537 | 0.096077 | -0.014298 | -0.205821 | 0.102727 | -1.378497 | 0.241109 | 0.046687 | 0.137557 | -0.591771 | 0.029661 | 0.056285 | 0.622292 |
| 32.86111 | -0.609119 | 0.086089 | 1.552464 | -2.285268 | 0.052913 | 0.650515 | -0.422334 | 0.174808 | -0.633989 | 0.270287 | 0.166816 | 0.781297 | 1.757593 | -0.403333 | 0.141649 | -0.017332 | 0.619578 | 0.356031 | 0.034888 | 0.08822 | -0.343337 | -0.000091 | -1.133562 | 0.471724 |
| 32.875 | -1.334616 | -0.161067 | 1.481608 | -4.840815 | 0.159962 | -0.046543 | -1.255607 | 0.082614 | -0.687372 | 0.281201 | 1.948849 | 1.29126 | 3.23601 | -0.779857 | 0.695232 | -0.183179 | 1.373901 | 0.417558 | 0.010182 | -0.028781 | -0.17358 | 0.013199 | -2.012223 | -0.09169 |
| 32.88889 | -0.532141 | -1.059286 | -5.363998 | -3.73193 | 0.136931 | -0.475853 | -2.241027 | -0.037102 | 0.256858 | 0.101108 | 1.277182 | -1.752846 | 1.496752 | -0.869212 | 0.524636 | -0.2263 | 0.931122 | 0.261167 | 0.027736 | -0.031997 | 0.392251 | 0.102767 | -3.062241 | -0.119326 |
| 32.90278 | -0.368114 | -0.554619 | 4.799106 | -1.827165 | -0.030343 | -0.520828 | -1.501635 | -0.096313 | 0.966193 | 0.042085 | 1.402075 | -0.658886 | -0.050835 | -0.702684 | 0.387728 | -0.195272 | -0.183571 | -0.235973 | 0.064648 | 0.002948 | -0.050784 | 0.115387 | -2.47815 | -0.099882 |
| 32.91667 | -0.357037 | 0.3794 | 6.053349 | 0.48599 | -0.227982 | 0.887389 | 1.32574 | 0.052389 | 0.016015 | 0.020906 | 4.640154 | 0.835755 | -0.492392 | -0.677604 | 0.09323 | -0.150982 | -0.541261 | -0.599475 | 0.024283 | 0.0494 | -0.995902 | 0.016829 | 0.392696 | -0.063955 |
| 32.93056 | 0.888982 | 1.345948 | 1.539946 | 2.70565 | -0.465038 | 1.778485 | 1.505964 | 0.11119 | -1.857588 | -0.195807 | 0.005993 | -0.700451 | -0.81293 | -0.537682 | -1.127603 | -0.126097 | -1.385174 | -0.610274 | -0.063549 | 0.063002 | -1.002906 | -0.051296 | 1.10392 | -0.375302 |
| 32.94444 | -0.446553 | 0.638715 | -1.4109 | 0.580959 | -1.442383 | -0.181306 | -0.27057 | 0.071573 | -2.290275 | -0.305204 | -2.770315 | -0.365519 | -1.345947 | -0.465906 | -1.880395 | -0.127226 | -3.709713 | -0.656076 | -0.121162 | 0.048622 | -0.161629 | -0.033617 | 0.536166 | -0.163111 |
| 32.95833 | -2.681344 | 0.087069 | -4.608891 | -1.232808 | -1.024383 | -2.107557 | -3.051238 | 0.001261 | -3.747823 | 0.096885 | -3.47417 | -0.130182 | -2.85847 | -0.358874 | -1.995589 | -0.135814 | -5.787379 | -0.953442 | -0.155525 | -0.01212 | 0.495051 | -0.036042 | 0.018848 | -0.005654 |
| 32.97222 | -2.831742 | -0.360711 | -2.656212 | -2.549719 | 0.579145 | -1.568131 | -4.317457 | 0.036294 | -1.885571 | 0.578534 | -4.663258 | -0.820778 | -3.332611 | 0.089151 | -1.14546 | -0.088897 | -5.080158 | -1.022205 | -0.179285 | 0.019192 | -0.765699 | -0.103741 | -1.683188 | -0.631003 |
| 32.98611 | -2.701939 | -0.43648 | -3.214143 | -5.541018 | 1.053342 | -0.632262 | -4.674674 | -0.124733 | -0.698234 | -0.214069 | -6.778782 | -3.529923 | -3.333024 | 0.146975 | -0.850186 | -0.091107 | -6.038737 | -1.402611 | -0.143288 | -0.012449 | -2.679516 | -0.200244 | -1.600015 | -1.546904 |
| 33 | -4.648982 | -0.734477 | -3.678552 | -6.663883 | -2.038421 | -0.66929 | -2.825169 | -0.164796 | -1.542356 | -1.712305 | -5.547009 | -5.068871 | -4.415358 | 0.54661 | -1.563702 | -0.23397 | -7.548258 | -1.901212 | -0.212282 | -0.127184 | -3.940559 | -0.305642 | -1.086305 | -2.525931 |
| 33.01389 | -4.076609 | -2.744771 | -4.215662 | -6.360433 | -3.319644 | -0.749171 | -0.25766 | -0.229956 | -3.225936 | -2.591574 | -2.851719 | -4.290393 | -3.750197 | 0.450459 | -2.692665 | -0.441536 | -5.618163 | -1.952953 | -0.230372 | 0.098609 | -3.530393 | -0.398177 | -1.284432 | -3.099958 |
| 33.02778 | -3.31837 | -2.538399 | -4.358902 | -5.909851 | -5.092112 | -1.199221 | -0.108981 | -0.265034 | -5.180331 | -2.844459 | -1.177413 | -4.2093 | -3.371881 | 0.014068 | -3.497101 | -0.666828 | -3.424266 | -1.824 | -0.200543 | 0.057652 | -2.360977 | -0.410828 | -1.434516 | -2.899402 |
| 33.04167 | -5.245131 | -1.998819 | -3.637218 | -5.310366 | -7.665409 | -2.385223 | -1.995001 | -0.04611 | -3.497382 | -2.147021 | -2.589471 | -4.035213 | -4.101985 | -0.561727 | -2.905375 | -0.697967 | -3.867367 | -1.526567 | -0.18556 | 0.110418 | -1.807257 | -0.644256 | -2.168989 | -2.967645 |
| 33.05556 | -6.738184 | -0.659019 | -2.409325 | -5.653601 | -6.368758 | -1.764649 | -3.125321 | 0.076073 | -0.327653 | -1.590444 | -4.011474 | -3.346911 | -4.488197 | -0.67744 | -2.127919 | -0.511557 | -5.81443 | -1.537176 | -0.200459 | 0.234643 | -1.690629 | -0.614103 | -2.631654 | -2.265147 |
| 33.06944 | -6.363632 | 0.660994 | -1.323837 | -2.829653 | -5.185352 | -0.924845 | -3.70324 | -0.137661 | -0.126426 | -0.916755 | -4.420346 | -3.014286 | -2.616873 | -0.396237 | -1.015839 | -0.294776 | -3.460981 | -1.202071 | -0.208414 | 0.192114 | -1.419116 | -0.441574 | -1.921154 | -1.505076 |
| 33.08333 | -1.98496 | 0.470474 | -0.514407 | -3.767185 | -2.071456 | -0.651143 | -4.372741 | -0.49464 | -1.503021 | -0.933937 | -3.806427 | -2.671198 | -0.484148 | -0.222484 | -0.025102 | -0.105124 | -1.279349 | -0.69497 | -0.210137 | 0.11291 | -1.30788 | -0.287789 | -0.962648 | -0.884997 |
| 33.09722 | 1.231869 | -0.604345 | -0.553684 | -4.030852 | -1.725232 | -1.310498 | -4.050223 | -0.696067 | -2.441444 | -1.185639 | -2.341157 | -2.127095 | 0.406464 | -0.113457 | 0.15626 | 0.02489 | -0.539382 | -0.245147 | -0.097811 | 0.044692 | -1.003954 | -0.141536 | -0.364849 | -0.68518 |
| 33.11111 | 0.597404 | -1.285421 | -1.059688 | -2.619853 | -2.613315 | -1.90676 | -2.846668 | -0.875235 | -2.629832 | -1.874641 | -1.178119 | -1.802198 | 0.133004 | -0.133259 | -0.257749 | -0.028162 | -0.360812 | -0.22969 | -0.039682 | -0.000586 | -0.744915 | 0.024051 | 0.122166 | -0.611076 |
| 33.125 | -1.914879 | -1.364997 | -1.982251 | -2.64426 | -3.221035 | -2.085925 | -2.082906 | -1.515974 | -2.383171 | -2.309652 | -1.456109 | -1.831012 | -0.339226 | -0.09768 | -0.426515 | -0.131846 | -0.500332 | -0.375351 | -0.024453 | -0.022486 | -0.757904 | 0.02811 | -0.019065 | -0.369525 |
| 33.13889 | -3.83846 | -1.369751 | -2.556749 | -4.781088 | -3.761689 | -1.808314 | -2.817467 | -2.700117 | -2.261793 | -2.006708 | -1.702803 | -2.2144 | -0.635036 | -0.043633 | -0.312711 | -0.143847 | -0.920027 | -0.50942 | -0.020179 | -0.034844 | -0.753287 | -0.042448 | -0.323966 | -0.28167 |
| 33.15278 | -3.578559 | -1.601233 | -2.330824 | -6.528672 | -3.259952 | -1.02199 | -2.994847 | -2.498078 | -2.788331 | -1.868212 | -1.596177 | -2.44102 | -0.72634 | -0.041289 | -0.16912 | -0.149861 | -1.447376 | -0.664702 | -0.000239 | -0.050972 | -0.659441 | -0.116982 | -0.585614 | -0.388831 |
| 33.16667 | -2.57103 | -1.880145 | -1.974728 | -6.786843 | -2.794284 | -0.616122 | -2.58717 | -1.827743 | -2.580303 | -2.299684 | -1.634638 | -2.002915 | -0.789171 | -0.063425 | -0.197917 | -0.174988 | -1.984147 | -0.844423 | 0.011602 | -0.092633 | -0.492391 | -0.184797 | -0.754542 | -0.447632 |
| 33.18056 | -1.844705 | -2.111623 | -1.95429 | -6.67434 | -2.958473 | -1.280361 | -1.536122 | -1.247572 | -2.271964 | -2.433685 | -1.619687 | -1.714176 | -0.884328 | -0.089014 | -0.395301 | -0.175274 | -2.550303 | -1.005137 | -0.020062 | -0.177744 | -0.390672 | -0.26153 | -0.792759 | -0.455321 |
| 33.19444 | -1.468892 | -2.140982 | -1.685605 | -5.380626 | -2.89178 | -1.764264 | -1.394076 | -0.757379 | -1.84966 | -2.390988 | -1.53698 | -2.098984 | -1.00319 | -0.125225 | -0.678124 | -0.166709 | -3.135581 | -1.077779 | -0.089359 | -0.211821 | -0.375413 | -0.320724 | -0.936779 | -0.546534 |
| 33.20833 | -1.428648 | -1.404776 | -1.21359 | -4.795458 | -2.329522 | -0.947663 | -1.55175 | -0.26109 | -0.497227 | -2.148224 | -1.201873 | -2.193707 | -1.276881 | -0.197034 | -0.85434 | -0.22311 | -3.936353 | -1.229871 | -0.165681 | -0.274829 | -0.489244 | -0.331975 | -1.217011 | -0.875066 |
| 33.22222 | -1.102907 | -0.508263 | -0.725594 | -4.144174 | -1.747086 | -0.448595 | -0.862252 | 0.018388 | 0.174708 | -1.317647 | -0.4824 | -1.033834 | -1.727907 | -0.393736 | -1.003624 | -0.38169 | -4.74134 | -1.631434 | -0.241452 | -0.384554 | -0.651264 | -0.31916 | -1.44107 | -1.397709 |
| 33.23611 | -0.412004 | -0.078971 | -0.362361 | -2.183025 | -1.375364 | -0.060312 | -0.039248 | -0.166075 | 0.126942 | -0.716991 | 0.058857 | -0.395431 | -2.059979 | -0.635733 | -0.95009 | -0.497027 | -5.412789 | -2.158079 | -0.306925 | -0.533534 | -0.631123 | -0.38005 | -1.536497 | -1.764471 |
| 33.25 | 0.331132 | -0.24392 | -0.007812 | -0.471105 | -0.797815 | 0.014945 | 0.084879 | -0.234159 | 0.014595 | -0.201426 | 0.261511 | -0.482754 | -1.83522 | -0.638646 | -0.544052 | -0.437868 | -5.668261 | -2.049054 | -0.313425 | -0.604739 | -0.374679 | -0.353688 | -1.746412 | -1.570102 |
| 33.26389 | 0.352352 | -0.410997 | 0.294814 | -0.06926 | -0.229359 | 0.30316 | -0.380584 | 0.130237 | 0.227329 | -0.071831 | 0.134146 | -0.740881 | -1.321665 | -0.501033 | -0.076235 | -0.308728 | -6.113124 | -1.10321 | -0.278437 | -0.647445 | -0.094282 | -0.202147 | -1.753682 | -1.178422 |
| 33.27778 | -0.192811 | -0.088019 | 0.563851 | -0.191564 | -0.456354 | 0.470247 | -0.268201 | 0.275816 | 0.24211 | -0.492841 | -0.115114 | -0.472145 | -0.940286 | -0.501972 | -0.182277 | -0.309696 | -5.861309 | -0.581942 | -0.275873 | -0.720012 | -0.004634 | -0.085328 | -1.44913 | -1.063378 |
| 33.29167 | -0.525129 | 0.297094 | 0.702042 | 0.019174 | -1.390855 | 0.236696 | -0.038816 | 0.004557 | -0.117406 | -0.708485 | -0.183428 | -0.429951 | -0.70419 | -0.582443 | -0.513854 | -0.313053 | -4.996191 | -0.460394 | -0.266311 | -0.739366 | 0.011463 | -0.061664 | -1.128812 | -1.166748 |
| 33.30556 | -0.43666 | 0.327143 | 0.814992 | 0.661233 | -1.633898 | -0.231566 | 0.380788 | -0.286188 | -0.278593 | 0.118319 | -0.025444 | -0.177976 | -0.44214 | -0.523441 | -0.582462 | -0.226695 | -3.968675 | -0.294048 | -0.20355 | -0.605079 | 0.038646 | 0.081435 | -0.740053 | -1.018893 |
| 33.31944 | -0.19013 | 0.332452 | 0.813199 | 1.367855 | -0.632124 | -0.234422 | -0.403539 | -0.297632 | 0.096133 | 0.906824 | -0.204502 | 0.207455 | -0.136937 | -0.298344 | -0.375646 | -0.113824 | -3.139209 | 0.111259 | -0.160699 | -0.475326 | 0.018878 | 0.141054 | -0.460639 | -0.535174 |
| 33.33333 | -0.142524 | 0.399643 | 0.496251 | 1.436932 | 0.152468 | 0.35669 | -0.205132 | -0.298026 | 0.248878 | 0.773449 | -0.504691 | 0.340466 | -0.012926 | -0.122397 | -0.236511 | -0.152495 | -2.291573 | 0.403468 | -0.203337 | -0.467352 | -0.167215 | 0.029366 | -0.346165 | -0.07717 |
| 33.34722 | -0.250238 | 0.425758 | -0.113945 | 0.876587 | 0.036258 | 0.264554 | 0.0451 | -0.372173 | -0.129707 | 0.457941 | -0.228754 | 0.09389 | -0.009647 | -0.05261 | -0.280489 | -0.255053 | -1.762679 | 0.343388 | -0.225236 | -0.481857 | -0.381654 | -0.084509 | -0.207542 | 0.133213 |
| 33.36111 | -0.057713 | 0.740363 | -0.249225 | 0.954201 | 0.327475 | -0.0719 | -0.097254 | -0.183127 | 0.008893 | 0.234039 | 0.408771 | 0.378135 | 0.059595 | 0.037219 | -0.316736 | -0.261948 | -0.93858 | 0.373032 | -0.162613 | -0.333069 | -0.430866 | -0.012985 | 0.089025 | 0.211119 |
| 33.375 | 0.522695 | 1.308343 | 0.356513 | 2.223719 | 1.739555 | 0.103894 | -0.181552 | 0.235078 | 0.687546 | 0.379126 | 1.009797 | 1.219858 | 0.270962 | 0.059503 | -0.071278 | -0.14891 | 0.356294 | 0.597095 | -0.087778 | -0.124885 | -0.142914 | 0.074157 | 0.481204 | 0.323999 |
| 33.38889 | 1.123244 | 1.219302 | 1.163874 | 3.770168 | 3.023768 | 0.959086 | 0.128961 | 0.538499 | 1.158708 | 1.064281 | 1.487546 | 1.056342 | 0.396929 | 0.052865 | 0.414642 | -0.05905 | 1.591985 | 0.538921 | -0.087179 | 0.017422 | 0.198498 | 0.083932 | 0.692455 | 0.478094 |
| 33.40278 | 1.554477 | 0.931957 | 1.653707 | 4.385827 | 3.600179 | 1.383319 | 0.714377 | 0.73121 | 1.633081 | 1.905347 | 1.828091 | 0.867209 | 0.469517 | 0.076783 | 0.837006 | 0.010054 | 2.785976 | 0.37186 | -0.081707 | 0.141147 | 0.429653 | 0.099592 | 0.802233 | 0.690685 |
| 33.41667 | 1.931775 | 1.208977 | 1.80647 | 5.448194 | 4.218263 | 1.617914 | 1.014262 | 0.888581 | 2.676645 | 3.341362 | 2.008236 | 1.503302 | 0.67673 | 0.188788 | 1.078193 | 0.093729 | 4.395346 | 0.560573 | -0.00048 | 0.369037 | 0.678107 | 0.160263 | 1.003098 | 1.047716 |
| 33.43056 | 2.238261 | 1.734423 | 2.011319 | 6.408446 | 4.579977 | 2.166456 | 1.475276 | 1.062693 | 4.043966 | 4.36147 | 2.391724 | 2.044966 | 0.961466 | 0.271937 | 1.415689 | 0.123896 | 6.17963 | 1.023865 | 0.073045 | 0.634592 | 0.8979 | 0.210322 | 1.324718 | 1.454365 |
| 33.44444 | 2.736623 | 2.194144 | 2.5993 | 7.325935 | 4.299567 | 2.167752 | 1.872401 | 1.090147 | 4.727249 | 4.647803 | 3.021455 | 2.382656 | 1.368648 | 0.355974 | 2.142739 | 0.196956 | 7.499606 | 1.576815 | 0.115449 | 0.688043 | 1.097006 | 0.22679 | 1.742028 | 1.80855 |
| 33.45833 | 3.752661 | 2.657243 | 3.459783 | 9.402517 | 5.271982 | 1.983049 | 2.386858 | 1.595852 | 4.779675 | 4.839663 | 3.602584 | 3.0332 | 1.946547 | 0.55951 | 3.144122 | 0.447119 | 8.384365 | 2.1848 | 0.189161 | 0.777084 | 1.654843 | 0.283059 | 2.292577 | 2.257146 |
| 33.47222 | 4.922236 | 2.55507 | 4.76083 | 11.766929 | 6.589439 | 2.496341 | 3.241978 | 2.288446 | 5.367482 | 5.466471 | 4.186958 | 3.661236 | 2.237662 | 0.644285 | 4.154918 | 0.725567 | 9.69548 | 2.980608 | 0.175621 | 0.978051 | 2.004528 | 0.409899 | 2.950535 | 2.731827 |
| 33.48611 | 6.185398 | 2.513212 | 6.810797 | 12.869355 | 8.111052 | 3.831973 | 4.802667 | 2.060717 | 5.478525 | 5.580939 | 5.353035 | 4.026828 | 2.41583 | 0.657853 | 4.105937 | 0.843984 | 9.861947 | 2.679867 | 0.150663 | 1.459995 | 1.792919 | 0.561525 | 3.164849 | 3.039295 |
| 33.5 | 7.442852 | 3.312882 | 9.116152 | 12.345293 | 7.205383 | 4.332201 | 5.778678 | 1.13374 | 6.275848 | 5.591389 | 6.683101 | 4.861922 | 2.773559 | 0.726154 | 3.811669 | 0.736594 | 9.068247 | 2.483211 | 0.199993 | 1.807467 | 2.84519 | 0.675062 | 2.752225 | 2.845343 |
| 33.51389 | 7.339545 | 3.311498 | 10.271342 | 12.432139 | 6.412989 | 3.815243 | 6.437211 | 0.361889 | 5.192267 | 3.377129 | 7.247674 | 6.643436 | 2.899018 | 0.697224 | 3.810384 | 0.448635 | 9.361711 | 2.297993 | 0.288835 | 1.48483 | 2.880892 | 0.630031 | 2.116078 | 2.377745 |
| 33.52778 | 7.425263 | 2.945523 | 8.684891 | 11.065594 | 3.514723 | 3.104771 | 6.231784 | 0.059393 | 2.886693 | 1.452272 | 7.406308 | 7.745569 | 2.505828 | 0.442099 | 3.825395 | 0.393488 | 9.12908 | 1.439435 | 0.186494 | 1.041363 | 3.244728 | 0.413636 | 1.731233 | 1.339753 |
| 33.54167 | 6.051915 | 1.412461 | 7.394569 | 8.903562 | 2.160834 | 1.697567 | 6.375211 | -0.202081 | 1.301255 | 0.739296 | 6.155089 | 6.126152 | 2.534845 | 0.327366 | 3.655863 | 0.557217 | 6.972818 | 0.697578 | 0.072595 | 0.94183 | 2.916554 | 0.112945 | 2.494941 | 1.540964 |
| 33.55556 | 2.258995 | 0.703635 | 4.017391 | 4.015478 | 0.836386 | 1.007984 | 4.44828 | -0.383559 | 0.334129 | 0.857106 | 1.36514 | 3.524342 | 2.901458 | 0.544175 | 2.828103 | 0.655075 | 6.830358 | 0.577446 | 0.097722 | 0.588056 | 2.389847 | 0.092148 | 2.130709 | 0.706535 |
| 33.56944 | -0.906702 | 0.112246 | 2.860942 | -3.886543 | -2.569868 | 1.113494 | 3.87983 | -0.356982 | 2.228716 | 0.456302 | -1.502421 | 2.694467 | 1.594697 | 0.597641 | 0.881472 | 0.472248 | 1.822423 | 0.53316 | 0.189147 | 0.041751 | 1.832981 | 0.142796 | 1.489447 | 0.429999 |
| 33.58333 | -0.336697 | -0.111267 | 4.016083 | -3.040813 | -4.151087 | 1.965848 | 4.041691 | -0.2901 | 3.343776 | 0.431951 | 0.501014 | 3.158776 | 1.773494 | 0.461755 | 0.139883 | 0.351195 | -0.389357 | 0.993661 | 0.157444 | 0.025245 | 1.046747 | 0.162062 | 1.442206 | -0.201867 |
| 33.59722 | 1.63342 | 0.154759 | 5.081098 | 0.201465 | -0.106488 | 3.869802 | 2.542744 | -0.139234 | 2.589939 | 0.541346 | 3.109944 | 2.106668 | 3.124513 | 0.273914 | 0.942543 | 0.244514 | 0.848249 | 1.075654 | 0.101613 | 0.054999 | 1.217473 | 0.121394 | 2.076876 | -0.692267 |
| 33.61111 | 4.166528 | 0.488404 | 6.972746 | 3.181608 | 4.917127 | 3.031792 | 2.215034 | 0.180733 | 0.778907 | 0.58003 | 5.480409 | 2.801865 | 4.62827 | 0.109739 | 1.79437 | 0.200644 | 1.928608 | 0.547378 | 0.061357 | 0.030932 | 1.591904 | 0.033234 | 0.97768 | -0.162498 |
| 33.625 | 2.984917 | 0.632735 | 4.000499 | 3.772173 | 2.583811 | 2.021075 | 2.650253 | 0.095047 | 0.638725 | 0.052654 | 7.891821 | 4.347765 | 3.292432 | 0.244953 | 1.615035 | 0.186674 | 2.555839 | 0.373438 | 0.007411 | 0.005953 | 1.044633 | -0.052283 | 1.464827 | 0.369979 |
| 33.63889 | 2.269133 | 0.595825 | 1.775301 | 3.462287 | 0.333476 | 0.767676 | 3.340636 | -0.07765 | 3.372271 | 0.352133 | 7.889718 | 3.542476 | 2.825085 | 0.450585 | 1.153066 | 0.226616 | 2.590376 | 0.628753 | 0.05576 | -0.06652 | 0.138149 | -0.063009 | 1.696778 | 0.392607 |
| 33.65278 | 3.31644 | 0.510981 | -0.285196 | 2.837655 | 0.149427 | -0.008839 | 4.176201 | -0.25846 | 3.015356 | 0.648101 | 3.353833 | 2.690196 | 1.413735 | 0.636924 | 0.333046 | 0.276078 | 2.560376 | 0.881679 | 0.041655 | -0.21553 | 0.43906 | -0.020119 | 1.830716 | 0.196859 |
| 33.66667 | 3.116949 | 0.377216 | -1.880343 | 1.946335 | 0.023229 | 1.025982 | 2.26687 | 0.178957 | 1.464846 | 0.605003 | -2.173543 | 0.555364 | 1.507323 | 0.637276 | -0.421547 | 0.109045 | 0.954692 | 0.766925 | 0.073367 | -0.064841 | 1.02652 | 0.016096 | 0.718408 | 0.049738 |
| 33.68056 | 2.298511 | 0.111455 | -1.85195 | -1.800327 | -0.00662 | 0.873528 | 1.307442 | 0.531153 | -0.024914 | 0.398968 | -1.594094 | -0.496522 | 0.185598 | 0.519035 | -0.376678 | -0.078009 | -0.281332 | 0.498057 | 0.197667 | 0.12087 | 1.104884 | 0.059911 | -1.37175 | 0.009649 |
| 33.69444 | 0.444667 | -0.266215 | -2.940273 | -6.272024 | 0.092501 | -1.322971 | 0.130266 | 0.274948 | -1.635011 | 0.064876 | -1.608568 | -1.349283 | -0.980742 | 0.269048 | 0.730225 | -0.114549 | -2.863426 | 0.269148 | 0.254436 | 0.131478 | 0.50609 | 0.070858 | -2.236318 | -0.280589 |
| 33.70833 | -0.424721 | -0.345648 | -3.837264 | -6.786058 | 0.167292 | -2.523919 | -1.634364 | -0.025949 | -1.663321 | -0.166065 | 0.360797 | -1.137726 | -0.000099 | -0.100771 | 1.064732 | -0.095009 | -4.447872 | -0.000695 | 0.139987 | 0.122649 | -0.190331 | 0.030691 | -3.61295 | -0.478985 |
| 33.72222 | -0.37921 | -0.006544 | 0.855288 | -0.787152 | 0.061849 | -1.713654 | -1.572705 | 0.106913 | -0.451477 | -0.233032 | 1.850625 | -0.427317 | -1.333093 | -0.516139 | 0.925798 | -0.073062 | -4.311268 | -0.464793 | 0.020022 | 0.103847 | -0.770104 | 0.009683 | -2.151828 | -0.861447 |
| 33.73611 | -0.533613 | 0.383106 | 3.500229 | 1.983621 | -0.061746 | 0.738292 | -0.02937 | 0.284982 | 0.578404 | -0.145855 | 1.02844 | 0.792836 | 0.002125 | -0.758594 | 0.281659 | 0.03538 | -1.67245 | -0.517119 | -0.059639 | 0.088776 | -0.839061 | 0.04256 | 0.891533 | 0.05197 |
| 33.75 | -0.752957 | 0.334822 | 3.218404 | 0.763109 | -0.053105 | 2.835245 | 1.80197 | 0.396911 | 0.533016 | 0.074117 | -3.417808 | 0.844922 | 1.279522 | -1.002832 | -0.679809 | 0.161593 | 1.287806 | -0.202121 | -0.193392 | 0.070175 | -0.539563 | 0.070267 | 3.494565 | 0.89621 |
| 33.76389 | 0.639897 | -0.054403 | 3.019175 | -2.718646 | -0.361616 | 1.058103 | 0.805891 | 0.347342 | -0.557609 | 0.122165 | -3.009024 | -0.448962 | 2.669292 | -0.684481 | -1.000178 | 0.219757 | 3.885928 | 0.073457 | -0.258506 | -0.008889 | 0.362512 | 0.041565 | 1.497072 | 0.448518 |
| 33.77778 | 0.390821 | -0.413167 | -3.073098 | -1.078696 | -1.370265 | -0.285783 | -1.587141 | 0.11632 | -1.006062 | -0.036829 | -3.781659 | -2.714519 | 3.154682 | -0.060796 | -0.365569 | 0.176327 | 4.223817 | -0.179358 | -0.186142 | -0.193979 | 1.059561 | 0.012854 | 0.341753 | 0.18834 |
| 33.79167 | -2.339825 | -0.53344 | -3.800599 | -0.79549 | -2.599531 | -0.117979 | -3.840302 | -0.032036 | -0.349569 | 0.054926 | -3.551311 | -2.201583 | 2.361636 | 0.447737 | 0.366188 | 0.119008 | 1.835609 | 0.001188 | -0.103788 | -0.092074 | 0.332417 | 0.042637 | -0.996358 | 0.176323 |
| 33.80556 | -1.036039 | -0.112828 | 0.37099 | -2.447583 | 0.855712 | 0.019104 | -2.579208 | 0.261243 | 1.056996 | 0.383193 | 0.385942 | -1.447584 | 2.623333 | 0.796981 | 0.613947 | 0.100594 | 0.55286 | 0.037401 | -0.031665 | 0.023018 | -0.004492 | 0.078116 | -2.505274 | -0.346335 |
| 33.81944 | 2.114211 | 0.233687 | 1.824809 | -4.599011 | 2.877647 | -0.794872 | -0.436903 | 0.250495 | 1.526813 | 0.377096 | 2.857647 | 0.144839 | 1.786327 | 0.784731 | 0.926835 | 0.02853 | 0.211226 | -0.132691 | 0.0231 | 0.011785 | -0.058767 | 0.094743 | -3.897876 | -0.72031 |
| 33.83333 | -0.452163 | 0.004134 | 0.384224 | -1.438677 | 0.318063 | 0.12465 | 2.48028 | 0.127987 | 0.337056 | 0.268345 | -0.394335 | -0.342012 | -0.82858 | 0.482369 | 0.464683 | -0.068633 | 0.249263 | -0.093386 | -0.002083 | 0.000677 | 0.177441 | 0.044929 | -1.165671 | -0.842306 |
| 33.84722 | -0.457092 | -0.12297 | -0.047225 | 2.61952 | -0.048765 | 0.390936 | 2.830046 | -0.105873 | -1.221413 | 0.09535 | -1.133029 | -1.046248 | -2.819936 | 0.057192 | -0.727814 | -0.154333 | -0.258383 | 0.058566 | -0.009654 | 0.000307 | -0.068921 | -0.021371 | 1.802707 | -0.741093 |
| 33.86111 | 0.198509 | 0.473221 | 0.653753 | 4.79903 | -0.432717 | 0.893192 | 2.543788 | -0.260119 | -1.003741 | 0.008535 | -2.540076 | -1.348701 | -3.768589 | -0.452531 | -1.409141 | -0.177948 | -1.63356 | 0.127081 | 0.017852 | 0.055571 | 0.048079 | -0.045815 | 2.801131 | -0.012488 |
| 33.875 | -1.085234 | 0.559786 | -0.425505 | 3.392457 | -0.534101 | 0.744821 | 1.319059 | -0.181187 | 1.118412 | 0.153874 | -1.641248 | -0.379474 | -3.493479 | -0.880069 | -0.237968 | -0.160123 | -4.48209 | 0.262817 | 0.032294 | 0.102915 | -0.426662 | -0.038864 | 0.352613 | 0.232534 |
| 33.88889 | -0.283437 | 0.005394 | -0.550588 | -0.615125 | -0.104588 | -0.841103 | -1.275263 | -0.070426 | 1.878213 | 0.410313 | 2.767329 | 1.535607 | -3.086347 | -0.894419 | 1.001777 | -0.153525 | -6.420253 | 0.354754 | 0.055705 | 0.043694 | -0.425407 | 0.013579 | 0.721849 | 0.396019 |
| 33.90278 | 2.601646 | -0.560619 | -2.945597 | -2.376396 | 0.315558 | -0.700188 | -3.388659 | -0.105566 | 0.456937 | 0.340568 | 4.067344 | 2.690622 | -4.009921 | -0.61516 | 0.451032 | -0.109343 | -5.752697 | -0.140892 | 0.062731 | -0.188629 | -0.405461 | 0.057259 | 0.221486 | -0.01574 |
| 33.91667 | 2.024209 | -0.572346 | -1.895008 | -3.551798 | 0.258549 | -0.422464 | -3.096617 | -0.139868 | -1.366958 | -0.07667 | 4.415519 | 3.090013 | -4.180725 | -0.465016 | -0.316114 | -0.027064 | -6.064765 | -0.273928 | 0.058332 | -0.169829 | -0.377973 | 0.040865 | -2.170224 | -0.44994 |
| 33.93056 | -1.869664 | -0.183963 | -3.057367 | -4.770179 | -0.494683 | -0.5905 | -1.968407 | -0.00912 | -3.024693 | -0.160255 | 1.714235 | 0.472896 | -3.217431 | -0.482384 | -1.923611 | 0.012019 | -7.763025 | -0.230255 | 0.028818 | -0.137004 | -0.647695 | -0.00743 | -1.769858 | -0.786198 |
| 33.94444 | -6.93521 | 0.063501 | -2.695289 | -7.608455 | -2.894 | -0.795304 | 0.257393 | -0.155573 | -1.673119 | 0.143432 | -2.186569 | -2.356594 | -2.456527 | -0.599997 | -2.576736 | -0.000015 | -7.529983 | -0.952916 | -0.032795 | 0.022974 | -0.796302 | -0.045344 | -0.73054 | -1.190288 |
| 33.95833 | -7.992774 | -0.567562 | -2.386816 | -2.834718 | -6.454348 | 0.635942 | 0.053641 | -0.622747 | -0.331505 | -0.066211 | -4.698013 | -3.33653 | -2.055554 | -0.603878 | -3.524301 | -0.034324 | -7.524702 | -1.180958 | -0.067275 | 0.175343 | -0.689189 | -0.078115 | -1.058123 | -0.982033 |
| 33.97222 | -4.655201 | -0.209625 | -0.601154 | -0.230085 | -6.321352 | -0.370136 | -3.461723 | -0.525404 | -0.448445 | -1.699426 | -2.465183 | -1.950682 | -2.068809 | -0.460516 | 0.358457 | -0.047757 | -9.242657 | -1.973636 | -0.02704 | 0.17716 | -0.648366 | -0.0746 | -0.537278 | 0.293234 |
| 33.98611 | -0.862926 | 0.645421 | -0.973914 | 2.82241 | -3.396504 | -1.031362 | -4.225663 | -0.330099 | -2.129705 | -1.677488 | -3.274129 | -0.50028 | -1.819234 | -0.31228 | -0.681616 | -0.115452 | -6.818925 | -1.78211 | -0.022625 | -0.073313 | -1.031274 | -0.084943 | -0.680362 | -0.073855 |
| 34 | -2.376698 | -0.088907 | -2.619244 | -0.660188 | -2.61022 | 0.178206 | -2.283085 | -0.166791 | -4.197346 | -3.208827 | -5.493144 | -2.748451 | -1.506899 | -0.11725 | -2.723544 | -0.231395 | -2.681816 | -1.082532 | -0.096297 | -0.839407 | -1.431911 | -0.18951 | -1.021266 | -1.241294 |
| 34.01389 | -5.42531 | -0.574158 | -7.234855 | -10.939932 | -5.126591 | -1.8642 | -4.896164 | -0.182673 | -4.285556 | -2.411239 | -3.789499 | -4.609375 | -1.565999 | 0.208156 | -3.077427 | -0.322257 | 1.20851 | -0.791043 | -0.22901 | -1.201097 | -2.934972 | -0.366817 | -1.688031 | -2.84892 |
| 34.02778 | -6.147991 | -2.369105 | -7.221142 | -10.640735 | -3.940947 | -1.418862 | -1.902219 | -0.312355 | -2.971519 | -2.35728 | -1.81607 | -5.327994 | -1.455377 | 0.146471 | -3.654662 | -0.383006 | 2.586549 | -1.348101 | -0.215838 | -0.528931 | -3.056559 | -0.584957 | -2.276366 | -4.190571 |
| 34.04167 | -5.90131 | -4.105726 | -5.589414 | -8.45475 | -7.85324 | -2.606155 | -2.566829 | -0.411007 | -2.971829 | -4.495137 | -1.940606 | -3.647109 | -1.67026 | -0.32507 | -3.101259 | -0.421028 | -0.709789 | -2.017296 | -0.266399 | -0.579104 | -1.912454 | -0.577839 | -1.964955 | -4.228446 |
| 34.05556 | -5.348485 | -2.071871 | -5.682909 | -6.581678 | -9.568914 | -3.199525 | -4.36036 | -1.225484 | -4.512121 | -6.317714 | -3.756366 | -2.559914 | -1.52195 | -0.939898 | -1.965198 | -0.375133 | -6.022764 | -2.293307 | -0.30913 | -0.195358 | -0.913471 | -0.784244 | -1.828334 | -4.186618 |
| 34.06944 | -5.113862 | 1.039134 | -5.316003 | -2.525838 | -7.648251 | -1.149247 | -3.803801 | -1.182973 | -5.651226 | -8.20979 | -4.710172 | -2.237367 | -0.871672 | -1.435033 | -0.698838 | -0.274166 | -4.795026 | -2.395167 | -0.326033 | -0.347424 | -0.256056 | -1.012771 | -1.552859 | -3.162962 |
| 34.08333 | -4.105573 | 1.886067 | -2.927544 | -1.623532 | -4.261073 | -0.493409 | -2.62317 | -1.305733 | -4.317191 | -7.129806 | -4.113421 | -2.446979 | -0.497023 | -1.080125 | -0.116227 | -0.279488 | -0.988282 | -1.752077 | -0.263133 | -0.323005 | -0.458345 | -1.065025 | -0.960176 | -2.168925 |
| 34.09722 | -2.948341 | 0.922427 | -1.744158 | -2.419934 | -3.127653 | -1.646862 | -0.89242 | -1.200556 | -2.328874 | -4.829664 | -2.392825 | -2.53636 | -0.425318 | -0.847653 | -0.448345 | -0.272181 | 0.594957 | -1.507671 | -0.223796 | -0.411244 | -0.655639 | -0.705468 | -0.671719 | -1.668568 |
| 34.11111 | -2.00856 | -0.007308 | -1.815805 | -1.39927 | -3.610371 | -1.425805 | -0.035114 | -1.199059 | -0.470179 | -1.503474 | -0.386638 | -0.728603 | -0.495852 | -0.678258 | -1.113804 | -0.253757 | 1.602386 | -1.080302 | -0.174566 | -0.403645 | -0.541299 | -0.284571 | -0.526429 | -1.292712 |
| 34.125 | -1.39821 | -2.077678 | -2.018589 | -1.027355 | -3.769366 | -1.517755 | -2.740644 | -0.68176 | 0.177737 | -0.314248 | -0.238231 | -1.649321 | -0.417709 | -0.462335 | -1.280045 | -0.198432 | 1.438526 | -0.578655 | -0.086708 | -0.222045 | -0.405345 | 0.001885 | -0.538156 | -0.933031 |
| 34.13889 | -1.785416 | -2.35958 | -2.613522 | -4.170809 | -5.179168 | -2.169476 | -2.887811 | -1.36398 | -0.999206 | -2.119919 | -0.733195 | -1.862346 | -0.369315 | -0.363375 | -1.023029 | -0.139449 | -0.163086 | -0.375673 | -0.067664 | -0.09253 | -0.383385 | -0.011475 | -0.648457 | -0.81648 |
| 34.15278 | -1.887227 | -2.185185 | -2.329199 | -8.11308 | -6.992094 | -1.497861 | -1.552022 | -2.247153 | -2.207639 | -4.345502 | -0.667728 | -1.930484 | -0.504916 | -0.377343 | -0.725802 | -0.133061 | -1.636256 | -0.46257 | -0.095043 | -0.153149 | -0.531606 | -0.213272 | -0.786202 | -0.866214 |
| 34.16667 | -1.670477 | -2.474301 | -2.141296 | -9.117946 | -8.116386 | -1.625297 | -0.935953 | -3.366827 | -2.361917 | -4.486383 | -1.137316 | -2.780784 | -0.621093 | -0.281618 | -0.475064 | -0.131867 | -2.161871 | -0.490726 | -0.083934 | -0.162087 | -0.626152 | -0.281051 | -0.848399 | -0.759884 |
| 34.18056 | -1.575362 | -2.543776 | -1.87924 | -8.4493 | -6.734571 | -2.042625 | -1.49249 | -3.156819 | -1.325546 | -2.438774 | -0.574914 | -1.629696 | -0.70396 | -0.098741 | -0.228957 | -0.113142 | -2.231619 | -0.453969 | -0.076527 | -0.174239 | -0.729671 | -0.255496 | -0.961762 | -0.739415 |
| 34.19444 | -1.720715 | -0.964299 | -1.489554 | -6.2418 | -4.73448 | -0.928031 | -0.656401 | -1.969837 | -0.3045 | -0.295791 | 0.184682 | -0.431553 | -0.78631 | -0.057278 | -0.265647 | -0.109261 | -2.325379 | -0.555781 | -0.120679 | -0.173809 | -0.930116 | -0.292764 | -1.061632 | -0.928033 |
| 34.20833 | -1.380255 | -0.822934 | -0.767834 | -4.262742 | -3.392951 | 0.422826 | 1.424728 | -0.970135 | -0.247705 | 0.359331 | 0.041991 | -0.147864 | -0.849182 | -0.178454 | -0.691739 | -0.188408 | -2.311784 | -0.832921 | -0.1752 | -0.143575 | -1.147729 | -0.358472 | -1.104898 | -1.245357 |
| 34.22222 | -0.81824 | -0.110015 | -0.195628 | -2.517539 | -2.526533 | 0.632938 | 0.847548 | -0.678242 | -0.418505 | -0.180401 | -0.669843 | -0.536608 | -0.848494 | -0.402465 | -1.032736 | -0.292966 | -2.273981 | -1.010917 | -0.195259 | -0.141342 | -1.174233 | -0.370446 | -1.126369 | -1.616092 |
| 34.23611 | -0.596489 | -0.428952 | 0.233838 | -0.961991 | -1.041553 | 0.171456 | -0.506504 | -0.564041 | -0.061488 | -0.22548 | -0.356613 | -0.799823 | -0.820376 | -0.654698 | -0.887923 | -0.266424 | -2.31823 | -0.796694 | -0.191497 | -0.188604 | -0.809296 | -0.285498 | -1.070171 | -1.75722 |
| 34.25 | -0.587043 | -0.207187 | 0.893711 | 0.039586 | 0.696076 | 1.053059 | 1.117362 | -0.11623 | 0.645169 | 1.093229 | 0.0509 | -0.869167 | -0.799201 | -0.691316 | -0.043392 | -0.191854 | -2.628873 | -0.45409 | -0.197786 | -0.218523 | -0.177745 | -0.062851 | -0.931212 | -1.571662 |
| 34.26389 | -0.510257 | -0.627929 | 1.328886 | 0.655074 | 1.429982 | 2.326729 | 3.320251 | 0.378671 | 1.201625 | 2.409753 | -0.390408 | -0.988143 | -0.832978 | -0.532045 | 0.582328 | -0.159309 | -3.153358 | -0.409471 | -0.267835 | -0.210755 | 0.434213 | 0.151228 | -0.871576 | -1.325145 |
| 34.27778 | -0.345997 | 0.142196 | 1.143419 | 1.604191 | 1.193258 | 1.490452 | 1.729822 | 0.384596 | 1.027207 | 1.86187 | -0.755034 | -0.431158 | -0.743143 | -0.358202 | 0.450992 | -0.203676 | -3.369604 | -0.430011 | -0.304629 | -0.189461 | 0.747775 | 0.251208 | -0.872665 | -1.019725 |
| 34.29167 | -0.094482 | 0.287435 | 0.951763 | 2.383312 | 0.885663 | 0.045356 | -0.337516 | 0.032955 | 0.742037 | 0.883476 | -0.154537 | -0.124733 | -0.530741 | -0.294989 | 0.231414 | -0.17015 | -2.773833 | -0.28934 | -0.288279 | -0.16122 | 0.543887 | 0.342144 | -0.695857 | -0.752385 |
| 34.30556 | 0.055541 | -0.123949 | 1.072262 | 2.377214 | 0.624554 | -0.074651 | -0.099522 | -0.375712 | 0.280802 | 0.497105 | 0.409715 | 0.02364 | -0.344052 | -0.341376 | 0.25062 | -0.115539 | -2.280227 | -0.200459 | -0.27455 | -0.146582 | 0.650159 | 0.317203 | -0.235016 | -0.636498 |
| 34.31944 | 0.090189 | -0.543832 | 1.236347 | 2.236231 | 1.035639 | 0.979373 | 1.303666 | -0.364085 | 0.4127 | 1.485847 | 0.264652 | 0.187606 | -0.242606 | -0.30414 | 0.42157 | -0.052569 | -1.688994 | -0.178189 | -0.26343 | -0.130909 | 0.960144 | 0.305661 | 0.217914 | -0.491402 |
| 34.33333 | 0.278042 | 0.19445 | 1.360895 | 2.489957 | 2.308887 | 1.739829 | 1.06813 | -0.074517 | 0.963501 | 3.679031 | 0.039309 | -0.089669 | -0.026428 | -0.014846 | 0.565825 | 0.013744 | -0.47679 | 0.1832 | -0.20714 | -0.071258 | 1.352116 | 0.72657 | 0.318084 | -0.092615 |
| 34.34722 | 0.798066 | -0.334552 | 1.289598 | 2.50396 | 2.983972 | 1.934075 | 0.030308 | 0.111405 | 1.351938 | 5.156805 | -0.07266 | -0.853872 | 0.157256 | 0.357271 | 0.516741 | -0.030856 | 0.744906 | 0.70114 | -0.155647 | -0.024789 | 1.929563 | 1.188798 | 0.187094 | 0.494834 |
| 34.36111 | 1.224604 | -0.439706 | 1.322033 | 2.660326 | 3.366276 | 2.40607 | 1.22606 | 0.305084 | 1.513337 | 4.559923 | 0.412398 | -0.325855 | 0.130567 | 0.570775 | 0.371034 | -0.09817 | 1.757324 | 0.934604 | -0.109571 | 0.049719 | 1.750662 | 0.898378 | 0.133541 | 1.089859 |
| 34.375 | 1.417264 | -0.305416 | 1.654378 | 3.4929 | 4.659107 | 2.432154 | 2.819143 | 0.712573 | 1.848277 | 4.372352 | 1.174978 | 0.734638 | 0.062687 | 0.758531 | 0.328901 | -0.064387 | 3.105203 | 0.991239 | -0.055902 | 0.190798 | 1.752213 | 0.792985 | 0.386352 | 1.610092 |
| 34.38889 | 1.731948 | 0.885916 | 1.684424 | 4.583547 | 5.130031 | 2.973127 | 2.861622 | 0.994123 | 2.938508 | 4.890483 | 1.570971 | 1.834089 | 0.041488 | 0.758486 | 0.531715 | -0.035236 | 4.594415 | 1.313763 | -0.013794 | 0.263818 | 1.697063 | 0.800021 | 0.853563 | 1.863261 |
| 34.40278 | 2.341412 | 0.501605 | 1.679427 | 5.219432 | 5.539925 | 3.317706 | 1.647468 | 1.192014 | 4.772223 | 6.137316 | 1.961724 | 1.123367 | 0.106603 | 0.586949 | 0.822733 | -0.038701 | 5.400439 | 1.593244 | 0.061626 | 0.332279 | 1.871869 | 0.773086 | 1.097937 | 1.903433 |
| 34.41667 | 2.813575 | 0.011667 | 2.148975 | 5.649637 | 8.227091 | 3.759285 | 2.188897 | 1.702764 | 5.342188 | 6.56207 | 2.317569 | 0.849569 | 0.254539 | 0.522623 | 1.293977 | 0.015256 | 6.690959 | 1.769144 | 0.208626 | 0.401598 | 2.270263 | 0.673772 | 1.225911 | 2.263074 |
| 34.43056 | 2.887935 | 0.403221 | 3.182314 | 5.964872 | 9.984619 | 5.514275 | 3.69682 | 2.141092 | 4.922469 | 4.342597 | 2.728511 | 1.315701 | 0.431501 | 0.553113 | 1.847487 | 0.110674 | 7.972866 | 1.895776 | 0.318428 | 0.460189 | 2.814412 | 0.64998 | 1.500831 | 2.766563 |
| 34.44444 | 3.492002 | 1.260315 | 4.229669 | 6.961927 | 11.65891 | 6.38184 | 3.824213 | 2.503537 | 3.782742 | 3.928069 | 2.918281 | 2.146694 | 0.674274 | 0.702041 | 2.362038 | 0.225045 | 7.742467 | 2.173947 | 0.403759 | 0.504584 | 2.536804 | 0.569573 | 1.940921 | 3.004392 |
| 34.45833 | 4.596876 | 2.443138 | 4.988757 | 8.648617 | 13.876318 | 5.214001 | 1.896396 | 2.249293 | 4.432439 | 4.486635 | 3.532664 | 3.191122 | 1.118918 | 0.838404 | 2.904381 | 0.420881 | 8.386709 | 2.631428 | 0.452945 | 0.626546 | 2.182923 | 0.523861 | 2.592128 | 3.096896 |
| 34.47222 | 5.207768 | 2.566194 | 5.140842 | 11.60304 | 14.452399 | 3.640157 | 2.072088 | 2.656995 | 4.946499 | 5.377902 | 4.480878 | 3.790045 | 1.567538 | 0.62496 | 3.740756 | 0.58715 | 9.606558 | 3.066023 | 0.364126 | 0.533982 | 2.952006 | 0.449407 | 3.063063 | 3.579294 |
| 34.48611 | 6.339014 | 2.579567 | 5.586253 | 15.475538 | 13.309842 | 4.062406 | 3.646704 | 2.503854 | 5.388739 | 5.703968 | 4.94634 | 3.846114 | 2.01583 | 0.537021 | 4.441465 | 0.662195 | 10.131896 | 3.365091 | 0.24238 | 0.466318 | 3.041693 | 0.516048 | 2.615263 | 3.064789 |
| 34.5 | 7.069332 | 3.069783 | 7.323684 | 17.633719 | 11.831805 | 4.030106 | 4.897066 | 1.807524 | 4.341658 | 4.744884 | 5.236675 | 4.910426 | 2.342628 | 0.455064 | 4.515264 | 0.538418 | 9.288696 | 3.319043 | 0.292106 | 0.287017 | 2.957048 | 0.505397 | 2.065899 | 2.324158 |
| 34.51389 | 7.716849 | 3.385062 | 10.277889 | 16.668899 | 10.435665 | 2.883945 | 3.75533 | 1.249307 | 3.206322 | 3.878272 | 4.62623 | 5.343529 | 2.664195 | 0.488365 | 3.910553 | 0.420861 | 10.027079 | 2.882852 | 0.309061 | -0.086068 | 2.875937 | 0.356646 | 2.594248 | 1.47929 |
| 34.52778 | 7.752661 | 2.991809 | 10.528261 | 17.884446 | 6.576662 | 2.579889 | 2.365233 | 0.177129 | 3.219328 | 3.65736 | 4.002314 | 5.283767 | 2.533005 | 0.58855 | 3.427767 | 0.386331 | 9.929866 | 1.35883 | -0.033617 | -0.136106 | 2.900863 | 0.184307 | 4.04043 | 1.061251 |
| 34.54167 | 6.03391 | 1.888308 | 9.009687 | 12.293233 | 2.335165 | 2.63477 | 2.501054 | -0.319536 | 3.437466 | 3.467516 | 4.300381 | 5.311351 | 2.321124 | 0.458823 | 2.799278 | 0.329479 | 8.091431 | 1.385536 | -0.261762 | -0.11636 | 1.859778 | 0.007171 | 3.616477 | 0.629277 |
| 34.55556 | -0.312759 | 1.246821 | 4.649968 | 2.893961 | -1.946002 | 2.43 | 2.868035 | -0.400284 | 3.127969 | 0.806587 | 2.431687 | 3.790423 | 2.441044 | 0.433437 | 1.007735 | 0.273952 | 5.460738 | 0.345105 | -0.156501 | -0.129651 | 1.125669 | -0.133685 | 2.367068 | 0.163995 |
| 34.56944 | -4.481012 | 0.417462 | 0.517914 | -3.252786 | -6.674088 | 1.685106 | 2.402753 | -0.274313 | 2.090967 | -2.185464 | -0.393411 | 0.782976 | 2.093492 | 0.727876 | -0.590588 | 0.214824 | 0.172842 | -0.041876 | 0.216264 | 0.003643 | 0.227063 | 0.10259 | -0.609097 | -0.451634 |
| 34.58333 | -1.454172 | -0.274543 | -0.106987 | -2.064729 | -3.300681 | 1.791429 | 2.458624 | 0.016009 | 2.556226 | -1.43154 | -0.333165 | 0.047277 | 0.991193 | 0.553118 | -1.784665 | 0.183818 | -6.649048 | 0.119957 | 0.388008 | 0.111513 | 0.07382 | 0.193508 | -2.394738 | -0.932579 |
| 34.59722 | 4.312819 | 0.11491 | 2.521828 | 0.239128 | 2.04385 | 1.714069 | 3.13038 | 0.249566 | 2.306188 | 0.435931 | 1.180305 | 1.044133 | 0.937393 | 0.316156 | -0.408977 | 0.104636 | -4.081842 | 0.566262 | 0.181896 | 0.0855 | 0.756942 | 0.114057 | -1.447508 | -0.928053 |
| 34.61111 | 9.616622 | 0.435839 | 3.654405 | 1.863944 | 7.930979 | 2.282203 | 1.979181 | 0.086189 | 0.020991 | 1.492867 | 2.809006 | 3.014202 | 1.701948 | -0.023412 | 1.255378 | 0.075153 | -1.418155 | 0.685002 | -0.078095 | -0.031223 | 0.454605 | 0.003985 | -0.713998 | 0.192627 |
| 34.625 | 6.021401 | 0.648204 | 5.097362 | 4.68328 | 5.414451 | 1.968001 | 0.448411 | -0.069282 | -0.079406 | 0.387534 | 4.267022 | 3.550038 | 2.093125 | 0.333292 | 1.47676 | 0.145136 | 1.67244 | 0.715153 | -0.085681 | -0.139177 | 0.657507 | -0.023901 | 0.138665 | 1.239991 |
| 34.63889 | 2.233335 | 0.941255 | 2.455026 | 5.638679 | 0.277523 | 2.890762 | 0.829999 | -0.28481 | 1.873272 | 0.002566 | 4.300569 | 3.556527 | 2.010776 | 0.883643 | 1.178749 | 0.254813 | 4.738875 | 0.469953 | -0.018526 | -0.181951 | 0.795839 | 0.026294 | 0.491055 | 0.883089 |
| 34.65278 | 3.194664 | 0.916153 | 0.813761 | 6.06948 | 0.199709 | 2.450685 | 1.623846 | 0.224382 | 2.373246 | 0.570889 | 2.416497 | 3.072936 | 2.759134 | 0.739524 | 0.38243 | 0.241391 | 5.783011 | 0.286939 | 0.003591 | -0.548656 | 0.748155 | 0.0574 | 1.095652 | 0.584706 |
| 34.66667 | 2.23655 | 0.793837 | 0.185604 | 4.866465 | 0.235488 | 0.6828 | 1.810641 | 0.397123 | 1.59416 | 0.836608 | -0.347913 | 1.713591 | 2.813124 | 0.581887 | -0.181901 | 0.17799 | 4.075894 | 0.361955 | 0.087979 | -0.110583 | 0.638245 | 0.030113 | 1.27343 | 0.343828 |
| 34.68056 | 0.915661 | 0.543875 | 1.251812 | 0.29453 | 0.189549 | 0.046498 | 1.328652 | 0.502274 | 0.819041 | 0.651188 | 0.320628 | 0.768561 | 0.227916 | 0.398109 | -0.627969 | 0.192097 | 0.27818 | 0.344181 | 0.165536 | 0.530505 | 0.30265 | -0.017377 | 0.920262 | 0.024277 |
| 34.69444 | 0.947753 | 0.201877 | 0.985418 | -3.361527 | 0.181691 | -0.601912 | 0.32951 | 0.440508 | -0.317601 | 0.28112 | -0.604254 | -0.907322 | -2.105 | 0.108173 | 0.010468 | 0.135533 | -2.988333 | 0.091227 | 0.119694 | 0.14932 | 0.614434 | 0.001261 | -0.612857 | -0.346535 |
| 34.70833 | 1.410762 | -0.002013 | -1.259926 | -3.943408 | 0.247325 | -1.936796 | -1.3011 | 0.643387 | -0.985164 | 0.195846 | -2.181018 | -1.429237 | -1.032518 | -0.157275 | 1.188212 | -0.002377 | -4.373657 | -0.049985 | 0.060144 | 0.037707 | 0.965391 | 0.066539 | -3.120148 | -0.87674 |
| 34.72222 | 0.435407 | -0.045485 | -1.330925 | -2.75056 | 0.2212 | -1.417546 | -1.939514 | 0.404781 | -0.13857 | 0.245806 | -1.224441 | 0.384355 | -2.002815 | -0.554933 | 1.526271 | -0.114406 | -3.578489 | 0.06205 | 0.041734 | 0.033288 | 0.805996 | 0.041715 | -2.473633 | 0.083248 |
| 34.73611 | -0.994046 | -0.124931 | -1.842639 | -1.613599 | 0.020927 | -0.121268 | -0.92607 | 0.141564 | 0.53012 | 0.217822 | 0.979854 | 0.346457 | -0.092298 | -0.845501 | 0.77162 | -0.175372 | -3.194285 | 0.224124 | 0.019971 | -0.021715 | 0.475453 | -0.021298 | 1.374804 | 0.798009 |
| 34.75 | -1.113839 | -0.122748 | -2.783056 | 0.469255 | 0.019027 | 0.170865 | 0.637397 | 0.109988 | 0.479551 | -0.31362 | 0.340547 | -0.782765 | 1.034603 | -0.873444 | -0.070403 | -0.222605 | 1.103382 | 0.486583 | 0.003814 | -0.08624 | -0.05668 | -0.010624 | 5.671047 | 1.301296 |
| 34.76389 | -3.765016 | 0.170909 | 0.720551 | 2.764507 | 0.228548 | -0.084036 | 1.914676 | 0.273915 | 0.54499 | -0.522407 | 0.37095 | 0.054291 | 1.216555 | -0.867056 | -0.570716 | -0.146845 | 3.715145 | 0.230572 | 0.011505 | -0.065039 | -0.401275 | 0.047811 | 2.552817 | 0.047927 |
| 34.77778 | -6.697086 | 0.387861 | 4.849234 | 1.070169 | 0.132299 | 0.396018 | 0.992514 | 0.527163 | -0.42088 | 0.385813 | -0.35093 | 0.067137 | 0.386323 | -0.366686 | -0.41089 | -0.026377 | 3.73974 | -0.078368 | 0.011349 | -0.023967 | -0.413605 | 0.041632 | 2.135897 | -0.034816 |
| 34.79167 | -6.437823 | 0.170095 | 3.126419 | -1.725759 | -0.889927 | 0.655478 | -0.936489 | 0.480844 | -0.788198 | 0.723367 | -0.911433 | -0.658285 | -0.102987 | 0.172277 | 0.272753 | 0.09181 | 0.233572 | -0.107699 | -0.046446 | -0.055561 | -0.181804 | 0.002378 | 1.462932 | 0.149608 |
| 34.80556 | -6.559371 | -0.182864 | -0.074564 | -1.8498 | -2.07294 | 0.818152 | -2.087936 | 0.330588 | -0.675863 | 0.01311 | -2.942398 | -1.308146 | -0.481527 | 0.654138 | 0.791998 | 0.230789 | -2.040692 | -0.357422 | -0.119846 | -0.008674 | 0.752094 | -0.001406 | -0.368398 | 0.184111 |
| 34.81944 | -2.558874 | -1.259901 | 4.306597 | -1.317449 | -3.166777 | -0.551663 | -0.947898 | 0.315757 | -0.131301 | -0.093918 | -3.748296 | -1.478229 | -2.228997 | 0.7118 | 0.835461 | 0.299743 | -2.903334 | -0.43452 | -0.145846 | -0.029612 | 0.98929 | 0.039259 | -3.352491 | 0.121553 |
| 34.83333 | 3.185501 | -1.795787 | 5.576214 | -3.093422 | 1.474983 | 0.6753 | -0.311044 | 0.259692 | 0.301164 | -0.237362 | 0.062204 | -0.495947 | -1.116281 | 0.643084 | 0.10338 | 0.255617 | -1.585247 | -0.346866 | -0.090841 | -0.011141 | 0.051538 | 0.091898 | -2.867591 | -0.267096 |
| 34.84722 | 1.917247 | 0.5916 | 2.696733 | -5.184155 | 1.500462 | 0.186667 | 0.137117 | 0.079664 | 0.879879 | -0.074041 | 0.56789 | 1.468107 | -0.150888 | 0.363879 | -0.438254 | 0.128029 | 0.119163 | -0.304289 | 0.024381 | 0.005124 | -0.820787 | 0.116618 | -2.820728 | -1.102638 |
| 34.86111 | -1.459303 | 2.517528 | 0.850309 | -6.067123 | -0.137755 | -0.500614 | 0.163855 | -0.10109 | 1.108231 | 0.52156 | 0.788174 | 1.892799 | 0.466776 | -0.054546 | -0.771875 | -0.066057 | 0.366361 | -0.227742 | 0.114554 | -0.006223 | -0.984679 | 0.100402 | -1.244574 | -1.333223 |
| 34.875 | -3.008933 | 1.311704 | -2.965481 | -0.061976 | -1.863093 | -0.074572 | 0.130765 | -0.117423 | 1.111871 | 0.793167 | -1.235871 | -0.430097 | 0.533588 | -0.438288 | -0.32057 | -0.286006 | -0.542361 | 0.005351 | 0.110848 | 0.047318 | -0.465443 | 0.04907 | 0.954081 | -1.410189 |
| 34.88889 | -0.850206 | 0.534101 | 2.779103 | 8.175807 | -1.53012 | -1.248181 | 0.207162 | -0.051163 | 0.356407 | 0.079916 | -2.04975 | -1.105855 | 0.771135 | -0.914074 | 0.564611 | -0.419572 | -0.332583 | 0.227842 | -0.018839 | 0.05151 | 0.527396 | -0.007956 | 1.168344 | -0.138954 |
| 34.90278 | -0.349246 | 0.215393 | 3.315557 | 9.794901 | -1.088994 | -1.686581 | -0.524323 | -0.189534 | -0.715663 | -0.310872 | -1.704218 | -1.216514 | 0.034072 | -1.029912 | 1.001592 | -0.376622 | -0.379818 | 0.046924 | -0.053989 | 0.049553 | 0.59542 | -0.025963 | -0.37147 | 1.055953 |
| 34.91667 | -2.121374 | -0.16402 | 1.630016 | 3.039323 | -1.858056 | -1.069734 | -1.147956 | -0.220747 | -1.039597 | -0.026809 | -2.924027 | -2.523313 | -1.18823 | -0.827687 | 0.193082 | -0.223293 | -1.758789 | -0.382808 | -0.080313 | 0.06154 | 0.388228 | -0.03963 | -1.010751 | 1.246235 |
| 34.93056 | -1.393177 | -0.888452 | -0.519157 | -6.936255 | -4.740074 | -0.895018 | -1.385645 | -0.033602 | -1.352356 | 0.007279 | -3.885222 | -6.18895 | -2.393629 | -0.994202 | -0.615444 | -0.015493 | -3.145999 | -0.793718 | -0.086473 | 0.026816 | -0.745931 | 0.034444 | -0.504389 | 0.642594 |
| 34.94444 | -4.988978 | -0.729428 | -0.746456 | -7.708859 | -3.95591 | -1.335044 | -0.612829 | 0.167844 | -2.07839 | -0.834629 | -3.653781 | -4.2441 | -2.144272 | -1.082499 | -0.16547 | 0.180036 | -3.737756 | -1.582515 | -0.027581 | 0.106974 | -1.236566 | 0.116842 | -0.84991 | 0.109925 |
| 34.95833 | -3.757109 | -0.070915 | -2.520807 | -0.608432 | -2.508125 | -0.802385 | 0.641324 | -0.018796 | -2.954666 | -1.354867 | -1.845402 | -1.1843 | -1.934443 | -0.898365 | -0.273033 | 0.141981 | -6.294449 | -1.318322 | 0.038265 | 0.132967 | -0.976139 | 0.311453 | -1.753187 | -0.921463 |
| 34.97222 | -1.455282 | -0.411108 | -3.586335 | -0.542023 | -2.949446 | -1.626649 | 1.009579 | -0.197173 | -2.302121 | -1.387454 | -0.232728 | 1.741912 | -2.292629 | -0.788906 | -1.257834 | -0.095102 | -7.367055 | -0.752013 | -0.054687 | 0.171299 | -0.56459 | -0.051933 | -0.237355 | -1.872959 |
| 34.98611 | -2.436988 | -0.547757 | -2.523009 | -2.101353 | -6.722958 | -2.553703 | -1.234749 | -0.02567 | -2.362616 | -2.027288 | -0.765467 | 1.404771 | -1.397544 | -0.58126 | -1.861025 | -0.231835 | -7.391982 | 0.27605 | 0.095572 | 0.155643 | -0.770329 | -0.143252 | 0.335097 | -2.260412 |
| 35 | -6.486114 | -0.641783 | -2.351138 | -7.748713 | -10.628005 | -1.956755 | -2.605361 | 0.146426 | -3.460111 | -2.711097 | -0.846427 | -0.464045 | -0.615717 | -0.681632 | -2.284102 | -0.161045 | -4.649912 | 1.59365 | 0.017354 | 0.184245 | -0.593836 | 0.073242 | -0.44467 | -4.405277 |
| 35.01389 | -4.555269 | -0.105484 | -2.186618 | -7.170214 | -9.645176 | -1.756291 | -2.518026 | -0.082655 | -5.603548 | -3.282242 | -3.271307 | -3.114628 | -0.10982 | -0.121145 | -2.722657 | -0.055893 | -2.096918 | -0.008615 | -0.131466 | 0.113314 | -1.018641 | 0.14448 | -2.083975 | -1.511357 |
| 35.02778 | -5.076327 | -1.068184 | -1.272991 | -5.673801 | -6.754857 | -2.440881 | -5.020897 | -0.458498 | -5.906226 | -4.380906 | -4.582805 | -4.797068 | -1.397195 | 0.231946 | -2.932072 | -0.029156 | -1.671801 | -1.040898 | -0.068376 | 0.095425 | -1.728668 | 0.18853 | -1.839065 | 1.119718 |
| 35.04167 | -4.697424 | -0.809085 | -2.265497 | -7.29923 | -6.679026 | -2.256726 | -3.590766 | -1.23045 | -7.637071 | -4.803579 | -4.821026 | -2.838591 | -2.24501 | 0.253047 | -3.084774 | 0.038226 | -4.387082 | -1.852229 | -0.075824 | 0.261795 | -3.654481 | -0.395921 | -2.40461 | -0.457887 |
| 35.05556 | -3.13722 | -1.486448 | -3.898386 | -11.740685 | -7.731335 | -3.202353 | -1.15601 | -1.945918 | -8.022607 | -5.557122 | -4.760396 | -0.922054 | -3.006366 | -0.045117 | -2.914115 | -0.073303 | -6.5701 | -2.895567 | -0.12182 | 0.290249 | -4.742895 | -2.375143 | -1.326608 | -1.762023 |
| 35.06944 | -2.456956 | -0.673655 | -4.350772 | -2.475048 | -10.092448 | -3.43716 | -1.697101 | -2.307539 | -7.137428 | -7.028651 | -3.98997 | -0.685586 | -3.200021 | -0.552529 | -2.776028 | -0.226645 | -5.219172 | -3.554049 | -0.446209 | 0.294686 | -3.149999 | -2.449927 | -1.69322 | -2.222244 |
| 35.08333 | -1.733488 | 0.165813 | -2.881616 | -0.839743 | -8.672613 | -5.635137 | -2.196702 | -2.477152 | -5.408976 | -6.367795 | -3.30517 | -1.261298 | -2.467332 | -0.697228 | -2.453208 | -0.380745 | -3.381229 | -3.025573 | -0.409173 | -0.415959 | -1.348337 | -1.78849 | -0.838759 | -2.03947 |
| 35.09722 | -2.142175 | -0.652129 | -1.244022 | -3.681 | -6.527674 | -7.181109 | -3.326903 | -3.877815 | -4.072357 | -4.954051 | -2.026988 | -1.466496 | -1.519082 | -0.588339 | -2.005339 | -0.356803 | -2.105329 | -2.164296 | -0.264742 | -0.591643 | -1.044519 | -1.365307 | -0.465246 | -2.087573 |
| 35.11111 | -1.187112 | -1.099163 | -0.470621 | -3.942048 | -6.670084 | -3.116648 | -0.81645 | -3.264688 | -1.636513 | -1.965113 | -1.365056 | -1.635003 | -0.961442 | -0.469579 | -1.656951 | -0.282734 | -1.294576 | -1.874391 | -0.199359 | -0.705527 | -1.035415 | -1.091895 | -0.432367 | -2.181073 |
| 35.125 | -1.103075 | -1.045171 | -0.536363 | -2.492829 | -8.385714 | -0.169775 | 1.748145 | -2.298425 | 0.44328 | 0.466195 | -1.638994 | -1.578019 | -0.870161 | -0.389281 | -1.373214 | -0.207545 | -0.93673 | -1.634527 | -0.209533 | -0.519957 | -0.86706 | -0.839974 | -0.631073 | -1.767994 |
| 35.13889 | -2.949672 | -0.593689 | -1.639681 | -2.867353 | -10.129417 | -0.812928 | -0.140036 | -1.940982 | 0.453404 | 0.901896 | -2.198947 | -1.490331 | -0.767241 | -0.413545 | -1.02842 | -0.161976 | -0.963191 | -1.330435 | -0.176347 | -0.32979 | -0.347013 | -0.48979 | -0.780611 | -1.660722 |
| 35.15278 | -5.586315 | 0.180541 | -2.417824 | -4.423702 | -10.71269 | -3.041185 | -2.34347 | -1.952704 | -1.461063 | -0.603269 | -2.704351 | -1.356482 | -0.368988 | -0.385493 | -0.709151 | -0.12428 | -1.104879 | -1.070964 | -0.098384 | -0.267445 | 0.072024 | -0.187064 | -0.829434 | -1.783021 |
| 35.16667 | -4.671043 | -0.274193 | -2.1503 | -5.471911 | -7.339485 | -5.222365 | -3.313295 | -1.993644 | -3.401896 | -2.938984 | -3.243636 | -1.810629 | -0.088036 | -0.399872 | -0.624456 | -0.168165 | -1.430456 | -0.900113 | -0.053903 | -0.295082 | 0.155977 | -0.173054 | -1.007057 | -1.76464 |
| 35.18056 | -2.730335 | 0.095624 | -0.86087 | -4.946195 | -4.636583 | -5.312199 | -2.549055 | -2.070182 | -5.054017 | -5.159429 | -1.707037 | -0.808886 | 0.065383 | -0.395996 | -0.831761 | -0.264363 | -1.820099 | -0.932115 | -0.057193 | -0.313998 | -0.084402 | -0.279408 | -1.331344 | -1.742045 |
| 35.19444 | -0.703331 | 0.856127 | 0.228209 | -3.213766 | -3.058307 | -6.261896 | -2.79792 | -1.692609 | -4.535847 | -5.749854 | 0.109637 | 0.463215 | 0.302693 | -0.488476 | -1.181038 | -0.332341 | -2.097612 | -1.22284 | -0.086124 | -0.311228 | -0.569712 | -0.446255 | -1.610807 | -2.192483 |
| 35.20833 | -0.312197 | 0.511777 | -0.2951 | -2.140259 | -1.21339 | -3.452143 | -1.542053 | -1.176237 | -1.140477 | -4.673644 | 1.558244 | 0.804604 | 0.692609 | -0.607184 | -1.467736 | -0.375965 | -2.193502 | -1.881496 | -0.154662 | -0.335839 | -1.231474 | -0.751005 | -1.42961 | -2.544273 |
| 35.22222 | 0.266309 | -1.410027 | -2.167566 | -2.525528 | 1.005157 | -0.924629 | -0.657579 | -0.797795 | -0.7271 | -4.165328 | 2.119492 | -0.598722 | 1.065561 | -0.777929 | -1.487356 | -0.465434 | -2.340583 | -2.644051 | -0.24837 | -0.37956 | -1.891329 | -1.131568 | -1.171235 | -2.455754 |
| 35.23611 | 0.471973 | -2.530703 | -3.355933 | -2.271805 | 3.227293 | 0.287281 | -0.672953 | -0.21648 | -0.615527 | -3.847734 | 1.280347 | -1.667607 | 1.193861 | -0.764024 | -1.137364 | -0.511679 | -2.667233 | -2.720841 | -0.323168 | -0.452889 | -1.837981 | -1.166656 | -1.114076 | -2.395902 |
| 35.25 | 0.904 | -0.360472 | -1.121445 | -0.673785 | 5.535188 | 2.175843 | 0.415458 | 0.017938 | 0.030268 | -1.989973 | -0.223089 | -1.061707 | 1.489585 | -0.745551 | -0.638733 | -0.39347 | -2.981008 | -2.263322 | -0.32625 | -0.533368 | -1.008921 | -0.704348 | -1.088409 | -2.118215 |
| 35.26389 | 0.831559 | 2.269569 | 1.049875 | 0.8278 | 6.051236 | 3.911479 | 1.481711 | -0.010851 | 0.982148 | 0.590162 | -0.490797 | 0.533553 | 2.093738 | -0.850256 | -0.239426 | -0.243685 | -2.912221 | -1.807039 | -0.35844 | -0.602083 | -0.138119 | -0.079335 | -0.824432 | -2.06473 |
| 35.27778 | 0.741054 | 2.002512 | 1.34334 | 1.307296 | 5.462184 | 2.598983 | 0.936552 | 0.040878 | 1.453411 | 2.898371 | 0.078692 | 1.84077 | 2.332122 | -0.805631 | -0.015329 | -0.136494 | -2.802245 | -1.385037 | -0.347898 | -0.607318 | 0.351364 | 0.320941 | -0.495313 | -1.760548 |
| 35.29167 | 1.178132 | 1.105041 | 0.577507 | 1.328918 | 5.347593 | 1.12667 | 0.092669 | 0.255457 | 1.36258 | 2.985948 | 0.720813 | 2.001959 | 2.560928 | -0.682968 | 0.168645 | -0.092718 | -2.631749 | -0.952457 | -0.310704 | -0.525718 | 0.65674 | 0.558165 | -0.365328 | -1.357239 |
| 35.30556 | 0.75179 | 1.292592 | 0.483808 | 1.693989 | 5.643313 | 1.452967 | 0.183317 | 0.239153 | 1.110288 | 2.106356 | 1.024937 | 1.348901 | 2.061818 | -0.581954 | 0.524998 | -0.001294 | -2.07258 | -0.476338 | -0.295701 | -0.467749 | 0.902445 | 0.802169 | -0.307712 | -1.186643 |
| 35.31944 | 0.164501 | 1.074711 | 1.08111 | 1.926437 | 3.948462 | 3.208701 | 0.872151 | 0.240192 | 0.821764 | 1.820635 | 1.143504 | 0.627993 | 1.46063 | -0.434611 | 0.858098 | 0.117565 | -1.801404 | -0.164527 | -0.251176 | -0.45091 | 1.04423 | 0.95256 | -0.212781 | -1.030388 |
| 35.33333 | -0.196886 | 0.063162 | 1.265028 | 1.880595 | 0.988674 | 2.688939 | 1.392769 | 0.239893 | 0.877867 | 2.945806 | 1.174625 | 0.44893 | 0.702391 | -0.30714 | 0.773182 | 0.124665 | -2.058121 | 0.023777 | -0.240854 | -0.479762 | 0.889658 | 0.820578 | -0.035647 | -0.646779 |
| 35.34722 | 1.808294 | -0.488362 | 1.068166 | 2.249346 | -0.383014 | 1.788468 | 2.362421 | 0.483455 | 1.925763 | 5.656186 | 1.363645 | 1.714685 | 0.559557 | -0.075043 | 0.410673 | 0.126563 | -1.966555 | 0.35555 | -0.19878 | -0.471657 | 0.856274 | 0.817196 | 0.199076 | -0.145485 |
| 35.36111 | 2.979553 | 0.726868 | 1.379322 | 3.428929 | 0.424659 | 1.930506 | 3.20137 | 0.917169 | 5.321105 | 9.394029 | 1.615788 | 2.402786 | 0.597422 | 0.25923 | 0.236399 | 0.130376 | -1.077397 | 0.875219 | -0.127934 | -0.33017 | 1.318466 | 1.119563 | 0.465745 | 0.351365 |
| 35.375 | 1.9371 | 0.871809 | 2.295297 | 4.882919 | 2.144349 | 2.896887 | 3.61971 | 0.93698 | 6.574774 | 11.01742 | 2.177648 | 1.88896 | 0.757828 | 0.458532 | 0.22603 | 0.076186 | -0.021154 | 1.363492 | -0.074996 | -0.237622 | 1.873896 | 1.462504 | 0.529337 | 0.703872 |
| 35.38889 | 1.214524 | -0.083355 | 2.973318 | 5.526812 | 3.935882 | 3.280439 | 3.479412 | 0.926338 | 6.362319 | 8.729176 | 2.869968 | 1.204154 | 0.551373 | 0.499735 | 0.225812 | 0.053805 | 0.762093 | 1.6815 | -0.045988 | -0.222007 | 1.941603 | 1.681638 | 0.495827 | 0.846629 |
| 35.40278 | 1.159851 | -0.250515 | 2.926757 | 5.663672 | 6.265708 | 3.982332 | 3.697119 | 1.090739 | 6.113284 | 6.878589 | 3.323283 | 1.815529 | 0.550251 | 0.527466 | 0.380484 | 0.184999 | 1.350072 | 1.726806 | 0.009261 | -0.142943 | 2.333017 | 1.744253 | 0.696245 | 0.932138 |
| 35.41667 | 2.551768 | -0.076279 | 3.516415 | 6.805346 | 8.144073 | 6.005668 | 3.942784 | 1.301835 | 6.403589 | 6.273416 | 3.403501 | 2.7208 | 0.734738 | 0.667999 | 0.799756 | 0.347185 | 1.992635 | 1.960656 | 0.031041 | -0.011935 | 3.002636 | 2.053223 | 1.04225 | 1.258072 |
| 35.43056 | 3.778103 | 0.881464 | 4.894243 | 8.547219 | 8.84272 | 5.848008 | 3.787299 | 1.749952 | 6.491421 | 4.928018 | 3.078321 | 2.572521 | 0.697306 | 0.826662 | 1.418198 | 0.40147 | 2.316962 | 2.392388 | 0.074077 | 0.100712 | 3.47271 | 1.806513 | 1.28139 | 1.585471 |
| 35.44444 | 4.831945 | 1.397703 | 6.299531 | 9.176794 | 10.170688 | 7.367127 | 3.463931 | 2.397906 | 8.976927 | 5.622466 | 3.443977 | 2.046085 | 0.777108 | 1.077613 | 2.039028 | 0.505163 | 2.536652 | 2.888397 | 0.182392 | 0.18112 | 4.151429 | 1.269115 | 1.445448 | 1.943308 |
| 35.45833 | 5.269968 | 1.135546 | 5.642711 | 9.790499 | 13.044838 | 6.514789 | 2.987158 | 3.014496 | 10.636134 | 4.746597 | 3.797354 | 1.55746 | 0.957494 | 1.400744 | 2.686379 | 0.531097 | 3.133819 | 3.412146 | 0.223842 | 0.296209 | 3.785322 | 1.117827 | 1.795826 | 2.425813 |
| 35.47222 | 5.562937 | 0.529145 | 4.986488 | 12.150038 | 14.382027 | 6.161202 | 2.579455 | 3.322763 | 10.591604 | 4.08476 | 3.562479 | 1.638 | 0.788897 | 1.412515 | 2.968565 | 0.440229 | 4.102649 | 3.694804 | 0.18212 | 0.385813 | 2.924442 | 0.70267 | 2.374194 | 3.300101 |
| 35.48611 | 6.777326 | 0.584332 | 5.237543 | 13.944503 | 14.715514 | 6.467645 | 2.352929 | 3.059258 | 10.430667 | 3.989097 | 3.320728 | 1.859452 | 0.584442 | 1.124973 | 2.89317 | 0.293394 | 5.198632 | 3.938398 | 0.135224 | 0.359019 | 1.526043 | 0.493443 | 2.915971 | 3.891504 |
| 35.5 | 8.401698 | 0.726698 | 6.685645 | 16.555028 | 14.638426 | 6.945193 | 2.210616 | 2.479611 | 10.232736 | 4.465354 | 3.078805 | 2.097251 | 0.527237 | 0.71433 | 2.78168 | 0.24542 | 5.536989 | 3.019527 | 0.112458 | 0.353298 | 1.972356 | 0.42925 | 2.910195 | 4.085661 |
| 35.51389 | 8.807531 | 1.551887 | 6.325408 | 16.788541 | 15.65886 | 7.384045 | 2.681853 | 2.154536 | 6.480108 | 4.183001 | 4.088468 | 4.110777 | 0.79235 | 0.372139 | 2.900464 | 0.444218 | 5.601594 | 2.956234 | 0.24459 | 0.302806 | 2.337989 | 0.431902 | 2.64095 | 3.462409 |
| 35.52778 | 9.431636 | 3.035124 | 3.861725 | 14.171281 | 15.723666 | 7.158972 | 3.050806 | 2.824139 | 3.735064 | 2.637544 | 4.249288 | 5.451739 | 0.789474 | 0.634476 | 3.224202 | 0.517807 | 5.632229 | 2.255694 | 0.168404 | 0.368723 | 2.27721 | 0.430229 | 2.782912 | 2.572185 |

| | | | | | | | | | | | | | | | | | | | | | | | | |
|---|---|---|---|---|---|---|---|---|---|---|---|---|---|---|---|---|---|---|---|---|---|---|---|---|
| 35.54167 | 6.030593 | 1.221898 | 2.690277 | 12.4606 | 9.973306 | 4.911765 | 2.8936 | 2.94295 | 2.504565 | 2.364674 | 2.774964 | 3.74764 | 0.746942 | 0.619242 | 3.141183 | 0.451195 | 5.279998 | 0.919545 | -0.17559 | 0.345559 | 2.24315 | 0.315519 | 1.872055 | 1.948046 |
| 35.55556 | 0.788297 | -0.625226 | 3.142239 | 7.544052 | 0.994409 | 2.904695 | 2.892793 | 1.493278 | 0.937706 | 1.24793 | 0.91416 | 0.566445 | 0.835406 | 0.851702 | 2.35938 | 0.183266 | 5.67224 | -0.317213 | -0.223596 | 0.613317 | 1.742512 | 0.100295 | 1.135984 | 1.733859 |
| 35.56944 | -2.462303 | -0.824006 | 4.992676 | 2.991095 | -7.016514 | 0.841296 | 3.004103 | -0.005093 | -0.501784 | -2.285719 | 0.382784 | -0.246272 | 0.900282 | 0.938551 | 1.415754 | -0.059835 | 3.364506 | -1.716373 | -0.100234 | 0.288436 | 1.48357 | 0.196973 | 0.827793 | 0.803638 |
| 35.58333 | 0.254558 | -0.349043 | 4.609932 | 4.556094 | -3.11882 | 0.541689 | 2.109162 | -0.487629 | 0.293917 | -2.345171 | 0.597967 | 0.98953 | 0.648463 | 0.864932 | 0.15032 | -0.111991 | 2.2977 | -1.423915 | -0.0148 | 0.154947 | -0.12926 | 0.245154 | -1.058835 | 0.560426 |
| 35.59722 | 1.697546 | 0.094638 | 3.159387 | 8.628527 | 3.542908 | 2.236811 | 1.389342 | -0.425179 | 1.476433 | 0.388663 | 2.023734 | 2.351805 | 0.551854 | 1.009328 | -0.572089 | 0.044935 | 1.672906 | -0.16836 | 0.216335 | 0.006503 | -0.048972 | -0.176566 | -2.45793 | 0.417644 |
| 35.61111 | 4.814689 | 1.086964 | -1.100788 | 10.957242 | 10.502211 | 2.270631 | 0.692856 | -0.269635 | 1.469072 | 2.146273 | 1.75982 | 2.8177 | 0.300522 | 0.643694 | -0.528897 | 0.124748 | 1.050451 | 0.676785 | 0.178018 | 0.016475 | -0.025626 | 0.505934 | -1.102779 | 0.140248 |
| 35.625 | 7.517315 | 1.108492 | 0.336314 | 6.824996 | 6.710276 | 1.680827 | 0.79102 | 0.123891 | 2.100238 | 1.417343 | 0.543283 | 2.260501 | 0.290705 | 0.319551 | 0.465724 | 0.149571 | 2.406146 | 0.539775 | -0.075641 | -0.042385 | 0.525323 | 0.844274 | 1.100099 | -0.069674 |
| 35.63889 | 6.98235 | 0.30318 | 2.469084 | 4.767318 | 2.017827 | 1.657796 | 2.344558 | -0.075666 | 2.050262 | 0.075792 | -1.093463 | 0.169182 | 0.218537 | 0.199844 | 1.081208 | 0.279727 | 3.99258 | 0.475875 | -0.045964 | -0.077196 | 0.232066 | 0.287317 | 1.456001 | -0.026961 |
| 35.65278 | 2.492127 | 0.431264 | -0.20173 | 4.648688 | 0.512245 | 2.051728 | 2.797977 | -0.224992 | 1.928126 | 0.358611 | 0.055713 | -1.048298 | 1.389676 | 0.367951 | 0.602126 | 0.428958 | 4.457055 | 0.204787 | 0.068587 | -0.067264 | 0.82553 | 0.16463 | 1.364041 | 0.179417 |
| 35.66667 | 1.243004 | 0.393929 | -4.608258 | 4.516221 | 0.033979 | 1.8148 | 1.04793 | -0.231443 | 1.513225 | 0.454991 | 0.168896 | -1.789943 | 1.007662 | 0.466572 | -0.564782 | 0.432192 | 1.074175 | 0.240329 | 0.252945 | 0.0179 | 0.46758 | 0.205805 | -0.0433 | 0.377137 |
| 35.68056 | -0.550125 | 0.344777 | -3.592922 | 2.428926 | 0.072271 | 1.027901 | 0.569252 | -0.092918 | 0.937873 | 0.239474 | -2.114623 | -0.702045 | 0.426298 | 0.336952 | -0.256097 | 0.355944 | -1.654649 | 0.171916 | 0.372689 | 0.035085 | 0.545936 | 0.231224 | 0.314015 | 0.455032 |
| 35.69444 | 0.780013 | 0.247979 | -3.363318 | -0.291144 | 0.176157 | -0.212439 | -0.003988 | 0.044118 | 0.220906 | 0.040613 | 0.987932 | 2.585379 | -0.264031 | 0.172462 | 1.011657 | 0.173748 | -0.234832 | -0.175273 | 0.250735 | 0.047357 | 0.087437 | 0.211327 | -1.105881 | 0.13102 |
| 35.70833 | 1.516635 | -0.014009 | 0.767749 | -3.467602 | 0.123427 | -1.331352 | -0.453263 | 0.141509 | -0.223894 | -0.052796 | 2.490372 | 3.317829 | -0.703865 | -0.029749 | 2.076949 | -0.019849 | -1.175236 | -0.520882 | 0.128195 | 0.187168 | 0.189756 | 0.13878 | -2.394462 | -0.153084 |
| 35.72222 | 0.347472 | -0.175753 | 5.563811 | -3.057924 | 0.047004 | -0.988453 | -1.340458 | 0.371957 | -0.05592 | -0.076419 | 2.106653 | 1.633098 | -0.705217 | -0.19225 | 1.30208 | -0.165647 | -2.428 | -0.346157 | 0.048228 | -0.304607 | 0.330315 | 0.066254 | -2.70698 | -0.514004 |
| 35.73611 | -1.737405 | -0.07956 | -1.681661 | -1.224948 | 0.014516 | -0.524823 | -0.932944 | 0.447566 | 0.302324 | 0.000647 | -0.836555 | -1.592232 | -0.059223 | -0.429724 | 0.887448 | -0.226916 | -2.19513 | 0.004995 | -0.065619 | -0.250203 | 0.349456 | 0.004194 | -2.405545 | -0.615999 |
| 35.75 | -1.345059 | 0.250141 | -4.144422 | -0.08155 | -0.026192 | -0.554138 | 0.454535 | 0.26603 | 0.241571 | 0.17499 | -4.393084 | -4.227333 | 0.42751 | -0.22483 | 0.654408 | -0.166702 | -1.25691 | 0.560561 | -0.104809 | 0.003489 | 0.27711 | -0.080306 | -2.352633 | -0.366819 |
| 35.76389 | -2.285177 | 0.510908 | -5.850765 | -2.655371 | -0.398754 | 0.181414 | 1.404718 | 0.08694 | 0.112561 | -0.144462 | -5.10438 | -3.055295 | 0.362657 | -0.4644 | 0.181192 | -0.063202 | -0.40577 | 0.490476 | -0.06115 | 0.091489 | 0.752724 | -0.080755 | -1.918879 | 0.087331 |
| 35.77778 | -4.977542 | 0.117726 | -3.009611 | -9.250293 | -2.599694 | -0.419015 | 0.41407 | 0.118882 | -0.273604 | -1.381666 | -0.196824 | -0.748813 | 0.211933 | -0.567894 | -0.414565 | 0.000287 | -1.966751 | 0.352523 | 0.098673 | 0.07979 | 1.09087 | -0.00945 | 1.326757 | 0.735045 |
| 35.79167 | -5.217359 | -0.194485 | -2.827253 | -6.815724 | -5.776679 | 0.578612 | -0.866849 | 0.420368 | -0.423125 | -0.621314 | -0.960672 | 1.906033 | -0.152967 | -0.228233 | -0.388988 | 0.032242 | -1.233366 | -0.032476 | 0.167728 | 0.115042 | 1.245448 | 0.057202 | 2.386956 | 0.265314 |
| 35.80556 | -2.173372 | -0.189297 | -0.197906 | -0.957035 | -4.383965 | 0.868059 | -1.949989 | 0.680338 | -0.543738 | 0.526886 | 0.346763 | 2.874614 | 0.083239 | 0.147979 | 0.109626 | 0.044633 | -2.905661 | -0.047947 | 0.13615 | -0.022426 | 0.925174 | 0.054741 | 2.486065 | 0.169778 |
| 35.81944 | 1.926484 | 0.054079 | 1.92059 | -0.200949 | -2.019777 | 1.066775 | -2.077312 | 0.481652 | 0.279764 | 1.040584 | -0.90389 | 1.467771 | 0.37166 | 0.525852 | 0.74284 | 0.085961 | -2.044418 | 0.081425 | 0.024103 | -0.148916 | 0.013518 | -0.010903 | 3.655151 | 0.155443 |
| 35.83333 | -1.852803 | 0.220452 | 1.806204 | -0.538539 | -1.79411 | -0.558215 | -1.139916 | -0.028904 | 0.606143 | 1.105939 | -2.445161 | -0.617049 | 0.644507 | 0.619499 | 0.364715 | 0.153685 | 1.000763 | 0.108702 | -0.097105 | -0.085545 | -0.455939 | -0.079448 | 2.138968 | 0.152126 |
| 35.84722 | -2.419615 | 0.225442 | 0.818757 | -1.420014 | -2.630299 | 0.392274 | 0.54021 | -0.052708 | 0.896936 | 0.030795 | -4.61606 | -3.366807 | 1.066269 | 0.472974 | 0.453119 | 0.237067 | 2.787363 | -0.03642 | -0.184203 | -0.060681 | -0.634462 | -0.126839 | 1.171704 | 0.177219 |
| 35.86111 | -4.232132 | -0.304386 | 0.958075 | -8.388697 | -0.627413 | -0.50241 | 1.886263 | -0.064465 | 0.69252 | 0.106276 | -0.508195 | -1.719938 | 0.693641 | 0.284945 | 0.408252 | 0.281466 | 3.212763 | -0.070855 | -0.179459 | -0.04274 | -0.426606 | -0.141355 | 0.408186 | 0.167328 |
| 35.875 | -2.458161 | -0.376664 | -0.158131 | -11.482464 | 0.056022 | -0.871702 | 1.326331 | -0.071266 | 0.173923 | -0.003906 | -1.272878 | 2.778631 | 0.182994 | 0.054762 | -0.576576 | 0.198407 | 2.316928 | -0.198857 | -0.119109 | -0.002106 | -0.084949 | -0.123515 | -0.454233 | 0.063058 |
| 35.88889 | -2.456644 | -0.005747 | 2.423615 | -1.667243 | -1.183589 | -0.903718 | -0.167009 | 0.014444 | -0.292604 | -0.136276 | 5.719684 | 3.066137 | -0.305752 | -0.198865 | -0.61633 | 0.062408 | 0.78071 | -0.773553 | 0.006599 | 0.022229 | -0.544289 | -0.057159 | -1.182006 | -0.273508 |
| 35.90278 | -0.606829 | 0.390859 | 3.491884 | 9.640333 | -4.307295 | -0.410191 | -0.575591 | 0.152403 | -1.171687 | -0.244082 | 3.198013 | 1.424875 | -0.523193 | -0.500796 | -0.799239 | -0.054547 | 0.086654 | -0.85068 | 0.070238 | 0.049184 | -0.28178 | 0.03006 | 0.425883 | -0.551967 |
| 35.91667 | 3.007036 | 0.338814 | -1.656432 | 2.853938 | -5.470427 | -2.241001 | -0.667476 | 0.241337 | -1.931077 | -0.494445 | -0.078259 | 0.874392 | -0.368423 | -0.456081 | -1.133833 | -0.146354 | 1.092628 | -0.202454 | -0.007395 | 0.018377 | -0.387302 | 0.076623 | 1.190636 | -0.84719 |
| 35.93056 | 2.108921 | 0.009683 | -5.82573 | -8.853258 | -7.008219 | -4.915505 | 0.92847 | -0.763976 | -0.651154 | 0.149161 | -2.439832 | -0.428127 | -0.302477 | -0.948842 | -0.641836 | -0.1706 | 2.98257 | 0.646227 | -0.094891 | 0.053616 | -0.111961 | 0.064598 | -1.017765 | -3.570828 |
| 35.94444 | 0.396217 | 0.034206 | -6.449878 | -13.401205 | -3.671669 | -2.189648 | 0.736182 | 0.021689 | 1.918305 | 0.525216 | -4.370747 | -3.224522 | -0.246027 | -1.940397 | 1.247934 | -0.162374 | 4.304121 | 0.65701 | -0.152591 | 0.154248 | -0.649334 | 0.05598 | -2.371831 | -5.168179 |
| 35.95833 | -0.624858 | 0.083189 | -6.389644 | -9.035798 | 2.036453 | -0.145478 | 1.096854 | 0.861465 | 2.133052 | -0.784706 | -3.934332 | -3.357966 | -0.350216 | -1.037612 | 1.776987 | -0.184843 | 3.916154 | 0.047264 | -0.058104 | 0.121712 | -0.664119 | -0.051165 | -2.26888 | -6.263359 |
| 35.97222 | -1.405607 | 0.12375 | -5.630864 | -2.665995 | 1.291041 | -1.284909 | 0.022628 | 0.057581 | -0.239004 | -1.760632 | -2.500185 | -1.584187 | -0.504521 | -0.593118 | -0.087928 | -0.251194 | 0.842836 | -1.225041 | 0.102611 | -0.029691 | -0.611572 | -0.21384 | -3.286786 | -4.108005 |
| 35.98611 | -1.962012 | 0.100908 | -4.474613 | 0.61044 | -2.32084 | -1.351153 | -0.82237 | 0.20386 | -2.445668 | -0.562736 | -3.874412 | -0.297878 | -0.635363 | -0.977231 | -2.620901 | -0.419847 | -2.667384 | -1.63145 | 0.103484 | 0.005779 | -0.757895 | -0.098655 | -3.317977 | -3.443184 |
| 36 | -2.898162 | -0.127616 | -4.442843 | -0.15079 | -5.031704 | -1.181004 | -0.729472 | 0.260545 | -5.276547 | -1.761455 | -6.83203 | -2.381121 | -0.678159 | -1.271199 | -4.04456 | -0.7132 | -5.84141 | -2.703389 | -0.029668 | 0.163505 | -0.473434 | -0.186249 | -2.590538 | -3.081906 |
| 36.01389 | 0.569065 | -0.528019 | -7.624273 | -1.560307 | -4.169373 | -0.910192 | -1.263151 | 0.115972 | -6.892288 | -3.230713 | -7.312022 | -5.819232 | -0.429805 | -1.204264 | -4.068899 | -0.750551 | -6.749563 | -2.529252 | -0.302588 | 0.182978 | -0.082861 | -0.436714 | -1.369484 | -2.571644 |
| 36.02778 | -3.585217 | -2.534084 | -7.990191 | -4.895212 | -5.105079 | -0.689988 | -1.74435 | -0.396559 | -5.828336 | -3.444148 | -2.417967 | -7.214855 | -0.286075 | -0.992529 | -1.81124 | -0.228471 | -6.810942 | -0.586304 | -0.444856 | -0.033096 | -0.184336 | -0.094453 | -1.364382 | -2.381782 |
| 36.04167 | -5.92641 | -3.856176 | -7.990964 | -8.50436 | -8.629668 | -1.782924 | -3.941422 | -2.514131 | -5.621685 | -3.862184 | -2.642338 | -6.329058 | -0.433673 | -2.2151 | -0.738732 | 0.295317 | -7.164964 | 0.021223 | -0.375588 | -0.065099 | -0.067053 | -0.235221 | -1.206062 | -2.669943 |
| 36.05556 | -3.576976 | -3.744283 | -6.73276 | -8.525802 | -8.219198 | -4.308824 | -5.151725 | -1.258807 | -6.5454 | -4.800472 | -4.118885 | -4.018006 | -0.574175 | -4.246369 | -1.765505 | 0.13205 | -5.610951 | -0.563507 | -0.313375 | 0.018396 | -0.760669 | -0.631435 | -1.440754 | -3.408804 |
| 36.06944 | -1.502945 | -1.252791 | -5.63188 | -6.387061 | -11.879145 | -5.282933 | -4.926324 | -0.653045 | -2.25256 | -7.313512 | -3.490775 | -2.779546 | -0.603219 | -3.409886 | -2.750671 | -0.261362 | -4.033061 | -1.365255 | -0.234221 | -0.336778 | -2.204004 | -0.793939 | -1.429136 | -2.858473 |
| 36.08333 | -0.574633 | -2.610747 | -4.34406 | -4.552449 | -10.279026 | -4.697302 | -2.122189 | -2.384616 | -5.864885 | -5.952552 | -2.821663 | -4.61852 | -0.571808 | -2.736457 | -2.983442 | -0.518454 | -2.828062 | -1.401611 | -0.164439 | -0.718995 | -2.689769 | -0.616623 | -1.501204 | -2.43129 |
| 36.09722 | -1.282465 | -4.585003 | -3.727691 | -9.658095 | -8.683766 | -1.730042 | 0.603069 | -4.58617 | -1.399375 | -3.271732 | -1.462705 | -4.301926 | -0.456276 | -2.217052 | -2.590971 | -0.596261 | -2.046248 | -1.127879 | -0.116384 | -0.980884 | -2.412275 | -0.565605 | -1.244785 | -1.872312 |
| 36.11111 | -3.744907 | -1.025444 | -4.185793 | -8.240793 | -8.90742 | -1.799046 | -0.213896 | -4.970502 | -0.2334 | -2.409311 | -0.534966 | -0.390421 | -0.402803 | -1.998829 | -1.981624 | -0.512495 | -1.681146 | -1.286798 | -0.13561 | -0.973819 | -2.465474 | -0.558641 | -0.852905 | -1.45986 |
| 36.125 | -6.609324 | -0.891958 | -3.388074 | -4.670706 | -12.018735 | -4.51902 | -2.860862 | -5.468316 | -1.861721 | -4.012185 | -0.802696 | -0.644144 | -0.37754 | -1.959371 | -1.381532 | -0.398541 | -1.629644 | -1.433734 | -0.172529 | -0.835399 | -2.652987 | -0.361719 | -0.658953 | -1.314115 |
| 36.13889 | -3.155778 | -1.592739 | -2.586587 | -4.558802 | -9.042544 | -4.740113 | -4.471855 | -5.196031 | -2.449213 | -6.165042 | -1.316116 | -1.417347 | -0.352504 | -1.94719 | -0.766612 | -0.19473 | -1.572519 | -1.385604 | -0.189776 | -0.630567 | -2.674336 | -0.37361 | -0.570274 | -1.214241 |
| 36.15278 | -1.845686 | -2.482875 | -2.285776 | -5.872018 | -6.684353 | -2.874543 | -3.286221 | -3.931347 | -1.879871 | -6.893928 | -2.217443 | -2.566106 | -0.436358 | -1.883748 | -0.311028 | -0.10671 | -1.513232 | -1.357504 | -0.183377 | -0.561542 | -2.721944 | -0.459752 | -0.790886 | -1.080721 |
| 36.16667 | -2.635622 | -3.481583 | -1.550571 | -6.681227 | -4.853047 | -2.654924 | -3.322041 | -2.734584 | -3.594678 | -6.949348 | -2.353157 | -3.227362 | -0.579513 | -1.667008 | -0.098476 | -0.145614 | -1.575745 | -1.482205 | -0.15965 | -0.552638 | -2.635469 | -0.485631 | -0.952639 | -0.977719 |
| 36.18056 | -6.022002 | -4.640194 | -1.274278 | -6.181538 | -5.882797 | -3.79415 | -3.042964 | -1.531336 | -4.656935 | -8.412268 | -1.541557 | -3.034429 | -0.662991 | -1.294987 | 0.023622 | -0.272625 | -1.834297 | -1.674922 | -0.185276 | -0.538978 | -2.656476 | -0.523646 | -0.769874 | -0.816208 |
| 36.19444 | -7.529345 | -2.645397 | -2.281507 | -4.293695 | -4.849775 | -4.778755 | -4.742464 | -0.310082 | -2.888259 | -9.822358 | -1.00246 | -2.111474 | -0.645256 | -0.984931 | 0.142698 | -0.404688 | -2.112016 | -1.874637 | -0.218163 | -0.556041 | -2.899618 | -0.598111 | -0.628216 | -0.774236 |
| 36.20833 | -2.23197 | 1.282378 | -1.427285 | -3.476688 | -1.666084 | -3.599104 | -4.301456 | 0.760693 | 0.877322 | -5.542013 | -0.785042 | -0.933197 | -0.602503 | -1.136851 | 0.101562 | -0.496714 | -2.257235 | -1.982033 | -0.252623 | -0.56446 | -3.196642 | -0.748033 | -0.609575 | -0.875327 |
| 36.22222 | 0.295365 | 5.837517 | 0.254508 | -2.451189 | 1.299438 | -1.386826 | -2.264222 | 1.553985 | 2.494975 | -1.075119 | -0.39115 | -0.061859 | -0.536997 | -1.913908 | -0.147652 | -0.468006 | -2.258587 | -2.110992 | -0.213434 | -0.662058 | -4.045093 | -0.984352 | -0.837972 | -1.153204 |
| 36.23611 | 0.469997 | 4.330021 | 1.341378 | -0.889753 | 3.295617 | -0.624587 | -1.379489 | 1.8629 | 2.018013 | 1.507808 | -0.017021 | 0.328514 | -0.482462 | -2.757694 | -0.39783 | -0.396261 | -2.34373 | -2.572973 | -0.124336 | -0.707794 | -5.643286 | -1.161067 | -0.908268 | -1.301592 |
| 36.25 | -0.832921 | 2.513589 | 1.582322 | 0.839375 | 4.150721 | 0.791805 | -0.674983 | 1.829823 | 1.346987 | 2.048566 | 0.29634 | 0.458228 | -0.509009 | -2.871103 | -0.473457 | -0.432681 | -2.623922 | -2.84927 | -0.088266 | -0.838335 | -5.605399 | -1.046014 | -0.68129 | -1.117109 |
| 36.26389 | -1.389716 | 1.268372 | 1.326867 | 2.385668 | 5.543566 | 1.78634 | 0.297041 | 1.719957 | 1.607542 | 3.06087 | 0.539372 | 0.822886 | -0.550545 | -2.21602 | -0.442235 | -0.478068 | -2.794414 | -2.2092 | -0.093522 | -0.946681 | -3.763497 | -0.700498 | -0.511918 | -0.897879 |
| 36.27778 | -0.384473 | 1.433429 | 1.459301 | 3.420147 | 6.715318 | 2.955245 | 1.286353 | 1.679342 | 3.387725 | 4.509508 | 0.531317 | 1.203679 | -0.574076 | -1.370047 | -0.567493 | -0.485985 | -2.820124 | -1.369815 | -0.142077 | -0.905064 | -2.312705 | -0.276834 | -0.478559 | -0.619545 |
| 36.29167 | 0.694878 | 3.436137 | 2.24304 | 3.639525 | 6.754889 | 4.021887 | 2.015384 | 1.281036 | 5.944601 | 6.664046 | 0.528803 | 1.701442 | -0.569026 | -0.71989 | -0.872968 | -0.486964 | -2.869818 | -1.16545 | -0.2369 | -0.782035 | -1.332298 | -0.404298 | -0.47909 | -0.457523 |
| 36.30556 | 1.625445 | 5.495797 | 2.644452 | 3.541869 | 6.660169 | 4.162281 | 1.901125 | 1.148321 | 8.542822 | 9.206262 | 0.764586 | 2.673839 | -0.526659 | -0.354757 | -1.086726 | -0.523035 | -2.759891 | -1.250981 | -0.296509 | -0.533221 | -0.866436 | -0.698215 | -0.464613 | -0.475899 |
| 36.31944 | 2.225173 | 5.739314 | 2.101805 | 2.97275 | 7.256931 | 3.81489 | 2.203877 | 1.247492 | 9.749356 | 10.487877 | 1.356421 | 3.222328 | -0.385434 | 0.090993 | -0.949431 | -0.444905 | -2.450001 | -1.156586 | -0.298565 | -0.391473 | -0.35493 | -0.902938 | -0.401932 | -0.41143 |
| 36.33333 | 1.756738 | 3.427431 | 1.566929 | 2.106519 | 7.19736 | 3.564319 | 2.497798 | 1.24434 | 9.008017 | 9.195886 | 2.168866 | 4.376946 | -0.203238 | 0.963736 | -0.573071 | -0.245207 | -1.998245 | -0.69271 | -0.259191 | -0.457049 | 0.047885 | -0.525907 | -0.329897 | -0.141728 |
| 36.34722 | 0.348433 | 2.616184 | 1.983703 | 1.788349 | 5.677677 | 2.852118 | 1.81422 | 1.141498 | 7.600497 | 7.79913 | 2.54174 | 5.364489 | -0.064207 | 2.164873 | -0.187938 | -0.027604 | -1.462757 | 0.006933 | -0.170063 | -0.51977 | 1.284907 | 0.377688 | -0.217374 | 0.22788 |
| 36.36111 | 0.445418 | 3.020578 | 2.871026 | 2.45579 | 4.991387 | 2.367843 | 0.992289 | 0.833419 | 7.207415 | 7.449472 | 2.460169 | 5.517534 | -0.020741 | 2.825937 | 0.119222 | 0.170742 | -1.069516 | 0.614687 | -0.05997 | -0.415396 | 2.87788 | 1.603989 | -0.11623 | 0.535626 |
| 36.375 | 2.1524 | 2.760065 | 3.287056 | 3.250486 | 4.97084 | 2.969551 | 1.373919 | 0.727966 | 7.9708 | 8.669322 | 2.207158 | 4.227258 | 0.013846 | 2.763107 | 0.313627 | 0.290699 | -0.754284 | 1.06009 | 0.051471 | -0.258215 | 4.335094 | 2.468738 | -0.066812 | 0.979079 |
| 36.38889 | 3.864402 | 2.268805 | 2.711596 | 3.973247 | 4.691111 | 4.436199 | 2.477293 | 0.73015 | 9.882545 | 8.809694 | 2.100218 | 3.334453 | 0.063025 | 2.629613 | 0.45056 | 0.318345 | -0.440898 | 1.456776 | 0.144857 | -0.088474 | 4.666263 | 2.695119 | 0.106335 | 1.263762 |
| 36.40278 | 3.928771 | 1.917375 | 2.719422 | 4.669797 | 5.456147 | 5.24663 | 2.878228 | 1.165577 | 9.427875 | 8.244515 | 2.557687 | 3.839166 | 0.131768 | 2.820341 | 0.615935 | 0.304012 | -0.136239 | 1.822051 | 0.178869 | 0.074049 | 4.820013 | 2.67841 | 0.382316 | 1.407388 |
| 36.41667 | 3.403334 | 2.01455 | 3.800867 | 5.492499 | 4.40575 | 5.2458 | 2.814024 | 1.367691 | 11.562741 | 9.260115 | 3.590139 | 4.873745 | 0.220255 | 3.040455 | 0.773557 | 0.292015 | 0.149343 | 2.023043 | 0.122253 | 0.115589 | 5.562379 | 2.728904 | 0.617627 | 1.571076 |
| 36.43056 | 3.476974 | 3.406083 | 5.011464 | 7.062219 | 3.816368 | 4.049822 | 2.297756 | 1.399683 | 13.984301 | 8.444597 | 2.069142 | 0.233183 | 0.282226 | 3.466036 | 1.067985 | 0.310933 | 0.503159 | 2.223065 | 0.089802 | 0.119049 | 6.071223 | 2.621213 | 0.834054 | 1.609046 |
| 36.44444 | 4.127793 | 4.080082 | 5.437254 | 8.737086 | 4.819809 | 4.405511 | 2.531878 | 1.655449 | 13.161837 | 7.825979 | 2.617871 | 2.167252 | 0.354326 | 3.401872 | 1.692929 | 0.393456 | 0.965517 | 2.492474 | 0.048933 | 0.164157 | 6.243001 | 2.335784 | 1.126836 | 1.670801 |
| 36.45833 | 5.571157 | 2.613024 | 4.835848 | 9.104826 | 7.373548 | 4.952335 | 2.935628 | 2.173287 | 12.665299 | 6.956279 | 3.194837 | 3.388986 | 0.434424 | 2.955907 | 2.449682 | 0.38362 | 1.615666 | 2.63907 | 0.041034 | 0.232642 | 5.117671 | 2.170206 | 1.462966 | 1.780973 |
| 36.47222 | 6.998771 | 0.482894 | 4.792042 | 9.660069 | 10.919392 | 4.342876 | 3.782439 | 2.676795 | 10.471919 | 6.663755 | 3.313012 | 3.146624 | 0.591105 | 3.067888 | 3.318857 | 0.333083 | 2.517478 | 2.772763 | 0.030622 | 0.404421 | 4.288341 | 1.982941 | 1.987674 | 2.383261 |
| 36.48611 | 9.296412 | 1.108625 | 5.748812 | 11.661758 | 13.951514 | 5.068513 | 3.052389 | 2.718364 | 9.71071 | 4.89345 | 3.164367 | 3.127386 | 0.760033 | 2.669845 | 4.508788 | 0.346489 | 3.627722 | 2.866534 | -0.029074 | 0.556568 | 2.952735 | 0.837405 | 2.520264 | 3.810259 |
| 36.5 | 10.144066 | 3.474686 | 6.883476 | 13.069167 | 14.387571 | 4.704714 | 1.658419 | 3.005888 | 8.341782 | 3.392657 | 3.411435 | 2.619066 | 0.651078 | 1.561568 | 5.32231 | 0.316223 | 4.418682 | 2.539158 | -0.068792 | 0.812999 | 2.867176 | 0.474608 | 2.634858 | 4.377393 |
| 36.51389 | 6.532341 | 4.38953 | 6.824105 | 13.787446 | 13.483698 | 5.623127 | 1.287971 | 3.026171 | 5.423423 | 2.446711 | 2.418901 | 3.865689 | 0.64001 | 1.258748 | 5.144292 | 0.319678 | 4.194666 | 2.842603 | 0.057216 | 0.825845 | 2.437503 | 0.302149 | 2.673985 | 4.506123 |
| 36.52778 | 3.432652 | 2.56131 | 6.71883 | 14.473211 | 14.746655 | 4.149156 | 0.963612 | 2.766352 | 3.507733 | 3.104748 | 2.612622 | 3.560934 | 0.609114 | 1.289084 | 4.843875 | 0.426765 | 3.866258 | 2.233625 | 0.112171 | 0.633052 | 2.667104 | 0.021261 | 2.713684 | 4.09736 |
| 36.54167 | 3.302743 | 0.745187 | 5.026281 | 15.311868 | 15.4564 | 3.27213 | 0.278817 | 2.379786 | 2.864867 | 3.729973 | 1.236856 | 0.727575 | 0.594561 | 1.025205 | 3.73857 | 0.575121 | 4.928807 | 1.282279 | -0.037743 | 0.497836 | 1.835524 | -0.117552 | 2.848387 | 3.597045 |
| 36.55556 | -0.436699 | -1.782185 | 3.1203 | 6.095163 | 6.780005 | 2.242311 | 0.598482 | 1.512726 | 2.983926 | 3.053024 | 0.571966 | -0.258759 | 0.358322 | 0.466922 | 2.821 | 0.398841 | 8.932787 | 1.271699 | -0.068821 | 0.429015 | 0.911322 | 0.07641 | 2.219756 | 1.537964 |
| 36.56944 | -1.883405 | -1.173401 | 2.798605 | -3.857592 | -4.6251 | 1.833049 | 1.794003 | 1.447923 | 1.14566 | 0.511697 | 0.51382 | 1.150791 | 0.491871 | 0.562172 | 2.079667 | 0.027349 | 6.855965 | 1.956791 | 0.110543 | 0.256992 | 1.091304 | 0.416809 | 1.594292 | -0.464559 |
| 36.58333 | -0.834844 | -0.517616 | 3.287601 | -1.103417 | -4.817489 | 2.441592 | 2.793071 | 1.145636 | 0.602558 | -0.113293 | 1.167652 | 1.889529 | 0.702741 | -0.335193 | 1.117606 | -0.106585 | 5.56117 | 2.831426 | 0.116251 | 0.249266 | 1.761302 | 0.410079 | 1.924018 | -1.909413 |
| 36.59722 | -0.122776 | 0.203631 | 4.128061 | 9.253055 | 2.815631 | 4.008245 | 2.162659 | 1.651694 | 1.398273 | 1.146172 | 2.479043 | 2.580825 | 0.603073 | 0.295077 | 0.241494 | 0.002794 | 6.152939 | 2.628166 | 0.003704 | 0.386703 | 1.549644 | 0.021359 | 2.382788 | -0.155149 |
| 36.61111 | 4.15609 | 0.700808 | 2.629833 | 16.593876 | 7.744871 | 4.884197 | 2.224948 | 1.281764 | 1.928743 | 1.676887 | 2.508682 | 2.431241 | 0.370976 | 0.555492 | -0.396581 | 0.232762 | 7.021584 | 2.188692 | -0.134004 | 0.253854 | 1.164089 | 0.235384 | 2.625055 | 2.307718 |
| 36.625 | 6.729353 | 0.830158 | 4.614993 | 14.548597 | 8.73163 | 4.216309 | 2.110254 | 0.02803 | 1.526642 | 0.707396 | 1.596798 | 1.507452 | 0.381208 | 0.586431 | -0.221868 | 0.287596 | 6.40036 | 1.134147 | -0.032321 | -0.054412 | 1.035921 | -0.355428 | 1.518489 | 3.384748 |
| 36.63889 | 8.063788 | 0.566306 | 2.705795 | 6.580896 | 7.191631 | 2.306583 | 1.681084 | 0.231554 | 1.079392 | -0.005997 | 1.735434 | 0.710014 | 0.648793 | 0.634697 | 0.203644 | 0.164241 | 5.749948 | 0.434186 | 0.055347 | 0.005019 | 0.827592 | -0.2751 | 0.688895 | 2.232747 |
| 36.65278 | 6.241303 | 0.406739 | -0.247785 | 1.85737 | 2.103166 | 0.03217 | 1.752883 | 0.567762 | 0.892009 | 0.106301 | 1.482494 | 1.570006 | 0.806512 | 0.485326 | 0.822161 | 0.145334 | 5.162787 | -0.024291 | 0.146144 | 0.042032 | 0.669031 | -0.023024 | 0.160102 | 1.238776 |
| 36.66667 | 3.715308 | 0.207724 | -2.12057 | -2.091762 | 0.943535 | -0.653046 | 2.76773 | 0.449493 | 0.970251 | 0.29547 | 1.307663 | 1.287984 | 1.144557 | 0.579682 | 0.583764 | 0.233148 | 4.213636 | -1.378154 | -0.022423 | 0.112968 | -0.01468 | 0.105865 | -0.299178 | 0.400476 |
| 36.68056 | 1.09737 | -0.108639 | -0.833358 | -3.27062 | 0.166066 | 0.008479 | 3.358491 | 0.116348 | 1.364996 | 0.35266 | 0.662894 | 0.864666 | 1.151621 | 0.436505 | 0.026575 | 0.302297 | 4.739727 | -1.715153 | 0.041475 | 0.135422 | 0.132202 | 0.025336 | -0.618751 | 0.305299 |
| 36.69444 | -1.610649 | -0.175255 | 3.366751 | -5.597255 | 0.198883 | 1.133655 | 2.367848 | -0.129647 | 1.49364 | 0.227873 | 0.01084 | 1.342767 | 0.918234 | 0.147408 | -0.089802 | 0.270308 | 3.922984 | 3.734304 | 0.263594 | 0.242711 | 0.53858 | 0.083855 | -1.429154 | 0.131334 |
| 36.70833 | -0.644253 | 0.588872 | 0.963933 | -1.716011 | 0.471251 | 1.892295 | 1.61798 | -0.110576 | 1.201934 | 0.22898 | -0.261703 | 0.358829 | 0.666663 | -0.085331 | -0.37606 | 0.161273 | 2.465238 | 0.962727 | 0.122434 | 0.244983 | 0.381552 | 0.062249 | -1.655405 | -0.315361 |
| 36.72222 | 0.037806 | 0.907695 | 2.582441 | 3.611343 | 0.499181 | 1.166623 | 0.321993 | -0.035849 | 0.567761 | 0.152971 | -0.278517 | -0.576014 | 0.455448 | -0.408179 | -1.009488 | 0.036079 | -0.13395 | 0.20878 | 0.042838 | 0.085488 | 0.080748 | 0.035016 | -1.35083 | -0.900782 |
| 36.73611 | -0.276329 | 0.435705 | 5.009445 | 6.069352 | -0.187081 | -0.362039 | -1.530183 | 0.144897 | 0.078814 | 0.151155 | -0.028549 | -0.625613 | 0.237101 | -0.219276 | -0.829349 | -0.105308 | -1.784247 | 0.207725 | 0.073502 | -0.139023 | -1.920065 | -0.011489 | -2.887212 | -1.516359 |
| 36.75 | -1.436076 | 0.121552 | 2.405897 | -1.52471 | -2.994884 | -1.109485 | -2.987951 | 0.280001 | 0.30224 | 0.233842 | -0.146043 | -0.179816 | 0.024347 | -0.216322 | -0.442641 | -0.212489 | -1.878164 | -0.453626 | -0.007424 | -0.150043 | -3.184118 | 0.020479 | -2.31487 | -1.475209 |
| 36.76389 | -3.372079 | -0.094986 | 0.619474 | -11.390757 | -4.785346 | -0.418793 | -2.250848 | 0.384147 | 0.484188 | 0.298722 | -0.241858 | 0.178637 | -0.05149 | -0.209145 | -0.184646 | -0.240312 | -2.437809 | -1.249865 | -0.118014 | -0.092493 | -1.943238 | 0.102279 | -1.382527 | -0.436739 |
| 36.77778 | -3.180805 | -0.512245 | -3.19881 | -13.757829 | -3.829328 | -0.040182 | -0.218883 | 0.44266 | 0.011891 | 0.21539 | -0.084802 | 0.010799 | 0.585787 | -0.211526 | -0.021626 | -0.108342 | -0.791717 | 0.083266 | -0.045481 | -0.134158 | 0.031743 | 0.166551 | 1.401022 | 0.722289 |
| 36.79167 | -1.192769 | -1.185931 | -5.891966 | -10.479632 | -1.639543 | 0.057079 | 1.44779 | 0.004724 | -0.488979 | 0.11526 | 0.536331 | 0.741289 | 1.158822 | -0.131701 | -0.054034 | 0.091135 | 0.515684 | 1.344369 | 0.101243 | -0.131264 | 2.01233 | 0.026318 | 3.518173 | 1.025024 |
| 36.80556 | -0.814367 | 0.384002 | 0.186879 | 0.678017 | 1.70981 | -0.392151 | 1.559671 | 0.098101 | -0.291005 | 0.003116 | 2.020172 | 1.943311 | 1.242951 | 0.067698 | -0.570194 | 0.184092 | 1.509994 | 1.117676 | -0.055017 | -0.061064 | 2.137085 | 0.006397 | 1.530635 | 0.271149 |
| 36.81944 | -0.209828 | 1.741593 | 2.33329 | 10.608387 | 0.508509 | -0.194065 | 0.875406 | 0.404058 | 0.106939 | -0.009777 | 3.095428 | 2.256429 | 0.805325 | 0.65217 | -0.169923 | 0.144063 | 2.113995 | 0.151743 | -0.160126 | 0.00641 | 1.525697 | 0.026144 | 0.315046 | 0.009746 |
| 36.83333 | -0.309617 | 1.430601 | 1.754016 | 10.831341 | -1.052691 | -0.271703 | -0.374351 | 0.497087 | 0.433347 | 0.102922 | 1.698324 | 2.082094 | 0.274755 | 0.682639 | 0.357817 | 0.093378 | 1.385505 | 0.028665 | -0.053074 | 0.078718 | 0.402891 | 0.100618 | -1.02449 | 0.207229 |
| 36.84722 | 0.68819 | 0.401445 | -0.009027 | 3.05234 | -2.084475 | -2.369786 | -1.406095 | 0.43756 | 0.605511 | 0.156282 | -0.936479 | 0.700385 | -0.209482 | 0.357604 | 0.998765 | 0.05205 | -1.201679 | 0.2913 | 0.154661 | 0.11189 | 0.390108 | 0.222509 | -1.519849 | 0.30147 |
| 36.86111 | 1.42514 | 0.326701 | -1.571028 | -4.661691 | -5.467261 | -0.776841 | 0.687519 | 0.217977 | 0.624717 | 0.001452 | -3.108467 | -1.881351 | -0.357532 | 0.074873 | 1.506555 | -0.031884 | -3.803581 | 0.796183 | 0.325303 | 0.025188 | -0.024106 | 0.174774 | -2.256444 | -0.244497 |
| 36.875 | -0.448997 | -1.441588 | 1.857856 | -5.267691 | -5.71211 | 3.933202 | 1.588153 | -0.418998 | 0.488451 | -0.107209 | -1.593011 | -1.890019 | -0.309708 | -0.156995 | 1.659763 | -0.020178 | -4.154697 | -1.596103 | -0.007815 | -0.111856 | 0.291023 | 0.083667 | 1.966296 | -1.56288 |
| 36.88889 | -1.863061 | -1.196118 | 3.533926 | -6.212812 | -5.24177 | 3.961708 | 0.417038 | 0.063894 | 0.01321 | 0.02428 | -0.303968 | -1.449729 | -0.147331 | -0.436709 | 0.431547 | 0.108698 | -3.242045 | -2.796204 | 0.070707 | -0.095011 | 0.218888 | 0.1985 | -0.701995 | -1.164361 |
| 36.90278 | -1.163475 | 0.293807 | 4.760498 | 7.062587 | -1.91822 | 0.890655 | 0.096631 | 0.608537 | -0.027008 | 0.159951 | 0.762126 | 0.181934 | -0.258724 | -0.539995 | -0.870412 | 0.229305 | -1.575511 | 0.989582 | 0.463387 | 0.004255 | 0.179284 | 0.287281 | -1.696453 | 0.2309 |
| 36.91667 | -1.47385 | 1.453538 | 4.684551 | 9.086679 | -2.034822 | -2.137481 | 1.063776 | 0.993874 | -0.280384 | 0.092899 | 0.382344 | 1.119113 | -0.298182 | -0.41042 | -2.792281 | 0.137395 | -0.240189 | 1.35728 | 0.121793 | 0.016537 | -2.305274 | 0.765109 | -1.92969 | 0.889867 |
| 36.93056 | -0.38863 | 2.582207 | 2.695323 | 9.053431 | 1.150023 | -1.505456 | 1.001961 | 0.591193 | -0.726855 | -0.40222 | -0.238541 | 0.426709 | -0.298636 | -0.365503 | -2.381026 | -0.055494 | -0.021321 | 0.876582 | 0.264304 | -0.001541 | -1.937587 | 0.962572 | -3.38682 | -1.021839 |
| 36.94444 | 0.33643 | -0.018516 | 1.901129 | 6.436638 | 4.085718 | -0.209169 | 1.487331 | 0.458844 | -1.764403 | -0.622574 | -1.82901 | -1.184305 | -0.271477 | -0.521312 | -0.286165 | -0.136562 | -1.088652 | 0.139524 | 0.182095 | 0.017134 | 1.328773 | 0.832698 | -3.347311 | -2.294869 |
| 36.95833 | -1.112466 | -0.062428 | -0.134143 | -1.730901 | 1.891494 | 0.152284 | 0.554679 | 0.207325 | -1.480766 | -0.812976 | 1.672334 | -0.528625 | -0.365934 | -0.593173 | 1.175181 | -0.220007 | -3.09416 | -1.240903 | -0.08004 | 0.005544 | 0.676652 | 0.118982 | -1.357241 | -2.393291 |
| 36.97222 | -5.971319 | -0.269944 | 0.79034 | -6.606186 | -5.483235 | -1.134904 | -0.647897 | 0.051438 | -1.121137 | -0.670022 | -0.421721 | 0.204303 | -0.493689 | -0.612855 | 0.613706 | -0.436377 | -5.021915 | -1.663346 | -0.226332 | 0.055377 | -3.41308 | -1.176633 | 0.887518 | -2.301768 |
| 36.98611 | -5.996639 | -0.514997 | -0.75217 | -7.707737 | -9.647119 | -1.817171 | -1.134003 | -0.409106 | -2.228604 | -0.271881 | -4.076144 | -0.431758 | -0.467198 | -0.768477 | -1.132678 | -0.498915 | -6.051441 | -3.71827 | -0.439211 | -0.199818 | -3.220874 | -2.170585 | -0.200638 | -1.042575 |
| 37 | -3.927318 | -1.616802 | -4.711916 | -6.331035 | -6.890866 | -3.277013 | -0.976233 | -0.426326 | -2.148414 | -0.450718 | -5.281378 | -1.503521 | -0.443682 | -0.278299 | -3.651198 | -0.176506 | -6.21812 | -4.472944 | -0.311919 | -0.146242 | -2.122263 | -0.733686 | -2.010316 | -0.682945 |
| 37.01389 | -2.259744 | -2.108845 | -2.975023 | -2.619774 | -5.426974 | 0.016228 | -1.394017 | -1.766776 | -1.515085 | -0.818757 | -8.378587 | -7.614667 | -0.40777 | 0.213991 | -3.805189 | 0.10404 | -5.659891 | -2.485159 | 0.172809 | -0.280511 | -1.975957 | 1.380755 | -3.702431 | -0.155803 |
| 37.02778 | -2.869407 | -1.522099 | -2.760463 | 0.515628 | -6.137448 | 2.470687 | -0.857117 | -1.821605 | -3.991366 | -1.326179 | -2.556291 | -8.503756 | -0.459881 | 0.094826 | -2.167081 | 0.088195 | -5.306019 | -0.634536 | 0.360287 | -0.586479 | -3.142057 | 1.09443 | -4.19537 | 0.24609 |
| 37.04167 | -5.558698 | -1.708973 | -5.214064 | -1.484315 | -7.703061 | 0.29025 | -2.140454 | -1.701759 | -7.671713 | -3.059886 | -3.111203 | -3.700819 | -0.351514 | -0.274818 | -0.935954 | -0.051033 | -5.813312 | -0.770199 | 0.094149 | -0.239602 | -2.894397 | -0.097169 | -2.567268 | -0.279891 |
| 37.05556 | -7.217383 | -2.540835 | -5.324612 | -1.786584 | -12.454594 | 0.475441 | -4.55147 | -2.106993 | -8.522431 | -7.977074 | -2.692992 | -2.327972 | -0.373858 | -1.062478 | -1.427165 | -0.172082 | -5.885362 | -1.569068 | -0.465091 | -0.04336 | -4.923935 | -1.325363 | -1.691459 | -1.631974 |
| 37.06944 | -0.596965 | 1.48271 | -2.815095 | -3.669388 | -10.82199 | -1.374465 | -6.45272 | -2.988164 | -9.38472 | -11.660547 | -5.104119 | -1.141548 | -0.398343 | -1.535077 | -2.181842 | -0.299159 | -4.869504 | -1.853514 | -0.599262 | -0.290783 | -3.468763 | -2.149628 | -0.930629 | -2.24332 |
| 37.08333 | 2.255465 | 1.989931 | -3.746552 | -4.780396 | -5.998751 | -3.706102 | -3.158617 | -2.71223 | -8.847892 | -11.776679 | -4.850785 | 1.762583 | -0.349817 | -1.623839 | -2.247492 | -0.421303 | -3.097563 | -1.77916 | -0.656864 | -0.779539 | -1.867074 | -3.389112 | -0.371871 | -2.420503 |
| 37.09722 | 5.292822 | 0.13949 | -1.982233 | -6.216342 | -3.349154 | -1.297175 | 0.901259 | -1.22146 | -3.844876 | -5.399487 | 0.850628 | 2.137161 | -0.357527 | -2.103095 | -2.229458 | -0.387956 | -1.743263 | -1.653145 | -0.673739 | -0.713089 | -1.35694 | -3.165585 | -0.195963 | -1.991106 |
| 37.11111 | 10.908661 | -1.888265 | -0.203493 | -5.844652 | -3.946063 | 0.353303 | 1.671853 | -0.861772 | -1.085696 | -2.070759 | 6.858262 | 2.145432 | -0.475528 | -2.214446 | -2.405848 | -0.280335 | -1.153258 | -1.572934 | -0.772571 | -0.59912 | -1.725256 | -3.5567 | -0.447868 | -1.848702 |
| 37.125 | 6.659374 | -0.583471 | -0.884385 | -5.03087 | -2.340632 | 0.201685 | 1.096165 | -2.201559 | -1.404815 | -2.176798 | 3.930167 | 1.772833 | -0.548894 | -2.751775 | -2.55852 | -0.20436 | -0.925847 | -1.6923 | -0.431443 | -0.583297 | -1.773519 | -3.260592 | -0.928551 | -1.946654 |
| 37.13889 | -2.048368 | -3.197128 | -1.446565 | -2.181774 | -3.641105 | -1.215225 | -0.009117 | -2.472512 | -2.542097 | -3.422269 | -2.916251 | -2.774739 | -0.706458 | -2.383139 | -2.308008 | -0.219577 | -0.873589 | -1.747987 | -0.602016 | -0.53939 | -1.415702 | -2.447063 | -1.090782 | -1.960545 |
| 37.15278 | -7.500062 | -5.097644 | -1.675219 | -0.980142 | -6.458161 | -1.261399 | -1.950836 | -1.532439 | -3.581112 | -3.613283 | -9.036558 | -6.604147 | -0.793006 | -2.106365 | -1.652016 | -0.229014 | -0.920341 | -1.593194 | -0.579282 | -0.345145 | -0.975567 | -1.569958 | -0.886664 | -1.552005 |
| 37.16667 | -10.825921 | -2.152482 | -1.385022 | -0.898118 | -7.523564 | -1.357278 | -3.330395 | -0.950237 | -4.279992 | -3.166199 | -9.244964 | -6.414211 | -0.814528 | -1.717939 | -1.236435 | -0.222885 | -0.899018 | -1.326289 | -0.441203 | -0.301711 | -0.646142 | -0.910801 | -0.74046 | -0.978175 |
| 37.18056 | -10.198017 | -3.688971 | -1.822437 | -1.562796 | -5.92909 | -2.250191 | -4.288764 | -0.29369 | -4.46845 | -3.113952 | -5.2298 | -6.100156 | -0.779189 | -1.2716 | -1.326086 | -0.210315 | -0.758309 | -1.115739 | -0.293523 | -0.239041 | -0.298512 | -0.347496 | -0.674697 | -0.569031 |
| 37.19444 | -8.639977 | -5.501918 | -2.209236 | -3.329146 | -5.198738 | -4.165771 | -4.074499 | 0.296201 | -5.464438 | -4.789649 | -2.442968 | -6.180932 | -0.730356 | -1.043919 | -1.73478 | -0.261471 | -0.681661 | -1.010046 | -0.201445 | -0.129213 | -0.069495 | 0.064375 | -0.830695 | -0.593057 |
| 37.20833 | -7.465562 | -6.520685 | -2.593981 | -4.341027 | -3.513131 | -4.282662 | -3.852978 | 0.679583 | -6.678139 | -8.348914 | -1.693107 | -5.012417 | -0.615376 | -1.224914 | -2.029884 | -0.307394 | -0.825672 | -1.101992 | -0.139677 | 0.005942 | -0.459595 | 0.118307 | -0.938531 | -0.83366 |
| 37.22222 | -3.6332 | -6.155498 | -2.190382 | -3.861676 | -1.667925 | -1.717231 | -4.547824 | 0.845387 | -5.349565 | -9.205125 | -1.798644 | -3.41313 | -0.446083 | -1.743187 | -2.105223 | -0.376376 | -1.134022 | -1.347957 | -0.057409 | 0.067234 | -1.650197 | -0.356061 | -0.764822 | -0.976037 |
| 37.23611 | -0.755171 | -3.933692 | -0.545383 | -1.149474 | -0.478872 | -0.218672 | -2.397737 | 0.43591 | -2.675292 | -5.955118 | -1.274822 | -1.523486 | -0.390803 | -2.689582 | -1.72852 | -0.43903 | -1.497246 | -1.632409 | -0.066884 | -0.056986 | -3.3731 | -1.508976 | -0.353385 | -0.86711 |
| 37.25 | -0.273419 | -2.557008 | 0.929917 | 1.289039 | 0.766393 | 0.263519 | -0.23614 | -0.102953 | -1.355049 | -3.745165 | 0.231616 | 0.217865 | -0.427279 | -3.606168 | -1.194802 | -0.400271 | -1.79503 | -1.642523 | -0.21233 | -0.25513 | -4.504678 | -3.104615 | -0.118824 | -0.775721 |
| 37.26389 | -1.467785 | -1.972018 | 1.484545 | 1.938457 | 0.639609 | 0.928775 | 1.057083 | -0.732884 | 0.069024 | -1.223329 | 3.057652 | 1.362675 | -0.413655 | -3.458909 | -0.62996 | -0.322756 | -1.892283 | -1.685253 | -0.374204 | -0.383351 | -4.746806 | -4.375307 | -0.164743 | -0.697178 |
| 37.27778 | -2.162444 | -1.566468 | 0.911733 | 0.677502 | -0.304724 | 1.767361 | 1.928516 | -1.231286 | 1.575975 | 2.012356 | 5.299618 | 1.178589 | -0.414366 | -3.196446 | -0.238099 | -0.28775 | -2.014005 | -2.141511 | -0.53338 | -0.472507 | -3.774943 | -5.028528 | -0.364606 | -0.749137 |
| 37.29167 | -1.64206 | -1.174421 | 1.106553 | -0.591832 | -0.145559 | 2.617821 | 3.230998 | -1.080356 | 2.369525 | 4.230239 | 6.318614 | 1.578167 | -0.433846 | -3.358463 | -0.189412 | -0.287761 | -2.259161 | -2.337775 | -1.099772 | -0.556844 | -3.304858 | -5.896146 | -0.478207 | -0.642043 |
| 37.30556 | -0.224313 | 1.788484 | 3.02221 | -0.278163 | 1.309355 | 3.498123 | 4.531113 | -0.518673 | 2.906444 | 4.276692 | 5.918891 | 3.299802 | -0.4376 | -3.367056 | -0.307811 | -0.259606 | -2.428195 | -2.379978 | -1.274397 | -0.586693 | -2.612311 | -6.243743 | -0.390402 | -0.551594 |
| 37.31944 | 1.922072 | 6.585716 | 4.844998 | 1.149814 | 3.195996 | 3.549017 | 4.56818 | -0.169916 | 4.348354 | 5.389365 | 5.285973 | 5.918134 | -0.336541 | -2.551608 | -0.244048 | -0.226819 | -2.268566 | -1.721471 | -1.082744 | -0.540844 | -1.982254 | -3.822198 | -0.186254 | -0.403417 |
| 37.33333 | 4.511636 | 7.760336 | 4.537158 | 2.644112 | 5.114664 | 3.63168 | 3.611671 | -0.02303 | 6.578819 | 7.183218 | 6.283122 | 6.723336 | -0.239883 | -1.404453 | 0.188046 | -0.191775 | -1.85953 | -0.911357 | -0.572825 | -0.396983 | -0.938277 | -0.938798 | -0.149237 | -0.280331 |
| 37.34722 | 6.115004 | 7.293158 | 2.715206 | 3.39525 | 5.768779 | 3.607056 | 3.154033 | -0.05186 | 7.882853 | 9.441715 | 7.58125 | 4.986606 | -0.190633 | -0.034833 | 0.929742 | -0.142513 | -1.351684 | -0.12143 | 0.026045 | -0.259345 | 0.279467 | 1.902065 | -0.343858 | -0.176826 |
| 37.36111 | 4.283535 | 6.701768 | 1.431727 | 3.310313 | 4.715492 | 3.832948 | 3.770037 | 0.147699 | 8.405856 | 9.829811 | 7.85358 | 3.847292 | -0.127487 | 1.08663 | 1.849073 | -0.058454 | -0.890873 | 0.615624 | 0.469657 | -0.163479 | 1.76766 | 4.68061 | -0.496562 | -0.082284 |
| 37.375 | 3.386959 | 6.344815 | 2.168981 | 3.824423 | 4.703995 | 4.59138 | 4.631043 | 0.414833 | 10.291477 | 10.748027 | 5.275031 | 3.514089 | -0.026901 | 1.709408 | 2.600229 | 0.014294 | -0.489239 | 1.144898 | 0.720661 | -0.100003 | 3.195659 | 5.563183 | -0.280297 | 0.002265 |
| 37.38889 | 3.518358 | 5.683242 | 3.28122 | 4.757265 | 5.431044 | 4.72757 | 6.014329 | 0.570241 | 11.014623 | 9.874214 | 1.445493 | 2.153367 | 0.043366 | 1.763738 | 2.803872 | 0.166049 | -0.191024 | 1.361133 | 0.507398 | 0.000101 | 4.353816 | 4.674192 | 0.271073 | 0.078085 |
| 37.40278 | 4.314436 | 5.258055 | 3.815239 | 5.250099 | 6.518278 | 4.22403 | 6.735387 | 0.705056 | 10.327529 | 8.717961 | 0.943577 | 0.985574 | -0.021189 | 1.418588 | 2.850764 | 0.317581 | -0.35181 | 1.48346 | 0.01472 | 0.123199 | 4.418765 | 3.876572 | 0.706658 | 0.064721 |
| 37.41667 | 5.08644 | 2.146031 | 3.014197 | 6.211588 | 6.213856 | 4.347638 | 7.03621 | 0.999167 | 10.730634 | 9.046745 | 7.088169 | 5.468391 | -0.1369 | 1.249631 | 2.667515 | 0.261094 | 0.020404 | 1.698037 | -0.336771 | 0.172139 | 5.637163 | 3.705656 | 0.85216 | 0.086314 |
| 37.43056 | 6.84622 | 0.377658 | 2.151438 | 7.448324 | 7.761448 | 4.803751 | 6.967035 | 1.048856 | 11.548046 | 10.969494 | 5.530052 | 5.817864 | 0.20663 | 1.710168 | 2.147648 | 0.389951 | 1.025138 | 1.956989 | 0.002436 | 0.10321 | 6.089776 | 3.633248 | 1.069658 | 0.265415 |
| 37.44444 | 5.86368 | 1.401057 | -0.064785 | 7.856124 | 8.964239 | 4.792832 | 6.860494 | 0.678207 | 8.841215 | 8.677455 | 5.59781 | 4.938869 | 0.58787 | 2.156574 | 1.637461 | 0.514319 | 2.167455 | 2.211956 | 0.405168 | -0.036534 | 5.780023 | 3.438255 | 1.50188 | 0.553139 |
| 37.45833 | 5.37666 | 1.123651 | 3.120777 | 8.142138 | 11.059173 | 5.180555 | 7.415305 | 0.741118 | 7.496574 | 6.127265 | 1.612679 | 1.094932 | 0.543681 | 3.016959 | 1.830416 | 0.527926 | 2.3504 | 2.435148 | 0.629791 | 0.157457 | 3.512863 | 2.223592 | 1.780645 | 0.728589 |
| 37.47222 | 7.081021 | 0.738456 | 4.870246 | 8.531873 | 11.701188 | 5.258779 | 5.976244 | 0.829906 | 5.463271 | 2.774711 | -1.275806 | -0.796967 | 0.628803 | 3.208401 | 2.663323 | 0.315016 | 2.476748 | 2.69728 | 0.736261 | 0.219077 | 2.102586 | 1.23037 | 1.673424 | 0.980405 |
| 37.48611 | 4.976445 | 1.287315 | 3.072713 | 8.376624 | 9.374116 | 3.042009 | 3.151519 | 1.181787 | 4.455211 | 1.471579 | 4.862773 | 5.900024 | 0.694563 | 2.489427 | 3.42223 | 0.144047 | 3.06666 | 3.872561 | 0.196534 | 0.27039 | 3.556871 | 1.561535 | 1.570631 | 1.20921 |
| 37.5 | 1.087095 | 0.492924 | 2.835115 | 6.825706 | 9.853316 | 1.153136 | 1.030051 | 1.364646 | 3.258885 | 0.849949 | 8.726375 | 6.306583 | 0.653144 | 1.259468 | 3.665667 | 0.053055 | 3.641108 | 3.929058 | -0.096904 | 0.332063 | 3.815693 | 0.680371 | 1.466797 | 2.030529 |
| 37.51389 | 4.95161 | -0.28003 | 2.009071 | 5.026202 | 11.009099 | 2.585001 | 0.560512 | 1.099645 | 4.715029 | 1.054241 | 5.315394 | 2.523572 | 0.580537 | 1.478295 | 2.997588 | 0.014069 | 3.42266 | 2.385895 | 0.055485 | 0.476619 | 2.207426 | 0.689559 | 1.720063 | 2.445288 |
| 37.52778 | 3.974736 | -0.120125 | 5.6788 | 4.857346 | 10.51792 | 3.587195 | 0.594348 | 1.094603 | 5.065857 | 0.623726 | 2.268604 | 1.733053 | 0.53691 | 0.906115 | 2.630322 | 0.179123 | 3.192099 | -0.028652 | -0.077273 | 0.57856 | 2.614475 | 0.558613 | 2.309523 | 1.206295 |
| 37.54167 | 4.092623 | 1.095588 | 8.345871 | 8.044249 | 10.692969 | 5.576358 | 0.159297 | 1.128892 | 2.510674 | -0.212038 | 0.338001 | 2.604344 | 0.427186 | 0.663042 | 2.120619 | 0.456233 | 4.369352 | -1.104422 | -0.317583 | 0.732893 | 2.850471 | 0.380866 | 1.995384 | -0.384737 |
| 37.55556 | 6.663319 | -0.112433 | 3.783181 | 10.088531 | 6.390704 | 5.217866 | 0.024108 | 0.519492 | 1.280139 | -1.161964 | 3.609221 | 1.724215 | 0.265443 | -0.027276 | 0.921164 | 0.356492 | 5.677038 | 0.080478 | 0.030372 | 0.950639 | 0.977109 | -0.22574 | 1.063046 | -1.375199 |
| 37.56944 | 4.771857 | -0.251155 | 1.005004 | 5.968489 | -1.791819 | 3.345581 | 0.146937 | 0.476928 | 0.849404 | -0.551524 | 3.534717 | 0.752866 | 0.50458 | -0.993823 | -0.439327 | 0.121738 | 3.909546 | 1.316977 | 0.477025 | 0.918116 | 0.015068 | 0.176986 | -0.49932 | -1.575847 |
| 37.58333 | -0.193603 | -0.330489 | -1.488529 | 0.052292 | -6.474569 | 1.754288 | 0.384801 | 0.677451 | 0.9861 | 0.017979 | 2.726643 | 0.568982 | 0.837844 | -0.509731 | 0.721906 | -0.022643 | 2.220495 | 1.415171 | 0.631035 | 0.845243 | -2.255735 | 0.39138 | 0.120813 | -0.431583 |
| 37.59722 | -2.691433 | 0.065794 | 0.780669 | -3.63632 | -1.105588 | 0.861512 | 0.57636 | 0.761018 | 0.689938 | 0.20746 | 1.375893 | 0.685981 | 0.900386 | 0.791265 | 1.618142 | 0.059936 | 2.99452 | 1.468261 | 0.164652 | 1.012503 | -0.462946 | 0.717724 | 2.118042 | 0.092785 |
| 37.61111 | 2.354333 | -0.490612 | 2.83373 | -3.352602 | 3.292981 | 1.239662 | 0.721015 | 1.532963 | 0.995338 | 0.246602 | 2.082928 | 1.842206 | 0.807023 | 0.894183 | 1.755804 | 0.222432 | 4.745717 | 1.970248 | -0.269941 | 1.081703 | 2.008015 | -0.10356 | 1.8481 | 1.089441 |
| 37.625 | 4.064585 | -0.973661 | 2.217273 | 1.703099 | 6.32751 | 1.862394 | 1.113971 | 0.923409 | 0.822094 | 0.259986 | 1.997908 | 1.614769 | 0.642253 | 0.855114 | 1.147776 | 0.279734 | 6.266658 | 1.138992 | -0.206426 | 0.243766 | 3.167046 | -0.696139 | 1.861125 | 2.611631 |
| 37.63889 | 3.644055 | 1.271771 | 3.611599 | 4.450092 | 6.172426 | 1.664789 | 1.417784 | 1.230936 | 0.404171 | 0.237656 | 1.935009 | 1.259591 | 0.88293 | 0.334953 | -0.166468 | 0.112808 | 5.96951 | 0.94282 | -0.135906 | 0.044834 | 1.834916 | -0.377436 | 1.832565 | 1.751672 |
| 37.65278 | 3.957604 | 1.66238 | 3.629407 | 18.107154 | 5.431784 | 2.318239 | 1.533263 | 0.612051 | 0.355836 | 0.181382 | 1.949574 | 1.453369 | 1.280037 | 0.448358 | -1.028638 | -0.037823 | 4.781223 | 0.029843 | -0.05957 | 0.093843 | 0.495984 | -0.058238 | 0.383712 | 0.403237 |
| 37.66667 | 3.649354 | 1.09442 | -0.534504 | 2.828691 | 3.151197 | 2.132015 | 1.183563 | 0.235936 | 0.412091 | 0.135855 | 2.280934 | 1.722532 | 1.001506 | 0.380335 | -1.492477 | 0.270373 | 3.522767 | -1.1805 | 0.062887 | -0.011872 | -0.605142 | -0.056029 | -3.25603 | -0.700474 |
| 37.68056 | 2.975042 | 0.840892 | -3.858969 | 6.417991 | 1.509814 | 1.501133 | 0.527975 | 0.056541 | 0.491494 | 0.08771 | 1.258432 | 1.097823 | 0.754142 | -0.142049 | -1.325078 | 0.349921 | 3.349376 | -1.829458 | -0.077108 | -0.0479 | -2.719848 | -0.138355 | -0.212159 | -0.13399 |
| 37.69444 | 1.444113 | 0.254538 | -2.237924 | 7.425131 | 0.563296 | -0.121325 | 0.330015 | 0.146026 | 0.149502 | 0.004037 | 0.56752 | 0.33539 | 0.537517 | -0.729664 | 0.384461 | 0.03479 | 1.992174 | -2.044239 | -0.048541 | 0.025276 | -5.053542 | 0.046877 | -0.377231 | 0.28559 |
| 37.70833 | 0.701511 | 0.031349 | -0.402073 | 7.71308 | 1.563658 | -0.931268 | 0.528299 | 0.262869 | -0.056307 | -0.027583 | -0.398491 | 0.012263 | 0.4003 | -0.000414 | 0.759456 | -0.065558 | 0.111745 | 0.680891 | 0.094073 | 0.104163 | -0.665401 | 0.271622 | 0.253585 | 1.204947 |
| 37.72222 | 1.468958 | -0.001385 | -0.032543 | 3.576433 | 3.516465 | -0.68048 | 0.753904 | 0.148423 | 0.000996 | 0.007569 | -0.900342 | -0.139248 | 0.30395 | 1.232233 | 1.254034 | 0.051111 | -0.380421 | 3.486342 | 0.215561 | 0.05671 | 0.793108 | 0.369766 | -0.080609 | 1.696011 |
| 37.73611 | 2.975742 | 0.127827 | 1.170519 | 0.999806 | 5.082957 | -0.023573 | 0.773358 | 0.014467 | -0.138999 | 0.097654 | -0.556644 | -0.320737 | 0.462123 | 0.118486 | 1.392953 | 0.345909 | 0.06961 | 1.445904 | 0.129696 | -0.117869 | 0.52088 | 0.143634 | 1.706219 | 1.981195 |
| 37.75 | 0.534492 | 0.26336 | 1.491883 | -0.886337 | -0.443637 | 0.212009 | 0.43753 | 0.002237 | 0.343809 | 0.10735 | -0.904882 | -0.48363 | 0.641823 | -0.305128 | 1.100118 | 0.251115 | 0.401565 | 0.95666 | -0.000773 | -0.117982 | 0.698107 | -0.523635 | 1.514913 | 0.366242 |
| 37.76389 | -2.537636 | 0.129963 | 0.5614 | -4.727352 | -2.831339 | 0.258199 | -0.16244 | 0.093476 | 0.536144 | 0.202523 | 0.043788 | 0.362236 | 0.76417 | -0.30432 | -0.279817 | 0.159607 | 0.384765 | 0.187463 | 0.071468 | -0.150663 | 0.903118 | -1.321521 | -0.591622 | -0.487583 |
| 37.77778 | -1.603207 | -0.001892 | -1.150782 | -3.124835 | -4.651685 | 0.507796 | -0.113277 | 0.14395 | 0.679606 | 0.231542 | 0.977703 | 1.785692 | 0.513989 | -0.124914 | -1.264532 | 0.031141 | 0.005694 | 0.035851 | -0.027929 | -0.084885 | -0.55145 | -1.647822 | -0.736105 | -1.192141 |
| 37.79167 | -1.412963 | 0.041554 | -0.894851 | 2.793972 | -5.164947 | 1.996146 | -0.25602 | -0.147192 | 0.744449 | 0.14732 | 1.620124 | 1.321708 | 0.388282 | -0.355894 | 0.151404 | 0.009318 | -0.362441 | 0.108445 | 0.043229 | -0.117173 | -2.659257 | -0.815627 | 0.228988 | -1.255984 |
| 37.80556 | -1.600853 | 0.060624 | 1.799605 | 8.431194 | -6.227511 | 2.342628 | -0.321039 | -0.626773 | 0.190903 | 0.128041 | 1.850683 | 0.579948 | 0.478207 | -0.387274 | 1.029767 | 0.026377 | -0.747318 | -0.531794 | 0.181187 | -0.16479 | -2.899102 | 0.016334 | -0.684766 | -0.621406 |
| 37.81944 | -1.349145 | 0.095596 | 3.228534 | 3.211355 | -6.441853 | 1.47018 | -0.426967 | 0.012566 | -0.318593 | 0.143841 | 0.39003 | -0.506462 | 0.418928 | -0.495782 | 0.904915 | 0.080154 | -0.568952 | -1.592275 | 0.01697 | -0.177455 | -1.093565 | 0.139326 | 0.006882 | -0.88037 |
| 37.83333 | -1.700069 | 0.090003 | -0.466032 | -3.895937 | -3.23227 | 0.811106 | 0.047896 | 0.647049 | -0.327105 | 0.158516 | -0.702279 | -1.294287 | 0.155388 | 0.304926 | 0.325083 | 0.104211 | -0.65372 | -1.600626 | -0.225827 | -0.092545 | 1.913107 | 0.321319 | -0.45929 | -1.16745 |
| 37.84722 | -1.803843 | 0.124795 | -0.40542 | -8.128627 | 0.275592 | -0.009908 | 0.685714 | 0.767538 | 0.095727 | 0.229554 | -1.478493 | -0.835559 | 0.103277 | 1.020248 | -0.449595 | 0.063312 | -1.053939 | 2.962258 | -0.119516 | 0.007757 | 1.840624 | 0.421282 | -0.051556 | -0.278799 |
| 37.86111 | 0.320011 | 0.222734 | -1.973967 | -7.851544 | 3.549878 | -0.396697 | 0.907937 | 0.223415 | 0.600415 | 0.285619 | -1.662704 | -0.458229 | 0.233785 | 1.173554 | -1.209223 | -0.018092 | -1.723392 | 2.376669 | -0.00075 | 0.067622 | 2.507425 | 0.725961 | -0.174902 | 0.602873 |
| 37.875 | 2.087535 | -0.926842 | -0.85153 | -4.606991 | 4.412384 | -0.469634 | 0.743238 | 0.378792 | -0.123069 | 0.202817 | -1.296261 | -0.203884 | -0.121348 | 0.335582 | -0.081768 | 0.003963 | -2.765799 | 1.10947 | 0.073644 | -0.071133 | 0.845692 | 0.308642 | -1.423559 | 1.184505 |
| 37.88889 | 1.222354 | -0.079511 | 0.906921 | -6.992429 | 1.626254 | -0.269157 | 0.235732 | 0.10664 | -1.357367 | 0.051608 | -0.527449 | 0.502331 | -0.658321 | 0.118721 | 1.437818 | 0.089986 | -3.284456 | 0.767071 | 0.004184 | -0.114657 | 0.134509 | -0.039638 | 0.867897 | 0.900087 |
| 37.90278 | 0.094769 | 0.233907 | -3.816547 | 0.934487 | -2.816118 | -0.217476 | -0.510248 | -0.244684 | 0.644446 | -0.132899 | 0.324277 | 0.870914 | -0.536916 | 0.142684 | 1.363187 | 0.145053 | -3.481214 | -0.037131 | 0.054796 | 0.005226 | 0.263861 | -0.231059 | 0.108417 | 0.815049 |
| 37.91667 | -1.656476 | 0.658253 | -5.901762 | 1.092447 | -7.825596 | -0.807153 | -1.320056 | -0.713002 | 2.318434 | -0.140687 | 0.685206 | 1.442194 | -0.095829 | 0.006475 | 1.397159 | 0.199194 | -3.893729 | -0.744696 | 0.117738 | 0.04587 | -2.118325 | -0.939625 | 0.930926 | 0.56849 |
| 37.93056 | -0.495002 | 0.444901 | -4.986358 | 3.031868 | -6.719176 | -2.735454 | -2.355835 | -0.364861 | 0.85458 | 0.024493 | -0.083652 | 1.147781 | -0.078155 | -0.181574 | 0.014761 | 0.140428 | -3.892096 | -3.050758 | 0.029933 | -0.079762 | -2.075662 | -1.466658 | -0.511774 | -0.234334 |
| 37.94444 | 1.481634 | 0.480435 | 1.104529 | 1.823929 | -2.498247 | -0.353751 | -1.713277 | -0.788624 | -0.098348 | 0.370447 | -0.331481 | 0.213856 | -0.458049 | -0.225981 | -2.422609 | -0.150856 | -2.740311 | -5.373412 | 0.173274 | -0.314323 | -1.911352 | -1.001891 | -1.783429 | -1.937729 |
| 37.95833 | -0.736056 | -0.0142 | 3.309234 | 0.28956 | 1.368716 | 1.584012 | 0.530077 | -0.30351 | -0.837842 | 0.145809 | -1.660416 | -0.47943 | -0.117083 | -0.742145 | -2.976293 | -0.068381 | -2.186277 | -3.461976 | 0.020494 | -0.503455 | -1.399762 | -0.974801 | -1.782936 | -2.181889 |
| 37.97222 | -6.458668 | 0.218536 | 3.666823 | -1.178237 | 1.654184 | 0.167222 | 1.051268 | -0.082769 | -1.129162 | -0.920179 | -3.900727 | -0.600628 | 0.223664 | -0.909613 | -2.960841 | -0.374805 | -1.935842 | -1.546135 | 0.186922 | -0.624128 | -1.557163 | -0.069173 | -1.388548 | -2.345133 |
| 37.98611 | -8.214178 | 0.174469 | -2.141183 | -5.170638 | -1.014425 | -1.329518 | -0.186916 | -0.121952 | -0.854733 | 0.071692 | -5.036373 | -1.990643 | 0.15189 | -1.941163 | -0.6883 | -0.620893 | -2.241141 | -0.518653 | 0.13741 | -0.772796 | -1.112523 | 0.413786 | -0.952963 | -2.295095 |
| | | | | | | | | | | | | | -1.371459 | -3.143307 | 1.968568 | -0.481058 | -3.377959 | -0.823952 | 0.294442 | -1.125547 | -2.982957 | 0.355572 | -1.209261 | -1.989668 |

**Supplementary Table S2.** Raw motion data from: **(c)** 100 mM NaCl treatment in Rocket

| Time | RocketControl1 | RocketControl10 | RocketControl13 | RocketControl2 | RocketControl3 | RocketControl4 | RocketControl5 | RocketControl6 | RocketControl7 | RocketControl8 | RocketControl9 | Rocket100mMNaCl1 | Rocket100mMNaCl10 | Rocket100mMNaCl11 | Rocket100mMNaCl12 | Rocket100mMNaCl2 | Rocket100mMNaCl3 | Rocket100mMNaCl4 | Rocket100mMNaCl5 | Rocket100mMNaCl6 | Rocket100mMNaCl7 | Rocket100mMNaCl8 | Rocket100mMNaCl9 |
|---|---|---|---|---|---|---|---|---|---|---|---|---|---|---|---|---|---|---|---|---|---|---|---|
| 18 | 0.166493 | 0.378636 | 0.015637 | -0.17327 | 0.043714 | -0.086632 | -0.115775 | 0.756694 | 0.039343 | -0.242163 | 0.298942 | 0.335548 | -0.003429 | 1.59596 | -0.112659 | -0.313457 | -0.530449 | -0.137041 | -0.197136 | -0.478729 | -0.15085 | 0.067854 | -0.528794 |
| 18.01389 | 0.139426 | 0.444956 | -0.029992 | -0.237723 | 0.484612 | -0.033586 | -0.040617 | 1.131965 | 0.080063 | 0.03302 | 0.29075 | 0.662808 | 0.318743 | 1.885257 | 0.20486 | 0.069128 | -0.353208 | 0.066263 | -0.0216 | -0.226887 | 1.015836 | 0.420638 | -0.23878 |
| 18.02778 | 0.242086 | 0.452709 | -0.043286 | -0.144012 | 0.646378 | 0.040497 | 0.007332 | 1.008509 | 0.165363 | 0.660136 | 0.243998 | 0.775436 | 0.579937 | 2.076519 | 0.308942 | 0.412947 | -0.141955 | 0.262914 | 0.22873 | -0.101873 | 0.778332 | 0.700973 | 0.015878 |
| 18.04167 | 0.435931 | 0.436221 | 0.079771 | 0.001393 | 0.606069 | 0.063769 | 0.057499 | 0.725313 | 0.238986 | 0.817014 | 0.290485 | 0.742613 | 0.583089 | 2.071497 | 0.291831 | 0.603106 | 0.075638 | 0.429246 | 0.3673 | -0.053051 | -0.325824 | 0.787902 | 0.182942 |
| 18.05556 | 0.559504 | 0.395925 | 0.163722 | 0.057078 | 0.362366 | 0.034305 | 0.091891 | 0.554202 | 0.215026 | 0.665746 | 0.398803 | 0.764514 | 0.455703 | 1.924827 | 0.227037 | 0.618684 | 0.153453 | 0.461025 | 0.322587 | -0.069466 | -0.825317 | 0.731117 | 0.226124 |
| 18.06944 | 0.42882 | 0.28702 | 0.064199 | 0.034893 | -0.077437 | -0.054124 | 0.029673 | 0.41177 | 0.041042 | 0.508452 | 0.266713 | 0.592626 | 0.31823 | 1.67284 | 0.147818 | 0.586807 | -0.017941 | 0.300736 | 0.287118 | -0.096202 | 0.59783 | 0.641386 | 0.195562 |
| 18.08333 | 0.300893 | 0.248249 | -0.010308 | 0.042954 | -0.347673 | -0.133644 | -0.102173 | 0.425693 | -0.215849 | 0.429906 | 0.198256 | -0.111045 | -0.55652 | 0.891987 | -0.435116 | -0.05248 | -1.197873 | -0.385754 | -0.130575 | -0.634531 | 2.20138 | -0.140566 | -0.285734 |
| 18.09722 | 0.376748 | 0.277492 | 0.097198 | 0.114085 | -0.199533 | -0.171845 | -0.227412 | 0.82785 | -0.384485 | 0.418968 | 0.496468 | -1.506664 | -2.51786 | -0.587904 | -1.878567 | -1.639814 | -4.577074 | -1.885957 | -1.405596 | -2.268175 | 0.978184 | -1.930977 | -1.472078 |
| 18.11111 | 0.314705 | 0.168868 | 0.231944 | 0.205236 | 0.189948 | -0.172648 | -0.23052 | 0.925502 | -0.287551 | 0.450731 | 0.430866 | -2.226577 | -3.008049 | -1.101614 | -2.524029 | -2.287083 | -4.166055 | -2.309638 | -2.112774 | -2.769298 | -1.302988 | -2.672157 | -2.022659 |
| 18.125 | 0.291202 | 0.102614 | 0.19723 | 0.311677 | 0.764417 | 0.006269 | -0.063493 | 0.694203 | 0.112881 | 0.90346 | 0.310475 | -1.574425 | -1.319722 | 0.210827 | -1.219331 | -1.090984 | -1.750692 | -0.855062 | -1.041084 | -1.461981 | -2.310637 | -1.017716 | -1.25641 |
| 18.13889 | 0.449065 | 0.176344 | 0.166094 | 0.59215 | 1.341845 | 0.206371 | 0.086562 | 0.489993 | 0.424476 | 1.315387 | 0.471708 | -0.566532 | -0.080487 | 1.881972 | 0.433104 | -0.004376 | -0.392159 | 0.285509 | 0.064054 | -0.594103 | -2.710262 | 0.573564 | -0.404884 |
| 18.15278 | 0.449302 | 0.412416 | 0.078516 | 0.666789 | 1.173377 | 0.288476 | 0.091373 | 0.293168 | 0.415567 | 1.096191 | 0.550011 | 0.280352 | 0.478631 | 1.363038 | 1.067969 | 0.346163 | 0.17335 | 0.361255 | 0.508266 | -0.300573 | -2.51794 | 1.008923 | 0.046224 |
| 18.16667 | 0.444551 | 0.903186 | 0.049441 | 0.54019 | 0.651769 | 0.321184 | 0.056934 | 0.529626 | 0.256267 | 0.760718 | 0.51099 | 0.706921 | 0.8065 | 0.569407 | 0.95096 | 0.420388 | 0.249942 | 0.153306 | 0.465526 | -0.305358 | -1.924974 | 0.537364 | 0.246785 |
| 18.18056 | 0.59768 | 1.246476 | 0.187998 | 0.518465 | 0.201488 | 0.340956 | 0.072578 | 0.916655 | 0.225029 | 0.486561 | 0.549815 | 0.712752 | 0.926074 | 0.090165 | 0.817903 | 0.454839 | 0.159909 | 0.08908 | 0.369027 | -0.172914 | -1.037227 | 0.153538 | 0.330177 |
| 18.19444 | 0.732886 | 1.007053 | 0.417856 | 0.526194 | 0.043018 | 0.300684 | 0.072908 | 1.43104 | 0.279373 | 0.55871 | 0.714717 | 0.514708 | 0.890772 | 0.130061 | 1.037116 | 0.507908 | 0.021348 | 0.217568 | 0.399581 | 0.142107 | 0.759659 | 0.049491 | 0.391926 |
| 18.20833 | 0.816055 | 0.522874 | 0.429552 | 0.482567 | 0.10372 | 0.200063 | 0.013795 | 1.520745 | 0.192635 | 0.993709 | 0.825625 | 0.374572 | 0.616668 | 0.728949 | 1.394679 | 0.510479 | 0.028148 | 0.470649 | 0.429988 | 0.389261 | 2.071006 | 0.105616 | 0.452481 |
| 18.22222 | 0.785203 | 0.218246 | 0.28212 | 0.301332 | 0.466149 | 0.073067 | -0.04391 | 1.074883 | -0.022134 | 1.463541 | 0.69887 | 0.427476 | 0.377773 | 1.176083 | 1.655097 | 0.439057 | 0.233459 | 0.662476 | 0.386379 | 0.529653 | 1.311256 | 0.326612 | 0.45918 |
| 18.23611 | 0.715915 | 0.083824 | 0.131667 | 0.158997 | 0.780603 | 0.031557 | -0.039679 | 0.567085 | -0.154136 | 1.48864 | 0.441981 | 0.629069 | 0.549809 | 1.049554 | 1.639411 | 0.403206 | 0.512814 | 0.743639 | 0.32436 | 0.723683 | -0.171565 | 0.682537 | 0.481173 |
| 18.25 | 0.655272 | 0.012367 | 0.040015 | 0.117955 | 0.875486 | 0.039747 | 0.020244 | 0.119956 | -0.140871 | 1.289747 | 0.23687 | 0.770791 | 0.905853 | 0.73337 | 1.473836 | 0.449233 | 0.756281 | 0.67258 | 0.32147 | 0.841198 | -1.59019 | 0.982649 | 0.515059 |
| 18.26389 | 0.479491 | -0.052086 | -0.004598 | 0.052196 | 0.677936 | -0.001866 | 0.035882 | -0.336231 | -0.111664 | 1.032241 | 0.041692 | 0.715355 | 0.987107 | 0.462558 | 1.209831 | 0.473666 | 0.821436 | 0.490477 | 0.289804 | 0.846869 | -2.731197 | 1.000184 | 0.509274 |
| 18.27778 | 0.291216 | -0.062038 | -0.016038 | -0.028317 | -0.048773 | -0.097214 | -0.02684 | -0.595419 | -0.096352 | 0.70755 | -0.166532 | 0.516369 | 0.642167 | 0.172191 | 0.937517 | 0.420344 | 0.707932 | 0.283764 | 0.246861 | 0.720073 | -1.423969 | 0.640056 | 0.436148 |
| 18.29167 | 0.146791 | 0.024118 | -0.008858 | -0.035658 | -0.593482 | -0.145731 | -0.053701 | -0.520091 | 0.001612 | 0.41477 | -0.290434 | 0.320945 | 0.294728 | -0.177426 | 0.605109 | 0.397412 | 0.53039 | 0.179453 | 0.335046 | 0.566315 | 1.424592 | 0.296918 | 0.34492 |
| [illegible] | [illegible] | [illegible] | [illegible] | [illegible] | [illegible] | [illegible] | [illegible] | [illegible] | [illegible] | [illegible] | [illegible] | [illegible] | [illegible] | [illegible] | [illegible] | [illegible] | [illegible] | [illegible] | [illegible] | [illegible] | [illegible] | [illegible] | [illegible] |

| | | | | | | | | | | | | | | | | | | | | | | | |
|---|---|---|---|---|---|---|---|---|---|---|---|---|---|---|---|---|---|---|---|---|---|---|---|
| 21.26389 | -0.35834 | -0.214632 | 0.21297 | 0.009138 | -1.10774 | -0.287156 | -0.204077 | 0.388288 | 0.16854 | 0.716187 | 0.11788 | 0.412257 | -0.381166 | 1.484351 | 0.384066 | 0.158741 | 0.221552 | 0.096411 | -0.126411 | 0.007956 | 0.165873 | 0.200174 | 0.237099 |
| 21.27778 | -0.210164 | -0.273439 | 0.415941 | -0.140617 | 0.132287 | -0.351957 | -0.000893 | 0.911591 | -0.010496 | 0.962825 | -0.08305 | 0.261506 | -0.400342 | 2.344213 | 1.255693 | 0.166863 | -0.043589 | -0.08731 | 0.140241 | 0.009731 | 0.035505 | 0.137336 | 0.10122 |
| 21.29167 | 0.043278 | -0.252182 | 0.446692 | 0.064682 | 2.203683 | -0.33525 | 0.171929 | 0.566086 | -0.078467 | 1.118674 | -0.177859 | 0.269544 | -0.254167 | 1.957039 | 1.542861 | 0.016992 | -0.266274 | -0.157751 | 0.405987 | -0.021392 | -0.143255 | 0.08786 | -0.031962 |
| 21.30556 | 0.339533 | -0.464408 | 0.321543 | 0.32471 | 2.754958 | -0.006525 | 0.215215 | 0.018218 | 0.001877 | 1.028484 | -0.193321 | 0.224593 | -0.181677 | 0.832367 | 1.038867 | -0.142933 | -0.633367 | -0.19851 | 0.395174 | -0.05711 | -0.304134 | 0.108904 | -0.101589 |
| 21.31944 | 0.459729 | -0.771536 | 0.169991 | 0.324188 | 1.775291 | -0.034396 | 0.047155 | -0.490528 | 0.160012 | 0.66835 | -0.211817 | 0.037318 | -0.288144 | -0.078327 | 0.174166 | -0.156795 | -0.799479 | -0.348437 | 0.123972 | -0.074293 | -0.33641 | -0.054663 | -0.119781 |
| 21.33333 | 0.167563 | -0.798943 | -0.051512 | 0.138062 | 1.117061 | -0.044578 | -0.066997 | -0.788636 | 0.265555 | 0.10121 | -0.266263 | -0.219415 | -0.506043 | -1.098849 | -0.831018 | -0.111024 | -0.403312 | -0.565506 | -0.24102 | -0.090095 | -0.267912 | -0.099533 | -0.108537 |
| 21.34722 | -0.272199 | -0.478445 | -0.395933 | -0.089397 | 0.661203 | -0.074128 | -0.165201 | -0.761735 | 0.279556 | -0.549426 | -0.210943 | -0.459558 | -0.739 | -2.473881 | -1.639047 | -0.190861 | 0.291598 | -0.662984 | -0.509444 | -0.108719 | -0.208565 | -0.219888 | -0.115887 |
| 21.36111 | -0.460559 | -0.22686 | -0.437084 | -0.258336 | 0.011673 | -0.053048 | -0.181263 | -0.373123 | 0.282466 | -0.928998 | -0.163395 | -0.562212 | -0.939133 | -3.263129 | -1.842932 | -0.288134 | 0.763618 | -0.529053 | -0.463485 | -0.060649 | -0.203303 | -0.066305 | -0.109995 |
| 21.375 | -0.423607 | -0.193911 | -0.407355 | -0.268445 | -1.800122 | 0.133572 | -0.142513 | -0.21573 | 0.291861 | -0.97311 | -0.209975 | -0.509204 | -1.069192 | -2.119614 | -1.565973 | -0.262355 | 0.72515 | -0.305049 | -0.210684 | -0.01336 | -0.197283 | -0.065483 | -0.103133 |
| 21.38889 | -0.271477 | -0.293507 | -0.174631 | -0.287375 | -3.28471 | 0.136212 | -0.07548 | 0.016938 | 0.260864 | -0.802492 | -0.274383 | -0.387117 | -1.078098 | -0.628484 | -1.270382 | -0.137669 | 0.208394 | -0.270171 | -0.092683 | -0.134807 | -0.182538 | -0.095108 | -0.192757 |
| 21.40278 | -0.103207 | -0.447578 | -0.101471 | -0.382122 | -3.070238 | -0.122184 | -0.077879 | 0.294478 | 0.086892 | -0.677002 | -0.422039 | -0.269811 | -1.004352 | 0.418305 | -1.028021 | -0.174418 | -0.401064 | -0.454301 | -0.205163 | -0.315264 | -0.198353 | -0.185445 | -0.336878 |
| 21.41667 | 0.039719 | -0.61933 | -0.155028 | -0.510888 | -1.582937 | -0.398399 | -0.153861 | -0.010265 | -0.155607 | -0.565753 | -0.681141 | -0.293346 | -0.913874 | 1.036604 | -0.66343 | -0.428404 | -0.7476 | -0.638434 | -0.404386 | -0.362775 | -0.19636 | -0.307663 | -0.451108 |
| 21.43056 | 0.126618 | -0.804129 | -0.201261 | -0.519146 | 0.154049 | -0.33287 | -0.093619 | -0.750124 | -0.32533 | -0.419446 | -0.689593 | -0.291453 | -0.769006 | 1.092664 | -0.338546 | -0.648569 | -0.859724 | -0.6815 | -0.581913 | -0.259153 | -0.041586 | -0.277518 | -0.473302 |
| 21.44444 | 0.238535 | -1.133111 | -0.173329 | -0.456708 | 1.213619 | -0.093517 | -0.011206 | -1.219075 | -0.339646 | -0.142923 | -0.559341 | -0.114272 | -0.732459 | 0.474868 | -0.249949 | -0.787315 | -0.648944 | -0.567342 | -0.714497 | -0.094031 | 0.187223 | -0.154672 | -0.444836 |
| 21.45833 | 0.229669 | -1.415883 | -0.298886 | -0.336758 | 1.109175 | -0.046607 | -0.060664 | -0.940813 | -0.466359 | -0.016742 | -0.38457 | 0.045674 | -0.824283 | -0.617487 | -0.435943 | -0.843743 | -0.16791 | -0.383394 | -0.727371 | 0.022208 | 0.264081 | -0.060054 | -0.491962 |
| 21.47222 | 0.10505 | -1.451514 | -0.48372 | -0.276884 | 0.628913 | -0.149487 | -0.217423 | -0.679051 | -0.553654 | -0.116849 | -0.081334 | -0.082586 | -0.836491 | -1.702134 | -0.838155 | -0.761274 | 0.259025 | -0.175259 | -0.621515 | 0.022181 | 0.011512 | -0.069725 | -0.640892 |
| 21.48611 | -0.121344 | -1.133057 | -0.599774 | -0.365966 | 0.596675 | -0.277055 | -0.340241 | -0.499443 | -0.46071 | -0.341251 | 0.201586 | -0.38534 | -0.691463 | -2.512402 | -1.292573 | -0.630624 | 0.466294 | -0.134668 | -0.516854 | 0.008877 | -0.489138 | -0.071087 | -0.654787 |
| 21.5 | -0.409109 | -0.885706 | -0.74696 | -0.442339 | 0.474282 | -0.355574 | -0.44256 | -0.666728 | -0.459889 | -0.480956 | 0.205484 | -0.617511 | -0.601687 | -2.644184 | -1.612913 | -0.689167 | 0.495906 | -0.431016 | -0.525604 | -0.017011 | -0.754562 | -0.008426 | -0.524342 |
| 21.51389 | -0.582379 | -0.796534 | -0.993615 | -0.431632 | -0.151571 | -0.416539 | -0.575556 | -0.720214 | -0.556935 | -0.55571 | 0.033524 | -0.839162 | -0.738457 | -2.103037 | -1.572934 | -1.007576 | 0.354955 | -0.710902 | -0.671933 | -0.087231 | -0.631996 | -0.083554 | -0.455004 |
| 21.52778 | -0.561116 | -0.739395 | -0.824382 | -0.475176 | -0.21866 | -0.337436 | -0.571389 | -0.549384 | -0.552435 | -0.545942 | -0.155904 | -1.082109 | -0.710951 | -1.462904 | -1.384914 | -1.328524 | 0.053995 | -0.603561 | -0.645635 | -0.084529 | -0.403895 | -0.139579 | -0.464614 |
| 21.54167 | -0.479004 | -0.966849 | -0.711239 | -0.767421 | -0.007952 | -0.276175 | -0.447009 | -0.934989 | -0.803853 | -0.661084 | -0.279218 | -1.235965 | -0.635147 | -1.275461 | -1.168112 | -1.685096 | -0.378084 | -0.47759 | -0.629668 | 0.052929 | -0.295015 | -0.334345 | -0.444746 |
| 21.55556 | -0.356447 | -1.349305 | -0.997951 | -0.921501 | -0.701039 | -0.457207 | -0.396678 | -1.413689 | -0.92805 | -0.892437 | -0.387427 | -0.942404 | -0.795362 | -1.602692 | -1.172071 | -1.875005 | -0.617066 | -0.716285 | -0.700002 | 0.23886 | -0.231807 | -0.636556 | -0.493587 |
| 21.56944 | -0.271149 | -1.458188 | -1.230991 | -0.833521 | -1.254528 | -0.53697 | -0.48397 | -1.491738 | -1.023644 | -1.019352 | -0.438202 | -0.508202 | -0.877594 | -2.089123 | -1.104382 | -1.356893 | -0.626657 | -0.883913 | -0.671739 | 0.440418 | -0.085299 | -0.486575 | -0.51512 |
| 21.58333 | -0.23284 | -1.610613 | -1.257319 | -0.721536 | -1.525762 | -0.483816 | -0.479123 | -1.643178 | -1.009688 | -1.066704 | -0.490933 | -0.255153 | -0.603409 | -2.434897 | -0.865158 | -0.912131 | -0.772008 | -0.570128 | -0.585726 | 0.578125 | 0.190169 | -0.02292 | -0.330563 |
| 21.59722 | -0.321305 | -2.209584 | -1.326692 | -0.728329 | -1.586389 | -0.442806 | -0.588051 | -2.112257 | -0.916237 | -1.23294 | -0.66532 | 0.103044 | -0.160008 | -2.547645 | -0.47962 | -0.845517 | -0.733032 | -0.150902 | -0.156552 | 0.700164 | 0.537786 | 0.3527 | -0.020486 |
| 21.61111 | -0.523343 | -2.364689 | -1.566484 | -0.770272 | -1.643017 | -0.549781 | -0.792395 | -2.876447 | -0.877347 | -1.332321 | -1.027174 | 0.805095 | 0.010844 | -2.406163 | -0.183928 | -0.962305 | -0.088929 | -0.131883 | 0.412603 | 0.857071 | 0.810057 | 0.533929 | 0.23464 |
| 21.625 | -0.930413 | -1.75287 | -1.499499 | -0.672108 | -1.3002 | -0.670829 | -0.675866 | -3.144201 | -0.756099 | -1.116901 | -1.371756 | 1.30121 | -0.189563 | -1.887349 | -0.166449 | -0.801956 | 0.548774 | -0.384136 | 0.674677 | 0.878874 | 0.817491 | 0.429666 | 0.410633 |
| 21.63889 | -1.214694 | -1.268986 | -1.027107 | -0.402712 | -0.812441 | -0.642116 | -0.352801 | -1.938813 | -0.48656 | -0.802942 | -1.301373 | 1.14346 | -0.408509 | -1.209916 | -0.384307 | -0.656779 | 1.214996 | -0.312262 | 0.429944 | 0.706256 | 0.526816 | 0.051467 | 0.652414 |
| 21.65278 | -0.779416 | -0.923601 | -0.461032 | -0.101894 | -0.497726 | -0.373829 | -0.129368 | -0.784452 | -0.243202 | -0.770625 | -1.00578 | 0.53507 | -0.33764 | -1.1514 | -0.382623 | -0.448911 | 1.599048 | -0.257359 | 0.046252 | 0.474398 | 0.111882 | -0.180862 | 0.746662 |
| 21.66667 | -0.144284 | -0.661599 | -0.090583 | -0.04841 | -0.551806 | -0.061612 | 0.018634 | 0.039137 | 0.015595 | -0.75826 | -0.490836 | 0.075078 | -0.157834 | -1.168321 | -0.148124 | -0.025195 | 1.448327 | -0.357818 | -0.135901 | 0.326872 | -0.148134 | -0.320043 | 0.617145 |
| 21.68056 | 0.265359 | -0.504722 | 0.07017 | -0.228721 | -0.519402 | -0.018108 | 0.028336 | 0.580663 | 0.197913 | -0.756976 | -0.093033 | -0.046809 | 0.036017 | -1.134982 | 0.142448 | 0.325272 | 1.346092 | -0.35532 | -0.123911 | 0.248613 | -0.26607 | -0.298018 | 0.407396 |
| 21.69444 | 0.474893 | -0.41151 | 0.081735 | -0.384598 | -0.484417 | -0.258593 | -0.036723 | 0.637245 | 0.160967 | -0.661026 | 0.195669 | -0.193027 | -0.020834 | -1.030461 | 0.191766 | 0.409683 | 0.976195 | -0.29494 | -0.225418 | 0.160373 | -0.324077 | -0.312534 | 0.098595 |
| 21.70833 | 0.545991 | -0.373946 | 0.088903 | -0.232907 | -0.438507 | -0.270708 | -0.102948 | 0.412017 | -0.002412 | -0.321137 | 0.376345 | -0.403112 | -0.188747 | -0.909886 | 0.130965 | 0.308954 | 0.427026 | -0.220874 | -0.412772 | 0.082498 | -0.445335 | -0.328519 | -0.179833 |
| 21.72222 | 0.487464 | -0.324281 | 0.146253 | -0.218689 | -0.336842 | -0.093176 | -0.13846 | 0.246526 | -0.15906 | -0.081415 | 0.36252 | -0.556225 | -0.239151 | -0.670084 | 0.099352 | 0.261902 | 0.035477 | -0.224804 | -0.535872 | -0.036221 | -0.456245 | -0.33775 | -0.320957 |
| 21.73611 | 0.433391 | -0.228768 | 0.171357 | -0.340153 | -0.182032 | 0.00861 | -0.142412 | 0.195199 | -0.235107 | -0.045163 | 0.178438 | -0.725316 | -0.339516 | -0.428263 | -0.03361 | 0.187545 | -0.294749 | -0.445968 | -0.61743 | -0.349979 | -0.515508 | -0.48232 | -0.50576 |
| 21.75 | 0.285635 | -0.15084 | 0.109178 | -0.168735 | -0.041833 | -0.0353 | -0.157924 | 0.129205 | -0.368735 | -0.123315 | -0.001694 | -0.920504 | -0.537458 | -0.309655 | -0.324121 | -0.021251 | -0.551095 | -0.800114 | -0.771771 | -0.716289 | -0.705218 | -0.778154 | -0.712521 |
| 21.76389 | 0.091411 | -0.134444 | -0.011847 | 0.010038 | 0.079067 | 0.008192 | -0.210083 | 0.029173 | -0.532421 | -0.253837 | -0.154136 | -0.893163 | -0.48828 | -0.084806 | -0.399355 | -0.032526 | -0.610215 | -0.916504 | -0.758636 | -0.685033 | -0.645364 | -0.885924 | -0.666377 |
| 21.77778 | -0.049002 | -0.146281 | -0.183298 | -0.203184 | 0.199568 | 0.197596 | -0.196676 | -0.152669 | -0.586591 | -0.333018 | -0.243533 | -0.575645 | -0.12032 | 0.274834 | -0.152179 | 0.412587 | -0.28834 | -0.743849 | -0.473923 | -0.222697 | -0.386652 | -0.540033 | -0.334548 |
| 21.79167 | -0.135405 | -0.231651 | -0.423765 | -0.301141 | 0.252779 | 0.226728 | -0.144096 | -0.456979 | -0.483951 | -0.339143 | -0.300831 | -0.207247 | 0.155567 | 0.379465 | 0.035142 | 0.829643 | -0.037449 | -0.562007 | -0.185924 | 0.133328 | -0.201656 | -0.247495 | -0.132567 |
| 21.80556 | -0.201137 | -0.295674 | -0.59466 | -0.174794 | 0.178767 | 0.026059 | -0.133048 | -0.766598 | -0.352094 | -0.327697 | -0.331314 | -0.033643 | 0.164918 | 0.112633 | 0.016617 | 0.962837 | 0.279836 | -0.486971 | -0.062082 | 0.080161 | -0.203534 | -0.785773 | -0.166567 |
| 21.81944 | -0.246714 | -0.258882 | -0.604535 | -0.147133 | 0.072438 | -0.047631 | -0.147982 | -0.901543 | -0.194318 | -0.296341 | -0.300571 | -0.058089 | 0.062825 | -0.160304 | 0.011133 | 0.894979 | 0.545022 | -0.399028 | -0.03823 | -0.175771 | -0.299953 | -0.164365 | -0.18908 |
| 21.83333 | -0.421304 | -0.144942 | -0.513026 | -0.254533 | -0.006662 | -0.048621 | -0.13014 | -0.852292 | -0.028454 | -0.246178 | -0.20331 | -0.121674 | -0.110893 | -0.259292 | 0.036598 | 0.858294 | 0.528711 | -0.235909 | -0.08199 | -0.354127 | -0.446064 | -0.298685 | -0.216748 |
| 21.84722 | -0.686602 | -0.017525 | -0.408572 | -0.199712 | -0.084715 | -0.105856 | -0.066627 | -0.715308 | 0.107527 | -0.214431 | -0.081166 | -0.054947 | -0.274757 | -0.315423 | 0.084338 | 0.892402 | 0.229313 | 0.043066 | -0.091665 | -0.279657 | -0.515716 | -0.354011 | -0.243933 |
| 21.86111 | -0.889048 | 0.077764 | -0.273996 | -0.060469 | -0.171896 | -0.234521 | 0.005 | -0.562551 | 0.208125 | -0.157822 | 0.049186 | 0.198361 | -0.21618 | -0.216852 | 0.254685 | 1.012711 | 0.070238 | 0.421893 | 0.043812 | -0.045964 | -0.35201 | -0.257908 | -0.141838 |
| 21.875 | -0.683655 | 0.265784 | -0.097842 | -0.051307 | -0.185322 | -0.217856 | 0.069273 | -0.376788 | 0.278649 | -0.074583 | 0.150155 | 0.317339 | -0.117923 | -0.074805 | 0.376686 | 0.937386 | 0.07605 | 0.655205 | 0.172041 | 0.025939 | -0.223468 | -0.138548 | -0.089872 |
| 21.88889 | -0.224274 | 0.643483 | 0.042485 | -0.045391 | -0.110481 | -0.104707 | 0.149588 | -0.151085 | 0.32546 | -0.037812 | 0.182351 | 0.274206 | -0.086981 | 0.032944 | 0.470452 | 0.717607 | 0.03144 | 0.821426 | 0.360617 | -0.151059 | -0.275925 | 0.088362 | -0.098847 |
| 21.90278 | 0.267572 | 1.018254 | 0.162468 | 0.122451 | 0.009496 | 0.061569 | 0.268292 | 0.142183 | 0.414058 | 0.000567 | 0.218048 | 0.263195 | 0.05252 | 0.224684 | 0.60113 | 0.64812 | 0.116313 | 1.142878 | 0.116972 | -0.273114 | -0.311647 | 0.468868 | 0.027254 |
| 21.91667 | 0.720153 | 1.308553 | 0.275141 | 0.343501 | 0.216692 | 0.243722 | 0.398575 | 0.481467 | 0.576051 | 0.137578 | 0.3395 | 0.453498 | 0.323252 | 0.544165 | 0.763627 | 0.76052 | 0.501347 | 0.947312 | 0.170957 | -0.091791 | -0.219636 | 0.532889 | 0.08871 |
| 21.93056 | 1.136053 | 1.63577 | 0.287224 | 0.334932 | 0.469461 | 0.4287 | 0.403669 | 0.69138 | 0.795382 | 0.244537 | 0.429176 | 0.625206 | 0.509348 | 0.915942 | 0.95347 | 0.805195 | 1.152644 | 0.421024 | 0.107017 | 0.318285 | -0.032978 | 0.354121 | 0.033073 |
| 21.94444 | 1.162578 | 1.814743 | 0.273678 | 0.212393 | 0.685338 | 0.583346 | 0.361955 | 0.691408 | 0.96675 | 0.257292 | 0.486358 | 0.659445 | 0.645108 | 1.157718 | 1.125737 | 0.715886 | 1.529302 | 0.660203 | 0.419848 | 0.65128 | 0.170226 | 0.682239 | 0.061342 |
| 21.95833 | 0.79346 | 1.624133 | 0.322868 | 0.170543 | 0.792038 | 0.598511 | 0.373664 | 0.652241 | 1.006485 | 0.261143 | 0.575646 | 0.712007 | 0.91055 | 1.102973 | 1.250073 | 0.557389 | 1.384635 | 1.004513 | 0.583513 | 0.745899 | 0.326064 | 0.672501 | 0.19454 |
| 21.97222 | 0.526898 | 1.640889 | 0.490379 | 0.202521 | 0.741103 | 0.500482 | 0.388881 | 0.579299 | 1.068496 | 0.341784 | 0.739794 | 0.850898 | 1.317946 | 1.031785 | 1.255445 | 0.406621 | 0.821707 | 1.003133 | 0.691495 | 0.74428 | 0.41609 | 0.58339 | 0.323309 |
| 21.98611 | 0.472061 | 1.880791 | 0.734948 | 0.23631 | 0.606451 | 0.460425 | 0.409442 | 0.761976 | 1.155435 | 0.453069 | 0.999026 | 0.990091 | 1.738011 | 1.283621 | 1.145756 | 0.314693 | 0.022338 | 0.805346 | 0.73927 | 0.672676 | 0.395711 | 0.382942 | 0.400815 |
| 22 | 0.461221 | 2.142359 | 0.953076 | 0.23625 | 0.524365 | 0.326876 | 0.379146 | 1.116999 | 1.09143 | 0.472746 | 1.112059 | 0.984786 | 2.007967 | 1.844864 | 1.032406 | 0.34099 | -0.341523 | 0.558582 | 0.585121 | 0.642782 | 0.332288 | 0.157271 | 0.459777 |
| 22.01389 | 0.417264 | 2.498598 | 1.040796 | 0.266398 | 0.518737 | 0.204175 | 0.249814 | 1.414441 | 0.971656 | 0.428057 | 0.968359 | 1.114994 | 2.177236 | 2.573965 | 1.088729 | 0.599824 | -0.127554 | 0.628333 | 0.460281 | 0.776464 | 0.30793 | 0.013197 | 0.589201 |
| 22.02778 | 0.411942 | 2.664721 | 1.016192 | 0.417466 | 0.566964 | 0.326605 | 0.161661 | 1.343035 | 0.881155 | 0.392403 | 0.694903 | 1.205088 | 2.05746 | 3.176931 | 1.303736 | 0.925141 | 0.459963 | 1.056468 | 0.43069 | 0.982484 | 0.283143 | 0.004896 | 0.762583 |
| 22.04167 | 0.59141 | 2.896165 | 1.135876 | 0.718401 | 0.731016 | 0.694329 | 0.278979 | 1.289201 | 0.990128 | 0.647161 | 0.53603 | 0.968588 | 1.774086 | 3.218253 | 1.52825 | 1.188803 | 1.336437 | 1.395647 | 0.512504 | 0.98238 | 0.334901 | 0.043988 | 0.797354 |
| 22.05556 | 0.702297 | 3.12923 | 1.185613 | 0.931918 | 0.843756 | 1.089733 | 0.366709 | 1.46753 | 1.047152 | 0.864755 | 0.459161 | 0.594234 | 1.677193 | 3.025144 | 1.653242 | 1.279097 | 2.161092 | 1.448828 | 0.742318 | 0.929352 | 0.355779 | 0.113387 | 0.895225 |
| 22.06944 | 0.527972 | 2.43179 | 1.029499 | 0.776287 | 0.911901 | 0.9629 | 0.254326 | 1.471906 | 0.830657 | 0.894667 | 0.275065 | -0.061345 | 1.602097 | 2.762585 | 1.629097 | 1.037404 | 1.638974 | 1.251372 | 0.953237 | 0.796892 | 0.297946 | 0.15565 | 0.884318 |
| 22.08333 | 0.197117 | 1.657115 | 0.601838 | 0.373372 | 0.944104 | 0.553581 | 0.150794 | 1.395414 | 0.440005 | 0.7737 | 0.045439 | -0.658952 | 1.017704 | 2.146644 | 1.000611 | 0.002679 | 0.746417 | 0.565489 | 0.559667 | 0.240295 | 0.069339 | -0.12979 | 0.476983 |
| 22.09722 | -0.082896 | 1.376979 | 0.293639 | -0.088442 | 1.024795 | 0.306921 | 0.17778 | 1.722209 | -0.13841 | 0.738117 | -0.311156 | -1.192239 | -0.270112 | 0.666964 | -0.356198 | -1.69554 | -0.402146 | -0.6243 | -0.512168 | -0.934814 | -0.496966 | -1.103811 | -0.494037 |
| 22.11111 | -0.061749 | 0.917719 | -0.021687 | -0.248834 | 0.642654 | 0.215402 | 0.082624 | 1.677953 | -0.356016 | 0.698294 | -0.534485 | -1.132037 | -1.235119 | -0.492463 | -0.9773 | -1.904733 | -1.005395 | -0.936556 | -1.104148 | -1.513665 | -0.986216 | -1.56497 | -1.175311 |
| 22.125 | 0.243796 | 0.323778 | 0.045103 | 0.171159 | 0.279801 | 0.411378 | -0.026461 | 1.120937 | 0.055612 | 0.833715 | 0.014483 | -0.355276 | -0.982483 | -0.286975 | -0.066269 | 0.10545 | -0.436431 | 0.361233 | -0.504634 | -0.908153 | -0.634376 | -0.697204 | -0.768602 |
| 22.13889 | 0.489429 | 0.724197 | 0.470992 | 0.993548 | 0.472714 | 0.590763 | 0.050823 | 1.303746 | 0.561633 | 1.235817 | 0.357832 | 0.559174 | -0.249107 | 0.790918 | 1.169308 | 1.837385 | -0.420343 | 1.676513 | 0.486347 | -0.198357 | -0.111144 | 0.69208 | -0.117163 |
| 22.15278 | 0.553281 | 1.077768 | 0.915405 | 1.254041 | 0.870449 | 0.762035 | 0.311744 | 1.184982 | 0.773826 | 1.805421 | 0.39755 | 0.724637 | 0.713587 | 1.793617 | 1.66306 | 2.318677 | -1.212934 | 1.747639 | 1.382799 | 0.220893 | 0.33131 | 1.689121 | 0.265107 |
| 22.16667 | 0.621901 | 0.950869 | 1.346019 | 1.306924 | 1.451874 | 0.799878 | 0.723327 | 1.087591 | 0.743452 | 2.112517 | 0.436684 | 1.277623 | 1.83565 | 2.499316 | 1.808303 | 2.047707 | -0.820902 | 1.299007 | 1.886393 | 0.644678 | 0.753474 | 1.827799 | 0.545599 |
| 22.18056 | 0.627444 | 1.108653 | 1.776734 | 1.085335 | 2.192311 | 0.625554 | 0.824832 | 1.206381 | 0.656751 | 2.038848 | 0.368599 | 1.517653 | 2.374776 | 2.972429 | 1.926545 | 1.185859 | 0.396241 | 0.735626 | 1.952001 | 0.579554 | 1.032926 | 1.419782 | 0.860161 |
| 22.19444 | 0.484666 | 1.345102 | 1.936013 | 0.897803 | 2.556843 | 0.495359 | 0.643077 | 1.534093 | 0.628494 | 1.67853 | 0.322653 | 1.04719 | 1.820237 | 3.125386 | 1.555018 | 0.435572 | 1.32277 | 0.361407 | 1.609234 | 0.575061 | 1.035686 | 1.149204 | 1.066431 |
| 22.20833 | 0.307848 | 1.230481 | 1.602323 | 0.972593 | 2.299951 | 0.629907 | 0.536224 | 1.468716 | 0.37292 | 1.148731 | 0.504374 | 0.796618 | 1.28516 | 2.83758 | 0.981404 | -0.078628 | 1.097648 | 0.202286 | 1.212855 | 0.494629 | 0.929309 | 1.100414 | 1.061552 |
| 22.22222 | 0.180719 | 0.902011 | 0.977608 | 1.031103 | 1.43682 | 0.633314 | 0.378469 | 1.039432 | 0.175375 | 0.868511 | 0.836909 | 0.818647 | 0.884819 | 2.44323 | 0.765918 | -0.490096 | 0.417273 | 0.180284 | 0.823616 | 0.377481 | 0.89637 | 1.422865 | 0.941554 |
| 22.23611 | 0.022009 | 0.564047 | 0.276606 | 0.970359 | 0.463354 | 0.512554 | 0.282992 | 0.474775 | 0.121039 | 0.862068 | 0.578078 | 0.802647 | 0.47172 | 1.738963 | 0.767196 | -0.801176 | 0.325569 | 0.315762 | 0.383983 | 0.318669 | 0.881327 | 1.769136 | 0.81381 |
| 22.25 | -0.244164 | 0.332755 | -0.215056 | 0.707638 | -0.473779 | 0.126085 | 0.144436 | 0.038427 | 0.285798 | 0.935148 | 0.274269 | 0.634762 | -0.014905 | 0.686133 | 0.747154 | -0.953818 | 0.328159 | 0.495994 | 0.008633 | 0.289473 | 0.681036 | 1.721416 | 0.585163 |
| 22.26389 | -0.457528 | 0.134357 | -0.504578 | 0.533722 | -1.392194 | -0.073135 | 0.039793 | -0.151662 | 0.291089 | 0.923011 | 0.035864 | 0.23714 | -0.297037 | -0.199627 | 0.599428 | -0.657996 | -0.269389 | 0.539186 | -0.218908 | 0.218769 | 0.398338 | 1.688907 | 0.39717 |
| 22.27778 | -0.557338 | -0.156331 | -0.348418 | 0.645397 | -1.839238 | -0.054188 | 0.037223 | -0.043157 | 0.36423 | 1.005218 | -0.228556 | -0.121246 | -0.131452 | -0.443009 | 0.467824 | 0.015521 | -0.952686 | 0.373748 | -0.16053 | 0.167362 | 0.223015 | 1.222353 | 0.332342 |
| 22.29167 | -0.470891 | -0.459907 | 0.02061 | 0.728252 | -1.567134 | -0.084774 | 0.107801 | 0.141117 | 0.333376 | 1.137998 | -0.45315 | -0.310805 | 0.018025 | 0.168808 | 0.528855 | 0.731994 | -1.269601 | 0.119442 | 0.110516 | 0.1483 | 0.174265 | 0.976403 | 0.235144 |
| 22.30556 | -0.258367 | -0.660399 | 0.353844 | 0.67952 | -1.096239 | -0.201984 | 0.117539 | 0.366617 | 0.267126 | 1.116574 | -0.475677 | -0.295248 | 0.009492 | 0.624741 | 0.596015 | 1.115096 | -1.336146 | -0.007906 | 0.400213 | 0.204206 | 0.134727 | 0.998216 | 0.113231 |
| 22.31944 | -0.031779 | -0.665517 | 0.425014 | 0.499309 | -0.625273 | -0.144425 | 0.025602 | 0.264729 | 0.169578 | 0.773157 | -0.21566 | 0.080025 | -0.125731 | 0.754328 | 0.517825 | 0.743627 | -0.745865 | -0.073384 | 0.428 | 0.188761 | 0.012833 | 0.713813 | 0.084746 |
| 22.33333 | 0.027353 | -0.597426 | 0.258724 | 0.259527 | -0.171164 | -0.045216 | -0.160631 | -0.255986 | 0.061999 | 0.249633 | -0.124735 | 0.666447 | -0.235263 | 0.711135 | 0.349568 | 0.041352 | 0.391573 | -0.262556 | 0.168174 | 0.103377 | -0.115546 | 0.514599 | 0.194273 |
| 22.34722 | 0.057104 | -0.68163 | -0.024959 | 0.081554 | 0.541644 | -0.03906 | -0.235286 | -0.569065 | 0.098163 | -0.226097 | -0.036178 | 0.940068 | -0.226332 | 0.524358 | 0.088457 | -0.292696 | 1.56022 | -0.532886 | -0.144666 | -0.014439 | -0.172745 | 0.340129 | 0.3531 |
| 22.36111 | 0.071964 | -0.790156 | -0.35969 | -0.046995 | 1.007888 | -0.000241 | -0.170697 | -0.670076 | 0.059152 | -0.538692 | 0.152545 | 0.517084 | -0.157597 | 0.002976 | -0.191759 | -0.336453 | 1.486041 | -0.701939 | -0.376493 | -0.116573 | -0.231044 | 0.281336 | 0.381015 |
| 22.375 | 0.076039 | -0.807479 | -0.609494 | -0.148036 | 1.255346 | -0.082208 | -0.033429 | -0.872934 | 0.058701 | -0.60523 | 0.344921 | 0.089594 | -0.113706 | -0.501649 | -0.381856 | -0.256007 | 0.528786 | -0.632225 | -0.425442 | -0.070722 | -0.291528 | 0.210045 | 0.262445 |
| 22.38889 | 0.075182 | -0.695258 | -0.699837 | -0.152337 | 0.542221 | -0.306935 | -0.04605 | -0.788802 | -0.025378 | -0.55646 | 0.496482 | -0.05584 | -0.280575 | -0.573027 | -0.430086 | -0.182423 | 0.095743 | -0.332236 | -0.312988 | -0.070781 | -0.360427 | 0.053829 | 0.10887 |
| 22.40278 | -0.051397 | -0.568781 | -0.885237 | -0.105305 | -0.150323 | -0.442431 | -0.193607 | -0.533069 | -0.288262 | -0.547882 | 0.518641 | -0.319636 | -0.529036 | -0.276267 | -0.495671 | -0.14855 | -0.103084 | -0.088912 | -0.111502 | -0.191816 | -0.466334 | -0.147528 | -0.055099 |
| 22.41667 | -0.169359 | -0.336231 | -0.957195 | -0.102699 | -0.500447 | -0.343632 | -0.301309 | -0.306892 | -0.518723 | -0.634898 | 0.421542 | -0.608355 | -0.732981 | 0.110197 | -0.647956 | 0.005954 | -0.619991 | -0.005684 | 0.011283 | -0.280385 | -0.52082 | -0.384394 | -0.152579 |
| 22.43056 | -0.206424 | -0.470691 | -0.951371 | -0.176488 | -0.88663 | -0.227037 | -0.240534 | -0.261054 | -0.475319 | -0.717267 | 0.221866 | -0.772842 | -0.7731 | 0.366493 | -0.719033 | -0.070879 | -1.139857 | -0.102562 | -0.095532 | -0.323767 | -0.567775 | -0.433875 | -0.202202 |
| 22.44444 | -0.269328 | -0.909288 | -0.828661 | -0.265598 | -1.247488 | -0.170974 | -0.158072 | -0.517159 | -0.336156 | -0.788789 | -0.18617 | -0.759293 | -0.821493 | 0.234786 | -0.818989 | -0.531348 | -0.791529 | -0.304504 | -0.335894 | -0.38047 | -0.696374 | -0.305902 | -0.357152 |
| 22.45833 | -0.298534 | -1.195778 | -0.720228 | -0.326369 | -1.664752 | -0.191536 | -0.132489 | -0.934787 | -0.358614 | -0.705628 | -0.67341 | -0.522464 | -0.947293 | -0.243487 | -0.80273 | -0.901124 | -0.012265 | -0.45867 | -0.443212 | -0.395771 | -0.771907 | -0.078742 | -0.580377 |
| 22.47222 | -0.246172 | -1.345653 | -0.724102 | -0.281754 | -1.694687 | -0.293745 | -0.301839 | -1.210295 | -0.466804 | -0.62208 | -0.917621 | -0.22899 | -1.171355 | -0.840717 | -0.802982 | -0.994911 | 0.573408 | -0.54098 | -0.411369 | -0.314623 | -0.696326 | -0.005236 | -0.711697 |
| 22.48611 | -0.207254 | -1.36536 | -0.896018 | -0.235739 | -1.149792 | -0.441201 | -0.536357 | -1.303286 | -0.655167 | -0.562253 | -0.947322 | -0.307456 | -1.490728 | -1.461593 | -0.914674 | -0.951933 | 0.568658 | -0.689959 | -0.510641 | -0.438527 | -0.666203 | -0.203841 | -0.761924 |
| 22.5 | -0.243005 | -1.376803 | -0.904396 | -0.299766 | -0.523433 | -0.56178 | -0.616491 | -1.229098 | -0.726158 | -0.599042 | -0.882544 | -0.767387 | -1.553874 | -2.041781 | -1.036445 | -0.942754 | 0.474269 | -0.822852 | -0.611498 | -0.703268 | -0.650117 | -0.536928 | -0.880896 |
| 22.51389 | -0.227086 | -1.669935 | -0.830476 | -0.410587 | -0.077666 | -0.497876 | -0.538009 | -0.999049 | -0.794522 | -0.706478 | -0.595655 | -1.137283 | -1.381582 | -2.370904 | -1.154395 | -1.004157 | 0.257295 | -0.848852 | -0.652956 | -0.662514 | -0.538061 | -0.841945 | -0.926466 |
| 22.52778 | -0.105502 | -1.810685 | -0.680736 | -0.604776 | 0.112854 | -0.31131 | -0.43824 | -0.664576 | -0.796145 | -0.641108 | -0.213892 | -1.071191 | -1.118224 | -2.432116 | -1.149134 | -1.170384 | 0.151444 | -0.695133 | -0.67978 | -0.276424 | -0.280182 | -0.831789 | -0.80258 |
| 22.54167 | -0.116963 | -1.635824 | -0.515576 | -0.950679 | 0.071847 | -0.327167 | -0.394418 | -0.339339 | -0.79065 | -0.545359 | -0.033667 | -0.903736 | -1.079885 | -2.057697 | -1.205626 | -1.412338 | -0.070474 | -0.588974 | -0.796183 | -0.024658 | -0.171253 | -0.998237 | -0.806471 |
| 22.55556 | -0.264196 | -1.400545 | -0.543192 | -1.117657 | -0.190426 | -0.694813 | -0.519023 | -0.463093 | -0.902219 | -0.530079 | -0.050921 | -0.839072 | -1.155994 | -1.677691 | -1.39512 | -1.718503 | -0.534561 | -0.59427 | -0.964905 | -0.17601 | -0.317774 | -1.056634 | -1.065548 |
| 22.56944 | -0.371257 | -1.63844 | -0.753181 | -1.180171 | -0.448931 | -1.085678 | -0.772782 | -0.979238 | -1.11484 | -0.595991 | -0.101807 | -0.994045 | -1.207193 | -1.742508 | -1.483094 | -1.941375 | -1.428488 | -0.746002 | -1.21667 | -0.469217 | -0.441279 | -0.838439 | -1.260176 |
| 22.58333 | -0.400804 | -2.1722 | -0.977991 | -1.163141 | -0.712443 | -1.149088 | -0.961764 | -1.423427 | -1.247738 | -0.734817 | -0.159419 | -1.07606 | -1.458407 | -2.172406 | -1.584999 | -2.036147 | -2.34389 | -1.055907 | -1.486195 | -0.721178 | -0.556127 | -0.909291 | -1.34432 |
| 22.59722 | -0.446658 | -2.942506 | -1.377331 | -1.261813 | -1.122404 | -1.172834 | -1.046778 | -2.142134 | -1.296318 | -0.863268 | -0.472114 | -1.081384 | -1.908091 | -2.769403 | -2.151165 | -2.229181 | -2.474877 | -1.455681 | -1.630947 | -0.945517 | -0.718842 | -1.280578 | -1.527095 |
| 22.61111 | -0.484227 | -2.813795 | -1.61849 | -1.404008 | -1.457733 | -1.365345 | -1.063449 | -3.148596 | -1.266019 | -0.595547 | -0.993239 | -0.604429 | -1.939305 | -3.065704 | -2.545658 | -2.211401 | -1.729358 | -1.589051 | -1.349095 | -0.821959 | -0.56524 | -1.578212 | -1.42627 |
| 22.625 | -0.678474 | -2.190477 | -1.508152 | -1.522656 | -1.367661 | -1.278085 | -1.047328 | -3.125209 | -0.959167 | -0.109895 | -1.276868 | 0.131394 | -1.497501 | -2.498675 | -2.208467 | -1.944427 | -0.351812 | -1.314996 | -0.704529 | -0.456813 | -0.122686 | -1.487728 | -0.795475 |
| 22.63889 | -0.769862 | -1.840352 | -0.812954 | -0.685432 | -1.104804 | -0.817657 | -0.707247 | -1.569373 | -0.50752 | -0.09321 | -1.382169 | 0.297896 | -1.222113 | -1.750821 | -1.80994 | -1.57134 | 0.819696 | -0.863962 | -0.330686 | -0.405448 | 0.147141 | -1.237042 | -0.232284 |
| 22.65278 | -0.521595 | -1.146936 | -0.211434 | 0.277088 | -0.873877 | -0.314806 | -0.379992 | -0.500694 | -0.132114 | -0.359326 | -1.004676 | 0.048606 | -0.869762 | -1.426432 | -1.391927 | -1.202356 | 1.798516 | -0.56412 | -0.232188 | -0.54761 | 0.235293 | -1.04157 | -0.035299 |
| 22.66667 | -0.198065 | -0.804023 | -0.01709 | 0.532708 | -0.713281 | -0.138351 | -0.195693 | -0.059945 | 0.029614 | -0.517003 | -0.688195 | -0.116026 | -0.478188 | -1.078099 | -0.942797 | -0.660985 | 1.690701 | -0.419032 | -0.102616 | -0.444114 | 0.295755 | -0.803971 | -0.009547 |
| 22.68056 | -0.075333 | -0.717732 | -0.040391 | -0.149328 | -0.678392 | -0.13298 | -0.086214 | 0.040183 | 0.048434 | -0.513387 | -0.552316 | -0.070619 | -0.157972 | -0.846518 | -0.509647 | -0.080639 | 1.111737 | -0.337546 | 0.012087 | -0.174531 | 0.288523 | -0.646518 | 0.003786 |
| 22.69444 | -0.005631 | -0.754103 | -0.113568 | -0.746348 | -0.689158 | -0.132439 | -0.021665 | 0.050999 | 0.035033 | -0.522274 | -0.455423 | 0.037759 | 0.061004 | -0.627256 | -0.181488 | 0.290085 | 0.63542 | -0.335806 | 0.006744 | 0.065395 | 0.192686 | -0.599743 | -0.045181 |
| 22.70833 | 0.071301 | -0.711775 | -0.17035 | -0.941498 | -0.669958 | -0.291257 | -0.016429 | 0.034115 | 0.007475 | -0.613024 | -0.287188 | -0.069814 | -0.032913 | -0.516792 | -0.08064 | 0.144324 | 0.159621 | -0.60118 | -0.201212 | -0.007095 | -0.075388 | -0.866803 | -0.283319 |
| 22.72222 | 0.086011 | -0.54217 | -0.151283 | -0.815408 | -0.679278 | -0.402259 | -0.037707 | 0.101772 | -0.03994 | -0.707776 | -0.068933 | -0.38934 | -0.401262 | -0.563432 | -0.10162 | -0.222243 | -0.509703 | -0.967647 | -0.524467 | -0.455387 | -0.449598 | -1.249007 | -0.628412 |
| 22.73611 | 0.014005 | -0.36894 | -0.060423 | -0.808744 | -0.630062 | -0.235143 | -0.074052 | 0.249984 | -0.113459 | -0.687985 | 0.061138 | -0.640064 | -0.676225 | -0.483689 | -0.042102 | -0.417477 | -1.105847 | -0.994326 | -0.651039 | -0.934195 | -0.621484 | -1.395539 | -0.814409 |
| 22.75 | -0.075895 | -0.248593 | 0.054444 | -0.6361 | -0.51335 | -0.026561 | -0.122798 | 0.321557 | -0.189912 | -0.602401 | 0.141055 | -0.631087 | -0.641638 | -0.243042 | 0.146313 | -0.320001 | -1.18656 | -0.689692 | -0.511506 | -1.128742 | -0.481221 | -1.131617 | -0.763849 |
| 22.76389 | -0.121653 | -0.22797 | 0.097747 | -0.291431 | -0.434228 | 0.011839 | -0.17191 | 0.303002 | -0.237647 | -0.531519 | 0.196397 | -0.523735 | -0.489253 | -0.088041 | 0.326809 | -0.069688 | -0.760076 | -0.417149 | -0.300995 | -0.991126 | -0.220921 | -0.744875 | -0.565119 |
| 22.77778 | -0.135156 | -0.249907 | 0.103777 | -0.072025 | -0.365453 | 0.019788 | -0.187094 | 0.214816 | -0.277329 | -0.480577 | 0.274334 | -0.550275 | -0.519908 | -0.064624 | 0.283076 | 0.120846 | -0.397771 | -0.428626 | -0.252471 | -0.716306 | -0.097765 | -0.592384 | -0.3574 |
| 22.79167 | -0.119516 | -0.193351 | 0.094456 | -0.08155 | -0.252901 | 0.115951 | -0.151551 | 0.051371 | -0.294224 | -0.329901 | 0.258984 | -0.751674 | -0.673483 | 0.043693 | 0.052842 | 0.215172 | -0.265886 | -0.528631 | -0.339379 | -0.479097 | -0.154767 | -0.655903 | -0.19255 |
| 22.80556 | -0.026985 | -0.1402 | 0.017654 | -0.163024 | -0.070847 | 0.193633 | -0.119673 | -0.185763 | -0.289479 | -0.124985 | 0.205464 | -0.716554 | -0.624807 | 0.262986 | -0.231487 | 0.27442 | -0.149846 | -0.460202 | -0.336616 | -0.339432 | -0.261485 | -0.648634 | -0.083329 |
| 22.81944 | 0.019831 | -0.169737 | -0.081491 | -0.213966 | 0.106668 | 0.209198 | -0.128015 | -0.374348 | -0.271226 | -0.037871 | 0.174327 | -0.515348 | -0.421878 | 0.346554 | -0.538231 | 0.397278 | 0.076513 | -0.26508 | -0.271176 | -0.279909 | -0.376546 | -0.615968 | -0.064953 |
| 22.83333 | 0.005117 | -0.152889 | -0.104811 | -0.298204 | 0.214046 | 0.177434 | -0.146533 | -0.424365 | -0.23339 | -0.043183 | 0.217728 | -0.28342 | -0.267006 | 0.268206 | -0.784645 | 0.584484 | 0.258961 | -0.128791 | -0.242717 | -0.278911 | -0.505061 | -0.637548 | -0.121677 |
| 22.84722 | -0.020853 | -0.063813 | -0.112577 | -0.377469 | 0.172092 | 0.090563 | -0.120432 | -0.300016 | -0.237148 | -0.069608 | 0.285166 | -0.240486 | -0.210975 | 0.123768 | -0.720324 | 0.708962 | 0.31417 | -0.079386 | -0.257477 | -0.200643 | -0.560069 | -0.770603 | -0.159911 |
| 22.86111 | 0.013949 | -0.023709 | -0.136951 | -0.43778 | 0.074439 | -0.001745 | -0.017552 | -0.123111 | -0.228658 | -0.12616 | 0.314514 | -0.188851 | -0.157504 | -0.025819 | -0.32592 | 0.824904 | 0.278424 | -0.030263 | -0.151052 | 0.047184 | -0.44642 | -0.6668 | -0.115318 |
| 22.875 | -0.172383 | -0.032394 | -0.083523 | -0.549215 | -0.007711 | -0.021659 | 0.090638 | -0.036053 | -0.077167 | -0.216578 | 0.307101 | 0.020787 | -0.026146 | -0.089984 | 0.130955 | 0.92677 | 0.168271 | 0.040026 | 0.089624 | 0.355465 | -0.36544 | -0.333577 | 0.00329 |
| 22.88889 | -0.41416 | -0.003792 | 0.032322 | -0.54192 | -0.058584 | 0.010659 | 0.134121 | -0.027819 | 0.171499 | -0.254355 | 0.292344 | 0.432324 | 0.134252 | 0.023998 | 0.532358 | 1.04354 | 0.08501 | 0.129952 | 0.351558 | 0.518341 | -0.83523 | -0.112236 | 0.167344 |
| 22.90278 | -0.494944 | 0.173859 | 0.097768 | -0.334385 | -0.055448 | 0.074934 | 0.151122 | -0.030369 | 0.369272 | -0.231572 | 0.302903 | 0.883931 | 0.267111 | 0.306521 | 0.929865 | 1.075638 | 0.123073 | 0.25439 | 0.552013 | 0.478069 | -1.904036 | 0.032532 | 0.311917 |
| 22.91667 | -0.362044 | 0.470521 | 0.150647 | 0.042742 | -0.009778 | 0.184223 | 0.211471 | 0.019693 | 0.510633 | -0.145659 | 0.354568 | 1.200033 | 0.37171 | 0.673499 | 1.137673 | 1.007675 | 0.324514 | 0.406343 | 0.66456 | 0.338618 | -1.853245 | 0.154355 | 0.358188 |
| 22.93056 | -0.143273 | 0.739543 | 0.228634 | 0.263919 | 0.057688 | 0.288986 | 0.283306 | 0.112127 | 0.608017 | -0.009033 | 0.441037 | 1.322769 | 0.431302 | 1.001349 | 1.129636 | 0.885573 | 0.657406 | 0.560369 | 0.640848 | 0.257877 | -0.588292 | 0.222491 | 0.307443 |
| 22.94444 | 0.13094 | 1.039302 | 0.325386 | 0.288553 | 0.159955 | 0.330652 | 0.35225 | 0.258657 | 0.678424 | 0.153971 | 0.547399 | 1.32552 | 0.440395 | 1.122851 | 0.974579 | 0.797179 | 1.096471 | 0.677647 | 0.512313 | 0.280429 | 0.037845 | 0.239515 | 0.213231 |
| 22.95833 | 0.492727 | 1.394573 | 0.42544 | 0.303168 | 0.320418 | 0.385828 | 0.428794 | 0.477438 | 0.796085 | 0.267009 | 0.645948 | 1.215252 | 0.328879 | 0.837605 | 0.743741 | 0.675793 | 1.358783 | 0.736694 | 0.341057 | 0.303487 | 0.035838 | 0.164003 | 0.09803 |
| 22.97222 | 0.768872 | 1.726246 | 0.51511 | 0.409506 | 0.484971 | 0.490828 | 0.505388 | 0.701614 | 0.933087 | 0.295856 | 0.702398 | 1.088311 | 0.188722 | 0.502847 | 0.595555 | 0.512655 | 1.238579 | 0.790373 | 0.194079 | 0.304877 | -0.057641 | 0.109991 | 0.013447 |
| 22.98611 | 0.908704 | 1.986232 | 0.583032 | 0.504775 | 0.620186 | 0.612327 | 0.574829 | 0.850089 | 1.00779 | 0.307325 | 0.719351 | 1.084203 | 0.314056 | 0.425664 | 0.690023 | 0.38744 | 1.030372 | 0.848878 | 0.192012 | 0.394282 | 0.008597 | 0.404247 | 0.024908 |
| 23 | 0.951248 | 2.199215 | 0.648121 | 0.565741 | 0.647657 | 0.632826 | 0.531839 | 0.981953 | 0.971313 | 0.281692 | 0.680369 | 1.119792 | 0.80216 | 0.438125 | 0.922964 | 0.342109 | 0.792105 | 0.872028 | 0.253267 | 0.570754 | 0.174395 | 1.122612 | 0.058491 |
| 23.01389 | 0.845793 | 2.501969 | 0.720689 | 0.561393 | 0.590287 | 0.542931 | 0.37064 | 1.070258 | 0.783053 | 0.142271 | 0.579033 | 1.123195 | 1.142102 | 0.336245 | 1.114863 | 0.401238 | 0.65644 | 0.913639 | 0.327343 | 0.697107 | 0.308415 | 0.86134 | 0.023835 |
| 23.02778 | 0.599547 | 2.925178 | 0.952245 | 0.697314 | 0.675917 | 0.580783 | 0.251597 | 1.334673 | 0.814911 | 0.152606 | 0.602283 | 1.171461 | 1.366444 | 0.38632 | 1.279811 | 0.564459 | 0.615499 | 1.034905 | 0.464382 | 0.777483 | 0.44966 | 0.640808 | 0.055087 |
| 23.04167 | 0.633474 | 3.48158 | 1.196431 | 1.100462 | 0.84179 | 0.842656 | 0.404863 | 1.607554 | 1.143355 | 0.546604 | 0.740051 | 1.034907 | 1.65094 | 0.613218 | 1.387176 | 0.815603 | 0.533513 | 1.166129 | 0.651422 | 0.840702 | 0.675092 | 1.1933 | 0.189977 |
| 23.05556 | 0.719466 | 3.317958 | 1.277145 | 1.360215 | 0.832819 | 0.977681 | 0.605169 | 1.684824 | 1.230966 | 0.912183 | 0.729057 | 1.017493 | 1.79648 | 0.800535 | 1.384033 | 0.914137 | 0.322222 | 1.137909 | 0.872317 | 0.911021 | 0.786631 | 1.806092 | 0.268227 |
| 23.06944 | 0.710255 | 2.745808 | 1.29259 | 1.36581 | 0.745315 | 0.913822 | 0.600964 | 1.536007 | 1.119066 | 0.961111 | 0.61085 | 0.838504 | 1.740973 | 0.880511 | 1.128746 | 0.792932 | -0.038166 | 0.94295 | 1.012007 | 0.796222 | 0.688299 | 1.004191 | 0.240573 |
| 23.08333 | 0.558662 | 2.053138 | 1.44022 | 1.033461 | 0.87101 | 0.864325 | 0.512483 | 1.584557 | 0.799643 | 0.923231 | 0.449958 | 0.151721 | 1.110988 | 0.452537 | 0.382515 | -0.076251 | -0.569277 | 0.246343 | 0.474065 | 0.026142 | 0.332605 | 0.245149 | -0.210075 |
| 23.09722 | 0.285878 | 1.439591 | 1.248689 | 0.342186 | 1.036975 | 0.654088 | 0.312481 | 2.057175 | 0.26182 | 0.767498 | 0.223987 | -0.918393 | -0.237548 | -0.733277 | -0.925222 | -1.565875 | -1.245443 | -0.862067 | -0.705711 | -1.382864 | -0.444176 | -0.906754 | -1.220126 |
| 23.11111 | 0.141366 | 1.020122 | 0.737733 | -0.160921 | 0.572866 | 0.378366 | 0.144301 | 1.922428 | -0.059805 | 0.619748 | -0.011464 | -1.200118 | -1.020586 | -1.15085 | -1.244083 | -1.579617 | -1.332118 | -1.070725 | -1.115548 | -1.891986 | -0.905104 | -1.385871 | -1.672566 |
| 23.125 | 0.397581 | 1.071941 | 0.467021 | 0.243032 | 0.197184 | 0.594522 | 0.118705 | 1.54437 | 0.216724 | 0.856123 | 0.090144 | -0.372359 | -0.492242 | -0.186166 | -0.023496 | 0.157686 | -0.535875 | 0.103553 | -0.311734 | -0.996811 | -0.568332 | -0.253643 | -1.020258 |
| 23.13889 | 1.011288 | 1.804275 | 0.590654 | 1.157047 | 0.456615 | 1.311108 | 0.200186 | 1.242941 | 0.796419 | 1.39272 | 0.465929 | 0.512159 | 0.247471 | 0.104196 | 1.537753 | 1.481749 | 0.19254 | 1.455494 | 0.572028 | -0.202068 | -0.006171 | 0.917456 | -0.266093 |
| 23.15278 | 0.992895 | 0.923588 | 0.770405 | 1.797486 | 0.876969 | 1.503957 | 0.283391 | 0.729224 | 0.989184 | 1.705399 | 0.222416 | 0.85784 | 0.98403 | -1.286702 | 2.100304 | 1.879274 | 0.417232 | 1.87901 | 1.196033 | 0.245666 | 0.441436 | 1.253512 | 0.247181 |
| 23.16667 | 0.305184 | 0.583935 | 0.887756 | 1.444152 | 1.274326 | 1.178607 | 0.54992 | 0.730652 | 1.064903 | 1.803615 | 0.007351 | 1.000892 | 2.122452 | -3.258413 | 2.206554 | 2.13438 | 0.466873 | 1.737453 | 1.804449 | 0.641816 | 0.818099 | 1.450427 | 0.64326 |
| 23.18056 | -0.203687 | 0.998012 | 1.072904 | 1.04744 | 1.661533 | 0.801605 | 0.850008 | 1.322028 | 0.87014 | 1.916316 | 0.077082 | 1.127977 | 2.889805 | -2.455181 | 2.257267 | 1.372839 | 0.34596 | 1.42499 | 1.795984 | 1.03573 | 0.981163 | 1.594257 | 1.031712 |
| 23.19444 | -0.371829 | 1.73895 | 1.269969 | 0.692117 | 1.728874 | 0.633311 | 1.085619 | 2.019624 | 0.75891 | 1.773287 | 0.235699 | 0.953253 | 2.621706 | 0.68425 | 1.890655 | 0.646634 | 0.31187 | 0.751555 | 1.562165 | 1.103583 | 1.048073 | 1.537268 | 1.350665 |
| 23.20833 | -0.284415 | 2.330057 | 1.272012 | 0.670188 | 1.647613 | 0.604181 | 1.050499 | 1.997865 | 0.471306 | 1.536293 | 0.221166 | 0.680426 | 2.160529 | 2.592601 | 1.260092 | 0.64405 | 0.305619 | 0.157672 | 1.238306 | 0.683497 | 1.061241 | 1.45536 | 1.426446 |
| 23.22222 | 0.303115 | 2.10723 | 1.103909 | 1.063421 | 1.302713 | 0.687782 | 0.794312 | 1.633436 | 0.166878 | 1.31842 | 0.113355 | 0.428595 | 1.824516 | 3.417272 | 0.775378 | -0.485585 | 0.337124 | -0.115211 | 0.792528 | 0.311651 | 0.968477 | 1.600057 | 1.382851 |
| 23.23611 | 1.054535 | 1.186262 | 0.871404 | 1.613026 | 0.881015 | 0.836443 | 0.517687 | 0.913071 | -0.032035 | 1.285858 | 0.456489 | 0.388399 | 1.298794 | 3.593809 | 0.478597 | -0.761798 | 0.369187 | -0.089375 | 0.36097 | 0.199979 | 0.738237 | 1.52557 | 1.263187 |
| 23.25 | 0.852894 | 0.563094 | 0.67644 | 1.351481 | 0.591403 | 0.833563 | 0.268925 | 0.164169 | -0.051229 | 1.300003 | 0.750045 | 0.447123 | 0.474821 | 3.183297 | 0.29982 | -0.720091 | 0.219575 | -0.050472 | 0.017982 | 0.089789 | 0.470186 | 1.456876 | 1.009083 |
| 23.26389 | 0.469049 | -0.012789 | 0.532452 | 1.011284 | 0.326628 | 0.481748 | 0.025774 | -0.303936 | 0.129639 | 1.490032 | 0.693711 | 0.379056 | -0.3858 | 2.630512 | 0.062057 | -0.359487 | -0.282886 | -0.160676 | -0.160188 | -0.0154 | 0.222546 | 1.258554 | 0.686148 |
| 23.27778 | 0.093725 | -0.386727 | 0.396783 | 0.585922 | 0.135017 | 0.179753 | -0.090436 | -0.313542 | 0.401838 | 1.615831 | 0.397134 | 0.163087 | -0.986527 | 2.324273 | -0.055315 | 0.324115 | -0.697727 | -0.26298 | -0.200441 | -0.022363 | 0.142887 | 0.769599 | 0.378227 |
| 23.29167 | -0.391453 | -0.580373 | 0.285404 | -0.025666 | -0.153373 | -0.11071 | -0.123448 | -0.11476 | 0.423453 | 1.44313 | 0.069008 | -0.112837 | -1.309136 | 2.092276 | -0.108459 | 0.807205 | -0.650439 | -0.218715 | -0.186644 | -0.034299 | 0.073579 | 0.857627 | 0.131046 |
| 23.30556 | -0.830846 | -0.622096 | 0.059886 | -0.239028 | -0.514881 | -0.242101 | -0.073546 | -0.057231 | 0.244652 | 1.076971 | -0.164718 | -0.405072 | -1.305618 | 1.778159 | -0.180324 | 0.901031 | -0.687292 | 0.054313 | -0.145439 | -0.075931 | -0.039452 | 1.195683 | -0.123776 |
| 23.31944 | -1.092347 | -0.467673 | -0.252224 | -0.009851 | -0.679613 | -0.173384 | 0.028494 | -0.132245 | 0.112409 | 0.748987 | -0.374515 | -0.621606 | -1.067259 | 1.098998 | -0.223527 | 0.549276 | -0.315177 | 0.452764 | -0.062345 | -0.102446 | -0.075431 | 0.854801 | -0.223059 |
| 23.33333 | -0.809868 | -0.322203 | -0.549424 | 0.265834 | -0.688468 | -0.044795 | 0.12602 | -0.415026 | -0.110323 | 0.50593 | -0.572919 | -0.497282 | -0.827783 | 0.176934 | -0.269877 | 0.039685 | 0.19523 | 0.438608 | -0.038921 | -0.119366 | -0.061358 | 0.140409 | -0.16581 |
| 23.34722 | -0.295794 | -0.328034 | -0.876467 | 0.048414 | -0.682212 | 0.028835 | 0.135939 | -0.654272 | -0.392977 | 0.327031 | -0.588567 | -0.199404 | -0.725857 | -0.778635 | -0.423146 | -0.291267 | 0.492692 | -0.092654 | -0.090426 | -0.105623 | -0.084033 | -0.272226 | -0.047346 |
| 23.36111 | 0.171921 | -0.459917 | -0.952218 | -0.351413 | -0.620452 | 0.009515 | 0.086531 | -0.881092 | -0.424198 | 0.237766 | -0.340489 | -0.077906 | -0.592264 | -1.502486 | -0.583607 | -0.298336 | 0.428529 | -0.662883 | -0.135937 | -0.088269 | -0.072466 | -0.388911 | 0.1163 |
| 23.375 | 0.368944 | -0.70647 | -0.77203 | -0.481356 | -0.500132 | -0.117035 | -0.014593 | -1.071917 | -0.26896 | 0.118883 | -0.076671 | -0.277895 | -0.541953 | -1.818783 | -0.653579 | -0.160691 | 0.080081 | -0.917436 | -0.218131 | -0.171878 | -0.052675 | -0.385176 | 0.222968 |
| 23.38889 | 0.478665 | -0.878366 | -0.542362 | -0.459373 | -0.398284 | -0.191091 | -0.116946 | -1.193335 | -0.258547 | -0.158383 | 0.134872 | -0.644288 | -0.629336 | -1.928276 | -0.643553 | -0.087752 | -0.216251 | -0.882584 | -0.307308 | -0.312485 | -0.147736 | -0.450888 | 0.117175 |
| 23.40278 | 0.611651 | -0.845733 | -0.33705 | -0.391335 | -0.355793 | -0.042523 | -0.110601 | -1.266024 | -0.212441 | -0.459269 | 0.127276 | -0.830588 | -0.730912 | -1.313758 | -0.563479 | -0.077715 | -0.373892 | -0.717401 | -0.295842 | -0.420056 | -0.284342 | -0.438195 | -0.091503 |
| 23.41667 | 0.559286 | -0.570148 | -0.222579 | -0.274957 | -0.355297 | 0.079217 | -0.098117 | -0.676429 | -0.160922 | -0.599012 | -0.075901 | -0.594514 | -0.748054 | -0.252668 | -0.44892 | -0.173918 | -0.823385 | -0.502826 | -0.241727 | -0.482614 | -0.374427 | -0.424284 | -0.210167 |
| 23.43056 | 0.260069 | -0.335924 | -0.269786 | -0.188102 | -0.387261 | -0.008186 | -0.214974 | -0.022467 | -0.199896 | -0.742599 | -0.279121 | -0.271025 | -0.859765 | 0.551765 | -0.400945 | -0.338458 | -1.212405 | -0.267153 | -0.257426 | -0.604548 | -0.416299 | -0.509955 | -0.25572 |
| 23.44444 | -0.099689 | -0.284854 | -0.299447 | -0.331493 | -0.438667 | -0.207997 | -0.317037 | 0.25473 | -0.318699 | -0.802607 | -0.368819 | -0.133675 | -1.058356 | 0.83924 | -0.407502 | -0.478717 | -0.835799 | -0.172048 | -0.390572 | -0.744733 | -0.403549 | -0.596816 | -0.345475 |
| 23.45833 | -0.364473 | -0.406881 | -0.501334 | -0.554813 | -0.483765 | -0.325209 | -0.367679 | 0.086635 | -0.466946 | -0.715694 | -0.249672 | -0.261192 | -1.128854 | 0.797915 | -0.47007 | -0.590388 | -0.234152 | -0.274213 | -0.554701 | -0.913158 | -0.399728 | -0.47655 | -0.511792 |
| 23.47222 | -0.528793 | -0.820653 | -0.851167 | -0.609203 | -0.524234 | -0.419673 | -0.315745 | -0.405545 | -0.686355 | -0.657832 | -0.088946 | -0.559724 | -1.081343 | 0.651728 | -0.617757 | -0.787618 | 0.365988 | -0.486324 | -0.711372 | -1.056957 | -0.398926 | -0.360837 | -0.710038 |
| 23.48611 | -0.770985 | -1.273735 | -1.220481 | -0.528383 | -0.572464 | -0.462244 | -0.393811 | -0.934533 | -0.916037 | -0.724776 | -0.053747 | -0.878362 | -1.094643 | 0.335956 | -0.803607 | -1.029533 | 0.491878 | -0.678307 | -0.744536 | -1.417232 | -0.369252 | -0.556151 | -0.848588 |
| 23.5 | -0.597567 | -1.620442 | -1.456598 | -0.543331 | -0.633563 | -0.456248 | -0.686456 | -1.061696 | -0.964733 | -0.764171 | -0.291452 | -1.042821 | -1.277531 | -0.007761 | -0.957092 | -1.189889 | 0.268908 | -0.708756 | -0.623686 | -1.354304 | -0.320534 | -1.476114 | -0.830965 |
| 23.51389 | -0.319183 | -1.565762 | -1.325148 | -0.589491 | -0.604139 | -0.373065 | -0.751099 | -0.751743 | -0.889782 | -0.491723 | -0.608733 | -0.824593 | -1.293683 | -0.219865 | -0.896905 | -1.105618 | 0.130623 | -0.549531 | -0.393004 | -0.871749 | -0.231433 | -2.297572 | -0.649313 |
| 23.52778 | -0.059437 | -1.298976 | -1.013493 | -0.554069 | -0.357178 | -0.301583 | -0.49509 | -0.487858 | -0.921333 | -0.09219 | -0.541814 | -0.581014 | -0.98595 | -0.362327 | -0.745598 | -0.812051 | 0.064393 | -0.327651 | -0.188727 | -0.494261 | -0.116249 | -1.705483 | -0.519993 |
| 23.54167 | -0.051025 | -1.333137 | -0.869236 | -0.629042 | -0.146816 | -0.405773 | -0.414377 | -0.606423 | -1.044871 | -0.063685 | -0.477443 | -0.527237 | -0.628362 | -0.710741 | -0.775629 | -0.66591 | -0.199892 | -0.271092 | -0.174139 | -0.375432 | -0.109896 | -1.612293 | -0.55939 |
| 23.55556 | -0.208546 | -1.767811 | -1.147437 | -0.920214 | -0.204147 | -0.75234 | -0.649631 | -1.025574 | -1.130677 | -0.266398 | -0.56657 | -0.633553 | -0.766649 | -1.091464 | -1.078416 | -0.986762 | -0.701563 | -0.434595 | -0.331237 | -0.436332 | -0.164531 | -2.199777 | -0.636781 |
| 23.56944 | -0.326823 | -2.10368 | -1.432982 | -0.984743 | -0.46716 | -1.022146 | -0.754615 | -1.379768 | -1.137862 | -0.28762 | -0.678254 | -0.620037 | -0.996191 | -1.069738 | -1.220102 | -1.236769 | -0.997393 | -0.69345 | -0.467872 | -0.425461 | -0.188894 | -1.975692 | -0.651583 |
| 23.58333 | -0.454406 | -1.892302 | -1.579208 | -1.062899 | -0.725668 | -1.043199 | -0.72547 | -1.572583 | -1.024827 | -0.414959 | -0.742099 | -0.435315 | -0.97421 | -0.792831 | -1.153202 | -0.966773 | -0.941081 | -0.628071 | -0.396994 | -0.269321 | -0.145063 | -1.237807 | -0.591573 |
| 23.59722 | -0.974537 | -2.170754 | -2.121682 | -1.654425 | -1.074192 | -1.28751 | -1.08096 | -2.193022 | -1.14691 | -0.873987 | -0.929205 | -0.145793 | -0.687399 | -0.809331 | -1.201575 | -0.941548 | -0.733435 | -0.322099 | -0.259726 | -0.068175 | -0.078253 | -0.823181 | -0.458812 |
| 23.61111 | -1.652755 | -2.926121 | -2.665421 | -2.142881 | -1.412744 | -1.608931 | -1.571132 | -2.954328 | -1.22928 | -1.238082 | -1.118219 | 0.128553 | -0.692847 | -0.908658 | -1.438874 | -1.315296 | -0.736515 | -0.280874 | -0.124096 | -0.042404 | -0.076617 | -0.69067 | -0.247226 |
| 23.625 | -1.99983 | -2.998893 | -2.530361 | -1.731614 | -1.224785 | -1.526159 | -1.362352 | -3.135506 | -0.822795 | -0.861158 | -1.206167 | 0.087243 | -1.017034 | -0.848529 | -1.736291 | -1.840839 | -0.789458 | -0.462347 | -0.112026 | -0.286831 | -0.150306 | -0.615456 | -0.002841 |
| 23.63889 | -1.737036 | -2.787008 | -1.610676 | -0.747247 | -0.878103 | -1.064727 | -0.835687 | -1.740326 | -0.316363 | -0.349188 | -1.194423 | -0.2431 | -1.106957 | -0.808058 | -2.125796 | -1.868058 | -0.301853 | -0.558582 | -0.158846 | -0.48261 | -0.31504 | -0.453549 | 0.058657 |
| 23.65278 | -0.927931 | -1.657713 | -0.528601 | 0.255021 | -0.535333 | -0.459041 | -0.376022 | -0.485774 | -0.016643 | -0.038815 | -1.065808 | -0.626781 | -0.839129 | -0.839555 | -1.979706 | -1.537634 | 0.144656 | -0.572462 | -0.271817 | -0.503827 | -0.482625 | -0.345182 | -0.021845 |
| 23.66667 | -0.499499 | -1.089271 | 0.054986 | 0.906802 | -0.369112 | -0.13082 | -0.227268 | 0.323904 | 0.224218 | 0.245036 | -0.789135 | -0.86543 | -0.500722 | -0.876712 | -1.492301 | -0.928811 | 0.021303 | -0.598133 | -0.349121 | -0.472427 | -0.549362 | -0.013887 | -0.134388 |
| 23.68056 | -0.475923 | -0.923564 | 0.117553 | 0.349361 | -0.349713 | -0.142999 | -0.301031 | 0.037786 | 0.405402 | 0.451487 | -0.539075 | -0.864843 | -0.285309 | -0.958145 | -1.246367 | -0.37625 | -0.56944 | -0.636321 | -0.359528 | -0.391806 | -0.581635 | -0.190319 | -0.291117 |
| 23.69444 | -0.108436 | -0.910332 | 0.028508 | -0.2274 | -0.279337 | -0.105501 | -0.219732 | -0.033557 | 0.342435 | 0.34404 | -0.344549 | -0.764438 | -0.141087 | -1.003082 | -1.063318 | -0.064617 | -0.806242 | -0.652947 | -0.395708 | -0.300316 | -0.649887 | -0.542609 | -0.468106 |
| 23.70833 | 0.191811 | -0.706384 | -0.00811 | -0.743431 | -0.152065 | -0.040445 | -0.107892 | -0.015446 | -0.02633 | 0.126529 | -0.195013 | -0.524184 | -0.010558 | -1.067782 | -0.94759 | 0.03539 | -0.456973 | -0.61088 | -0.480368 | -0.188724 | -0.673585 | -0.740208 | -0.509746 |
| 23.72222 | 0.223265 | -0.526217 | -0.041726 | -1.085568 | -0.040783 | -0.154136 | -0.10715 | 0.036796 | -0.374874 | 0.082083 | -0.091419 | -0.228826 | 0.159989 | -0.868609 | -0.873625 | 0.161419 | 0.024151 | -0.501737 | -0.442616 | -0.090031 | -0.630557 | -0.406497 | -0.340286 |
| 23.73611 | 0.150482 | -0.425758 | -0.009482 | -1.222464 | -0.018354 | -0.325174 | -0.157078 | 0.056888 | -0.437776 | 0.010898 | -0.035499 | -0.016252 | 0.330772 | -0.453586 | -0.693278 | 0.280758 | 0.397274 | -0.382304 | -0.288761 | -0.099916 | -0.479821 | -0.248593 | -0.12303 |
| 23.75 | 0.016298 | -0.413156 | 0.054609 | -1.068662 | -0.089634 | -0.315159 | -0.144303 | 0.085177 | -0.349624 | -0.14329 | 0.025563 | 0.040578 | 0.397157 | -0.264692 | -0.414229 | 0.134921 | 0.742584 | -0.340493 | -0.201306 | -0.20938 | -0.41549 | -0.111954 | -0.060215 |
| 23.76389 | -0.242166 | -0.402316 | 0.121068 | -0.632326 | -0.145332 | -0.185733 | -0.073206 | 0.056555 | -0.264894 | -0.274542 | 0.128353 | 0.009688 | 0.287269 | -0.377482 | -0.382027 | -0.101605 | 0.988602 | -0.339446 | -0.259461 | -0.17337 | -0.447515 | -0.030583 | -0.115535 |
| 23.77778 | -0.66277 | -0.351825 | 0.034941 | -0.293009 | -0.14944 | -0.097443 | -0.069109 | -0.218015 | -0.26855 | -0.369108 | 0.187946 | 0.148637 | 0.064149 | -0.510668 | -0.411305 | -0.169256 | 0.99116 | -0.281364 | -0.288763 | 0.268206 | -0.473986 | 0.021159 | -0.17232 |
| 23.79167 | -0.894911 | -0.278044 | -0.059073 | 0.033 | -0.116404 | -0.079994 | -0.117222 | -0.239418 | -0.313614 | -0.432504 | 0.178549 | -0.078348 | -0.117979 | -0.411963 | -0.41477 | -0.042981 | 0.607094 | -0.190422 | -0.276375 | -0.995851 | -0.486478 | 0.06087 | -0.219829 |
| 23.80556 | -0.793307 | -0.267941 | -0.010095 | 0.276003 | -0.119521 | -0.042663 | -0.159801 | -0.098007 | -0.345086 | -0.45491 | 0.168328 | -1.477145 | -0.313045 | -0.2615 | -0.476286 | 0.178411 | 0.266974 | -0.216379 | -0.355836 | -0.876793 | -0.570389 | -0.005807 | -0.352698 |
| 23.81944 | -0.330392 | -0.305133 | 0.107023 | 0.436055 | -0.213638 | 0.040615 | -0.173701 | 0.032349 | -0.316167 | -0.380401 | 0.17194 | -0.777484 | -0.488694 | -0.18706 | -0.578278 | 0.324184 | 0.013058 | -0.282182 | -0.36039 | -0.725069 | -0.646835 | -0.089225 | -0.441111 |
| 23.83333 | 0.198814 | -0.289295 | 0.174683 | 0.628831 | -0.272666 | 0.092166 | -0.19237 | 0.123335 | -0.257894 | -0.291469 | 0.181874 | -1.229275 | -0.458772 | -0.053172 | -0.561585 | 0.488296 | 0.011911 | -0.220001 | -0.179939 | -0.481157 | -0.59038 | 0.042761 | -0.303657 |
| 23.84722 | 0.539075 | -0.131529 | 0.127119 | 0.764114 | -0.180883 | 0.114358 | -0.179538 | 0.158417 | -0.189928 | -0.222825 | 0.166317 | -1.542311 | -0.330287 | 0.045143 | -0.571276 | 0.562188 | 0.022543 | -0.164002 | -0.023395 | -0.261839 | -0.571598 | 0.041476 | -0.13981 |
| 23.86111 | 0.795118 | 0.059579 | 0.022379 | 0.411996 | 0.019398 | 0.168822 | -0.108049 | 0.139137 | -0.140783 | -0.098255 | 0.151008 | -1.499887 | -0.176379 | -0.01174 | -0.724028 | 0.5743 | -0.106871 | -0.224258 | -0.040123 | -0.225736 | -0.612746 | -0.140501 | -0.124111 |
| 23.875 | 0.935784 | 0.167357 | -0.054831 | 0.03818 | 0.19953 | 0.191987 | -0.005057 | 0.081495 | -0.070062 | 0.067654 | 0.134564 | -1.310045 | 0.070556 | -0.041655 | -0.803372 | 0.72186 | -0.24595 | -0.219867 | -0.043995 | -0.34568 | -0.524751 | -0.238628 | -0.095939 |
| 23.88889 | 0.842773 | 0.199379 | -0.09097 | -0.234373 | 0.22356 | 0.163729 | 0.089545 | 0.045353 | 0.05068 | 0.18059 | 0.144835 | -0.681764 | 0.304811 | -0.018654 | -0.681044 | 0.868394 | -0.205944 | -0.148294 | -0.00181 | -0.590366 | -0.430049 | -0.189661 | -0.052393 |
| 23.90278 | 0.396801 | 0.292875 | -0.048384 | -0.353275 | 0.148399 | 0.094973 | 0.166017 | 0.087275 | 0.173879 | 0.203891 | 0.197318 | -0.072093 | 0.33554 | 0.063354 | -0.540145 | 0.874253 | -0.013861 | -0.145107 | -0.035425 | -0.582027 | -0.445496 | -0.110046 | -0.094286 |
| 23.91667 | 0.041783 | 0.543008 | 0.149669 | -0.238238 | 0.102009 | 0.09963 | 0.217713 | 0.209856 | 0.264169 | 0.206725 | 0.27779 | 0.525117 | 0.305626 | 0.243803 | -0.338598 | 0.904599 | 0.254497 | -0.13027 | -0.015291 | -0.181001 | -0.458568 | -0.065639 | -0.100979 |
| 23.93056 | -0.166668 | 0.858813 | 0.387974 | 0.038093 | 0.161012 | 0.217168 | 0.257893 | 0.353315 | 0.327418 | 0.220436 | 0.343507 | 0.855472 | 0.314298 | 0.442693 | -0.10848 | 0.886228 | 0.486108 | -0.084647 | 0.070544 | 0.034375 | -0.424856 | -0.020841 | -0.049504 |
| 23.94444 | -0.234079 | 1.054504 | 0.544451 | 0.318643 | 0.286048 | 0.352279 | 0.356954 | 0.482887 | 0.406374 | 0.16119 | 0.404067 | 0.977549 | 0.301708 | 0.488681 | 0.00579 | 0.764075 | 0.645022 | -0.082276 | 0.083227 | -0.069855 | -0.377863 | -0.062309 | -0.046788 |
| 23.95833 | 0.051713 | 1.131509 | 0.661741 | 0.522727 | 0.403796 | 0.496194 | 0.471857 | 0.649698 | 0.523677 | 0.156834 | 0.502473 | 0.916753 | 0.269772 | 0.474858 | 0.009881 | 0.577221 | 0.702304 | 0.045706 | 0.073253 | -0.251828 | -0.40957 | -0.154052 | -0.08508 |
| 23.97222 | 0.707185 | 1.26007 | 0.765076 | 0.631081 | 0.472474 | 0.534743 | 0.537921 | 0.878443 | 0.682121 | 0.164344 | 0.61159 | 0.820902 | 0.281243 | 0.56204 | 0.03499 | 0.376971 | 0.806268 | 0.135113 | 0.169985 | -0.341166 | -0.432221 | -0.172689 | 0.000539 |
| 23.98611 | 1.419135 | 1.538363 | 0.859409 | 0.667478 | 0.512245 | 0.537469 | 0.568907 | 1.09698 | 0.828987 | 0.162643 | 0.682779 | 0.784397 | 0.374677 | 0.76603 | 0.235485 | 0.29441 | 0.954519 | 0.394557 | 0.333235 | -0.293388 | -0.347318 | -0.08757 | 0.042937 |
| 24 | 1.616212 | 1.950147 | 0.87952 | 0.63329 | 0.517665 | 0.585435 | 0.534837 | 1.311707 | 0.932875 | 0.1409 | 0.72028 | 0.696051 | 0.441391 | 0.844687 | 0.464269 | 0.260405 | 1.034222 | 0.48344 | 0.364462 | -0.273757 | -0.351232 | 0.007975 | 0.071878 |
| 24.01389 | 1.17117 | 2.546121 | 0.826627 | 0.518978 | 0.522736 | 0.486713 | 0.397002 | 1.235802 | 0.920334 | 0.106875 | 0.685186 | 0.517343 | 0.370942 | 0.591674 | 0.679552 | 0.249641 | 0.907392 | 0.36443 | 0.171519 | -0.389104 | -0.518099 | -0.017572 | -0.028717 |
| 24.02778 | 0.779331 | 3.21204 | 0.922224 | 0.610422 | 0.644151 | 0.497185 | 0.382512 | 1.148539 | 0.975904 | 0.167986 | 0.671514 | 0.452711 | 0.300248 | 0.235773 | 0.912464 | 0.277129 | 0.610229 | 0.333845 | -0.00241 | -0.524166 | -0.610744 | -0.072747 | -0.149139 |
| 24.04167 | 0.791262 | 3.472954 | 1.178523 | 0.920333 | 0.849036 | 0.688904 | 0.508495 | 1.278667 | 1.256516 | 0.467588 | 0.746758 | 0.490238 | 0.495201 | 0.185069 | 1.220789 | 0.478402 | 0.395143 | 0.486386 | 0.006737 | -0.505947 | -0.463426 | 0.030826 | -0.106939 |
| 24.05556 | 0.579356 | 3.14833 | 1.455439 | 1.180806 | 0.978424 | 0.915658 | 0.686679 | 1.562708 | 1.489132 | 0.751539 | 0.867463 | 0.554145 | 0.727892 | 0.453846 | 1.468556 | 0.748128 | 0.38222 | 0.721508 | 0.175216 | -0.315294 | -0.240687 | 0.314873 | 0.032751 |
| 24.06944 | 0.051257 | 2.644922 | 1.631688 | 1.182846 | 0.897575 | 1.01659 | 0.704488 | 1.538202 | 1.265966 | 0.757845 | 0.832521 | 0.589083 | 0.899943 | 0.65132 | 1.62444 | 0.871598 | 0.377957 | 0.847567 | 0.400719 | -0.116163 | -0.114665 | 0.634409 | 0.092757 |
| 24.08333 | -0.471975 | 1.928412 | 1.55967 | 0.969659 | 1.02047 | 0.886284 | 0.593988 | 1.366005 | 0.820484 | 0.554961 | 0.698839 | 0.538003 | 0.668639 | 0.475142 | 1.352594 | 0.458222 | 0.46664 | 0.551964 | 0.298545 | 0.123466 | -0.145865 | 0.674324 | -0.111292 |
| 24.09722 | -0.531728 | 1.391806 | 1.076638 | 0.650221 | 1.250699 | 0.747029 | 0.529591 | 1.33678 | 0.254185 | 0.392781 | 0.736505 | 0.106996 | -0.103904 | -0.105039 | 0.409619 | -0.383051 | 0.452019 | -0.042927 | -0.205469 | 0.458079 | -0.454984 | 0.152743 | -0.555306 |
| 24.11111 | -0.047793 | 1.154127 | 0.534258 | 0.314537 | 0.79969 | 0.52986 | 0.329929 | 0.950098 | 0.081199 | 0.281729 | 0.585304 | -0.027826 | -0.638539 | -0.412655 | -0.169662 | -0.554546 | -0.023157 | -0.328728 | -0.594883 | 0.278235 | -0.935471 | -0.212977 | -0.907739 |
| 24.125 | 0.972237 | 1.883569 | 0.445016 | 0.519408 | 0.26451 | 0.616153 | 0.207027 | 0.894123 | 0.579533 | 0.324828 | 0.647839 | 0.799415 | -0.301914 | 0.338697 | 0.53997 | -0.207804 | -0.472301 | 0.234548 | -0.361854 | -0.047289 | -0.97964 | 0.281919 | -0.790915 |
| 24.13889 | 2.196071 | 2.931177 | 1.057574 | 1.541539 | 0.43495 | 1.253238 | 0.37834 | 1.806724 | 1.440075 | 0.527225 | 1.091407 | 2.489931 | 0.355133 | 1.28721 | 2.037895 | -0.63338 | -0.324778 | 1.392742 | 0.337775 | 0.00514 | -0.536586 | 1.029152 | -0.396697 |
| 24.15278 | 2.114455 | 1.698063 | 1.496304 | 2.087057 | 0.87241 | 1.388908 | 0.473347 | 1.595037 | 1.803338 | 0.757562 | 1.415708 | 2.433618 | 0.921916 | 1.321812 | 2.625764 | -0.378039 | -0.103295 | 1.970663 | 1.002573 | 0.258985 | -0.073762 | 1.635887 | -0.117775 |
| 24.16667 | 1.763117 | 0.908303 | 1.484337 | 1.671918 | 1.49085 | 1.228496 | 0.649059 | 1.06399 | 1.218011 | 1.046313 | 0.699558 | 1.417 | 1.636438 | 1.290794 | 2.601983 | 0.811231 | -0.120689 | 1.551343 | 1.486995 | 0.639375 | 0.172796 | 2.225922 | 0.01927 |
| 24.18056 | 1.068875 | 0.950536 | 1.264906 | 1.226744 | 2.163115 | 1.117874 | 1.078685 | 1.739035 | 0.575799 | 1.192677 | 0.440984 | 0.115757 | 2.201977 | 1.597683 | 2.331817 | 1.179777 | 0.053307 | 1.145476 | 1.498627 | 0.556139 | 0.227172 | 2.112837 | 0.164564 |
| 24.19444 | 0.205567 | 1.189232 | 1.005187 | 0.804026 | 2.430003 | 0.82846 | 1.324038 | 2.118369 | 0.243759 | 1.218966 | 0.274301 | -0.691746 | 2.323366 | 1.921731 | 1.852831 | 0.696178 | 0.466161 | 0.811511 | 1.200559 | -0.068853 | 0.163714 | 1.97272 | 0.33986 |
| 24.20833 | -0.532987 | 1.660695 | 0.671472 | 0.578665 | 2.222837 | 0.557203 | 1.294901 | 1.886205 | 0.097601 | 1.135768 | -0.034243 | -0.250525 | 1.972448 | 1.805148 | 1.27755 | 0.230162 | 1.035233 | 0.441679 | 0.920965 | -0.282145 | 0.133913 | 1.177094 | 0.535598 |
| 24.22222 | -0.722963 | 2.080778 | 0.501855 | 0.654704 | 1.788478 | 0.39803 | 1.228889 | 1.509084 | 0.164588 | 1.113274 | -0.297928 | 0.415995 | 1.465448 | 1.688118 | 0.979912 | -0.064619 | 1.436089 | 0.194932 | 0.720381 | -0.210878 | 0.191306 | 0.357983 | 0.677898 |
| 24.23611 | -0.302756 | 1.954782 | 0.360253 | 1.026385 | 1.395487 | 0.214394 | 1.043649 | 0.804157 | 0.247551 | 1.161626 | -0.231363 | 0.900806 | 1.110442 | 1.714227 | 0.945174 | -0.341413 | 1.173184 | 0.118176 | 0.389454 | -0.276976 | 0.198125 | 0.081852 | 0.645018 |
| 24.25 | 0.659572 | 1.081254 | 0.53989 | 1.400923 | 0.916507 | 0.246457 | 0.757639 | 0.075036 | 0.401339 | 1.210225 | 0.289653 | 1.017333 | 0.650611 | 1.249776 | 1.019225 | -0.447126 | 0.508383 | 0.120869 | 0.004625 | -0.291547 | 0.118632 | 0.03972 | 0.473206 |
| 24.26389 | 1.489112 | 0.123484 | 0.799834 | 1.42097 | 0.511035 | 0.485447 | 0.44143 | -0.358788 | 0.628444 | 1.325116 | 0.939447 | 0.705651 | 0.239166 | 0.584322 | 0.885161 | -0.254105 | 0.015403 | 0.173983 | -0.212722 | 0.224196 | 0.145279 | 0.030496 | 0.339991 |
| 24.27778 | 1.337523 | -0.414902 | 0.757736 | 1.064021 | 0.200934 | 0.643534 | 0.183845 | -0.667933 | 0.756026 | 1.371467 | 1.209042 | 0.358802 | -0.094653 | -0.10593 | 0.7648 | 0.046284 | -0.144712 | 0.197051 | -0.176022 | 1.337014 | 0.263384 | 0.093704 | 0.319402 |
| 24.29167 | 0.490484 | -0.651541 | 0.3848 | 0.582128 | -0.11372 | 0.562201 | -0.043523 | -0.875176 | 0.668506 | 1.329024 | 0.841846 | -0.234115 | -0.485685 | -0.631358 | 0.575399 | 0.31209 | -0.456579 | 0.20266 | 0.030134 | 1.921091 | 0.298847 | 0.446661 | 0.287689 |
| 24.30556 | -0.30077 | -0.745942 | 0.041488 | 0.147359 | -0.314255 | 0.311018 | -0.193243 | -0.828388 | 0.249955 | 1.191079 | 0.361598 | -0.860851 | -0.88055 | -0.59015 | 0.222397 | 0.521136 | -1.215295 | 0.27822 | 0.292259 | 0.834749 | 0.193032 | 0.877973 | 0.146874 |
| 24.31944 | -0.69954 | -0.407496 | -0.154344 | -0.092955 | -0.375062 | 0.091856 | -0.189597 | -0.450148 | -0.340363 | 0.979081 | -0.018221 | -0.790602 | -1.044588 | -0.048937 | -0.281163 | 0.564374 | -1.972496 | 0.500745 | 0.421815 | -0.252159 | 0.147259 | 1.042824 | 0.038074 |
| 24.33333 | -1.44363 | -0.210095 | -0.355429 | -0.179089 | -0.463095 | -0.079821 | -0.048269 | -0.144405 | -0.640249 | 0.744109 | -0.315158 | -0.188006 | -0.720221 | 0.571964 | -0.634193 | 0.408784 | -1.312643 | 0.703948 | 0.449309 | -0.550585 | 0.253916 | 0.666668 | 0.102076 |
| 24.34722 | -1.706189 | -0.216577 | -0.62446 | -0.133504 | -0.558556 | -0.183599 | 0.178106 | -0.07142 | -0.591488 | 0.45503 | -0.483943 | 0.219836 | -0.237094 | 0.80816 | -0.460308 | 0.156758 | 0.134675 | 0.603783 | 0.488588 | -0.411079 | 0.356139 | 0.37013 | 0.212677 |
| 24.36111 | -1.279528 | -0.512957 | -0.73827 | -0.007821 | -0.604378 | -0.236088 | 0.296497 | -0.253451 | -0.308732 | 0.199093 | -0.357506 | -0.004669 | 0.007439 | 0.56612 | -0.303541 | -0.235502 | 1.148966 | 0.213652 | 0.265957 | -0.100733 | 0.327269 | 0.083597 | 0.216402 |
| 24.375 | -0.684234 | -0.74424 | -0.669521 | -0.012925 | -0.52897 | -0.185469 | 0.285075 | -0.446021 | -0.109889 | 0.002234 | -0.233751 | -0.453489 | 0.030379 | 0.218538 | -0.441669 | -0.532776 | 0.774894 | -0.155006 | 0.02377 | 0.443214 | 0.276875 | -0.254653 | 0.16076 |
| 24.38889 | -0.182946 | -0.866756 | -0.507606 | -0.156804 | -0.356067 | -0.096711 | 0.192945 | -0.487354 | -0.011391 | -0.089458 | -0.170973 | -0.727966 | 0.001531 | 0.0262 | -0.676116 | -0.392467 | 1.068068 | -0.380365 | -0.042609 | 0.857397 | 0.285671 | -0.544719 | 0.109144 |
| 24.40278 | 0.081629 | -0.65348 | -0.387001 | -0.148935 | -0.275588 | 0.014038 | 0.051191 | -0.326928 | 0.072479 | -0.193649 | -0.13842 | -0.529286 | -0.038747 | -0.097453 | -0.589245 | -0.136077 | 1.332761 | -0.475946 | 0.002517 | 0.537179 | 0.355227 | -0.485151 | 0.070381 |
| 24.41667 | 0.285519 | -0.454939 | -0.41224 | -0.0119 | -0.280757 | -0.008994 | -0.072468 | -0.459902 | 0.023646 | -0.327572 | -0.180289 | -0.019206 | -0.17632 | -0.196273 | -0.223789 | 0.224776 | 1.357714 | -0.482099 | 0.056412 | -0.231792 | 0.432171 | -0.508125 | 0.056733 |
| 24.43056 | 0.268567 | -0.381482 | -0.505911 | 0.005762 | -0.453524 | -0.15621 | -0.238894 | -1.060539 | -0.147604 | -0.385827 | -0.324619 | 0.215856 | -0.405548 | -0.224366 | -0.024895 | 0.599003 | 0.662609 | -0.372621 | 0.082037 | -0.783392 | 0.401768 | -0.630666 | 0.041065 |
| 24.44444 | 0.119209 | -0.536454 | -0.515094 | -0.228643 | -0.659691 | -0.371773 | -0.32572 | -1.515473 | -0.339986 | -0.416392 | -0.497776 | -0.070103 | -0.436419 | -0.104732 | -0.162142 | 0.788629 | 0.396879 | -0.225544 | 0.035245 | -0.634175 | 0.268444 | -0.54714 | -0.008513 |
| 24.45833 | 0.37557 | -0.892165 | -0.527208 | -0.610743 | -0.760034 | -0.489009 | -0.30259 | -1.199651 | -0.557326 | -0.445409 | -0.589786 | -0.708122 | -0.321754 | 0.105516 | -0.386268 | 0.444322 | -0.819132 | -0.119261 | -0.018868 | -0.147078 | 0.172945 | -0.506188 | -0.073606 |
| 24.47222 | 0.199304 | -1.232538 | -0.674055 | -0.886747 | -0.71621 | -0.483144 | -0.197652 | -0.953161 | -0.747373 | -0.460666 | -0.627406 | -1.481145 | -0.312466 | 0.205625 | -0.48593 | -0.121415 | -1.753938 | -0.17588 | -0.065576 | 0.245019 | 0.181988 | -0.641371 | -0.054012 |
| 24.48611 | -0.216862 | -1.608172 | -0.809768 | -0.780451 | -0.622054 | -0.441757 | -0.051073 | -0.732735 | -0.827055 | -0.528432 | -0.709666 | -1.612179 | -0.440663 | 0.175881 | -0.461408 | -0.698419 | -1.553622 | -0.269145 | -0.080326 | 0.497252 | 0.193495 | -0.804787 | 0.019733 |
| 24.5 | -0.461973 | -1.837835 | -0.825731 | -0.501584 | -0.592339 | -0.368444 | -0.171669 | -0.548197 | -0.726101 | -0.45915 | -0.718429 | -1.413057 | -0.507114 | 0.158652 | -0.4855 | -1.006013 | -0.949101 | -0.333138 | -0.16981 | 0.563311 | 0.126868 | -0.764229 | 0.014492 |
| 24.51389 | -0.207603 | -1.913616 | -0.853962 | -0.392958 | -0.541815 | -0.261696 | -0.422879 | -0.501939 | -0.626828 | -0.305718 | -0.538227 | -1.243827 | -0.410843 | 0.226314 | -0.607538 | -0.668372 | -0.663381 | -0.251681 | -0.256698 | 0.629899 | 0.078669 | -0.913646 | -0.057583 |
| 24.52778 | -0.298945 | -1.729563 | -0.944755 | -0.474354 | -0.466615 | -0.238308 | -0.605983 | -0.711198 | -0.613121 | -0.027312 | -0.364581 | -1.177041 | -0.23072 | 0.389794 | -0.698775 | -0.453472 | -0.18584 | -0.108429 | -0.209668 | 0.88709 | 0.109127 | -1.569694 | -0.027965 |
| 24.54167 | -0.539116 | -1.655563 | -1.177557 | -0.772351 | -0.485042 | -0.531359 | -0.661237 | -1.107409 | -0.784144 | 0.066272 | -0.373461 | -1.135749 | -0.284638 | 0.34535 | -0.823589 | -0.565763 | -0.213193 | -0.188145 | -0.200517 | 0.702935 | 0.040322 | -1.989886 | -0.030725 |

| | | | | | | | | | | | | | | | | | | | | | | | |
|---|---|---|---|---|---|---|---|---|---|---|---|---|---|---|---|---|---|---|---|---|---|---|---|
| 24.55556 | -0.802282 | -1.861287 | -1.349006 | -1.033401 | -0.615814 | -0.91753 | -0.738014 | -1.324625 | -0.98352 | -0.037741 | -0.51065 | -1.263492 | -0.589764 | 0.117343 | -1.100054 | -1.032126 | -0.339716 | -0.446267 | -0.380007 | 0.313666 | -0.155453 | -1.925796 | -0.213495 |
| 24.56944 | -0.992005 | -2.438638 | -1.300243 | -1.083044 | -0.732372 | -0.991044 | -0.805242 | -1.449939 | -1.005156 | -0.163702 | -0.574105 | -1.328988 | -0.827057 | -0.108012 | -1.322919 | -1.217336 | -0.616063 | -0.650947 | -0.564929 | 0.096714 | -0.318325 | -1.872095 | -0.370285 |
| 24.58333 | -1.190784 | -2.838711 | -1.253514 | -1.371978 | -0.868507 | -1.042747 | -1.057708 | -1.600705 | -1.011718 | -0.287444 | -0.579254 | -1.277805 | -1.181799 | -0.658468 | -1.723598 | -1.412353 | -0.98144 | -0.861809 | -0.747397 | -0.299986 | -0.647863 | -1.906835 | -0.645572 |
| 24.59722 | -1.692408 | -3.224657 | -1.917473 | -2.067913 | -1.295946 | -1.234539 | -1.456777 | -2.553629 | -1.064958 | -0.419171 | -0.728324 | -1.273567 | -1.927849 | -1.847948 | -2.855215 | -2.078415 | -1.598571 | -1.375611 | -1.254721 | -0.738143 | -1.250897 | -2.345627 | -1.14112 |
| 24.61111 | -2.615186 | -3.665246 | -2.941581 | -2.561156 | -1.755995 | -1.547645 | -1.742899 | -3.826957 | -1.151454 | -0.476042 | -1.003474 | -0.992988 | -2.195886 | -2.455531 | -3.453891 | -2.432311 | -2.087615 | -1.463755 | -1.321354 | -0.273511 | -1.226173 | -2.172313 | -1.041949 |
| 24.625 | -2.691595 | -3.037135 | -2.777866 | -1.937239 | -1.559847 | -1.51615 | -1.549291 | -3.679317 | -0.835609 | -0.38325 | -1.097419 | -0.500415 | -1.56973 | -1.951567 | -2.812218 | -2.165568 | -1.79489 | -0.853038 | -0.673311 | 0.538494 | -0.4742 | -1.523818 | -0.240222 |
| 24.63889 | -1.785391 | -1.879028 | -1.501386 | -0.635229 | -1.160945 | -0.94697 | -0.789694 | -1.64453 | -0.279841 | -0.393228 | -0.964021 | -0.799747 | -1.083661 | -1.465498 | -2.299487 | -1.70637 | -1.567571 | -0.501366 | -0.29471 | 0.5506 | -0.149404 | -1.000797 | -0.223535 |
| 24.65278 | -0.988227 | -0.879914 | -0.327013 | 1.003782 | -0.821774 | -0.445209 | -0.252285 | -0.072237 | 0.071989 | -0.354226 | -0.745778 | -1.075372 | -1.024981 | -1.360255 | -2.075595 | -1.476551 | -1.414827 | -0.570399 | -0.407524 | 0.661307 | -0.34644 | -0.575979 | -0.699263 |
| 24.66667 | -0.202224 | -0.435103 | 0.20643 | 1.315943 | -0.509619 | 0.111951 | 0.03977 | 0.640309 | 0.254676 | -0.223979 | -0.477588 | -1.331358 | -1.038772 | -1.469948 | -1.895457 | -1.211614 | -1.461578 | -0.685887 | -0.519135 | 0.664586 | -0.607377 | -0.572 | -0.809304 |
| 24.68056 | 0.2376 | -0.22723 | 0.342173 | 1.244079 | -0.320501 | 0.405688 | 0.065385 | 0.510784 | 0.410907 | -0.246574 | -0.254424 | -1.226577 | -1.042934 | -1.476967 | -1.789009 | -0.898601 | -1.320278 | -0.750922 | -0.50002 | 0.454427 | -0.761974 | -0.683986 | -0.594192 |
| 24.69444 | 0.408618 | 0.016415 | 0.314185 | 0.893541 | -0.267825 | 0.232384 | -0.01957 | 0.574893 | 0.387332 | -0.280517 | -0.08914 | -0.977874 | -0.990619 | -1.195874 | -1.688987 | -0.566452 | -0.806483 | -0.759528 | -0.391551 | 0.329384 | -0.806349 | -0.401623 | -0.453268 |
| 24.70833 | 0.525517 | 0.126678 | 0.100461 | 0.157412 | -0.268911 | -0.027328 | -0.092135 | 0.436242 | 0.189246 | -0.208527 | 0.055453 | -0.792929 | -0.693432 | -0.964147 | -1.456633 | -0.270204 | -0.308248 | -0.690161 | -0.418669 | 0.172707 | -0.782849 | -0.255589 | -0.429428 |
| 24.72222 | 0.723784 | 0.021882 | -0.181524 | -0.556283 | -0.247036 | -0.10647 | -0.146825 | 0.177498 | -0.03986 | -0.129368 | 0.18047 | -0.589841 | -0.335416 | -0.941709 | -1.238805 | -0.062778 | 0.166513 | -0.556444 | -0.527366 | 0.102279 | -0.676126 | -0.296396 | -0.377905 |
| 24.73611 | 0.956623 | -0.102055 | -0.130357 | -0.947506 | -0.213776 | -0.071596 | -0.146199 | 0.145885 | -0.241977 | -0.201536 | 0.22548 | -0.216484 | -0.17368 | -0.945326 | -1.118851 | 0.136404 | 0.54506 | -0.261888 | -0.47927 | 0.090508 | -0.403037 | -0.243139 | -0.180106 |
| 24.75 | 0.691421 | -0.258719 | 0.40135 | -1.030269 | -0.167067 | -0.06821 | -0.101201 | 0.101015 | -0.412713 | -0.450651 | 0.085882 | 1.184217 | 0.554758 | -1.143622 | -1.13733 | 0.746558 | 1.207908 | 0.645525 | -0.150633 | 0.563097 | 0.317647 | -0.373605 | 0.454941 |
| 24.76389 | 0.208637 | -0.308561 | 0.70767 | -0.923885 | -0.164164 | -0.12541 | -0.033094 | -0.116809 | -0.59097 | -0.657789 | -0.158381 | 3.803094 | 1.88928 | -0.965937 | -0.34182 | 1.467717 | 2.415976 | 1.737256 | 0.235294 | 2.09062 | 1.387843 | -0.415726 | 1.681922 |
| 24.77778 | -0.139783 | -0.240677 | 0.44688 | -0.772239 | -0.499866 | -0.222317 | -0.105617 | -0.133395 | -0.549152 | -0.610569 | -0.111346 | 5.443736 | 2.345159 | -1.06848 | 1.289902 | 0.126406 | 1.83062 | 0.367158 | 0.791774 | 2.0483 | 1.30722 | -0.850829 | 2.073293 |
| 24.79167 | -0.692111 | -0.159134 | -0.396407 | -0.174936 | -1.553334 | -0.2186 | -0.117463 | -0.091107 | -0.324724 | -0.425914 | 0.137652 | 1.752144 | 0.873718 | -2.01935 | 0.652778 | -1.081472 | 1.759844 | -1.863679 | -0.369552 | -2.353005 | -0.342159 | -0.681549 | 0.069328 |
| 24.80556 | -1.483549 | -0.221168 | -0.421946 | 0.315496 | -1.548753 | 0.333662 | 0.166651 | -0.449526 | -0.192683 | -0.391562 | 0.205776 | -0.705922 | -0.549333 | -0.907618 | 0.298825 | -0.436742 | 0.520412 | -1.682453 | 0.035424 | -2.54506 | -0.959891 | -0.32062 | -0.641356 |
| 24.81944 | -0.99386 | -0.463388 | -0.571782 | 0.445365 | -0.609133 | -0.057011 | -0.234418 | -0.810706 | -0.194471 | -0.357539 | 0.085742 | -1.284584 | -0.994271 | -0.353562 | 0.170764 | 0.046916 | 0.15318 | -1.31433 | -0.084836 | -1.61499 | -0.597939 | -0.181819 | -0.952351 |
| 24.83333 | -0.631226 | -0.617266 | -0.756241 | 0.170404 | -0.181 | -0.18061 | -0.271583 | -0.828 | -0.236454 | -0.322684 | -0.024822 | -1.104824 | -0.984967 | -0.513489 | -0.057062 | 0.108824 | 0.409246 | -1.090271 | -0.275003 | -0.811991 | -0.56728 | -0.285748 | -1.409356 |
| 24.84722 | -0.447101 | -0.531822 | -0.559878 | -0.141864 | 0.068933 | -0.170474 | -0.176932 | -0.638656 | -0.228134 | -0.264022 | -0.091061 | -0.861489 | -0.780915 | -0.667978 | -0.289211 | 0.213408 | 0.349425 | -0.93937 | -0.395235 | -0.603257 | -0.534986 | -0.404783 | -1.655779 |
| 24.86111 | -0.284175 | -0.413 | -0.494555 | -0.334207 | 0.135902 | -0.217688 | -0.115862 | -0.481124 | -0.138864 | -0.314302 | -0.158056 | -0.477152 | -0.595315 | -0.708383 | -0.501266 | 0.336212 | 0.22657 | -0.871305 | -0.413865 | -0.571247 | -0.531165 | -0.457152 | -1.69659 |
| 24.875 | -0.030923 | -0.279947 | -0.359148 | -0.395614 | 0.115253 | -0.134423 | -0.021113 | -0.396462 | -0.048365 | -0.440431 | -0.214392 | -0.016891 | -0.514241 | -0.743643 | -0.638793 | 0.310078 | 0.086058 | -0.824004 | -0.435776 | -0.581018 | -0.581805 | -0.447129 | -1.443082 |
| 24.88889 | 0.320667 | 0.010144 | -0.197859 | -0.372463 | 0.179018 | -0.107256 | 0.110235 | -0.157162 | -0.038113 | -0.524563 | -0.198643 | 0.219557 | -0.496027 | -0.789158 | -0.735071 | 0.121017 | -0.129761 | -0.834631 | -0.516965 | -0.488258 | -0.675676 | -0.417717 | -1.072439 |
| 24.90278 | 0.66295 | 0.409556 | 0.055205 | -0.243603 | 0.331554 | 0.090835 | 0.222157 | 0.252769 | -0.045291 | -0.471585 | -0.068919 | -0.486181 | -0.770962 | -1.372715 | -1.268467 | -0.748681 | -0.336109 | -0.967095 | -0.781134 | 0.00471 | -0.823903 | -0.328243 | -1.112756 |
| 24.91667 | 1.015882 | 0.763373 | 0.317091 | 0.032235 | 0.454056 | 0.447476 | 0.320141 | 0.599087 | 0.031176 | -0.342772 | 0.121454 | -3.043614 | -2.013293 | -4.076649 | -3.120114 | -3.324674 | -1.140069 | -1.628896 | -1.74418 | 0.733002 | -0.918441 | -0.685298 | -2.086495 |
| 24.93056 | 1.11438 | 1.149811 | 0.461717 | 0.296208 | 0.525422 | 0.721361 | 0.296315 | 0.749027 | 0.152118 | -0.161361 | 0.313453 | -4.510914 | -2.286791 | -5.360752 | -3.026358 | -3.720154 | -1.936754 | -2.438195 | -1.941957 | 0.123247 | -0.767637 | -1.120847 | -2.341174 |
| 24.94444 | 1.022454 | 1.313524 | 0.646489 | 0.442364 | 0.612172 | 0.849406 | 0.122762 | 0.793702 | 0.275996 | 0.088974 | 0.478449 | -1.610974 | -0.771313 | -1.857474 | -0.9384 | -0.922795 | -0.715379 | -1.083396 | -0.676697 | -0.551079 | -0.54349 | -0.63415 | -0.905733 |
| 24.95833 | 1.235565 | 1.250585 | 0.810639 | 0.550882 | 0.634105 | 0.74657 | 0.094044 | 0.83031 | 0.414107 | 0.351656 | 0.603935 | 0.814662 | 0.011155 | 2.001279 | 0.405154 | 1.644752 | 0.545093 | 0.15612 | 0.2676 | -0.626686 | -0.389001 | 0.454023 | 0.376171 |
| 24.97222 | 1.304999 | 1.246987 | 0.98851 | 0.643493 | 0.715497 | 0.702453 | 0.422764 | 0.95369 | 0.550347 | 0.508635 | 0.61625 | 1.057764 | 0.288482 | 3.165582 | 0.561165 | 2.134585 | 0.870136 | 0.547628 | 0.474804 | -0.436565 | -0.271468 | 0.907283 | 0.629526 |
| 24.98611 | 1.062367 | 1.357743 | 1.147268 | 0.636411 | 0.862762 | 0.644639 | 0.639554 | 1.041573 | 0.615197 | 0.582278 | 0.642403 | 0.72637 | 0.45275 | 2.897987 | 0.437551 | 1.287231 | 0.544692 | 0.344466 | 0.26477 | -0.312983 | -0.241288 | 0.69538 | 0.399019 |
| 25 | 0.875005 | 1.45508 | 1.023421 | 0.554505 | 0.824788 | 0.567775 | 0.667527 | 0.989999 | 0.578868 | 0.543654 | 0.599129 | 0.406292 | 0.462666 | 2.262818 | 0.229179 | 0.776883 | 0.239865 | -0.031292 | -0.043477 | -0.354746 | -0.31158 | 0.250824 | 0.220289 |
| 25.01389 | 0.582451 | 1.625455 | 0.783108 | 0.558255 | 0.634615 | 0.364565 | 0.623179 | 0.906129 | 0.341583 | 0.312313 | 0.401995 | 0.076995 | 0.088991 | 1.250529 | -0.184936 | 0.409087 | -0.363139 | -0.467901 | -0.326391 | -0.646702 | -0.783963 | -0.186176 | 0.099539 |
| 25.02778 | 0.333252 | 1.943945 | 0.831998 | 0.611287 | 0.458563 | 0.288381 | 0.493528 | 0.727209 | 0.24125 | 0.119911 | 0.110925 | -0.05644 | -0.443661 | 0.876763 | -0.450715 | -0.032055 | -0.634163 | -0.589153 | -0.232951 | -0.774697 | -1.079641 | -0.324798 | 0.042496 |
| 25.04167 | 0.691646 | 2.201708 | 1.228497 | 0.925035 | 0.529729 | 0.667561 | 0.708481 | 0.9106 | 0.71385 | 0.391459 | 0.376025 | 0.154976 | -0.131006 | 1.101246 | -0.270799 | 0.069084 | -0.357334 | -0.285751 | 0.071542 | -0.567811 | -0.784643 | -0.126255 | 0.177919 |
| 25.05556 | 1.080224 | 2.703075 | 1.517563 | 1.217379 | 0.658924 | 1.089693 | 1.074473 | 1.246085 | 1.12507 | 0.68572 | 0.630441 | 0.456373 | 0.269391 | 1.420965 | -0.037131 | 0.296647 | -0.11183 | -0.110462 | 0.294222 | -0.425981 | -0.424156 | -0.001377 | 0.228429 |
| 25.06944 | 1.011532 | 2.305525 | 0.952409 | 1.192557 | 0.606055 | 1.100647 | 1.029573 | 1.363169 | 1.212553 | 0.744819 | 0.719915 | 0.699033 | 0.361862 | 1.586188 | 0.081659 | 0.340504 | -0.198718 | -0.116547 | 0.475308 | -0.498851 | -0.303656 | 0.011025 | 0.195835 |
| 25.08333 | 0.999299 | 1.739973 | 0.092239 | 1.18167 | 0.826294 | 0.906778 | 0.877912 | 1.862254 | 1.146016 | 0.890698 | 0.771335 | 0.753087 | 0.07192 | 1.677822 | -0.202396 | -0.092277 | -0.701255 | -0.597488 | 0.459431 | -1.105718 | -0.701945 | -0.332496 | -0.224737 |
| 25.09722 | 0.893075 | 1.713161 | 0.295664 | 0.677709 | 1.504173 | 0.841953 | 0.922613 | 2.651909 | 0.731583 | 0.874923 | 0.74589 | 0.392726 | -0.974897 | 1.452502 | -1.124267 | -0.983148 | -1.588437 | -1.719764 | 0.113673 | -2.321852 | -2.040951 | -1.291624 | -1.261883 |
| 25.11111 | 0.539718 | 1.489111 | 0.719437 | 0.21384 | 1.228377 | 0.467961 | 0.564947 | 1.918442 | 0.244679 | 0.715764 | 0.534784 | 0.422216 | -1.626596 | 0.97065 | -1.557452 | -1.093315 | -1.865735 | -2.090682 | -0.425306 | -2.395324 | -2.533547 | -1.733762 | -1.749858 |
| 25.125 | 0.80785 | 1.809106 | 1.335861 | 0.576633 | 0.593176 | 0.368164 | 0.349422 | 1.403521 | 0.549141 | 0.690574 | 0.471161 | 0.874085 | -0.920423 | 0.575469 | -0.998044 | 0.02742 | -1.045369 | -1.147849 | -0.399887 | -1.286186 | -1.500499 | -1.110428 | -1.135283 |
| 25.13889 | 1.917929 | 1.219969 | 2.060061 | 1.713471 | 0.743306 | 0.643207 | 0.492866 | 2.118541 | 1.471132 | 1.028875 | 1.037553 | 0.960221 | -0.173701 | 0.3886 | -0.395775 | 0.661813 | -0.451655 | -0.447408 | -0.193208 | -0.556069 | -0.799966 | -0.521776 | -0.560262 |
| 25.15278 | 1.894509 | 1.094845 | 2.247951 | 2.509018 | 0.975765 | 0.962479 | 0.612096 | 1.710607 | 1.940386 | 0.987551 | 1.347319 | 0.862073 | 0.19085 | 0.525409 | -0.032233 | 0.602581 | -0.203554 | -0.162417 | -0.057586 | -0.249363 | -0.590421 | -0.216307 | -0.340192 |
| 25.16667 | 1.096087 | 1.264549 | 1.671158 | 2.344446 | 1.238719 | 0.664137 | 0.474304 | 0.85118 | 1.449355 | 0.851107 | 1.162369 | 0.972104 | 0.407373 | 0.819645 | 0.270138 | 0.435894 | -0.031202 | 0.024054 | 0.075792 | -0.063845 | -0.403754 | 0.005202 | -0.266641 |
| 25.18056 | 0.834509 | 1.706854 | 1.065277 | 1.704997 | 1.668656 | 0.516975 | 0.664538 | 1.182987 | 0.859092 | 0.832307 | 0.80502 | 1.020852 | 0.453118 | 0.816422 | 0.431836 | 0.37692 | 0.108289 | 0.172223 | 0.150991 | 0.072183 | -0.178109 | 0.155738 | -0.134647 |
| 25.19444 | 0.20154 | 1.430837 | 0.9579 | 1.229012 | 1.913619 | 0.662221 | 1.068249 | 2.380379 | 0.507203 | 0.914296 | 0.340983 | 0.824835 | 0.385975 | 0.62871 | 0.407812 | 0.319159 | 0.138119 | 0.205133 | 0.124021 | 0.108598 | -0.112054 | 0.174526 | 0.280955 |
| 25.20833 | -1.169464 | 0.916935 | 0.673177 | 0.892619 | 2.035779 | 0.807085 | 1.086937 | 2.768324 | 0.372363 | 0.911064 | -0.254024 | 0.639189 | 0.259108 | 0.448939 | 0.332987 | 0.286533 | 0.101917 | 0.183924 | 0.074802 | 0.081995 | -0.12605 | 0.126654 | 0.770788 |
| 25.22222 | -1.199666 | 1.206586 | 0.609106 | 0.548709 | 2.089525 | 0.645507 | 1.037909 | 2.53156 | 0.18001 | 0.889375 | -0.446686 | 0.471346 | 0.146658 | 0.325548 | 0.243932 | 0.221929 | 0.053278 | 0.11501 | 0.065264 | 0.020763 | -0.124552 | 0.087808 | 0.571238 |
| 25.23611 | -0.375083 | 1.449403 | 0.779934 | 0.323726 | 2.159342 | 0.494541 | 0.97652 | 1.943145 | 0.07124 | 0.907718 | 0.037032 | 0.345691 | 0.084325 | 0.221346 | 0.149258 | 0.171002 | 0.01665 | 0.041556 | 0.055875 | -0.019252 | -0.11469 | 0.064182 | 0.126916 |
| 25.25 | 0.980452 | 1.202411 | 1.139791 | 0.176874 | 1.869961 | 0.387573 | 0.691999 | 1.049294 | 0.235032 | 0.804466 | 0.602764 | 0.252007 | 0.084878 | 0.127379 | 0.084978 | 0.115463 | -0.001823 | 0.044085 | 0.04337 | -0.02723 | -0.039338 | 0.036408 | -0.087363 |
| 25.26389 | 2.007117 | 0.690047 | 1.085326 | 0.184757 | 1.404883 | 0.297254 | 0.454355 | 0.364937 | 0.40711 | 0.82546 | 1.125586 | 0.113689 | 0.046214 | 0.014172 | -0.026851 | -0.038449 | -0.103416 | 0.017592 | -0.050622 | -0.128578 | -0.068035 | -0.068215 | -0.142888 |
| 25.27778 | 1.604392 | 0.327877 | 0.757729 | 0.317236 | 0.865433 | 0.292931 | 0.289027 | -0.373652 | 0.37207 | 0.864105 | 1.295758 | -0.137659 | -0.155501 | -0.217029 | -0.212919 | -0.338199 | -0.315675 | -0.157383 | -0.260799 | -0.355659 | -0.2978 | -0.275028 | -0.272401 |
| 25.29167 | 0.308927 | -0.19901 | 0.338439 | 0.533824 | 0.547237 | 0.090317 | 0.158074 | -1.29559 | 0.292676 | 0.594946 | 0.968533 | -0.368453 | -0.411366 | -0.473768 | -0.312193 | -0.569669 | -0.472349 | -0.322015 | -0.439265 | -0.506399 | -0.523944 | -0.454186 | -0.479286 |
| 25.30556 | -0.208332 | -0.424098 | -0.396373 | 0.676656 | 0.35548 | -0.10314 | 0.051802 | -1.800037 | 0.227476 | 0.338336 | 0.296952 | -0.392138 | -0.462833 | -0.519613 | -0.221114 | -0.597947 | -0.466955 | -0.326691 | -0.450956 | -0.498499 | -0.521504 | -0.471801 | -0.524455 |
| 25.31944 | -0.418107 | -0.093365 | -0.758121 | 0.61553 | 0.285043 | -0.150368 | -0.107178 | -1.716551 | 0.085726 | 0.400673 | -0.103955 | -0.213393 | -0.316151 | -0.335381 | 0.002752 | -0.484309 | -0.366633 | -0.196516 | -0.334674 | -0.402347 | -0.374936 | -0.352438 | -0.399293 |
| 25.33333 | -1.358054 | 0.183169 | -0.902357 | 0.445953 | 0.137136 | -0.115002 | 0.10166 | -1.094845 | -0.139775 | 0.616114 | -0.408359 | 0.078536 | -0.077067 | -0.069797 | 0.26399 | -0.252138 | -0.172024 | 0.024427 | -0.125574 | -0.179274 | -0.155375 | -0.126311 | -0.14307 |
| 25.34722 | -1.555231 | 0.315996 | -1.045964 | 0.192566 | -0.136125 | -0.018177 | 0.154539 | -0.463423 | -0.365509 | 0.780335 | -0.750164 | 0.310063 | 0.10839 | 0.089104 | 0.428635 | -0.053191 | 0.006034 | 0.187168 | 0.045678 | 0.011528 | 0.019139 | 0.071541 | 0.078945 |
| 25.36111 | -1.464488 | 0.246733 | -1.040071 | -0.091332 | -0.35235 | -0.019054 | 0.044485 | -0.169582 | -0.524761 | 0.6683 | -0.666911 | 0.299911 | 0.089372 | 0.015787 | 0.319096 | -0.070246 | -0.004353 | 0.130681 | 0.026356 | 0.001936 | 0.000943 | 0.101862 | 0.05059 |
| 25.375 | -0.827251 | -0.14671 | -0.738244 | -0.393502 | -0.444847 | -0.078722 | -0.116758 | -0.210708 | -0.593826 | 0.242099 | -0.393521 | 0.21656 | 0.001403 | -0.08097 | 0.170372 | -0.138795 | -0.068876 | 0.016433 | -0.049338 | -0.054917 | -0.095489 | 0.061019 | -0.105209 |
| 25.38889 | -0.174197 | -0.510812 | -0.341783 | -0.554852 | -0.451934 | -0.085346 | -0.133951 | -0.318907 | -0.503514 | -0.052188 | -0.15535 | 0.280258 | 0.065023 | -0.00379 | 0.225097 | -0.086069 | -0.017638 | 0.061411 | -0.001191 | -0.008628 | -0.075494 | 0.096866 | -0.120064 |
| 25.40278 | 0.227788 | -0.658994 | -0.09995 | -0.47247 | -0.407394 | -0.025395 | 0.006328 | -0.38116 | -0.419023 | -0.014979 | 0.106038 | 0.443886 | 0.285 | 0.201845 | 0.437128 | 0.035136 | 0.115254 | 0.216666 | 0.151738 | 0.115306 | 0.079175 | 0.224019 | 0.050687 |
| 25.41667 | 0.663114 | -0.843353 | 0.014795 | -0.355987 | -0.367923 | 0.019672 | 0.077916 | -0.563271 | -0.34809 | 0.052886 | 0.361899 | 0.572734 | 0.478723 | 0.346123 | 0.639481 | 0.13444 | 0.236034 | 0.348014 | 0.285239 | 0.236762 | 0.242941 | 0.353365 | 0.284723 |
| 25.43056 | 1.317356 | -1.020445 | 0.098599 | -0.266097 | -0.391621 | -0.002381 | -0.042478 | -0.717457 | -0.333848 | 0.07651 | 0.52269 | 0.544222 | 0.401138 | 0.235829 | 0.639658 | 0.136321 | 0.269209 | 0.34086 | 0.276936 | 0.287154 | 0.229037 | 0.332749 | 0.377514 |
| 25.44444 | 1.240839 | -1.125406 | 0.068388 | -0.250801 | -0.437018 | 0.059066 | -0.19865 | -0.779239 | -0.440099 | 0.010096 | 0.404722 | 0.397546 | 0.189072 | 0.07363 | 0.522169 | 0.060827 | 0.232454 | 0.248778 | 0.188455 | 0.247921 | 0.13094 | 0.221202 | 0.291598 |
| 25.45833 | 0.533866 | -1.049169 | -0.309613 | -0.374856 | -0.490937 | -0.233471 | -0.380531 | -0.741334 | -0.523599 | -0.157602 | -0.204146 | 0.390689 | 0.21526 | 0.145429 | 0.419341 | 0.041786 | 0.216275 | 0.240145 | 0.19737 | 0.215477 | 0.153962 | 0.219633 | 0.226604 |
| 25.47222 | 0.198215 | -1.041351 | -0.727507 | -0.513768 | -0.577098 | -0.410504 | -0.433337 | -0.717319 | -0.637548 | -0.364167 | -0.672781 | 0.451961 | 0.354872 | 0.344295 | 0.467533 | 0.111805 | 0.272816 | 0.307995 | 0.27414 | 0.269032 | 0.254575 | 0.306505 | 0.25371 |
| 25.48611 | -0.423614 | -1.042719 | -1.134115 | -0.721665 | -0.680051 | -0.5278 | -0.36206 | -0.768827 | -0.632263 | -0.507113 | -1.134439 | 0.455248 | 0.365654 | 0.466582 | 0.553205 | 0.15707 | 0.302174 | 0.327299 | 0.28365 | 0.295423 | 0.250314 | 0.317074 | 0.236188 |
| 25.5 | -0.732933 | -1.152349 | -1.641812 | -0.909686 | -0.588083 | -0.686711 | -0.376279 | -0.912624 | -0.585857 | -0.447457 | -1.070895 | 0.4609 | 0.350932 | 0.501115 | 0.541749 | 0.104499 | 0.200816 | 0.266333 | 0.218751 | 0.158505 | 0.169921 | 0.255651 | 0.157023 |
| 25.51389 | -1.158283 | -1.65092 | -2.069562 | -1.109662 | -0.404723 | -0.771766 | -0.524155 | -1.211603 | -0.679121 | -0.323678 | -1.015826 | 0.437869 | 0.290423 | 0.476371 | 0.502952 | -0.020059 | 0.042857 | 0.185444 | 0.10644 | -0.019735 | 0.03506 | 0.149204 | 0.030651 |
| 25.52778 | -1.698699 | -2.201298 | -2.268565 | -1.277419 | -0.463709 | -0.83147 | -0.833364 | -1.58317 | -1.092195 | -0.493932 | -0.885853 | 0.265835 | 0.014136 | 0.259441 | 0.429346 | -0.204267 | -0.095905 | 0.074966 | -0.070039 | -0.129567 | -0.212198 | -0.042162 | -0.194313 |
| 25.54167 | -1.323721 | -2.125587 | -1.948633 | -1.251663 | -0.710023 | -0.96021 | -1.202764 | -1.52788 | -1.335942 | -0.619313 | -0.751734 | 0.140287 | -0.150918 | 0.101998 | 0.372495 | -0.338792 | -0.15104 | 0.037613 | -0.155567 | -0.152612 | -0.261189 | -0.14326 | -0.306484 |
| 25.55556 | -1.048737 | -1.894226 | -1.526113 | -1.094872 | -0.836424 | -0.991394 | -1.150746 | -1.225922 | -1.138605 | -0.338537 | -0.499311 | 0.313362 | 0.11852 | 0.286835 | 0.471138 | -0.189395 | 0.002803 | 0.239616 | -0.012718 | -0.019285 | 0.059976 | 0.008095 | -0.09335 |
| 25.56944 | -1.205131 | -1.937275 | -0.608027 | -1.020599 | -0.792756 | -0.728927 | -0.796647 | -1.053778 | -1.014218 | -0.115619 | -0.188717 | 0.600469 | 0.47177 | 0.532388 | 0.890051 | 0.261306 | 0.484901 | 0.657354 | 0.23915 | 0.459376 | 0.483377 | 0.311491 | 0.331474 |
| 25.58333 | -1.872035 | -1.947389 | 0.434368 | -1.062713 | -0.769684 | -0.793632 | -0.639895 | -1.353809 | -1.139399 | -0.35561 | -0.217069 | 0.77973 | 0.704836 | 0.665713 | 1.580472 | 0.901634 | 1.29483 | 1.329249 | 0.685117 | 1.449078 | 0.984779 | 0.764477 | 0.903779 |
| 25.59722 | -2.14871 | -2.651288 | -0.606402 | -1.407208 | -1.025678 | -0.974192 | -1.2784 | -2.168421 | -1.233789 | -0.877607 | -0.680412 | 1.063146 | 0.939006 | 0.732452 | 1.842382 | 1.216753 | 1.698163 | 1.995986 | 1.183952 | 2.094484 | 1.562088 | 1.377954 | 1.508206 |
| 25.61111 | -2.68248 | -3.604031 | -2.134723 | -1.688851 | -1.275874 | -1.123596 | -1.707575 | -2.95152 | -0.984033 | -0.74495 | -1.159154 | 1.246851 | 1.319397 | 0.801149 | 1.565243 | 1.085056 | 1.511009 | 1.843763 | 1.217824 | 1.763831 | 1.760444 | 1.445996 | 1.510029 |
| 25.625 | -2.545315 | -2.614508 | -2.44983 | -1.506389 | -1.288218 | -1.268779 | -1.357765 | -3.107677 | -0.565153 | -0.680094 | -1.29681 | 1.061473 | 1.202938 | 0.546587 | 1.00602 | 0.797413 | 0.993775 | 1.270163 | 0.949605 | 1.010787 | 1.245374 | 1.023846 | 0.963459 |
| 25.63889 | -1.751488 | -1.445744 | -1.518487 | -0.892326 | -1.181558 | -0.95358 | -0.793067 | -1.761257 | -0.047404 | -0.674158 | -1.279671 | 0.855336 | 0.971732 | 0.351081 | 0.756592 | 0.692771 | 0.643883 | 0.867085 | 0.7799 | 0.638172 | 0.827925 | 0.68699 | 0.581105 |
| 25.65278 | -0.945291 | -0.816529 | -0.547026 | -0.01488 | -0.742286 | -0.731442 | -0.288678 | -0.206709 | 0.252221 | -0.539232 | -0.964236 | 0.737928 | 0.737304 | 0.263424 | 0.62051 | 0.636944 | 0.476669 | 0.698643 | 0.632161 | 0.481451 | 0.610138 | 0.434323 | 0.406396 |
| 25.66667 | -0.15813 | -0.764829 | 0.054046 | 0.399804 | -0.213669 | -0.347592 | 0.035537 | 0.694917 | 0.353902 | -0.476277 | -0.521495 | 0.625814 | 0.512073 | 0.227181 | 0.535697 | 0.530677 | 0.357795 | 0.623463 | 0.469589 | 0.396488 | 0.458187 | 0.260637 | 0.288079 |
| 25.68056 | 0.351359 | -0.981921 | 0.424561 | 0.440612 | 0.042845 | -0.006237 | 0.109955 | 0.974062 | 0.348927 | -0.532521 | -0.262056 | 0.554813 | 0.355546 | 0.290897 | 0.517875 | 0.459034 | 0.2703 | 0.606599 | 0.36747 | 0.349594 | 0.388144 | 0.148621 | 0.235816 |
| 25.69444 | 0.703272 | -0.812043 | 0.57162 | 0.321372 | 0.059594 | 0.303089 | 0.068254 | 1.18966 | 0.28864 | -0.515391 | -0.126636 | 0.553766 | 0.24514 | 0.363176 | 0.488588 | 0.435627 | 0.200886 | 0.473539 | 0.306823 | 0.24918 | 0.337744 | 0.062054 | 0.20583 |
| 25.70833 | 0.898592 | -0.291051 | 0.374855 | 0.251428 | -0.026983 | 0.529473 | 0.072567 | 0.810775 | 0.140531 | -0.298364 | -0.00054 | 0.631403 | 0.274145 | 0.521959 | 0.529928 | 0.474383 | 0.18806 | 0.447416 | 0.31866 | 0.215984 | 0.344157 | 0.084638 | 0.254728 |
| 25.72222 | 1.109503 | 0.153721 | -0.121406 | 0.302736 | -0.129174 | 0.546329 | 0.076702 | 0.098375 | -0.037983 | -0.054011 | 0.129897 | 0.724399 | 0.398174 | 0.76463 | 0.671243 | 0.542718 | 0.293331 | 0.546317 | 0.361346 | 0.364103 | 0.432294 | 0.225086 | 0.406593 |
| 25.73611 | 1.139284 | 0.379399 | -0.346354 | 0.262318 | -0.208777 | 0.384414 | 0.124275 | -0.452279 | -0.238581 | 0.069227 | 0.177143 | 0.789421 | 0.541538 | 0.959858 | 0.734156 | 0.600589 | 0.419501 | 0.739529 | 0.404275 | 0.465364 | 0.557772 | 0.382213 | 0.523747 |
| 25.75 | 0.962053 | 0.425826 | -0.080372 | -0.004592 | -0.27382 | 0.050608 | 0.21684 | -0.715603 | -0.391236 | -0.028418 | 0.143193 | 0.79892 | 0.536352 | 0.954431 | 0.609337 | 0.523101 | 0.469379 | 0.735272 | 0.388757 | 0.384957 | 0.565851 | 0.4333 | 0.47986 |
| 25.76389 | 0.577052 | 0.397252 | 0.153909 | -0.250998 | -0.27361 | -0.249542 | 0.233286 | -0.510554 | -0.416793 | -0.157233 | 0.080024 | 0.624971 | 0.235662 | 0.60264 | 0.385059 | 0.239515 | 0.297553 | 0.581068 | 0.188531 | 0.185043 | 0.349302 | 0.242466 | 0.218135 |
| 25.77778 | 0.121657 | 0.319578 | 0.236017 | -0.319322 | -0.260053 | -0.261081 | 0.128773 | -0.298765 | -0.385267 | -0.210599 | 0.008994 | 0.476341 | -0.087609 | 0.17382 | 0.234093 | 0.050701 | 0.057355 | 0.375296 | -0.050087 | -0.015361 | 0.143527 | 0.039295 | -0.0774 |
| 25.79167 | -0.117276 | 0.167459 | 0.167839 | -0.188353 | -0.222882 | -0.171722 | -0.012006 | -0.31534 | -0.344801 | -0.202841 | -0.070506 | 0.50267 | -0.133494 | -0.061003 | 0.391526 | 0.147571 | 0.010007 | 0.324188 | -0.126423 | -0.085393 | 0.121173 | 0.072249 | -0.191853 |
| 25.80556 | -0.232868 | -0.080446 | -0.033491 | -0.032792 | -0.166076 | -0.114275 | -0.179304 | -0.435411 | -0.284926 | -0.244737 | -0.140353 | 0.610211 | 0.008063 | -0.088651 | 0.783766 | 0.307102 | 0.132172 | 0.393427 | -0.01511 | -0.037505 | 0.240751 | 0.245725 | -0.170467 |
| 25.81944 | -0.665479 | -0.287676 | -0.085791 | 0.00821 | -0.083791 | -0.151727 | -0.321449 | -0.59241 | -0.199122 | -0.307918 | -0.165417 | 0.700626 | 0.130567 | -0.085073 | 0.849464 | 0.368934 | 0.196985 | 0.541063 | 0.127208 | 0.063234 | 0.359232 | 0.255057 | -0.129931 |
| 25.83333 | -0.698357 | -0.302292 | 0.017534 | -0.040796 | -0.071289 | -0.081091 | -0.331045 | -0.675783 | -0.119334 | -0.424985 | -0.160834 | 0.643062 | 0.101052 | -0.212347 | 0.526198 | 0.269738 | 0.112169 | 0.537357 | 0.141909 | 0.086645 | 0.31307 | 0.029626 | -0.18739 |
| 25.84722 | -0.343257 | -0.214118 | 0.002212 | -0.119294 | -0.097922 | 0.052429 | -0.18298 | -0.682479 | -0.075748 | -0.553684 | -0.181612 | 0.517514 | -0.051134 | -0.226265 | 0.157556 | 0.170281 | 0.076187 | 0.415379 | 0.071444 | -0.033029 | 0.186966 | -0.168939 | -0.318127 |
| 25.86111 | -0.256792 | -0.05979 | -0.022248 | -0.068252 | -0.098411 | 0.205354 | 0.032142 | -0.551888 | -0.046589 | -0.447244 | -0.169287 | 0.347551 | -0.221766 | -0.175129 | -0.075643 | 0.15127 | 0.120141 | 0.093487 | -0.111566 | -0.14126 | 0.034021 | -0.316342 | -0.44559 |
| 25.875 | -0.370824 | 0.132388 | -0.007353 | 0.201408 | 0.073118 | 0.310049 | 0.262328 | -0.342333 | 0.018452 | -0.07162 | -0.046108 | 0.145176 | -0.320159 | -0.112392 | -0.080861 | 0.064446 | 0.081046 | -0.022883 | -0.320837 | -0.169298 | -0.106342 | -0.453151 | -0.479936 |
| 25.88889 | -0.228749 | 0.294788 | -0.078555 | 0.404509 | 0.222132 | 0.346179 | 0.229625 | -0.247917 | 0.065054 | 0.241972 | 0.053012 | 0.193056 | -0.240417 | 0.079532 | 0.101478 | -0.03404 | -0.06714 | 0.044317 | -0.28603 | -0.15385 | -0.057599 | -0.395319 | -0.389131 |
| 25.90278 | -0.339059 | 0.4329 | 0.049887 | 0.372841 | 0.258722 | 0.286193 | 0.167375 | -0.156543 | 0.062815 | 0.281452 | 0.06011 | 0.499732 | -0.015521 | 0.295493 | 0.300589 | 0.011521 | -0.041976 | 0.1653 | -0.046783 | -0.086375 | 0.143379 | -0.11162 | -0.195403 |
| 25.91667 | -0.182045 | 0.690083 | 0.223929 | 0.370332 | 0.268144 | 0.203615 | 0.207451 | 0.183048 | 0.121832 | 0.291842 | 0.155434 | 0.75566 | 0.107746 | 0.28487 | 0.40165 | 0.180113 | 0.11557 | 0.169143 | 0.1303 | -0.061715 | 0.233157 | 0.106258 | -0.09991 |
| 25.93056 | 0.220906 | 1.052009 | 0.356778 | 0.419652 | 0.303794 | 0.239535 | 0.331769 | 0.664234 | 0.229992 | 0.325155 | 0.333507 | 0.779003 | 0.032289 | 0.040166 | 0.397797 | 0.258563 | 0.100032 | 0.063863 | 0.076425 | -0.115629 | 0.142165 | 0.165686 | -0.143609 |
| 25.94444 | 0.348082 | 1.332833 | 0.521997 | 0.482564 | 0.227809 | 0.353717 | 0.455403 | 1.021555 | 0.329985 | 0.230948 | 0.480803 | 0.688683 | -0.10824 | -0.248449 | 0.296278 | 0.030323 | -0.130798 | -0.062946 | -0.150514 | -0.218968 | 0.024151 | 0.126204 | -0.269779 |
| 25.95833 | 0.667137 | 1.579129 | 0.803273 | 0.526076 | 0.208037 | 0.546981 | 0.628067 | 1.295643 | 0.411933 | 0.072388 | 0.584501 | 0.64063 | -0.17185 | -0.280508 | 0.219912 | -0.230497 | -0.19672 | -0.097296 | -0.219895 | -0.267272 | 0.01133 | 0.0947 | -0.30448 |
| 25.97222 | 1.047423 | 1.828155 | 1.004497 | 0.563509 | 0.336939 | 0.761133 | 0.803096 | 1.362082 | 0.503495 | 0.008962 | 0.6685 | 0.700581 | -0.014061 | -0.058432 | 0.326341 | -0.146857 | -0.032448 | 0.038353 | 0.024556 | -0.172256 | 0.132014 | 0.177446 | -0.184208 |
| 25.98611 | 1.310791 | 1.977657 | 0.888414 | 0.644522 | 0.386285 | 0.888328 | 0.850515 | 1.291878 | 0.585169 | 0.022206 | 0.704104 | 0.685083 | 0.207502 | 0.09074 | 0.507927 | 0.200059 | 0.169214 | 0.200989 | 0.203134 | -0.025709 | 0.223973 | 0.289429 | -0.035461 |
| 26 | 1.724362 | 1.940887 | 0.564614 | 0.730989 | 0.270506 | 0.845602 | 0.748147 | 1.285052 | 0.616166 | 0.076574 | 0.619447 | 0.602646 | 0.219324 | 0.155346 | 0.648663 | 0.383342 | 0.140435 | 0.220168 | 0.093481 | -0.015171 | 0.251685 | 0.306126 | 0.069963 |
| 26.01389 | 2.148833 | 1.831615 | 0.433353 | 0.785585 | 0.160402 | 0.745783 | 0.667982 | 1.138295 | 0.409484 | 0.161238 | 0.34654 | 0.604153 | 0.176717 | 0.175273 | 0.701512 | 0.314664 | -0.003603 | 0.115607 | -0.03721 | -0.1077 | 0.155348 | 0.192432 | 0.050513 |
| 26.02778 | 2.111789 | 1.681936 | 0.746624 | 0.709014 | 0.361149 | 0.917012 | 0.716364 | 1.088158 | 0.138419 | 0.475815 | 0.221091 | 0.578985 | 0.088165 | 0.20557 | 0.678663 | 0.142947 | 0.061578 | 0.049742 | 0.005547 | -0.054531 | 0.054166 | 0.04547 | 0.085426 |
| 26.04167 | 2.185734 | 1.923965 | 1.4453 | 0.834289 | 0.73486 | 1.092893 | 0.910196 | 1.388905 | 0.384947 | 1.049758 | 0.558108 | 0.672257 | 0.106949 | 0.355963 | 0.655617 | 0.24874 | 0.227357 | 0.134004 | 0.12656 | 0.038424 | 0.148237 | 0.018547 | 0.273952 |
| 26.05556 | 2.082169 | 2.173036 | 1.985648 | 0.94046 | 0.966609 | 1.235854 | 1.031984 | 1.6029 | 0.636788 | 1.305268 | 0.75865 | 0.895051 | 0.144371 | 0.381692 | 0.524647 | 0.451321 | 0.189405 | 0.166109 | 0.126844 | 0.054999 | 0.261802 | 0.064113 | 0.304107 |
| 26.06944 | 1.705077 | 2.167028 | 2.179888 | 0.822752 | 0.877877 | 1.20196 | 1.011741 | 1.84419 | 0.850533 | 1.506647 | 0.677798 | 1.152468 | 0.539168 | 0.55205 | 0.365761 | 0.529744 | -0.161853 | 0.046839 | 0.181336 | -0.050959 | 0.420961 | 0.143352 | 0.246618 |
| 26.08333 | 1.262454 | 2.083714 | 1.311057 | 0.820739 | 0.822452 | 1.214853 | 1.089006 | 2.59757 | 1.162962 | 1.836309 | 0.596596 | 1.245259 | 0.627566 | 0.778214 | 0.2034 | 0.281625 | -0.569322 | -0.341041 | 0.210766 | -0.426848 | 0.401226 | -0.074296 | -0.071047 |
| 26.09722 | 0.703106 | 1.811881 | 1.019978 | 0.955105 | 1.032527 | 1.225582 | 1.022383 | 3.175804 | 1.08088 | 1.872066 | 0.522932 | 0.936871 | -0.21237 | 0.673838 | -0.078147 | -0.160403 | -1.206249 | -1.002648 | -0.205423 | -1.079031 | -0.200692 | -0.706824 | -0.691188 |
| 26.11111 | 0.201508 | 1.403529 | 1.126974 | 0.969775 | 0.681146 | 0.751343 | 0.27078 | 1.935248 | 0.61729 | 1.21141 | 0.460568 | 0.751058 | -1.260178 | 0.488356 | -0.159823 | 0.030803 | -1.080338 | -1.223543 | -0.698344 | -1.242587 | -0.964465 | -1.045319 | -1.030051 |
| 26.125 | 0.631163 | 1.590017 | 1.263223 | 1.365424 | 0.57717 | 0.726653 | 0.24602 | 1.554588 | 0.726765 | 0.911166 | 0.460307 | 1.055264 | -0.798456 | 0.729916 | 0.244207 | 0.831608 | -0.41663 | -0.693178 | -0.482052 | -0.679085 | -0.813694 | -0.485969 | -0.661966 |
| 26.13889 | 1.810755 | 2.021002 | 2.18166 | 2.007738 | 1.220449 | 1.001469 | 0.784651 | 1.650782 | 1.283285 | 0.826819 | 0.532231 | 1.570198 | 0.073991 | 1.060089 | 0.690809 | 1.266562 | -0.09427 | -0.101814 | 0.042676 | -0.135295 | -0.1766 | 0.249665 | -0.052209 |
| 26.15278 | 1.983627 | 1.515915 | 2.57641 | 2.104286 | 1.307557 | 1.236532 | 0.951726 | 1.193194 | 1.700944 | 0.646872 | 0.447977 | 1.519823 | 0.389515 | 1.257972 | 0.722332 | 1.169531 | -0.062322 | 0.185408 | 0.352522 | 0.09636 | 0.185698 | 0.486955 | 0.319454 |
| 26.16667 | 1.742882 | 1.16486 | 1.960715 | 1.644186 | 1.189285 | 0.816225 | 0.665405 | 0.21211 | 1.552359 | 0.664217 | 0.411989 | 1.228769 | 0.294317 | 1.099335 | 0.609808 | 0.65477 | -0.000046 | 0.310304 | 0.436753 | 0.198241 | 0.341661 | 0.443504 | 0.430426 |
| 26.18056 | 0.659068 | 1.175492 | 1.119218 | 1.239327 | 1.308054 | 0.499508 | 0.876168 | -0.086959 | 1.110964 | 0.937501 | 0.584962 | 1.01193 | 0.279128 | 0.872196 | 0.782313 | 0.171197 | 0.303108 | 0.449187 | 0.523217 | 0.301249 | 0.567913 | 0.392829 | 0.538174 |
| 26.19444 | -0.061939 | 1.511116 | 0.705951 | 0.852992 | 1.535658 | 0.300071 | 0.950434 | 0.816845 | 0.60177 | 1.145815 | 0.678837 | 0.930057 | 0.315615 | 0.549322 | 1.064432 | -0.124002 | 0.583363 | 0.511065 | 0.615276 | 0.33164 | 0.775721 | 0.36201 | 0.639381 |
| 26.20833 | -0.698045 | 1.790805 | 0.530735 | 0.550474 | 1.45761 | 0.030737 | 0.67291 | 1.924329 | 0.28697 | 1.116475 | 0.563955 | 0.684367 | 0.253792 | 0.236008 | 1.080604 | 0.264136 | 0.520022 | 0.429444 | 0.596182 | 0.25389 | 0.75389 | 0.18691 | 0.659571 |
| 26.22222 | -1.226131 | 1.781441 | 0.299497 | 0.29415 | 1.358472 | -0.108554 | 0.414202 | 2.241573 | 0.061097 | 1.077807 | 0.500222 | 0.418928 | 0.142557 | 0.152309 | 0.952966 | 0.857263 | 0.29189 | 0.330646 | 0.503399 | 0.192359 | 0.623906 | 0.025298 | 0.605515 |
| 26.23611 | -2.085674 | 1.611959 | 0.368302 | 0.16022 | 1.700395 | 0.033239 | 0.464642 | 2.357804 | 0.048744 | 0.937545 | 0.338241 | 0.325867 | 0.116508 | 0.51116 | 0.900249 | 1.34088 | 0.12453 | 0.279059 | 0.457824 | 0.169167 | 0.578011 | -0.0249 | 0.568671 |
| 26.25 | -1.389295 | 1.13558 | 0.60086 | 0.066696 | 1.83958 | -0.001796 | 0.411562 | 1.951952 | 0.140485 | 0.703145 | 0.251479 | 0.347652 | 0.097285 | 1.039226 | 0.916245 | 1.346889 | -0.040299 | 0.205542 | 0.407346 | 0.109145 | 0.569904 | -0.006113 | 0.526664 |
| 26.26389 | -0.000764 | 0.801516 | 0.961801 | 0.059163 | 1.36848 | 0.460553 | 0.393635 | 0.85141 | 0.439336 | 0.58259 | 0.348296 | 0.357871 | 0.011913 | 1.324092 | 0.900758 | 0.971401 | -0.158547 | 0.115466 | 0.393941 | 0.038401 | 0.503648 | -0.038401 | 0.466251 |
| 26.27778 | 1.200542 | 0.392721 | 1.091367 | 0.098627 | 0.862903 | 0.549814 | 0.10611 | -0.270588 | 0.714753 | 0.584659 | 0.567211 | 0.353921 | -0.083182 | 1.077759 | 0.803515 | 0.625236 | -0.101827 | 0.039614 | 0.41256 | -0.006985 | 0.436162 | -0.050478 | 0.392128 |
| 26.29167 | 1.678447 | -0.151566 | 0.9425 | 0.244743 | 0.395069 | 0.221272 | 0.083622 | -1.223925 | 0.744521 | 0.582501 | 0.494838 | 0.35508 | -0.081151 | 0.705581 | 0.662895 | 0.419756 | 0.031899 | 0.006028 | 0.447404 | -0.02151 | 0.452789 | 0.029943 | 0.335213 |
| 26.30556 | 1.736454 | -0.045631 | 0.614995 | 0.220605 | 0.00377 | -0.305126 | -0.031539 | -1.662082 | 0.553664 | 0.679757 | 0.41333 | 0.399305 | -0.07088 | 0.548747 | 0.586871 | 0.281041 | 0.13087 | 0.005723 | 0.394066 | -0.00644 | 0.43377 | 0.156312 | 0.265559 |
| 26.31944 | 1.155573 | 0.423636 | 0.213471 | -0.110753 | -0.307333 | -0.507632 | 0.000088 | -1.671946 | 0.320496 | 0.891694 | 0.157353 | 0.396979 | -0.096702 | 0.6179 | 0.475033 | 0.021731 | 0.112481 | -0.01229 | 0.268994 | -0.012209 | 0.364877 | 0.194732 | 0.159395 |
| 26.33333 | 0.424698 | 0.62539 | -0.385722 | -0.618464 | -0.592544 | -0.465975 | -0.103978 | -1.625687 | 0.059124 | 0.971136 | -0.050083 | 0.408 | -0.061751 | 0.718012 | 0.357411 | -0.25271 | -0.057869 | -0.057375 | 0.199598 | -0.054262 | 0.351283 | 0.165701 | -0.00233 |
| 26.34722 | -0.39507 | 0.255115 | -1.15618 | -1.155465 | -0.918666 | -0.408065 | -0.094426 | -1.243184 | -0.16261 | 0.784002 | -0.263988 | 0.421873 | 0.004036 | 0.609264 | 0.286743 | 0.027336 | -0.194982 | -0.065239 | 0.20788 | -0.071415 | 0.365586 | 0.093119 | -0.192249 |
| 26.36111 | -1.3086 | -0.326014 | -1.742581 | -1.443573 | -1.133985 | -0.65177 | -0.070999 | -0.873337 | -0.336897 | 0.557186 | -0.469921 | 0.420182 | 0.075434 | 0.312823 | 0.284624 | 0.590001 | -0.199961 | -0.022054 | 0.244369 | -0.035577 | 0.394061 | 0.039289 | -0.219003 |
| 26.375 | -1.724758 | -0.569931 | -1.790072 | -1.38122 | -1.088419 | -0.310012 | -0.14452 | -0.417706 | -0.375464 | 0.431067 | -0.399509 | 0.489004 | 0.111262 | 0.017901 | 0.30986 | 0.708226 | -0.118884 | 0.028353 | 0.268809 | 0.01432 | 0.484347 | 0.036515 | -0.064053 |
| 26.38889 | -1.331835 | -0.713304 | -1.326847 | -0.712364 | -0.93329 | 0.120339 | -0.032464 | -0.031241 | -0.235486 | 0.373407 | -0.246028 | 0.581558 | 0.140653 | -0.066482 | 0.272233 | 0.342868 | 0.067589 | 0.040083 | 0.253013 | 0.033461 | 0.569646 | 0.032895 | 0.092291 |
| 26.40278 | -0.53377 | -0.90364 | -0.701338 | -0.285635 | -0.674843 | 0.435856 | 0.076919 | 0.24893 | -0.154661 | 0.278189 | -0.11835 | 0.632057 | 0.255892 | 0.112124 | 0.150152 | 0.036779 | 0.217595 | -0.009257 | 0.216868 | -0.00129 | 0.536716 | -0.040092 | 0.182718 |
| 26.41667 | 0.05809 | -1.079033 | -0.365328 | -0.195229 | -0.436319 | 0.518285 | 0.146421 | 0.23504 | -0.1265 | 0.0881 | 0.061132 | 0.589848 | 0.375614 | 0.465258 | 0.104019 | -0.347263 | 0.204837 | -0.041057 | 0.205837 | -0.016875 | 0.484017 | -0.075912 | 0.156425 |
| 26.43056 | 0.118166 | -1.198979 | -0.479601 | -0.262065 | -0.532982 | 0.410988 | 0.075625 | 0.166213 | -0.197738 | -0.09748 | 0.097612 | 0.560317 | 0.390274 | 0.845138 | 0.14404 | -0.644142 | 0.11957 | 0.028695 | 0.278142 | 0.05807 | 0.517925 | -0.016827 | 0.096591 |
| 26.44444 | -0.012407 | -1.400293 | -0.872465 | -0.401884 | -0.798007 | 0.133799 | -0.214402 | 0.112324 | -0.358809 | -0.249494 | -0.11739 | 0.44131 | 0.086066 | 0.788464 | 0.171482 | -0.478443 | 0.065749 | 0.159926 | 0.371937 | 0.181703 | 0.462803 | 0.023446 | 0.040554 |
| 26.45833 | 0.145214 | -1.049162 | -0.949348 | -0.631689 | -1.10255 | -0.386364 | -0.45153 | -0.005421 | -0.425118 | -0.087501 | -0.387916 | 0.325907 | -0.196151 | 0.323631 | 0.063828 | 0.031001 | 0.05131 | 0.215154 | 0.342303 | 0.239968 | 0.371431 | 0.055345 | 0.013885 |
| 26.47222 | 0.581161 | -0.800699 | -0.816942 | -1.006107 | -1.450871 | -0.607593 | -0.359095 | -0.430688 | -0.554662 | 0.089444 | -0.494714 | 0.279523 | 0.108693 | 0.074656 | -0.178938 | 0.507751 | 0.053746 | 0.174538 | 0.268382 | 0.229779 | 0.511189 | 0.093805 | -0.026753 |
| 26.48611 | 0.556739 | -1.049746 | -1.081493 | -1.315121 | -1.625449 | -0.698677 | -0.131306 | -1.318667 | -0.834774 | -0.036507 | -0.627082 | 0.179248 | 0.138123 | 0.011276 | -0.400639 | 0.714063 | 0.098734 | 0.117891 | 0.226095 | 0.195763 | 0.602882 | 0.090592 | -0.105162 |
| 26.5 | 0.036473 | -1.57477 | -1.576838 | -1.499145 | -1.502636 | -0.481412 | -0.042828 | -1.511145 | -1.17645 | -0.370545 | -0.694675 | 0.061096 | 0.028599 | 0.049824 | -0.542214 | 0.468778 | 0.168332 | 0.069732 | 0.25444 | 0.162812 | 0.56907 | 0.03878 | -0.183379 |
| 26.51389 | -0.447895 | -1.768863 | -1.855856 | -1.546558 | -1.697329 | -0.811349 | -0.473118 | -1.348242 | -1.419605 | -0.437737 | -0.663711 | -0.002739 | 0.0673 | 0.142671 | -0.641502 | 0.022135 | 0.117804 | -0.013469 | 0.26902 | 0.120586 | 0.529789 | -0.021795 | -0.32071 |
| 26.52778 | -1.020793 | -1.553284 | -2.123186 | -1.514882 | -1.59002 | -1.204862 | -0.721501 | -1.186441 | -1.406427 | -0.174836 | -0.642718 | -0.072234 | 0.028805 | 0.218658 | -0.788528 | -0.468163 | -0.019705 | -0.14267 | 0.263514 | 0.077731 | 0.468993 | -0.10193 | -0.454793 |
| 26.54167 | -1.555036 | -1.447213 | -2.48905 | -1.411167 | -1.183097 | -1.56503 | -0.512851 | -1.220638 | -1.214627 | 0.039531 | -0.661137 | -0.145677 | -0.073143 | 0.156069 | -1.010336 | -0.786268 | -0.1062 | -0.214188 | 0.244457 | 0.064169 | 0.436996 | -0.141295 | -0.551682 |
| 26.55556 | -1.373189 | -1.534003 | -2.662254 | -1.239567 | -0.607387 | -1.516288 | -0.363651 | -1.213362 | -0.933505 | -0.040667 | -0.531838 | -0.181283 | -0.071989 | 0.046409 | -1.004833 | -0.548522 | -0.007864 | -0.105564 | 0.279497 | 0.137274 | 0.49262 | -0.109756 | -0.420901 |
| 26.56944 | -1.084287 | -1.756155 | -2.714111 | -1.289162 | -0.257223 | -1.087354 | -0.109397 | -1.223616 | -0.735899 | -0.523508 | -0.276271 | -0.275111 | 0.080252 | -0.033314 | -0.604096 | -0.097876 | 0.338631 | 0.18179 | 0.33631 | 0.311193 | 0.541069 | 0.016686 | -0.134641 |
| 26.58333 | -1.18699 | -2.272509 | -2.712607 | -1.460528 | -0.335164 | -0.647865 | -0.277782 | -1.329792 | -1.048164 | -1.162445 | -0.216277 | -0.740064 | -0.14185 | -0.476868 | -0.489592 | -0.360892 | 0.292283 | 0.24619 | 0.065299 | 0.207269 | 0.332576 | -0.201652 | -0.241892 |
| 26.59722 | -1.464553 | -2.750146 | -2.063752 | -1.619008 | -0.856265 | -0.708721 | -0.815585 | -2.185663 | -1.247156 | -1.961878 | -0.622813 | -1.4663 | -0.706758 | -1.550972 | -1.330827 | -1.441787 | -0.390169 | -0.277331 | -0.60157 | -0.286154 | -0.030247 | -1.080248 | -0.86905 |
| 26.61111 | -1.821304 | -2.620086 | -3.189374 | -1.950941 | -1.303045 | -1.200515 | -1.313853 | -3.069261 | -1.099931 | -2.043613 | -0.739975 | -1.379476 | -0.561553 | -1.793129 | -2.027725 | -1.790282 | -0.592347 | -0.653208 | -0.716 | -0.41282 | 0.261133 | -1.126631 | -1.045516 |
| 26.625 | -1.927353 | -2.206592 | -3.674274 | -1.457668 | -1.596426 | -1.459632 | -1.216887 | -3.052903 | -0.616617 | -1.814549 | -0.63942 | -0.687549 | 0.046461 | -1.259114 | -1.782479 | -1.031672 | -0.272931 | -0.498039 | -0.110021 | -0.040351 | 0.841244 | -0.8095 | -0.557226 |
| 26.63889 | -1.299916 | -1.757073 | -1.970033 | -0.84329 | -1.638477 | -1.542146 | -0.969025 | -1.708315 | -0.034809 | -0.650715 | -0.522767 | -0.340161 | 0.194145 | -0.790595 | -1.344006 | -0.344316 | 0.171554 | -0.339382 | 0.294702 | 0.183435 | 0.832728 | -0.815617 | -0.277512 |
| 26.65278 | -0.778552 | -1.012365 | -0.246859 | -0.294697 | -1.02794 | -1.181783 | -0.491027 | 0.47367 | 0.402004 | 0.428273 | -0.44074 | -0.388547 | -0.075538 | -0.324958 | -1.233226 | 0.062935 | 0.420074 | -0.418082 | 0.162512 | 0.074432 | 0.482534 | -0.716347 | -0.290776 |
| 26.66667 | -0.153641 | -0.674574 | 0.559744 | 0.060961 | -0.571537 | -0.510635 | -0.101154 | 1.870957 | 0.61275 | 0.534612 | -0.372868 | -0.414157 | -0.211497 | 0.007224 | -1.079097 | -0.197204 | 0.206125 | -0.499102 | -0.068494 | -0.067908 | 0.236451 | -0.251867 | -0.310114 |
| 26.68056 | 0.214182 | -0.936728 | 0.684464 | 0.134503 | -0.272207 | -0.105799 | -0.10142 | 1.584644 | 0.62077 | 0.008611 | -0.314632 | -0.441785 | -0.192465 | -0.144816 | -0.861782 | -0.630003 | -0.192911 | -0.531021 | -0.214086 | -0.151268 | 0.099192 | 0.133641 | -0.343799 |
| 26.69444 | 0.113124 | -1.162932 | 0.541775 | 0.121119 | -0.082852 | -0.134798 | 0.012094 | 1.23949 | 0.459105 | -0.729275 | -0.27603 | -0.375881 | -0.129033 | -0.383774 | -0.708697 | -0.868929 | -0.122448 | -0.51294 | -0.221162 | -0.172713 | 0.034898 | 0.207199 | -0.363099 |
| 26.70833 | -0.017868 | -1.020516 | 0.312459 | 0.165073 | 0.048398 | -0.345515 | 0.035857 | 0.670377 | 0.283315 | -1.113042 | -0.274632 | -0.151013 | 0.000883 | -0.238892 | -0.519395 | -0.61428 | 0.038712 | -0.527795 | -0.101059 | -0.147036 | 0.081474 | 0.237464 | -0.30413 |
| 26.72222 | 0.196914 | -0.727034 | 0.083462 | 0.262173 | 0.124082 | -0.610525 | -0.007691 | 0.197489 | 0.183113 | -1.065981 | -0.257186 | 0.03732 | 0.101314 | 0.0507 | -0.38044 | -0.269999 | -0.113542 | -0.540895 | -0.031511 | -0.105818 | 0.104494 | 0.357308 | -0.238306 |
| 26.73611 | 0.580479 | -0.567794 | -0.055998 | 0.324451 | 0.175072 | -0.682447 | 0.015466 | -0.247068 | 0.093747 | -0.810665 | -0.217716 | -0.023837 | 0.159191 | -0.094756 | -0.194564 | -0.245645 | -0.134771 | -0.485592 | -0.164564 | -0.089078 | -0.037674 | 0.448135 | -0.207642 |
| 26.75 | 0.813541 | -0.525662 | -0.204532 | 0.294995 | -0.021831 | -0.60158 | 0.045574 | -0.787571 | 0.072356 | -0.272455 | -0.162634 | -0.052176 | 0.240702 | -0.247188 | -0.115493 | -0.202377 | 0.018246 | -0.363801 | -0.236775 | -0.037383 | -0.07912 | 0.387574 | -0.124531 |
| 26.76389 | 0.86376 | -0.323753 | -0.467648 | 0.144681 | -0.24605 | -0.429703 | 0.010817 | -0.94736 | 0.068196 | 0.12102 | -0.102473 | 0.079666 | 0.285998 | -0.129705 | -0.300243 | -0.059959 | -0.092541 | -0.275556 | -0.050885 | 0.021081 | 0.106081 | 0.210814 | -0.056841 |
| 26.77778 | 0.660843 | -0.12267 | -0.530533 | -0.048802 | -0.436469 | -0.281369 | -0.028115 | -0.825117 | -0.016075 | 0.093901 | -0.066118 | 0.141192 | 0.291201 | -0.036174 | -0.558659 | 0.063173 | -0.197809 | -0.234393 | 0.133775 | 0.032459 | 0.202905 | -0.070465 | -0.064561 |
| 26.79167 | 0.472936 | -0.10157 | -0.444067 | -0.10178 | -0.68543 | -0.15319 | -0.075009 | -0.491552 | -0.132484 | -0.042331 | -0.037608 | 0.054268 | 0.320034 | -0.104355 | -0.640915 | 0.144333 | 0.035718 | -0.11319 | 0.082522 | 0.052955 | 0.105845 | -0.257566 | -0.096494 |
| 26.80556 | 0.47906 | -0.253169 | -0.454412 | -0.008206 | -0.6822 | -0.115814 | -0.145436 | -0.310759 | -0.201693 | -0.020437 | -0.017058 | 0.006555 | 0.269888 | -0.153568 | -0.601648 | 0.24811 | 0.253998 | 0.048679 | 0.008146 | 0.10114 | 0.024449 | -0.253836 | -0.085995 |
| 26.81944 | 0.522031 | -0.412005 | -0.443211 | 0.154967 | -0.163681 | -0.119476 | -0.23033 | -0.258852 | -0.275475 | 0.082723 | -0.043064 | 0.041343 | 0.146726 | -0.081594 | -0.490527 | 0.419597 | 0.203449 | 0.173414 | 0.071149 | 0.145222 | 0.071204 | -0.104256 | -0.038471 |
| 26.83333 | 0.438789 | -0.522745 | -0.302319 | 0.310799 | 0.328171 | 0.021258 | -0.311466 | -0.194083 | -0.324088 | 0.208854 | -0.0806 | 0.052901 | 0.011615 | -0.003545 | -0.442771 | 0.48364 | 0.123554 | 0.221633 | 0.113901 | 0.147513 | 0.132021 | 0.055418 | -0.006069 |
| 26.84722 | 0.084774 | -0.552154 | -0.203419 | 0.426766 | 0.465446 | 0.381121 | -0.328911 | -0.160517 | -0.310941 | 0.415694 | -0.116624 | 0.026944 | -0.145013 | -0.040322 | -0.375589 | 0.340931 | 0.147444 | 0.183119 | -0.001029 | 0.100114 | 0.089782 | 0.148426 | 0.018322 |
| 26.86111 | -0.448936 | -0.498366 | -0.190728 | 0.424664 | 0.313945 | 0.699624 | -0.26995 | -0.140382 | -0.27756 | 0.503515 | -0.135855 | 0.037817 | -0.266104 | -0.071629 | -0.267378 | 0.156468 | 0.132908 | 0.131341 | -0.0899 | 0.058653 | 0.037188 | 0.247936 | 0.022757 |
| 26.875 | -0.838924 | -0.386056 | -0.142905 | 0.351866 | 0.198658 | 0.580977 | -0.185356 | -0.158315 | -0.181777 | 0.236139 | -0.122318 | 0.056243 | -0.292841 | -0.125917 | -0.046469 | 0.115443 | 0.079808 | 0.10549 | -0.037709 | 0.049307 | 0.027815 | 0.374092 | 0.022411 |
| 26.88889 | -1.173368 | -0.225753 | -0.118334 | 0.280384 | 0.156466 | 0.333339 | -0.168762 | -0.142467 | -0.03606 | -0.168156 | -0.094642 | -0.035944 | -0.282135 | -0.267009 | 0.203471 | 0.1619 | 0.140899 | 0.098179 | 0.059875 | 0.021727 | -0.069269 | 0.373154 | 0.003584 |
| 26.90278 | -1.117256 | -0.015585 | -0.07388 | 0.213493 | 0.171258 | 0.05334 | -0.162148 | -0.139107 | 0.130897 | -0.3439 | -0.048403 | -0.065794 | -0.294197 | -0.347997 | 0.34737 | 0.162223 | 0.345387 | 0.067426 | 0.15387 | -0.032302 | -0.159817 | 0.270859 | -0.020342 |
| 26.91667 | -0.717562 | 0.259014 | -0.049717 | 0.179809 | 0.260871 | -0.135369 | -0.016336 | -0.039418 | 0.305692 | -0.198115 | 0.02924 | -0.001528 | -0.223913 | -0.319137 | 0.482158 | 0.095089 | 0.359795 | 0.052247 | 0.220399 | -0.056281 | -0.154734 | 0.166963 | -0.02506 |
| 26.93056 | -0.239393 | 0.637894 | 0.015558 | 0.196196 | 0.431889 | -0.146977 | 0.19171 | 0.193964 | 0.480497 | 0.056904 | 0.12572 | 0.031933 | -0.106065 | -0.276519 | 0.687738 | 0.082251 | 0.097719 | 0.070014 | 0.03877 | -0.056031 | -0.207719 | 0.076748 | -0.004858 |
| 26.94444 | 0.152553 | 1.088649 | 0.252964 | 0.246386 | 0.647999 | 0.008198 | 0.338511 | 0.413324 | 0.53891 | 0.304937 | 0.206478 | 0.186264 | -0.035906 | -0.225549 | 0.855925 | 0.088332 | -0.015432 | 0.104211 | -0.139618 | -0.0507 | -0.284721 | 0.073011 | 0.035466 |
| 26.95833 | 0.646027 | 1.377478 | 0.690822 | 0.346933 | 0.858314 | 0.484953 | 0.574465 | 0.643576 | 0.513643 | 0.510523 | 0.277142 | 0.475783 | 0.022326 | -0.155457 | 0.955566 | 0.095103 | 0.285882 | 0.12982 | -0.113611 | -0.016334 | -0.248324 | 0.263127 | 0.053799 |
| 26.97222 | 1.293158 | 1.637339 | 1.176068 | 0.528318 | 0.956024 | 1.17678 | 0.872485 | 0.913873 | 0.519815 | 0.69805 | 0.399822 | 0.735176 | 0.10566 | -0.003916 | 1.064392 | 0.167995 | 0.68624 | 0.132707 | 0.14729 | -0.008552 | -0.090432 | 0.473419 | 0.091031 |
| 26.98611 | 1.702167 | 1.92829 | 1.473142 | 0.650481 | 0.843784 | 1.500239 | 0.932132 | 1.114715 | 0.563808 | 0.731301 | 0.49859 | 0.886421 | 0.163532 | 0.088647 | 1.028659 | 0.416495 | 0.453665 | 0.091964 | 0.225625 | -0.035979 | 0.007953 | 0.519709 | 0.103289 |
| 27 | 1.876512 | 1.778146 | 1.524367 | 0.549304 | 0.52769 | 1.623577 | 0.94939 | 1.015022 | 0.490018 | 0.48473 | 0.485446 | 0.851068 | 0.127311 | 0.068069 | 0.83808 | 0.60817 | 0.036939 | -0.019398 | 0.084638 | -0.132184 | -0.140492 | 0.322867 | 0.050077 |
| 27.01389 | 1.540012 | 1.580254 | 1.516883 | 0.339805 | 0.207005 | 1.456713 | 0.759609 | 0.640649 | 0.318189 | 0.034429 | 0.327143 | 0.66035 | -0.133274 | -0.196617 | 0.545828 | 0.561328 | -0.12346 | -0.192056 | -0.11094 | -0.264129 | -0.526606 | 0.043318 | 0.01115 |
| 27.02778 | 0.836412 | 1.687597 | 1.864721 | 0.26535 | 0.264394 | 1.159484 | 0.682721 | 0.548236 | 0.144942 | -0.191373 | 0.103465 | 0.4669 | -0.210491 | -0.33139 | 0.337658 | 0.219872 | 0.05907 | -0.183994 | -0.185203 | -0.231094 | -0.534213 | -0.088384 | 0.071001 |
| 27.04167 | 1.118368 | 2.17731 | 2.283386 | 0.469232 | 0.723244 | 0.99365 | 0.779536 | 1.103675 | 0.57814 | 0.530417 | 0.32772 | 0.582248 | 0.100615 | -0.040297 | 0.325066 | 0.296349 | 0.257643 | -0.121126 | -0.017455 | -0.17748 | -0.137033 | -0.07758 | 0.1288 |
| 27.05556 | 1.212344 | 2.194358 | 2.810024 | 0.769295 | 1.148401 | 1.03238 | 0.984737 | 1.704386 | 1.063587 | 1.33647 | 0.558893 | 0.804405 | 0.406149 | 0.348867 | 0.430803 | 0.424228 | 0.09333 | -0.127003 | 0.151011 | -0.1864 | 0.09626 | -0.037347 | 0.104859 |
| 27.06944 | 1.16588 | 1.658563 | 2.665185 | 0.889077 | 1.275758 | 0.951418 | 0.914629 | 1.749169 | 1.206136 | 1.6664 | 0.571701 | 0.822945 | 0.553435 | 0.525938 | 0.590109 | 0.502557 | -0.236685 | -0.138584 | 0.209071 | -0.16168 | 0.170958 | 0.078594 | 0.183445 |
| 27.08333 | 0.899083 | 1.076981 | 2.857906 | 1.052905 | 1.557836 | 0.924228 | 0.824063 | 1.941621 | 1.288424 | 2.033582 | 0.58199 | 0.62967 | 0.26505 | 0.334608 | 0.215715 | 0.433482 | -0.589969 | -0.444816 | -0.076407 | -0.416572 | -0.100112 | -0.013735 | -0.103419 |
| 27.09722 | 0.957864 | 0.596721 | 3.352579 | 0.927336 | 1.956952 | 1.331034 | 1.056872 | 2.36716 | 1.019468 | 2.294269 | 0.673698 | 0.161164 | -0.575607 | -0.093579 | -0.521713 | 0.059086 | -0.90318 | -1.16302 | -0.738004 | -1.089158 | -0.854278 | -0.566229 | -0.889672 |
| 27.11111 | 0.482618 | 0.668508 | 2.17372 | 0.663315 | 1.38476 | 0.770469 | 0.631569 | 1.807914 | 0.435308 | 2.16184 | 0.437892 | -0.015897 | -0.977029 | -0.309795 | -0.76749 | 0.344002 | -0.989517 | -1.521524 | -1.141076 | -1.397381 | -1.387377 | -0.784613 | -1.29825 |
| 27.125 | 0.278838 | 1.768482 | 1.639201 | 1.133415 | 0.33501 | 0.819058 | 0.354155 | 1.188088 | 0.57897 | 2.347165 | 0.307578 | 0.516128 | -0.563947 | 0.009514 | -0.080723 | 1.530499 | -0.523406 | -0.935099 | -0.764029 | -0.920461 | -1.071511 | -0.030568 | -0.860127 |
| 27.13889 | 0.568884 | 3.128242 | 2.272951 | 2.199759 | -0.037213 | 1.341743 | 0.727459 | 1.661638 | 1.297497 | 2.719032 | 0.546896 | 0.954169 | 0.02216 | 0.390125 | 0.679429 | 2.004659 | 0.039691 | -0.183066 | -0.207834 | -0.363404 | -0.64592 | 0.485469 | -0.205482 |
| 27.15278 | 0.711297 | 2.535754 | 2.386064 | 2.610043 | -0.193616 | 1.152028 | 0.756019 | 1.649264 | 1.626199 | 2.062616 | 0.568255 | 0.978147 | 0.238404 | 0.789113 | 1.155404 | 1.971847 | 0.01952 | 0.184882 | 0.041539 | -0.089922 | -0.330029 | 0.274273 | 0.186249 |
| 27.16667 | 0.598017 | 1.622911 | 1.431795 | 1.970236 | -0.099503 | 0.894975 | 0.477217 | 0.396798 | 1.339798 | 1.189406 | 0.700596 | 0.75539 | 0.268342 | 0.988128 | 1.164736 | 0.913144 | -0.135363 | 0.245828 | 0.079148 | -0.024306 | -0.055581 | -0.056286 | 0.407891 |
| 27.18056 | 0.602363 | 0.984371 | 0.740175 | 0.998144 | 0.534635 | 0.578733 | 0.501124 | 0.123527 | 0.932541 | 0.494739 | 0.782403 | 0.340944 | 0.183466 | 0.74496 | 1.067782 | 0.44182 | -0.15899 | 0.21057 | -0.018234 | -0.046933 | 0.088058 | -0.206424 | 0.505556 |
| 27.19444 | 0.453722 | 0.646452 | 0.358297 | 0.140349 | 1.475987 | 0.422357 | 0.887741 | 0.69982 | 0.459025 | -0.227344 | 0.660241 | 0.125132 | 0.195298 | 0.425771 | 0.844475 | 0.239999 | 0.126976 | 0.189168 | -0.126453 | -0.058648 | 0.197786 | -0.341345 | 0.525784 |
| 27.20833 | 0.042182 | 0.437821 | 0.071678 | -0.382142 | 2.333403 | 0.362312 | 1.260573 | 1.203116 | 0.093858 | -0.370584 | 0.457065 | 0.162837 | 0.221337 | 0.332489 | 0.710274 | 0.112462 | 0.448618 | 0.206391 | -0.165009 | -0.054149 | 0.310482 | -0.298651 | 0.670106 |
| 27.22222 | -0.169917 | 0.586074 | 0.027579 | -0.359676 | 2.455841 | -0.255084 | 0.935954 | 1.335046 | -0.029216 | 0.572554 | 0.284205 | 0.163172 | 0.073049 | 0.310857 | 0.767142 | 0.002494 | 0.529694 | 0.178453 | -0.133671 | -0.048421 | 0.199236 | -0.127279 | 0.757695 |
| 27.23611 | -0.164704 | 0.838224 | 0.584111 | -0.093629 | 2.077972 | 0.000926 | 0.59732 | 1.28081 | 0.122185 | 1.917139 | 0.228692 | -0.055587 | -0.122153 | 0.300637 | 0.72759 | 0.391778 | 0.417666 | 0.144008 | -0.021784 | -0.046265 | 0.047938 | 0.24745 | 0.649483 |
| 27.25 | 0.065672 | 0.847447 | 1.390669 | 0.184952 | 1.721454 | 0.439196 | 0.223314 | 1.166936 | 0.332237 | 2.891966 | 0.32122 | -0.232096 | -0.216444 | 0.342687 | 0.658593 | 0.649226 | 0.192106 | 0.120161 | 0.129365 | -0.06648 | 0.101718 | 0.382986 | 0.435019 |
| 27.26389 | 0.511844 | 0.870539 | 1.759353 | 0.312076 | 1.172829 | 0.461089 | -0.359266 | 0.792043 | 0.46987 | 2.658337 | 0.380263 | -0.312172 | -0.289978 | 0.383775 | 0.561005 | 0.754642 | -0.074429 | 0.117196 | 0.212145 | -0.093751 | 0.147887 | 0.288109 | 0.31509 |
| 27.27778 | 0.968607 | 0.82699 | 1.815201 | 0.285247 | 0.425181 | 0.302993 | -0.301417 | 0.006206 | 0.489938 | 1.677231 | 0.376018 | -0.297503 | -0.336244 | 0.504659 | 0.405936 | 0.657724 | -0.276982 | 0.138742 | 0.203452 | -0.084271 | 0.080911 | 0.118344 | 0.19034 |
| 27.29167 | 1.322212 | 0.666234 | 1.624074 | 0.23451 | -0.368503 | 0.334023 | 0.21863 | -0.620333 | 0.394576 | 0.682193 | 0.283905 | -0.025704 | -0.352353 | 0.714071 | 0.295542 | 0.30267 | -0.191415 | 0.138909 | 0.150498 | -0.067454 | 0.099455 | 0.00242 | 0.096282 |
| 27.30556 | 0.728494 | 0.477113 | 1.275757 | 0.219603 | -0.886142 | 0.593193 | 0.673885 | -0.987357 | 0.211715 | 0.038348 | 0.230944 | 0.33781 | -0.389027 | 0.732554 | 0.355905 | -0.088951 | 0.024136 | 0.111002 | 0.067632 | -0.056967 | 0.173135 | -0.02886 | 0.082799 |
| 27.31944 | 0.445907 | 0.324287 | 0.615524 | 0.22844 | -0.870286 | 0.206394 | 0.466124 | -0.95406 | 0.040897 | -0.404606 | 0.217184 | 0.581612 | -0.324582 | 0.399075 | 0.426885 | -0.307331 | 0.168444 | 0.183322 | -0.00655 | -0.030196 | 0.069988 | -0.00602 | 0.137934 |
| 27.33333 | 0.192836 | 0.452721 | -0.332451 | 0.103272 | -0.484117 | -0.397974 | -0.061819 | -0.620226 | -0.156734 | -0.318276 | 0.259112 | 0.801816 | -0.181914 | -0.402873 | 0.445399 | -0.466538 | 0.23886 | 0.230308 | -0.051234 | 0.016501 | -0.013332 | 0.054718 | 0.157498 |
| 27.34722 | -0.303439 | 0.579929 | -1.113354 | -0.192737 | -0.161754 | -0.453177 | -0.437718 | -0.201336 | -0.255783 | 0.19471 | 0.183169 | 0.967423 | -0.042629 | -2.909144 | 0.436552 | -0.644304 | 0.263432 | 0.179277 | -0.02634 | 0.070113 | 0.049505 | 0.110653 | 0.142086 |
| 27.36111 | -0.943272 | 0.57239 | -1.659372 | -0.521182 | 0.136264 | -1.053595 | -0.429819 | 0.088484 | -0.254826 | 0.775715 | 0.094995 | 0.872945 | -0.040933 | -5.714338 | -0.257373 | -0.838162 | 0.117671 | 0.114249 | -0.007614 | 0.081495 | 0.099162 | 0.08133 | 0.041021 |
| 27.375 | -1.089575 | 0.316529 | -2.461863 | -0.718969 | 0.131121 | -1.424273 | -0.382139 | 0.144867 | -0.273306 | 0.973572 | 0.17637 | 0.483296 | -0.086471 | -3.83878 | -0.349371 | -1.111404 | -0.186531 | 0.01585 | -0.00587 | 0.049393 | 0.172559 | 0.017045 | -0.070518 |
| 27.38889 | -1.188534 | -0.088639 | -3.141383 | -0.682393 | -0.003376 | -1.284883 | -0.317922 | 0.036277 | -0.34733 | 0.632116 | 0.24044 | 0.016726 | -0.140005 | -1.529525 | -0.365483 | -0.541148 | -0.398982 | -0.039198 | -0.037046 | 0.041074 | 0.225516 | 0.025555 | -0.11257 |
| 27.40278 | -1.073735 | -0.439138 | -2.544307 | -0.628641 | -0.193089 | -0.550622 | -0.016164 | -0.001799 | -0.387799 | 0.273831 | 0.188971 | -0.3793 | -0.134453 | -0.446727 | -0.566735 | 0.44807 | -0.501574 | 0.013335 | -0.089025 | 0.049742 | 0.080489 | 0.034108 | -0.158471 |
| 27.41667 | -1.093157 | -0.470233 | -1.381627 | -0.545271 | -0.38178 | 0.198046 | 0.182003 | 0.069522 | -0.318361 | -0.031689 | 0.076875 | -0.537692 | -0.097656 | -0.043233 | -0.602432 | 0.528728 | -0.237651 | 0.04429 | -0.134296 | 0.033547 | -0.125483 | 0.020722 | -0.193142 |
| 27.43056 | -0.703601 | -0.109182 | -0.845569 | -0.318371 | -0.540496 | 0.936385 | 0.130932 | -0.049694 | -0.265586 | -0.397192 | -0.13568 | -0.501177 | -0.122929 | 0.17277 | -0.517901 | 0.188665 | 0.083662 | -0.010573 | -0.183839 | 0.006177 | -0.161741 | -0.010519 | -0.254539 |
| 27.44444 | -0.283216 | 0.219213 | -0.522897 | -0.273865 | -0.660679 | 1.293905 | -0.097783 | -0.339194 | -0.204595 | -0.79708 | -0.292538 | -0.442696 | -0.187709 | 0.252958 | -0.472368 | 0.078944 | 0.211037 | -0.07749 | -0.193256 | -0.009912 | -0.058698 | -0.034598 | -0.324205 |
| 27.45833 | 0.179879 | 0.005476 | -0.424449 | -0.522096 | -0.776824 | 1.264622 | -0.201497 | -0.639142 | -0.200845 | -0.870234 | -0.330426 | -0.490028 | -0.262488 | 0.314955 | -0.526494 | -0.081692 | 0.099648 | -0.072107 | -0.178447 | 0.006881 | 0.01482 | -0.047515 | -0.356158 |
| 27.47222 | 0.575529 | -0.612857 | -0.205414 | -0.858847 | -1.060475 | 0.746614 | -0.357221 | -0.871103 | -0.292991 | -0.473809 | -0.237966 | -0.531506 | -0.314144 | 0.314399 | -0.655602 | -0.391678 | -0.165283 | 0.00752 | -0.177274 | 0.035765 | -0.002465 | -0.084549 | -0.273758 |
| 27.48611 | 0.953766 | -1.21708 | -0.090803 | -1.081459 | -1.298207 | -0.119868 | -0.773993 | -1.148453 | -0.453282 | 0.028778 | -0.15595 | -0.294034 | -0.357962 | 0.176566 | -0.762386 | -0.90852 | -0.454817 | 0.090382 | -0.169339 | 0.061276 | 0.038012 | -0.167206 | -0.132298 |
| 27.5 | 1.016136 | -1.122044 | -0.14257 | -1.168296 | -1.258678 | -0.71506 | -1.210807 | -1.261484 | -0.63445 | 0.420659 | -0.18393 | 0.142287 | -0.416185 | -0.068518 | -0.915641 | -0.898955 | -0.585493 | 0.321018 | -0.171025 | 0.090059 | 0.095614 | -0.352797 | -0.086963 |
| 27.51389 | 0.746236 | -0.724628 | -0.404465 | -1.160578 | -0.982161 | -1.429288 | -1.080184 | -1.133615 | -0.679486 | 0.591978 | -0.149247 | 0.4009 | -0.415875 | -0.316849 | -1.170936 | -0.621643 | -0.3749 | 0.789808 | -0.18287 | 0.088311 | 0.056214 | -0.537576 | -0.206595 |
| 27.52778 | 0.338996 | -0.671858 | -0.923599 | -1.255329 | -0.847629 | -1.987337 | -0.408865 | -1.017706 | -0.732447 | 0.605636 | -0.147802 | 0.368622 | -0.406243 | -0.475299 | -1.255767 | -0.35091 | -0.139343 | 0.802786 | -0.196612 | 0.04607 | 0.004958 | -0.665841 | -0.400994 |
| 27.54167 | -0.062422 | -1.250752 | -1.809734 | -1.415763 | -0.829014 | -1.797787 | -0.01354 | -1.106802 | -0.944273 | 0.06595 | -0.167264 | 0.152959 | -0.323584 | -0.540263 | -1.165296 | -0.143096 | 0.023727 | 0.407637 | -0.223106 | 0.02367 | 0.014229 | -0.667368 | -0.553678 |
| 27.55556 | -0.420855 | -1.920047 | -2.434722 | -1.291838 | -0.750917 | -1.77477 | -0.35164 | -1.182489 | -0.821789 | -0.594088 | -0.104497 | 0.036245 | -0.056168 | -0.426749 | -0.944639 | 0.108134 | 0.125086 | 0.289186 | -0.146795 | 0.1218 | 0.122522 | -0.627931 | -0.429107 |
| 27.56944 | -0.616201 | -2.207679 | -3.194539 | -1.169126 | -0.745233 | -1.715235 | -0.594675 | -1.026955 | -0.802537 | -1.328896 | 0.08879 | 0.105135 | 0.252752 | -0.19902 | -0.7308 | 0.421398 | 0.345797 | 0.580958 | -0.013459 | 0.348229 | 0.276924 | -0.420952 | 0.018467 |
| 27.58333 | -1.147567 | -2.260749 | -4.424196 | -1.38197 | -0.939002 | -1.679932 | -0.585018 | -1.105854 | -1.547893 | -2.064345 | -0.096322 | 0.18557 | 0.37362 | -0.363406 | -0.614458 | 0.509854 | 0.270741 | 0.950937 | 0.005317 | 0.542977 | 0.359146 | -0.251792 | 0.361461 |
| 27.59722 | -1.546504 | -2.343005 | -5.259861 | -1.949855 | -1.365862 | -2.082554 | -0.988903 | -1.794886 | -2.090057 | -2.924647 | -0.651441 | -0.068982 | 0.240155 | -1.129279 | -0.639671 | -0.187464 | -0.201815 | 0.772722 | -0.010604 | 0.535851 | 0.371922 | -0.282322 | 0.306496 |
| 27.61111 | -1.730424 | -2.518981 | -6.340938 | -1.899902 | -1.531125 | -2.846427 | -1.628225 | -2.899014 | -1.566756 | -2.437398 | -0.763599 | -0.367172 | 0.131729 | -1.556062 | -0.69357 | -0.886065 | -0.277739 | 0.306478 | 0.184668 | 0.52694 | 0.509581 | -0.238594 | 0.12026 |
| 27.625 | -1.966364 | -2.393727 | -6.73195 | -1.60676 | -1.375546 | -2.488041 | -1.316152 | -3.411256 | -0.884609 | -2.093914 | -0.382167 | -0.624827 | 0.011951 | -1.703536 | -0.981336 | -0.873376 | -0.766619 | 0.047733 | 0.281346 | 0.434786 | 0.447821 | -0.241826 | -0.06243 |
| 27.63889 | -1.923201 | -1.441789 | -4.836844 | -1.198738 | -1.244118 | -1.729729 | -0.670509 | -1.912403 | -0.262753 | -0.79863 | -0.104242 | -0.857467 | -0.054394 | -1.309806 | -1.414681 | -0.811867 | -0.694232 | -0.053872 | 0.185762 | 0.279711 | 0.071587 | -0.452049 | -0.199995 |
| 27.65278 | -1.36152 | -0.460208 | -1.343942 | -0.492068 | -0.932852 | -0.700535 | 0.022931 | -0.135293 | 0.184793 | 0.22136 | -0.074325 | -0.77124 | -0.026529 | -0.67486 | -1.509816 | -0.223337 | 0.697141 | 0.11841 | 0.107107 | 0.188466 | -0.147052 | -0.383145 | -0.150168 |
| 27.66667 | -0.644161 | -0.140202 | 0.955794 | -0.15446 | -0.245057 | -0.112933 | 0.12267 | 1.08854 | 0.550045 | 0.618832 | -0.096986 | -0.29095 | 0.113832 | -0.017126 | -1.002707 | 0.267977 | 1.233295 | 0.298862 | 0.131931 | 0.18884 | -0.131694 | -0.09243 | 0.026895 |
| 27.68056 | -0.310386 | -0.235352 | 1.379776 | -0.171018 | 0.102697 | 0.163524 | -0.027337 | 1.329475 | 0.948369 | 0.227302 | -0.089414 | -0.11386 | 0.276715 | 0.192147 | -0.535841 | 0.121235 | 0.169875 | 0.281993 | 0.157934 | 0.182844 | -0.028627 | 0.236989 | 0.167947 |
| 27.69444 | -0.270299 | -0.3283 | 1.321261 | -0.232874 | -0.042877 | 0.463943 | 0.249994 | 1.745189 | 1.261823 | -0.50258 | -0.118636 | -0.116154 | 0.321517 | 0.228515 | -0.307544 | -0.145221 | -0.582258 | 0.147362 | 0.150386 | 0.147028 | 0.018885 | 0.288663 | 0.234317 |
| 27.70833 | -0.358292 | -0.535624 | 1.025311 | -0.29006 | -0.31148 | 0.774332 | 0.481478 | 1.333659 | 1.20761 | -0.945002 | -0.180095 | 0.128601 | 0.304432 | 0.498102 | -0.220207 | -0.171112 | -0.184539 | 0.077557 | 0.152883 | 0.154741 | 0.066726 | 0.37973 | 0.264009 |
| 27.72222 | 0.040737 | -0.790684 | 0.583139 | -0.283319 | -0.438936 | 1.050599 | 0.296291 | 0.564447 | 0.75464 | -1.247513 | -0.263186 | 0.63929 | 0.329154 | 0.726177 | -0.24326 | -0.052916 | 0.581733 | 0.054578 | 0.1695 | 0.187012 | 0.13804 | 0.519765 | 0.252929 |
| 27.73611 | 0.452495 | -0.948715 | 0.21916 | -0.137067 | -0.221688 | 1.119476 | 0.015197 | 0.0561 | 0.432888 | -1.401195 | -0.325296 | 0.956428 | 0.316867 | 0.516022 | -0.525605 | -0.087076 | 0.317006 | 0.048532 | 0.201709 | 0.187012 | 0.12535 | 0.605715 | 0.177553 |
| 27.75 | 0.755478 | -1.01813 | 0.069953 | -0.003982 | 0.223529 | 0.927286 | -0.209232 | -0.344602 | 0.19607 | -1.184982 | -0.373758 | 0.658469 | 0.274116 | 0.097488 | -0.874406 | -0.211393 | -0.552534 | 0.137934 | 0.166955 | 0.184987 | 0.062811 | 0.606708 | 0.056816 |
| 27.76389 | 0.832051 | -1.024135 | -0.094635 | 0.079562 | 0.396271 | 0.589659 | -0.175652 | -0.705608 | -0.036925 | -0.771882 | -0.363377 | 0.190822 | 0.241176 | -0.185812 | -0.900749 | -0.27162 | -0.516576 | 0.396953 | 0.104658 | 0.188777 | -0.026094 | 0.524357 | -0.005512 |
| 27.77778 | 0.853906 | -0.805259 | -0.509877 | 0.145393 | 0.337825 | 0.100843 | -0.118212 | -0.689312 | -0.19584 | -0.38123 | -0.268819 | -0.273492 | 0.184434 | -0.243706 | -0.617922 | -0.163751 | 0.221297 | 0.597352 | 0.013832 | 0.197039 | -0.107455 | 0.365166 | 0.057433 |
| 27.79167 | 0.935723 | -0.541966 | -0.906588 | 0.198502 | 0.312043 | -0.351096 | -0.141114 | -0.347909 | -0.346658 | -0.226562 | -0.198299 | -0.317694 | 0.026848 | -0.15501 | -0.553733 | -0.02061 | 0.391988 | 0.455486 | 0.02248 | 0.162231 | -0.059208 | 0.329039 | 0.107553 |
| 27.80556 | 0.953997 | -0.443817 | -0.830502 | 0.098097 | 0.181074 | -0.705695 | -0.179634 | -0.264003 | -0.525391 | -0.328817 | -0.209017 | -0.16312 | -0.150491 | -0.155059 | -0.664079 | -0.040029 | -0.059295 | 0.279219 | 0.044699 | 0.115077 | 0.020774 | 0.364678 | 0.062112 |
| 27.81944 | 0.750478 | -0.552989 | -0.809128 | 0.018518 | -0.107899 | -0.87928 | -0.184824 | -0.309641 | -0.550405 | -0.525775 | -0.267967 | -0.241652 | -0.391896 | -0.457688 | -0.682947 | -0.210278 | -0.534656 | 0.207858 | -0.105879 | -0.014748 | -0.143037 | 0.187212 | -0.098905 |
| 27.83333 | 0.446496 | -0.653807 | -0.748458 | 0.082017 | -0.585457 | -0.942205 | -0.027432 | -0.296791 | -0.374149 | -0.531873 | -0.331079 | -0.486709 | -0.695747 | -0.759209 | -0.623186 | -0.337952 | -0.587485 | 0.039678 | -0.313148 | -0.228725 | -0.410187 | 0.010705 | -0.303916 |

| | | | | | | | | | | | | | | | | | | | | | | | |
|---|---|---|---|---|---|---|---|---|---|---|---|---|---|---|---|---|---|---|---|---|---|---|---|
| 27.84722 | 0.345446 | -0.430345 | -0.650417 | 0.282724 | -1.661118 | -0.537298 | 0.369728 | -0.070469 | 0.033442 | -0.178862 | -0.270844 | -0.521644 | -0.592808 | -0.630976 | -0.384621 | -0.245052 | -0.287997 | -0.099514 | -0.327055 | -0.276283 | -0.387298 | 0.179058 | -0.245802 |
| 27.86111 | 0.198332 | -0.062841 | -0.703935 | 0.391481 | -1.485461 | -0.079325 | 0.733903 | 0.108684 | 0.328939 | 0.081144 | -0.117956 | -0.323128 | -0.22175 | -0.298725 | -0.143782 | -0.027853 | 0.121899 | -0.020368 | -0.215794 | -0.132995 | -0.213059 | 0.430947 | 0.001545 |
| 27.875 | -0.18718 | -0.023589 | -0.634583 | 0.31909 | 0.070861 | 0.04082 | 0.68898 | -0.023549 | 0.263991 | -0.131566 | -0.04919 | -0.160349 | 0.076084 | -0.174157 | -0.028462 | 0.17561 | 0.310051 | 0.135763 | -0.143159 | 0.000442 | -0.169461 | 0.52437 | 0.131822 |
| 27.88889 | -0.609497 | -0.074037 | -0.327214 | 0.268719 | 0.622925 | 0.010288 | 0.305792 | -0.229967 | 0.215085 | -0.386217 | -0.028002 | 0.035419 | 0.261876 | -0.152611 | 0.075784 | 0.362638 | 0.147662 | 0.163975 | -0.039986 | 0.054441 | -0.129558 | 0.54523 | 0.125301 |
| 27.90278 | -0.720116 | 0.089071 | -0.165579 | 0.26488 | 0.601074 | 0.028341 | -0.172375 | -0.317805 | 0.213529 | -0.545018 | -0.027174 | 0.071579 | 0.342579 | -0.085384 | 0.189986 | 0.420463 | 0.038256 | 0.124571 | -0.037487 | 0.016831 | -0.169626 | 0.479184 | 0.050174 |
| 27.91667 | -0.78346 | 0.328408 | -0.635316 | 0.198139 | 0.518989 | 0.150481 | -0.537887 | -0.319866 | 0.070867 | -0.633944 | -0.017238 | 0.036117 | 0.319708 | 0.036223 | 0.36891 | 0.369481 | 0.397154 | 0.121795 | -0.127457 | -0.056018 | -0.267133 | 0.346498 | 0.003006 |
| 27.93056 | -0.786591 | 0.607857 | -0.037642 | 0.260087 | 0.544463 | 0.917092 | -0.351028 | -0.071457 | -0.008041 | -0.494733 | 0.044941 | 0.199284 | 0.350099 | 0.233819 | 0.65649 | 0.407787 | 1.062074 | 0.165732 | -0.094372 | -0.065632 | -0.214965 | 0.378348 | 0.058047 |
| 27.94444 | -0.436168 | 0.887478 | 0.495633 | 0.480357 | 0.602797 | 1.041824 | -0.175592 | 0.33367 | 0.028469 | -0.290994 | 0.133688 | 0.402223 | 0.508179 | 0.349209 | 0.917139 | 0.505541 | 1.147825 | 0.26965 | -0.00817 | -0.030382 | -0.132565 | 0.433564 | 0.14719 |
| 27.95833 | -0.028738 | 1.149388 | 1.118818 | 0.673299 | 0.632779 | 0.978363 | -0.025824 | 0.716652 | 0.04057 | -0.070167 | 0.222729 | 0.601189 | 0.648459 | 0.501328 | 1.112509 | 0.589842 | 0.592312 | 0.350725 | 0.018396 | -0.014794 | -0.078834 | 0.392261 | 0.200052 |
| 27.97222 | 0.382373 | 1.534475 | 1.460563 | 0.812955 | 0.668249 | 0.838177 | 0.266563 | 1.071689 | 0.109084 | 0.396736 | 0.323417 | 0.781872 | 0.641311 | 0.566996 | 1.212992 | 0.572651 | 0.152924 | 0.308746 | 0.067464 | -0.023949 | -0.019638 | 0.4185 | 0.189159 |
| 27.98611 | 0.788116 | 2.029215 | 1.884953 | 0.967586 | 0.789066 | 0.65029 | 0.559102 | 1.327026 | 0.214425 | 0.886202 | 0.384117 | 0.899682 | 0.623361 | 0.559622 | 1.231129 | 0.511491 | 0.084141 | 0.248471 | 0.071343 | -0.064749 | -0.032471 | 0.413071 | 0.238873 |
| 28 | 1.053829 | 2.090471 | 2.362536 | 0.983593 | 0.95708 | 0.440556 | 0.572332 | 1.263351 | 0.222818 | 0.905018 | 0.306527 | 0.913886 | 0.318887 | 0.32542 | 1.224969 | 0.273471 | 0.326278 | 0.22164 | -0.090879 | -0.210989 | -0.219847 | 0.197444 | 0.23171 |
| 28.01389 | 0.546885 | 1.990537 | 2.479329 | 0.661528 | 0.95255 | 0.235281 | 0.348805 | 1.030044 | 0.093139 | 0.801661 | 0.045635 | 0.61797 | -0.320725 | -0.144608 | 0.973391 | -0.144678 | 0.598421 | 0.076571 | -0.367746 | -0.503346 | -0.730392 | -0.231754 | 0.003411 |
| 28.02778 | 0.608372 | 2.129527 | 2.708024 | 0.517557 | 0.923032 | 0.321787 | 0.193756 | 1.048279 | 0.03581 | 1.08835 | -0.114229 | 0.567263 | -0.589044 | -0.359794 | 0.997246 | -0.189647 | 0.885054 | 0.039439 | -0.468858 | -0.555 | -0.856514 | -0.432083 | -0.177768 |
| 28.04167 | 1.511095 | 2.569614 | 2.929769 | 0.708447 | 0.976228 | 0.874762 | 0.095597 | 1.409212 | 0.476783 | 1.729282 | 0.096078 | 0.548158 | -0.336477 | -0.084305 | 1.072599 | 0.113363 | 0.587866 | 0.236321 | -0.229713 | -0.308779 | -0.422033 | -0.202011 | -0.062614 |
| 28.05556 | 2.032509 | 2.601378 | 2.598179 | 0.932109 | 1.061552 | 0.968524 | -0.041977 | 1.431721 | 0.776785 | 2.03212 | 0.249021 | 0.582662 | -0.133247 | 0.207704 | 1.072297 | 0.353391 | -0.009696 | 0.25318 | 0.008834 | -0.209313 | -0.146714 | 0.088389 | 0.011356 |
| 28.06944 | 1.814425 | 2.378958 | 2.773918 | 0.952727 | 1.190033 | 0.276645 | -0.056887 | 1.305107 | 0.884594 | 1.886582 | 0.241173 | 0.498836 | -0.205959 | 0.418812 | 0.955397 | 0.513415 | -0.510183 | 0.223892 | 0.110482 | -0.218088 | -0.074549 | 0.277733 | -0.020383 |
| 28.08333 | 1.112543 | 1.784784 | 4.747168 | 1.037946 | 1.572644 | 0.280615 | -0.048273 | 1.618057 | 1.138376 | 2.057683 | 0.342793 | 0.180781 | -0.934316 | 0.438458 | 0.545613 | 0.336462 | -0.831998 | -0.06496 | -0.046524 | -0.507328 | -0.404485 | -0.030137 | -0.279459 |
| 28.09722 | 1.093274 | 0.974119 | 6.315462 | 0.798972 | 2.26613 | 0.308083 | -0.271765 | 2.044974 | 1.023704 | 3.057115 | 0.453832 | -0.351716 | -2.624679 | 0.220919 | -0.168683 | -0.250776 | -0.829001 | -0.688929 | -0.492532 | -1.061324 | -1.18166 | -0.810963 | -0.930067 |
| 28.11111 | 0.502783 | 0.545022 | 4.580704 | 0.507185 | 1.564791 | 0.278543 | -0.734109 | 1.858753 | 0.521603 | 2.541573 | 0.222357 | -0.514271 | -3.133428 | -0.163602 | -0.558867 | -0.445029 | -0.471991 | -1.035473 | -0.734263 | -1.342532 | -1.672358 | -1.287385 | -1.234059 |
| 28.125 | 0.05134 | 1.163245 | 2.003217 | 0.691941 | 0.207887 | 0.683429 | -1.196407 | 1.314134 | 0.410702 | 1.778856 | 0.063327 | -0.037183 | -1.744402 | -0.248716 | -0.044984 | 0.353021 | 0.055403 | -0.678539 | -0.417125 | -0.945984 | -1.248802 | -0.746824 | -0.937333 |
| 28.13889 | 0.18 | 2.589433 | 1.766161 | 1.578654 | 0.239175 | 1.959127 | 0.31168 | 1.511823 | 0.917885 | 2.042235 | 0.298619 | 0.631454 | -0.47135 | 0.201842 | 0.974183 | 1.168339 | 0.240617 | -0.051619 | -0.020477 | -0.400693 | -0.585174 | 0.000691 | -0.39546 |
| 28.15278 | 0.48582 | 2.72845 | 2.062651 | 2.017085 | 1.150762 | 2.167333 | 1.163541 | 1.300084 | 1.119284 | 2.363753 | 0.386567 | 1.115885 | 0.037492 | 0.814612 | 1.797573 | 0.818357 | 0.70093 | 0.416142 | 0.257346 | -0.106722 | -0.183188 | 0.305662 | -0.015159 |
| 28.16667 | 0.686011 | 2.627547 | 2.540403 | 2.157091 | 2.172048 | 1.115097 | 1.637815 | 0.605766 | 1.118567 | 2.492168 | 0.41505 | 1.257235 | 0.041567 | 0.972055 | 2.007518 | 0.24842 | 0.214474 | 0.710803 | 0.22828 | -0.003144 | -0.028268 | -0.052596 | 0.126299 |
| 28.18056 | 0.990463 | 1.878183 | 2.697349 | 1.685401 | 2.163448 | 0.74546 | 1.739319 | 0.302512 | 1.052133 | 1.964195 | 0.525283 | 1.256058 | 0.066759 | 0.632722 | 1.713192 | -1.004725 | -0.332859 | 0.988393 | 0.1824 | 0.03027 | 0.028153 | -0.081537 | 0.213639 |
| 28.19444 | 0.751346 | -0.3126 | 1.636767 | 0.884479 | 1.805036 | 0.180623 | 1.325916 | 0.2816 | 0.772511 | 0.997477 | 0.48511 | 1.174301 | 0.117162 | 0.35512 | 1.262865 | -1.354499 | -0.462302 | 0.965276 | 0.307409 | 0.092684 | 0.110183 | 0.038796 | 0.330215 |
| 28.20833 | 0.337122 | -0.674183 | 0.43894 | 0.182366 | 1.384828 | -0.888179 | 0.631576 | 0.255165 | 0.526708 | -0.019505 | 0.365734 | 1.043628 | 0.206377 | 0.382411 | 1.239865 | -2.434698 | -0.174637 | 0.640351 | 0.595016 | 0.216744 | 0.277469 | 0.22484 | 0.469934 |
| 28.22222 | 0.283969 | 0.053156 | 0.227312 | -0.120672 | 0.950754 | -1.049679 | 0.243917 | 0.283798 | 0.576278 | -0.294845 | 0.365299 | 0.611132 | 0.213847 | 0.160836 | 1.552561 | -1.467826 | 0.321796 | 0.35045 | 0.956863 | 0.280789 | 0.395302 | 0.34195 | 0.521211 |
| 28.23611 | 0.217788 | 1.171245 | 0.658054 | -0.225531 | 0.54372 | -0.46881 | 0.271922 | 0.395591 | 0.782877 | 0.442796 | 0.503909 | 0.51766 | 0.149871 | 0.058824 | 1.316113 | -0.563526 | 0.71146 | 0.150726 | 1.114264 | 0.228636 | 0.351265 | 0.330455 | 0.426103 |
| 28.25 | -0.040151 | 1.145788 | 0.549958 | -0.302732 | 0.277921 | 0.026963 | 0.831147 | 0.432536 | 0.629054 | 1.276157 | 0.572115 | 0.37063 | 0.083986 | 0.009987 | 1.030966 | -0.208532 | 0.937857 | 0.120627 | 1.061075 | 0.149431 | 0.237724 | 0.343886 | 0.299304 |
| 28.26389 | -0.393046 | 0.873536 | 0.157936 | -0.165455 | -0.365287 | 0.645005 | 1.320626 | 0.172053 | 0.290481 | 1.9237 | 0.53746 | 0.263759 | 0.002213 | -0.06296 | 0.579719 | -0.019996 | 1.244887 | 0.152738 | 0.912402 | 0.103615 | 0.177653 | 0.387713 | 0.212503 |
| 28.27778 | -0.614452 | 0.23721 | 0.012713 | 0.164786 | -0.808546 | 1.224991 | 0.770323 | -0.133105 | -0.001822 | 2.296845 | 0.413099 | 0.172681 | -0.007959 | 0.000437 | 0.1288 | 0.417178 | 0.654653 | 0.134223 | 0.748294 | 0.079994 | 0.183908 | 0.385061 | 0.19333 |
| 28.29167 | -0.334513 | -0.409343 | -0.274441 | 0.521917 | -1.588328 | 1.301493 | 0.38358 | -0.408797 | -0.037117 | 2.391997 | 0.315981 | -0.00821 | 0.023221 | 0.145203 | -0.180623 | 0.822001 | -0.062954 | 0.083643 | 0.598623 | 0.045804 | 0.161313 | 0.33406 | 0.19497 |
| 28.30556 | 0.315352 | -0.72479 | 0.798716 | 0.663464 | -0.690041 | 0.52926 | 0.728114 | -0.698729 | -0.083025 | 1.719108 | 0.237741 | -0.225781 | 0.08072 | 0.276926 | -0.170002 | 0.833737 | -0.722057 | 0.046027 | 0.432485 | 0.017538 | 0.110321 | 0.276218 | 0.192818 |
| 28.31944 | 1.090493 | -0.948735 | 1.310543 | 0.481415 | 0.684056 | 0.015305 | 0.307631 | -0.778875 | -0.212354 | 0.615203 | 0.172434 | -0.339182 | 0.169452 | 0.413136 | 0.110343 | 0.649863 | -0.799384 | 0.031163 | 0.272464 | 0.029129 | 0.117136 | 0.248002 | 0.144917 |
| 28.33333 | 1.039125 | 0.055526 | 1.18923 | 0.198222 | 1.80219 | -0.076534 | -0.419179 | -0.618373 | -0.217187 | -0.353428 | 0.06087 | -0.286072 | 0.205632 | 0.568233 | 0.431786 | 0.513452 | -0.758203 | 0.065448 | 0.151658 | 0.074269 | 0.183992 | 0.258775 | 0.104839 |
| 28.34722 | 0.867041 | 1.210618 | 0.775653 | -0.123081 | 2.121468 | -0.136743 | -0.922898 | -0.368831 | -0.140691 | -1.517229 | -0.006867 | -0.197496 | 0.242431 | 0.650135 | 0.581243 | 0.277909 | -0.310177 | 0.169904 | 0.114008 | 0.145883 | 0.255181 | 0.236875 | 0.121886 |
| 28.36111 | 0.381461 | 1.575942 | -0.141848 | -0.448132 | 1.023788 | -0.385629 | -1.138336 | -0.177334 | -0.070288 | -1.976801 | -0.043814 | -0.159779 | 0.301788 | 0.745136 | 0.756829 | 0.092065 | 0.514935 | 0.287118 | 0.133638 | 0.224853 | 0.313131 | 0.224006 | 0.176252 |
| 28.375 | -0.180643 | 0.775279 | -1.264234 | -0.705663 | 0.34798 | -1.493134 | -0.749177 | -0.053123 | -0.233829 | -1.507741 | -0.11332 | -0.094708 | 0.301871 | 0.865553 | 0.818797 | -0.141934 | 1.150838 | 0.291165 | 0.16102 | 0.271289 | 0.361278 | 0.230827 | 0.199675 |
| 28.38889 | -0.94393 | 0.062075 | -2.91901 | -0.67548 | -0.295569 | -1.482716 | -0.234587 | 0.187005 | -0.685436 | -0.343901 | -0.001966 | 0.022182 | 0.230925 | 0.766813 | 0.620448 | -0.338562 | 1.414861 | 0.170268 | 0.171499 | 0.283433 | 0.403339 | 0.277695 | 0.176564 |
| 28.40278 | -1.883664 | -0.754416 | -3.191425 | -0.538513 | -1.023304 | -0.566088 | -0.333525 | 0.497975 | -0.769425 | 0.801054 | 0.107844 | 0.169813 | 0.104785 | 0.504231 | 0.265373 | -0.424746 | 1.212858 | 0.009906 | 0.1883 | 0.292947 | 0.462933 | 0.368285 | 0.152944 |
| 28.41667 | -1.781522 | -1.122153 | -2.392831 | -0.398226 | -1.126504 | 0.394521 | -0.161149 | 0.536091 | -0.583884 | 1.455412 | 0.174958 | 0.286229 | 0.011468 | 0.195659 | -0.03668 | -0.378382 | 0.612038 | -0.095943 | 0.181445 | 0.298927 | 0.501875 | 0.419473 | 0.135742 |
| 28.43056 | -2.205563 | -0.556433 | -2.025889 | -0.224536 | -1.376602 | 1.002524 | 0.428565 | 0.176413 | -0.35607 | 1.106858 | 0.200905 | 0.161936 | -0.063722 | -0.054664 | -0.327777 | -0.285142 | -0.043417 | -0.103054 | 0.12274 | 0.255261 | 0.393696 | 0.338027 | 0.104532 |
| 28.44444 | -1.319692 | -0.089889 | -2.049334 | -0.031952 | -2.28991 | 0.78168 | 0.116192 | -0.364006 | -0.146691 | 0.440681 | 0.174194 | -0.028571 | -0.098418 | -0.210504 | -0.492922 | 0.049051 | -0.798578 | -0.077558 | 0.016068 | 0.182459 | 0.256725 | 0.231262 | 0.077199 |
| 28.45833 | -0.839366 | 0.21254 | -2.596825 | -0.003399 | -3.354052 | 0.310806 | -0.685313 | -0.872291 | -0.127189 | -0.534025 | -0.013733 | -0.106427 | -0.205184 | -0.173027 | -0.620627 | 0.269229 | -1.136405 | -0.10502 | -0.068929 | 0.122177 | 0.205509 | 0.148102 | 0.052027 |
| 28.47222 | -0.46831 | -0.170904 | -2.739597 | -0.308002 | -2.968136 | 0.28699 | -0.277939 | -1.112624 | -0.204836 | -1.248228 | -0.305153 | -0.163607 | -0.251089 | -0.050367 | -0.781129 | 0.384726 | -0.999985 | -0.141327 | -0.150045 | 0.078314 | 0.153223 | 0.021676 | 0.024926 |
| 28.48611 | -0.046328 | -0.818543 | -2.586154 | -0.676343 | -2.356564 | 0.206542 | 0.196996 | -1.224102 | -0.258438 | -1.730446 | -0.505648 | -0.214557 | -0.291713 | -0.008929 | -0.824719 | 0.232256 | -1.586038 | -0.130207 | -0.241945 | 0.047637 | 0.101449 | -0.162706 | -0.012563 |
| 28.5 | 0.377911 | -1.4576 | -2.452165 | -1.001289 | -2.01025 | 0.02338 | 0.186345 | -1.326459 | -0.387534 | -1.639848 | -0.624821 | -0.262959 | -0.420413 | -0.075705 | -0.773712 | -0.22989 | -1.062116 | -0.141468 | -0.312005 | 0.020626 | 0.081495 | -0.313392 | -0.070661 |
| 28.51389 | 0.395358 | -1.740048 | -2.050726 | -1.227214 | -1.787052 | -0.535064 | -0.265091 | -1.279984 | -0.608347 | -1.086565 | -0.539481 | -0.313011 | -0.469488 | -0.217751 | -0.639797 | -0.788381 | -0.064095 | -0.221575 | -0.374664 | -0.025091 | 0.074272 | -0.42735 | -0.170589 |
| 28.52778 | 0.206954 | -1.668562 | -1.615071 | -1.371897 | -1.493112 | -1.419089 | -1.345242 | -1.150457 | -0.806486 | -0.358734 | -0.402606 | -0.429876 | -0.404212 | -0.350048 | -0.547046 | -1.102089 | 0.491314 | -0.35958 | -0.469287 | -0.096164 | 0.028593 | -0.504191 | -0.271491 |
| 28.54167 | -0.418298 | -1.518641 | -1.362912 | -1.401864 | -1.486031 | -0.681005 | -1.367627 | -1.002146 | -0.979937 | 0.306134 | -0.273106 | -0.511651 | -0.349852 | -0.36351 | -0.611871 | -1.081732 | -0.066262 | -0.431447 | -0.53857 | -0.139817 | -0.000215 | -0.572335 | -0.31285 |
| 28.55556 | -0.891683 | -1.309675 | -1.517263 | -1.205453 | -1.931614 | -0.211058 | -1.239256 | -0.593256 | -0.991126 | 0.712885 | -0.138105 | -0.414486 | -0.215372 | -0.24596 | -0.803361 | -0.927483 | -0.705348 | -0.339787 | -0.486901 | -0.053287 | 0.100621 | -0.49845 | -0.190311 |
| 28.56944 | -0.905292 | -1.250385 | -2.734843 | -0.929884 | -1.168039 | -0.24127 | -0.762693 | -0.152244 | -0.842943 | 0.421513 | 0.080662 | -0.4331 | 0.137341 | -0.075446 | -0.930819 | -0.470668 | -0.394309 | -0.036173 | -0.390993 | 0.173444 | 0.271347 | -0.309209 | 0.094045 |
| 28.58333 | -1.197684 | -1.52363 | -5.779801 | -1.122409 | -1.827993 | -0.570427 | -0.088293 | -0.630933 | -1.204071 | -0.773127 | 0.129434 | -0.622342 | 0.248027 | -0.465484 | -1.216123 | -0.10499 | 0.271914 | 0.048194 | -0.476647 | 0.208381 | 0.230778 | -0.484374 | 0.14749 |
| 28.59722 | -1.549307 | -2.271322 | -7.736257 | -1.622579 | -2.31283 | -0.960596 | -0.110672 | -1.972146 | -1.303305 | -2.137497 | -0.150922 | -1.113134 | -0.349446 | -1.513403 | -2.006702 | -0.704154 | -0.002008 | -0.626763 | -0.843071 | -0.181101 | -0.117485 | -0.971046 | -0.30014 |
| 28.61111 | -1.947101 | -3.097011 | -8.344402 | -1.716538 | -2.52973 | -1.49772 | -0.4439 | -3.489187 | -0.843273 | -3.947111 | -0.278888 | -1.353277 | -0.853919 | -1.895457 | -2.529183 | -1.375393 | -0.535851 | -1.239812 | -0.708893 | -0.289889 | -0.066876 | -1.059806 | -0.591278 |
| 28.625 | -1.554902 | -3.035909 | -6.734237 | -1.488609 | -1.909777 | -1.355128 | -0.181779 | -3.913974 | -0.407908 | -3.835426 | -0.196506 | -1.074849 | -0.822744 | -1.358222 | -2.165989 | -2.004848 | -1.117542 | -1.291278 | -0.498315 | -0.097714 | 0.08782 | -0.8156 | -0.429412 |
| 28.63889 | -1.117538 | -1.891326 | -3.86179 | -1.260267 | -0.994483 | -1.012799 | -0.090705 | -2.154119 | -0.026925 | -2.331338 | -0.11515 | -1.323661 | -0.750751 | -1.54539 | -1.481925 | -1.690573 | -2.017165 | -0.951621 | -0.191421 | -0.025204 | -0.024158 | -0.88757 | -0.305432 |
| 28.65278 | -0.758273 | -0.99184 | -1.729696 | -1.057056 | -0.446916 | -0.170608 | 0.285831 | -0.357646 | 0.276983 | -0.546459 | -0.033469 | -1.005429 | -0.668287 | -1.163079 | -0.92913 | -0.767731 | -1.316192 | -0.54333 | 0.083722 | -0.040679 | -0.048457 | -0.559825 | -0.217428 |
| 28.66667 | -0.290759 | -0.588729 | -0.438231 | -0.665775 | 0.125505 | 0.303759 | 0.420442 | 0.719497 | 0.621131 | 0.24832 | 0.129056 | -0.187867 | -0.156244 | -0.475537 | -0.388726 | 0.351093 | 0.206636 | -0.202958 | 0.319834 | 0.063099 | 0.192768 | 0.00362 | 0.010645 |
| 28.68056 | -0.220844 | -0.618241 | 0.271049 | -0.348861 | 0.258349 | 0.19443 | 0.036493 | 0.680923 | 0.726006 | 0.05815 | 0.1983 | 0.289102 | 0.390983 | 0.297122 | 0.049822 | 1.0737 | 1.280365 | 0.098344 | 0.430608 | 0.318698 | 0.578425 | 0.863657 | 0.335233 |
| 28.69444 | -0.464475 | -0.605998 | 0.645825 | -0.248582 | -0.00503 | 0.21512 | -0.119758 | 0.613842 | 0.605508 | -0.467215 | 0.083122 | 0.339984 | 0.585137 | 0.75366 | 0.088311 | 0.815399 | 1.00438 | 0.272271 | 0.522291 | 0.495884 | 0.828037 | 0.958539 | 0.528099 |
| 28.70833 | -0.37945 | -0.509805 | 0.890166 | -0.138947 | -0.245571 | 0.40596 | -0.084292 | 0.592373 | 0.449762 | -0.622873 | -0.023011 | 0.319067 | 0.395803 | 0.647027 | -0.047995 | 0.25263 | 0.465825 | 0.318889 | 0.374817 | 0.468528 | 0.70608 | 0.695128 | 0.454212 |
| 28.72222 | -0.002419 | -0.332039 | 0.889143 | 0.099979 | -0.44331 | 0.369375 | -0.09239 | 0.419117 | 0.464189 | -0.606993 | -0.062717 | 0.390629 | 0.07762 | 0.381724 | -0.140687 | -0.216361 | 0.281005 | 0.25335 | 0.114537 | 0.29802 | 0.4182 | 0.33098 | 0.231965 |
| 28.73611 | 0.226532 | -0.24338 | 0.821923 | 0.353339 | -0.538105 | 0.008934 | -0.278618 | 0.173698 | 0.363813 | -0.581261 | -0.104808 | 0.424186 | -0.160395 | 0.140886 | -0.407395 | -0.550988 | 0.305367 | 0.057901 | -0.056051 | 0.11531 | 0.194085 | -0.012986 | 0.015569 |
| 28.75 | 0.404656 | -0.270769 | 0.813585 | 0.431952 | -0.447212 | -0.302815 | -0.481235 | 0.28374 | 0.156888 | -0.457153 | -0.184061 | 0.297039 | -0.328262 | -0.226391 | -0.859035 | -0.804916 | 0.113338 | -0.108644 | -0.209551 | -0.027139 | -0.055123 | -0.360332 | -0.160897 |
| 28.76389 | 0.366828 | -0.348698 | 0.686632 | 0.306927 | -0.365466 | -0.24037 | -0.406835 | 0.348742 | -0.040498 | -0.206179 | -0.276661 | -0.047992 | -0.549899 | -0.502537 | -1.098119 | -0.820143 | -0.556257 | -0.12821 | -0.367443 | -0.09042 | -0.274793 | -0.65118 | -0.258429 |
| 28.77778 | 0.297311 | -0.307611 | 0.634621 | 0.150952 | -0.264952 | 0.066639 | -0.076616 | 0.43193 | -0.073738 | 0.134204 | -0.313429 | -0.370279 | -0.734334 | -0.338114 | -0.920081 | -0.686582 | -1.129238 | -0.104696 | -0.307445 | -0.087383 | -0.211806 | -0.681483 | -0.247718 |
| 28.79167 | 0.77394 | -0.225696 | 0.992793 | 0.115982 | -0.073085 | 0.254119 | 0.239501 | 0.704189 | 0.018626 | 0.27148 | -0.295882 | -0.635628 | -0.790536 | -0.160291 | -0.878762 | -0.606093 | -1.135035 | -0.286659 | -0.158502 | -0.128917 | -0.058278 | -0.621856 | -0.282546 |
| 28.80556 | 1.646635 | -0.175569 | 1.487637 | 0.121072 | 0.083561 | 0.249037 | 0.539268 | 1.052413 | 0.050197 | 0.224933 | -0.318076 | -0.971643 | -1.038282 | -0.361034 | -1.096047 | -0.503149 | -0.833705 | -0.546927 | -0.25048 | -0.324579 | -0.177745 | -0.611432 | -0.457943 |
| 28.81944 | 1.770333 | -0.099528 | 1.47785 | 0.12646 | 0.056329 | 0.274039 | 0.937381 | 0.953164 | -0.076237 | 0.138874 | -0.371928 | -1.029484 | -0.902633 | -0.641856 | -1.057239 | -0.177206 | -0.831124 | -0.502726 | -0.430272 | -0.360929 | -0.322539 | -0.475559 | -0.514115 |
| 28.83333 | 1.443162 | -0.179385 | 0.768906 | 0.186891 | 0.01672 | 0.445374 | 1.155194 | 0.375402 | -0.282839 | 0.055594 | -0.402031 | -0.608969 | -0.522415 | -0.535532 | -0.661601 | 0.209469 | -0.87079 | -0.250767 | -0.317997 | -0.156873 | -0.144418 | -0.062351 | -0.317724 |
| 28.84722 | 0.757731 | -0.400101 | 0.003041 | 0.230017 | 0.045228 | 0.718962 | 0.991924 | -0.135168 | -0.279903 | -0.107063 | -0.378754 | 0.036248 | -0.273313 | -0.20498 | -0.401388 | 0.258161 | -0.512855 | -0.202637 | 0.032429 | 0.018149 | 0.230238 | 0.395443 | -0.091157 |
| 28.86111 | -0.065669 | -0.464833 | -0.452691 | 0.251953 | 0.119917 | 0.693569 | 0.655013 | -0.389365 | -0.102328 | -0.296977 | -0.303084 | 0.357533 | -0.283035 | -0.012508 | -0.320243 | 0.002997 | 0.10605 | -0.367083 | 0.27131 | 0.062243 | 0.46383 | 0.593411 | 0.042889 |
| 28.875 | -0.615989 | -0.375491 | -0.817205 | 0.212683 | 0.105669 | 0.45179 | 0.341889 | -0.494532 | -0.011985 | -0.296465 | -0.273363 | 0.350028 | -0.304234 | 0.131508 | -0.088194 | -0.093381 | 0.526804 | -0.335382 | 0.226818 | 0.104949 | 0.390519 | 0.495456 | 0.116009 |
| 28.88889 | -1.153262 | -0.198106 | -1.060917 | 0.148707 | 0.054018 | 0.277968 | 0.090292 | -0.424608 | 0.009892 | -0.162246 | -0.193253 | 0.275418 | -0.121041 | 0.257563 | 0.251538 | 0.068277 | 0.595927 | -0.081636 | 0.082455 | 0.150666 | 0.219839 | 0.306077 | 0.186024 |
| 28.90278 | -1.612298 | -0.027644 | -1.060328 | 0.111266 | 0.064189 | 0.193548 | -0.158337 | -0.201439 | 0.104098 | -0.034231 | -0.068587 | 0.289162 | 0.13349 | 0.25942 | 0.349786 | 0.233456 | 0.748546 | 0.100529 | 0.078663 | 0.156177 | 0.115868 | 0.160921 | 0.16234 |
| 28.91667 | -2.108855 | 0.185608 | -0.893936 | 0.09947 | 0.044158 | 0.042214 | -0.227889 | -0.027522 | 0.188 | 0.138813 | 0.002027 | 0.316341 | 0.179778 | 0.152619 | 0.271902 | 0.236774 | 1.076836 | 0.035598 | 0.18127 | 0.065332 | 0.089927 | 0.132196 | 0.047723 |
| 28.93056 | -1.576191 | 0.494063 | -0.313416 | 0.12672 | 0.25657 | 0.100587 | -0.057943 | 0.271562 | 0.189859 | 0.160709 | 0.051937 | 0.293154 | 0.103713 | 0.034424 | 0.246096 | 0.193006 | 1.18927 | -0.181632 | 0.199419 | -0.102828 | 0.061128 | 0.191866 | -0.073877 |
| 28.94444 | -0.683078 | 0.82766 | 0.353388 | 0.168508 | 0.592128 | 0.217866 | 0.187961 | 0.673992 | 0.09735 | 0.135917 | 0.092538 | 0.263999 | 0.20282 | 0.008226 | 0.468247 | 0.369787 | 0.952107 | -0.239038 | 0.086885 | -0.166295 | -0.044511 | 0.263956 | -0.068573 |
| 28.95833 | 0.08625 | 1.138878 | 0.714909 | 0.145721 | 0.633439 | 0.348984 | 0.362361 | 0.916236 | -0.031317 | 0.126667 | 0.096144 | 0.286026 | 0.504645 | 0.154272 | 0.725774 | 0.697249 | 0.942724 | -0.035654 | 0.073025 | -0.06778 | -0.120631 | 0.350868 | 0.032927 |
| 28.97222 | 0.316367 | 1.411537 | 0.982779 | 0.161809 | 0.56608 | 0.519188 | 0.549001 | 1.036148 | -0.026731 | 0.351727 | 0.104561 | 0.409596 | 0.562631 | 0.363596 | 0.863854 | 0.761811 | 1.251976 | 0.075677 | 0.056467 | -0.038188 | -0.069508 | 0.505909 | 0.075613 |
| 28.98611 | 0.729096 | 1.822077 | 1.315048 | 0.233613 | 0.678839 | 0.695092 | 0.708495 | 1.161012 | 0.064222 | 0.670711 | 0.145572 | 0.477214 | 0.440578 | 0.521404 | 0.866439 | 0.527818 | 1.597511 | -0.004761 | -0.083139 | -0.13793 | -0.042387 | 0.744842 | 0.035273 |
| 29 | 0.756571 | 2.079127 | 1.489923 | 0.177142 | 0.704249 | 0.749897 | 0.5617 | 0.79502 | 0.082497 | 0.772868 | 0.070774 | 0.476894 | 0.286021 | 0.534586 | 0.900468 | 0.298571 | 1.586951 | -0.142657 | -0.209334 | -0.282359 | -0.194754 | 0.945194 | -0.064678 |
| 29.01389 | 0.591359 | 2.161962 | 1.487369 | 0.016622 | 0.534792 | 0.587207 | 0.201602 | 0.324493 | -0.132508 | 0.697742 | -0.218762 | 0.35857 | 0.005921 | 0.316613 | 0.684896 | 0.113427 | 0.99844 | -0.28844 | -0.176093 | -0.423776 | -0.451848 | 0.817034 | -0.185561 |
| 29.02778 | 0.793048 | 2.262326 | 1.956073 | -0.072439 | 0.751598 | 0.297856 | 0.110862 | 0.281912 | -0.320974 | 0.795747 | -0.264534 | 0.273049 | 0.072936 | 0.201575 | 0.690957 | 0.164663 | 0.367854 | -0.094952 | 0.048668 | -0.336988 | -0.477185 | 0.672225 | -0.140309 |
| 29.04167 | 1.656263 | 2.276646 | 2.681382 | 0.235189 | 1.522021 | 0.410477 | 0.316787 | 0.721262 | 0.183317 | 1.334736 | 0.062601 | 0.754161 | 0.45706 | 0.366833 | 1.110894 | 0.453299 | 0.167719 | 0.259719 | 0.155549 | -0.098201 | -0.284087 | 0.593476 | 0.112659 |
| 29.05556 | 1.848737 | 2.358827 | 3.228139 | 0.500064 | 1.829701 | 0.554478 | 0.420335 | 1.057271 | 0.760079 | 1.936005 | 0.31182 | 0.989415 | 0.52694 | 0.513693 | 1.537024 | 0.620595 | 0.337273 | 0.386055 | 0.171471 | 0.003744 | -0.128571 | 0.676277 | 0.206348 |
| 29.06944 | 1.113538 | 2.489854 | 4.022026 | 0.590424 | 1.8926 | 0.499871 | 0.479713 | 1.316372 | 1.098514 | 2.041036 | 0.378883 | 0.87872 | 0.471541 | 0.616275 | 1.439895 | 0.690928 | 0.724046 | 0.278676 | 0.197778 | -0.054207 | 0.045667 | 0.750534 | 0.18489 |
| 29.08333 | 0.566833 | 2.470234 | 5.332055 | 0.652987 | 2.23145 | 0.447603 | 0.497172 | 1.513986 | 1.283262 | 2.148389 | 0.381177 | 0.540936 | 0.012895 | 0.504616 | 0.706058 | 0.308452 | 0.475244 | -0.275209 | -0.047612 | -0.495236 | -0.26099 | 0.412293 | -0.237307 |
| 29.09722 | 0.464918 | 2.413964 | 6.676884 | 0.57265 | 2.711523 | 0.326203 | 0.588917 | 1.699042 | 0.898624 | 2.656937 | 0.333669 | -0.162221 | -0.756067 | 0.100031 | -0.177182 | -0.525349 | -0.59289 | -1.062938 | -0.724584 | -1.238877 | -1.197115 | -0.491482 | -0.994937 |
| 29.11111 | 0.362741 | 1.714717 | 4.005189 | 0.462748 | 2.11065 | 0.504816 | 0.492035 | 0.920644 | 0.326291 | 1.893346 | 0.090975 | -0.489048 | -0.955037 | -0.206267 | -0.375539 | -0.559628 | -0.597036 | -1.171895 | -0.893721 | -1.27765 | -1.407688 | -0.81578 | -1.182118 |
| 29.125 | 0.432512 | 1.251409 | 1.649997 | 0.762022 | 1.117704 | 0.802758 | -0.145205 | 0.647602 | 0.446066 | 1.337122 | -0.040018 | -0.028867 | -0.320743 | 0.079701 | 0.27301 | 0.541305 | 0.787812 | -0.463302 | -0.292184 | -0.591712 | -0.612792 | 0.039973 | -0.611107 |
| 29.13889 | 0.827883 | 2.642347 | 2.254589 | 1.665271 | 1.051963 | 1.424727 | 0.710122 | 1.228019 | 1.096962 | 1.720582 | 0.109303 | 0.610787 | 0.344505 | 0.383283 | 1.118221 | 1.620269 | 2.198665 | 0.346161 | 0.225538 | -0.056161 | 0.04907 | 0.853384 | 0.012551 |
| 29.15278 | 1.222472 | 2.609878 | 2.720233 | 1.99097 | 1.39637 | 1.30015 | 1.219919 | 1.674623 | 1.263322 | 1.918794 | 0.241348 | 0.960996 | 0.773315 | 0.50556 | 1.915458 | 1.36743 | 1.581393 | 0.891506 | 0.452421 | 0.186186 | 0.294081 | 0.907913 | 0.408507 |
| 29.16667 | 1.334704 | 2.413233 | 1.913447 | 1.433001 | 1.920598 | 0.48975 | 1.237021 | 0.961589 | 1.184641 | 1.661151 | 0.277787 | 0.971147 | 1.005576 | 0.592821 | 2.188444 | 1.173234 | 0.368723 | 1.230911 | 0.364401 | 0.266213 | 0.33346 | 0.935551 | 0.586459 |
| 29.18056 | 1.276085 | 2.202278 | 1.926214 | 0.908183 | 2.417453 | 0.278142 | 0.702458 | 0.554094 | 1.103614 | 1.460624 | 0.238294 | 0.911163 | 0.885223 | 0.712801 | 1.962026 | 0.966209 | -0.671214 | 1.382845 | 0.278094 | 0.249499 | 0.308021 | 1.051721 | 0.544805 |
| 29.19444 | 1.208728 | 1.50853 | 2.61923 | 0.51288 | 2.621621 | 0.270076 | 0.244715 | 0.801592 | 1.147163 | 1.124744 | 0.220871 | 1.034768 | 0.649707 | 0.74901 | 1.745251 | 0.438383 | -1.480931 | 1.323668 | 0.298607 | 0.198419 | 0.285152 | 0.92161 | 0.471802 |
| 29.20833 | 1.180208 | 0.680875 | 2.621177 | 0.255634 | 2.634293 | 0.271955 | 0.552862 | 0.787047 | 1.176437 | 0.735641 | 0.228615 | 1.112939 | 0.417965 | 0.636521 | 1.483029 | -0.020771 | -0.953664 | 1.110622 | 0.404372 | 0.14878 | 0.248057 | 0.314442 | 0.353295 |
| 29.22222 | 0.946135 | -0.018439 | 1.742772 | 0.154764 | 1.592909 | 0.132824 | 0.782211 | 0.314621 | 0.996852 | 0.298877 | 0.213252 | 1.060401 | 0.235892 | 0.412362 | 1.31561 | -0.090149 | 0.080429 | 0.768703 | 0.55328 | 0.112556 | 0.187678 | 0.16042 | 0.264849 |
| 29.23611 | 0.513174 | -0.448889 | 1.28571 | 0.080912 | 0.624832 | -0.216297 | 0.320567 | -0.156281 | 0.818756 | -0.051309 | 0.255993 | 0.852846 | 0.096956 | 0.169794 | 1.250202 | 0.313551 | 1.294427 | 0.562023 | 0.680703 | 0.1306 | 0.194873 | 0.144578 | 0.215199 |
| 29.25 | 0.029415 | -0.385507 | 0.547848 | 0.022252 | 0.429257 | -1.049769 | 0.232627 | -0.436758 | 0.6991 | -0.293508 | 0.362108 | 0.741634 | -0.012587 | 0.07259 | 1.139808 | 0.440917 | 1.902855 | 0.49171 | 0.79157 | 0.16079 | 0.245609 | 0.178061 | 0.217189 |
| 29.26389 | -0.375167 | 0.192253 | -0.707111 | 0.00078 | -0.087609 | -0.539761 | -0.053109 | -0.630832 | 0.349325 | -0.100546 | 0.340262 | 0.670321 | -0.14448 | -0.016202 | 0.904878 | 0.498558 | 1.494152 | 0.336129 | 0.761027 | 0.155981 | 0.260064 | 0.206521 | 0.18747 |
| 29.27778 | -0.446974 | 0.555776 | -1.412551 | 0.067807 | -1.215374 | 0.07192 | 0.92991 | -0.589649 | 0.003083 | 0.237724 | 0.273373 | 0.63243 | -0.283608 | -0.172085 | 0.675135 | 0.557185 | 0.774726 | 0.236766 | 0.670239 | 0.174707 | 0.284975 | 0.232717 | 0.176776 |
| 29.29167 | -0.831722 | 0.888888 | -1.471459 | 0.170243 | -1.393342 | 0.660096 | 1.27153 | -0.290512 | -0.184313 | 0.673867 | 0.249451 | 0.60437 | -0.303806 | -0.167495 | 0.478021 | 0.729116 | 0.247088 | 0.260599 | 0.52478 | 0.198042 | 0.266459 | 0.177988 | 0.176361 |
| 29.30556 | -1.013874 | 1.19969 | -1.47319 | 0.095976 | -1.626402 | 1.218385 | 1.220654 | -0.04601 | -0.319632 | 0.759003 | 0.190345 | 0.602588 | -0.258587 | -0.077391 | 0.576677 | 0.726557 | -0.28968 | 0.266215 | 0.363895 | 0.152128 | 0.17375 | 0.158467 | 0.092692 |
| 29.31944 | -0.658194 | 0.703721 | -1.243806 | -0.114991 | -0.841076 | 0.922989 | 0.757334 | -0.107474 | -0.406444 | 0.641014 | 0.039993 | 0.629418 | -0.176326 | 0.167573 | 0.75797 | 0.647372 | -0.84182 | 0.267261 | 0.259946 | 0.1287 | 0.150801 | 0.230854 | 0.019701 |
| 29.33333 | -0.325297 | -0.134744 | -1.123965 | -0.251931 | -0.337992 | 0.176631 | 0.329547 | -0.231847 | -0.31696 | 0.552112 | -0.087626 | 0.495409 | -0.104026 | 0.264951 | 0.586203 | 0.663357 | -1.109055 | 0.287094 | 0.208938 | 0.130325 | 0.172215 | 0.253572 | -0.002662 |
| 29.34722 | -0.128881 | -0.554456 | -0.930058 | -0.178708 | -0.14259 | -0.058746 | -0.343213 | -0.223047 | -0.16555 | 0.42772 | -0.158643 | 0.270733 | -0.230553 | 0.089367 | 0.009066 | 0.614878 | -1.263793 | 0.108796 | 0.057202 | -0.035723 | 0.04562 | 0.047295 | -0.114402 |
| 29.36111 | -0.115957 | -0.485574 | -0.051192 | -0.165369 | -0.155113 | 0.016886 | -0.222563 | -0.154007 | -0.076126 | 0.295776 | -0.289096 | 0.037394 | -0.410894 | -0.166786 | -0.392024 | 0.230418 | -0.682823 | -0.211009 | -0.151928 | -0.295493 | -0.206547 | -0.225117 | -0.315885 |
| 29.375 | -0.140089 | 0.088206 | 1.023316 | -0.260134 | -0.327191 | -0.050546 | -0.564963 | -0.15267 | -0.121009 | 0.17197 | -0.491347 | -0.151787 | -0.281558 | -0.167142 | -0.488186 | -0.164696 | 0.302464 | -0.349207 | -0.199967 | -0.361896 | -0.296242 | -0.340633 | -0.378116 |
| 29.38889 | -0.223289 | 0.64471 | 0.925713 | -0.411603 | -0.539908 | -0.337509 | -0.84135 | -0.314545 | -0.238294 | 0.068424 | -0.377415 | -0.263076 | -0.058145 | 0.042599 | -0.482227 | -0.46337 | 1.339045 | -0.280937 | -0.118163 | -0.230275 | -0.198156 | -0.348025 | -0.283842 |
| 29.40278 | -0.434522 | 0.562509 | -0.664883 | -0.632909 | -0.830916 | -1.387633 | -1.692673 | -0.608525 | -0.521923 | -0.089975 | -0.209829 | -0.268056 | 0.045271 | 0.223598 | -0.510117 | -0.608036 | 1.678725 | -0.225311 | -0.056107 | -0.117792 | -0.10439 | -0.269441 | -0.184505 |
| 29.41667 | -0.660143 | -0.000495 | -1.52578 | -0.630308 | -0.837314 | -1.838819 | -1.721741 | -0.756782 | -0.7414 | -0.016094 | -0.009051 | -0.237643 | -0.039785 | 0.315923 | -0.442739 | -0.609361 | 1.112783 | -0.260534 | -0.059394 | -0.075489 | -0.058769 | -0.187004 | -0.121571 |
| 29.43056 | -0.859662 | -0.178726 | -1.390946 | -0.544411 | -0.782891 | -0.757332 | -1.156448 | -0.604823 | -0.786959 | 0.221705 | -0.030249 | -0.165644 | -0.184976 | 0.259994 | -0.302162 | -0.47853 | 0.390066 | -0.370122 | -0.127865 | -0.064128 | -0.033497 | -0.160321 | -0.120234 |
| 29.44444 | -0.834486 | -0.180783 | -1.000989 | -0.538216 | -0.827372 | -0.136123 | -0.884077 | -0.531343 | -0.748667 | 0.210208 | -0.116109 | -0.147039 | -0.352705 | 0.206255 | -0.144413 | -0.517005 | -0.093411 | -0.427146 | -0.161307 | -0.021819 | 0.003766 | -0.143049 | -0.108296 |
| 29.45833 | -0.746792 | -0.370869 | -1.10557 | -0.551454 | -0.993799 | 0.37181 | -0.68952 | -0.725231 | -0.810636 | -0.099957 | -0.157425 | -0.207547 | -0.50001 | 0.101409 | -0.147274 | -0.514445 | -0.5325 | -0.336913 | -0.132097 | 0.028042 | 0.055642 | -0.128165 | -0.06899 |
| 29.47222 | -0.645658 | -0.4617 | -1.764812 | -0.540068 | -1.507278 | 0.611076 | 0.694604 | -0.994758 | -0.772631 | -0.357182 | -0.121974 | -0.330092 | -0.650719 | 0.011862 | -0.300529 | -0.351546 | -1.004144 | -0.249844 | -0.102673 | 0.030983 | 0.110834 | -0.184556 | -0.019176 |
| 29.48611 | -0.555515 | -0.523315 | -2.246924 | -0.510208 | -1.998831 | 0.74844 | 1.208465 | -0.97395 | -0.59266 | -0.68441 | -0.113988 | -0.535211 | -0.800888 | -0.114332 | -0.51316 | -0.046678 | -1.450312 | -0.168753 | -0.131151 | 0.019311 | 0.126737 | -0.274398 | 0.016969 |
| 29.5 | -0.586095 | -0.533695 | -2.430701 | -0.544737 | -2.270943 | 0.476924 | 0.66061 | -0.756084 | -0.497437 | -0.941771 | -0.17898 | -0.726162 | -0.796359 | -0.344708 | -0.678745 | 0.087693 | -1.297922 | -0.133714 | -0.227691 | -0.008096 | 0.042032 | -0.384975 | -0.036713 |
| 29.51389 | -0.583915 | -0.749918 | -2.783 | -0.745208 | -2.166347 | 0.166007 | 0.656378 | -0.640229 | -0.581958 | -1.253718 | -0.303143 | -0.835471 | -0.601656 | -0.514571 | -0.814319 | -0.131524 | -1.902872 | -0.13284 | -0.323455 | -0.054902 | -0.037874 | -0.513936 | -0.147993 |
| 29.52778 | -0.623122 | -1.279377 | -3.363735 | -0.950693 | -2.503624 | -0.162456 | -0.019269 | -0.738525 | -0.720163 | -1.532384 | -0.39457 | -0.775718 | -0.499907 | -0.484559 | -0.887317 | -0.754211 | -1.373279 | -0.100767 | -0.343045 | -0.060002 | -0.013281 | -0.586216 | -0.189216 |
| 29.54167 | -0.618099 | -1.754247 | -3.414728 | -1.022139 | -2.923277 | -0.232964 | -0.821606 | -0.794079 | -0.617616 | -1.577086 | -0.378128 | -0.660454 | -0.420548 | -0.322253 | -0.943503 | -1.232582 | -0.801498 | -0.068983 | -0.287636 | -0.014058 | 0.07932 | -0.593776 | -0.155811 |
| 29.55556 | -0.589287 | -0.824836 | -2.89044 | -0.993411 | -2.461808 | -0.54422 | -1.424446 | -0.791511 | -0.438508 | -1.199318 | -0.250758 | -0.493209 | -0.248701 | -0.117201 | -0.963866 | -1.369492 | -0.760007 | 0.005793 | -0.20645 | 0.087474 | 0.212366 | -0.494659 | -0.041727 |
| 29.56944 | -0.410673 | -0.225201 | -2.97631 | -0.997566 | -1.95417 | -1.154952 | -1.577735 | -0.747005 | -0.530955 | -1.260261 | -0.024219 | -0.379795 | 0.065805 | -0.116852 | -0.809924 | -1.156985 | -0.935812 | 0.242105 | -0.193863 | 0.24512 | 0.330379 | -0.338184 | 0.15095 |
| 29.58333 | -0.429228 | -0.618844 | -3.808751 | -1.340731 | -2.028082 | -1.481396 | -1.586481 | -1.131544 | -1.090382 | -2.488422 | -0.025924 | -0.962071 | -0.099251 | -1.090939 | -1.248715 | -1.006211 | -1.019914 | 0.163164 | -0.51944 | 0.063576 | 0.087124 | -0.40478 | 0.014313 |
| 29.59722 | -0.669113 | -1.78644 | -4.743182 | -1.738683 | -2.724068 | -1.836964 | -1.918717 | -2.065228 | -1.241789 | -3.683138 | -0.44159 | -1.785245 | -1.109708 | -2.636008 | -2.576845 | -1.755368 | -1.117299 | -0.748676 | -1.5498 | -0.545979 | -0.7392 | -0.948816 | -0.827314 |
| 29.61111 | -0.830366 | -3.346548 | -6.01875 | -1.902991 | -3.132939 | -2.302197 | -2.032243 | -3.053308 | -0.94705 | -3.625124 | -0.629735 | -1.804784 | -1.633042 | -3.277097 | -3.481766 | -2.100032 | -1.002111 | -1.366052 | -1.517079 | -0.761706 | -0.605549 | -2.199773 | -1.116153 |
| 29.625 | -1.13893 | -3.419973 | -5.692365 | -1.370603 | -2.282291 | -1.816637 | -1.689903 | -3.216618 | -0.600734 | -3.145917 | -0.477608 | -1.344189 | -0.90495 | -3.052107 | -2.521265 | -1.542837 | -0.077441 | -1.030403 | -1.01412 | -0.22611 | 0.094567 | -2.52227 | -0.596275 |
| 29.63889 | -0.937457 | -1.889349 | -3.383668 | -0.805274 | -0.907338 | -1.299184 | -1.360224 | -1.771271 | -0.409446 | -3.080086 | -0.265518 | -1.125865 | -0.428073 | -2.236291 | -1.439305 | -0.683647 | 0.968398 | -0.687501 | -0.367625 | 0.105055 | 0.350756 | -1.769958 | -0.239714 |
| 29.65278 | -0.406095 | -0.80249 | -0.869137 | -0.549491 | 0.115529 | -0.212396 | -0.342654 | -0.115753 | -0.433879 | -2.367545 | -0.250699 | -0.857484 | -0.357642 | -1.173175 | -0.830604 | -0.136658 | 1.091378 | -0.557886 | 0.431757 | 0.028847 | 0.225417 | -0.341606 | -0.173633 |
| 29.66667 | -0.048989 | -0.581537 | 0.589572 | -0.380312 | 0.336729 | 0.641886 | 0.478687 | 0.982719 | -0.536649 | -1.384581 | -0.352456 | -0.3744 | -0.146566 | -0.739072 | -0.49304 | 0.168549 | 0.259841 | -0.482363 | 0.537557 | -0.093417 | 0.086594 | 1.272522 | -0.155386 |
| 29.68056 | 0.374769 | -0.646678 | 0.811722 | -0.22548 | 0.171204 | 0.506457 | 0.496505 | 0.546062 | -0.478347 | -0.558252 | -0.380624 | -0.192364 | 0.171782 | -0.581984 | -0.425761 | 0.332845 | -0.274449 | -0.479858 | 0.184673 | -0.111216 | 0.040998 | 1.920321 | -0.089694 |
| 29.69444 | 0.715182 | -0.41584 | 0.443125 | -0.054263 | -0.035573 | 0.588494 | 0.515479 | 0.404619 | -0.205133 | 0.1614 | -0.31696 | -0.389687 | 0.363888 | -0.532897 | -0.345719 | 0.165679 | -0.565797 | -0.408743 | -0.055552 | 0.003454 | -0.000157 | 1.281525 | 0.038089 |
| 29.70833 | 0.767298 | 0.042602 | 0.702298 | 0.029409 | -0.21514 | 0.920679 | 0.629946 | 0.337998 | 0.108949 | 0.436966 | -0.303948 | -0.517211 | 0.386281 | -0.682166 | -0.208819 | -0.187574 | -0.64924 | -0.342583 | -0.280729 | 0.102975 | -0.042497 | 0.69506 | 0.112155 |
| 29.72222 | 0.610281 | 0.377537 | 1.158064 | 0.016697 | -0.586785 | 0.904952 | 0.757102 | 0.146904 | 0.250477 | 0.197904 | -0.28972 | -0.417671 | 0.084641 | -0.632582 | -0.415631 | -0.467021 | -0.298125 | -0.464819 | -0.439463 | 0.057342 | 0.029192 | 0.309516 | 0.051194 |
| 29.73611 | 0.4134 | 0.232371 | 1.246913 | 0.10419 | -0.848049 | 0.35533 | 0.6031 | -0.388191 | 0.19243 | -0.185707 | -0.242667 | -0.360837 | -0.339133 | -0.383061 | -0.921373 | -0.502579 | -0.268807 | -0.723396 | -0.345419 | -0.111157 | 0.115928 | -0.192032 | -0.100076 |
| 29.75 | 0.219828 | -0.216399 | 1.352619 | 0.419559 | -0.750428 | -0.236202 | 0.30966 | -0.106739 | 0.187832 | -0.505348 | -0.142086 | -0.532673 | -0.505771 | -0.054601 | -1.100336 | -0.38405 | -0.775137 | -0.781529 | -0.367672 | -0.20722 | 0.099092 | -0.226049 | -0.175774 |
| 29.76389 | -0.006325 | -0.699609 | 2.03739 | 0.656677 | -0.583231 | -0.598786 | 0.046521 | 0.370299 | 0.21825 | -0.629219 | -0.101467 | -0.788198 | -0.478575 | -0.020646 | -0.981899 | -0.422663 | -0.837225 | -0.6552 | -0.651394 | -0.200561 | -0.014826 | -0.210006 | -0.191777 |
| 29.77778 | 0.061874 | -0.995884 | 1.794903 | 0.51339 | -0.092935 | -0.842937 | -0.060936 | 0.284583 | -0.008742 | -0.675831 | -0.143489 | -0.905667 | -0.514136 | -0.155951 | -1.016401 | -0.527823 | -0.003604 | -0.60068 | -0.443724 | -0.237067 | -0.054101 | -0.252548 | -0.238471 |
| 29.79167 | 0.037408 | -0.79236 | 0.992308 | 0.180943 | 0.546128 | -0.933928 | 0.194889 | 0.316188 | -0.3318 | -0.534697 | -0.177437 | -0.82328 | -0.419051 | -0.099716 | -1.022874 | -0.394514 | 0.378795 | -0.581519 | -0.11494 | -0.197306 | -0.064699 | -0.444293 | -0.215798 |
| 29.80556 | 0.045788 | -0.378496 | 0.465231 | -0.026878 | 0.876471 | -0.881828 | 0.550342 | 0.534314 | -0.464533 | -0.248992 | -0.143575 | -0.981258 | -0.231698 | -0.020926 | -1.024614 | -0.269716 | -0.008196 | -0.461769 | -0.181027 | -0.113737 | -0.173353 | -0.610905 | -0.187154 |
| 29.81944 | 0.730734 | -0.050551 | 0.472882 | 0.005814 | 0.670696 | -0.340701 | 0.275481 | 0.690908 | -0.365981 | -0.016868 | 0.114084 | -1.029924 | -0.193355 | -0.055881 | -1.077571 | -0.319155 | -0.866356 | -0.34716 | -0.301206 | -0.084741 | -0.31848 | -0.490507 | -0.214038 |
| 29.83333 | 1.097367 | -0.293654 | 0.700824 | 0.126912 | 0.301903 | 0.382768 | 0.089687 | 0.728012 | -0.29683 | -0.04921 | 0.058066 | -0.813217 | -0.294937 | 0.1185 | -0.994733 | -0.323198 | -1.038758 | -0.354528 | 0.053766 | -0.08539 | -0.262334 | -0.156309 | -0.189515 |
| 29.84722 | 0.931918 | -0.549361 | 0.896163 | 0.209636 | 0.054159 | 0.655775 | 0.175167 | 0.754092 | -0.331454 | -0.277583 | -0.392525 | -0.460224 | -0.318656 | 0.423819 | -0.72066 | -0.243491 | -0.240917 | -0.359013 | 0.395247 | -0.075522 | -0.093719 | 0.028724 | -0.089042 |
| 29.86111 | 0.455579 | -0.875525 | 0.58258 | 0.278119 | -0.126129 | 0.835544 | 0.37275 | 0.667171 | -0.277327 | -0.431515 | -0.135745 | -0.297022 | -0.20171 | 0.557648 | -0.37223 | -0.189908 | -0.011256 | -0.277381 | 0.125821 | -0.035043 | -0.051692 | -0.082071 | -0.001335 |
| 29.875 | -0.108219 | -1.37321 | -1.584296 | 0.136448 | 0.141633 | 0.935312 | 0.507878 | 0.135795 | -0.301903 | -0.577952 | -0.604419 | -0.257193 | -0.106721 | 0.322447 | -0.187916 | -0.229157 | -0.068168 | -0.228252 | -0.361711 | -0.076571 | -0.227381 | -0.387359 | -0.056046 |
| 29.88889 | -0.476279 | -1.918992 | -2.674484 | -0.062716 | 0.64103 | 0.603676 | 0.489807 | -1.405493 | -0.352287 | -0.756081 | -2.927724 | -0.526341 | -0.308414 | -0.198891 | -0.245916 | -0.480795 | -0.048755 | -0.334982 | -0.677843 | -0.260285 | -0.76937 | -0.69131 | -0.342575 |
| 29.90278 | -0.601729 | -1.166308 | -0.701667 | -0.091772 | 1.29717 | 0.635171 | -0.054386 | -2.330613 | -0.293403 | -0.669275 | -2.948094 | -1.100039 | -0.828732 | -0.780934 | -0.407545 | -0.929428 | 0.114802 | -0.666108 | -0.770668 | -0.603519 | -1.635732 | -0.972663 | -0.799437 |
| 29.91667 | -0.632115 | 0.123954 | 0.65807 | -0.09718 | 1.436827 | 0.644789 | -0.02615 | -2.07094 | -0.194019 | -0.450398 | -1.123742 | -1.018312 | -1.135379 | -0.815344 | -0.268226 | -0.922122 | 0.688527 | -0.847377 | -0.684527 | -0.82004 | -1.674591 | -0.913843 | -0.973103 |
| 29.93056 | -0.516024 | 1.276006 | 0.738918 | -0.086169 | 1.202827 | 0.711668 | 0.199813 | -1.259732 | -0.121124 | -0.15776 | -0.297787 | -0.42442 | -0.724043 | -0.489994 | 0.315565 | -0.268613 | 1.632053 | -0.410808 | -0.308463 | -0.564401 | -1.107236 | -0.393122 | -0.613545 |
| 29.94444 | -0.345075 | 2.34213 | 0.404926 | 0.024708 | 0.984157 | 0.469855 | 0.166251 | -0.600689 | -0.046977 | 0.27962 | -0.017854 | -0.026681 | -0.158186 | -0.235561 | 0.752617 | 0.273901 | 1.958852 | 0.060746 | 0.167882 | -0.244127 | -0.624362 | 0.313967 | -0.235707 |
| 29.95833 | -0.10375 | 2.94682 | -0.066899 | 0.205107 | 0.615888 | 0.238832 | 0.13597 | -0.357088 | 0.061531 | 0.668707 | 0.21553 | 0.175312 | 0.096469 | 0.044023 | 0.981834 | 0.436497 | 1.069467 | 0.163591 | 0.53326 | -0.127533 | -0.401922 | 0.639406 | -0.073449 |
| 29.97222 | 0.319494 | 3.041793 | -0.063545 | 0.318968 | 0.442736 | 0.016813 | -0.036654 | -0.122456 | 0.085604 | 1.024149 | 0.403792 | 0.237986 | 0.134078 | 0.178608 | 1.194887 | 0.56276 | 0.273446 | 0.198938 | 0.401022 | -0.125691 | -0.423373 | 0.389444 | -0.071018 |
| 29.98611 | 0.373704 | 2.804511 | 0.495112 | 0.291566 | 0.718567 | -0.229838 | -0.272483 | 0.276191 | -0.028996 | 1.262968 | 0.478781 | 0.3061 | -0.101582 | 0.285161 | 1.247067 | 0.452772 | 0.178864 | 0.117239 | 0.079146 | -0.293727 | -0.637882 | 0.232095 | -0.253399 |
| 30 | 0.178978 | 2.219621 | 1.39595 | 0.153635 | 0.594741 | -0.284074 | -0.398194 | 0.56636 | -0.113864 | 0.979761 | 0.457322 | 0.28884 | -0.379932 | 0.490969 | 1.22636 | 0.333291 | 0.439567 | -0.089961 | -0.203785 | -0.519319 | -0.9367 | 0.369262 | -0.488486 |
| 30.01389 | 0.085405 | 1.464802 | 2.269899 | 0.198274 | 1.053146 | -0.144643 | -0.330897 | 0.625201 | -0.250445 | 0.717084 | 0.324899 | 0.352865 | -0.526241 | 0.555109 | 1.133307 | 0.539894 | 0.388354 | -0.135306 | -0.222833 | -0.599233 | -1.212336 | 0.573702 | -0.574033 |
| 30.02778 | 0.348544 | 0.97494 | 3.017778 | 0.415285 | 1.83437 | 0.413127 | 0.197858 | 0.876826 | -0.32238 | 0.753016 | 0.237139 | 0.332841 | -0.343836 | 0.595202 | 1.173326 | 0.849213 | 0.237451 | 0.056005 | 0.063631 | -0.329849 | -0.968389 | 0.63491 | -0.357563 |
| 30.04167 | 1.201195 | 1.140322 | 3.042728 | 0.822996 | 2.113117 | 1.303966 | 0.984045 | 0.861273 | 0.1805 | 1.20634 | 0.554347 | 0.735881 | 0.078142 | 0.906046 | 1.411608 | 1.078756 | 0.258138 | 0.305034 | 0.476187 | -0.033277 | -0.662014 | 0.74182 | 0.021444 |
| 30.05556 | 1.822869 | 1.435233 | 2.7065 | 1.18884 | 2.061098 | 1.791038 | 1.584286 | 0.78624 | 0.568756 | 1.3862 | 0.787009 | 1.161526 | 0.465762 | 1.119336 | 1.648089 | 1.354134 | 0.495423 | 0.569233 | 0.578827 | 0.181204 | -0.589016 | 1.018901 | 0.290149 |
| 30.06944 | 2.07546 | 1.429681 | 2.775957 | 1.201737 | 1.814447 | 1.549578 | 1.518982 | 0.515912 | 0.694334 | 1.341208 | 0.811606 | 1.415458 | 0.750956 | 1.208937 | 1.74982 | 1.30783 | 0.882224 | 0.809605 | 0.569513 | 0.320935 | -0.505405 | 1.158677 | 0.475388 |
| 30.08333 | 1.865744 | 1.458993 | 4.321826 | 1.04492 | 1.283953 | 1.201639 | 1.042507 | 0.377363 | 0.888452 | 1.822677 | 0.926913 | 1.215424 | 0.758855 | 1.35325 | 1.506865 | 0.656127 | 0.996299 | 0.823276 | 0.499498 | 0.27869 | -0.307472 | 1.122704 | 0.501331 |
| 30.09722 | 1.159896 | 1.635235 | 5.600872 | 0.927068 | 1.499983 | 0.889061 | 0.731611 | 0.456874 | 0.761336 | 2.434982 | 0.967556 | 0.715372 | 0.413438 | 1.374436 | 1.008473 | 0.039636 | 0.467203 | 0.662541 | 0.227935 | -0.025411 | -0.392411 | 0.52826 | 0.192261 |
| 30.11111 | 0.721969 | 1.115104 | 4.551117 | 0.681484 | 0.790891 | 0.518603 | 0.270994 | 0.609826 | 0.388562 | 1.912858 | 0.663607 | 0.369872 | -0.113436 | 0.814152 | 0.590415 | 0.065081 | -0.128423 | 0.277468 | -0.083494 | -0.370936 | -0.700338 | -0.000402 | -0.338817 |
| 30.125 | 0.401598 | 0.757138 | 2.670169 | 0.794842 | 0.197667 | 0.361063 | 0.150102 | 0.401481 | 0.513193 | 1.444722 | 0.286989 | 0.267047 | -0.241244 | 0.371463 | 0.930853 | 1.059222 | 0.270085 | 0.193734 | -0.055506 | -0.360559 | -0.599799 | 0.219147 | -0.458405 |
| 30.13889 | 0.433751 | 1.009719 | 2.30848 | 1.52385 | 0.676317 | 0.338958 | 0.787293 | 0.505583 | 0.905773 | 1.475654 | 0.178525 | 0.980067 | 0.019871 | 0.416004 | 1.846299 | 2.658221 | 1.150263 | 0.52877 | 0.223515 | -0.112881 | -0.224199 | 0.86493 | -0.181166 |
| 30.15278 | 0.4092 | 1.344037 | 2.621689 | 1.974234 | 1.072392 | 0.419148 | 0.75035 | 0.759313 | 1.07541 | 0.207195 | 0.252187 | 1.018644 | 0.184246 | 0.234645 | 2.006148 | 3.162753 | 1.629676 | 0.827961 | 0.217956 | -0.016534 | -0.109491 | 0.877171 | 0.009031 |
| 30.16667 | 0.264613 | 1.574152 | 3.270027 | 1.511939 | 1.357282 | 0.42129 | 0.242083 | 0.894277 | 1.19595 | -0.047032 | 0.412285 | 0.849961 | 0.265682 | -0.064973 | 1.438276 | 1.760663 | 1.531974 | 0.8655 | 0.061807 | -0.042991 | -0.150948 | 0.580677 | -0.026021 |
| 30.18056 | 0.463575 | 1.84921 | 3.506579 | 1.058368 | 1.658589 | 0.854093 | 0.280679 | 1.024781 | 1.264666 | 0.587216 | 0.530078 | 0.832657 | 0.459604 | 0.067274 | 1.012216 | 1.044808 | 0.984587 | 0.959684 | -0.022459 | -0.058918 | -0.134363 | 0.52758 | -0.030955 |
| 30.19444 | 0.910656 | 1.808209 | 2.875643 | 0.686909 | 2.055301 | 1.40834 | 0.67282 | 1.122531 | 1.182125 | 1.412908 | 0.508724 | 0.860134 | 0.408913 | 0.256423 | 0.804697 | 0.683054 | 0.643163 | 0.895381 | -0.114988 | -0.140656 | -0.234607 | 0.41331 | -0.07982 |
| 30.20833 | 1.107615 | 1.139545 | 1.231672 | 0.26382 | 1.862266 | 1.294889 | 0.309837 | 0.806844 | 0.770026 | 1.020316 | 0.337744 | 0.574098 | -0.059932 | 0.152793 | 0.783505 | 0.130487 | 0.315346 | 0.51525 | -0.382866 | -0.415555 | -0.629839 | -0.13182 | -0.394489 |
| 30.22222 | 1.037433 | 0.605939 | 0.184323 | -0.101282 | 0.899163 | 0.682418 | 0.019383 | 0.396951 | 0.225159 | 0.301794 | 0.121318 | 0.039501 | -0.742992 | -0.242851 | 0.627346 | -0.359467 | -0.251014 | 0.154077 | -0.643768 | -0.702185 | -1.119304 | -0.575068 | -0.779339 |
| 30.23611 | 0.861042 | 0.477096 | 0.056781 | -0.201268 | 0.568167 | 0.391713 | -0.070274 | 0.026969 | 0.264264 | 0.206897 | 0.078583 | -0.327499 | -1.497965 | -0.766269 | 0.245429 | -0.544291 | -0.824314 | -0.072654 | -0.543998 | -0.788876 | -1.341606 | -0.852391 | -0.942511 |
| 30.25 | 0.824715 | 0.261329 | 0.770459 | -0.017606 | 0.299894 | 0.611158 | 0.069844 | -0.168397 | 0.556528 | 0.335019 | 0.181147 | -0.325427 | -2.419374 | -0.792233 | -0.367796 | -0.494635 | -0.747312 | -0.135479 | -0.219334 | -0.623313 | -1.137527 | -0.766923 | -0.769457 |
| 30.26389 | 0.579269 | 0.364226 | 0.497153 | 0.180375 | 0.028213 | 0.4647 | 0.022106 | -0.288363 | 0.82962 | 0.484491 | 0.225846 | -0.125527 | -2.908321 | -0.112632 | -0.930772 | -0.499167 | -0.353682 | 0.058244 | 0.115756 | -0.32374 | -0.724921 | -0.569784 | -0.405998 |
| 30.27778 | -0.027306 | 0.492371 | -1.573616 | 0.094212 | -0.527169 | -0.301305 | 0.055318 | -0.573047 | 0.64902 | 0.2473 | 0.125715 | -0.048808 | -2.924958 | -4.791686 | -0.266265 | -0.512483 | 0.065578 | 0.255747 | 0.300456 | -0.108789 | -0.391751 | -0.459817 | -0.166929 |
| 30.29167 | -0.682921 | 0.681207 | -3.292791 | 0.041496 | -0.730336 | -0.803242 | 0.157774 | -0.97287 | 0.506321 | -0.068608 | -0.000335 | 0.098143 | -1.72015 | -3.925857 | 0.345424 | -0.151431 | 0.1672 | 0.274378 | 0.365614 | 0.055635 | -0.126491 | -0.378404 | -0.000233 |
| 30.30556 | -1.345285 | 0.947678 | -2.877641 | 0.266217 | -0.237736 | -0.617483 | 0.453238 | -1.026729 | 0.60185 | 0.114939 | 0.067061 | 0.458761 | -0.825716 | -1.420675 | 0.701398 | 0.49691 | 0.334504 | 0.388429 | 0.476106 | 0.245657 | 0.155961 | -0.433809 | 0.228052 |
| 30.31944 | -1.450538 | 0.927175 | -1.533905 | 0.423129 | 0.269734 | -0.146956 | -0.022915 | -0.881493 | 0.705957 | 0.328295 | 0.080609 | 0.624431 | -0.521146 | 0.622934 | 0.987498 | 0.721254 | 0.28545 | 0.424695 | 0.444272 | 0.261291 | 0.190945 | -0.640654 | 0.23755 |
| 30.33333 | -1.067998 | 0.642025 | 0.210841 | 0.331907 | 0.736088 | 0.376369 | -0.48299 | -0.738911 | 0.74396 | 0.194422 | 0.040083 | 0.493171 | -0.189978 | 1.517539 | 1.145515 | 0.656288 | -0.006081 | 0.316603 | 0.274128 | 0.131037 | 0.053577 | -0.866423 | 0.070643 |
| 30.34722 | -0.939588 | 0.344766 | 1.201994 | 0.14172 | 1.187464 | 0.643652 | -0.205385 | -0.579501 | 0.685216 | 0.185091 | 0.012186 | 0.514186 | 0.559805 | 2.337784 | 1.38659 | 0.818674 | -0.387958 | 0.264617 | 0.249389 | 0.167031 | 0.127829 | -0.847931 | 0.119573 |
| 30.36111 | -0.786243 | 0.421615 | 0.383358 | -0.1252 | 0.978309 | 0.559515 | 0.433939 | -0.411986 | 0.488468 | 0.077938 | -0.005816 | 0.705706 | 1.282307 | 2.753591 | 1.336825 | 1.013472 | -0.524289 | 0.257229 | 0.323745 | 0.296264 | 0.307461 | -0.430335 | 0.296996 |
| 30.375 | -0.339925 | 0.38995 | -0.967902 | -0.370126 | 0.40466 | 0.508643 | 0.614595 | -0.264089 | 0.155579 | 0.120608 | -0.071047 | 0.755864 | 1.607105 | 2.673436 | 0.88711 | 1.055933 | -0.328918 | 0.181403 | 0.323835 | 0.31775 | 0.320811 | 0.131133 | 0.363783 |
| 30.38889 | -0.027607 | 0.281804 | -0.773179 | -0.455857 | -0.132206 | 0.567212 | 0.852642 | -0.053533 | -0.075282 | 0.338624 | -0.14279 | 0.677773 | 1.382673 | 2.062093 | 0.312035 | 0.928817 | 0.146321 | 0.065058 | 0.268978 | 0.255444 | 0.230827 | 0.32876 | 0.311572 |
| 30.40278 | 0.092957 | 0.409728 | 0.111552 | -0.518782 | -0.479795 | 0.382333 | 0.867569 | 0.130163 | -0.252258 | 0.45465 | -0.185155 | 0.51461 | 1.008988 | 1.463389 | -0.078623 | 0.632328 | 0.770525 | 0.066163 | 0.272932 | 0.214336 | 0.205076 | 0.521563 | 0.272608 |
| 30.41667 | 0.069076 | 0.638287 | 0.502726 | -0.641911 | -1.472706 | 0.221913 | 0.895743 | 0.19254 | -0.570032 | 0.420572 | -0.23292 | 0.449523 | 0.752951 | 1.251982 | -0.003288 | 0.157256 | 1.31538 | 0.198803 | 0.31893 | 0.22335 | 0.248728 | 0.61731 | 0.306163 |
| 30.43056 | -0.006523 | 0.677869 | -0.608036 | -0.688786 | -1.98813 | 0.122492 | 0.851164 | 0.247759 | -1.056384 | 0.293349 | -0.215209 | 0.358212 | 0.543375 | 1.156979 | 0.338153 | -0.368786 | 1.240069 | 0.345765 | 0.279132 | 0.210042 | 0.265635 | 0.499153 | 0.277983 |
| 30.44444 | -0.152747 | 0.661965 | -1.1796 | -0.556958 | -1.470118 | -0.038162 | 0.215491 | 0.257528 | -1.142474 | 0.186186 | -0.165787 | 0.236514 | 0.147333 | 0.588598 | 0.436637 | -0.56872 | 0.9036 | 0.346199 | 0.115983 | 0.107778 | 0.15684 | 0.361545 | 0.118118 |
| 30.45833 | -0.472472 | 0.579424 | -0.693537 | -0.438079 | -0.711869 | -0.692823 | -0.657906 | 0.139721 | -1.101883 | 0.276925 | -0.197764 | 0.128293 | -0.251535 | -0.376221 | 0.252152 | -0.591725 | 0.643639 | 0.21641 | 0.017589 | -0.0146 | -0.025387 | 0.097376 | 0.001018 |
| 30.47222 | -0.998163 | 0.300768 | -0.901517 | -0.5779 | -0.401246 | -1.033424 | -0.897233 | -0.353887 | -1.029251 | 0.156866 | -0.330537 | 0.083174 | -0.285992 | -0.695303 | 0.241923 | -0.645615 | 0.303689 | 0.140955 | 0.053942 | 0.021951 | -0.00907 | 0.01246 | -0.040379 |
| 30.48611 | -1.570253 | -0.124286 | -1.646722 | -0.783361 | -1.113736 | -0.797992 | -0.991302 | -0.892344 | -0.981166 | -0.306223 | -0.384545 | 0.124803 | 0.018794 | -0.434265 | 0.413946 | -0.57858 | -0.223799 | 0.1336 | 0.227721 | 0.234241 | 0.25469 | 0.20236 | 0.215203 |
| 30.5 | -1.831669 | -0.57085 | -2.296779 | -0.968437 | -1.327384 | -0.678358 | -0.835245 | -1.395834 | -0.957362 | -0.812029 | -0.41083 | 0.011082 | 0.301017 | -0.037352 | 0.573558 | -0.268035 | -0.729435 | 0.105333 | 0.364636 | 0.427285 | 0.478146 | 0.429432 | 0.445461 |
| 30.51389 | -1.579648 | -0.906932 | -2.568549 | -1.093225 | -1.380717 | -0.628236 | -0.975522 | -1.717994 | -1.117451 | -1.343909 | -0.435719 | -0.343589 | 0.365779 | 0.45379 | 0.429399 | -0.050209 | -1.294078 | 0.016503 | 0.343624 | 0.441673 | 0.481858 | 0.422579 | 0.468365 |
| 30.52778 | -1.25145 | -1.280205 | -2.329046 | -1.271992 | -1.090011 | -0.397109 | -0.921469 | -1.841434 | -1.019996 | -1.283503 | -0.435941 | -0.848375 | 0.248954 | 1.135718 | -0.16963 | -0.172905 | -1.93389 | -0.05785 | 0.209295 | 0.292841 | 0.317611 | 0.234952 | 0.307778 |
| 30.54167 | -1.229261 | -1.259413 | -1.798168 | -1.376008 | -1.44617 | -0.232018 | -0.827239 | -1.690691 | -0.904402 | -0.930976 | -0.379175 | -1.254174 | 0.039417 | 2.132815 | -0.813719 | -0.143558 | -2.020448 | -0.128054 | 0.050279 | 0.167142 | 0.1809 | 0.088106 | 0.146302 |
| 30.55556 | -1.604416 | -0.797407 | -1.762621 | -1.151968 | -1.768346 | -0.003758 | -0.877051 | -1.751802 | -0.718203 | -1.081263 | -0.301482 | -1.487391 | 0.003418 | 2.06256 | -0.776856 | -0.122264 | -1.945141 | -0.148807 | -0.066406 | 0.230714 | 0.255071 | 0.113116 | 0.139439 |
| 30.56944 | -1.603083 | -0.323192 | -3.288267 | -0.972598 | -1.371304 | 0.172633 | -0.892808 | -2.053736 | -0.531663 | -1.517413 | -0.115395 | -1.489884 | 0.251983 | 1.408237 | -0.405055 | -0.260763 | -1.581947 | 0.105651 | -0.248255 | 0.423823 | 0.528171 | 0.231193 | 0.448912 |
| 30.58333 | -1.190474 | -0.342751 | -6.307873 | -1.20589 | -1.619676 | -0.855904 | -1.30655 | -2.399602 | -0.530755 | -2.434559 | -0.065767 | -1.409608 | 0.572391 | 0.610655 | 0.186184 | -0.657284 | -1.080062 | 0.396212 | -0.613546 | 0.543159 | 0.759577 | 0.214089 | 0.82616 |
| 30.59722 | -1.175631 | -0.8772 | -8.200769 | -1.672953 | -2.487499 | -1.950749 | -2.090002 | -2.930966 | -0.774597 | -4.276879 | -0.262905 | -1.160486 | 0.70333 | -0.155953 | 0.577042 | -1.683172 | -0.425757 | 0.04965 | -0.843925 | 0.63636 | 0.828082 | 0.464641 | 0.918497 |
| 30.61111 | -1.631999 | -1.35127 | -6.289972 | -1.316631 | -2.579914 | -2.15428 | -2.210733 | -3.091827 | -0.706978 | -3.686579 | -0.279204 | -0.703618 | 0.95393 | 0.058446 | 0.541754 | -2.52414 | 0.329229 | -0.299509 | -0.547687 | 0.883687 | 1.042615 | 0.922133 | 0.913236 |
| 30.625 | -1.930331 | -1.796602 | -4.089796 | -0.595979 | -1.774443 | -1.972704 | -1.710311 | -2.667748 | -0.383015 | -3.350747 | -0.091763 | -0.271952 | 1.028401 | 0.392551 | -0.264484 | -1.996326 | 0.851616 | -0.10237 | 0.303792 | 0.992554 | 1.117479 | 0.763365 | 0.959508 |
| 30.63889 | -1.634569 | -2.381876 | -3.057206 | -0.195953 | -0.991438 | -1.389352 | -2.011659 | -1.32368 | -0.118879 | -3.292056 | 0.012146 | -0.549646 | 0.691594 | 0.074018 | -0.688834 | -1.297867 | 1.747607 | 0.294494 | 0.711726 | 0.826537 | 0.859324 | 0.213273 | 0.815058 |
| 30.65278 | -0.607666 | -2.254131 | -1.115524 | 0.111284 | -0.317727 | -1.050347 | -1.663827 | 0.254575 | 0.141128 | -3.462688 | -0.046427 | -0.58662 | 0.359239 | -0.417228 | -0.712911 | -0.636488 | 2.687303 | 0.384275 | 1.148723 | 0.620787 | 0.62845 | -0.463785 | 0.66052 |
| 30.66667 | -0.087165 | -1.336472 | 0.515054 | 0.304264 | 0.049973 | -0.938197 | -0.83079 | 0.831789 | 0.28253 | -2.616452 | -0.151328 | -0.330647 | 0.134539 | -0.807947 | -0.821049 | -0.142525 | 1.741321 | 0.394827 | 1.391112 | 0.522822 | 0.551699 | -0.663735 | 0.537311 |
| 30.68056 | 0.058366 | -0.856505 | 0.598841 | 0.381685 | 0.245652 | -0.471969 | -0.087101 | 1.114971 | 0.3049 | -1.202953 | -0.336039 | -0.070523 | -0.049748 | -1.035756 | -0.945786 | 0.160745 | 0.669533 | 0.304619 | 1.159923 | 0.447269 | 0.523124 | -0.780789 | 0.393716 |
| 30.69444 | -0.015092 | -0.55968 | 0.243731 | 0.395425 | 0.054867 | -0.003553 | 0.488658 | 0.778513 | 0.264296 | -0.320723 | -0.509218 | -0.120276 | -0.203286 | -1.379005 | -0.959445 | 0.216785 | 0.013533 | 0.036371 | 0.784039 | 0.292483 | 0.372823 | -0.907245 | 0.214818 |
| 30.70833 | 0.019695 | -0.182636 | 0.111738 | 0.374511 | -0.127915 | 0.584451 | 0.720631 | 0.27076 | 0.164254 | -0.068102 | -0.686663 | -0.304335 | -0.312594 | -1.377547 | -0.973007 | 0.248159 | -0.700005 | -0.277851 | 0.447463 | 0.113869 | 0.16477 | -1.235013 | 0.069509 |
| 30.72222 | 0.083674 | 0.161455 | 0.818219 | 0.423192 | -0.200816 | 1.098913 | 0.695575 | 0.167713 | 0.150868 | -0.198375 | -0.655073 | -0.298629 | -0.226331 | -0.995767 | -0.973121 | 0.015652 | -1.352763 | -0.586155 | 0.185626 | 0.029382 | 0.084467 | -0.929779 | -0.053957 |
| 30.73611 | 0.121353 | 0.240449 | 1.741286 | 0.577689 | -0.304494 | 1.548186 | 0.848444 | 0.138132 | 0.20617 | -0.544057 | -0.35191 | -0.308526 | -0.040156 | -0.817158 | -1.069224 | -0.260951 | -1.505019 | -0.793241 | -0.079687 | 0.010853 | 0.074124 | -0.70983 | -0.097098 |
| 30.75 | -0.144024 | 0.190186 | 1.940563 | 0.67401 | -0.081585 | 1.338133 | 1.01121 | -0.046013 | 0.270109 | -0.573586 | -0.19711 | -0.445678 | 0.132574 | -0.788982 | -1.116047 | -0.154516 | -1.218696 | -0.796411 | -0.399161 | 0.008489 | 0.045748 | -0.739301 | -0.044552 |
| 30.76389 | -0.494095 | 0.290311 | 1.095372 | 0.700456 | 0.09138 | 0.768005 | 0.805735 | -0.590996 | 0.192606 | -0.628336 | -0.186568 | -0.413213 | 0.19437 | -0.648526 | -1.087999 | -0.052393 | -0.934131 | -0.671348 | -0.668462 | 0.020004 | 0.052154 | -0.744697 | 0.023915 |
| 30.77778 | -0.637234 | 0.38536 | -0.493605 | 0.590873 | 0.017513 | 0.365979 | 0.585985 | -1.26346 | 0.109291 | -0.740375 | -0.201189 | -0.112882 | 0.251324 | -0.265851 | -0.940587 | -0.413823 | -0.244928 | -0.523095 | -0.667205 | 0.099289 | 0.153392 | -0.67407 | 0.051523 |
| 30.79167 | -0.115737 | 0.149813 | -1.583886 | 0.344704 | -0.181987 | -0.044264 | 0.274359 | -0.792136 | 0.074214 | -0.686497 | -0.251057 | 0.062528 | 0.274544 | 0.125027 | -0.729031 | -0.575692 | 0.509187 | -0.318511 | -0.581335 | 0.184271 | 0.252806 | -0.51705 | 0.061026 |
| 30.80556 | 0.583843 | 0.062617 | -1.144663 | 0.169859 | 0.151339 | -0.280121 | -0.102874 | 0.016019 | 0.028736 | -0.427367 | -0.276139 | 0.047334 | 0.151327 | 0.39073 | -0.743643 | -0.179095 | 0.895456 | -0.111491 | -0.405528 | 0.178461 | 0.217418 | -0.400095 | 0.038554 |
| 30.81944 | 1.194784 | 0.268107 | 0.514939 | 0.32717 | 0.318767 | -0.144065 | -0.237137 | 0.68212 | 0.04033 | 0.133597 | -0.231924 | -0.025832 | -0.065739 | 0.409391 | -1.074793 | 0.031434 | 0.951327 | -0.113177 | -0.167833 | 0.047431 | 0.068031 | -0.351933 | -0.087348 |
| 30.83333 | 1.78091 | 0.310232 | 1.955554 | 0.468812 | 0.3777 | 0.118475 | -0.345705 | 0.813971 | 0.021734 | 0.664598 | -0.208935 | -0.106611 | -0.198748 | 0.336616 | -1.383884 | -0.12949 | 0.538604 | -0.196795 | 0.107276 | -0.119658 | -0.09676 | -0.459299 | -0.263828 |
| 30.84722 | 1.513313 | -0.074336 | 1.36316 | 0.418347 | 0.389691 | 0.28842 | -0.273961 | 0.613318 | -0.056383 | 0.585817 | -0.214218 | -0.150659 | -0.22966 | 0.217806 | -1.243474 | -0.109038 | -0.336229 | -0.101756 | 0.405703 | -0.149925 | -0.136167 | -0.504943 | -0.285885 |
| 30.86111 | 1.119937 | -0.479472 | 0.289162 | 0.327594 | 0.521201 | 0.479803 | -0.145226 | 0.087569 | -0.064989 | -0.034882 | -0.222182 | -0.046742 | -0.126451 | 0.208821 | -0.9388 | 0.12663 | -1.185826 | 0.063611 | 0.774798 | -0.006991 | 0.004988 | -0.262123 | -0.115897 |
| 30.875 | 0.779176 | -0.458448 | -0.083741 | 0.222527 | 0.43601 | 0.65174 | 0.028718 | -0.153089 | -0.069911 | -0.836094 | -0.213537 | 0.22 | 0.136997 | 0.417194 | -0.764418 | 0.250723 | -0.846353 | 0.088263 | 1.094009 | 0.167687 | 0.197305 | 0.176636 | 0.021703 |
| 30.88889 | 0.337429 | -0.114993 | 0.00174 | 0.144789 | 0.175582 | 0.747289 | 0.281128 | -0.138369 | -0.035988 | -1.136133 | -0.146923 | 0.36584 | 0.24697 | 0.374734 | -0.566082 | 0.271595 | 0.253305 | -0.030578 | 0.919163 | 0.208278 | 0.265198 | 0.632525 | -0.008867 |
| 30.90278 | -0.551437 | 0.420811 | 0.258098 | 0.169363 | 0.358473 | 1.074164 | 0.587903 | -0.178231 | 0.033401 | -0.587077 | -0.049444 | 0.280658 | 0.102468 | -0.173329 | -0.170304 | 0.545643 | 1.547678 | -0.068356 | 0.421985 | 0.115771 | 0.174553 | 0.816029 | -0.098899 |
| 30.91667 | -1.464523 | 0.632503 | 0.522151 | 0.205564 | 0.607627 | 1.534879 | 0.937988 | -0.738904 | 0.048049 | 0.300128 | -0.031134 | 0.210817 | -0.044145 | -0.797017 | 0.395274 | 0.959173 | 2.377766 | 0.000381 | -0.053846 | 0.02776 | 0.068346 | 0.716738 | -0.111209 |
| 30.93056 | -1.184706 | 0.57349 | 0.497714 | 0.215906 | 0.952488 | 1.595555 | 1.26574 | -1.209063 | 0.02447 | 1.295451 | -0.11193 | 0.27975 | -0.069136 | -0.8596 | 0.940371 | 1.181996 | 2.227967 | 0.139941 | -0.365744 | -0.002716 | 0.029051 | 0.71654 | -0.02966 |
| 30.94444 | -0.067158 | 0.499179 | 0.457307 | 0.251724 | 1.305118 | 1.288225 | 1.370273 | -1.132507 | 0.014922 | 2.022865 | -0.137993 | 0.429741 | -0.079017 | -0.305237 | 1.327851 | 1.141245 | 1.899912 | 0.235804 | -0.386199 | -0.038916 | -0.002105 | 1.169738 | 0.004461 |
| 30.95833 | 1.126629 | 0.550689 | 1.103877 | 0.304141 | 1.326179 | 1.057785 | 1.229466 | -0.531451 | 0.053604 | 2.093389 | -0.042123 | 0.601226 | -0.018766 | 0.764595 | 1.588046 | 0.886436 | 1.709547 | 0.260212 | -0.112974 | -0.060163 | -0.025174 | 1.889515 | -0.059116 |
| 30.97222 | 1.845066 | 0.614928 | 1.942743 | 0.343357 | 1.116429 | 0.794689 | 0.970454 | 0.224502 | 0.135474 | 1.74671 | 0.052162 | 0.668886 | 0.087325 | 1.706176 | 1.682484 | 0.74771 | 1.533753 | 0.359939 | 0.296012 | -0.01894 | 0.014571 | 2.308512 | -0.073836 |
| 30.98611 | 2.072447 | 0.492425 | 2.131554 | 0.421166 | 1.189197 | 0.687951 | 0.814236 | 0.826949 | 0.197713 | 1.222867 | 0.055998 | 0.650302 | 0.186469 | 1.749788 | 1.724992 | 0.822077 | 1.219242 | 0.497026 | 0.678728 | -0.011437 | 0.010585 | 1.907055 | 0.009355 |
| 31 | 1.689937 | 0.734418 | 2.176872 | 0.550431 | 1.419038 | 0.698488 | 0.52686 | 0.890604 | 0.148382 | 1.164495 | 0.02727 | 0.468054 | 0.235759 | 1.40625 | 1.636636 | 0.679238 | 0.48684 | 0.391593 | 0.760232 | -0.132588 | -0.132557 | 1.257313 | -0.013385 |
| 31.01389 | 1.169496 | 0.957085 | 2.128407 | 0.556919 | 1.70079 | 0.714864 | 0.425445 | 0.498207 | -0.096535 | 1.114881 | -0.180107 | 0.258623 | 0.152257 | 0.861431 | 1.667068 | 0.497211 | -0.069297 | 0.245088 | 0.476843 | -0.361554 | -0.430204 | 1.171101 | -0.155665 |
| 31.02778 | 0.880634 | 0.565083 | 1.676105 | 0.381415 | 1.732263 | 0.480418 | 0.684099 | 0.157257 | -0.344966 | 1.230339 | -0.452916 | 0.230135 | 0.077424 | 0.472529 | 1.604975 | 0.571692 | -0.059008 | 0.261414 | 0.10142 | -0.423886 | -0.555082 | 1.172 | -0.189369 |
| 31.04167 | 0.927325 | 0.55844 | 1.8606 | 0.418463 | 1.529962 | 0.685914 | 1.142092 | 0.156288 | -0.122538 | 1.636779 | -0.247082 | 0.498746 | -0.020176 | 0.490868 | 1.511909 | 0.741179 | 0.454765 | 0.305632 | -0.135779 | -0.371091 | -0.459349 | 0.649554 | -0.097304 |
| 31.05556 | 0.78232 | 1.034527 | 2.636107 | 0.719875 | 1.573238 | 1.078019 | 1.784192 | 0.296321 | 0.214698 | 2.191436 | 0.078074 | 0.765151 | -0.171174 | 0.634301 | 1.506187 | 0.793375 | 1.179539 | 0.230523 | -0.214908 | -0.285589 | -0.343357 | 0.37398 | -0.108075 |
| 31.06944 | 0.330869 | 1.036387 | 2.896399 | 0.84355 | 1.59683 | 0.865653 | 2.157897 | 0.3062 | 0.28841 | 2.520647 | 0.156847 | 0.770061 | -0.289987 | 0.526787 | 1.269157 | 0.756716 | 1.705792 | 0.127957 | -0.28985 | -0.281399 | -0.320647 | 0.342489 | -0.166464 |
| 31.08333 | 0.292299 | 0.985644 | 3.275165 | 1.025932 | 1.644684 | 0.91072 | 2.396823 | 0.397304 | 0.244584 | 2.331914 | 0.190755 | 0.120835 | -0.727978 | -0.377191 | 0.362744 | 0.181281 | 1.400983 | -0.262194 | -0.705 | -0.675173 | -0.740583 | 0.177576 | -0.566269 |
| 31.09722 | 0.81154 | 0.855999 | 3.769627 | 1.082428 | 2.102237 | 0.86017 | 2.192005 | 0.35549 | 0.044925 | 2.253253 | 0.298165 | -0.970065 | -1.798735 | -2.033212 | -1.20179 | -1.062185 | -0.768812 | -1.09483 | -1.659011 | -1.629604 | -1.836949 | -0.715752 | -1.543714 |
| 31.11111 | 0.666876 | 0.937034 | 2.165189 | 0.693844 | 1.364439 | 0.688872 | 2.096683 | -0.069444 | -0.197409 | 1.501886 | 0.09113 | -1.515779 | -2.319836 | -3.884022 | -2.135631 | -1.615515 | -2.270196 | -1.500845 | -2.195111 | -2.266069 | -2.577774 | -1.56671 | -2.272039 |
| 31.125 | 0.667251 | 0.748542 | 0.707461 | 0.506267 | 0.769277 | 0.250196 | 1.139488 | -0.29794 | 0.025486 | 0.975209 | -0.149607 | -0.678664 | -1.61287 | -4.265815 | -1.349983 | -0.425006 | -1.823737 | -0.69948 | -1.375035 | -1.588887 | -1.769663 | -0.938515 | -1.719896 |

| | | | | | | | | | | | | | | | | | | | | | | | |
|---|---|---|---|---|---|---|---|---|---|---|---|---|---|---|---|---|---|---|---|---|---|---|---|
| 31.13889 | 0.588181 | 0.818789 | 0.548019 | 0.993179 | 0.636154 | 0.097517 | 0.687251 | 0.073469 | 0.590927 | 0.893807 | -0.069146 | 0.308343 | -0.884534 | -2.564389 | 0.1711 | 1.46291 | -0.834322 | 0.248428 | -0.499624 | -0.724997 | -0.778566 | -0.115963 | -0.780047 |
| 31.15278 | 0.187755 | 1.021298 | -0.017529 | 1.311528 | 0.30417 | 0.151824 | 0.516103 | 0.273028 | 0.847229 | 0.867309 | 0.141285 | 1.556985 | -0.574086 | -1.499885 | 0.989167 | 2.262325 | -0.22314 | 0.71256 | -0.168167 | -0.25557 | -0.259357 | 0.1524 | -0.261851 |
| 31.16667 | -0.023842 | 1.021049 | -0.637105 | 1.03952 | 0.598121 | 0.234712 | -0.637229 | 0.595904 | 0.695021 | 0.38495 | 0.273751 | 1.282362 | -0.606571 | -1.289853 | 1.028884 | 1.659703 | -0.068454 | 0.899374 | 0.051058 | -0.048783 | -0.04896 | -0.06391 | -0.116895 |
| 31.18056 | -0.257559 | 1.269975 | -0.580918 | 0.761767 | 0.732786 | -0.352278 | -1.399769 | 0.71161 | 0.666176 | 0.209855 | 0.371059 | 0.867784 | -0.511077 | -1.094026 | 0.879479 | 1.08691 | 0.472182 | 1.067099 | 0.136142 | 0.128917 | 0.139319 | -0.080296 | 0.027041 |
| 31.19444 | 0.407433 | 1.425228 | 0.759335 | 0.575126 | 0.879264 | -0.159609 | -0.801472 | 0.73842 | 0.736607 | 0.409539 | 0.490723 | 0.840435 | 0.480859 | -0.40587 | 0.759838 | 1.068188 | 1.056958 | 1.210652 | 0.359314 | 0.311554 | 0.346517 | 0.169616 | 0.350355 |
| 31.20833 | 0.994519 | 1.198265 | 2.025961 | 0.54425 | 1.399042 | 0.199958 | -0.390402 | 0.732178 | 0.734998 | 0.644317 | 0.529071 | 0.989266 | 1.546514 | 0.054322 | 0.88096 | 1.151932 | 1.640779 | 1.144495 | 0.540499 | 0.429567 | 0.466609 | 0.516531 | 0.623851 |
| 31.22222 | 0.60282 | 0.795756 | 1.817727 | 0.512389 | 1.391322 | 0.374742 | 0.093043 | 0.624547 | 0.594846 | 0.558854 | 0.449024 | 0.936177 | 1.628793 | 0.098042 | 1.306305 | 1.028839 | 1.906139 | 1.166333 | 0.648071 | 0.523895 | 0.5488 | 0.868556 | 0.708568 |
| 31.23611 | -0.026426 | 0.196372 | 0.762196 | 0.427469 | 0.893866 | 0.644979 | -0.025581 | 0.407434 | 0.50652 | 0.335148 | 0.343203 | 0.765593 | 1.243877 | 0.371315 | 1.485271 | 0.925304 | 1.804372 | 1.215166 | 0.729303 | 0.60337 | 0.606316 | 1.157635 | 0.7257 |
| 31.25 | -0.197569 | 0.022874 | -0.015927 | 0.277578 | 0.493848 | 1.110878 | 1.15455 | 0.152072 | 0.426256 | 0.133598 | 0.267819 | 0.717557 | 0.899077 | 0.754097 | 1.234688 | 0.69312 | 0.850042 | 1.109855 | 0.88371 | 0.596109 | 0.570208 | 1.229821 | 0.694244 |
| 31.26389 | -0.341465 | -0.055081 | -0.650288 | 0.108414 | 0.831725 | 1.267573 | 1.752657 | -0.058803 | 0.21562 | -0.029612 | 0.245477 | 0.743774 | 0.78724 | 0.994677 | 1.105004 | 0.367184 | 0.631512 | 1.043157 | 0.81357 | 0.451459 | 0.436764 | 0.729419 | 0.548838 |
| 31.27778 | -0.314073 | -0.177717 | -1.327351 | -0.007646 | 0.765077 | 1.037818 | 1.47 | -0.197829 | 0.12184 | -0.027496 | 0.281604 | 0.752075 | 0.845822 | 1.57432 | 1.26939 | 0.140369 | 0.653033 | 0.823804 | 0.60796 | 0.221119 | 0.24393 | 0.279079 | 0.316436 |
| 31.29167 | -0.267568 | -0.111459 | -1.894078 | -0.033162 | -0.181716 | 0.450623 | 0.937961 | -0.174789 | 0.16428 | 0.001726 | 0.346619 | 0.760492 | 0.758439 | 2.16993 | 1.514191 | 0.060428 | 0.224706 | 0.536174 | 0.421783 | 0.011783 | 0.033775 | -0.098543 | 0.101469 |
| 31.30556 | -0.899608 | -0.013394 | -2.200572 | -0.095195 | -0.53275 | -0.042611 | 1.018575 | -0.246584 | 0.18063 | -0.140494 | 0.28458 | 0.404115 | 0.420389 | 2.097426 | 1.152628 | 0.023608 | -0.638033 | 0.318701 | 0.231569 | -0.167292 | -0.159141 | -0.448099 | -0.097213 |
| 31.31944 | -1.311894 | 0.338167 | -1.783934 | -0.125075 | -0.95672 | -0.195282 | 0.438629 | -0.361006 | 0.249409 | -0.024127 | 0.203357 | -0.05966 | 0.177712 | 1.672408 | 0.19878 | -0.200454 | -1.01919 | 0.190278 | 0.103075 | -0.232547 | -0.229296 | -0.605238 | -0.193628 |
| 31.33333 | -1.154477 | 0.401196 | -0.647679 | -0.081378 | -0.801856 | 0.083533 | -0.243707 | -0.240456 | 0.409181 | 0.262573 | 0.238197 | -0.047162 | 0.245893 | 1.503906 | -0.385707 | -0.261179 | -1.20968 | 0.108159 | 0.053019 | -0.165852 | -0.157168 | -0.531365 | -0.132472 |
| 31.34722 | -1.338051 | 0.383913 | -0.30165 | -0.194511 | -0.824064 | 0.067434 | -0.006475 | 0.03151 | 0.382407 | 0.243096 | 0.262871 | 0.06334 | 0.401108 | 1.391922 | -0.477711 | -0.149988 | -0.459418 | 0.062183 | 0.019636 | -0.081925 | -0.077664 | -0.354452 | -0.048305 |
| 31.36111 | -1.619837 | 0.28035 | -1.415405 | -0.560319 | -1.077347 | -0.2321 | -0.022049 | 0.193352 | 0.168705 | -0.183874 | 0.19175 | -0.074037 | 0.458976 | 1.315146 | -0.401683 | 0.206694 | 0.352976 | 0.03166 | 0.044971 | -0.038457 | -0.048167 | -0.129965 | -0.034549 |
| 31.375 | -2.16959 | 0.242564 | -2.521379 | -0.740364 | -0.524124 | -0.417049 | -0.134787 | 0.217082 | -0.034656 | -0.72896 | 0.137849 | -0.163754 | 0.338028 | 1.362414 | -0.188168 | 0.623661 | 0.884282 | 0.015238 | 0.084383 | -0.034277 | -0.03891 | 0.061367 | -0.052264 |
| 31.38889 | -2.243132 | 0.064437 | -2.154753 | -0.675837 | 0.104341 | -0.298756 | -0.649893 | 0.220926 | -0.135277 | -0.833125 | 0.145544 | 0.37664 | 0.243634 | 1.33977 | 0.143349 | 0.901502 | 1.083367 | -0.011362 | 0.081552 | -0.045645 | -0.030078 | 0.041984 | -0.040809 |
| 31.40278 | -2.086958 | -0.02006 | -1.595698 | -0.604172 | 0.313286 | -0.40822 | -0.893253 | 0.204092 | -0.228574 | -0.930707 | 0.162529 | 0.397611 | 0.163785 | 1.027571 | 0.260917 | 0.836449 | 0.712179 | -0.007992 | 0.054511 | -0.037133 | -0.011791 | -0.092113 | 0.005137 |
| 31.41667 | -1.445017 | -0.378051 | -1.172018 | -0.618476 | 0.953672 | -0.202203 | -0.457524 | 0.106685 | -0.373869 | -1.232502 | 0.124553 | 0.448871 | 0.185239 | 0.642787 | 0.212875 | 0.51559 | 0.134071 | -0.103645 | 0.010806 | -0.014696 | 0.015822 | -0.215801 | 0.05048 |
| 31.43056 | -0.517143 | -0.870244 | -0.586077 | -0.638123 | 1.290695 | 0.24497 | 0.667797 | -0.260769 | -0.439928 | -1.686211 | 0.060519 | 0.424685 | 0.243857 | 0.404244 | 0.10673 | 0.091415 | -0.392823 | -0.092477 | -0.020765 | -0.005599 | 0.024198 | -0.323763 | 0.055919 |
| 31.44444 | 0.0936 | -0.770617 | -0.052192 | -0.636539 | 1.392977 | 0.41088 | 0.814875 | -0.550643 | -0.403312 | -2.217238 | 0.016945 | 0.357513 | 0.227472 | 0.201735 | -0.022295 | -0.208159 | 0.195172 | 0.008002 | -0.006357 | -0.013934 | 0.000918 | -0.383915 | 0.033664 |
| 31.45833 | 0.380631 | -0.102693 | 0.213325 | -0.620911 | 1.325082 | 0.270284 | 0.461697 | -0.789217 | -0.356317 | -2.138044 | 0.001949 | 0.270426 | 0.051998 | -0.161843 | -0.168221 | -0.351081 | -0.128807 | 0.044045 | -0.002522 | -0.039525 | -0.025423 | -0.132379 | -0.023842 |
| 31.47222 | 0.172818 | 0.509535 | -0.05578 | -0.662948 | 1.080446 | -0.009668 | -0.065307 | -0.7437 | -0.339449 | -1.748965 | 0.02289 | 0.179857 | -0.216778 | -0.510645 | -0.185629 | -0.406624 | -0.068151 | 0.049774 | 0.043795 | -0.059498 | -0.042233 | 0.200029 | -0.062774 |
| 31.48611 | -0.112759 | 0.058298 | -0.533932 | -0.804267 | 0.176469 | -0.317022 | 0.270255 | -0.68234 | -0.267502 | -1.21248 | 0.0599 | 0.060361 | -0.481911 | -0.619364 | 0.033275 | -0.532384 | -0.231229 | 0.105927 | 0.081116 | -0.042682 | -0.035523 | 0.383073 | -0.045041 |
| 31.5 | -0.15285 | -0.27478 | -0.807283 | -1.026004 | -0.589146 | -0.416463 | 0.550533 | -0.748806 | -0.126802 | -1.033889 | 0.079222 | -0.181597 | -0.69597 | -0.778083 | 0.490508 | -0.747118 | -0.901595 | 0.03682 | 0.02797 | -0.005062 | -0.012715 | 0.219635 | -0.014246 |
| 31.51389 | -0.308249 | -0.61886 | -1.252337 | -1.300951 | -1.656405 | -0.532944 | 0.889035 | -0.684435 | -0.101718 | -1.303844 | 0.054617 | -0.421753 | -0.850379 | -0.885821 | 1.000432 | -0.917293 | -1.355005 | -0.11188 | -0.077072 | 0.018027 | 0.014459 | -0.16584 | 0.003021 |
| 31.52778 | -0.364602 | -0.764496 | -1.879855 | -1.572747 | -2.029076 | -0.199522 | 1.139542 | -0.586752 | -0.223077 | -1.716816 | -0.038634 | -0.598788 | -0.834932 | -0.869271 | 1.367329 | -0.852792 | -1.267093 | -0.274165 | -0.120286 | 0.017031 | 0.030805 | -0.530119 | -0.0094 |
| 31.54167 | -0.733814 | -0.78304 | -2.133657 | -1.765973 | -2.532866 | -0.214306 | 0.57939 | -0.557075 | -0.356859 | -2.145992 | -0.138505 | -0.879926 | -0.788781 | -0.790957 | 1.163378 | -1.119841 | -1.284165 | -0.342067 | -0.078351 | 0.022342 | 0.033851 | -0.754282 | -0.037898 |
| 31.55556 | -1.031347 | -0.769163 | -2.363599 | -1.704536 | -2.630029 | -0.204012 | -0.382547 | -0.580039 | -0.385724 | -2.71657 | -0.13421 | -0.758681 | -0.650929 | -0.691301 | 0.607364 | -1.161325 | -1.375537 | -0.244666 | 0.033872 | 0.098895 | 0.104958 | -0.775021 | 0.029127 |
| 31.56944 | -0.732764 | -0.793811 | -3.216724 | -1.729372 | -2.680709 | 0.074497 | -1.086541 | -0.565243 | -0.271913 | -2.80714 | 0.033128 | -0.795742 | -0.50479 | -0.697675 | 0.261673 | -0.854821 | -1.652544 | 0.037519 | 0.104136 | 0.239144 | 0.278602 | -0.79618 | 0.211428 |
| 31.58333 | -0.454669 | -0.923277 | -4.367396 | -1.885839 | -2.391661 | 0.239255 | -1.080458 | -0.869902 | -0.288135 | -3.393057 | 0.061932 | -1.089949 | -0.572302 | -1.095003 | 0.078538 | -0.632487 | -1.948873 | 0.04188 | -0.148544 | 0.26773 | 0.430814 | -1.240226 | 0.286982 |
| 31.59722 | -0.773798 | -1.796013 | -5.098339 | -2.025999 | -2.926738 | -0.148256 | -1.397095 | -1.506106 | -0.554756 | -4.086318 | -0.319836 | -1.382416 | -1.051971 | -1.725284 | -0.077107 | -1.090922 | -1.553309 | -0.35053 | -0.774445 | 0.175522 | 0.309812 | -1.961785 | 0.108097 |
| 31.61111 | -0.888213 | -2.321888 | -6.01744 | -1.74828 | -3.460235 | -1.140135 | -2.293231 | -1.949428 | -0.766311 | -5.522859 | -0.632401 | -1.312467 | -1.255874 | -1.649362 | -0.206863 | -2.005543 | -1.081918 | -0.522476 | -1.261173 | 0.12746 | 0.161543 | -2.808726 | -0.045545 |
| 31.625 | -0.88044 | -2.13635 | -6.423853 | -1.57051 | -3.198045 | -1.255414 | -2.404856 | -2.248358 | -0.78682 | -4.716002 | -0.626715 | -1.005938 | -1.139206 | -1.229324 | -0.516679 | -2.396171 | -0.825309 | -0.367968 | -1.159312 | 0.137598 | 0.125897 | -2.81401 | -0.073495 |
| 31.63889 | -0.754059 | -1.58635 | -4.609182 | -1.082201 | -2.020859 | -0.624246 | -1.519677 | -2.023857 | -0.584951 | -3.157895 | -0.379676 | -0.782135 | -0.949092 | -0.496957 | -0.679239 | -1.994349 | -0.393926 | -0.082247 | -0.458528 | 0.109053 | 0.113294 | -2.500955 | -0.076937 |
| 31.65278 | -0.152339 | -0.772355 | -0.46886 | -0.201077 | 0.072719 | -0.370001 | -1.021291 | -0.488093 | -0.231276 | -2.669868 | -0.168869 | -0.67168 | -0.720788 | 0.19478 | -0.411537 | -1.2305 | 0.419215 | 0.189009 | 0.449184 | 0.116069 | 0.208689 | -1.264418 | -0.024041 |
| 31.66667 | 0.267053 | -0.194233 | 1.680475 | 0.564485 | 1.250591 | -0.564425 | -0.534252 | 0.733384 | 0.009947 | -2.87735 | -0.238879 | -0.668548 | -0.819156 | 0.20885 | -0.177666 | -0.214923 | 0.889837 | 0.416212 | 1.140158 | 0.226235 | 0.294648 | -0.433257 | 0.097268 |
| 31.68056 | 0.211802 | 0.189224 | 1.192937 | 0.524586 | 1.562627 | -0.688616 | -0.230734 | 1.367002 | 0.088379 | -1.830913 | -0.529256 | -0.670251 | -0.90597 | 0.023755 | -0.197621 | 0.428867 | 0.41616 | 0.379448 | 1.091178 | 0.301756 | 0.321112 | -0.144034 | 0.123638 |
| 31.69444 | 0.049595 | 0.538427 | 1.744335 | 0.374749 | 1.463194 | -0.245261 | 0.329954 | 2.01136 | -0.029628 | -0.846012 | -0.716706 | -0.584377 | -0.652364 | 0.053596 | -0.395726 | 0.082844 | -0.100802 | 0.261586 | 0.989946 | 0.309487 | 0.321319 | -0.029503 | 0.080573 |
| 31.70833 | 0.106057 | 0.517951 | 3.153112 | 0.279729 | 1.661476 | 0.372695 | 0.985768 | 2.40469 | -0.16817 | -0.084156 | -0.720356 | -0.606579 | -0.230858 | 0.256481 | -0.735565 | -0.23573 | -0.501791 | 0.157561 | 0.905063 | 0.28237 | 0.287553 | 0.040404 | 0.031749 |
| 31.72222 | -0.008527 | 0.552661 | 2.457923 | -0.030695 | 1.362522 | 0.875686 | 1.554107 | 1.603551 | -0.293754 | 0.583602 | -0.714958 | -0.647856 | 0.116021 | 0.160204 | -1.090347 | -0.160755 | -0.698086 | 0.026008 | 0.68866 | 0.261092 | 0.24567 | 0.047881 | 0.017276 |
| 31.73611 | -0.130446 | 0.467806 | 0.391828 | -0.270927 | 1.142136 | 0.894967 | 2.147434 | 0.094591 | -0.393938 | 1.059993 | -0.690622 | -0.508382 | 0.30473 | 0.102082 | -1.448006 | -0.136204 | -1.069191 | -0.198303 | 0.604355 | 0.29024 | 0.272656 | 0.082338 | 0.0361 |
| 31.75 | 0.08642 | 0.465885 | -0.82131 | -0.227639 | 0.755433 | 0.592316 | 1.964306 | -1.156555 | -0.399982 | 1.173366 | -0.606299 | -0.352842 | 0.259304 | -0.033912 | -1.731266 | -0.080483 | -0.914665 | -0.334727 | 0.593962 | 0.372864 | 0.3764 | 0.099592 | 0.159911 |
| 31.76389 | 0.650644 | 0.51794 | -0.119806 | 0.086495 | 0.555334 | 0.823564 | 1.628018 | -1.627277 | -0.210774 | 0.848142 | -0.370333 | -0.301676 | 0.202206 | -0.365656 | -1.681978 | 0.244799 | -0.890978 | -0.281664 | 0.408785 | 0.434128 | 0.488092 | -0.055777 | 0.260021 |
| 31.77778 | 1.241252 | 0.462086 | 0.349603 | 0.257232 | 0.201011 | 0.80088 | 1.524365 | -1.636611 | -0.170995 | 0.434788 | -0.163304 | -0.129412 | 0.459267 | -0.557016 | -1.380401 | 0.484504 | -0.547342 | -0.163112 | 0.147659 | 0.393947 | 0.583379 | -0.217571 | 0.302843 |
| 31.79167 | 1.516198 | 0.276896 | -0.695015 | 0.120903 | -0.327247 | 0.749501 | 0.933884 | -1.70784 | -0.329761 | -0.088309 | -0.147097 | -0.043409 | 0.832836 | -0.248203 | -0.94203 | 0.311981 | 0.391015 | -0.123056 | 0.065618 | 0.347722 | 0.60412 | -0.158538 | 0.314322 |
| 31.80556 | 1.55361 | 0.145679 | -1.683625 | 0.014676 | -0.572001 | 0.485378 | 0.241905 | -1.181734 | -0.494006 | -0.42223 | -0.187189 | -0.097781 | 0.737301 | 0.085698 | -0.641759 | -0.076549 | 1.158137 | -0.158969 | 0.018062 | 0.284841 | 0.468894 | -0.040957 | 0.24392 |
| 31.81944 | 1.034959 | 0.1255 | -1.220404 | 0.095757 | -0.253167 | 0.309194 | -0.324751 | -0.627259 | -0.600136 | -0.899597 | -0.196727 | -0.230492 | 0.21117 | 0.190323 | -0.588309 | -0.282101 | 1.133268 | -0.152359 | -0.054593 | 0.178462 | 0.264491 | -0.026617 | 0.10965 |
| 31.83333 | 0.441157 | 0.259668 | -0.107845 | 0.380179 | -0.052148 | 0.212281 | -0.756905 | 0.062653 | -0.653636 | -1.230994 | -0.086136 | -0.291836 | -0.197723 | 0.415612 | -0.651865 | -0.181554 | 0.774278 | -0.090895 | -0.046601 | 0.122537 | 0.137864 | -0.004829 | 0.031034 |
| 31.84722 | -0.191675 | 0.356397 | 0.51211 | 0.627326 | 0.156361 | 0.153094 | -0.577345 | 0.731217 | -0.721499 | -1.160524 | -0.036651 | -0.081406 | -0.217477 | 0.805503 | -0.552126 | -0.114689 | 0.530469 | -0.046137 | 0.135346 | 0.133763 | 0.126443 | 0.106097 | 0.042201 |
| 31.86111 | -0.436918 | 0.220143 | 0.76646 | 0.655324 | 0.497323 | 0.146801 | -0.123326 | 1.324777 | -0.407028 | -1.146394 | -0.091639 | 0.23027 | -0.014423 | 1.127856 | -0.205899 | -0.11995 | -0.274564 | -0.090752 | 0.429999 | 0.162716 | 0.171374 | 0.334898 | 0.085674 |
| 31.875 | -0.423438 | 0.16181 | 1.617698 | 0.69815 | 1.044624 | 0.362437 | 0.03434 | 1.862621 | -0.015078 | -1.130147 | -0.083523 | 0.40877 | 0.102045 | 1.203815 | 0.090273 | -0.040943 | -0.909038 | -0.147833 | 0.593776 | 0.154248 | 0.159149 | 0.528801 | 0.063053 |
| 31.88889 | -0.052385 | 0.20447 | 2.385465 | 0.665546 | 1.430006 | 0.662706 | 0.272199 | 2.192656 | -0.770757 | -0.972941 | -0.061525 | 0.260295 | 0.0876 | 0.78903 | -0.039263 | 0.059268 | -0.956772 | -0.291162 | 0.362472 | 0.023151 | 0.009311 | 0.541425 | -0.078072 |
| 31.90278 | 0.426573 | 0.403042 | 2.022692 | 0.55939 | 1.560735 | 0.651906 | 0.705375 | 2.182608 | -0.467688 | -0.792264 | -0.113791 | 0.021012 | 0.157419 | 0.170071 | -0.248215 | 0.135674 | -0.510327 | -0.42646 | 0.043152 | -0.153997 | -0.173738 | 0.612323 | -0.24914 |
| 31.91667 | 0.494185 | 0.793225 | 1.793492 | 0.533467 | 1.434919 | 0.571471 | 1.029295 | 1.760186 | 0.153112 | -0.557141 | -0.094277 | -0.012132 | 0.351562 | 0.09016 | -0.173824 | 0.215459 | 0.52395 | -0.443949 | -0.082431 | -0.219099 | -0.228518 | 0.888023 | -0.323279 |
| 31.93056 | 0.433846 | 1.005235 | 1.983257 | 0.668411 | 1.567862 | 0.351364 | 0.895229 | 1.436273 | 0.699273 | -0.092347 | 0.037479 | 0.174766 | 0.753637 | 0.239189 | 0.168311 | 0.344208 | 2.222477 | -0.279006 | -0.033697 | -0.150397 | -0.13465 | 0.990128 | -0.259478 |
| 31.94444 | 0.245341 | 0.915265 | 1.505451 | 0.594398 | 1.053754 | 0.182023 | 0.718423 | 0.377632 | 0.794379 | 0.380983 | 0.066195 | 0.326127 | 0.928002 | -0.036316 | 0.706359 | 0.561462 | 3.617198 | 0.074092 | 0.084397 | 0.003618 | 0.031957 | 0.729941 | -0.073621 |
| 31.95833 | 0.425006 | 0.666602 | -0.181535 | 0.337084 | 0.213568 | 0.666145 | 1.074169 | -0.883553 | 0.901947 | 0.77162 | 0.003633 | 0.402175 | 0.967514 | -0.226309 | 1.159262 | 0.787603 | 3.864806 | 0.368468 | 0.1787 | 0.144038 | 0.170064 | 0.492495 | 0.107342 |
| 31.97222 | 0.221849 | 0.434599 | -0.717332 | 0.328686 | -0.090162 | 0.896643 | 1.184956 | -1.685095 | 0.70905 | 1.249627 | 0.058815 | 0.498754 | 0.953487 | 0.077303 | 1.53178 | 0.917302 | 2.896691 | 0.45965 | 0.266508 | 0.185978 | 0.223543 | 0.467809 | 0.170888 |
| 31.98611 | 0.009951 | 0.463157 | -0.089492 | 0.429414 | -0.014642 | 0.922808 | 0.539985 | -1.381754 | 0.267605 | 1.622434 | 0.083336 | 0.447236 | 0.62842 | 0.143723 | 1.750853 | 0.916282 | 1.281412 | 0.386292 | 0.298217 | 0.110527 | 0.138037 | 0.488746 | 0.087212 |
| 32 | -0.449272 | 0.251973 | -0.082665 | 0.309926 | -0.219201 | 0.850377 | 0.17621 | -0.827802 | 0.206156 | 1.498019 | -0.150928 | 0.26921 | 0.118421 | -0.179438 | 1.633861 | 0.894507 | -0.076824 | 0.243041 | 0.170291 | -0.077725 | -0.109239 | 0.442472 | -0.138769 |
| 32.01389 | -0.466869 | 0.010368 | 0.433139 | 0.036133 | 0.235682 | 0.605848 | -0.073314 | -0.078942 | -0.428656 | 1.091805 | -0.470725 | 0.241539 | -0.29034 | -0.105014 | 1.054828 | 0.661799 | -1.263896 | 0.069718 | -0.015052 | -0.337576 | -0.467837 | 0.370328 | -0.423808 |
| 32.02778 | -0.12007 | -0.055525 | 2.306609 | 0.019227 | 0.916119 | 0.315438 | 0.535192 | 0.773364 | -1.140843 | 1.13214 | -0.400113 | 0.388995 | -0.587887 | 0.130647 | 0.702357 | 0.366738 | -1.633685 | -0.069197 | -0.245502 | -0.467775 | -0.568035 | 0.228337 | -0.548789 |
| 32.04167 | 0.02037 | 0.724424 | 3.87784 | 0.324752 | 1.688456 | 1.107153 | 2.074829 | 0.950336 | -0.976732 | 1.430683 | -0.084115 | 0.736148 | -0.709207 | 0.286606 | 0.507443 | 0.535891 | -0.825472 | -0.003347 | -0.367698 | -0.465953 | -0.502817 | 0.362473 | -0.514229 |
| 32.05556 | 0.545209 | 1.358067 | 3.586782 | 0.488172 | 1.837693 | 0.551258 | 3.250472 | 0.970767 | -0.311763 | 1.253961 | 0.020792 | 1.110347 | -0.664736 | 0.798379 | 0.302893 | 0.676766 | 0.379849 | 0.098137 | -0.266445 | -0.385214 | -0.424606 | 0.726379 | -0.437369 |
| 32.06944 | 0.81149 | 1.398293 | 3.418447 | 0.469546 | 1.516167 | -0.014418 | 3.01914 | 0.814568 | 0.601989 | 1.32935 | 0.023862 | 0.89435 | -0.54758 | 1.144936 | 0.034121 | 0.671922 | 1.592221 | 0.217947 | -0.123852 | -0.268977 | -0.28143 | 0.795617 | -0.333686 |
| 32.08333 | 1.051356 | 1.325057 | 6.089634 | 0.649021 | 1.646849 | 0.162413 | 1.840523 | 0.806465 | 0.336702 | 1.753022 | 0.246906 | -0.138918 | -0.930783 | 0.477017 | -0.688865 | -0.040571 | 1.899281 | -0.007386 | -0.520567 | -0.677121 | -0.668914 | -0.072277 | -0.686276 |
| 32.09722 | 1.551874 | 1.153925 | 8.168184 | 0.753563 | 1.304507 | -0.162296 | 0.585583 | 0.756872 | -0.766576 | 2.285238 | 0.334619 | -1.347526 | -1.94488 | -1.226307 | -2.037285 | -1.59049 | 0.012246 | -0.911113 | -1.556674 | -1.804182 | -1.866255 | -1.96243 | -1.740063 |
| 32.11111 | 1.546239 | 0.791969 | 5.191785 | 0.434473 | 0.580787 | 0.022621 | -0.088288 | 0.492735 | -0.893121 | 1.87714 | -0.075091 | -1.448038 | -2.607293 | -2.719789 | -2.492886 | -2.181932 | -1.20832 | -1.353297 | -1.906255 | -2.247191 | -2.364448 | -2.525054 | -2.197069 |
| 32.125 | 0.699889 | 0.410739 | 1.265933 | 0.352494 | 0.531233 | -0.226424 | -0.262906 | 0.345065 | 0.011819 | 1.3206 | -0.358722 | -0.196822 | -2.120545 | -3.077159 | -1.469032 | -0.881235 | -0.564893 | -0.665729 | -0.885209 | -1.25297 | -1.278671 | -1.176413 | -1.393282 |
| 32.13889 | 0.274346 | 0.624076 | 0.210666 | 0.879518 | 0.655731 | -0.309872 | 0.476458 | 0.367145 | 0.519512 | 0.838227 | -0.125635 | 1.107359 | -1.225853 | -2.559083 | -0.052079 | 0.727256 | 0.497893 | 0.06024 | -0.011305 | -0.373812 | -0.348375 | 0.131499 | -0.497324 |
| 32.15278 | -0.000198 | 0.974264 | 0.216675 | 1.266927 | 0.756658 | -0.695468 | 0.358251 | 0.164052 | 0.924578 | 0.924975 | 0.255727 | 1.527982 | -0.426728 | -1.723703 | 0.780154 | 1.924353 | 0.361847 | 0.545571 | 0.317372 | -0.005838 | 0.042915 | 0.751862 | -0.055127 |
| 32.16667 | -0.526925 | 1.150312 | -0.549439 | 0.954376 | 0.304672 | -1.404441 | -0.908609 | -0.331357 | 1.229847 | 0.606762 | 0.471023 | 1.182725 | -0.085043 | -1.215731 | 0.926198 | 2.070987 | -0.036576 | 0.699523 | 0.37707 | 0.062398 | 0.069331 | 1.031629 | 0.021898 |
| 32.18056 | -1.093117 | 0.941699 | -1.929321 | 0.588737 | -0.12821 | -1.656429 | -0.829427 | -0.632634 | 0.84251 | 0.368544 | 0.431854 | 0.661393 | 0.11268 | -1.044458 | 0.695986 | 2.099678 | -0.301479 | 0.720599 | 0.349258 | 0.047004 | 0.003354 | 1.471578 | -0.044293 |
| 32.19444 | -1.31993 | 0.820591 | -2.791046 | 0.444301 | -0.167095 | -1.599853 | 1.370991 | -0.612036 | 0.02779 | 0.243983 | 0.388948 | 0.57388 | 0.287449 | -0.944894 | 0.404813 | 2.625681 | -0.222253 | 0.822534 | 0.358562 | 0.065059 | 0.037022 | 2.087832 | -0.042697 |
| 32.20833 | -1.025702 | 0.757317 | -2.95863 | 0.457757 | 0.091392 | -0.594964 | 1.148862 | -0.367031 | 0.295475 | 0.204926 | 0.510317 | 0.369894 | 0.444821 | -0.783466 | 0.341036 | 2.695157 | 0.086264 | 0.903085 | 0.348957 | 0.12824 | 0.129519 | 1.969251 | 0.046862 |
| 32.22222 | -0.434712 | 0.619518 | -2.549176 | 0.567402 | 0.282792 | 0.045879 | 0.499134 | 0.13267 | 0.80081 | 0.056593 | 0.668466 | 0.354853 | 0.559253 | -0.421288 | 0.561671 | 2.236731 | 0.461932 | 0.895556 | 0.396497 | 0.212732 | 0.224638 | 1.292457 | 0.144267 |
| 32.23611 | -0.107972 | 0.406676 | -1.961307 | 0.61694 | 0.332435 | 0.159218 | -0.350145 | 0.7677 | 0.501682 | -0.110886 | 0.601621 | 0.519601 | 0.481178 | -0.022311 | 0.887725 | 1.906332 | 0.756475 | 0.909404 | 0.442359 | 0.272394 | 0.284227 | 0.927712 | 0.209283 |
| 32.25 | -0.072879 | 0.043001 | -1.377948 | 0.514328 | 0.315872 | 0.156257 | -1.62718 | 0.775391 | -0.04992 | -0.053662 | 0.412942 | 0.686804 | 0.34272 | 0.371168 | 1.028885 | 1.724825 | 1.032238 | 0.861314 | 0.422458 | 0.288911 | 0.298193 | 0.841974 | 0.234095 |
| 32.26389 | 0.04519 | -0.285755 | -0.915392 | 0.279912 | 0.425876 | -0.502608 | -2.296428 | 0.84807 | 0.013247 | -0.036737 | 0.240503 | 0.761559 | 0.317016 | 0.734497 | 0.745303 | 1.544588 | 1.148334 | 0.845233 | 0.371851 | 0.305146 | 0.307043 | 0.978553 | 0.259107 |
| 32.27778 | -0.028561 | -0.575189 | -0.673468 | 0.050219 | 0.560145 | -0.669778 | -1.319853 | 0.858541 | 0.315374 | -0.348136 | 0.135179 | 0.800867 | 0.313329 | 0.851986 | 0.500859 | 1.169409 | 0.887709 | 0.76028 | 0.383152 | 0.314616 | 0.322884 | 1.138069 | 0.28163 |
| 32.29167 | -0.333602 | -0.560854 | -0.377682 | -0.120301 | 0.603443 | -0.518964 | 0.856251 | 0.726412 | 0.440951 | -0.98006 | 0.097487 | 0.80075 | 0.262876 | 0.809096 | 0.413886 | 0.550466 | 0.380515 | 0.665799 | 0.415404 | 0.278227 | 0.304589 | 0.996177 | 0.258069 |
| 32.30556 | -0.534139 | -0.168453 | 0.16088 | -0.11351 | 0.671859 | -1.082348 | 2.726661 | 0.477221 | 0.843681 | -0.956644 | 0.108693 | 0.795513 | 0.121174 | 0.596351 | 0.34996 | -0.146372 | 0.229868 | 0.534386 | 0.398378 | 0.170075 | 0.209477 | 0.568803 | 0.15957 |
| 32.31944 | -0.355388 | 0.216334 | 0.693878 | 0.023941 | 0.252628 | -1.019202 | 2.532627 | 0.206782 | 1.066908 | -0.359663 | 0.099333 | 0.866345 | -0.103174 | 0.52964 | 0.249818 | -0.745408 | 0.021103 | 0.414643 | 0.38671 | 0.021986 | 0.053023 | 0.102771 | 0.008051 |
| 32.33333 | -0.34144 | 0.367974 | 0.190908 | 0.12063 | 0.326693 | -0.909787 | 1.503415 | -0.121936 | 0.392225 | 0.050101 | 0.013284 | 0.501108 | -0.306031 | 0.320122 | 0.21404 | -1.119957 | -0.322804 | 0.388681 | 0.405861 | -0.035346 | -0.02631 | -0.202715 | -0.064348 |
| 32.34722 | -0.270322 | 0.450265 | -0.607459 | 0.201109 | 0.408962 | -0.536606 | 0.906842 | -0.3587 | -0.252031 | 0.332226 | -0.046918 | 0.720666 | -0.367487 | -0.075867 | 0.332122 | -1.08875 | -0.008693 | 0.407452 | 0.395893 | 0.029055 | 0.028529 | -0.342961 | 0.002476 |
| 32.36111 | -0.19219 | 0.501816 | -0.565743 | 0.26338 | 0.175864 | 0.28572 | 1.775412 | -0.537564 | -0.039026 | 0.574807 | -0.03052 | 0.546354 | -0.250484 | -0.268015 | 0.56388 | -0.859099 | -0.070489 | 0.420846 | 0.364263 | 0.116652 | 0.121431 | -0.413421 | 0.10609 |
| 32.375 | -0.307465 | 0.573352 | -1.051848 | 0.199892 | -0.451256 | 0.605364 | 0.556192 | -0.871462 | 0.414746 | 0.644444 | -0.024156 | 0.133462 | -0.165161 | 0.220444 | 0.854559 | -0.575022 | -0.380934 | 0.447736 | 0.379217 | 0.181515 | 0.192957 | -0.338512 | 0.177668 |
| 32.38889 | -0.471789 | 0.548464 | -1.37225 | 0.036715 | -0.443406 | 0.835337 | -1.3044 | -0.938955 | 0.679394 | 0.344019 | -0.022818 | -0.26925 | -0.263055 | 1.142212 | 1.044308 | -0.157263 | -0.160881 | 0.48076 | 0.388372 | 0.216338 | 0.233143 | -0.178195 | 0.216459 |
| 32.40278 | -0.849074 | 0.258477 | -1.000815 | -0.209086 | 0.329693 | 1.155673 | -2.099029 | -0.844168 | 0.43678 | 0.0622 | 0.015457 | -0.525431 | -0.263479 | 1.832692 | 1.073286 | 0.308591 | -0.410831 | 0.458416 | 0.384571 | 0.232567 | 0.245558 | -0.060313 | 0.234226 |
| 32.41667 | -0.834556 | 0.030165 | -0.475999 | -0.435675 | 0.567591 | 1.677782 | -2.186539 | -0.493053 | -0.348638 | -0.103429 | 0.045416 | -0.611659 | -0.068071 | 1.912581 | 1.019392 | 0.689682 | -0.334619 | 0.40293 | 0.368389 | 0.242899 | 0.252439 | -0.043803 | 0.249592 |
| 32.43056 | -0.371116 | 0.032258 | 0.090419 | -0.428507 | 0.689363 | 2.094234 | -1.418879 | -0.096418 | -0.751154 | -0.210706 | 0.096391 | -0.574216 | 0.047836 | 1.237186 | 0.968171 | 0.825012 | -0.16778 | 0.357926 | 0.334904 | 0.254937 | 0.260638 | -0.010734 | 0.267938 |
| 32.44444 | -0.009325 | -0.15252 | 1.079997 | -0.236268 | 0.700799 | 1.304622 | -1.490333 | 0.554842 | -0.140084 | -0.271803 | 0.190001 | -0.379256 | -0.009646 | 0.697249 | 0.877084 | 0.812973 | -0.342504 | 0.31239 | 0.261748 | 0.243398 | 0.246721 | 0.076031 | 0.264443 |
| 32.45833 | -0.126817 | -0.494572 | 1.148513 | -0.084373 | 0.083666 | 0.326475 | -1.163171 | 1.059299 | 0.371537 | -0.623025 | 0.224922 | -0.176814 | -0.091091 | 0.274293 | 0.704188 | 0.588793 | -0.235734 | 0.234585 | 0.143533 | 0.194612 | 0.199302 | 0.148261 | 0.214146 |
| 32.47222 | -0.677857 | -0.690527 | 0.403363 | -0.018216 | -0.687649 | -0.503895 | -0.149109 | 1.207295 | 0.404596 | -1.191081 | 0.158834 | 0.112279 | -0.060162 | 0.048893 | 0.541077 | 0.33358 | -0.188099 | 0.144196 | 0.059528 | 0.160792 | 0.168744 | 0.191229 | 0.153469 |
| 32.48611 | -1.080928 | -0.611693 | -0.997734 | -0.001368 | -0.776071 | -0.867688 | -0.324232 | 1.10581 | 0.650996 | -1.67813 | 0.025443 | 0.411525 | 0.040154 | -0.035258 | 0.448231 | 0.188562 | -0.070228 | 0.058495 | 0.026325 | 0.153868 | 0.169599 | 0.14452 | 0.104945 |
| 32.5 | -1.582391 | -0.367524 | -2.145539 | -0.049884 | -0.465227 | -1.19647 | 0.322444 | 0.992174 | -0.19731 | -1.868515 | -0.062786 | 0.60722 | 0.161935 | -0.013708 | 0.428743 | 0.057197 | 0.05961 | 0.04273 | 0.048619 | 0.163971 | 0.193616 | 0.01469 | 0.106433 |
| 32.51389 | -2.248873 | -0.622783 | -2.012426 | -0.145498 | 0.133026 | -1.69805 | 0.838547 | 1.151858 | -1.103277 | -1.605328 | -0.048574 | 0.816816 | 0.217585 | 0.081252 | 0.45841 | -0.142535 | 0.443949 | 0.123772 | 0.079272 | 0.218058 | 0.253067 | -0.166545 | 0.190247 |
| 32.52778 | -1.747738 | -0.72624 | -1.539519 | -0.389246 | 0.345854 | -2.038964 | 0.674235 | 1.057439 | -0.672814 | -1.332432 | -0.056154 | 0.925337 | 0.216443 | 0.142048 | 0.450841 | -0.393095 | 0.664715 | 0.148882 | 0.1238 | 0.253526 | 0.281349 | -0.418605 | 0.249803 |
| 32.54167 | -1.207545 | -0.336104 | -1.46132 | -0.747438 | 0.117713 | -2.215716 | 0.262055 | 0.802672 | 0.285489 | -1.452677 | -0.176916 | 0.501382 | 0.170033 | 0.070674 | 0.321085 | -0.691115 | 0.16382 | 0.108698 | 0.123522 | 0.211153 | 0.236221 | -0.800385 | 0.208039 |
| 32.55556 | -0.88321 | 0.264216 | -2.588992 | -0.90708 | -1.029014 | -1.962957 | -0.197578 | 0.407871 | 0.841034 | -1.483008 | -0.220179 | 0.190484 | 0.209937 | 0.070509 | 0.202112 | -1.004649 | -0.062046 | 0.162009 | 0.173726 | 0.217047 | 0.2568 | -1.177884 | 0.226873 |
| 32.56944 | -1.206065 | 0.519588 | -5.305728 | -1.261956 | -2.05954 | -1.200406 | -0.438108 | -0.131679 | 0.13909 | -1.73754 | -0.15046 | -0.231436 | 0.449032 | 0.287936 | 0.19897 | -1.196002 | 0.235486 | 0.303753 | 0.334525 | 0.338363 | 0.420691 | -1.29021 | 0.375466 |
| 32.58333 | -1.487396 | -0.161551 | -7.375993 | -1.73845 | -2.695713 | -1.024681 | -0.700772 | -1.030097 | -1.561737 | -2.997323 | -0.284425 | -0.904386 | -0.036044 | 0.038485 | -0.208774 | -1.478589 | -0.255384 | 0.029555 | 0.011634 | 0.125261 | 0.255311 | -1.628253 | 0.119191 |
| 32.59722 | -1.957522 | -0.928038 | -6.995857 | -1.861002 | -2.665335 | -1.687934 | -1.717854 | -1.4795 | -2.574917 | -3.925236 | -0.664315 | -1.678074 | -1.237389 | -0.589334 | -1.232191 | -2.202747 | -1.632501 | -0.837357 | -0.727805 | -0.53401 | -0.490489 | -2.932924 | -0.633702 |
| 32.61111 | -1.482903 | -1.051842 | -6.479597 | -1.449931 | -3.407675 | -1.726779 | -3.123859 | -1.302906 | -1.881984 | -4.14528 | -0.760226 | -2.011308 | -1.670753 | -0.504456 | -1.708159 | -2.937644 | -2.365586 | -0.89031 | -1.002584 | -0.639208 | -0.627359 | -3.729674 | -0.752981 |
| 32.625 | -0.810704 | -1.000411 | -5.668708 | -0.937187 | -2.745634 | -0.909577 | -2.030598 | -1.702612 | -0.725828 | -4.752881 | -0.669441 | -1.878764 | -1.275517 | -0.57093 | -1.771551 | -3.263766 | -2.197393 | -0.590883 | -0.669026 | -0.231362 | -0.228069 | -2.984652 | -0.295515 |
| 32.63889 | -0.619831 | -0.726833 | -2.77058 | -0.569967 | -1.212939 | -0.736229 | -1.549269 | -1.425051 | 0.350311 | -3.657229 | -0.628249 | -1.585846 | -0.899943 | -1.138223 | -1.6192 | -2.579583 | -2.312474 | -0.41296 | -0.481652 | -0.051114 | -0.04411 | -2.370349 | -0.073318 |
| 32.65278 | -0.207058 | -0.287329 | 0.62743 | -0.196772 | -0.14655 | -0.213871 | -1.194401 | -1.002707 | 0.988118 | -2.71054 | -0.634748 | -0.908028 | -0.461484 | -1.245612 | -1.275056 | -1.385977 | -1.82084 | -0.165725 | -0.003022 | -0.015369 | 0.010844 | -1.468075 | -0.007997 |
| 32.66667 | -0.329355 | -0.294212 | 1.400776 | -0.174912 | 0.805992 | 0.15827 | -0.402375 | -0.219337 | 0.479603 | -2.648391 | -0.586911 | -0.34163 | -0.139748 | -1.125896 | -0.870723 | -0.542399 | -1.050601 | 0.152949 | 0.575005 | 0.082402 | 0.118117 | -0.515108 | 0.099212 |
| 32.68056 | -0.202373 | -0.425037 | 0.594729 | -0.389764 | 1.219186 | 0.359609 | 0.321558 | -0.057991 | 0.678582 | -2.542622 | -0.451881 | -0.181472 | 0.000972 | -1.187828 | -0.665745 | -0.124855 | -0.709385 | 0.268763 | 0.773508 | 0.185666 | 0.221102 | 0.262197 | 0.193659 |
| 32.69444 | 0.621842 | 0.25029 | 0.604715 | -0.254981 | 0.822516 | 0.774485 | 0.697401 | 0.207162 | 1.226991 | -1.292454 | -0.291913 | -0.248674 | -0.037296 | -1.312861 | -0.740371 | -0.121046 | -0.764403 | 0.266822 | 0.559009 | 0.260189 | 0.29784 | 0.451691 | 0.247814 |
| 32.70833 | 0.885301 | 0.629318 | 0.524956 | 0.0362 | 0.117471 | 1.033821 | 0.813597 | 0.660684 | 0.586787 | -0.480743 | -0.165413 | -0.351228 | 0.068629 | -1.464272 | -1.00495 | -0.425887 | -0.894089 | 0.230362 | 0.424934 | 0.250081 | 0.28831 | 0.449526 | 0.233898 |
| 32.72222 | 0.498631 | -0.094818 | 0.062322 | 0.076575 | -0.694816 | 0.773194 | 0.594429 | 1.070868 | 0.027332 | -0.180689 | -0.09734 | -0.410408 | 0.146842 | -1.493944 | -1.161584 | -0.804092 | -0.821685 | 0.135357 | 0.255345 | 0.153067 | 0.184591 | 0.454715 | 0.141892 |
| 32.73611 | 0.158592 | -0.841301 | 0.368877 | 0.110021 | -1.415792 | 0.415475 | 0.579887 | 1.3554 | -0.283328 | 0.282243 | -0.036972 | -0.438249 | 0.10209 | -1.3324 | -1.168658 | -0.762393 | -0.673634 | -0.022609 | 0.014035 | 0.046954 | 0.075763 | 0.334451 | 0.034077 |
| 32.75 | -0.023608 | -1.119598 | 1.972649 | 0.206181 | -1.240922 | 0.414161 | 1.013541 | 1.779224 | -0.176221 | 1.262115 | 0.026902 | -0.492495 | 0.048007 | -0.991939 | -1.172889 | -0.263561 | -0.530961 | -0.14126 | -0.153151 | 0.029176 | 0.0525 | 0.382506 | 0.015038 |
| 32.76389 | 0.073193 | -1.114076 | 3.760813 | 0.226141 | -1.084263 | 0.263885 | 0.93364 | 2.31884 | 0.11166 | 1.839293 | -0.003313 | -0.575756 | 0.035854 | -0.719356 | -1.174864 | 0.191556 | -0.003966 | -0.231769 | -0.226349 | 0.053443 | 0.064616 | 0.255029 | 0.04641 |
| 32.77778 | -0.277201 | -1.018842 | 3.95738 | -0.108927 | -1.059894 | 0.333231 | 0.897179 | 2.368437 | 0.598302 | 1.4378 | -0.069851 | -0.621644 | -0.015835 | -0.688314 | -1.23518 | 0.21629 | 1.078067 | -0.398917 | -0.203737 | 0.007625 | 0.013036 | 0.018102 | 0.00598 |
| 32.79167 | -0.565927 | -1.139625 | 1.595076 | -0.84557 | -0.972308 | 0.336328 | 1.072731 | 1.846446 | 0.344852 | 1.110976 | -0.133007 | -0.631611 | -0.100337 | -0.733507 | -1.211512 | -0.249786 | 1.591611 | -0.599717 | -0.161086 | -0.103867 | -0.080287 | -0.256154 | -0.120242 |
| 32.80556 | -0.066214 | -0.833112 | 0.4625 | -0.951888 | -0.771722 | 0.884054 | 1.441754 | 0.847026 | -0.646273 | 1.307565 | -0.129343 | -0.743715 | -0.242023 | -1.116475 | -1.119128 | -0.771954 | 1.007812 | -0.798071 | -0.1939 | -0.242119 | -0.197092 | -0.486314 | -0.27272 |
| 32.81944 | 0.770404 | -0.344059 | 1.047164 | -0.479222 | -0.605709 | 0.707057 | 1.4369 | -0.29752 | -0.996853 | 1.22422 | -0.041521 | -0.784664 | -0.328206 | -1.604535 | -1.03728 | -0.916582 | 0.277212 | -0.838541 | -0.244002 | -0.334043 | -0.29138 | -0.610046 | -0.378083 |
| 32.83333 | 1.330634 | -0.052709 | 0.115407 | -0.242631 | -0.730359 | 0.692243 | 1.357525 | -1.504577 | -1.272823 | 0.611132 | 0.011891 | -0.577171 | -0.302414 | -2.216348 | -0.859649 | -0.65587 | -0.184121 | -0.689707 | -0.143792 | -0.256463 | -0.2166 | -0.63448 | -0.304479 |
| 32.84722 | 1.963788 | -0.060355 | -1.558527 | -0.313966 | -0.355063 | 0.217339 | 1.143098 | -2.016572 | -1.42101 | -0.029976 | -0.064911 | -0.282048 | -0.209733 | -3.857936 | -0.555895 | -0.430967 | -0.516138 | -0.368444 | 0.134392 | -0.022342 | 0.022951 | -0.658857 | -0.053405 |
| 32.86111 | 2.116421 | -0.096235 | -0.656243 | -0.018773 | 0.453274 | -0.100673 | 0.63879 | -1.192102 | 0.532792 | -0.663116 | 0.004586 | 0.005001 | -0.013523 | -3.445843 | -0.307645 | -0.326796 | -1.105868 | -0.070962 | 0.338514 | 0.18188 | 0.257837 | -0.702878 | 0.182492 |
| 32.875 | 1.964561 | 0.122831 | 0.655267 | 0.401626 | 1.45034 | 0.394601 | -0.450208 | -1.318702 | 1.348128 | -1.248789 | 0.038432 | 0.215062 | 0.264941 | -1.180288 | -0.161777 | -0.407845 | -1.549043 | 0.057692 | 0.355921 | 0.254269 | 0.341224 | -0.562083 | 0.282702 |
| 32.88889 | 1.421841 | 0.413469 | 0.675072 | 0.346906 | 2.113937 | 0.873261 | -1.961412 | -0.540742 | -0.148065 | -2.008504 | -0.08596 | 0.284361 | 0.442311 | 0.077546 | -0.011459 | -0.410492 | -0.828985 | 0.118892 | 0.323405 | 0.263167 | 0.335986 | -0.364777 | 0.288005 |
| 32.90278 | 0.398137 | 0.553906 | -0.135148 | 0.055137 | 2.640433 | 0.184224 | -1.318845 | 0.048998 | -0.772121 | -2.408481 | -0.193559 | 0.329145 | 0.534507 | 0.657221 | 0.136638 | -0.331769 | 0.343966 | 0.166763 | 0.323364 | 0.250671 | 0.327657 | -0.189703 | 0.269047 |
| 32.91667 | -0.441564 | 0.75736 | -0.503947 | -0.131538 | 2.759577 | -0.09035 | -0.871696 | 0.39211 | -0.168918 | -2.423922 | -0.149078 | 0.373675 | 0.602666 | 0.855566 | 0.256197 | -0.10423 | 1.242609 | 0.1183 | 0.301888 | 0.193746 | 0.264515 | -0.013893 | 0.218365 |
| 32.93056 | -0.463895 | 1.063486 | -0.059328 | -0.092088 | 2.24629 | -0.809661 | -0.245334 | 0.494695 | 0.168928 | -1.997937 | -0.053963 | 0.34932 | 0.535573 | 0.752489 | 0.360533 | 0.064218 | 1.735348 | -0.047329 | 0.228067 | 0.087529 | 0.099414 | 0.252178 | 0.119656 |
| 32.94444 | 0.35465 | 1.364369 | 0.882078 | 0.133928 | 1.231151 | -0.165349 | 0.128175 | 0.538291 | 0.255497 | -1.052325 | 0.027841 | 0.322376 | 0.412925 | 0.740587 | 0.420261 | 0.073926 | 2.087372 | -0.170451 | 0.109076 | -0.042984 | -0.046581 | 0.590665 | -0.04604 |
| 32.95833 | 0.158152 | 1.47264 | 1.681184 | 0.38756 | -0.114086 | 0.072469 | 0.305115 | 0.845875 | -0.247978 | -0.127436 | -0.009686 | 0.400637 | 0.377779 | 1.131462 | 0.451762 | 0.063337 | 2.871193 | -0.205834 | 0.046368 | -0.106694 | -0.044337 | 0.74275 | -0.1254 |
| 32.97222 | -0.024858 | 1.346019 | 1.624544 | 0.402183 | -0.61039 | 0.445135 | 0.585797 | 1.363233 | -0.932319 | 0.720973 | -0.027709 | 0.474516 | 0.438188 | 1.665875 | 0.708561 | 0.06984 | 2.907266 | -0.125557 | 0.134259 | -0.044135 | 0.013805 | 0.791589 | -0.057136 |
| 32.98611 | 0.501111 | 1.265562 | 1.894039 | 0.3173 | -0.501555 | 0.716031 | 1.209979 | 1.429411 | -1.40599 | 1.311986 | 0.048403 | 0.498284 | 0.427414 | 1.759313 | 1.099674 | 0.064848 | 2.023685 | 0.000736 | 0.186145 | 0.028703 | 0.039247 | 0.792961 | 0.016961 |
| 33 | 0.920558 | 1.050619 | 2.652649 | 0.321279 | -0.172457 | 0.688134 | 1.666619 | 1.988566 | -1.255895 | 1.34726 | 0.113843 | 0.463414 | 0.311683 | 1.232609 | 1.134742 | 0.141439 | 0.987269 | 0.036864 | 0.145037 | -0.047322 | -0.044203 | 0.855944 | -0.050831 |
| 33.01389 | 0.944286 | 0.796345 | 2.467119 | 0.133537 | -0.115351 | 0.38011 | 1.50732 | 1.614955 | -1.202623 | 0.903146 | -0.047003 | 0.399237 | 0.140286 | 0.755339 | 0.9986 | 0.350962 | 0.252195 | -0.051479 | 0.10386 | -0.187012 | -0.295592 | 1.02175 | -0.211018 |
| 33.02778 | 0.232642 | 0.730113 | 1.921733 | 0.002582 | -0.198371 | 0.609255 | 1.024012 | 0.235234 | -1.921582 | 0.888547 | -0.099098 | 0.492672 | 0.088259 | 1.201504 | 0.697691 | 0.237679 | -0.39965 | 0.028734 | 0.107569 | -0.136255 | -0.271593 | 1.046229 | -0.221037 |
| 33.04167 | 0.545441 | 1.139926 | 2.334352 | 0.185617 | 0.785642 | 0.596066 | 0.979753 | -0.350151 | -1.334751 | 1.747174 | 0.158637 | 0.696619 | 0.137186 | 1.762074 | 0.451949 | 0.447602 | -0.954524 | 0.121637 | 0.026317 | -0.152837 | -0.179221 | 0.789457 | -0.239592 |
| 33.05556 | 0.765095 | 1.563895 | 1.856394 | 0.274419 | 2.001428 | 1.84748 | -0.068273 | -0.879063 | 0.606812 | 2.131076 | 0.241197 | 0.910106 | 0.063614 | 1.621167 | 0.226049 | 0.576026 | -1.248913 | 0.009497 | -0.152505 | -0.306504 | -0.330346 | 0.606555 | -0.380383 |
| 33.06944 | 0.509801 | 1.605408 | 1.032308 | 0.142027 | 2.111288 | 0.506937 | -0.860393 | -1.421474 | 1.579289 | 2.738911 | 0.125453 | 0.784446 | 0.031268 | 1.423933 | 0.056043 | 0.54391 | -0.986891 | -0.100807 | -0.196848 | -0.392101 | -0.409289 | 0.629689 | -0.450457 |
| 33.08333 | 0.8754 | 1.449017 | 2.222624 | 0.256069 | 1.845607 | 0.506953 | 1.018901 | -1.372899 | 1.008083 | 3.199818 | 0.194899 | 0.020184 | -0.319317 | 0.824679 | -0.554646 | 0.04626 | -0.857442 | -0.555162 | -0.572904 | -0.85547 | -0.865382 | 0.058424 | -0.876859 |
| 33.09722 | 0.615979 | 1.101651 | 3.114812 | 0.357238 | 1.471194 | 1.016205 | 1.560123 | -1.286458 | -0.415536 | 3.053912 | 0.282012 | -0.976251 | -1.076227 | -0.281215 | -1.857301 | -0.988922 | -1.052889 | -1.497798 | -1.292555 | -1.788181 | -1.870244 | -1.039511 | -1.729744 |
| 33.11111 | 0.451077 | 1.266202 | 1.891214 | 0.155228 | 0.232641 | 1.022887 | 0.80642 | -0.745677 | -2.211739 | 2.387523 | 0.077598 | -0.896142 | -1.126435 | -0.52511 | -2.849957 | -1.128797 | -0.657 | -1.609233 | -1.14582 | -1.733954 | -1.835029 | -1.297868 | -1.71494 |
| 33.125 | 0.108423 | 1.41798 | 0.659079 | -0.168579 | -0.967576 | -0.351132 | -0.16866 | 0.071223 | -2.120688 | 1.675989 | -0.125081 | 0.168743 | -0.154329 | 0.514316 | -2.525962 | 0.40519 | 0.818403 | -0.503813 | -0.264573 | -0.614038 | -0.634106 | -0.2874 | -0.719556 |
| 33.13889 | -0.176456 | 2.195864 | 1.117465 | -0.227626 | -0.83153 | -0.841508 | -0.427936 | 0.835218 | -0.826418 | 1.84274 | -0.090926 | 0.944203 | 0.808415 | 1.520892 | -1.311607 | 1.69006 | 2.045202 | 0.472084 | 0.413948 | 0.129045 | 0.157224 | 0.921095 | 0.057266 |
| 33.15278 | -0.423165 | 2.254639 | 1.269714 | -0.003585 | -1.048433 | -0.013174 | 0.003065 | 0.8154 | -0.318127 | 1.379645 | 0.069837 | 1.228605 | 1.096985 | 1.641165 | -0.464207 | 2.063958 | 1.596713 | 0.803851 | 0.510785 | 0.299569 | 0.349452 | 1.529878 | 0.26891 |
| 33.16667 | -0.447248 | 1.557622 | 0.110069 | 0.774171 | -1.368771 | 0.385372 | 0.477036 | 0.739565 | -0.506409 | 0.739823 | 0.141601 | 1.597707 | 1.018783 | 1.346521 | -0.033628 | 1.817042 | 1.191374 | 0.792329 | 0.616302 | 0.243777 | 0.286348 | 1.283224 | 0.201941 |
| 33.18056 | -0.179486 | 1.036923 | 0.015247 | 1.133999 | 0.07461 | 0.566603 | 0.492689 | 0.533597 | 0.175219 | 0.226418 | 0.19069 | 1.296254 | 0.658588 | 0.956646 | -0.054647 | 1.56554 | 0.83368 | 0.715234 | 0.60315 | 0.132096 | 0.150081 | 1.186192 | 0.054501 |
| 33.19444 | -0.287867 | 0.760241 | -0.694107 | 0.922458 | -0.067389 | 0.706454 | 1.258246 | 0.323872 | 1.875669 | 0.189363 | 0.318953 | 1.07028 | 0.347335 | 0.693817 | -0.375732 | 1.215536 | 0.657096 | 0.591722 | 0.431163 | 0.037924 | 0.040005 | 1.343739 | -0.057895 |
| 33.20833 | -0.693777 | 0.500026 | -0.456312 | -0.254004 | -0.66876 | 0.466 | 1.782249 | 0.284636 | 2.605337 | 0.293319 | 0.393293 | 0.645585 | 0.155104 | 0.507183 | -0.704006 | 1.074357 | 0.30998 | 0.494892 | 0.245289 | -0.020841 | -0.008131 | 1.389849 | -0.088472 |
| 33.22222 | -0.651747 | 0.439572 | -0.071726 | -0.368481 | -0.821577 | 0.203609 | 1.305977 | 0.150714 | 0.854608 | 0.21285 | 0.262384 | 0.481547 | 0.040985 | 0.236446 | -0.901969 | 1.327755 | 0.551236 | 0.401893 | 0.089903 | -0.060319 | -0.048988 | 1.206162 | -0.118269 |
| 33.23611 | -0.386539 | 0.883067 | 0.168619 | 0.36381 | -0.564127 | 0.076419 | 0.533098 | -0.08947 | 0.757112 | 0.026695 | 0.195712 | 0.36207 | 0.044755 | 0.14585 | -1.109082 | 1.250554 | 0.259678 | 0.309847 | -0.010869 | -0.076722 | -0.061758 | 1.058874 | -0.153695 |
| 33.25 | -0.161468 | 1.552601 | 0.30964 | 0.946635 | 0.392477 | 0.19514 | -0.366802 | -0.285448 | 1.126617 | 0.098625 | 0.200514 | 0.257304 | 0.072507 | 0.183865 | -1.236111 | 0.906913 | -0.019067 | 0.203913 | -0.007802 | -0.114627 | -0.079406 | 1.020377 | -0.185097 |
| 33.26389 | 0.151303 | 1.685231 | 0.401671 | 1.136856 | 1.226511 | -0.060447 | -0.599924 | -0.424608 | 0.629623 | 0.061857 | 0.177419 | -0.090153 | -0.056995 | -0.000834 | -1.155958 | 0.624135 | -0.188248 | 0.098857 | 0.026713 | -0.188126 | -0.180262 | 1.258107 | -0.253013 |
| 33.27778 | -0.078555 | 1.222512 | 0.215778 | 0.620241 | 1.268044 | -0.741234 | -0.779037 | -0.730421 | 0.447619 | 0.12554 | 0.108201 | -0.073024 | -0.186462 | -0.235543 | -1.004742 | 0.624388 | -0.317703 | 0.112043 | 0.043338 | -0.206393 | -0.239786 | 1.518471 | -0.288762 |
| 33.29167 | -0.521497 | 0.608953 | 0.287288 | -0.065746 | 1.123544 | -0.996599 | -0.893571 | -0.886893 | 0.397513 | 0.112792 | 0.038463 | 0.123461 | -0.112517 | -0.387448 | -0.846349 | 0.73122 | -0.21054 | 0.211515 | 0.093536 | -0.140207 | -0.161643 | 1.365987 | -0.212506 |
| 33.30556 | -0.473373 | 0.132769 | -0.257135 | -0.560245 | 1.559034 | -0.635339 | -0.907255 | -1.0739 | -0.130341 | 0.076147 | -0.00411 | 0.341174 | 0.062498 | -0.517349 | -0.698755 | 0.492909 | -0.301055 | 0.260762 | 0.135996 | -0.082804 | -0.070276 | 1.075027 | -0.117976 |
| 33.31944 | -0.61724 | -0.153992 | -2.134056 | -0.646536 | 0.477705 | -0.600308 | -0.374228 | -1.329367 | -0.408668 | 0.036181 | 0.00111 | 0.462977 | 0.156904 | -0.648937 | -0.609264 | 0.09687 | -0.735672 | 0.225871 | 0.100333 | -0.103439 | -0.074404 | 0.802319 | -0.115076 |
| 33.33333 | -0.404731 | -0.269773 | -2.6964 | -0.573639 | 1.502214 | -0.083245 | -0.417676 | -0.875791 | -0.295062 | -0.126371 | 0.01124 | 0.3619 | 0.132778 | -0.554863 | -0.643093 | -0.070032 | -0.866605 | 0.175329 | 0.04012 | -0.147742 | -0.139366 | 0.527728 | -0.17589 |
| 33.34722 | 0.020625 | -0.336633 | -1.936105 | -0.317194 | 1.127143 | 0.650265 | 0.131072 | -0.14059 | 0.467849 | -0.315517 | 0.017904 | 0.352561 | 0.085826 | -0.200553 | -0.672365 | 0.041842 | -0.452559 | 0.207917 | 0.080853 | -0.126943 | -0.146788 | 0.342967 | -0.195754 |
| 33.36111 | 0.204384 | -0.153906 | -0.535064 | 0.084622 | 1.202185 | 1.096279 | 1.127611 | 0.249152 | 1.26326 | -0.362441 | 0.032584 | 0.236808 | 0.121246 | 0.151805 | -0.528828 | 0.218138 | 0.568456 | 0.332934 | 0.144559 | -0.046893 | -0.063358 | 0.284982 | -0.140521 |
| 33.375 | 0.798288 | 0.095312 | 0.284711 | 0.627883 | 0.283571 | 1.360288 | 1.721826 | 0.519413 | 0.681601 | -0.244135 | 0.045616 | -0.019519 | 0.142556 | 0.299636 | -0.226699 | 0.213021 | 0.976862 | 0.401609 | 0.268813 | 0.022879 | 0.021199 | 0.167765 | -0.074113 |
| 33.38889 | 1.244906 | 0.131444 | 0.606056 | 0.566127 | -1.086714 | 1.438586 | 0.976521 | 0.79182 | -0.147204 | -0.198872 | 0.054327 | -0.21403 | 0.080365 | 0.275938 | -0.028886 | 0.109517 | 0.54322 | 0.407314 | 0.257041 | 0.031304 | 0.0548 | 0.001771 | -0.060525 |
| 33.40278 | 1.252647 | 0.112683 | 0.932316 | 0.208323 | -1.288501 | 0.991844 | 0.120932 | 1.001841 | -0.592115 | -0.2785 | 0.060255 | -0.312879 | -0.127466 | 0.173708 | 0.131775 | -0.076999 | 0.520218 | 0.326456 | 0.17034 | -0.028284 | 0.018332 | -0.15509 | -0.107006 |
| 33.41667 | 0.682285 | 0.276992 | 0.657862 | -0.034963 | -0.735605 | 0.746162 | -0.512581 | 1.147919 | -0.613191 | -0.47075 | -0.016817 | -0.682918 | -0.405643 | -0.062781 | 0.24591 | 0.017422 | 0.440234 | 0.219074 | 0.050779 | -0.095261 | -0.083642 | -0.182707 | -0.170651 |
| 33.43056 | 0.353962 | 0.460195 | 0.672636 | -0.660068 | -1.673695 | -0.364986 | -0.574818 | 0.901407 | -0.362898 | -0.401401 | -0.051607 | -0.744135 | -0.580472 | -0.30074 | 0.383243 | 0.421258 | 0.274933 | 0.15488 | -0.020887 | -0.092952 | -0.130732 | -0.160273 | -0.179476 |
| 33.44444 | 0.040348 | 0.552029 | 0.752132 | -1.041668 | -2.977697 | -0.346935 | -0.265838 | 1.124071 | -0.440979 | -0.125918 | 0.06482 | -0.718552 | -0.493622 | -0.429785 | 0.554574 | 0.686698 | 0.189154 | 0.187279 | -0.014135 | -0.030536 | -0.063437 | -0.1692 | -0.121125 |
| 33.45833 | 0.007904 | 0.563294 | 0.861815 | -1.087967 | -2.072928 | -0.616933 | -0.518038 | 1.290801 | -1.176371 | 0.028121 | 0.071101 | -0.737295 | -0.316036 | -0.546282 | 0.798804 | 0.480942 | 0.096127 | 0.182576 | 0.032127 | 0.015321 | 0.007053 | -0.221412 | -0.072892 |
| 33.47222 | -0.098447 | 0.298106 | 0.054413 | -1.074743 | -0.726906 | -1.266215 | -1.006139 | 1.201639 | -2.395369 | -0.154287 | -0.119505 | -0.615836 | -0.212153 | -0.6275 | 0.976426 | 0.031586 | -0.400148 | 0.118427 | 0.104241 | 0.002371 | 0.005645 | -0.312898 | -0.07298 |
| 33.48611 | -0.082777 | -0.056389 | -0.743204 | -0.824486 | -0.095592 | -1.773574 | -0.631113 | 0.593861 | -2.924032 | -0.272573 | -0.227481 | -0.33841 | -0.273507 | -0.843895 | 1.129107 | -0.363609 | -0.64813 | 0.015955 | 0.0912 | -0.046609 | -0.051412 | -0.358923 | -0.118163 |
| 33.5 | 0.111295 | -0.082996 | 0.410842 | -0.524798 | 1.291348 | -1.453613 | -0.103045 | 0.04517 | -2.00359 | -0.090303 | -0.075311 | -0.170455 | -0.415372 | -1.205706 | 1.23331 | -0.657892 | -0.743851 | -0.04714 | -0.063798 | -0.091913 | -0.109227 | -0.406186 | -0.163465 |
| 33.51389 | 0.297718 | -0.084811 | 1.529752 | -0.524079 | 2.104473 | -1.227787 | 0.6558 | -0.384403 | -0.38183 | -0.055438 | 0.037181 | -0.244462 | -0.446409 | -1.484888 | 1.164438 | -0.839904 | -0.443296 | -0.025864 | -0.166146 | -0.081412 | -0.095509 | -0.378554 | -0.144692 |
| 33.52778 | 0.319553 | -0.243178 | 0.40608 | -0.615332 | 0.649189 | -1.045901 | 0.912361 | -1.352818 | 1.594243 | -0.473118 | -0.080127 | -0.459658 | -0.324133 | -1.320834 | 1.103125 | -0.69023 | -0.435239 | 0.041139 | -0.136398 | -0.006656 | -0.006187 | -0.262541 | -0.051909 |
| 33.54167 | -0.173629 | -0.483713 | -1.976582 | -0.463941 | 0.685497 | -0.775465 | 0.482405 | -1.50686 | 2.933905 | -1.36752 | -0.184476 | -0.658351 | -0.209954 | -0.794409 | 1.022897 | -0.467684 | -0.459282 | 0.081686 | -0.084846 | 0.087568 | 0.097277 | -0.162674 | 0.071885 |
| 33.55556 | -0.893174 | -0.155458 | -4.001429 | -0.416402 | 0.98964 | -0.45122 | 0.434924 | -0.812462 | 1.436113 | -2.341889 | -0.003429 | -0.660671 | -0.025935 | -0.161857 | 1.142791 | -0.280585 | -0.259967 | 0.162234 | -0.033809 | 0.228197 | 0.245971 | -0.083507 | 0.265353 |
| 33.56944 | -1.089799 | 0.01027 | -4.985117 | -0.374541 | 0.225745 | -0.266112 | 0.749632 | -0.673177 | -0.014465 | -3.886929 | 0.321815 | -0.526146 | 0.29476 | 0.407585 | 1.514276 | -0.593327 | -0.371434 | 0.346056 | 0.146112 | 0.452073 | 0.518898 | 0.011282 | 0.531524 |
| 33.58333 | -0.12755 | -0.480111 | -5.822556 | -0.719928 | -0.635276 | -0.348822 | 0.824376 | -0.616319 | -1.246367 | -5.562281 | 0.282135 | -0.734234 | 0.227851 | 0.292863 | 1.501818 | -1.816265 | -0.572395 | 0.160604 | 0.057406 | 0.486968 | 0.618367 | -0.193757 | 0.522435 |
| 33.59722 | -1.315691 | -1.454305 | -5.656646 | -1.321584 | -1.421805 | -0.742514 | 0.168312 | -1.221073 | -0.14165 | -5.715207 | -0.120878 | -1.084777 | -0.414247 | -0.698187 | 0.930376 | -2.491611 | -1.016775 | -0.351134 | -0.492299 | 0.159443 | 0.165105 | -0.808131 | 0.069804 |
| 33.61111 | -2.952824 | -2.106027 | -4.708975 | -1.448603 | -1.243878 | -0.956142 | -0.727567 | -1.364474 | 2.207868 | -4.869885 | -0.345162 | -1.170518 | -0.81637 | -1.499926 | 0.674088 | -3.131824 | -1.644233 | -0.710725 | -0.65192 | -0.091507 | -0.057626 | -1.462645 | -0.227278 |
| 33.625 | -3.352144 | -2.134164 | -3.977766 | -0.901655 | -1.278213 | -0.855716 | -0.598521 | -1.440824 | 2.512909 | -4.809222 | -0.442356 | -1.008167 | -0.554226 | -1.091831 | 0.852838 | -3.270658 | -1.837906 | -0.54412 | -0.363849 | -0.0773 | -0.000936 | -1.748401 | -0.164462 |
| 33.63889 | -1.745825 | -1.471625 | -4.655365 | -0.248975 | -2.89012 | -0.854958 | 0.280365 | -1.094741 | 1.490966 | -4.762559 | -0.339414 | -0.733288 | -0.563123 | -0.696178 | 1.220893 | -2.444305 | -1.933921 | -0.367061 | -0.17041 | -0.037653 | -0.00308 | -1.649531 | -0.040044 |
| 33.65278 | -0.334924 | -0.530069 | -3.529861 | 0.225363 | -3.06192 | -1.336721 | 0.848906 | -0.65725 | 0.674919 | -5.278151 | -0.056543 | -0.396531 | -0.838078 | -0.75858 | 1.321984 | -1.77819 | -1.61775 | -0.304433 | -0.000024 | -0.046591 | -0.050944 | -1.398814 | -0.012495 |
| 33.66667 | 0.233718 | -0.206859 | -1.976602 | 0.269245 | -2.406319 | -0.751417 | 0.680534 | -0.196172 | 0.388857 | -5.444855 | 0.018789 | -0.035749 | -0.826231 | -0.983921 | 1.049833 | -1.48065 | -0.939375 | -0.235696 | 0.107092 | -0.052946 | -0.070194 | -0.919161 | -0.05046 |
| 33.68056 | 0.141925 | -0.635749 | -1.087199 | 0.160544 | -1.872234 | -0.37989 | 0.49026 | 0.032284 | 0.062051 | -3.708335 | -0.155817 | -0.00239 | -0.799467 | -1.847429 | 0.694561 | -1.156708 | -0.553747 | -0.199962 | 0.056035 | -0.061436 | -0.114015 | -0.72312 | -0.095398 |
| 33.69444 | -0.031171 | -0.538395 | -0.341392 | -0.089459 | -1.194331 | 0.315078 | 1.099978 | 0.410945 | 0.108949 | -2.335596 | -0.264316 | -0.212416 | -0.799087 | -2.91919 | 0.729253 | -1.337146 | -0.712957 | -0.167515 | -0.014136 | -0.055407 | -0.11576 | -0.499813 | -0.099942 |
| 33.70833 | 0.093328 | 0.062566 | 0.700818 | -0.118607 | 0.435391 | 0.258 | 1.086539 | 0.975668 | 0.256887 | -1.221716 | -0.11722 | -0.195858 | -0.690844 | -3.168787 | 0.524698 | -1.451893 | -0.624129 | -0.136656 | 0.011648 | -0.037019 | -0.031756 | -0.122159 | -0.066327 |
| 33.72222 | 0.320686 | 0.454334 | 1.754481 | 0.234373 | 2.313765 | 0.330436 | 0.602474 | 1.442779 | -0.179497 | -0.200289 | 0.024429 | -0.138207 | -0.521678 | -1.803623 | 0.325357 | -1.174146 | -0.317898 | -0.083216 | 0.110091 | -0.011924 | 0.041173 | 0.026368 | -0.024277 |
| 33.73611 | 0.348134 | 0.303182 | 2.607252 | 0.463725 | 3.39858 | 0.505123 | 0.086152 | 1.749563 | -0.979828 | 1.08784 | 0.083684 | -0.206274 | -0.392573 | -0.914317 | 0.216243 | -0.77524 | -0.050358 | 0.007203 | 0.134238 | 0.021371 | 0.053208 | -0.320257 | 0.007038 |
| 33.75 | 0.626819 | -0.203513 | 3.020876 | 0.386314 | 3.416207 | 0.239142 | -0.059796 | 1.96687 | -1.458896 | 2.766784 | 0.062964 | -0.187735 | -0.174285 | -0.096184 | 0.357736 | -0.633339 | 0.466381 | 0.124035 | 0.140086 | 0.091724 | 0.110585 | -0.663479 | 0.055553 |
| 33.76389 | 0.812329 | -0.856301 | 3.56491 | 0.142052 | 4.789252 | -0.197683 | -0.300287 | 1.867786 | -0.268983 | 4.301125 | 0.070239 | -0.117309 | 0.120311 | 0.664606 | 0.606967 | -0.592231 | 1.226838 | 0.19342 | 0.183021 | 0.167724 | 0.199195 | -0.690386 | 0.126056 |
| 33.77778 | 0.708286 | -1.375884 | 3.895651 | 0.082366 | 4.797903 | -0.677715 | -0.368898 | 1.515366 | 2.09565 | 4.606522 | 0.104018 | -0.224571 | 0.266724 | 1.226896 | 0.613878 | -0.558326 | 1.625879 | 0.132759 | 0.166617 | 0.182793 | 0.212656 | -0.530345 | 0.152668 |
| 33.79167 | 0.417828 | -1.725314 | 3.49016 | -0.027855 | 4.303809 | -0.717353 | -0.212679 | 1.271317 | 1.678596 | 3.647138 | 0.099977 | -0.367759 | 0.27302 | 1.656374 | 0.566841 | -0.18241 | 1.441715 | 0.033862 | 0.122335 | 0.223273 | 0.212918 | -0.463294 | 0.211638 |
| 33.80556 | 0.272366 | -1.697704 | 2.475531 | -0.160238 | 3.240791 | -0.267646 | -0.077229 | 0.901947 | 0.610944 | 2.105728 | 0.080953 | -0.401915 | 0.283785 | 2.030759 | 0.632502 | 0.173748 | 1.252084 | -0.014249 | 0.031742 | 0.269842 | 0.232476 | -0.371688 | 0.289741 |
| 33.81944 | 0.249447 | -1.344144 | 1.153829 | -0.124262 | 1.617609 | 0.375916 | 0.508733 | 0.671303 | -0.632229 | 0.693467 | 0.074662 | -0.231167 | 0.263375 | 2.297553 | 0.532251 | 0.158098 | 0.999835 | -0.185026 | -0.118774 | 0.126154 | 0.167687 | -0.219223 | 0.140803 |
| 33.83333 | -0.091423 | -0.833608 | -0.431611 | 0.010549 | -0.079768 | 1.099655 | 0.982499 | 0.436122 | -2.49588 | -0.769469 | 0.033172 | -0.112642 | 0.173673 | 1.863466 | 0.376684 | -0.055276 | 0.534146 | -0.340384 | -0.194746 | -0.063543 | 0.045282 | -0.109762 | -0.069177 |
| 33.84722 | -0.492817 | -0.362283 | -0.901654 | 0.133443 | -1.280655 | 1.367119 | 1.076981 | 0.414456 | -2.62285 | -2.130468 | 0.083408 | -0.055585 | 0.028261 | 1.297595 | 0.434982 | -0.004922 | 0.086834 | -0.228812 | -0.031487 | 0.021355 | 0.114056 | -0.017781 | 0.008964 |
| 33.86111 | -0.624585 | 0.069645 | -0.907607 | 0.288208 | -2.44665 | 0.947945 | 0.321094 | 0.506364 | -0.418989 | -2.703348 | 0.122079 | 0.133677 | -0.296577 | 0.957293 | 0.713015 | 0.263669 | -0.419328 | 0.081817 | 0.298883 | 0.268972 | 0.35762 | 0.24694 | 0.283639 |
| 33.875 | -0.605514 | 0.349268 | -1.923038 | 0.294811 | -3.474041 | 0.674677 | -0.057688 | 0.579962 | -0.156559 | -3.206291 | 0.173355 | 0.410274 | -0.282999 | 0.763772 | 0.96112 | 0.399447 | -0.916441 | 0.341234 | 0.607824 | 0.515683 | 0.60315 | 0.555436 | 0.533178 |
| 33.88889 | -0.814366 | 0.364815 | -2.74858 | -0.410541 | -2.522586 | 0.701266 | -0.016453 | 0.58989 | -0.904676 | -4.253253 | 0.164683 | 0.723784 | 0.500739 | 0.662657 | 1.284899 | 0.648246 | -0.770131 | 0.625982 | 0.913923 | 0.821999 | 0.919294 | 0.848755 | 0.8166 |
| 33.90278 | -0.55804 | 0.08823 | -2.185807 | -1.351509 | -1.34986 | 0.356298 | -0.233522 | 0.559053 | -1.011541 | -5.430051 | 0.084568 | 0.842125 | 0.662919 | 0.352172 | 1.324985 | 0.716432 | -0.44682 | 0.66025 | 0.97909 | 0.86121 | 0.979554 | 0.885697 | 0.867634 |
| 33.91667 | -0.070277 | -0.24219 | -1.248152 | -1.752882 | -0.72142 | 0.27387 | -0.301946 | 0.129877 | -0.187875 | -4.310756 | -0.143021 | 0.341029 | -0.032995 | -0.601303 | 0.728558 | 0.201983 | -0.37751 | 0.02444 | 0.398071 | 0.216304 | 0.288153 | 0.383213 | 0.232483 |
| 33.93056 | 0.662043 | -0.193252 | -0.114927 | -0.308791 | 0.248186 | 0.907621 | 0.14941 | -0.126056 | 0.355077 | -2.056713 | -0.264532 | -0.329067 | -0.803752 | -1.397192 | -0.034106 | -0.427035 | -0.044502 | -0.752706 | -0.326534 | -0.565869 | -0.532135 | -0.166755 | -0.58999 |
| 33.94444 | 1.647568 | 0.293588 | 1.307587 | 1.850096 | 1.636948 | 0.828344 | 0.519137 | -0.109117 | -0.176484 | -0.454804 | -0.039093 | -0.532782 | -0.856203 | -1.440614 | -0.56504 | -0.640424 | 0.308366 | -1.007867 | -0.594256 | -0.865058 | -0.82353 | -0.217345 | -0.92949 |
| 33.95833 | 1.34011 | 0.605369 | 2.503413 | 2.870773 | 2.259786 | -0.952112 | -0.379595 | -0.10218 | -0.911016 | 0.893209 | -0.058646 | -0.451505 | -0.360082 | -1.121702 | -0.832055 | -0.751065 | 0.686661 | -0.899911 | -0.518877 | -0.852425 | -0.81591 | -0.004485 | -0.920368 |
| 33.97222 | 1.031551 | 0.645644 | 3.408304 | 1.942143 | 1.565468 | -0.676412 | -0.026266 | 0.022812 | -0.804224 | 2.09741 | 0.039926 | -0.330111 | 0.137811 | -0.526184 | -0.918607 | -0.726394 | 0.646987 | -0.757289 | -0.392601 | -0.80667 | -0.799425 | -0.132866 | -0.869964 |
| 33.98611 | 0.770462 | 0.704346 | 4.124921 | 0.787272 | 2.781319 | -0.467143 | -0.203853 | 0.548827 | -0.097761 | 3.415491 | 0.1411 | -0.143552 | 0.429795 | 0.248619 | -0.87853 | -0.530888 | 0.431213 | -0.672669 | -0.357184 | -0.719602 | -0.763466 | -0.284268 | -0.790429 |
| 34 | -0.430391 | 0.711198 | 4.103278 | 0.424091 | 3.571529 | -0.119307 | -0.273062 | 0.986254 | 0.626132 | 3.59321 | 0.134518 | 0.030742 | 0.396377 | 0.852297 | -0.815647 | -0.213534 | 0.686219 | -0.546179 | -0.310833 | -0.609712 | -0.713814 | -0.23427 | -0.692175 |
| 34.01389 | -0.56771 | 0.653546 | 2.991861 | 0.142596 | 2.593473 | -0.109806 | -0.008398 | 1.210316 | 0.247466 | 3.353267 | -0.056699 | 0.149408 | 0.198045 | 0.880823 | -0.723297 | 0.074478 | 1.136299 | -0.473993 | -0.236144 | -0.646107 | -0.814589 | -0.034124 | -0.719001 |
| 34.02778 | 1.687713 | 0.598197 | 3.302526 | 0.591471 | 1.010686 | 0.141043 | 0.193125 | 1.328413 | -0.681914 | 2.998387 | 0.128048 | 0.408462 | -0.042021 | 0.501955 | -0.487128 | 0.079104 | 1.313291 | -0.331283 | -0.061825 | -0.551203 | -0.666448 | 0.428637 | -0.610089 |
| 34.04167 | 2.174158 | 1.079897 | 2.542803 | 1.245212 | 0.596614 | -0.366086 | 0.161468 | 1.090028 | -0.574542 | 3.31059 | 0.508764 | 0.738578 | -0.08895 | 0.131529 | -0.038955 | 0.47884 | 1.200991 | -0.042404 | 0.207955 | -0.312774 | -0.335668 | 0.679246 | -0.365729 |
| 34.05556 | 1.933504 | 1.614232 | -0.133198 | 1.04044 | 0.573179 | -0.587905 | 0.540201 | 0.610555 | 0.134274 | 4.015406 | 0.628005 | 0.940688 | -0.241939 | -0.012102 | 0.277485 | 0.749794 | 1.005099 | 0.142979 | 0.390934 | -0.131227 | -0.171949 | 0.711749 | -0.183594 |
| 34.06944 | 0.553062 | 1.754676 | -0.476546 | 0.471637 | 0.277094 | -0.192175 | 0.884021 | 0.067393 | 0.387691 | 4.556641 | 0.552779 | 0.951883 | -0.317972 | 0.352419 | 0.593609 | 0.876546 | 0.500643 | 0.311483 | 0.472898 | 0.028701 | 0.002898 | 0.629745 | -0.008506 |
| 34.08333 | 0.534408 | 1.727551 | 1.775749 | 0.334525 | 0.579343 | 0.170502 | 0.600015 | -0.149014 | 0.446554 | 4.263775 | 0.643858 | 0.727008 | -0.339707 | 0.405003 | 0.622657 | 0.665538 | 0.440134 | 0.27626 | 0.322892 | -0.07996 | -0.085506 | 0.30526 | -0.042303 |
| 34.09722 | 0.421985 | 1.81914 | 4.415522 | 0.42411 | 1.59438 | -0.113334 | -0.230216 | -0.399356 | 0.523509 | 3.722882 | 0.656554 | -0.002464 | -0.874337 | -1.184132 | -0.165932 | -0.134258 | -0.500982 | -0.261642 | -0.335913 | -0.729757 | -0.715659 | -0.69016 | -0.616497 |
| 34.11111 | 0.255043 | 2.072588 | 3.850655 | 0.342926 | 1.61897 | 0.237799 | -0.715958 | -0.791893 | 0.028848 | 2.98148 | 0.125779 | -0.844883 | -1.761229 | -3.525785 | -1.649848 | -1.232317 | -2.511661 | -1.028837 | -1.111017 | -1.427963 | -1.432995 | -2.331086 | -1.358926 |
| 34.125 | 0.852038 | 2.353891 | 2.293992 | 0.388246 | 1.499231 | 0.418328 | -0.682035 | -0.446413 | -0.595657 | 1.484972 | -0.168567 | -0.945955 | -1.874876 | -3.162795 | -2.806587 | -1.160569 | -2.330516 | -0.997345 | -1.101548 | -1.272532 | -1.27393 | -4.17449 | -1.32105 |
| 34.13889 | 0.873455 | 2.638097 | 1.605688 | 0.698163 | 1.193211 | -0.251218 | -0.033994 | 0.744248 | -0.492244 | 0.771329 | -0.056427 | 0.245157 | -0.425979 | -1.487003 | -3.10138 | 0.374352 | -1.185434 | -0.095526 | -0.348338 | -0.537333 | -0.496578 | -4.164699 | -0.618269 |
| 34.15278 | 0.115047 | 2.279961 | 0.767672 | 0.973617 | 0.655455 | -0.977559 | 0.45865 | 1.586549 | -0.032058 | -0.047226 | 0.228727 | 1.128655 | -0.191039 | -2.594023 | -2.744671 | 1.33724 | -1.033287 | 0.605993 | 0.1412 | -0.102995 | -0.089108 | -2.449052 | -0.17847 |
| 34.16667 | -0.382688 | 1.84559 | 0.399443 | 0.862193 | -0.113986 | -0.388929 | 0.820551 | 1.680098 | -0.123961 | -0.296228 | 0.46496 | 1.30649 | -0.396282 | -2.298512 | -1.79264 | 1.043101 | -0.960444 | 0.651967 | 0.152842 | -0.042609 | -0.052568 | -1.974016 | -0.126431 |
| 34.18056 | -0.093709 | 0.445461 | 0.023276 | 0.317243 | -0.313459 | 0.609256 | 1.443755 | 1.651004 | -0.301892 | -0.285642 | 0.69551 | 1.450155 | -0.383529 | -1.377631 | -0.635113 | 1.212504 | -0.312016 | 0.52485 | 0.397303 | -0.005054 | -0.015991 | -1.590924 | -0.051264 |
| 34.19444 | -0.280403 | -0.564364 | 0.858022 | 0.537782 | -0.469683 | 0.554585 | 1.124381 | 1.446072 | -0.004798 | -0.219074 | 0.869157 | 1.160197 | -0.283101 | -0.919138 | 0.192628 | 1.526166 | 0.862344 | 0.568133 | 0.404643 | 0.075259 | 0.062429 | -0.653607 | -0.178738 |
| 34.20833 | -0.165201 | -0.861248 | 1.223387 | 0.685947 | -0.235455 | 0.508297 | 0.607466 | 1.124399 | 0.021701 | -0.112835 | 0.933376 | 1.122667 | -0.131781 | -0.563599 | 0.597666 | 1.760263 | 2.460356 | 0.678079 | 0.493131 | 0.159473 | 0.141985 | 0.317398 | -0.163725 |
| 34.22222 | -0.293565 | -0.286778 | 1.39946 | 0.772703 | -0.041398 | 0.272223 | 0.180599 | 0.770757 | -0.276738 | 0.134085 | 0.960058 | 1.335907 | -0.07301 | -0.298612 | 0.77119 | 1.972551 | 2.870152 | 0.763026 | 0.536845 | 0.202589 | 0.197157 | 1.127469 | -0.272829 |
| 34.23611 | 0.199006 | 0.78396 | 2.132532 | 1.455227 | -0.095355 | -0.069894 | -0.136514 | 0.284228 | -0.512619 | 0.363993 | 0.884427 | 1.452477 | -0.150027 | -0.157293 | 0.761548 | 2.326559 | 2.27585 | 0.81308 | 0.500183 | 0.246254 | 0.252782 | 1.410617 | -0.462375 |
| 34.25 | 0.098391 | 1.942751 | 2.95544 | 1.55861 | -0.284842 | -0.498391 | 0.076376 | -0.00148 | 0.038788 | 0.393382 | 0.690155 | 1.649938 | -0.127105 | -0.14005 | 0.602128 | 2.542598 | 1.749871 | 0.91402 | 0.577535 | 0.368906 | 0.390143 | 1.684718 | -0.419753 |
| 34.26389 | -0.123764 | 2.661412 | 2.46249 | 1.412922 | -0.377415 | 1.114252 | 0.149315 | -0.067645 | 0.835523 | 0.356345 | 0.490228 | 2.060501 | 0.058136 | -0.243501 | 0.460864 | 2.545734 | 0.944937 | 0.920772 | 0.68835 | 0.489311 | 0.532024 | 2.295413 | -0.150903 |
| 34.27778 | -0.739076 | 2.593191 | 0.997088 | 1.360766 | -0.52642 | 2.093093 | 0.489228 | 0.07043 | 0.636177 | 0.23562 | 0.2988 | 2.115485 | 0.206316 | -0.404857 | 0.400586 | 2.367764 | 1.368279 | 0.837646 | 0.755786 | 0.537028 | 0.577041 | 2.554208 | 0.176173 |
| 34.29167 | -0.867116 | 2.017831 | 0.413252 | 1.446033 | -0.522286 | 0.801223 | 0.418761 | -0.028363 | 0.271546 | 0.332389 | 0.123876 | 1.39841 | 0.378347 | -0.438008 | 0.341035 | 1.930107 | 1.162003 | 0.822597 | 0.769726 | 0.560142 | 0.583298 | 2.482803 | 0.444676 |
| 34.30556 | -0.369379 | 1.364574 | -0.210266 | 2.007181 | -0.101012 | -0.76117 | 0.028154 | -0.361066 | 1.156603 | 0.334203 | -0.008126 | 1.030584 | 0.433165 | -0.478792 | 0.245396 | 1.221902 | 0.741158 | 0.856541 | 0.781365 | 0.572326 | 0.582255 | 2.640733 | 0.749049 |
| 34.31944 | -0.24778 | 0.772534 | -1.802683 | 2.315629 | 0.331965 | -1.269126 | -0.288914 | -0.52012 | 0.555608 | 0.295497 | -0.15424 | 0.791376 | 0.413346 | -0.471254 | 0.130993 | 0.510168 | 0.335824 | 0.86583 | 0.798036 | 0.539384 | 0.53887 | 2.366669 | 1.048661 |
| 34.33333 | -1.010607 | 0.312449 | -3.610414 | 1.417242 | 0.771693 | -2.660811 | -0.498004 | -0.443141 | -0.532894 | 0.179489 | -0.278341 | 0.708962 | 0.666303 | -0.519473 | 0.074697 | 0.221615 | -0.062517 | 0.814095 | 0.72743 | 0.462838 | 0.471045 | 1.802672 | 1.217959 |
| 34.34722 | -1.021008 | 0.158744 | -3.212779 | 1.310811 | 0.817228 | -2.209827 | -0.376625 | -0.567378 | 0.844752 | 0.152015 | -0.242878 | 0.587551 | 0.716661 | -0.668169 | -0.021867 | 0.356935 | -0.157589 | 0.551059 | 0.585182 | 0.325821 | 0.354337 | 1.808968 | 1.103504 |
| 34.36111 | -1.273352 | 0.313167 | -2.105193 | 1.676744 | 1.300393 | -1.794876 | 0.050567 | -0.558047 | 0.835245 | 0.115735 | -0.145705 | 0.42444 | 0.419018 | -0.86509 | -0.127582 | 0.318192 | -0.445655 | 0.117968 | 0.335306 | 0.144855 | 0.163046 | 1.151557 | 0.223533 |
| 34.375 | -1.629798 | 0.438379 | -1.622601 | 0.867354 | 1.461444 | -0.98451 | 0.611841 | -0.324667 | -0.98776 | -0.064387 | -0.213261 | 0.24001 | 0.206837 | -1.020563 | -0.313103 | -0.063754 | -0.755272 | -0.175011 | 0.110613 | 0.019316 | 0.008419 | 0.801419 | -0.034556 |
| 34.38889 | -1.729075 | 0.445197 | -0.697132 | 0.261284 | 1.65656 | -0.59111 | 1.813875 | 0.060705 | -0.858319 | -0.219505 | -0.31546 | 0.019148 | 0.143113 | -1.037469 | -0.574615 | -0.367836 | -1.515319 | -0.234733 | 0.022684 | -0.019834 | -0.035897 | 0.886366 | -0.048489 |
| 34.40278 | -2.633933 | 0.581985 | 0.398874 | 0.164663 | 0.949285 | -0.370507 | 1.808754 | 0.558259 | -0.072444 | -0.208701 | -0.187851 | -0.199144 | 0.021964 | -1.002324 | -0.889606 | -0.603563 | -2.424127 | -0.143443 | 0.012357 | -0.033801 | -0.041994 | 0.684475 | -0.068595 |
| 34.41667 | -2.820233 | 0.870249 | 1.212192 | 0.331882 | 0.493084 | -0.427334 | 0.805686 | 0.805233 | -0.630615 | -0.213128 | -0.07112 | -0.425507 | -0.095716 | -0.903744 | -1.115908 | -0.560593 | -2.233816 | -0.023612 | 0.022667 | -0.047592 | -0.048754 | 0.396081 | -0.075653 |

| | | | | | | | | | | | | | | | | | | | | | | | |
|---|---|---|---|---|---|---|---|---|---|---|---|---|---|---|---|---|---|---|---|---|---|---|---|
| 34.43056 | -3.050915 | 1.033581 | 1.492721 | 0.353537 | 0.355623 | -0.180331 | 0.415489 | 0.856984 | -0.698558 | -0.281301 | 0.009112 | -0.602333 | -0.181432 | -0.818896 | -1.188304 | -0.013791 | -1.421235 | 0.075518 | 0.060042 | -0.037678 | -0.036181 | 0.24767 | -0.10867 |
| 34.44444 | -2.356016 | 0.97558 | 0.940501 | 0.16235 | -0.941725 | 0.456896 | 0.065068 | 0.803848 | 0.384362 | -0.515358 | 0.039488 | -0.698241 | -0.25083 | -0.67262 | -1.029355 | 0.586113 | -0.405641 | 0.101592 | 0.088526 | -0.002087 | -0.005495 | 0.171612 | -0.150565 |
| 34.45833 | -1.109587 | 0.399391 | 0.211951 | -0.090339 | -1.581868 | 0.51685 | 0.335807 | 0.722602 | 0.502625 | -0.840042 | -0.108951 | -0.71834 | -0.265514 | -0.488462 | -0.735948 | 0.896972 | 0.322021 | 0.090555 | 0.092174 | 0.034213 | 0.031954 | 0.042004 | -0.099442 |
| 34.47222 | -0.309833 | 0.340479 | -0.390697 | -0.158065 | -1.237167 | 0.529303 | 0.31322 | 0.772773 | -0.271777 | -0.619117 | -0.223153 | -0.857606 | -0.273566 | -0.340719 | -0.439127 | 0.612695 | 0.596529 | 0.108193 | 0.106833 | 0.048151 | 0.060482 | -0.087931 | -0.03831 |
| 34.48611 | 0.137926 | 0.208536 | -0.27699 | -0.14818 | -1.151312 | 0.241335 | -0.22303 | 0.7447 | -0.338108 | -0.272641 | -0.247496 | -1.054815 | -0.304804 | -0.255796 | -0.218986 | 0.167341 | 0.392293 | 0.091821 | 0.129085 | 0.051632 | 0.074454 | -0.154668 | -0.003175 |
| 34.5 | 0.235391 | 0.001568 | -0.402386 | -0.18599 | -0.440909 | -0.071134 | -0.463259 | 0.591764 | -0.702923 | 0.003835 | -0.286573 | -1.263377 | -0.357501 | -0.244003 | -0.089316 | -0.201154 | 0.31194 | 0.01057 | 0.099275 | 0.034478 | 0.056902 | -0.201002 | -0.024057 |
| 34.51389 | -0.303883 | -0.234719 | -1.284672 | -0.410892 | 0.183609 | -0.57775 | -0.557377 | 0.190247 | -0.936174 | -0.04931 | -0.427345 | -1.378001 | -0.476669 | -0.303766 | -0.027043 | -0.338451 | 0.214837 | -0.133167 | -0.014247 | -0.016122 | 0.001934 | -0.239951 | -0.092996 |
| 34.52778 | -0.900182 | -0.476326 | -2.469782 | -0.554921 | 0.62963 | -0.74617 | -0.828842 | -0.376254 | -0.214135 | -0.172027 | -0.400947 | -1.421902 | -0.524468 | -0.317339 | 0.07003 | -0.454105 | 0.1837 | -0.295429 | -0.222 | -0.036619 | -0.022551 | -0.251264 | -0.138284 |
| 34.54167 | -1.29033 | -1.073995 | -2.704953 | -0.505575 | 0.71521 | -0.690026 | -0.767269 | -0.708353 | -0.515445 | -0.11915 | -0.135964 | -1.181331 | -0.490558 | -0.24635 | 0.20636 | -0.772306 | 0.655991 | -0.43753 | -0.35801 | 0.007512 | 0.025842 | -0.256709 | -0.12195 |
| 34.55556 | -0.950436 | -0.458355 | -2.61589 | -0.530705 | 1.675951 | -0.516968 | -0.59494 | -0.71059 | -0.891642 | -0.279926 | -0.002508 | -0.805897 | -0.302686 | -0.062765 | 0.379176 | -1.034947 | 0.486094 | -0.432609 | -0.290136 | 0.12659 | 0.1495 | -0.191447 | -0.005883 |
| 34.56944 | 0.201218 | 0.394609 | -3.433383 | -0.610095 | 2.50554 | -0.418041 | -0.824169 | -0.552974 | -0.100329 | -1.096598 | 0.126814 | -0.696718 | -0.033832 | 0.15923 | 0.617423 | -1.018885 | 0.251726 | -0.047895 | -0.041056 | 0.308522 | 0.326769 | -0.119816 | 0.171625 |
| 34.58333 | 0.351481 | 0.527826 | -4.00071 | -0.833986 | 2.55931 | -0.295589 | -0.873353 | -0.671055 | 0.345603 | -2.749936 | 0.071184 | -0.794366 | -0.004697 | 0.072474 | 0.583058 | -1.326987 | -0.132069 | 0.014287 | 0.098205 | 0.330868 | 0.341924 | -0.244356 | 0.075954 |
| 34.59722 | -0.125741 | -0.612435 | -3.88601 | -1.251746 | 1.358569 | -0.623466 | -1.113782 | -1.432769 | 0.027449 | -4.605221 | -0.493151 | -0.744526 | -0.307471 | -0.27459 | 0.242116 | -2.117299 | -0.820006 | -0.482842 | -0.274164 | -0.002515 | 0.067821 | -0.457909 | -0.355048 |
| 34.61111 | -0.249002 | -1.586443 | -2.992924 | -1.218383 | 0.241905 | -0.449525 | -0.645795 | -2.266549 | -0.105981 | -5.199663 | -0.727747 | -0.690739 | -0.364675 | -0.361545 | 0.446371 | -2.516767 | -1.261119 | -0.655718 | -0.491183 | -0.080752 | 0.053553 | -0.418576 | -0.469119 |
| 34.625 | -0.215071 | -2.807608 | -2.445887 | -0.613735 | 0.166821 | 0.09607 | -0.185738 | -2.128948 | 0.167204 | -5.365285 | -0.465077 | -0.825895 | -0.06236 | -0.066496 | 1.827989 | -2.148392 | -1.027951 | -0.633013 | -0.27371 | 0.148432 | 0.236798 | -0.198461 | -0.185486 |
| 34.63889 | 0.065591 | -2.40534 | -1.981493 | -0.135759 | 0.89306 | 0.192874 | -0.138967 | -1.618366 | 0.158071 | -4.916449 | -0.287269 | -0.802518 | 0.207802 | 0.114154 | 3.485526 | -1.577488 | -0.672986 | -0.549589 | 0.070337 | 0.255163 | 0.298525 | -0.092694 | -0.094727 |
| 34.65278 | -0.45386 | -2.093445 | -1.418664 | 0.33639 | 1.818316 | 0.605057 | 0.364261 | -1.222703 | 0.0667 | -5.094481 | -0.004864 | -0.468917 | 0.47334 | 0.069862 | 3.417874 | -1.146491 | -0.447291 | -0.407468 | 0.313591 | 0.190728 | 0.21725 | -0.121446 | -0.274048 |
| 34.66667 | -0.809261 | -1.762645 | -1.378338 | 0.661811 | 0.026878 | 0.470088 | 0.248214 | -0.995569 | 0.176433 | -5.792014 | 0.177568 | -0.085104 | 0.712131 | -0.010148 | 2.097247 | -0.783838 | -0.275092 | -0.155882 | 0.428115 | 0.13706 | 0.143718 | -0.15238 | -0.399164 |
| 34.68056 | -0.693319 | -1.467082 | -1.180267 | 0.633923 | -0.800756 | 0.222405 | -0.173397 | -0.710129 | 0.506043 | -5.002223 | 0.125656 | -0.163499 | 0.63151 | -0.036513 | 1.362049 | -0.711031 | -0.58392 | 0.011022 | 0.353988 | 0.137156 | 0.12876 | -0.22071 | -0.361666 |
| 34.69444 | -0.240506 | -0.759777 | -0.653867 | 0.569721 | -0.716761 | 0.115219 | -0.209802 | -0.211957 | 0.620934 | -3.027002 | 0.089408 | -0.304995 | 0.507234 | 0.01178 | 1.149426 | -0.704135 | -0.899606 | 0.157355 | 0.364632 | 0.217702 | 0.220031 | -0.166826 | -0.224913 |
| 34.70833 | 0.70362 | -0.238988 | -0.454616 | 0.336327 | -0.401829 | 0.119837 | -0.406357 | 0.381272 | 0.514095 | -1.809124 | 0.076176 | 0.073728 | 0.545326 | 0.110224 | 1.121189 | -0.769219 | -0.8233 | 0.27544 | 0.543015 | 0.345105 | 0.392637 | 0.02423 | -0.063455 |
| 34.72222 | 1.548773 | 0.049517 | -0.606564 | 0.05347 | -0.509941 | -0.358911 | -0.356999 | 0.771878 | 0.376333 | -0.527589 | -0.020952 | 0.536197 | 0.512683 | 0.22387 | 1.189091 | -0.762468 | -0.576368 | 0.309754 | 0.662898 | 0.424251 | 0.490676 | 0.145643 | 0.085037 |
| 34.73611 | 1.276485 | 0.402132 | -0.534309 | -0.002322 | -1.083273 | -0.924468 | -0.877336 | 0.536432 | 0.003665 | 0.907048 | -0.020184 | 0.550417 | 0.437893 | 0.320193 | 1.267372 | -0.641445 | -0.57912 | 0.308419 | 0.680117 | 0.456422 | 0.487962 | 0.184422 | 0.199764 |
| 34.75 | 1.082719 | 0.598374 | -0.441117 | 0.174679 | -1.392051 | -0.747125 | -1.318981 | 0.265432 | -0.466348 | 2.794635 | -0.026773 | 0.244019 | 0.350892 | 0.330699 | 1.303962 | -0.160538 | -0.825315 | 0.341184 | 0.58417 | 0.46702 | 0.427501 | 0.179091 | 0.196556 |
| 34.76389 | 1.192505 | 0.548728 | 0.707573 | 0.488282 | -0.945257 | -0.44005 | -1.297458 | 0.413976 | -0.563346 | 4.1274 | 0.270621 | -0.027326 | 0.235999 | 0.228868 | 1.036007 | 0.334637 | -0.8839 | 0.332464 | 0.349095 | 0.405719 | 0.324438 | 0.181401 | 0.104704 |
| 34.77778 | 1.021455 | 0.269005 | 1.576538 | 0.748097 | -0.812755 | -0.31263 | -1.075076 | 0.768043 | -0.394536 | 3.750419 | 0.543219 | -0.143392 | 0.08796 | 0.038556 | 0.777342 | 0.474264 | -0.384623 | 0.124718 | 0.076337 | 0.184343 | 0.208202 | 0.213406 | -0.017512 |
| 34.79167 | 0.338585 | -0.28571 | 0.950595 | 0.5692 | -0.943299 | -0.299231 | -0.784073 | 0.819408 | -0.175018 | 2.348135 | 0.535662 | -0.299772 | -0.126882 | -0.227295 | 0.516356 | 0.248616 | 0.07763 | -0.186227 | -0.224858 | -0.105996 | -0.005549 | 0.062913 | -0.205108 |
| 34.80556 | -0.144212 | -0.863216 | 0.120037 | 0.388474 | -0.803728 | 0.034873 | -0.579718 | 0.76532 | 0.140133 | 0.513929 | 0.413595 | -0.562514 | -0.292567 | -0.438525 | 0.281489 | 0.007454 | 0.326979 | -0.458246 | -0.416136 | -0.281155 | -0.218351 | -0.114876 | -0.405276 |
| 34.81944 | -0.048391 | -1.099959 | -0.345963 | 0.311491 | -0.845474 | 0.845686 | -0.032112 | 0.617973 | 0.401637 | -1.29645 | 0.345043 | -0.54287 | -0.156365 | -0.370808 | 0.21252 | -0.043733 | 0.419589 | -0.398486 | -0.355619 | -0.221223 | -0.189361 | -0.054881 | -0.3761 |
| 34.83333 | 0.297046 | -1.316967 | -0.819795 | 0.31139 | -0.792304 | 1.272734 | 0.559376 | 0.579367 | 0.525283 | -3.044396 | 0.277263 | -0.131923 | 0.137409 | -0.10114 | 0.297497 | 0.089264 | 0.891635 | -0.098939 | -0.109696 | 0.02163 | 0.064063 | 0.275701 | -0.130658 |
| 34.84722 | -0.100974 | -1.575664 | -0.740804 | 0.289502 | -0.651748 | 0.308381 | 0.762099 | 0.528012 | 0.432455 | -4.099544 | 0.246025 | 0.284797 | 0.341151 | 0.1179 | 0.314351 | 0.292208 | 0.98046 | 0.149109 | 0.138839 | 0.231982 | 0.285644 | 0.647432 | 0.065785 |
| 34.86111 | -0.714434 | -1.561475 | -0.984961 | 0.25507 | -0.403471 | -0.187607 | 0.983352 | 0.54943 | 0.205644 | -4.216076 | 0.191177 | 0.411796 | 0.355166 | 0.167027 | 0.23304 | 0.411584 | 0.453622 | 0.138475 | 0.211536 | 0.256088 | 0.309855 | 0.869785 | 0.039309 |
| 34.875 | -1.406375 | -1.080409 | -0.994521 | 0.165141 | -0.473403 | -0.92128 | 0.763957 | 0.445458 | -0.181378 | -4.402669 | 0.033125 | 0.306069 | 0.262799 | 0.098098 | 0.032438 | 0.312526 | 0.259961 | -0.06738 | 0.187454 | 0.177068 | 0.228367 | 0.933995 | -0.103067 |
| 34.88889 | -1.475932 | -0.657002 | -1.065467 | -0.042011 | -1.661461 | -2.642313 | 0.457663 | 0.017823 | -0.432377 | -4.258015 | 0.035092 | 0.216478 | 0.300062 | 0.105641 | 0.026232 | 0.169489 | 0.86144 | -0.130221 | 0.254631 | 0.202121 | 0.259044 | 0.853553 | -0.109984 |
| 34.90278 | -0.920687 | -0.169119 | -1.607468 | -0.324879 | -2.367655 | -1.623784 | 0.300701 | -0.107063 | -0.53598 | -3.356935 | 0.055252 | 0.174542 | 0.388374 | 0.155056 | 0.034107 | 0.161433 | 0.983573 | -0.014952 | 0.415084 | 0.333151 | 0.339513 | 0.802664 | -0.003204 |
| 34.91667 | 0.685513 | 0.275048 | -0.843315 | -0.065512 | -0.900654 | -0.3099 | -0.52355 | 0.150751 | -0.504539 | -2.295411 | 0.15943 | 0.152111 | 0.323988 | 0.049147 | -0.204002 | 0.039311 | 0.727083 | 0.035179 | 0.454765 | 0.311562 | 0.270425 | 0.780275 | -0.020562 |
| 34.93056 | 2.232445 | 0.532772 | 0.568506 | 0.248825 | 0.385047 | 1.130644 | -0.54815 | 0.17904 | -0.195911 | -1.20164 | 0.299113 | 0.381444 | 0.261449 | -0.038128 | -0.346309 | -0.256011 | 0.529074 | -0.038928 | 0.395089 | 0.089102 | 0.134666 | 0.896779 | -0.094321 |
| 34.94444 | 2.271619 | 0.63634 | 1.37288 | 0.468986 | 1.051808 | 1.552403 | -0.139271 | 0.03243 | 0.217616 | -0.405239 | 0.288168 | 0.674274 | 0.289469 | 0.023501 | -0.330488 | -0.307228 | 0.304842 | 0.014596 | 0.406868 | -0.004028 | 0.133584 | 1.034079 | -0.071829 |
| 34.95833 | 1.990229 | 0.709768 | 1.657371 | 0.616793 | 0.619931 | 1.129773 | 0.124827 | -0.086435 | 0.460383 | 0.472306 | 0.218789 | 0.639525 | 0.279255 | 0.003093 | -0.412715 | -0.098392 | 0.376995 | 0.061957 | 0.379401 | 0.028478 | 0.140787 | 0.940634 | -0.053718 |
| 34.97222 | 1.105038 | 0.547627 | 1.044561 | 0.613763 | -0.809426 | -0.028691 | 0.171249 | -0.045684 | 0.391001 | 1.301427 | 0.4289 | 0.527026 | 0.184895 | -0.117689 | -0.509098 | 0.071945 | 0.326579 | 0.006748 | 0.25041 | -0.013507 | 0.025183 | 0.724666 | -0.16196 |
| 34.98611 | 0.086862 | 0.231149 | -0.522331 | 0.565716 | -1.843722 | -0.927068 | -1.235262 | 0.169087 | 0.220825 | 1.642789 | 0.50017 | 0.662416 | 0.16345 | -0.139121 | -0.483267 | 0.191943 | -0.323506 | 0.063228 | 0.293633 | 0.003387 | 0.015329 | 0.57023 | -0.212448 |
| 35 | -0.287179 | 0.028761 | -1.615819 | -0.163472 | -1.191043 | -0.53218 | -0.858989 | 0.168378 | -0.139332 | 1.203023 | 0.355331 | 0.834818 | 0.146733 | -0.098413 | -0.297068 | 0.382997 | -0.582086 | 0.306824 | 0.354255 | 0.053806 | 0.05179 | 0.535402 | -0.197323 |
| 35.01389 | -0.982112 | -0.022348 | -1.708144 | -0.854363 | -1.318541 | -0.085482 | -0.253416 | -0.13643 | -0.677931 | 0.851009 | -0.045549 | 0.687619 | 0.009898 | -0.187486 | -0.144146 | 0.454988 | -1.555778 | 0.385214 | 0.333724 | -0.01188 | -0.082988 | 0.501903 | -0.233815 |
| 35.02778 | -1.597828 | 0.431694 | -0.808046 | -1.011107 | -1.153277 | 0.482154 | 0.467446 | 0.255596 | -0.80134 | 0.767756 | -0.299733 | 0.632551 | -0.016676 | -0.320109 | -0.041068 | 0.19282 | -2.946297 | 0.500763 | 0.345388 | 0.027391 | -0.04181 | 0.497424 | -0.144355 |
| 35.04167 | -1.280072 | 1.644618 | 1.312488 | -0.623467 | -0.956143 | 1.073095 | 0.583796 | 1.076702 | -0.532001 | 1.964071 | -0.059688 | 0.795478 | 0.064361 | -0.179677 | 0.080675 | 0.337641 | -1.384157 | 0.466597 | 0.381773 | 0.107528 | 0.107502 | 0.547971 | -0.013189 |
| 35.05556 | -0.458671 | 2.356909 | 2.314527 | -0.306975 | -0.958333 | 0.951504 | 0.740397 | 1.192694 | -0.309162 | 2.407215 | -0.028606 | 1.027326 | 0.120934 | -0.098915 | 0.079891 | 0.491091 | 0.139228 | 0.32562 | 0.458051 | 0.092117 | 0.125867 | 0.588667 | 0.0231 |
| 35.06944 | 0.53364 | 2.193339 | 2.773987 | -0.38455 | -0.436625 | 0.741957 | 0.982924 | 1.124502 | -0.092942 | 2.270164 | -0.008592 | 1.005998 | 0.086924 | -0.170467 | -0.019346 | 0.528745 | 1.576892 | 0.147912 | 0.502059 | 0.025183 | 0.060528 | 0.523329 | -0.054284 |
| 35.08333 | 0.921881 | 1.903699 | 4.027774 | 0.183604 | 0.139188 | 1.069507 | 0.927812 | 1.060401 | 0.19683 | 2.918597 | 0.417509 | 0.35117 | -0.470225 | -0.723999 | -0.479521 | 0.065835 | 2.05202 | -0.447108 | -0.061345 | -0.534507 | -0.506386 | -0.112725 | -0.634291 |
| 35.09722 | 0.755102 | 1.819883 | 5.853621 | 0.35884 | 0.726352 | 2.1912 | 0.730411 | 0.709604 | 0.146068 | 2.885484 | 0.910095 | -0.819016 | -1.514099 | -1.714485 | -1.333595 | -0.971858 | 1.837956 | -1.5034 | -1.241954 | -1.672602 | -1.680453 | -1.315521 | -1.696641 |
| 35.11111 | 0.511342 | 1.882746 | 5.424206 | 0.12938 | 0.824322 | 1.854743 | 0.237742 | 0.252688 | -0.073725 | 3.378531 | 0.633131 | -1.181452 | -1.947939 | -2.05316 | -1.720727 | -1.227378 | 1.176609 | -1.735898 | -1.563071 | -1.986899 | -2.060303 | -1.910568 | -2.018257 |
| 35.125 | 0.171335 | 2.684498 | 2.223656 | 0.221306 | 0.599685 | 1.206169 | -0.161897 | 0.37223 | -0.01807 | 3.321346 | -0.219675 | -0.814086 | -1.597242 | -1.661045 | -1.473215 | -0.330162 | 1.181494 | -1.026594 | -0.984492 | -1.373021 | -1.443827 | -1.432777 | -1.521918 |
| 35.13889 | 0.233408 | 3.760472 | 1.321539 | 0.266355 | 0.918658 | 0.978411 | -0.342494 | 0.857876 | 0.003528 | 3.433366 | -0.625469 | -0.61513 | -1.365776 | -1.421768 | -1.315111 | 0.619038 | 1.488763 | -0.734658 | -0.658538 | -1.149254 | -1.206978 | -1.031267 | -1.281143 |
| 35.15278 | 0.356012 | 4.38098 | 1.480911 | 0.265852 | 1.49673 | -0.0936 | -0.087669 | 0.776451 | -0.073093 | 3.707738 | -0.629742 | -0.106389 | -1.137088 | -1.206834 | -1.046129 | 0.690815 | 1.051156 | -0.653276 | -0.466956 | -1.006285 | -1.049172 | -0.779929 | -1.104386 |
| 35.16667 | 0.514983 | 4.297686 | 1.425629 | 0.119019 | 1.243417 | 0.088462 | -0.116976 | 0.708327 | -0.329753 | 3.342905 | -0.574268 | 0.450734 | -0.725243 | -0.804084 | -0.569113 | 0.49196 | 0.166074 | -0.287059 | -0.229835 | -0.643159 | -0.685284 | -0.450197 | -0.733245 |
| 35.18056 | 0.703747 | 3.498644 | 1.10752 | 0.096412 | 0.858316 | 0.873451 | 0.255571 | -0.111548 | -0.470859 | 2.659771 | -0.403928 | 0.912154 | -0.433181 | -0.474221 | -0.122422 | -0.114612 | -0.033392 | -0.00866 | -0.044754 | -0.364147 | -0.402757 | -0.134783 | -0.434911 |
| 35.19444 | 0.652608 | 1.771998 | 0.850112 | 0.237147 | 0.887583 | 0.98873 | 0.93476 | -0.840806 | -0.519471 | 1.621634 | 0.099773 | 1.011322 | -0.284161 | -0.280494 | 0.072383 | -0.291796 | -0.090838 | 0.099576 | 0.124456 | -0.213486 | -0.237603 | 0.252657 | -0.261888 |
| 35.20833 | 0.258013 | -0.042718 | -0.171104 | 0.330623 | 1.124441 | 1.315562 | 1.302032 | -1.359296 | -0.554834 | 0.914098 | 0.61266 | 1.078033 | -0.185173 | -0.112092 | 0.049957 | 0.085925 | 0.474368 | 0.158234 | 0.289293 | -0.057405 | -0.070354 | 0.701167 | -0.096709 |
| 35.22222 | -0.345088 | -1.825713 | -1.466455 | 0.313283 | 0.481791 | 1.534459 | 0.351349 | -1.901072 | -0.436908 | 0.411596 | 0.857462 | 1.102345 | -0.172963 | -0.070145 | 0.000759 | 0.282731 | 0.866402 | 0.297898 | 0.281736 | -0.00263 | -0.019408 | 0.898256 | -0.040671 |
| 35.23611 | -0.845657 | -2.228981 | -2.161457 | 0.137725 | -0.057067 | 1.502156 | -0.221345 | -2.020177 | -0.344909 | 0.030841 | 0.548417 | 0.973463 | -0.300765 | -0.204263 | -0.128346 | 0.105063 | 0.809402 | 0.316755 | 0.086176 | -0.13084 | -0.174186 | 0.666924 | -0.168846 |
| 35.25 | -0.68366 | -0.352605 | -2.029183 | 0.060384 | 0.352594 | 0.801504 | -0.370282 | -1.099636 | -0.038685 | -0.210649 | 0.349732 | 0.830371 | -0.466725 | -0.384495 | -0.330518 | 0.012457 | 0.66386 | 0.085118 | -0.114012 | -0.296403 | -0.390562 | 0.345485 | -0.336644 |
| 35.26389 | -0.119607 | 2.135005 | -0.820674 | 0.006585 | 1.077347 | 0.079287 | -0.441311 | -0.259024 | 0.384777 | -0.616002 | 0.300452 | 0.562546 | -0.464832 | -0.408338 | -0.441886 | -0.133601 | 0.483268 | -0.073774 | -0.181302 | -0.33221 | -0.453778 | 0.343264 | -0.360612 |
| 35.27778 | 0.62023 | 4.564067 | 0.517912 | -0.040174 | 1.251199 | -0.198705 | -0.460309 | 0.474717 | 0.667016 | -0.63708 | 0.223144 | 0.40503 | -0.352941 | -0.293487 | -0.452957 | 0.055534 | 0.751221 | -0.120915 | -0.060139 | -0.279935 | -0.396391 | 0.556526 | -0.296882 |
| 35.29167 | 0.917377 | 5.261475 | 1.682185 | -0.09565 | 1.209277 | 0.168914 | -0.167348 | 0.718251 | 0.556941 | -0.519943 | 0.146787 | 0.224831 | -0.423918 | -0.286906 | -0.595801 | 0.182877 | 1.056906 | -0.224326 | 0.009382 | -0.293009 | -0.38168 | 0.655636 | -0.337016 |
| 35.30556 | 0.968738 | 3.872432 | 2.044445 | -0.025394 | 1.017869 | 0.308342 | -0.310381 | 0.841089 | 0.287843 | 0.092193 | -0.020876 | 0.355653 | -0.509649 | -0.265907 | -0.781529 | 0.381391 | 1.205374 | -0.23631 | 0.013201 | -0.271324 | -0.336926 | 0.52987 | -0.346027 |
| 35.31944 | 1.072081 | 2.699387 | 1.613771 | 0.116455 | 0.373249 | -0.13601 | -0.109115 | 1.210281 | 0.101972 | 0.52211 | -0.161605 | 0.673663 | -0.349721 | -0.088165 | -0.838985 | 0.680293 | 1.531585 | -0.016702 | 0.186955 | -0.104171 | -0.129519 | 0.449575 | -0.155893 |
| 35.33333 | 1.012259 | 1.51477 | 1.318733 | 0.307415 | 0.728545 | 0.528949 | -0.032208 | 1.368053 | 0.130143 | 0.705192 | -0.063391 | 0.948395 | -0.104462 | 0.081423 | -0.878927 | 0.809917 | 1.476726 | 0.228655 | 0.385835 | 0.089115 | 0.100958 | 0.474537 | 0.066889 |
| 35.34722 | 0.655525 | 0.074092 | 1.197746 | 0.728606 | 0.806853 | 0.520964 | 0.107039 | 1.382416 | 0.254463 | 0.846796 | -0.070321 | 0.9425 | 0.092656 | 0.185358 | -0.865331 | 0.686616 | 1.208594 | 0.368626 | 0.500209 | 0.249148 | 0.278557 | 0.681911 | 0.22163 |
| 35.36111 | 0.182393 | -1.511851 | 1.397046 | 0.698022 | 0.959837 | 0.821795 | 0.479355 | 1.658605 | 0.292808 | 0.83632 | -0.20521 | 0.967036 | 0.235244 | 0.316498 | -0.765252 | 0.682227 | 0.935181 | 0.372173 | 0.648062 | 0.412467 | 0.449213 | 1.030683 | 0.362712 |
| 35.375 | -0.136584 | -3.080972 | 1.411309 | 0.838478 | 1.317206 | 0.474211 | 1.173093 | 1.079379 | 0.243898 | 1.397209 | -0.051509 | 1.096584 | 0.316746 | 0.454395 | -0.522115 | 0.734374 | 0.87001 | 0.345892 | 0.73802 | 0.531943 | 0.577875 | 1.327174 | 0.474451 |
| 35.38889 | -0.125027 | -5.212407 | 0.641795 | 0.894434 | 1.511917 | 0.382129 | 1.183763 | 0.212631 | 0.200176 | 1.670566 | 0.106172 | 1.134509 | 0.287367 | 0.471879 | -0.360301 | 0.694013 | 0.765134 | 0.322504 | 0.637184 | 0.519453 | 0.57923 | 1.35855 | 0.471233 |
| 35.40278 | -0.053 | -4.268348 | -0.379343 | 0.893468 | 0.744908 | -0.076414 | 0.35952 | -0.433718 | 0.121229 | 1.614273 | 0.137508 | 0.81124 | 0.166339 | 0.361896 | -0.360799 | 0.482556 | 0.503076 | 0.189888 | 0.406496 | 0.376503 | 0.438831 | 1.015078 | 0.326569 |
| 35.41667 | -0.117719 | -3.528015 | -0.969163 | 0.894991 | -0.151733 | -0.580612 | 0.404268 | -1.021077 | 0.017259 | 1.004393 | 0.03102 | 0.319608 | 0.014955 | 0.203391 | -0.438222 | 0.282662 | 0.07506 | -0.006819 | 0.231005 | 0.205366 | 0.242028 | 0.493745 | 0.150794 |
| 35.43056 | 0.079111 | -0.940342 | -1.239875 | 1.032439 | 0.634058 | -0.561188 | 0.423087 | -1.29369 | -0.03395 | 0.565883 | -0.125163 | -0.130857 | 0.032927 | 0.11445 | -0.432324 | 0.117722 | -0.161009 | -0.047689 | 0.183715 | 0.134289 | 0.148974 | 0.035562 | 0.089699 |
| 35.44444 | 0.5886 | -0.352092 | -1.00221 | 1.018269 | 1.709586 | -0.639682 | 0.240533 | -1.378194 | 0.053579 | 0.267026 | -0.220058 | -0.358418 | 0.137933 | 0.085324 | -0.362547 | 0.063767 | -0.352313 | 0.019777 | 0.195181 | 0.138346 | 0.155531 | -0.218688 | 0.095705 |
| 35.45833 | 0.336143 | -0.347871 | -1.91906 | 0.00367 | 1.832408 | -0.273881 | -0.001626 | -1.566419 | -0.120942 | -0.937949 | -0.039494 | -0.454698 | 0.139758 | -0.001525 | -0.322429 | -0.095474 | -0.708216 | 0.077768 | 0.080625 | 0.091437 | 0.106227 | -0.368098 | 0.039149 |
| 35.47222 | -0.385837 | -2.567791 | -1.453267 | -0.488724 | 1.75577 | -0.308616 | -0.029292 | -1.602984 | -0.236465 | -2.157473 | -0.046947 | -0.503372 | 0.039139 | -0.112465 | -0.352466 | -0.479262 | -0.804638 | 0.014329 | -0.107374 | -0.012491 | -0.001053 | -0.568971 | -0.067758 |
| 35.48611 | 0.089439 | -2.603056 | -0.995038 | -0.654038 | 0.084005 | 0.099661 | -0.851331 | -1.547435 | -0.45798 | -1.560082 | -0.113514 | -0.607878 | -0.077391 | -0.187156 | -0.396762 | -0.47163 | -0.80136 | -0.127915 | -0.222331 | -0.112041 | -0.090412 | -0.781653 | -0.158373 |
| 35.5 | 1.18878 | -1.260512 | -0.600857 | -0.451423 | 0.337621 | 0.565421 | -1.176673 | -1.125804 | -0.605378 | -0.922214 | -0.242734 | -0.76189 | -0.190464 | -0.2436 | -0.441505 | -0.351508 | -0.508499 | -0.320339 | -0.244958 | -0.169516 | -0.164168 | -0.971101 | -0.21855 |
| 35.51389 | 1.737786 | 0.122374 | -0.482736 | -0.333615 | 0.961711 | 0.723274 | -0.748451 | -0.438044 | -0.616611 | -1.049055 | -0.32967 | -0.906359 | -0.217038 | -0.263583 | -0.43116 | -0.083569 | -0.751085 | -0.488982 | -0.130595 | -0.162725 | -0.176001 | -1.080258 | -0.223685 |
| 35.52778 | 0.716936 | 1.249837 | -0.91795 | -0.370512 | 1.264424 | 0.637676 | -0.13533 | 0.276951 | -0.633875 | -1.218508 | -0.551638 | -0.790241 | -0.137951 | -0.213122 | -0.338088 | 0.116379 | -0.953922 | -0.534229 | -0.062784 | -0.107591 | -0.10747 | -1.102386 | -0.15349 |
| 35.54167 | 0.310753 | 1.795987 | -2.243053 | -0.430198 | 0.961807 | 0.458824 | 0.176136 | 0.786255 | -0.657243 | -1.311185 | -0.654294 | -0.672834 | -0.16383 | -0.133199 | -0.197083 | 0.182678 | -0.93045 | -0.449484 | -0.08871 | -0.052959 | -0.039521 | -1.051228 | -0.060453 |
| 35.55556 | 0.210599 | 2.499549 | -2.589463 | -0.489046 | 0.842471 | 0.267581 | -0.507197 | 0.905675 | -0.640042 | -0.708668 | -0.418717 | -0.560106 | -0.206227 | -0.001301 | 0.005427 | 0.181757 | -0.89268 | -0.216208 | -0.19681 | 0.01687 | 0.052546 | -0.866553 | 0.058269 |
| 35.56944 | 1.775472 | 3.757314 | -1.515628 | -0.367432 | 0.919139 | 0.520005 | -0.09866 | 1.123881 | -0.328112 | -0.461686 | -0.076658 | -0.404613 | -0.099503 | 0.20519 | 0.276545 | 0.253757 | -0.57374 | 0.015057 | -0.17216 | 0.144698 | 0.247001 | -0.582863 | 0.217157 |
| 35.58333 | 0.847689 | 4.993116 | -0.93866 | -0.623518 | 0.601168 | 0.484135 | 0.115222 | 0.987929 | -0.154381 | -1.130321 | -0.090611 | -0.437491 | 0.072258 | 0.276098 | 0.582257 | -0.192331 | -0.611789 | 0.04571 | -0.053988 | 0.316639 | 0.468384 | -0.482389 | 0.315085 |
| 35.59722 | -0.599744 | 4.00524 | -0.885345 | -0.97205 | -1.894083 | 0.162808 | 0.160243 | -0.138529 | -0.60375 | -1.535057 | -0.285972 | -0.476962 | 0.274689 | 0.376165 | 1.050219 | -0.886867 | -0.968965 | 0.025088 | 0.029851 | 0.523121 | 0.602468 | -0.519267 | 0.487372 |
| 35.61111 | -1.177453 | 1.017309 | -1.394418 | -1.095237 | -3.847247 | -0.090435 | -0.179155 | -1.43164 | -1.259255 | -1.055535 | -0.290818 | -0.165218 | 0.549098 | 0.62631 | 1.763611 | -1.368509 | -1.517918 | 0.0905 | -0.046036 | 0.694692 | 0.783268 | -0.409946 | 0.700534 |
| 35.625 | -0.100396 | 0.267309 | -2.737712 | -1.016784 | -4.145725 | -0.087814 | -0.252096 | -2.385372 | -1.111814 | -0.628274 | -0.087422 | 0.051365 | 0.613703 | 0.686926 | 2.39694 | -1.066795 | -1.707408 | 0.029355 | -0.217187 | 0.679396 | 0.765639 | -0.334416 | 0.708638 |
| 35.63889 | 0.475439 | 0.526937 | -2.282468 | -0.775441 | -4.090774 | -0.133691 | 0.239279 | -2.799231 | -0.449779 | -1.452674 | -0.009895 | -0.109016 | 0.424709 | 0.518891 | 2.453672 | -0.584436 | -0.99847 | -0.060821 | -0.195316 | 0.511347 | 0.549504 | -0.447785 | 0.518466 |
| 35.65278 | 0.044086 | 0.855694 | -0.158968 | -0.236515 | -3.715876 | -0.190878 | 0.445527 | -1.926093 | 0.288672 | -2.141041 | -0.073616 | -0.2431 | 0.294234 | 0.286992 | 1.838239 | -0.278041 | -0.244364 | -0.234776 | 0.097329 | 0.308783 | 0.314115 | -0.48066 | 0.312365 |
| 35.66667 | -0.179926 | -0.447682 | 0.127052 | 0.327266 | -2.602987 | -0.195561 | -0.243728 | -0.982256 | 0.929239 | -2.124906 | 0.060424 | 0.054674 | 0.293235 | 0.177372 | 1.050929 | -0.124253 | 0.107201 | -0.380349 | 0.365613 | 0.207857 | 0.209788 | -0.265364 | 0.215356 |
| 35.68056 | -0.340319 | -0.338546 | -0.134971 | 0.511516 | -1.290415 | -0.107242 | -0.318166 | -0.331944 | 1.065646 | -1.111591 | 0.084832 | 0.316875 | 0.418739 | 0.224398 | 0.515062 | 0.109257 | -0.127313 | -0.386773 | 0.436397 | 0.258482 | 0.276902 | 0.064198 | 0.26386 |
| 35.69444 | -0.508294 | 0.356078 | 0.18646 | 0.299662 | -1.23348 | 0.273833 | 0.244563 | 0.634352 | 0.459408 | -0.24749 | 0.078792 | 0.300829 | 0.539048 | 0.340878 | 0.339051 | 0.169773 | -0.506733 | -0.15513 | 0.469981 | 0.364699 | 0.40786 | 0.299438 | 0.370591 |
| 35.70833 | -0.957211 | 0.797373 | 0.220987 | -0.202975 | -1.935168 | 0.69939 | 0.534642 | 1.899773 | -0.243877 | 0.219149 | -0.039613 | 0.250158 | 0.546291 | 0.404332 | 0.324737 | 0.14686 | -0.641533 | 0.140368 | 0.611798 | 0.417676 | 0.452206 | 0.339802 | 0.424294 |
| 35.72222 | -1.203958 | 0.918738 | 0.288123 | -0.678862 | -1.515695 | 0.98329 | 0.87834 | 2.240918 | -0.896705 | 0.533228 | -0.15758 | 0.276233 | 0.464406 | 0.374615 | 0.289537 | 0.01052 | -0.702511 | 0.266247 | 0.8156 | 0.391125 | 0.389904 | 0.182251 | 0.39095 |
| 35.73611 | -1.104547 | 0.837681 | 1.488698 | -0.626666 | -0.019086 | 0.817581 | 0.658406 | 1.916359 | -0.925693 | 1.498524 | -0.125377 | 0.215073 | 0.306508 | 0.276845 | 0.207196 | -0.143848 | -0.914716 | 0.229495 | 0.84818 | 0.332638 | 0.300596 | -0.012083 | 0.306076 |
| 35.75 | -0.946411 | 0.540974 | 2.104738 | -0.565927 | 2.065567 | 0.34498 | 0.081333 | 1.193752 | -0.92034 | 2.286107 | 0.048335 | 0.068412 | 0.155346 | 0.122448 | 0.079805 | -0.429358 | -1.350215 | 0.087199 | 0.589146 | 0.2277 | 0.19633 | -0.143685 | 0.185466 |
| 35.76389 | -1.013765 | 0.605149 | 1.043917 | -0.542762 | 3.730842 | -0.085333 | -0.702455 | 0.117187 | -1.312407 | 2.62302 | 0.210436 | -0.036717 | 0.09619 | 0.025281 | -0.049278 | -0.773965 | -1.499343 | -0.03633 | 0.11073 | 0.066789 | 0.082818 | -0.161591 | 0.078157 |
| 35.77778 | -1.532783 | 0.892267 | 0.038744 | -0.391841 | 3.532197 | -0.496146 | -0.871915 | -0.968173 | -1.638055 | 2.660944 | 0.366738 | -0.194439 | 0.035114 | -0.026832 | -0.160332 | -0.987711 | -0.919106 | -0.080126 | -0.17545 | -0.009676 | 0.025373 | -0.247228 | -0.000193 |
| 35.79167 | -2.270935 | 1.351753 | -0.517258 | -0.249067 | 2.695936 | -0.561483 | -0.647838 | -1.868141 | -0.970292 | 2.761489 | 0.591433 | -0.392695 | -0.027364 | -0.075671 | -0.237285 | -0.823183 | -0.159212 | -0.022255 | -0.130156 | 0.036101 | 0.01866 | -0.315338 | -0.022289 |
| 35.80556 | -1.567429 | 1.452029 | -1.346081 | -0.28301 | 2.446114 | -0.536928 | 0.197348 | -2.501413 | 0.259579 | 2.779831 | 0.572107 | -0.295509 | 0.102005 | 0.015735 | -0.102137 | -0.585213 | -0.077921 | 0.048854 | -0.013941 | 0.088817 | 0.0885 | -0.260869 | 0.068956 |
| 35.81944 | -0.772453 | 1.484661 | -1.125703 | -0.298807 | 2.18889 | -0.27153 | 0.979392 | -2.529924 | 1.396428 | 2.632533 | 0.399312 | 0.038041 | 0.359059 | 0.181086 | 0.168692 | -0.459204 | -0.218155 | 0.004455 | 0.060711 | 0.098562 | 0.178101 | -0.035417 | 0.17807 |
| 35.83333 | -0.503903 | 1.120553 | 0.674511 | -0.134166 | 1.869572 | 0.288991 | 1.164409 | -1.631468 | 1.42392 | 2.385891 | 0.228846 | 0.136981 | 0.435245 | 0.19402 | 0.276663 | -0.172568 | 0.502073 | -0.001333 | 0.188237 | 0.114984 | 0.197914 | 0.053139 | 0.172267 |
| 35.84722 | -0.911178 | 0.562951 | 2.093628 | -0.047898 | 1.773136 | 0.865201 | 0.97428 | -0.557246 | 1.085018 | 2.014637 | 0.000783 | 0.078338 | 0.401826 | 0.105988 | 0.205754 | 0.193868 | 1.176131 | 0.1092 | 0.33998 | 0.161993 | 0.168398 | 0.093929 | 0.13147 |
| 35.86111 | -0.534131 | -0.049249 | 1.764229 | 0.085555 | 1.114973 | 1.145443 | 0.52674 | 0.181002 | 0.525022 | 1.191449 | 0.069225 | 0.093806 | 0.329104 | 0.045865 | 0.092391 | 0.179308 | 1.133599 | 0.113615 | 0.352207 | 0.073875 | 0.099592 | 0.113017 | 0.064392 |
| 35.875 | -0.211276 | -0.486195 | 0.844363 | 0.184597 | 0.758454 | 1.255778 | 0.150685 | 0.91603 | -0.380236 | 0.032592 | 0.170039 | 0.021832 | 0.175448 | -0.033087 | -0.084672 | -0.035416 | 0.377316 | 0.110294 | 0.256322 | -0.092955 | -0.015216 | 0.026158 | -0.056283 |
| 35.88889 | -0.82894 | -0.716695 | 0.635711 | 0.258628 | 0.515836 | 1.302622 | -0.112443 | 1.848648 | -1.229424 | -0.876116 | 0.423472 | -0.076554 | 0.138475 | -0.050511 | -0.205571 | -0.048138 | 0.242509 | 0.377756 | 0.266156 | -0.0695 | -0.000412 | 0.033845 | -0.056466 |
| 35.90278 | -1.127701 | -0.610555 | 0.699545 | 0.305993 | 0.434402 | 1.045363 | -0.444726 | 2.651085 | -1.778468 | -1.427723 | 0.458365 | 0.03631 | 0.324179 | 0.128565 | -0.126676 | 0.162645 | 0.398796 | 0.629684 | 0.379479 | 0.152937 | 0.203068 | 0.322464 | 0.135361 |
| 35.91667 | -0.895514 | -0.295845 | 0.132808 | 0.29627 | 0.282454 | 0.4842 | -0.004868 | 2.277533 | -1.096614 | -1.53947 | 0.717355 | 0.263174 | 0.473554 | 0.305515 | -0.015355 | 0.358244 | 0.459453 | 0.642068 | 0.482476 | 0.317453 | 0.368929 | 0.638371 | 0.298983 |
| 35.93056 | -1.189564 | 0.233169 | -0.539313 | 0.153722 | -0.282888 | 0.163079 | 0.003663 | 1.293058 | 0.13939 | -0.922519 | 0.751158 | 0.369762 | 0.408289 | 0.279544 | -0.095839 | 0.346307 | -0.185762 | 0.70437 | 0.461909 | 0.284988 | 0.315951 | 0.759528 | 0.271897 |
| 35.94444 | -1.426958 | 0.922903 | -0.013021 | 0.197811 | -1.447011 | -0.476419 | -0.503138 | 0.59331 | 1.28201 | 0.227937 | 0.813593 | 0.378279 | 0.271152 | 0.148567 | -0.284809 | 0.240348 | -0.589852 | 0.822695 | 0.381237 | 0.173189 | 0.195979 | 0.752774 | 0.164269 |
| 35.95833 | -1.656876 | 1.526868 | 0.618005 | 0.344252 | -2.518699 | -0.552112 | 0.015271 | 0.03443 | 1.566288 | 1.206684 | 0.479381 | 0.356997 | 0.123561 | -0.001331 | -0.480658 | 0.145141 | -0.195702 | 0.71384 | 0.349541 | 0.036755 | 0.074413 | 0.649916 | 0.039346 |
| 35.97222 | -1.21675 | 1.864209 | 0.616841 | 0.41717 | -3.927978 | -0.402374 | 0.666781 | -0.385754 | 1.283821 | 1.890214 | 0.045517 | 0.277995 | -0.023215 | -0.150208 | -0.568322 | 0.039553 | 0.787041 | 0.605652 | 0.330983 | -0.112391 | -0.100809 | 0.562299 | -0.108314 |
| 35.98611 | -1.180697 | 1.773573 | 0.344399 | 0.406969 | -3.740992 | -0.687633 | 1.283296 | -0.605308 | 0.856118 | 2.511462 | -0.198409 | 0.23796 | -0.095547 | -0.207285 | -0.559372 | 0.02052 | 1.242042 | 0.863478 | 0.435627 | -0.172304 | -0.197495 | 0.485632 | -0.17766 |
| 36 | -0.511455 | 1.357734 | -0.641141 | 0.147134 | -1.211891 | -0.255944 | 1.189255 | -0.859142 | 0.230174 | 2.592288 | -0.310917 | 0.167032 | -0.164709 | -0.175036 | -0.543704 | 0.043816 | 1.303383 | 0.958116 | 0.485709 | -0.18367 | -0.230251 | 0.383664 | -0.203299 |
| 36.01389 | 0.828628 | 0.684221 | -1.982184 | -0.191177 | 0.713754 | -0.270277 | 0.203808 | -1.399043 | -0.630979 | 1.263911 | -0.192242 | 0.039958 | -0.266313 | -0.264466 | -0.580226 | 0.020979 | 1.279701 | 0.86379 | 0.400158 | -0.287905 | -0.375337 | 0.330966 | -0.262638 |
| 36.02778 | 0.56648 | 0.196473 | -2.271462 | -0.449931 | 0.713261 | -0.800565 | -0.046354 | -1.655842 | -1.190353 | 0.48344 | -0.094532 | 0.231154 | -0.177782 | -0.4072 | -0.459627 | -0.008754 | 1.788129 | 0.926606 | 0.473259 | -0.240142 | -0.2717 | 0.375485 | -0.23981 |
| 36.04167 | 1.245608 | 0.865567 | -1.3453 | -0.440609 | 1.843681 | -0.928629 | 0.625083 | -0.937754 | -1.385639 | 0.380901 | -0.067818 | 0.646348 | 0.040179 | -0.175165 | -0.181504 | 0.177085 | 2.285944 | 0.994078 | 0.56106 | -0.050572 | -0.019777 | 0.399184 | -0.073255 |
| 36.05556 | 1.251798 | 2.143701 | 1.818908 | -0.344714 | 2.866312 | 0.039204 | 0.630943 | 0.075435 | -0.836758 | 0.33653 | 0.014748 | 0.899501 | 0.147471 | -0.002433 | -0.085777 | 0.31838 | 1.684917 | 0.790369 | 0.519699 | 0.033945 | 0.055008 | 0.305292 | 0.031762 |
| 36.06944 | 2.110954 | 3.009374 | 5.097865 | -0.045025 | 3.264316 | 0.282619 | 0.229453 | 1.009046 | -0.121635 | 0.483432 | 0.144739 | 0.732986 | 0.072963 | -0.09901 | -0.214854 | 0.229119 | 0.845256 | 0.450851 | 0.307894 | -0.047891 | -0.018867 | 0.035075 | -0.053137 |
| 36.08333 | 1.520512 | 2.963553 | 5.534535 | 0.027869 | 2.585387 | 1.40151 | 0.54684 | 1.320018 | 0.225483 | -0.114882 | 0.083936 | -0.145117 | -0.734323 | -0.809511 | -0.94525 | -0.433413 | -0.673998 | -0.367383 | -0.508173 | -0.731054 | -0.687674 | -0.723087 | -0.728246 |
| 36.09722 | 1.260746 | 2.362861 | 3.405591 | -0.13591 | 1.249536 | 1.595272 | 0.890868 | 0.987351 | -0.013686 | -1.93738 | -0.226126 | -1.736466 | -2.489607 | -2.136775 | -2.34253 | -1.702702 | -2.506997 | -1.964061 | -2.062121 | -2.201878 | -2.264555 | -2.099974 | -2.149011 |
| 36.11111 | 0.18479 | 1.784917 | 0.892412 | 0.038704 | 0.931291 | 0.978206 | 0.266672 | 0.392591 | -0.347699 | -2.541225 | -0.338229 | -2.360469 | -3.61925 | -2.659471 | -2.907373 | -2.247194 | -4.180815 | -2.456358 | -2.511497 | -2.813837 | -3.077787 | -2.824265 | -2.803082 |
| 36.125 | -0.978176 | 1.662605 | -0.368104 | 0.522184 | -0.237379 | -0.266412 | -0.456602 | 0.291827 | 0.373702 | 0.087499 | -0.268942 | -1.440126 | -2.703648 | -1.75458 | -1.895384 | -0.992164 | -2.458268 | -1.202227 | -1.188028 | -1.628925 | -1.816192 | -2.106466 | -1.77993 |
| 36.13889 | -1.056119 | 2.432782 | 0.113228 | 0.978178 | -1.093592 | -0.708559 | -0.256989 | 1.007535 | 0.635944 | 2.218501 | -0.030723 | -0.29267 | -1.280559 | -0.636925 | -0.641829 | 1.453221 | -1.30996 | -0.369523 | 0.090789 | -0.586391 | -0.671212 | -1.179092 | -0.660689 |
| 36.15278 | 0.321213 | 2.558912 | 2.167251 | 0.994453 | -0.733173 | -0.700789 | 0.460792 | 2.353978 | 0.131082 | 2.590202 | 0.260196 | 0.420807 | -0.187447 | -0.079395 | 0.020074 | 3.108606 | -0.8725 | 0.079228 | 0.792072 | -0.094262 | -0.079369 | -0.537366 | -0.092042 |
| 36.16667 | 0.821117 | 2.208692 | 1.541315 | 0.808082 | 0.048188 | -0.21934 | 0.978632 | 2.554322 | -0.013697 | 1.449647 | 0.405894 | 0.965683 | 0.483359 | 0.07705 | 0.231863 | 2.176581 | -0.287004 | 0.267099 | 0.662809 | 0.08664 | 0.144232 | -0.035582 | 0.101262 |
| 36.18056 | -0.759506 | 1.782779 | 0.493107 | 0.642979 | -0.036233 | -0.821072 | 1.132243 | 1.572286 | 0.11185 | 0.799588 | 0.167668 | 1.349296 | 0.713825 | 0.220602 | 0.397335 | 0.767647 | -0.12825 | 0.518628 | 0.334803 | 0.237255 | 0.267651 | 0.541418 | 0.252943 |
| 36.19444 | 0.823332 | 1.397428 | -0.05269 | 0.593355 | -0.085121 | -0.097964 | 1.417247 | 0.646975 | 0.000981 | 0.539312 | -0.046684 | 1.53302 | 0.709272 | 0.443743 | 0.66014 | -0.094235 | 0.167675 | 0.911896 | 0.20034 | 0.437267 | 0.462938 | 1.124538 | 0.465191 |
| 36.20833 | 1.835553 | 0.947266 | 0.245899 | 0.530783 | -0.217937 | 1.423208 | 1.08237 | 0.076217 | -0.136034 | 0.350865 | -0.102362 | 1.608903 | 0.576959 | 0.590005 | 0.831863 | -0.187953 | 0.895798 | 1.136162 | 0.263819 | 0.549071 | 0.621732 | 1.41974 | 0.59867 |
| 36.22222 | 2.18573 | 0.425821 | -0.266202 | 0.187238 | -0.805713 | 1.834688 | 0.716324 | -0.288075 | -0.15354 | 0.101104 | 0.006858 | 1.404003 | 0.332639 | 0.580564 | 0.787034 | 0.050679 | 2.134857 | 0.994145 | 0.289402 | 0.509482 | 0.566557 | 1.415908 | 0.568977 |
| 36.23611 | 2.596278 | -0.012326 | -1.64474 | 0.456631 | -0.37327 | 1.305287 | 0.677731 | -0.404717 | -0.116195 | -0.142851 | 0.142438 | 1.220757 | 0.171082 | 0.540939 | 0.703187 | 0.486049 | 2.982778 | 0.918664 | 0.31686 | 0.466826 | 0.481175 | 1.215107 | 0.510331 |
| 36.25 | 2.727413 | -0.192332 | -2.468071 | 1.942771 | -0.925448 | 0.575603 | -0.557328 | -0.313553 | 0.312196 | -0.239768 | 0.488585 | 0.980472 | 0.136725 | 0.505919 | 0.601874 | 0.95915 | 2.978403 | 1.0536 | 0.414956 | 0.432832 | 0.45878 | 1.003067 | 0.459643 |
| 36.26389 | 2.253894 | -0.214269 | -1.706156 | 2.877752 | -1.356195 | 0.360129 | -1.685072 | -0.267866 | 0.611612 | -0.337974 | 0.729528 | 0.687856 | 0.027312 | 0.351246 | 0.420536 | 1.46436 | 2.613622 | 0.958289 | 0.473489 | 0.274988 | 0.31135 | 0.842895 | 0.301582 |
| 36.27778 | 2.209373 | -0.558271 | -1.495039 | 2.443317 | -1.913958 | 0.489761 | -1.780053 | -0.287058 | 0.373945 | -0.522829 | 0.768764 | 0.380018 | -0.154325 | 0.10819 | 0.235186 | 1.263532 | 2.237395 | 0.644419 | 0.529542 | 0.057991 | 0.050611 | 0.833477 | 0.077961 |
| 36.29167 | 2.060548 | -0.196122 | -0.71642 | 2.188762 | -1.96428 | 0.348747 | -1.178383 | -0.18163 | 0.168853 | -0.55592 | 0.73815 | 0.32902 | -0.200577 | -0.038808 | 0.183154 | 1.147638 | 2.232051 | 0.505342 | 0.608844 | -0.042827 | -0.087195 | 1.015195 | -0.046842 |
| 36.30556 | 1.027792 | 1.722144 | -0.242642 | 1.666297 | -2.191131 | 0.447438 | -0.299239 | 0.011957 | 0.148383 | -0.477634 | 0.450404 | 0.489374 | -0.025512 | -0.008918 | 0.274637 | 1.187423 | 1.797939 | 0.527508 | 0.651617 | -0.006254 | -0.035504 | 1.052738 | -0.019884 |
| 36.31944 | 1.093378 | 2.424236 | 0.265067 | 2.664838 | -1.223339 | 0.448303 | 0.344095 | 0.085003 | 0.207576 | 0.343118 | 0.455595 | 0.657014 | 0.184114 | 0.067838 | 0.377383 | 1.322946 | 0.738482 | 0.506196 | 0.620117 | 0.03042 | 0.031165 | 0.908332 | 0.02835 |
| 36.33333 | 1.715823 | 2.149737 | 0.250253 | 3.069469 | -0.66476 | 0.228603 | 0.425529 | -0.105956 | 0.231173 | 1.073834 | 0.26248 | 0.755973 | 0.237679 | 0.047385 | 0.336478 | 1.267781 | 0.600422 | 0.336547 | 0.508553 | -0.01841 | -0.004752 | 0.634488 | -0.009856 |
| 36.34722 | 2.371758 | 1.239017 | -0.33095 | 4.365094 | -0.59151 | -0.606737 | -0.042082 | -0.440576 | 0.24511 | 1.00532 | -0.145651 | 0.71195 | 0.220207 | -0.0182 | 0.24361 | 0.825979 | 0.867717 | 0.161311 | 0.408697 | -0.065209 | -0.063947 | 0.407691 | -0.065373 |
| 36.36111 | 1.227992 | -0.199105 | -0.943274 | 5.580174 | -0.420604 | -1.075088 | -0.044406 | -0.747161 | 0.281121 | 0.829283 | -0.422757 | 0.696138 | 0.232479 | 0.002342 | 0.242708 | 0.209758 | -0.086099 | 0.058134 | 0.349739 | -0.018316 | -0.03634 | 0.29006 | -0.034752 |
| 36.375 | 0.284185 | -1.477313 | -1.351096 | 5.539836 | 0.075385 | -0.54684 | 0.407562 | -0.793078 | 0.266786 | 0.625966 | -0.308515 | 0.706307 | 0.218497 | 0.080654 | 0.295382 | -0.569112 | -0.397745 | -0.006957 | 0.29668 | 0.052955 | 0.035411 | 0.242345 | 0.034534 |
| 36.38889 | 0.183624 | -2.12078 | -1.302598 | 4.91711 | 0.106207 | -0.472433 | 0.538639 | -0.686621 | 0.057343 | 0.564442 | -0.119626 | 0.624198 | 0.10571 | 0.103485 | 0.277142 | -1.231384 | -0.324999 | -0.160617 | 0.114222 | 0.056082 | 0.050159 | 0.196645 | 0.047787 |
| 36.40278 | -0.171715 | -1.259431 | -1.442018 | 5.062367 | -0.398986 | 0.227662 | 0.599512 | -0.688755 | -0.38036 | 0.421904 | -0.075995 | 0.456041 | -0.01089 | 0.083223 | 0.143357 | -1.244582 | 0.008357 | -0.286344 | -0.027243 | 0.030876 | 0.038926 | 0.128164 | 0.023709 |
| 36.41667 | -0.605371 | -0.397463 | -1.078927 | 4.305061 | -0.741976 | 0.671742 | 0.717744 | -0.553258 | -0.799128 | 0.314154 | -0.076142 | 0.309788 | -0.030727 | 0.082471 | 0.006489 | -0.777232 | 0.232999 | -0.278516 | -0.069402 | 0.044612 | 0.054954 | 0.08072 | 0.025717 |
| 36.43056 | -0.792789 | 0.578548 | -0.459896 | 3.008206 | -0.521077 | 1.094414 | 0.382379 | -0.30107 | -0.888656 | 0.140761 | 0.036462 | 0.207481 | 0.019401 | 0.075627 | -0.031922 | 0.165946 | -0.036192 | -0.202869 | -0.055452 | 0.063576 | 0.065646 | 0.038876 | 0.036222 |
| 36.44444 | -0.376528 | 1.291043 | 0.409511 | 1.972669 | -0.229656 | 1.41792 | 0.278587 | -0.15323 | -0.635573 | -0.120713 | 0.164743 | 0.101984 | -0.018998 | 0.054567 | -0.052866 | 0.896956 | -0.344046 | -0.138512 | -0.091693 | 0.048035 | 0.05066 | -0.027179 | 0.0208 |
| 36.45833 | 0.156007 | 1.321031 | 0.891469 | 1.528466 | 0.153463 | 1.245479 | 0.699046 | 0.016168 | -0.24104 | -0.457099 | 0.096275 | -0.026497 | -0.10327 | 0.022313 | -0.098333 | 0.781139 | -0.833375 | -0.134022 | -0.204235 | 0.007493 | 0.015707 | -0.144034 | -0.014939 |
| 36.47222 | 0.502573 | 0.636057 | 1.184505 | 1.037756 | 0.752054 | 0.713651 | 0.696805 | 0.291113 | 0.160854 | -0.797753 | -0.149963 | -0.139571 | -0.14398 | -0.019961 | -0.139449 | 0.37646 | -1.269288 | -0.20169 | -0.367954 | -0.036547 | -0.031115 | -0.262928 | -0.052293 |
| 36.48611 | 0.636249 | -0.15631 | 1.228013 | 0.734664 | 1.257867 | 0.170012 | 0.336632 | 0.436499 | 0.438833 | -1.542091 | 0.015577 | -0.268786 | -0.170172 | -0.050353 | -0.154116 | -0.073745 | -1.240326 | -0.30677 | -0.501267 | -0.076219 | -0.076273 | -0.373336 | -0.087319 |
| 36.5 | 1.305796 | -0.740509 | 1.169653 | 0.47821 | 1.031648 | -0.235091 | 0.038172 | 0.33815 | 0.457323 | -2.411344 | 0.132751 | -0.356294 | -0.185458 | -0.073839 | -0.184059 | -0.500401 | -0.468666 | -0.408602 | -0.544324 | -0.111827 | -0.114839 | -0.446366 | -0.120391 |
| 36.51389 | 1.60661 | -0.659226 | 1.046694 | 0.249317 | 0.553711 | -0.559532 | -0.027913 | 0.094424 | 0.320918 | -1.594349 | 0.110281 | -0.614355 | -0.279779 | -0.098407 | -0.259375 | -0.977638 | -0.010955 | -0.57033 | -0.437852 | -0.1165 | -0.120966 | -0.516418 | -0.138116 |
| 36.52778 | 1.518552 | -0.019046 | 1.126288 | 0.125571 | 0.29231 | -0.275655 | -0.304311 | 0.033506 | 0.249952 | -0.317534 | 0.264316 | -0.941655 | -0.333391 | -0.080082 | -0.326014 | -0.957993 | -0.514592 | -0.539633 | -0.27021 | -0.079966 | -0.081207 | -0.530016 | -0.112476 |
| 36.54167 | 1.168433 | 0.781676 | 0.899592 | -0.016706 | 0.129337 | 0.489029 | 0.104369 | -0.181072 | 0.046271 | 0.602739 | 0.31771 | -0.696239 | -0.276111 | -0.012049 | -0.331199 | -1.407425 | -0.968888 | -0.387518 | -0.218807 | -0.028729 | -0.020209 | -0.505564 | -0.054605 |
| 36.55556 | 1.654423 | 1.20662 | 0.245716 | -0.181602 | -0.106448 | 1.389209 | 0.764197 | -0.668976 | -0.234406 | 1.300342 | 0.356073 | -0.334391 | -0.098642 | 0.145331 | -0.270022 | -1.472189 | -1.227708 | -0.259675 | -0.268953 | 0.059862 | 0.079341 | -0.464706 | 0.059291 |
| 36.56944 | 2.450445 | 1.358489 | -0.638449 | -0.14137 | -0.222782 | 2.085178 | 1.435132 | -1.04805 | -0.332386 | 1.717526 | 0.26149 | -0.036872 | 0.191005 | 0.355943 | -0.157105 | -0.533712 | -1.020114 | -0.212459 | -0.420952 | 0.186263 | 0.229144 | -0.389624 | 0.21914 |
| 36.58333 | 2.653411 | 1.40044 | -1.535883 | -0.082482 | -0.373524 | 2.22969 | 1.919596 | -1.464915 | -0.381779 | 0.950657 | 0.220738 | -0.046254 | 0.235015 | 0.252809 | -0.202212 | -0.042419 | -0.91116 | -0.443795 | -0.554243 | 0.171255 | 0.281093 | -0.524205 | 0.176066 |
| 36.59722 | 0.151768 | 1.42678 | -3.54736 | -0.555825 | -0.24233 | 1.672902 | 1.502027 | -2.213958 | -0.826869 | -0.332489 | -0.16376 | -0.239411 | -0.109224 | -0.077979 | -0.481774 | -0.897586 | -1.068458 | -1.186058 | -0.607199 | -0.02358 | 0.086215 | -0.840362 | -0.024014 |
| 36.61111 | -1.228527 | 1.006206 | -4.41831 | -1.050398 | -0.292227 | 0.792607 | 0.812861 | -3.193665 | -0.862818 | -1.345342 | -0.74028 | -0.35296 | -0.260929 | -0.16817 | -0.736595 | -1.453253 | -1.505377 | -1.570715 | -0.652689 | -0.077543 | -0.023448 | -0.957731 | -0.113928 |
| 36.625 | -1.486859 | 0.626055 | -3.571661 | -0.96263 | -0.027014 | 0.345578 | 0.1928 | -3.452644 | -0.945155 | -1.718646 | -0.604238 | -0.615091 | -0.070512 | -0.13433 | -0.941186 | -1.191632 | -1.902335 | -1.628837 | -0.699327 | -0.008879 | -0.020268 | -0.971164 | -0.104271 |
| 36.63889 | -0.573906 | 0.269057 | -2.41027 | -0.589631 | 0.044708 | -0.343474 | -0.416564 | -3.816145 | -1.118834 | -1.879077 | -0.057861 | -0.829584 | -0.001855 | -0.114484 | -1.007751 | -0.156067 | -1.549013 | -1.593679 | -0.788399 | 0.041235 | -0.005514 | -0.966152 | -0.043753 |
| 36.65278 | 0.383711 | -0.949211 | -0.89261 | -0.161896 | 0.02591 | -0.074452 | -0.026902 | -1.876825 | -0.68724 | -0.953288 | 0.510211 | -0.320932 | 0.241771 | 0.017297 | -0.731372 | 0.728465 | -0.767671 | -1.246696 | -0.841542 | 0.028461 | 0.021498 | -0.809131 | 0.029845 |
| 36.66667 | 1.397033 | -2.879864 | 0.780038 | 0.151423 | -0.028291 | 0.39985 | 0.309613 | -0.260742 | 0.562957 | -0.441889 | 0.696573 | 0.139172 | 0.392765 | 0.128784 | -0.534612 | 1.566556 | -0.356014 | -0.758202 | -0.552743 | -0.013376 | 0.034813 | -0.636648 | 0.051193 |
| 36.68056 | 1.311827 | -3.944812 | 0.861853 | -0.180036 | -0.058682 | 0.978527 | -0.100948 | 0.699737 | 1.51043 | -0.890159 | 0.326486 | 0.115323 | 0.243869 | 0.092885 | -0.67535 | 1.30463 | -0.552213 | -0.386582 | -0.257269 | -0.017411 | 0.029488 | -0.568684 | 0.005321 |
| 36.69444 | 0.261243 | -1.952888 | 1.276664 | -0.841483 | -0.067412 | 1.193695 | 0.16932 | 1.889188 | 2.235082 | -1.664566 | 0.004751 | 0.025707 | 0.215622 | 0.059998 | -0.703211 | 0.483681 | -0.718855 | 0.080238 | -0.089907 | 0.023852 | 0.054203 | -0.451242 | 0.018857 |
| 36.70833 | -0.377168 | -0.323317 | 2.268855 | -1.200809 | -0.176901 | 1.036259 | 0.947395 | 3.339849 | 2.136022 | -2.618366 | 0.06866 | 0.092115 | 0.309455 | 0.093563 | -0.649365 | -0.21156 | -1.088308 | 0.547473 | 0.008044 | 0.066656 | 0.094484 | -0.296089 | 0.062971 |
| 36.72222 | -0.87861 | 0.868336 | 2.510003 | -1.113069 | -0.430842 | 0.696058 | 1.302158 | 3.554643 | 2.303897 | -2.659041 | 0.00945 | 0.242343 | 0.362099 | 0.137183 | -0.593616 | -0.709122 | -1.321302 | 0.669285 | 0.172312 | 0.110972 | 0.137435 | -0.238804 | 0.0997 |
| 36.73611 | -1.406536 | 1.788499 | 1.54657 | -0.921624 | -0.403178 | 0.121467 | 0.748719 | 3.45177 | 2.205551 | -1.527366 | -0.194196 | 0.382613 | 0.365432 | 0.215375 | -0.407044 | -0.863562 | -1.016879 | 0.53248 | 0.487808 | 0.185418 | 0.216781 | -0.246774 | 0.161894 |
| 36.75 | -2.221966 | 2.240119 | 0.100268 | -0.478197 | -0.118106 | -0.945027 | 0.142523 | 3.221259 | 1.591033 | -0.412241 | -0.414114 | 0.448132 | 0.409154 | 0.336465 | -0.233151 | -0.586576 | -0.629837 | 0.166597 | 0.867644 | 0.285212 | 0.328266 | -0.269765 | 0.268965 |
| 36.76389 | -2.510499 | 3.248266 | 0.00636 | 0.022287 | 0.201551 | -2.016086 | -0.05212 | 2.896593 | 0.266859 | 1.011559 | -0.572863 | 0.419562 | 0.446312 | 0.418778 | -0.09575 | -0.587445 | -0.416588 | -0.14662 | 1.059401 | 0.347449 | 0.397897 | -0.238374 | 0.343048 |
| 36.77778 | -2.283506 | 3.978182 | 1.388372 | 0.379048 | 0.269216 | -1.905167 | -0.50326 | 2.441941 | -0.916149 | 3.1054 | -0.66282 | 0.342516 | 0.398635 | 0.385567 | -0.106597 | -0.999121 | -0.876474 | -0.420037 | 0.986463 | 0.325996 | 0.372646 | -0.182973 | 0.314388 |
| 36.79167 | -2.52902 | 2.985294 | 1.135269 | 0.219509 | 0.173296 | -1.08702 | -0.553444 | 0.934344 | -2.060506 | 4.028562 | -0.277719 | 0.245244 | 0.353644 | 0.313332 | -0.178305 | -0.978134 | -1.301745 | -0.640383 | 0.829024 | 0.271448 | 0.306007 | -0.108551 | 0.247256 |
| 36.80556 | -3.376165 | 1.630331 | -1.251079 | -0.716802 | 0.251183 | -0.477184 | -0.233543 | -0.588126 | -2.165434 | 2.454662 | 0.082731 | 0.188562 | 0.366914 | 0.273727 | -0.245537 | -0.392513 | -1.60927 | -0.643785 | 0.70931 | 0.23512 | 0.259572 | -0.089161 | 0.206187 |
| 36.81944 | -2.59594 | 0.707274 | -2.720769 | -2.10669 | 0.270358 | -0.248246 | -0.058003 | -1.709268 | -1.274412 | 0.587757 | 0.348837 | 0.127417 | 0.340647 | 0.239917 | -0.282016 | 0.194832 | -1.800144 | -0.501985 | 0.620567 | 0.199534 | 0.22035 | -0.081894 | 0.174758 |
| 36.83333 | -1.085053 | -0.079777 | -1.022562 | -1.425325 | 0.012988 | 0.240168 | 0.267685 | -1.681193 | 0.491271 | 0.386967 | 0.352191 | 0.116681 | 0.251055 | 0.194808 | -0.313454 | 0.465748 | -1.754978 | -0.400166 | 0.492122 | 0.156174 | 0.174985 | -0.049042 | 0.13997 |
| 36.84722 | 0.131116 | -0.575202 | 0.35661 | -1.088453 | -0.095944 | 0.829338 | 0.395109 | -1.364477 | 2.120068 | 0.246349 | 0.654855 | 0.091509 | 0.213324 | 0.150843 | -0.32292 | 0.444161 | -1.295138 | -0.272177 | 0.297726 | 0.111826 | 0.129846 | -0.02442 | 0.104632 |
| 36.86111 | -0.231105 | -0.769811 | -0.735135 | -1.158267 | 0.074576 | 1.02466 | 0.519816 | -1.228608 | 2.611405 | -0.564571 | 0.509941 | 0.018356 | 0.22409 | 0.09447 | -0.336012 | 0.367703 | -0.284214 | 0.066455 | 0.118949 | 0.063611 | 0.078814 | -0.013511 | 0.064418 |
| 36.875 | -0.655459 | -0.918719 | -0.336717 | -1.326144 | 0.124884 | 0.761578 | 0.369986 | -1.078144 | 1.993849 | -1.201566 | 0.24895 | -0.065164 | 0.232863 | 0.053169 | -0.325601 | 0.280504 | 0.781839 | 0.313799 | 0.038053 | 0.039068 | 0.049515 | -0.025104 | 0.049828 |
| 36.88889 | -0.15919 | -0.817022 | 0.461432 | -1.255094 | 0.125871 | 0.860954 | 0.182158 | -0.82909 | 1.364185 | -1.523962 | 0.332171 | -0.070345 | 0.246759 | 0.06321 | -0.254146 | 0.30214 | 1.171695 | 0.382256 | 0.064183 | 0.050209 | 0.064891 | -0.006979 | 0.076667 |
| 36.90278 | 0.724746 | -0.331592 | 0.028639 | -0.701962 | 0.371835 | 0.880626 | 0.042989 | -0.433107 | 0.475625 | -2.180386 | 0.289389 | 0.06059 | 0.350036 | 0.127096 | -0.153658 | 0.235478 | 1.092496 | 0.288845 | 0.184951 | 0.098866 | 0.118842 | 0.036088 | 0.135306 |
| 36.91667 | 1.803965 | 0.257945 | -0.118155 | -0.003423 | 0.45127 | 1.225151 | 0.126639 | -0.042276 | -0.540234 | -2.108551 | 0.081379 | 0.237887 | 0.577856 | 0.221496 | -0.043817 | 0.088537 | 1.363215 | 0.123402 | 0.33438 | 0.166237 | 0.187394 | 0.090648 | 0.192794 |
| 36.93056 | 1.49952 | 0.88419 | -0.24839 | 0.330776 | 0.240647 | 1.112907 | -0.032255 | 0.131294 | -1.511483 | -1.31443 | -0.101222 | 0.499544 | 0.750187 | 0.337159 | 0.102658 | 0.12258 | 1.858595 | 0.098448 | 0.54835 | 0.228052 | 0.251635 | 0.181775 | 0.237333 |
| 36.94444 | -0.262288 | 1.459302 | 0.179478 | 0.170208 | 0.058743 | 0.39841 | -0.290579 | 0.063129 | -1.975848 | 0.281861 | 0.033738 | 0.77812 | 0.785013 | 0.440366 | 0.251693 | 0.434619 | 2.581647 | 0.378117 | 0.787232 | 0.314673 | 0.338514 | 0.324046 | 0.301481 |
| 36.95833 | -1.508862 | 1.56071 | 0.496536 | 0.256693 | 0.147582 | -0.755612 | -0.186792 | -0.112492 | -2.076676 | 1.78752 | 0.215608 | 0.85784 | 0.694633 | 0.43729 | 0.295408 | 0.76943 | 2.874635 | 0.845967 | 0.940689 | 0.345841 | 0.377805 | 0.422943 | 0.329701 |
| 36.97222 | -1.389232 | 1.214813 | 0.02972 | 0.400309 | 0.12929 | -0.945504 | 0.02998 | -0.172335 | -0.806692 | 2.265361 | 0.227586 | 0.602037 | 0.466257 | 0.235596 | 0.145282 | 0.968997 | 2.027425 | 1.051219 | 0.851644 | 0.175677 | 0.218549 | 0.323605 | 0.166763 |
| 36.98611 | -0.30663 | 0.822021 | -0.255849 | -0.153557 | 0.361044 | -1.021892 | -0.118322 | -0.001231 | 0.85917 | 1.57697 | 0.425608 | 0.10903 | 0.122661 | -0.169086 | -0.184423 | 1.096721 | 0.58145 | 0.946564 | 0.602723 | -0.179757 | -0.141453 | 0.027024 | -0.196347 |
| 37 | -0.45684 | 0.500371 | 0.618613 | -0.81612 | 0.378234 | -0.970598 | -0.475207 | 0.15975 | 1.694316 | 1.137464 | 0.640705 | -0.264556 | -0.129352 | -0.47558 | -0.454335 | 0.831397 | -0.852852 | 0.768369 | 0.374998 | -0.443431 | -0.46356 | -0.138907 | -0.478644 |
| 37.01389 | -0.61129 | 0.220914 | 1.251039 | -1.217843 | -0.059019 | -0.342083 | 0.178988 | 0.271042 | 0.635284 | 0.567451 | 0.248493 | -0.368749 | -0.10926 | -0.517969 | -0.402298 | 0.015311 | -1.135778 | 0.673295 | 0.482258 | -0.461951 | -0.568884 | 0.011039 | -0.47028 |
| 37.02778 | 0.021371 | 0.529277 | 1.518283 | -1.332839 | -0.330943 | 0.21732 | 0.817035 | 0.541289 | 0.095821 | 0.085122 | -0.109307 | -0.082581 | 0.049809 | -0.328922 | -0.132113 | -0.729496 | -1.11547 | 0.850442 | 0.611685 | -0.277206 | -0.322028 | 0.235086 | -0.329294 |
| 37.04167 | 1.083433 | 1.67738 | 1.684519 | 0.203317 | -0.330456 | 0.644386 | 1.371725 | 0.815003 | 0.316281 | 0.304062 | -0.174031 | 0.255391 | -0.04533 | -0.166199 | 0.046901 | -0.465224 | -1.681453 | 0.875245 | 0.443741 | -0.161904 | -0.136823 | 0.273788 | -0.207941 |
| 37.05556 | 1.770797 | 2.752191 | 1.060543 | 0.468276 | -0.032482 | 0.375884 | 1.305042 | 0.773811 | 0.055729 | 0.105089 | -0.12418 | 0.410849 | -0.351882 | -0.148409 | 0.056923 | 0.368178 | -2.206454 | 0.800996 | 0.239569 | -0.171632 | -0.170156 | 0.193013 | -0.186684 |
| 37.06944 | 2.178377 | 2.897413 | 0.007521 | -0.107369 | 0.481847 | 0.225654 | 0.635574 | 0.712838 | -0.33109 | -0.765482 | -0.20464 | 0.290124 | -0.396818 | -0.100632 | 0.073753 | 0.822634 | -1.180322 | 0.704744 | 0.144058 | -0.145595 | -0.142906 | 0.158158 | -0.143255 |
| 37.08333 | 2.060768 | 2.81253 | -0.689979 | -0.244926 | 0.668684 | -0.513347 | 0.116606 | 0.749663 | -0.59255 | -1.34826 | 0.036912 | -0.165851 | -0.434027 | -0.346458 | -0.174349 | -0.162002 | -0.656638 | 0.217388 | -0.225559 | -0.433646 | -0.424971 | -0.167482 | -0.405718 |
| 37.09722 | 1.776911 | 3.020531 | -0.963793 | -0.179701 | 0.512589 | -0.344182 | 0.172226 | 0.55279 | -1.085475 | -2.127831 | 0.254316 | -1.212885 | -1.043722 | -1.184348 | -0.963149 | -2.343573 | -1.40924 | -0.709295 | -1.112644 | -1.320063 | -1.384034 | -1.000266 | -1.263007 |
| 37.11111 | 1.062566 | 2.030062 | -1.510229 | 0.890781 | 0.154669 | 0.774447 | 0.358541 | 0.024915 | -1.723284 | -2.248756 | 0.378487 | -1.804518 | -1.533119 | -1.660135 | -1.418481 | -2.534722 | -2.141895 | -0.907185 | -1.308857 | -1.741769 | -1.894635 | -1.395168 | -1.715941 |
| 37.125 | 0.004779 | 0.427299 | -1.533731 | 1.593616 | -0.179116 | 1.072625 | -0.011343 | 0.049369 | -0.152258 | -0.226714 | 0.376962 | -0.902496 | -1.032467 | -1.001898 | -0.787463 | -0.551669 | -0.897842 | -0.038133 | -0.235938 | -0.896524 | -0.988252 | -0.588743 | -0.964253 |
| 37.13889 | -0.705885 | 0.806488 | -0.395428 | 0.989259 | 0.095599 | 0.29016 | 0.169288 | 0.813602 | 1.827436 | 1.798573 | 0.83212 | 0.227023 | -0.006768 | -0.001121 | 0.216643 | 1.226387 | 0.866342 | 0.624217 | 1.307193 | 0.042188 | 0.036333 | 0.401959 | 0.005779 |
| 37.15278 | -1.005627 | 1.644387 | 0.427432 | 0.817818 | 0.207632 | 0.582384 | 0.587618 | 1.971469 | 1.905634 | 1.756681 | 1.128262 | 0.838536 | 0.725791 | 0.494747 | 0.815924 | 1.26057 | 1.591033 | 0.957951 | 1.75265 | 0.409497 | 0.455312 | 0.735252 | 0.432157 |
| 37.16667 | -0.498929 | 1.969664 | 0.517136 | 1.127427 | -0.047737 | -0.687762 | 0.760239 | 2.37535 | 1.56768 | 1.200049 | 1.431298 | 1.049856 | 1.091939 | 0.483165 | 0.9915 | 2.482791 | 2.204778 | 1.064454 | 1.223002 | 0.389816 | 0.419226 | 0.679906 | 0.402287 |
| 37.18056 | -0.507884 | 1.445378 | 0.242683 | 1.538033 | 0.009551 | -1.242451 | 0.488937 | 1.73256 | 1.126432 | 0.778642 | 1.107727 | 1.208167 | 1.088941 | 0.351956 | 1.024499 | 2.942255 | 1.804271 | 0.750702 | 0.716875 | 0.318412 | 0.322689 | 0.64389 | 0.292855 |
| 37.19444 | -1.012023 | 1.344737 | -0.175211 | 1.448039 | 0.460264 | -0.899355 | -0.376539 | 1.019649 | 0.719237 | 0.45361 | 0.66331 | 1.151939 | 0.651876 | 0.282888 | 0.851147 | 1.749831 | 1.722303 | 0.487548 | 0.32205 | 0.260162 | 0.267816 | 0.570971 | 0.239794 |
| 37.20833 | 0.564879 | 1.248037 | 0.121886 | 0.959208 | 1.225684 | -0.764442 | -0.775322 | 0.455654 | 0.480261 | 0.233443 | 0.354518 | 0.841679 | -0.000718 | 0.203381 | 0.569007 | 0.080819 | 2.107386 | 0.328389 | 0.052206 | 0.151414 | 0.161849 | 0.357779 | 0.159946 |
| 37.22222 | -0.297149 | 0.649129 | 0.685197 | 0.34396 | 1.847105 | 0.108566 | -0.998728 | -0.064399 | 0.213145 | 0.062429 | 0.120376 | 0.530147 | -0.324123 | 0.074748 | 0.326896 | -1.242347 | 1.507199 | 0.165423 | -0.036097 | 0.029552 | 0.022705 | 0.118177 | 0.037953 |
| 37.23611 | 0.022362 | 0.047613 | 0.491434 | 0.256366 | 2.450917 | 0.290879 | -0.623133 | -0.394835 | 0.053003 | -0.092428 | 0.003353 | 0.411954 | -0.210187 | 0.02994 | 0.280159 | -1.896372 | 1.139601 | 0.110587 | 0.058455 | 0.020747 | -0.004673 | 0.0563 | 0.004036 |
| 37.25 | 0.083536 | -0.352129 | -0.516316 | 0.502195 | 2.916892 | 0.292027 | -0.257248 | -0.61263 | -0.068367 | -0.455067 | -0.057506 | 0.48978 | 0.069395 | 0.125787 | 0.409205 | -2.002534 | 1.762237 | 0.584389 | 0.127021 | 0.129948 | 0.107582 | 0.201211 | 0.099917 |
| 37.26389 | -2.149052 | -0.828708 | -0.768552 | 0.295776 | 3.007467 | 0.689643 | 0.052672 | -1.022646 | -0.367863 | -0.7072 | 0.241203 | 0.540544 | 0.27413 | 0.230075 | 0.523618 | -2.118083 | 2.166399 | 0.978741 | 0.267818 | 0.226305 | 0.21145 | 0.320121 | 0.196961 |
| 37.27778 | -1.267456 | -1.387856 | 0.387884 | 0.1179 | 2.529564 | 0.594357 | 0.388062 | -0.626072 | -0.51741 | -1.36877 | 0.19639 | 0.488453 | 0.360065 | 0.248973 | 0.521904 | -1.378641 | 2.006885 | 0.805691 | 0.404232 | 0.236152 | 0.22448 | 0.309369 | 0.220684 |
| 37.29167 | 1.544674 | -1.594521 | -0.890442 | -0.074547 | 1.709721 | 0.255693 | 0.846299 | 0.202369 | -0.632549 | -1.884071 | 0.290416 | 0.392305 | 0.44452 | 0.206401 | 0.451392 | 0.292811 | 1.709124 | 0.424505 | 0.451569 | 0.181848 | 0.173648 | 0.244861 | 0.180872 |
| 37.30556 | 1.446479 | -0.670532 | -2.174857 | -0.047115 | 0.466601 | -0.444692 | 0.599348 | 0.570294 | -0.099281 | -2.666496 | 0.036387 | 0.379236 | 0.550658 | 0.129028 | 0.317281 | 1.58486 | 1.384203 | 0.159887 | 0.406335 | 0.101626 | 0.092038 | 0.168185 | 0.1087 |
| 37.31944 | 0.739023 | 0.766679 | -2.032617 | -0.006314 | -0.480393 | -0.546123 | 0.325914 | 0.684837 | 0.44715 | -2.616262 | -0.003351 | 0.328062 | 0.610512 | 0.054902 | 0.187524 | 2.3413 | 1.087036 | 0.239593 | 0.288234 | 0.02313 | 0.015592 | 0.030975 | 0.032162 |
| 37.33333 | 0.646013 | 1.446729 | -1.613476 | 0.013373 | -1.532865 | -0.293995 | -0.192907 | 0.257134 | 0.670498 | 0.473045 | 0.251791 | 0.296809 | 0.346597 | -0.012218 | 0.094849 | 2.042732 | 0.446693 | 0.448465 | 0.200873 | -0.049882 | -0.05098 | -0.089446 | -0.030089 |
| 37.34722 | 0.754077 | 1.726216 | -1.06392 | -0.027198 | -2.310169 | -0.388314 | 0.129768 | -0.402217 | 0.533101 | 4.677277 | 0.355957 | 0.240246 | -0.142372 | -0.088141 | 0.028336 | 1.898348 | -0.529339 | 0.360421 | 0.164582 | -0.104007 | -0.113762 | -0.217448 | -0.089215 |
| 37.36111 | 0.513753 | 1.936219 | -0.278468 | -0.222278 | -2.415423 | -1.582612 | 0.012304 | -0.154193 | 0.239955 | 7.134308 | 0.282069 | 0.242514 | -0.585346 | -0.110718 | 0.005827 | 1.544712 | -1.069072 | 0.158438 | 0.18824 | -0.089922 | -0.11158 | -0.292929 | -0.097288 |
| 37.375 | 0.192815 | 1.658023 | 0.551747 | -0.303477 | -2.061318 | -0.781506 | 0.0028 | 0.473856 | 0.092409 | 6.091225 | 0.133237 | 0.323389 | -0.712092 | -0.033755 | 0.043101 | 1.541039 | -0.512839 | 0.130723 | 0.271229 | -0.008113 | -0.030212 | -0.290271 | -0.031609 |
| 37.38889 | 0.129076 | 1.215189 | 1.215101 | -0.211116 | -0.996616 | -0.236689 | -0.089124 | 1.207065 | 0.094307 | 4.087738 | -0.101366 | 0.394271 | -0.506637 | 0.055694 | 0.087479 | 1.633891 | -0.107434 | 0.131138 | 0.245772 | 0.063197 | 0.049487 | -0.257207 | 0.035464 |
| 37.40278 | 0.290551 | 0.719606 | 1.211253 | -0.209678 | -0.171196 | 0.020313 | -0.057883 | 1.824079 | 0.13454 | 3.137734 | -0.133726 | 0.447726 | -0.098561 | 0.085651 | 0.098267 | 1.309619 | -0.80996 | 0.057874 | 0.154938 | 0.084334 | 0.08648 | -0.262402 | 0.055513 |
| 37.41667 | 0.264207 | 0.673728 | -0.218778 | -0.350484 | 0.020268 | -0.020957 | 0.118602 | 2.148819 | 0.202959 | 2.91731 | -0.338212 | 0.52049 | 0.311211 | 0.065725 | 0.079481 | 0.251695 | -1.378482 | -0.078646 | 0.065257 | 0.078598 | 0.095567 | -0.283293 | 0.048683 |
| 37.43056 | 0.412367 | 0.51028 | -2.047746 | -0.434648 | 0.704565 | 0.179519 | 0.449849 | 2.565732 | 0.089098 | 1.836694 | -0.165349 | 0.516438 | 0.587833 | 0.037159 | 0.060041 | -0.999221 | -0.980129 | -0.153458 | 0.040469 | 0.063488 | 0.087503 | -0.276939 | 0.037435 |
| 37.44444 | 0.546201 | -0.394766 | -3.018937 | -0.438455 | 1.600797 | 0.516829 | 0.281725 | 3.174199 | -0.093299 | 0.341897 | 0.054597 | 0.290439 | 0.656346 | 0.032656 | 0.057498 | -2.098861 | -1.875455 | -0.144256 | 0.123363 | 0.056319 | 0.080103 | -0.232477 | 0.028852 |
| 37.45833 | 0.456527 | -1.015447 | -1.693658 | -0.069676 | 1.030785 | 0.346892 | 0.551687 | 3.198193 | -0.517508 | 0.566852 | 0.297288 | -0.030551 | 0.684883 | 0.048298 | 0.086859 | -2.013758 | -1.349902 | -0.112761 | 0.156056 | 0.07166 | 0.087212 | -0.183516 | 0.037093 |
| 37.47222 | 0.414156 | 0.288148 | -0.073008 | 0.058249 | 2.131129 | 0.386972 | 1.574012 | 2.812202 | -1.355884 | 0.375099 | 0.43735 | -0.208482 | 0.670984 | 0.071525 | 0.109086 | -2.687484 | -0.628149 | -0.164219 | 0.030257 | 0.087199 | 0.090494 | -0.159275 | 0.053719 |
| 37.48611 | 0.344933 | 1.617267 | 1.559816 | -0.155541 | 0.663927 | 0.001119 | 1.29017 | 3.302899 | -1.142427 | 0.218084 | 0.288295 | -0.189346 | 0.534498 | 0.071883 | 0.11091 | -2.467841 | -0.22617 | -0.341954 | -0.14846 | 0.090263 | 0.089641 | -0.133295 | 0.060435 |
| 37.5 | 0.260552 | 1.264371 | 1.826075 | -0.312303 | 0.098247 | -0.265891 | 1.269683 | 3.642715 | -0.598266 | -0.066301 | 0.212323 | -0.116481 | 0.389272 | 0.040076 | 0.07967 | -0.917904 | -0.074119 | -0.500169 | -0.296112 | 0.087043 | 0.097671 | -0.085726 | 0.052908 |
| 37.51389 | 0.058984 | 0.489363 | 1.640766 | -0.045237 | 0.086445 | -0.393788 | 1.34691 | 2.632647 | -0.053579 | -0.280743 | 0.297485 | -0.130516 | 0.253606 | 0.001465 | 0.037842 | 0.973232 | 0.591427 | -0.591241 | -0.255011 | 0.067236 | 0.087128 | -0.055062 | 0.034535 |
| 37.52778 | -0.022448 | 0.546633 | 1.214006 | 0.055439 | 0.249503 | -0.209644 | 1.233725 | 1.942711 | 0.505668 | -1.013869 | 0.268292 | -0.112531 | 0.091294 | -0.031525 | 0.001198 | 1.35409 | 1.023813 | -0.663707 | -0.354388 | 0.018291 | 0.032721 | -0.088119 | -0.005406 |
| 37.54167 | -0.124648 | 1.260584 | 0.268782 | -0.397931 | 0.344898 | 0.084414 | 0.617259 | 1.245556 | 0.936713 | -1.509228 | 0.112281 | -0.083884 | -0.088452 | -0.066964 | -0.022571 | 0.565276 | 1.231532 | -0.847446 | -0.601001 | -0.021966 | -0.018535 | -0.121725 | -0.039801 |
| 37.55556 | -0.206083 | 1.613862 | -0.951084 | -0.789245 | 0.203378 | 0.367246 | 0.834842 | 0.300222 | 1.04141 | -1.068141 | -0.126084 | -0.1605 | -0.081334 | -0.018281 | 0.033513 | 0.342464 | 0.874879 | -1.020734 | -0.669204 | 0.019133 | 0.029203 | -0.090837 | 0.032308 |
| 37.56944 | 0.13428 | 1.116122 | -1.414825 | -0.710668 | -0.185616 | 0.901635 | 1.524615 | -0.501663 | 1.085696 | -0.736846 | -0.227936 | -0.254641 | 0.060582 | 0.114157 | 0.125326 | 0.479787 | 0.34582 | -1.194398 | -0.542068 | 0.124214 | 0.174438 | -0.069311 | 0.180996 |
| 37.58333 | 0.128767 | -0.054665 | -1.451526 | -1.040819 | 0.036415 | 0.761288 | 1.139346 | -1.297765 | 0.998405 | -2.126742 | 0.134759 | -0.357089 | -0.163673 | -0.001334 | -0.128817 | -0.055297 | -0.758436 | -1.774314 | -0.660201 | 0.043561 | 0.143481 | -0.387572 | 0.035646 |
| 37.59722 | 0.100816 | -0.086588 | -1.764941 | -1.636742 | -0.625294 | 0.264543 | -0.011668 | -1.789237 | -0.037089 | -2.4787 | 0.183184 | -0.726705 | -0.621362 | -0.33547 | -0.676356 | -1.900347 | -1.736019 | -2.276378 | -1.066446 | -0.252312 | -0.177276 | -0.984437 | -0.280094 |
| 37.61111 | -0.103573 | 0.268053 | -1.843792 | -1.832796 | -1.171338 | -0.135124 | -0.674797 | -1.826933 | -0.891184 | -2.534323 | -0.26987 | -0.891219 | -0.695872 | -0.417804 | -0.91119 | -3.316513 | -2.107699 | -2.583486 | -1.334602 | -0.278207 | -0.234293 | -1.159366 | -0.30669 |
| 37.625 | -0.27751 | -0.02615 | -2.155576 | -1.645413 | -0.72603 | -0.602004 | -0.37127 | -1.52762 | -1.417356 | -3.081596 | -0.362774 | -0.825014 | -0.349677 | -0.230069 | -0.809833 | -1.798883 | -1.461223 | -1.812383 | -1.225849 | -0.070062 | -0.054959 | -0.834995 | -0.101267 |
| 37.63889 | -0.2604 | -0.110096 | -2.548726 | -1.223637 | -0.105205 | -0.579865 | -0.299578 | -1.259558 | -1.74634 | -4.701154 | -0.048056 | -0.53327 | 0.036471 | -0.1166 | -0.675063 | 0.363933 | -0.647925 | -1.301797 | -0.930457 | 0.071322 | 0.07569 | -0.427206 | 0.045175 |
| 37.65278 | 0.074703 | 0.379394 | -1.363825 | -0.618255 | -0.198492 | -0.37226 | -0.417215 | -0.610713 | -1.076533 | -3.30147 | 0.168586 | -0.120119 | 0.388615 | -0.052433 | -0.565295 | 2.726854 | 0.070856 | -0.792864 | -0.369173 | 0.131559 | 0.140116 | -0.04459 | 0.108786 |
| 37.66667 | 0.448149 | 0.938289 | -0.564879 | -0.502865 | -0.659973 | -0.23264 | -0.540077 | 0.044617 | 0.278693 | -1.411608 | -0.00658 | 0.235223 | 0.638779 | 0.028932 | -0.448098 | 3.275848 | 0.843632 | -0.16402 | 0.128746 | 0.17098 | 0.182587 | 0.205253 | 0.150497 |
| 37.68056 | 0.739917 | 0.738227 | -1.440858 | -1.018174 | -0.983597 | -0.025549 | -0.54169 | -0.227729 | 0.744693 | 0.282874 | -0.1189 | 0.200512 | 0.677271 | 0.100592 | -0.406479 | 1.329168 | 1.388945 | 0.189798 | 0.341923 | 0.18333 | 0.201763 | 0.296239 | 0.165695 |
| 37.69444 | 0.788062 | 1.063315 | -1.89601 | -1.636055 | -1.068112 | 0.142247 | -0.363675 | -0.81122 | 0.747951 | 1.163096 | 0.276255 | 0.130766 | 0.638269 | 0.156985 | -0.378422 | -0.469521 | 1.225544 | 0.472905 | 0.410467 | 0.217591 | 0.24143 | 0.333161 | 0.205543 |
| 37.70833 | 0.639696 | 1.108462 | -2.105878 | -1.747283 | -1.308528 | 0.03239 | -0.740733 | -1.114873 | 0.733756 | 1.936194 | 0.451746 | 0.427263 | 0.567606 | 0.262752 | -0.278127 | -1.427451 | 0.688549 | 0.992934 | 0.521957 | 0.316961 | 0.343536 | 0.368629 | 0.309294 |

| | | | | | | | | | | | | | | | | | | | | | | | |
|---|---|---|---|---|---|---|---|---|---|---|---|---|---|---|---|---|---|---|---|---|---|---|---|
| 37.72222 | 0.704287 | 1.056741 | -1.735125 | -1.468085 | -1.227055 | -0.301235 | -1.09474 | -0.502023 | 1.056409 | 2.299014 | 0.133145 | 0.709088 | 0.52563 | 0.345674 | -0.166443 | -1.094338 | 0.392827 | 1.765563 | 0.640687 | 0.395267 | 0.434356 | 0.302878 | 0.400188 |
| 37.73611 | 0.622262 | 0.608438 | -0.288165 | -0.481084 | -0.3278 | 0.104725 | -0.463625 | 0.429273 | 1.421017 | 2.435724 | -0.161801 | 0.522919 | 0.374771 | 0.31028 | -0.17298 | -0.602917 | 0.482948 | 2.084614 | 0.722783 | 0.348269 | 0.387892 | 0.062854 | 0.360211 |
| 37.75 | 0.401624 | 0.345692 | 0.900827 | 0.238028 | 0.173067 | 0.453813 | 0.316451 | 0.574709 | 1.336215 | 2.348775 | -0.031422 | 0.223025 | 0.156223 | 0.183603 | -0.250205 | -0.457493 | 0.399713 | 1.559662 | 0.887596 | 0.250376 | 0.275023 | -0.239424 | 0.251707 |
| 37.76389 | 0.142811 | -0.347262 | 0.516838 | 0.728538 | 0.353263 | 0.501575 | 0.736236 | 0.033144 | 0.628043 | 1.910719 | 0.091521 | 0.153298 | 0.026654 | 0.117949 | -0.219248 | -0.470083 | -0.068626 | 0.658121 | 0.981457 | 0.213248 | 0.226092 | -0.329056 | 0.204543 |
| 37.77778 | -0.094774 | -0.735984 | -0.06158 | 1.030019 | 0.187211 | 0.435566 | 0.802121 | -0.45005 | -0.255504 | 1.313935 | 0.027875 | 0.144538 | -0.016098 | 0.097218 | -0.168755 | -0.284279 | 0.114038 | -0.162902 | 0.815787 | 0.203482 | 0.214372 | -0.270882 | 0.198018 |
| 37.79167 | -0.242824 | -0.677516 | -0.322158 | 1.070732 | 0.195932 | 0.457709 | 0.640765 | -0.508482 | -1.186976 | 1.003936 | 0.145606 | 0.048767 | -0.070796 | 0.055558 | -0.145387 | 0.303531 | 0.23751 | -0.77984 | 0.713744 | 0.18608 | 0.201127 | -0.207781 | 0.180379 |
| 37.80556 | -0.356153 | -0.15784 | -0.140504 | 1.037954 | 0.267185 | 0.499961 | 0.494666 | -0.258743 | -1.874547 | 1.513763 | 0.299766 | 0.019775 | -0.020008 | 0.070419 | -0.097462 | 0.696705 | 0.403758 | -1.089418 | 0.654235 | 0.18873 | 0.208241 | -0.074695 | 0.175332 |
| 37.81944 | -0.277654 | 0.130549 | 0.125397 | 0.725525 | 0.270794 | 0.318942 | 0.542134 | -0.162523 | -2.259781 | 2.203624 | 0.459491 | 0.223528 | 0.030734 | 0.094199 | -0.049685 | 0.733042 | 0.371662 | -1.263737 | 0.674293 | 0.196375 | 0.214695 | 0.083525 | 0.176615 |
| 37.83333 | -0.241637 | 0.092825 | 0.267146 | 0.302427 | 0.121189 | 0.018787 | 0.550963 | -0.202816 | -2.46554 | 2.724979 | 0.65394 | 0.266129 | 0.06047 | 0.068049 | -0.069859 | 0.698687 | 0.096048 | -1.171365 | 0.630735 | 0.169513 | 0.186113 | 0.215119 | 0.147889 |
| 37.84722 | -0.176223 | 0.205712 | 0.324343 | -0.005747 | -0.127678 | -0.136495 | 0.406362 | -0.038736 | -1.325673 | 3.013492 | 0.699978 | 0.010501 | 0.029606 | 0.011879 | -0.112841 | 0.671549 | -0.258236 | -0.919435 | 0.474778 | 0.105361 | 0.125625 | 0.250412 | 0.086829 |
| 37.86111 | 0.145991 | 0.74007 | 1.093529 | 0.145596 | -0.202299 | -0.015636 | 0.188098 | -0.083574 | 0.325522 | 3.442811 | 0.788804 | -0.100155 | 0.026921 | -0.020449 | -0.129274 | 0.480209 | -0.511944 | -0.716821 | 0.280901 | 0.050332 | 0.07275 | 0.319756 | 0.03881 |
| 37.875 | 0.450304 | 1.408129 | 2.190729 | 0.359819 | 0.160314 | 0.224845 | 0.169375 | -0.090022 | 1.996958 | 3.515195 | 0.675261 | 0.094672 | 0.126756 | 0.023351 | -0.085717 | 0.159199 | -0.466783 | -0.374252 | 0.13337 | 0.073621 | 0.09463 | 0.356288 | 0.05341 |
| 37.88889 | 0.385798 | 0.944759 | 1.592377 | 0.428516 | 0.736072 | 0.281959 | 0.071124 | 0.048516 | 2.891325 | 1.391764 | 0.72212 | 0.212115 | 0.279555 | 0.139376 | 0.01441 | -0.051597 | 0.12259 | 0.06044 | 0.077243 | 0.175216 | 0.207418 | 0.389026 | 0.141521 |
| 37.90278 | 0.277644 | 0.059105 | 0.733339 | 0.48826 | 0.987384 | 0.264622 | 0.042114 | 0.293121 | 2.485835 | -0.708908 | 0.705982 | 0.06277 | 0.362701 | 0.174172 | 0.081433 | -0.148552 | 0.91917 | 0.434128 | 0.053075 | 0.211746 | 0.257208 | 0.358837 | 0.181174 |
| 37.91667 | 0.216113 | -0.707629 | 0.492513 | 0.562754 | 1.240466 | 0.303177 | 0.077193 | 0.512232 | 1.574336 | -2.498696 | 0.421782 | -0.075098 | 0.332725 | 0.077977 | 0.054889 | -0.090781 | 1.148977 | 0.56059 | 0.073371 | 0.130072 | 0.165342 | 0.254615 | 0.10087 |
| 37.93056 | 0.201766 | -1.2098 | 0.484908 | 0.605901 | 1.356229 | 0.331685 | -0.171538 | 0.640909 | 0.468425 | -3.669721 | 0.212065 | 0.077499 | 0.362629 | 0.060116 | 0.076117 | 0.101291 | 1.775721 | 0.620743 | 0.250668 | 0.096246 | 0.113517 | 0.25676 | 0.060791 |
| 37.94444 | 0.008783 | -1.209058 | 0.765527 | 0.376509 | 1.233897 | 0.11388 | -0.308549 | 0.708193 | -0.398262 | -3.763201 | 0.229201 | 0.303051 | 0.45308 | 0.122346 | 0.161007 | 0.492446 | 1.802375 | 0.732365 | 0.503244 | 0.125378 | 0.142984 | 0.360126 | 0.102071 |
| 37.95833 | -0.223711 | -0.894903 | 2.036085 | -0.00053 | 0.683506 | 0.171216 | -0.025038 | 0.750198 | -1.151384 | -2.744686 | 0.134099 | 0.35976 | 0.446475 | 0.110601 | 0.193398 | 0.865905 | 1.740073 | 0.798798 | 0.656029 | 0.102672 | 0.114907 | 0.433946 | 0.09161 |
| 37.97222 | -0.360112 | -0.35349 | 3.427377 | -0.204612 | -0.185839 | 0.709577 | 0.740264 | 0.826583 | -1.363379 | -0.618083 | -0.023691 | 0.458043 | 0.359965 | 0.079597 | 0.2172 | 1.208203 | 1.492934 | 0.822746 | 0.773256 | 0.06882 | 0.072506 | 0.480434 | 0.062478 |
| 37.98611 | -0.401233 | 0.282021 | 4.035176 | -0.189485 | 0.062176 | 0.826947 | 0.881521 | 0.847082 | -0.960856 | 1.239748 | 0.238498 | 0.560309 | 0.253436 | 0.010208 | 0.202097 | 1.794178 | 1.114008 | 0.768801 | 0.85674 | 0.014991 | 0.017544 | 0.497793 | 0.007995 |
| 38 | -0.512666 | 0.475769 | 4.197951 | -0.349511 | 0.144026 | 0.580991 | 0.47454 | 0.65965 | -0.170983 | 2.84139 | 0.738972 | 0.350896 | -0.000867 | -0.205047 | 0.091624 | 1.513397 | 0.526564 | 0.743742 | 0.792402 | -0.134616 | -0.167346 | 0.43775 | -0.137167 |
| 38.01389 | -0.545161 | 0.651847 | 3.487469 | -0.535989 | -0.12204 | 0.012728 | -0.316215 | 0.326405 | 0.570331 | 5.251423 | 0.723719 | 0.210311 | -0.16786 | -0.337259 | 0.012084 | 0.949614 | 0.348281 | 0.99776 | 0.708882 | -0.249378 | -0.348693 | 0.481026 | -0.208014 |
| 38.02778 | -0.296501 | 0.083254 | 3.615083 | -0.535155 | 0.081246 | -0.759467 | -0.902524 | 0.375559 | 1.470193 | 4.346416 | 0.376561 | 0.470622 | -0.034231 | -0.162892 | 0.03386 | 0.452737 | 0.732766 | 1.507931 | 0.791108 | -0.125127 | -0.151212 | 0.596382 | -0.145214 |
| 38.04167 | -0.116733 | 1.175962 | 4.425417 | -0.183948 | 0.726474 | -0.552858 | -0.467864 | 0.938082 | 1.940387 | 3.117285 | 0.612238 | 0.631678 | 0.092412 | 0.021429 | 0.132665 | -0.24638 | 0.758341 | 1.798045 | 0.791895 | 0.013761 | 0.035978 | 0.661653 | -0.04312 |
| 38.05556 | 0.035175 | 1.275027 | 3.983336 | 0.234313 | 0.585749 | -0.275923 | 0.110434 | 1.486586 | 1.590763 | 1.890498 | 0.628713 | 0.534879 | -0.175472 | 0.046879 | 0.208927 | -0.312092 | 0.585985 | 1.723259 | 0.662893 | 0.040048 | 0.061479 | 0.625619 | 0.012153 |
| 38.06944 | 0.055561 | 0.592573 | 3.004614 | 0.395675 | 0.338219 | -0.083677 | 0.486668 | 1.782946 | 0.771683 | 0.60337 | 0.465219 | 0.172775 | -0.806888 | -0.018019 | 0.174909 | -0.297102 | 0.913872 | 1.191756 | 0.353283 | -0.014351 | 0.025869 | 0.526573 | -0.070692 |
| 38.08333 | 0.096262 | -0.15274 | 1.626622 | 0.274694 | 0.406767 | 0.487639 | 0.467352 | 1.939809 | 0.264911 | -0.124078 | 0.27767 | -0.535832 | -1.671338 | -0.671162 | -0.396888 | -0.734572 | 1.040683 | -0.077795 | -0.554187 | -0.648971 | -0.583758 | -0.253204 | -0.769423 |
| 38.09722 | 0.233341 | -0.100673 | 1.179035 | 0.108807 | 0.553682 | 0.337661 | -0.333435 | 2.260567 | -0.074251 | -0.659221 | 0.063938 | -1.551672 | -2.617718 | -1.906883 | -1.488402 | -1.738508 | -0.105404 | -1.645853 | -2.113475 | -2.015181 | -1.998896 | -1.407362 | -1.976365 |
| 38.11111 | 0.284537 | 0.601064 | 0.65166 | 0.034186 | 0.087737 | -0.235077 | -1.777311 | 1.434824 | -0.498102 | -0.37048 | -0.06608 | -1.709123 | -2.670115 | -2.294543 | -1.851073 | -1.752782 | -0.508028 | -1.545673 | -2.203977 | -2.391391 | -2.455657 | -1.582275 | -2.369649 |
| 38.125 | 0.379888 | 1.538298 | 0.31575 | 0.220965 | -0.024474 | -0.476693 | -1.657624 | 0.557995 | -0.265905 | 0.418619 | 0.233116 | -0.716915 | -1.62324 | -1.449397 | -1.094738 | -0.001169 | 0.145021 | 0.043398 | -0.617518 | -1.277109 | -1.323719 | -0.631369 | -1.380816 |
| 38.13889 | 0.632012 | 1.820823 | 0.673394 | 0.426739 | -0.09897 | -0.216484 | -1.163617 | 0.414624 | 0.392615 | 0.736157 | 0.814491 | 0.122764 | -0.733512 | -0.615378 | -0.307994 | 1.710316 | 0.918024 | 0.816559 | 0.525057 | -0.485096 | -0.517925 | 0.121474 | -0.553721 |
| 38.15278 | 0.81674 | 1.92844 | 1.049583 | 0.497337 | -0.047499 | -0.123578 | -0.995444 | 0.855936 | 0.693741 | 0.260391 | 0.933391 | 0.363018 | -0.337355 | -0.198292 | 0.149787 | 2.54692 | 1.46725 | 0.504716 | 1.120503 | -0.171725 | -0.197875 | 0.424581 | -0.18945 |
| 38.16667 | 0.765677 | 2.305184 | 1.318384 | 0.418343 | 0.329595 | -0.138832 | -1.258273 | 1.562656 | 0.396928 | 0.309192 | 0.547103 | 0.490376 | -0.198579 | -0.034143 | 0.34746 | 3.862565 | 1.129675 | 0.747641 | 1.039081 | -0.086652 | -0.112325 | 0.392456 | -0.084626 |
| 38.18056 | 0.668041 | 2.00323 | 1.500865 | 0.60584 | 0.70011 | -0.277675 | -2.252389 | 1.727873 | 0.056854 | 0.509548 | 0.744617 | 0.525268 | -0.200295 | -0.023191 | 0.354429 | 4.817293 | 0.585519 | 0.53957 | 1.124889 | -0.080916 | -0.126556 | 0.234048 | -0.09926 |
| 38.19444 | 0.662352 | 0.970764 | 2.005344 | 0.796161 | 0.49611 | -0.648241 | -2.528594 | 1.04487 | 0.056198 | 0.576305 | 0.85699 | 0.447089 | -0.2471 | -0.043355 | 0.318036 | 3.781211 | 0.428364 | 0.218992 | 0.856293 | -0.075345 | -0.14206 | 0.201321 | -0.120283 |
| 38.20833 | 0.724079 | 0.428125 | 2.273743 | 0.6966 | 0.434499 | -0.279592 | -0.754272 | 1.106353 | 0.236124 | 0.514553 | 0.655416 | 0.414266 | -0.274917 | -0.055621 | 0.350077 | 2.387501 | 0.343383 | -0.104603 | 0.265893 | -0.069473 | -0.130018 | 0.163441 | -0.111002 |
| 38.22222 | 0.646784 | -0.084233 | 1.599838 | 0.509698 | 0.428712 | -0.355661 | 0.19036 | 0.763944 | 0.206688 | 0.317865 | 0.446329 | 0.407677 | -0.279636 | -0.057334 | 0.381715 | 0.730742 | -0.100565 | -0.578112 | -0.109201 | -0.066883 | -0.115191 | 0.128529 | -0.103186 |
| 38.23611 | 0.345471 | -0.227995 | 0.869761 | 0.250459 | 0.345057 | -0.565669 | 0.267226 | 0.530221 | 0.10604 | 0.176754 | 0.374917 | 0.435892 | -0.2935 | -0.049027 | 0.39441 | -0.555423 | -0.253692 | -0.903654 | -0.448605 | -0.068164 | -0.109535 | 0.07957 | -0.109288 |
| 38.25 | 0.05677 | 0.223789 | 0.71156 | 0.249821 | -0.074171 | -0.79473 | -0.700384 | 0.798653 | 0.109414 | 0.038399 | 0.252031 | 0.486408 | -0.306306 | -0.041572 | 0.404635 | -1.965025 | -0.884179 | -0.972056 | -0.426828 | -0.074909 | -0.102162 | 0.016577 | -0.115135 |
| 38.26389 | -0.243673 | 0.373971 | 0.557895 | 0.423821 | 0.003438 | -1.07076 | -0.717751 | 0.577659 | 0.140562 | 0.008571 | 0.110382 | 0.478945 | -0.273645 | -0.022328 | 0.3779 | -3.854851 | -0.595777 | -0.196082 | -0.26796 | -0.062834 | -0.075062 | -0.056688 | -0.097902 |
| 38.27778 | -0.488792 | 0.588374 | 0.298945 | 0.562654 | 0.119679 | -0.828918 | -0.240423 | -0.480034 | 0.186999 | -0.115252 | -0.003691 | 0.461019 | -0.182503 | 0.021964 | 0.335929 | -5.009202 | -0.015978 | 0.281611 | -0.188429 | -0.021203 | -0.022678 | -0.086627 | -0.050488 |
| 38.29167 | -0.615981 | 0.86803 | 0.117436 | 0.634475 | 0.306803 | -0.458394 | 0.573266 | -0.286592 | 0.219834 | -0.197933 | -0.014962 | 0.447861 | -0.072239 | 0.066098 | 0.310117 | -4.132344 | 0.632904 | 0.011859 | -0.409811 | 0.024219 | 0.033299 | -0.027815 | 0.00708 |
| 38.30556 | -0.635484 | 0.884214 | 0.137532 | 0.611615 | 0.35471 | 0.004702 | 1.118393 | 0.295404 | 0.14783 | -0.219082 | 0.127928 | 0.361926 | -0.017511 | 0.073963 | 0.275939 | -2.921997 | 1.40983 | -0.235728 | -0.412671 | 0.042354 | 0.053196 | 0.020665 | 0.02721 |
| 38.31944 | -0.350504 | 0.850437 | 0.223969 | 0.507009 | 0.167751 | 0.588312 | 1.266235 | 0.782011 | 0.014742 | -0.476552 | 0.352477 | 0.257616 | -0.002338 | 0.061697 | 0.240749 | -1.535376 | 1.716325 | 0.108124 | -0.258038 | 0.043747 | 0.050759 | 0.087394 | 0.031209 |
| 38.33333 | -0.004809 | 0.531715 | -0.03133 | 0.310153 | 0.05204 | 0.927596 | 1.088983 | 1.220611 | -0.202075 | -0.687659 | 0.269968 | 0.171009 | 0.025714 | 0.073213 | 0.270079 | -0.044819 | 1.444741 | 0.664718 | -0.012785 | 0.053242 | 0.063717 | 0.152495 | 0.047416 |
| 38.34722 | 0.433933 | -0.496723 | -0.513364 | 0.269219 | -0.048307 | 0.983529 | 0.765725 | 1.277151 | -0.528363 | -0.960853 | -0.080007 | 0.107359 | 0.067825 | 0.082793 | 0.331292 | 1.537943 | 2.171811 | 0.911308 | 0.364094 | 0.063071 | 0.076272 | 0.163352 | 0.059352 |
| 38.36111 | 0.672603 | -1.200104 | -0.595757 | 0.255605 | -0.292837 | 0.65987 | 0.48979 | 1.362082 | -0.705948 | -0.82734 | -0.056271 | 0.068934 | 0.127233 | 0.058553 | 0.357848 | 2.932675 | 2.279053 | 0.914804 | 0.764031 | 0.053685 | 0.070252 | 0.114175 | 0.048563 |
| 38.375 | 0.511616 | -0.989141 | -0.293874 | 0.201447 | -0.31233 | 0.575329 | 0.270597 | 1.348513 | -0.738369 | -0.142743 | 0.139135 | 0.050076 | 0.212847 | 0.009028 | 0.351161 | 3.573249 | 2.262497 | 0.90836 | 0.971788 | 0.030978 | 0.058221 | 0.09484 | 0.025902 |
| 38.38889 | 0.425051 | -0.459216 | 0.093038 | 0.145437 | -0.119519 | 0.341096 | 0.12835 | 1.288909 | -0.675292 | 0.640562 | 0.162169 | 0.106052 | 0.157707 | -0.055244 | 0.309871 | 3.717297 | 2.559525 | 0.805306 | 0.957459 | 0.003252 | 0.032675 | 0.097489 | -0.015661 |
| 38.40278 | 0.395995 | 0.436961 | -0.050561 | 0.111933 | 0.222012 | -0.092932 | -0.028822 | 1.189741 | -0.515482 | 1.245889 | 0.272317 | 0.25404 | -0.634813 | -0.095132 | 0.244189 | 3.552064 | 2.431018 | 0.544213 | 0.757893 | -0.034717 | -0.021099 | 0.094724 | -0.074258 |
| 38.41667 | 0.396511 | 1.312652 | -0.336503 | 0.106662 | 0.021105 | -1.109461 | -0.201876 | 0.935783 | -0.287873 | 0.87546 | 0.143055 | 0.327796 | -1.213847 | -0.063707 | 0.174312 | 3.191863 | 1.565576 | 0.279255 | 0.440373 | -0.076389 | -0.081726 | 0.003251 | -0.12658 |
| 38.43056 | 0.412704 | 2.045094 | -0.319541 | 0.152897 | 0.137841 | -1.919932 | 0.413903 | 0.838415 | -0.123343 | 0.329069 | -0.22391 | 0.274523 | -1.339693 | -0.055847 | 0.116839 | 2.650294 | -1.39834 | 0.105584 | 0.2163 | -0.110535 | -0.124911 | -0.122141 | -0.161932 |
| 38.44444 | 0.461226 | 2.153546 | -0.15973 | 0.156771 | 0.226827 | -2.160775 | 1.094847 | 0.849797 | 0.002819 | 0.258068 | -0.195814 | 0.184778 | -1.391265 | -0.090222 | 0.091723 | 2.197831 | -1.689575 | -0.0124 | 0.135854 | -0.119468 | -0.128897 | -0.230905 | -0.160285 |
| 38.45833 | 0.361643 | 2.104078 | -0.188487 | 0.14434 | 0.050131 | -2.092521 | 1.586305 | 1.018999 | 0.115761 | 0.208638 | 0.007905 | 0.060124 | -1.280701 | -0.063914 | 0.110085 | 1.588196 | -0.879756 | -0.073676 | 0.073984 | -0.075824 | -0.071516 | -0.201646 | -0.096294 |
| 38.47222 | 0.124374 | 1.826455 | 0.067794 | 0.177139 | 0.156624 | -1.741985 | 1.760209 | 1.415969 | 0.224596 | 0.115126 | 0.035194 | 0.001271 | -0.966779 | 0.060157 | 0.209389 | 0.330754 | -0.753422 | -0.182931 | -0.010044 | 0.039803 | 0.061734 | -0.065441 | 0.040483 |
| 38.48611 | -0.150486 | 1.127172 | 0.255249 | 0.175857 | 0.492083 | -1.477619 | 1.32948 | 1.771783 | 0.175397 | 0.02861 | -0.060437 | 0.016998 | -0.447311 | 0.244961 | 0.349214 | -1.475481 | -0.227259 | -0.432534 | 0.034971 | 0.207717 | 0.240638 | 0.158338 | 0.223681 |
| 38.5 | -0.368984 | 0.190492 | 0.219635 | 0.192487 | 0.503686 | -1.476062 | 0.574113 | 1.757235 | -0.052479 | -0.077981 | -0.166188 | 0.081271 | -0.105629 | 0.403752 | 0.460499 | -3.016411 | -0.516581 | -0.777338 | -0.035501 | 0.355183 | 0.384881 | 0.352497 | 0.371835 |
| 38.51389 | -0.514006 | -0.476693 | 0.134455 | 0.130277 | 0.159928 | -1.62156 | -0.100721 | 1.876284 | -0.394868 | -0.266956 | -0.156278 | 0.045605 | 0.017015 | 0.474918 | 0.479758 | -4.076448 | -0.39523 | -0.897656 | -0.279202 | 0.422215 | 0.447144 | 0.474559 | 0.438528 |
| 38.52778 | -0.56976 | -0.6746 | 0.061906 | -0.041412 | -0.340218 | -1.756382 | -0.679911 | 2.045563 | -0.749999 | -0.485749 | -0.239392 | -0.10663 | 0.015096 | 0.448728 | 0.415591 | -4.706743 | -0.183769 | -1.184438 | -0.036689 | 0.398908 | 0.420659 | 0.541191 | 0.417056 |
| 38.54167 | -0.627377 | -0.561329 | -0.258345 | -0.289186 | -0.718106 | -2.088101 | -0.970468 | 2.282476 | -1.307651 | -0.690312 | -0.58552 | -0.021909 | 0.013788 | 0.372244 | 0.2663 | -2.980712 | -0.093864 | -1.813528 | 0.132379 | 0.331234 | 0.346537 | 0.566032 | 0.348127 |
| 38.55556 | -0.755534 | -0.527261 | -1.40061 | -0.395318 | -0.513802 | -1.850801 | -1.236939 | 2.210573 | -2.406737 | -0.463404 | -0.763749 | 0.285595 | 0.177581 | 0.384585 | 0.388469 | -1.979155 | -0.013335 | -1.733878 | 0.101097 | 0.335534 | 0.351929 | 0.584571 | 0.369849 |
| 38.56944 | -0.569405 | -0.912066 | -1.718273 | -0.221049 | 0.002983 | -0.808733 | -0.587371 | 2.320431 | -2.717854 | -0.007456 | -0.498808 | 0.582089 | 0.434396 | 0.520983 | 0.544044 | -1.318894 | 0.039376 | -1.389809 | -0.067026 | 0.430292 | 0.489762 | 0.607746 | 0.520246 |
