## Supplemental Table S2d for "Leaf movements as a quantitative metric for early stress detection"

**Supplementary Table S2.** Raw motion data from: **(d)** 100 mM KCl treatment under dimG$^{day/night}$ and RGB$^{day}$/dimG$^{night}$ with their respective controls.

| Time | RGBControl1 | RGBControl2 | RGBControl3 | RGBControl4 | RGBControl5 | RGBControl6 | RGBControl7 | RGBControl8 | RGBControl9 | dimGControl1 | dimGControl2 | dimGControl3 | dimGControl4 | dimGControl5 | dimGControl6 | dimGControl7 | dimGControl8 | dimGControl9 |
|---|---|---|---|---|---|---|---|---|---|---|---|---|---|---|---|---|---|---|
| 24.00694 | -0.329619 | -1.522532 | -1.194629 | -0.950083 | -0.299372 | -1.593943 | -0.329171 | 0.084531 | -0.426306 | -0.521689 | -1.889034 | -1.392723 | -1.091552 | -0.342768 | -1.576081 | -0.374496 | -0.096819 | -0.741038 |
| 24.02083 | -0.182888 | -1.029178 | -0.795705 | -0.550584 | -0.265909 | -0.805003 | -0.255711 | 0.024104 | -0.230874 | -0.238862 | -1.350883 | -1.076383 | -0.633803 | -0.224004 | -0.940478 | -0.273078 | 0.101338 | -0.393197 |
| 24.03472 | 0.084475 | -0.568318 | -0.365886 | -0.123747 | -0.111088 | -0.057082 | -0.027511 | -0.06597 | -0.020736 | -0.070358 | -0.8412 | -0.5981 | -0.231807 | -0.105076 | -0.237654 | -0.061933 | 0.065978 | -0.101413 |
| 24.04861 | 0.559122 | -0.165689 | -0.135156 | 0.386158 | 0.008925 | 0.368591 | 0.156402 | -0.190097 | 0.041936 | 0.27139 | -0.468568 | -0.174662 | 0.119035 | -0.038562 | 0.136617 | 0.153373 | -0.085343 | 0.08013 |
| 24.0625 | 1.07447 | 0.355972 | -0.014746 | 0.842416 | 0.165868 | 0.678065 | 0.15575 | -0.400421 | -0.154678 | 0.839268 | 0.017055 | -0.145389 | 0.440502 | -0.034951 | 0.413334 | 0.170784 | -0.392866 | -0.039085 |
| 24.07639 | 1.838242 | 1.528063 | 0.22447 | 1.367131 | 0.83429 | 1.318769 | 0.225072 | -0.253539 | -0.092186 | 1.78478 | 1.267036 | -0.298294 | 0.952348 | 0.407546 | 1.047033 | 0.158933 | -0.617251 | -0.357237 |
| 24.09028 | 2.675608 | 2.997102 | 0.245818 | 1.864147 | 1.917017 | 2.277775 | 0.487711 | 0.323246 | 0.464785 | 3.009239 | 3.074012 | -0.122316 | 1.649915 | 1.44551 | 2.009328 | 0.41883 | -0.281363 | -0.159963 |
| 24.10417 | 2.316379 | 2.737055 | 0.171586 | 1.850151 | 2.041176 | 2.971455 | 0.505664 | 0.808175 | 1.252712 | 2.78679 | 3.0579 | 0.25126 | 1.905341 | 1.763352 | 2.123693 | 0.531716 | 0.413213 | 0.675821 |
| 24.11806 | 1.031037 | 1.123699 | 0.305707 | 1.213923 | 1.056346 | 2.172413 | 0.371435 | 0.711632 | 1.455397 | 1.387258 | 1.350063 | 0.491065 | 1.506838 | 1.081911 | 2.218813 | 0.437322 | 0.807487 | 1.618184 |
| 24.13194 | 0.313906 | 0.287936 | 0.407853 | 0.561913 | 0.317282 | 1.056025 | 0.327277 | 0.336987 | 0.802229 | 0.590878 | 0.480252 | 0.524411 | 0.929142 | 0.518562 | 1.603648 | 0.418497 | 0.638054 | 1.655934 |
| 24.14583 | 0.226764 | 0.229979 | 0.257145 | 0.10495 | 0.120488 | 0.700708 | 0.199305 | 0.173137 | 0.465563 | 0.379665 | 0.346715 | 0.391261 | 0.324001 | 0.158618 | 0.970586 | 0.287381 | 0.262651 | 0.778298 |
| 24.15972 | 0.244096 | 0.223632 | -0.017054 | -0.244202 | 0.08926 | 0.606629 | 0.002697 | 0.129926 | 0.243133 | 0.157645 | 0.093264 | 0.068146 | -0.130151 | 0.035675 | 0.549508 | 0.029214 | 0.018447 | 0.253908 |
| 24.17361 | 0.412447 | 0.187211 | -0.116037 | -0.250777 | 0.128523 | 0.480472 | -0.047585 | 0.088214 | 0.090026 | 0.06533 | -0.05507 | -0.162467 | -0.34433 | 0.11879 | 0.424241 | -0.128712 | 0.022892 | 0.031546 |
| 24.1875 | 0.6723 | 0.214973 | 0.046864 | 0.064518 | 0.193239 | 0.326954 | 0.076026 | 0.05788 | 0.140708 | 0.349325 | 0.041217 | -0.058781 | -0.174086 | 0.218824 | 0.328955 | -0.026199 | 0.10235 | 0.07497 |
| 24.20139 | 0.699515 | 0.24164 | 0.259552 | 0.331023 | 0.16583 | 0.072186 | 0.196862 | 0.048021 | 0.305003 | 0.736299 | 0.210027 | 0.173969 | 0.216591 | 0.210062 | 0.192857 | 0.168742 | 0.114195 | 0.243271 |
| 24.21528 | 0.450814 | 0.173529 | 0.190811 | 0.295696 | 0.089712 | -0.301479 | 0.137709 | 0.063659 | 0.373378 | 0.743418 | 0.222018 | 0.255856 | 0.369319 | 0.089152 | -0.130377 | 0.182989 | 0.022838 | 0.328037 |
| 24.22917 | 0.009068 | -0.003472 | -0.109976 | 0.014931 | 0.01532 | -0.62152 | -0.034519 | 0.042926 | 0.275711 | 0.321633 | 0.082409 | 0.050102 | 0.168013 | 0.013977 | -0.41883 | 0.015934 | -0.025781 | 0.254087 |
| 24.24306 | -0.406812 | -0.177383 | -0.396399 | -0.334039 | -0.121169 | -0.787143 | -0.24329 | -0.013832 | 0.097752 | -0.205164 | -0.069442 | -0.215477 | -0.154689 | -0.002263 | -0.655818 | -0.134465 | 0.00834 | 0.188967 |
| 24.25694 | -0.529054 | -0.191009 | -0.390884 | -0.476095 | -0.228325 | -0.677261 | -0.355978 | 0.004045 | 0.043417 | -0.423627 | -0.061474 | -0.334032 | -0.375688 | -0.10873 | -0.604059 | -0.247956 | 0.061312 | 0.190902 |
| 24.27083 | -0.229489 | -0.049525 | -0.096879 | -0.239687 | -0.192737 | -0.579346 | -0.272638 | 0.095027 | 0.168848 | -0.333799 | 0.003216 | -0.228233 | -0.378385 | -0.228268 | -0.58434 | -0.352022 | 0.057781 | 0.214466 |
| 24.28472 | 0.29947 | 0.129161 | 0.207642 | 0.172378 | -0.137361 | -0.276099 | -0.062079 | 0.12092 | 0.28528 | -0.057354 | 0.042645 | -0.019316 | -0.164653 | -0.190067 | -0.460382 | -0.256747 | 0.054729 | 0.231753 |
| 24.29861 | 0.729468 | 0.232354 | 0.336468 | 0.579241 | -0.082947 | 0.155542 | 0.122942 | 0.060944 | 0.243737 | 0.387719 | 0.084829 | 0.264968 | 0.233527 | -0.081105 | -0.154897 | 0.005301 | 0.105965 | 0.246206 |
| 24.3125 | 0.828339 | 0.226694 | 0.287108 | 0.718048 | 0.004713 | 0.463313 | 0.234491 | 0.046104 | 0.123483 | 0.889692 | 0.229923 | 0.374672 | 0.701216 | 0.013743 | 0.296511 | 0.229568 | 0.121556 | 0.221156 |
| 24.32639 | 0.53164 | 0.105156 | 0.096034 | 0.588428 | 0.023005 | 0.573677 | 0.240418 | 0.135821 | 0.066328 | 0.947634 | 0.224996 | 0.219131 | 0.844938 | 0.061884 | 0.562895 | 0.281114 | 0.101761 | 0.110536 |
| 24.34028 | 0.415305 | -0.080454 | 0.042021 | 0.528851 | -0.037212 | 0.718056 | 0.250461 | 0.434717 | 0.25411 | 0.410034 | -0.107413 | 0.020044 | 0.597733 | 0.108397 | 0.541775 | 0.262073 | 0.226503 | 0.021276 |
| 24.35417 | 0.849783 | -0.050713 | 0.411441 | 0.76982 | -0.067905 | 0.397357 | 0.324851 | 0.970932 | 0.821592 | 0.536114 | -0.149822 | 0.169644 | 0.633513 | 0.045757 | 0.673571 | 0.330931 | 0.749792 | 0.391433 |
| 24.36806 | 0.963525 | 0.267773 | 0.873207 | 0.499829 | -0.030526 | 0.196851 | 0.437899 | 0.937995 | 1.128372 | 1.294981 | 0.301183 | 0.662627 | 0.696385 | 0.024942 | 0.4138 | 0.454692 | 1.247468 | 1.115971 |
| 24.38194 | 1.262091 | 0.513885 | 0.919304 | 0.684634 | 0.024777 | -0.076456 | 0.491642 | 0.568649 | 0.89877 | 1.447591 | 0.636736 | 0.885245 | 0.55308 | 0.019995 | 0.142919 | 0.529859 | 0.977333 | 1.182314 |
| 24.39583 | 0.897547 | 0.4083 | 0.503844 | 0.687286 | 0.021167 | -0.426802 | 0.377954 | 0.183446 | 0.400848 | 1.328212 | 0.571237 | 0.63992 | 0.666902 | 0.027009 | -0.178507 | 0.44735 | 0.437165 | 0.663797 |
| 24.40972 | 0.12906 | 0.103205 | 0.009483 | 0.294332 | -0.00366 | -0.61753 | 0.139795 | -0.080096 | -0.044777 | 0.469889 | 0.143441 | 0.193111 | 0.557936 | 0.057492 | -0.602103 | 0.218454 | 0.089569 | 0.13773 |
| 24.42361 | -0.068006 | 0.160175 | -0.151532 | -0.148334 | -0.001794 | -0.265985 | 0.025706 | -0.166156 | -0.123975 | -0.026929 | 0.077451 | -0.073277 | 0.176597 | 0.090022 | -0.533271 | 0.076472 | -0.017243 | -0.113654 |
| 24.4375 | 0.901685 | 1.501212 | -0.137486 | -0.445135 | -0.110466 | 0.655626 | 0.203964 | -0.121619 | 0.657823 | 0.757167 | 1.034228 | -0.128013 | -0.258046 | 0.112483 | 0.208879 | 0.104091 | 0.079407 | 0.30796 |
| 24.45139 | 2.896407 | 3.614088 | 0.014195 | -0.446458 | 0.021292 | 1.627179 | 0.567985 | 0.454496 | 2.026799 | 2.174009 | 2.322397 | 0.045715 | -0.345346 | 0.005506 | 1.0888 | 0.184791 | 0.738205 | 1.161313 |
| 24.46528 | 3.932692 | 4.142684 | 0.881251 | 0.136111 | 0.049351 | 1.805691 | 0.606826 | 2.383045 | 2.552956 | 2.803076 | 2.782147 | 1.001321 | 0.212431 | 0.080155 | 1.0824 | 0.519617 | 2.439106 | 1.90795 |
| 24.47917 | 3.255 | 3.085943 | 2.404454 | 1.431045 | 0.322986 | 1.403878 | 1.121616 | 5.28167 | 3.011769 | 2.653778 | 2.619539 | 2.489366 | 1.378386 | 0.359825 | 0.949884 | 1.112099 | 4.933286 | 2.729504 |
| 24.49306 | 2.121942 | 2.168577 | 2.4099 | 1.877731 | 0.459767 | 0.263473 | 0.92637 | 6.267035 | 3.499927 | 2.015287 | 2.102757 | 2.412343 | 1.871836 | 0.462461 | 0.203807 | 0.912852 | 6.232429 | 3.461202 |
| 24.5 | 1.525606 | 1.690997 | 1.525858 | 1.423272 | 0.475325 | -0.121538 | 0.555734 | 4.964117 | 2.774143 | 1.525606 | 1.690997 | 1.525858 | 1.423272 | 0.475325 | -0.121538 | 0.555734 | 4.964117 | 2.774143 |
| 24.51389 | 0.537652 | 1.260037 | 0.966465 | 0.499618 | 0.466485 | -0.377309 | 0.360036 | 2.357789 | 1.433037 | 0.537652 | 1.260037 | 0.966465 | 0.499618 | 0.466485 | -0.377309 | 0.360036 | 2.357789 | 1.433037 |
| 24.52778 | -0.883405 | 0.572073 | 0.372952 | -0.316832 | 0.446338 | -0.743161 | 0.102781 | 0.484741 | 0.35433 | -0.883405 | 0.572073 | 0.372952 | -0.316832 | 0.446338 | -0.743161 | 0.102781 | 0.484741 | 0.35433 |
| 24.54167 | -1.943064 | 0.155805 | -0.120018 | -0.954545 | 0.356849 | -1.119547 | -0.219792 | -1.051494 | -0.699155 | -1.943064 | 0.155805 | -0.120018 | -0.954545 | 0.356849 | -1.119547 | -0.219792 | -1.051494 | -0.699155 |
| 24.55556 | -1.932192 | 0.369407 | -0.487188 | -1.354264 | 0.393169 | -0.953283 | -0.367953 | -2.668877 | -1.519867 | -1.932192 | 0.369407 | -0.487188 | -1.354264 | 0.393169 | -0.953283 | -0.367953 | -2.668877 | -1.519867 |
| 24.56944 | -0.921547 | 0.692274 | -0.486429 | -1.394641 | 0.578955 | -0.178942 | -0.288072 | -4.627571 | -2.312634 | -0.921547 | 0.692274 | -0.486429 | -1.394641 | 0.578955 | -0.178942 | -0.288072 | -4.627571 | -2.312634 |
| 24.58333 | 0.142799 | 0.776588 | -0.1605 | -0.552872 | 0.939982 | 0.550465 | -0.027916 | -5.501867 | -2.36356 | 0.142799 | 0.776588 | -0.1605 | -0.552872 | 0.939982 | 0.550465 | -0.027916 | -5.501867 | -2.36356 |
| 24.59722 | 1.519901 | 0.790106 | 0.289722 | 0.463682 | 1.095528 | 0.613386 | 0.036542 | -4.512274 | -1.38781 | 1.519901 | 0.790106 | 0.289722 | 0.463682 | 1.095528 | 0.613386 | 0.036542 | -4.512274 | -1.38781 |
| 24.61111 | 2.203613 | 0.875554 | 0.630331 | 0.577917 | 0.764265 | 0.2768 | 0.222334 | -1.435806 | -0.20158 | 2.203613 | 0.875554 | 0.630331 | 0.577917 | 0.764265 | 0.2768 | 0.222334 | -1.435806 | -0.20158 |
| 24.625 | 1.415069 | 0.9299 | 0.598957 | 0.051062 | 0.278078 | -0.197207 | 0.116637 | 1.613226 | 0.649089 | 1.415069 | 0.9299 | 0.598957 | 0.051062 | 0.278078 | -0.197207 | 0.116637 | 1.613226 | 0.649089 |
| 24.63889 | 0.43634 | 0.684975 | 0.69447 | -0.384895 | -0.040187 | -0.448567 | 0.126677 | 3.848174 | 1.277044 | 0.43634 | 0.684975 | 0.69447 | -0.384895 | -0.040187 | -0.448567 | 0.126677 | 3.848174 | 1.277044 |
| 24.65278 | -0.518623 | 0.228651 | 1.032723 | -0.58803 | -0.189693 | -0.666996 | 0.202312 | 4.569364 | 1.871749 | -0.518623 | 0.228651 | 1.032723 | -0.58803 | -0.189693 | -0.666996 | 0.202312 | 4.569364 | 1.871749 |
| 24.66667 | -1.666626 | -0.142365 | 0.647083 | -0.661312 | -0.213005 | -1.098458 | -0.26703 | 4.14809 | 2.034364 | -1.666626 | -0.142365 | 0.647083 | -0.661312 | -0.213005 | -1.098458 | -0.26703 | 4.14809 | 2.034364 |
| 24.68056 | -2.43343 | -0.133698 | 0.394991 | -0.751083 | -0.209116 | -0.857436 | -0.228182 | 1.453157 | 0.904435 | -2.43343 | -0.133698 | 0.394991 | -0.751083 | -0.209116 | -0.857436 | -0.228182 | 1.453157 | 0.904435 |
| 24.69444 | -1.685317 | 0.581765 | 0.109847 | -0.788639 | 0.012598 | 0.059013 | 0.096581 | -1.281025 | -0.346931 | -1.685317 | 0.581765 | 0.109847 | -0.788639 | 0.012598 | 0.059013 | 0.096581 | -1.281025 | -0.346931 |
| 24.70833 | -0.91891 | 1.230609 | -0.115318 | -0.487126 | 0.076428 | 0.771863 | 0.449873 | -3.818686 | -1.680594 | -0.91891 | 1.230609 | -0.115318 | -0.487126 | 0.076428 | 0.771863 | 0.449873 | -3.818686 | -1.680594 |
| [illegible] | | | | | | | | | | | | | | | | | | |

| Time | RGBStress1 | RGBStress2 | RGBStress3 | RGBStress4 | RGBStress5 | RGBStress6 | RGBStress7 | RGBStress8 | RGBStress9 | RGBStress10 | dimGStress1 | dimGStress2 | dimGStress3 | dimGStress4 | dimGStress5 | dimGStress6 | dimGStress7 | dimGStress8 | dimGStress9 | dimGStress10 |
|---|---|---|---|---|---|---|---|---|---|---|---|---|---|---|---|---|---|---|---|---|
| 24 | -2.159448 | -1.08029 | -0.290082 | -0.248771 | -0.140491 | -0.23776 | -0.637132 | 0.014491 | -0.096975 | -1.634475 | -3.289443 | -1.319695 | -0.759037 | -0.320991 | -0.26295 | -0.276093 | -0.795027 | -0.197411 | 0.151016 | -1.279097 |
| 24.01389 | -0.92932 | -0.443798 | -0.163099 | -0.142001 | -0.13051 | -0.248656 | -0.322919 | -0.025512 | -0.095398 | -1.282105 | -1.982778 | -0.845516 | -0.323452 | -0.149662 | -0.170274 | -0.290734 | -0.428551 | -0.114674 | 0.004824 | -1.283453 |
| 24.02778 | -0.308167 | -0.150791 | 0.141787 | 0.091377 | 0.008086 | -0.273498 | -0.173908 | 0.030784 | -0.046329 | -0.816621 | -0.493348 | -0.23695 | 0.051291 | -0.069163 | -0.031882 | -0.282483 | -0.163006 | 0.066746 | -0.155596 | -0.821581 |
| 24.04167 | -0.393779 | -0.376637 | 0.238034 | 0.672711 | 0.423888 | -0.256118 | -0.053958 | 0.1048 | 0.028816 | -0.749655 | 0.020227 | -0.086233 | 0.370545 | 0.002025 | 0.029053 | -0.380421 | -0.181739 | 0.056393 | -0.34931 | -0.432068 |
| 24.05556 | -1.379117 | -1.428882 | -0.404508 | 0.347012 | 0.646569 | -1.323279 | -0.587804 | -0.655127 | -0.855967 | -1.460895 | -0.669323 | -1.01 | -0.180684 | -0.64276 | -0.168823 | -1.603289 | -1.036819 | -0.808912 | -0.886768 | -0.937422 |
| 24.06944 | -2.923172 | -2.745241 | -2.292642 | -2.974059 | -0.862892 | -5.000655 | -3.086477 | -2.304281 | -2.409792 | -3.165173 | -3.130448 | -3.164918 | -2.331743 | -2.856621 | -0.860338 | -5.12048 | -3.278867 | -2.806457 | -2.399389 | -2.701773 |
| 24.08333 | -2.194534 | -2.018984 | -2.594558 | -3.954 | -2.264225 | -4.610614 | -3.142473 | -1.94945 | -1.468377 | -2.715459 | -3.26364 | -2.940436 | -3.485135 | -4.236913 | -1.655769 | -4.145098 | -2.922388 | -3.039524 | -2.286402 | -3.082708 |
| 24.09722 | 0.123932 | -0.145547 | -0.455883 | -0.777061 | -0.825816 | 0.087898 | 0.060208 | -0.340899 | 0.032925 | -0.252746 | -0.454755 | -0.571441 | -1.125795 | -2.08442 | -0.906495 | -1.704291 | -1.130388 | -1.194429 | -1.047482 | -0.665866 |
| 24.11111 | 0.931431 | 0.650007 | 0.232753 | 1.182957 | 0.215872 | 2.916416 | 1.78242 | 0.745251 | 0.85918 | 0.997786 | 1.176376 | 0.677241 | 0.016611 | 0.688188 | 0.119089 | 1.78854 | 1.102097 | 0.747808 | 0.561928 | 1.030888 |
| 24.125 | 0.457231 | 0.296695 | -0.057827 | 0.745012 | 0.512235 | 1.156171 | 0.695466 | 0.453458 | 0.229962 | 0.681985 | 0.900659 | 0.424404 | -0.05041 | 1.197227 | 0.451618 | 2.131942 | 1.170038 | 0.781214 | 0.590803 | 0.938333 |
| 24.13889 | 0.008442 | -0.066484 | -0.339467 | 0.218953 | 0.297547 | -0.395353 | -0.331013 | -0.087099 | -0.274516 | 0.118317 | -0.137983 | -0.323091 | -0.701254 | 0.068703 | 0.073802 | -0.878974 | -0.565921 | -0.336447 | -0.542705 | 0.088782 |
| 24.15278 | -0.164522 | -0.177536 | -0.349336 | 0.271844 | 0.048083 | -0.892294 | -0.662649 | -0.294822 | -0.424885 | -0.152565 | -0.65464 | -0.610809 | -0.709636 | -0.636383 | -0.167461 | -2.44341 | -1.502995 | -0.695233 | -1.046084 | -0.481318 |
| 24.16667 | -0.257262 | -0.2434 | -0.276223 | 0.597382 | -0.090938 | -0.703197 | -0.670369 | -0.314496 | -0.464821 | -0.303096 | -0.533154 | -0.433451 | -0.503425 | -0.449234 | -0.09931 | -1.58035 | -0.93479 | -0.435599 | -0.760042 | -0.485338 |
| 24.18056 | -0.293743 | -0.295394 | -0.18906 | 0.754339 | -0.142063 | -0.828909 | -0.805119 | -0.313542 | -0.424744 | -0.461094 | -0.265807 | -0.210879 | -0.433822 | 0.21101 | -0.056286 | -0.285265 | -0.326061 | -0.26958 | -0.465528 | -0.347839 |
| 24.19444 | -0.261678 | -0.293244 | -0.243535 | 0.672949 | -0.14665 | -1.180136 | -0.930792 | -0.33428 | -0.179928 | -0.547128 | -0.108387 | -0.113853 | -0.48529 | 0.677855 | -0.100905 | 0.187372 | -0.195804 | -0.266894 | -0.359446 | -0.324409 |
| 24.20833 | -0.169787 | -0.232987 | -0.353605 | 0.534904 | -0.129008 | -1.232342 | -0.819323 | -0.32967 | 0.331399 | -0.592132 | -0.17443 | -0.186344 | -0.611782 | 0.638116 | -0.181487 | -0.030129 | -0.349439 | -0.374872 | -0.148647 | -0.422242 |
| 24.22222 | -0.10549 | -0.187698 | -0.427395 | 0.348064 | -0.136133 | -0.969797 | -0.595396 | -0.316284 | 0.830922 | -0.651484 | -0.258716 | -0.269155 | -0.610933 | 0.522317 | -0.221207 | -0.376051 | -0.424768 | -0.433404 | 0.294418 | -0.500907 |
| 24.23611 | -0.139978 | -0.21191 | -0.398072 | 0.102526 | -0.154475 | -0.851893 | -0.478131 | -0.271286 | 0.702696 | -0.709539 | -0.184706 | -0.218239 | -0.489343 | 0.447367 | -0.127698 | -0.498807 | -0.35184 | -0.344923 | 0.750024 | -0.457399 |
| 24.25 | -0.208962 | -0.230238 | -0.209829 | -0.087857 | -0.146304 | -0.815442 | -0.437789 | -0.210844 | 0.269372 | -0.636899 | -0.062435 | -0.144993 | -0.166873 | 0.327591 | -0.055924 | -0.506293 | -0.28069 | -0.248698 | 0.673466 | -0.428929 |
| 24.26389 | -0.233236 | -0.190422 | -0.023918 | -0.148651 | -0.11748 | -0.723508 | -0.376024 | -0.198863 | 0.043549 | -0.478764 | -0.178312 | -0.191929 | 0.025177 | 0.059103 | -0.109147 | -0.547096 | -0.29668 | -0.271714 | 0.408213 | -0.477699 |
| 24.27778 | -0.258567 | -0.183602 | -0.00094 | -0.17265 | -0.124531 | -0.614626 | -0.325338 | -0.203402 | -0.051983 | -0.390476 | -0.342216 | -0.248494 | 0.007797 | -0.098622 | -0.203935 | -0.627821 | -0.32203 | -0.336394 | 0.147029 | -0.413297 |
| 24.29167 | -0.315444 | -0.24959 | 0.016112 | -0.246816 | -0.151135 | -0.591619 | -0.322993 | -0.19772 | -0.111275 | -0.404795 | -0.298436 | -0.194476 | 0.078666 | -0.102553 | -0.152641 | -0.541201 | -0.255907 | -0.24419 | 0.148427 | -0.312532 |
| 24.30556 | -0.329855 | -0.244699 | 0.077351 | -0.229938 | -0.120786 | -0.500567 | -0.254577 | -0.132961 | -0.061444 | -0.347872 | -0.137944 | -0.08016 | 0.283947 | 0.030655 | -0.023222 | -0.315643 | -0.124631 | -0.034618 | 0.28081 | -0.148411 |
| 24.31944 | -0.208108 | -0.127186 | 0.169132 | -0.043095 | -0.038364 | -0.238778 | -0.105615 | -0.029528 | 0.054438 | -0.175111 | -0.120867 | -0.022356 | 0.300039 | 0.155973 | -0.021836 | -0.11789 | -0.032133 | 0.0358 | 0.415779 | -0.014146 |
| 24.33333 | 0.001082 | 0.014319 | 0.270962 | 0.247378 | 0.030784 | 0.187926 | 0.116792 | 0.080281 | 0.241554 | 0.053579 | -0.064465 | 0.056658 | 0.304269 | 0.329595 | 0.004428 | 0.170827 | 0.105581 | 0.104036 | 0.537621 | 0.16777 |
| 24.34722 | 0.270693 | 0.20271 | 0.446492 | 0.50861 | 0.114988 | 0.686757 | 0.419035 | 0.25176 | 0.596288 | 0.308982 | 0.271116 | 0.265913 | 0.655471 | 0.691368 | 0.183016 | 0.711007 | 0.371525 | 0.328474 | 0.77359 | 0.456568 |
| 24.36111 | 0.433589 | 0.368256 | 0.560952 | 0.567016 | 0.21814 | 0.907823 | 0.58898 | 0.435831 | 0.709628 | 0.46304 | 0.566185 | 0.433502 | 0.921658 | 0.888346 | 0.310756 | 1.047339 | 0.548114 | 0.537034 | 1.010608 | 0.598073 |
| 24.375 | 0.425808 | 0.426247 | 0.46797 | 0.494801 | 0.279891 | 0.864901 | 0.595985 | 0.43191 | 0.539218 | 0.42744 | 0.537281 | 0.426486 | 0.746508 | 0.664052 | 0.302204 | 0.913484 | 0.484887 | 0.594499 | 0.775359 | 0.465163 |
| 24.38889 | 0.39563 | 0.395693 | 0.422009 | 0.361752 | 0.280391 | 0.716223 | 0.513565 | 0.392015 | 0.312413 | 0.297195 | 0.398178 | 0.357082 | 0.488441 | 0.40323 | 0.226075 | 0.624968 | 0.39671 | 0.420496 | 0.529931 | 0.328556 |
| 24.40278 | 0.279038 | 0.256148 | 0.346262 | 0.190629 | 0.1906 | 0.487853 | 0.355481 | 0.292375 | 0.162228 | 0.146026 | 0.350918 | 0.319922 | 0.402203 | 0.277565 | 0.195586 | 0.478198 | 0.369495 | 0.315969 | 0.376504 | 0.267585 |
| 24.41667 | 0.147859 | 0.160676 | 0.131789 | 0.178777 | 0.062908 | 0.339519 | 0.25666 | 0.129375 | 0.29206 | 0.161425 | 0.295642 | 0.279015 | 0.3595 | 0.208815 | 0.197008 | 0.38588 | 0.30426 | 0.280857 | 0.201709 | 0.208321 |
| 24.43056 | 0.187783 | 0.287257 | -0.402472 | 0.680681 | -0.107923 | 0.635181 | 0.482103 | -0.012012 | 1.258413 | 0.665727 | 0.229761 | 0.287375 | 0.286065 | 0.190395 | 0.158551 | 0.365422 | 0.273138 | 0.243656 | 0.143638 | 0.18649 |
| 24.44444 | 0.433108 | 0.897877 | -1.125758 | 1.45345 | -0.333002 | 1.527022 | 1.201519 | -0.03709 | 2.969536 | 1.601341 | 0.327289 | 0.516058 | 0.380979 | 0.398246 | 0.259991 | 0.825213 | 0.545633 | 0.529408 | 0.422688 | 0.394628 |
| 24.45833 | 1.159967 | 1.892243 | -0.591807 | 2.403334 | -0.25965 | 2.56635 | 2.108846 | 0.270961 | 3.809884 | 2.107074 | 0.692759 | 0.872469 | 0.845971 | 1.059366 | 0.593258 | 2.500295 | 1.392541 | 1.196641 | 1.226733 | 0.908465 |
| 24.47222 | 1.357258 | 1.701388 | 0.736631 | 2.351565 | 0.378948 | 4.962287 | 3.036031 | 1.222568 | 3.282169 | 1.973209 | 0.90598 | 0.957011 | 1.224545 | 1.637707 | 0.741172 | 4.465517 | 2.588582 | 1.550034 | 1.96714 | 1.280044 |
| 24.48611 | 0.840832 | 0.802239 | 1.270706 | 1.745905 | 0.579373 | 3.639989 | 2.357129 | 1.132045 | 1.421115 | 1.113467 | 0.759124 | 0.677523 | 1.333258 | 1.643135 | 0.614535 | 3.478757 | 2.236093 | 1.137325 | 1.215518 | 0.994337 |
| 24.5 | 0.520417 | 0.370409 | 1.02378 | 0.920104 | 0.428795 | 1.671408 | 1.242251 | 0.570816 | 0.139002 | 0.568375 | 0.520417 | 0.370409 | 1.02378 | 0.920104 | 0.428795 | 1.671408 | 1.242251 | 0.570816 | 0.139002 | 0.568375 |
| 24.51389 | 0.357801 | 0.157898 | 0.610702 | 0.531319 | 0.27988 | 0.835207 | 0.730603 | 0.289143 | -0.066997 | 0.250671 | 0.357801 | 0.157898 | 0.610702 | 0.531319 | 0.27988 | 0.835207 | 0.730603 | 0.289143 | -0.066997 | 0.250671 |
| 24.52778 | 0.246174 | 0.043521 | 0.346581 | 0.294029 | 0.170319 | 0.487789 | 0.475872 | 0.181017 | 0.578478 | 0.079249 | 0.246174 | 0.043521 | 0.346581 | 0.294029 | 0.170319 | 0.487789 | 0.475872 | 0.181017 | 0.578478 | 0.079249 |
| 24.54167 | 0.166576 | 0.034681 | 0.218648 | 0.155559 | 0.096741 | 0.297234 | 0.321942 | 0.162905 | 1.001951 | 0.037514 | 0.166576 | 0.034681 | 0.218648 | 0.155559 | 0.096741 | 0.297234 | 0.321942 | 0.162905 | 1.001951 | 0.037514 |
| 24.55556 | 0.115846 | 0.100504 | 0.180549 | 0.079757 | 0.067094 | 0.218008 | 0.269628 | 0.197664 | 0.898526 | 0.10188 | 0.115846 | 0.100504 | 0.180549 | 0.079757 | 0.067094 | 0.218008 | 0.269628 | 0.197664 | 0.898526 | 0.10188 |
| 24.56944 | 0.107163 | 0.200601 | 0.169814 | 0.051481 | 0.077643 | 0.190393 | 0.281127 | 0.227879 | 0.717357 | 0.169923 | 0.107163 | 0.200601 | 0.169814 | 0.051481 | 0.077643 | 0.190393 | 0.281127 | 0.227879 | 0.717357 | 0.169923 |
| 24.58333 | 0.157789 | 0.297447 | 0.142365 | 0.063538 | 0.090664 | 0.174106 | 0.297556 | 0.243452 | 0.570666 | 0.204502 | 0.157789 | 0.297447 | 0.142365 | 0.063538 | 0.090664 | 0.174106 | 0.297556 | 0.243452 | 0.570666 | 0.204502 |
| 24.59722 | 0.216488 | 0.303366 | 0.101559 | 0.081081 | 0.082486 | 0.162898 | 0.289601 | 0.238555 | 0.423541 | 0.191476 | 0.216488 | 0.303366 | 0.101559 | 0.081081 | 0.082486 | 0.162898 | 0.289601 | 0.238555 | 0.423541 | 0.191476 |
| 24.61111 | 0.232435 | 0.182017 | 0.056672 | 0.075445 | 0.056256 | 0.151615 | 0.253277 | 0.171699 | 0.275314 | 0.127615 | 0.232435 | 0.182017 | 0.056672 | 0.075445 | 0.056256 | 0.151615 | 0.253277 | 0.171699 | 0.275314 | 0.127615 |
| 24.625 | 0.21173 | 0.052629 | 0.016867 | 0.057942 | 0.022117 | 0.146599 | 0.220675 | 0.095237 | 0.198775 | 0.062887 | 0.21173 | 0.052629 | 0.016867 | 0.057942 | 0.022117 | 0.146599 | 0.220675 | 0.095237 | 0.198775 | 0.062887 |
| 24.63889 | 0.187796 | 0.024667 | 0.055497 | 0.052976 | 0.012391 | 0.114422 | 0.206655 | 0.046594 | 0.233142 | 0.068149 | 0.187796 | 0.024667 | 0.055497 | 0.052976 | 0.012391 | 0.114422 | 0.206655 | 0.046594 | 0.233142 | 0.068149 |
| 24.65278 | 0.172221 | 0.124788 | 0.412889 | 0.049534 | 0.021217 | 0.051517 | 0.201927 | 0.029934 | 0.290813 | 0.124817 | 0.172221 | 0.124788 | 0.412889 | 0.049534 | 0.021217 | 0.051517 | 0.201927 | 0.029934 | 0.290813 | 0.124817 |
| 24.66667 | 0.161405 | 0.240844 | 1.374875 | 0.047281 | 0.031551 | -0.000435 | 0.191305 | 0.036611 | 0.281318 | 0.174575 | 0.161405 | 0.240844 | 1.374875 | 0.047281 | 0.031551 | -0.000435 | 0.191305 | 0.036611 | 0.281318 | 0.174575 |
| 24.68056 | 0.149166 | 0.253219 | 0.703858 | 0.050865 | 0.04831 | -0.018431 | 0.162035 | 0.061107 | 0.197801 | 0.175664 | 0.149166 | 0.253219 | 0.703858 | 0.050865 | 0.04831 | -0.018431 | 0.162035 | 0.061107 | 0.197801 | 0.175664 |
| 24.69444 | 0.159741 | 0.100108 | 0.277003 | 0.062944 | 0.085675 | 0.011756 | 0.139551 | 0.096353 | 0.090509 | 0.157341 | 0.159741 | 0.100108 | 0.277003 | 0.062944 | 0.085675 | 0.011756 | 0.139551 | 0.096353 | 0.090509 | 0.157341 |
| 24.70833 | 0.193227 | -0.053743 | 0.311613 | 0.086064 | 0.109724 | 0.07921 | 0.130536 | 0.122306 | -0.005973 | 0.140067 | 0.193227 | -0.053743 | 0.311613 | 0.086064 | 0.109724 | 0.07921 | 0.130536 | 0.122306 | -0.005973 | 0.140067 |
| [illegible] | | | | | | | | | | | | | | | | | | | | |

| | | | | | | | | | | | | | | | | | | |
|---|---|---|---|---|---|---|---|---|---|---|---|---|---|---|---|---|---|---|
| 27.91667 | -3.314543 | -3.288426 | -0.848906 | -0.745722 | -3.523937 | -2.081808 | -0.498755 | -2.099341 | -0.970299 | 0.680223 | 1.005556 | 0.793166 | 1.0318 | -0.824287 | 0.250266 | 1.249415 | -0.066191 | 0.666957 |
| 27.93056 | -4.609087 | -4.317601 | -3.061116 | -2.866471 | -3.69095 | -2.763683 | -2.149409 | -2.4482 | -2.839393 | 0.587255 | 0.792676 | 0.571599 | 1.187366 | -0.568363 | 0.384195 | 1.500819 | -0.154745 | 1.180288 |
| 27.94444 | -3.967624 | -4.413093 | -4.533519 | -3.347795 | -1.622659 | -1.666674 | -2.121043 | -1.707179 | -2.956017 | 0.135828 | 0.13267 | -0.038756 | 0.359288 | -0.626981 | 0.251028 | 1.002116 | -0.372262 | 1.086955 |
| 27.95833 | -3.094169 | -3.99789 | -4.090831 | -2.554443 | -0.384 | -0.626474 | -1.292532 | -0.9367 | -2.090033 | -1.046826 | -1.103736 | -1.448983 | -1.578705 | -1.13737 | -0.300442 | -0.58468 | -0.819148 | -0.481769 |
| 27.97222 | -1.744963 | -2.794669 | -1.933507 | -1.038103 | 0.214277 | -0.190155 | -0.508385 | -0.215245 | -1.791064 | -2.653205 | -2.880936 | -3.467189 | -3.45442 | -1.224591 | -0.861279 | -2.009994 | -0.98408 | -2.374364 |
| 27.98611 | -0.569341 | -1.169954 | -0.381811 | -0.024256 | 0.406998 | 0.013359 | -0.248255 | 0.379108 | -1.663432 | -3.287517 | -3.98202 | -4.831407 | -3.606819 | -0.506761 | -0.852376 | -1.831277 | -0.94658 | -2.624288 |
| 28.00694 | 0.190855 | 0.233013 | 0.654817 | 0.7398 | 0.374134 | 0.091307 | -0.175815 | 0.579216 | -0.971735 | -2.258389 | -3.666614 | -3.194539 | -1.950275 | 0.024538 | -0.387555 | -0.810013 | -0.404111 | -2.049625 |
| 28.02083 | 0.78316 | 1.399021 | 1.490466 | 1.127361 | 0.376858 | 0.137393 | -0.14995 | 0.535508 | -0.159773 | -1.09866 | -2.078637 | -1.184482 | -0.693682 | 0.335309 | -0.022438 | -0.286641 | 0.035837 | -1.989425 |
| 28.03472 | 1.216096 | 2.170291 | 2.092192 | 1.326164 | 0.390858 | 0.166847 | -0.118612 | 0.390905 | 0.428626 | -0.170597 | -0.464023 | 0.108454 | 0.350821 | 0.465071 | 0.120129 | -0.086486 | 0.341065 | -1.572303 |
| 28.04861 | 1.411034 | 2.403382 | 2.169026 | 1.565977 | 0.300904 | 0.051586 | 0.002277 | 0.101059 | 0.38692 | 0.423364 | 0.732271 | 1.139951 | 1.126588 | 0.345883 | 0.066507 | -0.041258 | 0.274198 | -0.778679 |
| 28.0625 | 1.462975 | 2.47167 | 1.695058 | 1.593153 | 0.114644 | 0.001429 | 0.406425 | -0.253315 | 0.051305 | 0.88619 | 1.533517 | 1.826691 | 1.51655 | 0.178528 | -0.100843 | -0.094569 | 0.074248 | -0.141885 |
| 28.07639 | 1.457934 | 2.442418 | 1.253075 | 1.269002 | 0.09157 | 0.207999 | 0.613052 | -0.288925 | -0.056579 | 1.269515 | 2.379079 | 2.329993 | 1.798095 | 0.20953 | -0.150246 | -0.184095 | 0.061717 | 0.315781 |
| 28.09028 | 1.050617 | 1.791518 | 1.033495 | 0.855866 | 0.141132 | 0.500671 | 0.807807 | 0.021276 | 0.45399 | 1.492178 | 3.025705 | 2.121893 | 1.862459 | 0.34387 | 0.034857 | -0.004345 | 0.05781 | 0.399734 |
| 28.10417 | 0.772208 | 1.095392 | 0.928186 | 0.694147 | 0.204645 | 0.533518 | 0.860436 | 0.328285 | 1.368093 | 1.581638 | 2.910088 | 1.547105 | 1.638773 | 0.313346 | 0.26778 | 0.410094 | -0.078835 | 0.288958 |
| 28.11806 | 0.694055 | 0.787227 | 0.875244 | 0.714362 | 0.175295 | 0.434881 | 0.850078 | 0.421147 | 1.806296 | 1.37912 | 2.312567 | 1.210423 | 1.115032 | 0.234042 | 0.447417 | 0.680919 | -0.046038 | 0.355839 |
| 28.13194 | 0.61292 | 0.470538 | 0.852402 | 0.637581 | 0.07229 | 0.356577 | 0.692472 | 0.275645 | 1.470735 | 0.976575 | 1.547007 | 1.034427 | 0.689907 | 0.174972 | 0.594059 | 0.862434 | 0.143197 | 1.050843 |
| 28.14583 | 0.498817 | 0.199163 | 0.915314 | 0.600007 | 0.080117 | 0.35939 | 0.737614 | 0.05477 | 0.984953 | 0.734069 | 1.118781 | 0.889994 | 0.403057 | 0.156735 | 0.513736 | 0.796665 | 0.266313 | 1.770608 |
| 28.15972 | 0.458841 | 0.140337 | 0.97314 | 0.565977 | 0.195406 | 0.418469 | 0.891716 | -0.086121 | 0.875738 | 0.646677 | 0.78845 | 0.759968 | 0.298659 | 0.021106 | 0.385364 | 0.675683 | 0.252299 | 1.793856 |
| 28.17361 | 0.535266 | 0.263493 | 1.021108 | 0.475827 | 0.216104 | 0.425222 | 0.877872 | -0.001527 | 1.036102 | 0.55817 | 0.446613 | 0.854171 | 0.2599 | 0.044852 | 0.345537 | 0.661689 | 0.197932 | 1.315816 |
| 28.1875 | 0.677596 | 0.467107 | 0.9743 | 0.281164 | 0.170504 | 0.340671 | 0.689012 | 0.194085 | 1.108386 | 0.523005 | 0.258231 | 1.029716 | 0.363558 | 0.169232 | 0.361783 | 0.781681 | 0.064034 | 1.134323 |
| 28.20139 | 0.716912 | 0.599259 | 0.711523 | 0.072697 | 0.140871 | 0.218322 | 0.487434 | 0.284727 | 0.817582 | 0.533597 | 0.218787 | 1.121845 | 0.435437 | 0.267834 | 0.356526 | 0.80632 | -0.033296 | 1.073524 |
| 28.21528 | 0.516307 | 0.434141 | 0.40096 | -0.065657 | 0.112258 | 0.117867 | 0.323627 | 0.230034 | 0.12696 | 0.574951 | 0.31474 | 1.077478 | 0.323254 | 0.274896 | 0.313519 | 0.707175 | 0.021414 | 1.12809 |
| 28.22917 | 0.177996 | -0.019243 | 0.151051 | -0.127599 | 0.096798 | 0.0749 | 0.172575 | 0.022043 | -0.60194 | 0.63256 | 0.499361 | 0.8596 | 0.136614 | 0.210328 | 0.238744 | 0.510792 | 0.151667 | 0.982391 |
| 28.24306 | -0.054532 | -0.318779 | -0.039926 | 0.032054 | 0.097744 | 0.029789 | 0.09245 | -0.209758 | -1.086827 | 0.613252 | 0.56242 | 0.534113 | -0.01291 | 0.168317 | 0.16294 | 0.348771 | 0.24677 | 0.478636 |
| 28.25694 | 0.038336 | -0.209432 | -0.057947 | 0.406408 | 0.148669 | -0.004435 | 0.164915 | -0.290668 | -0.836488 | 0.374915 | 0.250772 | 0.278829 | -0.018833 | 0.170415 | 0.120352 | 0.274844 | 0.172484 | -0.204139 |
| 28.27083 | 0.385865 | 0.136692 | 0.206664 | 0.856647 | 0.216614 | -0.002653 | 0.286668 | -0.203758 | -0.102667 | 0.030839 | -0.240712 | 0.065007 | 0.118963 | 0.135168 | 0.048872 | 0.18888 | -0.023934 | -0.817749 |
| 28.28472 | 0.701068 | 0.434641 | 0.61784 | 1.1426 | 0.214869 | 0.037832 | 0.420343 | 0.002798 | 0.591623 | -0.099251 | -0.446597 | -0.066594 | 0.419587 | 0.136896 | -0.038778 | 0.112825 | -0.224238 | -1.056179 |
| 28.29861 | 0.852718 | 0.668749 | 0.97743 | 1.041545 | 0.097965 | 0.109502 | 0.551049 | 0.180743 | 0.664094 | 0.136798 | -0.167082 | -0.025397 | 0.781979 | 0.190121 | -0.052319 | 0.142373 | -0.276435 | -0.678935 |
| 28.3125 | 0.830522 | 0.818667 | 1.097574 | 0.638975 | 0.003547 | 0.183834 | 0.521736 | 0.139603 | 0.147543 | 0.458234 | 0.245271 | 0.267812 | 1.080932 | 0.246369 | 0.007842 | 0.3178 | -0.122932 | 0.112551 |
| 28.32639 | 0.485962 | 0.474529 | 0.809083 | 0.090147 | -0.048315 | 0.220436 | 0.239549 | -0.069748 | -0.633489 | 0.804361 | 0.691055 | 0.753311 | 1.090073 | 0.241888 | 0.106522 | 0.518821 | 0.156682 | 0.719575 |
| 28.34028 | -0.119744 | -0.199196 | 0.210398 | -0.385201 | -0.091307 | 0.167133 | -0.090344 | -0.336291 | -1.364677 | 1.028117 | 1.084752 | 1.144264 | 0.723448 | 0.192938 | 0.142463 | 0.554829 | 0.272838 | 0.680219 |
| 28.35417 | -0.564058 | -0.807823 | -0.373652 | -0.490511 | -0.187127 | 0.120102 | -0.225574 | -0.499369 | -1.631797 | 0.882763 | 1.011361 | 1.2055 | 0.184752 | 0.091575 | 0.105466 | 0.32889 | 0.041053 | 0.079498 |
| 28.36806 | -0.595145 | -0.971926 | -0.694725 | -0.110123 | -0.214399 | 0.124141 | -0.125083 | -0.476473 | -1.066264 | 0.271682 | 0.263385 | 0.649486 | -0.318723 | -0.016001 | 0.062689 | -0.039092 | -0.247701 | -0.714543 |
| 28.38194 | -0.248641 | -0.686621 | -0.402219 | 0.378298 | -0.150696 | 0.128417 | 0.133549 | -0.196624 | -0.277739 | -0.442153 | -0.629849 | -0.045405 | -0.691368 | -0.077635 | 0.064238 | -0.24746 | -0.42317 | -1.343476 |
| 28.39583 | 0.16791 | -0.250888 | 0.257477 | 0.672403 | -0.013173 | 0.207146 | 0.293907 | 0.158329 | 0.245353 | -0.827745 | -1.141726 | -0.633431 | -0.555638 | -0.089642 | 0.073438 | -0.243052 | -0.441875 | -1.277154 |
| 28.40972 | 1.306766 | 0.614725 | 1.049145 | 1.031555 | 0.40936 | 0.873325 | 0.535532 | 0.564013 | 0.382442 | -0.607406 | -1.023276 | -0.75715 | -0.0446 | -0.040005 | 0.093947 | -0.014744 | -0.273326 | -0.634895 |
| 28.42361 | 3.024827 | 2.112838 | 1.832442 | 1.611422 | 1.311746 | 2.146109 | 1.200385 | 0.834932 | 0.975712 | -0.158244 | -0.602373 | -0.20401 | 0.555281 | 0.058575 | 0.13192 | 0.169351 | -0.014643 | 0.09605 |
| 28.4375 | 3.689931 | 3.077387 | 2.269512 | 1.760582 | 2.143589 | 2.682549 | 2.284885 | 1.682939 | 2.168808 | 0.238262 | -0.075953 | 0.586538 | 0.938 | 0.089853 | 0.128821 | 0.231973 | 0.166903 | 0.299086 |
| 28.45139 | 2.2893 | 1.734652 | 2.713897 | 3.040575 | 2.272509 | 1.523366 | 2.399536 | 2.105024 | 3.039499 | 0.597534 | 0.411545 | 1.406496 | 1.325608 | 0.245473 | 0.255641 | 0.595218 | 0.455641 | 0.404722 |
| 28.46528 | 1.148178 | 1.266624 | 3.227402 | 4.88356 | 1.888993 | 0.974381 | 2.6982 | 1.372728 | 3.133592 | 1.087284 | 0.806529 | 2.03521 | 2.305665 | 0.849572 | 0.606681 | 1.597312 | 1.032751 | 1.392608 |
| 28.47917 | 1.271886 | 1.50301 | 3.376044 | 5.23976 | 1.956232 | 1.034128 | 2.683489 | 0.88246 | 3.044784 | 1.176169 | 1.017711 | 2.523272 | 3.383499 | 1.631862 | 0.829012 | 2.325789 | 1.434806 | 2.370525 |
| 28.49306 | 1.25603 | 1.230316 | 2.264274 | 2.979341 | 1.605252 | 0.916453 | 2.089383 | 0.454177 | 1.655747 | 0.968354 | 1.146848 | 3.175224 | 4.953343 | 1.771955 | 0.886435 | 2.701939 | 1.280572 | 3.00264 |
| 28.5 | 0.983605 | 0.658843 | 1.344749 | 1.172588 | 0.794644 | 0.647626 | 1.475095 | -0.093488 | 0.298802 | 1.271886 | 1.50301 | 3.376044 | 5.23976 | 1.956232 | 1.034128 | 2.683489 | 0.88246 | 3.044784 |
| 28.51389 | 0.801713 | 0.173982 | 0.470436 | -0.070033 | 0.09529 | 0.568019 | 1.026922 | -0.436633 | -0.633579 | 1.25603 | 1.230316 | 2.264274 | 2.979341 | 1.605252 | 0.916453 | 2.089383 | 0.454177 | 1.655747 |
| 28.52778 | 0.884415 | -0.144176 | -0.538458 | -1.217854 | -0.309021 | 0.735866 | 0.577094 | -0.509275 | -0.664644 | 0.983605 | 0.658843 | 1.344749 | 1.172588 | 0.794644 | 0.647626 | 1.475095 | -0.093488 | 0.298802 |
| 28.54167 | 0.809337 | -0.42739 | -1.270187 | -1.583753 | -0.523849 | 0.921849 | 0.525646 | -0.158793 | -0.520186 | 0.801713 | 0.173982 | 0.470436 | -0.070033 | 0.09529 | 0.568019 | 1.026922 | -0.436633 | -0.633579 |
| 28.55556 | 0.73749 | -0.668998 | -1.149983 | -1.085478 | -0.769135 | 0.909113 | 0.552201 | -0.006578 | -0.628528 | 0.884415 | -0.144176 | -0.538458 | -1.217854 | -0.309021 | 0.735866 | 0.577094 | -0.509275 | -0.664644 |
| 28.56944 | 0.737016 | -0.642539 | -0.314371 | -0.502102 | -0.929233 | 0.646202 | 0.448192 | 0.015803 | -0.701779 | 0.809337 | -0.42739 | -1.270187 | -1.583753 | -0.523849 | 0.921849 | 0.525646 | -0.158793 | -0.520186 |
| 28.58333 | 0.526584 | -0.530525 | 0.353063 | -0.153491 | -0.999385 | 0.174528 | 0.139041 | -0.008036 | -0.736787 | 0.73749 | -0.668998 | -1.149983 | -1.085478 | -0.769135 | 0.909113 | 0.552201 | -0.006578 | -0.628528 |
| 28.59722 | 0.039736 | -0.384953 | 0.781745 | -0.238068 | -0.818759 | -0.459691 | -0.414831 | 0.069768 | -1.022284 | 0.737016 | -0.642539 | -0.314371 | -0.502102 | -0.929233 | 0.646202 | 0.448192 | 0.015803 | -0.701779 |
| 28.61111 | -0.580508 | -0.30805 | 0.775272 | -0.779566 | -0.402863 | -1.132826 | -1.205564 | 0.139529 | -1.224993 | 0.526584 | -0.530525 | 0.353063 | -0.153491 | -0.999385 | 0.174528 | 0.139041 | -0.008036 | -0.736787 |
| 28.625 | -1.382672 | -0.261537 | 0.603998 | -1.392204 | 0.046273 | -1.553377 | -1.882259 | 0.115206 | -0.978838 | 0.039736 | -0.384953 | 0.781745 | -0.238068 | -0.818759 | -0.459691 | -0.414831 | 0.069768 | -1.022284 |
| 28.63889 | -1.336162 | -0.228505 | 0.145835 | -1.069605 | 0.433047 | -1.398226 | -1.724223 | 0.036816 | -0.455962 | -0.580508 | -0.30805 | 0.775272 | -0.779566 | -0.402863 | -1.132826 | -1.205564 | 0.139529 | -1.224993 |
| 28.65278 | -0.694642 | -0.216766 | -0.386982 | -0.010468 | 1.091496 | -0.582963 | -0.91518 | -0.124927 | -0.030775 | -1.382672 | -0.261537 | 0.603998 | -1.392204 | 0.046273 | -1.553377 | -1.882259 | 0.115206 | -0.978838 |
| 28.66667 | -0.290625 | -0.252326 | -0.477556 | 0.623601 | 1.582717 | -0.036229 | -0.025807 | -0.185288 | -0.098974 | -1.336162 | -0.228505 | 0.145835 | -1.069605 | 0.433047 | -1.398226 | -1.724223 | 0.036816 | -0.455962 |
| 28.68056 | 0.068056 | -0.301599 | -0.808123 | 0.861159 | 1.408593 | 0.256584 | 0.703447 | -0.162913 | -0.779082 | -0.694642 | -0.216766 | -0.386982 | -0.010468 | 1.091496 | -0.582963 | -0.91518 | -0.124927 | -0.030775 |
| 28.69444 | 0.353442 | -0.386215 | -1.777355 | 0.478917 | 1.094014 | 0.287477 | 1.182162 | -0.2577 | -0.854086 | -0.290625 | -0.252326 | -0.477556 | 0.623601 | 1.582717 | -0.036229 | -0.025807 | -0.185288 | -0.098974 |
| 28.70833 | 0.574233 | -0.438924 | -2.308801 | -0.025899 | 0.512425 | 0.144386 | 1.421652 | -0.394686 | -0.672275 | 0.068056 | -0.301599 | -0.808123 | 0.861159 | 1.408593 | 0.256584 | 0.703447 | -0.162913 | -0.779082 |
| 28.72222 | 0.578617 | -0.675849 | -0.554937 | -0.547419 | -0.224422 | -0.080126 | 1.221011 | -0.335936 | -0.364166 | 0.353442 | -0.386215 | -1.777355 | 0.478917 | 1.094014 | 0.287477 | 1.182162 | -0.2577 | -0.854086 |
| 28.73611 | 0.668052 | -0.909843 | 0.607786 | -0.840219 | -0.989493 | -0.203902 | 0.88083 | -0.075186 | -0.247094 | 0.574233 | -0.438924 | -2.308801 | -0.025899 | 0.512425 | 0.144386 | 1.421652 | -0.394686 | -0.672275 |
| 28.75 | 0.427882 | -0.925256 | 1.066932 | -0.819018 | -1.512968 | -0.272368 | 0.265687 | 0.18621 | -0.478756 | 0.578617 | -0.675849 | -0.554937 | -0.547419 | -0.224422 | -0.080126 | 1.221011 | -0.335936 | -0.364166 |
| 28.76389 | -0.206851 | -0.746962 | 1.356553 | -0.781614 | -1.363418 | -0.316735 | -0.36344 | 0.399745 | -0.785394 | 0.668052 | -0.909843 | 0.607786 | -0.840219 | -0.989493 | -0.203902 | 0.88083 | -0.075186 | -0.247094 |
| 28.77778 | -0.942637 | -0.806041 | 1.25314 | -1.326866 | -1.242898 | -0.380722 | -1.399981 | 0.413728 | -1.411313 | 0.427882 | -0.925256 | 1.066932 | -0.819018 | -1.512968 | -0.272368 | 0.265687 | 0.18621 | -0.478756 |
| 28.79167 | -1.878863 | -1.089315 | -0.153802 | -1.906105 | -1.326042 | -0.502389 | -3.024841 | 0.22697 | -1.825651 | -0.206851 | -0.746962 | 1.356553 | -0.781614 | -1.363418 | -0.316735 | -0.36344 | 0.399745 | -0.785394 |
| 28.80556 | -2.205945 | -1.25252 | -1.484919 | -2.188891 | -0.997837 | -0.55929 | -3.430702 | -0.144482 | -2.215109 | -0.942637 | -0.806041 | 1.25314 | -1.326866 | -1.242898 | -0.380722 | -1.399981 | 0.413728 | -1.411313 |
| 28.81944 | -1.707065 | -1.170472 | -2.841811 | -1.834783 | -0.610192 | -0.476837 | -2.052511 | -0.788477 | -2.321658 | -1.878863 | -1.089315 | -0.153802 | -1.906105 | -1.326042 | -0.502389 | -3.024841 | 0.22697 | -1.825651 |
| 28.83333 | -0.953657 | -0.917127 | -2.909722 | -1.349206 | -0.395252 | -0.485394 | -0.836816 | -1.029466 | -2.025338 | -2.205945 | -1.25252 | -1.484919 | -2.188891 | -0.997837 | -0.55929 | -3.430702 | -0.144482 | -2.215109 |
| 28.84722 | -0.172873 | -0.527561 | -1.733338 | -1.032057 | -0.374906 | -0.384001 | -0.135049 | -1.055908 | -1.353893 | -1.707065 | -1.170472 | -2.841811 | -1.834783 | -0.610192 | -0.476837 | -2.052511 | -0.788477 | -2.321658 |
| 28.86111 | 0.440124 | -0.141992 | -0.264291 | -0.400486 | -0.265644 | -0.167326 | 0.24066 | -0.855518 | -0.641001 | -0.953657 | -0.917127 | -2.909722 | -1.349206 | -0.395252 | -0.485394 | -0.836816 | -1.029466 | -2.025338 |
| 28.875 | 0.674729 | 0.12867 | 0.966608 | 0.282355 | 0.067307 | 0.014373 | 0.438662 | -0.499503 | 0.192544 | -0.172873 | -0.527561 | -1.733338 | -1.032057 | -0.374906 | -0.384001 | -0.135049 | -1.055908 | -1.353893 |
| 28.88889 | 0.501776 | -0.025867 | 2.243323 | 0.691505 | 0.407958 | 0.102349 | 0.58798 | -0.284344 | 0.968186 | 0.440124 | -0.141992 | -0.264291 | -0.400486 | -0.265644 | -0.167326 | 0.24066 | -0.855518 | -0.641001 |
| 28.90278 | -1.019385 | -1.316337 | 3.327473 | 0.617276 | -0.041685 | -0.474301 | 0.696047 | -1.196541 | 1.252799 | 0.674729 | 0.12867 | 0.966608 | 0.282355 | 0.067307 | 0.014373 | 0.438662 | -0.499503 | 0.192544 |
| 28.91667 | -3.414032 | -3.658917 | 2.284869 | -0.693816 | -1.635306 | -1.909776 | 0.027816 | -3.415843 | -0.460488 | 0.657234 | 0.152272 | 2.167837 | 0.691371 | 0.500955 | 0.197006 | 0.569068 | -0.125905 | 1.028338 |
| 28.93056 | -5.448471 | -6.296367 | -1.053016 | -3.101074 | -2.729579 | -2.851147 | -1.318556 | -4.113577 | -2.758896 | 0.298546 | 0.038277 | 2.976806 | 0.790728 | 0.649507 | 0.320924 | 0.637979 | 0.155308 | 1.846581 |
| 28.94444 | -5.564733 | -7.098067 | -4.531489 | -3.54966 | -2.447257 | -2.836706 | -2.368857 | -3.546024 | -4.096658 | -0.446003 | -0.508874 | 2.50858 | 0.006429 | 0.196962 | 0.131673 | 0.318047 | -0.209039 | 1.526117 |
| 28.95833 | -3.537733 | -6.085636 | -5.45526 | -2.911119 | -1.504583 | -1.635581 | -1.705328 | -2.526095 | -3.564185 | -1.87541 | -2.090741 | -0.18431 | -1.937032 | -0.886854 | -0.382474 | -0.579225 | -1.501251 | -0.780318 |
| 28.97222 | -1.989594 | -4.414883 | -4.486528 | -1.711885 | -0.506957 | -0.719767 | -0.967885 | -1.03093 | -3.069671 | -3.595988 | -4.460326 | -3.175328 | -3.827675 | -1.94166 | -0.768697 | -1.331158 | -3.061728 | -3.331895 |
| 28.98611 | -0.527638 | -2.074262 | -2.959623 | -0.588575 | 0.188901 | -0.124029 | -0.483168 | -0.542116 | -2.575601 | -3.271101 | -5.274288 | -5.231009 | -3.726616 | -1.735801 | -0.783998 | -1.165019 | -2.220122 | -3.78285 |
| 29.00694 | 0.679548 | -0.120221 | -1.730397 | 0.111749 | 0.60938 | 0.285308 | -0.028631 | -0.140359 | -1.650171 | -2.429108 | -4.596521 | -4.372211 | -2.394688 | -0.779301 | -0.806245 | -0.696867 | -1.410652 | -2.944066 |
| 29.02083 | 1.616854 | 1.038845 | -0.832817 | 0.490351 | 0.815252 | 0.55525 | 0.41264 | 0.322532 | -0.555033 | -1.355191 | -3.061772 | -3.66743 | -1.298133 | -0.098813 | -0.453282 | -0.414429 | -0.842555 | -2.545395 |
| 29.03472 | 2.507811 | 1.746767 | 0.053233 | 0.7279 | 0.937275 | 0.582049 | 0.641039 | 0.836225 | 0.079802 | -0.079614 | -1.130189 | -2.697742 | -0.28693 | 0.346897 | -0.047719 | -0.18665 | -0.357172 | -2.146532 |
| 29.04861 | 2.932631 | 2.042734 | 0.878744 | 0.768617 | 0.938772 | 0.317337 | 0.688806 | 1.235549 | 0.197339 | 0.893812 | 0.379397 | -1.690213 | 0.346228 | 0.619217 | 0.215417 | 0.057881 | -0.040455 | -1.412435 |
| 29.0625 | 2.587766 | 1.545682 | 1.620588 | 0.734599 | 0.819516 | -0.091157 | 0.691043 | 1.32438 | 0.329835 | 1.442331 | 1.078486 | -0.642842 | 0.764061 | 0.684213 | 0.263789 | 0.360107 | 0.216279 | -0.716639 |
| 29.07639 | 1.968142 | 1.014899 | 2.793492 | 0.83542 | 0.547525 | 0.037611 | 0.828758 | 1.384328 | 0.322169 | 1.837036 | 1.449651 | 0.281955 | 1.155366 | 0.80586 | 0.031302 | 0.518198 | 0.759018 | -0.264651 |
| [illegible] | | | | | | | | | | | | | | | | | | |

[illegible]

| | | | | | | | | | | |
|---|---|---|---|---|---|---|---|---|---|---|
| 31.86111 | -0.097505 | -0.889404 | -0.527328 | -0.060573 | -0.085025 | -0.553742 | -0.765039 | -0.747772 | -0.934211 | -1.962007 |
| 31.875 | 0.648721 | -0.233384 | 0.112249 | 0.53866 | 0.135399 | -0.113589 | -0.113049 | -0.404165 | -0.105814 | -1.106744 |
| 31.88889 | 0.825064 | 0.434607 | 0.693554 | 0.944945 | 0.200249 | 0.011178 | 0.538034 | -0.401409 | 0.714419 | -0.097505 |
| 31.90278 | -0.812494 | -0.000938 | 1.216709 | 0.560781 | -0.297046 | -0.585844 | 0.735957 | -1.403515 | 1.203594 | 0.648721 |
| 31.91667 | -4.292531 | -2.012739 | 0.225649 | -1.933099 | -1.593623 | -2.00965 | -0.222096 | -3.298859 | 0.154903 | 1.075597 |
| 31.93056 | -6.366915 | -4.484189 | -2.248302 | -4.191982 | -2.192499 | -3.263809 | -2.048868 | -4.250741 | -2.283175 | 1.221614 |
| 31.94444 | -5.805382 | -4.896633 | -3.083016 | -4.587446 | -2.321565 | -3.649367 | -3.222429 | -2.645395 | -3.636879 | 0.618379 |
| 31.95833 | -3.653289 | -3.841461 | -2.557984 | -3.182717 | -1.894934 | -3.07221 | -2.567424 | -1.053392 | -3.067927 | -0.874282 |
| 31.97222 | -2.56249 | -3.517948 | -2.621087 | -2.029343 | -1.402335 | -2.618124 | -2.318676 | -0.569114 | -2.683201 | -2.476581 |
| 31.98611 | -1.978421 | -3.497291 | -2.483326 | -1.218384 | -1.258779 | -2.018952 | -1.673413 | -0.419107 | -1.631155 | -3.248408 |
| 32.00694 | -0.848571 | -2.40825 | -1.652987 | -0.402945 | -0.86102 | -1.050615 | -0.655406 | -0.086026 | -0.417558 | -3.10062 |
| 32.02083 | 0.143367 | -1.255808 | -0.703425 | 0.09804 | -0.283999 | 0.05987 | 0.050014 | 0.15713 | 0.499219 | -2.427111 |
| 32.03472 | 0.509758 | -0.76842 | -0.109724 | 0.309777 | 0.179957 | 0.940954 | 0.60539 | 0.10542 | 1.028577 | -1.565746 |
| 32.04861 | 0.715739 | -0.480013 | 0.307136 | 0.421973 | 0.488427 | 1.584992 | 0.939499 | 0.098001 | 1.300592 | -0.505206 |
| 32.0625 | 1.175647 | 0.099732 | 0.59776 | 0.545237 | 0.692538 | 1.891661 | 0.94055 | 0.249781 | 1.412344 | 0.508571 |
| 32.07639 | 1.775276 | 0.756926 | 0.592521 | 0.586956 | 0.878946 | 2.177868 | 0.952269 | 0.496901 | 1.566754 | 1.114929 |
| 32.09028 | 2.003821 | 1.124418 | 0.658077 | 0.402599 | 0.904614 | 2.279829 | 1.037451 | 0.729712 | 1.855889 | 1.406104 |
| 32.10417 | 2.003957 | 1.233178 | 0.879044 | 0.417671 | 0.877795 | 2.085259 | 1.155804 | 0.920109 | 2.300208 | 1.50972 |
| 32.11806 | 2.221602 | 1.590404 | 0.961355 | 0.699577 | 0.831224 | 1.649288 | 1.113465 | 1.278194 | 2.955525 | 1.734263 |
| 32.13194 | 2.172915 | 1.806024 | 0.859672 | 0.857483 | 0.624389 | 1.303 | 1.085698 | 1.270557 | 2.879193 | 2.05354 |
| 32.14583 | 1.760374 | 1.614231 | 0.691577 | 0.813246 | 0.473792 | 1.049324 | 1.236337 | 0.950712 | 2.851581 | 2.023842 |
| 32.15972 | 1.6629 | 1.707442 | 0.641028 | 0.703307 | 0.46949 | 0.978665 | 1.592323 | 0.776644 | 2.895265 | 1.827123 |
| 32.17361 | 1.479023 | 1.869137 | 0.665281 | 0.459821 | 0.484372 | 0.857035 | 1.758995 | 0.715907 | 2.566221 | 1.760273 |
| 32.1875 | 1.054521 | 1.744919 | 0.803407 | 0.076021 | 0.413134 | 0.537387 | 1.875411 | 0.478504 | 2.013534 | 1.523438 |
| 32.20139 | 0.849113 | 1.760955 | 1.114244 | -0.078859 | 0.398204 | 0.276961 | 2.061647 | 0.269972 | 1.578761 | 1.268294 |
| 32.21528 | 0.897607 | 1.602033 | 1.381296 | 0.050453 | 0.487254 | 0.254391 | 1.893021 | 0.24555 | 1.247558 | 1.149327 |
| 32.22917 | 1.088615 | 1.423838 | 1.49985 | 0.223675 | 0.609513 | 0.550965 | 1.805888 | 0.306331 | 0.955733 | 0.918925 |

| | | | | | | | | | | | | | |
|---|---|---|---|---|---|---|---|---|---|---|---|---|---|
| 35.80556 | -1.272396 | -0.44943 | -2.10291 | -0.144694 | -3.140116 | -2.504893 | -1.214167 | -1.791967 | -4.981229 | -1.019101 | -1.240537 | -1.346906 | 0.058391 |
| 35.81944 | -1.270978 | 0.141263 | -1.820181 | -0.359896 | -2.869884 | -2.20343 | -1.326779 | -1.611591 | -4.305671 | -1.172284 | -0.939065 | -1.925155 | 0.012844 |
| 35.83333 | -1.404292 | 0.664989 | -1.901437 | -0.493667 | -3.272227 | -1.958663 | -1.223234 | -1.580564 | -3.2411 | -1.272396 | -0.44943 | -2.10291 | -0.144694 |
| 35.84722 | -1.222378 | 1.110246 | -1.709256 | -0.36302 | -3.515272 | -2.189921 | -0.957154 | -1.545294 | -2.284918 | -1.270978 | 0.141263 | -1.820181 | -0.359896 |
| 35.86111 | -0.74211 | 1.362622 | -1.354914 | -0.090491 | -2.829686 | -1.941676 | -0.611555 | -1.388103 | -1.581833 | -1.404292 | 0.664989 | -1.901437 | -0.493667 |
| 35.875 | -0.024462 | 1.495451 | -0.762803 | 0.110085 | -1.731577 | -1.222553 | -0.237799 | -0.611087 | -1.196996 | -1.222378 | 1.110246 | -1.709256 | -0.36302 |
| 35.88889 | 0.604135 | 1.565301 | -0.085799 | 0.145301 | -0.412879 | -0.520635 | -0.173947 | 0.505714 | -0.930775 | -0.74211 | 1.362622 | -1.354914 | -0.090491 |
| 35.90278 | -0.292551 | 0.353842 | 0.272919 | -0.368102 | 0.189692 | -0.435954 | -1.730181 | 1.546551 | -1.648261 | -0.024462 | 1.495451 | -0.762803 | 0.110085 |
| 35.91667 | -3.005717 | -3.984617 | -0.636065 | -1.641497 | -1.579812 | -1.900109 | -4.716266 | 1.421214 | -5.182344 | 0.788081 | 1.750072 | -0.054745 | 0.227175 |
| 35.93056 | -5.318892 | -8.143591 | -2.668817 | -2.05456 | -4.373431 | -3.865042 | -5.440134 | -1.095564 | -8.296648 | 1.266411 | 1.946798 | 0.537817 | 0.298147 |
| 35.94444 | -5.112944 | -5.63039 | -2.407674 | -1.084656 | -5.469473 | -4.267728 | -3.334625 | -4.708707 | -7.175763 | 0.481787 | 0.917664 | 0.48047 | -0.096255 |
| 35.95833 | -2.615951 | -2.114639 | -1.032701 | -0.114865 | -3.95224 | -2.954807 | -1.431295 | -6.281677 | -4.899505 | -1.934433 | -1.606703 | -0.873011 | -1.181904 |
| 35.97222 | -1.065832 | -1.101405 | -0.427873 | 0.470707 | -3.020937 | -1.9807 | -0.525011 | -5.429275 | -3.542412 | -3.733692 | -3.289812 | -3.129167 | -1.467261 |
| 35.98611 | -0.466143 | -0.987185 | -0.173077 | 0.498787 | -2.774601 | -1.379674 | -0.128314 | -3.0103 | -2.830941 | -2.749072 | -2.138331 | -2.405347 | -0.718593 |
| 36.00694 | -0.171364 | -1.049756 | 0.01766 | 0.349954 | -2.118152 | -0.990512 | 0.028513 | -0.777851 | -1.529555 | -1.439768 | -1.158961 | -1.373345 | 0.132489 |
| 36.02083 | 0.063796 | -1.036352 | 0.106387 | 0.30404 | -1.643148 | -0.834556 | 0.07071 | 0.959109 | 0.009448 | -0.815955 | -0.760601 | -1.051374 | 0.416075 |
| 36.03472 | 0.227554 | -0.979567 | 0.209214 | 0.340879 | -1.479001 | -0.71013 | 0.114182 | 2.480444 | 1.33919 | -0.318072 | -0.789737 | -0.826416 | 0.396317 |
| 36.04861 | 0.383624 | -1.066835 | 0.351665 | 0.389367 | -0.839836 | -0.487235 | 0.230374 | 3.869347 | 1.967152 | -0.052556 | -1.025306 | -0.633216 | 0.29592 |
| 36.0625 | 0.60075 | -1.273804 | 0.529757 | 0.497021 | 0.056743 | -0.236891 | 0.360123 | 5.118152 | 1.940827 | 0.10236 | -1.165493 | -0.372423 | 0.285921 |
| 36.07639 | 0.949104 | -1.240868 | 0.707789 | 0.619872 | 1.026945 | 0.033904 | 0.475315 | 6.267906 | 1.877532 | 0.263476 | -1.085451 | -0.032513 | 0.390514 |
| 36.09028 | 1.490381 | -0.856053 | 0.901488 | 0.729833 | 1.929286 | 0.413264 | 0.597389 | 5.97577 | 1.952023 | 0.454059 | -1.147197 | 0.321586 | 0.48509 |
| 36.10417 | 1.964037 | -0.291603 | 1.100087 | 0.884678 | 2.762142 | 0.962852 | 0.640117 | 5.537392 | 1.71053 | 0.832386 | -1.094891 | 0.635784 | 0.559023 |
| 36.11806 | 2.263434 | 0.165218 | 1.39003 | 1.013991 | 3.752224 | 1.427295 | 0.533889 | 5.251575 | 1.601536 | 1.384142 | -0.790112 | 0.931771 | 0.686812 |
| 36.13194 | 2.323279 | 0.600882 | 1.671601 | 1.090773 | 4.98498 | 2.232282 | 0.46694 | 4.204025 | 1.452553 | 1.845789 | -0.330988 | 1.267196 | 0.796683 |
| 36.14583 | 2.215358 | 0.912754 | 1.768666 | 1.070222 | 5.304168 | 2.214227 | 0.507969 | 3.552166 | 1.554494 | 2.176145 | 0.049412 | 1.681613 | 0.898866 |
| 36.15972 | 2.158642 | 1.069743 | 1.687158 | 1.035384 | 5.014682 | 1.800337 | 0.544934 | 3.61049 | 1.996188 | 2.299938 | 0.316673 | 2.13622 | 0.977 |
| 36.17361 | 2.112734 | 1.110016 | 1.457465 | 1.003617 | 4.592551 | 0.928005 | 0.580578 | 3.214235 | 2.086983 | 2.176075 | 0.57523 | 2.553259 | 1.057438 |
| 36.1875 | 2.061857 | 1.050088 | 1.261839 | 1.046849 | 4.539167 | 0.788784 | 0.670453 | 2.678734 | 1.841045 | 2.148574 | 0.969412 | 2.525988 | 1.0833 |
| 36.20139 | 1.997056 | 0.946822 | 1.088705 | 1.112907 | 4.384943 | 0.972101 | 0.724788 | 2.420356 | 1.58178 | 2.298072 | 1.296381 | 2.284543 | 1.097706 |
| 36.21528 | 2.080199 | 0.913138 | 0.974728 | 1.207542 | 3.725311 | 0.868384 | 0.756864 | 2.105662 | 1.347659 | 2.254519 | 1.296755 | 1.725681 | 1.072479 |
| 36.22917 | 1.890004 | 0.961382 | 0.940858 | 1.297788 | 2.767391 | 0.410138 | 0.775888 | 1.230597 | 0.655113 | 2.193917 | 0.989699 | 1.292191 | 1.061743 |
| 36.24306 | 1.693563 | 1.098379 | 0.980194 | 1.338667 | 1.702831 | -0.054147 | 0.829085 | 0.236237 | 0.129397 | 2.136007 | 0.761982 | 0.999288 | 1.114252 |
| 36.25694 | 1.565402 | 1.292663 | 1.037548 | 1.296585 | 0.735805 | -0.253387 | 0.928763 | -0.491551 | -0.089185 | 2.329864 | 0.858147 | 0.908494 | 1.245797 |
| 36.27083 | 1.36574 | 1.369494 | 1.040674 | 1.21399 | -0.057157 | -0.327258 | 1.01014 | -1.189327 | -0.296922 | 2.147967 | 1.166838 | 0.915818 | 1.372821 |
| 36.28472 | 1.131532 | 1.395787 | 0.99779 | 1.181457 | -0.612817 | -0.425312 | 1.071966 | -1.82511 | -0.457083 | 1.914614 | 1.274796 | 0.816012 | 1.402124 |
| 36.29861 | 0.996911 | 1.385377 | 1.0539 | 1.148522 | -1.000932 | -0.681188 | 1.159623 | -2.162483 | -0.429195 | 1.59335 | 1.255138 | 0.705503 | 1.369933 |
| 36.3125 | 0.931328 | 1.215503 | 1.122012 | 1.079559 | -1.005539 | -0.915429 | 1.150471 | -1.953324 | -0.396252 | 1.302232 | 1.248909 | 0.69758 | 1.322456 |
| 36.32639 | 0.90873 | 0.960505 | 1.128625 | 1.016304 | -0.944904 | -0.951983 | 1.070822 | -1.56759 | -0.346213 | 1.073864 | 1.293413 | 0.740062 | 1.290336 |
| 36.34028 | 0.980579 | 0.933482 | 1.110545 | 1.058467 | -1.125197 | -0.856743 | 1.064712 | -1.491608 | -0.500897 | 1.002347 | 1.263354 | 0.825905 | 1.234811 |
| 36.35417 | 1.093104 | 0.90755 | 1.037226 | 1.112368 | -1.528403 | -0.853719 | 1.104452 | -1.77254 | -0.810255 | 0.948067 | 1.05839 | 0.826798 | 1.144666 |
| 36.36806 | 0.953494 | 0.644696 | 0.949768 | 1.067228 | -1.845667 | -0.997935 | 1.110046 | -1.988654 | -1.117796 | 0.923398 | 0.808404 | 0.7979 | 1.095528 |
| 36.38194 | 0.500219 | 0.249266 | 0.89824 | 1.013786 | -1.913882 | -1.133871 | 1.027668 | -2.024763 | -1.284848 | 0.940075 | 0.646099 | 0.790872 | 1.112301 |
| 36.39583 | 0.553801 | 0.386378 | 1.005546 | 1.023175 | -1.749534 | -1.174356 | 1.091773 | -1.941829 | -1.070828 | 1.159461 | 0.570474 | 0.799955 | 1.137035 |
| 36.40972 | 1.403025 | 2.167789 | 1.907553 | 1.184379 | -0.721547 | -1.126659 | 1.987459 | -1.575588 | -0.160522 | 1.025888 | 0.472109 | 0.777397 | 1.11542 |
| 36.42361 | 3.143656 | 5.739313 | 3.655559 | 1.728462 | 0.437915 | -0.831547 | 3.667536 | -0.995684 | 1.080509 | 0.605573 | 0.334621 | 0.71017 | 1.072074 |
| 36.4375 | 4.576068 | 8.216363 | 3.260317 | 2.904949 | 2.612438 | 0.474499 | 5.013772 | -1.336987 | 2.178671 | 0.616806 | 0.395362 | 0.654701 | 1.160851 |
| 36.45139 | 4.880778 | 6.368798 | 2.186514 | 2.744871 | 3.381463 | 1.391277 | 4.260299 | -0.115506 | 2.251761 | 1.310942 | 1.315021 | 0.993533 | 1.408642 |

[illegible]
