## Supplemental Table S3 for "Leaf movements as a quantitative metric for early stress detection"

**Supplementary Table S3.** Mean, SEM, fold changes, *p*-values (t-test), and percentage differences in motion obtained from the statistical comparison of 12 h integrated motion following stress imposition with the corresponding time point on day before stress imposition in **(a)** different stress treatments in lettuce, **(b)** salt stress applied at different locations on lettuce, **(c)** within location replication of different stresses, **(d)** 100 mM NaCl treatment on different CEA crops and **(e)** 100 mM KCl treatment under $dimG^{day/night}$ and $RGB^{day/}dimG^{night}$ with their respective non-stressed plants.

**Supplementary Table S3.** Mean, SEM, fold changes, p-values (t-test), and percentage differences in motion obtained from the statistical comparison of 12 h integrated motion following stress imposition with the corresponding time point on day before stress imposition in (a) different stress treatments in lettuce with their respective non-stressed plants.

UWA Exp 6

| | NutrientWithdrawal Mean | NutrientWithdrawal SEM | 150 mM Salt Mean | 150 mM Salt SEM | Control Mean | Control SEM | Hypoxia Mean | Hypoxia SEM | Days post stress | | NutrientWithdrawal T.test P values | NutrientWithdrawal Fold Change | NutrientWithdrawal % difference | 150 mM Salt T.test P values | 150 mM Salt Fold Change | 150 mM Salt % difference | Control T.test P values | Control Fold Change | Control % difference | Hypoxia T.test P values | Hypoxia Fold Change | Hypoxia % difference |
|---|---|---|---|---|---|---|---|---|---|---|---|---|---|---|---|---|---|---|---|---|---|---|
| 18.25 | 0.16 | 0.04 | 0.118777 | 0.02 | 0.090107047 | 0.015 | 0 | 0 | | | | | | | | | | | | | | |
| 18.75 | 0.2 | 0.05 | 0 | 0 | 0.141664774 | 0.024 | 0.29026 | 0.05 | | | | | | | | | | | | | | |
| 19.25 | 0.2 | 0.04 | 0.1412295 | 0.03 | 0.152279659 | 0.034 | 0.17479 | 0.03 | | | | | | | | | | | | | | |
| 19.75 | 0.17 | 0.04 | 0.1023291 | 0.02 | 0.230856693 | 0.042 | 0.24402 | 0.04 | | | | | | | | | | | | | | |
| 20.25 | 0.2 | 0.04 | 0.1167376 | 0.02 | 0.227019048 | 0.038 | 0.24338 | 0.05 | | | | | | | | | | | | | | |
| 20.75 | 0.22 | 0.06 | 0.1300423 | 0.03 | 0.236131282 | 0.037 | 0.33472 | 0.08 | | | | | | | | | | | | | | |
| 21.25 | 0.17 | 0.04 | 0.1224617 | 0.02 | 0.235601568 | 0.036 | 0.31662 | 0.08 | | | | | | | | | | | | | | |
| 21.75 | 0.25 | 0.06 | 0.1731307 | 0.03 | 0.303081559 | 0.052 | 0.36715 | 0.06 | | | | | | | | | | | | | | |
| 22.25 | 0.21 | 0.04 | 0.1460789 | 0.02 | 0.233935282 | 0.046 | 0.32811 | 0.06 | | | | | | | | | | | | | | |
| 22.75 | 0.28 | 0.05 | 0.2229279 | 0.04 | 0.344511852 | 0.071 | 0.45753 | 0.08 | | | | | | | | | | | | | | |
| 23.25 | 0.22 | 0.04 | 0.1712063 | 0.02 | 0.255595184 | 0.058 | 0.34364 | 0.05 | | | | | | | | | | | | | | |
| 23.75 | 0.34 | 0.05 | 0.2315979 | 0.03 | 0.348365955 | 0.077 | 0.5601 | 0.12 | | | | | | | | | | | | | | |
| 24.25 | 0.34 | 0.03 | 0.1825857 | 0.02 | 0.296511194 | 0.046 | 0.30999 | 0.03 | 1 | Day t | 0.02 | 0.64 | 56.6 | 0.72 | 0.94 | 6.65 | 0.58 | 0.86 | 16 | 0.58 | 1.11 | -9.8 |
| 24.75 | 0.34 | 0.05 | 0.1982692 | 0.03 | 0.379826099 | 0.076 | 0.34274 | 0.05 | | Night | 0.96 | 1.01 | -1 | 0.45 | 1.17 | -14 | 0.78 | 0.92 | 9.03 | 0.11 | 1.63 | -39 |
| 25.25 | 0.25 | 0.03 | 0.1835851 | 0.03 | 0.236855392 | 0.034 | 0.28969 | 0.03 | 2 | Day t | 0.49 | 0.86 | 16.1 | 0.74 | 0.93 | 7.23 | 0.78 | 1.08 | -7.3 | 0.37 | 1.19 | -16 |
| 25.75 | 0.35 | 0.06 | 0.3017326 | 0.06 | 0.325977952 | 0.053 | 0.43509 | 0.08 | | Night | 0.84 | 0.96 | 4.39 | 0.31 | 0.77 | 30.3 | 0.81 | 1.07 | -6.4 | 0.4 | 1.29 | -22 |
| 26.25 | 0.23 | 0.04 | 0.2550416 | 0.05 | 0.227430252 | 0.036 | 0.3247 | 0.08 | 3 | Day t | 0.87 | 0.96 | 4.03 | 0.14 | 0.67 | 49 | 0.68 | 1.12 | -11 | 0.84 | 1.06 | -5.5 |
| 26.75 | 0.29 | 0.05 | 0.3489906 | 0.07 | 0.308258013 | 0.051 | 0.46328 | 0.09 | | Night | 0.52 | 1.16 | -14 | 0.16 | 0.66 | 50.7 | 0.67 | 1.13 | -12 | 0.53 | 1.21 | -17 |
| 27.25 | 0.16 | 0.03 | 0.1541735 | 0.02 | 0.331332982 | 0.049 | 0.37615 | 0.08 | 4 | Day t | 0.21 | 1.35 | -26 | 0.63 | 1.11 | -9.9 | 0.33 | 0.77 | 29.6 | 0.73 | 0.91 | 9.46 |
| 27.75 | 0.22 | 0.04 | 0.118928 | 0.02 | 0.316784164 | 0.042 | 0.46464 | 0.13 | | Night | 0.05 | 1.57 | -36 | 0.01 | 1.95 | -49 | 0.72 | 1.1 | -9.1 | 0.6 | 1.21 | -17 |
| 28.25 | 0.09 | 0.01 | 0.1318805 | 0.03 | 0.24353074 | 0.036 | 0.4757 | 0.08 | 5 | Day t | 0 | 2.4 | -58 | 0.31 | 1.3 | -23 | 0.86 | 1.05 | -4.7 | 0.17 | 0.72 | 38.4 |
| 28.75 | 0.12 | 0.02 | 0.1327597 | 0.02 | 0.270499644 | 0.051 | 0.54313 | 0.14 | | Night | 0 | 2.8 | -64 | 0.02 | 1.74 | -43 | 0.41 | 1.29 | -22 | 0.93 | 1.03 | -3 |
| 29.25 | 0.07 | 0.01 | 0.078003 | 0.02 | 0.260599754 | 0.043 | 0.40731 | 0.07 | 6 | Day t | 0 | 3.2 | -69 | 0 | 2.19 | -54 | 0.94 | 0.98 | 1.96 | 0.45 | 0.84 | 18.5 |
| 29.75 | 0.09 | 0.01 | 0.0980248 | 0.02 | 0.336032656 | 0.068 | 0.6446 | 0.11 | | Night | 0 | 3.76 | -73 | 0 | 2.36 | -58 | 0.91 | 1.04 | -3.5 | 0.61 | 0.87 | 15.1 |
| 30.25 | 0.05 | 0.01 | 0.0629815 | 0.01 | 0.342626046 | 0.044 | 0.38376 | 0.04 | 7 | Day t | 0 | 4.11 | -76 | 0 | 2.72 | -63 | 0.24 | 0.75 | 34.1 | 0.53 | 0.9 | 11.7 |
| 30.75 | 0.08 | 0.01 | 0.1370039 | 0.03 | 0.372965447 | 0.056 | 0.64123 | 0.1 | | Night | 0 | 4.21 | -76 | 0.04 | 1.69 | -41 | 0.8 | 0.93 | 7.06 | 0.61 | 0.87 | 14.5 |
| 31.25 | 0.14 | 0.05 | 0.1102205 | 0.01 | 0.374937211 | 0.053 | 0.38142 | 0.05 | 8 | Day t | 0.21 | 1.55 | -35 | 0.04 | 1.55 | -36 | 0.14 | 0.68 | 46.7 | 0.59 | 0.9 | 11 |
| 31.75 | 0.06 | 0.01 | 0.1258118 | 0.02 | 0.360670977 | 0.063 | 0.4221 | 0.07 | | Night | 0 | 6.16 | -84 | 0.01 | 1.84 | -46 | 0.9 | 0.97 | 3.53 | 0.34 | 1.33 | -25 |
| 32.25 | 0.04 | 0.01 | 0.0763151 | 0.01 | 0.318152177 | 0.039 | 0.47306 | 0.06 | | | | | | | | | | | | | | |
| 32.75 | 0.06 | 0.01 | 0.1301464 | 0.02 | 0.492685595 | 0.104 | 0.56681 | 0.08 | | | | | | | | | | | | | | |
| 33.25 | 0.04 | 0.01 | 0.0733253 | 0.01 | 0.255992729 | 0.037 | 0.49177 | 0.05 | | | | | | | | | | | | | | |
| 33.75 | 0.06 | 0.01 | 0.1227115 | 0.02 | 0.558186015 | 0.115 | 0.66861 | 0.13 | | | | | | | | | | | | | | |
| 34.25 | 0.08 | 0.02 | 0.0839674 | 0.01 | 0.300391576 | 0.057 | 0.35149 | 0.06 | | | | | | | | | | | | | | |
| 34.75 | 0.07 | 0.01 | 0.1287911 | 0.02 | 0.403314527 | 0.055 | 0.51772 | 0.08 | | | | | | | | | | | | | | |
| 35.25 | 0.08 | 0.02 | 0.1216649 | 0.01 | 0.477663174 | 0.07 | 0.59985 | 0.08 | | | | | | | | | | | | | | |
| 35.75 | 0.07 | 0.01 | 0.1623435 | 0.02 | 0.459689344 | 0.085 | 0.67303 | 0.09 | | | | | | | | | | | | | | |
| 36.25 | 0.07 | 0.02 | 0.1120274 | 0.02 | 0.519850322 | 0.08 | 0.4757 | 0.08 | | | | | | | | | | | | | | |
| 36.75 | 0.26 | 0.06 | 0.1821373 | 0.03 | 0.463268611 | 0.091 | 0.6266 | 0.13 | | | | | | | | | | | | | | |
| 37.25 | 0.2 | 0.04 | 0.1035211 | 0.02 | 0.466761964 | 0.047 | 0.52782 | 0.09 | | | | | | | | | | | | | | |
| 37.75 | 0.26 | 0.09 | 0.1432368 | 0.02 | 0.329186913 | 0.052 | 0.56192 | 0.1 | | | | | | | | | | | | | | |

AU Exp4

| | 100 mM NaCl-SaltB Mean | 100 mM NaCl-SaltB SEM | ControlB Mean | ControlB SEM | Days post stress | | 100 mM NaCl T.test P values | 100 mM NaCl Fold Change | 100 mM NaCl % difference | Control T.test P values | Control Fold Change | Control % difference |
|---|---|---|---|---|---|---|---|---|---|---|---|---|
| 18.3 | 0.11771 | 0.02412 | 0.28664 | 0.02508 | | | | | | | | |
| 18.8 | 0.06925 | 0.01922 | 0.17432 | 0.01319 | | | | | | | | |
| 19.3 | 0.07241 | 0.02411 | 0.14678 | 0.0141 | | | | | | | | |
| 19.8 | 0.08495 | 0.0283 | 0.16259 | 0.02098 | | | | | | | | |
| 20.3 | 0.0639 | 0.01286 | 0.13791 | 0.00894 | | | | | | | | |
| 20.8 | 0.13953 | 0.05096 | 0.13975 | 0.02658 | | | | | | | | |
| 21.3 | 0.04198 | 0.00853 | 0.08607 | 0.00971 | | | | | | | | |
| 21.8 | 0.14389 | 0.0497 | 0.16798 | 0.03309 | | | | | | | | |
| 22.3 | 0.05379 | 0.00853 | 0.10143 | 0.01065 | | | | | | | | |
| 22.8 | 0.16591 | 0.05131 | 0.24761 | 0.03568 | | | | | | | | |
| 23.3 | 0.07986 | 0.0199 | 0.166 | 0.01246 | | | | | | | | |
| 23.8 | 0.20918 | 0.05911 | 0.25635 | 0.03753 | | | | | | | | |
| 24.3 | 0.17088 | 0.04125 | 0.17811 | 0.01544 | 1 | Day t | 0.06 | 0.97 | 2.99 | 0.55 | 0.93 | 7.29 |
| 24.8 | 0.07919 | 0.01548 | 0.33909 | 0.04679 | | Night | 0.05 | 1.01 | -0.9 | 0.19 | 0.76 | 32.3 |
| 25.3 | 0.0752 | 0.01856 | 0.19291 | 0.01411 | 2 | Day t | 0.87 | 2.21 | -55 | 0.17 | 0.86 | 16.2 |
| 25.8 | 0.10295 | 0.02898 | 0.34367 | 0.04761 | | Night | 0.12 | 0.78 | 28.9 | 0.17 | 0.75 | 34.1 |
| 26.3 | 0.06891 | 0.01572 | 0.21197 | 0.03435 | 3 | Day t | 0.67 | 2.41 | -58 | 0.23 | 0.78 | 27.7 |
| 26.8 | 0.12784 | 0.02809 | 0.42746 | 0.05771 | | Night | 0.23 | 0.62 | 60.1 | 0.02 | 0.6 | 66.7 |
| 27.3 | 0.07645 | 0.02181 | 0.18427 | 0.02036 | 4 | Day t | 0.91 | 2.17 | -54 | 0.46 | 0.9 | 11 |
| 27.8 | 0.16032 | 0.034 | 0.34796 | 0.02402 | | Night | 0.48 | 0.5 | 101 | 0.06 | 0.74 | 35.7 |
| 28.3 | 0.08996 | 0.02091 | 0.25185 | 0.03692 | 5 | Day t | 0.73 | 1.84 | -46 | 0.04 | 0.66 | 51.7 |
| 28.8 | 0.19618 | 0.04222 | 0.48124 | 0.05992 | | Night | 0.86 | 0.41 | 146 | 0.01 | 0.53 | 87.7 |
| 29.3 | 0.0857 | 0.01848 | 0.41883 | 0.05465 | 6 | Day t | 0.83 | 1.94 | -48 | 0 | 0.4 | 152 |
| 29.8 | 0.21916 | 0.03921 | 0.49069 | 0.06301 | | Night | 0.89 | 0.36 | 174 | 0.01 | 0.52 | 91.4 |
| 30.3 | 0.0904 | 0.01885 | 0.30143 | 0.0291 | 7 | Day t | 0.71 | 1.84 | -46 | 0 | 0.55 | 81.6 |
| 30.8 | 0.19065 | 0.03012 | 0.58884 | 0.0877 | | Night | 0.78 | 0.42 | 139 | 0 | 0.44 | 130 |
| 31.3 | 0.08407 | 0.01963 | 0.38267 | 0.04486 | 8 | Day t | 0.88 | 1.97 | -49 | 0 | 0.43 | 131 |
| 31.8 | 0.20626 | 0.0351 | 0.65946 | 0.09452 | | Night | 0.97 | 0.39 | 158 | 0 | 0.39 | 157 |
| 32.3 | 0.12496 | 0.0289 | 0.40658 | 0.07854 | | | | | | | | |
| 32.8 | 0.20315 | 0.04229 | 0.59615 | 0.10487 | | | | | | | | |
| 33.3 | 0.12776 | 0.03945 | 0.30439 | 0.06359 | | | | | | | | |
| 33.8 | 0.19235 | 0.05078 | 0.56479 | 0.0914 | | | | | | | | |
| 34.3 | 0.13368 | 0.03551 | 0.348 | 0.04521 | | | | | | | | |
| 34.8 | 0.216 | 0.05947 | 0.41045 | 0.05696 | | | | | | | | |
| 35.3 | 0.14669 | 0.02685 | 0.37076 | 0.05382 | | | | | | | | |
| 35.8 | 0.21818 | 0.06049 | 0.49047 | 0.07938 | | | | | | | | |
| 36.3 | 0.16857 | 0.02963 | 0.34757 | 0.06193 | | | | | | | | |
| 36.8 | 0.26687 | 0.06685 | 0.51139 | 0.09737 | | | | | | | | |
| 37.3 | 0.15976 | 0.02791 | 0.34615 | 0.0572 | | | | | | | | |
| 37.8 | 0.22572 | 0.0513 | 0.4508 | 0.10614 | | | | | | | | |

AU Exp6

| | Control Mean | Control SEM | Boron Mean | Boron SEM | 100mMKCl Mean | 100mMKCl SEM | Days post stress | | Control T.test P values | Control Fold Change | Control % difference | Boron T.test P values | Boron Fold Change | Boron % difference | 100 mM KCl T.test P values | 100 mM KCl Fold Change | 100 mM KCl % difference |
|---|---|---|---|---|---|---|---|---|---|---|---|---|---|---|---|---|---|
| 18.3 | 0.13004 | 0.02467 | 0.3225144 | 0.06 | 0.22 | 0.03 | | | | | | | | | | | |
| 18.8 | 0.12688 | 0.03609 | 0.34495926 | 0.06 | 0.14 | 0.02 | | | | | | | | | | | |
| 19.3 | 0.11189 | 0.03405 | 0.24179394 | 0.03 | 0.16 | 0.02 | | | | | | | | | | | |
| 19.8 | 0.17401 | 0.04266 | 0.44367463 | 0.1 | 0.18 | 0.03 | | | | | | | | | | | |
| 20.3 | 0.1308 | 0.02699 | 0.24395526 | 0.04 | 0.2 | 0.03 | | | | | | | | | | | |
| 20.8 | 0.25388 | 0.05794 | 0.58200885 | 0.11 | 0.19 | 0.04 | | | | | | | | | | | |
| 21.3 | 0.202 | 0.03245 | 0.33649215 | 0.02 | 0.15 | 0.02 | | | | | | | | | | | |
| 21.8 | 0.36322 | 0.08786 | 0.52916662 | 0.05 | 0.24 | 0.03 | | | | | | | | | | | |
| 22.3 | 0.2367 | 0.05688 | 0.38203014 | 0.07 | 0.17 | 0.03 | | | | | | | | | | | |
| 22.8 | 0.37049 | 0.06988 | 0.78687946 | 0.12 | 0.29 | 0.04 | | | | | | | | | | | |
| 23.3 | 0.25019 | 0.05255 | 0.5531477 | 0.15 | 0.28 | 0.05 | | | | | | | | | | | |
| 23.8 | 0.46098 | 0.10196 | 0.81309276 | 0.13 | 0.34 | 0.05 | | | | | | | | | | | |
| 24.3 | 0.25684 | 0.03719 | 0.50573213 | 0.07 | 0.3 | 0.03 | 24.3 | Day t | 0.92 | 0.97 | 2.66 | 0.78 | 1.09 | -8.6 | 0.65 | 0.91 | 9.54 |
| 24.8 | 0.47204 | 0.08936 | 0.61646605 | 0.12 | 0.17 | 0.02 | 24.8 | Night | 0.94 | 0.98 | 2.4 | 0.29 | 1.32 | -24 | 0.01 | 2.06 | -51 |
| 25.3 | 0.21813 | 0.03586 | 0.34897162 | 0.04 | 0.19 | 0.02 | 25.3 | Day t | 0.62 | 1.15 | -13 | 0.21 | 1.59 | -37 | 0.1 | 1.47 | -32 |
| 25.8 | 0.36892 | 0.03746 | 0.49230613 | 0.09 | 0.29 | 0.02 | 25.8 | Night | 0.41 | 1.25 | -20 | 0.06 | 1.65 | -39 | 0.37 | 1.19 | -16 |
| 26.3 | 0.24103 | 0.03713 | 0.36713093 | 0.06 | 0.16 | 0.01 | 26.3 | Day t | 0.89 | 1.04 | -3.7 | 0.27 | 1.51 | -34 | 0.03 | 1.72 | -42 |
| 26.8 | 0.34928 | 0.04876 | 0.40711186 | 0.06 | 0.33 | 0.03 | 26.8 | Night | 0.34 | 1.32 | -24 | 0.01 | 2 | -50 | 0.83 | 1.04 | -3.8 |
| 27.3 | 0.31686 | 0.03418 | 0.4703487 | 0.07 | 0.24 | 0.02 | 27.3 | Day t | 0.3 | 0.79 | 26.7 | 0.62 | 1.18 | -15 | 0.46 | 1.17 | -15 |
| 27.8 | 0.38914 | 0.05832 | 0.40773936 | 0.08 | 0.26 | 0.04 | 27.8 | Night | 0.55 | 1.18 | -16 | 0.02 | 1.99 | -50 | 0.24 | 1.29 | -23 |
| 28.3 | 0.33847 | 0.06456 | 0.4116868 | 0.05 | 0.24 | 0.02 | 28.3 | Day t | 0.3 | 0.74 | 35.3 | 0.39 | 1.34 | -26 | 0.45 | 1.18 | -15 |
| 28.8 | 0.43992 | 0.05617 | 0.5180265 | 0.07 | 0.24 | 0.04 | 28.8 | Night | 0.86 | 1.05 | -4.6 | 0.06 | 1.57 | -36 | 0.14 | 1.41 | -29 |
| 29.3 | 0.41447 | 0.07477 | 0.59822349 | 0.07 | 0.22 | 0.03 | 29.3 | Day t | 0.09 | 0.6 | 65.7 | 0.79 | 0.92 | 8.15 | 0.35 | 1.24 | -20 |
| 29.8 | 0.40263 | 0.05525 | 0.63011306 | 0.08 | 0.27 | 0.05 | 29.8 | Night | 0.62 | 1.14 | -13 | 0.24 | 1.29 | -23 | 0.32 | 1.27 | -21 |
| 30.3 | 0.53599 | 0.06876 | 0.96418695 | 0.06 | 0.23 | 0.03 | 30.3 | Day t | 0 | 0.47 | 114 | 0.02 | 0.57 | 74.3 | 0.46 | 1.18 | -15 |
| 30.8 | 0.49408 | 0.06315 | 0.73674071 | 0.11 | 0.24 | 0.04 | 30.8 | Night | 0.79 | 0.93 | 7.18 | 0.66 | 1.1 | -9.4 | 0.14 | 1.42 | -30 |
| 31.3 | 0.36485 | 0.06473 | 0.59966174 | 0.06 | 0.19 | 0.02 | 31.3 | Day t | 0.19 | 0.69 | 45.8 | 0.78 | 0.92 | 8.41 | 0.09 | 1.49 | -33 |
| 31.8 | 0.49871 | 0.04998 | 0.77998551 | 0.15 | 0.24 | 0.04 | 31.8 | Night | 0.74 | 0.92 | 8.18 | 0.87 | 1.04 | -4.1 | 0.16 | 1.42 | -30 |
| 32.3 | 0.51353 | 0.0676 | 0.55178011 | 0.06 | 0.2 | 0.02 | | | | | | | | | | | |
| 32.8 | 0.50919 | 0.06243 | 0.6990216 | 0.1 | 0.24 | 0.04 | | | | | | | | | | | |
| 33.3 | 0.49629 | 0.07162 | 0.6039511 | 0.07 | 0.22 | 0.04 | | | | | | | | | | | |
| 33.8 | 0.61621 | 0.1002 | 0.84823295 | 0.09 | 0.22 | 0.04 | | | | | | | | | | | |
| 34.3 | 0.52638 | 0.08361 | 0.76751192 | 0.14 | 0.28 | 0.03 | | | | | | | | | | | |
| 34.8 | 0.72228 | 0.08989 | 0.88929064 | 0.17 | 0.23 | 0.04 | | | | | | | | | | | |
| 35.3 | 0.6731 | 0.09103 | 0.5758611 | 0.11 | 0.24 | 0.04 | | | | | | | | | | | |
| 35.8 | 0.45622 | 0.09471 | 0.68897824 | 0.12 | 0.23 | 0.04 | | | | | | | | | | | |
| 36.3 | NaN | NaN | 0.9193308 | 0.1 | 0.39 | 0.04 | | | | | | | | | | | |
| 36.8 | NaN | NaN | 0.69086662 | 0.13 | 0.31 | 0.04 | | | | | | | | | | | |
| 37.3 | NaN | NaN | 0.05831028 | 0.01 | 0.03 | 0.01 | | | | | | | | | | | |

UoC Exp8

| | Control Mean | Control SEM | Water withdrawal Mean | Water withdrawal SEM | Days post stress | | Control T.test P values | Control Fold Change | Control % difference | Water withdrawal T.test P values | Water withdrawal Fold Change | Water withdrawal % difference |
|---|---|---|---|---|---|---|---|---|---|---|---|---|
| 18.3 | 0.2047 | 0.05 | 0.12 | 0.02 | | | | | | | | |
| 18.8 | 0.2654 | 0.04 | 0.18 | 0.03 | | | | | | | | |
| 19.3 | 0.2383 | 0.03 | 0.2 | 0.03 | | | | | | | | |
| 19.8 | 0.3655 | 0.06 | 0.19 | 0.02 | | | | | | | | |
| 20.3 | 0.2365 | 0.03 | 0.17 | 0.02 | | | | | | | | |
| 20.8 | 0.4238 | 0.06 | 0.27 | 0.05 | | | | | | | | |
| 21.3 | 0.298 | 0.04 | 0.23 | 0.03 | | | | | | | | |
| 21.8 | 0.5502 | 0.06 | 0.41 | 0.06 | | | | | | | | |
| 22.3 | 0.3731 | 0.04 | 0.33 | 0.04 | | | | | | | | |
| 22.8 | 0.7357 | 0.13 | 0.58 | 0.08 | | | | | | | | |
| 23.3 | 0.4577 | 0.05 | 0.41 | 0.04 | | | | | | | | |
| 23.8 | 0.7333 | 0.13 | 0.71 | 0.09 | | | | | | | | |
| 24.3 | 0.6089 | 0.06 | 0.55 | 0.05 | 1 | Day t | 0.06 | 0.75 | 33 | 0.06 | 0.75 | 33.6 |
| 24.8 | 0.8082 | 0.12 | 0.76 | 0.11 | | Night | 0.67 | 0.91 | 10.2 | 0.72 | 0.93 | 7.69 |
| 25.3 | 0.7157 | 0.08 | 0.56 | 0.06 | 2 | Day t | 0.01 | 0.64 | 56.4 | 0.07 | 0.74 | 36 |
| 25.8 | 1.0438 | 0.1 | 0.8 | 0.1 | | Night | 0.06 | 0.7 | 42.3 | 0.52 | 0.89 | 12.4 |
| 26.3 | 0.7414 | 0.11 | 0.5 | 0.07 | 3 | Day t | 0.02 | 0.62 | 62 | 0.27 | 0.82 | 21.4 |
| 26.8 | 1.0246 | 0.1 | 0.84 | 0.11 | | Night | 0.09 | 0.72 | 39.7 | 0.38 | 0.84 | 18.5 |
| 27.3 | 0.6664 | 0.12 | 0.42 | 0.06 | 4 | Day t | 0.11 | 0.69 | 45.6 | 0.88 | 0.97 | 2.68 |
| 27.8 | 0.9873 | 0.16 | 0.74 | 0.09 | | Night | 0.22 | 0.74 | 34.6 | 0.82 | 0.96 | 4.37 |
| 28.3 | 0.5252 | 0.06 | 0.43 | 0.05 | 5 | Day t | 0.38 | 0.87 | 14.7 | 0.78 | 0.96 | 4.54 |
| 28.8 | 0.975 | 0.17 | 0.73 | 0.1 | | Night | 0.26 | 0.75 | 33 | 0.87 | 0.97 | 3.29 |
| 29.3 | 0.5346 | 0.05 | 0.91 | 0.06 | 6 | Day t | 0.29 | 0.86 | 16.8 | 0 | 0.45 | 120 |
| 29.8 | 0.9638 | 0.18 | 0.19 | 0.07 | | Night | 0.3 | 0.76 | 31.4 | 0 | 3.74 | -73 |
| 30.3 | 0.5481 | 0.09 | 0.88 | 0.05 | 7 | Day t | 0.39 | 0.83 | 19.8 | 0 | 0.47 | 113 |
| 30.8 | 0.8283 | 0.13 | 0.56 | 0.05 | | Night | 0.6 | 0.89 | 13 | 0.18 | 1.26 | -21 |
| 31.3 | 0.6268 | 0.09 | 0.44 | 0.04 | 8 | Day t | 0.1 | 0.73 | 36.9 | 0.66 | 0.94 | 6.39 |
| 31.8 | 0.8438 | 0.17 | 0.66 | 0.08 | | Night | 0.61 | 0.87 | 15.1 | 0.67 | 1.08 | -7.5 |
| 32.3 | 0.7371 | 0.12 | 0.43 | 0.04 | | | | | | | | |
| 32.8 | 0.9858 | 0.24 | 0.77 | 0.1 | | | | | | | | |
| 33.3 | 0.7342 | 0.14 | 0.42 | 0.04 | | | | | | | | |
| 33.8 | 1.0597 | 0.28 | 0.87 | 0.1 | | | | | | | | |
| 34.3 | 0.6859 | 0.13 | 0.51 | 0.06 | | | | | | | | |
| 34.8 | 0.8071 | 0.18 | 0.88 | 0.17 | | | | | | | | |
| 35.3 | 0.799 | 0.24 | 0.52 | 0.1 | | | | | | | | |
| 35.8 | 1.014 | 0.31 | 0.86 | 0.18 | | | | | | | | |
| 36.3 | 0.7421 | 0.16 | 0.63 | 0.08 | | | | | | | | |
| 36.8 | 0.5956 | 0.07 | 0.88 | 0.16 | | | | | | | | |

**Fold change = (Average area under the curve before stress Day 23) / (Average area under the curve after stress)**

**Supplementary Table S3.** Mean, SEM, fold changes, *p* -values (t-test), and percentage differences in motion obtained from the statistical comparison of 12 h integrated motion following stress imposition with the corresponding time point on day before stress imposition in **(b)** salt stress applied at different locations on lettuce with their respective non-stressed plants.

**UWA Exp6**

| Day | 150 mM Salt Mean | 150 mM Salt SEM | Control Mean | Control SEM | Days post stress | | 150 mM Salt T.test P values | 150 mM Salt Fold Change | 150 mM Salt % difference | Control T.test P values | Control Fold Change | Control % difference |
|---|---|---|---|---|---|---|---|---|---|---|---|---|
| 18 | 0.1188 | 0 | 0.09011 | 0 | | | | | | | | |
| 19 | 0 | 0 | 0.14166 | 0 | | | | | | | | |
| 19 | 0.1412 | 0 | 0.15228 | 0 | | | | | | | | |
| 20 | 0.1023 | 0 | 0.23086 | 0 | | | | | | | | |
| 20 | 0.1167 | 0 | 0.22702 | 0 | | | | | | | | |
| 21 | 0.13 | 0 | 0.23613 | 0 | | | | | | | | |
| 21 | 0.1225 | 0 | 0.2356 | 0 | | | | | | | | |
| 22 | 0.1731 | 0 | 0.30308 | 0.1 | | | | | | | | |
| 22 | 0.1461 | 0 | 0.23394 | 0 | | | | | | | | |
| 23 | 0.2229 | 0 | 0.34451 | 0.1 | | | | | | | | |
| 23 | 0.1712 | 0 | 0.2556 | 0.1 | | | | | | | | |
| 24 | 0.2316 | 0 | 0.34837 | 0.1 | | | | | | | | |
| 24 | 0.1826 | 0 | 0.29651 | 0 | 1 | Day t | 0.7 | 0.9 | 6.6 | 0.6 | 0.9 | 16 |
| 25 | 0.1983 | 0 | 0.37983 | 0.1 | | Night | 0.5 | 1.2 | -14 | 0.8 | 0.9 | 9 |
| 25 | 0.1836 | 0 | 0.23686 | 0 | 2 | Day t | 0.7 | 0.9 | 7.2 | 0.8 | 1.1 | -7.3 |
| 26 | 0.3017 | 0.1 | 0.32598 | 0.1 | | Night | 0.3 | 0.8 | 30 | 0.8 | 1.1 | -6.4 |
| 26 | 0.255 | 0 | 0.22743 | 0 | 3 | Day t | 0.1 | 0.7 | 49 | 0.7 | 1.1 | -11 |
| 27 | 0.349 | 0.1 | 0.30826 | 0.1 | | Night | 0.2 | 0.7 | 51 | 0.7 | 1.1 | -12 |
| 27 | 0.1542 | 0 | 0.33133 | 0 | 4 | Day t | 0.6 | 1.1 | -9.9 | 0.3 | 0.8 | 30 |
| 28 | 0.1189 | 0 | 0.31678 | 0 | | Night | 0 | 1.9 | -49 | 0.7 | 1.1 | -9.1 |
| 28 | 0.1319 | 0 | 0.24353 | 0 | 5 | Day t | 0.3 | 1.3 | -23 | 0.9 | 1 | -4.7 |
| 29 | 0.1328 | 0 | 0.2705 | 0.1 | | Night | 0 | 1.7 | -43 | 0.4 | 1.3 | -22 |
| 29 | 0.078 | 0 | 0.2606 | 0 | 6 | Day t | 0 | 2.2 | -54 | 0.9 | 1 | 2 |
| 30 | 0.098 | 0 | 0.33603 | 0.1 | | Night | 0 | 2.4 | -58 | 0.9 | 1 | -3.5 |
| 30 | 0.063 | 0 | 0.34263 | 0 | 7 | Day t | 0 | 2.7 | -63 | 0.2 | 0.7 | 34 |
| 31 | 0.137 | 0 | 0.37297 | 0.1 | | Night | 0 | 1.7 | -41 | 0.8 | 0.9 | 7.1 |
| 31 | 0.1102 | 0 | 0.37494 | 0.1 | 8 | Day t | 0 | 1.6 | -36 | 0.1 | 0.7 | 47 |
| 32 | 0.1258 | 0 | 0.36067 | 0.1 | | Night | 0 | 1.8 | -46 | 0.9 | 1 | 3.5 |
| 32 | 0.0763 | 0 | 0.31815 | 0 | | | | | | | | |
| 33 | 0.1301 | 0 | 0.49269 | 0.1 | | | | | | | | |
| 33 | 0.0733 | 0 | 0.25599 | 0 | | | | | | | | |
| 34 | 0.1227 | 0 | 0.55819 | 0.1 | | | | | | | | |
| 34 | 0.084 | 0 | 0.30039 | 0.1 | | | | | | | | |
| 35 | 0.1288 | 0 | 0.40331 | 0.1 | | | | | | | | |
| 35 | 0.1217 | 0 | 0.47766 | 0.1 | | | | | | | | |
| 36 | 0.1623 | 0 | 0.45969 | 0.1 | | | | | | | | |
| 36 | 0.112 | 0 | 0.51985 | 0.1 | | | | | | | | |
| 37 | 0.1821 | 0 | 0.46327 | 0.1 | | | | | | | | |
| 37 | 0.1035 | 0 | 0.46676 | 0 | | | | | | | | |
| 38 | 0.1432 | 0 | 0.32919 | 0.1 | | | | | | | | |

**AU Exp4**

| | 100 mM NaCl-SaltB Mean | 100 mM NaCl-SaltB SEM | ControlB Mean | ControlB SEM | Days post stress | | 100 mM NaCl T.test P values | 100 mM NaCl Fold Change | 100 mM NaCl % difference | Control T.test P values | Control Fold Change | Control % difference |
|---|---|---|---|---|---|---|---|---|---|---|---|---|
| 18.25 | 0.117706 | 0.024124 | 0.286636 | 0.025076 | | | | | | | | |
| 18.75 | 0.069245 | 0.019218 | 0.174318 | 0.013193 | | | | | | | | |
| 19.25 | 0.072408 | 0.024109 | 0.146775 | 0.014099 | | | | | | | | |
| 19.75 | 0.084946 | 0.028301 | 0.162586 | 0.020976 | | | | | | | | |
| 20.25 | 0.063897 | 0.01286 | 0.137907 | 0.008943 | | | | | | | | |
| 20.75 | 0.139534 | 0.050963 | 0.139748 | 0.02658 | | | | | | | | |
| 21.25 | 0.041976 | 0.008526 | 0.086066 | 0.009714 | | | | | | | | |
| 21.75 | 0.143891 | 0.049702 | 0.167985 | 0.033091 | | | | | | | | |
| 22.25 | 0.053787 | 0.008535 | 0.101432 | 0.010649 | | | | | | | | |
| 22.75 | 0.165913 | 0.051311 | 0.247605 | 0.035679 | | | | | | | | |
| 23.25 | 0.079865 | 0.019903 | 0.166004 | 0.012461 | | | | | | | | |
| 23.75 | 0.209183 | 0.059111 | 0.256353 | 0.037533 | | | | | | | | |
| 24.25 | 0.170878 | 0.041253 | 0.178108 | 0.015442 | 1 | Day t | 0.1 | 1 | 2.99 | 0.6 | 0.9 | 7.29 |
| 24.75 | 0.079185 | 0.015483 | 0.339086 | 0.046791 | | Night | 0 | 1 | -0.9 | 0.2 | 0.8 | 32.3 |
| 25.25 | 0.075198 | 0.018563 | 0.19291 | 0.014112 | 2 | Day t | 0.9 | 2.2 | -55 | 0.2 | 0.9 | 16.2 |
| 25.75 | 0.102951 | 0.028984 | 0.343675 | 0.047614 | | Night | 0.1 | 0.8 | 28.9 | 0.2 | 0.7 | 34.1 |
| 26.25 | 0.068915 | 0.015722 | 0.211969 | 0.034347 | 3 | Day t | 0.7 | 2.4 | -58 | 0.2 | 0.8 | 27.7 |
| 26.75 | 0.127841 | 0.02809 | 0.427464 | 0.05771 | | Night | 0.2 | 0.6 | 60.1 | 0 | 0.6 | 66.7 |
| 27.25 | 0.076445 | 0.021807 | 0.184268 | 0.020356 | 4 | Day t | 0.9 | 2.2 | -54 | 0.5 | 0.9 | 11 |
| 27.75 | 0.160316 | 0.034 | 0.347963 | 0.024024 | | Night | 0.5 | 0.5 | 101 | 0.1 | 0.7 | 35.7 |
| 28.25 | 0.089961 | 0.020905 | 0.251852 | 0.036922 | 5 | Day t | 0.7 | 1.8 | -46 | 0 | 0.7 | 51.7 |
| 28.75 | 0.19618 | 0.042218 | 0.481242 | 0.05992 | | Night | 0.9 | 0.4 | 146 | 0 | 0.5 | 87.7 |
| 29.25 | 0.085699 | 0.018483 | 0.418832 | 0.05465 | 6 | Day t | 0.8 | 1.9 | -48 | 0 | 0.4 | 152 |
| 29.75 | 0.219161 | 0.039209 | 0.490692 | 0.063008 | | Night | 0.9 | 0.4 | 174 | 0 | 0.5 | 91.4 |
| 30.25 | 0.090397 | 0.018849 | 0.30143 | 0.029102 | 7 | Day t | 0.7 | 1.8 | -46 | 0 | 0.6 | 81.6 |
| 30.75 | 0.190645 | 0.030118 | 0.588837 | 0.087704 | | Night | 0.8 | 0.4 | 139 | 0 | 0.4 | 130 |
| 31.25 | 0.084067 | 0.019634 | 0.38267 | 0.044856 | 8 | Day t | 0.9 | 2 | -49 | 0 | 0.4 | 131 |
| 31.75 | 0.206263 | 0.0351 | 0.659464 | 0.094518 | | Night | 1 | 0.4 | 158 | 0 | 0.4 | 157 |
| 32.25 | 0.12496 | 0.028898 | 0.406576 | 0.078538 | | | | | | | | |
| 32.75 | 0.203151 | 0.042292 | 0.596152 | 0.104873 | | | | | | | | |
| 33.25 | 0.127761 | 0.039448 | 0.304389 | 0.063588 | | | | | | | | |
| 33.75 | 0.192349 | 0.050785 | 0.564786 | 0.091397 | | | | | | | | |
| 34.25 | 0.13368 | 0.035515 | 0.347998 | 0.045212 | | | | | | | | |
| 34.75 | 0.215997 | 0.059473 | 0.410447 | 0.056962 | | | | | | | | |
| 35.25 | 0.146688 | 0.026853 | 0.370764 | 0.053818 | | | | | | | | |
| 35.75 | 0.21818 | 0.060494 | 0.490475 | 0.079382 | | | | | | | | |
| 36.25 | 0.16857 | 0.029633 | 0.347566 | 0.061926 | | | | | | | | |
| 36.75 | 0.266866 | 0.066849 | 0.511388 | 0.097368 | | | | | | | | |
| 37.25 | 0.159759 | 0.027906 | 0.346153 | 0.057195 | | | | | | | | |
| 37.75 | 0.225722 | 0.051297 | 0.450798 | 0.106142 | | | | | | | | |

**UoC**

| Day | 100 mM NaCl Mean | 100 mM NaCl SEM | Control Mean | Control SEM | Days post stress | | 100 mM NaCl T.test P values | 100 mM NaCl Fold Change | 100 mM NaCl % difference | Control T.test P values | Control Fold Change | Control % difference |
|---|---|---|---|---|---|---|---|---|---|---|---|---|
| 18 | 0.17583 | 0.04057 | 0.165057 | 0.018919 | | | | | | | | |
| 18.5 | 0.22536 | 0.0508 | 0.204699 | 0.02056 | | | | | | | | |
| 19 | 0.18339 | 0.03692 | 0.178901 | 0.018554 | | | | | | | | |
| 19.5 | 0.26074 | 0.05087 | 0.246288 | 0.025906 | | | | | | | | |
| 20 | 0.23494 | 0.04364 | 0.206623 | 0.019536 | | | | | | | | |
| 20.5 | 0.31393 | 0.05067 | 0.262578 | 0.022771 | | | | | | | | |
| 21 | 0.28532 | 0.04178 | 0.26295 | 0.015168 | | | | | | | | |
| 21.5 | 0.31662 | 0.05396 | 0.239447 | 0.024262 | | | | | | | | |
| 22 | 0 | 0 | 0 | 0 | | | | | | | | |
| 22.5 | 0 | 0 | 0 | 0 | | | | | | | | |
| 23 | 0.4569 | 0.04775 | 0.341018 | 0.029489 | | | | | | | | |
| 23.5 | 0.62189 | 0.06932 | 0.375953 | 0.032146 | | | | | | | | |
| 24 | 0.3042 | 0.05324 | 0.294771 | 0.019894 | 1 | Day t | 0.05 | 1.5 | -33 | 0.21 | 1.16 | -14 |
| 24.5 | 0.36375 | 0.05168 | 0.398429 | 0.026941 | | Night | 0.01 | 1.71 | -42 | 0.6 | 0.94 | 5.98 |
| 25 | 0.29605 | 0.03396 | 0.340238 | 0.02217 | 2 | Day t | 0.01 | 1.54 | -35 | 0.98 | 1 | -0.2 |
| 25.5 | 0.36116 | 0.0336 | 0.403624 | 0.023413 | | Night | 0 | 1.72 | -42 | 0.5 | 0.93 | 7.36 |
| 26 | 0.32213 | 0.03701 | 0.336681 | 0.028197 | 3 | Day t | 0.04 | 1.42 | -29 | 0.92 | 1.01 | -1.3 |
| 26.5 | 0.48983 | 0.03254 | 0.409809 | 0.015446 | | Night | 0.1 | 1.27 | -21 | 0.36 | 0.92 | 9.01 |
| 27 | 0.31719 | 0.03024 | 0.35157 | 0.037889 | 4 | Day t | 0.02 | 1.44 | -31 | 0.83 | 0.97 | 3.09 |
| 27.5 | 0.55596 | 0.04202 | 0.417855 | 0.034948 | | Night | 0.43 | 1.12 | -11 | 0.39 | 0.9 | 11.1 |
| 28 | 0.31472 | 0.03449 | 0.34543 | 0.043163 | 5 | Day t | 0.03 | 1.45 | -31 | 0.93 | 0.99 | 1.29 |
| 28.5 | 0.54374 | 0.04059 | 0.40244 | 0.044324 | | Night | 0.34 | 1.14 | -13 | 0.64 | 0.93 | 7.05 |
| 29 | 0.3108 | 0.03234 | 0.361089 | 0.0497 | 6 | Day t | 0.02 | 1.47 | -32 | 0.73 | 0.94 | 5.89 |
| 29.5 | 0.55945 | 0.04696 | 0.421101 | 0.048123 | | Night | 0.46 | 1.11 | -10 | 0.45 | 0.89 | 12 |
| 30 | 0.38912 | 0.03992 | 0.43889 | 0.033862 | 7 | Day t | 0.29 | 1.17 | -15 | 0.04 | 0.78 | 28.7 |
| 30.5 | 0.51711 | 0.04456 | 0.490995 | 0.053907 | | Night | 0.22 | 1.2 | -17 | 0.09 | 0.77 | 30.6 |
| 31 | 0.38448 | 0.02627 | 0.425789 | 0.04509 | 8 | Day t | 0.2 | 1.19 | -16 | 0.14 | 0.8 | 24.9 |
| 31.5 | 0.55605 | 0.04811 | 0.688073 | 0.09624 | | Night | 0.44 | 1.12 | -11 | 0.01 | 0.55 | 83 |
| 32 | 0.30575 | 0.02478 | 0.436522 | 0.03957 | | | | | | | | |
| 32.5 | 0.45843 | 0.03506 | 0.638012 | 0.094242 | | | | | | | | |
| 33 | 0.3255 | 0.03405 | 0.392104 | 0.054648 | | | | | | | | |
| 33.5 | 0.38726 | 0.0402 | 0.522457 | 0.087866 | | | | | | | | |
| 34 | 0.35266 | 0.02811 | 0.382992 | 0.051159 | | | | | | | | |
| 34.5 | 0.42998 | 0.05541 | 0.469587 | 0.079179 | | | | | | | | |
| 35 | 0.37301 | 0.03552 | 0.335078 | 0.056346 | | | | | | | | |
| 35.5 | 0.47121 | 0.06442 | 0.448936 | 0.06627 | | | | | | | | |
| 36 | 0.4479 | 0.04658 | 0.278903 | 0.037368 | | | | | | | | |
| 36.5 | 0.52611 | 0.05642 | 0.357144 | 0.059536 | | | | | | | | |
| 37 | 0.43669 | 0.04384 | 0.226021 | 0.044838 | | | | | | | | |
| 37.5 | 0.33638 | 0.03735 | 0.164622 | 0.033278 | | | | | | | | |

**Fold change = (Average area under the curve before stress Day 23) / (Average area under the curve after stress)**

**Supplementary Table S3.** Mean, SEM, fold changes, p-values (t-test), and percentage differences in motion obtained from the statistical comparison of 12 h integrated motion following stress imposition with the corresponding time point on day before stress imposition in (c) within location replication of different stresses with their respective non-stressed plants.

| Time | UWA Exp4 | UWA Exp6 | UWA Exp4 | UWA Exp6 | UWA Exp4 | UWA Exp6 | | Days post stress | | UWA Exp4 | | UWA Exp6 | | | UWA Exp4 | | | UWA Exp6 | | | UWA Exp4 | | | UWA Exp6 | | |
|---|---|---|---|---|---|---|---|---|---|---|---|---|---|---|---|---|---|---|---|---|---|---|---|---|---|---|
| | Control | Control | Hypoxia | Hypoxia | NutrientWithdrawal | NutrientWithdrawal | | | | NutrientWithdrawal | | NutrientWithdrawal | | | Control | | | Control | | | Hypoxia | | | Hypoxia | | |
| | | | | | | | | | | T.test P values | Fold Change | T.test P values | Fold Change | % difference | T.test P values | Fold Change | | T.test P values | Fold Change | % difference | T.test P values | Fold Change | | T.test P values | Fold Change | % difference |
| 18 | 0.2 | 0.09 | 0.24 | 0 | 0.23 | 0.16 | | | | | | | | | | | | | | | | | | | | |
| 19 | 0.15 | 0.14 | 0.22 | 0.29 | 0.16 | 0.2 | | | | | | | | | | | | | | | | | | | | |
| 19 | 0.31 | 0.15 | 0.4 | 0.17 | 0.3 | 0.2 | | | | | | | | | | | | | | | | | | | | |
| 20 | 0.18 | 0.23 | 0.21 | 0.24 | 0.17 | 0.17 | | | | | | | | | | | | | | | | | | | | |
| 20 | 0.19 | 0.23 | 0.19 | 0.24 | 0.1 | 0.2 | | | | | | | | | | | | | | | | | | | | |
| 21 | 0.19 | 0.24 | 0.27 | 0.33 | 0.2 | 0.22 | | | | | | | | | | | | | | | | | | | | |
| 21 | 0.14 | 0.24 | 0.2 | 0.32 | 0.1 | 0.17 | | | | | | | | | | | | | | | | | | | | |
| 22 | 0.28 | 0.3 | 0.37 | 0.37 | 0.19 | 0.25 | | | | | | | | | | | | | | | | | | | | |
| 22 | 0.17 | 0.23 | 0.21 | 0.33 | 0.14 | 0.21 | | | | | | | | | | | | | | | | | | | | |
| 23 | 0.27 | 0.34 | 0.33 | 0.46 | 0.23 | 0.28 | | | | | | | | | | | | | | | | | | | | |
| 23 | 0.22 | 0.26 | 0.27 | 0.34 | 0.17 | 0.22 | | | | | | | | | | | | | | | | | | | | |
| 24 | 0.36 | 0.35 | 0.35 | 0.56 | 0.27 | 0.34 | | | | | | | | | | | | | | | | | | | | |
| 24 | 0.23 | 0.3 | 0.25 | 0.31 | 0.35 | 0.34 | | 1 | Day t | 0 | 0.49442 | 0 | 0.6 | 57 | 0.7 | 0.9 | | 0.6 | 0.9 | 16 | 0.8 | 1.1 | | 0.6 | 1.1 | -9.8 |
| 25 | 0.37 | 0.38 | 0.31 | 0.34 | 0.32 | 0.34 | | | Night | 0.6 | 0.86 | 1 | 1 | -1 | 0.9 | 1 | | 0.8 | 0.9 | 9 | 0.6 | 1.1 | | 0.1 | 1.6 | -39 |
| 25 | 0.21 | 0.24 | 0.22 | 0.29 | 0.19 | 0.25 | | 2 | Day t | 0.7 | 0.90398 | 0.5 | 0.9 | 16 | 0.8 | 1 | | 0.8 | 1.1 | -7.3 | 0.4 | 1.2 | | 0.4 | 1.2 | -16 |
| 26 | 0.36 | 0.33 | 0.33 | 0.44 | 0.31 | 0.35 | | | Night | 0.7 | 0.88052 | 0.8 | 1 | 4.4 | 1 | 1 | | 0.8 | 1.1 | -6.4 | 0.8 | 1.1 | | 0.4 | 1.3 | -22 |
| 26 | 0.19 | 0.23 | 0.28 | 0.32 | 0.25 | 0.23 | | 3 | Day t | 0.2 | 0.69512 | 0.9 | 1 | 4 | 0.5 | 1.2 | | 0.7 | 1.1 | -11 | 0.8 | 0.9 | | 0.8 | 1.1 | -5.5 |
| 27 | 0.31 | 0.31 | 0.35 | 0.46 | 0 | 0.29 | | | Night | 0 | | 0.5 | 1.2 | -14 | 0.5 | 1.2 | | 0.7 | 1.1 | -12 | 1 | 1 | | 0.5 | 1.2 | -17 |
| 27 | 0.17 | 0.33 | 0.33 | 0.38 | 0 | 0.16 | | 4 | Day t | 0 | | 0.2 | 1.4 | -26 | 0.3 | 1.3 | | 0.3 | 0.8 | 30 | 0.4 | 0.8 | | 0.7 | 0.9 | 9.5 |
| 28 | 0.3 | 0.32 | 0.49 | 0.46 | 0 | 0.22 | | | Night | 0 | | 0.1 | 1.6 | -36 | 0.5 | 1.2 | | 0.7 | 1.1 | -9.1 | 0.2 | 0.7 | | 0.6 | 1.2 | -17 |
| 28 | 0.19 | 0.24 | 0.42 | 0.48 | 0 | 0.09 | | 5 | Day t | 0 | | 0 | 2.4 | -58 | 0.4 | 1.1 | | 0.9 | 1 | -4.7 | 0.3 | 0.6 | | 0.2 | 0.7 | 38 |
| 29 | 0.24 | 0.27 | 0.46 | 0.54 | 0 | 0.12 | | | Night | 0 | | 0 | 2.8 | -64 | 0.1 | 1.5 | | 0.4 | 1.3 | -22 | 0.3 | 0.8 | | 0.9 | 1 | -3 |
| 29 | 0.21 | 0.26 | 0.38 | 0.41 | 0 | 0.07 | | 6 | Day t | 0 | | 0 | 3.2 | -69 | 0.8 | 1.1 | | 0.9 | 1 | 2 | 0.3 | 0.7 | | 0.4 | 0.8 | 19 |
| 30 | 0.31 | 0.34 | 0.39 | 0.64 | 0 | 0.09 | | | Night | 0 | | 0 | 3.8 | -73 | 0.4 | 1.2 | | 0.9 | 1 | -3.5 | 0.7 | 0.9 | | 0.6 | 0.9 | 15 |
| 30 | 0.47 | 0.34 | 0.32 | 0.38 | 0.23 | 0.05 | | 7 | Day t | 0.4 | 0.75949 | 0 | 4.1 | -76 | 0 | 0.5 | | 0.2 | 0.7 | 34 | 0.4 | 0.8 | | 0.5 | 0.9 | 12 |
| 31 | 0.26 | 0.37 | 0.33 | 0.64 | 0.1 | 0.08 | | | Night | 0 | 2.66041 | 0 | 4.2 | -76 | 0.2 | 1.4 | | 0.8 | 0.9 | 7.1 | 0.8 | 1.1 | | 0.6 | 0.9 | 14 |
| 31 | 0.27 | 0.37 | 0.31 | 0.38 | 0.09 | 0.14 | | 8 | Day t | 0.1 | 1.95955 | 0.2 | 1.5 | -35 | 0.5 | 0.8 | | 0.1 | 0.7 | 47 | 0.5 | 0.9 | | 0.6 | 0.9 | 11 |
| 32 | 0.44 | 0.36 | 0.4 | 0.42 | 0.11 | 0.06 | | | Night | 0 | 2.44767 | 0 | 6.2 | -84 | 0.5 | 0.8 | | 0.9 | 1 | 3.5 | 0.7 | 0.9 | | 0.3 | 1.3 | -25 |
| 32 | 0.33 | 0.32 | 0.29 | 0.47 | 0.1 | 0.04 | | | | | | | | | | | | | | | | | | | | |
| 33 | 0.42 | 0.49 | 0.42 | 0.57 | 0.11 | 0.06 | | | | | | | | | | | | | | | | | | | | |
| 33 | 0.36 | 0.26 | 0.28 | 0.49 | 0.08 | 0.04 | | | | | | | | | | | | | | | | | | | | |
| 34 | 0.34 | 0.56 | 0.55 | 0.67 | 0.1 | 0.06 | | | | | | | | | | | | | | | | | | | | |
| 34 | 0.33 | 0.3 | 0.32 | 0.35 | 0.07 | 0.08 | | | | | | | | | | | | | | | | | | | | |
| 35 | 0.32 | 0.4 | 0.44 | 0.52 | 0.1 | 0.07 | | | | | | | | | | | | | | | | | | | | |
| 35 | 0.3 | 0.48 | 0.36 | 0.6 | 0.07 | 0.08 | | | | | | | | | | | | | | | | | | | | |
| 36 | 0.26 | 0.46 | 0.44 | 0.67 | 0.1 | 0.07 | | | | | | | | | | | | | | | | | | | | |
| 36 | 0.25 | 0.52 | 0.39 | 0.48 | 0.07 | 0.07 | | | | | | | | | | | | | | | | | | | | |
| 37 | 0.24 | 0.46 | 0.36 | 0.63 | 0.05 | 0.26 | | | | | | | | | | | | | | | | | | | | |
| 37 | 0.3 | 0.47 | 0.62 | 0.53 | 0.05 | 0.2 | | | | | | | | | | | | | | | | | | | | |
| 38 | 0.28 | 0.33 | 0.56 | 0.56 | 0.05 | 0.26 | | | | | | | | | | | | | | | | | | | | |
| | | 0.09 | | 0.11 | | 0.09 | | | | | | | | | | | | | | | | | | | | |
| | | 0 | | 0 | | 0 | | | | | | | | | | | | | | | | | | | | |

| Time | AU Exp4 | AU Exp5 | AU Exp5 | AU Exp4 | AU Exp5 | AU Exp5 | AU Exp6 | | Days post stress | | AU Exp4 | | | AU Exp5 | | | AU Exp5 | | | AU Exp4 | | | AU Exp5 | | | AU Exp5 | | | AU Exp6 | | |
|---|---|---|---|---|---|---|---|---|---|---|---|---|---|---|---|---|---|---|---|---|---|---|---|---|---|---|---|---|---|---|---|
| | 100 mM NaCl | 100 mM NaCl | 100 mM NaCl | Control | Control | Control | Control | | | | 100 mM NaCl | | | 100 mM NaCl | | | 100 mM NaCl | | | Control | | | Control | | | Control | | | Control | | |
| | | | | | | | | | | | T.test P values | Fold Change | % difference | T.test P values | Fold Change | % difference | T.test P values | Fold Change | % difference | T.test P values | Fold Change | % difference | T.test P values | Fold Change | % difference | T.test P values | Fold Change | % difference | T.test P values | Fold Change | % difference |
| 18.25 | 0.1 | 0.2 | 0.2 | 0.3 | 0.2 | 0.1 | 0.1 | | | | | | | | | | | | | | | | | | | | | | | | |
| 18.75 | 0.1 | 0.2 | 0.1 | 0.2 | 0.1 | 0.1 | 0.1 | | | | | | | | | | | | | | | | | | | | | | | | |
| 19.25 | 0.1 | 0.1 | 0.2 | 0.1 | 0.3 | 0.1 | 0.1 | | | | | | | | | | | | | | | | | | | | | | | | |
| 19.75 | 0.1 | 0.3 | 0.2 | 0.2 | 0.3 | 0.2 | 0.2 | | | | | | | | | | | | | | | | | | | | | | | | |
| 20.25 | 0.1 | 0.2 | 0.2 | 0.1 | 0.2 | 0.1 | 0.1 | | | | | | | | | | | | | | | | | | | | | | | | |
| 20.75 | 0.1 | 0.3 | 0.3 | 0.1 | 0.4 | 0.3 | 0.3 | | | | | | | | | | | | | | | | | | | | | | | | |
| 21.25 | 0 | 0.2 | 0.2 | 0.1 | 0.2 | 0.1 | 0.2 | | | | | | | | | | | | | | | | | | | | | | | | |
| 21.75 | 0.1 | 0.5 | 0.3 | 0.2 | 0.5 | 0.4 | 0.4 | | | | | | | | | | | | | | | | | | | | | | | | |
| 22.25 | 0.1 | 0.2 | 0.2 | 0.1 | 0.2 | 0.1 | 0.2 | | | | | | | | | | | | | | | | | | | | | | | | |
| 22.75 | 0.2 | 0.5 | 0.4 | 0.2 | 0.4 | 0.4 | 0.4 | | | | | | | | | | | | | | | | | | | | | | | | |
| 23.25 | 0.1 | 0.2 | 0.2 | 0.2 | 0.2 | 0.2 | 0.3 | | | | | | | | | | | | | | | | | | | | | | | | |
| 23.75 | 0.2 | 0.5 | 0.4 | 0.3 | 0.4 | 0.4 | 0.5 | | | | | | | | | | | | | | | | | | | | | | | | |
| 24.25 | 0.2 | 0.3 | 0.2 | 0.2 | 0.4 | 0.3 | 0.3 | | 1 | Day t | 0.1 | 0.5 | 114 | 0 | 0.6 | 56 | 1 | 1 | 0.1 | 0.6 | 0.9 | 7.29 | 0 | 0.6 | 77.8 | 0 | 0.6 | 67.5 | 0.9 | 1 | 2.66 |
| 24.75 | 0.1 | 0.1 | 0.1 | 0.3 | 0.5 | 0.5 | 0.5 | | | Night | 0 | 2.6 | -62 | 0 | 4.4 | -77 | 0 | 4.1 | -76 | 0.2 | 0.8 | 32.3 | 0.5 | 0.9 | 12.1 | 0.2 | 0.8 | 30.2 | 0.9 | 1 | 2.4 |
| 25.25 | 0.1 | 0.2 | 0.2 | 0.2 | 0.3 | 0.3 | 0.2 | | 2 | Day t | 0.9 | 1.1 | -5.8 | 0.9 | 1 | 3.4 | 0.5 | 0.8 | 18 | 0.2 | 0.9 | 16.2 | 0 | 0.6 | 62.6 | 0 | 0.5 | 107 | 0.6 | 1.1 | -13 |
| 25.75 | 0.1 | 0.5 | 0.4 | 0.3 | 0.5 | 0.6 | 0.4 | | | Night | 0.1 | 2 | -51 | 0.9 | 1 | -4.3 | 0.8 | 1 | 5 | 0.2 | 0.7 | 34.1 | 0.1 | 0.7 | 33.4 | 0.1 | 0.6 | 53.9 | 0.4 | 1.2 | -20 |
| 26.25 | 0.1 | 0.3 | 0.3 | 0.2 | 0.3 | 0.3 | 0.2 | | 3 | Day t | 0.7 | 1.2 | -14 | 0 | 0.6 | 60 | 0.1 | 0.7 | 51 | 0.2 | 0.8 | 27.7 | 0.1 | 0.6 | 55.4 | 0 | 0.5 | 85.3 | 0.9 | 1 | -3.7 |
| 26.75 | 0.1 | 0.4 | 0.4 | 0.4 | 0.5 | 0.5 | 0.3 | | | Night | 0.2 | 1.6 | -39 | 0.3 | 1.2 | -18 | 0.8 | 1.1 | -5.8 | 0 | 0.6 | 66.7 | 0.1 | 0.7 | 36.7 | 0.2 | 0.8 | 27.6 | 0.3 | 1.3 | -24 |
| 27.25 | 0.1 | 0.2 | 0.3 | 0.2 | 0.3 | 0.3 | 0.3 | | 4 | Day t | 0.9 | 1 | -4.3 | 0.2 | 0.7 | 35 | 0 | 0.7 | 49 | 0.5 | 0.9 | 11 | 0.1 | 0.7 | 50.8 | 0 | 0.5 | 90.2 | 0.3 | 0.8 | 26.7 |
| 27.75 | 0.2 | 0.4 | 0.4 | 0.3 | 0.6 | 0.5 | 0.4 | | | Night | 0.5 | 1.3 | -23 | 0.4 | 1.1 | -13 | 0.9 | 1 | 2 | 0.1 | 0.7 | 35.7 | 0.1 | 0.7 | 46.8 | 0.3 | 0.8 | 21.1 | 0.5 | 1.2 | -16 |
| 28.25 | 0.1 | 0.2 | 0.3 | 0.3 | 0.4 | 0.4 | 0.3 | | 5 | Day t | 0.7 | 0.9 | 12.6 | 0.2 | 0.7 | 35 | 0 | 0.6 | 56 | 0 | 0.7 | 51.7 | 0 | 0.6 | 67.5 | 0 | 0.4 | 148 | 0.3 | 0.7 | 35.3 |
| 28.75 | 0.2 | 0.4 | 0.5 | 0.5 | 0.7 | 0.7 | 0.4 | | | Night | 0.9 | 1.1 | -6.2 | 0.3 | 1.2 | -15 | 0.5 | 0.9 | 16 | 0 | 0.5 | 87.7 | 0.1 | 0.6 | 64 | 0.1 | 0.6 | 68.5 | 0.9 | 1 | -4.6 |
| 29.25 | 0.1 | 0.3 | 0.3 | 0.4 | 0.4 | 0.6 | 0.4 | | 6 | Day t | 0.8 | 0.9 | 7.31 | 0.1 | 0.7 | 48 | 0.1 | 0.6 | 61 | 0 | 0.4 | 152 | 0 | 0.5 | 108 | 0 | 0.3 | 279 | 0.1 | 0.6 | 65.7 |
| 29.75 | 0.2 | 0.4 | 0.4 | 0.5 | 0.7 | 0.8 | 0.4 | | | Night | 0.9 | 1 | 4.77 | 0.2 | 1.3 | -20 | 0.9 | 1 | 4.1 | 0 | 0.5 | 91.4 | 0.1 | 0.5 | 86.3 | 0 | 0.5 | 103 | 0.6 | 1.1 | -13 |
| 30.25 | 0.1 | 0.3 | 0.3 | 0.3 | 0.6 | 0.6 | 0.5 | | 7 | Day t | 0.7 | 0.9 | 13.2 | 0.1 | 0.7 | 41 | 0.1 | 0.6 | 55 | 0 | 0.6 | 81.6 | 0 | 0.4 | 172 | 0 | 0.3 | 292 | 0 | 0.5 | 114 |
| 30.75 | 0.2 | 0.4 | 0.4 | 0.6 | 0.7 | 0.7 | 0.5 | | | Night | 0.8 | 1.1 | -8.9 | 0.1 | 1.4 | -27 | 0.7 | 0.9 | 9.7 | 0 | 0.4 | 130 | 0 | 0.6 | 75.4 | 0 | 0.6 | 60.9 | 0.8 | 0.9 | 7.18 |
| 31.25 | 0.1 | 0.2 | 0.3 | 0.4 | 0.4 | 0.4 | 0.4 | | 8 | Day t | 0.9 | 1 | 5.26 | 0.1 | 0.8 | 31 | 0.2 | 0.7 | 43 | 0 | 0.4 | 131 | 0 | 0.5 | 99 | 0 | 0.4 | 159 | 0.2 | 0.7 | 45.8 |
| 31.75 | 0.2 | 0.4 | 0.4 | 0.7 | 0.7 | 0.6 | 0.5 | | | Night | 1 | 1 | -1.4 | 0.2 | 1.3 | -21 | 0.8 | 0.9 | 5.5 | 0 | 0.4 | 157 | 0 | 0.5 | 85.3 | 0.2 | 0.7 | 43.9 | 0.7 | 0.9 | 8.18 |
| 32.25 | 0.1 | 0.3 | 0.4 | 0.4 | 0.5 | 0.6 | 0.5 | | | | | | | | | | | | | | | | | | | | | | | | |
| 32.75 | 0.2 | 0.4 | 0.5 | 0.6 | 0.7 | 0.6 | 0.5 | | | | | | | | | | | | | | | | | | | | | | | | |
| 33.25 | 0.1 | 0.3 | 0.4 | 0.3 | 0.3 | 0.6 | 0.5 | | | | | | | | | | | | | | | | | | | | | | | | |
| 33.75 | 0.2 | 0.5 | 0.4 | 0.6 | 0.7 | 0.8 | 0.6 | | | | | | | | | | | | | | | | | | | | | | | | |
| 34.25 | 0.1 | 0.4 | 0.3 | 0.3 | 0.3 | 0.4 | 0.5 | | | | | | | | | | | | | | | | | | | | | | | | |
| 34.75 | 0.2 | 0.5 | 0.3 | 0.4 | 0.6 | 0.4 | 0.7 | | | | | | | | | | | | | | | | | | | | | | | | |
| 35.25 | 0.1 | 0.4 | 0.3 | 0.4 | 0.4 | 0.4 | 0.7 | | | | | | | | | | | | | | | | | | | | | | | | |
| 35.75 | 0.2 | 0.5 | 0.3 | 0.5 | 0.5 | 0.5 | 0.5 | | | | | | | | | | | | | | | | | | | | | | | | |
| 36.25 | 0.2 | 0.5 | 0.3 | 0.3 | 0.4 | 0.5 | ### | | | | | | | | | | | | | | | | | | | | | | | | |
| 36.75 | 0.3 | 0.4 | 0.3 | 0.5 | 0.7 | 0.7 | ### | | | | | | | | | | | | | | | | | | | | | | | | |
| 37.25 | 0.2 | 0.5 | 0.3 | 0.3 | 0.6 | 0.5 | ### | | | | | | | | | | | | | | | | | | | | | | | | |
| 37.75 | 0.2 | 0.2 | 0.2 | 0.5 | 0.4 | 0.3 | 0 | | | | | | | | | | | | | | | | | | | | | | | | |

**Fold change = (Average area under the curve before stress Day 23) / (Average area under the curve after stress)**

**Supplementary Table S3**. Mean, SEM, fold changes, p-values (t-test), and percentage differences in motion obtained from the statistical comparison of 12 h integrated motion following stress imposition with the corresponding time point on day before stress imposition in (d) 100 mM NaCl treatment on different CEA crops with their respective non-stressed plants.

UWA Exp9

| | Radish 100 mM NaCl | | RadishControl | | MizunaControl | | Mizuna 100 mM NaCl | | | Days post stress | | Radish 100 mM NaCl | | | RadishControl | | | MizunaControl | | | Mizuna 100 mM NaCl | | |
|---|---|---|---|---|---|---|---|---|---|---|---|---|---|---|---|---|---|---|---|---|---|---|---|
| | Mean | SEM | Mean | SEM | Mean | SEM | Mean | SEM | | | | T.test P values | Fold Change | % difference | T.test P values | Fold Change | % difference | T.test P values | Fold Change | % difference | T.test P values | Fold Change | % difference |
| 18.25 | 0.323 | 0.068 | 0.119283145 | 0.016 | 0.157 | 0.04 | 0.08 | 0.021 | | | | | | | | | | | | | | | |
| 18.75 | 0.356 | 0.079 | 0.17579421 | 0.02 | 0.147 | 0.03 | 0.096 | 0.018 | | | | | | | | | | | | | | | |
| 19.25 | 0.401 | 0.052 | 0.135651417 | 0.012 | 0.168 | 0.046 | 0.094 | 0.016 | | | | | | | | | | | | | | | |
| 19.75 | 0.467 | 0.081 | 0.269386476 | 0.033 | 0.262 | 0.038 | 0.171 | 0.027 | | | | | | | | | | | | | | | |
| 20.25 | 0.418 | 0.046 | 0.282104509 | 0.041 | 0.279 | 0.042 | 0.202 | 0.025 | | | | | | | | | | | | | | | |
| 20.75 | 0.493 | 0.072 | 0.374753028 | 0.043 | 0.341 | 0.045 | 0.182 | 0.027 | | | | | | | | | | | | | | | |
| 21.25 | 0.432 | 0.054 | 0.236802367 | 0.035 | 0.424 | 0.043 | 0.28 | 0.036 | | | | | | | | | | | | | | | |
| 21.75 | 0.451 | 0.043 | 0.465149133 | 0.069 | 0.392 | 0.047 | 0.277 | 0.042 | | | | | | | | | | | | | | | |
| 22.25 | 0.357 | 0.038 | 0.274977715 | 0.041 | 0.348 | 0.041 | 0.326 | 0.051 | | | | | | | | | | | | | | | |
| 22.75 | 0.491 | 0.04 | 0.434450094 | 0.055 | 0.361 | 0.039 | 0.334 | 0.051 | | | | | | | | | | | | | | | |
| 23.25 | 0.379 | 0.039 | 0.291631804 | 0.048 | 0.331 | 0.025 | 0.343 | 0.045 | | | | | | | | | | | | | | | |
| 23.75 | 0.467 | 0.04 | 0.422143897 | 0.043 | 0.391 | 0.037 | 0.383 | 0.051 | | | | | | | | | | | | | | | |
| 24.25 | 0.41 | 0.034 | 0.229216616 | 0.033 | 0.344 | 0.032 | 0.564 | 0.073 | 23.75 | 1 | Day to D | 0.567 | 0.926 | 8.014 | 0.293 | 1.272 | -21.4 | 0.74 | 0.96 | 4.163 | 0.018 | 0.608 | 64.43 |
| 24.75 | 0.108 | 0.009 | 0.490080325 | 0.061 | 0.39 | 0.055 | 0.18 | 0.025 | 24.25 | | Night to | 1E-08 | 4.323 | -76.87 | 0.372 | 0.861 | 16.09 | 0.992 | 1.002 | -0.162 | 0.002 | 2.124 | -52.92 |
| 25.25 | 0.151 | 0.016 | 0.346789535 | 0.082 | 0.386 | 0.042 | 0.205 | 0.025 | 24.75 | 2 | Day to D | 2E-05 | 2.515 | -60.24 | 0.566 | 0.841 | 18.91 | 0.265 | 0.855 | 16.9 | 0.014 | 1.676 | -40.33 |
| 25.75 | 0.262 | 0.072 | 0.50143753 | 0.052 | 0.425 | 0.058 | 0.342 | 0.03 | 25.25 | | Night to | 0.021 | 1.783 | -43.93 | 0.253 | 0.842 | 18.78 | 0.624 | 0.919 | 8.8 | 0.493 | 1.12 | -10.72 |
| 26.25 | 0.3 | 0.037 | 0.247445798 | 0.022 | 0.43 | 0.05 | 0.312 | 0.031 | 25.75 | 3 | Day to D | 0.16 | 1.263 | -20.79 | 0.408 | 1.179 | -15.15 | 0.09 | 0.769 | 29.96 | 0.583 | 1.098 | -8.931 |
| 26.75 | 0.352 | 0.056 | 0.62748038 | 0.081 | 0.387 | 0.049 | 0.324 | 0.026 | 26.25 | | Night to | 0.112 | 1.324 | -24.48 | 0.036 | 0.673 | 48.64 | 0.951 | 1.01 | -0.977 | 0.311 | 1.183 | -15.49 |
| 27.25 | 0.388 | 0.035 | 0.311647206 | 0.037 | 0.389 | 0.055 | 0.375 | 0.034 | 26.75 | 4 | Day to D | 0.87 | 0.978 | 2.294 | 0.742 | 0.936 | 6.863 | 0.347 | 0.85 | 17.61 | 0.572 | 0.913 | 9.488 |
| 27.75 | 0.434 | 0.04 | 0.648283335 | 0.081 | 0.46 | 0.058 | 0.323 | 0.029 | 27.25 | | Night to | 0.572 | 1.075 | -6.953 | 0.022 | 0.651 | 53.57 | 0.329 | 0.85 | 17.59 | 0.32 | 1.184 | -15.55 |
| 28.25 | 0.477 | 0.055 | 0.38106546 | 0.044 | 0.39 | 0.046 | 0.401 | 0.049 | 27.75 | 5 | Day to D | 0.163 | 0.796 | 25.68 | 0.182 | 0.765 | 30.67 | 0.267 | 0.847 | 18.01 | 0.391 | 0.854 | 17.05 |
| 28.75 | 0.503 | 0.049 | 0.641001015 | 0.095 | 0.436 | 0.058 | 0.391 | 0.049 | 28.25 | | Night to | 0.572 | 0.928 | 7.771 | 0.047 | 0.659 | 51.84 | 0.52 | 0.896 | 11.56 | 0.91 | 0.979 | 2.111 |
| 29.25 | 0.425 | 0.065 | 0.394560132 | 0.069 | 0.543 | 0.09 | 0.47 | 0.071 | 28.75 | 6 | Day to D | 0.55 | 0.892 | 12.17 | 0.234 | 0.739 | 35.29 | 0.033 | 0.608 | 64.39 | 0.147 | 0.73 | 37 |
| 29.75 | 0.556 | 0.042 | 0.65325558 | 0.085 | 0.419 | 0.058 | 0.444 | 0.069 | 29.25 | | Night to | 0.139 | 0.839 | 19.23 | 0.025 | 0.646 | 54.75 | 0.687 | 0.933 | 7.227 | 0.485 | 0.862 | 15.96 |
| 30.25 | 0.421 | 0.08 | 0.39422607 | 0.064 | 0.6 | 0.054 | 0.495 | 0.073 | 29.75 | 7 | Day to D | 0.646 | 0.902 | 10.91 | 0.214 | 0.74 | 35.18 | 2E-04 | 0.551 | 81.44 | 0.09 | 0.692 | 44.45 |
| 30.75 | 0.63 | 0.112 | 0.738160513 | 0.08 | 0.452 | 0.058 | 0.481 | 0.088 | 30.25 | | Night to | 0.185 | 0.741 | 34.97 | 0.002 | 0.572 | 74.86 | 0.39 | 0.866 | 15.54 | 0.342 | 0.796 | 25.68 |
| 31.25 | 0.486 | 0.109 | 0.525771098 | 0.087 | 0.646 | 0.057 | 0.448 | 0.084 | 30.75 | 8 | Day to D | 0.368 | 0.781 | 28.05 | 0.027 | 0.555 | 80.29 | 4E-05 | 0.511 | 95.54 | 0.285 | 0.766 | 30.62 |
| 31.75 | 0.746 | 0.146 | 0.752744181 | 0.117 | 0.587 | 0.097 | 0.54 | 0.119 | 31.25 | | Night to | 0.079 | 0.626 | 59.79 | 0.015 | 0.561 | 78.31 | 0.073 | 0.666 | 50.09 | 0.239 | 0.709 | 40.99 |
| 32.25 | 0.604 | 0.149 | 0.54862746 | 0.128 | 0.68 | 0.079 | 0.573 | 0.111 | | | | | | | | | | | | | | | |
| 32.75 | 0.759 | 0.15 | 0.846538728 | 0.095 | 0.832 | 0.116 | 0.576 | 0.109 | | | | | | | | | | | | | | | |
| 33.25 | 0.712 | 0.217 | 0.630919008 | 0.133 | 0.836 | 0.065 | 0.641 | 0.13 | | | | | | | | | | | | | | | |
| 33.75 | 1.014 | 0.195 | 0.886238343 | 0.097 | 0.991 | 0.089 | 0.67 | 0.159 | | | | | | | | | | | | | | | |
| 34.25 | 0.716 | 0.154 | 0.805989857 | 0.116 | 1.051 | 0.13 | 0.717 | 0.13 | | | | | | | | | | | | | | | |
| 34.75 | 1.011 | 0.179 | 0.758782789 | 0.091 | 1.041 | 0.093 | 0.538 | 0.091 | | | | | | | | | | | | | | | |
| 35.25 | 0.52 | 0.082 | 0.738909022 | 0.133 | 0.97 | 0.09 | 0.617 | 0.107 | | | | | | | | | | | | | | | |
| 35.75 | 0.967 | 0.105 | 0.818527084 | 0.087 | 0.956 | 0.096 | 0.56 | 0.078 | | | | | | | | | | | | | | | |
| 36.25 | 0.47 | 0.073 | 0.477219023 | 0.055 | 0.658 | 0.076 | 0.514 | 0.075 | | | | | | | | | | | | | | | |

AU Exp8

| | RocketControl | | Rocket100mMNaCl | | | Days post stress | | RocketControl | | | Rocket100 mM NaCl | | |
|---|---|---|---|---|---|---|---|---|---|---|---|---|---|
| | Mean | SEM | Mean | SEM | | | | T.test P values | Fold Change | % difference | T.test P values | Fold Change | % difference |
| 18.25 | 0.083 | 0.013 | 0.241 | 0.035 | | | | | | | | | |
| 18.75 | 0.211 | 0.041 | 0.341 | 0.033 | | | | | | | | | |
| 19.25 | 0.167 | 0.025 | 0.305 | 0.039 | | | | | | | | | |
| 19.75 | 0.177 | 0.031 | 0.46 | 0.027 | | | | | | | | | |
| 20.25 | 0.231 | 0.034 | 0.395 | 0.041 | | | | | | | | | |
| 20.75 | 0.283 | 0.047 | 0.264 | 0.05 | | | | | | | | | |
| 21.25 | 0.217 | 0.039 | 0.28 | 0.026 | | | | | | | | | |
| 21.75 | 0.311 | 0.04 | 0.377 | 0.038 | | | | | | | | | |
| 22.25 | 0.221 | 0.034 | 0.274 | 0.018 | | | | | | | | | |
| 22.75 | 0.326 | 0.033 | 0.341 | 0.045 | | | | | | | | | |
| 23.25 | 0.234 | 0.035 | 0.199 | 0.023 | | | | | | | | | |
| 23.75 | 0.37 | 0.037 | 0.321 | 0.036 | | | | | | | | | |
| 24.25 | 0.262 | 0.028 | 0.416 | 0.038 | | 1 | Day to D | 0.541 | 0.893 | 11.98 | 7E-05 | 0.479 | 108.8 |
| 24.75 | 0.379 | 0.041 | 0.168 | 0.01 | | | Night to | 0.871 | 0.976 | 2.471 | 0.234 | 1.909 | -47.63 |
| 25.25 | 0.281 | 0.036 | 0.16 | 0.017 | | 2 | Day to D | 0.359 | 0.832 | 20.17 | 0.184 | 1.248 | -19.89 |
| 25.75 | 0.385 | 0.042 | 0.175 | 0.027 | | | Night to | 0.792 | 0.961 | 4.048 | 0.511 | 1.83 | -45.37 |
| 26.25 | 0.282 | 0.029 | 0.128 | 0.015 | | 3 | Day to D | 0.309 | 0.83 | 20.49 | 0.017 | 1.554 | -35.65 |
| 26.75 | 0.36 | 0.033 | 0.162 | 0.033 | | | Night to | 0.839 | 1.029 | -2.788 | 0.37 | 1.975 | -49.36 |
| 27.25 | 0.287 | 0.037 | 0.17 | 0.021 | | 4 | Day to D | 0.312 | 0.816 | 22.5 | 0.363 | 1.171 | -14.6 |
| 27.75 | 0.399 | 0.052 | 0.198 | 0.031 | | | Night to | 0.654 | 0.927 | 7.893 | 0.986 | 1.617 | -38.16 |
| 28.25 | 0.264 | 0.036 | 0.2 | 0.023 | | 5 | Day to D | 0.565 | 0.888 | 12.65 | 0.989 | 0.998 | 0.223 |
| 28.75 | 0.39 | 0.045 | 0.25 | 0.039 | | | Night to | 0.734 | 0.949 | 5.42 | 0.271 | 1.283 | -22.09 |
| 29.25 | 0.304 | 0.034 | 0.231 | 0.024 | | 6 | Day to D | 0.168 | 0.771 | 29.78 | 0.36 | 0.864 | 15.77 |
| 29.75 | 0.358 | 0.033 | 0.273 | 0.037 | | | Night to | 0.811 | 1.034 | -3.277 | 0.102 | 1.173 | -14.77 |
| 30.25 | 0.287 | 0.048 | 0.285 | 0.044 | | 7 | Day to D | 0.387 | 0.815 | 22.65 | 0.097 | 0.698 | 43.36 |
| 30.75 | 0.324 | 0.05 | 0.256 | 0.04 | | | Night to | 0.467 | 1.144 | -12.55 | 0.228 | 1.252 | -20.13 |
| 31.25 | 0.316 | 0.049 | 0.253 | 0.031 | | 8 | Day to D | 0.189 | 0.741 | 35.04 | 0.178 | 0.786 | 27.27 |
| 31.75 | 0.341 | 0.041 | 0.251 | 0.042 | | | Night to | 0.609 | 1.084 | -7.782 | 0.288 | 1.278 | -21.77 |
| 32.25 | 0.356 | 0.06 | 0.262 | 0.048 | | | | | | | | | |
| 32.75 | 0.342 | 0.037 | 0.214 | 0.038 | | | | | | | | | |
| 33.25 | 0.435 | 0.126 | 0.267 | 0.036 | | | | | | | | | |
| 33.75 | 0.323 | 0.028 | 0.283 | 0.049 | | | | | | | | | |
| 34.25 | 0.368 | 0.098 | 0.214 | 0.024 | | | | | | | | | |
| 34.75 | 0.377 | 0.094 | 0.2 | 0.027 | | | | | | | | | |
| 35.25 | 0.413 | 0.077 | 0.226 | 0.023 | | | | | | | | | |
| 35.75 | 0.384 | 0.068 | 0.211 | 0.042 | | | | | | | | | |
| 36.25 | 0.4 | 0.074 | 0.225 | 0.039 | | | | | | | | | |
| 36.75 | 0.387 | 0.065 | 0.234 | 0.063 | | | | | | | | | |
| 37.25 | 0.323 | 0.079 | 0.233 | 0.032 | | | | | | | | | |
| 37.75 | 0.276 | 0.043 | 0.23 | 0.09 | | | | | | | | | |

AU Exp9

| | AmaranthControl | | Amaranth100mMNaCl | | Tomato100mMNaCl | | TomatoControl | | | Days post stress | | AmaranthControl | | | Amaranth100 mM NaCl | | | Tomato100 mM NaCl | | | TomatoControl | | |
|---|---|---|---|---|---|---|---|---|---|---|---|---|---|---|---|---|---|---|---|---|---|---|---|
| | Mean | SEM | Mean | SEM | Mean | SEM | Mean | SEM | | | | T.test P values | Fold Change | % difference | T.test P values | Fold Change | % difference | T.test P values | Fold Change | % difference | T.test P values | Fold Change | % difference |
| 18.25 | 0.085 | 0.008 | 0.179 | 0.006 | 0.092 | 0.01 | 0.182 | 0.017 | | | | | | | | | | | | | | | |
| 18.75 | 0.058 | 0.005 | 0.162 | 0.006 | 0.09 | 0.017 | 0.172 | 0.03 | | | | | | | | | | | | | | | |
| 19.25 | 0.04 | 0.004 | 0.111 | 0.005 | 0.041 | 0.006 | 0.112 | 0.009 | | | | | | | | | | | | | | | |
| 19.75 | 0.046 | 0.008 | 0.12 | 0.008 | 0.165 | 0.035 | 0.153 | 0.032 | | | | | | | | | | | | | | | |
| 20.25 | 0.041 | 0.009 | 0.127 | 0.005 | 0.142 | 0.03 | 0.121 | 0.01 | | | | | | | | | | | | | | | |
| 20.75 | 0.06 | 0.009 | 0.138 | 0.013 | 0.217 | 0.041 | 0.232 | 0.032 | | | | | | | | | | | | | | | |
| 21.25 | 0.076 | 0.02 | 0.138 | 0.014 | 0.258 | 0.036 | 0.259 | 0.029 | | | | | | | | | | | | | | | |
| 21.75 | 0.082 | 0.017 | 0.286 | 0.013 | 0.256 | 0.038 | 0.264 | 0.027 | | | | | | | | | | | | | | | |
| 22.25 | 0.104 | 0.024 | 0.27 | 0.02 | 0.253 | 0.029 | 0.16 | 0.011 | | | | | | | | | | | | | | | |
| 22.75 | 0.089 | 0.018 | 0.15 | 0.019 | 0.335 | 0.039 | 0.294 | 0.032 | | | | | | | | | | | | | | | |
| 23.25 | 0.14 | 0.03 | 0.332 | 0.044 | 0.362 | 0.034 | 0.298 | 0.026 | | | | | | | | | | | | | | | |
| 23.75 | 0.113 | 0.018 | 0.2 | 0.036 | 0.407 | 0.043 | 0.43 | 0.039 | | | | | | | | | | | | | | | |
| 24.25 | 0.169 | 0.032 | 0.218 | 0.023 | 0.382 | 0.039 | 0.456 | 0.029 | | | Day to D | 0.518402 | 0.83 | 20.46 | 0.033 | 1.528 | -34.54 | 0.708 | 0.948 | 5.456 | 0.00053639 | 0.654 | 52.82677705 |
| 24.75 | 0.159 | 0.029 | 0.142 | 0.019 | 0.349 | 0.049 | 0.499 | 0.04 | | | Night to | 0.192445 | 0.712 | 40.36 | 9E-04 | 1.411 | -29.11 | 0.824 | 1.166 | -14.27 | 0.23315853 | 0.863 | 15.93528694 |
| 25.25 | 0.227 | 0.044 | 0.178 | 0.02 | 0.419 | 0.033 | 0.569 | 0.032 | | | Day to D | 0.116973 | 0.617 | 61.96 | 0.005 | 1.868 | -46.47 | 0.248 | 0.865 | 15.54 | 1.4284E-06 | 0.524 | 90.88944369 |
| 25.75 | 0.201 | 0.04 | 0.197 | 0.018 | 0.462 | 0.051 | 0.542 | 0.043 | | | Night to | 0.054765 | 0.561 | 78.22 | 0.011 | 1.016 | -1.563 | 0.122 | 0.882 | 13.38 | 0.06784813 | 0.794 | 25.92283379 |
| 26.25 | 0.314 | 0.046 | 0.297 | 0.036 | 0.401 | 0.033 | 0.624 | 0.025 | | | Day to D | 0.004653 | 0.447 | 123.7 | 0.546 | 1.119 | -10.61 | 0.43 | 0.904 | 10.61 | 7.0575E-09 | 0.478 | 109.1929289 |
| 26.75 | 0.282 | 0.049 | 0.21 | 0.042 | 0.549 | 0.084 | 0.703 | 0.052 | | | Night to | 0.003627 | 0.401 | 149.4 | 0.061 | 0.954 | 4.777 | 0.055 | 0.742 | 34.76 | 0.00036175 | 0.612 | 63.49084725 |
| 27.25 | 0.374 | 0.059 | 0.284 | 0.037 | 0.467 | 0.035 | 0.656 | 0.023 | | | Day to D | 0.001969 | 0.375 | 166.4 | 0.418 | 1.169 | -14.44 | 0.046 | 0.776 | 28.84 | 6.7706E-10 | 0.454 | 120.1874904 |
| 27.75 | 0.32 | 0.055 | 0.196 | 0.034 | 0.579 | 0.071 | 0.761 | 0.067 | | | Night to | 0.001769 | 0.354 | 182.9 | 0.025 | 1.02 | -1.948 | 0.013 | 0.703 | 42.15 | 0.00032918 | 0.565 | 76.97102742 |
| 28.25 | 0.41 | 0.055 | 0.309 | 0.051 | 0.562 | 0.046 | 0.757 | 0.033 | | | Day to D | 0.000272 | 0.342 | 192.2 | 0.73 | 1.077 | -7.117 | 0.003 | 0.645 | 54.97 | 2.7876E-10 | 0.394 | 153.8460908 |
| 28.75 | 0.347 | 0.043 | 0.219 | 0.046 | 0.572 | 0.081 | 0.894 | 0.063 | | | Night to | 5.44E-05 | 0.326 | 207.2 | 0.092 | 0.915 | 9.273 | 0.028 | 0.712 | 40.5 | 2.8926E-06 | 0.481 | 107.8414675 |
| 29.25 | 0.497 | 0.044 | 0.323 | 0.056 | 0.619 | 0.064 | 0.822 | 0.039 | | | Day to D | 9.76E-07 | 0.282 | 254.8 | 0.901 | 1.028 | -2.694 | 0.002 | 0.585 | 70.85 | 1.631E-10 | 0.363 | 175.8318737 |
| 29.75 | 0.428 | 0.048 | 0.28 | 0.041 | 0.612 | 0.075 | 0.941 | 0.086 | | | Night to | 3.64E-06 | 0.264 | 279.1 | 0.397 | 0.715 | 39.9 | 0.007 | 0.665 | 50.32 | 1.9079E-05 | 0.457 | 118.742186 |
| 30.25 | 0.524 | 0.047 | 0.334 | 0.05 | 0.587 | 0.076 | 0.815 | 0.042 | | | Day to D | 5.98E-07 | 0.268 | 273.5 | 0.977 | 0.994 | 0.586 | 0.015 | 0.618 | 61.9 | 5.1056E-10 | 0.366 | 173.3564732 |
| 30.75 | 0.449 | 0.048 | 0.262 | 0.059 | 0.703 | 0.103 | 1.02 | 0.078 | | | Night to | 1.52E-06 | 0.252 | 297.5 | 0.354 | 0.764 | 30.85 | 0.006 | 0.58 | 72.55 | 9.071E-07 | 0.422 | 137.2242741 |
| 31.25 | 0.562 | 0.042 | 0.377 | 0.059 | 0.714 | 0.114 | 0.971 | 0.039 | | | Day to D | 3.64E-08 | 0.25 | 300.6 | 0.555 | 0.883 | 13.3 | 0.008 | 0.508 | 96.88 | 1.081E-12 | 0.307 | 225.6092262 |
| 31.75 | 0.479 | 0.059 | 0.302 | 0.067 | 0.765 | 0.112 | 1.048 | 0.1 | | | Night to | 5.85E-06 | 0.236 | 324.2 | 0.712 | 0.663 | 50.89 | 0.003 | 0.532 | 87.8 | 8.7655E-06 | 0.411 | 143.577532 |
| 32.25 | 0.565 | 0.062 | 0.346 | 0.064 | 0.718 | 0.11 | 1.096 | 0.04 | | | | | | | | | | | | | | | |
| 32.75 | 0.51 | 0.05 | 0.354 | 0.085 | 0.818 | 0.132 | 1.16 | 0.092 | | | | | | | | | | | | | | | |
| 33.25 | 0.645 | 0.07 | 0.348 | 0.066 | 0.748 | 0.115 | 1.339 | 0.091 | | | | | | | | | | | | | | | |
| 33.75 | 0.577 | 0.08 | 0.417 | 0.111 | 0.934 | 0.187 | 1.46 | 0.096 | | | | | | | | | | | | | | | |
| 34.25 | 0.773 | 0.094 | 0.376 | 0.094 | 0.818 | 0.098 | 1.531 | 0.153 | | | | | | | | | | | | | | | |
| 34.75 | 0.685 | 0.068 | 0.426 | 0.108 | 0.782 | 0.132 | 1.495 | 0.13 | | | | | | | | | | | | | | | |
| 35.25 | 0.858 | 0.093 | 0.363 | 0.094 | 0.776 | 0.059 | 1.878 | 0.184 | | | | | | | | | | | | | | | |
| 35.75 | 0.671 | 0.071 | 0.385 | 0.108 | 0.742 | 0.092 | 1.809 | 0.182 | | | | | | | | | | | | | | | |
| 36.25 | 0.832 | 0.101 | 0.406 | 0.109 | 0.853 | 0.113 | 2.251 | 0.22 | | | | | | | | | | | | | | | |
| 36.75 | 0.838 | 0.085 | 0.443 | 0.142 | 0.844 | 0.098 | 1.864 | 0.179 | | | | | | | | | | | | | | | |
| 37.25 | 1.005 | 0.118 | 0.32 | 0.082 | 0.843 | 0.112 | 2.349 | 0.137 | | | | | | | | | | | | | | | |
| 37.75 | 0.603 | 0.076 | 0.329 | 0.097 | 0.686 | 0.066 | 1.171 | 0.163 | | | | | | | | | | | | | | | |

**Fold change = (Average area under the curve before stress Day 23) / (Average area under the curve after stress)**

**Supplementary Table S3.** Mean, SEM, fold changes, p-values (t-test), and percentage differences in motion obtained from the statistical comparison of 12 h integrated motion following stress imposition with the corresponding time point on day before stress imposition in (e) 100 mM KCl treatment under dimGday/night and RGBday/dimGnight with their respective non-stressed plants.

| AU Exp6 | | | | | | | | | |
|---|---|---|---|---|---|---|---|---|---|
| | RGB day DimG nightControl | | RGB day DimG night100mMI | | | DimG day/nightControl | | DimG day/night100mMKCl | |
| Time | Control_avg | Control_SEM | Stress_avg | Stress_SEM | Time | Control_avg | Control_SEM | Stress_avg | Stress_SEM |
| 24.25 | 0.294992 | 0.039737 | 0.315581 | 0.040209 | 24.25 | 0.282785 | 0.036177 | 0.319694 | 0.03379 |
| 24.75 | 0.459401 | 0.072917 | 0.147584 | 0.016045 | 24.75 | 0.443698 | 0.071679 | 0.123638 | 0.011141 |
| 25.25 | 0.23578 | 0.0273 | 0.231973 | 0.021909 | 25.25 | 0.213664 | 0.029433 | 0.216678 | 0.015992 |
| 25.75 | 0.405326 | 0.040558 | 0.277746 | 0.022054 | 25.75 | 0.363398 | 0.038414 | 0.248912 | 0.020884 |
| 26.25 | 0.274913 | 0.034498 | 0.218482 | 0.019763 | 26.25 | 0.236013 | 0.034406 | 0.178196 | 0.011853 |
| 26.75 | 0.355655 | 0.036824 | 0.330541 | 0.029009 | 26.75 | 0.315781 | 0.040355 | 0.302305 | 0.026609 |
| 27.25 | 0.351154 | 0.032598 | 0.242296 | 0.02189 | 27.25 | 0.326986 | 0.029859 | 0.219863 | 0.022484 |
| 27.75 | 0.436372 | 0.034949 | 0.31713 | 0.037172 | 27.75 | 0.383562 | 0.046517 | 0.274554 | 0.034024 |
| 28.25 | 0.388426 | 0.048747 | 0.219976 | 0.021584 | 28.25 | 0.316914 | 0.053371 | 0.186983 | 0.018388 |
| 28.75 | 0.519214 | 0.045898 | 0.319488 | 0.044827 | 28.75 | 0.445124 | 0.047135 | 0.267268 | 0.040313 |
| 29.25 | 0.44029 | 0.047427 | 0.229663 | 0.02335 | 29.25 | 0.365385 | 0.060856 | 0.19302 | 0.023982 |
| 29.75 | 0.523225 | 0.061312 | 0.299816 | 0.038565 | 29.75 | 0.432207 | 0.055057 | 0.248743 | 0.041225 |
| 30.25 | 0.612681 | 0.05864 | 0.265925 | 0.028169 | 30.25 | 0.479009 | 0.058617 | 0.221646 | 0.031795 |
| 30.75 | 0.512037 | 0.052663 | 0.296167 | 0.028967 | 30.75 | 0.47046 | 0.057381 | 0.235892 | 0.029253 |
| 31.25 | 0.426246 | 0.074725 | 0.257335 | 0.026236 | 31.25 | 0.354515 | 0.066877 | 0.179219 | 0.020354 |
| 31.75 | 0.590981 | 0.045632 | 0.269131 | 0.034559 | 31.75 | 0.504965 | 0.046228 | 0.231921 | 0.034015 |
| 32.25 | 0.556607 | 0.079624 | 0.231664 | 0.023772 | 32.25 | 0.486063 | 0.067145 | 0.182526 | 0.02846 |
| 32.75 | 0.617429 | 0.052714 | 0.29097 | 0.029412 | 32.75 | 0.499728 | 0.060064 | 0.239995 | 0.035172 |
| 33.25 | 0.547465 | 0.076196 | 0.300506 | 0.037209 | 33.25 | 0.454618 | 0.064769 | 0.207032 | 0.03763 |
| 33.75 | 0.784073 | 0.153698 | 0.295219 | 0.025771 | 33.75 | 0.640639 | 0.111545 | 0.24245 | 0.032115 |
| 34.25 | 0.609826 | 0.087939 | 0.35318 | 0.036278 | 34.25 | 0.487667 | 0.074205 | 0.237661 | 0.028404 |
| 34.75 | 0.648811 | 0.111442 | 0.336104 | 0.047329 | 34.75 | 0.63282 | 0.101255 | 0.250952 | 0.046117 |
| 35.25 | 0.551209 | 0.057719 | 0.326595 | 0.047716 | 35.25 | 0.443877 | 0.057202 | 0.199133 | 0.035861 |
| 35.75 | 0.688257 | 0.092694 | 0.309018 | 0.044845 | 35.75 | 0.614497 | 0.08289 | 0.260324 | 0.040462 |
| 36.25 | 0.728389 | 0.08782 | 0.343153 | 0.054693 | 36.25 | 0.604204 | 0.068798 | 0.258772 | 0.037092 |
| 36.75 | 0.756643 | 0.112057 | 0.338225 | 0.050542 | 36.75 | 0.674706 | 0.093924 | 0.294955 | 0.040746 |

**NO calculations were done**
