## Supplemental Table S4 for "Leaf movements as a quantitative metric for early stress detection"

**Supplementary Table S4.** Mean, SEM, fold changes, *p*-values (t-test), and percentage differences in motion obtained from the statistical comparison of day-today motion following stress imposition in **(a)** different stress treatments in lettuce, **(b)** salt stress applied at different locations on lettuce, **(c)** within location replication of different stresses and **(d)** 100 mM NaCl treatment on different CEA crops, with their respective non-stressed plants.

**Supplementary Table S4.** Mean, SEM, fold changes, p-values (t-test), and percentage differences in motion obtained from the statistical comparison of day-today motion following stress imposition in (a) different stress treatments in lettuce with their respective non-stressed plants.

UWA Exp6

| Day | NutrientWithdrawal | | 150 mM Salt | | Control | | Hypoxia | | Days post stress | NutrientWithdrawal | | | 150 mM Salt | | | Control | | | Hypoxia | | |
|---|---|---|---|---|---|---|---|---|---|---|---|---|---|---|---|---|---|---|---|---|---|
| | Mean | SEM | Mean | SEM | Mean | SEM | Mean | SEM | | T.test P values | Fold Change | % difference | T.test P values | Fold Change | % difference | T.test P values | Fold Change | % difference | T.test P values | Fold Change | % difference |
| 18.5 | 0.3661 | 0.0871 | 0.1188 | 0.0175 | 0.2361 | 0.0395 | 0.2903 | 0.0456 | | | | | | | | | | | | | |
| 19.5 | 0.3669 | 0.0754 | 0.2468 | 0.0405 | 0.4 | 0.0737 | 0.4252 | 0.0696 | | | | | | | | | | | | | |
| 20.5 | 0.4222 | 0.0899 | 0.251 | 0.0493 | 0.4737 | 0.0676 | 0.584 | 0.1323 | | | | | | | | | | | | | |
| 21.5 | 0.4246 | 0.0942 | 0.2982 | 0.0475 | 0.5474 | 0.0861 | 0.691 | 0.1402 | | | | | | | | | | | | | |
| 22.5 | 0.501 | 0.0904 | 0.374 | 0.0599 | 0.5854 | 0.1152 | 0.7961 | 0.1394 | | | | | | | | | | | | | |
| 23.5 | 0.5651 | 0.0811 | 0.4071 | 0.0567 | 0.6126 | 0.1341 | 0.919 | 0.1715 | | | | | | | | | | | | | |
| 24.5 | 0.6832 | 0.0782 | 0.3844 | 0.0418 | 0.6861 | 0.1196 | 0.662 | 0.0709 | 1 | 0.3058 | 0.827 | 20.915 | 0.7505 | 1.059 | -5.57 | 0.6865 | 0.8929 | 11.996 | 0.18 | 1.3882 | -27.96 |
| 25.5 | 0.6146 | 0.0907 | 0.493 | 0.085 | 0.5769 | 0.0892 | 0.7305 | 0.1087 | 2 | 0.6877 | 0.9194 | 8.7704 | 0.4096 | 0.8258 | 21.092 | 0.8263 | 1.062 | -5.839 | 0.3635 | 1.258 | -20.51 |
| 26.5 | 0.5242 | 0.0938 | 0.6124 | 0.1185 | 0.5487 | 0.0869 | 0.8004 | 0.1702 | 3 | 0.7449 | 1.078 | -7.233 | 0.1324 | 0.6648 | 50.416 | 0.6931 | 1.1164 | -10.43 | 0.6284 | 1.1482 | -12.91 |
| 27.5 | 0.382 | 0.0597 | 0.2742 | 0.0425 | 0.6587 | 0.0876 | 0.8512 | 0.2123 | 4 | 0.0828 | 1.4791 | -32.39 | 0.074 | 1.4845 | -32.64 | 0.776 | 0.93 | 7.5281 | 0.806 | 1.0797 | -7.38 |
| 28.5 | 0.2157 | 0.0242 | 0.2761 | 0.0503 | 0.5221 | 0.0846 | 1.046 | 0.2158 | 5 | 0.0004 | 2.6201 | -61.83 | 0.0978 | 1.4745 | -32.18 | 0.5736 | 1.1735 | -14.78 | 0.6494 | 0.8785 | 13.825 |
| 29.5 | 0.1597 | 0.0144 | 0.1789 | 0.034 | 0.6072 | 0.1061 | 1.0734 | 0.1793 | 6 | 6E-05 | 3.5386 | -71.74 | 0.0023 | 2.2751 | -56.05 | 0.975 | 1.0089 | -0.883 | 0.5402 | 0.8562 | 16.802 |
| 30.5 | 0.1352 | 0.0126 | 0.2018 | 0.0363 | 0.7271 | 0.0971 | 1.0454 | 0.1312 | 7 | 3E-05 | 4.1792 | -76.07 | 0.0059 | 2.0177 | -50.44 | 0.4964 | 0.8425 | 18.689 | 0.5643 | 0.8791 | 13.753 |
| 31.5 | 0.1967 | 0.0482 | 0.2398 | 0.0243 | 0.7542 | 0.1154 | 0.8193 | 0.12 | 8 | 0.0008 | 2.8723 | -65.18 | 0.0127 | 1.6979 | -41.1 | 0.4319 | 0.8122 | 23.116 | 0.6384 | 1.1217 | -10.85 |
| 32.5 | 0.0997 | 0.0114 | 0.2113 | 0.029 | 0.8313 | 0.138 | 1.0645 | 0.1402 | 9 | 1E-05 | 5.665 | -82.35 | 0.0055 | 1.927 | -48.1 | 0.268 | 0.737 | 35.693 | 0.5181 | 0.8633 | 15.836 |
| 33.5 | 0.1027 | 0.0116 | 0.2013 | 0.023 | 0.8362 | 0.1505 | 1.1897 | 0.173 | 10 | 1E-05 | 5.5001 | -81.82 | 0.0028 | 2.0223 | -50.55 | 0.2793 | 0.7326 | 36.494 | 0.2785 | 0.7725 | 29.454 |
| 34.5 | 0.1534 | 0.0226 | 0.219 | 0.0313 | 0.7207 | 0.1009 | 0.8896 | 0.1335 | 11 | 7E-05 | 3.6832 | -72.85 | 0.0082 | 1.8589 | -46.2 | 0.5263 | 0.8501 | 17.636 | 0.8935 | 1.0331 | -3.202 |
| 35.5 | 0.1531 | 0.0276 | 0.2975 | 0.0301 | 0.9549 | 0.1406 | 1.2958 | 0.1464 | 12 | 8E-05 | 3.6903 | -72.9 | 0.1018 | 1.3682 | -26.91 | 0.092 | 0.6416 | 55.867 | 0.1089 | 0.7092 | 41.006 |
| 36.5 | 0.3299 | 0.0662 | 0.2995 | 0.0378 | 0.9988 | 0.163 | 1.1373 | 0.1838 | 13 | 0.0351 | 1.7129 | -41.62 | 0.1286 | 1.3592 | -26.43 | 0.0809 | 0.6134 | 63.029 | 0.3946 | 0.808 | 23.758 |
| 37.5 | 0.4675 | 0.1207 | 0.2507 | 0.0341 | 0.8102 | 0.0873 | 1.1129 | 0.1812 | 14 | 0.5095 | 1.2086 | -17.26 | 0.0273 | 1.6238 | -38.42 | 0.23 | 0.7562 | 32.245 | 0.4452 | 0.8257 | 21.104 |
| 38.5 | 0.0895 | 0.0324 | 0.0297 | 0.0068 | 0.0883 | 0.0142 | 0.1111 | 0.0199 | | | | | | | | | | | | | |

AU Exp4

| Day | 100 mM NaCl | | Control | | Days post stress | 100 mM NaCl | | | Control | | |
|---|---|---|---|---|---|---|---|---|---|---|---|
| | Mean | SEM | Mean | SEM | | T.test P values | Fold Change | % difference | T.test P values | Fold Change | % difference |
| 18.5 | 0.1919 | 0.0428 | 0.4764 | 0.0293 | | | | | | | |
| 19.5 | 0.1608 | 0.0503 | 0.312 | 0.033 | | | | | | | |
| 20.5 | 0.2097 | 0.0644 | 0.2822 | 0.0261 | | | | | | | |
| 21.5 | 0.1942 | 0.059 | 0.2589 | 0.0374 | | | | | | | |
| 22.5 | 0.2258 | 0.0595 | 0.3544 | 0.039 | | | | | | | |
| 23.5 | 0.2948 | 0.0751 | 0.4269 | 0.0423 | | | | | | | |
| 24.5 | 0.2614 | 0.0603 | 0.5346 | 0.0508 | 1 | 0.7334 | 1.1274 | -11.3 | 0.1226 | 0.7985 | 25.231 |
| 25.5 | 0.1815 | 0.0449 | 0.5472 | 0.0464 | 2 | 0.2117 | 1.6242 | -38.43 | 0.0734 | 0.7801 | 28.187 |
| 26.5 | 0.2019 | 0.0437 | 0.6468 | 0.07 | 3 | 0.2992 | 1.46 | -31.51 | 0.0162 | 0.6601 | 51.499 |
| 27.5 | 0.2425 | 0.0536 | 0.5427 | 0.0348 | 4 | 0.5778 | 1.2156 | -17.74 | 0.0504 | 0.7866 | 27.127 |
| 28.5 | 0.2926 | 0.0645 | 0.7425 | 0.093 | 5 | 0.9827 | 1.0074 | -0.738 | 0.007 | 0.575 | 73.915 |
| 29.5 | 0.3131 | 0.0595 | 0.92 | 0.0934 | 6 | 0.8504 | 0.9415 | 6.2148 | 0.0002 | 0.464 | 115.5 |
| 30.5 | 0.289 | 0.047 | 0.9017 | 0.1019 | 7 | 0.9493 | 1.0198 | -1.938 | 0.0005 | 0.4734 | 111.22 |
| 31.5 | 0.2973 | 0.052 | 1.0613 | 0.1133 | 8 | 0.9781 | 0.9914 | 0.8643 | 8E-05 | 0.4023 | 148.6 |
| 32.5 | 0.3336 | 0.0713 | 1.0162 | 0.1414 | 9 | 0.7115 | 0.8834 | 13.195 | 0.0011 | 0.4201 | 138.03 |
| 33.5 | 0.326 | 0.089 | 0.8819 | 0.1211 | 10 | 0.7915 | 0.9042 | 10.598 | 0.0027 | 0.4841 | 106.59 |
| 34.5 | 0.3566 | 0.0928 | 0.7675 | 0.0946 | 11 | 0.6108 | 0.8266 | 20.977 | 0.0046 | 0.5562 | 79.783 |
| 35.5 | 0.3715 | 0.083 | 0.8705 | 0.1162 | 12 | 0.5015 | 0.7934 | 26.034 | 0.0025 | 0.4904 | 103.9 |
| 36.5 | 0.4418 | 0.0833 | 0.8676 | 0.1496 | 13 | 0.2062 | 0.6672 | 49.88 | 0.0119 | 0.492 | 103.24 |
| 37.5 | 0.3903 | 0.0662 | 0.8033 | 0.1549 | 14 | 0.3526 | 0.7553 | 32.405 | 0.0323 | 0.5314 | 88.168 |

AU Exp6

| Day | Control | | Boron | | 100 mM KCl | | Days post stress | Control | | | Boron | | | 100 mM KCl | | |
|---|---|---|---|---|---|---|---|---|---|---|---|---|---|---|---|---|
| | Mean | SEM | Mean | SEM | Mean | SEM | | T.test P values | Fold Change | % difference | T.test P values | Fold Change | % difference | T.test P values | Fold Change | % difference |
| 18.5 | 0.3297 | 0.0848 | 0.6944 | 0.1279 | 0.3203 | 0.0475 | | | | | | | | | | |
| 19.5 | 0.4486 | 0.0905 | 0.705 | 0.1307 | 0.4426 | 0.0697 | | | | | | | | | | |
| 20.5 | 0.62 | 0.1238 | 0.8539 | 0.1498 | 0.3757 | 0.0558 | | | | | | | | | | |
| 21.5 | 0.6576 | 0.1227 | 0.8911 | 0.0796 | 0.4676 | 0.0706 | | | | | | | | | | |
| 22.5 | 0.7769 | 0.1471 | 1.1857 | 0.1648 | 0.6054 | 0.089 | | | | | | | | | | |
| 23.5 | 0.7921 | 0.0934 | 1.3787 | 0.2635 | 0.4583 | 0.0451 | | | | | | | | | | |
| 24.5 | 0.6165 | 0.0576 | 1.1444 | 0.1882 | 0.4835 | 0.0345 | 1 | 0.1534 | 1.2199 | -18.03 | 0.4798 | 1.2048 | -17 | 0.6618 | 0.9478 | 5.5057 |
| 25.5 | 0.5481 | 0.0715 | 0.8542 | 0.1111 | 0.499 | 0.0364 | 2 | 0.0707 | 1.2864 | -22.27 | 0.0853 | 1.6141 | -38.05 | 0.491 | 0.9183 | 8.8928 |
| 26.5 | 0.6981 | 0.0606 | 0.7779 | 0.1144 | 0.5116 | 0.0556 | 3 | 0.4389 | 0.9705 | 3.039 | 0.0528 | 1.7724 | -43.58 | 0.4656 | 0.8958 | 11.637 |
| 27.5 | 0.7746 | 0.1044 | 0.8856 | 0.1372 | 0.4687 | 0.0566 | 4 | 0.9078 | 0.9178 | 8.962 | 0.1164 | 1.5568 | -35.77 | 0.8877 | 0.9779 | 2.261 |
| 28.5 | 0.8295 | 0.1168 | 0.9386 | 0.1014 | 0.457 | 0.0638 | 5 | 0.8172 | 0.8882 | 12.582 | 0.1386 | 1.469 | -31.93 | 0.9869 | 1.0028 | -0.283 |
| 29.5 | 0.9839 | 0.1151 | 1.2381 | 0.136 | 0.4735 | 0.0604 | 6 | 0.2428 | 0.793 | 26.097 | 0.6418 | 1.1136 | -10.2 | 0.8422 | 0.9679 | 3.3209 |
| 30.5 | 0.8267 | 0.0972 | 1.7054 | 0.1571 | 0.4268 | 0.0519 | 7 | 0.8124 | 1.0012 | -0.121 | 0.3028 | 0.8085 | 23.693 | 0.6519 | 1.0739 | -6.883 |
| 31.5 | 0.9469 | 0.0953 | 1.3922 | 0.1907 | 0.4383 | 0.0648 | 8 | 0.2924 | 0.8155 | 22.626 | 0.9675 | 0.9903 | 0.9753 | 0.803 | 1.0456 | -4.364 |
| 32.5 | 1.1781 | 0.1532 | 1.2585 | 0.1367 | 0.4655 | 0.0721 | 9 | 0.062 | 0.6535 | 53.021 | 0.6909 | 1.0955 | -8.718 | 0.9329 | 0.9844 | 1.5826 |
| 33.5 | 1.1924 | 0.1784 | 1.4666 | 0.1427 | 0.5039 | 0.0661 | 10 | 0.0819 | 0.6021 | 66.096 | 0.773 | 0.9401 | 6.3764 | 0.5759 | 0.9095 | 9.9474 |
| 34.5 | 1.0949 | 0.1419 | 1.6692 | 0.2916 | 0.4759 | 0.0773 | 11 | 0.115 | 0.6728 | 48.625 | 0.4705 | 0.826 | 21.07 | 0.8464 | 0.9631 | 3.836 |
| 35.5 | 1.3314 | 0.1593 | 1.2782 | 0.1813 | 0.5722 | 0.0757 | 12 | 0.0155 | 0.5586 | 79.004 | 0.7574 | 1.0787 | -7.293 | 0.2123 | 0.8009 | 24.863 |

UoC Exp8

| Day | Control | | Water withdrawal | | Days post stress | Control | | | Water withdrawal | | |
|---|---|---|---|---|---|---|---|---|---|---|---|
| | Mean | SEM | Mean | SEM | | T.test P values | Fold Change | % difference | T.test P values | Fold Change | % difference |
| 18.5 | 0.4864 | 0.0833 | 0.3167 | 0.0486 | | | | | | | |
| 19.5 | 0.6222 | 0.0918 | 0.3954 | 0.0556 | | | | | | | |
| 20.5 | 0.6814 | 0.0932 | 0.4581 | 0.0787 | | | | | | | |
| 21.5 | 0.8795 | 0.0928 | 0.6612 | 0.0856 | | | | | | | |
| 22.5 | 1.1488 | 0.1653 | 0.9454 | 0.116 | | | | | | | |
| 23.5 | 1.2262 | 0.1715 | 1.1585 | 0.1332 | | | | | | | |
| 24.5 | 1.4696 | 0.1744 | 1.3568 | 0.1612 | 1 | 0.3303 | 0.8343 | 19.855 | 0.3531 | 0.8538 | 17.123 |
| 25.5 | 1.8197 | 0.1798 | 1.4133 | 0.1546 | 2 | 0.0259 | 0.6738 | 48.405 | 0.2249 | 0.8197 | 22 |
| 26.5 | 1.8199 | 0.1877 | 1.3881 | 0.1653 | 3 | 0.029 | 0.6738 | 48.418 | 0.2912 | 0.8346 | 19.824 |
| 27.5 | 1.7241 | 0.2604 | 1.2201 | 0.1465 | 4 | 0.1245 | 0.7112 | 40.604 | 0.7587 | 0.9495 | 5.3178 |
| 28.5 | 1.574 | 0.2119 | 1.21 | 0.1467 | 5 | 0.2152 | 0.779 | 28.367 | 0.7973 | 0.9574 | 4.4467 |
| 29.5 | 1.5572 | 0.2253 | 1.1009 | 0.0523 | 6 | 0.2548 | 0.7874 | 27 | 0.6914 | 1.0523 | -4.97 |
| 30.5 | 1.427 | 0.1957 | 1.5066 | 0.0855 | 7 | 0.4485 | 0.8593 | 16.376 | 0.0387 | 0.7689 | 30.052 |
| 31.5 | 1.5119 | 0.238 | 1.129 | 0.1109 | 8 | 0.3406 | 0.811 | 23.299 | 0.8667 | 1.0261 | -2.541 |
| 32.5 | 1.7672 | 0.3492 | 1.2611 | 0.1284 | 9 | 0.1782 | 0.6938 | 44.124 | 0.5849 | 0.9186 | 8.856 |
| 33.5 | 1.8357 | 0.4066 | 1.3496 | 0.1239 | 10 | 0.1811 | 0.668 | 49.705 | 0.3049 | 0.8584 | 16.501 |
| 34.5 | 1.5332 | 0.2871 | 1.4405 | 0.2244 | 11 | 0.3684 | 0.7997 | 25.04 | 0.2916 | 0.8042 | 24.347 |
| 35.5 | 1.8539 | 0.5548 | 1.4175 | 0.263 | 12 | 0.2914 | 0.6614 | 51.19 | 0.3891 | 0.8172 | 22.363 |
| 36.5 | 1.3818 | 0.2267 | 1.562 | 0.2317 | 13 | 0.5897 | 0.8874 | 12.688 | 0.1453 | 0.7417 | 34.833 |

**Fold change = (Average area under the curve before stress Day 23) / (Average area under the curve after stress)**

**Supplementary Table S4**. Mean, SEM, fold changes, p-values (t-test), and percentage differences in motion obtained from the statistical comparison of day-today motion following stress imposition in (b) salt stress applied at different locations on lettuce with their respective non-stressed plants.

**UWA Exp6**

| Day | 150 mM Salt | | Control | | Days post stress | 150 mM Salt | | | Control | | |
|---|---|---|---|---|---|---|---|---|---|---|---|
| | Mean | SEM | Mean | SEM | | T.test P values | Fold Change | % difference | T.test P values | Fold Change | % difference |
| 18.5 | 0.1188 | 0.0175 | 0.2361 | 0.0395 | | | | | | | |
| 19.5 | 0.2468 | 0.0405 | 0.4 | 0.0737 | | | | | | | |
| 20.5 | 0.251 | 0.0493 | 0.4737 | 0.0676 | | | | | | | |
| 21.5 | 0.2982 | 0.0475 | 0.5474 | 0.0861 | | | | | | | |
| 22.5 | 0.374 | 0.0599 | 0.5854 | 0.1152 | | | | | | | |
| 23.5 | 0.4071 | 0.0567 | 0.6126 | 0.1341 | | | | | | | |
| 24.5 | 0.3844 | 0.0418 | 0.6861 | 0.1196 | 1 | 0.7505 | 1.059 | -5.57 | 0.6865 | 0.8929 | 11.996 |
| 25.5 | 0.493 | 0.085 | 0.5769 | 0.0892 | 2 | 0.4096 | 0.8258 | 21.092 | 0.8263 | 1.062 | -5.839 |
| 26.5 | 0.6124 | 0.1185 | 0.5487 | 0.0869 | 3 | 0.1324 | 0.6648 | 50.416 | 0.6931 | 1.1164 | -10.43 |
| 27.5 | 0.2742 | 0.0425 | 0.6587 | 0.0876 | 4 | 0.074 | 1.4845 | -32.64 | 0.776 | 0.93 | 7.5281 |
| 28.5 | 0.2761 | 0.0503 | 0.5221 | 0.0846 | 5 | 0.0978 | 1.4745 | -32.18 | 0.5736 | 1.1735 | -14.78 |
| 29.5 | 0.1789 | 0.034 | 0.6072 | 0.1061 | 6 | 0.0023 | 2.2751 | -56.05 | 0.975 | 1.0089 | -0.883 |
| 30.5 | 0.2018 | 0.0363 | 0.7271 | 0.0971 | 7 | 0.0059 | 2.0177 | -50.44 | 0.4964 | 0.8425 | 18.689 |
| 31.5 | 0.2398 | 0.0243 | 0.7542 | 0.1154 | 8 | 0.0127 | 1.6979 | -41.1 | 0.4319 | 0.8122 | 23.116 |
| 32.5 | 0.2113 | 0.029 | 0.8313 | 0.138 | 9 | 0.0055 | 1.927 | -48.1 | 0.268 | 0.737 | 35.693 |
| 33.5 | 0.2013 | 0.023 | 0.8362 | 0.1505 | 10 | 0.0028 | 2.0223 | -50.55 | 0.2793 | 0.7326 | 36.494 |
| 34.5 | 0.219 | 0.0313 | 0.7207 | 0.1009 | 11 | 0.0082 | 1.8589 | -46.2 | 0.5263 | 0.8501 | 17.636 |
| 35.5 | 0.2975 | 0.0301 | 0.9549 | 0.1406 | 12 | 0.1018 | 1.3682 | -26.91 | 0.092 | 0.6416 | 55.867 |
| 36.5 | 0.2995 | 0.0378 | 0.9988 | 0.163 | 13 | 0.1286 | 1.3592 | -26.43 | 0.0809 | 0.6134 | 63.029 |
| 37.5 | 0.2507 | 0.0341 | 0.8102 | 0.0873 | 14 | 0.0273 | 1.6238 | -38.42 | 0.23 | 0.7562 | 32.245 |
| 38.5 | 0.0297 | 0.0068 | 0.0883 | 0.0142 | | | | | | | |

**AU Exp4**

| Day | 100 mM NaCl | | Control | | Days post stress | 100 mM NaCl | | | Control | | |
|---|---|---|---|---|---|---|---|---|---|---|---|
| | Mean | SEM | Mean | SEM | | T.test P values | Fold Change | % difference | T.test P values | Fold Change | % difference |
| 18.5 | 0.1919 | 0.0428 | 0.4764 | 0.0293 | | | | | | | |
| 19.5 | 0.1608 | 0.0503 | 0.312 | 0.033 | | | | | | | |
| 20.5 | 0.2097 | 0.0644 | 0.2822 | 0.0261 | | | | | | | |
| 21.5 | 0.1942 | 0.059 | 0.2589 | 0.0374 | | | | | | | |
| 22.5 | 0.2258 | 0.0595 | 0.3544 | 0.039 | | | | | | | |
| 23.5 | 0.2948 | 0.0751 | 0.4269 | 0.0423 | | | | | | | |
| 24.5 | 0.2614 | 0.0603 | 0.5346 | 0.0508 | 1 | 0.7334 | 1.1274 | -11.3 | 0.1226 | 0.7985 | 25.231 |
| 25.5 | 0.1815 | 0.0449 | 0.5472 | 0.0464 | 2 | 0.2117 | 1.6242 | -38.43 | 0.0734 | 0.7801 | 28.187 |
| 26.5 | 0.2019 | 0.0437 | 0.6468 | 0.07 | 3 | 0.2992 | 1.46 | -31.51 | 0.0162 | 0.6601 | 51.499 |
| 27.5 | 0.2425 | 0.0536 | 0.5427 | 0.0348 | 4 | 0.5778 | 1.2156 | -17.74 | 0.0504 | 0.7866 | 27.127 |
| 28.5 | 0.2926 | 0.0645 | 0.7425 | 0.093 | 5 | 0.9827 | 1.0074 | -0.738 | 0.007 | 0.575 | 73.915 |
| 29.5 | 0.3131 | 0.0595 | 0.92 | 0.0934 | 6 | 0.8504 | 0.9415 | 6.2148 | 0.0002 | 0.464 | 115.5 |
| 30.5 | 0.289 | 0.047 | 0.9017 | 0.1019 | 7 | 0.9493 | 1.0198 | -1.938 | 0.0005 | 0.4734 | 111.22 |
| 31.5 | 0.2973 | 0.052 | 1.0613 | 0.1133 | 8 | 0.9781 | 0.9914 | 0.8643 | 8E-05 | 0.4023 | 148.6 |
| 32.5 | 0.3336 | 0.0713 | 1.0162 | 0.1414 | 9 | 0.7115 | 0.8834 | 13.195 | 0.0011 | 0.4201 | 138.03 |
| 33.5 | 0.326 | 0.089 | 0.8819 | 0.1211 | 10 | 0.7915 | 0.9042 | 10.598 | 0.0027 | 0.4841 | 106.59 |
| 34.5 | 0.3566 | 0.0928 | 0.7675 | 0.0946 | 11 | 0.6108 | 0.8266 | 20.977 | 0.0046 | 0.5562 | 79.783 |
| 35.5 | 0.3715 | 0.083 | 0.8705 | 0.1162 | 12 | 0.5015 | 0.7934 | 26.034 | 0.0025 | 0.4904 | 103.9 |
| 36.5 | 0.4418 | 0.0833 | 0.8676 | 0.1496 | 13 | 0.2062 | 0.6672 | 49.88 | 0.0119 | 0.492 | 103.24 |
| 37.5 | 0.3903 | 0.0662 | 0.8033 | 0.1549 | 14 | 0.3526 | 0.7553 | 32.405 | 0.0323 | 0.5314 | 88.168 |

**UoC Exp1**

| Day | Control | | 100 mM NaCl | | Days post stress | Control | | | 100 mM NaCl | | |
|---|---|---|---|---|---|---|---|---|---|---|---|
| | Mean | SEM | Mean | SEM | | T.test P values | Fold Change | % difference | T.test P values | Fold Change | % difference |
| 18.5 | 0.3799 | 0.0393 | 0.4072 | 0.1014 | | | | | | | |
| 19.5 | 0.4333 | 0.0442 | 0.4543 | 0.098 | | | | | | | |
| 20.5 | 0.4782 | 0.041 | 0.5633 | 0.1049 | | | | | | | |
| 21.5 | 0.5163 | 0.0392 | 0.6206 | 0.1052 | | | | | | | |
| 22.5 | 0 | 0 | 0 | 0 | | | | | | | |
| 23.5 | 0.7261 | 0.051 | 1.0895 | 0.1281 | | | | | | | |
| 24.5 | 0.7043 | 0.0421 | 0.7011 | 0.1109 | 1 | 0.7457 | 1.031 | -3.006 | 0.0198 | 1.5539 | -35.65 |
| 25.5 | 0.7606 | 0.0373 | 0.6777 | 0.0733 | 2 | 0.3318 | 0.926 | 7.9923 | 0.8473 | 1.0346 | -3.344 |
| 26.5 | 0.7634 | 0.04 | 0.8368 | 0.0757 | 3 | 0.9591 | 0.9963 | 0.3745 | 0.1104 | 0.8098 | 23.482 |
| 27.5 | 0.7881 | 0.0642 | 0.9066 | 0.0809 | 4 | 0.7484 | 0.9687 | 3.2362 | 0.4944 | 0.923 | 8.3371 |
| 28.5 | 0.7701 | 0.0791 | 0.8914 | 0.0802 | 5 | 0.8617 | 1.0234 | -2.289 | 0.8841 | 1.0171 | -1.679 |
| 29.5 | 0.8079 | 0.0778 | 0.9049 | 0.0852 | 6 | 0.7377 | 0.9532 | 4.9094 | 0.8996 | 0.9851 | 1.5172 |
| 30.5 | 0.9575 | 0.0795 | 0.9417 | 0.0827 | 7 | 0.1974 | 0.8438 | 18.515 | 0.7356 | 0.9609 | 4.0647 |
| 31.5 | 1.1496 | 0.1428 | 0.9796 | 0.0784 | 8 | 0.257 | 0.8329 | 20.067 | 0.7164 | 0.9612 | 4.033 |
| 32.5 | 1.1114 | 0.1116 | 0.7912 | 0.0628 | 9 | 0.8356 | 1.0344 | -3.326 | 0.0511 | 1.2381 | -19.23 |
| 33.5 | 0.9371 | 0.1241 | 0.7324 | 0.0808 | 10 | 0.3118 | 1.186 | -15.68 | 0.5317 | 1.0804 | -7.441 |
| 34.5 | 0.8739 | 0.1118 | 0.8041 | 0.0872 | 11 | 0.7101 | 1.0723 | -6.742 | 0.5123 | 0.9108 | 9.7938 |
| 35.5 | 0.8056 | 0.1183 | 0.8741 | 0.1084 | 12 | 0.6804 | 1.0848 | -7.814 | 0.5843 | 0.9199 | 8.7025 |
| 36.5 | 0.6515 | 0.0966 | 1.007 | 0.1133 | 13 | 0.3279 | 1.2366 | -19.13 | 0.3598 | 0.868 | 15.205 |

**Fold change = (Average area under the curve before stress Day 23) / (Average area under the curve after stress)**

**Supplementary Table S4.** Mean, SEM, fold changes, *p* -values (t-test), and percentage differences in motion obtained from the statistical comparison of day-today motion following stress imposition in **(c)** within location replication of different stresses with their respective non-stressed plants.

| | UWA E | UWA E | UWA E | UWA E | UWA E | UWA Exp6 | | | UWA Exp4 | | | UWA Exp6 | | | UWA Exp4 | | | UWA Exp6 | | | UWA Exp4 | | | UWA Exp6 | | |
|---|---|---|---|---|---|---|---|---|---|---|---|---|---|---|---|---|---|---|---|---|---|---|---|---|---|---|
| | Control | Control | Hypoxia | Hypoxia | NutrientWithdrawal | NutrientWithdrawal | | Days post stress | Control | | | Control | | | Hypoxia | | | Hypoxia | | | NutrientWithdrawal | | | NutrientWithdrawal | | |
| | | | | | | | | | T.test P values | Fold Change | % difference | T.test P values | Fold Change | % difference | T.test P values | Fold Change | % difference | T.test P values | Fold Change | % difference | T.test P values | Fold Change | % difference | T.test P values | Fold Change | % difference |
| 18.5 | 0.37 | 0.24 | 0.49 | 0.29 | 0.41 | 0.37 | | | | | | | | | | | | | | | | | | | | |
| 19.5 | 0.5 | 0.4 | 0.62 | 0.43 | 0.49 | 0.37 | | | | | | | | | | | | | | | | | | | | |
| 20.5 | 0.39 | 0.47 | 0.47 | 0.58 | 0.31 | 0.42 | | | | | | | | | | | | | | | | | | | | |
| 21.5 | 0.44 | 0.55 | 0.6 | 0.69 | 0.31 | 0.42 | | | | | | | | | | | | | | | | | | | | |
| 22.5 | 0.46 | 0.59 | 0.57 | 0.8 | 0.38 | 0.5 | | | | | | | | | | | | | | | | | | | | |
| 23.5 | 0.6 | 0.61 | 0.64 | 0.92 | 0.46 | 0.57 | | | | | | | | | | | | | | | | | | | | |
| 24.5 | 0.62 | 0.69 | 0.58 | 0.66 | 0.68 | 0.68 | | 1 | 0.86 | 0.97 | 3.23 | 0.69 | 0.89 | 12 | 0.63 | 1.1 | -9.2 | 0.18 | 1.39 | -28 | 0.12 | 0.67 | 48.2 | 0.31 | 0.83 | 20.9 |
| 25.5 | 0.59 | 0.58 | 0.57 | 0.73 | 0.52 | 0.61 | | 2 | 0.95 | 1.01 | -1.1 | 0.83 | 1.06 | -5.8 | 0.6 | 1.12 | -10 | 0.36 | 1.26 | -21 | 0.65 | 0.88 | 13.4 | 0.69 | 0.92 | 8.77 |
| 26.5 | 0.51 | 0.55 | 0.66 | 0.8 | 0.3 | 0.52 | | 3 | 0.49 | 1.17 | -15 | 0.69 | 1.12 | -10 | 0.89 | 0.97 | 3.1 | 0.63 | 1.15 | -13 | 0.16 | 1.51 | -34 | 0.74 | 1.08 | -7.2 |
| 27.5 | 0.49 | 0.66 | 0.85 | 0.85 | | 0.38 | | 4 | 0.36 | 1.22 | -18 | 0.78 | 0.93 | 7.53 | 0.28 | 0.75 | 33.2 | 0.81 | 1.08 | -7.4 | | | | 0.08 | 1.48 | -32 |
| 28.5 | 0.45 | 0.52 | 0.89 | 1.05 | | 0.22 | | 5 | 0.09 | 1.35 | -26 | 0.57 | 1.17 | -15 | 0.33 | 0.71 | 40.3 | 0.65 | 0.88 | 13.8 | | | | 0 | 2.62 | -62 |
| 29.5 | 0.53 | 0.61 | 0.79 | 1.07 | | 0.16 | | 6 | 0.44 | 1.14 | -12 | 0.98 | 1.01 | -0.9 | 0.46 | 0.81 | 23.9 | 0.54 | 0.86 | 16.8 | | | | 0 | 3.54 | -72 |
| 30.5 | 0.74 | 0.73 | 0.67 | 1.05 | 0.33 | 0.14 | | 7 | 0.26 | 0.81 | 23.6 | 0.5 | 0.84 | 18.7 | 0.85 | 0.95 | 4.81 | 0.56 | 0.88 | 13.8 | 0.28 | 1.39 | -28 | 0 | 4.18 | -76 |
| 31.5 | 0.73 | 0.75 | 0.73 | 0.82 | 0.2 | 0.2 | | 8 | 0.51 | 0.83 | 20.5 | 0.43 | 0.81 | 23.1 | 0.63 | 0.87 | 14.8 | 0.64 | 1.12 | -11 | 0.02 | 2.3 | -56 | 0 | 2.87 | -65 |
| 32.5 | 0.77 | 0.83 | 0.73 | 1.06 | 0.21 | 0.1 | | 9 | 0.19 | 0.78 | 28.7 | 0.27 | 0.74 | 35.7 | 0.58 | 0.87 | 15.2 | 0.52 | 0.86 | 15.8 | 0.02 | 2.18 | -54 | 0 | 5.67 | -82 |
| 33.5 | 0.73 | 0.84 | 0.84 | 1.19 | 0.19 | 0.1 | | 10 | 0.33 | 0.83 | 20.6 | 0.28 | 0.73 | 36.5 | 0.35 | 0.76 | 32.4 | 0.28 | 0.77 | 29.5 | 0.01 | 2.47 | -59 | 0 | 5.5 | -82 |
| 34.5 | 0.67 | 0.72 | 0.78 | 0.89 | 0.17 | 0.15 | | 11 | 0.61 | 0.9 | 11.4 | 0.53 | 0.85 | 17.6 | 0.34 | 0.82 | 22.1 | 0.89 | 1.03 | -3.2 | 0.01 | 2.66 | -62 | 0 | 3.68 | -73 |
| 35.5 | 0.57 | 0.95 | 0.83 | 1.3 | 0.17 | 0.15 | | 12 | 0.78 | 1.05 | -4.7 | 0.09 | 0.64 | 55.9 | 0.19 | 0.77 | 30.2 | 0.11 | 0.71 | 41 | 0.01 | 2.65 | -62 | 0 | 3.69 | -73 |
| 36.5 | 0.49 | 1 | 0.76 | 1.14 | 0.12 | 0.33 | | 13 | 0.25 | 1.23 | -19 | 0.08 | 0.61 | 63 | 0.33 | 0.83 | 19.9 | 0.39 | 0.81 | 23.8 | 0 | 3.81 | -74 | 0.04 | 1.71 | -42 |
| 37.5 | 0.6 | 0.81 | 1.21 | 1.11 | 0.09 | 0.47 | | 14 | 0.95 | 1.01 | -0.9 | 0.23 | 0.76 | 32.2 | 0 | 0.53 | 90.3 | 0.45 | 0.83 | 21.1 | 0 | 5 | -80 | 0.51 | 1.21 | -17 |

| AU Ex | AU Ex | AU Ex | AU Ex | AU Ex | AU Ex | AU Exp5 | | | AU Exp4 | | | AU Exp5 | | | AU Exp5 | | | AU Exp4 | | | AU Exp5 | | | AU Exp5 | | |
|---|---|---|---|---|---|---|---|---|---|---|---|---|---|---|---|---|---|---|---|---|---|---|---|---|---|---|
| | ControlB | ControlA | ControlB | 100 mM NaCl-SaltB | 100 mM NaCl-SaltA | 100 mM NaCl-SaltB | | Days post stress | ControlB | | | ControlA | | | ControlB | | | 100 mM NaCl-SaltB | | | 100 mM NaCl-SaltA | | | 100 mM NaCl-SaltB | | |
| | | | | | | | | | T.test P values | Fold Change | % difference | T.test P values | Fold Change | % difference | T.test P values | Fold Change | % difference | T.test P values | Fold Change | % difference | T.test P values | Fold Change | % difference | T.test P values | Fold Change | % difference |
| 18.5 | 0.48 | 0.36 | 0.27 | 0.19 | 0.42 | 0.3 | | | | | | | | | | | | | | | | | | | | |
| 19.5 | 0.31 | 0.51 | 0.3 | 0.16 | 0.38 | 0.37 | | | | | | | | | | | | | | | | | | | | |
| 20.5 | 0.28 | 0.63 | 0.44 | 0.21 | 0.52 | 0.48 | | | | | | | | | | | | | | | | | | | | |
| 21.5 | 0.26 | 0.74 | 0.52 | 0.19 | 0.66 | 0.6 | | | | | | | | | | | | | | | | | | | | |
| 22.5 | 0.35 | 0.68 | 0.55 | 0.23 | 0.69 | 0.56 | | | | | | | | | | | | | | | | | | | | |
| 23.5 | 0.43 | 0.64 | 0.59 | 0.29 | 0.69 | 0.59 | | | | | | | | | | | | | | | | | | | | |
| 24.5 | 0.53 | 0.85 | 0.82 | 0.26 | 0.4 | 0.28 | | 1 | 0.12 | 0.8 | 25.2 | 0.02 | 0.75 | 33.3 | 0.09 | 0.72 | 39 | 0.73 | 1.13 | -11 | 0.01 | 1.7 | -41 | 0 | 2.1 | -52 |
| 25.5 | 0.55 | 0.9 | 0.98 | 0.18 | 0.67 | 0.63 | | 2 | 0.07 | 0.78 | 28.2 | 0.03 | 0.71 | 40.8 | 0.04 | 0.6 | 66.1 | 0.21 | 1.62 | -38 | 0.87 | 1.04 | -3.4 | 0.71 | 0.93 | 7.7 |
| 26.5 | 0.65 | 0.89 | 0.84 | 0.2 | 0.71 | 0.65 | | 3 | 0.02 | 0.66 | 51.5 | 0.08 | 0.71 | 40.3 | 0.07 | 0.71 | 41.8 | 0.3 | 1.46 | -32 | 0.9 | 0.98 | 2.2 | 0.64 | 0.9 | 11.7 |
| 27.5 | 0.54 | 0.93 | 0.81 | 0.24 | 0.69 | 0.69 | | 4 | 0.05 | 0.79 | 27.1 | 0.09 | 0.69 | 45.4 | 0.09 | 0.72 | 38.2 | 0.58 | 1.22 | -18 | 0.97 | 1.01 | -0.5 | 0.36 | 0.85 | 17 |
| 28.5 | 0.74 | 1.03 | 1.1 | 0.29 | 0.68 | 0.75 | | 5 | 0.01 | 0.57 | 73.9 | 0.04 | 0.62 | 61.8 | 0.02 | 0.53 | 87.3 | 0.98 | 1.01 | -0.7 | 0.89 | 1.02 | -2 | 0.21 | 0.78 | 28.4 |
| 29.5 | 0.92 | 1.21 | 1.46 | 0.31 | 0.68 | 0.72 | | 6 | 0 | 0.46 | 116 | 0.04 | 0.53 | 90 | 0 | 0.4 | 147 | 0.85 | 0.94 | 6.21 | 0.9 | 1.02 | -2 | 0.44 | 0.82 | 22.4 |
| 30.5 | 0.9 | 1.3 | 1.31 | 0.29 | 0.62 | 0.73 | | 7 | 0 | 0.47 | 111 | 0.03 | 0.49 | 104 | 0 | 0.45 | 122 | 0.95 | 1.02 | -1.9 | 0.53 | 1.11 | -10 | 0.42 | 0.81 | 24.1 |
| 31.5 | 1.06 | 1.19 | 1.02 | 0.3 | 0.64 | 0.69 | | 8 | 0 | 0.4 | 149 | 0.02 | 0.54 | 86.3 | 0.01 | 0.58 | 73 | 0.98 | 0.99 | 0.86 | 0.62 | 1.08 | -7.4 | 0.47 | 0.85 | 17.4 |
| 32.5 | 1.02 | 1.18 | 1.18 | 0.33 | 0.76 | 0.88 | | 9 | 0 | 0.42 | 138 | 0.03 | 0.54 | 84.7 | 0.02 | 0.5 | 99.9 | 0.71 | 0.88 | 13.2 | 0.53 | 0.91 | 10.1 | 0.09 | 0.66 | 50.5 |
| 33.5 | 0.88 | 0.95 | 1.41 | 0.33 | 0.84 | 0.81 | | 10 | 0 | 0.48 | 107 | 0.03 | 0.67 | 49.8 | 0.02 | 0.42 | 139 | 0.79 | 0.9 | 10.6 | 0.23 | 0.82 | 22 | 0.2 | 0.73 | 37.9 |
| 34.5 | 0.77 | 0.9 | 0.84 | 0.36 | 0.88 | 0.67 | | 11 | 0 | 0.56 | 79.8 | 0.07 | 0.71 | 41.3 | 0.19 | 0.7 | 43.1 | 0.61 | 0.83 | 21 | 0.2 | 0.79 | 27.4 | 0.57 | 0.88 | 13.9 |
| 35.5 | 0.87 | 0.95 | 0.96 | 0.37 | 0.98 | 0.64 | | 12 | 0 | 0.49 | 104 | 0.17 | 0.67 | 49.5 | 0.08 | 0.62 | 62.4 | 0.5 | 0.79 | 26 | 0.11 | 0.71 | 41.2 | 0.64 | 0.92 | 9.03 |
| 36.5 | 0.87 | 1.06 | 1.17 | 0.44 | 0.9 | 0.6 | | 13 | 0.01 | 0.49 | 103 | 0.07 | 0.6 | 66.6 | 0.01 | 0.5 | 98.6 | 0.21 | 0.67 | 49.9 | 0.22 | 0.77 | 30.3 | 0.9 | 0.98 | 2 |
| 37.5 | 0.8 | 1.09 | 0.83 | 0.39 | 0.67 | 0.53 | | 14 | 0.03 | 0.53 | 88.2 | 0.11 | 0.59 | 70.7 | 0.13 | 0.71 | 41.5 | 0.35 | 0.76 | 32.4 | 0.88 | 1.03 | -2.5 | 0.59 | 1.1 | -9.2 |

**Fold change = (Average area under the curve before stress Day 23) / (Average area under the curve after stress)**

**Supplementary Table S4.** Mean, SEM, fold changes, p-values (t-test), and percentage differences in motion obtained from the statistical comparison of day-today motion following stress imposition in (d) 100 mM NaCl treatment on different CEA crops with their respective non-stressed plants.

UWA Exp9

| Average Area under the cuve | | Radish 100 mM NaCl | | RadishControl | | MizunaControl | | Mizuna 100 mM NaCl | | Days post stress | Radish 100 mM NaCl | | | RadishControl | | | MizunaControl | | | Mizuna 100 mM NaCl | | |
|---|---|---|---|---|---|---|---|---|---|---|---|---|---|---|---|---|---|---|---|---|---|---|
| | | Mean | SEM | Mean | SEM | Mean | SEM | | | | T.test P values | Fold Change | % difference | T.test P values | Fold Change | % difference | T.test P values | Fold Change | % difference | T.test P values | Fold Change | % difference |
| | 18.5 | 0.6824 | 0.1633 | 0.3022 | 0.034 | 0.3182 | 0.0668 | 0.1838 | 0.0407 | | | | | | | | | | | | | |
| | 19.5 | 0.8756 | 0.1472 | 0.4168 | 0.0385 | 0.4408 | 0.0828 | 0.2722 | 0.0387 | | | | | | | | | | | | | |
| | 20.5 | 0.9236 | 0.1113 | 0.6698 | 0.0734 | 0.6359 | 0.0812 | 0.3923 | 0.052 | | | | | | | | | | | | | |
| | 21.5 | 0.895 | 0.1055 | 0.7179 | 0.1036 | 0.8322 | 0.0874 | 0.5686 | 0.0768 | | | | | | | | | | | | | |
| | 22.5 | 0.8686 | 0.0828 | 0.7212 | 0.09 | 0.7223 | 0.0719 | 0.6749 | 0.102 | | | | | | | | | | | | | |
| | 23.5 | 0.8598 | 0.083 | 0.7274 | 0.0846 | 0.7358 | 0.0585 | 0.7412 | 0.0945 | | | | | | | | | | | | | |
| | 24.5 | 0.5219 | 0.046 | 0.7331 | 0.0882 | 0.7501 | 0.0694 | 0.7629 | 0.0848 | 1 | 0.0005 | 1.6475 | -39.3 | 0.9636 | 0.9923 | 0.7747 | 0.8768 | 0.981 | 1.9348 | 0.866 | 0.9716 | 2.9266 |
| | 25.5 | 0.4187 | 0.0983 | 0.8628 | 0.0974 | 0.8293 | 0.0963 | 0.558 | 0.0481 | 2 | 0.0007 | 2.0537 | -51.31 | 0.3053 | 0.8431 | 18.613 | 0.4155 | 0.8873 | 12.705 | 0.0981 | 1.3284 | -24.72 |
| | 26.5 | 0.6594 | 0.1025 | 0.8921 | 0.0979 | 0.8319 | 0.0893 | 0.6494 | 0.0525 | 3 | 0.0931 | 1.304 | -23.31 | 0.2163 | 0.8154 | 22.641 | 0.378 | 0.8845 | 13.055 | 0.4047 | 1.1415 | -12.39 |
| | 27.5 | 0.8382 | 0.0747 | 0.9758 | 0.113 | 0.8609 | 0.087 | 0.7144 | 0.0581 | 4 | 0.8252 | 1.0258 | -2.515 | 0.0924 | 0.7455 | 34.139 | 0.2455 | 0.8547 | 17.001 | 0.811 | 1.0376 | -3.624 |
| | 28.5 | 0.9971 | 0.1034 | 1.034 | 0.1273 | 0.8424 | 0.1024 | 0.8061 | 0.0946 | 5 | 0.2446 | 0.8623 | 15.964 | 0.0574 | 0.7035 | 42.143 | 0.3757 | 0.8735 | 14.488 | 0.6326 | 0.9196 | 8.7453 |
| | 29.5 | 0.999 | 0.1028 | 1.0663 | 0.1483 | 0.9773 | 0.1216 | 0.9257 | 0.1367 | 6 | 0.2368 | 0.8607 | 16.187 | 0.0599 | 0.6822 | 46.581 | 0.0874 | 0.753 | 32.811 | 0.279 | 0.8007 | 24.887 |
| | 30.5 | 1.0717 | 0.2119 | 1.1572 | 0.1404 | 1.0741 | 0.0959 | 0.9951 | 0.1598 | 7 | 0.2942 | 0.8023 | 24.636 | 0.0156 | 0.6286 | 59.081 | 0.0064 | 0.6851 | 45.974 | 0.1855 | 0.7449 | 34.245 |
| | 31.5 | 1.2616 | 0.2958 | 1.2988 | 0.1881 | 1.2646 | 0.138 | 0.9995 | 0.2055 | 8 | 0.1453 | 0.6816 | 46.72 | 0.0111 | 0.5601 | 78.55 | 0.0019 | 0.5818 | 71.866 | 0.2659 | 0.7416 | 34.837 |
| | 32.5 | 1.3892 | 0.3416 | 1.4297 | 0.2167 | 1.5491 | 0.1892 | 1.1762 | 0.2208 | 9 | 0.096 | 0.619 | 61.558 | 0.0063 | 0.5088 | 96.547 | 0.0005 | 0.475 | 110.53 | 0.0838 | 0.6302 | 58.684 |
| | 33.5 | 1.7584 | 0.4548 | 1.5518 | 0.2074 | 1.8643 | 0.0905 | 1.3375 | 0.2737 | 10 | 0.0352 | 0.489 | 104.51 | 0.0013 | 0.4688 | 113.32 | 5E-10 | 0.3947 | 153.36 | 0.0515 | 0.5542 | 80.44 |
| | 34.5 | 1.7535 | 0.3756 | 1.5991 | 0.1828 | 2.1462 | 0.2009 | 1.2801 | 0.2196 | 11 | 0.0136 | 0.4903 | 103.94 | 0.0003 | 0.4549 | 119.82 | 9E-07 | 0.3429 | 191.67 | 0.0345 | 0.579 | 72.7 |
| | 35.5 | 1.5089 | 0.2124 | 1.5948 | 0.1859 | 1.9837 | 0.173 | 1.2026 | 0.1767 | 12 | 0.0034 | 0.5699 | 75.479 | 0.0003 | 0.4561 | 119.24 | 7E-07 | 0.3709 | 169.58 | 0.0312 | 0.6164 | 62.241 |
| | 36.5 | 0.4697 | 0.0847 | 0.4772 | 0.0548 | 0.6583 | 0.0758 | 0.5136 | 0.0751 | 13 | 0.001 | 1.8307 | -45.38 | 0.0212 | 1.5243 | -34.4 | 0.4272 | 1.1177 | -10.53 | 0.0727 | 1.4431 | -30.7 |

AU Exp8

| Average Area under the cuve | Day | RocketControl | | Rocket100 mM NaCl | | Days post stress | RocketControl | | | Rocket100 mM NaCl | | |
|---|---|---|---|---|---|---|---|---|---|---|---|---|
| | | Mean | SEM | Mean | SEM | | T.test P values | Fold Change | % difference | T.test P values | Fold Change | % difference |
| | 18.5 | 0.3526 | 0.0516 | 0.774 | 0.0599 | | | | | | | |
| | 19.5 | 0.5211 | 0.0763 | 0.6716 | 0.087 | | | | | | | |
| | 20.5 | 0.5349 | 0.0723 | 0.665 | 0.06 | | | | | | | |
| | 21.5 | 0.557 | 0.0656 | 0.6222 | 0.0532 | | | | | | | |
| | 22.5 | 0.6181 | 0.072 | 0.5294 | 0.0517 | | | | | | | |
| | 23.5 | 0.6552 | 0.0696 | 0.5936 | 0.0446 | | | | | | | |
| | 24.5 | 0.6817 | 0.073 | 0.3426 | 0.0399 | 1 | 0.795 | 0.9611 | 4.0522 | 0.0004 | 1.7328 | -42.29 |
| | 25.5 | 0.6581 | 0.0591 | 0.2987 | 0.0419 | 2 | 0.9751 | 0.9956 | 0.4397 | 8E-05 | 1.9873 | -49.68 |
| | 26.5 | 0.6983 | 0.0847 | 0.3758 | 0.0471 | 3 | 0.698 | 0.9382 | 6.587 | 0.0028 | 1.5797 | -36.7 |
| | 27.5 | 0.6666 | 0.0714 | 0.4568 | 0.0616 | 4 | 0.91 | 0.9829 | 1.7425 | 0.0857 | 1.2996 | -23.05 |
| | 28.5 | 0.6707 | 0.0547 | 0.5122 | 0.0535 | 5 | 0.8627 | 0.9769 | 2.3676 | 0.255 | 1.1589 | -13.71 |
| | 29.5 | 0.6178 | 0.0934 | 0.5577 | 0.0768 | 6 | 0.7516 | 1.0605 | -5.708 | 0.6897 | 1.0644 | -6.051 |
| | 30.5 | 0.6641 | 0.082 | 0.5175 | 0.0562 | 7 | 0.9349 | 0.9866 | 1.358 | 0.2999 | 1.1472 | -12.83 |
| | 31.5 | 0.7076 | 0.0932 | 0.4873 | 0.079 | 8 | 0.6572 | 0.926 | 7.9971 | 0.2538 | 1.2182 | -17.91 |
| | 32.5 | 0.7686 | 0.1397 | 0.5705 | 0.0745 | 9 | 0.476 | 0.8525 | 17.307 | 0.7926 | 1.0405 | -3.892 |
| | 33.5 | 0.7585 | 0.1689 | 0.4299 | 0.0486 | 10 | 0.578 | 0.8638 | 15.767 | 0.0211 | 1.3808 | -27.58 |
| | 34.5 | 0.807 | 0.105 | 0.4543 | 0.0601 | 11 | 0.2421 | 0.8118 | 23.179 | 0.076 | 1.3068 | -23.48 |
| | 35.5 | 0.7949 | 0.119 | 0.4671 | 0.0975 | 12 | 0.3229 | 0.8242 | 21.325 | 0.2506 | 1.2708 | -21.31 |
| | 36.5 | 0.6066 | 0.0808 | 0.4737 | 0.1137 | 13 | 0.6538 | 1.08 | -7.411 | 0.3366 | 1.2533 | -20.21 |

AU Exp9

| Average Area under the cuve | | AmaranthControl | | Amaranth100 mM NaCl | | Tomato100 mM NaCl | | TomatoControl | | Days post stress | AmaranthControl | | | Amaranth100 mM NaCl | | | Tomato100 mM NaCl | | | TomatoControl | | |
|---|---|---|---|---|---|---|---|---|---|---|---|---|---|---|---|---|---|---|---|---|---|---|
| | | Mean | SEM | Mean | SEM | Mean | SEM | Mean | SEM | | T.test P values | Fold Change | % difference | T.test P values | Fold Change | % difference | T.test P values | Fold Change | % difference | T.test P values | Fold Change | % difference |
| | 18.5 | 0.0872 | 0.0116 | 0.236 | 0.0105 | 0.2078 | 0.0371 | 0.2689 | 0.0384 | | | | | | | | | | | | | |
| | 19.5 | 0.1022 | 0.0179 | 0.2729 | 0.0128 | 0.3711 | 0.071 | 0.3609 | 0.0408 | | | | | | | | | | | | | |
| | 20.5 | 0.1596 | 0.0356 | 0.4326 | 0.0195 | 0.5329 | 0.0759 | 0.5404 | 0.0549 | | | | | | | | | | | | | |
| | 21.5 | 0.1947 | 0.0397 | 0.4294 | 0.0347 | 0.6118 | 0.0706 | 0.4737 | 0.0384 | | | | | | | | | | | | | |
| | 22.5 | 0.2562 | 0.0462 | 0.5416 | 0.0788 | 0.7907 | 0.0771 | 0.7466 | 0.064 | | | | | | | | | | | | | |
| | 23.5 | 0.3327 | 0.0589 | 0.3638 | 0.0376 | 0.7523 | 0.0854 | 0.9768 | 0.0643 | | | | | | | | | | | | | |
| | 24.5 | 0.433 | 0.0829 | 0.3785 | 0.0337 | 0.8945 | 0.0834 | 1.1463 | 0.0671 | 1 | 0.3348 | 0.7684 | 30.133 | 0.7744 | 0.9612 | 4.0335 | 0.2489 | 0.841 | 18.905 | 0.0819 | 0.8522 | 17.35 |
| | 25.5 | 0.6008 | 0.0947 | 0.5177 | 0.0733 | 0.9601 | 0.1161 | 1.3364 | 0.0702 | 2 | 0.0251 | 0.5538 | 80.557 | 0.0783 | 0.7028 | 42.283 | 0.1663 | 0.7835 | 27.632 | 0.001 | 0.7309 | 36.813 |
| | 26.5 | 0.7002 | 0.1139 | 0.488 | 0.0694 | 1.0566 | 0.106 | 1.4304 | 0.0773 | 3 | 0.009 | 0.4752 | 110.42 | 0.1327 | 0.7455 | 34.143 | 0.0383 | 0.712 | 40.455 | 0.0002 | 0.6829 | 46.433 |
| | 27.5 | 0.7642 | 0.0973 | 0.5337 | 0.0944 | 1.1506 | 0.1275 | 1.6648 | 0.0801 | 4 | 0.001 | 0.4354 | 129.66 | 0.1117 | 0.6816 | 46.705 | 0.0183 | 0.6538 | 52.95 | 1E-06 | 0.5868 | 70.429 |
| | 28.5 | 0.9338 | 0.0809 | 0.6104 | 0.0952 | 1.2447 | 0.1381 | 1.7802 | 0.0918 | 5 | 5E-06 | 0.3563 | 180.62 | 0.0269 | 0.596 | 67.778 | 0.0072 | 0.6044 | 65.464 | 3E-07 | 0.5487 | 82.249 |
| | 29.5 | 0.9849 | 0.0862 | 0.6067 | 0.1054 | 1.31 | 0.1829 | 1.8594 | 0.085 | 6 | 3E-06 | 0.3379 | 195.98 | 0.0436 | 0.5997 | 66.758 | 0.0128 | 0.5742 | 74.144 | 3E-08 | 0.5253 | 90.359 |
| | 30.5 | 1.0578 | 0.0928 | 0.6864 | 0.124 | 1.5041 | 0.2284 | 2.0361 | 0.1142 | 7 | 1E-06 | 0.3146 | 217.89 | 0.0228 | 0.5301 | 88.656 | 0.0064 | 0.5002 | 99.936 | 5E-08 | 0.4798 | 108.44 |
| | 31.5 | 1.0924 | 0.1046 | 0.7105 | 0.1468 | 1.563 | 0.236 | 2.2761 | 0.1069 | 8 | 2E-06 | 0.3046 | 228.29 | 0.0345 | 0.5121 | 95.283 | 0.0046 | 0.4813 | 107.77 | 6E-10 | 0.4292 | 133.01 |
| | 32.5 | 1.2386 | 0.1301 | 0.7751 | 0.1748 | 1.7064 | 0.2952 | 2.8359 | 0.1766 | 9 | 2E-06 | 0.2686 | 272.25 | 0.0336 | 0.4694 | 113.05 | 0.0061 | 0.4409 | 126.83 | 1E-09 | 0.3444 | 190.32 |
| | 33.5 | 1.4831 | 0.1551 | 0.8098 | 0.1999 | 1.615 | 0.2285 | 3.0537 | 0.2725 | 10 | 6E-07 | 0.2244 | 345.72 | 0.0417 | 0.4493 | 122.58 | 0.0024 | 0.4658 | 114.68 | 2E-07 | 0.3199 | 212.62 |
| | 34.5 | 1.5493 | 0.1512 | 0.761 | 0.2006 | 1.537 | 0.1326 | 3.7264 | 0.3017 | 11 | 2E-07 | 0.2148 | 365.6 | 0.0674 | 0.4781 | 109.18 | 1E-04 | 0.4894 | 104.31 | 9E-09 | 0.2621 | 281.48 |
| | 35.5 | 1.689 | 0.1389 | 0.8605 | 0.246 | 1.7252 | 0.1668 | 4.1627 | 0.3221 | 12 | 8E-09 | 0.197 | 407.61 | 0.0613 | 0.4228 | 136.51 | 6E-05 | 0.436 | 129.33 | 2E-09 | 0.2347 | 326.15 |
| | 36.5 | 1.6344 | 0.1584 | 0.6612 | 0.1703 | 1.5523 | 0.1598 | 3.569 | 0.2192 | 13 | 1E-07 | 0.2036 | 391.19 | 0.1054 | 0.5503 | 81.727 | 0.0003 | 0.4846 | 106.35 | 1E-10 | 0.2737 | 265.38 |

**Fold change = (Average area under the curve before stress Day 23) / (Average area under the curve after stress)**
