## Supplemental Table S5 for "Leaf movements as a quantitative metric for early stress detection"

**Supplementary Table S5.** The end point root and shoot fresh weights, dry weights and chlorophyll content data of **(a)** different stress treatments in lettuce, **(b)** salt stress applied at different locations on lettuce, **(c)** within location replication of different stresses and **(d)** 100 mM NaCl treatment on different CEA crops, with their respective non-stressed plants.

**S5a Endpoint physiological data for different stress treatments on lettuce, grouped with their respecive controls.**

| Treatment | FW roots (g) | FW shoots (g) | DW roots (g) | DW shoots (g) | Ave. chlorophyll content (SPAD units) |
|---|---|---|---|---|---|
| Control | 3.1 | 24 | 0.1 | 1.2 | |
| Control | 2.9 | 19 | 0.1 | 1 | 17 |
| Control | 3 | 20 | 0.1 | 1 | 18 |
| Control | 1.8 | 16 | 0.1 | 0.8 | |
| Control | 3.7 | 27 | 0.1 | 1.2 | |
| Control | 2.3 | 20 | 0.1 | 1 | |
| Control | 4.2 | 27 | 0.2 | 1.3 | 11 |
| Control | 5.2 | 31 | 0.2 | 1.6 | 17 |
| Control | 1.9 | 16 | 0.1 | 0.8 | |
| Control | 3.5 | 24 | 0.1 | 1.1 | 20 |
| Control | 2.5 | 21 | 0.1 | 0.9 | 15 |
| Control | 1 | 10 | 0 | 0.5 | |
| Boron | 3.6 | 18 | 0.2 | 0.9 | 15 |
| Boron | 1.5 | 17 | 0.1 | 0.8 | |
| Boron | 3.1 | 23 | 0.1 | 1.2 | |
| Boron | 1.2 | 11 | 0 | 0.6 | 14 |
| Boron | 2.3 | 18 | 0.1 | 0.9 | |
| Boron | 2.6 | 23 | 0.1 | 1.1 | |
| Boron | 1.4 | 14 | 0 | 0.7 | 17 |
| Boron | 2.9 | 22 | 0.1 | 1.1 | 16 |
| Boron | 1.1 | 12 | 0.1 | 0.6 | |
| Boron | 2.2 | 18 | 0.1 | 0.8 | |
| Boron | 2.8 | 19 | 0.1 | 1 | 15 |
| Boron | 2.1 | 20 | 0.1 | 0.9 | 15 |
| 100mM KCl | 5.3 | 13 | 0.2 | 0.8 | |
| 100mM KCl | 1.7 | 7.5 | 0.1 | 0.5 | |
| 100mM KCl | 3.5 | 10 | 0.2 | 0.7 | 24 |
| 100mM KCl | 1.7 | 7.2 | 0.1 | 0.4 | |
| 100mM KCl | 2.7 | 8.4 | 0.1 | 0.6 | 22 |
| 100mM KCl | 2.8 | 7.3 | 0.1 | 0.5 | |
| 100mM KCl | 1.6 | 8.5 | 0.1 | 0.7 | 28 |
| 100mM KCl | 3 | 8.5 | 0.1 | 0.6 | 15 |

| Treatment | FW roots (g) | FW shoots (g) | DW roots (g) | DW shoots (g) | Ave. chlorophyll content (SPAD units) |
|---|---|---|---|---|---|
| Hypoxia | 1.837 | 14.752 | 0.104 | 0.731 | |
| Hypoxia | 1.445 | 10.916 | 0.074 | 0.559 | 13.2 |
| Hypoxia | 2..428 | 16.547 | 0.126 | 0.85 | 11.3 |
| Hypoxia | 2.568 | 20.663 | 0.138 | 1.015 | |
| Hypoxia | 1.513 | 10.541 | 0.078 | 0.525 | 14.4 |
| Hypoxia | 3.166 | 23.646 | 0.157 | 1.188 | |
| Hypoxia | 1.537 | 10.653 | 0 | 0.533 | 15.4 |
| Hypoxia | 2.029 | 13.246 | 0.103 | 0.696 | 14.06 |
| Hypoxia | 1.900 | 12.692 | 0 | 0.6 | |
| Hypoxia | 2.246 | 14.735 | 0 | 0.803 | |
| Hypoxia | 2.754 | 18.680 | 0 | 0.954 | |
| Hypoxia | 2.719 | 22.101 | 0.154 | 1.243 | 14.7 |
| Control | 4.126 | 19.053 | 0.215 | 1.13 | 20 |
| Control | 3.063 | 21.184 | 0.161 | 1.044 | 14.3 |
| Control | 2.386 | 14.645 | 0 | 0.746 | 14.8 |
| Control | 3.153 | 16.972 | 0 | 0.871 | |
| Control | 2.627 | 15.687 | 0.118 | 0.753 | 16 |
| Control | 2.589 | 14.238 | 0 | 0.729 | |
| Control | 1.724 | 11.452 | 0.132 | 0.636 | 16.3 |
| Control | 3.879 | 21.859 | 0.217 | 1.104 | |
| Control | 2.256 | 13.863 | 0 | 0.718 | |
| Control | 1.837 | 13.859 | 0.12 | 0.692 | |
| Control | 3.235 | 14.682 | 0.175 | 0.834 | 17 |
| Control | 3.374 | 21.399 | 0.192 | 1.141 | |
| 150mM NaCl | | dead | 0 | 0.147 | |
| 150mM NaCl | 0.816 | 1.736 | 0.05 | 0.256 | 24.1 |
| 150mM NaCl | 0.746 | 2.734 | 0.052 | 0.32 | 20.8 |
| 150mM NaCl | 1.324 | 3.885 | 0 | 0.475 | 22.2 |
| 150mM NaCl | 1.911 | 6.093 | 0.13 | 0.296 | |
| 150mM NaCl | 1.296 | 3.632 | 0 | 0.4 | |
| 150mM NaCl | 1.426 | 4.952 | 0.091 | 0.305 | 24.7 |
| 150mM NaCl | 1.234 | 3.894 | 0 | 0.254 | 21.3 |

| Treatment | FW roots (g) | FW shoots (g) | DW roots (g) | DW shoots (g) | Ave. chlorophyll content (SPAD units) |
|---|---|---|---|---|---|
| 100 mM NaCl | 1.1 | 3.5 | 0.1 | 0.3 | 20 |
| 100 mM NaCl | 1 | 3.1 | 0 | 0.2 | 19 |
| 100 mM NaCl | 1.1 | 3.4 | 0.1 | 0.3 | 23 |
| 100 mM NaCl | 1.6 | 4.8 | 0.1 | 0.3 | 21 |
| 100 mM NaCl | 0.7 | 2.7 | 0 | 0.2 | 22 |
| 100 mM NaCl | 1.2 | 4 | 0 | 0.2 | 22 |
| 100 mM NaCl | 1.1 | 4.2 | 0.1 | 0.3 | 24 |
| 100 mM NaCl | 1.5 | 4.2 | 0.1 | 0.3 | 25 |
| 100 mM NaCl | 1 | 3 | 0 | 0.2 | 21 |
| 100 mM NaCl | 0.7 | 2.2 | 0 | 0.2 | 22 |
| 100 mM NaCl | 1.1 | 3.9 | 0 | 0.3 | 26 |
| 100 mM NaCl | 1.7 | 5.3 | 0.1 | 0.4 | 23 |
| Control | 0.4 | 1.9 | 0 | 0.1 | 13 |
| Control | 2.2 | 11 | 0.1 | 0.6 | 15 |
| Control | 1.7 | 8.3 | 0.1 | 0.5 | 15 |
| Control | 1.7 | 9.6 | 0.1 | 0.5 | 15 |
| Control | 3.3 | 15 | 0.1 | 0.9 | 18 |
| Control | 0.7 | 3.8 | 0 | 0.2 | 15 |
| Control | 1 | 5.2 | 0 | 0.3 | 15 |
| Control | 1.7 | 9.1 | 0.1 | 0.5 | 13 |
| Control | 1.5 | 7.5 | 0.1 | 0.4 | 18 |
| Control | 1.4 | 6 | 0.1 | 0.3 | 18 |
| Control | 0.7 | 4.6 | 0 | 0.3 | 15 |
| Control | 2.3 | 11 | 0.1 | 0.5 | 15 |

| | FW roots (g) | FW shoots (g) | DW roots (g) | DW shoots (g) | Ave. chlorophyll content (SPAD units) |
|---|---|---|---|---|---|
| Control | 1.2 | 12 | 0.1 | 0.5 | 114 |
| Control | 1.9 | 15 | 0.1 | 0.8 | 133 |
| Control | 1.8 | 16 | 0.1 | 0.8 | 78 |
| Control | 1 | 9.1 | 0 | 0.4 | 189 |
| Control | 2 | 16 | 0.1 | 0.8 | 156 |
| Control | 1.9 | 15 | 0.1 | 0.7 | 118 |
| Control | 1.6 | 12 | 0.1 | 0.5 | 134 |
| Control | 1.2 | 10 | 0 | 0.4 | 190 |
| Control | 1.6 | 15 | 0.7 | 0.8 | 157 |
| Control | 1.2 | 11 | 0.5 | 0.5 | 130 |
| Control | 1.5 | 10 | 0.1 | 0.5 | 101 |
| Control | 1.5 | 12 | 0.1 | 0.6 | 151 |
| Water withdrawal | 0.9 | 8.2 | 0.1 | 0.3 | 205 |
| Water withdrawal | 0.7 | 5.3 | 0 | 0.2 | 157 |
| Water withdrawal | 1.5 | 10 | 0.1 | 0.6 | 169 |
| Water withdrawal | 1 | 7.8 | 0.1 | 0.4 | 194 |
| Water withdrawal | 1.2 | 9.3 | 0.1 | 0.5 | 178 |
| Water withdrawal | 1.3 | 11 | 0.1 | 0.5 | 159 |
| Water withdrawal | 1 | 10 | 0.1 | 0.5 | 179 |
| Water withdrawal | 1.5 | 9.9 | 0.1 | 0.6 | 213 |
| Water withdrawal | 1.7 | 14 | 0.1 | 0.7 | 189 |
| Water withdrawal | 0.9 | 7.9 | 0.1 | 0.6 | 162 |
| Water withdrawal | 1.1 | 8 | 0.1 | 0.4 | 149 |
| Water withdrawal | 0.9 | 6.9 | 0 | 0.3 | 189 |

| | | | | | |
|---|---|---|---|---|---|
| 100mM KCl | 2.2 | 9.8 | 0.1 | 0.6 | |
| 100mM KCl | 2.8 | 7.8 | 0.1 | 0.4 | 21 |
| 100mM KCl | 1.5 | 4.5 | 0.1 | 0.3 | |
| 100mM KCl | 1.1 | 3.7 | 0.1 | 0.2 | 18 |

| | | | | | |
|---|---|---|---|---|---|
| 150mM NaCl | 1.074 | 3.320 | 0.07 | 0.354 | |
| 150mM NaCl | 0.836 | 3.737 | 0.06 | 0.55 | 24.5 |
| 150mM NaCl | 1.840 | 6.645 | 0 | 0.334 | |
| 150mM NaCl | 1.204 | 3.862 | 0.085 | 0 | |
| NO nutrients | 0.873 | 1.173 | 0 | 0.112 | |
| NO nutrients | 1.272 | 2.808 | 0.119 | 0.249 | 11.7 |
| NO nutrients | 1.672 | 1.932 | 0.133 | 0.215 | |
| NO nutrients | 1.601 | 2.402 | 0.112 | 0.227 | 13 |
| NO nutrients | 1.56 | 2.406 | 0 | 0.215 | 12.3 |
| NO nutrients | 1.615 | 2.186 | 0.15 | 0.216 | |
| NO nutrients | 1.702 | 2.845 | 0.168 | 0.299 | 12.5 |
| NO nutrients | 1.360 | 1.957 | 0.112 | 0.182 | 12.4 |
| NO nutrients | 1.002 | 1.973 | 0 | 0.152 | |
| NO nutrients | 0.795 | 1.018 | 0.074 | 0.107 | |
| NO nutrients | 1.757 | 2.959 | 0.153 | 0.228 | 11.4 |
| NO nutrients | 1.849 | 3.127 | 0 | 0.295 | |

**S5b Endpoint physiological data for Control and NaCl treatments conducted in different locations.**

| UoC | | | | | |
|---|---|---|---|---|---|
| **Treatment** | **FW roots (g)** | **FW shoots (g)** | **DW roots (g)** | **DW shoots (g)** | **Ave. chlorophyll content (units)** |
| 100mM NaCl | 1.1 | 5.4 | 0.1 | 0.4 | 257 |
| 100mM NaCl | 0.9 | 5.4 | 0.1 | 0.4 | 282 |
| 100mM NaCl | 0.9 | 7.1 | 0.1 | 0.5 | 366 |
| 100mM NaCl | 1.5 | 5.8 | 0.1 | 0.4 | 316 |
| 100mM NaCl | 1.8 | 9.8 | 0.1 | 0.7 | 313 |
| 100mM NaCl | 1.1 | 5.8 | 0.1 | 0.4 | 192 |
| 100mM NaCl | 1.5 | 9.5 | 0.1 | 0.6 | 160 |
| 100mM NaCl | 0.9 | 4.1 | 0 | 0.2 | 236 |
| 100mM NaCl | 1.3 | 5.9 | 0.1 | 0.4 | 290 |
| 100mM NaCl | 1.2 | 8.9 | 0.1 | 0.6 | 236 |
| 100mM NaCl | 0.7 | 4.2 | 0 | 0.3 | 203 |
| 100mM NaCl | 1.3 | 8.4 | 0.1 | 0.6 | 301 |
| Control | 1.3 | 9.6 | 0.1 | 0.5 | 278 |
| Control | 0.7 | 5.1 | 0 | 0.3 | 319 |
| Control | 2 | 20 | 0.1 | 1 | 278 |
| Control | 3.2 | 25 | 0.1 | 1.3 | 300 |
| Control | 2 | 16 | 0.1 | 0.8 | 375 |
| Control | 3.9 | 23 | 0.2 | 1.1 | 319 |
| Control | 1.9 | 18 | 0.1 | 0.9 | 344 |
| Control | 2.3 | 17 | 0.1 | 0.9 | 265 |

| AU | | | | | |
|---|---|---|---|---|---|
| **Treatment** | **FW roots (g)** | **FW shoots (g)** | **DW roots (g)** | **DW shoots (g)** | **Ave. chlorophyll content (SPAD units)** |
| 100 mM NaCl | 1.1 | 3.5 | 0 | 0 | 20 |
| 100 mM NaCl | 1 | 3.1 | 0 | 0 | 19 |
| 100 mM NaCl | 1.1 | 3.4 | 0 | 0 | 23 |
| 100 mM NaCl | 1.6 | 4.8 | 0 | 0 | 21 |
| 100 mM NaCl | 0.7 | 2.7 | 0 | 0 | 22 |
| 100 mM NaCl | 1.2 | 4 | 0 | 0 | 22 |
| 100 mM NaCl | 1.1 | 4.2 | 0 | 0 | 24 |
| 100 mM NaCl | 1.5 | 4.2 | 0 | 0 | 25 |
| 100 mM NaCl | 1 | 3 | 0 | 0 | 21 |
| 100 mM NaCl | 0.7 | 2.2 | 0 | 0 | 22 |
| 100 mM NaCl | 1.1 | 3.9 | 0 | 0 | 26 |
| 100 mM NaCl | 1.7 | 5.3 | 0 | 0 | 23 |
| Control | 0.4 | 1.9 | 0 | 0 | 13 |
| Control | 2.2 | 11 | 0 | 1 | 15 |
| Control | 1.7 | 8.3 | 0 | 0 | 15 |
| Control | 1.7 | 9.6 | 0 | 0 | 15 |
| Control | 3.3 | 15 | 0 | 1 | 18 |
| Control | 0.7 | 3.8 | 0 | 0 | 15 |
| Control | 1 | 5.2 | 0 | 0 | 15 |
| Control | 1.7 | 9.1 | 0 | 0 | 13 |

| UWA | | | | | |
|---|---|---|---|---|---|
| **Treatment** | **FW roots (g)** | **FW shoots (g)** | **DW roots (g)** | **DW shoots (g)** | **Ave. chlorophyll content (SPAD units)** |
| 150mM NaCl | | dead | 0 | 0.15 | |
| 150mM NaCl | 0.82 | 1.736 | 0.05 | 0.26 | 24.1 |
| 150mM NaCl | 0.75 | 2.734 | 0.05 | 0.32 | 20.8 |
| 150mM NaCl | 1.32 | 3.885 | 0 | 0.48 | 22.2 |
| 150mM NaCl | 1.91 | 6.093 | 0.13 | 0.3 | |
| 150mM NaCl | 1.3 | 3.632 | 0 | 0.4 | |
| 150mM NaCl | 1.43 | 4.952 | 0.09 | 0.31 | 24.7 |
| 150mM NaCl | 1.234 | 3.894 | 0 | 0.25 | 21.3 |
| 150mM NaCl | 1.074 | 3.320 | 0.07 | 0.35 | |
| 150mM NaCl | 0.836 | 3.737 | 0.06 | 0.55 | 24.5 |
| 150mM NaCl | 1.840 | 6.645 | 0 | 0.33 | |
| 150mM NaCl | 1.204 | 3.862 | 0.09 | 0 | |
| Control | 4.13 | 19.053 | 0.22 | 1.13 | 20 |
| Control | 3.06 | 21.184 | 0.16 | 1.04 | 14.3 |
| Control | 2.39 | 14.645 | 0 | 0.75 | 14.8 |
| Control | 3.15 | 16.972 | 0 | 0.87 | |
| Control | 2.63 | 15.687 | 0.12 | 0.75 | 16 |
| Control | 2.59 | 14.238 | 0 | 0.73 | |
| Control | 1.72 | 11.452 | 0.13 | 0.64 | 16.3 |
| Control | 3.879 | 21.859 | 0.22 | 1.1 | |

| | | | | | |
|---|---|---|---|---|---|
| Control | 2.4 | 19 | 0.1 | 1 | 339 |
| Control | 1.7 | 13 | 0.1 | 0.6 | 305 |
| Control | 2.8 | 17 | 0.1 | 0.8 | 306 |
| Control | 2.1 | 17 | 0.1 | 0.9 | 259 |

| | | | | | |
|---|---|---|---|---|---|
| Control | 1.5 | 7.5 | 0 | 0 | 18 |
| Control | 1.4 | 6 | 0 | 0 | 18 |
| Control | 0.7 | 4.6 | 0 | 0 | 15 |
| Control | 2.3 | 11 | 0 | 1 | 15 |

| | | | | | |
|---|---|---|---|---|---|
| Control | 2.256 | 13.863 | 0 | 0.72 | |
| Control | 1.837 | 13.859 | 0.12 | 0.69 | |
| Control | 3.235 | 14.682 | 0.18 | 0.83 | 17 |
| Control | 3.374 | 21.399 | 0.19 | 1.14 | |

**S5c Endpoint physiological data for replicates of different stress treatments on lettuce conducted at the same location.**

| Experiment | Treatment | FW roots (g) | FW shoots (g) | DW roots (g) | DW shoots (g) | Ave. chlorophyll content (SPAD units) | | Experiment | Treatment | FW roots (g) | FW shoots (g) | DW roots (g) | DW shoots (g) | Ave. chlorophyll content (SPAD units) | Ave. stomatal cond. (mol m-2 s-1) |
|---|---|---|---|---|---|---|---|---|---|---|---|---|---|---|---|
| UWA Exp4 | Hypoxia | 2.907 | 14.432 | 0.1225 | 0.7303 | | | AU Exp5 | 100mM NaCl1 | 1.3408 | 5.1593 | 0.064 | 0.343 | 23.375 | 0.106 |
| UWA Exp4 | Hypoxia | 2.068 | 10.752 | 0 | 0.6309 | | | AU Exp5 | 100mM NaCl1 | 0.055 | 0.2183 | 0.002 | 0.023 | | |
| UWA Exp4 | Hypoxia | 1.959 | 9.825 | 0.1007 | 0.5205 | | | AU Exp5 | 100mM NaCl1 | 0.8397 | 3.0335 | 0.036 | 0.21 | 21.85 | 0.118 |
| UWA Exp4 | Hypoxia | 3.034 | 17.709 | 0 | 0.9301 | 17.175 | | AU Exp5 | 100mM NaCl1 | 2.4063 | 8.4155 | 0.082 | 0.528 | 23.225 | 0.128 |
| UWA Exp4 | Hypoxia | 1.701 | 9.608 | 0.1187 | 0.5504 | 17.725 | | AU Exp5 | 100mM NaCl1 | 1.3944 | 5.3014 | 0.065 | 0.37 | | |
| UWA Exp4 | Hypoxia | 2.68 | 15.793 | 0 | 0.9206 | | | AU Exp5 | 100mM NaCl1 | 1.5469 | 5.5789 | 0.07 | 0.382 | | |
| UWA Exp4 | Hypoxia | 1.48 | 8.942 | 0.0803 | 0.5536 | | | AU Exp5 | 100mM NaCl1 | 1.7263 | 6.7309 | 0.08 | 0.47 | 25.7 | 0.130 |
| UWA Exp4 | Hypoxia | 5.134 | 28.187 | 0.3002 | 1.5669 | 12.175 | | AU Exp5 | 100mM NaCl1 | 1.2009 | 5.3 | 0.06 | 0.353 | 22.55 | 0.097 |
| UWA Exp4 | Hypoxia | 1.373 | 8.592 | 0 | 0.5109 | 13.55 | | AU Exp5 | 100mM NaCl1 | 0.8846 | 3.1964 | 0.041 | 0.209 | | |
| UWA Exp4 | Hypoxia | 2.258 | 13.244 | 0.1189 | 0.7801 | | | AU Exp5 | 100mM NaCl1 | 1.321 | 4.3577 | 0.06 | 0.298 | | |
| UWA Exp4 | Hypoxia | 3.906 | 22.963 | 0 | 1.2121 | | | AU Exp5 | 100mM NaCl1 | 1.1322 | 4.0986 | 0.051 | 0.257 | | |
| UWA Exp4 | Hypoxia | 2.605 | 14.640 | 0 | 0.8408 | 13.25 | | AU Exp5 | 100mM NaCl1 | 1.6186 | 5.8969 | 0.066 | 0.394 | 22.75 | 0.099 |
| UWA Exp4 | Control | 4.12 | 23.877 | 0 | 1.2455 | 14.975 | | AU Exp5 | Control1 | 1.8481 | 7.7423 | 0.075 | 0.358 | 12.375 | 1.025 |
| UWA Exp4 | Control | 2.589 | 17.642 | 0.1289 | 0.9108 | | | AU Exp5 | Control1 | 1.3166 | 8.5844 | 0.05 | 0.354 | | |
| UWA Exp4 | Control | 3.269 | 17.879 | 0 | 0.9209 | | | AU Exp5 | Control1 | 2.2939 | 10.617 | 0.08 | 0.605 | 13.85 | 0.930 |
| UWA Exp4 | Control | 3.802 | 14.673 | 0.1435 | 0.8106 | 15.275 | | AU Exp5 | Control1 | 0.3978 | 2.0063 | 0.018 | 0.093 | | |
| UWA Exp4 | Control | 2.374 | 17.772 | 0 | 0.9485 | 16.975 | | AU Exp5 | Control1 | 0.881 | 5.4236 | 0.034 | 0.263 | 16.725 | 0.845 |
| UWA Exp4 | Control | 3.374 | 20.143 | 0.1982 | 1.2884 | 19.7 | | AU Exp5 | Control1 | 0.4689 | 1.985 | 0.019 | 0.115 | | |
| UWA Exp4 | Control | 2.618 | 15.993 | 0 | 0.8382 | 14.025 | | AU Exp5 | Control1 | 1.3024 | 6.5707 | 0.054 | 0.373 | 13.125 | 0.514 |
| UWA Exp4 | Control | 2.108 | 14.389 | 0.1112 | 0.6763 | 22.1 | | AU Exp5 | Control1 | 2.8768 | 13.267 | 0.113 | 0.806 | 16.2 | 0.579 |
| UWA Exp4 | Control | 2.902 | 19.584 | 0.1705 | 1.0944 | | | AU Exp5 | Control1 | 1.8277 | 8.7322 | 0.065 | 0.476 | | |
| UWA Exp4 | Control | 2.330 | 15.662 | 0 | 0.7473 | | | AU Exp5 | Control1 | 1.2961 | 7.9063 | 0.046 | 0.431 | | |
| UWA Exp4 | Control | 5.893 | 32.524 | 0.3304 | 1.8341 | | | AU Exp5 | Control1 | 2.2268 | 12.499 | 0.083 | 0.644 | | |
| UWA Exp4 | Control | 3.523 | 26.073 | 0 | 1.4042 | | | AU Exp5 | Control1 | 1.272 | 7.8259 | 0.043 | 0.371 | 14.05 | 0.354 |
| UWA Exp4 | NO nutrients | 1.571 | 2.155 | 0.1499 | 0.2721 | | | AU Exp5 | Control2 | 0.7797 | 4.962 | 0.029 | 0.299 | 12.825 | 0.363 |
| UWA Exp4 | NO nutrients | 1.909 | 2.956 | 0.1877 | 0.3266 | | | AU Exp5 | Control2 | 2.2298 | 13.007 | 0.062 | 0.702 | | |
| UWA Exp4 | NO nutrients | 1.202 | 1.506 | 0 | 0.2197 | | | AU Exp5 | Control2 | 2.739 | 11.379 | 0.071 | 0.654 | 16.775 | |
| UWA Exp4 | NO nutrients | 2.081 | 2.906 | 0 | 0.3898 | | | AU Exp5 | Control2 | 1.2103 | 7.2493 | 0.033 | 0.362 | | |
| UWA Exp4 | NO nutrients | 0.903 | 1.5 | 0 | 0.191 | | | AU Exp5 | Control2 | 0.7589 | 4.093 | 0.023 | 0.189 | 8.375 | 0.156 |
| UWA Exp4 | NO nutrients | 1.001 | 1.603 | 0.1003 | 0.1979 | 11.75 | | AU Exp5 | Control2 | 1.4721 | 9.3942 | 0.048 | 0.541 | | 0.233 |
| UWA Exp4 | NO nutrients | 1.344 | 1.786 | 0 | 0.2301 | 12.375 | | AU Exp5 | Control2 | 1.6947 | 13.179 | 0.057 | 0.736 | 15.45 | |
| UWA Exp4 | NO nutrients | 1.323 | 1.863 | 0.1297 | 0.2028 | 13.975 | | AU Exp5 | Control2 | 2.6091 | 13.689 | 0.074 | 0.737 | 16.8 | |
| UWA Exp4 | NO nutrients | 2.644 | 3.492 | 0.2763 | 0.4379 | | | AU Exp5 | Control2 | 0.2492 | 1.1939 | 0.008 | 0.072 | | |
| UWA Exp4 | NO nutrients | 0.926 | 1.532 | 0 | 0.2845 | 14.1 | | AU Exp5 | Control2 | 2.4867 | 9.6834 | 0.07 | 0.528 | | 0.173 |
| UWA Exp4 | NO nutrients | 1.291 | 1.928 | 0.1275 | 0.2473 | 13.25 | | AU Exp5 | Control2 | 1.3189 | 6.7054 | 0.043 | 0.355 | | 0.336 |
| UWA Exp4 | NO nutrients | 0.822 | 1.148 | 0 | 0.1801 | 12.7 | | AU Exp5 | Control2 | 2.1985 | 12.367 | 0.065 | 0.664 | 17.2 | 0.212 |
| UWA Exp6 | Hypoxia | 1.837 | 14.752 | 0.104 | 0.731 | | | AU Exp5 | 100mM NaCl | 0.8495 | 3.7372 | 0.037 | 0.251 | | |
| UWA Exp6 | Hypoxia | 1.445 | 10.916 | 0.074 | 0.559 | 13.2 | | AU Exp5 | 100mM NaCl | 1.0222 | 5.036 | 0.038 | 0.332 | 24.25 | 0.190 |
| UWA Exp6 | Hypoxia | 2..428 | 16.547 | 0.126 | 0.85 | 11.3 | | AU Exp5 | 100mM NaCl | 0.9664 | 5.0261 | 0.042 | 0.349 | 24.6 | 0.234 |
| UWA Exp6 | Hypoxia | 2.568 | 20.663 | 0.138 | 1.015 | | | AU Exp5 | 100mM NaCl | 1.0248 | 3.868 | 0.045 | 0.254 | | 0.139 |
| UWA Exp6 | Hypoxia | 1.513 | 10.541 | 0.078 | 0.525 | 14.4 | | AU Exp5 | 100mM NaCl | 1.5479 | 5.3488 | 0.059 | 0.345 | | |
| UWA Exp6 | Hypoxia | 3.166 | 23.646 | 0.157 | 1.188 | | | AU Exp5 | 100mM NaCl | 0.7225 | 3.0824 | 0.031 | 0.212 | 25.125 | |
| UWA Exp6 | Hypoxia | 1.537 | 10.653 | 0 | 0.533 | 15.4 | | AU Exp5 | 100mM NaCl | 1.857 | 6.531 | 0.079 | 0.446 | 24.425 | 0.253 |
| UWA Exp6 | Hypoxia | 2.029 | 13.246 | 0.103 | 0.696 | 14.06 | | AU Exp5 | 100mM NaCl | 0.717 | 2.6778 | 0.032 | 0.167 | | |
| UWA Exp6 | Hypoxia | 1.900 | 12.692 | 0 | 0.6 | | | AU Exp5 | 100mM NaCl | 0.6957 | 3.6367 | 0.037 | 0.241 | | |
| UWA Exp6 | Hypoxia | 2.246 | 14.735 | 0 | 0.803 | | | AU Exp5 | 100mM NaCl | 1.5656 | 4.5846 | 0.064 | 0.318 | 25.5 | 0.013 |
| UWA Exp6 | Hypoxia | 2.754 | 18.680 | 0 | 0.954 | | | AU Exp5 | 100mM NaCl | 0.3327 | 1.9375 | 0.016 | 0.146 | 18.6 | 0.015 |
| UWA Exp6 | Hypoxia | 2.719 | 22.101 | 0.154 | 1.243 | 14.7 | | AU Exp5 | 100mM NaCl | 1.5709 | 6.034 | 0.069 | 0.363 | | |
| UWA Exp6 | Control | 4.126 | 19.053 | 0.215 | 1.13 | 20 | | AU Exp6 | Control1 | 3.1488 | 23.945 | 0.13 | 1.178 | | |
| UWA Exp6 | Control | 3.063 | 21.184 | 0.161 | 1.044 | 14.3 | | AU Exp6 | Control1 | 2.8794 | 18.944 | 0.106 | 0.976 | 17 | 0.3103 |

| | | | | | | |
|---|---|---|---|---|---|---|
| UWA Exp6 | Control | 2.386 | 14.645 | 0 | 0.746 | 14.8 |
| UWA Exp6 | Control | 3.153 | 16.972 | 0 | 0.871 | |
| UWA Exp6 | Control | 2.627 | 15.687 | 0.118 | 0.753 | 16 |
| UWA Exp6 | Control | 2.589 | 14.238 | 0 | 0.729 | |
| UWA Exp6 | Control | 1.724 | 11.452 | 0.132 | 0.636 | 16.3 |
| UWA Exp6 | Control | 3.879 | 21.859 | 0.217 | 1.104 | |
| UWA Exp6 | Control | 2.256 | 13.863 | 0 | 0.718 | |
| UWA Exp6 | Control | 1.837 | 13.859 | 0.12 | 0.692 | |
| UWA Exp6 | Control | 3.235 | 14.682 | 0.175 | 0.834 | 17 |
| UWA Exp6 | Control | 3.374 | 21.399 | 0.192 | 1.141 | |
| UWA Exp6 | NO nutrients | 0.873 | 1.173 | 0 | 0.112 | |
| UWA Exp6 | NO nutrients | 1.272 | 2.808 | 0.119 | 0.249 | 11.7 |
| UWA Exp6 | NO nutrients | 1.672 | 1.932 | 0.133 | 0.215 | |
| UWA Exp6 | NO nutrients | 1.601 | 2.402 | 0.112 | 0.227 | 13 |
| UWA Exp6 | NO nutrients | 1.56 | 2.406 | 0 | 0.215 | 12.3 |
| UWA Exp6 | NO nutrients | 1.615 | 2.186 | 0.15 | 0.216 | |
| UWA Exp6 | NO nutrients | 1.702 | 2.845 | 0.168 | 0.299 | 12.5 |
| UWA Exp6 | NO nutrients | 1.360 | 1.957 | 0.112 | 0.182 | 12.4 |
| UWA Exp6 | NO nutrients | 1.002 | 1.973 | 0 | 0.152 | |
| UWA Exp6 | NO nutrients | 0.795 | 1.018 | 0.074 | 0.107 | |
| UWA Exp6 | NO nutrients | 1.757 | 2.959 | 0.153 | 0.228 | 11.4 |
| UWA Exp6 | NO nutrients | 1.849 | 3.127 | 0 | 0.295 | |

| | | | | | | | |
|---|---|---|---|---|---|---|---|
| AU Exp6 | Control1 | 2.9921 | 19.729 | 0.117 | 1.01 | 18.175 | 0.1487 |
| AU Exp6 | Control1 | 1.8352 | 15.977 | 0.06 | 0.788 | | |
| AU Exp6 | Control1 | 3.6697 | 26.526 | 0.129 | 1.234 | | |
| AU Exp6 | Control1 | 2.2818 | 20.214 | 0.07 | 0.958 | | |
| AU Exp6 | Control1 | 4.179 | 27.403 | 0.156 | 1.277 | 11.05 | 0.223 |
| AU Exp6 | Control1 | 5.1552 | 31.315 | 0.189 | 1.621 | 16.925 | 0.2652 |
| AU Exp6 | Control1 | 1.8969 | 15.621 | 0.078 | 0.795 | | |
| AU Exp6 | Control1 | 3.4671 | 23.945 | 0.141 | 1.112 | 20.475 | 0.218 |
| AU Exp6 | Control1 | 2.5183 | 21.403 | 0.09 | 0.938 | 15.05 | 0.2004 |
| AU Exp6 | Control1 | 1.0048 | 10.132 | 0.04 | 0.482 | | |
| AU Exp4 | 100 mM NaCl | 1.102 | 3.539 | 0.071 | 0.262 | 20.3 | 0.0792 |
| AU Exp4 | 100 mM NaCl | 1.018 | 3.061 | 0.044 | 0.213 | 19.1 | |
| AU Exp4 | 100 mM NaCl | 1.069 | 3.43 | 0.052 | 0.257 | 22.85 | |
| AU Exp4 | 100 mM NaCl | 1.643 | 4.821 | 0.063 | 0.334 | 20.825 | 0.1731 |
| AU Exp4 | 100 mM NaCl | 0.746 | 2.69 | 0.035 | 0.202 | 21.95 | 0.2318 |
| AU Exp4 | 100 mM NaCl | 1.17 | 4.022 | 0.047 | 0.249 | 22.025 | |
| AU Exp4 | 100 mM NaCl | 1.14 | 4.243 | 0.051 | 0.277 | 24.4 | |
| AU Exp4 | 100 mM NaCl | 1.508 | 4.186 | 0.065 | 0.294 | 25.425 | 0.2591 |
| AU Exp4 | 100 mM NaCl | 0.987 | 3.007 | 0.041 | 0.219 | 21.175 | |
| AU Exp4 | 100 mM NaCl | 0.723 | 2.232 | 0.028 | 0.153 | 21.8 | 0.0397 |
| AU Exp4 | 100 mM NaCl | 1.125 | 3.866 | 0.043 | 0.293 | 25.775 | |
| AU Exp4 | 100 mM NaCl | 1.741 | 5.316 | 0.062 | 0.365 | 22.6 | 0.0117 |
| AU Exp4 | Control | 0.415 | 1.911 | 0.012 | 0.103 | 13.25 | 0.1906 |
| AU Exp4 | Control | 2.199 | 11.009 | 0.094 | 0.608 | 15.3 | |
| AU Exp4 | Control | 1.718 | 8.255 | 0.064 | 0.451 | 15.025 | |
| AU Exp4 | Control | 1.734 | 9.583 | 0.066 | 0.479 | 14.75 | 0.253 |
| AU Exp4 | Control | 3.275 | 14.894 | 0.128 | 0.856 | 18.25 | |
| AU Exp4 | Control | 0.682 | 3.788 | 0.026 | 0.228 | 15.225 | |
| AU Exp4 | Control | 1.004 | 5.207 | 0.036 | 0.283 | 14.775 | |
| AU Exp4 | Control | 1.652 | 9.102 | 0.067 | 0.457 | 12.95 | 0.4396 |
| AU Exp4 | Control | 1.499 | 7.523 | 0.058 | 0.389 | 17.725 | 0.3712 |
| AU Exp4 | Control | 1.357 | 5.952 | 0.055 | 0.331 | 18.325 | 0.192 |
| AU Exp4 | Control | 0.747 | 4.56 | 0.026 | 0.271 | 14.5 | |
| AU Exp4 | Control | 2.304 | 10.587 | 0.079 | 0.549 | 14.725 | 0.2918 |

**S5d Endpoint physiological data for 100 mM NaCl stress treatment on different crop species and their respective controls.**

| Treatment | FW roots (g) | FW shoots (g) | DW roots (g) | DW shoots (g) | Ave. chlorophyll content (SPAD units) |
|---|---|---|---|---|---|
| Amaranth 100mM NaCl | 0.4301 | 0.6366 | 0.025 | 0.048 | 16.375 |
| Amaranth 100mM NaCl | 0.2227 | 0.3411 | 0.017 | 0.024 | 15.7 |
| Amaranth 100mM NaCl | | 0 | | 0 | |
| Amaranth 100mM NaCl | | 0 | | 0 | |
| Amaranth 100mM NaCl | 0.5546 | 0.9338 | 0.025 | 0.067 | 16.3 |
| Amaranth 100mM NaCl | | 0 | | 0 | |
| Amaranth 100mM NaCl | | 0 | | 0 | |
| Amaranth 100mM NaCl | | 0 | | 0 | |
| Amaranth 100mM NaCl | | 0 | | 0 | |
| Amaranth 100mM NaCl | 0.6834 | 1.3233 | 0.028 | 0.105 | 15.6 |
| Amaranth 100mM NaCl | | 0 | | 0 | |
| Amaranth 100mM NaCl | 0.2411 | 0.3789 | 0.015 | 0.036 | 15.425 |
| Amaranth control | 0.2373 | 0.7911 | 0.007 | 0.065 | 18.8 |
| Amaranth control | 0.0647 | 0.224 | 0.004 | 0.028 | |
| Amaranth control | 0.8969 | 3.0755 | 0.04 | 0.241 | 20.8 |
| Amaranth control | 1.1743 | 3.787 | 0.063 | 0.302 | 21.35 |

| | | | | | |
|---|---|---|---|---|---|
| Amaranth control | 0.5954 | 1.782 | 0.03 | 0.133 | 18.775 |
| Amaranth control | 0.5129 | 1.8513 | 0.023 | 0.15 | 20.75 |
| Amaranth control | 0.7494 | 1.0049 | 0.03 | 0.068 | 22.4 |
| Amaranth control | 0.9649 | 3.9205 | 0.048 | 0.289 | 21.275 |
| Amaranth control | 1.038 | 3.4399 | 0.052 | 0.268 | 20.9 |
| Amaranth control | 0.4526 | 1.628 | 0.011 | 0.117 | 19.7 |
| Amaranth control | 0.2577 | 0.8888 | 0.008 | 0.062 | 19.025 |
| Amaranth control | 0.2942 | 0.3536 | 0.017 | 0.044 | 17.7 |
| Mizuna 100mM NaCl | 0.909 | 15.118 | 69.8 | 890 | |
| Mizuna 100mM NaCl | 0.777 | 13.785 | 54.6 | 854 | 32.9 |
| Mizuna 100mM NaCl | 0.648 | 8.839 | | 564 | 28.7 |
| Mizuna 100mM NaCl | 0.827 | 11.175 | | 693 | |
| Mizuna 100mM NaCl | dead | dead | dead | dead | |
| Mizuna 100mM NaCl | 0.765 | 10.371 | 55.7 | 646 | |
| Mizuna 100mM NaCl | 0.655 | 8.707 | | 613 | |
| Mizuna 100mM NaCl | 0.985 | 10.354 | | 697 | 33.3 |
| Mizuna 100mM NaCl | 1.680 | 23.441 | 110.2 | 1371 | 36 |
| Mizuna 100mM NaCl | 0.376 | 4.947 | 30.1 | 298 | |
| Mizuna 100mM NaCl | 0.905 | 10.387 | 49.6 | 724 | 35.8 |
| Mizuna 100mM NaCl | 0.828 | 16.701 | 71.4 | 1052 | 37.6 |
| Mizuna control | 1.183 | 26.39 | 104.5 | 1498 | |
| Mizuna control | 0.678 | 18.014 | | 1039 | 22.5 |
| Mizuna control | 0.279 | 6.279 | | 354 | 25.5 |
| Mizuna control | 0.907 | 28.013 | | 1389 | |
| Mizuna control | 0.784 | 19.027 | | 1032 | |
| Mizuna control | 1.717 | 25.287 | 153.8 | 1399 | 27.5 |
| Mizuna control | 0.997 | 22.496 | 105.6 | 1169 | 29.2 |
| Mizuna control | 0.779 | 12.660 | 93.1 | 690 | 28.7 |
| Mizuna control | 3.280 | 56.745 | 47.9 | 2806 | 28.7 |
| Mizuna control | 0.364 | 7.885 | 258.1 | 448 | |
| Mizuna control | 0.414 | 4.188 | 50.3 | 291 | |
| Mizuna control | 0.791 | 17.320 | 79.2 | 946 | |
| Radish 100mM NaCl | 4.28 | 7.733 | 300.8 | 474.9 | 29.3 |

| | | | | | |
|---|---|---|---|---|---|
| Radish 100mM NaCl | 1.324 | 6.022 | 99 | 460.3 | 24.4 |
| Radish 100mM NaCl | dead | dead | dead | | |
| Radish 100mM NaCl | 4.211 | 9.84 | 261 | 585 | |
| Radish 100mM NaCl | 3.78 | 11.676 | | 642 | |
| Radish 100mM NaCl | 7.898 | 6.755 | | 401 | |
| Radish 100mM NaCl | 6.08 | 8.54 | | 540 | 34.3 |
| Radish 100mM NaCl | 6.952 | 19.238 | 333 | 1115 | 32.5 |
| Radish 100mM NaCl | 5.654 | 13.551 | 364 | 839 | 35.8 |
| Radish 100mM NaCl | 5.338 | 7.475 | 419 | 451 | |
| Radish 100mM NaCl | 5.280 | 8.604 | 311.9 | 458 | |
| Radish 100mM NaCl | 5.313 | 8.405 | | 477 | 31 |
| Radish control | 16.758 | 15.182 | 825.9 | 814 | |
| Radish control | 8.299 | 19.026 | 486.2 | 1020 | 28.9 |
| Radish control | 14.352 | 10.186 | 810.8 | 675 | 30.3 |
| Radish control | 9.553 | 12.125 | 594 | 685 | |
| Radish control | 11.463 | 12.899 | | 703 | |
| Radish control | 1.953 | 7.988 | 127 | 421 | |
| Radish control | 13.398 | 15.939 | 729.5 | 860 | 32.5 |
| Radish control | 22.944 | 16.577 | 1090 | 937 | 23.6 |
| Radish control | 18.657 | 16.363 | | 888 | 28.3 |
| Radish control | 12.542 | 10.697 | | 608 | |
| Radish control | 13.168 | 15.393 | | 855 | 32.88 |
| Radish control | 16.273 | 10.582 | 954 | 608 | |
| Rocket 100mM NaCl | 1.3067 | 2.7562 | 0.068 | 0.199 | 36.65 |
| Rocket 100mM NaCl | 0.9783 | 1.5802 | 0.033 | 0.125 | 28.325 |
| Rocket 100mM NaCl | 0.4567 | 3.1611 | 0.031 | 0.227 | 37.125 |
| Rocket 100mM NaCl | 1.2091 | 2.9486 | 0.038 | 0.226 | 41.9 |
| Rocket 100mM NaCl | 0.9947 | 3.0657 | 0.054 | 0.236 | 36.65 |
| Rocket 100mM NaCl | - | 0 | | | |
| Rocket 100mM NaCl | - | 0 | | | |
| Rocket 100mM NaCl | 0.3612 | 1.5942 | 0.025 | 0.167 | |
| Rocket 100mM NaCl | - | 0 | | | |
| Rocket 100mM NaCl | 0.7261 | 4.1255 | 0.061 | 0.306 | 34.15 |

| | | | | | |
|---|---|---|---|---|---|
| Rocket 100mM NaCl | - | 0 | | | |
| Rocket 100mM NaCl | 0.5841 | 2.2953 | 0.046 | 0.198 | 23.7 |
| Rocket control | 0.4468 | 2.8346 | 0.026 | 0.185 | 28.75 |
| Rocket control | 0.4879 | 3.8282 | 0.03 | 0.245 | 24.1 |
| Rocket control | 1.3671 | 10.1063 | 0.088 | 0.614 | 31.95 |
| Rocket control | 2.2034 | 7.5019 | 0.114 | 0.498 | 28.85 |
| Rocket control | 0.8187 | 4.8119 | 0.04 | 0.312 | 27.45 |
| Rocket control | 0.5468 | 4.746 | 0.042 | 0.48 | 22.35 |
| Rocket control | 0.5238 | 3.6051 | 0.036 | 0.207 | 32.075 |
| Rocket control | 0.5667 | 4.0145 | 0.033 | 0.23 | 32.7 |
| Rocket control | 0.483 | 4.45 | 0.037 | 0.302 | 30 |
| Rocket control | 0.3806 | 2.6702 | 0.025 | 0.194 | 23.55 |
| Rocket control | 1.8811 | 7.5099 | 0.111 | 0.461 | 25.5 |
| Rocket control | 1.0248 | 3.0607 | 0.062 | 0.207 | 25.725 |
| Tomato 100mM NaCl | 5.1305 | 10.739 | 0.193 | 0.833 | 33.8 |
| Tomato 100mM NaCl | 3.2354 | 7.1264 | 0.08 | 0.5 | 33.5 |
| Tomato 100mM NaCl | 5.246 | 9.2333 | 0.153 | 0.686 | 32.1 |
| Tomato 100mM NaCl | 2.9045 | 7.6162 | 0.106 | 0.508 | 31.275 |
| Tomato 100mM NaCl | 3.6469 | 9.9156 | 0.117 | 0.705 | 32.975 |
| Tomato 100mM NaCl | 4.872 | 11.5432 | 0.165 | 0.82 | 33.7 |
| Tomato 100mM NaCl | 1.5629 | 4.4052 | 0.052 | 0.281 | 31.5 |
| Tomato 100mM NaCl | | 0 | | | |
| Tomato 100mM NaCl | 3.2536 | 7.3972 | 0.143 | 0.488 | 33.475 |
| Tomato 100mM NaCl | 7.7804 | 16.5056 | 0.32 | 1.15 | 37.225 |
| Tomato 100mM NaCl | 1.8079 | 5.0257 | 0.078 | 0.361 | 28.95 |
| Tomato 100mM NaCl | 3.6425 | 9.0945 | 0.131 | 0.581 | 30.625 |
| Tomato control | 3.5825 | 19.7333 | 0.131 | 1.245 | 25.675 |
| Tomato control | 2.4116 | 10.7582 | 0.045 | 0.645 | 23.3 |
| Tomato control | 3.9264 | 10.7602 | 0.14 | 0.67 | 24.1 |
| Tomato control | 4.3073 | 17.2636 | 0.175 | 1.065 | 26.925 |
| Tomato control | 2.7257 | 11.4657 | 0.1 | 0.63 | 25.925 |
| Tomato control | 1.353 | 7.207 | 0.057 | 0.301 | 27.1 |
| Tomato control | 1.824 | 13.3705 | 0.072 | 0.789 | 27.55 |

| | | | | | |
|---|---|---|---|---|---|
| Tomato control | 2.1756 | 9.9106 | 0.091 | 0.59 | 22.95 |
| Tomato control | 2.6887 | 13.0171 | 0.112 | 0.786 | 23.975 |
| Tomato control | 4.4059 | 12.2485 | 0.12 | 0.783 | 23.7 |
| Tomato control | 1.8864 | 12.5962 | 0.07 | 0.73 | 21.95 |
| Tomato control | 2.4237 | 18.7286 | 0.095 | 1.119 | 24.7 |
