## Supplemental Table S6a for "Leaf movements as a quantitative metric for early stress detection"

**Supplementary Table S6.** 1 hour integrated motion data in **(a)** lettuce subjected to different stresses and their respective controls

Supplementary Table S6. 1 hour integrated motion data in (a) lettuce subjected to 100 mM NaCl

| IntervalMid | Control1 | Control2 | Control3 | Control4 | Control5 | Control6 | Control7 | Control8 | Control9 | Stress1 | Stress2 | Stress3 | Stress4 | Stress5 | Stress6 | Stress7 | Stress8 | Stress9 | Stress10 |
|---|---|---|---|---|---|---|---|---|---|---|---|---|---|---|---|---|---|---|---|
| 18.1111111 | 0.047205 | 0.044148 | 0.006314 | 0.006792 | 0.034329 | 0.003874 | 0.105912 | 0.010314 | 0.019421 | 0.054998 | 0.092849 | 0.010092 | 0.003612 | 0.008821 | 0.002732 | 0.004761 | 0.050216 | 0.045465 | 0.053755 |
| 18.125 | 0.058611 | 0.056889 | 0.012386 | 0.008116 | 0.052933 | 0.007093 | 0.084439 | 0.014114 | 0.026171 | 0.054221 | 0.079762 | 0.011981 | 0.004293 | 0.006347 | 0.004088 | 0.002296 | 0.073028 | 0.047602 | 0.042734 |
| 18.1388889 | 0.037157 | 0.040094 | 0.014749 | 0.0121 | 0.037336 | 0.010373 | 0.040689 | 0.011139 | 0.01748 | 0.030036 | 0.043177 | 0.009348 | 0.003176 | 0.004033 | 0.004899 | 0.001466 | 0.053348 | 0.029144 | 0.018869 |
| 18.1527778 | 0.01654 | 0.035798 | 0.010602 | 0.032521 | 0.015578 | 0.008486 | 0.019612 | 0.013256 | 0.007041 | 0.011966 | 0.040411 | 0.005833 | 0.003086 | 0.004958 | 0.003918 | 0.002406 | 0.024305 | 0.01222 | 0.004426 |
| 18.1666667 | 0.015979 | 0.056482 | 0.005497 | 0.063218 | 0.014876 | 0.004363 | 0.031525 | 0.021365 | 0.002646 | 0.007261 | 0.066104 | 0.003343 | 0.004086 | 0.006041 | 0.002849 | 0.003309 | 0.011402 | 0.005892 | 0.000646 |
| 18.1805556 | 0.014106 | 0.05387 | 0.004104 | 0.065056 | 0.014516 | 0.003273 | 0.044104 | 0.018145 | 0.002622 | 0.004654 | 0.060621 | 0.00235 | 0.004196 | 0.006456 | 0.002569 | 0.00342 | 0.006355 | 0.004499 | 0.000423 |
| 18.1944444 | 0.010243 | 0.026132 | 0.007326 | 0.034178 | 0.007606 | 0.0047 | 0.036823 | 0.008742 | 0.005318 | 0.001749 | 0.031247 | 0.002772 | 0.003687 | 0.007086 | 0.002871 | 0.002898 | 0.004465 | 0.004552 | 0.001103 |
| 18.2083333 | 0.01298 | 0.011088 | 0.010032 | 0.010022 | 0.007549 | 0.008502 | 0.022163 | 0.009535 | 0.00859 | 0.000896 | 0.014591 | 0.003684 | 0.003398 | 0.006696 | 0.003358 | 0.002347 | 0.00388 | 0.004534 | 0.001613 |
| 18.2222222 | 0.015981 | 0.010947 | 0.009928 | 0.005899 | 0.010764 | 0.012813 | 0.010272 | 0.013499 | 0.011773 | 0.001225 | 0.00943 | 0.004325 | 0.003427 | 0.005611 | 0.003773 | 0.00218 | 0.003646 | 0.004112 | 0.001462 |
| 18.2361111 | 0.016446 | 0.011908 | 0.00895 | 0.007275 | 0.011652 | 0.014995 | 0.005844 | 0.013536 | 0.01408 | 0.001473 | 0.008013 | 0.004339 | 0.003466 | 0.00521 | 0.003893 | 0.002323 | 0.003334 | 0.003303 | 0.000897 |
| 18.25 | 0.01531 | 0.010788 | 0.007425 | 0.007814 | 0.010826 | 0.014896 | 0.008479 | 0.011996 | 0.013718 | 0.001834 | 0.007994 | 0.003806 | 0.003364 | 0.005472 | 0.003754 | 0.002245 | 0.002749 | 0.002061 | 0.000571 |
| 18.2638889 | 0.014176 | 0.00964 | 0.006277 | 0.008138 | 0.009477 | 0.014421 | 0.012293 | 0.010882 | 0.011616 | 0.002582 | 0.008081 | 0.003328 | 0.003524 | 0.005814 | 0.003765 | 0.002087 | 0.002528 | 0.00104 | 0.000955 |
| 18.2777778 | 0.014539 | 0.01046 | 0.007018 | 0.009212 | 0.009676 | 0.016706 | 0.014523 | 0.011949 | 0.010659 | 0.003861 | 0.007349 | 0.003384 | 0.00401 | 0.005669 | 0.003892 | 0.002315 | 0.003001 | 0.00102 | 0.001267 |
| 18.2916667 | 0.015201 | 0.01239 | 0.008457 | 0.010496 | 0.010681 | 0.022414 | 0.014874 | 0.014302 | 0.011024 | 0.005186 | 0.005575 | 0.003783 | 0.004394 | 0.004872 | 0.003841 | 0.002817 | 0.00335 | 0.001918 | 0.000962 |
| 18.3055556 | 0.014504 | 0.013078 | 0.008475 | 0.010748 | 0.010098 | 0.030228 | 0.012646 | 0.016837 | 0.01076 | 0.005451 | 0.003052 | 0.003776 | 0.004384 | 0.004252 | 0.003565 | 0.003008 | 0.002675 | 0.00281 | 0.000592 |
| 18.3194444 | 0.013608 | 0.012694 | 0.007748 | 0.011176 | 0.00808 | 0.036391 | 0.010072 | 0.021071 | 0.010237 | 0.004436 | 0.00154 | 0.002918 | 0.003807 | 0.004318 | 0.003078 | 0.00244 | 0.001316 | 0.00251 | 0.000515 |
| 18.3333333 | 0.014876 | 0.013702 | 0.008638 | 0.013648 | 0.006012 | 0.037014 | 0.010514 | 0.027618 | 0.011566 | 0.002939 | 0.002561 | 0.002053 | 0.003073 | 0.004524 | 0.002694 | 0.001729 | 0.000565 | 0.001343 | 0.000607 |
| 18.3472222 | 0.018121 | 0.016618 | 0.011539 | 0.016811 | 0.003861 | 0.033919 | 0.013713 | 0.032402 | 0.0147 | 0.001668 | 0.004594 | 0.002177 | 0.002987 | 0.004468 | 0.002747 | 0.001853 | 0.001104 | 0.000657 | 0.001149 |
| 18.3611111 | 0.020967 | 0.019364 | 0.01456 | 0.018196 | 0.002193 | 0.031038 | 0.01657 | 0.031865 | 0.017462 | 0.001541 | 0.005572 | 0.002873 | 0.003393 | 0.00422 | 0.002929 | 0.002556 | 0.002463 | 0.000409 | 0.002202 |
| 18.375 | 0.022258 | 0.021019 | 0.01627 | 0.018333 | 0.002143 | 0.029384 | 0.017708 | 0.028205 | 0.018638 | 0.002425 | 0.005319 | 0.003139 | 0.003464 | 0.003904 | 0.002734 | 0.002959 | 0.003624 | 0.000171 | 0.002921 |
| 18.3888889 | 0.022835 | 0.022466 | 0.016848 | 0.019153 | 0.003831 | 0.028469 | 0.017914 | 0.02512 | 0.018918 | 0.002934 | 0.004546 | 0.002721 | 0.003109 | 0.004147 | 0.002334 | 0.002916 | 0.004224 | 0.000271 | 0.003113 |
| 18.4027778 | 0.022996 | 0.023609 | 0.01671 | 0.020196 | 0.006501 | 0.027377 | 0.017463 | 0.023177 | 0.018671 | 0.002751 | 0.00363 | 0.001786 | 0.002784 | 0.005652 | 0.002352 | 0.002748 | 0.004539 | 0.000811 | 0.003229 |
| 18.4166667 | 0.021387 | 0.022828 | 0.015352 | 0.019022 | 0.008758 | 0.024584 | 0.01514 | 0.020721 | 0.016972 | 0.002387 | 0.00245 | 0.001234 | 0.002613 | 0.008094 | 0.002885 | 0.002568 | 0.004705 | 0.001382 | 0.003156 |
| 18.4305556 | 0.017363 | 0.019557 | 0.012207 | 0.014838 | 0.010702 | 0.019721 | 0.010309 | 0.016942 | 0.013368 | 0.002073 | 0.001125 | 0.001794 | 0.002456 | 0.010423 | 0.003834 | 0.002325 | 0.00489 | 0.001667 | 0.002775 |
| 18.4444444 | 0.012176 | 0.015421 | 0.007811 | 0.009417 | 0.013816 | 0.014317 | 0.00483 | 0.012671 | 0.00876 | 0.001895 | 0.00068 | 0.002109 | 0.00213 | 0.012847 | 0.005503 | 0.001967 | 0.005347 | 0.001848 | 0.002375 |
| 18.4583333 | 0.007237 | 0.012406 | 0.003679 | 0.004495 | 0.019457 | 0.009555 | 0.001823 | 0.008683 | 0.004327 | 0.001824 | 0.00195 | 0.001363 | 0.001673 | 0.013542 | 0.007275 | 0.001613 | 0.006236 | 0.002311 | 0.002063 |
| 18.4722222 | 0.003577 | 0.011762 | 0.00167 | 0.001467 | 0.026872 | 0.006272 | 0.002202 | 0.005231 | 0.001439 | 0.001698 | 0.003971 | 0.000559 | 0.001415 | 0.012686 | 0.007504 | 0.001455 | 0.007473 | 0.00299 | 0.00172 |
| 18.4861111 | 0.002372 | 0.014299 | 0.001255 | 0.001002 | 0.033085 | 0.005758 | 0.002759 | 0.003094 | 0.000859 | 0.001504 | 0.005207 | 0.000235 | 0.001491 | 0.014777 | 0.006166 | 0.001442 | 0.008655 | 0.003343 | 0.001396 |
| 18.5 | 0.003957 | 0.019103 | 0.001438 | 0.002583 | 0.037462 | 0.008068 | 0.002254 | 0.003242 | 0.002229 | 0.00144 | 0.005276 | 0.000285 | 0.001773 | 0.015984 | 0.005003 | 0.00151 | 0.009394 | 0.003217 | 0.001262 |
| 18.5138889 | 0.006873 | 0.02298 | 0.003271 | 0.005668 | 0.040847 | 0.011391 | 0.003199 | 0.00513 | 0.004994 | 0.001471 | 0.004536 | 0.000688 | 0.002083 | 0.01284 | 0.004753 | 0.001676 | 0.009519 | 0.00305 | 0.001237 |
| 18.5277778 | 0.009232 | 0.023559 | 0.005641 | 0.008758 | 0.042809 | 0.014063 | 0.005261 | 0.007137 | 0.007746 | 0.001397 | 0.003807 | 0.001363 | 0.002363 | 0.0097 | 0.004963 | 0.001867 | 0.009125 | 0.003249 | 0.001133 |
| 18.5416667 | 0.01097 | 0.021875 | 0.007515 | 0.011198 | 0.044031 | 0.016208 | 0.007175 | 0.008795 | 0.009889 | 0.001341 | 0.003918 | 0.002248 | 0.002704 | 0.007422 | 0.005493 | 0.002117 | 0.00847 | 0.00392 | 0.001145 |
| 18.5555556 | 0.013741 | 0.021309 | 0.01044 | 0.014441 | 0.045654 | 0.019074 | 0.009836 | 0.011045 | 0.012727 | 0.001314 | 0.004121 | 0.003285 | 0.003124 | 0.004265 | 0.006263 | 0.00246 | 0.007786 | 0.004789 | 0.001228 |
| 18.5694444 | 0.019362 | 0.025025 | 0.015678 | 0.020535 | 0.047649 | 0.023994 | 0.011955 | 0.013891 | 0.017429 | 0.002126 | 0.004558 | 0.004273 | 0.003776 | 0.002989 | 0.006721 | 0.0027 | 0.007534 | 0.005218 | 0.002686 |
| 18.5833333 | 0.028427 | 0.034025 | 0.023232 | 0.029893 | 0.050508 | 0.031611 | 0.010694 | 0.015764 | 0.023869 | 0.007397 | 0.012694 | 0.00504 | 0.005127 | 0.008985 | 0.00674 | 0.002771 | 0.008194 | 0.004732 | 0.011011 |
| 18.5972222 | 0.036803 | 0.043612 | 0.030263 | 0.035125 | 0.052475 | 0.038383 | 0.008416 | 0.016794 | 0.029096 | 0.017408 | 0.045859 | 0.0055 | 0.006837 | 0.013776 | 0.006807 | 0.003068 | 0.009261 | 0.003794 | 0.025396 |
| 18.6111111 | 0.036414 | 0.044188 | 0.031328 | 0.025152 | 0.048004 | 0.036977 | 0.009597 | 0.017353 | 0.028226 | 0.022791 | 0.087821 | 0.00558 | 0.007263 | 0.008869 | 0.006498 | 0.003773 | 0.009339 | 0.003352 | 0.031078 |
| 18.625 | 0.025369 | 0.032646 | 0.023847 | 0.012208 | 0.035867 | 0.025857 | 0.011365 | 0.015128 | 0.020066 | 0.017611 | 0.090569 | 0.004878 | 0.005448 | 0.00415 | 0.0048 | 0.003944 | 0.007684 | 0.00318 | 0.022405 |
| 18.6388889 | 0.011292 | 0.017043 | 0.012193 | 0.012753 | 0.022361 | 0.012265 | 0.008771 | 0.009555 | 0.009462 | 0.008742 | 0.059818 | 0.003406 | 0.002624 | 0.002723 | 0.00229 | 0.00296 | 0.00519 | 0.002635 | 0.01163 |
| 18.6527778 | 0.003272 | 0.006064 | 0.003681 | 0.014209 | 0.013212 | 0.003741 | 0.004013 | 0.004078 | 0.002771 | 0.00328 | 0.032853 | 0.002381 | 0.000809 | 0.001191 | 0.000596 | 0.001999 | 0.003444 | 0.002392 | 0.005747 |
| 18.6666667 | 0.002859 | 0.001613 | 0.000769 | 0.00982 | 0.008626 | 0.001503 | 0.001823 | 0.002068 | 0.00131 | 0.001891 | 0.017876 | 0.002282 | 0.000477 | 0.000859 | 0.000117 | 0.001838 | 0.002802 | 0.002847 | 0.002671 |
| 18.6805556 | 0.00431 | 0.000923 | 0.00042 | 0.006024 | 0.00656 | 0.001599 | 0.002272 | 0.002432 | 0.00124 | 0.003052 | 0.011054 | 0.002262 | 0.000755 | 0.001109 | 0.000146 | 0.001777 | 0.002402 | 0.00322 | 0.000971 |
| 18.6944444 | 0.004533 | 0.000719 | 0.000419 | 0.002976 | 0.006255 | 0.001404 | 0.002334 | 0.001986 | 0.000927 | 0.004938 | 0.008483 | 0.002056 | 0.001128 | 0.000886 | 0.000279 | 0.001519 | 0.001719 | 0.003364 | 0.000313 |
| 18.7083333 | 0.004365 | 0.000405 | 0.000914 | 0.001467 | 0.007243 | 0.001957 | 0.001601 | 0.001104 | 0.001652 | 0.006142 | 0.005873 | 0.00217 | 0.001626 | 0.001088 | 0.00055 | 0.001484 | 0.001304 | 0.003747 | 0.000381 |
| 18.7222222 | 0.003844 | 0.000967 | 0.002091 | 0.002388 | 0.008612 | 0.00376 | 0.002296 | 0.001712 | 0.003324 | 0.00595 | 0.002947 | 0.002973 | 0.002224 | 0.004601 | 0.000972 | 0.001845 | 0.001951 | 0.004456 | 0.000873 |
| 18.7361111 | 0.00291 | 0.002112 | 0.003549 | 0.004692 | 0.009687 | 0.00516 | 0.004774 | 0.003152 | 0.004863 | 0.004112 | 0.001637 | 0.003871 | 0.002481 | 0.010421 | 0.001382 | 0.002054 | 0.003005 | 0.005098 | 0.001203 |
| 18.75 | 0.002698 | 0.0026 | 0.004509 | 0.005938 | 0.009918 | 0.005306 | 0.007004 | 0.004173 | 0.005512 | 0.002361 | 0.001321 | 0.003973 | 0.002241 | 0.011648 | 0.001707 | 0.001769 | 0.003257 | 0.005177 | 0.00092 |
| 18.7638889 | 0.002693 | 0.002992 | 0.00539 | 0.006489 | 0.010106 | 0.005609 | 0.008355 | 0.005308 | 0.006068 | 0.003204 | 0.001204 | 0.003598 | 0.002062 | 0.00663 | 0.001943 | 0.001701 | 0.002898 | 0.004915 | 0.000572 |
| 18.7777778 | 0.002819 | 0.005533 | 0.00759 | 0.008455 | 0.012187 | 0.007974 | 0.010547 | 0.007796 | 0.008449 | 0.005689 | 0.00166 | 0.003344 | 0.002104 | 0.001861 | 0.001888 | 0.002124 | 0.002575 | 0.004805 | 0.000574 |
| 18.7916667 | 0.005566 | 0.010179 | 0.011214 | 0.012653 | 0.016523 | 0.012522 | 0.014517 | 0.011824 | 0.012898 | 0.007365 | 0.002179 | 0.003227 | 0.002147 | 0.000833 | 0.001639 | 0.002501 | 0.002552 | 0.004825 | 0.000619 |
| 18.8055556 | 0.008167 | 0.013235 | 0.014062 | 0.016435 | 0.020149 | 0.016213 | 0.017932 | 0.015034 | 0.016462 | 0.007449 | 0.002191 | 0.003063 | 0.002223 | 0.001493 | 0.001425 | 0.00252 | 0.002844 | 0.004631 | 0.000519 |
| 18.8194444 | 0.006342 | 0.011999 | 0.014212 | 0.016787 | 0.020086 | 0.016304 | 0.018098 | 0.01507 | 0.016333 | 0.005995 | 0.001918 | 0.002604 | 0.002357 | 0.002622 | 0.001026 | 0.002082 | 0.003081 | 0.004002 | 0.000317 |
| 18.8333333 | 0.003199 | 0.00939 | 0.013806 | 0.015561 | 0.018518 | 0.015112 | 0.016954 | 0.014049 | 0.01481 | 0.003865 | 0.001904 | 0.002095 | 0.002453 | 0.00385 | 0.000452 | 0.001573 | 0.003121 | 0.003414 | 0.000183 |
| 18.8472222 | 0.002693 | 0.009048 | 0.015213 | 0.015854 | 0.018672 | 0.015364 | 0.017309 | 0.014703 | 0.015172 | 0.002413 | 0.002237 | 0.002071 | 0.002363 | 0.004086 | 0.000196 | 0.001612 | 0.003254 | 0.003482 | 0.000148 |
| 18.8611111 | 0.003415 | 0.009714 | 0.016667 | 0.016676 | 0.019248 | 0.015582 | 0.017917 | 0.015665 | 0.016216 | 0.002203 | 0.002899 | 0.002609 | 0.002161 | 0.00375 | 0.00045 | 0.00224 | 0.003611 | 0.004052 | 0.00029 |
| 18.875 | 0.003051 | 0.009292 | 0.016221 | 0.016261 | 0.018443 | 0.01445 | 0.017255 | 0.015154 | 0.015905 | 0.002516 | 0.003674 | 0.003122 | 0.001909 | 0.003913 | 0.000779 | 0.002755 | 0.003724 | 0.004394 | 0.000768 |
| 18.8888889 | 0.0018 | 0.008128 | 0.014479 | 0.014915 | 0.016789 | 0.012906 | 0.015981 | 0.013648 | 0.014408 | 0.002332 | 0.003754 | 0.002922 | 0.001438 | 0.004351 | 0.000721 | 0.002418 | 0.003144 | 0.003971 | 0.001042 |
| 18.9027778 | 0.001048 | 0.007313 | 0.013066 | 0.013517 | 0.015278 | 0.011935 | 0.01482 | 0.012279 | 0.012917 | 0.001802 | 0.003111 | 0.002327 | 0.000794 | 0.004321 | 0.000473 | 0.001615 | 0.002316 | 0.003214 | 0.000794 |
| 18.9166667 | 0.001633 | 0.007421 | 0.013095 | 0.012702 | 0.014717 | 0.011735 | 0.014131 | 0.011754 | 0.01244 | 0.001616 | 0.002627 | 0.002105 | 0.000272 | 0.003905 | 0.000364 | 0.001248 | 0.001947 | 0.002887 | 0.000734 |
| 18.9305556 | 0.003028 | 0.008113 | 0.014232 | 0.012577 | 0.015036 | 0.011818 | 0.01433 | 0.012013 | 0.012871 | 0.001758 | 0.002856 | 0.002306 | 0.000172 | 0.003922 | 0.000392 | 0.001447 | 0.002238 | 0.003019 | 0.001493 |
| 18.9444444 | 0.003745 | 0.008365 | 0.01502 | 0.012221 | 0.014761 | 0.011331 | 0.014991 | 0.011891 | 0.012997 | 0.001528 | 0.003417 | 0.002295 | 0.000271 | 0.004179 | 0.000379 | 0.001507 | 0.002607 | 0.002937 | 0.002419 |
| 18.9583333 | 0.003006 | 0.00755 | 0.014602 | 0.010969 | 0.013115 | 0.010051 | 0.01515 | 0.010578 | 0.012145 | 0.001002 | 0.00348 | 0.001594 | 0.000399 | 0.003603 | 0.000351 | 0.00098 | 0.002291 | 0.002111 | 0.002455 |
| 18.9722222 | 0.001806 | 0.006293 | 0.013651 | 0.009614 | 0.011382 | 0.008757 | 0.014885 | 0.00902 | 0.011197 | 0.000897 | 0.003408 | 0.000921 | 0.000705 | 0.00255 | 0.000448 | 0.000419 | 0.001699 | 0.001334 | 0.002162 |
| 18.9861111 | 0.001065 | 0.005095 | 0.01272 | 0.008661 | 0.010308 | 0.007761 | 0.014745 | 0.007941 | 0.010688 | 0.00075 | 0.004139 | 0.001135 | 0.000709 | 0.002397 | 0.000378 | 0.000394 | 0.001625 | 0.001524 | 0.002701 |
| 19 | 0.001352 | 0.003707 | 0.011561 | 0.007713 | 0.009195 | 0.006685 | 0.014566 | 0.006955 | 0.010024 | 0.000422 | 0.005631 | 0.001793 | 0.000421 | 0.003996 | 0.000212 | 0.000811 | 0.001823 | 0.002115 | 0.003793 |
| 19.0138889 | 0.003199 | 0.002062 | 0.009929 | 0.006482 | 0.007645 | 0.005448 | 0.013808 | 0.005779 | 0.008855 | 0.000601 | 0.007338 | 0.002083 | 0.000458 | 0.007858 | 0.000173 | 0.001075 | 0.001733 | 0.002279 | 0.00448 |
| 19.0277778 | 0.005571 | 0.000988 | 0.007782 | 0.00485 | 0.005655 | 0.003855 | 0.012141 | 0.004337 | 0.00723 | 0.001135 | 0.008913 | 0.001831 | 0.000745 | 0.011848 | 0.000107 | 0.000871 | 0.001461 | 0.00196 | 0.004484 |
| 19.0416667 | 0.006957 | 0.001154 | 0.005782 | 0.003183 | 0.003616 | 0.001978 | 0.010124 | 0.002832 | 0.005658 | 0.001771 | 0.009823 | 0.001373 | 0.000826 | 0.011134 | 8.28E-05 | 0.000448 | 0.001321 | 0.001613 | 0.004102 |
| 19.0555556 | 0.007185 | 0.001746 | 0.004599 | 0.002118 | 0.001858 | 0.001214 | 0.009021 | 0.001526 | 0.004597 | 0.002728 | 0.009711 | 0.001069 | 0.000601 | 0.006478 | 0.00025 | 0.000191 | 0.001172 | 0.00169 | 0.003839 |
| 19.0694444 | 0.008008 | 0.003734 | 0.003652 | 0.002062 | 0.002434 | 0.003622 | 0.010671 | 0.001429 | 0.003277 | 0.00437 | 0.011788 | 0.000902 | 0.000737 | 0.003036 | 0.000647 | 0.000267 | 0.000782 | 0.002389 | 0.006209 |
| 19.0833333 | 0.010681 | 0.009748 | 0.002174 | 0.003978 | 0.009103 | 0.010423 | 0.02181 | 0.004271 | 0.0041 | 0.008163 | 0.02226 | 0.000966 | 0.001909 | 0.00184 | 0.001357 | 0.00089 | 0.000822 | 0.003764 | 0.016836 |
| 19.0972222 | 0.009733 | 0.015759 | 0.002053 | 0.009577 | 0.018224 | 0.017597 | 0.045998 | 0.008088 | 0.009485 | 0.016467 | 0.047097 | 0.001792 | 0.003151 | 0.00199 | 0.001904 | 0.002539 | 0.00165 | 0.005616 | 0.034131 |
| 19.1111111 | 0.009436 | 0.013389 | 0.005699 | 0.015924 | 0.019364 | 0.016405 | 0.05609 | 0.008294 | 0.012896 | 0.026726 | 0.074571 | 0.002674 | 0.002651 | 0.002031 | 0.001999 | 0.004153 | 0.001697 | 0.007922 | 0.04188 |
| 19.125 | 0.018822 | 0.008504 | 0.011332 | 0.014457 | 0.010995 | 0.008921 | 0.039306 | 0.005569 | 0.009902 | 0.028236 | 0.073917 | 0.002239 | 0.001172 | 0.001587 | 0.003021 | 0.003748 | 0.001235 | 0.010453 | 0.031875 |
| 19.1388889 | 0.025278 | 0.01049 | 0.014534 | 0.006986 | 0.003396 | 0.005384 | 0.033141 | 0.003713 | 0.005304 | 0.017272 | 0.046055 | 0.001038 | 0.000626 | 0.002381 | 0.004889 | 0.001847 | 0.002012 | 0.011343 | 0.017291 |
| 19.1527778 | 0.02173 | 0.013079 | 0.012832 | 0.003053 | 0.00214 | 0.00554 | 0.042087 | 0.002742 | 0.002692 | 0.005954 | 0.021774 | 0.000507 | 0.00082 | 0.003604 | 0.005408 | 0.000655 | 0.003108 | 0.00956 | 0.00935 |
| 19.1666667 | 0.016053 | 0.011802 | 0.008076 | 0.002118 | 0.003193 | 0.005062 | 0.03721 | 0.001463 | 0.001403 | 0.002585 | 0.010554 | 0.000885 | 0.001199 | 0.004235 | 0.004586 | 0.000704 | 0.003572 | 0.006824 | 0.006638 |
| 19.1805556 | 0.011248 | 0.009555 | 0.003696 | 0.001496 | 0.003866 | 0.004026 | 0.021076 | 0.000658 | 0.000599 | 0.0031 | 0.005554 | 0.001474 | 0.001516 | 0.004367 | 0.00385 | 0.001034 | 0.003574 | 0.004769 | 0.005387 |
| 19.1944444 | 0.00787 | 0.007739 | 0.001241 | 0.001522 | 0.004222 | 0.002844 | 0.009171 | 0.000698 | 0.000286 | 0.003563 | 0.003042 | 0.001815 | 0.001373 | 0.00381 | 0.003237 | 0.001122 | 0.003155 | 0.00367 | 0.004122 |
| 19.2083333 | 0.005818 | 0.006395 | 0.000707 | 0.001212 | 0.003869 | 0.001751 | 0.004921 | 0.000745 | 0.000683 | 0.003705 | 0.001507 | 0.001826 | 0.000933 | 0.003124 | 0.002323 | 0.000973 | 0.002618 | 0.003042 | 0.002942 |
| 19.2222222 | 0.004043 | 0.00476 | 0.001463 | 0.001213 | 0.002654 | 0.001849 | 0.004949 | 0.000969 | 0.001939 | 0.003549 | 0.000585 | 0.001654 | 0.000637 | 0.002657 | 0.001501 | 0.000762 | 0.002325 | 0.002606 | 0.00233 |
| 19.2361111 | 0.002292 | 0.003139 | 0.002366 | 0.002103 | 0.001251 | 0.003119 | 0.005507 | 0.001509 | 0.003261 | 0.003059 | 0.000233 | 0.001665 | 0.000736 | 0.002362 | 0.001276 | 0.000727 | 0.002439 | 0.002564 | 0.002324 |
| 19.25 | 0.001456 | 0.002697 | 0.002271 | 0.002266 | 0.000878 | 0.003707 | 0.004818 | 0.001318 | 0.003369 | 0.002517 | 0.000275 | 0.00192 | 0.000994 | 0.002257 | 0.001364 | 0.000848 | 0.002675 | 0.00281 | 0.002468 |
| 19.2638889 | 0.001609 | 0.003027 | 0.001427 | 0.001204 | 0.001587 | 0.003352 | 0.003412 | 0.000757 | 0.002501 | 0.002278 | 0.000802 | 0.002222 | 0.000926 | 0.002153 | 0.001267 | 0.000938 | 0.002529 | 0.002995 | 0.002508 |

| | | | | | | | | | | | | | | | | | | | |
|---|---|---|---|---|---|---|---|---|---|---|---|---|---|---|---|---|---|---|---|
| 19.2777778 | 0.00139 | 0.00253 | 0.001221 | 0.000633 | 0.001956 | 0.003448 | 0.002748 | 0.000617 | 0.002226 | 0.002182 | 0.00163 | 0.002184 | 0.000728 | 0.001989 | 0.00102 | 0.00099 | 0.002123 | 0.002948 | 0.00257 |
| 19.2916667 | 0.001302 | 0.001659 | 0.002473 | 0.001856 | 0.001567 | 0.004608 | 0.003717 | 0.001185 | 0.003483 | 0.001807 | 0.001696 | 0.001575 | 0.000938 | 0.002085 | 0.000864 | 0.000999 | 0.002016 | 0.002705 | 0.002588 |
| 19.3055556 | 0.003132 | 0.002317 | 0.004839 | 0.004181 | 0.001833 | 0.006563 | 0.005913 | 0.003072 | 0.005767 | 0.001529 | 0.001029 | 0.000852 | 0.001114 | 0.002807 | 0.000696 | 0.000876 | 0.00202 | 0.002112 | 0.002155 |
| 19.3194444 | 0.005177 | 0.003604 | 0.006684 | 0.005478 | 0.002595 | 0.008116 | 0.007511 | 0.004748 | 0.007242 | 0.001866 | 0.000915 | 0.00069 | 0.000703 | 0.005001 | 0.000538 | 0.001009 | 0.001827 | 0.001106 | 0.001381 |
| 19.3333333 | 0.00423 | 0.003298 | 0.005593 | 0.004092 | 0.002562 | 0.007339 | 0.006525 | 0.003832 | 0.005773 | 0.001738 | 0.000971 | 0.001 | 0.000713 | 0.008776 | 0.001125 | 0.001262 | 0.002269 | 0.000837 | 0.001421 |
| 19.3472222 | 0.002579 | 0.003118 | 0.003188 | 0.002419 | 0.003183 | 0.004079 | 0.003385 | 0.002473 | 0.003104 | 0.001362 | 0.00056 | 0.001682 | 0.001789 | 0.011214 | 0.002394 | 0.001713 | 0.003466 | 0.002235 | 0.002672 |
| 19.3611111 | 0.003342 | 0.004359 | 0.002759 | 0.002867 | 0.004579 | 0.001133 | 0.000851 | 0.002975 | 0.002374 | 0.002024 | 0.000602 | 0.002537 | 0.002295 | 0.009797 | 0.002672 | 0.00259 | 0.003696 | 0.003683 | 0.003477 |
| 19.375 | 0.003426 | 0.004291 | 0.002303 | 0.003083 | 0.00386 | 0.000738 | 0.000446 | 0.002597 | 0.00191 | 0.002285 | 0.00107 | 0.002315 | 0.001368 | 0.006562 | 0.001497 | 0.002393 | 0.002507 | 0.003432 | 0.002812 |
| 19.3888889 | 0.001766 | 0.002783 | 0.001004 | 0.002586 | 0.001596 | 0.001821 | 0.001262 | 0.001029 | 0.000871 | 0.001529 | 0.001206 | 0.001215 | 0.00044 | 0.004981 | 0.000506 | 0.001127 | 0.001523 | 0.002215 | 0.001771 |
| 19.4027778 | 0.001118 | 0.002235 | 0.000572 | 0.003225 | 0.000885 | 0.002184 | 0.00183 | 0.000548 | 0.000617 | 0.000782 | 0.001326 | 0.000495 | 0.000268 | 0.005146 | 0.000678 | 0.000414 | 0.001538 | 0.001324 | 0.001297 |
| 19.4166667 | 0.002548 | 0.003665 | 0.001197 | 0.005419 | 0.00246 | 0.001683 | 0.001494 | 0.00144 | 0.001326 | 0.000377 | 0.002084 | 0.000416 | 0.000407 | 0.00427 | 0.001352 | 0.000348 | 0.001944 | 0.001195 | 0.001273 |
| 19.4305556 | 0.005751 | 0.006903 | 0.003584 | 0.009524 | 0.004812 | 0.001712 | 0.001359 | 0.003627 | 0.003526 | 0.000274 | 0.003349 | 0.000686 | 0.000688 | 0.002003 | 0.001413 | 0.00047 | 0.002046 | 0.001787 | 0.001274 |
| 19.4444444 | 0.009065 | 0.010332 | 0.006551 | 0.014811 | 0.005742 | 0.001749 | 0.001947 | 0.005831 | 0.005959 | 0.000485 | 0.004997 | 0.000656 | 0.000945 | 0.001141 | 0.000933 | 0.000658 | 0.001789 | 0.002448 | 0.001036 |
| 19.4583333 | 0.009262 | 0.010767 | 0.006378 | 0.017291 | 0.004801 | 0.002909 | 0.002877 | 0.005083 | 0.005317 | 0.001579 | 0.007542 | 0.000608 | 0.000822 | 0.001974 | 0.001005 | 0.00072 | 0.001433 | 0.002273 | 0.000702 |
| 19.4722222 | 0.007121 | 0.007974 | 0.005638 | 0.012419 | 0.006793 | 0.009128 | 0.007223 | 0.004817 | 0.005031 | 0.003618 | 0.011897 | 0.001404 | 0.00064 | 0.002271 | 0.001607 | 0.001116 | 0.000939 | 0.001452 | 0.001007 |
| 19.4861111 | 0.00909 | 0.008311 | 0.010799 | 0.006791 | 0.01469 | 0.018673 | 0.016042 | 0.011349 | 0.01131 | 0.005315 | 0.019163 | 0.002139 | 0.000863 | 0.001673 | 0.001756 | 0.001727 | 0.000437 | 0.000965 | 0.002051 |
| 19.5 | 0.015561 | 0.014077 | 0.018576 | 0.010029 | 0.022588 | 0.026125 | 0.02406 | 0.020042 | 0.019632 | 0.005366 | 0.028742 | 0.001573 | 0.000997 | 0.002029 | 0.001143 | 0.00142 | 0.000365 | 0.000805 | 0.002504 |
| 19.5138889 | 0.020747 | 0.019513 | 0.023098 | 0.016692 | 0.02567 | 0.029716 | 0.027754 | 0.024896 | 0.024208 | 0.00445 | 0.03597 | 0.000523 | 0.000864 | 0.005537 | 0.000465 | 0.000617 | 0.00055 | 0.000634 | 0.002116 |
| 19.5277778 | 0.022459 | 0.02146 | 0.024298 | 0.019386 | 0.02457 | 0.030006 | 0.027601 | 0.025945 | 0.025106 | 0.004205 | 0.037408 | 0.000377 | 0.001101 | 0.012569 | 0.000495 | 0.000467 | 0.000592 | 0.000518 | 0.002036 |
| 19.5416667 | 0.020918 | 0.020052 | 0.022811 | 0.017996 | 0.020686 | 0.02734 | 0.024601 | 0.023524 | 0.022698 | 0.004675 | 0.035654 | 0.001119 | 0.001597 | 0.014847 | 0.001105 | 0.001018 | 0.000792 | 0.000593 | 0.002615 |
| 19.5555556 | 0.01696 | 0.016101 | 0.019304 | 0.014152 | 0.015515 | 0.022536 | 0.020241 | 0.018927 | 0.018228 | 0.004993 | 0.032639 | 0.001702 | 0.001488 | 0.008556 | 0.001504 | 0.001553 | 0.000998 | 0.000939 | 0.003357 |
| 19.5694444 | 0.012014 | 0.010965 | 0.014596 | 0.009162 | 0.009704 | 0.016839 | 0.015895 | 0.013952 | 0.013172 | 0.005771 | 0.029879 | 0.001348 | 0.001156 | 0.003782 | 0.0012 | 0.001445 | 0.000912 | 0.001335 | 0.004065 |
| 19.5833333 | 0.006378 | 0.006377 | 0.008381 | 0.005383 | 0.00574 | 0.010747 | 0.011185 | 0.008461 | 0.007494 | 0.00892 | 0.032939 | 0.001091 | 0.001885 | 0.002795 | 0.001116 | 0.000823 | 0.001553 | 0.002054 | 0.005831 |
| 19.5972222 | 0.00197 | 0.005402 | 0.004881 | 0.005799 | 0.00736 | 0.00535 | 0.006096 | 0.004586 | 0.004702 | 0.01343 | 0.041414 | 0.002018 | 0.003036 | 0.002949 | 0.00193 | 0.000454 | 0.003297 | 0.003025 | 0.010846 |
| 19.6111111 | 0.002388 | 0.005825 | 0.007016 | 0.006338 | 0.010394 | 0.005711 | 0.00254 | 0.004675 | 0.006757 | 0.014516 | 0.047045 | 0.002745 | 0.003243 | 0.004815 | 0.002217 | 0.000653 | 0.004446 | 0.003102 | 0.019995 |
| 19.625 | 0.008315 | 0.005332 | 0.007098 | 0.004956 | 0.00777 | 0.018542 | 0.002541 | 0.004719 | 0.006544 | 0.010612 | 0.046009 | 0.002312 | 0.00261 | 0.006061 | 0.001543 | 0.000915 | 0.003693 | 0.001997 | 0.028734 |
| 19.6388889 | 0.016568 | 0.008339 | 0.003957 | 0.006902 | 0.003814 | 0.03958 | 0.006129 | 0.004843 | 0.003549 | 0.005708 | 0.038332 | 0.001397 | 0.001542 | 0.005801 | 0.001645 | 0.000947 | 0.001902 | 0.000767 | 0.030005 |
| 19.6527778 | 0.02314 | 0.013918 | 0.004765 | 0.010836 | 0.005574 | 0.051552 | 0.0112 | 0.008654 | 0.004148 | 0.003136 | 0.027133 | 0.000713 | 0.000659 | 0.004171 | 0.002721 | 0.000664 | 0.000838 | 0.000266 | 0.024897 |
| 19.6666667 | 0.027073 | 0.017434 | 0.009519 | 0.012466 | 0.009809 | 0.046408 | 0.01775 | 0.013274 | 0.007393 | 0.00259 | 0.018973 | 0.000579 | 0.000499 | 0.002248 | 0.00314 | 0.000316 | 0.000831 | 0.00033 | 0.020399 |
| 19.6805556 | 0.028567 | 0.017689 | 0.01393 | 0.011945 | 0.011693 | 0.03342 | 0.026411 | 0.015777 | 0.009616 | 0.002507 | 0.014966 | 0.000609 | 0.000817 | 0.001637 | 0.002716 | 0.000358 | 0.000936 | 0.000347 | 0.018321 |
| 19.6944444 | 0.028011 | 0.015866 | 0.01596 | 0.010934 | 0.01157 | 0.022404 | 0.034822 | 0.01717 | 0.0107 | 0.002489 | 0.012662 | 0.000682 | 0.001702 | 0.001693 | 0.002611 | 0.000776 | 0.001163 | 0.000355 | 0.01676 |
| 19.7083333 | 0.02706 | 0.014522 | 0.016258 | 0.010882 | 0.01178 | 0.017113 | 0.039845 | 0.018804 | 0.012122 | 0.002832 | 0.011376 | 0.001059 | 0.002604 | 0.002221 | 0.003027 | 0.00127 | 0.00184 | 0.00075 | 0.015197 |
| 19.7222222 | 0.025681 | 0.01365 | 0.015033 | 0.011 | 0.012386 | 0.016148 | 0.039954 | 0.019431 | 0.013007 | 0.002855 | 0.009385 | 0.001313 | 0.002446 | 0.002391 | 0.003266 | 0.001105 | 0.00253 | 0.001014 | 0.013711 |
| 19.7361111 | 0.023005 | 0.011742 | 0.011013 | 0.009607 | 0.011221 | 0.014703 | 0.035495 | 0.017014 | 0.011203 | 0.002159 | 0.005726 | 0.00134 | 0.001803 | 0.002444 | 0.003076 | 0.000451 | 0.002887 | 0.000839 | 0.012325 |
| 19.75 | 0.019335 | 0.009113 | 0.00558 | 0.006997 | 0.008202 | 0.010658 | 0.027179 | 0.012181 | 0.007197 | 0.001531 | 0.004214 | 0.001442 | 0.001476 | 0.002928 | 0.002509 | 0.000551 | 0.002833 | 0.000806 | 0.011176 |
| 19.7638889 | 0.015441 | 0.007138 | 0.002937 | 0.004855 | 0.00518 | 0.005594 | 0.017237 | 0.007586 | 0.003286 | 0.001347 | 0.005808 | 0.001296 | 0.001119 | 0.002045 | 0.001736 | 0.001471 | 0.002632 | 0.000762 | 0.010326 |
| 19.7777778 | 0.012833 | 0.00685 | 0.003295 | 0.004575 | 0.004079 | 0.002 | 0.009079 | 0.005579 | 0.001407 | 0.001623 | 0.0057 | 0.001166 | 0.000633 | 0.000623 | 0.001326 | 0.002528 | 0.002818 | 0.00054 | 0.009708 |
| 19.7916667 | 0.012787 | 0.008141 | 0.002971 | 0.006246 | 0.005695 | 0.001185 | 0.003802 | 0.00626 | 0.002258 | 0.002354 | 0.003638 | 0.001507 | 0.000296 | 0.000403 | 0.001488 | 0.003256 | 0.003251 | 0.000549 | 0.008772 |
| 19.8055556 | 0.013725 | 0.009302 | 0.001624 | 0.008256 | 0.008059 | 0.00161 | 0.002309 | 0.007571 | 0.004339 | 0.002904 | 0.00364 | 0.001569 | 0.000286 | 0.001255 | 0.002053 | 0.003284 | 0.003793 | 0.000672 | 0.007185 |
| 19.8194444 | 0.012184 | 0.00827 | 0.00079 | 0.008349 | 0.007695 | 0.00173 | 0.005542 | 0.007004 | 0.004747 | 0.002618 | 0.005319 | 0.000914 | 0.000646 | 0.002367 | 0.002615 | 0.002871 | 0.004903 | 0.000755 | 0.005491 |
| 19.8333333 | 0.007462 | 0.005066 | 0.000643 | 0.006915 | 0.004298 | 0.002011 | 0.009996 | 0.00423 | 0.002858 | 0.001517 | 0.005149 | 0.000272 | 0.0008 | 0.002264 | 0.002419 | 0.002388 | 0.006192 | 0.00074 | 0.004037 |
| 19.8472222 | 0.00314 | 0.00268 | 0.000945 | 0.01015 | 0.00158 | 0.002806 | 0.010832 | 0.001544 | 0.001326 | 0.000539 | 0.002819 | 0.000362 | 0.000676 | 0.001026 | 0.001743 | 0.002145 | 0.006934 | 0.000638 | 0.003073 |
| 19.8611111 | 0.00214 | 0.003392 | 0.001445 | 0.023385 | 0.002175 | 0.005072 | 0.007703 | 0.001233 | 0.001984 | 0.000542 | 0.001223 | 0.00103 | 0.001028 | 0.000256 | 0.001523 | 0.002352 | 0.007513 | 0.000887 | 0.002869 |
| 19.875 | 0.002569 | 0.004704 | 0.001858 | 0.037256 | 0.004021 | 0.007134 | 0.004973 | 0.002356 | 0.002888 | 0.000807 | 0.002028 | 0.00126 | 0.001402 | 0.000265 | 0.001431 | 0.002253 | 0.008602 | 0.001164 | 0.002709 |
| 19.8888889 | 0.002038 | 0.003469 | 0.002841 | 0.035813 | 0.003697 | 0.00516 | 0.00599 | 0.002563 | 0.002494 | 0.001235 | 0.004062 | 0.000699 | 0.000973 | 0.001014 | 0.00087 | 0.001412 | 0.010069 | 0.00086 | 0.001935 |
| 19.9027778 | 0.001879 | 0.001656 | 0.005023 | 0.02134 | 0.001687 | 0.002282 | 0.00996 | 0.003182 | 0.002985 | 0.002779 | 0.006402 | 0.000187 | 0.000404 | 0.002644 | 0.000267 | 0.00065 | 0.011175 | 0.000496 | 0.000953 |
| 19.9166667 | 0.002844 | 0.001158 | 0.006167 | 0.009192 | 0.000493 | 0.002837 | 0.014999 | 0.004718 | 0.004504 | 0.003901 | 0.008999 | 0.000228 | 0.000333 | 0.003853 | 0.000205 | 0.00053 | 0.012245 | 0.000554 | 0.000305 |
| 19.9305556 | 0.003268 | 0.001208 | 0.005459 | 0.004647 | 0.000589 | 0.004251 | 0.01935 | 0.004812 | 0.004613 | 0.003023 | 0.010648 | 0.000628 | 0.000535 | 0.003795 | 0.000677 | 0.00082 | 0.015368 | 0.000842 | 0.000189 |
| 19.9444444 | 0.002685 | 0.002189 | 0.004743 | 0.003218 | 0.001168 | 0.003757 | 0.021116 | 0.003444 | 0.003327 | 0.002601 | 0.00984 | 0.000858 | 0.000618 | 0.003442 | 0.001045 | 0.000974 | 0.020939 | 0.001016 | 0.000263 |
| 19.9583333 | 0.002073 | 0.004196 | 0.003841 | 0.002953 | 0.002616 | 0.003185 | 0.018971 | 0.002279 | 0.00225 | 0.004232 | 0.006619 | 0.000616 | 0.000367 | 0.00357 | 0.000913 | 0.000747 | 0.025381 | 0.000945 | 0.000402 |
| 19.9722222 | 0.003043 | 0.006924 | 0.002312 | 0.005404 | 0.005738 | 0.005572 | 0.014327 | 0.003733 | 0.002954 | 0.00598 | 0.003945 | 0.00032 | 0.000118 | 0.003978 | 0.001157 | 0.000425 | 0.026537 | 0.000933 | 0.001239 |
| 19.9861111 | 0.005612 | 0.009516 | 0.001686 | 0.010563 | 0.009255 | 0.008689 | 0.010956 | 0.009907 | 0.005279 | 0.006808 | 0.005589 | 0.000234 | 0.000139 | 0.004213 | 0.002271 | 0.000255 | 0.026405 | 0.000909 | 0.002553 |
| 20 | 0.006715 | 0.010621 | 0.002241 | 0.017473 | 0.011479 | 0.007926 | 0.011755 | 0.018846 | 0.006986 | 0.006622 | 0.011689 | 0.000213 | 0.000472 | 0.003964 | 0.003167 | 0.000424 | 0.02605 | 0.000561 | 0.003423 |
| 20.0138889 | 0.004763 | 0.009751 | 0.002686 | 0.024662 | 0.012084 | 0.004538 | 0.016251 | 0.024196 | 0.007048 | 0.005642 | 0.018847 | 0.000391 | 0.000997 | 0.003014 | 0.003185 | 0.000972 | 0.02498 | 0.000274 | 0.003881 |
| 20.0277778 | 0.004229 | 0.006497 | 0.00211 | 0.026783 | 0.010775 | 0.003754 | 0.021481 | 0.021525 | 0.005527 | 0.005397 | 0.024762 | 0.000679 | 0.001062 | 0.001875 | 0.002508 | 0.001354 | 0.024322 | 0.000231 | 0.004595 |
| 20.0416667 | 0.009248 | 0.00348 | 0.001696 | 0.020837 | 0.007456 | 0.007045 | 0.025264 | 0.01353 | 0.003229 | 0.006868 | 0.029125 | 0.000543 | 0.000558 | 0.001696 | 0.0015 | 0.001111 | 0.027166 | 0.000164 | 0.006006 |
| 20.0555556 | 0.014805 | 0.003361 | 0.002548 | 0.013166 | 0.004242 | 0.009852 | 0.025011 | 0.006386 | 0.001633 | 0.009197 | 0.030886 | 0.00024 | 0.000211 | 0.002704 | 0.000837 | 0.00073 | 0.034453 | 0.000151 | 0.008152 |
| 20.0694444 | 0.012821 | 0.004824 | 0.004258 | 0.010403 | 0.005798 | 0.008143 | 0.016814 | 0.005183 | 0.003314 | 0.011 | 0.029666 | 0.000436 | 0.000441 | 0.004662 | 0.000673 | 0.000767 | 0.037649 | 0.000386 | 0.010368 |
| 20.0833333 | 0.008904 | 0.011298 | 0.011589 | 0.012171 | 0.015619 | 0.008908 | 0.011258 | 0.011568 | 0.012752 | 0.012684 | 0.027991 | 0.00123 | 0.001513 | 0.007715 | 0.000496 | 0.001031 | 0.025867 | 0.001214 | 0.012011 |
| 20.0972222 | 0.012971 | 0.02405 | 0.026209 | 0.013945 | 0.029704 | 0.018884 | 0.021776 | 0.020589 | 0.027952 | 0.015791 | 0.028609 | 0.002451 | 0.003128 | 0.00956 | 0.000214 | 0.000958 | 0.015542 | 0.002293 | 0.01356 |
| 20.1111111 | 0.01611 | 0.031781 | 0.036608 | 0.01167 | 0.03689 | 0.02749 | 0.033904 | 0.022562 | 0.037335 | 0.018263 | 0.028013 | 0.003326 | 0.003642 | 0.007352 | 0.000503 | 0.000724 | 0.028015 | 0.002374 | 0.015397 |
| 20.125 | 0.010573 | 0.02752 | 0.034051 | 0.008028 | 0.031523 | 0.025617 | 0.032107 | 0.015166 | 0.033871 | 0.014286 | 0.019019 | 0.002837 | 0.002439 | 0.005664 | 0.001752 | 0.000852 | 0.043446 | 0.001622 | 0.01595 |
| 20.1388889 | 0.003764 | 0.017531 | 0.024171 | 0.007991 | 0.01984 | 0.018251 | 0.022276 | 0.007052 | 0.023402 | 0.007611 | 0.010192 | 0.001528 | 0.001509 | 0.007903 | 0.00313 | 0.001523 | 0.041263 | 0.001579 | 0.013188 |
| 20.1527778 | 0.001103 | 0.008781 | 0.015097 | 0.009331 | 0.00934 | 0.011203 | 0.012852 | 0.003655 | 0.013849 | 0.006766 | 0.011507 | 0.001028 | 0.001919 | 0.008 | 0.00373 | 0.002852 | 0.03079 | 0.002401 | 0.008091 |
| 20.1666667 | 0.001529 | 0.003375 | 0.008587 | 0.008393 | 0.003289 | 0.005641 | 0.00603 | 0.002341 | 0.007289 | 0.007317 | 0.012274 | 0.001438 | 0.002462 | 0.004328 | 0.003714 | 0.004042 | 0.022326 | 0.00293 | 0.003849 |
| 20.1805556 | 0.002127 | 0.001405 | 0.004432 | 0.006524 | 0.001903 | 0.001922 | 0.002025 | 0.001262 | 0.003768 | 0.004586 | 0.007437 | 0.001639 | 0.002438 | 0.001773 | 0.003454 | 0.004297 | 0.016686 | 0.002668 | 0.001861 |
| 20.1944444 | 0.001906 | 0.001308 | 0.002643 | 0.00563 | 0.001983 | 0.000532 | 0.000788 | 0.001072 | 0.002736 | 0.001724 | 0.00315 | 0.001354 | 0.002041 | 0.001131 | 0.003104 | 0.003689 | 0.012309 | 0.002052 | 0.001178 |
| 20.2083333 | 0.001883 | 0.001657 | 0.001931 | 0.004985 | 0.002025 | 0.001181 | 0.000598 | 0.0012 | 0.002238 | 0.000612 | 0.001894 | 0.001075 | 0.001735 | 0.001088 | 0.002875 | 0.002605 | 0.008789 | 0.001695 | 0.000926 |
| 20.2222222 | 0.002892 | 0.002642 | 0.001354 | 0.003388 | 0.003053 | 0.003512 | 0.000737 | 0.001117 | 0.001508 | 0.000793 | 0.003074 | 0.001204 | 0.001972 | 0.001526 | 0.002861 | 0.001651 | 0.00627 | 0.001736 | 0.001078 |
| 20.2361111 | 0.004659 | 0.004241 | 0.001629 | 0.001769 | 0.004867 | 0.006452 | 0.002029 | 0.001791 | 0.001836 | 0.001406 | 0.004763 | 0.001727 | 0.002618 | 0.001791 | 0.003036 | 0.001233 | 0.004661 | 0.001989 | 0.001536 |
| 20.25 | 0.006348 | 0.005846 | 0.002913 | 0.001908 | 0.006545 | 0.008772 | 0.003888 | 0.003348 | 0.003364 | 0.002029 | 0.005816 | 0.002236 | 0.003149 | 0.002306 | 0.003238 | 0.00121 | 0.003786 | 0.002342 | 0.001867 |
| 20.2638889 | 0.007764 | 0.007313 | 0.004546 | 0.003786 | 0.007814 | 0.010715 | 0.005478 | 0.005082 | 0.004956 | 0.002339 | 0.005944 | 0.002353 | 0.003231 | 0.003477 | 0.003154 | 0.001195 | 0.003231 | 0.002574 | 0.001821 |
| 20.2777778 | 0.008959 | 0.008589 | 0.00628 | 0.006014 | 0.008738 | 0.0125 | 0.006729 | 0.006734 | 0.006188 | 0.002273 | 0.005586 | 0.002148 | 0.003039 | 0.003814 | 0.002745 | 0.001141 | 0.002548 | 0.002515 | 0.001645 |
| 20.2916667 | 0.009246 | 0.008864 | 0.007227 | 0.00706 | 0.008755 | 0.01273 | 0.006918 | 0.007327 | 0.006554 | 0.00212 | 0.005327 | 0.001962 | 0.002922 | 0.00292 | 0.002341 | 0.001262 | 0.001755 | 0.002376 | 0.001709 |
| 20.3055556 | 0.008626 | 0.00797 | 0.006885 | 0.006799 | 0.007864 | 0.011188 | 0.006094 | 0.006475 | 0.005967 | 0.002073 | 0.005191 | 0.001934 | 0.002873 | 0.003811 | 0.002183 | 0.001483 | 0.001108 | 0.002392 | 0.002016 |
| 20.3194444 | 0.008577 | 0.007279 | 0.006668 | 0.006751 | 0.007442 | 0.010322 | 0.006007 | 0.006041 | 0.00581 | 0.002109 | 0.00496 | 0.001892 | 0.002633 | 0.006792 | 0.002064 | 0.001549 | 0.000696 | 0.002462 | 0.00218 |
| 20.3333333 | 0.009405 | 0.007455 | 0.007385 | 0.007077 | 0.00793 | 0.011172 | 0.007103 | 0.007044 | 0.006577 | 0.002116 | 0.004629 | 0.001685 | 0.002169 | 0.006406 | 0.002048 | 0.001394 | 0.000442 | 0.002317 | 0.001944 |
| 20.3472222 | 0.00958 | 0.00758 | 0.007786 | 0.006393 | 0.008122 | 0.011844 | 0.007603 | 0.007975 | 0.006898 | 0.002081 | 0.004416 | 0.001479 | 0.001825 | 0.003396 | 0.001915 | 0.00116 | 0.000322 | 0.001928 | 0.001648 |
| 20.3611111 | 0.008571 | 0.007217 | 0.007143 | 0.004929 | 0.007698 | 0.011755 | 0.006915 | 0.008015 | 0.006283 | 0.001988 | 0.004334 | 0.001436 | 0.001821 | 0.003172 | 0.001972 | 0.001004 | 0.00036 | 0.001617 | 0.001703 |
| 20.375 | 0.007545 | 0.006974 | 0.00647 | 0.00452 | 0.007813 | 0.013136 | 0.006464 | 0.008275 | 0.005933 | 0.001841 | 0.004292 | 0.001544 | 0.00201 | 0.004976 | 0.002169 | 0.000976 | 0.000672 | 0.001646 | 0.002006 |
| 20.3888889 | 0.007292 | 0.007112 | 0.006453 | 0.00564 | 0.008773 | 0.016409 | 0.006885 | 0.00915 | 0.006376 | 0.001849 | 0.004313 | 0.001802 | 0.002197 | 0.007572 | 0.002359 | 0.001101 | 0.001383 | 0.001831 | 0.002233 |
| 20.4027778 | 0.007772 | 0.007584 | 0.006837 | 0.007198 | 0.00967 | 0.018769 | 0.00757 | 0.009904 | 0.007017 | 0.001925 | 0.004022 | 0.001933 | 0.002123 | 0.00963 | 0.002336 | 0.001233 | 0.002101 | 0.001801 | 0.002108 |
| 20.4166667 | 0.008595 | 0.00827 | 0.00727 | 0.008032 | 0.009849 | 0.018512 | 0.007956 | 0.010119 | 0.007335 | 0.001807 | 0.003184 | 0.001756 | 0.001762 | 0.007215 | 0.00205 | 0.001262 | 0.002366 | 0.001628 | 0.001692 |
| 20.4305556 | 0.0087 | 0.00833 | 0.00705 | 0.006993 | 0.008959 | 0.016055 | 0.007394 | 0.009446 | 0.006789 | 0.001632 | 0.002424 | 0.001638 | 0.001561 | 0.00503 | 0.001771 | 0.001321 | 0.002334 | 0.001622 | 0.001295 |
| 20.4444444 | 0.007695 | 0.007479 | 0.005931 | 0.004239 | 0.007305 | 0.012337 | 0.005891 | 0.008051 | 0.005392 | 0.001428 | 0.002072 | 0.001665 | 0.001568 | 0.007824 | 0.001595 | 0.001373 | 0.002122 | 0.001721 | 0.000905 |
| 20.4583333 | 0.006981 | 0.00705 | 0.005174 | 0.002031 | 0.006361 | 0.009103 | 0.004851 | 0.007341 | 0.004487 | 0.001038 | 0.001849 | 0.001541 | 0.001369 | 0.007679 | 0.001478 | 0.001283 | 0.001621 | 0.001717 | 0.000479 |

| | | | | | | | | | | | | | | | | | | | |
|---|---|---|---|---|---|---|---|---|---|---|---|---|---|---|---|---|---|---|---|
| 20.4722222 | 0.007658 | 0.007946 | 0.005969 | 0.001976 | 0.007129 | 0.00802 | 0.005182 | 0.008219 | 0.005193 | 0.000603 | 0.001873 | 0.001328 | 0.000978 | 0.004256 | 0.001479 | 0.001179 | 0.001063 | 0.001615 | 0.000236 |
| 20.4861111 | 0.008584 | 0.009031 | 0.007452 | 0.002889 | 0.008448 | 0.008379 | 0.005576 | 0.009489 | 0.006529 | 0.000331 | 0.002135 | 0.001192 | 0.000696 | 0.003734 | 0.001463 | 0.001167 | 0.00062 | 0.001485 | 0.000475 |
| 20.5 | 0.007972 | 0.008805 | 0.007734 | 0.002836 | 0.008176 | 0.008049 | 0.004356 | 0.009451 | 0.006513 | 0.000476 | 0.001743 | 0.000984 | 0.000508 | 0.003908 | 0.001217 | 0.001066 | 0.000438 | 0.001326 | 0.001268 |
| 20.5138889 | 0.005995 | 0.007582 | 0.006674 | 0.00203 | 0.005431 | 0.006654 | 0.002168 | 0.008258 | 0.005003 | 0.001244 | 0.00152 | 0.000669 | 0.000363 | 0.004634 | 0.00093 | 0.000827 | 0.000789 | 0.001201 | 0.002088 |
| 20.5277778 | 0.004109 | 0.006549 | 0.005735 | 0.001515 | 0.00248 | 0.005315 | 0.000898 | 0.007166 | 0.003578 | 0.002299 | 0.003661 | 0.00058 | 0.000557 | 0.005215 | 0.001031 | 0.000776 | 0.001387 | 0.001266 | 0.002528 |
| 20.5416667 | 0.00284 | 0.005825 | 0.005438 | 0.001011 | 0.001878 | 0.004398 | 0.001217 | 0.006452 | 0.002887 | 0.00345 | 0.007727 | 0.000778 | 0.001067 | 0.00418 | 0.001435 | 0.001023 | 0.001999 | 0.001408 | 0.003096 |
| 20.5555556 | 0.001638 | 0.005006 | 0.005156 | 0.000821 | 0.004368 | 0.003249 | 0.002493 | 0.005659 | 0.002486 | 0.005238 | 0.01253 | 0.000913 | 0.001222 | 0.007992 | 0.001581 | 0.001236 | 0.003134 | 0.001314 | 0.004242 |
| 20.5694444 | 0.001565 | 0.004521 | 0.005185 | 0.002876 | 0.00956 | 0.002778 | 0.003697 | 0.005047 | 0.00265 | 0.008511 | 0.020019 | 0.001161 | 0.00096 | 0.015987 | 0.00147 | 0.001438 | 0.006305 | 0.000833 | 0.006097 |
| 20.5833333 | 0.005044 | 0.005794 | 0.007835 | 0.007516 | 0.015155 | 0.006442 | 0.004662 | 0.00614 | 0.004841 | 0.014434 | 0.034587 | 0.002128 | 0.001304 | 0.017525 | 0.001768 | 0.002362 | 0.015744 | 0.000691 | 0.010098 |
| 20.5972222 | 0.01173 | 0.009233 | 0.013904 | 0.009414 | 0.020563 | 0.013321 | 0.006896 | 0.010123 | 0.009553 | 0.021076 | 0.048696 | 0.003502 | 0.002718 | 0.012711 | 0.002567 | 0.004427 | 0.03437 | 0.00142 | 0.016576 |
| 20.6111111 | 0.017884 | 0.011903 | 0.018622 | 0.009326 | 0.024119 | 0.016866 | 0.011159 | 0.014204 | 0.01377 | 0.021891 | 0.043794 | 0.004211 | 0.004222 | 0.008765 | 0.002871 | 0.006116 | 0.057488 | 0.001545 | 0.021764 |
| 20.625 | 0.021075 | 0.009232 | 0.016664 | 0.01244 | 0.021708 | 0.016139 | 0.014442 | 0.012229 | 0.013912 | 0.015923 | 0.02361 | 0.003816 | 0.004363 | 0.007576 | 0.002023 | 0.00518 | 0.077794 | 0.000932 | 0.023247 |
| 20.6388889 | 0.022477 | 0.005245 | 0.009402 | 0.012438 | 0.014222 | 0.015791 | 0.013431 | 0.008232 | 0.010527 | 0.01052 | 0.013481 | 0.002612 | 0.002763 | 0.008791 | 0.001334 | 0.002468 | 0.090881 | 0.000853 | 0.023275 |
| 20.6527778 | 0.023206 | 0.007625 | 0.003594 | 0.007127 | 0.006443 | 0.016029 | 0.009957 | 0.012705 | 0.006155 | 0.009372 | 0.016284 | 0.001366 | 0.001094 | 0.009896 | 0.001715 | 0.00073 | 0.086628 | 0.00068 | 0.023276 |
| 20.6666667 | 0.023433 | 0.012425 | 0.003044 | 0.002886 | 0.001979 | 0.014281 | 0.00928 | 0.021559 | 0.003261 | 0.010464 | 0.016453 | 0.000723 | 0.000919 | 0.007724 | 0.002344 | 0.000472 | 0.061705 | 0.000247 | 0.022736 |
| 20.6805556 | 0.024272 | 0.014557 | 0.004666 | 0.001419 | 0.0009 | 0.011122 | 0.013424 | 0.025795 | 0.004065 | 0.012571 | 0.021483 | 0.000441 | 0.001605 | 0.003621 | 0.003251 | 0.000736 | 0.034711 | 0.000183 | 0.021554 |
| 20.6944444 | 0.026426 | 0.015103 | 0.004934 | 0.0017 | 0.000798 | 0.008764 | 0.0173 | 0.024756 | 0.007303 | 0.0159 | 0.034215 | 0.00044 | 0.001884 | 0.000992 | 0.004248 | 0.000838 | 0.017445 | 0.000536 | 0.019507 |
| 20.7083333 | 0.027783 | 0.01482 | 0.003953 | 0.003261 | 0.000836 | 0.008443 | 0.014362 | 0.020798 | 0.009474 | 0.019754 | 0.033706 | 0.000827 | 0.001308 | 0.001528 | 0.004238 | 0.000519 | 0.008759 | 0.001012 | 0.016523 |
| 20.7222222 | 0.025727 | 0.013545 | 0.003023 | 0.003821 | 0.00093 | 0.010159 | 0.008773 | 0.016334 | 0.008814 | 0.021241 | 0.018402 | 0.000918 | 0.000586 | 0.004338 | 0.003162 | 0.00033 | 0.011085 | 0.000902 | 0.012643 |
| 20.7361111 | 0.020486 | 0.011815 | 0.002307 | 0.002964 | 0.001078 | 0.012573 | 0.010559 | 0.012969 | 0.006267 | 0.017959 | 0.008729 | 0.00055 | 0.000578 | 0.004881 | 0.001669 | 0.000552 | 0.020947 | 0.000448 | 0.007838 |
| 20.75 | 0.014073 | 0.010489 | 0.001774 | 0.003776 | 0.002038 | 0.013917 | 0.019528 | 0.010892 | 0.003824 | 0.011817 | 0.016892 | 0.000385 | 0.00097 | 0.002294 | 0.000656 | 0.000739 | 0.027567 | 0.000431 | 0.003961 |
| 20.7638889 | 0.007665 | 0.008424 | 0.002881 | 0.00488 | 0.002616 | 0.012658 | 0.029691 | 0.008795 | 0.002594 | 0.005907 | 0.034874 | 0.000418 | 0.001147 | 0.001593 | 0.000651 | 0.000617 | 0.027099 | 0.000486 | 0.003718 |
| 20.7777778 | 0.003048 | 0.004735 | 0.005412 | 0.003652 | 0.001826 | 0.009195 | 0.035939 | 0.005875 | 0.003484 | 0.002186 | 0.044603 | 0.0005 | 0.000978 | 0.002353 | 0.001374 | 0.000495 | 0.022716 | 0.000438 | 0.005821 |
| 20.7916667 | 0.002147 | 0.001823 | 0.006327 | 0.001758 | 0.001249 | 0.006542 | 0.032386 | 0.003179 | 0.005124 | 0.001046 | 0.035551 | 0.000766 | 0.00056 | 0.00197 | 0.002281 | 0.000627 | 0.015977 | 0.000597 | 0.006642 |
| 20.8055556 | 0.003167 | 0.001139 | 0.004277 | 0.001125 | 0.001061 | 0.006834 | 0.019747 | 0.001872 | 0.004123 | 0.001584 | 0.018113 | 0.000815 | 0.000206 | 0.001704 | 0.002283 | 0.000564 | 0.008561 | 0.000597 | 0.0049 |
| 20.8194444 | 0.00388 | 0.000837 | 0.002325 | 0.001659 | 0.000705 | 0.008775 | 0.013371 | 0.001733 | 0.001963 | 0.003938 | 0.012736 | 0.000545 | 8.76E-05 | 0.001994 | 0.001277 | 0.000458 | 0.006124 | 0.000404 | 0.003193 |
| 20.8333333 | 0.004837 | 0.000475 | 0.002183 | 0.002468 | 0.000603 | 0.008596 | 0.021365 | 0.001693 | 0.001485 | 0.006404 | 0.020794 | 0.000489 | 0.000192 | 0.003794 | 0.000596 | 0.00089 | 0.010116 | 0.000416 | 0.005094 |
| 20.8472222 | 0.006712 | 0.000804 | 0.001997 | 0.003024 | 0.000384 | 0.005013 | 0.02827 | 0.001828 | 0.001358 | 0.006629 | 0.026308 | 0.00058 | 0.00048 | 0.005603 | 0.000762 | 0.001274 | 0.013206 | 0.000413 | 0.008445 |
| 20.8611111 | 0.008574 | 0.001179 | 0.002133 | 0.003354 | 0.000435 | 0.001686 | 0.022941 | 0.003192 | 0.001397 | 0.004385 | 0.019048 | 0.000533 | 0.000509 | 0.004201 | 0.001356 | 0.000948 | 0.010868 | 0.000346 | 0.008174 |
| 20.875 | 0.009264 | 0.001015 | 0.002853 | 0.00312 | 0.001374 | 0.001787 | 0.014655 | 0.005007 | 0.001922 | 0.002626 | 0.012542 | 0.000597 | 0.000291 | 0.001627 | 0.00161 | 0.000507 | 0.005812 | 0.000517 | 0.005048 |
| 20.8888889 | 0.008928 | 0.001068 | 0.003573 | 0.002069 | 0.002047 | 0.003399 | 0.01852 | 0.004304 | 0.00227 | 0.005079 | 0.025081 | 0.000683 | 0.00032 | 0.000513 | 0.001124 | 0.000498 | 0.003676 | 0.0006 | 0.004702 |
| 20.9027778 | 0.00831 | 0.001605 | 0.004612 | 0.000956 | 0.001546 | 0.003134 | 0.031224 | 0.003015 | 0.002611 | 0.00934 | 0.042262 | 0.000542 | 0.000416 | 0.000312 | 0.000782 | 0.00044 | 0.007102 | 0.000524 | 0.008912 |
| 20.9166667 | 0.007765 | 0.00139 | 0.004755 | 0.001064 | 0.001032 | 0.001411 | 0.034004 | 0.0053 | 0.002135 | 0.010793 | 0.039761 | 0.000577 | 0.000333 | 0.000324 | 0.00107 | 0.000279 | 0.013295 | 0.000815 | 0.013991 |
| 20.9305556 | 0.007826 | 0.000664 | 0.003246 | 0.00299 | 0.001708 | 0.000611 | 0.023164 | 0.007835 | 0.001306 | 0.009043 | 0.022526 | 0.000916 | 0.00019 | 0.000628 | 0.001173 | 0.000274 | 0.019217 | 0.001144 | 0.016888 |
| 20.9444444 | 0.008804 | 0.000428 | 0.001504 | 0.004994 | 0.00306 | 0.001025 | 0.011266 | 0.0075 | 0.001607 | 0.006475 | 0.01223 | 0.001134 | 0.000174 | 0.000972 | 0.000798 | 0.000281 | 0.021659 | 0.000867 | 0.017452 |
| 20.9583333 | 0.010418 | 0.000928 | 0.000679 | 0.004588 | 0.003602 | 0.001576 | 0.005622 | 0.005491 | 0.00188 | 0.005639 | 0.009614 | 0.001013 | 0.000612 | 0.000989 | 0.000747 | 0.000451 | 0.018328 | 0.000358 | 0.017462 |
| 20.9722222 | 0.012947 | 0.002763 | 0.000656 | 0.003334 | 0.002408 | 0.003298 | 0.004429 | 0.003311 | 0.002428 | 0.007839 | 0.010157 | 0.000669 | 0.001633 | 0.000977 | 0.001334 | 0.000704 | 0.010908 | 0.000199 | 0.018521 |
| 20.9861111 | 0.01668 | 0.00534 | 0.001518 | 0.005727 | 0.00112 | 0.007582 | 0.005891 | 0.003229 | 0.004812 | 0.012465 | 0.022253 | 0.000497 | 0.002778 | 0.00122 | 0.001831 | 0.000749 | 0.004138 | 0.000338 | 0.020292 |
| 21 | 0.020597 | 0.007385 | 0.003038 | 0.010878 | 0.001314 | 0.012066 | 0.009386 | 0.004567 | 0.007273 | 0.017026 | 0.037142 | 0.000478 | 0.003334 | 0.001373 | 0.002185 | 0.000777 | 0.002248 | 0.000509 | 0.021553 |
| 21.0138889 | 0.022753 | 0.008187 | 0.003842 | 0.012827 | 0.001533 | 0.013596 | 0.012775 | 0.003802 | 0.007651 | 0.018803 | 0.039017 | 0.000279 | 0.002773 | 0.001352 | 0.002993 | 0.000978 | 0.005345 | 0.000639 | 0.021319 |
| 21.0277778 | 0.0223 | 0.00756 | 0.00299 | 0.01 | 0.001121 | 0.012393 | 0.013937 | 0.002811 | 0.005835 | 0.016657 | 0.027756 | 0.000167 | 0.001446 | 0.001528 | 0.003995 | 0.001084 | 0.010502 | 0.000882 | 0.019237 |
| 21.0416667 | 0.020301 | 0.006307 | 0.001524 | 0.006875 | 0.001331 | 0.011172 | 0.012552 | 0.005778 | 0.003264 | 0.011409 | 0.013786 | 0.000458 | 0.000493 | 0.001789 | 0.004472 | 0.000793 | 0.014363 | 0.001412 | 0.016358 |
| 21.0555556 | 0.017366 | 0.004989 | 0.001276 | 0.006038 | 0.002415 | 0.011507 | 0.008882 | 0.010341 | 0.001604 | 0.006027 | 0.005591 | 0.000863 | 0.000356 | 0.001805 | 0.004254 | 0.000402 | 0.01722 | 0.001889 | 0.014386 |
| 21.0694444 | 0.011754 | 0.003797 | 0.003299 | 0.006395 | 0.004822 | 0.013447 | 0.006442 | 0.010225 | 0.002791 | 0.003241 | 0.002843 | 0.001135 | 0.000841 | 0.001737 | 0.003452 | 0.000348 | 0.02136 | 0.001596 | 0.013561 |
| 21.0833333 | 0.007119 | 0.005782 | 0.008404 | 0.012536 | 0.010521 | 0.018424 | 0.012251 | 0.007953 | 0.008676 | 0.003159 | 0.002144 | 0.001658 | 0.002012 | 0.001996 | 0.002047 | 0.000387 | 0.02651 | 0.001094 | 0.013163 |
| 21.0972222 | 0.009417 | 0.011976 | 0.015499 | 0.025119 | 0.019735 | 0.023828 | 0.024214 | 0.011496 | 0.016812 | 0.004183 | 0.001763 | 0.002633 | 0.00344 | 0.002637 | 0.001236 | 0.000697 | 0.032356 | 0.001433 | 0.012182 |
| 21.1111111 | 0.012376 | 0.015753 | 0.020125 | 0.029814 | 0.025681 | 0.019304 | 0.029453 | 0.015851 | 0.021607 | 0.004662 | 0.001191 | 0.003378 | 0.003826 | 0.002766 | 0.001762 | 0.00158 | 0.035197 | 0.001788 | 0.009603 |
| 21.125 | 0.009693 | 0.013273 | 0.018443 | 0.02071 | 0.021573 | 0.00935 | 0.022514 | 0.012641 | 0.019755 | 0.004398 | 0.000453 | 0.003058 | 0.002585 | 0.002019 | 0.001738 | 0.0018 | 0.027229 | 0.001551 | 0.006296 |
| 21.1388889 | 0.005287 | 0.007676 | 0.011858 | 0.009273 | 0.012066 | 0.006977 | 0.0113 | 0.006656 | 0.012898 | 0.003931 | 9.05E-05 | 0.00186 | 0.001001 | 0.001755 | 0.001261 | 0.000995 | 0.012882 | 0.001563 | 0.003848 |
| 21.1527778 | 0.002596 | 0.003 | 0.005384 | 0.003784 | 0.004814 | 0.006393 | 0.003693 | 0.00534 | 0.006209 | 0.003022 | 8.50E-05 | 0.000812 | 0.000279 | 0.002111 | 0.001742 | 0.000416 | 0.005031 | 0.001863 | 0.002521 |
| 21.1666667 | 0.001673 | 0.001023 | 0.002324 | 0.002339 | 0.001809 | 0.003765 | 0.000951 | 0.00519 | 0.003106 | 0.001597 | 0.000176 | 0.000334 | 8.97E-05 | 0.001907 | 0.00212 | 0.00033 | 0.003356 | 0.001927 | 0.002019 |
| 21.1805556 | 0.002006 | 0.001068 | 0.002551 | 0.001389 | 0.001859 | 0.004479 | 0.0014 | 0.003141 | 0.003196 | 0.000549 | 0.000247 | 0.000262 | 0.000189 | 0.001552 | 0.001863 | 0.000478 | 0.002429 | 0.001904 | 0.002142 |
| 21.1944444 | 0.002618 | 0.0021 | 0.003978 | 0.000541 | 0.002722 | 0.007111 | 0.002776 | 0.002626 | 0.004293 | 0.000354 | 0.000379 | 0.000566 | 0.00055 | 0.001786 | 0.001602 | 0.000791 | 0.003229 | 0.002024 | 0.002577 |
| 21.2083333 | 0.002707 | 0.002581 | 0.004677 | 0.000399 | 0.003039 | 0.006858 | 0.003648 | 0.002727 | 0.004682 | 0.000625 | 0.000689 | 0.000948 | 0.000729 | 0.00202 | 0.001528 | 0.000957 | 0.005126 | 0.002068 | 0.002986 |
| 21.2222222 | 0.002403 | 0.002005 | 0.00427 | 0.000401 | 0.002782 | 0.003523 | 0.003878 | 0.001738 | 0.00422 | 0.000992 | 0.000817 | 0.000983 | 0.000637 | 0.001602 | 0.001561 | 0.000818 | 0.005622 | 0.001749 | 0.0032 |
| 21.2361111 | 0.002003 | 0.001448 | 0.003524 | 0.000773 | 0.002245 | 0.001616 | 0.003552 | 0.001241 | 0.003616 | 0.001273 | 0.000637 | 0.000662 | 0.000405 | 0.00096 | 0.001599 | 0.000557 | 0.004626 | 0.001231 | 0.00323 |
| 21.25 | 0.002104 | 0.001895 | 0.003634 | 0.001957 | 0.001868 | 0.004433 | 0.003103 | 0.002506 | 0.003811 | 0.001712 | 0.000594 | 0.000426 | 0.00013 | 0.000957 | 0.001805 | 0.000513 | 0.003324 | 0.001081 | 0.003348 |
| 21.2638889 | 0.003259 | 0.003404 | 0.004893 | 0.002114 | 0.002236 | 0.009238 | 0.00349 | 0.004823 | 0.005021 | 0.002582 | 0.000891 | 0.00065 | 0.000246 | 0.001516 | 0.002234 | 0.000842 | 0.002668 | 0.001456 | 0.003723 |
| 21.2777778 | 0.004796 | 0.004711 | 0.00596 | 0.001854 | 0.003444 | 0.009472 | 0.005078 | 0.00574 | 0.00617 | 0.003396 | 0.001159 | 0.001052 | 0.000876 | 0.001858 | 0.002451 | 0.001262 | 0.002578 | 0.001801 | 0.003982 |
| 21.2916667 | 0.005568 | 0.004816 | 0.005815 | 0.003711 | 0.004737 | 0.005101 | 0.007138 | 0.004346 | 0.006341 | 0.00349 | 0.001084 | 0.00117 | 0.001352 | 0.001823 | 0.002249 | 0.001428 | 0.002496 | 0.001772 | 0.003765 |
| 21.3055556 | 0.00482 | 0.003865 | 0.004792 | 0.005392 | 0.004521 | 0.002008 | 0.007896 | 0.002444 | 0.005298 | 0.002799 | 0.000685 | 0.00099 | 0.001114 | 0.001469 | 0.001783 | 0.001247 | 0.002294 | 0.001521 | 0.003151 |
| 21.3194444 | 0.002691 | 0.002301 | 0.0033 | 0.004239 | 0.002731 | 0.002563 | 0.006297 | 0.001209 | 0.003067 | 0.001787 | 0.000227 | 0.000844 | 0.000555 | 0.000809 | 0.001088 | 0.000951 | 0.002243 | 0.001225 | 0.002563 |
| 21.3333333 | 0.00082 | 0.001034 | 0.001709 | 0.00269 | 0.001443 | 0.003848 | 0.00418 | 0.000706 | 0.001011 | 0.000961 | 0.000187 | 0.001058 | 0.000517 | 0.000817 | 0.000407 | 0.001011 | 0.002489 | 0.001141 | 0.002515 |
| 21.3472222 | 0.000382 | 0.000702 | 0.000854 | 0.003284 | 0.00109 | 0.002886 | 0.00373 | 0.000913 | 0.00067 | 0.000623 | 0.00041 | 0.00139 | 0.000954 | 0.001773 | 0.000185 | 0.001297 | 0.002725 | 0.001288 | 0.002786 |
| 21.3611111 | 0.000604 | 0.000542 | 0.000939 | 0.004486 | 0.0006 | 0.001493 | 0.004292 | 0.001019 | 0.001508 | 0.001132 | 0.000556 | 0.001354 | 0.001142 | 0.002125 | 0.000509 | 0.001172 | 0.002571 | 0.001265 | 0.002557 |
| 21.375 | 0.000637 | 0.000409 | 0.001169 | 0.004527 | 0.000299 | 0.001398 | 0.00433 | 0.001032 | 0.002007 | 0.002489 | 0.000996 | 0.00116 | 0.001044 | 0.001336 | 0.001088 | 0.000699 | 0.0021 | 0.001039 | 0.001726 |
| 21.3888889 | 0.000889 | 0.001102 | 0.001066 | 0.004137 | 0.00095 | 0.001269 | 0.003511 | 0.001924 | 0.001469 | 0.004007 | 0.001611 | 0.001171 | 0.00096 | 0.000761 | 0.001887 | 0.00041 | 0.001549 | 0.000914 | 0.000833 |
| 21.4027778 | 0.00156 | 0.002379 | 0.001161 | 0.004221 | 0.002104 | 0.000997 | 0.002377 | 0.003448 | 0.00073 | 0.005296 | 0.001809 | 0.001283 | 0.000826 | 0.000669 | 0.003393 | 0.000459 | 0.001069 | 0.000928 | 0.000309 |
| 21.4166667 | 0.001862 | 0.003005 | 0.002205 | 0.00481 | 0.002575 | 0.000933 | 0.001823 | 0.004292 | 0.000906 | 0.006585 | 0.001545 | 0.001299 | 0.000662 | 0.000667 | 0.00514 | 0.000634 | 0.000798 | 0.000893 | 0.00039 |
| 21.4305556 | 0.001662 | 0.002478 | 0.003979 | 0.005175 | 0.002167 | 0.000621 | 0.00192 | 0.003916 | 0.001815 | 0.008202 | 0.001016 | 0.000996 | 0.000468 | 0.000947 | 0.005356 | 0.000547 | 0.000572 | 0.000611 | 0.001032 |
| 21.4444444 | 0.001189 | 0.001464 | 0.005994 | 0.005209 | 0.001556 | 0.000718 | 0.0023 | 0.00282 | 0.002851 | 0.009889 | 0.0012 | 0.000446 | 0.000242 | 0.001094 | 0.00386 | 0.000262 | 0.000332 | 0.000296 | 0.00187 |
| 21.4583333 | 0.000831 | 0.00073 | 0.008258 | 0.00651 | 0.000888 | 0.001966 | 0.003496 | 0.001487 | 0.004209 | 0.01089 | 0.00238 | 0.000232 | 0.000174 | 0.001667 | 0.002257 | 0.00017 | 0.000267 | 0.000208 | 0.00229 |
| 21.4722222 | 0.001197 | 0.000658 | 0.010117 | 0.009587 | 0.000401 | 0.003678 | 0.005911 | 0.000695 | 0.006035 | 0.011075 | 0.002503 | 0.000572 | 0.000321 | 0.00252 | 0.001135 | 0.000279 | 0.000255 | 0.000261 | 0.002119 |
| 21.4861111 | 0.001703 | 0.000835 | 0.010364 | 0.012829 | 0.000294 | 0.004186 | 0.009202 | 0.000711 | 0.007312 | 0.010819 | 0.002312 | 0.000986 | 0.000488 | 0.002382 | 0.000471 | 0.00049 | 0.000254 | 0.000475 | 0.002003 |
| 21.5 | 0.002168 | 0.000656 | 0.008867 | 0.016047 | 0.00022 | 0.003151 | 0.012668 | 0.000778 | 0.007583 | 0.010263 | 0.003913 | 0.001099 | 0.000371 | 0.002502 | 0.000362 | 0.000651 | 0.000266 | 0.000625 | 0.002481 |
| 21.5138889 | 0.003255 | 0.000373 | 0.006749 | 0.019244 | 0.000247 | 0.002467 | 0.015308 | 0.001092 | 0.007688 | 0.009732 | 0.005095 | 0.000944 | 0.000177 | 0.003389 | 0.000597 | 0.00077 | 0.000184 | 0.000531 | 0.003341 |
| 21.5277778 | 0.004428 | 0.000319 | 0.004822 | 0.01927 | 0.000582 | 0.002086 | 0.01659 | 0.0022 | 0.007232 | 0.009979 | 0.00445 | 0.000929 | 0.000397 | 0.003911 | 0.000774 | 0.001151 | 0.000132 | 0.000632 | 0.003868 |
| 21.5416667 | 0.004673 | 0.000551 | 0.002746 | 0.014614 | 0.001555 | 0.001648 | 0.015542 | 0.003604 | 0.005147 | 0.010876 | 0.00471 | 0.001999 | 0.001002 | 0.004134 | 0.000824 | 0.001959 | 0.000261 | 0.001425 | 0.003465 |
| 21.5555556 | 0.004114 | 0.001016 | 0.000994 | 0.0081 | 0.003145 | 0.003956 | 0.011581 | 0.005388 | 0.002566 | 0.010997 | 0.00996 | 0.003449 | 0.001512 | 0.003275 | 0.001371 | 0.002813 | 0.001242 | 0.002614 | 0.003339 |
| 21.5694444 | 0.00405 | 0.001151 | 0.000805 | 0.004629 | 0.004842 | 0.011829 | 0.007944 | 0.00821 | 0.001654 | 0.009854 | 0.023037 | 0.002947 | 0.001679 | 0.001611 | 0.002278 | 0.003151 | 0.006891 | 0.002849 | 0.008875 |
| 21.5833333 | 0.00623 | 0.00074 | 0.003059 | 0.009333 | 0.008846 | 0.022109 | 0.007673 | 0.010024 | 0.001909 | 0.008652 | 0.047515 | 0.001953 | 0.002135 | 0.002189 | 0.002677 | 0.003309 | 0.025119 | 0.001865 | 0.027737 |
| 21.5972222 | 0.011165 | 0.00046 | 0.008804 | 0.020393 | 0.016341 | 0.023881 | 0.00792 | 0.007381 | 0.003981 | 0.007742 | 0.070798 | 0.003078 | 0.003426 | 0.005312 | 0.002146 | 0.003797 | 0.052695 | 0.001145 | 0.053421 |
| 21.6111111 | 0.014822 | 0.000664 | 0.015012 | 0.028092 | 0.022587 | 0.015402 | 0.006286 | 0.003778 | 0.010053 | 0.005387 | 0.069044 | 0.004594 | 0.004455 | 0.007842 | 0.001425 | 0.003778 | 0.06423 | 0.000966 | 0.061996 |
| 21.625 | 0.013333 | 0.001006 | 0.016551 | 0.026137 | 0.022782 | 0.00935 | 0.006171 | 0.003441 | 0.016069 | 0.002494 | 0.047592 | 0.004516 | 0.003813 | 0.007193 | 0.001659 | 0.002503 | 0.04888 | 0.000932 | 0.045872 |
| 21.6388889 | 0.009842 | 0.002254 | 0.014113 | 0.017757 | 0.017099 | 0.009083 | 0.008006 | 0.005571 | 0.017883 | 0.001955 | 0.023994 | 0.002934 | 0.002177 | 0.00423 | 0.002286 | 0.001574 | 0.027336 | 0.000878 | 0.023279 |
| 21.6527778 | 0.008398 | 0.004992 | 0.011467 | 0.010696 | 0.0097 | 0.007542 | 0.00739 | 0.01097 | 0.016529 | 0.002426 | 0.011684 | 0.001172 | 0.001207 | 0.002192 | 0.002277 | 0.002132 | 0.012685 | 0.000603 | 0.009021 |

| | | | | | | | | | | | | | | | | | | | |
|---|---|---|---|---|---|---|---|---|---|---|---|---|---|---|---|---|---|---|---|
| 21.6666667 | 0.010097 | 0.007629 | 0.009441 | 0.008124 | 0.004564 | 0.003934 | 0.006225 | 0.017535 | 0.01261 | 0.00197 | 0.020573 | 0.000407 | 0.001031 | 0.001548 | 0.00207 | 0.002689 | 0.007703 | 0.000539 | 0.004452 |
| 21.6805556 | 0.014702 | 0.008973 | 0.006122 | 0.006509 | 0.002142 | 0.00234 | 0.011953 | 0.022548 | 0.006688 | 0.002383 | 0.03625 | 0.00033 | 0.000814 | 0.001823 | 0.002015 | 0.002533 | 0.012943 | 0.000469 | 0.005475 |
| 21.6944444 | 0.018604 | 0.008467 | 0.003834 | 0.003756 | 0.001304 | 0.00495 | 0.024909 | 0.025252 | 0.003115 | 0.00521 | 0.033792 | 0.000412 | 0.000499 | 0.002759 | 0.00169 | 0.00221 | 0.016373 | 0.000351 | 0.007798 |
| 21.7083333 | 0.017237 | 0.005902 | 0.006713 | 0.001942 | 0.001241 | 0.01041 | 0.037209 | 0.023563 | 0.004708 | 0.008452 | 0.024126 | 0.00057 | 0.000382 | 0.002809 | 0.001545 | 0.00199 | 0.01164 | 0.000421 | 0.009613 |
| 21.7222222 | 0.010931 | 0.002817 | 0.010275 | 0.001661 | 0.001105 | 0.016932 | 0.040999 | 0.018118 | 0.007548 | 0.010068 | 0.033034 | 0.000943 | 0.000564 | 0.002156 | 0.002364 | 0.001958 | 0.009282 | 0.000469 | 0.007839 |
| 21.7361111 | 0.005774 | 0.001242 | 0.007992 | 0.001784 | 0.000793 | 0.022547 | 0.031727 | 0.011956 | 0.007643 | 0.010223 | 0.041126 | 0.00123 | 0.000976 | 0.00184 | 0.003306 | 0.001846 | 0.016227 | 0.000824 | 0.004677 |
| 21.75 | 0.007437 | 0.001227 | 0.004857 | 0.001536 | 0.000926 | 0.023811 | 0.019789 | 0.006804 | 0.005092 | 0.009485 | 0.03164 | 0.001132 | 0.001108 | 0.001767 | 0.002622 | 0.001355 | 0.024082 | 0.001386 | 0.0051 |
| 21.7638889 | 0.013755 | 0.001496 | 0.007937 | 0.001483 | 0.001052 | 0.019264 | 0.025455 | 0.003175 | 0.004154 | 0.008047 | 0.019537 | 0.001201 | 0.000849 | 0.001736 | 0.001652 | 0.000976 | 0.024316 | 0.001559 | 0.004943 |
| 21.7777778 | 0.018005 | 0.001895 | 0.012117 | 0.003347 | 0.000898 | 0.011306 | 0.043408 | 0.002111 | 0.006956 | 0.005742 | 0.011461 | 0.00196 | 0.000816 | 0.002331 | 0.002607 | 0.001298 | 0.016685 | 0.001327 | 0.004813 |
| 21.7916667 | 0.01666 | 0.002165 | 0.011865 | 0.006071 | 0.001242 | 0.004806 | 0.04994 | 0.002924 | 0.008431 | 0.002997 | 0.01201 | 0.002323 | 0.000921 | 0.002755 | 0.00367 | 0.001404 | 0.008046 | 0.001216 | 0.007504 |
| 21.8055556 | 0.010438 | 0.002158 | 0.00773 | 0.006095 | 0.001432 | 0.002639 | 0.0385 | 0.002728 | 0.005773 | 0.001644 | 0.022515 | 0.001443 | 0.000639 | 0.003342 | 0.002691 | 0.001111 | 0.006141 | 0.001522 | 0.009086 |
| 21.8194444 | 0.006717 | 0.00353 | 0.003915 | 0.003506 | 0.001433 | 0.002791 | 0.020085 | 0.001758 | 0.002215 | 0.002933 | 0.028611 | 0.000561 | 0.000239 | 0.004961 | 0.001534 | 0.001577 | 0.009104 | 0.001946 | 0.008588 |
| 21.8333333 | 0.009497 | 0.005914 | 0.004495 | 0.001459 | 0.001947 | 0.00297 | 0.012071 | 0.001594 | 0.00074 | 0.005646 | 0.021846 | 0.000312 | 0.000135 | 0.00452 | 0.002444 | 0.002198 | 0.012279 | 0.002039 | 0.007058 |
| 21.8472222 | 0.010163 | 0.006416 | 0.006294 | 0.000973 | 0.001955 | 0.00296 | 0.019661 | 0.002556 | 0.001056 | 0.007319 | 0.011725 | 0.000233 | 0.000168 | 0.00234 | 0.003713 | 0.001827 | 0.016183 | 0.001751 | 0.00493 |
| 21.8611111 | 0.006556 | 0.00472 | 0.005881 | 0.001494 | 0.001322 | 0.002266 | 0.02703 | 0.00596 | 0.001927 | 0.006396 | 0.010727 | 0.000187 | 0.000187 | 0.002516 | 0.003393 | 0.000986 | 0.022014 | 0.001528 | 0.006607 |
| 21.875 | 0.006779 | 0.002548 | 0.004175 | 0.001841 | 0.000767 | 0.001153 | 0.023167 | 0.011125 | 0.003212 | 0.003522 | 0.016823 | 0.000292 | 0.000223 | 0.005316 | 0.002012 | 0.000794 | 0.02694 | 0.001686 | 0.015123 |
| 21.8888889 | 0.012085 | 0.001761 | 0.004641 | 0.001652 | 0.000416 | 0.000768 | 0.012323 | 0.015726 | 0.004627 | 0.00217 | 0.016786 | 0.000453 | 0.000501 | 0.006882 | 0.001599 | 0.001396 | 0.026657 | 0.002004 | 0.024409 |
| 21.9027778 | 0.016951 | 0.004725 | 0.00804 | 0.001916 | 0.000584 | 0.001108 | 0.004026 | 0.018261 | 0.005047 | 0.005116 | 0.011542 | 0.000402 | 0.001253 | 0.005295 | 0.002783 | 0.002019 | 0.020792 | 0.002207 | 0.02709 |
| 21.9166667 | 0.018744 | 0.010364 | 0.009771 | 0.003568 | 0.001328 | 0.001342 | 0.002459 | 0.017916 | 0.004053 | 0.010632 | 0.015825 | 0.000352 | 0.001724 | 0.004015 | 0.002756 | 0.002014 | 0.013471 | 0.002454 | 0.020783 |
| 21.9305556 | 0.019089 | 0.012682 | 0.008113 | 0.006713 | 0.001451 | 0.001083 | 0.00427 | 0.015757 | 0.003013 | 0.015156 | 0.030656 | 0.000399 | 0.001416 | 0.003985 | 0.001543 | 0.001569 | 0.010489 | 0.00279 | 0.01018 |
| 21.9444444 | 0.019814 | 0.009255 | 0.005511 | 0.008876 | 0.000918 | 0.001448 | 0.00954 | 0.014608 | 0.00344 | 0.015183 | 0.043831 | 0.000363 | 0.001356 | 0.002594 | 0.001163 | 0.001268 | 0.014624 | 0.002939 | 0.00315 |
| 21.9583333 | 0.020902 | 0.004435 | 0.003894 | 0.008081 | 0.001014 | 0.003552 | 0.01794 | 0.017185 | 0.005282 | 0.010174 | 0.046951 | 0.00025 | 0.00183 | 0.000962 | 0.001632 | 0.001064 | 0.022861 | 0.00294 | 0.003275 |
| 21.9722222 | 0.021631 | 0.001539 | 0.003722 | 0.007198 | 0.001055 | 0.005863 | 0.023267 | 0.023789 | 0.007189 | 0.004944 | 0.0379 | 0.000293 | 0.001653 | 0.000456 | 0.00432 | 0.000667 | 0.030084 | 0.003202 | 0.010332 |
| 21.9861111 | 0.021628 | 0.000633 | 0.004548 | 0.009999 | 0.000741 | 0.006744 | 0.022356 | 0.031589 | 0.007956 | 0.004458 | 0.021502 | 0.000532 | 0.0011 | 0.000626 | 0.009082 | 0.000424 | 0.033913 | 0.003728 | 0.02006 |
| 22 | 0.020498 | 0.00094 | 0.005476 | 0.015536 | 0.001131 | 0.006368 | 0.01898 | 0.036251 | 0.007499 | 0.008684 | 0.008231 | 0.00058 | 0.001801 | 0.001553 | 0.012742 | 0.000845 | 0.034219 | 0.003838 | 0.027024 |
| 22.0138889 | 0.018322 | 0.001395 | 0.005724 | 0.018602 | 0.001672 | 0.005356 | 0.018321 | 0.035713 | 0.006541 | 0.014262 | 0.005909 | 0.000453 | 0.003444 | 0.002439 | 0.013128 | 0.001795 | 0.03026 | 0.003447 | 0.029456 |
| 22.0277778 | 0.016623 | 0.002454 | 0.005841 | 0.016314 | 0.001426 | 0.004825 | 0.022946 | 0.03262 | 0.006117 | 0.01866 | 0.012876 | 0.000351 | 0.004169 | 0.002468 | 0.010672 | 0.002561 | 0.022081 | 0.003254 | 0.028794 |
| 22.0416667 | 0.016818 | 0.004819 | 0.006357 | 0.011948 | 0.001065 | 0.005343 | 0.02975 | 0.030464 | 0.006497 | 0.02086 | 0.020314 | 0.000287 | 0.003336 | 0.001886 | 0.007368 | 0.002508 | 0.013463 | 0.003477 | 0.026168 |
| 22.0555556 | 0.017078 | 0.006795 | 0.006019 | 0.008564 | 0.001167 | 0.005915 | 0.030453 | 0.029183 | 0.006256 | 0.019868 | 0.020923 | 0.000433 | 0.001879 | 0.001189 | 0.004746 | 0.001781 | 0.008313 | 0.003679 | 0.02093 |
| 22.0694444 | 0.013995 | 0.005945 | 0.004675 | 0.006079 | 0.00306 | 0.005359 | 0.020295 | 0.025335 | 0.004842 | 0.015148 | 0.014856 | 0.000734 | 0.001061 | 0.000603 | 0.00264 | 0.001164 | 0.006877 | 0.003498 | 0.013566 |
| 22.0833333 | 0.008094 | 0.004346 | 0.006884 | 0.00776 | 0.009339 | 0.00325 | 0.011378 | 0.015471 | 0.006607 | 0.008702 | 0.008223 | 0.001755 | 0.00157 | 0.000638 | 0.001465 | 0.001638 | 0.009025 | 0.002861 | 0.007161 |
| 22.0972222 | 0.00477 | 0.005609 | 0.013953 | 0.014195 | 0.016836 | 0.001402 | 0.015696 | 0.007329 | 0.013241 | 0.003992 | 0.004813 | 0.003702 | 0.002432 | 0.002159 | 0.001931 | 0.002947 | 0.014784 | 0.001794 | 0.004473 |
| 22.1111111 | 0.005292 | 0.006599 | 0.017736 | 0.01632 | 0.018972 | 0.001207 | 0.020301 | 0.00853 | 0.016787 | 0.002487 | 0.003806 | 0.0045 | 0.002071 | 0.004574 | 0.002019 | 0.00319 | 0.018682 | 0.001445 | 0.004433 |
| 22.125 | 0.005556 | 0.004606 | 0.014168 | 0.011528 | 0.015634 | 0.002164 | 0.015264 | 0.010125 | 0.013176 | 0.002692 | 0.00527 | 0.003323 | 0.001003 | 0.005479 | 0.001055 | 0.002064 | 0.01382 | 0.002716 | 0.00409 |
| 22.1388889 | 0.005615 | 0.002842 | 0.008845 | 0.00831 | 0.011145 | 0.004807 | 0.00876 | 0.007387 | 0.008194 | 0.002992 | 0.007231 | 0.001706 | 0.000475 | 0.003658 | 0.000652 | 0.001669 | 0.007078 | 0.004321 | 0.002855 |
| 22.1527778 | 0.006615 | 0.002803 | 0.006031 | 0.009798 | 0.00801 | 0.007117 | 0.006202 | 0.004278 | 0.006078 | 0.002939 | 0.005619 | 0.000672 | 0.000305 | 0.001691 | 0.001027 | 0.002545 | 0.006063 | 0.005219 | 0.00187 |
| 22.1666667 | 0.007084 | 0.00312 | 0.005287 | 0.010518 | 0.00613 | 0.007022 | 0.005855 | 0.002231 | 0.005939 | 0.002064 | 0.002316 | 0.000362 | 0.000159 | 0.001305 | 0.0015 | 0.003006 | 0.00457 | 0.00553 | 0.001334 |
| 22.1805556 | 0.006799 | 0.003589 | 0.00559 | 0.00789 | 0.005238 | 0.00652 | 0.005763 | 0.002084 | 0.006373 | 0.001412 | 0.0008 | 0.000505 | 0.000252 | 0.001356 | 0.001654 | 0.002642 | 0.001809 | 0.005618 | 0.000844 |
| 22.1944444 | 0.006094 | 0.003952 | 0.005944 | 0.004814 | 0.005024 | 0.005967 | 0.005392 | 0.003054 | 0.006445 | 0.002046 | 0.000657 | 0.000728 | 0.000447 | 0.003696 | 0.001469 | 0.00216 | 0.001557 | 0.005604 | 0.000446 |
| 22.2083333 | 0.00463 | 0.00314 | 0.005126 | 0.002673 | 0.004225 | 0.003961 | 0.00412 | 0.00312 | 0.005174 | 0.002222 | 0.000708 | 0.000777 | 0.000546 | 0.006755 | 0.001222 | 0.001742 | 0.002556 | 0.004947 | 0.000876 |
| 22.2222222 | 0.002462 | 0.001669 | 0.003303 | 0.001503 | 0.002581 | 0.001535 | 0.002095 | 0.001932 | 0.002894 | 0.001221 | 0.000708 | 0.00064 | 0.001018 | 0.005149 | 0.001159 | 0.001421 | 0.003897 | 0.003747 | 0.002749 |
| 22.2361111 | 0.001165 | 0.001253 | 0.001488 | 0.002169 | 0.001088 | 0.000365 | 0.000837 | 0.001191 | 0.001134 | 0.000407 | 0.000976 | 0.000613 | 0.001802 | 0.002154 | 0.001246 | 0.001227 | 0.004871 | 0.002895 | 0.004568 |
| 22.25 | 0.001896 | 0.002151 | 0.000523 | 0.003903 | 0.000426 | 0.000155 | 0.001192 | 0.001578 | 0.000704 | 0.000316 | 0.001374 | 0.000659 | 0.001947 | 0.002417 | 0.00124 | 0.001037 | 0.005089 | 0.00242 | 0.004786 |
| 22.2638889 | 0.003105 | 0.002954 | 0.000472 | 0.004752 | 0.000526 | 0.000153 | 0.001894 | 0.001646 | 0.00079 | 0.000415 | 0.00311 | 0.000471 | 0.001452 | 0.003578 | 0.001102 | 0.000759 | 0.005134 | 0.001833 | 0.003968 |
| 22.2777778 | 0.003064 | 0.002656 | 0.000564 | 0.003508 | 0.0006 | 0.00054 | 0.001634 | 0.001104 | 0.000719 | 0.000492 | 0.007205 | 0.000226 | 0.00097 | 0.003373 | 0.000959 | 0.000606 | 0.004905 | 0.001365 | 0.002497 |
| 22.2916667 | 0.001796 | 0.001503 | 0.000778 | 0.002096 | 0.000391 | 0.001298 | 0.001046 | 0.001087 | 0.000932 | 0.000968 | 0.010179 | 0.000232 | 0.000534 | 0.002468 | 0.000794 | 0.000754 | 0.003814 | 0.001514 | 0.001547 |
| 22.3055556 | 0.000806 | 0.000706 | 0.001282 | 0.003602 | 0.000439 | 0.001555 | 0.001484 | 0.001427 | 0.001444 | 0.001516 | 0.007746 | 0.000657 | 0.000264 | 0.001426 | 0.000696 | 0.001175 | 0.002094 | 0.002275 | 0.002712 |
| 22.3194444 | 0.001371 | 0.000988 | 0.001828 | 0.007022 | 0.000853 | 0.001748 | 0.002768 | 0.001483 | 0.001995 | 0.001749 | 0.004136 | 0.001446 | 0.000339 | 0.000692 | 0.000799 | 0.001695 | 0.000739 | 0.003112 | 0.004109 |
| 22.3333333 | 0.002968 | 0.002266 | 0.002903 | 0.010029 | 0.001672 | 0.002402 | 0.0042 | 0.002246 | 0.003305 | 0.001987 | 0.004121 | 0.001786 | 0.000437 | 0.000851 | 0.000873 | 0.001805 | 0.000434 | 0.003235 | 0.003178 |
| 22.3472222 | 0.004741 | 0.004441 | 0.00483 | 0.012035 | 0.003041 | 0.003474 | 0.005512 | 0.004452 | 0.005712 | 0.00225 | 0.003554 | 0.00115 | 0.000591 | 0.001823 | 0.000682 | 0.001268 | 0.000886 | 0.002466 | 0.001609 |
| 22.3611111 | 0.00596 | 0.006191 | 0.006488 | 0.0137 | 0.004249 | 0.004172 | 0.00621 | 0.006472 | 0.007812 | 0.001913 | 0.002132 | 0.000357 | 0.00077 | 0.00276 | 0.000437 | 0.000623 | 0.00107 | 0.001639 | 0.001479 |
| 22.375 | 0.006018 | 0.006246 | 0.00669 | 0.015101 | 0.004051 | 0.002634 | 0.005841 | 0.007066 | 0.008315 | 0.001324 | 0.003226 | 0.000204 | 0.0006 | 0.002876 | 0.000424 | 0.00033 | 0.00104 | 0.001442 | 0.001368 |
| 22.3888889 | 0.005175 | 0.005314 | 0.005677 | 0.015522 | 0.002414 | 0.001077 | 0.004572 | 0.0071 | 0.007646 | 0.00204 | 0.004928 | 0.000424 | 0.000289 | 0.002808 | 0.000525 | 0.000211 | 0.001378 | 0.001718 | 0.000823 |
| 22.4027778 | 0.004167 | 0.003963 | 0.004256 | 0.015765 | 0.000806 | 0.001701 | 0.003482 | 0.007022 | 0.006691 | 0.004344 | 0.004786 | 0.000654 | 0.000259 | 0.002545 | 0.000578 | 0.000101 | 0.001196 | 0.001954 | 0.000545 |
| 22.4166667 | 0.003428 | 0.002311 | 0.003283 | 0.016508 | 0.000315 | 0.003247 | 0.004111 | 0.006881 | 0.005967 | 0.007521 | 0.004411 | 0.001047 | 0.000499 | 0.001562 | 0.000713 | 0.000189 | 0.000757 | 0.001909 | 0.001258 |
| 22.4305556 | 0.002778 | 0.001484 | 0.003607 | 0.015857 | 0.00036 | 0.00346 | 0.007729 | 0.007996 | 0.005796 | 0.010644 | 0.005512 | 0.001421 | 0.000551 | 0.000731 | 0.000776 | 0.000411 | 0.001146 | 0.001764 | 0.003649 |
| 22.4444444 | 0.00178 | 0.00178 | 0.00479 | 0.013057 | 0.000462 | 0.003144 | 0.013718 | 0.010575 | 0.005914 | 0.012498 | 0.006384 | 0.001177 | 0.000464 | 0.000519 | 0.000511 | 0.000368 | 0.001512 | 0.001647 | 0.006275 |
| 22.4583333 | 0.000978 | 0.001631 | 0.005314 | 0.011301 | 0.001137 | 0.004039 | 0.018289 | 0.012555 | 0.005694 | 0.012951 | 0.005132 | 0.000918 | 0.000822 | 0.000671 | 0.000247 | 0.000214 | 0.001237 | 0.001726 | 0.007487 |
| 22.4722222 | 0.001561 | 0.001447 | 0.004545 | 0.012492 | 0.002323 | 0.003339 | 0.017627 | 0.013031 | 0.004881 | 0.012798 | 0.003228 | 0.001983 | 0.001996 | 0.001716 | 0.000337 | 0.000628 | 0.001553 | 0.00228 | 0.007624 |
| 22.4861111 | 0.003642 | 0.002733 | 0.002711 | 0.014288 | 0.003707 | 0.001973 | 0.01211 | 0.013173 | 0.003488 | 0.013554 | 0.004461 | 0.003519 | 0.003766 | 0.004326 | 0.000483 | 0.001515 | 0.002922 | 0.002606 | 0.007973 |
| 22.5 | 0.007183 | 0.003452 | 0.001728 | 0.01133 | 0.005793 | 0.002254 | 0.005916 | 0.011949 | 0.002826 | 0.016701 | 0.008959 | 0.003253 | 0.004828 | 0.007241 | 0.000669 | 0.001814 | 0.00408 | 0.002678 | 0.008647 |
| 22.5138889 | 0.011125 | 0.0093 | 0.003024 | 0.00797 | 0.007181 | 0.003601 | 0.005469 | 0.007379 | 0.003951 | 0.019741 | 0.011979 | 0.002227 | 0.004414 | 0.008949 | 0.001439 | 0.001182 | 0.00422 | 0.012555 | 0.008274 |
| 22.5277778 | 0.010692 | 0.016714 | 0.003397 | 0.010212 | 0.009001 | 0.014149 | 0.012574 | 0.003217 | 0.00425 | 0.018401 | 0.010078 | 0.001997 | 0.003563 | 0.007867 | 0.002372 | 0.00064 | 0.003482 | 0.022451 | 0.006514 |
| 22.5416667 | 0.006043 | 0.011287 | 0.003151 | 0.011349 | 0.012513 | 0.026657 | 0.016432 | 0.006138 | 0.004503 | 0.013176 | 0.005645 | 0.003386 | 0.003113 | 0.00442 | 0.002123 | 0.00097 | 0.00301 | 0.013581 | 0.004148 |
| 22.5555556 | 0.004564 | 0.004784 | 0.005246 | 0.012154 | 0.011526 | 0.021284 | 0.009877 | 0.013664 | 0.006846 | 0.007742 | 0.002858 | 0.007584 | 0.003 | 0.002169 | 0.001977 | 0.001604 | 0.003861 | 0.004305 | 0.002509 |
| 22.5694444 | 0.008173 | 0.007538 | 0.0067 | 0.01278 | 0.006206 | 0.010408 | 0.005108 | 0.016316 | 0.00719 | 0.004724 | 0.005805 | 0.01082 | 0.002895 | 0.001902 | 0.003413 | 0.001619 | 0.005598 | 0.003 | 0.003859 |
| 22.5833333 | 0.014482 | 0.012548 | 0.005969 | 0.008705 | 0.002331 | 0.009627 | 0.012796 | 0.01461 | 0.005809 | 0.005075 | 0.029924 | 0.009701 | 0.001989 | 0.00301 | 0.003602 | 0.000986 | 0.014925 | 0.001997 | 0.014502 |
| 22.5972222 | 0.0203 | 0.015202 | 0.004404 | 0.005969 | 0.001279 | 0.013213 | 0.02518 | 0.015926 | 0.006667 | 0.007044 | 0.077253 | 0.00598 | 0.000661 | 0.003228 | 0.003178 | 0.00077 | 0.032555 | 0.002863 | 0.0347 |
| 22.6111111 | 0.021248 | 0.012615 | 0.002375 | 0.005398 | 0.002592 | 0.01294 | 0.030687 | 0.020573 | 0.008201 | 0.007216 | 0.106304 | 0.002393 | 0.00064 | 0.002498 | 0.005003 | 0.001507 | 0.039512 | 0.003021 | 0.04839 |
| 22.625 | 0.018342 | 0.007675 | 0.00096 | 0.00533 | 0.003837 | 0.007577 | 0.0287 | 0.02436 | 0.007675 | 0.005242 | 0.08882 | 0.000556 | 0.001405 | 0.003867 | 0.006312 | 0.001686 | 0.030823 | 0.002426 | 0.046611 |
| 22.6388889 | 0.015761 | 0.005389 | 0.00068 | 0.006688 | 0.003641 | 0.002879 | 0.02285 | 0.023826 | 0.004918 | 0.003921 | 0.049218 | 0.000218 | 0.0012 | 0.004414 | 0.004589 | 0.001071 | 0.019331 | 0.002737 | 0.03508 |
| 22.6527778 | 0.014289 | 0.005222 | 0.001383 | 0.006368 | 0.003445 | 0.00233 | 0.013521 | 0.018308 | 0.00316 | 0.005173 | 0.020618 | 0.000705 | 0.000585 | 0.002707 | 0.002187 | 0.000865 | 0.009889 | 0.002465 | 0.020615 |
| 22.6666667 | 0.013771 | 0.004891 | 0.002587 | 0.007171 | 0.003609 | 0.004328 | 0.007834 | 0.012245 | 0.005145 | 0.008689 | 0.015274 | 0.001411 | 0.000384 | 0.00192 | 0.001362 | 0.000786 | 0.005349 | 0.001425 | 0.010213 |
| 22.6805556 | 0.015237 | 0.004485 | 0.002471 | 0.009508 | 0.003039 | 0.004803 | 0.011211 | 0.010112 | 0.007223 | 0.01151 | 0.02544 | 0.001344 | 0.000278 | 0.002055 | 0.001311 | 0.000945 | 0.005406 | 0.00082 | 0.006894 |
| 22.6944444 | 0.017526 | 0.003815 | 0.002282 | 0.008698 | 0.003544 | 0.002849 | 0.013302 | 0.012058 | 0.005636 | 0.009715 | 0.035501 | 0.000705 | 0.000205 | 0.001955 | 0.002438 | 0.001654 | 0.007167 | 0.000949 | 0.007228 |
| 22.7083333 | 0.018061 | 0.002365 | 0.005629 | 0.005165 | 0.006179 | 0.001556 | 0.011379 | 0.016822 | 0.003086 | 0.005182 | 0.038438 | 0.000364 | 0.00025 | 0.001595 | 0.005389 | 0.002467 | 0.014073 | 0.001272 | 0.007833 |
| 22.7222222 | 0.016456 | 0.001701 | 0.010866 | 0.003904 | 0.006511 | 0.002259 | 0.019542 | 0.023204 | 0.003459 | 0.004083 | 0.036464 | 0.000271 | 0.000688 | 0.001065 | 0.008139 | 0.002954 | 0.020022 | 0.002557 | 0.007948 |
| 22.7361111 | 0.013963 | 0.003495 | 0.013017 | 0.007627 | 0.003288 | 0.002841 | 0.038361 | 0.028024 | 0.005774 | 0.006061 | 0.03215 | 0.000235 | 0.001409 | 0.001005 | 0.008708 | 0.002839 | 0.015748 | 0.006378 | 0.006888 |
| 22.75 | 0.011039 | 0.008244 | 0.0092 | 0.00859 | 0.001114 | 0.002554 | 0.048339 | 0.026917 | 0.006927 | 0.00634 | 0.022935 | 0.000241 | 0.001606 | 0.001178 | 0.00692 | 0.002218 | 0.00794 | 0.011554 | 0.004519 |
| 22.7638889 | 0.006726 | 0.012774 | 0.005529 | 0.007065 | 0.002085 | 0.004426 | 0.038215 | 0.019224 | 0.00575 | 0.004258 | 0.014332 | 0.000446 | 0.001259 | 0.001052 | 0.003984 | 0.00142 | 0.007666 | 0.015773 | 0.003579 |
| 22.7777778 | 0.003375 | 0.011821 | 0.008698 | 0.011975 | 0.004797 | 0.009428 | 0.022212 | 0.010117 | 0.003366 | 0.003344 | 0.019587 | 0.001213 | 0.001009 | 0.001291 | 0.002944 | 0.001256 | 0.01542 | 0.017367 | 0.005406 |
| 22.7916667 | 0.004201 | 0.012244 | 0.012665 | 0.018226 | 0.007413 | 0.013843 | 0.023899 | 0.007683 | 0.002346 | 0.007032 | 0.032084 | 0.002245 | 0.000872 | 0.001421 | 0.005008 | 0.002084 | 0.022383 | 0.015192 | 0.006966 |
| 22.8055556 | 0.00652 | 0.021069 | 0.012055 | 0.019886 | 0.008309 | 0.015012 | 0.041635 | 0.012618 | 0.002919 | 0.012644 | 0.033046 | 0.002465 | 0.001215 | 0.000871 | 0.006425 | 0.003051 | 0.021302 | 0.00938 | 0.006526 |
| 22.8194444 | 0.008271 | 0.028705 | 0.008307 | 0.01881 | 0.008177 | 0.014049 | 0.054339 | 0.016121 | 0.003153 | 0.012622 | 0.020912 | 0.001622 | 0.002186 | 0.000374 | 0.004902 | 0.003418 | 0.012532 | 0.00488 | 0.00464 |
| 22.8333333 | 0.009867 | 0.030118 | 0.004563 | 0.013494 | 0.010608 | 0.014122 | 0.051902 | 0.013128 | 0.002673 | 0.007303 | 0.014754 | 0.000639 | 0.002368 | 0.000284 | 0.002486 | 0.003091 | 0.005913 | 0.006653 | 0.002276 |
| 22.8472222 | 0.009428 | 0.02993 | 0.004527 | 0.007691 | 0.01825 | 0.01724 | 0.035146 | 0.006689 | 0.001705 | 0.005289 | 0.023914 | 0.000208 | 0.001342 | 0.000324 | 0.001684 | 0.002198 | 0.010716 | 0.010662 | 0.001307 |

| | | | | | | | | | | | | | | | | | | | |
|---|---|---|---|---|---|---|---|---|---|---|---|---|---|---|---|---|---|---|---|
| 22.8611111 | 0.005898 | 0.029271 | 0.006988 | 0.00847 | 0.027427 | 0.021584 | 0.017256 | 0.002279 | 0.000765 | 0.008521 | 0.034348 | 0.000179 | 0.000826 | 0.000754 | 0.00239 | 0.001059 | 0.02002 | 0.011682 | 0.001825 |
| 22.875 | 0.002011 | 0.026119 | 0.006819 | 0.009751 | 0.031102 | 0.02433 | 0.013649 | 0.00434 | 0.000608 | 0.012677 | 0.03612 | 0.000285 | 0.002105 | 0.002056 | 0.002267 | 0.00029 | 0.019149 | 0.009902 | 0.003397 |
| 22.8888889 | 0.001411 | 0.018938 | 0.004133 | 0.006318 | 0.028366 | 0.025128 | 0.017027 | 0.01375 | 0.001067 | 0.015565 | 0.031756 | 0.000731 | 0.004268 | 0.003263 | 0.002014 | 0.000274 | 0.012152 | 0.007484 | 0.009791 |
| 22.9027778 | 0.004497 | 0.010251 | 0.003694 | 0.004642 | 0.02248 | 0.024504 | 0.013246 | 0.026206 | 0.002959 | 0.015996 | 0.02711 | 0.001577 | 0.005517 | 0.003506 | 0.004581 | 0.000965 | 0.015529 | 0.00566 | 0.022176 |
| 22.9166667 | 0.008868 | 0.007991 | 0.006201 | 0.008143 | 0.016216 | 0.023276 | 0.0085 | 0.036295 | 0.006949 | 0.01292 | 0.025491 | 0.002619 | 0.005127 | 0.002975 | 0.008787 | 0.002239 | 0.027417 | 0.004527 | 0.033311 |
| 22.9305556 | 0.01263 | 0.015237 | 0.009171 | 0.012291 | 0.014231 | 0.024341 | 0.014961 | 0.04216 | 0.011155 | 0.00717 | 0.025667 | 0.003829 | 0.003898 | 0.002012 | 0.011169 | 0.003759 | 0.035797 | 0.004563 | 0.036101 |
| 22.9444444 | 0.015169 | 0.024151 | 0.011467 | 0.013262 | 0.020026 | 0.02923 | 0.026835 | 0.0444 | 0.014152 | 0.002549 | 0.02654 | 0.005281 | 0.002853 | 0.001451 | 0.011229 | 0.005225 | 0.03698 | 0.006282 | 0.031104 |
| 22.9583333 | 0.016148 | 0.027837 | 0.012509 | 0.012465 | 0.031282 | 0.03468 | 0.032417 | 0.043493 | 0.015829 | 0.00169 | 0.028935 | 0.006438 | 0.002032 | 0.001913 | 0.011014 | 0.006442 | 0.034096 | 0.009632 | 0.023777 |
| 22.9722222 | 0.015671 | 0.024369 | 0.01202 | 0.013115 | 0.042497 | 0.037383 | 0.027888 | 0.040679 | 0.015767 | 0.004103 | 0.032237 | 0.006438 | 0.001124 | 0.002731 | 0.011322 | 0.007021 | 0.031347 | 0.013164 | 0.019082 |
| 22.9861111 | 0.014349 | 0.014431 | 0.01055 | 0.01527 | 0.048973 | 0.036432 | 0.015609 | 0.038692 | 0.014234 | 0.009852 | 0.031736 | 0.005597 | 0.000374 | 0.003012 | 0.011879 | 0.006477 | 0.029072 | 0.014109 | 0.019035 |
| 23 | 0.012564 | 0.006205 | 0.008788 | 0.016528 | 0.048222 | 0.032355 | 0.004865 | 0.038412 | 0.012273 | 0.016159 | 0.023845 | 0.004837 | 0.000136 | 0.002683 | 0.012726 | 0.00434 | 0.025247 | 0.011262 | 0.02244 |
| 23.0138889 | 0.010655 | 0.005041 | 0.00752 | 0.016256 | 0.041057 | 0.027666 | 0.003083 | 0.037311 | 0.011186 | 0.019919 | 0.012521 | 0.00449 | 0.000259 | 0.001741 | 0.013069 | 0.002081 | 0.018867 | 0.006657 | 0.025811 |
| 23.0277778 | 0.009482 | 0.005654 | 0.006779 | 0.014703 | 0.030955 | 0.023816 | 0.009231 | 0.033104 | 0.011636 | 0.021852 | 0.004331 | 0.004427 | 0.000321 | 0.000958 | 0.012002 | 0.001434 | 0.011448 | 0.002871 | 0.025616 |
| 23.0416667 | 0.0098 | 0.009018 | 0.005385 | 0.011795 | 0.021515 | 0.020279 | 0.019694 | 0.026644 | 0.012223 | 0.02162 | 0.001243 | 0.004339 | 0.000169 | 0.00082 | 0.009717 | 0.001427 | 0.005719 | 0.000805 | 0.021683 |
| 23.0555556 | 0.010922 | 0.017946 | 0.002895 | 0.00777 | 0.01487 | 0.016941 | 0.029649 | 0.019978 | 0.011282 | 0.018054 | 0.002363 | 0.003927 | 7.32E-05 | 0.001368 | 0.006961 | 0.002009 | 0.003323 | 0.000573 | 0.016765 |
| 23.0694444 | 0.01038 | 0.026387 | 0.001671 | 0.004334 | 0.014736 | 0.01436 | 0.033216 | 0.012676 | 0.009764 | 0.011685 | 0.00713 | 0.002946 | 8.87E-05 | 0.003844 | 0.004075 | 0.002849 | 0.004926 | 0.004059 | 0.012783 |
| 23.0833333 | 0.0072 | 0.030009 | 0.004881 | 0.005348 | 0.027318 | 0.012756 | 0.025497 | 0.007621 | 0.010133 | 0.005376 | 0.015498 | 0.00156 | 0.00039 | 0.006344 | 0.002475 | 0.00294 | 0.010421 | 0.016328 | 0.01032 |
| 23.0972222 | 0.006653 | 0.025034 | 0.011312 | 0.010424 | 0.042238 | 0.01069 | 0.013179 | 0.009779 | 0.010408 | 0.002271 | 0.023162 | 0.000583 | 0.001272 | 0.005039 | 0.003468 | 0.00421 | 0.015308 | 0.033448 | 0.009325 |
| 23.1111111 | 0.011599 | 0.015304 | 0.014852 | 0.012277 | 0.034841 | 0.008138 | 0.009117 | 0.012231 | 0.008338 | 0.002155 | 0.0209 | 0.000341 | 0.001957 | 0.002148 | 0.004183 | 0.005695 | 0.013555 | 0.037283 | 0.008866 |
| 23.125 | 0.017945 | 0.011678 | 0.011074 | 0.008761 | 0.013904 | 0.012634 | 0.009572 | 0.010315 | 0.00904 | 0.003903 | 0.011313 | 0.000257 | 0.00188 | 0.001605 | 0.002447 | 0.004217 | 0.008073 | 0.023633 | 0.007365 |
| 23.1388889 | 0.024054 | 0.009781 | 0.005251 | 0.005231 | 0.004684 | 0.029769 | 0.006304 | 0.012302 | 0.012382 | 0.011201 | 0.006514 | 0.000164 | 0.002411 | 0.003382 | 0.000805 | 0.001557 | 0.006337 | 0.010008 | 0.004372 |
| 23.1527778 | 0.023627 | 0.007078 | 0.003955 | 0.004218 | 0.00376 | 0.034192 | 0.002157 | 0.013605 | 0.011058 | 0.016175 | 0.0056 | 0.000693 | 0.003313 | 0.006001 | 0.001497 | 0.000895 | 0.005987 | 0.005334 | 0.002199 |
| 23.1666667 | 0.014482 | 0.012533 | 0.003738 | 0.005415 | 0.003021 | 0.028088 | 0.002006 | 0.010938 | 0.005597 | 0.013825 | 0.00421 | 0.001874 | 0.00269 | 0.006181 | 0.003523 | 0.002037 | 0.004848 | 0.003632 | 0.002261 |
| 23.1805556 | 0.008636 | 0.019767 | 0.00229 | 0.005698 | 0.002089 | 0.039268 | 0.003725 | 0.012606 | 0.002364 | 0.012708 | 0.005519 | 0.002435 | 0.00174 | 0.003368 | 0.004243 | 0.002608 | 0.006235 | 0.002661 | 0.002245 |
| 23.1944444 | 0.008208 | 0.016613 | 0.00293 | 0.003314 | 0.001732 | 0.036197 | 0.003435 | 0.010785 | 0.003476 | 0.008207 | 0.009205 | 0.0023 | 0.001384 | 0.00133 | 0.003087 | 0.001818 | 0.005866 | 0.004705 | 0.001024 |
| 23.2083333 | 0.007398 | 0.008517 | 0.005795 | 0.001631 | 0.003496 | 0.014419 | 0.003941 | 0.004765 | 0.006399 | 0.00347 | 0.011528 | 0.002651 | 0.000922 | 0.001446 | 0.001835 | 0.001061 | 0.003189 | 0.007862 | 0.000599 |
| 23.2222222 | 0.00595 | 0.00484 | 0.008167 | 0.001581 | 0.006106 | 0.008933 | 0.006774 | 0.003896 | 0.008657 | 0.004584 | 0.009778 | 0.00338 | 0.001286 | 0.002341 | 0.001747 | 0.001181 | 0.003852 | 0.010219 | 0.001939 |
| 23.2361111 | 0.003585 | 0.005666 | 0.008445 | 0.002351 | 0.007533 | 0.016368 | 0.008719 | 0.004465 | 0.00881 | 0.005561 | 0.006771 | 0.003955 | 0.00254 | 0.003158 | 0.002791 | 0.001901 | 0.006478 | 0.013004 | 0.003879 |
| 23.25 | 0.001519 | 0.00844 | 0.006651 | 0.002446 | 0.006104 | 0.022093 | 0.00866 | 0.003846 | 0.006695 | 0.004072 | 0.007278 | 0.004158 | 0.002814 | 0.003549 | 0.003211 | 0.002106 | 0.006841 | 0.018091 | 0.004381 |
| 23.2638889 | 0.001482 | 0.010633 | 0.004119 | 0.001366 | 0.002875 | 0.020221 | 0.008499 | 0.004036 | 0.004434 | 0.001753 | 0.006966 | 0.003949 | 0.001728 | 0.003109 | 0.00199 | 0.001406 | 0.004282 | 0.024173 | 0.002755 |
| 23.2777778 | 0.003783 | 0.009474 | 0.003481 | 0.001744 | 0.001836 | 0.012653 | 0.01166 | 0.005267 | 0.004772 | 0.001083 | 0.005268 | 0.003915 | 0.000726 | 0.002467 | 0.000759 | 0.000833 | 0.001995 | 0.025814 | 0.001124 |
| 23.2916667 | 0.007892 | 0.00537 | 0.00617 | 0.004269 | 0.00495 | 0.008377 | 0.019357 | 0.007369 | 0.007835 | 0.002776 | 0.005819 | 0.004136 | 0.000515 | 0.002365 | 0.000555 | 0.000583 | 0.002292 | 0.021554 | 0.001009 |
| 23.3055556 | 0.009804 | 0.002437 | 0.010329 | 0.006679 | 0.009837 | 0.011758 | 0.026382 | 0.010269 | 0.011056 | 0.006028 | 0.004953 | 0.00391 | 0.000922 | 0.002436 | 0.000847 | 0.000473 | 0.003436 | 0.016127 | 0.001444 |
| 23.3194444 | 0.008427 | 0.002116 | 0.013326 | 0.007902 | 0.012871 | 0.015949 | 0.02703 | 0.012706 | 0.012616 | 0.007302 | 0.00229 | 0.002837 | 0.001253 | 0.002172 | 0.001012 | 0.000533 | 0.003274 | 0.013038 | 0.001559 |
| 23.3333333 | 0.00764 | 0.002472 | 0.01458 | 0.009067 | 0.013411 | 0.015398 | 0.02256 | 0.012845 | 0.012653 | 0.004997 | 0.0012 | 0.001345 | 0.001308 | 0.001857 | 0.000974 | 0.000491 | 0.002066 | 0.012096 | 0.001506 |
| 23.3472222 | 0.009191 | 0.003941 | 0.015035 | 0.010562 | 0.012803 | 0.013419 | 0.017562 | 0.010943 | 0.012207 | 0.002673 | 0.00129 | 0.00083 | 0.001899 | 0.002367 | 0.001415 | 0.00113 | 0.00131 | 0.011546 | 0.001847 |
| 23.3611111 | 0.01144 | 0.007031 | 0.015205 | 0.011102 | 0.011882 | 0.01307 | 0.014472 | 0.009247 | 0.011891 | 0.001752 | 0.001665 | 0.001904 | 0.002935 | 0.003371 | 0.002565 | 0.002932 | 0.00119 | 0.010037 | 0.002527 |
| 23.375 | 0.011265 | 0.008883 | 0.014248 | 0.008664 | 0.011161 | 0.013467 | 0.012833 | 0.008792 | 0.010966 | 0.000898 | 0.003369 | 0.002707 | 0.002967 | 0.00301 | 0.003025 | 0.003932 | 0.001043 | 0.008019 | 0.002373 |
| 23.3888889 | 0.007037 | 0.007278 | 0.011587 | 0.00538 | 0.010094 | 0.014575 | 0.011268 | 0.008636 | 0.008851 | 0.001134 | 0.005077 | 0.001878 | 0.001757 | 0.001359 | 0.001974 | 0.00279 | 0.001011 | 0.006401 | 0.001239 |
| 23.4027778 | 0.003251 | 0.004016 | 0.008419 | 0.005703 | 0.007685 | 0.016367 | 0.00927 | 0.007583 | 0.006707 | 0.003469 | 0.004898 | 0.000922 | 0.000939 | 0.000454 | 0.000981 | 0.001307 | 0.000861 | 0.005579 | 0.000597 |
| 23.4166667 | 0.003552 | 0.001867 | 0.006048 | 0.006875 | 0.004252 | 0.016004 | 0.00676 | 0.005342 | 0.005523 | 0.006096 | 0.005553 | 0.000732 | 0.00124 | 0.000685 | 0.0012 | 0.001341 | 0.000581 | 0.006138 | 0.001489 |
| 23.4305556 | 0.004157 | 0.000939 | 0.004426 | 0.004872 | 0.002894 | 0.012475 | 0.003691 | 0.003003 | 0.004446 | 0.007556 | 0.009128 | 0.000561 | 0.001762 | 0.000889 | 0.001661 | 0.002248 | 0.001055 | 0.007594 | 0.004211 |
| 23.4444444 | 0.00353 | 0.000771 | 0.002927 | 0.002365 | 0.00573 | 0.008537 | 0.001499 | 0.002006 | 0.002996 | 0.008235 | 0.012543 | 0.000635 | 0.001553 | 0.00088 | 0.001259 | 0.002306 | 0.002419 | 0.008087 | 0.008389 |
| 23.4583333 | 0.004028 | 0.002542 | 0.002057 | 0.002924 | 0.009817 | 0.005253 | 0.001545 | 0.004193 | 0.003458 | 0.009095 | 0.012726 | 0.001513 | 0.000912 | 0.001521 | 0.000792 | 0.001629 | 0.00374 | 0.007684 | 0.012684 |
| 23.4722222 | 0.006595 | 0.007074 | 0.0033 | 0.006693 | 0.012493 | 0.002661 | 0.002511 | 0.010388 | 0.006636 | 0.010742 | 0.010115 | 0.002479 | 0.000569 | 0.002368 | 0.001135 | 0.001729 | 0.00392 | 0.008478 | 0.015398 |
| 23.4861111 | 0.010276 | 0.012949 | 0.005695 | 0.011863 | 0.013762 | 0.001683 | 0.003171 | 0.017767 | 0.008818 | 0.012017 | 0.006093 | 0.002094 | 0.000876 | 0.002024 | 0.001228 | 0.002016 | 0.002638 | 0.010598 | 0.015015 |
| 23.5 | 0.013049 | 0.017418 | 0.006668 | 0.016179 | 0.014283 | 0.001607 | 0.003893 | 0.023039 | 0.007741 | 0.011594 | 0.002464 | 0.001126 | 0.001996 | 0.000957 | 0.000945 | 0.001386 | 0.001184 | 0.012211 | 0.011964 |
| 23.5138889 | 0.013839 | 0.01877 | 0.006157 | 0.01788 | 0.015276 | 0.00247 | 0.005775 | 0.026124 | 0.005438 | 0.01094 | 0.001188 | 0.000902 | 0.002635 | 0.000553 | 0.000891 | 0.001174 | 0.001441 | 0.012773 | 0.009328 |
| 23.5277778 | 0.012541 | 0.017831 | 0.005715 | 0.016432 | 0.01641 | 0.003708 | 0.010255 | 0.02827 | 0.004675 | 0.0116 | 0.001395 | 0.000862 | 0.001872 | 0.000764 | 0.001107 | 0.002265 | 0.002856 | 0.013113 | 0.009088 |
| 23.5416667 | 0.009833 | 0.016508 | 0.006092 | 0.013004 | 0.015809 | 0.002843 | 0.017108 | 0.028257 | 0.005766 | 0.012628 | 0.002248 | 0.001229 | 0.001116 | 0.000936 | 0.001807 | 0.003218 | 0.003136 | 0.013281 | 0.010342 |
| 23.5555556 | 0.00712 | 0.015133 | 0.006578 | 0.009288 | 0.013208 | 0.001493 | 0.023084 | 0.023769 | 0.006712 | 0.013258 | 0.003834 | 0.001975 | 0.001196 | 0.00069 | 0.002134 | 0.00306 | 0.00225 | 0.013069 | 0.0121 |
| 23.5694444 | 0.00428 | 0.013383 | 0.005238 | 0.005339 | 0.010024 | 0.004536 | 0.024822 | 0.015476 | 0.005508 | 0.015378 | 0.003966 | 0.002475 | 0.001082 | 0.00038 | 0.001817 | 0.002444 | 0.003334 | 0.015121 | 0.01486 |
| 23.5833333 | 0.003843 | 0.011902 | 0.00396 | 0.002802 | 0.008261 | 0.015111 | 0.023628 | 0.007113 | 0.0031 | 0.022223 | 0.004819 | 0.002532 | 0.000727 | 0.000929 | 0.002164 | 0.001923 | 0.010114 | 0.024436 | 0.02067 |
| 23.5972222 | 0.008566 | 0.010802 | 0.005779 | 0.002731 | 0.011143 | 0.024809 | 0.021662 | 0.002497 | 0.001639 | 0.030065 | 0.009879 | 0.002755 | 0.001179 | 0.001813 | 0.002899 | 0.001465 | 0.021535 | 0.039765 | 0.029369 |
| 23.6111111 | 0.011336 | 0.008178 | 0.006163 | 0.004786 | 0.015647 | 0.019941 | 0.018238 | 0.002008 | 0.001593 | 0.028736 | 0.014209 | 0.004527 | 0.001842 | 0.001534 | 0.002526 | 0.001324 | 0.031055 | 0.052117 | 0.03411 |
| 23.625 | 0.00746 | 0.004255 | 0.003541 | 0.009491 | 0.014094 | 0.008669 | 0.013935 | 0.004315 | 0.002194 | 0.01905 | 0.015385 | 0.006818 | 0.001591 | 0.000622 | 0.001703 | 0.002535 | 0.033889 | 0.053004 | 0.030553 |
| 23.6388889 | 0.004143 | 0.002507 | 0.002095 | 0.01144 | 0.008499 | 0.005012 | 0.009944 | 0.009418 | 0.004327 | 0.010209 | 0.018504 | 0.006476 | 0.000848 | 0.000233 | 0.00132 | 0.004762 | 0.030306 | 0.038687 | 0.022477 |
| 23.6527778 | 0.00622 | 0.002089 | 0.001611 | 0.007275 | 0.007398 | 0.004503 | 0.006427 | 0.016584 | 0.009591 | 0.004745 | 0.024153 | 0.003352 | 0.000493 | 0.000158 | 0.00102 | 0.005929 | 0.023807 | 0.017577 | 0.013834 |
| 23.6666667 | 0.01137 | 0.002464 | 0.002656 | 0.002604 | 0.011829 | 0.002474 | 0.003336 | 0.024553 | 0.012974 | 0.002693 | 0.032198 | 0.000842 | 0.000594 | 0.000475 | 0.001452 | 0.005538 | 0.018327 | 0.006343 | 0.008784 |
| 23.6805556 | 0.01766 | 0.006377 | 0.006967 | 0.001514 | 0.013279 | 0.001102 | 0.001083 | 0.031266 | 0.009954 | 0.00323 | 0.049302 | 0.000444 | 0.001235 | 0.001264 | 0.003076 | 0.004183 | 0.014416 | 0.008295 | 0.011404 |
| 23.6944444 | 0.022136 | 0.010535 | 0.011893 | 0.002357 | 0.008308 | 0.000774 | 0.001527 | 0.034327 | 0.005768 | 0.002956 | 0.076173 | 0.000645 | 0.002099 | 0.00151 | 0.004649 | 0.002391 | 0.011651 | 0.014168 | 0.020505 |
| 23.7083333 | 0.021224 | 0.011389 | 0.013652 | 0.002178 | 0.005725 | 0.000747 | 0.007853 | 0.034687 | 0.007435 | 0.002492 | 0.096073 | 0.000646 | 0.002197 | 0.001098 | 0.005228 | 0.00147 | 0.011689 | 0.018828 | 0.030433 |
| 23.7222222 | 0.016699 | 0.013051 | 0.011003 | 0.002603 | 0.010853 | 0.000831 | 0.02058 | 0.034403 | 0.012924 | 0.005486 | 0.090758 | 0.001263 | 0.002401 | 0.002347 | 0.005557 | 0.001612 | 0.012873 | 0.019124 | 0.036001 |
| 23.7361111 | 0.012934 | 0.020273 | 0.006439 | 0.007546 | 0.019186 | 0.001245 | 0.031657 | 0.032576 | 0.015978 | 0.010106 | 0.062095 | 0.002243 | 0.003941 | 0.005992 | 0.006214 | 0.001405 | 0.010073 | 0.014124 | 0.034662 |
| 23.75 | 0.011087 | 0.030424 | 0.005156 | 0.0149 | 0.025444 | 0.002254 | 0.031892 | 0.027416 | 0.015187 | 0.010721 | 0.032759 | 0.002068 | 0.005265 | 0.007757 | 0.006167 | 0.000792 | 0.004754 | 0.014776 | 0.026117 |
| 23.7638889 | 0.010142 | 0.035333 | 0.007191 | 0.018716 | 0.023141 | 0.002442 | 0.022742 | 0.019898 | 0.011375 | 0.006764 | 0.02971 | 0.001307 | 0.004782 | 0.00492 | 0.004441 | 0.001275 | 0.002165 | 0.024226 | 0.013891 |
| 23.7777778 | 0.010074 | 0.030163 | 0.00644 | 0.015948 | 0.012546 | 0.001882 | 0.011696 | 0.012269 | 0.006058 | 0.004369 | 0.052735 | 0.001397 | 0.003162 | 0.001752 | 0.001979 | 0.003321 | 0.002131 | 0.025569 | 0.005641 |
| 23.7916667 | 0.010578 | 0.01735 | 0.004282 | 0.009198 | 0.003866 | 0.00235 | 0.005286 | 0.007324 | 0.002659 | 0.005186 | 0.076179 | 0.001414 | 0.001819 | 0.001753 | 0.000525 | 0.005107 | 0.002349 | 0.015141 | 0.005666 |
| 23.8055556 | 0.008867 | 0.008821 | 0.006407 | 0.007314 | 0.002101 | 0.003255 | 0.007402 | 0.009744 | 0.00347 | 0.004434 | 0.081921 | 0.000998 | 0.002059 | 0.003211 | 0.000553 | 0.004819 | 0.003202 | 0.01052 | 0.008832 |
| 23.8194444 | 0.005447 | 0.012575 | 0.009465 | 0.01347 | 0.002953 | 0.003709 | 0.013826 | 0.019172 | 0.006704 | 0.003813 | 0.070381 | 0.001125 | 0.003456 | 0.00392 | 0.001435 | 0.002872 | 0.0063 | 0.017469 | 0.009178 |
| 23.8333333 | 0.005506 | 0.017668 | 0.008341 | 0.018628 | 0.003761 | 0.002919 | 0.015574 | 0.02698 | 0.009166 | 0.004438 | 0.055579 | 0.001441 | 0.003504 | 0.002974 | 0.002073 | 0.001257 | 0.009688 | 0.022143 | 0.006129 |
| 23.8472222 | 0.009002 | 0.015255 | 0.005173 | 0.01626 | 0.007409 | 0.001414 | 0.011073 | 0.02719 | 0.009602 | 0.004553 | 0.046425 | 0.001286 | 0.001742 | 0.001866 | 0.001849 | 0.001588 | 0.01159 | 0.015548 | 0.002521 |
| 23.8611111 | 0.011816 | 0.009796 | 0.005304 | 0.009293 | 0.012633 | 0.000778 | 0.006563 | 0.021277 | 0.008407 | 0.0099 | 0.039451 | 0.001285 | 0.000955 | 0.001524 | 0.002032 | 0.003629 | 0.013671 | 0.00627 | 0.001184 |
| 23.875 | 0.011199 | 0.004885 | 0.008944 | 0.003447 | 0.016881 | 0.000857 | 0.006358 | 0.012606 | 0.005626 | 0.019407 | 0.032972 | 0.001956 | 0.002077 | 0.00096 | 0.00326 | 0.005819 | 0.017885 | 0.00324 | 0.002908 |
| 23.8888889 | 0.006853 | 0.001563 | 0.012223 | 0.001754 | 0.020534 | 0.001497 | 0.008839 | 0.005809 | 0.002843 | 0.020943 | 0.026606 | 0.002445 | 0.002678 | 0.000648 | 0.003805 | 0.006353 | 0.023779 | 0.005199 | 0.008644 |
| 23.9027778 | 0.002961 | 0.002648 | 0.014485 | 0.003521 | 0.023988 | 0.00295 | 0.013279 | 0.00698 | 0.001879 | 0.012465 | 0.017156 | 0.002606 | 0.002042 | 0.001063 | 0.003674 | 0.004925 | 0.027555 | 0.010899 | 0.018178 |
| 23.9166667 | 0.003808 | 0.010273 | 0.016315 | 0.006763 | 0.028921 | 0.005228 | 0.020821 | 0.017667 | 0.0036 | 0.005329 | 0.007212 | 0.003679 | 0.00197 | 0.002622 | 0.005273 | 0.003453 | 0.027422 | 0.017251 | 0.027946 |
| 23.9305556 | 0.008639 | 0.021771 | 0.016588 | 0.00977 | 0.036086 | 0.007434 | 0.028499 | 0.031861 | 0.009198 | 0.004161 | 0.003021 | 0.005584 | 0.002537 | 0.004972 | 0.008708 | 0.003601 | 0.026193 | 0.020558 | 0.033409 |
| 23.9444444 | 0.014113 | 0.031229 | 0.014823 | 0.011844 | 0.044395 | 0.008442 | 0.033076 | 0.042705 | 0.015471 | 0.007617 | 0.006058 | 0.00699 | 0.002155 | 0.006359 | 0.011871 | 0.005187 | 0.026819 | 0.022667 | 0.033498 |
| 23.9583333 | 0.018827 | 0.035796 | 0.012757 | 0.01403 | 0.052368 | 0.008901 | 0.035508 | 0.048262 | 0.01951 | 0.01979 | 0.015203 | 0.007636 | 0.000993 | 0.00634 | 0.013919 | 0.007241 | 0.028522 | 0.026293 | 0.030553 |
| 23.9722222 | 0.02314 | 0.03675 | 0.012376 | 0.016783 | 0.056324 | 0.009752 | 0.035947 | 0.048233 | 0.021861 | 0.03359 | 0.025901 | 0.007473 | 0.001164 | 0.00541 | 0.014518 | 0.008731 | 0.027883 | 0.030088 | 0.027066 |
| 23.9861111 | 0.026202 | 0.035794 | 0.012832 | 0.017634 | 0.053757 | 0.010474 | 0.031358 | 0.043258 | 0.022114 | 0.032918 | 0.031182 | 0.00618 | 0.002749 | 0.003852 | 0.012799 | 0.00844 | 0.022727 | 0.030147 | 0.025387 |
| 24 | 0.026456 | 0.033005 | 0.012367 | 0.015326 | 0.047906 | 0.010712 | 0.024331 | 0.037876 | 0.020624 | 0.018699 | 0.026502 | 0.005597 | 0.003612 | 0.002955 | 0.009512 | 0.005952 | 0.014651 | 0.026321 | 0.027074 |
| 24.0138889 | 0.023931 | 0.028339 | 0.011189 | 0.012811 | 0.042879 | 0.011301 | 0.022905 | 0.035563 | 0.018797 | 0.008214 | 0.015124 | 0.007202 | 0.003349 | 0.003823 | 0.006914 | 0.002905 | 0.007783 | 0.021557 | 0.030087 |
| 24.0277778 | 0.019753 | 0.023807 | 0.00997 | 0.011992 | 0.040023 | 0.012469 | 0.026602 | 0.034229 | 0.016764 | 0.006542 | 0.009018 | 0.008284 | 0.00229 | 0.004944 | 0.005484 | 0.00162 | 0.003986 | 0.017307 | 0.030327 |
| 24.0416667 | 0.015986 | 0.022243 | 0.008777 | 0.011533 | 0.039491 | 0.013496 | 0.027929 | 0.031364 | 0.014507 | 0.010135 | 0.009742 | 0.006725 | 0.001144 | 0.00471 | 0.004161 | 0.002217 | 0.001963 | 0.013511 | 0.026848 |

| | | | | | | | | | | | | | | | | | | | |
|---|---|---|---|---|---|---|---|---|---|---|---|---|---|---|---|---|---|---|---|
| 24.0555556 | 0.014995 | 0.024737 | 0.007815 | 0.011284 | 0.039408 | 0.014441 | 0.025767 | 0.027367 | 0.013463 | 0.02271 | 0.007767 | 0.004507 | 0.000833 | 0.00378 | 0.003284 | 0.003034 | 0.001496 | 0.010996 | 0.021843 |
| 24.0694444 | 0.015331 | 0.029843 | 0.006893 | 0.012294 | 0.035157 | 0.017723 | 0.024141 | 0.022206 | 0.013982 | 0.033833 | 0.006036 | 0.003204 | 0.000757 | 0.002778 | 0.003069 | 0.002826 | 0.005013 | 0.012713 | 0.017534 |
| 24.0833333 | 0.012455 | 0.034973 | 0.004896 | 0.013231 | 0.024712 | 0.027373 | 0.024814 | 0.015239 | 0.013397 | 0.030963 | 0.011544 | 0.002533 | 0.000556 | 0.001826 | 0.002286 | 0.002567 | 0.013692 | 0.021628 | 0.014916 |
| 24.0972222 | 0.006823 | 0.035861 | 0.002148 | 0.011823 | 0.012889 | 0.03806 | 0.027365 | 0.007999 | 0.009385 | 0.020378 | 0.019993 | 0.002857 | 0.001076 | 0.001863 | 0.00178 | 0.004432 | 0.021746 | 0.031689 | 0.013353 |
| 24.1111111 | 0.004648 | 0.029566 | 0.001195 | 0.008265 | 0.007058 | 0.033009 | 0.029202 | 0.002935 | 0.004387 | 0.011593 | 0.026668 | 0.002877 | 0.002642 | 0.00218 | 0.002657 | 0.006173 | 0.021013 | 0.030821 | 0.011425 |
| 24.125 | 0.006199 | 0.019705 | 0.002667 | 0.005645 | 0.007474 | 0.015421 | 0.026816 | 0.002584 | 0.002747 | 0.007365 | 0.029931 | 0.001859 | 0.003281 | 0.001794 | 0.003098 | 0.004427 | 0.01378 | 0.020963 | 0.009439 |
| 24.1388889 | 0.006352 | 0.010206 | 0.003902 | 0.00747 | 0.006765 | 0.004705 | 0.020711 | 0.007309 | 0.004241 | 0.005284 | 0.026326 | 0.001382 | 0.002064 | 0.0022 | 0.002472 | 0.001634 | 0.009114 | 0.013778 | 0.008151 |
| 24.1527778 | 0.007465 | 0.005627 | 0.003381 | 0.012307 | 0.003033 | 0.002047 | 0.01463 | 0.010923 | 0.005407 | 0.003936 | 0.0153 | 0.001346 | 0.000764 | 0.003415 | 0.001917 | 0.000736 | 0.00809 | 0.01112 | 0.006479 |
| 24.1666667 | 0.010439 | 0.008182 | 0.002623 | 0.014579 | 0.001061 | 0.001436 | 0.010547 | 0.008139 | 0.004908 | 0.00555 | 0.008292 | 0.001633 | 0.000589 | 0.003974 | 0.001596 | 0.001091 | 0.005558 | 0.009318 | 0.004324 |
| 24.1805556 | 0.008459 | 0.009009 | 0.003716 | 0.012167 | 0.002478 | 0.002732 | 0.008562 | 0.00345 | 0.004256 | 0.009665 | 0.01359 | 0.001754 | 0.001209 | 0.0039 | 0.001116 | 0.001964 | 0.003513 | 0.007845 | 0.002731 |
| 24.1944444 | 0.005512 | 0.004652 | 0.007205 | 0.007643 | 0.004952 | 0.008712 | 0.008206 | 0.001452 | 0.005199 | 0.012939 | 0.020388 | 0.001921 | 0.001489 | 0.00368 | 0.001022 | 0.003313 | 0.007164 | 0.005696 | 0.001871 |
| 24.2083333 | 0.007277 | 0.001552 | 0.009628 | 0.003984 | 0.004878 | 0.017857 | 0.007107 | 0.000865 | 0.0064 | 0.017417 | 0.022444 | 0.002518 | 0.001399 | 0.00264 | 0.001533 | 0.003989 | 0.01321 | 0.003913 | 0.002582 |
| 24.2222222 | 0.006936 | 0.002792 | 0.006852 | 0.003463 | 0.004549 | 0.02056 | 0.004649 | 0.001562 | 0.004771 | 0.017339 | 0.025094 | 0.00226 | 0.00206 | 0.001471 | 0.001768 | 0.002729 | 0.015021 | 0.005667 | 0.004181 |
| 24.2361111 | 0.005001 | 0.00621 | 0.002399 | 0.005279 | 0.007411 | 0.01376 | 0.004054 | 0.002594 | 0.002154 | 0.009181 | 0.027378 | 0.001556 | 0.002405 | 0.001326 | 0.002463 | 0.000922 | 0.010344 | 0.007277 | 0.00387 |
| 24.25 | 0.005754 | 0.008212 | 0.002618 | 0.005259 | 0.007888 | 0.005304 | 0.00522 | 0.002319 | 0.001342 | 0.009113 | 0.023086 | 0.002371 | 0.00265 | 0.002533 | 0.003266 | 0.00171 | 0.010613 | 0.006912 | 0.007072 |
| 24.2638889 | 0.005841 | 0.007916 | 0.009403 | 0.005741 | 0.004925 | 0.001875 | 0.004655 | 0.004193 | 0.00179 | 0.022546 | 0.034836 | 0.00801 | 0.006604 | 0.007558 | 0.008398 | 0.005244 | 0.03517 | 0.016082 | 0.027126 |
| 24.2777778 | 0.008978 | 0.01509 | 0.019036 | 0.017237 | 0.002825 | 0.003654 | 0.007031 | 0.012793 | 0.003722 | 0.050821 | 0.079876 | 0.016241 | 0.012538 | 0.014915 | 0.021303 | 0.007544 | 0.07814 | 0.050617 | 0.060007 |
| 24.2916667 | 0.017865 | 0.032966 | 0.022089 | 0.037216 | 0.001932 | 0.006639 | 0.01705 | 0.02446 | 0.005954 | 0.083321 | 0.098685 | 0.017099 | 0.012592 | 0.018042 | 0.029704 | 0.007148 | 0.094513 | 0.089287 | 0.072169 |
| 24.3055556 | 0.02198 | 0.039442 | 0.01575 | 0.044141 | 0.002667 | 0.007115 | 0.024288 | 0.027791 | 0.006833 | 0.07612 | 0.05748 | 0.009245 | 0.006502 | 0.014307 | 0.021486 | 0.006313 | 0.060382 | 0.080261 | 0.045228 |
| 24.3194444 | 0.017323 | 0.026556 | 0.008134 | 0.03159 | 0.004487 | 0.006516 | 0.022242 | 0.02044 | 0.006428 | 0.034229 | 0.018769 | 0.003198 | 0.002142 | 0.007972 | 0.008305 | 0.004836 | 0.02077 | 0.035184 | 0.015463 |
| 24.3333333 | 0.010877 | 0.012733 | 0.005291 | 0.016763 | 0.005316 | 0.007906 | 0.018311 | 0.01281 | 0.005485 | 0.010536 | 0.014708 | 0.003371 | 0.001148 | 0.004018 | 0.002503 | 0.002342 | 0.01042 | 0.008958 | 0.010097 |
| 24.3472222 | 0.006423 | 0.006812 | 0.006967 | 0.009497 | 0.0049 | 0.009923 | 0.017521 | 0.010143 | 0.005778 | 0.00798 | 0.014377 | 0.003981 | 0.001388 | 0.003364 | 0.002504 | 0.001082 | 0.009882 | 0.005949 | 0.008754 |
| 24.3611111 | 0.004584 | 0.005451 | 0.010581 | 0.006052 | 0.004214 | 0.009842 | 0.018548 | 0.008426 | 0.008222 | 0.006021 | 0.007179 | 0.00249 | 0.003163 | 0.004258 | 0.004366 | 0.001659 | 0.005528 | 0.006094 | 0.003195 |
| 24.375 | 0.004527 | 0.005617 | 0.012313 | 0.004248 | 0.003052 | 0.008549 | 0.018198 | 0.005049 | 0.010271 | 0.00412 | 0.002365 | 0.001341 | 0.004687 | 0.00403 | 0.004558 | 0.002024 | 0.003231 | 0.004884 | 0.001585 |
| 24.3888889 | 0.003665 | 0.004633 | 0.009597 | 0.006849 | 0.002995 | 0.007832 | 0.01445 | 0.002808 | 0.008661 | 0.003738 | 0.000847 | 0.000831 | 0.004635 | 0.002299 | 0.002672 | 0.001229 | 0.004963 | 0.002822 | 0.001619 |
| 24.4027778 | 0.004218 | 0.003425 | 0.005111 | 0.013801 | 0.006613 | 0.006831 | 0.00815 | 0.003782 | 0.004693 | 0.002981 | 0.000715 | 0.000363 | 0.003876 | 0.000935 | 0.001238 | 0.000648 | 0.007159 | 0.000953 | 0.001934 |
| 24.4166667 | 0.010176 | 0.00612 | 0.004241 | 0.020402 | 0.011568 | 0.004734 | 0.003841 | 0.006138 | 0.003193 | 0.001837 | 0.001053 | 0.000265 | 0.002806 | 0.000531 | 0.000964 | 0.000888 | 0.007248 | 0.0005 | 0.003199 |
| 24.4305556 | 0.017653 | 0.010515 | 0.006875 | 0.022981 | 0.013847 | 0.003306 | 0.004123 | 0.00814 | 0.005107 | 0.002134 | 0.001523 | 0.000724 | 0.001857 | 0.000545 | 0.000752 | 0.000917 | 0.005663 | 0.001441 | 0.004244 |
| 24.4444444 | 0.01942 | 0.009682 | 0.007318 | 0.020581 | 0.011886 | 0.003635 | 0.00422 | 0.008353 | 0.005389 | 0.002489 | 0.001818 | 0.001237 | 0.001452 | 0.0007 | 0.000817 | 0.000791 | 0.004219 | 0.002856 | 0.004659 |
| 24.4583333 | 0.013148 | 0.006203 | 0.004626 | 0.015298 | 0.007671 | 0.006861 | 0.002638 | 0.006612 | 0.003901 | 0.001814 | 0.001856 | 0.000923 | 0.001144 | 0.000746 | 0.001564 | 0.00125 | 0.003425 | 0.003945 | 0.004599 |
| 24.4722222 | 0.006368 | 0.008253 | 0.004589 | 0.009266 | 0.004829 | 0.014337 | 0.003378 | 0.00408 | 0.006115 | 0.003285 | 0.001601 | 0.000497 | 0.001031 | 0.00095 | 0.001793 | 0.001388 | 0.002503 | 0.004851 | 0.003254 |
| 24.4861111 | 0.006586 | 0.01289 | 0.010283 | 0.005583 | 0.003591 | 0.022938 | 0.006426 | 0.00446 | 0.011789 | 0.010281 | 0.00163 | 0.00075 | 0.001383 | 0.001357 | 0.001368 | 0.000992 | 0.002067 | 0.005529 | 0.001337 |
| 24.5 | 0.006814 | 0.011215 | 0.016373 | 0.005735 | 0.003155 | 0.026762 | 0.007727 | 0.008603 | 0.015039 | 0.020973 | 0.002006 | 0.001359 | 0.001379 | 0.001115 | 0.001134 | 0.001137 | 0.002512 | 0.005545 | 0.000834 |
| 24.5138889 | 0.00671 | 0.008843 | 0.014747 | 0.007501 | 0.005434 | 0.021677 | 0.007291 | 0.008837 | 0.010982 | 0.031589 | 0.003157 | 0.001547 | 0.001956 | 0.000843 | 0.001112 | 0.001184 | 0.00356 | 0.005636 | 0.001756 |
| 24.5277778 | 0.011908 | 0.014581 | 0.007979 | 0.013537 | 0.01082 | 0.01081 | 0.010675 | 0.007416 | 0.005784 | 0.039396 | 0.008126 | 0.00108 | 0.004186 | 0.001153 | 0.001182 | 0.00059 | 0.004904 | 0.006103 | 0.002839 |
| 24.5416667 | 0.012799 | 0.019092 | 0.005971 | 0.021594 | 0.01653 | 0.004396 | 0.016246 | 0.011977 | 0.007815 | 0.042014 | 0.024156 | 0.000706 | 0.005732 | 0.001879 | 0.00083 | 0.000332 | 0.005645 | 0.005991 | 0.00279 |
| 24.5555556 | 0.009774 | 0.019233 | 0.010187 | 0.025401 | 0.021356 | 0.007618 | 0.019744 | 0.019554 | 0.013205 | 0.041535 | 0.045932 | 0.000911 | 0.005108 | 0.002913 | 0.000876 | 0.000915 | 0.006862 | 0.00612 | 0.002523 |
| 24.5694444 | 0.01609 | 0.026051 | 0.016629 | 0.027059 | 0.026053 | 0.016069 | 0.025921 | 0.027936 | 0.01907 | 0.046168 | 0.057458 | 0.002186 | 0.003819 | 0.003721 | 0.002274 | 0.002576 | 0.010972 | 0.009164 | 0.004775 |
| 24.5833333 | 0.029958 | 0.041826 | 0.020965 | 0.027391 | 0.030319 | 0.024395 | 0.039314 | 0.035572 | 0.026211 | 0.063564 | 0.0655 | 0.004015 | 0.002507 | 0.004473 | 0.003378 | 0.004789 | 0.019677 | 0.016006 | 0.010077 |
| 24.5972222 | 0.036152 | 0.053487 | 0.020925 | 0.023772 | 0.034322 | 0.028065 | 0.055862 | 0.038923 | 0.030807 | 0.085204 | 0.075993 | 0.005226 | 0.001831 | 0.004688 | 0.003009 | 0.005951 | 0.030975 | 0.022979 | 0.014874 |
| 24.6111111 | 0.029871 | 0.048769 | 0.02081 | 0.01867 | 0.039689 | 0.023605 | 0.066638 | 0.040562 | 0.028461 | 0.084552 | 0.070662 | 0.006011 | 0.002229 | 0.003625 | 0.002254 | 0.005521 | 0.035362 | 0.024853 | 0.015006 |
| 24.625 | 0.021482 | 0.033359 | 0.021883 | 0.016649 | 0.039774 | 0.015125 | 0.064431 | 0.04654 | 0.021192 | 0.05623 | 0.047078 | 0.006704 | 0.001949 | 0.001963 | 0.002391 | 0.004303 | 0.027522 | 0.021445 | 0.01146 |
| 24.6388889 | 0.016091 | 0.018443 | 0.017564 | 0.017342 | 0.025315 | 0.010119 | 0.047483 | 0.055103 | 0.013494 | 0.026784 | 0.02501 | 0.006203 | 0.000856 | 0.000827 | 0.003336 | 0.003194 | 0.015885 | 0.015697 | 0.008009 |
| 24.6527778 | 0.010856 | 0.007611 | 0.009655 | 0.016013 | 0.012989 | 0.010311 | 0.02387 | 0.056639 | 0.006995 | 0.01153 | 0.013331 | 0.004435 | 0.000413 | 0.000619 | 0.00409 | 0.002638 | 0.010628 | 0.009781 | 0.006176 |
| 24.6666667 | 0.006861 | 0.001811 | 0.006503 | 0.010219 | 0.020201 | 0.013016 | 0.010179 | 0.046434 | 0.004591 | 0.005941 | 0.00813 | 0.003107 | 0.000836 | 0.001368 | 0.004133 | 0.002525 | 0.015688 | 0.006079 | 0.005812 |
| 24.6805556 | 0.008327 | 0.000791 | 0.006248 | 0.004186 | 0.029466 | 0.01519 | 0.007644 | 0.031798 | 0.005818 | 0.004407 | 0.005799 | 0.002713 | 0.001453 | 0.002797 | 0.00359 | 0.002443 | 0.023618 | 0.00483 | 0.006516 |
| 24.6944444 | 0.015556 | 0.003342 | 0.009831 | 0.003196 | 0.022668 | 0.016321 | 0.011199 | 0.024558 | 0.005328 | 0.004078 | 0.004545 | 0.002588 | 0.001769 | 0.003895 | 0.002657 | 0.002083 | 0.02172 | 0.004192 | 0.007446 |
| 24.7083333 | 0.024591 | 0.010518 | 0.02201 | 0.00689 | 0.015384 | 0.01865 | 0.030242 | 0.027372 | 0.006617 | 0.003763 | 0.003315 | 0.002475 | 0.001691 | 0.004544 | 0.001636 | 0.001544 | 0.013015 | 0.003397 | 0.007615 |
| 24.7222222 | 0.030133 | 0.022925 | 0.032634 | 0.010697 | 0.028249 | 0.021617 | 0.052351 | 0.032147 | 0.01563 | 0.003163 | 0.001945 | 0.002487 | 0.001419 | 0.005618 | 0.00112 | 0.00122 | 0.007446 | 0.003143 | 0.007185 |
| 24.7361111 | 0.02902 | 0.037658 | 0.029776 | 0.011678 | 0.046895 | 0.022151 | 0.051516 | 0.032885 | 0.025657 | 0.002684 | 0.001545 | 0.002657 | 0.001404 | 0.007206 | 0.001473 | 0.001286 | 0.005841 | 0.004514 | 0.006892 |
| 24.75 | 0.022409 | 0.047717 | 0.016337 | 0.010973 | 0.048708 | 0.019385 | 0.029495 | 0.03363 | 0.030338 | 0.002739 | 0.001837 | 0.002906 | 0.001681 | 0.00845 | 0.002143 | 0.001508 | 0.00527 | 0.006888 | 0.006672 |
| 24.7638889 | 0.014727 | 0.045606 | 0.005703 | 0.011961 | 0.032778 | 0.014004 | 0.011715 | 0.038933 | 0.030226 | 0.003236 | 0.001539 | 0.003109 | 0.001828 | 0.008511 | 0.002282 | 0.001638 | 0.004269 | 0.008091 | 0.006474 |
| 24.7777778 | 0.010632 | 0.032939 | 0.004119 | 0.016043 | 0.016017 | 0.007535 | 0.013052 | 0.045089 | 0.025847 | 0.00376 | 0.001372 | 0.003004 | 0.001625 | 0.007202 | 0.001737 | 0.001583 | 0.003199 | 0.00745 | 0.006479 |
| 24.7916667 | 0.011578 | 0.019391 | 0.003884 | 0.019852 | 0.014029 | 0.00286 | 0.017359 | 0.04526 | 0.018569 | 0.004352 | 0.001754 | 0.002552 | 0.001291 | 0.005229 | 0.001095 | 0.001432 | 0.00298 | 0.006265 | 0.006402 |
| 24.8055556 | 0.015882 | 0.009933 | 0.003777 | 0.019905 | 0.020208 | 0.001378 | 0.012529 | 0.036597 | 0.010671 | 0.005661 | 0.00204 | 0.002302 | 0.001295 | 0.0038 | 0.001126 | 0.001592 | 0.003805 | 0.005657 | 0.006182 |
| 24.8194444 | 0.019634 | 0.005093 | 0.00811 | 0.015003 | 0.022486 | 0.001392 | 0.009494 | 0.020868 | 0.004342 | 0.006958 | 0.002412 | 0.002359 | 0.001701 | 0.003316 | 0.001809 | 0.002147 | 0.00477 | 0.005702 | 0.006118 |
| 24.8333333 | 0.018042 | 0.007158 | 0.011887 | 0.009543 | 0.01999 | 0.001709 | 0.010059 | 0.012935 | 0.001675 | 0.006422 | 0.002608 | 0.00181 | 0.001727 | 0.00307 | 0.002035 | 0.002335 | 0.004582 | 0.005657 | 0.005759 |
| 24.8472222 | 0.010791 | 0.013977 | 0.009334 | 0.010985 | 0.014661 | 0.002239 | 0.011801 | 0.026565 | 0.002614 | 0.004286 | 0.002016 | 0.000939 | 0.001033 | 0.002931 | 0.001297 | 0.001652 | 0.003157 | 0.004934 | 0.004565 |
| 24.8611111 | 0.004308 | 0.017169 | 0.005179 | 0.014176 | 0.008624 | 0.002143 | 0.025641 | 0.046064 | 0.004086 | 0.002558 | 0.001365 | 0.000614 | 0.000659 | 0.003603 | 0.000755 | 0.000941 | 0.002004 | 0.004221 | 0.00337 |
| 24.875 | 0.004085 | 0.011779 | 0.006981 | 0.010717 | 0.007842 | 0.002224 | 0.044908 | 0.05121 | 0.004093 | 0.002186 | 0.001517 | 0.000616 | 0.001167 | 0.004531 | 0.001133 | 0.001052 | 0.002025 | 0.00437 | 0.00306 |
| 24.8888889 | 0.008292 | 0.007144 | 0.012291 | 0.006383 | 0.011964 | 0.003159 | 0.04739 | 0.04192 | 0.003365 | 0.002175 | 0.001773 | 0.000868 | 0.001512 | 0.004309 | 0.001438 | 0.001323 | 0.002268 | 0.00479 | 0.002944 |
| 24.9027778 | 0.01109 | 0.012002 | 0.015597 | 0.007341 | 0.01216 | 0.003067 | 0.030346 | 0.026726 | 0.002319 | 0.001638 | 0.001548 | 0.001021 | 0.001224 | 0.003328 | 0.001168 | 0.001082 | 0.0021 | 0.004632 | 0.002347 |
| 24.9166667 | 0.009088 | 0.015373 | 0.016078 | 0.009239 | 0.008226 | 0.002014 | 0.016177 | 0.01321 | 0.001761 | 0.001193 | 0.001513 | 0.001086 | 0.001086 | 0.002661 | 0.001052 | 0.000883 | 0.002031 | 0.004183 | 0.001797 |
| 24.9305556 | 0.00599 | 0.010737 | 0.015582 | 0.009018 | 0.010681 | 0.003376 | 0.011914 | 0.008255 | 0.001818 | 0.001161 | 0.001779 | 0.001253 | 0.001085 | 0.001876 | 0.001132 | 0.000975 | 0.001992 | 0.003865 | 0.001485 |
| 24.9444444 | 0.007483 | 0.009265 | 0.01607 | 0.009987 | 0.023888 | 0.009156 | 0.012209 | 0.014923 | 0.002724 | 0.000885 | 0.001824 | 0.001079 | 0.00076 | 0.000875 | 0.00103 | 0.000864 | 0.001635 | 0.00356 | 0.00107 |
| 24.9583333 | 0.012936 | 0.015698 | 0.017095 | 0.013696 | 0.04013 | 0.016373 | 0.024794 | 0.027462 | 0.007797 | 0.000578 | 0.001969 | 0.000981 | 0.000685 | 0.000663 | 0.001319 | 0.000756 | 0.001492 | 0.003603 | 0.001087 |
| 24.9722222 | 0.018225 | 0.023252 | 0.018329 | 0.018779 | 0.050616 | 0.0212 | 0.038977 | 0.038437 | 0.016664 | 0.000773 | 0.002297 | 0.001332 | 0.001058 | 0.000941 | 0.002084 | 0.000946 | 0.001652 | 0.004172 | 0.001854 |
| 24.9861111 | 0.02127 | 0.029105 | 0.019786 | 0.02323 | 0.052494 | 0.022902 | 0.04206 | 0.045296 | 0.023737 | 0.000863 | 0.001881 | 0.001182 | 0.000933 | 0.00071 | 0.002133 | 0.00089 | 0.001246 | 0.004153 | 0.002359 |
| 25 | 0.021463 | 0.033003 | 0.018641 | 0.024129 | 0.045931 | 0.020908 | 0.036247 | 0.047522 | 0.023987 | 0.000557 | 0.001169 | 0.000579 | 0.000527 | 0.000647 | 0.001599 | 0.000472 | 0.000655 | 0.003343 | 0.002598 |
| 25.0138889 | 0.019674 | 0.033823 | 0.013583 | 0.021248 | 0.033209 | 0.016133 | 0.023682 | 0.045577 | 0.018529 | 0.000558 | 0.001445 | 0.000762 | 0.000842 | 0.001528 | 0.001735 | 0.000262 | 0.000923 | 0.00296 | 0.003496 |
| 25.0277778 | 0.0198 | 0.033722 | 0.009297 | 0.017481 | 0.020248 | 0.01235 | 0.011712 | 0.042722 | 0.013577 | 0.000754 | 0.001787 | 0.001205 | 0.001193 | 0.00257 | 0.00208 | 0.00032 | 0.001289 | 0.002774 | 0.004282 |
| 25.0416667 | 0.023799 | 0.033389 | 0.00918 | 0.014638 | 0.012338 | 0.010446 | 0.008823 | 0.038976 | 0.01179 | 0.000495 | 0.001169 | 0.001033 | 0.000896 | 0.003166 | 0.001941 | 0.000261 | 0.000981 | 0.002041 | 0.004098 |
| 25.0555556 | 0.028158 | 0.029961 | 0.009971 | 0.013248 | 0.010685 | 0.008513 | 0.016044 | 0.02998 | 0.011309 | 0.000201 | 0.000513 | 0.000805 | 0.000736 | 0.003462 | 0.001916 | 0.000291 | 0.000517 | 0.001313 | 0.003523 |
| 25.0694444 | 0.02784 | 0.021977 | 0.00717 | 0.011801 | 0.011688 | 0.0055 | 0.025754 | 0.016567 | 0.009223 | 0.000979 | 0.00199 | 0.000657 | 0.001267 | 0.003861 | 0.001978 | 0.00056 | 0.001028 | 0.001575 | 0.002839 |
| 25.0833333 | 0.02003 | 0.011776 | 0.006778 | 0.009657 | 0.009448 | 0.007154 | 0.02594 | 0.011168 | 0.008097 | 0.004586 | 0.01012 | 0.000466 | 0.002971 | 0.005637 | 0.001554 | 0.000921 | 0.005412 | 0.00499 | 0.004847 |
| 25.0972222 | 0.00898 | 0.004551 | 0.016388 | 0.012445 | 0.007117 | 0.01931 | 0.016131 | 0.018132 | 0.014158 | 0.011802 | 0.02761 | 0.000931 | 0.005535 | 0.00861 | 0.002179 | 0.001322 | 0.01736 | 0.014602 | 0.012071 |
| 25.1111111 | 0.005257 | 0.00461 | 0.025526 | 0.019559 | 0.014915 | 0.031129 | 0.010823 | 0.019906 | 0.020822 | 0.01778 | 0.041778 | 0.001932 | 0.006234 | 0.009091 | 0.002737 | 0.001449 | 0.029209 | 0.025585 | 0.017211 |
| 25.125 | 0.00959 | 0.011403 | 0.024283 | 0.021209 | 0.029091 | 0.030581 | 0.013469 | 0.011051 | 0.017014 | 0.014655 | 0.032669 | 0.002252 | 0.003919 | 0.005703 | 0.002396 | 0.001623 | 0.024583 | 0.022451 | 0.012912 |
| 25.1388889 | 0.010817 | 0.018093 | 0.017431 | 0.014783 | 0.033241 | 0.020737 | 0.013906 | 0.004705 | 0.008648 | 0.007784 | 0.01495 | 0.001871 | 0.001316 | 0.002133 | 0.000963 | 0.002136 | 0.013281 | 0.011215 | 0.007146 |
| 25.1527778 | 0.008155 | 0.01649 | 0.010367 | 0.007956 | 0.024272 | 0.009545 | 0.008833 | 0.003592 | 0.006692 | 0.008335 | 0.012548 | 0.001441 | 0.000385 | 0.000614 | 0.000386 | 0.002134 | 0.012305 | 0.010288 | 0.007074 |
| 25.1666667 | 0.01167 | 0.010357 | 0.005092 | 0.007211 | 0.013782 | 0.003049 | 0.003652 | 0.002976 | 0.00861 | 0.012481 | 0.013798 | 0.000949 | 0.000492 | 0.000273 | 0.000705 | 0.001263 | 0.011854 | 0.011206 | 0.006689 |
| 25.1805556 | 0.021226 | 0.011001 | 0.003107 | 0.0078 | 0.008021 | 0.001658 | 0.002108 | 0.003227 | 0.007974 | 0.014376 | 0.010825 | 0.000572 | 0.000598 | 0.000488 | 0.000999 | 0.000457 | 0.007455 | 0.007123 | 0.004157 |
| 25.1944444 | 0.027508 | 0.016889 | 0.004656 | 0.004896 | 0.004678 | 0.00105 | 0.001607 | 0.004535 | 0.005138 | 0.015234 | 0.009399 | 0.000646 | 0.000391 | 0.000844 | 0.000766 | 0.00036 | 0.004905 | 0.004266 | 0.002782 |
| 25.2083333 | 0.027327 | 0.019201 | 0.008187 | 0.002813 | 0.00306 | 0.000627 | 0.00135 | 0.004449 | 0.002575 | 0.016003 | 0.009734 | 0.001029 | 0.000226 | 0.000786 | 0.000454 | 0.000566 | 0.004116 | 0.003901 | 0.00258 |
| 25.2222222 | 0.024417 | 0.017086 | 0.011382 | 0.003807 | 0.004566 | 0.001544 | 0.003735 | 0.003224 | 0.001977 | 0.01739 | 0.011732 | 0.001531 | 0.0004 | 0.000672 | 0.000653 | 0.000651 | 0.004823 | 0.005041 | 0.00299 |
| 25.2361111 | 0.021616 | 0.013962 | 0.012027 | 0.004888 | 0.006627 | 0.002507 | 0.007717 | 0.002508 | 0.003238 | 0.018976 | 0.014548 | 0.002052 | 0.00076 | 0.001033 | 0.001358 | 0.000688 | 0.006972 | 0.005933 | 0.003729 |

| | | | | | | | | | | | | | | | | | | | |
|---|---|---|---|---|---|---|---|---|---|---|---|---|---|---|---|---|---|---|---|
| 25.25 | 0.019272 | 0.011575 | 0.009604 | 0.004669 | 0.007075 | 0.002228 | 0.010362 | 0.002757 | 0.004292 | 0.018634 | 0.015868 | 0.002083 | 0.00093 | 0.001538 | 0.002037 | 0.000624 | 0.008967 | 0.005221 | 0.004279 |
| 25.2638889 | 0.017398 | 0.010067 | 0.005792 | 0.004029 | 0.007554 | 0.001349 | 0.010272 | 0.003853 | 0.004217 | 0.015387 | 0.014559 | 0.001277 | 0.000687 | 0.001763 | 0.002127 | 0.000396 | 0.008839 | 0.003247 | 0.004087 |
| 25.2777778 | 0.015861 | 0.009073 | 0.002823 | 0.004125 | 0.008217 | 0.001587 | 0.008354 | 0.005166 | 0.003342 | 0.010162 | 0.011907 | 0.000757 | 0.00069 | 0.001528 | 0.002123 | 0.000496 | 0.006535 | 0.001488 | 0.002783 |
| 25.2916667 | 0.014263 | 0.008135 | 0.001568 | 0.005384 | 0.006235 | 0.002862 | 0.006169 | 0.005583 | 0.002383 | 0.004862 | 0.009721 | 0.001669 | 0.001899 | 0.000859 | 0.003046 | 0.001282 | 0.003788 | 0.000959 | 0.001371 |
| 25.3055556 | 0.012395 | 0.006604 | 0.00108 | 0.006672 | 0.004055 | 0.004015 | 0.004218 | 0.004346 | 0.002593 | 0.001522 | 0.010085 | 0.002663 | 0.003153 | 0.000627 | 0.004164 | 0.001849 | 0.002962 | 0.00094 | 0.001173 |
| 25.3194444 | 0.009794 | 0.003985 | 0.001212 | 0.006825 | 0.005492 | 0.004684 | 0.002472 | 0.002242 | 0.003969 | 0.001189 | 0.012807 | 0.002127 | 0.002549 | 0.001725 | 0.00355 | 0.001543 | 0.004291 | 0.000785 | 0.00227 |
| 25.3333333 | 0.005871 | 0.002374 | 0.002489 | 0.00608 | 0.005304 | 0.004807 | 0.002739 | 0.000992 | 0.004287 | 0.002187 | 0.013434 | 0.001124 | 0.001144 | 0.002956 | 0.001851 | 0.001181 | 0.004206 | 0.000948 | 0.003604 |
| 25.3472222 | 0.002404 | 0.004268 | 0.003324 | 0.005415 | 0.003676 | 0.004061 | 0.00599 | 0.001109 | 0.002657 | 0.002207 | 0.00991 | 0.000897 | 0.000937 | 0.002412 | 0.001391 | 0.001115 | 0.002963 | 0.001916 | 0.002941 |
| 25.3611111 | 0.001441 | 0.007176 | 0.002589 | 0.005416 | 0.005613 | 0.002789 | 0.007827 | 0.002711 | 0.001197 | 0.002318 | 0.007317 | 0.000901 | 0.001887 | 0.000958 | 0.001736 | 0.001215 | 0.003972 | 0.003797 | 0.001233 |
| 25.375 | 0.003436 | 0.006989 | 0.002439 | 0.006724 | 0.006958 | 0.001733 | 0.005492 | 0.006262 | 0.002059 | 0.004908 | 0.010921 | 0.001299 | 0.002834 | 0.000875 | 0.001225 | 0.001068 | 0.004245 | 0.004007 | 0.000953 |
| 25.3888889 | 0.00842 | 0.005318 | 0.004203 | 0.011059 | 0.005091 | 0.00403 | 0.003648 | 0.011277 | 0.005059 | 0.010378 | 0.013046 | 0.003303 | 0.002609 | 0.002367 | 0.000843 | 0.000515 | 0.003043 | 0.003379 | 0.001838 |
| 25.4027778 | 0.013839 | 0.007498 | 0.006202 | 0.018872 | 0.00393 | 0.012503 | 0.005519 | 0.016837 | 0.008166 | 0.01299 | 0.009877 | 0.005386 | 0.001398 | 0.004249 | 0.001925 | 0.000316 | 0.004042 | 0.005524 | 0.002856 |
| 25.4166667 | 0.01812 | 0.011819 | 0.007972 | 0.026572 | 0.003606 | 0.019638 | 0.007961 | 0.021062 | 0.010655 | 0.009232 | 0.008774 | 0.004558 | 0.000467 | 0.005709 | 0.002863 | 0.000389 | 0.005057 | 0.006784 | 0.003041 |
| 25.4305556 | 0.02226 | 0.013158 | 0.010111 | 0.029872 | 0.002757 | 0.017735 | 0.008102 | 0.021848 | 0.012056 | 0.0047 | 0.006428 | 0.001994 | 0.000395 | 0.005484 | 0.00215 | 0.000289 | 0.003491 | 0.004361 | 0.002764 |
| 25.4444444 | 0.026981 | 0.01354 | 0.012527 | 0.028478 | 0.004953 | 0.011696 | 0.005846 | 0.019456 | 0.011369 | 0.001995 | 0.002535 | 0.000511 | 0.001019 | 0.003263 | 0.001186 | 0.00018 | 0.00136 | 0.00194 | 0.002877 |
| 25.4583333 | 0.033151 | 0.017118 | 0.014951 | 0.025605 | 0.012211 | 0.007784 | 0.003443 | 0.01635 | 0.010428 | 0.000592 | 0.001582 | 0.000203 | 0.001705 | 0.001661 | 0.001311 | 0.000377 | 0.000404 | 0.001543 | 0.002954 |
| 25.4722222 | 0.037311 | 0.021466 | 0.016433 | 0.023444 | 0.02202 | 0.006428 | 0.00372 | 0.015051 | 0.012245 | 0.000329 | 0.001382 | 0.000137 | 0.001655 | 0.001618 | 0.001764 | 0.000641 | 0.00085 | 0.002052 | 0.002939 |
| 25.4861111 | 0.034094 | 0.020736 | 0.016665 | 0.021566 | 0.030209 | 0.006941 | 0.006252 | 0.015558 | 0.01651 | 0.000608 | 0.001358 | 0.000196 | 0.00107 | 0.001297 | 0.001537 | 0.000608 | 0.002692 | 0.002884 | 0.003837 |
| 25.5 | 0.025395 | 0.01512 | 0.016621 | 0.020319 | 0.033472 | 0.008344 | 0.008874 | 0.015757 | 0.020096 | 0.001587 | 0.001971 | 0.000531 | 0.00089 | 0.000767 | 0.000898 | 0.000658 | 0.004919 | 0.004077 | 0.005933 |
| 25.5138889 | 0.015532 | 0.008542 | 0.017023 | 0.02149 | 0.030416 | 0.009144 | 0.011207 | 0.014592 | 0.020832 | 0.002714 | 0.002701 | 0.000814 | 0.000872 | 0.000576 | 0.000684 | 0.000889 | 0.006039 | 0.005002 | 0.007798 |
| 25.5277778 | 0.00704 | 0.003285 | 0.018284 | 0.02535 | 0.02226 | 0.009111 | 0.014717 | 0.013371 | 0.018891 | 0.003236 | 0.003154 | 0.000704 | 0.000563 | 0.000537 | 0.001114 | 0.000863 | 0.005654 | 0.005117 | 0.008045 |
| 25.5416667 | 0.003407 | 0.001309 | 0.019916 | 0.028536 | 0.014352 | 0.009065 | 0.019996 | 0.014517 | 0.015901 | 0.002851 | 0.003057 | 0.000507 | 0.000473 | 0.000844 | 0.00171 | 0.000532 | 0.004505 | 0.004532 | 0.006623 |
| 25.5555556 | 0.005374 | 0.002934 | 0.019683 | 0.026539 | 0.012023 | 0.009348 | 0.024965 | 0.018563 | 0.013722 | 0.002153 | 0.002697 | 0.000596 | 0.000511 | 0.000859 | 0.001682 | 0.000244 | 0.003713 | 0.004068 | 0.004491 |
| 25.5694444 | 0.009399 | 0.007965 | 0.017095 | 0.019797 | 0.014787 | 0.009562 | 0.027814 | 0.02413 | 0.014542 | 0.002807 | 0.003556 | 0.000561 | 0.000482 | 0.000531 | 0.000924 | 0.000245 | 0.003963 | 0.006429 | 0.004469 |
| 25.5833333 | 0.012898 | 0.020493 | 0.016612 | 0.013636 | 0.015426 | 0.008306 | 0.031194 | 0.031021 | 0.022491 | 0.006126 | 0.006977 | 0.000379 | 0.00154 | 0.000519 | 0.000619 | 0.000378 | 0.005289 | 0.016051 | 0.009966 |
| 25.5972222 | 0.01505 | 0.040473 | 0.022139 | 0.011509 | 0.025078 | 0.006768 | 0.037482 | 0.036761 | 0.035309 | 0.010879 | 0.011447 | 0.000405 | 0.00404 | 0.000807 | 0.001391 | 0.000609 | 0.00714 | 0.032459 | 0.019419 |
| 25.6111111 | 0.014164 | 0.052423 | 0.029538 | 0.011652 | 0.066123 | 0.008166 | 0.043786 | 0.036042 | 0.038733 | 0.013595 | 0.012912 | 0.000402 | 0.005831 | 0.001306 | 0.001917 | 0.001067 | 0.008053 | 0.046376 | 0.024333 |
| 25.625 | 0.010964 | 0.046862 | 0.028243 | 0.010867 | 0.097221 | 0.008197 | 0.045106 | 0.030198 | 0.026895 | 0.012421 | 0.00966 | 0.000314 | 0.005365 | 0.001657 | 0.001262 | 0.001295 | 0.006832 | 0.050482 | 0.019587 |
| 25.6388889 | 0.009236 | 0.034675 | 0.018712 | 0.007394 | 0.076492 | 0.004874 | 0.034995 | 0.025711 | 0.012323 | 0.009001 | 0.004523 | 0.000437 | 0.003858 | 0.001656 | 0.000525 | 0.00116 | 0.004453 | 0.044908 | 0.010314 |
| 25.6527778 | 0.010433 | 0.024384 | 0.016705 | 0.003584 | 0.044418 | 0.004911 | 0.019919 | 0.023991 | 0.007791 | 0.005648 | 0.001554 | 0.000838 | 0.002549 | 0.001251 | 0.000782 | 0.000923 | 0.002934 | 0.032317 | 0.004349 |
| 25.6666667 | 0.013544 | 0.016619 | 0.020857 | 0.003188 | 0.030893 | 0.00816 | 0.017665 | 0.022861 | 0.011631 | 0.003161 | 0.00112 | 0.001062 | 0.001547 | 0.001235 | 0.002091 | 0.000701 | 0.002847 | 0.018442 | 0.003439 |
| 25.6805556 | 0.017094 | 0.011536 | 0.017603 | 0.003751 | 0.028123 | 0.009708 | 0.022598 | 0.022707 | 0.012559 | 0.001866 | 0.001098 | 0.001321 | 0.001086 | 0.00222 | 0.003745 | 0.000682 | 0.003044 | 0.008732 | 0.003828 |
| 25.6944444 | 0.018715 | 0.007015 | 0.01863 | 0.002757 | 0.028277 | 0.008725 | 0.018899 | 0.026619 | 0.008467 | 0.00268 | 0.001002 | 0.001787 | 0.000844 | 0.00252 | 0.004316 | 0.000614 | 0.002215 | 0.003652 | 0.005223 |
| 25.7083333 | 0.016759 | 0.003823 | 0.033633 | 0.002256 | 0.024551 | 0.007137 | 0.01946 | 0.036144 | 0.007258 | 0.004847 | 0.001413 | 0.0014 | 0.000413 | 0.001593 | 0.003549 | 0.000454 | 0.000878 | 0.002471 | 0.007474 |
| 25.7222222 | 0.012206 | 0.005692 | 0.041349 | 0.002512 | 0.017505 | 0.006254 | 0.035076 | 0.045964 | 0.00904 | 0.004753 | 0.003562 | 0.000647 | 0.000212 | 0.000687 | 0.002499 | 0.001099 | 0.001043 | 0.004069 | 0.006523 |
| 25.7361111 | 0.006678 | 0.010471 | 0.027505 | 0.002134 | 0.011596 | 0.004716 | 0.043634 | 0.048321 | 0.007248 | 0.003661 | 0.006042 | 0.000746 | 0.000366 | 0.000511 | 0.001575 | 0.00254 | 0.003693 | 0.006021 | 0.003151 |
| 25.75 | 0.00268 | 0.011702 | 0.015173 | 0.003539 | 0.009178 | 0.002259 | 0.030594 | 0.043952 | 0.003852 | 0.007671 | 0.005221 | 0.001146 | 0.000744 | 0.001097 | 0.001425 | 0.002978 | 0.007631 | 0.007817 | 0.001414 |
| 25.7638889 | 0.002039 | 0.00915 | 0.022488 | 0.008612 | 0.011167 | 0.001413 | 0.021152 | 0.039155 | 0.002763 | 0.015909 | 0.003246 | 0.001144 | 0.001067 | 0.001614 | 0.002371 | 0.001843 | 0.009606 | 0.009754 | 0.001716 |
| 25.7777778 | 0.002063 | 0.011947 | 0.028909 | 0.012106 | 0.017063 | 0.002916 | 0.035117 | 0.033751 | 0.004851 | 0.021185 | 0.004445 | 0.000924 | 0.000941 | 0.001259 | 0.003042 | 0.000894 | 0.007181 | 0.010763 | 0.001814 |
| 25.7916667 | 0.001444 | 0.023671 | 0.019524 | 0.009243 | 0.02162 | 0.004493 | 0.046929 | 0.023421 | 0.0121 | 0.019435 | 0.006865 | 0.000608 | 0.000889 | 0.000813 | 0.002865 | 0.001314 | 0.003228 | 0.010143 | 0.002018 |
| 25.8055556 | 0.004179 | 0.035851 | 0.010924 | 0.004255 | 0.019092 | 0.005369 | 0.033204 | 0.011717 | 0.017692 | 0.012693 | 0.007357 | 0.000467 | 0.001061 | 0.001369 | 0.002214 | 0.001942 | 0.001456 | 0.007753 | 0.004422 |
| 25.8194444 | 0.012515 | 0.040139 | 0.015874 | 0.002009 | 0.011327 | 0.006282 | 0.01869 | 0.009543 | 0.014395 | 0.006981 | 0.005692 | 0.000796 | 0.000868 | 0.001709 | 0.001325 | 0.001372 | 0.001044 | 0.004369 | 0.007953 |
| 25.8333333 | 0.020236 | 0.033874 | 0.018447 | 0.003485 | 0.004776 | 0.005844 | 0.026248 | 0.020605 | 0.00814 | 0.00945 | 0.003562 | 0.000879 | 0.000512 | 0.001267 | 0.000763 | 0.000531 | 0.001376 | 0.002527 | 0.009625 |
| 25.8472222 | 0.020363 | 0.020943 | 0.0141 | 0.00889 | 0.001707 | 0.003354 | 0.029542 | 0.035499 | 0.007025 | 0.019016 | 0.003521 | 0.000456 | 0.000573 | 0.001239 | 0.000752 | 0.000841 | 0.002653 | 0.002342 | 0.006906 |
| 25.8611111 | 0.01339 | 0.009861 | 0.021271 | 0.013331 | 0.001037 | 0.001945 | 0.018727 | 0.044418 | 0.009047 | 0.025284 | 0.004687 | 0.000261 | 0.001127 | 0.001502 | 0.001154 | 0.001762 | 0.003599 | 0.002504 | 0.005087 |
| 25.875 | 0.006547 | 0.008166 | 0.034609 | 0.013596 | 0.001773 | 0.004174 | 0.018374 | 0.044809 | 0.009621 | 0.021852 | 0.004841 | 0.000744 | 0.001491 | 0.00177 | 0.001385 | 0.001666 | 0.003495 | 0.005238 | 0.012035 |
| 25.8888889 | 0.004931 | 0.015301 | 0.034762 | 0.012943 | 0.001962 | 0.009214 | 0.03018 | 0.038158 | 0.00918 | 0.012973 | 0.004574 | 0.001563 | 0.001459 | 0.002058 | 0.001164 | 0.000975 | 0.002394 | 0.010585 | 0.022161 |
| 25.9027778 | 0.004119 | 0.023939 | 0.022445 | 0.013698 | 0.001915 | 0.014708 | 0.038951 | 0.028514 | 0.009019 | 0.010867 | 0.006482 | 0.002113 | 0.001571 | 0.00227 | 0.001102 | 0.001547 | 0.001319 | 0.014503 | 0.026396 |
| 25.9166667 | 0.002924 | 0.031231 | 0.012017 | 0.014408 | 0.004483 | 0.018619 | 0.041281 | 0.020886 | 0.01008 | 0.017628 | 0.012804 | 0.003095 | 0.001505 | 0.002648 | 0.001009 | 0.003067 | 0.002021 | 0.01571 | 0.024678 |
| 25.9305556 | 0.006063 | 0.03629 | 0.012088 | 0.012266 | 0.008975 | 0.021047 | 0.040096 | 0.019126 | 0.011658 | 0.019394 | 0.019763 | 0.004516 | 0.001112 | 0.002624 | 0.000676 | 0.00378 | 0.004529 | 0.015338 | 0.020197 |
| 25.9444444 | 0.015296 | 0.038475 | 0.018714 | 0.009071 | 0.012511 | 0.023885 | 0.035677 | 0.026118 | 0.012625 | 0.013051 | 0.019705 | 0.00502 | 0.000921 | 0.001713 | 0.000448 | 0.002883 | 0.007964 | 0.01392 | 0.014842 |
| 25.9583333 | 0.032098 | 0.043614 | 0.024728 | 0.010538 | 0.015394 | 0.027567 | 0.031782 | 0.040975 | 0.013786 | 0.013957 | 0.01235 | 0.005301 | 0.000742 | 0.000755 | 0.000554 | 0.00159 | 0.012628 | 0.012691 | 0.010772 |
| 25.9722222 | 0.050352 | 0.05075 | 0.02662 | 0.017844 | 0.018628 | 0.029012 | 0.032014 | 0.0556 | 0.015296 | 0.027212 | 0.005234 | 0.005944 | 0.000662 | 0.000313 | 0.000964 | 0.001077 | 0.018618 | 0.012749 | 0.009345 |
| 25.9861111 | 0.055026 | 0.04888 | 0.023854 | 0.025914 | 0.020684 | 0.024998 | 0.031759 | 0.06294 | 0.017627 | 0.038985 | 0.002538 | 0.005586 | 0.001113 | 0.000241 | 0.001144 | 0.000928 | 0.022618 | 0.013379 | 0.009773 |
| 26 | 0.041748 | 0.037125 | 0.018342 | 0.029798 | 0.020936 | 0.017025 | 0.02597 | 0.063471 | 0.022065 | 0.041524 | 0.002504 | 0.004074 | 0.001386 | 0.000593 | 0.001599 | 0.001235 | 0.019823 | 0.013679 | 0.010216 |
| 26.0138889 | 0.021142 | 0.023926 | 0.011714 | 0.027449 | 0.020603 | 0.010357 | 0.018614 | 0.057405 | 0.026807 | 0.036315 | 0.004827 | 0.00253 | 0.000975 | 0.001459 | 0.002531 | 0.002231 | 0.011119 | 0.013397 | 0.00956 |
| 26.0277778 | 0.008488 | 0.014803 | 0.005967 | 0.020824 | 0.020332 | 0.008895 | 0.016326 | 0.045024 | 0.028614 | 0.023322 | 0.008748 | 0.001569 | 0.000566 | 0.002733 | 0.002462 | 0.002983 | 0.00393 | 0.012103 | 0.008663 |
| 26.0416667 | 0.005575 | 0.01142 | 0.003419 | 0.015029 | 0.019134 | 0.011793 | 0.019727 | 0.031156 | 0.025344 | 0.010772 | 0.010902 | 0.001237 | 0.000608 | 0.003615 | 0.001252 | 0.002728 | 0.001752 | 0.009529 | 0.008164 |
| 26.0555556 | 0.005679 | 0.011832 | 0.004181 | 0.011878 | 0.015699 | 0.014557 | 0.022292 | 0.020041 | 0.017208 | 0.00825 | 0.010754 | 0.001279 | 0.000883 | 0.003197 | 0.000428 | 0.001587 | 0.002489 | 0.006564 | 0.008097 |
| 26.0694444 | 0.011666 | 0.013813 | 0.006509 | 0.010212 | 0.010761 | 0.013491 | 0.018004 | 0.013317 | 0.008977 | 0.009022 | 0.010873 | 0.001267 | 0.001365 | 0.001994 | 0.000363 | 0.000993 | 0.006386 | 0.005561 | 0.009307 |
| 26.0833333 | 0.021757 | 0.017847 | 0.007251 | 0.009032 | 0.006325 | 0.008594 | 0.009576 | 0.010033 | 0.004357 | 0.006858 | 0.014534 | 0.000997 | 0.002909 | 0.002507 | 0.001318 | 0.002629 | 0.012409 | 0.009361 | 0.012854 |
| 26.0972222 | 0.031384 | 0.023812 | 0.005009 | 0.008863 | 0.005764 | 0.004804 | 0.00405 | 0.009663 | 0.002021 | 0.008259 | 0.021799 | 0.000984 | 0.005397 | 0.005514 | 0.003617 | 0.005078 | 0.017345 | 0.01793 | 0.018193 |
| 26.1111111 | 0.043062 | 0.029381 | 0.002757 | 0.01127 | 0.008688 | 0.00421 | 0.003246 | 0.014266 | 0.001475 | 0.014634 | 0.026726 | 0.001322 | 0.006359 | 0.006824 | 0.005317 | 0.00502 | 0.017613 | 0.02519 | 0.022293 |
| 26.125 | 0.055666 | 0.032415 | 0.002344 | 0.012089 | 0.009286 | 0.005108 | 0.004617 | 0.021248 | 0.004019 | 0.018224 | 0.024428 | 0.001091 | 0.004713 | 0.004237 | 0.005059 | 0.002695 | 0.010892 | 0.02518 | 0.021825 |
| 26.1388889 | 0.060601 | 0.031489 | 0.002326 | 0.007515 | 0.006814 | 0.007394 | 0.005325 | 0.023605 | 0.008023 | 0.016501 | 0.018139 | 0.00051 | 0.002528 | 0.001658 | 0.003551 | 0.001524 | 0.004091 | 0.018857 | 0.015529 |
| 26.1527778 | 0.051265 | 0.026113 | 0.001721 | 0.003216 | 0.00388 | 0.007323 | 0.004114 | 0.020259 | 0.010492 | 0.012145 | 0.012237 | 0.000369 | 0.001392 | 0.001448 | 0.001941 | 0.002213 | 0.003096 | 0.009871 | 0.007809 |
| 26.1666667 | 0.032009 | 0.018388 | 0.0009 | 0.004605 | 0.002643 | 0.004554 | 0.002797 | 0.015193 | 0.010971 | 0.006587 | 0.006397 | 0.000332 | 0.00093 | 0.001613 | 0.001589 | 0.002769 | 0.003168 | 0.004546 | 0.004659 |
| 26.1805556 | 0.014815 | 0.010741 | 0.000809 | 0.008626 | 0.004811 | 0.00319 | 0.001941 | 0.009486 | 0.010108 | 0.003436 | 0.002576 | 0.000164 | 0.000597 | 0.001175 | 0.002149 | 0.00225 | 0.002127 | 0.006918 | 0.003865 |
| 26.1944444 | 0.010346 | 0.005429 | 0.002398 | 0.008872 | 0.007523 | 0.003347 | 0.00232 | 0.004531 | 0.00814 | 0.00441 | 0.002914 | 9.56E-05 | 0.000376 | 0.000685 | 0.001889 | 0.001143 | 0.001646 | 0.010776 | 0.002039 |
| 26.2083333 | 0.016506 | 0.005487 | 0.006054 | 0.005585 | 0.007894 | 0.003613 | 0.006875 | 0.00388 | 0.0055 | 0.004926 | 0.003642 | 0.00023 | 0.000244 | 0.000387 | 0.000904 | 0.000293 | 0.001185 | 0.01106 | 0.00102 |
| 26.2222222 | 0.022301 | 0.009667 | 0.010194 | 0.00552 | 0.005654 | 0.00342 | 0.014457 | 0.006908 | 0.003059 | 0.003072 | 0.002409 | 0.000433 | 0.00012 | 0.000345 | 0.000307 | 9.56E-05 | 0.000965 | 0.009518 | 0.001144 |
| 26.2361111 | 0.024546 | 0.013642 | 0.013548 | 0.010118 | 0.003572 | 0.002353 | 0.020842 | 0.010445 | 0.003045 | 0.00116 | 0.000949 | 0.000433 | 0.000191 | 0.000352 | 0.000387 | 0.000603 | 0.001242 | 0.007985 | 0.001562 |
| 26.25 | 0.026203 | 0.016425 | 0.016432 | 0.012861 | 0.005722 | 0.001851 | 0.023356 | 0.012711 | 0.006478 | 0.000283 | 0.000665 | 0.000346 | 0.000427 | 0.000304 | 0.000781 | 0.001623 | 0.001113 | 0.006325 | 0.003554 |
| 26.2638889 | 0.028498 | 0.019066 | 0.018338 | 0.009985 | 0.010292 | 0.003767 | 0.021454 | 0.012768 | 0.010528 | 0.00017 | 0.001361 | 0.000357 | 0.000511 | 0.000355 | 0.000849 | 0.002149 | 0.000845 | 0.003847 | 0.006512 |
| 26.2777778 | 0.03138 | 0.021516 | 0.018055 | 0.004679 | 0.013411 | 0.005448 | 0.016474 | 0.011008 | 0.012202 | 0.000202 | 0.001698 | 0.000487 | 0.000644 | 0.000603 | 0.000502 | 0.001468 | 0.00071 | 0.001519 | 0.007514 |
| 26.2916667 | 0.033761 | 0.02318 | 0.015086 | 0.001474 | 0.014551 | 0.004649 | 0.010275 | 0.009042 | 0.010991 | 0.000154 | 0.001261 | 0.000798 | 0.000916 | 0.000781 | 0.000193 | 0.000821 | 0.000946 | 0.000518 | 0.005981 |
| 26.3055556 | 0.034169 | 0.023411 | 0.010644 | 0.00263 | 0.01398 | 0.003338 | 0.004989 | 0.007655 | 0.007922 | 0.000123 | 0.001035 | 0.001042 | 0.001076 | 0.000877 | 0.000199 | 0.001446 | 0.002602 | 0.000426 | 0.003591 |
| 26.3194444 | 0.032008 | 0.020529 | 0.006525 | 0.006397 | 0.011333 | 0.003577 | 0.002964 | 0.005584 | 0.00415 | 0.000669 | 0.000943 | 0.000866 | 0.001127 | 0.001008 | 0.00036 | 0.002494 | 0.004521 | 0.000546 | 0.001707 |
| 26.3333333 | 0.027288 | 0.014017 | 0.003238 | 0.007687 | 0.006961 | 0.005052 | 0.003303 | 0.003016 | 0.001813 | 0.002506 | 0.000609 | 0.000653 | 0.001052 | 0.000885 | 0.000339 | 0.002855 | 0.004821 | 0.001181 | 0.000897 |
| 26.3472222 | 0.021289 | 0.007509 | 0.001132 | 0.004754 | 0.003311 | 0.006466 | 0.002788 | 0.001805 | 0.001974 | 0.004598 | 0.000509 | 0.000833 | 0.000876 | 0.000575 | 0.000164 | 0.002459 | 0.003845 | 0.003064 | 0.001139 |
| 26.3611111 | 0.016649 | 0.004637 | 0.000457 | 0.002217 | 0.002962 | 0.007354 | 0.0015 | 0.001918 | 0.001831 | 0.004915 | 0.00046 | 0.000767 | 0.000815 | 0.000517 | 0.000251 | 0.001816 | 0.002939 | 0.005174 | 0.002559 |
| 26.375 | 0.014223 | 0.00425 | 0.000434 | 0.005474 | 0.005115 | 0.007705 | 0.001374 | 0.003834 | 0.001059 | 0.003536 | 0.001153 | 0.000322 | 0.000856 | 0.000609 | 0.000644 | 0.001413 | 0.002286 | 0.005475 | 0.004498 |
| 26.3888889 | 0.012618 | 0.003897 | 0.00094 | 0.014432 | 0.00714 | 0.008387 | 0.002159 | 0.00772 | 0.0019 | 0.002585 | 0.00383 | 0.000139 | 0.000657 | 0.000463 | 0.000723 | 0.001191 | 0.00151 | 0.004034 | 0.003985 |
| 26.4027778 | 0.011158 | 0.003945 | 0.002681 | 0.022049 | 0.007984 | 0.011197 | 0.003078 | 0.011938 | 0.003828 | 0.003477 | 0.007737 | 0.000262 | 0.000305 | 0.000208 | 0.000372 | 0.000948 | 0.001406 | 0.004303 | 0.00263 |
| 26.4166667 | 0.010404 | 0.005355 | 0.005992 | 0.022375 | 0.007542 | 0.016191 | 0.004119 | 0.016959 | 0.005786 | 0.005621 | 0.010436 | 0.00035 | 0.00029 | 0.000247 | 0.000269 | 0.000747 | 0.002072 | 0.00756 | 0.004298 |
| 26.4305556 | 0.011578 | 0.007617 | 0.010326 | 0.018425 | 0.006135 | 0.020537 | 0.005252 | 0.024108 | 0.007615 | 0.007791 | 0.010424 | 0.00033 | 0.000751 | 0.000701 | 0.000764 | 0.000632 | 0.002425 | 0.009607 | 0.006163 |

| | | | | | | | | | | | | | | | | | | | |
|---|---|---|---|---|---|---|---|---|---|---|---|---|---|---|---|---|---|---|---|
| 26.4444444 | 0.015905 | 0.010374 | 0.01438 | 0.017195 | 0.004813 | 0.022609 | 0.007056 | 0.031667 | 0.009303 | 0.009393 | 0.009445 | 0.000469 | 0.001461 | 0.001214 | 0.001525 | 0.000429 | 0.002811 | 0.008145 | 0.00631 |
| 26.4583333 | 0.022837 | 0.013888 | 0.016725 | 0.020512 | 0.004396 | 0.023921 | 0.010239 | 0.036624 | 0.010782 | 0.010388 | 0.010072 | 0.000587 | 0.002229 | 0.001296 | 0.001913 | 0.000332 | 0.004029 | 0.005195 | 0.006639 |
| 26.4722222 | 0.029986 | 0.017447 | 0.01683 | 0.024246 | 0.004437 | 0.025015 | 0.013892 | 0.038542 | 0.011538 | 0.010701 | 0.01229 | 0.000448 | 0.003127 | 0.000852 | 0.001673 | 0.000826 | 0.005713 | 0.003083 | 0.007171 |
| 26.4861111 | 0.034133 | 0.019459 | 0.015907 | 0.024787 | 0.003485 | 0.024777 | 0.015761 | 0.038229 | 0.011124 | 0.010013 | 0.014436 | 0.000292 | 0.003863 | 0.000316 | 0.001335 | 0.001633 | 0.007521 | 0.002608 | 0.0066 |
| 26.5 | 0.033209 | 0.019358 | 0.01603 | 0.022439 | 0.00167 | 0.022365 | 0.014134 | 0.035495 | 0.009902 | 0.008601 | 0.015008 | 0.000359 | 0.003785 | 0.000268 | 0.001515 | 0.002043 | 0.009704 | 0.003101 | 0.004967 |
| 26.5138889 | 0.028363 | 0.017854 | 0.018455 | 0.019542 | 0.000507 | 0.018195 | 0.010082 | 0.030523 | 0.009502 | 0.007811 | 0.013639 | 0.000505 | 0.002961 | 0.00079 | 0.001926 | 0.001958 | 0.012263 | 0.003843 | 0.003489 |
| 26.5277778 | 0.022853 | 0.015375 | 0.021945 | 0.018162 | 0.000397 | 0.014029 | 0.007616 | 0.024935 | 0.012205 | 0.008529 | 0.011759 | 0.000686 | 0.002229 | 0.001431 | 0.001893 | 0.001706 | 0.014468 | 0.004585 | 0.002621 |
| 26.5416667 | 0.018622 | 0.011723 | 0.022324 | 0.01801 | 0.001276 | 0.010686 | 0.008806 | 0.019607 | 0.017079 | 0.009625 | 0.010796 | 0.000827 | 0.002045 | 0.002079 | 0.001692 | 0.001232 | 0.015069 | 0.004863 | 0.002071 |
| 26.5555556 | 0.014497 | 0.006721 | 0.016331 | 0.016985 | 0.003684 | 0.007451 | 0.011553 | 0.013304 | 0.019636 | 0.009523 | 0.010539 | 0.000634 | 0.002201 | 0.002955 | 0.002148 | 0.000674 | 0.014185 | 0.004784 | 0.00217 |
| 26.5694444 | 0.00912 | 0.004063 | 0.007485 | 0.015296 | 0.007916 | 0.004773 | 0.015566 | 0.006873 | 0.019234 | 0.009232 | 0.010962 | 0.000414 | 0.002714 | 0.003904 | 0.003198 | 0.000775 | 0.014221 | 0.006703 | 0.004357 |
| 26.5833333 | 0.004234 | 0.008381 | 0.006162 | 0.015459 | 0.016873 | 0.005634 | 0.023949 | 0.004293 | 0.022792 | 0.011614 | 0.013614 | 0.000732 | 0.004075 | 0.004869 | 0.004292 | 0.001711 | 0.017562 | 0.014165 | 0.010294 |
| 26.5972222 | 0.002119 | 0.013612 | 0.019266 | 0.016434 | 0.027302 | 0.008542 | 0.036009 | 0.008926 | 0.034651 | 0.017606 | 0.019031 | 0.00225 | 0.005778 | 0.005632 | 0.005342 | 0.002568 | 0.022199 | 0.026427 | 0.019585 |
| 26.6111111 | 0.003347 | 0.014788 | 0.039765 | 0.016436 | 0.024451 | 0.007535 | 0.04396 | 0.019736 | 0.045021 | 0.023654 | 0.023148 | 0.005132 | 0.006204 | 0.005038 | 0.005911 | 0.00228 | 0.022366 | 0.034264 | 0.027878 |
| 26.625 | 0.008771 | 0.022367 | 0.049733 | 0.01658 | 0.010566 | 0.004939 | 0.039901 | 0.029973 | 0.043166 | 0.024615 | 0.019858 | 0.008051 | 0.005188 | 0.00295 | 0.004982 | 0.001176 | 0.01779 | 0.031999 | 0.029744 |
| 26.6388889 | 0.015271 | 0.034829 | 0.037083 | 0.015653 | 0.002849 | 0.004844 | 0.023939 | 0.033127 | 0.031972 | 0.021246 | 0.010696 | 0.008378 | 0.003302 | 0.00215 | 0.00309 | 0.000894 | 0.01306 | 0.025565 | 0.024536 |
| 26.6527778 | 0.016764 | 0.034036 | 0.019515 | 0.012549 | 0.003093 | 0.004167 | 0.011584 | 0.029274 | 0.017203 | 0.016557 | 0.006187 | 0.005942 | 0.001371 | 0.004047 | 0.002827 | 0.00175 | 0.008822 | 0.019805 | 0.015819 |
| 26.6666667 | 0.013837 | 0.023438 | 0.020966 | 0.009402 | 0.005029 | 0.00373 | 0.015431 | 0.024623 | 0.008128 | 0.009649 | 0.009721 | 0.003725 | 0.000519 | 0.006299 | 0.004686 | 0.002167 | 0.004797 | 0.014538 | 0.007245 |
| 26.6805556 | 0.012711 | 0.020806 | 0.028956 | 0.008234 | 0.006947 | 0.004682 | 0.021042 | 0.026661 | 0.010826 | 0.003112 | 0.012056 | 0.003132 | 0.000411 | 0.007937 | 0.006586 | 0.001817 | 0.00316 | 0.009046 | 0.003426 |
| 26.6944444 | 0.016779 | 0.029118 | 0.022431 | 0.008956 | 0.008268 | 0.005587 | 0.017906 | 0.037118 | 0.012526 | 0.002979 | 0.007804 | 0.003409 | 0.000666 | 0.009167 | 0.00744 | 0.001466 | 0.002616 | 0.004324 | 0.006122 |
| 26.7083333 | 0.024052 | 0.039735 | 0.018786 | 0.009803 | 0.008247 | 0.010734 | 0.021849 | 0.051715 | 0.009014 | 0.008666 | 0.003277 | 0.003089 | 0.00082 | 0.008691 | 0.006685 | 0.001007 | 0.002523 | 0.002806 | 0.010512 |
| 26.7222222 | 0.031201 | 0.043435 | 0.041442 | 0.010056 | 0.005936 | 0.019423 | 0.037375 | 0.065094 | 0.012409 | 0.013945 | 0.003881 | 0.001765 | 0.00084 | 0.006669 | 0.004587 | 0.000777 | 0.006302 | 0.002694 | 0.011941 |
| 26.7361111 | 0.033301 | 0.034535 | 0.06551 | 0.010353 | 0.00333 | 0.023639 | 0.043965 | 0.068239 | 0.019807 | 0.014205 | 0.005514 | 0.000632 | 0.001296 | 0.004393 | 0.002303 | 0.001413 | 0.011957 | 0.002852 | 0.008877 |
| 26.75 | 0.026094 | 0.020838 | 0.05874 | 0.010936 | 0.003853 | 0.020517 | 0.031842 | 0.053987 | 0.0175 | 0.009742 | 0.004324 | 0.000405 | 0.001338 | 0.002685 | 0.001439 | 0.00217 | 0.013237 | 0.004816 | 0.00749 |
| 26.7638889 | 0.013098 | 0.021026 | 0.034265 | 0.012757 | 0.005189 | 0.013087 | 0.017085 | 0.028826 | 0.007516 | 0.006811 | 0.003379 | 0.000512 | 0.00097 | 0.002705 | 0.002749 | 0.002146 | 0.008306 | 0.004703 | 0.013346 |
| 26.7777778 | 0.006358 | 0.032155 | 0.02787 | 0.014969 | 0.007167 | 0.006069 | 0.02278 | 0.016351 | 0.001616 | 0.008144 | 0.005038 | 0.000304 | 0.001321 | 0.003846 | 0.004135 | 0.001536 | 0.003896 | 0.003463 | 0.017867 |
| 26.7916667 | 0.011465 | 0.034213 | 0.044652 | 0.015193 | 0.019861 | 0.002243 | 0.047806 | 0.026035 | 0.001131 | 0.009723 | 0.006491 | 0.000403 | 0.001976 | 0.004123 | 0.004976 | 0.001101 | 0.005412 | 0.005535 | 0.015083 |
| 26.8055556 | 0.013282 | 0.02153 | 0.043681 | 0.011013 | 0.027222 | 0.000782 | 0.049675 | 0.026791 | 0.001122 | 0.007772 | 0.004843 | 0.000702 | 0.001644 | 0.002742 | 0.004385 | 0.000841 | 0.005934 | 0.005502 | 0.008265 |
| 26.8194444 | 0.005068 | 0.006054 | 0.015775 | 0.003673 | 0.011885 | 0.000178 | 0.018491 | 0.009437 | 0.000396 | 0.002749 | 0.001549 | 0.000349 | 0.000546 | 0.000811 | 0.001627 | 0.000321 | 0.002172 | 0.001843 | 0.002141 |
| 26.8333333 | 0 | 0 | 0 | 0 | 0 | 0 | 0 | 0 | 0 | 0 | 0 | 0 | 0 | 0 | 0 | 0 | 0 | 0 | 0 |
| 26.8472222 | 0 | 0 | 0 | 0 | 0 | 0 | 0 | 0 | 0 | 0 | 0 | 0 | 0 | 0 | 0 | 0 | 0 | 0 | 0 |
| 26.8611111 | 0 | 0 | 0 | 0 | 0 | 0 | 0 | 0 | 0 | 0 | 0 | 0 | 0 | 0 | 0 | 0 | 0 | 0 | 0 |
| 26.875 | 0 | 0 | 0 | 0 | 0 | 0 | 0 | 0 | 0 | 0 | 0 | 0 | 0 | 0 | 0 | 0 | 0 | 0 | 0 |
| 26.8888889 | 0.004887 | 0.001797 | 0.012905 | 0.009249 | 0.002056 | 0.003143 | 0.010388 | 0.010203 | 0.008016 | 0.001463 | 0.003569 | 0.000768 | 0.002291 | 0.000517 | 0.002121 | 0.001238 | 0.001815 | 0.003017 | 0.005283 |
| 26.9027778 | 0.015323 | 0.009177 | 0.031273 | 0.022343 | 0.005367 | 0.007901 | 0.033319 | 0.023189 | 0.02479 | 0.005031 | 0.012334 | 0.001933 | 0.004587 | 0.001607 | 0.005058 | 0.003301 | 0.004857 | 0.006085 | 0.013129 |
| 26.9166667 | 0.02195 | 0.017414 | 0.024524 | 0.022515 | 0.006714 | 0.008357 | 0.05326 | 0.021167 | 0.032392 | 0.010069 | 0.0193 | 0.003468 | 0.002463 | 0.00185 | 0.004336 | 0.003349 | 0.005557 | 0.005529 | 0.011424 |
| 26.9305556 | 0.02144 | 0.016121 | 0.011423 | 0.022061 | 0.007405 | 0.008144 | 0.058641 | 0.02647 | 0.022967 | 0.013704 | 0.019208 | 0.006813 | 0.000674 | 0.001134 | 0.002697 | 0.00194 | 0.004231 | 0.009809 | 0.008892 |
| 26.9444444 | 0.014183 | 0.008788 | 0.018508 | 0.02665 | 0.00894 | 0.011351 | 0.041418 | 0.047961 | 0.011695 | 0.011307 | 0.013187 | 0.009189 | 0.001045 | 0.000606 | 0.002658 | 0.001171 | 0.003465 | 0.018373 | 0.015539 |
| 26.9583333 | 0.008678 | 0.008618 | 0.032361 | 0.026519 | 0.01264 | 0.015364 | 0.015971 | 0.062639 | 0.01561 | 0.011777 | 0.006453 | 0.008281 | 0.000795 | 0.000667 | 0.002374 | 0.001798 | 0.00623 | 0.023458 | 0.021983 |
| 26.9722222 | 0.016029 | 0.021644 | 0.040148 | 0.022974 | 0.018419 | 0.015997 | 0.003838 | 0.063465 | 0.028144 | 0.024417 | 0.004021 | 0.00625 | 0.000545 | 0.001474 | 0.001583 | 0.003648 | 0.011952 | 0.024633 | 0.021945 |
| 26.9861111 | 0.026827 | 0.043401 | 0.040851 | 0.021152 | 0.023681 | 0.014078 | 0.001926 | 0.060727 | 0.036886 | 0.03863 | 0.005236 | 0.005193 | 0.001012 | 0.002527 | 0.000863 | 0.005681 | 0.017026 | 0.023175 | 0.0185 |
| 27 | 0.028538 | 0.064082 | 0.035507 | 0.024085 | 0.025366 | 0.015659 | 0.002801 | 0.063206 | 0.040668 | 0.042691 | 0.0072 | 0.005565 | 0.002094 | 0.002995 | 0.000635 | 0.007092 | 0.020262 | 0.021238 | 0.017264 |
| 27.0138889 | 0.022645 | 0.074493 | 0.025695 | 0.030331 | 0.021432 | 0.021602 | 0.006647 | 0.067758 | 0.040668 | 0.037251 | 0.009766 | 0.005823 | 0.003264 | 0.002602 | 0.000933 | 0.007058 | 0.022097 | 0.020399 | 0.020393 |
| 27.0277778 | 0.017342 | 0.071561 | 0.015035 | 0.037868 | 0.013972 | 0.026796 | 0.012144 | 0.070441 | 0.039051 | 0.029718 | 0.014506 | 0.004855 | 0.0037 | 0.0018 | 0.000951 | 0.005578 | 0.023003 | 0.019797 | 0.024355 |
| 27.0416667 | 0.017907 | 0.058515 | 0.006806 | 0.043368 | 0.007441 | 0.026759 | 0.0147 | 0.068258 | 0.034998 | 0.024949 | 0.019877 | 0.003205 | 0.003341 | 0.001008 | 0.000549 | 0.003724 | 0.021689 | 0.01824 | 0.023842 |
| 27.0555556 | 0.022459 | 0.040556 | 0.00273 | 0.041182 | 0.006038 | 0.021942 | 0.012662 | 0.056879 | 0.026906 | 0.022621 | 0.022326 | 0.00151 | 0.002874 | 0.000498 | 0.000465 | 0.002244 | 0.017941 | 0.015759 | 0.018346 |
| 27.0694444 | 0.027461 | 0.025293 | 0.003314 | 0.032321 | 0.008136 | 0.016448 | 0.008587 | 0.039941 | 0.017734 | 0.022132 | 0.022686 | 0.000509 | 0.002536 | 0.00082 | 0.000589 | 0.001251 | 0.01467 | 0.014062 | 0.014083 |
| 27.0833333 | 0.030835 | 0.021551 | 0.006984 | 0.023295 | 0.00739 | 0.013002 | 0.00504 | 0.025395 | 0.01123 | 0.024038 | 0.02631 | 0.000299 | 0.003213 | 0.002795 | 0.00069 | 0.001258 | 0.013597 | 0.015806 | 0.016005 |
| 27.0972222 | 0.032679 | 0.033303 | 0.011266 | 0.017226 | 0.003499 | 0.013307 | 0.004747 | 0.016015 | 0.008151 | 0.027487 | 0.034978 | 0.000203 | 0.005441 | 0.006096 | 0.001119 | 0.002502 | 0.013936 | 0.020637 | 0.020699 |
| 27.1111111 | 0.035629 | 0.051171 | 0.01154 | 0.013575 | 0.00279 | 0.014887 | 0.009265 | 0.012175 | 0.006237 | 0.027483 | 0.042683 | 0.000305 | 0.006886 | 0.007194 | 0.001857 | 0.002846 | 0.014337 | 0.023931 | 0.020716 |
| 27.125 | 0.040736 | 0.050553 | 0.009396 | 0.010383 | 0.008635 | 0.011197 | 0.014821 | 0.01288 | 0.005812 | 0.020426 | 0.041973 | 0.000516 | 0.005694 | 0.004499 | 0.001766 | 0.001835 | 0.012762 | 0.021838 | 0.014778 |
| 27.1388889 | 0.045455 | 0.029026 | 0.014508 | 0.00751 | 0.015632 | 0.007679 | 0.017342 | 0.013301 | 0.010281 | 0.01077 | 0.031118 | 0.000714 | 0.003455 | 0.001976 | 0.00099 | 0.001458 | 0.007642 | 0.015265 | 0.007953 |
| 27.1527778 | 0.044301 | 0.014479 | 0.022315 | 0.005967 | 0.017182 | 0.0118 | 0.015519 | 0.009391 | 0.014518 | 0.005177 | 0.015315 | 0.001301 | 0.001886 | 0.001809 | 0.0012 | 0.001119 | 0.002812 | 0.007825 | 0.003437 |
| 27.1666667 | 0.033769 | 0.014206 | 0.021308 | 0.004805 | 0.014616 | 0.017207 | 0.010863 | 0.005365 | 0.011781 | 0.004453 | 0.005227 | 0.001369 | 0.000905 | 0.002015 | 0.002361 | 0.00044 | 0.00187 | 0.002732 | 0.00122 |
| 27.1805556 | 0.018473 | 0.014524 | 0.014625 | 0.003149 | 0.012337 | 0.019679 | 0.007346 | 0.006802 | 0.006095 | 0.005204 | 0.004899 | 0.00081 | 0.000423 | 0.001819 | 0.003144 | 0.000551 | 0.001677 | 0.000809 | 0.000601 |
| 27.1944444 | 0.007043 | 0.010352 | 0.010838 | 0.002731 | 0.01165 | 0.02028 | 0.007088 | 0.009523 | 0.004111 | 0.005414 | 0.005823 | 0.000769 | 0.000531 | 0.00173 | 0.003049 | 0.001249 | 0.00096 | 0.000557 | 0.000582 |
| 27.2083333 | 0.003428 | 0.006368 | 0.011571 | 0.004266 | 0.011686 | 0.018127 | 0.008347 | 0.00953 | 0.006164 | 0.004987 | 0.003749 | 0.000916 | 0.000698 | 0.001692 | 0.002469 | 0.001774 | 0.001426 | 0.000968 | 0.000509 |
| 27.2222222 | 0.004298 | 0.006374 | 0.013044 | 0.004802 | 0.010781 | 0.013337 | 0.008365 | 0.008916 | 0.007074 | 0.003711 | 0.002114 | 0.000773 | 0.000569 | 0.001574 | 0.001871 | 0.001844 | 0.002525 | 0.001613 | 0.000331 |
| 27.2361111 | 0.005065 | 0.008943 | 0.012513 | 0.003182 | 0.008543 | 0.008162 | 0.006636 | 0.008422 | 0.004805 | 0.002134 | 0.002216 | 0.000916 | 0.000492 | 0.001488 | 0.001601 | 0.001715 | 0.002639 | 0.001901 | 0.000159 |
| 27.25 | 0.004945 | 0.011708 | 0.010887 | 0.002646 | 0.006431 | 0.005295 | 0.004893 | 0.006792 | 0.003571 | 0.001829 | 0.00269 | 0.001655 | 0.000805 | 0.001625 | 0.001837 | 0.001821 | 0.001477 | 0.002038 | 0.000183 |
| 27.2638889 | 0.004727 | 0.013059 | 0.009897 | 0.004716 | 0.0052 | 0.006611 | 0.005 | 0.003892 | 0.004195 | 0.002702 | 0.002081 | 0.002297 | 0.001128 | 0.001768 | 0.002222 | 0.002195 | 0.00041 | 0.002524 | 0.000504 |
| 27.2777778 | 0.005003 | 0.012253 | 0.009273 | 0.005931 | 0.004498 | 0.011118 | 0.006747 | 0.002586 | 0.00333 | 0.003031 | 0.001764 | 0.002389 | 0.000912 | 0.001646 | 0.002249 | 0.00265 | 0.000433 | 0.003142 | 0.000873 |
| 27.2916667 | 0.005956 | 0.00998 | 0.007883 | 0.004709 | 0.003823 | 0.014152 | 0.007798 | 0.004589 | 0.001331 | 0.002732 | 0.003829 | 0.002713 | 0.000407 | 0.00144 | 0.001919 | 0.003194 | 0.001952 | 0.003335 | 0.001021 |
| 27.3055556 | 0.006798 | 0.007663 | 0.005768 | 0.003921 | 0.002928 | 0.01228 | 0.006285 | 0.005964 | 0.001143 | 0.003432 | 0.007035 | 0.00378 | 0.000181 | 0.001366 | 0.001506 | 0.003441 | 0.005263 | 0.003426 | 0.000989 |
| 27.3194444 | 0.006654 | 0.00518 | 0.003814 | 0.00539 | 0.001943 | 0.006799 | 0.00341 | 0.004401 | 0.003406 | 0.005011 | 0.00975 | 0.004726 | 0.000378 | 0.001412 | 0.001317 | 0.002838 | 0.009106 | 0.003829 | 0.000914 |
| 27.3333333 | 0.005743 | 0.003116 | 0.002558 | 0.00769 | 0.001477 | 0.002663 | 0.002263 | 0.003104 | 0.006266 | 0.004386 | 0.01156 | 0.004612 | 0.000718 | 0.001464 | 0.001453 | 0.001689 | 0.010702 | 0.003504 | 0.000774 |
| 27.3472222 | 0.004908 | 0.003661 | 0.001972 | 0.008476 | 0.002292 | 0.001799 | 0.003183 | 0.002753 | 0.007937 | 0.003308 | 0.01216 | 0.003496 | 0.000749 | 0.001317 | 0.001474 | 0.000674 | 0.008776 | 0.002124 | 0.000498 |
| 27.3611111 | 0.004373 | 0.005114 | 0.001788 | 0.006855 | 0.003575 | 0.002646 | 0.003561 | 0.001637 | 0.007966 | 0.005686 | 0.011909 | 0.002141 | 0.000514 | 0.000986 | 0.001028 | 0.000168 | 0.005594 | 0.001329 | 0.000249 |
| 27.375 | 0.004044 | 0.003965 | 0.001517 | 0.003936 | 0.003807 | 0.00584 | 0.002325 | 0.000606 | 0.006987 | 0.007509 | 0.011177 | 0.001213 | 0.000398 | 0.000719 | 0.000481 | 0.000109 | 0.003943 | 0.001716 | 0.000286 |
| 27.3888889 | 0.004027 | 0.001512 | 0.001101 | 0.002581 | 0.003437 | 0.009137 | 0.000816 | 0.000329 | 0.005867 | 0.006369 | 0.009389 | 0.00067 | 0.000307 | 0.000503 | 0.000213 | 0.000209 | 0.003269 | 0.002673 | 0.000581 |
| 27.4027778 | 0.004081 | 0.000837 | 0.001731 | 0.003539 | 0.003928 | 0.009008 | 0.000325 | 0.000891 | 0.004772 | 0.005599 | 0.00617 | 0.000482 | 0.000189 | 0.000289 | 0.000156 | 0.000343 | 0.002116 | 0.00386 | 0.00093 |
| 27.4166667 | 0.00395 | 0.001961 | 0.004737 | 0.005197 | 0.005403 | 0.00789 | 0.000565 | 0.003463 | 0.003371 | 0.006917 | 0.003089 | 0.000739 | 0.000193 | 0.000255 | 0.000244 | 0.000371 | 0.001552 | 0.004692 | 0.001183 |
| 27.4305556 | 0.003644 | 0.002445 | 0.009712 | 0.009112 | 0.006274 | 0.010733 | 0.001393 | 0.009058 | 0.001915 | 0.009556 | 0.002812 | 0.000854 | 0.000237 | 0.000265 | 0.000417 | 0.00026 | 0.002273 | 0.004905 | 0.00139 |
| 27.4444444 | 0.003329 | 0.003037 | 0.014474 | 0.016339 | 0.005219 | 0.018257 | 0.003248 | 0.015819 | 0.002088 | 0.012281 | 0.005899 | 0.00053 | 0.000311 | 0.000298 | 0.000562 | 0.000259 | 0.003065 | 0.005013 | 0.001653 |
| 27.4583333 | 0.004177 | 0.006196 | 0.017199 | 0.022558 | 0.003629 | 0.02593 | 0.005014 | 0.019366 | 0.004974 | 0.014643 | 0.010271 | 0.000495 | 0.00038 | 0.000535 | 0.000776 | 0.000347 | 0.003962 | 0.005571 | 0.001961 |
| 27.4722222 | 0.007691 | 0.008999 | 0.018155 | 0.024566 | 0.003313 | 0.0305 | 0.00574 | 0.018727 | 0.008606 | 0.016075 | 0.014296 | 0.001512 | 0.000485 | 0.000664 | 0.001251 | 0.00028 | 0.005632 | 0.006798 | 0.001906 |
| 27.4861111 | 0.013303 | 0.009755 | 0.019326 | 0.024114 | 0.003359 | 0.032537 | 0.005857 | 0.01731 | 0.010433 | 0.016082 | 0.01726 | 0.003132 | 0.00048 | 0.000718 | 0.001661 | 0.000172 | 0.007604 | 0.008199 | 0.001509 |
| 27.5 | 0.019252 | 0.010777 | 0.022027 | 0.022473 | 0.002523 | 0.031182 | 0.005299 | 0.016812 | 0.010019 | 0.015997 | 0.018611 | 0.004235 | 0.000357 | 0.001066 | 0.001465 | 0.000116 | 0.009085 | 0.009307 | 0.001578 |
| 27.5138889 | 0.024939 | 0.013639 | 0.025208 | 0.019092 | 0.00147 | 0.025169 | 0.003882 | 0.016646 | 0.008588 | 0.01737 | 0.018552 | 0.004511 | 0.000633 | 0.001629 | 0.000852 | 0.000236 | 0.009403 | 0.010047 | 0.001717 |
| 27.5277778 | 0.030399 | 0.018702 | 0.026835 | 0.015813 | 0.001725 | 0.018884 | 0.003623 | 0.017722 | 0.008067 | 0.018299 | 0.017836 | 0.004062 | 0.001291 | 0.002213 | 0.000603 | 0.000915 | 0.0084 | 0.009545 | 0.002046 |
| 27.5416667 | 0.034387 | 0.023645 | 0.025089 | 0.015269 | 0.002967 | 0.017885 | 0.007406 | 0.02063 | 0.009952 | 0.01599 | 0.017222 | 0.003184 | 0.001824 | 0.002297 | 0.000768 | 0.001574 | 0.006819 | 0.007625 | 0.003403 |
| 27.5555556 | 0.034411 | 0.023999 | 0.018513 | 0.01683 | 0.002945 | 0.021193 | 0.014005 | 0.022678 | 0.013471 | 0.012102 | 0.017609 | 0.002756 | 0.001745 | 0.001551 | 0.000656 | 0.001477 | 0.00597 | 0.006385 | 0.005012 |
| 27.5694444 | 0.030453 | 0.017847 | 0.009183 | 0.020331 | 0.0043 | 0.023746 | 0.019141 | 0.021709 | 0.016757 | 0.011251 | 0.019974 | 0.002964 | 0.002064 | 0.00067 | 0.000761 | 0.001577 | 0.006293 | 0.008691 | 0.006753 |
| 27.5833333 | 0.027342 | 0.009694 | 0.003584 | 0.028549 | 0.016019 | 0.023021 | 0.023047 | 0.0202 | 0.022058 | 0.01673 | 0.026181 | 0.003185 | 0.004854 | 0.000941 | 0.00263 | 0.002489 | 0.007394 | 0.017046 | 0.009899 |
| 27.5972222 | 0.029696 | 0.006892 | 0.009028 | 0.037084 | 0.035826 | 0.020252 | 0.029143 | 0.019376 | 0.034244 | 0.029127 | 0.037372 | 0.004499 | 0.008983 | 0.002666 | 0.005532 | 0.003203 | 0.010738 | 0.029349 | 0.016632 |
| 27.6111111 | 0.035088 | 0.013033 | 0.027236 | 0.036184 | 0.042204 | 0.016192 | 0.037084 | 0.017338 | 0.051815 | 0.039593 | 0.048462 | 0.008451 | 0.010406 | 0.003505 | 0.006435 | 0.003119 | 0.018485 | 0.036595 | 0.027226 |
| 27.625 | 0.036274 | 0.024517 | 0.043775 | 0.027003 | 0.029755 | 0.009595 | 0.04403 | 0.014847 | 0.063614 | 0.033917 | 0.051208 | 0.013089 | 0.008039 | 0.003104 | 0.004573 | 0.002551 | 0.028368 | 0.033782 | 0.036235 |

| | | | | | | | | | | | | | | | | | | | |
|---|---|---|---|---|---|---|---|---|---|---|---|---|---|---|---|---|---|---|---|
| 27.6388889 | 0.028298 | 0.032297 | 0.039539 | 0.016592 | 0.018394 | 0.005044 | 0.044503 | 0.011244 | 0.057973 | 0.016475 | 0.04222 | 0.014934 | 0.004769 | 0.004623 | 0.002669 | 0.001661 | 0.032969 | 0.025859 | 0.037633 |
| 27.6527778 | 0.014524 | 0.030093 | 0.022897 | 0.008168 | 0.015538 | 0.006409 | 0.033879 | 0.005496 | 0.037216 | 0.008006 | 0.024965 | 0.013375 | 0.002498 | 0.00765 | 0.003434 | 0.000702 | 0.028137 | 0.018563 | 0.032341 |
| 27.6666667 | 0.005808 | 0.020392 | 0.01949 | 0.004237 | 0.014702 | 0.007009 | 0.017639 | 0.002783 | 0.015536 | 0.011324 | 0.011107 | 0.010325 | 0.001608 | 0.009187 | 0.005948 | 0.000287 | 0.01826 | 0.014161 | 0.025297 |
| 27.6805556 | 0.004618 | 0.012377 | 0.028698 | 0.00487 | 0.011781 | 0.004498 | 0.010659 | 0.007305 | 0.003791 | 0.012481 | 0.008256 | 0.007127 | 0.001893 | 0.008639 | 0.007373 | 0.00043 | 0.009919 | 0.013606 | 0.017731 |
| 27.6944444 | 0.005112 | 0.01351 | 0.028163 | 0.007965 | 0.006463 | 0.004597 | 0.016739 | 0.017339 | 0.001468 | 0.016136 | 0.007965 | 0.004823 | 0.001981 | 0.007392 | 0.007246 | 0.000646 | 0.007037 | 0.015483 | 0.009604 |
| 27.7083333 | 0.008396 | 0.022442 | 0.020001 | 0.011697 | 0.002006 | 0.011152 | 0.018847 | 0.029282 | 0.002737 | 0.029984 | 0.00468 | 0.004777 | 0.002321 | 0.006698 | 0.006631 | 0.0009 | 0.008442 | 0.017282 | 0.004789 |
| 27.7222222 | 0.013778 | 0.031655 | 0.026584 | 0.011862 | 0.000697 | 0.021628 | 0.012944 | 0.039669 | 0.006101 | 0.03906 | 0.002439 | 0.005477 | 0.003486 | 0.005852 | 0.00541 | 0.001055 | 0.008408 | 0.017897 | 0.005641 |
| 27.7361111 | 0.014812 | 0.036135 | 0.043726 | 0.00732 | 0.002204 | 0.029148 | 0.014894 | 0.046929 | 0.010326 | 0.033011 | 0.001543 | 0.004057 | 0.003496 | 0.003875 | 0.003389 | 0.000719 | 0.005538 | 0.015715 | 0.007928 |
| 27.75 | 0.013341 | 0.033997 | 0.048825 | 0.00343 | 0.005869 | 0.028492 | 0.027473 | 0.051383 | 0.012537 | 0.021011 | 0.001599 | 0.002065 | 0.001877 | 0.001931 | 0.001656 | 0.000439 | 0.00495 | 0.010383 | 0.006455 |
| 27.7638889 | 0.018288 | 0.025298 | 0.034059 | 0.003637 | 0.010116 | 0.018233 | 0.034692 | 0.050105 | 0.010337 | 0.015897 | 0.004602 | 0.002925 | 0.000896 | 0.0018 | 0.001133 | 0.000901 | 0.007867 | 0.004966 | 0.003535 |
| 27.7777778 | 0.031523 | 0.013956 | 0.017979 | 0.007634 | 0.011319 | 0.009468 | 0.02449 | 0.040981 | 0.006201 | 0.015498 | 0.01062 | 0.006495 | 0.001607 | 0.002851 | 0.00123 | 0.001546 | 0.009871 | 0.001866 | 0.00324 |
| 27.7916667 | 0.04015 | 0.010015 | 0.020433 | 0.012529 | 0.00791 | 0.014805 | 0.014233 | 0.027691 | 0.004408 | 0.009817 | 0.016029 | 0.010188 | 0.002495 | 0.002751 | 0.001172 | 0.001997 | 0.008889 | 0.001126 | 0.002912 |
| 27.8055556 | 0.031796 | 0.01924 | 0.029342 | 0.013848 | 0.006379 | 0.025196 | 0.023518 | 0.014221 | 0.004872 | 0.003051 | 0.015468 | 0.010486 | 0.002073 | 0.001833 | 0.001862 | 0.002545 | 0.006005 | 0.001676 | 0.001773 |
| 27.8194444 | 0.017585 | 0.027449 | 0.0295 | 0.010195 | 0.012023 | 0.023778 | 0.039526 | 0.009066 | 0.008728 | 0.001501 | 0.009568 | 0.006485 | 0.0013 | 0.002831 | 0.003472 | 0.003074 | 0.003403 | 0.001825 | 0.002292 |
| 27.8333333 | 0.016459 | 0.022469 | 0.023761 | 0.006804 | 0.017515 | 0.012845 | 0.045723 | 0.018818 | 0.011233 | 0.003533 | 0.006145 | 0.003278 | 0.001412 | 0.005467 | 0.004189 | 0.003306 | 0.004244 | 0.001384 | 0.002786 |
| 27.8472222 | 0.027402 | 0.011831 | 0.019372 | 0.010545 | 0.015891 | 0.006475 | 0.040716 | 0.034138 | 0.008475 | 0.009324 | 0.006002 | 0.003589 | 0.001667 | 0.006835 | 0.003747 | 0.002751 | 0.009697 | 0.001246 | 0.002052 |
| 27.8611111 | 0.038076 | 0.00699 | 0.02023 | 0.016535 | 0.011038 | 0.005086 | 0.029356 | 0.040943 | 0.008737 | 0.018715 | 0.003883 | 0.003319 | 0.001974 | 0.0064 | 0.002953 | 0.001526 | 0.016115 | 0.001944 | 0.002006 |
| 27.875 | 0.043981 | 0.012334 | 0.024884 | 0.014951 | 0.007493 | 0.005269 | 0.017591 | 0.035714 | 0.017146 | 0.028068 | 0.001356 | 0.00227 | 0.001832 | 0.005236 | 0.001785 | 0.000686 | 0.017823 | 0.005542 | 0.003747 |
| 27.8888889 | 0.047006 | 0.024777 | 0.027366 | 0.008147 | 0.004675 | 0.008972 | 0.00863 | 0.027395 | 0.028765 | 0.031129 | 0.000745 | 0.004005 | 0.001322 | 0.003146 | 0.000691 | 0.000959 | 0.012614 | 0.012493 | 0.007894 |
| 27.9027778 | 0.044385 | 0.033837 | 0.020747 | 0.005081 | 0.004152 | 0.014314 | 0.00482 | 0.023786 | 0.035372 | 0.023101 | 0.001025 | 0.006997 | 0.002314 | 0.001606 | 0.000888 | 0.001808 | 0.006519 | 0.01894 | 0.013631 |
| 27.9166667 | 0.032839 | 0.031257 | 0.010469 | 0.010385 | 0.008621 | 0.021523 | 0.007461 | 0.023928 | 0.032189 | 0.010772 | 0.001788 | 0.00846 | 0.004038 | 0.002777 | 0.002595 | 0.001971 | 0.006888 | 0.021844 | 0.016588 |
| 27.9305556 | 0.01781 | 0.021631 | 0.005944 | 0.022253 | 0.014941 | 0.031476 | 0.01031 | 0.026837 | 0.02355 | 0.004338 | 0.004984 | 0.009263 | 0.004683 | 0.004942 | 0.003527 | 0.001631 | 0.011219 | 0.022075 | 0.013785 |
| 27.9444444 | 0.007216 | 0.013482 | 0.006765 | 0.032625 | 0.019406 | 0.040161 | 0.008986 | 0.033418 | 0.018395 | 0.006388 | 0.012856 | 0.01062 | 0.004092 | 0.005783 | 0.00244 | 0.001422 | 0.014198 | 0.022042 | 0.009063 |
| 27.9583333 | 0.004959 | 0.011018 | 0.014751 | 0.032252 | 0.021872 | 0.043447 | 0.011046 | 0.041721 | 0.021829 | 0.015371 | 0.021838 | 0.011226 | 0.002535 | 0.005375 | 0.001863 | 0.001434 | 0.015877 | 0.023186 | 0.010364 |
| 27.9722222 | 0.008325 | 0.016024 | 0.029142 | 0.020932 | 0.021735 | 0.041229 | 0.018031 | 0.045589 | 0.03148 | 0.024789 | 0.024949 | 0.009577 | 0.000917 | 0.004179 | 0.002878 | 0.002496 | 0.017706 | 0.025154 | 0.019961 |
| 27.9861111 | 0.00895 | 0.027805 | 0.039284 | 0.009876 | 0.018762 | 0.03717 | 0.02049 | 0.039823 | 0.038312 | 0.029607 | 0.019391 | 0.006811 | 0.000338 | 0.002237 | 0.003448 | 0.004551 | 0.018875 | 0.026052 | 0.03007 |
| 28 | 0.006147 | 0.042635 | 0.037354 | 0.01034 | 0.015229 | 0.033843 | 0.01665 | 0.02696 | 0.036686 | 0.030624 | 0.01015 | 0.005464 | 0.00045 | 0.000675 | 0.003118 | 0.005908 | 0.017454 | 0.024594 | 0.032783 |
| 28.0138889 | 0.006939 | 0.051803 | 0.025261 | 0.022552 | 0.011403 | 0.028878 | 0.012101 | 0.015528 | 0.028919 | 0.028665 | 0.005323 | 0.005882 | 0.000522 | 0.000375 | 0.002634 | 0.005544 | 0.013785 | 0.021439 | 0.028995 |
| 28.0277778 | 0.009959 | 0.046681 | 0.011617 | 0.038024 | 0.006017 | 0.019983 | 0.009147 | 0.012036 | 0.018655 | 0.023631 | 0.006892 | 0.006053 | 0.000613 | 0.000501 | 0.00216 | 0.004115 | 0.010811 | 0.017401 | 0.023443 |
| 28.0416667 | 0.008195 | 0.031716 | 0.003843 | 0.046274 | 0.001994 | 0.01033 | 0.007441 | 0.015914 | 0.009284 | 0.016725 | 0.009592 | 0.005065 | 0.000687 | 0.000734 | 0.002022 | 0.002981 | 0.01054 | 0.012809 | 0.018524 |
| 28.0555556 | 0.005284 | 0.018632 | 0.002963 | 0.040219 | 0.00169 | 0.003936 | 0.006381 | 0.020648 | 0.004312 | 0.010254 | 0.007979 | 0.003481 | 0.000502 | 0.001059 | 0.002081 | 0.002288 | 0.01142 | 0.008707 | 0.015263 |
| 28.0694444 | 0.007902 | 0.012839 | 0.005245 | 0.02471 | 0.00257 | 0.000993 | 0.005092 | 0.020282 | 0.004544 | 0.006953 | 0.003649 | 0.001865 | 0.000333 | 0.001409 | 0.001804 | 0.001454 | 0.010971 | 0.007309 | 0.015153 |
| 28.0833333 | 0.013378 | 0.017017 | 0.00809 | 0.012124 | 0.002998 | 0.001683 | 0.003604 | 0.013633 | 0.008231 | 0.011423 | 0.004967 | 0.000939 | 0.000622 | 0.003557 | 0.001095 | 0.001255 | 0.010634 | 0.011827 | 0.019883 |
| 28.0972222 | 0.019903 | 0.036611 | 0.013174 | 0.009608 | 0.002955 | 0.007772 | 0.003602 | 0.005785 | 0.015086 | 0.024401 | 0.014911 | 0.001148 | 0.001386 | 0.007412 | 0.000806 | 0.002188 | 0.01387 | 0.022327 | 0.026901 |
| 28.1111111 | 0.030393 | 0.066941 | 0.019185 | 0.013375 | 0.001867 | 0.016863 | 0.004706 | 0.004627 | 0.023466 | 0.035335 | 0.023353 | 0.003006 | 0.002038 | 0.008511 | 0.000951 | 0.003026 | 0.019556 | 0.030604 | 0.028733 |
| 28.125 | 0.04416 | 0.080682 | 0.016388 | 0.013089 | 0.000571 | 0.021701 | 0.004957 | 0.01332 | 0.024681 | 0.03351 | 0.020857 | 0.005517 | 0.001973 | 0.00553 | 0.001287 | 0.002818 | 0.021984 | 0.0289 | 0.022452 |
| 28.1388889 | 0.054063 | 0.057596 | 0.00992 | 0.007456 | 0.001389 | 0.017847 | 0.004855 | 0.027712 | 0.016654 | 0.021872 | 0.012213 | 0.005474 | 0.001379 | 0.003345 | 0.002635 | 0.001811 | 0.018103 | 0.02081 | 0.013595 |
| 28.1527778 | 0.053682 | 0.027336 | 0.014158 | 0.006572 | 0.005579 | 0.011322 | 0.004532 | 0.036961 | 0.011631 | 0.010759 | 0.006575 | 0.003008 | 0.000794 | 0.003777 | 0.00357 | 0.000741 | 0.011252 | 0.013301 | 0.008078 |
| 28.1666667 | 0.044297 | 0.024207 | 0.019248 | 0.012319 | 0.011512 | 0.014035 | 0.002828 | 0.033071 | 0.012447 | 0.006823 | 0.004881 | 0.001535 | 0.000685 | 0.003601 | 0.002887 | 0.000198 | 0.005787 | 0.008027 | 0.005833 |
| 28.1805556 | 0.0306 | 0.034299 | 0.016113 | 0.012483 | 0.016877 | 0.027066 | 0.001083 | 0.018624 | 0.011268 | 0.006502 | 0.003758 | 0.001617 | 0.000987 | 0.002206 | 0.001836 | 8.63E-05 | 0.002414 | 0.003986 | 0.003813 |
| 28.1944444 | 0.016523 | 0.040781 | 0.014015 | 0.009285 | 0.020973 | 0.04197 | 0.001956 | 0.008159 | 0.010528 | 0.004587 | 0.002147 | 0.001763 | 0.001171 | 0.001188 | 0.001057 | 9.07E-05 | 0.001275 | 0.001573 | 0.001548 |
| 28.2083333 | 0.006455 | 0.044796 | 0.018292 | 0.014614 | 0.023332 | 0.053334 | 0.004985 | 0.010505 | 0.014787 | 0.004312 | 0.001557 | 0.002182 | 0.001026 | 0.000828 | 0.00048 | 0.000123 | 0.003236 | 0.001034 | 0.000991 |
| 28.2222222 | 0.002532 | 0.04372 | 0.024302 | 0.020211 | 0.02284 | 0.058678 | 0.007468 | 0.01446 | 0.021046 | 0.006816 | 0.00404 | 0.003339 | 0.000736 | 0.00072 | 0.000255 | 0.000242 | 0.007433 | 0.001617 | 0.002852 |
| 28.2361111 | 0.002708 | 0.036003 | 0.025255 | 0.018075 | 0.0188 | 0.057797 | 0.007999 | 0.013605 | 0.024864 | 0.008885 | 0.008914 | 0.004 | 0.00074 | 0.000758 | 0.000588 | 0.000375 | 0.010404 | 0.003822 | 0.005336 |
| 28.25 | 0.004922 | 0.025926 | 0.019122 | 0.011564 | 0.012974 | 0.048652 | 0.006572 | 0.012098 | 0.023825 | 0.008831 | 0.012198 | 0.003205 | 0.001179 | 0.00095 | 0.001143 | 0.000313 | 0.009438 | 0.005824 | 0.006075 |
| 28.2638889 | 0.008844 | 0.01662 | 0.009873 | 0.00616 | 0.008151 | 0.03144 | 0.003813 | 0.010486 | 0.017798 | 0.006494 | 0.011394 | 0.001975 | 0.00161 | 0.001098 | 0.001354 | 0.000257 | 0.006633 | 0.005107 | 0.005097 |
| 28.2777778 | 0.012828 | 0.009128 | 0.003235 | 0.005709 | 0.004759 | 0.016646 | 0.001298 | 0.008333 | 0.00906 | 0.003722 | 0.008179 | 0.00138 | 0.001811 | 0.001087 | 0.00144 | 0.000317 | 0.005816 | 0.003493 | 0.004774 |
| 28.2916667 | 0.014672 | 0.005573 | 0.001665 | 0.005518 | 0.002821 | 0.011659 | 0.000826 | 0.006527 | 0.002984 | 0.002501 | 0.004784 | 0.000963 | 0.001906 | 0.00093 | 0.001866 | 0.000298 | 0.007714 | 0.003918 | 0.006171 |
| 28.3055556 | 0.014658 | 0.006574 | 0.005415 | 0.00419 | 0.003301 | 0.014072 | 0.002637 | 0.004825 | 0.001946 | 0.003299 | 0.003022 | 0.000972 | 0.001838 | 0.00082 | 0.002521 | 0.000268 | 0.009204 | 0.006212 | 0.007896 |
| 28.3194444 | 0.01436 | 0.009358 | 0.013079 | 0.003582 | 0.005956 | 0.018373 | 0.005312 | 0.002969 | 0.003713 | 0.005916 | 0.005316 | 0.002176 | 0.001601 | 0.000992 | 0.003079 | 0.000373 | 0.00673 | 0.007826 | 0.008571 |
| 28.3333333 | 0.01413 | 0.010536 | 0.019049 | 0.002839 | 0.008933 | 0.020231 | 0.007152 | 0.002084 | 0.006935 | 0.009782 | 0.010653 | 0.002958 | 0.001459 | 0.00127 | 0.003086 | 0.000453 | 0.00375 | 0.007611 | 0.007859 |
| 28.3472222 | 0.013232 | 0.008998 | 0.017919 | 0.003329 | 0.010261 | 0.018731 | 0.007416 | 0.002577 | 0.008476 | 0.013074 | 0.014352 | 0.002254 | 0.001536 | 0.001371 | 0.002334 | 0.000298 | 0.004992 | 0.006963 | 0.005995 |
| 28.3611111 | 0.011644 | 0.006024 | 0.01125 | 0.006738 | 0.009184 | 0.014629 | 0.00637 | 0.003229 | 0.006468 | 0.013052 | 0.013343 | 0.002119 | 0.001619 | 0.001263 | 0.0015 | 0.000116 | 0.005813 | 0.006487 | 0.003688 |
| 28.375 | 0.010784 | 0.003647 | 0.005113 | 0.012595 | 0.00641 | 0.010406 | 0.004773 | 0.002943 | 0.003735 | 0.00833 | 0.008588 | 0.003467 | 0.001425 | 0.000985 | 0.00129 | 0.000255 | 0.003698 | 0.004886 | 0.001767 |
| 28.3888889 | 0.011765 | 0.002901 | 0.003871 | 0.014197 | 0.003382 | 0.010189 | 0.003416 | 0.002456 | 0.002941 | 0.004438 | 0.003773 | 0.004541 | 0.001063 | 0.000657 | 0.001392 | 0.000645 | 0.002745 | 0.002609 | 0.000963 |
| 28.4027778 | 0.013324 | 0.003024 | 0.007874 | 0.008586 | 0.001227 | 0.015143 | 0.002224 | 0.003057 | 0.004149 | 0.007127 | 0.00138 | 0.004488 | 0.000758 | 0.000391 | 0.001123 | 0.000733 | 0.00316 | 0.002163 | 0.00146 |
| 28.4166667 | 0.013684 | 0.002604 | 0.013553 | 0.003483 | 0.000484 | 0.021514 | 0.001417 | 0.005251 | 0.005384 | 0.010961 | 0.000708 | 0.004199 | 0.000515 | 0.000328 | 0.000825 | 0.000394 | 0.003174 | 0.003846 | 0.002206 |
| 28.4305556 | 0.013174 | 0.001767 | 0.016213 | 0.005533 | 0.000668 | 0.024232 | 0.001503 | 0.009695 | 0.005321 | 0.009575 | 0.000586 | 0.004215 | 0.000674 | 0.000863 | 0.000792 | 0.00047 | 0.002835 | 0.005124 | 0.001826 |
| 28.4444444 | 0.012969 | 0.001717 | 0.014844 | 0.01326 | 0.000809 | 0.022126 | 0.001791 | 0.016649 | 0.004841 | 0.005086 | 0.001053 | 0.004278 | 0.001322 | 0.001691 | 0.000693 | 0.001333 | 0.002468 | 0.004567 | 0.001591 |
| 28.4583333 | 0.013088 | 0.002819 | 0.01304 | 0.02018 | 0.001403 | 0.018669 | 0.002828 | 0.024201 | 0.005745 | 0.001899 | 0.002832 | 0.004649 | 0.00147 | 0.001628 | 0.000866 | 0.00185 | 0.002661 | 0.003592 | 0.003934 |
| 28.4722222 | 0.013052 | 0.004804 | 0.015131 | 0.022675 | 0.003026 | 0.017091 | 0.004978 | 0.029213 | 0.009196 | 0.002617 | 0.006135 | 0.005812 | 0.000873 | 0.000761 | 0.001758 | 0.001282 | 0.003656 | 0.004033 | 0.008124 |
| 28.4861111 | 0.013856 | 0.008082 | 0.022321 | 0.022194 | 0.004468 | 0.017869 | 0.005808 | 0.030409 | 0.014751 | 0.008474 | 0.009735 | 0.007329 | 0.000417 | 0.000469 | 0.002845 | 0.000723 | 0.005018 | 0.005881 | 0.012205 |
| 28.5 | 0.01729 | 0.012663 | 0.030678 | 0.021236 | 0.003948 | 0.02027 | 0.003972 | 0.029491 | 0.020271 | 0.016899 | 0.012426 | 0.008178 | 0.000517 | 0.001422 | 0.002682 | 0.000682 | 0.006523 | 0.007852 | 0.015028 |
| 28.5138889 | 0.022887 | 0.016864 | 0.033802 | 0.020567 | 0.001875 | 0.024192 | 0.001363 | 0.028983 | 0.024073 | 0.02247 | 0.013641 | 0.007497 | 0.001068 | 0.00293 | 0.001341 | 0.00057 | 0.007772 | 0.0086 | 0.01604 |
| 28.5277778 | 0.027455 | 0.018852 | 0.029981 | 0.020061 | 0.001328 | 0.030171 | 0.001295 | 0.029118 | 0.02592 | 0.021526 | 0.013121 | 0.00549 | 0.001167 | 0.003368 | 0.000621 | 0.000389 | 0.008063 | 0.007612 | 0.016214 |
| 28.5416667 | 0.027578 | 0.017661 | 0.022763 | 0.019332 | 0.003524 | 0.035225 | 0.00417 | 0.027149 | 0.024793 | 0.015811 | 0.011516 | 0.003591 | 0.000649 | 0.002519 | 0.000945 | 0.000283 | 0.007036 | 0.005819 | 0.016144 |
| 28.5555556 | 0.022344 | 0.01225 | 0.014886 | 0.017896 | 0.00579 | 0.032236 | 0.006217 | 0.021403 | 0.01847 | 0.011292 | 0.00998 | 0.002567 | 0.000416 | 0.001968 | 0.001046 | 0.00027 | 0.005168 | 0.005053 | 0.014692 |
| 28.5694444 | 0.016241 | 0.005618 | 0.010694 | 0.020334 | 0.006071 | 0.020843 | 0.005435 | 0.015374 | 0.00922 | 0.013257 | 0.011218 | 0.002077 | 0.000789 | 0.002536 | 0.000839 | 0.000261 | 0.004095 | 0.007716 | 0.013333 |
| 28.5833333 | 0.019813 | 0.006009 | 0.019659 | 0.038291 | 0.006921 | 0.012131 | 0.004845 | 0.016255 | 0.006776 | 0.026298 | 0.020468 | 0.002114 | 0.002139 | 0.004045 | 0.001521 | 0.000371 | 0.005481 | 0.016845 | 0.017164 |
| 28.5972222 | 0.039214 | 0.018864 | 0.04364 | 0.068183 | 0.012919 | 0.01374 | 0.008408 | 0.02828 | 0.019027 | 0.049246 | 0.036985 | 0.002848 | 0.003773 | 0.005094 | 0.003412 | 0.000604 | 0.008635 | 0.030778 | 0.028708 |
| 28.6111111 | 0.063279 | 0.038813 | 0.068743 | 0.083671 | 0.020342 | 0.021464 | 0.015845 | 0.04457 | 0.042603 | 0.069841 | 0.049709 | 0.004414 | 0.004038 | 0.004121 | 0.003901 | 0.000745 | 0.012836 | 0.040723 | 0.042019 |
| 28.625 | 0.071261 | 0.048121 | 0.075708 | 0.071846 | 0.019761 | 0.027954 | 0.023092 | 0.049343 | 0.06683 | 0.07192 | 0.050337 | 0.007131 | 0.002995 | 0.003939 | 0.002582 | 0.001424 | 0.020135 | 0.041855 | 0.047328 |
| 28.6388889 | 0.056084 | 0.03864 | 0.055259 | 0.043071 | 0.012855 | 0.02484 | 0.026168 | 0.036506 | 0.075639 | 0.049752 | 0.040823 | 0.010965 | 0.00225 | 0.007114 | 0.002619 | 0.002615 | 0.028106 | 0.035987 | 0.040945 |
| 28.6527778 | 0.030307 | 0.021281 | 0.029241 | 0.019825 | 0.008209 | 0.013679 | 0.023406 | 0.018677 | 0.055319 | 0.026867 | 0.026845 | 0.013283 | 0.002691 | 0.010217 | 0.002392 | 0.002903 | 0.029802 | 0.025223 | 0.026358 |
| 28.6666667 | 0.014869 | 0.012181 | 0.028943 | 0.01723 | 0.008204 | 0.011021 | 0.016161 | 0.012398 | 0.027273 | 0.032081 | 0.012752 | 0.011059 | 0.003201 | 0.010639 | 0.00103 | 0.002391 | 0.024635 | 0.014433 | 0.01127 |
| 28.6805556 | 0.016553 | 0.016635 | 0.04454 | 0.016747 | 0.010154 | 0.022797 | 0.008591 | 0.017311 | 0.023751 | 0.045022 | 0.005831 | 0.006456 | 0.002679 | 0.009723 | 0.000694 | 0.002206 | 0.017454 | 0.010232 | 0.002662 |
| 28.6944444 | 0.018681 | 0.021997 | 0.045835 | 0.013985 | 0.010596 | 0.032328 | 0.0054 | 0.017972 | 0.032038 | 0.032232 | 0.009952 | 0.002926 | 0.002226 | 0.008446 | 0.001896 | 0.001936 | 0.011116 | 0.015159 | 0.000552 |
| 28.7083333 | 0.012747 | 0.018884 | 0.02976 | 0.027099 | 0.008761 | 0.025618 | 0.00746 | 0.012063 | 0.026151 | 0.017382 | 0.016116 | 0.000952 | 0.001865 | 0.006341 | 0.004446 | 0.001332 | 0.00611 | 0.024145 | 0.000328 |
| 28.7222222 | 0.008367 | 0.019638 | 0.023244 | 0.038079 | 0.008146 | 0.023107 | 0.01043 | 0.012975 | 0.016985 | 0.025817 | 0.016002 | 0.000537 | 0.000974 | 0.003665 | 0.006704 | 0.00128 | 0.002618 | 0.027266 | 0.000255 |
| 28.7361111 | 0.008404 | 0.030651 | 0.03458 | 0.02779 | 0.010081 | 0.042311 | 0.009151 | 0.021516 | 0.026256 | 0.032256 | 0.011168 | 0.000932 | 0.000742 | 0.001746 | 0.0071 | 0.001705 | 0.00168 | 0.020479 | 0.001493 |
| 28.75 | 0.007187 | 0.034977 | 0.038634 | 0.014132 | 0.010596 | 0.060935 | 0.005762 | 0.025862 | 0.039194 | 0.022062 | 0.006899 | 0.001599 | 0.001445 | 0.001127 | 0.005883 | 0.001871 | 0.003676 | 0.010204 | 0.004696 |
| 28.7638889 | 0.006666 | 0.028382 | 0.031197 | 0.015839 | 0.008452 | 0.057411 | 0.006377 | 0.024268 | 0.034417 | 0.009413 | 0.006845 | 0.00397 | 0.002304 | 0.000938 | 0.004369 | 0.001423 | 0.005503 | 0.005941 | 0.007121 |
| 28.7777778 | 0.010293 | 0.019531 | 0.021531 | 0.025816 | 0.006961 | 0.034772 | 0.009449 | 0.02438 | 0.018858 | 0.004216 | 0.011327 | 0.00776 | 0.002029 | 0.000952 | 0.003007 | 0.00088 | 0.004782 | 0.00836 | 0.006643 |
| 28.7916667 | 0.017906 | 0.012322 | 0.012752 | 0.022917 | 0.00835 | 0.017997 | 0.009152 | 0.026783 | 0.011334 | 0.004412 | 0.015399 | 0.01005 | 0.000762 | 0.001074 | 0.001829 | 0.001563 | 0.00289 | 0.010856 | 0.00784 |
| 28.8055556 | 0.033371 | 0.008841 | 0.010545 | 0.016269 | 0.011766 | 0.025125 | 0.006065 | 0.023845 | 0.017912 | 0.004101 | 0.013823 | 0.008615 | 0.000245 | 0.001078 | 0.001721 | 0.003266 | 0.002051 | 0.010596 | 0.011571 |
| 28.8194444 | 0.046986 | 0.014053 | 0.011751 | 0.028016 | 0.014047 | 0.039506 | 0.005674 | 0.016211 | 0.019866 | 0.0041 | 0.008909 | 0.00496 | 0.00048 | 0.001179 | 0.002607 | 0.004349 | 0.002381 | 0.007578 | 0.012526 |

| | | | | | | | | | | | | | | | | | | | |
|---|---|---|---|---|---|---|---|---|---|---|---|---|---|---|---|---|---|---|---|
| 28.8333333 | 0.039164 | 0.026861 | 0.013566 | 0.036642 | 0.012925 | 0.041384 | 0.007561 | 0.014876 | 0.012586 | 0.00439 | 0.004913 | 0.003628 | 0.001023 | 0.001178 | 0.003232 | 0.004219 | 0.002261 | 0.004527 | 0.010779 |
| 28.8472222 | 0.022329 | 0.026371 | 0.016332 | 0.023582 | 0.009862 | 0.027792 | 0.006895 | 0.02283 | 0.013042 | 0.003772 | 0.002348 | 0.004954 | 0.002041 | 0.001284 | 0.002453 | 0.003568 | 0.002241 | 0.005032 | 0.008641 |
| 28.8611111 | 0.026168 | 0.013535 | 0.023686 | 0.01198 | 0.007922 | 0.014078 | 0.004439 | 0.029921 | 0.017299 | 0.005794 | 0.001083 | 0.004675 | 0.002975 | 0.001576 | 0.001609 | 0.002781 | 0.003846 | 0.0051 | 0.006753 |
| 28.875 | 0.045543 | 0.009308 | 0.041967 | 0.017952 | 0.007214 | 0.014535 | 0.004261 | 0.025068 | 0.016151 | 0.010398 | 0.001485 | 0.003116 | 0.003894 | 0.002808 | 0.002959 | 0.001684 | 0.005426 | 0.003876 | 0.008058 |
| 28.8888889 | 0.057682 | 0.019285 | 0.066523 | 0.028389 | 0.004926 | 0.015683 | 0.00579 | 0.017982 | 0.024279 | 0.016209 | 0.003811 | 0.004647 | 0.005802 | 0.003586 | 0.005112 | 0.00108 | 0.004393 | 0.006513 | 0.015707 |
| 28.9027778 | 0.059566 | 0.051168 | 0.080279 | 0.029589 | 0.003334 | 0.020719 | 0.006151 | 0.03131 | 0.047882 | 0.0251 | 0.008066 | 0.009252 | 0.007922 | 0.002936 | 0.006497 | 0.002012 | 0.004242 | 0.011381 | 0.0274 |
| 28.9166667 | 0.055771 | 0.096517 | 0.069755 | 0.022933 | 0.007289 | 0.047461 | 0.00582 | 0.060509 | 0.066507 | 0.035623 | 0.01289 | 0.01367 | 0.00832 | 0.003522 | 0.007633 | 0.004078 | 0.009042 | 0.017093 | 0.034195 |
| 28.9305556 | 0.049626 | 0.125795 | 0.045994 | 0.016609 | 0.014446 | 0.07619 | 0.006677 | 0.080704 | 0.066969 | 0.04297 | 0.016317 | 0.017428 | 0.006894 | 0.004945 | 0.008275 | 0.006557 | 0.015113 | 0.025445 | 0.031119 |
| 28.9444444 | 0.045376 | 0.121662 | 0.029806 | 0.017779 | 0.0199 | 0.082514 | 0.008919 | 0.080881 | 0.058196 | 0.044318 | 0.017331 | 0.021123 | 0.005166 | 0.005701 | 0.007192 | 0.008486 | 0.01914 | 0.033288 | 0.025413 |
| 28.9583333 | 0.044089 | 0.093329 | 0.028393 | 0.029404 | 0.02154 | 0.072349 | 0.010932 | 0.069297 | 0.052174 | 0.039742 | 0.016197 | 0.022427 | 0.004311 | 0.006395 | 0.005239 | 0.008706 | 0.022355 | 0.035191 | 0.024948 |
| 28.9722222 | 0.045671 | 0.061725 | 0.03614 | 0.051295 | 0.01904 | 0.059509 | 0.011275 | 0.059531 | 0.049731 | 0.030848 | 0.013968 | 0.019623 | 0.004636 | 0.007494 | 0.00501 | 0.007706 | 0.025485 | 0.030881 | 0.028619 |
| 28.9861111 | 0.046319 | 0.036079 | 0.042481 | 0.076645 | 0.013648 | 0.047761 | 0.009115 | 0.055217 | 0.046187 | 0.020571 | 0.011446 | 0.014882 | 0.005827 | 0.008615 | 0.006454 | 0.006597 | 0.027477 | 0.02529 | 0.032035 |
| 29 | 0.040872 | 0.01786 | 0.040601 | 0.085988 | 0.008438 | 0.037289 | 0.004851 | 0.051664 | 0.040858 | 0.013248 | 0.008712 | 0.010835 | 0.006656 | 0.009206 | 0.007577 | 0.005445 | 0.027754 | 0.022159 | 0.03396 |
| 29.0138889 | 0.030537 | 0.009053 | 0.031216 | 0.066615 | 0.006495 | 0.029183 | 0.001708 | 0.045387 | 0.035152 | 0.012381 | 0.005962 | 0.008572 | 0.006354 | 0.009151 | 0.0078 | 0.004228 | 0.026467 | 0.022204 | 0.034981 |
| 29.0277778 | 0.019557 | 0.010542 | 0.020315 | 0.033529 | 0.007197 | 0.021443 | 0.001078 | 0.034846 | 0.028408 | 0.017251 | 0.00374 | 0.007488 | 0.005501 | 0.008665 | 0.007732 | 0.0034 | 0.022419 | 0.023633 | 0.033069 |
| 29.0416667 | 0.010046 | 0.018047 | 0.012619 | 0.010012 | 0.006934 | 0.012175 | 0.001438 | 0.019978 | 0.01936 | 0.021467 | 0.002338 | 0.005624 | 0.004404 | 0.007943 | 0.007332 | 0.002981 | 0.015104 | 0.023187 | 0.025926 |
| 29.0555556 | 0.003797 | 0.023588 | 0.008274 | 0.002818 | 0.004204 | 0.005365 | 0.002678 | 0.008164 | 0.009453 | 0.018483 | 0.002162 | 0.002861 | 0.00332 | 0.006489 | 0.00637 | 0.002605 | 0.007725 | 0.019887 | 0.016198 |
| 29.0694444 | 0.002301 | 0.025556 | 0.007033 | 0.007288 | 0.001532 | 0.003115 | 0.005466 | 0.00393 | 0.003081 | 0.012513 | 0.005533 | 0.001327 | 0.002837 | 0.003999 | 0.004865 | 0.002354 | 0.004782 | 0.017106 | 0.010647 |
| 29.0833333 | 0.00731 | 0.032794 | 0.012522 | 0.02288 | 0.001848 | 0.007342 | 0.010236 | 0.006037 | 0.006012 | 0.01822 | 0.016688 | 0.001441 | 0.003051 | 0.003395 | 0.002671 | 0.002508 | 0.008029 | 0.020389 | 0.014166 |
| 29.0972222 | 0.0179 | 0.048711 | 0.026267 | 0.04513 | 0.005881 | 0.023972 | 0.015366 | 0.016283 | 0.020091 | 0.03846 | 0.031598 | 0.001429 | 0.003276 | 0.00589 | 0.000977 | 0.002622 | 0.013142 | 0.028708 | 0.022173 |
| 29.1111111 | 0.026729 | 0.062283 | 0.041293 | 0.059589 | 0.014088 | 0.048119 | 0.017766 | 0.028211 | 0.038697 | 0.050955 | 0.03488 | 0.001563 | 0.002845 | 0.006265 | 0.000805 | 0.002258 | 0.013461 | 0.031607 | 0.022729 |
| 29.125 | 0.033397 | 0.069645 | 0.055345 | 0.061164 | 0.029046 | 0.07636 | 0.020038 | 0.039416 | 0.058163 | 0.04041 | 0.022682 | 0.003893 | 0.00293 | 0.006798 | 0.002306 | 0.0025 | 0.00988 | 0.024915 | 0.013039 |
| 29.1388889 | 0.044187 | 0.078737 | 0.073563 | 0.055905 | 0.052993 | 0.112478 | 0.026715 | 0.054527 | 0.077422 | 0.022174 | 0.009956 | 0.006982 | 0.004242 | 0.012265 | 0.005165 | 0.0037 | 0.007227 | 0.015865 | 0.004422 |
| 29.1527778 | 0.053337 | 0.081713 | 0.085236 | 0.046212 | 0.076961 | 0.139716 | 0.032657 | 0.065034 | 0.082092 | 0.01253 | 0.006517 | 0.007084 | 0.004501 | 0.015021 | 0.006305 | 0.004019 | 0.00536 | 0.008177 | 0.002584 |
| 29.1666667 | 0.0494 | 0.064165 | 0.072646 | 0.034891 | 0.08034 | 0.128331 | 0.030137 | 0.061608 | 0.061877 | 0.009757 | 0.008686 | 0.004405 | 0.00296 | 0.011206 | 0.00431 | 0.002923 | 0.00307 | 0.003373 | 0.002152 |
| 29.1805556 | 0.034772 | 0.035434 | 0.042574 | 0.0285 | 0.057379 | 0.082943 | 0.022472 | 0.048384 | 0.032099 | 0.008486 | 0.010327 | 0.002275 | 0.001756 | 0.006784 | 0.001878 | 0.001885 | 0.001461 | 0.003307 | 0.001605 |
| 29.1944444 | 0.021325 | 0.014476 | 0.017311 | 0.027625 | 0.027437 | 0.038633 | 0.018627 | 0.034849 | 0.011982 | 0.007319 | 0.008943 | 0.001889 | 0.001588 | 0.005124 | 0.000943 | 0.001469 | 0.000862 | 0.004135 | 0.001735 |
| 29.2083333 | 0.014404 | 0.00646 | 0.010199 | 0.024682 | 0.01085 | 0.015673 | 0.020061 | 0.024871 | 0.00385 | 0.006037 | 0.005706 | 0.002144 | 0.001538 | 0.005111 | 0.000914 | 0.001187 | 0.0006 | 0.004468 | 0.001345 |
| 29.2222222 | 0.009952 | 0.010433 | 0.018492 | 0.016049 | 0.01359 | 0.019643 | 0.020678 | 0.016635 | 0.004125 | 0.004114 | 0.002728 | 0.002363 | 0.001418 | 0.005496 | 0.00069 | 0.001067 | 0.000362 | 0.006287 | 0.000674 |
| 29.2361111 | 0.005562 | 0.024302 | 0.031678 | 0.012102 | 0.02712 | 0.046942 | 0.016031 | 0.009079 | 0.012263 | 0.002722 | 0.001347 | 0.002418 | 0.001339 | 0.005893 | 0.000449 | 0.001222 | 0.000549 | 0.008504 | 0.000358 |
| 29.25 | 0.005257 | 0.041374 | 0.044676 | 0.016665 | 0.037789 | 0.081629 | 0.008095 | 0.006612 | 0.027331 | 0.004256 | 0.001893 | 0.001941 | 0.00095 | 0.005463 | 0.000652 | 0.001154 | 0.001558 | 0.009162 | 0.000755 |
| 29.2638889 | 0.008583 | 0.053901 | 0.054944 | 0.016246 | 0.041071 | 0.10285 | 0.002709 | 0.011379 | 0.045989 | 0.007619 | 0.002915 | 0.001111 | 0.000455 | 0.004089 | 0.0007 | 0.000743 | 0.002314 | 0.007543 | 0.001732 |
| 29.2777778 | 0.008521 | 0.057366 | 0.056386 | 0.009266 | 0.039878 | 0.101084 | 0.002068 | 0.015866 | 0.059149 | 0.009444 | 0.002847 | 0.000699 | 0.000409 | 0.003033 | 0.00049 | 0.000531 | 0.001732 | 0.004104 | 0.002188 |
| 29.2916667 | 0.006328 | 0.053309 | 0.04619 | 0.007237 | 0.034547 | 0.088663 | 0.004924 | 0.014262 | 0.059309 | 0.008856 | 0.001847 | 0.000848 | 0.000639 | 0.003152 | 0.000508 | 0.000666 | 0.00069 | 0.002049 | 0.001488 |
| 29.3055556 | 0.009786 | 0.043858 | 0.030417 | 0.011757 | 0.024109 | 0.075585 | 0.010521 | 0.008786 | 0.048241 | 0.00683 | 0.000935 | 0.001219 | 0.000887 | 0.004176 | 0.000472 | 0.000906 | 0.000322 | 0.004131 | 0.000813 |
| 29.3194444 | 0.016422 | 0.030016 | 0.015938 | 0.016521 | 0.011973 | 0.059113 | 0.015349 | 0.00585 | 0.030542 | 0.003972 | 0.000777 | 0.00163 | 0.001237 | 0.005292 | 0.000447 | 0.001125 | 0.000483 | 0.007851 | 0.001368 |
| 29.3333333 | 0.019011 | 0.017226 | 0.006096 | 0.017329 | 0.005539 | 0.03854 | 0.016272 | 0.007824 | 0.014075 | 0.001771 | 0.000861 | 0.001804 | 0.001497 | 0.00565 | 0.000769 | 0.001139 | 0.000715 | 0.009742 | 0.002384 |
| 29.3472222 | 0.016896 | 0.008419 | 0.002514 | 0.01421 | 0.006458 | 0.018424 | 0.014825 | 0.009702 | 0.008824 | 0.001585 | 0.000722 | 0.001449 | 0.001307 | 0.004948 | 0.001237 | 0.000818 | 0.000808 | 0.00893 | 0.002595 |
| 29.3611111 | 0.01348 | 0.004902 | 0.002608 | 0.01033 | 0.008251 | 0.007655 | 0.013936 | 0.008776 | 0.01389 | 0.002427 | 0.000732 | 0.000925 | 0.000825 | 0.003769 | 0.001591 | 0.000562 | 0.000885 | 0.006392 | 0.001909 |
| 29.375 | 0.010923 | 0.010192 | 0.003863 | 0.009247 | 0.008897 | 0.009435 | 0.014645 | 0.006604 | 0.019151 | 0.003447 | 0.000677 | 0.000939 | 0.000473 | 0.00303 | 0.001666 | 0.000896 | 0.00129 | 0.004024 | 0.00116 |
| 29.3888889 | 0.008831 | 0.020469 | 0.006748 | 0.011753 | 0.010191 | 0.013707 | 0.015287 | 0.004956 | 0.020073 | 0.004139 | 0.000643 | 0.001332 | 0.000343 | 0.002598 | 0.001173 | 0.001444 | 0.001535 | 0.002667 | 0.000999 |
| 29.4027778 | 0.006868 | 0.029131 | 0.009195 | 0.014108 | 0.011654 | 0.013437 | 0.013816 | 0.003944 | 0.017699 | 0.003248 | 0.001297 | 0.001347 | 0.000439 | 0.001661 | 0.000694 | 0.001411 | 0.001202 | 0.001701 | 0.001174 |
| 29.4166667 | 0.006356 | 0.03425 | 0.008096 | 0.011389 | 0.013396 | 0.009586 | 0.01155 | 0.003616 | 0.014332 | 0.002336 | 0.002315 | 0.000841 | 0.00086 | 0.000667 | 0.001159 | 0.000802 | 0.001425 | 0.000894 | 0.001244 |
| 29.4305556 | 0.008769 | 0.036023 | 0.004373 | 0.006085 | 0.016501 | 0.005421 | 0.012431 | 0.006104 | 0.012884 | 0.004435 | 0.002904 | 0.000778 | 0.001702 | 0.000482 | 0.00194 | 0.000393 | 0.002925 | 0.000738 | 0.000992 |
| 29.4444444 | 0.012061 | 0.032615 | 0.00269 | 0.004755 | 0.019341 | 0.003208 | 0.016547 | 0.010553 | 0.012276 | 0.00798 | 0.002648 | 0.002359 | 0.002813 | 0.000779 | 0.002207 | 0.000498 | 0.004705 | 0.000902 | 0.000856 |
| 29.4583333 | 0.010922 | 0.020826 | 0.006402 | 0.006785 | 0.017393 | 0.003099 | 0.018752 | 0.011904 | 0.00946 | 0.009983 | 0.002154 | 0.004838 | 0.003633 | 0.000921 | 0.001839 | 0.000541 | 0.005705 | 0.000863 | 0.002432 |
| 29.4722222 | 0.006099 | 0.012493 | 0.012691 | 0.009275 | 0.010152 | 0.006498 | 0.01557 | 0.007902 | 0.009862 | 0.01093 | 0.002662 | 0.006134 | 0.00404 | 0.00089 | 0.001104 | 0.000386 | 0.006093 | 0.001513 | 0.007693 |
| 29.4861111 | 0.003253 | 0.019237 | 0.016414 | 0.014524 | 0.003854 | 0.01127 | 0.008554 | 0.004322 | 0.018432 | 0.012218 | 0.00402 | 0.005928 | 0.00446 | 0.001059 | 0.000858 | 0.000432 | 0.006623 | 0.003833 | 0.01623 |
| 29.5 | 0.001991 | 0.028527 | 0.015371 | 0.023899 | 0.00232 | 0.013971 | 0.003269 | 0.006193 | 0.028272 | 0.013985 | 0.005141 | 0.005758 | 0.00471 | 0.001419 | 0.001419 | 0.000828 | 0.007246 | 0.007077 | 0.022064 |
| 29.5138889 | 0.001281 | 0.028338 | 0.011147 | 0.031019 | 0.002065 | 0.015558 | 0.002015 | 0.008181 | 0.034168 | 0.016036 | 0.005839 | 0.006357 | 0.004306 | 0.00153 | 0.002044 | 0.001342 | 0.007616 | 0.009525 | 0.020471 |
| 29.5277778 | 0.003371 | 0.017612 | 0.008151 | 0.028225 | 0.001752 | 0.017432 | 0.002607 | 0.005482 | 0.03636 | 0.018037 | 0.00648 | 0.006668 | 0.003343 | 0.001384 | 0.001962 | 0.001477 | 0.007939 | 0.010246 | 0.015909 |
| 29.5416667 | 0.007147 | 0.008793 | 0.009284 | 0.017174 | 0.003092 | 0.019072 | 0.005484 | 0.001628 | 0.032883 | 0.019453 | 0.007122 | 0.006105 | 0.002343 | 0.001453 | 0.001139 | 0.001164 | 0.00802 | 0.009898 | 0.014788 |
| 29.5555556 | 0.008584 | 0.013326 | 0.012375 | 0.006282 | 0.004089 | 0.020207 | 0.008371 | 0.002061 | 0.022233 | 0.02057 | 0.008177 | 0.005666 | 0.002002 | 0.001976 | 0.000343 | 0.000947 | 0.007805 | 0.009976 | 0.017707 |
| 29.5694444 | 0.008619 | 0.016963 | 0.015225 | 0.006756 | 0.003983 | 0.021558 | 0.007699 | 0.008811 | 0.009695 | 0.023732 | 0.012696 | 0.006166 | 0.002289 | 0.002056 | 0.000186 | 0.001115 | 0.007871 | 0.012019 | 0.021842 |
| 29.5833333 | 0.017887 | 0.014254 | 0.022167 | 0.030449 | 0.005216 | 0.025083 | 0.004633 | 0.021559 | 0.004579 | 0.03512 | 0.025403 | 0.00834 | 0.002037 | 0.001685 | 0.000787 | 0.001517 | 0.009116 | 0.020513 | 0.027999 |
| 29.5972222 | 0.042249 | 0.020294 | 0.036161 | 0.0719 | 0.007829 | 0.030132 | 0.004538 | 0.038448 | 0.012498 | 0.054732 | 0.045113 | 0.012568 | 0.001069 | 0.001517 | 0.002116 | 0.00236 | 0.011783 | 0.03713 | 0.039929 |
| 29.6111111 | 0.065304 | 0.036114 | 0.051831 | 0.101453 | 0.007664 | 0.032349 | 0.011173 | 0.051894 | 0.033784 | 0.068278 | 0.058928 | 0.018319 | 0.001098 | 0.00226 | 0.003286 | 0.003776 | 0.013891 | 0.051677 | 0.054166 |
| 29.625 | 0.069542 | 0.049007 | 0.060715 | 0.098395 | 0.005303 | 0.03369 | 0.019021 | 0.052751 | 0.060838 | 0.067688 | 0.056238 | 0.023441 | 0.002874 | 0.005738 | 0.003972 | 0.005672 | 0.016099 | 0.055508 | 0.060478 |
| 29.6388889 | 0.05834 | 0.05097 | 0.056061 | 0.069404 | 0.005121 | 0.033977 | 0.021129 | 0.040289 | 0.071323 | 0.054856 | 0.04311 | 0.023469 | 0.004952 | 0.01016 | 0.004751 | 0.007732 | 0.021398 | 0.050101 | 0.054385 |
| 29.6527778 | 0.039664 | 0.038905 | 0.038519 | 0.034449 | 0.004134 | 0.024633 | 0.015779 | 0.021205 | 0.050707 | 0.031663 | 0.029207 | 0.017338 | 0.005406 | 0.011925 | 0.004396 | 0.007778 | 0.026149 | 0.037422 | 0.038525 |
| 29.6666667 | 0.018713 | 0.020134 | 0.019392 | 0.018021 | 0.003264 | 0.01214 | 0.009258 | 0.009307 | 0.023859 | 0.016634 | 0.017809 | 0.009552 | 0.004085 | 0.01134 | 0.002748 | 0.004647 | 0.025619 | 0.020449 | 0.018831 |
| 29.6805556 | 0.004545 | 0.011251 | 0.014251 | 0.021448 | 0.007351 | 0.008688 | 0.010982 | 0.01289 | 0.018919 | 0.024879 | 0.009553 | 0.004316 | 0.003247 | 0.011008 | 0.001928 | 0.001486 | 0.022262 | 0.008182 | 0.006019 |
| 29.6944444 | 0.002483 | 0.013555 | 0.022289 | 0.019167 | 0.010482 | 0.008356 | 0.01575 | 0.01907 | 0.026207 | 0.036453 | 0.005756 | 0.001834 | 0.004665 | 0.011589 | 0.002477 | 0.000996 | 0.019078 | 0.006207 | 0.002875 |
| 29.7083333 | 0.009226 | 0.014336 | 0.023945 | 0.016538 | 0.007607 | 0.005934 | 0.012756 | 0.015108 | 0.025525 | 0.031499 | 0.009324 | 0.00097 | 0.007465 | 0.011172 | 0.005252 | 0.001903 | 0.015773 | 0.009328 | 0.002328 |
| 29.7222222 | 0.020375 | 0.011305 | 0.017029 | 0.034438 | 0.007706 | 0.008796 | 0.012214 | 0.008867 | 0.017923 | 0.01839 | 0.016938 | 0.001352 | 0.00971 | 0.007928 | 0.010214 | 0.002076 | 0.011751 | 0.010894 | 0.00238 |
| 29.7361111 | 0.031634 | 0.016286 | 0.020863 | 0.051094 | 0.017551 | 0.020827 | 0.024822 | 0.012937 | 0.022914 | 0.016423 | 0.020426 | 0.002002 | 0.009151 | 0.003502 | 0.013523 | 0.001233 | 0.007491 | 0.007989 | 0.005335 |
| 29.75 | 0.037138 | 0.03245 | 0.033661 | 0.041321 | 0.027779 | 0.036483 | 0.035944 | 0.0234 | 0.046163 | 0.027274 | 0.01948 | 0.001734 | 0.005586 | 0.00226 | 0.013347 | 0.000394 | 0.003968 | 0.004893 | 0.011375 |
| 29.7638889 | 0.031991 | 0.043284 | 0.035474 | 0.020408 | 0.028447 | 0.045199 | 0.033698 | 0.027859 | 0.064457 | 0.032114 | 0.019062 | 0.001408 | 0.003652 | 0.004619 | 0.011762 | 0.000267 | 0.001699 | 0.008155 | 0.017322 |
| 29.7777778 | 0.018807 | 0.033503 | 0.023587 | 0.014049 | 0.020386 | 0.04034 | 0.021476 | 0.022311 | 0.057394 | 0.025786 | 0.019002 | 0.002357 | 0.005362 | 0.007701 | 0.010654 | 0.000805 | 0.000951 | 0.014229 | 0.020639 |
| 29.7916667 | 0.013091 | 0.012582 | 0.009663 | 0.022514 | 0.009955 | 0.027708 | 0.011939 | 0.013005 | 0.032798 | 0.017489 | 0.014485 | 0.00282 | 0.007097 | 0.009919 | 0.010588 | 0.00216 | 0.001071 | 0.017716 | 0.019242 |
| 29.8055556 | 0.019526 | 0.002417 | 0.003777 | 0.024887 | 0.003299 | 0.015844 | 0.012871 | 0.006119 | 0.018481 | 0.009494 | 0.007444 | 0.001995 | 0.007352 | 0.009514 | 0.010714 | 0.00401 | 0.001065 | 0.018698 | 0.012981 |
| 29.8194444 | 0.019688 | 0.002622 | 0.00399 | 0.024589 | 0.003197 | 0.007297 | 0.011416 | 0.004113 | 0.02476 | 0.0059 | 0.005191 | 0.002134 | 0.007123 | 0.006461 | 0.009848 | 0.005241 | 0.001492 | 0.017783 | 0.010417 |
| 29.8333333 | 0.02021 | 0.002286 | 0.008063 | 0.044896 | 0.007263 | 0.005172 | 0.008062 | 0.005571 | 0.036302 | 0.009253 | 0.006487 | 0.00411 | 0.006197 | 0.003678 | 0.007862 | 0.00537 | 0.002828 | 0.014645 | 0.0133 |
| 29.8472222 | 0.037378 | 0.008888 | 0.018638 | 0.062268 | 0.009386 | 0.012353 | 0.015994 | 0.004917 | 0.041944 | 0.009016 | 0.007673 | 0.0063 | 0.00404 | 0.002402 | 0.005478 | 0.004928 | 0.003996 | 0.009988 | 0.011439 |
| 29.8611111 | 0.048617 | 0.033166 | 0.02232 | 0.043695 | 0.006795 | 0.02285 | 0.026456 | 0.002509 | 0.036782 | 0.006775 | 0.008786 | 0.007307 | 0.001831 | 0.002877 | 0.003489 | 0.004635 | 0.004001 | 0.00578 | 0.010273 |
| 29.875 | 0.038713 | 0.045616 | 0.020155 | 0.024672 | 0.005632 | 0.029116 | 0.024967 | 0.001905 | 0.021701 | 0.011824 | 0.007995 | 0.007684 | 0.001101 | 0.004331 | 0.001936 | 0.004213 | 0.002898 | 0.003741 | 0.02037 |
| 29.8888889 | 0.023029 | 0.029799 | 0.037209 | 0.053365 | 0.011439 | 0.031018 | 0.015074 | 0.003416 | 0.009701 | 0.01343 | 0.005139 | 0.008877 | 0.003147 | 0.005316 | 0.001495 | 0.002983 | 0.002447 | 0.007071 | 0.033902 |
| 29.9027778 | 0.021834 | 0.016654 | 0.070725 | 0.104616 | 0.019169 | 0.030399 | 0.015422 | 0.003927 | 0.006564 | 0.011018 | 0.004108 | 0.011436 | 0.009122 | 0.005998 | 0.003029 | 0.001726 | 0.006268 | 0.017461 | 0.042608 |
| 29.9166667 | 0.037642 | 0.015213 | 0.097443 | 0.113824 | 0.023302 | 0.029104 | 0.026163 | 0.005579 | 0.00971 | 0.02057 | 0.005048 | 0.01413 | 0.016331 | 0.006642 | 0.005664 | 0.002166 | 0.013852 | 0.029828 | 0.044537 |
| 29.9305556 | 0.054271 | 0.028363 | 0.101198 | 0.068291 | 0.02334 | 0.032133 | 0.026785 | 0.015245 | 0.024837 | 0.037514 | 0.004483 | 0.015692 | 0.020245 | 0.007486 | 0.008818 | 0.004366 | 0.019815 | 0.038277 | 0.037205 |
| 29.9444444 | 0.061537 | 0.059794 | 0.076626 | 0.023405 | 0.020414 | 0.043469 | 0.015641 | 0.028587 | 0.049299 | 0.04566 | 0.00423 | 0.016468 | 0.0196 | 0.008576 | 0.011871 | 0.007197 | 0.021712 | 0.040642 | 0.023958 |
| 29.9583333 | 0.062013 | 0.099071 | 0.040661 | 0.011667 | 0.016552 | 0.056212 | 0.007516 | 0.039513 | 0.068438 | 0.041331 | 0.007502 | 0.016466 | 0.014706 | 0.009079 | 0.012711 | 0.010158 | 0.0197 | 0.03655 | 0.013208 |
| 29.9722222 | 0.061193 | 0.133425 | 0.015589 | 0.0209 | 0.012929 | 0.058687 | 0.006487 | 0.046803 | 0.074115 | 0.028763 | 0.012291 | 0.014759 | 0.007294 | 0.00824 | 0.010926 | 0.01263 | 0.01452 | 0.028418 | 0.009688 |
| 29.9861111 | 0.058074 | 0.140615 | 0.005058 | 0.046964 | 0.01099 | 0.050589 | 0.004734 | 0.047268 | 0.067835 | 0.013601 | 0.015477 | 0.011653 | 0.002321 | 0.00664 | 0.008714 | 0.013456 | 0.008924 | 0.020574 | 0.012869 |
| 30 | 0.051445 | 0.110933 | 0.003056 | 0.074554 | 0.011555 | 0.038461 | 0.00414 | 0.040441 | 0.055076 | 0.003844 | 0.016459 | 0.009048 | 0.001469 | 0.005626 | 0.007868 | 0.012379 | 0.005914 | 0.015563 | 0.017771 |
| 30.0138889 | 0.044008 | 0.063084 | 0.004207 | 0.083183 | 0.012343 | 0.026428 | 0.009144 | 0.030715 | 0.040105 | 0.002915 | 0.0153 | 0.007935 | 0.002823 | 0.005367 | 0.007986 | 0.01038 | 0.006504 | 0.01363 | 0.018034 |

| | | | | | | | | | | | | | | | | | | | |
|---|---|---|---|---|---|---|---|---|---|---|---|---|---|---|---|---|---|---|---|
| 30.0277778 | 0.035912 | 0.026454 | 0.004057 | 0.069115 | 0.010558 | 0.017554 | 0.010921 | 0.022692 | 0.026266 | 0.007751 | 0.011936 | 0.007251 | 0.00599 | 0.005075 | 0.007568 | 0.008263 | 0.009502 | 0.013617 | 0.012999 |
| 30.0416667 | 0.024822 | 0.016808 | 0.002366 | 0.041043 | 0.006226 | 0.013243 | 0.008902 | 0.017112 | 0.01701 | 0.015324 | 0.007565 | 0.00595 | 0.008797 | 0.004431 | 0.006448 | 0.006288 | 0.012682 | 0.013935 | 0.007414 |
| 30.0555556 | 0.012145 | 0.023452 | 0.001193 | 0.016148 | 0.003484 | 0.012317 | 0.014792 | 0.011994 | 0.012631 | 0.021994 | 0.004392 | 0.004098 | 0.008922 | 0.003603 | 0.005364 | 0.00467 | 0.014443 | 0.014291 | 0.00469 |
| 30.0694444 | 0.004142 | 0.025669 | 0.002336 | 0.004525 | 0.004267 | 0.014428 | 0.025918 | 0.007771 | 0.01158 | 0.027767 | 0.00727 | 0.002694 | 0.007539 | 0.002445 | 0.004175 | 0.003982 | 0.016018 | 0.017343 | 0.006063 |
| 30.0833333 | 0.004929 | 0.020623 | 0.010341 | 0.010379 | 0.005023 | 0.023 | 0.023626 | 0.009542 | 0.017855 | 0.039374 | 0.022467 | 0.003207 | 0.007565 | 0.001406 | 0.002569 | 0.004766 | 0.021779 | 0.028552 | 0.013065 |
| 30.0972222 | 0.013557 | 0.027069 | 0.027617 | 0.030695 | 0.01029 | 0.0428 | 0.015933 | 0.020207 | 0.036308 | 0.056292 | 0.045152 | 0.005526 | 0.009101 | 0.001161 | 0.00127 | 0.006062 | 0.031184 | 0.044983 | 0.022324 |
| 30.1111111 | 0.022649 | 0.048991 | 0.045734 | 0.045044 | 0.027085 | 0.06811 | 0.030379 | 0.033501 | 0.059353 | 0.061756 | 0.054549 | 0.00716 | 0.00843 | 0.002229 | 0.001451 | 0.006231 | 0.033631 | 0.049001 | 0.023906 |
| 30.125 | 0.023945 | 0.064193 | 0.049602 | 0.03901 | 0.045535 | 0.083492 | 0.050583 | 0.040214 | 0.068797 | 0.047344 | 0.040272 | 0.006626 | 0.005157 | 0.006055 | 0.003639 | 0.005617 | 0.024214 | 0.034479 | 0.015402 |
| 30.1388889 | 0.017172 | 0.062193 | 0.03602 | 0.025701 | 0.051714 | 0.081362 | 0.044958 | 0.038698 | 0.056373 | 0.027887 | 0.017922 | 0.004806 | 0.002678 | 0.010153 | 0.006516 | 0.005004 | 0.012326 | 0.018128 | 0.005962 |
| 30.1527778 | 0.009409 | 0.048221 | 0.01769 | 0.020123 | 0.042449 | 0.060457 | 0.024208 | 0.033044 | 0.031753 | 0.016277 | 0.006268 | 0.002658 | 0.00167 | 0.009833 | 0.006945 | 0.00375 | 0.005061 | 0.010255 | 0.001553 |
| 30.1666667 | 0.008503 | 0.028317 | 0.010444 | 0.019602 | 0.023426 | 0.032284 | 0.016796 | 0.024763 | 0.012192 | 0.010469 | 0.007288 | 0.001617 | 0.000879 | 0.006281 | 0.004503 | 0.002047 | 0.00168 | 0.006665 | 0.000664 |
| 30.1805556 | 0.012639 | 0.011515 | 0.016929 | 0.017072 | 0.010664 | 0.0215 | 0.022274 | 0.013628 | 0.006948 | 0.005374 | 0.009677 | 0.002129 | 0.00035 | 0.003341 | 0.003089 | 0.00114 | 0.000656 | 0.003881 | 0.000855 |
| 30.1944444 | 0.014313 | 0.004462 | 0.021963 | 0.011586 | 0.011825 | 0.026718 | 0.021141 | 0.005408 | 0.008879 | 0.001838 | 0.006483 | 0.00201 | 0.000365 | 0.002168 | 0.003929 | 0.001237 | 0.000406 | 0.002051 | 0.001074 |
| 30.2083333 | 0.011189 | 0.00311 | 0.017429 | 0.006024 | 0.016362 | 0.027303 | 0.012066 | 0.004743 | 0.010046 | 0.001522 | 0.002801 | 0.001055 | 0.000459 | 0.001923 | 0.004274 | 0.001608 | 0.00019 | 0.00113 | 0.001218 |
| 30.2222222 | 0.006816 | 0.002613 | 0.009118 | 0.003807 | 0.018841 | 0.022501 | 0.004315 | 0.006102 | 0.010693 | 0.002137 | 0.002304 | 0.000769 | 0.000326 | 0.001527 | 0.003244 | 0.001718 | 0.00046 | 0.001534 | 0.001245 |
| 30.2361111 | 0.006687 | 0.002839 | 0.002905 | 0.008802 | 0.021172 | 0.021521 | 0.002027 | 0.006212 | 0.012073 | 0.003026 | 0.00263 | 0.000688 | 0.000389 | 0.000796 | 0.002157 | 0.001325 | 0.001323 | 0.004738 | 0.00112 |
| 30.25 | 0.009298 | 0.00516 | 0.001625 | 0.020892 | 0.02381 | 0.026886 | 0.005664 | 0.007147 | 0.015307 | 0.005405 | 0.003845 | 0.000715 | 0.000989 | 0.000395 | 0.00219 | 0.000824 | 0.002348 | 0.009561 | 0.001406 |
| 30.2638889 | 0.007814 | 0.015698 | 0.00531 | 0.030603 | 0.025624 | 0.033157 | 0.011533 | 0.007905 | 0.021311 | 0.009234 | 0.004567 | 0.001403 | 0.001668 | 0.000505 | 0.00302 | 0.000844 | 0.003514 | 0.013 | 0.002112 |
| 30.2777778 | 0.006035 | 0.034304 | 0.012775 | 0.029787 | 0.025337 | 0.035349 | 0.013655 | 0.006908 | 0.029282 | 0.013061 | 0.002972 | 0.001875 | 0.001616 | 0.000647 | 0.003111 | 0.000801 | 0.004886 | 0.013785 | 0.002276 |
| 30.2916667 | 0.010479 | 0.051511 | 0.021662 | 0.020493 | 0.024516 | 0.038298 | 0.009973 | 0.005138 | 0.037312 | 0.014426 | 0.001314 | 0.001589 | 0.000851 | 0.000609 | 0.002047 | 0.000637 | 0.00536 | 0.012688 | 0.001747 |
| 30.3055556 | 0.014865 | 0.060313 | 0.028955 | 0.009977 | 0.024242 | 0.04238 | 0.00632 | 0.003169 | 0.043146 | 0.012506 | 0.001371 | 0.001134 | 0.000316 | 0.000387 | 0.000948 | 0.001106 | 0.004188 | 0.010937 | 0.001172 |
| 30.3194444 | 0.014601 | 0.059359 | 0.032595 | 0.00556 | 0.023106 | 0.041362 | 0.008309 | 0.002088 | 0.043609 | 0.008915 | 0.001773 | 0.000803 | 0.000604 | 0.000314 | 0.000567 | 0.00161 | 0.002429 | 0.008523 | 0.000716 |
| 30.3333333 | 0.011844 | 0.050672 | 0.0313 | 0.006332 | 0.019976 | 0.035383 | 0.009891 | 0.002646 | 0.036608 | 0.005003 | 0.002008 | 0.000503 | 0.001416 | 0.000656 | 0.000562 | 0.001726 | 0.001438 | 0.005049 | 0.000351 |
| 30.3472222 | 0.007378 | 0.037267 | 0.02537 | 0.005665 | 0.016056 | 0.028087 | 0.006804 | 0.002521 | 0.024949 | 0.002576 | 0.00429 | 0.000776 | 0.002136 | 0.000958 | 0.000589 | 0.001888 | 0.001076 | 0.002247 | 0.000237 |
| 30.3611111 | 0.003547 | 0.022986 | 0.016808 | 0.004535 | 0.011725 | 0.019753 | 0.003467 | 0.001544 | 0.012901 | 0.002875 | 0.006936 | 0.002066 | 0.002565 | 0.001112 | 0.00099 | 0.002125 | 0.000912 | 0.001623 | 0.000537 |
| 30.375 | 0.003208 | 0.011042 | 0.007878 | 0.009113 | 0.007011 | 0.01081 | 0.004432 | 0.003311 | 0.005976 | 0.003 | 0.006074 | 0.003033 | 0.003039 | 0.001243 | 0.001416 | 0.002059 | 0.00084 | 0.002032 | 0.001881 |
| 30.3888889 | 0.004061 | 0.005497 | 0.001914 | 0.018211 | 0.005546 | 0.006525 | 0.011593 | 0.008396 | 0.006873 | 0.002421 | 0.003995 | 0.00237 | 0.003303 | 0.001022 | 0.001016 | 0.001417 | 0.000975 | 0.003011 | 0.003765 |
| 30.4027778 | 0.003228 | 0.007417 | 0.001314 | 0.023718 | 0.007399 | 0.007428 | 0.018574 | 0.011315 | 0.00878 | 0.003005 | 0.004433 | 0.001056 | 0.002387 | 0.000698 | 0.001012 | 0.000606 | 0.001403 | 0.003101 | 0.003639 |
| 30.4166667 | 0.002434 | 0.011785 | 0.003082 | 0.021337 | 0.008172 | 0.006361 | 0.020679 | 0.00993 | 0.008854 | 0.002653 | 0.004119 | 0.000351 | 0.000927 | 0.000715 | 0.002258 | 0.000244 | 0.001115 | 0.002287 | 0.001853 |
| 30.4305556 | 0.002444 | 0.01568 | 0.003685 | 0.013509 | 0.008845 | 0.003578 | 0.020739 | 0.00856 | 0.011038 | 0.001074 | 0.001775 | 0.000193 | 0.000381 | 0.00074 | 0.002902 | 0.000409 | 0.000578 | 0.002819 | 0.001368 |
| 30.4444444 | 0.002241 | 0.017679 | 0.005409 | 0.006227 | 0.012124 | 0.003504 | 0.019912 | 0.009174 | 0.01477 | 0.000806 | 0.002167 | 0.000457 | 0.001119 | 0.000913 | 0.002162 | 0.000782 | 0.001278 | 0.003171 | 0.002122 |
| 30.4583333 | 0.003152 | 0.015017 | 0.008653 | 0.005489 | 0.015324 | 0.003543 | 0.016489 | 0.008468 | 0.015044 | 0.002839 | 0.00773 | 0.001889 | 0.003594 | 0.002437 | 0.002455 | 0.001402 | 0.003375 | 0.004183 | 0.00219 |
| 30.4722222 | 0.005494 | 0.008144 | 0.009003 | 0.011088 | 0.015193 | 0.002438 | 0.010533 | 0.005139 | 0.009377 | 0.006222 | 0.014726 | 0.004742 | 0.006923 | 0.005305 | 0.004845 | 0.002279 | 0.005816 | 0.006998 | 0.002102 |
| 30.4861111 | 0.005417 | 0.002886 | 0.005594 | 0.01613 | 0.013291 | 0.003349 | 0.006841 | 0.002177 | 0.003971 | 0.008126 | 0.016052 | 0.007004 | 0.008097 | 0.006938 | 0.006062 | 0.002537 | 0.006687 | 0.008359 | 0.002626 |
| 30.5 | 0.003837 | 0.004319 | 0.003219 | 0.015871 | 0.012524 | 0.005494 | 0.009974 | 0.002969 | 0.004459 | 0.007377 | 0.0101 | 0.007129 | 0.006076 | 0.005733 | 0.003992 | 0.001827 | 0.00525 | 0.007706 | 0.002242 |
| 30.5138889 | 0.006513 | 0.010079 | 0.005682 | 0.013561 | 0.013061 | 0.005368 | 0.017746 | 0.009211 | 0.006987 | 0.005921 | 0.004736 | 0.006486 | 0.003376 | 0.003377 | 0.001581 | 0.001117 | 0.002869 | 0.00715 | 0.001522 |
| 30.5277778 | 0.012365 | 0.013538 | 0.009729 | 0.014056 | 0.014465 | 0.003458 | 0.024052 | 0.016912 | 0.008187 | 0.005578 | 0.003599 | 0.006626 | 0.002599 | 0.001715 | 0.00133 | 0.001154 | 0.002164 | 0.007831 | 0.001869 |
| 30.5416667 | 0.015452 | 0.009944 | 0.010503 | 0.014399 | 0.01742 | 0.002832 | 0.02595 | 0.021404 | 0.008366 | 0.00703 | 0.004746 | 0.007463 | 0.003765 | 0.001175 | 0.00158 | 0.001611 | 0.005125 | 0.009217 | 0.002522 |
| 30.5555556 | 0.011329 | 0.007398 | 0.008638 | 0.010076 | 0.020731 | 0.003996 | 0.024134 | 0.021734 | 0.009145 | 0.010193 | 0.009891 | 0.008475 | 0.005045 | 0.001905 | 0.001076 | 0.002119 | 0.009931 | 0.010262 | 0.003339 |
| 30.5694444 | 0.007095 | 0.018635 | 0.009086 | 0.004145 | 0.020417 | 0.00665 | 0.019022 | 0.016159 | 0.014489 | 0.016594 | 0.016031 | 0.010034 | 0.004591 | 0.003547 | 0.000588 | 0.002924 | 0.012886 | 0.012485 | 0.005923 |
| 30.5833333 | 0.015884 | 0.040881 | 0.017556 | 0.009237 | 0.014739 | 0.014448 | 0.010734 | 0.012399 | 0.027612 | 0.031143 | 0.022162 | 0.01341 | 0.002423 | 0.004651 | 0.001117 | 0.004472 | 0.014533 | 0.021941 | 0.013346 |
| 30.5972222 | 0.036895 | 0.060872 | 0.035381 | 0.037583 | 0.007775 | 0.027468 | 0.00677 | 0.023158 | 0.046965 | 0.053229 | 0.032106 | 0.01875 | 0.001634 | 0.005237 | 0.001998 | 0.00788 | 0.018052 | 0.038192 | 0.028364 |
| 30.6111111 | 0.056295 | 0.069016 | 0.052288 | 0.079033 | 0.006956 | 0.038888 | 0.017427 | 0.039981 | 0.064756 | 0.06769 | 0.041775 | 0.023829 | 0.005349 | 0.007782 | 0.001915 | 0.013502 | 0.02134 | 0.047059 | 0.049275 |
| 30.625 | 0.060513 | 0.067025 | 0.056836 | 0.105233 | 0.010831 | 0.043983 | 0.034954 | 0.046001 | 0.07319 | 0.063597 | 0.042472 | 0.025818 | 0.01079 | 0.01153 | 0.002257 | 0.017592 | 0.019689 | 0.042824 | 0.065995 |
| 30.6388889 | 0.048577 | 0.056653 | 0.04821 | 0.094801 | 0.010748 | 0.038517 | 0.038344 | 0.037006 | 0.066665 | 0.049401 | 0.03125 | 0.022253 | 0.012567 | 0.01219 | 0.002832 | 0.01601 | 0.014111 | 0.033771 | 0.066981 |
| 30.6527778 | 0.027591 | 0.034806 | 0.028745 | 0.053334 | 0.007092 | 0.023025 | 0.022711 | 0.019911 | 0.043515 | 0.03487 | 0.016721 | 0.013247 | 0.010792 | 0.008812 | 0.001806 | 0.009644 | 0.008835 | 0.024887 | 0.052333 |
| 30.6666667 | 0.012389 | 0.017512 | 0.014393 | 0.021918 | 0.004388 | 0.01386 | 0.010653 | 0.009974 | 0.021666 | 0.02223 | 0.01179 | 0.004744 | 0.008813 | 0.005059 | 0.000878 | 0.003513 | 0.006981 | 0.018547 | 0.032682 |
| 30.6805556 | 0.010772 | 0.023416 | 0.021743 | 0.025486 | 0.007443 | 0.017677 | 0.017252 | 0.010457 | 0.022524 | 0.011089 | 0.013025 | 0.001231 | 0.00877 | 0.004592 | 0.001417 | 0.001239 | 0.008635 | 0.01583 | 0.018193 |
| 30.6944444 | 0.011584 | 0.039137 | 0.034125 | 0.032992 | 0.01708 | 0.01748 | 0.026151 | 0.009621 | 0.036405 | 0.006349 | 0.009046 | 0.000786 | 0.010276 | 0.006799 | 0.002294 | 0.001191 | 0.010386 | 0.014408 | 0.010831 |
| 30.7083333 | 0.017555 | 0.049672 | 0.028834 | 0.02531 | 0.021483 | 0.012713 | 0.022727 | 0.010556 | 0.042772 | 0.012194 | 0.004812 | 0.001041 | 0.010654 | 0.008529 | 0.002398 | 0.001249 | 0.00987 | 0.012608 | 0.007218 |
| 30.7222222 | 0.033311 | 0.05888 | 0.018022 | 0.034493 | 0.013874 | 0.018478 | 0.013483 | 0.020476 | 0.037484 | 0.017833 | 0.004256 | 0.002403 | 0.0091 | 0.008362 | 0.00266 | 0.001007 | 0.007547 | 0.010925 | 0.005397 |
| 30.7361111 | 0.044416 | 0.06061 | 0.02455 | 0.076162 | 0.008327 | 0.030967 | 0.012694 | 0.029871 | 0.02287 | 0.014002 | 0.004504 | 0.005282 | 0.007395 | 0.006949 | 0.004447 | 0.000557 | 0.004466 | 0.010076 | 0.004913 |
| 30.75 | 0.04375 | 0.05096 | 0.041838 | 0.09964 | 0.014645 | 0.038982 | 0.022826 | 0.030554 | 0.016766 | 0.012053 | 0.010625 | 0.007148 | 0.006834 | 0.005511 | 0.006004 | 0.000926 | 0.001975 | 0.011086 | 0.004242 |
| 30.7638889 | 0.034438 | 0.068822 | 0.053597 | 0.068041 | 0.019599 | 0.038087 | 0.030004 | 0.020595 | 0.039584 | 0.021907 | 0.022621 | 0.005557 | 0.007281 | 0.004884 | 0.006371 | 0.002169 | 0.001307 | 0.014143 | 0.006234 |
| 30.7777778 | 0.019589 | 0.108185 | 0.049909 | 0.030556 | 0.013458 | 0.026766 | 0.023365 | 0.011835 | 0.067572 | 0.030438 | 0.028536 | 0.004679 | 0.007264 | 0.005511 | 0.00756 | 0.002955 | 0.003172 | 0.014995 | 0.014784 |
| 30.7916667 | 0.012108 | 0.104911 | 0.032726 | 0.035629 | 0.009673 | 0.015621 | 0.013737 | 0.018409 | 0.065043 | 0.026776 | 0.022094 | 0.008705 | 0.005127 | 0.006788 | 0.010224 | 0.002464 | 0.006337 | 0.009981 | 0.022119 |
| 30.8055556 | 0.019757 | 0.06106 | 0.030095 | 0.069559 | 0.022673 | 0.021632 | 0.015632 | 0.027955 | 0.036151 | 0.015991 | 0.012844 | 0.011565 | 0.002938 | 0.006989 | 0.011873 | 0.001662 | 0.007262 | 0.006277 | 0.019986 |
| 30.8194444 | 0.028214 | 0.025219 | 0.04788 | 0.103039 | 0.038309 | 0.038611 | 0.018458 | 0.028056 | 0.016431 | 0.006715 | 0.009911 | 0.008768 | 0.003727 | 0.006273 | 0.0109 | 0.001732 | 0.00496 | 0.009575 | 0.011483 |
| 30.8333333 | 0.028319 | 0.020266 | 0.058389 | 0.097513 | 0.036362 | 0.038777 | 0.01242 | 0.02252 | 0.027835 | 0.0043 | 0.011488 | 0.004241 | 0.005253 | 0.006138 | 0.008693 | 0.001573 | 0.002034 | 0.012187 | 0.006414 |
| 30.8472222 | 0.025652 | 0.035082 | 0.05354 | 0.061681 | 0.020627 | 0.02119 | 0.006905 | 0.018777 | 0.051277 | 0.00701 | 0.013227 | 0.002986 | 0.005257 | 0.005138 | 0.006152 | 0.00087 | 0.000812 | 0.010042 | 0.007201 |
| 30.8611111 | 0.023446 | 0.056382 | 0.038352 | 0.063958 | 0.008823 | 0.012856 | 0.006641 | 0.021998 | 0.063694 | 0.011021 | 0.011737 | 0.006695 | 0.005862 | 0.003763 | 0.003184 | 0.000658 | 0.002357 | 0.00591 | 0.007281 |
| 30.875 | 0.018574 | 0.077606 | 0.019043 | 0.080797 | 0.008743 | 0.01158 | 0.007215 | 0.026493 | 0.062765 | 0.014474 | 0.007206 | 0.012995 | 0.008494 | 0.00577 | 0.00147 | 0.001047 | 0.006736 | 0.002876 | 0.00717 |
| 30.8888889 | 0.010142 | 0.099904 | 0.006433 | 0.052687 | 0.008856 | 0.010851 | 0.007044 | 0.023505 | 0.057493 | 0.015444 | 0.005089 | 0.018463 | 0.010781 | 0.008909 | 0.001877 | 0.002813 | 0.01277 | 0.002862 | 0.008103 |
| 30.9027778 | 0.00431 | 0.124914 | 0.008033 | 0.027612 | 0.010887 | 0.025764 | 0.010287 | 0.018018 | 0.055666 | 0.013443 | 0.005171 | 0.019588 | 0.011724 | 0.009075 | 0.002327 | 0.006144 | 0.019612 | 0.003505 | 0.008191 |
| 30.9166667 | 0.006393 | 0.13465 | 0.029036 | 0.057103 | 0.029562 | 0.050942 | 0.020449 | 0.020413 | 0.055507 | 0.011013 | 0.004534 | 0.015706 | 0.013624 | 0.007393 | 0.002253 | 0.009839 | 0.026139 | 0.005291 | 0.011111 |
| 30.9305556 | 0.014291 | 0.116473 | 0.054181 | 0.11167 | 0.05257 | 0.065101 | 0.03125 | 0.029345 | 0.049515 | 0.010643 | 0.006469 | 0.010555 | 0.018025 | 0.006489 | 0.002719 | 0.011764 | 0.028957 | 0.009454 | 0.018813 |
| 30.9444444 | 0.023707 | 0.081289 | 0.059688 | 0.143538 | 0.057989 | 0.058914 | 0.031727 | 0.036966 | 0.034865 | 0.012588 | 0.009521 | 0.006991 | 0.023168 | 0.008077 | 0.00361 | 0.010528 | 0.025893 | 0.012585 | 0.027586 |
| 30.9583333 | 0.034649 | 0.046977 | 0.049093 | 0.13403 | 0.044557 | 0.042616 | 0.021991 | 0.041477 | 0.021279 | 0.014718 | 0.009116 | 0.004916 | 0.023756 | 0.011269 | 0.00414 | 0.007267 | 0.018383 | 0.013775 | 0.033542 |
| 30.9722222 | 0.044591 | 0.020848 | 0.036994 | 0.095286 | 0.023307 | 0.030471 | 0.013653 | 0.043503 | 0.020205 | 0.014783 | 0.006331 | 0.003661 | 0.018183 | 0.01281 | 0.003995 | 0.003856 | 0.009918 | 0.015318 | 0.0336 |
| 30.9861111 | 0.046329 | 0.007333 | 0.028411 | 0.051003 | 0.009018 | 0.027528 | 0.015063 | 0.039693 | 0.033194 | 0.013093 | 0.004555 | 0.003221 | 0.01072 | 0.011263 | 0.004385 | 0.00206 | 0.004493 | 0.018621 | 0.027819 |
| 31 | 0.039181 | 0.00751 | 0.025282 | 0.020746 | 0.006925 | 0.033963 | 0.019932 | 0.030331 | 0.047475 | 0.010342 | 0.00423 | 0.0035 | 0.004644 | 0.008254 | 0.006525 | 0.002314 | 0.003093 | 0.023401 | 0.020705 |
| 31.0138889 | 0.028709 | 0.01224 | 0.02666 | 0.010964 | 0.006246 | 0.04299 | 0.019779 | 0.02005 | 0.050022 | 0.006642 | 0.002913 | 0.004287 | 0.001161 | 0.006001 | 0.009442 | 0.003337 | 0.003497 | 0.028033 | 0.0153 |
| 31.0277778 | 0.018828 | 0.017438 | 0.028159 | 0.013911 | 0.004943 | 0.043074 | 0.015822 | 0.011071 | 0.042143 | 0.0035 | 0.000972 | 0.005335 | 0.000269 | 0.005123 | 0.011205 | 0.0041 | 0.00374 | 0.031189 | 0.011411 |
| 31.0416667 | 0.011006 | 0.025044 | 0.025233 | 0.019462 | 0.006107 | 0.029253 | 0.011221 | 0.004518 | 0.027941 | 0.00244 | 0.00051 | 0.004631 | 0.000441 | 0.00477 | 0.01102 | 0.004095 | 0.00311 | 0.031416 | 0.008227 |
| 31.0555556 | 0.005726 | 0.03309 | 0.01583 | 0.020203 | 0.007404 | 0.016054 | 0.014134 | 0.002629 | 0.013878 | 0.003178 | 0.001374 | 0.002938 | 0.000419 | 0.003967 | 0.009074 | 0.002912 | 0.001902 | 0.027061 | 0.005464 |
| 31.0694444 | 0.005764 | 0.037206 | 0.010321 | 0.013107 | 0.014551 | 0.020674 | 0.028698 | 0.00579 | 0.011324 | 0.006527 | 0.00332 | 0.003419 | 0.000196 | 0.00231 | 0.005955 | 0.001233 | 0.001219 | 0.0213 | 0.004712 |
| 31.0833333 | 0.009916 | 0.033096 | 0.012647 | 0.007837 | 0.024949 | 0.026495 | 0.042125 | 0.011077 | 0.016989 | 0.016265 | 0.009317 | 0.003447 | 0.00016 | 0.000699 | 0.003235 | 0.000886 | 0.001282 | 0.024005 | 0.008469 |
| 31.0972222 | 0.012683 | 0.022828 | 0.018489 | 0.012091 | 0.024248 | 0.026685 | 0.037906 | 0.014385 | 0.017647 | 0.030926 | 0.021893 | 0.002717 | 0.000304 | 0.001622 | 0.002731 | 0.002735 | 0.00251 | 0.038018 | 0.014734 |
| 31.1111111 | 0.021852 | 0.02489 | 0.03995 | 0.01476 | 0.030066 | 0.049627 | 0.026311 | 0.011661 | 0.023339 | 0.038942 | 0.033614 | 0.00463 | 0.000453 | 0.006681 | 0.004787 | 0.005147 | 0.006036 | 0.046282 | 0.016563 |
| 31.125 | 0.034444 | 0.042845 | 0.065275 | 0.009842 | 0.056767 | 0.084605 | 0.028867 | 0.007379 | 0.042649 | 0.033526 | 0.032696 | 0.006014 | 0.000467 | 0.011994 | 0.007863 | 0.005926 | 0.009519 | 0.036042 | 0.011919 |
| 31.1388889 | 0.028676 | 0.049799 | 0.063033 | 0.008428 | 0.066475 | 0.094486 | 0.028581 | 0.011512 | 0.054693 | 0.022109 | 0.019464 | 0.004473 | 0.000697 | 0.012425 | 0.00965 | 0.00505 | 0.009082 | 0.019152 | 0.005801 |
| 31.1527778 | 0.016595 | 0.038349 | 0.035593 | 0.017523 | 0.042699 | 0.072945 | 0.017307 | 0.021774 | 0.04792 | 0.013481 | 0.009593 | 0.002183 | 0.001535 | 0.009541 | 0.008864 | 0.003398 | 0.004999 | 0.007932 | 0.002279 |
| 31.1666667 | 0.022772 | 0.02131 | 0.016544 | 0.027168 | 0.021031 | 0.037798 | 0.017257 | 0.028285 | 0.029932 | 0.009222 | 0.013062 | 0.001031 | 0.002077 | 0.006855 | 0.005699 | 0.001714 | 0.002046 | 0.002707 | 0.001578 |
| 31.1805556 | 0.036683 | 0.008872 | 0.026065 | 0.026888 | 0.023762 | 0.020935 | 0.024475 | 0.026526 | 0.013477 | 0.007389 | 0.018971 | 0.001557 | 0.001496 | 0.005023 | 0.002689 | 0.001066 | 0.002562 | 0.001518 | 0.00151 |
| 31.1944444 | 0.044481 | 0.009233 | 0.045923 | 0.017611 | 0.035752 | 0.03275 | 0.024997 | 0.018079 | 0.010483 | 0.006168 | 0.01832 | 0.002014 | 0.000531 | 0.00385 | 0.002445 | 0.001296 | 0.003819 | 0.002162 | 0.000969 |
| 31.2083333 | 0.048072 | 0.01949 | 0.058401 | 0.007973 | 0.04452 | 0.051479 | 0.021598 | 0.008807 | 0.018594 | 0.005084 | 0.013851 | 0.001316 | 0.00022 | 0.003108 | 0.003351 | 0.001285 | 0.004677 | 0.002339 | 0.000784 |

| | | | | | | | | | | | | | | | | | | |
|---|---|---|---|---|---|---|---|---|---|---|---|---|---|---|---|---|---|---|
| 31.2222222 | 0.05099 | 0.030221 | 0.061912 | 0.006298 | 0.049246 | 0.06458 | 0.019218 | 0.005288 | 0.028083 | 0.004632 | 0.007843 | 0.000648 | 0.000476 | 0.002697 | 0.003152 | 0.000906 | 0.005545 | 0.001441 | 0.000802 |
| 31.2361111 | 0.052116 | 0.034765 | 0.05702 | 0.01098 | 0.047386 | 0.073109 | 0.018514 | 0.008241 | 0.033153 | 0.004962 | 0.004297 | 0.000863 | 0.00071 | 0.00261 | 0.002472 | 0.00064 | 0.005986 | 0.000575 | 0.000951 |
| 31.25 | 0.045417 | 0.031553 | 0.045607 | 0.015172 | 0.04227 | 0.073848 | 0.019821 | 0.012769 | 0.033233 | 0.004562 | 0.004444 | 0.00153 | 0.000886 | 0.002722 | 0.002611 | 0.000748 | 0.006433 | 0.000973 | 0.001749 |
| 31.2638889 | 0.032833 | 0.026351 | 0.032275 | 0.015254 | 0.039689 | 0.064075 | 0.023158 | 0.01576 | 0.032089 | 0.002807 | 0.004399 | 0.001461 | 0.000957 | 0.00292 | 0.003468 | 0.001156 | 0.006917 | 0.002403 | 0.002455 |
| 31.2777778 | 0.02143 | 0.024882 | 0.021852 | 0.012294 | 0.040255 | 0.050127 | 0.025549 | 0.01599 | 0.031312 | 0.001791 | 0.005881 | 0.001245 | 0.000666 | 0.003346 | 0.003707 | 0.0014 | 0.006121 | 0.003679 | 0.002338 |
| 31.2916667 | 0.011262 | 0.025908 | 0.014882 | 0.008892 | 0.039867 | 0.036718 | 0.023636 | 0.012644 | 0.028668 | 0.001868 | 0.008275 | 0.00211 | 0.000258 | 0.003825 | 0.002978 | 0.001174 | 0.004115 | 0.005139 | 0.001443 |
| 31.3055556 | 0.004749 | 0.026965 | 0.012192 | 0.006576 | 0.034798 | 0.023167 | 0.017945 | 0.007417 | 0.024336 | 0.001847 | 0.007474 | 0.00242 | 0.000224 | 0.003631 | 0.002051 | 0.000687 | 0.002297 | 0.007144 | 0.000614 |
| 31.3194444 | 0.004809 | 0.027909 | 0.014576 | 0.006372 | 0.024735 | 0.012021 | 0.01176 | 0.003043 | 0.021122 | 0.003783 | 0.010652 | 0.002095 | 0.0004 | 0.002633 | 0.00123 | 0.000923 | 0.002357 | 0.00739 | 0.000543 |
| 31.3333333 | 0.004999 | 0.028786 | 0.018964 | 0.00692 | 0.013124 | 0.006097 | 0.006465 | 0.001641 | 0.019712 | 0.007656 | 0.02038 | 0.002938 | 0.000495 | 0.001568 | 0.00095 | 0.002394 | 0.005298 | 0.004802 | 0.000726 |
| 31.3472222 | 0.006792 | 0.030126 | 0.022175 | 0.005282 | 0.00586 | 0.006057 | 0.003191 | 0.003552 | 0.019074 | 0.007508 | 0.020998 | 0.003394 | 0.000709 | 0.000978 | 0.001255 | 0.003328 | 0.007104 | 0.002146 | 0.001237 |
| 31.3611111 | 0.020295 | 0.03266 | 0.024712 | 0.002535 | 0.006015 | 0.012885 | 0.002356 | 0.005163 | 0.018584 | 0.005002 | 0.013069 | 0.002131 | 0.001276 | 0.000895 | 0.001223 | 0.002355 | 0.004653 | 0.001607 | 0.002083 |
| 31.375 | 0.039333 | 0.034797 | 0.0268 | 0.001524 | 0.013887 | 0.024676 | 0.002965 | 0.003968 | 0.017735 | 0.00595 | 0.01283 | 0.000834 | 0.001865 | 0.001091 | 0.001087 | 0.000837 | 0.001849 | 0.002774 | 0.001983 |
| 31.3888889 | 0.044805 | 0.03121 | 0.025654 | 0.001622 | 0.025229 | 0.033632 | 0.005482 | 0.003131 | 0.015893 | 0.005626 | 0.014906 | 0.000549 | 0.001633 | 0.001076 | 0.00128 | 0.000151 | 0.001722 | 0.004121 | 0.001197 |
| 31.4027778 | 0.029061 | 0.020095 | 0.019679 | 0.001529 | 0.027981 | 0.032145 | 0.007698 | 0.004326 | 0.012853 | 0.002355 | 0.009627 | 0.000831 | 0.000996 | 0.000785 | 0.001276 | 9.87E-05 | 0.00199 | 0.004103 | 0.001177 |
| 31.4166667 | 0.017969 | 0.010377 | 0.010461 | 0.002127 | 0.017645 | 0.020682 | 0.007565 | 0.005558 | 0.008873 | 0.00128 | 0.004383 | 0.000755 | 0.00109 | 0.001143 | 0.001315 | 0.000367 | 0.002109 | 0.00265 | 0.00154 |
| 31.4305556 | 0.031446 | 0.009461 | 0.00392 | 0.00413 | 0.009501 | 0.008886 | 0.005927 | 0.006985 | 0.004762 | 0.002483 | 0.00555 | 0.000277 | 0.001618 | 0.002522 | 0.001751 | 0.000825 | 0.003883 | 0.001546 | 0.001833 |
| 31.4444444 | 0.037253 | 0.009076 | 0.003326 | 0.006127 | 0.01593 | 0.003615 | 0.003383 | 0.009619 | 0.002634 | 0.003543 | 0.006462 | 0.000171 | 0.00168 | 0.003768 | 0.002035 | 0.001066 | 0.004905 | 0.002188 | 0.002212 |
| 31.4583333 | 0.026739 | 0.007382 | 0.006214 | 0.006347 | 0.022903 | 0.004009 | 0.001376 | 0.012667 | 0.00422 | 0.003624 | 0.005084 | 0.000282 | 0.001318 | 0.004036 | 0.001884 | 0.001124 | 0.003934 | 0.003053 | 0.002615 |
| 31.4722222 | 0.034728 | 0.014383 | 0.012535 | 0.005359 | 0.016574 | 0.009827 | 0.002165 | 0.015594 | 0.008738 | 0.002895 | 0.008076 | 0.000223 | 0.000987 | 0.003591 | 0.001697 | 0.001447 | 0.002877 | 0.002455 | 0.003214 |
| 31.4861111 | 0.053319 | 0.022591 | 0.020207 | 0.005025 | 0.008823 | 0.017396 | 0.007002 | 0.018158 | 0.014157 | 0.001823 | 0.012122 | 0.000178 | 0.000659 | 0.00255 | 0.001834 | 0.00181 | 0.001828 | 0.002223 | 0.003787 |
| 31.5 | 0.048666 | 0.020879 | 0.025535 | 0.005705 | 0.010693 | 0.020087 | 0.013029 | 0.018768 | 0.018832 | 0.000854 | 0.009967 | 0.000195 | 0.000384 | 0.001265 | 0.002374 | 0.001752 | 0.000737 | 0.004305 | 0.003936 |
| 31.5138889 | 0.026904 | 0.012662 | 0.027276 | 0.00681 | 0.011754 | 0.016286 | 0.015035 | 0.016321 | 0.021795 | 0.001611 | 0.004614 | 0.001095 | 0.000486 | 0.001235 | 0.003189 | 0.001673 | 0.00109 | 0.007914 | 0.003664 |
| 31.5277778 | 0.015233 | 0.004947 | 0.026128 | 0.010228 | 0.009301 | 0.008704 | 0.011313 | 0.011838 | 0.023278 | 0.005289 | 0.001554 | 0.003594 | 0.000985 | 0.003018 | 0.003795 | 0.001901 | 0.003654 | 0.011457 | 0.003048 |
| 31.5416667 | 0.013164 | 0.00321 | 0.02463 | 0.021762 | 0.014899 | 0.002844 | 0.006309 | 0.006869 | 0.024763 | 0.009716 | 0.003097 | 0.006113 | 0.001629 | 0.005108 | 0.003859 | 0.002048 | 0.006805 | 0.013617 | 0.002127 |
| 31.5555556 | 0.01552 | 0.009369 | 0.024264 | 0.039985 | 0.022897 | 0.002098 | 0.004976 | 0.003514 | 0.027487 | 0.012841 | 0.010887 | 0.0073 | 0.002269 | 0.006392 | 0.003502 | 0.002202 | 0.008702 | 0.014629 | 0.001323 |
| 31.5694444 | 0.027773 | 0.020465 | 0.026567 | 0.055095 | 0.019827 | 0.006475 | 0.004825 | 0.005321 | 0.03317 | 0.015823 | 0.021628 | 0.007423 | 0.002876 | 0.00685 | 0.002481 | 0.002886 | 0.008887 | 0.016966 | 0.002066 |
| 31.5833333 | 0.039398 | 0.034898 | 0.035777 | 0.074596 | 0.012654 | 0.017572 | 0.006277 | 0.015767 | 0.048558 | 0.023811 | 0.03058 | 0.007456 | 0.003673 | 0.007124 | 0.001534 | 0.004506 | 0.008728 | 0.025064 | 0.005928 |
| 31.5972222 | 0.044536 | 0.049297 | 0.055117 | 0.098434 | 0.015355 | 0.035285 | 0.011908 | 0.032317 | 0.073166 | 0.038046 | 0.036299 | 0.009175 | 0.005451 | 0.010505 | 0.001736 | 0.007536 | 0.01129 | 0.038863 | 0.012888 |
| 31.6111111 | 0.050848 | 0.058208 | 0.07294 | 0.097874 | 0.02933 | 0.053644 | 0.016735 | 0.042797 | 0.090911 | 0.04892 | 0.0393 | 0.013138 | 0.008937 | 0.019541 | 0.001757 | 0.012033 | 0.019619 | 0.048404 | 0.022664 |
| 31.625 | 0.056684 | 0.065711 | 0.076315 | 0.072984 | 0.047036 | 0.067233 | 0.019747 | 0.040787 | 0.096349 | 0.049847 | 0.040146 | 0.016555 | 0.01219 | 0.028324 | 0.00114 | 0.016974 | 0.029553 | 0.047587 | 0.034341 |
| 31.6388889 | 0.050344 | 0.069091 | 0.066383 | 0.048268 | 0.054259 | 0.068497 | 0.021318 | 0.03286 | 0.09413 | 0.044422 | 0.035232 | 0.015678 | 0.012294 | 0.028284 | 0.000992 | 0.019292 | 0.029889 | 0.04042 | 0.043022 |
| 31.6527778 | 0.028921 | 0.056045 | 0.044817 | 0.02716 | 0.043433 | 0.048311 | 0.018056 | 0.022954 | 0.078486 | 0.037236 | 0.021059 | 0.010878 | 0.01014 | 0.019744 | 0.000971 | 0.016875 | 0.018131 | 0.031542 | 0.044407 |
| 31.6666667 | 0.01244 | 0.031463 | 0.022541 | 0.014211 | 0.022784 | 0.024321 | 0.01175 | 0.012246 | 0.049829 | 0.030152 | 0.010481 | 0.005718 | 0.007906 | 0.009436 | 0.000667 | 0.011733 | 0.006989 | 0.025074 | 0.042051 |
| 31.6805556 | 0.016039 | 0.013032 | 0.019361 | 0.019688 | 0.011089 | 0.023824 | 0.007333 | 0.0052 | 0.023564 | 0.023541 | 0.016033 | 0.002168 | 0.00653 | 0.003101 | 0.000845 | 0.006764 | 0.005941 | 0.022102 | 0.042117 |
| 31.6944444 | 0.025213 | 0.009714 | 0.035869 | 0.033912 | 0.018325 | 0.041341 | 0.004517 | 0.004726 | 0.014701 | 0.016772 | 0.025093 | 0.000669 | 0.007719 | 0.001663 | 0.002342 | 0.003579 | 0.006947 | 0.019143 | 0.043387 |
| 31.7083333 | 0.022734 | 0.016504 | 0.047797 | 0.033665 | 0.029243 | 0.057828 | 0.002856 | 0.009186 | 0.020964 | 0.01017 | 0.023108 | 0.000624 | 0.012034 | 0.001827 | 0.00455 | 0.002156 | 0.005305 | 0.013781 | 0.040319 |
| 31.7222222 | 0.015526 | 0.027813 | 0.038332 | 0.032236 | 0.030808 | 0.061969 | 0.005283 | 0.014075 | 0.029057 | 0.00528 | 0.013189 | 0.000617 | 0.014044 | 0.002995 | 0.005629 | 0.001268 | 0.006892 | 0.007212 | 0.031046 |
| 31.7361111 | 0.021898 | 0.045538 | 0.018381 | 0.064971 | 0.023585 | 0.046844 | 0.009685 | 0.015723 | 0.038095 | 0.002264 | 0.008222 | 0.000493 | 0.009691 | 0.006558 | 0.004158 | 0.00063 | 0.009982 | 0.00303 | 0.017317 |
| 31.75 | 0.036203 | 0.064952 | 0.00845 | 0.10204 | 0.01375 | 0.027682 | 0.010953 | 0.013216 | 0.046471 | 0.000986 | 0.008195 | 0.000664 | 0.004028 | 0.009903 | 0.002342 | 0.000428 | 0.009356 | 0.003181 | 0.008271 |
| 31.7638889 | 0.038207 | 0.075361 | 0.012863 | 0.097888 | 0.012079 | 0.0313 | 0.00779 | 0.010087 | 0.039721 | 0.001633 | 0.008797 | 0.001187 | 0.001888 | 0.010563 | 0.003085 | 0.000529 | 0.005601 | 0.005342 | 0.01149 |
| 31.7777778 | 0.028234 | 0.059944 | 0.02033 | 0.062802 | 0.015589 | 0.050639 | 0.007097 | 0.010561 | 0.027129 | 0.003327 | 0.017929 | 0.002594 | 0.003119 | 0.00978 | 0.004089 | 0.000704 | 0.004159 | 0.005448 | 0.018908 |
| 31.7916667 | 0.01745 | 0.042831 | 0.028034 | 0.031946 | 0.012963 | 0.060199 | 0.013071 | 0.010916 | 0.039552 | 0.00528 | 0.03027 | 0.004979 | 0.006336 | 0.008826 | 0.003068 | 0.000814 | 0.007318 | 0.004237 | 0.022765 |
| 31.8055556 | 0.009775 | 0.073273 | 0.032143 | 0.032644 | 0.012438 | 0.052275 | 0.015278 | 0.006453 | 0.065249 | 0.007072 | 0.031379 | 0.006428 | 0.009644 | 0.007189 | 0.001467 | 0.001747 | 0.009151 | 0.006027 | 0.023321 |
| 31.8194444 | 0.010109 | 0.111307 | 0.025883 | 0.062319 | 0.024408 | 0.032928 | 0.01225 | 0.003358 | 0.069009 | 0.008043 | 0.023149 | 0.004776 | 0.011254 | 0.004232 | 0.001032 | 0.003428 | 0.006666 | 0.009419 | 0.020364 |
| 31.8333333 | 0.015111 | 0.106282 | 0.023656 | 0.09407 | 0.037056 | 0.022562 | 0.017812 | 0.00607 | 0.051086 | 0.007749 | 0.01779 | 0.002681 | 0.010872 | 0.001986 | 0.001879 | 0.004137 | 0.003542 | 0.010144 | 0.01378 |
| 31.8472222 | 0.011401 | 0.07746 | 0.034318 | 0.096343 | 0.046256 | 0.032536 | 0.033225 | 0.010688 | 0.028491 | 0.006009 | 0.020821 | 0.003387 | 0.009068 | 0.002753 | 0.002994 | 0.003195 | 0.002977 | 0.008217 | 0.007057 |
| 31.8611111 | 0.005987 | 0.044194 | 0.043207 | 0.06092 | 0.051549 | 0.046285 | 0.046872 | 0.017271 | 0.021605 | 0.003835 | 0.027346 | 0.003732 | 0.005737 | 0.004649 | 0.003706 | 0.002083 | 0.004313 | 0.006468 | 0.005743 |
| 31.875 | 0.005768 | 0.02455 | 0.046159 | 0.039996 | 0.040318 | 0.052227 | 0.052208 | 0.026981 | 0.040166 | 0.004644 | 0.03008 | 0.0026 | 0.005218 | 0.004869 | 0.003836 | 0.002974 | 0.005626 | 0.007731 | 0.007905 |
| 31.8888889 | 0.006964 | 0.040378 | 0.050357 | 0.073272 | 0.019559 | 0.04615 | 0.04892 | 0.036587 | 0.065744 | 0.010154 | 0.026523 | 0.003213 | 0.01238 | 0.00307 | 0.003364 | 0.005439 | 0.008061 | 0.012307 | 0.006358 |
| 31.9027778 | 0.015842 | 0.080856 | 0.056102 | 0.104088 | 0.006543 | 0.034721 | 0.038386 | 0.04122 | 0.080515 | 0.017662 | 0.019197 | 0.004682 | 0.02179 | 0.001064 | 0.002206 | 0.007392 | 0.01304 | 0.01817 | 0.004988 |
| 31.9166667 | 0.032271 | 0.113434 | 0.063905 | 0.077542 | 0.007308 | 0.039775 | 0.02443 | 0.039006 | 0.080047 | 0.023338 | 0.013545 | 0.004354 | 0.026477 | 0.001059 | 0.001447 | 0.008612 | 0.018943 | 0.02286 | 0.011687 |
| 31.9305556 | 0.047798 | 0.122615 | 0.0761 | 0.041786 | 0.021696 | 0.059488 | 0.011951 | 0.033959 | 0.068801 | 0.025316 | 0.015024 | 0.002737 | 0.025865 | 0.003545 | 0.002173 | 0.009723 | 0.02284 | 0.024067 | 0.020777 |
| 31.9444444 | 0.052158 | 0.115527 | 0.090183 | 0.058262 | 0.042621 | 0.081004 | 0.004117 | 0.03023 | 0.05833 | 0.02489 | 0.023208 | 0.002114 | 0.021729 | 0.008224 | 0.002931 | 0.009532 | 0.023686 | 0.021402 | 0.02487 |
| 31.9583333 | 0.042791 | 0.096753 | 0.095142 | 0.105623 | 0.060113 | 0.106257 | 0.001506 | 0.029933 | 0.054413 | 0.023671 | 0.030004 | 0.003094 | 0.015837 | 0.014289 | 0.002775 | 0.007243 | 0.02308 | 0.017237 | 0.02535 |
| 31.9722222 | 0.030334 | 0.070798 | 0.081833 | 0.136975 | 0.065886 | 0.119184 | 0.001919 | 0.033953 | 0.051953 | 0.021347 | 0.026787 | 0.004653 | 0.010089 | 0.019571 | 0.001975 | 0.004869 | 0.022268 | 0.013675 | 0.024313 |
| 31.9861111 | 0.01979 | 0.048038 | 0.055945 | 0.135676 | 0.057717 | 0.098698 | 0.00247 | 0.038877 | 0.04516 | 0.017551 | 0.014925 | 0.00633 | 0.006319 | 0.022014 | 0.001092 | 0.003863 | 0.020794 | 0.010412 | 0.021619 |
| 32 | 0.011453 | 0.031667 | 0.028816 | 0.106374 | 0.039705 | 0.058289 | 0.002144 | 0.039739 | 0.034045 | 0.01399 | 0.006776 | 0.007487 | 0.005399 | 0.021211 | 0.001474 | 0.003551 | 0.017531 | 0.007872 | 0.017747 |
| 32.0138889 | 0.007466 | 0.020649 | 0.009924 | 0.063737 | 0.021211 | 0.027067 | 0.001489 | 0.035065 | 0.019335 | 0.012681 | 0.006297 | 0.007362 | 0.007943 | 0.01803 | 0.00323 | 0.003391 | 0.012577 | 0.007713 | 0.01398 |
| 32.0277778 | 0.006588 | 0.014007 | 0.002953 | 0.029162 | 0.01775 | 0.020844 | 0.000865 | 0.026925 | 0.009466 | 0.013017 | 0.006219 | 0.005767 | 0.011927 | 0.013195 | 0.005122 | 0.003235 | 0.007936 | 0.009522 | 0.010993 |
| 32.0416667 | 0.006773 | 0.009177 | 0.002539 | 0.012074 | 0.03017 | 0.027422 | 0.000694 | 0.018112 | 0.013122 | 0.013772 | 0.007908 | 0.003271 | 0.011416 | 0.007906 | 0.00617 | 0.00249 | 0.005382 | 0.01086 | 0.008467 |
| 32.0555556 | 0.006938 | 0.005212 | 0.005227 | 0.005896 | 0.041068 | 0.028425 | 0.001991 | 0.010209 | 0.020763 | 0.014204 | 0.011532 | 0.001788 | 0.006018 | 0.004708 | 0.006203 | 0.001258 | 0.004863 | 0.010921 | 0.006569 |
| 32.0694444 | 0.00744 | 0.004935 | 0.009694 | 0.004832 | 0.040118 | 0.023894 | 0.004115 | 0.004721 | 0.020381 | 0.014566 | 0.011522 | 0.001451 | 0.002224 | 0.005274 | 0.005812 | 0.000902 | 0.005056 | 0.013994 | 0.007808 |
| 32.0833333 | 0.012183 | 0.006244 | 0.01256 | 0.00798 | 0.026947 | 0.019489 | 0.00421 | 0.003141 | 0.015653 | 0.019604 | 0.016684 | 0.001658 | 0.002407 | 0.00811 | 0.006039 | 0.003248 | 0.006539 | 0.026822 | 0.01594 |
| 32.0972222 | 0.026464 | 0.008621 | 0.025555 | 0.016642 | 0.022083 | 0.031658 | 0.003966 | 0.00598 | 0.023653 | 0.030302 | 0.037995 | 0.003724 | 0.004456 | 0.010581 | 0.007567 | 0.008401 | 0.011422 | 0.045983 | 0.028715 |
| 32.1111111 | 0.047279 | 0.019263 | 0.057455 | 0.026516 | 0.042719 | 0.069104 | 0.009807 | 0.011371 | 0.047188 | 0.034916 | 0.060852 | 0.006043 | 0.006009 | 0.012387 | 0.009645 | 0.012587 | 0.017933 | 0.052902 | 0.03588 |
| 32.125 | 0.064508 | 0.034698 | 0.084558 | 0.027343 | 0.060985 | 0.10552 | 0.017447 | 0.015334 | 0.065709 | 0.025925 | 0.061923 | 0.006386 | 0.005723 | 0.013638 | 0.010051 | 0.012366 | 0.021825 | 0.039613 | 0.03252 |
| 32.1388889 | 0.074918 | 0.041599 | 0.087792 | 0.016691 | 0.053854 | 0.117041 | 0.015785 | 0.015049 | 0.068641 | 0.01241 | 0.041527 | 0.005099 | 0.004051 | 0.012924 | 0.009075 | 0.008681 | 0.021342 | 0.01979 | 0.022432 |
| 32.1527778 | 0.072839 | 0.033125 | 0.071922 | 0.006047 | 0.03178 | 0.103541 | 0.008633 | 0.011464 | 0.061191 | 0.006581 | 0.019916 | 0.0028 | 0.002019 | 0.010058 | 0.006843 | 0.004564 | 0.016236 | 0.007928 | 0.010521 |
| 32.1666667 | 0.050451 | 0.016788 | 0.043693 | 0.00213 | 0.012637 | 0.074717 | 0.00686 | 0.008307 | 0.047031 | 0.008491 | 0.01532 | 0.000959 | 0.000656 | 0.00658 | 0.004434 | 0.00182 | 0.008489 | 0.007478 | 0.003572 |
| 32.1805556 | 0.024954 | 0.009028 | 0.024044 | 0.00151 | 0.006953 | 0.041031 | 0.01002 | 0.005994 | 0.027518 | 0.008911 | 0.024342 | 0.000483 | 0.000232 | 0.00342 | 0.002885 | 0.00102 | 0.00491 | 0.010486 | 0.003786 |
| 32.1944444 | 0.02382 | 0.014188 | 0.035993 | 0.001801 | 0.010678 | 0.019153 | 0.013063 | 0.004997 | 0.012065 | 0.007089 | 0.032278 | 0.000526 | 0.00012 | 0.001456 | 0.001579 | 0.00218 | 0.008666 | 0.010541 | 0.005073 |
| 32.2083333 | 0.044765 | 0.022615 | 0.064238 | 0.003344 | 0.017106 | 0.025229 | 0.015394 | 0.009365 | 0.010267 | 0.007363 | 0.035367 | 0.000538 | 0.000135 | 0.001049 | 0.000789 | 0.003892 | 0.013874 | 0.008799 | 0.004593 |
| 32.2222222 | 0.065299 | 0.02903 | 0.08342 | 0.006194 | 0.024493 | 0.050171 | 0.017982 | 0.016894 | 0.016088 | 0.006745 | 0.034964 | 0.000506 | 0.000367 | 0.001227 | 0.000851 | 0.005073 | 0.015682 | 0.006752 | 0.004269 |
| 32.2361111 | 0.072769 | 0.033231 | 0.086787 | 0.007202 | 0.030137 | 0.076479 | 0.022222 | 0.021234 | 0.022031 | 0.003953 | 0.030904 | 0.000537 | 0.000557 | 0.002248 | 0.001266 | 0.006019 | 0.013886 | 0.005169 | 0.004735 |
| 32.25 | 0.067409 | 0.030767 | 0.073801 | 0.005331 | 0.032927 | 0.094348 | 0.027948 | 0.020141 | 0.027703 | 0.00315 | 0.022906 | 0.001228 | 0.000507 | 0.004019 | 0.002128 | 0.006574 | 0.010127 | 0.004713 | 0.00502 |
| 32.2638889 | 0.060231 | 0.023851 | 0.054024 | 0.005508 | 0.036177 | 0.099142 | 0.030611 | 0.015272 | 0.032631 | 0.00452 | 0.01497 | 0.002077 | 0.000346 | 0.00504 | 0.003788 | 0.006577 | 0.008201 | 0.005647 | 0.003705 |
| 32.2777778 | 0.060474 | 0.021069 | 0.036822 | 0.009082 | 0.038156 | 0.095491 | 0.026973 | 0.010076 | 0.036359 | 0.004105 | 0.013524 | 0.001833 | 0.000327 | 0.005395 | 0.005749 | 0.006625 | 0.009962 | 0.006286 | 0.002437 |
| 32.2916667 | 0.059708 | 0.022122 | 0.021369 | 0.0111 | 0.033362 | 0.087352 | 0.019006 | 0.006708 | 0.039295 | 0.00253 | 0.016529 | 0.001574 | 0.000538 | 0.005852 | 0.006689 | 0.006604 | 0.012142 | 0.004318 | 0.004721 |
| 32.3055556 | 0.053862 | 0.02499 | 0.012963 | 0.008305 | 0.023217 | 0.074194 | 0.011475 | 0.00596 | 0.037651 | 0.003199 | 0.018704 | 0.002548 | 0.000743 | 0.005594 | 0.00579 | 0.005531 | 0.011309 | 0.00211 | 0.009478 |
| 32.3194444 | 0.050859 | 0.032178 | 0.017613 | 0.003921 | 0.012617 | 0.056051 | 0.008517 | 0.008112 | 0.030529 | 0.004552 | 0.017835 | 0.002729 | 0.000661 | 0.004269 | 0.003388 | 0.003447 | 0.007069 | 0.002377 | 0.013107 |
| 32.3333333 | 0.046241 | 0.038476 | 0.026149 | 0.002139 | 0.006081 | 0.034093 | 0.00778 | 0.011004 | 0.023685 | 0.003642 | 0.011626 | 0.001938 | 0.000814 | 0.003206 | 0.00138 | 0.002782 | 0.004109 | 0.002644 | 0.013893 |
| 32.3472222 | 0.031274 | 0.034028 | 0.026218 | 0.002299 | 0.007921 | 0.017123 | 0.006631 | 0.011063 | 0.017039 | 0.003555 | 0.00689 | 0.002428 | 0.001665 | 0.002713 | 0.001023 | 0.004174 | 0.006505 | 0.001499 | 0.012076 |
| 32.3611111 | 0.015349 | 0.02 | 0.018502 | 0.004008 | 0.015714 | 0.015869 | 0.008694 | 0.007588 | 0.01128 | 0.007321 | 0.01198 | 0.003045 | 0.002397 | 0.002125 | 0.000999 | 0.003775 | 0.009438 | 0.000485 | 0.00919 |
| 32.375 | 0.012224 | 0.007847 | 0.010622 | 0.006419 | 0.020195 | 0.021615 | 0.012088 | 0.003637 | 0.008758 | 0.010064 | 0.01774 | 0.002365 | 0.002342 | 0.00123 | 0.001116 | 0.0026 | 0.00871 | 0.000375 | 0.00685 |
| 32.3888889 | 0.015475 | 0.002936 | 0.007306 | 0.006604 | 0.014936 | 0.016769 | 0.010098 | 0.002062 | 0.007508 | 0.007532 | 0.013913 | 0.00146 | 0.001643 | 0.000679 | 0.001804 | 0.004657 | 0.005823 | 0.000606 | 0.005495 |
| 32.4027778 | 0.0119 | 0.002532 | 0.010168 | 0.00408 | 0.007236 | 0.010129 | 0.005065 | 0.001765 | 0.007156 | 0.005503 | 0.008102 | 0.000913 | 0.000802 | 0.001058 | 0.001814 | 0.007028 | 0.002929 | 0.001141 | 0.004722 |

| | | | | | | | | | | | | | | | | | | | |
|---|---|---|---|---|---|---|---|---|---|---|---|---|---|---|---|---|---|---|---|
| 32.4166667 | 0.00828 | 0.005996 | 0.016805 | 0.002106 | 0.007038 | 0.017307 | 0.003682 | 0.002604 | 0.008199 | 0.008369 | 0.01044 | 0.000478 | 0.000542 | 0.001579 | 0.001087 | 0.006877 | 0.001221 | 0.00243 | 0.003777 |
| 32.4305556 | 0.015658 | 0.012141 | 0.021894 | 0.003579 | 0.009202 | 0.026134 | 0.005341 | 0.007477 | 0.009475 | 0.009076 | 0.014017 | 0.000269 | 0.000878 | 0.001932 | 0.000923 | 0.005601 | 0.000566 | 0.003391 | 0.002532 |
| 32.4444444 | 0.023943 | 0.015941 | 0.021877 | 0.005763 | 0.006955 | 0.024039 | 0.006062 | 0.014379 | 0.009772 | 0.005287 | 0.009997 | 0.000545 | 0.001084 | 0.002558 | 0.001179 | 0.004608 | 0.000352 | 0.002986 | 0.002469 |
| 32.4583333 | 0.020763 | 0.012887 | 0.0168 | 0.004745 | 0.004513 | 0.014066 | 0.004721 | 0.017062 | 0.008893 | 0.003568 | 0.003778 | 0.000838 | 0.000926 | 0.003379 | 0.000982 | 0.004539 | 0.001162 | 0.00234 | 0.003143 |
| 32.4722222 | 0.010691 | 0.006892 | 0.011026 | 0.002621 | 0.006828 | 0.005678 | 0.002816 | 0.013612 | 0.008069 | 0.00441 | 0.003144 | 0.00141 | 0.00092 | 0.004374 | 0.001117 | 0.005467 | 0.004485 | 0.003329 | 0.002662 |
| 32.4861111 | 0.004058 | 0.004005 | 0.011715 | 0.002866 | 0.009296 | 0.005841 | 0.003262 | 0.009673 | 0.009825 | 0.004109 | 0.007129 | 0.00303 | 0.00152 | 0.005729 | 0.002551 | 0.006763 | 0.009501 | 0.006291 | 0.002487 |
| 32.5 | 0.006286 | 0.004555 | 0.020879 | 0.00563 | 0.007074 | 0.014962 | 0.006689 | 0.010499 | 0.014599 | 0.004886 | 0.010919 | 0.004556 | 0.002255 | 0.007237 | 0.004207 | 0.00724 | 0.012951 | 0.009698 | 0.003956 |
| 32.5138889 | 0.015478 | 0.010068 | 0.030602 | 0.009145 | 0.00526 | 0.031694 | 0.010022 | 0.013382 | 0.018306 | 0.007972 | 0.01389 | 0.005094 | 0.002002 | 0.008101 | 0.004788 | 0.006386 | 0.013478 | 0.011455 | 0.004867 |
| 32.5277778 | 0.02249 | 0.018838 | 0.032263 | 0.009542 | 0.007415 | 0.04635 | 0.010938 | 0.014702 | 0.01745 | 0.008241 | 0.016626 | 0.004711 | 0.001081 | 0.007466 | 0.00423 | 0.004699 | 0.01203 | 0.010634 | 0.00388 |
| 32.5416667 | 0.022983 | 0.026142 | 0.026352 | 0.006704 | 0.008745 | 0.050359 | 0.009968 | 0.015307 | 0.014013 | 0.004923 | 0.016651 | 0.003456 | 0.000628 | 0.005831 | 0.00311 | 0.00288 | 0.009437 | 0.008178 | 0.00283 |
| 32.5555556 | 0.020284 | 0.029101 | 0.01907 | 0.004631 | 0.006238 | 0.043315 | 0.007853 | 0.015178 | 0.011914 | 0.00244 | 0.012737 | 0.00285 | 0.000945 | 0.005271 | 0.002239 | 0.0018 | 0.007241 | 0.006673 | 0.003362 |
| 32.5694444 | 0.017553 | 0.02766 | 0.017666 | 0.009271 | 0.003334 | 0.030025 | 0.004704 | 0.013565 | 0.01472 | 0.003727 | 0.009164 | 0.00399 | 0.002216 | 0.006505 | 0.00174 | 0.002492 | 0.007895 | 0.008988 | 0.005671 |
| 32.5833333 | 0.01857 | 0.028611 | 0.03089 | 0.040047 | 0.002908 | 0.023322 | 0.003101 | 0.014232 | 0.02694 | 0.010696 | 0.012089 | 0.005649 | 0.003642 | 0.008631 | 0.001288 | 0.005348 | 0.012492 | 0.016393 | 0.010219 |
| 32.5972222 | 0.025091 | 0.037685 | 0.059276 | 0.097898 | 0.005088 | 0.035612 | 0.006394 | 0.021994 | 0.046962 | 0.021713 | 0.023951 | 0.006994 | 0.003935 | 0.012808 | 0.001636 | 0.009322 | 0.02123 | 0.027951 | 0.016668 |
| 32.6111111 | 0.038536 | 0.053686 | 0.083932 | 0.145729 | 0.016292 | 0.061389 | 0.010075 | 0.030686 | 0.061139 | 0.029965 | 0.043076 | 0.009071 | 0.003675 | 0.023417 | 0.002549 | 0.012931 | 0.034871 | 0.038982 | 0.022174 |
| 32.625 | 0.057333 | 0.075323 | 0.090317 | 0.156304 | 0.039741 | 0.081099 | 0.010451 | 0.030892 | 0.060003 | 0.033243 | 0.060165 | 0.01205 | 0.004309 | 0.038055 | 0.003696 | 0.015175 | 0.045535 | 0.043903 | 0.022993 |
| 32.6388889 | 0.063078 | 0.090179 | 0.081795 | 0.127815 | 0.059839 | 0.082608 | 0.013844 | 0.025608 | 0.051357 | 0.035276 | 0.06214 | 0.013195 | 0.005272 | 0.043294 | 0.005229 | 0.014748 | 0.040579 | 0.041695 | 0.018204 |
| 32.6527778 | 0.043516 | 0.081356 | 0.060253 | 0.082765 | 0.058867 | 0.06374 | 0.019235 | 0.019898 | 0.045427 | 0.037276 | 0.044003 | 0.010541 | 0.005948 | 0.034276 | 0.005024 | 0.011345 | 0.02267 | 0.033864 | 0.011312 |
| 32.6666667 | 0.018084 | 0.057813 | 0.031548 | 0.046734 | 0.038389 | 0.032705 | 0.019129 | 0.01509 | 0.043389 | 0.037418 | 0.023832 | 0.005978 | 0.007807 | 0.020679 | 0.002896 | 0.006547 | 0.010773 | 0.023281 | 0.00747 |
| 32.6805556 | 0.007164 | 0.036265 | 0.011812 | 0.02277 | 0.017254 | 0.014788 | 0.015388 | 0.012535 | 0.042416 | 0.03347 | 0.027217 | 0.002321 | 0.010849 | 0.010361 | 0.00132 | 0.003092 | 0.016538 | 0.014577 | 0.009347 |
| 32.6944444 | 0.008729 | 0.019947 | 0.008566 | 0.017708 | 0.011401 | 0.024641 | 0.011639 | 0.010761 | 0.038904 | 0.026426 | 0.036641 | 0.001124 | 0.012875 | 0.004286 | 0.001642 | 0.002986 | 0.027538 | 0.010007 | 0.015806 |
| 32.7083333 | 0.012732 | 0.009445 | 0.013263 | 0.035038 | 0.017584 | 0.046483 | 0.00688 | 0.007741 | 0.031379 | 0.020113 | 0.028501 | 0.002291 | 0.012609 | 0.001489 | 0.003299 | 0.004295 | 0.028119 | 0.008252 | 0.021509 |
| 32.7222222 | 0.013506 | 0.010721 | 0.016855 | 0.06278 | 0.021564 | 0.062779 | 0.00414 | 0.006709 | 0.020714 | 0.017248 | 0.021345 | 0.003871 | 0.010237 | 0.000681 | 0.005058 | 0.003514 | 0.0174 | 0.007085 | 0.021274 |
| 32.7361111 | 0.009292 | 0.026545 | 0.018376 | 0.085412 | 0.018851 | 0.06492 | 0.006773 | 0.010737 | 0.012098 | 0.017722 | 0.019395 | 0.004137 | 0.006529 | 0.0005 | 0.006263 | 0.001743 | 0.006507 | 0.005123 | 0.01677 |
| 32.75 | 0.007951 | 0.046545 | 0.017386 | 0.072175 | 0.014451 | 0.051765 | 0.009517 | 0.016477 | 0.014166 | 0.017541 | 0.014588 | 0.003229 | 0.002954 | 0.00092 | 0.006394 | 0.001986 | 0.003715 | 0.002647 | 0.011859 |
| 32.7638889 | 0.01854 | 0.061958 | 0.012378 | 0.04481 | 0.010071 | 0.029369 | 0.009395 | 0.02041 | 0.022853 | 0.013651 | 0.016892 | 0.002298 | 0.001471 | 0.00243 | 0.005139 | 0.003525 | 0.00454 | 0.001092 | 0.006828 |
| 32.7777778 | 0.027047 | 0.076713 | 0.006676 | 0.069156 | 0.008784 | 0.009734 | 0.009927 | 0.020656 | 0.032232 | 0.008357 | 0.017246 | 0.002339 | 0.002605 | 0.004788 | 0.003055 | 0.005518 | 0.006181 | 0.000823 | 0.004001 |
| 32.7916667 | 0.019234 | 0.083025 | 0.004878 | 0.11943 | 0.01501 | 0.006614 | 0.012416 | 0.016873 | 0.041391 | 0.004465 | 0.009325 | 0.003759 | 0.004628 | 0.006705 | 0.002027 | 0.00813 | 0.009385 | 0.002074 | 0.006252 |
| 32.8055556 | 0.007872 | 0.062451 | 0.006255 | 0.124932 | 0.022217 | 0.020045 | 0.014779 | 0.012558 | 0.036673 | 0.002582 | 0.006611 | 0.00584 | 0.005824 | 0.006625 | 0.002771 | 0.010142 | 0.010041 | 0.004726 | 0.009634 |
| 32.8194444 | 0.007902 | 0.034649 | 0.009213 | 0.085083 | 0.020903 | 0.036711 | 0.015468 | 0.009367 | 0.021199 | 0.00348 | 0.010644 | 0.006893 | 0.005342 | 0.00505 | 0.003328 | 0.009417 | 0.007837 | 0.004968 | 0.010313 |
| 32.8333333 | 0.011999 | 0.040516 | 0.013373 | 0.039537 | 0.013002 | 0.043391 | 0.012577 | 0.00597 | 0.018906 | 0.006432 | 0.018888 | 0.005749 | 0.003276 | 0.003154 | 0.003129 | 0.005903 | 0.010509 | 0.004171 | 0.008961 |
| 32.8472222 | 0.009309 | 0.075067 | 0.013976 | 0.02189 | 0.005278 | 0.033744 | 0.008288 | 0.002831 | 0.027273 | 0.00887 | 0.032134 | 0.0033 | 0.001236 | 0.002466 | 0.002979 | 0.004202 | 0.018394 | 0.007889 | 0.006114 |
| 32.8611111 | 0.006314 | 0.095387 | 0.011591 | 0.043223 | 0.00349 | 0.029528 | 0.009153 | 0.002502 | 0.025579 | 0.008445 | 0.046513 | 0.001364 | 0.000408 | 0.005091 | 0.002233 | 0.006208 | 0.023307 | 0.012618 | 0.003465 |
| 32.875 | 0.008118 | 0.084139 | 0.017559 | 0.084331 | 0.009801 | 0.052177 | 0.011287 | 0.006987 | 0.014723 | 0.005387 | 0.05668 | 0.000504 | 0.000493 | 0.009721 | 0.001573 | 0.008077 | 0.022473 | 0.013995 | 0.004143 |
| 32.8888889 | 0.010323 | 0.053944 | 0.031965 | 0.129019 | 0.018429 | 0.07016 | 0.008087 | 0.014467 | 0.011898 | 0.0034 | 0.059872 | 0.0005 | 0.000858 | 0.013542 | 0.002518 | 0.008662 | 0.019036 | 0.014235 | 0.00704 |
| 32.9027778 | 0.010413 | 0.030575 | 0.04539 | 0.153309 | 0.022816 | 0.06469 | 0.006524 | 0.020853 | 0.018264 | 0.002962 | 0.054799 | 0.00096 | 0.000831 | 0.015268 | 0.004222 | 0.010749 | 0.016882 | 0.015666 | 0.008074 |
| 32.9166667 | 0.009088 | 0.041033 | 0.060724 | 0.125414 | 0.022294 | 0.055717 | 0.012612 | 0.023605 | 0.023319 | 0.003746 | 0.043232 | 0.001352 | 0.001017 | 0.013604 | 0.005825 | 0.015527 | 0.018291 | 0.017398 | 0.005649 |
| 32.9305556 | 0.00925 | 0.071356 | 0.079923 | 0.067534 | 0.021032 | 0.059843 | 0.018797 | 0.023686 | 0.025332 | 0.007763 | 0.032698 | 0.001568 | 0.002488 | 0.008959 | 0.006704 | 0.02011 | 0.022394 | 0.017606 | 0.00408 |
| 32.9444444 | 0.015489 | 0.092122 | 0.092006 | 0.029659 | 0.023701 | 0.084163 | 0.01993 | 0.023286 | 0.028971 | 0.012588 | 0.028702 | 0.001797 | 0.003393 | 0.004756 | 0.00573 | 0.020505 | 0.025554 | 0.016697 | 0.008808 |
| 32.9583333 | 0.029592 | 0.098743 | 0.091704 | 0.019409 | 0.030206 | 0.113554 | 0.018976 | 0.022582 | 0.036679 | 0.015939 | 0.025976 | 0.001825 | 0.00223 | 0.003801 | 0.003369 | 0.016481 | 0.024642 | 0.01564 | 0.016709 |
| 32.9722222 | 0.044984 | 0.093634 | 0.082634 | 0.019608 | 0.037144 | 0.123657 | 0.01696 | 0.022119 | 0.046074 | 0.01825 | 0.01654 | 0.001793 | 0.001058 | 0.005704 | 0.002146 | 0.011817 | 0.01997 | 0.0139 | 0.021219 |
| 32.9861111 | 0.051178 | 0.078421 | 0.066487 | 0.022188 | 0.039988 | 0.108887 | 0.011812 | 0.02361 | 0.053146 | 0.019262 | 0.007703 | 0.002468 | 0.001166 | 0.008425 | 0.003356 | 0.009545 | 0.01464 | 0.011119 | 0.02031 |
| 33 | 0.040909 | 0.056858 | 0.045504 | 0.025678 | 0.03333 | 0.079637 | 0.006316 | 0.026551 | 0.053974 | 0.018948 | 0.00865 | 0.003492 | 0.001291 | 0.010591 | 0.004611 | 0.009225 | 0.012121 | 0.008237 | 0.016181 |
| 33.0138889 | 0.022491 | 0.034146 | 0.024316 | 0.028737 | 0.018673 | 0.045591 | 0.006787 | 0.027406 | 0.046413 | 0.017208 | 0.010455 | 0.003685 | 0.000819 | 0.012105 | 0.004142 | 0.008962 | 0.013393 | 0.007101 | 0.011667 |
| 33.0277778 | 0.016658 | 0.015623 | 0.010226 | 0.027137 | 0.0091 | 0.020893 | 0.011652 | 0.022684 | 0.032464 | 0.014047 | 0.006962 | 0.002701 | 0.00054 | 0.012921 | 0.002927 | 0.007573 | 0.015928 | 0.008572 | 0.007448 |
| 33.0416667 | 0.025563 | 0.006551 | 0.008116 | 0.019524 | 0.011302 | 0.019107 | 0.016265 | 0.013081 | 0.016492 | 0.010269 | 0.005993 | 0.001389 | 0.001073 | 0.012093 | 0.002043 | 0.005087 | 0.016134 | 0.010732 | 0.003678 |
| 33.0555556 | 0.033407 | 0.008013 | 0.010323 | 0.009819 | 0.016052 | 0.027511 | 0.019638 | 0.005422 | 0.007753 | 0.007667 | 0.013815 | 0.000702 | 0.001582 | 0.009597 | 0.001308 | 0.002621 | 0.012899 | 0.010759 | 0.001466 |
| 33.0694444 | 0.031707 | 0.011014 | 0.008062 | 0.003204 | 0.015173 | 0.022524 | 0.020591 | 0.003984 | 0.010425 | 0.009618 | 0.028044 | 0.000714 | 0.001557 | 0.006971 | 0.00076 | 0.001979 | 0.009273 | 0.010306 | 0.001847 |
| 33.0833333 | 0.022653 | 0.007889 | 0.010147 | 0.00394 | 0.012402 | 0.016645 | 0.017173 | 0.004181 | 0.011045 | 0.020322 | 0.047472 | 0.001509 | 0.002639 | 0.005393 | 0.000886 | 0.004977 | 0.010463 | 0.018294 | 0.006479 |
| 33.0972222 | 0.024626 | 0.006176 | 0.029978 | 0.011988 | 0.017417 | 0.032558 | 0.01033 | 0.005396 | 0.010899 | 0.036949 | 0.071327 | 0.002891 | 0.005134 | 0.005535 | 0.001256 | 0.010956 | 0.01751 | 0.037525 | 0.014521 |
| 33.1111111 | 0.048533 | 0.01485 | 0.060443 | 0.017208 | 0.024361 | 0.051276 | 0.005977 | 0.008293 | 0.023311 | 0.044439 | 0.08235 | 0.003843 | 0.006673 | 0.007886 | 0.002357 | 0.014843 | 0.023849 | 0.050898 | 0.018013 |
| 33.125 | 0.072264 | 0.026491 | 0.081116 | 0.011703 | 0.021516 | 0.05415 | 0.004658 | 0.007401 | 0.033968 | 0.035821 | 0.067633 | 0.004323 | 0.006359 | 0.010798 | 0.004569 | 0.013084 | 0.023458 | 0.044583 | 0.012777 |
| 33.1388889 | 0.077211 | 0.030121 | 0.085198 | 0.004646 | 0.014909 | 0.047056 | 0.004075 | 0.004311 | 0.029766 | 0.024121 | 0.04416 | 0.004805 | 0.005359 | 0.010488 | 0.005835 | 0.008465 | 0.017771 | 0.028238 | 0.005367 |
| 33.1527778 | 0.06187 | 0.022777 | 0.077956 | 0.003022 | 0.01071 | 0.034633 | 0.007385 | 0.004717 | 0.020412 | 0.016803 | 0.025356 | 0.004048 | 0.004086 | 0.006578 | 0.004398 | 0.004463 | 0.01046 | 0.014181 | 0.001774 |
| 33.1666667 | 0.033254 | 0.011673 | 0.059608 | 0.002377 | 0.006313 | 0.019547 | 0.012331 | 0.007153 | 0.01205 | 0.010244 | 0.01281 | 0.0025 | 0.002414 | 0.002682 | 0.002636 | 0.00208 | 0.005048 | 0.010271 | 0.001878 |
| 33.1805556 | 0.013445 | 0.008603 | 0.034205 | 0.002005 | 0.002958 | 0.007862 | 0.015425 | 0.007137 | 0.006548 | 0.007378 | 0.012134 | 0.00231 | 0.000928 | 0.000914 | 0.003378 | 0.001792 | 0.0041 | 0.018282 | 0.003047 |
| 33.1944444 | 0.014516 | 0.012995 | 0.015144 | 0.002356 | 0.00216 | 0.00455 | 0.016765 | 0.004404 | 0.008774 | 0.009467 | 0.017468 | 0.002327 | 0.000285 | 0.000637 | 0.004741 | 0.00246 | 0.004586 | 0.026714 | 0.003522 |
| 33.2083333 | 0.021055 | 0.017113 | 0.012752 | 0.002968 | 0.00169 | 0.007021 | 0.016666 | 0.003324 | 0.015457 | 0.010262 | 0.019739 | 0.001422 | 0.000173 | 0.000812 | 0.004628 | 0.002402 | 0.003575 | 0.027794 | 0.003538 |
| 33.2222222 | 0.021511 | 0.018937 | 0.024084 | 0.006278 | 0.003996 | 0.012121 | 0.016793 | 0.005775 | 0.018142 | 0.008438 | 0.020968 | 0.000648 | 0.000129 | 0.001508 | 0.003258 | 0.001697 | 0.002931 | 0.023857 | 0.003051 |
| 33.2361111 | 0.020935 | 0.019744 | 0.039848 | 0.011648 | 0.011288 | 0.019636 | 0.017889 | 0.008373 | 0.01632 | 0.006474 | 0.023523 | 0.000431 | 0.00019 | 0.002386 | 0.001468 | 0.001082 | 0.004166 | 0.019743 | 0.002098 |
| 33.25 | 0.02396 | 0.021456 | 0.056942 | 0.015955 | 0.018906 | 0.025866 | 0.0205 | 0.009826 | 0.015226 | 0.005709 | 0.025197 | 0.000395 | 0.000515 | 0.002243 | 0.000467 | 0.001283 | 0.006453 | 0.017808 | 0.002305 |
| 33.2638889 | 0.031292 | 0.025798 | 0.073082 | 0.017315 | 0.021857 | 0.026047 | 0.023836 | 0.011229 | 0.01826 | 0.007686 | 0.024647 | 0.000365 | 0.000881 | 0.001503 | 0.000329 | 0.002333 | 0.008307 | 0.01642 | 0.003538 |
| 33.2777778 | 0.041311 | 0.030403 | 0.081611 | 0.014247 | 0.019105 | 0.020533 | 0.023853 | 0.011944 | 0.024557 | 0.011107 | 0.023397 | 0.000654 | 0.000765 | 0.001961 | 0.000464 | 0.002991 | 0.008958 | 0.01354 | 0.003449 |
| 33.2916667 | 0.050982 | 0.03168 | 0.078979 | 0.008113 | 0.011992 | 0.013974 | 0.018335 | 0.010943 | 0.028531 | 0.011441 | 0.02387 | 0.001175 | 0.000717 | 0.003624 | 0.000565 | 0.002375 | 0.008331 | 0.007997 | 0.001925 |
| 33.3055556 | 0.057459 | 0.030023 | 0.070081 | 0.003239 | 0.005283 | 0.012841 | 0.010184 | 0.008474 | 0.025446 | 0.007276 | 0.02609 | 0.001543 | 0.00158 | 0.005153 | 0.000876 | 0.001392 | 0.006603 | 0.004647 | 0.001401 |
| 33.3194444 | 0.058946 | 0.026525 | 0.060058 | 0.002077 | 0.004217 | 0.019362 | 0.004162 | 0.005877 | 0.017378 | 0.002779 | 0.028589 | 0.001649 | 0.00261 | 0.005785 | 0.00175 | 0.00147 | 0.00476 | 0.008577 | 0.002946 |
| 33.3333333 | 0.055968 | 0.020662 | 0.048493 | 0.003426 | 0.006095 | 0.027056 | 0.001848 | 0.003555 | 0.009635 | 0.001617 | 0.030398 | 0.001481 | 0.002785 | 0.005384 | 0.001973 | 0.002245 | 0.0035 | 0.01335 | 0.004584 |
| 33.3472222 | 0.049987 | 0.013518 | 0.036459 | 0.00519 | 0.004923 | 0.029522 | 0.001917 | 0.002063 | 0.0064 | 0.00209 | 0.029623 | 0.001179 | 0.002132 | 0.004268 | 0.00129 | 0.002184 | 0.002366 | 0.011499 | 0.00415 |
| 33.3611111 | 0.041211 | 0.007425 | 0.026714 | 0.006643 | 0.002444 | 0.026924 | 0.004052 | 0.00251 | 0.008462 | 0.00145 | 0.026621 | 0.000943 | 0.001101 | 0.003004 | 0.00135 | 0.001491 | 0.001931 | 0.006741 | 0.002335 |
| 33.375 | 0.029941 | 0.003312 | 0.020326 | 0.007592 | 0.002681 | 0.020952 | 0.006438 | 0.00384 | 0.011741 | 0.000516 | 0.025051 | 0.000644 | 0.000725 | 0.002426 | 0.002506 | 0.001896 | 0.00355 | 0.006286 | 0.000995 |
| 33.3888889 | 0.019077 | 0.001123 | 0.01774 | 0.00687 | 0.00302 | 0.013225 | 0.006623 | 0.00441 | 0.012033 | 0.000701 | 0.025256 | 0.000413 | 0.001482 | 0.002581 | 0.003972 | 0.003449 | 0.006364 | 0.008122 | 0.000993 |
| 33.4027778 | 0.011529 | 0.000663 | 0.017907 | 0.004558 | 0.001649 | 0.007044 | 0.005382 | 0.00451 | 0.009552 | 0.002268 | 0.0246 | 0.000544 | 0.001965 | 0.002679 | 0.005385 | 0.004915 | 0.008538 | 0.00676 | 0.001508 |
| 33.4166667 | 0.007344 | 0.001669 | 0.019886 | 0.004294 | 0.000619 | 0.003933 | 0.004014 | 0.005003 | 0.006502 | 0.004961 | 0.022722 | 0.00081 | 0.001408 | 0.002016 | 0.006451 | 0.006062 | 0.009526 | 0.003558 | 0.001268 |
| 33.4305556 | 0.006033 | 0.003662 | 0.022708 | 0.007612 | 0.000453 | 0.002886 | 0.003119 | 0.006024 | 0.004495 | 0.006762 | 0.01765 | 0.001069 | 0.000819 | 0.001237 | 0.006032 | 0.006858 | 0.008529 | 0.002658 | 0.000971 |
| 33.4444444 | 0.00724 | 0.006141 | 0.024285 | 0.011099 | 0.00114 | 0.003411 | 0.004016 | 0.007929 | 0.005355 | 0.005704 | 0.010229 | 0.001228 | 0.000765 | 0.001425 | 0.003684 | 0.006933 | 0.005244 | 0.003401 | 0.001257 |
| 33.4583333 | 0.009501 | 0.009029 | 0.023176 | 0.013365 | 0.004082 | 0.006525 | 0.007812 | 0.01175 | 0.010022 | 0.003073 | 0.00765 | 0.001262 | 0.000951 | 0.00185 | 0.001216 | 0.007013 | 0.001917 | 0.003299 | 0.001468 |
| 33.4722222 | 0.010543 | 0.011908 | 0.019402 | 0.01602 | 0.0092 | 0.013155 | 0.01334 | 0.016807 | 0.015788 | 0.001485 | 0.00863 | 0.00123 | 0.001213 | 0.00214 | 0.000342 | 0.00829 | 0.000775 | 0.004449 | 0.00166 |
| 33.4861111 | 0.009067 | 0.012818 | 0.015247 | 0.018332 | 0.012875 | 0.020316 | 0.016402 | 0.019323 | 0.018375 | 0.001431 | 0.006279 | 0.000992 | 0.001388 | 0.002852 | 0.000525 | 0.010278 | 0.001505 | 0.007455 | 0.003085 |
| 33.5 | 0.006693 | 0.010232 | 0.014793 | 0.017619 | 0.012533 | 0.024119 | 0.013857 | 0.016975 | 0.017447 | 0.002489 | 0.005317 | 0.000889 | 0.001091 | 0.003815 | 0.000751 | 0.011711 | 0.003144 | 0.008926 | 0.004852 |
| 33.5138889 | 0.006342 | 0.006278 | 0.01955 | 0.013773 | 0.011074 | 0.025907 | 0.00856 | 0.013205 | 0.01668 | 0.004637 | 0.012345 | 0.00098 | 0.000556 | 0.004531 | 0.001018 | 0.012036 | 0.005806 | 0.008295 | 0.004474 |
| 33.5277778 | 0.009368 | 0.004291 | 0.025952 | 0.009454 | 0.011563 | 0.028673 | 0.005302 | 0.012154 | 0.018021 | 0.007003 | 0.020663 | 0.000687 | 0.000311 | 0.004776 | 0.001143 | 0.01126 | 0.008808 | 0.006921 | 0.002969 |
| 33.5416667 | 0.01497 | 0.004703 | 0.029355 | 0.006177 | 0.012323 | 0.032251 | 0.004853 | 0.013234 | 0.019135 | 0.007828 | 0.022114 | 0.000219 | 0.000285 | 0.004533 | 0.000839 | 0.009931 | 0.010323 | 0.004671 | 0.00302 |
| 33.5555556 | 0.020841 | 0.006066 | 0.028025 | 0.003805 | 0.011313 | 0.035827 | 0.005499 | 0.013839 | 0.017707 | 0.006425 | 0.018049 | 0.000204 | 0.000268 | 0.003843 | 0.000769 | 0.008921 | 0.010109 | 0.002305 | 0.004294 |
| 33.5694444 | 0.025855 | 0.007775 | 0.024154 | 0.002623 | 0.010426 | 0.042719 | 0.004908 | 0.012721 | 0.016254 | 0.005039 | 0.016962 | 0.000539 | 0.00023 | 0.003083 | 0.001287 | 0.009485 | 0.010991 | 0.002688 | 0.005951 |
| 33.5833333 | 0.032989 | 0.011977 | 0.024551 | 0.004622 | 0.014991 | 0.065282 | 0.004175 | 0.011974 | 0.020092 | 0.006858 | 0.02776 | 0.000748 | 0.000911 | 0.002863 | 0.002069 | 0.012862 | 0.016871 | 0.008768 | 0.010269 |
| 33.5972222 | 0.041134 | 0.021209 | 0.034111 | 0.012843 | 0.025836 | 0.107441 | 0.006642 | 0.014725 | 0.029466 | 0.011834 | 0.04776 | 0.001715 | 0.003347 | 0.005334 | 0.003605 | 0.017974 | 0.026342 | 0.019387 | 0.017603 |

| | | | | | | | | | | | | | | | | | | | |
|---|---|---|---|---|---|---|---|---|---|---|---|---|---|---|---|---|---|---|---|
| 33.6111111 | 0.042052 | 0.032756 | 0.051969 | 0.026822 | 0.036774 | 0.143409 | 0.00822 | 0.019441 | 0.041512 | 0.016002 | 0.061993 | 0.005116 | 0.007046 | 0.012418 | 0.004943 | 0.0227 | 0.034906 | 0.027784 | 0.02393 |
| 33.625 | 0.038368 | 0.041394 | 0.072076 | 0.044527 | 0.042742 | 0.15629 | 0.006777 | 0.021017 | 0.061327 | 0.01623 | 0.065651 | 0.00991 | 0.009335 | 0.021785 | 0.003771 | 0.025194 | 0.044786 | 0.030367 | 0.025815 |
| 33.6388889 | 0.037651 | 0.047311 | 0.079324 | 0.06455 | 0.043106 | 0.149256 | 0.008158 | 0.020066 | 0.080614 | 0.012944 | 0.063848 | 0.012633 | 0.007413 | 0.026874 | 0.001404 | 0.022259 | 0.050207 | 0.029944 | 0.022377 |
| 33.6527778 | 0.033404 | 0.052545 | 0.065312 | 0.07871 | 0.037981 | 0.116434 | 0.010758 | 0.019292 | 0.074167 | 0.007527 | 0.052048 | 0.01079 | 0.003682 | 0.024148 | 0.000783 | 0.013312 | 0.039296 | 0.027897 | 0.015381 |
| 33.6666667 | 0.021089 | 0.05771 | 0.041025 | 0.079566 | 0.02854 | 0.068401 | 0.00876 | 0.015803 | 0.042134 | 0.00283 | 0.028472 | 0.005593 | 0.002541 | 0.018517 | 0.001734 | 0.00515 | 0.01934 | 0.024818 | 0.008909 |
| 33.6805556 | 0.009177 | 0.067833 | 0.019351 | 0.074551 | 0.023878 | 0.032859 | 0.004519 | 0.010572 | 0.021221 | 0.00192 | 0.013468 | 0.001897 | 0.002425 | 0.01483 | 0.002682 | 0.002865 | 0.008204 | 0.02228 | 0.005766 |
| 33.6944444 | 0.005982 | 0.080171 | 0.009969 | 0.074821 | 0.028667 | 0.018499 | 0.002761 | 0.007221 | 0.028731 | 0.005131 | 0.023165 | 0.001607 | 0.002722 | 0.013554 | 0.003401 | 0.002342 | 0.01219 | 0.019578 | 0.004606 |
| 33.7083333 | 0.008701 | 0.081444 | 0.016319 | 0.073791 | 0.030696 | 0.01262 | 0.002462 | 0.004548 | 0.032022 | 0.010475 | 0.036958 | 0.002096 | 0.005573 | 0.013128 | 0.004051 | 0.001109 | 0.02193 | 0.015645 | 0.003134 |
| 33.7222222 | 0.007464 | 0.071223 | 0.024554 | 0.055295 | 0.022044 | 0.009334 | 0.001932 | 0.003271 | 0.019253 | 0.013103 | 0.040446 | 0.00261 | 0.007628 | 0.010432 | 0.004595 | 0.001304 | 0.029155 | 0.010438 | 0.001467 |
| 33.7361111 | 0.004179 | 0.058471 | 0.022799 | 0.032123 | 0.011125 | 0.013382 | 0.002127 | 0.00388 | 0.010342 | 0.010644 | 0.041676 | 0.003399 | 0.006275 | 0.005533 | 0.004803 | 0.003187 | 0.031699 | 0.00549 | 0.00119 |
| 33.75 | 0.006409 | 0.043229 | 0.013864 | 0.042122 | 0.005006 | 0.021336 | 0.001867 | 0.004252 | 0.011469 | 0.006059 | 0.035567 | 0.003381 | 0.004318 | 0.002299 | 0.003693 | 0.00537 | 0.029544 | 0.004518 | 0.003245 |
| 33.7638889 | 0.009138 | 0.026063 | 0.005168 | 0.082017 | 0.002154 | 0.027631 | 0.001379 | 0.005634 | 0.017671 | 0.002374 | 0.019235 | 0.002815 | 0.003647 | 0.001674 | 0.001818 | 0.005824 | 0.023298 | 0.008051 | 0.006642 |
| 33.7777778 | 0.007081 | 0.02556 | 0.002305 | 0.109298 | 0.001138 | 0.033072 | 0.001716 | 0.006551 | 0.020674 | 0.002702 | 0.008683 | 0.002683 | 0.003307 | 0.001439 | 0.000985 | 0.003722 | 0.013182 | 0.012152 | 0.009083 |
| 33.7916667 | 0.004844 | 0.050128 | 0.00449 | 0.110932 | 0.002209 | 0.038293 | 0.001833 | 0.004654 | 0.016374 | 0.008638 | 0.011073 | 0.002736 | 0.002136 | 0.000971 | 0.001073 | 0.001428 | 0.008422 | 0.014195 | 0.009201 |
| 33.8055556 | 0.008348 | 0.080618 | 0.006826 | 0.093243 | 0.004736 | 0.037048 | 0.003032 | 0.002763 | 0.009064 | 0.013808 | 0.012487 | 0.002421 | 0.000797 | 0.002552 | 0.001148 | 0.000723 | 0.01569 | 0.014777 | 0.008101 |
| 33.8194444 | 0.011306 | 0.103128 | 0.006864 | 0.058162 | 0.008263 | 0.026461 | 0.006736 | 0.004351 | 0.008789 | 0.010582 | 0.007111 | 0.002219 | 0.000331 | 0.007424 | 0.001033 | 0.000655 | 0.024542 | 0.014955 | 0.006796 |
| 33.8333333 | 0.008039 | 0.111002 | 0.010969 | 0.023669 | 0.013927 | 0.013353 | 0.012663 | 0.01003 | 0.018395 | 0.005884 | 0.003573 | 0.002995 | 0.00062 | 0.011664 | 0.000709 | 0.000482 | 0.024422 | 0.013855 | 0.004266 |
| 33.8472222 | 0.007472 | 0.095672 | 0.022933 | 0.005966 | 0.022479 | 0.009367 | 0.019805 | 0.017968 | 0.029601 | 0.009927 | 0.008753 | 0.004598 | 0.00115 | 0.01116 | 0.000462 | 0.00073 | 0.015663 | 0.010317 | 0.00242 |
| 33.8611111 | 0.009243 | 0.062906 | 0.032017 | 0.001451 | 0.030926 | 0.017318 | 0.026426 | 0.025024 | 0.035201 | 0.017709 | 0.023168 | 0.00639 | 0.001073 | 0.007288 | 0.000797 | 0.002473 | 0.008887 | 0.00532 | 0.004411 |
| 33.875 | 0.008337 | 0.033659 | 0.03106 | 0.002578 | 0.036278 | 0.022648 | 0.030785 | 0.028372 | 0.033137 | 0.021768 | 0.041935 | 0.007435 | 0.00073 | 0.007714 | 0.001461 | 0.005137 | 0.015274 | 0.001616 | 0.008204 |
| 33.8888889 | 0.012783 | 0.019382 | 0.026255 | 0.011344 | 0.040194 | 0.018719 | 0.030971 | 0.027496 | 0.026617 | 0.020323 | 0.058483 | 0.006367 | 0.001164 | 0.017439 | 0.001586 | 0.007083 | 0.028499 | 0.00085 | 0.011164 |
| 33.9027778 | 0.027314 | 0.01485 | 0.024382 | 0.030539 | 0.044348 | 0.025926 | 0.027116 | 0.02316 | 0.022955 | 0.015108 | 0.069431 | 0.003723 | 0.001718 | 0.030894 | 0.001099 | 0.009026 | 0.03693 | 0.002752 | 0.012679 |
| 33.9166667 | 0.04469 | 0.013139 | 0.026445 | 0.042389 | 0.047838 | 0.060578 | 0.022231 | 0.018201 | 0.027104 | 0.011605 | 0.074254 | 0.001925 | 0.001781 | 0.037862 | 0.000741 | 0.011914 | 0.038949 | 0.00702 | 0.012167 |
| 33.9305556 | 0.053541 | 0.011748 | 0.034422 | 0.031755 | 0.049809 | 0.110926 | 0.020035 | 0.017325 | 0.037546 | 0.011819 | 0.069604 | 0.001817 | 0.001982 | 0.032434 | 0.000978 | 0.013982 | 0.035498 | 0.012235 | 0.009946 |
| 33.9444444 | 0.048475 | 0.010002 | 0.048212 | 0.0269 | 0.048286 | 0.147893 | 0.021501 | 0.022431 | 0.048122 | 0.012631 | 0.053042 | 0.002067 | 0.002166 | 0.019426 | 0.001495 | 0.013191 | 0.027114 | 0.015552 | 0.007299 |
| 33.9583333 | 0.037726 | 0.00876 | 0.061067 | 0.052136 | 0.041793 | 0.148191 | 0.022822 | 0.030203 | 0.052684 | 0.011796 | 0.02914 | 0.001632 | 0.001584 | 0.008005 | 0.001709 | 0.009976 | 0.016902 | 0.015399 | 0.00573 |
| 33.9722222 | 0.031243 | 0.01129 | 0.071013 | 0.082062 | 0.035112 | 0.123816 | 0.021814 | 0.037044 | 0.051026 | 0.009148 | 0.011682 | 0.000871 | 0.000922 | 0.002614 | 0.00153 | 0.00693 | 0.009206 | 0.013398 | 0.006095 |
| 33.9861111 | 0.028175 | 0.018162 | 0.078318 | 0.089106 | 0.035151 | 0.100168 | 0.020303 | 0.040774 | 0.04562 | 0.004992 | 0.009066 | 0.000353 | 0.00108 | 0.003077 | 0.001051 | 0.005492 | 0.006339 | 0.012208 | 0.007676 |
| 34 | 0.025462 | 0.02616 | 0.077564 | 0.071823 | 0.040697 | 0.082127 | 0.018206 | 0.039116 | 0.036447 | 0.001976 | 0.011838 | 0.000157 | 0.001732 | 0.005792 | 0.000566 | 0.004866 | 0.00688 | 0.011892 | 0.008499 |
| 34.0138889 | 0.025115 | 0.033203 | 0.064563 | 0.043546 | 0.042913 | 0.064645 | 0.014618 | 0.03208 | 0.025565 | 0.001328 | 0.013057 | 0.00046 | 0.001897 | 0.006945 | 0.000808 | 0.004028 | 0.007729 | 0.011744 | 0.006932 |
| 34.0277778 | 0.027033 | 0.036812 | 0.042607 | 0.019353 | 0.035461 | 0.048231 | 0.009449 | 0.021009 | 0.015938 | 0.001138 | 0.014857 | 0.001445 | 0.001268 | 0.005888 | 0.001851 | 0.00254 | 0.007963 | 0.012246 | 0.00373 |
| 34.0416667 | 0.025549 | 0.033222 | 0.019827 | 0.007944 | 0.020592 | 0.037975 | 0.004717 | 0.009194 | 0.007662 | 0.001385 | 0.016391 | 0.002527 | 0.000744 | 0.004577 | 0.002616 | 0.00109 | 0.008468 | 0.012945 | 0.001775 |
| 34.0555556 | 0.019097 | 0.0246 | 0.007494 | 0.006812 | 0.00833 | 0.033429 | 0.003056 | 0.002312 | 0.003192 | 0.00175 | 0.014071 | 0.002936 | 0.00082 | 0.004465 | 0.002478 | 0.000622 | 0.008997 | 0.012101 | 0.00147 |
| 34.0694444 | 0.014432 | 0.019967 | 0.007388 | 0.014061 | 0.008569 | 0.034346 | 0.003544 | 0.005208 | 0.004253 | 0.003767 | 0.012298 | 0.002386 | 0.002099 | 0.005729 | 0.00156 | 0.001434 | 0.010844 | 0.011093 | 0.002482 |
| 34.0833333 | 0.024257 | 0.028393 | 0.011153 | 0.036827 | 0.031975 | 0.050304 | 0.008017 | 0.020545 | 0.014822 | 0.015378 | 0.028637 | 0.001939 | 0.005632 | 0.008604 | 0.001023 | 0.005221 | 0.018851 | 0.01752 | 0.008189 |
| 34.0972222 | 0.055236 | 0.048904 | 0.022787 | 0.063195 | 0.072289 | 0.078767 | 0.016345 | 0.041909 | 0.038286 | 0.038347 | 0.062775 | 0.003496 | 0.009345 | 0.012706 | 0.001294 | 0.011932 | 0.030698 | 0.033693 | 0.017675 |
| 34.1111111 | 0.094262 | 0.066225 | 0.039721 | 0.065524 | 0.083971 | 0.092815 | 0.018702 | 0.051319 | 0.055644 | 0.055165 | 0.078003 | 0.006044 | 0.009415 | 0.017219 | 0.001206 | 0.01591 | 0.033937 | 0.047513 | 0.022512 |
| 34.125 | 0.119152 | 0.070983 | 0.04766 | 0.039069 | 0.052693 | 0.081707 | 0.011337 | 0.039411 | 0.050156 | 0.048398 | 0.058875 | 0.006253 | 0.006313 | 0.019162 | 0.001234 | 0.013472 | 0.024976 | 0.048799 | 0.0191 |
| 34.1388889 | 0.120299 | 0.066501 | 0.046133 | 0.016117 | 0.020509 | 0.063215 | 0.003781 | 0.020055 | 0.034241 | 0.025239 | 0.033925 | 0.003852 | 0.003063 | 0.014654 | 0.001333 | 0.00775 | 0.01313 | 0.039005 | 0.011871 |
| 34.1527778 | 0.099749 | 0.051532 | 0.03984 | 0.017324 | 0.007013 | 0.045197 | 0.003839 | 0.012757 | 0.023345 | 0.009827 | 0.019453 | 0.002887 | 0.001205 | 0.006786 | 0.001094 | 0.003315 | 0.005729 | 0.022421 | 0.005207 |
| 34.1666667 | 0.062066 | 0.02852 | 0.030296 | 0.021433 | 0.003536 | 0.023936 | 0.010079 | 0.016994 | 0.01646 | 0.013259 | 0.010907 | 0.005087 | 0.001059 | 0.001998 | 0.001585 | 0.002271 | 0.005083 | 0.010338 | 0.001797 |
| 34.1805556 | 0.030159 | 0.017815 | 0.01955 | 0.015114 | 0.004228 | 0.01029 | 0.016782 | 0.019144 | 0.009749 | 0.020361 | 0.005785 | 0.007504 | 0.001758 | 0.00145 | 0.002261 | 0.00354 | 0.008426 | 0.012525 | 0.001651 |
| 34.1944444 | 0.031879 | 0.027757 | 0.010103 | 0.00712 | 0.004368 | 0.012999 | 0.019744 | 0.017252 | 0.008409 | 0.018512 | 0.002856 | 0.008042 | 0.002456 | 0.002914 | 0.002235 | 0.004577 | 0.009263 | 0.020657 | 0.002654 |
| 34.2083333 | 0.058301 | 0.039547 | 0.004508 | 0.004468 | 0.002203 | 0.019705 | 0.019044 | 0.017221 | 0.015753 | 0.010737 | 0.001131 | 0.006766 | 0.002886 | 0.004909 | 0.002126 | 0.004609 | 0.006396 | 0.025272 | 0.004088 |
| 34.2222222 | 0.077945 | 0.042559 | 0.004605 | 0.008519 | 0.003497 | 0.023136 | 0.018021 | 0.020721 | 0.028297 | 0.004523 | 0.001033 | 0.004734 | 0.002608 | 0.007116 | 0.002136 | 0.004172 | 0.004151 | 0.027019 | 0.004541 |
| 34.2361111 | 0.076174 | 0.039371 | 0.010057 | 0.016245 | 0.01126 | 0.02519 | 0.020804 | 0.024868 | 0.043628 | 0.003791 | 0.003027 | 0.002984 | 0.001556 | 0.009208 | 0.001816 | 0.003559 | 0.004414 | 0.028547 | 0.002776 |
| 34.25 | 0.060594 | 0.033104 | 0.019326 | 0.023114 | 0.02064 | 0.026413 | 0.027434 | 0.026113 | 0.061269 | 0.003878 | 0.005779 | 0.001608 | 0.000745 | 0.010433 | 0.001448 | 0.003209 | 0.005073 | 0.030171 | 0.001168 |
| 34.2638889 | 0.043709 | 0.026902 | 0.03029 | 0.024425 | 0.02896 | 0.030197 | 0.031284 | 0.024366 | 0.07335 | 0.004435 | 0.008058 | 0.000726 | 0.000615 | 0.009953 | 0.001582 | 0.003229 | 0.004468 | 0.029884 | 0.001881 |
| 34.2777778 | 0.031381 | 0.020559 | 0.039154 | 0.018715 | 0.03311 | 0.034598 | 0.026652 | 0.020701 | 0.069428 | 0.008896 | 0.010724 | 0.000673 | 0.000554 | 0.007544 | 0.00168 | 0.003007 | 0.003613 | 0.025855 | 0.002909 |
| 34.2916667 | 0.023637 | 0.014425 | 0.042029 | 0.010008 | 0.028943 | 0.033048 | 0.016922 | 0.015499 | 0.052855 | 0.012613 | 0.014599 | 0.001413 | 0.000539 | 0.004085 | 0.001082 | 0.001938 | 0.003775 | 0.018683 | 0.002441 |
| 34.3055556 | 0.017428 | 0.009703 | 0.037804 | 0.003385 | 0.019164 | 0.024376 | 0.009127 | 0.009865 | 0.031937 | 0.011252 | 0.017548 | 0.002656 | 0.000887 | 0.002106 | 0.000338 | 0.001295 | 0.004619 | 0.011411 | 0.002321 |
| 34.3194444 | 0.010548 | 0.006189 | 0.030095 | 0.001607 | 0.0091 | 0.012865 | 0.004381 | 0.005323 | 0.015394 | 0.006751 | 0.017278 | 0.003495 | 0.001293 | 0.003144 | 0.000178 | 0.002403 | 0.005164 | 0.005757 | 0.003315 |
| 34.3333333 | 0.004266 | 0.004361 | 0.023655 | 0.003951 | 0.004003 | 0.007234 | 0.001824 | 0.002461 | 0.013994 | 0.002613 | 0.014655 | 0.003112 | 0.001115 | 0.004791 | 0.000518 | 0.003368 | 0.004728 | 0.002797 | 0.004289 |
| 34.3472222 | 0.002798 | 0.004172 | 0.019801 | 0.005128 | 0.006412 | 0.009988 | 0.002575 | 0.002882 | 0.024802 | 0.002175 | 0.012356 | 0.002036 | 0.001074 | 0.004719 | 0.000687 | 0.002952 | 0.004096 | 0.003964 | 0.005919 |
| 34.3611111 | 0.008725 | 0.003779 | 0.017868 | 0.005151 | 0.011575 | 0.012437 | 0.005205 | 0.007399 | 0.030937 | 0.0076 | 0.01346 | 0.001092 | 0.001211 | 0.003221 | 0.000858 | 0.001891 | 0.005457 | 0.006606 | 0.007814 |
| 34.375 | 0.018324 | 0.002692 | 0.017923 | 0.009507 | 0.014413 | 0.010571 | 0.006712 | 0.010995 | 0.021449 | 0.015481 | 0.018328 | 0.000648 | 0.001511 | 0.002398 | 0.001882 | 0.00103 | 0.009208 | 0.005625 | 0.006977 |
| 34.3888889 | 0.024272 | 0.002223 | 0.019377 | 0.013743 | 0.012742 | 0.006802 | 0.005104 | 0.008432 | 0.011699 | 0.017428 | 0.022597 | 0.000675 | 0.002486 | 0.002747 | 0.002738 | 0.001081 | 0.012075 | 0.003285 | 0.003589 |
| 34.4027778 | 0.022699 | 0.002693 | 0.019723 | 0.011126 | 0.007175 | 0.004482 | 0.00294 | 0.004469 | 0.018493 | 0.011337 | 0.02254 | 0.000793 | 0.002474 | 0.002001 | 0.002233 | 0.001393 | 0.011153 | 0.003768 | 0.001503 |
| 34.4166667 | 0.016366 | 0.003441 | 0.017825 | 0.005486 | 0.003514 | 0.007925 | 0.00399 | 0.005442 | 0.028919 | 0.004532 | 0.019099 | 0.001033 | 0.001708 | 0.000595 | 0.001517 | 0.001515 | 0.007499 | 0.003637 | 0.001582 |
| 34.4305556 | 0.009817 | 0.004111 | 0.015134 | 0.003222 | 0.004992 | 0.015311 | 0.006029 | 0.007347 | 0.029351 | 0.003431 | 0.01513 | 0.001909 | 0.002017 | 0.000263 | 0.001559 | 0.001516 | 0.003693 | 0.002552 | 0.001402 |
| 34.4444444 | 0.004602 | 0.004084 | 0.012875 | 0.00331 | 0.007087 | 0.02045 | 0.007003 | 0.006272 | 0.021056 | 0.007439 | 0.01285 | 0.003115 | 0.00284 | 0.000422 | 0.001768 | 0.001017 | 0.001255 | 0.002808 | 0.000686 |
| 34.4583333 | 0.001696 | 0.003185 | 0.011965 | 0.002819 | 0.006932 | 0.022533 | 0.007 | 0.004454 | 0.010908 | 0.011362 | 0.012378 | 0.003257 | 0.002958 | 0.000827 | 0.001939 | 0.000724 | 0.000539 | 0.002873 | 0.000562 |
| 34.4722222 | 0.003177 | 0.002656 | 0.014237 | 0.003783 | 0.00644 | 0.025253 | 0.005901 | 0.004105 | 0.007391 | 0.011173 | 0.013153 | 0.002036 | 0.002212 | 0.002039 | 0.002137 | 0.001586 | 0.000853 | 0.002462 | 0.000657 |
| 34.4861111 | 0.007906 | 0.003472 | 0.019609 | 0.006149 | 0.007509 | 0.029491 | 0.004733 | 0.005897 | 0.012206 | 0.007893 | 0.016207 | 0.001161 | 0.001203 | 0.0029 | 0.001802 | 0.002559 | 0.001531 | 0.001786 | 0.000655 |
| 34.5 | 0.011352 | 0.005146 | 0.024256 | 0.005783 | 0.009738 | 0.030145 | 0.004344 | 0.009557 | 0.019546 | 0.005096 | 0.021701 | 0.00123 | 0.000938 | 0.00249 | 0.001027 | 0.002387 | 0.003524 | 0.001943 | 0.001382 |
| 34.5138889 | 0.011355 | 0.006312 | 0.024705 | 0.004139 | 0.010331 | 0.021998 | 0.004275 | 0.013693 | 0.022126 | 0.004921 | 0.026843 | 0.001417 | 0.002003 | 0.002309 | 0.000853 | 0.001892 | 0.007863 | 0.004552 | 0.003766 |
| 34.5277778 | 0.008574 | 0.005221 | 0.021401 | 0.006379 | 0.007648 | 0.010838 | 0.003216 | 0.015987 | 0.017362 | 0.007948 | 0.030582 | 0.002243 | 0.00339 | 0.002931 | 0.001487 | 0.001889 | 0.012783 | 0.007269 | 0.006562 |
| 34.5416667 | 0.005631 | 0.00336 | 0.016909 | 0.009084 | 0.004125 | 0.007941 | 0.002149 | 0.01423 | 0.008912 | 0.012646 | 0.034011 | 0.003084 | 0.003572 | 0.002878 | 0.001856 | 0.001702 | 0.01512 | 0.007295 | 0.007119 |
| 34.5555556 | 0.00682 | 0.004005 | 0.014342 | 0.006597 | 0.002578 | 0.010447 | 0.002998 | 0.00858 | 0.003469 | 0.015917 | 0.035622 | 0.002273 | 0.002697 | 0.001726 | 0.001287 | 0.001279 | 0.014115 | 0.005208 | 0.005484 |
| 34.5694444 | 0.008677 | 0.004388 | 0.018402 | 0.002168 | 0.004632 | 0.014496 | 0.004157 | 0.003206 | 0.003899 | 0.016742 | 0.03725 | 0.000899 | 0.002188 | 0.000812 | 0.000789 | 0.002207 | 0.013188 | 0.00446 | 0.00504 |
| 34.5833333 | 0.008566 | 0.003884 | 0.032181 | 0.000316 | 0.013864 | 0.03859 | 0.004529 | 0.002294 | 0.012787 | 0.017128 | 0.047156 | 0.000606 | 0.00298 | 0.000764 | 0.001556 | 0.005925 | 0.017201 | 0.009453 | 0.009815 |
| 34.5972222 | 0.015104 | 0.006111 | 0.049136 | 0.001147 | 0.027061 | 0.087176 | 0.00514 | 0.006734 | 0.032859 | 0.0203 | 0.065484 | 0.002672 | 0.005547 | 0.001236 | 0.002824 | 0.012657 | 0.027029 | 0.019904 | 0.020362 |
| 34.6111111 | 0.028265 | 0.008879 | 0.058125 | 0.008112 | 0.035804 | 0.121978 | 0.005839 | 0.015735 | 0.058477 | 0.027769 | 0.074403 | 0.00842 | 0.009415 | 0.004976 | 0.00295 | 0.022407 | 0.037346 | 0.029433 | 0.030967 |
| 34.625 | 0.040506 | 0.012031 | 0.062108 | 0.021364 | 0.042658 | 0.119655 | 0.007956 | 0.026581 | 0.079528 | 0.033657 | 0.065494 | 0.014396 | 0.01249 | 0.015759 | 0.001751 | 0.031536 | 0.044815 | 0.032633 | 0.034127 |
| 34.6388889 | 0.047728 | 0.017387 | 0.067524 | 0.030261 | 0.0523 | 0.106208 | 0.012892 | 0.033394 | 0.087139 | 0.028516 | 0.053442 | 0.01445 | 0.012669 | 0.026594 | 0.000595 | 0.032692 | 0.049841 | 0.029536 | 0.027587 |
| 34.6527778 | 0.046195 | 0.021746 | 0.065901 | 0.030988 | 0.056847 | 0.095788 | 0.016921 | 0.031762 | 0.077489 | 0.015347 | 0.045238 | 0.008446 | 0.009304 | 0.028001 | 0.000349 | 0.025598 | 0.047699 | 0.023507 | 0.016576 |
| 34.6666667 | 0.038451 | 0.022783 | 0.05326 | 0.029254 | 0.048779 | 0.071564 | 0.016858 | 0.023326 | 0.051561 | 0.009298 | 0.036343 | 0.003793 | 0.004496 | 0.023439 | 0.000647 | 0.01728 | 0.036408 | 0.018338 | 0.009116 |
| 34.6805556 | 0.030029 | 0.022012 | 0.037959 | 0.029397 | 0.033871 | 0.03643 | 0.014637 | 0.012659 | 0.026374 | 0.015098 | 0.023875 | 0.004782 | 0.001473 | 0.018597 | 0.000913 | 0.011561 | 0.020929 | 0.015368 | 0.007211 |
| 34.6944444 | 0.019916 | 0.021037 | 0.025403 | 0.037144 | 0.021564 | 0.011035 | 0.013135 | 0.006481 | 0.021367 | 0.021427 | 0.014156 | 0.006952 | 0.000889 | 0.014797 | 0.000831 | 0.008078 | 0.01019 | 0.013171 | 0.007515 |
| 34.7083333 | 0.014332 | 0.019158 | 0.015032 | 0.051462 | 0.014808 | 0.003528 | 0.013446 | 0.006665 | 0.02824 | 0.019316 | 0.019535 | 0.006824 | 0.00185 | 0.01233 | 0.000628 | 0.005841 | 0.012523 | 0.010518 | 0.007272 |
| 34.7222222 | 0.021397 | 0.016764 | 0.007354 | 0.05822 | 0.010544 | 0.004799 | 0.012926 | 0.005778 | 0.032265 | 0.011097 | 0.036891 | 0.005288 | 0.003078 | 0.010326 | 0.000715 | 0.003542 | 0.023287 | 0.00721 | 0.005387 |
| 34.7361111 | 0.023012 | 0.016614 | 0.006589 | 0.047734 | 0.00861 | 0.004554 | 0.009135 | 0.006203 | 0.028911 | 0.007173 | 0.050674 | 0.003658 | 0.002721 | 0.008408 | 0.000819 | 0.001869 | 0.032573 | 0.004414 | 0.002975 |
| 34.75 | 0.017669 | 0.019843 | 0.010124 | 0.029355 | 0.008814 | 0.003632 | 0.00425 | 0.013714 | 0.019692 | 0.011626 | 0.05523 | 0.002599 | 0.001661 | 0.006962 | 0.000664 | 0.002725 | 0.0373 | 0.00568 | 0.001895 |
| 34.7638889 | 0.027686 | 0.022722 | 0.008628 | 0.028826 | 0.005749 | 0.003097 | 0.00182 | 0.018863 | 0.010773 | 0.015934 | 0.053001 | 0.002158 | 0.001414 | 0.00668 | 0.000471 | 0.004894 | 0.035144 | 0.011554 | 0.002758 |
| 34.7777778 | 0.041498 | 0.021209 | 0.004968 | 0.05043 | 0.001452 | 0.005628 | 0.003384 | 0.015809 | 0.010582 | 0.014644 | 0.03922 | 0.001809 | 0.001749 | 0.008667 | 0.000547 | 0.005347 | 0.023217 | 0.016846 | 0.005138 |
| 34.7916667 | 0.040482 | 0.016663 | 0.008715 | 0.06619 | 0.002732 | 0.012947 | 0.007665 | 0.010383 | 0.022772 | 0.009596 | 0.020563 | 0.001357 | 0.003111 | 0.013225 | 0.000905 | 0.003284 | 0.010634 | 0.01757 | 0.006743 |

| | | | | | | | | | | | | | | | | | | | |
|---|---|---|---|---|---|---|---|---|---|---|---|---|---|---|---|---|---|---|---|
| 34.8055556 | 0.026619 | 0.010383 | 0.018461 | 0.059836 | 0.009397 | 0.016959 | 0.012597 | 0.006071 | 0.037473 | 0.004086 | 0.017226 | 0.001559 | 0.003849 | 0.018148 | 0.001102 | 0.00133 | 0.009091 | 0.01465 | 0.005107 |
| 34.8194444 | 0.01567 | 0.005862 | 0.02486 | 0.042708 | 0.014395 | 0.01468 | 0.016237 | 0.005383 | 0.043899 | 0.001916 | 0.028657 | 0.002519 | 0.002718 | 0.019817 | 0.001033 | 0.001476 | 0.01467 | 0.010865 | 0.00223 |
| 34.8333333 | 0.020645 | 0.008286 | 0.018566 | 0.030117 | 0.014556 | 0.012496 | 0.016675 | 0.008406 | 0.041014 | 0.004184 | 0.037953 | 0.003582 | 0.002082 | 0.017038 | 0.00099 | 0.002259 | 0.020015 | 0.00733 | 0.001367 |
| 34.8472222 | 0.025157 | 0.013661 | 0.011523 | 0.019393 | 0.015398 | 0.016442 | 0.013256 | 0.008378 | 0.032125 | 0.008108 | 0.036072 | 0.004459 | 0.002181 | 0.012422 | 0.001074 | 0.002797 | 0.023114 | 0.003872 | 0.001553 |
| 34.8611111 | 0.017813 | 0.01656 | 0.022611 | 0.008878 | 0.019623 | 0.026774 | 0.007139 | 0.007532 | 0.020395 | 0.011329 | 0.022867 | 0.004938 | 0.001763 | 0.008748 | 0.001215 | 0.003943 | 0.021935 | 0.001536 | 0.001273 |
| 34.875 | 0.013074 | 0.014845 | 0.048788 | 0.006641 | 0.02207 | 0.037745 | 0.002893 | 0.011256 | 0.010882 | 0.013273 | 0.01619 | 0.004776 | 0.001602 | 0.006069 | 0.001295 | 0.005545 | 0.014061 | 0.001675 | 0.001468 |
| 34.8888889 | 0.011849 | 0.009674 | 0.074075 | 0.010803 | 0.016813 | 0.042741 | 0.002817 | 0.011834 | 0.010893 | 0.013289 | 0.034372 | 0.00396 | 0.001464 | 0.004023 | 0.001552 | 0.006573 | 0.009564 | 0.003529 | 0.003023 |
| 34.9027778 | 0.008788 | 0.007455 | 0.072984 | 0.013738 | 0.009006 | 0.041078 | 0.002571 | 0.008254 | 0.020019 | 0.011259 | 0.060174 | 0.002576 | 0.00094 | 0.005765 | 0.002123 | 0.00527 | 0.022233 | 0.006587 | 0.004841 |
| 34.9166667 | 0.010064 | 0.007419 | 0.043619 | 0.012451 | 0.009569 | 0.040061 | 0.001168 | 0.011551 | 0.02905 | 0.008779 | 0.070522 | 0.001126 | 0.000962 | 0.014153 | 0.002481 | 0.003573 | 0.043341 | 0.010602 | 0.005443 |
| 34.9305556 | 0.012653 | 0.005292 | 0.015809 | 0.00818 | 0.016821 | 0.045876 | 0.001212 | 0.024268 | 0.032635 | 0.006721 | 0.066597 | 0.000428 | 0.001342 | 0.026044 | 0.002322 | 0.005818 | 0.056815 | 0.012948 | 0.005073 |
| 34.9444444 | 0.010074 | 0.003685 | 0.005805 | 0.007385 | 0.022187 | 0.055644 | 0.003214 | 0.038365 | 0.032313 | 0.004544 | 0.05808 | 0.000536 | 0.001364 | 0.035851 | 0.001978 | 0.009608 | 0.058084 | 0.010658 | 0.004235 |
| 34.9583333 | 0.00999 | 0.003084 | 0.012825 | 0.011366 | 0.0243 | 0.061161 | 0.006581 | 0.045067 | 0.032967 | 0.003623 | 0.045616 | 0.001048 | 0.001902 | 0.039292 | 0.002143 | 0.010796 | 0.050221 | 0.006561 | 0.002594 |
| 34.9722222 | 0.020458 | 0.002212 | 0.032882 | 0.010948 | 0.02633 | 0.059275 | 0.009676 | 0.039923 | 0.035553 | 0.004683 | 0.028039 | 0.002019 | 0.003479 | 0.03322 | 0.002709 | 0.010306 | 0.037586 | 0.007197 | 0.001439 |
| 34.9861111 | 0.032576 | 0.003624 | 0.053919 | 0.005413 | 0.029222 | 0.054043 | 0.011814 | 0.023663 | 0.034307 | 0.006143 | 0.012659 | 0.003089 | 0.004264 | 0.020571 | 0.002398 | 0.010216 | 0.024213 | 0.012034 | 0.001459 |
| 35 | 0.039301 | 0.007935 | 0.063142 | 0.002293 | 0.031646 | 0.04968 | 0.013469 | 0.009487 | 0.028051 | 0.006758 | 0.007291 | 0.003592 | 0.003519 | 0.00961 | 0.001148 | 0.009489 | 0.013746 | 0.016537 | 0.001219 |
| 35.0138889 | 0.039022 | 0.011576 | 0.059768 | 0.002666 | 0.030663 | 0.042168 | 0.015328 | 0.009363 | 0.019889 | 0.005472 | 0.011086 | 0.003599 | 0.002649 | 0.00381 | 0.000264 | 0.006157 | 0.006595 | 0.018547 | 0.001056 |
| 35.0277778 | 0.029481 | 0.010992 | 0.049873 | 0.002716 | 0.023414 | 0.024914 | 0.017787 | 0.014215 | 0.010608 | 0.003145 | 0.015715 | 0.00382 | 0.002712 | 0.002072 | 8.67E-05 | 0.002808 | 0.004089 | 0.016927 | 0.001011 |
| 35.0416667 | 0.015843 | 0.006267 | 0.035468 | 0.001724 | 0.012329 | 0.011527 | 0.020683 | 0.01419 | 0.006014 | 0.003326 | 0.017424 | 0.004904 | 0.003536 | 0.002187 | 0.000389 | 0.002851 | 0.007361 | 0.012724 | 0.001102 |
| 35.0555556 | 0.009967 | 0.003769 | 0.019526 | 0.0024 | 0.007204 | 0.016983 | 0.02376 | 0.010244 | 0.013123 | 0.006924 | 0.01691 | 0.006089 | 0.00399 | 0.001646 | 0.001221 | 0.004392 | 0.011628 | 0.009546 | 0.002416 |
| 35.0694444 | 0.01516 | 0.010027 | 0.014597 | 0.006184 | 0.010455 | 0.028077 | 0.02703 | 0.007172 | 0.02799 | 0.007963 | 0.01339 | 0.005674 | 0.002931 | 0.001014 | 0.002182 | 0.00384 | 0.010319 | 0.011772 | 0.003194 |
| 35.0833333 | 0.016025 | 0.02419 | 0.020363 | 0.013168 | 0.011905 | 0.026726 | 0.031921 | 0.006471 | 0.040516 | 0.010931 | 0.020588 | 0.003806 | 0.002325 | 0.002338 | 0.003003 | 0.003985 | 0.01044 | 0.023013 | 0.006416 |
| 35.0972222 | 0.020481 | 0.047861 | 0.019007 | 0.027154 | 0.015533 | 0.015714 | 0.035029 | 0.012343 | 0.040113 | 0.029179 | 0.046419 | 0.004161 | 0.005906 | 0.005303 | 0.004125 | 0.009664 | 0.024253 | 0.03937 | 0.016897 |
| 35.1111111 | 0.048332 | 0.077966 | 0.017715 | 0.040828 | 0.028475 | 0.01186 | 0.028614 | 0.025626 | 0.026003 | 0.048809 | 0.060288 | 0.00791 | 0.0126 | 0.008268 | 0.005467 | 0.015879 | 0.039181 | 0.048041 | 0.027593 |
| 35.125 | 0.07001 | 0.094007 | 0.031221 | 0.037069 | 0.03863 | 0.014268 | 0.014744 | 0.02844 | 0.010696 | 0.04542 | 0.047755 | 0.009777 | 0.01849 | 0.010458 | 0.006161 | 0.016238 | 0.037482 | 0.043521 | 0.02864 |
| 35.1388889 | 0.063663 | 0.083077 | 0.045327 | 0.02279 | 0.040732 | 0.01024 | 0.004414 | 0.018153 | 0.004801 | 0.025172 | 0.032094 | 0.006963 | 0.018748 | 0.010318 | 0.005176 | 0.011811 | 0.023969 | 0.031226 | 0.018955 |
| 35.1527778 | 0.050435 | 0.056412 | 0.04803 | 0.013217 | 0.036528 | 0.006812 | 0.007514 | 0.008362 | 0.012102 | 0.015697 | 0.021959 | 0.005671 | 0.012889 | 0.006586 | 0.003547 | 0.006431 | 0.01111 | 0.017113 | 0.009877 |
| 35.1666667 | 0.035118 | 0.028789 | 0.040303 | 0.01066 | 0.022843 | 0.012003 | 0.021866 | 0.007297 | 0.017416 | 0.027055 | 0.012402 | 0.010842 | 0.01168 | 0.002766 | 0.003665 | 0.004923 | 0.007229 | 0.011647 | 0.011569 |
| 35.1805556 | 0.019489 | 0.016953 | 0.024958 | 0.018155 | 0.010151 | 0.018434 | 0.027844 | 0.022186 | 0.010277 | 0.038665 | 0.00817 | 0.017323 | 0.017514 | 0.002119 | 0.004469 | 0.007979 | 0.011861 | 0.018809 | 0.015847 |
| 35.1944444 | 0.020283 | 0.033309 | 0.011212 | 0.028113 | 0.01062 | 0.020641 | 0.016492 | 0.039706 | 0.008603 | 0.035416 | 0.009642 | 0.020074 | 0.019147 | 0.002466 | 0.003796 | 0.010651 | 0.015577 | 0.026606 | 0.013045 |
| 35.2083333 | 0.035558 | 0.055825 | 0.008335 | 0.024901 | 0.01478 | 0.019978 | 0.004074 | 0.040167 | 0.017035 | 0.024585 | 0.009143 | 0.018785 | 0.014721 | 0.002048 | 0.002473 | 0.010422 | 0.013289 | 0.028306 | 0.006741 |
| 35.2222222 | 0.049719 | 0.065465 | 0.014749 | 0.013192 | 0.01102 | 0.018261 | 0.00343 | 0.032332 | 0.026713 | 0.013671 | 0.006043 | 0.014045 | 0.009841 | 0.001552 | 0.002263 | 0.008486 | 0.008174 | 0.025758 | 0.002959 |
| 35.2361111 | 0.05714 | 0.070667 | 0.02298 | 0.005666 | 0.005844 | 0.016107 | 0.010993 | 0.031672 | 0.037447 | 0.006803 | 0.003927 | 0.008137 | 0.00657 | 0.001545 | 0.002844 | 0.006248 | 0.004847 | 0.02116 | 0.002516 |
| 35.25 | 0.057973 | 0.075605 | 0.029555 | 0.002958 | 0.007312 | 0.013111 | 0.017648 | 0.037834 | 0.043237 | 0.007182 | 0.003558 | 0.004106 | 0.005556 | 0.0023 | 0.002877 | 0.005013 | 0.004961 | 0.018067 | 0.003549 |
| 35.2638889 | 0.058577 | 0.078151 | 0.035867 | 0.003871 | 0.010121 | 0.010763 | 0.019256 | 0.047117 | 0.035009 | 0.007612 | 0.003265 | 0.003408 | 0.005747 | 0.003261 | 0.002512 | 0.005 | 0.006055 | 0.017073 | 0.004166 |
| 35.2777778 | 0.066951 | 0.075489 | 0.042373 | 0.009871 | 0.010662 | 0.012379 | 0.018434 | 0.056104 | 0.031435 | 0.005425 | 0.002313 | 0.00477 | 0.00441 | 0.003167 | 0.002293 | 0.004764 | 0.005644 | 0.015709 | 0.003407 |
| 35.2916667 | 0.078224 | 0.066235 | 0.045052 | 0.017313 | 0.012653 | 0.017756 | 0.017486 | 0.05597 | 0.046996 | 0.008072 | 0.002389 | 0.005656 | 0.002219 | 0.001921 | 0.001707 | 0.003153 | 0.004516 | 0.01373 | 0.001738 |
| 35.3055556 | 0.079041 | 0.052776 | 0.040789 | 0.019657 | 0.017649 | 0.023785 | 0.015878 | 0.042934 | 0.057204 | 0.012613 | 0.003194 | 0.0042 | 0.001805 | 0.000729 | 0.000781 | 0.00197 | 0.003676 | 0.011813 | 0.001529 |
| 35.3194444 | 0.060486 | 0.035804 | 0.031032 | 0.015112 | 0.020425 | 0.028109 | 0.013153 | 0.02397 | 0.046454 | 0.010133 | 0.002607 | 0.003332 | 0.002507 | 0.000568 | 0.00049 | 0.003138 | 0.002365 | 0.008696 | 0.004553 |
| 35.3333333 | 0.032869 | 0.017537 | 0.021171 | 0.008556 | 0.014704 | 0.02707 | 0.009493 | 0.01051 | 0.026159 | 0.006089 | 0.002289 | 0.006476 | 0.003503 | 0.001583 | 0.00096 | 0.004703 | 0.000942 | 0.004798 | 0.008081 |
| 35.3472222 | 0.013777 | 0.00704 | 0.016608 | 0.007031 | 0.005946 | 0.021193 | 0.005349 | 0.008879 | 0.017227 | 0.008819 | 0.004726 | 0.009336 | 0.004818 | 0.003069 | 0.00148 | 0.004181 | 0.000921 | 0.002887 | 0.008552 |
| 35.3611111 | 0.007363 | 0.004734 | 0.017633 | 0.009016 | 0.002347 | 0.018506 | 0.002489 | 0.011599 | 0.023924 | 0.009853 | 0.007841 | 0.008064 | 0.005419 | 0.003871 | 0.0013 | 0.002544 | 0.003307 | 0.002454 | 0.006759 |
| 35.375 | 0.007539 | 0.003237 | 0.019991 | 0.008813 | 0.002408 | 0.021873 | 0.001336 | 0.011442 | 0.027354 | 0.010074 | 0.009565 | 0.004389 | 0.004852 | 0.00319 | 0.001004 | 0.002033 | 0.008118 | 0.001877 | 0.004498 |
| 35.3888889 | 0.006551 | 0.002044 | 0.020129 | 0.008367 | 0.002509 | 0.026523 | 0.001555 | 0.012521 | 0.019761 | 0.019855 | 0.010236 | 0.001887 | 0.003558 | 0.001571 | 0.001357 | 0.001839 | 0.012587 | 0.002333 | 0.002409 |
| 35.4027778 | 0.005255 | 0.003156 | 0.018169 | 0.011028 | 0.001815 | 0.026979 | 0.002897 | 0.01839 | 0.010284 | 0.029176 | 0.011165 | 0.002269 | 0.001934 | 0.000514 | 0.001405 | 0.001197 | 0.013244 | 0.003521 | 0.001625 |
| 35.4166667 | 0.008717 | 0.008124 | 0.015465 | 0.015808 | 0.001143 | 0.019328 | 0.00278 | 0.025416 | 0.007116 | 0.026692 | 0.012913 | 0.003138 | 0.00114 | 0.000768 | 0.00101 | 0.00162 | 0.010078 | 0.003945 | 0.001834 |
| 35.4305556 | 0.011903 | 0.011298 | 0.011907 | 0.020516 | 0.001239 | 0.008794 | 0.001344 | 0.0293 | 0.007406 | 0.015229 | 0.013626 | 0.002371 | 0.002069 | 0.001669 | 0.001179 | 0.002589 | 0.005262 | 0.003179 | 0.002148 |
| 35.4444444 | 0.009881 | 0.018773 | 0.007067 | 0.021613 | 0.001162 | 0.003399 | 0.001117 | 0.028012 | 0.005081 | 0.00629 | 0.013103 | 0.000939 | 0.003256 | 0.001891 | 0.001387 | 0.002249 | 0.001656 | 0.001614 | 0.003568 |
| 35.4583333 | 0.011378 | 0.016992 | 0.0032 | 0.015302 | 0.001206 | 0.003685 | 0.001602 | 0.018559 | 0.002145 | 0.005347 | 0.01295 | 0.00054 | 0.003692 | 0.001164 | 0.001556 | 0.001775 | 0.000495 | 0.000497 | 0.006548 |
| 35.4722222 | 0.023149 | 0.010188 | 0.004369 | 0.006411 | 0.002523 | 0.008931 | 0.003025 | 0.009683 | 0.004731 | 0.006362 | 0.013456 | 0.001905 | 0.004118 | 0.001005 | 0.002476 | 0.002957 | 0.000307 | 0.000387 | 0.009498 |
| 35.4861111 | 0.035912 | 0.006096 | 0.010023 | 0.00373 | 0.004237 | 0.014423 | 0.005895 | 0.011255 | 0.015889 | 0.01177 | 0.015245 | 0.004917 | 0.005164 | 0.00218 | 0.003208 | 0.004251 | 0.000388 | 0.0013 | 0.010499 |
| 35.5 | 0.040012 | 0.008344 | 0.014676 | 0.007794 | 0.004697 | 0.014517 | 0.007033 | 0.012806 | 0.031386 | 0.021909 | 0.017183 | 0.007386 | 0.00636 | 0.003205 | 0.002841 | 0.004118 | 0.000856 | 0.002874 | 0.008529 |
| 35.5138889 | 0.036661 | 0.013911 | 0.016473 | 0.01335 | 0.003789 | 0.01201 | 0.005723 | 0.009972 | 0.045183 | 0.024732 | 0.016758 | 0.006731 | 0.005829 | 0.003094 | 0.001965 | 0.002705 | 0.001348 | 0.003742 | 0.004416 |
| 35.5277778 | 0.030859 | 0.021056 | 0.018824 | 0.01388 | 0.002998 | 0.012805 | 0.005172 | 0.009734 | 0.050941 | 0.016098 | 0.015339 | 0.003974 | 0.003212 | 0.002936 | 0.001494 | 0.001134 | 0.001837 | 0.003733 | 0.001087 |
| 35.5416667 | 0.022801 | 0.026224 | 0.02324 | 0.008679 | 0.003761 | 0.018487 | 0.006072 | 0.012072 | 0.044352 | 0.006311 | 0.016216 | 0.001838 | 0.001643 | 0.003689 | 0.001477 | 0.00109 | 0.002852 | 0.004144 | 0.000572 |
| 35.5555556 | 0.012511 | 0.025148 | 0.026195 | 0.004739 | 0.005742 | 0.025007 | 0.006671 | 0.014535 | 0.030014 | 0.003198 | 0.02056 | 0.001064 | 0.002998 | 0.004683 | 0.00187 | 0.002541 | 0.004649 | 0.005495 | 0.001804 |
| 35.5694444 | 0.005651 | 0.019288 | 0.025351 | 0.008712 | 0.007328 | 0.029265 | 0.005051 | 0.0171 | 0.014892 | 0.004554 | 0.026911 | 0.001236 | 0.004934 | 0.004975 | 0.001781 | 0.003938 | 0.008293 | 0.008132 | 0.003065 |
| 35.5833333 | 0.009991 | 0.013264 | 0.025445 | 0.021693 | 0.008251 | 0.03663 | 0.004465 | 0.021229 | 0.004783 | 0.012462 | 0.035408 | 0.003087 | 0.005864 | 0.004946 | 0.001322 | 0.00522 | 0.017487 | 0.014017 | 0.005453 |
| 35.5972222 | 0.029582 | 0.009011 | 0.031899 | 0.035601 | 0.00904 | 0.053677 | 0.00978 | 0.030068 | 0.006413 | 0.027339 | 0.047358 | 0.008714 | 0.007572 | 0.005732 | 0.001779 | 0.008756 | 0.031683 | 0.023408 | 0.011183 |
| 35.6111111 | 0.057873 | 0.006402 | 0.039638 | 0.042601 | 0.007812 | 0.06974 | 0.014076 | 0.051815 | 0.022593 | 0.037341 | 0.062745 | 0.017664 | 0.00978 | 0.008394 | 0.002238 | 0.01601 | 0.042895 | 0.030437 | 0.018932 |
| 35.625 | 0.085644 | 0.008627 | 0.043336 | 0.047163 | 0.004441 | 0.069694 | 0.010005 | 0.086639 | 0.048943 | 0.03659 | 0.075602 | 0.023272 | 0.00904 | 0.012469 | 0.002346 | 0.023752 | 0.049288 | 0.030637 | 0.024301 |
| 35.6388889 | 0.096691 | 0.014093 | 0.050485 | 0.045223 | 0.0049 | 0.059652 | 0.004886 | 0.099918 | 0.072616 | 0.029171 | 0.076289 | 0.019944 | 0.00565 | 0.014776 | 0.001944 | 0.025577 | 0.05285 | 0.026973 | 0.022912 |
| 35.6527778 | 0.074729 | 0.014355 | 0.056824 | 0.029647 | 0.01206 | 0.049913 | 0.005385 | 0.069301 | 0.077326 | 0.017894 | 0.067284 | 0.011134 | 0.002729 | 0.0134 | 0.000986 | 0.019702 | 0.050634 | 0.0233 | 0.013941 |
| 35.6666667 | 0.036658 | 0.009012 | 0.047293 | 0.012641 | 0.020616 | 0.040159 | 0.006111 | 0.028583 | 0.061189 | 0.009402 | 0.055626 | 0.007104 | 0.001012 | 0.010602 | 0.00128 | 0.011347 | 0.044909 | 0.021731 | 0.006434 |
| 35.6805556 | 0.020244 | 0.005175 | 0.027655 | 0.004761 | 0.02614 | 0.027419 | 0.003627 | 0.011901 | 0.037162 | 0.011089 | 0.041079 | 0.011786 | 0.001202 | 0.008833 | 0.003507 | 0.005098 | 0.039719 | 0.022962 | 0.007221 |
| 35.6944444 | 0.031716 | 0.005857 | 0.01137 | 0.005391 | 0.030256 | 0.017403 | 0.001508 | 0.019133 | 0.017758 | 0.01864 | 0.022534 | 0.016261 | 0.003392 | 0.007953 | 0.005326 | 0.001709 | 0.032803 | 0.023813 | 0.007982 |
| 35.7083333 | 0.043845 | 0.00966 | 0.003768 | 0.010678 | 0.035223 | 0.022506 | 0.000914 | 0.033528 | 0.013685 | 0.023208 | 0.012108 | 0.015196 | 0.004758 | 0.006883 | 0.004031 | 0.000484 | 0.021637 | 0.020773 | 0.004299 |
| 35.7222222 | 0.046298 | 0.01382 | 0.00494 | 0.018004 | 0.0402 | 0.034326 | 0.000804 | 0.045944 | 0.021261 | 0.022412 | 0.024067 | 0.01192 | 0.00356 | 0.004947 | 0.001702 | 0.000431 | 0.012368 | 0.014817 | 0.00225 |
| 35.7361111 | 0.039596 | 0.011627 | 0.009794 | 0.026711 | 0.043498 | 0.033024 | 0.001559 | 0.053781 | 0.022847 | 0.016837 | 0.047833 | 0.008924 | 0.001452 | 0.003043 | 0.001071 | 0.001305 | 0.016121 | 0.009327 | 0.002877 |
| 35.75 | 0.02635 | 0.009632 | 0.013554 | 0.03283 | 0.043127 | 0.020515 | 0.002126 | 0.048721 | 0.01602 | 0.009512 | 0.065219 | 0.00698 | 0.001849 | 0.002995 | 0.001623 | 0.00349 | 0.03075 | 0.00644 | 0.00266 |
| 35.7638889 | 0.027189 | 0.018859 | 0.014221 | 0.031444 | 0.03748 | 0.008997 | 0.00168 | 0.028049 | 0.010893 | 0.007737 | 0.073464 | 0.00602 | 0.005994 | 0.005 | 0.003126 | 0.00589 | 0.04283 | 0.005454 | 0.002758 |
| 35.7777778 | 0.045008 | 0.029038 | 0.009842 | 0.026678 | 0.028284 | 0.003164 | 0.002057 | 0.010312 | 0.010221 | 0.01234 | 0.073165 | 0.004607 | 0.010078 | 0.007863 | 0.003933 | 0.006332 | 0.046286 | 0.004559 | 0.005133 |
| 35.7916667 | 0.059351 | 0.030336 | 0.007348 | 0.023735 | 0.019972 | 0.004785 | 0.002858 | 0.009432 | 0.008077 | 0.016495 | 0.063074 | 0.002789 | 0.009648 | 0.010315 | 0.002893 | 0.004526 | 0.044463 | 0.002843 | 0.006442 |
| 35.8055556 | 0.054122 | 0.02303 | 0.014817 | 0.019645 | 0.015979 | 0.012699 | 0.002524 | 0.011932 | 0.007471 | 0.016037 | 0.049685 | 0.002188 | 0.00547 | 0.011569 | 0.001588 | 0.002348 | 0.039762 | 0.002243 | 0.005192 |
| 35.8194444 | 0.029692 | 0.013008 | 0.025605 | 0.014388 | 0.01596 | 0.022765 | 0.003053 | 0.008609 | 0.017224 | 0.011442 | 0.037444 | 0.002142 | 0.002021 | 0.01133 | 0.001237 | 0.002002 | 0.031633 | 0.005215 | 0.00503 |
| 35.8333333 | 0.016665 | 0.012889 | 0.032898 | 0.014201 | 0.017282 | 0.030284 | 0.005379 | 0.014884 | 0.031014 | 0.005826 | 0.023892 | 0.002793 | 0.001853 | 0.009384 | 0.001802 | 0.003366 | 0.022351 | 0.00978 | 0.007464 |
| 35.8472222 | 0.034585 | 0.025222 | 0.031579 | 0.025727 | 0.018699 | 0.032777 | 0.007136 | 0.04971 | 0.039966 | 0.004614 | 0.01308 | 0.00498 | 0.004068 | 0.006193 | 0.002753 | 0.004413 | 0.014673 | 0.013381 | 0.009899 |
| 35.8611111 | 0.060011 | 0.038794 | 0.020503 | 0.039678 | 0.019672 | 0.031836 | 0.00746 | 0.079997 | 0.041523 | 0.009946 | 0.013777 | 0.006489 | 0.006536 | 0.003073 | 0.003772 | 0.004178 | 0.007626 | 0.016255 | 0.00892 |
| 35.875 | 0.076233 | 0.045326 | 0.010816 | 0.037695 | 0.019579 | 0.027476 | 0.006305 | 0.072792 | 0.034333 | 0.016654 | 0.021237 | 0.006047 | 0.007403 | 0.001609 | 0.004313 | 0.003097 | 0.003353 | 0.018449 | 0.004808 |
| 35.8888889 | 0.08086 | 0.045704 | 0.012897 | 0.030326 | 0.018962 | 0.01973 | 0.003693 | 0.066464 | 0.019922 | 0.020808 | 0.020628 | 0.005074 | 0.006293 | 0.00188 | 0.003456 | 0.002556 | 0.004151 | 0.018738 | 0.001873 |
| 35.9027778 | 0.071353 | 0.041074 | 0.023722 | 0.036556 | 0.018411 | 0.012948 | 0.002148 | 0.079783 | 0.011106 | 0.021404 | 0.013248 | 0.004549 | 0.004371 | 0.001809 | 0.001654 | 0.003919 | 0.00419 | 0.017363 | 0.002431 |
| 35.9166667 | 0.053054 | 0.029181 | 0.033638 | 0.048005 | 0.01685 | 0.007978 | 0.002736 | 0.085213 | 0.019686 | 0.017997 | 0.014944 | 0.004167 | 0.002865 | 0.001401 | 0.001069 | 0.006972 | 0.004329 | 0.015639 | 0.005744 |
| 35.9305556 | 0.032656 | 0.016463 | 0.039347 | 0.053151 | 0.012547 | 0.005716 | 0.002961 | 0.073641 | 0.035115 | 0.013421 | 0.028845 | 0.00311 | 0.002147 | 0.00266 | 0.002657 | 0.009887 | 0.011841 | 0.013438 | 0.009217 |
| 35.9444444 | 0.019343 | 0.014966 | 0.042876 | 0.05407 | 0.00687 | 0.0099 | 0.002107 | 0.061122 | 0.044751 | 0.010697 | 0.042577 | 0.001543 | 0.001505 | 0.004945 | 0.004692 | 0.010369 | 0.023072 | 0.010399 | 0.010348 |
| 35.9583333 | 0.016303 | 0.025375 | 0.045427 | 0.053969 | 0.005284 | 0.016191 | 0.00112 | 0.059969 | 0.046697 | 0.008769 | 0.047503 | 0.00078 | 0.001571 | 0.006005 | 0.005064 | 0.007322 | 0.031036 | 0.007368 | 0.008521 |
| 35.9722222 | 0.018509 | 0.037709 | 0.044523 | 0.04999 | 0.009972 | 0.019555 | 0.001188 | 0.069504 | 0.042979 | 0.005992 | 0.043745 | 0.001289 | 0.003456 | 0.00477 | 0.004295 | 0.003851 | 0.033472 | 0.004639 | 0.004888 |
| 35.9861111 | 0.019029 | 0.038312 | 0.039555 | 0.038042 | 0.017646 | 0.022402 | 0.00317 | 0.083823 | 0.036941 | 0.006335 | 0.03455 | 0.002597 | 0.004602 | 0.002609 | 0.004033 | 0.003843 | 0.031399 | 0.002018 | 0.001981 |

| | | | | | | | | | | | | | | | | | | | |
|---|---|---|---|---|---|---|---|---|---|---|---|---|---|---|---|---|---|---|---|
| 36 | 0.013439 | 0.024017 | 0.032068 | 0.020795 | 0.024553 | 0.026873 | 0.005227 | 0.097112 | 0.030371 | 0.012577 | 0.024314 | 0.004233 | 0.003151 | 0.002064 | 0.004042 | 0.005583 | 0.028111 | 0.000581 | 0.001188 |
| 36.0138889 | 0.011135 | 0.016029 | 0.024787 | 0.008633 | 0.026453 | 0.029402 | 0.006704 | 0.108982 | 0.022978 | 0.019278 | 0.017238 | 0.005683 | 0.001278 | 0.002989 | 0.003682 | 0.00643 | 0.02461 | 0.000795 | 0.001297 |
| 36.0277778 | 0.018607 | 0.028078 | 0.020414 | 0.0085 | 0.021446 | 0.025693 | 0.009587 | 0.101845 | 0.013842 | 0.020703 | 0.014051 | 0.006529 | 0.000918 | 0.003329 | 0.00323 | 0.006341 | 0.020109 | 0.001278 | 0.001875 |
| 36.0416667 | 0.025936 | 0.043934 | 0.016445 | 0.014481 | 0.012228 | 0.016246 | 0.014107 | 0.059988 | 0.010783 | 0.015654 | 0.011603 | 0.006863 | 0.001432 | 0.00254 | 0.002656 | 0.005289 | 0.014985 | 0.001684 | 0.004104 |
| 36.0555556 | 0.023728 | 0.048796 | 0.01109 | 0.020483 | 0.005856 | 0.007865 | 0.019025 | 0.02903 | 0.020521 | 0.008057 | 0.007056 | 0.007371 | 0.001778 | 0.001499 | 0.002125 | 0.00391 | 0.01078 | 0.003695 | 0.007152 |
| 36.0694444 | 0.018029 | 0.039038 | 0.006957 | 0.026595 | 0.005378 | 0.004998 | 0.023223 | 0.051743 | 0.033754 | 0.007378 | 0.004676 | 0.00757 | 0.002122 | 0.000943 | 0.002278 | 0.002561 | 0.011057 | 0.008676 | 0.007827 |
| 36.0833333 | 0.030073 | 0.025838 | 0.012191 | 0.026083 | 0.007111 | 0.006546 | 0.024167 | 0.08079 | 0.037023 | 0.021759 | 0.015775 | 0.006227 | 0.003939 | 0.002355 | 0.002867 | 0.003344 | 0.020356 | 0.018859 | 0.009043 |
| 36.0972222 | 0.063553 | 0.03195 | 0.031691 | 0.019274 | 0.01327 | 0.013477 | 0.018818 | 0.063606 | 0.024696 | 0.047708 | 0.038923 | 0.00761 | 0.010206 | 0.005991 | 0.003645 | 0.009603 | 0.033808 | 0.033083 | 0.019598 |
| 36.1111111 | 0.08844 | 0.050141 | 0.049256 | 0.016127 | 0.021217 | 0.020724 | 0.009895 | 0.045892 | 0.010076 | 0.068288 | 0.051911 | 0.013242 | 0.019648 | 0.007487 | 0.005177 | 0.01646 | 0.035795 | 0.041431 | 0.032542 |
| 36.125 | 0.086708 | 0.058025 | 0.04632 | 0.013392 | 0.022475 | 0.021666 | 0.005293 | 0.070317 | 0.007443 | 0.068453 | 0.042717 | 0.015274 | 0.024836 | 0.004957 | 0.005761 | 0.016105 | 0.022932 | 0.038146 | 0.032575 |
| 36.1388889 | 0.065913 | 0.051686 | 0.029157 | 0.010867 | 0.020211 | 0.017397 | 0.004906 | 0.088641 | 0.009722 | 0.049652 | 0.023513 | 0.010725 | 0.020464 | 0.002246 | 0.004204 | 0.00976 | 0.009141 | 0.028392 | 0.020262 |
| 36.1527778 | 0.033904 | 0.031098 | 0.012607 | 0.014036 | 0.019812 | 0.009639 | 0.004456 | 0.070678 | 0.010675 | 0.025228 | 0.011208 | 0.006171 | 0.012199 | 0.002176 | 0.004001 | 0.004607 | 0.004512 | 0.017527 | 0.010618 |
| 36.1666667 | 0.013028 | 0.011761 | 0.004089 | 0.018279 | 0.019577 | 0.00347 | 0.005758 | 0.039661 | 0.010809 | 0.012722 | 0.013854 | 0.008493 | 0.011671 | 0.004048 | 0.005894 | 0.005847 | 0.006122 | 0.008677 | 0.014509 |
| 36.1805556 | 0.016529 | 0.009102 | 0.004282 | 0.019542 | 0.015053 | 0.002448 | 0.007488 | 0.017719 | 0.008774 | 0.017594 | 0.02292 | 0.014118 | 0.017124 | 0.005115 | 0.005464 | 0.011439 | 0.005868 | 0.005976 | 0.023626 |
| 36.1944444 | 0.023976 | 0.012898 | 0.007449 | 0.017012 | 0.007351 | 0.005558 | 0.006017 | 0.016759 | 0.005023 | 0.025171 | 0.023209 | 0.015289 | 0.018329 | 0.004139 | 0.002428 | 0.014695 | 0.002773 | 0.008702 | 0.026275 |
| 36.2083333 | 0.028352 | 0.0176 | 0.009145 | 0.010783 | 0.002707 | 0.01213 | 0.003162 | 0.03071 | 0.003969 | 0.026402 | 0.013732 | 0.012698 | 0.015043 | 0.002432 | 0.000928 | 0.012864 | 0.000923 | 0.010583 | 0.022247 |
| 36.2222222 | 0.039423 | 0.023774 | 0.009861 | 0.00496 | 0.001594 | 0.016381 | 0.001882 | 0.043579 | 0.009059 | 0.024162 | 0.007803 | 0.010996 | 0.012032 | 0.001065 | 0.001179 | 0.008328 | 0.00107 | 0.010386 | 0.016621 |
| 36.2361111 | 0.0548 | 0.029246 | 0.01124 | 0.003904 | 0.001158 | 0.017604 | 0.003131 | 0.050598 | 0.019373 | 0.020858 | 0.010284 | 0.012041 | 0.011441 | 0.000469 | 0.001283 | 0.00506 | 0.001883 | 0.01146 | 0.012969 |
| 36.25 | 0.065934 | 0.035202 | 0.013114 | 0.007064 | 0.001081 | 0.01939 | 0.007266 | 0.055094 | 0.034583 | 0.017671 | 0.01029 | 0.014979 | 0.013277 | 0.000881 | 0.001688 | 0.005752 | 0.002484 | 0.015442 | 0.01284 |
| 36.2638889 | 0.070804 | 0.042498 | 0.015128 | 0.009149 | 0.001851 | 0.022528 | 0.013025 | 0.056918 | 0.051758 | 0.013837 | 0.005727 | 0.017861 | 0.015301 | 0.001571 | 0.003 | 0.008775 | 0.002666 | 0.021572 | 0.015199 |
| 36.2777778 | 0.073008 | 0.049348 | 0.017988 | 0.006622 | 0.003435 | 0.028285 | 0.017158 | 0.053197 | 0.064271 | 0.009137 | 0.003246 | 0.018142 | 0.014405 | 0.001704 | 0.004107 | 0.009953 | 0.002566 | 0.025307 | 0.016263 |
| 36.2916667 | 0.072178 | 0.052378 | 0.021412 | 0.003038 | 0.005071 | 0.030772 | 0.018061 | 0.046252 | 0.069788 | 0.005427 | 0.002512 | 0.014324 | 0.010903 | 0.00182 | 0.003531 | 0.007527 | 0.002644 | 0.022763 | 0.013967 |
| 36.3055556 | 0.068039 | 0.052003 | 0.026329 | 0.002144 | 0.005987 | 0.024847 | 0.016254 | 0.041408 | 0.068806 | 0.00442 | 0.002271 | 0.007689 | 0.007706 | 0.002353 | 0.00223 | 0.003717 | 0.003148 | 0.016839 | 0.008342 |
| 36.3194444 | 0.062681 | 0.051075 | 0.031609 | 0.001638 | 0.006044 | 0.014104 | 0.011382 | 0.036761 | 0.05815 | 0.005056 | 0.002377 | 0.003669 | 0.004736 | 0.00219 | 0.001843 | 0.002417 | 0.003392 | 0.011562 | 0.005508 |
| 36.3333333 | 0.056712 | 0.04937 | 0.032937 | 0.001443 | 0.007299 | 0.007181 | 0.005807 | 0.027703 | 0.03867 | 0.003961 | 0.00258 | 0.005924 | 0.003881 | 0.001653 | 0.001634 | 0.004054 | 0.002414 | 0.007096 | 0.011714 |
| 36.3472222 | 0.054823 | 0.046942 | 0.032763 | 0.005876 | 0.013564 | 0.012876 | 0.007296 | 0.016764 | 0.022222 | 0.002094 | 0.00306 | 0.008787 | 0.006735 | 0.002035 | 0.001652 | 0.004544 | 0.001272 | 0.003724 | 0.019998 |
| 36.3611111 | 0.05669 | 0.044188 | 0.033341 | 0.013891 | 0.022496 | 0.026773 | 0.016339 | 0.009231 | 0.016023 | 0.001669 | 0.003064 | 0.007014 | 0.007126 | 0.002131 | 0.00231 | 0.003576 | 0.001026 | 0.003014 | 0.020766 |
| 36.375 | 0.05222 | 0.035981 | 0.028552 | 0.016042 | 0.024547 | 0.033441 | 0.02108 | 0.005523 | 0.012558 | 0.003642 | 0.004302 | 0.00556 | 0.005978 | 0.002157 | 0.001862 | 0.005445 | 0.0009 | 0.005185 | 0.012859 |
| 36.3888889 | 0.036443 | 0.020693 | 0.017787 | 0.010216 | 0.016119 | 0.028676 | 0.015952 | 0.004669 | 0.006348 | 0.0112 | 0.007242 | 0.007511 | 0.007918 | 0.003365 | 0.001273 | 0.00954 | 0.001451 | 0.008719 | 0.005774 |
| 36.4027778 | 0.019176 | 0.012165 | 0.010569 | 0.006329 | 0.007061 | 0.023212 | 0.009463 | 0.004748 | 0.004369 | 0.02189 | 0.008595 | 0.007407 | 0.007454 | 0.004208 | 0.003327 | 0.011927 | 0.003397 | 0.012169 | 0.00753 |
| 36.4166667 | 0.011857 | 0.015133 | 0.010067 | 0.005786 | 0.005946 | 0.022051 | 0.008039 | 0.003147 | 0.008193 | 0.029888 | 0.009268 | 0.003973 | 0.003691 | 0.004076 | 0.006854 | 0.011734 | 0.005732 | 0.013796 | 0.01097 |
| 36.4305556 | 0.009956 | 0.013721 | 0.008672 | 0.00389 | 0.007227 | 0.019203 | 0.006956 | 0.002488 | 0.007832 | 0.033098 | 0.012538 | 0.001536 | 0.001497 | 0.003545 | 0.007828 | 0.009344 | 0.006217 | 0.011612 | 0.00959 |
| 36.4444444 | 0.009163 | 0.006473 | 0.004782 | 0.00255 | 0.006326 | 0.013071 | 0.005465 | 0.005437 | 0.003492 | 0.030413 | 0.01673 | 0.001269 | 0.001537 | 0.002677 | 0.005018 | 0.005454 | 0.004107 | 0.00656 | 0.006688 |
| 36.4583333 | 0.011189 | 0.003894 | 0.003182 | 0.002305 | 0.006416 | 0.009584 | 0.007989 | 0.008166 | 0.002046 | 0.021477 | 0.017619 | 0.001528 | 0.003735 | 0.001961 | 0.001721 | 0.002095 | 0.00173 | 0.002645 | 0.006154 |
| 36.4722222 | 0.011468 | 0.005069 | 0.002862 | 0.001758 | 0.006981 | 0.013157 | 0.009331 | 0.006828 | 0.003189 | 0.010907 | 0.01424 | 0.003682 | 0.007444 | 0.001677 | 0.000573 | 0.000843 | 0.000884 | 0.001757 | 0.008141 |
| 36.4861111 | 0.00969 | 0.006421 | 0.003837 | 0.002153 | 0.005561 | 0.023106 | 0.007283 | 0.007367 | 0.008064 | 0.005102 | 0.009266 | 0.008798 | 0.009887 | 0.00133 | 0.001135 | 0.001494 | 0.001478 | 0.001685 | 0.010685 |
| 36.5 | 0.009624 | 0.008086 | 0.009558 | 0.003487 | 0.003477 | 0.032578 | 0.006945 | 0.016204 | 0.016549 | 0.004919 | 0.005761 | 0.012658 | 0.00948 | 0.000724 | 0.002151 | 0.00338 | 0.003179 | 0.00183 | 0.011588 |
| 36.5138889 | 0.011681 | 0.008513 | 0.019164 | 0.005052 | 0.002151 | 0.033068 | 0.007565 | 0.027924 | 0.024599 | 0.010515 | 0.005277 | 0.012285 | 0.007413 | 0.000702 | 0.00281 | 0.006441 | 0.005628 | 0.003416 | 0.009922 |
| 36.5277778 | 0.015609 | 0.005699 | 0.02665 | 0.006564 | 0.001373 | 0.024047 | 0.005854 | 0.036019 | 0.029992 | 0.02012 | 0.008332 | 0.009639 | 0.00599 | 0.001438 | 0.002749 | 0.009631 | 0.007418 | 0.005096 | 0.007264 |
| 36.5416667 | 0.01938 | 0.002281 | 0.025524 | 0.007482 | 0.001069 | 0.013353 | 0.00348 | 0.039541 | 0.031009 | 0.025747 | 0.01262 | 0.007151 | 0.00563 | 0.001482 | 0.001875 | 0.010357 | 0.006979 | 0.005123 | 0.005311 |
| 36.5555556 | 0.019497 | 0.001384 | 0.017788 | 0.007826 | 0.001029 | 0.013224 | 0.004078 | 0.037993 | 0.027595 | 0.022681 | 0.013701 | 0.005656 | 0.005161 | 0.000785 | 0.001154 | 0.008458 | 0.005415 | 0.003711 | 0.004084 |
| 36.5694444 | 0.01716 | 0.001791 | 0.010587 | 0.008357 | 0.001414 | 0.02111 | 0.007645 | 0.034195 | 0.022329 | 0.01557 | 0.013699 | 0.006419 | 0.00555 | 0.000514 | 0.000928 | 0.007751 | 0.007375 | 0.003612 | 0.0042 |
| 36.5833333 | 0.022385 | 0.001968 | 0.015858 | 0.009546 | 0.002272 | 0.018646 | 0.013621 | 0.038156 | 0.01723 | 0.014009 | 0.020064 | 0.010575 | 0.008001 | 0.000827 | 0.000415 | 0.011333 | 0.017684 | 0.008226 | 0.006932 |
| 36.5972222 | 0.04276 | 0.003073 | 0.041851 | 0.010219 | 0.003039 | 0.01117 | 0.020226 | 0.057906 | 0.014267 | 0.022218 | 0.036517 | 0.017263 | 0.011013 | 0.001726 | 0.000306 | 0.018061 | 0.035688 | 0.016752 | 0.011639 |
| 36.6111111 | 0.070851 | 0.007987 | 0.072013 | 0.011547 | 0.006375 | 0.019309 | 0.020168 | 0.09487 | 0.019769 | 0.031446 | 0.056318 | 0.024735 | 0.012779 | 0.003416 | 0.000579 | 0.023912 | 0.051394 | 0.024047 | 0.016441 |
| 36.625 | 0.08966 | 0.017356 | 0.084986 | 0.017549 | 0.014918 | 0.03681 | 0.013368 | 0.148264 | 0.038969 | 0.031663 | 0.068569 | 0.029665 | 0.013098 | 0.00836 | 0.00036 | 0.02553 | 0.058063 | 0.026793 | 0.01999 |
| 36.6388889 | 0.086024 | 0.025345 | 0.077907 | 0.025499 | 0.025246 | 0.043834 | 0.010522 | 0.186422 | 0.065216 | 0.025642 | 0.075171 | 0.02705 | 0.012042 | 0.016125 | 0.000469 | 0.022648 | 0.058841 | 0.026096 | 0.020032 |
| 36.6527778 | 0.060253 | 0.025729 | 0.060808 | 0.027882 | 0.033934 | 0.039064 | 0.012541 | 0.166683 | 0.07962 | 0.019369 | 0.082021 | 0.016715 | 0.008479 | 0.020076 | 0.00129 | 0.016565 | 0.056077 | 0.024243 | 0.01477 |
| 36.6666667 | 0.031044 | 0.020586 | 0.044705 | 0.023903 | 0.039219 | 0.031248 | 0.011119 | 0.107675 | 0.07275 | 0.011751 | 0.086939 | 0.008029 | 0.004041 | 0.018096 | 0.001693 | 0.008882 | 0.054503 | 0.023757 | 0.007744 |
| 36.6805556 | 0.021854 | 0.016486 | 0.03158 | 0.01782 | 0.040218 | 0.0229 | 0.005797 | 0.055978 | 0.059728 | 0.006761 | 0.085252 | 0.009008 | 0.002702 | 0.014111 | 0.001145 | 0.003395 | 0.056779 | 0.026299 | 0.005592 |
| 36.6944444 | 0.033361 | 0.015761 | 0.018884 | 0.012083 | 0.040975 | 0.014279 | 0.002474 | 0.027002 | 0.054148 | 0.011142 | 0.073141 | 0.015122 | 0.003048 | 0.010548 | 0.000489 | 0.002499 | 0.056718 | 0.029576 | 0.008369 |
| 36.7083333 | 0.046081 | 0.015581 | 0.009792 | 0.006983 | 0.044462 | 0.010522 | 0.00226 | 0.012264 | 0.051424 | 0.019663 | 0.049606 | 0.018148 | 0.002345 | 0.006824 | 0.000506 | 0.003678 | 0.049305 | 0.030025 | 0.009876 |
| 36.7222222 | 0.049868 | 0.013306 | 0.010595 | 0.00288 | 0.047379 | 0.017736 | 0.002414 | 0.007885 | 0.041752 | 0.026783 | 0.028522 | 0.015344 | 0.001825 | 0.003196 | 0.000969 | 0.004181 | 0.034982 | 0.026948 | 0.008039 |
| 36.7361111 | 0.049983 | 0.008667 | 0.017635 | 0.000902 | 0.046102 | 0.03348 | 0.002761 | 0.019548 | 0.023784 | 0.028496 | 0.034621 | 0.009658 | 0.001826 | 0.001467 | 0.001187 | 0.003348 | 0.018417 | 0.020897 | 0.004601 |
| 36.75 | 0.051775 | 0.007565 | 0.024601 | 0.00069 | 0.040024 | 0.047643 | 0.003006 | 0.041332 | 0.014881 | 0.018548 | 0.066072 | 0.005705 | 0.001788 | 0.001137 | 0.000848 | 0.003266 | 0.012186 | 0.014157 | 0.001955 |
| 36.7638889 | 0.051522 | 0.013821 | 0.035487 | 0.002577 | 0.032087 | 0.050233 | 0.004321 | 0.059199 | 0.027281 | 0.009341 | 0.094995 | 0.004428 | 0.003449 | 0.001396 | 0.000411 | 0.005858 | 0.023876 | 0.009632 | 0.001927 |
| 36.7777778 | 0.037422 | 0.01767 | 0.044089 | 0.009047 | 0.027386 | 0.039828 | 0.008487 | 0.062476 | 0.045367 | 0.015749 | 0.104546 | 0.003221 | 0.006096 | 0.003242 | 0.000377 | 0.008414 | 0.04342 | 0.007041 | 0.004855 |
| 36.7916667 | 0.025491 | 0.013924 | 0.036046 | 0.018833 | 0.025607 | 0.023268 | 0.012188 | 0.053692 | 0.053268 | 0.026948 | 0.096999 | 0.002562 | 0.005735 | 0.006889 | 0.000754 | 0.008553 | 0.060828 | 0.004591 | 0.009904 |
| 36.8055556 | 0.042516 | 0.009434 | 0.018358 | 0.028688 | 0.0235 | 0.009317 | 0.012426 | 0.045761 | 0.04859 | 0.029923 | 0.081632 | 0.005121 | 0.003157 | 0.011185 | 0.001674 | 0.006703 | 0.071593 | 0.002541 | 0.013185 |
| 36.8194444 | 0.062502 | 0.006125 | 0.014226 | 0.036019 | 0.02209 | 0.005922 | 0.00967 | 0.042829 | 0.032965 | 0.025835 | 0.063005 | 0.007492 | 0.001675 | 0.01407 | 0.002838 | 0.004113 | 0.071232 | 0.00358 | 0.012702 |
| 36.8333333 | 0.062643 | 0.003987 | 0.032821 | 0.037103 | 0.022753 | 0.015291 | 0.005207 | 0.036598 | 0.016264 | 0.019337 | 0.041454 | 0.005781 | 0.001669 | 0.014146 | 0.003213 | 0.004202 | 0.060274 | 0.008502 | 0.011086 |
| 36.8472222 | 0.054584 | 0.007552 | 0.051154 | 0.031255 | 0.023723 | 0.029339 | 0.002704 | 0.024404 | 0.012933 | 0.011354 | 0.02188 | 0.003319 | 0.002698 | 0.011832 | 0.002461 | 0.008514 | 0.044698 | 0.01448 | 0.010881 |
| 36.8611111 | 0.036064 | 0.013968 | 0.047537 | 0.021893 | 0.022971 | 0.036425 | 0.005084 | 0.018703 | 0.024802 | 0.005782 | 0.009254 | 0.004576 | 0.002955 | 0.007726 | 0.00131 | 0.012702 | 0.030745 | 0.018888 | 0.011897 |
| 36.875 | 0.018201 | 0.017108 | 0.031524 | 0.011883 | 0.020197 | 0.035942 | 0.009027 | 0.02587 | 0.041561 | 0.009311 | 0.003209 | 0.008579 | 0.001607 | 0.004592 | 0.000534 | 0.014005 | 0.02126 | 0.020945 | 0.013 |
| 36.8888889 | 0.018167 | 0.016823 | 0.017805 | 0.005471 | 0.016642 | 0.033918 | 0.010532 | 0.037158 | 0.051176 | 0.02093 | 0.001207 | 0.012982 | 0.00085 | 0.006272 | 0.000298 | 0.014636 | 0.015857 | 0.021123 | 0.014297 |
| 36.9027778 | 0.02611 | 0.013962 | 0.010027 | 0.003867 | 0.013606 | 0.030754 | 0.008948 | 0.046377 | 0.043286 | 0.032759 | 0.00149 | 0.016962 | 0.002109 | 0.010131 | 0.000346 | 0.016446 | 0.013025 | 0.020774 | 0.014907 |
| 36.9166667 | 0.031044 | 0.00902 | 0.014014 | 0.005033 | 0.011079 | 0.024957 | 0.004999 | 0.059399 | 0.024865 | 0.037163 | 0.00369 | 0.01949 | 0.004592 | 0.010719 | 0.000704 | 0.018291 | 0.012065 | 0.020543 | 0.013521 |
| 36.9305556 | 0.031768 | 0.009033 | 0.031658 | 0.010458 | 0.007665 | 0.017393 | 0.002028 | 0.075117 | 0.021129 | 0.03457 | 0.006068 | 0.019266 | 0.005983 | 0.007265 | 0.001434 | 0.018432 | 0.012303 | 0.020274 | 0.011101 |
| 36.9444444 | 0.04075 | 0.019645 | 0.047995 | 0.022821 | 0.004986 | 0.0104 | 0.002249 | 0.080307 | 0.035361 | 0.028121 | 0.006169 | 0.014679 | 0.004985 | 0.003374 | 0.001965 | 0.015957 | 0.013023 | 0.020399 | 0.008544 |
| 36.9583333 | 0.058647 | 0.031364 | 0.044266 | 0.032916 | 0.007691 | 0.008984 | 0.003196 | 0.071802 | 0.047045 | 0.01722 | 0.00381 | 0.007634 | 0.003032 | 0.002713 | 0.001873 | 0.009866 | 0.013981 | 0.021367 | 0.005318 |
| 36.9722222 | 0.065826 | 0.032451 | 0.026692 | 0.042597 | 0.01472 | 0.014797 | 0.004507 | 0.057171 | 0.050066 | 0.009296 | 0.00246 | 0.004519 | 0.002662 | 0.003922 | 0.001172 | 0.004506 | 0.015206 | 0.021782 | 0.00438 |
| 36.9861111 | 0.054247 | 0.023051 | 0.017235 | 0.04713 | 0.02117 | 0.021418 | 0.007545 | 0.048896 | 0.051431 | 0.014099 | 0.004824 | 0.006287 | 0.00416 | 0.004085 | 0.000764 | 0.004444 | 0.017004 | 0.019449 | 0.008106 |
| 37 | 0.03067 | 0.011244 | 0.013341 | 0.039757 | 0.025218 | 0.022351 | 0.010647 | 0.052601 | 0.052924 | 0.025344 | 0.007507 | 0.007925 | 0.005498 | 0.00324 | 0.001373 | 0.005869 | 0.018979 | 0.014601 | 0.011142 |
| 37.0138889 | 0.013701 | 0.00429 | 0.007293 | 0.027583 | 0.027589 | 0.016315 | 0.011264 | 0.059076 | 0.048967 | 0.031956 | 0.007455 | 0.007952 | 0.006365 | 0.00263 | 0.001826 | 0.00604 | 0.02031 | 0.009344 | 0.008969 |
| 37.0277778 | 0.015332 | 0.005772 | 0.005858 | 0.020946 | 0.028611 | 0.008762 | 0.008387 | 0.064196 | 0.038349 | 0.03033 | 0.005584 | 0.007548 | 0.007416 | 0.002492 | 0.001502 | 0.006086 | 0.020631 | 0.005316 | 0.005296 |
| 37.0416667 | 0.026644 | 0.013599 | 0.008095 | 0.018155 | 0.026223 | 0.006485 | 0.004136 | 0.070885 | 0.026609 | 0.022979 | 0.003451 | 0.007459 | 0.008748 | 0.002173 | 0.001011 | 0.00641 | 0.018769 | 0.002787 | 0.005547 |
| 37.0555556 | 0.034854 | 0.022545 | 0.008455 | 0.015674 | 0.019219 | 0.007953 | 0.002231 | 0.075672 | 0.016252 | 0.014171 | 0.00186 | 0.008215 | 0.009954 | 0.001443 | 0.000681 | 0.006815 | 0.014094 | 0.00139 | 0.007573 |
| 37.0694444 | 0.037032 | 0.028608 | 0.007279 | 0.013539 | 0.013791 | 0.007837 | 0.003226 | 0.074241 | 0.00773 | 0.009593 | 0.0041 | 0.007868 | 0.009593 | 0.001973 | 0.000917 | 0.005394 | 0.011209 | 0.001764 | 0.008564 |
| 37.0833333 | 0.028763 | 0.02383 | 0.00495 | 0.012612 | 0.019465 | 0.005149 | 0.004299 | 0.073705 | 0.004357 | 0.02231 | 0.017019 | 0.007694 | 0.00793 | 0.006523 | 0.001713 | 0.005492 | 0.019644 | 0.006979 | 0.013743 |
| 37.0972222 | 0.022244 | 0.015796 | 0.004163 | 0.017916 | 0.032984 | 0.003293 | 0.00852 | 0.079818 | 0.007426 | 0.049566 | 0.035925 | 0.014667 | 0.010518 | 0.012748 | 0.002067 | 0.013205 | 0.037595 | 0.017196 | 0.025449 |
| 37.1111111 | 0.039581 | 0.022993 | 0.009779 | 0.028275 | 0.039879 | 0.003428 | 0.017187 | 0.080412 | 0.011883 | 0.062498 | 0.041237 | 0.025757 | 0.017517 | 0.013475 | 0.001857 | 0.02143 | 0.044764 | 0.022569 | 0.033786 |
| 37.125 | 0.057219 | 0.034988 | 0.017349 | 0.028561 | 0.037491 | 0.006012 | 0.022393 | 0.070289 | 0.013387 | 0.054819 | 0.027063 | 0.028899 | 0.018787 | 0.007802 | 0.002698 | 0.020269 | 0.03223 | 0.016802 | 0.028981 |
| 37.1388889 | 0.049839 | 0.037055 | 0.021058 | 0.01679 | 0.034502 | 0.013092 | 0.023269 | 0.063693 | 0.011746 | 0.042875 | 0.011293 | 0.020895 | 0.012415 | 0.003343 | 0.004322 | 0.011611 | 0.014548 | 0.007572 | 0.015459 |
| 37.1527778 | 0.03048 | 0.033383 | 0.022489 | 0.011991 | 0.031694 | 0.01719 | 0.020937 | 0.065123 | 0.00766 | 0.027508 | 0.007975 | 0.010784 | 0.008341 | 0.003789 | 0.004874 | 0.006866 | 0.005264 | 0.003893 | 0.00626 |
| 37.1666667 | 0.014453 | 0.028614 | 0.023267 | 0.017616 | 0.023911 | 0.011922 | 0.012681 | 0.063865 | 0.006046 | 0.013462 | 0.013249 | 0.009295 | 0.010952 | 0.005244 | 0.005908 | 0.013453 | 0.00548 | 0.005969 | 0.008107 |
| 37.1805556 | 0.006807 | 0.023435 | 0.022208 | 0.016607 | 0.012635 | 0.006657 | 0.004288 | 0.055085 | 0.010097 | 0.012319 | 0.017939 | 0.015218 | 0.015918 | 0.005046 | 0.007323 | 0.022723 | 0.009703 | 0.007972 | 0.016414 |

| | | | | | | | | | | | | | | | | | | | |
|---|---|---|---|---|---|---|---|---|---|---|---|---|---|---|---|---|---|---|---|
| 37.1944444 | 0.007159 | 0.018169 | 0.01707 | 0.008993 | 0.007443 | 0.009136 | 0.001424 | 0.043308 | 0.013676 | 0.020522 | 0.018843 | 0.019105 | 0.019847 | 0.003849 | 0.005498 | 0.025387 | 0.012282 | 0.007337 | 0.023912 |
| 37.2083333 | 0.011879 | 0.01291 | 0.009153 | 0.007254 | 0.011601 | 0.016405 | 0.002267 | 0.030056 | 0.011836 | 0.02634 | 0.016691 | 0.018551 | 0.020336 | 0.002714 | 0.002672 | 0.021728 | 0.011454 | 0.005915 | 0.025871 |
| 37.2222222 | 0.019667 | 0.00921 | 0.007377 | 0.007021 | 0.018813 | 0.027687 | 0.007876 | 0.019183 | 0.008593 | 0.02305 | 0.012115 | 0.016844 | 0.01715 | 0.002755 | 0.002398 | 0.014773 | 0.009449 | 0.005361 | 0.022788 |
| 37.2361111 | 0.031449 | 0.013133 | 0.014663 | 0.005837 | 0.024472 | 0.040308 | 0.018377 | 0.02533 | 0.011962 | 0.013751 | 0.006999 | 0.01516 | 0.013278 | 0.004527 | 0.002668 | 0.00857 | 0.0083 | 0.006281 | 0.019786 |
| 37.25 | 0.045416 | 0.025382 | 0.020858 | 0.009006 | 0.024269 | 0.044856 | 0.027149 | 0.049145 | 0.020088 | 0.006331 | 0.003293 | 0.013966 | 0.011606 | 0.006919 | 0.002441 | 0.0066 | 0.008191 | 0.008194 | 0.018109 |
| 37.2638889 | 0.064067 | 0.041419 | 0.021795 | 0.01251 | 0.018509 | 0.040168 | 0.031006 | 0.072787 | 0.025035 | 0.005251 | 0.001927 | 0.013494 | 0.012181 | 0.007868 | 0.003203 | 0.008178 | 0.007953 | 0.00988 | 0.017046 |
| 37.2777778 | 0.090174 | 0.056373 | 0.022185 | 0.014527 | 0.013219 | 0.034376 | 0.032526 | 0.085669 | 0.026241 | 0.010548 | 0.003091 | 0.012546 | 0.01168 | 0.006026 | 0.005062 | 0.008646 | 0.006387 | 0.010188 | 0.017127 |
| 37.2916667 | 0.112629 | 0.06387 | 0.023549 | 0.017108 | 0.00998 | 0.032565 | 0.032028 | 0.08876 | 0.025381 | 0.016902 | 0.004446 | 0.009526 | 0.007494 | 0.003161 | 0.006609 | 0.005359 | 0.003727 | 0.008959 | 0.016867 |
| 37.3055556 | 0.118091 | 0.063485 | 0.024952 | 0.018645 | 0.007762 | 0.034455 | 0.028428 | 0.086281 | 0.024217 | 0.014255 | 0.003476 | 0.004961 | 0.003239 | 0.00241 | 0.006264 | 0.002216 | 0.001324 | 0.006648 | 0.014159 |
| 37.3194444 | 0.107061 | 0.058054 | 0.025485 | 0.017184 | 0.00622 | 0.036083 | 0.021872 | 0.078897 | 0.024623 | 0.009085 | 0.002549 | 0.002282 | 0.002945 | 0.00373 | 0.004067 | 0.002162 | 0.0004 | 0.003825 | 0.008539 |
| 37.3333333 | 0.086319 | 0.049515 | 0.022818 | 0.011748 | 0.00422 | 0.033572 | 0.012071 | 0.064296 | 0.025028 | 0.015294 | 0.004862 | 0.002571 | 0.003392 | 0.00495 | 0.001816 | 0.002544 | 0.000758 | 0.002258 | 0.005639 |
| 37.3472222 | 0.058749 | 0.038088 | 0.015684 | 0.006087 | 0.002541 | 0.027461 | 0.004743 | 0.041956 | 0.023964 | 0.024296 | 0.008116 | 0.003129 | 0.0024 | 0.00451 | 0.000942 | 0.00196 | 0.001137 | 0.00191 | 0.009525 |
| 37.3611111 | 0.028324 | 0.023893 | 0.007793 | 0.005627 | 0.00312 | 0.021313 | 0.005097 | 0.019533 | 0.02258 | 0.0238 | 0.007486 | 0.002901 | 0.002329 | 0.002935 | 0.001029 | 0.002392 | 0.001385 | 0.001755 | 0.013828 |
| 37.375 | 0.00788 | 0.012505 | 0.003746 | 0.007655 | 0.004082 | 0.017518 | 0.007064 | 0.009986 | 0.021976 | 0.014364 | 0.004738 | 0.004002 | 0.004706 | 0.002268 | 0.00101 | 0.005163 | 0.002504 | 0.003423 | 0.012608 |
| 37.3888889 | 0.003371 | 0.007485 | 0.002805 | 0.007196 | 0.003252 | 0.015624 | 0.006141 | 0.011618 | 0.021494 | 0.01022 | 0.006215 | 0.00744 | 0.006974 | 0.002324 | 0.001491 | 0.008234 | 0.00345 | 0.00629 | 0.007768 |
| 37.4027778 | 0.006429 | 0.006332 | 0.001879 | 0.004855 | 0.001628 | 0.014494 | 0.00367 | 0.011405 | 0.020363 | 0.021692 | 0.010181 | 0.011459 | 0.00671 | 0.001821 | 0.002909 | 0.009262 | 0.002863 | 0.008674 | 0.005196 |
| 37.4166667 | 0.009983 | 0.005839 | 0.001383 | 0.003415 | 0.00122 | 0.014253 | 0.002653 | 0.007025 | 0.018979 | 0.03287 | 0.009847 | 0.014046 | 0.004947 | 0.001437 | 0.003681 | 0.008458 | 0.001906 | 0.010529 | 0.004827 |
| 37.4305556 | 0.011373 | 0.004005 | 0.003376 | 0.004797 | 0.002728 | 0.015242 | 0.004656 | 0.00346 | 0.017827 | 0.031291 | 0.006051 | 0.013009 | 0.003524 | 0.001094 | 0.002805 | 0.006998 | 0.001602 | 0.010729 | 0.003571 |
| 37.4444444 | 0.011727 | 0.002474 | 0.006594 | 0.00837 | 0.003742 | 0.015791 | 0.005453 | 0.001752 | 0.016424 | 0.023833 | 0.005835 | 0.008269 | 0.003173 | 0.000451 | 0.001238 | 0.005399 | 0.001329 | 0.008236 | 0.003693 |
| 37.4583333 | 0.011931 | 0.004242 | 0.008351 | 0.011513 | 0.00271 | 0.014688 | 0.004162 | 0.000762 | 0.014526 | 0.017214 | 0.008326 | 0.004984 | 0.003218 | 0.000272 | 0.00062 | 0.004177 | 0.001015 | 0.004904 | 0.004827 |
| 37.4722222 | 0.010921 | 0.007275 | 0.009062 | 0.012602 | 0.001309 | 0.013858 | 0.005551 | 0.000861 | 0.012833 | 0.014178 | 0.007262 | 0.004875 | 0.004445 | 0.001115 | 0.001388 | 0.004495 | 0.000903 | 0.004126 | 0.004626 |
| 37.4861111 | 0.011819 | 0.007775 | 0.010077 | 0.011539 | 0.001218 | 0.014251 | 0.009174 | 0.003429 | 0.01141 | 0.016831 | 0.006075 | 0.004283 | 0.007629 | 0.002219 | 0.001746 | 0.005156 | 0.000711 | 0.005569 | 0.006403 |
| 37.5 | 0.016759 | 0.005473 | 0.01186 | 0.008605 | 0.00325 | 0.014845 | 0.009899 | 0.00884 | 0.010122 | 0.021046 | 0.007848 | 0.0051 | 0.010081 | 0.001913 | 0.001194 | 0.00368 | 0.00061 | 0.006169 | 0.011412 |
| 37.5138889 | 0.021931 | 0.002521 | 0.015352 | 0.004517 | 0.006895 | 0.016176 | 0.006208 | 0.015818 | 0.009698 | 0.019429 | 0.006829 | 0.009681 | 0.009776 | 0.000996 | 0.00094 | 0.001555 | 0.000746 | 0.005614 | 0.014951 |
| 37.5277778 | 0.023261 | 0.000902 | 0.01971 | 0.00151 | 0.01033 | 0.017908 | 0.00201 | 0.021459 | 0.009956 | 0.013186 | 0.004592 | 0.013874 | 0.006839 | 0.000809 | 0.000718 | 0.001009 | 0.001004 | 0.004938 | 0.014172 |
| 37.5416667 | 0.02031 | 0.001264 | 0.022224 | 0.001133 | 0.011151 | 0.018034 | 0.001449 | 0.023968 | 0.010376 | 0.007057 | 0.007505 | 0.01412 | 0.003205 | 0.00094 | 0.000442 | 0.001413 | 0.001645 | 0.003694 | 0.009398 |
| 37.5555556 | 0.014182 | 0.00308 | 0.022399 | 0.001265 | 0.008221 | 0.01655 | 0.004398 | 0.023746 | 0.011595 | 0.002969 | 0.011512 | 0.011509 | 0.001625 | 0.000942 | 0.000826 | 0.002664 | 0.002365 | 0.002372 | 0.003946 |
| 37.5694444 | 0.007097 | 0.00573 | 0.021574 | 0.00177 | 0.004927 | 0.013503 | 0.00975 | 0.022512 | 0.013387 | 0.005842 | 0.010059 | 0.008846 | 0.00202 | 0.000712 | 0.001305 | 0.004861 | 0.002497 | 0.002608 | 0.001471 |
| 37.5833333 | 0.003622 | 0.00773 | 0.020633 | 0.003854 | 0.007014 | 0.007752 | 0.015967 | 0.024929 | 0.015253 | 0.021971 | 0.006374 | 0.009836 | 0.004252 | 0.001272 | 0.002195 | 0.007741 | 0.007553 | 0.006244 | 0.001825 |
| 37.5972222 | 0.009006 | 0.006464 | 0.021331 | 0.005158 | 0.015256 | 0.00262 | 0.016776 | 0.034481 | 0.018011 | 0.045815 | 0.008729 | 0.018865 | 0.00945 | 0.002108 | 0.004829 | 0.012166 | 0.025761 | 0.014362 | 0.004554 |
| 37.6111111 | 0.029035 | 0.00846 | 0.024782 | 0.009342 | 0.023151 | 0.00223 | 0.009745 | 0.049682 | 0.023388 | 0.059696 | 0.017103 | 0.033088 | 0.017872 | 0.002291 | 0.00675 | 0.017072 | 0.049577 | 0.023179 | 0.008448 |
| 37.625 | 0.068726 | 0.023299 | 0.02982 | 0.018181 | 0.024688 | 0.00575 | 0.004298 | 0.067088 | 0.031889 | 0.060131 | 0.022013 | 0.040445 | 0.024521 | 0.005016 | 0.00584 | 0.019241 | 0.064491 | 0.026587 | 0.011472 |
| 37.6388889 | 0.099133 | 0.035925 | 0.034824 | 0.023174 | 0.024062 | 0.01127 | 0.007131 | 0.081294 | 0.04205 | 0.054686 | 0.01747 | 0.034335 | 0.022177 | 0.010434 | 0.003638 | 0.017135 | 0.067402 | 0.02386 | 0.01259 |
| 37.6527778 | 0.086347 | 0.03075 | 0.036041 | 0.020379 | 0.027986 | 0.016296 | 0.012303 | 0.08683 | 0.050586 | 0.038923 | 0.009169 | 0.020641 | 0.012345 | 0.013833 | 0.001596 | 0.011732 | 0.062349 | 0.019021 | 0.010571 |
| 37.6666667 | 0.051535 | 0.020112 | 0.03114 | 0.0125 | 0.031677 | 0.017671 | 0.012002 | 0.082577 | 0.052626 | 0.019324 | 0.007812 | 0.008828 | 0.005317 | 0.014321 | 0.000484 | 0.00639 | 0.056701 | 0.015988 | 0.006262 |
| 37.6805556 | 0.025195 | 0.012163 | 0.02642 | 0.008649 | 0.028835 | 0.014846 | 0.006585 | 0.073452 | 0.046938 | 0.014355 | 0.016308 | 0.004929 | 0.004648 | 0.014117 | 0.000519 | 0.004178 | 0.054796 | 0.017297 | 0.002882 |
| 37.6944444 | 0.019872 | 0.005425 | 0.028647 | 0.013045 | 0.022066 | 0.012514 | 0.003396 | 0.067122 | 0.037919 | 0.020814 | 0.027166 | 0.006887 | 0.003436 | 0.013877 | 0.002042 | 0.004965 | 0.05393 | 0.021159 | 0.002602 |
| 37.7083333 | 0.031199 | 0.003559 | 0.032848 | 0.017617 | 0.017706 | 0.01383 | 0.005695 | 0.065478 | 0.030833 | 0.025171 | 0.030851 | 0.008371 | 0.001833 | 0.013037 | 0.004702 | 0.006777 | 0.048482 | 0.023143 | 0.004639 |
| 37.7222222 | 0.042085 | 0.006391 | 0.030217 | 0.020215 | 0.018281 | 0.016081 | 0.008324 | 0.060826 | 0.027781 | 0.025952 | 0.023238 | 0.008107 | 0.003358 | 0.011321 | 0.006199 | 0.008464 | 0.03532 | 0.022097 | 0.007092 |
| 37.7361111 | 0.050347 | 0.011494 | 0.020949 | 0.023216 | 0.019561 | 0.017849 | 0.008202 | 0.042201 | 0.025272 | 0.02314 | 0.013936 | 0.00778 | 0.005385 | 0.008792 | 0.005858 | 0.009905 | 0.019393 | 0.019135 | 0.008274 |
| 37.75 | 0.055718 | 0.018873 | 0.016338 | 0.025359 | 0.017948 | 0.01857 | 0.009134 | 0.027037 | 0.018567 | 0.018093 | 0.018505 | 0.00757 | 0.005516 | 0.005442 | 0.005712 | 0.009811 | 0.014858 | 0.015669 | 0.006385 |
| 37.7638889 | 0.051065 | 0.024339 | 0.022033 | 0.023787 | 0.013944 | 0.017201 | 0.014114 | 0.048273 | 0.019831 | 0.016339 | 0.032956 | 0.007648 | 0.003438 | 0.004357 | 0.006773 | 0.007658 | 0.027579 | 0.012224 | 0.002727 |
| 37.7777778 | 0.037355 | 0.023428 | 0.022354 | 0.017902 | 0.009315 | 0.014988 | 0.020091 | 0.092665 | 0.035957 | 0.017053 | 0.041858 | 0.007615 | 0.001718 | 0.007737 | 0.007641 | 0.005698 | 0.045696 | 0.008759 | 0.000721 |
| 37.7916667 | 0.020722 | 0.018561 | 0.015121 | 0.012005 | 0.006468 | 0.013304 | 0.022686 | 0.121356 | 0.045558 | 0.014346 | 0.039709 | 0.006298 | 0.002507 | 0.01067 | 0.006672 | 0.00511 | 0.059378 | 0.00613 | 0.00117 |
| 37.8055556 | 0.010076 | 0.01209 | 0.015477 | 0.007428 | 0.006807 | 0.012087 | 0.02187 | 0.120092 | 0.041603 | 0.009147 | 0.029555 | 0.003694 | 0.003048 | 0.008557 | 0.0045 | 0.004462 | 0.066293 | 0.004408 | 0.002556 |
| 37.8194444 | 0.011592 | 0.010584 | 0.014936 | 0.00372 | 0.008353 | 0.0116 | 0.019685 | 0.097795 | 0.036852 | 0.009624 | 0.017833 | 0.001715 | 0.002276 | 0.00579 | 0.003244 | 0.002658 | 0.065335 | 0.002826 | 0.003549 |
| 37.8333333 | 0.017863 | 0.016711 | 0.006312 | 0.002436 | 0.007302 | 0.012092 | 0.016466 | 0.06952 | 0.033582 | 0.014761 | 0.009366 | 0.002552 | 0.002898 | 0.009355 | 0.003989 | 0.001431 | 0.058386 | 0.002963 | 0.004091 |
| 37.8472222 | 0.019757 | 0.018153 | 0.002074 | 0.002854 | 0.004791 | 0.01321 | 0.012371 | 0.045181 | 0.026538 | 0.017483 | 0.005053 | 0.004597 | 0.004454 | 0.016186 | 0.006072 | 0.002429 | 0.049439 | 0.006566 | 0.004784 |
| 37.8611111 | 0.018011 | 0.011412 | 0.002997 | 0.00565 | 0.00599 | 0.015411 | 0.007718 | 0.025313 | 0.015106 | 0.017644 | 0.003201 | 0.004697 | 0.00451 | 0.018663 | 0.008229 | 0.003944 | 0.041152 | 0.011943 | 0.006752 |
| 37.875 | 0.015438 | 0.006471 | 0.003816 | 0.011809 | 0.009955 | 0.018496 | 0.003231 | 0.011473 | 0.00529 | 0.016947 | 0.001818 | 0.003347 | 0.003563 | 0.015607 | 0.009509 | 0.004334 | 0.032871 | 0.016872 | 0.010429 |
| 37.8888889 | 0.00999 | 0.008865 | 0.006513 | 0.017525 | 0.011268 | 0.019984 | 0.001394 | 0.007091 | 0.002416 | 0.012854 | 0.000851 | 0.003983 | 0.003358 | 0.010589 | 0.008818 | 0.003492 | 0.022567 | 0.019814 | 0.014749 |
| 37.9027778 | 0.007558 | 0.014047 | 0.008737 | 0.021391 | 0.009949 | 0.01752 | 0.004181 | 0.01494 | 0.004459 | 0.008259 | 0.000834 | 0.00811 | 0.003764 | 0.005746 | 0.006328 | 0.002238 | 0.012722 | 0.02032 | 0.018155 |
| 37.9166667 | 0.013413 | 0.016836 | 0.008168 | 0.026945 | 0.010129 | 0.010534 | 0.012517 | 0.041658 | 0.006935 | 0.012672 | 0.001894 | 0.013127 | 0.003361 | 0.002962 | 0.004721 | 0.002211 | 0.008775 | 0.020062 | 0.019645 |
| 37.9305556 | 0.020221 | 0.018027 | 0.009026 | 0.035195 | 0.01589 | 0.006032 | 0.027324 | 0.082801 | 0.009705 | 0.023393 | 0.00421 | 0.016218 | 0.00203 | 0.00461 | 0.005824 | 0.004038 | 0.011539 | 0.021344 | 0.019132 |
| 37.9444444 | 0.025464 | 0.017973 | 0.013856 | 0.04382 | 0.030646 | 0.010967 | 0.045058 | 0.12423 | 0.013815 | 0.030002 | 0.006087 | 0.016375 | 0.001515 | 0.008139 | 0.007521 | 0.006483 | 0.016689 | 0.024789 | 0.015785 |
| 37.9583333 | 0.026484 | 0.015541 | 0.016333 | 0.049236 | 0.045771 | 0.017079 | 0.056517 | 0.147016 | 0.016344 | 0.031448 | 0.005443 | 0.013318 | 0.002424 | 0.009498 | 0.006422 | 0.007243 | 0.019927 | 0.028432 | 0.009257 |
| 37.9722222 | 0.019679 | 0.014876 | 0.013253 | 0.048855 | 0.047914 | 0.017817 | 0.058265 | 0.145653 | 0.012758 | 0.031476 | 0.00305 | 0.007925 | 0.002795 | 0.008306 | 0.00367 | 0.005353 | 0.019784 | 0.029519 | 0.004329 |
| 37.9861111 | 0.013187 | 0.0213 | 0.015379 | 0.045121 | 0.042814 | 0.016389 | 0.055553 | 0.139704 | 0.006143 | 0.029683 | 0.002053 | 0.004105 | 0.001692 | 0.005969 | 0.003096 | 0.002812 | 0.01671 | 0.025416 | 0.005514 |
| 38 | 0.016987 | 0.032624 | 0.024909 | 0.041261 | 0.037848 | 0.012573 | 0.049962 | 0.134172 | 0.002372 | 0.023421 | 0.00379 | 0.00542 | 0.000577 | 0.003614 | 0.004253 | 0.00249 | 0.012749 | 0.016056 | 0.009326 |
| 38.0138889 | 0.0266 | 0.040265 | 0.029628 | 0.036713 | 0.029709 | 0.006727 | 0.039526 | 0.119283 | 0.001179 | 0.013626 | 0.005293 | 0.009567 | 0.000258 | 0.001862 | 0.004419 | 0.004224 | 0.010613 | 0.006682 | 0.01125 |
| 38.0277778 | 0.024192 | 0.036022 | 0.025715 | 0.030365 | 0.020017 | 0.004054 | 0.02608 | 0.097096 | 0.002077 | 0.007243 | 0.004701 | 0.012718 | 0.000866 | 0.001091 | 0.003201 | 0.00552 | 0.010797 | 0.002566 | 0.010792 |
| 38.0416667 | 0.013817 | 0.022164 | 0.017448 | 0.021438 | 0.01131 | 0.006996 | 0.013626 | 0.072054 | 0.005949 | 0.009964 | 0.00347 | 0.014502 | 0.003187 | 0.001796 | 0.001528 | 0.005763 | 0.011499 | 0.00286 | 0.008775 |
| 38.0555556 | 0.011851 | 0.010238 | 0.00966 | 0.011646 | 0.005562 | 0.013004 | 0.010843 | 0.044651 | 0.011826 | 0.017099 | 0.003014 | 0.015447 | 0.007041 | 0.003328 | 0.000479 | 0.0057 | 0.011772 | 0.003957 | 0.005984 |
| 38.0694445 | 0.013263 | 0.006656 | 0.009458 | 0.009602 | 0.004035 | 0.019139 | 0.018991 | 0.020733 | 0.017398 | 0.016527 | 0.004632 | 0.013545 | 0.009534 | 0.003284 | 0.001546 | 0.004544 | 0.012808 | 0.003839 | 0.004294 |

Supplementary Table S6. 1 hour integrated motion data in (a) lettuce subjected to 150 mM NaCl gradual stress

| IntervalMid | Control1 | Control2 | Control3 | Control4 | Control5 | Control6 | Control7 | Control8 | Control9 | Control10 | Control11 | Control12 | Stress1 | Stress2 | Stress3 | Stress4 | Stress5 | Stress6 | Stress7 | Stress8 | Stress9 | Stress10 | Stress11 | Stress12 |
|---|---|---|---|---|---|---|---|---|---|---|---|---|---|---|---|---|---|---|---|---|---|---|---|---|
| 18.11111111 | 0.027509 | 0.020401 | 0.016219 | 0.003804 | 0.014822 | 0.001633 | 0.016574 | 0.001892 | 0.047122 | 0.007611 | 0.006356 | 0.018381 | 0.018472 | 0.038272 | 0.00236 | 0.001908 | 0.002628 | 0.001672 | 0.001815 | 0.000376 | 0.029014 | 0.027527 | 0.019592 | 0.002826 |
| 18.125 | 0.013083 | 0.011243 | 0.010006 | 0.002767 | 0.006354 | 0.001563 | 0.005995 | 0.000962 | 0.034442 | 0.005058 | 0.004698 | 0.008786 | 0.007936 | 0.024067 | 0.000752 | 0.00276 | 0.00095 | 0.002184 | 0.001538 | 0.000844 | 0.013722 | 0.016637 | 0.009563 | 0.002459 |
| 18.13888889 | 0.005518 | 0.004912 | 0.008158 | 0.002457 | 0.002307 | 0.001823 | 0.001458 | 0.000744 | 0.019448 | 0.00263 | 0.004386 | 0.004156 | 0.002877 | 0.012067 | 0.000375 | 0.003429 | 0.000854 | 0.002629 | 0.001224 | 0.001263 | 0.00421 | 0.006744 | 0.003992 | 0.002452 |
| 18.15277778 | 0.002549 | 0.001898 | 0.00781 | 0.002686 | 0.000722 | 0.002619 | 0.001203 | 0.001098 | 0.011505 | 0.001029 | 0.004964 | 0.00212 | 0.001215 | 0.006794 | 0.000294 | 0.00319 | 0.000799 | 0.002908 | 0.000784 | 0.00143 | 0.001005 | 0.001606 | 0.00153 | 0.002345 |
| 18.16666667 | 0.00168 | 0.000881 | 0.00572 | 0.002929 | 0.000536 | 0.003628 | 0.001215 | 0.001138 | 0.007587 | 0.000342 | 0.005835 | 0.001573 | 0.001308 | 0.005373 | 0.000197 | 0.002485 | 0.000649 | 0.002844 | 0.000544 | 0.001565 | 0.000563 | 0.000378 | 0.001127 | 0.002232 |
| 18.18055556 | 0.001584 | 0.000858 | 0.00323 | 0.002746 | 0.000496 | 0.004031 | 0.000731 | 0.000753 | 0.006393 | 0.000272 | 0.006312 | 0.001846 | 0.001944 | 0.005806 | 0.000278 | 0.002017 | 0.000557 | 0.002544 | 0.00058 | 0.001756 | 0.000478 | 0.000341 | 0.00169 | 0.002138 |
| 18.19444444 | 0.001539 | 0.001271 | 0.002268 | 0.002269 | 0.000468 | 0.003506 | 0.000341 | 0.000646 | 0.006053 | 0.000546 | 0.006117 | 0.002003 | 0.002497 | 0.006638 | 0.000453 | 0.0018 | 0.000473 | 0.00227 | 0.000602 | 0.002036 | 0.000331 | 0.000441 | 0.002141 | 0.002049 |
| 18.20833333 | 0.001194 | 0.001779 | 0.001988 | 0.001937 | 0.000511 | 0.002643 | 0.000318 | 0.000741 | 0.00521 | 0.000899 | 0.005476 | 0.001496 | 0.002892 | 0.007166 | 0.000682 | 0.001637 | 0.000417 | 0.002043 | 0.000487 | 0.002529 | 0.000291 | 0.000472 | 0.00188 | 0.001851 |
| 18.22222222 | 0.000674 | 0.00212 | 0.001633 | 0.001894 | 0.000385 | 0.002029 | 0.000363 | 0.000566 | 0.003965 | 0.000969 | 0.004989 | 0.000681 | 0.00311 | 0.007318 | 0.00097 | 0.001527 | 0.000429 | 0.001802 | 0.000354 | 0.003291 | 0.0003 | 0.000349 | 0.001155 | 0.001772 |
| 18.23611111 | 0.0003 | 0.002026 | 0.001302 | 0.001931 | 0.000312 | 0.001736 | 0.000291 | 0.000254 | 0.003036 | 0.000736 | 0.005165 | 0.000206 | 0.003105 | 0.007246 | 0.001274 | 0.001488 | 0.000458 | 0.001617 | 0.000324 | 0.004322 | 0.000206 | 0.000439 | 0.000667 | 0.002172 |
| 18.25 | 0.000207 | 0.001395 | 0.000945 | 0.002008 | 0.000277 | 0.001742 | 0.00025 | 0.000118 | 0.002589 | 0.000447 | 0.005971 | 0.000159 | 0.003205 | 0.007096 | 0.001504 | 0.001415 | 0.000408 | 0.001567 | 0.000325 | 0.005756 | 0.000113 | 0.000665 | 0.000642 | 0.002717 |
| 18.26388889 | 0.000345 | 0.000801 | 0.000547 | 0.002153 | 0.000229 | 0.001928 | 0.000361 | 0.000149 | 0.002399 | 0.000434 | 0.007106 | 0.000239 | 0.003689 | 0.006877 | 0.001543 | 0.001162 | 0.00028 | 0.001592 | 0.000233 | 0.007487 | 0.000154 | 0.000649 | 0.000802 | 0.00301 |
| 18.27777778 | 0.00038 | 0.000848 | 0.000479 | 0.002041 | 0.000261 | 0.001989 | 0.000452 | 0.000181 | 0.002493 | 0.000586 | 0.008557 | 0.000356 | 0.004208 | 0.006745 | 0.00133 | 0.000751 | 0.00022 | 0.001528 | 0.000153 | 0.008815 | 0.000233 | 0.000518 | 0.001046 | 0.003236 |
| 18.29166667 | 0.000236 | 0.001326 | 0.000921 | 0.001652 | 0.000289 | 0.001897 | 0.000363 | 0.000236 | 0.002937 | 0.000525 | 0.010316 | 0.000458 | 0.004344 | 0.006416 | 0.000959 | 0.000379 | 0.000254 | 0.00133 | 0.000177 | 0.009003 | 0.000213 | 0.000623 | 0.001344 | 0.003397 |
| 18.30555556 | 0.000218 | 0.00186 | 0.001368 | 0.001602 | 0.000434 | 0.001953 | 0.000321 | 0.000287 | 0.003291 | 0.000597 | 0.012379 | 0.000361 | 0.004047 | 0.005445 | 0.000628 | 0.000237 | 0.000227 | 0.001159 | 0.000157 | 0.008118 | 0.000144 | 0.000921 | 0.001446 | 0.0032 |
| 18.31944444 | 0.000315 | 0.002421 | 0.001461 | 0.002063 | 0.000635 | 0.002113 | 0.000273 | 0.000251 | 0.003193 | 0.001208 | 0.015541 | 0.000145 | 0.003535 | 0.004156 | 0.000457 | 0.000391 | 0.000214 | 0.001131 | 8.6694412 | 0.007039 | 0.000123 | 0.001113 | 0.001305 | 0.002773 |
| 18.33333333 | 0.000419 | 0.00268 | 0.001506 | 0.002441 | 0.000753 | 0.002027 | 0.000241 | 0.000183 | 0.002913 | 0.001879 | 0.019489 | 0.00012 | 0.003038 | 0.003171 | 0.000413 | 0.000753 | 0.000425 | 0.00112 | 7.7250006 | 0.006461 | 0.000125 | 0.001052 | 0.001147 | 0.002567 |
| 18.34722222 | 0.000423 | 0.002122 | 0.001741 | 0.002435 | 0.000728 | 0.001768 | 0.000384 | 0.000135 | 0.002777 | 0.001926 | 0.021779 | 0.000348 | 0.002626 | 0.002626 | 0.000398 | 0.001142 | 0.001352 | 0.001008 | 6.5784727 | 0.006396 | 0.000123 | 0.001047 | 0.00105 | 0.002666 |
| 18.36111111 | 0.000321 | 0.001288 | 0.002231 | 0.002285 | 0.000673 | 0.001571 | 0.000463 | 0.000143 | 0.003032 | 0.001465 | 0.02073 | 0.000644 | 0.002287 | 0.002287 | 0.000331 | 0.001381 | 0.00378 | 0.000889 | 3.4472224 | 0.006758 | 0.000176 | 0.001456 | 0.001006 | 0.00268 |
| 18.375 | 0.000413 | 0.001337 | 0.002892 | 0.001984 | 0.00105 | 0.0013 | 0.000566 | 0.000316 | 0.003788 | 0.001237 | 0.016755 | 0.000874 | 0.001986 | 0.00208 | 0.00022 | 0.001404 | 0.006309 | 0.000817 | 4.6854170 | 0.007542 | 0.000228 | 0.001919 | 0.001107 | 0.002505 |
| 18.38888889 | 0.000687 | 0.002206 | 0.003268 | 0.001605 | 0.00174 | 0.000957 | 0.000683 | 0.000662 | 0.004464 | 0.00141 | 0.012032 | 0.00096 | 0.001705 | 0.001996 | 0.000188 | 0.001309 | 0.006201 | 0.000762 | 0.000148 | 0.008745 | 0.000264 | 0.001929 | 0.00142 | 0.002565 |
| 18.40277778 | 0.000816 | 0.002899 | 0.003149 | 0.001537 | 0.002148 | 0.000854 | 0.000477 | 0.00084 | 0.004539 | 0.001635 | 0.008757 | 0.00094 | 0.001531 | 0.001915 | 0.000434 | 0.001277 | 0.004021 | 0.000757 | 0.000306 | 0.010444 | 0.000458 | 0.001712 | 0.00185 | 0.002971 |
| 18.41666667 | 0.000745 | 0.003051 | 0.002853 | 0.001808 | 0.002112 | 0.001043 | 0.00021 | 0.000712 | 0.004437 | 0.001956 | 0.007187 | 0.000805 | 0.001548 | 0.001779 | 0.001115 | 0.001455 | 0.002342 | 0.00085 | 0.000366 | 0.012298 | 0.000494 | 0.00181 | 0.002169 | 0.003337 |
| 18.43055556 | 0.000664 | 0.003092 | 0.002776 | 0.00203 | 0.001905 | 0.001264 | 0.000305 | 0.000567 | 0.004774 | 0.002581 | 0.006603 | 0.000545 | 0.001225 | 0.001249 | 0.00133 | 0.00126 | 0.001401 | 0.000722 | 0.000241 | 0.01005 | 0.000217 | 0.001583 | 0.001699 | 0.002597 |
| 18.44444444 | 0.00069 | 0.003075 | 0.002878 | 0.001965 | 0.001606 | 0.001456 | 0.000791 | 0.000556 | 0.005452 | 0.003175 | 0.006567 | 0.000692 | 0.000423 | 0.000399 | 0.000541 | 0.000456 | 0.000459 | 0.000259 | 6.7437456 | 0.003465 | 2.7652760 | 0.000586 | 0.000566 | 0.000871 |
| 18.45833333 | 0.000623 | 0.002276 | 0.002524 | 0.001995 | 0.001243 | 0.002 | 0.001271 | 0.000465 | 0.005777 | 0.002974 | 0.007111 | 0.001766 | 0 | 0 | 0 | 0 | 0 | 0 | 0 | 0 | 0 | 0 | 0 | 0 |
| 18.47222222 | 0.000329 | 0.001081 | 0.001511 | 0.002506 | 0.001037 | 0.002893 | 0.001681 | 0.000223 | 0.005734 | 0.00214 | 0.008053 | 0.003063 | 0 | 0 | 0 | 0 | 0 | 0 | 0 | 0 | 0 | 0 | 0 | 0 |
| 18.48611111 | 0.000917 | 0.002991 | 0.001305 | 0.002887 | 0.002706 | 0.003028 | 0.005025 | 0.00113 | 0.008361 | 0.003081 | 0.008078 | 0.002921 | 0 | 0 | 0 | 0 | 0 | 0 | 0 | 0 | 0 | 0 | 0 | 0 |
| 18.5 | 0.004121 | 0.012844 | 0.00435 | 0.002149 | 0.010639 | 0.002247 | 0.017661 | 0.008158 | 0.018499 | 0.008941 | 0.005836 | 0.002251 | 0 | 0 | 0 | 0 | 0 | 0 | 0 | 0 | 0 | 0 | 0 | 0 |
| 18.51388889 | 0.008602 | 0.027775 | 0.009099 | 0.000789 | 0.023844 | 0.002752 | 0.035863 | 0.019158 | 0.033815 | 0.020186 | 0.003004 | 0.003432 | 0 | 0 | 0 | 0 | 0 | 0 | 0 | 0 | 0 | 0 | 0 | 0 |
| 18.52777778 | 0.009987 | 0.034299 | 0.008565 | 0.000531 | 0.030896 | 0.005091 | 0.041448 | 0.019845 | 0.041909 | 0.03064 | 0.002623 | 0.00484 | 0 | 0 | 0 | 0 | 0 | 0 | 0 | 0 | 0 | 0 | 0 | 0 |
| 18.54166667 | 0.007575 | 0.026309 | 0.003364 | 0.001123 | 0.026361 | 0.006485 | 0.029974 | 0.010741 | 0.036777 | 0.032841 | 0.004137 | 0.004664 | 0 | 0 | 0 | 0 | 0 | 0 | 0 | 0 | 0 | 0 | 0 | 0 |
| 18.55555556 | 0.005622 | 0.0141 | 0.000738 | 0.001461 | 0.019363 | 0.00604 | 0.016778 | 0.005641 | 0.027076 | 0.028897 | 0.005314 | 0.003931 | 0 | 0 | 0 | 0 | 0 | 0 | 0 | 0 | 0 | 0 | 0 | 0 |
| 18.56944444 | 0.006343 | 0.00697 | 0.00047 | 0.002266 | 0.017004 | 0.005409 | 0.010224 | 0.005324 | 0.021172 | 0.025313 | 0.00578 | 0.002811 | 0 | 0 | 0 | 0 | 0 | 0 | 0 | 0 | 0 | 0 | 0 | 0 |
| 18.58333333 | 0.008314 | 0.00444 | 0.000523 | 0.003359 | 0.017521 | 0.005016 | 0.008195 | 0.004922 | 0.020109 | 0.023594 | 0.005918 | 0.001356 | 0 | 0 | 0 | 0 | 0 | 0 | 0 | 0 | 0 | 0 | 0 | 0 |
| 18.59722222 | 0.010371 | 0.004153 | 0.001222 | 0.003745 | 0.018363 | 0.004655 | 0.007836 | 0.003969 | 0.022517 | 0.022147 | 0.005821 | 0.000572 | 0 | 0 | 0 | 0 | 0 | 0 | 0 | 0 | 0 | 0 | 0 | 0 |
| 18.61111111 | 0.012067 | 0.005284 | 0.002331 | 0.003604 | 0.018482 | 0.00438 | 0.006977 | 0.003025 | 0.026445 | 0.020637 | 0.005809 | 0.000424 | 0 | 0 | 0 | 0 | 0 | 0 | 0 | 0 | 0 | 0 | 0 | 0 |
| 18.625 | 0.013018 | 0.007275 | 0.00373 | 0.003345 | 0.017448 | 0.003979 | 0.004911 | 0.002798 | 0.030444 | 0.019178 | 0.006603 | 0.000586 | 0 | 0 | 0 | 0 | 0 | 0 | 0 | 0 | 0 | 0 | 0 | 0 |
| 18.63888889 | 0.012829 | 0.009684 | 0.005034 | 0.003226 | 0.015579 | 0.003228 | 0.002435 | 0.004022 | 0.033958 | 0.017202 | 0.00817 | 0.00129 | 0 | 0 | 0 | 0 | 0 | 0 | 0 | 0 | 0 | 0 | 0 | 0 |
| 18.65277778 | 0.011287 | 0.011858 | 0.005728 | 0.003321 | 0.012992 | 0.002168 | 0.001219 | 0.006198 | 0.036188 | 0.01509 | 0.00923 | 0.001907 | 0 | 0 | 0 | 0 | 0 | 0 | 0 | 0 | 0 | 0 | 0 | 0 |
| 18.66666667 | 0.008969 | 0.012817 | 0.005402 | 0.003321 | 0.0097 | 0.001347 | 0.00167 | 0.007249 | 0.035946 | 0.013441 | 0.009096 | 0.001943 | 0 | 0 | 0 | 0 | 0 | 0 | 0 | 0 | 0 | 0 | 0 | 0 |
| 18.68055556 | 0.006544 | 0.01224 | 0.004168 | 0.002859 | 0.006128 | 0.001189 | 0.002078 | 0.006084 | 0.032696 | 0.011879 | 0.008455 | 0.001783 | 0 | 0 | 0 | 0 | 0 | 0 | 0 | 0 | 0 | 0 | 0 | 0 |
| 18.69444444 | 0.004358 | 0.010841 | 0.002738 | 0.002397 | 0.003193 | 0.001415 | 0.001626 | 0.004114 | 0.027626 | 0.009635 | 0.007916 | 0.001571 | 0 | 0 | 0 | 0 | 0 | 0 | 0 | 0 | 0 | 0 | 0 | 0 |
| 18.70833333 | 0.002383 | 0.009589 | 0.001623 | 0.002549 | 0.001405 | 0.001604 | 0.001059 | 0.002984 | 0.022519 | 0.006578 | 0.007545 | 0.001083 | 0 | 0 | 0 | 0 | 0 | 0 | 0 | 0 | 0 | 0 | 0 | 0 |
| 18.72222222 | 0.001088 | 0.008787 | 0.000883 | 0.002958 | 0.001192 | 0.001674 | 0.000793 | 0.002553 | 0.017883 | 0.003463 | 0.007322 | 0.000453 | 0 | 0 | 0 | 0 | 0 | 0 | 0 | 0 | 0 | 0 | 0 | 0 |
| 18.73611111 | 0.001158 | 0.008504 | 0.000511 | 0.003169 | 0.003139 | 0.001758 | 0.000533 | 0.00223 | 0.013648 | 0.001486 | 0.007256 | 0.000234 | 0 | 0 | 0 | 0 | 0 | 0 | 0 | 0 | 0 | 0 | 0 | 0 |
| 18.75 | 0.001785 | 0.00919 | 0.000413 | 0.002973 | 0.005826 | 0.001863 | 0.0003 | 0.002031 | 0.010034 | 0.001452 | 0.00728 | 0.000456 | 0 | 0 | 0 | 0 | 0 | 0 | 0 | 0 | 0 | 0 | 0 | 0 |
| 18.76388889 | 0.002081 | 0.010401 | 0.000433 | 0.002536 | 0.006978 | 0.002152 | 0.000371 | 0.002106 | 0.007522 | 0.002307 | 0.007154 | 0.000669 | 0 | 0 | 0 | 0 | 0 | 0 | 0 | 0 | 0 | 0 | 0 | 0 |
| 18.77777778 | 0.001899 | 0.010535 | 0.000493 | 0.0023 | 0.005857 | 0.003691 | 0.000395 | 0.002485 | 0.006534 | 0.002255 | 0.006705 | 0.000754 | 0 | 0 | 0 | 0 | 0 | 0 | 0 | 0 | 0 | 0 | 0 | 0 |
| 18.79166667 | 0.00125 | 0.008974 | 0.000551 | 0.00249 | 0.003651 | 0.006712 | 0.000248 | 0.002875 | 0.00649 | 0.002479 | 0.006233 | 0.000827 | 0 | 0 | 0 | 0 | 0 | 0 | 0 | 0 | 0 | 0 | 0 | 0 |
| 18.80555556 | 0.000543 | 0.006932 | 0.000633 | 0.003349 | 0.002069 | 0.008883 | 0.000394 | 0.002935 | 0.006269 | 0.00468 | 0.006464 | 0.000713 | 0 | 0 | 0 | 0 | 0 | 0 | 0 | 0 | 0 | 0 | 0 | 0 |
| 18.81944444 | 0.00036 | 0.005647 | 0.000643 | 0.00449 | 0.001685 | 0.008886 | 0.001101 | 0.002649 | 0.005557 | 0.006722 | 0.008296 | 0.000403 | 0 | 0 | 0 | 0 | 0 | 0 | 0 | 0 | 0 | 0 | 0 | 0 |
| 18.83333333 | 0.001039 | 0.005083 | 0.000419 | 0.004775 | 0.002462 | 0.007894 | 0.002269 | 0.002067 | 0.00449 | 0.006407 | 0.012181 | 0.000206 | 0 | 0 | 0 | 0 | 0 | 0 | 0 | 0 | 0 | 0 | 0 | 0 |
| 18.84722222 | 0.002292 | 0.005057 | 0.00019 | 0.004077 | 0.00446 | 0.006782 | 0.003705 | 0.001258 | 0.002856 | 0.003888 | 0.015757 | 0.000227 | 0 | 0 | 0 | 0 | 0 | 0 | 0 | 0 | 0 | 0 | 0 | 0 |
| 18.86111111 | 0.0036 | 0.005732 | 0.000118 | 0.003311 | 0.007557 | 0.005884 | 0.005368 | 0.000554 | 0.001637 | 0.002434 | 0.015377 | 0.000455 | 0 | 0 | 0 | 0 | 0 | 0 | 0 | 0 | 0 | 0 | 0 | 0 |
| 18.875 | 0.004822 | 0.006283 | 8.9152784 | 0.002723 | 0.010796 | 0.005298 | 0.007598 | 0.00039 | 0.0023 | 0.004733 | 0.011756 | 0.000498 | 0 | 0 | 0 | 0 | 0 | 0 | 0 | 0 | 0 | 0 | 0 | 0 |
| 18.88888889 | 0.006199 | 0.005664 | 0.000116 | 0.002076 | 0.0127 | 0.004643 | 0.010484 | 0.00073 | 0.004036 | 0.00665 | 0.008427 | 0.000413 | 0 | 0 | 0 | 0 | 0 | 0 | 0 | 0 | 0 | 0 | 0 | 0 |
| 18.90277778 | 0.008237 | 0.004431 | 0.000356 | 0.001569 | 0.012817 | 0.003757 | 0.013278 | 0.00107 | 0.005877 | 0.004651 | 0.006865 | 0.000785 | 0 | 0 | 0 | 0 | 0 | 0 | 0 | 0 | 0 | 0 | 0 | 0 |
| 18.91666667 | 0.011003 | 0.00403 | 0.000851 | 0.001334 | 0.011787 | 0.002984 | 0.015634 | 0.001112 | 0.00788 | 0.001781 | 0.006596 | 0.001257 | 0 | 0 | 0 | 0 | 0 | 0 | 0 | 0 | 0 | 0 | 0 | 0 |
| 18.93055556 | 0.013441 | 0.005096 | 0.001219 | 0.001161 | 0.010273 | 0.002553 | 0.017723 | 0.000964 | 0.009403 | 0.000828 | 0.006853 | 0.001519 | 0 | 0 | 0 | 0 | 0 | 0 | 0 | 0 | 0 | 0 | 0 | 0 |
| 18.94444444 | 0.014617 | 0.006086 | 0.001232 | 0.00097 | 0.008694 | 0.002394 | 0.019483 | 0.000655 | 0.009713 | 0.001794 | 0.00716 | 0.00188 | 0 | 0 | 0 | 0 | 0 | 0 | 0 | 0 | 0 | 0 | 0 | 0 |
| 18.95833333 | 0.014309 | 0.0048 | 0.001282 | 0.000893 | 0.007119 | 0.002351 | 0.02023 | 0.000414 | 0.009186 | 0.005112 | 0.007484 | 0.002315 | 0 | 0 | 0 | 0 | 0 | 0 | 0 | 0 | 0 | 0 | 0 | 0 |
| 18.97222222 | 0.01227 | 0.002096 | 0.00158 | 0.000942 | 0.005648 | 0.002284 | 0.01902 | 0.000492 | 0.008827 | 0.009555 | 0.008075 | 0.002557 | 0 | 0 | 0 | 0 | 0 | 0 | 0 | 0 | 0 | 0 | 0 | 0 |
| 18.98611111 | 0.009454 | 0.001149 | 0.002286 | 0.00098 | 0.005035 | 0.001824 | 0.016853 | 0.000705 | 0.009948 | 0.013086 | 0.008767 | 0.00318 | 0 | 0 | 0 | 0 | 0 | 0 | 0 | 0 | 0 | 0 | 0 | 0 |
| 19 | 0.007945 | 0.005767 | 0.00428 | 0.0012 | 0.006472 | 0.001773 | 0.017528 | 0.001431 | 0.015711 | 0.018493 | 0.008124 | 0.006797 | 0 | 0 | 0 | 0 | 0 | 0 | 0 | 0 | 0 | 0 | 0 | 0 |
| 19.01388889 | 0.00851 | 0.01853 | 0.008534 | 0.002771 | 0.011062 | 0.005384 | 0.024898 | 0.002812 | 0.030583 | 0.031864 | 0.00646 | 0.016616 | 0 | 0 | 0 | 0 | 0 | 0 | 0 | 0 | 0 | 0 | 0 | 0 |
| 19.02777778 | 0.009403 | 0.031468 | 0.01442 | 0.00565 | 0.015641 | 0.013172 | 0.038025 | 0.009872 | 0.055654 | 0.049627 | 0.009929 | 0.031727 | 0 | 0 | 0 | 0 | 0 | 0 | 0 | 0 | 0 | 0 | 0 | 0 |
| 19.04166667 | 0.008901 | 0.028165 | 0.0152 | 0.007593 | 0.012974 | 0.017859 | 0.043914 | 0.018667 | 0.064282 | 0.05507 | 0.020982 | 0.03778 | 0 | 0 | 0 | 0 | 0 | 0 | 0 | 0 | 0 | 0 | 0 | 0 |
| 19.05555556 | 0.007418 | 0.015622 | 0.010546 | 0.007981 | 0.00739 | 0.014553 | 0.030278 | 0.017235 | 0.047496 | 0.037714 | 0.031666 | 0.026817 | 0 | 0 | 0 | 0 | 0 | 0 | 0 | 0 | 0 | 0 | 0 | 0 |
| 19.06944444 | 0.005817 | 0.012649 | 0.008272 | 0.007662 | 0.00633 | 0.009108 | 0.014852 | 0.013219 | 0.042283 | 0.01693 | 0.03646 | 0.016849 | 0 | 0 | 0 | 0 | 0 | 0 | 0 | 0 | 0 | 0 | 0 | 0 |
| 19.08333333 | 0.004379 | 0.012164 | 0.006137 | 0.006931 | 0.004684 | 0.006297 | 0.013518 | 0.010701 | 0.041769 | 0.013698 | 0.036319 | 0.014544 | 0 | 0 | 0 | 0 | 0 | 0 | 0 | 0 | 0 | 0 | 0 | 0 |
| 19.09722222 | 0.003128 | 0.007375 | 0.002789 | 0.005945 | 0.001899 | 0.004903 | 0.012441 | 0.004947 | 0.025306 | 0.015167 | 0.031943 | 0.009972 | 0 | 0 | 0 | 0 | 0 | 0 | 0 | 0 | 0 | 0 | 0 | 0 |
| 19.11111111 | 0.00197 | 0.003981 | 0.0011 | 0.004625 | 0.001229 | 0.003958 | 0.006968 | 0.001429 | 0.010403 | 0.010648 | 0.025032 | 0.004306 | 0 | 0 | 0 | 0 | 0 | 0 | 0 | 0 | 0 | 0 | 0 | 0 |
| 19.125 | 0.00099 | 0.00264 | 0.00036 | 0.002987 | 0.000928 | 0.00329 | 0.003156 | 0.001396 | 0.004277 | 0.005809 | 0.01899 | 0.001309 | 0 | 0 | 0 | 0 | 0 | 0 | 0 | 0 | 0 | 0 | 0 | 0 |
| 19.13888889 | 0.00038 | 0.002246 | 0.000121 | 0.001554 | 0.000432 | 0.002695 | 0.00113 | 0.001247 | 0.001908 | 0.002931 | 0.015279 | 0.000328 | 0 | 0 | 0 | 0 | 0 | 0 | 0 | 0 | 0 | 0 | 0 | 0 |
| 19.15277778 | 0.000196 | 0.002491 | 0.000182 | 0.000891 | 0.000188 | 0.002104 | 0.000367 | 0.000632 | 0.001052 | 0.001824 | 0.012674 | 0.000573 | 0 | 0 | 0 | 0 | 0 | 0 | 0 | 0 | 0 | 0 | 0 | 0 |
| 19.16666667 | 0.000215 | 0.002288 | 0.000687 | 0.000789 | 0.000366 | 0.001665 | 0.001427 | 0.000341 | 0.002322 | 0.001295 | 0.009984 | 0.001446 | 0 | 0 | 0 | 0 | 0 | 0 | 0 | 0 | 0 | 0 | 0 | 0 |
| 19.18055556 | 0.00042 | 0.001157 | 0.001633 | 0.000742 | 0.001756 | 0.001798 | 0.005362 | 0.000447 | 0.007649 | 0.001227 | 0.007808 | 0.001799 | 0 | 0 | 0 | 0 | 0 | 0 | 0 | 0 | 0 | 0 | 0 | 0 |
| 19.19444444 | 0.000751 | 0.001312 | 0.001974 | 0.001317 | 0.002872 | 0.002534 | 0.007676 | 0.000669 | 0.010098 | 0.001662 | 0.006581 | 0.00141 | 0 | 0 | 0 | 0 | 0 | 0 | 0 | 0 | 0 | 0 | 0 | 0 |
| 19.20833333 | 0.001664 | 0.004896 | 0.001494 | 0.002909 | 0.003109 | 0.002725 | 0.007042 | 0.000904 | 0.009336 | 0.003359 | 0.00494 | 0.001067 | 0 | 0 | 0 | 0 | 0 | 0 | 0 | 0 | 0 | 0 | 0 | 0 |
| 19.22222222 | 0.003683 | 0.009579 | 0.001414 | 0.003326 | 0.004457 | 0.001779 | 0.008919 | 0.00101 | 0.013551 | 0.007585 | 0.002454 | 0.001245 | 0 | 0 | 0 | 0 | 0 | 0 | 0 | 0 | 0 | 0 | 0 | 0 |
| 19.23611111 | 0.004897 | 0.011065 | 0.001608 | 0.001666 | 0.004734 | 0.000846 | 0.009838 | 0.001124 | 0.014415 | 0.00903 | 0.000791 | 0.002225 | 0 | 0 | 0 | 0 | 0 | 0 | 0 | 0 | 0 | 0 | 0 | 0 |
| 19.25 | 0.00404 | 0.008949 | 0.001443 | 0.000417 | 0.002987 | 0.000591 | 0.00572 | 0.000997 | 0.007661 | 0.005707 | 0.000371 | 0.002613 | 0 | 0 | 0 | 0 | 0 | 0 | 0 | 0 | 0 | 0 | 0 | 0 |
| 19.26388889 | 0.002482 | 0.005325 | 0.001313 | 0.000378 | 0.001307 | 0.000758 | 0.001778 | 0.000505 | 0.002073 | 0.00366 | 0.00032 | 0.00181 | 4.4104170 | 0.000252 | 8.3118062 | 0.000206 | 0.000228 | 0.000983 | 0.000229 | 0.000152 | 0.000357 | 0.000175 | 0.000588 | 0.001609 |
| 19.27777778 | 0.00161 | 0.002009 | 0.00114 | 0.00051 | 0.000624 | 0.000917 | 0.000734 | 0.000354 | 0.000602 | 0.003157 | 0.000479 | 0.001049 | 0.00011 | 0.000727 | 0.000345 | 0.000563 | 0.000722 | 0.002429 | 0.000704 | 0.000443 | 0.000895 | 0.000641 | 0.001509 | 0.004441 |
| 19.29166667 | 0.00189 | 0.001079 | 0.000828 | 0.000537 | 0.001169 | 0.00071 | 0.001042 | 0.000875 | 0.001672 | 0.002457 | 0.000519 | 0.001086 | 0.000196 | 0.000962 | 0.000659 | 0.000562 | 0.001219 | 0.001924 | 0.000835 | 0.00046 | 0.001076 | 0.000962 | 0.001394 | 0.004544 |
| 19.30555556 | 0.002904 | 0.003163 | 0.000911 | 0.000804 | 0.002114 | 0.000543 | 0.002526 | 0.001824 | 0.003532 | 0.003863 | 0.000437 | 0.001861 | 0.000469 | 0.001059 | 0.000843 | 0.000311 | 0.001904 | 0.000856 | 0.000495 | 0.000302 | 0.001652 | 0.000814 | 0.001165 | 0.002383 |
| 19.31944444 | 0.00373 | 0.006041 | 0.001096 | 0.001097 | 0.002553 | 0.000581 | 0.004164 | 0.00237 | 0.004892 | 0.006361 | 0.000557 | 0.002506 | 0.000841 | 0.001141 | 0.000927 | 0.000256 | 0.002757 | 0.001373 | 0.000164 | 0.000415 | 0.002529 | 0.000801 | 0.001912 | 0.001518 |
| 19.33333333 | 0.003849 | 0.007312 | 0.000746 | 0.000957 | 0.002438 | 0.000412 | 0.004616 | 0.002185 | 0.005337 | 0.007053 | 0.000587 | 0.002466 | 0.000996 | 0.000923 | 0.000973 | 0.000322 | 0.003256 | 0.002411 | 8.8750007 | 0.000637 | 0.002352 | 0.001319 | 0.002473 | 0.002336 |
| 19.34722222 | 0.003574 | 0.006953 | 0.00025 | 0.00074 | 0.002338 | 0.000159 | 0.004485 | 0.001878 | 0.005304 | 0.006032 | 0.00061 | 0.001981 | 0.000861 | 0.000457 | 0.000981 | 0.000313 | 0.003255 | 0.003096 | 0.000176 | 0.000706 | 0.00128 | 0.00179 | 0.002338 | 0.002757 |
| 19.36111111 | 0.003385 | 0.006049 | 8.6604173 | 0.00072 | 0.00245 | 9.4631952 | 0.004515 | 0.001732 | 0.005237 | 0.004615 | 0.000851 | 0.001644 | 0.000645 | 0.000247 | 0.001032 | 0.00026 | 0.003131 | 0.003837 | 0.000246 | 0.000668 | 0.000574 | 0.001852 | 0.002268 | 0.0018 |
| 19.375 | 0.003532 | 0.005372 | 9.9638896 | 0.000767 | 0.002663 | 0.000193 | 0.005016 | 0.001627 | 0.005566 | 0.003866 | 0.001179 | 0.001886 | 0.000624 | 0.000646 | 0.00112 | 0.000234 | 0.0033 | 0.005134 | 0.000238 | 0.000633 | 0.000884 | 0.001629 | 0.003183 | 0.000755 |
| 19.38888889 | 0.004061 | 0.005299 | 0.000161 | 0.000893 | 0.003017 | 0.00041 | 0.005884 | 0.001495 | 0.006641 | 0.004139 | 0.001543 | 0.002303 | 0.000799 | 0.001385 | 0.001071 | 0.000239 | 0.003836 | 0.006795 | 0.000235 | 0.000712 | 0.001808 | 0.001426 | 0.005342 | 0.000599 |
| 19.40277778 | 0.004816 | 0.005313 | 0.000254 | 0.001006 | 0.003359 | 0.000598 | 0.006498 | 0.001329 | 0.007932 | 0.004916 | 0.001909 | 0.002217 | 0.000932 | 0.001945 | 0.000837 | 0.000268 | 0.004498 | 0.007855 | 0.000324 | 0.000927 | 0.002442 | 0.001718 | 0.007833 | 0.001252 |
| 19.41666667 | 0.005773 | 0.004778 | 0.000273 | 0.000898 | 0.003457 | 0.000652 | 0.006564 | 0.001225 | 0.008603 | 0.005498 | 0.00226 | 0.001753 | 0.000962 | 0.002116 | 0.000515 | 0.000335 | 0.005163 | 0.00787 | 0.000387 | 0.001162 | 0.002381 | 0.002564 | 0.009592 | 0.002561 |
| 19.43055556 | 0.007112 | 0.004045 | 0.000306 | 0.000661 | 0.003406 | 0.00069 | 0.006668 | 0.001309 | 0.008858 | 0.005711 | 0.002585 | 0.00143 | 0.001017 | 0.002058 | 0.000249 | 0.000471 | 0.005983 | 0.007731 | 0.000312 | 0.001301 | 0.001927 | 0.003436 | 0.010442 | 0.004058 |
| 19.44444444 | 0.008526 | 0.003127 | 0.000402 | 0.000411 | 0.002954 | 0.000577 | 0.007152 | 0.001343 | 0.009584 | 0.00542 | 0.002659 | 0.00116 | 0.001221 | 0.002023 | 0.000292 | 0.000716 | 0.006808 | 0.008295 | 0.000338 | 0.001382 | 0.001547 | 0.003811 | 0.010836 | 0.004913 |
| 19.45833333 | 0.009184 | 0.002063 | 0.000359 | 0.000246 | 0.001784 | 0.000532 | 0.007656 | 0.001091 | 0.011116 | 0.00425 | 0.002272 | 0.001686 | 0.001746 | 0.002249 | 0.0007 | 0.001239 | 0.00727 | 0.009545 | 0.000692 | 0.001653 | 0.001686 | 0.003679 | 0.011215 | 0.0048 |
| 19.47222222 | 0.008924 | 0.00144 | 0.000197 | 0.000311 | 0.000666 | 0.00092 | 0.008228 | 0.000696 | 0.013192 | 0.002631 | 0.001821 | 0.002993 | 0.002542 | 0.002829 | 0.000928 | 0.002106 | 0.0074 | 0.010766 | 0.001243 | 0.002333 | 0.002524 | 0.003464 | 0.012383 | 0.004255 |
| 19.48611111 | 0.009492 | 0.002975 | 0.000581 | 0.000367 | 0.002349 | 0.001295 | 0.01102 | 0.000479 | 0.018272 | 0.00316 | 0.00259 | 0.005633 | 0.003116 | 0.003279 | 0.000708 | 0.002696 | 0.007378 | 0.010777 | 0.001451 | 0.003171 | 0.00383 | 0.003177 | 0.016275 | 0.00328 |
| 19.5 | 0.014689 | 0.010541 | 0.002524 | 0.000243 | 0.009804 | 0.002853 | 0.021952 | 0.001024 | 0.0343 | 0.008907 | 0.006725 | 0.023269 | 0.002959 | 0.003079 | 0.00043 | 0.002179 | 0.007088 | 0.012204 | 0.001207 | 0.003743 | 0.005691 | 0.002288 | 0.027236 | 0.001751 |
| 19.51388889 | 0.024317 | 0.022378 | 0.005915 | 0.00049 | 0.020276 | 0.007062 | 0.035052 | 0.002161 | 0.062995 | 0.018801 | 0.017182 | 0.07972 | 0.002111 | 0.00273 | 0.0007 | 0.001079 | 0.006395 | 0.018825 | 0.001828 | 0.00421 | 0.007939 | 0.00131 | 0.046041 | 0.00209 |
| 19.52777778 | 0.030079 | 0.028708 | 0.008885 | 0.001563 | 0.026146 | 0.011995 | 0.036559 | 0.002898 | 0.084569 | 0.027907 | 0.03897 | 0.159135 | 0.001109 | 0.002806 | 0.002182 | 0.000522 | 0.005218 | 0.02182 | 0.004016 | 0.005115 | 0.008515 | 0.001153 | 0.059758 | 0.005797 |
| 19.54166667 | 0.027925 | 0.025543 | 0.010467 | 0.006019 | 0.025099 | 0.014715 | 0.028963 | 0.003285 | 0.080805 | 0.03272 | 0.064744 | 0.193585 | 0.000646 | 0.003302 | 0.003876 | 0.000828 | 0.003526 | 0.01424 | 0.005678 | 0.007545 | 0.006444 | 0.001529 | 0.056113 | 0.009104 |
| 19.55555556 | 0.024351 | 0.019738 | 0.012862 | 0.016866 | 0.022658 | 0.016529 | 0.01976 | 0.003815 | 0.063654 | 0.033422 | 0.078826 | 0.161824 | 0.000827 | 0.004318 | 0.003679 | 0.002473 | 0.002676 | 0.006388 | 0.004353 | 0.012912 | 0.004304 | 0.002852 | 0.040862 | 0.008807 |
| 19.56944444 | 0.022481 | 0.016762 | 0.017856 | 0.029727 | 0.022514 | 0.019702 | 0.013068 | 0.00403 | 0.050114 | 0.03102 | 0.103995 | 0.110904 | 0.001619 | 0.005455 | 0.001901 | 0.005095 | 0.003462 | 0.004288 | 0.001909 | 0.019143 | 0.003764 | 0.004914 | 0.025555 | 0.006655 |
| 19.58333333 | 0.018458 | 0.015856 | 0.023071 | 0.036272 | 0.022618 | 0.021899 | 0.009552 | 0.003335 | 0.041242 | 0.024871 | 0.135244 | 0.071345 | 0.003369 | 0.005685 | 0.000636 | 0.008484 | 0.003749 | 0.003983 | 0.00141 | 0.021503 | 0.004052 | 0.005784 | 0.015855 | 0.005038 |
| 19.59722222 | 0.01142 | 0.016458 | 0.026255 | 0.033382 | 0.020558 | 0.021895 | 0.006594 | 0.002116 | 0.035199 | 0.016546 | 0.116901 | 0.045848 | 0.005668 | 0.004631 | 0.000475 | 0.017351 | 0.002293 | 0.003071 | 0.001824 | 0.019775 | 0.004957 | 0.005773 | 0.01275 | 0.005473 |
| 19.61111111 | 0.005121 | 0.019315 | 0.026703 | 0.025816 | 0.01573 | 0.020299 | 0.00452 | 0.001273 | 0.030259 | 0.01075 | 0.076278 | 0.029879 | 0.007936 | 0.002982 | 0.000803 | 0.035236 | 0.000814 | 0.001843 | 0.002062 | 0.017039 | 0.006754 | 0.00642 | 0.014365 | 0.00906 |
| 19.625 | 0.004543 | 0.023414 | 0.023699 | 0.019873 | 0.008684 | 0.018067 | 0.003945 | 0.001302 | 0.023622 | 0.009316 | 0.070273 | 0.017991 | 0.009707 | 0.001624 | 0.001711 | 0.053916 | 0.000751 | 0.001393 | 0.00264 | 0.014118 | 0.008689 | 0.00759 | 0.017401 | 0.014684 |
| 19.63888889 | 0.012102 | 0.026504 | 0.01919 | 0.015883 | 0.004177 | 0.017665 | 0.003905 | 0.001992 | 0.015437 | 0.010848 | 0.086697 | 0.009769 | 0.010274 | 0.001005 | 0.003687 | 0.056514 | 0.002089 | 0.002112 | 0.003311 | 0.011019 | 0.009163 | 0.008089 | 0.01878 | 0.018797 |
| 19.65277778 | 0.02298 | 0.027392 | 0.015552 | 0.01239 | 0.004889 | 0.018678 | 0.003565 | 0.0026 | 0.008746 | 0.013807 | 0.098049 | 0.004429 | 0.009486 | 0.00108 | 0.006496 | 0.040842 | 0.004137 | 0.00347 | 0.003378 | 0.008349 | 0.007431 | 0.007207 | 0.017282 | 0.018212 |
| 19.66666667 | 0.026785 | 0.026532 | 0.01304 | 0.010112 | 0.004743 | 0.018106 | 0.003395 | 0.002752 | 0.005653 | 0.016346 | 0.093303 | 0.0018 | 0.007926 | 0.001841 | 0.009285 | 0.02438 | 0.00527 | 0.004776 | 0.002669 | 0.006073 | 0.004625 | 0.005597 | 0.012884 | 0.012975 |
| 19.68055556 | 0.01901 | 0.025521 | 0.011004 | 0.008901 | 0.003365 | 0.015509 | 0.003789 | 0.002656 | 0.004843 | 0.015815 | 0.075021 | 0.001405 | 0.006216 | 0.002862 | 0.011303 | 0.015403 | 0.004765 | 0.005966 | 0.001789 | 0.004028 | 0.002245 | 0.004168 | 0.006994 | 0.006305 |
| 19.69444444 | 0.00918 | 0.025406 | 0.008742 | 0.007801 | 0.004712 | 0.013516 | 0.004115 | 0.002301 | 0.003811 | 0.010754 | 0.05404 | 0.001369 | 0.004528 | 0.003138 | 0.012554 | 0.010304 | 0.003356 | 0.007121 | 0.001218 | 0.002311 | 0.000799 | 0.003065 | 0.004139 | 0.00326 |

| | | | | | | | | | | | | | | | | | | | | | | | | |
|---|---|---|---|---|---|---|---|---|---|---|---|---|---|---|---|---|---|---|---|---|---|---|---|---|
| 19.70833333 | 0.008313 | 0.025243 | 0.006212 | 0.006635 | 0.005871 | 0.013314 | 0.003741 | 0.00156 | 0.001999 | 0.005535 | 0.038206 | 0.002089 | 0.003109 | 0.002466 | 0.013329 | 0.007352 | 0.001721 | 0.008022 | 0.000867 | 0.000989 | 0.000264 | 0.002042 | 0.005734 | 0.005442 |
| 19.72222222 | 0.014452 | 0.023597 | 0.004046 | 0.005338 | 0.005647 | 0.013459 | 0.002846 | 0.000844 | 0.00083 | 0.005263 | 0.02846 | 0.003471 | 0.00227 | 0.001406 | 0.013234 | 0.005962 | 0.00101 | 0.008398 | 0.000522 | 0.000253 | 0.000567 | 0.001333 | 0.006596 | 0.008003 |
| 19.73611111 | 0.018142 | 0.020617 | 0.002849 | 0.003767 | 0.005554 | 0.012625 | 0.001913 | 0.000603 | 0.000741 | 0.007268 | 0.022363 | 0.005411 | 0.00194 | 0.000736 | 0.012162 | 0.005135 | 0.001966 | 0.007643 | 0.000215 | 0.000308 | 0.00145 | 0.001661 | 0.004208 | 0.00625 |
| 19.75 | 0.013052 | 0.017636 | 0.002691 | 0.002199 | 0.004898 | 0.010718 | 0.001322 | 0.00072 | 0.000659 | 0.006346 | 0.017581 | 0.007612 | 0.00175 | 0.000732 | 0.010524 | 0.003398 | 0.003211 | 0.005481 | 6.6868060 | 0.001053 | 0.002295 | 0.003098 | 0.002188 | 0.004004 |
| 19.76388889 | 0.007733 | 0.015452 | 0.003273 | 0.001258 | 0.00317 | 0.008304 | 0.001492 | 0.000913 | 0.000403 | 0.004915 | 0.01347 | 0.008294 | 0.001258 | 0.000869 | 0.00863 | 0.001474 | 0.002935 | 0.002864 | 9.0798618 | 0.001812 | 0.002355 | 0.00463 | 0.003225 | 0.006277 |
| 19.77777778 | 0.011976 | 0.013724 | 0.004038 | 0.001294 | 0.001445 | 0.005997 | 0.002426 | 0.001028 | 0.0004 | 0.008388 | 0.010824 | 0.007567 | 0.000791 | 0.000671 | 0.006622 | 0.001046 | 0.002044 | 0.001078 | 0.000226 | 0.001881 | 0.001447 | 0.005473 | 0.004178 | 0.008733 |
| 19.79166667 | 0.015627 | 0.011359 | 0.004538 | 0.001716 | 0.000926 | 0.00443 | 0.003331 | 0.001003 | 0.000683 | 0.013057 | 0.009617 | 0.006475 | 0.001188 | 0.000282 | 0.004474 | 0.001414 | 0.00285 | 0.000766 | 0.00041 | 0.001217 | 0.000713 | 0.005453 | 0.003649 | 0.007438 |
| 19.80555556 | 0.011085 | 0.0081 | 0.004575 | 0.001791 | 0.001263 | 0.003903 | 0.003552 | 0.000812 | 0.001851 | 0.013229 | 0.008633 | 0.004733 | 0.002189 | 0.00024 | 0.002638 | 0.001772 | 0.004542 | 0.001422 | 0.000655 | 0.000632 | 0.000944 | 0.00422 | 0.00515 | 0.004605 |
| 19.81944444 | 0.005758 | 0.005427 | 0.003972 | 0.001812 | 0.001359 | 0.003748 | 0.003275 | 0.000461 | 0.004672 | 0.009068 | 0.007396 | 0.002973 | 0.002888 | 0.000785 | 0.001581 | 0.001551 | 0.005005 | 0.001972 | 0.000903 | 0.000867 | 0.001234 | 0.002212 | 0.007821 | 0.00377 |
| 19.83333333 | 0.006559 | 0.003766 | 0.002928 | 0.001999 | 0.000935 | 0.002681 | 0.002646 | 0.000213 | 0.008341 | 0.005487 | 0.006513 | 0.004038 | 0.002904 | 0.001431 | 0.000977 | 0.001154 | 0.004207 | 0.002009 | 0.000972 | 0.001327 | 0.001181 | 0.0012 | 0.007901 | 0.004773 |
| 19.84722222 | 0.010921 | 0.002216 | 0.001965 | 0.001664 | 0.000724 | 0.00186 | 0.001601 | 0.000349 | 0.011483 | 0.0082 | 0.005972 | 0.009213 | 0.00246 | 0.00147 | 0.000527 | 0.001881 | 0.002703 | 0.001606 | 0.000823 | 0.001109 | 0.001282 | 0.001871 | 0.005738 | 0.003796 |
| 19.86111111 | 0.011267 | 0.001861 | 0.001429 | 0.000986 | 0.001387 | 0.003603 | 0.001273 | 0.000622 | 0.013709 | 0.015006 | 0.004864 | 0.017833 | 0.002116 | 0.001134 | 0.000476 | 0.002534 | 0.001446 | 0.000876 | 0.000545 | 0.000792 | 0.001548 | 0.002878 | 0.00418 | 0.002166 |
| 19.875 | 0.006489 | 0.003666 | 0.001393 | 0.000916 | 0.001909 | 0.006422 | 0.002988 | 0.000625 | 0.015358 | 0.019203 | 0.003153 | 0.028075 | 0.002195 | 0.001126 | 0.000853 | 0.001793 | 0.001882 | 0.000294 | 0.000289 | 0.001094 | 0.001761 | 0.003472 | 0.004844 | 0.002913 |
| 19.88888889 | 0.001899 | 0.006648 | 0.001232 | 0.001148 | 0.002036 | 0.00773 | 0.005598 | 0.000395 | 0.01716 | 0.017382 | 0.001489 | 0.035134 | 0.002798 | 0.001008 | 0.001123 | 0.001057 | 0.004293 | 0.000301 | 0.000232 | 0.001163 | 0.002365 | 0.00433 | 0.008727 | 0.00316 |
| 19.90277778 | 0.002593 | 0.011015 | 0.001776 | 0.001062 | 0.004279 | 0.007047 | 0.007027 | 0.000325 | 0.020056 | 0.011474 | 0.000761 | 0.03619 | 0.003989 | 0.000615 | 0.00084 | 0.001267 | 0.007765 | 0.001189 | 0.000656 | 0.000904 | 0.00404 | 0.005931 | 0.016977 | 0.002306 |
| 19.91666667 | 0.0104 | 0.016912 | 0.0051 | 0.000703 | 0.009405 | 0.005332 | 0.007433 | 0.000525 | 0.024093 | 0.006881 | 0.001383 | 0.035044 | 0.005811 | 0.000457 | 0.000467 | 0.001146 | 0.010566 | 0.003178 | 0.001651 | 0.001622 | 0.006674 | 0.007891 | 0.028577 | 0.003671 |
| 19.93055556 | 0.022688 | 0.022635 | 0.010334 | 0.00053 | 0.015287 | 0.004035 | 0.009016 | 0.000816 | 0.026665 | 0.007098 | 0.002104 | 0.036441 | 0.008461 | 0.000567 | 0.000593 | 0.00077 | 0.011168 | 0.005535 | 0.002667 | 0.003531 | 0.009147 | 0.009568 | 0.039063 | 0.006689 |
| 19.94444444 | 0.033148 | 0.026936 | 0.014613 | 0.000739 | 0.019579 | 0.003593 | 0.013301 | 0.001292 | 0.025182 | 0.011204 | 0.002187 | 0.041893 | 0.011594 | 0.000929 | 0.000771 | 0.001261 | 0.010547 | 0.007105 | 0.003327 | 0.005227 | 0.010313 | 0.010699 | 0.042936 | 0.01009 |
| 19.95833333 | 0.036111 | 0.029101 | 0.017219 | 0.000754 | 0.019021 | 0.003503 | 0.018733 | 0.001813 | 0.021266 | 0.016439 | 0.002348 | 0.049821 | 0.013776 | 0.000841 | 0.00063 | 0.001853 | 0.009827 | 0.007495 | 0.003706 | 0.005229 | 0.010305 | 0.011982 | 0.038168 | 0.014659 |
| 19.97222222 | 0.032355 | 0.028205 | 0.019685 | 0.000881 | 0.014189 | 0.003349 | 0.021845 | 0.001971 | 0.018219 | 0.021195 | 0.003247 | 0.054038 | 0.013793 | 0.000387 | 0.000449 | 0.00162 | 0.008066 | 0.006913 | 0.003698 | 0.003482 | 0.009492 | 0.013517 | 0.028022 | 0.019174 |
| 19.98611111 | 0.026921 | 0.024694 | 0.021418 | 0.001615 | 0.010304 | 0.003449 | 0.021394 | 0.00172 | 0.016802 | 0.024619 | 0.005471 | 0.049355 | 0.011653 | 0.000446 | 0.000471 | 0.00123 | 0.005117 | 0.00558 | 0.002771 | 0.001542 | 0.00761 | 0.013076 | 0.01787 | 0.020456 |
| 20 | 0.022396 | 0.020859 | 0.021174 | 0.002107 | 0.008373 | 0.005421 | 0.019806 | 0.001239 | 0.015799 | 0.029224 | 0.010075 | 0.042086 | 0.00812 | 0.000734 | 0.001232 | 0.001781 | 0.002697 | 0.003373 | 0.002146 | 0.00078 | 0.004657 | 0.009039 | 0.011844 | 0.017254 |
| 20.01388889 | 0.018426 | 0.018507 | 0.021351 | 0.001625 | 0.00621 | 0.013864 | 0.019594 | 0.001032 | 0.014672 | 0.042468 | 0.019636 | 0.044585 | 0.004276 | 0.001031 | 0.003043 | 0.002176 | 0.002357 | 0.001574 | 0.003986 | 0.003763 | 0.001973 | 0.004384 | 0.013847 | 0.014053 |
| 20.02777778 | 0.014919 | 0.016871 | 0.024277 | 0.000847 | 0.003337 | 0.028206 | 0.020821 | 0.000962 | 0.014114 | 0.058554 | 0.043089 | 0.062643 | 0.001545 | 0.001042 | 0.004918 | 0.006817 | 0.002958 | 0.001582 | 0.006551 | 0.011923 | 0.001413 | 0.003174 | 0.0261 | 0.016439 |
| 20.04166667 | 0.013258 | 0.014684 | 0.026054 | 0.003009 | 0.0012 | 0.033131 | 0.020925 | 0.002809 | 0.014382 | 0.055026 | 0.074329 | 0.077702 | 0.000478 | 0.001108 | 0.005362 | 0.024096 | 0.002304 | 0.001725 | 0.006754 | 0.017119 | 0.002307 | 0.003935 | 0.037162 | 0.019437 |
| 20.05555556 | 0.013199 | 0.011505 | 0.021991 | 0.007982 | 0.001614 | 0.023069 | 0.015647 | 0.006531 | 0.014228 | 0.031293 | 0.077623 | 0.057471 | 0.000671 | 0.002468 | 0.003566 | 0.038592 | 0.001796 | 0.002538 | 0.004679 | 0.013455 | 0.002232 | 0.003347 | 0.029404 | 0.014004 |
| 20.06944444 | 0.012131 | 0.007505 | 0.015026 | 0.009657 | 0.00343 | 0.012213 | 0.007442 | 0.007101 | 0.012085 | 0.016149 | 0.044715 | 0.026055 | 0.001788 | 0.003998 | 0.001686 | 0.030428 | 0.003195 | 0.008031 | 0.002268 | 0.007749 | 0.001156 | 0.001699 | 0.012792 | 0.006152 |
| 20.08333333 | 0.008219 | 0.004289 | 0.009664 | 0.007183 | 0.004059 | 0.006818 | 0.00395 | 0.004871 | 0.007756 | 0.016945 | 0.01384 | 0.02246 | 0.002722 | 0.00417 | 0.001935 | 0.012603 | 0.00379 | 0.014007 | 0.000998 | 0.004896 | 0.000784 | 0.000738 | 0.007677 | 0.003589 |
| 20.09722222 | 0.004651 | 0.002832 | 0.006483 | 0.004666 | 0.003529 | 0.004151 | 0.004318 | 0.003078 | 0.004629 | 0.014231 | 0.004712 | 0.025355 | 0.002694 | 0.003643 | 0.002491 | 0.003355 | 0.00221 | 0.012663 | 0.001049 | 0.003575 | 0.001188 | 0.00064 | 0.009803 | 0.002416 |
| 20.11111111 | 0.006283 | 0.002027 | 0.004574 | 0.003275 | 0.003257 | 0.00293 | 0.003353 | 0.002063 | 0.00703 | 0.006036 | 0.003455 | 0.019554 | 0.002242 | 0.003267 | 0.002249 | 0.002306 | 0.000661 | 0.007163 | 0.001568 | 0.003101 | 0.001402 | 0.000561 | 0.010959 | 0.001869 |
| 20.125 | 0.011168 | 0.001053 | 0.003337 | 0.002567 | 0.003421 | 0.002158 | 0.001725 | 0.00183 | 0.012533 | 0.002076 | 0.001685 | 0.013849 | 0.001797 | 0.00311 | 0.002039 | 0.002889 | 0.000239 | 0.003118 | 0.001773 | 0.003211 | 0.001246 | 0.000491 | 0.010943 | 0.002909 |
| 20.13888889 | 0.014587 | 0.000522 | 0.002535 | 0.002179 | 0.003189 | 0.001562 | 0.001574 | 0.001867 | 0.014841 | 0.002656 | 0.001203 | 0.009431 | 0.001338 | 0.002843 | 0.001988 | 0.002596 | 0.000739 | 0.001049 | 0.001657 | 0.003387 | 0.001097 | 0.000692 | 0.009519 | 0.004147 |
| 20.15277778 | 0.015368 | 0.000605 | 0.001844 | 0.001762 | 0.002084 | 0.001199 | 0.001472 | 0.001434 | 0.012755 | 0.004165 | 0.001223 | 0.007026 | 0.000834 | 0.002426 | 0.002018 | 0.001919 | 0.001664 | 0.000445 | 0.001415 | 0.003465 | 0.001034 | 0.000935 | 0.008019 | 0.004041 |
| 20.16666667 | 0.014589 | 0.000572 | 0.001111 | 0.001146 | 0.000978 | 0.001002 | 0.000841 | 0.00071 | 0.010375 | 0.005192 | 0.000887 | 0.006647 | 0.000605 | 0.003183 | 0.002419 | 0.001207 | 0.002064 | 0.000604 | 0.001216 | 0.003595 | 0.001496 | 0.001425 | 0.007937 | 0.002735 |
| 20.18055556 | 0.013298 | 0.000312 | 0.001149 | 0.00054 | 0.000732 | 0.00076 | 0.00191 | 0.000409 | 0.009721 | 0.006414 | 0.000947 | 0.005272 | 0.001794 | 0.009699 | 0.004518 | 0.002672 | 0.002449 | 0.002019 | 0.001789 | 0.004844 | 0.005836 | 0.004179 | 0.007946 | 0.002801 |
| 20.19444444 | 0.012054 | 0.000422 | 0.002562 | 0.000437 | 0.000695 | 0.000458 | 0.005586 | 0.00035 | 0.008609 | 0.007506 | 0.00167 | 0.003152 | 0.004481 | 0.023243 | 0.008421 | 0.006983 | 0.005171 | 0.004453 | 0.003313 | 0.008106 | 0.014762 | 0.010166 | 0.006072 | 0.004479 |
| 20.20833333 | 0.011175 | 0.001392 | 0.003795 | 0.001139 | 0.000692 | 0.00037 | 0.007262 | 0.000249 | 0.009989 | 0.005998 | 0.003151 | 0.005095 | 0.004794 | 0.025559 | 0.009123 | 0.007919 | 0.006776 | 0.004354 | 0.003316 | 0.009044 | 0.016038 | 0.011748 | 0.004773 | 0.004984 |
| 20.22222222 | 0.010408 | 0.003346 | 0.003502 | 0.002289 | 0.001408 | 0.000597 | 0.007555 | 0.000245 | 0.018303 | 0.004131 | 0.007161 | 0.012981 | 0.003577 | 0.018541 | 0.006117 | 0.007299 | 0.00701 | 0.003183 | 0.002352 | 0.00634 | 0.0127 | 0.011006 | 0.007681 | 0.005619 |
| 20.23611111 | 0.008979 | 0.005329 | 0.002707 | 0.002595 | 0.002334 | 0.000716 | 0.00973 | 0.000259 | 0.02531 | 0.005105 | 0.012514 | 0.021852 | 0.004499 | 0.020944 | 0.004789 | 0.011691 | 0.010061 | 0.004675 | 0.002999 | 0.005095 | 0.01701 | 0.015459 | 0.011702 | 0.009405 |
| 20.25 | 0.006233 | 0.005843 | 0.003 | 0.001899 | 0.002727 | 0.000492 | 0.007513 | 0.000496 | 0.022849 | 0.00418 | 0.013565 | 0.023373 | 0.003836 | 0.016735 | 0.003769 | 0.013309 | 0.00954 | 0.004897 | 0.002628 | 0.004019 | 0.015254 | 0.014179 | 0.011715 | 0.009435 |
| 20.26388889 | 0.003012 | 0.005044 | 0.00445 | 0.001566 | 0.00301 | 0.000354 | 0.002622 | 0.001426 | 0.017155 | 0.00135 | 0.010266 | 0.01917 | 0.001817 | 0.004958 | 0.002061 | 0.009199 | 0.004817 | 0.00266 | 0.000862 | 0.001585 | 0.005873 | 0.006637 | 0.009349 | 0.00406 |
| 20.27777778 | 0.00103 | 0.004402 | 0.005803 | 0.001673 | 0.003431 | 0.000463 | 0.001752 | 0.002506 | 0.015199 | 0.000892 | 0.006964 | 0.015179 | 0.001782 | 0.00101 | 0.001757 | 0.005876 | 0.002689 | 0.001443 | 0.000155 | 0.000332 | 0.001771 | 0.00269 | 0.008262 | 0.001318 |
| 20.29166667 | 0.001203 | 0.004676 | 0.006192 | 0.001497 | 0.003632 | 0.000482 | 0.003548 | 0.002663 | 0.01705 | 0.002343 | 0.005616 | 0.012602 | 0.002275 | 0.000698 | 0.001608 | 0.003874 | 0.00298 | 0.001324 | 0.000188 | 0.000296 | 0.001746 | 0.001603 | 0.008432 | 0.001896 |
| 20.30555556 | 0.003901 | 0.005479 | 0.005937 | 0.001118 | 0.00406 | 0.000386 | 0.00591 | 0.002168 | 0.021534 | 0.004706 | 0.006484 | 0.012879 | 0.002479 | 0.000522 | 0.000922 | 0.002172 | 0.0035 | 0.001345 | 0.00035 | 0.000721 | 0.002385 | 0.001599 | 0.008856 | 0.002445 |
| 20.31944444 | 0.008762 | 0.006061 | 0.005853 | 0.000906 | 0.005303 | 0.000413 | 0.008139 | 0.002023 | 0.02599 | 0.007314 | 0.010215 | 0.015748 | 0.002739 | 0.000513 | 0.000446 | 0.001191 | 0.003551 | 0.00142 | 0.000487 | 0.001801 | 0.00338 | 0.002532 | 0.009172 | 0.002209 |
| 20.33333333 | 0.013808 | 0.006103 | 0.006286 | 0.001015 | 0.006605 | 0.00062 | 0.010158 | 0.002408 | 0.027235 | 0.009214 | 0.019078 | 0.01863 | 0.003157 | 0.000508 | 0.000334 | 0.001087 | 0.003643 | 0.001513 | 0.000525 | 0.003439 | 0.00497 | 0.0038 | 0.009495 | 0.00192 |
| 20.34722222 | 0.016592 | 0.006026 | 0.007025 | 0.001283 | 0.006829 | 0.000936 | 0.012644 | 0.002781 | 0.025172 | 0.010056 | 0.030907 | 0.020316 | 0.00392 | 0.000995 | 0.000267 | 0.001837 | 0.004656 | 0.001589 | 0.000428 | 0.004201 | 0.006663 | 0.004399 | 0.010231 | 0.002352 |
| 20.36111111 | 0.016804 | 0.006554 | 0.007782 | 0.001326 | 0.006035 | 0.001319 | 0.015837 | 0.002827 | 0.021788 | 0.010151 | 0.03755 | 0.021495 | 0.005006 | 0.002288 | 0.000141 | 0.002937 | 0.00657 | 0.001916 | 0.000209 | 0.003467 | 0.007669 | 0.003921 | 0.01145 | 0.003322 |
| 20.375 | 0.016042 | 0.007948 | 0.008355 | 0.001093 | 0.005265 | 0.001815 | 0.018185 | 0.002733 | 0.020094 | 0.010245 | 0.036393 | 0.02304 | 0.006068 | 0.003981 | 8.7493062 | 0.003725 | 0.008061 | 0.002791 | 7.4416672 | 0.002447 | 0.007994 | 0.003026 | 0.013247 | 0.003995 |
| 20.38888889 | 0.015325 | 0.009655 | 0.008694 | 0.001184 | 0.005441 | 0.0025 | 0.018567 | 0.002741 | 0.022589 | 0.010754 | 0.032653 | 0.025237 | 0.007011 | 0.005222 | 0.000154 | 0.004056 | 0.008084 | 0.003983 | 0.000174 | 0.001816 | 0.008149 | 0.002782 | 0.015322 | 0.003942 |
| 20.40277778 | 0.01447 | 0.01056 | 0.008887 | 0.002129 | 0.006544 | 0.003118 | 0.017575 | 0.002774 | 0.028051 | 0.011637 | 0.029469 | 0.027631 | 0.008108 | 0.005374 | 0.000369 | 0.004066 | 0.007315 | 0.004914 | 0.000519 | 0.001333 | 0.008431 | 0.003833 | 0.016766 | 0.003753 |
| 20.41666667 | 0.012893 | 0.010257 | 0.009475 | 0.003781 | 0.007632 | 0.00334 | 0.016391 | 0.002608 | 0.032584 | 0.013427 | 0.026953 | 0.029936 | 0.009202 | 0.004977 | 0.000597 | 0.003748 | 0.007384 | 0.005291 | 0.000923 | 0.00093 | 0.008773 | 0.005832 | 0.017415 | 0.004755 |
| 20.43055556 | 0.010145 | 0.009856 | 0.010977 | 0.005506 | 0.008306 | 0.003592 | 0.016378 | 0.002458 | 0.034223 | 0.016629 | 0.026293 | 0.032514 | 0.009865 | 0.006291 | 0.000495 | 0.003386 | 0.009119 | 0.005403 | 0.001081 | 0.000635 | 0.009343 | 0.007597 | 0.017837 | 0.007393 |
| 20.44444444 | 0.006415 | 0.010422 | 0.012396 | 0.006822 | 0.008731 | 0.004693 | 0.01755 | 0.002683 | 0.033424 | 0.019865 | 0.02681 | 0.034916 | 0.010123 | 0.012222 | 0.000362 | 0.004234 | 0.011962 | 0.005321 | 0.00126 | 0.00054 | 0.011121 | 0.008361 | 0.018097 | 0.00971 |
| 20.45833333 | 0.00284 | 0.011125 | 0.012116 | 0.007081 | 0.008274 | 0.006761 | 0.017927 | 0.003054 | 0.030429 | 0.020583 | 0.025812 | 0.035748 | 0.010067 | 0.02001 | 0.000732 | 0.007127 | 0.014282 | 0.004664 | 0.00221 | 0.000671 | 0.014302 | 0.008659 | 0.018292 | 0.009625 |
| 20.47222222 | 0.000868 | 0.011486 | 0.010376 | 0.005699 | 0.007374 | 0.00877 | 0.015146 | 0.002929 | 0.025712 | 0.017397 | 0.024066 | 0.035247 | 0.009886 | 0.021202 | 0.001089 | 0.010577 | 0.014412 | 0.003978 | 0.003704 | 0.000714 | 0.017211 | 0.00963 | 0.019494 | 0.00801 |
| 20.48611111 | 0.002465 | 0.015755 | 0.010217 | 0.004328 | 0.010303 | 0.011112 | 0.009647 | 0.002493 | 0.021857 | 0.013183 | 0.025623 | 0.039536 | 0.010352 | 0.015869 | 0.000843 | 0.012413 | 0.012156 | 0.003842 | 0.004223 | 0.000912 | 0.017764 | 0.012223 | 0.024862 | 0.005971 |
| 20.5 | 0.00981 | 0.030358 | 0.015947 | 0.005459 | 0.022737 | 0.017398 | 0.005083 | 0.002874 | 0.022726 | 0.018708 | 0.033576 | 0.058617 | 0.012108 | 0.013353 | 0.000362 | 0.014023 | 0.008632 | 0.003489 | 0.003039 | 0.0031 | 0.016111 | 0.016441 | 0.04039 | 0.00383 |
| 20.51388889 | 0.024863 | 0.052386 | 0.027415 | 0.008611 | 0.038183 | 0.029249 | 0.004424 | 0.004137 | 0.027565 | 0.040824 | 0.046142 | 0.08599 | 0.013664 | 0.017205 | 0.000684 | 0.016382 | 0.004942 | 0.002802 | 0.00155 | 0.008234 | 0.014763 | 0.019717 | 0.063382 | 0.003323 |
| 20.52777778 | 0.041205 | 0.067138 | 0.036768 | 0.010951 | 0.040645 | 0.040187 | 0.006451 | 0.004922 | 0.029955 | 0.06604 | 0.054941 | 0.093517 | 0.011911 | 0.02397 | 0.0024 | 0.012945 | 0.002394 | 0.002198 | 0.001033 | 0.013985 | 0.014362 | 0.019118 | 0.076132 | 0.003666 |
| 20.54166667 | 0.042898 | 0.068819 | 0.038337 | 0.010779 | 0.031829 | 0.039585 | 0.006415 | 0.004912 | 0.027857 | 0.074041 | 0.05418 | 0.072375 | 0.007365 | 0.029826 | 0.004028 | 0.005348 | 0.001801 | 0.001571 | 0.001818 | 0.019812 | 0.012797 | 0.016814 | 0.066774 | 0.002361 |
| 20.55555556 | 0.029175 | 0.0691 | 0.035438 | 0.008952 | 0.028662 | 0.028453 | 0.003793 | 0.004838 | 0.026789 | 0.06164 | 0.047201 | 0.044795 | 0.004322 | 0.031311 | 0.003525 | 0.002517 | 0.002094 | 0.001843 | 0.003908 | 0.027649 | 0.009132 | 0.016828 | 0.041846 | 0.000696 |
| 20.56944444 | 0.013554 | 0.073984 | 0.031021 | 0.007072 | 0.043894 | 0.017898 | 0.00373 | 0.004331 | 0.027461 | 0.037445 | 0.037871 | 0.02428 | 0.004119 | 0.027158 | 0.001583 | 0.001964 | 0.002889 | 0.003296 | 0.005843 | 0.032494 | 0.004657 | 0.017591 | 0.020031 | 0.000722 |
| 20.58333333 | 0.00693 | 0.073713 | 0.023696 | 0.006513 | 0.06246 | 0.011306 | 0.006976 | 0.003019 | 0.022321 | 0.016447 | 0.027615 | 0.014142 | 0.004607 | 0.020301 | 0.00038 | 0.001253 | 0.003711 | 0.004177 | 0.006416 | 0.027632 | 0.002208 | 0.015885 | 0.017104 | 0.002575 |
| 20.59722222 | 0.013897 | 0.060862 | 0.014623 | 0.008458 | 0.053634 | 0.006461 | 0.008944 | 0.001826 | 0.011476 | 0.008477 | 0.018995 | 0.020955 | 0.004664 | 0.014839 | 0.000296 | 0.001314 | 0.002853 | 0.003651 | 0.005896 | 0.016043 | 0.003166 | 0.012493 | 0.025769 | 0.00553 |
| 20.61111111 | 0.023226 | 0.041517 | 0.007338 | 0.012079 | 0.0267 | 0.004343 | 0.006529 | 0.001261 | 0.005907 | 0.008477 | 0.012984 | 0.035864 | 0.004223 | 0.012461 | 0.000622 | 0.001065 | 0.001366 | 0.002497 | 0.00462 | 0.006278 | 0.004901 | 0.009363 | 0.026318 | 0.008578 |
| 20.625 | 0.019632 | 0.025222 | 0.003749 | 0.014817 | 0.009275 | 0.006998 | 0.004062 | 0.00102 | 0.009348 | 0.017231 | 0.008234 | 0.039619 | 0.002943 | 0.01276 | 0.001072 | 0.00067 | 0.000787 | 0.001317 | 0.003075 | 0.002408 | 0.004489 | 0.007146 | 0.015811 | 0.010989 |
| 20.63888889 | 0.013551 | 0.015139 | 0.004741 | 0.015466 | 0.006116 | 0.013875 | 0.006938 | 0.001153 | 0.014368 | 0.031935 | 0.004423 | 0.027489 | 0.001614 | 0.014637 | 0.001386 | 0.000621 | 0.001162 | 0.000542 | 0.002274 | 0.002547 | 0.003937 | 0.006253 | 0.010123 | 0.011663 |
| 20.65277778 | 0.025086 | 0.009446 | 0.009067 | 0.014279 | 0.00893 | 0.020604 | 0.009635 | 0.001625 | 0.016044 | 0.040189 | 0.003403 | 0.011352 | 0.001832 | 0.016043 | 0.001376 | 0.001021 | 0.002936 | 0.000566 | 0.002816 | 0.005607 | 0.007734 | 0.007234 | 0.016452 | 0.010129 |
| 20.66666667 | 0.041423 | 0.006029 | 0.012742 | 0.011356 | 0.008123 | 0.020307 | 0.006611 | 0.001914 | 0.013837 | 0.034513 | 0.005095 | 0.003892 | 0.003529 | 0.015152 | 0.001313 | 0.001278 | 0.004495 | 0.001145 | 0.003873 | 0.009559 | 0.013142 | 0.009654 | 0.023755 | 0.007517 |
| 20.68055556 | 0.042271 | 0.004086 | 0.013148 | 0.007787 | 0.006883 | 0.011657 | 0.003474 | 0.001645 | 0.008699 | 0.020396 | 0.005665 | 0.004688 | 0.005267 | 0.01214 | 0.00141 | 0.001044 | 0.003865 | 0.001708 | 0.004094 | 0.008995 | 0.013453 | 0.011023 | 0.022817 | 0.004324 |
| 20.69444444 | 0.030535 | 0.002666 | 0.011244 | 0.005407 | 0.015054 | 0.004545 | 0.005925 | 0.00107 | 0.007263 | 0.017708 | 0.00345 | 0.005022 | 0.005866 | 0.008354 | 0.001461 | 0.000706 | 0.001914 | 0.002069 | 0.003301 | 0.010345 | 0.008166 | 0.009159 | 0.013912 | 0.00218 |
| 20.70833333 | 0.016285 | 0.002163 | 0.008783 | 0.005089 | 0.027235 | 0.005155 | 0.010128 | 0.000537 | 0.012502 | 0.027099 | 0.002146 | 0.002464 | 0.004936 | 0.004899 | 0.001372 | 0.000656 | 0.000716 | 0.002594 | 0.002065 | 0.025114 | 0.004409 | 0.004971 | 0.00658 | 0.001971 |
| 20.72222222 | 0.011763 | 0.004492 | 0.006838 | 0.006537 | 0.030461 | 0.007522 | 0.010787 | 0.00023 | 0.016177 | 0.026624 | 0.005075 | 0.000663 | 0.003141 | 0.002567 | 0.001279 | 0.001045 | 0.000647 | 0.003626 | 0.001035 | 0.039676 | 0.006732 | 0.002011 | 0.005719 | 0.002487 |
| 20.73611111 | 0.020503 | 0.008954 | 0.005938 | 0.008298 | 0.022066 | 0.008315 | 0.008347 | 0.000219 | 0.012649 | 0.015034 | 0.011554 | 0.000651 | 0.001656 | 0.002116 | 0.001233 | 0.001864 | 0.000586 | 0.004946 | 0.000901 | 0.039522 | 0.00985 | 0.00195 | 0.006223 | 0.004947 |
| 20.75 | 0.03322 | 0.012814 | 0.004859 | 0.008821 | 0.011669 | 0.007656 | 0.004933 | 0.000221 | 0.007214 | 0.009627 | 0.018825 | 0.002899 | 0.001032 | 0.003464 | 0.001212 | 0.002686 | 0.001429 | 0.006167 | 0.001432 | 0.034301 | 0.01024 | 0.001842 | 0.009775 | 0.007433 |
| 20.76388889 | 0.03937 | 0.014058 | 0.002978 | 0.007883 | 0.009469 | 0.00563 | 0.003235 | 0.000162 | 0.007731 | 0.012523 | 0.021684 | 0.007329 | 0.001021 | 0.005758 | 0.00112 | 0.003167 | 0.005016 | 0.006654 | 0.001485 | 0.029952 | 0.009077 | 0.001641 | 0.01624 | 0.006892 |
| 20.77777778 | 0.028812 | 0.011974 | 0.002089 | 0.006512 | 0.012081 | 0.008269 | 0.006478 | 0.000331 | 0.012185 | 0.012695 | 0.01757 | 0.008886 | 0.001269 | 0.007916 | 0.000871 | 0.00333 | 0.009165 | 0.00535 | 0.001042 | 0.02315 | 0.005951 | 0.004037 | 0.018824 | 0.0049 |
| 20.79166667 | 0.015357 | 0.008045 | 0.002692 | 0.005684 | 0.011682 | 0.016842 | 0.010795 | 0.000724 | 0.012647 | 0.010081 | 0.009965 | 0.006952 | 0.001644 | 0.00819 | 0.000592 | 0.003067 | 0.008955 | 0.003133 | 0.00103 | 0.015424 | 0.003776 | 0.006888 | 0.014091 | 0.003034 |
| 20.80555556 | 0.023573 | 0.004558 | 0.003436 | 0.005177 | 0.009663 | 0.020975 | 0.009069 | 0.000854 | 0.007983 | 0.01687 | 0.003736 | 0.009547 | 0.002176 | 0.005493 | 0.000486 | 0.002371 | 0.005068 | 0.002827 | 0.001114 | 0.00986 | 0.005242 | 0.007041 | 0.008754 | 0.002696 |
| 20.81944444 | 0.039758 | 0.00232 | 0.003638 | 0.004241 | 0.009452 | 0.015831 | 0.004656 | 0.000556 | 0.004734 | 0.031629 | 0.002227 | 0.018169 | 0.002761 | 0.002995 | 0.000596 | 0.001731 | 0.003153 | 0.00444 | 0.000766 | 0.006191 | 0.0056 | 0.004703 | 0.012748 | 0.003966 |
| 20.83333333 | 0.039406 | 0.001214 | 0.003288 | 0.002944 | 0.009628 | 0.007589 | 0.004462 | 0.000544 | 0.005451 | 0.035473 | 0.005625 | 0.024578 | 0.003103 | 0.004452 | 0.000845 | 0.001457 | 0.004347 | 0.004621 | 0.000504 | 0.003559 | 0.003563 | 0.003623 | 0.020308 | 0.003614 |
| 20.84722222 | 0.024399 | 0.00132 | 0.002704 | 0.001995 | 0.008233 | 0.003528 | 0.006285 | 0.001198 | 0.004755 | 0.022347 | 0.010138 | 0.022407 | 0.00323 | 0.007603 | 0.001113 | 0.001278 | 0.005933 | 0.003141 | 0.000919 | 0.002829 | 0.003916 | 0.0061 | 0.019741 | 0.002375 |
| 20.86111111 | 0.011886 | 0.002631 | 0.002478 | 0.001432 | 0.005436 | 0.005463 | 0.006206 | 0.00146 | 0.003284 | 0.0089 | 0.01143 | 0.013577 | 0.003497 | 0.009021 | 0.001293 | 0.000995 | 0.006653 | 0.003538 | 0.001991 | 0.004288 | 0.007302 | 0.008829 | 0.011679 | 0.003436 |
| 20.875 | 0.014754 | 0.005188 | 0.003315 | 0.001301 | 0.003202 | 0.008918 | 0.005141 | 0.000897 | 0.004305 | 0.005094 | 0.009365 | 0.006883 | 0.004214 | 0.008512 | 0.001224 | 0.000767 | 0.005226 | 0.005914 | 0.003301 | 0.006107 | 0.010167 | 0.010326 | 0.007163 | 0.004075 |
| 20.88888889 | 0.029761 | 0.008759 | 0.005602 | 0.002662 | 0.003478 | 0.011268 | 0.004803 | 0.000637 | 0.004054 | 0.005677 | 0.006619 | 0.008892 | 0.005453 | 0.00712 | 0.0008 | 0.000546 | 0.003442 | 0.007468 | 0.004632 | 0.006336 | 0.010517 | 0.011946 | 0.010048 | 0.002877 |
| 20.90277778 | 0.046243 | 0.012106 | 0.009162 | 0.004898 | 0.005745 | 0.012101 | 0.005778 | 0.000751 | 0.002215 | 0.011597 | 0.005515 | 0.016999 | 0.006913 | 0.006137 | 0.000358 | 0.000383 | 0.005311 | 0.007681 | 0.005541 | 0.005087 | 0.009715 | 0.013007 | 0.01127 | 0.003227 |
| 20.91666667 | 0.05458 | 0.014081 | 0.0126 | 0.005449 | 0.007786 | 0.010051 | 0.007271 | 0.000739 | 0.001807 | 0.022582 | 0.006758 | 0.023796 | 0.008179 | 0.00702 | 0.000248 | 0.000539 | 0.009026 | 0.007514 | 0.005324 | 0.003983 | 0.010123 | 0.012134 | 0.007564 | 0.005339 |
| 20.93055556 | 0.05149 | 0.016017 | 0.014144 | 0.003808 | 0.008603 | 0.006228 | 0.008331 | 0.001474 | 0.004461 | 0.0307 | 0.009214 | 0.025296 | 0.009243 | 0.011371 | 0.000249 | 0.00073 | 0.011867 | 0.007002 | 0.004158 | 0.004002 | 0.012703 | 0.009943 | 0.006907 | 0.005406 |
| 20.94444444 | 0.040981 | 0.020735 | 0.014991 | 0.002086 | 0.008965 | 0.003177 | 0.009069 | 0.003789 | 0.012662 | 0.029599 | 0.012094 | 0.022764 | 0.01102 | 0.01896 | 0.00016 | 0.000657 | 0.014742 | 0.005508 | 0.002916 | 0.005097 | 0.016242 | 0.008887 | 0.013157 | 0.002978 |
| 20.95833333 | 0.02503 | 0.02767 | 0.017803 | 0.001792 | 0.00957 | 0.001796 | 0.009655 | 0.006708 | 0.0241 | 0.018467 | 0.01619 | 0.019438 | 0.014557 | 0.026543 | 0.000138 | 0.000608 | 0.018363 | 0.003411 | 0.002282 | 0.006403 | 0.018513 | 0.010116 | 0.01948 | 0.001326 |
| 20.97222222 | 0.011973 | 0.032598 | 0.022489 | 0.002096 | 0.010084 | 0.001627 | 0.009528 | 0.007737 | 0.03307 | 0.011613 | 0.021065 | 0.016972 | 0.018574 | 0.030521 | 0.000264 | 0.000823 | 0.020191 | 0.001946 | 0.00235 | 0.006726 | 0.018102 | 0.012207 | 0.020124 | 0.002626 |
| 20.98611111 | 0.011007 | 0.032258 | 0.02653 | 0.001839 | 0.009664 | 0.002179 | 0.008551 | 0.005433 | 0.035299 | 0.016933 | 0.024512 | 0.015662 | 0.019479 | 0.029639 | 0.000345 | 0.000952 | 0.01603 | 0.001128 | 0.002466 | 0.005981 | 0.014876 | 0.012419 | 0.016678 | 0.00705 |
| 21 | 0.011744 | 0.028266 | 0.028816 | 0.001862 | 0.008699 | 0.003434 | 0.009253 | 0.002197 | 0.032271 | 0.016904 | 0.025991 | 0.018762 | 0.015985 | 0.02603 | 0.00023 | 0.000769 | 0.007981 | 0.001062 | 0.001927 | 0.006143 | 0.01014 | 0.010005 | 0.015527 | 0.012304 |
| 21.01388889 | 0.010054 | 0.02539 | 0.030242 | 0.002832 | 0.007602 | 0.005903 | 0.014864 | 0.001098 | 0.031411 | 0.016784 | 0.02691 | 0.030882 | 0.011525 | 0.022401 | 0.000208 | 0.000687 | 0.003901 | 0.002284 | 0.001258 | 0.010042 | 0.006777 | 0.007804 | 0.024721 | 0.018111 |
| 21.02777778 | 0.018917 | 0.026603 | 0.030819 | 0.002817 | 0.005651 | 0.010161 | 0.023824 | 0.001602 | 0.034677 | 0.037632 | 0.02859 | 0.051627 | 0.009336 | 0.018729 | 0.000695 | 0.001295 | 0.006827 | 0.003424 | 0.001317 | 0.016709 | 0.006608 | 0.008178 | 0.047182 | 0.02503 |
| 21.04166667 | 0.037081 | 0.029663 | 0.031072 | 0.001554 | 0.004091 | 0.014355 | 0.031199 | 0.001812 | 0.035601 | 0.056805 | 0.03005 | 0.069592 | 0.008526 | 0.014795 | 0.001671 | 0.001718 | 0.012743 | 0.00347 | 0.001942 | 0.021019 | 0.008061 | 0.009898 | 0.068886 | 0.029121 |
| 21.05555556 | 0.047752 | 0.028647 | 0.030347 | 0.00185 | 0.005365 | 0.01436 | 0.032356 | 0.001526 | 0.030007 | 0.046463 | 0.027176 | 0.068002 | 0.007897 | 0.011606 | 0.001931 | 0.001554 | 0.017397 | 0.003096 | 0.002621 | 0.019649 | 0.008732 | 0.009312 | 0.064669 | 0.025597 |
| 21.06944444 | 0.039223 | 0.021648 | 0.025143 | 0.003529 | 0.00806 | 0.009848 | 0.025017 | 0.001817 | 0.02073 | 0.023434 | 0.018861 | 0.046347 | 0.00796 | 0.008897 | 0.001361 | 0.001967 | 0.017601 | 0.002826 | 0.002647 | 0.013986 | 0.007586 | 0.005764 | 0.034731 | 0.014749 |
| 21.08333333 | 0.019498 | 0.011832 | 0.015985 | 0.003411 | 0.008576 | 0.005013 | 0.013294 | 0.002952 | 0.010769 | 0.015401 | 0.010674 | 0.020881 | 0.008262 | 0.00589 | 0.001377 | 0.001925 | 0.01128 | 0.002607 | 0.001662 | 0.007746 | 0.004482 | 0.002925 | 0.01176 | 0.005613 |
| 21.09722222 | 0.008693 | 0.004259 | 0.00781 | 0.002927 | 0.006712 | 0.002418 | 0.004415 | 0.004133 | 0.004544 | 0.018354 | 0.006763 | 0.005354 | 0.008046 | 0.003369 | 0.001389 | 0.000898 | 0.005555 | 0.002309 | 0.000616 | 0.003407 | 0.001965 | 0.002437 | 0.011403 | 0.003863 |
| 21.11111111 | 0.011296 | 0.001796 | 0.003017 | 0.005952 | 0.005181 | 0.001378 | 0.001977 | 0.004838 | 0.00369 | 0.014929 | 0.005494 | 0.001282 | 0.007465 | 0.002439 | 0.000974 | 0.000265 | 0.007803 | 0.001581 | 0.000399 | 0.002329 | 0.002189 | 0.003765 | 0.016268 | 0.003701 |
| 21.125 | 0.014535 | 0.001471 | 0.00093 | 0.010378 | 0.005399 | 0.000967 | 0.004018 | 0.004534 | 0.00299 | 0.008316 | 0.004848 | 0.002195 | 0.006434 | 0.003041 | 0.000824 | 0.000128 | 0.011903 | 0.000653 | 0.000884 | 0.004666 | 0.002846 | 0.005814 | 0.014939 | 0.003727 |
| 21.13888889 | 0.012965 | 0.001313 | 0.000698 | 0.011936 | 0.005915 | 0.001028 | 0.005421 | 0.00276 | 0.002214 | 0.009311 | 0.005008 | 0.003995 | 0.004869 | 0.00384 | 0.000747 | 7.7868061 | 0.01163 | 0.000489 | 0.001796 | 0.008205 | 0.002357 | 0.006699 | 0.010376 | 0.006183 |
| 21.15277778 | 0.010424 | 0.002188 | 0.001455 | 0.008993 | 0.004503 | 0.001204 | 0.004056 | 0.001375 | 0.003108 | 0.01513 | 0.005958 | 0.004177 | 0.003353 | 0.00345 | 0.000555 | 8.9909729 | 0.008371 | 0.000909 | 0.002671 | 0.010281 | 0.001483 | 0.006583 | 0.005999 | 0.006249 |
| 21.16666667 | 0.008845 | 0.003261 | 0.002241 | 0.00458 | 0.002226 | 0.001155 | 0.002293 | 0.001337 | 0.003136 | 0.018769 | 0.006762 | 0.003359 | 0.00229 | 0.00192 | 0.000459 | 0.000234 | 0.004669 | 0.001283 | 0.002979 | 0.009665 | 0.000798 | 0.006883 | 0.003934 | 0.005483 |
| 21.18055556 | 0.007768 | 0.004228 | 0.00303 | 0.002262 | 0.001184 | 0.001211 | 0.001924 | 0.001467 | 0.002313 | 0.017998 | 0.007361 | 0.004251 | 0.001693 | 0.000615 | 0.000556 | 0.000622 | 0.002382 | 0.001382 | 0.002704 | 0.00735 | 0.000813 | 0.007879 | 0.004525 | 0.008359 |
| 21.19444444 | 0.006716 | 0.005188 | 0.004217 | 0.002224 | 0.00095 | 0.001592 | 0.001676 | 0.002191 | 0.004624 | 0.011958 | 0.008211 | 0.003885 | 0.00175 | 0.000844 | 0.000685 | 0.001206 | 0.00229 | 0.001041 | 0.002337 | 0.005104 | 0.001075 | 0.008805 | 0.007602 | 0.00757 |
| 21.20833333 | 0.007401 | 0.004132 | 0.004723 | 0.00293 | 0.000992 | 0.001611 | 0.00183 | 0.002725 | 0.012445 | 0.007815 | 0.006998 | 0.003035 | 0.002076 | 0.002229 | 0.000836 | 0.001356 | 0.004648 | 0.000573 | 0.002207 | 0.003472 | 0.001331 | 0.00824 | 0.015468 | 0.003974 |
| 21.22222222 | 0.011516 | 0.002648 | 0.006218 | 0.0032 | 0.001128 | 0.001712 | 0.004068 | 0.002038 | 0.021849 | 0.012625 | 0.003985 | 0.006165 | 0.001612 | 0.0031 | 0.00102 | 0.000984 | 0.009275 | 0.000652 | 0.002076 | 0.00247 | 0.002592 | 0.005906 | 0.026156 | 0.00474 |
| 21.23611111 | 0.016282 | 0.003833 | 0.009141 | 0.004479 | 0.00178 | 0.002491 | 0.005547 | 0.002039 | 0.024497 | 0.015358 | 0.002266 | 0.009633 | 0.000955 | 0.00257 | 0.000919 | 0.000856 | 0.014095 | 0.001506 | 0.001863 | 0.002212 | 0.004022 | 0.003915 | 0.028438 | 0.00482 |
| 21.25 | 0.016598 | 0.004604 | 0.008749 | 0.005359 | 0.003223 | 0.002461 | 0.003775 | 0.002919 | 0.018592 | 0.010796 | 0.001918 | 0.007821 | 0.000975 | 0.001466 | 0.000522 | 0.000776 | 0.016297 | 0.002706 | 0.001763 | 0.002471 | 0.004111 | 0.003515 | 0.019597 | 0.003156 |
| 21.26388889 | 0.010963 | 0.002961 | 0.006131 | 0.005198 | 0.004632 | 0.001625 | 0.001406 | 0.002971 | 0.012621 | 0.01101 | 0.00304 | 0.003517 | 0.001021 | 0.000932 | 0.000221 | 0.000398 | 0.014915 | 0.003381 | 0.00155 | 0.002886 | 0.003688 | 0.003132 | 0.01067 | 0.003357 |
| 21.27777778 | 0.005786 | 0.001552 | 0.005192 | 0.005355 | 0.004967 | 0.001377 | 0.001555 | 0.002701 | 0.01456 | 0.014491 | 0.003515 | 0.003467 | 0.001158 | 0.000721 | 9.5722229 | 0.000131 | 0.011418 | 0.002545 | 0.001016 | 0.003145 | 0.003791 | 0.001898 | 0.007864 | 0.002646 |
| 21.29166667 | 0.006543 | 0.001296 | 0.0058 | 0.005322 | 0.00425 | 0.001802 | 0.004574 | 0.002331 | 0.024503 | 0.013348 | 0.002304 | 0.010651 | 0.001227 | 0.000861 | 8.0708339 | 0.000151 | 0.008895 | 0.001616 | 0.000807 | 0.002942 | 0.004254 | 0.001254 | 0.00884 | 0.001189 |
| 21.30555556 | 0.00768 | 0.001365 | 0.007189 | 0.005434 | 0.003615 | 0.002359 | 0.008316 | 0.00242 | 0.034947 | 0.016328 | 0.002034 | 0.02103 | 0.001553 | 0.00239 | 0.000171 | 0.000321 | 0.009097 | 0.002686 | 0.001259 | 0.002276 | 0.004747 | 0.001886 | 0.0102 | 0.000846 |
| 21.31944444 | 0.00516 | 0.002821 | 0.008996 | 0.005725 | 0.003536 | 0.002732 | 0.011009 | 0.004814 | 0.037352 | 0.023957 | 0.002754 | 0.024573 | 0.002961 | 0.004838 | 0.000218 | 0.000403 | 0.010456 | 0.004396 | 0.001878 | 0.001686 | 0.005288 | 0.003278 | 0.009546 | 0.000942 |

| | | | | | | | | | | | | | | | | | | | | | | | | |
|---|---|---|---|---|---|---|---|---|---|---|---|---|---|---|---|---|---|---|---|---|---|---|---|---|
| 21.33333333 | 0.002584 | 0.00559 | 0.011511 | 0.005646 | 0.003967 | 0.003146 | 0.013001 | 0.007154 | 0.03237 | 0.021762 | 0.003218 | 0.019177 | 0.004136 | 0.006721 | 0.000143 | 0.000326 | 0.010836 | 0.005079 | 0.002546 | 0.001803 | 0.006017 | 0.005537 | 0.006836 | 0.001939 |
| 21.34722222 | 0.003917 | 0.007561 | 0.015093 | 0.00534 | 0.004567 | 0.004021 | 0.015276 | 0.005975 | 0.02711 | 0.011619 | 0.00367 | 0.01527 | 0.004261 | 0.007391 | 0.000108 | 0.000316 | 0.009973 | 0.004872 | 0.003303 | 0.002641 | 0.006906 | 0.007889 | 0.004513 | 0.003664 |
| 21.36111111 | 0.009276 | 0.008046 | 0.018901 | 0.005777 | 0.004938 | 0.005224 | 0.018343 | 0.002805 | 0.024684 | 0.005011 | 0.004638 | 0.021584 | 0.004262 | 0.006921 | 0.000245 | 0.000484 | 0.008414 | 0.004202 | 0.003875 | 0.003537 | 0.007818 | 0.008313 | 0.004391 | 0.00547 |
| 21.375 | 0.015393 | 0.007828 | 0.020958 | 0.007353 | 0.005191 | 0.006549 | 0.021602 | 0.001189 | 0.023798 | 0.010047 | 0.005795 | 0.035439 | 0.005271 | 0.006329 | 0.000422 | 0.000936 | 0.007326 | 0.003729 | 0.004142 | 0.003801 | 0.008657 | 0.006479 | 0.005668 | 0.006529 |
| 21.38888889 | 0.019662 | 0.008017 | 0.020354 | 0.009125 | 0.005318 | 0.008101 | 0.023425 | 0.002746 | 0.022846 | 0.030688 | 0.006768 | 0.045048 | 0.007506 | 0.007029 | 0.000445 | 0.001856 | 0.008011 | 0.004046 | 0.004202 | 0.003404 | 0.00927 | 0.004567 | 0.006937 | 0.005686 |
| 21.40277778 | 0.021646 | 0.009133 | 0.018374 | 0.009995 | 0.005155 | 0.009195 | 0.021799 | 0.005517 | 0.020586 | 0.049218 | 0.007731 | 0.042181 | 0.009741 | 0.009043 | 0.000396 | 0.002973 | 0.010321 | 0.004928 | 0.004204 | 0.002807 | 0.00978 | 0.004798 | 0.008323 | 0.003447 |
| 21.41666667 | 0.022552 | 0.011226 | 0.016032 | 0.009755 | 0.005121 | 0.009433 | 0.016833 | 0.006453 | 0.017112 | 0.046724 | 0.009062 | 0.032471 | 0.010565 | 0.010939 | 0.000383 | 0.003749 | 0.013 | 0.00575 | 0.004318 | 0.002652 | 0.01086 | 0.007436 | 0.010988 | 0.002205 |
| 21.43055556 | 0.024124 | 0.01383 | 0.014005 | 0.008793 | 0.005705 | 0.009399 | 0.011678 | 0.005755 | 0.014548 | 0.029079 | 0.010902 | 0.02708 | 0.009973 | 0.011685 | 0.000329 | 0.004097 | 0.014848 | 0.006798 | 0.004503 | 0.003538 | 0.01271 | 0.01093 | 0.01456 | 0.004171 |
| 21.44444444 | 0.026903 | 0.015686 | 0.013702 | 0.006926 | 0.006462 | 0.008996 | 0.008579 | 0.005016 | 0.014226 | 0.011823 | 0.012671 | 0.030224 | 0.009467 | 0.011907 | 0.000325 | 0.004459 | 0.015279 | 0.008742 | 0.004725 | 0.005255 | 0.01435 | 0.01372 | 0.016914 | 0.008532 |
| 21.45833333 | 0.02897 | 0.015455 | 0.014245 | 0.004408 | 0.00647 | 0.007888 | 0.006822 | 0.004335 | 0.014752 | 0.00605 | 0.013259 | 0.03764 | 0.010181 | 0.012343 | 0.000635 | 0.005347 | 0.015001 | 0.010075 | 0.00519 | 0.006841 | 0.015173 | 0.015348 | 0.017731 | 0.012234 |
| 21.47222222 | 0.026586 | 0.014372 | 0.013459 | 0.002844 | 0.006074 | 0.007069 | 0.004514 | 0.003461 | 0.013913 | 0.014371 | 0.012681 | 0.039899 | 0.012081 | 0.012203 | 0.00094 | 0.006525 | 0.01483 | 0.00801 | 0.006014 | 0.008028 | 0.015856 | 0.015546 | 0.018746 | 0.013412 |
| 21.48611111 | 0.018335 | 0.017684 | 0.013564 | 0.004361 | 0.008398 | 0.008715 | 0.003534 | 0.003334 | 0.011163 | 0.030466 | 0.014614 | 0.036257 | 0.013886 | 0.011508 | 0.0007 | 0.007024 | 0.013672 | 0.004373 | 0.007359 | 0.010329 | 0.016994 | 0.015087 | 0.022388 | 0.011924 |
| 21.5 | 0.009142 | 0.031872 | 0.021383 | 0.011254 | 0.019099 | 0.015269 | 0.006051 | 0.004837 | 0.010235 | 0.046761 | 0.026769 | 0.044464 | 0.01425 | 0.01384 | 0.000284 | 0.006507 | 0.010365 | 0.003142 | 0.009412 | 0.018131 | 0.018847 | 0.016745 | 0.033498 | 0.007916 |
| 21.51388889 | 0.006142 | 0.055907 | 0.037408 | 0.020653 | 0.034044 | 0.024025 | 0.007029 | 0.006795 | 0.016398 | 0.065723 | 0.048635 | 0.072822 | 0.013306 | 0.023129 | 0.00054 | 0.00535 | 0.006682 | 0.002831 | 0.009764 | 0.030286 | 0.019632 | 0.021131 | 0.052336 | 0.003957 |
| 21.52777778 | 0.013346 | 0.071495 | 0.05011 | 0.023051 | 0.038589 | 0.025098 | 0.006548 | 0.007386 | 0.029163 | 0.084505 | 0.05925 | 0.092797 | 0.011291 | 0.036243 | 0.001361 | 0.003667 | 0.004464 | 0.002424 | 0.007437 | 0.033453 | 0.016281 | 0.027332 | 0.066676 | 0.005188 |
| 21.54166667 | 0.029411 | 0.066868 | 0.050607 | 0.016082 | 0.033485 | 0.016331 | 0.00719 | 0.006631 | 0.043018 | 0.084706 | 0.048034 | 0.081147 | 0.008208 | 0.043129 | 0.002214 | 0.002407 | 0.002696 | 0.003356 | 0.006925 | 0.023644 | 0.011002 | 0.033676 | 0.065026 | 0.010737 |
| 21.55555556 | 0.0434 | 0.0558 | 0.042068 | 0.00817 | 0.030223 | 0.006843 | 0.005996 | 0.005934 | 0.048925 | 0.061285 | 0.032408 | 0.054537 | 0.005219 | 0.039296 | 0.002508 | 0.002686 | 0.001274 | 0.003619 | 0.008492 | 0.012917 | 0.007093 | 0.03441 | 0.047952 | 0.011805 |
| 21.56944444 | 0.041849 | 0.047382 | 0.030595 | 0.004083 | 0.028548 | 0.003616 | 0.008457 | 0.005725 | 0.040915 | 0.033676 | 0.02154 | 0.030234 | 0.003196 | 0.029505 | 0.001579 | 0.002575 | 0.000801 | 0.002509 | 0.007912 | 0.006492 | 0.004439 | 0.02742 | 0.024289 | 0.007544 |
| 21.58333333 | 0.026945 | 0.038589 | 0.019303 | 0.001935 | 0.02414 | 0.007353 | 0.016888 | 0.005088 | 0.025963 | 0.032925 | 0.011153 | 0.017367 | 0.002123 | 0.017214 | 0.000509 | 0.001703 | 0.001512 | 0.002647 | 0.004711 | 0.003153 | 0.004711 | 0.01679 | 0.01204 | 0.007089 |
| 21.59722222 | 0.012153 | 0.028145 | 0.010026 | 0.000587 | 0.017811 | 0.013673 | 0.022327 | 0.003668 | 0.012755 | 0.060231 | 0.005199 | 0.024217 | 0.002765 | 0.007401 | 0.000296 | 0.002012 | 0.003644 | 0.004142 | 0.002353 | 0.00398 | 0.007638 | 0.008402 | 0.020268 | 0.011601 |
| 21.61111111 | 0.011328 | 0.015579 | 0.00462 | 0.000818 | 0.011378 | 0.016741 | 0.0181 | 0.002222 | 0.007766 | 0.065956 | 0.007721 | 0.033503 | 0.00467 | 0.004174 | 0.000367 | 0.0023 | 0.005317 | 0.004279 | 0.002628 | 0.007169 | 0.0092 | 0.006463 | 0.033891 | 0.013051 |
| 21.625 | 0.02082 | 0.008333 | 0.004479 | 0.00323 | 0.005807 | 0.013066 | 0.011905 | 0.002028 | 0.012556 | 0.042172 | 0.010347 | 0.025086 | 0.006424 | 0.006781 | 0.000399 | 0.001742 | 0.004929 | 0.00315 | 0.00278 | 0.008278 | 0.007613 | 0.006323 | 0.036501 | 0.010518 |
| 21.63888889 | 0.022591 | 0.01308 | 0.006824 | 0.006371 | 0.003881 | 0.007645 | 0.015616 | 0.002891 | 0.014835 | 0.038964 | 0.007862 | 0.012405 | 0.007261 | 0.015896 | 0.000524 | 0.001096 | 0.002934 | 0.00384 | 0.001709 | 0.005751 | 0.004672 | 0.00516 | 0.023946 | 0.013162 |
| 21.65277778 | 0.018446 | 0.021688 | 0.006501 | 0.007656 | 0.006614 | 0.009677 | 0.019916 | 0.002923 | 0.008986 | 0.053023 | 0.007891 | 0.008015 | 0.006688 | 0.024569 | 0.000731 | 0.001079 | 0.001034 | 0.006555 | 0.001222 | 0.004337 | 0.004328 | 0.008572 | 0.011736 | 0.019663 |
| 21.66666667 | 0.01566 | 0.025787 | 0.004292 | 0.006387 | 0.009776 | 0.01727 | 0.013237 | 0.001843 | 0.00342 | 0.045596 | 0.013978 | 0.007306 | 0.005175 | 0.024263 | 0.000862 | 0.002064 | 0.000523 | 0.007993 | 0.001603 | 0.007607 | 0.007293 | 0.014847 | 0.014301 | 0.018292 |
| 21.68055556 | 0.022791 | 0.024547 | 0.004703 | 0.003874 | 0.009681 | 0.021521 | 0.006976 | 0.000982 | 0.002451 | 0.023292 | 0.018198 | 0.014028 | 0.003452 | 0.016502 | 0.000915 | 0.002943 | 0.001238 | 0.006736 | 0.001557 | 0.010897 | 0.008757 | 0.016419 | 0.019574 | 0.010825 |
| 21.69444444 | 0.043721 | 0.019744 | 0.006411 | 0.001677 | 0.006165 | 0.018316 | 0.01334 | 0.000815 | 0.003679 | 0.017095 | 0.015856 | 0.025366 | 0.002092 | 0.008786 | 0.001 | 0.003024 | 0.002773 | 0.003956 | 0.000935 | 0.010495 | 0.005928 | 0.0116 | 0.019134 | 0.009671 |
| 21.70833333 | 0.053206 | 0.013641 | 0.006345 | 0.000929 | 0.003055 | 0.010462 | 0.02325 | 0.001026 | 0.004542 | 0.034312 | 0.010382 | 0.029884 | 0.001685 | 0.009165 | 0.001087 | 0.002617 | 0.003516 | 0.00249 | 0.000463 | 0.007618 | 0.002896 | 0.006031 | 0.017385 | 0.014371 |
| 21.72222222 | 0.037024 | 0.007907 | 0.005411 | 0.001345 | 0.003732 | 0.004952 | 0.0228 | 0.001252 | 0.005692 | 0.052328 | 0.005273 | 0.021595 | 0.002187 | 0.016716 | 0.00101 | 0.001864 | 0.002973 | 0.004226 | 0.000287 | 0.004222 | 0.003799 | 0.005595 | 0.015178 | 0.012523 |
| 21.73611111 | 0.015035 | 0.004799 | 0.003825 | 0.001317 | 0.005871 | 0.006479 | 0.013399 | 0.001001 | 0.008556 | 0.046878 | 0.003459 | 0.012337 | 0.00257 | 0.021874 | 0.00079 | 0.000948 | 0.003573 | 0.00639 | 0.000384 | 0.003415 | 0.006427 | 0.008642 | 0.009958 | 0.007278 |
| 21.75 | 0.007716 | 0.004953 | 0.002474 | 0.001454 | 0.00689 | 0.012794 | 0.006391 | 0.000434 | 0.009374 | 0.027621 | 0.005695 | 0.019012 | 0.001954 | 0.018877 | 0.000657 | 0.000582 | 0.00477 | 0.007335 | 0.000574 | 0.006337 | 0.007863 | 0.011111 | 0.005126 | 0.009744 |
| 21.76388889 | 0.014309 | 0.007812 | 0.004224 | 0.002996 | 0.006393 | 0.017394 | 0.004083 | 0.000338 | 0.006425 | 0.025565 | 0.008918 | 0.029824 | 0.001104 | 0.010951 | 0.000678 | 0.001308 | 0.003727 | 0.007885 | 0.001136 | 0.009074 | 0.007742 | 0.011333 | 0.005566 | 0.01136 |
| 21.77777778 | 0.023792 | 0.013213 | 0.008765 | 0.004313 | 0.004035 | 0.015832 | 0.003536 | 0.000617 | 0.003246 | 0.035499 | 0.010143 | 0.026709 | 0.001784 | 0.004253 | 0.00068 | 0.002389 | 0.001706 | 0.006676 | 0.002377 | 0.008122 | 0.00621 | 0.008235 | 0.005502 | 0.007091 |
| 21.79166667 | 0.033029 | 0.019901 | 0.01333 | 0.004191 | 0.0022 | 0.010038 | 0.006829 | 0.000734 | 0.00259 | 0.029678 | 0.007684 | 0.016687 | 0.004616 | 0.002223 | 0.000588 | 0.002663 | 0.000837 | 0.003962 | 0.003826 | 0.00464 | 0.003793 | 0.004944 | 0.003582 | 0.007687 |
| 21.80555556 | 0.041428 | 0.027111 | 0.015704 | 0.00318 | 0.003721 | 0.005391 | 0.013732 | 0.001315 | 0.007885 | 0.017078 | 0.004859 | 0.018917 | 0.008422 | 0.006014 | 0.000532 | 0.002042 | 0.001245 | 0.002542 | 0.005043 | 0.00284 | 0.001996 | 0.006049 | 0.004684 | 0.013573 |
| 21.81944444 | 0.044589 | 0.03421 | 0.015434 | 0.001998 | 0.006916 | 0.003036 | 0.020339 | 0.003182 | 0.020153 | 0.024239 | 0.006259 | 0.030524 | 0.011836 | 0.014703 | 0.000539 | 0.001184 | 0.003737 | 0.00266 | 0.005909 | 0.004101 | 0.002314 | 0.007703 | 0.005548 | 0.01602 |
| 21.83333333 | 0.039565 | 0.036958 | 0.014352 | 0.001672 | 0.009792 | 0.002639 | 0.020909 | 0.006302 | 0.033789 | 0.047832 | 0.007499 | 0.0329 | 0.014397 | 0.02396 | 0.000507 | 0.000537 | 0.007861 | 0.004374 | 0.006323 | 0.004778 | 0.003283 | 0.006198 | 0.004418 | 0.0116 |
| 21.84722222 | 0.028009 | 0.033726 | 0.014374 | 0.002198 | 0.012394 | 0.003749 | 0.015917 | 0.010116 | 0.040362 | 0.063106 | 0.005571 | 0.02537 | 0.01541 | 0.02845 | 0.000531 | 0.000809 | 0.011572 | 0.0074 | 0.006838 | 0.003073 | 0.002665 | 0.00722 | 0.006004 | 0.005291 |
| 21.86111111 | 0.017101 | 0.031718 | 0.016328 | 0.002156 | 0.015019 | 0.004651 | 0.01429 | 0.01275 | 0.037238 | 0.057005 | 0.006741 | 0.020231 | 0.014519 | 0.027378 | 0.00078 | 0.002512 | 0.014441 | 0.009903 | 0.008576 | 0.002329 | 0.001746 | 0.013584 | 0.010878 | 0.002859 |
| 21.875 | 0.014723 | 0.036295 | 0.019987 | 0.001251 | 0.017738 | 0.008327 | 0.020196 | 0.012413 | 0.030769 | 0.03673 | 0.013872 | 0.022873 | 0.013744 | 0.024152 | 0.001093 | 0.005113 | 0.01667 | 0.013147 | 0.01104 | 0.005744 | 0.003236 | 0.020644 | 0.015418 | 0.007373 |
| 21.88888889 | 0.022084 | 0.043459 | 0.024404 | 0.000621 | 0.019987 | 0.014359 | 0.029706 | 0.009575 | 0.028379 | 0.018087 | 0.022082 | 0.029959 | 0.015146 | 0.01999 | 0.001186 | 0.008072 | 0.017831 | 0.017775 | 0.012735 | 0.011684 | 0.006766 | 0.026098 | 0.015654 | 0.021249 |
| 21.90277778 | 0.03267 | 0.047339 | 0.028239 | 0.001053 | 0.021098 | 0.016308 | 0.037053 | 0.00609 | 0.032114 | 0.010679 | 0.027872 | 0.03738 | 0.018252 | 0.015451 | 0.001057 | 0.010938 | 0.01793 | 0.02075 | 0.013532 | 0.016506 | 0.010346 | 0.029985 | 0.00987 | 0.035966 |
| 21.91666667 | 0.038403 | 0.047284 | 0.029512 | 0.002351 | 0.01948 | 0.013145 | 0.037455 | 0.004448 | 0.038207 | 0.010603 | 0.031447 | 0.043759 | 0.021726 | 0.013921 | 0.000924 | 0.012474 | 0.018043 | 0.020559 | 0.014444 | 0.017144 | 0.012397 | 0.031576 | 0.004933 | 0.037388 |
| 21.93055556 | 0.036284 | 0.043567 | 0.027044 | 0.003798 | 0.014567 | 0.008888 | 0.028605 | 0.006407 | 0.040616 | 0.019269 | 0.032113 | 0.04722 | 0.024691 | 0.017854 | 0.000787 | 0.011521 | 0.018785 | 0.018364 | 0.015506 | 0.012648 | 0.011622 | 0.029854 | 0.006856 | 0.023872 |
| 21.94444444 | 0.030528 | 0.035329 | 0.022134 | 0.004982 | 0.008803 | 0.005958 | 0.014921 | 0.010478 | 0.035788 | 0.042445 | 0.028477 | 0.043093 | 0.026108 | 0.026707 | 0.000462 | 0.008641 | 0.018821 | 0.015439 | 0.015192 | 0.006036 | 0.008386 | 0.023488 | 0.010814 | 0.010899 |
| 21.95833333 | 0.027186 | 0.025875 | 0.017427 | 0.005557 | 0.004868 | 0.004167 | 0.006455 | 0.01255 | 0.025635 | 0.067883 | 0.022021 | 0.03196 | 0.025701 | 0.036932 | 0.000241 | 0.005776 | 0.017659 | 0.012868 | 0.013073 | 0.002035 | 0.004701 | 0.014423 | 0.011823 | 0.009176 |
| 21.97222222 | 0.026862 | 0.019423 | 0.014056 | 0.004643 | 0.003588 | 0.003588 | 0.005041 | 0.010587 | 0.015788 | 0.075922 | 0.016062 | 0.021034 | 0.023748 | 0.042305 | 0.000526 | 0.004768 | 0.016166 | 0.011296 | 0.010084 | 0.001632 | 0.002409 | 0.007821 | 0.009573 | 0.008923 |
| 21.98611111 | 0.02517 | 0.016463 | 0.011996 | 0.002386 | 0.003617 | 0.004291 | 0.003789 | 0.006496 | 0.010034 | 0.060641 | 0.012425 | 0.01308 | 0.019707 | 0.039319 | 0.000773 | 0.005276 | 0.014214 | 0.010014 | 0.007248 | 0.001906 | 0.001876 | 0.006109 | 0.005461 | 0.009489 |
| 22 | 0.021756 | 0.015842 | 0.013103 | 0.001624 | 0.00357 | 0.005653 | 0.003035 | 0.00306 | 0.010266 | 0.036682 | 0.013694 | 0.008522 | 0.014313 | 0.030511 | 0.000863 | 0.004907 | 0.012171 | 0.008336 | 0.005596 | 0.004698 | 0.002842 | 0.007766 | 0.005841 | 0.018753 |
| 22.01388889 | 0.022445 | 0.016718 | 0.020095 | 0.006786 | 0.003166 | 0.008232 | 0.005972 | 0.001564 | 0.016236 | 0.025624 | 0.026073 | 0.01139 | 0.010064 | 0.021617 | 0.001729 | 0.002903 | 0.012298 | 0.007201 | 0.005651 | 0.014193 | 0.005755 | 0.011498 | 0.018287 | 0.030464 |
| 22.02777778 | 0.028963 | 0.018569 | 0.031297 | 0.016815 | 0.002055 | 0.012587 | 0.009933 | 0.002022 | 0.022308 | 0.036476 | 0.05031 | 0.024313 | 0.009011 | 0.016963 | 0.003315 | 0.001341 | 0.017738 | 0.00831 | 0.006538 | 0.023546 | 0.010669 | 0.016502 | 0.040898 | 0.039509 |
| 22.04166667 | 0.036297 | 0.023028 | 0.039511 | 0.021727 | 0.000733 | 0.015745 | 0.013426 | 0.005035 | 0.022142 | 0.052555 | 0.063988 | 0.040934 | 0.010802 | 0.015746 | 0.004391 | 0.001375 | 0.03028 | 0.010431 | 0.005635 | 0.022177 | 0.01456 | 0.019066 | 0.055293 | 0.043033 |
| 22.05555556 | 0.037924 | 0.02961 | 0.037195 | 0.014849 | 0.000633 | 0.014499 | 0.017246 | 0.008082 | 0.016843 | 0.047933 | 0.04435 | 0.049519 | 0.0124 | 0.014514 | 0.00356 | 0.00247 | 0.042114 | 0.009236 | 0.002706 | 0.013533 | 0.014785 | 0.015652 | 0.04428 | 0.034156 |
| 22.06944444 | 0.029618 | 0.030974 | 0.023208 | 0.007844 | 0.001636 | 0.010402 | 0.01947 | 0.007301 | 0.011903 | 0.02616 | 0.016494 | 0.043359 | 0.011151 | 0.011217 | 0.001615 | 0.003701 | 0.039497 | 0.00494 | 0.000909 | 0.006794 | 0.012834 | 0.009263 | 0.020714 | 0.017139 |
| 22.08333333 | 0.015331 | 0.023353 | 0.009816 | 0.011635 | 0.00207 | 0.006365 | 0.017163 | 0.008581 | 0.010077 | 0.018688 | 0.009646 | 0.025923 | 0.007879 | 0.006486 | 0.000648 | 0.003386 | 0.022666 | 0.001737 | 0.001145 | 0.003351 | 0.009374 | 0.005808 | 0.006641 | 0.008111 |
| 22.09722222 | 0.005582 | 0.011876 | 0.007125 | 0.015947 | 0.001443 | 0.003434 | 0.011572 | 0.015365 | 0.009697 | 0.038843 | 0.009673 | 0.015107 | 0.005337 | 0.004968 | 0.000525 | 0.001791 | 0.008679 | 0.00092 | 0.002402 | 0.00146 | 0.004631 | 0.006432 | 0.004422 | 0.010332 |
| 22.11111111 | 0.004217 | 0.004981 | 0.007495 | 0.013494 | 0.000874 | 0.001783 | 0.007771 | 0.019537 | 0.007715 | 0.065422 | 0.005667 | 0.018422 | 0.004751 | 0.009123 | 0.000493 | 0.000717 | 0.007418 | 0.001496 | 0.003956 | 0.001032 | 0.0015 | 0.00806 | 0.00588 | 0.010409 |
| 22.125 | 0.004489 | 0.004706 | 0.005091 | 0.00864 | 0.001107 | 0.001227 | 0.008933 | 0.017143 | 0.004743 | 0.077931 | 0.002446 | 0.019226 | 0.005743 | 0.014323 | 0.0005 | 0.000666 | 0.010092 | 0.001906 | 0.004824 | 0.001862 | 0.000747 | 0.00789 | 0.005035 | 0.00636 |
| 22.13888889 | 0.002824 | 0.004647 | 0.002645 | 0.00449 | 0.001401 | 0.001221 | 0.011084 | 0.010043 | 0.002918 | 0.072358 | 0.001109 | 0.011022 | 0.007657 | 0.016818 | 0.000478 | 0.001093 | 0.009837 | 0.00137 | 0.004757 | 0.003417 | 0.000583 | 0.005403 | 0.002521 | 0.006622 |
| 22.15277778 | 0.002337 | 0.003445 | 0.001453 | 0.002021 | 0.001053 | 0.00111 | 0.009152 | 0.004896 | 0.002458 | 0.054937 | 0.001008 | 0.004931 | 0.009496 | 0.016917 | 0.000402 | 0.001511 | 0.008637 | 0.000704 | 0.004055 | 0.005292 | 0.000379 | 0.002512 | 0.001489 | 0.009992 |
| 22.16666667 | 0.002417 | 0.004043 | 0.00101 | 0.000851 | 0.001081 | 0.000861 | 0.007366 | 0.007023 | 0.002791 | 0.033633 | 0.001294 | 0.005297 | 0.009592 | 0.015049 | 0.000242 | 0.001866 | 0.007842 | 0.00072 | 0.002543 | 0.006344 | 0.000373 | 0.001226 | 0.001891 | 0.009848 |
| 22.18055556 | 0.003358 | 0.005634 | 0.00061 | 0.000315 | 0.002158 | 0.001003 | 0.008335 | 0.009782 | 0.003344 | 0.017143 | 0.002303 | 0.006072 | 0.007145 | 0.010821 | 7.9111109 | 0.001999 | 0.007413 | 0.001243 | 0.001605 | 0.005637 | 0.000921 | 0.001075 | 0.00195 | 0.006413 |
| 22.19444444 | 0.007735 | 0.006456 | 0.000279 | 0.000352 | 0.002767 | 0.001508 | 0.007152 | 0.007214 | 0.003658 | 0.016808 | 0.003845 | 0.00787 | 0.004049 | 0.005754 | 4.0256940 | 0.001707 | 0.007292 | 0.001963 | 0.003113 | 0.003597 | 0.00151 | 0.001194 | 0.001235 | 0.005034 |
| 22.20833333 | 0.011767 | 0.005647 | 0.000149 | 0.000978 | 0.001921 | 0.001671 | 0.003626 | 0.002746 | 0.003586 | 0.027502 | 0.004639 | 0.011568 | 0.003311 | 0.003842 | 9.9729174 | 0.001099 | 0.007386 | 0.002863 | 0.005382 | 0.001597 | 0.001278 | 0.001604 | 0.00055 | 0.004538 |
| 22.22222222 | 0.012025 | 0.003278 | 0.000192 | 0.001448 | 0.001031 | 0.00126 | 0.001993 | 0.001317 | 0.003623 | 0.029898 | 0.004245 | 0.009753 | 0.005508 | 0.005355 | 0.000132 | 0.000576 | 0.007203 | 0.003468 | 0.006013 | 0.000937 | 0.000669 | 0.001575 | 0.000596 | 0.004084 |
| 22.23611111 | 0.009018 | 0.001446 | 0.000294 | 0.00112 | 0.001009 | 0.000766 | 0.003404 | 0.003471 | 0.004145 | 0.019515 | 0.003333 | 0.006942 | 0.008051 | 0.006805 | 7.9305561 | 0.000602 | 0.006205 | 0.003268 | 0.004911 | 0.001587 | 0.000396 | 0.001218 | 0.000884 | 0.006744 |
| 22.25 | 0.004868 | 0.001651 | 0.000232 | 0.000515 | 0.000853 | 0.000585 | 0.004486 | 0.006462 | 0.004573 | 0.010968 | 0.002717 | 0.011229 | 0.007694 | 0.007278 | 4.4520836 | 0.001324 | 0.004378 | 0.002603 | 0.003218 | 0.002546 | 0.000482 | 0.001832 | 0.00064 | 0.007872 |
| 22.26388889 | 0.003368 | 0.002244 | 0.000207 | 0.000561 | 0.000582 | 0.000581 | 0.003729 | 0.006506 | 0.004138 | 0.008217 | 0.002579 | 0.014095 | 0.004584 | 0.006874 | 0.00011 | 0.002131 | 0.002559 | 0.002172 | 0.001886 | 0.003188 | 0.000894 | 0.002873 | 0.000286 | 0.004837 |
| 22.27777778 | 0.004394 | 0.002221 | 0.000704 | 0.001511 | 0.000942 | 0.000477 | 0.002422 | 0.004942 | 0.002844 | 0.007461 | 0.002476 | 0.009351 | 0.00235 | 0.004826 | 0.000213 | 0.002026 | 0.001679 | 0.001904 | 0.000992 | 0.003215 | 0.0019 | 0.002751 | 0.000488 | 0.002333 |
| 22.29166667 | 0.004491 | 0.002154 | 0.002064 | 0.002811 | 0.001275 | 0.00031 | 0.002221 | 0.006442 | 0.001383 | 0.011853 | 0.001757 | 0.005162 | 0.002098 | 0.004265 | 0.000233 | 0.001331 | 0.002172 | 0.001256 | 0.000413 | 0.002604 | 0.003217 | 0.001679 | 0.000875 | 0.002305 |
| 22.30555556 | 0.003619 | 0.001814 | 0.004214 | 0.003862 | 0.000966 | 0.000174 | 0.005168 | 0.007677 | 0.000972 | 0.013149 | 0.001611 | 0.006052 | 0.003221 | 0.007963 | 0.000164 | 0.001472 | 0.003871 | 0.001 | 0.000247 | 0.001736 | 0.003401 | 0.001307 | 0.001111 | 0.003918 |
| 22.31944444 | 0.006149 | 0.001419 | 0.006498 | 0.00484 | 0.000757 | 0.000194 | 0.009671 | 0.005126 | 0.002518 | 0.011425 | 0.00397 | 0.006934 | 0.00539 | 0.011751 | 0.000151 | 0.002304 | 0.005856 | 0.002281 | 0.000769 | 0.000881 | 0.002426 | 0.001913 | 0.001936 | 0.004587 |
| 22.33333333 | 0.014186 | 0.002502 | 0.008586 | 0.006039 | 0.001414 | 0.000677 | 0.013117 | 0.00257 | 0.005752 | 0.022575 | 0.006922 | 0.008057 | 0.005091 | 0.012951 | 0.000227 | 0.002675 | 0.007151 | 0.004471 | 0.002211 | 0.000483 | 0.002887 | 0.002525 | 0.003951 | 0.004408 |
| 22.34722222 | 0.022876 | 0.005196 | 0.01046 | 0.007284 | 0.002364 | 0.00156 | 0.015018 | 0.003391 | 0.009879 | 0.039585 | 0.007458 | 0.013437 | 0.004012 | 0.01463 | 0.00028 | 0.002414 | 0.007626 | 0.00663 | 0.004261 | 0.000851 | 0.005146 | 0.003047 | 0.006874 | 0.008016 |
| 22.36111111 | 0.024959 | 0.006862 | 0.011595 | 0.008126 | 0.002552 | 0.002227 | 0.015001 | 0.008761 | 0.013623 | 0.040151 | 0.005695 | 0.021301 | 0.007638 | 0.01885 | 0.000245 | 0.001921 | 0.007825 | 0.00841 | 0.005553 | 0.001298 | 0.006507 | 0.00362 | 0.00979 | 0.013689 |
| 22.375 | 0.01933 | 0.006177 | 0.012287 | 0.008633 | 0.002293 | 0.002607 | 0.014013 | 0.015843 | 0.016026 | 0.024347 | 0.004116 | 0.027226 | 0.011961 | 0.0231 | 0.000198 | 0.0018 | 0.008544 | 0.00966 | 0.005221 | 0.001587 | 0.005991 | 0.003862 | 0.011746 | 0.016124 |
| 22.38888889 | 0.012638 | 0.004918 | 0.013729 | 0.009457 | 0.002972 | 0.003551 | 0.014441 | 0.017623 | 0.01738 | 0.010561 | 0.005168 | 0.027584 | 0.011789 | 0.024356 | 0.000276 | 0.002536 | 0.010226 | 0.010858 | 0.004437 | 0.002293 | 0.00473 | 0.003335 | 0.012602 | 0.015435 |
| 22.40277778 | 0.01067 | 0.004249 | 0.015345 | 0.010061 | 0.00418 | 0.005092 | 0.016384 | 0.011361 | 0.018167 | 0.012386 | 0.008657 | 0.023076 | 0.008441 | 0.023894 | 0.000427 | 0.003648 | 0.012038 | 0.012491 | 0.004532 | 0.003311 | 0.003595 | 0.002274 | 0.012786 | 0.013605 |
| 22.41666667 | 0.014309 | 0.004139 | 0.015679 | 0.00898 | 0.004098 | 0.006273 | 0.018204 | 0.0041 | 0.018999 | 0.032786 | 0.011856 | 0.018649 | 0.004815 | 0.023777 | 0.00053 | 0.004248 | 0.01303 | 0.014565 | 0.005803 | 0.004064 | 0.00363 | 0.00253 | 0.012927 | 0.011456 |
| 22.43055556 | 0.021905 | 0.004893 | 0.014704 | 0.006193 | 0.002586 | 0.006845 | 0.019543 | 0.003299 | 0.021127 | 0.053656 | 0.012388 | 0.018035 | 0.003469 | 0.023104 | 0.000538 | 0.00422 | 0.013543 | 0.016363 | 0.007814 | 0.004455 | 0.005966 | 0.005848 | 0.013823 | 0.009747 |
| 22.44444444 | 0.029707 | 0.0062 | 0.013791 | 0.003284 | 0.001067 | 0.00716 | 0.019724 | 0.00893 | 0.024632 | 0.051441 | 0.011044 | 0.020188 | 0.006092 | 0.020079 | 0.000434 | 0.004321 | 0.014201 | 0.016437 | 0.009175 | 0.004742 | 0.01004 | 0.011028 | 0.015378 | 0.009476 |
| 22.45833333 | 0.032826 | 0.006843 | 0.01453 | 0.002027 | 0.000668 | 0.007486 | 0.017677 | 0.016709 | 0.027597 | 0.028861 | 0.01039 | 0.02141 | 0.011977 | 0.014459 | 0.000433 | 0.005441 | 0.015591 | 0.014652 | 0.009136 | 0.005232 | 0.014274 | 0.015481 | 0.016955 | 0.01076 |
| 22.47222222 | 0.028766 | 0.006527 | 0.016591 | 0.002477 | 0.001202 | 0.007775 | 0.01477 | 0.018643 | 0.029679 | 0.013259 | 0.011852 | 0.019968 | 0.017397 | 0.008213 | 0.000459 | 0.006949 | 0.017585 | 0.012141 | 0.00889 | 0.006102 | 0.017512 | 0.018233 | 0.018967 | 0.012261 |
| 22.48611111 | 0.020755 | 0.007554 | 0.01931 | 0.004259 | 0.002034 | 0.008687 | 0.013104 | 0.012752 | 0.03747 | 0.012531 | 0.017062 | 0.018752 | 0.019669 | 0.006503 | 0.000373 | 0.006673 | 0.019113 | 0.009376 | 0.009556 | 0.007469 | 0.019358 | 0.019729 | 0.022195 | 0.012212 |
| 22.5 | 0.017323 | 0.013872 | 0.026519 | 0.010877 | 0.003663 | 0.012695 | 0.0139 | 0.007032 | 0.061975 | 0.025555 | 0.028864 | 0.024627 | 0.020868 | 0.015908 | 0.000387 | 0.004577 | 0.020204 | 0.006592 | 0.012803 | 0.012583 | 0.02204 | 0.021715 | 0.029465 | 0.011592 |
| 22.51388889 | 0.024794 | 0.028026 | 0.041306 | 0.023568 | 0.00579 | 0.019214 | 0.019146 | 0.009092 | 0.087738 | 0.071624 | 0.041345 | 0.040521 | 0.022964 | 0.037922 | 0.000514 | 0.004318 | 0.021629 | 0.005847 | 0.019422 | 0.022943 | 0.026231 | 0.026507 | 0.041951 | 0.013487 |
| 22.52777778 | 0.037621 | 0.04187 | 0.052625 | 0.032345 | 0.005455 | 0.022023 | 0.030875 | 0.017448 | 0.081234 | 0.119059 | 0.040993 | 0.058068 | 0.022493 | 0.063041 | 0.000974 | 0.007012 | 0.02145 | 0.010167 | 0.022638 | 0.027769 | 0.026263 | 0.033246 | 0.048838 | 0.018584 |
| 22.54166667 | 0.045411 | 0.042978 | 0.051061 | 0.029573 | 0.002494 | 0.020233 | 0.037893 | 0.020206 | 0.052883 | 0.11608 | 0.029142 | 0.063438 | 0.017417 | 0.073896 | 0.00116 | 0.011129 | 0.017781 | 0.017164 | 0.01772 | 0.020658 | 0.021594 | 0.038653 | 0.041999 | 0.024003 |
| 22.55555556 | 0.042018 | 0.03476 | 0.046213 | 0.021399 | 0.00185 | 0.018928 | 0.025779 | 0.013577 | 0.033236 | 0.076412 | 0.018003 | 0.049971 | 0.012431 | 0.063154 | 0.001191 | 0.014246 | 0.01362 | 0.017751 | 0.011155 | 0.012526 | 0.017123 | 0.040733 | 0.029385 | 0.021769 |
| 22.56944444 | 0.026286 | 0.028618 | 0.042697 | 0.013443 | 0.004688 | 0.01662 | 0.014563 | 0.007072 | 0.020833 | 0.035836 | 0.01092 | 0.026036 | 0.010609 | 0.039567 | 0.001816 | 0.011639 | 0.011034 | 0.011304 | 0.007001 | 0.010615 | 0.01154 | 0.036449 | 0.017589 | 0.014719 |
| 22.58333333 | 0.01414 | 0.027222 | 0.035134 | 0.006949 | 0.006473 | 0.010819 | 0.024636 | 0.003847 | 0.009362 | 0.01937 | 0.006312 | 0.012675 | 0.010035 | 0.021175 | 0.001967 | 0.006223 | 0.008165 | 0.009881 | 0.004403 | 0.011842 | 0.006493 | 0.024068 | 0.008697 | 0.019853 |
| 22.59722222 | 0.021683 | 0.027106 | 0.024349 | 0.004888 | 0.005931 | 0.005318 | 0.038495 | 0.00427 | 0.005157 | 0.037393 | 0.004266 | 0.016098 | 0.009643 | 0.022062 | 0.002142 | 0.00596 | 0.004226 | 0.016142 | 0.004284 | 0.013001 | 0.006667 | 0.010834 | 0.008699 | 0.032704 |
| 22.61111111 | 0.036312 | 0.028276 | 0.013672 | 0.005581 | 0.00604 | 0.003633 | 0.038932 | 0.011647 | 0.009502 | 0.06426 | 0.003703 | 0.015437 | 0.010402 | 0.034556 | 0.00258 | 0.008819 | 0.001279 | 0.020071 | 0.00406 | 0.012453 | 0.007299 | 0.005825 | 0.015838 | 0.033573 |
| 22.625 | 0.0392 | 0.032261 | 0.005845 | 0.004453 | 0.00761 | 0.004399 | 0.024629 | 0.016057 | 0.014105 | 0.056527 | 0.002854 | 0.011482 | 0.012638 | 0.03741 | 0.002021 | 0.007287 | 0.000554 | 0.0165 | 0.003227 | 0.010098 | 0.004775 | 0.008998 | 0.019108 | 0.020622 |
| 22.63888889 | 0.025956 | 0.037873 | 0.004004 | 0.004079 | 0.008727 | 0.003994 | 0.015359 | 0.011974 | 0.011381 | 0.037369 | 0.002169 | 0.02268 | 0.014942 | 0.025965 | 0.001947 | 0.003315 | 0.000685 | 0.008942 | 0.004942 | 0.007977 | 0.003646 | 0.014319 | 0.015653 | 0.011362 |
| 22.65277778 | 0.01546 | 0.042903 | 0.007612 | 0.007224 | 0.008285 | 0.002037 | 0.031583 | 0.014059 | 0.004988 | 0.057359 | 0.002338 | 0.038419 | 0.014974 | 0.025466 | 0.003711 | 0.002809 | 0.001059 | 0.004708 | 0.006088 | 0.007761 | 0.005777 | 0.018588 | 0.01051 | 0.018762 |
| 22.66666667 | 0.027978 | 0.044713 | 0.012559 | 0.010255 | 0.006297 | 0.000785 | 0.044637 | 0.020156 | 0.002771 | 0.077215 | 0.003704 | 0.04134 | 0.011459 | 0.044995 | 0.005935 | 0.004434 | 0.001483 | 0.005819 | 0.004917 | 0.007834 | 0.007734 | 0.01997 | 0.006684 | 0.028744 |
| 22.68055556 | 0.045099 | 0.042227 | 0.014673 | 0.011368 | 0.003325 | 0.000914 | 0.028255 | 0.015942 | 0.004418 | 0.058841 | 0.006741 | 0.027872 | 0.005953 | 0.056871 | 0.007157 | 0.00565 | 0.001721 | 0.007571 | 0.003502 | 0.005802 | 0.007044 | 0.015422 | 0.00561 | 0.024587 |
| 22.69444444 | 0.047451 | 0.036026 | 0.010762 | 0.010855 | 0.001144 | 0.00135 | 0.012967 | 0.006801 | 0.005011 | 0.027862 | 0.010332 | 0.013153 | 0.002796 | 0.04412 | 0.006414 | 0.006989 | 0.002728 | 0.008762 | 0.002243 | 0.003893 | 0.004509 | 0.008031 | 0.010401 | 0.013252 |
| 22.70833333 | 0.040559 | 0.02652 | 0.00425 | 0.007755 | 0.000542 | 0.001389 | 0.025119 | 0.002251 | 0.005688 | 0.012679 | 0.012413 | 0.014187 | 0.00429 | 0.023594 | 0.004007 | 0.008752 | 0.004322 | 0.009834 | 0.002623 | 0.006421 | 0.002853 | 0.004353 | 0.020187 | 0.010469 |
| 22.72222222 | 0.027688 | 0.014658 | 0.001991 | 0.00432 | 0.000855 | 0.002154 | 0.048183 | 0.003187 | 0.010915 | 0.021072 | 0.011672 | 0.022284 | 0.0079 | 0.021414 | 0.003085 | 0.00813 | 0.004729 | 0.00799 | 0.004362 | 0.01123 | 0.002587 | 0.005649 | 0.026292 | 0.01749 |
| 22.73611111 | 0.016455 | 0.006337 | 0.003339 | 0.00463 | 0.001897 | 0.004421 | 0.054628 | 0.007793 | 0.016736 | 0.042934 | 0.008178 | 0.029851 | 0.009998 | 0.036427 | 0.004734 | 0.005937 | 0.003407 | 0.005162 | 0.003876 | 0.01256 | 0.003275 | 0.005917 | 0.022765 | 0.021201 |
| 22.75 | 0.021796 | 0.006596 | 0.003882 | 0.007085 | 0.002677 | 0.006129 | 0.034745 | 0.015078 | 0.017356 | 0.066524 | 0.0039 | 0.03488 | 0.007509 | 0.04895 | 0.005351 | 0.008245 | 0.001956 | 0.006955 | 0.002248 | 0.009124 | 0.005816 | 0.003235 | 0.01256 | 0.014937 |
| 22.76388889 | 0.037456 | 0.011948 | 0.002653 | 0.006517 | 0.002096 | 0.005752 | 0.017134 | 0.018899 | 0.013687 | 0.073838 | 0.002269 | 0.027028 | 0.004702 | 0.04803 | 0.003418 | 0.01265 | 0.002063 | 0.011708 | 0.002814 | 0.004572 | 0.006174 | 0.001827 | 0.005715 | 0.007554 |
| 22.77777778 | 0.045178 | 0.01712 | 0.002209 | 0.003844 | 0.001514 | 0.003733 | 0.0231 | 0.013091 | 0.009089 | 0.050681 | 0.004186 | 0.016372 | 0.007519 | 0.032792 | 0.00202 | 0.011509 | 0.003498 | 0.014679 | 0.003178 | 0.002684 | 0.005841 | 0.003364 | 0.008318 | 0.008097 |
| 22.79166667 | 0.041598 | 0.019597 | 0.003953 | 0.004223 | 0.002131 | 0.00269 | 0.024958 | 0.006829 | 0.005697 | 0.026836 | 0.006888 | 0.022756 | 0.010007 | 0.015291 | 0.002848 | 0.005897 | 0.005016 | 0.014874 | 0.002303 | 0.002148 | 0.01082 | 0.009637 | 0.01314 | 0.010747 |
| 22.80555556 | 0.028504 | 0.019308 | 0.006411 | 0.006757 | 0.002059 | 0.004541 | 0.015594 | 0.010793 | 0.003836 | 0.038643 | 0.007804 | 0.030554 | 0.006782 | 0.010047 | 0.00348 | 0.001659 | 0.005893 | 0.012442 | 0.00307 | 0.001865 | 0.016386 | 0.018022 | 0.013872 | 0.008478 |
| 22.81944444 | 0.012949 | 0.017921 | 0.009506 | 0.006852 | 0.001325 | 0.00702 | 0.020772 | 0.016956 | 0.003441 | 0.0571 | 0.006633 | 0.020126 | 0.004235 | 0.013059 | 0.002722 | 0.000511 | 0.006449 | 0.008002 | 0.005677 | 0.004185 | 0.017328 | 0.023755 | 0.013167 | 0.008657 |
| 22.83333333 | 0.00504 | 0.01768 | 0.013782 | 0.004787 | 0.001411 | 0.008836 | 0.042241 | 0.013555 | 0.004108 | 0.042664 | 0.00512 | 0.009958 | 0.009232 | 0.011422 | 0.003201 | 0.001257 | 0.007293 | 0.003587 | 0.008374 | 0.009055 | 0.014522 | 0.026648 | 0.013474 | 0.01842 |
| 22.84722222 | 0.004693 | 0.020635 | 0.017978 | 0.005276 | 0.001265 | 0.010009 | 0.05351 | 0.010422 | 0.007924 | 0.021207 | 0.005526 | 0.017652 | 0.018552 | 0.009626 | 0.005835 | 0.00279 | 0.008525 | 0.001225 | 0.009956 | 0.014038 | 0.009787 | 0.027611 | 0.014152 | 0.027752 |
| 22.86111111 | 0.010769 | 0.026732 | 0.021801 | 0.008623 | 0.001729 | 0.009728 | 0.039724 | 0.018621 | 0.017218 | 0.031852 | 0.009154 | 0.031623 | 0.026169 | 0.021808 | 0.00739 | 0.003886 | 0.010238 | 0.001651 | 0.009945 | 0.016969 | 0.005577 | 0.026347 | 0.014725 | 0.02475 |
| 22.875 | 0.022181 | 0.03355 | 0.026101 | 0.009603 | 0.004217 | 0.008601 | 0.017848 | 0.025645 | 0.029265 | 0.069445 | 0.014274 | 0.03779 | 0.029243 | 0.044441 | 0.006329 | 0.004642 | 0.012087 | 0.004187 | 0.009361 | 0.018871 | 0.005794 | 0.023806 | 0.015771 | 0.013134 |
| 22.88888889 | 0.039441 | 0.037381 | 0.030096 | 0.006464 | 0.006386 | 0.007897 | 0.005547 | 0.020609 | 0.03821 | 0.103013 | 0.016112 | 0.034697 | 0.028163 | 0.063975 | 0.004281 | 0.005761 | 0.013081 | 0.007473 | 0.009951 | 0.02147 | 0.011134 | 0.023814 | 0.01722 | 0.00392 |
| 22.90277778 | 0.052014 | 0.035922 | 0.032798 | 0.003039 | 0.006083 | 0.007686 | 0.003738 | 0.012443 | 0.03842 | 0.10386 | 0.013189 | 0.027272 | 0.02581 | 0.070978 | 0.002222 | 0.006256 | 0.012907 | 0.011088 | 0.011723 | 0.021817 | 0.016371 | 0.028391 | 0.018424 | 0.00257 |
| 22.91666667 | 0.044113 | 0.029605 | 0.034541 | 0.002409 | 0.004333 | 0.007732 | 0.007906 | 0.007484 | 0.032148 | 0.074376 | 0.009379 | 0.019912 | 0.024281 | 0.060416 | 0.001013 | 0.005426 | 0.012259 | 0.014326 | 0.012693 | 0.017167 | 0.016853 | 0.036087 | 0.019071 | 0.008315 |
| 22.93055556 | 0.02534 | 0.018394 | 0.034581 | 0.002633 | 0.002772 | 0.007966 | 0.016365 | 0.005256 | 0.02701 | 0.036497 | 0.007747 | 0.014281 | 0.02548 | 0.037849 | 0.000775 | 0.004826 | 0.011625 | 0.014904 | 0.011869 | 0.010055 | 0.012545 | 0.043137 | 0.018764 | 0.018893 |
| 22.94444444 | 0.011001 | 0.008207 | 0.031128 | 0.001728 | 0.002002 | 0.007922 | 0.024941 | 0.007067 | 0.025223 | 0.012826 | 0.008212 | 0.012149 | 0.029316 | 0.016346 | 0.000905 | 0.006596 | 0.011017 | 0.012775 | 0.010311 | 0.005174 | 0.006347 | 0.044814 | 0.017067 | 0.028991 |

| | | | | | | | | | | | | | | | | | | | | | | | | |
|---|---|---|---|---|---|---|---|---|---|---|---|---|---|---|---|---|---|---|---|---|---|---|---|---|
| 22.95833333 | 0.004484 | 0.006621 | 0.024741 | 0.001106 | 0.00197 | 0.007517 | 0.025835 | 0.008242 | 0.024107 | 0.01066 | 0.010254 | 0.013876 | 0.032438 | 0.005213 | 0.001783 | 0.009879 | 0.010067 | 0.010466 | 0.009022 | 0.004753 | 0.002252 | 0.038516 | 0.014256 | 0.031984 |
| 22.97222222 | 0.003141 | 0.007908 | 0.01765 | 0.001684 | 0.00229 | 0.007178 | 0.01663 | 0.006594 | 0.021316 | 0.013128 | 0.012739 | 0.016276 | 0.029882 | 0.003074 | 0.002346 | 0.011322 | 0.008116 | 0.008181 | 0.00804 | 0.006802 | 0.001231 | 0.026833 | 0.010956 | 0.025544 |
| 22.98611111 | 0.005184 | 0.005627 | 0.012294 | 0.00299 | 0.002725 | 0.006918 | 0.008627 | 0.004688 | 0.018404 | 0.022139 | 0.014137 | 0.016761 | 0.020398 | 0.0028 | 0.001583 | 0.009283 | 0.005457 | 0.004661 | 0.007136 | 0.008303 | 0.001842 | 0.016495 | 0.008628 | 0.014253 |
| 23 | 0.012288 | 0.004576 | 0.011762 | 0.005929 | 0.003474 | 0.006638 | 0.009523 | 0.00541 | 0.018607 | 0.046169 | 0.016126 | 0.016267 | 0.010733 | 0.004192 | 0.001102 | 0.005386 | 0.004214 | 0.001883 | 0.006636 | 0.01053 | 0.005084 | 0.011887 | 0.010443 | 0.007733 |
| 23.01388889 | 0.023295 | 0.009687 | 0.016832 | 0.009385 | 0.004435 | 0.007114 | 0.009408 | 0.009207 | 0.022652 | 0.073286 | 0.024473 | 0.020254 | 0.007629 | 0.009935 | 0.002118 | 0.002612 | 0.006698 | 0.001508 | 0.007255 | 0.018209 | 0.011609 | 0.012438 | 0.02005 | 0.012829 |
| 23.02777778 | 0.036554 | 0.01581 | 0.026303 | 0.00965 | 0.00484 | 0.009597 | 0.010575 | 0.013482 | 0.027014 | 0.099441 | 0.041528 | 0.03497 | 0.011635 | 0.019101 | 0.003714 | 0.002348 | 0.014327 | 0.001613 | 0.008736 | 0.028826 | 0.019955 | 0.015538 | 0.037823 | 0.026642 |
| 23.04166667 | 0.050477 | 0.018505 | 0.038074 | 0.007931 | 0.004627 | 0.012652 | 0.023302 | 0.023681 | 0.025997 | 0.12538 | 0.056949 | 0.057189 | 0.020434 | 0.029304 | 0.004589 | 0.002961 | 0.026647 | 0.002513 | 0.008445 | 0.030958 | 0.027123 | 0.018655 | 0.0525 | 0.036311 |
| 23.05555556 | 0.056895 | 0.021447 | 0.042799 | 0.007223 | 0.004264 | 0.012405 | 0.037991 | 0.047008 | 0.01964 | 0.131461 | 0.056757 | 0.07146 | 0.027865 | 0.035835 | 0.00384 | 0.002491 | 0.036751 | 0.005529 | 0.004994 | 0.021792 | 0.029405 | 0.018014 | 0.048648 | 0.033627 |
| 23.06944444 | 0.047071 | 0.024271 | 0.031708 | 0.005644 | 0.0032 | 0.008939 | 0.038734 | 0.069334 | 0.013651 | 0.099629 | 0.041124 | 0.064392 | 0.024589 | 0.032699 | 0.00186 | 0.001851 | 0.036255 | 0.008506 | 0.001899 | 0.010219 | 0.025168 | 0.012056 | 0.029134 | 0.024359 |
| 23.08333333 | 0.025097 | 0.021898 | 0.016186 | 0.002981 | 0.001811 | 0.006047 | 0.023588 | 0.062569 | 0.011053 | 0.048678 | 0.021395 | 0.040266 | 0.013892 | 0.019533 | 0.000406 | 0.001942 | 0.027012 | 0.008255 | 0.00132 | 0.003817 | 0.015832 | 0.008294 | 0.011943 | 0.015099 |
| 23.09722222 | 0.01008 | 0.01478 | 0.01329 | 0.002129 | 0.002371 | 0.004766 | 0.01079 | 0.034128 | 0.010897 | 0.023083 | 0.012117 | 0.018529 | 0.009779 | 0.009115 | 5.4284726 | 0.001891 | 0.016597 | 0.004693 | 0.00123 | 0.00356 | 0.006312 | 0.011009 | 0.006572 | 0.011984 |
| 23.11111111 | 0.011162 | 0.007657 | 0.018365 | 0.002283 | 0.004798 | 0.003731 | 0.012963 | 0.022267 | 0.012198 | 0.036613 | 0.016775 | 0.013984 | 0.012168 | 0.011963 | 0.0001 | 0.001629 | 0.008832 | 0.002071 | 0.00066 | 0.004845 | 0.001435 | 0.01369 | 0.006682 | 0.01659 |
| 23.125 | 0.013999 | 0.003326 | 0.019025 | 0.002022 | 0.005695 | 0.002705 | 0.016481 | 0.032339 | 0.013621 | 0.057784 | 0.021514 | 0.020023 | 0.011723 | 0.020033 | 0.000109 | 0.0016 | 0.004337 | 0.002636 | 0.000233 | 0.004377 | 0.000553 | 0.013159 | 0.005066 | 0.019704 |
| 23.13888889 | 0.01194 | 0.001809 | 0.014842 | 0.00165 | 0.004011 | 0.002009 | 0.013602 | 0.041603 | 0.012466 | 0.067739 | 0.019846 | 0.022828 | 0.007377 | 0.023199 | 0.000102 | 0.001757 | 0.002198 | 0.003847 | 0.000449 | 0.002806 | 0.001545 | 0.010457 | 0.002406 | 0.018866 |
| 23.15277778 | 0.010255 | 0.001458 | 0.009535 | 0.00104 | 0.003261 | 0.001596 | 0.011457 | 0.04276 | 0.00846 | 0.070578 | 0.016482 | 0.018745 | 0.003506 | 0.021124 | 0.000188 | 0.001834 | 0.001273 | 0.004221 | 0.00131 | 0.001461 | 0.004312 | 0.006351 | 0.000895 | 0.017439 |
| 23.16666667 | 0.009724 | 0.001983 | 0.004559 | 0.000437 | 0.00403 | 0.001472 | 0.013694 | 0.039398 | 0.00467 | 0.067845 | 0.0144 | 0.011422 | 0.003305 | 0.01734 | 0.000304 | 0.001472 | 0.001401 | 0.004345 | 0.00201 | 0.001613 | 0.006531 | 0.005332 | 0.000931 | 0.015545 |
| 23.18055556 | 0.009844 | 0.003995 | 0.001317 | 0.000191 | 0.003571 | 0.001443 | 0.017918 | 0.034759 | 0.003056 | 0.062001 | 0.013309 | 0.005803 | 0.005051 | 0.014317 | 0.000249 | 0.000869 | 0.00208 | 0.00438 | 0.001856 | 0.003203 | 0.006335 | 0.009073 | 0.002058 | 0.013677 |
| 23.19444444 | 0.011138 | 0.005727 | 0.000309 | 0.000198 | 0.004214 | 0.001115 | 0.01821 | 0.031098 | 0.002585 | 0.055843 | 0.011749 | 0.003235 | 0.004904 | 0.01225 | 0.000136 | 0.000893 | 0.002484 | 0.004157 | 0.001193 | 0.004778 | 0.005134 | 0.011537 | 0.003304 | 0.01287 |
| 23.20833333 | 0.013853 | 0.005592 | 0.000324 | 0.000365 | 0.006773 | 0.000625 | 0.011606 | 0.028117 | 0.001762 | 0.047595 | 0.009215 | 0.001729 | 0.002996 | 0.01112 | 0.000173 | 0.001794 | 0.00247 | 0.00372 | 0.001271 | 0.005672 | 0.00379 | 0.008851 | 0.003747 | 0.012494 |
| 23.22222222 | 0.016048 | 0.004275 | 0.000288 | 0.000562 | 0.006328 | 0.000422 | 0.006799 | 0.025194 | 0.000805 | 0.035521 | 0.006424 | 0.000505 | 0.001795 | 0.010748 | 0.000191 | 0.00279 | 0.002281 | 0.003185 | 0.002699 | 0.006 | 0.002178 | 0.005048 | 0.003129 | 0.010329 |
| 23.23611111 | 0.015616 | 0.002875 | 0.000202 | 0.00052 | 0.003255 | 0.000574 | 0.013221 | 0.02165 | 0.001133 | 0.02118 | 0.004044 | 0.000402 | 0.002022 | 0.009931 | 9.5715285 | 0.003206 | 0.00234 | 0.002503 | 0.004593 | 0.005423 | 0.000861 | 0.004824 | 0.001968 | 0.006015 |
| 23.25 | 0.01273 | 0.001965 | 0.000737 | 0.000636 | 0.001679 | 0.00101 | 0.021892 | 0.01649 | 0.002576 | 0.009778 | 0.00196 | 0.001588 | 0.00436 | 0.007833 | 6.3326393 | 0.002969 | 0.002934 | 0.00178 | 0.00585 | 0.003984 | 0.000538 | 0.004697 | 0.001126 | 0.002794 |
| 23.26388889 | 0.008677 | 0.001523 | 0.002154 | 0.001649 | 0.001249 | 0.00144 | 0.020804 | 0.00923 | 0.002557 | 0.007379 | 0.000839 | 0.00405 | 0.008065 | 0.005291 | 0.000185 | 0.002265 | 0.003935 | 0.001164 | 0.005772 | 0.002737 | 0.001214 | 0.003697 | 0.000932 | 0.00353 |
| 23.27777778 | 0.004722 | 0.001359 | 0.003299 | 0.002881 | 0.000754 | 0.001258 | 0.011596 | 0.003558 | 0.001049 | 0.011618 | 0.0011 | 0.008513 | 0.008075 | 0.003269 | 0.000353 | 0.001325 | 0.005134 | 0.000645 | 0.004442 | 0.002364 | 0.002381 | 0.005872 | 0.001077 | 0.006017 |
| 23.29166667 | 0.002095 | 0.001288 | 0.002769 | 0.003006 | 0.000513 | 0.001013 | 0.004314 | 0.002944 | 0.000337 | 0.015352 | 0.001822 | 0.014701 | 0.006238 | 0.001884 | 0.000404 | 0.000516 | 0.006328 | 0.000326 | 0.002962 | 0.002538 | 0.003313 | 0.006799 | 0.001163 | 0.006254 |
| 23.30555556 | 0.00139 | 0.001151 | 0.001449 | 0.001868 | 0.000433 | 0.001387 | 0.003419 | 0.004863 | 0.000805 | 0.014102 | 0.00202 | 0.018863 | 0.008994 | 0.001425 | 0.000352 | 0.000157 | 0.007037 | 0.000502 | 0.002142 | 0.00271 | 0.003469 | 0.004251 | 0.000873 | 0.004684 |
| 23.31944444 | 0.002249 | 0.001068 | 0.001127 | 0.000825 | 0.000471 | 0.001421 | 0.002965 | 0.00566 | 0.002276 | 0.009465 | 0.001649 | 0.018222 | 0.01189 | 0.002573 | 0.000377 | 0.000334 | 0.007223 | 0.001387 | 0.001661 | 0.002522 | 0.002997 | 0.003351 | 0.000433 | 0.005111 |
| 23.33333333 | 0.00421 | 0.001014 | 0.001631 | 0.000771 | 0.000752 | 0.000764 | 0.002021 | 0.005057 | 0.004418 | 0.009397 | 0.000951 | 0.015353 | 0.009532 | 0.005366 | 0.000606 | 0.001002 | 0.00774 | 0.002773 | 0.001189 | 0.001915 | 0.002552 | 0.004346 | 0.000259 | 0.005966 |
| 23.34722222 | 0.007248 | 0.000874 | 0.002533 | 0.000837 | 0.001437 | 0.000254 | 0.004398 | 0.007214 | 0.006198 | 0.014688 | 0.000558 | 0.013718 | 0.004951 | 0.008679 | 0.001152 | 0.001888 | 0.008953 | 0.004289 | 0.001718 | 0.001197 | 0.002749 | 0.004694 | 0.000551 | 0.004494 |
| 23.36111111 | 0.011052 | 0.00093 | 0.003998 | 0.000941 | 0.001599 | 0.000294 | 0.009333 | 0.012773 | 0.006911 | 0.018593 | 0.001098 | 0.014816 | 0.002257 | 0.012473 | 0.00208 | 0.002838 | 0.010081 | 0.006042 | 0.003721 | 0.0006 | 0.003967 | 0.004853 | 0.0019 | 0.003672 |
| 23.375 | 0.014842 | 0.001561 | 0.005598 | 0.001926 | 0.001729 | 0.000756 | 0.014365 | 0.015262 | 0.006982 | 0.020637 | 0.001825 | 0.018612 | 0.002369 | 0.018769 | 0.003086 | 0.003855 | 0.010518 | 0.008214 | 0.00537 | 0.000406 | 0.0057 | 0.00592 | 0.004304 | 0.003623 |
| 23.38888889 | 0.018318 | 0.002519 | 0.00706 | 0.003535 | 0.003924 | 0.002567 | 0.017638 | 0.012531 | 0.00731 | 0.025139 | 0.002116 | 0.023198 | 0.005736 | 0.026258 | 0.003823 | 0.004598 | 0.010565 | 0.010479 | 0.00517 | 0.000786 | 0.006701 | 0.007151 | 0.006815 | 0.005354 |
| 23.40277778 | 0.021235 | 0.003146 | 0.008354 | 0.004923 | 0.006147 | 0.00556 | 0.017462 | 0.00826 | 0.007769 | 0.031452 | 0.002675 | 0.025551 | 0.011468 | 0.02848 | 0.004253 | 0.004868 | 0.010695 | 0.012352 | 0.003742 | 0.001391 | 0.0066 | 0.006996 | 0.008804 | 0.012289 |
| 23.41666667 | 0.022355 | 0.003062 | 0.009094 | 0.005518 | 0.005124 | 0.005469 | 0.013972 | 0.00513 | 0.007746 | 0.033992 | 0.004129 | 0.023559 | 0.01461 | 0.022755 | 0.004535 | 0.004974 | 0.010776 | 0.013188 | 0.002166 | 0.002045 | 0.006174 | 0.00541 | 0.010132 | 0.018126 |
| 23.43055556 | 0.021761 | 0.002213 | 0.009736 | 0.005026 | 0.002994 | 0.003017 | 0.010616 | 0.003636 | 0.007857 | 0.032298 | 0.00633 | 0.01831 | 0.012308 | 0.014309 | 0.004797 | 0.005263 | 0.010521 | 0.013078 | 0.001019 | 0.002613 | 0.006543 | 0.004242 | 0.010659 | 0.015637 |
| 23.44444444 | 0.021902 | 0.001078 | 0.011566 | 0.00405 | 0.003189 | 0.003523 | 0.010662 | 0.00302 | 0.0098 | 0.032547 | 0.008313 | 0.013565 | 0.007216 | 0.008935 | 0.004741 | 0.006278 | 0.011142 | 0.013368 | 0.001221 | 0.003071 | 0.008842 | 0.006423 | 0.010849 | 0.008441 |
| 23.45833333 | 0.0239 | 0.000381 | 0.013899 | 0.003867 | 0.003505 | 0.005714 | 0.013121 | 0.004194 | 0.013803 | 0.037624 | 0.008756 | 0.011339 | 0.003786 | 0.00824 | 0.004222 | 0.008639 | 0.01434 | 0.01456 | 0.003504 | 0.003738 | 0.012699 | 0.013368 | 0.011891 | 0.004246 |
| 23.47222222 | 0.026152 | 0.001076 | 0.01399 | 0.004596 | 0.003199 | 0.006716 | 0.013957 | 0.007684 | 0.017944 | 0.043447 | 0.008146 | 0.011163 | 0.004876 | 0.01123 | 0.003779 | 0.011411 | 0.019323 | 0.015878 | 0.006521 | 0.004806 | 0.015842 | 0.021753 | 0.014332 | 0.006474 |
| 23.48611111 | 0.028615 | 0.005131 | 0.012247 | 0.005785 | 0.005743 | 0.007331 | 0.012978 | 0.010536 | 0.025458 | 0.046637 | 0.010274 | 0.01586 | 0.011103 | 0.017377 | 0.004351 | 0.012662 | 0.023299 | 0.016916 | 0.008712 | 0.008165 | 0.018568 | 0.027886 | 0.019204 | 0.013461 |
| 23.5 | 0.036722 | 0.0152 | 0.015183 | 0.009446 | 0.009202 | 0.010716 | 0.016227 | 0.013738 | 0.046193 | 0.055848 | 0.019937 | 0.03156 | 0.021893 | 0.028334 | 0.00637 | 0.011374 | 0.026127 | 0.018827 | 0.012206 | 0.019315 | 0.028408 | 0.033434 | 0.032818 | 0.022229 |
| 23.51388889 | 0.049458 | 0.030583 | 0.025394 | 0.015834 | 0.009575 | 0.017172 | 0.025709 | 0.021301 | 0.073154 | 0.087491 | 0.03356 | 0.054046 | 0.033688 | 0.045077 | 0.009009 | 0.007166 | 0.028516 | 0.022446 | 0.01644 | 0.039771 | 0.04801 | 0.040176 | 0.060085 | 0.028206 |
| 23.52777778 | 0.057255 | 0.04054 | 0.032536 | 0.019501 | 0.007026 | 0.020985 | 0.037088 | 0.029866 | 0.082343 | 0.136952 | 0.039197 | 0.065282 | 0.038377 | 0.061602 | 0.011421 | 0.003542 | 0.025512 | 0.023633 | 0.015792 | 0.053629 | 0.061814 | 0.04452 | 0.077056 | 0.026393 |
| 23.54166667 | 0.05861 | 0.035435 | 0.029116 | 0.018463 | 0.003409 | 0.020049 | 0.039618 | 0.032954 | 0.07331 | 0.159745 | 0.034217 | 0.057798 | 0.032197 | 0.069434 | 0.011813 | 0.002838 | 0.017265 | 0.0172 | 0.010322 | 0.046892 | 0.05807 | 0.044144 | 0.062811 | 0.018399 |
| 23.55555556 | 0.051418 | 0.02569 | 0.023289 | 0.016173 | 0.000991 | 0.016423 | 0.026243 | 0.028561 | 0.06352 | 0.125257 | 0.025215 | 0.042633 | 0.021569 | 0.066194 | 0.009086 | 0.002967 | 0.012185 | 0.009441 | 0.005723 | 0.031204 | 0.042493 | 0.039502 | 0.040187 | 0.009369 |
| 23.56944444 | 0.031989 | 0.023621 | 0.024252 | 0.013708 | 0.000328 | 0.011364 | 0.019743 | 0.017935 | 0.054879 | 0.065899 | 0.015558 | 0.027153 | 0.012772 | 0.051493 | 0.00638 | 0.004106 | 0.010792 | 0.010226 | 0.00415 | 0.019303 | 0.02254 | 0.028654 | 0.026507 | 0.004619 |
| 23.58333333 | 0.014102 | 0.022895 | 0.03079 | 0.009823 | 0.000938 | 0.011809 | 0.032911 | 0.007955 | 0.043304 | 0.031446 | 0.007715 | 0.012644 | 0.006715 | 0.028385 | 0.004987 | 0.004082 | 0.008816 | 0.01534 | 0.003039 | 0.011674 | 0.011675 | 0.013941 | 0.018694 | 0.005127 |
| 23.59722222 | 0.012815 | 0.019261 | 0.034552 | 0.005013 | 0.003342 | 0.017239 | 0.04003 | 0.00603 | 0.035972 | 0.045771 | 0.004283 | 0.005647 | 0.003772 | 0.015137 | 0.003243 | 0.002438 | 0.006644 | 0.016014 | 0.001748 | 0.008652 | 0.012588 | 0.005308 | 0.016111 | 0.004703 |
| 23.61111111 | 0.022841 | 0.018536 | 0.029104 | 0.0022 | 0.005973 | 0.015268 | 0.028208 | 0.008268 | 0.038257 | 0.076896 | 0.004859 | 0.010901 | 0.004684 | 0.01978 | 0.002277 | 0.002844 | 0.005676 | 0.011185 | 0.002388 | 0.009311 | 0.009668 | 0.00454 | 0.016283 | 0.004615 |
| 23.625 | 0.031701 | 0.020595 | 0.018357 | 0.002239 | 0.006126 | 0.00788 | 0.017296 | 0.007959 | 0.040178 | 0.072325 | 0.007069 | 0.024224 | 0.009249 | 0.023233 | 0.004337 | 0.004783 | 0.005775 | 0.005887 | 0.005024 | 0.011098 | 0.00607 | 0.005472 | 0.015763 | 0.009711 |
| 23.63888889 | 0.033679 | 0.021175 | 0.009871 | 0.003131 | 0.003779 | 0.005211 | 0.024507 | 0.005353 | 0.036306 | 0.043404 | 0.009221 | 0.037765 | 0.01518 | 0.017004 | 0.007773 | 0.005688 | 0.006265 | 0.005845 | 0.006968 | 0.01303 | 0.009614 | 0.010962 | 0.012555 | 0.016212 |
| 23.65277778 | 0.02614 | 0.020028 | 0.004535 | 0.003127 | 0.002019 | 0.006188 | 0.037224 | 0.004157 | 0.03249 | 0.039456 | 0.010987 | 0.038636 | 0.018471 | 0.010898 | 0.009018 | 0.004266 | 0.005513 | 0.010077 | 0.006207 | 0.015941 | 0.013819 | 0.022456 | 0.007735 | 0.017286 |
| 23.66666667 | 0.01363 | 0.020233 | 0.001496 | 0.002741 | 0.002553 | 0.005387 | 0.037424 | 0.007995 | 0.028453 | 0.06826 | 0.012134 | 0.021922 | 0.017028 | 0.014746 | 0.007846 | 0.002602 | 0.003454 | 0.014186 | 0.003521 | 0.018894 | 0.012214 | 0.030896 | 0.007437 | 0.011966 |
| 23.68055556 | 0.009942 | 0.022373 | 0.000704 | 0.004728 | 0.002836 | 0.002665 | 0.024394 | 0.014388 | 0.022236 | 0.086026 | 0.012275 | 0.008973 | 0.01203 | 0.025464 | 0.005139 | 0.003949 | 0.003549 | 0.016365 | 0.001178 | 0.016038 | 0.008832 | 0.029504 | 0.013498 | 0.009844 |
| 23.69444444 | 0.018691 | 0.02175 | 0.001987 | 0.008492 | 0.002593 | 0.001174 | 0.012827 | 0.017345 | 0.013907 | 0.063704 | 0.010994 | 0.012251 | 0.006195 | 0.030443 | 0.002964 | 0.007417 | 0.006583 | 0.015167 | 0.000708 | 0.008199 | 0.013196 | 0.018672 | 0.019114 | 0.014894 |
| 23.70833333 | 0.026622 | 0.01525 | 0.005614 | 0.010209 | 0.004594 | 0.001596 | 0.018633 | 0.015418 | 0.009357 | 0.033511 | 0.007822 | 0.018079 | 0.003905 | 0.026272 | 0.002887 | 0.009927 | 0.008896 | 0.009893 | 0.001222 | 0.0052 | 0.018069 | 0.010688 | 0.019179 | 0.016447 |
| 23.72222222 | 0.021556 | 0.006783 | 0.009719 | 0.008875 | 0.007766 | 0.002036 | 0.03442 | 0.009907 | 0.017235 | 0.043031 | 0.004189 | 0.021171 | 0.006705 | 0.01611 | 0.002747 | 0.010482 | 0.007403 | 0.006609 | 0.00131 | 0.007621 | 0.012442 | 0.018759 | 0.013432 | 0.010164 |
| 23.73611111 | 0.012157 | 0.002568 | 0.012486 | 0.006331 | 0.008066 | 0.003551 | 0.03774 | 0.005702 | 0.0286 | 0.082107 | 0.003151 | 0.021978 | 0.010219 | 0.009063 | 0.002135 | 0.008246 | 0.004374 | 0.009482 | 0.001846 | 0.008153 | 0.004862 | 0.031534 | 0.00605 | 0.007376 |
| 23.75 | 0.014593 | 0.005243 | 0.013102 | 0.003843 | 0.005512 | 0.008302 | 0.022841 | 0.00913 | 0.029865 | 0.111581 | 0.004964 | 0.014848 | 0.010402 | 0.014946 | 0.002526 | 0.005119 | 0.004448 | 0.01242 | 0.004001 | 0.005443 | 0.005466 | 0.028976 | 0.002022 | 0.01464 |
| 23.76388889 | 0.025884 | 0.014013 | 0.01166 | 0.001868 | 0.005042 | 0.013302 | 0.00955 | 0.017728 | 0.023656 | 0.116434 | 0.00702 | 0.009289 | 0.006995 | 0.027657 | 0.003432 | 0.006295 | 0.005828 | 0.012697 | 0.006139 | 0.003309 | 0.00741 | 0.017097 | 0.001923 | 0.018213 |
| 23.77777778 | 0.036599 | 0.024745 | 0.009548 | 0.000766 | 0.007893 | 0.014079 | 0.009127 | 0.02292 | 0.017739 | 0.092984 | 0.006876 | 0.011022 | 0.005204 | 0.035679 | 0.003464 | 0.010088 | 0.004461 | 0.013378 | 0.005918 | 0.004366 | 0.006928 | 0.015867 | 0.004684 | 0.012884 |
| 23.79166667 | 0.040764 | 0.030425 | 0.009174 | 0.000464 | 0.008813 | 0.009917 | 0.008454 | 0.020757 | 0.010768 | 0.049625 | 0.004394 | 0.009886 | 0.008455 | 0.035598 | 0.00256 | 0.011709 | 0.003167 | 0.01553 | 0.003524 | 0.007531 | 0.012946 | 0.019772 | 0.007927 | 0.011866 |
| 23.80555556 | 0.032729 | 0.027629 | 0.011753 | 0.001343 | 0.006089 | 0.005028 | 0.005766 | 0.013972 | 0.00481 | 0.020108 | 0.002742 | 0.0099 | 0.011085 | 0.030168 | 0.001655 | 0.010712 | 0.005595 | 0.01564 | 0.001686 | 0.011331 | 0.025048 | 0.016491 | 0.007754 | 0.015127 |
| 23.81944444 | 0.017164 | 0.021095 | 0.01503 | 0.003239 | 0.00556 | 0.004264 | 0.012526 | 0.006925 | 0.00346 | 0.013368 | 0.002771 | 0.016001 | 0.008172 | 0.023748 | 0.001122 | 0.007853 | 0.008775 | 0.010113 | 0.00249 | 0.013007 | 0.029642 | 0.021366 | 0.004688 | 0.014783 |
| 23.83333333 | 0.005514 | 0.01764 | 0.01758 | 0.00462 | 0.008257 | 0.007564 | 0.026428 | 0.00271 | 0.002677 | 0.009172 | 0.001988 | 0.028039 | 0.005659 | 0.017515 | 0.001273 | 0.004035 | 0.0096 | 0.004363 | 0.005229 | 0.01006 | 0.021387 | 0.043159 | 0.003665 | 0.011509 |
| 23.84722222 | 0.003435 | 0.018313 | 0.020374 | 0.004456 | 0.00721 | 0.011465 | 0.034509 | 0.0015 | 0.00388 | 0.012469 | 0.001459 | 0.043358 | 0.012431 | 0.012221 | 0.002972 | 0.001083 | 0.008867 | 0.003422 | 0.008348 | 0.006325 | 0.010823 | 0.060502 | 0.00791 | 0.008345 |
| 23.86111111 | 0.009556 | 0.020364 | 0.023082 | 0.002786 | 0.00506 | 0.013153 | 0.029838 | 0.00177 | 0.010322 | 0.033731 | 0.003839 | 0.049672 | 0.026308 | 0.009829 | 0.006576 | 0.000557 | 0.008833 | 0.003106 | 0.010964 | 0.00604 | 0.006221 | 0.057791 | 0.017261 | 0.007664 |
| 23.875 | 0.020938 | 0.023104 | 0.023964 | 0.001713 | 0.007674 | 0.012661 | 0.018196 | 0.003347 | 0.020727 | 0.06733 | 0.008876 | 0.041418 | 0.039022 | 0.010983 | 0.010214 | 0.002038 | 0.011044 | 0.00186 | 0.012218 | 0.00955 | 0.008252 | 0.043551 | 0.029123 | 0.011494 |
| 23.88888889 | 0.03322 | 0.027365 | 0.02239 | 0.003179 | 0.009914 | 0.013011 | 0.007774 | 0.005641 | 0.029712 | 0.104312 | 0.014416 | 0.023251 | 0.043798 | 0.015249 | 0.011239 | 0.004052 | 0.015542 | 0.002784 | 0.011225 | 0.016124 | 0.016611 | 0.032237 | 0.038797 | 0.018756 |
| 23.90277778 | 0.043713 | 0.030733 | 0.019537 | 0.004589 | 0.007415 | 0.015534 | 0.002133 | 0.010797 | 0.032586 | 0.137776 | 0.018102 | 0.007376 | 0.038755 | 0.021885 | 0.009618 | 0.005306 | 0.021706 | 0.005518 | 0.00869 | 0.023489 | 0.027815 | 0.029306 | 0.041522 | 0.024729 |
| 23.91666667 | 0.053222 | 0.028792 | 0.017021 | 0.003479 | 0.005597 | 0.017827 | 0.001482 | 0.016722 | 0.032675 | 0.142068 | 0.01907 | 0.002293 | 0.02895 | 0.029874 | 0.007202 | 0.005044 | 0.027103 | 0.00848 | 0.006344 | 0.026959 | 0.033935 | 0.028945 | 0.036435 | 0.025219 |
| 23.93055556 | 0.053952 | 0.021805 | 0.015642 | 0.002613 | 0.009974 | 0.017893 | 0.003901 | 0.013856 | 0.032412 | 0.099353 | 0.017133 | 0.004996 | 0.019852 | 0.03517 | 0.004909 | 0.003498 | 0.027417 | 0.011628 | 0.004712 | 0.023563 | 0.029425 | 0.024509 | 0.026873 | 0.019831 |
| 23.94444444 | 0.038092 | 0.013767 | 0.015604 | 0.003732 | 0.013256 | 0.014963 | 0.008092 | 0.006451 | 0.028846 | 0.046444 | 0.011728 | 0.012817 | 0.01365 | 0.032553 | 0.002563 | 0.001627 | 0.020773 | 0.014427 | 0.003178 | 0.015633 | 0.017256 | 0.016294 | 0.016953 | 0.011075 |
| 23.95833333 | 0.017708 | 0.006914 | 0.015548 | 0.003685 | 0.009233 | 0.009822 | 0.011014 | 0.005188 | 0.022634 | 0.027965 | 0.005959 | 0.021567 | 0.010763 | 0.024071 | 0.000973 | 0.00064 | 0.010981 | 0.015689 | 0.001741 | 0.007706 | 0.007026 | 0.008004 | 0.008713 | 0.005377 |
| 23.97222222 | 0.009396 | 0.002981 | 0.013044 | 0.002264 | 0.004972 | 0.005086 | 0.009738 | 0.00607 | 0.018416 | 0.040583 | 0.004723 | 0.029367 | 0.010906 | 0.015666 | 0.000859 | 0.001568 | 0.003839 | 0.015304 | 0.001454 | 0.002601 | 0.003997 | 0.004484 | 0.003484 | 0.005582 |
| 23.98611111 | 0.009512 | 0.00318 | 0.009422 | 0.002677 | 0.006572 | 0.00259 | 0.006505 | 0.005457 | 0.019401 | 0.047674 | 0.005831 | 0.039385 | 0.012937 | 0.008838 | 0.001539 | 0.003981 | 0.001263 | 0.013522 | 0.002799 | 0.001243 | 0.003632 | 0.00914 | 0.001893 | 0.004801 |
| 24 | 0.008967 | 0.004557 | 0.010101 | 0.004778 | 0.009911 | 0.003603 | 0.006415 | 0.009015 | 0.025057 | 0.035512 | 0.005822 | 0.048798 | 0.015435 | 0.003984 | 0.002119 | 0.005769 | 0.002764 | 0.010217 | 0.004968 | 0.004638 | 0.005802 | 0.020061 | 0.003713 | 0.004407 |
| 24.01388889 | 0.014241 | 0.003853 | 0.017057 | 0.006396 | 0.009226 | 0.009584 | 0.011951 | 0.015447 | 0.030373 | 0.035498 | 0.010396 | 0.052762 | 0.017916 | 0.004571 | 0.002269 | 0.006416 | 0.008619 | 0.00687 | 0.00692 | 0.014648 | 0.015313 | 0.032252 | 0.013113 | 0.013598 |
| 24.02777778 | 0.030446 | 0.004418 | 0.025834 | 0.006095 | 0.008292 | 0.019918 | 0.020678 | 0.016903 | 0.029189 | 0.068568 | 0.024114 | 0.057618 | 0.02198 | 0.013225 | 0.00334 | 0.008032 | 0.015934 | 0.006072 | 0.008083 | 0.028762 | 0.028585 | 0.041821 | 0.031689 | 0.030312 |
| 24.04166667 | 0.051776 | 0.009833 | 0.032509 | 0.00544 | 0.013696 | 0.028069 | 0.02974 | 0.017933 | 0.021423 | 0.09741 | 0.038705 | 0.073091 | 0.029407 | 0.030366 | 0.006099 | 0.011061 | 0.019935 | 0.008434 | 0.007997 | 0.032945 | 0.034362 | 0.045906 | 0.046417 | 0.04312 |
| 24.05555556 | 0.065411 | 0.01638 | 0.035556 | 0.007949 | 0.015626 | 0.02705 | 0.034761 | 0.03061 | 0.0144 | 0.082025 | 0.0461 | 0.085302 | 0.03503 | 0.049092 | 0.008724 | 0.011746 | 0.020003 | 0.012098 | 0.006529 | 0.020285 | 0.029238 | 0.041866 | 0.043302 | 0.044488 |
| 24.06944444 | 0.058956 | 0.018872 | 0.03283 | 0.011075 | 0.00958 | 0.019935 | 0.028202 | 0.044559 | 0.013185 | 0.040633 | 0.046272 | 0.078354 | 0.029358 | 0.051339 | 0.009201 | 0.007888 | 0.01761 | 0.013864 | 0.004777 | 0.006069 | 0.022607 | 0.028514 | 0.027222 | 0.035117 |
| 24.08333333 | 0.037368 | 0.015196 | 0.023627 | 0.008951 | 0.00868 | 0.01347 | 0.015742 | 0.040107 | 0.015387 | 0.011549 | 0.037023 | 0.058238 | 0.016466 | 0.030654 | 0.006387 | 0.004315 | 0.012659 | 0.011061 | 0.003452 | 0.000999 | 0.017084 | 0.015379 | 0.012183 | 0.019352 |
| 24.09722222 | 0.016506 | 0.007709 | 0.012371 | 0.004855 | 0.009696 | 0.008871 | 0.013841 | 0.021806 | 0.016755 | 0.002809 | 0.020096 | 0.031526 | 0.011755 | 0.013521 | 0.004234 | 0.005446 | 0.006618 | 0.00589 | 0.001867 | 0.001697 | 0.011268 | 0.015533 | 0.006676 | 0.010054 |
| 24.11111111 | 0.007171 | 0.002868 | 0.007065 | 0.005346 | 0.00706 | 0.005286 | 0.020294 | 0.011047 | 0.014961 | 0.002476 | 0.009047 | 0.011834 | 0.016429 | 0.019454 | 0.007255 | 0.007901 | 0.003024 | 0.004142 | 0.000522 | 0.00416 | 0.010289 | 0.023317 | 0.008352 | 0.014261 |
| 24.125 | 0.007128 | 0.002672 | 0.007617 | 0.007575 | 0.007511 | 0.003017 | 0.021857 | 0.017266 | 0.010335 | 0.002611 | 0.009405 | 0.010017 | 0.018985 | 0.029853 | 0.010829 | 0.007179 | 0.002395 | 0.006089 | 0.000192 | 0.004248 | 0.011097 | 0.027969 | 0.007997 | 0.01963 |
| 24.13888889 | 0.008178 | 0.002584 | 0.007397 | 0.006589 | 0.007621 | 0.002361 | 0.017727 | 0.024851 | 0.006377 | 0.001724 | 0.011167 | 0.015679 | 0.015549 | 0.02903 | 0.010889 | 0.004367 | 0.002488 | 0.007228 | 0.000498 | 0.002082 | 0.007688 | 0.028583 | 0.004369 | 0.020162 |
| 24.15277778 | 0.007836 | 0.001516 | 0.009703 | 0.003672 | 0.003981 | 0.002453 | 0.012617 | 0.023844 | 0.005332 | 0.001948 | 0.012024 | 0.019887 | 0.008782 | 0.021176 | 0.010345 | 0.00598 | 0.004676 | 0.005644 | 0.002156 | 0.001546 | 0.007739 | 0.027018 | 0.001743 | 0.024209 |
| 24.16666667 | 0.007245 | 0.002249 | 0.011858 | 0.002517 | 0.003053 | 0.003457 | 0.016549 | 0.027115 | 0.00665 | 0.014002 | 0.019415 | 0.023541 | 0.004314 | 0.023705 | 0.012031 | 0.016119 | 0.013506 | 0.003748 | 0.003586 | 0.002628 | 0.018257 | 0.02267 | 0.002225 | 0.031716 |
| 24.18055556 | 0.013877 | 0.005057 | 0.014294 | 0.007689 | 0.007254 | 0.0066 | 0.035077 | 0.039945 | 0.013055 | 0.046811 | 0.034292 | 0.03491 | 0.007973 | 0.038859 | 0.010887 | 0.024486 | 0.024475 | 0.003477 | 0.012239 | 0.005306 | 0.030618 | 0.016554 | 0.010934 | 0.033665 |
| 24.19444444 | 0.019258 | 0.006622 | 0.018699 | 0.012827 | 0.011366 | 0.008709 | 0.043547 | 0.04278 | 0.018799 | 0.07654 | 0.038414 | 0.053805 | 0.017161 | 0.036337 | 0.009751 | 0.023669 | 0.029095 | 0.002629 | 0.032838 | 0.011958 | 0.034624 | 0.017673 | 0.028845 | 0.041762 |
| 24.20833333 | 0.018737 | 0.005679 | 0.018679 | 0.010567 | 0.009206 | 0.007445 | 0.036286 | 0.032459 | 0.024683 | 0.062568 | 0.023578 | 0.056386 | 0.022518 | 0.015241 | 0.015545 | 0.02531 | 0.023867 | 0.001696 | 0.039428 | 0.018306 | 0.028612 | 0.025171 | 0.042392 | 0.053755 |
| 24.22222222 | 0.027056 | 0.005223 | 0.020213 | 0.00752 | 0.00495 | 0.005067 | 0.037667 | 0.029724 | 0.039082 | 0.022338 | 0.011227 | 0.039999 | 0.019473 | 0.006157 | 0.017002 | 0.035359 | 0.011225 | 0.002849 | 0.022554 | 0.017704 | 0.016499 | 0.025071 | 0.042468 | 0.042214 |
| 24.23611111 | 0.033394 | 0.005816 | 0.015244 | 0.004448 | 0.004055 | 0.003248 | 0.035575 | 0.025766 | 0.038673 | 0.010023 | 0.008437 | 0.034266 | 0.01334 | 0.011301 | 0.010582 | 0.03995 | 0.002606 | 0.004578 | 0.006978 | 0.013703 | 0.007578 | 0.016178 | 0.033284 | 0.017996 |
| 24.25 | 0.033626 | 0.006039 | 0.005205 | 0.001545 | 0.003166 | 0.003172 | 0.021965 | 0.012034 | 0.018012 | 0.02004 | 0.007752 | 0.041242 | 0.007963 | 0.012861 | 0.004474 | 0.027409 | 0.001924 | 0.005589 | 0.003264 | 0.011512 | 0.00739 | 0.009224 | 0.019643 | 0.005576 |
| 24.26388889 | 0.03674 | 0.005367 | 0.003145 | 0.00205 | 0.001825 | 0.003552 | 0.016156 | 0.004868 | 0.00524 | 0.020285 | 0.006701 | 0.040824 | 0.003441 | 0.007139 | 0.001309 | 0.011212 | 0.003291 | 0.005986 | 0.004217 | 0.010459 | 0.010028 | 0.007084 | 0.007548 | 0.004504 |
| 24.27777778 | 0.037862 | 0.004089 | 0.003878 | 0.003948 | 0.001549 | 0.003377 | 0.015627 | 0.003048 | 0.003916 | 0.011751 | 0.005143 | 0.032011 | 0.000995 | 0.002137 | 0.000811 | 0.007435 | 0.003946 | 0.005681 | 0.003965 | 0.009623 | 0.008953 | 0.006654 | 0.001928 | 0.00581 |
| 24.29166667 | 0.036461 | 0.003566 | 0.005273 | 0.004898 | 0.002255 | 0.004749 | 0.010097 | 0.001361 | 0.003576 | 0.006609 | 0.005985 | 0.023184 | 0.000614 | 0.001609 | 0.000862 | 0.01159 | 0.003683 | 0.004379 | 0.002093 | 0.009158 | 0.005535 | 0.00692 | 0.00206 | 0.007467 |
| 24.30555556 | 0.036163 | 0.004453 | 0.00762 | 0.003504 | 0.004191 | 0.007676 | 0.005251 | 0.003141 | 0.002402 | 0.008616 | 0.009446 | 0.01707 | 0.001002 | 0.002765 | 0.000526 | 0.016004 | 0.002653 | 0.002435 | 0.001054 | 0.00852 | 0.003296 | 0.007298 | 0.004129 | 0.010631 |
| 24.31944444 | 0.034646 | 0.005369 | 0.008425 | 0.002062 | 0.006532 | 0.00896 | 0.007383 | 0.005957 | 0.001579 | 0.012667 | 0.011984 | 0.011786 | 0.001109 | 0.002616 | 0.000346 | 0.014317 | 0.001552 | 0.001157 | 0.001055 | 0.007044 | 0.002447 | 0.006791 | 0.004167 | 0.010604 |
| 24.33333333 | 0.027973 | 0.004796 | 0.00683 | 0.002969 | 0.007444 | 0.007149 | 0.009988 | 0.005811 | 0.001178 | 0.012947 | 0.010874 | 0.006639 | 0.000949 | 0.001381 | 0.000444 | 0.006807 | 0.001612 | 0.000894 | 0.000892 | 0.005268 | 0.001849 | 0.005327 | 0.001991 | 0.006546 |
| 24.34722222 | 0.018845 | 0.003216 | 0.004349 | 0.003435 | 0.005674 | 0.004429 | 0.007276 | 0.003028 | 0.001058 | 0.008351 | 0.007186 | 0.005654 | 0.001042 | 0.000853 | 0.000426 | 0.001994 | 0.002186 | 0.000681 | 0.000918 | 0.003824 | 0.001711 | 0.003698 | 0.000393 | 0.003682 |
| 24.36111111 | 0.011289 | 0.001829 | 0.002614 | 0.002197 | 0.002877 | 0.003031 | 0.002768 | 0.001892 | 0.001097 | 0.005105 | 0.004352 | 0.008142 | 0.001479 | 0.001332 | 0.000233 | 0.001525 | 0.001999 | 0.000426 | 0.001315 | 0.002231 | 0.002657 | 0.002086 | 0.000252 | 0.004195 |
| 24.375 | 0.006549 | 0.000947 | 0.002555 | 0.001645 | 0.001726 | 0.003834 | 0.001864 | 0.005523 | 0.002612 | 0.00766 | 0.005471 | 0.007005 | 0.002053 | 0.002124 | 0.000154 | 0.003969 | 0.001476 | 0.000547 | 0.001778 | 0.001526 | 0.004556 | 0.001447 | 0.001174 | 0.00651 |
| 24.38888889 | 0.005497 | 0.000634 | 0.00441 | 0.002046 | 0.001699 | 0.006982 | 0.006623 | 0.014106 | 0.006831 | 0.008153 | 0.010533 | 0.004983 | 0.002691 | 0.003269 | 0.000302 | 0.008713 | 0.001389 | 0.000797 | 0.002645 | 0.003326 | 0.006646 | 0.003372 | 0.003517 | 0.009057 |
| 24.40277778 | 0.009056 | 0.000798 | 0.007408 | 0.00396 | 0.002504 | 0.011589 | 0.015324 | 0.024099 | 0.009854 | 0.00947 | 0.015922 | 0.009253 | 0.003241 | 0.004785 | 0.000626 | 0.013576 | 0.001558 | 0.001103 | 0.004152 | 0.005665 | 0.007633 | 0.006672 | 0.006496 | 0.010391 |
| 24.41666667 | 0.016389 | 0.001514 | 0.010369 | 0.007443 | 0.005148 | 0.015893 | 0.023026 | 0.028887 | 0.008803 | 0.022776 | 0.018279 | 0.014583 | 0.003347 | 0.005891 | 0.000936 | 0.015534 | 0.001622 | 0.001511 | 0.005406 | 0.006207 | 0.006745 | 0.008759 | 0.007814 | 0.009499 |
| 24.43055556 | 0.02446 | 0.002977 | 0.012301 | 0.009398 | 0.006969 | 0.017893 | 0.024536 | 0.025637 | 0.006205 | 0.04074 | 0.01752 | 0.016172 | 0.00298 | 0.00596 | 0.000983 | 0.01236 | 0.001617 | 0.001822 | 0.005238 | 0.005275 | 0.004947 | 0.008422 | 0.006499 | 0.007075 |
| 24.44444444 | 0.030036 | 0.004318 | 0.01277 | 0.008552 | 0.005322 | 0.016711 | 0.019984 | 0.017589 | 0.005282 | 0.052811 | 0.015033 | 0.016918 | 0.00286 | 0.005645 | 0.000725 | 0.007416 | 0.001817 | 0.002114 | 0.00426 | 0.004515 | 0.004126 | 0.006237 | 0.004505 | 0.00558 |
| 24.45833333 | 0.031372 | 0.004739 | 0.011931 | 0.006304 | 0.002982 | 0.01415 | 0.013332 | 0.008656 | 0.007468 | 0.057829 | 0.01196 | 0.018245 | 0.00364 | 0.005969 | 0.000403 | 0.006308 | 0.002661 | 0.002582 | 0.004005 | 0.005339 | 0.005261 | 0.003707 | 0.004239 | 0.006568 |
| 24.47222222 | 0.028277 | 0.00435 | 0.01044 | 0.004223 | 0.003025 | 0.012863 | 0.00824 | 0.002735 | 0.012261 | 0.054757 | 0.009628 | 0.019434 | 0.005024 | 0.006985 | 0.000364 | 0.009175 | 0.004167 | 0.003223 | 0.004643 | 0.007828 | 0.007565 | 0.001837 | 0.0063 | 0.008664 |
| 24.48611111 | 0.02418 | 0.004796 | 0.011403 | 0.004799 | 0.00293 | 0.015352 | 0.008535 | 0.001513 | 0.019305 | 0.0447 | 0.010312 | 0.020518 | 0.006566 | 0.009061 | 0.001385 | 0.013945 | 0.005878 | 0.004312 | 0.005819 | 0.01155 | 0.00944 | 0.001409 | 0.010264 | 0.009881 |
| 24.5 | 0.027047 | 0.009054 | 0.021774 | 0.009626 | 0.002865 | 0.025258 | 0.017606 | 0.00382 | 0.030718 | 0.03948 | 0.017034 | 0.025305 | 0.009327 | 0.01478 | 0.004202 | 0.021231 | 0.00798 | 0.006784 | 0.007794 | 0.017957 | 0.01047 | 0.003543 | 0.016825 | 0.010865 |
| 24.51388889 | 0.038913 | 0.016615 | 0.040842 | 0.015937 | 0.006164 | 0.040989 | 0.03267 | 0.010389 | 0.043502 | 0.05434 | 0.029821 | 0.036614 | 0.014968 | 0.025295 | 0.007863 | 0.028117 | 0.011167 | 0.011845 | 0.010078 | 0.031397 | 0.012209 | 0.008604 | 0.027546 | 0.015046 |
| 24.52777778 | 0.053453 | 0.020462 | 0.050983 | 0.018964 | 0.010098 | 0.049343 | 0.043926 | 0.018548 | 0.04705 | 0.08992 | 0.042361 | 0.04621 | 0.022221 | 0.034682 | 0.009275 | 0.026888 | 0.01498 | 0.017153 | 0.011056 | 0.050325 | 0.015515 | 0.015497 | 0.038432 | 0.022448 |
| 24.54166667 | 0.071393 | 0.015775 | 0.043884 | 0.018236 | 0.01126 | 0.043127 | 0.042631 | 0.022994 | 0.050386 | 0.123386 | 0.046496 | 0.044006 | 0.026653 | 0.036826 | 0.006836 | 0.016581 | 0.016382 | 0.017639 | 0.010083 | 0.060355 | 0.018241 | 0.020733 | 0.038226 | 0.025025 |
| 24.55555556 | 0.083523 | 0.009558 | 0.035012 | 0.01691 | 0.009493 | 0.032204 | 0.028436 | 0.024768 | 0.068676 | 0.130728 | 0.039517 | 0.033414 | 0.024998 | 0.033815 | 0.004056 | 0.006194 | 0.01434 | 0.013694 | 0.008083 | 0.051349 | 0.017498 | 0.021782 | 0.025477 | 0.018311 |
| 24.56944444 | 0.068465 | 0.009611 | 0.034737 | 0.015721 | 0.005459 | 0.023973 | 0.014738 | 0.024345 | 0.079283 | 0.108311 | 0.024591 | 0.022704 | 0.018547 | 0.028858 | 0.003981 | 0.002041 | 0.011915 | 0.010224 | 0.005394 | 0.031913 | 0.013069 | 0.020302 | 0.011529 | 0.008997 |

| | | | | | | | | | | | | | | | | | | | | | | | | |
|---|---|---|---|---|---|---|---|---|---|---|---|---|---|---|---|---|---|---|---|---|---|---|---|---|
| 24.58333333 | 0.035124 | 0.01256 | 0.037801 | 0.013887 | 0.002413 | 0.017508 | 0.016654 | 0.018131 | 0.062329 | 0.0628 | 0.011434 | 0.013436 | 0.01154 | 0.022346 | 0.004833 | 0.001157 | 0.008959 | 0.009542 | 0.00272 | 0.014674 | 0.007602 | 0.016318 | 0.005294 | 0.00435 |
| 24.59722222 | 0.014211 | 0.013157 | 0.036603 | 0.012872 | 0.002193 | 0.012312 | 0.027865 | 0.009033 | 0.034238 | 0.030719 | 0.006202 | 0.006677 | 0.007882 | 0.015873 | 0.004971 | 0.000891 | 0.004592 | 0.009746 | 0.001419 | 0.007506 | 0.003414 | 0.009099 | 0.005368 | 0.005628 |
| 24.61111111 | 0.01735 | 0.011881 | 0.030526 | 0.016008 | 0.001797 | 0.008728 | 0.032764 | 0.003345 | 0.02109 | 0.042274 | 0.006552 | 0.005447 | 0.008616 | 0.01129 | 0.005044 | 0.00198 | 0.002015 | 0.008544 | 0.001306 | 0.009721 | 0.001541 | 0.005178 | 0.006925 | 0.009193 |
| 24.625 | 0.030453 | 0.0106 | 0.026006 | 0.021484 | 0.003147 | 0.0057 | 0.026264 | 0.001502 | 0.030546 | 0.071239 | 0.013703 | 0.007765 | 0.010363 | 0.00898 | 0.005504 | 0.003609 | 0.003686 | 0.006191 | 0.002272 | 0.008954 | 0.003785 | 0.010154 | 0.013095 | 0.010394 |
| 24.63888889 | 0.037477 | 0.009693 | 0.024084 | 0.022266 | 0.007669 | 0.002874 | 0.015895 | 0.001309 | 0.041356 | 0.092848 | 0.022492 | 0.007475 | 0.009404 | 0.008049 | 0.005709 | 0.003823 | 0.008454 | 0.003324 | 0.004031 | 0.005887 | 0.010721 | 0.019335 | 0.018573 | 0.00818 |
| 24.65277778 | 0.036784 | 0.009326 | 0.021049 | 0.019001 | 0.009318 | 0.001305 | 0.018979 | 0.003521 | 0.039965 | 0.096588 | 0.023861 | 0.008388 | 0.006224 | 0.006927 | 0.005229 | 0.002228 | 0.012129 | 0.001729 | 0.005035 | 0.009666 | 0.018209 | 0.026235 | 0.013942 | 0.004748 |
| 24.66666667 | 0.031492 | 0.010179 | 0.017519 | 0.014915 | 0.006799 | 0.000635 | 0.034196 | 0.007237 | 0.032971 | 0.06757 | 0.020309 | 0.017903 | 0.004303 | 0.004892 | 0.004767 | 0.000762 | 0.011249 | 0.002835 | 0.004701 | 0.014029 | 0.0202 | 0.025606 | 0.006582 | 0.002295 |
| 24.68055556 | 0.02186 | 0.012904 | 0.013429 | 0.009114 | 0.003692 | 0.000919 | 0.038889 | 0.007419 | 0.023466 | 0.050786 | 0.014278 | 0.027101 | 0.005952 | 0.002991 | 0.005027 | 0.000832 | 0.006623 | 0.003598 | 0.004158 | 0.014365 | 0.016494 | 0.017144 | 0.006552 | 0.002277 |
| 24.69444444 | 0.018221 | 0.017665 | 0.007874 | 0.006264 | 0.003343 | 0.001409 | 0.0247 | 0.00777 | 0.011847 | 0.092068 | 0.009099 | 0.025591 | 0.009754 | 0.003606 | 0.005671 | 0.001544 | 0.002497 | 0.003179 | 0.004464 | 0.013063 | 0.011501 | 0.015019 | 0.010971 | 0.003798 |
| 24.70833333 | 0.029686 | 0.024749 | 0.005871 | 0.009218 | 0.005279 | 0.001149 | 0.013698 | 0.012962 | 0.004625 | 0.127929 | 0.008693 | 0.01496 | 0.012848 | 0.008665 | 0.006091 | 0.001449 | 0.001116 | 0.005036 | 0.005528 | 0.008662 | 0.006625 | 0.028345 | 0.010055 | 0.003692 |
| 24.72222222 | 0.044605 | 0.032538 | 0.009956 | 0.011957 | 0.004771 | 0.001501 | 0.02534 | 0.016328 | 0.013493 | 0.115674 | 0.007529 | 0.005076 | 0.014082 | 0.017233 | 0.006158 | 0.000733 | 0.002162 | 0.008968 | 0.006257 | 0.005645 | 0.005213 | 0.04054 | 0.006612 | 0.003563 |
| 24.73611111 | 0.052273 | 0.035177 | 0.014908 | 0.011158 | 0.003059 | 0.00245 | 0.042002 | 0.013466 | 0.029061 | 0.083403 | 0.006344 | 0.002202 | 0.013246 | 0.025583 | 0.006058 | 0.000356 | 0.006684 | 0.011642 | 0.005659 | 0.010735 | 0.013488 | 0.034339 | 0.010169 | 0.005968 |
| 24.75 | 0.047152 | 0.02916 | 0.017321 | 0.009659 | 0.004144 | 0.002792 | 0.046006 | 0.006937 | 0.024262 | 0.046305 | 0.009271 | 0.003408 | 0.009835 | 0.029651 | 0.005887 | 0.00034 | 0.012314 | 0.011532 | 0.004923 | 0.016098 | 0.022182 | 0.018504 | 0.015945 | 0.007738 |
| 24.76388889 | 0.027633 | 0.017391 | 0.018064 | 0.009378 | 0.004909 | 0.003337 | 0.039052 | 0.002665 | 0.013505 | 0.031796 | 0.010877 | 0.009636 | 0.004994 | 0.027016 | 0.005381 | 0.000703 | 0.013719 | 0.009415 | 0.005115 | 0.011905 | 0.017236 | 0.013128 | 0.015373 | 0.006096 |
| 24.77777778 | 0.015998 | 0.007561 | 0.017781 | 0.010149 | 0.003778 | 0.004959 | 0.022607 | 0.003866 | 0.025568 | 0.059366 | 0.008258 | 0.020962 | 0.002688 | 0.018843 | 0.004396 | 0.001519 | 0.00901 | 0.007305 | 0.004684 | 0.007239 | 0.011961 | 0.020023 | 0.00921 | 0.00411 |
| 24.79166667 | 0.027244 | 0.005913 | 0.016345 | 0.013085 | 0.005683 | 0.005323 | 0.009701 | 0.007291 | 0.04232 | 0.099504 | 0.007896 | 0.0305 | 0.004606 | 0.009419 | 0.003047 | 0.003229 | 0.00517 | 0.006293 | 0.00274 | 0.010506 | 0.018092 | 0.027013 | 0.006004 | 0.005199 |
| 24.80555556 | 0.040499 | 0.009229 | 0.014199 | 0.018365 | 0.009094 | 0.003239 | 0.009831 | 0.010082 | 0.040443 | 0.121962 | 0.01333 | 0.031016 | 0.007735 | 0.006148 | 0.001584 | 0.005522 | 0.009353 | 0.005289 | 0.001469 | 0.011382 | 0.019692 | 0.030279 | 0.009773 | 0.00537 |
| 24.81944444 | 0.039816 | 0.012385 | 0.01166 | 0.020406 | 0.007311 | 0.00193 | 0.009131 | 0.011906 | 0.028374 | 0.130293 | 0.015106 | 0.022378 | 0.009982 | 0.013314 | 0.000578 | 0.005833 | 0.016311 | 0.003196 | 0.002847 | 0.007164 | 0.012346 | 0.03105 | 0.011947 | 0.002902 |
| 24.83333333 | 0.033211 | 0.014336 | 0.009553 | 0.015017 | 0.004337 | 0.00347 | 0.006786 | 0.012397 | 0.020932 | 0.132391 | 0.008534 | 0.012269 | 0.011547 | 0.024514 | 0.000398 | 0.0039 | 0.018145 | 0.002044 | 0.005606 | 0.008326 | 0.014473 | 0.024494 | 0.008137 | 0.001491 |
| 24.84722222 | 0.031356 | 0.0148 | 0.008855 | 0.010528 | 0.005977 | 0.006055 | 0.014406 | 0.010412 | 0.018704 | 0.120701 | 0.002377 | 0.010603 | 0.013407 | 0.032407 | 0.000879 | 0.002062 | 0.013068 | 0.003962 | 0.006692 | 0.016657 | 0.033832 | 0.014843 | 0.004748 | 0.002724 |
| 24.86111111 | 0.038701 | 0.014588 | 0.010079 | 0.015965 | 0.006972 | 0.008061 | 0.026972 | 0.006112 | 0.016093 | 0.093045 | 0.001195 | 0.019101 | 0.016327 | 0.035485 | 0.001826 | 0.00106 | 0.005872 | 0.007147 | 0.004415 | 0.022116 | 0.049994 | 0.016536 | 0.003836 | 0.0079 |
| 24.875 | 0.051573 | 0.015417 | 0.014488 | 0.02765 | 0.004028 | 0.00914 | 0.034088 | 0.003042 | 0.013071 | 0.059485 | 0.003449 | 0.021585 | 0.018385 | 0.034476 | 0.003186 | 0.000713 | 0.002544 | 0.010277 | 0.001785 | 0.017775 | 0.042475 | 0.023057 | 0.004025 | 0.016529 |
| 24.88888889 | 0.061105 | 0.017752 | 0.02245 | 0.037336 | 0.001664 | 0.009687 | 0.032625 | 0.00512 | 0.013106 | 0.029944 | 0.008952 | 0.015916 | 0.016981 | 0.031201 | 0.004906 | 0.002909 | 0.004596 | 0.013183 | 0.001168 | 0.009964 | 0.023045 | 0.018453 | 0.008487 | 0.025794 |
| 24.90277778 | 0.065004 | 0.020057 | 0.029899 | 0.040157 | 0.001725 | 0.010604 | 0.025033 | 0.010561 | 0.017674 | 0.010587 | 0.016248 | 0.021814 | 0.013471 | 0.027614 | 0.006527 | 0.006739 | 0.010094 | 0.01391 | 0.00171 | 0.00704 | 0.013108 | 0.010251 | 0.014583 | 0.031334 |
| 24.91666667 | 0.064206 | 0.020852 | 0.03169 | 0.036446 | 0.001499 | 0.012125 | 0.014116 | 0.014959 | 0.023971 | 0.002358 | 0.023721 | 0.036635 | 0.011839 | 0.024059 | 0.007114 | 0.006229 | 0.015272 | 0.011718 | 0.004071 | 0.009664 | 0.015089 | 0.011794 | 0.015508 | 0.028288 |
| 24.93055556 | 0.054846 | 0.019389 | 0.028842 | 0.027266 | 0.000542 | 0.014181 | 0.005811 | 0.019807 | 0.02732 | 0.000798 | 0.029898 | 0.045332 | 0.013443 | 0.019976 | 0.006591 | 0.003618 | 0.016504 | 0.008894 | 0.007305 | 0.014756 | 0.022509 | 0.017821 | 0.010342 | 0.016733 |
| 24.94444444 | 0.03544 | 0.01463 | 0.024576 | 0.01422 | 0.000345 | 0.016222 | 0.004573 | 0.028901 | 0.026082 | 0.002659 | 0.029407 | 0.047983 | 0.016657 | 0.016227 | 0.005534 | 0.003064 | 0.013044 | 0.007823 | 0.009569 | 0.018943 | 0.029959 | 0.020654 | 0.00434 | 0.006624 |
| 24.95833333 | 0.016514 | 0.007971 | 0.019786 | 0.005574 | 0.000825 | 0.01747 | 0.004862 | 0.038496 | 0.021519 | 0.010436 | 0.019018 | 0.04643 | 0.018434 | 0.014332 | 0.003949 | 0.002661 | 0.007132 | 0.008642 | 0.010247 | 0.018409 | 0.029357 | 0.020059 | 0.001779 | 0.004878 |
| 24.97222222 | 0.011222 | 0.00343 | 0.016147 | 0.004539 | 0.001005 | 0.016386 | 0.003711 | 0.039869 | 0.015186 | 0.024391 | 0.009455 | 0.037284 | 0.015832 | 0.014113 | 0.002026 | 0.003375 | 0.003171 | 0.009942 | 0.008718 | 0.012767 | 0.017682 | 0.015792 | 0.001482 | 0.004674 |
| 24.98611111 | 0.013251 | 0.002541 | 0.015432 | 0.006251 | 0.000567 | 0.012178 | 0.005854 | 0.031442 | 0.008875 | 0.038865 | 0.01011 | 0.02118 | 0.010346 | 0.014377 | 0.001033 | 0.005054 | 0.002787 | 0.010162 | 0.005464 | 0.00684 | 0.006641 | 0.009449 | 0.003699 | 0.002298 |
| 25 | 0.011851 | 0.00251 | 0.017788 | 0.009525 | 0.001036 | 0.008613 | 0.012912 | 0.020683 | 0.005785 | 0.048492 | 0.010974 | 0.009597 | 0.007695 | 0.014724 | 0.001618 | 0.00806 | 0.003601 | 0.008755 | 0.002197 | 0.005785 | 0.00472 | 0.006225 | 0.010377 | 0.002428 |
| 25.01388889 | 0.016181 | 0.002587 | 0.022759 | 0.014535 | 0.003566 | 0.010649 | 0.022749 | 0.014789 | 0.009224 | 0.052904 | 0.013166 | 0.007414 | 0.011985 | 0.015983 | 0.001957 | 0.015835 | 0.006218 | 0.007117 | 0.001817 | 0.012742 | 0.010284 | 0.005734 | 0.02209 | 0.009479 |
| 25.02777778 | 0.034241 | 0.00587 | 0.02806 | 0.019001 | 0.005451 | 0.018541 | 0.033476 | 0.015137 | 0.020089 | 0.055015 | 0.029102 | 0.011979 | 0.021662 | 0.020781 | 0.001703 | 0.026943 | 0.010808 | 0.006672 | 0.005273 | 0.024049 | 0.023611 | 0.004521 | 0.037013 | 0.022045 |
| 25.04166667 | 0.058219 | 0.01217 | 0.030562 | 0.02312 | 0.003971 | 0.02818 | 0.042872 | 0.020367 | 0.02689 | 0.054659 | 0.049054 | 0.027349 | 0.03059 | 0.032649 | 0.002938 | 0.030673 | 0.017813 | 0.006495 | 0.009474 | 0.028987 | 0.037924 | 0.006712 | 0.045511 | 0.029181 |
| 25.05555556 | 0.074861 | 0.019375 | 0.030202 | 0.026299 | 0.001399 | 0.033685 | 0.045101 | 0.026004 | 0.020132 | 0.048206 | 0.057883 | 0.04407 | 0.032538 | 0.05003 | 0.004929 | 0.022267 | 0.02123 | 0.005559 | 0.010554 | 0.022932 | 0.03758 | 0.012109 | 0.035852 | 0.021875 |
| 25.06944444 | 0.072648 | 0.022905 | 0.027357 | 0.022693 | 0.000555 | 0.031409 | 0.033478 | 0.022727 | 0.018042 | 0.034029 | 0.052547 | 0.047826 | 0.024677 | 0.060036 | 0.005663 | 0.016803 | 0.017876 | 0.004906 | 0.007115 | 0.011714 | 0.026786 | 0.01132 | 0.018033 | 0.011048 |
| 25.08333333 | 0.051845 | 0.017565 | 0.020556 | 0.012322 | 0.001325 | 0.022583 | 0.016558 | 0.010914 | 0.029799 | 0.018016 | 0.037012 | 0.039266 | 0.013092 | 0.050221 | 0.005108 | 0.019165 | 0.021733 | 0.004617 | 0.004074 | 0.006229 | 0.03084 | 0.00871 | 0.011282 | 0.012799 |
| 25.09722222 | 0.025525 | 0.008537 | 0.011094 | 0.005767 | 0.003136 | 0.01146 | 0.011194 | 0.002465 | 0.036236 | 0.013209 | 0.020769 | 0.023994 | 0.00779 | 0.026183 | 0.003537 | 0.016878 | 0.02947 | 0.003122 | 0.005047 | 0.011131 | 0.049208 | 0.015298 | 0.012447 | 0.021174 |
| 25.11111111 | 0.012848 | 0.004724 | 0.004422 | 0.008211 | 0.005943 | 0.004159 | 0.015328 | 0.000692 | 0.02791 | 0.020113 | 0.016568 | 0.010399 | 0.007319 | 0.008649 | 0.001801 | 0.010071 | 0.025982 | 0.001376 | 0.005041 | 0.018257 | 0.058865 | 0.022691 | 0.011662 | 0.02108 |
| 25.125 | 0.016699 | 0.005063 | 0.003505 | 0.011666 | 0.008056 | 0.00325 | 0.016638 | 0.001079 | 0.013426 | 0.023549 | 0.023472 | 0.00577 | 0.005783 | 0.006402 | 0.001267 | 0.004969 | 0.015388 | 0.001096 | 0.002604 | 0.018706 | 0.051291 | 0.022804 | 0.007574 | 0.012367 |
| 25.13888889 | 0.019962 | 0.004355 | 0.00443 | 0.010245 | 0.007061 | 0.004166 | 0.013539 | 0.001559 | 0.005107 | 0.017186 | 0.025259 | 0.007376 | 0.004033 | 0.006913 | 0.001461 | 0.003671 | 0.006463 | 0.001249 | 0.001362 | 0.011443 | 0.032532 | 0.019078 | 0.003593 | 0.007708 |
| 25.15277778 | 0.015773 | 0.002627 | 0.003932 | 0.005795 | 0.00411 | 0.003212 | 0.010129 | 0.001462 | 0.006506 | 0.007529 | 0.018815 | 0.008495 | 0.002974 | 0.003892 | 0.001116 | 0.006299 | 0.002248 | 0.001082 | 0.001739 | 0.005176 | 0.015208 | 0.014223 | 0.003705 | 0.011099 |
| 25.16666667 | 0.008687 | 0.001617 | 0.003121 | 0.002732 | 0.003389 | 0.001687 | 0.00811 | 0.000947 | 0.009182 | 0.001566 | 0.013285 | 0.008428 | 0.001909 | 0.001391 | 0.000556 | 0.008242 | 0.001832 | 0.000916 | 0.002252 | 0.007473 | 0.010982 | 0.009362 | 0.005943 | 0.013016 |
| 25.18055556 | 0.003123 | 0.001459 | 0.002748 | 0.003415 | 0.005662 | 0.001365 | 0.006975 | 0.001216 | 0.006947 | 0.000694 | 0.01257 | 0.009829 | 0.001769 | 0.000546 | 0.000611 | 0.006663 | 0.001763 | 0.000717 | 0.002182 | 0.012359 | 0.018213 | 0.004794 | 0.006881 | 0.008543 |
| 25.19444444 | 0.00199 | 0.001685 | 0.002731 | 0.004152 | 0.00582 | 0.001409 | 0.005982 | 0.003192 | 0.004341 | 0.002548 | 0.015177 | 0.013585 | 0.002518 | 0.000439 | 0.001011 | 0.003356 | 0.001418 | 0.00074 | 0.001411 | 0.011625 | 0.020764 | 0.002624 | 0.005795 | 0.004464 |
| 25.20833333 | 0.005562 | 0.001779 | 0.002985 | 0.002566 | 0.003957 | 0.00146 | 0.004796 | 0.006355 | 0.005717 | 0.005034 | 0.017662 | 0.018912 | 0.00251 | 0.00074 | 0.001182 | 0.000922 | 0.00187 | 0.001244 | 0.000801 | 0.00647 | 0.013666 | 0.004696 | 0.00323 | 0.006706 |
| 25.22222222 | 0.012155 | 0.001625 | 0.00309 | 0.000841 | 0.004503 | 0.003173 | 0.002918 | 0.009345 | 0.006642 | 0.006694 | 0.018052 | 0.025083 | 0.001521 | 0.001975 | 0.00111 | 0.000402 | 0.002529 | 0.001943 | 0.001108 | 0.003263 | 0.009023 | 0.008117 | 0.001673 | 0.009495 |
| 25.23611111 | 0.017616 | 0.001413 | 0.00265 | 0.001166 | 0.004336 | 0.005924 | 0.001644 | 0.012047 | 0.004993 | 0.007121 | 0.01676 | 0.031132 | 0.000958 | 0.003703 | 0.000995 | 0.001052 | 0.002577 | 0.002624 | 0.001129 | 0.002572 | 0.007731 | 0.00953 | 0.003258 | 0.007214 |
| 25.25 | 0.018519 | 0.00116 | 0.001749 | 0.003892 | 0.002894 | 0.006709 | 0.003041 | 0.015318 | 0.003513 | 0.007064 | 0.014687 | 0.035506 | 0.001497 | 0.005147 | 0.000896 | 0.001989 | 0.002095 | 0.003274 | 0.000596 | 0.00211 | 0.005343 | 0.009067 | 0.005904 | 0.00339 |
| 25.26388889 | 0.014995 | 0.000711 | 0.000934 | 0.006656 | 0.003891 | 0.004518 | 0.005936 | 0.018108 | 0.003514 | 0.006637 | 0.012552 | 0.036616 | 0.002519 | 0.006406 | 0.000874 | 0.002432 | 0.001523 | 0.003475 | 0.000383 | 0.002913 | 0.005575 | 0.008223 | 0.006278 | 0.001452 |
| 25.27777778 | 0.009453 | 0.000478 | 0.000717 | 0.006769 | 0.004296 | 0.002045 | 0.007372 | 0.0184 | 0.00528 | 0.005568 | 0.010715 | 0.033169 | 0.004018 | 0.008075 | 0.000978 | 0.001832 | 0.000957 | 0.002986 | 0.000537 | 0.003133 | 0.005918 | 0.007071 | 0.003957 | 0.002708 |
| 25.29166667 | 0.004495 | 0.000914 | 0.000894 | 0.004459 | 0.003914 | 0.001165 | 0.005982 | 0.016215 | 0.00684 | 0.004528 | 0.009429 | 0.026546 | 0.006249 | 0.010296 | 0.001208 | 0.000949 | 0.000533 | 0.002164 | 0.00117 | 0.001941 | 0.003492 | 0.004693 | 0.001987 | 0.007693 |
| 25.30555556 | 0.0028 | 0.001563 | 0.001003 | 0.003329 | 0.005635 | 0.00127 | 0.003057 | 0.013855 | 0.00656 | 0.004348 | 0.008621 | 0.019818 | 0.008307 | 0.012458 | 0.001576 | 0.000908 | 0.000621 | 0.001426 | 0.001956 | 0.001811 | 0.001658 | 0.001993 | 0.001907 | 0.011225 |
| 25.31944444 | 0.00504 | 0.001859 | 0.001061 | 0.005475 | 0.006112 | 0.003026 | 0.001811 | 0.013621 | 0.007243 | 0.00512 | 0.007654 | 0.013836 | 0.008884 | 0.014026 | 0.001789 | 0.001417 | 0.000949 | 0.001072 | 0.002124 | 0.002312 | 0.001159 | 0.000646 | 0.001696 | 0.009066 |
| 25.33333333 | 0.007906 | 0.001916 | 0.001296 | 0.008291 | 0.004257 | 0.006515 | 0.003425 | 0.015197 | 0.010918 | 0.005957 | 0.006667 | 0.008007 | 0.007725 | 0.014666 | 0.001491 | 0.001629 | 0.001122 | 0.001185 | 0.002011 | 0.002603 | 0.001255 | 0.000558 | 0.001046 | 0.004421 |
| 25.34722222 | 0.008926 | 0.001874 | 0.001834 | 0.009622 | 0.003095 | 0.009833 | 0.005521 | 0.014823 | 0.013681 | 0.005895 | 0.006211 | 0.004404 | 0.00648 | 0.01419 | 0.000826 | 0.001397 | 0.002675 | 0.00155 | 0.002079 | 0.003732 | 0.004014 | 0.000608 | 0.001389 | 0.002391 |
| 25.36111111 | 0.008318 | 0.001637 | 0.002784 | 0.009644 | 0.003535 | 0.011523 | 0.006737 | 0.009648 | 0.011517 | 0.005164 | 0.006284 | 0.005728 | 0.007441 | 0.013444 | 0.000407 | 0.001287 | 0.006739 | 0.00198 | 0.001989 | 0.0066 | 0.012688 | 0.001074 | 0.003201 | 0.005397 |
| 25.375 | 0.007634 | 0.001602 | 0.004226 | 0.009039 | 0.004652 | 0.010752 | 0.008096 | 0.003712 | 0.006269 | 0.004488 | 0.006947 | 0.008549 | 0.010679 | 0.013577 | 0.000618 | 0.001708 | 0.010134 | 0.002531 | 0.001604 | 0.010512 | 0.024321 | 0.003039 | 0.005168 | 0.011956 |
| 25.38888889 | 0.007857 | 0.001988 | 0.005792 | 0.008415 | 0.005285 | 0.008 | 0.0102 | 0.001126 | 0.003106 | 0.004748 | 0.008539 | 0.007702 | 0.013892 | 0.014548 | 0.001401 | 0.002446 | 0.009551 | 0.003331 | 0.001322 | 0.012952 | 0.030475 | 0.005934 | 0.006555 | 0.018058 |
| 25.40277778 | 0.007274 | 0.002005 | 0.006937 | 0.008524 | 0.005319 | 0.006241 | 0.012728 | 0.001136 | 0.004767 | 0.007472 | 0.011531 | 0.004839 | 0.015399 | 0.015444 | 0.002553 | 0.003363 | 0.006544 | 0.004463 | 0.00159 | 0.01222 | 0.026726 | 0.007733 | 0.008945 | 0.020188 |
| 25.41666667 | 0.004954 | 0.001477 | 0.007568 | 0.010308 | 0.005237 | 0.007951 | 0.014644 | 0.004121 | 0.010137 | 0.01201 | 0.014753 | 0.005205 | 0.015235 | 0.01517 | 0.003615 | 0.00437 | 0.004662 | 0.006059 | 0.002306 | 0.009858 | 0.017778 | 0.007384 | 0.013049 | 0.017863 |
| 25.43055556 | 0.005412 | 0.001567 | 0.007922 | 0.013443 | 0.0049 | 0.012077 | 0.014414 | 0.009483 | 0.015469 | 0.014465 | 0.014945 | 0.008885 | 0.014183 | 0.013189 | 0.003981 | 0.00508 | 0.005164 | 0.007421 | 0.00279 | 0.008972 | 0.012698 | 0.005903 | 0.015899 | 0.013452 |
| 25.44444444 | 0.01196 | 0.002319 | 0.008266 | 0.016318 | 0.004239 | 0.015094 | 0.012401 | 0.013644 | 0.016419 | 0.013869 | 0.011322 | 0.011911 | 0.013648 | 0.010424 | 0.003271 | 0.005468 | 0.006833 | 0.007224 | 0.002844 | 0.011486 | 0.016029 | 0.005005 | 0.013559 | 0.010001 |
| 25.45833333 | 0.019726 | 0.002518 | 0.008399 | 0.017335 | 0.003817 | 0.016097 | 0.010658 | 0.015149 | 0.011897 | 0.013412 | 0.007472 | 0.013713 | 0.014766 | 0.008886 | 0.001887 | 0.006184 | 0.00773 | 0.005902 | 0.003304 | 0.016145 | 0.024075 | 0.006423 | 0.007284 | 0.009026 |
| 25.47222222 | 0.023777 | 0.002221 | 0.007985 | 0.0166 | 0.004007 | 0.017186 | 0.01059 | 0.014018 | 0.009232 | 0.01503 | 0.007413 | 0.016647 | 0.017016 | 0.009929 | 0.001029 | 0.007229 | 0.007182 | 0.00533 | 0.004638 | 0.019271 | 0.027127 | 0.011351 | 0.003333 | 0.01028 |
| 25.48611111 | 0.026106 | 0.002764 | 0.00795 | 0.017129 | 0.00526 | 0.017758 | 0.013671 | 0.010714 | 0.02187 | 0.021458 | 0.012359 | 0.021849 | 0.019049 | 0.014036 | 0.002045 | 0.008942 | 0.006003 | 0.006893 | 0.006151 | 0.019764 | 0.019502 | 0.016533 | 0.002707 | 0.012515 |
| 25.5 | 0.033686 | 0.005427 | 0.010373 | 0.023412 | 0.008181 | 0.021484 | 0.022805 | 0.006581 | 0.033471 | 0.039285 | 0.022815 | 0.031928 | 0.021492 | 0.022298 | 0.005526 | 0.014052 | 0.005013 | 0.011187 | 0.006814 | 0.023847 | 0.007826 | 0.017874 | 0.005458 | 0.015132 |
| 25.51388889 | 0.050926 | 0.009556 | 0.015404 | 0.0344 | 0.011595 | 0.035132 | 0.036526 | 0.004244 | 0.030012 | 0.065683 | 0.039341 | 0.047852 | 0.027894 | 0.035276 | 0.009453 | 0.022414 | 0.00579 | 0.019057 | 0.006651 | 0.036089 | 0.005641 | 0.022071 | 0.017628 | 0.021788 |
| 25.52777778 | 0.071675 | 0.011759 | 0.019864 | 0.039442 | 0.014061 | 0.047228 | 0.045241 | 0.003832 | 0.033489 | 0.082876 | 0.055965 | 0.059557 | 0.038051 | 0.050297 | 0.010109 | 0.025919 | 0.010121 | 0.028665 | 0.005766 | 0.047833 | 0.019031 | 0.038443 | 0.03498 | 0.032185 |
| 25.54166667 | 0.088641 | 0.010515 | 0.020104 | 0.033652 | 0.015055 | 0.043777 | 0.043904 | 0.00353 | 0.042301 | 0.08734 | 0.061217 | 0.056333 | 0.044109 | 0.06208 | 0.008718 | 0.01932 | 0.016047 | 0.033112 | 0.004453 | 0.051698 | 0.035782 | 0.059374 | 0.045019 | 0.036663 |
| 25.55555556 | 0.105625 | 0.008128 | 0.017757 | 0.025851 | 0.013233 | 0.037804 | 0.035384 | 0.006077 | 0.05544 | 0.092234 | 0.051149 | 0.043012 | 0.040783 | 0.068878 | 0.009451 | 0.009681 | 0.01951 | 0.02975 | 0.004235 | 0.048194 | 0.041486 | 0.064351 | 0.044031 | 0.031503 |
| 25.56944444 | 0.113877 | 0.006086 | 0.016066 | 0.020887 | 0.009078 | 0.035838 | 0.022309 | 0.009894 | 0.078942 | 0.097272 | 0.032499 | 0.031023 | 0.030746 | 0.073732 | 0.01079 | 0.004342 | 0.020026 | 0.02217 | 0.004261 | 0.03493 | 0.034355 | 0.045049 | 0.034798 | 0.019705 |
| 25.58333333 | 0.096174 | 0.004479 | 0.012725 | 0.01623 | 0.004621 | 0.026889 | 0.010254 | 0.009314 | 0.085739 | 0.089676 | 0.017459 | 0.02316 | 0.01688 | 0.076971 | 0.00923 | 0.002361 | 0.015755 | 0.012705 | 0.002909 | 0.019251 | 0.020461 | 0.023718 | 0.020608 | 0.011216 |
| 25.59722222 | 0.062496 | 0.003483 | 0.00857 | 0.011002 | 0.001861 | 0.0157 | 0.007486 | 0.006047 | 0.061268 | 0.061605 | 0.01852 | 0.016564 | 0.009505 | 0.073111 | 0.005147 | 0.002034 | 0.008648 | 0.005105 | 0.002437 | 0.019919 | 0.015831 | 0.027897 | 0.009849 | 0.016562 |
| 25.61111111 | 0.03088 | 0.002646 | 0.009441 | 0.008486 | 0.00125 | 0.010381 | 0.014596 | 0.005827 | 0.032193 | 0.031972 | 0.029092 | 0.010144 | 0.01542 | 0.055607 | 0.002238 | 0.002793 | 0.005636 | 0.002087 | 0.003418 | 0.032042 | 0.031065 | 0.052845 | 0.009911 | 0.023501 |
| 25.625 | 0.013918 | 0.001505 | 0.014838 | 0.012202 | 0.001258 | 0.008573 | 0.0223 | 0.008234 | 0.023301 | 0.023791 | 0.026036 | 0.010009 | 0.019346 | 0.028425 | 0.002549 | 0.002797 | 0.004306 | 0.001563 | 0.003321 | 0.035457 | 0.054013 | 0.068668 | 0.011573 | 0.017522 |
| 25.63888889 | 0.017939 | 0.000571 | 0.019665 | 0.016574 | 0.003139 | 0.005787 | 0.021286 | 0.009479 | 0.022784 | 0.030005 | 0.017854 | 0.019323 | 0.012354 | 0.012291 | 0.004578 | 0.002217 | 0.002076 | 0.001771 | 0.004138 | 0.026619 | 0.054155 | 0.058736 | 0.008966 | 0.010908 |
| 25.65277778 | 0.030434 | 0.001074 | 0.021749 | 0.013197 | 0.008142 | 0.003119 | 0.014385 | 0.011361 | 0.019171 | 0.04383 | 0.029226 | 0.026953 | 0.008249 | 0.019287 | 0.006542 | 0.002839 | 0.002894 | 0.002327 | 0.007215 | 0.01442 | 0.035456 | 0.0335 | 0.011303 | 0.015316 |
| 25.66666667 | 0.042107 | 0.002541 | 0.020971 | 0.006317 | 0.01314 | 0.004309 | 0.01695 | 0.016789 | 0.017069 | 0.055887 | 0.042754 | 0.022504 | 0.016949 | 0.037403 | 0.005658 | 0.003703 | 0.008444 | 0.002725 | 0.006986 | 0.01456 | 0.041311 | 0.019771 | 0.01652 | 0.018376 |
| 25.68055556 | 0.055233 | 0.002776 | 0.020005 | 0.005135 | 0.014396 | 0.007985 | 0.028448 | 0.017679 | 0.012836 | 0.045797 | 0.034418 | 0.013292 | 0.026587 | 0.055149 | 0.005602 | 0.002895 | 0.015823 | 0.002922 | 0.003305 | 0.026438 | 0.063534 | 0.033265 | 0.015646 | 0.012381 |
| 25.69444444 | 0.060176 | 0.001873 | 0.01952 | 0.008051 | 0.01125 | 0.011558 | 0.030433 | 0.01169 | 0.009019 | 0.023894 | 0.016121 | 0.013801 | 0.026348 | 0.069144 | 0.011371 | 0.00197 | 0.018979 | 0.003649 | 0.002127 | 0.038496 | 0.066397 | 0.059403 | 0.009193 | 0.008307 |
| 25.70833333 | 0.043487 | 0.001793 | 0.016096 | 0.010769 | 0.006009 | 0.011008 | 0.019702 | 0.010729 | 0.01356 | 0.022872 | 0.008601 | 0.020235 | 0.018471 | 0.076385 | 0.015919 | 0.002012 | 0.012967 | 0.00419 | 0.002059 | 0.042892 | 0.049063 | 0.072563 | 0.003787 | 0.011857 |
| 25.72222222 | 0.017409 | 0.00207 | 0.011231 | 0.01293 | 0.002432 | 0.006973 | 0.009949 | 0.013295 | 0.022619 | 0.04878 | 0.018786 | 0.02105 | 0.008736 | 0.07325 | 0.015668 | 0.001446 | 0.006795 | 0.004144 | 0.001209 | 0.028829 | 0.026257 | 0.055382 | 0.004157 | 0.011965 |
| 25.73611111 | 0.005903 | 0.001418 | 0.007445 | 0.013305 | 0.00201 | 0.006607 | 0.011337 | 0.016056 | 0.026557 | 0.076021 | 0.034789 | 0.015627 | 0.002926 | 0.055737 | 0.013481 | 0.00113 | 0.010184 | 0.004997 | 0.001459 | 0.01453 | 0.021815 | 0.027939 | 0.007872 | 0.010021 |
| 25.75 | 0.010579 | 0.001017 | 0.004705 | 0.009816 | 0.003605 | 0.00936 | 0.019006 | 0.018503 | 0.020894 | 0.082803 | 0.035304 | 0.008892 | 0.002532 | 0.028451 | 0.008253 | 0.001402 | 0.014216 | 0.006921 | 0.001733 | 0.019776 | 0.043914 | 0.024534 | 0.009586 | 0.018006 |
| 25.76388889 | 0.019064 | 0.001106 | 0.005006 | 0.006336 | 0.005686 | 0.009093 | 0.024818 | 0.012574 | 0.011514 | 0.06877 | 0.021042 | 0.006488 | 0.00386 | 0.009558 | 0.004154 | 0.001382 | 0.01138 | 0.008734 | 0.001559 | 0.028979 | 0.066675 | 0.041195 | 0.007358 | 0.02677 |
| 25.77777778 | 0.026467 | 0.00098 | 0.008512 | 0.010576 | 0.005032 | 0.005379 | 0.027751 | 0.005014 | 0.00564 | 0.040923 | 0.015061 | 0.00862 | 0.005831 | 0.006882 | 0.006095 | 0.002561 | 0.006513 | 0.00969 | 0.001972 | 0.029973 | 0.069658 | 0.054364 | 0.007328 | 0.023621 |
| 25.79166667 | 0.029625 | 0.001898 | 0.01246 | 0.019892 | 0.00245 | 0.002726 | 0.027238 | 0.00486 | 0.003808 | 0.02583 | 0.021584 | 0.011598 | 0.013635 | 0.006972 | 0.008897 | 0.00491 | 0.005166 | 0.009538 | 0.00185 | 0.022424 | 0.051856 | 0.055877 | 0.011934 | 0.015628 |
| 25.80555556 | 0.02265 | 0.003847 | 0.015037 | 0.025807 | 0.00271 | 0.002504 | 0.022643 | 0.005415 | 0.002721 | 0.045986 | 0.02106 | 0.015956 | 0.025775 | 0.004474 | 0.008165 | 0.007461 | 0.005044 | 0.00863 | 0.00104 | 0.011015 | 0.028979 | 0.048411 | 0.014091 | 0.019176 |
| 25.81944444 | 0.015167 | 0.00504 | 0.016071 | 0.026405 | 0.00568 | 0.003316 | 0.016043 | 0.003008 | 0.001778 | 0.075401 | 0.013135 | 0.023614 | 0.036217 | 0.006325 | 0.006269 | 0.010331 | 0.006125 | 0.007542 | 0.001318 | 0.004525 | 0.028247 | 0.034901 | 0.011546 | 0.03058 |
| 25.83333333 | 0.025416 | 0.005201 | 0.015624 | 0.024701 | 0.007648 | 0.00473 | 0.009532 | 0.001264 | 0.002097 | 0.091119 | 0.014405 | 0.033278 | 0.042118 | 0.013522 | 0.00639 | 0.01278 | 0.011926 | 0.007384 | 0.002987 | 0.005197 | 0.045458 | 0.02149 | 0.006901 | 0.0341 |
| 25.84722222 | 0.046095 | 0.005787 | 0.012838 | 0.024321 | 0.005677 | 0.005876 | 0.004806 | 0.002146 | 0.003861 | 0.09036 | 0.023494 | 0.041792 | 0.041883 | 0.021956 | 0.008412 | 0.013734 | 0.016988 | 0.009518 | 0.006227 | 0.006068 | 0.050214 | 0.011765 | 0.005702 | 0.030813 |
| 25.86111111 | 0.05692 | 0.007874 | 0.009257 | 0.027314 | 0.003549 | 0.01058 | 0.002905 | 0.006769 | 0.010233 | 0.069614 | 0.028816 | 0.044842 | 0.036336 | 0.028953 | 0.009581 | 0.013155 | 0.016823 | 0.013294 | 0.010772 | 0.004786 | 0.031997 | 0.00571 | 0.012812 | 0.030054 |
| 25.875 | 0.054395 | 0.010227 | 0.008523 | 0.031089 | 0.005931 | 0.021062 | 0.002105 | 0.016076 | 0.02253 | 0.040971 | 0.025918 | 0.039577 | 0.027518 | 0.0317 | 0.008131 | 0.011427 | 0.013436 | 0.015741 | 0.014831 | 0.008375 | 0.014306 | 0.005692 | 0.022912 | 0.034039 |
| 25.88888889 | 0.046571 | 0.010359 | 0.01136 | 0.030381 | 0.009984 | 0.0292 | 0.001404 | 0.02769 | 0.036807 | 0.02219 | 0.017406 | 0.029732 | 0.018664 | 0.028597 | 0.004634 | 0.008635 | 0.009464 | 0.014475 | 0.016753 | 0.014735 | 0.016885 | 0.011987 | 0.028217 | 0.037182 |
| 25.90277778 | 0.039796 | 0.007673 | 0.01724 | 0.026349 | 0.012291 | 0.030354 | 0.001258 | 0.033035 | 0.04885 | 0.017766 | 0.010145 | 0.022209 | 0.013429 | 0.023198 | 0.001847 | 0.005882 | 0.007131 | 0.010722 | 0.018124 | 0.019273 | 0.028506 | 0.017885 | 0.027069 | 0.034217 |
| 25.91666667 | 0.037049 | 0.00388 | 0.02468 | 0.024337 | 0.011052 | 0.029563 | 0.002891 | 0.026655 | 0.053523 | 0.020855 | 0.010483 | 0.018313 | 0.012229 | 0.019605 | 0.001412 | 0.004566 | 0.006379 | 0.007709 | 0.019891 | 0.02435 | 0.042519 | 0.017116 | 0.021911 | 0.023514 |
| 25.93055556 | 0.034103 | 0.001575 | 0.026939 | 0.023923 | 0.00661 | 0.023742 | 0.008322 | 0.016407 | 0.045846 | 0.024615 | 0.017577 | 0.014438 | 0.011818 | 0.018068 | 0.001675 | 0.004399 | 0.004873 | 0.006807 | 0.016887 | 0.030008 | 0.059493 | 0.010734 | 0.015494 | 0.011705 |
| 25.94444444 | 0.023921 | 0.001592 | 0.020523 | 0.02238 | 0.002382 | 0.010776 | 0.015439 | 0.008947 | 0.028557 | 0.029035 | 0.024099 | 0.008216 | 0.008695 | 0.01666 | 0.001202 | 0.004372 | 0.002822 | 0.007059 | 0.009118 | 0.031871 | 0.068782 | 0.008142 | 0.009486 | 0.008741 |
| 25.95833333 | 0.012015 | 0.001921 | 0.011702 | 0.019313 | 0.000846 | 0.00516 | 0.019241 | 0.003863 | 0.014003 | 0.033424 | 0.024882 | 0.004557 | 0.005015 | 0.014539 | 0.000729 | 0.004143 | 0.004623 | 0.007534 | 0.00431 | 0.02785 | 0.062971 | 0.013047 | 0.005433 | 0.009526 |
| 25.97222222 | 0.005597 | 0.002188 | 0.005428 | 0.015141 | 0.001363 | 0.009855 | 0.016398 | 0.002093 | 0.010953 | 0.030762 | 0.019362 | 0.008657 | 0.00546 | 0.012012 | 0.001119 | 0.003635 | 0.011627 | 0.007271 | 0.003906 | 0.022051 | 0.045923 | 0.015826 | 0.004137 | 0.007314 |
| 25.98611111 | 0.004603 | 0.002598 | 0.002875 | 0.011427 | 0.002353 | 0.013906 | 0.008475 | 0.003964 | 0.017785 | 0.021765 | 0.011195 | 0.014663 | 0.007233 | 0.009397 | 0.001952 | 0.003155 | 0.01882 | 0.005276 | 0.003175 | 0.018244 | 0.02859 | 0.011606 | 0.005814 | 0.004766 |
| 26 | 0.010263 | 0.002502 | 0.005889 | 0.011026 | 0.005474 | 0.015991 | 0.002443 | 0.007442 | 0.029843 | 0.016884 | 0.007832 | 0.014828 | 0.006162 | 0.007054 | 0.002921 | 0.00447 | 0.022637 | 0.002564 | 0.001819 | 0.016412 | 0.01994 | 0.005786 | 0.010967 | 0.00385 |
| 26.01388889 | 0.02428 | 0.003335 | 0.012743 | 0.014627 | 0.012107 | 0.018178 | 0.002041 | 0.010321 | 0.042691 | 0.017987 | 0.013329 | 0.009489 | 0.007954 | 0.004585 | 0.003575 | 0.009222 | 0.024574 | 0.001621 | 0.002358 | 0.014959 | 0.02178 | 0.004167 | 0.01783 | 0.007391 |
| 26.02777778 | 0.045477 | 0.005202 | 0.017194 | 0.019546 | 0.018011 | 0.020819 | 0.005085 | 0.011652 | 0.054282 | 0.019199 | 0.021788 | 0.008737 | 0.018973 | 0.003078 | 0.002968 | 0.015618 | 0.025669 | 0.003717 | 0.004922 | 0.013116 | 0.032369 | 0.006437 | 0.022329 | 0.015297 |
| 26.04166667 | 0.069664 | 0.008412 | 0.015215 | 0.025538 | 0.019249 | 0.027086 | 0.011797 | 0.009879 | 0.061862 | 0.020043 | 0.026035 | 0.021099 | 0.034346 | 0.008077 | 0.003432 | 0.019879 | 0.025824 | 0.008034 | 0.010033 | 0.013381 | 0.050832 | 0.00739 | 0.023314 | 0.021288 |
| 26.05555556 | 0.08564 | 0.014888 | 0.010601 | 0.032549 | 0.015252 | 0.035621 | 0.021868 | 0.005995 | 0.05923 | 0.021911 | 0.030781 | 0.039212 | 0.042881 | 0.022024 | 0.008643 | 0.019957 | 0.023696 | 0.012697 | 0.01663 | 0.017225 | 0.074971 | 0.004714 | 0.023716 | 0.020975 |
| 26.06944444 | 0.0791 | 0.021172 | 0.009153 | 0.033659 | 0.009486 | 0.038315 | 0.026264 | 0.003211 | 0.04783 | 0.022625 | 0.037028 | 0.048818 | 0.035971 | 0.034172 | 0.014084 | 0.015224 | 0.015926 | 0.015196 | 0.018748 | 0.019429 | 0.090705 | 0.001625 | 0.024497 | 0.015951 |
| 26.08333333 | 0.051965 | 0.021505 | 0.00977 | 0.023744 | 0.009449 | 0.031425 | 0.01976 | 0.002001 | 0.035388 | 0.02287 | 0.029262 | 0.042349 | 0.019895 | 0.031546 | 0.01304 | 0.008064 | 0.008866 | 0.014059 | 0.012956 | 0.014861 | 0.078908 | 0.000657 | 0.022059 | 0.009152 |
| 26.09722222 | 0.022958 | 0.016101 | 0.008233 | 0.011195 | 0.014005 | 0.01799 | 0.009636 | 0.003164 | 0.021829 | 0.028741 | 0.018878 | 0.025316 | 0.01333 | 0.017557 | 0.00826 | 0.003787 | 0.013029 | 0.009692 | 0.00688 | 0.006942 | 0.04501 | 0.001664 | 0.014257 | 0.006789 |
| 26.11111111 | 0.009042 | 0.008784 | 0.004671 | 0.007284 | 0.014841 | 0.007427 | 0.003167 | 0.005869 | 0.01182 | 0.03534 | 0.034282 | 0.010283 | 0.018482 | 0.005753 | 0.004349 | 0.002626 | 0.020903 | 0.005414 | 0.009419 | 0.002319 | 0.031026 | 0.006876 | 0.007497 | 0.014643 |
| 26.125 | 0.010412 | 0.003802 | 0.003184 | 0.009096 | 0.011409 | 0.005135 | 0.001619 | 0.006981 | 0.014589 | 0.026858 | 0.056779 | 0.003023 | 0.021237 | 0.001523 | 0.001877 | 0.001529 | 0.020045 | 0.004972 | 0.016292 | 0.001549 | 0.05852 | 0.017663 | 0.009861 | 0.024163 |
| 26.13888889 | 0.013359 | 0.003512 | 0.004528 | 0.00878 | 0.007003 | 0.004945 | 0.003396 | 0.005873 | 0.024313 | 0.01164 | 0.058106 | 0.0019 | 0.018375 | 0.001307 | 0.000586 | 0.000844 | 0.012949 | 0.005738 | 0.019201 | 0.001223 | 0.096599 | 0.03022 | 0.017612 | 0.025419 |
| 26.15277778 | 0.011079 | 0.004798 | 0.004355 | 0.00626 | 0.003539 | 0.006861 | 0.005846 | 0.004046 | 0.030336 | 0.007574 | 0.042005 | 0.002551 | 0.014817 | 0.001644 | 0.000195 | 0.000865 | 0.006857 | 0.003983 | 0.017548 | 0.002383 | 0.117141 | 0.040369 | 0.020797 | 0.020101 |
| 26.16666667 | 0.006101 | 0.004542 | 0.002753 | 0.003394 | 0.003368 | 0.013509 | 0.006944 | 0.002731 | 0.026631 | 0.008237 | 0.020614 | 0.002191 | 0.011328 | 0.001734 | 0.000234 | 0.000812 | 0.003219 | 0.002766 | 0.014039 | 0.006273 | 0.116131 | 0.045571 | 0.015118 | 0.011397 |
| 26.18055556 | 0.002338 | 0.003532 | 0.0031 | 0.001196 | 0.005184 | 0.017477 | 0.006386 | 0.002425 | 0.014567 | 0.006834 | 0.00726 | 0.001395 | 0.007025 | 0.00344 | 0.000268 | 0.000573 | 0.001078 | 0.004974 | 0.010361 | 0.009489 | 0.100595 | 0.040685 | 0.007112 | 0.004646 |
| 26.19444444 | 0.001414 | 0.002829 | 0.00349 | 0.000962 | 0.00549 | 0.013519 | 0.005437 | 0.003792 | 0.005563 | 0.005393 | 0.004921 | 0.001449 | 0.003678 | 0.007403 | 0.000277 | 0.000424 | 0.000306 | 0.006914 | 0.007281 | 0.00993 | 0.07799 | 0.026075 | 0.004419 | 0.003002 |

| | | | | | | | | | | | | | | | | | | | | | | | | |
|---|---|---|---|---|---|---|---|---|---|---|---|---|---|---|---|---|---|---|---|---|---|---|---|---|
| 26.20833333 | 0.001513 | 0.002648 | 0.002702 | 0.002677 | 0.003637 | 0.0067 | 0.005167 | 0.005059 | 0.005378 | 0.003507 | 0.006168 | 0.002491 | 0.0044 | 0.011318 | 0.000305 | 0.00093 | 0.000629 | 0.005029 | 0.004736 | 0.007825 | 0.052113 | 0.021161 | 0.003559 | 0.004178 |
| 26.22222222 | 0.003 | 0.0029 | 0.00346 | 0.005055 | 0.001753 | 0.003781 | 0.005101 | 0.003746 | 0.007341 | 0.001579 | 0.012492 | 0.003613 | 0.007581 | 0.014464 | 0.000229 | 0.002055 | 0.001557 | 0.002805 | 0.00237 | 0.004285 | 0.027406 | 0.033921 | 0.00312 | 0.010135 |
| 26.23611111 | 0.00517 | 0.003054 | 0.005881 | 0.006457 | 0.001693 | 0.004518 | 0.004209 | 0.002474 | 0.005587 | 0.000678 | 0.02025 | 0.004539 | 0.00814 | 0.018114 | 0.000172 | 0.003369 | 0.002461 | 0.004139 | 0.001162 | 0.0017 | 0.013477 | 0.042236 | 0.006857 | 0.017684 |
| 26.25 | 0.004799 | 0.002824 | 0.0078 | 0.006198 | 0.002342 | 0.004772 | 0.002402 | 0.002978 | 0.002498 | 0.000467 | 0.017106 | 0.005415 | 0.004928 | 0.022504 | 0.000321 | 0.004215 | 0.002444 | 0.006121 | 0.001828 | 0.001158 | 0.017926 | 0.036324 | 0.012217 | 0.01947 |
| 26.26388889 | 0.004654 | 0.002419 | 0.008313 | 0.004844 | 0.002419 | 0.004148 | 0.001331 | 0.00253 | 0.001295 | 0.000371 | 0.010021 | 0.005988 | 0.001951 | 0.026962 | 0.00058 | 0.003879 | 0.001534 | 0.006016 | 0.002721 | 0.001269 | 0.02669 | 0.02452 | 0.014123 | 0.015029 |
| 26.27777778 | 0.007717 | 0.001997 | 0.007897 | 0.003336 | 0.002171 | 0.004766 | 0.002121 | 0.003345 | 0.002114 | 0.000387 | 0.011834 | 0.006249 | 0.001703 | 0.03091 | 0.000745 | 0.002816 | 0.000987 | 0.004655 | 0.002221 | 0.001877 | 0.022789 | 0.012127 | 0.010415 | 0.007772 |
| 26.29166667 | 0.009397 | 0.001661 | 0.007436 | 0.00202 | 0.001707 | 0.006695 | 0.00316 | 0.007045 | 0.005762 | 0.000706 | 0.012331 | 0.006547 | 0.003516 | 0.032812 | 0.000749 | 0.002416 | 0.000874 | 0.003229 | 0.001723 | 0.004157 | 0.012988 | 0.006489 | 0.004295 | 0.002434 |
| 26.30555556 | 0.006874 | 0.001347 | 0.007197 | 0.001193 | 0.001006 | 0.008614 | 0.003335 | 0.00855 | 0.009529 | 0.001161 | 0.007962 | 0.006974 | 0.0062 | 0.031936 | 0.000661 | 0.00344 | 0.000843 | 0.003193 | 0.003682 | 0.007326 | 0.011079 | 0.009609 | 0.001802 | 0.002 |
| 26.31944444 | 0.003123 | 0.000911 | 0.006631 | 0.001376 | 0.000828 | 0.009619 | 0.003132 | 0.005563 | 0.008209 | 0.001409 | 0.009351 | 0.00739 | 0.006348 | 0.029507 | 0.000475 | 0.005055 | 0.001743 | 0.005539 | 0.006645 | 0.009626 | 0.011698 | 0.012995 | 0.002278 | 0.005009 |
| 26.33333333 | 0.001912 | 0.000699 | 0.005492 | 0.001406 | 0.002159 | 0.010134 | 0.002702 | 0.003417 | 0.006564 | 0.002054 | 0.012661 | 0.007987 | 0.004606 | 0.026867 | 0.000291 | 0.005799 | 0.003312 | 0.009763 | 0.007817 | 0.010265 | 0.008735 | 0.012781 | 0.002297 | 0.008147 |
| 26.34722222 | 0.00382 | 0.001276 | 0.004129 | 0.001519 | 0.003881 | 0.011579 | 0.002257 | 0.00619 | 0.010704 | 0.004238 | 0.011428 | 0.009182 | 0.006161 | 0.024622 | 0.00036 | 0.005347 | 0.004433 | 0.013922 | 0.007207 | 0.009946 | 0.00813 | 0.011301 | 0.002933 | 0.009471 |
| 26.36111111 | 0.005835 | 0.002339 | 0.003041 | 0.003883 | 0.003448 | 0.014067 | 0.002183 | 0.009924 | 0.013925 | 0.008434 | 0.007811 | 0.01081 | 0.009365 | 0.022537 | 0.000666 | 0.004636 | 0.004888 | 0.015813 | 0.006239 | 0.01044 | 0.0095 | 0.011074 | 0.004458 | 0.008688 |
| 26.375 | 0.006448 | 0.003109 | 0.002802 | 0.007357 | 0.002146 | 0.015598 | 0.001811 | 0.010596 | 0.011542 | 0.012495 | 0.004622 | 0.01141 | 0.00942 | 0.019983 | 0.000792 | 0.004774 | 0.004696 | 0.015056 | 0.00691 | 0.013216 | 0.017188 | 0.014201 | 0.006031 | 0.006832 |
| 26.38888889 | 0.007627 | 0.003498 | 0.003818 | 0.010469 | 0.002989 | 0.014755 | 0.001702 | 0.007106 | 0.006427 | 0.013096 | 0.003855 | 0.010294 | 0.007168 | 0.01713 | 0.000511 | 0.005979 | 0.003645 | 0.013978 | 0.009 | 0.017979 | 0.034341 | 0.020438 | 0.008391 | 0.005363 |
| 26.40277778 | 0.01189 | 0.003716 | 0.00527 | 0.014183 | 0.00441 | 0.012638 | 0.004 | 0.003464 | 0.003644 | 0.010962 | 0.007062 | 0.009374 | 0.006543 | 0.014915 | 0.00017 | 0.007394 | 0.002108 | 0.014511 | 0.008937 | 0.021725 | 0.042985 | 0.026674 | 0.011678 | 0.00558 |
| 26.41666667 | 0.019037 | 0.003659 | 0.005814 | 0.018542 | 0.005051 | 0.011731 | 0.007444 | 0.003512 | 0.006469 | 0.009971 | 0.012319 | 0.009706 | 0.009273 | 0.013807 | 0.000132 | 0.007691 | 0.001105 | 0.015205 | 0.006129 | 0.021648 | 0.033622 | 0.028681 | 0.013947 | 0.008215 |
| 26.43055556 | 0.025612 | 0.003248 | 0.005593 | 0.02148 | 0.005918 | 0.013535 | 0.009474 | 0.003663 | 0.010702 | 0.012209 | 0.016431 | 0.010053 | 0.013881 | 0.013169 | 0.000207 | 0.006387 | 0.001218 | 0.013937 | 0.003267 | 0.019415 | 0.021816 | 0.02468 | 0.013452 | 0.012891 |
| 26.44444444 | 0.027889 | 0.002523 | 0.005291 | 0.022033 | 0.006404 | 0.016944 | 0.010015 | 0.003056 | 0.015446 | 0.016928 | 0.019132 | 0.010018 | 0.017907 | 0.012383 | 0.000829 | 0.005867 | 0.002167 | 0.011512 | 0.001359 | 0.019095 | 0.023611 | 0.016273 | 0.011295 | 0.017818 |
| 26.45833333 | 0.025132 | 0.001569 | 0.005104 | 0.022151 | 0.005166 | 0.018973 | 0.009852 | 0.003851 | 0.021545 | 0.020479 | 0.018591 | 0.010403 | 0.020181 | 0.012298 | 0.003407 | 0.009856 | 0.003455 | 0.010064 | 0.000848 | 0.021816 | 0.036724 | 0.007781 | 0.009237 | 0.020113 |
| 26.47222222 | 0.020672 | 0.00096 | 0.005304 | 0.023358 | 0.00285 | 0.01816 | 0.009723 | 0.006279 | 0.019613 | 0.020038 | 0.011881 | 0.0113 | 0.021701 | 0.013839 | 0.006941 | 0.015841 | 0.004828 | 0.011064 | 0.002656 | 0.024274 | 0.043208 | 0.004151 | 0.008022 | 0.018809 |
| 26.48611111 | 0.019199 | 0.001542 | 0.006694 | 0.026703 | 0.002173 | 0.016466 | 0.010548 | 0.00973 | 0.013712 | 0.01981 | 0.005302 | 0.01371 | 0.024605 | 0.017249 | 0.009299 | 0.018026 | 0.005944 | 0.014931 | 0.006158 | 0.02434 | 0.034331 | 0.006115 | 0.008583 | 0.016193 |
| 26.5 | 0.025824 | 0.003745 | 0.010088 | 0.036867 | 0.004546 | 0.017486 | 0.014458 | 0.013282 | 0.020488 | 0.028444 | 0.012888 | 0.021222 | 0.031951 | 0.024852 | 0.012116 | 0.019613 | 0.006333 | 0.021069 | 0.008243 | 0.026184 | 0.022954 | 0.009842 | 0.012847 | 0.015985 |
| 26.51388889 | 0.04503 | 0.007873 | 0.014633 | 0.053474 | 0.008174 | 0.022178 | 0.021727 | 0.020393 | 0.029454 | 0.050928 | 0.040024 | 0.033028 | 0.04707 | 0.037874 | 0.015897 | 0.024922 | 0.005892 | 0.02982 | 0.007805 | 0.033873 | 0.024287 | 0.014526 | 0.023146 | 0.020918 |
| 26.52777778 | 0.069782 | 0.010923 | 0.017137 | 0.061026 | 0.01061 | 0.025283 | 0.027285 | 0.036335 | 0.023155 | 0.082063 | 0.073983 | 0.0411 | 0.064565 | 0.049507 | 0.016455 | 0.028091 | 0.004873 | 0.036193 | 0.006477 | 0.044522 | 0.038465 | 0.022373 | 0.038085 | 0.028077 |
| 26.54166667 | 0.085465 | 0.0103 | 0.015488 | 0.051302 | 0.011923 | 0.022772 | 0.026999 | 0.051175 | 0.013095 | 0.113902 | 0.090069 | 0.03942 | 0.069735 | 0.052587 | 0.014097 | 0.023578 | 0.004357 | 0.032547 | 0.004058 | 0.050795 | 0.050702 | 0.029871 | 0.047634 | 0.030263 |
| 26.55555556 | 0.090202 | 0.009952 | 0.012482 | 0.038052 | 0.012195 | 0.01719 | 0.02321 | 0.049139 | 0.013545 | 0.13647 | 0.076212 | 0.029876 | 0.058407 | 0.050905 | 0.013842 | 0.015461 | 0.006074 | 0.024912 | 0.001347 | 0.043914 | 0.047296 | 0.033459 | 0.044314 | 0.024812 |
| 26.56944444 | 0.088202 | 0.011277 | 0.011788 | 0.030087 | 0.009666 | 0.009866 | 0.01896 | 0.035435 | 0.021262 | 0.138718 | 0.045328 | 0.019059 | 0.03982 | 0.053937 | 0.016092 | 0.009617 | 0.00896 | 0.020457 | 0.000197 | 0.025629 | 0.030118 | 0.03275 | 0.033597 | 0.014833 |
| 26.58333333 | 0.080029 | 0.01127 | 0.013141 | 0.021312 | 0.005233 | 0.005101 | 0.014063 | 0.020501 | 0.029129 | 0.122689 | 0.021822 | 0.010864 | 0.020561 | 0.065008 | 0.016843 | 0.005434 | 0.009691 | 0.014165 | 0.000803 | 0.018681 | 0.017385 | 0.025797 | 0.020934 | 0.006986 |
| 26.59722222 | 0.067765 | 0.009297 | 0.013765 | 0.01083 | 0.001883 | 0.007279 | 0.008209 | 0.010492 | 0.028855 | 0.090439 | 0.023313 | 0.005476 | 0.012047 | 0.076366 | 0.013082 | 0.002086 | 0.007617 | 0.005771 | 0.00184 | 0.031699 | 0.031743 | 0.013975 | 0.011252 | 0.00824 |
| 26.61111111 | 0.052196 | 0.006309 | 0.012376 | 0.005484 | 0.000445 | 0.00994 | 0.004171 | 0.010514 | 0.018908 | 0.047626 | 0.041365 | 0.003601 | 0.021014 | 0.074606 | 0.00737 | 0.001066 | 0.004731 | 0.003563 | 0.001575 | 0.041026 | 0.065312 | 0.007169 | 0.013667 | 0.016921 |
| 26.625 | 0.034152 | 0.003306 | 0.008587 | 0.005523 | 0.000223 | 0.007954 | 0.004761 | 0.015878 | 0.010484 | 0.028209 | 0.05012 | 0.007075 | 0.031056 | 0.054414 | 0.006195 | 0.001041 | 0.00273 | 0.009331 | 0.001594 | 0.034623 | 0.080966 | 0.012225 | 0.021801 | 0.02234 |
| 26.63888889 | 0.01816 | 0.001536 | 0.003316 | 0.006421 | 0.000978 | 0.004185 | 0.005251 | 0.024142 | 0.012196 | 0.047431 | 0.03504 | 0.015982 | 0.030943 | 0.02965 | 0.010733 | 0.001226 | 0.002139 | 0.015427 | 0.003998 | 0.019725 | 0.058819 | 0.020725 | 0.020028 | 0.017728 |
| 26.65277778 | 0.014367 | 0.001986 | 0.000446 | 0.007421 | 0.003008 | 0.002205 | 0.004804 | 0.028434 | 0.014372 | 0.075067 | 0.022846 | 0.026581 | 0.023798 | 0.029298 | 0.014939 | 0.001777 | 0.002893 | 0.014465 | 0.007542 | 0.011638 | 0.035686 | 0.024392 | 0.017751 | 0.010259 |
| 26.66666667 | 0.027482 | 0.003725 | 0.000769 | 0.007077 | 0.006708 | 0.0022 | 0.009983 | 0.018268 | 0.011627 | 0.095576 | 0.040715 | 0.033983 | 0.018162 | 0.062705 | 0.013982 | 0.001569 | 0.00498 | 0.008518 | 0.009537 | 0.023228 | 0.059898 | 0.023086 | 0.028507 | 0.006691 |
| 26.68055556 | 0.044635 | 0.005097 | 0.002019 | 0.009047 | 0.011688 | 0.002163 | 0.017098 | 0.00559 | 0.009376 | 0.101019 | 0.057825 | 0.03396 | 0.0299 | 0.104392 | 0.008648 | 0.000868 | 0.008469 | 0.004552 | 0.008527 | 0.042322 | 0.093555 | 0.018604 | 0.038516 | 0.003991 |
| 26.69444444 | 0.051899 | 0.005915 | 0.004439 | 0.018677 | 0.015016 | 0.005243 | 0.01758 | 0.003496 | 0.00867 | 0.07672 | 0.044012 | 0.024538 | 0.054526 | 0.128055 | 0.006204 | 0.000577 | 0.012279 | 0.003146 | 0.005321 | 0.047165 | 0.076353 | 0.013578 | 0.034923 | 0.002257 |
| 26.70833333 | 0.047785 | 0.006815 | 0.009165 | 0.028576 | 0.013476 | 0.015578 | 0.010606 | 0.005575 | 0.011586 | 0.039212 | 0.022853 | 0.012639 | 0.06354 | 0.120542 | 0.011243 | 0.000563 | 0.014277 | 0.003346 | 0.002415 | 0.029833 | 0.038057 | 0.009894 | 0.020627 | 0.003397 |
| 26.72222222 | 0.036284 | 0.00783 | 0.011274 | 0.030889 | 0.007844 | 0.024913 | 0.005815 | 0.005083 | 0.01012 | 0.013901 | 0.028652 | 0.010014 | 0.04918 | 0.084954 | 0.017557 | 0.00084 | 0.014103 | 0.007976 | 0.001743 | 0.015109 | 0.040747 | 0.008485 | 0.010213 | 0.006018 |
| 26.73611111 | 0.027155 | 0.008283 | 0.00881 | 0.025017 | 0.005494 | 0.025402 | 0.010778 | 0.003473 | 0.005631 | 0.003726 | 0.057073 | 0.017979 | 0.026212 | 0.040827 | 0.019496 | 0.001723 | 0.011984 | 0.01645 | 0.002439 | 0.021702 | 0.074416 | 0.008887 | 0.016055 | 0.008325 |
| 26.75 | 0.027312 | 0.007838 | 0.006192 | 0.014098 | 0.00987 | 0.020406 | 0.018862 | 0.003346 | 0.007772 | 0.000971 | 0.082081 | 0.026211 | 0.011705 | 0.016846 | 0.017027 | 0.00321 | 0.007244 | 0.023126 | 0.003241 | 0.028971 | 0.089775 | 0.010014 | 0.02688 | 0.00841 |
| 26.76388889 | 0.025357 | 0.006799 | 0.00339 | 0.008413 | 0.012994 | 0.012618 | 0.023457 | 0.003326 | 0.008486 | 0.000681 | 0.087562 | 0.026711 | 0.014826 | 0.023743 | 0.012352 | 0.004903 | 0.00473 | 0.025171 | 0.003552 | 0.026191 | 0.071147 | 0.010954 | 0.027676 | 0.005993 |
| 26.77777778 | 0.014156 | 0.005787 | 0.00111 | 0.014823 | 0.010812 | 0.005557 | 0.023487 | 0.003435 | 0.007547 | 0.001468 | 0.068967 | 0.020194 | 0.025514 | 0.032394 | 0.007515 | 0.005145 | 0.008149 | 0.024901 | 0.002679 | 0.02021 | 0.038537 | 0.011381 | 0.018611 | 0.003591 |
| 26.79166667 | 0.005526 | 0.004965 | 0.001043 | 0.024307 | 0.006137 | 0.002781 | 0.018457 | 0.00602 | 0.011692 | 0.00266 | 0.037073 | 0.010745 | 0.033351 | 0.023942 | 0.00375 | 0.003571 | 0.013255 | 0.023127 | 0.001584 | 0.016289 | 0.025319 | 0.011309 | 0.008046 | 0.003948 |
| 26.80555556 | 0.004127 | 0.003807 | 0.003642 | 0.025944 | 0.002577 | 0.002291 | 0.010919 | 0.008025 | 0.013677 | 0.00345 | 0.019526 | 0.005925 | 0.037845 | 0.01034 | 0.003263 | 0.003657 | 0.018175 | 0.019115 | 0.002002 | 0.019198 | 0.02468 | 0.010703 | 0.002025 | 0.007163 |
| 26.81944444 | 0.006168 | 0.002279 | 0.008573 | 0.019364 | 0.002904 | 0.005309 | 0.005041 | 0.008488 | 0.009666 | 0.004229 | 0.023407 | 0.010148 | 0.0408 | 0.004191 | 0.005772 | 0.007018 | 0.020941 | 0.015318 | 0.003836 | 0.027119 | 0.022592 | 0.010448 | 0.001637 | 0.013094 |
| 26.83333333 | 0.015307 | 0.001149 | 0.013452 | 0.011403 | 0.0048 | 0.014374 | 0.002622 | 0.007961 | 0.005752 | 0.005775 | 0.025671 | 0.018279 | 0.043472 | 0.006931 | 0.009426 | 0.011079 | 0.01786 | 0.014861 | 0.006064 | 0.033029 | 0.046653 | 0.011119 | 0.005917 | 0.022504 |
| 26.84722222 | 0.028665 | 0.001099 | 0.017744 | 0.010764 | 0.004294 | 0.024787 | 0.003797 | 0.010413 | 0.007492 | 0.007718 | 0.016208 | 0.024522 | 0.046009 | 0.01297 | 0.014659 | 0.014926 | 0.010692 | 0.019108 | 0.008491 | 0.032121 | 0.08728 | 0.012373 | 0.011362 | 0.030342 |
| 26.86111111 | 0.041533 | 0.001817 | 0.021844 | 0.020781 | 0.004117 | 0.028964 | 0.007969 | 0.022506 | 0.016835 | 0.010035 | 0.011938 | 0.026914 | 0.048191 | 0.02006 | 0.021194 | 0.019072 | 0.004657 | 0.027055 | 0.010317 | 0.025542 | 0.103515 | 0.013542 | 0.01428 | 0.029091 |
| 26.875 | 0.05375 | 0.002053 | 0.024325 | 0.036192 | 0.008856 | 0.024352 | 0.012746 | 0.034459 | 0.029304 | 0.013401 | 0.028987 | 0.02712 | 0.047202 | 0.028571 | 0.02556 | 0.021635 | 0.002831 | 0.033422 | 0.009644 | 0.020345 | 0.089012 | 0.013013 | 0.015229 | 0.022185 |
| 26.88888889 | 0.056989 | 0.001348 | 0.023062 | 0.047938 | 0.013776 | 0.017601 | 0.015058 | 0.036281 | 0.038602 | 0.017957 | 0.053545 | 0.027706 | 0.041187 | 0.038757 | 0.025541 | 0.021074 | 0.004427 | 0.033641 | 0.007637 | 0.020782 | 0.062718 | 0.01033 | 0.016616 | 0.018056 |
| 26.90277778 | 0.047623 | 0.000988 | 0.01858 | 0.051372 | 0.012691 | 0.016954 | 0.01383 | 0.029703 | 0.042107 | 0.021486 | 0.060998 | 0.027422 | 0.032982 | 0.049844 | 0.023003 | 0.019051 | 0.008172 | 0.029624 | 0.007339 | 0.025322 | 0.041782 | 0.007155 | 0.017633 | 0.016814 |
| 26.91666667 | 0.0333 | 0.001497 | 0.013448 | 0.047414 | 0.007831 | 0.021307 | 0.009923 | 0.018616 | 0.039574 | 0.019685 | 0.042608 | 0.021985 | 0.024815 | 0.057844 | 0.019506 | 0.017394 | 0.012085 | 0.025692 | 0.006997 | 0.029033 | 0.026695 | 0.005858 | 0.016548 | 0.014131 |
| 26.93055556 | 0.02254 | 0.001913 | 0.009025 | 0.037389 | 0.007259 | 0.023828 | 0.004805 | 0.008576 | 0.030002 | 0.012614 | 0.020464 | 0.011957 | 0.015216 | 0.059397 | 0.014685 | 0.01545 | 0.011605 | 0.022375 | 0.004549 | 0.026108 | 0.016852 | 0.007753 | 0.013035 | 0.009287 |
| 26.94444444 | 0.017685 | 0.00259 | 0.00547 | 0.023074 | 0.012339 | 0.0208 | 0.001243 | 0.003174 | 0.016072 | 0.006496 | 0.018353 | 0.004216 | 0.007657 | 0.054312 | 0.009459 | 0.012366 | 0.007649 | 0.017332 | 0.002607 | 0.017894 | 0.014433 | 0.011577 | 0.008377 | 0.004437 |
| 26.95833333 | 0.015197 | 0.003898 | 0.00362 | 0.009752 | 0.012962 | 0.013354 | 0.000856 | 0.001699 | 0.010081 | 0.005908 | 0.020459 | 0.00251 | 0.007936 | 0.044101 | 0.005978 | 0.009089 | 0.008626 | 0.010494 | 0.002574 | 0.010319 | 0.016694 | 0.014757 | 0.004355 | 0.001804 |
| 26.97222222 | 0.012355 | 0.003936 | 0.005563 | 0.00347 | 0.008183 | 0.006011 | 0.002019 | 0.00342 | 0.016218 | 0.009142 | 0.012654 | 0.003954 | 0.011722 | 0.032545 | 0.004592 | 0.006163 | 0.014791 | 0.004702 | 0.003044 | 0.005742 | 0.01935 | 0.014953 | 0.001916 | 0.002366 |
| 26.98611111 | 0.009833 | 0.002102 | 0.00898 | 0.0027 | 0.008189 | 0.002366 | 0.002146 | 0.00821 | 0.020199 | 0.012415 | 0.007802 | 0.005221 | 0.011518 | 0.023853 | 0.004295 | 0.003186 | 0.016034 | 0.001774 | 0.003404 | 0.003842 | 0.021084 | 0.011731 | 0.001094 | 0.003501 |
| 27 | 0.010276 | 0.000546 | 0.007683 | 0.004941 | 0.013358 | 0.005236 | 0.002363 | 0.009024 | 0.015368 | 0.013831 | 0.015465 | 0.004241 | 0.007734 | 0.020433 | 0.004721 | 0.002104 | 0.010075 | 0.00233 | 0.004287 | 0.004314 | 0.026612 | 0.006598 | 0.00181 | 0.003661 |
| 27.01388889 | 0.017635 | 0.000304 | 0.005946 | 0.012301 | 0.017721 | 0.016328 | 0.005007 | 0.007129 | 0.018427 | 0.012991 | 0.030987 | 0.003354 | 0.008076 | 0.022368 | 0.005402 | 0.006557 | 0.009064 | 0.007283 | 0.005691 | 0.007262 | 0.038791 | 0.003498 | 0.005399 | 0.006801 |
| 27.02777778 | 0.030349 | 0.000989 | 0.009632 | 0.020585 | 0.018895 | 0.027923 | 0.007472 | 0.00958 | 0.042131 | 0.011077 | 0.046868 | 0.006654 | 0.016867 | 0.026596 | 0.005516 | 0.015625 | 0.021069 | 0.015049 | 0.006854 | 0.012154 | 0.054944 | 0.004508 | 0.011372 | 0.013859 |
| 27.04166667 | 0.042183 | 0.002724 | 0.010789 | 0.027293 | 0.0176 | 0.031765 | 0.009273 | 0.011593 | 0.07718 | 0.011332 | 0.054493 | 0.010459 | 0.029326 | 0.030117 | 0.006254 | 0.022331 | 0.035334 | 0.023645 | 0.008268 | 0.01591 | 0.070869 | 0.004685 | 0.01763 | 0.020176 |
| 27.05555556 | 0.053225 | 0.008244 | 0.006623 | 0.03894 | 0.016834 | 0.030126 | 0.012336 | 0.011344 | 0.113516 | 0.014903 | 0.048997 | 0.011508 | 0.037943 | 0.03471 | 0.009831 | 0.021797 | 0.041595 | 0.033114 | 0.011273 | 0.01795 | 0.078228 | 0.002477 | 0.02273 | 0.022353 |
| 27.06944444 | 0.061624 | 0.016819 | 0.003097 | 0.052949 | 0.01513 | 0.025393 | 0.013502 | 0.00944 | 0.126376 | 0.017473 | 0.036178 | 0.010947 | 0.036018 | 0.04194 | 0.014377 | 0.015653 | 0.03231 | 0.036054 | 0.014768 | 0.020201 | 0.067242 | 0.001615 | 0.023989 | 0.018248 |
| 27.08333333 | 0.056842 | 0.020484 | 0.001921 | 0.054453 | 0.009649 | 0.016115 | 0.010281 | 0.006608 | 0.085002 | 0.015021 | 0.021856 | 0.008357 | 0.022543 | 0.044203 | 0.015641 | 0.008879 | 0.022244 | 0.024761 | 0.016029 | 0.018493 | 0.041761 | 0.002138 | 0.020073 | 0.009505 |
| 27.09722222 | 0.037104 | 0.017034 | 0.003094 | 0.04042 | 0.006335 | 0.007897 | 0.005017 | 0.008711 | 0.038138 | 0.009248 | 0.011207 | 0.004448 | 0.010652 | 0.035408 | 0.012996 | 0.007669 | 0.035076 | 0.011052 | 0.014007 | 0.010275 | 0.018382 | 0.002921 | 0.013266 | 0.004224 |
| 27.11111111 | 0.016592 | 0.01109 | 0.005846 | 0.023218 | 0.008292 | 0.008231 | 0.001851 | 0.015442 | 0.04761 | 0.004334 | 0.012387 | 0.002014 | 0.011903 | 0.022785 | 0.008755 | 0.012083 | 0.044709 | 0.009067 | 0.009401 | 0.003393 | 0.009926 | 0.003907 | 0.006417 | 0.006152 |
| 27.125 | 0.00865 | 0.005457 | 0.009056 | 0.012015 | 0.009608 | 0.012698 | 0.001913 | 0.017268 | 0.075934 | 0.00179 | 0.025426 | 0.001229 | 0.017797 | 0.015556 | 0.004977 | 0.013663 | 0.032893 | 0.014298 | 0.004423 | 0.002087 | 0.01496 | 0.00465 | 0.002943 | 0.010686 |
| 27.13888889 | 0.010819 | 0.002312 | 0.013079 | 0.007438 | 0.008759 | 0.013722 | 0.002069 | 0.011878 | 0.080636 | 0.000971 | 0.03949 | 0.001224 | 0.018966 | 0.014796 | 0.002618 | 0.008699 | 0.017 | 0.017878 | 0.002151 | 0.001979 | 0.025249 | 0.004754 | 0.005349 | 0.014453 |
| 27.15277778 | 0.011164 | 0.002899 | 0.015962 | 0.006636 | 0.007956 | 0.010765 | 0.001098 | 0.006854 | 0.070557 | 0.000867 | 0.045575 | 0.002173 | 0.016796 | 0.015255 | 0.001559 | 0.004047 | 0.008389 | 0.018651 | 0.003489 | 0.001408 | 0.036448 | 0.003811 | 0.010875 | 0.017189 |
| 27.16666667 | 0.00708 | 0.005057 | 0.013291 | 0.006672 | 0.007291 | 0.006011 | 0.000305 | 0.007325 | 0.054658 | 0.001027 | 0.043895 | 0.00252 | 0.014156 | 0.012656 | 0.001038 | 0.005374 | 0.009617 | 0.018351 | 0.005983 | 0.001637 | 0.044943 | 0.002231 | 0.016549 | 0.018212 |
| 27.18055556 | 0.005031 | 0.006889 | 0.007242 | 0.006396 | 0.006342 | 0.00232 | 0.001092 | 0.009348 | 0.03931 | 0.002093 | 0.0375 | 0.002441 | 0.011394 | 0.008148 | 0.000521 | 0.009511 | 0.016324 | 0.016903 | 0.007129 | 0.002767 | 0.046662 | 0.001449 | 0.020132 | 0.015647 |
| 27.19444444 | 0.00556 | 0.009571 | 0.005644 | 0.008051 | 0.005989 | 0.002759 | 0.007839 | 0.014544 | 0.052924 | 0.012063 | 0.026721 | 0.008156 | 0.008045 | 0.004657 | 0.000152 | 0.012875 | 0.01912 | 0.012459 | 0.006334 | 0.003022 | 0.035 | 0.00168 | 0.017459 | 0.011646 |
| 27.20833333 | 0.008098 | 0.010644 | 0.008605 | 0.013663 | 0.006089 | 0.00467 | 0.019076 | 0.030212 | 0.085629 | 0.028353 | 0.01384 | 0.018191 | 0.011579 | 0.007256 | 0.000383 | 0.018746 | 0.017783 | 0.009336 | 0.010881 | 0.004723 | 0.029057 | 0.003611 | 0.02449 | 0.021283 |
| 27.22222222 | 0.015533 | 0.008371 | 0.009588 | 0.019653 | 0.004537 | 0.006417 | 0.029615 | 0.039225 | 0.071016 | 0.033318 | 0.007147 | 0.02546 | 0.029089 | 0.022345 | 0.00187 | 0.030166 | 0.029441 | 0.015336 | 0.029506 | 0.01157 | 0.060229 | 0.007268 | 0.055043 | 0.05062 |
| 27.23611111 | 0.034315 | 0.011838 | 0.006641 | 0.034785 | 0.002583 | 0.010746 | 0.049633 | 0.048198 | 0.028351 | 0.044471 | 0.012122 | 0.043863 | 0.044219 | 0.038891 | 0.004593 | 0.038131 | 0.044212 | 0.021816 | 0.044858 | 0.016154 | 0.089874 | 0.00752 | 0.074727 | 0.067244 |
| 27.25 | 0.046434 | 0.01617 | 0.005278 | 0.051446 | 0.003821 | 0.014131 | 0.064983 | 0.066446 | 0.034362 | 0.068987 | 0.017055 | 0.061939 | 0.03781 | 0.032263 | 0.006574 | 0.03089 | 0.033287 | 0.016799 | 0.032904 | 0.012512 | 0.069207 | 0.004742 | 0.058822 | 0.048203 |
| 27.26388889 | 0.030743 | 0.010853 | 0.008732 | 0.040398 | 0.006439 | 0.011641 | 0.046406 | 0.056139 | 0.055107 | 0.060813 | 0.013625 | 0.046635 | 0.019704 | 0.013895 | 0.005656 | 0.015396 | 0.011599 | 0.007969 | 0.010789 | 0.006501 | 0.030095 | 0.004176 | 0.03309 | 0.020415 |
| 27.27777778 | 0.013574 | 0.006848 | 0.012119 | 0.016846 | 0.005101 | 0.007558 | 0.022395 | 0.026838 | 0.045232 | 0.030118 | 0.012019 | 0.025224 | 0.00738 | 0.008676 | 0.002824 | 0.006249 | 0.002425 | 0.004565 | 0.002943 | 0.00285 | 0.010466 | 0.003542 | 0.018624 | 0.00738 |
| 27.29166667 | 0.00774 | 0.007189 | 0.010912 | 0.00498 | 0.001648 | 0.007181 | 0.017023 | 0.01569 | 0.037895 | 0.016751 | 0.01152 | 0.020951 | 0.003309 | 0.006532 | 0.000806 | 0.00468 | 0.000801 | 0.004187 | 0.002374 | 0.001388 | 0.005302 | 0.001639 | 0.010181 | 0.003149 |
| 27.30555556 | 0.004605 | 0.005535 | 0.006276 | 0.001649 | 0.000777 | 0.006292 | 0.013413 | 0.0155 | 0.029386 | 0.014889 | 0.008565 | 0.015957 | 0.002734 | 0.002195 | 0.000271 | 0.005725 | 0.000388 | 0.003675 | 0.00185 | 0.001188 | 0.005741 | 0.000856 | 0.004219 | 0.001242 |
| 27.31944444 | 0.003951 | 0.00298 | 0.003904 | 0.002149 | 0.00189 | 0.002975 | 0.014278 | 0.011244 | 0.021431 | 0.025746 | 0.011621 | 0.0152 | 0.00244 | 0.001905 | 0.000226 | 0.006917 | 0.000257 | 0.002942 | 0.002642 | 0.001278 | 0.00772 | 0.001033 | 0.003356 | 0.001348 |
| 27.33333333 | 0.003483 | 0.001487 | 0.003853 | 0.004599 | 0.003231 | 0.000938 | 0.02049 | 0.004414 | 0.032393 | 0.036239 | 0.019486 | 0.021317 | 0.001789 | 0.002351 | 0.000142 | 0.006923 | 0.000211 | 0.002048 | 0.003605 | 0.001207 | 0.007839 | 0.001335 | 0.004247 | 0.001721 |
| 27.34722222 | 0.004946 | 0.001488 | 0.005993 | 0.006408 | 0.004068 | 0.001194 | 0.017022 | 0.001048 | 0.032646 | 0.025853 | 0.021247 | 0.018876 | 0.000972 | 0.001766 | 7.3631950 | 0.00557 | 0.000338 | 0.001078 | 0.004187 | 0.000971 | 0.007002 | 0.001451 | 0.003789 | 0.001217 |
| 27.36111111 | 0.007195 | 0.002339 | 0.015131 | 0.005429 | 0.003795 | 0.003021 | 0.010894 | 0.001162 | 0.018701 | 0.013701 | 0.014102 | 0.013854 | 0.000448 | 0.001003 | 0.000118 | 0.003665 | 0.000481 | 0.00039 | 0.004516 | 0.000802 | 0.006207 | 0.001389 | 0.002935 | 0.001294 |
| 27.375 | 0.008145 | 0.003126 | 0.023397 | 0.003733 | 0.003421 | 0.005011 | 0.01001 | 0.005074 | 0.014919 | 0.011421 | 0.009612 | 0.012161 | 0.000441 | 0.000617 | 0.000225 | 0.002311 | 0.00043 | 0.000195 | 0.004549 | 0.000983 | 0.005032 | 0.001011 | 0.002182 | 0.003009 |
| 27.38888889 | 0.009754 | 0.002436 | 0.020035 | 0.003396 | 0.004199 | 0.00664 | 0.009332 | 0.011857 | 0.01326 | 0.011575 | 0.014155 | 0.012223 | 0.000909 | 0.000874 | 0.000292 | 0.001923 | 0.000256 | 0.000342 | 0.004836 | 0.001256 | 0.004609 | 0.000481 | 0.001965 | 0.004433 |
| 27.40277778 | 0.011717 | 0.001226 | 0.011041 | 0.00376 | 0.004025 | 0.008828 | 0.008636 | 0.013968 | 0.005158 | 0.013985 | 0.016694 | 0.017614 | 0.002951 | 0.003406 | 0.00058 | 0.001564 | 0.00042 | 0.001256 | 0.007754 | 0.001134 | 0.008419 | 0.000391 | 0.004218 | 0.004283 |
| 27.41666667 | 0.009363 | 0.002537 | 0.006427 | 0.004129 | 0.002725 | 0.011984 | 0.011098 | 0.009842 | 0.005987 | 0.016102 | 0.01188 | 0.019839 | 0.005546 | 0.009383 | 0.001161 | 0.001363 | 0.001017 | 0.003344 | 0.013772 | 0.001226 | 0.015619 | 0.000736 | 0.009013 | 0.005072 |
| 27.43055556 | 0.004141 | 0.005185 | 0.006909 | 0.007733 | 0.00405 | 0.015078 | 0.021404 | 0.005999 | 0.015796 | 0.021409 | 0.009168 | 0.021272 | 0.005043 | 0.013611 | 0.001415 | 0.001946 | 0.001342 | 0.004682 | 0.017096 | 0.001826 | 0.017792 | 0.000938 | 0.011129 | 0.005363 |
| 27.44444444 | 0.005632 | 0.004911 | 0.008549 | 0.016082 | 0.007112 | 0.015907 | 0.02579 | 0.005903 | 0.018358 | 0.022998 | 0.01241 | 0.021488 | 0.003864 | 0.010342 | 0.000974 | 0.001899 | 0.00114 | 0.003447 | 0.012319 | 0.001917 | 0.011783 | 0.000742 | 0.007687 | 0.005002 |
| 27.45833333 | 0.014276 | 0.00244 | 0.008764 | 0.024094 | 0.007777 | 0.012882 | 0.014613 | 0.010203 | 0.01907 | 0.013264 | 0.01256 | 0.013831 | 0.005918 | 0.004599 | 0.000605 | 0.00114 | 0.001325 | 0.001416 | 0.005247 | 0.001546 | 0.007173 | 0.000606 | 0.004487 | 0.008939 |
| 27.47222222 | 0.018199 | 0.001237 | 0.008905 | 0.02459 | 0.006184 | 0.009569 | 0.007992 | 0.011089 | 0.022651 | 0.01439 | 0.007095 | 0.011841 | 0.007091 | 0.002736 | 0.000622 | 0.001097 | 0.001697 | 0.000593 | 0.002444 | 0.001662 | 0.007055 | 0.00076 | 0.004169 | 0.011956 |
| 27.48611111 | 0.011557 | 0.003003 | 0.010427 | 0.018626 | 0.005229 | 0.012593 | 0.021833 | 0.009638 | 0.035199 | 0.044677 | 0.003688 | 0.028131 | 0.005077 | 0.003219 | 0.002099 | 0.001653 | 0.001281 | 0.000996 | 0.003132 | 0.002631 | 0.006826 | 0.000831 | 0.004003 | 0.008783 |
| 27.5 | 0.008421 | 0.006092 | 0.012277 | 0.025381 | 0.006211 | 0.02233 | 0.043342 | 0.011769 | 0.065347 | 0.087695 | 0.008925 | 0.052322 | 0.004515 | 0.005784 | 0.009509 | 0.001786 | 0.000524 | 0.002538 | 0.007092 | 0.004104 | 0.009315 | 0.001679 | 0.005921 | 0.007317 |
| 27.51388889 | 0.02714 | 0.006598 | 0.014585 | 0.05248 | 0.008666 | 0.034392 | 0.047278 | 0.015732 | 0.075423 | 0.108778 | 0.025445 | 0.058081 | 0.005545 | 0.008271 | 0.015397 | 0.002085 | 0.000967 | 0.003587 | 0.008474 | 0.004516 | 0.00999 | 0.004238 | 0.008435 | 0.008344 |
| 27.52777778 | 0.057545 | 0.004757 | 0.018033 | 0.071167 | 0.009523 | 0.041444 | 0.033847 | 0.030204 | 0.045377 | 0.101588 | 0.045514 | 0.045643 | 0.004839 | 0.006103 | 0.009869 | 0.004457 | 0.002852 | 0.002498 | 0.007122 | 0.00287 | 0.008799 | 0.00654 | 0.006628 | 0.007364 |
| 27.54166667 | 0.079304 | 0.003858 | 0.021089 | 0.066174 | 0.006926 | 0.038617 | 0.023123 | 0.047023 | 0.013453 | 0.087714 | 0.055138 | 0.032262 | 0.004472 | 0.002548 | 0.003509 | 0.007063 | 0.003781 | 0.000751 | 0.010155 | 0.001252 | 0.013282 | 0.00594 | 0.003138 | 0.00813 |
| 27.55555556 | 0.089725 | 0.004852 | 0.021435 | 0.055223 | 0.004435 | 0.031064 | 0.02012 | 0.048854 | 0.005088 | 0.081949 | 0.046847 | 0.02083 | 0.004936 | 0.001623 | 0.002735 | 0.005914 | 0.002488 | 0.000372 | 0.010882 | 0.002039 | 0.012837 | 0.003409 | 0.002 | 0.007394 |
| 27.56944444 | 0.083396 | 0.005992 | 0.018016 | 0.045614 | 0.003622 | 0.02191 | 0.018588 | 0.035775 | 0.010808 | 0.077965 | 0.026533 | 0.013888 | 0.004793 | 0.001906 | 0.002182 | 0.002985 | 0.001067 | 0.000914 | 0.005786 | 0.004139 | 0.005234 | 0.002254 | 0.002623 | 0.003248 |
| 27.58333333 | 0.061424 | 0.0062 | 0.012313 | 0.035489 | 0.003162 | 0.014555 | 0.017637 | 0.018941 | 0.018216 | 0.067388 | 0.015598 | 0.014615 | 0.005591 | 0.002061 | 0.000972 | 0.002705 | 0.000725 | 0.000905 | 0.001632 | 0.00518 | 0.001139 | 0.002525 | 0.003174 | 0.001288 |
| 27.59722222 | 0.037028 | 0.005811 | 0.006431 | 0.028523 | 0.004083 | 0.014155 | 0.018389 | 0.015761 | 0.022934 | 0.056186 | 0.024969 | 0.01854 | 0.007535 | 0.001644 | 0.000289 | 0.004166 | 0.000671 | 0.000462 | 0.000352 | 0.004777 | 0.001213 | 0.001894 | 0.002982 | 0.000841 |
| 27.61111111 | 0.017217 | 0.004359 | 0.002151 | 0.021371 | 0.005043 | 0.014458 | 0.018057 | 0.02604 | 0.025645 | 0.047371 | 0.038037 | 0.019183 | 0.009402 | 0.001054 | 0.00015 | 0.005127 | 0.001032 | 0.000358 | 0.000268 | 0.003788 | 0.002491 | 0.001249 | 0.002108 | 0.000619 |
| 27.625 | 0.00639 | 0.002153 | 0.002685 | 0.011254 | 0.003811 | 0.010637 | 0.01381 | 0.030636 | 0.029193 | 0.042738 | 0.038286 | 0.01702 | 0.010424 | 0.00152 | 0.000128 | 0.005343 | 0.001844 | 0.000577 | 0.00028 | 0.002867 | 0.003856 | 0.002244 | 0.001803 | 0.001329 |
| 27.63888889 | 0.003453 | 0.000761 | 0.006814 | 0.006687 | 0.003652 | 0.007316 | 0.007496 | 0.021858 | 0.031246 | 0.042617 | 0.023638 | 0.014432 | 0.010144 | 0.002818 | 8.4888895 | 0.005022 | 0.002356 | 0.000873 | 0.000296 | 0.002349 | 0.004741 | 0.003225 | 0.004141 | 0.0022 |
| 27.65277778 | 0.003883 | 0.000595 | 0.007493 | 0.010775 | 0.005224 | 0.00493 | 0.004584 | 0.018639 | 0.025728 | 0.035702 | 0.013512 | 0.010054 | 0.008688 | 0.003871 | 6.9333338 | 0.004737 | 0.002437 | 0.001231 | 0.000366 | 0.002069 | 0.004683 | 0.00302 | 0.007607 | 0.00249 |
| 27.66666667 | 0.005269 | 0.000861 | 0.006301 | 0.013597 | 0.004418 | 0.002035 | 0.006925 | 0.031161 | 0.014156 | 0.021706 | 0.027285 | 0.009216 | 0.006664 | 0.004346 | 0.000142 | 0.004535 | 0.002383 | 0.001619 | 0.00062 | 0.0016 | 0.003784 | 0.002495 | 0.008379 | 0.002364 |
| 27.68055556 | 0.008531 | 0.001045 | 0.007984 | 0.009734 | 0.004024 | 0.001498 | 0.008446 | 0.039049 | 0.005801 | 0.022393 | 0.044681 | 0.017833 | 0.00471 | 0.0042 | 0.000271 | 0.003958 | 0.002313 | 0.001873 | 0.001128 | 0.001016 | 0.002566 | 0.001997 | 0.006219 | 0.002199 |
| 27.69444444 | 0.016087 | 0.0017 | 0.007154 | 0.005765 | 0.008436 | 0.003239 | 0.006364 | 0.030752 | 0.007679 | 0.047928 | 0.039322 | 0.028731 | 0.003302 | 0.0033 | 0.000432 | 0.003161 | 0.002115 | 0.001797 | 0.001686 | 0.000693 | 0.001515 | 0.001376 | 0.003717 | 0.002023 |
| 27.70833333 | 0.018008 | 0.002555 | 0.004285 | 0.010622 | 0.01395 | 0.004722 | 0.007062 | 0.018743 | 0.020948 | 0.072093 | 0.021502 | 0.031537 | 0.002369 | 0.002033 | 0.00063 | 0.00241 | 0.001698 | 0.001322 | 0.002258 | 0.000657 | 0.000782 | 0.000816 | 0.002081 | 0.001825 |
| 27.72222222 | 0.011305 | 0.002716 | 0.005549 | 0.020245 | 0.016328 | 0.008258 | 0.013265 | 0.021391 | 0.034668 | 0.069993 | 0.018169 | 0.026135 | 0.001541 | 0.001003 | 0.000817 | 0.001732 | 0.001194 | 0.000663 | 0.002691 | 0.000748 | 0.000693 | 0.000475 | 0.001062 | 0.001776 |
| 27.73611111 | 0.011548 | 0.002781 | 0.008734 | 0.024928 | 0.013505 | 0.014685 | 0.018271 | 0.034676 | 0.038025 | 0.051067 | 0.032017 | 0.019058 | 0.000771 | 0.000521 | 0.000965 | 0.00135 | 0.000764 | 0.000299 | 0.002841 | 0.000957 | 0.001465 | 0.00025 | 0.000578 | 0.001722 |
| 27.75 | 0.017991 | 0.00326 | 0.010945 | 0.026969 | 0.007309 | 0.015879 | 0.017663 | 0.030956 | 0.028561 | 0.030629 | 0.046338 | 0.011864 | 0.000351 | 0.000539 | 0.001105 | 0.001425 | 0.000398 | 0.000404 | 0.002808 | 0.001336 | 0.002557 | 0.000185 | 0.000411 | 0.001512 |
| 27.76388889 | 0.020182 | 0.003728 | 0.014262 | 0.02616 | 0.004185 | 0.00929 | 0.013204 | 0.018265 | 0.015648 | 0.013815 | 0.053073 | 0.005928 | 0.000496 | 0.00086 | 0.001257 | 0.001816 | 0.000199 | 0.000619 | 0.002683 | 0.001827 | 0.003465 | 0.000444 | 0.000302 | 0.00137 |
| 27.77777778 | 0.015779 | 0.004233 | 0.017851 | 0.017617 | 0.005239 | 0.003194 | 0.007583 | 0.021557 | 0.015487 | 0.005164 | 0.049027 | 0.00333 | 0.001037 | 0.001349 | 0.00134 | 0.002304 | 0.000319 | 0.00068 | 0.002623 | 0.002248 | 0.004187 | 0.000906 | 0.000289 | 0.001159 |
| 27.79166667 | 0.008286 | 0.004702 | 0.017523 | 0.009389 | 0.00601 | 0.001157 | 0.006265 | 0.022341 | 0.019325 | 0.003765 | 0.03617 | 0.004375 | 0.001681 | 0.001792 | 0.001274 | 0.002798 | 0.000542 | 0.000545 | 0.002681 | 0.002492 | 0.004679 | 0.001231 | 0.000448 | 0.000659 |
| 27.80555556 | 0.00744 | 0.004452 | 0.015396 | 0.012303 | 0.005508 | 0.001054 | 0.013748 | 0.015637 | 0.014487 | 0.005248 | 0.023733 | 0.010829 | 0.002175 | 0.002127 | 0.001078 | 0.003195 | 0.000633 | 0.00033 | 0.002651 | 0.002646 | 0.004958 | 0.001385 | 0.000695 | 0.000227 |
| 27.81944444 | 0.01507 | 0.003735 | 0.01643 | 0.022967 | 0.007281 | 0.001694 | 0.023893 | 0.018369 | 0.00746 | 0.006911 | 0.015038 | 0.015408 | 0.002308 | 0.002598 | 0.000818 | 0.003256 | 0.000631 | 0.00035 | 0.002544 | 0.002919 | 0.005376 | 0.001606 | 0.000756 | 0.000177 |

| | | | | | | | | | | | | | | | | | | | | | | | | |
|---|---|---|---|---|---|---|---|---|---|---|---|---|---|---|---|---|---|---|---|---|---|---|---|---|
| 27.83333333 | 0.024112 | 0.003387 | 0.019899 | 0.031804 | 0.011942 | 0.005113 | 0.029566 | 0.027694 | 0.006536 | 0.005852 | 0.008539 | 0.011142 | 0.002153 | 0.003191 | 0.000667 | 0.002773 | 0.000487 | 0.000784 | 0.002555 | 0.003362 | 0.006244 | 0.001962 | 0.00069 | 0.00062 |
| 27.84722222 | 0.031549 | 0.004016 | 0.02279 | 0.034295 | 0.014405 | 0.013024 | 0.03118 | 0.038161 | 0.014124 | 0.005625 | 0.003963 | 0.005267 | 0.001982 | 0.003379 | 0.000829 | 0.001895 | 0.00033 | 0.001504 | 0.002629 | 0.003777 | 0.007512 | 0.002321 | 0.001066 | 0.00131 |
| 27.86111111 | 0.039054 | 0.00574 | 0.02333 | 0.03095 | 0.01187 | 0.020498 | 0.028663 | 0.038367 | 0.02238 | 0.01123 | 0.002274 | 0.006655 | 0.00174 | 0.003036 | 0.001369 | 0.001105 | 0.000277 | 0.00215 | 0.002598 | 0.003946 | 0.008757 | 0.002615 | 0.001848 | 0.001609 |
| 27.875 | 0.048198 | 0.007348 | 0.021093 | 0.026728 | 0.008515 | 0.023542 | 0.02335 | 0.02403 | 0.025819 | 0.017932 | 0.003543 | 0.013891 | 0.001239 | 0.002799 | 0.002162 | 0.000697 | 0.000198 | 0.002401 | 0.002424 | 0.003722 | 0.009469 | 0.002853 | 0.002287 | 0.001377 |
| 27.88888889 | 0.057059 | 0.007647 | 0.017137 | 0.027336 | 0.008482 | 0.022528 | 0.019554 | 0.009801 | 0.025551 | 0.020096 | 0.004345 | 0.02048 | 0.000881 | 0.002765 | 0.002663 | 0.000699 | 0.00019 | 0.002265 | 0.002277 | 0.003334 | 0.009239 | 0.003079 | 0.002298 | 0.001105 |
| 27.90277778 | 0.061471 | 0.007024 | 0.013177 | 0.031582 | 0.010739 | 0.01947 | 0.017012 | 0.00535 | 0.021111 | 0.019588 | 0.003093 | 0.022671 | 0.001258 | 0.002487 | 0.002352 | 0.001065 | 0.000342 | 0.002141 | 0.002448 | 0.003293 | 0.008923 | 0.003292 | 0.002462 | 0.001289 |
| 27.91666667 | 0.060672 | 0.005973 | 0.009928 | 0.032155 | 0.012515 | 0.017212 | 0.01298 | 0.008982 | 0.015513 | 0.01893 | 0.001999 | 0.017814 | 0.002288 | 0.001734 | 0.001498 | 0.001543 | 0.000586 | 0.002332 | 0.002921 | 0.003762 | 0.010439 | 0.003325 | 0.003056 | 0.001881 |
| 27.93055556 | 0.054663 | 0.003972 | 0.006051 | 0.02577 | 0.011667 | 0.014229 | 0.007858 | 0.014815 | 0.013068 | 0.015549 | 0.001363 | 0.011287 | 0.003407 | 0.000999 | 0.00076 | 0.001872 | 0.000812 | 0.002723 | 0.003356 | 0.004505 | 0.013733 | 0.003077 | 0.0038 | 0.002684 |
| 27.94444444 | 0.042506 | 0.002763 | 0.002834 | 0.016187 | 0.007336 | 0.008094 | 0.005701 | 0.018137 | 0.01283 | 0.008607 | 0.002435 | 0.013546 | 0.004173 | 0.001124 | 0.000561 | 0.002058 | 0.001079 | 0.003264 | 0.003528 | 0.005278 | 0.017021 | 0.00268 | 0.004385 | 0.003475 |
| 27.95833333 | 0.02624 | 0.004129 | 0.001891 | 0.008605 | 0.00436 | 0.002245 | 0.005705 | 0.017736 | 0.009587 | 0.005899 | 0.007398 | 0.017972 | 0.00434 | 0.001677 | 0.000817 | 0.002044 | 0.001386 | 0.003851 | 0.003479 | 0.006218 | 0.019508 | 0.002301 | 0.004917 | 0.003908 |
| 27.97222222 | 0.015283 | 0.004376 | 0.002854 | 0.007274 | 0.007529 | 0.00267 | 0.009225 | 0.016259 | 0.009091 | 0.023918 | 0.013975 | 0.021632 | 0.003779 | 0.001755 | 0.000887 | 0.001631 | 0.001446 | 0.004099 | 0.00348 | 0.007474 | 0.021348 | 0.002074 | 0.005547 | 0.004056 |
| 27.98611111 | 0.018924 | 0.004296 | 0.008132 | 0.01086 | 0.010902 | 0.010131 | 0.027633 | 0.023725 | 0.022326 | 0.073494 | 0.018655 | 0.045282 | 0.00305 | 0.002516 | 0.000854 | 0.001276 | 0.001386 | 0.003475 | 0.002786 | 0.008393 | 0.019834 | 0.002471 | 0.004913 | 0.003599 |
| 28 | 0.02497 | 0.008828 | 0.015561 | 0.014268 | 0.007799 | 0.019498 | 0.050642 | 0.03686 | 0.030646 | 0.12354 | 0.015936 | 0.079213 | 0.007083 | 0.00989 | 0.001963 | 0.003892 | 0.004668 | 0.004003 | 0.005044 | 0.007768 | 0.018954 | 0.0059 | 0.005082 | 0.007017 |
| 28.01388889 | 0.02251 | 0.010688 | 0.01927 | 0.018225 | 0.004244 | 0.025005 | 0.048713 | 0.030831 | 0.022169 | 0.121702 | 0.012452 | 0.077071 | 0.02312 | 0.031781 | 0.004584 | 0.016612 | 0.016987 | 0.015636 | 0.019727 | 0.004958 | 0.042275 | 0.017815 | 0.016336 | 0.02061 |
| 28.02777778 | 0.025492 | 0.005327 | 0.019723 | 0.022961 | 0.005546 | 0.024829 | 0.023867 | 0.015418 | 0.016675 | 0.079669 | 0.019908 | 0.040471 | 0.045067 | 0.058908 | 0.007297 | 0.043965 | 0.033177 | 0.038804 | 0.041601 | 0.002804 | 0.092036 | 0.03642 | 0.029553 | 0.032099 |
| 28.04166667 | 0.03437 | 0.005595 | 0.019752 | 0.02935 | 0.00845 | 0.021493 | 0.008144 | 0.016708 | 0.013233 | 0.056028 | 0.025509 | 0.018988 | 0.05481 | 0.063745 | 0.007817 | 0.067788 | 0.039305 | 0.044621 | 0.044043 | 0.004476 | 0.121041 | 0.045848 | 0.023079 | 0.027534 |
| 28.05555556 | 0.035432 | 0.015888 | 0.018004 | 0.034709 | 0.012373 | 0.017684 | 0.011754 | 0.022015 | 0.006049 | 0.055268 | 0.019056 | 0.022643 | 0.04527 | 0.040745 | 0.005357 | 0.059576 | 0.032151 | 0.02496 | 0.023986 | 0.007212 | 0.098593 | 0.036742 | 0.009406 | 0.015612 |
| 28.06944444 | 0.028961 | 0.023617 | 0.01284 | 0.028088 | 0.014558 | 0.013784 | 0.01651 | 0.021368 | 0.002936 | 0.039865 | 0.012512 | 0.022116 | 0.024743 | 0.016069 | 0.002175 | 0.029039 | 0.018967 | 0.011191 | 0.008102 | 0.00731 | 0.050409 | 0.019715 | 0.005893 | 0.006761 |
| 28.08333333 | 0.017139 | 0.02132 | 0.00633 | 0.015783 | 0.011124 | 0.008548 | 0.012221 | 0.016533 | 0.002992 | 0.021959 | 0.013278 | 0.012186 | 0.009704 | 0.007523 | 0.000535 | 0.011464 | 0.008226 | 0.010841 | 0.003469 | 0.004917 | 0.017969 | 0.00744 | 0.005424 | 0.003462 |
| 28.09722222 | 0.006666 | 0.012135 | 0.00266 | 0.011903 | 0.005971 | 0.003822 | 0.00698 | 0.009679 | 0.00533 | 0.034174 | 0.014261 | 0.01009 | 0.008994 | 0.009183 | 0.000223 | 0.012077 | 0.00504 | 0.010559 | 0.003402 | 0.00226 | 0.013904 | 0.002354 | 0.002888 | 0.004426 |
| 28.11111111 | 0.005433 | 0.004658 | 0.005577 | 0.012664 | 0.00392 | 0.003349 | 0.008047 | 0.010206 | 0.011439 | 0.053515 | 0.009601 | 0.014426 | 0.012948 | 0.011835 | 0.000373 | 0.014066 | 0.007008 | 0.009244 | 0.004776 | 0.00137 | 0.019626 | 0.0024 | 0.000834 | 0.005452 |
| 28.125 | 0.008556 | 0.003369 | 0.013941 | 0.011334 | 0.003522 | 0.006186 | 0.008304 | 0.016986 | 0.017236 | 0.043224 | 0.006533 | 0.012726 | 0.013438 | 0.012933 | 0.000561 | 0.01282 | 0.008993 | 0.008612 | 0.003801 | 0.001854 | 0.021337 | 0.004286 | 0.000305 | 0.005249 |
| 28.13888889 | 0.011612 | 0.004096 | 0.02294 | 0.008428 | 0.00266 | 0.009498 | 0.004138 | 0.020093 | 0.018572 | 0.019174 | 0.010446 | 0.008408 | 0.011439 | 0.011622 | 0.000443 | 0.011292 | 0.00941 | 0.008101 | 0.00191 | 0.001932 | 0.020731 | 0.005656 | 0.000302 | 0.004614 |
| 28.15277778 | 0.012662 | 0.005065 | 0.02834 | 0.004907 | 0.002784 | 0.011474 | 0.004036 | 0.014048 | 0.01781 | 0.012988 | 0.013255 | 0.005949 | 0.00895 | 0.007891 | 0.000198 | 0.009884 | 0.009362 | 0.007796 | 0.00156 | 0.001348 | 0.019921 | 0.006522 | 0.000217 | 0.004238 |
| 28.16666667 | 0.010652 | 0.005876 | 0.02687 | 0.002647 | 0.003533 | 0.011603 | 0.006868 | 0.005828 | 0.016333 | 0.015867 | 0.011717 | 0.002776 | 0.007423 | 0.004168 | 0.000163 | 0.008525 | 0.009887 | 0.008284 | 0.002007 | 0.001002 | 0.018942 | 0.007176 | 0.000173 | 0.004438 |
| 28.18055556 | 0.00633 | 0.00501 | 0.019583 | 0.002891 | 0.003377 | 0.010366 | 0.007054 | 0.004873 | 0.013242 | 0.013749 | 0.009955 | 0.002562 | 0.006744 | 0.003107 | 0.000322 | 0.007344 | 0.010526 | 0.009385 | 0.00341 | 0.001256 | 0.017976 | 0.007635 | 0.000167 | 0.004645 |
| 28.19444444 | 0.001814 | 0.00325 | 0.012175 | 0.002949 | 0.002619 | 0.007913 | 0.005263 | 0.005674 | 0.009671 | 0.011598 | 0.00877 | 0.004443 | 0.006165 | 0.004237 | 0.000507 | 0.006359 | 0.010392 | 0.010171 | 0.00504 | 0.00192 | 0.017045 | 0.007516 | 0.00028 | 0.004331 |
| 28.20833333 | 0.001254 | 0.002504 | 0.008678 | 0.00243 | 0.002141 | 0.00563 | 0.004517 | 0.004087 | 0.007349 | 0.01615 | 0.008631 | 0.004001 | 0.005164 | 0.005392 | 0.000442 | 0.00566 | 0.009446 | 0.010209 | 0.005566 | 0.002631 | 0.01587 | 0.006591 | 0.00056 | 0.003658 |
| 28.22222222 | 0.003971 | 0.002832 | 0.010005 | 0.00411 | 0.001663 | 0.004674 | 0.006683 | 0.006319 | 0.006349 | 0.021758 | 0.009812 | 0.003996 | 0.004008 | 0.005666 | 0.000255 | 0.00581 | 0.008189 | 0.00994 | 0.005582 | 0.003004 | 0.014609 | 0.005446 | 0.000835 | 0.003068 |
| 28.23611111 | 0.005566 | 0.00272 | 0.01455 | 0.005998 | 0.001803 | 0.004317 | 0.006602 | 0.010695 | 0.006051 | 0.016671 | 0.010647 | 0.003794 | 0.003513 | 0.005293 | 0.000301 | 0.006636 | 0.007525 | 0.009889 | 0.005564 | 0.002868 | 0.013547 | 0.004681 | 0.00104 | 0.002813 |
| 28.25 | 0.005555 | 0.005438 | 0.017888 | 0.005716 | 0.004294 | 0.003616 | 0.007412 | 0.009527 | 0.008535 | 0.02029 | 0.007899 | 0.007899 | 0.003709 | 0.004832 | 0.000422 | 0.006743 | 0.008071 | 0.010052 | 0.005319 | 0.002442 | 0.012425 | 0.004389 | 0.001235 | 0.002702 |
| 28.26388889 | 0.007451 | 0.011918 | 0.015275 | 0.006891 | 0.007813 | 0.002652 | 0.016337 | 0.00601 | 0.015346 | 0.049921 | 0.004393 | 0.021095 | 0.003705 | 0.004571 | 0.000415 | 0.005841 | 0.009233 | 0.010095 | 0.004822 | 0.00199 | 0.010913 | 0.004348 | 0.00149 | 0.002478 |
| 28.27777778 | 0.008463 | 0.014509 | 0.009463 | 0.007762 | 0.007737 | 0.002198 | 0.01961 | 0.006024 | 0.018224 | 0.061315 | 0.003649 | 0.025107 | 0.002947 | 0.004379 | 0.000341 | 0.004805 | 0.009119 | 0.009576 | 0.004252 | 0.001566 | 0.009371 | 0.003999 | 0.001819 | 0.002019 |
| 28.29166667 | 0.007284 | 0.010103 | 0.009056 | 0.008011 | 0.005157 | 0.002487 | 0.015903 | 0.006883 | 0.014345 | 0.047159 | 0.00442 | 0.020222 | 0.001951 | 0.004066 | 0.000236 | 0.003951 | 0.006922 | 0.008365 | 0.003528 | 0.001213 | 0.007881 | 0.003047 | 0.002149 | 0.001194 |
| 28.30555556 | 0.009738 | 0.005242 | 0.01247 | 0.015324 | 0.004867 | 0.002618 | 0.021565 | 0.009719 | 0.01114 | 0.051808 | 0.007232 | 0.026446 | 0.001476 | 0.003615 | 0.000139 | 0.003173 | 0.00408 | 0.007001 | 0.002695 | 0.001167 | 0.006019 | 0.001875 | 0.002546 | 0.000576 |
| 28.31944444 | 0.010593 | 0.004487 | 0.010878 | 0.020135 | 0.005953 | 0.00232 | 0.021789 | 0.013053 | 0.00766 | 0.050321 | 0.007748 | 0.026705 | 0.001371 | 0.003259 | 0.000151 | 0.002398 | 0.001862 | 0.005901 | 0.001933 | 0.001539 | 0.003916 | 0.000916 | 0.002913 | 0.000816 |
| 28.33333333 | 0.009265 | 0.007323 | 0.00484 | 0.013565 | 0.009076 | 0.003197 | 0.013393 | 0.009816 | 0.004713 | 0.039892 | 0.007694 | 0.020062 | 0.001082 | 0.003271 | 0.000253 | 0.001661 | 0.000684 | 0.005208 | 0.001296 | 0.002001 | 0.002128 | 0.000367 | 0.002854 | 0.001378 |
| 28.34722222 | 0.010649 | 0.010597 | 0.002203 | 0.006945 | 0.012384 | 0.00389 | 0.014299 | 0.003504 | 0.004682 | 0.048927 | 0.009756 | 0.023647 | 0.000614 | 0.003471 | 0.000264 | 0.001063 | 0.000314 | 0.004916 | 0.000862 | 0.002174 | 0.001006 | 0.000385 | 0.002452 | 0.001681 |
| 28.36111111 | 0.00944 | 0.008593 | 0.004991 | 0.00623 | 0.009352 | 0.003052 | 0.014173 | 0.003094 | 0.004202 | 0.042533 | 0.008201 | 0.02116 | 0.000371 | 0.003249 | 0.000158 | 0.000669 | 0.0003 | 0.004892 | 0.000885 | 0.002063 | 0.000734 | 0.000778 | 0.002144 | 0.001536 |
| 28.375 | 0.008751 | 0.00326 | 0.008598 | 0.009472 | 0.003541 | 0.003746 | 0.009885 | 0.009123 | 0.002824 | 0.022285 | 0.0072 | 0.013734 | 0.000637 | 0.002587 | 6.8875005 | 0.000569 | 0.000396 | 0.005052 | 0.001416 | 0.002033 | 0.001282 | 0.000995 | 0.001897 | 0.000964 |
| 28.38888889 | 0.009375 | 0.002102 | 0.010074 | 0.01239 | 0.002256 | 0.005798 | 0.008664 | 0.013816 | 0.00248 | 0.018716 | 0.007771 | 0.010917 | 0.001336 | 0.002359 | 7.9895839 | 0.000732 | 0.000304 | 0.005182 | 0.00205 | 0.002285 | 0.002164 | 0.001012 | 0.001602 | 0.000607 |
| 28.40277778 | 0.006745 | 0.004915 | 0.00947 | 0.013325 | 0.003916 | 0.006904 | 0.007642 | 0.013315 | 0.003501 | 0.022066 | 0.004552 | 0.007731 | 0.00234 | 0.00316 | 0.000231 | 0.000984 | 0.000129 | 0.004998 | 0.002363 | 0.002504 | 0.002817 | 0.001479 | 0.001632 | 0.001044 |
| 28.41666667 | 0.006789 | 0.006702 | 0.008206 | 0.018251 | 0.004513 | 0.008014 | 0.006001 | 0.011928 | 0.005568 | 0.018338 | 0.00203 | 0.005187 | 0.003616 | 0.004492 | 0.000487 | 0.001468 | 0.00012 | 0.004593 | 0.002569 | 0.002533 | 0.003241 | 0.002518 | 0.002045 | 0.001737 |
| 28.43055556 | 0.009859 | 0.007075 | 0.007188 | 0.022937 | 0.004716 | 0.010292 | 0.006427 | 0.011589 | 0.008517 | 0.013403 | 0.002572 | 0.005151 | 0.004657 | 0.005225 | 0.000688 | 0.002486 | 0.000282 | 0.004504 | 0.003306 | 0.002806 | 0.003913 | 0.003315 | 0.002133 | 0.00157 |
| 28.44444444 | 0.009587 | 0.00788 | 0.006103 | 0.01722 | 0.006002 | 0.01277 | 0.012284 | 0.010531 | 0.011963 | 0.022042 | 0.004322 | 0.010443 | 0.00475 | 0.004999 | 0.000651 | 0.004173 | 0.000324 | 0.004981 | 0.004543 | 0.00351 | 0.004941 | 0.003499 | 0.001584 | 0.000705 |
| 28.45833333 | 0.007541 | 0.007335 | 0.006388 | 0.007004 | 0.005523 | 0.013722 | 0.016215 | 0.009784 | 0.012199 | 0.029544 | 0.005643 | 0.014165 | 0.004178 | 0.004745 | 0.00042 | 0.006683 | 0.000314 | 0.005677 | 0.005532 | 0.004291 | 0.005917 | 0.003557 | 0.000884 | 0.000153 |
| 28.47222222 | 0.006906 | 0.004947 | 0.007635 | 0.002453 | 0.00247 | 0.01274 | 0.013513 | 0.009994 | 0.009131 | 0.024181 | 0.003664 | 0.010243 | 0.003983 | 0.005372 | 0.00032 | 0.009211 | 0.000782 | 0.006075 | 0.005811 | 0.005041 | 0.007187 | 0.00346 | 0.00054 | 0.000238 |
| 28.48611111 | 0.006707 | 0.004832 | 0.006653 | 0.005228 | 0.0034 | 0.011208 | 0.016564 | 0.009047 | 0.012657 | 0.032692 | 0.002044 | 0.012072 | 0.005481 | 0.007754 | 0.000458 | 0.010606 | 0.002463 | 0.006463 | 0.005395 | 0.006742 | 0.012488 | 0.002538 | 0.001202 | 0.001386 |
| 28.5 | 0.00962 | 0.009553 | 0.006126 | 0.017722 | 0.01072 | 0.012622 | 0.030338 | 0.008922 | 0.02525 | 0.062735 | 0.003873 | 0.024237 | 0.011245 | 0.013954 | 0.00052 | 0.012776 | 0.007792 | 0.008271 | 0.004445 | 0.011428 | 0.030351 | 0.002875 | 0.002898 | 0.004564 |
| 28.51388889 | 0.024199 | 0.014062 | 0.013865 | 0.045763 | 0.017719 | 0.02158 | 0.039808 | 0.01694 | 0.02654 | 0.082203 | 0.012238 | 0.03031 | 0.021643 | 0.02444 | 0.000739 | 0.017642 | 0.018141 | 0.012613 | 0.003953 | 0.018951 | 0.063608 | 0.005568 | 0.00453 | 0.011076 |
| 28.52777778 | 0.050774 | 0.011438 | 0.030575 | 0.073636 | 0.019107 | 0.037721 | 0.033871 | 0.036892 | 0.025415 | 0.073615 | 0.024962 | 0.02246 | 0.028891 | 0.033977 | 0.001475 | 0.021386 | 0.028114 | 0.016496 | 0.004546 | 0.024606 | 0.095784 | 0.006239 | 0.007211 | 0.020222 |
| 28.54166667 | 0.071165 | 0.004802 | 0.042452 | 0.066914 | 0.016401 | 0.048448 | 0.023097 | 0.058747 | 0.046981 | 0.052835 | 0.025566 | 0.011217 | 0.026422 | 0.037849 | 0.00236 | 0.01977 | 0.031727 | 0.015392 | 0.006407 | 0.025817 | 0.109692 | 0.003835 | 0.013425 | 0.028146 |
| 28.55555556 | 0.070817 | 0.00227 | 0.035082 | 0.035201 | 0.014475 | 0.04305 | 0.021143 | 0.063329 | 0.055949 | 0.03435 | 0.013254 | 0.006046 | 0.019649 | 0.037031 | 0.002682 | 0.015152 | 0.031133 | 0.01139 | 0.009962 | 0.02641 | 0.108782 | 0.002427 | 0.021047 | 0.031706 |
| 28.56944444 | 0.057646 | 0.003784 | 0.017848 | 0.019614 | 0.014596 | 0.030201 | 0.024782 | 0.047914 | 0.034149 | 0.018512 | 0.004637 | 0.006653 | 0.015509 | 0.033416 | 0.001869 | 0.012073 | 0.030658 | 0.009124 | 0.013065 | 0.028651 | 0.100348 | 0.003919 | 0.025932 | 0.030393 |
| 28.58333333 | 0.042108 | 0.004592 | 0.011048 | 0.021439 | 0.014153 | 0.020658 | 0.025125 | 0.025726 | 0.020236 | 0.012814 | 0.003559 | 0.009347 | 0.013673 | 0.028257 | 0.001052 | 0.010928 | 0.029938 | 0.008736 | 0.013551 | 0.029843 | 0.086453 | 0.006623 | 0.029918 | 0.026996 |
| 28.59722222 | 0.026664 | 0.003883 | 0.01326 | 0.018839 | 0.012081 | 0.015365 | 0.020349 | 0.01299 | 0.024149 | 0.024235 | 0.004059 | 0.010685 | 0.011645 | 0.022415 | 0.001361 | 0.010039 | 0.027211 | 0.008566 | 0.012392 | 0.028075 | 0.070708 | 0.008384 | 0.03442 | 0.024473 |
| 28.61111111 | 0.01467 | 0.003208 | 0.012711 | 0.011824 | 0.008192 | 0.01211 | 0.013427 | 0.017692 | 0.020404 | 0.033731 | 0.005478 | 0.009818 | 0.00891 | 0.016866 | 0.002017 | 0.009241 | 0.023076 | 0.008012 | 0.011633 | 0.024375 | 0.056124 | 0.008529 | 0.035567 | 0.022193 |
| 28.625 | 0.007916 | 0.002975 | 0.008561 | 0.010849 | 0.003937 | 0.00843 | 0.007162 | 0.024591 | 0.011681 | 0.024759 | 0.009517 | 0.009005 | 0.006086 | 0.012898 | 0.002264 | 0.008653 | 0.018396 | 0.007118 | 0.012056 | 0.020741 | 0.043973 | 0.007161 | 0.03337 | 0.018689 |
| 28.63888889 | 0.005135 | 0.002649 | 0.004228 | 0.015227 | 0.002417 | 0.00542 | 0.005639 | 0.018808 | 0.014885 | 0.012782 | 0.013587 | 0.008866 | 0.003934 | 0.01089 | 0.001835 | 0.007707 | 0.013774 | 0.006279 | 0.012543 | 0.018226 | 0.035026 | 0.005062 | 0.029752 | 0.014567 |
| 28.65277778 | 0.005762 | 0.00199 | 0.001643 | 0.01998 | 0.002703 | 0.005316 | 0.011961 | 0.010298 | 0.019041 | 0.015025 | 0.011255 | 0.007776 | 0.003361 | 0.010087 | 0.000942 | 0.006559 | 0.010286 | 0.00601 | 0.01262 | 0.01659 | 0.029377 | 0.003128 | 0.025405 | 0.011735 |
| 28.66666667 | 0.008953 | 0.001417 | 0.001387 | 0.022207 | 0.002924 | 0.007459 | 0.019276 | 0.016722 | 0.013705 | 0.021039 | 0.007578 | 0.005048 | 0.004716 | 0.009373 | 0.000375 | 0.005981 | 0.008147 | 0.006203 | 0.012122 | 0.015446 | 0.026518 | 0.001607 | 0.021901 | 0.010731 |
| 28.68055556 | 0.011804 | 0.001341 | 0.003439 | 0.020008 | 0.003515 | 0.009914 | 0.020365 | 0.039058 | 0.008012 | 0.017091 | 0.011985 | 0.003167 | 0.007103 | 0.01128 | 0.000576 | 0.005865 | 0.007409 | 0.006292 | 0.010434 | 0.014505 | 0.025504 | 0.000962 | 0.019598 | 0.010729 |
| 28.69444444 | 0.010251 | 0.001487 | 0.007981 | 0.015463 | 0.003677 | 0.011856 | 0.016933 | 0.062642 | 0.010584 | 0.009907 | 0.015184 | 0.00536 | 0.008919 | 0.019755 | 0.001668 | 0.005582 | 0.007686 | 0.005944 | 0.008382 | 0.013677 | 0.024261 | 0.001389 | 0.018029 | 0.010816 |
| 28.70833333 | 0.006648 | 0.001215 | 0.009581 | 0.010549 | 0.002468 | 0.013062 | 0.010684 | 0.074289 | 0.014367 | 0.011522 | 0.009984 | 0.009992 | 0.009184 | 0.028369 | 0.003312 | 0.005002 | 0.007691 | 0.005147 | 0.006757 | 0.013077 | 0.02217 | 0.002028 | 0.01649 | 0.010212 |
| 28.72222222 | 0.009052 | 0.001052 | 0.008438 | 0.006949 | 0.001795 | 0.012736 | 0.007942 | 0.068231 | 0.012017 | 0.020332 | 0.003409 | 0.013082 | 0.007936 | 0.025731 | 0.004505 | 0.004148 | 0.007006 | 0.003882 | 0.005098 | 0.012674 | 0.020172 | 0.002415 | 0.014791 | 0.008919 |
| 28.73611111 | 0.013457 | 0.001427 | 0.012145 | 0.007635 | 0.003318 | 0.010557 | 0.014507 | 0.041822 | 0.006106 | 0.029196 | 0.000881 | 0.012166 | 0.00563 | 0.014653 | 0.00433 | 0.002984 | 0.006261 | 0.002272 | 0.003292 | 0.012207 | 0.018511 | 0.0027 | 0.013507 | 0.007601 |
| 28.75 | 0.010351 | 0.001238 | 0.016169 | 0.008417 | 0.005364 | 0.007633 | 0.021529 | 0.02167 | 0.017757 | 0.036234 | 0.00124 | 0.007558 | 0.002842 | 0.006206 | 0.002975 | 0.001745 | 0.005948 | 0.001005 | 0.002152 | 0.011258 | 0.017325 | 0.002759 | 0.012423 | 0.006672 |
| 28.76388889 | 0.007478 | 0.000578 | 0.017801 | 0.010243 | 0.007907 | 0.004558 | 0.023068 | 0.035953 | 0.033384 | 0.039443 | 0.002629 | 0.003082 | 0.001208 | 0.003008 | 0.002645 | 0.000889 | 0.006078 | 0.000647 | 0.002016 | 0.009668 | 0.01628 | 0.002132 | 0.011089 | 0.006137 |
| 28.77777778 | 0.013221 | 0.000446 | 0.018389 | 0.018391 | 0.011701 | 0.003346 | 0.021479 | 0.058147 | 0.010745 | 0.036084 | 0.005547 | 0.001736 | 0.001683 | 0.001973 | 0.004174 | 0.000922 | 0.006116 | 0.000656 | 0.002251 | 0.007847 | 0.014625 | 0.001083 | 0.009857 | 0.005883 |
| 28.79166667 | 0.01821 | 0.000818 | 0.014861 | 0.023034 | 0.013855 | 0.006752 | 0.016818 | 0.060339 | 0.016942 | 0.02793 | 0.006262 | 0.002289 | 0.002638 | 0.00169 | 0.004943 | 0.001786 | 0.00563 | 0.00071 | 0.001978 | 0.006139 | 0.012192 | 0.000417 | 0.008996 | 0.005732 |
| 28.80555556 | 0.013946 | 0.001573 | 0.010643 | 0.015887 | 0.011254 | 0.011253 | 0.009465 | 0.049974 | 0.015107 | 0.021949 | 0.00528 | 0.002221 | 0.002498 | 0.001807 | 0.003858 | 0.002288 | 0.004777 | 0.000849 | 0.001144 | 0.004708 | 0.009279 | 0.000455 | 0.008342 | 0.005283 |
| 28.81944444 | 0.009654 | 0.002633 | 0.009372 | 0.008895 | 0.006649 | 0.012373 | 0.003502 | 0.039761 | 0.008594 | 0.023353 | 0.009168 | 0.001657 | 0.001513 | 0.001471 | 0.00242 | 0.00197 | 0.00396 | 0.000979 | 0.000418 | 0.003683 | 0.006268 | 0.001302 | 0.007856 | 0.004486 |
| 28.83333333 | 0.017897 | 0.003774 | 0.008411 | 0.012484 | 0.006352 | 0.011297 | 0.002444 | 0.033287 | 0.006811 | 0.033264 | 0.01432 | 0.002761 | 0.001104 | 0.001446 | 0.001393 | 0.002894 | 0.003532 | 0.002134 | 0.00041 | 0.002974 | 0.003767 | 0.002264 | 0.007529 | 0.004003 |
| 28.84722222 | 0.032121 | 0.005125 | 0.006652 | 0.016889 | 0.011941 | 0.010697 | 0.005386 | 0.03051 | 0.010707 | 0.049322 | 0.014441 | 0.005946 | 0.001521 | 0.003095 | 0.000824 | 0.006424 | 0.003545 | 0.004665 | 0.001071 | 0.002712 | 0.002486 | 0.002471 | 0.0071 | 0.004118 |
| 28.86111111 | 0.04047 | 0.007269 | 0.006916 | 0.014056 | 0.018352 | 0.010901 | 0.009937 | 0.031799 | 0.011651 | 0.062342 | 0.010693 | 0.00841 | 0.001478 | 0.005287 | 0.000968 | 0.011217 | 0.003731 | 0.007318 | 0.002052 | 0.002898 | 0.002416 | 0.002002 | 0.006056 | 0.004473 |
| 28.875 | 0.040122 | 0.009812 | 0.008737 | 0.010253 | 0.020003 | 0.010699 | 0.013767 | 0.034275 | 0.028954 | 0.062937 | 0.007199 | 0.007687 | 0.000819 | 0.006625 | 0.001687 | 0.015688 | 0.003792 | 0.008908 | 0.002669 | 0.002973 | 0.003252 | 0.001221 | 0.004334 | 0.004652 |
| 28.88888889 | 0.032736 | 0.012383 | 0.008242 | 0.012765 | 0.017466 | 0.009926 | 0.015313 | 0.034024 | 0.010212 | 0.050621 | 0.005178 | 0.00432 | 0.000538 | 0.006873 | 0.002237 | 0.017953 | 0.003756 | 0.008977 | 0.002413 | 0.002798 | 0.005051 | 0.000518 | 0.002865 | 0.004482 |
| 28.90277778 | 0.02572 | 0.013656 | 0.004977 | 0.019389 | 0.013496 | 0.009103 | 0.014638 | 0.029881 | 0.010368 | 0.03039 | 0.003643 | 0.001885 | 0.000885 | 0.006724 | 0.002174 | 0.015962 | 0.003821 | 0.007765 | 0.00161 | 0.002799 | 0.00754 | 0.000193 | 0.002608 | 0.004151 |
| 28.91666667 | 0.02442 | 0.009997 | 0.001818 | 0.024559 | 0.008652 | 0.007515 | 0.012791 | 0.022965 | 0.008737 | 0.013971 | 0.00508 | 0.002774 | 0.001512 | 0.007863 | 0.001725 | 0.010531 | 0.003918 | 0.00606 | 0.000857 | 0.002828 | 0.00982 | 0.000157 | 0.003353 | 0.003734 |
| 28.93055556 | 0.02495 | 0.004939 | 0.000728 | 0.025606 | 0.00422 | 0.004725 | 0.010574 | 0.014623 | 0.017104 | 0.012729 | 0.010413 | 0.005111 | 0.00253 | 0.010118 | 0.001245 | 0.004743 | 0.003807 | 0.005256 | 0.000568 | 0.00234 | 0.01083 | 0.000342 | 0.004238 | 0.003073 |
| 28.94444444 | 0.022238 | 0.004881 | 0.001029 | 0.022036 | 0.002708 | 0.001968 | 0.008455 | 0.006866 | 0.031094 | 0.020943 | 0.016184 | 0.007342 | 0.003579 | 0.010809 | 0.000831 | 0.001486 | 0.003423 | 0.005949 | 0.000728 | 0.001465 | 0.010661 | 0.000671 | 0.004743 | 0.002318 |
| 28.95833333 | 0.01558 | 0.007273 | 0.001148 | 0.016985 | 0.004394 | 0.000655 | 0.006781 | 0.002478 | 0.038384 | 0.024127 | 0.018368 | 0.00944 | 0.003689 | 0.009243 | 0.000466 | 0.000814 | 0.002937 | 0.00743 | 0.000971 | 0.000729 | 0.009845 | 0.000889 | 0.004677 | 0.001812 |
| 28.97222222 | 0.009314 | 0.008066 | 0.000714 | 0.016067 | 0.006696 | 0.000685 | 0.007183 | 0.001188 | 0.035372 | 0.018209 | 0.015404 | 0.010259 | 0.003006 | 0.007507 | 0.000207 | 0.001169 | 0.002502 | 0.008825 | 0.001196 | 0.00053 | 0.008114 | 0.001051 | 0.004049 | 0.001582 |
| 28.98611111 | 0.012289 | 0.006289 | 0.000691 | 0.022264 | 0.006901 | 0.002602 | 0.011233 | 0.002113 | 0.021924 | 0.022302 | 0.008847 | 0.007984 | 0.0026 | 0.006786 | 0.000369 | 0.002506 | 0.002238 | 0.009383 | 0.001202 | 0.000943 | 0.005733 | 0.001284 | 0.003231 | 0.001594 |
| 29 | 0.025283 | 0.004842 | 0.003042 | 0.030334 | 0.004242 | 0.0077 | 0.017656 | 0.007527 | 0.013652 | 0.056366 | 0.005358 | 0.00894 | 0.002085 | 0.007752 | 0.001756 | 0.003368 | 0.001703 | 0.009159 | 0.001023 | 0.001953 | 0.003311 | 0.002613 | 0.002571 | 0.001445 |
| 29.01388889 | 0.036646 | 0.005774 | 0.008451 | 0.037649 | 0.002651 | 0.012883 | 0.02315 | 0.014796 | 0.026881 | 0.105138 | 0.011602 | 0.021385 | 0.005257 | 0.012334 | 0.004408 | 0.00248 | 0.002881 | 0.01029 | 0.001949 | 0.005929 | 0.005732 | 0.007289 | 0.006217 | 0.00133 |
| 29.02777778 | 0.041112 | 0.005887 | 0.013101 | 0.043228 | 0.004322 | 0.013989 | 0.023808 | 0.021048 | 0.043775 | 0.132212 | 0.021574 | 0.039085 | 0.009124 | 0.020276 | 0.006195 | 0.00115 | 0.012037 | 0.014052 | 0.002792 | 0.016156 | 0.02019 | 0.015453 | 0.018493 | 0.002296 |
| 29.04166667 | 0.043589 | 0.006957 | 0.012659 | 0.03806 | 0.00451 | 0.014089 | 0.019599 | 0.028696 | 0.046474 | 0.109731 | 0.024692 | 0.045899 | 0.011347 | 0.024408 | 0.005884 | 0.000711 | 0.024196 | 0.015534 | 0.0038 | 0.023316 | 0.034982 | 0.022218 | 0.027962 | 0.005641 |
| 29.05555556 | 0.042895 | 0.00955 | 0.007678 | 0.022321 | 0.002682 | 0.016133 | 0.012999 | 0.033796 | 0.037571 | 0.062149 | 0.020631 | 0.031119 | 0.010872 | 0.01791 | 0.004627 | 0.001526 | 0.021632 | 0.01059 | 0.006143 | 0.016909 | 0.028776 | 0.022008 | 0.021166 | 0.009339 |
| 29.06944444 | 0.035078 | 0.007981 | 0.004689 | 0.010785 | 0.003249 | 0.014917 | 0.006226 | 0.027019 | 0.022355 | 0.05212 | 0.016179 | 0.015952 | 0.007083 | 0.009134 | 0.003223 | 0.002948 | 0.009813 | 0.005853 | 0.005646 | 0.008341 | 0.013668 | 0.015811 | 0.010192 | 0.007609 |
| 29.08333333 | 0.022544 | 0.003377 | 0.008248 | 0.012127 | 0.00619 | 0.008523 | 0.003284 | 0.016061 | 0.018308 | 0.079913 | 0.01414 | 0.024371 | 0.003922 | 0.008896 | 0.001917 | 0.006045 | 0.006947 | 0.006442 | 0.003744 | 0.006719 | 0.011087 | 0.009072 | 0.009826 | 0.003541 |
| 29.09722222 | 0.011046 | 0.001808 | 0.012234 | 0.0171 | 0.008211 | 0.00369 | 0.004422 | 0.015914 | 0.032226 | 0.097292 | 0.012215 | 0.038439 | 0.006152 | 0.01249 | 0.00094 | 0.010522 | 0.008233 | 0.008028 | 0.0036 | 0.006248 | 0.012567 | 0.004741 | 0.01236 | 0.003358 |
| 29.11111111 | 0.007053 | 0.004568 | 0.012042 | 0.017084 | 0.007609 | 0.004683 | 0.005756 | 0.02353 | 0.040794 | 0.093236 | 0.008933 | 0.040015 | 0.009207 | 0.013865 | 0.00045 | 0.01158 | 0.007502 | 0.008535 | 0.002778 | 0.004786 | 0.010488 | 0.00222 | 0.009936 | 0.004296 |
| 29.125 | 0.011303 | 0.008624 | 0.011865 | 0.015699 | 0.00518 | 0.007784 | 0.006314 | 0.030365 | 0.029302 | 0.082295 | 0.00521 | 0.033148 | 0.008509 | 0.013604 | 0.000506 | 0.008637 | 0.007241 | 0.008711 | 0.001281 | 0.003726 | 0.008493 | 0.001241 | 0.006522 | 0.004239 |
| 29.13888889 | 0.017749 | 0.011236 | 0.014621 | 0.015586 | 0.002653 | 0.009304 | 0.00673 | 0.030307 | 0.012387 | 0.067828 | 0.002391 | 0.02302 | 0.00534 | 0.012406 | 0.000954 | 0.005291 | 0.007066 | 0.007728 | 0.000583 | 0.003297 | 0.007656 | 0.001941 | 0.005295 | 0.004199 |
| 29.15277778 | 0.021043 | 0.012495 | 0.018307 | 0.01575 | 0.001383 | 0.009182 | 0.006776 | 0.02287 | 0.004823 | 0.048333 | 0.001412 | 0.012803 | 0.002384 | 0.010656 | 0.001393 | 0.002788 | 0.006515 | 0.005275 | 0.000494 | 0.003411 | 0.008113 | 0.002943 | 0.004501 | 0.004142 |
| 29.16666667 | 0.021492 | 0.012832 | 0.019481 | 0.016142 | 0.00134 | 0.008484 | 0.006666 | 0.013655 | 0.003699 | 0.026271 | 0.002609 | 0.005374 | 0.001776 | 0.008559 | 0.001433 | 0.001223 | 0.005811 | 0.00255 | 0.000398 | 0.003848 | 0.009255 | 0.003509 | 0.00279 | 0.003901 |
| 29.18055556 | 0.020501 | 0.01124 | 0.016976 | 0.017162 | 0.001793 | 0.008553 | 0.006995 | 0.007006 | 0.006762 | 0.011225 | 0.004757 | 0.002767 | 0.003137 | 0.006204 | 0.001215 | 0.000916 | 0.005162 | 0.001258 | 0.000261 | 0.004516 | 0.010748 | 0.003683 | 0.002111 | 0.003597 |
| 29.19444444 | 0.016502 | 0.008043 | 0.013867 | 0.018915 | 0.002285 | 0.009458 | 0.00778 | 0.00327 | 0.012555 | 0.01132 | 0.007152 | 0.003218 | 0.004334 | 0.00424 | 0.001025 | 0.001636 | 0.004701 | 0.001896 | 0.000288 | 0.00525 | 0.012421 | 0.003178 | 0.003877 | 0.00342 |
| 29.20833333 | 0.010889 | 0.004568 | 0.012528 | 0.020573 | 0.002216 | 0.010353 | 0.008279 | 0.001333 | 0.01716 | 0.017794 | 0.00976 | 0.002727 | 0.004608 | 0.003229 | 0.000851 | 0.002431 | 0.004522 | 0.00323 | 0.000572 | 0.005866 | 0.01359 | 0.002146 | 0.006495 | 0.003251 |
| 29.22222222 | 0.005688 | 0.002034 | 0.011456 | 0.02051 | 0.001401 | 0.010517 | 0.007649 | 0.002568 | 0.021601 | 0.018858 | 0.011746 | 0.001677 | 0.004519 | 0.002982 | 0.000669 | 0.002887 | 0.00451 | 0.003757 | 0.001115 | 0.006524 | 0.013716 | 0.001286 | 0.00761 | 0.003079 |
| 29.23611111 | 0.002759 | 0.001489 | 0.009874 | 0.017759 | 0.000608 | 0.010028 | 0.005697 | 0.006594 | 0.025853 | 0.013221 | 0.012015 | 0.002513 | 0.004202 | 0.00298 | 0.000527 | 0.00309 | 0.004601 | 0.003475 | 0.00163 | 0.007162 | 0.013168 | 0.000849 | 0.00645 | 0.002959 |
| 29.25 | 0.004508 | 0.002079 | 0.00842 | 0.012826 | 0.000308 | 0.009362 | 0.003002 | 0.009616 | 0.026933 | 0.006323 | 0.010484 | 0.004132 | 0.003553 | 0.00298 | 0.00041 | 0.002877 | 0.004879 | 0.003339 | 0.001841 | 0.007514 | 0.012905 | 0.000552 | 0.00405 | 0.002841 |
| 29.26388889 | 0.008411 | 0.001985 | 0.00688 | 0.007055 | 0.000775 | 0.008152 | 0.001487 | 0.008434 | 0.023708 | 0.004244 | 0.008388 | 0.004828 | 0.002683 | 0.002718 | 0.000406 | 0.001981 | 0.00504 | 0.003493 | 0.001895 | 0.008146 | 0.014388 | 0.000599 | 0.002107 | 0.002909 |
| 29.27777778 | 0.010704 | 0.001835 | 0.005035 | 0.00242 | 0.002441 | 0.005873 | 0.002155 | 0.005841 | 0.016741 | 0.008433 | 0.006569 | 0.004568 | 0.001627 | 0.001997 | 0.000454 | 0.000965 | 0.004185 | 0.00286 | 0.001746 | 0.009434 | 0.01805 | 0.001272 | 0.0014 | 0.00289 |
| 29.29166667 | 0.010783 | 0.002415 | 0.003734 | 0.001365 | 0.004132 | 0.003291 | 0.002333 | 0.006055 | 0.008417 | 0.014881 | 0.00535 | 0.004653 | 0.000711 | 0.001915 | 0.000375 | 0.000802 | 0.002262 | 0.001689 | 0.001172 | 0.010531 | 0.021102 | 0.002267 | 0.001583 | 0.002224 |
| 29.30555556 | 0.009941 | 0.002578 | 0.003274 | 0.004823 | 0.003948 | 0.00151 | 0.001942 | 0.00606 | 0.004187 | 0.017042 | 0.005279 | 0.006756 | 0.000549 | 0.002471 | 0.000284 | 0.001039 | 0.000728 | 0.00132 | 0.000823 | 0.010725 | 0.020461 | 0.002907 | 0.002485 | 0.001466 |
| 29.31944444 | 0.00923 | 0.002715 | 0.002905 | 0.011584 | 0.002138 | 0.000592 | 0.00373 | 0.006877 | 0.007044 | 0.011856 | 0.006452 | 0.010405 | 0.001153 | 0.002156 | 0.000217 | 0.000945 | 0.00053 | 0.001167 | 0.001065 | 0.010134 | 0.017174 | 0.002643 | 0.003749 | 0.001388 |
| 29.33333333 | 0.007721 | 0.003747 | 0.002894 | 0.017822 | 0.000621 | 0.000569 | 0.006142 | 0.014184 | 0.014375 | 0.007832 | 0.007973 | 0.013024 | 0.002186 | 0.00253 | 0.000149 | 0.000596 | 0.001085 | 0.001113 | 0.001021 | 0.009327 | 0.014349 | 0.001725 | 0.004306 | 0.001577 |
| 29.34722222 | 0.004593 | 0.004522 | 0.00399 | 0.019012 | 0.000198 | 0.002087 | 0.006958 | 0.021625 | 0.021241 | 0.0128 | 0.008696 | 0.012914 | 0.003503 | 0.005205 | 0.000177 | 0.00025 | 0.001666 | 0.001814 | 0.000654 | 0.008966 | 0.012924 | 0.000856 | 0.003826 | 0.001533 |
| 29.36111111 | 0.003692 | 0.003349 | 0.005746 | 0.015432 | 0.000558 | 0.00451 | 0.006455 | 0.02113 | 0.023396 | 0.017742 | 0.00845 | 0.010343 | 0.004819 | 0.008162 | 0.000216 | 0.000172 | 0.001868 | 0.002304 | 0.000713 | 0.009179 | 0.012274 | 0.000515 | 0.002778 | 0.001452 |
| 29.375 | 0.007461 | 0.002027 | 0.006966 | 0.011075 | 0.001311 | 0.005739 | 0.005211 | 0.015027 | 0.020065 | 0.014655 | 0.007889 | 0.00614 | 0.005804 | 0.00912 | 0.000173 | 0.000629 | 0.001449 | 0.002073 | 0.000803 | 0.009228 | 0.011873 | 0.000629 | 0.001588 | 0.001332 |
| 29.38888889 | 0.011407 | 0.002794 | 0.00707 | 0.008783 | 0.001554 | 0.005194 | 0.00417 | 0.009033 | 0.014552 | 0.007741 | 0.007432 | 0.002648 | 0.006465 | 0.007439 | 0.00016 | 0.001832 | 0.000796 | 0.001851 | 0.000447 | 0.008608 | 0.011567 | 0.000704 | 0.000802 | 0.000993 |
| 29.40277778 | 0.012875 | 0.00394 | 0.006517 | 0.010118 | 0.001161 | 0.004294 | 0.004623 | 0.006769 | 0.010127 | 0.003825 | 0.007228 | 0.00132 | 0.006748 | 0.004233 | 0.000333 | 0.003333 | 0.00039 | 0.002007 | 0.000109 | 0.007753 | 0.011248 | 0.00077 | 0.000678 | 0.000657 |
| 29.41666667 | 0.01396 | 0.004215 | 0.005292 | 0.014279 | 0.00125 | 0.004316 | 0.007288 | 0.009388 | 0.007335 | 0.003742 | 0.007534 | 0.001535 | 0.006288 | 0.001606 | 0.000586 | 0.003997 | 0.000303 | 0.002187 | 3.1909724 | 0.006962 | 0.010466 | 0.001022 | 0.00082 | 0.00046 |
| 29.43055556 | 0.016216 | 0.003863 | 0.003522 | 0.018491 | 0.002572 | 0.005813 | 0.011761 | 0.015338 | 0.005766 | 0.003233 | 0.007985 | 0.003705 | 0.005018 | 0.000511 | 0.00059 | 0.003302 | 0.000457 | 0.002134 | 1.5979167 | 0.006138 | 0.008856 | 0.00099 | 0.000767 | 0.000337 |
| 29.44444444 | 0.018715 | 0.002818 | 0.002007 | 0.020126 | 0.004862 | 0.0083 | 0.016296 | 0.021549 | 0.006155 | 0.002237 | 0.007658 | 0.006057 | 0.003649 | 0.000322 | 0.00041 | 0.002344 | 0.000502 | 0.002087 | 0.000236 | 0.00523 | 0.006967 | 0.000754 | 0.000459 | 0.000383 |

| | | | | | | | | | | | | | | | | | | | | | | | | |
|---|---|---|---|---|---|---|---|---|---|---|---|---|---|---|---|---|---|---|---|---|---|---|---|---|
| 29.45833333 | 0.020133 | 0.001737 | 0.001331 | 0.017895 | 0.007077 | 0.011219 | 0.019178 | 0.026196 | 0.009306 | 0.003708 | 0.006745 | 0.005074 | 0.00294 | 0.000521 | 0.000452 | 0.002761 | 0.000518 | 0.002286 | 0.001095 | 0.004323 | 0.005921 | 0.001185 | 0.000213 | 0.000692 |
| 29.47222222 | 0.021741 | 0.00262 | 0.002198 | 0.016522 | 0.006893 | 0.016147 | 0.021221 | 0.031358 | 0.013807 | 0.008495 | 0.007043 | 0.002855 | 0.002944 | 0.001346 | 0.000512 | 0.004673 | 0.000737 | 0.00264 | 0.002239 | 0.003876 | 0.005878 | 0.001788 | 0.000178 | 0.001039 |
| 29.48611111 | 0.02658 | 0.006898 | 0.004761 | 0.024871 | 0.006169 | 0.026705 | 0.023794 | 0.041074 | 0.02114 | 0.020073 | 0.010252 | 0.002434 | 0.003707 | 0.003363 | 0.000279 | 0.006403 | 0.000864 | 0.002864 | 0.002717 | 0.004719 | 0.00648 | 0.001904 | 0.000594 | 0.00091 |
| 29.5 | 0.03679 | 0.011146 | 0.008604 | 0.040647 | 0.010686 | 0.037886 | 0.02369 | 0.056003 | 0.038026 | 0.043433 | 0.01511 | 0.003087 | 0.006131 | 0.007425 | 0.000326 | 0.00678 | 0.001392 | 0.002938 | 0.002479 | 0.007807 | 0.008455 | 0.003459 | 0.002477 | 0.000799 |
| 29.51388889 | 0.048521 | 0.010474 | 0.012866 | 0.047223 | 0.018778 | 0.03633 | 0.01636 | 0.071961 | 0.062103 | 0.069907 | 0.020204 | 0.005997 | 0.010079 | 0.014341 | 0.001194 | 0.006976 | 0.002638 | 0.002782 | 0.002557 | 0.012202 | 0.011848 | 0.006336 | 0.005935 | 0.001333 |
| 29.52777778 | 0.056244 | 0.007284 | 0.017124 | 0.035969 | 0.024232 | 0.023955 | 0.010532 | 0.084977 | 0.090369 | 0.084544 | 0.026011 | 0.009738 | 0.012652 | 0.022662 | 0.00236 | 0.008232 | 0.002828 | 0.002181 | 0.003456 | 0.014162 | 0.013526 | 0.00585 | 0.008619 | 0.002515 |
| 29.54166667 | 0.063586 | 0.006633 | 0.024081 | 0.021026 | 0.024635 | 0.015708 | 0.01271 | 0.092148 | 0.11147 | 0.081195 | 0.029154 | 0.010162 | 0.011688 | 0.028388 | 0.002612 | 0.009179 | 0.001391 | 0.002093 | 0.004498 | 0.01218 | 0.01173 | 0.003303 | 0.008979 | 0.00491 |
| 29.55555556 | 0.069058 | 0.007683 | 0.030016 | 0.018892 | 0.021598 | 0.016395 | 0.011298 | 0.084904 | 0.093682 | 0.060781 | 0.027083 | 0.008129 | 0.008885 | 0.029401 | 0.002102 | 0.008919 | 0.001073 | 0.002407 | 0.005093 | 0.009942 | 0.009385 | 0.003707 | 0.009086 | 0.007446 |
| 29.56944444 | 0.06401 | 0.007596 | 0.026541 | 0.024722 | 0.01735 | 0.019917 | 0.008213 | 0.063285 | 0.051399 | 0.037251 | 0.023077 | 0.006408 | 0.006464 | 0.026862 | 0.00204 | 0.007766 | 0.002902 | 0.001655 | 0.004992 | 0.010185 | 0.008645 | 0.004554 | 0.010049 | 0.008183 |
| 29.58333333 | 0.04965 | 0.006502 | 0.017678 | 0.027899 | 0.011538 | 0.020808 | 0.012581 | 0.039627 | 0.044638 | 0.021524 | 0.018679 | 0.004365 | 0.003912 | 0.021184 | 0.001749 | 0.004898 | 0.004468 | 0.000841 | 0.003723 | 0.011193 | 0.008832 | 0.003129 | 0.009614 | 0.007234 |
| 29.59722222 | 0.031181 | 0.004996 | 0.009323 | 0.029989 | 0.006786 | 0.017556 | 0.017349 | 0.020558 | 0.069512 | 0.013215 | 0.013908 | 0.004806 | 0.001995 | 0.011908 | 0.001528 | 0.00342 | 0.004264 | 0.000697 | 0.001755 | 0.01113 | 0.009268 | 0.001932 | 0.006593 | 0.005543 |
| 29.61111111 | 0.013777 | 0.002871 | 0.004326 | 0.03261 | 0.008832 | 0.011426 | 0.017728 | 0.016238 | 0.068179 | 0.011044 | 0.011337 | 0.011927 | 0.002659 | 0.005144 | 0.003254 | 0.007244 | 0.002687 | 0.000979 | 0.000746 | 0.009462 | 0.009732 | 0.002141 | 0.003263 | 0.003629 |
| 29.625 | 0.006389 | 0.00105 | 0.004377 | 0.032453 | 0.014606 | 0.005735 | 0.015281 | 0.032066 | 0.036384 | 0.011548 | 0.011756 | 0.01895 | 0.004152 | 0.006414 | 0.005433 | 0.013066 | 0.000928 | 0.002492 | 0.001141 | 0.006771 | 0.010052 | 0.001881 | 0.002308 | 0.001916 |
| 29.63888889 | 0.010573 | 0.000578 | 0.003756 | 0.02946 | 0.015516 | 0.002449 | 0.010117 | 0.052385 | 0.016729 | 0.01972 | 0.01206 | 0.018942 | 0.004392 | 0.010331 | 0.005633 | 0.016768 | 0.000309 | 0.004169 | 0.001591 | 0.004888 | 0.010484 | 0.001969 | 0.002753 | 0.000827 |
| 29.65277778 | 0.015508 | 0.001205 | 0.002047 | 0.025254 | 0.011328 | 0.001495 | 0.005431 | 0.065714 | 0.02393 | 0.034453 | 0.009549 | 0.013284 | 0.003761 | 0.012383 | 0.004246 | 0.018627 | 0.000819 | 0.004421 | 0.001445 | 0.004235 | 0.011478 | 0.003657 | 0.002373 | 0.000344 |
| 29.66666667 | 0.015513 | 0.002556 | 0.003147 | 0.020861 | 0.006949 | 0.00244 | 0.004948 | 0.0599 | 0.034705 | 0.039262 | 0.005271 | 0.007573 | 0.002989 | 0.012919 | 0.002842 | 0.019916 | 0.001476 | 0.003698 | 0.001056 | 0.004216 | 0.01223 | 0.004586 | 0.00163 | 0.000184 |
| 29.68055556 | 0.013123 | 0.003887 | 0.005193 | 0.017246 | 0.004152 | 0.003113 | 0.005501 | 0.037295 | 0.030364 | 0.02838 | 0.003267 | 0.008087 | 0.00213 | 0.011735 | 0.00187 | 0.019389 | 0.001881 | 0.00255 | 0.00061 | 0.004935 | 0.011935 | 0.003434 | 0.002352 | 0.000562 |
| 29.69444444 | 0.008356 | 0.004353 | 0.006765 | 0.013538 | 0.003147 | 0.002613 | 0.003636 | 0.042842 | 0.019313 | 0.016283 | 0.004626 | 0.012415 | 0.001254 | 0.009404 | 0.000982 | 0.015474 | 0.002412 | 0.001462 | 0.000295 | 0.00595 | 0.011194 | 0.001599 | 0.004427 | 0.001798 |
| 29.70833333 | 0.004308 | 0.005568 | 0.00812 | 0.009119 | 0.00425 | 0.002746 | 0.001818 | 0.085962 | 0.020656 | 0.018394 | 0.005233 | 0.012066 | 0.001077 | 0.007819 | 0.000345 | 0.010532 | 0.002658 | 0.001165 | 0.000724 | 0.005359 | 0.00933 | 0.000816 | 0.004957 | 0.002945 |
| 29.72222222 | 0.00557 | 0.008355 | 0.009024 | 0.005867 | 0.005461 | 0.004541 | 0.002659 | 0.12156 | 0.031088 | 0.027216 | 0.003908 | 0.008277 | 0.00151 | 0.008211 | 0.000188 | 0.007568 | 0.001817 | 0.00105 | 0.002209 | 0.003507 | 0.005312 | 0.001641 | 0.003682 | 0.002826 |
| 29.73611111 | 0.007193 | 0.009055 | 0.009168 | 0.005391 | 0.004747 | 0.008743 | 0.004651 | 0.129989 | 0.034733 | 0.028713 | 0.005541 | 0.011217 | 0.001458 | 0.009575 | 0.000309 | 0.005635 | 0.000742 | 0.000551 | 0.003477 | 0.003472 | 0.002118 | 0.002972 | 0.004421 | 0.002066 |
| 29.75 | 0.004763 | 0.005745 | 0.00788 | 0.006907 | 0.003563 | 0.013596 | 0.00476 | 0.116997 | 0.032954 | 0.025029 | 0.012205 | 0.022149 | 0.00152 | 0.010126 | 0.000697 | 0.002971 | 0.000419 | 0.000296 | 0.003284 | 0.004968 | 0.002161 | 0.00323 | 0.00739 | 0.001863 |
| 29.76388889 | 0.003752 | 0.003505 | 0.006094 | 0.008827 | 0.003085 | 0.015944 | 0.003277 | 0.084598 | 0.030195 | 0.021272 | 0.018806 | 0.031878 | 0.002715 | 0.009135 | 0.001754 | 0.000824 | 0.000544 | 0.000859 | 0.002397 | 0.00578 | 0.002988 | 0.002578 | 0.009154 | 0.002132 |
| 29.77777778 | 0.008664 | 0.005522 | 0.008181 | 0.010451 | 0.002499 | 0.014205 | 0.004025 | 0.043201 | 0.024082 | 0.018116 | 0.020779 | 0.033543 | 0.00364 | 0.007482 | 0.002967 | 0.00035 | 0.000637 | 0.002266 | 0.001841 | 0.005634 | 0.002905 | 0.001996 | 0.008396 | 0.002124 |
| 29.79166667 | 0.014109 | 0.007251 | 0.014009 | 0.010769 | 0.001437 | 0.00854 | 0.007828 | 0.018818 | 0.013438 | 0.017688 | 0.018981 | 0.02702 | 0.003108 | 0.007173 | 0.003117 | 0.000567 | 0.000571 | 0.00387 | 0.001643 | 0.005055 | 0.002461 | 0.001803 | 0.006305 | 0.00142 |
| 29.80555556 | 0.014524 | 0.006142 | 0.01593 | 0.008256 | 0.000552 | 0.003787 | 0.009906 | 0.031056 | 0.004441 | 0.022354 | 0.015537 | 0.018213 | 0.001735 | 0.009311 | 0.001957 | 0.000746 | 0.000417 | 0.005098 | 0.001696 | 0.004481 | 0.001975 | 0.001897 | 0.004455 | 0.000609 |
| 29.81944444 | 0.011078 | 0.004604 | 0.013185 | 0.004605 | 0.000361 | 0.003918 | 0.008747 | 0.063083 | 0.006611 | 0.033565 | 0.011166 | 0.015078 | 0.001095 | 0.012709 | 0.000995 | 0.00085 | 0.00033 | 0.005446 | 0.00192 | 0.00386 | 0.001245 | 0.001859 | 0.003304 | 0.000331 |
| 29.83333333 | 0.011582 | 0.004913 | 0.010829 | 0.004156 | 0.000377 | 0.007831 | 0.008286 | 0.082998 | 0.016242 | 0.052247 | 0.006659 | 0.019149 | 0.002112 | 0.01523 | 0.001284 | 0.000878 | 0.00039 | 0.004252 | 0.002408 | 0.003332 | 0.000485 | 0.001387 | 0.003193 | 0.000334 |
| 29.84722222 | 0.021151 | 0.006316 | 0.009798 | 0.007333 | 0.001346 | 0.013614 | 0.010165 | 0.083794 | 0.018366 | 0.070848 | 0.003086 | 0.024564 | 0.003713 | 0.015712 | 0.001675 | 0.000818 | 0.000427 | 0.002148 | 0.003324 | 0.003853 | 0.001418 | 0.0007 | 0.005117 | 0.000314 |
| 29.86111111 | 0.030111 | 0.005866 | 0.010531 | 0.01092 | 0.004746 | 0.019894 | 0.013625 | 0.076073 | 0.011516 | 0.077885 | 0.001841 | 0.02574 | 0.004375 | 0.01443 | 0.001267 | 0.000934 | 0.000414 | 0.000979 | 0.004128 | 0.006353 | 0.00548 | 0.000435 | 0.008954 | 0.000209 |
| 29.875 | 0.032206 | 0.004313 | 0.01406 | 0.014648 | 0.010012 | 0.025151 | 0.014545 | 0.071154 | 0.008577 | 0.073072 | 0.004873 | 0.022408 | 0.003784 | 0.01192 | 0.000879 | 0.001925 | 0.000465 | 0.000777 | 0.003999 | 0.010849 | 0.012212 | 0.000637 | 0.013016 | 0.000185 |
| 29.88888889 | 0.030193 | 0.004289 | 0.019555 | 0.020913 | 0.014372 | 0.027493 | 0.011789 | 0.069066 | 0.011037 | 0.059778 | 0.012824 | 0.015084 | 0.003262 | 0.008843 | 0.001291 | 0.003617 | 0.000834 | 0.001088 | 0.003446 | 0.016928 | 0.020183 | 0.000881 | 0.016258 | 0.000447 |
| 29.90277778 | 0.024414 | 0.004693 | 0.021314 | 0.029535 | 0.014695 | 0.025584 | 0.013606 | 0.06217 | 0.011011 | 0.039446 | 0.023296 | 0.01097 | 0.003829 | 0.006638 | 0.00164 | 0.005027 | 0.001692 | 0.002787 | 0.003382 | 0.023385 | 0.028603 | 0.001589 | 0.019227 | 0.001291 |
| 29.91666667 | 0.017405 | 0.004299 | 0.013871 | 0.037234 | 0.010997 | 0.019095 | 0.022854 | 0.056181 | 0.009432 | 0.018608 | 0.031142 | 0.019181 | 0.004241 | 0.00547 | 0.001476 | 0.00634 | 0.001878 | 0.004774 | 0.00326 | 0.0274 | 0.035209 | 0.002724 | 0.02174 | 0.002624 |
| 29.93055556 | 0.011933 | 0.004622 | 0.007969 | 0.036004 | 0.006352 | 0.009872 | 0.031244 | 0.051433 | 0.007261 | 0.009272 | 0.027796 | 0.027651 | 0.003403 | 0.003982 | 0.001421 | 0.007976 | 0.0012 | 0.005246 | 0.002644 | 0.027606 | 0.036906 | 0.003225 | 0.022233 | 0.003831 |
| 29.94444444 | 0.007482 | 0.005135 | 0.014305 | 0.026839 | 0.005048 | 0.005063 | 0.02788 | 0.037799 | 0.004599 | 0.014616 | 0.013862 | 0.023841 | 0.002257 | 0.00203 | 0.001415 | 0.00915 | 0.001122 | 0.004676 | 0.001991 | 0.025377 | 0.034266 | 0.002744 | 0.020545 | 0.004772 |
| 29.95833333 | 0.00421 | 0.003817 | 0.023439 | 0.022004 | 0.007973 | 0.009114 | 0.017073 | 0.019735 | 0.006455 | 0.021582 | 0.004322 | 0.015219 | 0.001561 | 0.000833 | 0.00115 | 0.009623 | 0.00167 | 0.004191 | 0.001635 | 0.022229 | 0.029625 | 0.00176 | 0.017291 | 0.005063 |
| 29.97222222 | 0.003343 | 0.002335 | 0.028363 | 0.021568 | 0.010051 | 0.014874 | 0.012897 | 0.009494 | 0.012057 | 0.017576 | 0.004322 | 0.008893 | 0.001312 | 0.000782 | 0.000669 | 0.009301 | 0.001967 | 0.00402 | 0.001772 | 0.019066 | 0.025039 | 0.001586 | 0.012707 | 0.004349 |
| 29.98611111 | 0.005901 | 0.00251 | 0.028747 | 0.019584 | 0.007088 | 0.013881 | 0.011643 | 0.007938 | 0.013688 | 0.015857 | 0.005632 | 0.007199 | 0.001458 | 0.000882 | 0.000459 | 0.007249 | 0.001778 | 0.003756 | 0.002758 | 0.016276 | 0.022128 | 0.002615 | 0.008519 | 0.003566 |
| 30 | 0.014742 | 0.004291 | 0.019514 | 0.016321 | 0.004009 | 0.009406 | 0.009381 | 0.014782 | 0.012036 | 0.041444 | 0.00882 | 0.016179 | 0.001819 | 0.002025 | 0.001699 | 0.004383 | 0.002476 | 0.002969 | 0.004115 | 0.013455 | 0.020155 | 0.002615 | 0.006769 | 0.003777 |
| 30.01388889 | 0.028929 | 0.008657 | 0.010692 | 0.018278 | 0.006489 | 0.012052 | 0.009178 | 0.034937 | 0.018705 | 0.083982 | 0.018741 | 0.033261 | 0.002922 | 0.006529 | 0.004396 | 0.003569 | 0.006175 | 0.002721 | 0.004816 | 0.009647 | 0.01605 | 0.003993 | 0.008432 | 0.004545 |
| 30.02777778 | 0.040528 | 0.01521 | 0.014311 | 0.027758 | 0.011051 | 0.021815 | 0.009845 | 0.052331 | 0.028982 | 0.103929 | 0.027051 | 0.044535 | 0.005974 | 0.013321 | 0.005091 | 0.005115 | 0.014078 | 0.004747 | 0.00472 | 0.005858 | 0.00921 | 0.01145 | 0.013391 | 0.003835 |
| 30.04166667 | 0.047724 | 0.023012 | 0.01773 | 0.03769 | 0.014774 | 0.02964 | 0.013446 | 0.052118 | 0.028412 | 0.083435 | 0.026472 | 0.043194 | 0.011019 | 0.017162 | 0.002588 | 0.005315 | 0.0202 | 0.007234 | 0.005418 | 0.006142 | 0.005172 | 0.017988 | 0.017661 | 0.002082 |
| 30.05555556 | 0.05207 | 0.029175 | 0.012947 | 0.043023 | 0.01635 | 0.030386 | 0.014831 | 0.046708 | 0.020617 | 0.046826 | 0.02158 | 0.030591 | 0.014982 | 0.014444 | 0.000811 | 0.004541 | 0.015485 | 0.006334 | 0.006388 | 0.011041 | 0.007942 | 0.014824 | 0.016382 | 0.002777 |
| 30.06944444 | 0.048983 | 0.026194 | 0.006682 | 0.041158 | 0.013935 | 0.025379 | 0.008889 | 0.045145 | 0.013016 | 0.020946 | 0.015615 | 0.016647 | 0.012493 | 0.00821 | 0.000654 | 0.005557 | 0.006791 | 0.002891 | 0.004758 | 0.014377 | 0.011149 | 0.007131 | 0.010811 | 0.003757 |
| 30.08333333 | 0.035256 | 0.015409 | 0.004753 | 0.032707 | 0.009576 | 0.01962 | 0.00294 | 0.040596 | 0.006788 | 0.008244 | 0.008091 | 0.017097 | 0.007375 | 0.006478 | 0.000549 | 0.004982 | 0.00333 | 0.000847 | 0.002426 | 0.012435 | 0.00857 | 0.003846 | 0.005048 | 0.002409 |
| 30.09722222 | 0.017479 | 0.012112 | 0.007827 | 0.022235 | 0.005389 | 0.014505 | 0.001221 | 0.028426 | 0.005328 | 0.00732 | 0.003226 | 0.030058 | 0.007519 | 0.012189 | 0.001326 | 0.002929 | 0.001815 | 0.001009 | 0.001761 | 0.007999 | 0.004126 | 0.005142 | 0.002205 | 0.001923 |
| 30.11111111 | 0.010073 | 0.01652 | 0.011286 | 0.012458 | 0.003103 | 0.008858 | 0.000829 | 0.014679 | 0.009479 | 0.018889 | 0.004037 | 0.040111 | 0.008003 | 0.017304 | 0.002776 | 0.003682 | 0.000749 | 0.001817 | 0.001782 | 0.004302 | 0.002408 | 0.006979 | 0.00322 | 0.002436 |
| 30.125 | 0.016301 | 0.016737 | 0.013592 | 0.007266 | 0.00462 | 0.004113 | 0.001716 | 0.006846 | 0.013535 | 0.043437 | 0.0061 | 0.043193 | 0.004741 | 0.015714 | 0.002913 | 0.005697 | 0.000538 | 0.001943 | 0.001502 | 0.002005 | 0.002392 | 0.00723 | 0.004584 | 0.002785 |
| 30.13888889 | 0.023156 | 0.01559 | 0.015473 | 0.011568 | 0.007795 | 0.002637 | 0.006653 | 0.005054 | 0.015057 | 0.067185 | 0.006271 | 0.043364 | 0.001844 | 0.009497 | 0.001699 | 0.006223 | 0.000309 | 0.001384 | 0.00078 | 0.000834 | 0.002306 | 0.005376 | 0.003346 | 0.002891 |
| 30.15277778 | 0.022357 | 0.01437 | 0.016231 | 0.020683 | 0.00956 | 0.003523 | 0.013497 | 0.004114 | 0.016629 | 0.067453 | 0.006143 | 0.039885 | 0.001122 | 0.005285 | 0.000628 | 0.00569 | 0.000179 | 0.001691 | 0.000355 | 0.000629 | 0.002987 | 0.002545 | 0.00213 | 0.002183 |
| 30.16666667 | 0.016332 | 0.012626 | 0.016049 | 0.026272 | 0.009814 | 0.004101 | 0.015991 | 0.002996 | 0.019382 | 0.056211 | 0.008706 | 0.030113 | 0.001357 | 0.007793 | 0.000194 | 0.004589 | 0.000231 | 0.003415 | 0.000496 | 0.000605 | 0.005035 | 0.000659 | 0.003613 | 0.001146 |
| 30.18055556 | 0.009588 | 0.013734 | 0.014855 | 0.027901 | 0.009169 | 0.004059 | 0.013021 | 0.005062 | 0.023737 | 0.049389 | 0.014577 | 0.015515 | 0.001847 | 0.011827 | 0.00016 | 0.003166 | 0.000542 | 0.005201 | 0.000783 | 0.000607 | 0.006884 | 0.000313 | 0.005116 | 0.000551 |
| 30.19444444 | 0.00835 | 0.013599 | 0.012829 | 0.025491 | 0.007898 | 0.004136 | 0.012202 | 0.010826 | 0.028154 | 0.037131 | 0.019071 | 0.006944 | 0.00305 | 0.012259 | 0.000443 | 0.002269 | 0.00101 | 0.005873 | 0.000893 | 0.001198 | 0.007424 | 0.000517 | 0.004615 | 0.000422 |
| 30.20833333 | 0.016475 | 0.010536 | 0.013342 | 0.01587 | 0.006483 | 0.004362 | 0.014767 | 0.019441 | 0.029593 | 0.018582 | 0.019437 | 0.008915 | 0.004894 | 0.011531 | 0.000912 | 0.002055 | 0.001524 | 0.005173 | 0.000872 | 0.001807 | 0.007216 | 0.000644 | 0.003244 | 0.000228 |
| 30.22222222 | 0.027824 | 0.008218 | 0.018124 | 0.006359 | 0.005278 | 0.00551 | 0.012909 | 0.030705 | 0.029385 | 0.005247 | 0.02004 | 0.010734 | 0.006292 | 0.011917 | 0.001144 | 0.002209 | 0.001672 | 0.003539 | 0.000904 | 0.001618 | 0.006735 | 0.000936 | 0.001932 | 6.96319500150024e-05 |
| 30.23611111 | 0.033001 | 0.006719 | 0.024454 | 0.005208 | 0.004345 | 0.008787 | 0.007768 | 0.042864 | 0.030295 | 0.002616 | 0.020755 | 0.007483 | 0.005897 | 0.012705 | 0.000831 | 0.002437 | 0.001219 | 0.001958 | 0.000947 | 0.000977 | 0.005655 | 0.001017 | 0.001262 | 0.000188 |
| 30.25 | 0.028482 | 0.004534 | 0.028795 | 0.00597 | 0.003565 | 0.013067 | 0.004765 | 0.049796 | 0.030222 | 0.007103 | 0.018643 | 0.003836 | 0.003638 | 0.012885 | 0.00059 | 0.0023 | 0.000682 | 0.001071 | 0.000943 | 0.000823 | 0.003727 | 0.00136 | 0.001247 | 0.000392 |
| 30.26388889 | 0.018205 | 0.002824 | 0.028279 | 0.003573 | 0.002556 | 0.01515 | 0.003712 | 0.045841 | 0.027272 | 0.013359 | 0.016062 | 0.003152 | 0.002556 | 0.011844 | 0.001112 | 0.001554 | 0.000537 | 0.000758 | 0.001023 | 0.001144 | 0.001664 | 0.002895 | 0.001156 | 0.000457 |
| 30.27777778 | 0.008261 | 0.004421 | 0.022612 | 0.001802 | 0.001395 | 0.013313 | 0.00287 | 0.03457 | 0.023179 | 0.014968 | 0.013861 | 0.002913 | 0.004725 | 0.009116 | 0.001974 | 0.001039 | 0.000682 | 0.000679 | 0.001171 | 0.001224 | 0.000681 | 0.004294 | 0.000776 | 0.000658 |
| 30.29166667 | 0.003502 | 0.009489 | 0.014345 | 0.003176 | 0.000595 | 0.008101 | 0.001835 | 0.022611 | 0.017911 | 0.011476 | 0.010847 | 0.002757 | 0.007351 | 0.004961 | 0.002467 | 0.001922 | 0.000926 | 0.000501 | 0.001255 | 0.000878 | 0.000679 | 0.004272 | 0.000798 | 0.001011 |
| 30.30555556 | 0.003959 | 0.017305 | 0.006738 | 0.005609 | 0.000382 | 0.004521 | 0.002118 | 0.011393 | 0.011064 | 0.013332 | 0.007237 | 0.006186 | 0.007623 | 0.002163 | 0.002185 | 0.003727 | 0.001099 | 0.000453 | 0.001154 | 0.000499 | 0.000749 | 0.003025 | 0.001209 | 0.000872 |
| 30.31944444 | 0.003396 | 0.028156 | 0.003018 | 0.006773 | 0.000581 | 0.006498 | 0.004487 | 0.004542 | 0.005184 | 0.021609 | 0.004855 | 0.012148 | 0.005279 | 0.002531 | 0.001266 | 0.005043 | 0.000894 | 0.001142 | 0.000732 | 0.000434 | 0.000709 | 0.001508 | 0.001297 | 0.000606 |
| 30.33333333 | 0.001922 | 0.035199 | 0.002963 | 0.006756 | 0.000613 | 0.007803 | 0.005914 | 0.003811 | 0.003687 | 0.023545 | 0.005905 | 0.015937 | 0.002817 | 0.003491 | 0.000444 | 0.005317 | 0.000565 | 0.002412 | 0.0003 | 0.000438 | 0.000543 | 0.000776 | 0.000975 | 0.000844 |
| 30.34722222 | 0.004557 | 0.034272 | 0.002487 | 0.005926 | 0.000853 | 0.005729 | 0.004975 | 0.005314 | 0.007984 | 0.015332 | 0.009896 | 0.013391 | 0.003007 | 0.003381 | 0.000264 | 0.004601 | 0.000592 | 0.003695 | 0.00017 | 0.00063 | 0.000358 | 0.000727 | 0.000576 | 0.000973 |
| 30.36111111 | 0.010338 | 0.032056 | 0.001744 | 0.004493 | 0.002029 | 0.006291 | 0.005731 | 0.011411 | 0.013361 | 0.009895 | 0.013999 | 0.007052 | 0.004375 | 0.003357 | 0.000606 | 0.003001 | 0.000665 | 0.004581 | 0.000365 | 0.00165 | 0.000685 | 0.00141 | 0.000369 | 0.000642 |
| 30.375 | 0.0133 | 0.031626 | 0.002927 | 0.003153 | 0.003304 | 0.009784 | 0.009523 | 0.021623 | 0.016407 | 0.015158 | 0.015536 | 0.003547 | 0.003833 | 0.004863 | 0.001054 | 0.001465 | 0.000492 | 0.004871 | 0.001104 | 0.00329 | 0.001763 | 0.002944 | 0.000327 | 0.00044 |
| 30.38888889 | 0.010956 | 0.02892 | 0.004234 | 0.002395 | 0.004108 | 0.011818 | 0.012998 | 0.029212 | 0.021134 | 0.021963 | 0.014835 | 0.002823 | 0.002462 | 0.008177 | 0.001509 | 0.001256 | 0.000313 | 0.004624 | 0.002042 | 0.004838 | 0.002882 | 0.00404 | 0.000297 | 0.00074 |
| 30.40277778 | 0.007404 | 0.020463 | 0.004351 | 0.002462 | 0.004608 | 0.011752 | 0.013436 | 0.032845 | 0.027524 | 0.023385 | 0.014665 | 0.003665 | 0.003671 | 0.011534 | 0.002135 | 0.002471 | 0.000471 | 0.004088 | 0.002644 | 0.005845 | 0.003144 | 0.004068 | 0.000484 | 0.001153 |
| 30.41666667 | 0.005361 | 0.009803 | 0.004164 | 0.003757 | 0.004451 | 0.011288 | 0.009692 | 0.035292 | 0.031239 | 0.017899 | 0.015369 | 0.007676 | 0.006522 | 0.013106 | 0.002848 | 0.003994 | 0.00093 | 0.003638 | 0.002963 | 0.006186 | 0.002818 | 0.004238 | 0.000818 | 0.001173 |
| 30.43055556 | 0.006065 | 0.004729 | 0.004482 | 0.006119 | 0.003312 | 0.01096 | 0.008794 | 0.036796 | 0.030898 | 0.011628 | 0.015161 | 0.010887 | 0.007866 | 0.012876 | 0.003298 | 0.004767 | 0.00143 | 0.003485 | 0.003181 | 0.006014 | 0.002774 | 0.004943 | 0.00089 | 0.00089 |
| 30.44444444 | 0.010212 | 0.006211 | 0.004865 | 0.008317 | 0.001964 | 0.010037 | 0.016463 | 0.036041 | 0.027678 | 0.014149 | 0.013962 | 0.008903 | 0.006795 | 0.011811 | 0.002689 | 0.005436 | 0.002078 | 0.003576 | 0.003569 | 0.00555 | 0.004011 | 0.00456 | 0.000801 | 0.000854 |
| 30.45833333 | 0.015591 | 0.009268 | 0.005293 | 0.008647 | 0.001769 | 0.008723 | 0.02091 | 0.03528 | 0.024795 | 0.015317 | 0.013638 | 0.004996 | 0.004018 | 0.010641 | 0.001676 | 0.007509 | 0.003038 | 0.003857 | 0.004525 | 0.005226 | 0.007377 | 0.003037 | 0.000935 | 0.00148 |
| 30.47222222 | 0.020906 | 0.013951 | 0.007187 | 0.007794 | 0.001778 | 0.010196 | 0.016788 | 0.038938 | 0.026621 | 0.013742 | 0.015508 | 0.004491 | 0.00149 | 0.00931 | 0.001314 | 0.010071 | 0.003954 | 0.004157 | 0.005851 | 0.005553 | 0.010958 | 0.002114 | 0.001279 | 0.002418 |
| 30.48611111 | 0.029123 | 0.024613 | 0.012767 | 0.011021 | 0.002224 | 0.020201 | 0.022669 | 0.05091 | 0.037005 | 0.028153 | 0.020478 | 0.00819 | 0.000663 | 0.008031 | 0.001752 | 0.010626 | 0.004212 | 0.004127 | 0.006423 | 0.006847 | 0.011037 | 0.002036 | 0.001722 | 0.002562 |
| 30.5 | 0.041483 | 0.044206 | 0.022747 | 0.024267 | 0.004928 | 0.041135 | 0.048618 | 0.064264 | 0.053517 | 0.051967 | 0.02612 | 0.017899 | 0.0013 | 0.008063 | 0.003108 | 0.009108 | 0.004242 | 0.00386 | 0.005913 | 0.008771 | 0.007837 | 0.003203 | 0.003033 | 0.001605 |
| 30.51388889 | 0.042888 | 0.070235 | 0.034001 | 0.037222 | 0.006565 | 0.056224 | 0.069807 | 0.068142 | 0.07694 | 0.064949 | 0.026801 | 0.036081 | 0.002942 | 0.010631 | 0.00387 | 0.006929 | 0.005013 | 0.003539 | 0.005032 | 0.009843 | 0.004214 | 0.005375 | 0.006293 | 0.000592 |
| 30.52777778 | 0.038993 | 0.087833 | 0.040006 | 0.031887 | 0.005997 | 0.05392 | 0.067654 | 0.067516 | 0.110715 | 0.062392 | 0.02143 | 0.051802 | 0.005374 | 0.015278 | 0.005281 | 0.004376 | 0.005475 | 0.002728 | 0.003275 | 0.008712 | 0.002085 | 0.00559 | 0.010839 | 0.000427 |
| 30.54166667 | 0.048568 | 0.08678 | 0.037414 | 0.020089 | 0.006504 | 0.048262 | 0.05516 | 0.073501 | 0.13723 | 0.049454 | 0.016487 | 0.049333 | 0.007787 | 0.018657 | 0.008246 | 0.00253 | 0.004563 | 0.002975 | 0.001077 | 0.006497 | 0.001612 | 0.003198 | 0.014639 | 0.001786 |
| 30.55555556 | 0.052962 | 0.074217 | 0.03201 | 0.021602 | 0.007482 | 0.041716 | 0.041202 | 0.083527 | 0.126634 | 0.037198 | 0.018527 | 0.031836 | 0.008821 | 0.017999 | 0.008867 | 0.003218 | 0.004085 | 0.00429 | 0.000896 | 0.005995 | 0.00166 | 0.001058 | 0.016218 | 0.004449 |
| 30.56944444 | 0.045957 | 0.057206 | 0.026179 | 0.029724 | 0.007033 | 0.027555 | 0.024745 | 0.085539 | 0.073872 | 0.027894 | 0.02369 | 0.013946 | 0.007784 | 0.014051 | 0.005958 | 0.005135 | 0.004793 | 0.003341 | 0.002504 | 0.007186 | 0.00215 | 0.000379 | 0.014558 | 0.005902 |
| 30.58333333 | 0.036798 | 0.040228 | 0.015773 | 0.034144 | 0.005918 | 0.01248 | 0.011237 | 0.08054 | 0.033567 | 0.018656 | 0.025061 | 0.006249 | 0.004622 | 0.008143 | 0.003414 | 0.005148 | 0.004728 | 0.001558 | 0.00342 | 0.007814 | 0.002148 | 0.000548 | 0.009691 | 0.004258 |
| 30.59722222 | 0.022838 | 0.026244 | 0.007304 | 0.03461 | 0.00573 | 0.005358 | 0.007515 | 0.074243 | 0.045035 | 0.011269 | 0.02095 | 0.011128 | 0.002202 | 0.004457 | 0.004827 | 0.003212 | 0.003059 | 0.001744 | 0.003101 | 0.007686 | 0.001658 | 0.001079 | 0.004506 | 0.002035 |
| 30.61111111 | 0.016095 | 0.013826 | 0.009415 | 0.032774 | 0.006149 | 0.00442 | 0.01105 | 0.065461 | 0.069131 | 0.005941 | 0.01315 | 0.019555 | 0.003248 | 0.007378 | 0.008746 | 0.003011 | 0.001559 | 0.002437 | 0.002679 | 0.007629 | 0.002147 | 0.001326 | 0.002573 | 0.002014 |
| 30.625 | 0.021442 | 0.005969 | 0.015151 | 0.029152 | 0.006084 | 0.005854 | 0.01563 | 0.053745 | 0.07372 | 0.003227 | 0.009692 | 0.022898 | 0.00532 | 0.013197 | 0.011656 | 0.006254 | 0.001708 | 0.001961 | 0.002432 | 0.007552 | 0.002868 | 0.001221 | 0.003428 | 0.00278 |
| 30.63888889 | 0.019468 | 0.007197 | 0.019234 | 0.024356 | 0.005779 | 0.007263 | 0.022475 | 0.035934 | 0.058852 | 0.002757 | 0.015339 | 0.021902 | 0.00659 | 0.017028 | 0.011847 | 0.010118 | 0.002246 | 0.00106 | 0.001792 | 0.007055 | 0.002882 | 0.001363 | 0.003797 | 0.002715 |
| 30.65277778 | 0.010615 | 0.013249 | 0.025056 | 0.020293 | 0.005566 | 0.004944 | 0.032284 | 0.018607 | 0.034451 | 0.002077 | 0.021974 | 0.018141 | 0.007495 | 0.018126 | 0.010049 | 0.012554 | 0.001772 | 0.001156 | 0.000925 | 0.006306 | 0.00197 | 0.001956 | 0.002405 | 0.001855 |
| 30.66666667 | 0.00967 | 0.016559 | 0.026138 | 0.017196 | 0.005488 | 0.00234 | 0.032207 | 0.01647 | 0.01687 | 0.001365 | 0.020559 | 0.010394 | 0.007919 | 0.017019 | 0.008286 | 0.013417 | 0.000837 | 0.00211 | 0.000497 | 0.006133 | 0.001496 | 0.002643 | 0.001178 | 0.001015 |
| 30.68055556 | 0.015102 | 0.016912 | 0.017019 | 0.014421 | 0.00622 | 0.00322 | 0.025541 | 0.022888 | 0.008811 | 0.002623 | 0.012104 | 0.005674 | 0.007945 | 0.015357 | 0.007166 | 0.01251 | 0.000422 | 0.00279 | 0.000679 | 0.007093 | 0.003954 | 0.002991 | 0.001997 | 0.001423 |
| 30.69444444 | 0.022712 | 0.01722 | 0.012429 | 0.012645 | 0.007851 | 0.006642 | 0.03188 | 0.02563 | 0.006075 | 0.005433 | 0.005548 | 0.01084 | 0.007661 | 0.014614 | 0.006168 | 0.010146 | 0.000577 | 0.002738 | 0.001227 | 0.008956 | 0.009282 | 0.003301 | 0.00384 | 0.002818 |
| 30.70833333 | 0.022501 | 0.016491 | 0.021604 | 0.012796 | 0.008901 | 0.009772 | 0.037191 | 0.029782 | 0.005303 | 0.008548 | 0.004933 | 0.018766 | 0.007082 | 0.015078 | 0.004854 | 0.007609 | 0.001039 | 0.002651 | 0.001717 | 0.010928 | 0.015382 | 0.004769 | 0.005034 | 0.003889 |
| 30.72222222 | 0.01618 | 0.013498 | 0.028854 | 0.013805 | 0.008094 | 0.008378 | 0.023448 | 0.031977 | 0.007655 | 0.012483 | 0.007387 | 0.022228 | 0.006475 | 0.016583 | 0.003614 | 0.006427 | 0.001286 | 0.003082 | 0.001609 | 0.011981 | 0.019777 | 0.007806 | 0.005199 | 0.004046 |
| 30.73611111 | 0.018125 | 0.009216 | 0.024508 | 0.013877 | 0.0059 | 0.008435 | 0.010033 | 0.024439 | 0.017428 | 0.016775 | 0.009567 | 0.019777 | 0.005957 | 0.018527 | 0.003231 | 0.006939 | 0.000947 | 0.003623 | 0.001217 | 0.011728 | 0.020833 | 0.010931 | 0.00501 | 0.003681 |
| 30.75 | 0.022148 | 0.006124 | 0.016004 | 0.013177 | 0.003691 | 0.01439 | 0.010438 | 0.024438 | 0.024564 | 0.01884 | 0.007755 | 0.012646 | 0.005437 | 0.020384 | 0.003386 | 0.008956 | 0.001008 | 0.003851 | 0.001826 | 0.010353 | 0.018607 | 0.012874 | 0.004811 | 0.003114 |
| 30.76388889 | 0.023987 | 0.004438 | 0.009505 | 0.011497 | 0.002199 | 0.013719 | 0.011622 | 0.041427 | 0.020693 | 0.017744 | 0.006 | 0.011641 | 0.005069 | 0.022556 | 0.003597 | 0.012429 | 0.002237 | 0.003844 | 0.003556 | 0.007564 | 0.013776 | 0.013862 | 0.003732 | 0.002284 |
| 30.77777778 | 0.031724 | 0.003112 | 0.011337 | 0.009112 | 0.001223 | 0.008319 | 0.009219 | 0.053291 | 0.012731 | 0.016665 | 0.01027 | 0.024381 | 0.005278 | 0.025051 | 0.003726 | 0.016156 | 0.004072 | 0.003679 | 0.005165 | 0.004291 | 0.007468 | 0.014293 | 0.001912 | 0.001328 |
| 30.79166667 | 0.043363 | 0.003441 | 0.020241 | 0.007704 | 0.000579 | 0.008979 | 0.012909 | 0.045494 | 0.00691 | 0.01708 | 0.015967 | 0.040327 | 0.006228 | 0.026873 | 0.003497 | 0.017498 | 0.005773 | 0.003206 | 0.005699 | 0.00351 | 0.003163 | 0.013909 | 0.000793 | 0.000617 |
| 30.80555556 | 0.044735 | 0.00381 | 0.024038 | 0.007634 | 0.001022 | 0.010708 | 0.030684 | 0.02622 | 0.005693 | 0.018007 | 0.016987 | 0.049913 | 0.007585 | 0.027424 | 0.003102 | 0.015473 | 0.006535 | 0.002257 | 0.004963 | 0.005717 | 0.003613 | 0.012151 | 0.000853 | 0.000454 |
| 30.81944444 | 0.029623 | 0.003102 | 0.017831 | 0.007325 | 0.002694 | 0.01052 | 0.052211 | 0.010771 | 0.009192 | 0.022048 | 0.012341 | 0.052392 | 0.008996 | 0.026844 | 0.002803 | 0.01257 | 0.005804 | 0.001074 | 0.004004 | 0.007874 | 0.006747 | 0.009132 | 0.001806 | 0.000491 |
| 30.83333333 | 0.015796 | 0.00423 | 0.007454 | 0.005421 | 0.004488 | 0.009743 | 0.05916 | 0.009802 | 0.015121 | 0.030802 | 0.007126 | 0.048403 | 0.010109 | 0.025251 | 0.00237 | 0.011153 | 0.004384 | 0.000324 | 0.004006 | 0.008878 | 0.010249 | 0.00564 | 0.003916 | 0.000827 |
| 30.84722222 | 0.021008 | 0.007011 | 0.001868 | 0.002737 | 0.006277 | 0.008826 | 0.055148 | 0.023354 | 0.022192 | 0.042739 | 0.008264 | 0.041661 | 0.01024 | 0.022612 | 0.001459 | 0.010696 | 0.003262 | 0.000323 | 0.004807 | 0.010138 | 0.014226 | 0.002923 | 0.007249 | 0.002225 |
| 30.86111111 | 0.040313 | 0.009825 | 0.001744 | 0.001838 | 0.008319 | 0.010232 | 0.049857 | 0.04486 | 0.028826 | 0.055131 | 0.012952 | 0.035735 | 0.00943 | 0.019141 | 0.00079 | 0.009436 | 0.002857 | 0.000725 | 0.005759 | 0.012569 | 0.020104 | 0.00275 | 0.010643 | 0.003966 |
| 30.875 | 0.062885 | 0.01529 | 0.002565 | 0.003871 | 0.00948 | 0.014067 | 0.040411 | 0.062298 | 0.030862 | 0.063837 | 0.014853 | 0.030391 | 0.008623 | 0.01534 | 0.00095 | 0.007194 | 0.003358 | 0.000851 | 0.006491 | 0.016064 | 0.027752 | 0.005213 | 0.0129 | 0.004907 |
| 30.88888889 | 0.074849 | 0.024399 | 0.002387 | 0.005953 | 0.007995 | 0.017174 | 0.025887 | 0.065523 | 0.027791 | 0.065506 | 0.01442 | 0.024285 | 0.008151 | 0.011385 | 0.000976 | 0.004872 | 0.004184 | 0.000516 | 0.006506 | 0.019534 | 0.033554 | 0.00856 | 0.013905 | 0.004982 |
| 30.90277778 | 0.065792 | 0.034483 | 0.001582 | 0.006216 | 0.004225 | 0.017408 | 0.015105 | 0.062922 | 0.02333 | 0.059695 | 0.013295 | 0.017104 | 0.007372 | 0.007033 | 0.000623 | 0.002631 | 0.004221 | 0.000222 | 0.005505 | 0.021355 | 0.033863 | 0.011879 | 0.014144 | 0.004351 |
| 30.91666667 | 0.046086 | 0.043126 | 0.002116 | 0.005682 | 0.001705 | 0.014671 | 0.01187 | 0.067282 | 0.017159 | 0.04661 | 0.012655 | 0.009805 | 0.005788 | 0.003426 | 0.000558 | 0.001506 | 0.003191 | 0.000288 | 0.004079 | 0.021198 | 0.027479 | 0.015143 | 0.013875 | 0.003278 |
| 30.93055556 | 0.027177 | 0.04708 | 0.003965 | 0.004976 | 0.002715 | 0.010212 | 0.011008 | 0.072974 | 0.010703 | 0.026164 | 0.013746 | 0.009 | 0.003801 | 0.002793 | 0.000844 | 0.002385 | 0.00192 | 0.000664 | 0.002574 | 0.020266 | 0.018305 | 0.017726 | 0.012739 | 0.002303 |
| 30.94444444 | 0.013356 | 0.04603 | 0.006381 | 0.004619 | 0.005401 | 0.006492 | 0.008089 | 0.068736 | 0.013338 | 0.012583 | 0.011774 | 0.018019 | 0.002077 | 0.004674 | 0.001216 | 0.003552 | 0.00113 | 0.001063 | 0.001111 | 0.019247 | 0.011583 | 0.01851 | 0.010961 | 0.001692 |
| 30.95833333 | 0.007183 | 0.040852 | 0.008928 | 0.004101 | 0.007553 | 0.004331 | 0.004389 | 0.053995 | 0.027325 | 0.019623 | 0.005373 | 0.028545 | 0.001002 | 0.006837 | 0.001548 | 0.003941 | 0.000712 | 0.001407 | 0.000334 | 0.01822 | 0.008196 | 0.017588 | 0.009498 | 0.001138 |
| 30.97222222 | 0.008521 | 0.030182 | 0.010337 | 0.002923 | 0.008139 | 0.002579 | 0.003571 | 0.036402 | 0.038845 | 0.033504 | 0.001689 | 0.032303 | 0.001203 | 0.008789 | 0.002078 | 0.004372 | 0.000439 | 0.002036 | 0.000199 | 0.016787 | 0.007112 | 0.015905 | 0.007984 | 0.000586 |
| 30.98611111 | 0.009787 | 0.017477 | 0.009031 | 0.003218 | 0.006237 | 0.002194 | 0.004087 | 0.026182 | 0.032612 | 0.034299 | 0.004123 | 0.024992 | 0.00341 | 0.010784 | 0.0039 | 0.005266 | 0.00105 | 0.00282 | 0.001064 | 0.014551 | 0.007628 | 0.013831 | 0.005414 | 0.000651 |
| 31 | 0.010124 | 0.010091 | 0.005403 | 0.006504 | 0.003215 | 0.006009 | 0.007822 | 0.031289 | 0.024829 | 0.02853 | 0.012467 | 0.016491 | 0.007351 | 0.015357 | 0.008341 | 0.007998 | 0.003815 | 0.002807 | 0.003917 | 0.01181 | 0.010026 | 0.011027 | 0.002552 | 0.0015 |
| 31.01388889 | 0.019976 | 0.007958 | 0.003192 | 0.010038 | 0.002355 | 0.01338 | 0.019913 | 0.050393 | 0.045727 | 0.042737 | 0.026742 | 0.027712 | 0.012428 | 0.030131 | 0.014121 | 0.017143 | 0.009133 | 0.001718 | 0.007152 | 0.009309 | 0.013426 | 0.007285 | 0.00201 | 0.002133 |
| 31.02777778 | 0.033905 | 0.007057 | 0.004581 | 0.011205 | 0.004564 | 0.021372 | 0.03251 | 0.070822 | 0.079851 | 0.070767 | 0.04628 | 0.050626 | 0.017093 | 0.053808 | 0.016326 | 0.02836 | 0.01542 | 0.000657 | 0.008382 | 0.008475 | 0.014329 | 0.008732 | 0.006087 | 0.0018 |
| 31.04166667 | 0.04239 | 0.010795 | 0.006461 | 0.011767 | 0.008557 | 0.029227 | 0.035202 | 0.080634 | 0.086952 | 0.076314 | 0.061667 | 0.056992 | 0.017185 | 0.06498 | 0.012138 | 0.026189 | 0.018943 | 0.000423 | 0.008225 | 0.010398 | 0.011602 | 0.019108 | 0.012209 | 0.001556 |
| 31.05555556 | 0.044107 | 0.023564 | 0.005296 | 0.013392 | 0.012707 | 0.031438 | 0.034773 | 0.077936 | 0.0619 | 0.05728 | 0.064804 | 0.046387 | 0.011276 | 0.049529 | 0.005713 | 0.014129 | 0.017046 | 0.000648 | 0.007395 | 0.014471 | 0.009103 | 0.026536 | 0.014384 | 0.002535 |
| 31.06944444 | 0.038318 | 0.042651 | 0.003223 | 0.0152 | 0.015256 | 0.024298 | 0.036497 | 0.066792 | 0.032949 | 0.039314 | 0.052232 | 0.036169 | 0.004628 | 0.022924 | 0.002533 | 0.008157 | 0.01152 | 0.001407 | 0.00564 | 0.017633 | 0.009217 | 0.021282 | 0.011207 | 0.003381 |

| | | | | | | | | | | | | | | | | | | | | | | | | |
|---|---|---|---|---|---|---|---|---|---|---|---|---|---|---|---|---|---|---|---|---|---|---|---|---|
| 31.08333333 | 0.028404 | 0.057983 | 0.00354 | 0.016022 | 0.015879 | 0.016423 | 0.03054 | 0.05083 | 0.015046 | 0.030173 | 0.029122 | 0.028115 | 0.002086 | 0.006516 | 0.002096 | 0.006721 | 0.005572 | 0.003038 | 0.003579 | 0.01686 | 0.009241 | 0.009709 | 0.005997 | 0.002493 |
| 31.09722222 | 0.017725 | 0.060589 | 0.00423 | 0.015572 | 0.015072 | 0.012416 | 0.018308 | 0.03252 | 0.00836 | 0.022882 | 0.023547 | 0.016484 | 0.003276 | 0.0033 | 0.001944 | 0.003865 | 0.002377 | 0.004445 | 0.001955 | 0.012443 | 0.006529 | 0.002802 | 0.002285 | 0.001598 |
| 31.11111111 | 0.008684 | 0.051362 | 0.004282 | 0.014677 | 0.012851 | 0.009763 | 0.011935 | 0.017157 | 0.016507 | 0.014831 | 0.040478 | 0.007898 | 0.006153 | 0.003659 | 0.001712 | 0.001553 | 0.003227 | 0.004701 | 0.001511 | 0.006876 | 0.004177 | 0.002225 | 0.00137 | 0.002051 |
| 31.125 | 0.003415 | 0.037934 | 0.005256 | 0.013784 | 0.009666 | 0.008548 | 0.014011 | 0.007636 | 0.028558 | 0.007455 | 0.049747 | 0.010703 | 0.008606 | 0.003957 | 0.001112 | 0.001362 | 0.003774 | 0.004156 | 0.001486 | 0.00294 | 0.004855 | 0.002448 | 0.001548 | 0.002058 |
| 31.13888889 | 0.001283 | 0.027827 | 0.006832 | 0.012704 | 0.006475 | 0.008711 | 0.018253 | 0.003305 | 0.029417 | 0.002466 | 0.036369 | 0.018687 | 0.008367 | 0.007558 | 0.00056 | 0.00158 | 0.002692 | 0.00343 | 0.001079 | 0.001641 | 0.004747 | 0.001442 | 0.001425 | 0.001144 |
| 31.15277778 | 0.002827 | 0.02119 | 0.008336 | 0.010339 | 0.003507 | 0.008768 | 0.016784 | 0.004053 | 0.019752 | 0.000716 | 0.015658 | 0.025042 | 0.005204 | 0.009923 | 0.000839 | 0.000976 | 0.003664 | 0.003045 | 0.001158 | 0.001949 | 0.002941 | 0.000631 | 0.001008 | 0.00077 |
| 31.16666667 | 0.008839 | 0.013553 | 0.009352 | 0.006646 | 0.001892 | 0.007267 | 0.010758 | 0.007576 | 0.010156 | 0.002968 | 0.005387 | 0.029678 | 0.002248 | 0.007248 | 0.002475 | 0.000383 | 0.005938 | 0.003251 | 0.001274 | 0.002453 | 0.001596 | 0.000675 | 0.000557 | 0.001316 |
| 31.18055556 | 0.017617 | 0.011978 | 0.009631 | 0.005156 | 0.003271 | 0.004058 | 0.011946 | 0.010257 | 0.00873 | 0.012076 | 0.012038 | 0.032299 | 0.001836 | 0.007465 | 0.004333 | 0.000699 | 0.006935 | 0.004067 | 0.000819 | 0.002136 | 0.002071 | 0.002578 | 0.000422 | 0.002013 |
| 31.19444444 | 0.026122 | 0.026704 | 0.010064 | 0.006065 | 0.006085 | 0.002259 | 0.023381 | 0.009464 | 0.009 | 0.028372 | 0.029122 | 0.037128 | 0.001952 | 0.01406 | 0.004128 | 0.001748 | 0.006964 | 0.005417 | 0.000488 | 0.001102 | 0.0054 | 0.00808 | 0.000373 | 0.00234 |
| 31.20833333 | 0.032245 | 0.050272 | 0.012032 | 0.005903 | 0.007739 | 0.003814 | 0.033237 | 0.00541 | 0.00739 | 0.047397 | 0.032325 | 0.045325 | 0.001253 | 0.020192 | 0.002388 | 0.002568 | 0.007009 | 0.006742 | 0.00042 | 0.000269 | 0.008355 | 0.014261 | 0.000242 | 0.002237 |
| 31.22222222 | 0.035662 | 0.060346 | 0.014899 | 0.005581 | 0.00814 | 0.006273 | 0.035485 | 0.003267 | 0.016273 | 0.065005 | 0.022626 | 0.051557 | 0.001371 | 0.019476 | 0.001538 | 0.002767 | 0.007176 | 0.006585 | 0.00061 | 8.5451395 | 0.007365 | 0.014471 | 0.000428 | 0.001832 |
| 31.23611111 | 0.03739 | 0.052726 | 0.01656 | 0.006363 | 0.008305 | 0.007956 | 0.034768 | 0.006658 | 0.043855 | 0.073235 | 0.032903 | 0.054379 | 0.002753 | 0.011561 | 0.001722 | 0.002545 | 0.007164 | 0.004416 | 0.001109 | 0.000161 | 0.003823 | 0.008933 | 0.001133 | 0.001312 |
| 31.25 | 0.039119 | 0.045333 | 0.016632 | 0.006835 | 0.008129 | 0.008566 | 0.035267 | 0.012394 | 0.07255 | 0.064646 | 0.056225 | 0.050242 | 0.003827 | 0.005985 | 0.001483 | 0.001974 | 0.007013 | 0.001949 | 0.001265 | 0.000148 | 0.001354 | 0.005851 | 0.001563 | 0.000763 |
| 31.26388889 | 0.039265 | 0.040853 | 0.015846 | 0.007221 | 0.006645 | 0.007671 | 0.035509 | 0.015959 | 0.071015 | 0.043062 | 0.057006 | 0.037485 | 0.00338 | 0.009649 | 0.001386 | 0.001181 | 0.007016 | 0.000631 | 0.000851 | 0.000214 | 0.001147 | 0.007573 | 0.001283 | 0.000473 |
| 31.27777778 | 0.036449 | 0.034582 | 0.014583 | 0.007963 | 0.003863 | 0.005409 | 0.02961 | 0.013893 | 0.041136 | 0.027123 | 0.035908 | 0.020926 | 0.001865 | 0.014909 | 0.00185 | 0.000579 | 0.007439 | 0.00045 | 0.000723 | 0.000391 | 0.001695 | 0.008383 | 0.001711 | 0.00155 |
| 31.29166667 | 0.032465 | 0.026281 | 0.013218 | 0.007284 | 0.001804 | 0.003044 | 0.016606 | 0.008082 | 0.024977 | 0.031194 | 0.015096 | 0.00927 | 0.00099 | 0.01433 | 0.007267 | 0.004142 | 0.007188 | 0.00314 | 0.004295 | 0.000374 | 0.004712 | 0.006906 | 0.004942 | 0.011005 |
| 31.30555556 | 0.027547 | 0.019063 | 0.011964 | 0.004153 | 0.00171 | 0.003675 | 0.005202 | 0.004723 | 0.037017 | 0.032948 | 0.00492 | 0.007954 | 0.002107 | 0.013617 | 0.02291 | 0.010903 | 0.017721 | 0.006449 | 0.010713 | 0.001716 | 0.024977 | 0.009636 | 0.011353 | 0.035146 |
| 31.31944444 | 0.022133 | 0.015419 | 0.011129 | 0.001457 | 0.002903 | 0.007853 | 0.00114 | 0.003204 | 0.043503 | 0.019173 | 0.003926 | 0.008288 | 0.003538 | 0.019779 | 0.029905 | 0.011328 | 0.031895 | 0.006232 | 0.010289 | 0.003248 | 0.040858 | 0.014382 | 0.013617 | 0.046122 |
| 31.33333333 | 0.018942 | 0.012408 | 0.01111 | 0.001568 | 0.004145 | 0.011488 | 0.002316 | 0.002142 | 0.028598 | 0.008199 | 0.008837 | 0.005298 | 0.003461 | 0.021405 | 0.023389 | 0.009264 | 0.031849 | 0.006197 | 0.005836 | 0.002562 | 0.033838 | 0.013673 | 0.010843 | 0.039947 |
| 31.34722222 | 0.019901 | 0.015106 | 0.012195 | 0.002814 | 0.003914 | 0.011657 | 0.006886 | 0.003678 | 0.014393 | 0.003622 | 0.014899 | 0.003076 | 0.0037 | 0.011764 | 0.023649 | 0.01245 | 0.025775 | 0.006618 | 0.006975 | 0.003437 | 0.028412 | 0.013204 | 0.009721 | 0.043319 |
| 31.36111111 | 0.022902 | 0.024164 | 0.013827 | 0.003487 | 0.002721 | 0.009093 | 0.010258 | 0.005582 | 0.018588 | 0.001583 | 0.016092 | 0.003988 | 0.00389 | 0.006058 | 0.019656 | 0.012359 | 0.013138 | 0.004529 | 0.007158 | 0.004381 | 0.016101 | 0.014801 | 0.006455 | 0.033471 |
| 31.375 | 0.023651 | 0.030195 | 0.014167 | 0.003471 | 0.00162 | 0.005536 | 0.00877 | 0.005124 | 0.024509 | 0.003921 | 0.011901 | 0.006587 | 0.002501 | 0.007471 | 0.006943 | 0.00576 | 0.002752 | 0.002457 | 0.003672 | 0.002334 | 0.002349 | 0.013768 | 0.001864 | 0.010756 |
| 31.38888889 | 0.020488 | 0.034377 | 0.012543 | 0.003214 | 0.000758 | 0.002518 | 0.006062 | 0.002972 | 0.01603 | 0.013614 | 0.00596 | 0.006135 | 0.001618 | 0.005921 | 0.001398 | 0.00117 | 0.002361 | 0.002023 | 0.002436 | 0.001356 | 0.000965 | 0.011619 | 0.000722 | 0.002171 |
| 31.40277778 | 0.014728 | 0.040025 | 0.009869 | 0.002836 | 0.000661 | 0.001819 | 0.005717 | 0.001426 | 0.008066 | 0.028958 | 0.004029 | 0.005168 | 0.002183 | 0.002655 | 0.002285 | 0.000619 | 0.003279 | 0.002191 | 0.002769 | 0.001913 | 0.000778 | 0.009671 | 0.001715 | 0.001973 |
| 31.41666667 | 0.009036 | 0.044331 | 0.00738 | 0.002104 | 0.001032 | 0.003827 | 0.007874 | 0.002006 | 0.01249 | 0.041788 | 0.007797 | 0.007614 | 0.003148 | 0.001819 | 0.005498 | 0.00133 | 0.004417 | 0.002971 | 0.003192 | 0.002927 | 0.00069 | 0.007583 | 0.0034 | 0.002132 |
| 31.43055556 | 0.00551 | 0.047923 | 0.006606 | 0.001609 | 0.001135 | 0.006315 | 0.010119 | 0.003467 | 0.018439 | 0.044065 | 0.012737 | 0.008207 | 0.003712 | 0.002568 | 0.008923 | 0.002417 | 0.005127 | 0.00448 | 0.003772 | 0.004635 | 0.001093 | 0.005945 | 0.004539 | 0.001769 |
| 31.44444444 | 0.004838 | 0.052939 | 0.007942 | 0.001794 | 0.001006 | 0.007122 | 0.009512 | 0.004699 | 0.016441 | 0.033345 | 0.013358 | 0.005032 | 0.003243 | 0.005906 | 0.009651 | 0.003335 | 0.004707 | 0.005643 | 0.00373 | 0.00572 | 0.001126 | 0.005026 | 0.004555 | 0.001581 |
| 31.45833333 | 0.006302 | 0.057342 | 0.009273 | 0.002093 | 0.000837 | 0.006632 | 0.006267 | 0.004673 | 0.010184 | 0.01782 | 0.007657 | 0.003644 | 0.002122 | 0.010921 | 0.007638 | 0.003809 | 0.003108 | 0.006058 | 0.003181 | 0.005686 | 0.000631 | 0.004197 | 0.003662 | 0.002205 |
| 31.47222222 | 0.010999 | 0.05086 | 0.010028 | 0.002687 | 0.001286 | 0.008161 | 0.006422 | 0.005749 | 0.009088 | 0.010808 | 0.002447 | 0.00605 | 0.001499 | 0.013747 | 0.005145 | 0.004141 | 0.001738 | 0.006162 | 0.003163 | 0.005466 | 0.000291 | 0.003603 | 0.002769 | 0.003059 |
| 31.48611111 | 0.022904 | 0.037589 | 0.013055 | 0.008236 | 0.004588 | 0.016844 | 0.021485 | 0.01753 | 0.017533 | 0.02447 | 0.00428 | 0.011268 | 0.001736 | 0.013745 | 0.004113 | 0.004634 | 0.002626 | 0.006036 | 0.004085 | 0.005831 | 0.00093 | 0.004232 | 0.002922 | 0.002823 |
| 31.5 | 0.04161 | 0.046454 | 0.020265 | 0.022883 | 0.011134 | 0.031169 | 0.055305 | 0.043976 | 0.027749 | 0.056115 | 0.014833 | 0.020795 | 0.00234 | 0.012964 | 0.00454 | 0.004722 | 0.006431 | 0.005477 | 0.00562 | 0.00718 | 0.004111 | 0.00644 | 0.005092 | 0.001764 |
| 31.51388889 | 0.058772 | 0.07218 | 0.030424 | 0.036345 | 0.016314 | 0.038918 | 0.085036 | 0.068186 | 0.030385 | 0.079987 | 0.034003 | 0.0348 | 0.002985 | 0.013749 | 0.006163 | 0.004065 | 0.011435 | 0.00457 | 0.007582 | 0.008873 | 0.009601 | 0.008705 | 0.009412 | 0.001733 |
| 31.52777778 | 0.062719 | 0.081815 | 0.037689 | 0.033677 | 0.01715 | 0.032301 | 0.084989 | 0.071706 | 0.02776 | 0.081783 | 0.048494 | 0.05036 | 0.003732 | 0.017666 | 0.008639 | 0.003263 | 0.014633 | 0.003312 | 0.008942 | 0.008382 | 0.01327 | 0.008424 | 0.01429 | 0.002913 |
| 31.54166667 | 0.048674 | 0.077196 | 0.03633 | 0.020303 | 0.016822 | 0.02275 | 0.06872 | 0.065349 | 0.040023 | 0.076627 | 0.044309 | 0.059874 | 0.004518 | 0.022054 | 0.011068 | 0.003616 | 0.013471 | 0.001905 | 0.008429 | 0.005759 | 0.012479 | 0.00714 | 0.017531 | 0.004038 |
| 31.55555556 | 0.027094 | 0.074058 | 0.028435 | 0.011271 | 0.017014 | 0.025971 | 0.070434 | 0.066528 | 0.066232 | 0.068696 | 0.027956 | 0.053235 | 0.005007 | 0.02167 | 0.012701 | 0.006257 | 0.009238 | 0.001449 | 0.007414 | 0.004801 | 0.0104 | 0.008937 | 0.017876 | 0.004815 |
| 31.56944444 | 0.021589 | 0.079175 | 0.019015 | 0.008578 | 0.015545 | 0.034906 | 0.088894 | 0.071818 | 0.077241 | 0.051576 | 0.015211 | 0.03326 | 0.005142 | 0.016698 | 0.012835 | 0.009576 | 0.006402 | 0.002228 | 0.007872 | 0.006273 | 0.009739 | 0.01122 | 0.016103 | 0.004913 |
| 31.58333333 | 0.03527 | 0.086051 | 0.009632 | 0.008264 | 0.012059 | 0.037869 | 0.086051 | 0.073624 | 0.068338 | 0.033135 | 0.012953 | 0.014079 | 0.004946 | 0.011332 | 0.010746 | 0.010858 | 0.005271 | 0.002769 | 0.008604 | 0.007906 | 0.009071 | 0.008942 | 0.014436 | 0.003831 |
| 31.59722222 | 0.043386 | 0.079616 | 0.004645 | 0.009361 | 0.008189 | 0.034034 | 0.051983 | 0.068222 | 0.049607 | 0.018985 | 0.012249 | 0.007071 | 0.004086 | 0.006941 | 0.006268 | 0.010257 | 0.003438 | 0.002146 | 0.007999 | 0.009128 | 0.007665 | 0.004884 | 0.013272 | 0.002126 |
| 31.61111111 | 0.039871 | 0.061556 | 0.00805 | 0.011462 | 0.004847 | 0.02028 | 0.024581 | 0.054839 | 0.027529 | 0.009007 | 0.006678 | 0.013794 | 0.002515 | 0.0037 | 0.003849 | 0.009343 | 0.001498 | 0.001105 | 0.006503 | 0.009953 | 0.006847 | 0.004567 | 0.010188 | 0.001361 |
| 31.625 | 0.030889 | 0.04238 | 0.01597 | 0.013499 | 0.00253 | 0.009668 | 0.032495 | 0.040031 | 0.01891 | 0.006974 | 0.004103 | 0.021441 | 0.001057 | 0.003772 | 0.006821 | 0.008403 | 0.000919 | 0.000679 | 0.004643 | 0.009903 | 0.007223 | 0.006297 | 0.004991 | 0.002107 |
| 31.63888889 | 0.018439 | 0.024391 | 0.024056 | 0.013638 | 0.002324 | 0.012287 | 0.060941 | 0.029455 | 0.028013 | 0.01322 | 0.006784 | 0.023228 | 0.000491 | 0.007399 | 0.01059 | 0.007327 | 0.001774 | 0.001025 | 0.002574 | 0.009254 | 0.00812 | 0.006068 | 0.002165 | 0.002831 |
| 31.65277778 | 0.008572 | 0.012448 | 0.030543 | 0.01094 | 0.003393 | 0.013479 | 0.078373 | 0.022623 | 0.035006 | 0.021618 | 0.011726 | 0.02463 | 0.00036 | 0.011622 | 0.012399 | 0.006168 | 0.003557 | 0.001781 | 0.001124 | 0.0089 | 0.008711 | 0.005484 | 0.003246 | 0.002497 |
| 31.66666667 | 0.006463 | 0.015619 | 0.031414 | 0.006307 | 0.003649 | 0.008981 | 0.073441 | 0.017546 | 0.03027 | 0.02946 | 0.02087 | 0.025457 | 0.000246 | 0.014278 | 0.013437 | 0.005109 | 0.005226 | 0.002731 | 0.001064 | 0.009347 | 0.008455 | 0.006691 | 0.005142 | 0.001437 |
| 31.68055556 | 0.016969 | 0.029578 | 0.022176 | 0.003223 | 0.003122 | 0.004301 | 0.065665 | 0.014151 | 0.020941 | 0.036765 | 0.028181 | 0.02225 | 0.000353 | 0.014397 | 0.013777 | 0.004266 | 0.005935 | 0.00374 | 0.00165 | 0.010477 | 0.007579 | 0.007528 | 0.006013 | 0.000648 |
| 31.69444444 | 0.037962 | 0.041237 | 0.0097 | 0.003547 | 0.002091 | 0.001779 | 0.070069 | 0.013655 | 0.014007 | 0.042515 | 0.02946 | 0.013489 | 0.001111 | 0.011448 | 0.013434 | 0.003418 | 0.004716 | 0.004558 | 0.001453 | 0.012004 | 0.006618 | 0.005194 | 0.005869 | 0.000783 |
| 31.70833333 | 0.042441 | 0.045702 | 0.004071 | 0.004171 | 0.001331 | 0.002516 | 0.07483 | 0.017115 | 0.009772 | 0.047423 | 0.031609 | 0.005274 | 0.002892 | 0.006477 | 0.011678 | 0.00233 | 0.003134 | 0.00488 | 0.000988 | 0.013461 | 0.005519 | 0.002437 | 0.00521 | 0.001071 |
| 31.72222222 | 0.022448 | 0.052159 | 0.002546 | 0.005887 | 0.002076 | 0.004538 | 0.068961 | 0.021717 | 0.00773 | 0.052997 | 0.03318 | 0.003567 | 0.005016 | 0.002285 | 0.008414 | 0.001598 | 0.004573 | 0.004485 | 0.001604 | 0.014256 | 0.004091 | 0.002966 | 0.003984 | 0.000858 |
| 31.73611111 | 0.012153 | 0.06609 | 0.000886 | 0.010117 | 0.002484 | 0.005785 | 0.055424 | 0.022348 | 0.01135 | 0.058552 | 0.025337 | 0.003699 | 0.006704 | 0.000906 | 0.005003 | 0.002464 | 0.007295 | 0.003368 | 0.002716 | 0.013975 | 0.002604 | 0.004099 | 0.002715 | 0.000542 |
| 31.75 | 0.017175 | 0.078202 | 9.7076396 | 0.011665 | 0.001525 | 0.005936 | 0.041154 | 0.019014 | 0.020147 | 0.063554 | 0.017219 | 0.003056 | 0.007644 | 0.002271 | 0.002214 | 0.004527 | 0.008385 | 0.00185 | 0.003562 | 0.012445 | 0.002984 | 0.003308 | 0.002394 | 0.000779 |
| 31.76388889 | 0.017927 | 0.081929 | 0.000706 | 0.007978 | 0.000963 | 0.006041 | 0.029769 | 0.013916 | 0.028327 | 0.066116 | 0.016751 | 0.002967 | 0.00788 | 0.005946 | 0.00054 | 0.006576 | 0.007649 | 0.001202 | 0.00428 | 0.010283 | 0.006141 | 0.002384 | 0.002379 | 0.001612 |
| 31.77777778 | 0.012004 | 0.080729 | 0.002283 | 0.004091 | 0.000784 | 0.006325 | 0.022445 | 0.008523 | 0.030916 | 0.06206 | 0.01155 | 0.005918 | 0.007884 | 0.011629 | 0.000717 | 0.008192 | 0.005122 | 0.002345 | 0.00493 | 0.008363 | 0.009055 | 0.003825 | 0.002036 | 0.002406 |
| 31.79166667 | 0.008885 | 0.078706 | 0.003682 | 0.00476 | 0.000459 | 0.005966 | 0.020432 | 0.004317 | 0.028069 | 0.056552 | 0.00355 | 0.015266 | 0.007918 | 0.018363 | 0.002933 | 0.009125 | 0.002477 | 0.003885 | 0.005468 | 0.006614 | 0.009426 | 0.007861 | 0.001582 | 0.002624 |
| 31.80555556 | 0.00788 | 0.071018 | 0.00436 | 0.006667 | 0.000468 | 0.004029 | 0.022424 | 0.002278 | 0.021333 | 0.058715 | 0.002481 | 0.028471 | 0.00809 | 0.024349 | 0.005981 | 0.009705 | 0.002194 | 0.004456 | 0.005803 | 0.004648 | 0.008344 | 0.013818 | 0.001266 | 0.002247 |
| 31.81944444 | 0.005202 | 0.054934 | 0.006835 | 0.004979 | 0.000817 | 0.001878 | 0.025903 | 0.002326 | 0.01252 | 0.067883 | 0.004136 | 0.042101 | 0.008508 | 0.027459 | 0.008151 | 0.010498 | 0.003711 | 0.004395 | 0.005947 | 0.00247 | 0.007487 | 0.020933 | 0.002145 | 0.001371 |
| 31.83333333 | 0.005222 | 0.037729 | 0.013552 | 0.002472 | 0.001413 | 0.002229 | 0.031587 | 0.002912 | 0.005862 | 0.074365 | 0.006445 | 0.057234 | 0.008987 | 0.027003 | 0.009053 | 0.011154 | 0.005468 | 0.004377 | 0.005782 | 0.001458 | 0.007598 | 0.024977 | 0.004502 | 0.0007 |
| 31.84722222 | 0.006897 | 0.026439 | 0.020036 | 0.00198 | 0.001578 | 0.004834 | 0.041201 | 0.004343 | 0.007152 | 0.07096 | 0.01123 | 0.068561 | 0.009168 | 0.024152 | 0.00872 | 0.010821 | 0.007181 | 0.005044 | 0.005607 | 0.002871 | 0.008876 | 0.021754 | 0.006344 | 0.000791 |
| 31.86111111 | 0.007588 | 0.02239 | 0.020697 | 0.002147 | 0.000908 | 0.008461 | 0.052322 | 0.00827 | 0.01267 | 0.060596 | 0.010996 | 0.067197 | 0.00927 | 0.020285 | 0.006817 | 0.00955 | 0.008524 | 0.007049 | 0.006227 | 0.005146 | 0.010753 | 0.01337 | 0.006448 | 0.00084 |
| 31.875 | 0.007672 | 0.023484 | 0.018364 | 0.003764 | 0.000297 | 0.011797 | 0.057601 | 0.014876 | 0.011231 | 0.045645 | 0.004535 | 0.057642 | 0.009546 | 0.01649 | 0.004186 | 0.008593 | 0.009772 | 0.010171 | 0.007298 | 0.006761 | 0.012334 | 0.005528 | 0.005757 | 0.000867 |
| 31.88888889 | 0.007919 | 0.026526 | 0.016973 | 0.006484 | 0.000557 | 0.012999 | 0.054434 | 0.022452 | 0.005345 | 0.027739 | 0.002311 | 0.047093 | 0.009697 | 0.013436 | 0.002157 | 0.008493 | 0.011178 | 0.012729 | 0.007915 | 0.00809 | 0.013558 | 0.001316 | 0.005436 | 0.001386 |
| 31.90277778 | 0.008009 | 0.02598 | 0.015539 | 0.007908 | 0.001712 | 0.011945 | 0.046841 | 0.027419 | 0.003915 | 0.01345 | 0.005214 | 0.035242 | 0.009303 | 0.011412 | 0.001105 | 0.008243 | 0.011771 | 0.013008 | 0.00775 | 0.009096 | 0.014864 | 0.000845 | 0.005934 | 0.002094 |
| 31.91666667 | 0.008739 | 0.020536 | 0.012215 | 0.007421 | 0.002938 | 0.009032 | 0.037755 | 0.026908 | 0.005645 | 0.005411 | 0.008365 | 0.023272 | 0.007892 | 0.010049 | 0.000632 | 0.006805 | 0.010746 | 0.011222 | 0.006821 | 0.008761 | 0.015824 | 0.003194 | 0.007235 | 0.002649 |
| 31.93055556 | 0.012602 | 0.013204 | 0.007256 | 0.005249 | 0.003666 | 0.005203 | 0.028998 | 0.021518 | 0.006906 | 0.002301 | 0.011206 | 0.012988 | 0.005682 | 0.009096 | 0.000545 | 0.004293 | 0.008347 | 0.009019 | 0.005576 | 0.007131 | 0.015954 | 0.007505 | 0.008929 | 0.002587 |
| 31.94444444 | 0.019176 | 0.009395 | 0.005502 | 0.0026 | 0.004116 | 0.004241 | 0.021502 | 0.013865 | 0.008081 | 0.003577 | 0.014035 | 0.00636 | 0.003463 | 0.008517 | 0.000718 | 0.002063 | 0.005763 | 0.007568 | 0.004409 | 0.006067 | 0.015688 | 0.013149 | 0.010068 | 0.001846 |
| 31.95833333 | 0.02559 | 0.01613 | 0.009961 | 0.001113 | 0.004463 | 0.007572 | 0.015592 | 0.006422 | 0.01002 | 0.005696 | 0.016197 | 0.006852 | 0.00178 | 0.007006 | 0.000594 | 0.001805 | 0.003583 | 0.006961 | 0.003528 | 0.00651 | 0.015012 | 0.017351 | 0.009656 | 0.001083 |
| 31.97222222 | 0.030093 | 0.026913 | 0.014993 | 0.002267 | 0.00456 | 0.010714 | 0.012788 | 0.00272 | 0.010936 | 0.006671 | 0.013303 | 0.011085 | 0.001261 | 0.00388 | 0.000246 | 0.003276 | 0.0018 | 0.00681 | 0.002994 | 0.007113 | 0.013629 | 0.016769 | 0.00785 | 0.000817 |
| 31.98611111 | 0.028329 | 0.027039 | 0.013871 | 0.010914 | 0.00489 | 0.008951 | 0.016577 | 0.006682 | 0.008897 | 0.016302 | 0.009606 | 0.010267 | 0.001982 | 0.001222 | 0.000179 | 0.004634 | 0.000897 | 0.006889 | 0.003055 | 0.006759 | 0.012258 | 0.012306 | 0.005522 | 0.001014 |
| 32 | 0.017879 | 0.019001 | 0.009114 | 0.027524 | 0.005855 | 0.005784 | 0.025892 | 0.016693 | 0.011335 | 0.043729 | 0.01888 | 0.008921 | 0.002783 | 0.002453 | 0.000906 | 0.005436 | 0.00238 | 0.007534 | 0.004647 | 0.005607 | 0.013605 | 0.00856 | 0.004069 | 0.001801 |
| 32.01388889 | 0.010008 | 0.018967 | 0.009405 | 0.038889 | 0.005911 | 0.007514 | 0.033373 | 0.025277 | 0.023348 | 0.080638 | 0.037262 | 0.020663 | 0.00236 | 0.009321 | 0.002952 | 0.006249 | 0.009891 | 0.00906 | 0.008113 | 0.004173 | 0.019454 | 0.010489 | 0.005618 | 0.003288 |
| 32.02777778 | 0.013429 | 0.028599 | 0.014572 | 0.037525 | 0.004775 | 0.011177 | 0.034584 | 0.028413 | 0.036574 | 0.096982 | 0.046527 | 0.03878 | 0.00199 | 0.018534 | 0.00549 | 0.005869 | 0.022908 | 0.00954 | 0.011974 | 0.00366 | 0.02463 | 0.019302 | 0.009072 | 0.004039 |
| 32.04166667 | 0.021169 | 0.036853 | 0.017335 | 0.031996 | 0.005615 | 0.012497 | 0.031026 | 0.027542 | 0.042638 | 0.068056 | 0.041546 | 0.045777 | 0.005053 | 0.023858 | 0.005505 | 0.005197 | 0.032402 | 0.007255 | 0.013456 | 0.006094 | 0.022085 | 0.028725 | 0.010342 | 0.002756 |
| 32.05555556 | 0.022626 | 0.041122 | 0.015704 | 0.026097 | 0.009096 | 0.011056 | 0.027444 | 0.024417 | 0.038095 | 0.031672 | 0.03423 | 0.03829 | 0.010896 | 0.023306 | 0.003199 | 0.008463 | 0.032413 | 0.005382 | 0.011821 | 0.011223 | 0.015259 | 0.031988 | 0.009565 | 0.001326 |
| 32.06944444 | 0.015683 | 0.043516 | 0.011056 | 0.015609 | 0.011003 | 0.006768 | 0.023049 | 0.019359 | 0.026374 | 0.022765 | 0.035467 | 0.022438 | 0.014827 | 0.018456 | 0.001888 | 0.012908 | 0.025181 | 0.005733 | 0.0088 | 0.014013 | 0.011242 | 0.028971 | 0.009153 | 0.001569 |
| 32.08333333 | 0.01363 | 0.044484 | 0.006095 | 0.00843 | 0.009044 | 0.003493 | 0.014439 | 0.011591 | 0.014468 | 0.016166 | 0.042025 | 0.009465 | 0.014319 | 0.010662 | 0.001808 | 0.011952 | 0.014436 | 0.004964 | 0.005016 | 0.010678 | 0.008163 | 0.021075 | 0.007792 | 0.001872 |
| 32.09722222 | 0.021665 | 0.042912 | 0.005269 | 0.012002 | 0.005829 | 0.00519 | 0.006764 | 0.005926 | 0.007313 | 0.008071 | 0.037709 | 0.006821 | 0.01048 | 0.003896 | 0.002631 | 0.006614 | 0.009856 | 0.002515 | 0.002632 | 0.004857 | 0.003687 | 0.010576 | 0.004545 | 0.001308 |
| 32.11111111 | 0.028426 | 0.036876 | 0.007726 | 0.016385 | 0.004125 | 0.009195 | 0.005006 | 0.007827 | 0.007078 | 0.008217 | 0.020252 | 0.008693 | 0.00577 | 0.001513 | 0.003284 | 0.002095 | 0.016142 | 0.000984 | 0.003229 | 0.001914 | 0.000894 | 0.00465 | 0.001616 | 0.001234 |
| 32.125 | 0.030821 | 0.029043 | 0.009334 | 0.015665 | 0.00387 | 0.011529 | 0.005379 | 0.012088 | 0.00963 | 0.008145 | 0.008405 | 0.008654 | 0.002312 | 0.00118 | 0.002477 | 0.000664 | 0.020698 | 0.000726 | 0.00357 | 0.001533 | 0.000781 | 0.00607 | 0.000423 | 0.001425 |
| 32.13888889 | 0.033905 | 0.022832 | 0.011584 | 0.013164 | 0.003741 | 0.01064 | 0.003581 | 0.013291 | 0.010096 | 0.005744 | 0.007521 | 0.007993 | 0.001 | 0.001644 | 0.001404 | 0.001117 | 0.015793 | 0.000679 | 0.002079 | 0.00101 | 0.001338 | 0.008508 | 0.0003 | 0.001175 |
| 32.15277778 | 0.037261 | 0.016542 | 0.016149 | 0.011049 | 0.003026 | 0.006407 | 0.001309 | 0.011889 | 0.008122 | 0.003763 | 0.007037 | 0.008749 | 0.00116 | 0.004153 | 0.001052 | 0.001711 | 0.008086 | 0.000509 | 0.00068 | 0.00033 | 0.001767 | 0.008457 | 0.000746 | 0.001328 |
| 32.16666667 | 0.038332 | 0.009416 | 0.019499 | 0.01017 | 0.001807 | 0.002555 | 0.001957 | 0.008819 | 0.005044 | 0.005401 | 0.005057 | 0.010646 | 0.001528 | 0.006878 | 0.001124 | 0.001485 | 0.005382 | 0.000701 | 0.000518 | 0.000207 | 0.002534 | 0.006836 | 0.001622 | 0.00162 |
| 32.18055556 | 0.039643 | 0.008158 | 0.019532 | 0.0106 | 0.001255 | 0.002319 | 0.006349 | 0.005213 | 0.004354 | 0.009932 | 0.007538 | 0.013965 | 0.00147 | 0.007082 | 0.00109 | 0.000875 | 0.00709 | 0.001409 | 0.001462 | 0.000293 | 0.002775 | 0.004119 | 0.002001 | 0.001149 |
| 32.19444444 | 0.04296 | 0.015909 | 0.018343 | 0.012082 | 0.002673 | 0.003204 | 0.01272 | 0.002657 | 0.007456 | 0.013373 | 0.014 | 0.019734 | 0.001232 | 0.004729 | 0.000961 | 0.000962 | 0.008515 | 0.001865 | 0.003126 | 0.000636 | 0.001892 | 0.002809 | 0.00133 | 0.000629 |
| 32.20833333 | 0.043003 | 0.02502 | 0.017313 | 0.013797 | 0.005094 | 0.002392 | 0.01801 | 0.002333 | 0.011009 | 0.01404 | 0.01887 | 0.026337 | 0.001409 | 0.002094 | 0.000898 | 0.001324 | 0.009308 | 0.00165 | 0.004039 | 0.001424 | 0.001016 | 0.004055 | 0.000516 | 0.000617 |
| 32.22222222 | 0.035368 | 0.030272 | 0.016019 | 0.014607 | 0.006884 | 0.002063 | 0.020094 | 0.0043 | 0.013917 | 0.012613 | 0.023432 | 0.029742 | 0.002204 | 0.00105 | 0.000733 | 0.001075 | 0.010371 | 0.001048 | 0.003094 | 0.002117 | 0.001392 | 0.004546 | 0.000585 | 0.000572 |
| 32.23611111 | 0.025817 | 0.031739 | 0.016008 | 0.014039 | 0.007546 | 0.005136 | 0.020103 | 0.00799 | 0.01616 | 0.011469 | 0.030981 | 0.029421 | 0.003266 | 0.000919 | 0.000425 | 0.000638 | 0.010804 | 0.000523 | 0.002834 | 0.0025 | 0.00281 | 0.003085 | 0.001363 | 0.000536 |
| 32.25 | 0.017534 | 0.030231 | 0.018467 | 0.012632 | 0.006937 | 0.009097 | 0.020057 | 0.011992 | 0.015919 | 0.012286 | 0.034946 | 0.028434 | 0.004038 | 0.000527 | 0.000239 | 0.000575 | 0.009762 | 0.000532 | 0.005336 | 0.003007 | 0.002841 | 0.001843 | 0.002176 | 0.000722 |
| 32.26388889 | 0.009227 | 0.0286 | 0.020502 | 0.010957 | 0.005503 | 0.010264 | 0.019744 | 0.013731 | 0.012354 | 0.014675 | 0.02944 | 0.02701 | 0.004034 | 0.000379 | 0.000221 | 0.000765 | 0.007411 | 0.001099 | 0.007712 | 0.003949 | 0.001338 | 0.001787 | 0.002828 | 0.001009 |
| 32.27777778 | 0.00375 | 0.02751 | 0.018996 | 0.009437 | 0.004009 | 0.009102 | 0.018846 | 0.012739 | 0.008712 | 0.017764 | 0.016764 | 0.024509 | 0.003165 | 0.001118 | 0.000193 | 0.001508 | 0.004291 | 0.001684 | 0.007389 | 0.005325 | 0.00067 | 0.002053 | 0.003221 | 0.00153 |
| 32.29166667 | 0.003274 | 0.025356 | 0.014171 | 0.008314 | 0.00285 | 0.008386 | 0.017492 | 0.01022 | 0.008 | 0.020322 | 0.007707 | 0.021489 | 0.001899 | 0.002442 | 0.000207 | 0.002703 | 0.001774 | 0.001738 | 0.005153 | 0.006288 | 0.001742 | 0.001947 | 0.003255 | 0.002098 |
| 32.30555556 | 0.007365 | 0.022581 | 0.008713 | 0.007755 | 0.002154 | 0.010174 | 0.016801 | 0.006552 | 0.008412 | 0.021456 | 0.009364 | 0.017619 | 0.001051 | 0.004175 | 0.000373 | 0.003278 | 0.0012 | 0.001452 | 0.002492 | 0.00569 | 0.00339 | 0.001687 | 0.003003 | 0.002395 |
| 32.31944444 | 0.012584 | 0.019067 | 0.005591 | 0.007867 | 0.001612 | 0.014481 | 0.018365 | 0.006052 | 0.00825 | 0.021084 | 0.010246 | 0.012707 | 0.001556 | 0.006314 | 0.000577 | 0.002691 | 0.001669 | 0.001457 | 0.000738 | 0.003697 | 0.003765 | 0.001499 | 0.002249 | 0.002422 |
| 32.33333333 | 0.016326 | 0.014429 | 0.005978 | 0.008267 | 0.001088 | 0.020159 | 0.020589 | 0.010423 | 0.006899 | 0.019255 | 0.007606 | 0.007283 | 0.003502 | 0.00826 | 0.000765 | 0.001542 | 0.001646 | 0.001787 | 0.00035 | 0.001683 | 0.003529 | 0.002579 | 0.00112 | 0.002083 |
| 32.34722222 | 0.017699 | 0.009959 | 0.008204 | 0.008397 | 0.001018 | 0.025023 | 0.019695 | 0.012513 | 0.004638 | 0.016441 | 0.010011 | 0.003331 | 0.005748 | 0.009279 | 0.000828 | 0.001142 | 0.001283 | 0.001908 | 0.000678 | 0.001191 | 0.003586 | 0.005343 | 0.000344 | 0.001348 |
| 32.36111111 | 0.016816 | 0.006325 | 0.009264 | 0.007331 | 0.0012 | 0.026674 | 0.014798 | 0.010259 | 0.003065 | 0.013293 | 0.013872 | 0.00268 | 0.007126 | 0.009571 | 0.000614 | 0.00188 | 0.002012 | 0.001689 | 0.001053 | 0.002015 | 0.002872 | 0.007476 | 0.000472 | 0.000568 |
| 32.375 | 0.014115 | 0.003978 | 0.008078 | 0.004369 | 0.000933 | 0.024206 | 0.008319 | 0.008988 | 0.002686 | 0.009665 | 0.014686 | 0.003806 | 0.007217 | 0.009506 | 0.000575 | 0.002804 | 0.004021 | 0.001611 | 0.000937 | 0.00244 | 0.001995 | 0.007254 | 0.001596 | 0.000175 |
| 32.38888889 | 0.010076 | 0.003776 | 0.005924 | 0.001674 | 0.000429 | 0.018935 | 0.003981 | 0.010854 | 0.003453 | 0.005402 | 0.013462 | 0.003864 | 0.006409 | 0.008462 | 0.001049 | 0.003185 | 0.005503 | 0.001905 | 0.000724 | 0.001797 | 0.00209 | 0.004609 | 0.003161 | 0.000185 |
| 32.40277778 | 0.007854 | 0.007103 | 0.005423 | 0.001436 | 0.000312 | 0.014814 | 0.003325 | 0.010965 | 0.005851 | 0.002092 | 0.014574 | 0.002385 | 0.005319 | 0.006073 | 0.001499 | 0.002894 | 0.005405 | 0.002074 | 0.000884 | 0.000879 | 0.002139 | 0.002444 | 0.003869 | 0.000209 |
| 32.41666667 | 0.01027 | 0.014208 | 0.008482 | 0.001611 | 0.000322 | 0.01427 | 0.003018 | 0.008875 | 0.010307 | 0.000798 | 0.022433 | 0.001168 | 0.004347 | 0.003575 | 0.001629 | 0.002271 | 0.00389 | 0.001763 | 0.000845 | 0.000952 | 0.001989 | 0.003621 | 0.00291 | 0.000172 |
| 32.43055556 | 0.016566 | 0.021029 | 0.013393 | 0.001217 | 0.000521 | 0.015154 | 0.00305 | 0.011506 | 0.016411 | 0.000687 | 0.034125 | 0.001317 | 0.003987 | 0.002984 | 0.001696 | 0.001563 | 0.00247 | 0.001435 | 0.001175 | 0.002304 | 0.002754 | 0.004968 | 0.001584 | 0.000179 |
| 32.44444444 | 0.024167 | 0.023969 | 0.017926 | 0.00109 | 0.001883 | 0.014515 | 0.00596 | 0.016067 | 0.020066 | 0.001297 | 0.035334 | 0.001437 | 0.004477 | 0.005109 | 0.001643 | 0.001104 | 0.003614 | 0.001681 | 0.002948 | 0.003813 | 0.004019 | 0.004163 | 0.001514 | 0.000294 |
| 32.45833333 | 0.030814 | 0.023078 | 0.021745 | 0.000626 | 0.00402 | 0.011938 | 0.009563 | 0.015384 | 0.016601 | 0.002165 | 0.02237 | 0.001383 | 0.005406 | 0.008915 | 0.001423 | 0.001709 | 0.006781 | 0.002552 | 0.005413 | 0.004747 | 0.004981 | 0.005072 | 0.002343 | 0.000951 |
| 32.47222222 | 0.035882 | 0.018364 | 0.024318 | 0.001437 | 0.004837 | 0.010236 | 0.01454 | 0.010997 | 0.011562 | 0.005334 | 0.014808 | 0.002058 | 0.006078 | 0.012755 | 0.001603 | 0.002821 | 0.009109 | 0.00361 | 0.007039 | 0.005198 | 0.005576 | 0.009285 | 0.003236 | 0.001844 |
| 32.48611111 | 0.040544 | 0.012775 | 0.02602 | 0.007409 | 0.003706 | 0.012659 | 0.025716 | 0.010574 | 0.016251 | 0.01487 | 0.017569 | 0.003429 | 0.005951 | 0.015231 | 0.002476 | 0.003058 | 0.009705 | 0.004326 | 0.007495 | 0.005212 | 0.006296 | 0.0132 | 0.004362 | 0.001731 |
| 32.5 | 0.045707 | 0.016451 | 0.027116 | 0.019398 | 0.003539 | 0.017766 | 0.04751 | 0.021196 | 0.030314 | 0.026233 | 0.024303 | 0.010283 | 0.005266 | 0.015536 | 0.003325 | 0.002426 | 0.010148 | 0.004569 | 0.007988 | 0.004584 | 0.009263 | 0.015275 | 0.006129 | 0.00156 |
| 32.51388889 | 0.053012 | 0.032312 | 0.026689 | 0.030645 | 0.004455 | 0.019103 | 0.075327 | 0.040726 | 0.038093 | 0.031039 | 0.044381 | 0.027363 | 0.005538 | 0.015189 | 0.003136 | 0.002017 | 0.012269 | 0.004382 | 0.010069 | 0.003028 | 0.014721 | 0.017364 | 0.008598 | 0.0026 |
| 32.52777778 | 0.061559 | 0.049265 | 0.027992 | 0.033893 | 0.003289 | 0.013369 | 0.093319 | 0.060742 | 0.03073 | 0.033191 | 0.074142 | 0.051517 | 0.007793 | 0.018241 | 0.001945 | 0.002067 | 0.015934 | 0.003456 | 0.014284 | 0.002109 | 0.017538 | 0.020271 | 0.010779 | 0.002795 |
| 32.54166667 | 0.065187 | 0.068342 | 0.032381 | 0.03221 | 0.002052 | 0.005805 | 0.100595 | 0.074364 | 0.019381 | 0.042658 | 0.083894 | 0.068074 | 0.010364 | 0.026469 | 0.001457 | 0.002275 | 0.018746 | 0.002151 | 0.019674 | 0.002112 | 0.014594 | 0.022413 | 0.011102 | 0.001704 |
| 32.55555556 | 0.05939 | 0.099661 | 0.03216 | 0.029794 | 0.00293 | 0.003133 | 0.107716 | 0.076059 | 0.020164 | 0.063794 | 0.059147 | 0.065802 | 0.011209 | 0.034708 | 0.003308 | 0.00372 | 0.018843 | 0.00174 | 0.024035 | 0.002336 | 0.010177 | 0.02217 | 0.009953 | 0.001882 |
| 32.56944444 | 0.049525 | 0.117109 | 0.024033 | 0.024127 | 0.003586 | 0.004368 | 0.113527 | 0.067937 | 0.034042 | 0.08414 | 0.023669 | 0.055347 | 0.010413 | 0.035697 | 0.005968 | 0.006448 | 0.016748 | 0.00176 | 0.025145 | 0.004837 | 0.008158 | 0.018426 | 0.008835 | 0.003391 |
| 32.58333333 | 0.037306 | 0.099145 | 0.013442 | 0.01742 | 0.00356 | 0.005631 | 0.110768 | 0.054318 | 0.046843 | 0.087306 | 0.013729 | 0.044978 | 0.008637 | 0.029978 | 0.006363 | 0.008315 | 0.013158 | 0.001741 | 0.022615 | 0.009044 | 0.007072 | 0.011001 | 0.007358 | 0.003866 |
| 32.59722222 | 0.023427 | 0.078586 | 0.009435 | 0.016482 | 0.003645 | 0.007089 | 0.115547 | 0.037792 | 0.05198 | 0.081119 | 0.029426 | 0.033301 | 0.006426 | 0.022646 | 0.004468 | 0.008312 | 0.008569 | 0.002914 | 0.017816 | 0.012526 | 0.005064 | 0.005112 | 0.004448 | 0.002724 |
| 32.61111111 | 0.03129 | 0.072411 | 0.010535 | 0.019186 | 0.00261 | 0.011384 | 0.137616 | 0.032403 | 0.049411 | 0.072585 | 0.034218 | 0.022815 | 0.004727 | 0.014532 | 0.002242 | 0.006846 | 0.004192 | 0.003947 | 0.011475 | 0.014268 | 0.002586 | 0.005756 | 0.001668 | 0.001322 |
| 32.625 | 0.059571 | 0.065859 | 0.007029 | 0.020613 | 0.001715 | 0.017651 | 0.131628 | 0.043978 | 0.038314 | 0.053339 | 0.021209 | 0.01384 | 0.004195 | 0.007487 | 0.001291 | 0.004879 | 0.002169 | 0.003102 | 0.006351 | 0.013838 | 0.000903 | 0.009971 | 0.000704 | 0.000583 |
| 32.63888889 | 0.072601 | 0.056704 | 0.002676 | 0.021304 | 0.002591 | 0.019774 | 0.084726 | 0.051265 | 0.023869 | 0.030068 | 0.013739 | 0.006291 | 0.004193 | 0.007137 | 0.002192 | 0.003501 | 0.003533 | 0.001527 | 0.007588 | 0.01249 | 0.000627 | 0.013274 | 0.000647 | 0.000451 |
| 32.65277778 | 0.064865 | 0.047805 | 0.001802 | 0.019386 | 0.003226 | 0.017239 | 0.042234 | 0.039734 | 0.01127 | 0.027303 | 0.016071 | 0.003736 | 0.003601 | 0.01371 | 0.003733 | 0.002139 | 0.006396 | 0.00045 | 0.013771 | 0.011258 | 0.000893 | 0.014282 | 0.0008 | 0.000833 |
| 32.66666667 | 0.049909 | 0.039415 | 0.002508 | 0.012602 | 0.00249 | 0.013205 | 0.041458 | 0.021706 | 0.00391 | 0.048741 | 0.024122 | 0.008368 | 0.002733 | 0.020312 | 0.004664 | 0.000642 | 0.009327 | 0.000697 | 0.016697 | 0.010071 | 0.001884 | 0.013546 | 0.001701 | 0.001425 |
| 32.68055556 | 0.032782 | 0.032954 | 0.003617 | 0.006184 | 0.002383 | 0.008063 | 0.074767 | 0.010404 | 0.004209 | 0.066834 | 0.03944 | 0.017551 | 0.002927 | 0.022818 | 0.004796 | 0.000287 | 0.011583 | 0.002429 | 0.0119 | 0.009256 | 0.004807 | 0.011655 | 0.003561 | 0.001277 |
| 32.69444444 | 0.020026 | 0.026954 | 0.004505 | 0.006563 | 0.004355 | 0.003809 | 0.096701 | 0.007336 | 0.009783 | 0.066062 | 0.053831 | 0.0265 | 0.004071 | 0.022564 | 0.004395 | 0.000956 | 0.011966 | 0.004138 | 0.005703 | 0.009167 | 0.008315 | 0.008689 | 0.00488 | 0.000528 |

| | | | | | | | | | | | | | | | | | | | | | | | | |
|---|---|---|---|---|---|---|---|---|---|---|---|---|---|---|---|---|---|---|---|---|---|---|---|---|
| 32.70833333 | 0.021699 | 0.016963 | 0.005268 | 0.010811 | 0.006559 | 0.002548 | 0.09409 | 0.010225 | 0.01509 | 0.059676 | 0.056207 | 0.030036 | 0.004496 | 0.0195 | 0.003911 | 0.001853 | 0.009928 | 0.00482 | 0.004748 | 0.009718 | 0.010232 | 0.004848 | 0.003859 | 0.000451 |
| 32.72222222 | 0.032306 | 0.007305 | 0.005423 | 0.011448 | 0.007166 | 0.003245 | 0.078357 | 0.016726 | 0.015849 | 0.056321 | 0.044762 | 0.026009 | 0.003537 | 0.012376 | 0.003501 | 0.002844 | 0.005848 | 0.004878 | 0.005041 | 0.01038 | 0.010116 | 0.002919 | 0.002104 | 0.001777 |
| 32.73611111 | 0.036136 | 0.007122 | 0.005025 | 0.007986 | 0.006745 | 0.002977 | 0.061484 | 0.023587 | 0.011233 | 0.051702 | 0.026007 | 0.019148 | 0.002017 | 0.007032 | 0.002848 | 0.003644 | 0.003633 | 0.004578 | 0.00322 | 0.010528 | 0.008357 | 0.00583 | 0.00221 | 0.004292 |
| 32.75 | 0.03462 | 0.015973 | 0.00466 | 0.006258 | 0.007085 | 0.001995 | 0.057619 | 0.026878 | 0.005177 | 0.048046 | 0.017119 | 0.015924 | 0.000796 | 0.010498 | 0.001795 | 0.003596 | 0.006253 | 0.003794 | 0.00172 | 0.010037 | 0.005697 | 0.01027 | 0.00322 | 0.007432 |
| 32.76388889 | 0.033014 | 0.031287 | 0.005246 | 0.008774 | 0.008929 | 0.001988 | 0.065531 | 0.026542 | 0.00292 | 0.048245 | 0.017025 | 0.018279 | 0.000317 | 0.01677 | 0.001417 | 0.002497 | 0.009048 | 0.002372 | 0.001155 | 0.009288 | 0.003355 | 0.01221 | 0.003597 | 0.010351 |
| 32.77777778 | 0.031917 | 0.048189 | 0.00618 | 0.012659 | 0.011287 | 0.002286 | 0.078017 | 0.024739 | 0.003819 | 0.051048 | 0.017447 | 0.023153 | 0.000536 | 0.014979 | 0.002301 | 0.002527 | 0.007523 | 0.001289 | 0.001909 | 0.008505 | 0.002169 | 0.010317 | 0.003139 | 0.011907 |
| 32.79166667 | 0.031292 | 0.057301 | 0.006325 | 0.015695 | 0.014176 | 0.003632 | 0.089911 | 0.020578 | 0.003945 | 0.054188 | 0.025706 | 0.025726 | 0.000949 | 0.010072 | 0.002852 | 0.005438 | 0.004299 | 0.001608 | 0.006029 | 0.007654 | 0.001944 | 0.006608 | 0.001909 | 0.012295 |
| 32.80555556 | 0.028428 | 0.058053 | 0.007107 | 0.018209 | 0.019063 | 0.005 | 0.102052 | 0.014556 | 0.002813 | 0.053123 | 0.034458 | 0.023511 | 0.001414 | 0.015304 | 0.002186 | 0.008838 | 0.004053 | 0.002595 | 0.012427 | 0.006812 | 0.001805 | 0.007645 | 0.000853 | 0.012829 |
| 32.81944444 | 0.02628 | 0.054431 | 0.010127 | 0.020697 | 0.024711 | 0.003565 | 0.121207 | 0.012279 | 0.002822 | 0.046759 | 0.035789 | 0.017724 | 0.002686 | 0.026301 | 0.001072 | 0.010948 | 0.005366 | 0.003617 | 0.016985 | 0.005748 | 0.001454 | 0.013607 | 0.000954 | 0.01347 |
| 32.83333333 | 0.027778 | 0.047938 | 0.016701 | 0.022189 | 0.025624 | 0.002077 | 0.138443 | 0.015858 | 0.003144 | 0.039961 | 0.03546 | 0.012597 | 0.005121 | 0.031312 | 0.000352 | 0.012164 | 0.004583 | 0.004623 | 0.016959 | 0.004131 | 0.001928 | 0.016968 | 0.001635 | 0.012939 |
| 32.84722222 | 0.030095 | 0.045568 | 0.026853 | 0.022297 | 0.020208 | 0.004031 | 0.143903 | 0.019294 | 0.003372 | 0.037186 | 0.038073 | 0.01299 | 0.00752 | 0.028681 | 0.000252 | 0.012194 | 0.005049 | 0.004905 | 0.013299 | 0.002232 | 0.003396 | 0.014719 | 0.001699 | 0.010938 |
| 32.86111111 | 0.029955 | 0.056019 | 0.036834 | 0.021692 | 0.013803 | 0.007747 | 0.150939 | 0.017395 | 0.008003 | 0.039458 | 0.042791 | 0.019141 | 0.008678 | 0.022251 | 0.000612 | 0.011008 | 0.008451 | 0.004296 | 0.009053 | 0.001423 | 0.004997 | 0.009765 | 0.001519 | 0.00883 |
| 32.875 | 0.023112 | 0.073649 | 0.043728 | 0.02139 | 0.010057 | 0.011287 | 0.1647 | 0.010961 | 0.015531 | 0.044817 | 0.043416 | 0.026487 | 0.008873 | 0.016476 | 0.001819 | 0.009055 | 0.008872 | 0.00403 | 0.006814 | 0.003015 | 0.00718 | 0.005662 | 0.00265 | 0.0078 |
| 32.88888889 | 0.012491 | 0.081124 | 0.04458 | 0.021547 | 0.008467 | 0.013771 | 0.177218 | 0.004472 | 0.016675 | 0.045573 | 0.038082 | 0.030891 | 0.008889 | 0.013755 | 0.003752 | 0.006686 | 0.005277 | 0.004832 | 0.00756 | 0.006358 | 0.01048 | 0.003593 | 0.005636 | 0.007908 |
| 32.90277778 | 0.006001 | 0.071321 | 0.038132 | 0.019884 | 0.007132 | 0.014045 | 0.181055 | 0.002446 | 0.009965 | 0.036549 | 0.026758 | 0.031423 | 0.008809 | 0.013018 | 0.005093 | 0.004689 | 0.003694 | 0.005845 | 0.010213 | 0.009884 | 0.014221 | 0.002821 | 0.009246 | 0.008859 |
| 32.91666667 | 0.004607 | 0.054823 | 0.02656 | 0.016153 | 0.004586 | 0.012511 | 0.156743 | 0.00451 | 0.004251 | 0.022895 | 0.014054 | 0.02857 | 0.008179 | 0.01145 | 0.005344 | 0.003712 | 0.005374 | 0.005858 | 0.011952 | 0.012464 | 0.017478 | 0.002677 | 0.011805 | 0.010154 |
| 32.93055556 | 0.007251 | 0.041424 | 0.013133 | 0.012928 | 0.002487 | 0.009821 | 0.106255 | 0.006546 | 0.00255 | 0.011346 | 0.016568 | 0.022805 | 0.006694 | 0.007807 | 0.004937 | 0.003371 | 0.005917 | 0.004696 | 0.01034 | 0.013284 | 0.01934 | 0.002841 | 0.012597 | 0.010678 |
| 32.94444444 | 0.012735 | 0.028992 | 0.005372 | 0.010392 | 0.002803 | 0.006312 | 0.064174 | 0.005152 | 0.002427 | 0.004256 | 0.03239 | 0.015205 | 0.004445 | 0.003918 | 0.003799 | 0.002952 | 0.003843 | 0.003208 | 0.006117 | 0.012059 | 0.019394 | 0.003306 | 0.011853 | 0.0093 |
| 32.95833333 | 0.017937 | 0.015239 | 0.003959 | 0.006675 | 0.003052 | 0.003829 | 0.038933 | 0.003602 | 0.006012 | 0.001607 | 0.044404 | 0.007495 | 0.002228 | 0.002891 | 0.002119 | 0.001967 | 0.001423 | 0.001969 | 0.002209 | 0.00943 | 0.018061 | 0.003796 | 0.010134 | 0.006557 |
| 32.97222222 | 0.021719 | 0.006091 | 0.004984 | 0.003046 | 0.003129 | 0.003971 | 0.018234 | 0.005376 | 0.011402 | 0.001893 | 0.046845 | 0.002786 | 0.001593 | 0.004284 | 0.001165 | 0.001482 | 0.000426 | 0.001024 | 0.000701 | 0.00638 | 0.015713 | 0.003345 | 0.007943 | 0.003775 |
| 32.98611111 | 0.019073 | 0.00488 | 0.012347 | 0.005147 | 0.00762 | 0.00661 | 0.004876 | 0.008979 | 0.010174 | 0.008532 | 0.03283 | 0.003806 | 0.003026 | 0.005228 | 0.002387 | 0.002793 | 0.00069 | 0.000661 | 0.001681 | 0.003467 | 0.013773 | 0.001866 | 0.005756 | 0.001667 |
| 33 | 0.011705 | 0.009538 | 0.02659 | 0.015157 | 0.014302 | 0.008753 | 0.003899 | 0.020727 | 0.009122 | 0.027648 | 0.021838 | 0.009512 | 0.004814 | 0.004651 | 0.004957 | 0.003802 | 0.002501 | 0.001933 | 0.004392 | 0.001257 | 0.017071 | 0.001732 | 0.00456 | 0.001158 |
| 33.01388889 | 0.009666 | 0.018136 | 0.040811 | 0.023567 | 0.017895 | 0.006193 | 0.014335 | 0.043517 | 0.023928 | 0.049113 | 0.030545 | 0.017407 | 0.004347 | 0.006286 | 0.006107 | 0.002636 | 0.008855 | 0.005694 | 0.008537 | 0.000408 | 0.030294 | 0.006875 | 0.005577 | 0.003428 |
| 33.02777778 | 0.011327 | 0.026038 | 0.051137 | 0.020952 | 0.019688 | 0.002487 | 0.023396 | 0.064168 | 0.045463 | 0.052603 | 0.099161 | 0.026941 | 0.003339 | 0.012635 | 0.005347 | 0.001369 | 0.01974 | 0.009792 | 0.014249 | 0.001876 | 0.045634 | 0.017346 | 0.008676 | 0.007505 |
| 33.04166667 | 0.015811 | 0.033197 | 0.058945 | 0.01143 | 0.020405 | 0.001961 | 0.017433 | 0.068107 | 0.05535 | 0.03425 | 0.194291 | 0.03312 | 0.006485 | 0.017471 | 0.003615 | 0.002401 | 0.028242 | 0.010022 | 0.020117 | 0.00746 | 0.046088 | 0.025391 | 0.011748 | 0.010219 |
| 33.05555556 | 0.022625 | 0.042865 | 0.05838 | 0.005475 | 0.01622 | 0.0026 | 0.006917 | 0.051592 | 0.049284 | 0.0134 | 0.17365 | 0.027895 | 0.012271 | 0.016781 | 0.00262 | 0.005362 | 0.029213 | 0.006572 | 0.02228 | 0.015944 | 0.030678 | 0.025045 | 0.013354 | 0.010523 |
| 33.06944444 | 0.025585 | 0.052348 | 0.042184 | 0.007076 | 0.010675 | 0.005708 | 0.004946 | 0.026624 | 0.030928 | 0.005356 | 0.086405 | 0.014517 | 0.015052 | 0.013112 | 0.003029 | 0.007417 | 0.024128 | 0.003547 | 0.018136 | 0.020806 | 0.014936 | 0.019248 | 0.012639 | 0.008859 |
| 33.08333333 | 0.022675 | 0.056133 | 0.022016 | 0.010131 | 0.012374 | 0.011233 | 0.007114 | 0.014327 | 0.012755 | 0.00345 | 0.036521 | 0.007408 | 0.011942 | 0.007904 | 0.003195 | 0.005624 | 0.015721 | 0.00212 | 0.009774 | 0.01711 | 0.006165 | 0.011518 | 0.008574 | 0.005251 |
| 33.09722222 | 0.015055 | 0.052639 | 0.016306 | 0.00993 | 0.01526 | 0.013675 | 0.007539 | 0.018485 | 0.005867 | 0.002602 | 0.014415 | 0.012529 | 0.005673 | 0.003075 | 0.004038 | 0.003454 | 0.007254 | 0.001145 | 0.003488 | 0.008575 | 0.003379 | 0.004909 | 0.005901 | 0.002433 |
| 33.11111111 | 0.006548 | 0.046187 | 0.021157 | 0.008224 | 0.010748 | 0.012099 | 0.006362 | 0.023371 | 0.006085 | 0.005377 | 0.01058 | 0.021582 | 0.001953 | 0.000877 | 0.004963 | 0.005258 | 0.00368 | 0.000417 | 0.003065 | 0.003273 | 0.005978 | 0.00301 | 0.007516 | 0.002142 |
| 33.125 | 0.002469 | 0.04098 | 0.018145 | 0.007164 | 0.006518 | 0.010052 | 0.004495 | 0.021364 | 0.004745 | 0.009325 | 0.012427 | 0.028708 | 0.001693 | 0.000976 | 0.003965 | 0.006887 | 0.005933 | 0.000164 | 0.005912 | 0.00252 | 0.009064 | 0.003155 | 0.007851 | 0.001827 |
| 33.13888889 | 0.00322 | 0.037923 | 0.007515 | 0.007568 | 0.007915 | 0.009391 | 0.002596 | 0.014985 | 0.002102 | 0.011135 | 0.013415 | 0.029649 | 0.001757 | 0.002603 | 0.002691 | 0.005023 | 0.009385 | 0.000178 | 0.008852 | 0.00195 | 0.008941 | 0.00217 | 0.005497 | 0.001742 |
| 33.15277778 | 0.006557 | 0.035159 | 0.003924 | 0.009146 | 0.008363 | 0.010789 | 0.001315 | 0.007649 | 0.001174 | 0.010226 | 0.015933 | 0.022069 | 0.001031 | 0.004853 | 0.002729 | 0.002336 | 0.010458 | 0.000139 | 0.01004 | 0.00127 | 0.006643 | 0.001597 | 0.002865 | 0.003644 |
| 33.16666667 | 0.01061 | 0.030704 | 0.010765 | 0.010951 | 0.005599 | 0.014227 | 0.000883 | 0.005094 | 0.001568 | 0.007582 | 0.022882 | 0.01373 | 0.000727 | 0.006367 | 0.003578 | 0.001307 | 0.008762 | 7.9263895 | 0.008879 | 0.001942 | 0.003403 | 0.001698 | 0.001313 | 0.005069 |
| 33.18055556 | 0.010071 | 0.025412 | 0.019691 | 0.012227 | 0.003254 | 0.017873 | 0.001651 | 0.008605 | 0.003007 | 0.00476 | 0.032359 | 0.009807 | 0.001788 | 0.006114 | 0.003944 | 0.001082 | 0.005716 | 8.4215246 | 0.006857 | 0.002364 | 0.001477 | 0.001214 | 0.001643 | 0.004141 |
| 33.19444444 | 0.004974 | 0.020593 | 0.024694 | 0.012672 | 0.0043 | 0.019229 | 0.003552 | 0.010352 | 0.005386 | 0.002423 | 0.031014 | 0.008332 | 0.003523 | 0.003997 | 0.003124 | 0.000837 | 0.004169 | 0.000289 | 0.004968 | 0.00167 | 0.002261 | 0.000807 | 0.002997 | 0.002415 |
| 33.20833333 | 0.003295 | 0.015971 | 0.026567 | 0.012262 | 0.008924 | 0.016466 | 0.005754 | 0.007569 | 0.00749 | 0.001331 | 0.018933 | 0.00758 | 0.004533 | 0.001674 | 0.001657 | 0.001325 | 0.004228 | 0.001058 | 0.00306 | 0.000967 | 0.003808 | 0.001246 | 0.005173 | 0.001362 |
| 33.22222222 | 0.009109 | 0.011688 | 0.026335 | 0.011056 | 0.012584 | 0.009876 | 0.006488 | 0.007615 | 0.007939 | 0.001323 | 0.011277 | 0.007038 | 0.003902 | 0.000709 | 0.000888 | 0.001985 | 0.003115 | 0.002186 | 0.00134 | 0.000741 | 0.004508 | 0.001722 | 0.007324 | 0.000964 |
| 33.23611111 | 0.016453 | 0.008636 | 0.023072 | 0.0092 | 0.010956 | 0.005899 | 0.004852 | 0.013482 | 0.00631 | 0.000937 | 0.008248 | 0.006641 | 0.002368 | 0.000529 | 0.001124 | 0.002103 | 0.001931 | 0.002569 | 0.000429 | 0.000514 | 0.004464 | 0.001661 | 0.007338 | 0.000912 |
| 33.25 | 0.018709 | 0.007188 | 0.015385 | 0.007384 | 0.006864 | 0.006752 | 0.00403 | 0.019871 | 0.00443 | 0.000382 | 0.005956 | 0.006706 | 0.002083 | 0.000505 | 0.001218 | 0.001657 | 0.003266 | 0.001931 | 0.000476 | 0.000408 | 0.004036 | 0.001254 | 0.004669 | 0.000997 |
| 33.26388889 | 0.017205 | 0.007302 | 0.009883 | 0.00619 | 0.006924 | 0.005655 | 0.004193 | 0.022413 | 0.004621 | 0.000295 | 0.005048 | 0.007342 | 0.003083 | 0.001357 | 0.000895 | 0.000983 | 0.005444 | 0.001149 | 0.001836 | 0.0006 | 0.003822 | 0.001661 | 0.002516 | 0.000827 |
| 33.27777778 | 0.015748 | 0.008597 | 0.016001 | 0.005422 | 0.009637 | 0.004265 | 0.003775 | 0.02134 | 0.00712 | 0.001277 | 0.003507 | 0.008174 | 0.00353 | 0.00329 | 0.000765 | 0.00073 | 0.00675 | 0.000791 | 0.004749 | 0.000872 | 0.004218 | 0.003572 | 0.003807 | 0.000464 |
| 33.29166667 | 0.015043 | 0.009907 | 0.023685 | 0.004852 | 0.008435 | 0.00604 | 0.005971 | 0.01923 | 0.009399 | 0.003872 | 0.001615 | 0.008396 | 0.003194 | 0.005633 | 0.001256 | 0.000591 | 0.007418 | 0.000806 | 0.007833 | 0.001768 | 0.004915 | 0.005236 | 0.005685 | 0.000366 |
| 33.30555556 | 0.013627 | 0.010065 | 0.017712 | 0.004417 | 0.004403 | 0.00724 | 0.008723 | 0.017327 | 0.009649 | 0.005906 | 0.001084 | 0.007217 | 0.00303 | 0.007331 | 0.002657 | 0.000481 | 0.007726 | 0.000965 | 0.009671 | 0.003357 | 0.004945 | 0.005093 | 0.00503 | 0.000403 |
| 33.31944444 | 0.011772 | 0.009414 | 0.009363 | 0.003901 | 0.002295 | 0.007163 | 0.008155 | 0.015408 | 0.009239 | 0.005391 | 0.00099 | 0.004629 | 0.003141 | 0.007228 | 0.004647 | 0.001379 | 0.008015 | 0.001127 | 0.009844 | 0.004783 | 0.003876 | 0.003893 | 0.00287 | 0.000273 |
| 33.33333333 | 0.010035 | 0.00956 | 0.012835 | 0.003109 | 0.002145 | 0.007443 | 0.006536 | 0.014335 | 0.007963 | 0.003834 | 0.00068 | 0.002333 | 0.003028 | 0.005493 | 0.006393 | 0.003228 | 0.007832 | 0.001076 | 0.008213 | 0.00533 | 0.002119 | 0.003307 | 0.001432 | 0.000291 |
| 33.34722222 | 0.007379 | 0.011043 | 0.014097 | 0.001968 | 0.003871 | 0.007784 | 0.006031 | 0.014051 | 0.004745 | 0.003536 | 0.000541 | 0.00251 | 0.003041 | 0.003678 | 0.00657 | 0.004946 | 0.006453 | 0.000776 | 0.005327 | 0.00458 | 0.001127 | 0.003749 | 0.001329 | 0.000903 |
| 33.36111111 | 0.00443 | 0.012013 | 0.008309 | 0.001159 | 0.007728 | 0.007757 | 0.006219 | 0.013609 | 0.002249 | 0.00407 | 0.00055 | 0.003981 | 0.003687 | 0.002458 | 0.004231 | 0.006208 | 0.004847 | 0.000755 | 0.002402 | 0.003231 | 0.001489 | 0.004336 | 0.001966 | 0.001433 |
| 33.375 | 0.003825 | 0.011248 | 0.005588 | 0.001159 | 0.010146 | 0.008718 | 0.008323 | 0.013787 | 0.001618 | 0.003476 | 0.001347 | 0.003834 | 0.00425 | 0.00131 | 0.002081 | 0.007153 | 0.004655 | 0.001051 | 0.00107 | 0.00282 | 0.001314 | 0.004483 | 0.002563 | 0.001128 |
| 33.38888889 | 0.005885 | 0.010238 | 0.006015 | 0.002099 | 0.008598 | 0.011781 | 0.013421 | 0.015575 | 0.001489 | 0.003361 | 0.002911 | 0.00244 | 0.004189 | 0.000922 | 0.00294 | 0.0072 | 0.005593 | 0.0012 | 0.002069 | 0.003357 | 0.000513 | 0.003344 | 0.002292 | 0.00064 |
| 33.40277778 | 0.00774 | 0.010324 | 0.01166 | 0.004348 | 0.00586 | 0.015344 | 0.018401 | 0.018759 | 0.003268 | 0.005846 | 0.004817 | 0.002717 | 0.004086 | 0.001842 | 0.00447 | 0.005875 | 0.006361 | 0.001355 | 0.00383 | 0.003499 | 0.000485 | 0.002081 | 0.001665 | 0.000508 |
| 33.41666667 | 0.007367 | 0.010742 | 0.020085 | 0.006729 | 0.007453 | 0.017288 | 0.019602 | 0.023047 | 0.006772 | 0.008007 | 0.008097 | 0.005031 | 0.00412 | 0.00223 | 0.004115 | 0.004143 | 0.006364 | 0.001416 | 0.004504 | 0.002937 | 0.000771 | 0.003066 | 0.001171 | 0.000571 |
| 33.43055556 | 0.005723 | 0.011417 | 0.024056 | 0.00798 | 0.012488 | 0.017018 | 0.016117 | 0.027099 | 0.008828 | 0.007413 | 0.013027 | 0.006553 | 0.003858 | 0.001508 | 0.002361 | 0.00267 | 0.005496 | 0.001071 | 0.004394 | 0.002115 | 0.001174 | 0.005005 | 0.000976 | 0.000465 |
| 33.44444444 | 0.003977 | 0.014178 | 0.022958 | 0.007922 | 0.0127 | 0.014859 | 0.010622 | 0.028732 | 0.00763 | 0.004551 | 0.017798 | 0.005185 | 0.00361 | 0.001535 | 0.001031 | 0.001529 | 0.003964 | 0.00073 | 0.004735 | 0.001256 | 0.002378 | 0.006275 | 0.001777 | 0.000224 |
| 33.45833333 | 0.002282 | 0.01881 | 0.019583 | 0.006719 | 0.008174 | 0.012274 | 0.006751 | 0.027568 | 0.004486 | 0.001887 | 0.021611 | 0.002946 | 0.003895 | 0.002972 | 0.001059 | 0.001459 | 0.002188 | 0.000888 | 0.005498 | 0.000559 | 0.003865 | 0.006305 | 0.003017 | 0.000213 |
| 33.47222222 | 0.002697 | 0.020755 | 0.016667 | 0.005772 | 0.00737 | 0.011515 | 0.006743 | 0.024984 | 0.003282 | 0.002741 | 0.027913 | 0.001997 | 0.004478 | 0.004422 | 0.00142 | 0.002377 | 0.0012 | 0.001682 | 0.006275 | 0.000288 | 0.004608 | 0.004953 | 0.003202 | 0.00034 |
| 33.48611111 | 0.009864 | 0.016116 | 0.0151 | 0.00745 | 0.008152 | 0.013729 | 0.010759 | 0.021467 | 0.00662 | 0.008815 | 0.040863 | 0.003682 | 0.004763 | 0.004842 | 0.002784 | 0.003726 | 0.001723 | 0.003024 | 0.00751 | 0.000291 | 0.005579 | 0.004027 | 0.002214 | 0.000568 |
| 33.5 | 0.024703 | 0.016454 | 0.016454 | 0.011624 | 0.007194 | 0.017032 | 0.015702 | 0.017461 | 0.0122 | 0.016782 | 0.055762 | 0.011184 | 0.00456 | 0.004774 | 0.006123 | 0.005809 | 0.002484 | 0.004631 | 0.009508 | 0.000768 | 0.00942 | 0.004946 | 0.001815 | 0.001446 |
| 33.51388889 | 0.040046 | 0.029833 | 0.024167 | 0.015259 | 0.00793 | 0.017356 | 0.017504 | 0.017702 | 0.015707 | 0.019798 | 0.066754 | 0.023194 | 0.004912 | 0.004463 | 0.008223 | 0.007731 | 0.003 | 0.005516 | 0.01223 | 0.002095 | 0.015362 | 0.007602 | 0.004252 | 0.003079 |
| 33.52777778 | 0.048506 | 0.037449 | 0.040297 | 0.018331 | 0.007139 | 0.015118 | 0.015025 | 0.026477 | 0.018189 | 0.021059 | 0.075757 | 0.037199 | 0.006644 | 0.003663 | 0.00658 | 0.007758 | 0.005938 | 0.004293 | 0.016171 | 0.002755 | 0.017762 | 0.009535 | 0.009407 | 0.004569 |
| 33.54166667 | 0.050939 | 0.03207 | 0.059987 | 0.024366 | 0.005918 | 0.014921 | 0.0103 | 0.039128 | 0.020692 | 0.027119 | 0.090304 | 0.051517 | 0.008396 | 0.005615 | 0.003835 | 0.007728 | 0.013387 | 0.001823 | 0.021952 | 0.002042 | 0.015374 | 0.009365 | 0.013133 | 0.005373 |
| 33.55555556 | 0.049099 | 0.030236 | 0.070931 | 0.031769 | 0.006839 | 0.016764 | 0.006025 | 0.047769 | 0.018503 | 0.030201 | 0.131549 | 0.060506 | 0.00868 | 0.012402 | 0.003051 | 0.009849 | 0.022579 | 0.000688 | 0.027284 | 0.003308 | 0.013802 | 0.010373 | 0.012283 | 0.00615 |
| 33.56944444 | 0.039464 | 0.041409 | 0.06935 | 0.032195 | 0.007224 | 0.017738 | 0.005112 | 0.048265 | 0.012449 | 0.022896 | 0.192377 | 0.061681 | 0.007722 | 0.017558 | 0.003017 | 0.011334 | 0.028015 | 0.000714 | 0.028939 | 0.008525 | 0.014974 | 0.011844 | 0.00882 | 0.007198 |
| 33.58333333 | 0.025144 | 0.061206 | 0.063979 | 0.024043 | 0.008747 | 0.017223 | 0.013762 | 0.038332 | 0.009363 | 0.013693 | 0.211919 | 0.058913 | 0.005743 | 0.014701 | 0.002051 | 0.009722 | 0.027082 | 0.000757 | 0.026905 | 0.014344 | 0.015942 | 0.009074 | 0.005603 | 0.007544 |
| 33.59722222 | 0.015194 | 0.078312 | 0.058214 | 0.014126 | 0.009515 | 0.016755 | 0.031887 | 0.02143 | 0.009972 | 0.00989 | 0.161329 | 0.05359 | 0.002981 | 0.007383 | 0.001643 | 0.006987 | 0.021961 | 0.00118 | 0.022052 | 0.017821 | 0.015577 | 0.004246 | 0.002937 | 0.00689 |
| 33.61111111 | 0.012866 | 0.088735 | 0.049842 | 0.007959 | 0.008699 | 0.018064 | 0.045484 | 0.007984 | 0.011777 | 0.007992 | 0.080456 | 0.04291 | 0.001368 | 0.003957 | 0.002532 | 0.00558 | 0.016289 | 0.00163 | 0.014625 | 0.018434 | 0.013951 | 0.002971 | 0.002034 | 0.00535 |
| 33.625 | 0.015421 | 0.097946 | 0.039823 | 0.007301 | 0.007051 | 0.01979 | 0.046105 | 0.00359 | 0.015263 | 0.00585 | 0.03274 | 0.026596 | 0.001699 | 0.005906 | 0.003471 | 0.005555 | 0.011315 | 0.001788 | 0.007826 | 0.016827 | 0.010338 | 0.005677 | 0.0043 | 0.003258 |
| 33.63888889 | 0.01886 | 0.103786 | 0.030899 | 0.009498 | 0.003969 | 0.019519 | 0.041216 | 0.003444 | 0.019275 | 0.005477 | 0.075116 | 0.012505 | 0.001982 | 0.007895 | 0.003243 | 0.006627 | 0.006832 | 0.001471 | 0.0078 | 0.014814 | 0.005553 | 0.008848 | 0.006317 | 0.002835 |
| 33.65277778 | 0.022143 | 0.101656 | 0.02444 | 0.010411 | 0.002826 | 0.017691 | 0.040546 | 0.004373 | 0.020825 | 0.007507 | 0.160922 | 0.011384 | 0.001383 | 0.007471 | 0.002173 | 0.00965 | 0.003223 | 0.001533 | 0.014018 | 0.012908 | 0.001986 | 0.0097 | 0.00546 | 0.004808 |
| 33.66666667 | 0.025483 | 0.095233 | 0.018704 | 0.007447 | 0.004697 | 0.01542 | 0.044781 | 0.008184 | 0.017171 | 0.008805 | 0.170524 | 0.022074 | 0.001245 | 0.005684 | 0.002359 | 0.014043 | 0.002182 | 0.003431 | 0.018368 | 0.010446 | 0.000811 | 0.008059 | 0.003861 | 0.007279 |
| 33.68055556 | 0.027437 | 0.089218 | 0.012373 | 0.003469 | 0.009979 | 0.013286 | 0.050497 | 0.010594 | 0.011131 | 0.00664 | 0.109377 | 0.034004 | 0.002031 | 0.003504 | 0.00429 | 0.017451 | 0.003822 | 0.005676 | 0.016744 | 0.007894 | 0.000934 | 0.004847 | 0.002482 | 0.008485 |
| 33.69444444 | 0.025906 | 0.081087 | 0.008386 | 0.002935 | 0.017216 | 0.012371 | 0.054342 | 0.007311 | 0.014356 | 0.00489 | 0.060227 | 0.039625 | 0.002631 | 0.002624 | 0.006371 | 0.018061 | 0.004583 | 0.006356 | 0.011721 | 0.006538 | 0.001419 | 0.002278 | 0.001489 | 0.007628 |
| 33.70833333 | 0.01903 | 0.061495 | 0.007502 | 0.004007 | 0.023084 | 0.012181 | 0.051557 | 0.003663 | 0.02285 | 0.006634 | 0.033198 | 0.039194 | 0.002385 | 0.003733 | 0.007256 | 0.015713 | 0.003136 | 0.005387 | 0.005931 | 0.006955 | 0.003226 | 0.002341 | 0.001885 | 0.00564 |
| 33.72222222 | 0.009216 | 0.031491 | 0.005374 | 0.003801 | 0.026461 | 0.01143 | 0.041052 | 0.004072 | 0.023604 | 0.009037 | 0.019321 | 0.036477 | 0.001817 | 0.00532 | 0.006299 | 0.011708 | 0.002254 | 0.003395 | 0.001565 | 0.008696 | 0.005872 | 0.002697 | 0.002422 | 0.004005 |
| 33.73611111 | 0.00542 | 0.014027 | 0.003427 | 0.002451 | 0.027686 | 0.01039 | 0.028999 | 0.005867 | 0.020136 | 0.010917 | 0.037847 | 0.032137 | 0.001575 | 0.006292 | 0.003927 | 0.007268 | 0.003743 | 0.001426 | 0.001502 | 0.011474 | 0.007666 | 0.001869 | 0.002579 | 0.003665 |
| 33.75 | 0.011583 | 0.021014 | 0.006975 | 0.001717 | 0.029283 | 0.009424 | 0.021487 | 0.007388 | 0.016986 | 0.012668 | 0.075205 | 0.028422 | 0.00143 | 0.007013 | 0.001794 | 0.003493 | 0.005618 | 0.000372 | 0.005908 | 0.015238 | 0.008616 | 0.001872 | 0.003356 | 0.004946 |
| 33.76388889 | 0.020076 | 0.030909 | 0.014576 | 0.002789 | 0.033198 | 0.008169 | 0.016663 | 0.009384 | 0.016161 | 0.01406 | 0.10886 | 0.030537 | 0.001023 | 0.008064 | 0.001439 | 0.002062 | 0.004783 | 0.000158 | 0.011928 | 0.018476 | 0.00873 | 0.002771 | 0.003935 | 0.007038 |
| 33.77777778 | 0.024174 | 0.028302 | 0.025364 | 0.005366 | 0.038421 | 0.006663 | 0.010698 | 0.010879 | 0.016391 | 0.015458 | 0.13022 | 0.038266 | 0.000873 | 0.008621 | 0.0025 | 0.002626 | 0.002972 | 0.000215 | 0.015843 | 0.01912 | 0.006998 | 0.003111 | 0.00351 | 0.008872 |
| 33.79166667 | 0.027928 | 0.023256 | 0.041305 | 0.008514 | 0.045293 | 0.005196 | 0.005925 | 0.010059 | 0.014533 | 0.016949 | 0.175351 | 0.045939 | 0.001002 | 0.007398 | 0.003455 | 0.002784 | 0.003811 | 0.000384 | 0.015105 | 0.01675 | 0.004022 | 0.00277 | 0.00265 | 0.009202 |
| 33.80555556 | 0.041636 | 0.024172 | 0.059811 | 0.010153 | 0.055279 | 0.003169 | 0.007063 | 0.007108 | 0.013574 | 0.018427 | 0.238734 | 0.049507 | 0.000915 | 0.004962 | 0.003525 | 0.002563 | 0.00502 | 0.000471 | 0.010449 | 0.011984 | 0.003031 | 0.002571 | 0.001682 | 0.008156 |
| 33.81944444 | 0.066092 | 0.027762 | 0.075192 | 0.009508 | 0.065501 | 0.0027 | 0.013105 | 0.00471 | 0.012547 | 0.019139 | 0.216628 | 0.048424 | 0.001229 | 0.002734 | 0.002681 | 0.00407 | 0.003914 | 0.000581 | 0.005364 | 0.006217 | 0.004646 | 0.003084 | 0.001351 | 0.007805 |
| 33.83333333 | 0.084924 | 0.031189 | 0.0831 | 0.008432 | 0.06817 | 0.005763 | 0.020523 | 0.005586 | 0.007943 | 0.018326 | 0.125259 | 0.043071 | 0.002648 | 0.001217 | 0.001438 | 0.007487 | 0.004093 | 0.001125 | 0.002085 | 0.003246 | 0.005694 | 0.004165 | 0.002372 | 0.007888 |
| 33.84722222 | 0.086777 | 0.036279 | 0.08483 | 0.007507 | 0.058542 | 0.008582 | 0.027718 | 0.009333 | 0.005024 | 0.016695 | 0.064453 | 0.034767 | 0.004746 | 0.000691 | 0.000729 | 0.011098 | 0.008648 | 0.001905 | 0.000574 | 0.004696 | 0.005314 | 0.005313 | 0.00385 | 0.007108 |
| 33.86111111 | 0.074981 | 0.044379 | 0.08297 | 0.005887 | 0.046733 | 0.00925 | 0.032667 | 0.013328 | 0.009874 | 0.015229 | 0.089674 | 0.025688 | 0.00683 | 0.001777 | 0.000592 | 0.013892 | 0.014968 | 0.002607 | 0.000599 | 0.007412 | 0.004898 | 0.005988 | 0.005453 | 0.006289 |
| 33.875 | 0.061492 | 0.053751 | 0.076744 | 0.004543 | 0.044487 | 0.010304 | 0.034298 | 0.016009 | 0.016232 | 0.014074 | 0.172004 | 0.015895 | 0.008595 | 0.004604 | 0.000895 | 0.015672 | 0.020453 | 0.003089 | 0.002134 | 0.009739 | 0.005123 | 0.006138 | 0.00732 | 0.006261 |
| 33.88888889 | 0.055995 | 0.060469 | 0.069011 | 0.004453 | 0.043881 | 0.01288 | 0.038147 | 0.017387 | 0.018555 | 0.013173 | 0.200504 | 0.00821 | 0.009797 | 0.008589 | 0.002361 | 0.01666 | 0.024819 | 0.00332 | 0.004487 | 0.011587 | 0.006246 | 0.005849 | 0.008756 | 0.006547 |
| 33.90277778 | 0.052003 | 0.060081 | 0.06596 | 0.004587 | 0.031646 | 0.016232 | 0.050892 | 0.017715 | 0.020664 | 0.012723 | 0.149914 | 0.008276 | 0.010052 | 0.012663 | 0.004581 | 0.016819 | 0.02613 | 0.003319 | 0.005841 | 0.011805 | 0.008168 | 0.004617 | 0.0091 | 0.006681 |
| 33.91666667 | 0.046603 | 0.051273 | 0.070142 | 0.004218 | 0.015236 | 0.019828 | 0.067788 | 0.01696 | 0.026719 | 0.011653 | 0.096912 | 0.013603 | 0.008895 | 0.015114 | 0.006142 | 0.015419 | 0.022471 | 0.00286 | 0.00518 | 0.009961 | 0.010346 | 0.002594 | 0.00848 | 0.006651 |
| 33.93055556 | 0.040305 | 0.03552 | 0.076751 | 0.003509 | 0.008419 | 0.023124 | 0.077133 | 0.015466 | 0.03673 | 0.008442 | 0.073968 | 0.019085 | 0.006243 | 0.015005 | 0.006474 | 0.013093 | 0.016397 | 0.002034 | 0.003745 | 0.007264 | 0.011884 | 0.001605 | 0.00754 | 0.006478 |
| 33.94444444 | 0.030143 | 0.018878 | 0.075112 | 0.00241 | 0.00635 | 0.025787 | 0.075949 | 0.014031 | 0.045897 | 0.004288 | 0.052998 | 0.024101 | 0.003204 | 0.012985 | 0.005742 | 0.010822 | 0.012324 | 0.001471 | 0.003221 | 0.005231 | 0.01229 | 0.002301 | 0.007291 | 0.006424 |
| 33.95833333 | 0.021062 | 0.011728 | 0.059567 | 0.004008 | 0.003384 | 0.026339 | 0.065086 | 0.012838 | 0.051173 | 0.003837 | 0.027879 | 0.026686 | 0.001255 | 0.009787 | 0.004108 | 0.00826 | 0.010505 | 0.001373 | 0.003459 | 0.004097 | 0.011324 | 0.003284 | 0.007708 | 0.00641 |
| 33.97222222 | 0.01642 | 0.009989 | 0.038836 | 0.011945 | 0.003336 | 0.025347 | 0.044935 | 0.01068 | 0.049619 | 0.006415 | 0.017483 | 0.02444 | 0.000633 | 0.005492 | 0.002569 | 0.005125 | 0.008391 | 0.001169 | 0.003082 | 0.003207 | 0.00886 | 0.004653 | 0.007301 | 0.00587 |
| 33.98611111 | 0.017777 | 0.00816 | 0.029714 | 0.020103 | 0.004102 | 0.026333 | 0.031242 | 0.006518 | 0.033004 | 0.009785 | 0.027456 | 0.015996 | 0.000757 | 0.003046 | 0.003487 | 0.004148 | 0.005463 | 0.000924 | 0.002135 | 0.002146 | 0.007463 | 0.006557 | 0.005753 | 0.004471 |
| 34 | 0.02806 | 0.016543 | 0.035723 | 0.01794 | 0.009675 | 0.025634 | 0.039336 | 0.002907 | 0.018008 | 0.022124 | 0.045504 | 0.008802 | 0.000884 | 0.005474 | 0.006829 | 0.006439 | 0.004175 | 0.001954 | 0.001925 | 0.001412 | 0.013156 | 0.006777 | 0.004235 | 0.00345 |
| 34.01388889 | 0.040794 | 0.032753 | 0.044047 | 0.008475 | 0.027182 | 0.018995 | 0.058962 | 0.00201 | 0.029386 | 0.042304 | 0.05556 | 0.013401 | 0.000585 | 0.009465 | 0.009093 | 0.006713 | 0.007866 | 0.005048 | 0.003317 | 0.002078 | 0.030861 | 0.005391 | 0.004953 | 0.005181 |
| 34.02777778 | 0.045072 | 0.047739 | 0.047935 | 0.003008 | 0.042664 | 0.011161 | 0.068549 | 0.002754 | 0.049457 | 0.052346 | 0.049485 | 0.022262 | 0.000819 | 0.010718 | 0.008464 | 0.004219 | 0.016541 | 0.008672 | 0.008184 | 0.004796 | 0.052625 | 0.007139 | 0.009255 | 0.009715 |
| 34.04166667 | 0.04304 | 0.063206 | 0.052982 | 0.003242 | 0.036558 | 0.008934 | 0.06135 | 0.00575 | 0.057708 | 0.04426 | 0.031112 | 0.02333 | 0.002941 | 0.007654 | 0.005394 | 0.004768 | 0.024351 | 0.009205 | 0.016538 | 0.010133 | 0.056503 | 0.012516 | 0.014516 | 0.013722 |
| 34.05555556 | 0.043792 | 0.078416 | 0.061037 | 0.004859 | 0.024225 | 0.011519 | 0.047001 | 0.007637 | 0.057264 | 0.027739 | 0.014593 | 0.017336 | 0.006262 | 0.003733 | 0.00312 | 0.009789 | 0.025724 | 0.005507 | 0.023025 | 0.016373 | 0.036939 | 0.01539 | 0.01659 | 0.014039 |
| 34.06944444 | 0.045514 | 0.086309 | 0.063398 | 0.005222 | 0.035127 | 0.012097 | 0.034841 | 0.008118 | 0.048892 | 0.013673 | 0.009423 | 0.010198 | 0.007691 | 0.003224 | 0.003607 | 0.014501 | 0.021816 | 0.001531 | 0.021289 | 0.018432 | 0.01542 | 0.013425 | 0.014508 | 0.009391 |
| 34.08333333 | 0.044752 | 0.083241 | 0.053893 | 0.003841 | 0.05206 | 0.008626 | 0.025928 | 0.01385 | 0.033854 | 0.005622 | 0.008036 | 0.0051 | 0.005665 | 0.002985 | 0.003043 | 0.013806 | 0.014714 | 0.000331 | 0.011901 | 0.014812 | 0.005179 | 0.008994 | 0.009083 | 0.005558 |
| 34.09722222 | 0.04276 | 0.074636 | 0.03876 | 0.001858 | 0.048455 | 0.003767 | 0.01849 | 0.018803 | 0.019659 | 0.002167 | 0.005507 | 0.002504 | 0.002504 | 0.001341 | 0.001439 | 0.007887 | 0.007344 | 0.000747 | 0.005384 | 0.00937 | 0.003308 | 0.004262 | 0.005202 | 0.008488 |
| 34.11111111 | 0.037242 | 0.065979 | 0.026938 | 0.000819 | 0.033348 | 0.00116 | 0.012886 | 0.014516 | 0.009128 | 0.001827 | 0.004738 | 0.002922 | 0.001057 | 0.000523 | 0.00129 | 0.00327 | 0.004941 | 0.000868 | 0.006677 | 0.00506 | 0.007499 | 0.001746 | 0.007157 | 0.012292 |
| 34.125 | 0.030072 | 0.057259 | 0.02179 | 0.001371 | 0.021709 | 0.000933 | 0.009371 | 0.00664 | 0.003731 | 0.003675 | 0.00451 | 0.00531 | 0.001504 | 0.000821 | 0.001357 | 0.003282 | 0.005694 | 0.000934 | 0.009407 | 0.002224 | 0.012955 | 0.001899 | 0.010527 | 0.012181 |
| 34.13888889 | 0.02421 | 0.047569 | 0.024214 | 0.002649 | 0.015144 | 0.00077 | 0.008242 | 0.003327 | 0.004744 | 0.00527 | 0.008334 | 0.006921 | 0.002368 | 0.00206 | 0.001002 | 0.003627 | 0.004979 | 0.001886 | 0.010954 | 0.000984 | 0.014431 | 0.002131 | 0.012032 | 0.00867 |
| 34.15277778 | 0.016455 | 0.038968 | 0.028748 | 0.004073 | 0.010287 | 0.001145 | 0.008519 | 0.004659 | 0.005856 | 0.004712 | 0.019279 | 0.007206 | 0.00298 | 0.003406 | 0.000796 | 0.003961 | 0.002993 | 0.002857 | 0.012402 | 0.000756 | 0.011578 | 0.001694 | 0.011447 | 0.004081 |
| 34.16666667 | 0.007204 | 0.0339 | 0.024594 | 0.006508 | 0.01063 | 0.003113 | 0.006559 | 0.006627 | 0.004272 | 0.002647 | 0.034101 | 0.007423 | 0.003122 | 0.003899 | 0.001011 | 0.005526 | 0.002014 | 0.003143 | 0.013498 | 0.000781 | 0.006368 | 0.00179 | 0.008903 | 0.002272 |
| 34.18055556 | 0.003344 | 0.032253 | 0.012573 | 0.009994 | 0.014309 | 0.006038 | 0.003268 | 0.007606 | 0.004167 | 0.001091 | 0.047479 | 0.007769 | 0.002405 | 0.003491 | 0.001655 | 0.004981 | 0.002514 | 0.00302 | 0.014149 | 0.001417 | 0.003563 | 0.002339 | 0.006411 | 0.002013 |
| 34.19444444 | 0.005713 | 0.032456 | 0.005258 | 0.012742 | 0.012374 | 0.008517 | 0.002015 | 0.00757 | 0.007697 | 0.000723 | 0.054567 | 0.007805 | 0.001293 | 0.002222 | 0.002603 | 0.002474 | 0.002761 | 0.002556 | 0.013693 | 0.001865 | 0.005831 | 0.002219 | 0.004697 | 0.001192 |
| 34.20833333 | 0.007772 | 0.033033 | 0.006434 | 0.013843 | 0.011667 | 0.010673 | 0.002332 | 0.007343 | 0.012121 | 0.000739 | 0.056099 | 0.007877 | 0.000875 | 0.001431 | 0.003548 | 0.001122 | 0.002885 | 0.001868 | 0.011505 | 0.001594 | 0.008896 | 0.00124 | 0.002971 | 0.000543 |
| 34.22222222 | 0.008629 | 0.032415 | 0.008997 | 0.013883 | 0.020018 | 0.012775 | 0.003747 | 0.008338 | 0.01265 | 0.000826 | 0.0559 | 0.008461 | 0.000978 | 0.002185 | 0.003822 | 0.000977 | 0.004029 | 0.001057 | 0.008496 | 0.001656 | 0.009661 | 0.000444 | 0.001273 | 0.00045 |
| 34.23611111 | 0.009874 | 0.029364 | 0.008582 | 0.013961 | 0.028828 | 0.014003 | 0.004611 | 0.010127 | 0.007954 | 0.001363 | 0.055089 | 0.00878 | 0.001132 | 0.003274 | 0.003019 | 0.001059 | 0.005548 | 0.000418 | 0.006734 | 0.002174 | 0.010018 | 0.00034 | 0.00061 | 0.001029 |
| 34.25 | 0.010572 | 0.024571 | 0.005687 | 0.01391 | 0.031511 | 0.013658 | 0.003819 | 0.010511 | 0.003785 | 0.001837 | 0.053573 | 0.008003 | 0.002284 | 0.003995 | 0.001715 | 0.002225 | 0.006615 | 0.000291 | 0.006603 | 0.002294 | 0.012494 | 0.000465 | 0.000857 | 0.002263 |
| 34.26388889 | 0.010735 | 0.019726 | 0.002638 | 0.01194 | 0.025596 | 0.01221 | 0.003404 | 0.008754 | 0.003496 | 0.001963 | 0.052275 | 0.006757 | 0.004291 | 0.004646 | 0.000949 | 0.004493 | 0.007209 | 0.000296 | 0.00642 | 0.001697 | 0.017249 | 0.000527 | 0.001202 | 0.003385 |
| 34.27777778 | 0.011545 | 0.016753 | 0.001891 | 0.007872 | 0.016822 | 0.010877 | 0.003389 | 0.006741 | 0.002863 | 0.001975 | 0.050871 | 0.005664 | 0.005747 | 0.005327 | 0.001288 | 0.005611 | 0.006911 | 0.000403 | 0.005139 | 0.000927 | 0.021228 | 0.000952 | 0.002327 | 0.003227 |
| 34.29166667 | 0.014723 | 0.017365 | 0.00306 | 0.003737 | 0.021211 | 0.010118 | 0.002752 | 0.006509 | 0.002429 | 0.001418 | 0.048274 | 0.00393 | 0.00625 | 0.005914 | 0.00245 | 0.004406 | 0.005549 | 0.000945 | 0.003 | 0.00075 | 0.02071 | 0.00179 | 0.004422 | 0.00193 |
| 34.30555556 | 0.01925 | 0.021525 | 0.003214 | 0.002071 | 0.036514 | 0.00991 | 0.001834 | 0.008244 | 0.004567 | 0.000959 | 0.043891 | 0.002307 | 0.006002 | 0.005698 | 0.003444 | 0.002189 | 0.004184 | 0.001313 | 0.002333 | 0.001087 | 0.015663 | 0.001868 | 0.005225 | 0.001246 |
| 34.31944444 | 0.0219 | 0.027615 | 0.001871 | 0.003304 | 0.045912 | 0.010079 | 0.000943 | 0.0106 | 0.006449 | 0.001866 | 0.037803 | 0.003132 | 0.00519 | 0.004554 | 0.003236 | 0.000656 | 0.003634 | 0.001126 | 0.00482 | 0.001092 | 0.010353 | 0.001253 | 0.003638 | 0.001531 |

| | | | | | | | | | | | | | | | | | | | | | | | | |
|---|---|---|---|---|---|---|---|---|---|---|---|---|---|---|---|---|---|---|---|---|---|---|---|---|
| 34.33333333 | 0.022126 | 0.033755 | 0.001111 | 0.004975 | 0.043451 | 0.010526 | 0.000404 | 0.01219 | 0.005527 | 0.003432 | 0.031342 | 0.00579 | 0.003976 | 0.003498 | 0.00227 | 0.000303 | 0.003813 | 0.000673 | 0.007408 | 0.000723 | 0.008032 | 0.001726 | 0.002373 | 0.001483 |
| 34.34722222 | 0.019845 | 0.038201 | 0.001058 | 0.004871 | 0.031281 | 0.01135 | 0.000473 | 0.013128 | 0.00308 | 0.004556 | 0.026149 | 0.008378 | 0.002749 | 0.002956 | 0.002562 | 0.000473 | 0.004417 | 0.000476 | 0.006986 | 0.000863 | 0.008083 | 0.003613 | 0.003862 | 0.000953 |
| 34.36111111 | 0.014642 | 0.039994 | 0.001941 | 0.002827 | 0.015065 | 0.012 | 0.000778 | 0.014691 | 0.002893 | 0.004896 | 0.023662 | 0.010482 | 0.002313 | 0.002721 | 0.00405 | 0.000666 | 0.00453 | 0.000735 | 0.004515 | 0.001663 | 0.008185 | 0.005269 | 0.005522 | 0.000617 |
| 34.375 | 0.008904 | 0.040348 | 0.004455 | 0.000803 | 0.003794 | 0.012088 | 0.001177 | 0.017404 | 0.006003 | 0.004548 | 0.021377 | 0.012279 | 0.00294 | 0.002828 | 0.004148 | 0.001598 | 0.003057 | 0.000686 | 0.003708 | 0.002346 | 0.007084 | 0.005101 | 0.00456 | 0.000642 |
| 34.38888889 | 0.004987 | 0.040651 | 0.007032 | 0.000687 | 0.003808 | 0.011912 | 0.00214 | 0.020329 | 0.00951 | 0.004201 | 0.017448 | 0.013261 | 0.003427 | 0.002887 | 0.002536 | 0.003389 | 0.001446 | 0.000249 | 0.005798 | 0.002711 | 0.005374 | 0.00316 | 0.002328 | 0.001007 |
| 34.40277778 | 0.003492 | 0.040658 | 0.007987 | 0.002639 | 0.012603 | 0.012173 | 0.004349 | 0.02167 | 0.010192 | 0.003885 | 0.014802 | 0.012642 | 0.002646 | 0.002476 | 0.002023 | 0.00495 | 0.001989 | 5.5305559 | 0.007182 | 0.002993 | 0.004028 | 0.002291 | 0.000848 | 0.001373 |
| 34.41666667 | 0.003345 | 0.040036 | 0.007563 | 0.004554 | 0.017455 | 0.013502 | 0.007908 | 0.021008 | 0.008713 | 0.003039 | 0.014574 | 0.011345 | 0.001667 | 0.00212 | 0.003869 | 0.006121 | 0.004239 | 0.000341 | 0.006391 | 0.002835 | 0.002756 | 0.004301 | 0.000508 | 0.00191 |
| 34.43055556 | 0.003241 | 0.038807 | 0.006944 | 0.003928 | 0.013107 | 0.014585 | 0.011952 | 0.02008 | 0.009753 | 0.001905 | 0.016556 | 0.011317 | 0.001984 | 0.00248 | 0.005944 | 0.007233 | 0.00646 | 0.001159 | 0.004796 | 0.002087 | 0.00141 | 0.006484 | 0.000503 | 0.002865 |
| 34.44444444 | 0.003911 | 0.037676 | 0.007448 | 0.001923 | 0.011952 | 0.013693 | 0.015755 | 0.01985 | 0.01501 | 0.001572 | 0.019896 | 0.0133 | 0.003083 | 0.003072 | 0.006082 | 0.00752 | 0.007052 | 0.00198 | 0.002654 | 0.001514 | 0.000614 | 0.006317 | 0.000503 | 0.003085 |
| 34.45833333 | 0.003828 | 0.037098 | 0.009948 | 0.000826 | 0.019254 | 0.011667 | 0.019002 | 0.019616 | 0.021196 | 0.001764 | 0.026758 | 0.015512 | 0.003878 | 0.003125 | 0.004252 | 0.006327 | 0.006044 | 0.002314 | 0.001044 | 0.001599 | 0.000524 | 0.004238 | 0.000578 | 0.002079 |
| 34.47222222 | 0.003104 | 0.035214 | 0.011721 | 0.000684 | 0.025727 | 0.01002 | 0.021206 | 0.01893 | 0.022946 | 0.002524 | 0.039177 | 0.015655 | 0.004525 | 0.003069 | 0.002193 | 0.004872 | 0.005655 | 0.002549 | 0.00073 | 0.001561 | 0.000694 | 0.002331 | 0.001 | 0.001451 |
| 34.48611111 | 0.00874 | 0.028379 | 0.009129 | 0.001324 | 0.029084 | 0.009705 | 0.021184 | 0.018937 | 0.015944 | 0.007513 | 0.056102 | 0.012489 | 0.005122 | 0.003163 | 0.001787 | 0.005107 | 0.007347 | 0.0033 | 0.001409 | 0.001057 | 0.001745 | 0.002684 | 0.002267 | 0.002514 |
| 34.5 | 0.022074 | 0.015842 | 0.004644 | 0.004292 | 0.037588 | 0.01038 | 0.01672 | 0.019459 | 0.016129 | 0.017828 | 0.07164 | 0.008392 | 0.004738 | 0.002367 | 0.002832 | 0.007677 | 0.010219 | 0.004974 | 0.003896 | 0.001018 | 0.00569 | 0.005379 | 0.003231 | 0.004645 |
| 34.51388889 | 0.032531 | 0.007922 | 0.002693 | 0.010607 | 0.058513 | 0.010661 | 0.010957 | 0.019236 | 0.042457 | 0.026895 | 0.078812 | 0.01076 | 0.003289 | 0.001102 | 0.003288 | 0.010901 | 0.013049 | 0.007013 | 0.008305 | 0.001202 | 0.010283 | 0.008695 | 0.004864 | 0.006679 |
| 34.52777778 | 0.033073 | 0.013492 | 0.00579 | 0.01871 | 0.085144 | 0.0115 | 0.013592 | 0.023153 | 0.068662 | 0.027932 | 0.085405 | 0.021255 | 0.001968 | 0.000942 | 0.00233 | 0.011699 | 0.015808 | 0.007177 | 0.013725 | 0.000853 | 0.011012 | 0.010886 | 0.010928 | 0.008637 |
| 34.54166667 | 0.02927 | 0.026248 | 0.016332 | 0.02487 | 0.089209 | 0.01488 | 0.023173 | 0.03544 | 0.066517 | 0.021992 | 0.09315 | 0.03183 | 0.001277 | 0.003274 | 0.002515 | 0.010658 | 0.018811 | 0.004784 | 0.020739 | 0.00091 | 0.010243 | 0.011097 | 0.020266 | 0.011117 |
| 34.55555556 | 0.027825 | 0.038337 | 0.031146 | 0.024511 | 0.054669 | 0.019072 | 0.025993 | 0.048283 | 0.043707 | 0.015049 | 0.089623 | 0.037612 | 0.001019 | 0.008706 | 0.005776 | 0.012091 | 0.02329 | 0.002889 | 0.027424 | 0.002796 | 0.013649 | 0.009897 | 0.0253 | 0.012889 |
| 34.56944444 | 0.029543 | 0.045529 | 0.041882 | 0.017637 | 0.02605 | 0.020391 | 0.017585 | 0.054112 | 0.024272 | 0.011469 | 0.075832 | 0.039557 | 0.00098 | 0.013924 | 0.009332 | 0.015731 | 0.028705 | 0.002975 | 0.029077 | 0.006092 | 0.019916 | 0.008635 | 0.021632 | 0.011558 |
| 34.58333333 | 0.032629 | 0.046955 | 0.045753 | 0.009754 | 0.040353 | 0.019288 | 0.007458 | 0.054861 | 0.027422 | 0.013284 | 0.058965 | 0.03905 | 0.000843 | 0.014702 | 0.009926 | 0.016987 | 0.030997 | 0.00275 | 0.026305 | 0.008839 | 0.023192 | 0.007989 | 0.012281 | 0.007171 |
| 34.59722222 | 0.034468 | 0.04696 | 0.047828 | 0.005671 | 0.058904 | 0.017722 | 0.003433 | 0.054289 | 0.033871 | 0.018604 | 0.040722 | 0.03794 | 0.000967 | 0.011124 | 0.007859 | 0.015003 | 0.029221 | 0.001568 | 0.021917 | 0.010119 | 0.022258 | 0.007301 | 0.006602 | 0.002866 |
| 34.61111111 | 0.034568 | 0.048652 | 0.053395 | 0.006707 | 0.048343 | 0.016093 | 0.003166 | 0.05493 | 0.024305 | 0.021573 | 0.024343 | 0.036515 | 0.001517 | 0.006485 | 0.004905 | 0.012808 | 0.02598 | 0.001329 | 0.016151 | 0.010454 | 0.019347 | 0.005505 | 0.010428 | 0.000864 |
| 34.625 | 0.03335 | 0.050416 | 0.060225 | 0.010969 | 0.024406 | 0.014896 | 0.004958 | 0.055314 | 0.010146 | 0.019315 | 0.012213 | 0.032999 | 0.001772 | 0.003252 | 0.002147 | 0.012455 | 0.022013 | 0.002348 | 0.008881 | 0.010506 | 0.015211 | 0.003133 | 0.014669 | 0.001104 |
| 34.63888889 | 0.030988 | 0.048975 | 0.059894 | 0.013623 | 0.011274 | 0.014501 | 0.009217 | 0.050554 | 0.004118 | 0.014252 | 0.004928 | 0.026188 | 0.001487 | 0.001729 | 0.000654 | 0.013096 | 0.01568 | 0.003101 | 0.004996 | 0.010612 | 0.01024 | 0.001404 | 0.012449 | 0.003287 |
| 34.65277778 | 0.029529 | 0.046163 | 0.051079 | 0.011801 | 0.009938 | 0.014465 | 0.012415 | 0.039981 | 0.011129 | 0.010043 | 0.003579 | 0.017143 | 0.001103 | 0.001002 | 0.000851 | 0.012752 | 0.009154 | 0.002627 | 0.009852 | 0.011267 | 0.005142 | 0.000604 | 0.007151 | 0.005583 |
| 34.66666667 | 0.030004 | 0.046732 | 0.041214 | 0.00783 | 0.008889 | 0.013988 | 0.012623 | 0.026949 | 0.0273 | 0.007369 | 0.006446 | 0.008501 | 0.001191 | 0.001148 | 0.001693 | 0.011027 | 0.00925 | 0.001498 | 0.017536 | 0.012461 | 0.002186 | 0.000436 | 0.002536 | 0.005797 |
| 34.68055556 | 0.031535 | 0.048647 | 0.03436 | 0.004138 | 0.01609 | 0.013801 | 0.011989 | 0.013519 | 0.044483 | 0.005002 | 0.009283 | 0.005335 | 0.001711 | 0.003322 | 0.002027 | 0.008221 | 0.015108 | 0.001076 | 0.019829 | 0.013338 | 0.00252 | 0.000712 | 0.001131 | 0.00395 |
| 34.69444444 | 0.032635 | 0.046 | 0.029933 | 0.003898 | 0.043307 | 0.014219 | 0.012458 | 0.004777 | 0.055437 | 0.004354 | 0.009796 | 0.009303 | 0.001999 | 0.006704 | 0.001414 | 0.004451 | 0.018975 | 0.001466 | 0.01538 | 0.012924 | 0.002534 | 0.001221 | 0.002446 | 0.001839 |
| 34.70833333 | 0.033186 | 0.037678 | 0.026574 | 0.009685 | 0.077167 | 0.0143 | 0.01262 | 0.003909 | 0.054991 | 0.007317 | 0.007464 | 0.014908 | 0.002024 | 0.009542 | 0.00097 | 0.002208 | 0.018211 | 0.001301 | 0.00882 | 0.010957 | 0.001751 | 0.00186 | 0.003672 | 0.001635 |
| 34.72222222 | 0.034368 | 0.027139 | 0.023295 | 0.016244 | 0.092732 | 0.013846 | 0.009021 | 0.004136 | 0.046402 | 0.011669 | 0.007021 | 0.020096 | 0.002239 | 0.011119 | 0.002103 | 0.003898 | 0.014432 | 0.001354 | 0.007567 | 0.008566 | 0.00289 | 0.003223 | 0.003333 | 0.003904 |
| 34.73611111 | 0.036987 | 0.018592 | 0.018948 | 0.018162 | 0.082413 | 0.012672 | 0.006586 | 0.002552 | 0.039179 | 0.014371 | 0.019161 | 0.02736 | 0.002423 | 0.011703 | 0.004253 | 0.008051 | 0.009405 | 0.003651 | 0.009665 | 0.007216 | 0.004144 | 0.005518 | 0.001998 | 0.006805 |
| 34.75 | 0.041202 | 0.014165 | 0.012743 | 0.016826 | 0.057406 | 0.011116 | 0.013051 | 0.002773 | 0.037887 | 0.015411 | 0.038534 | 0.03486 | 0.002106 | 0.01187 | 0.006618 | 0.012329 | 0.005812 | 0.006832 | 0.008281 | 0.007694 | 0.003423 | 0.007921 | 0.001591 | 0.008737 |
| 34.76388889 | 0.045267 | 0.011842 | 0.007141 | 0.015444 | 0.033814 | 0.010461 | 0.021595 | 0.004716 | 0.03775 | 0.017203 | 0.044561 | 0.039219 | 0.001646 | 0.012607 | 0.008715 | 0.01538 | 0.00625 | 0.008379 | 0.008754 | 0.009426 | 0.00338 | 0.009618 | 0.003449 | 0.010097 |
| 34.77777778 | 0.046814 | 0.007654 | 0.005204 | 0.016034 | 0.023761 | 0.011512 | 0.024773 | 0.005483 | 0.032283 | 0.021068 | 0.035641 | 0.041221 | 0.001604 | 0.014437 | 0.009869 | 0.01655 | 0.011328 | 0.007854 | 0.017437 | 0.01115 | 0.005408 | 0.009712 | 0.006261 | 0.010686 |
| 34.79166667 | 0.044423 | 0.006206 | 0.005589 | 0.018183 | 0.030029 | 0.013013 | 0.02602 | 0.003972 | 0.023035 | 0.024441 | 0.030034 | 0.041186 | 0.001575 | 0.015488 | 0.010253 | 0.015598 | 0.019028 | 0.005665 | 0.027173 | 0.012246 | 0.006205 | 0.008044 | 0.007685 | 0.010125 |
| 34.80555556 | 0.037568 | 0.014491 | 0.005371 | 0.02028 | 0.044747 | 0.013486 | 0.02824 | 0.002651 | 0.01681 | 0.024834 | 0.034184 | 0.039269 | 0.001186 | 0.014614 | 0.01024 | 0.011943 | 0.025491 | 0.00296 | 0.030491 | 0.011888 | 0.003915 | 0.005681 | 0.006681 | 0.00943 |
| 34.81944444 | 0.028241 | 0.026969 | 0.005186 | 0.021833 | 0.052088 | 0.012765 | 0.029364 | 0.004779 | 0.012987 | 0.023076 | 0.038386 | 0.03721 | 0.001159 | 0.012292 | 0.009943 | 0.006721 | 0.027435 | 0.002084 | 0.027457 | 0.009735 | 0.001122 | 0.00346 | 0.00411 | 0.009486 |
| 34.83333333 | 0.019606 | 0.036649 | 0.006264 | 0.022804 | 0.048741 | 0.009878 | 0.027543 | 0.011287 | 0.007858 | 0.021783 | 0.037569 | 0.035059 | 0.001888 | 0.007971 | 0.009225 | 0.004374 | 0.024612 | 0.003332 | 0.020811 | 0.006951 | 0.000787 | 0.001725 | 0.001959 | 0.009992 |
| 34.84722222 | 0.011517 | 0.041986 | 0.007169 | 0.023595 | 0.049968 | 0.006067 | 0.024383 | 0.021234 | 0.005909 | 0.021922 | 0.031558 | 0.030922 | 0.002452 | 0.004323 | 0.007776 | 0.006577 | 0.020213 | 0.004379 | 0.014196 | 0.004179 | 0.002982 | 0.001009 | 0.00179 | 0.009868 |
| 34.86111111 | 0.006454 | 0.043565 | 0.006526 | 0.024373 | 0.062708 | 0.003998 | 0.022402 | 0.027842 | 0.010074 | 0.021958 | 0.021246 | 0.023449 | 0.001804 | 0.005514 | 0.005838 | 0.008845 | 0.016813 | 0.003727 | 0.010168 | 0.002061 | 0.006132 | 0.001383 | 0.00316 | 0.008496 |
| 34.875 | 0.009776 | 0.045353 | 0.005567 | 0.024105 | 0.08256 | 0.003344 | 0.02174 | 0.022931 | 0.017108 | 0.020687 | 0.010567 | 0.013353 | 0.00127 | 0.008401 | 0.004214 | 0.008312 | 0.013598 | 0.002273 | 0.008834 | 0.002202 | 0.008747 | 0.001416 | 0.004345 | 0.0064 |
| 34.88888889 | 0.018131 | 0.047174 | 0.00643 | 0.021594 | 0.099993 | 0.003246 | 0.019883 | 0.011207 | 0.025038 | 0.019142 | 0.006219 | 0.006259 | 0.002519 | 0.008436 | 0.003418 | 0.006299 | 0.009371 | 0.002094 | 0.008291 | 0.004123 | 0.00989 | 0.000986 | 0.00382 | 0.00438 |
| 34.90277778 | 0.023913 | 0.043926 | 0.00918 | 0.016382 | 0.103459 | 0.003937 | 0.014851 | 0.003438 | 0.032689 | 0.019288 | 0.011891 | 0.007469 | 0.004087 | 0.006336 | 0.003149 | 0.003708 | 0.004594 | 0.00297 | 0.006689 | 0.005771 | 0.009046 | 0.001082 | 0.001961 | 0.002946 |
| 34.91666667 | 0.024694 | 0.036441 | 0.012048 | 0.00945 | 0.092466 | 0.006079 | 0.007753 | 0.001906 | 0.038148 | 0.019225 | 0.017321 | 0.012022 | 0.004656 | 0.004077 | 0.003138 | 0.001832 | 0.001668 | 0.003165 | 0.003883 | 0.00687 | 0.00648 | 0.001065 | 0.000727 | 0.002392 |
| 34.93055556 | 0.021262 | 0.029208 | 0.012543 | 0.00372 | 0.075552 | 0.009908 | 0.004023 | 0.00355 | 0.042154 | 0.014019 | 0.015089 | 0.014534 | 0.004617 | 0.002671 | 0.003336 | 0.001886 | 0.002157 | 0.0025 | 0.002693 | 0.007724 | 0.003578 | 0.000706 | 0.000676 | 0.002546 |
| 34.94444444 | 0.016182 | 0.022163 | 0.00979 | 0.002241 | 0.058931 | 0.012887 | 0.006512 | 0.006993 | 0.044945 | 0.007396 | 0.012729 | 0.015123 | 0.004252 | 0.001938 | 0.003311 | 0.002151 | 0.003626 | 0.001418 | 0.00537 | 0.008163 | 0.002534 | 0.000745 | 0.000756 | 0.003032 |
| 34.95833333 | 0.012438 | 0.014624 | 0.007022 | 0.004926 | 0.043163 | 0.011345 | 0.01045 | 0.0103 | 0.045559 | 0.006053 | 0.018214 | 0.012928 | 0.003493 | 0.00127 | 0.002687 | 0.001729 | 0.00416 | 0.000691 | 0.00928 | 0.007899 | 0.004502 | 0.000987 | 0.000592 | 0.003279 |
| 34.97222222 | 0.013177 | 0.016451 | 0.007122 | 0.008241 | 0.031387 | 0.006903 | 0.012353 | 0.00966 | 0.043793 | 0.008128 | 0.037263 | 0.009654 | 0.002284 | 0.001216 | 0.00208 | 0.002948 | 0.004418 | 0.000969 | 0.011785 | 0.006175 | 0.007634 | 0.001158 | 0.000923 | 0.002837 |
| 34.98611111 | 0.019045 | 0.029827 | 0.007555 | 0.007354 | 0.03158 | 0.005935 | 0.010629 | 0.010565 | 0.034067 | 0.018543 | 0.066805 | 0.012204 | 0.001506 | 0.002648 | 0.003843 | 0.006741 | 0.004815 | 0.001618 | 0.012224 | 0.003349 | 0.010872 | 0.002159 | 0.002837 | 0.002491 |
| 35 | 0.023208 | 0.031277 | 0.0065 | 0.003919 | 0.047607 | 0.008827 | 0.00648 | 0.01832 | 0.019197 | 0.044846 | 0.085931 | 0.019401 | 0.002481 | 0.004359 | 0.00823 | 0.009154 | 0.004139 | 0.001522 | 0.009417 | 0.002466 | 0.017096 | 0.003072 | 0.004663 | 0.004057 |
| 35.01388889 | 0.019625 | 0.018758 | 0.006991 | 0.003148 | 0.067385 | 0.007904 | 0.006768 | 0.021865 | 0.013094 | 0.07378 | 0.078643 | 0.020369 | 0.004324 | 0.004133 | 0.010822 | 0.007211 | 0.004235 | 0.002298 | 0.006442 | 0.006759 | 0.03011 | 0.00413 | 0.004977 | 0.007873 |
| 35.02777778 | 0.010543 | 0.016584 | 0.008203 | 0.005817 | 0.071577 | 0.005238 | 0.012024 | 0.015876 | 0.014243 | 0.088978 | 0.052997 | 0.013075 | 0.004457 | 0.002951 | 0.009552 | 0.00755 | 0.010898 | 0.00524 | 0.010782 | 0.013309 | 0.044897 | 0.008547 | 0.007581 | 0.011159 |
| 35.04166667 | 0.003036 | 0.025281 | 0.006157 | 0.010806 | 0.057795 | 0.008245 | 0.017425 | 0.008171 | 0.017106 | 0.09432 | 0.025733 | 0.00525 | 0.00318 | 0.003868 | 0.006806 | 0.016876 | 0.020743 | 0.005604 | 0.020175 | 0.01616 | 0.047039 | 0.012571 | 0.009944 | 0.011466 |
| 35.05555556 | 0.000751 | 0.030689 | 0.002395 | 0.013254 | 0.035614 | 0.012257 | 0.020455 | 0.0038 | 0.02244 | 0.093013 | 0.013093 | 0.001647 | 0.003832 | 0.005976 | 0.003784 | 0.028642 | 0.025018 | 0.003983 | 0.025411 | 0.016132 | 0.034446 | 0.011678 | 0.008061 | 0.009376 |
| 35.06944444 | 0.001145 | 0.02963 | 0.000635 | 0.011472 | 0.014915 | 0.013185 | 0.019635 | 0.00277 | 0.023904 | 0.078422 | 0.017688 | 0.001946 | 0.006633 | 0.00606 | 0.001424 | 0.032201 | 0.022866 | 0.005185 | 0.023797 | 0.015954 | 0.019243 | 0.009628 | 0.006757 | 0.00622 |
| 35.08333333 | 0.003362 | 0.02712 | 0.000997 | 0.010128 | 0.003827 | 0.013037 | 0.016567 | 0.002823 | 0.019099 | 0.053228 | 0.022984 | 0.004016 | 0.007918 | 0.00364 | 0.000668 | 0.025087 | 0.016445 | 0.006335 | 0.015593 | 0.013892 | 0.00989 | 0.00791 | 0.006432 | 0.003988 |
| 35.09722222 | 0.008307 | 0.027746 | 0.002899 | 0.010213 | 0.001113 | 0.013119 | 0.013645 | 0.002429 | 0.01328 | 0.027949 | 0.021795 | 0.004408 | 0.006397 | 0.001923 | 0.000595 | 0.012768 | 0.009037 | 0.005897 | 0.007405 | 0.009238 | 0.010521 | 0.004587 | 0.006244 | 0.005477 |
| 35.11111111 | 0.013209 | 0.031537 | 0.006041 | 0.009722 | 0.00286 | 0.013391 | 0.010326 | 0.003247 | 0.009708 | 0.010501 | 0.018272 | 0.00296 | 0.00405 | 0.003089 | 0.000955 | 0.004618 | 0.00698 | 0.006103 | 0.007079 | 0.004215 | 0.017066 | 0.002128 | 0.007977 | 0.009663 |
| 35.125 | 0.015473 | 0.034111 | 0.008295 | 0.008413 | 0.009717 | 0.012764 | 0.005623 | 0.00694 | 0.008427 | 0.004908 | 0.014654 | 0.002817 | 0.002482 | 0.00427 | 0.001322 | 0.004343 | 0.01024 | 0.007216 | 0.010181 | 0.001397 | 0.019192 | 0.002327 | 0.009107 | 0.012934 |
| 35.13888889 | 0.015202 | 0.031821 | 0.008369 | 0.007058 | 0.020585 | 0.011133 | 0.002283 | 0.011526 | 0.007714 | 0.010724 | 0.012982 | 0.004259 | 0.001815 | 0.003318 | 0.000982 | 0.006046 | 0.011658 | 0.00828 | 0.011333 | 0.000925 | 0.015796 | 0.002247 | 0.008826 | 0.013731 |
| 35.15277778 | 0.013706 | 0.027314 | 0.006957 | 0.006958 | 0.032048 | 0.009793 | 0.002757 | 0.014568 | 0.006266 | 0.023682 | 0.014758 | 0.005791 | 0.001438 | 0.001555 | 0.000691 | 0.006262 | 0.008933 | 0.008178 | 0.011676 | 0.000857 | 0.012163 | 0.001094 | 0.008575 | 0.012257 |
| 35.16666667 | 0.012583 | 0.025282 | 0.005473 | 0.008336 | 0.043151 | 0.00952 | 0.00378 | 0.015464 | 0.004816 | 0.036988 | 0.019259 | 0.006584 | 0.00091 | 0.000449 | 0.000646 | 0.005976 | 0.004747 | 0.006801 | 0.01253 | 0.000975 | 0.009941 | 0.00076 | 0.009165 | 0.009611 |
| 35.18055556 | 0.01256 | 0.025972 | 0.005454 | 0.010404 | 0.053702 | 0.009409 | 0.002712 | 0.014729 | 0.005113 | 0.043848 | 0.024239 | 0.006495 | 0.000592 | 0.000197 | 0.000721 | 0.004381 | 0.001943 | 0.005262 | 0.012961 | 0.001307 | 0.007727 | 0.001438 | 0.010853 | 0.007007 |
| 35.19444444 | 0.013624 | 0.026772 | 0.007454 | 0.013387 | 0.062125 | 0.009261 | 0.001685 | 0.012593 | 0.007804 | 0.04262 | 0.027294 | 0.005991 | 0.00079 | 0.000194 | 0.00158 | 0.00215 | 0.001959 | 0.004521 | 0.011233 | 0.001501 | 0.005232 | 0.002229 | 0.012545 | 0.005567 |
| 35.20833333 | 0.014632 | 0.026633 | 0.010593 | 0.016137 | 0.070052 | 0.010251 | 0.002466 | 0.009441 | 0.010458 | 0.036603 | 0.02757 | 0.005723 | 0.001161 | 0.000223 | 0.002691 | 0.001521 | 0.003392 | 0.004395 | 0.007725 | 0.001867 | 0.003892 | 0.002729 | 0.013089 | 0.005592 |
| 35.22222222 | 0.014033 | 0.024774 | 0.013537 | 0.018079 | 0.076662 | 0.012351 | 0.00308 | 0.007572 | 0.010185 | 0.02966 | 0.025769 | 0.005474 | 0.001694 | 0.000389 | 0.003173 | 0.002216 | 0.004592 | 0.003866 | 0.004616 | 0.002283 | 0.003867 | 0.00296 | 0.011361 | 0.006352 |
| 35.23611111 | 0.012098 | 0.019487 | 0.015577 | 0.020233 | 0.077995 | 0.014191 | 0.002151 | 0.008389 | 0.008763 | 0.024516 | 0.022874 | 0.00477 | 0.002502 | 0.000495 | 0.003197 | 0.002995 | 0.005293 | 0.002335 | 0.003238 | 0.002668 | 0.003309 | 0.00312 | 0.00711 | 0.007149 |
| 35.25 | 0.010589 | 0.011783 | 0.016304 | 0.021613 | 0.070863 | 0.014718 | 0.001255 | 0.009801 | 0.008971 | 0.022559 | 0.02046 | 0.003802 | 0.003211 | 0.000602 | 0.003115 | 0.003264 | 0.005967 | 0.001088 | 0.003593 | 0.003315 | 0.002002 | 0.003313 | 0.003504 | 0.007927 |
| 35.26388889 | 0.009592 | 0.005409 | 0.01474 | 0.021563 | 0.053194 | 0.013837 | 0.001755 | 0.00915 | 0.010199 | 0.02403 | 0.018842 | 0.002436 | 0.003429 | 0.000874 | 0.00334 | 0.003155 | 0.007053 | 0.001176 | 0.005289 | 0.004094 | 0.002344 | 0.003276 | 0.003431 | 0.008897 |
| 35.27777778 | 0.008082 | 0.002262 | 0.01071 | 0.021186 | 0.034042 | 0.012134 | 0.003625 | 0.00646 | 0.010327 | 0.029412 | 0.018673 | 0.001498 | 0.003443 | 0.000818 | 0.003856 | 0.003475 | 0.008338 | 0.001206 | 0.007262 | 0.004112 | 0.005224 | 0.003137 | 0.004805 | 0.00923 |
| 35.29166667 | 0.007091 | 0.001229 | 0.006288 | 0.020919 | 0.02319 | 0.010276 | 0.005843 | 0.003914 | 0.00963 | 0.038243 | 0.02167 | 0.001495 | 0.003821 | 0.000823 | 0.003952 | 0.004173 | 0.009552 | 0.000791 | 0.007184 | 0.003002 | 0.008906 | 0.003645 | 0.005446 | 0.008315 |
| 35.30555556 | 0.008085 | 0.001303 | 0.003327 | 0.018424 | 0.014687 | 0.008681 | 0.007032 | 0.002919 | 0.010308 | 0.046319 | 0.026818 | 0.001086 | 0.004452 | 0.001721 | 0.00326 | 0.004545 | 0.010555 | 0.001205 | 0.004385 | 0.001505 | 0.010919 | 0.00508 | 0.005133 | 0.007104 |
| 35.31944444 | 0.010026 | 0.002605 | 0.001832 | 0.012324 | 0.008347 | 0.007687 | 0.006811 | 0.003068 | 0.012206 | 0.048333 | 0.03142 | 0.000573 | 0.004914 | 0.002618 | 0.002254 | 0.003776 | 0.010771 | 0.002437 | 0.002334 | 0.0007 | 0.009539 | 0.006466 | 0.003307 | 0.006249 |
| 35.33333333 | 0.011454 | 0.005199 | 0.001918 | 0.007006 | 0.009785 | 0.007596 | 0.005894 | 0.003812 | 0.013891 | 0.04333 | 0.033852 | 0.001172 | 0.00508 | 0.002475 | 0.001216 | 0.002377 | 0.010469 | 0.00369 | 0.002733 | 0.001092 | 0.005583 | 0.006448 | 0.001678 | 0.004969 |
| 35.34722222 | 0.012357 | 0.008156 | 0.003598 | 0.005016 | 0.01026 | 0.007887 | 0.005084 | 0.004534 | 0.015064 | 0.035457 | 0.033318 | 0.002399 | 0.004946 | 0.001693 | 0.000384 | 0.002366 | 0.010397 | 0.004566 | 0.002761 | 0.001991 | 0.002687 | 0.004958 | 0.002451 | 0.002767 |
| 35.36111111 | 0.012798 | 0.01061 | 0.005576 | 0.003273 | 0.009773 | 0.008044 | 0.004787 | 0.004444 | 0.016998 | 0.028737 | 0.031028 | 0.00328 | 0.004643 | 0.00147 | 0.000109 | 0.003441 | 0.009908 | 0.004977 | 0.001774 | 0.002582 | 0.003486 | 0.003179 | 0.004041 | 0.001338 |
| 35.375 | 0.01298 | 0.012855 | 0.006296 | 0.00138 | 0.013888 | 0.008763 | 0.004257 | 0.004477 | 0.020771 | 0.024943 | 0.029492 | 0.003647 | 0.004388 | 0.001684 | 0.000328 | 0.00393 | 0.008273 | 0.005041 | 0.001881 | 0.002635 | 0.006242 | 0.00225 | 0.004935 | 0.001629 |
| 35.38888889 | 0.013615 | 0.015207 | 0.005452 | 0.000957 | 0.014656 | 0.010372 | 0.002701 | 0.006873 | 0.023752 | 0.02434 | 0.029875 | 0.003868 | 0.003994 | 0.001349 | 0.001039 | 0.004314 | 0.005622 | 0.004778 | 0.00321 | 0.00218 | 0.008106 | 0.002487 | 0.005636 | 0.001565 |
| 35.40277778 | 0.015374 | 0.017817 | 0.004485 | 0.001007 | 0.008631 | 0.011695 | 0.001114 | 0.011877 | 0.023975 | 0.02495 | 0.032254 | 0.004026 | 0.003344 | 0.001163 | 0.001896 | 0.005322 | 0.002895 | 0.003931 | 0.004194 | 0.001527 | 0.008435 | 0.003313 | 0.006094 | 0.001203 |
| 35.41666667 | 0.018113 | 0.020604 | 0.004227 | 0.000881 | 0.004028 | 0.011788 | 0.000905 | 0.016452 | 0.021556 | 0.024362 | 0.036374 | 0.003518 | 0.003108 | 0.001794 | 0.002357 | 0.006593 | 0.002047 | 0.002957 | 0.004644 | 0.001299 | 0.008056 | 0.004004 | 0.005933 | 0.002508 |
| 35.43055556 | 0.020606 | 0.023323 | 0.003374 | 0.000915 | 0.007954 | 0.010133 | 0.001577 | 0.018386 | 0.014265 | 0.02208 | 0.043223 | 0.003788 | 0.003729 | 0.002682 | 0.002358 | 0.007943 | 0.003368 | 0.002794 | 0.005481 | 0.001612 | 0.007828 | 0.004329 | 0.005368 | 0.004713 |
| 35.44444444 | 0.022603 | 0.025184 | 0.0022 | 0.001459 | 0.021254 | 0.007356 | 0.001572 | 0.018687 | 0.006248 | 0.019959 | 0.051802 | 0.005774 | 0.004244 | 0.003274 | 0.001683 | 0.009002 | 0.004194 | 0.002871 | 0.006148 | 0.001555 | 0.007627 | 0.004021 | 0.004707 | 0.006075 |
| 35.45833333 | 0.025469 | 0.021847 | 0.002787 | 0.00538 | 0.039548 | 0.007177 | 0.001479 | 0.016584 | 0.009012 | 0.02003 | 0.051431 | 0.010323 | 0.003762 | 0.003323 | 0.001181 | 0.009197 | 0.003073 | 0.002147 | 0.006047 | 0.000848 | 0.007226 | 0.002865 | 0.003936 | 0.00602 |
| 35.47222222 | 0.031435 | 0.01178 | 0.006712 | 0.014476 | 0.057438 | 0.010249 | 0.002434 | 0.013822 | 0.020527 | 0.01972 | 0.041415 | 0.018835 | 0.00316 | 0.00339 | 0.001461 | 0.009013 | 0.00158 | 0.001357 | 0.006616 | 0.00048 | 0.007311 | 0.001966 | 0.003301 | 0.005449 |
| 35.48611111 | 0.040137 | 0.007811 | 0.017928 | 0.019751 | 0.080235 | 0.009774 | 0.002539 | 0.022639 | 0.030638 | 0.016681 | 0.050509 | 0.02288 | 0.003084 | 0.003577 | 0.001711 | 0.009244 | 0.001726 | 0.001508 | 0.009157 | 0.001174 | 0.009182 | 0.002661 | 0.002684 | 0.005455 |
| 35.5 | 0.044376 | 0.035468 | 0.040866 | 0.024056 | 0.120198 | 0.023246 | 0.001332 | 0.050822 | 0.077258 | 0.013976 | 0.112737 | 0.043172 | 0.00278 | 0.00277 | 0.002726 | 0.010279 | 0.00416 | 0.002838 | 0.013438 | 0.002999 | 0.01404 | 0.004618 | 0.002565 | 0.005836 |
| 35.51388889 | 0.039255 | 0.150805 | 0.080378 | 0.057291 | 0.170838 | 0.058178 | 0.008805 | 0.064776 | 0.154073 | 0.016471 | 0.216195 | 0.096382 | 0.00316 | 0.00233 | 0.003245 | 0.013899 | 0.007616 | 0.005767 | 0.018399 | 0.005907 | 0.020601 | 0.005785 | 0.007122 | 0.007372 |
| 35.52777778 | 0.035828 | 0.215106 | 0.092615 | 0.076463 | 0.164191 | 0.058628 | 0.032853 | 0.06078 | 0.145527 | 0.024564 | 0.242475 | 0.111631 | 0.006015 | 0.00436 | 0.010633 | 0.024914 | 0.007335 | 0.008595 | 0.02218 | 0.006891 | 0.023064 | 0.010517 | 0.016657 | 0.015822 |
| 35.54166667 | 0.036786 | 0.174449 | 0.047095 | 0.052421 | 0.110052 | 0.05741 | 0.052731 | 0.055779 | 0.112483 | 0.058993 | 0.174082 | 0.098894 | 0.008903 | 0.012362 | 0.035031 | 0.037621 | 0.013803 | 0.008675 | 0.023546 | 0.009033 | 0.027928 | 0.032845 | 0.032159 | 0.03895 |
| 35.55555556 | 0.031766 | 0.183224 | 0.013786 | 0.064869 | 0.082873 | 0.089557 | 0.050018 | 0.03606 | 0.167523 | 0.099997 | 0.128742 | 0.111968 | 0.008243 | 0.017847 | 0.042752 | 0.03685 | 0.027986 | 0.007448 | 0.019891 | 0.012127 | 0.031116 | 0.046422 | 0.042615 | 0.048251 |
| 35.56944444 | 0.028356 | 0.136758 | 0.020517 | 0.092444 | 0.063635 | 0.072766 | 0.039888 | 0.025318 | 0.169026 | 0.082456 | 0.094606 | 0.085036 | 0.010577 | 0.019396 | 0.036275 | 0.025845 | 0.040521 | 0.006689 | 0.01462 | 0.018083 | 0.02998 | 0.041376 | 0.029022 | 0.024216 |
| 35.58333333 | 0.032019 | 0.048212 | 0.033834 | 0.073086 | 0.054467 | 0.032825 | 0.033634 | 0.029439 | 0.074099 | 0.036319 | 0.065858 | 0.029739 | 0.01354 | 0.021154 | 0.049455 | 0.020367 | 0.049035 | 0.005558 | 0.014485 | 0.028528 | 0.03289 | 0.045628 | 0.011192 | 0.010213 |
| 35.59722222 | 0.035947 | 0.02799 | 0.042815 | 0.037769 | 0.047221 | 0.026163 | 0.028247 | 0.036124 | 0.013894 | 0.013629 | 0.068781 | 0.00956 | 0.006868 | 0.011822 | 0.043958 | 0.016673 | 0.04156 | 0.005585 | 0.013791 | 0.025966 | 0.024154 | 0.034853 | 0.00887 | 0.012897 |
| 35.61111111 | 0.037874 | 0.030395 | 0.041656 | 0.015347 | 0.040055 | 0.027923 | 0.023473 | 0.040603 | 0.005256 | 0.00803 | 0.062447 | 0.007288 | 0.00058 | 0.003723 | 0.017471 | 0.007516 | 0.03442 | 0.007455 | 0.008561 | 0.016264 | 0.013691 | 0.011576 | 0.012693 | 0.007474 |
| 35.625 | 0.037926 | 0.039043 | 0.03216 | 0.011265 | 0.068271 | 0.025094 | 0.01974 | 0.037057 | 0.008812 | 0.00774 | 0.040827 | 0.004829 | 0.000173 | 0.003585 | 0.005268 | 0.003078 | 0.036891 | 0.007944 | 0.004983 | 0.012897 | 0.012892 | 0.002671 | 0.016977 | 0.001915 |
| 35.63888889 | 0.036537 | 0.044165 | 0.023443 | 0.019568 | 0.104685 | 0.020447 | 0.015999 | 0.032019 | 0.013552 | 0.005518 | 0.024146 | 0.004149 | 0.000117 | 0.002065 | 0.003022 | 0.004198 | 0.0362 | 0.007033 | 0.007385 | 0.011146 | 0.010087 | 0.000991 | 0.020199 | 0.001522 |
| 35.65277778 | 0.034643 | 0.043205 | 0.019068 | 0.029 | 0.11759 | 0.01917 | 0.010933 | 0.027833 | 0.012781 | 0.005191 | 0.020416 | 0.004193 | 0.000129 | 0.001349 | 0.002941 | 0.003828 | 0.031972 | 0.006931 | 0.013969 | 0.009612 | 0.005945 | 0.00181 | 0.021601 | 0.002692 |
| 35.66666667 | 0.033131 | 0.045897 | 0.016564 | 0.034876 | 0.108187 | 0.021407 | 0.007186 | 0.027092 | 0.012186 | 0.012239 | 0.02629 | 0.003321 | 0.000161 | 0.003002 | 0.003182 | 0.002444 | 0.023254 | 0.007138 | 0.021233 | 0.0087 | 0.004467 | 0.003945 | 0.020897 | 0.003508 |
| 35.68055556 | 0.030823 | 0.067077 | 0.013714 | 0.036599 | 0.09372 | 0.023996 | 0.006888 | 0.025702 | 0.015155 | 0.022807 | 0.033053 | 0.004344 | 0.000249 | 0.007169 | 0.003213 | 0.002439 | 0.011898 | 0.006134 | 0.025416 | 0.008988 | 0.006372 | 0.006425 | 0.016748 | 0.003425 |
| 35.69444444 | 0.026967 | 0.102393 | 0.011152 | 0.033245 | 0.072052 | 0.025095 | 0.00657 | 0.021705 | 0.013788 | 0.031189 | 0.039644 | 0.005678 | 0.000886 | 0.011892 | 0.002572 | 0.004246 | 0.005457 | 0.003993 | 0.02307 | 0.009962 | 0.007417 | 0.008693 | 0.009938 | 0.003374 |
| 35.70833333 | 0.023704 | 0.123464 | 0.00996 | 0.021962 | 0.037749 | 0.025454 | 0.004883 | 0.019933 | 0.011018 | 0.034054 | 0.046565 | 0.004839 | 0.002402 | 0.014962 | 0.001669 | 0.006144 | 0.00699 | 0.002095 | 0.015357 | 0.010353 | 0.005447 | 0.010152 | 0.006924 | 0.003062 |
| 35.72222222 | 0.024289 | 0.116454 | 0.010062 | 0.011467 | 0.019067 | 0.025632 | 0.004117 | 0.020553 | 0.011394 | 0.030266 | 0.048135 | 0.00652 | 0.004464 | 0.016245 | 0.002081 | 0.007456 | 0.008693 | 0.001121 | 0.007308 | 0.00993 | 0.002517 | 0.010792 | 0.010007 | 0.001865 |
| 35.73611111 | 0.028651 | 0.098019 | 0.01147 | 0.014831 | 0.028914 | 0.026402 | 0.004439 | 0.021038 | 0.009988 | 0.02276 | 0.045146 | 0.012721 | 0.006137 | 0.017316 | 0.004001 | 0.008116 | 0.007504 | 0.001007 | 0.004609 | 0.009392 | 0.001311 | 0.01125 | 0.01248 | 0.001055 |
| 35.75 | 0.033237 | 0.083841 | 0.013097 | 0.025306 | 0.039348 | 0.026962 | 0.005963 | 0.017967 | 0.007754 | 0.014711 | 0.038177 | 0.021095 | 0.006783 | 0.018917 | 0.006041 | 0.008124 | 0.007545 | 0.001654 | 0.007206 | 0.009421 | 0.001988 | 0.011167 | 0.010982 | 0.001866 |
| 35.76388889 | 0.035995 | 0.073134 | 0.013473 | 0.029134 | 0.02935 | 0.025837 | 0.008283 | 0.010431 | 0.009669 | 0.008783 | 0.027001 | 0.027643 | 0.006428 | 0.020864 | 0.00749 | 0.008294 | 0.009966 | 0.003038 | 0.008681 | 0.010386 | 0.002246 | 0.01097 | 0.006691 | 0.002916 |
| 35.77777778 | 0.035409 | 0.056314 | 0.013574 | 0.019874 | 0.017457 | 0.026307 | 0.00824 | 0.005031 | 0.011749 | 0.00896 | 0.021728 | 0.027087 | 0.005545 | 0.023578 | 0.008258 | 0.009787 | 0.015082 | 0.005007 | 0.006693 | 0.011525 | 0.003264 | 0.011309 | 0.003458 | 0.003571 |
| 35.79166667 | 0.033449 | 0.036937 | 0.01415 | 0.010177 | 0.029155 | 0.028942 | 0.005764 | 0.004421 | 0.011907 | 0.015207 | 0.023912 | 0.022532 | 0.004883 | 0.026185 | 0.008258 | 0.012527 | 0.023583 | 0.006533 | 0.007579 | 0.011578 | 0.007683 | 0.011554 | 0.004837 | 0.004846 |
| 35.80555556 | 0.035837 | 0.028448 | 0.014201 | 0.014041 | 0.051166 | 0.027206 | 0.003281 | 0.008925 | 0.013782 | 0.022867 | 0.023257 | 0.019254 | 0.004859 | 0.026843 | 0.00773 | 0.015285 | 0.03256 | 0.006244 | 0.01412 | 0.009995 | 0.013166 | 0.011139 | 0.007546 | 0.006687 |
| 35.81944444 | 0.04261 | 0.031715 | 0.013785 | 0.01964 | 0.062975 | 0.018288 | 0.002548 | 0.017501 | 0.015631 | 0.027589 | 0.018528 | 0.016919 | 0.005675 | 0.025843 | 0.00716 | 0.017042 | 0.036777 | 0.00418 | 0.02004 | 0.006819 | 0.017744 | 0.010173 | 0.008187 | 0.007852 |
| 35.83333333 | 0.047384 | 0.038059 | 0.014695 | 0.019599 | 0.072021 | 0.008297 | 0.005363 | 0.022137 | 0.016237 | 0.027838 | 0.0155 | 0.015292 | 0.007108 | 0.024008 | 0.006912 | 0.017191 | 0.03411 | 0.002307 | 0.022648 | 0.003614 | 0.021766 | 0.00903 | 0.007108 | 0.008025 |
| 35.84722222 | 0.045154 | 0.037027 | 0.017475 | 0.021575 | 0.08598 | 0.003379 | 0.010842 | 0.021957 | 0.017881 | 0.025703 | 0.015118 | 0.013103 | 0.008061 | 0.021303 | 0.006757 | 0.014921 | 0.027157 | 0.002808 | 0.023699 | 0.002905 | 0.02407 | 0.007349 | 0.004449 | 0.007936 |
| 35.86111111 | 0.038172 | 0.024808 | 0.02013 | 0.026802 | 0.099426 | 0.002627 | 0.015978 | 0.020949 | 0.017815 | 0.023712 | 0.01506 | 0.008044 | 0.007617 | 0.01806 | 0.006385 | 0.009554 | 0.019373 | 0.004871 | 0.023445 | 0.004351 | 0.024173 | 0.005018 | 0.001701 | 0.007589 |
| 35.875 | 0.032208 | 0.010117 | 0.020976 | 0.027267 | 0.106361 | 0.002427 | 0.017482 | 0.020735 | 0.014488 | 0.023009 | 0.012934 | 0.003696 | 0.005878 | 0.014814 | 0.006164 | 0.004951 | 0.011784 | 0.005655 | 0.021698 | 0.006151 | 0.023391 | 0.002617 | 0.00086 | 0.006797 |
| 35.88888889 | 0.02912 | 0.002572 | 0.019465 | 0.021112 | 0.105482 | 0.002272 | 0.014232 | 0.021345 | 0.013511 | 0.022797 | 0.007454 | 0.003266 | 0.00339 | 0.011782 | 0.005662 | 0.00614 | 0.005532 | 0.004482 | 0.017709 | 0.008451 | 0.022025 | 0.001788 | 0.001015 | 0.004997 |
| 35.90277778 | 0.026717 | 0.001003 | 0.016375 | 0.017203 | 0.099293 | 0.002218 | 0.0094 | 0.020171 | 0.0134 | 0.021181 | 0.002378 | 0.003986 | 0.001277 | 0.008612 | 0.004298 | 0.010423 | 0.002649 | 0.002791 | 0.012326 | 0.011534 | 0.018866 | 0.003542 | 0.000891 | 0.002387 |
| 35.91666667 | 0.023222 | 0.000813 | 0.013652 | 0.016923 | 0.089634 | 0.002923 | 0.005086 | 0.01824 | 0.009423 | 0.0184 | 0.001306 | 0.004902 | 0.000421 | 0.00591 | 0.003093 | 0.01363 | 0.00203 | 0.003018 | 0.007715 | 0.014787 | 0.014386 | 0.004846 | 0.001371 | 0.000656 |
| 35.93055556 | 0.019995 | 0.000741 | 0.013179 | 0.013457 | 0.07193 | 0.006232 | 0.003614 | 0.017923 | 0.005984 | 0.015272 | 0.001622 | 0.006398 | 0.000597 | 0.004418 | 0.002855 | 0.014634 | 0.001104 | 0.005739 | 0.004068 | 0.017118 | 0.010065 | 0.003399 | 0.002774 | 0.000673 |
| 35.94444444 | 0.019056 | 0.00156 | 0.015656 | 0.007336 | 0.047852 | 0.011955 | 0.007886 | 0.017129 | 0.004413 | 0.013088 | 0.001083 | 0.007585 | 0.001497 | 0.003387 | 0.002864 | 0.013325 | 0.000897 | 0.007312 | 0.003153 | 0.016817 | 0.00549 | 0.001343 | 0.004579 | 0.001783 |

| | | | | | | | | | | | | | | | | | | | | | | | | |
|---|---|---|---|---|---|---|---|---|---|---|---|---|---|---|---|---|---|---|---|---|---|---|---|---|
| 35.95833333 | 0.019548 | 0.005063 | 0.018613 | 0.004708 | 0.031044 | 0.017769 | 0.01409 | 0.014033 | 0.005845 | 0.014436 | 0.001547 | 0.006624 | 0.002346 | 0.002144 | 0.002301 | 0.012544 | 0.002854 | 0.005341 | 0.007371 | 0.013486 | 0.002581 | 0.000863 | 0.00648 | 0.002681 |
| 35.97222222 | 0.018084 | 0.009612 | 0.015862 | 0.006428 | 0.031581 | 0.01818 | 0.017622 | 0.013625 | 0.016502 | 0.022222 | 0.007362 | 0.013841 | 0.002573 | 0.000987 | 0.001447 | 0.014363 | 0.005703 | 0.002626 | 0.013518 | 0.009232 | 0.003428 | 0.001516 | 0.007717 | 0.0032 |
| 35.98611111 | 0.014307 | 0.009017 | 0.010242 | 0.006475 | 0.047483 | 0.012105 | 0.016893 | 0.021473 | 0.030526 | 0.040341 | 0.019678 | 0.038936 | 0.002167 | 0.000477 | 0.000743 | 0.016746 | 0.007036 | 0.002416 | 0.01715 | 0.006482 | 0.00357 | 0.002039 | 0.007589 | 0.004414 |
| 36 | 0.017639 | 0.013984 | 0.008028 | 0.010365 | 0.065901 | 0.008581 | 0.011659 | 0.02967 | 0.028321 | 0.061872 | 0.022322 | 0.045423 | 0.001798 | 0.002917 | 0.001039 | 0.017593 | 0.006229 | 0.003417 | 0.015296 | 0.006695 | 0.005198 | 0.001825 | 0.005349 | 0.007179 |
| 36.01388889 | 0.031269 | 0.040683 | 0.007724 | 0.018686 | 0.076918 | 0.009536 | 0.009161 | 0.028213 | 0.014858 | 0.07141 | 0.016823 | 0.033825 | 0.003189 | 0.012749 | 0.004095 | 0.017833 | 0.010296 | 0.003103 | 0.010517 | 0.01048 | 0.017857 | 0.001633 | 0.004182 | 0.011546 |
| 36.02777778 | 0.041005 | 0.062499 | 0.007228 | 0.018099 | 0.09132 | 0.010064 | 0.012095 | 0.026579 | 0.01117 | 0.081939 | 0.01863 | 0.036647 | 0.005132 | 0.026028 | 0.008875 | 0.01876 | 0.022983 | 0.001931 | 0.012055 | 0.015728 | 0.037075 | 0.002425 | 0.008657 | 0.015799 |
| 36.04166667 | 0.039328 | 0.050381 | 0.005533 | 0.016956 | 0.117062 | 0.015529 | 0.012218 | 0.033813 | 0.018721 | 0.113093 | 0.015476 | 0.047321 | 0.004658 | 0.030181 | 0.010391 | 0.018206 | 0.032481 | 0.002321 | 0.018934 | 0.0184 | 0.046255 | 0.002753 | 0.01444 | 0.01515 |
| 36.05555556 | 0.034799 | 0.032784 | 0.004526 | 0.020213 | 0.127255 | 0.019308 | 0.009655 | 0.034793 | 0.02431 | 0.138087 | 0.013559 | 0.05178 | 0.004074 | 0.02388 | 0.006517 | 0.013564 | 0.029405 | 0.004958 | 0.020944 | 0.017303 | 0.036434 | 0.002325 | 0.015494 | 0.008394 |
| 36.06944444 | 0.037458 | 0.04482 | 0.005758 | 0.019991 | 0.115631 | 0.011021 | 0.009498 | 0.022439 | 0.024951 | 0.124533 | 0.031612 | 0.047239 | 0.005644 | 0.017015 | 0.001957 | 0.0081 | 0.018561 | 0.007342 | 0.015182 | 0.013723 | 0.018714 | 0.003284 | 0.012611 | 0.002933 |
| 36.08333333 | 0.042108 | 0.062442 | 0.006892 | 0.018759 | 0.097682 | 0.005938 | 0.009386 | 0.012342 | 0.025605 | 0.089501 | 0.047782 | 0.049232 | 0.00589 | 0.013018 | 0.000714 | 0.005418 | 0.008142 | 0.007978 | 0.007522 | 0.00818 | 0.014 | 0.004934 | 0.008426 | 0.003154 |
| 36.09722222 | 0.033716 | 0.046676 | 0.006453 | 0.019249 | 0.06439 | 0.019729 | 0.006208 | 0.009358 | 0.02095 | 0.068299 | 0.034709 | 0.051882 | 0.003608 | 0.0091 | 0.001384 | 0.003921 | 0.003179 | 0.008293 | 0.005078 | 0.003498 | 0.025904 | 0.004982 | 0.008746 | 0.005692 |
| 36.11111111 | 0.017191 | 0.017816 | 0.007133 | 0.020934 | 0.029967 | 0.040794 | 0.002701 | 0.006407 | 0.023407 | 0.063286 | 0.011605 | 0.056178 | 0.001475 | 0.004818 | 0.001952 | 0.004015 | 0.00432 | 0.009136 | 0.007165 | 0.003129 | 0.039497 | 0.00355 | 0.016493 | 0.007897 |
| 36.125 | 0.009527 | 0.007737 | 0.006283 | 0.021672 | 0.01343 | 0.042747 | 0.001817 | 0.004542 | 0.029913 | 0.058095 | 0.003011 | 0.053709 | 0.000743 | 0.002269 | 0.001886 | 0.007064 | 0.007177 | 0.009164 | 0.008414 | 0.004682 | 0.043484 | 0.002224 | 0.022729 | 0.009923 |
| 36.13888889 | 0.011568 | 0.012092 | 0.003803 | 0.019257 | 0.005802 | 0.026803 | 0.002077 | 0.009593 | 0.022291 | 0.052453 | 0.005909 | 0.030735 | 0.000906 | 0.001718 | 0.001395 | 0.01092 | 0.007721 | 0.007315 | 0.008009 | 0.004559 | 0.037086 | 0.001428 | 0.0211 | 0.010937 |
| 36.15277778 | 0.015538 | 0.042544 | 0.002847 | 0.016526 | 0.007709 | 0.016738 | 0.001934 | 0.023432 | 0.018996 | 0.045329 | 0.022955 | 0.034679 | 0.001309 | 0.001364 | 0.000792 | 0.014912 | 0.005467 | 0.004488 | 0.007918 | 0.002886 | 0.024964 | 0.000869 | 0.015461 | 0.009609 |
| 36.16666667 | 0.015506 | 0.092956 | 0.004212 | 0.01684 | 0.022513 | 0.012934 | 0.0046 | 0.043559 | 0.028747 | 0.030541 | 0.046473 | 0.084431 | 0.001504 | 0.000688 | 0.000378 | 0.01833 | 0.002913 | 0.002198 | 0.009213 | 0.002082 | 0.012343 | 0.000621 | 0.011594 | 0.007188 |
| 36.18055556 | 0.009703 | 0.106023 | 0.008999 | 0.01926 | 0.026968 | 0.011237 | 0.013362 | 0.056417 | 0.028141 | 0.02179 | 0.051349 | 0.103311 | 0.0011 | 0.00062 | 0.00033 | 0.018664 | 0.002606 | 0.001122 | 0.011686 | 0.002986 | 0.004059 | 0.001035 | 0.011656 | 0.005731 |
| 36.19444444 | 0.005673 | 0.070023 | 0.012416 | 0.021799 | 0.020026 | 0.023053 | 0.022041 | 0.051992 | 0.022123 | 0.028014 | 0.037154 | 0.060121 | 0.00057 | 0.00205 | 0.000248 | 0.015481 | 0.002941 | 0.001488 | 0.013717 | 0.003463 | 0.001498 | 0.002859 | 0.013184 | 0.005818 |
| 36.20833333 | 0.009107 | 0.03048 | 0.012997 | 0.022498 | 0.031169 | 0.03646 | 0.023374 | 0.038791 | 0.030138 | 0.033558 | 0.020348 | 0.027928 | 0.000803 | 0.004599 | 0.000333 | 0.009816 | 0.002343 | 0.003296 | 0.013577 | 0.002869 | 0.00223 | 0.005327 | 0.01287 | 0.005943 |
| 36.22222222 | 0.01816 | 0.02519 | 0.013894 | 0.019447 | 0.052432 | 0.037033 | 0.021007 | 0.023815 | 0.041267 | 0.024165 | 0.010296 | 0.049396 | 0.00149 | 0.006398 | 0.001101 | 0.004984 | 0.00208 | 0.005622 | 0.010974 | 0.002467 | 0.004595 | 0.0069 | 0.009533 | 0.004617 |
| 36.23611111 | 0.026923 | 0.045831 | 0.016025 | 0.014623 | 0.064312 | 0.036022 | 0.019634 | 0.012583 | 0.042373 | 0.01016 | 0.008909 | 0.082525 | 0.001775 | 0.006537 | 0.002123 | 0.005336 | 0.002958 | 0.006803 | 0.006794 | 0.002938 | 0.005604 | 0.007154 | 0.005199 | 0.002562 |
| 36.25 | 0.025952 | 0.048544 | 0.020075 | 0.015765 | 0.059867 | 0.033507 | 0.015891 | 0.019327 | 0.038184 | 0.004995 | 0.015903 | 0.075341 | 0.001751 | 0.00609 | 0.002274 | 0.009791 | 0.004021 | 0.006158 | 0.00281 | 0.003766 | 0.004549 | 0.006837 | 0.00324 | 0.002072 |
| 36.26388889 | 0.016887 | 0.040096 | 0.024349 | 0.021879 | 0.039558 | 0.020139 | 0.008278 | 0.040267 | 0.03982 | 0.004588 | 0.031229 | 0.057634 | 0.002387 | 0.006251 | 0.001557 | 0.01343 | 0.003734 | 0.004681 | 0.000976 | 0.003759 | 0.004361 | 0.007319 | 0.004775 | 0.004504 |
| 36.27777778 | 0.012064 | 0.036799 | 0.02288 | 0.023427 | 0.028794 | 0.011009 | 0.002351 | 0.047142 | 0.041173 | 0.008745 | 0.036043 | 0.051537 | 0.004132 | 0.007577 | 0.00134 | 0.013429 | 0.002029 | 0.003268 | 0.0009 | 0.002772 | 0.005462 | 0.00834 | 0.007203 | 0.007713 |
| 36.29166667 | 0.015821 | 0.038109 | 0.014175 | 0.020631 | 0.047649 | 0.015324 | 0.001125 | 0.030009 | 0.030887 | 0.017343 | 0.021302 | 0.049238 | 0.005922 | 0.009456 | 0.001588 | 0.008856 | 0.000633 | 0.001902 | 0.001518 | 0.002026 | 0.00599 | 0.008444 | 0.006877 | 0.008738 |
| 36.30555556 | 0.019316 | 0.052542 | 0.005851 | 0.019822 | 0.076857 | 0.022332 | 0.002171 | 0.012293 | 0.016371 | 0.028068 | 0.008829 | 0.067319 | 0.006579 | 0.010669 | 0.001223 | 0.004392 | 0.000569 | 0.001007 | 0.003566 | 0.001982 | 0.005633 | 0.007621 | 0.003768 | 0.007328 |
| 36.31944444 | 0.013413 | 0.04523 | 0.003009 | 0.020705 | 0.084823 | 0.026643 | 0.003816 | 0.012936 | 0.010582 | 0.044935 | 0.008513 | 0.063115 | 0.00643 | 0.011481 | 0.000576 | 0.005039 | 0.00159 | 0.001589 | 0.006038 | 0.001604 | 0.005633 | 0.006836 | 0.001641 | 0.00449 |
| 36.33333333 | 0.005838 | 0.016285 | 0.003092 | 0.019278 | 0.066362 | 0.021631 | 0.004947 | 0.022466 | 0.012083 | 0.064379 | 0.013283 | 0.027656 | 0.006315 | 0.012166 | 0.00033 | 0.006072 | 0.003243 | 0.003484 | 0.006746 | 0.00073 | 0.006083 | 0.006386 | 0.002412 | 0.003165 |
| 36.34722222 | 0.008338 | 0.009879 | 0.006032 | 0.014817 | 0.037584 | 0.010779 | 0.004891 | 0.030004 | 0.014222 | 0.079561 | 0.019196 | 0.014962 | 0.006281 | 0.011422 | 0.000381 | 0.004285 | 0.004708 | 0.005478 | 0.005721 | 0.000226 | 0.006919 | 0.005729 | 0.003828 | 0.005548 |
| 36.36111111 | 0.018661 | 0.036849 | 0.013957 | 0.012313 | 0.02546 | 0.00628 | 0.00477 | 0.041391 | 0.023539 | 0.09809 | 0.029976 | 0.045236 | 0.006122 | 0.008895 | 0.00069 | 0.003752 | 0.005552 | 0.006975 | 0.003924 | 9.1979174 | 0.008497 | 0.004183 | 0.004089 | 0.007999 |
| 36.375 | 0.024601 | 0.066234 | 0.02033 | 0.015389 | 0.041921 | 0.010915 | 0.004078 | 0.05427 | 0.037499 | 0.112996 | 0.039768 | 0.085295 | 0.005752 | 0.005459 | 0.001278 | 0.006357 | 0.006182 | 0.007973 | 0.001977 | 0.000215 | 0.010507 | 0.00211 | 0.003118 | 0.008252 |
| 36.38888889 | 0.017115 | 0.056415 | 0.015643 | 0.0174 | 0.047055 | 0.019436 | 0.002673 | 0.048988 | 0.033635 | 0.095126 | 0.031202 | 0.080876 | 0.005159 | 0.00228 | 0.001906 | 0.009331 | 0.006923 | 0.008296 | 0.000751 | 0.000759 | 0.012374 | 0.000882 | 0.002316 | 0.00774 |
| 36.40277778 | 0.006594 | 0.032567 | 0.005778 | 0.019452 | 0.025865 | 0.032151 | 0.002954 | 0.026185 | 0.01993 | 0.053858 | 0.01553 | 0.065029 | 0.004511 | 0.000612 | 0.00243 | 0.011041 | 0.007684 | 0.007552 | 0.000631 | 0.001453 | 0.013256 | 0.001258 | 0.004045 | 0.006589 |
| 36.41666667 | 0.004031 | 0.023054 | 0.002655 | 0.022941 | 0.014122 | 0.041488 | 0.004143 | 0.013201 | 0.01574 | 0.028015 | 0.012097 | 0.06953 | 0.004056 | 0.000328 | 0.002801 | 0.011729 | 0.008489 | 0.005934 | 0.000715 | 0.001856 | 0.012613 | 0.002845 | 0.007378 | 0.00483 |
| 36.43055556 | 0.007074 | 0.01929 | 0.006664 | 0.017323 | 0.011925 | 0.034396 | 0.003874 | 0.018199 | 0.011764 | 0.026669 | 0.011541 | 0.050749 | 0.004042 | 0.000537 | 0.002855 | 0.011323 | 0.009185 | 0.004373 | 0.000662 | 0.001738 | 0.010905 | 0.004718 | 0.009001 | 0.003102 |
| 36.44444444 | 0.012968 | 0.033352 | 0.013083 | 0.007119 | 0.006808 | 0.021095 | 0.002955 | 0.02599 | 0.009932 | 0.031194 | 0.011643 | 0.019491 | 0.004403 | 0.000759 | 0.002162 | 0.010464 | 0.009115 | 0.003483 | 0.000863 | 0.001224 | 0.0092 | 0.005564 | 0.007548 | 0.00156 |
| 36.45833333 | 0.016134 | 0.055067 | 0.015918 | 0.00224 | 0.005167 | 0.014343 | 0.003573 | 0.026752 | 0.015438 | 0.027114 | 0.019201 | 0.015009 | 0.004824 | 0.00072 | 0.00132 | 0.009888 | 0.008096 | 0.003549 | 0.001277 | 0.000669 | 0.008433 | 0.004817 | 0.004067 | 0.000768 |
| 36.47222222 | 0.01671 | 0.072414 | 0.011379 | 0.002294 | 0.005849 | 0.010925 | 0.007581 | 0.022185 | 0.025624 | 0.020173 | 0.033996 | 0.034333 | 0.005126 | 0.000486 | 0.001218 | 0.009331 | 0.007028 | 0.005015 | 0.001918 | 0.000306 | 0.008769 | 0.003633 | 0.001849 | 0.001452 |
| 36.48611111 | 0.017725 | 0.089469 | 0.007896 | 0.004649 | 0.007714 | 0.006844 | 0.012663 | 0.017778 | 0.038459 | 0.014988 | 0.055398 | 0.059222 | 0.00527 | 0.000749 | 0.001595 | 0.007977 | 0.007794 | 0.008138 | 0.00337 | 0.000596 | 0.010494 | 0.003596 | 0.002191 | 0.002783 |
| 36.5 | 0.016946 | 0.092394 | 0.012765 | 0.004698 | 0.018584 | 0.005185 | 0.014915 | 0.017611 | 0.043532 | 0.021662 | 0.072257 | 0.060887 | 0.004864 | 0.00161 | 0.00299 | 0.006132 | 0.011834 | 0.012149 | 0.006422 | 0.00211 | 0.014744 | 0.005125 | 0.003014 | 0.00367 |
| 36.51388889 | 0.01614 | 0.082146 | 0.017019 | 0.002763 | 0.032287 | 0.007581 | 0.016925 | 0.025779 | 0.034895 | 0.078178 | 0.075467 | 0.033592 | 0.003442 | 0.001831 | 0.004495 | 0.005481 | 0.017806 | 0.014521 | 0.009878 | 0.004177 | 0.020409 | 0.006542 | 0.007587 | 0.004041 |
| 36.52777778 | 0.013629 | 0.064201 | 0.013825 | 0.006437 | 0.044202 | 0.015614 | 0.025303 | 0.043208 | 0.018012 | 0.194737 | 0.071172 | 0.012181 | 0.002005 | 0.001884 | 0.003706 | 0.007382 | 0.023203 | 0.013231 | 0.0109 | 0.005695 | 0.023345 | 0.005973 | 0.019108 | 0.004097 |
| 36.54166667 | 0.010417 | 0.034461 | 0.010031 | 0.02239 | 0.072991 | 0.02954 | 0.037261 | 0.059422 | 0.007674 | 0.272163 | 0.06663 | 0.014902 | 0.002062 | 0.00291 | 0.001886 | 0.010925 | 0.028169 | 0.009614 | 0.010742 | 0.007127 | 0.0226 | 0.004652 | 0.030539 | 0.00472 |
| 36.55555556 | 0.012734 | 0.010607 | 0.01168 | 0.041281 | 0.120396 | 0.040102 | 0.037888 | 0.060971 | 0.010722 | 0.220336 | 0.060111 | 0.021176 | 0.002935 | 0.003588 | 0.001207 | 0.012723 | 0.033385 | 0.006127 | 0.013789 | 0.009579 | 0.021616 | 0.00475 | 0.034078 | 0.006557 |
| 36.56944444 | 0.014668 | 0.010325 | 0.016076 | 0.04504 | 0.151314 | 0.036776 | 0.024081 | 0.051509 | 0.015326 | 0.109374 | 0.053525 | 0.015097 | 0.002775 | 0.003664 | 0.000969 | 0.010184 | 0.036164 | 0.003508 | 0.01885 | 0.012715 | 0.021581 | 0.005395 | 0.029668 | 0.007641 |
| 36.58333333 | 0.015027 | 0.016935 | 0.02079 | 0.036608 | 0.137301 | 0.026331 | 0.011485 | 0.044542 | 0.014161 | 0.043899 | 0.05097 | 0.005615 | 0.001517 | 0.004661 | 0.000818 | 0.005003 | 0.034037 | 0.003019 | 0.021398 | 0.015264 | 0.020484 | 0.004698 | 0.021465 | 0.006499 |
| 36.59722222 | 0.023178 | 0.017532 | 0.026303 | 0.029379 | 0.092517 | 0.023526 | 0.007894 | 0.042393 | 0.010096 | 0.046128 | 0.042748 | 0.004436 | 0.000571 | 0.006429 | 0.001382 | 0.001645 | 0.0285 | 0.00534 | 0.022008 | 0.01656 | 0.017417 | 0.002691 | 0.01213 | 0.004616 |
| 36.61111111 | 0.035783 | 0.027468 | 0.029669 | 0.025185 | 0.043568 | 0.027742 | 0.007188 | 0.039752 | 0.005289 | 0.072252 | 0.028518 | 0.009508 | 0.000372 | 0.007611 | 0.002124 | 0.001347 | 0.023857 | 0.007806 | 0.023402 | 0.016777 | 0.01322 | 0.001208 | 0.006218 | 0.003382 |
| 36.625 | 0.040965 | 0.050073 | 0.027314 | 0.018973 | 0.015657 | 0.031218 | 0.005612 | 0.034588 | 0.003196 | 0.07682 | 0.019471 | 0.014296 | 0.000427 | 0.007903 | 0.002277 | 0.001455 | 0.022719 | 0.008556 | 0.02567 | 0.016336 | 0.008978 | 0.001425 | 0.009048 | 0.003292 |
| 36.63888889 | 0.036263 | 0.070069 | 0.021094 | 0.00992 | 0.01661 | 0.030932 | 0.004073 | 0.028203 | 0.00541 | 0.062223 | 0.017228 | 0.014835 | 0.000843 | 0.007216 | 0.001685 | 0.001375 | 0.022938 | 0.007704 | 0.027637 | 0.015504 | 0.005279 | 0.0019 | 0.01586 | 0.004563 |
| 36.65277778 | 0.02609 | 0.077057 | 0.015375 | 0.007585 | 0.02631 | 0.026101 | 0.003674 | 0.02349 | 0.008714 | 0.053754 | 0.019942 | 0.011026 | 0.001641 | 0.004737 | 0.001068 | 0.002824 | 0.020738 | 0.005919 | 0.028038 | 0.014162 | 0.003455 | 0.001628 | 0.020565 | 0.006598 |
| 36.66666667 | 0.014768 | 0.075317 | 0.011487 | 0.018347 | 0.042998 | 0.019458 | 0.003521 | 0.020099 | 0.011089 | 0.068832 | 0.022619 | 0.008453 | 0.002413 | 0.002172 | 0.00153 | 0.005308 | 0.013116 | 0.00458 | 0.025663 | 0.012004 | 0.00634 | 0.001597 | 0.023878 | 0.007659 |
| 36.68055556 | 0.008508 | 0.079179 | 0.009117 | 0.031325 | 0.073818 | 0.014905 | 0.002713 | 0.014132 | 0.012722 | 0.10698 | 0.019504 | 0.011909 | 0.002793 | 0.001854 | 0.0028 | 0.006591 | 0.008185 | 0.00414 | 0.020371 | 0.009574 | 0.01353 | 0.002631 | 0.02525 | 0.006494 |
| 36.69444444 | 0.01072 | 0.09479 | 0.007946 | 0.032964 | 0.103055 | 0.012922 | 0.004257 | 0.007408 | 0.013363 | 0.152699 | 0.012342 | 0.020782 | 0.002292 | 0.001837 | 0.003796 | 0.005576 | 0.016405 | 0.00388 | 0.013405 | 0.008765 | 0.020292 | 0.00431 | 0.022677 | 0.00417 |
| 36.70833333 | 0.017912 | 0.115313 | 0.006836 | 0.021484 | 0.110965 | 0.01185 | 0.008992 | 0.005651 | 0.012583 | 0.18509 | 0.006644 | 0.029553 | 0.001541 | 0.002138 | 0.003454 | 0.003511 | 0.030377 | 0.003808 | 0.007123 | 0.010345 | 0.023469 | 0.006145 | 0.017348 | 0.002159 |
| 36.72222222 | 0.02363 | 0.123951 | 0.006987 | 0.007769 | 0.097833 | 0.011638 | 0.012635 | 0.006723 | 0.01077 | 0.185522 | 0.007259 | 0.030674 | 0.002076 | 0.005753 | 0.002395 | 0.003522 | 0.040658 | 0.004732 | 0.005693 | 0.012309 | 0.023674 | 0.008221 | 0.011754 | 0.001782 |
| 36.73611111 | 0.02556 | 0.111661 | 0.009916 | 0.003174 | 0.070515 | 0.014053 | 0.011588 | 0.004811 | 0.009456 | 0.160325 | 0.013695 | 0.024187 | 0.003544 | 0.012185 | 0.003068 | 0.006286 | 0.044745 | 0.00645 | 0.009683 | 0.012627 | 0.023177 | 0.010393 | 0.006421 | 0.003386 |
| 36.75 | 0.028002 | 0.087844 | 0.012172 | 0.005879 | 0.04265 | 0.018113 | 0.006807 | 0.002686 | 0.00981 | 0.128812 | 0.018169 | 0.018781 | 0.004523 | 0.019697 | 0.005407 | 0.00958 | 0.042349 | 0.008177 | 0.013423 | 0.010806 | 0.022726 | 0.01287 | 0.003879 | 0.005345 |
| 36.76388889 | 0.031001 | 0.063546 | 0.012615 | 0.009118 | 0.032061 | 0.022215 | 0.003288 | 0.005561 | 0.010725 | 0.104785 | 0.018957 | 0.016054 | 0.004895 | 0.026558 | 0.007272 | 0.012686 | 0.035749 | 0.009435 | 0.014558 | 0.007666 | 0.022112 | 0.01542 | 0.005745 | 0.006878 |
| 36.77777778 | 0.029427 | 0.041924 | 0.014297 | 0.009412 | 0.032097 | 0.023077 | 0.003501 | 0.01209 | 0.010978 | 0.107388 | 0.023198 | 0.010787 | 0.005333 | 0.031744 | 0.008008 | 0.015695 | 0.027438 | 0.00945 | 0.01605 | 0.004462 | 0.022772 | 0.016916 | 0.006477 | 0.007821 |
| 36.79166667 | 0.024511 | 0.027325 | 0.016329 | 0.00797 | 0.032386 | 0.019213 | 0.004025 | 0.01938 | 0.012201 | 0.114722 | 0.029968 | 0.0052 | 0.00618 | 0.034545 | 0.007964 | 0.018076 | 0.020456 | 0.007828 | 0.018913 | 0.002327 | 0.025728 | 0.017209 | 0.005002 | 0.008662 |
| 36.80555556 | 0.021111 | 0.021545 | 0.016105 | 0.007165 | 0.030035 | 0.015654 | 0.003154 | 0.025726 | 0.015388 | 0.088817 | 0.029741 | 0.0048 | 0.007057 | 0.034598 | 0.007005 | 0.019329 | 0.015133 | 0.005226 | 0.022822 | 0.002757 | 0.029265 | 0.016248 | 0.007015 | 0.010179 |
| 36.81944444 | 0.019283 | 0.017743 | 0.015134 | 0.006526 | 0.023365 | 0.013583 | 0.004249 | 0.02937 | 0.018237 | 0.044609 | 0.020153 | 0.008239 | 0.007639 | 0.033285 | 0.005079 | 0.019349 | 0.010927 | 0.002534 | 0.027532 | 0.00633 | 0.030279 | 0.01379 | 0.011204 | 0.010754 |
| 36.83333333 | 0.017256 | 0.013388 | 0.01586 | 0.006833 | 0.016881 | 0.010507 | 0.008793 | 0.029481 | 0.020037 | 0.014613 | 0.008851 | 0.011394 | 0.00807 | 0.030207 | 0.003114 | 0.019014 | 0.009269 | 0.001027 | 0.032563 | 0.012407 | 0.027658 | 0.01084 | 0.012093 | 0.009143 |
| 36.84722222 | 0.014173 | 0.010431 | 0.016306 | 0.008713 | 0.013079 | 0.007692 | 0.012761 | 0.026422 | 0.021852 | 0.007427 | 0.004737 | 0.013266 | 0.008224 | 0.02478 | 0.002012 | 0.018492 | 0.009192 | 0.001347 | 0.036342 | 0.019636 | 0.023166 | 0.008239 | 0.008986 | 0.007821 |
| 36.86111111 | 0.01003 | 0.007101 | 0.01289 | 0.009888 | 0.012157 | 0.007278 | 0.012118 | 0.02252 | 0.022717 | 0.012482 | 0.008402 | 0.013724 | 0.007567 | 0.017989 | 0.001458 | 0.016915 | 0.00758 | 0.002787 | 0.034991 | 0.025879 | 0.018409 | 0.005684 | 0.005229 | 0.008231 |
| 36.875 | 0.010266 | 0.00531 | 0.009448 | 0.008706 | 0.017046 | 0.008196 | 0.008412 | 0.020848 | 0.022995 | 0.012816 | 0.012374 | 0.011597 | 0.006197 | 0.011393 | 0.000927 | 0.014555 | 0.004377 | 0.00465 | 0.028302 | 0.02946 | 0.013355 | 0.003363 | 0.004567 | 0.008474 |
| 36.88888889 | 0.017327 | 0.010552 | 0.012196 | 0.005412 | 0.027322 | 0.007621 | 0.005931 | 0.021556 | 0.024402 | 0.007026 | 0.015038 | 0.007402 | 0.005077 | 0.006898 | 0.000689 | 0.01175 | 0.003428 | 0.005347 | 0.020453 | 0.029865 | 0.008267 | 0.00223 | 0.006338 | 0.008191 |
| 36.90277778 | 0.023925 | 0.020967 | 0.018139 | 0.003216 | 0.035245 | 0.004801 | 0.006502 | 0.023718 | 0.026248 | 0.005783 | 0.021508 | 0.00352 | 0.004576 | 0.005271 | 0.000965 | 0.008011 | 0.005778 | 0.003654 | 0.014439 | 0.027293 | 0.004579 | 0.002312 | 0.006981 | 0.00775 |
| 36.91666667 | 0.024829 | 0.030782 | 0.022432 | 0.003853 | 0.03699 | 0.002792 | 0.007828 | 0.026931 | 0.027864 | 0.013155 | 0.030788 | 0.001456 | 0.004263 | 0.005408 | 0.001503 | 0.003861 | 0.008445 | 0.002544 | 0.009901 | 0.022352 | 0.003422 | 0.002806 | 0.005792 | 0.006861 |
| 36.93055556 | 0.021495 | 0.035999 | 0.024792 | 0.003593 | 0.036758 | 0.003659 | 0.006741 | 0.029009 | 0.030882 | 0.024284 | 0.035727 | 0.002315 | 0.003868 | 0.005167 | 0.00193 | 0.001607 | 0.01035 | 0.004401 | 0.005807 | 0.016186 | 0.004507 | 0.00293 | 0.004407 | 0.006169 |
| 36.94444444 | 0.017609 | 0.035645 | 0.025477 | 0.003668 | 0.034825 | 0.006688 | 0.004022 | 0.027853 | 0.034525 | 0.028009 | 0.033921 | 0.004811 | 0.003227 | 0.003864 | 0.001935 | 0.002295 | 0.012087 | 0.005879 | 0.002721 | 0.010586 | 0.005721 | 0.002181 | 0.003771 | 0.005645 |
| 36.95833333 | 0.015615 | 0.028782 | 0.023404 | 0.008745 | 0.024719 | 0.013369 | 0.002901 | 0.022595 | 0.034579 | 0.02134 | 0.031376 | 0.008787 | 0.002384 | 0.00251 | 0.001599 | 0.003972 | 0.013702 | 0.004536 | 0.001067 | 0.006971 | 0.005025 | 0.001048 | 0.003801 | 0.004798 |
| 36.97222222 | 0.01669 | 0.015425 | 0.017446 | 0.015292 | 0.010597 | 0.022765 | 0.003413 | 0.015012 | 0.029813 | 0.015904 | 0.032084 | 0.020529 | 0.001645 | 0.001782 | 0.001278 | 0.005395 | 0.015016 | 0.002443 | 0.000738 | 0.005011 | 0.002906 | 0.000464 | 0.004685 | 0.003885 |
| 36.98611111 | 0.021382 | 0.006909 | 0.010717 | 0.015818 | 0.009076 | 0.023045 | 0.003658 | 0.012479 | 0.025319 | 0.015659 | 0.028446 | 0.030524 | 0.001544 | 0.001904 | 0.001201 | 0.007083 | 0.015567 | 0.002472 | 0.001719 | 0.003913 | 0.001189 | 0.00101 | 0.00701 | 0.003401 |
| 37 | 0.027877 | 0.015143 | 0.008288 | 0.009694 | 0.029174 | 0.014333 | 0.002981 | 0.018842 | 0.02684 | 0.027922 | 0.015557 | 0.022186 | 0.004619 | 0.007698 | 0.003025 | 0.010457 | 0.011806 | 0.004706 | 0.00242 | 0.004581 | 0.002455 | 0.002617 | 0.009297 | 0.005047 |
| 37.01388889 | 0.034272 | 0.029199 | 0.009469 | 0.004583 | 0.057055 | 0.013291 | 0.004696 | 0.023944 | 0.029764 | 0.058162 | 0.006561 | 0.013287 | 0.012964 | 0.028399 | 0.008674 | 0.015666 | 0.00913 | 0.006091 | 0.004998 | 0.008526 | 0.010367 | 0.00374 | 0.00886 | 0.010602 |
| 37.02777778 | 0.040121 | 0.02565 | 0.009641 | 0.008804 | 0.073001 | 0.022186 | 0.013359 | 0.017871 | 0.024035 | 0.075387 | 0.012053 | 0.029327 | 0.020896 | 0.054724 | 0.015174 | 0.019423 | 0.016385 | 0.005092 | 0.012973 | 0.013701 | 0.023198 | 0.002956 | 0.01157 | 0.017287 |
| 37.04166667 | 0.042195 | 0.013331 | 0.011559 | 0.017699 | 0.098626 | 0.031158 | 0.020731 | 0.009604 | 0.014155 | 0.071118 | 0.015898 | 0.046935 | 0.020244 | 0.058591 | 0.016504 | 0.020151 | 0.022107 | 0.002857 | 0.019552 | 0.014915 | 0.029419 | 0.001895 | 0.01967 | 0.018808 |
| 37.05555556 | 0.034065 | 0.035473 | 0.023022 | 0.019421 | 0.127875 | 0.037897 | 0.015931 | 0.021645 | 0.016171 | 0.08742 | 0.02923 | 0.05147 | 0.012927 | 0.037693 | 0.011444 | 0.018554 | 0.017366 | 0.001417 | 0.017763 | 0.011043 | 0.021729 | 0.002776 | 0.020246 | 0.013614 |
| 37.06944444 | 0.019265 | 0.077786 | 0.035826 | 0.015619 | 0.113447 | 0.051052 | 0.00663 | 0.044447 | 0.025347 | 0.118045 | 0.065504 | 0.059962 | 0.006682 | 0.017867 | 0.004906 | 0.014371 | 0.008803 | 0.002426 | 0.009888 | 0.005679 | 0.010813 | 0.004662 | 0.011532 | 0.006417 |
| 37.08333333 | 0.011874 | 0.07101 | 0.034664 | 0.017749 | 0.069357 | 0.049658 | 0.005808 | 0.044088 | 0.024047 | 0.095857 | 0.064341 | 0.060278 | 0.003842 | 0.010557 | 0.001492 | 0.008872 | 0.003406 | 0.004792 | 0.003796 | 0.002632 | 0.00972 | 0.005601 | 0.008663 | 0.002016 |
| 37.09722222 | 0.013214 | 0.031108 | 0.024326 | 0.027683 | 0.028507 | 0.02223 | 0.013642 | 0.032797 | 0.016343 | 0.043891 | 0.022588 | 0.055603 | 0.002113 | 0.007937 | 0.001619 | 0.004696 | 0.003611 | 0.006698 | 0.00347 | 0.003199 | 0.016172 | 0.004662 | 0.019014 | 0.001499 |
| 37.11111111 | 0.012584 | 0.018591 | 0.017746 | 0.029447 | 0.009264 | 0.01013 | 0.020805 | 0.026504 | 0.010554 | 0.023556 | 0.008074 | 0.04676 | 0.001047 | 0.005064 | 0.003102 | 0.002354 | 0.00558 | 0.009017 | 0.004284 | 0.004978 | 0.022735 | 0.0024 | 0.032028 | 0.002495 |
| 37.125 | 0.00995 | 0.023922 | 0.014362 | 0.020167 | 0.009949 | 0.016194 | 0.022459 | 0.015416 | 0.007293 | 0.014011 | 0.013482 | 0.026413 | 0.001323 | 0.003854 | 0.003752 | 0.001318 | 0.006259 | 0.011953 | 0.003714 | 0.005504 | 0.026881 | 0.001072 | 0.035331 | 0.002837 |
| 37.13888889 | 0.010746 | 0.028143 | 0.009834 | 0.013568 | 0.009702 | 0.020071 | 0.020127 | 0.007559 | 0.006671 | 0.005161 | 0.007384 | 0.014341 | 0.001864 | 0.005552 | 0.003246 | 0.0012 | 0.006274 | 0.012338 | 0.002992 | 0.004723 | 0.027773 | 0.001495 | 0.02841 | 0.002211 |
| 37.15277778 | 0.01003 | 0.046321 | 0.011483 | 0.011215 | 0.013228 | 0.027531 | 0.01377 | 0.01437 | 0.009543 | 0.019933 | 0.014193 | 0.020784 | 0.002125 | 0.007257 | 0.00242 | 0.002681 | 0.00616 | 0.008742 | 0.002349 | 0.003797 | 0.024315 | 0.001924 | 0.016627 | 0.001236 |
| 37.16666667 | 0.005632 | 0.074652 | 0.021648 | 0.011687 | 0.023228 | 0.032067 | 0.008985 | 0.031728 | 0.019665 | 0.051368 | 0.045397 | 0.044755 | 0.002156 | 0.005832 | 0.001547 | 0.007197 | 0.005662 | 0.004286 | 0.00214 | 0.003239 | 0.017183 | 0.001599 | 0.006552 | 0.00086 |
| 37.18055556 | 0.00265 | 0.086831 | 0.028964 | 0.013218 | 0.025683 | 0.026639 | 0.011817 | 0.04357 | 0.030222 | 0.064892 | 0.069016 | 0.062311 | 0.001446 | 0.003117 | 0.000683 | 0.010961 | 0.004513 | 0.002847 | 0.003146 | 0.002947 | 0.009923 | 0.001388 | 0.002081 | 0.001078 |
| 37.19444444 | 0.002882 | 0.076567 | 0.027677 | 0.009911 | 0.018149 | 0.016328 | 0.017052 | 0.041863 | 0.031448 | 0.049718 | 0.068715 | 0.052957 | 0.000887 | 0.002146 | 0.000573 | 0.009654 | 0.003015 | 0.004944 | 0.005426 | 0.002682 | 0.005225 | 0.001297 | 0.002357 | 0.001315 |
| 37.20833333 | 0.008688 | 0.050222 | 0.019426 | 0.009182 | 0.021209 | 0.006074 | 0.016493 | 0.029009 | 0.023599 | 0.03339 | 0.045091 | 0.035923 | 0.001696 | 0.002245 | 0.001998 | 0.005199 | 0.001605 | 0.00723 | 0.007593 | 0.001774 | 0.002829 | 0.000877 | 0.005002 | 0.001578 |
| 37.22222222 | 0.015096 | 0.024734 | 0.011994 | 0.015961 | 0.039319 | 0.001985 | 0.011454 | 0.015716 | 0.014171 | 0.030369 | 0.017449 | 0.028215 | 0.00276 | 0.001779 | 0.004362 | 0.003014 | 0.000919 | 0.006793 | 0.008051 | 0.001042 | 0.002225 | 0.000485 | 0.005858 | 0.00199 |
| 37.23611111 | 0.013181 | 0.026544 | 0.016871 | 0.015785 | 0.036215 | 0.004411 | 0.011595 | 0.018428 | 0.01583 | 0.031818 | 0.025441 | 0.029361 | 0.002902 | 0.002422 | 0.005769 | 0.006165 | 0.001453 | 0.005201 | 0.006934 | 0.001784 | 0.003366 | 0.000405 | 0.004202 | 0.002266 |
| 37.25 | 0.007577 | 0.041427 | 0.027445 | 0.008209 | 0.014896 | 0.00818 | 0.015353 | 0.029508 | 0.024818 | 0.032913 | 0.054445 | 0.034657 | 0.002371 | 0.006091 | 0.005249 | 0.011873 | 0.002478 | 0.005707 | 0.005643 | 0.003298 | 0.004825 | 0.000456 | 0.004512 | 0.00209 |
| 37.26388889 | 0.004193 | 0.032712 | 0.027491 | 0.005585 | 0.017555 | 0.007943 | 0.01179 | 0.029752 | 0.021723 | 0.026614 | 0.055656 | 0.028888 | 0.001686 | 0.010863 | 0.003641 | 0.01594 | 0.002907 | 0.007716 | 0.005036 | 0.004623 | 0.004601 | 0.000657 | 0.006757 | 0.001397 |
| 37.27777778 | 0.003675 | 0.017324 | 0.023333 | 0.005637 | 0.034169 | 0.011994 | 0.005291 | 0.024508 | 0.013395 | 0.020685 | 0.047291 | 0.019266 | 0.00134 | 0.013811 | 0.001868 | 0.018056 | 0.002205 | 0.008035 | 0.005226 | 0.005248 | 0.003024 | 0.000671 | 0.006614 | 0.000805 |
| 37.29166667 | 0.005646 | 0.023303 | 0.025237 | 0.003218 | 0.033468 | 0.025451 | 0.003644 | 0.023876 | 0.015028 | 0.025659 | 0.059674 | 0.02154 | 0.001561 | 0.013752 | 0.001006 | 0.02035 | 0.001228 | 0.006152 | 0.005836 | 0.004829 | 0.002069 | 0.000373 | 0.003952 | 0.000996 |
| 37.30555556 | 0.007704 | 0.02501 | 0.0222 | 0.004923 | 0.030507 | 0.035437 | 0.006062 | 0.018937 | 0.015355 | 0.028445 | 0.060472 | 0.026177 | 0.002285 | 0.011633 | 0.001759 | 0.021477 | 0.001277 | 0.00399 | 0.006278 | 0.003943 | 0.002294 | 0.000556 | 0.002792 | 0.00134 |
| 37.31944444 | 0.010106 | 0.011249 | 0.011938 | 0.011259 | 0.033607 | 0.042726 | 0.014008 | 0.008079 | 0.010817 | 0.021928 | 0.041926 | 0.026079 | 0.003256 | 0.009334 | 0.002695 | 0.019105 | 0.001477 | 0.002876 | 0.00626 | 0.003113 | 0.003021 | 0.002088 | 0.003901 | 0.001012 |
| 37.33333333 | 0.010545 | 0.010823 | 0.008402 | 0.01184 | 0.025985 | 0.048578 | 0.032153 | 0.004258 | 0.007586 | 0.016895 | 0.040819 | 0.020079 | 0.004243 | 0.007934 | 0.002611 | 0.013945 | 0.000977 | 0.003183 | 0.005682 | 0.002122 | 0.003654 | 0.005141 | 0.004122 | 0.000617 |
| 37.34722222 | 0.008344 | 0.024679 | 0.011223 | 0.007088 | 0.018538 | 0.050163 | 0.063288 | 0.008703 | 0.003888 | 0.017267 | 0.052638 | 0.01025 | 0.004861 | 0.007692 | 0.001858 | 0.007718 | 0.000654 | 0.004264 | 0.004463 | 0.0011 | 0.003863 | 0.00757 | 0.002682 | 0.000968 |
| 37.36111111 | 0.008603 | 0.031262 | 0.01233 | 0.005217 | 0.019893 | 0.049566 | 0.081508 | 0.015149 | 0.002169 | 0.014214 | 0.054579 | 0.003656 | 0.004631 | 0.008449 | 0.000922 | 0.003069 | 0.000774 | 0.005074 | 0.003067 | 0.000671 | 0.003603 | 0.00714 | 0.001603 | 0.001496 |
| 37.375 | 0.008537 | 0.03431 | 0.01512 | 0.005414 | 0.027531 | 0.039067 | 0.061397 | 0.023764 | 0.005719 | 0.016233 | 0.059866 | 0.010211 | 0.003731 | 0.009671 | 0.000265 | 0.001207 | 0.000822 | 0.005938 | 0.003023 | 0.000553 | 0.003646 | 0.005159 | 0.00249 | 0.001265 |
| 37.38888889 | 0.006733 | 0.045088 | 0.024119 | 0.006449 | 0.046236 | 0.020673 | 0.029644 | 0.031803 | 0.015526 | 0.03439 | 0.080414 | 0.0351 | 0.002776 | 0.010675 | 0.00041 | 0.001414 | 0.001418 | 0.007688 | 0.005067 | 0.000607 | 0.005256 | 0.004031 | 0.005443 | 0.000754 |
| 37.40277778 | 0.00636 | 0.049034 | 0.030188 | 0.00846 | 0.059837 | 0.009603 | 0.012389 | 0.033032 | 0.022348 | 0.044744 | 0.091564 | 0.050973 | 0.002027 | 0.010969 | 0.001369 | 0.003774 | 0.003188 | 0.010019 | 0.007239 | 0.001338 | 0.008067 | 0.003638 | 0.008818 | 0.000655 |
| 37.41666667 | 0.0065 | 0.031964 | 0.021719 | 0.009638 | 0.043629 | 0.005673 | 0.005215 | 0.025519 | 0.015753 | 0.033176 | 0.067879 | 0.04171 | 0.001445 | 0.010429 | 0.002459 | 0.006928 | 0.004853 | 0.011957 | 0.007416 | 0.002066 | 0.010436 | 0.002545 | 0.010564 | 0.000615 |
| 37.43055556 | 0.010601 | 0.018482 | 0.010362 | 0.015794 | 0.024209 | 0.013349 | 0.006238 | 0.013151 | 0.011742 | 0.030088 | 0.034422 | 0.045658 | 0.001073 | 0.009186 | 0.003133 | 0.00845 | 0.00526 | 0.012578 | 0.006201 | 0.002308 | 0.011706 | 0.001069 | 0.01024 | 0.000547 |
| 37.44444444 | 0.014663 | 0.023728 | 0.008829 | 0.024322 | 0.032078 | 0.034669 | 0.009386 | 0.004636 | 0.020008 | 0.040373 | 0.02113 | 0.067877 | 0.001128 | 0.007216 | 0.003298 | 0.008047 | 0.004976 | 0.011935 | 0.005559 | 0.002522 | 0.012076 | 0.000626 | 0.008846 | 0.000448 |
| 37.45833333 | 0.013537 | 0.026373 | 0.012374 | 0.021148 | 0.034993 | 0.044559 | 0.006451 | 0.008749 | 0.021399 | 0.036246 | 0.029522 | 0.059244 | 0.001791 | 0.004855 | 0.002854 | 0.007735 | 0.004865 | 0.01096 | 0.00617 | 0.002789 | 0.011343 | 0.001274 | 0.007494 | 0.000287 |
| 37.47222222 | 0.021302 | 0.032356 | 0.023874 | 0.015046 | 0.029745 | 0.062614 | 0.00402 | 0.027454 | 0.021307 | 0.033452 | 0.069704 | 0.042696 | 0.0029 | 0.003106 | 0.002163 | 0.009702 | 0.004813 | 0.010368 | 0.007092 | 0.002992 | 0.009693 | 0.002015 | 0.007369 | 0.000407 |
| 37.48611111 | 0.042396 | 0.04451 | 0.036798 | 0.018304 | 0.027898 | 0.099497 | 0.004971 | 0.048855 | 0.036109 | 0.045789 | 0.112461 | 0.059124 | 0.003865 | 0.002378 | 0.00265 | 0.014174 | 0.005078 | 0.010717 | 0.007842 | 0.003203 | 0.009288 | 0.002364 | 0.00961 | 0.001457 |
| 37.5 | 0.059415 | 0.038232 | 0.035 | 0.017769 | 0.033626 | 0.089527 | 0.00456 | 0.058136 | 0.043799 | 0.041193 | 0.104103 | 0.060144 | 0.003907 | 0.001833 | 0.005269 | 0.019623 | 0.006919 | 0.01211 | 0.009655 | 0.003372 | 0.01298 | 0.002125 | 0.014843 | 0.003504 |
| 37.51388889 | 0.063318 | 0.022766 | 0.024867 | 0.01689 | 0.068735 | 0.039134 | 0.005556 | 0.053591 | 0.03221 | 0.018844 | 0.058008 | 0.030462 | 0.00261 | 0.000991 | 0.007884 | 0.023008 | 0.009378 | 0.013773 | 0.012528 | 0.002634 | 0.019541 | 0.00169 | 0.02327 | 0.004873 |
| 37.52777778 | 0.056959 | 0.019043 | 0.019137 | 0.025571 | 0.100885 | 0.013564 | 0.01135 | 0.04483 | 0.022665 | 0.005919 | 0.022503 | 0.008756 | 0.001463 | 0.000878 | 0.007071 | 0.021625 | 0.009734 | 0.014947 | 0.013477 | 0.001476 | 0.022992 | 0.002212 | 0.033095 | 0.00399 |
| 37.54166667 | 0.043477 | 0.022748 | 0.018521 | 0.032284 | 0.109832 | 0.018155 | 0.021645 | 0.04269 | 0.02564 | 0.004155 | 0.010677 | 0.002578 | 0.001754 | 0.001747 | 0.003577 | 0.016288 | 0.009104 | 0.014866 | 0.012195 | 0.00141 | 0.021397 | 0.002357 | 0.037405 | 0.001992 |
| 37.55555556 | 0.029537 | 0.021979 | 0.018935 | 0.030931 | 0.108326 | 0.028325 | 0.034526 | 0.04895 | 0.034386 | 0.004753 | 0.011904 | 0.001621 | 0.001759 | 0.00195 | 0.00151 | 0.009724 | 0.011573 | 0.012712 | 0.012636 | 0.003287 | 0.019517 | 0.002093 | 0.031608 | 0.001435 |
| 37.56944444 | 0.020822 | 0.015239 | 0.017103 | 0.026721 | 0.096525 | 0.028614 | 0.044109 | 0.056298 | 0.038863 | 0.004466 | 0.018299 | 0.002872 | 0.000825 | 0.001254 | 0.001869 | 0.004679 | 0.016438 | 0.008102 | 0.015402 | 0.006859 | 0.019268 | 0.003923 | 0.022 | 0.002239 |

| | | | | | | | | | | | | | | | | | | | | | | | | |
|---|---|---|---|---|---|---|---|---|---|---|---|---|---|---|---|---|---|---|---|---|---|---|---|---|
| 37.58333333 | 0.015707 | 0.013981 | 0.013692 | 0.025957 | 0.084132 | 0.023023 | 0.045642 | 0.057566 | 0.033608 | 0.004421 | 0.024966 | 0.007197 | 0.000423 | 0.001068 | 0.002056 | 0.003326 | 0.019369 | 0.004767 | 0.017293 | 0.0091 | 0.017648 | 0.005821 | 0.014744 | 0.002609 |
| 37.59722222 | 0.012753 | 0.021134 | 0.013295 | 0.025621 | 0.068676 | 0.021045 | 0.042663 | 0.053498 | 0.025469 | 0.005081 | 0.033153 | 0.009039 | 0.000928 | 0.001178 | 0.001215 | 0.004499 | 0.019667 | 0.007513 | 0.017694 | 0.008998 | 0.013915 | 0.005748 | 0.009039 | 0.002279 |
| 37.61111111 | 0.011903 | 0.022469 | 0.016901 | 0.020757 | 0.045911 | 0.024923 | 0.040066 | 0.049486 | 0.023055 | 0.003463 | 0.041698 | 0.005378 | 0.001521 | 0.001215 | 0.00075 | 0.004903 | 0.019236 | 0.013192 | 0.018956 | 0.008486 | 0.010399 | 0.004642 | 0.004789 | 0.002062 |
| 37.625 | 0.00877 | 0.016335 | 0.019374 | 0.018143 | 0.030816 | 0.028222 | 0.036272 | 0.04606 | 0.024084 | 0.001995 | 0.043463 | 0.002168 | 0.001536 | 0.002067 | 0.001049 | 0.003288 | 0.018831 | 0.016531 | 0.02215 | 0.008095 | 0.008661 | 0.00276 | 0.003284 | 0.002677 |
| 37.63888889 | 0.00443 | 0.010937 | 0.019 | 0.020528 | 0.027096 | 0.026327 | 0.03064 | 0.041718 | 0.023819 | 0.00284 | 0.040662 | 0.001821 | 0.001205 | 0.003766 | 0.001495 | 0.00152 | 0.018365 | 0.016303 | 0.02558 | 0.007104 | 0.008316 | 0.001262 | 0.00377 | 0.004059 |
| 37.65277778 | 0.003928 | 0.006501 | 0.020877 | 0.018102 | 0.022049 | 0.029612 | 0.026698 | 0.041224 | 0.025955 | 0.00426 | 0.044239 | 0.003277 | 0.001038 | 0.005637 | 0.001671 | 0.002537 | 0.018299 | 0.01347 | 0.027091 | 0.004973 | 0.007911 | 0.001551 | 0.003707 | 0.00556 |
| 37.66666667 | 0.005559 | 0.003597 | 0.025152 | 0.009609 | 0.02137 | 0.038007 | 0.026353 | 0.045541 | 0.030953 | 0.005554 | 0.0549 | 0.006176 | 0.000974 | 0.006873 | 0.001204 | 0.007525 | 0.019263 | 0.009824 | 0.026457 | 0.0024 | 0.006271 | 0.002765 | 0.002743 | 0.006867 |
| 37.68055556 | 0.006127 | 0.005294 | 0.025243 | 0.003825 | 0.031801 | 0.035139 | 0.026537 | 0.04548 | 0.0299 | 0.005474 | 0.061918 | 0.006161 | 0.000647 | 0.007414 | 0.000772 | 0.013966 | 0.020345 | 0.006542 | 0.025155 | 0.000771 | 0.003542 | 0.003787 | 0.003551 | 0.00808 |
| 37.69444444 | 0.004509 | 0.012109 | 0.021375 | 0.002344 | 0.042565 | 0.024424 | 0.024676 | 0.039278 | 0.022801 | 0.004318 | 0.061421 | 0.00356 | 0.000264 | 0.008978 | 0.001199 | 0.017076 | 0.018664 | 0.004399 | 0.022193 | 0.000391 | 0.002713 | 0.004813 | 0.006077 | 0.00919 |
| 37.70833333 | 0.002197 | 0.018419 | 0.019377 | 0.002192 | 0.052412 | 0.019804 | 0.022828 | 0.034696 | 0.018391 | 0.002592 | 0.059203 | 0.003359 | 0.000347 | 0.011991 | 0.00153 | 0.014751 | 0.013601 | 0.003849 | 0.015397 | 0.000501 | 0.005721 | 0.0061 | 0.008157 | 0.009367 |
| 37.72222222 | 0.001753 | 0.022157 | 0.01892 | 0.00321 | 0.061305 | 0.017043 | 0.022363 | 0.032284 | 0.016373 | 0.001185 | 0.056007 | 0.005894 | 0.000756 | 0.014996 | 0.001191 | 0.009112 | 0.008197 | 0.004504 | 0.00769 | 0.001338 | 0.009606 | 0.007289 | 0.01032 | 0.007814 |
| 37.73611111 | 0.003395 | 0.021659 | 0.018713 | 0.006378 | 0.066621 | 0.013751 | 0.021648 | 0.030917 | 0.015473 | 0.002513 | 0.051528 | 0.008073 | 0.001233 | 0.017047 | 0.000657 | 0.004083 | 0.004592 | 0.005736 | 0.005159 | 0.003464 | 0.011933 | 0.008271 | 0.014161 | 0.00493 |
| 37.75 | 0.004312 | 0.0137 | 0.020788 | 0.009138 | 0.071437 | 0.016845 | 0.019966 | 0.034935 | 0.019345 | 0.007169 | 0.049853 | 0.00704 | 0.001997 | 0.01884 | 0.000512 | 0.002443 | 0.005005 | 0.007234 | 0.008852 | 0.006879 | 0.013302 | 0.009654 | 0.019244 | 0.002412 |
| 37.76388889 | 0.003085 | 0.006581 | 0.023779 | 0.009797 | 0.069505 | 0.021513 | 0.018313 | 0.041777 | 0.023773 | 0.010871 | 0.048031 | 0.004738 | 0.003082 | 0.021916 | 0.001189 | 0.003807 | 0.010212 | 0.008563 | 0.014077 | 0.011024 | 0.014754 | 0.011053 | 0.023031 | 0.002136 |
| 37.77777778 | 0.00189 | 0.006735 | 0.025366 | 0.009903 | 0.052612 | 0.019425 | 0.016916 | 0.044558 | 0.022421 | 0.009086 | 0.041365 | 0.004651 | 0.004059 | 0.024512 | 0.002213 | 0.006605 | 0.01647 | 0.008692 | 0.017791 | 0.013443 | 0.016263 | 0.011648 | 0.022872 | 0.003562 |
| 37.79166667 | 0.002355 | 0.009197 | 0.025611 | 0.010174 | 0.027913 | 0.015345 | 0.016254 | 0.043858 | 0.018142 | 0.00421 | 0.033217 | 0.005931 | 0.004236 | 0.024276 | 0.002871 | 0.00991 | 0.020799 | 0.006775 | 0.019974 | 0.011832 | 0.016309 | 0.011937 | 0.019505 | 0.004625 |
| 37.80555556 | 0.003722 | 0.010024 | 0.024796 | 0.010927 | 0.010769 | 0.014989 | 0.016401 | 0.043111 | 0.015326 | 0.001658 | 0.027671 | 0.006633 | 0.003393 | 0.022278 | 0.002853 | 0.012049 | 0.022042 | 0.00355 | 0.020946 | 0.007701 | 0.014544 | 0.012094 | 0.016126 | 0.004743 |
| 37.81944444 | 0.004517 | 0.010959 | 0.023564 | 0.012083 | 0.006663 | 0.017105 | 0.016753 | 0.042741 | 0.014115 | 0.002249 | 0.023296 | 0.005899 | 0.002348 | 0.020044 | 0.002339 | 0.011213 | 0.02069 | 0.00133 | 0.021026 | 0.003947 | 0.012136 | 0.011719 | 0.013939 | 0.004761 |
| 37.83333333 | 0.004133 | 0.014751 | 0.022941 | 0.013391 | 0.005983 | 0.019437 | 0.018089 | 0.043265 | 0.014944 | 0.002945 | 0.018408 | 0.003653 | 0.002083 | 0.018173 | 0.001833 | 0.008126 | 0.019132 | 0.000911 | 0.020569 | 0.002063 | 0.009271 | 0.011107 | 0.011457 | 0.005131 |
| 37.84722222 | 0.002677 | 0.019835 | 0.023313 | 0.014568 | 0.005079 | 0.020477 | 0.02031 | 0.044607 | 0.017821 | 0.003127 | 0.012461 | 0.001424 | 0.002601 | 0.017054 | 0.001372 | 0.004434 | 0.018582 | 0.001153 | 0.019667 | 0.002165 | 0.006687 | 0.010396 | 0.007786 | 0.005391 |
| 37.86111111 | 0.001826 | 0.022508 | 0.023918 | 0.014684 | 0.009506 | 0.018791 | 0.020993 | 0.044332 | 0.020283 | 0.003647 | 0.006485 | 0.000498 | 0.003286 | 0.017801 | 0.000852 | 0.002623 | 0.018603 | 0.002419 | 0.017925 | 0.00228 | 0.005346 | 0.009462 | 0.003957 | 0.005282 |
| 37.875 | 0.002634 | 0.022309 | 0.023589 | 0.013642 | 0.016002 | 0.014304 | 0.018356 | 0.041354 | 0.019701 | 0.003872 | 0.00514 | 0.000383 | 0.004008 | 0.0201 | 0.00058 | 0.004683 | 0.018134 | 0.00496 | 0.015092 | 0.001481 | 0.004459 | 0.00857 | 0.002508 | 0.004635 |
| 37.88888889 | 0.004469 | 0.021079 | 0.02254 | 0.012662 | 0.019757 | 0.010823 | 0.013533 | 0.038322 | 0.01654 | 0.003395 | 0.007151 | 0.000623 | 0.004964 | 0.021465 | 0.000522 | 0.007326 | 0.015334 | 0.007575 | 0.011262 | 0.001193 | 0.003054 | 0.007684 | 0.004955 | 0.003312 |
| 37.90277778 | 0.006996 | 0.02005 | 0.021883 | 0.012187 | 0.022179 | 0.009372 | 0.008516 | 0.036377 | 0.014196 | 0.002601 | 0.008334 | 0.001446 | 0.005858 | 0.021065 | 0.00075 | 0.007721 | 0.008908 | 0.009323 | 0.006894 | 0.001895 | 0.001556 | 0.006319 | 0.008579 | 0.002111 |
| 37.91666667 | 0.009225 | 0.019242 | 0.021656 | 0.012156 | 0.026004 | 0.006627 | 0.004686 | 0.03322 | 0.013407 | 0.001826 | 0.010058 | 0.002428 | 0.006301 | 0.019445 | 0.001724 | 0.006846 | 0.003804 | 0.009373 | 0.003306 | 0.002846 | 0.001061 | 0.004865 | 0.010511 | 0.001639 |
| 37.93055556 | 0.010343 | 0.017339 | 0.021532 | 0.012866 | 0.030508 | 0.003048 | 0.003779 | 0.029194 | 0.013356 | 0.001456 | 0.012575 | 0.002403 | 0.006476 | 0.016851 | 0.003309 | 0.005388 | 0.004335 | 0.007401 | 0.001258 | 0.003724 | 0.001279 | 0.003934 | 0.010361 | 0.001588 |
| 37.94444444 | 0.01063 | 0.013766 | 0.021766 | 0.015131 | 0.028928 | 0.001732 | 0.006689 | 0.025625 | 0.016454 | 0.002035 | 0.014317 | 0.001346 | 0.006463 | 0.013868 | 0.00495 | 0.003516 | 0.006516 | 0.004333 | 0.000732 | 0.004322 | 0.00123 | 0.003111 | 0.008538 | 0.001553 |
| 37.95833333 | 0.011135 | 0.008919 | 0.022249 | 0.019223 | 0.017831 | 0.004798 | 0.012114 | 0.022317 | 0.022623 | 0.003288 | 0.015489 | 0.000489 | 0.005913 | 0.010858 | 0.005859 | 0.002129 | 0.007438 | 0.002247 | 0.001615 | 0.004098 | 0.000892 | 0.00219 | 0.006034 | 0.00136 |
| 37.97222222 | 0.012382 | 0.004823 | 0.022393 | 0.022898 | 0.008087 | 0.01152 | 0.017209 | 0.019057 | 0.026857 | 0.004291 | 0.017196 | 0.000305 | 0.004795 | 0.008386 | 0.005109 | 0.002621 | 0.007632 | 0.002726 | 0.002763 | 0.002989 | 0.000682 | 0.001491 | 0.004409 | 0.000964 |
| 37.98611111 | 0.012758 | 0.003692 | 0.021129 | 0.022248 | 0.00599 | 0.016083 | 0.018478 | 0.016946 | 0.025427 | 0.005238 | 0.020802 | 0.001512 | 0.003241 | 0.006368 | 0.003134 | 0.004806 | 0.00677 | 0.004472 | 0.002866 | 0.001881 | 0.000674 | 0.001044 | 0.004712 | 0.000839 |

Supplementary Table S6. 1 hour integrated motion data in (a) lettuce subjected to 100 mM Boron

| IntervalMid | Control_1 | Control_2 | Control_3 | Control_4 | Control_5 | Control_6 | Control_7 | Control_8 | Control_9 | Stress1 | Stress2 | Stress3 | Stress4 | Stress5 | Stress6 | Stress7 | Stress8 | Stress9 |
|---|---|---|---|---|---|---|---|---|---|---|---|---|---|---|---|---|---|---|
| 18.1111111 | 0.088445 | 0.063452 | 0.009109 | 0.025994 | 0.040796 | 0.035389 | 0.011188 | 0.035538 | 0.086846 | 0.147044 | 0.021065 | 0.255035 | 0.151093 | 0.085775 | 0.120818 | 0.08544 | 0.068111 | 0.010188 |
| 18.125 | 0.058518 | 0.032657 | 0.005873 | 0.019293 | 0.030365 | 0.022632 | 0.012877 | 0.022934 | 0.052316 | 0.107914 | 0.014153 | 0.119739 | 0.104406 | 0.0432 | 0.067548 | 0.038679 | 0.03608 | 0.018517 |
| 18.1388889 | 0.020591 | 0.021001 | 0.006079 | 0.011408 | 0.015467 | 0.009679 | 0.007911 | 0.02416 | 0.041711 | 0.044232 | 0.016932 | 0.057917 | 0.045866 | 0.019998 | 0.035264 | 0.034646 | 0.039199 | 0.030712 |
| 18.1527778 | 0.005512 | 0.025243 | 0.006489 | 0.008184 | 0.0143 | 0.011171 | 0.002568 | 0.028883 | 0.054189 | 0.028921 | 0.018659 | 0.07552 | 0.037211 | 0.020483 | 0.035343 | 0.042288 | 0.043435 | 0.026662 |
| 18.1666667 | 0.004611 | 0.021333 | 0.003649 | 0.004569 | 0.012374 | 0.013106 | 0.000518 | 0.021173 | 0.050023 | 0.036828 | 0.012081 | 0.088023 | 0.04742 | 0.022668 | 0.032831 | 0.031369 | 0.02875 | 0.011584 |
| 18.1805556 | 0.004644 | 0.01046 | 0.001063 | 0.001444 | 0.005346 | 0.008212 | 0.000844 | 0.010345 | 0.031553 | 0.030816 | 0.005911 | 0.06551 | 0.039495 | 0.024693 | 0.025016 | 0.015645 | 0.013125 | 0.008489 |
| 18.1944444 | 0.003717 | 0.003601 | 0.0003 | 0.000303 | 0.001219 | 0.003884 | 0.002138 | 0.003818 | 0.016136 | 0.017258 | 0.004826 | 0.029492 | 0.021425 | 0.023719 | 0.017281 | 0.005833 | 0.004553 | 0.009509 |
| 18.2083333 | 0.003314 | 0.001912 | 0.000547 | 0.000333 | 0.000182 | 0.002131 | 0.003194 | 0.001196 | 0.006951 | 0.006704 | 0.006605 | 0.00942 | 0.007451 | 0.014001 | 0.008552 | 0.006429 | 0.004548 | 0.007164 |
| 18.2222222 | 0.004009 | 0.001607 | 0.001158 | 0.001361 | 0.000331 | 0.001382 | 0.003007 | 0.000634 | 0.00332 | 0.004491 | 0.00898 | 0.010934 | 0.003314 | 0.007122 | 0.006136 | 0.012658 | 0.01044 | 0.008182 |
| 18.2361111 | 0.005575 | 0.001514 | 0.001583 | 0.001989 | 0.000971 | 0.001603 | 0.005364 | 0.001195 | 0.004506 | 0.009579 | 0.014797 | 0.017774 | 0.006081 | 0.01153 | 0.012946 | 0.024819 | 0.024898 | 0.017471 |
| 18.25 | 0.006256 | 0.001931 | 0.00625 | 0.008202 | 0.00137 | 0.002499 | 0.014152 | 0.001863 | 0.004928 | 0.015203 | 0.022193 | 0.015824 | 0.008029 | 0.018042 | 0.02054 | 0.038904 | 0.043129 | 0.02883 |
| 18.2638889 | 0.004295 | 0.003755 | 0.011542 | 0.018131 | 0.001165 | 0.002914 | 0.016832 | 0.001763 | 0.00278 | 0.015814 | 0.02249 | 0.009651 | 0.006518 | 0.017012 | 0.019829 | 0.040606 | 0.047056 | 0.027626 |
| 18.2777778 | 0.002474 | 0.005578 | 0.007267 | 0.014876 | 0.001029 | 0.003111 | 0.007925 | 0.00186 | 0.003165 | 0.013695 | 0.014629 | 0.017415 | 0.00763 | 0.013016 | 0.014535 | 0.030818 | 0.034263 | 0.014844 |
| 18.2916667 | 0.002923 | 0.005819 | 0.001846 | 0.00521 | 0.000808 | 0.003224 | 0.002209 | 0.002371 | 0.006977 | 0.011614 | 0.006457 | 0.03089 | 0.011073 | 0.011983 | 0.011896 | 0.021097 | 0.019333 | 0.006267 |
| 18.3055556 | 0.002867 | 0.004061 | 0.001166 | 0.002109 | 0.000374 | 0.002573 | 0.001903 | 0.001877 | 0.008721 | 0.007678 | 0.002969 | 0.026424 | 0.00888 | 0.008969 | 0.00885 | 0.012335 | 0.009492 | 0.005641 |
| 18.3194444 | 0.002278 | 0.002386 | 0.000711 | 0.001289 | 0.000643 | 0.001727 | 0.00152 | 0.000915 | 0.006301 | 0.004134 | 0.003924 | 0.016842 | 0.004712 | 0.004665 | 0.004384 | 0.006241 | 0.005913 | 0.005985 |
| 18.3333333 | 0.002305 | 0.002294 | 0.00049 | 0.000665 | 0.00172 | 0.0017 | 0.001658 | 0.001071 | 0.004595 | 0.004448 | 0.005754 | 0.020994 | 0.00577 | 0.004976 | 0.003296 | 0.006773 | 0.007041 | 0.004762 |
| 18.3472222 | 0.003282 | 0.002848 | 0.00074 | 0.000953 | 0.0023 | 0.002118 | 0.002 | 0.002511 | 0.008075 | 0.006239 | 0.00565 | 0.029483 | 0.007428 | 0.00557 | 0.004249 | 0.008688 | 0.006775 | 0.002608 |
| 18.3611111 | 0.004288 | 0.004263 | 0.00072 | 0.001286 | 0.001696 | 0.001974 | 0.001693 | 0.003782 | 0.012963 | 0.005807 | 0.003808 | 0.029187 | 0.006265 | 0.00318 | 0.003539 | 0.006768 | 0.004037 | 0.001474 |
| 18.375 | 0.003202 | 0.004181 | 0.000382 | 0.001018 | 0.001012 | 0.001365 | 0.001124 | 0.003 | 0.011832 | 0.004074 | 0.001847 | 0.017308 | 0.00494 | 0.001171 | 0.00273 | 0.004977 | 0.002856 | 0.002595 |
| 18.3888889 | 0.001895 | 0.002394 | 0.00027 | 0.000519 | 0.001203 | 0.001053 | 0.001243 | 0.001246 | 0.00625 | 0.003908 | 0.001107 | 0.006577 | 0.004456 | 0.001102 | 0.003289 | 0.007063 | 0.004031 | 0.002992 |
| 18.4027778 | 0.002626 | 0.002553 | 0.000187 | 0.000272 | 0.001451 | 0.000864 | 0.001507 | 0.000609 | 0.003114 | 0.002973 | 0.000902 | 0.007126 | 0.004161 | 0.001959 | 0.002526 | 0.007583 | 0.004738 | 0.001345 |
| 18.4166667 | 0.003086 | 0.0034 | 0.000122 | 0.000185 | 0.000997 | 0.000678 | 0.001051 | 0.001213 | 0.002451 | 0.000973 | 0.000519 | 0.009176 | 0.005379 | 0.002818 | 0.000841 | 0.005792 | 0.003451 | 0.001581 |
| 18.4305556 | 0.002742 | 0.00362 | 0.000244 | 0.000274 | 0.000471 | 0.000903 | 0.000639 | 0.002682 | 0.002617 | 0.000941 | 0.000517 | 0.010732 | 0.008765 | 0.004372 | 0.001036 | 0.004459 | 0.001932 | 0.003159 |
| 18.4444444 | 0.003973 | 0.00475 | 0.000461 | 0.000773 | 0.000257 | 0.000922 | 0.001016 | 0.003752 | 0.003513 | 0.003477 | 0.001313 | 0.022958 | 0.00989 | 0.005968 | 0.003111 | 0.005135 | 0.002248 | 0.003393 |
| 18.4583333 | 0.004009 | 0.004043 | 0.000499 | 0.001025 | 0.000266 | 0.000426 | 0.001404 | 0.002877 | 0.002945 | 0.006371 | 0.002451 | 0.035248 | 0.00689 | 0.005306 | 0.004679 | 0.008214 | 0.00338 | 0.003083 |
| 18.4722222 | 0.002704 | 0.002476 | 0.000375 | 0.000835 | 0.000397 | 0.000264 | 0.001133 | 0.001165 | 0.001986 | 0.005277 | 0.00299 | 0.027774 | 0.004691 | 0.003455 | 0.003523 | 0.007116 | 0.003859 | 0.002745 |
| 18.4861111 | 0.002545 | 0.002307 | 0.0006 | 0.00094 | 0.00068 | 0.000733 | 0.00062 | 0.000781 | 0.001692 | 0.001891 | 0.002218 | 0.011628 | 0.005176 | 0.002496 | 0.00147 | 0.002594 | 0.003372 | 0.002599 |
| 18.5 | 0.003086 | 0.002815 | 0.001167 | 0.001226 | 0.001697 | 0.001763 | 0.000506 | 0.002508 | 0.002215 | 0.000678 | 0.001747 | 0.008909 | 0.005124 | 0.001742 | 0.000715 | 0.000941 | 0.00189 | 0.002905 |
| 18.5138889 | 0.005479 | 0.005523 | 0.002381 | 0.002263 | 0.003577 | 0.003955 | 0.001369 | 0.00587 | 0.005763 | 0.00053 | 0.003237 | 0.018862 | 0.004704 | 0.002003 | 0.001157 | 0.002068 | 0.000728 | 0.005349 |
| 18.5277778 | 0.007177 | 0.007946 | 0.004155 | 0.004486 | 0.005451 | 0.006985 | 0.003052 | 0.008695 | 0.009795 | 0.000521 | 0.004397 | 0.02516 | 0.006094 | 0.004013 | 0.002462 | 0.003429 | 0.000763 | 0.007364 |
| 18.5416667 | 0.006554 | 0.007868 | 0.004932 | 0.005801 | 0.006412 | 0.008598 | 0.00424 | 0.009206 | 0.010723 | 0.001125 | 0.003365 | 0.018768 | 0.00833 | 0.00497 | 0.003402 | 0.002954 | 0.001387 | 0.005191 |
| 18.5555556 | 0.008791 | 0.010495 | 0.004018 | 0.004683 | 0.006083 | 0.007603 | 0.003791 | 0.009127 | 0.012454 | 0.005239 | 0.002421 | 0.021971 | 0.010894 | 0.006158 | 0.006921 | 0.007026 | 0.004349 | 0.002382 |
| 18.5694444 | 0.020039 | 0.0235 | 0.002264 | 0.002379 | 0.004796 | 0.005624 | 0.002913 | 0.012219 | 0.02078 | 0.025127 | 0.004445 | 0.071014 | 0.017177 | 0.01542 | 0.021479 | 0.028962 | 0.014294 | 0.00455 |
| 18.5833333 | 0.050575 | 0.041805 | 0.001822 | 0.001172 | 0.00501 | 0.006639 | 0.002367 | 0.018968 | 0.038134 | 0.074803 | 0.008264 | 0.162612 | 0.039669 | 0.043515 | 0.050536 | 0.069936 | 0.031886 | 0.010718 |
| 18.5972222 | 0.092246 | 0.056949 | 0.003672 | 0.003032 | 0.00842 | 0.012635 | 0.003248 | 0.022235 | 0.057189 | 0.142052 | 0.008427 | 0.238035 | 0.080539 | 0.080193 | 0.081744 | 0.099392 | 0.043748 | 0.015918 |
| 18.6111111 | 0.101402 | 0.059885 | 0.004671 | 0.008823 | 0.013675 | 0.016759 | 0.006132 | 0.015645 | 0.055957 | 0.169398 | 0.005618 | 0.227473 | 0.104454 | 0.087248 | 0.091224 | 0.083166 | 0.033769 | 0.019219 |
| 18.625 | 0.070484 | 0.04187 | 0.002937 | 0.013792 | 0.017638 | 0.012907 | 0.006236 | 0.006261 | 0.034294 | 0.127026 | 0.004742 | 0.148068 | 0.091138 | 0.058347 | 0.072625 | 0.040231 | 0.013637 | 0.018879 |
| 18.6388889 | 0.039848 | 0.02034 | 0.000984 | 0.012704 | 0.01664 | 0.006641 | 0.00338 | 0.002515 | 0.017255 | 0.065434 | 0.004639 | 0.08119 | 0.066125 | 0.027394 | 0.044518 | 0.011093 | 0.003907 | 0.012128 |
| 18.6527778 | 0.021997 | 0.008555 | 0.000518 | 0.009207 | 0.011863 | 0.003782 | 0.001634 | 0.001953 | 0.008869 | 0.028997 | 0.006077 | 0.045861 | 0.046512 | 0.012301 | 0.02249 | 0.00319 | 0.004481 | 0.007066 |
| 18.6666667 | 0.011712 | 0.002345 | 0.000763 | 0.00673 | 0.007581 | 0.003264 | 0.000916 | 0.001279 | 0.004166 | 0.015986 | 0.008548 | 0.022666 | 0.030363 | 0.008947 | 0.011235 | 0.008083 | 0.009879 | 0.009466 |
| 18.6805556 | 0.010665 | 0.001022 | 0.000772 | 0.004637 | 0.005781 | 0.003405 | 0.000457 | 0.001331 | 0.006458 | 0.015334 | 0.008105 | 0.013281 | 0.016068 | 0.011991 | 0.012906 | 0.020108 | 0.015621 | 0.009023 |
| 18.6944444 | 0.012447 | 0.001397 | 0.000604 | 0.003763 | 0.005973 | 0.003582 | 0.000394 | 0.002128 | 0.012576 | 0.017851 | 0.006008 | 0.012581 | 0.007427 | 0.018096 | 0.022127 | 0.031494 | 0.018848 | 0.003409 |
| 18.7083333 | 0.008942 | 0.001932 | 0.000695 | 0.00446 | 0.006029 | 0.002627 | 0.000546 | 0.001972 | 0.016431 | 0.016347 | 0.004779 | 0.010539 | 0.008664 | 0.023675 | 0.025852 | 0.033598 | 0.020036 | 0.001858 |
| 18.7222222 | 0.005531 | 0.004209 | 0.000746 | 0.005334 | 0.004523 | 0.001483 | 0.000708 | 0.001693 | 0.015984 | 0.009523 | 0.004248 | 0.007466 | 0.013431 | 0.026701 | 0.020171 | 0.026995 | 0.018044 | 0.003769 |
| 18.7361111 | 0.008995 | 0.006254 | 0.000685 | 0.005158 | 0.002205 | 0.001968 | 0.000848 | 0.001879 | 0.011096 | 0.004399 | 0.003678 | 0.007115 | 0.013416 | 0.027346 | 0.011496 | 0.020453 | 0.013042 | 0.003752 |
| 18.75 | 0.010758 | 0.005329 | 0.001178 | 0.003878 | 0.000801 | 0.002432 | 0.000887 | 0.001907 | 0.008145 | 0.005898 | 0.003714 | 0.010687 | 0.009752 | 0.023851 | 0.007375 | 0.015312 | 0.007822 | 0.005702 |
| 18.7638889 | 0.006726 | 0.002575 | 0.001851 | 0.002676 | 0.00067 | 0.001699 | 0.000711 | 0.002072 | 0.013952 | 0.007328 | 0.004333 | 0.008997 | 0.009023 | 0.016176 | 0.009915 | 0.007875 | 0.003869 | 0.01125 |
| 18.7777778 | 0.003585 | 0.001272 | 0.001608 | 0.002679 | 0.000822 | 0.00088 | 0.000504 | 0.001923 | 0.018988 | 0.005509 | 0.004188 | 0.007194 | 0.010088 | 0.010184 | 0.009747 | 0.00279 | 0.001492 | 0.012217 |
| 18.7916667 | 0.00236 | 0.002981 | 0.000834 | 0.002754 | 0.000695 | 0.000864 | 0.000614 | 0.001742 | 0.013973 | 0.006906 | 0.003413 | 0.016791 | 0.007674 | 0.009041 | 0.00685 | 0.004169 | 0.000811 | 0.007203 |
| 18.8055556 | 0.001898 | 0.00664 | 0.000728 | 0.00191 | 0.000441 | 0.001665 | 0.000658 | 0.001509 | 0.005609 | 0.010019 | 0.003262 | 0.024165 | 0.003832 | 0.009861 | 0.008136 | 0.006431 | 0.001067 | 0.003702 |
| 18.8194444 | 0.003721 | 0.009762 | 0.001253 | 0.001799 | 0.000567 | 0.002375 | 0.000412 | 0.001769 | 0.001949 | 0.009754 | 0.003858 | 0.019055 | 0.002234 | 0.010365 | 0.0083 | 0.004784 | 0.001711 | 0.003984 |
| 18.8333333 | 0.00539 | 0.010544 | 0.001559 | 0.002402 | 0.001183 | 0.002444 | 0.000285 | 0.002543 | 0.004806 | 0.008251 | 0.004372 | 0.012432 | 0.00371 | 0.010966 | 0.00664 | 0.003384 | 0.002337 | 0.003669 |
| 18.8472222 | 0.005036 | 0.008405 | 0.001125 | 0.002031 | 0.001542 | 0.001834 | 0.000569 | 0.002711 | 0.012367 | 0.007779 | 0.004399 | 0.013826 | 0.005458 | 0.011726 | 0.006893 | 0.007934 | 0.003101 | 0.001743 |
| 18.8611111 | 0.006313 | 0.004701 | 0.000583 | 0.001963 | 0.001178 | 0.000976 | 0.001287 | 0.003514 | 0.018962 | 0.007365 | 0.004166 | 0.021356 | 0.006066 | 0.011382 | 0.006251 | 0.016522 | 0.004454 | 0.001979 |
| 18.875 | 0.00875 | 0.002607 | 0.000635 | 0.002898 | 0.000531 | 0.000782 | 0.002181 | 0.004373 | 0.017709 | 0.006601 | 0.003954 | 0.032466 | 0.010367 | 0.009321 | 0.003613 | 0.022638 | 0.005477 | 0.00463 |
| 18.8888889 | 0.007603 | 0.003729 | 0.00057 | 0.002805 | 0.000249 | 0.001379 | 0.002666 | 0.003242 | 0.010911 | 0.005762 | 0.003811 | 0.039309 | 0.015646 | 0.005954 | 0.002706 | 0.021824 | 0.004356 | 0.006722 |
| 18.9027778 | 0.004241 | 0.007068 | 0.000465 | 0.001716 | 0.000274 | 0.002016 | 0.002171 | 0.002382 | 0.009095 | 0.004174 | 0.0039 | 0.030876 | 0.01261 | 0.003228 | 0.005754 | 0.013644 | 0.00313 | 0.008662 |
| 18.9166667 | 0.003303 | 0.010181 | 0.001039 | 0.001433 | 0.000218 | 0.002099 | 0.001074 | 0.003621 | 0.011646 | 0.002015 | 0.004598 | 0.024764 | 0.009308 | 0.004328 | 0.009437 | 0.006496 | 0.004408 | 0.010843 |
| 18.9305556 | 0.003235 | 0.011498 | 0.001417 | 0.001803 | 0.000325 | 0.001395 | 0.000418 | 0.003825 | 0.010003 | 0.000816 | 0.005098 | 0.033867 | 0.018632 | 0.009384 | 0.008085 | 0.00768 | 0.006398 | 0.009498 |
| 18.9444444 | 0.00363 | 0.01131 | 0.001288 | 0.002097 | 0.001019 | 0.000712 | 0.000434 | 0.002436 | 0.009608 | 0.001584 | 0.004256 | 0.034726 | 0.031937 | 0.014481 | 0.003647 | 0.013327 | 0.010494 | 0.005383 |
| 18.9583333 | 0.008313 | 0.01093 | 0.0018 | 0.002196 | 0.001846 | 0.000828 | 0.000618 | 0.003263 | 0.016037 | 0.004129 | 0.002899 | 0.023976 | 0.036757 | 0.015319 | 0.002997 | 0.022036 | 0.017877 | 0.00339 |
| 18.9722222 | 0.013912 | 0.010741 | 0.002505 | 0.001907 | 0.002122 | 0.000809 | 0.000875 | 0.006009 | 0.020927 | 0.006926 | 0.001976 | 0.014425 | 0.032571 | 0.011105 | 0.006532 | 0.031866 | 0.023086 | 0.003296 |
| 18.9861111 | 0.014649 | 0.012167 | 0.00213 | 0.002165 | 0.002031 | 0.000473 | 0.001008 | 0.006079 | 0.020537 | 0.009103 | 0.001635 | 0.010451 | 0.025742 | 0.005534 | 0.010305 | 0.035237 | 0.023803 | 0.004685 |
| 19 | 0.013156 | 0.015452 | 0.001664 | 0.003362 | 0.002373 | 0.001126 | 0.001073 | 0.003341 | 0.020098 | 0.012572 | 0.002748 | 0.016748 | 0.02275 | 0.003183 | 0.011416 | 0.031687 | 0.024203 | 0.008612 |
| 19.0138889 | 0.014789 | 0.016454 | 0.002916 | 0.004342 | 0.003838 | 0.003257 | 0.001748 | 0.002245 | 0.022334 | 0.01851 | 0.004811 | 0.023797 | 0.023702 | 0.00487 | 0.012762 | 0.028191 | 0.02608 | 0.012931 |
| 19.0277778 | 0.016955 | 0.015766 | 0.005392 | 0.005144 | 0.005488 | 0.005248 | 0.002759 | 0.003092 | 0.022738 | 0.02441 | 0.005146 | 0.01835 | 0.024858 | 0.006752 | 0.018819 | 0.027343 | 0.026008 | 0.014041 |
| 19.0416667 | 0.016311 | 0.017638 | 0.006885 | 0.005327 | 0.005492 | 0.005049 | 0.002827 | 0.003847 | 0.017375 | 0.028026 | 0.003018 | 0.018179 | 0.025064 | 0.005369 | 0.028649 | 0.024568 | 0.021005 | 0.010048 |
| 19.0555556 | 0.018064 | 0.019112 | 0.006797 | 0.003592 | 0.004536 | 0.003236 | 0.002204 | 0.006679 | 0.012168 | 0.030702 | 0.002113 | 0.052607 | 0.026122 | 0.004237 | 0.040103 | 0.018932 | 0.01541 | 0.005421 |
| 19.0694444 | 0.025499 | 0.015183 | 0.006498 | 0.002728 | 0.005163 | 0.002566 | 0.002257 | 0.013451 | 0.018685 | 0.040885 | 0.008458 | 0.116134 | 0.033005 | 0.010965 | 0.055777 | 0.022992 | 0.018806 | 0.008109 |
| 19.0833333 | 0.038281 | 0.016351 | 0.007152 | 0.007254 | 0.008298 | 0.0063 | 0.002953 | 0.02079 | 0.041304 | 0.076685 | 0.023174 | 0.192144 | 0.057124 | 0.033368 | 0.076984 | 0.05248 | 0.033943 | 0.018634 |
| 19.0972222 | 0.051441 | 0.027611 | 0.008483 | 0.016411 | 0.012955 | 0.013006 | 0.005401 | 0.022897 | 0.060824 | 0.131538 | 0.035589 | 0.232611 | 0.085603 | 0.062386 | 0.087973 | 0.085568 | 0.04492 | 0.024816 |
| 19.1111111 | 0.047146 | 0.026164 | 0.00787 | 0.021439 | 0.016545 | 0.015095 | 0.008575 | 0.015687 | 0.051113 | 0.134903 | 0.029417 | 0.169894 | 0.073805 | 0.059714 | 0.066362 | 0.07103 | 0.034463 | 0.01962 |
| 19.125 | 0.024122 | 0.013317 | 0.00459 | 0.014764 | 0.014503 | 0.009877 | 0.00774 | 0.008771 | 0.027235 | 0.070441 | 0.018252 | 0.065552 | 0.035856 | 0.031034 | 0.033478 | 0.033589 | 0.024792 | 0.015234 |
| 19.1388889 | 0.006278 | 0.012861 | 0.002779 | 0.004784 | 0.008092 | 0.003795 | 0.004175 | 0.009211 | 0.019285 | 0.022015 | 0.025856 | 0.01831 | 0.023673 | 0.018089 | 0.024702 | 0.028088 | 0.03246 | 0.013735 |
| 19.1527778 | 0.002009 | 0.018451 | 0.002577 | 0.001656 | 0.004539 | 0.001328 | 0.001902 | 0.006895 | 0.016091 | 0.014069 | 0.030906 | 0.010584 | 0.022525 | 0.013813 | 0.023514 | 0.027391 | 0.030493 | 0.009854 |
| 19.1666667 | 0.003361 | 0.024718 | 0.002594 | 0.001577 | 0.004043 | 0.000883 | 0.0014 | 0.003674 | 0.010885 | 0.019337 | 0.017904 | 0.021444 | 0.012043 | 0.007697 | 0.014509 | 0.01274 | 0.015825 | 0.012193 |
| 19.1805556 | 0.005155 | 0.03148 | 0.003699 | 0.001359 | 0.002975 | 0.001126 | 0.002017 | 0.005048 | 0.013758 | 0.024667 | 0.00577 | 0.042276 | 0.005083 | 0.004711 | 0.008403 | 0.003922 | 0.008719 | 0.016261 |
| 19.1944444 | 0.004637 | 0.026937 | 0.003182 | 0.001543 | 0.001793 | 0.001272 | 0.001931 | 0.004792 | 0.011853 | 0.017324 | 0.00192 | 0.037237 | 0.002801 | 0.00223 | 0.005626 | 0.002757 | 0.00802 | 0.010781 |
| 19.2083333 | 0.002686 | 0.016679 | 0.001215 | 0.001235 | 0.001513 | 0.001059 | 0.00122 | 0.001946 | 0.004021 | 0.005298 | 0.001094 | 0.014014 | 0.001877 | 0.001435 | 0.003318 | 0.003849 | 0.005295 | 0.003706 |
| 19.2222222 | 0.001549 | 0.010139 | 0.000308 | 0.001 | 0.001385 | 0.000994 | 0.001101 | 0.000752 | 0.001751 | 0.001506 | 0.001063 | 0.007124 | 0.00178 | 0.00143 | 0.002764 | 0.005926 | 0.002546 | 0.002414 |

| | | | | | | | | | | | | | | | | | | |
|---|---|---|---|---|---|---|---|---|---|---|---|---|---|---|---|---|---|---|
| 19.2361111 | 0.001466 | 0.00675 | 0.000182 | 0.001058 | 0.000998 | 0.000876 | 0.001074 | 0.001252 | 0.002081 | 0.001977 | 0.001729 | 0.013011 | 0.001831 | 0.001006 | 0.003501 | 0.009307 | 0.003108 | 0.002355 |
| 19.25 | 0.001593 | 0.007281 | 0.000222 | 0.000746 | 0.00089 | 0.00061 | 0.000859 | 0.001461 | 0.002022 | 0.001768 | 0.002296 | 0.011794 | 0.00343 | 0.000828 | 0.003311 | 0.011616 | 0.004845 | 0.002106 |
| 19.2638889 | 0.001383 | 0.008862 | 0.0002 | 0.000431 | 0.00101 | 0.000297 | 0.000528 | 0.00102 | 0.001677 | 0.001311 | 0.001661 | 0.005153 | 0.004963 | 0.001088 | 0.002343 | 0.010328 | 0.004228 | 0.002126 |
| 19.2777778 | 0.001765 | 0.006861 | 0.000323 | 0.000569 | 0.000944 | 8.7680562 | 0.000284 | 0.000502 | 0.000833 | 0.001541 | 0.000912 | 0.003415 | 0.004087 | 0.002677 | 0.002172 | 0.006675 | 0.002639 | 0.001923 |
| 19.2916667 | 0.003328 | 0.002875 | 0.000758 | 0.0008 | 0.000606 | 8.4972229 | 0.000504 | 0.000361 | 0.000572 | 0.003785 | 0.001535 | 0.007621 | 0.002938 | 0.005217 | 0.00374 | 0.003307 | 0.002924 | 0.002328 |
| 19.3055556 | 0.004816 | 0.000569 | 0.001204 | 0.000781 | 0.000474 | 0.000172 | 0.000916 | 0.000342 | 0.001362 | 0.007535 | 0.002769 | 0.012226 | 0.00301 | 0.007074 | 0.007268 | 0.001448 | 0.00282 | 0.002401 |
| 19.3194444 | 0.004432 | 0.001888 | 0.001026 | 0.000791 | 0.000865 | 0.000257 | 0.000854 | 0.000372 | 0.001674 | 0.008945 | 0.004 | 0.014886 | 0.005585 | 0.007352 | 0.008646 | 0.001699 | 0.002836 | 0.005714 |
| 19.3333333 | 0.002513 | 0.007055 | 0.000407 | 0.00119 | 0.001332 | 0.000522 | 0.000704 | 0.000556 | 0.00152 | 0.008641 | 0.006557 | 0.0156 | 0.011162 | 0.007981 | 0.007621 | 0.003831 | 0.005917 | 0.014127 |
| 19.3472222 | 0.001565 | 0.011864 | 0.000144 | 0.001332 | 0.001515 | 0.000937 | 0.001048 | 0.000709 | 0.002296 | 0.012041 | 0.010779 | 0.01586 | 0.014384 | 0.011705 | 0.011081 | 0.009338 | 0.011038 | 0.019331 |
| 19.3611111 | 0.001393 | 0.013838 | 0.000505 | 0.000871 | 0.001184 | 0.001101 | 0.00123 | 0.000628 | 0.002987 | 0.015864 | 0.012931 | 0.015423 | 0.011614 | 0.015069 | 0.016204 | 0.0158 | 0.014422 | 0.016983 |
| 19.375 | 0.001025 | 0.016175 | 0.001353 | 0.000464 | 0.00058 | 0.000829 | 0.00095 | 0.000689 | 0.003404 | 0.01295 | 0.01003 | 0.013637 | 0.006455 | 0.012677 | 0.013928 | 0.014258 | 0.013621 | 0.012209 |
| 19.3888889 | 0.001825 | 0.020657 | 0.002151 | 0.000482 | 0.000396 | 0.000396 | 0.000838 | 0.00111 | 0.004903 | 0.006873 | 0.005276 | 0.014041 | 0.002761 | 0.006642 | 0.007154 | 0.006999 | 0.01021 | 0.008797 |
| 19.4027778 | 0.004328 | 0.025802 | 0.00251 | 0.000934 | 0.000677 | 0.000258 | 0.001094 | 0.002054 | 0.007725 | 0.004991 | 0.003513 | 0.012465 | 0.001181 | 0.003035 | 0.00485 | 0.003555 | 0.007419 | 0.006758 |
| 19.4166667 | 0.007224 | 0.028748 | 0.002691 | 0.001355 | 0.001122 | 0.000526 | 0.001362 | 0.003416 | 0.010683 | 0.006871 | 0.004883 | 0.005925 | 0.001351 | 0.00346 | 0.007794 | 0.002869 | 0.006058 | 0.006419 |
| 19.4305556 | 0.007399 | 0.027602 | 0.002795 | 0.001251 | 0.00122 | 0.000713 | 0.001417 | 0.003766 | 0.011598 | 0.008153 | 0.006235 | 0.001884 | 0.002558 | 0.005131 | 0.010606 | 0.003454 | 0.005106 | 0.007474 |
| 19.4444444 | 0.004317 | 0.025293 | 0.002581 | 0.000799 | 0.000933 | 0.000647 | 0.001336 | 0.00247 | 0.010483 | 0.007103 | 0.005611 | 0.002989 | 0.003095 | 0.005268 | 0.010101 | 0.00866 | 0.004092 | 0.007761 |
| 19.4583333 | 0.001821 | 0.025959 | 0.002472 | 0.000928 | 0.00106 | 0.000923 | 0.001028 | 0.00127 | 0.010152 | 0.004561 | 0.003418 | 0.005108 | 0.003827 | 0.003767 | 0.007328 | 0.015339 | 0.002865 | 0.005543 |
| 19.4722222 | 0.001616 | 0.027792 | 0.00308 | 0.002479 | 0.001576 | 0.001732 | 0.000972 | 0.00132 | 0.01153 | 0.002632 | 0.00223 | 0.008283 | 0.007857 | 0.003927 | 0.004208 | 0.01966 | 0.002964 | 0.003577 |
| 19.4861111 | 0.00193 | 0.026602 | 0.003712 | 0.004172 | 0.001893 | 0.00236 | 0.001418 | 0.002111 | 0.012593 | 0.003687 | 0.003306 | 0.018598 | 0.012243 | 0.007246 | 0.003631 | 0.023386 | 0.006027 | 0.003898 |
| 19.5 | 0.002427 | 0.024431 | 0.003857 | 0.004578 | 0.00256 | 0.002813 | 0.001621 | 0.003615 | 0.012968 | 0.006308 | 0.004644 | 0.030515 | 0.014214 | 0.010474 | 0.005811 | 0.026448 | 0.010912 | 0.004739 |
| 19.5138889 | 0.004645 | 0.025252 | 0.004447 | 0.005371 | 0.004491 | 0.004471 | 0.002411 | 0.005872 | 0.013943 | 0.006127 | 0.005811 | 0.026275 | 0.017779 | 0.011598 | 0.005117 | 0.024972 | 0.016633 | 0.007856 |
| 19.5277778 | 0.007506 | 0.02678 | 0.00556 | 0.007681 | 0.007048 | 0.007262 | 0.004378 | 0.007966 | 0.015214 | 0.003975 | 0.007716 | 0.009998 | 0.021966 | 0.011852 | 0.002099 | 0.019618 | 0.021796 | 0.012712 |
| 19.5416667 | 0.008773 | 0.026273 | 0.005962 | 0.008976 | 0.00826 | 0.00872 | 0.005606 | 0.009041 | 0.016311 | 0.00493 | 0.009449 | 0.008105 | 0.020683 | 0.013129 | 0.003891 | 0.016488 | 0.023559 | 0.013437 |
| 19.5555556 | 0.010514 | 0.027516 | 0.005286 | 0.006717 | 0.007106 | 0.007447 | 0.004381 | 0.009559 | 0.018683 | 0.011521 | 0.009922 | 0.029228 | 0.014305 | 0.015419 | 0.011917 | 0.018286 | 0.021122 | 0.008969 |
| 19.5694444 | 0.018221 | 0.034267 | 0.005043 | 0.004084 | 0.005239 | 0.004942 | 0.002707 | 0.011039 | 0.02648 | 0.030223 | 0.008376 | 0.063919 | 0.009081 | 0.018218 | 0.023014 | 0.028852 | 0.017692 | 0.004835 |
| 19.5833333 | 0.037575 | 0.042709 | 0.006582 | 0.003482 | 0.004903 | 0.00476 | 0.00229 | 0.013054 | 0.048004 | 0.078694 | 0.00544 | 0.112057 | 0.018643 | 0.028137 | 0.036265 | 0.061133 | 0.021858 | 0.006065 |
| 19.5972222 | 0.058988 | 0.037327 | 0.00778 | 0.005422 | 0.008173 | 0.009588 | 0.003881 | 0.012772 | 0.068264 | 0.146534 | 0.003431 | 0.146347 | 0.051945 | 0.045068 | 0.048687 | 0.089053 | 0.029158 | 0.010897 |
| 19.6111111 | 0.061582 | 0.019706 | 0.005488 | 0.010173 | 0.014864 | 0.016252 | 0.008549 | 0.009753 | 0.057873 | 0.163481 | 0.003961 | 0.1102 | 0.07444 | 0.049363 | 0.055449 | 0.06776 | 0.025848 | 0.011925 |
| 19.625 | 0.045558 | 0.014204 | 0.002308 | 0.011809 | 0.017676 | 0.018084 | 0.011675 | 0.00699 | 0.029059 | 0.106824 | 0.008279 | 0.044848 | 0.053394 | 0.035245 | 0.060614 | 0.025427 | 0.020219 | 0.008041 |
| 19.6388889 | 0.026327 | 0.025247 | 0.002392 | 0.009741 | 0.013306 | 0.013937 | 0.010587 | 0.006548 | 0.010962 | 0.047112 | 0.014657 | 0.032762 | 0.02114 | 0.019493 | 0.06261 | 0.011566 | 0.019161 | 0.007188 |
| 19.6527778 | 0.014834 | 0.040742 | 0.003771 | 0.007047 | 0.00745 | 0.008469 | 0.007965 | 0.006717 | 0.005856 | 0.022465 | 0.01811 | 0.050756 | 0.011367 | 0.010604 | 0.051711 | 0.028411 | 0.013528 | 0.010217 |
| 19.6666667 | 0.012858 | 0.048239 | 0.003632 | 0.003713 | 0.003313 | 0.005004 | 0.005356 | 0.005158 | 0.005087 | 0.016062 | 0.017095 | 0.054685 | 0.011041 | 0.008989 | 0.034903 | 0.046084 | 0.006368 | 0.011431 |
| 19.6805556 | 0.012795 | 0.042294 | 0.002514 | 0.001532 | 0.001197 | 0.003493 | 0.003431 | 0.003635 | 0.005008 | 0.013514 | 0.013519 | 0.040502 | 0.013316 | 0.013431 | 0.024997 | 0.036003 | 0.005323 | 0.010733 |
| 19.6944444 | 0.009226 | 0.027475 | 0.002581 | 0.000835 | 0.000657 | 0.002776 | 0.002186 | 0.005102 | 0.006111 | 0.011364 | 0.008567 | 0.038336 | 0.032466 | 0.015803 | 0.020339 | 0.020942 | 0.004743 | 0.009351 |
| 19.7083333 | 0.004505 | 0.012385 | 0.00363 | 0.000758 | 0.001007 | 0.001832 | 0.001234 | 0.00823 | 0.013281 | 0.011313 | 0.003997 | 0.074208 | 0.060514 | 0.010823 | 0.012765 | 0.031439 | 0.004042 | 0.006113 |
| 19.7222222 | 0.002553 | 0.005812 | 0.003077 | 0.001069 | 0.002538 | 0.00111 | 0.000502 | 0.009261 | 0.026832 | 0.015236 | 0.00264 | 0.097157 | 0.070882 | 0.005341 | 0.006959 | 0.040558 | 0.006496 | 0.002897 |
| 19.7361111 | 0.00336 | 0.007988 | 0.002746 | 0.000912 | 0.004104 | 0.001362 | 0.000254 | 0.006363 | 0.039901 | 0.020975 | 0.003909 | 0.07456 | 0.053762 | 0.00417 | 0.012429 | 0.027391 | 0.006253 | 0.002833 |
| 19.75 | 0.005933 | 0.011052 | 0.005046 | 0.000711 | 0.003362 | 0.001948 | 0.000434 | 0.003009 | 0.042144 | 0.031062 | 0.004192 | 0.069801 | 0.043804 | 0.004002 | 0.022784 | 0.016035 | 0.007324 | 0.003635 |
| 19.7638889 | 0.009076 | 0.011145 | 0.006772 | 0.001387 | 0.002301 | 0.001998 | 0.00088 | 0.0028 | 0.026733 | 0.040624 | 0.002555 | 0.099138 | 0.060249 | 0.005011 | 0.028585 | 0.027757 | 0.017363 | 0.002463 |
| 19.7777778 | 0.008316 | 0.014348 | 0.005665 | 0.00163 | 0.004271 | 0.001564 | 0.001602 | 0.002912 | 0.011142 | 0.03238 | 0.000895 | 0.096423 | 0.0676 | 0.006233 | 0.028581 | 0.046126 | 0.029605 | 0.001942 |
| 19.7916667 | 0.00636 | 0.021755 | 0.003233 | 0.001602 | 0.006688 | 0.001531 | 0.001799 | 0.001798 | 0.012924 | 0.016926 | 0.000701 | 0.054655 | 0.053685 | 0.004528 | 0.025705 | 0.044925 | 0.033935 | 0.005347 |
| 19.8055556 | 0.007193 | 0.022993 | 0.002101 | 0.00205 | 0.006665 | 0.001414 | 0.001225 | 0.00122 | 0.020303 | 0.013281 | 0.002011 | 0.031754 | 0.033954 | 0.003862 | 0.023893 | 0.02698 | 0.026846 | 0.011528 |
| 19.8194444 | 0.006225 | 0.015212 | 0.003315 | 0.00155 | 0.004778 | 0.001216 | 0.001123 | 0.001338 | 0.0202 | 0.017779 | 0.003706 | 0.041391 | 0.01565 | 0.00875 | 0.024524 | 0.014243 | 0.014006 | 0.015461 |
| 19.8333333 | 0.004884 | 0.010244 | 0.003407 | 0.000995 | 0.002839 | 0.001944 | 0.001664 | 0.001243 | 0.012566 | 0.021757 | 0.004174 | 0.037056 | 0.006431 | 0.014182 | 0.024122 | 0.011737 | 0.006891 | 0.01395 |
| 19.8472222 | 0.005873 | 0.014678 | 0.002088 | 0.001297 | 0.002804 | 0.00261 | 0.00184 | 0.001114 | 0.005691 | 0.020409 | 0.002905 | 0.019851 | 0.010931 | 0.012109 | 0.017751 | 0.009803 | 0.007343 | 0.009989 |
| 19.8611111 | 0.006216 | 0.0206 | 0.002539 | 0.001098 | 0.003854 | 0.002435 | 0.001272 | 0.002455 | 0.007136 | 0.013105 | 0.002195 | 0.022578 | 0.024343 | 0.006026 | 0.008372 | 0.012597 | 0.006916 | 0.00674 |
| 19.875 | 0.005228 | 0.022891 | 0.003545 | 0.001475 | 0.003696 | 0.001572 | 0.000654 | 0.004183 | 0.012125 | 0.004964 | 0.003412 | 0.02439 | 0.045947 | 0.004367 | 0.003038 | 0.018771 | 0.0072 | 0.003511 |
| 19.8888889 | 0.004694 | 0.023834 | 0.003934 | 0.00395 | 0.002245 | 0.000877 | 0.000626 | 0.003369 | 0.011133 | 0.001695 | 0.004482 | 0.014264 | 0.061956 | 0.004198 | 0.002954 | 0.019987 | 0.012939 | 0.001089 |
| 19.9027778 | 0.006064 | 0.018695 | 0.004295 | 0.006288 | 0.00312 | 0.001144 | 0.000969 | 0.001823 | 0.013222 | 0.003801 | 0.003702 | 0.015722 | 0.045073 | 0.003822 | 0.007144 | 0.020172 | 0.017387 | 0.001101 |
| 19.9166667 | 0.010885 | 0.011742 | 0.003657 | 0.005387 | 0.002837 | 0.001318 | 0.001199 | 0.003512 | 0.032719 | 0.009677 | 0.002823 | 0.034343 | 0.029355 | 0.01599 | 0.014367 | 0.025419 | 0.016989 | 0.003819 |
| 19.9305556 | 0.015373 | 0.015367 | 0.002015 | 0.003738 | 0.006224 | 0.001535 | 0.001122 | 0.006465 | 0.048755 | 0.016232 | 0.00512 | 0.05744 | 0.061337 | 0.015996 | 0.023389 | 0.035814 | 0.017821 | 0.010523 |
| 19.9444444 | 0.012739 | 0.023221 | 0.000946 | 0.006398 | 0.009159 | 0.003093 | 0.001561 | 0.007412 | 0.042663 | 0.020716 | 0.0092 | 0.068056 | 0.101979 | 0.023967 | 0.033753 | 0.045053 | 0.023844 | 0.020413 |
| 19.9583333 | 0.008773 | 0.030018 | 0.000964 | 0.011075 | 0.010801 | 0.005446 | 0.002763 | 0.008659 | 0.02668 | 0.02392 | 0.010567 | 0.055926 | 0.106906 | 0.026487 | 0.040114 | 0.045432 | 0.031779 | 0.027868 |
| 19.9722222 | 0.011227 | 0.03574 | 0.001955 | 0.01183 | 0.01113 | 0.006711 | 0.003098 | 0.012964 | 0.013592 | 0.026516 | 0.008218 | 0.03285 | 0.085017 | 0.017522 | 0.037419 | 0.036521 | 0.035239 | 0.026887 |
| 19.9861111 | 0.016864 | 0.033247 | 0.003192 | 0.009074 | 0.010395 | 0.006082 | 0.001882 | 0.018531 | 0.00928 | 0.027004 | 0.004444 | 0.017854 | 0.055962 | 0.006521 | 0.028901 | 0.023903 | 0.028535 | 0.01785 |
| 20 | 0.02221 | 0.024007 | 0.003642 | 0.007169 | 0.009588 | 0.00579 | 0.000697 | 0.021079 | 0.012382 | 0.025635 | 0.002233 | 0.020495 | 0.028325 | 0.003769 | 0.022523 | 0.014333 | 0.014459 | 0.008283 |
| 20.0138889 | 0.02522 | 0.016306 | 0.003735 | 0.009029 | 0.009548 | 0.007892 | 0.001428 | 0.019102 | 0.015499 | 0.026768 | 0.001751 | 0.028169 | 0.011851 | 0.005624 | 0.022443 | 0.012388 | 0.004527 | 0.005051 |
| 20.0277778 | 0.024441 | 0.013039 | 0.004001 | 0.013717 | 0.010249 | 0.010713 | 0.003922 | 0.015441 | 0.016024 | 0.031628 | 0.001318 | 0.025119 | 0.009036 | 0.00794 | 0.023753 | 0.01561 | 0.002385 | 0.007116 |
| 20.0416667 | 0.022739 | 0.01399 | 0.003486 | 0.015481 | 0.011058 | 0.011289 | 0.005492 | 0.012641 | 0.01548 | 0.034775 | 0.001293 | 0.032014 | 0.013494 | 0.008136 | 0.020874 | 0.016993 | 0.002385 | 0.008143 |
| 20.0555556 | 0.023499 | 0.019985 | 0.001896 | 0.011106 | 0.012094 | 0.010005 | 0.004386 | 0.012503 | 0.014553 | 0.036328 | 0.001516 | 0.069659 | 0.033698 | 0.008368 | 0.019879 | 0.015766 | 0.003168 | 0.006612 |
| 20.0694444 | 0.026998 | 0.033737 | 0.000801 | 0.005137 | 0.013691 | 0.010218 | 0.002027 | 0.017081 | 0.017969 | 0.048289 | 0.003629 | 0.12347 | 0.058137 | 0.011683 | 0.03035 | 0.017055 | 0.007041 | 0.0049 |
| 20.0833333 | 0.031134 | 0.055511 | 0.001248 | 0.004209 | 0.015552 | 0.013364 | 0.001368 | 0.024422 | 0.032715 | 0.082437 | 0.008408 | 0.160933 | 0.06649 | 0.015369 | 0.044978 | 0.019852 | 0.008408 | 0.009215 |
| 20.0972222 | 0.036255 | 0.070222 | 0.002822 | 0.010333 | 0.018061 | 0.016413 | 0.003547 | 0.027693 | 0.050906 | 0.123445 | 0.00885 | 0.1531 | 0.067248 | 0.014726 | 0.044343 | 0.016469 | 0.00704 | 0.020329 |
| 20.1111111 | 0.03389 | 0.057423 | 0.004036 | 0.016496 | 0.018047 | 0.014852 | 0.005319 | 0.022421 | 0.053951 | 0.114196 | 0.004573 | 0.104789 | 0.061423 | 0.008939 | 0.026454 | 0.007573 | 0.011669 | 0.023296 |
| 20.125 | 0.017643 | 0.028167 | 0.004121 | 0.0144 | 0.01118 | 0.008996 | 0.003952 | 0.013501 | 0.040968 | 0.055802 | 0.00283 | 0.058055 | 0.043395 | 0.005617 | 0.012557 | 0.002972 | 0.017281 | 0.014699 |
| 20.1388889 | 0.004178 | 0.013618 | 0.004266 | 0.007183 | 0.00355 | 0.003393 | 0.001583 | 0.008414 | 0.025649 | 0.025521 | 0.002926 | 0.03791 | 0.025205 | 0.006565 | 0.010618 | 0.003369 | 0.017879 | 0.006174 |
| 20.1527778 | 0.001604 | 0.015564 | 0.003681 | 0.002945 | 0.002198 | 0.001699 | 0.001287 | 0.00559 | 0.012975 | 0.028963 | 0.002508 | 0.026879 | 0.013156 | 0.00565 | 0.008948 | 0.003755 | 0.016301 | 0.003623 |
| 20.1666667 | 0.001787 | 0.014011 | 0.00239 | 0.003174 | 0.003987 | 0.003726 | 0.003311 | 0.003155 | 0.004695 | 0.022476 | 0.002366 | 0.012211 | 0.007308 | 0.003926 | 0.006432 | 0.003891 | 0.013995 | 0.00614 |
| 20.1805556 | 0.002005 | 0.007521 | 0.002253 | 0.003688 | 0.004437 | 0.005033 | 0.004534 | 0.0029 | 0.002359 | 0.01048 | 0.002496 | 0.005553 | 0.008587 | 0.005101 | 0.007033 | 0.003527 | 0.010745 | 0.007956 |
| 20.1944444 | 0.002389 | 0.004396 | 0.001658 | 0.002347 | 0.002964 | 0.003463 | 0.003012 | 0.002187 | 0.001161 | 0.007019 | 0.00403 | 0.012636 | 0.009754 | 0.007225 | 0.009581 | 0.003745 | 0.008472 | 0.005432 |
| 20.2083333 | 0.002958 | 0.004673 | 0.000803 | 0.001135 | 0.001776 | 0.00177 | 0.00123 | 0.001458 | 0.001154 | 0.004903 | 0.007289 | 0.018195 | 0.01052 | 0.008765 | 0.011136 | 0.007637 | 0.010733 | 0.003133 |
| 20.2222222 | 0.003914 | 0.007643 | 0.001483 | 0.001044 | 0.001981 | 0.001316 | 0.000972 | 0.002551 | 0.003097 | 0.004319 | 0.010084 | 0.011445 | 0.014616 | 0.009492 | 0.009109 | 0.01257 | 0.015521 | 0.005492 |
| 20.2361111 | 0.003761 | 0.008561 | 0.002209 | 0.001056 | 0.001831 | 0.000951 | 0.001086 | 0.002554 | 0.002878 | 0.006975 | 0.011708 | 0.006459 | 0.018465 | 0.009543 | 0.008006 | 0.014234 | 0.018462 | 0.009526 |
| 20.25 | 0.003089 | 0.005302 | 0.00163 | 0.000755 | 0.001021 | 0.000602 | 0.000975 | 0.001235 | 0.001219 | 0.008184 | 0.012423 | 0.01247 | 0.018317 | 0.009524 | 0.010677 | 0.013637 | 0.018267 | 0.01084 |
| 20.2638889 | 0.002718 | 0.002912 | 0.00079 | 0.000663 | 0.000594 | 0.000614 | 0.001092 | 0.000761 | 0.001138 | 0.006394 | 0.011401 | 0.019152 | 0.013966 | 0.00828 | 0.011641 | 0.012072 | 0.013817 | 0.008941 |
| 20.2777778 | 0.002605 | 0.002639 | 0.000518 | 0.000848 | 0.000596 | 0.000514 | 0.001315 | 0.001444 | 0.001872 | 0.0039 | 0.008875 | 0.019094 | 0.008052 | 0.005163 | 0.008222 | 0.008401 | 0.00753 | 0.005634 |
| 20.2916667 | 0.003392 | 0.002851 | 0.000514 | 0.000708 | 0.000593 | 0.000438 | 0.001653 | 0.002601 | 0.00197 | 0.003255 | 0.006076 | 0.011316 | 0.004418 | 0.002597 | 0.004185 | 0.003708 | 0.004557 | 0.002674 |
| 20.3055556 | 0.003002 | 0.003716 | 0.000812 | 0.000337 | 0.0007 | 0.000787 | 0.001955 | 0.002634 | 0.002496 | 0.002527 | 0.004673 | 0.006215 | 0.005322 | 0.002129 | 0.002548 | 0.002184 | 0.004656 | 0.001848 |
| 20.3194444 | 0.001612 | 0.006223 | 0.001194 | 0.000325 | 0.001388 | 0.001241 | 0.001613 | 0.002553 | 0.005028 | 0.001315 | 0.00473 | 0.009639 | 0.009849 | 0.001527 | 0.001843 | 0.00386 | 0.006266 | 0.003583 |
| 20.3333333 | 0.0018 | 0.008349 | 0.004062 | 0.000888 | 0.002195 | 0.001635 | 0.001211 | 0.004705 | 0.008722 | 0.00198 | 0.004041 | 0.011276 | 0.014776 | 0.001095 | 0.003642 | 0.004246 | 0.006841 | 0.00534 |
| 20.3472222 | 0.002585 | 0.007688 | 0.00213 | 0.001978 | 0.002442 | 0.002194 | 0.001657 | 0.006776 | 0.010829 | 0.003608 | 0.003156 | 0.008141 | 0.016572 | 0.002444 | 0.009392 | 0.003025 | 0.003853 | 0.005628 |
| 20.3611111 | 0.002045 | 0.005255 | 0.001892 | 0.002332 | 0.002015 | 0.002072 | 0.002022 | 0.005474 | 0.010414 | 0.004906 | 0.002607 | 0.00646 | 0.014978 | 0.003244 | 0.011582 | 0.00265 | 0.001056 | 0.005705 |
| 20.375 | 0.001568 | 0.003327 | 0.001763 | 0.001813 | 0.002107 | 0.001614 | 0.001736 | 0.003218 | 0.009367 | 0.010633 | 0.002079 | 0.011284 | 0.013505 | 0.002514 | 0.007836 | 0.004034 | 0.000902 | 0.006454 |

| | | | | | | | | | | | | | | | | | | |
|---|---|---|---|---|---|---|---|---|---|---|---|---|---|---|---|---|---|---|
| 20.3888889 | 0.002608 | 0.003447 | 0.002348 | 0.001551 | 0.002487 | 0.001846 | 0.001841 | 0.002719 | 0.0103 | 0.020338 | 0.003167 | 0.025568 | 0.015311 | 0.00284 | 0.005591 | 0.007246 | 0.001558 | 0.006453 |
| 20.4027778 | 0.002802 | 0.006828 | 0.002813 | 0.002452 | 0.002615 | 0.003245 | 0.003283 | 0.003393 | 0.014663 | 0.023455 | 0.004017 | 0.033988 | 0.020132 | 0.004619 | 0.005266 | 0.009212 | 0.002954 | 0.005401 |
| 20.4166667 | 0.002078 | 0.012561 | 0.004282 | 0.004871 | 0.004115 | 0.006015 | 0.005165 | 0.006119 | 0.020334 | 0.016298 | 0.003109 | 0.02482 | 0.024472 | 0.006354 | 0.00316 | 0.008693 | 0.006226 | 0.007038 |
| 20.4305556 | 0.00325 | 0.016508 | 0.005662 | 0.0061 | 0.005485 | 0.007224 | 0.004712 | 0.008565 | 0.022925 | 0.009061 | 0.003071 | 0.013802 | 0.024652 | 0.007887 | 0.002024 | 0.008524 | 0.009786 | 0.010714 |
| 20.4444444 | 0.005879 | 0.01738 | 0.005407 | 0.005764 | 0.005782 | 0.006775 | 0.002917 | 0.008613 | 0.022597 | 0.008884 | 0.004474 | 0.014008 | 0.020393 | 0.008263 | 0.002985 | 0.008997 | 0.010792 | 0.011105 |
| 20.4583333 | 0.008764 | 0.019325 | 0.004814 | 0.006633 | 0.006417 | 0.007327 | 0.002605 | 0.0095 | 0.024549 | 0.01308 | 0.005938 | 0.019908 | 0.016207 | 0.006948 | 0.004228 | 0.011309 | 0.012129 | 0.009631 |
| 20.4722222 | 0.011261 | 0.023968 | 0.004833 | 0.008597 | 0.007057 | 0.008341 | 0.003105 | 0.012973 | 0.030074 | 0.018706 | 0.007619 | 0.022201 | 0.015955 | 0.005974 | 0.004781 | 0.018902 | 0.019472 | 0.01105 |
| 20.4861111 | 0.011383 | 0.025726 | 0.005223 | 0.010027 | 0.007437 | 0.00867 | 0.003241 | 0.015872 | 0.033251 | 0.022974 | 0.009292 | 0.017646 | 0.016611 | 0.006801 | 0.004849 | 0.028016 | 0.028302 | 0.014499 |
| 20.5 | 0.009666 | 0.020747 | 0.005584 | 0.01088 | 0.007897 | 0.008302 | 0.00333 | 0.014722 | 0.029805 | 0.025013 | 0.009893 | 0.014244 | 0.013969 | 0.008629 | 0.006079 | 0.032154 | 0.029507 | 0.015485 |
| 20.5138889 | 0.010189 | 0.014442 | 0.005909 | 0.011808 | 0.008492 | 0.007873 | 0.003797 | 0.01108 | 0.023695 | 0.026808 | 0.009291 | 0.017555 | 0.011792 | 0.009696 | 0.007008 | 0.03294 | 0.023925 | 0.01286 |
| 20.5277778 | 0.013183 | 0.013114 | 0.006062 | 0.012228 | 0.009278 | 0.007886 | 0.004481 | 0.009018 | 0.019687 | 0.026142 | 0.007542 | 0.016629 | 0.015312 | 0.008974 | 0.005012 | 0.032555 | 0.017101 | 0.010502 |
| 20.5416667 | 0.014962 | 0.016321 | 0.00626 | 0.012085 | 0.010247 | 0.008389 | 0.005283 | 0.010218 | 0.01651 | 0.02279 | 0.006696 | 0.009803 | 0.020817 | 0.007825 | 0.003532 | 0.030285 | 0.01288 | 0.011247 |
| 20.5555556 | 0.016189 | 0.021991 | 0.006975 | 0.011743 | 0.010498 | 0.009197 | 0.005829 | 0.012178 | 0.013956 | 0.026791 | 0.01154 | 0.019743 | 0.021523 | 0.010026 | 0.012363 | 0.03313 | 0.015666 | 0.014238 |
| 20.5694444 | 0.02411 | 0.03123 | 0.00762 | 0.00978 | 0.009669 | 0.010314 | 0.005595 | 0.012642 | 0.018819 | 0.048348 | 0.024846 | 0.06631 | 0.022295 | 0.017168 | 0.034298 | 0.050104 | 0.028347 | 0.018602 |
| 20.5833333 | 0.047871 | 0.048554 | 0.007643 | 0.009325 | 0.012879 | 0.013338 | 0.008159 | 0.014384 | 0.039049 | 0.091132 | 0.041754 | 0.139224 | 0.043254 | 0.030323 | 0.060848 | 0.078516 | 0.046135 | 0.024397 |
| 20.5972222 | 0.078157 | 0.069301 | 0.007758 | 0.017681 | 0.024693 | 0.020869 | 0.018436 | 0.019303 | 0.075849 | 0.136487 | 0.05101 | 0.158729 | 0.078635 | 0.047203 | 0.082454 | 0.09515 | 0.055663 | 0.028177 |
| 20.6111111 | 0.080915 | 0.071402 | 0.007468 | 0.027057 | 0.032908 | 0.026845 | 0.027416 | 0.022896 | 0.099866 | 0.13503 | 0.044301 | 0.13662 | 0.086273 | 0.052234 | 0.086603 | 0.077862 | 0.050286 | 0.024422 |
| 20.625 | 0.049453 | 0.047665 | 0.004995 | 0.024517 | 0.027155 | 0.023045 | 0.024181 | 0.019202 | 0.076059 | 0.081928 | 0.026734 | 0.157024 | 0.059028 | 0.035274 | 0.068236 | 0.041435 | 0.037853 | 0.014983 |
| 20.6388889 | 0.019577 | 0.020993 | 0.00402 | 0.015733 | 0.017155 | 0.014792 | 0.016358 | 0.01117 | 0.038737 | 0.032149 | 0.013608 | 0.147327 | 0.027314 | 0.016254 | 0.039602 | 0.022442 | 0.023376 | 0.006938 |
| 20.6527778 | 0.012152 | 0.010898 | 0.008583 | 0.008983 | 0.009196 | 0.008388 | 0.011594 | 0.009301 | 0.038605 | 0.013987 | 0.011677 | 0.096067 | 0.011127 | 0.011746 | 0.01902 | 0.036566 | 0.010348 | 0.003633 |
| 20.6666667 | 0.017005 | 0.01175 | 0.012382 | 0.004072 | 0.003173 | 0.004511 | 0.008336 | 0.011954 | 0.059876 | 0.01885 | 0.016008 | 0.077055 | 0.009321 | 0.013838 | 0.018232 | 0.058725 | 0.004242 | 0.006568 |
| 20.6805556 | 0.019337 | 0.011094 | 0.009284 | 0.001585 | 0.000946 | 0.004239 | 0.005953 | 0.010233 | 0.065279 | 0.02966 | 0.020561 | 0.06305 | 0.009448 | 0.024327 | 0.028087 | 0.056762 | 0.003825 | 0.013781 |
| 20.6944444 | 0.014845 | 0.015589 | 0.004231 | 0.002963 | 0.001326 | 0.005936 | 0.00378 | 0.01007 | 0.04504 | 0.031501 | 0.021351 | 0.038384 | 0.009021 | 0.041433 | 0.028725 | 0.034184 | 0.008278 | 0.01965 |
| 20.7083333 | 0.016696 | 0.031115 | 0.003967 | 0.007642 | 0.002076 | 0.005413 | 0.002703 | 0.01687 | 0.023426 | 0.020518 | 0.01691 | 0.037803 | 0.013079 | 0.046042 | 0.018245 | 0.028843 | 0.015906 | 0.02346 |
| 20.7222222 | 0.030235 | 0.040905 | 0.006399 | 0.012327 | 0.001808 | 0.00362 | 0.004716 | 0.021712 | 0.028141 | 0.011975 | 0.009378 | 0.063217 | 0.023728 | 0.030506 | 0.012693 | 0.055571 | 0.023222 | 0.024074 |
| 20.7361111 | 0.031754 | 0.029217 | 0.006354 | 0.01268 | 0.001693 | 0.004471 | 0.006303 | 0.018261 | 0.043119 | 0.016634 | 0.003493 | 0.095726 | 0.032867 | 0.012416 | 0.016335 | 0.076358 | 0.025848 | 0.016964 |
| 20.75 | 0.018112 | 0.014253 | 0.004415 | 0.009251 | 0.003605 | 0.005397 | 0.004399 | 0.010864 | 0.037868 | 0.022324 | 0.001795 | 0.099155 | 0.033442 | 0.00635 | 0.016207 | 0.062017 | 0.020394 | 0.007847 |
| 20.7638889 | 0.011827 | 0.016486 | 0.004403 | 0.005479 | 0.005412 | 0.003637 | 0.002657 | 0.008708 | 0.020652 | 0.025086 | 0.002646 | 0.072864 | 0.03821 | 0.00648 | 0.00905 | 0.034404 | 0.019081 | 0.004004 |
| 20.7777778 | 0.011756 | 0.017071 | 0.005619 | 0.002586 | 0.004548 | 0.002001 | 0.003219 | 0.011556 | 0.01909 | 0.035871 | 0.003077 | 0.090446 | 0.055787 | 0.010711 | 0.00363 | 0.03171 | 0.029476 | 0.002138 |
| 20.7916667 | 0.009064 | 0.010958 | 0.004194 | 0.001452 | 0.002841 | 0.003223 | 0.003985 | 0.010402 | 0.03713 | 0.047407 | 0.002529 | 0.134596 | 0.063606 | 0.013823 | 0.002891 | 0.0528 | 0.036072 | 0.001171 |
| 20.8055556 | 0.00622 | 0.014918 | 0.002145 | 0.001646 | 0.00342 | 0.005896 | 0.004667 | 0.008453 | 0.054128 | 0.041685 | 0.003533 | 0.133448 | 0.052094 | 0.010188 | 0.002486 | 0.072244 | 0.033331 | 0.00208 |
| 20.8194444 | 0.003886 | 0.022687 | 0.00183 | 0.002597 | 0.005416 | 0.006172 | 0.004842 | 0.009164 | 0.05054 | 0.023489 | 0.005938 | 0.089636 | 0.040394 | 0.008578 | 0.002212 | 0.076468 | 0.030212 | 0.005636 |
| 20.8333333 | 0.001749 | 0.022425 | 0.002414 | 0.006559 | 0.006267 | 0.004041 | 0.003443 | 0.006644 | 0.028671 | 0.008703 | 0.005969 | 0.05328 | 0.029094 | 0.012899 | 0.00524 | 0.061968 | 0.029251 | 0.01361 |
| 20.8472222 | 0.001031 | 0.013919 | 0.004811 | 0.012347 | 0.005448 | 0.003169 | 0.00245 | 0.005491 | 0.010989 | 0.006225 | 0.003911 | 0.070304 | 0.019404 | 0.017604 | 0.010668 | 0.037456 | 0.027244 | 0.02102 |
| 20.8611111 | 0.001165 | 0.00866 | 0.005192 | 0.013735 | 0.003416 | 0.002969 | 0.002091 | 0.009006 | 0.008659 | 0.014458 | 0.006126 | 0.108552 | 0.028791 | 0.019776 | 0.015547 | 0.01939 | 0.018781 | 0.020394 |
| 20.875 | 0.000912 | 0.014247 | 0.003226 | 0.009082 | 0.001393 | 0.00224 | 0.002209 | 0.009098 | 0.009418 | 0.028258 | 0.019267 | 0.107907 | 0.04298 | 0.019401 | 0.016069 | 0.021814 | 0.010841 | 0.011932 |
| 20.8888889 | 0.000848 | 0.017641 | 0.004355 | 0.003456 | 0.000685 | 0.003028 | 0.004029 | 0.005469 | 0.005587 | 0.03073 | 0.039473 | 0.066081 | 0.032578 | 0.01834 | 0.012834 | 0.039281 | 0.013589 | 0.005726 |
| 20.9027778 | 0.002243 | 0.012642 | 0.004875 | 0.002419 | 0.001443 | 0.004231 | 0.004423 | 0.004391 | 0.003564 | 0.024325 | 0.043239 | 0.036394 | 0.014404 | 0.018789 | 0.013534 | 0.047093 | 0.014342 | 0.007706 |
| 20.9166667 | 0.003622 | 0.010523 | 0.004076 | 0.006755 | 0.003605 | 0.00354 | 0.002689 | 0.004944 | 0.010646 | 0.038727 | 0.028938 | 0.059678 | 0.019111 | 0.019313 | 0.021648 | 0.032344 | 0.011412 | 0.008498 |
| 20.9305556 | 0.008446 | 0.010636 | 0.007627 | 0.011675 | 0.006791 | 0.002563 | 0.002244 | 0.008009 | 0.026929 | 0.061072 | 0.033576 | 0.100448 | 0.044368 | 0.018545 | 0.032807 | 0.021685 | 0.019486 | 0.007236 |
| 20.9444444 | 0.022057 | 0.012933 | 0.012145 | 0.014005 | 0.009454 | 0.004161 | 0.003094 | 0.016674 | 0.04368 | 0.065477 | 0.061424 | 0.118252 | 0.077234 | 0.019202 | 0.041778 | 0.038786 | 0.028718 | 0.014731 |
| 20.9583333 | 0.036141 | 0.029165 | 0.012766 | 0.014903 | 0.010872 | 0.007473 | 0.004115 | 0.024811 | 0.049963 | 0.059425 | 0.079459 | 0.109453 | 0.102608 | 0.02458 | 0.043932 | 0.05533 | 0.026777 | 0.024264 |
| 20.9722222 | 0.040363 | 0.05009 | 0.010396 | 0.01486 | 0.011939 | 0.009245 | 0.004399 | 0.026898 | 0.043374 | 0.055207 | 0.075004 | 0.081535 | 0.102793 | 0.034118 | 0.039576 | 0.04998 | 0.021712 | 0.024516 |
| 20.9861111 | 0.033848 | 0.057502 | 0.006133 | 0.013286 | 0.012749 | 0.007822 | 0.003084 | 0.023503 | 0.0316 | 0.055465 | 0.057136 | 0.05322 | 0.07702 | 0.041489 | 0.036165 | 0.035372 | 0.023361 | 0.018602 |
| 21 | 0.021893 | 0.047675 | 0.002937 | 0.010783 | 0.012924 | 0.00421 | 0.00155 | 0.017544 | 0.02284 | 0.052577 | 0.034322 | 0.033639 | 0.040318 | 0.037729 | 0.034745 | 0.022201 | 0.028136 | 0.011709 |
| 21.0138889 | 0.013043 | 0.028028 | 0.002396 | 0.008434 | 0.012719 | 0.001351 | 0.001431 | 0.012723 | 0.018299 | 0.037756 | 0.015931 | 0.0181 | 0.018743 | 0.022752 | 0.028383 | 0.012736 | 0.02468 | 0.005472 |
| 21.0277778 | 0.014645 | 0.012551 | 0.003456 | 0.007925 | 0.012319 | 0.002088 | 0.001592 | 0.011908 | 0.016508 | 0.017822 | 0.010852 | 0.010545 | 0.021262 | 0.008843 | 0.019827 | 0.005804 | 0.014976 | 0.001662 |
| 21.0416667 | 0.026207 | 0.009508 | 0.006836 | 0.011141 | 0.012242 | 0.005613 | 0.001199 | 0.015397 | 0.017958 | 0.009267 | 0.012206 | 0.013476 | 0.021122 | 0.006378 | 0.018064 | 0.001801 | 0.01311 | 0.001913 |
| 21.0555556 | 0.03795 | 0.015407 | 0.009456 | 0.015242 | 0.011931 | 0.007673 | 0.002266 | 0.019324 | 0.021799 | 0.018139 | 0.009966 | 0.014752 | 0.012186 | 0.01403 | 0.025276 | 0.00119 | 0.018321 | 0.004989 |
| 21.0694444 | 0.04315 | 0.025906 | 0.011509 | 0.015895 | 0.010959 | 0.008159 | 0.005373 | 0.02222 | 0.028774 | 0.037939 | 0.014216 | 0.026288 | 0.01595 | 0.028096 | 0.042499 | 0.008103 | 0.018048 | 0.009268 |
| 21.0833333 | 0.045437 | 0.043062 | 0.0135 | 0.015326 | 0.012651 | 0.012932 | 0.007195 | 0.029411 | 0.044225 | 0.062164 | 0.035428 | 0.063038 | 0.038143 | 0.044986 | 0.06412 | 0.028045 | 0.021777 | 0.014781 |
| 21.0972222 | 0.049157 | 0.063238 | 0.01674 | 0.016837 | 0.017958 | 0.020779 | 0.008933 | 0.042442 | 0.065249 | 0.081032 | 0.056413 | 0.089886 | 0.065184 | 0.053176 | 0.06957 | 0.050923 | 0.038668 | 0.020962 |
| 21.1111111 | 0.046685 | 0.071392 | 0.018551 | 0.017203 | 0.020178 | 0.021626 | 0.012264 | 0.050687 | 0.070785 | 0.078565 | 0.054655 | 0.082128 | 0.079674 | 0.045313 | 0.0509 | 0.056892 | 0.047888 | 0.027111 |
| 21.125 | 0.033157 | 0.059392 | 0.01501 | 0.013501 | 0.015388 | 0.013571 | 0.011065 | 0.043929 | 0.04878 | 0.055992 | 0.036261 | 0.061269 | 0.080413 | 0.030828 | 0.02794 | 0.039745 | 0.035293 | 0.028583 |
| 21.1388889 | 0.020962 | 0.038278 | 0.008019 | 0.008751 | 0.008985 | 0.005545 | 0.005353 | 0.025409 | 0.020919 | 0.031011 | 0.020959 | 0.048314 | 0.073441 | 0.020678 | 0.015657 | 0.018614 | 0.019366 | 0.018892 |
| 21.1527778 | 0.015884 | 0.020938 | 0.005132 | 0.005104 | 0.005813 | 0.003133 | 0.00179 | 0.012019 | 0.010573 | 0.012894 | 0.014622 | 0.041767 | 0.054 | 0.014824 | 0.009443 | 0.012138 | 0.02041 | 0.009257 |
| 21.1666667 | 0.010678 | 0.011057 | 0.007085 | 0.002555 | 0.004522 | 0.002965 | 0.002543 | 0.013221 | 0.01299 | 0.004242 | 0.013147 | 0.033264 | 0.025898 | 0.011068 | 0.004553 | 0.010047 | 0.025551 | 0.011504 |
| 21.1805556 | 0.005302 | 0.007186 | 0.007542 | 0.001808 | 0.002995 | 0.002821 | 0.005883 | 0.015129 | 0.012542 | 0.003911 | 0.012151 | 0.021509 | 0.00841 | 0.008917 | 0.001808 | 0.006591 | 0.018078 | 0.013464 |
| 21.1944444 | 0.004645 | 0.004518 | 0.005466 | 0.001842 | 0.001847 | 0.00393 | 0.008987 | 0.009155 | 0.007393 | 0.006041 | 0.009226 | 0.013474 | 0.007064 | 0.005944 | 0.002217 | 0.010103 | 0.0071 | 0.008266 |
| 21.2083333 | 0.005727 | 0.0018 | 0.003083 | 0.001193 | 0.001815 | 0.005257 | 0.009674 | 0.004975 | 0.006695 | 0.00829 | 0.005086 | 0.015282 | 0.009433 | 0.00271 | 0.004752 | 0.014316 | 0.003572 | 0.003202 |
| 21.2222222 | 0.00388 | 0.00122 | 0.001429 | 0.00065 | 0.001764 | 0.005745 | 0.008176 | 0.006767 | 0.01065 | 0.009625 | 0.001911 | 0.014774 | 0.009864 | 0.001549 | 0.004868 | 0.0133 | 0.004919 | 0.0016 |
| 21.2361111 | 0.002055 | 0.00159 | 0.001056 | 0.00086 | 0.001206 | 0.005981 | 0.006723 | 0.007348 | 0.011561 | 0.007232 | 0.000999 | 0.012336 | 0.010448 | 0.001712 | 0.003713 | 0.010406 | 0.005451 | 0.001469 |
| 21.25 | 0.003803 | 0.00182 | 0.001512 | 0.000857 | 0.001118 | 0.006062 | 0.006308 | 0.004976 | 0.007675 | 0.003569 | 0.002356 | 0.018672 | 0.011155 | 0.002491 | 0.005271 | 0.006626 | 0.004874 | 0.000812 |
| 21.2638889 | 0.006653 | 0.003299 | 0.002548 | 0.00079 | 0.001494 | 0.0055 | 0.005764 | 0.005342 | 0.005505 | 0.003326 | 0.005141 | 0.018564 | 0.009915 | 0.002594 | 0.005949 | 0.003538 | 0.006725 | 0.000862 |
| 21.2777778 | 0.007907 | 0.005302 | 0.002877 | 0.001306 | 0.001836 | 0.004526 | 0.004593 | 0.008278 | 0.007197 | 0.005295 | 0.006951 | 0.01309 | 0.006186 | 0.001442 | 0.005796 | 0.002693 | 0.009358 | 0.002864 |
| 21.2916667 | 0.007918 | 0.00451 | 0.001826 | 0.001277 | 0.002191 | 0.003599 | 0.004152 | 0.009138 | 0.006041 | 0.007729 | 0.005089 | 0.012472 | 0.003581 | 0.00173 | 0.006252 | 0.002297 | 0.006954 | 0.005331 |
| 21.3055556 | 0.007649 | 0.002286 | 0.000693 | 0.000512 | 0.002287 | 0.002873 | 0.004338 | 0.007044 | 0.002723 | 0.009255 | 0.001677 | 0.008549 | 0.003299 | 0.004061 | 0.00445 | 0.0019 | 0.00279 | 0.006248 |
| 21.3194444 | 0.008101 | 0.001961 | 0.000168 | 0.0007 | 0.002096 | 0.002693 | 0.003241 | 0.003884 | 0.001963 | 0.009662 | 0.000922 | 0.005464 | 0.003382 | 0.006092 | 0.002166 | 0.003072 | 0.003724 | 0.007737 |
| 21.3333333 | 0.008372 | 0.004023 | 0.0007 | 0.001649 | 0.002318 | 0.003776 | 0.001293 | 0.001535 | 0.002476 | 0.012414 | 0.00351 | 0.006888 | 0.004478 | 0.008869 | 0.00296 | 0.008342 | 0.009735 | 0.013663 |
| 21.3472222 | 0.006469 | 0.006851 | 0.002212 | 0.001973 | 0.003605 | 0.005596 | 0.001279 | 0.001119 | 0.002862 | 0.01696 | 0.007767 | 0.007452 | 0.00658 | 0.013404 | 0.007492 | 0.01597 | 0.015776 | 0.021494 |
| 21.3611111 | 0.005577 | 0.007792 | 0.002604 | 0.002816 | 0.003802 | 0.006068 | 0.00414 | 0.001638 | 0.00389 | 0.019786 | 0.011899 | 0.013139 | 0.009875 | 0.018877 | 0.016719 | 0.020138 | 0.016482 | 0.026442 |
| 21.375 | 0.007554 | 0.0077 | 0.001543 | 0.004594 | 0.004164 | 0.005484 | 0.007717 | 0.002261 | 0.005761 | 0.023137 | 0.017864 | 0.029621 | 0.018939 | 0.02811 | 0.03193 | 0.023264 | 0.016021 | 0.032147 |
| 21.3888889 | 0.007741 | 0.009011 | 0.001203 | 0.005416 | 0.00521 | 0.00716 | 0.010168 | 0.004468 | 0.005952 | 0.030605 | 0.027983 | 0.045196 | 0.032778 | 0.040877 | 0.047499 | 0.031949 | 0.023793 | 0.043176 |
| 21.4027778 | 0.008155 | 0.013097 | 0.002493 | 0.0054 | 0.006746 | 0.011598 | 0.012041 | 0.008012 | 0.005629 | 0.037825 | 0.037572 | 0.047886 | 0.040396 | 0.048768 | 0.054497 | 0.041956 | 0.035866 | 0.053282 |
| 21.4166667 | 0.011604 | 0.014882 | 0.004527 | 0.005013 | 0.007223 | 0.015018 | 0.013533 | 0.009354 | 0.005986 | 0.037795 | 0.039028 | 0.041168 | 0.035732 | 0.044883 | 0.051257 | 0.042218 | 0.038352 | 0.05331 |
| 21.4305556 | 0.013291 | 0.011454 | 0.004914 | 0.003659 | 0.006137 | 0.015153 | 0.013401 | 0.007668 | 0.005007 | 0.02874 | 0.030005 | 0.029538 | 0.025239 | 0.031086 | 0.039934 | 0.030106 | 0.027538 | 0.042198 |
| 21.4444444 | 0.013601 | 0.011841 | 0.003412 | 0.002894 | 0.005104 | 0.014192 | 0.012033 | 0.008503 | 0.007235 | 0.016192 | 0.015826 | 0.017059 | 0.015407 | 0.014993 | 0.02298 | 0.013825 | 0.013213 | 0.024778 |
| 21.4583333 | 0.016171 | 0.021448 | 0.003084 | 0.005419 | 0.005944 | 0.014878 | 0.011943 | 0.016725 | 0.015686 | 0.013346 | 0.006114 | 0.014337 | 0.008798 | 0.004934 | 0.01047 | 0.00465 | 0.004684 | 0.01255 |
| 21.4722222 | 0.020627 | 0.034702 | 0.005754 | 0.010577 | 0.008165 | 0.016693 | 0.013463 | 0.027187 | 0.023324 | 0.023378 | 0.005618 | 0.020327 | 0.009677 | 0.003377 | 0.012314 | 0.006163 | 0.004088 | 0.015282 |
| 21.4861111 | 0.02563 | 0.04306 | 0.009456 | 0.013421 | 0.009282 | 0.017748 | 0.014397 | 0.029509 | 0.021761 | 0.032993 | 0.010527 | 0.024681 | 0.014089 | 0.005559 | 0.023296 | 0.013546 | 0.010188 | 0.024189 |
| 21.5 | 0.028618 | 0.040721 | 0.01013 | 0.011278 | 0.008341 | 0.017558 | 0.013721 | 0.020451 | 0.014366 | 0.035417 | 0.015661 | 0.02356 | 0.011152 | 0.007201 | 0.032096 | 0.021621 | 0.017758 | 0.026689 |
| 21.5138889 | 0.028714 | 0.03041 | 0.006742 | 0.007494 | 0.007252 | 0.016192 | 0.011781 | 0.00922 | 0.013318 | 0.032033 | 0.017184 | 0.017183 | 0.005745 | 0.006779 | 0.030604 | 0.020992 | 0.015935 | 0.021673 |
| 21.5277778 | 0.02994 | 0.02235 | 0.002872 | 0.005869 | 0.007886 | 0.014376 | 0.010146 | 0.004536 | 0.014048 | 0.024203 | 0.014264 | 0.01308 | 0.005001 | 0.004157 | 0.018095 | 0.012279 | 0.009927 | 0.012023 |

| | | | | | | | | | | | | | | | | | | |
|---|---|---|---|---|---|---|---|---|---|---|---|---|---|---|---|---|---|---|
| 21.5416667 | 0.034952 | 0.023541 | 0.002525 | 0.007031 | 0.010116 | 0.01395 | 0.011063 | 0.008486 | 0.013765 | 0.019242 | 0.012317 | 0.011743 | 0.007799 | 0.001443 | 0.00588 | 0.006799 | 0.010715 | 0.00544 |
| 21.5555556 | 0.042522 | 0.034136 | 0.006734 | 0.009297 | 0.012297 | 0.01532 | 0.014212 | 0.022792 | 0.028516 | 0.025203 | 0.016224 | 0.0135 | 0.019137 | 0.001926 | 0.007599 | 0.008939 | 0.013382 | 0.005978 |
| 21.5694444 | 0.049892 | 0.048782 | 0.013922 | 0.010889 | 0.013642 | 0.016882 | 0.016997 | 0.035612 | 0.053972 | 0.043885 | 0.026108 | 0.036865 | 0.039069 | 0.009217 | 0.029638 | 0.023436 | 0.023615 | 0.007543 |
| 21.5833333 | 0.062303 | 0.063757 | 0.019598 | 0.016535 | 0.018736 | 0.017677 | 0.018704 | 0.034171 | 0.082503 | 0.075818 | 0.045765 | 0.090677 | 0.074981 | 0.025759 | 0.069308 | 0.050271 | 0.03928 | 0.012012 |
| 21.5972222 | 0.082486 | 0.078969 | 0.018443 | 0.034227 | 0.031938 | 0.01814 | 0.021443 | 0.026004 | 0.112374 | 0.104764 | 0.073251 | 0.155785 | 0.131094 | 0.044801 | 0.102294 | 0.079491 | 0.041967 | 0.019445 |
| 21.6111111 | 0.086986 | 0.082707 | 0.012054 | 0.050367 | 0.040225 | 0.014809 | 0.020271 | 0.020379 | 0.117522 | 0.096303 | 0.083442 | 0.186207 | 0.161339 | 0.049711 | 0.090738 | 0.089507 | 0.028275 | 0.018872 |
| 21.625 | 0.062008 | 0.062023 | 0.005697 | 0.043583 | 0.032485 | 0.009003 | 0.012981 | 0.015923 | 0.087571 | 0.052192 | 0.061058 | 0.152494 | 0.125835 | 0.033158 | 0.05064 | 0.066672 | 0.01518 | 0.013511 |
| 21.6388889 | 0.032934 | 0.034609 | 0.003709 | 0.022356 | 0.020582 | 0.010005 | 0.011985 | 0.010323 | 0.044542 | 0.023985 | 0.029587 | 0.081792 | 0.065796 | 0.016105 | 0.045248 | 0.033044 | 0.01486 | 0.017417 |
| 21.6527778 | 0.028441 | 0.032839 | 0.009098 | 0.008082 | 0.012993 | 0.014909 | 0.017435 | 0.008515 | 0.022067 | 0.032061 | 0.010843 | 0.028412 | 0.047199 | 0.01388 | 0.069915 | 0.018222 | 0.01499 | 0.024805 |
| 21.6666667 | 0.043217 | 0.05137 | 0.015297 | 0.005859 | 0.007901 | 0.01505 | 0.016996 | 0.009459 | 0.037542 | 0.037416 | 0.006661 | 0.013627 | 0.07004 | 0.012149 | 0.070591 | 0.023283 | 0.017193 | 0.019318 |
| 21.6805556 | 0.047564 | 0.054742 | 0.012689 | 0.007492 | 0.004297 | 0.009543 | 0.010132 | 0.008818 | 0.062426 | 0.021721 | 0.017525 | 0.010717 | 0.084809 | 0.009082 | 0.046666 | 0.032565 | 0.03 | 0.012263 |
| 21.6944444 | 0.03423 | 0.03501 | 0.008971 | 0.008931 | 0.003287 | 0.004793 | 0.00616 | 0.015661 | 0.06712 | 0.005657 | 0.03436 | 0.009121 | 0.071699 | 0.018102 | 0.025214 | 0.03758 | 0.037631 | 0.020593 |
| 21.7083333 | 0.02859 | 0.028501 | 0.015147 | 0.009614 | 0.005935 | 0.006221 | 0.007286 | 0.02903 | 0.052254 | 0.002802 | 0.041796 | 0.020366 | 0.044319 | 0.036372 | 0.013271 | 0.038648 | 0.026846 | 0.027211 |
| 21.7222222 | 0.03847 | 0.05265 | 0.020382 | 0.007724 | 0.008982 | 0.007811 | 0.0059 | 0.031589 | 0.028568 | 0.007773 | 0.035131 | 0.032216 | 0.019009 | 0.04425 | 0.008186 | 0.037761 | 0.01573 | 0.019637 |
| 21.7361111 | 0.041258 | 0.070861 | 0.015824 | 0.005598 | 0.009036 | 0.005561 | 0.002531 | 0.018857 | 0.013505 | 0.008577 | 0.020493 | 0.027086 | 0.006117 | 0.03202 | 0.005938 | 0.033543 | 0.02768 | 0.016549 |
| 21.75 | 0.026663 | 0.057149 | 0.00848 | 0.004149 | 0.005662 | 0.00339 | 0.003018 | 0.009193 | 0.018961 | 0.010502 | 0.008843 | 0.020863 | 0.003912 | 0.015351 | 0.00365 | 0.025674 | 0.041792 | 0.022535 |
| 21.7638889 | 0.018307 | 0.033799 | 0.005098 | 0.002354 | 0.002198 | 0.003327 | 0.005303 | 0.014858 | 0.033865 | 0.022451 | 0.007742 | 0.027819 | 0.006803 | 0.011711 | 0.002714 | 0.030083 | 0.031135 | 0.021211 |
| 21.7777778 | 0.035529 | 0.034294 | 0.007348 | 0.001217 | 0.001762 | 0.003907 | 0.005191 | 0.022227 | 0.042785 | 0.026042 | 0.013498 | 0.025617 | 0.010806 | 0.018043 | 0.002777 | 0.055235 | 0.019175 | 0.011377 |
| 21.7916667 | 0.05539 | 0.05071 | 0.008402 | 0.001001 | 0.003406 | 0.003114 | 0.003272 | 0.019204 | 0.039403 | 0.019875 | 0.014833 | 0.016606 | 0.015201 | 0.02202 | 0.00255 | 0.074709 | 0.024653 | 0.006442 |
| 21.8055556 | 0.05357 | 0.053076 | 0.006241 | 0.00245 | 0.00521 | 0.001757 | 0.003387 | 0.013191 | 0.024302 | 0.026248 | 0.010503 | 0.013948 | 0.019819 | 0.018692 | 0.002525 | 0.06446 | 0.02374 | 0.011152 |
| 21.8194444 | 0.031902 | 0.035188 | 0.00653 | 0.004539 | 0.00474 | 0.001837 | 0.006004 | 0.017259 | 0.011088 | 0.030655 | 0.010706 | 0.01617 | 0.021553 | 0.010823 | 0.004768 | 0.038682 | 0.016315 | 0.01601 |
| 21.8333333 | 0.012815 | 0.017478 | 0.006981 | 0.004527 | 0.00239 | 0.00446 | 0.007872 | 0.026854 | 0.014038 | 0.022322 | 0.016401 | 0.037261 | 0.021024 | 0.005074 | 0.015522 | 0.037106 | 0.029901 | 0.014714 |
| 21.8472222 | 0.008475 | 0.014746 | 0.007194 | 0.005714 | 0.001385 | 0.008944 | 0.007494 | 0.024059 | 0.031588 | 0.025125 | 0.019929 | 0.0597 | 0.020569 | 0.005721 | 0.033573 | 0.064252 | 0.059804 | 0.017662 |
| 21.8611111 | 0.011442 | 0.016843 | 0.009471 | 0.012457 | 0.002239 | 0.011268 | 0.004947 | 0.01435 | 0.051428 | 0.041278 | 0.018094 | 0.054244 | 0.01878 | 0.008746 | 0.047471 | 0.079349 | 0.069406 | 0.028722 |
| 21.875 | 0.027527 | 0.026321 | 0.010429 | 0.019971 | 0.003604 | 0.008438 | 0.002951 | 0.017054 | 0.064804 | 0.055469 | 0.01274 | 0.036269 | 0.017187 | 0.007577 | 0.04826 | 0.066046 | 0.042778 | 0.031272 |
| 21.8888889 | 0.052353 | 0.046189 | 0.007334 | 0.021153 | 0.004245 | 0.003797 | 0.003817 | 0.02463 | 0.068319 | 0.057889 | 0.009596 | 0.028392 | 0.019138 | 0.008253 | 0.037093 | 0.041351 | 0.016889 | 0.020529 |
| 21.9027778 | 0.065894 | 0.055925 | 0.003702 | 0.017007 | 0.00331 | 0.003186 | 0.00585 | 0.020484 | 0.057183 | 0.046024 | 0.012638 | 0.031145 | 0.02196 | 0.019023 | 0.023358 | 0.018622 | 0.016964 | 0.008662 |
| 21.9166667 | 0.058816 | 0.046467 | 0.00377 | 0.011531 | 0.001551 | 0.007068 | 0.008474 | 0.011462 | 0.033211 | 0.03299 | 0.01848 | 0.040856 | 0.02409 | 0.030919 | 0.018283 | 0.005163 | 0.016368 | 0.003931 |
| 21.9305556 | 0.04115 | 0.02973 | 0.004206 | 0.005902 | 0.000825 | 0.011725 | 0.010478 | 0.006978 | 0.013271 | 0.029184 | 0.021075 | 0.05461 | 0.031016 | 0.033798 | 0.026385 | 0.005486 | 0.01457 | 0.008679 |
| 21.9444444 | 0.030674 | 0.017667 | 0.005815 | 0.003123 | 0.002304 | 0.014433 | 0.010039 | 0.008924 | 0.006886 | 0.033512 | 0.020126 | 0.063453 | 0.043644 | 0.028565 | 0.039719 | 0.015833 | 0.035933 | 0.019764 |
| 21.9583333 | 0.033718 | 0.013865 | 0.011961 | 0.006571 | 0.006762 | 0.014416 | 0.008587 | 0.021787 | 0.007502 | 0.041767 | 0.020895 | 0.061308 | 0.051209 | 0.019921 | 0.045867 | 0.026913 | 0.061963 | 0.028432 |
| 21.9722222 | 0.046292 | 0.016754 | 0.017745 | 0.013329 | 0.013839 | 0.012924 | 0.007352 | 0.036193 | 0.015825 | 0.047841 | 0.022794 | 0.051624 | 0.045678 | 0.012204 | 0.042241 | 0.029493 | 0.05938 | 0.029737 |
| 21.9861111 | 0.061836 | 0.02601 | 0.019914 | 0.017195 | 0.020346 | 0.011873 | 0.006149 | 0.040957 | 0.030371 | 0.047274 | 0.022483 | 0.044087 | 0.032106 | 0.006694 | 0.036532 | 0.021636 | 0.034566 | 0.025148 |
| 22 | 0.072349 | 0.039707 | 0.019386 | 0.015318 | 0.022447 | 0.011348 | 0.005044 | 0.03665 | 0.038629 | 0.03964 | 0.027217 | 0.044297 | 0.020889 | 0.003467 | 0.031373 | 0.011441 | 0.023 | 0.015274 |
| 22.0138889 | 0.070435 | 0.050449 | 0.016659 | 0.011082 | 0.019625 | 0.010168 | 0.004586 | 0.025331 | 0.033863 | 0.028299 | 0.039147 | 0.047107 | 0.018772 | 0.004013 | 0.023851 | 0.007575 | 0.028468 | 0.006748 |
| 22.0277778 | 0.058935 | 0.054703 | 0.011763 | 0.0088 | 0.015571 | 0.008029 | 0.005011 | 0.01211 | 0.020616 | 0.01925 | 0.037503 | 0.041467 | 0.022216 | 0.005265 | 0.015022 | 0.008888 | 0.023836 | 0.005414 |
| 22.0416667 | 0.046425 | 0.053194 | 0.00625 | 0.007821 | 0.012926 | 0.005372 | 0.005139 | 0.004409 | 0.009435 | 0.017098 | 0.018741 | 0.023228 | 0.022793 | 0.003398 | 0.008984 | 0.007575 | 0.019075 | 0.006394 |
| 22.0555556 | 0.0374 | 0.044938 | 0.003231 | 0.006804 | 0.010814 | 0.00309 | 0.00371 | 0.007343 | 0.006411 | 0.021178 | 0.006765 | 0.01245 | 0.020996 | 0.002328 | 0.011448 | 0.006765 | 0.034324 | 0.012498 |
| 22.0694444 | 0.037476 | 0.036471 | 0.003515 | 0.006969 | 0.009391 | 0.003715 | 0.001811 | 0.021847 | 0.006898 | 0.029312 | 0.006108 | 0.033726 | 0.022949 | 0.005777 | 0.025883 | 0.017406 | 0.047487 | 0.026726 |
| 22.0833333 | 0.046877 | 0.041731 | 0.00877 | 0.00961 | 0.010266 | 0.008985 | 0.002575 | 0.044934 | 0.012805 | 0.042096 | 0.009344 | 0.074595 | 0.032291 | 0.013237 | 0.049387 | 0.04126 | 0.051709 | 0.040885 |
| 22.0972222 | 0.049631 | 0.058153 | 0.019599 | 0.013444 | 0.012684 | 0.015659 | 0.006958 | 0.069763 | 0.034846 | 0.056372 | 0.011746 | 0.108244 | 0.051551 | 0.019602 | 0.07219 | 0.074265 | 0.073095 | 0.054113 |
| 22.1111111 | 0.034225 | 0.068262 | 0.028582 | 0.014843 | 0.012783 | 0.017961 | 0.011279 | 0.082292 | 0.053268 | 0.058768 | 0.009188 | 0.122334 | 0.064921 | 0.017123 | 0.078311 | 0.09937 | 0.110528 | 0.066023 |
| 22.125 | 0.015885 | 0.064044 | 0.02891 | 0.012736 | 0.008773 | 0.014042 | 0.012064 | 0.068585 | 0.040692 | 0.039963 | 0.004611 | 0.097065 | 0.050743 | 0.008586 | 0.057028 | 0.088931 | 0.118319 | 0.058268 |
| 22.1388889 | 0.011061 | 0.047375 | 0.022289 | 0.009635 | 0.003762 | 0.007889 | 0.009467 | 0.038395 | 0.015883 | 0.015748 | 0.002663 | 0.05215 | 0.02266 | 0.003825 | 0.025268 | 0.047845 | 0.073573 | 0.030173 |
| 22.1527778 | 0.010954 | 0.026923 | 0.014419 | 0.006591 | 0.001064 | 0.003981 | 0.005269 | 0.014726 | 0.004211 | 0.004295 | 0.003213 | 0.041593 | 0.005804 | 0.005629 | 0.009583 | 0.024462 | 0.039465 | 0.014407 |
| 22.1666667 | 0.00981 | 0.012249 | 0.008344 | 0.003302 | 0.000543 | 0.00215 | 0.002864 | 0.006476 | 0.001517 | 0.002569 | 0.00371 | 0.054248 | 0.002451 | 0.009536 | 0.006423 | 0.035486 | 0.05944 | 0.018058 |
| 22.1805556 | 0.009633 | 0.01006 | 0.004186 | 0.001674 | 0.001285 | 0.002066 | 0.003915 | 0.013189 | 0.004631 | 0.004031 | 0.00564 | 0.055927 | 0.002718 | 0.009476 | 0.003588 | 0.045078 | 0.084275 | 0.021406 |
| 22.1944444 | 0.006521 | 0.020337 | 0.001828 | 0.002921 | 0.003066 | 0.004131 | 0.005483 | 0.028713 | 0.015492 | 0.004616 | 0.009709 | 0.045088 | 0.001841 | 0.006371 | 0.002773 | 0.039213 | 0.082117 | 0.020729 |
| 22.2083333 | 0.005161 | 0.028565 | 0.002414 | 0.004682 | 0.004295 | 0.005016 | 0.00584 | 0.042078 | 0.027164 | 0.003166 | 0.012658 | 0.025181 | 0.001059 | 0.004627 | 0.005386 | 0.029963 | 0.071514 | 0.02129 |
| 22.2222222 | 0.010479 | 0.030639 | 0.007205 | 0.004733 | 0.004349 | 0.003769 | 0.005067 | 0.049099 | 0.034614 | 0.005239 | 0.013392 | 0.01321 | 0.001685 | 0.003832 | 0.010805 | 0.023307 | 0.068512 | 0.025686 |
| 22.2361111 | 0.02068 | 0.035827 | 0.01294 | 0.003592 | 0.004173 | 0.003106 | 0.003139 | 0.050799 | 0.0383 | 0.013389 | 0.013765 | 0.020839 | 0.005248 | 0.002609 | 0.017959 | 0.016284 | 0.067269 | 0.033767 |
| 22.25 | 0.029696 | 0.042094 | 0.014855 | 0.002979 | 0.003496 | 0.003277 | 0.001553 | 0.04545 | 0.037186 | 0.021904 | 0.014794 | 0.022198 | 0.012589 | 0.001444 | 0.022885 | 0.010206 | 0.049586 | 0.039504 |
| 22.2638889 | 0.030755 | 0.042802 | 0.012604 | 0.003472 | 0.002235 | 0.003347 | 0.0015 | 0.032705 | 0.030058 | 0.025627 | 0.015752 | 0.015853 | 0.021702 | 0.001369 | 0.022704 | 0.01193 | 0.024769 | 0.03757 |
| 22.2777778 | 0.027972 | 0.041754 | 0.010537 | 0.004519 | 0.001906 | 0.004003 | 0.001911 | 0.01969 | 0.020115 | 0.023759 | 0.015556 | 0.021278 | 0.028738 | 0.001523 | 0.018934 | 0.012657 | 0.016203 | 0.030517 |
| 22.2916667 | 0.02666 | 0.042239 | 0.011787 | 0.005609 | 0.003111 | 0.005901 | 0.001686 | 0.013594 | 0.012225 | 0.017581 | 0.013899 | 0.021483 | 0.029799 | 0.001441 | 0.0146 | 0.012843 | 0.015626 | 0.023408 |
| 22.3055556 | 0.021724 | 0.033498 | 0.012808 | 0.005895 | 0.003763 | 0.00684 | 0.001099 | 0.012393 | 0.007389 | 0.010408 | 0.011665 | 0.012009 | 0.023814 | 0.002388 | 0.010612 | 0.021467 | 0.017644 | 0.016889 |
| 22.3194444 | 0.012685 | 0.021341 | 0.009121 | 0.00511 | 0.002546 | 0.004777 | 0.000538 | 0.009501 | 0.006018 | 0.005141 | 0.00967 | 0.006054 | 0.014557 | 0.00283 | 0.007313 | 0.021504 | 0.023177 | 0.011379 |
| 22.3333333 | 0.009094 | 0.026782 | 0.006127 | 0.004017 | 0.001358 | 0.003247 | 0.000256 | 0.006341 | 0.007306 | 0.003784 | 0.008023 | 0.008402 | 0.006462 | 0.001785 | 0.008721 | 0.01644 | 0.021533 | 0.013348 |
| 22.3472222 | 0.009095 | 0.037114 | 0.009296 | 0.002489 | 0.00123 | 0.004813 | 0.00023 | 0.010407 | 0.008941 | 0.005221 | 0.007688 | 0.023146 | 0.001865 | 0.001311 | 0.015351 | 0.02569 | 0.021897 | 0.022765 |
| 22.3611111 | 0.007144 | 0.028627 | 0.011398 | 0.001248 | 0.002015 | 0.005288 | 0.000364 | 0.01908 | 0.012149 | 0.005413 | 0.009805 | 0.038396 | 0.000878 | 0.001486 | 0.019306 | 0.026352 | 0.02297 | 0.027126 |
| 22.375 | 0.00823 | 0.013675 | 0.00754 | 0.001068 | 0.003719 | 0.003111 | 0.000615 | 0.022422 | 0.017884 | 0.003741 | 0.013416 | 0.03744 | 0.001946 | 0.001087 | 0.01671 | 0.016133 | 0.017677 | 0.021087 |
| 22.3888889 | 0.012353 | 0.011964 | 0.003402 | 0.001403 | 0.004493 | 0.00196 | 0.000877 | 0.01559 | 0.020364 | 0.001958 | 0.016735 | 0.021425 | 0.003571 | 0.000506 | 0.010773 | 0.018524 | 0.020772 | 0.011251 |
| 22.4027778 | 0.0171 | 0.015668 | 0.003451 | 0.002489 | 0.003944 | 0.003177 | 0.000868 | 0.007269 | 0.015342 | 0.001436 | 0.019572 | 0.009602 | 0.003776 | 0.000526 | 0.005122 | 0.017177 | 0.025104 | 0.004445 |
| 22.4166667 | 0.023419 | 0.015065 | 0.004986 | 0.003359 | 0.003917 | 0.004808 | 0.000412 | 0.00853 | 0.015701 | 0.002864 | 0.02079 | 0.011271 | 0.002908 | 0.000921 | 0.002636 | 0.009473 | 0.017876 | 0.002897 |
| 22.4305556 | 0.028265 | 0.013099 | 0.00558 | 0.003656 | 0.004665 | 0.005728 | 0.000476 | 0.013737 | 0.029875 | 0.005334 | 0.018595 | 0.016403 | 0.004921 | 0.001699 | 0.003541 | 0.010665 | 0.010644 | 0.005122 |
| 22.4444444 | 0.029368 | 0.014933 | 0.006525 | 0.004654 | 0.004924 | 0.006338 | 0.001577 | 0.014887 | 0.037964 | 0.010862 | 0.015645 | 0.019933 | 0.012238 | 0.002341 | 0.00659 | 0.012432 | 0.011932 | 0.010081 |
| 22.4583333 | 0.026492 | 0.020153 | 0.007522 | 0.006762 | 0.003707 | 0.006228 | 0.002626 | 0.014173 | 0.025003 | 0.02263 | 0.015987 | 0.039685 | 0.021843 | 0.002868 | 0.010971 | 0.018006 | 0.026585 | 0.016742 |
| 22.4722222 | 0.018705 | 0.021161 | 0.006853 | 0.009169 | 0.001897 | 0.005125 | 0.002835 | 0.016053 | 0.011725 | 0.036436 | 0.019806 | 0.061317 | 0.027515 | 0.005745 | 0.013946 | 0.029812 | 0.051302 | 0.020799 |
| 22.4861111 | 0.011008 | 0.019896 | 0.00565 | 0.010201 | 0.001861 | 0.004986 | 0.002847 | 0.019122 | 0.013042 | 0.041946 | 0.024505 | 0.059994 | 0.026431 | 0.009404 | 0.01228 | 0.028386 | 0.055567 | 0.017988 |
| 22.5 | 0.009565 | 0.024717 | 0.006759 | 0.009593 | 0.004017 | 0.006933 | 0.0038 | 0.020332 | 0.013658 | 0.034312 | 0.026874 | 0.036871 | 0.02136 | 0.009842 | 0.006752 | 0.015197 | 0.03356 | 0.009573 |
| 22.5138889 | 0.011432 | 0.030296 | 0.009439 | 0.008437 | 0.006136 | 0.008555 | 0.005058 | 0.021946 | 0.017369 | 0.020429 | 0.026203 | 0.019984 | 0.0172 | 0.008036 | 0.003182 | 0.006993 | 0.020562 | 0.002865 |
| 22.5277778 | 0.009249 | 0.024303 | 0.009576 | 0.007814 | 0.007649 | 0.007631 | 0.004845 | 0.023134 | 0.032356 | 0.009923 | 0.025384 | 0.015818 | 0.016091 | 0.007811 | 0.006708 | 0.007749 | 0.018809 | 0.004284 |
| 22.5416667 | 0.004182 | 0.010749 | 0.006339 | 0.007556 | 0.00983 | 0.006031 | 0.003594 | 0.016841 | 0.034613 | 0.006495 | 0.025229 | 0.020742 | 0.01528 | 0.00987 | 0.015302 | 0.020343 | 0.017067 | 0.011818 |
| 22.5555556 | 0.007346 | 0.011291 | 0.004915 | 0.005498 | 0.009582 | 0.007042 | 0.002943 | 0.007943 | 0.019678 | 0.011964 | 0.023319 | 0.038409 | 0.011222 | 0.011782 | 0.023714 | 0.037276 | 0.029313 | 0.018554 |
| 22.5694444 | 0.021001 | 0.035576 | 0.009865 | 0.002729 | 0.005602 | 0.010182 | 0.003564 | 0.005261 | 0.017129 | 0.029332 | 0.019799 | 0.056412 | 0.006482 | 0.014371 | 0.034445 | 0.043619 | 0.045783 | 0.023022 |
| 22.5833333 | 0.033292 | 0.070834 | 0.022042 | 0.004604 | 0.007161 | 0.013926 | 0.008677 | 0.010624 | 0.021926 | 0.057152 | 0.015013 | 0.064356 | 0.009049 | 0.027339 | 0.063511 | 0.046045 | 0.047745 | 0.031395 |
| 22.5972222 | 0.042565 | 0.099578 | 0.038077 | 0.016348 | 0.019131 | 0.020358 | 0.020894 | 0.028233 | 0.033317 | 0.081833 | 0.008207 | 0.072109 | 0.025162 | 0.060037 | 0.120631 | 0.059817 | 0.043713 | 0.047845 |
| 22.6111111 | 0.049021 | 0.099589 | 0.046188 | 0.033017 | 0.029832 | 0.025589 | 0.03006 | 0.042706 | 0.068739 | 0.084023 | 0.004836 | 0.098805 | 0.051377 | 0.092854 | 0.162924 | 0.076556 | 0.044191 | 0.060425 |
| 22.625 | 0.04051 | 0.064004 | 0.037167 | 0.037343 | 0.028175 | 0.02099 | 0.024573 | 0.032058 | 0.083618 | 0.0584 | 0.01121 | 0.128559 | 0.06461 | 0.090171 | 0.135714 | 0.079204 | 0.042331 | 0.051803 |
| 22.6388889 | 0.020702 | 0.029038 | 0.01986 | 0.023983 | 0.019922 | 0.010968 | 0.012623 | 0.015026 | 0.056298 | 0.032336 | 0.019626 | 0.100573 | 0.045379 | 0.052847 | 0.071551 | 0.056802 | 0.03172 | 0.030095 |
| 22.6527778 | 0.010647 | 0.02978 | 0.013779 | 0.011186 | 0.014271 | 0.008674 | 0.010397 | 0.020382 | 0.026776 | 0.037718 | 0.017315 | 0.059366 | 0.024974 | 0.028648 | 0.051174 | 0.032304 | 0.03339 | 0.02519 |
| 22.6666667 | 0.017643 | 0.050125 | 0.021874 | 0.011936 | 0.010921 | 0.015868 | 0.017713 | 0.037185 | 0.022227 | 0.048722 | 0.007966 | 0.088951 | 0.034243 | 0.038248 | 0.083873 | 0.045454 | 0.051436 | 0.047575 |
| 22.6805556 | 0.026049 | 0.053536 | 0.026672 | 0.014521 | 0.007402 | 0.019482 | 0.021084 | 0.046876 | 0.044434 | 0.049026 | 0.004926 | 0.105532 | 0.042034 | 0.043403 | 0.110287 | 0.073815 | 0.047601 | 0.068761 |

| | | | | | | | | | | | | | | | | | | |
|---|---|---|---|---|---|---|---|---|---|---|---|---|---|---|---|---|---|---|
| 22.6944444 | 0.023037 | 0.033494 | 0.01887 | 0.011518 | 0.004014 | 0.013387 | 0.014305 | 0.038501 | 0.060972 | 0.079932 | 0.008456 | 0.072798 | 0.025866 | 0.026872 | 0.097456 | 0.079201 | 0.033322 | 0.060317 |
| 22.7083333 | 0.014847 | 0.021822 | 0.010467 | 0.012825 | 0.00287 | 0.008852 | 0.008209 | 0.029847 | 0.044633 | 0.120045 | 0.01007 | 0.088564 | 0.01019 | 0.016826 | 0.05691 | 0.067094 | 0.044795 | 0.035648 |
| 22.7222222 | 0.014611 | 0.036112 | 0.015175 | 0.017086 | 0.004698 | 0.009736 | 0.01089 | 0.051464 | 0.031656 | 0.108547 | 0.011725 | 0.127442 | 0.012655 | 0.028224 | 0.02867 | 0.056217 | 0.049652 | 0.030768 |
| 22.7361111 | 0.019468 | 0.045521 | 0.022845 | 0.013394 | 0.006529 | 0.007596 | 0.011649 | 0.062508 | 0.038708 | 0.061506 | 0.020325 | 0.0972 | 0.016543 | 0.035702 | 0.025063 | 0.05358 | 0.035396 | 0.043343 |
| 22.75 | 0.015321 | 0.029976 | 0.01845 | 0.008005 | 0.00629 | 0.006189 | 0.006867 | 0.042386 | 0.031722 | 0.047283 | 0.030292 | 0.071797 | 0.014637 | 0.026555 | 0.02296 | 0.052435 | 0.044066 | 0.03695 |
| 22.7638889 | 0.007788 | 0.017163 | 0.011208 | 0.012708 | 0.005248 | 0.012688 | 0.006421 | 0.040567 | 0.025239 | 0.079157 | 0.02505 | 0.127515 | 0.021614 | 0.023257 | 0.034003 | 0.045922 | 0.064993 | 0.018556 |
| 22.7777778 | 0.008501 | 0.0307 | 0.012616 | 0.021974 | 0.005272 | 0.019411 | 0.012279 | 0.05978 | 0.035569 | 0.094987 | 0.012784 | 0.152795 | 0.028287 | 0.028318 | 0.054695 | 0.034425 | 0.054088 | 0.019254 |
| 22.7916667 | 0.011284 | 0.050668 | 0.015962 | 0.022454 | 0.00675 | 0.017057 | 0.017623 | 0.060671 | 0.033454 | 0.07026 | 0.014597 | 0.093363 | 0.020701 | 0.021771 | 0.059049 | 0.023202 | 0.031367 | 0.033042 |
| 22.8055556 | 0.010081 | 0.051051 | 0.012574 | 0.013078 | 0.008714 | 0.009138 | 0.017569 | 0.037403 | 0.02031 | 0.038876 | 0.017575 | 0.058816 | 0.01587 | 0.010587 | 0.04036 | 0.023259 | 0.028163 | 0.038131 |
| 22.8194444 | 0.007214 | 0.031765 | 0.005848 | 0.006992 | 0.009645 | 0.004626 | 0.01241 | 0.024478 | 0.022076 | 0.026012 | 0.013363 | 0.08628 | 0.030808 | 0.011806 | 0.026339 | 0.035769 | 0.024056 | 0.028159 |
| 22.8333333 | 0.00557 | 0.018319 | 0.002987 | 0.00956 | 0.007976 | 0.003766 | 0.006123 | 0.034373 | 0.031551 | 0.021962 | 0.017451 | 0.087716 | 0.03761 | 0.017036 | 0.042202 | 0.034231 | 0.023783 | 0.022467 |
| 22.8472222 | 0.005973 | 0.019528 | 0.005179 | 0.011151 | 0.004252 | 0.004607 | 0.003313 | 0.031792 | 0.027799 | 0.021817 | 0.023531 | 0.079703 | 0.026155 | 0.013701 | 0.046831 | 0.030778 | 0.052782 | 0.023642 |
| 22.8611111 | 0.00699 | 0.018336 | 0.014067 | 0.007911 | 0.001702 | 0.009095 | 0.002921 | 0.023143 | 0.028961 | 0.033001 | 0.020383 | 0.134942 | 0.033927 | 0.009874 | 0.029939 | 0.068672 | 0.082087 | 0.022073 |
| 22.875 | 0.010084 | 0.025734 | 0.025968 | 0.007235 | 0.001368 | 0.013762 | 0.002092 | 0.045288 | 0.05665 | 0.038358 | 0.021584 | 0.178856 | 0.058064 | 0.016605 | 0.036491 | 0.124358 | 0.07795 | 0.042643 |
| 22.8888889 | 0.01743 | 0.052413 | 0.032154 | 0.01314 | 0.001351 | 0.014497 | 0.002319 | 0.06869 | 0.08249 | 0.032344 | 0.02514 | 0.138241 | 0.061169 | 0.023263 | 0.064533 | 0.15092 | 0.050383 | 0.083166 |
| 22.9027778 | 0.02485 | 0.077439 | 0.033464 | 0.018925 | 0.001587 | 0.012603 | 0.004648 | 0.057329 | 0.08058 | 0.026652 | 0.01865 | 0.068156 | 0.035878 | 0.021635 | 0.077266 | 0.142489 | 0.021863 | 0.10003 |
| 22.9166667 | 0.026103 | 0.081917 | 0.032178 | 0.019101 | 0.002819 | 0.010482 | 0.00732 | 0.031654 | 0.063418 | 0.02421 | 0.021352 | 0.032378 | 0.015218 | 0.019546 | 0.061932 | 0.115805 | 0.007817 | 0.08234 |
| 22.9305556 | 0.020184 | 0.068018 | 0.027121 | 0.015887 | 0.00503 | 0.010161 | 0.008974 | 0.019851 | 0.048643 | 0.028084 | 0.04765 | 0.028831 | 0.012967 | 0.022307 | 0.035981 | 0.073067 | 0.007522 | 0.055984 |
| 22.9444444 | 0.012059 | 0.048635 | 0.020692 | 0.013984 | 0.008528 | 0.012061 | 0.008913 | 0.025118 | 0.038591 | 0.040653 | 0.074139 | 0.05137 | 0.013451 | 0.027934 | 0.020169 | 0.027414 | 0.022637 | 0.038809 |
| 22.9583333 | 0.007468 | 0.028985 | 0.018469 | 0.01713 | 0.012375 | 0.014714 | 0.007903 | 0.038233 | 0.029648 | 0.052871 | 0.07382 | 0.091031 | 0.025163 | 0.032007 | 0.019678 | 0.009497 | 0.04039 | 0.031615 |
| 22.9722222 | 0.006977 | 0.013856 | 0.021701 | 0.024151 | 0.015076 | 0.017806 | 0.00803 | 0.050948 | 0.023172 | 0.052617 | 0.050047 | 0.115605 | 0.055578 | 0.030753 | 0.025828 | 0.020736 | 0.033996 | 0.030222 |
| 22.9861111 | 0.008685 | 0.009767 | 0.026173 | 0.029313 | 0.016443 | 0.020767 | 0.008883 | 0.054031 | 0.017888 | 0.040291 | 0.02332 | 0.114367 | 0.086255 | 0.025106 | 0.026566 | 0.044842 | 0.01924 | 0.027829 |
| 23 | 0.01061 | 0.01012 | 0.026448 | 0.027553 | 0.016344 | 0.020017 | 0.008064 | 0.040816 | 0.011317 | 0.021747 | 0.01162 | 0.095109 | 0.090388 | 0.019926 | 0.018332 | 0.057109 | 0.013456 | 0.017895 |
| 23.0138889 | 0.01009 | 0.009121 | 0.020199 | 0.019777 | 0.013893 | 0.014525 | 0.005 | 0.020904 | 0.009681 | 0.008577 | 0.018186 | 0.064134 | 0.061349 | 0.017779 | 0.013613 | 0.039733 | 0.01122 | 0.009977 |
| 23.0277778 | 0.00651 | 0.008714 | 0.011214 | 0.010439 | 0.008947 | 0.007799 | 0.002095 | 0.012753 | 0.012789 | 0.007601 | 0.024174 | 0.037492 | 0.026095 | 0.019138 | 0.01788 | 0.020205 | 0.011127 | 0.014335 |
| 23.0416667 | 0.004541 | 0.006735 | 0.004377 | 0.00472 | 0.00391 | 0.003516 | 0.001193 | 0.01502 | 0.011313 | 0.006966 | 0.018048 | 0.043685 | 0.009662 | 0.024119 | 0.016669 | 0.027714 | 0.013043 | 0.015986 |
| 23.0555556 | 0.005465 | 0.008109 | 0.001829 | 0.005505 | 0.001176 | 0.004039 | 0.001021 | 0.012511 | 0.010872 | 0.008149 | 0.010907 | 0.071734 | 0.011948 | 0.029371 | 0.018695 | 0.03895 | 0.012459 | 0.012346 |
| 23.0694444 | 0.007326 | 0.024075 | 0.002855 | 0.007009 | 0.000425 | 0.009255 | 0.001114 | 0.012482 | 0.018999 | 0.028247 | 0.017102 | 0.070725 | 0.023163 | 0.029711 | 0.042166 | 0.029824 | 0.012561 | 0.020313 |
| 23.0833333 | 0.014159 | 0.050439 | 0.010044 | 0.00633 | 0.000888 | 0.016464 | 0.004871 | 0.028246 | 0.03049 | 0.065888 | 0.037666 | 0.073321 | 0.041846 | 0.028988 | 0.06917 | 0.029783 | 0.033043 | 0.036841 |
| 23.0972222 | 0.022263 | 0.072961 | 0.025239 | 0.009862 | 0.002242 | 0.022552 | 0.014494 | 0.048367 | 0.040387 | 0.100629 | 0.06267 | 0.144602 | 0.065864 | 0.032806 | 0.075436 | 0.068881 | 0.07982 | 0.046574 |
| 23.1111111 | 0.022011 | 0.078937 | 0.037384 | 0.016566 | 0.003988 | 0.024806 | 0.025401 | 0.055101 | 0.039716 | 0.10976 | 0.078973 | 0.221268 | 0.074602 | 0.03095 | 0.056898 | 0.101059 | 0.130289 | 0.041866 |
| 23.125 | 0.016229 | 0.066206 | 0.036079 | 0.01987 | 0.005136 | 0.019405 | 0.029113 | 0.042843 | 0.024041 | 0.084058 | 0.072086 | 0.209053 | 0.053567 | 0.018451 | 0.027903 | 0.079563 | 0.153981 | 0.025094 |
| 23.1388889 | 0.015959 | 0.049856 | 0.025622 | 0.020258 | 0.005657 | 0.009016 | 0.022861 | 0.024674 | 0.01006 | 0.042013 | 0.043054 | 0.123134 | 0.022676 | 0.008961 | 0.009448 | 0.035906 | 0.139493 | 0.009176 |
| 23.1527778 | 0.022301 | 0.041101 | 0.015613 | 0.019235 | 0.006462 | 0.002775 | 0.013104 | 0.017424 | 0.007105 | 0.013728 | 0.018863 | 0.046437 | 0.006226 | 0.006668 | 0.005152 | 0.012451 | 0.101435 | 0.005199 |
| 23.1666667 | 0.023558 | 0.035063 | 0.009054 | 0.014692 | 0.006552 | 0.00147 | 0.006138 | 0.018253 | 0.004732 | 0.008925 | 0.019296 | 0.031099 | 0.002312 | 0.007136 | 0.005369 | 0.010436 | 0.058209 | 0.008911 |
| 23.1805556 | 0.014823 | 0.025049 | 0.00429 | 0.007418 | 0.004773 | 0.002712 | 0.002745 | 0.016075 | 0.001301 | 0.016197 | 0.030573 | 0.076548 | 0.001273 | 0.008971 | 0.004423 | 0.014195 | 0.026928 | 0.010872 |
| 23.1944444 | 0.005338 | 0.012487 | 0.001914 | 0.003226 | 0.002908 | 0.007401 | 0.00137 | 0.009729 | 0.001237 | 0.022338 | 0.036098 | 0.144247 | 0.001187 | 0.007031 | 0.002791 | 0.01602 | 0.022185 | 0.009999 |
| 23.2083333 | 0.003346 | 0.005291 | 0.002697 | 0.004522 | 0.004127 | 0.013919 | 0.001423 | 0.004897 | 0.00417 | 0.02287 | 0.035405 | 0.18219 | 0.001901 | 0.003731 | 0.001927 | 0.014682 | 0.03517 | 0.009454 |
| 23.2222222 | 0.007037 | 0.006752 | 0.004455 | 0.007071 | 0.007439 | 0.017818 | 0.002764 | 0.006123 | 0.009106 | 0.017961 | 0.030675 | 0.177988 | 0.00308 | 0.004137 | 0.003034 | 0.010171 | 0.047349 | 0.009904 |
| 23.2361111 | 0.010278 | 0.008036 | 0.004911 | 0.007679 | 0.008493 | 0.015436 | 0.00482 | 0.012882 | 0.015187 | 0.010616 | 0.023184 | 0.14565 | 0.004284 | 0.003982 | 0.009013 | 0.010429 | 0.053178 | 0.009678 |
| 23.25 | 0.009569 | 0.004089 | 0.004575 | 0.007289 | 0.007115 | 0.00842 | 0.007072 | 0.022876 | 0.023047 | 0.005742 | 0.016348 | 0.094275 | 0.004909 | 0.00353 | 0.014199 | 0.025008 | 0.056831 | 0.008035 |
| 23.2638889 | 0.008401 | 0.009266 | 0.006424 | 0.008567 | 0.006419 | 0.002613 | 0.008199 | 0.035624 | 0.031609 | 0.005782 | 0.014585 | 0.048423 | 0.004074 | 0.004134 | 0.012542 | 0.045303 | 0.062241 | 0.00903 |
| 23.2777778 | 0.010337 | 0.030688 | 0.011278 | 0.011617 | 0.006109 | 0.001431 | 0.007522 | 0.045758 | 0.037313 | 0.011404 | 0.016938 | 0.027545 | 0.003268 | 0.006093 | 0.011943 | 0.057395 | 0.064923 | 0.012708 |
| 23.2916667 | 0.012725 | 0.049258 | 0.015853 | 0.013531 | 0.004315 | 0.003395 | 0.00642 | 0.048033 | 0.038942 | 0.018896 | 0.018572 | 0.017762 | 0.005827 | 0.013052 | 0.014649 | 0.059939 | 0.062109 | 0.013153 |
| 23.3055556 | 0.013436 | 0.054098 | 0.017371 | 0.013142 | 0.001998 | 0.005934 | 0.006686 | 0.045715 | 0.037308 | 0.023037 | 0.017534 | 0.011332 | 0.011001 | 0.021281 | 0.016401 | 0.057698 | 0.059639 | 0.012451 |
| 23.3194444 | 0.012544 | 0.050501 | 0.01582 | 0.011228 | 0.001407 | 0.007771 | 0.007935 | 0.041064 | 0.032648 | 0.024127 | 0.014564 | 0.011133 | 0.016653 | 0.025655 | 0.019652 | 0.051469 | 0.058359 | 0.014796 |
| 23.3333333 | 0.011007 | 0.041693 | 0.013195 | 0.009034 | 0.002505 | 0.007982 | 0.008223 | 0.031796 | 0.02687 | 0.024589 | 0.011076 | 0.024261 | 0.020831 | 0.02735 | 0.024564 | 0.038245 | 0.05441 | 0.017055 |
| 23.3472222 | 0.009216 | 0.031434 | 0.011084 | 0.007491 | 0.002507 | 0.006098 | 0.007113 | 0.018717 | 0.021398 | 0.021633 | 0.007766 | 0.050912 | 0.019397 | 0.026624 | 0.024378 | 0.021854 | 0.043459 | 0.01348 |
| 23.3611111 | 0.00566 | 0.019208 | 0.007987 | 0.005567 | 0.001846 | 0.00323 | 0.005812 | 0.010725 | 0.017154 | 0.013978 | 0.004664 | 0.05025 | 0.012876 | 0.020745 | 0.016374 | 0.019979 | 0.039898 | 0.00961 |
| 23.375 | 0.001956 | 0.010067 | 0.004402 | 0.00272 | 0.002816 | 0.001594 | 0.004643 | 0.017751 | 0.021073 | 0.011818 | 0.003446 | 0.043515 | 0.012022 | 0.010816 | 0.012162 | 0.037589 | 0.05794 | 0.015699 |
| 23.3888889 | 0.001273 | 0.013136 | 0.003752 | 0.000754 | 0.003426 | 0.001507 | 0.002837 | 0.033733 | 0.030218 | 0.018701 | 0.005596 | 0.082785 | 0.018928 | 0.004008 | 0.017054 | 0.047657 | 0.067858 | 0.022962 |
| 23.4027778 | 0.003118 | 0.01826 | 0.004666 | 0.000766 | 0.002319 | 0.001556 | 0.001004 | 0.042403 | 0.028696 | 0.021192 | 0.009112 | 0.097793 | 0.019976 | 0.003114 | 0.018164 | 0.032457 | 0.051425 | 0.020682 |
| 23.4166667 | 0.005497 | 0.013356 | 0.004101 | 0.002225 | 0.00147 | 0.002411 | 0.000414 | 0.037491 | 0.016757 | 0.013278 | 0.012034 | 0.068909 | 0.011971 | 0.00582 | 0.010673 | 0.018719 | 0.025941 | 0.013759 |
| 23.4305556 | 0.007655 | 0.00933 | 0.002497 | 0.004682 | 0.001404 | 0.004555 | 0.000908 | 0.022301 | 0.011634 | 0.00802 | 0.015959 | 0.074268 | 0.007777 | 0.010638 | 0.005906 | 0.028724 | 0.009906 | 0.008061 |
| 23.4444444 | 0.010594 | 0.020227 | 0.002353 | 0.007717 | 0.002266 | 0.006503 | 0.001892 | 0.01361 | 0.020275 | 0.01259 | 0.02064 | 0.076735 | 0.013841 | 0.013301 | 0.011166 | 0.039134 | 0.006582 | 0.003922 |
| 23.4583333 | 0.014374 | 0.035453 | 0.005029 | 0.010474 | 0.004836 | 0.0076 | 0.003261 | 0.027258 | 0.031534 | 0.016656 | 0.021432 | 0.05784 | 0.021361 | 0.012429 | 0.018294 | 0.03135 | 0.008031 | 0.001691 |
| 23.4722222 | 0.016472 | 0.03775 | 0.008355 | 0.011827 | 0.006628 | 0.007992 | 0.004971 | 0.048278 | 0.034425 | 0.01704 | 0.016748 | 0.073704 | 0.02335 | 0.012453 | 0.019458 | 0.015189 | 0.015562 | 0.002192 |
| 23.4861111 | 0.014889 | 0.024339 | 0.009926 | 0.01103 | 0.005633 | 0.007674 | 0.006197 | 0.056815 | 0.02804 | 0.019429 | 0.011223 | 0.077024 | 0.022043 | 0.014987 | 0.016942 | 0.004378 | 0.024461 | 0.004107 |
| 23.5 | 0.011798 | 0.009691 | 0.010254 | 0.00944 | 0.003571 | 0.006908 | 0.006517 | 0.05192 | 0.019174 | 0.025594 | 0.009673 | 0.078438 | 0.021958 | 0.016867 | 0.015198 | 0.004091 | 0.024695 | 0.006152 |
| 23.5138889 | 0.012291 | 0.004657 | 0.011094 | 0.009437 | 0.002133 | 0.006813 | 0.007157 | 0.039545 | 0.014291 | 0.037273 | 0.013167 | 0.132717 | 0.025704 | 0.015948 | 0.017391 | 0.014483 | 0.016832 | 0.008887 |
| 23.5277778 | 0.017303 | 0.004971 | 0.011937 | 0.010792 | 0.001274 | 0.007975 | 0.008602 | 0.024685 | 0.011385 | 0.045475 | 0.019951 | 0.145158 | 0.030488 | 0.013962 | 0.023404 | 0.035684 | 0.014254 | 0.012469 |
| 23.5416667 | 0.020941 | 0.010073 | 0.010689 | 0.010989 | 0.000717 | 0.008822 | 0.009308 | 0.013671 | 0.007908 | 0.033532 | 0.024411 | 0.09339 | 0.029574 | 0.011947 | 0.027198 | 0.05166 | 0.021874 | 0.01392 |
| 23.5555556 | 0.019302 | 0.020882 | 0.007167 | 0.010381 | 0.001606 | 0.00948 | 0.007735 | 0.012608 | 0.007711 | 0.016527 | 0.022294 | 0.090867 | 0.019803 | 0.010321 | 0.025383 | 0.042307 | 0.02202 | 0.010657 |
| 23.5694444 | 0.017093 | 0.043628 | 0.00534 | 0.012232 | 0.005658 | 0.013292 | 0.00426 | 0.013846 | 0.01167 | 0.01883 | 0.01923 | 0.123821 | 0.01112 | 0.012304 | 0.027362 | 0.019711 | 0.01101 | 0.007081 |
| 23.5833333 | 0.022457 | 0.084589 | 0.01371 | 0.016147 | 0.010774 | 0.019591 | 0.001616 | 0.023136 | 0.016536 | 0.040325 | 0.023288 | 0.101069 | 0.017267 | 0.023197 | 0.043958 | 0.015265 | 0.00649 | 0.011524 |
| 23.5972222 | 0.036807 | 0.127882 | 0.034525 | 0.021959 | 0.011754 | 0.024057 | 0.005144 | 0.057673 | 0.03319 | 0.07591 | 0.033298 | 0.08519 | 0.043551 | 0.048517 | 0.073589 | 0.044478 | 0.019118 | 0.025356 |
| 23.6111111 | 0.054013 | 0.140521 | 0.053707 | 0.030772 | 0.010991 | 0.024349 | 0.012664 | 0.090929 | 0.060811 | 0.10803 | 0.040714 | 0.139686 | 0.081653 | 0.077653 | 0.102855 | 0.091923 | 0.041723 | 0.03891 |
| 23.625 | 0.061921 | 0.113137 | 0.055837 | 0.034972 | 0.015605 | 0.018736 | 0.014163 | 0.08852 | 0.071481 | 0.110857 | 0.037225 | 0.154408 | 0.098416 | 0.082333 | 0.105764 | 0.113417 | 0.053528 | 0.03856 |
| 23.6388889 | 0.048617 | 0.067777 | 0.041776 | 0.028476 | 0.023573 | 0.010201 | 0.008579 | 0.057374 | 0.05895 | 0.075014 | 0.021877 | 0.09026 | 0.068491 | 0.053511 | 0.070631 | 0.084873 | 0.040105 | 0.023722 |
| 23.6527778 | 0.025895 | 0.035075 | 0.02145 | 0.015523 | 0.027965 | 0.009188 | 0.005089 | 0.03096 | 0.034034 | 0.03857 | 0.011584 | 0.063685 | 0.034389 | 0.025433 | 0.032365 | 0.040468 | 0.019553 | 0.009231 |
| 23.6666667 | 0.020435 | 0.038204 | 0.009421 | 0.008226 | 0.027356 | 0.01819 | 0.009437 | 0.038844 | 0.018105 | 0.043249 | 0.019791 | 0.11982 | 0.045935 | 0.027351 | 0.026612 | 0.028588 | 0.020021 | 0.005367 |
| 23.6805556 | 0.032018 | 0.05982 | 0.01278 | 0.014205 | 0.024821 | 0.025094 | 0.015051 | 0.070048 | 0.030437 | 0.064708 | 0.032133 | 0.183526 | 0.077571 | 0.034713 | 0.043726 | 0.053181 | 0.035782 | 0.010083 |
| 23.6944444 | 0.033526 | 0.067304 | 0.018842 | 0.021116 | 0.022721 | 0.019259 | 0.013722 | 0.096954 | 0.052058 | 0.069318 | 0.032272 | 0.184481 | 0.092258 | 0.024579 | 0.059917 | 0.083941 | 0.045646 | 0.017966 |
| 23.7083333 | 0.023768 | 0.047547 | 0.016996 | 0.016583 | 0.020027 | 0.011947 | 0.007438 | 0.097621 | 0.053624 | 0.048701 | 0.022997 | 0.122513 | 0.077851 | 0.021261 | 0.061931 | 0.085743 | 0.041255 | 0.023784 |
| 23.7222222 | 0.028056 | 0.030769 | 0.010322 | 0.010726 | 0.014208 | 0.014565 | 0.004557 | 0.067015 | 0.033115 | 0.033166 | 0.026792 | 0.093935 | 0.052357 | 0.036279 | 0.041002 | 0.054405 | 0.024971 | 0.019697 |
| 23.7361111 | 0.04187 | 0.054114 | 0.010174 | 0.017827 | 0.007254 | 0.014445 | 0.007207 | 0.053096 | 0.021788 | 0.057859 | 0.049799 | 0.153798 | 0.060904 | 0.039634 | 0.022475 | 0.036178 | 0.009584 | 0.01464 |
| 23.75 | 0.042232 | 0.080595 | 0.019945 | 0.02488 | 0.0059 | 0.007715 | 0.009759 | 0.071938 | 0.030983 | 0.078999 | 0.057593 | 0.181233 | 0.087923 | 0.024439 | 0.038728 | 0.040687 | 0.003247 | 0.027526 |
| 23.7638889 | 0.027873 | 0.073548 | 0.029951 | 0.016875 | 0.011913 | 0.005547 | 0.00921 | 0.06842 | 0.030103 | 0.054972 | 0.038246 | 0.120894 | 0.07765 | 0.011452 | 0.062774 | 0.030643 | 0.004826 | 0.040653 |
| 23.7777778 | 0.014185 | 0.044671 | 0.032636 | 0.008949 | 0.019133 | 0.010945 | 0.00599 | 0.042493 | 0.023179 | 0.02657 | 0.029634 | 0.097977 | 0.042517 | 0.009278 | 0.057085 | 0.018687 | 0.014688 | 0.029035 |
| 23.7916667 | 0.012363 | 0.01983 | 0.024636 | 0.013017 | 0.021092 | 0.014946 | 0.002658 | 0.045024 | 0.037092 | 0.032391 | 0.047813 | 0.151491 | 0.032944 | 0.015496 | 0.033685 | 0.031157 | 0.024833 | 0.011781 |
| 23.8055556 | 0.013662 | 0.019656 | 0.014339 | 0.01878 | 0.015954 | 0.010168 | 0.002222 | 0.076324 | 0.058539 | 0.053572 | 0.06421 | 0.190197 | 0.05453 | 0.018416 | 0.033038 | 0.055323 | 0.028461 | 0.012103 |
| 23.8194444 | 0.009892 | 0.032964 | 0.015348 | 0.016376 | 0.007896 | 0.003808 | 0.003709 | 0.104308 | 0.070528 | 0.065988 | 0.05942 | 0.170892 | 0.08085 | 0.011847 | 0.062548 | 0.066534 | 0.026359 | 0.024507 |
| 23.8333333 | 0.010215 | 0.045703 | 0.022228 | 0.010645 | 0.004027 | 0.002431 | 0.00477 | 0.10352 | 0.062002 | 0.06043 | 0.036904 | 0.102145 | 0.087522 | 0.006088 | 0.086526 | 0.044998 | 0.021217 | 0.035003 |

| | | | | | | | | | | | | | | | | | | |
|---|---|---|---|---|---|---|---|---|---|---|---|---|---|---|---|---|---|---|
| 23.8472222 | 0.014223 | 0.050484 | 0.021017 | 0.012422 | 0.004939 | 0.004499 | 0.006534 | 0.066573 | 0.034114 | 0.036141 | 0.016703 | 0.033219 | 0.063985 | 0.006258 | 0.079422 | 0.027235 | 0.015999 | 0.034185 |
| 23.8611111 | 0.011626 | 0.035762 | 0.012335 | 0.016061 | 0.0052 | 0.013049 | 0.009717 | 0.041172 | 0.018639 | 0.020046 | 0.016186 | 0.020502 | 0.035153 | 0.012605 | 0.045656 | 0.045393 | 0.010226 | 0.022245 |
| 23.875 | 0.007928 | 0.020571 | 0.008877 | 0.012262 | 0.003587 | 0.026547 | 0.01033 | 0.060491 | 0.023688 | 0.030304 | 0.021702 | 0.053767 | 0.035961 | 0.029521 | 0.020017 | 0.049392 | 0.006394 | 0.010663 |
| 23.8888889 | 0.013441 | 0.025752 | 0.014752 | 0.010083 | 0.004424 | 0.032973 | 0.00638 | 0.07301 | 0.022085 | 0.036896 | 0.016005 | 0.089335 | 0.050224 | 0.051581 | 0.021502 | 0.032872 | 0.009566 | 0.00897 |
| 23.9027778 | 0.02036 | 0.026197 | 0.017056 | 0.016017 | 0.009665 | 0.027454 | 0.002342 | 0.048677 | 0.015855 | 0.029387 | 0.013932 | 0.098412 | 0.04094 | 0.065571 | 0.021662 | 0.049819 | 0.028883 | 0.009465 |
| 23.9166667 | 0.018312 | 0.016475 | 0.014668 | 0.023149 | 0.014924 | 0.016486 | 0.001302 | 0.034453 | 0.027598 | 0.039631 | 0.034724 | 0.082146 | 0.030798 | 0.060288 | 0.019769 | 0.091032 | 0.055921 | 0.009325 |
| 23.9305556 | 0.011069 | 0.02359 | 0.023161 | 0.026651 | 0.016411 | 0.008913 | 0.001303 | 0.062472 | 0.0486 | 0.07207 | 0.061309 | 0.061768 | 0.055972 | 0.039802 | 0.033836 | 0.10828 | 0.068288 | 0.017934 |
| 23.9444444 | 0.005228 | 0.045467 | 0.038728 | 0.026027 | 0.015719 | 0.008661 | 0.002336 | 0.091701 | 0.055565 | 0.099212 | 0.072932 | 0.050694 | 0.086812 | 0.020718 | 0.046191 | 0.093262 | 0.067581 | 0.029272 |
| 23.9583333 | 0.002709 | 0.065544 | 0.047236 | 0.021252 | 0.015105 | 0.01241 | 0.006 | 0.085919 | 0.042455 | 0.100138 | 0.068439 | 0.042029 | 0.09357 | 0.012157 | 0.045439 | 0.068446 | 0.063204 | 0.033094 |
| 23.9722222 | 0.003096 | 0.075447 | 0.047746 | 0.013921 | 0.014313 | 0.01334 | 0.009708 | 0.059008 | 0.023459 | 0.07301 | 0.047261 | 0.025988 | 0.082487 | 0.013317 | 0.037799 | 0.048383 | 0.049734 | 0.028278 |
| 23.9861111 | 0.003772 | 0.06819 | 0.042337 | 0.006504 | 0.012393 | 0.008334 | 0.010226 | 0.02963 | 0.010712 | 0.036306 | 0.023396 | 0.01914 | 0.054708 | 0.0167 | 0.023715 | 0.029377 | 0.025998 | 0.016611 |
| 24 | 0.003601 | 0.045947 | 0.033958 | 0.001954 | 0.009856 | 0.002554 | 0.009117 | 0.010992 | 0.00491 | 0.020283 | 0.018187 | 0.039713 | 0.027061 | 0.015486 | 0.011445 | 0.015029 | 0.010347 | 0.009769 |
| 24.0138889 | 0.002882 | 0.022353 | 0.023396 | 0.00054 | 0.007931 | 0.002087 | 0.007566 | 0.008683 | 0.005672 | 0.028968 | 0.024684 | 0.065321 | 0.0252 | 0.011912 | 0.01233 | 0.018582 | 0.015853 | 0.015426 |
| 24.0277778 | 0.001627 | 0.013304 | 0.012846 | 0.001923 | 0.006408 | 0.006869 | 0.004714 | 0.016201 | 0.010391 | 0.032642 | 0.022257 | 0.071315 | 0.035465 | 0.010122 | 0.018013 | 0.035929 | 0.031255 | 0.022411 |
| 24.0416667 | 0.002134 | 0.020982 | 0.012881 | 0.006903 | 0.005041 | 0.013411 | 0.004095 | 0.025156 | 0.012733 | 0.020513 | 0.013101 | 0.056659 | 0.031979 | 0.008765 | 0.01688 | 0.051816 | 0.042921 | 0.020599 |
| 24.0555556 | 0.00472 | 0.025594 | 0.023266 | 0.012153 | 0.005915 | 0.019689 | 0.008439 | 0.031413 | 0.011038 | 0.018482 | 0.013867 | 0.031162 | 0.018997 | 0.007293 | 0.012358 | 0.05249 | 0.041842 | 0.013498 |
| 24.0694444 | 0.010002 | 0.023267 | 0.030043 | 0.012723 | 0.010118 | 0.028437 | 0.012024 | 0.028313 | 0.007667 | 0.046743 | 0.026167 | 0.019719 | 0.019317 | 0.01003 | 0.019398 | 0.037449 | 0.032433 | 0.014347 |
| 24.0833333 | 0.022652 | 0.035049 | 0.02358 | 0.013333 | 0.012998 | 0.039828 | 0.009713 | 0.021743 | 0.009159 | 0.088568 | 0.040235 | 0.047033 | 0.043612 | 0.028056 | 0.042802 | 0.038405 | 0.042863 | 0.027555 |
| 24.0972222 | 0.038052 | 0.057558 | 0.018605 | 0.023225 | 0.011725 | 0.047425 | 0.008456 | 0.027455 | 0.021167 | 0.113951 | 0.051026 | 0.096559 | 0.074413 | 0.070829 | 0.063001 | 0.071367 | 0.076174 | 0.039474 |
| 24.1111111 | 0.042374 | 0.067019 | 0.029614 | 0.036639 | 0.0101 | 0.050373 | 0.012545 | 0.032921 | 0.033598 | 0.108003 | 0.05163 | 0.115177 | 0.082724 | 0.111128 | 0.064335 | 0.092094 | 0.097587 | 0.040344 |
| 24.125 | 0.032139 | 0.061497 | 0.03949 | 0.039548 | 0.009727 | 0.050132 | 0.013942 | 0.022116 | 0.033185 | 0.074689 | 0.03806 | 0.081431 | 0.059905 | 0.104964 | 0.051849 | 0.069568 | 0.092203 | 0.034114 |
| 24.1388889 | 0.017351 | 0.050943 | 0.037359 | 0.030485 | 0.008738 | 0.041885 | 0.010819 | 0.007785 | 0.022336 | 0.036722 | 0.020754 | 0.032472 | 0.027493 | 0.061136 | 0.036529 | 0.034461 | 0.076804 | 0.029618 |
| 24.1527778 | 0.007429 | 0.037722 | 0.028775 | 0.017993 | 0.006221 | 0.025658 | 0.006824 | 0.002538 | 0.011311 | 0.014733 | 0.008825 | 0.006968 | 0.008464 | 0.024497 | 0.022855 | 0.019251 | 0.060744 | 0.027445 |
| 24.1666667 | 0.004521 | 0.024318 | 0.016995 | 0.008448 | 0.003283 | 0.010781 | 0.003822 | 0.002213 | 0.004696 | 0.007115 | 0.005332 | 0.004295 | 0.005603 | 0.016819 | 0.012738 | 0.016773 | 0.037386 | 0.018793 |
| 24.1805556 | 0.010086 | 0.012468 | 0.007589 | 0.006322 | 0.001508 | 0.006418 | 0.003746 | 0.004372 | 0.002089 | 0.008573 | 0.011778 | 0.008159 | 0.009574 | 0.029085 | 0.013373 | 0.01139 | 0.015302 | 0.009431 |
| 24.1944444 | 0.025935 | 0.01229 | 0.005304 | 0.013558 | 0.003583 | 0.013961 | 0.004541 | 0.010336 | 0.00358 | 0.01787 | 0.020863 | 0.013574 | 0.015767 | 0.044736 | 0.023718 | 0.008431 | 0.006708 | 0.009173 |
| 24.2083333 | 0.051514 | 0.039063 | 0.006002 | 0.027744 | 0.015941 | 0.031366 | 0.006302 | 0.013255 | 0.006344 | 0.023362 | 0.024659 | 0.023248 | 0.027919 | 0.06122 | 0.032831 | 0.0152 | 0.006363 | 0.009358 |
| 24.2222222 | 0.073542 | 0.072729 | 0.005515 | 0.042761 | 0.035152 | 0.049926 | 0.010613 | 0.011064 | 0.009396 | 0.016162 | 0.020895 | 0.037079 | 0.039265 | 0.067635 | 0.028482 | 0.020996 | 0.007616 | 0.006907 |
| 24.2361111 | 0.069237 | 0.073194 | 0.007749 | 0.048385 | 0.042043 | 0.058858 | 0.01333 | 0.013301 | 0.021735 | 0.008931 | 0.011868 | 0.055295 | 0.034755 | 0.050283 | 0.015581 | 0.015388 | 0.007925 | 0.008029 |
| 24.25 | 0.042724 | 0.043321 | 0.012207 | 0.040612 | 0.030873 | 0.056701 | 0.012673 | 0.018516 | 0.038668 | 0.009385 | 0.005215 | 0.07741 | 0.021633 | 0.032343 | 0.017469 | 0.010483 | 0.015411 | 0.017048 |
| 24.2638889 | 0.020477 | 0.018453 | 0.013411 | 0.025619 | 0.015817 | 0.044421 | 0.010845 | 0.016293 | 0.039641 | 0.011629 | 0.008575 | 0.092888 | 0.019023 | 0.03483 | 0.031086 | 0.019525 | 0.034744 | 0.032619 |
| 24.2777778 | 0.010471 | 0.008798 | 0.009549 | 0.011856 | 0.006052 | 0.028433 | 0.007101 | 0.008207 | 0.024072 | 0.018905 | 0.01893 | 0.093023 | 0.024497 | 0.04553 | 0.03757 | 0.032966 | 0.049047 | 0.043133 |
| 24.2916667 | 0.00528 | 0.004086 | 0.004792 | 0.006449 | 0.002422 | 0.017318 | 0.003295 | 0.002239 | 0.00915 | 0.024632 | 0.023726 | 0.083059 | 0.025431 | 0.047044 | 0.035644 | 0.038152 | 0.042782 | 0.041816 |
| 24.3055556 | 0.004428 | 0.001699 | 0.003138 | 0.006895 | 0.003417 | 0.011993 | 0.002172 | 0.001157 | 0.002722 | 0.023277 | 0.022135 | 0.07221 | 0.023968 | 0.039143 | 0.032807 | 0.036421 | 0.027915 | 0.035228 |
| 24.3194444 | 0.010419 | 0.002413 | 0.003153 | 0.006313 | 0.005323 | 0.008854 | 0.00243 | 0.002374 | 0.00295 | 0.023674 | 0.022432 | 0.065305 | 0.027524 | 0.032423 | 0.032999 | 0.035111 | 0.026105 | 0.032002 |
| 24.3333333 | 0.017815 | 0.004745 | 0.004601 | 0.006782 | 0.005056 | 0.005868 | 0.003796 | 0.002455 | 0.006177 | 0.025263 | 0.022816 | 0.060009 | 0.030207 | 0.028862 | 0.031971 | 0.035861 | 0.03642 | 0.031163 |
| 24.3472222 | 0.017672 | 0.005114 | 0.00511 | 0.0078 | 0.002794 | 0.006059 | 0.003824 | 0.001289 | 0.00801 | 0.021027 | 0.016786 | 0.050104 | 0.02094 | 0.022723 | 0.023864 | 0.032786 | 0.039495 | 0.025194 |
| 24.3611111 | 0.011055 | 0.003169 | 0.003969 | 0.005972 | 0.000829 | 0.011277 | 0.002426 | 0.000575 | 0.007119 | 0.011155 | 0.007254 | 0.031825 | 0.010291 | 0.012322 | 0.011837 | 0.023069 | 0.026202 | 0.013531 |
| 24.375 | 0.008025 | 0.001964 | 0.00566 | 0.005924 | 0.000884 | 0.016212 | 0.0037 | 0.000721 | 0.005715 | 0.003523 | 0.002029 | 0.015844 | 0.011559 | 0.005445 | 0.004242 | 0.011984 | 0.015361 | 0.005738 |
| 24.3888889 | 0.009627 | 0.001358 | 0.007721 | 0.00892 | 0.003111 | 0.017002 | 0.006822 | 0.001311 | 0.005453 | 0.001558 | 0.002027 | 0.020519 | 0.011146 | 0.007788 | 0.002159 | 0.005566 | 0.018732 | 0.005745 |
| 24.4027778 | 0.007976 | 0.000768 | 0.005624 | 0.009008 | 0.005245 | 0.015508 | 0.008394 | 0.001608 | 0.005914 | 0.001971 | 0.003921 | 0.043717 | 0.004733 | 0.015063 | 0.001946 | 0.002987 | 0.020699 | 0.007408 |
| 24.4166667 | 0.005807 | 0.001204 | 0.003693 | 0.006536 | 0.004788 | 0.011528 | 0.006047 | 0.001897 | 0.006418 | 0.002027 | 0.006475 | 0.068017 | 0.003048 | 0.020373 | 0.001794 | 0.001336 | 0.014592 | 0.007326 |
| 24.4305556 | 0.011962 | 0.003071 | 0.006416 | 0.009256 | 0.002542 | 0.007408 | 0.003451 | 0.002696 | 0.006565 | 0.002969 | 0.009999 | 0.074323 | 0.00376 | 0.016346 | 0.001687 | 0.000895 | 0.008709 | 0.005974 |
| 24.4444444 | 0.02163 | 0.005345 | 0.008566 | 0.017228 | 0.001184 | 0.009103 | 0.005177 | 0.003131 | 0.005549 | 0.004448 | 0.010546 | 0.05032 | 0.004688 | 0.00846 | 0.003351 | 0.00164 | 0.006185 | 0.006993 |
| 24.4583333 | 0.022187 | 0.005468 | 0.005785 | 0.020756 | 0.001708 | 0.013639 | 0.007319 | 0.00383 | 0.003219 | 0.011094 | 0.013142 | 0.030334 | 0.01519 | 0.006257 | 0.014021 | 0.007108 | 0.011794 | 0.019242 |
| 24.4722222 | 0.015999 | 0.004095 | 0.002978 | 0.018569 | 0.002253 | 0.016111 | 0.00789 | 0.009059 | 0.003781 | 0.035766 | 0.033795 | 0.057617 | 0.038263 | 0.011704 | 0.038424 | 0.020424 | 0.034332 | 0.045582 |
| 24.4861111 | 0.019286 | 0.004918 | 0.007097 | 0.017786 | 0.001561 | 0.015991 | 0.009574 | 0.02065 | 0.013334 | 0.063623 | 0.059651 | 0.094116 | 0.055927 | 0.024546 | 0.060207 | 0.030367 | 0.055755 | 0.068386 |
| 24.5 | 0.031762 | 0.009645 | 0.016529 | 0.017912 | 0.000803 | 0.011417 | 0.012293 | 0.02932 | 0.026428 | 0.062421 | 0.058178 | 0.080503 | 0.04798 | 0.027774 | 0.056755 | 0.025677 | 0.04787 | 0.065619 |
| 24.5138889 | 0.038322 | 0.014902 | 0.021341 | 0.017149 | 0.000638 | 0.006098 | 0.013623 | 0.025273 | 0.028781 | 0.034405 | 0.031578 | 0.045605 | 0.023592 | 0.015108 | 0.032138 | 0.014163 | 0.022094 | 0.042 |
| 24.5277778 | 0.031763 | 0.013352 | 0.016376 | 0.016978 | 0.000913 | 0.007653 | 0.01141 | 0.013481 | 0.018386 | 0.013074 | 0.011591 | 0.05669 | 0.009451 | 0.005956 | 0.010493 | 0.007003 | 0.008942 | 0.018369 |
| 24.5416667 | 0.015937 | 0.006497 | 0.007635 | 0.013607 | 0.001611 | 0.013305 | 0.006672 | 0.0044 | 0.007312 | 0.010862 | 0.008382 | 0.104459 | 0.011036 | 0.006474 | 0.004367 | 0.003572 | 0.014816 | 0.006979 |
| 24.5555556 | 0.008895 | 0.009254 | 0.003248 | 0.008119 | 0.002431 | 0.013038 | 0.003302 | 0.001413 | 0.004674 | 0.011156 | 0.009715 | 0.117892 | 0.01277 | 0.009979 | 0.010789 | 0.004101 | 0.01968 | 0.009464 |
| 24.5694444 | 0.0258 | 0.03103 | 0.002604 | 0.007209 | 0.002226 | 0.014165 | 0.00326 | 0.006349 | 0.013131 | 0.01351 | 0.015024 | 0.082824 | 0.016418 | 0.023324 | 0.027203 | 0.014939 | 0.021254 | 0.021588 |
| 24.5833333 | 0.054918 | 0.058758 | 0.008478 | 0.008064 | 0.001414 | 0.024089 | 0.006898 | 0.027743 | 0.031518 | 0.031253 | 0.029983 | 0.095839 | 0.03688 | 0.048152 | 0.048265 | 0.040313 | 0.042166 | 0.038505 |
| 24.5972222 | 0.072458 | 0.07296 | 0.031512 | 0.014921 | 0.00365 | 0.029191 | 0.016223 | 0.073262 | 0.053519 | 0.063158 | 0.046508 | 0.188902 | 0.059848 | 0.060641 | 0.057722 | 0.08546 | 0.074568 | 0.049848 |
| 24.6111111 | 0.070319 | 0.070305 | 0.058281 | 0.033618 | 0.008766 | 0.022125 | 0.025394 | 0.128737 | 0.075196 | 0.091524 | 0.048034 | 0.235703 | 0.068974 | 0.047357 | 0.048565 | 0.136983 | 0.097259 | 0.046774 |
| 24.625 | 0.057014 | 0.059139 | 0.061388 | 0.045454 | 0.012223 | 0.010271 | 0.024261 | 0.155293 | 0.086292 | 0.096096 | 0.032778 | 0.172226 | 0.066075 | 0.023785 | 0.033322 | 0.163099 | 0.100248 | 0.033078 |
| 24.6388889 | 0.038918 | 0.046839 | 0.044656 | 0.036236 | 0.013053 | 0.005724 | 0.016558 | 0.1286 | 0.072518 | 0.067781 | 0.017112 | 0.088279 | 0.045367 | 0.013717 | 0.018712 | 0.149098 | 0.071327 | 0.017548 |
| 24.6527778 | 0.024197 | 0.033216 | 0.026609 | 0.019023 | 0.012879 | 0.011245 | 0.009574 | 0.070586 | 0.041629 | 0.034269 | 0.015995 | 0.086208 | 0.023399 | 0.027822 | 0.010063 | 0.098591 | 0.034936 | 0.010776 |
| 24.6666667 | 0.029497 | 0.017778 | 0.012725 | 0.014499 | 0.011917 | 0.020717 | 0.005454 | 0.030408 | 0.019728 | 0.033872 | 0.026349 | 0.156331 | 0.027769 | 0.046187 | 0.013758 | 0.046904 | 0.032637 | 0.013553 |
| 24.6805556 | 0.04654 | 0.008702 | 0.00764 | 0.024862 | 0.010786 | 0.02733 | 0.006322 | 0.036504 | 0.022726 | 0.054345 | 0.03118 | 0.233168 | 0.045758 | 0.039801 | 0.018182 | 0.038154 | 0.053722 | 0.016475 |
| 24.6944444 | 0.046729 | 0.01102 | 0.010978 | 0.035123 | 0.011959 | 0.022257 | 0.008637 | 0.076506 | 0.042024 | 0.068667 | 0.023446 | 0.240366 | 0.056405 | 0.027174 | 0.016653 | 0.064199 | 0.067652 | 0.017963 |
| 24.7083333 | 0.027209 | 0.017573 | 0.011254 | 0.032614 | 0.017299 | 0.012928 | 0.00675 | 0.121013 | 0.059088 | 0.069678 | 0.012712 | 0.179289 | 0.055787 | 0.042841 | 0.015919 | 0.085401 | 0.066752 | 0.017903 |
| 24.7222222 | 0.018938 | 0.02108 | 0.007619 | 0.020584 | 0.024684 | 0.013148 | 0.002642 | 0.139886 | 0.058525 | 0.048412 | 0.009869 | 0.110009 | 0.039163 | 0.060111 | 0.015419 | 0.081768 | 0.052825 | 0.012301 |
| 24.7361111 | 0.037404 | 0.022447 | 0.009516 | 0.014293 | 0.027051 | 0.014264 | 0.002245 | 0.110849 | 0.037089 | 0.03303 | 0.01661 | 0.123355 | 0.020131 | 0.044269 | 0.01009 | 0.058882 | 0.031801 | 0.005722 |
| 24.75 | 0.050987 | 0.024105 | 0.014926 | 0.011601 | 0.020154 | 0.009474 | 0.004152 | 0.06248 | 0.016945 | 0.061223 | 0.028146 | 0.126166 | 0.023111 | 0.020772 | 0.007534 | 0.030388 | 0.013698 | 0.005454 |
| 24.7638889 | 0.037987 | 0.023752 | 0.017519 | 0.007395 | 0.009449 | 0.007776 | 0.004044 | 0.0591 | 0.019283 | 0.089147 | 0.03507 | 0.083162 | 0.03776 | 0.022029 | 0.017022 | 0.018859 | 0.006558 | 0.010395 |
| 24.7777778 | 0.019489 | 0.017559 | 0.020977 | 0.009784 | 0.003807 | 0.012232 | 0.003974 | 0.096382 | 0.035243 | 0.070574 | 0.027806 | 0.10006 | 0.046776 | 0.036675 | 0.028648 | 0.031916 | 0.006062 | 0.01838 |
| 24.7916667 | 0.021807 | 0.008921 | 0.02366 | 0.015432 | 0.004393 | 0.020007 | 0.005544 | 0.118993 | 0.048992 | 0.034103 | 0.016384 | 0.162378 | 0.048517 | 0.041507 | 0.030674 | 0.043644 | 0.003769 | 0.026728 |
| 24.8055556 | 0.043648 | 0.004494 | 0.018902 | 0.018484 | 0.005728 | 0.025843 | 0.006698 | 0.099435 | 0.047534 | 0.017499 | 0.011748 | 0.187343 | 0.037891 | 0.031172 | 0.024937 | 0.0393 | 0.00304 | 0.028371 |
| 24.8194444 | 0.057075 | 0.006886 | 0.010742 | 0.020501 | 0.004471 | 0.019947 | 0.005694 | 0.057885 | 0.029098 | 0.01977 | 0.008094 | 0.162433 | 0.020259 | 0.027393 | 0.015533 | 0.026726 | 0.008018 | 0.019122 |
| 24.8333333 | 0.046687 | 0.017554 | 0.005069 | 0.019552 | 0.002158 | 0.012134 | 0.00605 | 0.054402 | 0.02277 | 0.030061 | 0.005872 | 0.11819 | 0.016769 | 0.037333 | 0.010332 | 0.013572 | 0.013726 | 0.008833 |
| 24.8472222 | 0.024974 | 0.038228 | 0.005221 | 0.013765 | 0.001619 | 0.016278 | 0.009631 | 0.097841 | 0.041018 | 0.043869 | 0.008071 | 0.076129 | 0.033153 | 0.032383 | 0.017633 | 0.009569 | 0.016317 | 0.008284 |
| 24.8611111 | 0.007847 | 0.055786 | 0.010936 | 0.007575 | 0.001594 | 0.015678 | 0.009957 | 0.128972 | 0.053655 | 0.052655 | 0.01525 | 0.041556 | 0.046532 | 0.027329 | 0.026869 | 0.021964 | 0.018546 | 0.010365 |
| 24.875 | 0.002003 | 0.043834 | 0.017402 | 0.004464 | 0.001578 | 0.011292 | 0.009571 | 0.115378 | 0.044538 | 0.050115 | 0.020556 | 0.017847 | 0.040855 | 0.049201 | 0.024243 | 0.033893 | 0.021973 | 0.00842 |
| 24.8888889 | 0.002595 | 0.024745 | 0.020361 | 0.0043 | 0.003627 | 0.022037 | 0.015595 | 0.074534 | 0.028443 | 0.038711 | 0.015485 | 0.007411 | 0.0242 | 0.067813 | 0.01261 | 0.028633 | 0.023206 | 0.004784 |
| 24.9027778 | 0.006379 | 0.037106 | 0.016281 | 0.004861 | 0.006757 | 0.038497 | 0.022662 | 0.035657 | 0.017558 | 0.025786 | 0.00825 | 0.005707 | 0.009601 | 0.060321 | 0.003843 | 0.014996 | 0.022318 | 0.002296 |
| 24.9166667 | 0.018478 | 0.055714 | 0.009449 | 0.008358 | 0.009209 | 0.048082 | 0.023519 | 0.012566 | 0.010091 | 0.017046 | 0.006394 | 0.005445 | 0.003456 | 0.043443 | 0.003314 | 0.00453 | 0.023431 | 0.003604 |
| 24.9305556 | 0.043705 | 0.054668 | 0.008203 | 0.024565 | 0.011628 | 0.051059 | 0.017673 | 0.003418 | 0.004473 | 0.014328 | 0.011251 | 0.007558 | 0.002777 | 0.035735 | 0.004337 | 0.000821 | 0.024404 | 0.006622 |
| 24.9444444 | 0.077889 | 0.046645 | 0.009321 | 0.049545 | 0.018209 | 0.049206 | 0.00928 | 0.001624 | 0.002867 | 0.01135 | 0.026204 | 0.008521 | 0.00188 | 0.040938 | 0.003132 | 0.001779 | 0.01963 | 0.007783 |
| 24.9583333 | 0.100194 | 0.04109 | 0.008555 | 0.065576 | 0.030383 | 0.041078 | 0.00338 | 0.001982 | 0.002364 | 0.012432 | 0.038257 | 0.008744 | 0.00241 | 0.050628 | 0.004206 | 0.004957 | 0.010108 | 0.009459 |
| 24.9722222 | 0.096527 | 0.035665 | 0.01136 | 0.063095 | 0.041856 | 0.026123 | 0.002155 | 0.002679 | 0.002005 | 0.023407 | 0.033634 | 0.015171 | 0.004191 | 0.050205 | 0.007879 | 0.008533 | 0.003339 | 0.013368 |
| 24.9861111 | 0.075298 | 0.030065 | 0.017353 | 0.048537 | 0.046655 | 0.012286 | 0.00416 | 0.002652 | 0.003935 | 0.03127 | 0.019687 | 0.018731 | 0.005249 | 0.033047 | 0.00898 | 0.008981 | 0.002122 | 0.014569 |

| | | | | | | | | | | | | | | | | | | |
|---|---|---|---|---|---|---|---|---|---|---|---|---|---|---|---|---|---|---|
| 25 | 0.0482 | 0.025235 | 0.019317 | 0.03068 | 0.041016 | 0.010836 | 0.005481 | 0.003333 | 0.005989 | 0.029473 | 0.010221 | 0.014424 | 0.00911 | 0.022436 | 0.008904 | 0.005881 | 0.003154 | 0.012334 |
| 25.0138889 | 0.023807 | 0.019782 | 0.017075 | 0.017113 | 0.025144 | 0.02152 | 0.003849 | 0.005741 | 0.005755 | 0.022223 | 0.014633 | 0.012972 | 0.013498 | 0.037166 | 0.012418 | 0.004633 | 0.006918 | 0.011887 |
| 25.0277778 | 0.011934 | 0.014414 | 0.01514 | 0.017081 | 0.010721 | 0.030738 | 0.001984 | 0.006881 | 0.003565 | 0.014296 | 0.024851 | 0.017512 | 0.012634 | 0.054497 | 0.015862 | 0.010906 | 0.014644 | 0.013128 |
| 25.0416667 | 0.012495 | 0.010002 | 0.01565 | 0.022546 | 0.006787 | 0.03151 | 0.002604 | 0.005844 | 0.002178 | 0.008366 | 0.025398 | 0.021784 | 0.007633 | 0.054725 | 0.01283 | 0.021891 | 0.0223 | 0.011394 |
| 25.0555556 | 0.011448 | 0.007183 | 0.017468 | 0.02007 | 0.006443 | 0.028467 | 0.003716 | 0.004618 | 0.003773 | 0.005286 | 0.016216 | 0.020328 | 0.005978 | 0.041638 | 0.009704 | 0.029745 | 0.021469 | 0.010044 |
| 25.0694444 | 0.013298 | 0.011209 | 0.01896 | 0.012776 | 0.009175 | 0.019809 | 0.003471 | 0.004191 | 0.004871 | 0.007014 | 0.015584 | 0.020933 | 0.015325 | 0.025313 | 0.022593 | 0.025829 | 0.018928 | 0.022508 |
| 25.0833333 | 0.038531 | 0.024286 | 0.020817 | 0.019468 | 0.022634 | 0.01545 | 0.006159 | 0.006972 | 0.007974 | 0.022644 | 0.035197 | 0.041713 | 0.036768 | 0.031837 | 0.054621 | 0.025406 | 0.040753 | 0.049392 |
| 25.0972222 | 0.073707 | 0.033391 | 0.022111 | 0.047474 | 0.036061 | 0.032516 | 0.016082 | 0.012292 | 0.019541 | 0.055691 | 0.058141 | 0.068179 | 0.059045 | 0.076946 | 0.085965 | 0.047653 | 0.084601 | 0.071073 |
| 25.1111111 | 0.083426 | 0.028116 | 0.019974 | 0.064552 | 0.033646 | 0.055986 | 0.026709 | 0.012221 | 0.03143 | 0.081323 | 0.060241 | 0.064548 | 0.063424 | 0.121153 | 0.093451 | 0.058949 | 0.109329 | 0.069712 |
| 25.125 | 0.064054 | 0.018473 | 0.015221 | 0.05105 | 0.02082 | 0.067591 | 0.029017 | 0.006759 | 0.033076 | 0.079408 | 0.038233 | 0.035152 | 0.046698 | 0.117962 | 0.075897 | 0.03707 | 0.089612 | 0.047261 |
| 25.1388889 | 0.038555 | 0.014048 | 0.010205 | 0.029708 | 0.011359 | 0.061219 | 0.023203 | 0.004334 | 0.02515 | 0.057362 | 0.014643 | 0.011844 | 0.023347 | 0.075322 | 0.047692 | 0.017784 | 0.044659 | 0.021354 |
| 25.1527778 | 0.019396 | 0.010127 | 0.005446 | 0.015892 | 0.006704 | 0.037112 | 0.014399 | 0.005555 | 0.013798 | 0.028656 | 0.006924 | 0.006916 | 0.00977 | 0.031503 | 0.021085 | 0.019379 | 0.018343 | 0.009603 |
| 25.1666667 | 0.015624 | 0.007256 | 0.003924 | 0.0073 | 0.003437 | 0.01486 | 0.007409 | 0.005982 | 0.006274 | 0.014751 | 0.006355 | 0.01196 | 0.011159 | 0.010247 | 0.008647 | 0.023269 | 0.027176 | 0.013988 |
| 25.1805556 | 0.030847 | 0.013803 | 0.008036 | 0.004144 | 0.002408 | 0.009965 | 0.006649 | 0.005339 | 0.006984 | 0.027685 | 0.010885 | 0.018346 | 0.014627 | 0.009343 | 0.015407 | 0.018109 | 0.046765 | 0.019417 |
| 25.1944444 | 0.045002 | 0.024087 | 0.01324 | 0.007905 | 0.003845 | 0.014682 | 0.010814 | 0.005481 | 0.013492 | 0.041676 | 0.019883 | 0.019709 | 0.010308 | 0.010816 | 0.027398 | 0.011613 | 0.056547 | 0.017719 |
| 25.2083333 | 0.043503 | 0.025057 | 0.015144 | 0.015824 | 0.005885 | 0.017235 | 0.013867 | 0.00653 | 0.019201 | 0.038396 | 0.018303 | 0.013047 | 0.003837 | 0.013462 | 0.03336 | 0.009914 | 0.053957 | 0.013745 |
| 25.2222222 | 0.032662 | 0.016075 | 0.013728 | 0.023982 | 0.006523 | 0.015639 | 0.013683 | 0.006417 | 0.018353 | 0.033364 | 0.01317 | 0.008846 | 0.00364 | 0.031068 | 0.036902 | 0.009429 | 0.046289 | 0.014648 |
| 25.2361111 | 0.019398 | 0.006023 | 0.009115 | 0.028354 | 0.005463 | 0.012886 | 0.01042 | 0.004617 | 0.010499 | 0.035207 | 0.017808 | 0.015074 | 0.010415 | 0.050302 | 0.040657 | 0.013486 | 0.036295 | 0.021706 |
| 25.25 | 0.009767 | 0.001652 | 0.004624 | 0.027206 | 0.004436 | 0.013244 | 0.005436 | 0.003356 | 0.002748 | 0.035613 | 0.022872 | 0.023227 | 0.015186 | 0.054405 | 0.041512 | 0.017109 | 0.021562 | 0.026184 |
| 25.2638889 | 0.011776 | 0.002431 | 0.004168 | 0.024207 | 0.00612 | 0.01913 | 0.002484 | 0.003433 | 0.003075 | 0.031608 | 0.021648 | 0.024084 | 0.013003 | 0.049702 | 0.036838 | 0.010521 | 0.011426 | 0.021553 |
| 25.2777778 | 0.024856 | 0.004355 | 0.004266 | 0.022965 | 0.012136 | 0.027703 | 0.003948 | 0.00377 | 0.012298 | 0.028968 | 0.021213 | 0.019177 | 0.009793 | 0.045009 | 0.026585 | 0.003421 | 0.011056 | 0.012158 |
| 25.2916667 | 0.039771 | 0.00624 | 0.00461 | 0.023645 | 0.020029 | 0.034817 | 0.008338 | 0.00406 | 0.023224 | 0.032183 | 0.024527 | 0.015005 | 0.011136 | 0.042653 | 0.014713 | 0.002423 | 0.019316 | 0.006128 |
| 25.3055556 | 0.043977 | 0.008345 | 0.008015 | 0.023221 | 0.02296 | 0.037237 | 0.01159 | 0.004524 | 0.02709 | 0.036339 | 0.026145 | 0.014797 | 0.014338 | 0.03907 | 0.007487 | 0.006224 | 0.030201 | 0.006255 |
| 25.3194444 | 0.029315 | 0.0091 | 0.009415 | 0.01716 | 0.015582 | 0.032777 | 0.0106 | 0.0048 | 0.021936 | 0.03074 | 0.023366 | 0.017087 | 0.013233 | 0.029361 | 0.006022 | 0.014585 | 0.024866 | 0.009519 |
| 25.3333333 | 0.012845 | 0.008387 | 0.005827 | 0.01008 | 0.007728 | 0.024929 | 0.006015 | 0.004754 | 0.013111 | 0.016387 | 0.017643 | 0.018662 | 0.007842 | 0.015151 | 0.006147 | 0.019328 | 0.013765 | 0.011854 |
| 25.3472222 | 0.01081 | 0.006712 | 0.002632 | 0.010367 | 0.009246 | 0.017552 | 0.00256 | 0.004536 | 0.00655 | 0.00619 | 0.010227 | 0.018468 | 0.003984 | 0.004669 | 0.005725 | 0.015969 | 0.017281 | 0.010776 |
| 25.3611111 | 0.010064 | 0.004309 | 0.002699 | 0.010689 | 0.010177 | 0.010975 | 0.002013 | 0.004024 | 0.005147 | 0.004249 | 0.005399 | 0.016518 | 0.005246 | 0.002474 | 0.006025 | 0.008847 | 0.017257 | 0.006424 |
| 25.375 | 0.009214 | 0.003341 | 0.003373 | 0.007951 | 0.005962 | 0.005879 | 0.002822 | 0.003377 | 0.00815 | 0.005558 | 0.00661 | 0.012751 | 0.009279 | 0.006014 | 0.007237 | 0.002823 | 0.014085 | 0.002708 |
| 25.3888889 | 0.017236 | 0.003439 | 0.005963 | 0.011193 | 0.005134 | 0.002559 | 0.006439 | 0.003442 | 0.012382 | 0.010684 | 0.007351 | 0.009202 | 0.011051 | 0.010715 | 0.007251 | 0.002775 | 0.025749 | 0.002268 |
| 25.4027778 | 0.024285 | 0.004311 | 0.010287 | 0.015731 | 0.008561 | 0.001665 | 0.010927 | 0.004547 | 0.015107 | 0.016617 | 0.004943 | 0.008106 | 0.008327 | 0.012133 | 0.005423 | 0.009066 | 0.03433 | 0.002207 |
| 25.4166667 | 0.019664 | 0.006403 | 0.01013 | 0.012531 | 0.00869 | 0.004926 | 0.011549 | 0.005466 | 0.014317 | 0.014509 | 0.005051 | 0.009313 | 0.004189 | 0.009183 | 0.003839 | 0.016606 | 0.024473 | 0.001783 |
| 25.4305556 | 0.009175 | 0.005959 | 0.005892 | 0.00541 | 0.004982 | 0.010066 | 0.008329 | 0.005158 | 0.010361 | 0.008458 | 0.006842 | 0.009628 | 0.001953 | 0.005245 | 0.002686 | 0.018149 | 0.010052 | 0.002179 |
| 25.4444444 | 0.003721 | 0.003202 | 0.003559 | 0.002788 | 0.002658 | 0.011274 | 0.004698 | 0.004329 | 0.007608 | 0.006168 | 0.007267 | 0.006409 | 0.001089 | 0.003439 | 0.001516 | 0.011839 | 0.004928 | 0.003419 |
| 25.4583333 | 0.005228 | 0.001872 | 0.003275 | 0.00643 | 0.002365 | 0.007294 | 0.003451 | 0.003916 | 0.010048 | 0.008807 | 0.010454 | 0.003117 | 0.002525 | 0.003412 | 0.001361 | 0.005708 | 0.010801 | 0.006002 |
| 25.4722222 | 0.010404 | 0.001659 | 0.004087 | 0.011324 | 0.004401 | 0.002582 | 0.004619 | 0.003263 | 0.015297 | 0.02041 | 0.019494 | 0.002348 | 0.007825 | 0.004302 | 0.001807 | 0.005911 | 0.021245 | 0.007205 |
| 25.4861111 | 0.012275 | 0.001972 | 0.007702 | 0.012518 | 0.008664 | 0.000721 | 0.006321 | 0.001984 | 0.016117 | 0.034743 | 0.029225 | 0.002015 | 0.011622 | 0.004407 | 0.002354 | 0.010018 | 0.022673 | 0.006233 |
| 25.5 | 0.007816 | 0.002343 | 0.0109 | 0.00878 | 0.012321 | 0.001044 | 0.006785 | 0.000924 | 0.01029 | 0.038031 | 0.032644 | 0.002622 | 0.009418 | 0.003089 | 0.002484 | 0.013849 | 0.014266 | 0.004348 |
| 25.5138889 | 0.002636 | 0.003435 | 0.009377 | 0.004297 | 0.012647 | 0.003282 | 0.005669 | 0.001264 | 0.004151 | 0.027388 | 0.028243 | 0.004943 | 0.004621 | 0.002753 | 0.001574 | 0.01537 | 0.012938 | 0.002731 |
| 25.5277778 | 0.001666 | 0.006554 | 0.005539 | 0.002322 | 0.009048 | 0.008145 | 0.003661 | 0.003177 | 0.002421 | 0.013839 | 0.019897 | 0.007229 | 0.001912 | 0.003117 | 0.001283 | 0.01434 | 0.02261 | 0.003454 |
| 25.5416667 | 0.005212 | 0.00687 | 0.004059 | 0.002941 | 0.004278 | 0.012989 | 0.002223 | 0.004264 | 0.003158 | 0.008417 | 0.011926 | 0.008345 | 0.004403 | 0.006506 | 0.003819 | 0.011261 | 0.028798 | 0.004705 |
| 25.5555556 | 0.011336 | 0.00349 | 0.00384 | 0.00744 | 0.001594 | 0.016315 | 0.003164 | 0.003754 | 0.007398 | 0.009553 | 0.007359 | 0.009605 | 0.012209 | 0.017664 | 0.008443 | 0.008253 | 0.02207 | 0.004606 |
| 25.5694444 | 0.019221 | 0.004023 | 0.005491 | 0.012989 | 0.002648 | 0.020515 | 0.00712 | 0.003384 | 0.016585 | 0.019171 | 0.008966 | 0.015807 | 0.02072 | 0.035462 | 0.013357 | 0.008276 | 0.020305 | 0.004296 |
| 25.5833333 | 0.029902 | 0.012324 | 0.016092 | 0.017202 | 0.010112 | 0.027729 | 0.014486 | 0.003827 | 0.024315 | 0.037397 | 0.017231 | 0.034693 | 0.026201 | 0.060642 | 0.018904 | 0.012802 | 0.039849 | 0.005471 |
| 25.5972222 | 0.044443 | 0.02477 | 0.037616 | 0.025781 | 0.023389 | 0.0358 | 0.023544 | 0.004595 | 0.02606 | 0.053815 | 0.028921 | 0.074171 | 0.028752 | 0.090128 | 0.025334 | 0.020638 | 0.061092 | 0.009671 |
| 25.6111111 | 0.059293 | 0.037788 | 0.060052 | 0.039946 | 0.030498 | 0.03783 | 0.029368 | 0.008312 | 0.02371 | 0.062189 | 0.03655 | 0.125496 | 0.029173 | 0.09978 | 0.030103 | 0.026704 | 0.06745 | 0.014942 |
| 25.625 | 0.061406 | 0.049415 | 0.066896 | 0.047034 | 0.023083 | 0.031083 | 0.029535 | 0.019647 | 0.020581 | 0.060348 | 0.031032 | 0.155912 | 0.028599 | 0.074924 | 0.028967 | 0.026084 | 0.058514 | 0.017006 |
| 25.6388889 | 0.039711 | 0.050974 | 0.052427 | 0.034778 | 0.011656 | 0.018141 | 0.022478 | 0.034184 | 0.016287 | 0.04531 | 0.015654 | 0.139442 | 0.026433 | 0.036162 | 0.021516 | 0.021391 | 0.034559 | 0.017701 |
| 25.6527778 | 0.023673 | 0.038827 | 0.026886 | 0.021346 | 0.011446 | 0.010538 | 0.011979 | 0.042879 | 0.009851 | 0.023831 | 0.00551 | 0.082687 | 0.020395 | 0.01581 | 0.013271 | 0.019898 | 0.018506 | 0.02163 |
| 25.6666667 | 0.040873 | 0.023875 | 0.012705 | 0.030263 | 0.021592 | 0.018748 | 0.008519 | 0.045717 | 0.007265 | 0.012171 | 0.005864 | 0.03762 | 0.011014 | 0.025141 | 0.007705 | 0.024398 | 0.030651 | 0.028148 |
| 25.6805556 | 0.058215 | 0.012998 | 0.020532 | 0.044398 | 0.026563 | 0.028081 | 0.010802 | 0.046995 | 0.009226 | 0.01058 | 0.010157 | 0.040058 | 0.004183 | 0.042638 | 0.004313 | 0.030626 | 0.052038 | 0.034719 |
| 25.6944444 | 0.053707 | 0.008557 | 0.027102 | 0.047638 | 0.020509 | 0.027298 | 0.009088 | 0.045134 | 0.008401 | 0.012492 | 0.01365 | 0.058201 | 0.004442 | 0.049706 | 0.004896 | 0.032852 | 0.056189 | 0.038335 |
| 25.7083333 | 0.035531 | 0.01556 | 0.016894 | 0.039829 | 0.010795 | 0.024335 | 0.004029 | 0.03763 | 0.010616 | 0.020688 | 0.012831 | 0.076196 | 0.009243 | 0.042121 | 0.009118 | 0.031926 | 0.036734 | 0.034276 |
| 25.7222222 | 0.022416 | 0.029411 | 0.005399 | 0.02454 | 0.006824 | 0.019407 | 0.001864 | 0.02522 | 0.022361 | 0.030378 | 0.010125 | 0.097061 | 0.013722 | 0.028285 | 0.010846 | 0.027182 | 0.019943 | 0.020408 |
| 25.7361111 | 0.035882 | 0.040681 | 0.005508 | 0.021499 | 0.012111 | 0.010232 | 0.002096 | 0.021952 | 0.02972 | 0.029725 | 0.015811 | 0.100184 | 0.014409 | 0.032449 | 0.007734 | 0.021437 | 0.022476 | 0.009807 |
| 25.75 | 0.049949 | 0.043351 | 0.008201 | 0.028177 | 0.018067 | 0.006115 | 0.001679 | 0.038576 | 0.020679 | 0.024795 | 0.022587 | 0.077077 | 0.011012 | 0.055918 | 0.003545 | 0.019671 | 0.019988 | 0.01173 |
| 25.7638889 | 0.034909 | 0.033633 | 0.01061 | 0.022195 | 0.016442 | 0.007709 | 0.001543 | 0.056704 | 0.01073 | 0.037617 | 0.01572 | 0.047825 | 0.007566 | 0.067249 | 0.001663 | 0.014189 | 0.011299 | 0.012327 |
| 25.7777778 | 0.01863 | 0.017579 | 0.020958 | 0.012166 | 0.009024 | 0.005103 | 0.001704 | 0.061752 | 0.016972 | 0.059283 | 0.006231 | 0.057354 | 0.010117 | 0.049011 | 0.002531 | 0.010813 | 0.016169 | 0.014261 |
| 25.7916667 | 0.030971 | 0.012226 | 0.026193 | 0.01687 | 0.004503 | 0.002998 | 0.001384 | 0.052577 | 0.027876 | 0.063019 | 0.010864 | 0.091628 | 0.013592 | 0.034409 | 0.004268 | 0.017997 | 0.031839 | 0.031006 |
| 25.8055556 | 0.055405 | 0.025931 | 0.016605 | 0.030827 | 0.010698 | 0.010288 | 0.001618 | 0.032492 | 0.02991 | 0.042761 | 0.025046 | 0.073812 | 0.012439 | 0.047766 | 0.004602 | 0.022876 | 0.046381 | 0.052801 |
| 25.8194444 | 0.067874 | 0.043813 | 0.009231 | 0.042297 | 0.023854 | 0.020511 | 0.003368 | 0.014587 | 0.020794 | 0.026505 | 0.032955 | 0.039864 | 0.014565 | 0.058969 | 0.004054 | 0.019104 | 0.047883 | 0.063918 |
| 25.8333333 | 0.05521 | 0.049327 | 0.013613 | 0.04245 | 0.028243 | 0.02926 | 0.006269 | 0.009641 | 0.011892 | 0.051717 | 0.025463 | 0.061413 | 0.016578 | 0.048773 | 0.005709 | 0.010965 | 0.030707 | 0.049215 |
| 25.8472222 | 0.0292 | 0.040746 | 0.016482 | 0.030429 | 0.018012 | 0.039936 | 0.006951 | 0.014674 | 0.017731 | 0.092551 | 0.019561 | 0.088886 | 0.013123 | 0.029562 | 0.007814 | 0.006201 | 0.013195 | 0.020294 |
| 25.8611111 | 0.022349 | 0.022767 | 0.012303 | 0.017237 | 0.009522 | 0.043458 | 0.004645 | 0.016522 | 0.026971 | 0.091789 | 0.029455 | 0.089489 | 0.01686 | 0.014542 | 0.006842 | 0.007114 | 0.007872 | 0.008041 |
| 25.875 | 0.026622 | 0.011855 | 0.012397 | 0.010301 | 0.0137 | 0.03475 | 0.002782 | 0.010105 | 0.024922 | 0.052983 | 0.032989 | 0.089961 | 0.029017 | 0.008871 | 0.005614 | 0.007114 | 0.007762 | 0.006011 |
| 25.8888889 | 0.019853 | 0.022234 | 0.018838 | 0.011588 | 0.020237 | 0.024884 | 0.005781 | 0.005412 | 0.015499 | 0.037208 | 0.021113 | 0.088384 | 0.0371 | 0.010777 | 0.006785 | 0.005286 | 0.014017 | 0.002307 |
| 25.9027778 | 0.014448 | 0.041146 | 0.025449 | 0.017327 | 0.020386 | 0.023488 | 0.01692 | 0.009939 | 0.014744 | 0.06753 | 0.019702 | 0.073652 | 0.040363 | 0.022117 | 0.013315 | 0.003483 | 0.024773 | 0.002922 |
| 25.9166667 | 0.01813 | 0.047717 | 0.030544 | 0.019994 | 0.015593 | 0.027335 | 0.027826 | 0.016236 | 0.028709 | 0.097721 | 0.03844 | 0.049587 | 0.037793 | 0.039174 | 0.023915 | 0.003886 | 0.036483 | 0.006706 |
| 25.9305556 | 0.022324 | 0.036062 | 0.031288 | 0.018549 | 0.009076 | 0.026412 | 0.026374 | 0.015736 | 0.039447 | 0.098454 | 0.057416 | 0.025523 | 0.02624 | 0.048076 | 0.026722 | 0.005819 | 0.047589 | 0.008186 |
| 25.9444444 | 0.027128 | 0.017882 | 0.029113 | 0.016061 | 0.010042 | 0.022117 | 0.017526 | 0.00912 | 0.034153 | 0.079158 | 0.065357 | 0.009523 | 0.012773 | 0.046929 | 0.01797 | 0.01012 | 0.051344 | 0.005383 |
| 25.9583333 | 0.032495 | 0.006146 | 0.028023 | 0.015793 | 0.02628 | 0.020453 | 0.012209 | 0.003152 | 0.019204 | 0.049127 | 0.061523 | 0.003139 | 0.004829 | 0.041383 | 0.009132 | 0.016072 | 0.042945 | 0.002893 |
| 25.9722222 | 0.03213 | 0.002792 | 0.023234 | 0.018832 | 0.041146 | 0.018639 | 0.01148 | 0.001684 | 0.008106 | 0.022494 | 0.045623 | 0.002503 | 0.003027 | 0.031264 | 0.007784 | 0.01351 | 0.023874 | 0.004152 |
| 25.9861111 | 0.026055 | 0.004952 | 0.012552 | 0.02326 | 0.034288 | 0.012049 | 0.011884 | 0.00148 | 0.007394 | 0.009008 | 0.024811 | 0.004548 | 0.002989 | 0.018643 | 0.007092 | 0.008219 | 0.011409 | 0.005951 |
| 26 | 0.020773 | 0.008462 | 0.005478 | 0.026642 | 0.017863 | 0.007736 | 0.010108 | 0.001118 | 0.00892 | 0.005405 | 0.011044 | 0.00732 | 0.004245 | 0.008503 | 0.004475 | 0.012679 | 0.017225 | 0.00656 |
| 26.0138889 | 0.017562 | 0.010527 | 0.006869 | 0.024452 | 0.013222 | 0.015524 | 0.005797 | 0.002354 | 0.007024 | 0.006452 | 0.010932 | 0.007331 | 0.009648 | 0.006073 | 0.004597 | 0.017819 | 0.026613 | 0.007702 |
| 26.0277778 | 0.011353 | 0.010854 | 0.008721 | 0.015047 | 0.016233 | 0.02843 | 0.003388 | 0.004694 | 0.005296 | 0.005673 | 0.017467 | 0.00493 | 0.014938 | 0.01311 | 0.004227 | 0.015952 | 0.029673 | 0.010461 |
| 26.0416667 | 0.007515 | 0.010719 | 0.00641 | 0.007144 | 0.013933 | 0.035607 | 0.004752 | 0.00618 | 0.006365 | 0.003762 | 0.021473 | 0.006466 | 0.013548 | 0.022371 | 0.003228 | 0.013631 | 0.032908 | 0.015799 |
| 26.0555556 | 0.014808 | 0.011058 | 0.004135 | 0.00832 | 0.008676 | 0.035095 | 0.006049 | 0.005568 | 0.009348 | 0.005362 | 0.020437 | 0.018152 | 0.013464 | 0.026633 | 0.005808 | 0.015182 | 0.041748 | 0.02322 |
| 26.0694444 | 0.020585 | 0.009916 | 0.004255 | 0.010667 | 0.005751 | 0.026484 | 0.004456 | 0.00348 | 0.009384 | 0.00822 | 0.014053 | 0.03175 | 0.020263 | 0.020449 | 0.007499 | 0.014957 | 0.044982 | 0.025275 |
| 26.0833333 | 0.021449 | 0.007976 | 0.005292 | 0.012535 | 0.008806 | 0.018098 | 0.005056 | 0.003185 | 0.010843 | 0.014869 | 0.01482 | 0.032612 | 0.021241 | 0.017891 | 0.014429 | 0.023787 | 0.038609 | 0.021035 |
| 26.0972222 | 0.041313 | 0.011801 | 0.008831 | 0.027381 | 0.020684 | 0.030153 | 0.01446 | 0.007795 | 0.02847 | 0.034391 | 0.032158 | 0.036818 | 0.024385 | 0.036258 | 0.034422 | 0.056398 | 0.053924 | 0.026367 |
| 26.1111111 | 0.073686 | 0.019249 | 0.013433 | 0.047391 | 0.037071 | 0.053687 | 0.024401 | 0.015307 | 0.056425 | 0.05596 | 0.044892 | 0.059652 | 0.049073 | 0.047038 | 0.044612 | 0.075298 | 0.080635 | 0.040023 |
| 26.125 | 0.077492 | 0.02024 | 0.0149 | 0.052131 | 0.047997 | 0.055511 | 0.026241 | 0.018323 | 0.071951 | 0.05658 | 0.039115 | 0.076187 | 0.074957 | 0.029926 | 0.031003 | 0.052274 | 0.068458 | 0.0425 |
| 26.1388889 | 0.048219 | 0.014438 | 0.018403 | 0.046562 | 0.042298 | 0.035891 | 0.030936 | 0.013238 | 0.064388 | 0.03909 | 0.025372 | 0.067499 | 0.071822 | 0.009685 | 0.013533 | 0.021718 | 0.030871 | 0.032611 |

| | | | | | | | | | | | | | | | | | | |
|---|---|---|---|---|---|---|---|---|---|---|---|---|---|---|---|---|---|---|
| 26.1527778 | 0.018356 | 0.007789 | 0.023862 | 0.035272 | 0.024799 | 0.015262 | 0.030657 | 0.007042 | 0.037438 | 0.019847 | 0.013398 | 0.039384 | 0.045098 | 0.002622 | 0.005956 | 0.016192 | 0.008393 | 0.020054 |
| 26.1666667 | 0.006967 | 0.003981 | 0.018728 | 0.017007 | 0.018629 | 0.004681 | 0.013331 | 0.006886 | 0.017604 | 0.010904 | 0.010133 | 0.017663 | 0.018715 | 0.001848 | 0.003339 | 0.031701 | 0.005249 | 0.009237 |
| 26.1805556 | 0.011589 | 0.003873 | 0.007788 | 0.006512 | 0.029289 | 0.004925 | 0.004248 | 0.009409 | 0.025647 | 0.018738 | 0.014947 | 0.021186 | 0.008751 | 0.002191 | 0.002013 | 0.044244 | 0.010124 | 0.003767 |
| 26.1944444 | 0.021191 | 0.004069 | 0.006166 | 0.007219 | 0.036074 | 0.007775 | 0.01029 | 0.008978 | 0.039997 | 0.030484 | 0.019119 | 0.031839 | 0.012797 | 0.005651 | 0.002786 | 0.040056 | 0.016723 | 0.004863 |
| 26.2083333 | 0.028554 | 0.003055 | 0.00863 | 0.012218 | 0.030594 | 0.008285 | 0.015095 | 0.006272 | 0.041673 | 0.034801 | 0.019617 | 0.041663 | 0.02206 | 0.010858 | 0.003096 | 0.026282 | 0.025623 | 0.007201 |
| 26.2222222 | 0.030961 | 0.002836 | 0.007154 | 0.018024 | 0.022807 | 0.006709 | 0.015209 | 0.004317 | 0.034493 | 0.032689 | 0.017841 | 0.057478 | 0.035236 | 0.013801 | 0.00148 | 0.013585 | 0.041477 | 0.007389 |
| 26.2361111 | 0.028716 | 0.002316 | 0.004258 | 0.02026 | 0.019677 | 0.004115 | 0.012096 | 0.003782 | 0.024638 | 0.026639 | 0.014665 | 0.070102 | 0.049958 | 0.015027 | 0.001338 | 0.006772 | 0.061543 | 0.008003 |
| 26.25 | 0.025405 | 0.00141 | 0.003172 | 0.018589 | 0.018212 | 0.002534 | 0.008398 | 0.003447 | 0.017278 | 0.02347 | 0.011906 | 0.067503 | 0.058259 | 0.016502 | 0.003758 | 0.006047 | 0.074194 | 0.01001 |
| 26.2638889 | 0.022766 | 0.002878 | 0.005744 | 0.015538 | 0.013015 | 0.003819 | 0.007459 | 0.003065 | 0.01463 | 0.025458 | 0.010345 | 0.060637 | 0.054382 | 0.01704 | 0.005057 | 0.008872 | 0.076182 | 0.0117 |
| 26.2777778 | 0.021689 | 0.007327 | 0.011141 | 0.013525 | 0.006313 | 0.009202 | 0.011147 | 0.003436 | 0.016874 | 0.026215 | 0.009772 | 0.060363 | 0.040336 | 0.015751 | 0.004061 | 0.010888 | 0.068096 | 0.012778 |
| 26.2916667 | 0.022123 | 0.011704 | 0.015385 | 0.013293 | 0.003243 | 0.016533 | 0.016398 | 0.004435 | 0.022953 | 0.023096 | 0.010298 | 0.05204 | 0.023175 | 0.014973 | 0.003979 | 0.009609 | 0.050715 | 0.013867 |
| 26.3055556 | 0.021516 | 0.011987 | 0.015147 | 0.014155 | 0.005169 | 0.019879 | 0.018005 | 0.005235 | 0.029574 | 0.018136 | 0.012101 | 0.03215 | 0.009567 | 0.017091 | 0.006057 | 0.008239 | 0.028981 | 0.014312 |
| 26.3194444 | 0.018852 | 0.008918 | 0.01157 | 0.015439 | 0.008836 | 0.018022 | 0.015566 | 0.005613 | 0.032297 | 0.012729 | 0.014672 | 0.016357 | 0.003596 | 0.019728 | 0.007338 | 0.009755 | 0.01334 | 0.012795 |
| 26.3333333 | 0.016158 | 0.007119 | 0.007818 | 0.016068 | 0.009014 | 0.016099 | 0.011662 | 0.005838 | 0.029423 | 0.008932 | 0.016093 | 0.014328 | 0.004197 | 0.016915 | 0.006096 | 0.01099 | 0.01102 | 0.008889 |
| 26.3472222 | 0.014303 | 0.007595 | 0.005161 | 0.014832 | 0.005254 | 0.016303 | 0.007191 | 0.005582 | 0.0223 | 0.008735 | 0.014751 | 0.013959 | 0.005377 | 0.009706 | 0.007378 | 0.009549 | 0.010122 | 0.004279 |
| 26.3611111 | 0.012755 | 0.006698 | 0.003923 | 0.012202 | 0.002671 | 0.015883 | 0.003393 | 0.004304 | 0.014333 | 0.012571 | 0.011705 | 0.015157 | 0.008375 | 0.006436 | 0.013698 | 0.007697 | 0.009907 | 0.001708 |
| 26.375 | 0.011377 | 0.00415 | 0.004673 | 0.009755 | 0.003509 | 0.012733 | 0.002482 | 0.002526 | 0.010879 | 0.018251 | 0.009376 | 0.029255 | 0.014956 | 0.007827 | 0.01864 | 0.008223 | 0.024473 | 0.001929 |
| 26.3888889 | 0.008054 | 0.003755 | 0.005492 | 0.007312 | 0.004354 | 0.008108 | 0.004029 | 0.001509 | 0.014673 | 0.021065 | 0.008645 | 0.03546 | 0.016363 | 0.008819 | 0.019555 | 0.011633 | 0.043691 | 0.003252 |
| 26.4027778 | 0.005446 | 0.004137 | 0.004338 | 0.004665 | 0.003293 | 0.004516 | 0.004563 | 0.002147 | 0.019044 | 0.017675 | 0.008568 | 0.025133 | 0.010231 | 0.008198 | 0.019545 | 0.016069 | 0.046439 | 0.004073 |
| 26.4166667 | 0.009441 | 0.003407 | 0.003628 | 0.00483 | 0.003871 | 0.002499 | 0.003058 | 0.004026 | 0.014575 | 0.00964 | 0.007971 | 0.028079 | 0.008504 | 0.007181 | 0.019686 | 0.018221 | 0.029923 | 0.003994 |
| 26.4305556 | 0.013409 | 0.004984 | 0.00342 | 0.00763 | 0.008234 | 0.001729 | 0.001905 | 0.00555 | 0.008816 | 0.002691 | 0.006748 | 0.050862 | 0.015958 | 0.006412 | 0.019903 | 0.015476 | 0.018391 | 0.003 |
| 26.4444444 | 0.009852 | 0.008535 | 0.003893 | 0.007224 | 0.01106 | 0.002805 | 0.002205 | 0.005746 | 0.00863 | 0.001505 | 0.006 | 0.056527 | 0.019401 | 0.004531 | 0.019206 | 0.009878 | 0.021258 | 0.002601 |
| 26.4583333 | 0.007289 | 0.010059 | 0.009409 | 0.004785 | 0.008658 | 0.005519 | 0.004702 | 0.004663 | 0.009957 | 0.00569 | 0.006549 | 0.035223 | 0.013797 | 0.002589 | 0.014906 | 0.007026 | 0.020209 | 0.003957 |
| 26.4722222 | 0.01316 | 0.007212 | 0.016283 | 0.007454 | 0.004281 | 0.008885 | 0.009555 | 0.002619 | 0.019556 | 0.013367 | 0.00784 | 0.024538 | 0.012528 | 0.003375 | 0.008184 | 0.009532 | 0.028269 | 0.005219 |
| 26.4861111 | 0.018806 | 0.003667 | 0.017173 | 0.011946 | 0.002299 | 0.011615 | 0.014418 | 0.001145 | 0.031581 | 0.021493 | 0.009668 | 0.034316 | 0.017578 | 0.005476 | 0.005724 | 0.015643 | 0.055679 | 0.005147 |
| 26.5 | 0.016612 | 0.003006 | 0.01262 | 0.011619 | 0.003717 | 0.010771 | 0.017636 | 0.000944 | 0.029374 | 0.025197 | 0.014241 | 0.032456 | 0.016725 | 0.00799 | 0.010283 | 0.022692 | 0.06802 | 0.003957 |
| 26.5138889 | 0.011049 | 0.004176 | 0.008459 | 0.008267 | 0.006341 | 0.007758 | 0.018507 | 0.001902 | 0.01622 | 0.02284 | 0.023085 | 0.018451 | 0.009287 | 0.012365 | 0.017012 | 0.026466 | 0.047132 | 0.002126 |
| 26.5277778 | 0.007312 | 0.007721 | 0.00878 | 0.004859 | 0.008084 | 0.008745 | 0.016015 | 0.004429 | 0.006852 | 0.016995 | 0.031756 | 0.018148 | 0.003872 | 0.017391 | 0.020645 | 0.023214 | 0.024421 | 0.001374 |
| 26.5416667 | 0.00525 | 0.009313 | 0.011792 | 0.002403 | 0.00698 | 0.0121 | 0.010532 | 0.005805 | 0.004697 | 0.011242 | 0.033854 | 0.023497 | 0.003894 | 0.019062 | 0.019201 | 0.015899 | 0.028786 | 0.00286 |
| 26.5555556 | 0.004363 | 0.005859 | 0.01252 | 0.002356 | 0.004184 | 0.012622 | 0.004711 | 0.004013 | 0.007242 | 0.006777 | 0.028519 | 0.016642 | 0.004878 | 0.015374 | 0.013611 | 0.00936 | 0.043376 | 0.004383 |
| 26.5694444 | 0.007732 | 0.002778 | 0.009476 | 0.007238 | 0.002669 | 0.013128 | 0.003752 | 0.002224 | 0.020606 | 0.003349 | 0.020291 | 0.007973 | 0.003155 | 0.010235 | 0.006993 | 0.005112 | 0.040812 | 0.003544 |
| 26.5833333 | 0.021285 | 0.003123 | 0.009024 | 0.020849 | 0.003553 | 0.021628 | 0.013637 | 0.005291 | 0.038124 | 0.003328 | 0.018651 | 0.010278 | 0.002211 | 0.013563 | 0.002746 | 0.003085 | 0.028677 | 0.002083 |
| 26.5972222 | 0.046355 | 0.007653 | 0.021127 | 0.043767 | 0.007367 | 0.03749 | 0.033795 | 0.013541 | 0.045891 | 0.012301 | 0.030961 | 0.019095 | 0.006681 | 0.03268 | 0.002488 | 0.006748 | 0.038837 | 0.002388 |
| 26.6111111 | 0.068794 | 0.015778 | 0.044649 | 0.066905 | 0.00804 | 0.04776 | 0.055599 | 0.019523 | 0.045638 | 0.032592 | 0.046888 | 0.039733 | 0.016458 | 0.055969 | 0.006556 | 0.02035 | 0.071796 | 0.005389 |
| 26.625 | 0.075505 | 0.023111 | 0.063933 | 0.079096 | 0.007394 | 0.042857 | 0.068627 | 0.018766 | 0.040963 | 0.053248 | 0.051222 | 0.076306 | 0.029381 | 0.064404 | 0.014006 | 0.038316 | 0.091599 | 0.009692 |
| 26.6388889 | 0.065814 | 0.024279 | 0.062875 | 0.073967 | 0.017006 | 0.026509 | 0.06192 | 0.014979 | 0.029318 | 0.054436 | 0.039987 | 0.099383 | 0.037816 | 0.055518 | 0.019359 | 0.050958 | 0.084634 | 0.009413 |
| 26.6527778 | 0.043098 | 0.016507 | 0.03893 | 0.049268 | 0.031048 | 0.011427 | 0.035188 | 0.010132 | 0.01593 | 0.032225 | 0.024618 | 0.084427 | 0.031823 | 0.035217 | 0.01698 | 0.0481 | 0.06025 | 0.005525 |
| 26.6666667 | 0.019808 | 0.006195 | 0.018747 | 0.023999 | 0.0378 | 0.008462 | 0.019114 | 0.005327 | 0.014715 | 0.01737 | 0.026024 | 0.047336 | 0.017176 | 0.01653 | 0.009531 | 0.031897 | 0.03101 | 0.003981 |
| 26.6805556 | 0.009106 | 0.001028 | 0.025598 | 0.020287 | 0.037896 | 0.017199 | 0.034572 | 0.003369 | 0.027715 | 0.027128 | 0.037708 | 0.01663 | 0.00723 | 0.012543 | 0.003655 | 0.019151 | 0.016408 | 0.003711 |
| 26.6944444 | 0.012219 | 0.000351 | 0.039075 | 0.025969 | 0.034104 | 0.027707 | 0.053967 | 0.002803 | 0.037003 | 0.033746 | 0.036949 | 0.005564 | 0.005656 | 0.014374 | 0.001058 | 0.021361 | 0.016899 | 0.005516 |
| 26.7083333 | 0.019691 | 0.0012 | 0.039716 | 0.021981 | 0.0253 | 0.038506 | 0.049983 | 0.002111 | 0.026885 | 0.022334 | 0.022113 | 0.005255 | 0.007894 | 0.011752 | 0.000742 | 0.038452 | 0.028032 | 0.009892 |
| 26.7222222 | 0.02125 | 0.00168 | 0.034304 | 0.011554 | 0.014182 | 0.049624 | 0.030444 | 0.002155 | 0.014459 | 0.008248 | 0.01099 | 0.008599 | 0.010162 | 0.010245 | 0.001582 | 0.051786 | 0.048607 | 0.011895 |
| 26.7361111 | 0.014437 | 0.001336 | 0.022772 | 0.00707 | 0.006307 | 0.052994 | 0.022319 | 0.002424 | 0.020415 | 0.002799 | 0.011462 | 0.015733 | 0.014358 | 0.014892 | 0.00325 | 0.039357 | 0.055349 | 0.010708 |
| 26.75 | 0.013762 | 0.001502 | 0.015913 | 0.013923 | 0.007185 | 0.044626 | 0.035996 | 0.003543 | 0.025424 | 0.004393 | 0.009494 | 0.017849 | 0.021391 | 0.022228 | 0.007171 | 0.025968 | 0.035946 | 0.007128 |
| 26.7638889 | 0.030497 | 0.002374 | 0.027027 | 0.026407 | 0.017285 | 0.025757 | 0.04985 | 0.005354 | 0.015068 | 0.006059 | 0.004469 | 0.019393 | 0.026616 | 0.025045 | 0.012053 | 0.037631 | 0.023108 | 0.004615 |
| 26.7777778 | 0.044211 | 0.002392 | 0.029262 | 0.029899 | 0.025052 | 0.015833 | 0.038224 | 0.006326 | 0.007166 | 0.006564 | 0.003956 | 0.027572 | 0.024045 | 0.021177 | 0.014644 | 0.041307 | 0.03723 | 0.007783 |
| 26.7916667 | 0.034967 | 0.004967 | 0.013501 | 0.018149 | 0.023101 | 0.025934 | 0.019381 | 0.005522 | 0.011002 | 0.011864 | 0.004301 | 0.035209 | 0.014914 | 0.014417 | 0.013086 | 0.026911 | 0.042753 | 0.011188 |
| 26.8055556 | 0.016334 | 0.013051 | 0.005375 | 0.006846 | 0.014812 | 0.035161 | 0.02655 | 0.002991 | 0.01885 | 0.013215 | 0.003636 | 0.037474 | 0.011205 | 0.016522 | 0.008139 | 0.018919 | 0.0304 | 0.010941 |
| 26.8194444 | 0.00869 | 0.017473 | 0.009828 | 0.004368 | 0.006438 | 0.03448 | 0.04523 | 0.000887 | 0.023638 | 0.014117 | 0.006509 | 0.03179 | 0.01446 | 0.033725 | 0.0036 | 0.022447 | 0.028855 | 0.009502 |
| 26.8333333 | 0.009393 | 0.011523 | 0.015259 | 0.002848 | 0.004526 | 0.028823 | 0.049995 | 0.000459 | 0.022409 | 0.031067 | 0.016557 | 0.018061 | 0.012923 | 0.04772 | 0.002865 | 0.022303 | 0.034195 | 0.00715 |
| 26.8472222 | 0.01419 | 0.004351 | 0.014698 | 0.00529 | 0.011418 | 0.018582 | 0.036724 | 0.001362 | 0.016875 | 0.040903 | 0.025551 | 0.009308 | 0.007164 | 0.043229 | 0.006625 | 0.015054 | 0.031798 | 0.005174 |
| 26.8611111 | 0.030295 | 0.002978 | 0.012132 | 0.0179 | 0.020298 | 0.016942 | 0.016295 | 0.00341 | 0.009335 | 0.029466 | 0.024048 | 0.015158 | 0.004366 | 0.025376 | 0.011986 | 0.020152 | 0.024304 | 0.004069 |
| 26.875 | 0.045532 | 0.002968 | 0.014236 | 0.02733 | 0.020333 | 0.03236 | 0.003626 | 0.005207 | 0.004869 | 0.016283 | 0.017099 | 0.021331 | 0.005113 | 0.012474 | 0.016076 | 0.029775 | 0.017459 | 0.005254 |
| 26.8888889 | 0.037894 | 0.004156 | 0.017095 | 0.020209 | 0.012673 | 0.049548 | 0.000721 | 0.005268 | 0.007847 | 0.014619 | 0.020622 | 0.015998 | 0.00649 | 0.018235 | 0.020213 | 0.023205 | 0.020387 | 0.014401 |
| 26.9027778 | 0.020555 | 0.009306 | 0.012479 | 0.011799 | 0.00672 | 0.059735 | 0.001289 | 0.003614 | 0.010256 | 0.028361 | 0.034509 | 0.01038 | 0.005419 | 0.031602 | 0.022484 | 0.008641 | 0.026037 | 0.026967 |
| 26.9166667 | 0.014707 | 0.013929 | 0.005497 | 0.018087 | 0.004018 | 0.065922 | 0.002917 | 0.001687 | 0.009236 | 0.043053 | 0.033799 | 0.015361 | 0.00259 | 0.043067 | 0.018846 | 0.004983 | 0.018008 | 0.028734 |
| 26.9305556 | 0.015608 | 0.012266 | 0.004641 | 0.029092 | 0.006432 | 0.062632 | 0.008513 | 0.000886 | 0.013547 | 0.038055 | 0.018716 | 0.027329 | 0.00135 | 0.051949 | 0.017059 | 0.012948 | 0.010073 | 0.018859 |
| 26.9444444 | 0.014712 | 0.006714 | 0.005305 | 0.032169 | 0.016648 | 0.045358 | 0.018692 | 0.001195 | 0.023195 | 0.024067 | 0.020039 | 0.037829 | 0.003947 | 0.04873 | 0.022011 | 0.020486 | 0.015421 | 0.009088 |
| 26.9583333 | 0.012122 | 0.004478 | 0.007112 | 0.028299 | 0.02298 | 0.026862 | 0.029038 | 0.00293 | 0.029104 | 0.02978 | 0.037948 | 0.046385 | 0.012077 | 0.03308 | 0.023348 | 0.020562 | 0.020501 | 0.004526 |
| 26.9722222 | 0.011707 | 0.004595 | 0.012726 | 0.024104 | 0.017444 | 0.017042 | 0.034072 | 0.005365 | 0.027556 | 0.044083 | 0.048876 | 0.065824 | 0.02254 | 0.016397 | 0.017057 | 0.013701 | 0.016238 | 0.004486 |
| 26.9861111 | 0.015811 | 0.003819 | 0.015604 | 0.024719 | 0.008845 | 0.01053 | 0.030565 | 0.007316 | 0.022771 | 0.043573 | 0.04739 | 0.090768 | 0.023894 | 0.005939 | 0.00834 | 0.012186 | 0.017402 | 0.007861 |
| 27 | 0.020838 | 0.004498 | 0.011814 | 0.030344 | 0.006753 | 0.005256 | 0.0201 | 0.009508 | 0.018865 | 0.040802 | 0.036723 | 0.095226 | 0.013469 | 0.003229 | 0.002504 | 0.020555 | 0.029452 | 0.010673 |
| 27.0138889 | 0.019209 | 0.005228 | 0.006759 | 0.033033 | 0.012102 | 0.004323 | 0.010119 | 0.012275 | 0.014621 | 0.042187 | 0.019338 | 0.082838 | 0.005074 | 0.004994 | 0.002088 | 0.020472 | 0.027912 | 0.009748 |
| 27.0277778 | 0.011397 | 0.003971 | 0.006766 | 0.025694 | 0.017931 | 0.005892 | 0.007161 | 0.013939 | 0.00891 | 0.03334 | 0.008465 | 0.073168 | 0.007152 | 0.007791 | 0.00426 | 0.016901 | 0.015549 | 0.00667 |
| 27.0416667 | 0.008037 | 0.004543 | 0.008129 | 0.013034 | 0.019326 | 0.006479 | 0.009224 | 0.013461 | 0.007955 | 0.01793 | 0.012465 | 0.064981 | 0.015126 | 0.009995 | 0.006355 | 0.026181 | 0.016734 | 0.00539 |
| 27.0555556 | 0.011795 | 0.007748 | 0.005589 | 0.006546 | 0.017952 | 0.006681 | 0.01028 | 0.011176 | 0.014862 | 0.016092 | 0.019451 | 0.055268 | 0.021994 | 0.010369 | 0.006791 | 0.027978 | 0.026111 | 0.007707 |
| 27.0694444 | 0.013207 | 0.00863 | 0.003754 | 0.00719 | 0.01643 | 0.00682 | 0.008055 | 0.008596 | 0.018622 | 0.022884 | 0.017695 | 0.057894 | 0.019353 | 0.007357 | 0.007635 | 0.019163 | 0.026432 | 0.009389 |
| 27.0833333 | 0.011426 | 0.007917 | 0.008855 | 0.010099 | 0.018649 | 0.006955 | 0.011465 | 0.010013 | 0.016815 | 0.022242 | 0.013999 | 0.101157 | 0.020489 | 0.008061 | 0.020724 | 0.028374 | 0.025097 | 0.012831 |
| 27.0972222 | 0.018846 | 0.014774 | 0.021053 | 0.021685 | 0.022898 | 0.011462 | 0.027954 | 0.016467 | 0.027043 | 0.024976 | 0.021746 | 0.171405 | 0.040346 | 0.021199 | 0.043946 | 0.053127 | 0.042462 | 0.027648 |
| 27.1111111 | 0.035351 | 0.028364 | 0.032507 | 0.040262 | 0.019989 | 0.018825 | 0.041389 | 0.020214 | 0.049103 | 0.036247 | 0.030671 | 0.182983 | 0.049647 | 0.03402 | 0.051036 | 0.058885 | 0.056641 | 0.042642 |
| 27.125 | 0.043856 | 0.036013 | 0.034425 | 0.04699 | 0.010843 | 0.019891 | 0.036482 | 0.015098 | 0.062139 | 0.046045 | 0.026591 | 0.121263 | 0.033658 | 0.03251 | 0.038338 | 0.039554 | 0.037846 | 0.045106 |
| 27.1388889 | 0.037928 | 0.033973 | 0.027457 | 0.035121 | 0.004199 | 0.01424 | 0.021689 | 0.007872 | 0.057028 | 0.060755 | 0.018286 | 0.069821 | 0.016598 | 0.021094 | 0.027547 | 0.023217 | 0.011127 | 0.044015 |
| 27.1527778 | 0.024032 | 0.022935 | 0.016713 | 0.019054 | 0.003883 | 0.008219 | 0.010512 | 0.006057 | 0.043572 | 0.066088 | 0.013071 | 0.04918 | 0.008138 | 0.011885 | 0.019489 | 0.020616 | 0.009139 | 0.038235 |
| 27.1666667 | 0.011783 | 0.015642 | 0.008997 | 0.008537 | 0.009632 | 0.008695 | 0.005669 | 0.005606 | 0.039944 | 0.042038 | 0.007901 | 0.028473 | 0.005112 | 0.013513 | 0.011492 | 0.020531 | 0.023459 | 0.023473 |
| 27.1805556 | 0.012315 | 0.029595 | 0.009757 | 0.004127 | 0.020662 | 0.019845 | 0.00711 | 0.008206 | 0.039425 | 0.015097 | 0.006777 | 0.014415 | 0.005362 | 0.019859 | 0.010094 | 0.014767 | 0.028919 | 0.011217 |
| 27.1944444 | 0.025717 | 0.05269 | 0.015344 | 0.005326 | 0.033268 | 0.026671 | 0.013993 | 0.021383 | 0.026848 | 0.005929 | 0.01053 | 0.025052 | 0.00816 | 0.01815 | 0.010342 | 0.007924 | 0.020497 | 0.008981 |
| 27.2083333 | 0.028455 | 0.051162 | 0.030954 | 0.011695 | 0.035439 | 0.040366 | 0.017398 | 0.034244 | 0.011637 | 0.00341 | 0.01085 | 0.051294 | 0.017893 | 0.010847 | 0.013721 | 0.007164 | 0.012192 | 0.015633 |
| 27.2222222 | 0.024386 | 0.038958 | 0.038744 | 0.021293 | 0.025178 | 0.066042 | 0.012952 | 0.042534 | 0.009069 | 0.002248 | 0.010985 | 0.064833 | 0.025706 | 0.00823 | 0.019449 | 0.012693 | 0.010473 | 0.018264 |
| 27.2361111 | 0.03157 | 0.043295 | 0.017657 | 0.027115 | 0.015958 | 0.061008 | 0.009185 | 0.043235 | 0.026539 | 0.007121 | 0.010189 | 0.048067 | 0.018382 | 0.007083 | 0.015085 | 0.023749 | 0.019789 | 0.013863 |
| 27.25 | 0.026267 | 0.031416 | 0.005348 | 0.029115 | 0.01721 | 0.045808 | 0.015147 | 0.027614 | 0.050223 | 0.016323 | 0.006451 | 0.035892 | 0.007727 | 0.004548 | 0.006043 | 0.035462 | 0.042818 | 0.015406 |
| 27.2638889 | 0.012952 | 0.009681 | 0.010496 | 0.027782 | 0.02446 | 0.042069 | 0.023086 | 0.015125 | 0.059876 | 0.02065 | 0.006646 | 0.060057 | 0.004732 | 0.004263 | 0.002907 | 0.038773 | 0.06354 | 0.016038 |
| 27.2777778 | 0.008754 | 0.004457 | 0.01195 | 0.019279 | 0.023906 | 0.025732 | 0.022224 | 0.014797 | 0.058109 | 0.018151 | 0.010629 | 0.073722 | 0.010095 | 0.007543 | 0.007114 | 0.032115 | 0.07041 | 0.011976 |
| 27.2916667 | 0.008895 | 0.0094 | 0.019509 | 0.009457 | 0.016094 | 0.010788 | 0.019962 | 0.023166 | 0.066248 | 0.032083 | 0.029875 | 0.052679 | 0.038321 | 0.026089 | 0.028197 | 0.022232 | 0.066241 | 0.022201 |

| | | | | | | | | | | | | | | | | | | |
|---|---|---|---|---|---|---|---|---|---|---|---|---|---|---|---|---|---|---|
| 27.3055556 | 0.016027 | 0.02395 | 0.034745 | 0.004803 | 0.016851 | 0.009117 | 0.022652 | 0.03578 | 0.08712 | 0.072177 | 0.065856 | 0.047693 | 0.083002 | 0.061578 | 0.06386 | 0.024452 | 0.052104 | 0.049905 |
| 27.3194444 | 0.017096 | 0.02862 | 0.043737 | 0.008985 | 0.018932 | 0.011124 | 0.025685 | 0.035474 | 0.088834 | 0.090952 | 0.079814 | 0.064184 | 0.099583 | 0.077985 | 0.08071 | 0.032628 | 0.030926 | 0.063961 |
| 27.3333333 | 0.011086 | 0.017927 | 0.04213 | 0.023782 | 0.009523 | 0.010654 | 0.025671 | 0.022522 | 0.062469 | 0.057886 | 0.05156 | 0.051615 | 0.065226 | 0.052833 | 0.054705 | 0.028049 | 0.018238 | 0.045573 |
| 27.3472222 | 0.015215 | 0.015154 | 0.035156 | 0.033372 | 0.003924 | 0.007075 | 0.02111 | 0.010083 | 0.033887 | 0.020326 | 0.019798 | 0.02501 | 0.02443 | 0.025742 | 0.021753 | 0.028186 | 0.021864 | 0.026496 |
| 27.3611111 | 0.020284 | 0.018739 | 0.022538 | 0.023453 | 0.008197 | 0.003596 | 0.012887 | 0.003277 | 0.015275 | 0.01485 | 0.015334 | 0.019541 | 0.015839 | 0.024418 | 0.017946 | 0.039264 | 0.024782 | 0.025022 |
| 27.375 | 0.014495 | 0.013685 | 0.009545 | 0.010713 | 0.014243 | 0.002821 | 0.009301 | 0.00125 | 0.008165 | 0.017613 | 0.021036 | 0.033191 | 0.017897 | 0.024635 | 0.021848 | 0.042791 | 0.015961 | 0.019617 |
| 27.3888889 | 0.006485 | 0.006704 | 0.004737 | 0.010046 | 0.016476 | 0.003708 | 0.011183 | 0.00114 | 0.010206 | 0.014796 | 0.023275 | 0.06726 | 0.017462 | 0.019987 | 0.020852 | 0.040481 | 0.0087 | 0.010909 |
| 27.4027778 | 0.002994 | 0.00698 | 0.005082 | 0.011002 | 0.013582 | 0.003047 | 0.0101 | 0.002093 | 0.009448 | 0.013501 | 0.023741 | 0.095889 | 0.019837 | 0.022051 | 0.022484 | 0.040818 | 0.017847 | 0.013512 |
| 27.4166667 | 0.004054 | 0.007775 | 0.009696 | 0.014782 | 0.008705 | 0.003331 | 0.011206 | 0.005172 | 0.009067 | 0.014292 | 0.023405 | 0.09898 | 0.021857 | 0.028665 | 0.024749 | 0.042396 | 0.040308 | 0.022463 |
| 27.4305556 | 0.010559 | 0.008711 | 0.017041 | 0.025406 | 0.005316 | 0.007686 | 0.018561 | 0.008363 | 0.018392 | 0.015912 | 0.022167 | 0.089536 | 0.019817 | 0.03158 | 0.024397 | 0.040351 | 0.05793 | 0.026441 |
| 27.4444444 | 0.016427 | 0.012356 | 0.019321 | 0.029431 | 0.006941 | 0.010655 | 0.021561 | 0.008714 | 0.02748 | 0.016257 | 0.016439 | 0.079409 | 0.012486 | 0.025638 | 0.019553 | 0.032518 | 0.055282 | 0.021215 |
| 27.4583333 | 0.014352 | 0.010984 | 0.01521 | 0.022073 | 0.010016 | 0.008228 | 0.016019 | 0.006361 | 0.026074 | 0.014215 | 0.007594 | 0.072348 | 0.005117 | 0.015255 | 0.011341 | 0.02206 | 0.035689 | 0.011164 |
| 27.4722222 | 0.007445 | 0.005726 | 0.009769 | 0.011156 | 0.012597 | 0.003822 | 0.007942 | 0.003685 | 0.018117 | 0.010689 | 0.002108 | 0.061942 | 0.002899 | 0.007151 | 0.004835 | 0.014258 | 0.015753 | 0.00533 |
| 27.4861111 | 0.002191 | 0.003707 | 0.006809 | 0.005455 | 0.016589 | 0.00178 | 0.003946 | 0.002123 | 0.010597 | 0.007082 | 0.001968 | 0.045657 | 0.003773 | 0.003283 | 0.00348 | 0.010792 | 0.006675 | 0.005468 |
| 27.5 | 0.001788 | 0.004068 | 0.006691 | 0.005196 | 0.019842 | 0.002953 | 0.004308 | 0.001277 | 0.006095 | 0.005609 | 0.005475 | 0.026878 | 0.003455 | 0.004127 | 0.003328 | 0.011779 | 0.006789 | 0.004908 |
| 27.5138889 | 0.005103 | 0.003068 | 0.008264 | 0.00549 | 0.020563 | 0.004409 | 0.003787 | 0.001027 | 0.006287 | 0.008172 | 0.011505 | 0.012455 | 0.002556 | 0.008495 | 0.003375 | 0.018184 | 0.016168 | 0.004283 |
| 27.5277778 | 0.008553 | 0.001636 | 0.009899 | 0.008139 | 0.019287 | 0.003643 | 0.003489 | 0.00171 | 0.012052 | 0.012764 | 0.01671 | 0.007599 | 0.002771 | 0.010884 | 0.006376 | 0.027012 | 0.035239 | 0.006433 |
| 27.5416667 | 0.009143 | 0.001553 | 0.010339 | 0.01214 | 0.016182 | 0.001827 | 0.006306 | 0.00273 | 0.020464 | 0.015586 | 0.017852 | 0.00669 | 0.002372 | 0.007615 | 0.008702 | 0.031414 | 0.048034 | 0.006942 |
| 27.5555556 | 0.007786 | 0.001502 | 0.010797 | 0.012315 | 0.012585 | 0.001785 | 0.008687 | 0.003766 | 0.025623 | 0.015489 | 0.016778 | 0.007968 | 0.001467 | 0.003078 | 0.008655 | 0.03095 | 0.042818 | 0.004933 |
| 27.5694444 | 0.008907 | 0.001124 | 0.013804 | 0.011143 | 0.011253 | 0.004958 | 0.010342 | 0.006708 | 0.024309 | 0.013638 | 0.016879 | 0.011227 | 0.002636 | 0.0024 | 0.010392 | 0.031376 | 0.032643 | 0.004703 |
| 27.5833333 | 0.016247 | 0.001653 | 0.021571 | 0.018861 | 0.014112 | 0.015336 | 0.019855 | 0.014924 | 0.022993 | 0.011955 | 0.021548 | 0.009172 | 0.007522 | 0.007014 | 0.018044 | 0.035377 | 0.041283 | 0.006879 |
| 27.5972222 | 0.030524 | 0.002547 | 0.035115 | 0.041904 | 0.020601 | 0.037622 | 0.043844 | 0.026641 | 0.029373 | 0.012296 | 0.033147 | 0.004768 | 0.010909 | 0.016692 | 0.028594 | 0.040118 | 0.075225 | 0.009396 |
| 27.6111111 | 0.045686 | 0.004247 | 0.051436 | 0.072747 | 0.023437 | 0.063045 | 0.075786 | 0.031412 | 0.041127 | 0.018288 | 0.049663 | 0.010388 | 0.007571 | 0.024651 | 0.033376 | 0.040745 | 0.114619 | 0.011884 |
| 27.625 | 0.052316 | 0.007833 | 0.06287 | 0.092051 | 0.018545 | 0.073158 | 0.09755 | 0.026088 | 0.048123 | 0.02941 | 0.060792 | 0.028365 | 0.002837 | 0.024697 | 0.030593 | 0.037472 | 0.132862 | 0.013616 |
| 27.6388889 | 0.047692 | 0.010309 | 0.06134 | 0.081704 | 0.011631 | 0.062868 | 0.091472 | 0.017963 | 0.040639 | 0.033117 | 0.052266 | 0.044932 | 0.00187 | 0.018491 | 0.022483 | 0.033555 | 0.123376 | 0.011368 |
| 27.6527778 | 0.034618 | 0.009965 | 0.045897 | 0.046437 | 0.006573 | 0.043297 | 0.059722 | 0.00974 | 0.021618 | 0.022602 | 0.030277 | 0.038649 | 0.002456 | 0.010358 | 0.01181 | 0.028805 | 0.097861 | 0.00918 |
| 27.6666667 | 0.018503 | 0.00838 | 0.023666 | 0.020744 | 0.006307 | 0.025151 | 0.026772 | 0.00359 | 0.010345 | 0.010392 | 0.023616 | 0.024702 | 0.003073 | 0.007988 | 0.007901 | 0.023128 | 0.057305 | 0.014684 |
| 27.6805556 | 0.007673 | 0.007886 | 0.010454 | 0.026634 | 0.010362 | 0.011612 | 0.01447 | 0.001718 | 0.01719 | 0.008607 | 0.040512 | 0.038545 | 0.004744 | 0.013961 | 0.01596 | 0.015226 | 0.031077 | 0.020565 |
| 27.6944444 | 0.006145 | 0.008755 | 0.014449 | 0.04301 | 0.013125 | 0.005134 | 0.02076 | 0.001768 | 0.028256 | 0.014151 | 0.056246 | 0.066554 | 0.006614 | 0.017562 | 0.025157 | 0.009866 | 0.057811 | 0.01745 |
| 27.7083333 | 0.009949 | 0.009585 | 0.019671 | 0.047042 | 0.01369 | 0.008306 | 0.03041 | 0.002791 | 0.029196 | 0.019575 | 0.05558 | 0.076554 | 0.008566 | 0.011801 | 0.023551 | 0.010337 | 0.104458 | 0.009974 |
| 27.7222222 | 0.013048 | 0.010951 | 0.015516 | 0.037549 | 0.013952 | 0.019558 | 0.036783 | 0.004535 | 0.018464 | 0.020729 | 0.037429 | 0.062256 | 0.009817 | 0.006573 | 0.012866 | 0.010233 | 0.123432 | 0.007754 |
| 27.7361111 | 0.011249 | 0.010373 | 0.00842 | 0.020286 | 0.012114 | 0.037145 | 0.039246 | 0.004964 | 0.010574 | 0.014333 | 0.022797 | 0.034575 | 0.007511 | 0.007798 | 0.004712 | 0.014774 | 0.100842 | 0.01108 |
| 27.75 | 0.006054 | 0.008634 | 0.00372 | 0.00898 | 0.008664 | 0.056249 | 0.033149 | 0.003245 | 0.015744 | 0.010207 | 0.036859 | 0.020649 | 0.00428 | 0.006812 | 0.006081 | 0.018152 | 0.054912 | 0.011376 |
| 27.7638889 | 0.003416 | 0.011302 | 0.003573 | 0.010765 | 0.011132 | 0.065318 | 0.019933 | 0.002157 | 0.018044 | 0.020435 | 0.054785 | 0.042232 | 0.003871 | 0.002972 | 0.01031 | 0.013038 | 0.03586 | 0.013098 |
| 27.7777778 | 0.008357 | 0.01347 | 0.00757 | 0.014031 | 0.018144 | 0.056444 | 0.017157 | 0.004159 | 0.013487 | 0.034946 | 0.049576 | 0.075134 | 0.004675 | 0.002767 | 0.009644 | 0.01717 | 0.045446 | 0.022554 |
| 27.7916667 | 0.020166 | 0.008986 | 0.012081 | 0.014497 | 0.022934 | 0.035117 | 0.032866 | 0.00715 | 0.015643 | 0.041023 | 0.028255 | 0.075419 | 0.006753 | 0.006561 | 0.010599 | 0.031473 | 0.041267 | 0.02777 |
| 27.8055556 | 0.028035 | 0.003516 | 0.012043 | 0.012963 | 0.025429 | 0.016812 | 0.050274 | 0.007907 | 0.018974 | 0.033349 | 0.013625 | 0.047037 | 0.008757 | 0.011605 | 0.01919 | 0.03356 | 0.022265 | 0.024019 |
| 27.8194444 | 0.020541 | 0.001943 | 0.007147 | 0.011842 | 0.019479 | 0.014978 | 0.054145 | 0.005635 | 0.013124 | 0.023175 | 0.020181 | 0.043392 | 0.006363 | 0.015706 | 0.027475 | 0.021222 | 0.011089 | 0.019289 |
| 27.8333333 | 0.01061 | 0.002707 | 0.003495 | 0.021144 | 0.012293 | 0.028874 | 0.041802 | 0.00368 | 0.007045 | 0.029577 | 0.033286 | 0.077941 | 0.002705 | 0.018546 | 0.03093 | 0.017081 | 0.017824 | 0.015611 |
| 27.8472222 | 0.014868 | 0.005605 | 0.003092 | 0.034004 | 0.01941 | 0.042943 | 0.021054 | 0.0046 | 0.008685 | 0.037406 | 0.037533 | 0.102648 | 0.001939 | 0.017022 | 0.02884 | 0.020835 | 0.035341 | 0.009932 |
| 27.8611111 | 0.019824 | 0.009682 | 0.004492 | 0.036848 | 0.031934 | 0.045661 | 0.011673 | 0.005214 | 0.008745 | 0.029764 | 0.037664 | 0.088427 | 0.002915 | 0.009722 | 0.019507 | 0.016778 | 0.052993 | 0.00467 |
| 27.875 | 0.013985 | 0.011119 | 0.005133 | 0.029075 | 0.036967 | 0.038054 | 0.024233 | 0.003368 | 0.005328 | 0.023811 | 0.043314 | 0.054171 | 0.008804 | 0.002938 | 0.008401 | 0.015693 | 0.058261 | 0.00537 |
| 27.8888889 | 0.009848 | 0.007673 | 0.007159 | 0.016576 | 0.032596 | 0.022233 | 0.046558 | 0.001257 | 0.009047 | 0.026545 | 0.046334 | 0.030014 | 0.02048 | 0.001938 | 0.004659 | 0.022154 | 0.043175 | 0.012494 |
| 27.9027778 | 0.013934 | 0.003535 | 0.016141 | 0.0062 | 0.022372 | 0.012284 | 0.057476 | 0.000626 | 0.017935 | 0.020931 | 0.032967 | 0.020341 | 0.02707 | 0.004519 | 0.009295 | 0.01889 | 0.022851 | 0.022104 |
| 27.9166667 | 0.015136 | 0.002854 | 0.024558 | 0.00391 | 0.01246 | 0.025341 | 0.045919 | 0.001475 | 0.020781 | 0.011971 | 0.018662 | 0.014078 | 0.021026 | 0.005318 | 0.015022 | 0.008934 | 0.010576 | 0.027489 |
| 27.9305556 | 0.017767 | 0.005629 | 0.031051 | 0.013822 | 0.011064 | 0.050329 | 0.02668 | 0.003257 | 0.015066 | 0.016575 | 0.025785 | 0.00952 | 0.009145 | 0.004508 | 0.019645 | 0.009409 | 0.005188 | 0.025867 |
| 27.9444444 | 0.030049 | 0.014623 | 0.042289 | 0.030567 | 0.018208 | 0.06463 | 0.021631 | 0.003478 | 0.008367 | 0.022914 | 0.043181 | 0.011119 | 0.002213 | 0.003655 | 0.026381 | 0.01734 | 0.003287 | 0.019318 |
| 27.9583333 | 0.036768 | 0.027258 | 0.049973 | 0.041538 | 0.02371 | 0.058506 | 0.030575 | 0.003197 | 0.006613 | 0.017955 | 0.047037 | 0.018252 | 0.002521 | 0.003115 | 0.034691 | 0.01912 | 0.004621 | 0.010851 |
| 27.9722222 | 0.033315 | 0.035277 | 0.043513 | 0.044753 | 0.02174 | 0.042266 | 0.040838 | 0.006304 | 0.008805 | 0.009246 | 0.035115 | 0.025676 | 0.006 | 0.004452 | 0.037966 | 0.01248 | 0.010137 | 0.005199 |
| 27.9861111 | 0.02997 | 0.034489 | 0.024829 | 0.046343 | 0.013261 | 0.03037 | 0.041774 | 0.011408 | 0.014545 | 0.003875 | 0.018482 | 0.030468 | 0.008641 | 0.006636 | 0.029524 | 0.006414 | 0.015384 | 0.004797 |
| 28 | 0.024788 | 0.024408 | 0.011737 | 0.043625 | 0.006998 | 0.021872 | 0.030079 | 0.016038 | 0.019639 | 0.002751 | 0.01191 | 0.033115 | 0.008414 | 0.010151 | 0.014626 | 0.005429 | 0.013955 | 0.006005 |
| 28.0138889 | 0.01436 | 0.013021 | 0.017711 | 0.031195 | 0.010111 | 0.013143 | 0.014857 | 0.017931 | 0.01579 | 0.008416 | 0.022433 | 0.034423 | 0.006028 | 0.01462 | 0.005072 | 0.005617 | 0.009289 | 0.004826 |
| 28.0277778 | 0.008355 | 0.013044 | 0.027993 | 0.016842 | 0.017409 | 0.006038 | 0.010425 | 0.01436 | 0.008603 | 0.019535 | 0.033127 | 0.033129 | 0.003394 | 0.014698 | 0.002354 | 0.006004 | 0.012491 | 0.00242 |
| 28.0416667 | 0.01242 | 0.021663 | 0.028036 | 0.015394 | 0.02206 | 0.003818 | 0.020077 | 0.007639 | 0.009704 | 0.025444 | 0.030236 | 0.027784 | 0.001807 | 0.008619 | 0.001458 | 0.011193 | 0.026328 | 0.002289 |
| 28.0555556 | 0.017008 | 0.025558 | 0.022233 | 0.02516 | 0.021346 | 0.006597 | 0.03289 | 0.003658 | 0.019372 | 0.020771 | 0.019638 | 0.02116 | 0.002503 | 0.00329 | 0.001317 | 0.024435 | 0.041338 | 0.005894 |
| 28.0694444 | 0.013823 | 0.018914 | 0.013716 | 0.026152 | 0.017972 | 0.008817 | 0.03648 | 0.005194 | 0.028573 | 0.013738 | 0.01217 | 0.021995 | 0.003657 | 0.002572 | 0.003423 | 0.0349 | 0.040962 | 0.007439 |
| 28.0833333 | 0.013234 | 0.015012 | 0.01457 | 0.024199 | 0.020553 | 0.008241 | 0.028401 | 0.011933 | 0.026639 | 0.018222 | 0.018811 | 0.043927 | 0.009566 | 0.008355 | 0.016237 | 0.031224 | 0.032874 | 0.013776 |
| 28.0972222 | 0.033908 | 0.036257 | 0.044472 | 0.048411 | 0.028655 | 0.011897 | 0.029038 | 0.020796 | 0.030728 | 0.041035 | 0.044162 | 0.088261 | 0.030683 | 0.02748 | 0.048243 | 0.037697 | 0.046015 | 0.043353 |
| 28.1111111 | 0.06695 | 0.075331 | 0.091769 | 0.083989 | 0.028426 | 0.019968 | 0.044694 | 0.02593 | 0.054547 | 0.064688 | 0.070517 | 0.122652 | 0.053703 | 0.050971 | 0.082353 | 0.070015 | 0.066171 | 0.077752 |
| 28.125 | 0.079768 | 0.100775 | 0.113365 | 0.087627 | 0.015713 | 0.020511 | 0.045018 | 0.022787 | 0.067171 | 0.063401 | 0.06923 | 0.108634 | 0.052721 | 0.053914 | 0.083081 | 0.089398 | 0.061245 | 0.078979 |
| 28.1388889 | 0.061826 | 0.093013 | 0.086146 | 0.056952 | 0.006189 | 0.011458 | 0.025958 | 0.012435 | 0.060507 | 0.03964 | 0.041292 | 0.058691 | 0.032577 | 0.031822 | 0.046887 | 0.069 | 0.034395 | 0.047332 |
| 28.1527778 | 0.032127 | 0.057555 | 0.039389 | 0.025614 | 0.008057 | 0.003837 | 0.010207 | 0.005673 | 0.052783 | 0.020455 | 0.017885 | 0.021846 | 0.024959 | 0.015817 | 0.021827 | 0.040471 | 0.015824 | 0.028305 |
| 28.1666667 | 0.012939 | 0.025965 | 0.017648 | 0.017513 | 0.01119 | 0.002286 | 0.003478 | 0.00689 | 0.04106 | 0.022877 | 0.01695 | 0.017369 | 0.033575 | 0.023849 | 0.034287 | 0.048628 | 0.022153 | 0.046357 |
| 28.1805556 | 0.013219 | 0.024042 | 0.029271 | 0.028615 | 0.009273 | 0.002458 | 0.00183 | 0.006692 | 0.022719 | 0.033513 | 0.024902 | 0.026763 | 0.033448 | 0.036324 | 0.054824 | 0.072589 | 0.025558 | 0.064054 |
| 28.1944444 | 0.024064 | 0.042905 | 0.049468 | 0.041373 | 0.006337 | 0.002906 | 0.002878 | 0.003364 | 0.009571 | 0.033838 | 0.025451 | 0.031604 | 0.022983 | 0.040453 | 0.058869 | 0.067656 | 0.013746 | 0.056119 |
| 28.2083333 | 0.034149 | 0.064704 | 0.059782 | 0.048439 | 0.006538 | 0.003029 | 0.003244 | 0.001774 | 0.008147 | 0.024175 | 0.018963 | 0.031058 | 0.013591 | 0.041267 | 0.050099 | 0.046093 | 0.005954 | 0.039939 |
| 28.2222222 | 0.040524 | 0.078754 | 0.056395 | 0.049735 | 0.008407 | 0.003387 | 0.004187 | 0.001779 | 0.009751 | 0.011921 | 0.010624 | 0.031455 | 0.007341 | 0.041874 | 0.039871 | 0.026574 | 0.005769 | 0.026094 |
| 28.2361111 | 0.041907 | 0.077489 | 0.044629 | 0.043438 | 0.008365 | 0.007068 | 0.010455 | 0.001816 | 0.00926 | 0.003996 | 0.004609 | 0.036684 | 0.004645 | 0.038858 | 0.030209 | 0.011754 | 0.006019 | 0.013087 |
| 28.25 | 0.03692 | 0.063071 | 0.034739 | 0.031658 | 0.006642 | 0.012199 | 0.018294 | 0.002181 | 0.014246 | 0.002148 | 0.003303 | 0.046234 | 0.00575 | 0.030754 | 0.019531 | 0.005074 | 0.004795 | 0.00437 |
| 28.2638889 | 0.028238 | 0.045315 | 0.028953 | 0.020124 | 0.005144 | 0.014925 | 0.022239 | 0.004158 | 0.029362 | 0.002311 | 0.004663 | 0.057203 | 0.0086 | 0.021325 | 0.011315 | 0.007016 | 0.004014 | 0.002903 |
| 28.2777778 | 0.021468 | 0.031757 | 0.024822 | 0.012463 | 0.003539 | 0.013937 | 0.021746 | 0.006445 | 0.044347 | 0.004213 | 0.005541 | 0.067784 | 0.009663 | 0.011784 | 0.005592 | 0.018772 | 0.007632 | 0.005815 |
| 28.2916667 | 0.017956 | 0.021821 | 0.022667 | 0.008752 | 0.001693 | 0.011319 | 0.019512 | 0.006728 | 0.046348 | 0.007843 | 0.005045 | 0.075378 | 0.00772 | 0.005146 | 0.002562 | 0.030878 | 0.010897 | 0.007411 |
| 28.3055556 | 0.015875 | 0.013472 | 0.024292 | 0.008208 | 0.001945 | 0.009988 | 0.019311 | 0.004946 | 0.03861 | 0.010289 | 0.003682 | 0.07898 | 0.004485 | 0.005042 | 0.003556 | 0.032316 | 0.008697 | 0.005213 |
| 28.3194444 | 0.014846 | 0.008207 | 0.028024 | 0.009878 | 0.004522 | 0.0099 | 0.021051 | 0.002495 | 0.032347 | 0.011918 | 0.001874 | 0.07987 | 0.002915 | 0.006322 | 0.004417 | 0.027358 | 0.003925 | 0.002729 |
| 28.3333333 | 0.015036 | 0.007018 | 0.030214 | 0.010817 | 0.006804 | 0.009641 | 0.021538 | 0.001056 | 0.030621 | 0.013877 | 0.000527 | 0.075511 | 0.004736 | 0.005173 | 0.003115 | 0.024496 | 0.001105 | 0.001799 |
| 28.3472222 | 0.016084 | 0.009359 | 0.028725 | 0.008462 | 0.007139 | 0.008488 | 0.018968 | 0.001582 | 0.029945 | 0.014341 | 0.000962 | 0.063743 | 0.008946 | 0.006694 | 0.003155 | 0.025683 | 0.002537 | 0.001515 |
| 28.3611111 | 0.017037 | 0.013027 | 0.02313 | 0.004232 | 0.005999 | 0.006625 | 0.014427 | 0.003969 | 0.024802 | 0.013579 | 0.004412 | 0.048297 | 0.014 | 0.012914 | 0.006271 | 0.028729 | 0.009281 | 0.002643 |
| 28.375 | 0.015514 | 0.013021 | 0.015324 | 0.001259 | 0.004982 | 0.004757 | 0.0103 | 0.005678 | 0.014888 | 0.015371 | 0.01143 | 0.030791 | 0.018546 | 0.018931 | 0.011108 | 0.031447 | 0.018892 | 0.003699 |
| 28.3888889 | 0.00968 | 0.00906 | 0.008033 | 0.001177 | 0.004474 | 0.003142 | 0.007551 | 0.004276 | 0.011838 | 0.021173 | 0.020491 | 0.015403 | 0.022032 | 0.022945 | 0.017258 | 0.034609 | 0.026405 | 0.005057 |
| 28.4027778 | 0.003721 | 0.008186 | 0.003302 | 0.004697 | 0.004011 | 0.001784 | 0.005315 | 0.003087 | 0.02011 | 0.027863 | 0.028025 | 0.011993 | 0.026155 | 0.026789 | 0.024642 | 0.03831 | 0.029194 | 0.00963 |
| 28.4166667 | 0.002543 | 0.009035 | 0.001553 | 0.012084 | 0.00416 | 0.001241 | 0.003867 | 0.0052 | 0.025063 | 0.029754 | 0.029436 | 0.017536 | 0.029281 | 0.028338 | 0.029314 | 0.037085 | 0.026029 | 0.01326 |
| 28.4305556 | 0.005771 | 0.007125 | 0.002675 | 0.021281 | 0.005302 | 0.00105 | 0.004968 | 0.00625 | 0.017546 | 0.024184 | 0.022607 | 0.020393 | 0.025223 | 0.023602 | 0.026033 | 0.027891 | 0.017137 | 0.009806 |
| 28.4444444 | 0.0129 | 0.009366 | 0.009127 | 0.028013 | 0.006422 | 0.001212 | 0.009006 | 0.004715 | 0.011275 | 0.015394 | 0.012506 | 0.018082 | 0.015614 | 0.014679 | 0.016497 | 0.017404 | 0.00731 | 0.004328 |

| | | | | | | | | | | | | | | | | | | |
|---|---|---|---|---|---|---|---|---|---|---|---|---|---|---|---|---|---|---|
| 28.4583333 | 0.021494 | 0.018834 | 0.020269 | 0.02767 | 0.00641 | 0.002523 | 0.013266 | 0.004925 | 0.015499 | 0.009417 | 0.006441 | 0.013418 | 0.009575 | 0.007733 | 0.007737 | 0.013961 | 0.002692 | 0.003193 |
| 28.4722222 | 0.025996 | 0.026888 | 0.029495 | 0.018901 | 0.004995 | 0.003451 | 0.013593 | 0.005163 | 0.014997 | 0.008174 | 0.005466 | 0.009891 | 0.009152 | 0.004905 | 0.002974 | 0.017897 | 0.00378 | 0.002579 |
| 28.4861111 | 0.021287 | 0.023409 | 0.0292 | 0.009803 | 0.002723 | 0.002889 | 0.008692 | 0.004185 | 0.01079 | 0.010163 | 0.005637 | 0.010259 | 0.010749 | 0.003813 | 0.002327 | 0.023176 | 0.006557 | 0.001233 |
| 28.5 | 0.012974 | 0.015055 | 0.017707 | 0.010511 | 0.001397 | 0.002049 | 0.004545 | 0.006664 | 0.019806 | 0.013663 | 0.004083 | 0.011381 | 0.011998 | 0.002955 | 0.005312 | 0.024269 | 0.008162 | 0.00247 |
| 28.5138889 | 0.013776 | 0.018506 | 0.00954 | 0.015674 | 0.001812 | 0.001838 | 0.005396 | 0.010666 | 0.032491 | 0.017785 | 0.001899 | 0.008238 | 0.011669 | 0.004111 | 0.008373 | 0.020848 | 0.008744 | 0.004451 |
| 28.5277778 | 0.018785 | 0.027337 | 0.014371 | 0.012828 | 0.002062 | 0.002118 | 0.005197 | 0.010974 | 0.031477 | 0.021704 | 0.003753 | 0.006623 | 0.011104 | 0.005558 | 0.007467 | 0.019344 | 0.010105 | 0.005482 |
| 28.5416667 | 0.015283 | 0.026324 | 0.016332 | 0.008334 | 0.001585 | 0.002731 | 0.003069 | 0.006966 | 0.018354 | 0.024382 | 0.012302 | 0.014641 | 0.012685 | 0.008741 | 0.007841 | 0.025742 | 0.012489 | 0.009383 |
| 28.5555556 | 0.008071 | 0.016 | 0.012165 | 0.014536 | 0.001715 | 0.003379 | 0.004065 | 0.003261 | 0.00782 | 0.024383 | 0.023518 | 0.025755 | 0.015778 | 0.020263 | 0.016172 | 0.035475 | 0.013326 | 0.013454 |
| 28.5694444 | 0.008558 | 0.008096 | 0.01933 | 0.02609 | 0.003359 | 0.004481 | 0.008531 | 0.005584 | 0.007632 | 0.021346 | 0.029921 | 0.034459 | 0.017509 | 0.031092 | 0.022857 | 0.040596 | 0.012035 | 0.012223 |
| 28.5833333 | 0.017504 | 0.011844 | 0.037741 | 0.040934 | 0.009933 | 0.008658 | 0.02097 | 0.014659 | 0.017369 | 0.016552 | 0.029826 | 0.048442 | 0.015546 | 0.027523 | 0.019523 | 0.040812 | 0.01446 | 0.011044 |
| 28.5972222 | 0.027419 | 0.021127 | 0.055557 | 0.06472 | 0.024837 | 0.015958 | 0.04247 | 0.027472 | 0.038614 | 0.010886 | 0.031917 | 0.066942 | 0.011093 | 0.016475 | 0.017273 | 0.03912 | 0.023884 | 0.015801 |
| 28.6111111 | 0.030611 | 0.0277 | 0.071229 | 0.0974 | 0.04087 | 0.021883 | 0.062159 | 0.035993 | 0.063447 | 0.007087 | 0.044485 | 0.077524 | 0.011083 | 0.011244 | 0.02507 | 0.037758 | 0.040014 | 0.021489 |
| 28.625 | 0.03045 | 0.033433 | 0.085068 | 0.12868 | 0.049528 | 0.02525 | 0.072314 | 0.033878 | 0.07931 | 0.006669 | 0.056874 | 0.078177 | 0.012872 | 0.011418 | 0.033001 | 0.040415 | 0.058893 | 0.021074 |
| 28.6388889 | 0.033112 | 0.037383 | 0.084664 | 0.127863 | 0.050623 | 0.026883 | 0.070544 | 0.024303 | 0.074639 | 0.007363 | 0.04659 | 0.077536 | 0.009921 | 0.009982 | 0.029199 | 0.043433 | 0.06549 | 0.015002 |
| 28.6527778 | 0.033108 | 0.032101 | 0.064232 | 0.08591 | 0.041398 | 0.024407 | 0.057898 | 0.013085 | 0.046216 | 0.009234 | 0.02361 | 0.077986 | 0.010233 | 0.009362 | 0.016854 | 0.038497 | 0.062168 | 0.009066 |
| 28.6666667 | 0.027951 | 0.018903 | 0.037668 | 0.037462 | 0.022846 | 0.019304 | 0.042128 | 0.007485 | 0.020048 | 0.012602 | 0.021471 | 0.067994 | 0.016232 | 0.018545 | 0.01414 | 0.029088 | 0.072868 | 0.010854 |
| 28.6805556 | 0.024107 | 0.007993 | 0.019612 | 0.017573 | 0.008988 | 0.017497 | 0.028514 | 0.01025 | 0.01549 | 0.016349 | 0.039377 | 0.041299 | 0.022239 | 0.027454 | 0.026978 | 0.021237 | 0.093554 | 0.022705 |
| 28.6944444 | 0.023471 | 0.006179 | 0.019566 | 0.028399 | 0.008592 | 0.020567 | 0.018797 | 0.011208 | 0.017243 | 0.01662 | 0.053445 | 0.020358 | 0.027147 | 0.020735 | 0.033656 | 0.012611 | 0.101229 | 0.034295 |
| 28.7083333 | 0.022504 | 0.011583 | 0.029367 | 0.037992 | 0.014763 | 0.024227 | 0.015143 | 0.005788 | 0.016205 | 0.010644 | 0.052362 | 0.032542 | 0.029461 | 0.01396 | 0.022656 | 0.008395 | 0.087108 | 0.031464 |
| 28.7222222 | 0.020981 | 0.016722 | 0.026976 | 0.029561 | 0.020773 | 0.023516 | 0.014432 | 0.001304 | 0.017215 | 0.004673 | 0.036863 | 0.059185 | 0.027393 | 0.023731 | 0.016103 | 0.009143 | 0.051838 | 0.017272 |
| 28.7361111 | 0.019015 | 0.017254 | 0.014804 | 0.015578 | 0.025187 | 0.0165 | 0.011025 | 0.000321 | 0.019228 | 0.005062 | 0.016737 | 0.060081 | 0.023953 | 0.035142 | 0.028001 | 0.011579 | 0.031412 | 0.011116 |
| 28.75 | 0.012708 | 0.014504 | 0.012516 | 0.007272 | 0.026019 | 0.010104 | 0.007924 | 0.000706 | 0.022206 | 0.007669 | 0.010244 | 0.03392 | 0.022973 | 0.034037 | 0.037983 | 0.026912 | 0.054092 | 0.018713 |
| 28.7638889 | 0.00824 | 0.01117 | 0.018693 | 0.009786 | 0.021109 | 0.015463 | 0.015099 | 0.001994 | 0.027822 | 0.009336 | 0.021873 | 0.016137 | 0.021567 | 0.024584 | 0.028647 | 0.041864 | 0.078527 | 0.024807 |
| 28.7777778 | 0.01794 | 0.008768 | 0.020391 | 0.022149 | 0.011602 | 0.029713 | 0.032696 | 0.003222 | 0.03091 | 0.017629 | 0.034318 | 0.028236 | 0.015747 | 0.015058 | 0.014671 | 0.031879 | 0.072002 | 0.020554 |
| 28.7916667 | 0.032514 | 0.007359 | 0.014785 | 0.032178 | 0.006448 | 0.039151 | 0.046488 | 0.002825 | 0.025268 | 0.026797 | 0.037695 | 0.048791 | 0.008702 | 0.017461 | 0.020045 | 0.015375 | 0.048394 | 0.011351 |
| 28.8055556 | 0.032984 | 0.006495 | 0.008907 | 0.024596 | 0.013916 | 0.034255 | 0.043374 | 0.002179 | 0.013344 | 0.025688 | 0.029957 | 0.059427 | 0.005803 | 0.036597 | 0.040128 | 0.015951 | 0.028673 | 0.010012 |
| 28.8194444 | 0.020945 | 0.00635 | 0.009704 | 0.011904 | 0.029158 | 0.018058 | 0.024864 | 0.003277 | 0.004281 | 0.0237 | 0.017019 | 0.058331 | 0.00515 | 0.051339 | 0.054622 | 0.020674 | 0.02056 | 0.022804 |
| 28.8333333 | 0.009333 | 0.007104 | 0.014932 | 0.014714 | 0.039344 | 0.006333 | 0.011599 | 0.004572 | 0.006999 | 0.023956 | 0.013377 | 0.03951 | 0.004819 | 0.041528 | 0.055043 | 0.017724 | 0.014781 | 0.038988 |
| 28.8472222 | 0.005418 | 0.008623 | 0.026883 | 0.019617 | 0.038152 | 0.005812 | 0.018159 | 0.005339 | 0.017439 | 0.018794 | 0.015993 | 0.020841 | 0.009837 | 0.02341 | 0.044019 | 0.016984 | 0.009585 | 0.046786 |
| 28.8611111 | 0.009369 | 0.010507 | 0.046331 | 0.012812 | 0.028535 | 0.006777 | 0.031177 | 0.007451 | 0.021941 | 0.012543 | 0.015799 | 0.026385 | 0.023598 | 0.021504 | 0.025752 | 0.024619 | 0.008044 | 0.042263 |
| 28.875 | 0.014448 | 0.013472 | 0.048263 | 0.007487 | 0.016273 | 0.004558 | 0.036434 | 0.009604 | 0.017797 | 0.009559 | 0.018863 | 0.04682 | 0.037239 | 0.026922 | 0.011166 | 0.027019 | 0.012599 | 0.027191 |
| 28.8888889 | 0.016663 | 0.018753 | 0.027962 | 0.013618 | 0.013547 | 0.003532 | 0.032948 | 0.007929 | 0.011442 | 0.009129 | 0.030871 | 0.067376 | 0.035442 | 0.019227 | 0.011646 | 0.016121 | 0.028825 | 0.011255 |
| 28.9027778 | 0.016268 | 0.023755 | 0.019704 | 0.021159 | 0.025808 | 0.00528 | 0.022558 | 0.00467 | 0.009285 | 0.014946 | 0.044612 | 0.084239 | 0.020814 | 0.011459 | 0.014214 | 0.004916 | 0.044245 | 0.004914 |
| 28.9166667 | 0.012019 | 0.024356 | 0.02846 | 0.022638 | 0.037353 | 0.007398 | 0.012331 | 0.005884 | 0.013819 | 0.023581 | 0.04985 | 0.094455 | 0.011322 | 0.023422 | 0.017224 | 0.004269 | 0.040888 | 0.005828 |
| 28.9305556 | 0.01239 | 0.022397 | 0.034953 | 0.025758 | 0.038074 | 0.008934 | 0.016615 | 0.009718 | 0.024034 | 0.027111 | 0.044042 | 0.097893 | 0.009188 | 0.040842 | 0.034538 | 0.012024 | 0.023792 | 0.006277 |
| 28.9444444 | 0.027576 | 0.023947 | 0.027893 | 0.037093 | 0.035939 | 0.010976 | 0.042974 | 0.010098 | 0.037734 | 0.026399 | 0.035074 | 0.094384 | 0.005628 | 0.047489 | 0.055075 | 0.017336 | 0.024102 | 0.004602 |
| 28.9583333 | 0.04796 | 0.029425 | 0.02115 | 0.050889 | 0.033978 | 0.013505 | 0.075558 | 0.007029 | 0.05054 | 0.023982 | 0.030744 | 0.083061 | 0.004782 | 0.046011 | 0.068767 | 0.017638 | 0.052199 | 0.005893 |
| 28.9722222 | 0.05554 | 0.033089 | 0.041427 | 0.05638 | 0.027305 | 0.014568 | 0.082908 | 0.009058 | 0.059566 | 0.016895 | 0.027927 | 0.065992 | 0.013585 | 0.03872 | 0.075072 | 0.031061 | 0.068999 | 0.014431 |
| 28.9861111 | 0.045651 | 0.031324 | 0.069988 | 0.050053 | 0.018149 | 0.013877 | 0.058143 | 0.019103 | 0.061693 | 0.007433 | 0.022074 | 0.04648 | 0.031416 | 0.024531 | 0.068189 | 0.062448 | 0.058351 | 0.021986 |
| 29 | 0.0263 | 0.02453 | 0.072185 | 0.038648 | 0.012331 | 0.01272 | 0.026814 | 0.027106 | 0.053654 | 0.004825 | 0.015219 | 0.025808 | 0.043991 | 0.012759 | 0.043879 | 0.082907 | 0.039259 | 0.020094 |
| 29.0138889 | 0.01208 | 0.014682 | 0.046116 | 0.026485 | 0.009797 | 0.009866 | 0.009358 | 0.027756 | 0.03732 | 0.007749 | 0.008779 | 0.011764 | 0.035274 | 0.016344 | 0.024255 | 0.073357 | 0.019883 | 0.011321 |
| 29.0277778 | 0.011999 | 0.006529 | 0.02242 | 0.01469 | 0.00676 | 0.00509 | 0.007327 | 0.022684 | 0.019642 | 0.011699 | 0.008141 | 0.011764 | 0.027391 | 0.029753 | 0.034469 | 0.044095 | 0.00863 | 0.005607 |
| 29.0416667 | 0.016992 | 0.003831 | 0.030315 | 0.011504 | 0.006258 | 0.00273 | 0.011716 | 0.013753 | 0.014267 | 0.018275 | 0.017556 | 0.014959 | 0.041243 | 0.038858 | 0.049997 | 0.021377 | 0.01197 | 0.010886 |
| 29.0555556 | 0.015887 | 0.003274 | 0.057494 | 0.017054 | 0.011936 | 0.005065 | 0.01538 | 0.006296 | 0.028443 | 0.027016 | 0.026699 | 0.010696 | 0.05135 | 0.035946 | 0.044409 | 0.023978 | 0.023736 | 0.022246 |
| 29.0694444 | 0.011808 | 0.005123 | 0.07382 | 0.015828 | 0.013868 | 0.00674 | 0.015021 | 0.004483 | 0.043386 | 0.030917 | 0.023988 | 0.013107 | 0.037201 | 0.02418 | 0.028133 | 0.033661 | 0.027395 | 0.02552 |
| 29.0833333 | 0.021291 | 0.021853 | 0.056794 | 0.019032 | 0.013405 | 0.006714 | 0.01287 | 0.014407 | 0.039438 | 0.025052 | 0.019837 | 0.042921 | 0.024187 | 0.02774 | 0.032751 | 0.035632 | 0.027741 | 0.027392 |
| 29.0972222 | 0.054117 | 0.063546 | 0.042031 | 0.053529 | 0.027169 | 0.011565 | 0.019498 | 0.043564 | 0.044574 | 0.027938 | 0.040321 | 0.083013 | 0.042201 | 0.058728 | 0.073605 | 0.068046 | 0.053013 | 0.045189 |
| 29.1111111 | 0.085684 | 0.113095 | 0.081708 | 0.092493 | 0.04518 | 0.018777 | 0.030601 | 0.068367 | 0.077965 | 0.052299 | 0.087967 | 0.096068 | 0.059373 | 0.082081 | 0.10705 | 0.151946 | 0.094817 | 0.059084 |
| 29.125 | 0.087273 | 0.136149 | 0.125066 | 0.094969 | 0.043004 | 0.021826 | 0.030264 | 0.061893 | 0.096123 | 0.064424 | 0.124057 | 0.083861 | 0.040222 | 0.066364 | 0.092818 | 0.223867 | 0.120832 | 0.052544 |
| 29.1388889 | 0.065865 | 0.12173 | 0.12252 | 0.068154 | 0.023564 | 0.01979 | 0.020647 | 0.040861 | 0.084836 | 0.041957 | 0.108213 | 0.069381 | 0.015239 | 0.030875 | 0.052632 | 0.219664 | 0.114295 | 0.036455 |
| 29.1527778 | 0.036244 | 0.082293 | 0.100033 | 0.036652 | 0.009193 | 0.012226 | 0.011891 | 0.023979 | 0.070704 | 0.020137 | 0.055208 | 0.057119 | 0.011858 | 0.012119 | 0.021055 | 0.151291 | 0.076038 | 0.019941 |
| 29.1666667 | 0.016724 | 0.039594 | 0.074674 | 0.015404 | 0.009804 | 0.005307 | 0.005872 | 0.011093 | 0.057298 | 0.027821 | 0.025896 | 0.046594 | 0.017407 | 0.016682 | 0.012443 | 0.072615 | 0.035593 | 0.015472 |
| 29.1805556 | 0.022983 | 0.020607 | 0.046674 | 0.012107 | 0.015761 | 0.005155 | 0.004601 | 0.004544 | 0.0395 | 0.047134 | 0.046865 | 0.039628 | 0.020031 | 0.026004 | 0.02436 | 0.043598 | 0.028376 | 0.024912 |
| 29.1944444 | 0.038997 | 0.027681 | 0.022624 | 0.02104 | 0.019399 | 0.005377 | 0.009002 | 0.008556 | 0.0216 | 0.055712 | 0.08632 | 0.033043 | 0.022014 | 0.029546 | 0.04336 | 0.078286 | 0.050492 | 0.036008 |
| 29.2083333 | 0.050789 | 0.040297 | 0.015384 | 0.028552 | 0.022145 | 0.002512 | 0.012359 | 0.019542 | 0.008704 | 0.053652 | 0.114317 | 0.023204 | 0.027036 | 0.028424 | 0.059354 | 0.125629 | 0.072997 | 0.045295 |
| 29.2222222 | 0.057677 | 0.04599 | 0.029076 | 0.027362 | 0.023388 | 0.001743 | 0.010154 | 0.03128 | 0.003354 | 0.044019 | 0.117168 | 0.01197 | 0.035287 | 0.02494 | 0.066517 | 0.145523 | 0.076813 | 0.054982 |
| 29.2361111 | 0.056856 | 0.042939 | 0.056165 | 0.022857 | 0.020826 | 0.004273 | 0.008433 | 0.040561 | 0.00486 | 0.029344 | 0.098205 | 0.007728 | 0.048594 | 0.023888 | 0.07222 | 0.133081 | 0.059024 | 0.062715 |
| 29.25 | 0.04931 | 0.038202 | 0.087694 | 0.022141 | 0.016089 | 0.007147 | 0.013095 | 0.042575 | 0.008131 | 0.02141 | 0.077594 | 0.01484 | 0.063197 | 0.028758 | 0.082454 | 0.096346 | 0.035112 | 0.064021 |
| 29.2638889 | 0.037913 | 0.040843 | 0.10968 | 0.02094 | 0.011438 | 0.008396 | 0.021384 | 0.037639 | 0.011496 | 0.021736 | 0.060428 | 0.027963 | 0.063408 | 0.040479 | 0.090698 | 0.053366 | 0.019808 | 0.058676 |
| 29.2777778 | 0.026555 | 0.054414 | 0.1133 | 0.015675 | 0.006606 | 0.008114 | 0.028075 | 0.029728 | 0.016019 | 0.021022 | 0.041691 | 0.042193 | 0.045543 | 0.055592 | 0.094943 | 0.02223 | 0.010828 | 0.049323 |
| 29.2916667 | 0.018722 | 0.073485 | 0.10436 | 0.009417 | 0.002488 | 0.007808 | 0.026765 | 0.020658 | 0.02076 | 0.01654 | 0.022434 | 0.055415 | 0.024337 | 0.062774 | 0.08699 | 0.008848 | 0.005647 | 0.036315 |
| 29.3055556 | 0.014709 | 0.081826 | 0.089746 | 0.0043 | 0.000809 | 0.00825 | 0.017196 | 0.012293 | 0.02423 | 0.009513 | 0.00999 | 0.065885 | 0.009868 | 0.057981 | 0.065949 | 0.005428 | 0.005771 | 0.021707 |
| 29.3194444 | 0.012021 | 0.071015 | 0.066853 | 0.001786 | 0.001161 | 0.008703 | 0.008769 | 0.007965 | 0.025507 | 0.006389 | 0.0094 | 0.069777 | 0.003792 | 0.046842 | 0.0426 | 0.005905 | 0.009064 | 0.010228 |
| 29.3333333 | 0.010708 | 0.05212 | 0.038939 | 0.001731 | 0.001369 | 0.009567 | 0.005104 | 0.007655 | 0.027799 | 0.010235 | 0.012286 | 0.063473 | 0.004767 | 0.03185 | 0.020605 | 0.008321 | 0.011107 | 0.006601 |
| 29.3472222 | 0.013376 | 0.038743 | 0.019899 | 0.004566 | 0.001776 | 0.011767 | 0.003206 | 0.006917 | 0.034694 | 0.011067 | 0.011089 | 0.053243 | 0.005555 | 0.018069 | 0.006203 | 0.011755 | 0.012234 | 0.007868 |
| 29.3611111 | 0.017829 | 0.031907 | 0.01222 | 0.012397 | 0.003716 | 0.013493 | 0.002884 | 0.006805 | 0.041943 | 0.007443 | 0.012814 | 0.049042 | 0.003439 | 0.01193 | 0.003141 | 0.018725 | 0.015664 | 0.005995 |
| 29.375 | 0.019968 | 0.025646 | 0.007052 | 0.019877 | 0.004661 | 0.012491 | 0.004356 | 0.007066 | 0.04358 | 0.007514 | 0.017983 | 0.051372 | 0.002425 | 0.009183 | 0.005039 | 0.023952 | 0.01674 | 0.003303 |
| 29.3888889 | 0.019283 | 0.017925 | 0.002522 | 0.020775 | 0.003613 | 0.009312 | 0.00581 | 0.004263 | 0.040061 | 0.007882 | 0.014261 | 0.055435 | 0.004453 | 0.006503 | 0.00905 | 0.02078 | 0.010557 | 0.006518 |
| 29.4027778 | 0.017067 | 0.010859 | 0.001313 | 0.016047 | 0.002449 | 0.005955 | 0.006428 | 0.001678 | 0.034936 | 0.005005 | 0.007121 | 0.059164 | 0.011028 | 0.00851 | 0.0143 | 0.016335 | 0.003973 | 0.015283 |
| 29.4166667 | 0.014997 | 0.006072 | 0.001234 | 0.009128 | 0.001779 | 0.00395 | 0.006039 | 0.001476 | 0.030705 | 0.003729 | 0.007721 | 0.058443 | 0.017167 | 0.011973 | 0.017093 | 0.017373 | 0.002582 | 0.022653 |
| 29.4305556 | 0.01519 | 0.005646 | 0.000884 | 0.004583 | 0.001181 | 0.003998 | 0.005117 | 0.002176 | 0.029573 | 0.00699 | 0.00857 | 0.049054 | 0.013705 | 0.010414 | 0.014539 | 0.021849 | 0.004859 | 0.023603 |
| 29.4444444 | 0.017959 | 0.01067 | 0.001333 | 0.005126 | 0.002042 | 0.00468 | 0.00437 | 0.004234 | 0.031032 | 0.012792 | 0.010144 | 0.031892 | 0.006261 | 0.011381 | 0.008969 | 0.026077 | 0.00991 | 0.020743 |
| 29.4583333 | 0.019674 | 0.016887 | 0.002209 | 0.008223 | 0.00693 | 0.003892 | 0.003798 | 0.005707 | 0.031776 | 0.016432 | 0.017609 | 0.016686 | 0.004461 | 0.02041 | 0.007909 | 0.02786 | 0.012857 | 0.017167 |
| 29.4722222 | 0.016085 | 0.018677 | 0.004235 | 0.010836 | 0.013494 | 0.002057 | 0.00458 | 0.004626 | 0.030773 | 0.016581 | 0.02283 | 0.017093 | 0.006876 | 0.027839 | 0.012044 | 0.028103 | 0.011271 | 0.012768 |
| 29.4861111 | 0.00872 | 0.01554 | 0.009739 | 0.01336 | 0.016164 | 0.000967 | 0.0083 | 0.003024 | 0.028546 | 0.015022 | 0.019523 | 0.030776 | 0.016503 | 0.028107 | 0.016164 | 0.027872 | 0.007704 | 0.009643 |
| 29.5 | 0.004616 | 0.010354 | 0.018002 | 0.01587 | 0.013334 | 0.000764 | 0.014191 | 0.001815 | 0.025697 | 0.011728 | 0.012174 | 0.042244 | 0.028972 | 0.024002 | 0.017741 | 0.026711 | 0.00365 | 0.010749 |
| 29.5138889 | 0.004614 | 0.00575 | 0.023749 | 0.017335 | 0.008179 | 0.000642 | 0.018843 | 0.001368 | 0.022728 | 0.007966 | 0.012129 | 0.042734 | 0.028114 | 0.018563 | 0.016303 | 0.02511 | 0.001408 | 0.013329 |
| 29.5277778 | 0.003359 | 0.002885 | 0.022526 | 0.017419 | 0.004605 | 0.000835 | 0.018682 | 0.00253 | 0.018224 | 0.006305 | 0.01978 | 0.036001 | 0.016332 | 0.013713 | 0.010739 | 0.023248 | 0.002333 | 0.013418 |
| 29.5416667 | 0.002749 | 0.001349 | 0.01609 | 0.01582 | 0.003684 | 0.00096 | 0.013591 | 0.003369 | 0.01167 | 0.007388 | 0.021628 | 0.029376 | 0.013169 | 0.008267 | 0.00913 | 0.019725 | 0.005424 | 0.011288 |
| 29.5555556 | 0.006951 | 0.000665 | 0.010885 | 0.014291 | 0.003102 | 0.001016 | 0.008105 | 0.002541 | 0.007107 | 0.010424 | 0.01447 | 0.024085 | 0.02297 | 0.003148 | 0.019243 | 0.014503 | 0.008361 | 0.007732 |
| 29.5694444 | 0.014567 | 0.002189 | 0.012332 | 0.018003 | 0.005823 | 0.001579 | 0.00803 | 0.003074 | 0.010469 | 0.01578 | 0.012629 | 0.018824 | 0.023692 | 0.002272 | 0.023638 | 0.015072 | 0.01175 | 0.005212 |
| 29.5833333 | 0.023707 | 0.008058 | 0.022581 | 0.032172 | 0.020185 | 0.003962 | 0.017303 | 0.008679 | 0.026672 | 0.025215 | 0.027952 | 0.031484 | 0.019962 | 0.007296 | 0.019375 | 0.027717 | 0.017944 | 0.010426 |
| 29.5972222 | 0.03345 | 0.016161 | 0.041278 | 0.053211 | 0.043038 | 0.011343 | 0.03556 | 0.018097 | 0.050611 | 0.034704 | 0.057881 | 0.090211 | 0.039591 | 0.018947 | 0.033856 | 0.039936 | 0.018208 | 0.023122 |

| | | | | | | | | | | | | | | | | | | |
|---|---|---|---|---|---|---|---|---|---|---|---|---|---|---|---|---|---|---|
| 29.6111111 | 0.041425 | 0.021713 | 0.062512 | 0.069323 | 0.058 | 0.021053 | 0.055265 | 0.026081 | 0.065747 | 0.036437 | 0.089413 | 0.181028 | 0.063448 | 0.03589 | 0.067933 | 0.03794 | 0.013981 | 0.03289 |
| 29.625 | 0.046355 | 0.023995 | 0.081404 | 0.074408 | 0.056341 | 0.025256 | 0.065681 | 0.028285 | 0.07078 | 0.034939 | 0.103335 | 0.23109 | 0.058715 | 0.048748 | 0.08702 | 0.027804 | 0.024287 | 0.038779 |
| 29.6388889 | 0.045114 | 0.024466 | 0.089403 | 0.067789 | 0.04583 | 0.022625 | 0.060856 | 0.02673 | 0.07354 | 0.039505 | 0.090295 | 0.206166 | 0.035772 | 0.046409 | 0.074708 | 0.016421 | 0.038156 | 0.042189 |
| 29.6527778 | 0.033234 | 0.023599 | 0.069703 | 0.051155 | 0.036848 | 0.017896 | 0.045245 | 0.025022 | 0.063844 | 0.04201 | 0.060642 | 0.162355 | 0.020328 | 0.031368 | 0.050646 | 0.005541 | 0.032307 | 0.035619 |
| 29.6666667 | 0.016948 | 0.022809 | 0.032765 | 0.032294 | 0.034641 | 0.014123 | 0.030296 | 0.017976 | 0.038352 | 0.032648 | 0.029763 | 0.139168 | 0.012802 | 0.018859 | 0.027157 | 0.000816 | 0.015054 | 0.020513 |
| 29.6805556 | 0.009036 | 0.02202 | 0.016934 | 0.016519 | 0.036678 | 0.0121 | 0.021132 | 0.009735 | 0.015875 | 0.020077 | 0.015114 | 0.134599 | 0.005457 | 0.02227 | 0.014922 | 0.000647 | 0.006168 | 0.01075 |
| 29.6944444 | 0.012073 | 0.018649 | 0.034879 | 0.005682 | 0.035042 | 0.011923 | 0.017089 | 0.008587 | 0.007974 | 0.011925 | 0.025742 | 0.13704 | 0.005112 | 0.035222 | 0.027183 | 0.001681 | 0.012978 | 0.01515 |
| 29.7083333 | 0.017628 | 0.01139 | 0.057434 | 0.001962 | 0.027131 | 0.013713 | 0.017711 | 0.007162 | 0.008127 | 0.0061 | 0.043897 | 0.126112 | 0.020621 | 0.035926 | 0.046579 | 0.009165 | 0.032996 | 0.025977 |
| 29.7222222 | 0.021315 | 0.008877 | 0.062428 | 0.002787 | 0.015146 | 0.013669 | 0.019441 | 0.004682 | 0.010399 | 0.004371 | 0.054931 | 0.099929 | 0.039825 | 0.019803 | 0.050062 | 0.031513 | 0.056411 | 0.034527 |
| 29.7361111 | 0.022032 | 0.019595 | 0.052698 | 0.005513 | 0.008234 | 0.013187 | 0.018085 | 0.005577 | 0.012981 | 0.00674 | 0.053861 | 0.069634 | 0.036518 | 0.009805 | 0.035017 | 0.048433 | 0.07097 | 0.034264 |
| 29.75 | 0.015456 | 0.032475 | 0.03441 | 0.010174 | 0.014877 | 0.014588 | 0.012731 | 0.00694 | 0.01044 | 0.009905 | 0.037198 | 0.042811 | 0.02678 | 0.020746 | 0.028522 | 0.045171 | 0.071858 | 0.021566 |
| 29.7638889 | 0.009956 | 0.036272 | 0.027384 | 0.014402 | 0.026926 | 0.010148 | 0.006695 | 0.007091 | 0.005199 | 0.012711 | 0.017262 | 0.020953 | 0.036313 | 0.036637 | 0.049081 | 0.064599 | 0.059317 | 0.01372 |
| 29.7777778 | 0.014964 | 0.033606 | 0.04215 | 0.016236 | 0.036558 | 0.005147 | 0.005831 | 0.006345 | 0.005573 | 0.014819 | 0.005986 | 0.011901 | 0.043041 | 0.041796 | 0.064277 | 0.106378 | 0.037823 | 0.021732 |
| 29.7916667 | 0.019585 | 0.029884 | 0.056092 | 0.0171 | 0.043849 | 0.009645 | 0.010867 | 0.004956 | 0.010769 | 0.015531 | 0.005939 | 0.02255 | 0.033017 | 0.034834 | 0.049067 | 0.110716 | 0.022519 | 0.029897 |
| 29.8055556 | 0.016813 | 0.026847 | 0.056427 | 0.019135 | 0.043115 | 0.015464 | 0.017547 | 0.003048 | 0.016692 | 0.011152 | 0.01578 | 0.035275 | 0.018032 | 0.020294 | 0.025364 | 0.06744 | 0.027256 | 0.026127 |
| 29.8194444 | 0.010896 | 0.022008 | 0.048726 | 0.022261 | 0.028376 | 0.011447 | 0.024319 | 0.001283 | 0.018378 | 0.008197 | 0.028179 | 0.031365 | 0.007564 | 0.013711 | 0.021959 | 0.034597 | 0.040126 | 0.017474 |
| 29.8333333 | 0.006538 | 0.014946 | 0.034572 | 0.022289 | 0.014394 | 0.003188 | 0.029277 | 0.00042 | 0.013458 | 0.017843 | 0.033076 | 0.018813 | 0.008766 | 0.021172 | 0.03818 | 0.049688 | 0.039052 | 0.019377 |
| 29.8472222 | 0.00937 | 0.010385 | 0.018491 | 0.016361 | 0.019971 | 0.000457 | 0.028957 | 0.001219 | 0.009815 | 0.033202 | 0.024897 | 0.016991 | 0.026127 | 0.03047 | 0.061551 | 0.073635 | 0.02336 | 0.034926 |
| 29.8611111 | 0.019891 | 0.014261 | 0.01661 | 0.009504 | 0.033742 | 0.002037 | 0.024271 | 0.006344 | 0.013733 | 0.042414 | 0.019939 | 0.023042 | 0.050901 | 0.032518 | 0.084233 | 0.068968 | 0.009025 | 0.054971 |
| 29.875 | 0.034451 | 0.02668 | 0.034543 | 0.009813 | 0.038232 | 0.007525 | 0.017066 | 0.019102 | 0.020673 | 0.041249 | 0.03496 | 0.022186 | 0.068221 | 0.03138 | 0.097064 | 0.039945 | 0.005842 | 0.065441 |
| 29.8888889 | 0.047161 | 0.040283 | 0.056569 | 0.017338 | 0.033125 | 0.011852 | 0.008649 | 0.035899 | 0.023755 | 0.028624 | 0.055675 | 0.014551 | 0.070425 | 0.035143 | 0.095509 | 0.011209 | 0.008857 | 0.057615 |
| 29.9027778 | 0.051076 | 0.047673 | 0.068572 | 0.024978 | 0.021127 | 0.008597 | 0.002911 | 0.044789 | 0.024457 | 0.017797 | 0.063957 | 0.015463 | 0.052669 | 0.038723 | 0.08008 | 0.009519 | 0.01878 | 0.045021 |
| 29.9166667 | 0.044735 | 0.046743 | 0.064241 | 0.028327 | 0.012456 | 0.005533 | 0.001231 | 0.04081 | 0.026635 | 0.02601 | 0.058733 | 0.03373 | 0.024722 | 0.034046 | 0.056536 | 0.017881 | 0.038814 | 0.036902 |
| 29.9305556 | 0.029912 | 0.039856 | 0.049665 | 0.027425 | 0.019599 | 0.004849 | 0.002211 | 0.031301 | 0.024388 | 0.041049 | 0.050341 | 0.056002 | 0.007094 | 0.02371 | 0.034506 | 0.010528 | 0.060433 | 0.026917 |
| 29.9444444 | 0.01288 | 0.031661 | 0.033156 | 0.025718 | 0.034459 | 0.003523 | 0.006601 | 0.022514 | 0.014024 | 0.041735 | 0.044067 | 0.069265 | 0.00527 | 0.014258 | 0.019645 | 0.00696 | 0.065592 | 0.015413 |
| 29.9583333 | 0.004214 | 0.02499 | 0.019455 | 0.026778 | 0.042216 | 0.003403 | 0.01415 | 0.015701 | 0.007283 | 0.029744 | 0.041269 | 0.070901 | 0.009343 | 0.009801 | 0.011616 | 0.010146 | 0.041188 | 0.010809 |
| 29.9722222 | 0.005543 | 0.018512 | 0.014794 | 0.028987 | 0.044508 | 0.003192 | 0.020006 | 0.009052 | 0.010437 | 0.016118 | 0.043217 | 0.059183 | 0.014121 | 0.010555 | 0.00844 | 0.01159 | 0.015809 | 0.01501 |
| 29.9861111 | 0.009975 | 0.012091 | 0.01161 | 0.02739 | 0.049752 | 0.002176 | 0.017221 | 0.006066 | 0.014744 | 0.006975 | 0.038884 | 0.036561 | 0.020376 | 0.013043 | 0.009283 | 0.026644 | 0.01781 | 0.017035 |
| 30 | 0.013171 | 0.007195 | 0.010092 | 0.018828 | 0.048245 | 0.003117 | 0.01004 | 0.011311 | 0.017911 | 0.004313 | 0.024729 | 0.014671 | 0.025112 | 0.012686 | 0.013304 | 0.050368 | 0.033621 | 0.013019 |
| 30.0138889 | 0.013074 | 0.003473 | 0.022634 | 0.008289 | 0.031693 | 0.006071 | 0.011193 | 0.016241 | 0.020373 | 0.006778 | 0.022594 | 0.003505 | 0.025213 | 0.00837 | 0.018208 | 0.065944 | 0.048523 | 0.008885 |
| 30.0277778 | 0.008457 | 0.002208 | 0.040682 | 0.004874 | 0.014135 | 0.005888 | 0.018961 | 0.014932 | 0.018047 | 0.012358 | 0.04274 | 0.001375 | 0.02289 | 0.005567 | 0.019072 | 0.0713 | 0.057364 | 0.00709 |
| 30.0416667 | 0.003771 | 0.00445 | 0.049818 | 0.007986 | 0.00829 | 0.00339 | 0.02456 | 0.010565 | 0.012549 | 0.019433 | 0.061713 | 0.003605 | 0.022252 | 0.009472 | 0.013463 | 0.077156 | 0.056601 | 0.006364 |
| 30.0555556 | 0.003436 | 0.006731 | 0.047649 | 0.010509 | 0.011483 | 0.003724 | 0.026475 | 0.006557 | 0.008499 | 0.022174 | 0.060039 | 0.009631 | 0.022594 | 0.014485 | 0.005744 | 0.084774 | 0.048794 | 0.004315 |
| 30.0694444 | 0.007858 | 0.008837 | 0.033684 | 0.010854 | 0.011455 | 0.004059 | 0.022649 | 0.010477 | 0.014696 | 0.015233 | 0.040096 | 0.011393 | 0.018266 | 0.012411 | 0.010591 | 0.074735 | 0.031834 | 0.008519 |
| 30.0833333 | 0.024242 | 0.025491 | 0.025148 | 0.021454 | 0.015661 | 0.007842 | 0.017788 | 0.031483 | 0.047468 | 0.017685 | 0.025607 | 0.017618 | 0.027259 | 0.015847 | 0.053265 | 0.049716 | 0.03483 | 0.036539 |
| 30.0972222 | 0.057091 | 0.064441 | 0.053114 | 0.048918 | 0.03865 | 0.020303 | 0.028278 | 0.057129 | 0.100485 | 0.046692 | 0.046727 | 0.047232 | 0.062928 | 0.035909 | 0.116861 | 0.047243 | 0.084501 | 0.072111 |
| 30.1111111 | 0.086675 | 0.098355 | 0.109047 | 0.075689 | 0.056149 | 0.03092 | 0.056576 | 0.060016 | 0.132506 | 0.073727 | 0.092721 | 0.059955 | 0.079875 | 0.04578 | 0.129548 | 0.063985 | 0.11221 | 0.069084 |
| 30.125 | 0.090248 | 0.098605 | 0.143562 | 0.081625 | 0.048954 | 0.03274 | 0.079758 | 0.03916 | 0.121265 | 0.066653 | 0.114356 | 0.035375 | 0.05267 | 0.029772 | 0.075888 | 0.063132 | 0.076776 | 0.034153 |
| 30.1388889 | 0.074583 | 0.077296 | 0.131949 | 0.06072 | 0.03351 | 0.026764 | 0.073075 | 0.015632 | 0.088669 | 0.033465 | 0.098087 | 0.024256 | 0.020555 | 0.011121 | 0.022292 | 0.044022 | 0.03089 | 0.013097 |
| 30.1527778 | 0.057187 | 0.060442 | 0.094759 | 0.030837 | 0.01985 | 0.016285 | 0.039983 | 0.003398 | 0.062128 | 0.011941 | 0.071005 | 0.03748 | 0.011175 | 0.005729 | 0.005485 | 0.022436 | 0.008068 | 0.012839 |
| 30.1666667 | 0.038083 | 0.045609 | 0.052292 | 0.01485 | 0.008595 | 0.00769 | 0.020353 | 0.002164 | 0.047133 | 0.015868 | 0.040514 | 0.050663 | 0.011512 | 0.00733 | 0.005703 | 0.011854 | 0.004787 | 0.013972 |
| 30.1805556 | 0.02057 | 0.025438 | 0.020075 | 0.017101 | 0.004346 | 0.005598 | 0.035259 | 0.004211 | 0.034999 | 0.022574 | 0.0162 | 0.058705 | 0.007903 | 0.008975 | 0.005579 | 0.010711 | 0.009357 | 0.011986 |
| 30.1944444 | 0.022332 | 0.017589 | 0.010447 | 0.030545 | 0.008298 | 0.010153 | 0.055091 | 0.004024 | 0.019622 | 0.022144 | 0.013698 | 0.054167 | 0.003477 | 0.010316 | 0.00403 | 0.009726 | 0.009784 | 0.011283 |
| 30.2083333 | 0.041537 | 0.030989 | 0.012022 | 0.043161 | 0.015248 | 0.015723 | 0.061347 | 0.002498 | 0.01055 | 0.02001 | 0.021884 | 0.0397 | 0.002424 | 0.011744 | 0.004642 | 0.013058 | 0.00971 | 0.013205 |
| 30.2222222 | 0.055624 | 0.045972 | 0.014529 | 0.04691 | 0.01874 | 0.017811 | 0.060267 | 0.002253 | 0.016819 | 0.019011 | 0.027871 | 0.028117 | 0.003878 | 0.013748 | 0.008021 | 0.02234 | 0.017484 | 0.016429 |
| 30.2361111 | 0.05595 | 0.049259 | 0.018509 | 0.040385 | 0.019408 | 0.016304 | 0.05233 | 0.00211 | 0.023578 | 0.01989 | 0.034099 | 0.017515 | 0.005521 | 0.015577 | 0.011208 | 0.030918 | 0.0237 | 0.017103 |
| 30.25 | 0.056347 | 0.045323 | 0.023663 | 0.027237 | 0.022324 | 0.015714 | 0.037993 | 0.001905 | 0.022227 | 0.01952 | 0.044537 | 0.011933 | 0.00707 | 0.014117 | 0.012939 | 0.033759 | 0.022495 | 0.013176 |
| 30.2638889 | 0.062057 | 0.033547 | 0.030538 | 0.016566 | 0.025776 | 0.020947 | 0.020817 | 0.005558 | 0.027059 | 0.01736 | 0.055888 | 0.010509 | 0.01084 | 0.009067 | 0.017104 | 0.033427 | 0.023749 | 0.008706 |
| 30.2777778 | 0.062594 | 0.015355 | 0.040651 | 0.013469 | 0.026102 | 0.028072 | 0.007155 | 0.014415 | 0.040895 | 0.015374 | 0.066011 | 0.00653 | 0.01775 | 0.004638 | 0.029644 | 0.032503 | 0.034421 | 0.008781 |
| 30.2916667 | 0.060814 | 0.006476 | 0.055358 | 0.013642 | 0.023147 | 0.029043 | 0.003126 | 0.023754 | 0.050393 | 0.010286 | 0.074529 | 0.003885 | 0.023572 | 0.003349 | 0.047088 | 0.027307 | 0.047721 | 0.010505 |
| 30.3055556 | 0.060921 | 0.014653 | 0.072443 | 0.0184 | 0.018672 | 0.02481 | 0.006444 | 0.030812 | 0.050232 | 0.006079 | 0.073511 | 0.002765 | 0.025031 | 0.00678 | 0.062682 | 0.016298 | 0.044888 | 0.00766 |
| 30.3194444 | 0.056937 | 0.030486 | 0.084712 | 0.029612 | 0.014885 | 0.023486 | 0.013811 | 0.0357 | 0.044175 | 0.00534 | 0.060499 | 0.002366 | 0.031562 | 0.011532 | 0.073315 | 0.00739 | 0.028559 | 0.003358 |
| 30.3333333 | 0.048684 | 0.043732 | 0.086344 | 0.036306 | 0.011632 | 0.025166 | 0.023688 | 0.037082 | 0.038713 | 0.005291 | 0.045874 | 0.005249 | 0.046244 | 0.010217 | 0.074661 | 0.008253 | 0.024867 | 0.004048 |
| 30.3472222 | 0.035452 | 0.05085 | 0.077539 | 0.032127 | 0.009242 | 0.02432 | 0.030345 | 0.034816 | 0.039876 | 0.009767 | 0.031278 | 0.007996 | 0.042649 | 0.007022 | 0.058966 | 0.019113 | 0.036119 | 0.013792 |
| 30.3611111 | 0.018571 | 0.052397 | 0.064059 | 0.020913 | 0.008967 | 0.021607 | 0.029435 | 0.029623 | 0.051424 | 0.031407 | 0.024655 | 0.019455 | 0.03713 | 0.005581 | 0.041239 | 0.045238 | 0.05737 | 0.045754 |
| 30.375 | 0.007797 | 0.048794 | 0.055162 | 0.010625 | 0.011402 | 0.019768 | 0.026002 | 0.023309 | 0.069799 | 0.081216 | 0.052702 | 0.06097 | 0.081099 | 0.025205 | 0.062462 | 0.099327 | 0.097373 | 0.105262 |
| 30.3888889 | 0.007116 | 0.039839 | 0.052021 | 0.009699 | 0.016288 | 0.018703 | 0.028352 | 0.021185 | 0.079373 | 0.135759 | 0.111471 | 0.119135 | 0.146232 | 0.082164 | 0.118824 | 0.163283 | 0.145098 | 0.165396 |
| 30.4027778 | 0.014288 | 0.030059 | 0.04795 | 0.016987 | 0.021832 | 0.01798 | 0.035317 | 0.026725 | 0.077074 | 0.166138 | 0.164091 | 0.161756 | 0.181546 | 0.135471 | 0.163936 | 0.190219 | 0.176184 | 0.19718 |
| 30.4166667 | 0.023977 | 0.024915 | 0.038969 | 0.02308 | 0.026317 | 0.01707 | 0.03711 | 0.034054 | 0.071501 | 0.177333 | 0.19032 | 0.175619 | 0.189245 | 0.155259 | 0.18194 | 0.175914 | 0.185113 | 0.202955 |
| 30.4305556 | 0.027134 | 0.023099 | 0.023693 | 0.022447 | 0.028054 | 0.013189 | 0.027142 | 0.031928 | 0.053012 | 0.181786 | 0.198343 | 0.16371 | 0.183502 | 0.156223 | 0.185087 | 0.15465 | 0.17951 | 0.196476 |
| 30.4444444 | 0.021007 | 0.019333 | 0.01384 | 0.017241 | 0.025192 | 0.006523 | 0.012479 | 0.018979 | 0.026166 | 0.180871 | 0.198684 | 0.138001 | 0.169895 | 0.148732 | 0.179504 | 0.141052 | 0.165665 | 0.180339 |
| 30.4583333 | 0.013218 | 0.011842 | 0.01549 | 0.012482 | 0.018354 | 0.002314 | 0.003846 | 0.009298 | 0.012759 | 0.162554 | 0.180328 | 0.109052 | 0.148868 | 0.128922 | 0.158726 | 0.125975 | 0.144041 | 0.151786 |
| 30.4722222 | 0.014931 | 0.004962 | 0.013646 | 0.012831 | 0.010563 | 0.002734 | 0.003445 | 0.007657 | 0.010367 | 0.114559 | 0.128535 | 0.079784 | 0.112887 | 0.091462 | 0.117375 | 0.098829 | 0.110716 | 0.111888 |
| 30.4861111 | 0.029174 | 0.003344 | 0.006731 | 0.01987 | 0.005364 | 0.005893 | 0.009315 | 0.005366 | 0.015224 | 0.0531 | 0.06417 | 0.048975 | 0.063819 | 0.045574 | 0.066483 | 0.058786 | 0.069595 | 0.06859 |
| 30.5 | 0.047259 | 0.007251 | 0.004875 | 0.030578 | 0.004136 | 0.009173 | 0.016269 | 0.003099 | 0.028814 | 0.0142 | 0.036445 | 0.021756 | 0.024243 | 0.020071 | 0.026191 | 0.028815 | 0.032891 | 0.033543 |
| 30.5138889 | 0.057651 | 0.012841 | 0.006678 | 0.038943 | 0.004659 | 0.00977 | 0.018701 | 0.002433 | 0.038433 | 0.005694 | 0.04704 | 0.009847 | 0.007978 | 0.023505 | 0.009864 | 0.034802 | 0.011294 | 0.012722 |
| 30.5277778 | 0.055061 | 0.016126 | 0.006043 | 0.040091 | 0.003934 | 0.006808 | 0.016092 | 0.002974 | 0.030448 | 0.010558 | 0.044469 | 0.013782 | 0.006238 | 0.027679 | 0.012409 | 0.062695 | 0.005772 | 0.003699 |
| 30.5416667 | 0.043702 | 0.01763 | 0.003614 | 0.035153 | 0.003181 | 0.003112 | 0.011518 | 0.006923 | 0.017548 | 0.020993 | 0.024737 | 0.022877 | 0.007441 | 0.021248 | 0.017298 | 0.081114 | 0.005989 | 0.00146 |
| 30.5555556 | 0.032185 | 0.020942 | 0.003432 | 0.029644 | 0.00508 | 0.001468 | 0.00837 | 0.010197 | 0.021861 | 0.025747 | 0.019506 | 0.029309 | 0.009581 | 0.014172 | 0.017276 | 0.07164 | 0.004089 | 0.002012 |
| 30.5694444 | 0.025959 | 0.0262 | 0.006764 | 0.027609 | 0.006223 | 0.00123 | 0.008616 | 0.008228 | 0.030189 | 0.018605 | 0.020933 | 0.025852 | 0.009261 | 0.010648 | 0.013113 | 0.039013 | 0.002102 | 0.003473 |
| 30.5833333 | 0.028776 | 0.03051 | 0.015887 | 0.033024 | 0.012138 | 0.001068 | 0.013969 | 0.006212 | 0.024637 | 0.009226 | 0.01788 | 0.016548 | 0.007771 | 0.015393 | 0.008645 | 0.02061 | 0.008646 | 0.003998 |
| 30.5972222 | 0.044475 | 0.033418 | 0.03535 | 0.049478 | 0.035332 | 0.002376 | 0.027118 | 0.013185 | 0.023518 | 0.011044 | 0.031633 | 0.01481 | 0.008986 | 0.031354 | 0.017702 | 0.026207 | 0.029493 | 0.010758 |
| 30.6111111 | 0.067616 | 0.03682 | 0.067186 | 0.069933 | 0.062356 | 0.006974 | 0.047249 | 0.02832 | 0.039421 | 0.034103 | 0.078381 | 0.014656 | 0.021322 | 0.05251 | 0.050765 | 0.031662 | 0.050814 | 0.03116 |
| 30.625 | 0.082042 | 0.044044 | 0.099919 | 0.079464 | 0.070419 | 0.010434 | 0.06292 | 0.046065 | 0.055234 | 0.067527 | 0.129623 | 0.009166 | 0.051133 | 0.070821 | 0.086504 | 0.059225 | 0.053506 | 0.052039 |
| 30.6388889 | 0.079435 | 0.051385 | 0.109036 | 0.070047 | 0.063355 | 0.011553 | 0.061185 | 0.056623 | 0.059874 | 0.081209 | 0.133847 | 0.0084 | 0.069771 | 0.072454 | 0.089866 | 0.097211 | 0.038845 | 0.056016 |
| 30.6527778 | 0.061215 | 0.049282 | 0.087824 | 0.048741 | 0.055385 | 0.014853 | 0.044438 | 0.052044 | 0.055052 | 0.058702 | 0.084455 | 0.015304 | 0.057717 | 0.047868 | 0.062597 | 0.080724 | 0.020957 | 0.039652 |
| 30.6666667 | 0.03396 | 0.039738 | 0.054173 | 0.028111 | 0.051243 | 0.017685 | 0.023137 | 0.042393 | 0.047806 | 0.024815 | 0.035647 | 0.02421 | 0.032114 | 0.02149 | 0.030577 | 0.031566 | 0.01168 | 0.019604 |
| 30.6805556 | 0.02043 | 0.030975 | 0.024318 | 0.012549 | 0.051013 | 0.017859 | 0.009984 | 0.034643 | 0.03936 | 0.011007 | 0.031775 | 0.032749 | 0.020336 | 0.022426 | 0.018834 | 0.006118 | 0.014586 | 0.017838 |
| 30.6944444 | 0.025455 | 0.025217 | 0.011887 | 0.004074 | 0.050168 | 0.017016 | 0.010747 | 0.023998 | 0.025691 | 0.012673 | 0.058478 | 0.040473 | 0.027878 | 0.040226 | 0.038522 | 0.001906 | 0.027324 | 0.033809 |
| 30.7083333 | 0.030226 | 0.020372 | 0.022221 | 0.002302 | 0.043808 | 0.014122 | 0.011846 | 0.012484 | 0.019732 | 0.012158 | 0.090703 | 0.044661 | 0.040193 | 0.048744 | 0.064829 | 0.02233 | 0.039163 | 0.04232 |
| 30.7222222 | 0.032762 | 0.013151 | 0.04054 | 0.002911 | 0.031383 | 0.008182 | 0.006329 | 0.010935 | 0.026396 | 0.011646 | 0.126673 | 0.044138 | 0.061458 | 0.046711 | 0.078213 | 0.069041 | 0.037124 | 0.029387 |
| 30.7361111 | 0.032998 | 0.010919 | 0.050515 | 0.005581 | 0.016799 | 0.005147 | 0.002032 | 0.021081 | 0.031795 | 0.016849 | 0.154847 | 0.042293 | 0.079788 | 0.047812 | 0.085965 | 0.088955 | 0.031154 | 0.011789 |
| 30.75 | 0.027005 | 0.020219 | 0.054484 | 0.009157 | 0.012278 | 0.009972 | 0.001287 | 0.030158 | 0.034381 | 0.019002 | 0.143381 | 0.040171 | 0.065468 | 0.049648 | 0.077406 | 0.057828 | 0.037817 | 0.00511 |

| | | | | | | | | | | | | | | | | | | |
|---|---|---|---|---|---|---|---|---|---|---|---|---|---|---|---|---|---|---|
| 30.7638889 | 0.025511 | 0.028604 | 0.058758 | 0.01016 | 0.023028 | 0.015253 | 0.002158 | 0.032332 | 0.035024 | 0.014108 | 0.088978 | 0.033938 | 0.03439 | 0.042671 | 0.046506 | 0.022732 | 0.048112 | 0.010076 |
| 30.7777778 | 0.031075 | 0.027129 | 0.045297 | 0.007142 | 0.035073 | 0.015904 | 0.004167 | 0.031138 | 0.025036 | 0.008686 | 0.041901 | 0.02202 | 0.028102 | 0.031839 | 0.04062 | 0.018892 | 0.048998 | 0.024581 |
| 30.7916667 | 0.031591 | 0.018607 | 0.023399 | 0.005155 | 0.039457 | 0.016628 | 0.005328 | 0.030716 | 0.017057 | 0.008306 | 0.039578 | 0.009402 | 0.051125 | 0.021594 | 0.080358 | 0.035304 | 0.047515 | 0.047903 |
| 30.8055556 | 0.027784 | 0.009136 | 0.027636 | 0.011007 | 0.043446 | 0.017376 | 0.003816 | 0.034972 | 0.027334 | 0.011419 | 0.060098 | 0.003194 | 0.077966 | 0.011275 | 0.113823 | 0.050382 | 0.04522 | 0.066036 |
| 30.8194444 | 0.022012 | 0.006404 | 0.039211 | 0.020909 | 0.049357 | 0.011974 | 0.002509 | 0.040971 | 0.04405 | 0.010488 | 0.074469 | 0.002751 | 0.088212 | 0.009212 | 0.106626 | 0.058914 | 0.035644 | 0.061433 |
| 30.8333333 | 0.012781 | 0.011467 | 0.030078 | 0.029858 | 0.04818 | 0.004098 | 0.004688 | 0.039633 | 0.048576 | 0.007183 | 0.07383 | 0.002798 | 0.077043 | 0.018428 | 0.080592 | 0.072394 | 0.024184 | 0.040587 |
| 30.8472222 | 0.00736 | 0.018325 | 0.012444 | 0.035064 | 0.036083 | 0.001243 | 0.007496 | 0.030171 | 0.034332 | 0.013613 | 0.0563 | 0.003934 | 0.055339 | 0.02915 | 0.056983 | 0.080602 | 0.0144 | 0.020596 |
| 30.8611111 | 0.007368 | 0.021963 | 0.007154 | 0.031747 | 0.019454 | 0.001429 | 0.007143 | 0.021536 | 0.015103 | 0.023745 | 0.030157 | 0.004521 | 0.030868 | 0.02659 | 0.034695 | 0.060682 | 0.010148 | 0.016397 |
| 30.875 | 0.006783 | 0.021722 | 0.012647 | 0.019835 | 0.009158 | 0.001711 | 0.005344 | 0.016531 | 0.008669 | 0.022523 | 0.019989 | 0.0056 | 0.018747 | 0.016699 | 0.031623 | 0.030157 | 0.012635 | 0.029723 |
| 30.8888889 | 0.005565 | 0.018597 | 0.016514 | 0.010927 | 0.011479 | 0.002992 | 0.007264 | 0.012019 | 0.018651 | 0.013238 | 0.048101 | 0.014482 | 0.035961 | 0.020084 | 0.056649 | 0.01522 | 0.010892 | 0.042764 |
| 30.9027778 | 0.007327 | 0.014626 | 0.017984 | 0.015471 | 0.017886 | 0.008241 | 0.014452 | 0.007506 | 0.034803 | 0.006618 | 0.091282 | 0.026341 | 0.068055 | 0.026891 | 0.093589 | 0.008887 | 0.004299 | 0.037006 |
| 30.9166667 | 0.013371 | 0.014733 | 0.026875 | 0.026271 | 0.017445 | 0.016042 | 0.020726 | 0.004792 | 0.042633 | 0.011137 | 0.122226 | 0.031928 | 0.098134 | 0.021065 | 0.135296 | 0.003543 | 0.003624 | 0.019277 |
| 30.9305556 | 0.02002 | 0.023073 | 0.053621 | 0.035802 | 0.011665 | 0.017059 | 0.023882 | 0.003672 | 0.039011 | 0.026258 | 0.133445 | 0.028929 | 0.110459 | 0.012697 | 0.151394 | 0.011118 | 0.010966 | 0.010946 |
| 30.9444444 | 0.023355 | 0.036993 | 0.087748 | 0.042432 | 0.012714 | 0.009223 | 0.02899 | 0.002779 | 0.031296 | 0.043735 | 0.111033 | 0.017433 | 0.09632 | 0.011962 | 0.126789 | 0.048024 | 0.020772 | 0.008701 |
| 30.9583333 | 0.023467 | 0.046563 | 0.106071 | 0.044121 | 0.026674 | 0.002262 | 0.034759 | 0.003344 | 0.026868 | 0.058129 | 0.070504 | 0.008686 | 0.072652 | 0.01312 | 0.094988 | 0.097949 | 0.030521 | 0.00561 |
| 30.9722222 | 0.018855 | 0.044218 | 0.1035 | 0.041932 | 0.04288 | 0.001867 | 0.036734 | 0.006681 | 0.027153 | 0.062677 | 0.040369 | 0.011086 | 0.048885 | 0.009322 | 0.068644 | 0.119103 | 0.038302 | 0.006853 |
| 30.9861111 | 0.011882 | 0.032889 | 0.089465 | 0.037036 | 0.050747 | 0.006209 | 0.032629 | 0.013427 | 0.026789 | 0.054619 | 0.023884 | 0.012748 | 0.027248 | 0.006076 | 0.042875 | 0.118885 | 0.043142 | 0.007887 |
| 31 | 0.01361 | 0.01876 | 0.068907 | 0.026141 | 0.048804 | 0.010893 | 0.019829 | 0.020776 | 0.020823 | 0.038682 | 0.015452 | 0.008072 | 0.013736 | 0.009947 | 0.01978 | 0.126623 | 0.046364 | 0.005562 |
| 31.0138889 | 0.03026 | 0.008774 | 0.041517 | 0.014033 | 0.038444 | 0.011913 | 0.012334 | 0.022505 | 0.012364 | 0.021264 | 0.011149 | 0.008208 | 0.0144 | 0.011352 | 0.008107 | 0.121909 | 0.039453 | 0.006578 |
| 31.0277778 | 0.053701 | 0.006206 | 0.02195 | 0.014224 | 0.023098 | 0.009999 | 0.026323 | 0.017745 | 0.011103 | 0.016049 | 0.009747 | 0.019049 | 0.030434 | 0.008945 | 0.01736 | 0.079783 | 0.023977 | 0.014816 |
| 31.0416667 | 0.071142 | 0.007039 | 0.029252 | 0.021749 | 0.011225 | 0.007926 | 0.047445 | 0.01303 | 0.019494 | 0.026848 | 0.01877 | 0.033212 | 0.055504 | 0.009713 | 0.045816 | 0.037822 | 0.021763 | 0.025851 |
| 31.0555556 | 0.07491 | 0.009758 | 0.051142 | 0.024316 | 0.010664 | 0.006631 | 0.05661 | 0.013006 | 0.02729 | 0.036693 | 0.042304 | 0.043904 | 0.081936 | 0.00945 | 0.084901 | 0.039941 | 0.032646 | 0.03188 |
| 31.0694444 | 0.060596 | 0.010939 | 0.062447 | 0.022717 | 0.013022 | 0.008476 | 0.050485 | 0.015038 | 0.02575 | 0.032231 | 0.057565 | 0.045986 | 0.090618 | 0.008935 | 0.104957 | 0.057694 | 0.033251 | 0.024305 |
| 31.0833333 | 0.042795 | 0.011092 | 0.046837 | 0.018795 | 0.015841 | 0.016949 | 0.032143 | 0.019862 | 0.019927 | 0.024128 | 0.051136 | 0.032261 | 0.062265 | 0.020571 | 0.08362 | 0.051132 | 0.032251 | 0.022063 |
| 31.0972222 | 0.055513 | 0.018387 | 0.033006 | 0.023339 | 0.033125 | 0.026684 | 0.025185 | 0.026824 | 0.027197 | 0.033409 | 0.069988 | 0.019418 | 0.032444 | 0.052257 | 0.079014 | 0.053316 | 0.062028 | 0.050736 |
| 31.1111111 | 0.08647 | 0.026387 | 0.059174 | 0.041948 | 0.05316 | 0.024745 | 0.050566 | 0.02341 | 0.045089 | 0.04945 | 0.125222 | 0.021367 | 0.040171 | 0.076217 | 0.116296 | 0.08564 | 0.093444 | 0.079531 |
| 31.125 | 0.091028 | 0.027582 | 0.088771 | 0.052959 | 0.053501 | 0.012781 | 0.087438 | 0.013335 | 0.053747 | 0.047545 | 0.150193 | 0.018952 | 0.053831 | 0.069165 | 0.119398 | 0.095946 | 0.07615 | 0.077159 |
| 31.1388889 | 0.073075 | 0.026699 | 0.088007 | 0.048469 | 0.038189 | 0.006612 | 0.10531 | 0.011239 | 0.053128 | 0.032258 | 0.130886 | 0.010264 | 0.056839 | 0.052135 | 0.083042 | 0.072224 | 0.037201 | 0.061238 |
| 31.1527778 | 0.056391 | 0.026962 | 0.071253 | 0.03871 | 0.021872 | 0.008133 | 0.089953 | 0.012405 | 0.048886 | 0.017563 | 0.097684 | 0.007853 | 0.062508 | 0.043017 | 0.058974 | 0.049549 | 0.021961 | 0.053369 |
| 31.1666667 | 0.04811 | 0.032449 | 0.052733 | 0.025851 | 0.011857 | 0.009401 | 0.049355 | 0.011371 | 0.038531 | 0.01087 | 0.065446 | 0.014019 | 0.067763 | 0.041941 | 0.058839 | 0.043791 | 0.033729 | 0.057143 |
| 31.1805556 | 0.04421 | 0.036792 | 0.034514 | 0.012746 | 0.006157 | 0.011307 | 0.028255 | 0.010957 | 0.023771 | 0.019115 | 0.04844 | 0.031872 | 0.079573 | 0.047564 | 0.078521 | 0.067106 | 0.064809 | 0.072531 |
| 31.1944444 | 0.029564 | 0.025625 | 0.021702 | 0.010339 | 0.003766 | 0.017561 | 0.05704 | 0.012749 | 0.012802 | 0.033349 | 0.040822 | 0.052583 | 0.096519 | 0.047948 | 0.097728 | 0.112973 | 0.099209 | 0.09089 |
| 31.2083333 | 0.017251 | 0.014453 | 0.024682 | 0.023808 | 0.006635 | 0.025791 | 0.101528 | 0.01424 | 0.014596 | 0.029627 | 0.031166 | 0.054893 | 0.089315 | 0.037958 | 0.088766 | 0.134191 | 0.105318 | 0.086649 |
| 31.2222222 | 0.028791 | 0.023869 | 0.033384 | 0.036998 | 0.009954 | 0.028405 | 0.12028 | 0.010695 | 0.024166 | 0.017842 | 0.034022 | 0.033951 | 0.049902 | 0.041596 | 0.05143 | 0.102583 | 0.071607 | 0.050558 |
| 31.2361111 | 0.042027 | 0.036503 | 0.02918 | 0.035684 | 0.010071 | 0.022121 | 0.107926 | 0.004513 | 0.026175 | 0.024219 | 0.04931 | 0.014257 | 0.019564 | 0.059335 | 0.021677 | 0.053564 | 0.031778 | 0.019794 |
| 31.25 | 0.038031 | 0.036495 | 0.015345 | 0.026284 | 0.008311 | 0.011456 | 0.076947 | 0.001247 | 0.016524 | 0.033874 | 0.056068 | 0.015556 | 0.023734 | 0.059599 | 0.023606 | 0.02427 | 0.019514 | 0.022526 |
| 31.2638889 | 0.028257 | 0.034657 | 0.007178 | 0.018344 | 0.005036 | 0.004218 | 0.044648 | 0.001653 | 0.007305 | 0.03038 | 0.054188 | 0.028029 | 0.038022 | 0.041327 | 0.036945 | 0.012381 | 0.023805 | 0.033962 |
| 31.2777778 | 0.02522 | 0.041087 | 0.005992 | 0.014013 | 0.001779 | 0.002566 | 0.022515 | 0.004571 | 0.007658 | 0.024219 | 0.049272 | 0.038736 | 0.046948 | 0.027296 | 0.04352 | 0.010198 | 0.02853 | 0.03558 |
| 31.2916667 | 0.030427 | 0.049774 | 0.010241 | 0.013198 | 0.000958 | 0.002673 | 0.012581 | 0.006967 | 0.009052 | 0.028136 | 0.043517 | 0.04455 | 0.054281 | 0.028897 | 0.046482 | 0.015729 | 0.033397 | 0.032856 |
| 31.3055556 | 0.040243 | 0.054329 | 0.022984 | 0.014708 | 0.002668 | 0.004183 | 0.01157 | 0.005595 | 0.006564 | 0.041336 | 0.046818 | 0.044077 | 0.058808 | 0.041987 | 0.048557 | 0.023232 | 0.032645 | 0.029 |
| 31.3194444 | 0.053926 | 0.055889 | 0.037136 | 0.016752 | 0.005828 | 0.006351 | 0.016044 | 0.00274 | 0.008818 | 0.049905 | 0.060052 | 0.037673 | 0.057095 | 0.054203 | 0.048039 | 0.028517 | 0.02357 | 0.024617 |
| 31.3333333 | 0.067541 | 0.052025 | 0.043262 | 0.017908 | 0.00875 | 0.007472 | 0.022228 | 0.002416 | 0.017174 | 0.045315 | 0.066667 | 0.027461 | 0.048492 | 0.054405 | 0.042014 | 0.029144 | 0.011814 | 0.018433 |
| 31.3472222 | 0.073477 | 0.041829 | 0.039141 | 0.016407 | 0.009439 | 0.007592 | 0.024081 | 0.004358 | 0.021525 | 0.030482 | 0.052471 | 0.017119 | 0.03769 | 0.039652 | 0.031668 | 0.025269 | 0.003718 | 0.010157 |
| 31.3611111 | 0.069221 | 0.026813 | 0.027891 | 0.01246 | 0.008109 | 0.0068 | 0.018605 | 0.00545 | 0.016065 | 0.01362 | 0.026628 | 0.00965 | 0.030294 | 0.018724 | 0.02155 | 0.024361 | 0.004069 | 0.003575 |
| 31.375 | 0.055396 | 0.013805 | 0.017218 | 0.00805 | 0.0065 | 0.005166 | 0.009583 | 0.004052 | 0.007277 | 0.003088 | 0.009797 | 0.005755 | 0.027952 | 0.00503 | 0.015604 | 0.031535 | 0.011626 | 0.003391 |
| 31.3888889 | 0.033906 | 0.012073 | 0.012708 | 0.004123 | 0.005366 | 0.003411 | 0.003364 | 0.002297 | 0.00346 | 0.002018 | 0.009506 | 0.004203 | 0.027597 | 0.001367 | 0.014725 | 0.038994 | 0.018435 | 0.007975 |
| 31.4027778 | 0.014597 | 0.013601 | 0.013708 | 0.001906 | 0.004017 | 0.004215 | 0.001839 | 0.002915 | 0.003388 | 0.006997 | 0.018548 | 0.002754 | 0.027415 | 0.001589 | 0.016542 | 0.041649 | 0.018851 | 0.01138 |
| 31.4166667 | 0.007762 | 0.009712 | 0.018363 | 0.001776 | 0.002522 | 0.008402 | 0.002267 | 0.005396 | 0.002622 | 0.013814 | 0.02746 | 0.001141 | 0.029172 | 0.002514 | 0.019458 | 0.046876 | 0.01384 | 0.010901 |
| 31.4305556 | 0.008582 | 0.009848 | 0.024194 | 0.001848 | 0.001777 | 0.013108 | 0.005209 | 0.007942 | 0.002533 | 0.021459 | 0.035502 | 0.000771 | 0.032041 | 0.002529 | 0.023552 | 0.055018 | 0.008512 | 0.007928 |
| 31.4444444 | 0.008385 | 0.016663 | 0.028603 | 0.001541 | 0.001198 | 0.015483 | 0.009605 | 0.009106 | 0.00518 | 0.027601 | 0.044277 | 0.001761 | 0.032108 | 0.002711 | 0.026821 | 0.056001 | 0.006435 | 0.005477 |
| 31.4583333 | 0.009422 | 0.01653 | 0.030591 | 0.001135 | 0.000902 | 0.013572 | 0.008953 | 0.00818 | 0.00775 | 0.029753 | 0.050826 | 0.002967 | 0.027707 | 0.004795 | 0.026231 | 0.047239 | 0.005972 | 0.004781 |
| 31.4722222 | 0.015754 | 0.010608 | 0.02977 | 0.00207 | 0.002038 | 0.007864 | 0.007497 | 0.006449 | 0.00774 | 0.028691 | 0.052169 | 0.002577 | 0.020769 | 0.005294 | 0.021443 | 0.033697 | 0.004606 | 0.004154 |
| 31.4861111 | 0.02555 | 0.012175 | 0.026841 | 0.005317 | 0.003736 | 0.002697 | 0.012608 | 0.006015 | 0.006277 | 0.024641 | 0.046109 | 0.001 | 0.01431 | 0.002758 | 0.015693 | 0.020051 | 0.0028 | 0.004243 |
| 31.5 | 0.03121 | 0.014468 | 0.022594 | 0.007884 | 0.004532 | 0.002045 | 0.015373 | 0.007493 | 0.004941 | 0.019524 | 0.034771 | 0.001815 | 0.01048 | 0.001048 | 0.012674 | 0.009241 | 0.002737 | 0.007048 |
| 31.5138889 | 0.026961 | 0.008786 | 0.018234 | 0.006085 | 0.004016 | 0.005297 | 0.009575 | 0.009546 | 0.003926 | 0.015528 | 0.024945 | 0.005874 | 0.009283 | 0.003093 | 0.012855 | 0.005189 | 0.003422 | 0.009675 |
| 31.5277778 | 0.019288 | 0.006653 | 0.016502 | 0.004061 | 0.003796 | 0.006287 | 0.003905 | 0.009887 | 0.002523 | 0.0109 | 0.020297 | 0.007175 | 0.007418 | 0.007684 | 0.012452 | 0.012749 | 0.004985 | 0.008232 |
| 31.5416667 | 0.016462 | 0.015421 | 0.018022 | 0.007376 | 0.00648 | 0.004475 | 0.004786 | 0.007565 | 0.000925 | 0.006844 | 0.020383 | 0.006948 | 0.005564 | 0.009777 | 0.009288 | 0.028674 | 0.013104 | 0.009034 |
| 31.5555556 | 0.012983 | 0.019425 | 0.019766 | 0.011527 | 0.011562 | 0.003619 | 0.006259 | 0.004565 | 0.000965 | 0.006296 | 0.023703 | 0.012853 | 0.00832 | 0.007309 | 0.00905 | 0.043153 | 0.02534 | 0.016155 |
| 31.5694444 | 0.012705 | 0.016246 | 0.023006 | 0.014638 | 0.015486 | 0.00542 | 0.006743 | 0.003967 | 0.00515 | 0.011561 | 0.032229 | 0.016325 | 0.009807 | 0.007192 | 0.012856 | 0.043304 | 0.026357 | 0.017218 |
| 31.5833333 | 0.028889 | 0.020781 | 0.039759 | 0.021652 | 0.015392 | 0.012552 | 0.014629 | 0.006979 | 0.015966 | 0.029534 | 0.054796 | 0.016939 | 0.010856 | 0.018806 | 0.035179 | 0.031559 | 0.024014 | 0.022549 |
| 31.5972222 | 0.05177 | 0.023241 | 0.071432 | 0.03437 | 0.011304 | 0.019249 | 0.031655 | 0.010133 | 0.030438 | 0.062533 | 0.090677 | 0.036505 | 0.024618 | 0.043586 | 0.099867 | 0.040393 | 0.040627 | 0.050313 |
| 31.6111111 | 0.07111 | 0.018373 | 0.093337 | 0.0523 | 0.01292 | 0.020983 | 0.057492 | 0.008027 | 0.038625 | 0.09613 | 0.120486 | 0.070724 | 0.04929 | 0.074104 | 0.17677 | 0.077199 | 0.058846 | 0.074145 |
| 31.625 | 0.088225 | 0.019527 | 0.091071 | 0.069381 | 0.025975 | 0.018623 | 0.081585 | 0.004067 | 0.039405 | 0.117115 | 0.137384 | 0.088988 | 0.074982 | 0.10244 | 0.225812 | 0.110311 | 0.062027 | 0.076531 |
| 31.6388889 | 0.097141 | 0.022233 | 0.075866 | 0.074179 | 0.03536 | 0.011188 | 0.079865 | 0.003368 | 0.036648 | 0.117913 | 0.146103 | 0.081453 | 0.091596 | 0.124872 | 0.246055 | 0.12864 | 0.065218 | 0.073092 |
| 31.6527778 | 0.089862 | 0.015923 | 0.058218 | 0.062125 | 0.032919 | 0.004083 | 0.051291 | 0.003239 | 0.028882 | 0.088696 | 0.135197 | 0.059605 | 0.087585 | 0.127872 | 0.230344 | 0.136141 | 0.071283 | 0.066994 |
| 31.6666667 | 0.068308 | 0.007424 | 0.042368 | 0.039764 | 0.030976 | 0.003459 | 0.022061 | 0.004856 | 0.02509 | 0.042837 | 0.101051 | 0.034237 | 0.06252 | 0.100462 | 0.177096 | 0.126546 | 0.068778 | 0.052451 |
| 31.6805556 | 0.042511 | 0.006055 | 0.030914 | 0.018067 | 0.040271 | 0.00736 | 0.013817 | 0.010574 | 0.030367 | 0.014183 | 0.058307 | 0.014145 | 0.030035 | 0.059234 | 0.11369 | 0.099306 | 0.056288 | 0.034441 |
| 31.6944444 | 0.021845 | 0.008212 | 0.020425 | 0.005457 | 0.052821 | 0.0102 | 0.025219 | 0.016777 | 0.030194 | 0.012788 | 0.025401 | 0.004327 | 0.009888 | 0.024685 | 0.060277 | 0.073126 | 0.043801 | 0.024891 |
| 31.7083333 | 0.015988 | 0.008711 | 0.009099 | 0.002858 | 0.053018 | 0.008993 | 0.036798 | 0.019591 | 0.017578 | 0.013678 | 0.01441 | 0.003527 | 0.010692 | 0.010018 | 0.034985 | 0.068172 | 0.036817 | 0.025609 |
| 31.7222222 | 0.026834 | 0.007439 | 0.003793 | 0.004647 | 0.044433 | 0.005917 | 0.037423 | 0.016397 | 0.01176 | 0.009934 | 0.01334 | 0.008129 | 0.02229 | 0.019419 | 0.062215 | 0.069632 | 0.029701 | 0.022853 |
| 31.7361111 | 0.039259 | 0.007228 | 0.004519 | 0.008574 | 0.037918 | 0.003774 | 0.033934 | 0.010486 | 0.028835 | 0.013575 | 0.00769 | 0.011841 | 0.036134 | 0.035685 | 0.121425 | 0.051476 | 0.018134 | 0.014843 |
| 31.75 | 0.041113 | 0.011904 | 0.006578 | 0.011769 | 0.030974 | 0.005001 | 0.033774 | 0.01136 | 0.049709 | 0.017156 | 0.001955 | 0.011952 | 0.049222 | 0.036826 | 0.159531 | 0.02613 | 0.009665 | 0.016804 |
| 31.7638889 | 0.031694 | 0.017198 | 0.009104 | 0.0086 | 0.0194 | 0.010261 | 0.035623 | 0.019788 | 0.052758 | 0.014178 | 0.002333 | 0.010385 | 0.052434 | 0.022895 | 0.157206 | 0.023003 | 0.012538 | 0.030024 |
| 31.7777778 | 0.016873 | 0.016737 | 0.009027 | 0.005156 | 0.011504 | 0.013114 | 0.033717 | 0.026535 | 0.041946 | 0.008787 | 0.011121 | 0.008035 | 0.045201 | 0.009972 | 0.137237 | 0.042366 | 0.021659 | 0.039528 |
| 31.7916667 | 0.013626 | 0.014212 | 0.005151 | 0.009001 | 0.020352 | 0.011279 | 0.026145 | 0.025219 | 0.026383 | 0.005237 | 0.023954 | 0.005579 | 0.038834 | 0.005517 | 0.116052 | 0.058759 | 0.026794 | 0.035455 |
| 31.8055556 | 0.027443 | 0.015082 | 0.00195 | 0.015782 | 0.041546 | 0.009954 | 0.015771 | 0.016638 | 0.016321 | 0.008142 | 0.025856 | 0.004304 | 0.03418 | 0.00617 | 0.085199 | 0.060952 | 0.021785 | 0.021855 |
| 31.8194444 | 0.042419 | 0.018569 | 0.002462 | 0.019371 | 0.057588 | 0.011598 | 0.007907 | 0.007491 | 0.023271 | 0.017879 | 0.017374 | 0.005424 | 0.022337 | 0.006022 | 0.048769 | 0.055498 | 0.011261 | 0.011286 |
| 31.8333333 | 0.051554 | 0.020969 | 0.005779 | 0.019041 | 0.060011 | 0.017708 | 0.007229 | 0.005623 | 0.043149 | 0.0223 | 0.017243 | 0.008183 | 0.013652 | 0.00774 | 0.043062 | 0.044932 | 0.007361 | 0.013875 |
| 31.8472222 | 0.059355 | 0.018477 | 0.011085 | 0.014791 | 0.053103 | 0.02782 | 0.00993 | 0.014194 | 0.057746 | 0.014209 | 0.02518 | 0.009933 | 0.023369 | 0.015922 | 0.077072 | 0.03401 | 0.010947 | 0.025876 |
| 31.8611111 | 0.057455 | 0.012348 | 0.016064 | 0.006826 | 0.046042 | 0.03393 | 0.012677 | 0.024445 | 0.056144 | 0.00731 | 0.026002 | 0.007921 | 0.037116 | 0.021418 | 0.102499 | 0.023086 | 0.013625 | 0.033104 |
| 31.875 | 0.037172 | 0.0113 | 0.014223 | 0.00374 | 0.042797 | 0.033535 | 0.013933 | 0.026159 | 0.0406 | 0.010332 | 0.017978 | 0.005569 | 0.037763 | 0.016878 | 0.084581 | 0.013901 | 0.016681 | 0.028235 |
| 31.8888889 | 0.01646 | 0.014612 | 0.007179 | 0.00995 | 0.042212 | 0.035965 | 0.011834 | 0.02478 | 0.022947 | 0.016684 | 0.016519 | 0.009689 | 0.026499 | 0.010285 | 0.048517 | 0.0143 | 0.023566 | 0.017565 |
| 31.9027778 | 0.018791 | 0.014895 | 0.006615 | 0.019447 | 0.042602 | 0.045114 | 0.016216 | 0.025741 | 0.021188 | 0.022958 | 0.03082 | 0.016461 | 0.013294 | 0.009654 | 0.046933 | 0.015707 | 0.026734 | 0.010283 |

| | | | | | | | | | | | | | | | | | | |
|---|---|---|---|---|---|---|---|---|---|---|---|---|---|---|---|---|---|---|
| 31.9166667 | 0.038067 | 0.020334 | 0.01209 | 0.025512 | 0.040771 | 0.05346 | 0.02917 | 0.024253 | 0.033043 | 0.029465 | 0.047419 | 0.016844 | 0.009936 | 0.014307 | 0.090265 | 0.01269 | 0.02437 | 0.006651 |
| 31.9305556 | 0.054495 | 0.035422 | 0.017479 | 0.025404 | 0.036635 | 0.052916 | 0.039415 | 0.019884 | 0.043929 | 0.040114 | 0.059385 | 0.010633 | 0.017157 | 0.019586 | 0.140331 | 0.010586 | 0.023294 | 0.007297 |
| 31.9444444 | 0.062815 | 0.047633 | 0.025404 | 0.024208 | 0.033678 | 0.04627 | 0.046363 | 0.017727 | 0.052457 | 0.057388 | 0.076607 | 0.004758 | 0.030144 | 0.023211 | 0.166656 | 0.010441 | 0.02289 | 0.019367 |
| 31.9583333 | 0.066312 | 0.051257 | 0.037762 | 0.025319 | 0.031725 | 0.043143 | 0.056915 | 0.019457 | 0.058821 | 0.073129 | 0.102182 | 0.004687 | 0.044107 | 0.02882 | 0.166236 | 0.017121 | 0.018772 | 0.041532 |
| 31.9722222 | 0.065559 | 0.051072 | 0.051202 | 0.026633 | 0.028374 | 0.045929 | 0.068722 | 0.023809 | 0.059822 | 0.07946 | 0.121092 | 0.012082 | 0.051363 | 0.035509 | 0.146112 | 0.030545 | 0.011067 | 0.060738 |
| 31.9861111 | 0.060921 | 0.055333 | 0.058156 | 0.026383 | 0.023123 | 0.047146 | 0.069643 | 0.029322 | 0.053527 | 0.077296 | 0.114311 | 0.021893 | 0.051977 | 0.038364 | 0.11309 | 0.041375 | 0.00651 | 0.067311 |
| 32 | 0.050268 | 0.059377 | 0.051387 | 0.023438 | 0.01655 | 0.040212 | 0.056465 | 0.033222 | 0.044837 | 0.068203 | 0.089164 | 0.023459 | 0.047302 | 0.036552 | 0.077802 | 0.049146 | 0.010872 | 0.063657 |
| 32.0138889 | 0.029674 | 0.047477 | 0.033985 | 0.015358 | 0.009486 | 0.028064 | 0.038647 | 0.030746 | 0.037268 | 0.050062 | 0.064989 | 0.014553 | 0.035353 | 0.028878 | 0.049594 | 0.057413 | 0.018487 | 0.056175 |
| 32.0277778 | 0.013545 | 0.025879 | 0.016427 | 0.00895 | 0.004509 | 0.01547 | 0.021082 | 0.021599 | 0.023861 | 0.027151 | 0.041429 | 0.008818 | 0.021993 | 0.016017 | 0.03062 | 0.059514 | 0.021046 | 0.04625 |
| 32.0416667 | 0.017157 | 0.013659 | 0.009847 | 0.014973 | 0.004536 | 0.006073 | 0.010808 | 0.012452 | 0.013066 | 0.019256 | 0.022703 | 0.016303 | 0.013093 | 0.009421 | 0.01796 | 0.04872 | 0.015725 | 0.031121 |
| 32.0555556 | 0.027927 | 0.018153 | 0.018334 | 0.026505 | 0.007768 | 0.003139 | 0.016601 | 0.007522 | 0.021208 | 0.03415 | 0.024827 | 0.025728 | 0.007119 | 0.018046 | 0.009903 | 0.032518 | 0.00682 | 0.014973 |
| 32.0694444 | 0.028731 | 0.026575 | 0.027601 | 0.028414 | 0.007759 | 0.004598 | 0.02682 | 0.007011 | 0.034752 | 0.045536 | 0.032013 | 0.021141 | 0.005429 | 0.025887 | 0.006989 | 0.031206 | 0.009795 | 0.010644 |
| 32.0833333 | 0.023143 | 0.025144 | 0.028033 | 0.026992 | 0.008408 | 0.012257 | 0.024463 | 0.010457 | 0.032878 | 0.041852 | 0.038298 | 0.021977 | 0.015261 | 0.027838 | 0.018518 | 0.068725 | 0.04835 | 0.034727 |
| 32.0972222 | 0.033636 | 0.033595 | 0.041763 | 0.049541 | 0.020727 | 0.027703 | 0.028337 | 0.015408 | 0.037744 | 0.059273 | 0.078495 | 0.054589 | 0.033043 | 0.049462 | 0.048667 | 0.138402 | 0.116028 | 0.081933 |
| 32.1111111 | 0.063027 | 0.063821 | 0.070278 | 0.085319 | 0.034988 | 0.037936 | 0.054107 | 0.01887 | 0.066998 | 0.105266 | 0.125797 | 0.089742 | 0.03942 | 0.078637 | 0.065068 | 0.17299 | 0.149149 | 0.105088 |
| 32.125 | 0.083847 | 0.087082 | 0.079348 | 0.095183 | 0.03713 | 0.039542 | 0.069834 | 0.019255 | 0.084641 | 0.13272 | 0.125314 | 0.094648 | 0.029032 | 0.078222 | 0.04666 | 0.134147 | 0.122205 | 0.080655 |
| 32.1388889 | 0.082477 | 0.093206 | 0.073166 | 0.080481 | 0.032147 | 0.043809 | 0.06554 | 0.01587 | 0.076994 | 0.119966 | 0.097398 | 0.075959 | 0.014686 | 0.058354 | 0.020328 | 0.077954 | 0.086238 | 0.041163 |
| 32.1527778 | 0.066115 | 0.093413 | 0.070731 | 0.058086 | 0.027314 | 0.046797 | 0.05792 | 0.010526 | 0.058036 | 0.081633 | 0.077277 | 0.045872 | 0.005113 | 0.043916 | 0.008773 | 0.047897 | 0.062425 | 0.017392 |
| 32.1666667 | 0.04211 | 0.083499 | 0.060538 | 0.031306 | 0.021773 | 0.03754 | 0.041876 | 0.006019 | 0.029964 | 0.043344 | 0.06421 | 0.018879 | 0.005189 | 0.036109 | 0.007111 | 0.030764 | 0.038593 | 0.010473 |
| 32.1805556 | 0.021422 | 0.05913 | 0.037932 | 0.012805 | 0.013043 | 0.022202 | 0.01902 | 0.003986 | 0.011896 | 0.020036 | 0.046249 | 0.007957 | 0.009985 | 0.029903 | 0.010243 | 0.015341 | 0.030192 | 0.008744 |
| 32.1944444 | 0.018314 | 0.030086 | 0.016114 | 0.012952 | 0.007871 | 0.020419 | 0.007115 | 0.005852 | 0.0174 | 0.007976 | 0.023264 | 0.005519 | 0.010243 | 0.022672 | 0.016392 | 0.006431 | 0.041582 | 0.006263 |
| 32.2083333 | 0.028781 | 0.010466 | 0.006633 | 0.021003 | 0.012089 | 0.034373 | 0.0106 | 0.009542 | 0.029306 | 0.003364 | 0.011591 | 0.002458 | 0.007147 | 0.013905 | 0.01919 | 0.007812 | 0.047939 | 0.003495 |
| 32.2222222 | 0.037756 | 0.00394 | 0.009526 | 0.027346 | 0.018771 | 0.04556 | 0.01945 | 0.011561 | 0.035501 | 0.005876 | 0.02115 | 0.001809 | 0.006271 | 0.012747 | 0.016192 | 0.015508 | 0.051044 | 0.002265 |
| 32.2361111 | 0.042776 | 0.005682 | 0.014875 | 0.031881 | 0.023005 | 0.048709 | 0.026468 | 0.012785 | 0.038636 | 0.01447 | 0.036216 | 0.005986 | 0.010439 | 0.026327 | 0.009291 | 0.024174 | 0.058861 | 0.004871 |
| 32.25 | 0.048832 | 0.013184 | 0.016935 | 0.032854 | 0.024851 | 0.049435 | 0.028751 | 0.01578 | 0.04144 | 0.027244 | 0.036028 | 0.013075 | 0.018577 | 0.045546 | 0.005782 | 0.029999 | 0.058538 | 0.011339 |
| 32.2638889 | 0.054619 | 0.02408 | 0.017768 | 0.030235 | 0.02386 | 0.046827 | 0.027222 | 0.019458 | 0.047447 | 0.040401 | 0.025061 | 0.017502 | 0.02666 | 0.060102 | 0.011578 | 0.032851 | 0.050176 | 0.019762 |
| 32.2777778 | 0.055058 | 0.032114 | 0.019004 | 0.02801 | 0.020866 | 0.039402 | 0.02663 | 0.021932 | 0.05817 | 0.045393 | 0.02714 | 0.01568 | 0.030155 | 0.067558 | 0.020232 | 0.034935 | 0.042339 | 0.02662 |
| 32.2916667 | 0.051655 | 0.036324 | 0.01979 | 0.02756 | 0.017463 | 0.030025 | 0.028986 | 0.023956 | 0.067395 | 0.043337 | 0.050854 | 0.01341 | 0.027949 | 0.07266 | 0.026434 | 0.037757 | 0.036066 | 0.03049 |
| 32.3055556 | 0.047716 | 0.039336 | 0.019466 | 0.026508 | 0.014694 | 0.022659 | 0.035289 | 0.023468 | 0.070022 | 0.044575 | 0.074278 | 0.019245 | 0.020772 | 0.077424 | 0.030834 | 0.040522 | 0.03092 | 0.033484 |
| 32.3194444 | 0.042191 | 0.040129 | 0.018446 | 0.023054 | 0.012402 | 0.018411 | 0.046457 | 0.018506 | 0.067879 | 0.04862 | 0.075155 | 0.029012 | 0.011069 | 0.075621 | 0.031799 | 0.039995 | 0.026941 | 0.034075 |
| 32.3333333 | 0.036176 | 0.041271 | 0.016553 | 0.016323 | 0.011121 | 0.01619 | 0.057447 | 0.013466 | 0.063128 | 0.047636 | 0.058544 | 0.030456 | 0.007157 | 0.065301 | 0.028682 | 0.035234 | 0.022226 | 0.029961 |
| 32.3472222 | 0.031152 | 0.04594 | 0.016114 | 0.008517 | 0.011456 | 0.014093 | 0.064641 | 0.011639 | 0.058024 | 0.037645 | 0.035352 | 0.020588 | 0.012403 | 0.051405 | 0.023714 | 0.029097 | 0.016043 | 0.022418 |
| 32.3611111 | 0.027302 | 0.050765 | 0.021025 | 0.004062 | 0.012634 | 0.010368 | 0.069589 | 0.01122 | 0.052583 | 0.02271 | 0.015054 | 0.008498 | 0.018579 | 0.039438 | 0.018621 | 0.024471 | 0.010815 | 0.013788 |
| 32.375 | 0.027269 | 0.051991 | 0.028578 | 0.003259 | 0.013473 | 0.00684 | 0.070868 | 0.010221 | 0.044628 | 0.013839 | 0.007345 | 0.003854 | 0.017209 | 0.032762 | 0.01478 | 0.022728 | 0.00998 | 0.008397 |
| 32.3888889 | 0.030726 | 0.048858 | 0.032773 | 0.0048 | 0.013548 | 0.008123 | 0.068826 | 0.008351 | 0.03393 | 0.016548 | 0.006719 | 0.008213 | 0.012258 | 0.033713 | 0.013616 | 0.025171 | 0.015291 | 0.010666 |
| 32.4027778 | 0.035391 | 0.045256 | 0.032657 | 0.007861 | 0.013971 | 0.015135 | 0.067526 | 0.007348 | 0.023417 | 0.024066 | 0.008625 | 0.016043 | 0.014474 | 0.040438 | 0.010764 | 0.032058 | 0.024507 | 0.019634 |
| 32.4166667 | 0.040226 | 0.045992 | 0.030009 | 0.008958 | 0.015581 | 0.022041 | 0.065445 | 0.008611 | 0.017268 | 0.025299 | 0.011033 | 0.018321 | 0.017145 | 0.043147 | 0.012925 | 0.038116 | 0.032303 | 0.027753 |
| 32.4305556 | 0.040574 | 0.048599 | 0.027306 | 0.006555 | 0.015823 | 0.022561 | 0.057941 | 0.00941 | 0.014327 | 0.017182 | 0.009227 | 0.012863 | 0.01314 | 0.030086 | 0.029752 | 0.035121 | 0.032477 | 0.026365 |
| 32.4444444 | 0.034946 | 0.048222 | 0.026564 | 0.003644 | 0.012172 | 0.016947 | 0.044495 | 0.006816 | 0.009803 | 0.011294 | 0.007278 | 0.009264 | 0.012985 | 0.012258 | 0.042225 | 0.023201 | 0.024465 | 0.015901 |
| 32.4583333 | 0.03257 | 0.045974 | 0.027083 | 0.004125 | 0.008258 | 0.011489 | 0.027776 | 0.003665 | 0.004304 | 0.010626 | 0.008757 | 0.00894 | 0.015487 | 0.006691 | 0.036585 | 0.011422 | 0.016486 | 0.00781 |
| 32.4722222 | 0.041006 | 0.046503 | 0.027897 | 0.008985 | 0.00921 | 0.012153 | 0.012444 | 0.005158 | 0.003875 | 0.012976 | 0.01833 | 0.007504 | 0.011313 | 0.00855 | 0.022189 | 0.006192 | 0.014919 | 0.010765 |
| 32.4861111 | 0.052543 | 0.048737 | 0.029635 | 0.014567 | 0.012972 | 0.016971 | 0.003597 | 0.009357 | 0.009518 | 0.024645 | 0.038984 | 0.008187 | 0.00696 | 0.014703 | 0.008867 | 0.007502 | 0.017549 | 0.021171 |
| 32.5 | 0.053746 | 0.047105 | 0.030408 | 0.015749 | 0.013787 | 0.017699 | 0.002329 | 0.009264 | 0.014184 | 0.032874 | 0.056346 | 0.007367 | 0.007658 | 0.018635 | 0.00183 | 0.012535 | 0.01984 | 0.029908 |
| 32.5138889 | 0.043671 | 0.040329 | 0.023526 | 0.010276 | 0.010987 | 0.011719 | 0.008574 | 0.004813 | 0.012892 | 0.03098 | 0.057298 | 0.0034 | 0.008775 | 0.015636 | 0.00201 | 0.019546 | 0.02305 | 0.034812 |
| 32.5277778 | 0.030687 | 0.030481 | 0.012183 | 0.005401 | 0.00746 | 0.009405 | 0.021345 | 0.001886 | 0.007729 | 0.032272 | 0.048425 | 0.006174 | 0.014421 | 0.016836 | 0.010112 | 0.031222 | 0.032389 | 0.042844 |
| 32.5416667 | 0.021444 | 0.021693 | 0.007845 | 0.00736 | 0.004573 | 0.017478 | 0.034185 | 0.002016 | 0.003414 | 0.03614 | 0.041018 | 0.017392 | 0.027257 | 0.025057 | 0.022003 | 0.0459 | 0.044687 | 0.054229 |
| 32.5555556 | 0.020467 | 0.021235 | 0.008545 | 0.009079 | 0.00311 | 0.026578 | 0.040928 | 0.001978 | 0.003117 | 0.030599 | 0.034925 | 0.022547 | 0.033496 | 0.027783 | 0.024538 | 0.050785 | 0.045994 | 0.054739 |
| 32.5694444 | 0.026984 | 0.030565 | 0.008973 | 0.006773 | 0.004084 | 0.027237 | 0.038536 | 0.001781 | 0.003819 | 0.016826 | 0.030014 | 0.015684 | 0.023083 | 0.019385 | 0.015865 | 0.042566 | 0.033386 | 0.039325 |
| 32.5833333 | 0.037412 | 0.043409 | 0.021457 | 0.011074 | 0.008447 | 0.019067 | 0.026344 | 0.00417 | 0.007217 | 0.006479 | 0.039701 | 0.009167 | 0.007911 | 0.013443 | 0.012966 | 0.041061 | 0.02697 | 0.027027 |
| 32.5972222 | 0.05346 | 0.052164 | 0.048897 | 0.029438 | 0.01583 | 0.017678 | 0.025166 | 0.008693 | 0.020669 | 0.01354 | 0.065387 | 0.012654 | 0.010151 | 0.024685 | 0.024006 | 0.056734 | 0.036227 | 0.035013 |
| 32.6111111 | 0.072391 | 0.050772 | 0.066227 | 0.049992 | 0.020835 | 0.03279 | 0.055646 | 0.01453 | 0.037867 | 0.057048 | 0.085565 | 0.030488 | 0.034009 | 0.054714 | 0.031478 | 0.073679 | 0.042887 | 0.048189 |
| 32.625 | 0.084184 | 0.042415 | 0.067211 | 0.068099 | 0.020403 | 0.046219 | 0.095203 | 0.022514 | 0.050269 | 0.11486 | 0.100858 | 0.051379 | 0.055937 | 0.086645 | 0.02346 | 0.075801 | 0.033585 | 0.049254 |
| 32.6388889 | 0.084093 | 0.034118 | 0.068048 | 0.088514 | 0.020538 | 0.047378 | 0.114208 | 0.029969 | 0.056043 | 0.133052 | 0.116635 | 0.048531 | 0.056307 | 0.099275 | 0.025158 | 0.060925 | 0.020462 | 0.050542 |
| 32.6527778 | 0.073924 | 0.027488 | 0.069012 | 0.09716 | 0.024659 | 0.041672 | 0.107995 | 0.031797 | 0.050854 | 0.10224 | 0.110368 | 0.02582 | 0.044704 | 0.084405 | 0.057457 | 0.036439 | 0.014772 | 0.055435 |
| 32.6666667 | 0.057229 | 0.022126 | 0.064395 | 0.081496 | 0.030745 | 0.02928 | 0.086883 | 0.028573 | 0.037276 | 0.05846 | 0.077386 | 0.015175 | 0.036353 | 0.047778 | 0.100634 | 0.019822 | 0.011438 | 0.047988 |
| 32.6805556 | 0.037818 | 0.016135 | 0.056521 | 0.051565 | 0.039225 | 0.013873 | 0.066458 | 0.023703 | 0.025093 | 0.057133 | 0.038813 | 0.025226 | 0.031034 | 0.016 | 0.12904 | 0.023476 | 0.013851 | 0.026936 |
| 32.6944444 | 0.022582 | 0.010033 | 0.047327 | 0.02404 | 0.047599 | 0.00724 | 0.052544 | 0.018348 | 0.01573 | 0.101903 | 0.024069 | 0.045849 | 0.022878 | 0.012193 | 0.136707 | 0.028749 | 0.022768 | 0.012994 |
| 32.7083333 | 0.015808 | 0.006846 | 0.038259 | 0.010723 | 0.049695 | 0.006714 | 0.042264 | 0.014411 | 0.007525 | 0.124538 | 0.043534 | 0.067213 | 0.014011 | 0.02839 | 0.12225 | 0.021258 | 0.031422 | 0.01662 |
| 32.7222222 | 0.012396 | 0.005166 | 0.030119 | 0.012974 | 0.044163 | 0.005839 | 0.033736 | 0.011386 | 0.006921 | 0.107106 | 0.059968 | 0.072852 | 0.008157 | 0.041082 | 0.085416 | 0.013939 | 0.03908 | 0.02631 |
| 32.7361111 | 0.007531 | 0.003584 | 0.021824 | 0.018312 | 0.033857 | 0.007765 | 0.021489 | 0.007299 | 0.017243 | 0.083464 | 0.048578 | 0.055619 | 0.004771 | 0.043902 | 0.045052 | 0.011863 | 0.038709 | 0.030919 |
| 32.75 | 0.004971 | 0.004208 | 0.014366 | 0.018943 | 0.023148 | 0.007253 | 0.016545 | 0.003739 | 0.031276 | 0.05941 | 0.029733 | 0.028224 | 0.004776 | 0.04039 | 0.020308 | 0.010883 | 0.02757 | 0.025076 |
| 32.7638889 | 0.008712 | 0.006398 | 0.008143 | 0.017148 | 0.014492 | 0.008826 | 0.040476 | 0.001776 | 0.044358 | 0.030778 | 0.017478 | 0.00891 | 0.007805 | 0.027232 | 0.013323 | 0.017574 | 0.018931 | 0.015465 |
| 32.7777778 | 0.014441 | 0.007621 | 0.004908 | 0.017337 | 0.007939 | 0.026133 | 0.077322 | 0.000998 | 0.055069 | 0.013236 | 0.010427 | 0.002736 | 0.018027 | 0.013834 | 0.030861 | 0.030037 | 0.020864 | 0.01115 |
| 32.7916667 | 0.017335 | 0.007855 | 0.006807 | 0.019805 | 0.004779 | 0.052504 | 0.099148 | 0.002199 | 0.056315 | 0.020069 | 0.013542 | 0.001096 | 0.033164 | 0.01607 | 0.070435 | 0.032683 | 0.025129 | 0.010662 |
| 32.8055556 | 0.016132 | 0.008117 | 0.011395 | 0.01774 | 0.005626 | 0.067369 | 0.093525 | 0.006656 | 0.040567 | 0.037419 | 0.030711 | 0.002249 | 0.039286 | 0.03588 | 0.106923 | 0.029705 | 0.024861 | 0.012804 |
| 32.8194444 | 0.010524 | 0.009698 | 0.016002 | 0.012051 | 0.007974 | 0.059965 | 0.061777 | 0.012404 | 0.021482 | 0.051087 | 0.056237 | 0.007719 | 0.033348 | 0.059191 | 0.122166 | 0.03243 | 0.021853 | 0.016255 |
| 32.8333333 | 0.005526 | 0.012505 | 0.018884 | 0.014596 | 0.008026 | 0.039618 | 0.026467 | 0.013722 | 0.022606 | 0.057149 | 0.082227 | 0.01095 | 0.025392 | 0.073071 | 0.115015 | 0.034643 | 0.020087 | 0.018911 |
| 32.8472222 | 0.007132 | 0.01371 | 0.018617 | 0.028497 | 0.005608 | 0.019239 | 0.008259 | 0.008953 | 0.035657 | 0.050272 | 0.088863 | 0.008331 | 0.026149 | 0.079796 | 0.094387 | 0.027869 | 0.022283 | 0.021158 |
| 32.8611111 | 0.010654 | 0.013089 | 0.016262 | 0.042046 | 0.007395 | 0.010018 | 0.00672 | 0.005625 | 0.04503 | 0.031122 | 0.061633 | 0.007885 | 0.036992 | 0.069184 | 0.079519 | 0.017931 | 0.026995 | 0.020269 |
| 32.875 | 0.00978 | 0.013477 | 0.013469 | 0.045124 | 0.017898 | 0.01457 | 0.008903 | 0.008142 | 0.053226 | 0.022899 | 0.029401 | 0.01486 | 0.050879 | 0.049796 | 0.078494 | 0.012132 | 0.026871 | 0.012821 |
| 32.8888889 | 0.011024 | 0.013639 | 0.010471 | 0.033878 | 0.031666 | 0.020039 | 0.009276 | 0.009469 | 0.055035 | 0.040908 | 0.026412 | 0.023712 | 0.059337 | 0.063992 | 0.077479 | 0.008482 | 0.017911 | 0.006944 |
| 32.9027778 | 0.020261 | 0.011282 | 0.0063 | 0.022103 | 0.04313 | 0.019696 | 0.009285 | 0.005915 | 0.042372 | 0.063286 | 0.051779 | 0.030124 | 0.057422 | 0.089674 | 0.066022 | 0.007238 | 0.010195 | 0.013273 |
| 32.9166667 | 0.031287 | 0.008713 | 0.004305 | 0.032215 | 0.052376 | 0.012933 | 0.008642 | 0.003272 | 0.022934 | 0.076528 | 0.093634 | 0.032469 | 0.046787 | 0.084926 | 0.045236 | 0.01165 | 0.009366 | 0.024889 |
| 32.9305556 | 0.039521 | 0.00846 | 0.009368 | 0.053317 | 0.058739 | 0.007771 | 0.006275 | 0.005053 | 0.017344 | 0.078595 | 0.136274 | 0.029107 | 0.030801 | 0.069662 | 0.02405 | 0.016842 | 0.008532 | 0.0301 |
| 32.9444444 | 0.047146 | 0.010787 | 0.018198 | 0.063899 | 0.059356 | 0.015257 | 0.00688 | 0.006572 | 0.0292 | 0.06496 | 0.149202 | 0.020514 | 0.014438 | 0.056999 | 0.012945 | 0.017472 | 0.009187 | 0.027835 |
| 32.9583333 | 0.054508 | 0.015307 | 0.025085 | 0.063089 | 0.053125 | 0.027457 | 0.018549 | 0.00643 | 0.042809 | 0.042962 | 0.128788 | 0.010182 | 0.005618 | 0.041294 | 0.012405 | 0.025771 | 0.018017 | 0.019207 |
| 32.9722222 | 0.058783 | 0.020998 | 0.027115 | 0.056192 | 0.040622 | 0.035844 | 0.03596 | 0.010966 | 0.058149 | 0.02571 | 0.10667 | 0.003958 | 0.004077 | 0.036575 | 0.017116 | 0.044373 | 0.030494 | 0.010079 |
| 32.9861111 | 0.056843 | 0.025391 | 0.024214 | 0.045469 | 0.023435 | 0.043431 | 0.04828 | 0.018518 | 0.073264 | 0.022121 | 0.088817 | 0.003397 | 0.003893 | 0.061927 | 0.026404 | 0.052339 | 0.035797 | 0.009762 |
| 33 | 0.046797 | 0.025194 | 0.019108 | 0.032255 | 0.01088 | 0.049268 | 0.049443 | 0.021655 | 0.07963 | 0.028778 | 0.061773 | 0.006262 | 0.00454 | 0.095139 | 0.034207 | 0.040933 | 0.031375 | 0.015585 |
| 33.0138889 | 0.03091 | 0.020102 | 0.014555 | 0.019604 | 0.012771 | 0.046382 | 0.040251 | 0.021 | 0.080291 | 0.030565 | 0.030434 | 0.012607 | 0.006886 | 0.089665 | 0.029882 | 0.022174 | 0.023582 | 0.021167 |
| 33.0277778 | 0.014804 | 0.013504 | 0.009795 | 0.011058 | 0.021933 | 0.031963 | 0.02833 | 0.0179 | 0.068091 | 0.021885 | 0.017249 | 0.021606 | 0.010058 | 0.049571 | 0.017672 | 0.015823 | 0.015485 | 0.024001 |
| 33.0416667 | 0.007324 | 0.007453 | 0.007876 | 0.014347 | 0.028343 | 0.017142 | 0.015804 | 0.011382 | 0.041548 | 0.020606 | 0.035396 | 0.029657 | 0.013433 | 0.02627 | 0.008932 | 0.027576 | 0.016586 | 0.022473 |
| 33.0555556 | 0.012397 | 0.003128 | 0.015062 | 0.032166 | 0.032777 | 0.017537 | 0.011073 | 0.00526 | 0.032809 | 0.036141 | 0.066107 | 0.032133 | 0.014291 | 0.04887 | 0.009292 | 0.043123 | 0.033499 | 0.017399 |

| | | | | | | | | | | | | | | | | | | |
|---|---|---|---|---|---|---|---|---|---|---|---|---|---|---|---|---|---|---|
| 33.0694444 | 0.019861 | 0.002725 | 0.02182 | 0.048901 | 0.034577 | 0.024829 | 0.019545 | 0.003273 | 0.051151 | 0.048603 | 0.082822 | 0.026938 | 0.009609 | 0.079986 | 0.010171 | 0.046498 | 0.04278 | 0.013494 |
| 33.0833333 | 0.017972 | 0.009313 | 0.020254 | 0.04097 | 0.026447 | 0.02525 | 0.024418 | 0.003489 | 0.052538 | 0.038054 | 0.066771 | 0.018632 | 0.008601 | 0.067036 | 0.015579 | 0.034621 | 0.03397 | 0.023636 |
| 33.0972222 | 0.022839 | 0.030995 | 0.026961 | 0.029951 | 0.017851 | 0.03816 | 0.03508 | 0.007217 | 0.040466 | 0.031378 | 0.044436 | 0.020598 | 0.026532 | 0.050415 | 0.045265 | 0.032964 | 0.036564 | 0.050467 |
| 33.1111111 | 0.056555 | 0.070793 | 0.044937 | 0.055168 | 0.027597 | 0.069196 | 0.063218 | 0.017061 | 0.060324 | 0.053908 | 0.060323 | 0.031994 | 0.051325 | 0.094547 | 0.082911 | 0.051783 | 0.059773 | 0.070123 |
| 33.125 | 0.10074 | 0.108545 | 0.052098 | 0.086799 | 0.046412 | 0.085957 | 0.074018 | 0.025438 | 0.090707 | 0.07644 | 0.094337 | 0.031371 | 0.065403 | 0.146293 | 0.102008 | 0.065756 | 0.079475 | 0.069978 |
| 33.1388889 | 0.121296 | 0.123266 | 0.045929 | 0.087674 | 0.057743 | 0.082409 | 0.058519 | 0.026529 | 0.105092 | 0.085363 | 0.121442 | 0.024398 | 0.068217 | 0.153706 | 0.10156 | 0.068718 | 0.092891 | 0.065261 |
| 33.1527778 | 0.101118 | 0.118747 | 0.037874 | 0.067554 | 0.055768 | 0.075257 | 0.042317 | 0.023556 | 0.110537 | 0.078885 | 0.135831 | 0.023098 | 0.055043 | 0.134289 | 0.090333 | 0.058492 | 0.094379 | 0.061039 |
| 33.1666667 | 0.057405 | 0.101739 | 0.030798 | 0.042706 | 0.041197 | 0.070067 | 0.030617 | 0.020243 | 0.100015 | 0.051075 | 0.121185 | 0.019126 | 0.031543 | 0.095105 | 0.071148 | 0.031294 | 0.06408 | 0.047081 |
| 33.1805556 | 0.023006 | 0.073689 | 0.024957 | 0.021228 | 0.022017 | 0.06711 | 0.019667 | 0.016446 | 0.076521 | 0.021625 | 0.090794 | 0.018362 | 0.014771 | 0.048957 | 0.043449 | 0.012609 | 0.024813 | 0.027611 |
| 33.1944444 | 0.011348 | 0.03961 | 0.018978 | 0.010056 | 0.010059 | 0.06164 | 0.009718 | 0.009695 | 0.052953 | 0.009183 | 0.06577 | 0.023131 | 0.013758 | 0.017193 | 0.01979 | 0.011946 | 0.006708 | 0.02172 |
| 33.2083333 | 0.016373 | 0.015017 | 0.011705 | 0.013527 | 0.012622 | 0.050504 | 0.004199 | 0.003153 | 0.034086 | 0.011788 | 0.041923 | 0.027037 | 0.021552 | 0.004036 | 0.011117 | 0.012543 | 0.005207 | 0.027667 |
| 33.2222222 | 0.021993 | 0.011031 | 0.005476 | 0.024156 | 0.020811 | 0.035161 | 0.006819 | 0.000994 | 0.022395 | 0.019162 | 0.018974 | 0.032065 | 0.031425 | 0.003067 | 0.010304 | 0.011429 | 0.010988 | 0.029492 |
| 33.2361111 | 0.024527 | 0.014749 | 0.005138 | 0.03379 | 0.026771 | 0.021024 | 0.014255 | 0.001388 | 0.014885 | 0.027612 | 0.00961 | 0.036277 | 0.043453 | 0.009719 | 0.00967 | 0.014511 | 0.022132 | 0.032366 |
| 33.25 | 0.030958 | 0.018304 | 0.012525 | 0.043917 | 0.030514 | 0.021912 | 0.021753 | 0.004013 | 0.010729 | 0.036829 | 0.019382 | 0.033633 | 0.049022 | 0.020656 | 0.014296 | 0.023656 | 0.035474 | 0.045169 |
| 33.2638889 | 0.041598 | 0.021677 | 0.022517 | 0.05111 | 0.032265 | 0.041667 | 0.026863 | 0.008781 | 0.018547 | 0.045704 | 0.038302 | 0.023484 | 0.042298 | 0.031683 | 0.022777 | 0.033041 | 0.047905 | 0.054401 |
| 33.2777778 | 0.052429 | 0.02125 | 0.027976 | 0.047172 | 0.033839 | 0.062767 | 0.029309 | 0.013497 | 0.039686 | 0.053824 | 0.057195 | 0.013152 | 0.028638 | 0.038318 | 0.024239 | 0.03451 | 0.057839 | 0.045732 |
| 33.2916667 | 0.059458 | 0.017715 | 0.027877 | 0.036486 | 0.034305 | 0.073393 | 0.030119 | 0.017577 | 0.064104 | 0.061079 | 0.073854 | 0.009711 | 0.017314 | 0.042561 | 0.019015 | 0.027881 | 0.0612 | 0.026006 |
| 33.3055556 | 0.059045 | 0.010907 | 0.025514 | 0.027553 | 0.031001 | 0.074558 | 0.028676 | 0.020795 | 0.083886 | 0.068031 | 0.090733 | 0.015837 | 0.014211 | 0.053764 | 0.014897 | 0.023309 | 0.05898 | 0.009317 |
| 33.3194444 | 0.051521 | 0.005318 | 0.022079 | 0.020822 | 0.023642 | 0.067769 | 0.024789 | 0.023288 | 0.093384 | 0.073699 | 0.105248 | 0.02774 | 0.017364 | 0.066856 | 0.015018 | 0.025124 | 0.057026 | 0.002425 |
| 33.3333333 | 0.041613 | 0.005118 | 0.017359 | 0.013646 | 0.014946 | 0.057214 | 0.02014 | 0.026397 | 0.089026 | 0.076656 | 0.101937 | 0.032382 | 0.020124 | 0.063616 | 0.017761 | 0.028664 | 0.056302 | 0.00191 |
| 33.3472222 | 0.030552 | 0.005173 | 0.011195 | 0.007157 | 0.007229 | 0.045333 | 0.014798 | 0.029418 | 0.078186 | 0.075311 | 0.078859 | 0.021589 | 0.016838 | 0.046732 | 0.020109 | 0.03464 | 0.05103 | 0.002341 |
| 33.3611111 | 0.016629 | 0.006662 | 0.004836 | 0.005595 | 0.00273 | 0.028462 | 0.008043 | 0.028079 | 0.073195 | 0.065359 | 0.051607 | 0.009991 | 0.009835 | 0.033227 | 0.019662 | 0.039365 | 0.038623 | 0.004226 |
| 33.375 | 0.005689 | 0.008988 | 0.001424 | 0.007323 | 0.002471 | 0.013066 | 0.003705 | 0.022686 | 0.074293 | 0.050956 | 0.031144 | 0.012885 | 0.009327 | 0.03013 | 0.016444 | 0.04124 | 0.028236 | 0.008121 |
| 33.3888889 | 0.002956 | 0.011229 | 0.002726 | 0.006182 | 0.002818 | 0.008701 | 0.003763 | 0.018837 | 0.072333 | 0.043413 | 0.01911 | 0.01794 | 0.014671 | 0.038729 | 0.014053 | 0.045744 | 0.025426 | 0.010941 |
| 33.4027778 | 0.004345 | 0.019718 | 0.006816 | 0.003187 | 0.003452 | 0.008778 | 0.003815 | 0.01628 | 0.059948 | 0.043246 | 0.01416 | 0.01327 | 0.012746 | 0.051093 | 0.016411 | 0.048201 | 0.028294 | 0.009094 |
| 33.4166667 | 0.008238 | 0.026408 | 0.010189 | 0.003641 | 0.006809 | 0.00685 | 0.00336 | 0.012679 | 0.042017 | 0.041989 | 0.016905 | 0.010078 | 0.008627 | 0.056856 | 0.021133 | 0.044682 | 0.036344 | 0.008984 |
| 33.4305556 | 0.015738 | 0.02976 | 0.011833 | 0.007209 | 0.012101 | 0.004843 | 0.005049 | 0.010341 | 0.032377 | 0.033885 | 0.02525 | 0.015033 | 0.014116 | 0.058081 | 0.022705 | 0.040662 | 0.045287 | 0.018955 |
| 33.4444444 | 0.023737 | 0.038328 | 0.012151 | 0.010081 | 0.016841 | 0.006411 | 0.006204 | 0.009286 | 0.029483 | 0.020723 | 0.031879 | 0.017041 | 0.018793 | 0.058709 | 0.019657 | 0.032714 | 0.044399 | 0.030321 |
| 33.4583333 | 0.026371 | 0.042916 | 0.012283 | 0.009813 | 0.017799 | 0.015093 | 0.004543 | 0.007395 | 0.021353 | 0.009039 | 0.031761 | 0.012666 | 0.015007 | 0.055259 | 0.013854 | 0.019754 | 0.028851 | 0.034001 |
| 33.4722222 | 0.023912 | 0.037585 | 0.014859 | 0.008068 | 0.014743 | 0.027236 | 0.002852 | 0.005362 | 0.010855 | 0.003944 | 0.026852 | 0.008514 | 0.008116 | 0.045237 | 0.009485 | 0.015923 | 0.014735 | 0.030741 |
| 33.4861111 | 0.022849 | 0.031849 | 0.019863 | 0.007868 | 0.011031 | 0.035319 | 0.002243 | 0.004021 | 0.008366 | 0.005807 | 0.021774 | 0.007528 | 0.003411 | 0.034141 | 0.00853 | 0.022043 | 0.019471 | 0.021231 |
| 33.5 | 0.024436 | 0.029901 | 0.024285 | 0.008272 | 0.008167 | 0.03788 | 0.002865 | 0.002995 | 0.011004 | 0.01157 | 0.017648 | 0.009055 | 0.003259 | 0.028119 | 0.010077 | 0.020218 | 0.029902 | 0.010691 |
| 33.5138889 | 0.026105 | 0.028428 | 0.026462 | 0.008144 | 0.00596 | 0.035426 | 0.00576 | 0.00251 | 0.011045 | 0.01868 | 0.014069 | 0.009999 | 0.004813 | 0.027193 | 0.014437 | 0.011722 | 0.027534 | 0.008478 |
| 33.5277778 | 0.027471 | 0.026612 | 0.025934 | 0.007817 | 0.004671 | 0.031297 | 0.008051 | 0.002011 | 0.009078 | 0.026404 | 0.013136 | 0.00811 | 0.006183 | 0.028713 | 0.019024 | 0.014454 | 0.015402 | 0.013491 |
| 33.5416667 | 0.026979 | 0.022927 | 0.022132 | 0.005967 | 0.003256 | 0.033759 | 0.006015 | 0.001518 | 0.008641 | 0.033539 | 0.015885 | 0.00488 | 0.008842 | 0.030476 | 0.015699 | 0.024842 | 0.008384 | 0.014645 |
| 33.5555556 | 0.02215 | 0.015847 | 0.017239 | 0.002864 | 0.002253 | 0.042664 | 0.003091 | 0.002017 | 0.010818 | 0.037695 | 0.020646 | 0.004012 | 0.012486 | 0.030788 | 0.006174 | 0.0304 | 0.015228 | 0.009921 |
| 33.5694444 | 0.015555 | 0.014538 | 0.016412 | 0.003772 | 0.002595 | 0.04727 | 0.002422 | 0.002519 | 0.011174 | 0.041024 | 0.028581 | 0.004223 | 0.013472 | 0.035171 | 0.004555 | 0.031946 | 0.027132 | 0.009251 |
| 33.5833333 | 0.013595 | 0.032217 | 0.024321 | 0.014712 | 0.004142 | 0.036189 | 0.005816 | 0.005088 | 0.00991 | 0.051662 | 0.045519 | 0.007488 | 0.010735 | 0.056592 | 0.018736 | 0.038004 | 0.037954 | 0.022738 |
| 33.5972222 | 0.021952 | 0.065091 | 0.043663 | 0.034992 | 0.009924 | 0.018857 | 0.019129 | 0.012294 | 0.018836 | 0.070141 | 0.070448 | 0.020801 | 0.014761 | 0.096082 | 0.045741 | 0.049548 | 0.045349 | 0.049145 |
| 33.6111111 | 0.03993 | 0.085415 | 0.057007 | 0.056282 | 0.019286 | 0.018545 | 0.03805 | 0.019877 | 0.039018 | 0.088466 | 0.093615 | 0.041437 | 0.034954 | 0.131144 | 0.06633 | 0.053143 | 0.037268 | 0.076294 |
| 33.625 | 0.059225 | 0.078646 | 0.062806 | 0.070523 | 0.025097 | 0.031439 | 0.048452 | 0.02342 | 0.060007 | 0.108953 | 0.114128 | 0.062052 | 0.060784 | 0.141405 | 0.063787 | 0.050725 | 0.026454 | 0.098688 |
| 33.6388889 | 0.071894 | 0.067169 | 0.057896 | 0.072875 | 0.027209 | 0.039919 | 0.046136 | 0.023724 | 0.076035 | 0.138776 | 0.129367 | 0.072221 | 0.074618 | 0.129668 | 0.047636 | 0.057107 | 0.027199 | 0.113805 |
| 33.6527778 | 0.075706 | 0.067156 | 0.047247 | 0.064609 | 0.030525 | 0.043103 | 0.037843 | 0.022797 | 0.079026 | 0.154072 | 0.122942 | 0.064888 | 0.071174 | 0.100561 | 0.030075 | 0.059892 | 0.026859 | 0.113146 |
| 33.6666667 | 0.072651 | 0.070711 | 0.036636 | 0.053908 | 0.031951 | 0.044598 | 0.024669 | 0.020967 | 0.065874 | 0.121705 | 0.09681 | 0.045975 | 0.056131 | 0.058106 | 0.017182 | 0.042553 | 0.02156 | 0.101423 |
| 33.6805556 | 0.063019 | 0.071288 | 0.028215 | 0.043795 | 0.030507 | 0.041204 | 0.01105 | 0.01896 | 0.046814 | 0.059601 | 0.069784 | 0.025635 | 0.037071 | 0.02197 | 0.01503 | 0.023418 | 0.016683 | 0.085924 |
| 33.6944444 | 0.047227 | 0.068725 | 0.021603 | 0.032771 | 0.029864 | 0.035076 | 0.006447 | 0.018468 | 0.030232 | 0.024705 | 0.042439 | 0.021205 | 0.019302 | 0.008623 | 0.024856 | 0.026605 | 0.030461 | 0.060636 |
| 33.7083333 | 0.028939 | 0.05905 | 0.01666 | 0.02286 | 0.027689 | 0.031594 | 0.005709 | 0.017848 | 0.015949 | 0.043549 | 0.02268 | 0.038175 | 0.011437 | 0.010709 | 0.047617 | 0.040438 | 0.05933 | 0.030652 |
| 33.7222222 | 0.013734 | 0.037994 | 0.013132 | 0.014949 | 0.020314 | 0.028445 | 0.005159 | 0.012334 | 0.010459 | 0.082036 | 0.030093 | 0.056859 | 0.018939 | 0.014356 | 0.073858 | 0.042676 | 0.080064 | 0.022614 |
| 33.7361111 | 0.011494 | 0.018071 | 0.011636 | 0.009062 | 0.01216 | 0.020655 | 0.009546 | 0.008454 | 0.02189 | 0.104529 | 0.04707 | 0.062514 | 0.028346 | 0.016735 | 0.082961 | 0.035955 | 0.091475 | 0.04775 |
| 33.75 | 0.022366 | 0.017351 | 0.011274 | 0.009677 | 0.013584 | 0.0111 | 0.014251 | 0.016261 | 0.040276 | 0.108938 | 0.046664 | 0.052726 | 0.026608 | 0.015456 | 0.072531 | 0.025508 | 0.087909 | 0.075987 |
| 33.7638889 | 0.031657 | 0.030908 | 0.010021 | 0.015631 | 0.02365 | 0.009783 | 0.01451 | 0.030678 | 0.05702 | 0.096668 | 0.032696 | 0.03624 | 0.018961 | 0.017091 | 0.058298 | 0.019426 | 0.058487 | 0.079163 |
| 33.7777778 | 0.032179 | 0.042678 | 0.008195 | 0.018388 | 0.030424 | 0.016813 | 0.012241 | 0.044833 | 0.073645 | 0.065436 | 0.016846 | 0.025871 | 0.014121 | 0.032666 | 0.048372 | 0.030756 | 0.034174 | 0.062893 |
| 33.7916667 | 0.028525 | 0.046682 | 0.007714 | 0.014315 | 0.029499 | 0.021873 | 0.009143 | 0.053484 | 0.084932 | 0.034216 | 0.006054 | 0.021488 | 0.012816 | 0.046788 | 0.037045 | 0.04609 | 0.052793 | 0.044153 |
| 33.8055556 | 0.025 | 0.046966 | 0.009935 | 0.00848 | 0.023912 | 0.020645 | 0.008102 | 0.049598 | 0.083749 | 0.030549 | 0.004003 | 0.017851 | 0.012558 | 0.043706 | 0.020932 | 0.044368 | 0.086875 | 0.032765 |
| 33.8194444 | 0.020112 | 0.046333 | 0.015584 | 0.0114 | 0.016147 | 0.017286 | 0.011104 | 0.033957 | 0.067771 | 0.052146 | 0.009099 | 0.01129 | 0.014784 | 0.03426 | 0.020626 | 0.028813 | 0.095005 | 0.033776 |
| 33.8333333 | 0.014809 | 0.041457 | 0.023114 | 0.023686 | 0.008404 | 0.014121 | 0.015439 | 0.015985 | 0.03926 | 0.079138 | 0.019312 | 0.00585 | 0.020145 | 0.029367 | 0.049512 | 0.017171 | 0.068996 | 0.044723 |
| 33.8472222 | 0.014134 | 0.032546 | 0.029036 | 0.033582 | 0.003376 | 0.009317 | 0.017875 | 0.004731 | 0.024221 | 0.099756 | 0.03446 | 0.006891 | 0.021122 | 0.027432 | 0.081765 | 0.019374 | 0.035098 | 0.051906 |
| 33.8611111 | 0.01687 | 0.023921 | 0.031391 | 0.034539 | 0.003169 | 0.005725 | 0.017893 | 0.001445 | 0.04197 | 0.10172 | 0.051032 | 0.007134 | 0.013653 | 0.021673 | 0.096541 | 0.03286 | 0.036893 | 0.043154 |
| 33.875 | 0.02007 | 0.016838 | 0.029158 | 0.028083 | 0.009071 | 0.009082 | 0.015231 | 0.002474 | 0.063854 | 0.073524 | 0.055895 | 0.00538 | 0.009479 | 0.013435 | 0.102136 | 0.050138 | 0.075538 | 0.023126 |
| 33.8888889 | 0.027609 | 0.013072 | 0.021851 | 0.01669 | 0.021616 | 0.016839 | 0.009739 | 0.005072 | 0.066709 | 0.03527 | 0.041477 | 0.006942 | 0.01861 | 0.015428 | 0.103526 | 0.064849 | 0.120657 | 0.008889 |
| 33.9027778 | 0.046643 | 0.022478 | 0.012739 | 0.012985 | 0.03962 | 0.022279 | 0.00747 | 0.007027 | 0.055517 | 0.018683 | 0.022366 | 0.011193 | 0.029568 | 0.036276 | 0.0916 | 0.066685 | 0.150671 | 0.006475 |
| 33.9166667 | 0.074529 | 0.049534 | 0.011336 | 0.027494 | 0.059973 | 0.023199 | 0.012336 | 0.005522 | 0.033272 | 0.026357 | 0.01108 | 0.026445 | 0.027236 | 0.062135 | 0.059591 | 0.044515 | 0.140618 | 0.005463 |
| 33.9305556 | 0.09866 | 0.078586 | 0.020852 | 0.046861 | 0.080862 | 0.021029 | 0.015342 | 0.004296 | 0.018445 | 0.046315 | 0.008508 | 0.054162 | 0.015299 | 0.068141 | 0.033304 | 0.025827 | 0.088334 | 0.003248 |
| 33.9444444 | 0.107658 | 0.091876 | 0.030638 | 0.059897 | 0.1013 | 0.017531 | 0.011341 | 0.009203 | 0.034825 | 0.060939 | 0.017289 | 0.073166 | 0.005764 | 0.063625 | 0.044458 | 0.04512 | 0.059227 | 0.005975 |
| 33.9583333 | 0.102899 | 0.096029 | 0.034802 | 0.068701 | 0.113811 | 0.012955 | 0.005462 | 0.016081 | 0.070514 | 0.06329 | 0.034121 | 0.069411 | 0.00421 | 0.075717 | 0.063132 | 0.072428 | 0.094514 | 0.00912 |
| 33.9722222 | 0.094152 | 0.104896 | 0.036745 | 0.072075 | 0.110638 | 0.007159 | 0.002158 | 0.019429 | 0.103515 | 0.067561 | 0.05137 | 0.045409 | 0.009403 | 0.096879 | 0.050693 | 0.072235 | 0.138496 | 0.010214 |
| 33.9861111 | 0.08011 | 0.113628 | 0.036214 | 0.06748 | 0.092846 | 0.003768 | 0.002 | 0.019015 | 0.123458 | 0.071013 | 0.060206 | 0.022844 | 0.017047 | 0.104088 | 0.025266 | 0.055685 | 0.145832 | 0.012618 |
| 34 | 0.05942 | 0.110998 | 0.03019 | 0.055935 | 0.063906 | 0.00664 | 0.003341 | 0.016709 | 0.126441 | 0.063675 | 0.059665 | 0.023994 | 0.021898 | 0.099555 | 0.01763 | 0.039034 | 0.123599 | 0.018027 |
| 34.0138889 | 0.038754 | 0.096847 | 0.019307 | 0.038842 | 0.033507 | 0.01034 | 0.004239 | 0.014407 | 0.120023 | 0.049251 | 0.067565 | 0.03388 | 0.023493 | 0.098015 | 0.024829 | 0.026824 | 0.075122 | 0.034567 |
| 34.0277778 | 0.019699 | 0.080174 | 0.010198 | 0.02113 | 0.02679 | 0.007903 | 0.004619 | 0.011121 | 0.108257 | 0.028291 | 0.083918 | 0.034403 | 0.023126 | 0.096828 | 0.030093 | 0.017351 | 0.035996 | 0.055562 |
| 34.0416667 | 0.010214 | 0.06424 | 0.011305 | 0.017724 | 0.049954 | 0.006913 | 0.008504 | 0.006416 | 0.086614 | 0.020358 | 0.076688 | 0.03571 | 0.016798 | 0.078704 | 0.025389 | 0.01304 | 0.042348 | 0.056413 |
| 34.0555556 | 0.016144 | 0.04777 | 0.019862 | 0.032976 | 0.065307 | 0.017488 | 0.017359 | 0.004631 | 0.058549 | 0.046629 | 0.042398 | 0.045256 | 0.013795 | 0.046758 | 0.01597 | 0.019238 | 0.051189 | 0.033843 |
| 34.0694444 | 0.019506 | 0.037434 | 0.022045 | 0.046915 | 0.049394 | 0.026171 | 0.023458 | 0.004938 | 0.039413 | 0.076755 | 0.018711 | 0.046205 | 0.021656 | 0.030984 | 0.011106 | 0.0253 | 0.039965 | 0.012445 |
| 34.0833333 | 0.020951 | 0.048514 | 0.021221 | 0.039957 | 0.024919 | 0.027475 | 0.019455 | 0.007865 | 0.052352 | 0.064354 | 0.020397 | 0.037331 | 0.027279 | 0.037258 | 0.020781 | 0.023862 | 0.04789 | 0.016932 |
| 34.0972222 | 0.050048 | 0.088467 | 0.035775 | 0.039301 | 0.024566 | 0.037644 | 0.017214 | 0.020425 | 0.094253 | 0.041916 | 0.046274 | 0.052972 | 0.042756 | 0.073755 | 0.05862 | 0.037711 | 0.106131 | 0.054896 |
| 34.1111111 | 0.094497 | 0.13643 | 0.057344 | 0.069707 | 0.05111 | 0.049335 | 0.024719 | 0.033506 | 0.125974 | 0.058142 | 0.099609 | 0.086849 | 0.069698 | 0.137841 | 0.097014 | 0.072128 | 0.199318 | 0.101462 |
| 34.125 | 0.113386 | 0.152529 | 0.067067 | 0.084285 | 0.072623 | 0.050742 | 0.027095 | 0.035386 | 0.134032 | 0.077619 | 0.148639 | 0.095132 | 0.078419 | 0.159917 | 0.095727 | 0.090688 | 0.257526 | 0.112556 |
| 34.1388889 | 0.109139 | 0.129625 | 0.063872 | 0.064114 | 0.073351 | 0.047011 | 0.020857 | 0.031047 | 0.129938 | 0.081828 | 0.152131 | 0.07777 | 0.071879 | 0.127932 | 0.068143 | 0.082737 | 0.247135 | 0.084271 |
| 34.1527778 | 0.099147 | 0.100048 | 0.051285 | 0.042187 | 0.061565 | 0.040121 | 0.012895 | 0.027843 | 0.114745 | 0.084897 | 0.110858 | 0.060778 | 0.05943 | 0.093486 | 0.04613 | 0.068632 | 0.210202 | 0.05019 |
| 34.1666667 | 0.078695 | 0.079395 | 0.034974 | 0.026328 | 0.041458 | 0.032844 | 0.006022 | 0.025174 | 0.090941 | 0.082587 | 0.057579 | 0.045626 | 0.039853 | 0.069148 | 0.034408 | 0.049633 | 0.175786 | 0.031756 |
| 34.1805556 | 0.048755 | 0.062997 | 0.020936 | 0.012165 | 0.01943 | 0.029621 | 0.005582 | 0.020254 | 0.064771 | 0.071547 | 0.02145 | 0.028368 | 0.02005 | 0.04836 | 0.023616 | 0.027793 | 0.136094 | 0.030484 |
| 34.1944444 | 0.022888 | 0.047373 | 0.011292 | 0.004042 | 0.008004 | 0.029154 | 0.011681 | 0.013871 | 0.041665 | 0.052116 | 0.013212 | 0.028301 | 0.007693 | 0.02988 | 0.013242 | 0.022771 | 0.083771 | 0.035873 |
| 34.2083333 | 0.008792 | 0.03277 | 0.006937 | 0.005313 | 0.01232 | 0.026844 | 0.018083 | 0.007908 | 0.024449 | 0.029439 | 0.029357 | 0.054597 | 0.006919 | 0.015963 | 0.008141 | 0.037178 | 0.044232 | 0.035591 |

| | | | | | | | | | | | | | | | | | | |
|---|---|---|---|---|---|---|---|---|---|---|---|---|---|---|---|---|---|---|
| 34.2222222 | 0.008019 | 0.020433 | 0.009447 | 0.012181 | 0.021952 | 0.02051 | 0.022094 | 0.003837 | 0.011543 | 0.014195 | 0.057315 | 0.100732 | 0.010866 | 0.006895 | 0.009431 | 0.054116 | 0.052765 | 0.030423 |
| 34.2361111 | 0.020224 | 0.009924 | 0.016303 | 0.021186 | 0.023605 | 0.011172 | 0.022771 | 0.002209 | 0.004742 | 0.015777 | 0.079939 | 0.131091 | 0.010945 | 0.002488 | 0.014869 | 0.057996 | 0.088243 | 0.031164 |
| 34.25 | 0.037448 | 0.003613 | 0.022549 | 0.031714 | 0.017926 | 0.006838 | 0.020722 | 0.001767 | 0.007281 | 0.027755 | 0.079739 | 0.104016 | 0.012004 | 0.003802 | 0.021223 | 0.045044 | 0.112657 | 0.040545 |
| 34.2638889 | 0.048429 | 0.005176 | 0.028495 | 0.041328 | 0.017434 | 0.016157 | 0.021602 | 0.002547 | 0.01579 | 0.04385 | 0.067046 | 0.061299 | 0.018938 | 0.011885 | 0.024016 | 0.032301 | 0.13298 | 0.048932 |
| 34.2777778 | 0.049387 | 0.013489 | 0.033805 | 0.048654 | 0.0254 | 0.034988 | 0.028282 | 0.006079 | 0.026577 | 0.06409 | 0.064318 | 0.048152 | 0.025459 | 0.025013 | 0.020564 | 0.036609 | 0.15312 | 0.04771 |
| 34.2916667 | 0.043435 | 0.019731 | 0.03431 | 0.055286 | 0.033631 | 0.059223 | 0.03522 | 0.00901 | 0.035054 | 0.081109 | 0.06229 | 0.042369 | 0.027421 | 0.03955 | 0.014537 | 0.045565 | 0.143428 | 0.03749 |
| 34.3055556 | 0.033442 | 0.016591 | 0.029167 | 0.059744 | 0.034903 | 0.08387 | 0.035821 | 0.007787 | 0.035027 | 0.090721 | 0.05508 | 0.029343 | 0.026008 | 0.056048 | 0.009187 | 0.039969 | 0.102057 | 0.022014 |
| 34.3194444 | 0.02086 | 0.014262 | 0.022527 | 0.054966 | 0.025089 | 0.094612 | 0.030469 | 0.004692 | 0.03058 | 0.100767 | 0.054318 | 0.018825 | 0.024825 | 0.073928 | 0.005929 | 0.029479 | 0.065672 | 0.010161 |
| 34.3333333 | 0.012804 | 0.036667 | 0.017767 | 0.040103 | 0.017104 | 0.084542 | 0.021908 | 0.003063 | 0.031768 | 0.112653 | 0.052742 | 0.014078 | 0.02456 | 0.083338 | 0.005837 | 0.02734 | 0.048223 | 0.007747 |
| 34.3472222 | 0.015058 | 0.076872 | 0.013432 | 0.022968 | 0.028658 | 0.062635 | 0.01239 | 0.003967 | 0.04066 | 0.110576 | 0.039469 | 0.011282 | 0.017847 | 0.073075 | 0.006765 | 0.024287 | 0.031171 | 0.010341 |
| 34.3611111 | 0.016239 | 0.090417 | 0.007981 | 0.010846 | 0.044432 | 0.043001 | 0.006564 | 0.006495 | 0.050933 | 0.090616 | 0.024677 | 0.021392 | 0.008527 | 0.047373 | 0.008819 | 0.017527 | 0.01527 | 0.01697 |
| 34.375 | 0.009638 | 0.068998 | 0.005203 | 0.006461 | 0.045294 | 0.028939 | 0.006235 | 0.00881 | 0.055234 | 0.066853 | 0.019973 | 0.041752 | 0.006179 | 0.0246 | 0.008743 | 0.016429 | 0.012762 | 0.018823 |
| 34.3888889 | 0.005163 | 0.049232 | 0.00583 | 0.00516 | 0.0402 | 0.015769 | 0.007905 | 0.009682 | 0.053222 | 0.043393 | 0.021471 | 0.039133 | 0.005571 | 0.015444 | 0.006668 | 0.020766 | 0.010817 | 0.010733 |
| 34.4027778 | 0.004406 | 0.044934 | 0.005808 | 0.00593 | 0.037523 | 0.01008 | 0.009168 | 0.009258 | 0.050284 | 0.020779 | 0.020409 | 0.018488 | 0.004768 | 0.013746 | 0.008372 | 0.025684 | 0.00408 | 0.005019 |
| 34.4166667 | 0.002491 | 0.043922 | 0.005041 | 0.00898 | 0.034456 | 0.023392 | 0.009308 | 0.007853 | 0.04935 | 0.008473 | 0.016972 | 0.006873 | 0.007463 | 0.010859 | 0.008315 | 0.026308 | 0.001255 | 0.005074 |
| 34.4305556 | 0.000881 | 0.040099 | 0.004531 | 0.008563 | 0.03067 | 0.047643 | 0.007468 | 0.005212 | 0.051552 | 0.007836 | 0.01233 | 0.0102 | 0.011794 | 0.006349 | 0.005411 | 0.019504 | 0.004344 | 0.005941 |
| 34.4444444 | 0.003588 | 0.038319 | 0.003633 | 0.005899 | 0.028812 | 0.06513 | 0.004965 | 0.002564 | 0.056936 | 0.011751 | 0.008565 | 0.029876 | 0.015622 | 0.006308 | 0.008322 | 0.010372 | 0.012551 | 0.006631 |
| 34.4583333 | 0.012797 | 0.047098 | 0.003253 | 0.008366 | 0.032924 | 0.067763 | 0.004497 | 0.002263 | 0.064781 | 0.014772 | 0.011737 | 0.048127 | 0.017714 | 0.012564 | 0.016611 | 0.004176 | 0.029405 | 0.009618 |
| 34.4722222 | 0.02201 | 0.059925 | 0.005818 | 0.014941 | 0.040996 | 0.055136 | 0.00506 | 0.004033 | 0.070355 | 0.011493 | 0.01569 | 0.041818 | 0.01624 | 0.015708 | 0.0217 | 0.002936 | 0.064533 | 0.013085 |
| 34.4861111 | 0.019604 | 0.056615 | 0.009766 | 0.014837 | 0.039667 | 0.033042 | 0.005631 | 0.004556 | 0.065355 | 0.006455 | 0.013431 | 0.024252 | 0.010889 | 0.011914 | 0.019943 | 0.005013 | 0.114825 | 0.009957 |
| 34.5 | 0.008662 | 0.037506 | 0.010631 | 0.007249 | 0.029952 | 0.013878 | 0.009597 | 0.00259 | 0.054047 | 0.008753 | 0.00956 | 0.029033 | 0.005458 | 0.006375 | 0.015145 | 0.005941 | 0.14193 | 0.005487 |
| 34.5138889 | 0.003886 | 0.022725 | 0.007332 | 0.002607 | 0.030765 | 0.007638 | 0.014831 | 0.001247 | 0.046699 | 0.019093 | 0.007074 | 0.067067 | 0.004633 | 0.00315 | 0.010757 | 0.006515 | 0.116569 | 0.007631 |
| 34.5277778 | 0.008284 | 0.01766 | 0.004173 | 0.003861 | 0.042017 | 0.013053 | 0.015091 | 0.002999 | 0.039954 | 0.033479 | 0.004425 | 0.14256 | 0.007543 | 0.005088 | 0.006621 | 0.010626 | 0.075468 | 0.009284 |
| 34.5416667 | 0.011073 | 0.015091 | 0.004584 | 0.006271 | 0.044125 | 0.020888 | 0.011586 | 0.006035 | 0.029376 | 0.042109 | 0.002218 | 0.23494 | 0.008616 | 0.010082 | 0.003627 | 0.017314 | 0.066651 | 0.007236 |
| 34.5555556 | 0.007154 | 0.010857 | 0.005197 | 0.006977 | 0.030262 | 0.021945 | 0.010213 | 0.006801 | 0.020981 | 0.035936 | 0.002859 | 0.272574 | 0.0073 | 0.014736 | 0.004748 | 0.020148 | 0.081507 | 0.009442 |
| 34.5694444 | 0.006536 | 0.008965 | 0.008397 | 0.009485 | 0.012874 | 0.014503 | 0.014125 | 0.004988 | 0.020019 | 0.021345 | 0.009272 | 0.223954 | 0.005466 | 0.022254 | 0.012485 | 0.016331 | 0.077163 | 0.011377 |
| 34.5833333 | 0.017254 | 0.021295 | 0.022055 | 0.019257 | 0.007249 | 0.009713 | 0.022911 | 0.00285 | 0.029918 | 0.015168 | 0.025301 | 0.129345 | 0.005525 | 0.03981 | 0.031178 | 0.017263 | 0.046339 | 0.019003 |
| 34.5972222 | 0.035384 | 0.048346 | 0.044783 | 0.037831 | 0.017382 | 0.023771 | 0.035366 | 0.003733 | 0.053529 | 0.023706 | 0.047001 | 0.063919 | 0.014157 | 0.065054 | 0.057107 | 0.03489 | 0.037279 | 0.049731 |
| 34.6111111 | 0.056279 | 0.077542 | 0.069842 | 0.060764 | 0.036009 | 0.051245 | 0.049296 | 0.009828 | 0.081399 | 0.042713 | 0.061986 | 0.067156 | 0.02702 | 0.087048 | 0.071348 | 0.054303 | 0.075008 | 0.072702 |
| 34.625 | 0.076387 | 0.10168 | 0.088333 | 0.080178 | 0.052456 | 0.070184 | 0.060134 | 0.015883 | 0.101277 | 0.067084 | 0.062981 | 0.066324 | 0.033439 | 0.106704 | 0.068737 | 0.057986 | 0.115388 | 0.062118 |
| 34.6388889 | 0.087277 | 0.123964 | 0.088984 | 0.088833 | 0.059244 | 0.07497 | 0.062077 | 0.015764 | 0.112261 | 0.092617 | 0.054863 | 0.032819 | 0.032466 | 0.132509 | 0.070571 | 0.05787 | 0.134445 | 0.048534 |
| 34.6527778 | 0.081611 | 0.145191 | 0.071116 | 0.079818 | 0.056908 | 0.07546 | 0.052742 | 0.013306 | 0.118564 | 0.111077 | 0.048543 | 0.022309 | 0.029739 | 0.157269 | 0.087572 | 0.066238 | 0.168169 | 0.057431 |
| 34.6666667 | 0.062782 | 0.157149 | 0.045494 | 0.054027 | 0.048723 | 0.075922 | 0.036403 | 0.015011 | 0.122526 | 0.113654 | 0.045835 | 0.020659 | 0.030139 | 0.164127 | 0.102769 | 0.068143 | 0.203695 | 0.083215 |
| 34.6805556 | 0.044215 | 0.154283 | 0.021091 | 0.024843 | 0.036241 | 0.071012 | 0.021348 | 0.018701 | 0.124413 | 0.104127 | 0.04423 | 0.008587 | 0.033282 | 0.144216 | 0.093355 | 0.051366 | 0.178248 | 0.106011 |
| 34.6944444 | 0.035863 | 0.13682 | 0.006933 | 0.008974 | 0.025439 | 0.061342 | 0.015108 | 0.017709 | 0.120173 | 0.087784 | 0.047289 | 0.002377 | 0.033673 | 0.099885 | 0.061399 | 0.032685 | 0.10105 | 0.108943 |
| 34.7083333 | 0.031272 | 0.108401 | 0.006309 | 0.008873 | 0.020465 | 0.050469 | 0.018967 | 0.010573 | 0.099559 | 0.06376 | 0.052234 | 0.008709 | 0.023389 | 0.048012 | 0.029111 | 0.026315 | 0.065223 | 0.090979 |
| 34.7222222 | 0.01978 | 0.070543 | 0.010289 | 0.016208 | 0.014726 | 0.032379 | 0.026632 | 0.005336 | 0.062801 | 0.039146 | 0.048594 | 0.048973 | 0.010932 | 0.020391 | 0.015454 | 0.02705 | 0.097451 | 0.062867 |
| 34.7361111 | 0.01208 | 0.036693 | 0.015419 | 0.028793 | 0.014925 | 0.025158 | 0.030816 | 0.008673 | 0.031602 | 0.0312 | 0.032203 | 0.118042 | 0.011183 | 0.027938 | 0.02615 | 0.027338 | 0.113078 | 0.038609 |
| 34.75 | 0.022542 | 0.035573 | 0.022441 | 0.0358 | 0.032692 | 0.049831 | 0.032936 | 0.015761 | 0.032796 | 0.046152 | 0.016232 | 0.19412 | 0.018929 | 0.03875 | 0.036911 | 0.025009 | 0.08074 | 0.025671 |
| 34.7638889 | 0.038992 | 0.062254 | 0.026717 | 0.027267 | 0.054324 | 0.078899 | 0.037798 | 0.021954 | 0.064954 | 0.069375 | 0.013711 | 0.244214 | 0.02373 | 0.035374 | 0.028483 | 0.022711 | 0.039091 | 0.01752 |
| 34.7777778 | 0.045614 | 0.084835 | 0.025432 | 0.014422 | 0.068623 | 0.088093 | 0.04249 | 0.02635 | 0.098243 | 0.08493 | 0.015994 | 0.20859 | 0.024004 | 0.02862 | 0.014608 | 0.020471 | 0.011892 | 0.012076 |
| 34.7916667 | 0.042176 | 0.089953 | 0.020242 | 0.006496 | 0.072686 | 0.078719 | 0.044562 | 0.02628 | 0.113575 | 0.082791 | 0.011652 | 0.120121 | 0.020755 | 0.019428 | 0.006866 | 0.016948 | 0.008482 | 0.01993 |
| 34.8055556 | 0.03271 | 0.080504 | 0.013355 | 0.003909 | 0.061005 | 0.055765 | 0.046159 | 0.022469 | 0.116169 | 0.053324 | 0.009198 | 0.052447 | 0.012597 | 0.014022 | 0.007561 | 0.016269 | 0.013413 | 0.039852 |
| 34.8194444 | 0.020979 | 0.06367 | 0.008538 | 0.006946 | 0.038357 | 0.027836 | 0.047336 | 0.019076 | 0.114299 | 0.029527 | 0.019181 | 0.045657 | 0.005792 | 0.022009 | 0.015127 | 0.018747 | 0.016012 | 0.057453 |
| 34.8333333 | 0.011566 | 0.046016 | 0.013428 | 0.013942 | 0.016818 | 0.012042 | 0.047534 | 0.016266 | 0.10344 | 0.046351 | 0.031341 | 0.112795 | 0.006678 | 0.032086 | 0.020042 | 0.02066 | 0.035522 | 0.067357 |
| 34.8472222 | 0.005803 | 0.030631 | 0.02742 | 0.021664 | 0.005438 | 0.017022 | 0.046549 | 0.011884 | 0.078878 | 0.066804 | 0.034912 | 0.177034 | 0.007979 | 0.034187 | 0.018444 | 0.019243 | 0.065941 | 0.07889 |
| 34.8611111 | 0.002349 | 0.017799 | 0.038019 | 0.024712 | 0.00246 | 0.022212 | 0.043972 | 0.007957 | 0.049247 | 0.061845 | 0.031048 | 0.169117 | 0.007644 | 0.03102 | 0.014645 | 0.016188 | 0.084791 | 0.096483 |
| 34.875 | 0.000893 | 0.008191 | 0.037647 | 0.021759 | 0.002539 | 0.014519 | 0.04067 | 0.008639 | 0.024826 | 0.03934 | 0.023091 | 0.117807 | 0.009363 | 0.025078 | 0.010686 | 0.016909 | 0.078828 | 0.105154 |
| 34.8888889 | 0.000864 | 0.00459 | 0.030123 | 0.016264 | 0.006122 | 0.007884 | 0.036344 | 0.012928 | 0.010611 | 0.023111 | 0.014493 | 0.063996 | 0.010379 | 0.018425 | 0.005906 | 0.015107 | 0.053939 | 0.092266 |
| 34.9027778 | 0.001826 | 0.00614 | 0.02005 | 0.008975 | 0.013957 | 0.014289 | 0.029527 | 0.016159 | 0.009167 | 0.040733 | 0.009129 | 0.024085 | 0.009063 | 0.012209 | 0.003426 | 0.015738 | 0.055587 | 0.068407 |
| 34.9166667 | 0.002106 | 0.00671 | 0.010618 | 0.003797 | 0.025073 | 0.027624 | 0.020588 | 0.017462 | 0.01404 | 0.067998 | 0.007438 | 0.009193 | 0.009087 | 0.008217 | 0.007602 | 0.025031 | 0.085473 | 0.049639 |
| 34.9305556 | 0.002721 | 0.005483 | 0.010161 | 0.005653 | 0.035405 | 0.039589 | 0.010684 | 0.01913 | 0.01349 | 0.070747 | 0.006799 | 0.016005 | 0.009441 | 0.011004 | 0.015161 | 0.025808 | 0.102225 | 0.038693 |
| 34.9444444 | 0.008702 | 0.009687 | 0.020661 | 0.010591 | 0.040529 | 0.049809 | 0.004258 | 0.021598 | 0.011365 | 0.057833 | 0.005269 | 0.070471 | 0.00995 | 0.019186 | 0.020249 | 0.021915 | 0.107381 | 0.035517 |
| 34.9583333 | 0.018598 | 0.022219 | 0.031332 | 0.014189 | 0.046255 | 0.061237 | 0.004258 | 0.022792 | 0.022954 | 0.049746 | 0.006345 | 0.142905 | 0.01553 | 0.025773 | 0.02129 | 0.027237 | 0.119978 | 0.043303 |
| 34.9722222 | 0.026958 | 0.034605 | 0.035695 | 0.015372 | 0.059099 | 0.074932 | 0.005139 | 0.021173 | 0.045061 | 0.054367 | 0.017137 | 0.15816 | 0.020059 | 0.028972 | 0.020478 | 0.032847 | 0.140272 | 0.056999 |
| 34.9861111 | 0.015166 | 0.01952 | 0.01821 | 0.007549 | 0.033656 | 0.041037 | 0.002347 | 0.009805 | 0.028187 | 0.067531 | 0.030385 | 0.165579 | 0.018877 | 0.031032 | 0.019454 | 0.026247 | 0.134979 | 0.062961 |
| 35 | 0 | 0 | 0 | 0 | 0 | 0 | 0 | 0 | 0 | 0.084355 | 0.036611 | 0.229543 | 0.017771 | 0.032113 | 0.017316 | 0.012985 | 0.093083 | 0.056029 |
| 35.0138889 | 0 | 0 | 0 | 0 | 0 | 0 | 0 | 0 | 0 | 0.099821 | 0.035029 | 0.279202 | 0.018086 | 0.028858 | 0.014254 | 0.008982 | 0.064308 | 0.04729 |
| 35.0277778 | 0 | 0 | 0 | 0 | 0 | 0 | 0 | 0 | 0 | 0.103273 | 0.022195 | 0.223545 | 0.017806 | 0.018066 | 0.010698 | 0.011782 | 0.066433 | 0.047282 |
| 35.0416667 | 0 | 0 | 0 | 0 | 0 | 0 | 0 | 0 | 0 | 0.08627 | 0.011647 | 0.12359 | 0.016931 | 0.00848 | 0.00665 | 0.010642 | 0.061818 | 0.050675 |
| 35.0555556 | 0 | 0 | 0 | 0 | 0 | 0 | 0 | 0 | 0 | 0.054212 | 0.016434 | 0.074238 | 0.015332 | 0.008971 | 0.005412 | 0.006063 | 0.044583 | 0.048676 |
| 35.0694444 | 0 | 0 | 0 | 0 | 0 | 0 | 0 | 0 | 0 | 0.030252 | 0.022975 | 0.067583 | 0.015913 | 0.009597 | 0.012165 | 0.004033 | 0.043904 | 0.046431 |
| 35.0833333 | 0 | 0 | 0 | 0 | 0 | 0 | 0 | 0 | 0 | 0.037688 | 0.018367 | 0.079465 | 0.019683 | 0.007708 | 0.025062 | 0.006507 | 0.04459 | 0.057993 |
| 35.0972222 | 0 | 0 | 0 | 0 | 0 | 0 | 0 | 0 | 0 | 0.077065 | 0.013438 | 0.139863 | 0.021453 | 0.015505 | 0.034976 | 0.014802 | 0.053759 | 0.085566 |
| 35.1111111 | 0 | 0 | 0 | 0 | 0 | 0 | 0 | 0 | 0 | 0.11566 | 0.026412 | 0.203753 | 0.02338 | 0.030984 | 0.036304 | 0.031901 | 0.098969 | 0.108247 |
| 35.125 | 0 | 0 | 0 | 0 | 0 | 0 | 0 | 0 | 0 | 0.122778 | 0.043089 | 0.173388 | 0.027689 | 0.038494 | 0.030113 | 0.046413 | 0.128033 | 0.107254 |
| 35.1388889 | 0 | 0 | 0 | 0 | 0 | 0 | 0 | 0 | 0 | 0.108892 | 0.043296 | 0.0991 | 0.028101 | 0.032697 | 0.024603 | 0.045116 | 0.123363 | 0.098827 |
| 35.1527778 | 0 | 0 | 0 | 0 | 0 | 0 | 0 | 0 | 0 | 0.094258 | 0.033817 | 0.061898 | 0.025241 | 0.021226 | 0.025546 | 0.034804 | 0.123075 | 0.109097 |
| 35.1666667 | 0 | 0 | 0 | 0 | 0 | 0 | 0 | 0 | 0 | 0.082152 | 0.025151 | 0.061197 | 0.023995 | 0.010544 | 0.026664 | 0.028387 | 0.140753 | 0.122309 |
| 35.1805556 | 0 | 0 | 0 | 0 | 0 | 0 | 0 | 0 | 0 | 0.071241 | 0.018257 | 0.069583 | 0.023857 | 0.004013 | 0.022461 | 0.024321 | 0.165184 | 0.106225 |
| 35.1944444 | 0 | 0 | 0 | 0 | 0 | 0 | 0 | 0 | 0 | 0.062598 | 0.011837 | 0.069744 | 0.020678 | 0.003065 | 0.015428 | 0.017371 | 0.162022 | 0.070839 |
| 35.2083333 | 0 | 0 | 0 | 0 | 0 | 0 | 0 | 0 | 0 | 0.056401 | 0.007413 | 0.068701 | 0.012115 | 0.006443 | 0.011852 | 0.009789 | 0.133342 | 0.046855 |
| 35.2222222 | 0 | 0 | 0 | 0 | 0 | 0 | 0 | 0 | 0 | 0.051615 | 0.005742 | 0.074129 | 0.004694 | 0.011402 | 0.014842 | 0.004174 | 0.118755 | 0.042784 |
| 35.2361111 | 0 | 0 | 0 | 0 | 0 | 0 | 0 | 0 | 0 | 0.047973 | 0.005592 | 0.08035 | 0.003122 | 0.014263 | 0.020362 | 0.001918 | 0.117204 | 0.047837 |
| 35.25 | 0 | 0 | 0 | 0 | 0 | 0 | 0 | 0 | 0 | 0.042747 | 0.005664 | 0.078859 | 0.004197 | 0.012704 | 0.023371 | 0.001849 | 0.099758 | 0.048407 |
| 35.2638889 | 0 | 0 | 0 | 0 | 0 | 0 | 0 | 0 | 0 | 0.031431 | 0.009923 | 0.06969 | 0.007534 | 0.007877 | 0.022088 | 0.005973 | 0.067984 | 0.041066 |
| 35.2777778 | 0 | 0 | 0 | 0 | 0 | 0 | 0 | 0 | 0 | 0.016083 | 0.021012 | 0.055373 | 0.010871 | 0.006264 | 0.017276 | 0.019169 | 0.039073 | 0.029839 |
| 35.2916667 | 0 | 0 | 0 | 0 | 0 | 0 | 0 | 0 | 0 | 0.007211 | 0.033235 | 0.040403 | 0.008715 | 0.011301 | 0.011779 | 0.039461 | 0.031953 | 0.018442 |
| 35.3055556 | 0 | 0 | 0 | 0 | 0 | 0 | 0 | 0 | 0 | 0.008509 | 0.042519 | 0.033538 | 0.004332 | 0.017844 | 0.006757 | 0.045423 | 0.04892 | 0.00939 |
| 35.3194444 | 0 | 0 | 0 | 0 | 0 | 0 | 0 | 0 | 0 | 0.023272 | 0.048158 | 0.037918 | 0.003224 | 0.023645 | 0.005426 | 0.025392 | 0.062021 | 0.00736 |
| 35.3333333 | 0 | 0 | 0 | 0 | 0 | 0 | 0 | 0 | 0 | 0.053934 | 0.046758 | 0.0474 | 0.003151 | 0.028211 | 0.012834 | 0.005593 | 0.064227 | 0.016599 |
| 35.3472222 | 0 | 0 | 0 | 0 | 0 | 0 | 0 | 0 | 0 | 0.073177 | 0.038011 | 0.053606 | 0.00317 | 0.02774 | 0.023304 | 0.002934 | 0.056776 | 0.032768 |
| 35.3611111 | 0 | 0 | 0 | 0 | 0 | 0 | 0 | 0 | 0 | 0.060824 | 0.025867 | 0.051861 | 0.004507 | 0.022024 | 0.0264 | 0.005412 | 0.034285 | 0.040989 |

| | | | | | | | | | | | | | | | | | | |
|---|---|---|---|---|---|---|---|---|---|---|---|---|---|---|---|---|---|---|
| 35.375 | 0 | 0 | 0 | 0 | 0 | 0 | 0 | 0 | 0 | 0.037007 | 0.013215 | 0.042992 | 0.003978 | 0.015486 | 0.022099 | 0.003643 | 0.016317 | 0.029576 |
| 35.3888889 | 0 | 0 | 0 | 0 | 0 | 0 | 0 | 0 | 0 | 0.019896 | 0.006164 | 0.028735 | 0.002852 | 0.012683 | 0.017099 | 0.002155 | 0.009391 | 0.013564 |
| 35.4027778 | 0 | 0 | 0 | 0 | 0 | 0 | 0 | 0 | 0 | 0.009531 | 0.007024 | 0.01371 | 0.004664 | 0.012656 | 0.012136 | 0.003986 | 0.003527 | 0.010098 |
| 35.4166667 | 0 | 0 | 0 | 0 | 0 | 0 | 0 | 0 | 0 | 0.004416 | 0.008591 | 0.006718 | 0.005542 | 0.009358 | 0.007149 | 0.006926 | 0.00089 | 0.012377 |
| 35.4305556 | 0 | 0 | 0 | 0 | 0 | 0 | 0 | 0 | 0 | 0.002806 | 0.007462 | 0.008434 | 0.003331 | 0.004204 | 0.006272 | 0.013334 | 0.003803 | 0.012348 |
| 35.4444444 | 0 | 0 | 0 | 0 | 0 | 0 | 0 | 0 | 0 | 0.002985 | 0.00599 | 0.011313 | 0.003117 | 0.00334 | 0.008744 | 0.024572 | 0.014797 | 0.014051 |
| 35.4583333 | 0 | 0 | 0 | 0 | 0 | 0 | 0 | 0 | 0 | 0.007116 | 0.004738 | 0.012568 | 0.008972 | 0.008108 | 0.012419 | 0.032531 | 0.032328 | 0.025198 |
| 35.4722222 | 0 | 0 | 0 | 0 | 0 | 0 | 0 | 0 | 0 | 0.0159 | 0.005668 | 0.030228 | 0.027849 | 0.019327 | 0.02305 | 0.029615 | 0.046715 | 0.045092 |
| 35.4861111 | 0 | 0 | 0 | 0 | 0 | 0 | 0 | 0 | 0 | 0.02269 | 0.00751 | 0.066332 | 0.067328 | 0.028258 | 0.037576 | 0.017643 | 0.052473 | 0.06202 |
| 35.5 | 0 | 0 | 0 | 0 | 0 | 0 | 0 | 0 | 0 | 0.019546 | 0.00928 | 0.072473 | 0.087148 | 0.021247 | 0.037018 | 0.011873 | 0.055344 | 0.063188 |
| 35.5138889 | 0 | 0 | 0 | 0 | 0 | 0 | 0 | 0 | 0 | 0.010335 | 0.014668 | 0.039947 | 0.060982 | 0.00788 | 0.020502 | 0.020313 | 0.054563 | 0.045779 |
| 35.5277778 | 0 | 0 | 0 | 0 | 0 | 0 | 0 | 0 | 0 | 0.007333 | 0.017004 | 0.015209 | 0.038983 | 0.002823 | 0.008396 | 0.02541 | 0.040415 | 0.021883 |
| 35.5416667 | 0 | 0 | 0 | 0 | 0 | 0 | 0 | 0 | 0 | 0.011678 | 0.014083 | 0.011174 | 0.036952 | 0.002352 | 0.007883 | 0.018653 | 0.021403 | 0.006707 |
| 35.5555556 | 0 | 0 | 0 | 0 | 0 | 0 | 0 | 0 | 0 | 0.015698 | 0.012496 | 0.012218 | 0.028972 | 0.00315 | 0.010302 | 0.014734 | 0.020575 | 0.002823 |
| 35.5694444 | 0 | 0 | 0 | 0 | 0 | 0 | 0 | 0 | 0 | 0.014864 | 0.011994 | 0.014688 | 0.015143 | 0.003591 | 0.011244 | 0.022783 | 0.055867 | 0.013498 |
| 35.5833333 | 0 | 0 | 0 | 0 | 0 | 0 | 0 | 0 | 0 | 0.011823 | 0.014461 | 0.035257 | 0.008165 | 0.005208 | 0.012758 | 0.034833 | 0.101714 | 0.04366 |
| 35.5972222 | 0 | 0 | 0 | 0 | 0 | 0 | 0 | 0 | 0 | 0.021452 | 0.024085 | 0.075488 | 0.015575 | 0.016655 | 0.01199 | 0.035244 | 0.099111 | 0.063063 |
| 35.6111111 | 0 | 0 | 0 | 0 | 0 | 0 | 0 | 0 | 0 | 0.051786 | 0.037386 | 0.104878 | 0.036596 | 0.035493 | 0.010822 | 0.033403 | 0.052688 | 0.04272 |
| 35.625 | 0 | 0 | 0 | 0 | 0 | 0 | 0 | 0 | 0 | 0.092861 | 0.045881 | 0.114344 | 0.061199 | 0.050294 | 0.015405 | 0.050302 | 0.029468 | 0.019531 |
| 35.6388889 | 0 | 0 | 0 | 0 | 0 | 0 | 0 | 0 | 0 | 0.133347 | 0.047459 | 0.121585 | 0.074 | 0.058549 | 0.021538 | 0.061337 | 0.044054 | 0.023241 |
| 35.6527778 | 0 | 0 | 0 | 0 | 0 | 0 | 0 | 0 | 0 | 0.162004 | 0.047453 | 0.126547 | 0.069625 | 0.062969 | 0.027358 | 0.047542 | 0.052779 | 0.029177 |
| 35.6666667 | 0 | 0 | 0 | 0 | 0 | 0 | 0 | 0 | 0 | 0.174946 | 0.046089 | 0.121358 | 0.057105 | 0.062041 | 0.033505 | 0.026607 | 0.043149 | 0.022599 |
| 35.6805556 | 0 | 0 | 0 | 0 | 0 | 0 | 0 | 0 | 0 | 0.179213 | 0.041022 | 0.111202 | 0.049735 | 0.056178 | 0.037175 | 0.011704 | 0.025758 | 0.014068 |
| 35.6944444 | 0 | 0 | 0 | 0 | 0 | 0 | 0 | 0 | 0 | 0.178123 | 0.03651 | 0.101473 | 0.059415 | 0.051397 | 0.039801 | 0.00897 | 0.01387 | 0.016556 |
| 35.7083333 | 0 | 0 | 0 | 0 | 0 | 0 | 0 | 0 | 0 | 0.170506 | 0.035738 | 0.092849 | 0.079019 | 0.051502 | 0.044598 | 0.014511 | 0.022065 | 0.026074 |
| 35.7222222 | 0 | 0 | 0 | 0 | 0 | 0 | 0 | 0 | 0 | 0.149575 | 0.037166 | 0.082564 | 0.076895 | 0.052759 | 0.048514 | 0.017677 | 0.039526 | 0.033136 |
| 35.7361111 | 0 | 0 | 0 | 0 | 0 | 0 | 0 | 0 | 0 | 0.110737 | 0.038179 | 0.066144 | 0.048232 | 0.050537 | 0.045273 | 0.012641 | 0.054349 | 0.041267 |
| 35.75 | 0 | 0 | 0 | 0 | 0 | 0 | 0 | 0 | 0 | 0.062978 | 0.037537 | 0.044566 | 0.037865 | 0.04561 | 0.034266 | 0.006645 | 0.064568 | 0.050497 |
| 35.7638889 | 0 | 0 | 0 | 0 | 0 | 0 | 0 | 0 | 0 | 0.024811 | 0.03595 | 0.02239 | 0.060802 | 0.039644 | 0.020332 | 0.008978 | 0.064396 | 0.052456 |
| 35.7777778 | 0 | 0 | 0 | 0 | 0 | 0 | 0 | 0 | 0 | 0.008155 | 0.032781 | 0.010438 | 0.079022 | 0.032854 | 0.009657 | 0.013278 | 0.05384 | 0.043981 |
| 35.7916667 | 0 | 0 | 0 | 0 | 0 | 0 | 0 | 0 | 0 | 0.004313 | 0.028035 | 0.016569 | 0.071975 | 0.027161 | 0.005099 | 0.011517 | 0.04053 | 0.030405 |
| 35.8055556 | 0 | 0 | 0 | 0 | 0 | 0 | 0 | 0 | 0 | 0.003936 | 0.023685 | 0.030893 | 0.057311 | 0.023332 | 0.005565 | 0.005802 | 0.02736 | 0.016514 |
| 35.8194444 | 0 | 0 | 0 | 0 | 0 | 0 | 0 | 0 | 0 | 0.005395 | 0.021198 | 0.044557 | 0.049696 | 0.021062 | 0.008801 | 0.002622 | 0.014885 | 0.006146 |
| 35.8333333 | 0 | 0 | 0 | 0 | 0 | 0 | 0 | 0 | 0 | 0.005212 | 0.021894 | 0.056104 | 0.037553 | 0.021013 | 0.012714 | 0.003782 | 0.007334 | 0.003331 |
| 35.8472222 | 0 | 0 | 0 | 0 | 0 | 0 | 0 | 0 | 0 | 0.003112 | 0.026647 | 0.059699 | 0.022066 | 0.023359 | 0.017067 | 0.005923 | 0.003631 | 0.005949 |
| 35.8611111 | 0 | 0 | 0 | 0 | 0 | 0 | 0 | 0 | 0 | 0.002636 | 0.032902 | 0.05121 | 0.022253 | 0.027462 | 0.021741 | 0.006759 | 0.00264 | 0.00766 |
| 35.875 | 0 | 0 | 0 | 0 | 0 | 0 | 0 | 0 | 0 | 0.005719 | 0.034936 | 0.035914 | 0.039229 | 0.029897 | 0.024486 | 0.007571 | 0.00559 | 0.006028 |
| 35.8888889 | 0 | 0 | 0 | 0 | 0 | 0 | 0 | 0 | 0 | 0.008356 | 0.030848 | 0.02328 | 0.063869 | 0.027759 | 0.023807 | 0.010123 | 0.007444 | 0.003459 |
| 35.9027778 | 0 | 0 | 0 | 0 | 0 | 0 | 0 | 0 | 0 | 0.008392 | 0.023487 | 0.017985 | 0.090174 | 0.021996 | 0.021035 | 0.011757 | 0.006112 | 0.003493 |
| 35.9166667 | 0 | 0 | 0 | 0 | 0 | 0 | 0 | 0 | 0 | 0.017287 | 0.013568 | 0.016569 | 0.104072 | 0.015346 | 0.018239 | 0.009323 | 0.008934 | 0.007399 |
| 35.9305556 | 0 | 0 | 0 | 0 | 0 | 0 | 0 | 0 | 0 | 0.0468 | 0.012833 | 0.014247 | 0.093371 | 0.009223 | 0.015058 | 0.005413 | 0.018428 | 0.014863 |
| 35.9444444 | 0 | 0 | 0 | 0 | 0 | 0 | 0 | 0 | 0 | 0.088573 | 0.037686 | 0.012074 | 0.057476 | 0.006414 | 0.009916 | 0.005243 | 0.031017 | 0.025722 |
| 35.9583333 | 0 | 0 | 0 | 0 | 0 | 0 | 0 | 0 | 0 | 0.119342 | 0.072544 | 0.01326 | 0.034602 | 0.011963 | 0.008562 | 0.008401 | 0.046249 | 0.03759 |
| 35.9722222 | 0 | 0 | 0 | 0 | 0 | 0 | 0 | 0 | 0 | 0.126608 | 0.091478 | 0.018241 | 0.062107 | 0.022094 | 0.015541 | 0.009559 | 0.057268 | 0.046177 |
| 35.9861111 | 0 | 0 | 0 | 0 | 0 | 0 | 0 | 0 | 0 | 0.118797 | 0.094037 | 0.027181 | 0.099947 | 0.026462 | 0.021957 | 0.006287 | 0.055612 | 0.0467 |
| 36 | 0 | 0 | 0 | 0 | 0 | 0 | 0 | 0 | 0 | 0.104171 | 0.082304 | 0.039025 | 0.110825 | 0.02037 | 0.019841 | 0.003103 | 0.04245 | 0.03801 |
| 36.0138889 | 0 | 0 | 0 | 0 | 0 | 0 | 0 | 0 | 0 | 0.089577 | 0.051512 | 0.049142 | 0.096978 | 0.010738 | 0.011868 | 0.00624 | 0.024765 | 0.025439 |
| 36.0277778 | 0 | 0 | 0 | 0 | 0 | 0 | 0 | 0 | 0 | 0.079357 | 0.022662 | 0.054933 | 0.076491 | 0.007762 | 0.006463 | 0.010701 | 0.011155 | 0.014668 |
| 36.0416667 | 0 | 0 | 0 | 0 | 0 | 0 | 0 | 0 | 0 | 0.06426 | 0.02135 | 0.056258 | 0.077385 | 0.010492 | 0.005397 | 0.00916 | 0.006932 | 0.009585 |
| 36.0555556 | 0 | 0 | 0 | 0 | 0 | 0 | 0 | 0 | 0 | 0.043032 | 0.034982 | 0.055965 | 0.083161 | 0.014864 | 0.00628 | 0.009729 | 0.012305 | 0.011697 |
| 36.0694444 | 0 | 0 | 0 | 0 | 0 | 0 | 0 | 0 | 0 | 0.032728 | 0.032253 | 0.066276 | 0.073551 | 0.014243 | 0.015014 | 0.022009 | 0.027572 | 0.02277 |
| 36.0833333 | 0 | 0 | 0 | 0 | 0 | 0 | 0 | 0 | 0 | 0.056527 | 0.039718 | 0.124836 | 0.084799 | 0.023305 | 0.040251 | 0.054699 | 0.05948 | 0.054055 |
| 36.0972222 | 0 | 0 | 0 | 0 | 0 | 0 | 0 | 0 | 0 | 0.108603 | 0.095659 | 0.215489 | 0.139644 | 0.068373 | 0.074735 | 0.111342 | 0.105266 | 0.099343 |
| 36.1111111 | 0 | 0 | 0 | 0 | 0 | 0 | 0 | 0 | 0 | 0.134941 | 0.129121 | 0.238762 | 0.188087 | 0.108196 | 0.084748 | 0.146449 | 0.131264 | 0.107777 |
| 36.125 | 0 | 0 | 0 | 0 | 0 | 0 | 0 | 0 | 0 | 0.111673 | 0.097272 | 0.180831 | 0.180772 | 0.09709 | 0.056168 | 0.115674 | 0.111064 | 0.064227 |
| 36.1388889 | 0 | 0 | 0 | 0 | 0 | 0 | 0 | 0 | 0 | 0.082782 | 0.058366 | 0.116761 | 0.145692 | 0.063434 | 0.023359 | 0.056095 | 0.068228 | 0.0332 |
| 36.1527778 | 0 | 0 | 0 | 0 | 0 | 0 | 0 | 0 | 0 | 0.078622 | 0.040577 | 0.081866 | 0.126382 | 0.037866 | 0.00982 | 0.018166 | 0.035517 | 0.038182 |
| 36.1666667 | 0 | 0 | 0 | 0 | 0 | 0 | 0 | 0 | 0 | 0.082613 | 0.038428 | 0.07152 | 0.119544 | 0.021969 | 0.004714 | 0.006635 | 0.016199 | 0.038332 |
| 36.1805556 | 0 | 0 | 0 | 0 | 0 | 0 | 0 | 0 | 0 | 0.076418 | 0.040749 | 0.070868 | 0.106104 | 0.013943 | 0.003274 | 0.007616 | 0.009178 | 0.024926 |
| 36.1944444 | 0 | 0 | 0 | 0 | 0 | 0 | 0 | 0 | 0 | 0.063629 | 0.034583 | 0.066341 | 0.08201 | 0.012635 | 0.009202 | 0.010257 | 0.015449 | 0.01407 |
| 36.2083333 | 0 | 0 | 0 | 0 | 0 | 0 | 0 | 0 | 0 | 0.050241 | 0.024541 | 0.055899 | 0.056739 | 0.01414 | 0.015517 | 0.009378 | 0.019859 | 0.007821 |
| 36.2222222 | 0 | 0 | 0 | 0 | 0 | 0 | 0 | 0 | 0 | 0.034424 | 0.01666 | 0.044358 | 0.035815 | 0.013048 | 0.015365 | 0.006731 | 0.01318 | 0.004567 |
| 36.2361111 | 0 | 0 | 0 | 0 | 0 | 0 | 0 | 0 | 0 | 0.017801 | 0.009493 | 0.035518 | 0.018537 | 0.008937 | 0.013481 | 0.009725 | 0.004612 | 0.005046 |
| 36.25 | 0 | 0 | 0 | 0 | 0 | 0 | 0 | 0 | 0 | 0.00706 | 0.007009 | 0.031226 | 0.009351 | 0.005396 | 0.013244 | 0.016276 | 0.001811 | 0.006005 |
| 36.2638889 | 0 | 0 | 0 | 0 | 0 | 0 | 0 | 0 | 0 | 0.004306 | 0.012192 | 0.029842 | 0.015196 | 0.003589 | 0.012498 | 0.017363 | 0.004566 | 0.00421 |
| 36.2777778 | 0 | 0 | 0 | 0 | 0 | 0 | 0 | 0 | 0 | 0.004245 | 0.02022 | 0.02816 | 0.033089 | 0.002756 | 0.010387 | 0.011404 | 0.01117 | 0.002506 |
| 36.2916667 | 0 | 0 | 0 | 0 | 0 | 0 | 0 | 0 | 0 | 0.005058 | 0.030646 | 0.023365 | 0.053024 | 0.005247 | 0.007541 | 0.005975 | 0.022874 | 0.003356 |
| 36.3055556 | 0 | 0 | 0 | 0 | 0 | 0 | 0 | 0 | 0 | 0.00894 | 0.042935 | 0.013718 | 0.069943 | 0.011672 | 0.006887 | 0.009585 | 0.039714 | 0.005898 |
| 36.3194444 | 0 | 0 | 0 | 0 | 0 | 0 | 0 | 0 | 0 | 0.014906 | 0.048784 | 0.006557 | 0.081478 | 0.01848 | 0.010232 | 0.019263 | 0.054932 | 0.01049 |
| 36.3333333 | 0 | 0 | 0 | 0 | 0 | 0 | 0 | 0 | 0 | 0.020529 | 0.050025 | 0.006617 | 0.082431 | 0.023974 | 0.014576 | 0.026024 | 0.061144 | 0.015414 |
| 36.3472222 | 0 | 0 | 0 | 0 | 0 | 0 | 0 | 0 | 0 | 0.023978 | 0.05554 | 0.006362 | 0.070837 | 0.026937 | 0.016167 | 0.028062 | 0.060692 | 0.017321 |
| 36.3611111 | 0 | 0 | 0 | 0 | 0 | 0 | 0 | 0 | 0 | 0.012475 | 0.029946 | 0.002488 | 0.031147 | 0.013735 | 0.007994 | 0.014124 | 0.029938 | 0.008664 |
| 36.375 | 0 | 0 | 0 | 0 | 0 | 0 | 0 | 0 | 0 | 0 | 0 | 0 | 0 | 0 | 0 | 0 | 0 | 0 |
| 36.3888889 | 0 | 0 | 0 | 0 | 0 | 0 | 0 | 0 | 0 | 0 | 0 | 0 | 0 | 0 | 0 | 0 | 0 | 0 |
| 36.4027778 | 0 | 0 | 0 | 0 | 0 | 0 | 0 | 0 | 0 | 0 | 0 | 0 | 0 | 0 | 0 | 0 | 0 | 0 |
| 36.4166667 | 0 | 0 | 0 | 0 | 0 | 0 | 0 | 0 | 0 | 0 | 0 | 0 | 0 | 0 | 0 | 0 | 0 | 0 |
| 36.4305556 | 0 | 0 | 0 | 0 | 0 | 0 | 0 | 0 | 0 | 0 | 0 | 0 | 0 | 0 | 0 | 0 | 0 | 0 |
| 36.4444444 | 0 | 0 | 0 | 0 | 0 | 0 | 0 | 0 | 0 | 0 | 0 | 0 | 0 | 0 | 0 | 0 | 0 | 0 |
| 36.4583333 | 0 | 0 | 0 | 0 | 0 | 0 | 0 | 0 | 0 | 0 | 0 | 0 | 0 | 0 | 0 | 0 | 0 | 0 |
| 36.4722222 | 0 | 0 | 0 | 0 | 0 | 0 | 0 | 0 | 0 | 0 | 0 | 0 | 0 | 0 | 0 | 0 | 0 | 0 |
| 36.4861111 | 0 | 0 | 0 | 0 | 0 | 0 | 0 | 0 | 0 | 0 | 0 | 0 | 0 | 0 | 0 | 0 | 0 | 0 |
| 36.5 | 0 | 0 | 0 | 0 | 0 | 0 | 0 | 0 | 0 | 0 | 0 | 0 | 0 | 0 | 0 | 0 | 0 | 0 |
| 36.5138889 | 0 | 0 | 0 | 0 | 0 | 0 | 0 | 0 | 0 | 0 | 0 | 0 | 0 | 0 | 0 | 0 | 0 | 0 |

| | | | | | | | | | | | | | | | | | | |
|---|---|---|---|---|---|---|---|---|---|---|---|---|---|---|---|---|---|---|
| 36.5277778 | 0 | 0 | 0 | 0 | 0 | 0 | 0 | 0 | 0 | 0 | 0 | 0 | 0 | 0 | 0 | 0 | 0 | 0 |
| 36.5416667 | 0 | 0 | 0 | 0 | 0 | 0 | 0 | 0 | 0 | 0 | 0 | 0 | 0 | 0 | 0 | 0 | 0 | 0 |
| 36.5555556 | 0 | 0 | 0 | 0 | 0 | 0 | 0 | 0 | 0 | 0 | 0 | 0 | 0 | 0 | 0 | 0 | 0 | 0 |
| 36.5694444 | 0 | 0 | 0 | 0 | 0 | 0 | 0 | 0 | 0 | 0 | 0 | 0 | 0 | 0 | 0 | 0 | 0 | 0 |
| 36.5833333 | 0 | 0 | 0 | 0 | 0 | 0 | 0 | 0 | 0 | 0 | 0 | 0 | 0 | 0 | 0 | 0 | 0 | 0 |
| 36.5972222 | 0.045727 | 0.05748 | 0.025527 | 0.02436 | 0.021253 | 0.010056 | 0.026382 | 0.01053 | 0.013187 | 0 | 0 | 0 | 0 | 0 | 0 | 0 | 0 | 0 |
| 36.6111111 | 0.093022 | 0.119438 | 0.04858 | 0.046299 | 0.055568 | 0.025455 | 0.05621 | 0.018739 | 0.038849 | 0.056034 | 0.059513 | 0.038742 | 0.011674 | 0.025831 | 0.021221 | 0.049684 | 0.038746 | 0.032084 |
| 36.625 | 0.078739 | 0.103409 | 0.038658 | 0.036031 | 0.079102 | 0.028149 | 0.055126 | 0.023988 | 0.049411 | 0.118043 | 0.11586 | 0.063932 | 0.041197 | 0.048783 | 0.040054 | 0.083961 | 0.061626 | 0.05859 |
| 36.6388889 | 0.053167 | 0.06698 | 0.025591 | 0.024699 | 0.096367 | 0.023747 | 0.045802 | 0.030646 | 0.041458 | 0.117774 | 0.108234 | 0.046998 | 0.075056 | 0.046539 | 0.0375 | 0.061415 | 0.04519 | 0.051532 |
| 36.6527778 | 0.04522 | 0.046864 | 0.017385 | 0.024362 | 0.103787 | 0.023036 | 0.03849 | 0.031355 | 0.037805 | 0.102964 | 0.099739 | 0.054383 | 0.103735 | 0.0499 | 0.039398 | 0.056365 | 0.054364 | 0.052023 |
| 36.6666667 | 0.052033 | 0.044344 | 0.014517 | 0.036147 | 0.108391 | 0.027857 | 0.038489 | 0.03195 | 0.041974 | 0.090152 | 0.086313 | 0.078758 | 0.131203 | 0.053304 | 0.041275 | 0.065313 | 0.074199 | 0.051435 |
| 36.6805556 | 0.060045 | 0.047388 | 0.015 | 0.053941 | 0.110212 | 0.03216 | 0.046405 | 0.021997 | 0.037176 | 0.087086 | 0.062091 | 0.099499 | 0.155438 | 0.053088 | 0.037394 | 0.078586 | 0.089147 | 0.041164 |
| 36.6944444 | 0.062125 | 0.048057 | 0.017023 | 0.070943 | 0.100937 | 0.029689 | 0.053131 | 0.012645 | 0.020281 | 0.09048 | 0.03357 | 0.108204 | 0.161988 | 0.050763 | 0.026855 | 0.08444 | 0.089131 | 0.025135 |
| 36.7083333 | 0.058036 | 0.04424 | 0.02084 | 0.085314 | 0.088403 | 0.026525 | 0.055797 | 0.020828 | 0.010646 | 0.090474 | 0.015137 | 0.110655 | 0.146957 | 0.047973 | 0.017256 | 0.07499 | 0.071617 | 0.011909 |
| 36.7222222 | 0.050461 | 0.035281 | 0.027444 | 0.091867 | 0.072778 | 0.026593 | 0.059011 | 0.030372 | 0.019772 | 0.077611 | 0.016909 | 0.108623 | 0.111678 | 0.043373 | 0.020325 | 0.055425 | 0.045976 | 0.005155 |
| 36.7361111 | 0.040963 | 0.022322 | 0.03464 | 0.078856 | 0.04963 | 0.021828 | 0.051506 | 0.027439 | 0.028287 | 0.051178 | 0.032555 | 0.099597 | 0.061836 | 0.035464 | 0.032658 | 0.038956 | 0.023703 | 0.003332 |
| 36.75 | 0.028729 | 0.010971 | 0.035234 | 0.044601 | 0.025289 | 0.015031 | 0.03049 | 0.022784 | 0.020463 | 0.022974 | 0.048926 | 0.088512 | 0.034128 | 0.026081 | 0.040983 | 0.03169 | 0.009231 | 0.005837 |
| 36.7638889 | 0.014706 | 0.009441 | 0.027026 | 0.022251 | 0.015828 | 0.020315 | 0.023056 | 0.029652 | 0.007703 | 0.007417 | 0.061825 | 0.07672 | 0.054349 | 0.016434 | 0.041261 | 0.029144 | 0.004491 | 0.011658 |
| 36.7777778 | 0.006009 | 0.018232 | 0.018061 | 0.033045 | 0.032379 | 0.034757 | 0.035601 | 0.053018 | 0.002879 | 0.011737 | 0.068675 | 0.057425 | 0.095544 | 0.007633 | 0.036093 | 0.026078 | 0.008068 | 0.016948 |
| 36.7916667 | 0.006885 | 0.029364 | 0.014181 | 0.048355 | 0.064822 | 0.041289 | 0.038856 | 0.085142 | 0.006467 | 0.032877 | 0.066142 | 0.03199 | 0.142598 | 0.002306 | 0.02718 | 0.022022 | 0.012578 | 0.019347 |
| 36.8055556 | 0.011669 | 0.037294 | 0.015622 | 0.049561 | 0.093 | 0.038187 | 0.02383 | 0.104373 | 0.011525 | 0.05428 | 0.046252 | 0.022431 | 0.192951 | 0.000739 | 0.017189 | 0.017992 | 0.014083 | 0.019395 |
| 36.8194444 | 0.011781 | 0.039859 | 0.021808 | 0.04071 | 0.107187 | 0.036261 | 0.010692 | 0.089173 | 0.009016 | 0.057415 | 0.023307 | 0.039401 | 0.212815 | 0.003143 | 0.01144 | 0.013711 | 0.013903 | 0.018153 |
| 36.8333333 | 0.006744 | 0.036523 | 0.030288 | 0.027429 | 0.113219 | 0.039502 | 0.009312 | 0.053173 | 0.005437 | 0.046023 | 0.023381 | 0.059714 | 0.184396 | 0.008639 | 0.010376 | 0.008443 | 0.013311 | 0.015046 |
| 36.8472222 | 0.003214 | 0.029452 | 0.035297 | 0.015229 | 0.115803 | 0.046879 | 0.013661 | 0.044144 | 0.007793 | 0.03384 | 0.028729 | 0.066904 | 0.132297 | 0.011732 | 0.009324 | 0.003482 | 0.012077 | 0.009938 |
| 36.8611111 | 0.002504 | 0.022809 | 0.031994 | 0.008454 | 0.113118 | 0.050644 | 0.017086 | 0.073723 | 0.01069 | 0.023268 | 0.024052 | 0.062763 | 0.083114 | 0.009511 | 0.008098 | 0.001446 | 0.008967 | 0.006042 |
| 36.875 | 0.002414 | 0.019467 | 0.023126 | 0.012924 | 0.095075 | 0.047602 | 0.016979 | 0.099037 | 0.011054 | 0.015758 | 0.01783 | 0.052646 | 0.044718 | 0.007945 | 0.008067 | 0.00246 | 0.004696 | 0.005977 |
| 36.8888889 | 0.003743 | 0.018557 | 0.015213 | 0.026768 | 0.064225 | 0.042866 | 0.01294 | 0.097851 | 0.010136 | 0.01336 | 0.01107 | 0.043838 | 0.018635 | 0.014313 | 0.010029 | 0.005699 | 0.004042 | 0.007216 |
| 36.9027778 | 0.004503 | 0.018342 | 0.008726 | 0.041707 | 0.040179 | 0.040415 | 0.007804 | 0.085419 | 0.013208 | 0.016419 | 0.007701 | 0.03894 | 0.009882 | 0.021216 | 0.015734 | 0.009514 | 0.008771 | 0.006145 |
| 36.9166667 | 0.004902 | 0.019327 | 0.008698 | 0.052468 | 0.026416 | 0.040386 | 0.00777 | 0.068053 | 0.025368 | 0.026763 | 0.013985 | 0.038755 | 0.026856 | 0.021177 | 0.022664 | 0.008643 | 0.011962 | 0.004949 |
| 36.9305556 | 0.006757 | 0.023295 | 0.018547 | 0.057134 | 0.014753 | 0.037763 | 0.014194 | 0.04071 | 0.055492 | 0.040407 | 0.026151 | 0.043911 | 0.064137 | 0.02144 | 0.026501 | 0.005206 | 0.009285 | 0.00804 |
| 36.9444444 | 0.007045 | 0.029357 | 0.025496 | 0.055038 | 0.006773 | 0.032096 | 0.024367 | 0.027461 | 0.089516 | 0.049879 | 0.038362 | 0.049944 | 0.097768 | 0.025533 | 0.025942 | 0.007387 | 0.009091 | 0.014995 |
| 36.9583333 | 0.010124 | 0.033609 | 0.027088 | 0.046698 | 0.009261 | 0.02846 | 0.036508 | 0.053487 | 0.096478 | 0.054228 | 0.052405 | 0.051233 | 0.11433 | 0.028846 | 0.022648 | 0.012916 | 0.01661 | 0.021925 |
| 36.9722222 | 0.019379 | 0.032415 | 0.032191 | 0.035155 | 0.023524 | 0.033326 | 0.046482 | 0.090964 | 0.082504 | 0.056761 | 0.064226 | 0.047258 | 0.12008 | 0.027118 | 0.02118 | 0.015863 | 0.024761 | 0.026312 |
| 36.9861111 | 0.02584 | 0.025384 | 0.042615 | 0.025898 | 0.045019 | 0.047342 | 0.051303 | 0.116821 | 0.067188 | 0.061311 | 0.064617 | 0.04252 | 0.119055 | 0.019682 | 0.026816 | 0.014772 | 0.031042 | 0.02761 |
| 37 | 0.024615 | 0.01516 | 0.052211 | 0.02207 | 0.062198 | 0.056822 | 0.052175 | 0.141312 | 0.05351 | 0.068214 | 0.055559 | 0.039736 | 0.111179 | 0.010928 | 0.039191 | 0.01136 | 0.036215 | 0.025422 |
| 37.0138889 | 0.017514 | 0.007875 | 0.051496 | 0.021305 | 0.065303 | 0.051288 | 0.048622 | 0.161261 | 0.041967 | 0.070064 | 0.043667 | 0.034722 | 0.097052 | 0.009026 | 0.049249 | 0.007275 | 0.036516 | 0.020494 |
| 37.0277778 | 0.010492 | 0.010263 | 0.039367 | 0.020086 | 0.057526 | 0.037596 | 0.039738 | 0.157225 | 0.031589 | 0.058504 | 0.030736 | 0.023447 | 0.073929 | 0.01666 | 0.052183 | 0.004647 | 0.030534 | 0.014029 |
| 37.0416667 | 0.013107 | 0.021078 | 0.020904 | 0.017745 | 0.046021 | 0.02436 | 0.027426 | 0.114603 | 0.02081 | 0.033241 | 0.018009 | 0.010575 | 0.043145 | 0.02653 | 0.053568 | 0.006849 | 0.020858 | 0.007711 |
| 37.0555556 | 0.023576 | 0.032853 | 0.008014 | 0.014262 | 0.033619 | 0.014394 | 0.014642 | 0.062986 | 0.022479 | 0.014221 | 0.014619 | 0.004618 | 0.019682 | 0.034547 | 0.057243 | 0.010288 | 0.010753 | 0.004944 |
| 37.0694444 | 0.026302 | 0.030126 | 0.005708 | 0.009374 | 0.026323 | 0.011147 | 0.006793 | 0.052877 | 0.038417 | 0.012592 | 0.017017 | 0.015764 | 0.014111 | 0.035847 | 0.054516 | 0.012765 | 0.011272 | 0.012991 |
| 37.0833333 | 0.026415 | 0.032125 | 0.012403 | 0.005199 | 0.035575 | 0.021015 | 0.010262 | 0.052593 | 0.039963 | 0.040396 | 0.026725 | 0.057074 | 0.048044 | 0.033361 | 0.045923 | 0.035271 | 0.042628 | 0.057216 |
| 37.0972222 | 0.047252 | 0.066559 | 0.03126 | 0.003105 | 0.060027 | 0.041369 | 0.022468 | 0.052898 | 0.038153 | 0.107492 | 0.069151 | 0.110891 | 0.139517 | 0.051992 | 0.059136 | 0.087762 | 0.099022 | 0.14218 |
| 37.1111111 | 0.065835 | 0.080952 | 0.041559 | 0.006009 | 0.075026 | 0.052626 | 0.029694 | 0.098634 | 0.048074 | 0.146079 | 0.113588 | 0.125434 | 0.215794 | 0.081638 | 0.084728 | 0.12136 | 0.132677 | 0.185775 |
| 37.125 | 0.049223 | 0.045474 | 0.029082 | 0.018756 | 0.069744 | 0.044237 | 0.026638 | 0.133947 | 0.03524 | 0.111123 | 0.110247 | 0.091212 | 0.20733 | 0.08015 | 0.079603 | 0.099849 | 0.111218 | 0.133589 |
| 37.1388889 | 0.020106 | 0.010677 | 0.016028 | 0.034563 | 0.067869 | 0.027188 | 0.022564 | 0.117595 | 0.014198 | 0.058032 | 0.083907 | 0.054348 | 0.160562 | 0.051438 | 0.049106 | 0.054043 | 0.059798 | 0.05545 |
| 37.1527778 | 0.006959 | 0.003325 | 0.016848 | 0.04009 | 0.078453 | 0.012585 | 0.022388 | 0.089858 | 0.019184 | 0.041916 | 0.071755 | 0.041422 | 0.133454 | 0.024511 | 0.024558 | 0.020544 | 0.020651 | 0.014027 |
| 37.1666667 | 0.009953 | 0.010136 | 0.02313 | 0.034743 | 0.079346 | 0.004565 | 0.023042 | 0.069394 | 0.034059 | 0.054005 | 0.069965 | 0.042958 | 0.118878 | 0.009633 | 0.022743 | 0.008807 | 0.007832 | 0.0059 |
| 37.1805556 | 0.018735 | 0.020106 | 0.026567 | 0.028323 | 0.067744 | 0.002475 | 0.022853 | 0.05211 | 0.042049 | 0.058873 | 0.065859 | 0.045695 | 0.096908 | 0.006507 | 0.038514 | 0.009891 | 0.011719 | 0.014344 |
| 37.1944444 | 0.025593 | 0.029366 | 0.02785 | 0.024496 | 0.062819 | 0.002915 | 0.022894 | 0.036078 | 0.04382 | 0.051816 | 0.059037 | 0.043124 | 0.071871 | 0.009117 | 0.051516 | 0.011 | 0.01745 | 0.026756 |
| 37.2083333 | 0.027643 | 0.034778 | 0.029659 | 0.021625 | 0.065457 | 0.003984 | 0.022442 | 0.020181 | 0.039608 | 0.041132 | 0.05287 | 0.037588 | 0.052308 | 0.012586 | 0.054844 | 0.010118 | 0.017122 | 0.037742 |
| 37.2222222 | 0.023347 | 0.034488 | 0.030234 | 0.018927 | 0.063587 | 0.004279 | 0.019425 | 0.010472 | 0.026839 | 0.027097 | 0.045242 | 0.032184 | 0.037329 | 0.014823 | 0.04948 | 0.010749 | 0.01312 | 0.043942 |
| 37.2361111 | 0.014633 | 0.028069 | 0.024965 | 0.016618 | 0.051374 | 0.00306 | 0.014563 | 0.014399 | 0.016862 | 0.014309 | 0.032752 | 0.026138 | 0.021517 | 0.010958 | 0.031735 | 0.015559 | 0.008328 | 0.042295 |
| 37.25 | 0.007486 | 0.018448 | 0.015144 | 0.016426 | 0.03144 | 0.001373 | 0.00919 | 0.023082 | 0.026598 | 0.011486 | 0.017734 | 0.021139 | 0.012492 | 0.00692 | 0.018793 | 0.022363 | 0.006809 | 0.036133 |
| 37.2638889 | 0.007135 | 0.011567 | 0.006286 | 0.018324 | 0.013045 | 0.00043 | 0.004466 | 0.032234 | 0.048692 | 0.01633 | 0.007117 | 0.020612 | 0.018275 | 0.012293 | 0.032072 | 0.02674 | 0.011758 | 0.030868 |
| 37.2777778 | 0.011824 | 0.009819 | 0.002563 | 0.020178 | 0.005553 | 0.000162 | 0.002324 | 0.044935 | 0.068916 | 0.021951 | 0.005581 | 0.022321 | 0.027446 | 0.017993 | 0.052906 | 0.028814 | 0.01885 | 0.026116 |
| 37.2916667 | 0.01833 | 0.008494 | 0.004708 | 0.020715 | 0.009743 | 0.00051 | 0.0027 | 0.05749 | 0.084165 | 0.030102 | 0.01 | 0.019756 | 0.036471 | 0.015234 | 0.058146 | 0.02918 | 0.026714 | 0.018832 |
| 37.3055556 | 0.025133 | 0.00451 | 0.008852 | 0.019807 | 0.01734 | 0.001583 | 0.003406 | 0.064682 | 0.095541 | 0.041433 | 0.015178 | 0.011791 | 0.051532 | 0.008413 | 0.049008 | 0.028521 | 0.036538 | 0.009737 |
| 37.3194444 | 0.028679 | 0.001628 | 0.011549 | 0.017085 | 0.022272 | 0.002619 | 0.002869 | 0.060811 | 0.102688 | 0.053203 | 0.020765 | 0.007188 | 0.069602 | 0.006913 | 0.030915 | 0.029597 | 0.046542 | 0.006457 |
| 37.3333333 | 0.028011 | 0.001278 | 0.012057 | 0.01256 | 0.024179 | 0.002729 | 0.002051 | 0.05323 | 0.094535 | 0.061548 | 0.026761 | 0.011247 | 0.082493 | 0.014909 | 0.013501 | 0.032061 | 0.052859 | 0.013147 |
| 37.3472222 | 0.024108 | 0.003165 | 0.009943 | 0.007805 | 0.023305 | 0.00175 | 0.001872 | 0.056031 | 0.07081 | 0.061388 | 0.028637 | 0.014135 | 0.083088 | 0.022591 | 0.005277 | 0.033769 | 0.051916 | 0.020662 |
| 37.3611111 | 0.016306 | 0.00973 | 0.005817 | 0.006531 | 0.020269 | 0.000689 | 0.002731 | 0.069494 | 0.050066 | 0.051087 | 0.022133 | 0.009673 | 0.070957 | 0.022496 | 0.002513 | 0.035353 | 0.044808 | 0.019377 |
| 37.375 | 0.007474 | 0.01691 | 0.002693 | 0.011347 | 0.017118 | 0.000374 | 0.005019 | 0.086064 | 0.034822 | 0.035576 | 0.011046 | 0.003823 | 0.05458 | 0.019218 | 0.001831 | 0.036944 | 0.036496 | 0.012286 |
| 37.3888889 | 0.002503 | 0.017216 | 0.002958 | 0.019605 | 0.014875 | 0.000319 | 0.006593 | 0.093484 | 0.017296 | 0.025634 | 0.003571 | 0.002025 | 0.046036 | 0.019238 | 0.006338 | 0.036534 | 0.031478 | 0.008031 |
| 37.4027778 | 0.003255 | 0.010844 | 0.005454 | 0.026826 | 0.013152 | 0.00075 | 0.006511 | 0.085276 | 0.005171 | 0.028808 | 0.001942 | 0.00364 | 0.050356 | 0.021328 | 0.018857 | 0.03313 | 0.029178 | 0.006732 |
| 37.4166667 | 0.009657 | 0.00508 | 0.007721 | 0.031746 | 0.011443 | 0.00232 | 0.00603 | 0.066866 | 0.004379 | 0.039425 | 0.002875 | 0.004279 | 0.057498 | 0.022024 | 0.034021 | 0.029112 | 0.026492 | 0.006154 |
| 37.4305556 | 0.018078 | 0.004971 | 0.009065 | 0.035445 | 0.009245 | 0.003945 | 0.00462 | 0.049399 | 0.01285 | 0.044452 | 0.004091 | 0.002409 | 0.052168 | 0.01986 | 0.0416 | 0.025303 | 0.021692 | 0.008584 |
| 37.4444444 | 0.023408 | 0.00771 | 0.010129 | 0.038174 | 0.005652 | 0.003778 | 0.002621 | 0.042666 | 0.031807 | 0.040364 | 0.006638 | 0.001271 | 0.032116 | 0.012309 | 0.036337 | 0.020173 | 0.014849 | 0.010481 |
| 37.4583333 | 0.02429 | 0.009975 | 0.011069 | 0.03875 | 0.004681 | 0.001945 | 0.002852 | 0.044543 | 0.054846 | 0.034481 | 0.013284 | 0.003036 | 0.027543 | 0.005152 | 0.023172 | 0.016437 | 0.008113 | 0.007606 |
| 37.4722222 | 0.02299 | 0.01223 | 0.011911 | 0.037194 | 0.011453 | 0.000593 | 0.006112 | 0.046182 | 0.06856 | 0.033893 | 0.023164 | 0.005993 | 0.05229 | 0.003952 | 0.016889 | 0.017251 | 0.004345 | 0.004364 |
| 37.4861111 | 0.02148 | 0.014034 | 0.013426 | 0.036343 | 0.020441 | 0.000615 | 0.011141 | 0.041055 | 0.065649 | 0.039562 | 0.030259 | 0.007547 | 0.059321 | 0.008183 | 0.015303 | 0.020425 | 0.004763 | 0.005931 |
| 37.5 | 0.018876 | 0.012867 | 0.01567 | 0.03833 | 0.025765 | 0.000889 | 0.015879 | 0.025464 | 0.045108 | 0.045904 | 0.030594 | 0.007343 | 0.033891 | 0.016011 | 0.01305 | 0.021741 | 0.009107 | 0.010071 |
| 37.5138889 | 0.014155 | 0.008737 | 0.016962 | 0.040906 | 0.029511 | 0.000641 | 0.016768 | 0.015666 | 0.024448 | 0.048436 | 0.026733 | 0.007546 | 0.017242 | 0.023153 | 0.022647 | 0.020539 | 0.015591 | 0.013041 |
| 37.5277778 | 0.009101 | 0.005494 | 0.015958 | 0.039866 | 0.033855 | 0.00043 | 0.012449 | 0.030159 | 0.027276 | 0.046934 | 0.022074 | 0.009402 | 0.020159 | 0.027377 | 0.03604 | 0.019493 | 0.021135 | 0.014097 |
| 37.5416667 | 0.00547 | 0.005298 | 0.013415 | 0.035018 | 0.03691 | 0.000992 | 0.006813 | 0.053564 | 0.046589 | 0.041591 | 0.016065 | 0.010103 | 0.020061 | 0.027621 | 0.037488 | 0.019246 | 0.023336 | 0.010451 |
| 37.5555556 | 0.003325 | 0.006674 | 0.011464 | 0.030517 | 0.036439 | 0.001444 | 0.005696 | 0.071002 | 0.064361 | 0.033174 | 0.008142 | 0.008262 | 0.011287 | 0.023619 | 0.029615 | 0.018933 | 0.022502 | 0.004809 |
| 37.5694444 | 0.006628 | 0.013178 | 0.014326 | 0.029445 | 0.029739 | 0.003378 | 0.014389 | 0.083301 | 0.076353 | 0.031769 | 0.005856 | 0.015571 | 0.015557 | 0.018813 | 0.022783 | 0.023498 | 0.028201 | 0.007493 |
| 37.5833333 | 0.026232 | 0.037782 | 0.027397 | 0.033637 | 0.022411 | 0.012525 | 0.039959 | 0.079193 | 0.071099 | 0.059239 | 0.024935 | 0.053081 | 0.059282 | 0.01875 | 0.028273 | 0.04123 | 0.058649 | 0.028831 |
| 37.5972222 | 0.063407 | 0.080751 | 0.047807 | 0.042235 | 0.036965 | 0.025997 | 0.077367 | 0.055045 | 0.050903 | 0.12599 | 0.069702 | 0.110876 | 0.13147 | 0.02636 | 0.050275 | 0.068042 | 0.103965 | 0.06598 |
| 37.6111111 | 0.093087 | 0.112836 | 0.061798 | 0.046763 | 0.067652 | 0.030563 | 0.099588 | 0.047032 | 0.060125 | 0.190781 | 0.11486 | 0.135853 | 0.173252 | 0.037387 | 0.086691 | 0.080969 | 0.114381 | 0.090223 |
| 37.625 | 0.095516 | 0.112289 | 0.062925 | 0.039769 | 0.091905 | 0.02267 | 0.093566 | 0.067911 | 0.107801 | 0.198674 | 0.133286 | 0.109473 | 0.157682 | 0.043437 | 0.124619 | 0.066833 | 0.079098 | 0.08662 |
| 37.6388889 | 0.085967 | 0.094521 | 0.058336 | 0.026555 | 0.121789 | 0.01316 | 0.077376 | 0.077962 | 0.130991 | 0.16814 | 0.129575 | 0.072563 | 0.12807 | 0.045719 | 0.145004 | 0.039757 | 0.047903 | 0.078094 |
| 37.6527778 | 0.081432 | 0.081317 | 0.052215 | 0.019331 | 0.161772 | 0.010628 | 0.067218 | 0.062222 | 0.09894 | 0.146631 | 0.117397 | 0.051064 | 0.116312 | 0.047569 | 0.131123 | 0.016804 | 0.046763 | 0.080189 |
| 37.6666667 | 0.078446 | 0.07586 | 0.043829 | 0.022752 | 0.186079 | 0.017625 | 0.063623 | 0.0355 | 0.054006 | 0.133046 | 0.095633 | 0.039003 | 0.106289 | 0.038657 | 0.083493 | 0.005591 | 0.057314 | 0.081931 |

| | | | | | | | | | | | | | | | | | | |
|---|---|---|---|---|---|---|---|---|---|---|---|---|---|---|---|---|---|---|
| 37.6805556 | 0.071505 | 0.071356 | 0.034851 | 0.031897 | 0.175443 | 0.032612 | 0.061617 | 0.019037 | 0.030126 | 0.113975 | 0.062964 | 0.030582 | 0.08736 | 0.020121 | 0.034813 | 0.003078 | 0.060909 | 0.073988 |
| 37.6944444 | 0.060644 | 0.06309 | 0.027568 | 0.039605 | 0.144547 | 0.048073 | 0.058086 | 0.027053 | 0.035372 | 0.097618 | 0.032111 | 0.025783 | 0.07067 | 0.010403 | 0.016326 | 0.003731 | 0.053731 | 0.061196 |
| 37.7083333 | 0.047712 | 0.052982 | 0.023193 | 0.041143 | 0.116109 | 0.05768 | 0.053396 | 0.043765 | 0.069518 | 0.085547 | 0.01605 | 0.024195 | 0.058087 | 0.009012 | 0.022118 | 0.00648 | 0.040046 | 0.051739 |
| 37.7222222 | 0.037714 | 0.044478 | 0.020607 | 0.037374 | 0.093582 | 0.058367 | 0.048496 | 0.053491 | 0.103184 | 0.070416 | 0.012304 | 0.022904 | 0.040494 | 0.007802 | 0.021957 | 0.007231 | 0.025313 | 0.045735 |
| 37.7361111 | 0.032012 | 0.038829 | 0.017018 | 0.031569 | 0.077933 | 0.048929 | 0.043063 | 0.059577 | 0.113144 | 0.05292 | 0.011896 | 0.017752 | 0.022173 | 0.010124 | 0.01114 | 0.005171 | 0.012139 | 0.037925 |
| 37.75 | 0.026463 | 0.03652 | 0.013153 | 0.024756 | 0.06592 | 0.030354 | 0.036638 | 0.066617 | 0.10598 | 0.035595 | 0.010442 | 0.010674 | 0.025274 | 0.01275 | 0.008903 | 0.003488 | 0.006134 | 0.028016 |
| 37.7638889 | 0.020712 | 0.036457 | 0.012182 | 0.016932 | 0.046659 | 0.016629 | 0.030532 | 0.07736 | 0.092821 | 0.018966 | 0.009083 | 0.012609 | 0.066975 | 0.013252 | 0.017131 | 0.00315 | 0.010233 | 0.01996 |
| 37.7777778 | 0.013758 | 0.035198 | 0.013558 | 0.008904 | 0.027655 | 0.025171 | 0.026316 | 0.095881 | 0.083094 | 0.007138 | 0.012278 | 0.029341 | 0.161492 | 0.016466 | 0.031053 | 0.004413 | 0.014996 | 0.018228 |
| 37.7916667 | 0.005977 | 0.03058 | 0.013702 | 0.006748 | 0.032829 | 0.044866 | 0.022669 | 0.123251 | 0.075998 | 0.016928 | 0.021022 | 0.050684 | 0.293274 | 0.022736 | 0.047205 | 0.006581 | 0.013756 | 0.02415 |
| 37.8055556 | 0.002429 | 0.024776 | 0.009951 | 0.013243 | 0.060336 | 0.04848 | 0.015239 | 0.15154 | 0.065128 | 0.046114 | 0.032662 | 0.050417 | 0.350124 | 0.028603 | 0.05832 | 0.009383 | 0.008177 | 0.029147 |
| 37.8194444 | 0.003047 | 0.020848 | 0.010212 | 0.01842 | 0.086513 | 0.035091 | 0.010319 | 0.166371 | 0.049244 | 0.065627 | 0.045702 | 0.02648 | 0.270334 | 0.032149 | 0.062177 | 0.013166 | 0.004953 | 0.022398 |
| 37.8333333 | 0.004589 | 0.018839 | 0.02069 | 0.016904 | 0.105565 | 0.026205 | 0.017886 | 0.156019 | 0.029689 | 0.06787 | 0.053684 | 0.01404 | 0.15592 | 0.033536 | 0.056808 | 0.016455 | 0.011267 | 0.017699 |
| 37.8472222 | 0.004965 | 0.016397 | 0.028489 | 0.014361 | 0.127675 | 0.020682 | 0.026957 | 0.122793 | 0.013737 | 0.052017 | 0.047662 | 0.034626 | 0.088506 | 0.031049 | 0.03967 | 0.018528 | 0.022735 | 0.029952 |
| 37.8611111 | 0.00526 | 0.012275 | 0.026909 | 0.01399 | 0.147619 | 0.011045 | 0.027391 | 0.082462 | 0.005764 | 0.031811 | 0.027722 | 0.099197 | 0.066704 | 0.023687 | 0.019615 | 0.019476 | 0.031287 | 0.04442 |
| 37.875 | 0.00985 | 0.007473 | 0.023598 | 0.01091 | 0.15126 | 0.004928 | 0.024086 | 0.050278 | 0.006191 | 0.032699 | 0.01054 | 0.202852 | 0.070017 | 0.016252 | 0.012561 | 0.019822 | 0.038229 | 0.054594 |
| 37.8888889 | 0.015267 | 0.004286 | 0.022808 | 0.01178 | 0.141459 | 0.002667 | 0.016386 | 0.03488 | 0.019894 | 0.033343 | 0.008373 | 0.248118 | 0.069557 | 0.012009 | 0.022432 | 0.0212 | 0.045562 | 0.067897 |
| 37.9027778 | 0.017293 | 0.005985 | 0.023073 | 0.028082 | 0.125027 | 0.001192 | 0.007986 | 0.03726 | 0.04874 | 0.02106 | 0.010799 | 0.184825 | 0.066333 | 0.008328 | 0.037396 | 0.022732 | 0.050733 | 0.079867 |
| 37.9166667 | 0.019687 | 0.012989 | 0.022776 | 0.046495 | 0.101909 | 0.000837 | 0.006969 | 0.052051 | 0.083397 | 0.01943 | 0.015256 | 0.121331 | 0.077246 | 0.006964 | 0.04391 | 0.022469 | 0.051718 | 0.082933 |
| 37.9305556 | 0.024126 | 0.021804 | 0.02344 | 0.054809 | 0.078521 | 0.001676 | 0.006799 | 0.07371 | 0.113521 | 0.026427 | 0.028139 | 0.108679 | 0.095138 | 0.010625 | 0.040799 | 0.017991 | 0.051324 | 0.078941 |

Supplementary Table S6. 1 hour integrated motion data in (a) lettuce subjected to Hypoxia

| IntervalMid | Control1 | Control2 | Control3 | Control4 | Control5 | Control6 | Control7 | Control8 | Control9 | Control10 | Control11 | Control12 | Stress1 | Stress2 | Stress3 | Stress4 | Stress5 | Stress6 | Stress7 | Stress8 | Stress9 | Stress10 | Stress11 | Stress12 |
|---|---|---|---|---|---|---|---|---|---|---|---|---|---|---|---|---|---|---|---|---|---|---|---|---|
| 18.1111111 | 0.027509 | 0.020401 | 0.016219 | 0.003804 | 0.014822 | 0.001633 | 0.016574 | 0.001892 | 0.047122 | 0.007611 | 0.006356 | 0.018381 | 0 | 0 | 0 | 0 | 0 | 0 | 0 | 0 | 0 | 0 | 0 | 0 |
| 18.125 | 0.013083 | 0.011243 | 0.010006 | 0.002767 | 0.006354 | 0.001563 | 0.005995 | 0.000962 | 0.034442 | 0.005058 | 0.004698 | 0.008786 | 0 | 0 | 0 | 0 | 0 | 0 | 0 | 0 | 0 | 0 | 0 | 0 |
| 18.1388889 | 0.005518 | 0.004912 | 0.008158 | 0.002457 | 0.002307 | 0.001823 | 0.001458 | 0.000744 | 0.019448 | 0.00263 | 0.004386 | 0.004156 | 0 | 0 | 0 | 0 | 0 | 0 | 0 | 0 | 0 | 0 | 0 | 0 |
| 18.1527778 | 0.002549 | 0.001898 | 0.00781 | 0.002686 | 0.000722 | 0.002619 | 0.001203 | 0.001098 | 0.011505 | 0.001029 | 0.004964 | 0.00212 | 0 | 0 | 0 | 0 | 0 | 0 | 0 | 0 | 0 | 0 | 0 | 0 |
| 18.1666667 | 0.00168 | 0.000881 | 0.00572 | 0.002929 | 0.000536 | 0.003628 | 0.001215 | 0.001138 | 0.007587 | 0.000342 | 0.005835 | 0.001573 | 0 | 0 | 0 | 0 | 0 | 0 | 0 | 0 | 0 | 0 | 0 | 0 |
| 18.1805556 | 0.001584 | 0.000858 | 0.00323 | 0.002746 | 0.000496 | 0.004031 | 0.000731 | 0.000753 | 0.006393 | 0.000272 | 0.006312 | 0.001846 | 0 | 0 | 0 | 0 | 0 | 0 | 0 | 0 | 0 | 0 | 0 | 0 |
| 18.1944444 | 0.001539 | 0.001271 | 0.002268 | 0.002269 | 0.000468 | 0.003506 | 0.000341 | 0.000646 | 0.006053 | 0.000546 | 0.006117 | 0.002003 | 0 | 0 | 0 | 0 | 0 | 0 | 0 | 0 | 0 | 0 | 0 | 0 |
| 18.2083333 | 0.001194 | 0.001779 | 0.001988 | 0.001937 | 0.000511 | 0.002643 | 0.000318 | 0.000741 | 0.00521 | 0.000899 | 0.005476 | 0.001496 | 0 | 0 | 0 | 0 | 0 | 0 | 0 | 0 | 0 | 0 | 0 | 0 |
| 18.2222222 | 0.000674 | 0.00212 | 0.001633 | 0.001894 | 0.000385 | 0.002029 | 0.000363 | 0.000566 | 0.003965 | 0.000969 | 0.004989 | 0.000681 | 0 | 0 | 0 | 0 | 0 | 0 | 0 | 0 | 0 | 0 | 0 | 0 |
| 18.2361111 | 0.0003 | 0.002026 | 0.001302 | 0.001931 | 0.000312 | 0.001736 | 0.000291 | 0.000254 | 0.003036 | 0.000736 | 0.005165 | 0.000206 | 0 | 0 | 0 | 0 | 0 | 0 | 0 | 0 | 0 | 0 | 0 | 0 |
| 18.25 | 0.000207 | 0.001395 | 0.000945 | 0.002008 | 0.000277 | 0.001742 | 0.00025 | 0.000118 | 0.002589 | 0.000447 | 0.005971 | 0.000159 | 0 | 0 | 0 | 0 | 0 | 0 | 0 | 0 | 0 | 0 | 0 | 0 |
| 18.2638889 | 0.000345 | 0.000801 | 0.000547 | 0.002153 | 0.000229 | 0.001928 | 0.000361 | 0.000149 | 0.002399 | 0.000434 | 0.007106 | 0.000239 | 0 | 0 | 0 | 0 | 0 | 0 | 0 | 0 | 0 | 0 | 0 | 0 |
| 18.2777778 | 0.00038 | 0.000848 | 0.000479 | 0.002041 | 0.000261 | 0.001989 | 0.000452 | 0.000181 | 0.002493 | 0.000586 | 0.008557 | 0.000356 | 0 | 0 | 0 | 0 | 0 | 0 | 0 | 0 | 0 | 0 | 0 | 0 |
| 18.2916667 | 0.000236 | 0.001326 | 0.000921 | 0.001652 | 0.000289 | 0.001897 | 0.000363 | 0.000236 | 0.002937 | 0.000525 | 0.010316 | 0.000458 | 0 | 0 | 0 | 0 | 0 | 0 | 0 | 0 | 0 | 0 | 0 | 0 |
| 18.3055556 | 0.000218 | 0.00186 | 0.001368 | 0.001602 | 0.000434 | 0.001953 | 0.000321 | 0.000287 | 0.003291 | 0.000597 | 0.012379 | 0.000361 | 0 | 0 | 0 | 0 | 0 | 0 | 0 | 0 | 0 | 0 | 0 | 0 |
| 18.3194444 | 0.000315 | 0.002421 | 0.001461 | 0.002063 | 0.000635 | 0.002113 | 0.000273 | 0.000251 | 0.003193 | 0.001208 | 0.015541 | 0.000145 | 0 | 0 | 0 | 0 | 0 | 0 | 0 | 0 | 0 | 0 | 0 | 0 |
| 18.3333333 | 0.000419 | 0.00268 | 0.001506 | 0.002441 | 0.000753 | 0.002027 | 0.000241 | 0.000183 | 0.002913 | 0.001879 | 0.019489 | 0.00012 | 0 | 0 | 0 | 0 | 0 | 0 | 0 | 0 | 0 | 0 | 0 | 0 |
| 18.3472222 | 0.000423 | 0.002122 | 0.001741 | 0.002435 | 0.000728 | 0.001768 | 0.000384 | 0.000135 | 0.002777 | 0.001926 | 0.021779 | 0.000348 | 0 | 0 | 0 | 0 | 0 | 0 | 0 | 0 | 0 | 0 | 0 | 0 |
| 18.3611111 | 0.000321 | 0.001288 | 0.002231 | 0.002285 | 0.000673 | 0.001571 | 0.000463 | 0.000143 | 0.003032 | 0.001465 | 0.02073 | 0.000644 | 0 | 0 | 0 | 0 | 0 | 0 | 0 | 0 | 0 | 0 | 0 | 0 |
| 18.375 | 0.000413 | 0.001337 | 0.002892 | 0.001984 | 0.00105 | 0.0013 | 0.000566 | 0.000316 | 0.003788 | 0.001237 | 0.016755 | 0.000874 | 0 | 0 | 0 | 0 | 0 | 0 | 0 | 0 | 0 | 0 | 0 | 0 |
| 18.3888889 | 0.000687 | 0.002206 | 0.003268 | 0.001605 | 0.00174 | 0.000957 | 0.000683 | 0.000662 | 0.004464 | 0.00141 | 0.012032 | 0.00096 | 0 | 0 | 0 | 0 | 0 | 0 | 0 | 0 | 0 | 0 | 0 | 0 |
| 18.4027778 | 0.000816 | 0.002899 | 0.003149 | 0.001537 | 0.002148 | 0.000854 | 0.000477 | 0.00084 | 0.004539 | 0.001635 | 0.008757 | 0.00094 | 0 | 0 | 0 | 0 | 0 | 0 | 0 | 0 | 0 | 0 | 0 | 0 |
| 18.4166667 | 0.000745 | 0.003051 | 0.002853 | 0.001808 | 0.002112 | 0.001043 | 0.00021 | 0.000712 | 0.004437 | 0.001956 | 0.007187 | 0.000805 | 0 | 0 | 0 | 0 | 0 | 0 | 0 | 0 | 0 | 0 | 0 | 0 |
| 18.4305556 | 0.000664 | 0.003092 | 0.002776 | 0.00203 | 0.001905 | 0.001264 | 0.000305 | 0.000567 | 0.004774 | 0.002581 | 0.006603 | 0.000545 | 0 | 0 | 0 | 0 | 0 | 0 | 0 | 0 | 0 | 0 | 0 | 0 |
| 18.4444444 | 0.00069 | 0.003075 | 0.002878 | 0.001965 | 0.001606 | 0.001456 | 0.000791 | 0.000556 | 0.005452 | 0.003175 | 0.006567 | 0.000692 | 0 | 0 | 0 | 0 | 0 | 0 | 0 | 0 | 0 | 0 | 0 | 0 |
| 18.4583333 | 0.000623 | 0.002276 | 0.002524 | 0.001995 | 0.001243 | 0.002 | 0.001271 | 0.000465 | 0.005777 | 0.002974 | 0.007111 | 0.001766 | 0 | 0 | 0 | 0 | 0 | 0 | 0 | 0 | 0 | 0 | 0 | 0 |
| 18.4722222 | 0.000329 | 0.001081 | 0.001511 | 0.002506 | 0.001037 | 0.002893 | 0.001681 | 0.000223 | 0.005734 | 0.00214 | 0.008053 | 0.003063 | 0 | 0 | 0 | 0 | 0 | 0 | 0 | 0 | 0 | 0 | 0 | 0 |
| 18.4861111 | 0.000917 | 0.002991 | 0.001305 | 0.002887 | 0.002706 | 0.003028 | 0.005025 | 0.00113 | 0.008361 | 0.003081 | 0.008078 | 0.002921 | 0 | 0 | 0 | 0 | 0 | 0 | 0 | 0 | 0 | 0 | 0 | 0 |
| 18.5 | 0.004121 | 0.012844 | 0.00435 | 0.002149 | 0.010639 | 0.002247 | 0.017661 | 0.008158 | 0.018499 | 0.008941 | 0.005836 | 0.002251 | 0 | 0 | 0 | 0 | 0 | 0 | 0 | 0 | 0 | 0 | 0 | 0 |
| 18.5138889 | 0.008602 | 0.027775 | 0.009099 | 0.000789 | 0.023844 | 0.002752 | 0.035863 | 0.019158 | 0.033815 | 0.020186 | 0.003004 | 0.003432 | 0 | 0 | 0 | 0 | 0 | 0 | 0 | 0 | 0 | 0 | 0 | 0 |
| 18.5277778 | 0.009987 | 0.034299 | 0.008565 | 0.000531 | 0.030896 | 0.005091 | 0.041448 | 0.019845 | 0.041909 | 0.03064 | 0.002623 | 0.00484 | 0 | 0 | 0 | 0 | 0 | 0 | 0 | 0 | 0 | 0 | 0 | 0 |
| 18.5416667 | 0.007575 | 0.026309 | 0.003364 | 0.001123 | 0.026361 | 0.006485 | 0.029974 | 0.010741 | 0.036777 | 0.032841 | 0.004137 | 0.004664 | 0 | 0 | 0 | 0 | 0 | 0 | 0 | 0 | 0 | 0 | 0 | 0 |
| 18.5555556 | 0.005622 | 0.0141 | 0.000738 | 0.001461 | 0.019363 | 0.00604 | 0.016778 | 0.005641 | 0.027076 | 0.028897 | 0.005314 | 0.003931 | 0 | 0 | 0 | 0 | 0 | 0 | 0 | 0 | 0 | 0 | 0 | 0 |
| 18.5694444 | 0.006343 | 0.00697 | 0.00047 | 0.002266 | 0.017004 | 0.005409 | 0.010224 | 0.005324 | 0.021172 | 0.025313 | 0.00578 | 0.002811 | 0 | 0 | 0 | 0 | 0 | 0 | 0 | 0 | 0 | 0 | 0 | 0 |
| 18.5833333 | 0.008314 | 0.00444 | 0.000523 | 0.003359 | 0.017521 | 0.005016 | 0.008195 | 0.004922 | 0.020109 | 0.023594 | 0.005918 | 0.001356 | 0 | 0 | 0 | 0 | 0 | 0 | 0 | 0 | 0 | 0 | 0 | 0 |
| 18.5972222 | 0.010371 | 0.004153 | 0.001222 | 0.003745 | 0.018363 | 0.004655 | 0.007836 | 0.003969 | 0.022517 | 0.022147 | 0.005821 | 0.000572 | 0 | 0 | 0 | 0 | 0 | 0 | 0 | 0 | 0 | 0 | 0 | 0 |
| 18.6111111 | 0.012067 | 0.005284 | 0.002331 | 0.003604 | 0.018482 | 0.00438 | 0.006977 | 0.003025 | 0.026445 | 0.020637 | 0.005809 | 0.000424 | 0 | 0 | 0 | 0 | 0 | 0 | 0 | 0 | 0 | 0 | 0 | 0 |
| 18.625 | 0.013018 | 0.007275 | 0.00373 | 0.003345 | 0.017448 | 0.003979 | 0.004911 | 0.002798 | 0.030444 | 0.019178 | 0.006603 | 0.000586 | 0 | 0 | 0 | 0 | 0 | 0 | 0 | 0 | 0 | 0 | 0 | 0 |
| 18.6388889 | 0.012829 | 0.009684 | 0.005034 | 0.003226 | 0.015579 | 0.003228 | 0.002435 | 0.004022 | 0.033958 | 0.017202 | 0.00817 | 0.00129 | 0 | 0 | 0 | 0 | 0 | 0 | 0 | 0 | 0 | 0 | 0 | 0 |
| 18.6527778 | 0.011287 | 0.011858 | 0.005728 | 0.003321 | 0.012992 | 0.002168 | 0.001219 | 0.006198 | 0.036188 | 0.01509 | 0.00923 | 0.001907 | 0 | 0 | 0 | 0 | 0 | 0 | 0 | 0 | 0 | 0 | 0 | 0 |
| 18.6666667 | 0.008969 | 0.012817 | 0.005402 | 0.003321 | 0.0097 | 0.001347 | 0.00167 | 0.007249 | 0.035946 | 0.013441 | 0.009096 | 0.001943 | 0 | 0 | 0 | 0 | 0 | 0 | 0 | 0 | 0 | 0 | 0 | 0 |
| 18.6805556 | 0.006544 | 0.01224 | 0.004168 | 0.002859 | 0.006128 | 0.001189 | 0.002078 | 0.006084 | 0.032696 | 0.011879 | 0.008455 | 0.001783 | 0 | 0 | 0 | 0 | 0 | 0 | 0 | 0 | 0 | 0 | 0 | 0 |
| 18.6944444 | 0.004358 | 0.010841 | 0.002738 | 0.002397 | 0.003193 | 0.001415 | 0.001626 | 0.004114 | 0.027626 | 0.009635 | 0.007916 | 0.001571 | 0 | 0 | 0 | 0 | 0 | 0 | 0 | 0 | 0 | 0 | 0 | 0 |
| 18.7083333 | 0.002383 | 0.009589 | 0.001623 | 0.002549 | 0.001405 | 0.001604 | 0.001059 | 0.002984 | 0.022519 | 0.006578 | 0.007545 | 0.001083 | 0 | 0 | 0 | 0 | 0 | 0 | 0 | 0 | 0 | 0 | 0 | 0 |
| 18.7222222 | 0.001088 | 0.008787 | 0.000883 | 0.002958 | 0.001192 | 0.001674 | 0.000793 | 0.002553 | 0.017883 | 0.003463 | 0.007322 | 0.000453 | 0 | 0 | 0 | 0 | 0 | 0 | 0 | 0 | 0 | 0 | 0 | 0 |
| 18.7361111 | 0.001158 | 0.008504 | 0.000511 | 0.003169 | 0.003139 | 0.001758 | 0.000533 | 0.00223 | 0.013648 | 0.001486 | 0.007256 | 0.000234 | 0 | 0 | 0 | 0 | 0 | 0 | 0 | 0 | 0 | 0 | 0 | 0 |
| 18.75 | 0.001785 | 0.00919 | 0.000413 | 0.002973 | 0.005826 | 0.001863 | 0.0003 | 0.002031 | 0.010034 | 0.001452 | 0.00728 | 0.000456 | 0 | 0 | 0 | 0 | 0 | 0 | 0 | 0 | 0 | 0 | 0 | 0 |
| 18.7638889 | 0.002081 | 0.010401 | 0.000433 | 0.002536 | 0.006978 | 0.002152 | 0.000371 | 0.002106 | 0.007522 | 0.002307 | 0.007154 | 0.000669 | 0 | 0 | 0 | 0 | 0 | 0 | 0 | 0 | 0 | 0 | 0 | 0 |
| 18.7777778 | 0.001899 | 0.010535 | 0.000493 | 0.0023 | 0.005857 | 0.003691 | 0.000395 | 0.002485 | 0.006534 | 0.002255 | 0.006705 | 0.000754 | 0 | 0 | 0 | 0 | 0 | 0 | 0 | 0 | 0 | 0 | 0 | 0 |
| 18.7916667 | 0.00125 | 0.008974 | 0.000551 | 0.00249 | 0.003651 | 0.006712 | 0.000248 | 0.002875 | 0.00649 | 0.002479 | 0.006233 | 0.000827 | 0 | 0 | 0 | 0 | 0 | 0 | 0 | 0 | 0 | 0 | 0 | 0 |
| 18.8055556 | 0.000543 | 0.006932 | 0.000633 | 0.003349 | 0.002069 | 0.008883 | 0.000394 | 0.002935 | 0.006269 | 0.00468 | 0.006464 | 0.000713 | 0 | 0 | 0 | 0 | 0 | 0 | 0 | 0 | 0 | 0 | 0 | 0 |
| 18.8194444 | 0.00036 | 0.005647 | 0.000643 | 0.00449 | 0.001685 | 0.008886 | 0.001101 | 0.002649 | 0.005557 | 0.006722 | 0.008296 | 0.000403 | 0.002805 | 0.001055 | 0.000336 | 0.000842 | 0.005221 | 0.000236 | 0.000147 | 0.000637 | 0.00015 | 0.003751 | 0.000212 | 5.00555595599889e-05 |
| 18.8333333 | 0.001039 | 0.005083 | 0.000419 | 0.004775 | 0.002462 | 0.007894 | 0.002269 | 0.002067 | 0.00449 | 0.006407 | 0.012181 | 0.000206 | 0.005752 | 0.003226 | 0.000836 | 0.002617 | 0.015705 | 0.000623 | 0.000848 | 0.001346 | 0.000684 | 0.010669 | 0.000794 | 0.000572 |
| 18.8472222 | 0.002292 | 0.005057 | 0.00019 | 0.004077 | 0.00446 | 0.006782 | 0.003705 | 0.001258 | 0.002856 | 0.003888 | 0.015757 | 0.000227 | 0.005901 | 0.004242 | 0.000731 | 0.003687 | 0.01854 | 0.001048 | 0.001983 | 0.000784 | 0.001176 | 0.011195 | 0.00181 | 0.00198 |
| 18.8611111 | 0.0036 | 0.005732 | 0.000118 | 0.003311 | 0.007557 | 0.005884 | 0.005368 | 0.000554 | 0.001637 | 0.002434 | 0.015377 | 0.000455 | 0.009525 | 0.004278 | 0.000449 | 0.003748 | 0.011225 | 0.001934 | 0.002327 | 0.000755 | 0.000912 | 0.007537 | 0.003125 | 0.003785 |
| 18.875 | 0.004822 | 0.006283 | 8.9152784 | 0.002723 | 0.010796 | 0.005298 | 0.007598 | 0.00039 | 0.0023 | 0.004733 | 0.011756 | 0.000498 | 0.012392 | 0.005652 | 0.000952 | 0.003256 | 0.003844 | 0.002731 | 0.001396 | 0.003144 | 0.000701 | 0.010927 | 0.003921 | 0.005235 |
| 18.8888889 | 0.006199 | 0.005664 | 0.000116 | 0.002076 | 0.0127 | 0.004643 | 0.010484 | 0.00073 | 0.004036 | 0.00665 | 0.008427 | 0.000413 | 0.009181 | 0.009118 | 0.00221 | 0.002179 | 0.001318 | 0.002674 | 0.000595 | 0.006853 | 0.002004 | 0.020405 | 0.003771 | 0.006364 |
| 18.9027778 | 0.008237 | 0.004431 | 0.000356 | 0.001569 | 0.012817 | 0.003757 | 0.013278 | 0.00107 | 0.005877 | 0.004651 | 0.006865 | 0.000785 | 0.009474 | 0.013362 | 0.003188 | 0.000981 | 0.001376 | 0.002273 | 0.001359 | 0.009781 | 0.004411 | 0.025828 | 0.003316 | 0.007961 |
| 18.9166667 | 0.011003 | 0.00403 | 0.000851 | 0.001334 | 0.011787 | 0.002984 | 0.015634 | 0.001112 | 0.00788 | 0.001781 | 0.006596 | 0.001257 | 0.017003 | 0.016004 | 0.003235 | 0.000333 | 0.00163 | 0.002321 | 0.004303 | 0.010742 | 0.006406 | 0.022041 | 0.003701 | 0.010281 |
| 18.9305556 | 0.013441 | 0.005096 | 0.001219 | 0.001161 | 0.010273 | 0.002553 | 0.017723 | 0.000964 | 0.009403 | 0.000828 | 0.006853 | 0.001519 | 0.020645 | 0.016161 | 0.002851 | 0.00035 | 0.002236 | 0.002553 | 0.009511 | 0.010167 | 0.007104 | 0.012628 | 0.005386 | 0.012203 |
| 18.9444444 | 0.014617 | 0.006086 | 0.001232 | 0.00097 | 0.008694 | 0.002394 | 0.019483 | 0.000655 | 0.009713 | 0.001794 | 0.00716 | 0.00188 | 0.013486 | 0.015124 | 0.002523 | 0.000492 | 0.003216 | 0.001853 | 0.01595 | 0.009227 | 0.006443 | 0.005049 | 0.00759 | 0.012706 |
| 18.9583333 | 0.014309 | 0.0048 | 0.001282 | 0.000893 | 0.007119 | 0.002351 | 0.02023 | 0.000414 | 0.009186 | 0.005112 | 0.007484 | 0.002315 | 0.006448 | 0.013783 | 0.002395 | 0.000359 | 0.003975 | 0.000866 | 0.021715 | 0.008609 | 0.004906 | 0.00302 | 0.009469 | 0.0124 |
| 18.9722222 | 0.01227 | 0.002096 | 0.00158 | 0.000942 | 0.005648 | 0.002284 | 0.01902 | 0.000492 | 0.008827 | 0.009555 | 0.008075 | 0.002557 | 0.008103 | 0.01186 | 0.002693 | 0.00025 | 0.00399 | 0.00074 | 0.024201 | 0.008016 | 0.003341 | 0.002595 | 0.010423 | 0.013284 |
| 18.9861111 | 0.009454 | 0.001149 | 0.002286 | 0.00098 | 0.005035 | 0.001824 | 0.016853 | 0.000705 | 0.009948 | 0.013086 | 0.008767 | 0.00318 | 0.009244 | 0.009862 | 0.004606 | 0.001383 | 0.003043 | 0.002003 | 0.023293 | 0.00717 | 0.003059 | 0.003815 | 0.010747 | 0.016562 |
| 19 | 0.007945 | 0.005767 | 0.00428 | 0.0012 | 0.006472 | 0.001773 | 0.017528 | 0.001431 | 0.015711 | 0.018493 | 0.008124 | 0.006797 | 0.016229 | 0.009381 | 0.010134 | 0.006268 | 0.004578 | 0.0064 | 0.028016 | 0.006117 | 0.006175 | 0.013822 | 0.013562 | 0.020616 |
| 19.0138889 | 0.00851 | 0.01853 | 0.008534 | 0.002771 | 0.011062 | 0.005384 | 0.024898 | 0.002812 | 0.030583 | 0.031864 | 0.00646 | 0.016616 | 0.055914 | 0.010899 | 0.018819 | 0.017054 | 0.012317 | 0.013636 | 0.056377 | 0.005438 | 0.017433 | 0.034267 | 0.025042 | 0.024122 |
| 19.0277778 | 0.009403 | 0.031468 | 0.01442 | 0.00565 | 0.015641 | 0.013172 | 0.038025 | 0.009872 | 0.055654 | 0.049627 | 0.009929 | 0.031727 | 0.115865 | 0.012191 | 0.025813 | 0.029437 | 0.017971 | 0.019718 | 0.093362 | 0.004337 | 0.040489 | 0.060136 | 0.048697 | 0.03081 |
| 19.0416667 | 0.008901 | 0.028165 | 0.0152 | 0.007593 | 0.012974 | 0.017859 | 0.043914 | 0.018667 | 0.064282 | 0.05507 | 0.020982 | 0.03778 | 0.129318 | 0.011075 | 0.024788 | 0.02892 | 0.015374 | 0.017817 | 0.098284 | 0.007186 | 0.050975 | 0.072978 | 0.065087 | 0.035789 |
| 19.0555556 | 0.007418 | 0.015622 | 0.010546 | 0.007981 | 0.00739 | 0.014553 | 0.030278 | 0.017235 | 0.047496 | 0.037714 | 0.031666 | 0.026817 | 0.084784 | 0.008382 | 0.014955 | 0.016493 | 0.015253 | 0.009625 | 0.072792 | 0.012762 | 0.034319 | 0.052628 | 0.047999 | 0.027162 |
| 19.0694444 | 0.005817 | 0.012649 | 0.008272 | 0.007662 | 0.00633 | 0.009108 | 0.014852 | 0.013219 | 0.042283 | 0.01693 | 0.03646 | 0.016849 | 0.074291 | 0.006022 | 0.006555 | 0.011134 | 0.019309 | 0.006371 | 0.041399 | 0.01317 | 0.022592 | 0.026467 | 0.032512 | 0.015175 |
| 19.0833333 | 0.004379 | 0.012164 | 0.006137 | 0.006931 | 0.004684 | 0.006297 | 0.013518 | 0.010701 | 0.041769 | 0.013698 | 0.036319 | 0.014544 | 0.112498 | 0.004421 | 0.005973 | 0.011069 | 0.016757 | 0.00783 | 0.018779 | 0.014225 | 0.021755 | 0.023829 | 0.043492 | 0.010748 |
| 19.0972222 | 0.003128 | 0.007375 | 0.002789 | 0.005945 | 0.001899 | 0.004903 | 0.012441 | 0.004947 | 0.025306 | 0.015167 | 0.031943 | 0.009972 | 0.112456 | 0.002937 | 0.006315 | 0.007242 | 0.009776 | 0.007336 | 0.00933 | 0.016189 | 0.015149 | 0.023104 | 0.039065 | 0.006198 |
| 19.1111111 | 0.00197 | 0.003981 | 0.0011 | 0.004625 | 0.001229 | 0.003958 | 0.006968 | 0.001429 | 0.010403 | 0.010648 | 0.025032 | 0.004306 | 0.069084 | 0.001435 | 0.00436 | 0.003375 | 0.005894 | 0.004761 | 0.006516 | 0.014047 | 0.00694 | 0.014269 | 0.019343 | 0.001753 |
| 19.125 | 0.00099 | 0.00264 | 0.00036 | 0.002987 | 0.000928 | 0.00329 | 0.003156 | 0.001396 | 0.004277 | 0.005809 | 0.01899 | 0.001309 | 0.030576 | 0.000678 | 0.00262 | 0.001668 | 0.004513 | 0.002336 | 0.005413 | 0.010259 | 0.002554 | 0.007314 | 0.00814 | 0.000984 |
| 19.1388889 | 0.00038 | 0.002246 | 0.000121 | 0.001554 | 0.000432 | 0.002695 | 0.00113 | 0.001247 | 0.001908 | 0.002931 | 0.015279 | 0.000328 | 0.010075 | 0.000729 | 0.001747 | 0.001388 | 0.003307 | 0.00165 | 0.004621 | 0.006817 | 0.002176 | 0.003948 | 0.002834 | 0.000905 |
| 19.1527778 | 0.000196 | 0.002491 | 0.000182 | 0.000891 | 0.000188 | 0.002104 | 0.000367 | 0.000632 | 0.001052 | 0.001824 | 0.012674 | 0.000573 | 0.003448 | 0.000784 | 0.00122 | 0.00159 | 0.002926 | 0.001721 | 0.003792 | 0.005054 | 0.003662 | 0.003622 | 0.000767 | 0.000709 |
| 19.1666667 | 0.000215 | 0.002288 | 0.000687 | 0.000789 | 0.000366 | 0.001665 | 0.001427 | 0.000341 | 0.002322 | 0.001295 | 0.009984 | 0.001446 | 0.003974 | 0.001091 | 0.0014 | 0.001285 | 0.004911 | 0.001193 | 0.00281 | 0.005225 | 0.005647 | 0.003189 | 0.001018 | 0.002395 |
| 19.1805556 | 0.00042 | 0.001157 | 0.001633 | 0.000742 | 0.001756 | 0.001798 | 0.005362 | 0.000447 | 0.007649 | 0.001227 | 0.007808 | 0.001799 | 0.012113 | 0.002821 | 0.00303 | 0.000655 | 0.009108 | 0.001054 | 0.001738 | 0.006066 | 0.008104 | 0.002576 | 0.003396 | 0.005776 |
| 19.1944444 | 0.000751 | 0.001312 | 0.001974 | 0.001317 | 0.002872 | 0.002534 | 0.007676 | 0.000669 | 0.010098 | 0.001662 | 0.006581 | 0.00141 | 0.019174 | 0.005099 | 0.004138 | 0.000912 | 0.01185 | 0.001873 | 0.001347 | 0.004961 | 0.007286 | 0.005746 | 0.004194 | 0.006313 |
| 19.2083333 | 0.001664 | 0.004896 | 0.001494 | 0.002909 | 0.003109 | 0.002725 | 0.007042 | 0.000904 | 0.009336 | 0.003359 | 0.00494 | 0.001067 | 0.022104 | 0.004966 | 0.005526 | 0.00239 | 0.009128 | 0.002584 | 0.001337 | 0.002102 | 0.004991 | 0.015828 | 0.003755 | 0.005629 |
| 19.2222222 | 0.003683 | 0.009579 | 0.001414 | 0.003326 | 0.004457 | 0.001779 | 0.008919 | 0.00101 | 0.013551 | 0.007585 | 0.002454 | 0.001245 | 0.032554 | 0.002571 | 0.007557 | 0.003199 | 0.004198 | 0.002773 | 0.000856 | 0.001329 | 0.005917 | 0.027616 | 0.005863 | 0.007059 |
| 19.2361111 | 0.004897 | 0.011065 | 0.001608 | 0.001666 | 0.004734 | 0.000846 | 0.009838 | 0.001124 | 0.014415 | 0.00903 | 0.000791 | 0.002225 | 0.031629 | 0.001175 | 0.006089 | 0.002308 | 0.002909 | 0.002944 | 0.00056 | 0.003197 | 0.004886 | 0.028812 | 0.005771 | 0.00538 |
| 19.25 | 0.00404 | 0.008949 | 0.001443 | 0.000417 | 0.002987 | 0.000591 | 0.00572 | 0.000997 | 0.007661 | 0.005707 | 0.000371 | 0.002613 | 0.016644 | 0.001536 | 0.002527 | 0.001107 | 0.004313 | 0.002776 | 0.00064 | 0.004521 | 0.001952 | 0.020674 | 0.003087 | 0.00154 |
| 19.2638889 | 0.002482 | 0.005325 | 0.001313 | 0.000378 | 0.001307 | 0.000758 | 0.001778 | 0.000505 | 0.002073 | 0.00366 | 0.00032 | 0.00181 | 0.013739 | 0.001991 | 0.001004 | 0.000441 | 0.005108 | 0.001856 | 0.000559 | 0.004683 | 0.001337 | 0.01359 | 0.003214 | 0.000314 |
| 19.2777778 | 0.00161 | 0.002009 | 0.00114 | 0.00051 | 0.000624 | 0.000917 | 0.000734 | 0.000354 | 0.000602 | 0.003157 | 0.000479 | 0.001049 | 0.016279 | 0.0023 | 0.001301 | 0.000394 | 0.00544 | 0.000815 | 0.000365 | 0.00528 | 0.00112 | 0.010749 | 0.004229 | 0.000632 |
| 19.2916667 | 0.00189 | 0.001079 | 0.000828 | 0.000537 | 0.001169 | 0.00071 | 0.001042 | 0.000875 | 0.001672 | 0.002457 | 0.000519 | 0.001086 | 0.011438 | 0.003519 | 0.00127 | 0.001158 | 0.006126 | 0.001363 | 0.000394 | 0.00625 | 0.001071 | 0.011631 | 0.003099 | 0.001787 |
| 19.3055556 | 0.002904 | 0.003163 | 0.000911 | 0.000804 | 0.002114 | 0.000543 | 0.002526 | 0.001824 | 0.003532 | 0.003863 | 0.000437 | 0.001861 | 0.008785 | 0.005549 | 0.000883 | 0.002256 | 0.006442 | 0.004419 | 0.000854 | 0.006727 | 0.002428 | 0.014753 | 0.002194 | 0.003718 |
| 19.3194444 | 0.00373 | 0.006041 | 0.001096 | 0.001097 | 0.002553 | 0.000581 | 0.004164 | 0.00237 | 0.004892 | 0.006361 | 0.000557 | 0.002506 | 0.014071 | 0.00708 | 0.001279 | 0.002898 | 0.005521 | 0.007329 | 0.001471 | 0.006969 | 0.003643 | 0.01782 | 0.003597 | 0.005661 |
| 19.3333333 | 0.003849 | 0.007312 | 0.000746 | 0.000957 | 0.002438 | 0.000412 | 0.004616 | 0.002185 | 0.005337 | 0.007053 | 0.000587 | 0.002466 | 0.018067 | 0.007393 | 0.001549 | 0.002927 | 0.004 | 0.007558 | 0.001828 | 0.007294 | 0.003124 | 0.019486 | 0.004918 | 0.00691 |
| 19.3472222 | 0.003574 | 0.006953 | 0.00025 | 0.00074 | 0.002338 | 0.000159 | 0.004485 | 0.001878 | 0.005304 | 0.006032 | 0.00061 | 0.001981 | 0.016063 | 0.006982 | 0.001136 | 0.002798 | 0.00334 | 0.005857 | 0.002107 | 0.0071 | 0.0018 | 0.019863 | 0.004698 | 0.006958 |
| 19.3611111 | 0.003385 | 0.006049 | 8.6604173 | 0.00072 | 0.00245 | 9.4631952 | 0.004515 | 0.001732 | 0.005237 | 0.004615 | 0.000851 | 0.001644 | 0.012116 | 0.006656 | 0.000706 | 0.002883 | 0.003801 | 0.00386 | 0.00272 | 0.00581 | 0.001439 | 0.019755 | 0.00414 | 0.006162 |
| 19.375 | 0.003532 | 0.005372 | 9.9638896 | 0.000767 | 0.002663 | 0.000193 | 0.005016 | 0.001627 | 0.005566 | 0.003866 | 0.001179 | 0.001886 | 0.009714 | 0.007058 | 0.000584 | 0.00309 | 0.004378 | 0.002865 | 0.003613 | 0.003612 | 0.002991 | 0.02104 | 0.00413 | 0.005802 |
| 19.3888889 | 0.004061 | 0.005299 | 0.000161 | 0.000893 | 0.003017 | 0.00041 | 0.005884 | 0.001495 | 0.006641 | 0.004139 | 0.001543 | 0.002303 | 0.009866 | 0.008381 | 0.000645 | 0.003089 | 0.004084 | 0.003544 | 0.004353 | 0.002078 | 0.005671 | 0.025473 | 0.004592 | 0.006464 |
| 19.4027778 | 0.004816 | 0.005313 | 0.000254 | 0.001006 | 0.003359 | 0.000598 | 0.006498 | 0.001329 | 0.007932 | 0.004916 | 0.001909 | 0.002217 | 0.010773 | 0.009811 | 0.000671 | 0.002735 | 0.002722 | 0.005284 | 0.004606 | 0.003138 | 0.007482 | 0.032016 | 0.004651 | 0.007274 |
| 19.4166667 | 0.005773 | 0.004778 | 0.000273 | 0.000898 | 0.003457 | 0.000652 | 0.006564 | 0.001225 | 0.008603 | 0.005498 | 0.00226 | 0.001753 | 0.010632 | 0.010382 | 0.000627 | 0.002226 | 0.001111 | 0.006482 | 0.004613 | 0.005902 | 0.007094 | 0.037675 | 0.003836 | 0.007769 |
| 19.4305556 | 0.007112 | 0.004045 | 0.000306 | 0.000661 | 0.003406 | 0.00069 | 0.006668 | 0.001309 | 0.008858 | 0.005711 | 0.002585 | 0.00143 | 0.01018 | 0.010211 | 0.000502 | 0.002037 | 0.000232 | 0.006143 | 0.004932 | 0.007863 | 0.005026 | 0.04098 | 0.002891 | 0.008878 |
| 19.4444444 | 0.008526 | 0.003127 | 0.000402 | 0.000411 | 0.002954 | 0.000577 | 0.007152 | 0.001343 | 0.009584 | 0.00542 | 0.002659 | 0.00116 | 0.010579 | 0.010272 | 0.000695 | 0.002748 | 0.000284 | 0.005307 | 0.006213 | 0.008696 | 0.003016 | 0.044054 | 0.003114 | 0.011068 |
| 19.4583333 | 0.009184 | 0.002063 | 0.000359 | 0.000246 | 0.001784 | 0.000532 | 0.007656 | 0.001091 | 0.011116 | 0.00425 | 0.002272 | 0.001686 | 0.011401 | 0.011211 | 0.002289 | 0.004619 | 0.001017 | 0.005887 | 0.008965 | 0.010084 | 0.002606 | 0.048858 | 0.00511 | 0.013311 |
| 19.4722222 | 0.008924 | 0.00144 | 0.000197 | 0.000311 | 0.000666 | 0.00092 | 0.008228 | 0.000696 | 0.013192 | 0.002631 | 0.001821 | 0.002993 | 0.011605 | 0.012293 | 0.005307 | 0.00657 | 0.002439 | 0.007656 | 0.012564 | 0.012438 | 0.003636 | 0.051311 | 0.007562 | 0.013854 |
| 19.4861111 | 0.009492 | 0.002975 | 0.000581 | 0.000367 | 0.002349 | 0.001295 | 0.01102 | 0.000479 | 0.018272 | 0.00316 | 0.00259 | 0.005633 | 0.013346 | 0.013356 | 0.009066 | 0.007391 | 0.0084 | 0.008465 | 0.016815 | 0.014001 | 0.004222 | 0.048737 | 0.008406 | 0.01262 |
| 19.5 | 0.014689 | 0.010541 | 0.002524 | 0.000243 | 0.009804 | 0.002853 | 0.021952 | 0.001024 | 0.0343 | 0.008907 | 0.006725 | 0.023269 | 0.024047 | 0.018102 | 0.014905 | 0.008237 | 0.027026 | 0.008153 | 0.025582 | 0.014261 | 0.003452 | 0.048105 | 0.007877 | 0.01313 |
| 19.5138889 | 0.024317 | 0.022378 | 0.005915 | 0.00049 | 0.020276 | 0.007062 | 0.035052 | 0.002161 | 0.062995 | 0.018801 | 0.017182 | 0.07972 | 0.044796 | 0.030327 | 0.024045 | 0.009772 | 0.054081 | 0.008313 | 0.046452 | 0.015575 | 0.00197 | 0.051414 | 0.006775 | 0.016507 |
| 19.5277778 | 0.030079 | 0.028708 | 0.008885 | 0.001563 | 0.026146 | 0.011995 | 0.036559 | 0.002898 | 0.084569 | 0.027907 | 0.03897 | 0.159135 | 0.056697 | 0.042008 | 0.029774 | 0.00828 | 0.069455 | 0.00759 | 0.07305 | 0.018025 | 0.001936 | 0.045819 | 0.004231 | 0.017707 |
| 19.5416667 | 0.027925 | 0.025543 | 0.010467 | 0.006019 | 0.025099 | 0.014715 | 0.028963 | 0.003285 | 0.080805 | 0.03272 | 0.064744 | 0.193585 | 0.044988 | 0.040642 | 0.026015 | 0.003499 | 0.066353 | 0.004751 | 0.086638 | 0.018287 | 0.003758 | 0.026526 | 0.001446 | 0.014604 |
| 19.5555556 | 0.024351 | 0.019738 | 0.012862 | 0.016866 | 0.022658 | 0.016529 | 0.01976 | 0.003815 | 0.063654 | 0.033422 | 0.078826 | 0.161824 | 0.024071 | 0.029964 | 0.020426 | 0.001626 | 0.055977 | 0.003122 | 0.082861 | 0.016627 | 0.004683 | 0.009098 | 0.001821 | 0.010805 |
| 19.5694444 | 0.022481 | 0.016762 | 0.017856 | 0.029727 | 0.022514 | 0.019702 | 0.013068 | 0.00403 | 0.050114 | 0.03102 | 0.103995 | 0.110904 | 0.01041 | 0.020052 | 0.018824 | 0.003416 | 0.043313 | 0.004241 | 0.068309 | 0.015859 | 0.007209 | 0.002848 | 0.005734 | 0.008084 |
| 19.5833333 | 0.018458 | 0.015856 | 0.023071 | 0.036272 | 0.022618 | 0.021899 | 0.009552 | 0.003335 | 0.041242 | 0.024871 | 0.135244 | 0.071345 | 0.00453 | 0.012229 | 0.017462 | 0.004666 | 0.027652 | 0.005131 | 0.050248 | 0.016139 | 0.014018 | 0.003645 | 0.010683 | 0.005112 |
| 19.5972222 | 0.01142 | 0.016458 | 0.026255 | 0.033382 | 0.020558 | 0.021895 | 0.006594 | 0.002116 | 0.035199 | 0.016546 | 0.116901 | 0.045848 | 0.002884 | 0.00568 | 0.014784 | 0.004124 | 0.013536 | 0.004258 | 0.033892 | 0.015805 | 0.020004 | 0.008768 | 0.014933 | 0.002655 |
| 19.6111111 | 0.005121 | 0.019315 | 0.026703 | 0.025816 | 0.01573 | 0.020299 | 0.00452 | 0.001273 | 0.030259 | 0.01075 | 0.076278 | 0.029879 | 0.003304 | 0.001689 | 0.012379 | 0.002793 | 0.005629 | 0.003063 | 0.02007 | 0.013961 | 0.021099 | 0.012551 | 0.0175 | 0.001985 |
| 19.625 | 0.004543 | 0.023414 | 0.023699 | 0.019873 | 0.008684 | 0.018067 | 0.003945 | 0.001302 | 0.023622 | 0.009316 | 0.070273 | 0.017991 | 0.005809 | 0.000674 | 0.011299 | 0.001724 | 0.005014 | 0.002675 | 0.010283 | 0.012152 | 0.019488 | 0.009503 | 0.018415 | 0.00362 |
| 19.6388889 | 0.012102 | 0.026504 | 0.01919 | 0.015883 | 0.004177 | 0.017665 | 0.003905 | 0.001992 | 0.015437 | 0.010848 | 0.086697 | 0.009769 | 0.008955 | 0.000782 | 0.011694 | 0.001361 | 0.01215 | 0.002871 | 0.00802 | 0.012168 | 0.019273 | 0.005384 | 0.019298 | 0.010792 |
| 19.6527778 | 0.02298 | 0.027392 | 0.015552 | 0.01239 | 0.004889 | 0.018678 | 0.003565 | 0.0026 | 0.008746 | 0.013807 | 0.098049 | 0.004429 | 0.009521 | 0.000914 | 0.012844 | 0.00162 | 0.025485 | 0.002846 | 0.013071 | 0.013032 | 0.020361 | 0.006887 | 0.020613 | 0.02182 |
| 19.6666667 | 0.026785 | 0.026532 | 0.01304 | 0.010112 | 0.004743 | 0.018106 | 0.003395 | 0.002752 | 0.005653 | 0.016346 | 0.093303 | 0.0018 | 0.007381 | 0.001182 | 0.0131 | 0.002184 | 0.040494 | 0.002311 | 0.019135 | 0.012738 | 0.019286 | 0.009078 | 0.020712 | 0.028628 |
| 19.6805556 | 0.01901 | 0.025521 | 0.011004 | 0.008901 | 0.003365 | 0.015509 | 0.003789 | 0.002656 | 0.004843 | 0.015815 | 0.075021 | 0.001405 | 0.004123 | 0.001903 | 0.011319 | 0.002798 | 0.052622 | 0.001825 | 0.020049 | 0.010529 | 0.014664 | 0.006729 | 0.018305 | 0.027372 |
| 19.6944444 | 0.00918 | 0.025406 | 0.008742 | 0.007801 | 0.004712 | 0.013516 | 0.004115 | 0.002301 | 0.003811 | 0.010754 | 0.05404 | 0.001369 | 0.002486 | 0.004296 | 0.008423 | 0.003107 | 0.057681 | 0.001684 | 0.014532 | 0.006618 | 0.007837 | 0.004541 | 0.013997 | 0.020532 |

| | | | | | | | | | | | | | | | | | | | | | | | | |
|---|---|---|---|---|---|---|---|---|---|---|---|---|---|---|---|---|---|---|---|---|---|---|---|---|
| 19.7083333 | 0.008313 | 0.025243 | 0.006212 | 0.006635 | 0.005871 | 0.013314 | 0.003741 | 0.00156 | 0.001999 | 0.005535 | 0.038206 | 0.002089 | 0.005834 | 0.007016 | 0.006038 | 0.002818 | 0.054464 | 0.00154 | 0.006674 | 0.003321 | 0.002451 | 0.005833 | 0.009456 | 0.012396 |
| 19.7222222 | 0.014452 | 0.023597 | 0.004046 | 0.005338 | 0.005647 | 0.013459 | 0.002846 | 0.000844 | 0.00083 | 0.005263 | 0.02846 | 0.003471 | 0.01119 | 0.007642 | 0.005074 | 0.002296 | 0.047455 | 0.001387 | 0.001824 | 0.003263 | 0.001507 | 0.008912 | 0.005625 | 0.005862 |
| 19.7361111 | 0.018142 | 0.020617 | 0.002849 | 0.003767 | 0.005554 | 0.012625 | 0.001913 | 0.000603 | 0.000741 | 0.007268 | 0.022363 | 0.005411 | 0.010988 | 0.006357 | 0.005778 | 0.002125 | 0.041435 | 0.001741 | 0.000845 | 0.005068 | 0.004282 | 0.013973 | 0.002799 | 0.00203 |
| 19.75 | 0.013052 | 0.017636 | 0.002691 | 0.002199 | 0.004898 | 0.010718 | 0.001322 | 0.00072 | 0.000659 | 0.006346 | 0.017581 | 0.007612 | 0.007314 | 0.004179 | 0.007011 | 0.002033 | 0.036158 | 0.002307 | 0.001157 | 0.005772 | 0.008474 | 0.017226 | 0.002191 | 0.001238 |
| 19.7638889 | 0.007733 | 0.015452 | 0.003273 | 0.001258 | 0.00317 | 0.008304 | 0.001492 | 0.000913 | 0.000403 | 0.004915 | 0.01347 | 0.008294 | 0.007709 | 0.002136 | 0.007121 | 0.001397 | 0.02817 | 0.002044 | 0.002653 | 0.004181 | 0.011665 | 0.012908 | 0.003765 | 0.003143 |
| 19.7777778 | 0.011976 | 0.013724 | 0.004038 | 0.001294 | 0.001445 | 0.005997 | 0.002426 | 0.001028 | 0.0004 | 0.008388 | 0.010824 | 0.007567 | 0.008489 | 0.001245 | 0.006439 | 0.000537 | 0.01687 | 0.001274 | 0.005265 | 0.002386 | 0.012487 | 0.006282 | 0.005043 | 0.006135 |
| 19.7916667 | 0.015627 | 0.011359 | 0.004538 | 0.001716 | 0.000926 | 0.00443 | 0.003331 | 0.001003 | 0.000683 | 0.013057 | 0.009617 | 0.006475 | 0.005259 | 0.001398 | 0.006261 | 0.000142 | 0.006893 | 0.001638 | 0.005252 | 0.003417 | 0.011416 | 0.004338 | 0.004989 | 0.007937 |
| 19.8055556 | 0.011085 | 0.0081 | 0.004575 | 0.001791 | 0.001263 | 0.003903 | 0.003552 | 0.000812 | 0.001851 | 0.013229 | 0.008633 | 0.004733 | 0.003527 | 0.00123 | 0.006258 | 0.000256 | 0.004106 | 0.003141 | 0.002747 | 0.006382 | 0.009507 | 0.003612 | 0.004598 | 0.008141 |
| 19.8194444 | 0.005758 | 0.005427 | 0.003972 | 0.001812 | 0.001359 | 0.003748 | 0.003275 | 0.000461 | 0.004672 | 0.009068 | 0.007396 | 0.002973 | 0.00647 | 0.000875 | 0.005488 | 0.000856 | 0.009958 | 0.003917 | 0.001867 | 0.008698 | 0.007019 | 0.001572 | 0.00476 | 0.008477 |
| 19.8333333 | 0.006559 | 0.003766 | 0.002928 | 0.001999 | 0.000935 | 0.002681 | 0.002646 | 0.000213 | 0.008341 | 0.005487 | 0.006513 | 0.004038 | 0.011141 | 0.0018 | 0.004 | 0.002001 | 0.021749 | 0.003222 | 0.001899 | 0.008636 | 0.004133 | 0.001647 | 0.005524 | 0.009263 |
| 19.8472222 | 0.010921 | 0.002216 | 0.001965 | 0.001664 | 0.000724 | 0.00186 | 0.001601 | 0.000349 | 0.011483 | 0.0082 | 0.005972 | 0.009213 | 0.012973 | 0.004064 | 0.002176 | 0.00319 | 0.035637 | 0.002235 | 0.002337 | 0.005548 | 0.002251 | 0.003994 | 0.00596 | 0.00949 |
| 19.8611111 | 0.011267 | 0.001861 | 0.001429 | 0.000986 | 0.001387 | 0.003603 | 0.001273 | 0.000622 | 0.013709 | 0.015006 | 0.004864 | 0.017833 | 0.010191 | 0.007236 | 0.001254 | 0.003666 | 0.046218 | 0.002257 | 0.005286 | 0.003254 | 0.001674 | 0.005171 | 0.005368 | 0.009018 |
| 19.875 | 0.006489 | 0.003666 | 0.001393 | 0.000916 | 0.001909 | 0.006422 | 0.002988 | 0.000625 | 0.015358 | 0.019203 | 0.003153 | 0.028075 | 0.008554 | 0.010656 | 0.002494 | 0.003315 | 0.049566 | 0.003489 | 0.006764 | 0.005992 | 0.002024 | 0.006178 | 0.004558 | 0.008948 |
| 19.8888889 | 0.001899 | 0.006648 | 0.001232 | 0.001148 | 0.002036 | 0.00773 | 0.005598 | 0.000395 | 0.01716 | 0.017382 | 0.001489 | 0.035134 | 0.008229 | 0.012544 | 0.005455 | 0.002739 | 0.047118 | 0.005275 | 0.005166 | 0.010941 | 0.003444 | 0.012164 | 0.004859 | 0.011025 |
| 19.9027778 | 0.002593 | 0.011015 | 0.001776 | 0.001062 | 0.004279 | 0.007047 | 0.007027 | 0.000325 | 0.020056 | 0.011474 | 0.000761 | 0.03619 | 0.006338 | 0.011383 | 0.009644 | 0.002407 | 0.045681 | 0.007243 | 0.005239 | 0.014617 | 0.00409 | 0.02479 | 0.006831 | 0.015718 |
| 19.9166667 | 0.0104 | 0.016912 | 0.0051 | 0.000703 | 0.009405 | 0.005332 | 0.007433 | 0.000525 | 0.024093 | 0.006881 | 0.001383 | 0.035044 | 0.009799 | 0.008553 | 0.014693 | 0.002461 | 0.052154 | 0.009235 | 0.006113 | 0.017383 | 0.002768 | 0.040744 | 0.010474 | 0.021823 |
| 19.9305556 | 0.022688 | 0.022635 | 0.010334 | 0.00053 | 0.015287 | 0.004035 | 0.009016 | 0.000816 | 0.026665 | 0.007098 | 0.002104 | 0.036441 | 0.019629 | 0.008232 | 0.019303 | 0.002913 | 0.065723 | 0.010173 | 0.005566 | 0.019693 | 0.001557 | 0.054795 | 0.015144 | 0.029272 |
| 19.9444444 | 0.033148 | 0.026936 | 0.014613 | 0.000739 | 0.019579 | 0.003593 | 0.013301 | 0.001292 | 0.025182 | 0.011204 | 0.002187 | 0.041893 | 0.028566 | 0.012837 | 0.02178 | 0.003422 | 0.076676 | 0.009164 | 0.00632 | 0.021263 | 0.0014 | 0.062239 | 0.018992 | 0.037841 |
| 19.9583333 | 0.036111 | 0.029101 | 0.017219 | 0.000754 | 0.019021 | 0.003503 | 0.018733 | 0.001813 | 0.021266 | 0.016439 | 0.002348 | 0.049821 | 0.031337 | 0.020437 | 0.021771 | 0.003742 | 0.074355 | 0.007373 | 0.008848 | 0.021379 | 0.000931 | 0.060673 | 0.020235 | 0.044625 |
| 19.9722222 | 0.032355 | 0.028205 | 0.019685 | 0.000881 | 0.014189 | 0.003349 | 0.021845 | 0.001971 | 0.018219 | 0.021195 | 0.003247 | 0.054038 | 0.028066 | 0.025108 | 0.020479 | 0.004068 | 0.059064 | 0.006298 | 0.0093 | 0.018344 | 0.000388 | 0.050695 | 0.019342 | 0.046196 |
| 19.9861111 | 0.026921 | 0.024694 | 0.021418 | 0.001615 | 0.010304 | 0.003449 | 0.021394 | 0.00172 | 0.016802 | 0.024619 | 0.005471 | 0.049355 | 0.024817 | 0.02224 | 0.019432 | 0.004814 | 0.039816 | 0.005888 | 0.007955 | 0.010786 | 0.000374 | 0.036451 | 0.017705 | 0.0411 |
| 20 | 0.022396 | 0.020859 | 0.021174 | 0.002107 | 0.008373 | 0.005421 | 0.019806 | 0.001239 | 0.015799 | 0.029224 | 0.010075 | 0.042086 | 0.027472 | 0.015704 | 0.020394 | 0.006617 | 0.024187 | 0.006265 | 0.014959 | 0.003907 | 0.000637 | 0.026137 | 0.01604 | 0.032587 |
| 20.0138889 | 0.018426 | 0.018507 | 0.021351 | 0.001625 | 0.00621 | 0.013864 | 0.019594 | 0.001032 | 0.014672 | 0.042468 | 0.019636 | 0.044585 | 0.038958 | 0.011697 | 0.023652 | 0.009112 | 0.014092 | 0.007543 | 0.045495 | 0.002033 | 0.001152 | 0.027122 | 0.01641 | 0.025296 |
| 20.0277778 | 0.014919 | 0.016871 | 0.024277 | 0.000847 | 0.003337 | 0.028206 | 0.020821 | 0.000962 | 0.014114 | 0.058554 | 0.043089 | 0.062643 | 0.062785 | 0.010579 | 0.026774 | 0.009993 | 0.008215 | 0.008376 | 0.095119 | 0.001864 | 0.001116 | 0.038598 | 0.025143 | 0.020719 |
| 20.0416667 | 0.013258 | 0.014684 | 0.026054 | 0.003009 | 0.0012 | 0.033131 | 0.020925 | 0.002809 | 0.014382 | 0.055026 | 0.074329 | 0.077702 | 0.093849 | 0.011345 | 0.02575 | 0.007817 | 0.007005 | 0.008053 | 0.121545 | 0.00327 | 0.001488 | 0.049589 | 0.040939 | 0.019585 |
| 20.0555556 | 0.013199 | 0.011505 | 0.021991 | 0.007982 | 0.001614 | 0.023069 | 0.015647 | 0.006531 | 0.014228 | 0.031293 | 0.077623 | 0.057471 | 0.092569 | 0.014476 | 0.018549 | 0.004359 | 0.007217 | 0.006679 | 0.095025 | 0.008166 | 0.004778 | 0.048668 | 0.041449 | 0.021373 |
| 20.0694444 | 0.012131 | 0.007505 | 0.015026 | 0.009657 | 0.00343 | 0.012213 | 0.007442 | 0.007101 | 0.012085 | 0.016149 | 0.044715 | 0.026055 | 0.058748 | 0.016633 | 0.008652 | 0.001835 | 0.004461 | 0.004658 | 0.046172 | 0.014643 | 0.008896 | 0.037494 | 0.029379 | 0.021418 |
| 20.0833333 | 0.008219 | 0.004289 | 0.009664 | 0.007183 | 0.004059 | 0.006818 | 0.00395 | 0.004871 | 0.007756 | 0.016945 | 0.01384 | 0.02246 | 0.050841 | 0.012785 | 0.002948 | 0.000654 | 0.001562 | 0.005782 | 0.015455 | 0.018825 | 0.008851 | 0.025076 | 0.025174 | 0.014463 |
| 20.0972222 | 0.004651 | 0.002832 | 0.006483 | 0.004666 | 0.003529 | 0.004151 | 0.004318 | 0.003078 | 0.004629 | 0.014231 | 0.004712 | 0.025355 | 0.058802 | 0.007072 | 0.002991 | 0.000309 | 0.001145 | 0.010971 | 0.005441 | 0.01792 | 0.005215 | 0.015193 | 0.01714 | 0.007537 |
| 20.1111111 | 0.006283 | 0.002027 | 0.004574 | 0.003275 | 0.003257 | 0.00293 | 0.003353 | 0.002063 | 0.00703 | 0.006036 | 0.003455 | 0.019554 | 0.046457 | 0.007202 | 0.003726 | 0.000162 | 0.001773 | 0.012823 | 0.00373 | 0.012995 | 0.002863 | 0.007798 | 0.005492 | 0.009921 |
| 20.125 | 0.011168 | 0.001053 | 0.003337 | 0.002567 | 0.003421 | 0.002158 | 0.001725 | 0.00183 | 0.012533 | 0.002076 | 0.001685 | 0.013849 | 0.026859 | 0.010653 | 0.003613 | 0.000378 | 0.001826 | 0.008511 | 0.00411 | 0.007371 | 0.002807 | 0.003196 | 0.001597 | 0.013944 |
| 20.1388889 | 0.014587 | 0.000522 | 0.002535 | 0.002179 | 0.003189 | 0.001562 | 0.001574 | 0.001867 | 0.014841 | 0.002656 | 0.001203 | 0.009431 | 0.013101 | 0.012113 | 0.004689 | 0.001127 | 0.001368 | 0.003768 | 0.004623 | 0.003166 | 0.003128 | 0.001591 | 0.00084 | 0.012706 |
| 20.1527778 | 0.015368 | 0.000605 | 0.001844 | 0.001762 | 0.002084 | 0.001199 | 0.001472 | 0.001434 | 0.012755 | 0.004165 | 0.001223 | 0.007026 | 0.005607 | 0.012095 | 0.007907 | 0.001269 | 0.001347 | 0.002908 | 0.004604 | 0.00104 | 0.003101 | 0.001273 | 0.000578 | 0.009043 |
| 20.1666667 | 0.014589 | 0.000572 | 0.001111 | 0.001146 | 0.000978 | 0.001002 | 0.000841 | 0.00071 | 0.010375 | 0.005192 | 0.000887 | 0.006647 | 0.004033 | 0.01196 | 0.011569 | 0.001007 | 0.002749 | 0.004986 | 0.003703 | 0.00126 | 0.002105 | 0.00087 | 0.001264 | 0.005452 |
| 20.1805556 | 0.013298 | 0.000312 | 0.001149 | 0.00054 | 0.000732 | 0.00076 | 0.00191 | 0.000409 | 0.009721 | 0.006414 | 0.000947 | 0.005272 | 0.006241 | 0.013055 | 0.011689 | 0.001415 | 0.008155 | 0.006262 | 0.002559 | 0.003452 | 0.001065 | 0.001461 | 0.002981 | 0.002841 |
| 20.1944444 | 0.012054 | 0.000422 | 0.002562 | 0.000437 | 0.000695 | 0.000458 | 0.005586 | 0.00035 | 0.008609 | 0.007506 | 0.00167 | 0.003152 | 0.008107 | 0.016433 | 0.008359 | 0.001588 | 0.014921 | 0.005676 | 0.00285 | 0.006543 | 0.001855 | 0.002666 | 0.004429 | 0.002044 |
| 20.2083333 | 0.011175 | 0.001392 | 0.003795 | 0.001139 | 0.000692 | 0.00037 | 0.007262 | 0.000249 | 0.009989 | 0.005998 | 0.003151 | 0.005095 | 0.007502 | 0.017734 | 0.005824 | 0.001995 | 0.013461 | 0.003862 | 0.003503 | 0.008637 | 0.003871 | 0.004652 | 0.004459 | 0.002081 |
| 20.2222222 | 0.010408 | 0.003346 | 0.003502 | 0.002289 | 0.001408 | 0.000597 | 0.007555 | 0.000245 | 0.018303 | 0.004131 | 0.007161 | 0.012981 | 0.008624 | 0.013129 | 0.005904 | 0.003343 | 0.006701 | 0.002432 | 0.002687 | 0.008278 | 0.005502 | 0.010326 | 0.004878 | 0.002125 |
| 20.2361111 | 0.008979 | 0.005329 | 0.002707 | 0.002595 | 0.002334 | 0.000716 | 0.00973 | 0.000259 | 0.02531 | 0.005105 | 0.012514 | 0.021852 | 0.01263 | 0.009103 | 0.006498 | 0.003644 | 0.004584 | 0.003274 | 0.001653 | 0.006171 | 0.005532 | 0.016074 | 0.005474 | 0.004261 |
| 20.25 | 0.006233 | 0.005843 | 0.003 | 0.001899 | 0.002727 | 0.000492 | 0.007513 | 0.000496 | 0.022849 | 0.00418 | 0.013565 | 0.023373 | 0.012478 | 0.011248 | 0.006241 | 0.00273 | 0.006612 | 0.003569 | 0.001588 | 0.004157 | 0.004646 | 0.015971 | 0.003735 | 0.007701 |
| 20.2638889 | 0.003012 | 0.005044 | 0.00445 | 0.001566 | 0.00301 | 0.000354 | 0.002622 | 0.001426 | 0.017155 | 0.00135 | 0.010266 | 0.01917 | 0.008498 | 0.014651 | 0.006195 | 0.002211 | 0.011657 | 0.00232 | 0.002571 | 0.003135 | 0.00522 | 0.013501 | 0.001757 | 0.010904 |
| 20.2777778 | 0.00103 | 0.004402 | 0.005803 | 0.001673 | 0.003431 | 0.000463 | 0.001752 | 0.002506 | 0.015199 | 0.000892 | 0.006964 | 0.015179 | 0.006124 | 0.014641 | 0.00671 | 0.002236 | 0.017326 | 0.002898 | 0.003164 | 0.002792 | 0.00734 | 0.013135 | 0.001517 | 0.013891 |
| 20.2916667 | 0.001203 | 0.004676 | 0.006192 | 0.001497 | 0.003632 | 0.000482 | 0.003548 | 0.002663 | 0.01705 | 0.002343 | 0.005616 | 0.012602 | 0.005966 | 0.01227 | 0.007364 | 0.00232 | 0.020751 | 0.004877 | 0.002576 | 0.002814 | 0.008845 | 0.014018 | 0.001842 | 0.015887 |
| 20.3055556 | 0.003901 | 0.005479 | 0.005937 | 0.001118 | 0.00406 | 0.000386 | 0.00591 | 0.002168 | 0.021534 | 0.004706 | 0.006484 | 0.012879 | 0.007073 | 0.010128 | 0.00824 | 0.00233 | 0.022964 | 0.004718 | 0.001787 | 0.002484 | 0.00907 | 0.015488 | 0.002151 | 0.016346 |
| 20.3194444 | 0.008762 | 0.006061 | 0.005853 | 0.000906 | 0.005303 | 0.000413 | 0.008139 | 0.002023 | 0.02599 | 0.007314 | 0.010215 | 0.015748 | 0.00833 | 0.009643 | 0.009023 | 0.002378 | 0.0243 | 0.003036 | 0.00138 | 0.00211 | 0.010355 | 0.017757 | 0.002453 | 0.015336 |
| 20.3333333 | 0.013808 | 0.006103 | 0.006286 | 0.001015 | 0.006605 | 0.00062 | 0.010158 | 0.002408 | 0.027235 | 0.009214 | 0.019078 | 0.01863 | 0.008671 | 0.010633 | 0.008931 | 0.00254 | 0.024851 | 0.003418 | 0.001543 | 0.004053 | 0.014025 | 0.021425 | 0.002705 | 0.0139 |
| 20.3472222 | 0.016592 | 0.006026 | 0.007025 | 0.001283 | 0.006829 | 0.000936 | 0.012644 | 0.002781 | 0.025172 | 0.010056 | 0.030907 | 0.020316 | 0.008191 | 0.012986 | 0.008004 | 0.002789 | 0.02589 | 0.005167 | 0.001947 | 0.008681 | 0.017229 | 0.026849 | 0.002983 | 0.013684 |
| 20.3611111 | 0.016804 | 0.006554 | 0.007782 | 0.001326 | 0.006035 | 0.001319 | 0.015837 | 0.002827 | 0.021788 | 0.010151 | 0.03755 | 0.021495 | 0.007809 | 0.016865 | 0.007056 | 0.002986 | 0.030246 | 0.005502 | 0.002066 | 0.014276 | 0.017272 | 0.032526 | 0.003535 | 0.015767 |
| 20.375 | 0.016042 | 0.007948 | 0.008355 | 0.001093 | 0.005265 | 0.001815 | 0.018185 | 0.002733 | 0.020094 | 0.010245 | 0.036393 | 0.02304 | 0.008028 | 0.021026 | 0.006554 | 0.002763 | 0.039196 | 0.004407 | 0.001897 | 0.020524 | 0.015444 | 0.037192 | 0.004063 | 0.019874 |
| 20.3888889 | 0.015325 | 0.009655 | 0.008694 | 0.001184 | 0.005441 | 0.0025 | 0.018567 | 0.002741 | 0.022589 | 0.010754 | 0.032653 | 0.025237 | 0.009039 | 0.022955 | 0.006455 | 0.002134 | 0.049519 | 0.003049 | 0.001744 | 0.025801 | 0.014579 | 0.039935 | 0.004062 | 0.02512 |
| 20.4027778 | 0.01447 | 0.01056 | 0.008887 | 0.002129 | 0.006544 | 0.003118 | 0.017575 | 0.002774 | 0.028051 | 0.011637 | 0.029469 | 0.027631 | 0.011047 | 0.020615 | 0.006794 | 0.002153 | 0.055418 | 0.002019 | 0.001712 | 0.02523 | 0.015179 | 0.039885 | 0.00431 | 0.029421 |
| 20.4166667 | 0.012893 | 0.010257 | 0.009475 | 0.003781 | 0.007632 | 0.00334 | 0.016391 | 0.002608 | 0.032584 | 0.013427 | 0.026953 | 0.029936 | 0.013009 | 0.01437 | 0.007249 | 0.003374 | 0.054768 | 0.001724 | 0.001783 | 0.018541 | 0.015374 | 0.038659 | 0.005512 | 0.029412 |
| 20.4305556 | 0.010145 | 0.009856 | 0.010977 | 0.005506 | 0.008306 | 0.003592 | 0.016378 | 0.002458 | 0.034223 | 0.016629 | 0.026293 | 0.032514 | 0.01345 | 0.007019 | 0.007186 | 0.004649 | 0.050749 | 0.002697 | 0.00199 | 0.010552 | 0.013854 | 0.039012 | 0.006779 | 0.02463 |
| 20.4444444 | 0.006415 | 0.010422 | 0.012396 | 0.006822 | 0.008731 | 0.004693 | 0.01755 | 0.002683 | 0.033424 | 0.019865 | 0.02681 | 0.034916 | 0.013105 | 0.002064 | 0.007187 | 0.005457 | 0.046434 | 0.004786 | 0.002829 | 0.004182 | 0.011684 | 0.04036 | 0.007975 | 0.018581 |
| 20.4583333 | 0.00284 | 0.011125 | 0.012116 | 0.007081 | 0.008274 | 0.006761 | 0.017927 | 0.003054 | 0.030429 | 0.020583 | 0.025812 | 0.035748 | 0.013702 | 0.003692 | 0.008638 | 0.006788 | 0.042226 | 0.006841 | 0.004998 | 0.002248 | 0.011086 | 0.03954 | 0.010086 | 0.014853 |
| 20.4722222 | 0.000868 | 0.011486 | 0.010376 | 0.005699 | 0.007374 | 0.00877 | 0.015146 | 0.002929 | 0.025712 | 0.017397 | 0.024066 | 0.035247 | 0.014955 | 0.013479 | 0.010696 | 0.008358 | 0.03925 | 0.007525 | 0.007903 | 0.006863 | 0.012663 | 0.035594 | 0.012664 | 0.013723 |
| 20.4861111 | 0.002465 | 0.015755 | 0.010217 | 0.004328 | 0.010303 | 0.011112 | 0.009647 | 0.002493 | 0.021857 | 0.013183 | 0.025623 | 0.039536 | 0.014543 | 0.024726 | 0.010171 | 0.009151 | 0.047395 | 0.006787 | 0.014238 | 0.01608 | 0.014678 | 0.031568 | 0.015275 | 0.012881 |
| 20.5 | 0.00981 | 0.030358 | 0.015947 | 0.005459 | 0.022737 | 0.017398 | 0.005083 | 0.002874 | 0.022726 | 0.018708 | 0.033576 | 0.058617 | 0.01198 | 0.029113 | 0.006652 | 0.010398 | 0.087134 | 0.006963 | 0.040055 | 0.025885 | 0.015233 | 0.032299 | 0.020061 | 0.011674 |
| 20.5138889 | 0.024863 | 0.052386 | 0.027415 | 0.008611 | 0.038183 | 0.029249 | 0.004424 | 0.004137 | 0.027565 | 0.040824 | 0.046142 | 0.08599 | 0.008844 | 0.027131 | 0.002966 | 0.012932 | 0.158062 | 0.010528 | 0.092049 | 0.030917 | 0.013773 | 0.035761 | 0.028479 | 0.012252 |
| 20.5277778 | 0.041205 | 0.067138 | 0.036768 | 0.010951 | 0.040645 | 0.040187 | 0.006451 | 0.004922 | 0.029955 | 0.06604 | 0.054941 | 0.093517 | 0.006675 | 0.024663 | 0.000989 | 0.013588 | 0.207406 | 0.015783 | 0.122792 | 0.026647 | 0.009927 | 0.028777 | 0.039211 | 0.01571 |
| 20.5416667 | 0.042898 | 0.068819 | 0.038337 | 0.010779 | 0.031829 | 0.039585 | 0.006415 | 0.004912 | 0.027857 | 0.074041 | 0.05418 | 0.072375 | 0.005823 | 0.021557 | 0.000762 | 0.010507 | 0.188136 | 0.016924 | 0.096357 | 0.015992 | 0.006344 | 0.016113 | 0.050144 | 0.020139 |
| 20.5555556 | 0.029175 | 0.0691 | 0.035438 | 0.008952 | 0.028662 | 0.028453 | 0.003793 | 0.004838 | 0.026789 | 0.06164 | 0.047201 | 0.044795 | 0.004332 | 0.014753 | 0.002207 | 0.007801 | 0.127481 | 0.012473 | 0.054099 | 0.007758 | 0.009046 | 0.01649 | 0.054212 | 0.021548 |
| 20.5694444 | 0.013554 | 0.073984 | 0.031021 | 0.007072 | 0.043894 | 0.017898 | 0.00373 | 0.004331 | 0.027461 | 0.037445 | 0.037871 | 0.02428 | 0.002777 | 0.007363 | 0.005561 | 0.008283 | 0.071822 | 0.006723 | 0.030799 | 0.004488 | 0.016583 | 0.01976 | 0.046216 | 0.016616 |
| 20.5833333 | 0.00693 | 0.073713 | 0.023696 | 0.006513 | 0.06246 | 0.011306 | 0.006976 | 0.003019 | 0.022321 | 0.016447 | 0.027615 | 0.014142 | 0.004607 | 0.004126 | 0.008916 | 0.009091 | 0.038296 | 0.003203 | 0.019428 | 0.003773 | 0.020081 | 0.012476 | 0.034106 | 0.0095 |
| 20.5972222 | 0.013897 | 0.060862 | 0.014623 | 0.008458 | 0.053634 | 0.006461 | 0.008944 | 0.001826 | 0.011476 | 0.008477 | 0.018995 | 0.020955 | 0.007616 | 0.006808 | 0.010345 | 0.008346 | 0.048626 | 0.002258 | 0.010523 | 0.006631 | 0.016902 | 0.003891 | 0.025262 | 0.009834 |
| 20.6111111 | 0.023226 | 0.041517 | 0.007338 | 0.012079 | 0.0267 | 0.004343 | 0.006529 | 0.001261 | 0.005907 | 0.008477 | 0.012984 | 0.035864 | 0.005969 | 0.011328 | 0.010047 | 0.007076 | 0.102848 | 0.001936 | 0.004199 | 0.010271 | 0.012298 | 0.003421 | 0.017411 | 0.018591 |
| 20.625 | 0.019632 | 0.025222 | 0.003749 | 0.014817 | 0.009275 | 0.006998 | 0.004062 | 0.00102 | 0.009348 | 0.017231 | 0.008234 | 0.039619 | 0.003659 | 0.010997 | 0.008094 | 0.00517 | 0.133521 | 0.002404 | 0.003653 | 0.008356 | 0.010463 | 0.013197 | 0.010378 | 0.027022 |
| 20.6388889 | 0.013551 | 0.015139 | 0.004741 | 0.015466 | 0.006116 | 0.013875 | 0.006938 | 0.001153 | 0.014368 | 0.031935 | 0.004423 | 0.027489 | 0.006564 | 0.009178 | 0.004647 | 0.003086 | 0.093633 | 0.003667 | 0.008081 | 0.006124 | 0.011493 | 0.02973 | 0.011769 | 0.027728 |
| 20.6527778 | 0.025086 | 0.009446 | 0.009067 | 0.014279 | 0.00893 | 0.020604 | 0.009635 | 0.001625 | 0.016044 | 0.040189 | 0.003403 | 0.011352 | 0.009365 | 0.013594 | 0.002981 | 0.0029 | 0.058803 | 0.004241 | 0.014654 | 0.010332 | 0.013639 | 0.042858 | 0.022335 | 0.018724 |
| 20.6666667 | 0.041423 | 0.006029 | 0.012742 | 0.011356 | 0.008123 | 0.020307 | 0.006611 | 0.001914 | 0.013837 | 0.034513 | 0.005095 | 0.003892 | 0.007275 | 0.018945 | 0.00535 | 0.00407 | 0.088277 | 0.004 | 0.015584 | 0.014016 | 0.016058 | 0.044341 | 0.028071 | 0.009218 |
| 20.6805556 | 0.042271 | 0.004086 | 0.013148 | 0.007787 | 0.006883 | 0.011657 | 0.003474 | 0.001645 | 0.008699 | 0.020396 | 0.005665 | 0.004688 | 0.003686 | 0.015626 | 0.006324 | 0.004111 | 0.122643 | 0.002925 | 0.01016 | 0.013056 | 0.019294 | 0.034097 | 0.019769 | 0.009096 |
| 20.6944444 | 0.030535 | 0.002666 | 0.011244 | 0.005407 | 0.015054 | 0.004545 | 0.005925 | 0.00107 | 0.007263 | 0.017708 | 0.00345 | 0.005022 | 0.003822 | 0.00958 | 0.003966 | 0.002841 | 0.119495 | 0.002935 | 0.007648 | 0.010265 | 0.021467 | 0.018931 | 0.010234 | 0.012567 |
| 20.7083333 | 0.016285 | 0.002163 | 0.008783 | 0.005089 | 0.027235 | 0.005155 | 0.010128 | 0.000537 | 0.012502 | 0.027099 | 0.002146 | 0.002464 | 0.006798 | 0.012368 | 0.003974 | 0.001769 | 0.081161 | 0.005562 | 0.009697 | 0.006336 | 0.018692 | 0.014585 | 0.014549 | 0.013198 |
| 20.7222222 | 0.011763 | 0.004492 | 0.006838 | 0.006537 | 0.030461 | 0.007522 | 0.010787 | 0.00023 | 0.016177 | 0.026624 | 0.005075 | 0.000663 | 0.006688 | 0.019611 | 0.006708 | 0.001099 | 0.055284 | 0.007141 | 0.009505 | 0.004931 | 0.011779 | 0.027514 | 0.025616 | 0.011052 |
| 20.7361111 | 0.020503 | 0.008954 | 0.005938 | 0.008298 | 0.022066 | 0.008315 | 0.008347 | 0.000219 | 0.012649 | 0.015034 | 0.011554 | 0.000651 | 0.004353 | 0.019837 | 0.008248 | 0.000766 | 0.089359 | 0.004918 | 0.00615 | 0.008774 | 0.005845 | 0.042078 | 0.030401 | 0.008039 |
| 20.75 | 0.03322 | 0.012814 | 0.004859 | 0.008821 | 0.011669 | 0.007656 | 0.004933 | 0.000221 | 0.007214 | 0.009627 | 0.018825 | 0.002899 | 0.005112 | 0.012442 | 0.007148 | 0.000732 | 0.131872 | 0.002496 | 0.005353 | 0.012759 | 0.003222 | 0.044918 | 0.029072 | 0.011366 |
| 20.7638889 | 0.03937 | 0.014058 | 0.002978 | 0.007883 | 0.009469 | 0.00563 | 0.003235 | 0.000162 | 0.007731 | 0.012523 | 0.021684 | 0.007329 | 0.007287 | 0.009033 | 0.004236 | 0.000982 | 0.135668 | 0.002626 | 0.006624 | 0.012202 | 0.004741 | 0.034661 | 0.03094 | 0.023535 |
| 20.7777778 | 0.028812 | 0.011974 | 0.002089 | 0.006512 | 0.012081 | 0.008269 | 0.006478 | 0.000331 | 0.012185 | 0.012695 | 0.01757 | 0.008886 | 0.007363 | 0.0152 | 0.001826 | 0.002044 | 0.102358 | 0.00261 | 0.005453 | 0.007343 | 0.009458 | 0.018772 | 0.033275 | 0.034935 |
| 20.7916667 | 0.015357 | 0.008045 | 0.002692 | 0.005684 | 0.011682 | 0.016842 | 0.010795 | 0.000724 | 0.012647 | 0.010081 | 0.009965 | 0.006952 | 0.005175 | 0.021241 | 0.002059 | 0.00249 | 0.059175 | 0.001625 | 0.003754 | 0.004218 | 0.012627 | 0.013335 | 0.02334 | 0.036788 |
| 20.8055556 | 0.023573 | 0.004558 | 0.003436 | 0.005177 | 0.009663 | 0.020975 | 0.009069 | 0.000854 | 0.007983 | 0.01687 | 0.003736 | 0.009547 | 0.002842 | 0.020408 | 0.004626 | 0.002188 | 0.059594 | 0.002263 | 0.006079 | 0.006753 | 0.012152 | 0.021835 | 0.011439 | 0.029191 |
| 20.8194444 | 0.039758 | 0.00232 | 0.003638 | 0.004241 | 0.009452 | 0.015831 | 0.004656 | 0.000556 | 0.004734 | 0.031629 | 0.002227 | 0.018169 | 0.003727 | 0.012769 | 0.007324 | 0.00218 | 0.089764 | 0.004887 | 0.010816 | 0.007568 | 0.00906 | 0.030721 | 0.014615 | 0.016848 |
| 20.8333333 | 0.039406 | 0.001214 | 0.003288 | 0.002944 | 0.009628 | 0.007589 | 0.004462 | 0.000544 | 0.005451 | 0.035473 | 0.005625 | 0.024578 | 0.007144 | 0.006286 | 0.007913 | 0.004137 | 0.088667 | 0.007865 | 0.012435 | 0.004425 | 0.004803 | 0.032696 | 0.027397 | 0.00639 |
| 20.8472222 | 0.024399 | 0.00132 | 0.002704 | 0.001995 | 0.008233 | 0.003528 | 0.006285 | 0.001198 | 0.004755 | 0.022347 | 0.010138 | 0.022407 | 0.009611 | 0.008929 | 0.007004 | 0.00698 | 0.050806 | 0.010391 | 0.008816 | 0.004058 | 0.002784 | 0.029039 | 0.038377 | 0.001781 |
| 20.8611111 | 0.011886 | 0.002631 | 0.002478 | 0.001432 | 0.005436 | 0.005463 | 0.006206 | 0.00146 | 0.003284 | 0.0089 | 0.01143 | 0.013577 | 0.009835 | 0.012931 | 0.006938 | 0.00584 | 0.023998 | 0.012173 | 0.00715 | 0.007694 | 0.004673 | 0.021573 | 0.041506 | 0.002885 |
| 20.875 | 0.014754 | 0.005188 | 0.003315 | 0.001301 | 0.003202 | 0.008918 | 0.005141 | 0.000897 | 0.004305 | 0.005094 | 0.009365 | 0.006883 | 0.008281 | 0.012681 | 0.006815 | 0.003226 | 0.039052 | 0.012908 | 0.012633 | 0.011866 | 0.00539 | 0.013768 | 0.034913 | 0.009645 |
| 20.8888889 | 0.029761 | 0.008759 | 0.005602 | 0.002662 | 0.003478 | 0.011268 | 0.004803 | 0.000637 | 0.004054 | 0.005677 | 0.006619 | 0.008892 | 0.006537 | 0.01224 | 0.005859 | 0.0024 | 0.067791 | 0.012734 | 0.016214 | 0.014833 | 0.003583 | 0.009468 | 0.023265 | 0.019816 |
| 20.9027778 | 0.046243 | 0.012106 | 0.009162 | 0.004898 | 0.005745 | 0.012101 | 0.005778 | 0.000751 | 0.002215 | 0.011597 | 0.005515 | 0.016999 | 0.006321 | 0.01311 | 0.005372 | 0.002414 | 0.079281 | 0.011388 | 0.010999 | 0.017929 | 0.002901 | 0.009295 | 0.014633 | 0.028729 |
| 20.9166667 | 0.05458 | 0.014081 | 0.0126 | 0.005449 | 0.007786 | 0.010051 | 0.007271 | 0.000739 | 0.001807 | 0.022582 | 0.006758 | 0.023796 | 0.007643 | 0.014372 | 0.005261 | 0.003141 | 0.074209 | 0.008482 | 0.005195 | 0.020626 | 0.003222 | 0.010653 | 0.012384 | 0.033777 |
| 20.9305556 | 0.05149 | 0.016017 | 0.014144 | 0.003808 | 0.008603 | 0.006228 | 0.008331 | 0.001474 | 0.004461 | 0.0307 | 0.009214 | 0.025296 | 0.009385 | 0.016944 | 0.005279 | 0.003993 | 0.064064 | 0.005011 | 0.006378 | 0.018899 | 0.0059 | 0.011321 | 0.013378 | 0.035833 |
| 20.9444444 | 0.040981 | 0.020735 | 0.014991 | 0.002086 | 0.008965 | 0.003177 | 0.009069 | 0.003789 | 0.012662 | 0.029599 | 0.012094 | 0.022764 | 0.010818 | 0.022563 | 0.006482 | 0.004116 | 0.0584 | 0.003919 | 0.008844 | 0.012762 | 0.011203 | 0.011412 | 0.016388 | 0.037753 |
| 20.9583333 | 0.02503 | 0.02767 | 0.017803 | 0.001792 | 0.00957 | 0.001796 | 0.009655 | 0.006708 | 0.0241 | 0.018467 | 0.01619 | 0.019438 | 0.011687 | 0.028092 | 0.009566 | 0.003182 | 0.054932 | 0.00797 | 0.007339 | 0.006465 | 0.015928 | 0.012579 | 0.024284 | 0.040641 |
| 20.9722222 | 0.011973 | 0.032598 | 0.022489 | 0.002096 | 0.010084 | 0.001627 | 0.009528 | 0.007737 | 0.03307 | 0.011613 | 0.021065 | 0.016972 | 0.011202 | 0.025985 | 0.013709 | 0.001819 | 0.044794 | 0.016292 | 0.004299 | 0.002865 | 0.01636 | 0.015477 | 0.036426 | 0.042179 |
| 20.9861111 | 0.011007 | 0.032258 | 0.02653 | 0.001839 | 0.009664 | 0.002179 | 0.008551 | 0.005433 | 0.035299 | 0.016933 | 0.024512 | 0.015662 | 0.009344 | 0.014711 | 0.01736 | 0.001083 | 0.026499 | 0.02396 | 0.005419 | 0.002471 | 0.010695 | 0.018726 | 0.04614 | 0.039041 |
| 21 | 0.011744 | 0.028266 | 0.028816 | 0.001862 | 0.008699 | 0.003434 | 0.009253 | 0.002197 | 0.032271 | 0.016904 | 0.025991 | 0.018762 | 0.009519 | 0.003953 | 0.020275 | 0.001043 | 0.010961 | 0.027412 | 0.01659 | 0.005349 | 0.004318 | 0.021163 | 0.046368 | 0.032543 |
| 21.0138889 | 0.010054 | 0.02539 | 0.030242 | 0.002832 | 0.007602 | 0.005903 | 0.014864 | 0.001098 | 0.031411 | 0.016784 | 0.02691 | 0.030882 | 0.01573 | 0.003859 | 0.022745 | 0.001433 | 0.005503 | 0.032004 | 0.040184 | 0.011556 | 0.002196 | 0.023499 | 0.037267 | 0.028027 |
| 21.0277778 | 0.018917 | 0.026603 | 0.030819 | 0.002817 | 0.005651 | 0.010161 | 0.023824 | 0.001602 | 0.034677 | 0.037632 | 0.02859 | 0.051627 | 0.026313 | 0.013131 | 0.024788 | 0.0035 | 0.004829 | 0.04057 | 0.079225 | 0.020492 | 0.003349 | 0.02759 | 0.025555 | 0.027136 |
| 21.0416667 | 0.037081 | 0.029663 | 0.031072 | 0.001554 | 0.004091 | 0.014355 | 0.031199 | 0.001812 | 0.035601 | 0.056805 | 0.03005 | 0.069592 | 0.034626 | 0.022767 | 0.027687 | 0.00641 | 0.003913 | 0.048468 | 0.127441 | 0.031126 | 0.006483 | 0.035397 | 0.01748 | 0.029065 |
| 21.0555556 | 0.047752 | 0.028647 | 0.030347 | 0.00185 | 0.005365 | 0.01436 | 0.032356 | 0.001526 | 0.030007 | 0.046463 | 0.027176 | 0.068002 | 0.03554 | 0.026008 | 0.028233 | 0.006302 | 0.002754 | 0.050506 | 0.134749 | 0.040511 | 0.007004 | 0.043703 | 0.015127 | 0.033499 |
| 21.0694444 | 0.039223 | 0.021648 | 0.025143 | 0.003529 | 0.00806 | 0.009848 | 0.025017 | 0.001817 | 0.02073 | 0.023434 | 0.018861 | 0.046347 | 0.028192 | 0.020188 | 0.021063 | 0.003517 | 0.003372 | 0.037184 | 0.076662 | 0.041908 | 0.004291 | 0.043692 | 0.015463 | 0.035018 |
| 21.0833333 | 0.019498 | 0.011832 | 0.015985 | 0.003411 | 0.008576 | 0.005013 | 0.013294 | 0.002952 | 0.010769 | 0.015401 | 0.010674 | 0.020881 | 0.015289 | 0.01066 | 0.010908 | 0.001853 | 0.00585 | 0.018824 | 0.026472 | 0.032661 | 0.002813 | 0.032914 | 0.01449 | 0.026499 |
| 21.0972222 | 0.008693 | 0.004259 | 0.00781 | 0.002927 | 0.006712 | 0.002418 | 0.004415 | 0.004133 | 0.004544 | 0.018354 | 0.006763 | 0.005354 | 0.00614 | 0.006927 | 0.008621 | 0.001471 | 0.006245 | 0.020254 | 0.022915 | 0.01881 | 0.001927 | 0.018278 | 0.011812 | 0.01336 |
| 21.1111111 | 0.011296 | 0.001796 | 0.003017 | 0.005952 | 0.005181 | 0.001378 | 0.001977 | 0.004838 | 0.00369 | 0.014929 | 0.005494 | 0.001282 | 0.004499 | 0.006877 | 0.014188 | 0.001413 | 0.004762 | 0.036688 | 0.025903 | 0.008533 | 0.001398 | 0.007174 | 0.009323 | 0.007475 |
| 21.125 | 0.014535 | 0.001471 | 0.00093 | 0.010378 | 0.005399 | 0.000967 | 0.004018 | 0.004534 | 0.00299 | 0.008316 | 0.004848 | 0.002195 | 0.003982 | 0.005003 | 0.016232 | 0.001912 | 0.004039 | 0.047456 | 0.020095 | 0.007025 | 0.001243 | 0.001902 | 0.007598 | 0.008192 |
| 21.1388889 | 0.012965 | 0.001313 | 0.000698 | 0.011936 | 0.005915 | 0.001028 | 0.005421 | 0.00276 | 0.002214 | 0.009311 | 0.005008 | 0.003995 | 0.004632 | 0.004529 | 0.011767 | 0.002341 | 0.00308 | 0.047834 | 0.012659 | 0.011075 | 0.001647 | 0.000372 | 0.006027 | 0.008742 |
| 21.1527778 | 0.010424 | 0.002188 | 0.001455 | 0.008993 | 0.004503 | 0.001204 | 0.004056 | 0.001375 | 0.003108 | 0.01513 | 0.005958 | 0.004177 | 0.006887 | 0.007783 | 0.006961 | 0.002382 | 0.001526 | 0.040671 | 0.005811 | 0.014649 | 0.002507 | 0.000404 | 0.004327 | 0.007984 |
| 21.1666667 | 0.008845 | 0.003261 | 0.002241 | 0.00458 | 0.002226 | 0.001155 | 0.002293 | 0.001337 | 0.003136 | 0.018769 | 0.006762 | 0.003359 | 0.00774 | 0.01207 | 0.004379 | 0.001993 | 0.000504 | 0.02742 | 0.002065 | 0.015449 | 0.002247 | 0.000694 | 0.00265 | 0.005853 |
| 21.1805556 | 0.007768 | 0.004228 | 0.00303 | 0.002262 | 0.001184 | 0.001211 | 0.001924 | 0.001467 | 0.002313 | 0.017998 | 0.007361 | 0.004251 | 0.00741 | 0.014551 | 0.002725 | 0.001068 | 0.000491 | 0.012953 | 0.002268 | 0.012873 | 0.002426 | 0.000901 | 0.001723 | 0.002674 |
| 21.1944444 | 0.006716 | 0.005188 | 0.004217 | 0.002224 | 0.00095 | 0.001592 | 0.001676 | 0.002191 | 0.004624 | 0.011958 | 0.008211 | 0.003885 | 0.007033 | 0.014436 | 0.001644 | 0.000629 | 0.001072 | 0.005536 | 0.005761 | 0.009368 | 0.00368 | 0.00225 | 0.002073 | 0.002007 |
| 21.2083333 | 0.007401 | 0.004132 | 0.004723 | 0.00293 | 0.000992 | 0.001611 | 0.00183 | 0.002725 | 0.012445 | 0.007815 | 0.006998 | 0.003035 | 0.0053 | 0.013679 | 0.001502 | 0.001399 | 0.001572 | 0.007687 | 0.007644 | 0.008975 | 0.003225 | 0.003101 | 0.002711 | 0.007098 |
| 21.2222222 | 0.011516 | 0.002648 | 0.006218 | 0.0032 | 0.001128 | 0.001712 | 0.004068 | 0.002038 | 0.021849 | 0.012625 | 0.003985 | 0.006165 | 0.003716 | 0.015674 | 0.002139 | 0.002037 | 0.002237 | 0.010483 | 0.008925 | 0.012148 | 0.001471 | 0.00345 | 0.002172 | 0.015698 |
| 21.2361111 | 0.016282 | 0.003833 | 0.009141 | 0.004479 | 0.00178 | 0.002491 | 0.005547 | 0.002039 | 0.024497 | 0.015358 | 0.002266 | 0.009633 | 0.005094 | 0.019508 | 0.004538 | 0.001366 | 0.002644 | 0.009151 | 0.012188 | 0.014799 | 0.002137 | 0.006363 | 0.001551 | 0.020684 |
| 21.25 | 0.016598 | 0.004604 | 0.008749 | 0.005359 | 0.003223 | 0.002461 | 0.003775 | 0.002919 | 0.018592 | 0.010796 | 0.001918 | 0.007821 | 0.004826 | 0.020374 | 0.007466 | 0.000528 | 0.002093 | 0.008054 | 0.009495 | 0.013953 | 0.005498 | 0.00884 | 0.002262 | 0.019045 |
| 21.2638889 | 0.010963 | 0.002961 | 0.006131 | 0.005198 | 0.004632 | 0.001625 | 0.001406 | 0.002971 | 0.012621 | 0.01101 | 0.00304 | 0.003517 | 0.003695 | 0.01703 | 0.008392 | 0.00039 | 0.001705 | 0.009578 | 0.003244 | 0.010937 | 0.007578 | 0.008515 | 0.003378 | 0.015661 |
| 21.2777778 | 0.005786 | 0.001552 | 0.005192 | 0.005355 | 0.004967 | 0.001377 | 0.001555 | 0.002701 | 0.01456 | 0.014491 | 0.003515 | 0.003467 | 0.005653 | 0.012365 | 0.007336 | 0.00078 | 0.00251 | 0.011776 | 0.001389 | 0.007859 | 0.00635 | 0.007462 | 0.004046 | 0.015398 |
| 21.2916667 | 0.006543 | 0.001296 | 0.0058 | 0.005322 | 0.00425 | 0.001802 | 0.004574 | 0.002331 | 0.024503 | 0.013348 | 0.002304 | 0.010651 | 0.005447 | 0.009923 | 0.006337 | 0.001293 | 0.004709 | 0.011409 | 0.001277 | 0.004742 | 0.005005 | 0.007428 | 0.004854 | 0.017619 |
| 21.3055556 | 0.00768 | 0.001365 | 0.007189 | 0.005434 | 0.003615 | 0.002359 | 0.008316 | 0.00242 | 0.034947 | 0.016328 | 0.002034 | 0.02103 | 0.00415 | 0.010668 | 0.006892 | 0.001211 | 0.007714 | 0.007928 | 0.00088 | 0.001921 | 0.00624 | 0.009383 | 0.006693 | 0.020406 |
| 21.3194444 | 0.00516 | 0.002821 | 0.008996 | 0.005725 | 0.003536 | 0.002732 | 0.011009 | 0.004814 | 0.037352 | 0.023957 | 0.002754 | 0.024573 | 0.00763 | 0.012046 | 0.007814 | 0.001 | 0.009971 | 0.003722 | 0.001434 | 0.000853 | 0.008803 | 0.011959 | 0.009347 | 0.024069 |

| | | | | | | | | | | | | | | | | | | | | | | | | |
|---|---|---|---|---|---|---|---|---|---|---|---|---|---|---|---|---|---|---|---|---|---|---|---|---|
| 21.3333333 | 0.002584 | 0.00559 | 0.011511 | 0.005646 | 0.003967 | 0.003146 | 0.013001 | 0.007154 | 0.03237 | 0.021762 | 0.003218 | 0.019177 | 0.010735 | 0.012142 | 0.00864 | 0.000888 | 0.012848 | 0.002202 | 0.003141 | 0.001377 | 0.011512 | 0.013984 | 0.012315 | 0.029161 |
| 21.3472222 | 0.003917 | 0.007561 | 0.015093 | 0.00534 | 0.004567 | 0.004021 | 0.015276 | 0.005975 | 0.02711 | 0.011619 | 0.00367 | 0.01527 | 0.009249 | 0.011317 | 0.01051 | 0.000576 | 0.020326 | 0.005111 | 0.005824 | 0.002461 | 0.014112 | 0.016378 | 0.015893 | 0.03571 |
| 21.3611111 | 0.009276 | 0.008046 | 0.018901 | 0.005777 | 0.004938 | 0.005224 | 0.018343 | 0.002805 | 0.024684 | 0.005011 | 0.004638 | 0.021584 | 0.007034 | 0.010439 | 0.01319 | 0.000298 | 0.03186 | 0.010509 | 0.008834 | 0.003902 | 0.016337 | 0.018832 | 0.020033 | 0.043491 |
| 21.375 | 0.015393 | 0.007828 | 0.020958 | 0.007353 | 0.005191 | 0.006549 | 0.021602 | 0.001189 | 0.023798 | 0.010047 | 0.005795 | 0.035439 | 0.006721 | 0.009962 | 0.014512 | 0.000357 | 0.042302 | 0.014624 | 0.010869 | 0.006906 | 0.019304 | 0.019727 | 0.024073 | 0.050835 |
| 21.3888889 | 0.019662 | 0.008017 | 0.020354 | 0.009125 | 0.005318 | 0.008101 | 0.023425 | 0.002746 | 0.022846 | 0.030688 | 0.006768 | 0.045048 | 0.007372 | 0.009365 | 0.01298 | 0.000937 | 0.050187 | 0.014373 | 0.011829 | 0.012115 | 0.023383 | 0.01896 | 0.028118 | 0.053277 |
| 21.4027778 | 0.021646 | 0.009133 | 0.018374 | 0.009995 | 0.005155 | 0.009195 | 0.021799 | 0.005517 | 0.020586 | 0.049218 | 0.007731 | 0.042181 | 0.008674 | 0.008557 | 0.010784 | 0.001528 | 0.05891 | 0.01003 | 0.013384 | 0.017424 | 0.02701 | 0.018348 | 0.033367 | 0.045987 |
| 21.4166667 | 0.022552 | 0.011226 | 0.016032 | 0.009755 | 0.005121 | 0.009433 | 0.016833 | 0.006453 | 0.017112 | 0.046724 | 0.009062 | 0.032471 | 0.011845 | 0.009186 | 0.010568 | 0.00291 | 0.068659 | 0.005043 | 0.01735 | 0.020003 | 0.029083 | 0.020399 | 0.040097 | 0.030599 |
| 21.4305556 | 0.024124 | 0.01383 | 0.014005 | 0.008793 | 0.005705 | 0.009399 | 0.011678 | 0.005755 | 0.014548 | 0.029079 | 0.010902 | 0.02708 | 0.014884 | 0.012704 | 0.011229 | 0.005399 | 0.075253 | 0.002826 | 0.022278 | 0.019196 | 0.02957 | 0.024278 | 0.045615 | 0.016538 |
| 21.4444444 | 0.026903 | 0.015686 | 0.013702 | 0.006926 | 0.006462 | 0.008996 | 0.008579 | 0.005016 | 0.014226 | 0.011823 | 0.012671 | 0.030224 | 0.014516 | 0.018253 | 0.011446 | 0.007014 | 0.076793 | 0.004649 | 0.024286 | 0.016523 | 0.029078 | 0.027759 | 0.04663 | 0.012355 |
| 21.4583333 | 0.02897 | 0.015455 | 0.014245 | 0.004408 | 0.00647 | 0.007888 | 0.006822 | 0.004335 | 0.014752 | 0.00605 | 0.013259 | 0.03764 | 0.013022 | 0.023598 | 0.012119 | 0.008101 | 0.075482 | 0.007897 | 0.02315 | 0.013436 | 0.028171 | 0.03274 | 0.043892 | 0.018189 |
| 21.4722222 | 0.026586 | 0.014372 | 0.013459 | 0.002844 | 0.006074 | 0.007069 | 0.004514 | 0.003461 | 0.013913 | 0.014371 | 0.012681 | 0.039899 | 0.013437 | 0.026213 | 0.01298 | 0.009436 | 0.074296 | 0.008665 | 0.021344 | 0.010881 | 0.026135 | 0.041068 | 0.040192 | 0.027772 |
| 21.4861111 | 0.018335 | 0.017684 | 0.013564 | 0.004361 | 0.008398 | 0.008715 | 0.003534 | 0.003334 | 0.011163 | 0.030466 | 0.014614 | 0.036257 | 0.012437 | 0.024205 | 0.013432 | 0.009872 | 0.08469 | 0.007754 | 0.022431 | 0.010014 | 0.021768 | 0.050241 | 0.037199 | 0.035141 |
| 21.5 | 0.009142 | 0.031872 | 0.021383 | 0.011254 | 0.019099 | 0.015269 | 0.006051 | 0.004837 | 0.010235 | 0.046761 | 0.026769 | 0.044464 | 0.007558 | 0.021256 | 0.015513 | 0.010158 | 0.137742 | 0.010921 | 0.035067 | 0.012125 | 0.01661 | 0.061186 | 0.038129 | 0.04107 |
| 21.5138889 | 0.006142 | 0.055907 | 0.037408 | 0.020653 | 0.034044 | 0.024025 | 0.007029 | 0.006795 | 0.016398 | 0.065723 | 0.048635 | 0.072822 | 0.004355 | 0.024611 | 0.020916 | 0.011775 | 0.23671 | 0.020914 | 0.062907 | 0.015755 | 0.013478 | 0.070827 | 0.043172 | 0.04723 |
| 21.5277778 | 0.013346 | 0.071495 | 0.05011 | 0.023051 | 0.038589 | 0.025098 | 0.006548 | 0.007386 | 0.029163 | 0.084505 | 0.05925 | 0.092797 | 0.005613 | 0.03406 | 0.025744 | 0.012888 | 0.311308 | 0.028093 | 0.086021 | 0.01594 | 0.011393 | 0.060524 | 0.041821 | 0.043459 |
| 21.5416667 | 0.029411 | 0.066868 | 0.050607 | 0.016082 | 0.033485 | 0.016331 | 0.00719 | 0.006631 | 0.043018 | 0.084706 | 0.048034 | 0.081147 | 0.006074 | 0.035826 | 0.023912 | 0.011166 | 0.302354 | 0.023793 | 0.081633 | 0.012405 | 0.012236 | 0.032674 | 0.0264 | 0.026138 |
| 21.5555556 | 0.0434 | 0.0558 | 0.042068 | 0.00817 | 0.030223 | 0.006843 | 0.005996 | 0.005934 | 0.048925 | 0.061285 | 0.032408 | 0.054537 | 0.004596 | 0.02178 | 0.017527 | 0.008826 | 0.238145 | 0.015387 | 0.058484 | 0.010868 | 0.020008 | 0.013317 | 0.011864 | 0.014199 |
| 21.5694444 | 0.041849 | 0.047382 | 0.030595 | 0.004083 | 0.028548 | 0.003616 | 0.008457 | 0.005725 | 0.040915 | 0.033676 | 0.02154 | 0.030234 | 0.004292 | 0.009041 | 0.011588 | 0.008341 | 0.167114 | 0.009041 | 0.032975 | 0.011788 | 0.029395 | 0.006325 | 0.011084 | 0.015725 |
| 21.5833333 | 0.026945 | 0.038589 | 0.019303 | 0.001935 | 0.02414 | 0.007353 | 0.016888 | 0.005088 | 0.025963 | 0.032925 | 0.011153 | 0.017367 | 0.005612 | 0.012034 | 0.005925 | 0.00754 | 0.096355 | 0.005398 | 0.013744 | 0.010135 | 0.030343 | 0.003191 | 0.011532 | 0.020094 |
| 21.5972222 | 0.012153 | 0.028145 | 0.010026 | 0.000587 | 0.017811 | 0.013673 | 0.022327 | 0.003668 | 0.012755 | 0.060231 | 0.005199 | 0.024217 | 0.007111 | 0.022202 | 0.004053 | 0.004656 | 0.05424 | 0.008534 | 0.008689 | 0.006456 | 0.019711 | 0.003201 | 0.007349 | 0.025082 |
| 21.6111111 | 0.011328 | 0.015579 | 0.00462 | 0.000818 | 0.011378 | 0.016741 | 0.0181 | 0.002222 | 0.007766 | 0.065956 | 0.007721 | 0.033503 | 0.007045 | 0.025853 | 0.009191 | 0.001666 | 0.08563 | 0.015749 | 0.008154 | 0.00776 | 0.00834 | 0.006164 | 0.006822 | 0.031535 |
| 21.625 | 0.02082 | 0.008333 | 0.004479 | 0.00323 | 0.005807 | 0.013066 | 0.011905 | 0.002028 | 0.012556 | 0.042172 | 0.010347 | 0.025086 | 0.005173 | 0.017344 | 0.013078 | 0.000811 | 0.160826 | 0.01658 | 0.007759 | 0.013609 | 0.003989 | 0.011925 | 0.008351 | 0.033148 |
| 21.6388889 | 0.022591 | 0.01308 | 0.006824 | 0.006371 | 0.003881 | 0.007645 | 0.015616 | 0.002891 | 0.014835 | 0.038964 | 0.007862 | 0.012405 | 0.002896 | 0.009914 | 0.008959 | 0.002871 | 0.21481 | 0.009839 | 0.01669 | 0.013132 | 0.002686 | 0.018371 | 0.006396 | 0.023594 |
| 21.6527778 | 0.018446 | 0.021688 | 0.006501 | 0.007656 | 0.006614 | 0.009677 | 0.019916 | 0.002923 | 0.008986 | 0.053023 | 0.007891 | 0.008015 | 0.001474 | 0.014717 | 0.005445 | 0.005495 | 0.21145 | 0.005596 | 0.025312 | 0.009052 | 0.005815 | 0.021648 | 0.005317 | 0.013558 |
| 21.6666667 | 0.01566 | 0.025787 | 0.004292 | 0.006387 | 0.009776 | 0.01727 | 0.013237 | 0.001843 | 0.00342 | 0.045596 | 0.013978 | 0.007306 | 0.001283 | 0.017449 | 0.009823 | 0.004741 | 0.155711 | 0.007539 | 0.019979 | 0.014627 | 0.015003 | 0.02255 | 0.010041 | 0.019127 |
| 21.6805556 | 0.022791 | 0.024547 | 0.004703 | 0.003874 | 0.009681 | 0.021521 | 0.006976 | 0.000982 | 0.002451 | 0.023292 | 0.018198 | 0.014028 | 0.003512 | 0.010808 | 0.014071 | 0.002481 | 0.084569 | 0.009332 | 0.009774 | 0.025108 | 0.023681 | 0.025042 | 0.01588 | 0.029147 |
| 21.6944444 | 0.043721 | 0.019744 | 0.006411 | 0.001677 | 0.006165 | 0.018316 | 0.01334 | 0.000815 | 0.003679 | 0.017095 | 0.015856 | 0.025366 | 0.009974 | 0.00514 | 0.011449 | 0.002853 | 0.030989 | 0.006713 | 0.006793 | 0.026292 | 0.022437 | 0.029231 | 0.019651 | 0.026112 |
| 21.7083333 | 0.053206 | 0.013641 | 0.006345 | 0.000929 | 0.003055 | 0.010462 | 0.02325 | 0.001026 | 0.004542 | 0.034312 | 0.010382 | 0.029884 | 0.016498 | 0.006175 | 0.006353 | 0.004964 | 0.006523 | 0.003628 | 0.006302 | 0.01797 | 0.012018 | 0.029612 | 0.02253 | 0.015183 |
| 21.7222222 | 0.037024 | 0.007907 | 0.005411 | 0.001345 | 0.003732 | 0.004952 | 0.0228 | 0.001252 | 0.005692 | 0.052328 | 0.005273 | 0.021595 | 0.014321 | 0.009218 | 0.006478 | 0.005473 | 0.002607 | 0.005177 | 0.00702 | 0.009396 | 0.004552 | 0.021058 | 0.020518 | 0.007756 |
| 21.7361111 | 0.015035 | 0.004799 | 0.003825 | 0.001317 | 0.005871 | 0.006479 | 0.013399 | 0.001001 | 0.008556 | 0.046878 | 0.003459 | 0.012337 | 0.008186 | 0.007456 | 0.012821 | 0.004049 | 0.003316 | 0.009435 | 0.011354 | 0.004383 | 0.004887 | 0.011573 | 0.014533 | 0.005167 |
| 21.75 | 0.007716 | 0.004953 | 0.002474 | 0.001454 | 0.00689 | 0.012794 | 0.006391 | 0.000434 | 0.009374 | 0.027621 | 0.005695 | 0.019012 | 0.00881 | 0.006941 | 0.018909 | 0.003936 | 0.006249 | 0.01217 | 0.012134 | 0.003932 | 0.008687 | 0.012565 | 0.016934 | 0.007962 |
| 21.7638889 | 0.014309 | 0.007812 | 0.004224 | 0.002996 | 0.006393 | 0.017394 | 0.004083 | 0.000338 | 0.006425 | 0.025565 | 0.008918 | 0.029824 | 0.009858 | 0.01409 | 0.019164 | 0.006097 | 0.021383 | 0.010797 | 0.00781 | 0.008472 | 0.01196 | 0.017409 | 0.028338 | 0.018877 |
| 21.7777778 | 0.023792 | 0.013213 | 0.008765 | 0.004313 | 0.004035 | 0.015832 | 0.003536 | 0.000617 | 0.003246 | 0.035499 | 0.010143 | 0.026709 | 0.006476 | 0.018701 | 0.012051 | 0.007152 | 0.051112 | 0.00631 | 0.007207 | 0.0172 | 0.012513 | 0.015886 | 0.032532 | 0.035369 |
| 21.7916667 | 0.033029 | 0.019901 | 0.01333 | 0.004191 | 0.0022 | 0.010038 | 0.006829 | 0.000734 | 0.00259 | 0.029678 | 0.007684 | 0.016687 | 0.006912 | 0.013681 | 0.004917 | 0.005269 | 0.085597 | 0.003982 | 0.014071 | 0.028113 | 0.00874 | 0.008886 | 0.021328 | 0.052209 |
| 21.8055556 | 0.041428 | 0.027111 | 0.015704 | 0.00318 | 0.003721 | 0.005391 | 0.013732 | 0.001315 | 0.007885 | 0.017078 | 0.004859 | 0.018917 | 0.013724 | 0.007997 | 0.004855 | 0.004565 | 0.10416 | 0.004291 | 0.019504 | 0.035999 | 0.004243 | 0.003688 | 0.013302 | 0.063302 |
| 21.8194444 | 0.044589 | 0.03421 | 0.015434 | 0.001998 | 0.006916 | 0.003036 | 0.020339 | 0.003182 | 0.020153 | 0.024239 | 0.006259 | 0.030524 | 0.021337 | 0.012341 | 0.006518 | 0.007888 | 0.095278 | 0.003195 | 0.015219 | 0.036309 | 0.005057 | 0.004701 | 0.027958 | 0.062138 |
| 21.8333333 | 0.039565 | 0.036958 | 0.014352 | 0.001672 | 0.009792 | 0.002639 | 0.020909 | 0.006302 | 0.033789 | 0.047832 | 0.007499 | 0.0329 | 0.024178 | 0.026216 | 0.004699 | 0.011664 | 0.072736 | 0.002326 | 0.008426 | 0.027797 | 0.010448 | 0.009004 | 0.050107 | 0.047072 |
| 21.8472222 | 0.028009 | 0.033726 | 0.014374 | 0.002198 | 0.012394 | 0.003749 | 0.015917 | 0.010116 | 0.040362 | 0.063106 | 0.005571 | 0.02537 | 0.020844 | 0.040837 | 0.003534 | 0.012647 | 0.053076 | 0.005227 | 0.008659 | 0.014375 | 0.016739 | 0.014073 | 0.061151 | 0.026384 |
| 21.8611111 | 0.017101 | 0.031718 | 0.016328 | 0.002156 | 0.015019 | 0.004651 | 0.01429 | 0.01275 | 0.037238 | 0.057005 | 0.006741 | 0.020231 | 0.016288 | 0.045952 | 0.008828 | 0.011489 | 0.039432 | 0.011431 | 0.009441 | 0.004429 | 0.01617 | 0.019357 | 0.062545 | 0.01387 |
| 21.875 | 0.014723 | 0.036295 | 0.019987 | 0.001251 | 0.017738 | 0.008327 | 0.020196 | 0.012413 | 0.030769 | 0.03673 | 0.013872 | 0.022873 | 0.01552 | 0.040331 | 0.019015 | 0.009523 | 0.027604 | 0.019425 | 0.007392 | 0.001718 | 0.012934 | 0.023997 | 0.058265 | 0.016203 |
| 21.8888889 | 0.022084 | 0.043459 | 0.024404 | 0.000621 | 0.019987 | 0.014359 | 0.029706 | 0.009575 | 0.028379 | 0.018087 | 0.022082 | 0.029959 | 0.016369 | 0.033733 | 0.028529 | 0.007016 | 0.018902 | 0.023857 | 0.01059 | 0.003914 | 0.02286 | 0.026741 | 0.049967 | 0.02622 |
| 21.9027778 | 0.03267 | 0.047339 | 0.028239 | 0.001053 | 0.021098 | 0.016308 | 0.037053 | 0.00609 | 0.032114 | 0.010679 | 0.027872 | 0.03738 | 0.015304 | 0.031161 | 0.032708 | 0.005062 | 0.024514 | 0.019962 | 0.01751 | 0.009601 | 0.036211 | 0.026523 | 0.039123 | 0.032674 |
| 21.9166667 | 0.038403 | 0.047284 | 0.029512 | 0.002351 | 0.01948 | 0.013145 | 0.037455 | 0.004448 | 0.038207 | 0.010603 | 0.031447 | 0.043759 | 0.013797 | 0.029207 | 0.029444 | 0.00538 | 0.048601 | 0.012285 | 0.021929 | 0.015998 | 0.034548 | 0.023272 | 0.028174 | 0.034044 |
| 21.9305556 | 0.036284 | 0.043567 | 0.027044 | 0.003798 | 0.014567 | 0.008888 | 0.028605 | 0.006407 | 0.040616 | 0.019269 | 0.032113 | 0.04722 | 0.013178 | 0.024652 | 0.021298 | 0.007283 | 0.074838 | 0.006278 | 0.022416 | 0.019688 | 0.02006 | 0.017039 | 0.018336 | 0.034063 |
| 21.9444444 | 0.030528 | 0.035329 | 0.022134 | 0.004982 | 0.008803 | 0.005958 | 0.014921 | 0.010478 | 0.035788 | 0.042445 | 0.028477 | 0.043093 | 0.011814 | 0.018117 | 0.012957 | 0.008065 | 0.083696 | 0.002633 | 0.017721 | 0.020214 | 0.009067 | 0.009479 | 0.009521 | 0.033815 |
| 21.9583333 | 0.027186 | 0.025875 | 0.017427 | 0.005557 | 0.004868 | 0.004167 | 0.006455 | 0.01255 | 0.025635 | 0.067883 | 0.022021 | 0.03196 | 0.010761 | 0.013467 | 0.008224 | 0.006914 | 0.071647 | 0.001677 | 0.010091 | 0.0189 | 0.009144 | 0.004114 | 0.004757 | 0.032487 |
| 21.9722222 | 0.026862 | 0.019423 | 0.014056 | 0.004643 | 0.003588 | 0.003588 | 0.005041 | 0.010587 | 0.015788 | 0.075922 | 0.016062 | 0.021034 | 0.012932 | 0.013581 | 0.008624 | 0.005666 | 0.046691 | 0.00389 | 0.005954 | 0.017748 | 0.010413 | 0.001686 | 0.007312 | 0.029309 |
| 21.9861111 | 0.02517 | 0.016463 | 0.011996 | 0.002386 | 0.003617 | 0.004291 | 0.003789 | 0.006496 | 0.010034 | 0.060641 | 0.012425 | 0.01308 | 0.016237 | 0.016072 | 0.013367 | 0.005345 | 0.021918 | 0.008386 | 0.009675 | 0.016748 | 0.007935 | 0.001259 | 0.013475 | 0.025567 |
| 22 | 0.021756 | 0.015842 | 0.013103 | 0.001624 | 0.00357 | 0.005653 | 0.003035 | 0.00306 | 0.010266 | 0.036682 | 0.013694 | 0.008522 | 0.017862 | 0.018792 | 0.020887 | 0.005215 | 0.008585 | 0.014138 | 0.023475 | 0.014011 | 0.006436 | 0.001606 | 0.01659 | 0.024409 |
| 22.0138889 | 0.022445 | 0.016718 | 0.020095 | 0.006786 | 0.003166 | 0.008232 | 0.005972 | 0.001564 | 0.016236 | 0.025624 | 0.026073 | 0.01139 | 0.020561 | 0.02555 | 0.031203 | 0.004572 | 0.007454 | 0.021767 | 0.047492 | 0.010003 | 0.012986 | 0.003955 | 0.017087 | 0.027852 |
| 22.0277778 | 0.028963 | 0.018569 | 0.031297 | 0.016815 | 0.002055 | 0.012587 | 0.009933 | 0.002022 | 0.022308 | 0.036476 | 0.05031 | 0.024313 | 0.02674 | 0.041854 | 0.044661 | 0.003291 | 0.010256 | 0.032809 | 0.076048 | 0.008027 | 0.030032 | 0.012967 | 0.0203 | 0.033704 |
| 22.0416667 | 0.036297 | 0.023028 | 0.039511 | 0.021727 | 0.000733 | 0.015745 | 0.013426 | 0.005035 | 0.022142 | 0.052555 | 0.063988 | 0.040934 | 0.033196 | 0.064455 | 0.055671 | 0.002531 | 0.009664 | 0.045232 | 0.082368 | 0.011281 | 0.05591 | 0.025328 | 0.02413 | 0.037697 |
| 22.0555556 | 0.037924 | 0.02961 | 0.037195 | 0.014849 | 0.000633 | 0.014499 | 0.017246 | 0.008082 | 0.016843 | 0.047933 | 0.04435 | 0.049519 | 0.034904 | 0.073091 | 0.054829 | 0.00227 | 0.006401 | 0.04919 | 0.050852 | 0.02028 | 0.073769 | 0.033621 | 0.020536 | 0.035922 |
| 22.0694444 | 0.029618 | 0.030974 | 0.023208 | 0.007844 | 0.001636 | 0.010402 | 0.01947 | 0.007301 | 0.011903 | 0.02616 | 0.016494 | 0.043359 | 0.030069 | 0.048983 | 0.040429 | 0.001321 | 0.007661 | 0.036954 | 0.022465 | 0.029504 | 0.063974 | 0.034685 | 0.011258 | 0.027284 |
| 22.0833333 | 0.015331 | 0.023353 | 0.009816 | 0.011635 | 0.00207 | 0.006365 | 0.017163 | 0.008581 | 0.010077 | 0.018688 | 0.009646 | 0.025923 | 0.019597 | 0.025269 | 0.021122 | 0.001005 | 0.016043 | 0.017517 | 0.019588 | 0.031453 | 0.03919 | 0.027589 | 0.008721 | 0.016964 |
| 22.0972222 | 0.005582 | 0.011876 | 0.007125 | 0.015947 | 0.001443 | 0.003434 | 0.011572 | 0.015365 | 0.009697 | 0.038843 | 0.009673 | 0.015107 | 0.009814 | 0.037113 | 0.012308 | 0.00223 | 0.023942 | 0.007943 | 0.017005 | 0.025481 | 0.021638 | 0.015986 | 0.015137 | 0.01045 |
| 22.1111111 | 0.004217 | 0.004981 | 0.007495 | 0.013494 | 0.000874 | 0.001783 | 0.007771 | 0.019537 | 0.007715 | 0.065422 | 0.005667 | 0.018422 | 0.008367 | 0.049988 | 0.017993 | 0.003308 | 0.024696 | 0.010621 | 0.008849 | 0.016537 | 0.023895 | 0.006822 | 0.017964 | 0.008792 |
| 22.125 | 0.004489 | 0.004706 | 0.005091 | 0.00864 | 0.001107 | 0.001227 | 0.008933 | 0.017143 | 0.004743 | 0.077931 | 0.002446 | 0.019226 | 0.011521 | 0.039961 | 0.024578 | 0.002621 | 0.019116 | 0.014938 | 0.003622 | 0.008588 | 0.035655 | 0.003455 | 0.012932 | 0.010023 |
| 22.1388889 | 0.002824 | 0.004647 | 0.002645 | 0.00449 | 0.001401 | 0.001221 | 0.011084 | 0.010043 | 0.002918 | 0.072358 | 0.001109 | 0.011022 | 0.012665 | 0.024932 | 0.025825 | 0.001206 | 0.011967 | 0.016121 | 0.001732 | 0.003291 | 0.036342 | 0.002936 | 0.006509 | 0.009456 |
| 22.1527778 | 0.002337 | 0.003445 | 0.001453 | 0.002021 | 0.001053 | 0.00111 | 0.009152 | 0.004896 | 0.002458 | 0.054937 | 0.001008 | 0.004931 | 0.012878 | 0.015449 | 0.025294 | 0.000996 | 0.006905 | 0.015513 | 0.001129 | 0.00146 | 0.032177 | 0.002101 | 0.002466 | 0.006866 |
| 22.1666667 | 0.002417 | 0.004043 | 0.00101 | 0.000851 | 0.001081 | 0.000861 | 0.007366 | 0.007023 | 0.002791 | 0.033633 | 0.001294 | 0.005297 | 0.013566 | 0.011198 | 0.025098 | 0.001753 | 0.004259 | 0.014463 | 0.000636 | 0.002149 | 0.032704 | 0.001624 | 0.001782 | 0.007786 |
| 22.1805556 | 0.003358 | 0.005634 | 0.00061 | 0.000315 | 0.002158 | 0.001003 | 0.008335 | 0.009782 | 0.003344 | 0.017143 | 0.002303 | 0.006072 | 0.01244 | 0.008485 | 0.023901 | 0.001811 | 0.002997 | 0.014297 | 0.000819 | 0.002744 | 0.032378 | 0.001506 | 0.004996 | 0.009779 |
| 22.1944444 | 0.007735 | 0.006456 | 0.000279 | 0.000352 | 0.002767 | 0.001508 | 0.007152 | 0.007214 | 0.003658 | 0.016808 | 0.003845 | 0.00787 | 0.008436 | 0.005543 | 0.020671 | 0.001422 | 0.002252 | 0.015346 | 0.001963 | 0.002372 | 0.027192 | 0.001145 | 0.010164 | 0.007213 |
| 22.2083333 | 0.011767 | 0.005647 | 0.000149 | 0.000978 | 0.001921 | 0.001671 | 0.003626 | 0.002746 | 0.003586 | 0.027502 | 0.004639 | 0.011568 | 0.004513 | 0.002922 | 0.019281 | 0.001589 | 0.00164 | 0.017244 | 0.003169 | 0.001651 | 0.019339 | 0.001044 | 0.01166 | 0.004205 |
| 22.2222222 | 0.012025 | 0.003278 | 0.000192 | 0.001448 | 0.001031 | 0.00126 | 0.001993 | 0.001317 | 0.003623 | 0.029898 | 0.004245 | 0.009753 | 0.00264 | 0.003418 | 0.020494 | 0.001258 | 0.002011 | 0.019041 | 0.003376 | 0.000917 | 0.011058 | 0.000904 | 0.007146 | 0.003748 |
| 22.2361111 | 0.009018 | 0.001446 | 0.000294 | 0.00112 | 0.001009 | 0.000766 | 0.003404 | 0.003471 | 0.004145 | 0.019515 | 0.003333 | 0.006942 | 0.002142 | 0.009915 | 0.01979 | 0.000636 | 0.003796 | 0.019101 | 0.003351 | 0.000782 | 0.005591 | 0.000716 | 0.00224 | 0.004948 |
| 22.25 | 0.004868 | 0.001651 | 0.000232 | 0.000515 | 0.000853 | 0.000585 | 0.004486 | 0.006462 | 0.004573 | 0.010968 | 0.002717 | 0.011229 | 0.003534 | 0.017286 | 0.015455 | 0.00075 | 0.004424 | 0.016845 | 0.004452 | 0.001596 | 0.00603 | 0.000565 | 0.00118 | 0.006653 |
| 22.2638889 | 0.003368 | 0.002244 | 0.000207 | 0.000561 | 0.000582 | 0.000581 | 0.003729 | 0.006506 | 0.004138 | 0.008217 | 0.002579 | 0.014095 | 0.005467 | 0.016447 | 0.009732 | 0.000892 | 0.003383 | 0.012988 | 0.005838 | 0.00276 | 0.008092 | 0.000393 | 0.002339 | 0.005725 |
| 22.2777778 | 0.004394 | 0.002221 | 0.000704 | 0.001511 | 0.000942 | 0.000477 | 0.002422 | 0.004942 | 0.002844 | 0.007461 | 0.002476 | 0.009351 | 0.004778 | 0.008923 | 0.004735 | 0.000874 | 0.003782 | 0.008501 | 0.005916 | 0.003356 | 0.007099 | 0.000347 | 0.004125 | 0.004181 |
| 22.2916667 | 0.004491 | 0.002154 | 0.002064 | 0.002811 | 0.001275 | 0.00031 | 0.002221 | 0.006442 | 0.001383 | 0.011853 | 0.001757 | 0.005162 | 0.002256 | 0.003652 | 0.00224 | 0.001371 | 0.00486 | 0.004291 | 0.004872 | 0.002976 | 0.00404 | 0.000428 | 0.004312 | 0.00429 |
| 22.3055556 | 0.003619 | 0.001814 | 0.004214 | 0.003862 | 0.000966 | 0.000174 | 0.005168 | 0.007677 | 0.000972 | 0.013149 | 0.001611 | 0.006052 | 0.001519 | 0.003635 | 0.001957 | 0.002344 | 0.003708 | 0.001546 | 0.003121 | 0.00196 | 0.002096 | 0.00057 | 0.005439 | 0.007071 |
| 22.3194444 | 0.006149 | 0.001419 | 0.006498 | 0.00484 | 0.000757 | 0.000194 | 0.009671 | 0.005126 | 0.002518 | 0.011425 | 0.00397 | 0.006934 | 0.003293 | 0.006282 | 0.002128 | 0.00321 | 0.001624 | 0.000871 | 0.002695 | 0.001425 | 0.002778 | 0.000557 | 0.013844 | 0.01302 |
| 22.3333333 | 0.014186 | 0.002502 | 0.008586 | 0.006039 | 0.001414 | 0.000677 | 0.013117 | 0.00257 | 0.005752 | 0.022575 | 0.006922 | 0.008057 | 0.00551 | 0.006079 | 0.004073 | 0.003581 | 0.001754 | 0.001717 | 0.006607 | 0.002041 | 0.005528 | 0.000907 | 0.026677 | 0.016051 |
| 22.3472222 | 0.022876 | 0.005196 | 0.01046 | 0.007284 | 0.002364 | 0.00156 | 0.015018 | 0.003391 | 0.009879 | 0.039585 | 0.007458 | 0.013437 | 0.009187 | 0.005956 | 0.007676 | 0.004587 | 0.0046 | 0.003137 | 0.013188 | 0.002616 | 0.012316 | 0.001928 | 0.036462 | 0.015467 |
| 22.3611111 | 0.024959 | 0.006862 | 0.011595 | 0.008126 | 0.002552 | 0.002227 | 0.015001 | 0.008761 | 0.013623 | 0.040151 | 0.005695 | 0.021301 | 0.013754 | 0.011939 | 0.009538 | 0.006505 | 0.007018 | 0.003647 | 0.017961 | 0.002692 | 0.02112 | 0.002598 | 0.038108 | 0.016844 |
| 22.375 | 0.01933 | 0.006177 | 0.012287 | 0.008633 | 0.002293 | 0.002607 | 0.014013 | 0.015843 | 0.016026 | 0.024347 | 0.004116 | 0.027226 | 0.014716 | 0.016817 | 0.008348 | 0.007017 | 0.006436 | 0.003794 | 0.018492 | 0.004238 | 0.024603 | 0.00267 | 0.032154 | 0.019998 |
| 22.3888889 | 0.012638 | 0.004918 | 0.013729 | 0.009457 | 0.002972 | 0.003551 | 0.014441 | 0.017623 | 0.01738 | 0.010561 | 0.005168 | 0.027584 | 0.012835 | 0.016668 | 0.007086 | 0.005881 | 0.004381 | 0.005588 | 0.016785 | 0.008059 | 0.022126 | 0.003151 | 0.02637 | 0.021715 |
| 22.4027778 | 0.01067 | 0.004249 | 0.015345 | 0.010061 | 0.00418 | 0.005092 | 0.016384 | 0.011361 | 0.018167 | 0.012386 | 0.008657 | 0.023076 | 0.01133 | 0.016014 | 0.00951 | 0.005109 | 0.003122 | 0.009048 | 0.016251 | 0.012201 | 0.017926 | 0.004183 | 0.026477 | 0.021462 |
| 22.4166667 | 0.014309 | 0.004139 | 0.015679 | 0.00898 | 0.004098 | 0.006273 | 0.018204 | 0.0041 | 0.018999 | 0.032786 | 0.011856 | 0.018649 | 0.009732 | 0.017478 | 0.015627 | 0.005134 | 0.003023 | 0.013011 | 0.016652 | 0.014437 | 0.014398 | 0.005448 | 0.030201 | 0.021012 |
| 22.4305556 | 0.021905 | 0.004893 | 0.014704 | 0.006193 | 0.002586 | 0.006845 | 0.019543 | 0.003299 | 0.021127 | 0.053656 | 0.012388 | 0.018035 | 0.007115 | 0.022047 | 0.021348 | 0.005553 | 0.003716 | 0.016143 | 0.015523 | 0.014736 | 0.012911 | 0.006932 | 0.03205 | 0.021151 |
| 22.4444444 | 0.029707 | 0.0062 | 0.013791 | 0.003284 | 0.001067 | 0.00716 | 0.019724 | 0.00893 | 0.024632 | 0.051441 | 0.011044 | 0.020188 | 0.004654 | 0.027789 | 0.023138 | 0.006831 | 0.00577 | 0.017017 | 0.012416 | 0.014897 | 0.014742 | 0.009031 | 0.030529 | 0.021147 |
| 22.4583333 | 0.032826 | 0.006843 | 0.01453 | 0.002027 | 0.000668 | 0.007486 | 0.017677 | 0.016709 | 0.027597 | 0.028861 | 0.01039 | 0.02141 | 0.007044 | 0.03128 | 0.023671 | 0.009592 | 0.010208 | 0.016258 | 0.010019 | 0.016542 | 0.020882 | 0.012434 | 0.029346 | 0.021353 |
| 22.4722222 | 0.028766 | 0.006527 | 0.016591 | 0.002477 | 0.001202 | 0.007775 | 0.01477 | 0.018643 | 0.029679 | 0.013259 | 0.011852 | 0.019968 | 0.016147 | 0.030418 | 0.026449 | 0.012328 | 0.017261 | 0.015607 | 0.011045 | 0.019767 | 0.028362 | 0.0159 | 0.031261 | 0.023092 |
| 22.4861111 | 0.020755 | 0.007554 | 0.01931 | 0.004259 | 0.002034 | 0.008687 | 0.013104 | 0.012752 | 0.03747 | 0.012531 | 0.017062 | 0.018752 | 0.024371 | 0.02468 | 0.02955 | 0.012211 | 0.037041 | 0.015246 | 0.017966 | 0.022509 | 0.030297 | 0.018579 | 0.036707 | 0.028321 |
| 22.5 | 0.017323 | 0.013872 | 0.026519 | 0.010877 | 0.003663 | 0.012695 | 0.0139 | 0.007032 | 0.061975 | 0.025555 | 0.028864 | 0.024627 | 0.026501 | 0.019559 | 0.032994 | 0.010246 | 0.108539 | 0.015806 | 0.036756 | 0.023845 | 0.027427 | 0.026115 | 0.049388 | 0.045791 |
| 22.5138889 | 0.024794 | 0.028026 | 0.041306 | 0.023568 | 0.00579 | 0.019214 | 0.019146 | 0.009092 | 0.087738 | 0.071624 | 0.041345 | 0.040521 | 0.026321 | 0.024198 | 0.040985 | 0.009643 | 0.25113 | 0.018642 | 0.066267 | 0.024076 | 0.027483 | 0.042989 | 0.068858 | 0.081273 |
| 22.5277778 | 0.037621 | 0.04187 | 0.052625 | 0.032345 | 0.005455 | 0.022023 | 0.030875 | 0.017448 | 0.081234 | 0.119059 | 0.040993 | 0.058068 | 0.023984 | 0.044441 | 0.049092 | 0.011001 | 0.359687 | 0.020475 | 0.081621 | 0.022266 | 0.032994 | 0.060814 | 0.080369 | 0.105887 |
| 22.5416667 | 0.045411 | 0.042978 | 0.051061 | 0.029573 | 0.002494 | 0.020233 | 0.037893 | 0.020206 | 0.052883 | 0.11608 | 0.029142 | 0.063438 | 0.017249 | 0.072108 | 0.047307 | 0.012692 | 0.332304 | 0.01886 | 0.061459 | 0.018887 | 0.036736 | 0.066997 | 0.071359 | 0.090945 |
| 22.5555556 | 0.042018 | 0.03476 | 0.046213 | 0.021399 | 0.00185 | 0.018928 | 0.025779 | 0.013577 | 0.033236 | 0.076412 | 0.018003 | 0.049971 | 0.010192 | 0.082163 | 0.036846 | 0.012851 | 0.237508 | 0.017809 | 0.027705 | 0.016142 | 0.030382 | 0.061839 | 0.048246 | 0.059261 |
| 22.5694444 | 0.026286 | 0.028618 | 0.042697 | 0.013443 | 0.004688 | 0.01662 | 0.014563 | 0.007072 | 0.020833 | 0.035836 | 0.01092 | 0.026036 | 0.006954 | 0.063903 | 0.024167 | 0.010251 | 0.160628 | 0.016474 | 0.01994 | 0.013939 | 0.018058 | 0.050118 | 0.023613 | 0.035461 |
| 22.5833333 | 0.01414 | 0.027222 | 0.035134 | 0.006949 | 0.006473 | 0.010819 | 0.024636 | 0.003847 | 0.009362 | 0.01937 | 0.006312 | 0.012675 | 0.011463 | 0.036103 | 0.014328 | 0.006547 | 0.10085 | 0.010742 | 0.042339 | 0.009127 | 0.010698 | 0.03258 | 0.011346 | 0.017126 |
| 22.5972222 | 0.021683 | 0.027106 | 0.024349 | 0.004888 | 0.005931 | 0.005318 | 0.038495 | 0.00427 | 0.005157 | 0.037393 | 0.004266 | 0.016098 | 0.021105 | 0.027567 | 0.016659 | 0.006167 | 0.054904 | 0.006269 | 0.05381 | 0.004179 | 0.01639 | 0.016871 | 0.016386 | 0.006877 |
| 22.6111111 | 0.036312 | 0.028276 | 0.013672 | 0.005581 | 0.00604 | 0.003633 | 0.038932 | 0.011647 | 0.009502 | 0.06426 | 0.003703 | 0.015437 | 0.023975 | 0.03814 | 0.029912 | 0.007013 | 0.047722 | 0.009133 | 0.038364 | 0.005324 | 0.035178 | 0.00983 | 0.021886 | 0.006208 |
| 22.625 | 0.0392 | 0.032261 | 0.005845 | 0.004453 | 0.00761 | 0.004399 | 0.024629 | 0.016057 | 0.014105 | 0.056527 | 0.002854 | 0.011482 | 0.017471 | 0.035162 | 0.035623 | 0.005516 | 0.076173 | 0.01139 | 0.040565 | 0.008128 | 0.050684 | 0.010988 | 0.018521 | 0.005692 |
| 22.6388889 | 0.025956 | 0.037873 | 0.004004 | 0.004079 | 0.008727 | 0.003994 | 0.015359 | 0.011974 | 0.011381 | 0.037369 | 0.002169 | 0.02268 | 0.008832 | 0.021187 | 0.024807 | 0.006691 | 0.108447 | 0.008844 | 0.069503 | 0.008338 | 0.046173 | 0.016295 | 0.012704 | 0.004625 |
| 22.6527778 | 0.01546 | 0.042903 | 0.007612 | 0.007224 | 0.008285 | 0.002037 | 0.031583 | 0.014059 | 0.004988 | 0.057359 | 0.002338 | 0.038419 | 0.006989 | 0.024707 | 0.016875 | 0.01241 | 0.140546 | 0.010722 | 0.081658 | 0.011881 | 0.027684 | 0.020352 | 0.007086 | 0.00589 |
| 22.6666667 | 0.027978 | 0.044713 | 0.012559 | 0.010255 | 0.006297 | 0.000785 | 0.044637 | 0.020156 | 0.002771 | 0.077215 | 0.003704 | 0.04134 | 0.013852 | 0.037706 | 0.028476 | 0.016143 | 0.174577 | 0.018339 | 0.060843 | 0.015718 | 0.022799 | 0.020007 | 0.004024 | 0.005385 |
| 22.6805556 | 0.045099 | 0.042227 | 0.014673 | 0.011368 | 0.003325 | 0.000914 | 0.028255 | 0.015942 | 0.004418 | 0.058841 | 0.006741 | 0.027872 | 0.015645 | 0.039087 | 0.0413 | 0.013377 | 0.189258 | 0.024257 | 0.034344 | 0.012152 | 0.04291 | 0.013735 | 0.005915 | 0.004292 |
| 22.6944444 | 0.047451 | 0.036026 | 0.010762 | 0.010855 | 0.001144 | 0.00135 | 0.012967 | 0.006801 | 0.005011 | 0.027862 | 0.010332 | 0.013153 | 0.011342 | 0.025217 | 0.037013 | 0.006847 | 0.163486 | 0.024003 | 0.034221 | 0.006365 | 0.056207 | 0.009857 | 0.007378 | 0.005336 |
| 22.7083333 | 0.040559 | 0.02652 | 0.00425 | 0.007755 | 0.000542 | 0.001389 | 0.025119 | 0.002251 | 0.005688 | 0.012679 | 0.012413 | 0.014187 | 0.016458 | 0.011452 | 0.021927 | 0.003725 | 0.108849 | 0.016573 | 0.045235 | 0.00457 | 0.0436 | 0.013239 | 0.005947 | 0.007391 |
| 22.7222222 | 0.027688 | 0.014658 | 0.001991 | 0.00432 | 0.000855 | 0.002154 | 0.048183 | 0.003187 | 0.010915 | 0.021072 | 0.011672 | 0.022284 | 0.02604 | 0.013982 | 0.01083 | 0.00692 | 0.057888 | 0.008482 | 0.033696 | 0.004524 | 0.02356 | 0.011906 | 0.006491 | 0.007081 |
| 22.7361111 | 0.016455 | 0.006337 | 0.003339 | 0.00463 | 0.001897 | 0.004421 | 0.054628 | 0.007793 | 0.016736 | 0.042934 | 0.008178 | 0.029851 | 0.026389 | 0.025715 | 0.014797 | 0.011112 | 0.028691 | 0.008549 | 0.014801 | 0.003764 | 0.010488 | 0.005201 | 0.01077 | 0.00591 |
| 22.75 | 0.021796 | 0.006596 | 0.003882 | 0.007085 | 0.002677 | 0.006129 | 0.034745 | 0.015078 | 0.017356 | 0.066524 | 0.0039 | 0.03488 | 0.018269 | 0.035559 | 0.029573 | 0.010695 | 0.024218 | 0.014595 | 0.010745 | 0.003516 | 0.012262 | 0.00185 | 0.015734 | 0.011351 |
| 22.7638889 | 0.037456 | 0.011948 | 0.002653 | 0.006517 | 0.002096 | 0.005752 | 0.017134 | 0.018899 | 0.013687 | 0.073838 | 0.002269 | 0.027028 | 0.008665 | 0.038002 | 0.04205 | 0.006015 | 0.041187 | 0.01901 | 0.010087 | 0.004705 | 0.032382 | 0.001072 | 0.015572 | 0.0193 |
| 22.7777778 | 0.045178 | 0.01712 | 0.002209 | 0.003844 | 0.001514 | 0.003733 | 0.0231 | 0.013091 | 0.009089 | 0.050681 | 0.004186 | 0.016372 | 0.004447 | 0.028205 | 0.04554 | 0.002322 | 0.066062 | 0.019912 | 0.012758 | 0.00538 | 0.053715 | 0.002318 | 0.012698 | 0.019762 |
| 22.7916667 | 0.041598 | 0.019597 | 0.003953 | 0.004223 | 0.002131 | 0.00269 | 0.024958 | 0.006829 | 0.005697 | 0.026836 | 0.006888 | 0.022756 | 0.0064 | 0.014388 | 0.040599 | 0.001845 | 0.079432 | 0.018666 | 0.025058 | 0.005457 | 0.056509 | 0.006493 | 0.02015 | 0.01459 |
| 22.8055556 | 0.028504 | 0.019308 | 0.006411 | 0.006757 | 0.002059 | 0.004541 | 0.015594 | 0.010793 | 0.003836 | 0.038643 | 0.007804 | 0.030554 | 0.00809 | 0.01054 | 0.030352 | 0.001999 | 0.075933 | 0.016534 | 0.033734 | 0.008955 | 0.039202 | 0.009748 | 0.036876 | 0.017439 |
| 22.8194444 | 0.012949 | 0.017921 | 0.009506 | 0.006852 | 0.001325 | 0.00702 | 0.020772 | 0.016956 | 0.003441 | 0.0571 | 0.006633 | 0.020126 | 0.00626 | 0.013123 | 0.017105 | 0.003376 | 0.06994 | 0.014228 | 0.028876 | 0.017593 | 0.021614 | 0.008003 | 0.050632 | 0.030167 |
| 22.8333333 | 0.00504 | 0.01768 | 0.013782 | 0.004787 | 0.001411 | 0.008836 | 0.042241 | 0.013555 | 0.004108 | 0.042664 | 0.00512 | 0.009958 | 0.004913 | 0.010253 | 0.008002 | 0.008037 | 0.075674 | 0.010421 | 0.015945 | 0.024837 | 0.024828 | 0.003467 | 0.060619 | 0.038385 |
| 22.8472222 | 0.004693 | 0.020635 | 0.017978 | 0.005276 | 0.001265 | 0.010009 | 0.05351 | 0.010422 | 0.007924 | 0.021207 | 0.005526 | 0.017652 | 0.009023 | 0.00741 | 0.008443 | 0.014485 | 0.094373 | 0.005214 | 0.008555 | 0.020892 | 0.036844 | 0.000759 | 0.064785 | 0.036838 |
| 22.8611111 | 0.010769 | 0.026732 | 0.021801 | 0.008623 | 0.001729 | 0.009728 | 0.039724 | 0.018621 | 0.017218 | 0.031852 | 0.009154 | 0.031623 | 0.016407 | 0.014269 | 0.011079 | 0.020582 | 0.121511 | 0.002344 | 0.009325 | 0.010945 | 0.035833 | 0.001124 | 0.053232 | 0.030186 |
| 22.875 | 0.022181 | 0.03355 | 0.026101 | 0.009603 | 0.004217 | 0.008601 | 0.017848 | 0.025645 | 0.029265 | 0.069445 | 0.014274 | 0.03779 | 0.022897 | 0.026875 | 0.009798 | 0.025389 | 0.149172 | 0.00296 | 0.00912 | 0.006607 | 0.024113 | 0.005349 | 0.028488 | 0.02822 |
| 22.8888889 | 0.039441 | 0.037381 | 0.030096 | 0.006464 | 0.006386 | 0.007897 | 0.005547 | 0.020609 | 0.03821 | 0.103013 | 0.016112 | 0.034697 | 0.027691 | 0.038827 | 0.007181 | 0.02697 | 0.15463 | 0.003199 | 0.01197 | 0.00563 | 0.024684 | 0.013669 | 0.01017 | 0.037717 |
| 22.9027778 | 0.052014 | 0.035922 | 0.032798 | 0.003039 | 0.006083 | 0.007686 | 0.003738 | 0.012443 | 0.03842 | 0.10386 | 0.013189 | 0.027272 | 0.027138 | 0.045241 | 0.009556 | 0.023218 | 0.122091 | 0.002958 | 0.023829 | 0.006295 | 0.045127 | 0.021019 | 0.007351 | 0.050915 |
| 22.9166667 | 0.044113 | 0.029605 | 0.034541 | 0.002409 | 0.004333 | 0.007732 | 0.007906 | 0.007484 | 0.032148 | 0.074376 | 0.009379 | 0.019912 | 0.018955 | 0.043265 | 0.019849 | 0.01619 | 0.070716 | 0.005626 | 0.037854 | 0.011316 | 0.063517 | 0.022297 | 0.008017 | 0.059145 |
| 22.9305556 | 0.02534 | 0.018394 | 0.034581 | 0.002633 | 0.002772 | 0.007966 | 0.016365 | 0.005256 | 0.02701 | 0.036497 | 0.007747 | 0.014281 | 0.008721 | 0.036199 | 0.032494 | 0.010836 | 0.029454 | 0.009328 | 0.047045 | 0.017093 | 0.066858 | 0.018842 | 0.01127 | 0.063118 |
| 22.9444444 | 0.011001 | 0.008207 | 0.031128 | 0.001728 | 0.002002 | 0.007922 | 0.024941 | 0.007067 | 0.025223 | 0.012826 | 0.008212 | 0.012149 | 0.004974 | 0.028069 | 0.038541 | 0.008229 | 0.011172 | 0.009998 | 0.044954 | 0.020675 | 0.05533 | 0.013328 | 0.023229 | 0.06205 |

| | | | | | | | | | | | | | | | | | | | | | | | | |
|---|---|---|---|---|---|---|---|---|---|---|---|---|---|---|---|---|---|---|---|---|---|---|---|---|
| 22.9583333 | 0.004484 | 0.006621 | 0.024741 | 0.001106 | 0.00197 | 0.007517 | 0.025835 | 0.008242 | 0.024107 | 0.01066 | 0.010254 | 0.013876 | 0.010361 | 0.020155 | 0.032998 | 0.005801 | 0.008779 | 0.007242 | 0.029511 | 0.022631 | 0.033503 | 0.007105 | 0.034729 | 0.050626 |
| 22.9722222 | 0.003141 | 0.007908 | 0.01765 | 0.001684 | 0.00229 | 0.007178 | 0.01663 | 0.006594 | 0.021316 | 0.013128 | 0.012739 | 0.016276 | 0.017724 | 0.012959 | 0.020885 | 0.003214 | 0.008477 | 0.003666 | 0.012664 | 0.022446 | 0.014612 | 0.002632 | 0.036696 | 0.029407 |
| 22.9861111 | 0.005184 | 0.005627 | 0.012294 | 0.00299 | 0.002725 | 0.006918 | 0.008627 | 0.004688 | 0.018404 | 0.022139 | 0.014137 | 0.016761 | 0.017484 | 0.00812 | 0.012448 | 0.00162 | 0.007318 | 0.001498 | 0.006225 | 0.019406 | 0.005997 | 0.00463 | 0.029847 | 0.00996 |
| 23 | 0.012288 | 0.004576 | 0.011762 | 0.005929 | 0.003474 | 0.006638 | 0.009523 | 0.00541 | 0.018607 | 0.046169 | 0.016126 | 0.016267 | 0.010944 | 0.00932 | 0.011071 | 0.002207 | 0.006424 | 0.002834 | 0.013185 | 0.01594 | 0.00629 | 0.01551 | 0.020984 | 0.005474 |
| 23.0138889 | 0.023295 | 0.009687 | 0.016832 | 0.009385 | 0.004435 | 0.007114 | 0.009408 | 0.009207 | 0.022652 | 0.073286 | 0.024473 | 0.020254 | 0.010356 | 0.018745 | 0.012547 | 0.005479 | 0.008471 | 0.009177 | 0.035648 | 0.01491 | 0.019861 | 0.029257 | 0.018436 | 0.017263 |
| 23.0277778 | 0.036554 | 0.01581 | 0.026303 | 0.00965 | 0.00484 | 0.009597 | 0.010575 | 0.013482 | 0.027014 | 0.099441 | 0.041528 | 0.03497 | 0.019536 | 0.03461 | 0.015842 | 0.007599 | 0.014587 | 0.018567 | 0.064113 | 0.015845 | 0.049695 | 0.039272 | 0.025619 | 0.037614 |
| 23.0416667 | 0.050477 | 0.018505 | 0.038074 | 0.007931 | 0.004627 | 0.012652 | 0.023302 | 0.023681 | 0.025997 | 0.12538 | 0.056949 | 0.057189 | 0.029504 | 0.053401 | 0.025768 | 0.005448 | 0.019174 | 0.029094 | 0.078011 | 0.020409 | 0.081427 | 0.046777 | 0.036567 | 0.060687 |
| 23.0555556 | 0.056895 | 0.021447 | 0.042799 | 0.007223 | 0.004264 | 0.012405 | 0.037991 | 0.047008 | 0.01964 | 0.131461 | 0.056757 | 0.07146 | 0.034087 | 0.062671 | 0.040244 | 0.002315 | 0.01769 | 0.037559 | 0.068576 | 0.03187 | 0.10055 | 0.056729 | 0.041385 | 0.072899 |
| 23.0694444 | 0.047071 | 0.024271 | 0.031708 | 0.005644 | 0.0032 | 0.008939 | 0.038734 | 0.069334 | 0.013651 | 0.099629 | 0.041124 | 0.064392 | 0.03256 | 0.046243 | 0.045031 | 0.001615 | 0.012372 | 0.035841 | 0.04847 | 0.043904 | 0.099093 | 0.0637 | 0.03701 | 0.052636 |
| 23.0833333 | 0.025097 | 0.021898 | 0.016186 | 0.002981 | 0.001811 | 0.006047 | 0.023588 | 0.062569 | 0.011053 | 0.048678 | 0.021395 | 0.040266 | 0.025624 | 0.023417 | 0.031952 | 0.001971 | 0.012451 | 0.021637 | 0.029475 | 0.045353 | 0.070868 | 0.055635 | 0.02468 | 0.030688 |
| 23.0972222 | 0.01008 | 0.01478 | 0.01329 | 0.002129 | 0.002371 | 0.004766 | 0.01079 | 0.034128 | 0.010897 | 0.023083 | 0.012117 | 0.018529 | 0.015992 | 0.023939 | 0.015614 | 0.001766 | 0.018989 | 0.00908 | 0.016415 | 0.036128 | 0.034066 | 0.036449 | 0.013891 | 0.044756 |
| 23.1111111 | 0.011162 | 0.007657 | 0.018365 | 0.002283 | 0.004798 | 0.003731 | 0.012963 | 0.022267 | 0.012198 | 0.036613 | 0.016775 | 0.013984 | 0.007818 | 0.034387 | 0.014064 | 0.002151 | 0.020626 | 0.009372 | 0.021232 | 0.02485 | 0.02026 | 0.022652 | 0.019017 | 0.054189 |
| 23.125 | 0.013999 | 0.003326 | 0.019025 | 0.002022 | 0.005695 | 0.002705 | 0.016481 | 0.032339 | 0.013621 | 0.057784 | 0.021514 | 0.020023 | 0.004536 | 0.037409 | 0.021723 | 0.003382 | 0.015025 | 0.013764 | 0.039151 | 0.016931 | 0.030496 | 0.021558 | 0.032888 | 0.035167 |
| 23.1388889 | 0.01194 | 0.001809 | 0.014842 | 0.00165 | 0.004011 | 0.002009 | 0.013602 | 0.041603 | 0.012466 | 0.067739 | 0.019846 | 0.022828 | 0.003959 | 0.035415 | 0.023457 | 0.002965 | 0.00976 | 0.014084 | 0.048409 | 0.012643 | 0.038471 | 0.026142 | 0.036956 | 0.014617 |
| 23.1527778 | 0.010255 | 0.001458 | 0.009535 | 0.00104 | 0.003261 | 0.001596 | 0.011457 | 0.04276 | 0.00846 | 0.070578 | 0.016482 | 0.018745 | 0.002335 | 0.031328 | 0.018546 | 0.001055 | 0.007807 | 0.012471 | 0.04458 | 0.010658 | 0.034501 | 0.027397 | 0.029185 | 0.005153 |
| 23.1666667 | 0.009724 | 0.001983 | 0.004559 | 0.000437 | 0.00403 | 0.001472 | 0.013694 | 0.039398 | 0.00467 | 0.067845 | 0.0144 | 0.011422 | 0.002005 | 0.02796 | 0.014358 | 0.000578 | 0.007465 | 0.010747 | 0.037147 | 0.008942 | 0.028046 | 0.022924 | 0.017383 | 0.004467 |
| 23.1805556 | 0.009844 | 0.003995 | 0.001317 | 0.000191 | 0.003571 | 0.001443 | 0.017918 | 0.034759 | 0.003056 | 0.062001 | 0.013309 | 0.005803 | 0.004346 | 0.026211 | 0.014083 | 0.001638 | 0.007552 | 0.009293 | 0.030943 | 0.006405 | 0.025453 | 0.015729 | 0.00888 | 0.00783 |
| 23.1944444 | 0.011138 | 0.005727 | 0.000309 | 0.000198 | 0.004214 | 0.001115 | 0.01821 | 0.031098 | 0.002585 | 0.055843 | 0.011749 | 0.003235 | 0.006214 | 0.024596 | 0.015751 | 0.002493 | 0.008208 | 0.009167 | 0.026271 | 0.004098 | 0.02693 | 0.009887 | 0.010042 | 0.009641 |
| 23.2083333 | 0.013853 | 0.005592 | 0.000324 | 0.000365 | 0.006773 | 0.000625 | 0.011606 | 0.028117 | 0.001762 | 0.047595 | 0.009215 | 0.001729 | 0.005949 | 0.021421 | 0.016304 | 0.002038 | 0.009172 | 0.011217 | 0.022274 | 0.002943 | 0.028027 | 0.00624 | 0.016187 | 0.006821 |
| 23.2222222 | 0.016048 | 0.004275 | 0.000288 | 0.000562 | 0.006328 | 0.000422 | 0.006799 | 0.025194 | 0.000805 | 0.035521 | 0.006424 | 0.000505 | 0.006688 | 0.01641 | 0.01731 | 0.001489 | 0.009139 | 0.014175 | 0.020058 | 0.002192 | 0.026335 | 0.003852 | 0.015098 | 0.003407 |
| 23.2361111 | 0.015616 | 0.002875 | 0.000202 | 0.00052 | 0.003255 | 0.000574 | 0.013221 | 0.02165 | 0.001133 | 0.02118 | 0.004044 | 0.000402 | 0.011411 | 0.010866 | 0.02213 | 0.002394 | 0.007129 | 0.015823 | 0.02108 | 0.001388 | 0.02313 | 0.003345 | 0.008637 | 0.00258 |
| 23.25 | 0.01273 | 0.001965 | 0.000737 | 0.000636 | 0.001679 | 0.00101 | 0.021892 | 0.01649 | 0.002576 | 0.009778 | 0.00196 | 0.001588 | 0.015207 | 0.005587 | 0.027503 | 0.002627 | 0.004211 | 0.0162 | 0.020732 | 0.0012 | 0.017035 | 0.003182 | 0.005587 | 0.002735 |
| 23.2638889 | 0.008677 | 0.001523 | 0.002154 | 0.001649 | 0.001249 | 0.00144 | 0.020804 | 0.00923 | 0.002557 | 0.007379 | 0.000839 | 0.00405 | 0.013056 | 0.002184 | 0.027789 | 0.001164 | 0.001934 | 0.016179 | 0.01391 | 0.001208 | 0.010164 | 0.00222 | 0.005117 | 0.002893 |
| 23.2777778 | 0.004722 | 0.001359 | 0.003299 | 0.002881 | 0.000754 | 0.001258 | 0.011596 | 0.003558 | 0.001049 | 0.011618 | 0.0011 | 0.008513 | 0.008845 | 0.001599 | 0.023611 | 0.000652 | 0.00071 | 0.015058 | 0.007171 | 0.000929 | 0.010485 | 0.002377 | 0.007075 | 0.004477 |
| 23.2916667 | 0.002095 | 0.001288 | 0.002769 | 0.003006 | 0.000513 | 0.001013 | 0.004314 | 0.002944 | 0.000337 | 0.015352 | 0.001822 | 0.014701 | 0.006985 | 0.002441 | 0.018102 | 0.0014 | 0.000532 | 0.012171 | 0.008862 | 0.000728 | 0.015957 | 0.002484 | 0.010459 | 0.008819 |
| 23.3055556 | 0.00139 | 0.001151 | 0.001449 | 0.001868 | 0.000433 | 0.001387 | 0.003419 | 0.004863 | 0.000805 | 0.014102 | 0.00202 | 0.018863 | 0.006819 | 0.003263 | 0.013082 | 0.00164 | 0.001123 | 0.00847 | 0.013382 | 0.000893 | 0.018692 | 0.001648 | 0.008507 | 0.009717 |
| 23.3194444 | 0.002249 | 0.001068 | 0.001127 | 0.000825 | 0.000471 | 0.001421 | 0.002965 | 0.00566 | 0.002276 | 0.009465 | 0.001649 | 0.018222 | 0.006479 | 0.00303 | 0.009548 | 0.00142 | 0.002046 | 0.005293 | 0.012323 | 0.001522 | 0.016404 | 0.002041 | 0.006527 | 0.007272 |
| 23.3333333 | 0.00421 | 0.001014 | 0.001631 | 0.000771 | 0.000752 | 0.000764 | 0.002021 | 0.005057 | 0.004418 | 0.009397 | 0.000951 | 0.015353 | 0.004909 | 0.00282 | 0.006044 | 0.001397 | 0.003107 | 0.003408 | 0.007092 | 0.001949 | 0.010642 | 0.004209 | 0.013555 | 0.010355 |
| 23.3472222 | 0.007248 | 0.000874 | 0.002533 | 0.000837 | 0.001437 | 0.000254 | 0.004398 | 0.007214 | 0.006198 | 0.014688 | 0.000558 | 0.013718 | 0.003005 | 0.003642 | 0.0033 | 0.001412 | 0.003647 | 0.002794 | 0.005744 | 0.002069 | 0.006413 | 0.006222 | 0.023325 | 0.015686 |
| 23.3611111 | 0.011052 | 0.00093 | 0.003998 | 0.000941 | 0.001599 | 0.000294 | 0.009333 | 0.012773 | 0.006911 | 0.018593 | 0.001098 | 0.014816 | 0.003917 | 0.006672 | 0.007672 | 0.001958 | 0.002788 | 0.002667 | 0.012285 | 0.00352 | 0.009225 | 0.006082 | 0.029022 | 0.015433 |
| 23.375 | 0.014842 | 0.001561 | 0.005598 | 0.001926 | 0.001729 | 0.000756 | 0.014365 | 0.015262 | 0.006982 | 0.020637 | 0.001825 | 0.018612 | 0.008609 | 0.011391 | 0.018911 | 0.003785 | 0.001808 | 0.002864 | 0.022045 | 0.006517 | 0.016476 | 0.00396 | 0.028684 | 0.010385 |
| 23.3888889 | 0.018318 | 0.002519 | 0.00706 | 0.003535 | 0.003924 | 0.002567 | 0.017638 | 0.012531 | 0.00731 | 0.025139 | 0.002116 | 0.023198 | 0.011514 | 0.012116 | 0.0256 | 0.005348 | 0.00159 | 0.004296 | 0.027716 | 0.00828 | 0.018483 | 0.00231 | 0.02097 | 0.005235 |
| 23.4027778 | 0.021235 | 0.003146 | 0.008354 | 0.004923 | 0.006147 | 0.00556 | 0.017462 | 0.00826 | 0.007769 | 0.031452 | 0.002675 | 0.025551 | 0.00826 | 0.00778 | 0.021513 | 0.005249 | 0.002056 | 0.007119 | 0.024273 | 0.007114 | 0.01309 | 0.00293 | 0.010378 | 0.002972 |
| 23.4166667 | 0.022355 | 0.003062 | 0.009094 | 0.005518 | 0.005124 | 0.005469 | 0.013972 | 0.00513 | 0.007746 | 0.033992 | 0.004129 | 0.023559 | 0.003286 | 0.004366 | 0.013628 | 0.004566 | 0.004057 | 0.009891 | 0.01422 | 0.00475 | 0.007695 | 0.005009 | 0.004549 | 0.004174 |
| 23.4305556 | 0.021761 | 0.002213 | 0.009736 | 0.005026 | 0.002994 | 0.003017 | 0.010616 | 0.003636 | 0.007857 | 0.032298 | 0.00633 | 0.01831 | 0.002451 | 0.005995 | 0.011173 | 0.004931 | 0.005586 | 0.010999 | 0.007936 | 0.004094 | 0.008701 | 0.006071 | 0.008108 | 0.00694 |
| 23.4444444 | 0.021902 | 0.001078 | 0.011566 | 0.00405 | 0.003189 | 0.003523 | 0.010662 | 0.00302 | 0.0098 | 0.032547 | 0.008313 | 0.013565 | 0.006673 | 0.012274 | 0.015236 | 0.00688 | 0.004645 | 0.010612 | 0.010992 | 0.006447 | 0.015307 | 0.004432 | 0.019159 | 0.008318 |
| 23.4583333 | 0.0239 | 0.000381 | 0.013899 | 0.003867 | 0.003505 | 0.005714 | 0.013121 | 0.004194 | 0.013803 | 0.037624 | 0.008756 | 0.011339 | 0.011977 | 0.020272 | 0.019126 | 0.008988 | 0.004646 | 0.010703 | 0.016679 | 0.010119 | 0.019735 | 0.002435 | 0.029669 | 0.007513 |
| 23.4722222 | 0.026152 | 0.001076 | 0.01399 | 0.004596 | 0.003199 | 0.006716 | 0.013957 | 0.007684 | 0.017944 | 0.043447 | 0.008146 | 0.011163 | 0.013618 | 0.026377 | 0.018407 | 0.009889 | 0.008813 | 0.012313 | 0.018386 | 0.013452 | 0.016998 | 0.003844 | 0.032477 | 0.006369 |
| 23.4861111 | 0.028615 | 0.005131 | 0.012247 | 0.005785 | 0.005743 | 0.007331 | 0.012978 | 0.010536 | 0.025458 | 0.046637 | 0.010274 | 0.01586 | 0.014713 | 0.029549 | 0.019315 | 0.009979 | 0.017601 | 0.014877 | 0.022159 | 0.019438 | 0.012957 | 0.008328 | 0.029594 | 0.010158 |
| 23.5 | 0.036722 | 0.0152 | 0.015183 | 0.009446 | 0.009202 | 0.010716 | 0.016227 | 0.013738 | 0.046193 | 0.055848 | 0.019937 | 0.03156 | 0.023253 | 0.034688 | 0.029015 | 0.009955 | 0.041557 | 0.020388 | 0.039715 | 0.036878 | 0.015844 | 0.015129 | 0.033551 | 0.029232 |
| 23.5138889 | 0.049458 | 0.030583 | 0.025394 | 0.015834 | 0.009575 | 0.017172 | 0.025709 | 0.021301 | 0.073154 | 0.087491 | 0.03356 | 0.054046 | 0.036905 | 0.048024 | 0.041153 | 0.009273 | 0.106022 | 0.027991 | 0.071524 | 0.060683 | 0.025946 | 0.025963 | 0.055103 | 0.064771 |
| 23.5277778 | 0.057255 | 0.04054 | 0.032536 | 0.019501 | 0.007026 | 0.020985 | 0.037088 | 0.029866 | 0.082343 | 0.136952 | 0.039197 | 0.065282 | 0.041779 | 0.068381 | 0.044048 | 0.006938 | 0.19828 | 0.030109 | 0.100641 | 0.065964 | 0.037085 | 0.03797 | 0.083788 | 0.091601 |
| 23.5416667 | 0.05861 | 0.035435 | 0.029116 | 0.018463 | 0.003409 | 0.020049 | 0.039618 | 0.032954 | 0.07331 | 0.159745 | 0.034217 | 0.057798 | 0.033899 | 0.084355 | 0.040398 | 0.005498 | 0.259251 | 0.026357 | 0.10875 | 0.049999 | 0.04598 | 0.043959 | 0.093386 | 0.084857 |
| 23.5555556 | 0.051418 | 0.02569 | 0.023289 | 0.016173 | 0.000991 | 0.016423 | 0.026243 | 0.028561 | 0.06352 | 0.125257 | 0.025215 | 0.042633 | 0.022902 | 0.081722 | 0.035785 | 0.008306 | 0.270393 | 0.023279 | 0.086591 | 0.033723 | 0.048914 | 0.039551 | 0.070772 | 0.055708 |
| 23.5694444 | 0.031989 | 0.023621 | 0.024252 | 0.013708 | 0.000328 | 0.011364 | 0.019743 | 0.017935 | 0.054879 | 0.065899 | 0.015558 | 0.027153 | 0.012758 | 0.061236 | 0.02607 | 0.01399 | 0.244743 | 0.021111 | 0.048141 | 0.024227 | 0.038092 | 0.02491 | 0.036864 | 0.027677 |
| 23.5833333 | 0.014102 | 0.022895 | 0.03079 | 0.009823 | 0.000938 | 0.011809 | 0.032911 | 0.007955 | 0.043304 | 0.031446 | 0.007715 | 0.012644 | 0.006292 | 0.035106 | 0.013002 | 0.019219 | 0.195397 | 0.016361 | 0.039712 | 0.018343 | 0.01935 | 0.012377 | 0.028032 | 0.012822 |
| 23.5972222 | 0.012815 | 0.019261 | 0.034552 | 0.005013 | 0.003342 | 0.017239 | 0.04003 | 0.00603 | 0.035972 | 0.045771 | 0.004283 | 0.005647 | 0.007493 | 0.020368 | 0.007525 | 0.023479 | 0.149153 | 0.009647 | 0.07053 | 0.015176 | 0.011981 | 0.010618 | 0.045764 | 0.012191 |
| 23.6111111 | 0.022841 | 0.018536 | 0.029104 | 0.0022 | 0.005973 | 0.015268 | 0.028208 | 0.008268 | 0.038257 | 0.076896 | 0.004859 | 0.010901 | 0.009647 | 0.031512 | 0.01469 | 0.023885 | 0.114412 | 0.004395 | 0.083931 | 0.012078 | 0.019866 | 0.012834 | 0.06298 | 0.012774 |
| 23.625 | 0.031701 | 0.020595 | 0.018357 | 0.002239 | 0.006126 | 0.00788 | 0.017296 | 0.007959 | 0.040178 | 0.072325 | 0.007069 | 0.024224 | 0.007011 | 0.050955 | 0.02466 | 0.016489 | 0.083935 | 0.003938 | 0.057596 | 0.006919 | 0.028991 | 0.024299 | 0.063384 | 0.014462 |
| 23.6388889 | 0.033679 | 0.021175 | 0.009871 | 0.003131 | 0.003779 | 0.005211 | 0.024507 | 0.005353 | 0.036306 | 0.043404 | 0.009221 | 0.037765 | 0.004388 | 0.047744 | 0.029885 | 0.012134 | 0.052883 | 0.007347 | 0.03187 | 0.00324 | 0.034268 | 0.04793 | 0.041036 | 0.024936 |
| 23.6527778 | 0.02614 | 0.020028 | 0.004535 | 0.003127 | 0.002019 | 0.006188 | 0.037224 | 0.004157 | 0.03249 | 0.039456 | 0.010987 | 0.038636 | 0.007525 | 0.031005 | 0.028753 | 0.019173 | 0.053414 | 0.008828 | 0.024331 | 0.003331 | 0.03531 | 0.065182 | 0.019821 | 0.030545 |
| 23.6666667 | 0.01363 | 0.020233 | 0.001496 | 0.002741 | 0.002553 | 0.005387 | 0.037424 | 0.007995 | 0.028453 | 0.06826 | 0.012134 | 0.021922 | 0.015206 | 0.039737 | 0.018155 | 0.027635 | 0.119374 | 0.005888 | 0.02184 | 0.00421 | 0.024331 | 0.064462 | 0.027965 | 0.02079 |
| 23.6805556 | 0.009942 | 0.022373 | 0.000704 | 0.004728 | 0.002836 | 0.002665 | 0.024394 | 0.014388 | 0.022236 | 0.086026 | 0.012275 | 0.008973 | 0.022332 | 0.075103 | 0.009306 | 0.029568 | 0.204548 | 0.004184 | 0.031525 | 0.003422 | 0.011764 | 0.050519 | 0.048964 | 0.008275 |
| 23.6944444 | 0.018691 | 0.02175 | 0.001987 | 0.008492 | 0.002593 | 0.001174 | 0.012827 | 0.017345 | 0.013907 | 0.063704 | 0.010994 | 0.012251 | 0.023212 | 0.093387 | 0.016476 | 0.022916 | 0.246903 | 0.009515 | 0.053701 | 0.003019 | 0.01302 | 0.029308 | 0.05891 | 0.007263 |
| 23.7083333 | 0.026622 | 0.01525 | 0.005614 | 0.010209 | 0.004594 | 0.001596 | 0.018633 | 0.015418 | 0.009357 | 0.033511 | 0.007822 | 0.018079 | 0.015029 | 0.073347 | 0.03817 | 0.01344 | 0.232352 | 0.019178 | 0.060953 | 0.007674 | 0.016857 | 0.009761 | 0.050901 | 0.016471 |
| 23.7222222 | 0.021556 | 0.006783 | 0.009719 | 0.008875 | 0.007766 | 0.002036 | 0.03442 | 0.009907 | 0.017235 | 0.043031 | 0.004189 | 0.021171 | 0.007728 | 0.038102 | 0.054205 | 0.015816 | 0.176539 | 0.024947 | 0.044492 | 0.014746 | 0.015528 | 0.001857 | 0.031969 | 0.022731 |
| 23.7361111 | 0.012157 | 0.002568 | 0.012486 | 0.006331 | 0.008066 | 0.003551 | 0.03774 | 0.005702 | 0.0286 | 0.082107 | 0.003151 | 0.021978 | 0.010738 | 0.027237 | 0.044084 | 0.028824 | 0.096509 | 0.020922 | 0.026101 | 0.018593 | 0.012213 | 0.007483 | 0.016407 | 0.017874 |
| 23.75 | 0.014593 | 0.005243 | 0.013102 | 0.003843 | 0.005512 | 0.008302 | 0.022841 | 0.00913 | 0.029865 | 0.111581 | 0.004964 | 0.014848 | 0.016404 | 0.051551 | 0.020428 | 0.03415 | 0.046566 | 0.010935 | 0.029152 | 0.019303 | 0.007536 | 0.02417 | 0.011764 | 0.01448 |
| 23.7638889 | 0.025884 | 0.014013 | 0.01166 | 0.001868 | 0.005042 | 0.013302 | 0.00955 | 0.017728 | 0.023656 | 0.116434 | 0.00702 | 0.009289 | 0.016467 | 0.078408 | 0.008987 | 0.025996 | 0.073274 | 0.004724 | 0.049088 | 0.016632 | 0.007155 | 0.041157 | 0.013726 | 0.020462 |
| 23.7777778 | 0.036599 | 0.024745 | 0.009548 | 0.000766 | 0.007893 | 0.014079 | 0.009127 | 0.02292 | 0.017739 | 0.092984 | 0.006876 | 0.011022 | 0.012546 | 0.084254 | 0.016908 | 0.01467 | 0.119991 | 0.006365 | 0.06689 | 0.010926 | 0.014227 | 0.043892 | 0.015659 | 0.021977 |
| 23.7916667 | 0.040764 | 0.030425 | 0.009174 | 0.000464 | 0.008813 | 0.009917 | 0.008454 | 0.020757 | 0.010768 | 0.049625 | 0.004394 | 0.009886 | 0.008429 | 0.068243 | 0.030806 | 0.006937 | 0.122783 | 0.011578 | 0.078911 | 0.005422 | 0.02341 | 0.031717 | 0.021617 | 0.014251 |
| 23.8055556 | 0.032729 | 0.027629 | 0.011753 | 0.001343 | 0.006089 | 0.005028 | 0.005766 | 0.013972 | 0.00481 | 0.020108 | 0.002742 | 0.0099 | 0.005529 | 0.037104 | 0.03409 | 0.003498 | 0.076886 | 0.015569 | 0.082766 | 0.00389 | 0.028739 | 0.015745 | 0.033907 | 0.01263 |
| 23.8194444 | 0.017164 | 0.021095 | 0.01503 | 0.003239 | 0.00556 | 0.004264 | 0.012526 | 0.006925 | 0.00346 | 0.013368 | 0.002771 | 0.016001 | 0.004879 | 0.017692 | 0.027905 | 0.003568 | 0.043547 | 0.017765 | 0.075032 | 0.007788 | 0.029482 | 0.011371 | 0.046628 | 0.025509 |
| 23.8333333 | 0.005514 | 0.01764 | 0.01758 | 0.00462 | 0.008257 | 0.007564 | 0.026428 | 0.00271 | 0.002677 | 0.009172 | 0.001988 | 0.028039 | 0.006259 | 0.021516 | 0.023215 | 0.00493 | 0.07856 | 0.020506 | 0.056971 | 0.013113 | 0.026622 | 0.018506 | 0.051452 | 0.037892 |
| 23.8472222 | 0.003435 | 0.018313 | 0.020374 | 0.004456 | 0.00721 | 0.011465 | 0.034509 | 0.0015 | 0.00388 | 0.012469 | 0.001459 | 0.043358 | 0.006314 | 0.019878 | 0.024005 | 0.00922 | 0.14073 | 0.025474 | 0.030732 | 0.016767 | 0.018609 | 0.016723 | 0.043173 | 0.035994 |
| 23.8611111 | 0.009556 | 0.020364 | 0.023082 | 0.002786 | 0.00506 | 0.013153 | 0.029838 | 0.00177 | 0.010322 | 0.033731 | 0.003839 | 0.049672 | 0.005838 | 0.013851 | 0.025193 | 0.017561 | 0.196395 | 0.028736 | 0.010048 | 0.022316 | 0.008895 | 0.007 | 0.027299 | 0.027214 |
| 23.875 | 0.020938 | 0.023104 | 0.023964 | 0.001713 | 0.007674 | 0.012661 | 0.018196 | 0.003347 | 0.020727 | 0.06733 | 0.008876 | 0.041418 | 0.008355 | 0.027446 | 0.021514 | 0.025333 | 0.237198 | 0.025275 | 0.00589 | 0.030143 | 0.003908 | 0.002426 | 0.014886 | 0.024619 |
| 23.8888889 | 0.03322 | 0.027365 | 0.02239 | 0.003179 | 0.009914 | 0.013011 | 0.007774 | 0.005641 | 0.029712 | 0.104312 | 0.014416 | 0.023251 | 0.012279 | 0.047488 | 0.014373 | 0.02871 | 0.235408 | 0.016308 | 0.011215 | 0.033221 | 0.002868 | 0.002902 | 0.011832 | 0.030831 |
| 23.9027778 | 0.043713 | 0.030733 | 0.019537 | 0.004589 | 0.007415 | 0.015534 | 0.002133 | 0.010797 | 0.032586 | 0.137776 | 0.018102 | 0.007376 | 0.014307 | 0.057679 | 0.00717 | 0.027119 | 0.197017 | 0.008163 | 0.020595 | 0.03132 | 0.002229 | 0.009225 | 0.015485 | 0.040636 |
| 23.9166667 | 0.053222 | 0.028792 | 0.017021 | 0.003479 | 0.005597 | 0.017827 | 0.001482 | 0.016722 | 0.032675 | 0.142068 | 0.01907 | 0.002293 | 0.014032 | 0.058585 | 0.00337 | 0.01926 | 0.149143 | 0.007272 | 0.034093 | 0.02847 | 0.002909 | 0.021939 | 0.019363 | 0.051025 |
| 23.9305556 | 0.053952 | 0.021805 | 0.015642 | 0.002613 | 0.009974 | 0.017893 | 0.003901 | 0.013856 | 0.032412 | 0.099353 | 0.017133 | 0.004996 | 0.013116 | 0.052916 | 0.003107 | 0.008766 | 0.100259 | 0.012658 | 0.044494 | 0.022702 | 0.007579 | 0.03684 | 0.021123 | 0.056708 |
| 23.9444444 | 0.038092 | 0.013767 | 0.015604 | 0.003732 | 0.013256 | 0.014963 | 0.008092 | 0.006451 | 0.028846 | 0.046444 | 0.011728 | 0.012817 | 0.012128 | 0.043805 | 0.002595 | 0.004042 | 0.05543 | 0.016901 | 0.041478 | 0.014035 | 0.013023 | 0.045818 | 0.022084 | 0.049912 |
| 23.9583333 | 0.017708 | 0.006914 | 0.015548 | 0.003685 | 0.009233 | 0.009822 | 0.011014 | 0.005188 | 0.022634 | 0.027965 | 0.005959 | 0.021567 | 0.009763 | 0.033931 | 0.003638 | 0.004417 | 0.025379 | 0.017295 | 0.026411 | 0.00663 | 0.01477 | 0.043437 | 0.022544 | 0.035111 |
| 23.9722222 | 0.009396 | 0.002981 | 0.013044 | 0.002264 | 0.004972 | 0.005086 | 0.009738 | 0.00607 | 0.018416 | 0.040583 | 0.004723 | 0.029367 | 0.005576 | 0.026513 | 0.007116 | 0.005194 | 0.010047 | 0.01641 | 0.0124 | 0.002473 | 0.013453 | 0.033029 | 0.023484 | 0.022597 |
| 23.9861111 | 0.009512 | 0.00318 | 0.009422 | 0.002677 | 0.006572 | 0.00259 | 0.006505 | 0.005457 | 0.019401 | 0.047674 | 0.005831 | 0.039385 | 0.002037 | 0.022006 | 0.007686 | 0.004826 | 0.002861 | 0.014525 | 0.009318 | 0.000982 | 0.012367 | 0.022934 | 0.027447 | 0.016287 |
| 24 | 0.008967 | 0.004557 | 0.010101 | 0.004778 | 0.009911 | 0.003603 | 0.006415 | 0.009015 | 0.025057 | 0.035512 | 0.005822 | 0.048798 | 0.002692 | 0.019546 | 0.007135 | 0.00516 | 0.001281 | 0.009438 | 0.017911 | 0.00164 | 0.014606 | 0.022556 | 0.034483 | 0.018748 |
| 24.0138889 | 0.014241 | 0.003853 | 0.017057 | 0.006396 | 0.009226 | 0.009584 | 0.011951 | 0.015447 | 0.030373 | 0.035498 | 0.010396 | 0.052762 | 0.008704 | 0.022549 | 0.013447 | 0.012042 | 0.003083 | 0.00681 | 0.032158 | 0.004699 | 0.019527 | 0.034712 | 0.044067 | 0.03234 |
| 24.0277778 | 0.030446 | 0.004418 | 0.025834 | 0.006095 | 0.008292 | 0.019918 | 0.020678 | 0.016903 | 0.029189 | 0.068568 | 0.024114 | 0.057618 | 0.01774 | 0.036564 | 0.024184 | 0.022429 | 0.008339 | 0.013895 | 0.046839 | 0.009937 | 0.026738 | 0.052471 | 0.058139 | 0.055257 |
| 24.0416667 | 0.051776 | 0.009833 | 0.032509 | 0.00544 | 0.013696 | 0.028069 | 0.02974 | 0.017933 | 0.021423 | 0.09741 | 0.038705 | 0.073091 | 0.024121 | 0.062545 | 0.03731 | 0.024781 | 0.024534 | 0.027573 | 0.057822 | 0.014826 | 0.038125 | 0.067868 | 0.070755 | 0.077334 |
| 24.0555556 | 0.065411 | 0.01638 | 0.035556 | 0.007949 | 0.015626 | 0.02705 | 0.034761 | 0.03061 | 0.0144 | 0.082025 | 0.0461 | 0.085302 | 0.025681 | 0.08439 | 0.052534 | 0.017474 | 0.042325 | 0.041258 | 0.057502 | 0.01733 | 0.048902 | 0.077405 | 0.068956 | 0.083477 |
| 24.0694444 | 0.058956 | 0.018872 | 0.03283 | 0.011075 | 0.00958 | 0.019935 | 0.028202 | 0.044559 | 0.013185 | 0.040633 | 0.046272 | 0.078354 | 0.024831 | 0.074063 | 0.059878 | 0.010627 | 0.036914 | 0.044148 | 0.048318 | 0.020176 | 0.051993 | 0.07514 | 0.052993 | 0.065294 |
| 24.0833333 | 0.037368 | 0.015196 | 0.023627 | 0.008951 | 0.00868 | 0.01347 | 0.015742 | 0.040107 | 0.015387 | 0.011549 | 0.037023 | 0.058238 | 0.019774 | 0.041128 | 0.052696 | 0.008265 | 0.017789 | 0.030904 | 0.036753 | 0.024784 | 0.042374 | 0.053975 | 0.032446 | 0.038761 |
| 24.0972222 | 0.016506 | 0.007709 | 0.012371 | 0.004855 | 0.009696 | 0.008871 | 0.013841 | 0.021806 | 0.016755 | 0.002809 | 0.020096 | 0.031526 | 0.011239 | 0.030419 | 0.032243 | 0.008219 | 0.011661 | 0.014408 | 0.021637 | 0.026749 | 0.021877 | 0.027147 | 0.016307 | 0.037808 |
| 24.1111111 | 0.007171 | 0.002868 | 0.007065 | 0.005346 | 0.00706 | 0.005286 | 0.020294 | 0.011047 | 0.014961 | 0.002476 | 0.009047 | 0.011834 | 0.007358 | 0.046372 | 0.01474 | 0.015388 | 0.019625 | 0.009337 | 0.01521 | 0.022841 | 0.008915 | 0.017635 | 0.013832 | 0.059805 |
| 24.125 | 0.007128 | 0.002672 | 0.007617 | 0.007575 | 0.007511 | 0.003017 | 0.021857 | 0.017266 | 0.010335 | 0.002611 | 0.009405 | 0.010017 | 0.008501 | 0.056392 | 0.014219 | 0.024264 | 0.025733 | 0.013916 | 0.025389 | 0.016029 | 0.01074 | 0.019042 | 0.020061 | 0.071367 |
| 24.1388889 | 0.008178 | 0.002584 | 0.007397 | 0.006589 | 0.007621 | 0.002361 | 0.017727 | 0.024851 | 0.006377 | 0.001724 | 0.011167 | 0.015679 | 0.008657 | 0.050524 | 0.020832 | 0.019404 | 0.025675 | 0.01882 | 0.031688 | 0.012152 | 0.010376 | 0.013863 | 0.020124 | 0.061529 |
| 24.1527778 | 0.007836 | 0.001516 | 0.009703 | 0.003672 | 0.003981 | 0.002453 | 0.012617 | 0.023844 | 0.005332 | 0.001948 | 0.012024 | 0.019887 | 0.008614 | 0.047294 | 0.031123 | 0.009353 | 0.036138 | 0.021779 | 0.023529 | 0.012331 | 0.007884 | 0.009905 | 0.012689 | 0.035699 |
| 24.1666667 | 0.007245 | 0.002249 | 0.011858 | 0.002517 | 0.003053 | 0.003457 | 0.016549 | 0.027115 | 0.00665 | 0.014002 | 0.019415 | 0.023541 | 0.010286 | 0.05318 | 0.043884 | 0.007785 | 0.043109 | 0.023016 | 0.015175 | 0.012886 | 0.012974 | 0.019397 | 0.008002 | 0.013801 |
| 24.1805556 | 0.013877 | 0.005057 | 0.014294 | 0.007689 | 0.007254 | 0.0066 | 0.035077 | 0.039945 | 0.013055 | 0.046811 | 0.034292 | 0.03491 | 0.011308 | 0.048768 | 0.043803 | 0.006912 | 0.028647 | 0.020504 | 0.021627 | 0.010204 | 0.013069 | 0.040451 | 0.007307 | 0.009168 |
| 24.1944444 | 0.019258 | 0.006622 | 0.018699 | 0.012827 | 0.011366 | 0.008709 | 0.043547 | 0.04278 | 0.018799 | 0.07654 | 0.038414 | 0.053805 | 0.010921 | 0.030388 | 0.032319 | 0.004015 | 0.018473 | 0.01513 | 0.03822 | 0.005487 | 0.005307 | 0.060077 | 0.007474 | 0.006727 |
| 24.2083333 | 0.018737 | 0.005679 | 0.018679 | 0.010567 | 0.009206 | 0.007445 | 0.036286 | 0.032459 | 0.024683 | 0.062568 | 0.023578 | 0.056386 | 0.010307 | 0.015611 | 0.02278 | 0.004841 | 0.015311 | 0.01138 | 0.042444 | 0.002602 | 0.001599 | 0.051703 | 0.008499 | 0.00426 |
| 24.2222222 | 0.027056 | 0.005223 | 0.020213 | 0.00752 | 0.00495 | 0.005067 | 0.037667 | 0.029724 | 0.039082 | 0.022338 | 0.011227 | 0.039999 | 0.009006 | 0.01029 | 0.01607 | 0.006871 | 0.007921 | 0.009358 | 0.02773 | 0.00214 | 0.002203 | 0.020722 | 0.008705 | 0.007551 |
| 24.2361111 | 0.033394 | 0.005816 | 0.015244 | 0.004448 | 0.004055 | 0.003248 | 0.035575 | 0.025766 | 0.038673 | 0.010023 | 0.008437 | 0.034266 | 0.007362 | 0.008656 | 0.009785 | 0.005644 | 0.003299 | 0.006102 | 0.011534 | 0.003684 | 0.003192 | 0.007154 | 0.013532 | 0.01297 |
| 24.25 | 0.033626 | 0.006039 | 0.005205 | 0.001545 | 0.003166 | 0.003172 | 0.021965 | 0.012034 | 0.018012 | 0.02004 | 0.007752 | 0.041242 | 0.006997 | 0.006651 | 0.005253 | 0.003117 | 0.002781 | 0.00247 | 0.007238 | 0.005425 | 0.003374 | 0.013205 | 0.022088 | 0.019045 |
| 24.2638889 | 0.03674 | 0.005367 | 0.003145 | 0.00205 | 0.001825 | 0.003552 | 0.016156 | 0.004868 | 0.00524 | 0.020285 | 0.006701 | 0.040824 | 0.008138 | 0.003806 | 0.003384 | 0.003174 | 0.002903 | 0.000631 | 0.011503 | 0.005501 | 0.003565 | 0.016241 | 0.023748 | 0.021996 |
| 24.2777778 | 0.037862 | 0.004089 | 0.003878 | 0.003948 | 0.001549 | 0.003377 | 0.015627 | 0.003048 | 0.003916 | 0.011751 | 0.005143 | 0.032011 | 0.010218 | 0.001719 | 0.004591 | 0.003686 | 0.002154 | 0.001541 | 0.013845 | 0.003727 | 0.003791 | 0.012369 | 0.015715 | 0.016347 |
| 24.2916667 | 0.036461 | 0.003566 | 0.005273 | 0.004898 | 0.002255 | 0.004749 | 0.010097 | 0.001361 | 0.003576 | 0.006609 | 0.005985 | 0.023184 | 0.012375 | 0.001835 | 0.008142 | 0.002606 | 0.001595 | 0.004944 | 0.009702 | 0.002296 | 0.002547 | 0.01396 | 0.008051 | 0.009928 |
| 24.3055556 | 0.036163 | 0.004453 | 0.00762 | 0.003504 | 0.004191 | 0.007676 | 0.005251 | 0.003141 | 0.002402 | 0.008616 | 0.009446 | 0.01707 | 0.012997 | 0.003443 | 0.010479 | 0.002286 | 0.002114 | 0.008163 | 0.003898 | 0.002922 | 0.000884 | 0.026494 | 0.010367 | 0.014219 |
| 24.3194444 | 0.034646 | 0.005369 | 0.008425 | 0.002062 | 0.006532 | 0.00896 | 0.007383 | 0.005957 | 0.001579 | 0.012667 | 0.011984 | 0.011786 | 0.011043 | 0.004923 | 0.009358 | 0.003055 | 0.002627 | 0.008387 | 0.00288 | 0.002806 | 0.000624 | 0.034074 | 0.015148 | 0.019901 |
| 24.3333333 | 0.027973 | 0.004796 | 0.00683 | 0.002969 | 0.007444 | 0.007149 | 0.009988 | 0.005811 | 0.001178 | 0.012947 | 0.010874 | 0.006639 | 0.007583 | 0.006232 | 0.006956 | 0.00291 | 0.001974 | 0.005984 | 0.006352 | 0.001521 | 0.001456 | 0.024042 | 0.013972 | 0.01584 |
| 24.3472222 | 0.018845 | 0.003216 | 0.004349 | 0.003435 | 0.005674 | 0.004429 | 0.007276 | 0.003028 | 0.001058 | 0.008351 | 0.007186 | 0.005654 | 0.004818 | 0.007786 | 0.006086 | 0.001717 | 0.001047 | 0.003105 | 0.008732 | 0.001227 | 0.002009 | 0.011133 | 0.009688 | 0.006939 |
| 24.3611111 | 0.011289 | 0.001829 | 0.002614 | 0.002197 | 0.002877 | 0.003031 | 0.002768 | 0.001892 | 0.001097 | 0.005105 | 0.004352 | 0.008142 | 0.004398 | 0.008367 | 0.007017 | 0.000652 | 0.001137 | 0.001353 | 0.006644 | 0.002353 | 0.001824 | 0.009062 | 0.008553 | 0.003812 |
| 24.375 | 0.006549 | 0.000947 | 0.002555 | 0.001645 | 0.001726 | 0.003834 | 0.001864 | 0.005523 | 0.002612 | 0.00766 | 0.005471 | 0.007005 | 0.006539 | 0.00717 | 0.008665 | 0.000415 | 0.002123 | 0.0016 | 0.005348 | 0.004279 | 0.001967 | 0.009792 | 0.014143 | 0.010846 |
| 24.3888889 | 0.005497 | 0.000634 | 0.00441 | 0.002046 | 0.001699 | 0.006982 | 0.006623 | 0.014106 | 0.006831 | 0.008153 | 0.010533 | 0.004983 | 0.009662 | 0.006086 | 0.01056 | 0.001086 | 0.002605 | 0.003738 | 0.010376 | 0.006017 | 0.003647 | 0.013785 | 0.025664 | 0.021381 |
| 24.4027778 | 0.009056 | 0.000798 | 0.007408 | 0.00396 | 0.002504 | 0.011589 | 0.015324 | 0.024099 | 0.009854 | 0.00947 | 0.015922 | 0.009253 | 0.011686 | 0.007834 | 0.012993 | 0.002083 | 0.002107 | 0.006709 | 0.016905 | 0.007029 | 0.006154 | 0.022679 | 0.037625 | 0.025852 |
| 24.4166667 | 0.016389 | 0.001514 | 0.010369 | 0.007443 | 0.005148 | 0.015893 | 0.023026 | 0.028887 | 0.008803 | 0.022776 | 0.018279 | 0.014583 | 0.011689 | 0.012344 | 0.015652 | 0.003133 | 0.001623 | 0.009326 | 0.020578 | 0.007049 | 0.008385 | 0.024714 | 0.042435 | 0.0212 |
| 24.4305556 | 0.02446 | 0.002977 | 0.012301 | 0.009398 | 0.006969 | 0.017893 | 0.024536 | 0.025637 | 0.006205 | 0.04074 | 0.01752 | 0.016172 | 0.010442 | 0.017175 | 0.018023 | 0.004413 | 0.002495 | 0.011001 | 0.023903 | 0.006668 | 0.011078 | 0.016129 | 0.039265 | 0.011949 |
| 24.4444444 | 0.030036 | 0.004318 | 0.01277 | 0.008552 | 0.005322 | 0.016711 | 0.019984 | 0.017589 | 0.005282 | 0.052811 | 0.015033 | 0.016918 | 0.009894 | 0.020666 | 0.02067 | 0.006091 | 0.005334 | 0.011764 | 0.029346 | 0.007313 | 0.01467 | 0.007612 | 0.03413 | 0.004593 |
| 24.4583333 | 0.031372 | 0.004739 | 0.011931 | 0.006304 | 0.002982 | 0.01415 | 0.013332 | 0.008656 | 0.007468 | 0.057829 | 0.01196 | 0.018245 | 0.01107 | 0.021359 | 0.022837 | 0.00838 | 0.007058 | 0.0114 | 0.03357 | 0.0089 | 0.01695 | 0.007038 | 0.031614 | 0.002994 |
| 24.4722222 | 0.028277 | 0.00435 | 0.01044 | 0.004223 | 0.003025 | 0.012863 | 0.00824 | 0.002735 | 0.012261 | 0.054757 | 0.009628 | 0.019434 | 0.013729 | 0.019739 | 0.022855 | 0.011551 | 0.007738 | 0.010642 | 0.033747 | 0.010926 | 0.016415 | 0.007302 | 0.034941 | 0.008796 |
| 24.4861111 | 0.02418 | 0.004796 | 0.011403 | 0.004799 | 0.00293 | 0.015352 | 0.008535 | 0.001513 | 0.019305 | 0.0447 | 0.010312 | 0.020518 | 0.019683 | 0.021911 | 0.024862 | 0.018237 | 0.013451 | 0.013032 | 0.03922 | 0.016862 | 0.017283 | 0.007309 | 0.051224 | 0.022769 |
| 24.5 | 0.027047 | 0.009054 | 0.021774 | 0.009626 | 0.002865 | 0.025258 | 0.017606 | 0.00382 | 0.030718 | 0.03948 | 0.017034 | 0.025305 | 0.030634 | 0.035172 | 0.032649 | 0.029171 | 0.019582 | 0.02093 | 0.058815 | 0.029976 | 0.023383 | 0.01506 | 0.082253 | 0.041079 |
| 24.5138889 | 0.038913 | 0.016615 | 0.040842 | 0.015937 | 0.006164 | 0.040989 | 0.03267 | 0.010389 | 0.043502 | 0.05434 | 0.029821 | 0.036614 | 0.03961 | 0.056754 | 0.036714 | 0.03549 | 0.017831 | 0.028035 | 0.076969 | 0.041354 | 0.030408 | 0.023772 | 0.105927 | 0.052617 |
| 24.5277778 | 0.053453 | 0.020462 | 0.050983 | 0.018964 | 0.010098 | 0.049343 | 0.043926 | 0.018548 | 0.04705 | 0.08992 | 0.042361 | 0.04621 | 0.041063 | 0.073705 | 0.030776 | 0.031539 | 0.010804 | 0.027561 | 0.073538 | 0.041759 | 0.032681 | 0.024339 | 0.098413 | 0.050723 |
| 24.5416667 | 0.071393 | 0.015775 | 0.043884 | 0.018236 | 0.01126 | 0.043127 | 0.042631 | 0.022994 | 0.050386 | 0.123386 | 0.046496 | 0.044006 | 0.040797 | 0.078053 | 0.024515 | 0.02238 | 0.010009 | 0.022899 | 0.0553 | 0.039632 | 0.030194 | 0.01983 | 0.069468 | 0.040712 |
| 24.5555556 | 0.083523 | 0.009558 | 0.035012 | 0.01691 | 0.009493 | 0.032204 | 0.028436 | 0.024768 | 0.068676 | 0.130728 | 0.039517 | 0.033414 | 0.04015 | 0.070004 | 0.021552 | 0.011735 | 0.023939 | 0.018292 | 0.03659 | 0.04055 | 0.024937 | 0.012961 | 0.039926 | 0.028482 |
| 24.5694444 | 0.068465 | 0.009611 | 0.034737 | 0.015721 | 0.005459 | 0.023973 | 0.014738 | 0.024345 | 0.079283 | 0.108311 | 0.024591 | 0.022704 | 0.035157 | 0.054966 | 0.016893 | 0.004685 | 0.041912 | 0.014537 | 0.019145 | 0.038133 | 0.017922 | 0.009988 | 0.017637 | 0.017541 |

| | | | | | | | | | | | | | | | | | | | | | | | | |
|---|---|---|---|---|---|---|---|---|---|---|---|---|---|---|---|---|---|---|---|---|---|---|---|---|
| 24.5833333 | 0.035124 | 0.01256 | 0.037801 | 0.013887 | 0.002413 | 0.017508 | 0.016654 | 0.018131 | 0.062329 | 0.0628 | 0.011434 | 0.013436 | 0.02768 | 0.040092 | 0.011383 | 0.004636 | 0.054107 | 0.013004 | 0.008248 | 0.03295 | 0.011735 | 0.014028 | 0.005518 | 0.013086 |
| 24.5972222 | 0.014211 | 0.013157 | 0.036603 | 0.012872 | 0.002193 | 0.012312 | 0.027865 | 0.009033 | 0.034238 | 0.030719 | 0.006202 | 0.006677 | 0.021343 | 0.025229 | 0.007457 | 0.005484 | 0.062245 | 0.01355 | 0.008724 | 0.029368 | 0.007978 | 0.012068 | 0.002198 | 0.015895 |
| 24.6111111 | 0.01735 | 0.011881 | 0.030526 | 0.016008 | 0.001797 | 0.008728 | 0.032764 | 0.003345 | 0.02109 | 0.042274 | 0.006552 | 0.005447 | 0.01605 | 0.014329 | 0.00523 | 0.003633 | 0.060714 | 0.014127 | 0.010121 | 0.026199 | 0.004744 | 0.007518 | 0.003592 | 0.022756 |
| 24.625 | 0.030453 | 0.0106 | 0.026006 | 0.021484 | 0.003147 | 0.0057 | 0.026264 | 0.001502 | 0.030546 | 0.071239 | 0.013703 | 0.007765 | 0.009696 | 0.019473 | 0.004687 | 0.003226 | 0.045767 | 0.013815 | 0.006983 | 0.021142 | 0.002598 | 0.014797 | 0.007999 | 0.030904 |
| 24.6388889 | 0.037477 | 0.009693 | 0.024084 | 0.022266 | 0.007669 | 0.002874 | 0.015895 | 0.001309 | 0.041356 | 0.092848 | 0.022492 | 0.007475 | 0.004543 | 0.035892 | 0.004913 | 0.007328 | 0.029392 | 0.0127 | 0.00384 | 0.015028 | 0.003519 | 0.026765 | 0.012176 | 0.034963 |
| 24.6527778 | 0.036784 | 0.009326 | 0.021049 | 0.019001 | 0.009318 | 0.001305 | 0.018979 | 0.003521 | 0.039965 | 0.096588 | 0.023861 | 0.008388 | 0.003281 | 0.042948 | 0.003729 | 0.013035 | 0.02072 | 0.009809 | 0.004711 | 0.010341 | 0.004002 | 0.032549 | 0.012699 | 0.03049 |
| 24.6666667 | 0.031492 | 0.010179 | 0.017519 | 0.014915 | 0.006799 | 0.000635 | 0.034196 | 0.007237 | 0.032971 | 0.06757 | 0.020309 | 0.017903 | 0.002895 | 0.031734 | 0.002622 | 0.016671 | 0.014099 | 0.005846 | 0.012024 | 0.007702 | 0.003916 | 0.027887 | 0.011508 | 0.019684 |
| 24.6805556 | 0.02186 | 0.012904 | 0.013429 | 0.009114 | 0.003692 | 0.000919 | 0.038889 | 0.007419 | 0.023466 | 0.050786 | 0.014278 | 0.027101 | 0.001882 | 0.017165 | 0.005306 | 0.01796 | 0.01367 | 0.004633 | 0.022175 | 0.005882 | 0.008565 | 0.016689 | 0.011366 | 0.008903 |
| 24.6944444 | 0.018221 | 0.017665 | 0.007874 | 0.006264 | 0.003343 | 0.001409 | 0.0247 | 0.00777 | 0.011847 | 0.092068 | 0.009099 | 0.025591 | 0.001549 | 0.018633 | 0.008458 | 0.017588 | 0.030189 | 0.006195 | 0.03046 | 0.00387 | 0.015951 | 0.014258 | 0.012646 | 0.004226 |
| 24.7083333 | 0.029686 | 0.024749 | 0.005871 | 0.009218 | 0.005279 | 0.001149 | 0.013698 | 0.012962 | 0.004625 | 0.127929 | 0.008693 | 0.01496 | 0.002913 | 0.032628 | 0.00677 | 0.014375 | 0.051838 | 0.005892 | 0.034363 | 0.00368 | 0.020948 | 0.022419 | 0.01374 | 0.004276 |
| 24.7222222 | 0.044605 | 0.032538 | 0.009956 | 0.011957 | 0.004771 | 0.001501 | 0.02534 | 0.016328 | 0.013493 | 0.115674 | 0.007529 | 0.005076 | 0.007509 | 0.047817 | 0.003328 | 0.008016 | 0.063502 | 0.003698 | 0.030698 | 0.006137 | 0.022211 | 0.029043 | 0.011078 | 0.003383 |
| 24.7361111 | 0.052273 | 0.035177 | 0.014908 | 0.011158 | 0.003059 | 0.00245 | 0.042002 | 0.013466 | 0.029061 | 0.083403 | 0.006344 | 0.002202 | 0.014358 | 0.05447 | 0.002521 | 0.003344 | 0.065935 | 0.003622 | 0.019234 | 0.006819 | 0.018906 | 0.02652 | 0.007014 | 0.00256 |
| 24.75 | 0.047152 | 0.02916 | 0.017321 | 0.009659 | 0.004144 | 0.002792 | 0.046006 | 0.006937 | 0.024262 | 0.046305 | 0.009271 | 0.003408 | 0.018008 | 0.041101 | 0.002832 | 0.003272 | 0.061974 | 0.005011 | 0.008791 | 0.004955 | 0.011789 | 0.0201 | 0.006608 | 0.004071 |
| 24.7638889 | 0.027633 | 0.017391 | 0.018064 | 0.009378 | 0.004909 | 0.003337 | 0.039052 | 0.002665 | 0.013505 | 0.031796 | 0.010877 | 0.009636 | 0.014971 | 0.018632 | 0.003878 | 0.004372 | 0.051686 | 0.004463 | 0.00933 | 0.004142 | 0.005438 | 0.027242 | 0.005915 | 0.006337 |
| 24.7777778 | 0.015998 | 0.007561 | 0.017781 | 0.010149 | 0.003778 | 0.004959 | 0.022607 | 0.003866 | 0.025568 | 0.059366 | 0.008258 | 0.020962 | 0.007965 | 0.009831 | 0.006238 | 0.005507 | 0.036224 | 0.002341 | 0.018694 | 0.004634 | 0.004418 | 0.04036 | 0.006554 | 0.008953 |
| 24.7916667 | 0.027244 | 0.005913 | 0.016345 | 0.013085 | 0.005683 | 0.005323 | 0.009701 | 0.007291 | 0.04232 | 0.099504 | 0.007896 | 0.0305 | 0.004427 | 0.017543 | 0.007635 | 0.008541 | 0.021313 | 0.001276 | 0.028325 | 0.004078 | 0.007917 | 0.037736 | 0.015614 | 0.009709 |
| 24.8055556 | 0.040499 | 0.009229 | 0.014199 | 0.018365 | 0.009094 | 0.003239 | 0.009831 | 0.010082 | 0.040443 | 0.121962 | 0.01333 | 0.031016 | 0.008433 | 0.029432 | 0.006634 | 0.012321 | 0.015607 | 0.001979 | 0.033367 | 0.002777 | 0.011273 | 0.022498 | 0.025881 | 0.007576 |
| 24.8194444 | 0.039816 | 0.012385 | 0.01166 | 0.020406 | 0.007311 | 0.00193 | 0.009131 | 0.011906 | 0.028374 | 0.130293 | 0.015106 | 0.022378 | 0.014359 | 0.038584 | 0.004511 | 0.012794 | 0.020512 | 0.002421 | 0.032938 | 0.003244 | 0.013034 | 0.012492 | 0.028327 | 0.008151 |
| 24.8333333 | 0.033211 | 0.014336 | 0.009553 | 0.015017 | 0.004337 | 0.00347 | 0.006786 | 0.012397 | 0.020932 | 0.132391 | 0.008534 | 0.012269 | 0.015918 | 0.045323 | 0.006025 | 0.009873 | 0.026851 | 0.001964 | 0.028643 | 0.00457 | 0.01452 | 0.023006 | 0.022943 | 0.012596 |
| 24.8472222 | 0.031356 | 0.0148 | 0.008855 | 0.010528 | 0.005977 | 0.006055 | 0.014406 | 0.010412 | 0.018704 | 0.120701 | 0.002377 | 0.010603 | 0.01116 | 0.051391 | 0.013642 | 0.007866 | 0.028971 | 0.003234 | 0.023289 | 0.003805 | 0.015196 | 0.044603 | 0.012857 | 0.017049 |
| 24.8611111 | 0.038701 | 0.014588 | 0.010079 | 0.015965 | 0.006972 | 0.008061 | 0.026972 | 0.006112 | 0.016093 | 0.093045 | 0.001195 | 0.019101 | 0.004063 | 0.055726 | 0.020966 | 0.008091 | 0.032257 | 0.006828 | 0.018619 | 0.00303 | 0.013651 | 0.046358 | 0.004326 | 0.022576 |
| 24.875 | 0.051573 | 0.015417 | 0.014488 | 0.02765 | 0.004028 | 0.00914 | 0.034088 | 0.003042 | 0.013071 | 0.059485 | 0.003449 | 0.021585 | 0.001838 | 0.050204 | 0.022846 | 0.008491 | 0.041266 | 0.010509 | 0.015693 | 0.006398 | 0.009513 | 0.030351 | 0.002741 | 0.028659 |
| 24.8888889 | 0.061105 | 0.017752 | 0.02245 | 0.037336 | 0.001664 | 0.009687 | 0.032625 | 0.00512 | 0.013106 | 0.029944 | 0.008952 | 0.015916 | 0.006589 | 0.031331 | 0.021212 | 0.00742 | 0.054182 | 0.012506 | 0.015376 | 0.011569 | 0.005194 | 0.034142 | 0.0075 | 0.02628 |
| 24.9027778 | 0.065004 | 0.020057 | 0.029899 | 0.040157 | 0.001725 | 0.010604 | 0.025033 | 0.010561 | 0.017674 | 0.010587 | 0.016248 | 0.021814 | 0.016015 | 0.014091 | 0.018439 | 0.006527 | 0.066987 | 0.011675 | 0.016734 | 0.014444 | 0.003711 | 0.052613 | 0.01409 | 0.015807 |
| 24.9166667 | 0.064206 | 0.020852 | 0.03169 | 0.036446 | 0.001499 | 0.012125 | 0.014116 | 0.014959 | 0.023971 | 0.002358 | 0.023721 | 0.036635 | 0.025455 | 0.012281 | 0.015168 | 0.008796 | 0.072416 | 0.008573 | 0.017114 | 0.015964 | 0.00339 | 0.04912 | 0.019448 | 0.007436 |
| 24.9305556 | 0.054846 | 0.019389 | 0.028842 | 0.027266 | 0.000542 | 0.014181 | 0.005811 | 0.019807 | 0.02732 | 0.000798 | 0.029898 | 0.045332 | 0.02976 | 0.016947 | 0.011294 | 0.013004 | 0.06131 | 0.005888 | 0.01373 | 0.018009 | 0.002423 | 0.028118 | 0.023271 | 0.006942 |
| 24.9444444 | 0.03544 | 0.01463 | 0.024576 | 0.01422 | 0.000345 | 0.016222 | 0.004573 | 0.028901 | 0.026082 | 0.002659 | 0.029407 | 0.047983 | 0.025393 | 0.016426 | 0.009265 | 0.015251 | 0.034746 | 0.005242 | 0.007586 | 0.018212 | 0.002886 | 0.023028 | 0.024156 | 0.015432 |
| 24.9583333 | 0.016514 | 0.007971 | 0.019786 | 0.005574 | 0.000825 | 0.01747 | 0.004862 | 0.038496 | 0.021519 | 0.010436 | 0.019018 | 0.04643 | 0.015622 | 0.012166 | 0.011615 | 0.012813 | 0.018126 | 0.005251 | 0.002719 | 0.014351 | 0.005706 | 0.040909 | 0.020099 | 0.025497 |
| 24.9722222 | 0.011222 | 0.00343 | 0.016147 | 0.004539 | 0.001005 | 0.016386 | 0.003711 | 0.039869 | 0.015186 | 0.024391 | 0.009455 | 0.037284 | 0.01212 | 0.015304 | 0.01558 | 0.006392 | 0.025508 | 0.004838 | 0.001164 | 0.008754 | 0.009101 | 0.053681 | 0.012477 | 0.03229 |
| 24.9861111 | 0.013251 | 0.002541 | 0.015432 | 0.006251 | 0.000567 | 0.012178 | 0.005854 | 0.031442 | 0.008875 | 0.038865 | 0.01011 | 0.02118 | 0.01317 | 0.029993 | 0.017841 | 0.005667 | 0.02888 | 0.006279 | 0.004567 | 0.004636 | 0.011444 | 0.046416 | 0.006814 | 0.039204 |
| 25 | 0.011851 | 0.00251 | 0.017788 | 0.009525 | 0.001036 | 0.008613 | 0.012912 | 0.020683 | 0.005785 | 0.048492 | 0.010974 | 0.009597 | 0.012167 | 0.043804 | 0.017652 | 0.01877 | 0.01786 | 0.011729 | 0.014651 | 0.004916 | 0.013393 | 0.032749 | 0.00956 | 0.048421 |
| 25.0138889 | 0.016181 | 0.002587 | 0.022759 | 0.014535 | 0.003566 | 0.010649 | 0.022749 | 0.014789 | 0.009224 | 0.052904 | 0.013166 | 0.007414 | 0.01828 | 0.050589 | 0.016996 | 0.039251 | 0.019558 | 0.020242 | 0.03031 | 0.010054 | 0.016827 | 0.023167 | 0.02356 | 0.060734 |
| 25.0277778 | 0.034241 | 0.00587 | 0.02806 | 0.019001 | 0.005451 | 0.018541 | 0.033476 | 0.015137 | 0.020089 | 0.055015 | 0.029102 | 0.011979 | 0.028231 | 0.06051 | 0.025592 | 0.051021 | 0.040288 | 0.03106 | 0.046349 | 0.015745 | 0.022284 | 0.014124 | 0.044381 | 0.072896 |
| 25.0416667 | 0.058219 | 0.01217 | 0.030562 | 0.02312 | 0.003971 | 0.02818 | 0.042872 | 0.020367 | 0.02689 | 0.054659 | 0.049054 | 0.027349 | 0.027396 | 0.075353 | 0.048775 | 0.042195 | 0.051438 | 0.042114 | 0.053922 | 0.017156 | 0.025663 | 0.009046 | 0.054119 | 0.070386 |
| 25.0555556 | 0.074861 | 0.019375 | 0.030202 | 0.026299 | 0.001399 | 0.033685 | 0.045101 | 0.026004 | 0.020132 | 0.048206 | 0.057883 | 0.04407 | 0.015714 | 0.078027 | 0.065903 | 0.024139 | 0.038653 | 0.04494 | 0.050757 | 0.014006 | 0.023745 | 0.020191 | 0.040916 | 0.045641 |
| 25.0694444 | 0.072648 | 0.022905 | 0.027357 | 0.022693 | 0.000555 | 0.031409 | 0.033478 | 0.022727 | 0.018042 | 0.034029 | 0.052547 | 0.047826 | 0.007277 | 0.05842 | 0.054992 | 0.020974 | 0.034787 | 0.034613 | 0.040054 | 0.008564 | 0.018084 | 0.04419 | 0.019898 | 0.025879 |
| 25.0833333 | 0.051845 | 0.017565 | 0.020556 | 0.012322 | 0.001325 | 0.022583 | 0.016558 | 0.010914 | 0.029799 | 0.018016 | 0.037012 | 0.039266 | 0.008255 | 0.029388 | 0.027029 | 0.033166 | 0.051518 | 0.018472 | 0.023613 | 0.003845 | 0.011431 | 0.063274 | 0.012114 | 0.030521 |
| 25.0972222 | 0.025525 | 0.008537 | 0.011094 | 0.005767 | 0.003136 | 0.01146 | 0.011194 | 0.002465 | 0.036236 | 0.013209 | 0.020769 | 0.023994 | 0.008009 | 0.014791 | 0.009254 | 0.042555 | 0.050323 | 0.007003 | 0.014675 | 0.002117 | 0.010518 | 0.063689 | 0.022271 | 0.039801 |
| 25.1111111 | 0.012848 | 0.004724 | 0.004422 | 0.008211 | 0.005943 | 0.004159 | 0.015328 | 0.000692 | 0.02791 | 0.020113 | 0.016568 | 0.010399 | 0.003757 | 0.018095 | 0.009231 | 0.040775 | 0.025928 | 0.003634 | 0.019747 | 0.002456 | 0.0153 | 0.046145 | 0.035768 | 0.039178 |
| 25.125 | 0.016699 | 0.005063 | 0.003505 | 0.011666 | 0.008056 | 0.00325 | 0.016638 | 0.001079 | 0.013426 | 0.023549 | 0.023472 | 0.00577 | 0.001679 | 0.021218 | 0.011245 | 0.029121 | 0.010983 | 0.003326 | 0.02324 | 0.004784 | 0.01727 | 0.023194 | 0.036881 | 0.027977 |
| 25.1388889 | 0.019962 | 0.004355 | 0.00443 | 0.010245 | 0.007061 | 0.004166 | 0.013539 | 0.001559 | 0.005107 | 0.017186 | 0.025259 | 0.007376 | 0.002692 | 0.018147 | 0.007033 | 0.014331 | 0.014705 | 0.002479 | 0.019606 | 0.007779 | 0.015467 | 0.009697 | 0.026638 | 0.012673 |
| 25.1527778 | 0.015773 | 0.002627 | 0.003932 | 0.005795 | 0.00411 | 0.003212 | 0.010129 | 0.001462 | 0.006506 | 0.007529 | 0.018815 | 0.008495 | 0.003379 | 0.01476 | 0.002653 | 0.004677 | 0.016954 | 0.002446 | 0.015776 | 0.008785 | 0.013434 | 0.011746 | 0.014874 | 0.004096 |
| 25.1666667 | 0.008687 | 0.001617 | 0.003121 | 0.002732 | 0.003389 | 0.001687 | 0.00811 | 0.000947 | 0.009182 | 0.001566 | 0.013285 | 0.008428 | 0.002486 | 0.015033 | 0.001649 | 0.002264 | 0.010801 | 0.002886 | 0.013521 | 0.007664 | 0.012078 | 0.018499 | 0.007701 | 0.004267 |
| 25.1805556 | 0.003123 | 0.001459 | 0.002748 | 0.003415 | 0.005662 | 0.001365 | 0.006975 | 0.001216 | 0.006947 | 0.000694 | 0.01257 | 0.009829 | 0.002401 | 0.018797 | 0.00375 | 0.003989 | 0.007791 | 0.004997 | 0.011723 | 0.005386 | 0.011025 | 0.015379 | 0.005575 | 0.009419 |
| 25.1944444 | 0.00199 | 0.001685 | 0.002731 | 0.004152 | 0.00582 | 0.001409 | 0.005982 | 0.003192 | 0.004341 | 0.002548 | 0.015177 | 0.013585 | 0.003535 | 0.022043 | 0.010329 | 0.00668 | 0.006823 | 0.010443 | 0.009656 | 0.003491 | 0.00993 | 0.009672 | 0.006344 | 0.013439 |
| 25.2083333 | 0.005562 | 0.001779 | 0.002985 | 0.002566 | 0.003957 | 0.00146 | 0.004796 | 0.006355 | 0.005717 | 0.005034 | 0.017662 | 0.018912 | 0.004017 | 0.021805 | 0.019784 | 0.009047 | 0.005292 | 0.015441 | 0.007593 | 0.002917 | 0.009318 | 0.013998 | 0.006746 | 0.010978 |
| 25.2222222 | 0.012155 | 0.001625 | 0.00309 | 0.000841 | 0.004503 | 0.003173 | 0.002918 | 0.009345 | 0.006642 | 0.006694 | 0.018052 | 0.025083 | 0.0035 | 0.017415 | 0.026354 | 0.00932 | 0.008381 | 0.017037 | 0.005949 | 0.003734 | 0.009864 | 0.014575 | 0.004555 | 0.004879 |
| 25.2361111 | 0.017616 | 0.001413 | 0.00265 | 0.001166 | 0.004336 | 0.005924 | 0.001644 | 0.012047 | 0.004993 | 0.007121 | 0.01676 | 0.031132 | 0.00328 | 0.010044 | 0.028759 | 0.006521 | 0.010955 | 0.016888 | 0.004267 | 0.005687 | 0.010826 | 0.013004 | 0.002357 | 0.002095 |
| 25.25 | 0.018519 | 0.00116 | 0.001749 | 0.003892 | 0.002894 | 0.006709 | 0.003041 | 0.015318 | 0.003513 | 0.007064 | 0.014687 | 0.035506 | 0.004123 | 0.004223 | 0.029449 | 0.002833 | 0.008725 | 0.016974 | 0.002509 | 0.007372 | 0.011257 | 0.031865 | 0.002947 | 0.003764 |
| 25.2638889 | 0.014995 | 0.000711 | 0.000934 | 0.006656 | 0.003891 | 0.004518 | 0.005936 | 0.018108 | 0.003514 | 0.006637 | 0.012552 | 0.036616 | 0.006618 | 0.003068 | 0.028593 | 0.001561 | 0.009669 | 0.017549 | 0.001195 | 0.007187 | 0.01108 | 0.055575 | 0.003089 | 0.008041 |
| 25.2777778 | 0.009453 | 0.000478 | 0.000717 | 0.006769 | 0.004296 | 0.002045 | 0.007372 | 0.0184 | 0.00528 | 0.005568 | 0.010715 | 0.033169 | 0.010736 | 0.003342 | 0.026263 | 0.00304 | 0.013568 | 0.017538 | 0.000758 | 0.005623 | 0.010286 | 0.048374 | 0.003272 | 0.011293 |
| 25.2916667 | 0.004495 | 0.000914 | 0.000894 | 0.004459 | 0.003914 | 0.001165 | 0.005982 | 0.016215 | 0.00684 | 0.004528 | 0.009429 | 0.026546 | 0.013398 | 0.002297 | 0.024202 | 0.004433 | 0.011622 | 0.016352 | 0.001395 | 0.003802 | 0.009615 | 0.030499 | 0.00704 | 0.009406 |
| 25.3055556 | 0.0028 | 0.001563 | 0.001003 | 0.003329 | 0.005635 | 0.00127 | 0.003057 | 0.013855 | 0.00656 | 0.004348 | 0.008621 | 0.019818 | 0.011991 | 0.001701 | 0.02336 | 0.004739 | 0.006099 | 0.014808 | 0.001707 | 0.001978 | 0.010504 | 0.040729 | 0.011474 | 0.005841 |
| 25.3194444 | 0.00504 | 0.001859 | 0.001061 | 0.005475 | 0.006112 | 0.003026 | 0.001811 | 0.013621 | 0.007243 | 0.00512 | 0.007654 | 0.013836 | 0.008342 | 0.001847 | 0.022906 | 0.005445 | 0.005057 | 0.012899 | 0.001968 | 0.000761 | 0.012869 | 0.04632 | 0.01493 | 0.006742 |
| 25.3333333 | 0.007906 | 0.001916 | 0.001296 | 0.008291 | 0.004257 | 0.006515 | 0.003425 | 0.015197 | 0.010918 | 0.005957 | 0.006667 | 0.008007 | 0.005133 | 0.003595 | 0.020693 | 0.006463 | 0.007284 | 0.009932 | 0.00378 | 0.000866 | 0.014675 | 0.032939 | 0.018657 | 0.008684 |
| 25.3472222 | 0.008926 | 0.001874 | 0.001834 | 0.009622 | 0.003095 | 0.009833 | 0.005521 | 0.014823 | 0.013681 | 0.005895 | 0.006211 | 0.004404 | 0.003705 | 0.00967 | 0.01635 | 0.005595 | 0.007566 | 0.006812 | 0.005697 | 0.001239 | 0.014398 | 0.042096 | 0.022287 | 0.006145 |
| 25.3611111 | 0.008318 | 0.001637 | 0.002784 | 0.009644 | 0.003535 | 0.011523 | 0.006737 | 0.009648 | 0.011517 | 0.005164 | 0.006284 | 0.005728 | 0.004495 | 0.015766 | 0.011575 | 0.003 | 0.005065 | 0.005309 | 0.006932 | 0.001765 | 0.012537 | 0.060517 | 0.025179 | 0.003867 |
| 25.375 | 0.007634 | 0.001602 | 0.004226 | 0.009039 | 0.004652 | 0.010752 | 0.008096 | 0.003712 | 0.006269 | 0.004488 | 0.006947 | 0.008549 | 0.00602 | 0.015139 | 0.008222 | 0.001989 | 0.002683 | 0.005974 | 0.008267 | 0.004733 | 0.010493 | 0.046143 | 0.027145 | 0.00702 |
| 25.3888889 | 0.007857 | 0.001988 | 0.005792 | 0.008415 | 0.005285 | 0.008 | 0.0102 | 0.001126 | 0.003106 | 0.004748 | 0.008539 | 0.007702 | 0.006564 | 0.009309 | 0.007773 | 0.004258 | 0.003345 | 0.007903 | 0.009977 | 0.009611 | 0.009493 | 0.027305 | 0.02887 | 0.008412 |
| 25.4027778 | 0.007274 | 0.002005 | 0.006937 | 0.008524 | 0.005319 | 0.006241 | 0.012728 | 0.001136 | 0.004767 | 0.007472 | 0.011531 | 0.004839 | 0.005704 | 0.007106 | 0.009914 | 0.009263 | 0.006578 | 0.009661 | 0.012418 | 0.013311 | 0.010316 | 0.045315 | 0.030656 | 0.008503 |
| 25.4166667 | 0.004954 | 0.001477 | 0.007568 | 0.010308 | 0.005237 | 0.007951 | 0.014644 | 0.004121 | 0.010137 | 0.01201 | 0.014753 | 0.005205 | 0.003762 | 0.012164 | 0.012623 | 0.014792 | 0.009143 | 0.010052 | 0.015617 | 0.013404 | 0.012691 | 0.059994 | 0.03164 | 0.01624 |
| 25.4305556 | 0.005412 | 0.001567 | 0.007922 | 0.013443 | 0.0049 | 0.012077 | 0.014414 | 0.009483 | 0.015469 | 0.014465 | 0.014945 | 0.008885 | 0.002825 | 0.018578 | 0.015005 | 0.019021 | 0.007736 | 0.009548 | 0.019759 | 0.010515 | 0.015677 | 0.039289 | 0.031636 | 0.024239 |
| 25.4444444 | 0.01196 | 0.002319 | 0.008266 | 0.016318 | 0.004239 | 0.015094 | 0.012401 | 0.013644 | 0.016419 | 0.013869 | 0.011322 | 0.011911 | 0.005116 | 0.023294 | 0.017733 | 0.022614 | 0.004581 | 0.00991 | 0.024276 | 0.00836 | 0.018956 | 0.028089 | 0.031364 | 0.022081 |
| 25.4583333 | 0.019726 | 0.002518 | 0.008399 | 0.017335 | 0.003817 | 0.016097 | 0.010658 | 0.015149 | 0.011897 | 0.013412 | 0.007472 | 0.013713 | 0.009315 | 0.026969 | 0.019367 | 0.024609 | 0.003896 | 0.010965 | 0.026019 | 0.009268 | 0.022164 | 0.044268 | 0.030285 | 0.011887 |
| 25.4722222 | 0.023777 | 0.002221 | 0.007985 | 0.0166 | 0.004007 | 0.017186 | 0.01059 | 0.014018 | 0.009232 | 0.01503 | 0.007413 | 0.016647 | 0.013402 | 0.028598 | 0.018349 | 0.022815 | 0.003346 | 0.011405 | 0.024068 | 0.013118 | 0.023431 | 0.043064 | 0.02922 | 0.004437 |
| 25.4861111 | 0.026106 | 0.002764 | 0.00795 | 0.017129 | 0.00526 | 0.017758 | 0.013671 | 0.010714 | 0.02187 | 0.021458 | 0.012359 | 0.021849 | 0.02034 | 0.031409 | 0.018963 | 0.021983 | 0.00167 | 0.013628 | 0.023997 | 0.022252 | 0.021615 | 0.028006 | 0.034095 | 0.00735 |
| 25.5 | 0.033686 | 0.005427 | 0.010373 | 0.023412 | 0.008181 | 0.021484 | 0.022805 | 0.006581 | 0.033471 | 0.039285 | 0.022815 | 0.031928 | 0.032426 | 0.041446 | 0.023827 | 0.028008 | 0.001316 | 0.020536 | 0.028388 | 0.037194 | 0.0189 | 0.038225 | 0.047797 | 0.025429 |
| 25.5138889 | 0.050926 | 0.009556 | 0.015404 | 0.0344 | 0.011595 | 0.035132 | 0.036526 | 0.004244 | 0.030012 | 0.065683 | 0.039341 | 0.047852 | 0.046556 | 0.054589 | 0.026689 | 0.036039 | 0.002173 | 0.026403 | 0.028348 | 0.047071 | 0.0141 | 0.050755 | 0.0597 | 0.053766 |
| 25.5277778 | 0.071675 | 0.011759 | 0.019864 | 0.039442 | 0.014061 | 0.047228 | 0.045241 | 0.003832 | 0.033489 | 0.082876 | 0.055965 | 0.059557 | 0.06139 | 0.064431 | 0.025874 | 0.038624 | 0.003545 | 0.025736 | 0.018833 | 0.04475 | 0.006608 | 0.043423 | 0.058661 | 0.06692 |
| 25.5416667 | 0.088641 | 0.010515 | 0.020104 | 0.033652 | 0.015055 | 0.043777 | 0.043904 | 0.00353 | 0.042301 | 0.08734 | 0.061217 | 0.056333 | 0.071988 | 0.063167 | 0.026965 | 0.035165 | 0.00491 | 0.022555 | 0.011073 | 0.038674 | 0.002806 | 0.030988 | 0.047604 | 0.055748 |
| 25.5555556 | 0.105625 | 0.008128 | 0.017757 | 0.025851 | 0.013233 | 0.037804 | 0.035384 | 0.006077 | 0.05544 | 0.092234 | 0.051149 | 0.043012 | 0.067991 | 0.045225 | 0.02661 | 0.028272 | 0.004887 | 0.01792 | 0.011545 | 0.035026 | 0.002687 | 0.021049 | 0.03247 | 0.036392 |
| 25.5694444 | 0.113877 | 0.006086 | 0.016066 | 0.020887 | 0.009078 | 0.035838 | 0.022309 | 0.009894 | 0.078942 | 0.097272 | 0.032499 | 0.031023 | 0.051836 | 0.02537 | 0.019044 | 0.018369 | 0.007602 | 0.010125 | 0.011667 | 0.03053 | 0.001858 | 0.021618 | 0.017054 | 0.020031 |
| 25.5833333 | 0.096174 | 0.004479 | 0.012725 | 0.01623 | 0.004621 | 0.026889 | 0.010254 | 0.009314 | 0.085739 | 0.089676 | 0.017459 | 0.02316 | 0.033116 | 0.013487 | 0.009105 | 0.010463 | 0.017064 | 0.003336 | 0.006923 | 0.024636 | 0.000795 | 0.029735 | 0.007387 | 0.008688 |
| 25.5972222 | 0.062496 | 0.003483 | 0.00857 | 0.011002 | 0.001861 | 0.0157 | 0.007486 | 0.006047 | 0.061268 | 0.061605 | 0.01852 | 0.016564 | 0.017864 | 0.008684 | 0.003114 | 0.015978 | 0.031035 | 0.001971 | 0.005191 | 0.020696 | 0.00189 | 0.037468 | 0.004091 | 0.003639 |
| 25.6111111 | 0.03088 | 0.002646 | 0.009441 | 0.008486 | 0.00125 | 0.010381 | 0.014596 | 0.005827 | 0.032193 | 0.031972 | 0.029092 | 0.010144 | 0.017212 | 0.019072 | 0.002962 | 0.028925 | 0.041347 | 0.005787 | 0.011824 | 0.018543 | 0.005535 | 0.054882 | 0.002429 | 0.006335 |
| 25.625 | 0.013918 | 0.001505 | 0.014838 | 0.012202 | 0.001258 | 0.008573 | 0.0223 | 0.008234 | 0.023301 | 0.023791 | 0.026036 | 0.010009 | 0.032013 | 0.046232 | 0.008354 | 0.028163 | 0.035543 | 0.010588 | 0.018937 | 0.01627 | 0.007944 | 0.064299 | 0.001733 | 0.011926 |
| 25.6388889 | 0.017939 | 0.000571 | 0.019665 | 0.016574 | 0.003139 | 0.005787 | 0.021286 | 0.009479 | 0.022784 | 0.030005 | 0.017854 | 0.019323 | 0.044365 | 0.065763 | 0.017658 | 0.018078 | 0.017774 | 0.011901 | 0.019511 | 0.01366 | 0.00669 | 0.045775 | 0.003123 | 0.013925 |
| 25.6527778 | 0.030434 | 0.001074 | 0.021749 | 0.013197 | 0.008142 | 0.003119 | 0.014385 | 0.011361 | 0.019171 | 0.04383 | 0.029226 | 0.026953 | 0.040649 | 0.056941 | 0.028209 | 0.01859 | 0.004861 | 0.008928 | 0.01677 | 0.00899 | 0.004736 | 0.038416 | 0.008352 | 0.019138 |
| 25.6666667 | 0.042107 | 0.002541 | 0.020971 | 0.006317 | 0.01314 | 0.004309 | 0.01695 | 0.016789 | 0.017069 | 0.055887 | 0.042754 | 0.022504 | 0.02281 | 0.031554 | 0.034094 | 0.027564 | 0.002612 | 0.004738 | 0.012212 | 0.005712 | 0.005159 | 0.051802 | 0.015796 | 0.021792 |
| 25.6805556 | 0.055233 | 0.002776 | 0.020005 | 0.005135 | 0.014396 | 0.007985 | 0.028448 | 0.017679 | 0.012836 | 0.045797 | 0.034418 | 0.013292 | 0.013024 | 0.019403 | 0.035556 | 0.034786 | 0.004557 | 0.004441 | 0.010347 | 0.010102 | 0.005883 | 0.049367 | 0.01941 | 0.014122 |
| 25.6944444 | 0.060176 | 0.001873 | 0.01952 | 0.008051 | 0.01125 | 0.011558 | 0.030433 | 0.01169 | 0.009019 | 0.023894 | 0.016121 | 0.013801 | 0.03134 | 0.038771 | 0.042784 | 0.037237 | 0.009598 | 0.009635 | 0.018744 | 0.016469 | 0.004146 | 0.031255 | 0.024696 | 0.017639 |
| 25.7083333 | 0.043487 | 0.001793 | 0.016096 | 0.010769 | 0.006009 | 0.011008 | 0.019702 | 0.010729 | 0.01356 | 0.022872 | 0.008601 | 0.020235 | 0.057607 | 0.065376 | 0.050035 | 0.029099 | 0.022983 | 0.013773 | 0.028472 | 0.017461 | 0.003502 | 0.017475 | 0.034947 | 0.032209 |
| 25.7222222 | 0.017409 | 0.00207 | 0.011231 | 0.01293 | 0.002432 | 0.006973 | 0.009949 | 0.013295 | 0.022619 | 0.04878 | 0.018786 | 0.02105 | 0.06169 | 0.066744 | 0.037856 | 0.015595 | 0.042382 | 0.010191 | 0.028908 | 0.013654 | 0.005977 | 0.018667 | 0.03665 | 0.037323 |
| 25.7361111 | 0.005903 | 0.001418 | 0.007445 | 0.013305 | 0.00201 | 0.006607 | 0.011337 | 0.016056 | 0.026557 | 0.076021 | 0.034789 | 0.015627 | 0.044716 | 0.046357 | 0.018244 | 0.01229 | 0.06224 | 0.005857 | 0.023278 | 0.007587 | 0.006296 | 0.038255 | 0.027745 | 0.031102 |
| 25.75 | 0.010579 | 0.001017 | 0.004705 | 0.009816 | 0.003605 | 0.00936 | 0.019006 | 0.018503 | 0.020894 | 0.082803 | 0.035304 | 0.008892 | 0.023018 | 0.022941 | 0.020044 | 0.015225 | 0.063707 | 0.010071 | 0.018064 | 0.003863 | 0.003452 | 0.075989 | 0.017495 | 0.021658 |
| 25.7638889 | 0.019064 | 0.001106 | 0.005006 | 0.006336 | 0.005686 | 0.009093 | 0.024818 | 0.012574 | 0.011514 | 0.06877 | 0.021042 | 0.006488 | 0.012445 | 0.010259 | 0.029124 | 0.014808 | 0.03952 | 0.017 | 0.013333 | 0.003506 | 0.002415 | 0.110223 | 0.009761 | 0.01531 |
| 25.7777778 | 0.026467 | 0.00098 | 0.008512 | 0.010576 | 0.005032 | 0.005379 | 0.027751 | 0.005014 | 0.00564 | 0.040923 | 0.015061 | 0.00862 | 0.014467 | 0.014654 | 0.026353 | 0.009893 | 0.027649 | 0.018655 | 0.007003 | 0.003149 | 0.004345 | 0.099524 | 0.010136 | 0.01462 |
| 25.7916667 | 0.029625 | 0.001898 | 0.01246 | 0.019892 | 0.00245 | 0.002726 | 0.027238 | 0.00486 | 0.003808 | 0.02583 | 0.021584 | 0.011598 | 0.015206 | 0.03073 | 0.026652 | 0.005668 | 0.041609 | 0.014628 | 0.003742 | 0.004749 | 0.005428 | 0.059867 | 0.017599 | 0.020144 |
| 25.8055556 | 0.02265 | 0.003847 | 0.015037 | 0.025807 | 0.00271 | 0.002504 | 0.022643 | 0.005415 | 0.002721 | 0.045986 | 0.02106 | 0.015956 | 0.01787 | 0.047986 | 0.034803 | 0.014276 | 0.046409 | 0.007712 | 0.007117 | 0.008217 | 0.005054 | 0.06454 | 0.022857 | 0.021091 |
| 25.8194444 | 0.015167 | 0.00504 | 0.016071 | 0.026405 | 0.00568 | 0.003316 | 0.016043 | 0.003008 | 0.001778 | 0.075401 | 0.013135 | 0.023614 | 0.030461 | 0.057463 | 0.036936 | 0.033667 | 0.029679 | 0.002645 | 0.014851 | 0.009894 | 0.007976 | 0.110701 | 0.020644 | 0.018815 |
| 25.8333333 | 0.025416 | 0.005201 | 0.015624 | 0.024701 | 0.007648 | 0.00473 | 0.009532 | 0.001264 | 0.002097 | 0.091119 | 0.014405 | 0.033278 | 0.040236 | 0.061091 | 0.024807 | 0.044093 | 0.016027 | 0.0027 | 0.022413 | 0.009025 | 0.014016 | 0.123269 | 0.012447 | 0.028577 |
| 25.8472222 | 0.046095 | 0.005787 | 0.012838 | 0.024321 | 0.005677 | 0.005876 | 0.004806 | 0.002146 | 0.003861 | 0.09036 | 0.023494 | 0.041792 | 0.03591 | 0.066165 | 0.011208 | 0.037662 | 0.021809 | 0.006814 | 0.024979 | 0.006938 | 0.015515 | 0.084453 | 0.009279 | 0.036155 |
| 25.8611111 | 0.05692 | 0.007874 | 0.009257 | 0.027314 | 0.003549 | 0.01058 | 0.002905 | 0.006769 | 0.010233 | 0.069614 | 0.028816 | 0.044842 | 0.025296 | 0.062976 | 0.010611 | 0.025872 | 0.031036 | 0.011911 | 0.022446 | 0.00633 | 0.01052 | 0.043822 | 0.016352 | 0.026215 |
| 25.875 | 0.054395 | 0.010227 | 0.008523 | 0.031089 | 0.005931 | 0.021062 | 0.002105 | 0.016076 | 0.02253 | 0.040971 | 0.025918 | 0.039577 | 0.021493 | 0.046111 | 0.016707 | 0.019705 | 0.027386 | 0.016478 | 0.018705 | 0.008981 | 0.005219 | 0.050175 | 0.023324 | 0.015582 |
| 25.8888889 | 0.046571 | 0.010359 | 0.01136 | 0.030381 | 0.009984 | 0.0292 | 0.001404 | 0.02769 | 0.036807 | 0.02219 | 0.017406 | 0.029732 | 0.022673 | 0.028177 | 0.02123 | 0.021589 | 0.018633 | 0.020459 | 0.016366 | 0.015983 | 0.003267 | 0.082047 | 0.025497 | 0.022432 |
| 25.9027778 | 0.039796 | 0.007673 | 0.01724 | 0.026349 | 0.012291 | 0.030354 | 0.001258 | 0.033035 | 0.04885 | 0.017766 | 0.010145 | 0.022209 | 0.019462 | 0.015684 | 0.027583 | 0.024053 | 0.021943 | 0.022085 | 0.015586 | 0.024341 | 0.003851 | 0.095572 | 0.027368 | 0.037509 |
| 25.9166667 | 0.037049 | 0.00388 | 0.02468 | 0.024337 | 0.011052 | 0.029563 | 0.002891 | 0.026655 | 0.053523 | 0.020855 | 0.010483 | 0.018313 | 0.011641 | 0.00818 | 0.034163 | 0.017608 | 0.039436 | 0.018397 | 0.014115 | 0.028331 | 0.004659 | 0.086803 | 0.029166 | 0.043188 |
| 25.9305556 | 0.034103 | 0.001575 | 0.026939 | 0.023923 | 0.00661 | 0.023742 | 0.008322 | 0.016407 | 0.045846 | 0.024615 | 0.017577 | 0.014438 | 0.007541 | 0.008599 | 0.03122 | 0.012535 | 0.053597 | 0.010323 | 0.009041 | 0.027673 | 0.004593 | 0.05358 | 0.027952 | 0.039441 |
| 25.9444444 | 0.023921 | 0.001592 | 0.020523 | 0.02238 | 0.002382 | 0.010776 | 0.015439 | 0.008947 | 0.028557 | 0.029035 | 0.024099 | 0.008216 | 0.007236 | 0.015686 | 0.020419 | 0.01899 | 0.050512 | 0.004246 | 0.004079 | 0.025204 | 0.004375 | 0.035725 | 0.024426 | 0.032486 |
| 25.9583333 | 0.012015 | 0.001921 | 0.011702 | 0.019313 | 0.000846 | 0.00516 | 0.019241 | 0.003863 | 0.014003 | 0.033424 | 0.024882 | 0.004557 | 0.005564 | 0.021764 | 0.010493 | 0.023921 | 0.032603 | 0.004604 | 0.004252 | 0.022077 | 0.003331 | 0.074073 | 0.020242 | 0.022904 |
| 25.9722222 | 0.005597 | 0.002188 | 0.005428 | 0.015141 | 0.001363 | 0.009855 | 0.016398 | 0.002093 | 0.010953 | 0.030762 | 0.019362 | 0.008657 | 0.006041 | 0.022706 | 0.007073 | 0.023527 | 0.016476 | 0.006891 | 0.004901 | 0.018829 | 0.001459 | 0.113701 | 0.017104 | 0.012777 |
| 25.9861111 | 0.004603 | 0.002598 | 0.002875 | 0.011427 | 0.002353 | 0.013906 | 0.008475 | 0.003964 | 0.017785 | 0.021765 | 0.011195 | 0.014663 | 0.009338 | 0.017038 | 0.007491 | 0.020458 | 0.01892 | 0.00675 | 0.003321 | 0.016453 | 0.000858 | 0.096709 | 0.018584 | 0.008961 |
| 26 | 0.010263 | 0.002502 | 0.005889 | 0.011026 | 0.005474 | 0.015991 | 0.002443 | 0.007442 | 0.029843 | 0.016884 | 0.007832 | 0.014828 | 0.011538 | 0.011242 | 0.007263 | 0.015351 | 0.033626 | 0.009675 | 0.002389 | 0.014219 | 0.001613 | 0.05337 | 0.024791 | 0.013414 |
| 26.0138889 | 0.02428 | 0.003335 | 0.012743 | 0.014627 | 0.012107 | 0.018178 | 0.002041 | 0.010321 | 0.042691 | 0.017987 | 0.013329 | 0.009489 | 0.008835 | 0.018418 | 0.009134 | 0.019586 | 0.037558 | 0.020422 | 0.002349 | 0.009691 | 0.004459 | 0.042838 | 0.033387 | 0.021647 |
| 26.0277778 | 0.045477 | 0.005202 | 0.017194 | 0.019546 | 0.018011 | 0.020819 | 0.005085 | 0.011652 | 0.054282 | 0.019199 | 0.021788 | 0.008737 | 0.005099 | 0.034392 | 0.019575 | 0.02951 | 0.024927 | 0.030795 | 0.007964 | 0.006007 | 0.008162 | 0.068728 | 0.046 | 0.031988 |
| 26.0416667 | 0.069664 | 0.008412 | 0.015215 | 0.025538 | 0.019249 | 0.027086 | 0.011797 | 0.009879 | 0.061862 | 0.020043 | 0.026035 | 0.021099 | 0.006199 | 0.049138 | 0.043391 | 0.030691 | 0.016726 | 0.033018 | 0.023015 | 0.010395 | 0.007047 | 0.096221 | 0.063006 | 0.044783 |
| 26.0555556 | 0.08564 | 0.014888 | 0.010601 | 0.032549 | 0.015252 | 0.035621 | 0.021868 | 0.005995 | 0.05923 | 0.021911 | 0.030781 | 0.039212 | 0.007479 | 0.057461 | 0.062825 | 0.024063 | 0.023814 | 0.030156 | 0.035758 | 0.020312 | 0.002558 | 0.095712 | 0.071168 | 0.053731 |
| 26.0694444 | 0.0791 | 0.021172 | 0.009153 | 0.033659 | 0.009486 | 0.038315 | 0.026264 | 0.003211 | 0.04783 | 0.022625 | 0.037028 | 0.048818 | 0.008743 | 0.051277 | 0.057275 | 0.015061 | 0.030187 | 0.023882 | 0.034625 | 0.025561 | 0.000308 | 0.059405 | 0.059356 | 0.050483 |
| 26.0833333 | 0.051965 | 0.021505 | 0.00977 | 0.023744 | 0.009449 | 0.031425 | 0.01976 | 0.002001 | 0.035388 | 0.02287 | 0.029262 | 0.042349 | 0.012763 | 0.033869 | 0.036733 | 0.008036 | 0.028357 | 0.014541 | 0.023492 | 0.022151 | 0.000114 | 0.038153 | 0.035997 | 0.033989 |
| 26.0972222 | 0.022958 | 0.016101 | 0.008233 | 0.011195 | 0.014005 | 0.01799 | 0.009636 | 0.003164 | 0.021829 | 0.028741 | 0.018878 | 0.025316 | 0.014458 | 0.016275 | 0.017641 | 0.008501 | 0.026039 | 0.007312 | 0.011519 | 0.012537 | 0.000355 | 0.070147 | 0.014757 | 0.015019 |
| 26.1111111 | 0.009042 | 0.008784 | 0.004671 | 0.007284 | 0.014841 | 0.007427 | 0.003167 | 0.005869 | 0.01182 | 0.03534 | 0.034282 | 0.010283 | 0.010435 | 0.00523 | 0.007603 | 0.014361 | 0.025899 | 0.006402 | 0.003915 | 0.005114 | 0.000971 | 0.116968 | 0.004209 | 0.0078 |
| 26.125 | 0.010412 | 0.003802 | 0.003184 | 0.009096 | 0.011409 | 0.005135 | 0.001619 | 0.006981 | 0.014589 | 0.026858 | 0.056779 | 0.003023 | 0.004462 | 0.001606 | 0.00701 | 0.017483 | 0.0251 | 0.010334 | 0.001413 | 0.004889 | 0.00179 | 0.142134 | 0.00335 | 0.012142 |
| 26.1388889 | 0.013359 | 0.003512 | 0.004528 | 0.00878 | 0.007003 | 0.004945 | 0.003396 | 0.005873 | 0.024313 | 0.01164 | 0.058106 | 0.0019 | 0.00143 | 0.002039 | 0.007722 | 0.015759 | 0.021237 | 0.014319 | 0.001523 | 0.005578 | 0.002556 | 0.141089 | 0.0037 | 0.018485 |
| 26.1527778 | 0.011079 | 0.004798 | 0.004355 | 0.00626 | 0.003539 | 0.006861 | 0.005846 | 0.004046 | 0.030336 | 0.007574 | 0.042005 | 0.002551 | 0.001848 | 0.003891 | 0.005451 | 0.014225 | 0.014758 | 0.01488 | 0.001434 | 0.003734 | 0.003143 | 0.116982 | 0.00218 | 0.024513 |
| 26.1666667 | 0.006101 | 0.004542 | 0.002753 | 0.003394 | 0.003368 | 0.013509 | 0.006944 | 0.002731 | 0.026631 | 0.008237 | 0.020614 | 0.002191 | 0.002708 | 0.003963 | 0.002141 | 0.015161 | 0.00811 | 0.011949 | 0.001403 | 0.002084 | 0.003697 | 0.078167 | 0.001303 | 0.034022 |
| 26.1805556 | 0.002338 | 0.003532 | 0.0031 | 0.001196 | 0.005184 | 0.017477 | 0.006386 | 0.002425 | 0.014567 | 0.006834 | 0.00726 | 0.001395 | 0.002119 | 0.003496 | 0.002678 | 0.016048 | 0.004249 | 0.007494 | 0.002337 | 0.001594 | 0.004482 | 0.039214 | 0.003458 | 0.043622 |
| 26.1944444 | 0.001414 | 0.002829 | 0.00349 | 0.000962 | 0.00549 | 0.013519 | 0.005437 | 0.003792 | 0.005563 | 0.005393 | 0.004921 | 0.001449 | 0.001415 | 0.007333 | 0.008277 | 0.015888 | 0.004299 | 0.003533 | 0.002542 | 0.001682 | 0.004907 | 0.017454 | 0.009044 | 0.04477 |

| | | | | | | | | | | | | | | | | | | | | | | | | |
|---|---|---|---|---|---|---|---|---|---|---|---|---|---|---|---|---|---|---|---|---|---|---|---|---|
| 26.2083333 | 0.001513 | 0.002648 | 0.002702 | 0.002677 | 0.003637 | 0.0067 | 0.005167 | 0.005059 | 0.005378 | 0.003507 | 0.006168 | 0.002491 | 0.002319 | 0.01315 | 0.013254 | 0.015001 | 0.006954 | 0.002431 | 0.001913 | 0.003577 | 0.004081 | 0.0206 | 0.01447 | 0.036705 |
| 26.2222222 | 0.003 | 0.0029 | 0.00346 | 0.005055 | 0.001753 | 0.003781 | 0.005101 | 0.003746 | 0.007341 | 0.001579 | 0.012492 | 0.003613 | 0.003527 | 0.016068 | 0.01542 | 0.013486 | 0.009499 | 0.004689 | 0.002924 | 0.006871 | 0.002466 | 0.032519 | 0.016707 | 0.026572 |
| 26.2361111 | 0.00517 | 0.003054 | 0.005881 | 0.006457 | 0.001693 | 0.004518 | 0.004209 | 0.002474 | 0.005587 | 0.000678 | 0.02025 | 0.004539 | 0.003137 | 0.015406 | 0.016657 | 0.011915 | 0.008414 | 0.007288 | 0.005391 | 0.008163 | 0.001638 | 0.029564 | 0.017252 | 0.0217 |
| 26.25 | 0.004799 | 0.002824 | 0.0078 | 0.006198 | 0.002342 | 0.004772 | 0.002402 | 0.002978 | 0.002498 | 0.000467 | 0.017106 | 0.005415 | 0.002101 | 0.012774 | 0.017836 | 0.010426 | 0.005358 | 0.007961 | 0.007159 | 0.006414 | 0.002201 | 0.017641 | 0.016398 | 0.01954 |
| 26.2638889 | 0.004654 | 0.002419 | 0.008313 | 0.004844 | 0.002419 | 0.004148 | 0.001331 | 0.00253 | 0.001295 | 0.000371 | 0.010021 | 0.005988 | 0.002107 | 0.009948 | 0.018826 | 0.008322 | 0.006572 | 0.007059 | 0.007196 | 0.004382 | 0.00352 | 0.013121 | 0.013629 | 0.012975 |
| 26.2777778 | 0.007717 | 0.001997 | 0.007897 | 0.003336 | 0.002171 | 0.004766 | 0.002121 | 0.003345 | 0.002114 | 0.000387 | 0.011834 | 0.006249 | 0.002931 | 0.007601 | 0.019149 | 0.006057 | 0.011949 | 0.005952 | 0.005631 | 0.003395 | 0.004717 | 0.01727 | 0.009921 | 0.007034 |
| 26.2916667 | 0.009397 | 0.001661 | 0.007436 | 0.00202 | 0.001707 | 0.006695 | 0.00316 | 0.007045 | 0.005762 | 0.000706 | 0.012331 | 0.006547 | 0.003356 | 0.005737 | 0.018379 | 0.007593 | 0.015901 | 0.006059 | 0.003516 | 0.002668 | 0.005036 | 0.041436 | 0.005401 | 0.010645 |
| 26.3055556 | 0.006874 | 0.001347 | 0.007197 | 0.001193 | 0.001006 | 0.008614 | 0.003335 | 0.00855 | 0.009529 | 0.001161 | 0.007962 | 0.006974 | 0.00275 | 0.004726 | 0.016959 | 0.012283 | 0.015309 | 0.007896 | 0.00195 | 0.002491 | 0.004318 | 0.087969 | 0.003025 | 0.019419 |
| 26.3194444 | 0.003123 | 0.000911 | 0.006631 | 0.001376 | 0.000828 | 0.009619 | 0.003132 | 0.005563 | 0.008209 | 0.001409 | 0.009351 | 0.00739 | 0.001702 | 0.005245 | 0.016259 | 0.013942 | 0.01079 | 0.010996 | 0.001422 | 0.003971 | 0.003244 | 0.12318 | 0.005542 | 0.023514 |
| 26.3333333 | 0.001912 | 0.000699 | 0.005492 | 0.001406 | 0.002159 | 0.010134 | 0.002702 | 0.003417 | 0.006564 | 0.002054 | 0.012661 | 0.007987 | 0.001232 | 0.006935 | 0.016565 | 0.01115 | 0.005106 | 0.013791 | 0.002068 | 0.007189 | 0.002956 | 0.105018 | 0.008779 | 0.02062 |
| 26.3472222 | 0.00382 | 0.001276 | 0.004129 | 0.001519 | 0.003881 | 0.011579 | 0.002257 | 0.00619 | 0.010704 | 0.004238 | 0.011428 | 0.009182 | 0.001592 | 0.008076 | 0.016738 | 0.006515 | 0.00142 | 0.014005 | 0.00346 | 0.008811 | 0.00368 | 0.063305 | 0.008991 | 0.014572 |
| 26.3611111 | 0.005835 | 0.002339 | 0.003041 | 0.003883 | 0.003448 | 0.014067 | 0.002183 | 0.009924 | 0.013925 | 0.008434 | 0.007811 | 0.01081 | 0.003152 | 0.00717 | 0.015728 | 0.004541 | 0.001634 | 0.011452 | 0.004353 | 0.006683 | 0.004745 | 0.067576 | 0.007855 | 0.009996 |
| 26.375 | 0.006448 | 0.003109 | 0.002802 | 0.007357 | 0.002146 | 0.015598 | 0.001811 | 0.010596 | 0.011542 | 0.012495 | 0.004622 | 0.01141 | 0.005196 | 0.004172 | 0.013703 | 0.007161 | 0.004613 | 0.008817 | 0.003995 | 0.006005 | 0.005311 | 0.089021 | 0.006845 | 0.008884 |
| 26.3888889 | 0.007627 | 0.003498 | 0.003818 | 0.010469 | 0.002989 | 0.014755 | 0.001702 | 0.007106 | 0.006427 | 0.013096 | 0.003855 | 0.010294 | 0.005322 | 0.001863 | 0.013219 | 0.009436 | 0.007483 | 0.007886 | 0.003145 | 0.010563 | 0.005239 | 0.070646 | 0.004978 | 0.008611 |
| 26.4027778 | 0.01189 | 0.003716 | 0.00527 | 0.014183 | 0.00441 | 0.012638 | 0.004 | 0.003464 | 0.003644 | 0.010962 | 0.007062 | 0.009374 | 0.00318 | 0.002194 | 0.015954 | 0.008128 | 0.008082 | 0.008107 | 0.00288 | 0.012398 | 0.004898 | 0.040049 | 0.004518 | 0.005771 |
| 26.4166667 | 0.019037 | 0.003659 | 0.005814 | 0.018542 | 0.005051 | 0.011731 | 0.007444 | 0.003512 | 0.006469 | 0.009971 | 0.012319 | 0.009706 | 0.00139 | 0.002221 | 0.019497 | 0.005743 | 0.005562 | 0.008854 | 0.003819 | 0.008255 | 0.004461 | 0.043793 | 0.008731 | 0.004293 |
| 26.4305556 | 0.025612 | 0.003248 | 0.005593 | 0.02148 | 0.005918 | 0.013535 | 0.009474 | 0.003663 | 0.010702 | 0.012209 | 0.016431 | 0.010053 | 0.001314 | 0.002323 | 0.021535 | 0.005394 | 0.002468 | 0.009995 | 0.006238 | 0.007311 | 0.004763 | 0.063486 | 0.015761 | 0.011021 |
| 26.4444444 | 0.027889 | 0.002523 | 0.005291 | 0.022033 | 0.006404 | 0.016944 | 0.010015 | 0.003056 | 0.015446 | 0.016928 | 0.019132 | 0.010018 | 0.00295 | 0.006504 | 0.023034 | 0.007528 | 0.001492 | 0.011268 | 0.009336 | 0.012052 | 0.006478 | 0.053956 | 0.024168 | 0.023081 |
| 26.4583333 | 0.025132 | 0.001569 | 0.005104 | 0.022151 | 0.005166 | 0.018973 | 0.009852 | 0.003851 | 0.021545 | 0.020479 | 0.018591 | 0.010403 | 0.006121 | 0.012573 | 0.024946 | 0.010478 | 0.00169 | 0.011805 | 0.010603 | 0.014096 | 0.008416 | 0.033653 | 0.032956 | 0.031435 |
| 26.4722222 | 0.020672 | 0.00096 | 0.005304 | 0.023358 | 0.00285 | 0.01816 | 0.009723 | 0.006279 | 0.019613 | 0.020038 | 0.011881 | 0.0113 | 0.008574 | 0.017219 | 0.029106 | 0.013024 | 0.002073 | 0.011451 | 0.009549 | 0.008957 | 0.009175 | 0.050483 | 0.039234 | 0.032935 |
| 26.4861111 | 0.019199 | 0.001542 | 0.006694 | 0.026703 | 0.002173 | 0.016466 | 0.010548 | 0.00973 | 0.013712 | 0.01981 | 0.005302 | 0.01371 | 0.0105 | 0.024742 | 0.044766 | 0.019212 | 0.003819 | 0.016127 | 0.011439 | 0.003952 | 0.010832 | 0.092729 | 0.044449 | 0.032436 |
| 26.5 | 0.025824 | 0.003745 | 0.010088 | 0.036867 | 0.004546 | 0.017486 | 0.014458 | 0.013282 | 0.020488 | 0.028444 | 0.012888 | 0.021222 | 0.01596 | 0.038771 | 0.073694 | 0.030801 | 0.01091 | 0.034284 | 0.020431 | 0.008089 | 0.014974 | 0.118012 | 0.050184 | 0.032813 |
| 26.5138889 | 0.04503 | 0.007873 | 0.014633 | 0.053474 | 0.008174 | 0.022178 | 0.021727 | 0.020393 | 0.029454 | 0.050928 | 0.040024 | 0.033028 | 0.024409 | 0.052296 | 0.091655 | 0.039172 | 0.025349 | 0.058118 | 0.029731 | 0.018616 | 0.01705 | 0.114868 | 0.047651 | 0.029918 |
| 26.5277778 | 0.069782 | 0.010923 | 0.017137 | 0.061026 | 0.01061 | 0.025283 | 0.027285 | 0.036335 | 0.023155 | 0.082063 | 0.073983 | 0.0411 | 0.035813 | 0.062378 | 0.079284 | 0.03959 | 0.038427 | 0.067039 | 0.028368 | 0.027506 | 0.015659 | 0.099674 | 0.034779 | 0.022582 |
| 26.5416667 | 0.085465 | 0.0103 | 0.015488 | 0.051302 | 0.011923 | 0.022772 | 0.026999 | 0.051175 | 0.013095 | 0.113902 | 0.090069 | 0.03942 | 0.056409 | 0.066428 | 0.055169 | 0.037714 | 0.036381 | 0.057564 | 0.020649 | 0.034431 | 0.018618 | 0.081336 | 0.025286 | 0.016314 |
| 26.5555556 | 0.090202 | 0.009952 | 0.012482 | 0.038052 | 0.012195 | 0.01719 | 0.02321 | 0.049139 | 0.013545 | 0.13647 | 0.076212 | 0.029876 | 0.073088 | 0.056646 | 0.042025 | 0.033309 | 0.020513 | 0.039769 | 0.019458 | 0.040136 | 0.026703 | 0.051974 | 0.021222 | 0.010954 |
| 26.5694444 | 0.088202 | 0.011277 | 0.011788 | 0.030087 | 0.009666 | 0.009866 | 0.01896 | 0.035435 | 0.021262 | 0.138718 | 0.045328 | 0.019059 | 0.067339 | 0.040712 | 0.036064 | 0.022588 | 0.012688 | 0.020687 | 0.02302 | 0.040607 | 0.029187 | 0.027286 | 0.014931 | 0.00711 |
| 26.5833333 | 0.080029 | 0.01127 | 0.013141 | 0.021312 | 0.005233 | 0.005101 | 0.014063 | 0.020501 | 0.029129 | 0.122689 | 0.021822 | 0.010864 | 0.050327 | 0.027204 | 0.03175 | 0.012675 | 0.023919 | 0.01079 | 0.022134 | 0.03427 | 0.023605 | 0.037276 | 0.009515 | 0.013581 |
| 26.5972222 | 0.067765 | 0.009297 | 0.013765 | 0.01083 | 0.001883 | 0.007279 | 0.008209 | 0.010492 | 0.028855 | 0.090439 | 0.023313 | 0.005476 | 0.032878 | 0.015214 | 0.031607 | 0.008493 | 0.03722 | 0.012422 | 0.014067 | 0.023818 | 0.016119 | 0.071455 | 0.010632 | 0.027218 |
| 26.6111111 | 0.052196 | 0.006309 | 0.012376 | 0.005484 | 0.000445 | 0.00994 | 0.004171 | 0.010514 | 0.018908 | 0.047626 | 0.041365 | 0.003601 | 0.017017 | 0.007448 | 0.033414 | 0.009685 | 0.039645 | 0.011879 | 0.006568 | 0.013389 | 0.010901 | 0.10987 | 0.016514 | 0.031886 |
| 26.625 | 0.034152 | 0.003306 | 0.008587 | 0.005523 | 0.000223 | 0.007954 | 0.004761 | 0.015878 | 0.010484 | 0.028209 | 0.05012 | 0.007075 | 0.011513 | 0.009204 | 0.03092 | 0.017003 | 0.033928 | 0.008941 | 0.007337 | 0.006383 | 0.0095 | 0.13518 | 0.023606 | 0.023073 |
| 26.6388889 | 0.01816 | 0.001536 | 0.003316 | 0.006421 | 0.000978 | 0.004185 | 0.005251 | 0.024142 | 0.012196 | 0.047431 | 0.03504 | 0.015982 | 0.014852 | 0.018904 | 0.023341 | 0.02516 | 0.023819 | 0.014891 | 0.010927 | 0.002914 | 0.009931 | 0.101678 | 0.031057 | 0.02099 |
| 26.6527778 | 0.014367 | 0.001986 | 0.000446 | 0.007421 | 0.003008 | 0.002205 | 0.004804 | 0.028434 | 0.014372 | 0.075067 | 0.022846 | 0.026581 | 0.019162 | 0.029125 | 0.014739 | 0.026193 | 0.012612 | 0.024449 | 0.012385 | 0.001466 | 0.009515 | 0.056219 | 0.034091 | 0.034066 |
| 26.6666667 | 0.027482 | 0.003725 | 0.000769 | 0.007077 | 0.006708 | 0.0022 | 0.009983 | 0.018268 | 0.011627 | 0.095576 | 0.040715 | 0.033983 | 0.028473 | 0.037344 | 0.007247 | 0.018526 | 0.005544 | 0.031337 | 0.013035 | 0.001245 | 0.008272 | 0.076966 | 0.025975 | 0.037044 |
| 26.6805556 | 0.044635 | 0.005097 | 0.002019 | 0.009047 | 0.011688 | 0.002163 | 0.017098 | 0.00559 | 0.009376 | 0.101019 | 0.057825 | 0.03396 | 0.041264 | 0.045217 | 0.006329 | 0.012908 | 0.003687 | 0.033823 | 0.013777 | 0.001515 | 0.010323 | 0.106404 | 0.011716 | 0.020505 |
| 26.6944444 | 0.051899 | 0.005915 | 0.004439 | 0.018677 | 0.015016 | 0.005243 | 0.01758 | 0.003496 | 0.00867 | 0.07672 | 0.044012 | 0.024538 | 0.040534 | 0.047456 | 0.014373 | 0.023147 | 0.005802 | 0.024893 | 0.010714 | 0.002146 | 0.015608 | 0.094131 | 0.00352 | 0.007709 |
| 26.7083333 | 0.047785 | 0.006815 | 0.009165 | 0.028576 | 0.013476 | 0.015578 | 0.010606 | 0.005575 | 0.011586 | 0.039212 | 0.022853 | 0.012639 | 0.023961 | 0.03463 | 0.020884 | 0.038648 | 0.015221 | 0.013399 | 0.007551 | 0.00603 | 0.017873 | 0.056898 | 0.002702 | 0.011183 |
| 26.7222222 | 0.036284 | 0.00783 | 0.011274 | 0.030889 | 0.007844 | 0.024913 | 0.005815 | 0.005083 | 0.01012 | 0.013901 | 0.028652 | 0.010014 | 0.008623 | 0.018622 | 0.022445 | 0.036124 | 0.028235 | 0.019071 | 0.013635 | 0.0138 | 0.016553 | 0.028636 | 0.002635 | 0.019063 |
| 26.7361111 | 0.027155 | 0.008283 | 0.00881 | 0.025017 | 0.005494 | 0.025402 | 0.010778 | 0.003473 | 0.005631 | 0.003726 | 0.057073 | 0.017979 | 0.007358 | 0.022239 | 0.026912 | 0.021752 | 0.038187 | 0.035894 | 0.020011 | 0.018819 | 0.015723 | 0.044091 | 0.002251 | 0.021365 |
| 26.75 | 0.027312 | 0.007838 | 0.006192 | 0.014098 | 0.00987 | 0.020406 | 0.018862 | 0.003346 | 0.007772 | 0.000971 | 0.082081 | 0.026211 | 0.021311 | 0.041101 | 0.028752 | 0.022605 | 0.043584 | 0.041838 | 0.015002 | 0.019696 | 0.016899 | 0.082276 | 0.003774 | 0.021392 |
| 26.7638889 | 0.025357 | 0.006799 | 0.00339 | 0.008413 | 0.012994 | 0.012618 | 0.023457 | 0.003326 | 0.008486 | 0.000681 | 0.087562 | 0.026711 | 0.033845 | 0.050381 | 0.018707 | 0.039581 | 0.042277 | 0.035204 | 0.006305 | 0.019003 | 0.018383 | 0.114643 | 0.004549 | 0.027651 |
| 26.7777778 | 0.014156 | 0.005787 | 0.00111 | 0.014823 | 0.010812 | 0.005557 | 0.023487 | 0.003435 | 0.007547 | 0.001468 | 0.068967 | 0.020194 | 0.029442 | 0.040842 | 0.006884 | 0.049111 | 0.035246 | 0.0249 | 0.0052 | 0.013079 | 0.017236 | 0.125705 | 0.006811 | 0.037542 |
| 26.7916667 | 0.005526 | 0.004965 | 0.001043 | 0.024307 | 0.006137 | 0.002781 | 0.018457 | 0.00602 | 0.011692 | 0.00266 | 0.037073 | 0.010745 | 0.018014 | 0.024645 | 0.00408 | 0.044087 | 0.026851 | 0.012625 | 0.007685 | 0.00649 | 0.012019 | 0.093232 | 0.01816 | 0.041516 |
| 26.8055556 | 0.004127 | 0.003807 | 0.003642 | 0.025944 | 0.002577 | 0.002291 | 0.010919 | 0.008025 | 0.013677 | 0.00345 | 0.019526 | 0.005925 | 0.009015 | 0.011768 | 0.00744 | 0.031588 | 0.019925 | 0.003865 | 0.007668 | 0.007502 | 0.005694 | 0.046573 | 0.032757 | 0.038748 |
| 26.8194444 | 0.006168 | 0.002279 | 0.008573 | 0.019364 | 0.002904 | 0.005309 | 0.005041 | 0.008488 | 0.009666 | 0.004229 | 0.023407 | 0.010148 | 0.007214 | 0.007461 | 0.012797 | 0.018289 | 0.016799 | 0.005755 | 0.006213 | 0.01049 | 0.002764 | 0.037015 | 0.041146 | 0.02914 |
| 26.8333333 | 0.015307 | 0.001149 | 0.013452 | 0.011403 | 0.0048 | 0.014374 | 0.002622 | 0.007961 | 0.005752 | 0.005775 | 0.025671 | 0.018279 | 0.018804 | 0.013361 | 0.01617 | 0.010595 | 0.019694 | 0.015621 | 0.006713 | 0.008065 | 0.002912 | 0.068023 | 0.043435 | 0.016911 |
| 26.8472222 | 0.028665 | 0.001099 | 0.017744 | 0.010764 | 0.004294 | 0.024787 | 0.003797 | 0.010413 | 0.007492 | 0.007718 | 0.016208 | 0.024522 | 0.037663 | 0.021733 | 0.018038 | 0.01084 | 0.027049 | 0.026096 | 0.014014 | 0.006585 | 0.0052 | 0.111611 | 0.039977 | 0.009904 |
| 26.8611111 | 0.041533 | 0.001817 | 0.021844 | 0.020781 | 0.004117 | 0.028964 | 0.007969 | 0.022506 | 0.016835 | 0.010035 | 0.011938 | 0.026914 | 0.056291 | 0.027522 | 0.023931 | 0.0165 | 0.031516 | 0.032511 | 0.026904 | 0.014537 | 0.011584 | 0.115669 | 0.03033 | 0.007971 |
| 26.875 | 0.05375 | 0.002053 | 0.024325 | 0.036192 | 0.008856 | 0.024352 | 0.012746 | 0.034459 | 0.029304 | 0.013401 | 0.028987 | 0.02712 | 0.068121 | 0.034156 | 0.040402 | 0.022799 | 0.027524 | 0.033869 | 0.03518 | 0.026236 | 0.019011 | 0.076536 | 0.018895 | 0.008663 |
| 26.8888889 | 0.056989 | 0.001348 | 0.023062 | 0.047938 | 0.013776 | 0.017601 | 0.015058 | 0.036281 | 0.038602 | 0.017957 | 0.053545 | 0.027706 | 0.065257 | 0.048774 | 0.064432 | 0.027412 | 0.017983 | 0.031708 | 0.032358 | 0.033181 | 0.020518 | 0.06415 | 0.012183 | 0.008516 |
| 26.9027778 | 0.047623 | 0.000988 | 0.01858 | 0.051372 | 0.012691 | 0.016954 | 0.01383 | 0.029703 | 0.042107 | 0.021486 | 0.060998 | 0.027422 | 0.048986 | 0.064429 | 0.081715 | 0.032375 | 0.009532 | 0.027623 | 0.023379 | 0.033472 | 0.014901 | 0.07608 | 0.011547 | 0.006617 |
| 26.9166667 | 0.0333 | 0.001497 | 0.013448 | 0.047414 | 0.007831 | 0.021307 | 0.009923 | 0.018616 | 0.039574 | 0.019685 | 0.042608 | 0.021985 | 0.027974 | 0.06381 | 0.085731 | 0.042316 | 0.008922 | 0.022871 | 0.014481 | 0.02555 | 0.008168 | 0.078421 | 0.012279 | 0.008731 |
| 26.9305556 | 0.02254 | 0.001913 | 0.009025 | 0.037389 | 0.007259 | 0.023828 | 0.004805 | 0.008576 | 0.030002 | 0.012614 | 0.020464 | 0.011957 | 0.010999 | 0.049149 | 0.077767 | 0.04944 | 0.013973 | 0.019804 | 0.00742 | 0.014874 | 0.005701 | 0.072658 | 0.01047 | 0.014884 |
| 26.9444444 | 0.017685 | 0.00259 | 0.00547 | 0.023074 | 0.012339 | 0.0208 | 0.001243 | 0.003174 | 0.016072 | 0.006496 | 0.018353 | 0.004216 | 0.004002 | 0.03113 | 0.063021 | 0.036898 | 0.017178 | 0.019367 | 0.003373 | 0.014282 | 0.00496 | 0.045419 | 0.007564 | 0.01932 |
| 26.9583333 | 0.015197 | 0.003898 | 0.00362 | 0.009752 | 0.012962 | 0.013354 | 0.000856 | 0.001699 | 0.010081 | 0.005908 | 0.020459 | 0.00251 | 0.005794 | 0.014987 | 0.044803 | 0.01846 | 0.019063 | 0.018832 | 0.002632 | 0.022122 | 0.004062 | 0.026799 | 0.008867 | 0.020855 |
| 26.9722222 | 0.012355 | 0.003936 | 0.005563 | 0.00347 | 0.008183 | 0.006011 | 0.002019 | 0.00342 | 0.016218 | 0.009142 | 0.012654 | 0.003954 | 0.008543 | 0.00685 | 0.025845 | 0.017304 | 0.018108 | 0.016651 | 0.003313 | 0.025514 | 0.006623 | 0.052512 | 0.012163 | 0.019236 |
| 26.9861111 | 0.009833 | 0.002102 | 0.00898 | 0.0027 | 0.008189 | 0.002366 | 0.002146 | 0.00821 | 0.020199 | 0.012415 | 0.007802 | 0.005221 | 0.006634 | 0.005467 | 0.015572 | 0.017309 | 0.013086 | 0.015105 | 0.006732 | 0.019224 | 0.009813 | 0.084556 | 0.013711 | 0.013834 |
| 27 | 0.010276 | 0.000546 | 0.007683 | 0.004941 | 0.013358 | 0.005236 | 0.002363 | 0.009024 | 0.015368 | 0.013831 | 0.015465 | 0.004241 | 0.004788 | 0.007496 | 0.016972 | 0.015517 | 0.013733 | 0.015287 | 0.013556 | 0.010334 | 0.007536 | 0.070691 | 0.025931 | 0.014673 |
| 27.0138889 | 0.017635 | 0.000304 | 0.005946 | 0.012301 | 0.017721 | 0.016328 | 0.005007 | 0.007129 | 0.018427 | 0.012991 | 0.030987 | 0.003354 | 0.008839 | 0.014144 | 0.021399 | 0.029584 | 0.02276 | 0.01336 | 0.018108 | 0.011674 | 0.003798 | 0.039605 | 0.049925 | 0.028819 |
| 27.0277778 | 0.030349 | 0.000989 | 0.009632 | 0.020585 | 0.018895 | 0.027923 | 0.007472 | 0.00958 | 0.042131 | 0.011077 | 0.046868 | 0.006654 | 0.015572 | 0.023685 | 0.024778 | 0.043007 | 0.032124 | 0.007899 | 0.018953 | 0.020013 | 0.005648 | 0.05045 | 0.06663 | 0.044432 |
| 27.0416667 | 0.042183 | 0.002724 | 0.010789 | 0.027293 | 0.0176 | 0.031765 | 0.009273 | 0.011593 | 0.07718 | 0.011332 | 0.054493 | 0.010459 | 0.021172 | 0.035514 | 0.032851 | 0.045097 | 0.037816 | 0.004237 | 0.022013 | 0.022988 | 0.009918 | 0.085434 | 0.068523 | 0.051829 |
| 27.0555556 | 0.053225 | 0.008244 | 0.006623 | 0.03894 | 0.016834 | 0.030126 | 0.012336 | 0.011344 | 0.113516 | 0.014903 | 0.048997 | 0.011508 | 0.022781 | 0.045866 | 0.042488 | 0.046479 | 0.03566 | 0.003019 | 0.026593 | 0.023468 | 0.009296 | 0.111854 | 0.057196 | 0.050563 |
| 27.0694444 | 0.061624 | 0.016819 | 0.003097 | 0.052949 | 0.01513 | 0.025393 | 0.013502 | 0.00944 | 0.126376 | 0.017473 | 0.036178 | 0.010947 | 0.018244 | 0.046499 | 0.042579 | 0.044773 | 0.02331 | 0.00265 | 0.024841 | 0.026077 | 0.009494 | 0.111824 | 0.036133 | 0.03834 |
| 27.0833333 | 0.056842 | 0.020484 | 0.001921 | 0.054453 | 0.009649 | 0.016115 | 0.010281 | 0.006608 | 0.085002 | 0.015021 | 0.021856 | 0.008357 | 0.009908 | 0.037547 | 0.033735 | 0.035928 | 0.010404 | 0.00455 | 0.017198 | 0.024359 | 0.015445 | 0.074023 | 0.016123 | 0.019088 |
| 27.0972222 | 0.037104 | 0.017034 | 0.003094 | 0.04042 | 0.006335 | 0.007897 | 0.005017 | 0.008711 | 0.038138 | 0.009248 | 0.011207 | 0.004448 | 0.004526 | 0.026545 | 0.02342 | 0.025317 | 0.008264 | 0.004823 | 0.010714 | 0.01508 | 0.015842 | 0.036539 | 0.00761 | 0.007551 |
| 27.1111111 | 0.016592 | 0.01109 | 0.005846 | 0.023218 | 0.008292 | 0.008231 | 0.001851 | 0.015442 | 0.04761 | 0.004334 | 0.012387 | 0.002014 | 0.005904 | 0.019371 | 0.014984 | 0.017317 | 0.015277 | 0.00275 | 0.00785 | 0.007131 | 0.009307 | 0.039544 | 0.010567 | 0.007467 |
| 27.125 | 0.00865 | 0.005457 | 0.009056 | 0.012015 | 0.009608 | 0.012698 | 0.001913 | 0.017268 | 0.075934 | 0.00179 | 0.025426 | 0.001229 | 0.008536 | 0.017016 | 0.008763 | 0.011749 | 0.023196 | 0.002837 | 0.006865 | 0.008592 | 0.008219 | 0.067321 | 0.016452 | 0.008182 |
| 27.1388889 | 0.010819 | 0.002312 | 0.013079 | 0.007438 | 0.008759 | 0.013722 | 0.002069 | 0.011878 | 0.080636 | 0.000971 | 0.03949 | 0.001224 | 0.008398 | 0.017408 | 0.004575 | 0.007488 | 0.026845 | 0.003499 | 0.004991 | 0.014926 | 0.016526 | 0.090567 | 0.019375 | 0.005131 |
| 27.1527778 | 0.011164 | 0.002899 | 0.015962 | 0.006636 | 0.007956 | 0.010765 | 0.001098 | 0.006854 | 0.070557 | 0.000867 | 0.045575 | 0.002173 | 0.007091 | 0.017674 | 0.002097 | 0.003893 | 0.025914 | 0.002501 | 0.002906 | 0.019793 | 0.024678 | 0.104188 | 0.019395 | 0.002717 |
| 27.1666667 | 0.00708 | 0.005057 | 0.013291 | 0.006672 | 0.007291 | 0.006011 | 0.000305 | 0.007325 | 0.054658 | 0.001027 | 0.043895 | 0.00252 | 0.00595 | 0.015869 | 0.000956 | 0.001283 | 0.023115 | 0.002306 | 0.003047 | 0.021308 | 0.020546 | 0.10742 | 0.017785 | 0.004242 |
| 27.1805556 | 0.005031 | 0.006889 | 0.007242 | 0.006396 | 0.006342 | 0.00232 | 0.001092 | 0.009348 | 0.03931 | 0.002093 | 0.0375 | 0.002441 | 0.004603 | 0.010223 | 0.001203 | 0.00169 | 0.017278 | 0.003387 | 0.003397 | 0.018447 | 0.016908 | 0.093967 | 0.016143 | 0.005888 |
| 27.1944444 | 0.00556 | 0.009571 | 0.005644 | 0.008051 | 0.005989 | 0.002759 | 0.007839 | 0.014544 | 0.052924 | 0.012063 | 0.026721 | 0.008156 | 0.006903 | 0.012993 | 0.005461 | 0.011259 | 0.010921 | 0.005531 | 0.0034 | 0.011083 | 0.03347 | 0.06062 | 0.017035 | 0.003988 |
| 27.2083333 | 0.008098 | 0.010644 | 0.008605 | 0.013663 | 0.006089 | 0.00467 | 0.019076 | 0.030212 | 0.085629 | 0.028353 | 0.01384 | 0.018191 | 0.018114 | 0.023381 | 0.019587 | 0.030175 | 0.00742 | 0.015067 | 0.006289 | 0.005911 | 0.059297 | 0.040052 | 0.021957 | 0.006328 |
| 27.2222222 | 0.015533 | 0.008371 | 0.009588 | 0.019653 | 0.004537 | 0.006417 | 0.029615 | 0.039225 | 0.071016 | 0.033318 | 0.007147 | 0.02546 | 0.028599 | 0.033654 | 0.029274 | 0.037665 | 0.015345 | 0.024715 | 0.009218 | 0.012765 | 0.062316 | 0.068408 | 0.025851 | 0.019388 |
| 27.2361111 | 0.034315 | 0.011838 | 0.006641 | 0.034785 | 0.002583 | 0.010746 | 0.049633 | 0.048198 | 0.028351 | 0.044471 | 0.012122 | 0.043863 | 0.028207 | 0.058823 | 0.023439 | 0.043746 | 0.041276 | 0.02373 | 0.008557 | 0.020281 | 0.044532 | 0.10526 | 0.032434 | 0.025183 |
| 27.25 | 0.046434 | 0.01617 | 0.005278 | 0.051446 | 0.003821 | 0.014131 | 0.064983 | 0.066446 | 0.034362 | 0.068987 | 0.017055 | 0.061939 | 0.023866 | 0.072626 | 0.022864 | 0.058489 | 0.049196 | 0.018759 | 0.006088 | 0.016934 | 0.036049 | 0.092793 | 0.041935 | 0.014582 |
| 27.2638889 | 0.030743 | 0.010853 | 0.008732 | 0.040398 | 0.006439 | 0.011641 | 0.046406 | 0.056139 | 0.055107 | 0.060813 | 0.013625 | 0.046635 | 0.016934 | 0.049267 | 0.024676 | 0.046355 | 0.02805 | 0.010822 | 0.00381 | 0.012313 | 0.02487 | 0.048159 | 0.032626 | 0.005339 |
| 27.2777778 | 0.013574 | 0.006848 | 0.012119 | 0.016846 | 0.005101 | 0.007558 | 0.022395 | 0.026838 | 0.045232 | 0.030118 | 0.012019 | 0.025224 | 0.006373 | 0.02088 | 0.014853 | 0.020531 | 0.013678 | 0.00444 | 0.003191 | 0.007719 | 0.009444 | 0.02161 | 0.016302 | 0.003295 |
| 27.2916667 | 0.00774 | 0.007189 | 0.010912 | 0.00498 | 0.001648 | 0.007181 | 0.017023 | 0.01569 | 0.037895 | 0.016751 | 0.01152 | 0.020951 | 0.001268 | 0.010234 | 0.005077 | 0.011193 | 0.007849 | 0.006368 | 0.00296 | 0.003341 | 0.009488 | 0.026265 | 0.017206 | 0.00321 |
| 27.3055556 | 0.004605 | 0.005535 | 0.006276 | 0.001649 | 0.000777 | 0.006292 | 0.013413 | 0.0155 | 0.029386 | 0.014889 | 0.008565 | 0.015957 | 0.001828 | 0.007785 | 0.001651 | 0.012928 | 0.00306 | 0.009473 | 0.003642 | 0.002743 | 0.013708 | 0.04266 | 0.017752 | 0.003908 |
| 27.3194444 | 0.003951 | 0.00298 | 0.003904 | 0.002149 | 0.00189 | 0.002975 | 0.014278 | 0.011244 | 0.021431 | 0.025746 | 0.011621 | 0.0152 | 0.003179 | 0.004772 | 0.001473 | 0.019601 | 0.003573 | 0.007254 | 0.006011 | 0.002916 | 0.011235 | 0.052388 | 0.011337 | 0.006953 |
| 27.3333333 | 0.003483 | 0.001487 | 0.003853 | 0.004599 | 0.003231 | 0.000938 | 0.02049 | 0.004414 | 0.032393 | 0.036239 | 0.019486 | 0.021317 | 0.004076 | 0.004654 | 0.003635 | 0.025704 | 0.005936 | 0.004476 | 0.007763 | 0.00601 | 0.00595 | 0.049907 | 0.014119 | 0.011118 |
| 27.3472222 | 0.004946 | 0.001488 | 0.005993 | 0.006408 | 0.004068 | 0.001194 | 0.017022 | 0.001048 | 0.032646 | 0.025853 | 0.021247 | 0.018876 | 0.00369 | 0.009142 | 0.007578 | 0.029216 | 0.00656 | 0.007111 | 0.007059 | 0.013529 | 0.003979 | 0.03848 | 0.023876 | 0.013885 |
| 27.3611111 | 0.007195 | 0.002339 | 0.015131 | 0.005429 | 0.003795 | 0.003021 | 0.010894 | 0.001162 | 0.018701 | 0.013701 | 0.014102 | 0.013854 | 0.00247 | 0.01258 | 0.012248 | 0.031004 | 0.004546 | 0.00997 | 0.003876 | 0.019874 | 0.004517 | 0.024543 | 0.026824 | 0.015942 |
| 27.375 | 0.008145 | 0.003126 | 0.023397 | 0.003733 | 0.003421 | 0.005011 | 0.01001 | 0.005074 | 0.014919 | 0.011421 | 0.009612 | 0.012161 | 0.00197 | 0.009673 | 0.015838 | 0.030852 | 0.00282 | 0.008679 | 0.001754 | 0.019516 | 0.004366 | 0.012909 | 0.019035 | 0.01556 |
| 27.3888889 | 0.009754 | 0.002436 | 0.020035 | 0.003396 | 0.004199 | 0.00664 | 0.009332 | 0.011857 | 0.01326 | 0.011575 | 0.014155 | 0.012223 | 0.003035 | 0.006409 | 0.01712 | 0.028697 | 0.003191 | 0.004994 | 0.003253 | 0.012701 | 0.007314 | 0.006699 | 0.013396 | 0.011152 |
| 27.4027778 | 0.011717 | 0.001226 | 0.011041 | 0.00376 | 0.004025 | 0.008828 | 0.008636 | 0.013968 | 0.005158 | 0.013985 | 0.016694 | 0.017614 | 0.005296 | 0.006065 | 0.016742 | 0.027475 | 0.003018 | 0.002369 | 0.005922 | 0.006065 | 0.013716 | 0.011081 | 0.013878 | 0.006559 |
| 27.4166667 | 0.009363 | 0.002537 | 0.006427 | 0.004129 | 0.002725 | 0.011984 | 0.011098 | 0.009842 | 0.005987 | 0.016102 | 0.01188 | 0.019839 | 0.010932 | 0.00698 | 0.01551 | 0.028805 | 0.002947 | 0.00262 | 0.005325 | 0.006189 | 0.015837 | 0.025269 | 0.013012 | 0.007472 |
| 27.4305556 | 0.004141 | 0.005185 | 0.006909 | 0.007733 | 0.00405 | 0.015078 | 0.021404 | 0.005999 | 0.015796 | 0.021409 | 0.009168 | 0.021272 | 0.017494 | 0.01315 | 0.014934 | 0.02817 | 0.004818 | 0.004726 | 0.003248 | 0.012307 | 0.012458 | 0.035856 | 0.01965 | 0.013576 |
| 27.4444444 | 0.005632 | 0.004911 | 0.008549 | 0.016082 | 0.007112 | 0.015907 | 0.02579 | 0.005903 | 0.018358 | 0.022998 | 0.01241 | 0.021488 | 0.017463 | 0.018119 | 0.015505 | 0.023997 | 0.006537 | 0.008215 | 0.004115 | 0.019425 | 0.008337 | 0.032915 | 0.033895 | 0.018726 |
| 27.4583333 | 0.014276 | 0.00244 | 0.008764 | 0.024094 | 0.007777 | 0.012882 | 0.014613 | 0.010203 | 0.01907 | 0.013264 | 0.01256 | 0.013831 | 0.012291 | 0.017212 | 0.015122 | 0.020918 | 0.006615 | 0.009302 | 0.005261 | 0.023259 | 0.004735 | 0.022695 | 0.035927 | 0.017826 |
| 27.4722222 | 0.018199 | 0.001237 | 0.008905 | 0.02459 | 0.006184 | 0.009569 | 0.007992 | 0.011089 | 0.022651 | 0.01439 | 0.007095 | 0.011841 | 0.008036 | 0.01551 | 0.014176 | 0.02163 | 0.005417 | 0.006014 | 0.005095 | 0.021962 | 0.003443 | 0.021214 | 0.020467 | 0.012255 |
| 27.4861111 | 0.011557 | 0.003003 | 0.010427 | 0.018626 | 0.005229 | 0.012593 | 0.021833 | 0.009638 | 0.035199 | 0.044677 | 0.003688 | 0.028131 | 0.006173 | 0.016862 | 0.017955 | 0.030027 | 0.008796 | 0.004449 | 0.009966 | 0.020662 | 0.00873 | 0.045337 | 0.009309 | 0.010487 |
| 27.5 | 0.008421 | 0.006092 | 0.012277 | 0.025381 | 0.006211 | 0.02233 | 0.043342 | 0.011769 | 0.065347 | 0.087695 | 0.008925 | 0.052322 | 0.00507 | 0.024978 | 0.034309 | 0.046379 | 0.019843 | 0.013567 | 0.023343 | 0.024444 | 0.01926 | 0.096093 | 0.019321 | 0.017946 |
| 27.5138889 | 0.02714 | 0.006598 | 0.014585 | 0.05248 | 0.008666 | 0.034392 | 0.047278 | 0.015732 | 0.075423 | 0.108778 | 0.025445 | 0.058081 | 0.003768 | 0.034697 | 0.062155 | 0.059025 | 0.027216 | 0.031741 | 0.034296 | 0.028026 | 0.026602 | 0.150355 | 0.037932 | 0.029498 |
| 27.5277778 | 0.057545 | 0.004757 | 0.018033 | 0.071167 | 0.009523 | 0.041444 | 0.033847 | 0.030204 | 0.045377 | 0.101588 | 0.045514 | 0.045643 | 0.003615 | 0.036666 | 0.084031 | 0.061261 | 0.025666 | 0.045075 | 0.032648 | 0.026999 | 0.030244 | 0.184076 | 0.046847 | 0.03798 |
| 27.5416667 | 0.079304 | 0.003858 | 0.021089 | 0.066174 | 0.006926 | 0.038617 | 0.023123 | 0.047023 | 0.013453 | 0.087714 | 0.055138 | 0.032262 | 0.003272 | 0.032199 | 0.087583 | 0.062087 | 0.02402 | 0.04713 | 0.028688 | 0.025928 | 0.037142 | 0.187865 | 0.047961 | 0.03838 |
| 27.5555556 | 0.089725 | 0.004852 | 0.021435 | 0.055223 | 0.004435 | 0.031064 | 0.02012 | 0.048854 | 0.005088 | 0.081949 | 0.046847 | 0.02083 | 0.001989 | 0.027045 | 0.073233 | 0.062217 | 0.024445 | 0.038249 | 0.030667 | 0.027252 | 0.0439 | 0.165831 | 0.044922 | 0.026502 |
| 27.5694444 | 0.083396 | 0.005992 | 0.018016 | 0.045614 | 0.003622 | 0.02191 | 0.018588 | 0.035775 | 0.010808 | 0.077965 | 0.026533 | 0.013888 | 0.002074 | 0.022812 | 0.047648 | 0.050028 | 0.022877 | 0.022367 | 0.02964 | 0.028421 | 0.043026 | 0.132722 | 0.030357 | 0.0136 |
| 27.5833333 | 0.061424 | 0.0062 | 0.012313 | 0.035489 | 0.003162 | 0.014555 | 0.017637 | 0.018941 | 0.018216 | 0.067388 | 0.015598 | 0.014615 | 0.002996 | 0.020401 | 0.023133 | 0.030117 | 0.018896 | 0.009454 | 0.021198 | 0.026227 | 0.036688 | 0.094225 | 0.013309 | 0.014347 |
| 27.5972222 | 0.037028 | 0.005811 | 0.006431 | 0.028523 | 0.004083 | 0.014155 | 0.018389 | 0.015761 | 0.022934 | 0.056186 | 0.024969 | 0.01854 | 0.003835 | 0.020637 | 0.008951 | 0.017129 | 0.011875 | 0.005011 | 0.010555 | 0.019895 | 0.029485 | 0.052026 | 0.006132 | 0.021145 |
| 27.6111111 | 0.017217 | 0.004359 | 0.002151 | 0.021371 | 0.005043 | 0.014458 | 0.018057 | 0.02604 | 0.025645 | 0.047371 | 0.038037 | 0.019183 | 0.004717 | 0.022713 | 0.004678 | 0.017591 | 0.006157 | 0.005979 | 0.004307 | 0.013725 | 0.020188 | 0.038628 | 0.00496 | 0.024679 |
| 27.625 | 0.00639 | 0.002153 | 0.002685 | 0.011254 | 0.003811 | 0.010637 | 0.01381 | 0.030636 | 0.029193 | 0.042738 | 0.038286 | 0.01702 | 0.005806 | 0.025273 | 0.00989 | 0.024008 | 0.008109 | 0.004717 | 0.0043 | 0.011257 | 0.010618 | 0.073949 | 0.006045 | 0.025284 |
| 27.6388889 | 0.003453 | 0.000761 | 0.006814 | 0.006687 | 0.003652 | 0.007316 | 0.007496 | 0.021858 | 0.031246 | 0.042617 | 0.023638 | 0.014432 | 0.007273 | 0.027088 | 0.025123 | 0.023284 | 0.01407 | 0.003123 | 0.008104 | 0.01081 | 0.009987 | 0.126695 | 0.006665 | 0.02157 |
| 27.6527778 | 0.003883 | 0.000595 | 0.007493 | 0.010775 | 0.005224 | 0.00493 | 0.004584 | 0.018639 | 0.025728 | 0.035702 | 0.013512 | 0.010054 | 0.007968 | 0.026158 | 0.034734 | 0.018361 | 0.018089 | 0.007478 | 0.013536 | 0.009187 | 0.020711 | 0.178916 | 0.005633 | 0.012695 |
| 27.6666667 | 0.005269 | 0.000861 | 0.006301 | 0.013597 | 0.004418 | 0.002035 | 0.006925 | 0.031161 | 0.014156 | 0.021706 | 0.027285 | 0.009216 | 0.007325 | 0.020906 | 0.030399 | 0.016065 | 0.017622 | 0.014973 | 0.014856 | 0.00542 | 0.031693 | 0.210179 | 0.007521 | 0.005383 |
| 27.6805556 | 0.008531 | 0.001045 | 0.007984 | 0.009734 | 0.004024 | 0.001498 | 0.008446 | 0.039049 | 0.005801 | 0.022393 | 0.044681 | 0.017833 | 0.006462 | 0.012336 | 0.021108 | 0.011816 | 0.013741 | 0.021436 | 0.010212 | 0.0019 | 0.031212 | 0.173722 | 0.011513 | 0.006456 |
| 27.6944444 | 0.016087 | 0.0017 | 0.007154 | 0.005765 | 0.008436 | 0.003239 | 0.006364 | 0.030752 | 0.007679 | 0.047928 | 0.039322 | 0.028731 | 0.006433 | 0.005901 | 0.014178 | 0.008703 | 0.011676 | 0.021738 | 0.005074 | 0.002649 | 0.01888 | 0.096 | 0.009231 | 0.007863 |
| 27.7083333 | 0.018008 | 0.002555 | 0.004285 | 0.010622 | 0.01395 | 0.004722 | 0.007062 | 0.018743 | 0.020948 | 0.072093 | 0.021502 | 0.031537 | 0.0061 | 0.005887 | 0.019255 | 0.011273 | 0.014077 | 0.018095 | 0.003934 | 0.007207 | 0.008602 | 0.071015 | 0.006139 | 0.004513 |
| 27.7222222 | 0.011305 | 0.002716 | 0.005549 | 0.020245 | 0.016328 | 0.008258 | 0.013265 | 0.021391 | 0.034668 | 0.069993 | 0.018169 | 0.026135 | 0.004519 | 0.006835 | 0.033339 | 0.010024 | 0.017876 | 0.016586 | 0.005867 | 0.008531 | 0.009667 | 0.112694 | 0.006387 | 0.003446 |
| 27.7361111 | 0.011548 | 0.002781 | 0.008734 | 0.024928 | 0.013505 | 0.014685 | 0.018271 | 0.034676 | 0.038025 | 0.051067 | 0.032017 | 0.019058 | 0.003623 | 0.005838 | 0.033333 | 0.00611 | 0.01939 | 0.011192 | 0.006677 | 0.005227 | 0.010718 | 0.153554 | 0.004326 | 0.004257 |
| 27.75 | 0.017991 | 0.00326 | 0.010945 | 0.026969 | 0.007309 | 0.015879 | 0.017663 | 0.030956 | 0.028561 | 0.030629 | 0.046338 | 0.011864 | 0.004064 | 0.008504 | 0.024507 | 0.008983 | 0.018893 | 0.006209 | 0.007331 | 0.002705 | 0.007123 | 0.159286 | 0.002769 | 0.005121 |
| 27.7638889 | 0.020182 | 0.003728 | 0.014262 | 0.02616 | 0.004185 | 0.00929 | 0.013204 | 0.018265 | 0.015648 | 0.013815 | 0.053073 | 0.005928 | 0.00408 | 0.01319 | 0.033389 | 0.017605 | 0.01922 | 0.011148 | 0.010807 | 0.003563 | 0.006081 | 0.132589 | 0.008498 | 0.00911 |
| 27.7777778 | 0.015779 | 0.004233 | 0.017851 | 0.017617 | 0.005239 | 0.003194 | 0.007583 | 0.021557 | 0.015487 | 0.005164 | 0.049027 | 0.00333 | 0.004306 | 0.014168 | 0.047756 | 0.025695 | 0.020903 | 0.019874 | 0.014647 | 0.006748 | 0.006092 | 0.084426 | 0.024041 | 0.014242 |
| 27.7916667 | 0.008286 | 0.004702 | 0.017523 | 0.009389 | 0.00601 | 0.001157 | 0.006265 | 0.022341 | 0.019325 | 0.003765 | 0.03617 | 0.004375 | 0.005468 | 0.011634 | 0.043698 | 0.026954 | 0.021566 | 0.024283 | 0.016035 | 0.009781 | 0.00675 | 0.045432 | 0.037282 | 0.015103 |
| 27.8055556 | 0.00744 | 0.004452 | 0.015396 | 0.012303 | 0.005508 | 0.001054 | 0.013748 | 0.015637 | 0.014487 | 0.005248 | 0.023733 | 0.010829 | 0.004713 | 0.006946 | 0.028492 | 0.021255 | 0.01941 | 0.023161 | 0.015978 | 0.014078 | 0.01278 | 0.057012 | 0.034293 | 0.011191 |
| 27.8194444 | 0.01507 | 0.003735 | 0.01643 | 0.022967 | 0.007281 | 0.001694 | 0.023893 | 0.018369 | 0.00746 | 0.006911 | 0.015038 | 0.015408 | 0.003161 | 0.004543 | 0.01689 | 0.015156 | 0.014486 | 0.024343 | 0.016786 | 0.017089 | 0.02188 | 0.114207 | 0.022483 | 0.013495 |

| | | | | | | | | | | | | | | | | | | | | | | | | |
|---|---|---|---|---|---|---|---|---|---|---|---|---|---|---|---|---|---|---|---|---|---|---|---|---|
| 27.8333333 | 0.024112 | 0.003387 | 0.019899 | 0.031804 | 0.011942 | 0.005113 | 0.029566 | 0.027694 | 0.006536 | 0.005852 | 0.008539 | 0.011142 | 0.005196 | 0.009399 | 0.008723 | 0.012943 | 0.008925 | 0.028709 | 0.018177 | 0.01609 | 0.025345 | 0.16111 | 0.023351 | 0.028857 |
| 27.8472222 | 0.031549 | 0.004016 | 0.02279 | 0.034295 | 0.014405 | 0.013024 | 0.03118 | 0.038161 | 0.014124 | 0.005625 | 0.003963 | 0.005267 | 0.010119 | 0.0178 | 0.004917 | 0.013207 | 0.006094 | 0.02847 | 0.016633 | 0.014029 | 0.019813 | 0.173297 | 0.035761 | 0.046459 |
| 27.8611111 | 0.039054 | 0.00574 | 0.02333 | 0.03095 | 0.01187 | 0.020498 | 0.028663 | 0.038367 | 0.02238 | 0.01123 | 0.002274 | 0.006655 | 0.015732 | 0.025265 | 0.00814 | 0.015085 | 0.007541 | 0.02744 | 0.015973 | 0.009804 | 0.0107 | 0.170779 | 0.048788 | 0.053337 |
| 27.875 | 0.048198 | 0.007348 | 0.021093 | 0.026728 | 0.008515 | 0.023542 | 0.02335 | 0.02403 | 0.025819 | 0.017932 | 0.003543 | 0.013891 | 0.019833 | 0.029522 | 0.009615 | 0.019874 | 0.010965 | 0.031286 | 0.022196 | 0.005506 | 0.007238 | 0.156943 | 0.05739 | 0.044762 |
| 27.8888889 | 0.057059 | 0.007647 | 0.017137 | 0.027336 | 0.008482 | 0.022528 | 0.019554 | 0.009801 | 0.025551 | 0.020096 | 0.004345 | 0.02048 | 0.018839 | 0.028146 | 0.013172 | 0.025472 | 0.013551 | 0.035383 | 0.031034 | 0.005881 | 0.008798 | 0.120497 | 0.048017 | 0.029401 |
| 27.9027778 | 0.061471 | 0.007024 | 0.013177 | 0.031582 | 0.010739 | 0.01947 | 0.017012 | 0.00535 | 0.021111 | 0.019588 | 0.003093 | 0.022671 | 0.012561 | 0.022983 | 0.027701 | 0.028608 | 0.016185 | 0.032286 | 0.038659 | 0.008139 | 0.007347 | 0.067692 | 0.024713 | 0.016788 |
| 27.9166667 | 0.060672 | 0.005973 | 0.009928 | 0.032155 | 0.012515 | 0.017212 | 0.01298 | 0.008982 | 0.015513 | 0.01893 | 0.001999 | 0.017814 | 0.00594 | 0.016608 | 0.036074 | 0.028561 | 0.019549 | 0.020097 | 0.044568 | 0.006342 | 0.004294 | 0.033964 | 0.010314 | 0.01077 |
| 27.9305556 | 0.054663 | 0.003972 | 0.006051 | 0.02577 | 0.011667 | 0.014229 | 0.007858 | 0.014815 | 0.013068 | 0.015549 | 0.001363 | 0.011287 | 0.002773 | 0.009065 | 0.02684 | 0.023354 | 0.020079 | 0.009448 | 0.045656 | 0.002392 | 0.003111 | 0.053687 | 0.011489 | 0.014622 |
| 27.9444444 | 0.042506 | 0.002763 | 0.002834 | 0.016187 | 0.007336 | 0.008094 | 0.005701 | 0.018137 | 0.01283 | 0.008607 | 0.002435 | 0.013546 | 0.001571 | 0.003463 | 0.012446 | 0.013006 | 0.015078 | 0.010037 | 0.037715 | 0.003222 | 0.002474 | 0.094676 | 0.010884 | 0.018214 |
| 27.9583333 | 0.02624 | 0.004129 | 0.001891 | 0.008605 | 0.00436 | 0.002245 | 0.005705 | 0.017736 | 0.009587 | 0.005899 | 0.007398 | 0.017972 | 0.001436 | 0.002646 | 0.004841 | 0.007777 | 0.007376 | 0.014536 | 0.021525 | 0.00657 | 0.002216 | 0.105912 | 0.008042 | 0.013425 |
| 27.9722222 | 0.015283 | 0.004376 | 0.002854 | 0.007274 | 0.007529 | 0.00267 | 0.009225 | 0.016259 | 0.009091 | 0.023918 | 0.013975 | 0.021632 | 0.002342 | 0.003931 | 0.003459 | 0.011456 | 0.003308 | 0.013543 | 0.007739 | 0.006489 | 0.003139 | 0.081952 | 0.018216 | 0.011708 |
| 27.9861111 | 0.018924 | 0.004296 | 0.008132 | 0.01086 | 0.010902 | 0.010131 | 0.027633 | 0.023725 | 0.022326 | 0.073494 | 0.018655 | 0.045282 | 0.003566 | 0.009655 | 0.006948 | 0.01326 | 0.009712 | 0.012106 | 0.004402 | 0.004098 | 0.007614 | 0.051541 | 0.038029 | 0.020497 |
| 28 | 0.02497 | 0.008828 | 0.015561 | 0.014268 | 0.007799 | 0.019498 | 0.050642 | 0.03686 | 0.030646 | 0.12354 | 0.015936 | 0.079213 | 0.007942 | 0.022835 | 0.018951 | 0.018215 | 0.029119 | 0.024798 | 0.010261 | 0.006331 | 0.017918 | 0.057125 | 0.063351 | 0.034954 |
| 28.0138889 | 0.02251 | 0.010688 | 0.01927 | 0.018225 | 0.004244 | 0.025005 | 0.048713 | 0.030831 | 0.022169 | 0.121702 | 0.012452 | 0.077071 | 0.015676 | 0.037522 | 0.031254 | 0.032906 | 0.053271 | 0.052208 | 0.022482 | 0.016107 | 0.032257 | 0.087503 | 0.088336 | 0.056215 |
| 28.0277778 | 0.025492 | 0.005327 | 0.019723 | 0.022961 | 0.005546 | 0.024829 | 0.023867 | 0.015418 | 0.016675 | 0.079669 | 0.019908 | 0.040471 | 0.020674 | 0.050178 | 0.032134 | 0.045735 | 0.069257 | 0.068831 | 0.031192 | 0.024341 | 0.051747 | 0.102509 | 0.087102 | 0.078842 |
| 28.0416667 | 0.03437 | 0.005595 | 0.019752 | 0.02935 | 0.00845 | 0.021493 | 0.008144 | 0.016708 | 0.013233 | 0.056028 | 0.025509 | 0.018988 | 0.020312 | 0.058969 | 0.024007 | 0.055578 | 0.071522 | 0.058388 | 0.026395 | 0.026808 | 0.065986 | 0.106563 | 0.052033 | 0.079668 |
| 28.0555556 | 0.035432 | 0.015888 | 0.018004 | 0.034709 | 0.012373 | 0.017684 | 0.011754 | 0.022015 | 0.006049 | 0.055268 | 0.019056 | 0.022643 | 0.017051 | 0.05696 | 0.015285 | 0.065951 | 0.060746 | 0.033651 | 0.013594 | 0.026734 | 0.051627 | 0.119023 | 0.027924 | 0.054797 |
| 28.0694444 | 0.028961 | 0.023617 | 0.01284 | 0.028088 | 0.014558 | 0.013784 | 0.01651 | 0.021368 | 0.002936 | 0.039865 | 0.012512 | 0.022116 | 0.011685 | 0.042534 | 0.00895 | 0.064296 | 0.0414 | 0.025227 | 0.006878 | 0.021507 | 0.03432 | 0.131886 | 0.038734 | 0.045883 |
| 28.0833333 | 0.017139 | 0.02132 | 0.00633 | 0.015783 | 0.011124 | 0.008548 | 0.012221 | 0.016533 | 0.002992 | 0.021959 | 0.013278 | 0.012186 | 0.005865 | 0.024229 | 0.004525 | 0.044992 | 0.021012 | 0.056434 | 0.010106 | 0.013449 | 0.040684 | 0.114734 | 0.04838 | 0.073735 |
| 28.0972222 | 0.006666 | 0.012135 | 0.00266 | 0.011903 | 0.005971 | 0.003822 | 0.00698 | 0.009679 | 0.00533 | 0.034174 | 0.014261 | 0.01009 | 0.004894 | 0.012754 | 0.002488 | 0.023413 | 0.00887 | 0.09697 | 0.013786 | 0.014276 | 0.034207 | 0.067162 | 0.04229 | 0.091101 |
| 28.1111111 | 0.005433 | 0.004658 | 0.005577 | 0.012664 | 0.00392 | 0.003349 | 0.008047 | 0.010206 | 0.011439 | 0.053515 | 0.009601 | 0.014426 | 0.009222 | 0.011719 | 0.002208 | 0.010584 | 0.009206 | 0.099447 | 0.013178 | 0.023517 | 0.012033 | 0.031639 | 0.033792 | 0.072226 |
| 28.125 | 0.008556 | 0.003369 | 0.013941 | 0.011334 | 0.003522 | 0.006186 | 0.008304 | 0.016986 | 0.017236 | 0.043224 | 0.006533 | 0.012726 | 0.012731 | 0.017468 | 0.002738 | 0.005894 | 0.013497 | 0.067425 | 0.009582 | 0.026896 | 0.001973 | 0.036618 | 0.022198 | 0.045313 |
| 28.1388889 | 0.011612 | 0.004096 | 0.02294 | 0.008428 | 0.00266 | 0.009498 | 0.004138 | 0.020093 | 0.018572 | 0.019174 | 0.010446 | 0.008408 | 0.012291 | 0.025239 | 0.003238 | 0.006155 | 0.014428 | 0.032029 | 0.005019 | 0.019475 | 0.001011 | 0.062399 | 0.010791 | 0.025321 |
| 28.1527778 | 0.012662 | 0.005065 | 0.02834 | 0.004907 | 0.002784 | 0.011474 | 0.004036 | 0.014048 | 0.01781 | 0.012988 | 0.013255 | 0.005949 | 0.009692 | 0.031129 | 0.002249 | 0.007109 | 0.01453 | 0.012418 | 0.003661 | 0.009095 | 0.000953 | 0.082869 | 0.008121 | 0.011696 |
| 28.1666667 | 0.010652 | 0.005876 | 0.02687 | 0.002647 | 0.003533 | 0.011603 | 0.006868 | 0.005828 | 0.016333 | 0.015867 | 0.011717 | 0.002776 | 0.007396 | 0.031997 | 0.00117 | 0.00636 | 0.016415 | 0.012501 | 0.00586 | 0.004478 | 0.00101 | 0.096247 | 0.009375 | 0.007095 |
| 28.1805556 | 0.00633 | 0.00501 | 0.019583 | 0.002891 | 0.003377 | 0.010366 | 0.007054 | 0.004873 | 0.013242 | 0.013749 | 0.009955 | 0.002562 | 0.006129 | 0.026634 | 0.002231 | 0.003823 | 0.018994 | 0.020342 | 0.007923 | 0.006388 | 0.002397 | 0.105519 | 0.007447 | 0.005658 |
| 28.1944444 | 0.001814 | 0.00325 | 0.012175 | 0.002949 | 0.002619 | 0.007913 | 0.005263 | 0.005674 | 0.009671 | 0.011598 | 0.00877 | 0.004443 | 0.005152 | 0.018015 | 0.004447 | 0.00198 | 0.021465 | 0.021024 | 0.007682 | 0.0095 | 0.004418 | 0.109953 | 0.004441 | 0.004757 |
| 28.2083333 | 0.001254 | 0.002504 | 0.008678 | 0.00243 | 0.002141 | 0.00563 | 0.004517 | 0.004087 | 0.007349 | 0.01615 | 0.008631 | 0.004001 | 0.004501 | 0.011083 | 0.006488 | 0.003158 | 0.024974 | 0.015092 | 0.006786 | 0.010568 | 0.005951 | 0.10516 | 0.002955 | 0.008411 |
| 28.2222222 | 0.003971 | 0.002832 | 0.010005 | 0.00411 | 0.001663 | 0.004674 | 0.006683 | 0.006319 | 0.006349 | 0.021758 | 0.009812 | 0.003996 | 0.004582 | 0.007878 | 0.00828 | 0.005782 | 0.029269 | 0.01465 | 0.0072 | 0.009182 | 0.00672 | 0.090226 | 0.004963 | 0.010253 |
| 28.2361111 | 0.005566 | 0.00272 | 0.01455 | 0.005998 | 0.001803 | 0.004317 | 0.006602 | 0.010695 | 0.006051 | 0.016671 | 0.010647 | 0.003794 | 0.005296 | 0.007217 | 0.009446 | 0.008045 | 0.030827 | 0.018641 | 0.00761 | 0.005717 | 0.006567 | 0.071679 | 0.011271 | 0.006885 |
| 28.25 | 0.005555 | 0.005438 | 0.017888 | 0.005716 | 0.004294 | 0.003616 | 0.007412 | 0.009527 | 0.008535 | 0.02029 | 0.007899 | 0.007899 | 0.006343 | 0.006721 | 0.009534 | 0.010057 | 0.027156 | 0.013788 | 0.00712 | 0.002149 | 0.005161 | 0.053937 | 0.015647 | 0.005402 |
| 28.2638889 | 0.007451 | 0.011918 | 0.015275 | 0.006891 | 0.007813 | 0.002652 | 0.016337 | 0.00601 | 0.015346 | 0.049921 | 0.004393 | 0.021095 | 0.007084 | 0.005347 | 0.009465 | 0.011921 | 0.020661 | 0.00709 | 0.006201 | 0.000927 | 0.002991 | 0.039131 | 0.012161 | 0.007056 |
| 28.2777778 | 0.008463 | 0.014509 | 0.009463 | 0.007762 | 0.007737 | 0.002198 | 0.01961 | 0.006024 | 0.018224 | 0.061315 | 0.003649 | 0.025107 | 0.00739 | 0.004716 | 0.010239 | 0.014015 | 0.015033 | 0.011583 | 0.004705 | 0.001421 | 0.00173 | 0.027627 | 0.005913 | 0.006059 |
| 28.2916667 | 0.007284 | 0.010103 | 0.009056 | 0.008011 | 0.005157 | 0.002487 | 0.015903 | 0.006883 | 0.014345 | 0.047159 | 0.00442 | 0.020222 | 0.007445 | 0.006085 | 0.011321 | 0.01625 | 0.011154 | 0.018292 | 0.003696 | 0.001985 | 0.001642 | 0.01709 | 0.003846 | 0.003534 |
| 28.3055556 | 0.009738 | 0.005242 | 0.01247 | 0.015324 | 0.004867 | 0.002618 | 0.021565 | 0.009719 | 0.01114 | 0.051808 | 0.007232 | 0.026446 | 0.007147 | 0.00809 | 0.011658 | 0.01713 | 0.008007 | 0.014956 | 0.003883 | 0.004076 | 0.001576 | 0.009491 | 0.003704 | 0.003282 |
| 28.3194444 | 0.010593 | 0.004487 | 0.010878 | 0.020135 | 0.005953 | 0.00232 | 0.021789 | 0.013053 | 0.00766 | 0.050321 | 0.007748 | 0.026705 | 0.006511 | 0.009033 | 0.011189 | 0.016363 | 0.00546 | 0.008874 | 0.004052 | 0.008685 | 0.001542 | 0.011235 | 0.004966 | 0.006492 |
| 28.3333333 | 0.009265 | 0.007323 | 0.00484 | 0.013565 | 0.009076 | 0.003197 | 0.013393 | 0.009816 | 0.004713 | 0.039892 | 0.007694 | 0.020062 | 0.005906 | 0.009015 | 0.01102 | 0.01592 | 0.004478 | 0.009758 | 0.003962 | 0.014004 | 0.00354 | 0.020939 | 0.009813 | 0.014974 |
| 28.3472222 | 0.010649 | 0.010597 | 0.002203 | 0.006945 | 0.012384 | 0.00389 | 0.014299 | 0.003504 | 0.004682 | 0.048927 | 0.009756 | 0.023647 | 0.005614 | 0.008655 | 0.0117 | 0.017837 | 0.004976 | 0.009121 | 0.005 | 0.016384 | 0.007502 | 0.02766 | 0.014221 | 0.019507 |
| 28.3611111 | 0.00944 | 0.008593 | 0.004991 | 0.00623 | 0.009352 | 0.003052 | 0.014173 | 0.003094 | 0.004202 | 0.042533 | 0.008201 | 0.02116 | 0.005352 | 0.007053 | 0.012227 | 0.022247 | 0.00547 | 0.007423 | 0.006746 | 0.01297 | 0.008813 | 0.027308 | 0.015158 | 0.01522 |
| 28.375 | 0.008751 | 0.00326 | 0.008598 | 0.009472 | 0.003541 | 0.003746 | 0.009885 | 0.009123 | 0.002824 | 0.022285 | 0.0072 | 0.013734 | 0.004772 | 0.004426 | 0.01146 | 0.026319 | 0.004905 | 0.015537 | 0.006365 | 0.006484 | 0.005885 | 0.025282 | 0.014214 | 0.014221 |
| 28.3888889 | 0.009375 | 0.002102 | 0.010074 | 0.01239 | 0.002256 | 0.005798 | 0.008664 | 0.013816 | 0.00248 | 0.018716 | 0.007771 | 0.010917 | 0.003889 | 0.004015 | 0.009722 | 0.026895 | 0.003624 | 0.02667 | 0.003934 | 0.002948 | 0.003575 | 0.02458 | 0.012786 | 0.0172 |
| 28.4027778 | 0.006745 | 0.004915 | 0.00947 | 0.013325 | 0.003916 | 0.006904 | 0.007642 | 0.013315 | 0.003501 | 0.022066 | 0.004552 | 0.007731 | 0.00322 | 0.005806 | 0.008398 | 0.024444 | 0.002633 | 0.02939 | 0.002987 | 0.004284 | 0.004732 | 0.021705 | 0.011676 | 0.014157 |
| 28.4166667 | 0.006789 | 0.006702 | 0.008206 | 0.018251 | 0.004513 | 0.008014 | 0.006001 | 0.011928 | 0.005568 | 0.018338 | 0.00203 | 0.005187 | 0.003333 | 0.006164 | 0.008456 | 0.021569 | 0.002632 | 0.022182 | 0.00285 | 0.006314 | 0.005571 | 0.013162 | 0.012362 | 0.01276 |
| 28.4305556 | 0.009859 | 0.007075 | 0.007188 | 0.022937 | 0.004716 | 0.010292 | 0.006427 | 0.011589 | 0.008517 | 0.013403 | 0.002572 | 0.005151 | 0.004277 | 0.004141 | 0.009591 | 0.020841 | 0.003373 | 0.010694 | 0.00321 | 0.008964 | 0.007242 | 0.00601 | 0.015784 | 0.018609 |
| 28.4444444 | 0.009587 | 0.00788 | 0.006103 | 0.01722 | 0.006002 | 0.01277 | 0.012284 | 0.010531 | 0.011963 | 0.022042 | 0.004322 | 0.010443 | 0.005536 | 0.002982 | 0.011148 | 0.023372 | 0.003654 | 0.004089 | 0.005283 | 0.012267 | 0.013797 | 0.007988 | 0.02016 | 0.0171 |
| 28.4583333 | 0.007541 | 0.007335 | 0.006388 | 0.007004 | 0.005523 | 0.013722 | 0.016215 | 0.009784 | 0.012199 | 0.029544 | 0.005643 | 0.014165 | 0.006232 | 0.004799 | 0.013253 | 0.027517 | 0.00312 | 0.003964 | 0.005418 | 0.011159 | 0.016825 | 0.01363 | 0.021213 | 0.013244 |
| 28.4722222 | 0.006906 | 0.004947 | 0.007635 | 0.002453 | 0.00247 | 0.01274 | 0.013513 | 0.009994 | 0.009131 | 0.024181 | 0.003664 | 0.010243 | 0.006762 | 0.00508 | 0.015928 | 0.032488 | 0.006325 | 0.003269 | 0.005597 | 0.009253 | 0.012818 | 0.019397 | 0.021031 | 0.023585 |
| 28.4861111 | 0.006707 | 0.004832 | 0.006653 | 0.005228 | 0.0034 | 0.011208 | 0.016564 | 0.009047 | 0.012657 | 0.032692 | 0.002044 | 0.012072 | 0.00977 | 0.00783 | 0.019182 | 0.041657 | 0.023947 | 0.006492 | 0.00632 | 0.01228 | 0.013926 | 0.02886 | 0.03181 | 0.04207 |
| 28.5 | 0.00962 | 0.009553 | 0.006126 | 0.017722 | 0.01072 | 0.012622 | 0.030338 | 0.008922 | 0.02525 | 0.062735 | 0.003873 | 0.024237 | 0.016253 | 0.026941 | 0.027209 | 0.050565 | 0.052308 | 0.026752 | 0.009158 | 0.018072 | 0.028223 | 0.042638 | 0.05121 | 0.060665 |
| 28.5138889 | 0.024199 | 0.014062 | 0.013865 | 0.045763 | 0.017719 | 0.02158 | 0.039808 | 0.01694 | 0.02654 | 0.082203 | 0.012238 | 0.03031 | 0.025473 | 0.059542 | 0.059388 | 0.043371 | 0.071798 | 0.055892 | 0.021871 | 0.017169 | 0.057105 | 0.061873 | 0.050738 | 0.071354 |
| 28.5277778 | 0.050774 | 0.011438 | 0.030575 | 0.073636 | 0.019107 | 0.037721 | 0.033871 | 0.036892 | 0.025415 | 0.073615 | 0.024962 | 0.02246 | 0.031734 | 0.091013 | 0.157539 | 0.028977 | 0.08024 | 0.067027 | 0.03012 | 0.018591 | 0.173912 | 0.115198 | 0.038379 | 0.089347 |
| 28.5416667 | 0.071165 | 0.004802 | 0.042452 | 0.066914 | 0.016401 | 0.048448 | 0.023097 | 0.058747 | 0.046981 | 0.052835 | 0.025566 | 0.011217 | 0.027065 | 0.099221 | 0.197939 | 0.047199 | 0.068395 | 0.046893 | 0.071832 | 0.079832 | 0.247913 | 0.238956 | 0.032866 | 0.151209 |
| 28.5555556 | 0.070817 | 0.00227 | 0.035082 | 0.035201 | 0.014475 | 0.04305 | 0.021143 | 0.063329 | 0.055949 | 0.03435 | 0.013254 | 0.006046 | 0.017142 | 0.084224 | 0.083085 | 0.091574 | 0.060335 | 0.016226 | 0.167273 | 0.170479 | 0.111709 | 0.365665 | 0.052767 | 0.206052 |
| 28.5694444 | 0.057646 | 0.003784 | 0.017848 | 0.019614 | 0.014596 | 0.030201 | 0.024782 | 0.047914 | 0.034149 | 0.018512 | 0.004637 | 0.006653 | 0.016284 | 0.090034 | 0.035375 | 0.09004 | 0.066032 | 0.005628 | 0.174663 | 0.163035 | 0.044576 | 0.345983 | 0.085261 | 0.192408 |
| 28.5833333 | 0.042108 | 0.004592 | 0.011048 | 0.021439 | 0.014153 | 0.020658 | 0.025125 | 0.025726 | 0.020236 | 0.012814 | 0.003559 | 0.009347 | 0.018831 | 0.097799 | 0.085604 | 0.042301 | 0.042386 | 0.013608 | 0.082039 | 0.079924 | 0.092587 | 0.184896 | 0.060593 | 0.136092 |
| 28.5972222 | 0.026664 | 0.003883 | 0.01326 | 0.018839 | 0.012081 | 0.015365 | 0.020349 | 0.01299 | 0.024149 | 0.024235 | 0.004059 | 0.010685 | 0.012146 | 0.061545 | 0.067108 | 0.017698 | 0.016984 | 0.01654 | 0.033927 | 0.02892 | 0.053859 | 0.049539 | 0.015982 | 0.071699 |
| 28.6111111 | 0.01467 | 0.003208 | 0.012711 | 0.011824 | 0.008192 | 0.01211 | 0.013427 | 0.017692 | 0.020404 | 0.033731 | 0.005478 | 0.009818 | 0.007672 | 0.025431 | 0.027159 | 0.016113 | 0.009981 | 0.0121 | 0.023686 | 0.017987 | 0.012266 | 0.022205 | 0.005022 | 0.032721 |
| 28.625 | 0.007916 | 0.002975 | 0.008561 | 0.010849 | 0.003937 | 0.00843 | 0.007162 | 0.024591 | 0.011681 | 0.024759 | 0.009517 | 0.009005 | 0.012378 | 0.020474 | 0.026242 | 0.01189 | 0.009921 | 0.018703 | 0.0141 | 0.016315 | 0.014795 | 0.070148 | 0.007168 | 0.020982 |
| 28.6388889 | 0.005135 | 0.002649 | 0.004228 | 0.015227 | 0.002417 | 0.00542 | 0.005639 | 0.018808 | 0.014885 | 0.012782 | 0.013587 | 0.008866 | 0.018562 | 0.023255 | 0.023751 | 0.007809 | 0.018266 | 0.033791 | 0.023882 | 0.013727 | 0.013725 | 0.144821 | 0.013297 | 0.013522 |
| 28.6527778 | 0.005762 | 0.00199 | 0.001643 | 0.01998 | 0.002703 | 0.005316 | 0.011961 | 0.010298 | 0.019041 | 0.015025 | 0.011255 | 0.007776 | 0.020434 | 0.024156 | 0.012812 | 0.008865 | 0.030254 | 0.044463 | 0.037247 | 0.008606 | 0.010326 | 0.1932 | 0.018264 | 0.008349 |
| 28.6666667 | 0.008953 | 0.001417 | 0.001387 | 0.022207 | 0.002924 | 0.007459 | 0.019276 | 0.016722 | 0.013705 | 0.021039 | 0.007578 | 0.005048 | 0.014312 | 0.022648 | 0.015775 | 0.015678 | 0.042814 | 0.043942 | 0.034932 | 0.008267 | 0.0132 | 0.203393 | 0.015353 | 0.006433 |
| 28.6805556 | 0.011804 | 0.001341 | 0.003439 | 0.020008 | 0.003515 | 0.009914 | 0.020365 | 0.039058 | 0.008012 | 0.017091 | 0.011985 | 0.003167 | 0.007701 | 0.017853 | 0.020152 | 0.015767 | 0.045278 | 0.027016 | 0.01967 | 0.021541 | 0.015045 | 0.208553 | 0.008461 | 0.009708 |
| 28.6944444 | 0.010251 | 0.001487 | 0.007981 | 0.015463 | 0.003677 | 0.011856 | 0.016933 | 0.062642 | 0.010584 | 0.009907 | 0.015184 | 0.00536 | 0.010038 | 0.011356 | 0.015212 | 0.007722 | 0.037383 | 0.015363 | 0.009481 | 0.040459 | 0.009879 | 0.193545 | 0.008043 | 0.016817 |
| 28.7083333 | 0.006648 | 0.001215 | 0.009581 | 0.010549 | 0.002468 | 0.013062 | 0.010684 | 0.074289 | 0.014367 | 0.011522 | 0.009984 | 0.009992 | 0.016008 | 0.005865 | 0.010548 | 0.006934 | 0.028699 | 0.027011 | 0.017086 | 0.053131 | 0.006697 | 0.134202 | 0.008464 | 0.018482 |
| 28.7222222 | 0.009052 | 0.001052 | 0.008438 | 0.006949 | 0.001795 | 0.012736 | 0.007942 | 0.068231 | 0.012017 | 0.020332 | 0.003409 | 0.013082 | 0.015409 | 0.005401 | 0.011696 | 0.013754 | 0.021923 | 0.040489 | 0.034823 | 0.051552 | 0.010879 | 0.063216 | 0.006621 | 0.011164 |
| 28.7361111 | 0.013457 | 0.001427 | 0.012145 | 0.007635 | 0.003318 | 0.010557 | 0.014507 | 0.041822 | 0.006106 | 0.029196 | 0.000881 | 0.012166 | 0.009755 | 0.009717 | 0.017209 | 0.022159 | 0.013029 | 0.036647 | 0.046287 | 0.033514 | 0.011747 | 0.020086 | 0.006307 | 0.004546 |
| 28.75 | 0.010351 | 0.001238 | 0.016169 | 0.008417 | 0.005364 | 0.007633 | 0.021529 | 0.02167 | 0.001757 | 0.036234 | 0.00124 | 0.007558 | 0.00955 | 0.014179 | 0.027179 | 0.026911 | 0.006856 | 0.020309 | 0.041406 | 0.024353 | 0.00608 | 0.022991 | 0.004632 | 0.005375 |
| 28.7638889 | 0.007478 | 0.000578 | 0.017801 | 0.010243 | 0.007907 | 0.004558 | 0.023068 | 0.035953 | 0.003384 | 0.039443 | 0.002629 | 0.003082 | 0.014372 | 0.014872 | 0.039275 | 0.020918 | 0.00972 | 0.009676 | 0.025765 | 0.042232 | 0.003759 | 0.08132 | 0.005103 | 0.010858 |
| 28.7777778 | 0.013221 | 0.000446 | 0.018389 | 0.018391 | 0.011701 | 0.003346 | 0.021479 | 0.058147 | 0.010745 | 0.036084 | 0.005547 | 0.001736 | 0.016608 | 0.010036 | 0.038861 | 0.01088 | 0.012508 | 0.017266 | 0.012862 | 0.062756 | 0.00471 | 0.177686 | 0.013634 | 0.017479 |
| 28.7916667 | 0.01821 | 0.000818 | 0.014861 | 0.023034 | 0.013855 | 0.006752 | 0.016818 | 0.060339 | 0.016942 | 0.02793 | 0.006262 | 0.002289 | 0.017526 | 0.005708 | 0.024578 | 0.00979 | 0.00866 | 0.032963 | 0.012041 | 0.066173 | 0.005862 | 0.244457 | 0.024524 | 0.020594 |
| 28.8055556 | 0.013946 | 0.001573 | 0.010643 | 0.015887 | 0.011254 | 0.011253 | 0.009465 | 0.049974 | 0.015107 | 0.021949 | 0.00528 | 0.002221 | 0.019689 | 0.00897 | 0.018791 | 0.019202 | 0.003409 | 0.04041 | 0.018111 | 0.055813 | 0.006157 | 0.240668 | 0.028324 | 0.017639 |
| 28.8194444 | 0.009654 | 0.002633 | 0.009372 | 0.008895 | 0.006649 | 0.012373 | 0.003502 | 0.039761 | 0.008594 | 0.023353 | 0.009168 | 0.001657 | 0.02073 | 0.015248 | 0.027055 | 0.032282 | 0.00125 | 0.033074 | 0.01817 | 0.035679 | 0.005572 | 0.192218 | 0.023258 | 0.012866 |
| 28.8333333 | 0.017897 | 0.003774 | 0.008411 | 0.012484 | 0.006352 | 0.011297 | 0.002444 | 0.033287 | 0.006811 | 0.033264 | 0.01432 | 0.002761 | 0.017876 | 0.018381 | 0.028027 | 0.046604 | 0.001801 | 0.0178 | 0.0112 | 0.019318 | 0.008395 | 0.120757 | 0.016206 | 0.011552 |
| 28.8472222 | 0.032121 | 0.005125 | 0.006652 | 0.016889 | 0.011941 | 0.010697 | 0.005386 | 0.03051 | 0.010707 | 0.049322 | 0.014441 | 0.005946 | 0.011401 | 0.017814 | 0.017454 | 0.0613 | 0.00281 | 0.007695 | 0.010025 | 0.021858 | 0.016993 | 0.067573 | 0.010305 | 0.015028 |
| 28.8611111 | 0.04047 | 0.007269 | 0.006916 | 0.014056 | 0.018352 | 0.010901 | 0.009937 | 0.031799 | 0.011651 | 0.062342 | 0.010693 | 0.00841 | 0.006657 | 0.013826 | 0.009717 | 0.068181 | 0.002779 | 0.013266 | 0.01956 | 0.030904 | 0.021259 | 0.086246 | 0.00914 | 0.019707 |
| 28.875 | 0.040122 | 0.009812 | 0.008737 | 0.010253 | 0.020003 | 0.010699 | 0.013767 | 0.034275 | 0.008954 | 0.062937 | 0.007199 | 0.007687 | 0.009833 | 0.00767 | 0.006371 | 0.066493 | 0.002276 | 0.024006 | 0.023324 | 0.030657 | 0.014505 | 0.159753 | 0.018765 | 0.021339 |
| 28.8888889 | 0.032736 | 0.012383 | 0.008242 | 0.012765 | 0.017466 | 0.009926 | 0.015313 | 0.034024 | 0.010212 | 0.050621 | 0.005178 | 0.00432 | 0.018431 | 0.00482 | 0.003844 | 0.061661 | 0.004486 | 0.019626 | 0.01463 | 0.02056 | 0.012609 | 0.223091 | 0.031485 | 0.016896 |
| 28.9027778 | 0.02572 | 0.013656 | 0.004977 | 0.019389 | 0.013496 | 0.009103 | 0.014638 | 0.029881 | 0.010368 | 0.03039 | 0.003643 | 0.001885 | 0.025789 | 0.008487 | 0.002344 | 0.05169 | 0.008593 | 0.008167 | 0.008823 | 0.010211 | 0.019047 | 0.228892 | 0.038067 | 0.00964 |
| 28.9166667 | 0.02442 | 0.009997 | 0.001818 | 0.024559 | 0.008652 | 0.007515 | 0.012791 | 0.022965 | 0.008737 | 0.013971 | 0.00508 | 0.002774 | 0.028438 | 0.013703 | 0.00186 | 0.03911 | 0.010072 | 0.005737 | 0.013763 | 0.012042 | 0.017283 | 0.179332 | 0.035471 | 0.009099 |
| 28.9305556 | 0.02495 | 0.004939 | 0.000728 | 0.025606 | 0.00422 | 0.004725 | 0.010574 | 0.014623 | 0.017104 | 0.012729 | 0.010413 | 0.005111 | 0.025965 | 0.016957 | 0.003441 | 0.026729 | 0.009044 | 0.012678 | 0.018844 | 0.020058 | 0.009642 | 0.111475 | 0.022448 | 0.014363 |
| 28.9444444 | 0.022238 | 0.004881 | 0.001029 | 0.022036 | 0.002708 | 0.001968 | 0.008455 | 0.006866 | 0.031094 | 0.020943 | 0.016184 | 0.007342 | 0.022272 | 0.019162 | 0.007074 | 0.014286 | 0.009098 | 0.017434 | 0.01481 | 0.023867 | 0.010576 | 0.059162 | 0.011239 | 0.015596 |
| 28.9583333 | 0.01558 | 0.007273 | 0.001148 | 0.016985 | 0.004394 | 0.000655 | 0.006781 | 0.002478 | 0.038384 | 0.024127 | 0.018368 | 0.00944 | 0.020689 | 0.022138 | 0.008444 | 0.005822 | 0.010388 | 0.014012 | 0.007091 | 0.029486 | 0.019308 | 0.029119 | 0.013691 | 0.01196 |
| 28.9722222 | 0.009314 | 0.008066 | 0.000714 | 0.016067 | 0.006696 | 0.000685 | 0.007183 | 0.001188 | 0.035372 | 0.018209 | 0.015404 | 0.010259 | 0.020071 | 0.027524 | 0.006495 | 0.002835 | 0.008707 | 0.010723 | 0.006037 | 0.041097 | 0.022108 | 0.020547 | 0.016577 | 0.015538 |
| 28.9861111 | 0.012289 | 0.006289 | 0.000691 | 0.022264 | 0.006901 | 0.002602 | 0.011233 | 0.002113 | 0.021924 | 0.022302 | 0.008847 | 0.007984 | 0.020827 | 0.034587 | 0.010555 | 0.007739 | 0.007705 | 0.017985 | 0.014693 | 0.047451 | 0.015022 | 0.021012 | 0.01403 | 0.030646 |
| 29 | 0.025283 | 0.004842 | 0.003042 | 0.030334 | 0.004242 | 0.0077 | 0.017656 | 0.007527 | 0.013652 | 0.056366 | 0.005358 | 0.00894 | 0.024623 | 0.040909 | 0.024559 | 0.024765 | 0.016747 | 0.043619 | 0.027873 | 0.046453 | 0.014219 | 0.02179 | 0.025526 | 0.047039 |
| 29.0138889 | 0.036646 | 0.005774 | 0.008451 | 0.037649 | 0.002651 | 0.012883 | 0.02315 | 0.014796 | 0.026881 | 0.105138 | 0.011602 | 0.021385 | 0.028612 | 0.046339 | 0.037611 | 0.044668 | 0.032181 | 0.07989 | 0.035609 | 0.0446 | 0.028136 | 0.042345 | 0.056387 | 0.066893 |
| 29.0277778 | 0.041112 | 0.005887 | 0.013101 | 0.043228 | 0.004322 | 0.013989 | 0.023808 | 0.021048 | 0.043775 | 0.132212 | 0.021574 | 0.039085 | 0.029543 | 0.053254 | 0.040241 | 0.055269 | 0.046502 | 0.098549 | 0.030149 | 0.040532 | 0.049429 | 0.081344 | 0.082318 | 0.09175 |
| 29.0416667 | 0.043589 | 0.006957 | 0.012659 | 0.03806 | 0.00451 | 0.014089 | 0.019599 | 0.028696 | 0.046474 | 0.109731 | 0.024692 | 0.045899 | 0.026558 | 0.05824 | 0.03297 | 0.058883 | 0.059064 | 0.081746 | 0.018781 | 0.032111 | 0.069135 | 0.121296 | 0.079197 | 0.108155 |
| 29.0555556 | 0.042895 | 0.00955 | 0.007678 | 0.022321 | 0.002682 | 0.016133 | 0.012999 | 0.033796 | 0.037571 | 0.062149 | 0.020631 | 0.031119 | 0.021228 | 0.049758 | 0.022706 | 0.061088 | 0.064656 | 0.043209 | 0.022683 | 0.018549 | 0.071585 | 0.14388 | 0.051 | 0.097762 |
| 29.0694444 | 0.035078 | 0.007981 | 0.004689 | 0.010785 | 0.003249 | 0.014917 | 0.006226 | 0.027019 | 0.022355 | 0.05212 | 0.016179 | 0.015952 | 0.015425 | 0.029426 | 0.018151 | 0.056625 | 0.055584 | 0.020942 | 0.03541 | 0.008475 | 0.045853 | 0.125447 | 0.034315 | 0.063048 |
| 29.0833333 | 0.022544 | 0.003377 | 0.008248 | 0.012127 | 0.00619 | 0.008523 | 0.003284 | 0.016061 | 0.018308 | 0.079913 | 0.01414 | 0.024371 | 0.008431 | 0.021939 | 0.020242 | 0.040763 | 0.035777 | 0.034997 | 0.02964 | 0.011219 | 0.021036 | 0.074445 | 0.051383 | 0.05306 |
| 29.0972222 | 0.011046 | 0.001808 | 0.012234 | 0.0171 | 0.008211 | 0.00369 | 0.004422 | 0.015914 | 0.032226 | 0.097292 | 0.012215 | 0.038439 | 0.003254 | 0.031397 | 0.022387 | 0.023142 | 0.01622 | 0.061387 | 0.012576 | 0.016245 | 0.028219 | 0.028395 | 0.070701 | 0.07743 |
| 29.1111111 | 0.007053 | 0.004568 | 0.012042 | 0.017084 | 0.007609 | 0.004683 | 0.005756 | 0.02353 | 0.040794 | 0.093236 | 0.008933 | 0.040015 | 0.002737 | 0.032343 | 0.018821 | 0.013087 | 0.006243 | 0.080406 | 0.00351 | 0.012768 | 0.044061 | 0.00882 | 0.067511 | 0.097088 |
| 29.125 | 0.011303 | 0.008624 | 0.011865 | 0.015699 | 0.00518 | 0.007784 | 0.006314 | 0.030365 | 0.029302 | 0.082295 | 0.00521 | 0.033148 | 0.00318 | 0.018886 | 0.009669 | 0.010273 | 0.007879 | 0.093263 | 0.002151 | 0.007545 | 0.043329 | 0.00752 | 0.050319 | 0.096102 |
| 29.1388889 | 0.017749 | 0.011236 | 0.014621 | 0.015586 | 0.002653 | 0.009304 | 0.00673 | 0.030307 | 0.012387 | 0.067828 | 0.002391 | 0.02302 | 0.002095 | 0.007988 | 0.00318 | 0.010472 | 0.01269 | 0.096108 | 0.003321 | 0.007399 | 0.029934 | 0.007476 | 0.027833 | 0.070148 |
| 29.1527778 | 0.021043 | 0.012495 | 0.018307 | 0.01575 | 0.001383 | 0.009182 | 0.006776 | 0.02287 | 0.004823 | 0.048333 | 0.001412 | 0.012803 | 0.000969 | 0.009193 | 0.002088 | 0.010167 | 0.015635 | 0.079234 | 0.005117 | 0.009494 | 0.014503 | 0.004569 | 0.009948 | 0.039014 |
| 29.1666667 | 0.021492 | 0.012832 | 0.019481 | 0.016142 | 0.00134 | 0.008484 | 0.006666 | 0.013655 | 0.003699 | 0.026271 | 0.002609 | 0.005374 | 0.000622 | 0.015705 | 0.002412 | 0.008523 | 0.01898 | 0.048626 | 0.005536 | 0.011202 | 0.008122 | 0.005008 | 0.003298 | 0.021999 |
| 29.1805556 | 0.020501 | 0.01124 | 0.016976 | 0.017162 | 0.001793 | 0.008553 | 0.006995 | 0.007006 | 0.006762 | 0.011225 | 0.004757 | 0.002767 | 0.00099 | 0.018956 | 0.006122 | 0.005726 | 0.024443 | 0.022687 | 0.003946 | 0.012961 | 0.00938 | 0.012975 | 0.003293 | 0.011054 |
| 29.1944444 | 0.016502 | 0.008043 | 0.013867 | 0.018915 | 0.002285 | 0.009458 | 0.00778 | 0.00327 | 0.012555 | 0.01132 | 0.007152 | 0.003218 | 0.002935 | 0.017716 | 0.013 | 0.003149 | 0.031656 | 0.008822 | 0.002196 | 0.013782 | 0.008494 | 0.026901 | 0.003088 | 0.005228 |
| 29.2083333 | 0.010889 | 0.004568 | 0.012528 | 0.020573 | 0.002216 | 0.010353 | 0.008279 | 0.001333 | 0.01716 | 0.017794 | 0.00976 | 0.002727 | 0.00589 | 0.01296 | 0.018063 | 0.003116 | 0.038278 | 0.006885 | 0.001448 | 0.012713 | 0.005488 | 0.042544 | 0.002166 | 0.006557 |
| 29.2222222 | 0.005688 | 0.002034 | 0.011456 | 0.02051 | 0.001401 | 0.010517 | 0.007649 | 0.002568 | 0.021601 | 0.018858 | 0.011746 | 0.001677 | 0.008087 | 0.006503 | 0.019166 | 0.00497 | 0.0411 | 0.012241 | 0.003379 | 0.010405 | 0.005636 | 0.056596 | 0.002882 | 0.007311 |
| 29.2361111 | 0.002759 | 0.001489 | 0.009874 | 0.017759 | 0.000608 | 0.010028 | 0.005697 | 0.006594 | 0.025853 | 0.013221 | 0.012015 | 0.002513 | 0.008667 | 0.002047 | 0.016932 | 0.005973 | 0.038305 | 0.015272 | 0.007886 | 0.008279 | 0.008947 | 0.065422 | 0.004776 | 0.004673 |
| 29.25 | 0.004508 | 0.002079 | 0.00842 | 0.012826 | 0.000308 | 0.009362 | 0.003002 | 0.009616 | 0.026933 | 0.006323 | 0.010484 | 0.004132 | 0.007518 | 0.004344 | 0.013564 | 0.00531 | 0.031002 | 0.011182 | 0.010909 | 0.00789 | 0.011935 | 0.066617 | 0.006604 | 0.003676 |
| 29.2638889 | 0.008411 | 0.001985 | 0.00688 | 0.007055 | 0.000775 | 0.008152 | 0.001487 | 0.008434 | 0.023708 | 0.004244 | 0.008388 | 0.004828 | 0.00501 | 0.015038 | 0.010759 | 0.003445 | 0.022461 | 0.005797 | 0.010019 | 0.009364 | 0.012709 | 0.060835 | 0.008146 | 0.004883 |
| 29.2777778 | 0.010704 | 0.001835 | 0.005035 | 0.00242 | 0.002441 | 0.005873 | 0.002155 | 0.005841 | 0.016741 | 0.008433 | 0.006569 | 0.004568 | 0.002217 | 0.029056 | 0.009139 | 0.001587 | 0.015302 | 0.005726 | 0.006485 | 0.011117 | 0.012295 | 0.051248 | 0.009952 | 0.004595 |
| 29.2916667 | 0.010783 | 0.002415 | 0.003734 | 0.001365 | 0.004132 | 0.003291 | 0.002333 | 0.006055 | 0.008417 | 0.014881 | 0.00535 | 0.004653 | 0.000961 | 0.036243 | 0.009257 | 0.001941 | 0.010784 | 0.005942 | 0.003005 | 0.010542 | 0.012062 | 0.044613 | 0.012042 | 0.004438 |
| 29.3055556 | 0.009941 | 0.002578 | 0.003274 | 0.004823 | 0.003948 | 0.00151 | 0.001942 | 0.00606 | 0.004187 | 0.017042 | 0.005279 | 0.006756 | 0.002084 | 0.030387 | 0.010475 | 0.005939 | 0.00971 | 0.005922 | 0.001754 | 0.006595 | 0.012319 | 0.043639 | 0.014319 | 0.006442 |
| 29.3194444 | 0.00923 | 0.002715 | 0.002905 | 0.011584 | 0.002138 | 0.000592 | 0.00373 | 0.006877 | 0.007044 | 0.011856 | 0.006452 | 0.010405 | 0.004753 | 0.016538 | 0.011511 | 0.012416 | 0.011098 | 0.015376 | 0.00212 | 0.002747 | 0.012297 | 0.043349 | 0.017683 | 0.006614 |
| 29.3333333 | 0.007721 | 0.003747 | 0.002894 | 0.017822 | 0.000621 | 0.000569 | 0.006142 | 0.014184 | 0.014375 | 0.007832 | 0.007973 | 0.013024 | 0.007426 | 0.005751 | 0.011993 | 0.018615 | 0.012249 | 0.032641 | 0.003745 | 0.001546 | 0.011457 | 0.04163 | 0.02145 | 0.004406 |
| 29.3472222 | 0.004593 | 0.004522 | 0.00399 | 0.019012 | 0.000198 | 0.002087 | 0.006958 | 0.021625 | 0.021241 | 0.0128 | 0.008696 | 0.012914 | 0.008796 | 0.003965 | 0.011673 | 0.022814 | 0.011437 | 0.046869 | 0.006955 | 0.002155 | 0.010439 | 0.041317 | 0.023028 | 0.003909 |
| 29.3611111 | 0.003692 | 0.003349 | 0.005746 | 0.015432 | 0.000558 | 0.00451 | 0.006455 | 0.021133 | 0.023396 | 0.017742 | 0.00845 | 0.010343 | 0.008347 | 0.011318 | 0.010679 | 0.024697 | 0.008968 | 0.051351 | 0.007788 | 0.005639 | 0.009709 | 0.043078 | 0.021072 | 0.003762 |
| 29.375 | 0.007461 | 0.002027 | 0.006966 | 0.011075 | 0.001311 | 0.005739 | 0.005211 | 0.015027 | 0.020065 | 0.014655 | 0.007889 | 0.00614 | 0.006912 | 0.024545 | 0.010141 | 0.024314 | 0.005807 | 0.044006 | 0.004459 | 0.011945 | 0.009275 | 0.044116 | 0.015547 | 0.003681 |
| 29.3888889 | 0.011407 | 0.002794 | 0.00707 | 0.008783 | 0.001554 | 0.005194 | 0.00417 | 0.009033 | 0.014552 | 0.007741 | 0.007432 | 0.002648 | 0.006464 | 0.032247 | 0.010492 | 0.022704 | 0.002759 | 0.026209 | 0.002047 | 0.018601 | 0.009437 | 0.042209 | 0.007849 | 0.00817 |
| 29.4027778 | 0.012875 | 0.00394 | 0.006517 | 0.010118 | 0.001161 | 0.004294 | 0.004623 | 0.006769 | 0.010127 | 0.003825 | 0.007228 | 0.00132 | 0.007878 | 0.02711 | 0.0108 | 0.021758 | 0.001423 | 0.009126 | 0.003836 | 0.021994 | 0.009719 | 0.038146 | 0.002351 | 0.014297 |
| 29.4166667 | 0.01396 | 0.004215 | 0.005292 | 0.014279 | 0.00125 | 0.004316 | 0.007288 | 0.009388 | 0.007335 | 0.003742 | 0.007534 | 0.001535 | 0.010252 | 0.015697 | 0.010323 | 0.023171 | 0.002653 | 0.004386 | 0.007377 | 0.020518 | 0.01024 | 0.035149 | 0.004161 | 0.015472 |
| 29.4305556 | 0.016216 | 0.003863 | 0.003522 | 0.018491 | 0.002572 | 0.005813 | 0.011761 | 0.015338 | 0.005766 | 0.003233 | 0.007985 | 0.003705 | 0.012844 | 0.006602 | 0.009905 | 0.028307 | 0.004318 | 0.011151 | 0.007834 | 0.018862 | 0.012704 | 0.034917 | 0.013679 | 0.010914 |
| 29.4444444 | 0.018715 | 0.002818 | 0.002007 | 0.020126 | 0.004862 | 0.0083 | 0.016296 | 0.021549 | 0.006155 | 0.002237 | 0.007658 | 0.006057 | 0.015069 | 0.00256 | 0.01052 | 0.036332 | 0.004544 | 0.025527 | 0.005152 | 0.019991 | 0.016803 | 0.034083 | 0.024569 | 0.010106 |

| | | | | | | | | | | | | | | | | | | | | | | | | |
|---|---|---|---|---|---|---|---|---|---|---|---|---|---|---|---|---|---|---|---|---|---|---|---|---|
| 29.4583333 | 0.020133 | 0.001737 | 0.001331 | 0.017895 | 0.007077 | 0.011219 | 0.019178 | 0.026196 | 0.009306 | 0.003708 | 0.006745 | 0.005074 | 0.01589 | 0.002864 | 0.011157 | 0.043737 | 0.00378 | 0.040099 | 0.0029 | 0.021608 | 0.019748 | 0.029533 | 0.029567 | 0.018617 |
| 29.4722222 | 0.021741 | 0.00262 | 0.002198 | 0.016522 | 0.006893 | 0.016147 | 0.021221 | 0.031358 | 0.013807 | 0.008495 | 0.007043 | 0.002855 | 0.016985 | 0.005377 | 0.011312 | 0.049888 | 0.002837 | 0.039012 | 0.001343 | 0.022309 | 0.020284 | 0.024253 | 0.030124 | 0.025252 |
| 29.4861111 | 0.02658 | 0.006898 | 0.004761 | 0.024871 | 0.006169 | 0.026705 | 0.023794 | 0.041074 | 0.02114 | 0.020073 | 0.010252 | 0.002434 | 0.025622 | 0.008998 | 0.01432 | 0.05978 | 0.005667 | 0.033197 | 0.001987 | 0.0249 | 0.021051 | 0.023805 | 0.037042 | 0.028489 |
| 29.5 | 0.03679 | 0.011146 | 0.008604 | 0.040647 | 0.010686 | 0.037886 | 0.02369 | 0.056003 | 0.038026 | 0.043433 | 0.01511 | 0.003087 | 0.043341 | 0.023031 | 0.02331 | 0.071664 | 0.014592 | 0.060596 | 0.005551 | 0.03064 | 0.025837 | 0.030886 | 0.051444 | 0.043491 |
| 29.5138889 | 0.048521 | 0.010474 | 0.012866 | 0.047223 | 0.018778 | 0.03633 | 0.01636 | 0.071961 | 0.062103 | 0.069907 | 0.020204 | 0.005997 | 0.052944 | 0.04481 | 0.033678 | 0.071147 | 0.019343 | 0.112566 | 0.00624 | 0.03519 | 0.033779 | 0.045266 | 0.06264 | 0.069159 |
| 29.5277778 | 0.056244 | 0.007284 | 0.017124 | 0.035969 | 0.024232 | 0.023955 | 0.010532 | 0.084977 | 0.090369 | 0.084544 | 0.026011 | 0.009738 | 0.047996 | 0.048591 | 0.037673 | 0.06094 | 0.014031 | 0.138738 | 0.003541 | 0.036881 | 0.041496 | 0.074758 | 0.063403 | 0.078066 |
| 29.5416667 | 0.063586 | 0.006633 | 0.024081 | 0.021026 | 0.024635 | 0.015708 | 0.01271 | 0.092148 | 0.11147 | 0.081195 | 0.029154 | 0.010162 | 0.047277 | 0.036317 | 0.037278 | 0.064782 | 0.009547 | 0.121598 | 0.002531 | 0.038998 | 0.049566 | 0.123496 | 0.052228 | 0.06488 |
| 29.5555556 | 0.069058 | 0.007683 | 0.030016 | 0.018892 | 0.021598 | 0.016395 | 0.011298 | 0.084904 | 0.093682 | 0.060781 | 0.027083 | 0.008129 | 0.052894 | 0.032048 | 0.04486 | 0.07585 | 0.012369 | 0.085262 | 0.003866 | 0.040449 | 0.056553 | 0.1899 | 0.034898 | 0.050165 |
| 29.5694444 | 0.06401 | 0.007596 | 0.026541 | 0.024722 | 0.01735 | 0.019917 | 0.008213 | 0.063285 | 0.051399 | 0.037251 | 0.023077 | 0.006408 | 0.048776 | 0.034375 | 0.06201 | 0.07088 | 0.016172 | 0.05518 | 0.00636 | 0.035356 | 0.048501 | 0.259344 | 0.017424 | 0.033615 |
| 29.5833333 | 0.04965 | 0.006502 | 0.017678 | 0.027899 | 0.011538 | 0.020808 | 0.012581 | 0.039627 | 0.044638 | 0.021524 | 0.018679 | 0.004365 | 0.03214 | 0.032179 | 0.07878 | 0.056042 | 0.014881 | 0.039201 | 0.006581 | 0.021171 | 0.028818 | 0.25942 | 0.009639 | 0.014846 |
| 29.5972222 | 0.031181 | 0.004996 | 0.009323 | 0.029989 | 0.006786 | 0.017556 | 0.017349 | 0.020558 | 0.069512 | 0.013215 | 0.013908 | 0.004806 | 0.013752 | 0.024497 | 0.084029 | 0.042571 | 0.00988 | 0.026659 | 0.004036 | 0.012468 | 0.017176 | 0.176146 | 0.019998 | 0.008006 |
| 29.6111111 | 0.013777 | 0.002871 | 0.004326 | 0.03261 | 0.008832 | 0.011426 | 0.017728 | 0.016238 | 0.068179 | 0.011044 | 0.011337 | 0.011927 | 0.005697 | 0.014382 | 0.068885 | 0.031875 | 0.004704 | 0.014083 | 0.003034 | 0.023646 | 0.033265 | 0.098259 | 0.032741 | 0.017075 |
| 29.625 | 0.006389 | 0.00105 | 0.004377 | 0.032453 | 0.014606 | 0.005735 | 0.015281 | 0.032066 | 0.036384 | 0.011548 | 0.011756 | 0.01895 | 0.006223 | 0.008459 | 0.04354 | 0.023451 | 0.002766 | 0.018991 | 0.004177 | 0.037597 | 0.058845 | 0.053097 | 0.026953 | 0.031566 |
| 29.6388889 | 0.010573 | 0.000578 | 0.003756 | 0.02946 | 0.015516 | 0.002449 | 0.010117 | 0.052385 | 0.016729 | 0.01972 | 0.01206 | 0.018942 | 0.005619 | 0.00777 | 0.021741 | 0.014892 | 0.006757 | 0.044025 | 0.004114 | 0.035228 | 0.049817 | 0.02519 | 0.019203 | 0.04149 |
| 29.6527778 | 0.015508 | 0.001205 | 0.002047 | 0.025254 | 0.011328 | 0.001495 | 0.005431 | 0.065714 | 0.02393 | 0.034453 | 0.009549 | 0.013284 | 0.006539 | 0.010764 | 0.011689 | 0.010385 | 0.013881 | 0.068718 | 0.00265 | 0.020726 | 0.025914 | 0.026148 | 0.031492 | 0.036632 |
| 29.6666667 | 0.015513 | 0.002556 | 0.003147 | 0.020861 | 0.006949 | 0.00244 | 0.004948 | 0.0599 | 0.034705 | 0.039262 | 0.005271 | 0.007573 | 0.013008 | 0.023852 | 0.011247 | 0.017169 | 0.020674 | 0.068951 | 0.004638 | 0.011082 | 0.021399 | 0.055795 | 0.045734 | 0.022503 |
| 29.6805556 | 0.013123 | 0.003887 | 0.005193 | 0.017246 | 0.004152 | 0.003113 | 0.005501 | 0.037295 | 0.030364 | 0.02838 | 0.003267 | 0.008087 | 0.022423 | 0.040641 | 0.010458 | 0.032875 | 0.026068 | 0.040509 | 0.009213 | 0.014204 | 0.027042 | 0.097623 | 0.046816 | 0.01765 |
| 29.6944444 | 0.008356 | 0.004353 | 0.006765 | 0.013538 | 0.003147 | 0.002613 | 0.003636 | 0.042842 | 0.019313 | 0.016283 | 0.004626 | 0.012415 | 0.028318 | 0.044278 | 0.011584 | 0.051857 | 0.029081 | 0.019707 | 0.009597 | 0.019989 | 0.022548 | 0.152199 | 0.038058 | 0.014855 |
| 29.7083333 | 0.004308 | 0.005568 | 0.00812 | 0.009119 | 0.00425 | 0.002746 | 0.001818 | 0.085962 | 0.020656 | 0.018394 | 0.005233 | 0.012066 | 0.023434 | 0.032342 | 0.012504 | 0.057065 | 0.028917 | 0.02693 | 0.005879 | 0.024916 | 0.017402 | 0.215296 | 0.022432 | 0.008335 |
| 29.7222222 | 0.00557 | 0.008355 | 0.009024 | 0.005867 | 0.005461 | 0.004541 | 0.002659 | 0.12156 | 0.031088 | 0.027216 | 0.003908 | 0.008277 | 0.013753 | 0.01533 | 0.011444 | 0.037456 | 0.024526 | 0.036459 | 0.002654 | 0.036382 | 0.022599 | 0.238561 | 0.012315 | 0.007129 |
| 29.7361111 | 0.007193 | 0.009055 | 0.009168 | 0.005391 | 0.004747 | 0.008743 | 0.004651 | 0.129989 | 0.034733 | 0.028713 | 0.005541 | 0.011217 | 0.011504 | 0.00518 | 0.013334 | 0.029363 | 0.017854 | 0.040589 | 0.001867 | 0.050758 | 0.026289 | 0.207033 | 0.016115 | 0.013932 |
| 29.75 | 0.004763 | 0.005745 | 0.00788 | 0.006907 | 0.003563 | 0.013596 | 0.00476 | 0.116997 | 0.032954 | 0.025029 | 0.012205 | 0.022149 | 0.016085 | 0.005347 | 0.013573 | 0.054334 | 0.012609 | 0.045971 | 0.003088 | 0.054744 | 0.019416 | 0.155947 | 0.025236 | 0.021319 |
| 29.7638889 | 0.003752 | 0.003505 | 0.006094 | 0.008827 | 0.003085 | 0.015944 | 0.003277 | 0.084598 | 0.030195 | 0.021272 | 0.018806 | 0.031878 | 0.019421 | 0.011142 | 0.014262 | 0.079339 | 0.008942 | 0.043521 | 0.003763 | 0.04617 | 0.015485 | 0.095235 | 0.030734 | 0.025482 |
| 29.7777778 | 0.008664 | 0.005522 | 0.008181 | 0.010451 | 0.002499 | 0.014205 | 0.004025 | 0.043201 | 0.024082 | 0.018116 | 0.020779 | 0.033543 | 0.018338 | 0.023133 | 0.026178 | 0.077058 | 0.005323 | 0.028556 | 0.003311 | 0.027859 | 0.020253 | 0.046484 | 0.025039 | 0.035861 |
| 29.7916667 | 0.014109 | 0.007251 | 0.014009 | 0.010769 | 0.001437 | 0.00854 | 0.007828 | 0.018818 | 0.013438 | 0.017688 | 0.018981 | 0.02702 | 0.013142 | 0.04471 | 0.043467 | 0.05264 | 0.002093 | 0.011991 | 0.002897 | 0.01528 | 0.017294 | 0.05117 | 0.01607 | 0.045918 |
| 29.8055556 | 0.014524 | 0.006142 | 0.01593 | 0.008256 | 0.000552 | 0.003787 | 0.009906 | 0.031056 | 0.004441 | 0.022354 | 0.015537 | 0.018213 | 0.006249 | 0.063969 | 0.048294 | 0.026318 | 0.001173 | 0.012982 | 0.004027 | 0.023012 | 0.010674 | 0.090159 | 0.022059 | 0.041724 |
| 29.8194444 | 0.011078 | 0.004604 | 0.013185 | 0.004605 | 0.000361 | 0.003918 | 0.008747 | 0.063083 | 0.006611 | 0.033565 | 0.011166 | 0.015078 | 0.001719 | 0.061082 | 0.032725 | 0.011719 | 0.003194 | 0.037583 | 0.008185 | 0.035374 | 0.00966 | 0.111386 | 0.036748 | 0.027306 |
| 29.8333333 | 0.011582 | 0.004913 | 0.010829 | 0.004156 | 0.000377 | 0.007831 | 0.008286 | 0.082998 | 0.016242 | 0.052247 | 0.006659 | 0.019149 | 0.001774 | 0.035177 | 0.021519 | 0.013101 | 0.006358 | 0.0667 | 0.013812 | 0.038414 | 0.010229 | 0.100378 | 0.045677 | 0.021769 |
| 29.8472222 | 0.021151 | 0.006316 | 0.009798 | 0.007333 | 0.001346 | 0.013614 | 0.010165 | 0.083794 | 0.018366 | 0.070848 | 0.003086 | 0.024564 | 0.005306 | 0.016671 | 0.035165 | 0.0265 | 0.009368 | 0.08749 | 0.019983 | 0.034994 | 0.012933 | 0.070073 | 0.041397 | 0.03981 |
| 29.8611111 | 0.030111 | 0.005866 | 0.010531 | 0.01092 | 0.004746 | 0.019894 | 0.013625 | 0.076073 | 0.011516 | 0.077885 | 0.001841 | 0.02574 | 0.007715 | 0.027082 | 0.048363 | 0.045034 | 0.012612 | 0.103083 | 0.026765 | 0.027776 | 0.022165 | 0.036838 | 0.030192 | 0.06353 |
| 29.875 | 0.032206 | 0.004313 | 0.01406 | 0.014648 | 0.010012 | 0.025151 | 0.014545 | 0.071154 | 0.008577 | 0.073072 | 0.004873 | 0.022408 | 0.007146 | 0.041903 | 0.046454 | 0.066811 | 0.016609 | 0.102819 | 0.030484 | 0.019484 | 0.02317 | 0.029819 | 0.039231 | 0.066012 |
| 29.8888889 | 0.030193 | 0.004289 | 0.019555 | 0.020913 | 0.014372 | 0.027493 | 0.011789 | 0.069066 | 0.011037 | 0.059778 | 0.012824 | 0.015084 | 0.010319 | 0.038958 | 0.043477 | 0.085918 | 0.021926 | 0.074522 | 0.027362 | 0.016738 | 0.012695 | 0.068159 | 0.068862 | 0.042037 |
| 29.9027778 | 0.024414 | 0.004693 | 0.021314 | 0.029535 | 0.014695 | 0.025584 | 0.013606 | 0.06217 | 0.011011 | 0.039446 | 0.023296 | 0.01097 | 0.019682 | 0.02728 | 0.039459 | 0.093304 | 0.026732 | 0.043226 | 0.017487 | 0.01634 | 0.005038 | 0.112556 | 0.082357 | 0.020064 |
| 29.9166667 | 0.017405 | 0.004299 | 0.013871 | 0.037234 | 0.010997 | 0.019095 | 0.022854 | 0.056181 | 0.009432 | 0.018608 | 0.031142 | 0.019181 | 0.030208 | 0.017543 | 0.029376 | 0.089029 | 0.027831 | 0.037246 | 0.008542 | 0.014929 | 0.00325 | 0.121079 | 0.072384 | 0.024544 |
| 29.9305556 | 0.011933 | 0.004622 | 0.007969 | 0.036004 | 0.006352 | 0.009872 | 0.031244 | 0.051433 | 0.007261 | 0.009272 | 0.027796 | 0.027651 | 0.038213 | 0.009598 | 0.020421 | 0.082038 | 0.026166 | 0.044962 | 0.007264 | 0.011738 | 0.006312 | 0.104228 | 0.057744 | 0.035503 |
| 29.9444444 | 0.007482 | 0.005135 | 0.014305 | 0.026839 | 0.005048 | 0.005063 | 0.02788 | 0.037799 | 0.004599 | 0.014616 | 0.013862 | 0.023841 | 0.039624 | 0.006286 | 0.014249 | 0.074723 | 0.024071 | 0.057222 | 0.006502 | 0.007629 | 0.009249 | 0.071005 | 0.037916 | 0.046004 |
| 29.9583333 | 0.00421 | 0.003817 | 0.023439 | 0.022004 | 0.007973 | 0.009114 | 0.017073 | 0.019735 | 0.006455 | 0.021582 | 0.004322 | 0.015219 | 0.035148 | 0.014044 | 0.007075 | 0.060294 | 0.021462 | 0.080899 | 0.005067 | 0.012043 | 0.008304 | 0.028672 | 0.018926 | 0.0607 |
| 29.9722222 | 0.003343 | 0.002335 | 0.028363 | 0.021568 | 0.010051 | 0.014874 | 0.012897 | 0.009494 | 0.012057 | 0.017576 | 0.004322 | 0.008893 | 0.032216 | 0.027962 | 0.005819 | 0.041318 | 0.018783 | 0.101373 | 0.009731 | 0.022077 | 0.005862 | 0.006219 | 0.016861 | 0.066685 |
| 29.9861111 | 0.005901 | 0.00251 | 0.028747 | 0.019584 | 0.007088 | 0.013881 | 0.011643 | 0.007938 | 0.013688 | 0.015857 | 0.005632 | 0.007199 | 0.03662 | 0.042909 | 0.022303 | 0.028744 | 0.019308 | 0.084833 | 0.016814 | 0.02538 | 0.007276 | 0.005198 | 0.019584 | 0.065255 |
| 30 | 0.014742 | 0.004291 | 0.019514 | 0.016321 | 0.004009 | 0.009406 | 0.009381 | 0.014782 | 0.012036 | 0.041444 | 0.00882 | 0.016179 | 0.046129 | 0.054518 | 0.047718 | 0.025124 | 0.025874 | 0.054658 | 0.026301 | 0.017669 | 0.014895 | 0.016658 | 0.028278 | 0.070549 |
| 30.0138889 | 0.028929 | 0.008657 | 0.010692 | 0.018278 | 0.006489 | 0.012052 | 0.009178 | 0.034937 | 0.018705 | 0.083982 | 0.018741 | 0.033261 | 0.052317 | 0.059886 | 0.056823 | 0.023338 | 0.037434 | 0.077659 | 0.04193 | 0.010983 | 0.026185 | 0.045341 | 0.061946 | 0.088938 |
| 30.0277778 | 0.040528 | 0.01521 | 0.014311 | 0.027758 | 0.011051 | 0.021815 | 0.009845 | 0.052331 | 0.028982 | 0.103929 | 0.027051 | 0.044535 | 0.05063 | 0.067479 | 0.050813 | 0.017994 | 0.050244 | 0.138148 | 0.053859 | 0.026812 | 0.037204 | 0.082918 | 0.092101 | 0.099916 |
| 30.0416667 | 0.047724 | 0.023012 | 0.01773 | 0.03769 | 0.014774 | 0.02964 | 0.013446 | 0.052118 | 0.028412 | 0.083435 | 0.026472 | 0.043194 | 0.04606 | 0.078741 | 0.043169 | 0.014228 | 0.056132 | 0.17182 | 0.049355 | 0.058255 | 0.040185 | 0.119532 | 0.086768 | 0.097459 |
| 30.0555556 | 0.05207 | 0.029175 | 0.012947 | 0.043023 | 0.01635 | 0.030386 | 0.014831 | 0.046708 | 0.020617 | 0.046826 | 0.02158 | 0.030591 | 0.039124 | 0.077855 | 0.029163 | 0.015222 | 0.047466 | 0.15498 | 0.032672 | 0.067947 | 0.035539 | 0.141053 | 0.058245 | 0.1053 |
| 30.0694444 | 0.048983 | 0.026194 | 0.006682 | 0.041158 | 0.013935 | 0.025379 | 0.008889 | 0.045145 | 0.013016 | 0.020946 | 0.015615 | 0.016647 | 0.023943 | 0.049595 | 0.014398 | 0.016522 | 0.028697 | 0.10549 | 0.016217 | 0.047195 | 0.031508 | 0.121203 | 0.028174 | 0.09944 |
| 30.0833333 | 0.035256 | 0.015409 | 0.004753 | 0.032707 | 0.009576 | 0.01962 | 0.00294 | 0.040596 | 0.006788 | 0.008244 | 0.008091 | 0.017097 | 0.010933 | 0.02447 | 0.008118 | 0.016332 | 0.012717 | 0.060506 | 0.011969 | 0.023055 | 0.024052 | 0.06839 | 0.015015 | 0.063628 |
| 30.0972222 | 0.017479 | 0.012112 | 0.007827 | 0.022235 | 0.005389 | 0.014505 | 0.001221 | 0.028426 | 0.005328 | 0.00732 | 0.003226 | 0.030058 | 0.01057 | 0.035128 | 0.009676 | 0.014108 | 0.008175 | 0.03155 | 0.027348 | 0.014785 | 0.014041 | 0.028495 | 0.026576 | 0.054279 |
| 30.1111111 | 0.010073 | 0.01652 | 0.011286 | 0.012458 | 0.003103 | 0.008858 | 0.000829 | 0.014679 | 0.009479 | 0.018889 | 0.004037 | 0.040111 | 0.014101 | 0.051411 | 0.0201 | 0.009551 | 0.010288 | 0.013505 | 0.049176 | 0.019229 | 0.014862 | 0.014474 | 0.049886 | 0.08042 |
| 30.125 | 0.016301 | 0.016737 | 0.013592 | 0.007266 | 0.00462 | 0.004113 | 0.001716 | 0.006846 | 0.013535 | 0.043437 | 0.0061 | 0.043193 | 0.015258 | 0.050915 | 0.030497 | 0.004814 | 0.008578 | 0.010719 | 0.059361 | 0.024317 | 0.024098 | 0.009319 | 0.071149 | 0.089663 |
| 30.1388889 | 0.023156 | 0.01559 | 0.015473 | 0.011568 | 0.007795 | 0.002637 | 0.006653 | 0.005054 | 0.015057 | 0.067185 | 0.006271 | 0.043364 | 0.017571 | 0.043371 | 0.033502 | 0.003025 | 0.005118 | 0.018337 | 0.052634 | 0.033425 | 0.028217 | 0.00498 | 0.081117 | 0.077598 |
| 30.1527778 | 0.022357 | 0.01437 | 0.016231 | 0.020683 | 0.00956 | 0.003523 | 0.013497 | 0.004114 | 0.016629 | 0.067453 | 0.006143 | 0.039885 | 0.020729 | 0.034332 | 0.030591 | 0.004478 | 0.00542 | 0.026747 | 0.036772 | 0.051251 | 0.025564 | 0.00248 | 0.073371 | 0.065826 |
| 30.1666667 | 0.016332 | 0.012626 | 0.016049 | 0.026272 | 0.009814 | 0.004101 | 0.015991 | 0.002996 | 0.019382 | 0.056211 | 0.008706 | 0.030113 | 0.020037 | 0.02594 | 0.02119 | 0.006469 | 0.00527 | 0.038527 | 0.021329 | 0.055054 | 0.023341 | 0.00275 | 0.055097 | 0.053287 |
| 30.1805556 | 0.009588 | 0.013734 | 0.014855 | 0.027901 | 0.009169 | 0.004059 | 0.013021 | 0.005062 | 0.023737 | 0.049389 | 0.014577 | 0.015515 | 0.016939 | 0.018851 | 0.010948 | 0.008233 | 0.003386 | 0.053984 | 0.009013 | 0.033621 | 0.020833 | 0.003374 | 0.033718 | 0.038974 |
| 30.1944444 | 0.00835 | 0.013599 | 0.012829 | 0.025491 | 0.007898 | 0.004136 | 0.012202 | 0.010826 | 0.028154 | 0.037131 | 0.019071 | 0.006944 | 0.014996 | 0.013139 | 0.011425 | 0.010198 | 0.002621 | 0.064492 | 0.003964 | 0.011151 | 0.015351 | 0.006339 | 0.015927 | 0.028587 |
| 30.2083333 | 0.016475 | 0.010536 | 0.013342 | 0.01587 | 0.006483 | 0.004362 | 0.014767 | 0.019441 | 0.029593 | 0.018582 | 0.019437 | 0.008915 | 0.014338 | 0.008538 | 0.019998 | 0.0119 | 0.002899 | 0.064195 | 0.005155 | 0.003415 | 0.010136 | 0.011376 | 0.013162 | 0.023938 |
| 30.2222222 | 0.027824 | 0.008218 | 0.018124 | 0.006359 | 0.005278 | 0.00551 | 0.012909 | 0.030705 | 0.029385 | 0.005247 | 0.02004 | 0.010734 | 0.015686 | 0.005771 | 0.022489 | 0.012953 | 0.003872 | 0.054147 | 0.006952 | 0.003785 | 0.006078 | 0.012919 | 0.01838 | 0.021493 |
| 30.2361111 | 0.033001 | 0.006719 | 0.024454 | 0.005208 | 0.004345 | 0.008787 | 0.007768 | 0.042864 | 0.030295 | 0.002616 | 0.020755 | 0.007483 | 0.019107 | 0.005502 | 0.017271 | 0.012773 | 0.006174 | 0.041127 | 0.008198 | 0.006601 | 0.003459 | 0.01026 | 0.0163 | 0.017971 |
| 30.25 | 0.028482 | 0.004534 | 0.028795 | 0.00597 | 0.003565 | 0.013067 | 0.004765 | 0.049796 | 0.030222 | 0.007103 | 0.018643 | 0.003836 | 0.021748 | 0.006115 | 0.012842 | 0.011292 | 0.009565 | 0.028483 | 0.009186 | 0.009559 | 0.003681 | 0.007317 | 0.009929 | 0.014052 |
| 30.2638889 | 0.018205 | 0.002824 | 0.028279 | 0.003573 | 0.002556 | 0.01515 | 0.003712 | 0.045841 | 0.027272 | 0.013359 | 0.016062 | 0.003152 | 0.020823 | 0.00538 | 0.011555 | 0.00938 | 0.011777 | 0.016444 | 0.009406 | 0.009188 | 0.006312 | 0.006334 | 0.009658 | 0.010466 |
| 30.2777778 | 0.008261 | 0.004421 | 0.022612 | 0.001802 | 0.001395 | 0.013313 | 0.00287 | 0.03457 | 0.023179 | 0.014968 | 0.013861 | 0.002913 | 0.016401 | 0.003854 | 0.011321 | 0.007879 | 0.011917 | 0.007049 | 0.007797 | 0.005452 | 0.007739 | 0.006391 | 0.012414 | 0.00628 |
| 30.2916667 | 0.003502 | 0.009489 | 0.014345 | 0.003176 | 0.000595 | 0.008101 | 0.001835 | 0.022611 | 0.017911 | 0.011476 | 0.010847 | 0.002757 | 0.009875 | 0.005056 | 0.012102 | 0.007636 | 0.012203 | 0.00284 | 0.004232 | 0.002632 | 0.006364 | 0.006294 | 0.010057 | 0.00649 |
| 30.3055556 | 0.003959 | 0.017305 | 0.006738 | 0.005609 | 0.000382 | 0.004521 | 0.002118 | 0.011393 | 0.011064 | 0.013332 | 0.007237 | 0.006186 | 0.003751 | 0.007559 | 0.013068 | 0.008473 | 0.014972 | 0.005062 | 0.001362 | 0.003219 | 0.004328 | 0.005453 | 0.005955 | 0.01945 |
| 30.3194444 | 0.003396 | 0.028156 | 0.003018 | 0.006773 | 0.000581 | 0.006498 | 0.004487 | 0.004542 | 0.005184 | 0.021609 | 0.004855 | 0.012148 | 0.001728 | 0.006458 | 0.010811 | 0.008926 | 0.02003 | 0.01058 | 0.002123 | 0.003835 | 0.002778 | 0.0037 | 0.006243 | 0.038688 |
| 30.3333333 | 0.001922 | 0.035199 | 0.002963 | 0.006756 | 0.000613 | 0.007803 | 0.005914 | 0.003811 | 0.003687 | 0.023545 | 0.005905 | 0.015937 | 0.005377 | 0.007549 | 0.005419 | 0.007475 | 0.023125 | 0.013935 | 0.0056 | 0.004134 | 0.002872 | 0.003608 | 0.008165 | 0.045992 |
| 30.3472222 | 0.004557 | 0.034272 | 0.002487 | 0.005926 | 0.000853 | 0.005729 | 0.004975 | 0.005314 | 0.007984 | 0.015332 | 0.009896 | 0.013391 | 0.007483 | 0.015627 | 0.001618 | 0.005339 | 0.020841 | 0.011733 | 0.006844 | 0.008558 | 0.003793 | 0.00681 | 0.009604 | 0.031366 |
| 30.3611111 | 0.010338 | 0.032056 | 0.001744 | 0.004493 | 0.002029 | 0.006291 | 0.005731 | 0.011411 | 0.013361 | 0.009895 | 0.013999 | 0.007052 | 0.004154 | 0.018945 | 0.001712 | 0.005439 | 0.017402 | 0.007682 | 0.00546 | 0.016649 | 0.002628 | 0.01054 | 0.016492 | 0.020006 |
| 30.375 | 0.0133 | 0.031626 | 0.002927 | 0.003153 | 0.003304 | 0.009784 | 0.009523 | 0.021623 | 0.016407 | 0.015158 | 0.015536 | 0.003547 | 0.004274 | 0.011937 | 0.003658 | 0.007107 | 0.01644 | 0.00811 | 0.007704 | 0.022499 | 0.001407 | 0.011989 | 0.029542 | 0.038432 |
| 30.3888889 | 0.010956 | 0.02892 | 0.004234 | 0.002395 | 0.004108 | 0.011818 | 0.012998 | 0.029212 | 0.021134 | 0.021963 | 0.014835 | 0.002823 | 0.008277 | 0.005897 | 0.005147 | 0.008712 | 0.01513 | 0.009287 | 0.012493 | 0.023095 | 0.002713 | 0.010132 | 0.040008 | 0.060017 |
| 30.4027778 | 0.007404 | 0.020463 | 0.004351 | 0.002462 | 0.004608 | 0.011752 | 0.013436 | 0.032845 | 0.027524 | 0.023385 | 0.014665 | 0.003665 | 0.008926 | 0.007809 | 0.006563 | 0.010382 | 0.012275 | 0.007232 | 0.013188 | 0.020171 | 0.004694 | 0.00614 | 0.040185 | 0.057162 |
| 30.4166667 | 0.005361 | 0.009803 | 0.004164 | 0.003757 | 0.004451 | 0.011288 | 0.009692 | 0.035292 | 0.031239 | 0.017899 | 0.015369 | 0.007676 | 0.005512 | 0.014583 | 0.009179 | 0.011588 | 0.01002 | 0.008633 | 0.010142 | 0.016342 | 0.006244 | 0.002917 | 0.029313 | 0.038168 |
| 30.4305556 | 0.006065 | 0.004729 | 0.004482 | 0.006119 | 0.003312 | 0.01096 | 0.008794 | 0.036796 | 0.030898 | 0.011628 | 0.015161 | 0.010887 | 0.002537 | 0.021729 | 0.012997 | 0.011869 | 0.009299 | 0.014283 | 0.00599 | 0.012949 | 0.007651 | 0.002678 | 0.016259 | 0.018109 |
| 30.4444444 | 0.010212 | 0.006211 | 0.004865 | 0.008317 | 0.001964 | 0.010037 | 0.016463 | 0.036041 | 0.027678 | 0.014149 | 0.013962 | 0.008903 | 0.002659 | 0.025942 | 0.015925 | 0.011593 | 0.009377 | 0.016249 | 0.003631 | 0.011129 | 0.008844 | 0.005364 | 0.010835 | 0.006458 |
| 30.4583333 | 0.015591 | 0.009268 | 0.005293 | 0.008647 | 0.001769 | 0.008723 | 0.02091 | 0.03528 | 0.024795 | 0.015317 | 0.013638 | 0.004996 | 0.00318 | 0.026029 | 0.015954 | 0.010906 | 0.009712 | 0.011733 | 0.00512 | 0.013013 | 0.009115 | 0.008557 | 0.008321 | 0.003969 |
| 30.4722222 | 0.020906 | 0.013951 | 0.007187 | 0.007794 | 0.001778 | 0.010196 | 0.016788 | 0.038938 | 0.026621 | 0.013742 | 0.015508 | 0.004491 | 0.006435 | 0.019674 | 0.013827 | 0.010831 | 0.011765 | 0.008345 | 0.006714 | 0.019039 | 0.008048 | 0.012316 | 0.009203 | 0.00615 |
| 30.4861111 | 0.029123 | 0.024613 | 0.012767 | 0.011021 | 0.002224 | 0.020201 | 0.022669 | 0.05091 | 0.037005 | 0.028153 | 0.020478 | 0.00819 | 0.018232 | 0.0176 | 0.012172 | 0.01392 | 0.0191 | 0.014483 | 0.007604 | 0.028515 | 0.007628 | 0.024729 | 0.029542 | 0.016989 |
| 30.5 | 0.041483 | 0.044206 | 0.022747 | 0.024267 | 0.004928 | 0.041135 | 0.048618 | 0.064264 | 0.053517 | 0.051967 | 0.02612 | 0.017899 | 0.03738 | 0.0353 | 0.012337 | 0.021305 | 0.031975 | 0.024955 | 0.007519 | 0.036047 | 0.009094 | 0.057278 | 0.065192 | 0.0374 |
| 30.5138889 | 0.042888 | 0.070235 | 0.034001 | 0.037222 | 0.006565 | 0.056224 | 0.069807 | 0.068142 | 0.07694 | 0.064949 | 0.026801 | 0.036081 | 0.056117 | 0.051441 | 0.011633 | 0.030372 | 0.04175 | 0.032004 | 0.0056 | 0.032697 | 0.010661 | 0.096179 | 0.090641 | 0.05084 |
| 30.5277778 | 0.038993 | 0.087833 | 0.040006 | 0.031887 | 0.005997 | 0.05392 | 0.067654 | 0.067516 | 0.110715 | 0.062392 | 0.02143 | 0.051802 | 0.069361 | 0.048928 | 0.00841 | 0.044886 | 0.043508 | 0.041765 | 0.005986 | 0.020622 | 0.011705 | 0.111761 | 0.091903 | 0.051349 |
| 30.5416667 | 0.048568 | 0.08678 | 0.037414 | 0.020089 | 0.006504 | 0.048262 | 0.05516 | 0.073501 | 0.13723 | 0.049454 | 0.016487 | 0.049333 | 0.080788 | 0.044284 | 0.007512 | 0.075793 | 0.047993 | 0.060318 | 0.011598 | 0.01158 | 0.018525 | 0.110289 | 0.080094 | 0.051904 |
| 30.5555556 | 0.052962 | 0.074217 | 0.03201 | 0.021602 | 0.007482 | 0.041716 | 0.041202 | 0.083527 | 0.126634 | 0.037198 | 0.018527 | 0.031836 | 0.085676 | 0.040617 | 0.014406 | 0.110732 | 0.059092 | 0.073147 | 0.020524 | 0.01034 | 0.029021 | 0.113807 | 0.068883 | 0.053883 |
| 30.5694444 | 0.045957 | 0.057206 | 0.026179 | 0.029724 | 0.007033 | 0.027555 | 0.024745 | 0.085539 | 0.073872 | 0.027894 | 0.02369 | 0.013946 | 0.074971 | 0.029005 | 0.026493 | 0.120578 | 0.063904 | 0.067959 | 0.025206 | 0.011668 | 0.02808 | 0.120106 | 0.056765 | 0.048153 |
| 30.5833333 | 0.036798 | 0.040228 | 0.015773 | 0.034144 | 0.005918 | 0.01248 | 0.011237 | 0.08054 | 0.033567 | 0.018656 | 0.025061 | 0.006249 | 0.059051 | 0.013225 | 0.037435 | 0.08956 | 0.059055 | 0.051799 | 0.023134 | 0.013078 | 0.016265 | 0.123756 | 0.042461 | 0.032848 |
| 30.5972222 | 0.022838 | 0.026244 | 0.007304 | 0.03461 | 0.00573 | 0.005358 | 0.007515 | 0.074243 | 0.045035 | 0.011269 | 0.02095 | 0.011128 | 0.047746 | 0.003683 | 0.044523 | 0.044684 | 0.052472 | 0.036238 | 0.019265 | 0.013762 | 0.008786 | 0.123741 | 0.02702 | 0.014738 |
| 30.6111111 | 0.016095 | 0.013826 | 0.009415 | 0.032774 | 0.006149 | 0.00442 | 0.01105 | 0.065461 | 0.069131 | 0.005941 | 0.01315 | 0.019555 | 0.033492 | 0.005453 | 0.048018 | 0.032409 | 0.048725 | 0.028707 | 0.018931 | 0.010332 | 0.010764 | 0.114669 | 0.013478 | 0.007265 |
| 30.625 | 0.021442 | 0.005969 | 0.015151 | 0.029152 | 0.006084 | 0.005854 | 0.01563 | 0.053745 | 0.07372 | 0.003227 | 0.009692 | 0.022898 | 0.015702 | 0.020047 | 0.051026 | 0.042427 | 0.046316 | 0.02655 | 0.020166 | 0.004882 | 0.010659 | 0.09949 | 0.007647 | 0.013378 |
| 30.6388889 | 0.019468 | 0.007197 | 0.019234 | 0.024356 | 0.005779 | 0.007263 | 0.022475 | 0.035934 | 0.058852 | 0.002757 | 0.015339 | 0.021902 | 0.007397 | 0.037031 | 0.055555 | 0.050019 | 0.039033 | 0.02355 | 0.015053 | 0.001571 | 0.006878 | 0.082719 | 0.010067 | 0.024911 |
| 30.6527778 | 0.010615 | 0.013249 | 0.025056 | 0.020293 | 0.005566 | 0.004944 | 0.032284 | 0.018607 | 0.034451 | 0.002077 | 0.021974 | 0.018141 | 0.013751 | 0.038675 | 0.058812 | 0.057002 | 0.026539 | 0.014752 | 0.007567 | 0.002943 | 0.010115 | 0.059169 | 0.019763 | 0.039352 |
| 30.6666667 | 0.00967 | 0.016559 | 0.026138 | 0.017196 | 0.005488 | 0.00234 | 0.032207 | 0.01647 | 0.01687 | 0.001365 | 0.020559 | 0.010394 | 0.026352 | 0.02553 | 0.056964 | 0.05931 | 0.013751 | 0.009845 | 0.006198 | 0.010663 | 0.01413 | 0.029795 | 0.029628 | 0.049729 |
| 30.6805556 | 0.015102 | 0.016912 | 0.017019 | 0.014421 | 0.00622 | 0.00322 | 0.025541 | 0.022888 | 0.008811 | 0.002623 | 0.012104 | 0.005674 | 0.035946 | 0.02095 | 0.04972 | 0.053896 | 0.005276 | 0.01915 | 0.010118 | 0.019685 | 0.009761 | 0.015007 | 0.025824 | 0.044089 |
| 30.6944444 | 0.022712 | 0.01722 | 0.012429 | 0.012645 | 0.007851 | 0.006642 | 0.03188 | 0.02563 | 0.006075 | 0.005433 | 0.005548 | 0.01084 | 0.040371 | 0.039149 | 0.039498 | 0.043301 | 0.00506 | 0.02391 | 0.015201 | 0.022617 | 0.005354 | 0.025782 | 0.021231 | 0.025743 |
| 30.7083333 | 0.022501 | 0.016491 | 0.021604 | 0.012796 | 0.008901 | 0.009772 | 0.037191 | 0.029782 | 0.005303 | 0.008548 | 0.004933 | 0.018766 | 0.042565 | 0.061832 | 0.028374 | 0.027022 | 0.011116 | 0.01746 | 0.019378 | 0.020258 | 0.00855 | 0.046421 | 0.033122 | 0.010678 |
| 30.7222222 | 0.01618 | 0.013498 | 0.028854 | 0.013805 | 0.008094 | 0.008378 | 0.023448 | 0.031977 | 0.007655 | 0.012483 | 0.007387 | 0.022228 | 0.040078 | 0.068265 | 0.020047 | 0.013843 | 0.018161 | 0.011459 | 0.022412 | 0.018979 | 0.016734 | 0.065574 | 0.041131 | 0.01085 |
| 30.7361111 | 0.018125 | 0.009216 | 0.024508 | 0.013877 | 0.0059 | 0.008435 | 0.010033 | 0.024439 | 0.017428 | 0.016775 | 0.009567 | 0.019777 | 0.026754 | 0.055347 | 0.016597 | 0.015641 | 0.023853 | 0.006724 | 0.025992 | 0.020241 | 0.023434 | 0.076212 | 0.028892 | 0.022697 |
| 30.75 | 0.022148 | 0.006124 | 0.016004 | 0.013177 | 0.003691 | 0.01439 | 0.010438 | 0.024438 | 0.024564 | 0.01884 | 0.007755 | 0.012646 | 0.020819 | 0.033869 | 0.013458 | 0.020289 | 0.027672 | 0.004276 | 0.032122 | 0.01845 | 0.020974 | 0.075867 | 0.014255 | 0.030983 |
| 30.7638889 | 0.023987 | 0.004438 | 0.009505 | 0.011497 | 0.002199 | 0.013719 | 0.011622 | 0.041427 | 0.020693 | 0.017744 | 0.006 | 0.011641 | 0.042374 | 0.01582 | 0.012242 | 0.014752 | 0.028575 | 0.007519 | 0.037304 | 0.010897 | 0.016192 | 0.0741 | 0.015668 | 0.029667 |
| 30.7777778 | 0.031724 | 0.003112 | 0.011337 | 0.009112 | 0.001223 | 0.008319 | 0.009219 | 0.053291 | 0.012731 | 0.016665 | 0.01027 | 0.024381 | 0.070083 | 0.011973 | 0.019833 | 0.007725 | 0.02688 | 0.020609 | 0.036753 | 0.006093 | 0.018386 | 0.072325 | 0.028703 | 0.026975 |
| 30.7916667 | 0.043363 | 0.003441 | 0.020241 | 0.007704 | 0.000579 | 0.008979 | 0.012909 | 0.045494 | 0.00691 | 0.01708 | 0.015967 | 0.040327 | 0.083166 | 0.028153 | 0.029842 | 0.007794 | 0.023712 | 0.031011 | 0.031262 | 0.009062 | 0.024908 | 0.06785 | 0.043458 | 0.029833 |
| 30.8055556 | 0.044735 | 0.00381 | 0.024038 | 0.007634 | 0.001022 | 0.010708 | 0.030684 | 0.02622 | 0.005693 | 0.018007 | 0.016987 | 0.049913 | 0.086293 | 0.060014 | 0.02865 | 0.01496 | 0.019756 | 0.029825 | 0.024836 | 0.00929 | 0.028935 | 0.056797 | 0.054939 | 0.03643 |
| 30.8194444 | 0.029623 | 0.003102 | 0.017831 | 0.007325 | 0.002694 | 0.01052 | 0.052211 | 0.010771 | 0.009192 | 0.022048 | 0.012341 | 0.052392 | 0.081379 | 0.103462 | 0.019535 | 0.024044 | 0.017004 | 0.026611 | 0.02266 | 0.004935 | 0.02159 | 0.035016 | 0.064973 | 0.040514 |
| 30.8333333 | 0.015796 | 0.00423 | 0.007454 | 0.005421 | 0.004488 | 0.009743 | 0.05916 | 0.009802 | 0.015121 | 0.030802 | 0.007126 | 0.048403 | 0.071828 | 0.137297 | 0.017277 | 0.026392 | 0.016295 | 0.0191 | 0.020617 | 0.004118 | 0.014097 | 0.028631 | 0.07172 | 0.039404 |
| 30.8472222 | 0.021008 | 0.007011 | 0.001868 | 0.002737 | 0.006277 | 0.008826 | 0.055148 | 0.023354 | 0.022192 | 0.042739 | 0.008264 | 0.041661 | 0.059567 | 0.135426 | 0.019744 | 0.019423 | 0.014688 | 0.016631 | 0.014883 | 0.003838 | 0.019334 | 0.04879 | 0.068204 | 0.03992 |
| 30.8611111 | 0.040313 | 0.009825 | 0.001744 | 0.001838 | 0.008319 | 0.010232 | 0.049857 | 0.04486 | 0.028826 | 0.055131 | 0.012952 | 0.035735 | 0.041688 | 0.099257 | 0.020037 | 0.010477 | 0.00915 | 0.036019 | 0.018038 | 0.006044 | 0.021892 | 0.054245 | 0.058261 | 0.048897 |
| 30.875 | 0.062885 | 0.01529 | 0.002565 | 0.003871 | 0.00948 | 0.014067 | 0.040411 | 0.062298 | 0.030862 | 0.063837 | 0.014853 | 0.030391 | 0.022899 | 0.049747 | 0.023812 | 0.008202 | 0.004314 | 0.068955 | 0.029107 | 0.013271 | 0.017143 | 0.036093 | 0.040543 | 0.068415 |
| 30.8888889 | 0.074849 | 0.024399 | 0.002387 | 0.005953 | 0.007995 | 0.017174 | 0.025887 | 0.065523 | 0.027791 | 0.065506 | 0.01442 | 0.024285 | 0.010472 | 0.021178 | 0.030447 | 0.013136 | 0.005716 | 0.091102 | 0.029616 | 0.017084 | 0.018071 | 0.017744 | 0.01712 | 0.096736 |
| 30.9027778 | 0.065792 | 0.034483 | 0.001582 | 0.006216 | 0.004225 | 0.017408 | 0.015105 | 0.062922 | 0.02333 | 0.059695 | 0.013295 | 0.017104 | 0.009952 | 0.033088 | 0.031561 | 0.022814 | 0.00934 | 0.079369 | 0.020306 | 0.018313 | 0.020673 | 0.012533 | 0.011161 | 0.122083 |
| 30.9166667 | 0.046086 | 0.043126 | 0.002116 | 0.005682 | 0.001705 | 0.014671 | 0.01187 | 0.067282 | 0.017159 | 0.04661 | 0.012655 | 0.009805 | 0.014998 | 0.067941 | 0.02754 | 0.033664 | 0.011872 | 0.044978 | 0.014977 | 0.02024 | 0.014247 | 0.027629 | 0.029938 | 0.120109 |
| 30.9305556 | 0.027177 | 0.04708 | 0.003965 | 0.004976 | 0.002715 | 0.010212 | 0.011008 | 0.072974 | 0.010703 | 0.026164 | 0.013746 | 0.009 | 0.014472 | 0.099235 | 0.021065 | 0.037783 | 0.013562 | 0.020109 | 0.012167 | 0.020322 | 0.008925 | 0.050435 | 0.054822 | 0.075867 |
| 30.9444444 | 0.013356 | 0.04603 | 0.006381 | 0.004619 | 0.005401 | 0.006492 | 0.008089 | 0.068736 | 0.013338 | 0.012583 | 0.011774 | 0.018019 | 0.01206 | 0.109212 | 0.011765 | 0.031647 | 0.012812 | 0.008841 | 0.007329 | 0.019929 | 0.015601 | 0.064935 | 0.064769 | 0.043473 |
| 30.9583333 | 0.007183 | 0.040852 | 0.008928 | 0.004101 | 0.007553 | 0.004331 | 0.004389 | 0.053995 | 0.027325 | 0.019623 | 0.005373 | 0.028545 | 0.01102 | 0.094482 | 0.006512 | 0.019935 | 0.008703 | 0.004784 | 0.00614 | 0.017976 | 0.017992 | 0.057774 | 0.052221 | 0.071498 |
| 30.9722222 | 0.008521 | 0.030182 | 0.010337 | 0.002923 | 0.008139 | 0.002579 | 0.003571 | 0.036402 | 0.038845 | 0.033504 | 0.001689 | 0.032303 | 0.009881 | 0.062173 | 0.010002 | 0.009244 | 0.00628 | 0.004263 | 0.010413 | 0.011491 | 0.00944 | 0.040765 | 0.028228 | 0.11278 |
| 30.9861111 | 0.009787 | 0.017477 | 0.009031 | 0.003218 | 0.006237 | 0.002194 | 0.004087 | 0.026182 | 0.032612 | 0.034299 | 0.004123 | 0.024992 | 0.018512 | 0.03499 | 0.013566 | 0.004534 | 0.014049 | 0.006828 | 0.017219 | 0.006912 | 0.00488 | 0.046387 | 0.022047 | 0.109713 |
| 31 | 0.010124 | 0.010091 | 0.005403 | 0.006504 | 0.003215 | 0.006009 | 0.007822 | 0.031289 | 0.024829 | 0.02853 | 0.012467 | 0.016491 | 0.052915 | 0.045137 | 0.01397 | 0.007166 | 0.029702 | 0.02157 | 0.026916 | 0.008947 | 0.005796 | 0.069633 | 0.059261 | 0.074708 |
| 31.0138889 | 0.019976 | 0.007958 | 0.003192 | 0.010038 | 0.002355 | 0.01338 | 0.019913 | 0.050393 | 0.045727 | 0.042737 | 0.026742 | 0.027712 | 0.088264 | 0.090853 | 0.016676 | 0.01208 | 0.043208 | 0.043336 | 0.040379 | 0.012771 | 0.006235 | 0.096468 | 0.128097 | 0.083315 |
| 31.0277778 | 0.033905 | 0.007057 | 0.004581 | 0.011205 | 0.004564 | 0.021372 | 0.03251 | 0.070822 | 0.079851 | 0.070767 | 0.04628 | 0.050626 | 0.084371 | 0.137215 | 0.020823 | 0.013491 | 0.053399 | 0.051762 | 0.055992 | 0.0136 | 0.004947 | 0.128294 | 0.180906 | 0.127243 |
| 31.0416667 | 0.04239 | 0.010795 | 0.006461 | 0.011767 | 0.008557 | 0.029227 | 0.035202 | 0.080634 | 0.086952 | 0.076314 | 0.061667 | 0.056992 | 0.060465 | 0.155432 | 0.018659 | 0.014371 | 0.066995 | 0.038138 | 0.073734 | 0.012125 | 0.004768 | 0.155514 | 0.180121 | 0.136736 |
| 31.0555556 | 0.044107 | 0.023564 | 0.005296 | 0.013392 | 0.012707 | 0.031438 | 0.034773 | 0.077936 | 0.0619 | 0.05728 | 0.064804 | 0.046387 | 0.045859 | 0.13933 | 0.009868 | 0.018949 | 0.082205 | 0.019465 | 0.09121 | 0.012085 | 0.008825 | 0.161695 | 0.144377 | 0.124337 |
| 31.0694444 | 0.038318 | 0.042651 | 0.003223 | 0.0152 | 0.015256 | 0.024298 | 0.036497 | 0.066792 | 0.032949 | 0.039314 | 0.052232 | 0.036169 | 0.036272 | 0.102971 | 0.003842 | 0.020705 | 0.086486 | 0.00978 | 0.095086 | 0.014894 | 0.012842 | 0.139309 | 0.102218 | 0.105897 |

| | | | | | | | | | | | | | | | | | | | | | | | | |
|---|---|---|---|---|---|---|---|---|---|---|---|---|---|---|---|---|---|---|---|---|---|---|---|---|
| 31.0833333 | 0.028404 | 0.057983 | 0.00354 | 0.016022 | 0.015879 | 0.016423 | 0.03054 | 0.05083 | 0.015046 | 0.030173 | 0.029122 | 0.028115 | 0.022036 | 0.066049 | 0.001869 | 0.013519 | 0.074325 | 0.005973 | 0.069837 | 0.016139 | 0.018203 | 0.102227 | 0.056498 | 0.0798 |
| 31.0972222 | 0.017725 | 0.060589 | 0.00423 | 0.015572 | 0.015072 | 0.012416 | 0.018308 | 0.03252 | 0.00836 | 0.022882 | 0.023547 | 0.016484 | 0.009355 | 0.036716 | 0.001146 | 0.005178 | 0.05289 | 0.007824 | 0.034567 | 0.011612 | 0.025707 | 0.070119 | 0.024921 | 0.05581 |
| 31.1111111 | 0.008684 | 0.051362 | 0.004282 | 0.014677 | 0.012851 | 0.009763 | 0.011935 | 0.017157 | 0.016507 | 0.014831 | 0.040478 | 0.007898 | 0.007501 | 0.015925 | 0.002168 | 0.003133 | 0.032462 | 0.011389 | 0.025462 | 0.005534 | 0.025803 | 0.049054 | 0.020737 | 0.034456 |
| 31.125 | 0.003415 | 0.037934 | 0.005256 | 0.013784 | 0.009666 | 0.008548 | 0.014011 | 0.007636 | 0.028558 | 0.007455 | 0.049747 | 0.010703 | 0.009827 | 0.006973 | 0.004155 | 0.002705 | 0.019183 | 0.010132 | 0.031544 | 0.003091 | 0.019583 | 0.036166 | 0.019573 | 0.018632 |
| 31.1388889 | 0.001283 | 0.027827 | 0.006832 | 0.012704 | 0.006475 | 0.008711 | 0.018253 | 0.003305 | 0.029417 | 0.002466 | 0.036369 | 0.018687 | 0.00901 | 0.008338 | 0.006185 | 0.002262 | 0.013883 | 0.005717 | 0.027009 | 0.003189 | 0.012806 | 0.025809 | 0.011323 | 0.012251 |
| 31.1527778 | 0.002827 | 0.02119 | 0.008336 | 0.010339 | 0.003507 | 0.008768 | 0.016784 | 0.004053 | 0.019752 | 0.000716 | 0.015658 | 0.025042 | 0.005449 | 0.013464 | 0.007714 | 0.004522 | 0.012665 | 0.003556 | 0.016735 | 0.003224 | 0.006635 | 0.015347 | 0.006651 | 0.012903 |
| 31.1666667 | 0.008839 | 0.013553 | 0.009352 | 0.006646 | 0.001892 | 0.007267 | 0.010758 | 0.007576 | 0.010156 | 0.002968 | 0.005387 | 0.029678 | 0.00381 | 0.022582 | 0.008993 | 0.007362 | 0.009941 | 0.00577 | 0.012602 | 0.003019 | 0.00457 | 0.006793 | 0.005597 | 0.012072 |
| 31.1805556 | 0.017617 | 0.011978 | 0.009631 | 0.005156 | 0.003271 | 0.004058 | 0.011946 | 0.010257 | 0.00873 | 0.012076 | 0.012038 | 0.032299 | 0.005543 | 0.036242 | 0.011287 | 0.007068 | 0.005273 | 0.008457 | 0.015706 | 0.005398 | 0.007542 | 0.002859 | 0.005136 | 0.009674 |
| 31.1944444 | 0.026122 | 0.026704 | 0.010064 | 0.006065 | 0.006085 | 0.002259 | 0.023381 | 0.009464 | 0.009 | 0.028372 | 0.029122 | 0.037128 | 0.006261 | 0.050945 | 0.014864 | 0.006095 | 0.002879 | 0.00851 | 0.020455 | 0.01443 | 0.011813 | 0.003654 | 0.007092 | 0.015842 |
| 31.2083333 | 0.032245 | 0.050272 | 0.012032 | 0.005903 | 0.007739 | 0.003814 | 0.033237 | 0.00541 | 0.00739 | 0.047397 | 0.032325 | 0.045325 | 0.009978 | 0.064249 | 0.018499 | 0.012839 | 0.003664 | 0.007877 | 0.021954 | 0.027325 | 0.015332 | 0.00688 | 0.01638 | 0.02885 |
| 31.2222222 | 0.035662 | 0.060346 | 0.014899 | 0.005581 | 0.00814 | 0.006273 | 0.035485 | 0.003267 | 0.016273 | 0.065005 | 0.022626 | 0.051557 | 0.024119 | 0.072242 | 0.021311 | 0.026103 | 0.004619 | 0.009283 | 0.01766 | 0.037476 | 0.019705 | 0.009123 | 0.029298 | 0.041671 |
| 31.2361111 | 0.03739 | 0.052726 | 0.01656 | 0.006363 | 0.008305 | 0.007956 | 0.034768 | 0.006658 | 0.043855 | 0.073235 | 0.032903 | 0.054379 | 0.042821 | 0.075102 | 0.022731 | 0.036612 | 0.005012 | 0.013119 | 0.011873 | 0.043424 | 0.025188 | 0.008507 | 0.042445 | 0.052448 |
| 31.25 | 0.039119 | 0.045333 | 0.016632 | 0.006835 | 0.008129 | 0.008566 | 0.035267 | 0.012394 | 0.07255 | 0.064646 | 0.056225 | 0.050242 | 0.051403 | 0.07537 | 0.022369 | 0.040491 | 0.005828 | 0.017342 | 0.011679 | 0.047319 | 0.029828 | 0.005605 | 0.061352 | 0.056532 |
| 31.2638889 | 0.039265 | 0.040853 | 0.015846 | 0.007221 | 0.006645 | 0.007671 | 0.035509 | 0.015959 | 0.071015 | 0.043062 | 0.057006 | 0.037485 | 0.045823 | 0.070049 | 0.020771 | 0.038219 | 0.006316 | 0.019125 | 0.013966 | 0.050545 | 0.031499 | 0.002915 | 0.080457 | 0.051711 |
| 31.2777778 | 0.036449 | 0.034582 | 0.014583 | 0.007963 | 0.003863 | 0.005409 | 0.02961 | 0.013893 | 0.041136 | 0.027123 | 0.035908 | 0.020926 | 0.037175 | 0.057215 | 0.017572 | 0.032182 | 0.004839 | 0.018019 | 0.011008 | 0.048656 | 0.028529 | 0.002598 | 0.081657 | 0.040256 |
| 31.2916667 | 0.032465 | 0.026281 | 0.013218 | 0.007284 | 0.001804 | 0.003044 | 0.016606 | 0.008082 | 0.024977 | 0.031194 | 0.015096 | 0.00927 | 0.030033 | 0.040834 | 0.0125 | 0.025546 | 0.002786 | 0.01554 | 0.007565 | 0.037824 | 0.021603 | 0.002444 | 0.066555 | 0.025231 |
| 31.3055556 | 0.027547 | 0.019063 | 0.011964 | 0.004153 | 0.00171 | 0.003675 | 0.005202 | 0.004723 | 0.037017 | 0.032948 | 0.00492 | 0.007954 | 0.022702 | 0.026545 | 0.00758 | 0.020206 | 0.002564 | 0.012177 | 0.011761 | 0.026584 | 0.015004 | 0.001621 | 0.051013 | 0.016708 |
| 31.3194444 | 0.022133 | 0.015419 | 0.011129 | 0.001457 | 0.002903 | 0.007853 | 0.00114 | 0.003204 | 0.043503 | 0.019173 | 0.003926 | 0.008288 | 0.0174 | 0.016759 | 0.004716 | 0.01765 | 0.002805 | 0.007858 | 0.016671 | 0.022977 | 0.011974 | 0.002339 | 0.040723 | 0.024951 |
| 31.3333333 | 0.018942 | 0.012408 | 0.01111 | 0.001568 | 0.004145 | 0.011488 | 0.002316 | 0.002142 | 0.028598 | 0.008199 | 0.008837 | 0.005298 | 0.01499 | 0.009705 | 0.003378 | 0.01794 | 0.003047 | 0.003893 | 0.016231 | 0.022948 | 0.010279 | 0.004312 | 0.033166 | 0.040039 |
| 31.3472222 | 0.019901 | 0.015106 | 0.012195 | 0.002814 | 0.003914 | 0.011657 | 0.006886 | 0.003678 | 0.014393 | 0.003622 | 0.014899 | 0.003076 | 0.014027 | 0.004778 | 0.002087 | 0.018056 | 0.004782 | 0.001872 | 0.014563 | 0.020639 | 0.007143 | 0.005744 | 0.023622 | 0.041111 |
| 31.3611111 | 0.022902 | 0.024164 | 0.013827 | 0.003487 | 0.002721 | 0.009093 | 0.010258 | 0.005582 | 0.018588 | 0.001583 | 0.016092 | 0.003988 | 0.013385 | 0.003951 | 0.000922 | 0.015742 | 0.007228 | 0.001908 | 0.014815 | 0.014525 | 0.003638 | 0.006078 | 0.013347 | 0.026119 |
| 31.375 | 0.023651 | 0.030195 | 0.014167 | 0.003471 | 0.00162 | 0.005536 | 0.00877 | 0.005124 | 0.024509 | 0.003921 | 0.011901 | 0.006587 | 0.01237 | 0.005462 | 0.000632 | 0.011858 | 0.00895 | 0.001853 | 0.017542 | 0.008091 | 0.002052 | 0.007287 | 0.005966 | 0.011652 |
| 31.3888889 | 0.020488 | 0.034377 | 0.012543 | 0.003214 | 0.000758 | 0.002518 | 0.006062 | 0.002972 | 0.01603 | 0.013614 | 0.00596 | 0.006135 | 0.009589 | 0.006713 | 0.000844 | 0.008688 | 0.009575 | 0.001406 | 0.022175 | 0.006325 | 0.00283 | 0.009646 | 0.002748 | 0.008601 |
| 31.4027778 | 0.014728 | 0.040025 | 0.009869 | 0.002836 | 0.000661 | 0.001819 | 0.005717 | 0.001426 | 0.008066 | 0.028958 | 0.004029 | 0.005168 | 0.00542 | 0.008147 | 0.001008 | 0.00841 | 0.008647 | 0.00299 | 0.02661 | 0.008393 | 0.003879 | 0.010597 | 0.003269 | 0.012419 |
| 31.4166667 | 0.009036 | 0.044331 | 0.00738 | 0.002104 | 0.001032 | 0.003827 | 0.007874 | 0.002006 | 0.01249 | 0.041788 | 0.007797 | 0.007614 | 0.002392 | 0.008973 | 0.001032 | 0.010369 | 0.006164 | 0.006559 | 0.028073 | 0.010693 | 0.003758 | 0.010046 | 0.003506 | 0.013359 |
| 31.4305556 | 0.00551 | 0.047923 | 0.006606 | 0.001609 | 0.001135 | 0.006315 | 0.010119 | 0.003467 | 0.018439 | 0.044065 | 0.012737 | 0.008207 | 0.002293 | 0.007248 | 0.001043 | 0.012121 | 0.003309 | 0.009842 | 0.024584 | 0.011275 | 0.003133 | 0.009455 | 0.002136 | 0.008308 |
| 31.4444444 | 0.004838 | 0.052939 | 0.007942 | 0.001794 | 0.001006 | 0.007122 | 0.009512 | 0.004699 | 0.016441 | 0.033345 | 0.013358 | 0.005032 | 0.00532 | 0.005037 | 0.002529 | 0.012918 | 0.001706 | 0.009762 | 0.015631 | 0.009096 | 0.002352 | 0.008859 | 0.002225 | 0.004988 |
| 31.4583333 | 0.006302 | 0.057342 | 0.009273 | 0.002093 | 0.000837 | 0.006632 | 0.006267 | 0.004673 | 0.010184 | 0.01782 | 0.007657 | 0.003644 | 0.010995 | 0.008198 | 0.005473 | 0.013283 | 0.002661 | 0.006177 | 0.008215 | 0.006197 | 0.002962 | 0.006778 | 0.005928 | 0.010559 |
| 31.4722222 | 0.010999 | 0.05086 | 0.010028 | 0.002687 | 0.001286 | 0.008161 | 0.006422 | 0.005749 | 0.009088 | 0.010808 | 0.002447 | 0.00605 | 0.021172 | 0.02007 | 0.006562 | 0.013944 | 0.00755 | 0.005398 | 0.011804 | 0.005185 | 0.006817 | 0.004782 | 0.013673 | 0.02644 |
| 31.4861111 | 0.022904 | 0.037589 | 0.013055 | 0.008236 | 0.004588 | 0.016844 | 0.021485 | 0.01753 | 0.017533 | 0.02447 | 0.00428 | 0.011268 | 0.041147 | 0.043616 | 0.005329 | 0.017313 | 0.020751 | 0.010915 | 0.019894 | 0.007408 | 0.012948 | 0.009536 | 0.02904 | 0.062798 |
| 31.5 | 0.04161 | 0.046454 | 0.020265 | 0.022883 | 0.011134 | 0.031169 | 0.055305 | 0.043976 | 0.027749 | 0.056115 | 0.014833 | 0.020795 | 0.063995 | 0.081776 | 0.003907 | 0.025205 | 0.042782 | 0.016699 | 0.02225 | 0.015662 | 0.019818 | 0.022922 | 0.057281 | 0.107061 |
| 31.5138889 | 0.058772 | 0.07218 | 0.030424 | 0.036345 | 0.016314 | 0.038918 | 0.085036 | 0.068186 | 0.030385 | 0.079987 | 0.034003 | 0.0348 | 0.069837 | 0.116817 | 0.003293 | 0.032301 | 0.06319 | 0.016471 | 0.020709 | 0.026073 | 0.023971 | 0.043013 | 0.083014 | 0.117407 |
| 31.5277778 | 0.062719 | 0.081815 | 0.037689 | 0.033677 | 0.01715 | 0.032301 | 0.084989 | 0.071706 | 0.02776 | 0.081783 | 0.048494 | 0.05036 | 0.055943 | 0.126014 | 0.005287 | 0.03109 | 0.073316 | 0.016091 | 0.022943 | 0.027254 | 0.021998 | 0.070754 | 0.082554 | 0.095252 |
| 31.5416667 | 0.048674 | 0.077196 | 0.03633 | 0.020303 | 0.016822 | 0.02275 | 0.06872 | 0.065349 | 0.040023 | 0.076627 | 0.044309 | 0.059874 | 0.04528 | 0.112256 | 0.008341 | 0.024608 | 0.076354 | 0.022567 | 0.03408 | 0.020666 | 0.018442 | 0.094727 | 0.065932 | 0.073607 |
| 31.5555556 | 0.027094 | 0.074058 | 0.028435 | 0.011271 | 0.017014 | 0.025971 | 0.070434 | 0.066528 | 0.066232 | 0.068696 | 0.027956 | 0.053235 | 0.046092 | 0.08225 | 0.008619 | 0.020727 | 0.076193 | 0.028717 | 0.045022 | 0.015908 | 0.019338 | 0.108919 | 0.051679 | 0.056918 |
| 31.5694444 | 0.021589 | 0.079175 | 0.019015 | 0.008578 | 0.015545 | 0.034906 | 0.088894 | 0.071818 | 0.077241 | 0.051576 | 0.015211 | 0.03326 | 0.043376 | 0.046239 | 0.009375 | 0.02 | 0.068315 | 0.02981 | 0.048398 | 0.014127 | 0.023046 | 0.106828 | 0.039503 | 0.039111 |
| 31.5833333 | 0.03527 | 0.086051 | 0.009632 | 0.008264 | 0.012059 | 0.037869 | 0.086051 | 0.073624 | 0.068338 | 0.033135 | 0.012953 | 0.014079 | 0.03492 | 0.030898 | 0.015716 | 0.019242 | 0.05264 | 0.032979 | 0.048363 | 0.014338 | 0.022492 | 0.07751 | 0.033258 | 0.026465 |
| 31.5972222 | 0.043386 | 0.079616 | 0.004645 | 0.009361 | 0.008189 | 0.034034 | 0.051983 | 0.068222 | 0.049607 | 0.018985 | 0.012249 | 0.007071 | 0.022989 | 0.03808 | 0.02656 | 0.017034 | 0.036117 | 0.038445 | 0.043945 | 0.016981 | 0.01604 | 0.048177 | 0.035421 | 0.022781 |
| 31.6111111 | 0.039871 | 0.061556 | 0.00805 | 0.011462 | 0.004847 | 0.02028 | 0.024581 | 0.054839 | 0.027529 | 0.009007 | 0.006678 | 0.013794 | 0.013041 | 0.042993 | 0.035603 | 0.014493 | 0.021708 | 0.036436 | 0.032906 | 0.022613 | 0.011136 | 0.034744 | 0.033556 | 0.021327 |
| 31.625 | 0.030889 | 0.04238 | 0.01597 | 0.013499 | 0.00253 | 0.009668 | 0.032495 | 0.040031 | 0.01891 | 0.006974 | 0.004103 | 0.021441 | 0.015421 | 0.045544 | 0.037447 | 0.01311 | 0.010714 | 0.026799 | 0.020331 | 0.029079 | 0.008461 | 0.024502 | 0.022012 | 0.015821 |
| 31.6388889 | 0.018439 | 0.024391 | 0.024056 | 0.013638 | 0.002324 | 0.012287 | 0.060941 | 0.029455 | 0.028013 | 0.01322 | 0.006784 | 0.023228 | 0.025382 | 0.051845 | 0.03331 | 0.012401 | 0.004305 | 0.02015 | 0.012271 | 0.032687 | 0.004281 | 0.014779 | 0.011046 | 0.007693 |
| 31.6527778 | 0.008572 | 0.012448 | 0.030543 | 0.01094 | 0.003393 | 0.013479 | 0.078373 | 0.022623 | 0.035006 | 0.021618 | 0.011726 | 0.02463 | 0.032641 | 0.055142 | 0.027352 | 0.011202 | 0.002302 | 0.019529 | 0.007667 | 0.031339 | 0.001302 | 0.006457 | 0.00681 | 0.003459 |
| 31.6666667 | 0.006463 | 0.015619 | 0.031414 | 0.006307 | 0.003649 | 0.008981 | 0.073441 | 0.017546 | 0.03027 | 0.02946 | 0.02087 | 0.025457 | 0.036656 | 0.059669 | 0.022174 | 0.00935 | 0.002995 | 0.01759 | 0.006599 | 0.024828 | 0.001301 | 0.003445 | 0.005861 | 0.006948 |
| 31.6805556 | 0.016969 | 0.029578 | 0.022176 | 0.003223 | 0.003122 | 0.004301 | 0.065665 | 0.014151 | 0.020941 | 0.036765 | 0.028181 | 0.02225 | 0.040253 | 0.071845 | 0.0185 | 0.007804 | 0.003385 | 0.01149 | 0.012361 | 0.016552 | 0.002232 | 0.00763 | 0.007552 | 0.014423 |
| 31.6944444 | 0.037962 | 0.041237 | 0.0097 | 0.003547 | 0.002091 | 0.001779 | 0.070069 | 0.013655 | 0.014007 | 0.042515 | 0.02946 | 0.013489 | 0.040581 | 0.083401 | 0.015899 | 0.007073 | 0.00259 | 0.005818 | 0.018739 | 0.008622 | 0.002347 | 0.012367 | 0.014068 | 0.02123 |
| 31.7083333 | 0.042441 | 0.045702 | 0.004071 | 0.004171 | 0.001331 | 0.002516 | 0.07483 | 0.017115 | 0.009772 | 0.047423 | 0.031609 | 0.005274 | 0.034963 | 0.081583 | 0.012735 | 0.007196 | 0.001621 | 0.005879 | 0.018335 | 0.004302 | 0.002217 | 0.011978 | 0.024596 | 0.026708 |
| 31.7222222 | 0.022448 | 0.052159 | 0.002546 | 0.005887 | 0.002076 | 0.004538 | 0.068961 | 0.021717 | 0.00773 | 0.052997 | 0.03318 | 0.003567 | 0.025793 | 0.067007 | 0.009122 | 0.008698 | 0.001132 | 0.013577 | 0.011932 | 0.007402 | 0.002899 | 0.007137 | 0.038496 | 0.03109 |
| 31.7361111 | 0.012153 | 0.06609 | 0.000886 | 0.010117 | 0.002484 | 0.005785 | 0.055424 | 0.022348 | 0.01135 | 0.058552 | 0.025337 | 0.003699 | 0.01529 | 0.047924 | 0.007284 | 0.012329 | 0.00111 | 0.024067 | 0.005258 | 0.013461 | 0.004199 | 0.003314 | 0.050527 | 0.032619 |
| 31.75 | 0.017175 | 0.078202 | 9.7076396 | 0.011665 | 0.001525 | 0.005936 | 0.041154 | 0.019014 | 0.020147 | 0.063554 | 0.017219 | 0.003056 | 0.00857 | 0.029797 | 0.006353 | 0.017934 | 0.000976 | 0.027978 | 0.003756 | 0.016121 | 0.005954 | 0.005163 | 0.051384 | 0.029554 |
| 31.7638889 | 0.017927 | 0.081929 | 0.000706 | 0.007978 | 0.000963 | 0.006041 | 0.029769 | 0.013916 | 0.028327 | 0.066116 | 0.016751 | 0.002967 | 0.01223 | 0.016524 | 0.006149 | 0.023294 | 0.000682 | 0.021414 | 0.006262 | 0.013197 | 0.007752 | 0.008741 | 0.042988 | 0.02297 |
| 31.7777778 | 0.012004 | 0.080729 | 0.002283 | 0.004091 | 0.000784 | 0.006325 | 0.022445 | 0.008523 | 0.030916 | 0.06206 | 0.01155 | 0.005918 | 0.020475 | 0.009036 | 0.008542 | 0.025976 | 0.000587 | 0.010625 | 0.007917 | 0.00773 | 0.009664 | 0.008997 | 0.033905 | 0.01625 |
| 31.7916667 | 0.008885 | 0.078706 | 0.003682 | 0.00476 | 0.000459 | 0.005966 | 0.020432 | 0.004317 | 0.028069 | 0.056552 | 0.00355 | 0.015266 | 0.025067 | 0.004872 | 0.011824 | 0.025127 | 0.001202 | 0.004549 | 0.007239 | 0.005532 | 0.012055 | 0.007073 | 0.026157 | 0.012911 |
| 31.8055556 | 0.00788 | 0.071018 | 0.00436 | 0.006667 | 0.000468 | 0.004029 | 0.022424 | 0.002278 | 0.021333 | 0.058715 | 0.002481 | 0.028471 | 0.02561 | 0.003026 | 0.012691 | 0.021275 | 0.003733 | 0.00798 | 0.006476 | 0.005224 | 0.013198 | 0.004438 | 0.0159 | 0.014071 |
| 31.8194444 | 0.005202 | 0.054934 | 0.006835 | 0.004979 | 0.000817 | 0.001878 | 0.025903 | 0.002326 | 0.01252 | 0.067883 | 0.004136 | 0.042101 | 0.02188 | 0.00594 | 0.012066 | 0.015443 | 0.007896 | 0.016003 | 0.00512 | 0.004012 | 0.011783 | 0.003094 | 0.006359 | 0.017037 |
| 31.8333333 | 0.005222 | 0.037729 | 0.013552 | 0.002472 | 0.001413 | 0.002229 | 0.031587 | 0.002912 | 0.005862 | 0.074365 | 0.006445 | 0.057234 | 0.013017 | 0.016426 | 0.011236 | 0.009117 | 0.012848 | 0.019519 | 0.003581 | 0.003453 | 0.008997 | 0.007023 | 0.004174 | 0.018861 |
| 31.8472222 | 0.006897 | 0.026439 | 0.020036 | 0.00198 | 0.001578 | 0.004834 | 0.041201 | 0.004343 | 0.007152 | 0.07096 | 0.01123 | 0.068561 | 0.005323 | 0.032828 | 0.008029 | 0.00692 | 0.018532 | 0.01395 | 0.008046 | 0.00231 | 0.006481 | 0.015403 | 0.010371 | 0.021423 |
| 31.8611111 | 0.007588 | 0.02239 | 0.020697 | 0.002147 | 0.000908 | 0.008461 | 0.052322 | 0.00827 | 0.01267 | 0.060596 | 0.010996 | 0.067197 | 0.00319 | 0.049411 | 0.005099 | 0.010284 | 0.023602 | 0.008245 | 0.020687 | 0.000904 | 0.004553 | 0.024083 | 0.027491 | 0.030222 |
| 31.875 | 0.007672 | 0.023484 | 0.018364 | 0.003764 | 0.000297 | 0.011797 | 0.057601 | 0.014876 | 0.011231 | 0.045645 | 0.004535 | 0.057642 | 0.002237 | 0.059971 | 0.007131 | 0.012231 | 0.026451 | 0.007787 | 0.03797 | 0.000854 | 0.002782 | 0.029292 | 0.048833 | 0.046697 |
| 31.8888889 | 0.007919 | 0.026526 | 0.016973 | 0.006484 | 0.000557 | 0.012999 | 0.054434 | 0.022452 | 0.005345 | 0.027739 | 0.002311 | 0.047093 | 0.001678 | 0.059649 | 0.009747 | 0.008486 | 0.026675 | 0.003982 | 0.055571 | 0.003178 | 0.001364 | 0.028994 | 0.061604 | 0.062856 |
| 31.9027778 | 0.008009 | 0.02598 | 0.015539 | 0.007908 | 0.001712 | 0.011945 | 0.046841 | 0.027419 | 0.003915 | 0.01345 | 0.005214 | 0.035242 | 0.002671 | 0.050786 | 0.006928 | 0.003249 | 0.024658 | 0.001884 | 0.067115 | 0.009088 | 0.001503 | 0.025002 | 0.06173 | 0.070412 |
| 31.9166667 | 0.008739 | 0.020536 | 0.012215 | 0.007421 | 0.002938 | 0.009032 | 0.037755 | 0.026908 | 0.005645 | 0.005411 | 0.008365 | 0.023272 | 0.006412 | 0.042738 | 0.002911 | 0.002668 | 0.022675 | 0.005907 | 0.068262 | 0.015258 | 0.003357 | 0.019802 | 0.05718 | 0.070551 |
| 31.9305556 | 0.012602 | 0.013204 | 0.007256 | 0.005249 | 0.003666 | 0.005203 | 0.028998 | 0.021518 | 0.006906 | 0.002301 | 0.011206 | 0.012988 | 0.012549 | 0.038845 | 0.003368 | 0.006409 | 0.022532 | 0.010628 | 0.05747 | 0.016662 | 0.004771 | 0.014691 | 0.059172 | 0.069775 |
| 31.9444444 | 0.019176 | 0.009395 | 0.005502 | 0.0026 | 0.004116 | 0.004241 | 0.021502 | 0.013865 | 0.008081 | 0.003577 | 0.014035 | 0.00636 | 0.016519 | 0.031305 | 0.005444 | 0.008151 | 0.022466 | 0.013499 | 0.033968 | 0.012542 | 0.004086 | 0.009545 | 0.063137 | 0.073166 |
| 31.9583333 | 0.02559 | 0.01613 | 0.009961 | 0.001113 | 0.004463 | 0.007572 | 0.015592 | 0.006422 | 0.01002 | 0.005696 | 0.016197 | 0.006852 | 0.01729 | 0.01692 | 0.005689 | 0.005256 | 0.020927 | 0.013223 | 0.014553 | 0.007456 | 0.002244 | 0.004392 | 0.060626 | 0.079577 |
| 31.9722222 | 0.030093 | 0.026913 | 0.014993 | 0.002267 | 0.00456 | 0.010714 | 0.012788 | 0.00272 | 0.010936 | 0.006671 | 0.013303 | 0.011085 | 0.015863 | 0.006342 | 0.004086 | 0.001693 | 0.021354 | 0.011082 | 0.014583 | 0.005029 | 0.00163 | 0.005772 | 0.058337 | 0.088627 |
| 31.9861111 | 0.028329 | 0.027039 | 0.013871 | 0.010914 | 0.00489 | 0.008951 | 0.016577 | 0.006682 | 0.008897 | 0.016302 | 0.009606 | 0.010267 | 0.014531 | 0.016816 | 0.005828 | 0.003591 | 0.031035 | 0.008527 | 0.016527 | 0.006269 | 0.006049 | 0.029246 | 0.072612 | 0.114029 |
| 32 | 0.017879 | 0.019001 | 0.009114 | 0.027524 | 0.005855 | 0.005784 | 0.025892 | 0.016693 | 0.011335 | 0.043729 | 0.01888 | 0.008921 | 0.022786 | 0.054818 | 0.015444 | 0.01266 | 0.047483 | 0.008998 | 0.025984 | 0.010039 | 0.014467 | 0.063038 | 0.099939 | 0.153817 |
| 32.0138889 | 0.010008 | 0.018967 | 0.009405 | 0.038889 | 0.005911 | 0.007514 | 0.033373 | 0.025277 | 0.023348 | 0.080638 | 0.037262 | 0.020663 | 0.03825 | 0.094154 | 0.022468 | 0.020008 | 0.052959 | 0.01899 | 0.059635 | 0.012993 | 0.017812 | 0.080087 | 0.115307 | 0.175725 |
| 32.0277778 | 0.013429 | 0.028599 | 0.014572 | 0.037525 | 0.004775 | 0.011177 | 0.034584 | 0.028413 | 0.036574 | 0.096982 | 0.046527 | 0.03878 | 0.044191 | 0.104 | 0.023343 | 0.016527 | 0.039131 | 0.036354 | 0.084235 | 0.015953 | 0.012575 | 0.077034 | 0.104704 | 0.156734 |
| 32.0416667 | 0.021169 | 0.036853 | 0.017335 | 0.031996 | 0.005615 | 0.012497 | 0.031026 | 0.027542 | 0.042638 | 0.068056 | 0.041546 | 0.045777 | 0.037407 | 0.092388 | 0.025461 | 0.008466 | 0.022556 | 0.049277 | 0.080091 | 0.021213 | 0.005634 | 0.056403 | 0.07042 | 0.114986 |
| 32.0555556 | 0.022626 | 0.041122 | 0.015704 | 0.026097 | 0.009096 | 0.011056 | 0.027444 | 0.024417 | 0.038095 | 0.031672 | 0.03423 | 0.03829 | 0.02605 | 0.078032 | 0.022118 | 0.006964 | 0.021403 | 0.042787 | 0.067914 | 0.025879 | 0.003912 | 0.034471 | 0.030427 | 0.085132 |
| 32.0694444 | 0.015683 | 0.043516 | 0.011056 | 0.015609 | 0.011003 | 0.006768 | 0.023049 | 0.019359 | 0.026374 | 0.022765 | 0.035467 | 0.022438 | 0.025261 | 0.064177 | 0.012768 | 0.013729 | 0.029834 | 0.027186 | 0.061283 | 0.026867 | 0.006886 | 0.022933 | 0.006897 | 0.063574 |
| 32.0833333 | 0.01363 | 0.044484 | 0.006095 | 0.00843 | 0.009044 | 0.003493 | 0.014439 | 0.011591 | 0.014468 | 0.016166 | 0.042025 | 0.009465 | 0.037495 | 0.044559 | 0.008392 | 0.018594 | 0.032699 | 0.034884 | 0.047871 | 0.023235 | 0.009848 | 0.015524 | 0.002896 | 0.037807 |
| 32.0972222 | 0.021665 | 0.042912 | 0.005269 | 0.012002 | 0.005829 | 0.00519 | 0.006764 | 0.005926 | 0.007313 | 0.008071 | 0.037709 | 0.006821 | 0.041402 | 0.025783 | 0.009286 | 0.013031 | 0.028755 | 0.054575 | 0.025903 | 0.013706 | 0.010025 | 0.009997 | 0.005607 | 0.017151 |
| 32.1111111 | 0.028426 | 0.036876 | 0.007726 | 0.016385 | 0.004125 | 0.009195 | 0.005006 | 0.007827 | 0.007078 | 0.008217 | 0.020252 | 0.008693 | 0.035121 | 0.029498 | 0.010338 | 0.006889 | 0.022641 | 0.05096 | 0.018889 | 0.005717 | 0.007961 | 0.010473 | 0.007089 | 0.00609 |
| 32.125 | 0.030821 | 0.029043 | 0.009334 | 0.015665 | 0.00387 | 0.011529 | 0.005379 | 0.012088 | 0.00963 | 0.008145 | 0.008405 | 0.008654 | 0.031172 | 0.045079 | 0.010957 | 0.008422 | 0.016567 | 0.031027 | 0.026666 | 0.006519 | 0.005872 | 0.018312 | 0.008709 | 0.002889 |
| 32.1388889 | 0.033905 | 0.022832 | 0.011584 | 0.013164 | 0.003741 | 0.01064 | 0.003581 | 0.013291 | 0.010096 | 0.005744 | 0.007521 | 0.007993 | 0.029052 | 0.051255 | 0.011083 | 0.012344 | 0.011201 | 0.020152 | 0.033659 | 0.009566 | 0.005076 | 0.023585 | 0.009736 | 0.005284 |
| 32.1527778 | 0.037261 | 0.016542 | 0.016149 | 0.011049 | 0.003026 | 0.006407 | 0.001309 | 0.011889 | 0.008122 | 0.003763 | 0.007037 | 0.008749 | 0.026815 | 0.046484 | 0.008856 | 0.01492 | 0.00592 | 0.015087 | 0.038821 | 0.009052 | 0.00567 | 0.020901 | 0.008788 | 0.008973 |
| 32.1666667 | 0.038332 | 0.009416 | 0.019499 | 0.01017 | 0.001807 | 0.002555 | 0.001957 | 0.008819 | 0.005044 | 0.005401 | 0.005057 | 0.010646 | 0.025496 | 0.035522 | 0.005264 | 0.01561 | 0.002717 | 0.008231 | 0.046148 | 0.006044 | 0.007375 | 0.015085 | 0.008059 | 0.012107 |
| 32.1805556 | 0.039643 | 0.008158 | 0.019532 | 0.0106 | 0.001255 | 0.002319 | 0.006349 | 0.005213 | 0.004354 | 0.009932 | 0.007538 | 0.013965 | 0.027681 | 0.024554 | 0.003235 | 0.015577 | 0.003576 | 0.003822 | 0.056171 | 0.003553 | 0.00963 | 0.010247 | 0.00927 | 0.014943 |
| 32.1944444 | 0.04296 | 0.015909 | 0.018343 | 0.012082 | 0.002673 | 0.003204 | 0.01272 | 0.002657 | 0.007456 | 0.013373 | 0.014 | 0.019734 | 0.034942 | 0.01529 | 0.003527 | 0.016579 | 0.005907 | 0.00402 | 0.064859 | 0.00347 | 0.011654 | 0.007024 | 0.01245 | 0.016751 |
| 32.2083333 | 0.043003 | 0.02502 | 0.017313 | 0.013797 | 0.005094 | 0.002392 | 0.01801 | 0.002333 | 0.011009 | 0.01404 | 0.01887 | 0.026337 | 0.042509 | 0.009243 | 0.004671 | 0.018113 | 0.006838 | 0.00765 | 0.065837 | 0.005614 | 0.013254 | 0.004958 | 0.015456 | 0.017083 |
| 32.2222222 | 0.035368 | 0.030272 | 0.016019 | 0.014607 | 0.006884 | 0.002063 | 0.020094 | 0.0043 | 0.013917 | 0.012613 | 0.023432 | 0.029742 | 0.043824 | 0.006744 | 0.005445 | 0.018878 | 0.005927 | 0.00994 | 0.057315 | 0.010602 | 0.014917 | 0.004161 | 0.016622 | 0.015587 |
| 32.2361111 | 0.025817 | 0.031739 | 0.016008 | 0.014039 | 0.007546 | 0.005136 | 0.020103 | 0.00799 | 0.01616 | 0.011469 | 0.030981 | 0.029421 | 0.038768 | 0.005269 | 0.006233 | 0.018356 | 0.00362 | 0.01058 | 0.043347 | 0.019072 | 0.01648 | 0.004912 | 0.016911 | 0.012578 |
| 32.25 | 0.017534 | 0.030231 | 0.018467 | 0.012632 | 0.006937 | 0.009097 | 0.020057 | 0.011992 | 0.015919 | 0.012286 | 0.034946 | 0.028434 | 0.032328 | 0.003749 | 0.007499 | 0.017338 | 0.0015 | 0.012474 | 0.028836 | 0.027828 | 0.017183 | 0.006487 | 0.016822 | 0.010131 |
| 32.2638889 | 0.009227 | 0.0286 | 0.020502 | 0.010957 | 0.005503 | 0.010264 | 0.019744 | 0.013731 | 0.012354 | 0.014675 | 0.02944 | 0.02701 | 0.027528 | 0.004926 | 0.009495 | 0.016677 | 0.001077 | 0.014873 | 0.016609 | 0.032465 | 0.016772 | 0.007809 | 0.016712 | 0.011225 |
| 32.2777778 | 0.00375 | 0.02751 | 0.018996 | 0.009437 | 0.004009 | 0.009102 | 0.018846 | 0.012739 | 0.008712 | 0.017764 | 0.016764 | 0.024509 | 0.024776 | 0.00932 | 0.011923 | 0.016512 | 0.001774 | 0.015676 | 0.007761 | 0.033777 | 0.015322 | 0.007679 | 0.015718 | 0.017695 |
| 32.2916667 | 0.003274 | 0.025356 | 0.014171 | 0.008314 | 0.00285 | 0.008386 | 0.017492 | 0.01022 | 0.008 | 0.020322 | 0.007707 | 0.021489 | 0.023791 | 0.009878 | 0.013839 | 0.016576 | 0.002457 | 0.015654 | 0.005292 | 0.033994 | 0.013564 | 0.005822 | 0.012863 | 0.028724 |
| 32.3055556 | 0.007365 | 0.022581 | 0.008713 | 0.007755 | 0.002154 | 0.010174 | 0.016801 | 0.006552 | 0.008412 | 0.021456 | 0.009364 | 0.017619 | 0.023354 | 0.006683 | 0.015822 | 0.016835 | 0.003667 | 0.016344 | 0.006519 | 0.033171 | 0.012386 | 0.003201 | 0.010897 | 0.040738 |
| 32.3194444 | 0.012584 | 0.019067 | 0.005591 | 0.007867 | 0.001612 | 0.014481 | 0.018365 | 0.006052 | 0.00825 | 0.021084 | 0.010246 | 0.012707 | 0.024245 | 0.008237 | 0.018122 | 0.017448 | 0.007113 | 0.017157 | 0.007736 | 0.031572 | 0.011811 | 0.002604 | 0.012259 | 0.050229 |
| 32.3333333 | 0.016326 | 0.014429 | 0.005978 | 0.008267 | 0.001088 | 0.020159 | 0.020589 | 0.010423 | 0.006899 | 0.019255 | 0.007606 | 0.007283 | 0.0269 | 0.013984 | 0.019616 | 0.017523 | 0.01216 | 0.016911 | 0.012015 | 0.028566 | 0.011207 | 0.006748 | 0.015394 | 0.054588 |
| 32.3472222 | 0.017699 | 0.009959 | 0.008204 | 0.008397 | 0.001018 | 0.025023 | 0.019695 | 0.012513 | 0.004638 | 0.016441 | 0.010011 | 0.003331 | 0.027462 | 0.017366 | 0.020059 | 0.016736 | 0.015106 | 0.014485 | 0.012483 | 0.023671 | 0.009719 | 0.013469 | 0.018281 | 0.051209 |
| 32.3611111 | 0.016816 | 0.006325 | 0.009264 | 0.007331 | 0.0012 | 0.026674 | 0.014798 | 0.010259 | 0.003065 | 0.013293 | 0.013872 | 0.00268 | 0.022495 | 0.016433 | 0.018694 | 0.016256 | 0.014468 | 0.009812 | 0.00896 | 0.018845 | 0.00694 | 0.018139 | 0.020473 | 0.043292 |
| 32.375 | 0.014115 | 0.003978 | 0.008078 | 0.004369 | 0.000933 | 0.024206 | 0.008319 | 0.008988 | 0.002686 | 0.009665 | 0.014686 | 0.003806 | 0.014657 | 0.013503 | 0.015346 | 0.015581 | 0.012338 | 0.005425 | 0.014899 | 0.015589 | 0.003854 | 0.019069 | 0.022413 | 0.038773 |
| 32.3888889 | 0.010076 | 0.003776 | 0.005924 | 0.001674 | 0.000429 | 0.018935 | 0.003981 | 0.010854 | 0.003453 | 0.005402 | 0.013462 | 0.003864 | 0.008766 | 0.010173 | 0.011224 | 0.014186 | 0.010371 | 0.003252 | 0.026831 | 0.012615 | 0.001676 | 0.018936 | 0.024311 | 0.038263 |
| 32.4027778 | 0.007854 | 0.007103 | 0.005423 | 0.001436 | 0.000312 | 0.014814 | 0.003325 | 0.010965 | 0.005851 | 0.002092 | 0.014574 | 0.002385 | 0.005754 | 0.007364 | 0.006071 | 0.013361 | 0.008638 | 0.003847 | 0.033679 | 0.00918 | 0.000678 | 0.020812 | 0.025079 | 0.038124 |
| 32.4166667 | 0.01027 | 0.014208 | 0.008482 | 0.001611 | 0.000322 | 0.01427 | 0.003018 | 0.008875 | 0.010307 | 0.000798 | 0.022433 | 0.001168 | 0.006877 | 0.006872 | 0.002355 | 0.013886 | 0.007072 | 0.009049 | 0.031226 | 0.006581 | 0.000547 | 0.02286 | 0.024976 | 0.04046 |
| 32.4305556 | 0.016566 | 0.021029 | 0.013393 | 0.001217 | 0.000521 | 0.015154 | 0.00305 | 0.011506 | 0.016411 | 0.000687 | 0.034125 | 0.001317 | 0.01293 | 0.009109 | 0.004642 | 0.015561 | 0.005849 | 0.017619 | 0.019079 | 0.0063 | 0.000822 | 0.021606 | 0.025918 | 0.046525 |
| 32.4444444 | 0.024167 | 0.023969 | 0.017926 | 0.00109 | 0.001883 | 0.014515 | 0.00596 | 0.016067 | 0.020066 | 0.001297 | 0.035334 | 0.001437 | 0.021463 | 0.012492 | 0.01143 | 0.0175 | 0.005098 | 0.025134 | 0.010961 | 0.009405 | 0.001333 | 0.017984 | 0.028074 | 0.050347 |
| 32.4583333 | 0.030814 | 0.023078 | 0.021745 | 0.000626 | 0.00402 | 0.011938 | 0.009563 | 0.015384 | 0.016601 | 0.002165 | 0.02237 | 0.001383 | 0.028667 | 0.014579 | 0.017445 | 0.019028 | 0.00471 | 0.0306 | 0.018471 | 0.014313 | 0.001622 | 0.014246 | 0.030524 | 0.049454 |
| 32.4722222 | 0.035882 | 0.018364 | 0.024318 | 0.001437 | 0.004837 | 0.010236 | 0.01454 | 0.010997 | 0.011562 | 0.005334 | 0.014808 | 0.002058 | 0.032008 | 0.017166 | 0.020807 | 0.020797 | 0.003535 | 0.034181 | 0.031754 | 0.018838 | 0.003397 | 0.010368 | 0.037788 | 0.055451 |
| 32.4861111 | 0.040544 | 0.012775 | 0.02602 | 0.007409 | 0.003706 | 0.012659 | 0.025716 | 0.010574 | 0.016251 | 0.01487 | 0.017569 | 0.003429 | 0.038923 | 0.027756 | 0.023616 | 0.025759 | 0.006122 | 0.037297 | 0.043201 | 0.024554 | 0.010271 | 0.016637 | 0.065198 | 0.087645 |
| 32.5 | 0.045707 | 0.016451 | 0.027116 | 0.019398 | 0.003539 | 0.017766 | 0.04751 | 0.021196 | 0.030314 | 0.026233 | 0.024303 | 0.010283 | 0.059143 | 0.046301 | 0.027441 | 0.034626 | 0.020475 | 0.046144 | 0.060097 | 0.033855 | 0.020055 | 0.045233 | 0.108422 | 0.137803 |
| 32.5138889 | 0.053012 | 0.032312 | 0.026689 | 0.030645 | 0.004455 | 0.019103 | 0.075327 | 0.040726 | 0.038093 | 0.031039 | 0.044381 | 0.027363 | 0.079903 | 0.058066 | 0.027679 | 0.0426 | 0.039666 | 0.063791 | 0.084659 | 0.04313 | 0.024606 | 0.079861 | 0.126707 | 0.171856 |
| 32.5277778 | 0.061559 | 0.049265 | 0.027992 | 0.033893 | 0.003289 | 0.013369 | 0.093319 | 0.060742 | 0.03073 | 0.033191 | 0.074142 | 0.051517 | 0.084606 | 0.059204 | 0.023616 | 0.046505 | 0.051414 | 0.082831 | 0.114013 | 0.043371 | 0.024782 | 0.09742 | 0.10768 | 0.189389 |
| 32.5416667 | 0.065187 | 0.068342 | 0.032381 | 0.03221 | 0.002052 | 0.005805 | 0.100595 | 0.074364 | 0.019381 | 0.042658 | 0.083894 | 0.068074 | 0.083011 | 0.063813 | 0.021373 | 0.046198 | 0.058306 | 0.087611 | 0.13036 | 0.032041 | 0.029538 | 0.100761 | 0.083388 | 0.203194 |
| 32.5555556 | 0.05939 | 0.099661 | 0.03216 | 0.029794 | 0.00293 | 0.003133 | 0.107716 | 0.076059 | 0.020164 | 0.063794 | 0.059147 | 0.065802 | 0.081646 | 0.074299 | 0.022714 | 0.040711 | 0.065015 | 0.074501 | 0.110009 | 0.016084 | 0.035777 | 0.101451 | 0.069555 | 0.193263 |
| 32.5694444 | 0.049525 | 0.117109 | 0.024033 | 0.024127 | 0.003586 | 0.004368 | 0.113527 | 0.067937 | 0.034042 | 0.08414 | 0.023669 | 0.055347 | 0.074132 | 0.07864 | 0.022986 | 0.03315 | 0.068675 | 0.059391 | 0.079525 | 0.004923 | 0.036053 | 0.100948 | 0.057771 | 0.148929 |
| 32.5833333 | 0.037306 | 0.099145 | 0.013442 | 0.01742 | 0.00356 | 0.005631 | 0.110768 | 0.054318 | 0.046843 | 0.087306 | 0.013729 | 0.044978 | 0.065175 | 0.071717 | 0.014837 | 0.029991 | 0.070056 | 0.050006 | 0.073173 | 0.001888 | 0.034288 | 0.090407 | 0.045606 | 0.097408 |
| 32.5972222 | 0.023427 | 0.078586 | 0.009435 | 0.016482 | 0.003645 | 0.007089 | 0.115547 | 0.037792 | 0.05198 | 0.081119 | 0.029426 | 0.033301 | 0.046098 | 0.060387 | 0.005964 | 0.031547 | 0.071853 | 0.043735 | 0.080557 | 0.001596 | 0.032376 | 0.064986 | 0.040452 | 0.064834 |
| 32.6111111 | 0.03129 | 0.072411 | 0.010535 | 0.019186 | 0.00261 | 0.011384 | 0.137616 | 0.032403 | 0.049411 | 0.072585 | 0.034218 | 0.022815 | 0.019627 | 0.048937 | 0.007432 | 0.034758 | 0.07177 | 0.047133 | 0.075696 | 0.003762 | 0.022213 | 0.036702 | 0.043398 | 0.046715 |
| 32.625 | 0.059571 | 0.065859 | 0.007029 | 0.020613 | 0.001715 | 0.017651 | 0.131628 | 0.043978 | 0.038314 | 0.053339 | 0.021209 | 0.01384 | 0.011867 | 0.0338 | 0.014661 | 0.036271 | 0.06597 | 0.055284 | 0.052492 | 0.012804 | 0.010683 | 0.016241 | 0.042964 | 0.027565 |
| 32.6388889 | 0.072601 | 0.056704 | 0.002676 | 0.021304 | 0.002591 | 0.019774 | 0.084726 | 0.051265 | 0.023869 | 0.030068 | 0.013739 | 0.006291 | 0.013248 | 0.020216 | 0.020422 | 0.031645 | 0.053218 | 0.054239 | 0.030066 | 0.024411 | 0.013451 | 0.006753 | 0.033698 | 0.016346 |
| 32.6527778 | 0.064865 | 0.047805 | 0.001802 | 0.019386 | 0.003226 | 0.017239 | 0.042234 | 0.039734 | 0.01127 | 0.027303 | 0.016071 | 0.003736 | 0.019655 | 0.024435 | 0.021002 | 0.021311 | 0.035378 | 0.05109 | 0.018934 | 0.030591 | 0.01939 | 0.008914 | 0.026754 | 0.024738 |
| 32.6666667 | 0.049909 | 0.039415 | 0.002508 | 0.012602 | 0.00249 | 0.013205 | 0.041458 | 0.021706 | 0.00391 | 0.048741 | 0.024122 | 0.008368 | 0.035725 | 0.046436 | 0.019688 | 0.010631 | 0.017148 | 0.050089 | 0.015283 | 0.031981 | 0.019378 | 0.015665 | 0.023088 | 0.03702 |
| 32.6805556 | 0.032782 | 0.032954 | 0.003617 | 0.006184 | 0.002383 | 0.008063 | 0.074767 | 0.010404 | 0.004209 | 0.066834 | 0.03944 | 0.017551 | 0.038047 | 0.066098 | 0.021608 | 0.005791 | 0.008766 | 0.040098 | 0.022245 | 0.031354 | 0.022077 | 0.02088 | 0.014992 | 0.046567 |
| 32.6944444 | 0.020026 | 0.026954 | 0.004505 | 0.006563 | 0.004355 | 0.003809 | 0.096701 | 0.007336 | 0.009783 | 0.066062 | 0.053831 | 0.0265 | 0.022889 | 0.068095 | 0.024459 | 0.007243 | 0.012727 | 0.021397 | 0.038088 | 0.027443 | 0.025846 | 0.023381 | 0.011496 | 0.060789 |

| | | | | | | | | | | | | | | | | | | | | | | | | |
|---|---|---|---|---|---|---|---|---|---|---|---|---|---|---|---|---|---|---|---|---|---|---|---|---|
| 32.7083333 | 0.021699 | 0.016963 | 0.005268 | 0.010811 | 0.006559 | 0.002548 | 0.09409 | 0.010225 | 0.01509 | 0.059676 | 0.056207 | 0.030036 | 0.009893 | 0.057042 | 0.025194 | 0.009606 | 0.016365 | 0.011579 | 0.048012 | 0.019012 | 0.03017 | 0.022227 | 0.016067 | 0.085426 |
| 32.7222222 | 0.032306 | 0.007305 | 0.005423 | 0.011448 | 0.007166 | 0.003245 | 0.078357 | 0.016726 | 0.015849 | 0.056321 | 0.044762 | 0.026009 | 0.005004 | 0.043895 | 0.025827 | 0.009501 | 0.013341 | 0.012463 | 0.043597 | 0.009161 | 0.032125 | 0.01491 | 0.021768 | 0.121941 |
| 32.7361111 | 0.036136 | 0.007122 | 0.005025 | 0.007986 | 0.006745 | 0.002977 | 0.061484 | 0.023587 | 0.011233 | 0.051702 | 0.026007 | 0.019148 | 0.002566 | 0.028771 | 0.023409 | 0.007298 | 0.00718 | 0.009585 | 0.027346 | 0.005173 | 0.021863 | 0.010006 | 0.031752 | 0.147294 |
| 32.75 | 0.03462 | 0.015973 | 0.00466 | 0.006258 | 0.007085 | 0.001995 | 0.057619 | 0.026878 | 0.005177 | 0.048046 | 0.017119 | 0.015924 | 0.002978 | 0.01377 | 0.014952 | 0.004519 | 0.003415 | 0.00351 | 0.019964 | 0.004983 | 0.015565 | 0.019279 | 0.036722 | 0.140291 |
| 32.7638889 | 0.033014 | 0.031287 | 0.005246 | 0.008774 | 0.008929 | 0.001988 | 0.065531 | 0.026542 | 0.00292 | 0.048245 | 0.017025 | 0.018279 | 0.005788 | 0.006009 | 0.00682 | 0.003948 | 0.004657 | 0.001453 | 0.032878 | 0.007134 | 0.027958 | 0.026865 | 0.027917 | 0.113983 |
| 32.7777778 | 0.031917 | 0.048189 | 0.00618 | 0.012659 | 0.011287 | 0.002286 | 0.078017 | 0.024739 | 0.003819 | 0.051048 | 0.017447 | 0.023153 | 0.008527 | 0.00826 | 0.002603 | 0.007835 | 0.005154 | 0.002142 | 0.040558 | 0.018183 | 0.039702 | 0.019851 | 0.025624 | 0.081556 |
| 32.7916667 | 0.031292 | 0.057301 | 0.006325 | 0.015695 | 0.014176 | 0.003632 | 0.089911 | 0.020578 | 0.003945 | 0.054188 | 0.025706 | 0.025726 | 0.013047 | 0.015817 | 0.002856 | 0.013381 | 0.005788 | 0.002565 | 0.027565 | 0.031717 | 0.039276 | 0.01028 | 0.030865 | 0.044049 |
| 32.8055556 | 0.028428 | 0.058053 | 0.007107 | 0.018209 | 0.019063 | 0.005 | 0.102052 | 0.014556 | 0.002813 | 0.053123 | 0.034458 | 0.023511 | 0.012694 | 0.019107 | 0.005405 | 0.01837 | 0.011612 | 0.003782 | 0.011052 | 0.035262 | 0.03677 | 0.00628 | 0.02495 | 0.027285 |
| 32.8194444 | 0.02628 | 0.054431 | 0.010127 | 0.020697 | 0.024711 | 0.003565 | 0.121207 | 0.012279 | 0.002822 | 0.046759 | 0.035789 | 0.017724 | 0.00678 | 0.016032 | 0.006291 | 0.024089 | 0.01591 | 0.004557 | 0.00515 | 0.031694 | 0.037236 | 0.008356 | 0.016828 | 0.04089 |
| 32.8333333 | 0.027778 | 0.047938 | 0.016701 | 0.022189 | 0.025624 | 0.002077 | 0.138443 | 0.015858 | 0.003144 | 0.039961 | 0.03546 | 0.012597 | 0.004439 | 0.012193 | 0.004995 | 0.030328 | 0.013393 | 0.003646 | 0.006393 | 0.027762 | 0.034999 | 0.015632 | 0.018348 | 0.042798 |
| 32.8472222 | 0.030095 | 0.045568 | 0.026853 | 0.022297 | 0.020208 | 0.004031 | 0.143903 | 0.019294 | 0.003372 | 0.037186 | 0.038073 | 0.01299 | 0.00693 | 0.010012 | 0.003474 | 0.033679 | 0.011049 | 0.007253 | 0.013097 | 0.019934 | 0.02427 | 0.027068 | 0.020509 | 0.024227 |
| 32.8611111 | 0.029955 | 0.056019 | 0.036834 | 0.021692 | 0.013803 | 0.007747 | 0.150939 | 0.017395 | 0.008003 | 0.039458 | 0.042791 | 0.019141 | 0.010866 | 0.007935 | 0.002999 | 0.032193 | 0.017261 | 0.01315 | 0.024729 | 0.009574 | 0.016676 | 0.039989 | 0.013866 | 0.022049 |
| 32.875 | 0.023112 | 0.073649 | 0.043728 | 0.02139 | 0.010057 | 0.011287 | 0.1647 | 0.010961 | 0.015531 | 0.044817 | 0.043416 | 0.026487 | 0.013264 | 0.008932 | 0.004777 | 0.028099 | 0.030906 | 0.012146 | 0.034879 | 0.005244 | 0.026149 | 0.048698 | 0.008435 | 0.04228 |
| 32.8888889 | 0.012491 | 0.081124 | 0.04458 | 0.021547 | 0.008467 | 0.013771 | 0.177218 | 0.004472 | 0.016675 | 0.045573 | 0.038082 | 0.030891 | 0.016036 | 0.016476 | 0.018426 | 0.022251 | 0.044777 | 0.008036 | 0.041881 | 0.00845 | 0.04012 | 0.053161 | 0.014049 | 0.063055 |
| 32.9027778 | 0.006001 | 0.071321 | 0.038132 | 0.019884 | 0.007132 | 0.014045 | 0.181055 | 0.002446 | 0.009965 | 0.036549 | 0.026758 | 0.031423 | 0.017261 | 0.027611 | 0.036746 | 0.014859 | 0.051659 | 0.014603 | 0.047967 | 0.014702 | 0.043137 | 0.056662 | 0.021675 | 0.079644 |
| 32.9166667 | 0.004607 | 0.054823 | 0.02656 | 0.016153 | 0.004586 | 0.012511 | 0.156743 | 0.00451 | 0.004251 | 0.022895 | 0.014054 | 0.02857 | 0.014361 | 0.038965 | 0.037416 | 0.010241 | 0.048769 | 0.035538 | 0.054454 | 0.018971 | 0.039551 | 0.058711 | 0.025944 | 0.09192 |
| 32.9305556 | 0.007251 | 0.041424 | 0.013133 | 0.012928 | 0.002487 | 0.009821 | 0.106255 | 0.006546 | 0.00255 | 0.011346 | 0.016568 | 0.022805 | 0.009428 | 0.048629 | 0.021917 | 0.013681 | 0.04297 | 0.057552 | 0.067356 | 0.019426 | 0.035833 | 0.05698 | 0.02923 | 0.09296 |
| 32.9444444 | 0.012735 | 0.028992 | 0.005372 | 0.010392 | 0.002803 | 0.006312 | 0.064174 | 0.005152 | 0.002427 | 0.004256 | 0.03239 | 0.015205 | 0.008476 | 0.055172 | 0.013572 | 0.019429 | 0.042474 | 0.074888 | 0.091786 | 0.01564 | 0.02883 | 0.047444 | 0.033316 | 0.093044 |
| 32.9583333 | 0.017937 | 0.015239 | 0.003959 | 0.006675 | 0.003052 | 0.003829 | 0.038933 | 0.003602 | 0.006012 | 0.001607 | 0.044404 | 0.007495 | 0.010036 | 0.053618 | 0.027927 | 0.018032 | 0.047836 | 0.08161 | 0.122733 | 0.010374 | 0.017068 | 0.031669 | 0.043244 | 0.100466 |
| 32.9722222 | 0.021719 | 0.006091 | 0.004984 | 0.003046 | 0.003129 | 0.003971 | 0.018234 | 0.005376 | 0.011402 | 0.001893 | 0.046845 | 0.002786 | 0.009686 | 0.040584 | 0.05161 | 0.013506 | 0.056268 | 0.065772 | 0.142688 | 0.008325 | 0.007036 | 0.019671 | 0.052889 | 0.096743 |
| 32.9861111 | 0.019073 | 0.00488 | 0.012347 | 0.005147 | 0.00762 | 0.00661 | 0.004876 | 0.008979 | 0.010174 | 0.008532 | 0.03283 | 0.003806 | 0.017181 | 0.029757 | 0.057107 | 0.01949 | 0.064791 | 0.037924 | 0.135829 | 0.009183 | 0.008533 | 0.024553 | 0.057851 | 0.092797 |
| 33 | 0.011705 | 0.009538 | 0.02659 | 0.015157 | 0.014302 | 0.008753 | 0.003899 | 0.020727 | 0.009122 | 0.027648 | 0.021838 | 0.009512 | 0.032246 | 0.030838 | 0.040913 | 0.037006 | 0.06894 | 0.01679 | 0.102417 | 0.020674 | 0.021804 | 0.053181 | 0.069577 | 0.118792 |
| 33.0138889 | 0.009666 | 0.018136 | 0.040811 | 0.023567 | 0.017895 | 0.006193 | 0.014335 | 0.043517 | 0.023928 | 0.049113 | 0.030545 | 0.017407 | 0.037709 | 0.028195 | 0.033918 | 0.045228 | 0.066197 | 0.007576 | 0.062217 | 0.044811 | 0.032158 | 0.090189 | 0.080673 | 0.140587 |
| 33.0277778 | 0.011327 | 0.026038 | 0.051137 | 0.020952 | 0.019688 | 0.002487 | 0.023396 | 0.064168 | 0.045463 | 0.052603 | 0.099161 | 0.026941 | 0.026048 | 0.0203 | 0.042207 | 0.036127 | 0.059992 | 0.009103 | 0.050828 | 0.064695 | 0.026797 | 0.10714 | 0.071351 | 0.114018 |
| 33.0416667 | 0.015811 | 0.033197 | 0.058945 | 0.01143 | 0.020405 | 0.001961 | 0.017433 | 0.068107 | 0.05535 | 0.03425 | 0.194291 | 0.03312 | 0.010124 | 0.024602 | 0.039841 | 0.020195 | 0.057409 | 0.009788 | 0.080341 | 0.066753 | 0.012766 | 0.097368 | 0.045876 | 0.069263 |
| 33.0555556 | 0.022625 | 0.042865 | 0.05838 | 0.005475 | 0.01622 | 0.0026 | 0.006917 | 0.051592 | 0.049284 | 0.0134 | 0.17365 | 0.027895 | 0.001916 | 0.032595 | 0.024969 | 0.008241 | 0.05937 | 0.006086 | 0.115323 | 0.045313 | 0.008641 | 0.076905 | 0.031866 | 0.054991 |
| 33.0694444 | 0.025585 | 0.052348 | 0.042184 | 0.007076 | 0.010675 | 0.005708 | 0.004946 | 0.026624 | 0.030928 | 0.005356 | 0.086405 | 0.014517 | 0.00057 | 0.02773 | 0.018663 | 0.006172 | 0.05687 | 0.009982 | 0.128585 | 0.019562 | 0.015185 | 0.0542 | 0.037478 | 0.056388 |
| 33.0833333 | 0.022675 | 0.056133 | 0.022016 | 0.010131 | 0.012374 | 0.011233 | 0.007114 | 0.014327 | 0.012755 | 0.00345 | 0.036521 | 0.007408 | 0.002213 | 0.014594 | 0.02785 | 0.008364 | 0.046293 | 0.018863 | 0.12945 | 0.019402 | 0.019199 | 0.029756 | 0.041061 | 0.044081 |
| 33.0972222 | 0.015055 | 0.052639 | 0.016306 | 0.00993 | 0.01526 | 0.013675 | 0.007539 | 0.018485 | 0.005867 | 0.002602 | 0.014415 | 0.012529 | 0.004074 | 0.005524 | 0.034627 | 0.009821 | 0.033765 | 0.02242 | 0.116738 | 0.031578 | 0.019231 | 0.01898 | 0.030049 | 0.0246 |
| 33.1111111 | 0.006548 | 0.046187 | 0.021157 | 0.008224 | 0.010748 | 0.012099 | 0.006362 | 0.023371 | 0.006085 | 0.005377 | 0.01058 | 0.021582 | 0.004173 | 0.004906 | 0.031374 | 0.010592 | 0.022948 | 0.021355 | 0.091208 | 0.030658 | 0.020267 | 0.029955 | 0.01325 | 0.009958 |
| 33.125 | 0.002469 | 0.04098 | 0.018145 | 0.007164 | 0.006518 | 0.010052 | 0.004495 | 0.021364 | 0.004745 | 0.009325 | 0.012427 | 0.028708 | 0.002611 | 0.007046 | 0.02641 | 0.010029 | 0.014857 | 0.017166 | 0.068427 | 0.022526 | 0.022114 | 0.040558 | 0.004457 | 0.003373 |
| 33.1388889 | 0.00322 | 0.037923 | 0.007515 | 0.007568 | 0.007915 | 0.009391 | 0.002596 | 0.014985 | 0.002102 | 0.011135 | 0.013415 | 0.029649 | 0.001645 | 0.007251 | 0.023009 | 0.008073 | 0.010775 | 0.009373 | 0.053126 | 0.014867 | 0.021406 | 0.035615 | 0.005861 | 0.004161 |
| 33.1527778 | 0.006557 | 0.035159 | 0.003924 | 0.009146 | 0.008363 | 0.010789 | 0.001315 | 0.007649 | 0.001174 | 0.010226 | 0.015933 | 0.022069 | 0.001921 | 0.006541 | 0.020446 | 0.005885 | 0.009801 | 0.005003 | 0.04331 | 0.00794 | 0.016596 | 0.022263 | 0.008208 | 0.006934 |
| 33.1666667 | 0.01061 | 0.030704 | 0.010765 | 0.010951 | 0.005599 | 0.014227 | 0.000883 | 0.005094 | 0.001568 | 0.007582 | 0.022882 | 0.01373 | 0.002882 | 0.007699 | 0.017354 | 0.004767 | 0.009487 | 0.006755 | 0.035911 | 0.006465 | 0.010268 | 0.010343 | 0.00731 | 0.005198 |
| 33.1805556 | 0.010071 | 0.025412 | 0.019691 | 0.012227 | 0.003254 | 0.017873 | 0.001651 | 0.008605 | 0.003007 | 0.00476 | 0.032359 | 0.009807 | 0.004637 | 0.011731 | 0.012316 | 0.004978 | 0.008434 | 0.008175 | 0.02702 | 0.007343 | 0.005788 | 0.003346 | 0.007816 | 0.003201 |
| 33.1944444 | 0.004974 | 0.020593 | 0.024694 | 0.012672 | 0.0043 | 0.019229 | 0.003552 | 0.010352 | 0.005386 | 0.002423 | 0.031014 | 0.008332 | 0.006712 | 0.017235 | 0.005997 | 0.005754 | 0.006746 | 0.007185 | 0.014777 | 0.008502 | 0.005629 | 0.000805 | 0.01108 | 0.005437 |
| 33.2083333 | 0.003295 | 0.015971 | 0.026567 | 0.012262 | 0.008924 | 0.016466 | 0.005754 | 0.007569 | 0.00749 | 0.001331 | 0.018933 | 0.00758 | 0.008733 | 0.021945 | 0.002773 | 0.006805 | 0.004677 | 0.006319 | 0.007615 | 0.017093 | 0.009999 | 0.000691 | 0.013899 | 0.006691 |
| 33.2222222 | 0.009109 | 0.011688 | 0.026335 | 0.011056 | 0.012584 | 0.009876 | 0.006488 | 0.007615 | 0.007939 | 0.001323 | 0.011277 | 0.007038 | 0.011061 | 0.023173 | 0.005689 | 0.008672 | 0.002659 | 0.005255 | 0.013343 | 0.028035 | 0.016462 | 0.002027 | 0.014975 | 0.00439 |
| 33.2361111 | 0.016453 | 0.008636 | 0.023072 | 0.0092 | 0.010956 | 0.005899 | 0.004852 | 0.013482 | 0.00631 | 0.000937 | 0.008248 | 0.006641 | 0.012597 | 0.020279 | 0.012585 | 0.01071 | 0.001493 | 0.00302 | 0.023552 | 0.030251 | 0.02266 | 0.004945 | 0.016086 | 0.001819 |
| 33.25 | 0.018709 | 0.007188 | 0.015385 | 0.007384 | 0.006864 | 0.006752 | 0.00403 | 0.019871 | 0.00443 | 0.000382 | 0.005956 | 0.006706 | 0.012734 | 0.017091 | 0.019333 | 0.01161 | 0.001229 | 0.001184 | 0.029695 | 0.023856 | 0.027681 | 0.008048 | 0.018029 | 0.001404 |
| 33.2638889 | 0.017205 | 0.007302 | 0.009883 | 0.00619 | 0.006924 | 0.005655 | 0.004193 | 0.022413 | 0.004621 | 0.000295 | 0.005048 | 0.007342 | 0.015812 | 0.017514 | 0.022729 | 0.012145 | 0.001071 | 0.000972 | 0.032124 | 0.015652 | 0.030673 | 0.010061 | 0.019376 | 0.003405 |
| 33.2777778 | 0.015748 | 0.008597 | 0.016001 | 0.005422 | 0.009637 | 0.004265 | 0.003775 | 0.02134 | 0.00712 | 0.001277 | 0.003507 | 0.008174 | 0.022203 | 0.020416 | 0.021584 | 0.013661 | 0.000732 | 0.002289 | 0.032047 | 0.011049 | 0.03104 | 0.010474 | 0.019417 | 0.005905 |
| 33.2916667 | 0.015043 | 0.009907 | 0.023685 | 0.004852 | 0.008435 | 0.00604 | 0.005971 | 0.01923 | 0.009399 | 0.003872 | 0.001615 | 0.008396 | 0.026064 | 0.022394 | 0.017731 | 0.01538 | 0.00076 | 0.004534 | 0.02935 | 0.012395 | 0.029239 | 0.009623 | 0.018705 | 0.006688 |
| 33.3055556 | 0.013627 | 0.010065 | 0.017712 | 0.004417 | 0.004403 | 0.00724 | 0.008723 | 0.017327 | 0.009649 | 0.005906 | 0.001084 | 0.007217 | 0.029963 | 0.022551 | 0.014186 | 0.016486 | 0.001191 | 0.007161 | 0.023816 | 0.01865 | 0.026276 | 0.008557 | 0.017632 | 0.005425 |
| 33.3194444 | 0.011772 | 0.009414 | 0.009363 | 0.003901 | 0.002295 | 0.007163 | 0.008155 | 0.015408 | 0.009239 | 0.005391 | 0.00099 | 0.004629 | 0.035214 | 0.021309 | 0.012574 | 0.017368 | 0.001577 | 0.009534 | 0.017265 | 0.025901 | 0.023783 | 0.008242 | 0.016284 | 0.003554 |
| 33.3333333 | 0.010035 | 0.00956 | 0.012835 | 0.003109 | 0.002145 | 0.007443 | 0.006536 | 0.014335 | 0.007963 | 0.003834 | 0.00068 | 0.002333 | 0.035759 | 0.017666 | 0.013363 | 0.018002 | 0.001813 | 0.012221 | 0.012872 | 0.032934 | 0.022595 | 0.008357 | 0.014855 | 0.003121 |
| 33.3472222 | 0.007379 | 0.011043 | 0.014097 | 0.001968 | 0.003871 | 0.007784 | 0.006031 | 0.014051 | 0.004745 | 0.003536 | 0.000541 | 0.00251 | 0.032106 | 0.012777 | 0.015976 | 0.018057 | 0.001887 | 0.014213 | 0.010905 | 0.03689 | 0.022183 | 0.008342 | 0.014724 | 0.003665 |
| 33.3611111 | 0.00443 | 0.012013 | 0.008309 | 0.001159 | 0.007728 | 0.007757 | 0.006219 | 0.013609 | 0.002249 | 0.00407 | 0.00055 | 0.003981 | 0.0267 | 0.01152 | 0.01933 | 0.018322 | 0.001575 | 0.013157 | 0.008749 | 0.032639 | 0.024525 | 0.009522 | 0.016756 | 0.003549 |
| 33.375 | 0.003825 | 0.011248 | 0.005588 | 0.001159 | 0.010146 | 0.008718 | 0.008323 | 0.013787 | 0.001618 | 0.003476 | 0.001347 | 0.003834 | 0.022167 | 0.016691 | 0.023541 | 0.019774 | 0.001112 | 0.011991 | 0.007577 | 0.023486 | 0.032915 | 0.012664 | 0.020422 | 0.003937 |
| 33.3888889 | 0.005885 | 0.010238 | 0.006015 | 0.002099 | 0.008598 | 0.011781 | 0.013421 | 0.015575 | 0.001489 | 0.003361 | 0.002911 | 0.00244 | 0.021752 | 0.025273 | 0.028178 | 0.021339 | 0.000912 | 0.014212 | 0.011485 | 0.01776 | 0.043771 | 0.014725 | 0.026444 | 0.006135 |
| 33.4027778 | 0.00774 | 0.010324 | 0.01166 | 0.004348 | 0.00586 | 0.015344 | 0.018401 | 0.018759 | 0.003268 | 0.005846 | 0.004817 | 0.002717 | 0.024058 | 0.031133 | 0.031838 | 0.02166 | 0.00066 | 0.018948 | 0.022994 | 0.019348 | 0.048057 | 0.015486 | 0.033456 | 0.009958 |
| 33.4166667 | 0.007367 | 0.010742 | 0.020085 | 0.006729 | 0.007453 | 0.017288 | 0.019602 | 0.023047 | 0.006772 | 0.008007 | 0.008097 | 0.005031 | 0.02643 | 0.031631 | 0.034069 | 0.021062 | 0.000484 | 0.024274 | 0.042097 | 0.027533 | 0.042364 | 0.017126 | 0.038066 | 0.015771 |
| 33.4305556 | 0.005723 | 0.011417 | 0.024056 | 0.00798 | 0.012488 | 0.017018 | 0.016117 | 0.027099 | 0.008828 | 0.007413 | 0.013027 | 0.006553 | 0.026202 | 0.0284 | 0.034778 | 0.020112 | 0.000738 | 0.028512 | 0.059667 | 0.039018 | 0.031209 | 0.017014 | 0.038948 | 0.022242 |
| 33.4444444 | 0.003977 | 0.014178 | 0.022958 | 0.007922 | 0.0127 | 0.014859 | 0.010622 | 0.028732 | 0.00763 | 0.004551 | 0.017798 | 0.005185 | 0.022425 | 0.023206 | 0.033977 | 0.019032 | 0.000816 | 0.030209 | 0.060305 | 0.047828 | 0.020036 | 0.013743 | 0.035743 | 0.025233 |
| 33.4583333 | 0.002282 | 0.01881 | 0.019583 | 0.006719 | 0.008174 | 0.012274 | 0.006751 | 0.027568 | 0.004486 | 0.001887 | 0.021611 | 0.002946 | 0.018906 | 0.018845 | 0.032884 | 0.018165 | 0.000496 | 0.029371 | 0.043228 | 0.04866 | 0.012963 | 0.009599 | 0.031089 | 0.02347 |
| 33.4722222 | 0.002697 | 0.020755 | 0.016667 | 0.005772 | 0.00737 | 0.011515 | 0.006743 | 0.024984 | 0.003282 | 0.002741 | 0.027913 | 0.001997 | 0.021191 | 0.021208 | 0.033708 | 0.019025 | 0.000574 | 0.028149 | 0.031101 | 0.043281 | 0.014932 | 0.008649 | 0.030792 | 0.02708 |
| 33.4861111 | 0.009864 | 0.016116 | 0.0151 | 0.00745 | 0.008152 | 0.013729 | 0.010759 | 0.021467 | 0.00662 | 0.008815 | 0.040863 | 0.003682 | 0.032048 | 0.03355 | 0.036744 | 0.023478 | 0.003321 | 0.028103 | 0.0531 | 0.042405 | 0.030554 | 0.018063 | 0.042072 | 0.052256 |
| 33.5 | 0.024703 | 0.016454 | 0.016454 | 0.011624 | 0.007194 | 0.017032 | 0.015702 | 0.017461 | 0.0122 | 0.016782 | 0.055762 | 0.011184 | 0.053731 | 0.051256 | 0.03991 | 0.030357 | 0.011509 | 0.029082 | 0.080046 | 0.052516 | 0.052999 | 0.03932 | 0.066868 | 0.092619 |
| 33.5138889 | 0.040046 | 0.029833 | 0.024167 | 0.015259 | 0.00793 | 0.017356 | 0.017504 | 0.017702 | 0.015707 | 0.019798 | 0.066754 | 0.023194 | 0.077718 | 0.063106 | 0.04188 | 0.033578 | 0.022108 | 0.031151 | 0.063473 | 0.054203 | 0.068695 | 0.060725 | 0.094791 | 0.130153 |
| 33.5277778 | 0.048506 | 0.037449 | 0.040297 | 0.018331 | 0.007139 | 0.015118 | 0.015025 | 0.026477 | 0.018189 | 0.021059 | 0.075757 | 0.037199 | 0.081821 | 0.070822 | 0.047371 | 0.035286 | 0.030342 | 0.035276 | 0.044245 | 0.036329 | 0.08264 | 0.076191 | 0.117011 | 0.176655 |
| 33.5416667 | 0.050939 | 0.03207 | 0.059987 | 0.024366 | 0.005918 | 0.014921 | 0.0103 | 0.039128 | 0.020692 | 0.027119 | 0.090304 | 0.051517 | 0.071918 | 0.08359 | 0.043895 | 0.051446 | 0.038463 | 0.047719 | 0.040121 | 0.024684 | 0.107349 | 0.092356 | 0.134939 | 0.227252 |
| 33.5555556 | 0.049099 | 0.030236 | 0.070931 | 0.031769 | 0.006839 | 0.016764 | 0.006025 | 0.047769 | 0.018503 | 0.030201 | 0.131549 | 0.060506 | 0.064727 | 0.083831 | 0.026991 | 0.071285 | 0.045645 | 0.071139 | 0.034486 | 0.032388 | 0.126812 | 0.105624 | 0.143053 | 0.247465 |
| 33.5694444 | 0.039464 | 0.041409 | 0.06935 | 0.032195 | 0.007224 | 0.017738 | 0.005112 | 0.048265 | 0.012449 | 0.022896 | 0.192377 | 0.061681 | 0.052239 | 0.075062 | 0.020921 | 0.066529 | 0.044034 | 0.079731 | 0.046907 | 0.043541 | 0.120349 | 0.10292 | 0.127438 | 0.227506 |
| 33.5833333 | 0.025144 | 0.061206 | 0.063979 | 0.024043 | 0.008747 | 0.017223 | 0.013762 | 0.038332 | 0.009363 | 0.013693 | 0.211919 | 0.058913 | 0.039851 | 0.079095 | 0.017308 | 0.046671 | 0.033169 | 0.055955 | 0.053885 | 0.045539 | 0.106287 | 0.084905 | 0.082436 | 0.178481 |
| 33.5972222 | 0.015194 | 0.078312 | 0.058214 | 0.014126 | 0.009515 | 0.016755 | 0.031887 | 0.02143 | 0.009972 | 0.00989 | 0.161329 | 0.05359 | 0.032975 | 0.078909 | 0.008693 | 0.03384 | 0.021043 | 0.025347 | 0.045763 | 0.040083 | 0.107257 | 0.063158 | 0.034639 | 0.12724 |
| 33.6111111 | 0.012866 | 0.088735 | 0.049842 | 0.007959 | 0.008699 | 0.018064 | 0.045484 | 0.007984 | 0.011777 | 0.007992 | 0.080456 | 0.04291 | 0.023226 | 0.067753 | 0.014027 | 0.027609 | 0.012707 | 0.008844 | 0.045593 | 0.032541 | 0.106771 | 0.042594 | 0.017728 | 0.091927 |
| 33.625 | 0.015421 | 0.097946 | 0.039823 | 0.007301 | 0.007051 | 0.01979 | 0.046105 | 0.00359 | 0.015263 | 0.00585 | 0.03274 | 0.026596 | 0.019554 | 0.050651 | 0.033378 | 0.020306 | 0.008508 | 0.00363 | 0.037872 | 0.023822 | 0.075382 | 0.023658 | 0.01968 | 0.065031 |
| 33.6388889 | 0.01886 | 0.103786 | 0.030899 | 0.009498 | 0.003969 | 0.019519 | 0.041216 | 0.003444 | 0.019275 | 0.005477 | 0.075116 | 0.012505 | 0.01988 | 0.029711 | 0.052692 | 0.010717 | 0.007305 | 0.005418 | 0.026423 | 0.013037 | 0.040653 | 0.012581 | 0.015687 | 0.038752 |
| 33.6527778 | 0.022143 | 0.101656 | 0.02444 | 0.010411 | 0.002826 | 0.017691 | 0.040546 | 0.004373 | 0.020825 | 0.007507 | 0.160922 | 0.011384 | 0.011189 | 0.02796 | 0.063065 | 0.004818 | 0.007173 | 0.008623 | 0.036727 | 0.004732 | 0.050978 | 0.015619 | 0.007606 | 0.023824 |
| 33.6666667 | 0.025483 | 0.095233 | 0.018704 | 0.007447 | 0.004697 | 0.01542 | 0.044781 | 0.008184 | 0.017171 | 0.008805 | 0.170524 | 0.022074 | 0.006282 | 0.046837 | 0.066622 | 0.006063 | 0.006449 | 0.012625 | 0.048673 | 0.002184 | 0.080832 | 0.022834 | 0.004579 | 0.034685 |
| 33.6805556 | 0.027437 | 0.089218 | 0.012373 | 0.003469 | 0.009979 | 0.013286 | 0.050497 | 0.010594 | 0.011131 | 0.00664 | 0.109377 | 0.034004 | 0.01506 | 0.062066 | 0.066488 | 0.008183 | 0.004236 | 0.022625 | 0.043697 | 0.003224 | 0.084507 | 0.023117 | 0.003802 | 0.062614 |
| 33.6944444 | 0.025906 | 0.081087 | 0.008386 | 0.002935 | 0.017216 | 0.012371 | 0.054342 | 0.007311 | 0.014356 | 0.00489 | 0.060227 | 0.039625 | 0.023494 | 0.065807 | 0.059322 | 0.006819 | 0.00233 | 0.026199 | 0.034589 | 0.008213 | 0.063826 | 0.018443 | 0.002991 | 0.094334 |
| 33.7083333 | 0.01903 | 0.061495 | 0.007502 | 0.004007 | 0.023084 | 0.012181 | 0.051557 | 0.003663 | 0.02285 | 0.006634 | 0.033198 | 0.039194 | 0.021023 | 0.062962 | 0.039065 | 0.004196 | 0.002978 | 0.017082 | 0.036779 | 0.013349 | 0.045625 | 0.011923 | 0.004813 | 0.125812 |
| 33.7222222 | 0.009216 | 0.031491 | 0.005374 | 0.003801 | 0.026461 | 0.01143 | 0.041052 | 0.004072 | 0.023604 | 0.009037 | 0.019321 | 0.036477 | 0.01228 | 0.057978 | 0.021538 | 0.002792 | 0.004514 | 0.00847 | 0.05311 | 0.012269 | 0.044068 | 0.007549 | 0.007811 | 0.145016 |
| 33.7361111 | 0.00542 | 0.014027 | 0.003427 | 0.002451 | 0.027686 | 0.01039 | 0.028999 | 0.005867 | 0.020136 | 0.010917 | 0.037847 | 0.032137 | 0.004559 | 0.057414 | 0.031394 | 0.004425 | 0.00587 | 0.006968 | 0.075197 | 0.00797 | 0.052992 | 0.00797 | 0.00815 | 0.1398 |
| 33.75 | 0.011583 | 0.021014 | 0.006975 | 0.001717 | 0.029283 | 0.009424 | 0.021487 | 0.007388 | 0.016986 | 0.012668 | 0.075205 | 0.028422 | 0.001828 | 0.061507 | 0.057629 | 0.008218 | 0.007963 | 0.016047 | 0.085644 | 0.005724 | 0.062043 | 0.010443 | 0.005533 | 0.112372 |
| 33.7638889 | 0.020076 | 0.030909 | 0.014576 | 0.002789 | 0.033198 | 0.008169 | 0.016663 | 0.009384 | 0.016161 | 0.01406 | 0.10886 | 0.030537 | 0.00444 | 0.060286 | 0.075382 | 0.010539 | 0.010026 | 0.034926 | 0.071702 | 0.003641 | 0.061132 | 0.012249 | 0.006459 | 0.064824 |
| 33.7777778 | 0.024174 | 0.028302 | 0.025364 | 0.005366 | 0.038421 | 0.006663 | 0.010698 | 0.010879 | 0.016391 | 0.015458 | 0.13022 | 0.038266 | 0.008848 | 0.048542 | 0.079138 | 0.008059 | 0.011275 | 0.046207 | 0.047809 | 0.001437 | 0.045122 | 0.01124 | 0.013728 | 0.032395 |
| 33.7916667 | 0.027928 | 0.023256 | 0.041305 | 0.008514 | 0.045293 | 0.005196 | 0.005925 | 0.010059 | 0.014533 | 0.016949 | 0.175351 | 0.045939 | 0.008567 | 0.02759 | 0.077579 | 0.00596 | 0.013041 | 0.034429 | 0.030511 | 0.001454 | 0.021772 | 0.007377 | 0.021847 | 0.046744 |
| 33.8055556 | 0.041636 | 0.024172 | 0.059811 | 0.010153 | 0.055279 | 0.003169 | 0.007063 | 0.007108 | 0.013574 | 0.018427 | 0.238734 | 0.049507 | 0.006198 | 0.012237 | 0.068481 | 0.011443 | 0.015767 | 0.015082 | 0.022881 | 0.004979 | 0.008855 | 0.004606 | 0.027289 | 0.07618 |
| 33.8194444 | 0.066092 | 0.027762 | 0.075192 | 0.009508 | 0.065501 | 0.0027 | 0.013105 | 0.00471 | 0.012547 | 0.019139 | 0.216628 | 0.048424 | 0.006735 | 0.013379 | 0.043538 | 0.017918 | 0.017676 | 0.010437 | 0.021849 | 0.013551 | 0.011769 | 0.003652 | 0.034203 | 0.104554 |
| 33.8333333 | 0.084924 | 0.031189 | 0.0831 | 0.008432 | 0.06817 | 0.005763 | 0.020523 | 0.005586 | 0.007943 | 0.018326 | 0.125259 | 0.043071 | 0.008858 | 0.013804 | 0.023484 | 0.017275 | 0.018873 | 0.01729 | 0.023609 | 0.02334 | 0.015314 | 0.002788 | 0.043344 | 0.124414 |
| 33.8472222 | 0.086777 | 0.036279 | 0.08483 | 0.007507 | 0.058542 | 0.008582 | 0.027718 | 0.009333 | 0.005024 | 0.016695 | 0.064453 | 0.034767 | 0.019348 | 0.009404 | 0.037506 | 0.01083 | 0.021419 | 0.023187 | 0.035527 | 0.027843 | 0.014025 | 0.004003 | 0.048012 | 0.103419 |
| 33.8611111 | 0.074981 | 0.044379 | 0.08297 | 0.005887 | 0.046733 | 0.00925 | 0.032667 | 0.013328 | 0.009874 | 0.015229 | 0.089674 | 0.025688 | 0.036608 | 0.014489 | 0.077309 | 0.008633 | 0.026067 | 0.023341 | 0.084198 | 0.024747 | 0.025606 | 0.009496 | 0.045154 | 0.05813 |
| 33.875 | 0.061492 | 0.053751 | 0.076744 | 0.004543 | 0.044487 | 0.010304 | 0.034298 | 0.016009 | 0.016232 | 0.014074 | 0.172004 | 0.015895 | 0.047999 | 0.026044 | 0.107699 | 0.016773 | 0.030177 | 0.019125 | 0.140857 | 0.016315 | 0.050129 | 0.017136 | 0.038769 | 0.053771 |
| 33.8888889 | 0.055995 | 0.060469 | 0.069011 | 0.004453 | 0.043881 | 0.01288 | 0.038147 | 0.017387 | 0.018555 | 0.013173 | 0.200504 | 0.00821 | 0.051517 | 0.033007 | 0.106526 | 0.028324 | 0.029008 | 0.014791 | 0.145763 | 0.007347 | 0.060684 | 0.022666 | 0.035702 | 0.100175 |
| 33.9027778 | 0.052003 | 0.060081 | 0.06596 | 0.004587 | 0.031646 | 0.016232 | 0.050892 | 0.017715 | 0.020664 | 0.012723 | 0.149914 | 0.008276 | 0.05167 | 0.031342 | 0.077604 | 0.03631 | 0.023197 | 0.009877 | 0.136969 | 0.002712 | 0.049436 | 0.024343 | 0.037533 | 0.141435 |
| 33.9166667 | 0.046603 | 0.051273 | 0.070142 | 0.004218 | 0.015236 | 0.019828 | 0.067788 | 0.01696 | 0.026719 | 0.011653 | 0.096912 | 0.013603 | 0.051771 | 0.024562 | 0.041089 | 0.037619 | 0.018011 | 0.005414 | 0.149311 | 0.003781 | 0.030583 | 0.02443 | 0.041037 | 0.149533 |
| 33.9305556 | 0.040305 | 0.03552 | 0.076751 | 0.003509 | 0.008419 | 0.023124 | 0.077133 | 0.015466 | 0.03673 | 0.008442 | 0.073968 | 0.019085 | 0.052601 | 0.020228 | 0.01664 | 0.031379 | 0.014943 | 0.008214 | 0.14876 | 0.007151 | 0.017073 | 0.024997 | 0.044973 | 0.142795 |
| 33.9444444 | 0.030143 | 0.018878 | 0.075112 | 0.00241 | 0.00635 | 0.025787 | 0.075949 | 0.014031 | 0.045897 | 0.004288 | 0.052998 | 0.024101 | 0.053603 | 0.01713 | 0.007225 | 0.018919 | 0.012104 | 0.014159 | 0.128238 | 0.008191 | 0.009633 | 0.023697 | 0.04605 | 0.131047 |
| 33.9583333 | 0.021062 | 0.011728 | 0.059567 | 0.004008 | 0.003384 | 0.026339 | 0.065086 | 0.012838 | 0.051173 | 0.003837 | 0.027879 | 0.026686 | 0.056311 | 0.012091 | 0.009326 | 0.008217 | 0.009101 | 0.012577 | 0.108277 | 0.006017 | 0.007832 | 0.018041 | 0.039272 | 0.113912 |
| 33.9722222 | 0.01642 | 0.009989 | 0.038836 | 0.011945 | 0.003336 | 0.025347 | 0.044935 | 0.01068 | 0.049619 | 0.006415 | 0.017483 | 0.02444 | 0.065509 | 0.011866 | 0.022777 | 0.005325 | 0.009853 | 0.011853 | 0.108091 | 0.007686 | 0.014364 | 0.013915 | 0.030989 | 0.105649 |
| 33.9861111 | 0.017777 | 0.00816 | 0.029714 | 0.020103 | 0.004102 | 0.026333 | 0.031242 | 0.006518 | 0.033004 | 0.009785 | 0.027456 | 0.015996 | 0.081606 | 0.017484 | 0.028214 | 0.007217 | 0.019136 | 0.023139 | 0.129945 | 0.017624 | 0.020912 | 0.023991 | 0.037677 | 0.130835 |
| 34 | 0.02806 | 0.016543 | 0.035723 | 0.01794 | 0.009675 | 0.025634 | 0.039336 | 0.002907 | 0.018008 | 0.022124 | 0.045504 | 0.008802 | 0.080803 | 0.046313 | 0.030056 | 0.016443 | 0.034718 | 0.033511 | 0.14465 | 0.031056 | 0.038108 | 0.051187 | 0.068703 | 0.190072 |
| 34.0138889 | 0.040794 | 0.032753 | 0.044047 | 0.008475 | 0.027182 | 0.018995 | 0.058962 | 0.00201 | 0.029386 | 0.042304 | 0.05556 | 0.013401 | 0.053855 | 0.100779 | 0.063578 | 0.032703 | 0.043265 | 0.036957 | 0.144886 | 0.043782 | 0.07297 | 0.076299 | 0.10965 | 0.216708 |
| 34.0277778 | 0.045072 | 0.047739 | 0.047935 | 0.003008 | 0.042664 | 0.011161 | 0.068549 | 0.002754 | 0.049457 | 0.052346 | 0.049485 | 0.022262 | 0.028927 | 0.121869 | 0.094155 | 0.040865 | 0.038562 | 0.035401 | 0.134631 | 0.046112 | 0.085103 | 0.076354 | 0.123819 | 0.165988 |
| 34.0416667 | 0.04304 | 0.063206 | 0.052982 | 0.003242 | 0.036558 | 0.008934 | 0.06135 | 0.00575 | 0.057708 | 0.04426 | 0.031112 | 0.02333 | 0.016215 | 0.085771 | 0.06813 | 0.034203 | 0.028388 | 0.023827 | 0.085913 | 0.029949 | 0.053342 | 0.051475 | 0.090196 | 0.090615 |
| 34.0555556 | 0.043792 | 0.078416 | 0.061037 | 0.004859 | 0.024225 | 0.011519 | 0.047001 | 0.007637 | 0.057264 | 0.027739 | 0.014593 | 0.017336 | 0.014185 | 0.040986 | 0.027712 | 0.021841 | 0.015184 | 0.021248 | 0.034999 | 0.013136 | 0.023876 | 0.022169 | 0.03754 | 0.046234 |
| 34.0694444 | 0.045514 | 0.086309 | 0.063398 | 0.005222 | 0.035127 | 0.012097 | 0.034841 | 0.008118 | 0.048892 | 0.013673 | 0.009423 | 0.010198 | 0.021784 | 0.028545 | 0.023338 | 0.010426 | 0.004578 | 0.037411 | 0.031833 | 0.012979 | 0.028692 | 0.006045 | 0.009742 | 0.033358 |
| 34.0833333 | 0.044752 | 0.083241 | 0.053893 | 0.003841 | 0.05206 | 0.008626 | 0.025928 | 0.01385 | 0.033854 | 0.005622 | 0.008036 | 0.0051 | 0.027634 | 0.037517 | 0.03309 | 0.004757 | 0.003756 | 0.047462 | 0.038098 | 0.020194 | 0.038044 | 0.00567 | 0.006649 | 0.031272 |
| 34.0972222 | 0.04276 | 0.074636 | 0.03876 | 0.001858 | 0.048455 | 0.003767 | 0.01849 | 0.018803 | 0.019659 | 0.002167 | 0.005507 | 0.002504 | 0.0274 | 0.039778 | 0.039892 | 0.006996 | 0.007418 | 0.043797 | 0.029971 | 0.024438 | 0.034764 | 0.019537 | 0.005355 | 0.029513 |
| 34.1111111 | 0.037242 | 0.065979 | 0.026938 | 0.000819 | 0.033348 | 0.00116 | 0.012886 | 0.014516 | 0.009128 | 0.001827 | 0.004738 | 0.002922 | 0.023443 | 0.031359 | 0.044098 | 0.011687 | 0.009725 | 0.036181 | 0.020126 | 0.025329 | 0.027793 | 0.040277 | 0.002016 | 0.030104 |
| 34.125 | 0.030072 | 0.057259 | 0.02179 | 0.001371 | 0.021709 | 0.000933 | 0.009371 | 0.00664 | 0.003731 | 0.003675 | 0.00451 | 0.00531 | 0.018621 | 0.019782 | 0.039752 | 0.014186 | 0.010228 | 0.029874 | 0.01126 | 0.024863 | 0.019719 | 0.053787 | 0.001397 | 0.028886 |
| 34.1388889 | 0.02421 | 0.047569 | 0.024214 | 0.002649 | 0.015144 | 0.00077 | 0.008242 | 0.003327 | 0.004744 | 0.00527 | 0.008334 | 0.006921 | 0.015183 | 0.01059 | 0.030677 | 0.014155 | 0.009458 | 0.024945 | 0.004595 | 0.023224 | 0.01255 | 0.05505 | 0.003294 | 0.019201 |
| 34.1527778 | 0.016455 | 0.038968 | 0.028748 | 0.004073 | 0.010287 | 0.001145 | 0.008519 | 0.004659 | 0.005856 | 0.004712 | 0.019279 | 0.007206 | 0.01327 | 0.005753 | 0.021641 | 0.012064 | 0.007474 | 0.018928 | 0.001081 | 0.018675 | 0.007622 | 0.046053 | 0.00474 | 0.009308 |
| 34.1666667 | 0.007204 | 0.0339 | 0.024594 | 0.006508 | 0.01063 | 0.003113 | 0.006559 | 0.006627 | 0.004272 | 0.002647 | 0.034101 | 0.007423 | 0.012129 | 0.003672 | 0.013712 | 0.008624 | 0.004572 | 0.013319 | 0.000187 | 0.012014 | 0.005118 | 0.032547 | 0.004005 | 0.00705 |
| 34.1805556 | 0.003344 | 0.032253 | 0.012573 | 0.009994 | 0.014309 | 0.006038 | 0.003268 | 0.007606 | 0.004167 | 0.001091 | 0.047479 | 0.007769 | 0.011091 | 0.002472 | 0.00746 | 0.005138 | 0.002451 | 0.008447 | 0.000285 | 0.005636 | 0.005113 | 0.021513 | 0.002088 | 0.007444 |
| 34.1944444 | 0.005713 | 0.032456 | 0.005258 | 0.012742 | 0.012374 | 0.008517 | 0.002015 | 0.00757 | 0.007697 | 0.000723 | 0.054567 | 0.007805 | 0.009956 | 0.001278 | 0.003552 | 0.002437 | 0.002817 | 0.003927 | 0.000363 | 0.003771 | 0.006871 | 0.015938 | 0.001026 | 0.00549 |
| 34.2083333 | 0.007772 | 0.033033 | 0.006434 | 0.013843 | 0.011667 | 0.010673 | 0.002332 | 0.007343 | 0.012121 | 0.000739 | 0.056099 | 0.007877 | 0.009425 | 0.000561 | 0.002589 | 0.001737 | 0.004303 | 0.001312 | 0.001081 | 0.007066 | 0.009942 | 0.017196 | 0.001821 | 0.002854 |
| 34.2222222 | 0.008629 | 0.032415 | 0.008997 | 0.013883 | 0.020018 | 0.012775 | 0.003747 | 0.008338 | 0.01265 | 0.000826 | 0.0559 | 0.008461 | 0.009964 | 0.000889 | 0.004081 | 0.00307 | 0.005089 | 0.001058 | 0.0031 | 0.007503 | 0.013641 | 0.024048 | 0.003743 | 0.003054 |
| 34.2361111 | 0.009874 | 0.029364 | 0.008582 | 0.013961 | 0.028828 | 0.014003 | 0.004611 | 0.010127 | 0.007954 | 0.001363 | 0.055089 | 0.00878 | 0.010444 | 0.001102 | 0.005637 | 0.003791 | 0.006894 | 0.001367 | 0.003721 | 0.00328 | 0.015278 | 0.03161 | 0.004011 | 0.006361 |
| 34.25 | 0.010572 | 0.024571 | 0.005687 | 0.01391 | 0.031511 | 0.013658 | 0.003819 | 0.010511 | 0.003785 | 0.001837 | 0.053573 | 0.008003 | 0.009827 | 0.000959 | 0.005657 | 0.003186 | 0.010595 | 0.00094 | 0.002574 | 0.00072 | 0.013477 | 0.034812 | 0.002233 | 0.010755 |
| 34.2638889 | 0.010735 | 0.019726 | 0.002638 | 0.01194 | 0.025596 | 0.01221 | 0.003404 | 0.008754 | 0.003496 | 0.001963 | 0.052275 | 0.006757 | 0.009163 | 0.001134 | 0.005209 | 0.002241 | 0.012532 | 0.000549 | 0.002508 | 0.00113 | 0.010026 | 0.032112 | 0.001184 | 0.012666 |
| 34.2777778 | 0.011545 | 0.016753 | 0.001891 | 0.007872 | 0.016822 | 0.010877 | 0.003389 | 0.006741 | 0.002863 | 0.001975 | 0.050871 | 0.005664 | 0.009632 | 0.001216 | 0.005478 | 0.001281 | 0.010383 | 0.001067 | 0.002465 | 0.003512 | 0.007124 | 0.02496 | 0.002054 | 0.011098 |
| 34.2916667 | 0.014723 | 0.017365 | 0.00306 | 0.003737 | 0.021211 | 0.010118 | 0.002752 | 0.006509 | 0.002429 | 0.001418 | 0.048274 | 0.00393 | 0.010564 | 0.001491 | 0.005631 | 0.001333 | 0.005647 | 0.003049 | 0.001407 | 0.00637 | 0.005237 | 0.01636 | 0.003792 | 0.010036 |
| 34.3055556 | 0.01925 | 0.021525 | 0.003214 | 0.002071 | 0.036514 | 0.00991 | 0.001834 | 0.008244 | 0.004567 | 0.000959 | 0.043891 | 0.002307 | 0.010825 | 0.001417 | 0.004853 | 0.002743 | 0.002663 | 0.005524 | 0.000856 | 0.007484 | 0.004008 | 0.010206 | 0.004641 | 0.012343 |
| 34.3194444 | 0.0219 | 0.027615 | 0.001871 | 0.003304 | 0.045912 | 0.010079 | 0.000943 | 0.0106 | 0.006449 | 0.001866 | 0.037803 | 0.003132 | 0.009999 | 0.001457 | 0.003875 | 0.004456 | 0.003661 | 0.006258 | 0.001778 | 0.00696 | 0.003159 | 0.006442 | 0.003592 | 0.016906 |

| | | | | | | | | | | | | | | | | | | | | | | | | |
|---|---|---|---|---|---|---|---|---|---|---|---|---|---|---|---|---|---|---|---|---|---|---|---|---|
| 34.3333333 | 0.022126 | 0.033755 | 0.001111 | 0.004975 | 0.043451 | 0.010526 | 0.000404 | 0.01219 | 0.005527 | 0.003432 | 0.031342 | 0.00579 | 0.008354 | 0.003264 | 0.003359 | 0.005078 | 0.003959 | 0.00519 | 0.004389 | 0.005439 | 0.001974 | 0.004541 | 0.002197 | 0.021408 |
| 34.3472222 | 0.019845 | 0.038201 | 0.001058 | 0.004871 | 0.031281 | 0.01135 | 0.000473 | 0.013128 | 0.00308 | 0.004556 | 0.026149 | 0.008378 | 0.007065 | 0.00499 | 0.003408 | 0.004266 | 0.002725 | 0.004562 | 0.006715 | 0.003883 | 0.001085 | 0.004296 | 0.002366 | 0.023545 |
| 34.3611111 | 0.014642 | 0.039994 | 0.001941 | 0.002827 | 0.015065 | 0.012 | 0.000778 | 0.014691 | 0.002893 | 0.004896 | 0.023662 | 0.010482 | 0.007841 | 0.004217 | 0.004873 | 0.003613 | 0.003153 | 0.006086 | 0.006168 | 0.004219 | 0.002957 | 0.006639 | 0.003058 | 0.020308 |
| 34.375 | 0.008904 | 0.040348 | 0.004455 | 0.000803 | 0.003794 | 0.012088 | 0.001177 | 0.017404 | 0.006003 | 0.004548 | 0.021377 | 0.012279 | 0.011236 | 0.004356 | 0.009605 | 0.005083 | 0.003326 | 0.009673 | 0.007832 | 0.007278 | 0.009377 | 0.017664 | 0.005686 | 0.013397 |
| 34.3888889 | 0.004987 | 0.040651 | 0.007032 | 0.000687 | 0.003808 | 0.011912 | 0.00214 | 0.020329 | 0.00951 | 0.004201 | 0.017448 | 0.013261 | 0.015901 | 0.008888 | 0.018716 | 0.008516 | 0.004459 | 0.013949 | 0.01666 | 0.011817 | 0.018267 | 0.034039 | 0.011802 | 0.013151 |
| 34.4027778 | 0.003492 | 0.040658 | 0.007987 | 0.002639 | 0.012603 | 0.012173 | 0.004349 | 0.02167 | 0.010192 | 0.003885 | 0.014802 | 0.012642 | 0.020028 | 0.013277 | 0.030002 | 0.01189 | 0.009734 | 0.01691 | 0.025001 | 0.015229 | 0.024071 | 0.046405 | 0.017188 | 0.021306 |
| 34.4166667 | 0.003345 | 0.040036 | 0.007563 | 0.004554 | 0.017455 | 0.013502 | 0.007908 | 0.021008 | 0.008713 | 0.003039 | 0.014574 | 0.011345 | 0.022405 | 0.014202 | 0.038921 | 0.013813 | 0.014881 | 0.017844 | 0.026165 | 0.015924 | 0.024267 | 0.053533 | 0.018965 | 0.028241 |
| 34.4305556 | 0.003241 | 0.038807 | 0.006944 | 0.003928 | 0.013107 | 0.014585 | 0.011952 | 0.02008 | 0.009753 | 0.001905 | 0.016556 | 0.011317 | 0.023546 | 0.014006 | 0.043336 | 0.014318 | 0.015647 | 0.017286 | 0.023693 | 0.015144 | 0.022158 | 0.058196 | 0.017403 | 0.031676 |
| 34.4444444 | 0.003911 | 0.037676 | 0.007448 | 0.001923 | 0.011952 | 0.013693 | 0.015755 | 0.01985 | 0.01501 | 0.001572 | 0.019896 | 0.0133 | 0.025189 | 0.014098 | 0.043933 | 0.013811 | 0.013272 | 0.016019 | 0.022865 | 0.014682 | 0.021994 | 0.061731 | 0.013227 | 0.035197 |
| 34.4583333 | 0.003828 | 0.037098 | 0.009948 | 0.000826 | 0.019254 | 0.011667 | 0.019002 | 0.019616 | 0.021196 | 0.001764 | 0.026758 | 0.015512 | 0.027893 | 0.013781 | 0.042474 | 0.01281 | 0.010429 | 0.01532 | 0.024095 | 0.014145 | 0.024759 | 0.06455 | 0.008528 | 0.042802 |
| 34.4722222 | 0.003104 | 0.035214 | 0.011721 | 0.000684 | 0.025727 | 0.01002 | 0.021206 | 0.01893 | 0.022946 | 0.002524 | 0.039177 | 0.015655 | 0.031987 | 0.014506 | 0.041514 | 0.012728 | 0.008922 | 0.015818 | 0.02747 | 0.013232 | 0.030175 | 0.070066 | 0.008072 | 0.052758 |
| 34.4861111 | 0.00874 | 0.028379 | 0.009129 | 0.001324 | 0.029084 | 0.009705 | 0.021184 | 0.018937 | 0.015944 | 0.007513 | 0.056102 | 0.012489 | 0.038434 | 0.022036 | 0.043463 | 0.015166 | 0.011256 | 0.018212 | 0.040081 | 0.015034 | 0.039395 | 0.08617 | 0.019195 | 0.06201 |
| 34.5 | 0.022074 | 0.015842 | 0.004644 | 0.004292 | 0.037588 | 0.01038 | 0.01672 | 0.019459 | 0.016129 | 0.017828 | 0.07164 | 0.008392 | 0.048047 | 0.036795 | 0.049922 | 0.021781 | 0.01777 | 0.020121 | 0.061589 | 0.019931 | 0.048675 | 0.118268 | 0.040192 | 0.070942 |
| 34.5138889 | 0.032531 | 0.007922 | 0.002693 | 0.010607 | 0.058513 | 0.010661 | 0.010957 | 0.019236 | 0.042457 | 0.026895 | 0.078812 | 0.01076 | 0.055465 | 0.047276 | 0.054601 | 0.032818 | 0.023081 | 0.01542 | 0.072397 | 0.021245 | 0.048288 | 0.160938 | 0.055174 | 0.068891 |
| 34.5277778 | 0.033073 | 0.013492 | 0.00579 | 0.01871 | 0.085144 | 0.0115 | 0.013592 | 0.023153 | 0.068662 | 0.027932 | 0.085405 | 0.021255 | 0.053924 | 0.045952 | 0.048051 | 0.04114 | 0.022999 | 0.007642 | 0.062463 | 0.015803 | 0.038734 | 0.190827 | 0.056055 | 0.061428 |
| 34.5416667 | 0.02927 | 0.026248 | 0.016332 | 0.02487 | 0.089209 | 0.01488 | 0.023173 | 0.03544 | 0.066517 | 0.021992 | 0.09315 | 0.03183 | 0.047254 | 0.040046 | 0.034158 | 0.04141 | 0.020151 | 0.005461 | 0.052388 | 0.012164 | 0.032161 | 0.170353 | 0.051655 | 0.072751 |
| 34.5555556 | 0.027825 | 0.038337 | 0.031146 | 0.024511 | 0.054669 | 0.019072 | 0.025993 | 0.048283 | 0.043707 | 0.015049 | 0.089623 | 0.037612 | 0.041113 | 0.036269 | 0.025263 | 0.039736 | 0.018217 | 0.010084 | 0.060013 | 0.014817 | 0.032444 | 0.104542 | 0.051766 | 0.092108 |
| 34.5694444 | 0.029543 | 0.045529 | 0.041882 | 0.017637 | 0.02605 | 0.020391 | 0.017585 | 0.054112 | 0.024272 | 0.011469 | 0.075832 | 0.039557 | 0.036265 | 0.033015 | 0.026942 | 0.040552 | 0.01672 | 0.018876 | 0.075125 | 0.013449 | 0.034098 | 0.042964 | 0.055442 | 0.099061 |
| 34.5833333 | 0.032629 | 0.046955 | 0.045753 | 0.009754 | 0.040353 | 0.019288 | 0.007458 | 0.054861 | 0.027422 | 0.013284 | 0.058965 | 0.03905 | 0.032776 | 0.029716 | 0.036485 | 0.042776 | 0.014674 | 0.023711 | 0.08277 | 0.00797 | 0.037787 | 0.014916 | 0.05504 | 0.105077 |
| 34.5972222 | 0.034468 | 0.04696 | 0.047828 | 0.005671 | 0.058904 | 0.017722 | 0.003433 | 0.054289 | 0.033871 | 0.018604 | 0.040722 | 0.03794 | 0.030411 | 0.028411 | 0.048289 | 0.044116 | 0.012682 | 0.019667 | 0.08299 | 0.005311 | 0.049674 | 0.017759 | 0.052477 | 0.114811 |
| 34.6111111 | 0.034568 | 0.048652 | 0.053395 | 0.006707 | 0.048343 | 0.016093 | 0.003166 | 0.05493 | 0.024305 | 0.021573 | 0.024343 | 0.036515 | 0.028288 | 0.028919 | 0.057277 | 0.043626 | 0.011396 | 0.011071 | 0.083424 | 0.00559 | 0.06819 | 0.019919 | 0.051435 | 0.11088 |
| 34.625 | 0.03335 | 0.050416 | 0.060225 | 0.010969 | 0.024406 | 0.014896 | 0.004958 | 0.055314 | 0.010146 | 0.019315 | 0.012213 | 0.032999 | 0.026873 | 0.029242 | 0.063177 | 0.04456 | 0.01059 | 0.004556 | 0.088521 | 0.012269 | 0.081366 | 0.019435 | 0.048774 | 0.091674 |
| 34.6388889 | 0.030988 | 0.048975 | 0.059894 | 0.013623 | 0.011274 | 0.014501 | 0.009217 | 0.050554 | 0.004118 | 0.014252 | 0.004928 | 0.026188 | 0.026531 | 0.029743 | 0.067479 | 0.048413 | 0.009433 | 0.006228 | 0.094032 | 0.020431 | 0.080018 | 0.0433 | 0.046653 | 0.07231 |
| 34.6527778 | 0.029529 | 0.046163 | 0.051079 | 0.011801 | 0.009938 | 0.014465 | 0.012415 | 0.039981 | 0.011129 | 0.010043 | 0.003579 | 0.017143 | 0.026291 | 0.03181 | 0.068019 | 0.05096 | 0.00792 | 0.01585 | 0.088988 | 0.023405 | 0.066895 | 0.070502 | 0.041163 | 0.060608 |
| 34.6666667 | 0.030004 | 0.046732 | 0.041214 | 0.00783 | 0.008889 | 0.013988 | 0.012623 | 0.026949 | 0.0273 | 0.007369 | 0.006446 | 0.008501 | 0.025592 | 0.034451 | 0.062614 | 0.045697 | 0.006377 | 0.02714 | 0.070423 | 0.023803 | 0.049754 | 0.074007 | 0.026927 | 0.053436 |
| 34.6805556 | 0.031535 | 0.048647 | 0.03436 | 0.004138 | 0.01609 | 0.013801 | 0.011989 | 0.013519 | 0.044483 | 0.005002 | 0.009283 | 0.005335 | 0.022663 | 0.033898 | 0.051032 | 0.032445 | 0.004324 | 0.030603 | 0.048387 | 0.02307 | 0.03337 | 0.055495 | 0.024831 | 0.042363 |
| 34.6944444 | 0.032635 | 0.046 | 0.029933 | 0.003898 | 0.043307 | 0.014219 | 0.012458 | 0.004777 | 0.055437 | 0.004354 | 0.009796 | 0.009303 | 0.017514 | 0.026892 | 0.034693 | 0.018029 | 0.002194 | 0.022229 | 0.030937 | 0.021897 | 0.019545 | 0.029472 | 0.042236 | 0.025723 |
| 34.7083333 | 0.033186 | 0.037678 | 0.026574 | 0.009685 | 0.077167 | 0.0143 | 0.01262 | 0.003909 | 0.054991 | 0.007317 | 0.007464 | 0.014908 | 0.014206 | 0.01747 | 0.019494 | 0.008059 | 0.001471 | 0.01262 | 0.017103 | 0.022474 | 0.009281 | 0.021744 | 0.06012 | 0.011091 |
| 34.7222222 | 0.034368 | 0.027139 | 0.023295 | 0.016244 | 0.092732 | 0.013846 | 0.009021 | 0.004136 | 0.046402 | 0.011669 | 0.007021 | 0.020096 | 0.015173 | 0.010725 | 0.0113 | 0.00442 | 0.00227 | 0.010524 | 0.007031 | 0.025789 | 0.003721 | 0.038544 | 0.059409 | 0.004139 |
| 34.7361111 | 0.036987 | 0.018592 | 0.018948 | 0.018162 | 0.082413 | 0.012672 | 0.006586 | 0.002552 | 0.039179 | 0.014371 | 0.019161 | 0.02736 | 0.019145 | 0.005513 | 0.010384 | 0.006387 | 0.00368 | 0.008653 | 0.004155 | 0.027228 | 0.004157 | 0.063268 | 0.03421 | 0.003327 |
| 34.75 | 0.041202 | 0.014165 | 0.012743 | 0.016826 | 0.057406 | 0.011116 | 0.013051 | 0.002773 | 0.037887 | 0.015411 | 0.038534 | 0.03486 | 0.023368 | 0.001719 | 0.013501 | 0.010977 | 0.005254 | 0.006285 | 0.007164 | 0.023011 | 0.009674 | 0.081288 | 0.011541 | 0.003774 |
| 34.7638889 | 0.045267 | 0.011842 | 0.007141 | 0.015444 | 0.033814 | 0.010461 | 0.021595 | 0.004716 | 0.03775 | 0.017203 | 0.044561 | 0.039219 | 0.025374 | 0.000662 | 0.018711 | 0.013182 | 0.007311 | 0.010719 | 0.010868 | 0.016225 | 0.016155 | 0.082599 | 0.011243 | 0.005217 |
| 34.7777778 | 0.046814 | 0.007654 | 0.005204 | 0.016034 | 0.023761 | 0.011512 | 0.024773 | 0.005483 | 0.032283 | 0.021068 | 0.035641 | 0.041221 | 0.025359 | 0.001036 | 0.024827 | 0.009613 | 0.010065 | 0.014422 | 0.01355 | 0.010511 | 0.019863 | 0.073039 | 0.030371 | 0.009193 |
| 34.7916667 | 0.044423 | 0.006206 | 0.005589 | 0.018183 | 0.030029 | 0.013013 | 0.02602 | 0.003972 | 0.023035 | 0.024441 | 0.030034 | 0.041186 | 0.025668 | 0.002806 | 0.027056 | 0.008369 | 0.012569 | 0.012374 | 0.016265 | 0.010536 | 0.019903 | 0.06605 | 0.054994 | 0.015115 |
| 34.8055556 | 0.037568 | 0.014491 | 0.005371 | 0.02028 | 0.044747 | 0.013486 | 0.02824 | 0.002651 | 0.01681 | 0.024834 | 0.034184 | 0.039269 | 0.027376 | 0.005117 | 0.021609 | 0.015724 | 0.013616 | 0.013999 | 0.019305 | 0.014165 | 0.017206 | 0.06478 | 0.066235 | 0.023682 |
| 34.8194444 | 0.028241 | 0.026969 | 0.005186 | 0.021833 | 0.052088 | 0.012765 | 0.029364 | 0.004779 | 0.012987 | 0.023076 | 0.038386 | 0.03721 | 0.029405 | 0.005347 | 0.011529 | 0.02085 | 0.012723 | 0.021621 | 0.025298 | 0.015013 | 0.014055 | 0.072993 | 0.058038 | 0.033289 |
| 34.8333333 | 0.019606 | 0.036649 | 0.006264 | 0.022804 | 0.048741 | 0.009878 | 0.027543 | 0.011287 | 0.007858 | 0.021783 | 0.037569 | 0.035059 | 0.028638 | 0.004728 | 0.007756 | 0.019698 | 0.01041 | 0.025567 | 0.03438 | 0.015343 | 0.011725 | 0.093354 | 0.034364 | 0.042517 |
| 34.8472222 | 0.011517 | 0.041986 | 0.007169 | 0.023595 | 0.049968 | 0.006067 | 0.024383 | 0.021234 | 0.005909 | 0.021922 | 0.031558 | 0.030922 | 0.023076 | 0.005962 | 0.014509 | 0.019915 | 0.008859 | 0.022018 | 0.040292 | 0.015723 | 0.010356 | 0.112573 | 0.020654 | 0.053868 |
| 34.8611111 | 0.006454 | 0.043565 | 0.006526 | 0.024373 | 0.062708 | 0.003998 | 0.022402 | 0.027842 | 0.010074 | 0.021958 | 0.021246 | 0.023449 | 0.017216 | 0.008025 | 0.024149 | 0.02216 | 0.009286 | 0.015263 | 0.040131 | 0.014538 | 0.01258 | 0.117548 | 0.042302 | 0.065993 |
| 34.875 | 0.009776 | 0.045353 | 0.005567 | 0.024105 | 0.08256 | 0.003344 | 0.02174 | 0.022931 | 0.017108 | 0.020687 | 0.010567 | 0.013353 | 0.016703 | 0.010376 | 0.033427 | 0.023341 | 0.009694 | 0.008095 | 0.035783 | 0.013053 | 0.016563 | 0.110067 | 0.082582 | 0.070929 |
| 34.8888889 | 0.018131 | 0.047174 | 0.00643 | 0.021594 | 0.099993 | 0.003246 | 0.019883 | 0.011207 | 0.025038 | 0.019142 | 0.006219 | 0.006259 | 0.018105 | 0.014598 | 0.040836 | 0.023967 | 0.008586 | 0.004523 | 0.029848 | 0.012079 | 0.015193 | 0.094794 | 0.113417 | 0.071843 |
| 34.9027778 | 0.023913 | 0.043926 | 0.00918 | 0.016382 | 0.103459 | 0.003937 | 0.014851 | 0.003438 | 0.032689 | 0.019288 | 0.011891 | 0.007469 | 0.015207 | 0.019256 | 0.042406 | 0.024603 | 0.007872 | 0.005209 | 0.026112 | 0.011388 | 0.010276 | 0.072492 | 0.114809 | 0.079761 |
| 34.9166667 | 0.024694 | 0.036441 | 0.012048 | 0.00945 | 0.092466 | 0.006079 | 0.007753 | 0.001906 | 0.038148 | 0.019225 | 0.017321 | 0.012022 | 0.009539 | 0.021732 | 0.038156 | 0.024224 | 0.009221 | 0.005061 | 0.025136 | 0.011646 | 0.006876 | 0.046232 | 0.078923 | 0.093808 |
| 34.9305556 | 0.021262 | 0.029208 | 0.012543 | 0.00372 | 0.075552 | 0.009908 | 0.004023 | 0.00355 | 0.042154 | 0.014019 | 0.015089 | 0.014534 | 0.006668 | 0.021622 | 0.033059 | 0.022092 | 0.011157 | 0.004165 | 0.02488 | 0.015409 | 0.004581 | 0.024517 | 0.038318 | 0.103389 |
| 34.9444444 | 0.016182 | 0.022163 | 0.00979 | 0.002241 | 0.058931 | 0.012887 | 0.006512 | 0.006993 | 0.044945 | 0.007396 | 0.012729 | 0.015123 | 0.008883 | 0.019677 | 0.029493 | 0.01901 | 0.011571 | 0.003239 | 0.024038 | 0.022381 | 0.003428 | 0.011077 | 0.026201 | 0.097975 |
| 34.9583333 | 0.012438 | 0.014624 | 0.007022 | 0.004926 | 0.043163 | 0.011345 | 0.01045 | 0.0103 | 0.045559 | 0.006053 | 0.018214 | 0.012928 | 0.014965 | 0.013368 | 0.021548 | 0.014218 | 0.010096 | 0.00477 | 0.024988 | 0.027605 | 0.003984 | 0.004642 | 0.028191 | 0.079045 |
| 34.9722222 | 0.013177 | 0.016451 | 0.007122 | 0.008241 | 0.031387 | 0.006903 | 0.012353 | 0.00966 | 0.043793 | 0.008128 | 0.037263 | 0.009654 | 0.030424 | 0.010459 | 0.016191 | 0.010242 | 0.006998 | 0.012054 | 0.040867 | 0.026445 | 0.006481 | 0.004123 | 0.028986 | 0.052275 |
| 34.9861111 | 0.019045 | 0.029827 | 0.007555 | 0.007354 | 0.03158 | 0.005935 | 0.010629 | 0.010565 | 0.034067 | 0.018543 | 0.066805 | 0.012204 | 0.05165 | 0.015798 | 0.025689 | 0.010818 | 0.00435 | 0.015469 | 0.081255 | 0.019095 | 0.011903 | 0.006871 | 0.032742 | 0.027851 |
| 35 | 0.023208 | 0.031277 | 0.0065 | 0.003919 | 0.047607 | 0.008827 | 0.00648 | 0.01832 | 0.019197 | 0.044846 | 0.085931 | 0.019401 | 0.056901 | 0.01766 | 0.02956 | 0.012609 | 0.008785 | 0.009289 | 0.121722 | 0.009984 | 0.022014 | 0.019061 | 0.041307 | 0.029934 |
| 35.0138889 | 0.019625 | 0.018758 | 0.006991 | 0.003148 | 0.067385 | 0.007904 | 0.006768 | 0.021865 | 0.013094 | 0.07378 | 0.078643 | 0.020369 | 0.039089 | 0.020086 | 0.020745 | 0.01467 | 0.020182 | 0.002955 | 0.130059 | 0.004986 | 0.03621 | 0.041046 | 0.054467 | 0.053986 |
| 35.0277778 | 0.010543 | 0.016584 | 0.008203 | 0.005817 | 0.071577 | 0.005238 | 0.012024 | 0.015876 | 0.014243 | 0.088978 | 0.052997 | 0.013075 | 0.019142 | 0.025847 | 0.02319 | 0.012261 | 0.029803 | 0.001308 | 0.109183 | 0.004782 | 0.04602 | 0.063989 | 0.063384 | 0.062139 |
| 35.0416667 | 0.003036 | 0.025281 | 0.006157 | 0.010806 | 0.057795 | 0.008245 | 0.017425 | 0.008171 | 0.017106 | 0.09432 | 0.025733 | 0.00525 | 0.017467 | 0.025879 | 0.029793 | 0.005011 | 0.03349 | 0.002252 | 0.077739 | 0.006398 | 0.040177 | 0.084906 | 0.063642 | 0.054098 |
| 35.0555556 | 0.000751 | 0.030689 | 0.002395 | 0.013254 | 0.035614 | 0.012257 | 0.020455 | 0.0038 | 0.02244 | 0.093013 | 0.013093 | 0.001647 | 0.023888 | 0.021956 | 0.021823 | 0.002474 | 0.033651 | 0.006889 | 0.049265 | 0.011475 | 0.021533 | 0.082251 | 0.061072 | 0.052077 |
| 35.0694444 | 0.001145 | 0.02963 | 0.000635 | 0.011472 | 0.014915 | 0.013185 | 0.019635 | 0.00277 | 0.023904 | 0.078422 | 0.017688 | 0.001946 | 0.024042 | 0.018042 | 0.010906 | 0.004893 | 0.031498 | 0.013376 | 0.028527 | 0.014361 | 0.008912 | 0.051226 | 0.058022 | 0.047376 |
| 35.0833333 | 0.003362 | 0.02712 | 0.000997 | 0.010128 | 0.003827 | 0.013037 | 0.016567 | 0.002823 | 0.019099 | 0.053228 | 0.022984 | 0.004016 | 0.020799 | 0.014035 | 0.012797 | 0.010338 | 0.021248 | 0.017861 | 0.013022 | 0.017864 | 0.010756 | 0.023956 | 0.043889 | 0.030655 |
| 35.0972222 | 0.008307 | 0.027746 | 0.002899 | 0.010213 | 0.001113 | 0.013119 | 0.013645 | 0.002429 | 0.01328 | 0.027949 | 0.021795 | 0.004408 | 0.016937 | 0.010034 | 0.021153 | 0.017942 | 0.012522 | 0.021242 | 0.004543 | 0.036761 | 0.014123 | 0.022847 | 0.021189 | 0.012982 |
| 35.1111111 | 0.013209 | 0.031537 | 0.006041 | 0.009722 | 0.00286 | 0.013391 | 0.010326 | 0.003247 | 0.009708 | 0.010501 | 0.018272 | 0.00296 | 0.012288 | 0.007002 | 0.025838 | 0.024191 | 0.020428 | 0.023663 | 0.003995 | 0.054299 | 0.012044 | 0.044335 | 0.009779 | 0.006987 |
| 35.125 | 0.015473 | 0.034111 | 0.008295 | 0.008413 | 0.009717 | 0.012764 | 0.005623 | 0.00694 | 0.008427 | 0.004908 | 0.014654 | 0.002817 | 0.007391 | 0.005314 | 0.026341 | 0.027362 | 0.031835 | 0.024638 | 0.005121 | 0.043037 | 0.008064 | 0.068269 | 0.009473 | 0.014796 |
| 35.1388889 | 0.015202 | 0.031821 | 0.008369 | 0.007058 | 0.020585 | 0.011133 | 0.002283 | 0.011526 | 0.007714 | 0.010724 | 0.012982 | 0.004259 | 0.003594 | 0.004167 | 0.02579 | 0.028369 | 0.031351 | 0.024573 | 0.004674 | 0.024315 | 0.004195 | 0.086143 | 0.010049 | 0.022985 |
| 35.1527778 | 0.013706 | 0.027314 | 0.006957 | 0.006958 | 0.032048 | 0.009793 | 0.002757 | 0.014568 | 0.006266 | 0.023682 | 0.014758 | 0.005791 | 0.001707 | 0.003095 | 0.027207 | 0.030266 | 0.023309 | 0.02097 | 0.003967 | 0.037568 | 0.002515 | 0.100572 | 0.008954 | 0.0216 |
| 35.1666667 | 0.012583 | 0.025282 | 0.005473 | 0.008336 | 0.043151 | 0.00952 | 0.00378 | 0.015464 | 0.004816 | 0.036988 | 0.019259 | 0.006584 | 0.001723 | 0.002505 | 0.02964 | 0.033248 | 0.016483 | 0.016413 | 0.003259 | 0.055383 | 0.001857 | 0.109479 | 0.006792 | 0.015562 |
| 35.1805556 | 0.01256 | 0.025972 | 0.005454 | 0.010404 | 0.053702 | 0.009409 | 0.002712 | 0.014729 | 0.005113 | 0.043848 | 0.024239 | 0.006495 | 0.003677 | 0.002426 | 0.030985 | 0.034318 | 0.012181 | 0.015087 | 0.002152 | 0.041487 | 0.001051 | 0.108342 | 0.004941 | 0.009284 |
| 35.1944444 | 0.013624 | 0.026772 | 0.007454 | 0.013387 | 0.062125 | 0.009261 | 0.001685 | 0.012593 | 0.007804 | 0.04262 | 0.027294 | 0.005991 | 0.006922 | 0.002789 | 0.030486 | 0.031626 | 0.008308 | 0.013102 | 0.00189 | 0.019578 | 0.001337 | 0.095161 | 0.003229 | 0.007299 |
| 35.2083333 | 0.014632 | 0.026633 | 0.010593 | 0.016137 | 0.070052 | 0.010251 | 0.002466 | 0.009441 | 0.010458 | 0.036603 | 0.02757 | 0.005723 | 0.009913 | 0.003258 | 0.028332 | 0.025373 | 0.004339 | 0.008578 | 0.004038 | 0.008956 | 0.001533 | 0.079875 | 0.001928 | 0.012925 |
| 35.2222222 | 0.014033 | 0.024774 | 0.013537 | 0.018079 | 0.076662 | 0.012351 | 0.00308 | 0.007572 | 0.010185 | 0.02966 | 0.025769 | 0.005474 | 0.011797 | 0.003458 | 0.026394 | 0.017758 | 0.002051 | 0.004844 | 0.007165 | 0.012423 | 0.000786 | 0.076333 | 0.002782 | 0.023256 |
| 35.2361111 | 0.012098 | 0.019487 | 0.015577 | 0.020233 | 0.077995 | 0.014191 | 0.002151 | 0.008389 | 0.008763 | 0.024516 | 0.022874 | 0.00477 | 0.013454 | 0.003705 | 0.025841 | 0.010428 | 0.003153 | 0.00594 | 0.008793 | 0.020104 | 0.000849 | 0.08652 | 0.005928 | 0.033646 |
| 35.25 | 0.010589 | 0.011783 | 0.016304 | 0.021613 | 0.070863 | 0.014718 | 0.001255 | 0.009801 | 0.008971 | 0.022559 | 0.02046 | 0.003802 | 0.015558 | 0.004194 | 0.026045 | 0.004518 | 0.005292 | 0.012701 | 0.00856 | 0.014425 | 0.002125 | 0.095665 | 0.010786 | 0.039674 |
| 35.2638889 | 0.009592 | 0.005409 | 0.01474 | 0.021563 | 0.053194 | 0.013837 | 0.001755 | 0.00915 | 0.010199 | 0.02403 | 0.018842 | 0.002436 | 0.017058 | 0.004962 | 0.025707 | 0.001815 | 0.006728 | 0.020583 | 0.008526 | 0.008017 | 0.002422 | 0.087769 | 0.015999 | 0.041956 |
| 35.2777778 | 0.008082 | 0.002262 | 0.01071 | 0.021186 | 0.034042 | 0.012134 | 0.003625 | 0.00646 | 0.010327 | 0.029412 | 0.018673 | 0.001498 | 0.017959 | 0.005708 | 0.024226 | 0.002507 | 0.00719 | 0.025672 | 0.00925 | 0.015716 | 0.001327 | 0.067452 | 0.019271 | 0.042333 |
| 35.2916667 | 0.007091 | 0.001229 | 0.006288 | 0.020919 | 0.02319 | 0.010276 | 0.005843 | 0.003914 | 0.00963 | 0.038243 | 0.02167 | 0.001495 | 0.019708 | 0.005472 | 0.024146 | 0.004857 | 0.005704 | 0.028534 | 0.00802 | 0.029551 | 0.001587 | 0.045626 | 0.022637 | 0.043961 |
| 35.3055556 | 0.008085 | 0.001303 | 0.003327 | 0.018424 | 0.014687 | 0.008681 | 0.007032 | 0.002919 | 0.010308 | 0.046319 | 0.026818 | 0.001086 | 0.021928 | 0.004154 | 0.027463 | 0.007883 | 0.003023 | 0.03031 | 0.004409 | 0.038668 | 0.004152 | 0.024786 | 0.028905 | 0.050572 |
| 35.3194444 | 0.010026 | 0.002605 | 0.001832 | 0.012324 | 0.008347 | 0.007687 | 0.006811 | 0.003068 | 0.012206 | 0.048333 | 0.03142 | 0.000573 | 0.023214 | 0.003001 | 0.031196 | 0.011737 | 0.001534 | 0.030513 | 0.002171 | 0.042054 | 0.007415 | 0.012584 | 0.035086 | 0.053034 |
| 35.3333333 | 0.011454 | 0.005199 | 0.001918 | 0.007006 | 0.009785 | 0.007596 | 0.005894 | 0.003812 | 0.013891 | 0.04333 | 0.033852 | 0.001172 | 0.022566 | 0.002895 | 0.032676 | 0.015715 | 0.002005 | 0.027382 | 0.002395 | 0.0485 | 0.009396 | 0.017435 | 0.036975 | 0.044767 |
| 35.3472222 | 0.012357 | 0.008156 | 0.003598 | 0.005016 | 0.01026 | 0.007887 | 0.005084 | 0.004534 | 0.015064 | 0.035457 | 0.033318 | 0.002399 | 0.019774 | 0.004599 | 0.033162 | 0.018071 | 0.002357 | 0.01785 | 0.002028 | 0.066679 | 0.009515 | 0.029865 | 0.034376 | 0.032861 |
| 35.3611111 | 0.012798 | 0.01061 | 0.005576 | 0.003273 | 0.009773 | 0.008044 | 0.004787 | 0.004444 | 0.016998 | 0.028737 | 0.031028 | 0.00328 | 0.016278 | 0.009514 | 0.035256 | 0.01748 | 0.001693 | 0.006523 | 0.001093 | 0.092915 | 0.008417 | 0.036173 | 0.030033 | 0.01918 |
| 35.375 | 0.01298 | 0.012855 | 0.006296 | 0.00138 | 0.013888 | 0.008763 | 0.004257 | 0.004477 | 0.020771 | 0.024943 | 0.029492 | 0.003647 | 0.013653 | 0.016692 | 0.040059 | 0.014843 | 0.001955 | 0.003031 | 0.000984 | 0.113837 | 0.007391 | 0.033495 | 0.026587 | 0.012248 |
| 35.3888889 | 0.013615 | 0.015207 | 0.005452 | 0.000957 | 0.014656 | 0.010372 | 0.002701 | 0.006873 | 0.023752 | 0.02434 | 0.029875 | 0.003868 | 0.012423 | 0.021748 | 0.045677 | 0.01143 | 0.003505 | 0.007314 | 0.001553 | 0.114716 | 0.0078 | 0.024084 | 0.025829 | 0.020551 |
| 35.4027778 | 0.015374 | 0.017817 | 0.004485 | 0.001007 | 0.008631 | 0.011695 | 0.001114 | 0.011877 | 0.023975 | 0.02495 | 0.032254 | 0.004026 | 0.012294 | 0.021332 | 0.04871 | 0.008522 | 0.004456 | 0.014092 | 0.003361 | 0.09049 | 0.0099 | 0.012006 | 0.027352 | 0.02812 |
| 35.4166667 | 0.018113 | 0.020604 | 0.004227 | 0.000881 | 0.004028 | 0.011788 | 0.000905 | 0.016452 | 0.021556 | 0.024362 | 0.036374 | 0.003518 | 0.012583 | 0.017487 | 0.04791 | 0.007706 | 0.004504 | 0.019549 | 0.004421 | 0.054895 | 0.012358 | 0.006418 | 0.02842 | 0.026443 |
| 35.4305556 | 0.020606 | 0.023323 | 0.003374 | 0.000915 | 0.007954 | 0.010133 | 0.001577 | 0.018386 | 0.014265 | 0.02208 | 0.043223 | 0.003788 | 0.012655 | 0.014613 | 0.041168 | 0.008579 | 0.00493 | 0.023487 | 0.003115 | 0.026088 | 0.014743 | 0.015743 | 0.027332 | 0.020875 |
| 35.4444444 | 0.022603 | 0.025184 | 0.0022 | 0.001459 | 0.021254 | 0.007356 | 0.001572 | 0.018687 | 0.006248 | 0.019959 | 0.051802 | 0.005774 | 0.012955 | 0.012725 | 0.027214 | 0.012532 | 0.005705 | 0.028329 | 0.003298 | 0.020654 | 0.019468 | 0.036977 | 0.0241 | 0.015648 |
| 35.4583333 | 0.025469 | 0.021847 | 0.002787 | 0.00538 | 0.039548 | 0.007177 | 0.001479 | 0.016584 | 0.009012 | 0.02003 | 0.051431 | 0.010323 | 0.014238 | 0.008377 | 0.023958 | 0.019883 | 0.00591 | 0.031755 | 0.005087 | 0.031251 | 0.024657 | 0.06062 | 0.020871 | 0.017548 |
| 35.4722222 | 0.031435 | 0.01178 | 0.006712 | 0.014476 | 0.057438 | 0.010249 | 0.002434 | 0.013822 | 0.020527 | 0.01972 | 0.041415 | 0.018835 | 0.017864 | 0.005422 | 0.037067 | 0.022625 | 0.004523 | 0.029051 | 0.012433 | 0.027254 | 0.024825 | 0.072793 | 0.021854 | 0.022207 |
| 35.4861111 | 0.040137 | 0.007811 | 0.017928 | 0.019751 | 0.080235 | 0.009774 | 0.002539 | 0.022639 | 0.030638 | 0.016681 | 0.050509 | 0.02288 | 0.0308 | 0.013652 | 0.038063 | 0.019789 | 0.009391 | 0.019287 | 0.045119 | 0.019936 | 0.01739 | 0.069902 | 0.031007 | 0.028004 |
| 35.5 | 0.044376 | 0.035468 | 0.040866 | 0.024056 | 0.120198 | 0.023246 | 0.001332 | 0.050822 | 0.077258 | 0.013976 | 0.112737 | 0.043172 | 0.07015 | 0.037322 | 0.024116 | 0.027697 | 0.025135 | 0.027866 | 0.122415 | 0.02031 | 0.029683 | 0.068314 | 0.054634 | 0.042796 |
| 35.5138889 | 0.039255 | 0.150805 | 0.080378 | 0.057291 | 0.170838 | 0.058178 | 0.008805 | 0.064776 | 0.154073 | 0.016471 | 0.216195 | 0.096382 | 0.142619 | 0.048733 | 0.036612 | 0.06101 | 0.065166 | 0.097978 | 0.147078 | 0.00946 | 0.135016 | 0.133281 | 0.090257 | 0.101215 |
| 35.5277778 | 0.035828 | 0.215106 | 0.092615 | 0.076463 | 0.164191 | 0.058628 | 0.032853 | 0.06078 | 0.145527 | 0.024564 | 0.242475 | 0.111631 | 0.155839 | 0.040769 | 0.052253 | 0.078278 | 0.105178 | 0.142254 | 0.060164 | 0 | 0.19915 | 0.176972 | 0.108133 | 0.134274 |
| 35.5416667 | 0.036786 | 0.174449 | 0.047095 | 0.052421 | 0.110052 | 0.05741 | 0.052731 | 0.055779 | 0.112483 | 0.058993 | 0.174082 | 0.098894 | 0.126497 | 0.038191 | 0.024186 | 0.075115 | 0.095706 | 0.154291 | 0.141468 | 0 | 0.160702 | 0.186364 | 0.116104 | 0.059769 |
| 35.5555556 | 0.031766 | 0.183224 | 0.013786 | 0.064869 | 0.082873 | 0.089557 | 0.050018 | 0.03606 | 0.167523 | 0.099997 | 0.128742 | 0.111968 | 0.146868 | 0.031996 | 0.018028 | 0.120336 | 0.060234 | 0.198411 | 0.291619 | 0.019288 | 0.165049 | 0.250085 | 0.132461 | 0.01661 |
| 35.5694444 | 0.028356 | 0.136758 | 0.020517 | 0.092444 | 0.063635 | 0.072766 | 0.039888 | 0.025318 | 0.169026 | 0.082456 | 0.094606 | 0.085036 | 0.112407 | 0.027352 | 0.048072 | 0.163121 | 0.025228 | 0.150651 | 0.173427 | 0.040581 | 0.118809 | 0.196978 | 0.116926 | 0.050408 |
| 35.5833333 | 0.032019 | 0.048212 | 0.033834 | 0.073086 | 0.054467 | 0.032825 | 0.033634 | 0.029439 | 0.074099 | 0.036319 | 0.065858 | 0.029739 | 0.033567 | 0.032811 | 0.050395 | 0.143087 | 0.003762 | 0.057642 | 0.058661 | 0.026201 | 0.039286 | 0.084754 | 0.084952 | 0.067263 |
| 35.5972222 | 0.035947 | 0.02799 | 0.042815 | 0.037769 | 0.047221 | 0.026163 | 0.028247 | 0.036124 | 0.013894 | 0.013629 | 0.068781 | 0.00956 | 0.005969 | 0.038817 | 0.04516 | 0.110074 | 0.000576 | 0.027792 | 0.080158 | 0.013469 | 0.02685 | 0.049325 | 0.077823 | 0.06531 |
| 35.6111111 | 0.037874 | 0.030395 | 0.041656 | 0.015347 | 0.040055 | 0.027923 | 0.023473 | 0.040603 | 0.005256 | 0.00803 | 0.062447 | 0.007288 | 0.006994 | 0.045209 | 0.052239 | 0.089028 | 0.000483 | 0.016406 | 0.094842 | 0.02261 | 0.036323 | 0.032712 | 0.063731 | 0.065104 |
| 35.625 | 0.037926 | 0.039043 | 0.03216 | 0.011265 | 0.068271 | 0.025094 | 0.01974 | 0.037057 | 0.008812 | 0.00774 | 0.040827 | 0.004829 | 0.014918 | 0.057541 | 0.044894 | 0.078473 | 0.000827 | 0.009515 | 0.100966 | 0.038354 | 0.057268 | 0.025878 | 0.030618 | 0.06382 |
| 35.6388889 | 0.036537 | 0.044165 | 0.023443 | 0.019568 | 0.104685 | 0.020447 | 0.015999 | 0.032019 | 0.013552 | 0.005518 | 0.024146 | 0.004149 | 0.021041 | 0.07492 | 0.027238 | 0.074761 | 0.000803 | 0.021199 | 0.093561 | 0.040718 | 0.076082 | 0.018658 | 0.020352 | 0.060091 |
| 35.6527778 | 0.034643 | 0.043205 | 0.019068 | 0.029 | 0.11759 | 0.01917 | 0.010933 | 0.027833 | 0.012781 | 0.005191 | 0.020416 | 0.004193 | 0.023355 | 0.080926 | 0.014469 | 0.058836 | 0.000672 | 0.046067 | 0.07054 | 0.024976 | 0.078815 | 0.013072 | 0.036364 | 0.063293 |
| 35.6666667 | 0.033131 | 0.045897 | 0.016564 | 0.034876 | 0.108187 | 0.021407 | 0.007186 | 0.027092 | 0.012186 | 0.012239 | 0.02629 | 0.003321 | 0.023063 | 0.071255 | 0.011817 | 0.033324 | 0.000622 | 0.056069 | 0.041463 | 0.020248 | 0.068301 | 0.02568 | 0.046122 | 0.089624 |
| 35.6805556 | 0.030823 | 0.067077 | 0.013714 | 0.036599 | 0.09372 | 0.023996 | 0.006888 | 0.025702 | 0.015155 | 0.022807 | 0.033053 | 0.004344 | 0.022258 | 0.058096 | 0.023614 | 0.025909 | 0.00059 | 0.049209 | 0.019529 | 0.021289 | 0.053551 | 0.0563 | 0.051002 | 0.110953 |
| 35.6944444 | 0.026967 | 0.102393 | 0.011152 | 0.033245 | 0.072052 | 0.025095 | 0.00657 | 0.021705 | 0.013788 | 0.031189 | 0.039644 | 0.005678 | 0.02213 | 0.048227 | 0.036767 | 0.038138 | 0.000993 | 0.042362 | 0.013363 | 0.026287 | 0.037866 | 0.086367 | 0.068833 | 0.086313 |
| 35.7083333 | 0.023704 | 0.123464 | 0.00996 | 0.021962 | 0.037749 | 0.025454 | 0.004883 | 0.019933 | 0.011018 | 0.034054 | 0.046565 | 0.004839 | 0.022737 | 0.041135 | 0.031052 | 0.044433 | 0.001462 | 0.044751 | 0.0216 | 0.041207 | 0.024989 | 0.104068 | 0.085234 | 0.050878 |
| 35.7222222 | 0.024289 | 0.116454 | 0.010062 | 0.011467 | 0.019067 | 0.025632 | 0.004117 | 0.020553 | 0.011394 | 0.030266 | 0.048135 | 0.00652 | 0.02383 | 0.03442 | 0.016491 | 0.035437 | 0.001394 | 0.054389 | 0.035387 | 0.05382 | 0.021687 | 0.114536 | 0.07903 | 0.031485 |
| 35.7361111 | 0.028651 | 0.098019 | 0.01147 | 0.014831 | 0.028914 | 0.026402 | 0.004439 | 0.021038 | 0.009988 | 0.02276 | 0.045146 | 0.012721 | 0.024561 | 0.025761 | 0.016444 | 0.01935 | 0.00129 | 0.060672 | 0.045931 | 0.059298 | 0.028602 | 0.114977 | 0.057153 | 0.019422 |
| 35.75 | 0.033237 | 0.083841 | 0.013097 | 0.025306 | 0.039348 | 0.026962 | 0.005963 | 0.017967 | 0.007754 | 0.014711 | 0.038177 | 0.021095 | 0.023771 | 0.017159 | 0.023322 | 0.011473 | 0.001582 | 0.058345 | 0.050002 | 0.062689 | 0.038628 | 0.09271 | 0.031939 | 0.012651 |
| 35.7638889 | 0.035995 | 0.073134 | 0.013473 | 0.029134 | 0.02935 | 0.025837 | 0.008283 | 0.010431 | 0.009669 | 0.008783 | 0.027001 | 0.027643 | 0.021322 | 0.010667 | 0.024561 | 0.022119 | 0.00159 | 0.05005 | 0.049047 | 0.073229 | 0.044209 | 0.063411 | 0.024346 | 0.007173 |
| 35.7777778 | 0.035409 | 0.056314 | 0.013574 | 0.019874 | 0.017457 | 0.026307 | 0.00824 | 0.005031 | 0.011749 | 0.00896 | 0.021728 | 0.027087 | 0.018281 | 0.005874 | 0.02264 | 0.038343 | 0.000921 | 0.037764 | 0.041289 | 0.088578 | 0.041011 | 0.0529 | 0.045219 | 0.005224 |
| 35.7916667 | 0.033449 | 0.036937 | 0.01415 | 0.010177 | 0.029155 | 0.028942 | 0.005764 | 0.004421 | 0.011907 | 0.015207 | 0.023912 | 0.022532 | 0.01487 | 0.005517 | 0.019531 | 0.044248 | 0.000379 | 0.021097 | 0.024627 | 0.084377 | 0.028024 | 0.061107 | 0.074394 | 0.006092 |
| 35.8055556 | 0.035837 | 0.028448 | 0.014201 | 0.014041 | 0.051166 | 0.027206 | 0.003281 | 0.008925 | 0.013782 | 0.022867 | 0.023257 | 0.019254 | 0.011145 | 0.011722 | 0.01542 | 0.039642 | 0.001065 | 0.01186 | 0.009906 | 0.05315 | 0.014373 | 0.073813 | 0.093748 | 0.018662 |
| 35.8194444 | 0.04261 | 0.031715 | 0.013785 | 0.01964 | 0.062975 | 0.018288 | 0.002548 | 0.017501 | 0.015631 | 0.027589 | 0.018528 | 0.016919 | 0.008919 | 0.021476 | 0.010132 | 0.03228 | 0.002936 | 0.020841 | 0.006477 | 0.031368 | 0.01199 | 0.0737 | 0.099422 | 0.036485 |
| 35.8333333 | 0.047384 | 0.038059 | 0.014695 | 0.019599 | 0.072021 | 0.008297 | 0.005363 | 0.022137 | 0.016237 | 0.027838 | 0.0155 | 0.015292 | 0.010332 | 0.032719 | 0.004848 | 0.027576 | 0.004189 | 0.029147 | 0.007192 | 0.032227 | 0.014453 | 0.058441 | 0.09834 | 0.043006 |
| 35.8472222 | 0.045154 | 0.037027 | 0.017475 | 0.021575 | 0.08598 | 0.003379 | 0.010842 | 0.021957 | 0.017881 | 0.025703 | 0.015118 | 0.013103 | 0.013264 | 0.045619 | 0.005555 | 0.024065 | 0.003802 | 0.023578 | 0.009592 | 0.02909 | 0.013886 | 0.050348 | 0.093121 | 0.053006 |
| 35.8611111 | 0.038172 | 0.024808 | 0.02013 | 0.026802 | 0.099426 | 0.002627 | 0.015978 | 0.020949 | 0.017815 | 0.023712 | 0.01506 | 0.008044 | 0.011793 | 0.059668 | 0.020009 | 0.019257 | 0.003082 | 0.015299 | 0.016148 | 0.013797 | 0.016772 | 0.054007 | 0.077381 | 0.065781 |
| 35.875 | 0.032208 | 0.010117 | 0.020976 | 0.027267 | 0.106361 | 0.002427 | 0.017482 | 0.020735 | 0.014488 | 0.023009 | 0.012934 | 0.003696 | 0.007593 | 0.071803 | 0.042683 | 0.01489 | 0.003186 | 0.010397 | 0.021671 | 0.00406 | 0.021112 | 0.054873 | 0.047284 | 0.078726 |
| 35.8888889 | 0.02912 | 0.002572 | 0.019465 | 0.021112 | 0.105482 | 0.002272 | 0.014232 | 0.021345 | 0.013511 | 0.022797 | 0.007454 | 0.003266 | 0.008955 | 0.077515 | 0.053403 | 0.010309 | 0.004123 | 0.009486 | 0.019326 | 0.019582 | 0.019654 | 0.051859 | 0.022937 | 0.095474 |
| 35.9027778 | 0.026717 | 0.001003 | 0.016375 | 0.017203 | 0.099293 | 0.002218 | 0.0094 | 0.020171 | 0.0134 | 0.021181 | 0.002378 | 0.003986 | 0.016262 | 0.078612 | 0.048296 | 0.008118 | 0.006074 | 0.012038 | 0.011943 | 0.054568 | 0.013445 | 0.047781 | 0.028015 | 0.102877 |
| 35.9166667 | 0.023222 | 0.000813 | 0.013652 | 0.016923 | 0.089634 | 0.002923 | 0.005086 | 0.01824 | 0.009423 | 0.0184 | 0.001306 | 0.004902 | 0.023505 | 0.079318 | 0.041156 | 0.017627 | 0.008523 | 0.014213 | 0.008828 | 0.06342 | 0.007527 | 0.045747 | 0.044028 | 0.108441 |
| 35.9305556 | 0.019995 | 0.000741 | 0.013179 | 0.013457 | 0.07193 | 0.006232 | 0.003614 | 0.017923 | 0.005984 | 0.015272 | 0.001622 | 0.006398 | 0.027553 | 0.075202 | 0.033195 | 0.032747 | 0.009069 | 0.015653 | 0.009582 | 0.040451 | 0.006925 | 0.045757 | 0.052479 | 0.112455 |
| 35.9444444 | 0.019056 | 0.00156 | 0.015656 | 0.007336 | 0.047852 | 0.011955 | 0.007886 | 0.017129 | 0.004413 | 0.013088 | 0.001083 | 0.007585 | 0.027272 | 0.062493 | 0.023944 | 0.040606 | 0.006673 | 0.013344 | 0.014695 | 0.032583 | 0.007542 | 0.043878 | 0.057364 | 0.10188 |

| | | | | | | | | | | | | | | | | | | | | | | | | |
|---|---|---|---|---|---|---|---|---|---|---|---|---|---|---|---|---|---|---|---|---|---|---|---|---|
| 35.9583333 | 0.019548 | 0.005063 | 0.018613 | 0.004708 | 0.031044 | 0.017769 | 0.01409 | 0.014033 | 0.005845 | 0.014436 | 0.001547 | 0.006624 | 0.023373 | 0.04728 | 0.018607 | 0.041371 | 0.003691 | 0.008003 | 0.023756 | 0.039883 | 0.004692 | 0.039607 | 0.059845 | 0.088486 |
| 35.9722222 | 0.018084 | 0.009612 | 0.015862 | 0.006428 | 0.031581 | 0.01818 | 0.017622 | 0.013625 | 0.016502 | 0.022222 | 0.007362 | 0.013841 | 0.019362 | 0.039251 | 0.015916 | 0.039044 | 0.002781 | 0.007035 | 0.02095 | 0.036533 | 0.002668 | 0.033978 | 0.061969 | 0.078661 |
| 35.9861111 | 0.014307 | 0.009017 | 0.010242 | 0.006475 | 0.047483 | 0.012105 | 0.016893 | 0.021473 | 0.030526 | 0.040341 | 0.019678 | 0.038936 | 0.01757 | 0.043471 | 0.01384 | 0.039281 | 0.005047 | 0.007887 | 0.019811 | 0.025157 | 0.009329 | 0.035705 | 0.065881 | 0.067344 |
| 36 | 0.017639 | 0.013984 | 0.008028 | 0.010365 | 0.065901 | 0.008581 | 0.011659 | 0.02967 | 0.028321 | 0.061872 | 0.022322 | 0.045423 | 0.016901 | 0.052693 | 0.01101 | 0.054428 | 0.010299 | 0.012169 | 0.055706 | 0.014543 | 0.02966 | 0.056565 | 0.068065 | 0.066992 |
| 36.0138889 | 0.031269 | 0.040683 | 0.007724 | 0.018686 | 0.076918 | 0.009536 | 0.009161 | 0.028213 | 0.014858 | 0.07141 | 0.016823 | 0.033825 | 0.01208 | 0.053719 | 0.00728 | 0.083048 | 0.017865 | 0.020907 | 0.126554 | 0.009258 | 0.056519 | 0.093896 | 0.061858 | 0.090522 |
| 36.0277778 | 0.041005 | 0.062499 | 0.007228 | 0.018099 | 0.09132 | 0.010064 | 0.012095 | 0.026579 | 0.01117 | 0.081939 | 0.01863 | 0.036647 | 0.007488 | 0.043928 | 0.00437 | 0.096904 | 0.026368 | 0.020853 | 0.175659 | 0.020412 | 0.070637 | 0.118111 | 0.051081 | 0.102226 |
| 36.0416667 | 0.039328 | 0.050381 | 0.005533 | 0.016956 | 0.117062 | 0.015529 | 0.012218 | 0.033813 | 0.018721 | 0.113093 | 0.015476 | 0.047321 | 0.013887 | 0.031696 | 0.003738 | 0.082333 | 0.034752 | 0.012298 | 0.17355 | 0.057319 | 0.056522 | 0.115175 | 0.041634 | 0.070855 |
| 36.0555556 | 0.034799 | 0.032784 | 0.004526 | 0.020213 | 0.127255 | 0.019308 | 0.009655 | 0.034793 | 0.02431 | 0.138087 | 0.013559 | 0.05178 | 0.02659 | 0.023124 | 0.004296 | 0.058213 | 0.039676 | 0.00707 | 0.15583 | 0.070732 | 0.028468 | 0.103344 | 0.030516 | 0.037169 |
| 36.0694444 | 0.037458 | 0.04482 | 0.005758 | 0.019991 | 0.115631 | 0.011021 | 0.009498 | 0.022439 | 0.024951 | 0.124533 | 0.031612 | 0.047239 | 0.035854 | 0.017884 | 0.006621 | 0.039356 | 0.037979 | 0.008116 | 0.122277 | 0.031085 | 0.014686 | 0.08963 | 0.01912 | 0.033662 |
| 36.0833333 | 0.042108 | 0.062442 | 0.006892 | 0.018759 | 0.097682 | 0.005938 | 0.009386 | 0.012342 | 0.025605 | 0.089501 | 0.047782 | 0.049232 | 0.035664 | 0.014955 | 0.012616 | 0.024595 | 0.031168 | 0.013223 | 0.077093 | 0.00392 | 0.013709 | 0.074362 | 0.010499 | 0.038799 |
| 36.0972222 | 0.033716 | 0.046676 | 0.006453 | 0.019249 | 0.06439 | 0.019729 | 0.006208 | 0.009358 | 0.02095 | 0.068299 | 0.034709 | 0.051882 | 0.025666 | 0.013629 | 0.017088 | 0.014012 | 0.018393 | 0.015937 | 0.048308 | 0.011063 | 0.009585 | 0.057075 | 0.005486 | 0.02935 |
| 36.1111111 | 0.017191 | 0.017816 | 0.007133 | 0.020934 | 0.029967 | 0.040794 | 0.002701 | 0.006407 | 0.023407 | 0.063286 | 0.011605 | 0.056178 | 0.014349 | 0.011862 | 0.017809 | 0.015362 | 0.008295 | 0.014411 | 0.031468 | 0.024047 | 0.004823 | 0.038808 | 0.003106 | 0.01772 |
| 36.125 | 0.009527 | 0.007737 | 0.006283 | 0.021672 | 0.01343 | 0.042747 | 0.001817 | 0.004542 | 0.029913 | 0.058095 | 0.003011 | 0.053709 | 0.007993 | 0.008616 | 0.01807 | 0.025622 | 0.009173 | 0.013952 | 0.019884 | 0.028728 | 0.007637 | 0.024737 | 0.001737 | 0.017386 |
| 36.1388889 | 0.011568 | 0.012092 | 0.003803 | 0.019257 | 0.005802 | 0.026803 | 0.002077 | 0.009593 | 0.022291 | 0.052453 | 0.005909 | 0.030735 | 0.00526 | 0.00641 | 0.017561 | 0.032667 | 0.010726 | 0.018736 | 0.01442 | 0.020277 | 0.014482 | 0.014601 | 0.002284 | 0.019189 |
| 36.1527778 | 0.015538 | 0.042544 | 0.002847 | 0.016526 | 0.007709 | 0.016738 | 0.001934 | 0.023432 | 0.018996 | 0.045329 | 0.022955 | 0.034679 | 0.003066 | 0.006908 | 0.014859 | 0.035269 | 0.008226 | 0.023645 | 0.01361 | 0.010969 | 0.018774 | 0.008957 | 0.005593 | 0.018007 |
| 36.1666667 | 0.015506 | 0.092956 | 0.004212 | 0.01684 | 0.022513 | 0.012934 | 0.0046 | 0.043559 | 0.028747 | 0.030541 | 0.046473 | 0.084431 | 0.001137 | 0.007094 | 0.010958 | 0.037167 | 0.00568 | 0.019199 | 0.012149 | 0.009437 | 0.017515 | 0.011649 | 0.009656 | 0.012367 |
| 36.1805556 | 0.009703 | 0.106023 | 0.008999 | 0.01926 | 0.026968 | 0.011237 | 0.013362 | 0.056417 | 0.028141 | 0.02179 | 0.051349 | 0.103311 | 0.000675 | 0.005624 | 0.007368 | 0.03989 | 0.004671 | 0.012198 | 0.007504 | 0.013158 | 0.010447 | 0.025632 | 0.011205 | 0.008159 |
| 36.1944444 | 0.005673 | 0.070023 | 0.012416 | 0.021799 | 0.020026 | 0.023053 | 0.022041 | 0.051992 | 0.022123 | 0.028014 | 0.037154 | 0.060121 | 0.003678 | 0.004438 | 0.0043 | 0.043893 | 0.005691 | 0.011862 | 0.003539 | 0.018804 | 0.005447 | 0.049028 | 0.010021 | 0.014836 |
| 36.2083333 | 0.009107 | 0.03048 | 0.012997 | 0.022498 | 0.031169 | 0.03646 | 0.023374 | 0.038791 | 0.030138 | 0.033558 | 0.020348 | 0.027928 | 0.008188 | 0.003883 | 0.001941 | 0.048283 | 0.007299 | 0.014198 | 0.003711 | 0.024001 | 0.008898 | 0.076076 | 0.006332 | 0.027516 |
| 36.2222222 | 0.01816 | 0.02519 | 0.013894 | 0.019447 | 0.052432 | 0.037033 | 0.021007 | 0.023815 | 0.041267 | 0.024165 | 0.010296 | 0.049396 | 0.009474 | 0.00333 | 0.002171 | 0.052518 | 0.007773 | 0.015632 | 0.005428 | 0.02674 | 0.013901 | 0.10105 | 0.003776 | 0.033966 |
| 36.2361111 | 0.026923 | 0.045831 | 0.016025 | 0.014623 | 0.064312 | 0.036022 | 0.019634 | 0.012583 | 0.042373 | 0.01016 | 0.008909 | 0.082525 | 0.009221 | 0.0025 | 0.006269 | 0.057506 | 0.00853 | 0.017698 | 0.0062 | 0.030595 | 0.015088 | 0.121699 | 0.007113 | 0.027497 |
| 36.25 | 0.025952 | 0.048544 | 0.020075 | 0.015765 | 0.059867 | 0.033507 | 0.015891 | 0.019327 | 0.038184 | 0.004995 | 0.015903 | 0.075341 | 0.010406 | 0.001553 | 0.012129 | 0.063322 | 0.010973 | 0.021664 | 0.007188 | 0.038178 | 0.013297 | 0.13518 | 0.014135 | 0.027278 |
| 36.2638889 | 0.016887 | 0.040096 | 0.024349 | 0.021879 | 0.039558 | 0.020139 | 0.008278 | 0.040267 | 0.03982 | 0.004588 | 0.031229 | 0.057634 | 0.012197 | 0.001212 | 0.016167 | 0.066136 | 0.013544 | 0.025886 | 0.008555 | 0.043939 | 0.008494 | 0.133833 | 0.021722 | 0.047954 |
| 36.2777778 | 0.012064 | 0.036799 | 0.02288 | 0.023427 | 0.028794 | 0.011009 | 0.002351 | 0.047142 | 0.041173 | 0.008745 | 0.036043 | 0.051537 | 0.013516 | 0.001039 | 0.016538 | 0.063004 | 0.013854 | 0.027299 | 0.008869 | 0.043841 | 0.003216 | 0.116752 | 0.028109 | 0.070879 |
| 36.2916667 | 0.015821 | 0.038109 | 0.014175 | 0.020631 | 0.047649 | 0.015324 | 0.001125 | 0.030009 | 0.030887 | 0.017343 | 0.021302 | 0.049238 | 0.014692 | 0.000462 | 0.014002 | 0.0552 | 0.011042 | 0.0262 | 0.008364 | 0.04118 | 0.001017 | 0.090936 | 0.031798 | 0.087724 |
| 36.3055556 | 0.019316 | 0.052542 | 0.005851 | 0.019822 | 0.076857 | 0.022332 | 0.002171 | 0.012293 | 0.016371 | 0.028068 | 0.008829 | 0.067319 | 0.016206 | 0.000293 | 0.009435 | 0.043293 | 0.007385 | 0.025219 | 0.008262 | 0.04061 | 0.002364 | 0.061054 | 0.033434 | 0.089378 |
| 36.3194444 | 0.013413 | 0.04523 | 0.003009 | 0.020705 | 0.084823 | 0.026643 | 0.003816 | 0.012936 | 0.010582 | 0.044935 | 0.008513 | 0.063115 | 0.017435 | 0.000967 | 0.009847 | 0.027849 | 0.004141 | 0.02569 | 0.008095 | 0.04156 | 0.004539 | 0.032798 | 0.033667 | 0.075391 |
| 36.3333333 | 0.005838 | 0.016285 | 0.003092 | 0.019278 | 0.066362 | 0.021631 | 0.004947 | 0.022466 | 0.012083 | 0.064379 | 0.013283 | 0.027656 | 0.01811 | 0.002063 | 0.019209 | 0.0148 | 0.002632 | 0.024004 | 0.005996 | 0.040084 | 0.004863 | 0.024713 | 0.032106 | 0.074849 |
| 36.3472222 | 0.008338 | 0.009879 | 0.006032 | 0.014817 | 0.037584 | 0.010779 | 0.004891 | 0.030004 | 0.014222 | 0.079561 | 0.019196 | 0.014962 | 0.017954 | 0.00311 | 0.025405 | 0.014416 | 0.005175 | 0.015661 | 0.00394 | 0.037625 | 0.007289 | 0.042731 | 0.031 | 0.087823 |
| 36.3611111 | 0.018661 | 0.036849 | 0.013957 | 0.012313 | 0.02546 | 0.00628 | 0.00477 | 0.041391 | 0.023539 | 0.09809 | 0.029976 | 0.045236 | 0.01655 | 0.003682 | 0.020735 | 0.024842 | 0.009373 | 0.006313 | 0.005616 | 0.039115 | 0.012595 | 0.061334 | 0.034011 | 0.099152 |
| 36.375 | 0.024601 | 0.066234 | 0.02033 | 0.015389 | 0.041921 | 0.010915 | 0.004078 | 0.05427 | 0.037499 | 0.112996 | 0.039768 | 0.085295 | 0.014774 | 0.003085 | 0.01127 | 0.030998 | 0.011142 | 0.002382 | 0.008418 | 0.043947 | 0.012822 | 0.066865 | 0.038252 | 0.105623 |
| 36.3888889 | 0.017115 | 0.056415 | 0.015643 | 0.0174 | 0.047055 | 0.019436 | 0.002673 | 0.048988 | 0.033635 | 0.095126 | 0.031202 | 0.080876 | 0.013559 | 0.001716 | 0.006506 | 0.027049 | 0.010596 | 0.00139 | 0.009596 | 0.046318 | 0.008262 | 0.065415 | 0.037907 | 0.097047 |
| 36.4027778 | 0.006594 | 0.032567 | 0.005778 | 0.019452 | 0.025865 | 0.032151 | 0.002954 | 0.026185 | 0.01993 | 0.053858 | 0.01553 | 0.065029 | 0.013498 | 0.001018 | 0.008115 | 0.017871 | 0.009734 | 0.001606 | 0.01027 | 0.042254 | 0.005098 | 0.05798 | 0.032375 | 0.072797 |
| 36.4166667 | 0.004031 | 0.023054 | 0.002655 | 0.022941 | 0.014122 | 0.041488 | 0.004143 | 0.013201 | 0.01574 | 0.028015 | 0.012097 | 0.06953 | 0.017386 | 0.000777 | 0.009974 | 0.009143 | 0.008288 | 0.003379 | 0.010574 | 0.034148 | 0.003939 | 0.045063 | 0.024036 | 0.041913 |
| 36.4305556 | 0.007074 | 0.01929 | 0.006664 | 0.017323 | 0.011925 | 0.034396 | 0.003874 | 0.018199 | 0.011764 | 0.026669 | 0.011541 | 0.050749 | 0.027419 | 0.000664 | 0.008668 | 0.006022 | 0.006182 | 0.00433 | 0.010339 | 0.026502 | 0.003672 | 0.032556 | 0.015394 | 0.019875 |
| 36.4444444 | 0.012968 | 0.033352 | 0.013083 | 0.007119 | 0.006808 | 0.021095 | 0.002955 | 0.02599 | 0.009932 | 0.031194 | 0.011643 | 0.019491 | 0.038416 | 0.001173 | 0.00485 | 0.006969 | 0.004919 | 0.003047 | 0.009189 | 0.022606 | 0.004992 | 0.024258 | 0.00977 | 0.019846 |
| 36.4583333 | 0.016134 | 0.055067 | 0.015918 | 0.00224 | 0.005167 | 0.014343 | 0.003573 | 0.026752 | 0.015438 | 0.027114 | 0.019201 | 0.015009 | 0.043522 | 0.001382 | 0.003133 | 0.00741 | 0.004264 | 0.001589 | 0.006092 | 0.024956 | 0.007944 | 0.017774 | 0.009479 | 0.027448 |
| 36.4722222 | 0.01671 | 0.072414 | 0.011379 | 0.002294 | 0.005849 | 0.010925 | 0.007581 | 0.022185 | 0.025624 | 0.020173 | 0.033996 | 0.034333 | 0.044661 | 0.002306 | 0.005214 | 0.011545 | 0.002722 | 0.005005 | 0.006808 | 0.033148 | 0.00878 | 0.011811 | 0.014525 | 0.022805 |
| 36.4861111 | 0.017725 | 0.089469 | 0.007896 | 0.004649 | 0.007714 | 0.006844 | 0.012663 | 0.017778 | 0.038459 | 0.014988 | 0.055398 | 0.059222 | 0.048866 | 0.008136 | 0.00742 | 0.027266 | 0.002454 | 0.018551 | 0.021721 | 0.045609 | 0.007993 | 0.015826 | 0.026626 | 0.024466 |
| 36.5 | 0.016946 | 0.092394 | 0.012765 | 0.004698 | 0.018584 | 0.005185 | 0.014915 | 0.017611 | 0.043532 | 0.021662 | 0.072257 | 0.060887 | 0.056772 | 0.01793 | 0.015536 | 0.054888 | 0.008277 | 0.040766 | 0.048109 | 0.056091 | 0.011758 | 0.036285 | 0.045361 | 0.05027 |
| 36.5138889 | 0.01614 | 0.082146 | 0.017019 | 0.002763 | 0.032287 | 0.007581 | 0.016925 | 0.025779 | 0.034895 | 0.078178 | 0.075467 | 0.033592 | 0.058216 | 0.024828 | 0.03606 | 0.088622 | 0.015198 | 0.058879 | 0.067537 | 0.055332 | 0.015974 | 0.059237 | 0.058297 | 0.069307 |
| 36.5277778 | 0.013629 | 0.064201 | 0.013825 | 0.006437 | 0.044202 | 0.015614 | 0.025303 | 0.043208 | 0.018012 | 0.194737 | 0.071172 | 0.012181 | 0.04772 | 0.028851 | 0.064794 | 0.12215 | 0.014566 | 0.061994 | 0.070446 | 0.036938 | 0.01701 | 0.072437 | 0.051578 | 0.063167 |
| 36.5416667 | 0.010417 | 0.034461 | 0.010031 | 0.02239 | 0.072991 | 0.02954 | 0.037261 | 0.059422 | 0.007674 | 0.272163 | 0.06663 | 0.014902 | 0.035903 | 0.035402 | 0.09064 | 0.150769 | 0.01143 | 0.055951 | 0.070062 | 0.022748 | 0.02239 | 0.078469 | 0.03316 | 0.040638 |
| 36.5555556 | 0.012734 | 0.010607 | 0.01168 | 0.041281 | 0.120396 | 0.040102 | 0.037888 | 0.060971 | 0.010722 | 0.220336 | 0.060111 | 0.021176 | 0.033352 | 0.041609 | 0.101374 | 0.168798 | 0.011398 | 0.049883 | 0.076303 | 0.032374 | 0.032808 | 0.079537 | 0.027081 | 0.026653 |
| 36.5694444 | 0.014668 | 0.010325 | 0.016076 | 0.04504 | 0.151314 | 0.036776 | 0.024081 | 0.051509 | 0.015326 | 0.109374 | 0.053525 | 0.015097 | 0.037708 | 0.041975 | 0.106981 | 0.170683 | 0.009546 | 0.046265 | 0.081707 | 0.035659 | 0.039146 | 0.075127 | 0.038183 | 0.07276 |
| 36.5833333 | 0.015027 | 0.016935 | 0.02079 | 0.036608 | 0.137301 | 0.026331 | 0.011485 | 0.044542 | 0.014161 | 0.043899 | 0.05097 | 0.005615 | 0.04066 | 0.038874 | 0.116403 | 0.155283 | 0.004737 | 0.048113 | 0.080534 | 0.02037 | 0.036627 | 0.067055 | 0.049477 | 0.162539 |
| 36.5972222 | 0.023178 | 0.017532 | 0.026303 | 0.029379 | 0.092517 | 0.023526 | 0.007894 | 0.042393 | 0.010096 | 0.046128 | 0.042748 | 0.004436 | 0.03992 | 0.036738 | 0.110703 | 0.129164 | 0.002319 | 0.055938 | 0.074805 | 0.014295 | 0.031964 | 0.058363 | 0.054228 | 0.199084 |
| 36.6111111 | 0.035783 | 0.027468 | 0.029669 | 0.025185 | 0.043568 | 0.027742 | 0.007188 | 0.039752 | 0.005289 | 0.072252 | 0.028518 | 0.009508 | 0.038444 | 0.034413 | 0.088812 | 0.10102 | 0.002654 | 0.062831 | 0.066804 | 0.019147 | 0.032746 | 0.050125 | 0.055228 | 0.166416 |
| 36.625 | 0.040965 | 0.050073 | 0.027314 | 0.018973 | 0.015657 | 0.031218 | 0.005612 | 0.034588 | 0.003196 | 0.07682 | 0.019471 | 0.014296 | 0.037413 | 0.029809 | 0.067506 | 0.067587 | 0.003066 | 0.061243 | 0.055909 | 0.014998 | 0.036547 | 0.040456 | 0.040857 | 0.138476 |
| 36.6388889 | 0.036263 | 0.070069 | 0.021094 | 0.00992 | 0.01661 | 0.030932 | 0.004073 | 0.028203 | 0.00541 | 0.062223 | 0.017228 | 0.014835 | 0.038459 | 0.027079 | 0.043887 | 0.032405 | 0.003617 | 0.05353 | 0.042629 | 0.011606 | 0.035703 | 0.027655 | 0.022571 | 0.134336 |
| 36.6527778 | 0.02609 | 0.077057 | 0.015375 | 0.007585 | 0.02631 | 0.026101 | 0.003674 | 0.02349 | 0.008714 | 0.053754 | 0.019942 | 0.011026 | 0.041321 | 0.03033 | 0.018974 | 0.017993 | 0.004792 | 0.047434 | 0.030925 | 0.024225 | 0.026931 | 0.013377 | 0.025418 | 0.139039 |
| 36.6666667 | 0.014768 | 0.075317 | 0.011487 | 0.018347 | 0.042998 | 0.019458 | 0.003521 | 0.020099 | 0.011089 | 0.068832 | 0.022619 | 0.008453 | 0.041666 | 0.0376 | 0.00769 | 0.032844 | 0.006697 | 0.044883 | 0.020919 | 0.044906 | 0.01403 | 0.004244 | 0.024319 | 0.150766 |
| 36.6805556 | 0.008508 | 0.079179 | 0.009117 | 0.031325 | 0.073818 | 0.014905 | 0.002713 | 0.014132 | 0.012722 | 0.10698 | 0.019504 | 0.011909 | 0.036745 | 0.043632 | 0.011097 | 0.055746 | 0.008251 | 0.045512 | 0.011124 | 0.058867 | 0.006855 | 0.004419 | 0.010047 | 0.146733 |
| 36.6944444 | 0.01072 | 0.09479 | 0.007946 | 0.032964 | 0.103055 | 0.012922 | 0.004257 | 0.007408 | 0.013363 | 0.152699 | 0.012342 | 0.020782 | 0.027357 | 0.042392 | 0.020339 | 0.0682 | 0.007934 | 0.047206 | 0.006558 | 0.06322 | 0.010478 | 0.00932 | 0.00684 | 0.124205 |
| 36.7083333 | 0.017912 | 0.115313 | 0.006836 | 0.021484 | 0.110965 | 0.01185 | 0.008992 | 0.005651 | 0.012583 | 0.18509 | 0.006644 | 0.029553 | 0.016818 | 0.033627 | 0.027926 | 0.072497 | 0.006772 | 0.046459 | 0.011127 | 0.064784 | 0.014842 | 0.014531 | 0.013491 | 0.099025 |
| 36.7222222 | 0.02363 | 0.123951 | 0.006987 | 0.007769 | 0.097833 | 0.011638 | 0.012635 | 0.006723 | 0.01077 | 0.185522 | 0.007259 | 0.030674 | 0.008872 | 0.023198 | 0.029333 | 0.076576 | 0.006405 | 0.042694 | 0.018933 | 0.064784 | 0.01256 | 0.016715 | 0.018772 | 0.069102 |
| 36.7361111 | 0.02556 | 0.111661 | 0.009916 | 0.003174 | 0.070515 | 0.014053 | 0.011588 | 0.004811 | 0.009456 | 0.160325 | 0.013695 | 0.024187 | 0.004071 | 0.014915 | 0.027166 | 0.080108 | 0.006421 | 0.036306 | 0.021099 | 0.064132 | 0.007375 | 0.011854 | 0.016322 | 0.036631 |
| 36.75 | 0.028002 | 0.087844 | 0.012172 | 0.005879 | 0.04265 | 0.018113 | 0.006807 | 0.002686 | 0.00981 | 0.128812 | 0.018169 | 0.018781 | 0.001251 | 0.00996 | 0.022968 | 0.074403 | 0.005693 | 0.030528 | 0.015766 | 0.064595 | 0.003779 | 0.004415 | 0.012602 | 0.017517 |
| 36.7638889 | 0.031001 | 0.063546 | 0.012615 | 0.009118 | 0.032061 | 0.022215 | 0.003288 | 0.005561 | 0.010725 | 0.104785 | 0.018957 | 0.016054 | 0.000518 | 0.008039 | 0.02028 | 0.057157 | 0.004246 | 0.028047 | 0.00791 | 0.06701 | 0.003004 | 0.001618 | 0.019188 | 0.021757 |
| 36.7777778 | 0.029427 | 0.041924 | 0.014297 | 0.009412 | 0.032097 | 0.023077 | 0.003501 | 0.01209 | 0.010978 | 0.107388 | 0.023198 | 0.010787 | 0.000857 | 0.006505 | 0.019136 | 0.033519 | 0.003478 | 0.027864 | 0.004271 | 0.058127 | 0.002459 | 0.003089 | 0.033725 | 0.037762 |
| 36.7916667 | 0.024511 | 0.027325 | 0.016329 | 0.00797 | 0.032386 | 0.019213 | 0.004025 | 0.01938 | 0.012201 | 0.114722 | 0.029968 | 0.0052 | 0.001095 | 0.004383 | 0.013606 | 0.012356 | 0.003818 | 0.027232 | 0.006667 | 0.039703 | 0.001667 | 0.007432 | 0.046409 | 0.044717 |
| 36.8055556 | 0.021111 | 0.021545 | 0.016105 | 0.007165 | 0.030035 | 0.015654 | 0.003154 | 0.025726 | 0.015388 | 0.088817 | 0.029741 | 0.0048 | 0.002313 | 0.005634 | 0.008279 | 0.003186 | 0.004251 | 0.022771 | 0.007709 | 0.034561 | 0.002064 | 0.012686 | 0.049048 | 0.035334 |
| 36.8194444 | 0.019283 | 0.017743 | 0.015134 | 0.006526 | 0.023365 | 0.013583 | 0.004249 | 0.02937 | 0.018237 | 0.044609 | 0.020153 | 0.008239 | 0.004965 | 0.012937 | 0.010369 | 0.004749 | 0.004296 | 0.015253 | 0.00617 | 0.02712 | 0.003675 | 0.015773 | 0.042191 | 0.026656 |
| 36.8333333 | 0.017256 | 0.013388 | 0.01586 | 0.006833 | 0.016881 | 0.010507 | 0.008793 | 0.029481 | 0.020037 | 0.014613 | 0.008851 | 0.011394 | 0.00724 | 0.023154 | 0.022335 | 0.007476 | 0.004166 | 0.008423 | 0.011433 | 0.021748 | 0.004473 | 0.015169 | 0.034778 | 0.039103 |
| 36.8472222 | 0.014173 | 0.010431 | 0.016306 | 0.008713 | 0.013079 | 0.007692 | 0.012761 | 0.026422 | 0.021852 | 0.007427 | 0.004737 | 0.013266 | 0.009515 | 0.032802 | 0.040344 | 0.00789 | 0.003923 | 0.006537 | 0.022517 | 0.074565 | 0.003567 | 0.009793 | 0.033686 | 0.072649 |
| 36.8611111 | 0.01003 | 0.007101 | 0.01289 | 0.009888 | 0.012157 | 0.007278 | 0.012118 | 0.02252 | 0.022717 | 0.012482 | 0.008402 | 0.013724 | 0.013815 | 0.040073 | 0.054563 | 0.012796 | 0.00307 | 0.009809 | 0.029871 | 0.110057 | 0.002853 | 0.004495 | 0.037373 | 0.118366 |
| 36.875 | 0.010266 | 0.00531 | 0.009448 | 0.008706 | 0.017046 | 0.008196 | 0.008412 | 0.020848 | 0.022995 | 0.012816 | 0.012374 | 0.011597 | 0.019441 | 0.042621 | 0.059213 | 0.025662 | 0.003881 | 0.013172 | 0.030098 | 0.063321 | 0.002761 | 0.004841 | 0.038254 | 0.169844 |
| 36.8888889 | 0.017327 | 0.010552 | 0.012196 | 0.005412 | 0.027322 | 0.007621 | 0.005931 | 0.021556 | 0.024402 | 0.007026 | 0.015038 | 0.007402 | 0.024296 | 0.041374 | 0.051302 | 0.042427 | 0.008204 | 0.014882 | 0.027191 | 0.032924 | 0.004567 | 0.008668 | 0.03176 | 0.221015 |
| 36.9027778 | 0.023925 | 0.020967 | 0.018139 | 0.003216 | 0.035245 | 0.004801 | 0.006502 | 0.023718 | 0.026248 | 0.005783 | 0.021508 | 0.00352 | 0.027573 | 0.040535 | 0.031903 | 0.060766 | 0.011417 | 0.014845 | 0.02697 | 0.024539 | 0.008267 | 0.011778 | 0.019295 | 0.252779 |
| 36.9166667 | 0.024829 | 0.030782 | 0.022432 | 0.003853 | 0.03699 | 0.002792 | 0.007828 | 0.026931 | 0.027864 | 0.013155 | 0.030788 | 0.001456 | 0.029652 | 0.041379 | 0.012525 | 0.077222 | 0.010665 | 0.015286 | 0.028842 | 0.01616 | 0.012452 | 0.013423 | 0.008244 | 0.248654 |
| 36.9305556 | 0.021495 | 0.035999 | 0.024792 | 0.003593 | 0.036758 | 0.003659 | 0.006741 | 0.029009 | 0.030882 | 0.024284 | 0.035727 | 0.002315 | 0.029752 | 0.042189 | 0.002969 | 0.086569 | 0.008528 | 0.020476 | 0.027282 | 0.025959 | 0.017579 | 0.01461 | 0.006038 | 0.222214 |
| 36.9444444 | 0.017609 | 0.035645 | 0.025477 | 0.003668 | 0.034825 | 0.006688 | 0.004022 | 0.027853 | 0.034525 | 0.028009 | 0.033921 | 0.004811 | 0.025487 | 0.041767 | 0.002972 | 0.084907 | 0.006862 | 0.027906 | 0.02109 | 0.034434 | 0.022264 | 0.01432 | 0.009096 | 0.194902 |
| 36.9583333 | 0.015615 | 0.028782 | 0.023404 | 0.008745 | 0.024719 | 0.013369 | 0.002901 | 0.022595 | 0.034579 | 0.02134 | 0.031376 | 0.008787 | 0.017188 | 0.041142 | 0.009157 | 0.070156 | 0.00455 | 0.032823 | 0.014936 | 0.036683 | 0.023907 | 0.01488 | 0.010171 | 0.169827 |
| 36.9722222 | 0.01669 | 0.015425 | 0.017446 | 0.015292 | 0.010597 | 0.022765 | 0.003413 | 0.015012 | 0.029813 | 0.015904 | 0.032084 | 0.020529 | 0.012492 | 0.041604 | 0.015373 | 0.046247 | 0.003349 | 0.03208 | 0.015561 | 0.032837 | 0.019589 | 0.018758 | 0.008221 | 0.136173 |
| 36.9861111 | 0.021382 | 0.006909 | 0.010717 | 0.015818 | 0.009076 | 0.023045 | 0.003658 | 0.012479 | 0.025319 | 0.015659 | 0.028446 | 0.030524 | 0.018837 | 0.045115 | 0.017697 | 0.029217 | 0.006428 | 0.028536 | 0.030104 | 0.023991 | 0.013428 | 0.017048 | 0.009892 | 0.100505 |
| 37 | 0.027877 | 0.015143 | 0.008288 | 0.009694 | 0.029174 | 0.014333 | 0.002981 | 0.018842 | 0.02684 | 0.027922 | 0.015557 | 0.022186 | 0.03287 | 0.052345 | 0.015843 | 0.037912 | 0.017211 | 0.029183 | 0.057924 | 0.01421 | 0.017426 | 0.016297 | 0.019971 | 0.083177 |
| 37.0138889 | 0.034272 | 0.029199 | 0.009469 | 0.004583 | 0.057055 | 0.013291 | 0.004696 | 0.023944 | 0.029764 | 0.058162 | 0.006561 | 0.013287 | 0.040564 | 0.0566 | 0.010164 | 0.065138 | 0.059829 | 0.027532 | 0.082073 | 0.006084 | 0.026797 | 0.023627 | 0.032005 | 0.076537 |
| 37.0277778 | 0.040121 | 0.02565 | 0.009641 | 0.008804 | 0.073001 | 0.022186 | 0.013359 | 0.017871 | 0.024035 | 0.075387 | 0.012053 | 0.029327 | 0.030955 | 0.052914 | 0.006588 | 0.087697 | 0.180051 | 0.016543 | 0.084392 | 0.003635 | 0.025234 | 0.022656 | 0.040191 | 0.056289 |
| 37.0416667 | 0.042195 | 0.013331 | 0.011559 | 0.017699 | 0.098626 | 0.031158 | 0.020731 | 0.009604 | 0.014155 | 0.071118 | 0.015898 | 0.046935 | 0.01501 | 0.045129 | 0.008209 | 0.096118 | 0.319284 | 0.008775 | 0.065993 | 0.007171 | 0.014594 | 0.01576 | 0.03996 | 0.027181 |
| 37.0555556 | 0.034065 | 0.035473 | 0.023022 | 0.019421 | 0.127875 | 0.037897 | 0.015931 | 0.021645 | 0.016171 | 0.08742 | 0.02923 | 0.05147 | 0.011136 | 0.036817 | 0.010386 | 0.087322 | 0.351678 | 0.011367 | 0.040774 | 0.008773 | 0.007813 | 0.012531 | 0.028642 | 0.016877 |
| 37.0694444 | 0.019265 | 0.077786 | 0.035826 | 0.015619 | 0.113447 | 0.051052 | 0.00663 | 0.044447 | 0.025347 | 0.118045 | 0.065504 | 0.059962 | 0.017293 | 0.028361 | 0.009756 | 0.064207 | 0.291894 | 0.014924 | 0.021556 | 0.005939 | 0.009204 | 0.011301 | 0.014405 | 0.037143 |
| 37.0833333 | 0.011874 | 0.07101 | 0.034664 | 0.017749 | 0.069357 | 0.049658 | 0.005808 | 0.044088 | 0.024047 | 0.095857 | 0.064341 | 0.060278 | 0.022108 | 0.0206 | 0.008212 | 0.040996 | 0.210749 | 0.015435 | 0.013251 | 0.005964 | 0.013176 | 0.013672 | 0.005301 | 0.070516 |
| 37.0972222 | 0.013214 | 0.031108 | 0.024326 | 0.027683 | 0.028507 | 0.02223 | 0.013642 | 0.032797 | 0.016343 | 0.043891 | 0.022588 | 0.055603 | 0.022578 | 0.014527 | 0.007401 | 0.024448 | 0.140098 | 0.015095 | 0.01114 | 0.011657 | 0.014385 | 0.021393 | 0.002146 | 0.088441 |
| 37.1111111 | 0.012584 | 0.018591 | 0.017746 | 0.029447 | 0.009264 | 0.01013 | 0.020805 | 0.026504 | 0.010554 | 0.023556 | 0.008074 | 0.04676 | 0.021446 | 0.010387 | 0.006351 | 0.012526 | 0.097501 | 0.01677 | 0.010335 | 0.017862 | 0.013969 | 0.029852 | 0.004173 | 0.084378 |
| 37.125 | 0.00995 | 0.023922 | 0.014362 | 0.020167 | 0.009949 | 0.016194 | 0.022459 | 0.015416 | 0.007293 | 0.014011 | 0.013482 | 0.026413 | 0.020566 | 0.008631 | 0.005844 | 0.005704 | 0.075781 | 0.021092 | 0.01101 | 0.018304 | 0.013855 | 0.032682 | 0.010439 | 0.07188 |
| 37.1388889 | 0.010746 | 0.028143 | 0.009834 | 0.013568 | 0.009702 | 0.020071 | 0.020127 | 0.007559 | 0.006671 | 0.005161 | 0.007384 | 0.014341 | 0.018118 | 0.009246 | 0.006575 | 0.004991 | 0.067328 | 0.024292 | 0.013276 | 0.013472 | 0.01313 | 0.030421 | 0.018192 | 0.059061 |
| 37.1527778 | 0.01003 | 0.046321 | 0.011483 | 0.011215 | 0.013228 | 0.027531 | 0.01377 | 0.01437 | 0.009543 | 0.019933 | 0.014193 | 0.020784 | 0.012413 | 0.010528 | 0.00653 | 0.009625 | 0.069633 | 0.024206 | 0.014611 | 0.012933 | 0.010745 | 0.025196 | 0.023364 | 0.047266 |
| 37.1666667 | 0.005632 | 0.074652 | 0.021648 | 0.011687 | 0.023228 | 0.032067 | 0.008985 | 0.031728 | 0.019665 | 0.051368 | 0.045397 | 0.044755 | 0.006235 | 0.01111 | 0.004739 | 0.016093 | 0.078672 | 0.023177 | 0.013027 | 0.015821 | 0.007647 | 0.017868 | 0.023558 | 0.037656 |
| 37.1805556 | 0.00265 | 0.086831 | 0.028964 | 0.013218 | 0.025683 | 0.026639 | 0.011817 | 0.04357 | 0.030222 | 0.064892 | 0.069016 | 0.062311 | 0.00516 | 0.011527 | 0.002968 | 0.019751 | 0.087061 | 0.024123 | 0.00976 | 0.012134 | 0.008245 | 0.009187 | 0.020488 | 0.029691 |
| 37.1944444 | 0.002882 | 0.076567 | 0.027677 | 0.009911 | 0.018149 | 0.016328 | 0.017052 | 0.041863 | 0.031448 | 0.049718 | 0.068715 | 0.052957 | 0.01023 | 0.012209 | 0.004665 | 0.019218 | 0.087047 | 0.025836 | 0.007163 | 0.005652 | 0.016379 | 0.003266 | 0.017639 | 0.022446 |
| 37.2083333 | 0.008688 | 0.050222 | 0.019426 | 0.009182 | 0.021209 | 0.006074 | 0.016493 | 0.029009 | 0.023599 | 0.03339 | 0.045091 | 0.035923 | 0.016399 | 0.012407 | 0.013412 | 0.016495 | 0.076774 | 0.026451 | 0.005233 | 0.004758 | 0.029488 | 0.003603 | 0.016566 | 0.016112 |
| 37.2222222 | 0.015096 | 0.024734 | 0.011994 | 0.015961 | 0.039319 | 0.001985 | 0.011454 | 0.015716 | 0.014171 | 0.030369 | 0.017449 | 0.028215 | 0.01916 | 0.011231 | 0.027425 | 0.01254 | 0.060552 | 0.026289 | 0.003942 | 0.009471 | 0.03788 | 0.006836 | 0.016157 | 0.009683 |
| 37.2361111 | 0.013181 | 0.026544 | 0.016871 | 0.015785 | 0.036215 | 0.004411 | 0.011595 | 0.018428 | 0.01583 | 0.031818 | 0.025441 | 0.029361 | 0.017436 | 0.008557 | 0.037051 | 0.00756 | 0.043361 | 0.025901 | 0.004529 | 0.015537 | 0.033982 | 0.010875 | 0.014593 | 0.005382 |
| 37.25 | 0.007577 | 0.041427 | 0.027445 | 0.008209 | 0.014896 | 0.00818 | 0.015353 | 0.029508 | 0.024818 | 0.032913 | 0.054445 | 0.034657 | 0.013414 | 0.005324 | 0.033329 | 0.004018 | 0.02462 | 0.02548 | 0.006814 | 0.019227 | 0.022975 | 0.016337 | 0.011824 | 0.007826 |
| 37.2638889 | 0.004193 | 0.032712 | 0.027491 | 0.005585 | 0.017555 | 0.007943 | 0.01179 | 0.029752 | 0.021723 | 0.026614 | 0.055656 | 0.028888 | 0.00966 | 0.002732 | 0.018446 | 0.00533 | 0.013899 | 0.02602 | 0.009087 | 0.019804 | 0.014222 | 0.022475 | 0.008763 | 0.016302 |
| 37.2777778 | 0.003675 | 0.017324 | 0.023333 | 0.005637 | 0.034169 | 0.011994 | 0.005291 | 0.024508 | 0.013395 | 0.020685 | 0.047291 | 0.019266 | 0.005725 | 0.002362 | 0.00656 | 0.01162 | 0.025519 | 0.027813 | 0.010744 | 0.018665 | 0.009461 | 0.028767 | 0.005291 | 0.028705 |
| 37.2916667 | 0.005646 | 0.023303 | 0.025237 | 0.003218 | 0.033468 | 0.025451 | 0.003644 | 0.023876 | 0.015028 | 0.025659 | 0.059674 | 0.02154 | 0.004706 | 0.00434 | 0.005784 | 0.018927 | 0.047901 | 0.029813 | 0.012703 | 0.018202 | 0.005839 | 0.034335 | 0.00239 | 0.04248 |
| 37.3055556 | 0.007704 | 0.02501 | 0.0222 | 0.004923 | 0.030507 | 0.035437 | 0.006062 | 0.018937 | 0.015355 | 0.028445 | 0.060472 | 0.026177 | 0.011741 | 0.005955 | 0.008299 | 0.023756 | 0.062491 | 0.031663 | 0.013725 | 0.01881 | 0.002685 | 0.035734 | 0.001882 | 0.050557 |
| 37.3194444 | 0.010106 | 0.011249 | 0.011938 | 0.011259 | 0.033607 | 0.042726 | 0.014008 | 0.008079 | 0.010817 | 0.021928 | 0.041926 | 0.026079 | 0.022024 | 0.005673 | 0.00862 | 0.025995 | 0.06786 | 0.032233 | 0.012786 | 0.017583 | 0.001932 | 0.030481 | 0.002949 | 0.049981 |
| 37.3333333 | 0.010545 | 0.010823 | 0.008402 | 0.01184 | 0.025985 | 0.048578 | 0.032153 | 0.004258 | 0.007586 | 0.016895 | 0.040819 | 0.020079 | 0.025653 | 0.004582 | 0.008811 | 0.027504 | 0.072401 | 0.029831 | 0.011743 | 0.014084 | 0.004267 | 0.023337 | 0.002936 | 0.047904 |
| 37.3472222 | 0.008344 | 0.024679 | 0.011223 | 0.007088 | 0.018538 | 0.050163 | 0.063288 | 0.008703 | 0.003888 | 0.017267 | 0.052638 | 0.01025 | 0.020657 | 0.003879 | 0.010145 | 0.028781 | 0.07906 | 0.025242 | 0.011676 | 0.011735 | 0.007733 | 0.019955 | 0.003984 | 0.052416 |
| 37.3611111 | 0.008603 | 0.031262 | 0.01233 | 0.005217 | 0.019893 | 0.049566 | 0.081508 | 0.015149 | 0.002169 | 0.014214 | 0.054579 | 0.003656 | 0.014517 | 0.00415 | 0.012691 | 0.028923 | 0.082002 | 0.020639 | 0.01215 | 0.012448 | 0.00869 | 0.019171 | 0.012806 | 0.06867 |
| 37.375 | 0.008537 | 0.03431 | 0.01512 | 0.005414 | 0.027531 | 0.039067 | 0.061397 | 0.023764 | 0.005719 | 0.016233 | 0.059866 | 0.010211 | 0.014749 | 0.005476 | 0.016155 | 0.028163 | 0.081565 | 0.017029 | 0.013518 | 0.015198 | 0.006355 | 0.018126 | 0.030485 | 0.091875 |
| 37.3888889 | 0.006733 | 0.045088 | 0.024119 | 0.006449 | 0.046236 | 0.020673 | 0.029644 | 0.031803 | 0.015526 | 0.03439 | 0.080414 | 0.0351 | 0.020294 | 0.006947 | 0.020152 | 0.027874 | 0.08101 | 0.014501 | 0.015163 | 0.017519 | 0.005967 | 0.015964 | 0.046685 | 0.100393 |
| 37.4027778 | 0.00636 | 0.049034 | 0.030188 | 0.00846 | 0.059837 | 0.009603 | 0.012389 | 0.033032 | 0.022348 | 0.044744 | 0.091564 | 0.050973 | 0.024259 | 0.007287 | 0.024238 | 0.02755 | 0.078605 | 0.012985 | 0.014052 | 0.017587 | 0.008318 | 0.013196 | 0.046289 | 0.08686 |
| 37.4166667 | 0.0065 | 0.031964 | 0.021719 | 0.009638 | 0.043629 | 0.005673 | 0.005215 | 0.025519 | 0.015753 | 0.033176 | 0.067879 | 0.04171 | 0.023517 | 0.006478 | 0.027731 | 0.027524 | 0.072731 | 0.01229 | 0.009303 | 0.01574 | 0.006777 | 0.010736 | 0.03123 | 0.069716 |
| 37.4305556 | 0.010601 | 0.018482 | 0.010362 | 0.015794 | 0.024209 | 0.013349 | 0.006238 | 0.013151 | 0.011742 | 0.030088 | 0.034422 | 0.045658 | 0.020422 | 0.006237 | 0.030849 | 0.030758 | 0.065788 | 0.012126 | 0.004915 | 0.014669 | 0.003686 | 0.009264 | 0.017464 | 0.062051 |
| 37.4444444 | 0.014663 | 0.023728 | 0.008829 | 0.024322 | 0.032078 | 0.034669 | 0.009386 | 0.004636 | 0.020008 | 0.040373 | 0.02113 | 0.067877 | 0.017927 | 0.007711 | 0.035273 | 0.036237 | 0.06271 | 0.012402 | 0.004814 | 0.018046 | 0.004821 | 0.010598 | 0.009239 | 0.063194 |
| 37.4583333 | 0.013537 | 0.026373 | 0.012374 | 0.021148 | 0.034993 | 0.044559 | 0.006451 | 0.008749 | 0.021399 | 0.036246 | 0.029522 | 0.059244 | 0.017452 | 0.009469 | 0.040454 | 0.040234 | 0.061716 | 0.013008 | 0.00704 | 0.025025 | 0.004866 | 0.013793 | 0.004719 | 0.065456 |
| 37.4722222 | 0.021302 | 0.032356 | 0.023874 | 0.015046 | 0.029745 | 0.062614 | 0.00402 | 0.027454 | 0.021307 | 0.033452 | 0.069704 | 0.042696 | 0.020447 | 0.011701 | 0.044266 | 0.042783 | 0.058397 | 0.014756 | 0.009271 | 0.031379 | 0.003633 | 0.017101 | 0.008292 | 0.064181 |
| 37.4861111 | 0.042396 | 0.04451 | 0.036798 | 0.018304 | 0.027898 | 0.099497 | 0.004971 | 0.048855 | 0.036109 | 0.045789 | 0.112461 | 0.059124 | 0.030359 | 0.019053 | 0.049494 | 0.048738 | 0.061897 | 0.020398 | 0.018545 | 0.038389 | 0.01133 | 0.022977 | 0.022141 | 0.069485 |
| 37.5 | 0.059415 | 0.038232 | 0.035 | 0.017769 | 0.033626 | 0.089527 | 0.00456 | 0.058136 | 0.043799 | 0.041193 | 0.104103 | 0.060144 | 0.044441 | 0.030691 | 0.057811 | 0.059655 | 0.081909 | 0.030627 | 0.040237 | 0.04901 | 0.032457 | 0.031187 | 0.038846 | 0.096397 |
| 37.5138889 | 0.063318 | 0.022766 | 0.024867 | 0.01689 | 0.068735 | 0.039134 | 0.005556 | 0.053591 | 0.03221 | 0.018844 | 0.058008 | 0.030462 | 0.054068 | 0.036038 | 0.060094 | 0.069274 | 0.108794 | 0.038298 | 0.062174 | 0.051159 | 0.063234 | 0.038465 | 0.046775 | 0.120846 |
| 37.5277778 | 0.056959 | 0.019043 | 0.019137 | 0.025571 | 0.100885 | 0.013564 | 0.01135 | 0.04483 | 0.022665 | 0.005919 | 0.022503 | 0.008756 | 0.060363 | 0.032334 | 0.051787 | 0.078849 | 0.131823 | 0.035967 | 0.071277 | 0.045933 | 0.092257 | 0.044615 | 0.04275 | 0.110431 |
| 37.5416667 | 0.043477 | 0.022748 | 0.018521 | 0.032284 | 0.109832 | 0.018155 | 0.021645 | 0.04269 | 0.02564 | 0.004155 | 0.010677 | 0.002578 | 0.064779 | 0.029451 | 0.043518 | 0.101663 | 0.146244 | 0.029127 | 0.070108 | 0.063392 | 0.100342 | 0.050267 | 0.030902 | 0.084473 |
| 37.5555556 | 0.029537 | 0.021979 | 0.018935 | 0.030931 | 0.108326 | 0.028325 | 0.034526 | 0.04895 | 0.034386 | 0.004753 | 0.011904 | 0.001621 | 0.056866 | 0.031052 | 0.041509 | 0.131023 | 0.146305 | 0.026591 | 0.063734 | 0.068333 | 0.087713 | 0.054691 | 0.021154 | 0.061927 |
| 37.5694444 | 0.020822 | 0.015239 | 0.017103 | 0.026721 | 0.096525 | 0.028614 | 0.044109 | 0.056298 | 0.038863 | 0.004466 | 0.018299 | 0.002872 | 0.038181 | 0.032347 | 0.040092 | 0.148319 | 0.127943 | 0.027865 | 0.053166 | 0.030371 | 0.069436 | 0.05401 | 0.018081 | 0.051353 |

| | | | | | | | | | | | | | | | | | | | | | | | | |
|---|---|---|---|---|---|---|---|---|---|---|---|---|---|---|---|---|---|---|---|---|---|---|---|---|
| 37.5833333 | 0.015707 | 0.013981 | 0.013692 | 0.025957 | 0.084132 | 0.023023 | 0.045642 | 0.057566 | 0.033608 | 0.004421 | 0.024966 | 0.007197 | 0.0264 | 0.031063 | 0.036223 | 0.153945 | 0.092104 | 0.030465 | 0.041755 | 0.009236 | 0.043682 | 0.046896 | 0.014216 | 0.066451 |
| 37.5972222 | 0.012753 | 0.021134 | 0.013295 | 0.025621 | 0.068676 | 0.021045 | 0.042663 | 0.053498 | 0.025469 | 0.005081 | 0.033153 | 0.009039 | 0.026406 | 0.028164 | 0.03454 | 0.151704 | 0.056277 | 0.033736 | 0.034111 | 0.014902 | 0.018028 | 0.040201 | 0.008708 | 0.095114 |
| 37.6111111 | 0.011903 | 0.022469 | 0.016901 | 0.020757 | 0.045911 | 0.024923 | 0.040066 | 0.049486 | 0.023055 | 0.003463 | 0.041698 | 0.005378 | 0.028402 | 0.024964 | 0.039772 | 0.146698 | 0.038079 | 0.035077 | 0.030577 | 0.020679 | 0.011704 | 0.036496 | 0.004845 | 0.110824 |
| 37.625 | 0.00877 | 0.016335 | 0.019374 | 0.018143 | 0.030816 | 0.028222 | 0.036272 | 0.04606 | 0.024084 | 0.001995 | 0.043463 | 0.002168 | 0.024971 | 0.020298 | 0.04787 | 0.143128 | 0.034365 | 0.032836 | 0.026504 | 0.032956 | 0.024684 | 0.031667 | 0.00738 | 0.10769 |
| 37.6388889 | 0.00443 | 0.010937 | 0.019 | 0.020528 | 0.027096 | 0.026327 | 0.03064 | 0.041718 | 0.023819 | 0.00284 | 0.040662 | 0.001821 | 0.01668 | 0.013189 | 0.05059 | 0.128009 | 0.035179 | 0.02919 | 0.019081 | 0.041702 | 0.04649 | 0.025801 | 0.016485 | 0.098946 |
| 37.6527778 | 0.003928 | 0.006501 | 0.020877 | 0.018102 | 0.022049 | 0.029612 | 0.026698 | 0.041224 | 0.025955 | 0.00426 | 0.044239 | 0.003277 | 0.010517 | 0.007217 | 0.047428 | 0.09246 | 0.036932 | 0.026615 | 0.011448 | 0.032448 | 0.060269 | 0.025146 | 0.023555 | 0.095714 |
| 37.6666667 | 0.005559 | 0.003597 | 0.025152 | 0.009609 | 0.02137 | 0.038007 | 0.026353 | 0.045541 | 0.030953 | 0.005554 | 0.0549 | 0.006176 | 0.010939 | 0.004856 | 0.038553 | 0.049174 | 0.038345 | 0.024814 | 0.005681 | 0.033077 | 0.052702 | 0.032002 | 0.025561 | 0.09937 |
| 37.6805556 | 0.006127 | 0.005294 | 0.025243 | 0.003825 | 0.031801 | 0.035139 | 0.026537 | 0.04548 | 0.0299 | 0.005474 | 0.061918 | 0.006161 | 0.014097 | 0.004146 | 0.022404 | 0.024786 | 0.036792 | 0.021683 | 0.002162 | 0.067967 | 0.036694 | 0.040875 | 0.024396 | 0.093983 |
| 37.6944444 | 0.004509 | 0.012109 | 0.021375 | 0.002344 | 0.042565 | 0.024424 | 0.024676 | 0.039278 | 0.022801 | 0.004318 | 0.061421 | 0.00356 | 0.015232 | 0.003236 | 0.012056 | 0.035311 | 0.032636 | 0.016076 | 0.002676 | 0.078487 | 0.027846 | 0.046264 | 0.022282 | 0.066711 |
| 37.7083333 | 0.002197 | 0.018419 | 0.019377 | 0.002192 | 0.052412 | 0.019804 | 0.022828 | 0.034696 | 0.018391 | 0.002592 | 0.059203 | 0.003359 | 0.015107 | 0.002277 | 0.016572 | 0.057578 | 0.027118 | 0.010101 | 0.006397 | 0.038622 | 0.025831 | 0.047291 | 0.020224 | 0.031205 |
| 37.7222222 | 0.001753 | 0.022157 | 0.01892 | 0.00321 | 0.061305 | 0.017043 | 0.022363 | 0.032284 | 0.016373 | 0.001185 | 0.056007 | 0.005894 | 0.017005 | 0.001653 | 0.025359 | 0.067681 | 0.017733 | 0.006348 | 0.013434 | 0.021252 | 0.028494 | 0.045467 | 0.017716 | 0.014586 |
| 37.7361111 | 0.003395 | 0.021659 | 0.018713 | 0.006378 | 0.066621 | 0.013751 | 0.021648 | 0.030917 | 0.015473 | 0.002513 | 0.051528 | 0.008073 | 0.019894 | 0.001489 | 0.030969 | 0.069014 | 0.009214 | 0.00404 | 0.023256 | 0.036632 | 0.034778 | 0.039461 | 0.016441 | 0.023694 |
| 37.75 | 0.004312 | 0.0137 | 0.020788 | 0.009138 | 0.071437 | 0.016845 | 0.019966 | 0.034935 | 0.019345 | 0.007169 | 0.049853 | 0.00704 | 0.019823 | 0.002758 | 0.031516 | 0.068369 | 0.011646 | 0.002072 | 0.029558 | 0.050948 | 0.04119 | 0.028764 | 0.016598 | 0.050551 |
| 37.7638889 | 0.003085 | 0.006581 | 0.023779 | 0.009797 | 0.069505 | 0.021513 | 0.018313 | 0.041777 | 0.023773 | 0.010871 | 0.048031 | 0.004738 | 0.016359 | 0.005393 | 0.031527 | 0.070313 | 0.023171 | 0.001477 | 0.030499 | 0.062536 | 0.044865 | 0.01564 | 0.013874 | 0.098333 |
| 37.7777778 | 0.00189 | 0.006735 | 0.025366 | 0.009903 | 0.052612 | 0.019425 | 0.016916 | 0.044558 | 0.022421 | 0.009086 | 0.041365 | 0.004651 | 0.010474 | 0.007151 | 0.034145 | 0.075878 | 0.035748 | 0.003907 | 0.028996 | 0.072071 | 0.045918 | 0.007148 | 0.008623 | 0.139366 |
| 37.7916667 | 0.002355 | 0.009197 | 0.025611 | 0.010174 | 0.027913 | 0.015345 | 0.016254 | 0.043858 | 0.018142 | 0.00421 | 0.033217 | 0.005931 | 0.004961 | 0.007336 | 0.034353 | 0.082194 | 0.044419 | 0.010252 | 0.024133 | 0.085295 | 0.045941 | 0.008428 | 0.00619 | 0.136621 |
| 37.8055556 | 0.003722 | 0.010024 | 0.024796 | 0.010927 | 0.010769 | 0.014989 | 0.016401 | 0.043111 | 0.015326 | 0.001658 | 0.027671 | 0.006633 | 0.004318 | 0.007525 | 0.030657 | 0.088237 | 0.046281 | 0.019237 | 0.018247 | 0.092911 | 0.046772 | 0.011192 | 0.00762 | 0.115792 |
| 37.8194444 | 0.004517 | 0.010959 | 0.023564 | 0.012083 | 0.006663 | 0.017105 | 0.016753 | 0.042741 | 0.014115 | 0.002249 | 0.023296 | 0.005899 | 0.007714 | 0.008626 | 0.027934 | 0.096263 | 0.043299 | 0.027661 | 0.01638 | 0.078506 | 0.048678 | 0.0136 | 0.012474 | 0.10644 |
| 37.8333333 | 0.004133 | 0.014751 | 0.022941 | 0.013391 | 0.005983 | 0.019437 | 0.018089 | 0.043265 | 0.014944 | 0.002945 | 0.018408 | 0.003653 | 0.012174 | 0.010468 | 0.025946 | 0.103999 | 0.040186 | 0.0327 | 0.017013 | 0.054081 | 0.049874 | 0.020677 | 0.021176 | 0.107847 |
| 37.8472222 | 0.002677 | 0.019835 | 0.023313 | 0.014568 | 0.005079 | 0.020477 | 0.02031 | 0.044607 | 0.017821 | 0.003127 | 0.012461 | 0.001424 | 0.017313 | 0.013472 | 0.018086 | 0.107461 | 0.039152 | 0.033255 | 0.016462 | 0.03546 | 0.046351 | 0.026768 | 0.02904 | 0.110363 |
| 37.8611111 | 0.001826 | 0.022508 | 0.023918 | 0.014684 | 0.009506 | 0.018791 | 0.020993 | 0.044332 | 0.020283 | 0.003647 | 0.006485 | 0.000498 | 0.022326 | 0.01747 | 0.012553 | 0.109931 | 0.041924 | 0.02994 | 0.015053 | 0.022486 | 0.038896 | 0.025969 | 0.029587 | 0.091835 |
| 37.875 | 0.002634 | 0.022309 | 0.023589 | 0.013642 | 0.016002 | 0.014304 | 0.018356 | 0.041354 | 0.019701 | 0.003872 | 0.00514 | 0.000383 | 0.025659 | 0.01965 | 0.016778 | 0.111105 | 0.05151 | 0.024724 | 0.013717 | 0.012603 | 0.034631 | 0.019035 | 0.023553 | 0.061414 |
| 37.8888889 | 0.004469 | 0.021079 | 0.02254 | 0.012662 | 0.019757 | 0.010823 | 0.013533 | 0.038322 | 0.01654 | 0.003395 | 0.007151 | 0.000623 | 0.02589 | 0.018886 | 0.024214 | 0.109429 | 0.069074 | 0.01929 | 0.010407 | 0.005857 | 0.034282 | 0.010041 | 0.014869 | 0.046634 |
| 37.9027778 | 0.006996 | 0.02005 | 0.021883 | 0.012187 | 0.022179 | 0.009372 | 0.008516 | 0.036377 | 0.014196 | 0.002601 | 0.008334 | 0.001446 | 0.024202 | 0.018075 | 0.029321 | 0.106482 | 0.084858 | 0.013041 | 0.006015 | 0.003376 | 0.031967 | 0.004077 | 0.006346 | 0.044682 |
| 37.9166667 | 0.009225 | 0.019242 | 0.021656 | 0.012156 | 0.026004 | 0.006627 | 0.004686 | 0.03322 | 0.013407 | 0.001826 | 0.010058 | 0.002428 | 0.024006 | 0.020147 | 0.03045 | 0.095595 | 0.082779 | 0.00666 | 0.005547 | 0.003252 | 0.025938 | 0.004889 | 0.003754 | 0.039616 |
| 37.9305556 | 0.010343 | 0.017339 | 0.021532 | 0.012866 | 0.030508 | 0.003048 | 0.003779 | 0.029194 | 0.013356 | 0.001456 | 0.012575 | 0.002403 | 0.023381 | 0.023949 | 0.027712 | 0.078114 | 0.064248 | 0.002623 | 0.008686 | 0.003059 | 0.017914 | 0.01041 | 0.007049 | 0.027496 |
| 37.9444444 | 0.01063 | 0.013766 | 0.021766 | 0.015131 | 0.028928 | 0.001732 | 0.006689 | 0.025625 | 0.016454 | 0.002035 | 0.014317 | 0.001346 | 0.019399 | 0.025397 | 0.022395 | 0.064119 | 0.045295 | 0.00114 | 0.009672 | 0.003199 | 0.009703 | 0.015421 | 0.010691 | 0.013649 |
| 37.9583333 | 0.011135 | 0.008919 | 0.022249 | 0.019223 | 0.017831 | 0.004798 | 0.012114 | 0.022317 | 0.022623 | 0.003288 | 0.015489 | 0.000489 | 0.015912 | 0.025571 | 0.018118 | 0.049848 | 0.03257 | 0.000951 | 0.009752 | 0.003981 | 0.008673 | 0.022384 | 0.017707 | 0.008489 |
| 37.9722222 | 0.012382 | 0.004823 | 0.022393 | 0.022898 | 0.008087 | 0.01152 | 0.017209 | 0.019057 | 0.026857 | 0.004291 | 0.017196 | 0.000305 | 0.017584 | 0.029968 | 0.018356 | 0.02962 | 0.023708 | 0.000962 | 0.018333 | 0.005783 | 0.017666 | 0.035313 | 0.028072 | 0.015037 |
| 37.9861111 | 0.012758 | 0.003692 | 0.021129 | 0.022248 | 0.00599 | 0.016083 | 0.018478 | 0.016946 | 0.025427 | 0.005238 | 0.020802 | 0.001512 | 0.024722 | 0.04049 | 0.01956 | 0.013072 | 0.022696 | 0.00178 | 0.035993 | 0.01046 | 0.021134 | 0.040873 | 0.030684 | 0.028039 |
| 38 | 0.0115 | 0.003952 | 0.017294 | 0.016364 | 0.006999 | 0.013535 | 0.014147 | 0.016293 | 0.016511 | 0.009317 | 0.028864 | 0.008262 | 0.03204 | 0.051327 | 0.01565 | 0.012483 | 0.033132 | 0.004737 | 0.051895 | 0.012199 | 0.025779 | 0.027112 | 0.019944 | 0.064292 |
| 38.0138889 | 0.016416 | 0.005194 | 0.012581 | 0.009956 | 0.010774 | 0.014391 | 0.007278 | 0.015738 | 0.009842 | 0.018391 | 0.047756 | 0.023408 | 0.028833 | 0.052468 | 0.008548 | 0.018662 | 0.043622 | 0.009361 | 0.058462 | 0.007135 | 0.051965 | 0.013298 | 0.007285 | 0.108368 |
| 38.0277778 | 0.030758 | 0.007743 | 0.012838 | 0.00646 | 0.018097 | 0.022535 | 0.004188 | 0.014343 | 0.013815 | 0.022526 | 0.072951 | 0.034525 | 0.017505 | 0.042413 | 0.00517 | 0.015351 | 0.042014 | 0.012592 | 0.055196 | 0.002176 | 0.074453 | 0.013942 | 0.004973 | 0.118565 |
| 38.0416667 | 0.039629 | 0.007056 | 0.020006 | 0.007003 | 0.034487 | 0.022663 | 0.007686 | 0.012399 | 0.015445 | 0.017492 | 0.081017 | 0.02566 | 0.013866 | 0.029383 | 0.006985 | 0.008787 | 0.034393 | 0.012888 | 0.044182 | 0.001753 | 0.069717 | 0.018643 | 0.009924 | 0.104659 |
| 38.0555556 | 0.033131 | 0.008182 | 0.028221 | 0.00901 | 0.058797 | 0.021217 | 0.015895 | 0.012738 | 0.014107 | 0.016466 | 0.062258 | 0.009971 | 0.018029 | 0.019551 | 0.009264 | 0.008697 | 0.031719 | 0.011095 | 0.029394 | 0.004749 | 0.048739 | 0.022522 | 0.015444 | 0.081488 |
| 38.0694444 | 0.017676 | 0.01254 | 0.032836 | 0.007151 | 0.075797 | 0.028391 | 0.023386 | 0.014107 | 0.0203 | 0.020151 | 0.032187 | 0.006979 | 0.020496 | 0.013181 | 0.009988 | 0.009438 | 0.031758 | 0.007092 | 0.016903 | 0.008916 | 0.024823 | 0.024307 | 0.017571 | 0.060798 |
| 38.0833333 | 0.007332 | 0.012231 | 0.032697 | 0.004007 | 0.077698 | 0.027713 | 0.022894 | 0.011875 | 0.027838 | 0.015845 | 0.010133 | 0.00836 | 0.019999 | 0.008989 | 0.010209 | 0.007419 | 0.027944 | 0.002986 | 0.01036 | 0.012299 | 0.013213 | 0.022265 | 0.015811 | 0.057874 |
| 38.0972222 | 0.007235 | 0.011214 | 0.027362 | 0.004596 | 0.069828 | 0.018001 | 0.015661 | 0.006666 | 0.0326 | 0.011163 | 0.003597 | 0.011123 | 0.019272 | 0.006234 | 0.010239 | 0.005578 | 0.020708 | 0.001351 | 0.007746 | 0.014057 | 0.017782 | 0.018249 | 0.013155 | 0.061284 |
| 38.1111111 | 0.008516 | 0.010248 | 0.019868 | 0.008048 | 0.060426 | 0.014141 | 0.009067 | 0.002505 | 0.033135 | 0.01629 | 0.003498 | 0.018604 | 0.019715 | 0.004529 | 0.010263 | 0.00513 | 0.014422 | 0.001206 | 0.005437 | 0.014987 | 0.02406 | 0.013387 | 0.011009 | 0.0602 |
| 38.125 | 0.006764 | 0.012185 | 0.014465 | 0.00978 | 0.053431 | 0.011348 | 0.006098 | 0.003117 | 0.024513 | 0.01591 | 0.004745 | 0.017677 | 0.020025 | 0.004075 | 0.010701 | 0.005927 | 0.011386 | 0.001836 | 0.003259 | 0.015181 | 0.025546 | 0.009656 | 0.009336 | 0.055417 |

Supplementary Table S6. 1 hour integrated motion data in (a) lettuce subjected to Water withdrawal

| IntervalMid | Control1 | Control2 | Control3 | Control4 | Control5 | Control6 | Control7 | Control8 | Control9 | Control10 | Control11 | Control12 | Stress1 | Stress2 | Stress3 | Stress4 | Stress5 | Stress6 | Stress7 | Stress8 | Stress9 | Stress10 | Stress11 | Stress12 |
|---|---|---|---|---|---|---|---|---|---|---|---|---|---|---|---|---|---|---|---|---|---|---|---|---|
| 27.1111111 | 0.192183 | 0.057376 | 0.088221 | 0.079221 | 0.068732 | 0.100092 | 0.022325 | 0.093917 | 0.021177 | 0.002766 | 0.024844 | 0.043028 | 0.019381 | 0.009932 | 0.025022 | 0.022977 | 0.039936 | 0.089959 | 0.005127 | 0.050743 | 0.060342 | 0.046298 | 0.065619 | 0.022724 |
| 27.125 | 0.131176 | 0.017655 | 0.056364 | 0.0602 | 0.025398 | 0.07632 | 0.008911 | 0.068549 | 0.010427 | 0.002921 | 0.013994 | 0.017085 | 0.019303 | 0.003487 | 0.018486 | 0.019576 | 0.026015 | 0.052167 | 0.002329 | 0.022773 | 0.02765 | 0.020895 | 0.044665 | 0.015634 |
| 27.1388889 | 0.076561 | 0.01517 | 0.038747 | 0.047996 | 0.011772 | 0.054372 | 0.010082 | 0.048954 | 0.014132 | 0.005183 | 0.012421 | 0.009691 | 0.013579 | 0.002593 | 0.016087 | 0.020835 | 0.014162 | 0.023857 | 0.001464 | 0.011825 | 0.012962 | 0.007558 | 0.027592 | 0.013455 |
| 27.1527778 | 0.034507 | 0.021716 | 0.027774 | 0.03592 | 0.013552 | 0.032256 | 0.012375 | 0.034404 | 0.016833 | 0.010087 | 0.015791 | 0.005934 | 0.006022 | 0.001566 | 0.010134 | 0.016476 | 0.012099 | 0.00819 | 0.002067 | 0.021213 | 0.013432 | 0.007352 | 0.01407 | 0.009157 |
| 27.1666667 | 0.017978 | 0.025166 | 0.016522 | 0.023338 | 0.016287 | 0.015134 | 0.015497 | 0.02587 | 0.017722 | 0.014196 | 0.013902 | 0.004878 | 0.004059 | 0.002243 | 0.005714 | 0.010031 | 0.01893 | 0.007439 | 0.003482 | 0.039206 | 0.032377 | 0.012972 | 0.005763 | 0.013568 |
| 27.1805556 | 0.032329 | 0.027043 | 0.007428 | 0.011552 | 0.020085 | 0.005946 | 0.020664 | 0.022413 | 0.019695 | 0.014992 | 0.013821 | 0.011668 | 0.006503 | 0.003723 | 0.004349 | 0.010984 | 0.023567 | 0.011456 | 0.00558 | 0.050032 | 0.059414 | 0.013571 | 0.002016 | 0.027864 |
| 27.1944444 | 0.053749 | 0.024887 | 0.002626 | 0.005855 | 0.023674 | 0.003099 | 0.024208 | 0.022926 | 0.019393 | 0.012362 | 0.023405 | 0.020957 | 0.011387 | 0.00339 | 0.004892 | 0.018646 | 0.02195 | 0.012468 | 0.00967 | 0.047751 | 0.075709 | 0.008063 | 0.001494 | 0.030815 |
| 27.2083333 | 0.071681 | 0.019774 | 0.001688 | 0.013155 | 0.02627 | 0.003656 | 0.022842 | 0.024693 | 0.012467 | 0.007912 | 0.028892 | 0.025056 | 0.011063 | 0.003003 | 0.010498 | 0.02153 | 0.016993 | 0.010333 | 0.012748 | 0.035473 | 0.076075 | 0.009452 | 0.004652 | 0.019351 |
| 27.2222222 | 0.096724 | 0.013179 | 0.003646 | 0.02823 | 0.029431 | 0.0052 | 0.017615 | 0.024711 | 0.003892 | 0.004405 | 0.021313 | 0.023814 | 0.011183 | 0.004701 | 0.01764 | 0.013985 | 0.01117 | 0.005805 | 0.010257 | 0.022418 | 0.069779 | 0.020229 | 0.010898 | 0.019628 |
| 27.2361111 | 0.132983 | 0.008561 | 0.009798 | 0.035053 | 0.03012 | 0.004614 | 0.011808 | 0.02145 | 0.003127 | 0.00471 | 0.008985 | 0.02126 | 0.019612 | 0.005537 | 0.017808 | 0.006805 | 0.00587 | 0.002052 | 0.00503 | 0.015759 | 0.066497 | 0.030378 | 0.019174 | 0.031686 |
| 27.25 | 0.177898 | 0.011408 | 0.022279 | 0.031146 | 0.029503 | 0.00351 | 0.010434 | 0.014154 | 0.011455 | 0.00991 | 0.007101 | 0.02348 | 0.019378 | 0.005955 | 0.013272 | 0.006631 | 0.00356 | 0.002728 | 0.001963 | 0.017662 | 0.065814 | 0.039821 | 0.032476 | 0.026979 |
| 27.2638889 | 0.217208 | 0.022717 | 0.037063 | 0.026567 | 0.033348 | 0.006177 | 0.015529 | 0.006842 | 0.021572 | 0.01668 | 0.02331 | 0.033906 | 0.012749 | 0.00704 | 0.01241 | 0.008483 | 0.005022 | 0.009124 | 0.001983 | 0.024255 | 0.06154 | 0.047674 | 0.051533 | 0.01566 |
| 27.2777778 | 0.226036 | 0.036583 | 0.048176 | 0.022818 | 0.04106 | 0.009466 | 0.021558 | 0.004636 | 0.026749 | 0.01945 | 0.047717 | 0.049319 | 0.018991 | 0.006021 | 0.015783 | 0.015946 | 0.006839 | 0.01771 | 0.00492 | 0.02575 | 0.048891 | 0.046911 | 0.066555 | 0.023502 |
| 27.2916667 | 0.193007 | 0.046074 | 0.057647 | 0.016735 | 0.044678 | 0.011165 | 0.022214 | 0.004473 | 0.026368 | 0.016471 | 0.059202 | 0.06241 | 0.026814 | 0.004222 | 0.019063 | 0.026781 | 0.006696 | 0.02312 | 0.008929 | 0.017365 | 0.027012 | 0.040387 | 0.070669 | 0.029297 |
| 27.3055556 | 0.131801 | 0.048414 | 0.069071 | 0.010366 | 0.039004 | 0.014798 | 0.016187 | 0.002805 | 0.023676 | 0.010716 | 0.04952 | 0.070197 | 0.023516 | 0.005618 | 0.021921 | 0.030526 | 0.005996 | 0.026462 | 0.011098 | 0.008427 | 0.012258 | 0.034401 | 0.069544 | 0.018107 |
| 27.3194444 | 0.064247 | 0.041281 | 0.078984 | 0.006772 | 0.028642 | 0.020782 | 0.007904 | 0.001842 | 0.020572 | 0.005399 | 0.033603 | 0.076052 | 0.014318 | 0.008741 | 0.023438 | 0.023767 | 0.004759 | 0.029949 | 0.009943 | 0.005058 | 0.01489 | 0.030036 | 0.068398 | 0.009874 |
| 27.3333333 | 0.027641 | 0.02891 | 0.080437 | 0.005641 | 0.019969 | 0.024813 | 0.003625 | 0.002285 | 0.016859 | 0.00212 | 0.026719 | 0.078409 | 0.006765 | 0.00874 | 0.019584 | 0.012699 | 0.0028 | 0.030384 | 0.006153 | 0.004714 | 0.015152 | 0.024995 | 0.065528 | 0.008546 |
| 27.3472222 | 0.044478 | 0.02465 | 0.070679 | 0.005182 | 0.016543 | 0.024529 | 0.003278 | 0.002461 | 0.013631 | 0.001663 | 0.0309 | 0.072542 | 0.006677 | 0.004954 | 0.011802 | 0.007122 | 0.002045 | 0.026616 | 0.002333 | 0.007722 | 0.011706 | 0.017917 | 0.056411 | 0.009503 |
| 27.3611111 | 0.081946 | 0.030383 | 0.055163 | 0.00515 | 0.022064 | 0.022857 | 0.005924 | 0.002276 | 0.013192 | 0.003623 | 0.039741 | 0.063997 | 0.014192 | 0.00168 | 0.00675 | 0.00567 | 0.002056 | 0.02353 | 0.00142 | 0.009091 | 0.023394 | 0.010591 | 0.041565 | 0.018856 |
| 27.375 | 0.106762 | 0.035492 | 0.040571 | 0.006039 | 0.033539 | 0.022655 | 0.012298 | 0.002289 | 0.01521 | 0.006323 | 0.041083 | 0.057342 | 0.022131 | 0.000911 | 0.007604 | 0.006485 | 0.00307 | 0.024292 | 0.004296 | 0.005518 | 0.040624 | 0.004829 | 0.024759 | 0.028614 |
| 27.3888889 | 0.108117 | 0.033159 | 0.028741 | 0.007204 | 0.039615 | 0.024473 | 0.016193 | 0.002212 | 0.016627 | 0.008704 | 0.028385 | 0.048971 | 0.021443 | 0.00096 | 0.012617 | 0.01186 | 0.006032 | 0.02684 | 0.008373 | 0.003246 | 0.043285 | 0.001928 | 0.010581 | 0.027565 |
| 27.4027778 | 0.089988 | 0.025248 | 0.018605 | 0.007248 | 0.033959 | 0.02749 | 0.013029 | 0.002154 | 0.01514 | 0.010872 | 0.012747 | 0.038754 | 0.012464 | 0.00156 | 0.017841 | 0.015744 | 0.00852 | 0.028025 | 0.010325 | 0.005049 | 0.030218 | 0.003226 | 0.005258 | 0.020201 |
| 27.4166667 | 0.066811 | 0.017587 | 0.009702 | 0.005418 | 0.021415 | 0.029396 | 0.006854 | 0.002874 | 0.010398 | 0.012604 | 0.006982 | 0.028192 | 0.005077 | 0.002652 | 0.019951 | 0.013477 | 0.008637 | 0.026065 | 0.008958 | 0.006253 | 0.015456 | 0.009659 | 0.006283 | 0.012946 |
| 27.4305556 | 0.03872 | 0.014684 | 0.005976 | 0.002909 | 0.012175 | 0.029591 | 0.002733 | 0.00406 | 0.00465 | 0.013544 | 0.006219 | 0.017712 | 0.005129 | 0.003518 | 0.019536 | 0.008264 | 0.006342 | 0.023274 | 0.005191 | 0.004358 | 0.013034 | 0.020333 | 0.006474 | 0.00793 |
| 27.4444444 | 0.021207 | 0.01823 | 0.00821 | 0.002329 | 0.013003 | 0.029395 | 0.003166 | 0.004877 | 0.001755 | 0.01392 | 0.00519 | 0.008351 | 0.011787 | 0.003507 | 0.018865 | 0.003757 | 0.003335 | 0.023032 | 0.00191 | 0.003202 | 0.016439 | 0.030039 | 0.008714 | 0.005462 |
| 27.4583333 | 0.040536 | 0.026126 | 0.01003 | 0.005097 | 0.020688 | 0.02965 | 0.008809 | 0.006082 | 0.003415 | 0.013851 | 0.00861 | 0.002223 | 0.021187 | 0.002334 | 0.01882 | 0.001204 | 0.002139 | 0.026101 | 0.001295 | 0.004721 | 0.016569 | 0.033416 | 0.016934 | 0.006006 |
| 27.4722222 | 0.081441 | 0.033695 | 0.007646 | 0.010239 | 0.027408 | 0.031206 | 0.016203 | 0.008424 | 0.007777 | 0.013279 | 0.013976 | 0.002022 | 0.026188 | 0.001508 | 0.020303 | 0.001529 | 0.001973 | 0.033424 | 0.003402 | 0.007429 | 0.029791 | 0.03151 | 0.027032 | 0.01087 |
| 27.4861111 | 0.112223 | 0.035441 | 0.007568 | 0.015742 | 0.029327 | 0.034179 | 0.020528 | 0.010733 | 0.011871 | 0.012419 | 0.014038 | 0.007271 | 0.023841 | 0.002427 | 0.022741 | 0.005408 | 0.002866 | 0.043271 | 0.007492 | 0.011022 | 0.060503 | 0.027929 | 0.035459 | 0.019217 |
| 27.5 | 0.120434 | 0.029048 | 0.0221 | 0.020261 | 0.026541 | 0.037944 | 0.021846 | 0.012659 | 0.015165 | 0.01212 | 0.008461 | 0.014824 | 0.017214 | 0.004423 | 0.023151 | 0.010084 | 0.007123 | 0.049711 | 0.010195 | 0.015144 | 0.077507 | 0.023574 | 0.041903 | 0.025575 |
| 27.5138889 | 0.116234 | 0.018224 | 0.0536 | 0.022651 | 0.020891 | 0.042299 | 0.022293 | 0.015295 | 0.018993 | 0.013263 | 0.004102 | 0.021714 | 0.010105 | 0.006886 | 0.020336 | 0.013031 | 0.012607 | 0.049557 | 0.008972 | 0.018108 | 0.06076 | 0.01701 | 0.046818 | 0.024408 |
| 27.5277778 | 0.115001 | 0.013352 | 0.095367 | 0.026044 | 0.018749 | 0.048716 | 0.025463 | 0.019248 | 0.0285 | 0.016756 | 0.006747 | 0.031641 | 0.01051 | 0.013609 | 0.019816 | 0.017717 | 0.016462 | 0.050928 | 0.006463 | 0.020932 | 0.030191 | 0.008478 | 0.05172 | 0.021159 |
| 27.5416667 | 0.122443 | 0.031589 | 0.158921 | 0.052055 | 0.03465 | 0.07225 | 0.043254 | 0.037501 | 0.063178 | 0.025297 | 0.030664 | 0.061162 | 0.032744 | 0.035461 | 0.037457 | 0.031371 | 0.022134 | 0.078439 | 0.006791 | 0.0311 | 0.01186 | 0.004403 | 0.06552 | 0.03988 |
| 27.5555556 | 0.145231 | 0.075986 | 0.233244 | 0.112868 | 0.076277 | 0.128449 | 0.085111 | 0.091563 | 0.130613 | 0.039626 | 0.103394 | 0.112987 | 0.081453 | 0.079826 | 0.086608 | 0.053059 | 0.033144 | 0.142854 | 0.011472 | 0.060326 | 0.017044 | 0.013431 | 0.105582 | 0.097438 |
| 27.5694444 | 0.171136 | 0.10794 | 0.251298 | 0.161401 | 0.117062 | 0.187292 | 0.127565 | 0.163141 | 0.174829 | 0.053895 | 0.192737 | 0.146204 | 0.128581 | 0.123933 | 0.136173 | 0.071022 | 0.04409 | 0.202569 | 0.017575 | 0.109598 | 0.0455 | 0.035058 | 0.14394 | 0.152695 |
| 27.5833333 | 0.158873 | 0.100483 | 0.19592 | 0.155491 | 0.122321 | 0.196628 | 0.136697 | 0.205672 | 0.152501 | 0.062485 | 0.215269 | 0.141821 | 0.142702 | 0.133431 | 0.141116 | 0.076247 | 0.046567 | 0.208882 | 0.021989 | 0.160701 | 0.080222 | 0.058995 | 0.124465 | 0.159189 |
| 27.5972222 | 0.093771 | 0.078295 | 0.119086 | 0.117998 | 0.099191 | 0.153169 | 0.119126 | 0.202269 | 0.105086 | 0.062756 | 0.165583 | 0.124552 | 0.122767 | 0.113724 | 0.112704 | 0.068799 | 0.041006 | 0.167991 | 0.023738 | 0.184223 | 0.104904 | 0.074881 | 0.06426 | 0.131021 |
| 27.6111111 | 0.048802 | 0.056394 | 0.055563 | 0.075938 | 0.073253 | 0.094495 | 0.09523 | 0.162037 | 0.064167 | 0.051407 | 0.099865 | 0.090088 | 0.081374 | 0.087813 | 0.078939 | 0.053984 | 0.03343 | 0.113819 | 0.021121 | 0.166406 | 0.103727 | 0.07232 | 0.039299 | 0.090034 |
| 27.625 | 0.088779 | 0.030842 | 0.036713 | 0.03799 | 0.051184 | 0.044229 | 0.065929 | 0.10703 | 0.033444 | 0.029143 | 0.050836 | 0.046901 | 0.040452 | 0.059802 | 0.043485 | 0.036281 | 0.024675 | 0.063224 | 0.015199 | 0.124926 | 0.067008 | 0.045933 | 0.076013 | 0.045513 |
| 27.6388889 | 0.170931 | 0.015248 | 0.080878 | 0.021774 | 0.032458 | 0.015782 | 0.033207 | 0.063564 | 0.033435 | 0.013633 | 0.030759 | 0.033996 | 0.025623 | 0.03225 | 0.017211 | 0.019416 | 0.015786 | 0.026158 | 0.008412 | 0.078441 | 0.032635 | 0.016891 | 0.109201 | 0.023837 |
| 27.6527778 | 0.218357 | 0.021779 | 0.134478 | 0.02651 | 0.019326 | 0.019909 | 0.017351 | 0.042712 | 0.060643 | 0.016337 | 0.023247 | 0.044808 | 0.034207 | 0.013768 | 0.011668 | 0.009749 | 0.012069 | 0.012952 | 0.00267 | 0.042965 | 0.038903 | 0.007618 | 0.089932 | 0.035698 |
| 27.6666667 | 0.181597 | 0.032182 | 0.133504 | 0.021911 | 0.013368 | 0.040546 | 0.016698 | 0.039052 | 0.074386 | 0.016077 | 0.020401 | 0.044566 | 0.034389 | 0.007763 | 0.018756 | 0.013164 | 0.017324 | 0.014941 | 0.002204 | 0.033877 | 0.061412 | 0.0114 | 0.051613 | 0.042191 |
| 27.6805556 | 0.108484 | 0.024835 | 0.085607 | 0.010387 | 0.015128 | 0.05759 | 0.01632 | 0.040767 | 0.052747 | 0.009473 | 0.04028 | 0.029108 | 0.023094 | 0.008714 | 0.034289 | 0.026529 | 0.030734 | 0.01729 | 0.007642 | 0.041574 | 0.063245 | 0.017804 | 0.046522 | 0.027056 |
| 27.6944444 | 0.113485 | 0.016892 | 0.071865 | 0.011985 | 0.020676 | 0.062909 | 0.027618 | 0.035929 | 0.022381 | 0.013985 | 0.06061 | 0.026475 | 0.028031 | 0.007282 | 0.047728 | 0.036699 | 0.040558 | 0.020603 | 0.015391 | 0.042086 | 0.042897 | 0.019624 | 0.081631 | 0.024042 |
| 27.7083333 | 0.190393 | 0.03125 | 0.128371 | 0.018251 | 0.025494 | 0.05581 | 0.041827 | 0.026343 | 0.008964 | 0.022591 | 0.049522 | 0.043492 | 0.041439 | 0.00635 | 0.040969 | 0.035435 | 0.03552 | 0.034113 | 0.022104 | 0.030405 | 0.035716 | 0.014266 | 0.110886 | 0.039897 |
| 27.7222222 | 0.225584 | 0.050335 | 0.179087 | 0.02378 | 0.02906 | 0.050002 | 0.043406 | 0.036531 | 0.007344 | 0.025372 | 0.030013 | 0.047416 | 0.033871 | 0.015479 | 0.024015 | 0.025569 | 0.020419 | 0.060124 | 0.025415 | 0.033492 | 0.060935 | 0.015084 | 0.097366 | 0.043651 |
| 27.7361111 | 0.179319 | 0.049669 | 0.172535 | 0.029159 | 0.029639 | 0.045845 | 0.03724 | 0.077351 | 0.012831 | 0.023444 | 0.042507 | 0.027313 | 0.017515 | 0.028718 | 0.024453 | 0.013864 | 0.009484 | 0.089088 | 0.025575 | 0.064611 | 0.087059 | 0.028542 | 0.053351 | 0.028665 |
| 27.75 | 0.097543 | 0.0296 | 0.127795 | 0.029971 | 0.021498 | 0.03458 | 0.026569 | 0.120391 | 0.027035 | 0.018281 | 0.078875 | 0.009063 | 0.022058 | 0.036845 | 0.044033 | 0.005654 | 0.010459 | 0.097846 | 0.021698 | 0.101278 | 0.083763 | 0.038511 | 0.022745 | 0.013715 |
| 27.7638889 | 0.042325 | 0.012719 | 0.071564 | 0.027481 | 0.010995 | 0.040498 | 0.02529 | 0.135456 | 0.042907 | 0.012896 | 0.100043 | 0.010551 | 0.04302 | 0.038448 | 0.057799 | 0.004182 | 0.016698 | 0.069178 | 0.012527 | 0.126068 | 0.052159 | 0.032626 | 0.028088 | 0.009692 |
| 27.7777778 | 0.059099 | 0.014974 | 0.037959 | 0.028054 | 0.009664 | 0.075194 | 0.042216 | 0.118613 | 0.04839 | 0.010309 | 0.086003 | 0.026899 | 0.059848 | 0.032458 | 0.053672 | 0.008906 | 0.020388 | 0.037765 | 0.006945 | 0.129274 | 0.024648 | 0.017838 | 0.044861 | 0.021228 |
| 27.7916667 | 0.134937 | 0.022145 | 0.057222 | 0.034071 | 0.014131 | 0.105774 | 0.056994 | 0.076261 | 0.041369 | 0.013024 | 0.046421 | 0.034622 | 0.059773 | 0.018973 | 0.041426 | 0.015567 | 0.019824 | 0.049522 | 0.012532 | 0.105719 | 0.033276 | 0.00779 | 0.053083 | 0.036574 |
| 27.8055556 | 0.212214 | 0.019561 | 0.10657 | 0.034291 | 0.02399 | 0.109639 | 0.060153 | 0.043548 | 0.027596 | 0.019811 | 0.01466 | 0.023028 | 0.040655 | 0.009905 | 0.035364 | 0.020907 | 0.01373 | 0.088351 | 0.018943 | 0.067388 | 0.064906 | 0.006161 | 0.050907 | 0.034363 |
| 27.8194444 | 0.245045 | 0.012913 | 0.142089 | 0.021772 | 0.033328 | 0.08844 | 0.054684 | 0.070308 | 0.016544 | 0.022732 | 0.00646 | 0.015579 | 0.018669 | 0.015557 | 0.039159 | 0.022517 | 0.006735 | 0.099354 | 0.018727 | 0.033518 | 0.077254 | 0.009833 | 0.038678 | 0.018697 |
| 27.8333333 | 0.222979 | 0.014464 | 0.147504 | 0.00872 | 0.028625 | 0.050532 | 0.039593 | 0.137239 | 0.017297 | 0.016451 | 0.006341 | 0.021422 | 0.011404 | 0.023688 | 0.042764 | 0.019822 | 0.005532 | 0.066047 | 0.017082 | 0.023876 | 0.053618 | 0.020049 | 0.025095 | 0.012368 |
| 27.8472222 | 0.154654 | 0.018969 | 0.115648 | 0.005211 | 0.016951 | 0.027488 | 0.025478 | 0.172814 | 0.021318 | 0.008496 | 0.012487 | 0.021148 | 0.012397 | 0.024245 | 0.037031 | 0.013748 | 0.00656 | 0.030217 | 0.014873 | 0.035958 | 0.025101 | 0.034999 | 0.023685 | 0.015197 |
| 27.8611111 | 0.076686 | 0.014995 | 0.059154 | 0.013357 | 0.013697 | 0.052873 | 0.02082 | 0.143239 | 0.017071 | 0.008641 | 0.028386 | 0.015382 | 0.010465 | 0.020293 | 0.02348 | 0.007193 | 0.005654 | 0.023753 | 0.009926 | 0.042229 | 0.016192 | 0.040687 | 0.037169 | 0.015135 |
| 27.875 | 0.028782 | 0.016003 | 0.024539 | 0.032821 | 0.023176 | 0.112459 | 0.025286 | 0.076216 | 0.014977 | 0.012729 | 0.039932 | 0.02558 | 0.013892 | 0.013846 | 0.011133 | 0.003845 | 0.004799 | 0.036662 | 0.005013 | 0.029798 | 0.017525 | 0.026568 | 0.045324 | 0.02089 |
| 27.8888889 | 0.018892 | 0.035726 | 0.026157 | 0.053428 | 0.03855 | 0.160936 | 0.031826 | 0.033072 | 0.03106 | 0.015628 | 0.036707 | 0.051276 | 0.028834 | 0.007199 | 0.008977 | 0.007755 | 0.010157 | 0.051055 | 0.003117 | 0.016925 | 0.031706 | 0.008583 | 0.03155 | 0.03915 |
| 27.9027778 | 0.041546 | 0.060332 | 0.025375 | 0.065364 | 0.050552 | 0.1671 | 0.032531 | 0.046852 | 0.056854 | 0.019728 | 0.022582 | 0.067371 | 0.044342 | 0.005903 | 0.02009 | 0.023144 | 0.028936 | 0.069234 | 0.007232 | 0.024402 | 0.06074 | 0.003987 | 0.025447 | 0.0578 |
| 27.9166667 | 0.091947 | 0.073933 | 0.018583 | 0.074629 | 0.057667 | 0.148692 | 0.02894 | 0.092175 | 0.072289 | 0.025164 | 0.009169 | 0.058704 | 0.056466 | 0.012026 | 0.046494 | 0.043977 | 0.052029 | 0.101353 | 0.018065 | 0.044744 | 0.082198 | 0.010334 | 0.051232 | 0.064647 |
| 27.9305556 | 0.15004 | 0.071815 | 0.034686 | 0.088658 | 0.062698 | 0.135024 | 0.027006 | 0.139946 | 0.070574 | 0.030276 | 0.005515 | 0.036044 | 0.068298 | 0.021189 | 0.080485 | 0.0594 | 0.061771 | 0.148504 | 0.026481 | 0.063477 | 0.076861 | 0.020839 | 0.086648 | 0.057739 |
| 27.9444444 | 0.176842 | 0.055079 | 0.068227 | 0.103177 | 0.063188 | 0.116339 | 0.028661 | 0.181457 | 0.059615 | 0.033572 | 0.012332 | 0.017023 | 0.069978 | 0.030065 | 0.100581 | 0.069276 | 0.061532 | 0.182137 | 0.025761 | 0.077401 | 0.049305 | 0.029805 | 0.11503 | 0.043967 |
| 27.9583333 | 0.155883 | 0.02961 | 0.09344 | 0.109801 | 0.053177 | 0.074249 | 0.03054 | 0.210259 | 0.050153 | 0.031803 | 0.02247 | 0.008845 | 0.059809 | 0.035186 | 0.102226 | 0.078125 | 0.062617 | 0.171566 | 0.019763 | 0.08483 | 0.022253 | 0.032831 | 0.119547 | 0.032492 |
| 27.9722222 | 0.100907 | 0.012798 | 0.094963 | 0.101525 | 0.032906 | 0.040926 | 0.028673 | 0.208146 | 0.04664 | 0.023057 | 0.02861 | 0.009722 | 0.052694 | 0.032669 | 0.096409 | 0.07933 | 0.059809 | 0.141593 | 0.013682 | 0.083693 | 0.009109 | 0.028201 | 0.089438 | 0.026592 |
| 27.9861111 | 0.051616 | 0.016012 | 0.0705 | 0.076628 | 0.015256 | 0.064768 | 0.021906 | 0.16451 | 0.046319 | 0.011426 | 0.030304 | 0.015532 | 0.049222 | 0.024015 | 0.083238 | 0.063638 | 0.04532 | 0.115664 | 0.008458 | 0.077563 | 0.007251 | 0.017872 | 0.049747 | 0.021625 |
| 28 | 0.053801 | 0.021926 | 0.037928 | 0.041917 | 0.012541 | 0.116852 | 0.012075 | 0.103138 | 0.040775 | 0.003676 | 0.028668 | 0.020955 | 0.042446 | 0.01485 | 0.059819 | 0.037217 | 0.024743 | 0.080435 | 0.00411 | 0.073074 | 0.00972 | 0.007865 | 0.041331 | 0.013118 |
| 28.0138889 | 0.096333 | 0.019197 | 0.027374 | 0.017627 | 0.017122 | 0.145822 | 0.00506 | 0.051004 | 0.028204 | 0.001621 | 0.023366 | 0.019576 | 0.034419 | 0.010393 | 0.031517 | 0.014828 | 0.009719 | 0.040884 | 0.002308 | 0.066821 | 0.009556 | 0.003329 | 0.066279 | 0.007941 |
| 28.0277778 | 0.123068 | 0.013166 | 0.031689 | 0.011678 | 0.015456 | 0.145703 | 0.004522 | 0.02057 | 0.019479 | 0.002858 | 0.021553 | 0.015295 | 0.030491 | 0.01389 | 0.014158 | 0.006523 | 0.005868 | 0.023881 | 0.002654 | 0.056194 | 0.007736 | 0.004619 | 0.08168 | 0.008173 |
| 28.0416667 | 0.094043 | 0.01967 | 0.034164 | 0.020149 | 0.024617 | 0.104369 | 0.018538 | 0.017148 | 0.031594 | 0.011024 | 0.041836 | 0.027001 | 0.041818 | 0.028652 | 0.029083 | 0.019077 | 0.015659 | 0.023596 | 0.006931 | 0.055226 | 0.01979 | 0.01628 | 0.062695 | 0.021933 |
| 28.0555556 | 0.078 | 0.054825 | 0.064368 | 0.056832 | 0.073556 | 0.066209 | 0.064359 | 0.034886 | 0.074701 | 0.031437 | 0.089833 | 0.07452 | 0.080626 | 0.057096 | 0.081972 | 0.050753 | 0.038685 | 0.036732 | 0.017274 | 0.073809 | 0.058594 | 0.04384 | 0.057267 | 0.074541 |
| 28.0694444 | 0.148893 | 0.096192 | 0.123672 | 0.103165 | 0.139189 | 0.110486 | 0.123213 | 0.050519 | 0.119706 | 0.051815 | 0.128839 | 0.133813 | 0.12934 | 0.082431 | 0.140318 | 0.078036 | 0.060826 | 0.074819 | 0.025174 | 0.095757 | 0.107287 | 0.072977 | 0.101507 | 0.140831 |
| 28.0833333 | 0.201774 | 0.107439 | 0.16003 | 0.110871 | 0.166915 | 0.171878 | 0.141452 | 0.041849 | 0.117136 | 0.053488 | 0.119519 | 0.147612 | 0.139012 | 0.077736 | 0.150968 | 0.077054 | 0.066971 | 0.100428 | 0.023916 | 0.096767 | 0.128387 | 0.077181 | 0.124724 | 0.154318 |
| 28.0972222 | 0.160767 | 0.08749 | 0.148359 | 0.078836 | 0.14881 | 0.176397 | 0.113045 | 0.018505 | 0.073467 | 0.039026 | 0.075875 | 0.115479 | 0.096993 | 0.048651 | 0.102301 | 0.055032 | 0.059457 | 0.094227 | 0.019168 | 0.073451 | 0.110029 | 0.053569 | 0.09251 | 0.103616 |
| 28.1111111 | 0.089842 | 0.056938 | 0.110249 | 0.044638 | 0.119712 | 0.160003 | 0.079709 | 0.003608 | 0.031265 | 0.022187 | 0.033749 | 0.079274 | 0.043473 | 0.021212 | 0.045368 | 0.032596 | 0.050194 | 0.078157 | 0.014578 | 0.041472 | 0.076628 | 0.025701 | 0.052378 | 0.04592 |
| 28.125 | 0.037791 | 0.02949 | 0.071725 | 0.02506 | 0.092165 | 0.139744 | 0.056823 | 0.00104 | 0.012082 | 0.010133 | 0.014406 | 0.050146 | 0.013914 | 0.010026 | 0.020598 | 0.017081 | 0.042105 | 0.061025 | 0.008276 | 0.016377 | 0.048347 | 0.008804 | 0.028859 | 0.027686 |
| 28.1388889 | 0.011383 | 0.012523 | 0.039666 | 0.013595 | 0.053673 | 0.105832 | 0.034079 | 0.002794 | 0.015589 | 0.007802 | 0.016667 | 0.024679 | 0.010821 | 0.012512 | 0.01379 | 0.00711 | 0.028522 | 0.044429 | 0.003392 | 0.007471 | 0.027169 | 0.002596 | 0.016754 | 0.035916 |
| 28.1527778 | 0.003376 | 0.012145 | 0.017065 | 0.012034 | 0.025839 | 0.064306 | 0.025873 | 0.00456 | 0.024408 | 0.014702 | 0.022994 | 0.015839 | 0.017197 | 0.014018 | 0.0096 | 0.003061 | 0.012718 | 0.028217 | 0.003043 | 0.015301 | 0.012422 | 0.003873 | 0.010818 | 0.046248 |
| 28.1666667 | 0.003238 | 0.017051 | 0.006892 | 0.021648 | 0.034804 | 0.029163 | 0.0458 | 0.008593 | 0.025999 | 0.02366 | 0.031579 | 0.030219 | 0.025074 | 0.008703 | 0.012684 | 0.005491 | 0.003489 | 0.014173 | 0.003415 | 0.03428 | 0.007881 | 0.008472 | 0.013985 | 0.058144 |
| 28.1805556 | 0.003559 | 0.01595 | 0.004375 | 0.030366 | 0.052664 | 0.012807 | 0.06408 | 0.014374 | 0.022561 | 0.031225 | 0.044798 | 0.050489 | 0.033892 | 0.00398 | 0.026974 | 0.011282 | 0.000684 | 0.008954 | 0.002084 | 0.054558 | 0.012814 | 0.012129 | 0.020334 | 0.067599 |
| 28.1944444 | 0.002178 | 0.018885 | 0.005242 | 0.029074 | 0.055367 | 0.015346 | 0.062225 | 0.014662 | 0.018541 | 0.034698 | 0.051777 | 0.062399 | 0.037255 | 0.003392 | 0.042097 | 0.016829 | 0.000295 | 0.011834 | 0.002135 | 0.061336 | 0.01724 | 0.012008 | 0.018087 | 0.064775 |
| 28.2083333 | 0.001178 | 0.025833 | 0.008017 | 0.021323 | 0.04847 | 0.021913 | 0.053204 | 0.008925 | 0.017878 | 0.02964 | 0.045091 | 0.061979 | 0.032173 | 0.003507 | 0.039189 | 0.019951 | 0.000205 | 0.013801 | 0.005139 | 0.047498 | 0.016552 | 0.008249 | 0.009361 | 0.048467 |
| 28.2222222 | 0.001245 | 0.030751 | 0.008905 | 0.014768 | 0.037791 | 0.023345 | 0.043211 | 0.004247 | 0.022861 | 0.01859 | 0.027844 | 0.0499 | 0.021833 | 0.005563 | 0.023499 | 0.019472 | 0.000506 | 0.013199 | 0.00791 | 0.025768 | 0.013426 | 0.003829 | 0.003817 | 0.026058 |
| 28.2361111 | 0.003879 | 0.034571 | 0.005812 | 0.011629 | 0.02918 | 0.02132 | 0.032518 | 0.005017 | 0.03122 | 0.009068 | 0.013071 | 0.035407 | 0.010805 | 0.009975 | 0.018269 | 0.014289 | 0.001389 | 0.014029 | 0.01016 | 0.017848 | 0.011903 | 0.001919 | 0.003366 | 0.016381 |
| 28.25 | 0.014106 | 0.034221 | 0.002806 | 0.011552 | 0.023339 | 0.015252 | 0.022376 | 0.008178 | 0.037413 | 0.005451 | 0.011248 | 0.023046 | 0.005628 | 0.011533 | 0.022012 | 0.007354 | 0.002095 | 0.017386 | 0.013174 | 0.020682 | 0.017474 | 0.004436 | 0.008946 | 0.022253 |
| 28.2638889 | 0.032148 | 0.027604 | 0.003954 | 0.014556 | 0.016446 | 0.007678 | 0.014038 | 0.007682 | 0.038474 | 0.008874 | 0.013185 | 0.012052 | 0.008295 | 0.008315 | 0.019086 | 0.004753 | 0.002995 | 0.024768 | 0.014807 | 0.019776 | 0.03043 | 0.0117 | 0.025106 | 0.02026 |
| 28.2777778 | 0.052919 | 0.017034 | 0.007649 | 0.018543 | 0.009299 | 0.005229 | 0.009106 | 0.005867 | 0.036405 | 0.017423 | 0.01123 | 0.004513 | 0.009285 | 0.003822 | 0.021376 | 0.004826 | 0.005294 | 0.037191 | 0.013829 | 0.026769 | 0.043534 | 0.021371 | 0.045037 | 0.014677 |
| 28.2916667 | 0.07207 | 0.007709 | 0.011925 | 0.021847 | 0.005943 | 0.008868 | 0.008823 | 0.00969 | 0.034549 | 0.024912 | 0.015137 | 0.002407 | 0.008558 | 0.001447 | 0.037506 | 0.003838 | 0.008422 | 0.05196 | 0.010379 | 0.046566 | 0.051117 | 0.030231 | 0.056584 | 0.023841 |
| 28.3055556 | 0.087398 | 0.002765 | 0.016918 | 0.025159 | 0.009427 | 0.01345 | 0.013328 | 0.016273 | 0.033231 | 0.025572 | 0.022586 | 0.0057 | 0.01657 | 0.001236 | 0.043717 | 0.005269 | 0.010978 | 0.066168 | 0.0056 | 0.057952 | 0.051775 | 0.035126 | 0.06071 | 0.029908 |
| 28.3194444 | 0.093814 | 0.00275 | 0.021005 | 0.027038 | 0.018571 | 0.014909 | 0.019658 | 0.018339 | 0.029817 | 0.018947 | 0.019841 | 0.011459 | 0.025467 | 0.003029 | 0.028934 | 0.010598 | 0.012199 | 0.071159 | 0.002781 | 0.0492 | 0.044852 | 0.034637 | 0.061772 | 0.01911 |
| 28.3333333 | 0.084682 | 0.008903 | 0.022399 | 0.024191 | 0.026254 | 0.013502 | 0.022403 | 0.015744 | 0.023457 | 0.009985 | 0.011322 | 0.016021 | 0.021662 | 0.005692 | 0.018149 | 0.014583 | 0.011941 | 0.062689 | 0.002515 | 0.030017 | 0.034139 | 0.030259 | 0.058692 | 0.009217 |
| 28.3472222 | 0.063637 | 0.019746 | 0.022512 | 0.018467 | 0.02767 | 0.011716 | 0.020425 | 0.013301 | 0.016944 | 0.005654 | 0.009677 | 0.01777 | 0.01138 | 0.006228 | 0.024861 | 0.013148 | 0.010627 | 0.053994 | 0.002973 | 0.015555 | 0.026215 | 0.024832 | 0.055792 | 0.010708 |
| 28.3611111 | 0.039767 | 0.030428 | 0.022812 | 0.014441 | 0.024379 | 0.010752 | 0.017755 | 0.013298 | 0.012289 | 0.007773 | 0.010389 | 0.016043 | 0.007673 | 0.003891 | 0.025426 | 0.007597 | 0.008958 | 0.050081 | 0.005893 | 0.012944 | 0.022944 | 0.020216 | 0.057422 | 0.010392 |
| 28.375 | 0.01861 | 0.036413 | 0.02412 | 0.013222 | 0.019179 | 0.011449 | 0.016524 | 0.013138 | 0.009751 | 0.011408 | 0.008648 | 0.011994 | 0.008461 | 0.001404 | 0.017085 | 0.003271 | 0.00681 | 0.044506 | 0.010595 | 0.020191 | 0.022409 | 0.016428 | 0.05875 | 0.008881 |
| 28.3888889 | 0.007265 | 0.035071 | 0.027043 | 0.013239 | 0.014246 | 0.01459 | 0.015461 | 0.009865 | 0.009724 | 0.011944 | 0.012854 | 0.009178 | 0.007373 | 0.001194 | 0.022164 | 0.00316 | 0.004274 | 0.03671 | 0.01249 | 0.029477 | 0.022303 | 0.013294 | 0.05258 | 0.017665 |
| 28.4027778 | 0.010743 | 0.026551 | 0.029556 | 0.012883 | 0.011162 | 0.018331 | 0.013253 | 0.004722 | 0.011493 | 0.009319 | 0.019497 | 0.009887 | 0.011367 | 0.003176 | 0.030253 | 0.003343 | 0.005036 | 0.027085 | 0.011333 | 0.031752 | 0.02059 | 0.010539 | 0.035515 | 0.025593 |
| 28.4166667 | 0.024084 | 0.01512 | 0.029758 | 0.0117 | 0.010559 | 0.020953 | 0.010517 | 0.00131 | 0.013612 | 0.005995 | 0.016439 | 0.013213 | 0.023621 | 0.005278 | 0.0236 | 0.00411 | 0.014021 | 0.016555 | 0.010334 | 0.023279 | 0.016564 | 0.007628 | 0.018664 | 0.01959 |
| 28.4305556 | 0.039666 | 0.008422 | 0.028119 | 0.010555 | 0.013222 | 0.022851 | 0.009204 | 0.0012 | 0.015555 | 0.003932 | 0.009588 | 0.017786 | 0.030634 | 0.005185 | 0.020331 | 0.010327 | 0.026692 | 0.008857 | 0.01087 | 0.012228 | 0.011628 | 0.004144 | 0.017135 | 0.009706 |
| 28.4444444 | 0.052149 | 0.011375 | 0.026319 | 0.010467 | 0.019031 | 0.02412 | 0.010334 | 0.003967 | 0.016415 | 0.004092 | 0.010838 | 0.022078 | 0.021785 | 0.005797 | 0.031126 | 0.017866 | 0.032465 | 0.006409 | 0.011631 | 0.009214 | 0.007551 | 0.001966 | 0.02013 | 0.007684 |
| 28.4583333 | 0.059828 | 0.020941 | 0.024009 | 0.010533 | 0.0245 | 0.022761 | 0.012906 | 0.007492 | 0.014459 | 0.006703 | 0.013869 | 0.022631 | 0.01179 | 0.009051 | 0.031899 | 0.020751 | 0.030555 | 0.008075 | 0.011765 | 0.008532 | 0.00485 | 0.002633 | 0.016112 | 0.007926 |
| 28.4722222 | 0.062471 | 0.03081 | 0.01977 | 0.009376 | 0.025801 | 0.017929 | 0.015801 | 0.009486 | 0.010801 | 0.010417 | 0.010992 | 0.018167 | 0.014253 | 0.008521 | 0.023048 | 0.019605 | 0.026504 | 0.011077 | 0.01374 | 0.006354 | 0.006043 | 0.006286 | 0.019193 | 0.010333 |
| 28.4861111 | 0.054873 | 0.038118 | 0.016078 | 0.008651 | 0.023962 | 0.01347 | 0.019772 | 0.010275 | 0.010032 | 0.013435 | 0.011371 | 0.012657 | 0.013729 | 0.005156 | 0.03134 | 0.017273 | 0.022664 | 0.014487 | 0.019006 | 0.008461 | 0.007822 | 0.009314 | 0.033418 | 0.019425 |
| 28.5 | 0.035189 | 0.039155 | 0.015979 | 0.010453 | 0.020425 | 0.013387 | 0.023827 | 0.011002 | 0.01371 | 0.015435 | 0.018273 | 0.009531 | 0.008638 | 0.005967 | 0.046112 | 0.014816 | 0.018986 | 0.018022 | 0.023691 | 0.011955 | 0.005782 | 0.010162 | 0.042335 | 0.030011 |
| 28.5138889 | 0.017242 | 0.029566 | 0.018491 | 0.013666 | 0.013135 | 0.016652 | 0.023467 | 0.011189 | 0.017869 | 0.017008 | 0.018666 | 0.008959 | 0.014595 | 0.007481 | 0.038108 | 0.011602 | 0.015557 | 0.01992 | 0.023158 | 0.015558 | 0.007041 | 0.014192 | 0.038309 | 0.033416 |
| 28.5277778 | 0.024468 | 0.01601 | 0.02447 | 0.022229 | 0.006356 | 0.02269 | 0.018565 | 0.010804 | 0.022554 | 0.02054 | 0.01067 | 0.016187 | 0.02878 | 0.007444 | 0.019819 | 0.011502 | 0.015466 | 0.023818 | 0.018078 | 0.019933 | 0.016984 | 0.021703 | 0.034023 | 0.027785 |
| 28.5416667 | 0.087877 | 0.018261 | 0.0425 | 0.051676 | 0.01835 | 0.039082 | 0.022599 | 0.013706 | 0.03968 | 0.031441 | 0.006454 | 0.044077 | 0.048636 | 0.016124 | 0.012165 | 0.023806 | 0.024501 | 0.045761 | 0.013724 | 0.024325 | 0.03983 | 0.03859 | 0.052926 | 0.030074 |
| 28.5555556 | 0.216831 | 0.053415 | 0.079922 | 0.099936 | 0.062709 | 0.0779 | 0.052429 | 0.029522 | 0.074664 | 0.053259 | 0.021174 | 0.092702 | 0.076714 | 0.042676 | 0.021119 | 0.050072 | 0.043615 | 0.101048 | 0.012303 | 0.037223 | 0.077699 | 0.065742 | 0.0997 | 0.054395 |
| 28.5694444 | 0.343617 | 0.1023 | 0.114109 | 0.138948 | 0.110939 | 0.137501 | 0.097012 | 0.063006 | 0.103906 | 0.078183 | 0.059737 | 0.132289 | 0.097357 | 0.074825 | 0.064464 | 0.072984 | 0.059553 | 0.159335 | 0.012993 | 0.065537 | 0.117193 | 0.081431 | 0.124322 | 0.082528 |
| 28.5833333 | 0.37996 | 0.1233 | 0.118868 | 0.150191 | 0.121248 | 0.187244 | 0.119144 | 0.104008 | 0.108604 | 0.089714 | 0.099205 | 0.14673 | 0.09527 | 0.083443 | 0.117893 | 0.073535 | 0.058977 | 0.168786 | 0.016201 | 0.102445 | 0.134887 | 0.0759 | 0.089911 | 0.096942 |

| | | | | | | | | | | | | | | | | | | | | | | | | |
|---|---|---|---|---|---|---|---|---|---|---|---|---|---|---|---|---|---|---|---|---|---|---|---|---|
| 28.5972222 | 0.32474 | 0.110783 | 0.107151 | 0.128299 | 0.100556 | 0.198753 | 0.104649 | 0.142038 | 0.098859 | 0.080196 | 0.108321 | 0.142797 | 0.082104 | 0.062568 | 0.128383 | 0.057705 | 0.050077 | 0.134022 | 0.020614 | 0.129641 | 0.117601 | 0.060684 | 0.038411 | 0.10051 |
| 28.6111111 | 0.233642 | 0.085318 | 0.095077 | 0.086296 | 0.073699 | 0.181071 | 0.07116 | 0.166403 | 0.080886 | 0.057241 | 0.079816 | 0.112731 | 0.064876 | 0.037867 | 0.101658 | 0.043365 | 0.0438 | 0.088284 | 0.020817 | 0.131533 | 0.07376 | 0.040985 | 0.017295 | 0.077514 |
| 28.625 | 0.141173 | 0.057288 | 0.08038 | 0.046894 | 0.044748 | 0.15098 | 0.034623 | 0.15857 | 0.057703 | 0.031761 | 0.041096 | 0.064755 | 0.036361 | 0.022147 | 0.07259 | 0.033444 | 0.035388 | 0.045338 | 0.013132 | 0.114673 | 0.035629 | 0.023491 | 0.031062 | 0.044535 |
| 28.6388889 | 0.065971 | 0.02881 | 0.052633 | 0.02369 | 0.020044 | 0.119418 | 0.015364 | 0.122362 | 0.039123 | 0.01483 | 0.029773 | 0.029192 | 0.018142 | 0.011309 | 0.046978 | 0.023902 | 0.019957 | 0.022109 | 0.004615 | 0.09341 | 0.033855 | 0.024051 | 0.060292 | 0.055707 |
| 28.6527778 | 0.036743 | 0.012163 | 0.024646 | 0.029436 | 0.010509 | 0.09843 | 0.02229 | 0.093823 | 0.028161 | 0.014361 | 0.039364 | 0.017752 | 0.034097 | 0.010956 | 0.022058 | 0.016902 | 0.009201 | 0.027382 | 0.004735 | 0.0773 | 0.052258 | 0.036858 | 0.085702 | 0.095256 |
| 28.6666667 | 0.058332 | 0.018094 | 0.016651 | 0.044276 | 0.009123 | 0.08394 | 0.027275 | 0.091141 | 0.022112 | 0.019284 | 0.030452 | 0.013469 | 0.058353 | 0.021428 | 0.007346 | 0.015182 | 0.009918 | 0.033515 | 0.011557 | 0.079191 | 0.057717 | 0.037117 | 0.080147 | 0.114278 |
| 28.6805556 | 0.096048 | 0.034543 | 0.013528 | 0.037582 | 0.007052 | 0.062257 | 0.01966 | 0.099062 | 0.020757 | 0.017369 | 0.018794 | 0.009568 | 0.049127 | 0.031501 | 0.009803 | 0.016145 | 0.010216 | 0.021634 | 0.017973 | 0.09571 | 0.046005 | 0.023869 | 0.042121 | 0.094487 |
| 28.6944444 | 0.141809 | 0.042741 | 0.007592 | 0.021621 | 0.009611 | 0.03457 | 0.020562 | 0.106233 | 0.024758 | 0.01855 | 0.039391 | 0.017151 | 0.026668 | 0.033743 | 0.030741 | 0.015277 | 0.009174 | 0.007898 | 0.022188 | 0.105976 | 0.026163 | 0.023292 | 0.012404 | 0.056478 |
| 28.7083333 | 0.195778 | 0.032165 | 0.008156 | 0.015274 | 0.016143 | 0.029237 | 0.029921 | 0.102598 | 0.028928 | 0.026446 | 0.06634 | 0.029463 | 0.037211 | 0.024867 | 0.061058 | 0.014521 | 0.01133 | 0.004599 | 0.024033 | 0.10307 | 0.015316 | 0.045317 | 0.006587 | 0.053959 |
| 28.7222222 | 0.233345 | 0.023045 | 0.020246 | 0.021654 | 0.020564 | 0.055998 | 0.031921 | 0.073243 | 0.026973 | 0.027136 | 0.067367 | 0.032339 | 0.060538 | 0.011848 | 0.072729 | 0.02197 | 0.012884 | 0.008153 | 0.022632 | 0.089261 | 0.025684 | 0.069981 | 0.015074 | 0.089967 |
| 28.7361111 | 0.229727 | 0.038263 | 0.03796 | 0.040904 | 0.022852 | 0.08622 | 0.027568 | 0.040946 | 0.021861 | 0.019044 | 0.045599 | 0.023117 | 0.062435 | 0.005519 | 0.050174 | 0.036543 | 0.010643 | 0.017988 | 0.018656 | 0.060507 | 0.042775 | 0.079027 | 0.03122 | 0.112872 |
| 28.75 | 0.161958 | 0.05883 | 0.041721 | 0.061654 | 0.025167 | 0.106401 | 0.027403 | 0.049351 | 0.017679 | 0.011016 | 0.026838 | 0.017852 | 0.045881 | 0.008868 | 0.026447 | 0.044465 | 0.007956 | 0.028854 | 0.012212 | 0.034216 | 0.050464 | 0.069103 | 0.046056 | 0.096329 |
| 28.7638889 | 0.081486 | 0.059598 | 0.027541 | 0.066624 | 0.028322 | 0.130946 | 0.034525 | 0.089738 | 0.011178 | 0.007642 | 0.035918 | 0.024451 | 0.023459 | 0.019037 | 0.037713 | 0.037845 | 0.013315 | 0.032854 | 0.005347 | 0.048638 | 0.047946 | 0.046107 | 0.047727 | 0.053173 |
| 28.7777778 | 0.075876 | 0.038532 | 0.02287 | 0.050976 | 0.029505 | 0.152077 | 0.042305 | 0.122399 | 0.005092 | 0.009302 | 0.055644 | 0.027078 | 0.016748 | 0.02418 | 0.065057 | 0.02333 | 0.022898 | 0.035136 | 0.001839 | 0.106282 | 0.038816 | 0.022816 | 0.032824 | 0.023615 |
| 28.7916667 | 0.121681 | 0.01502 | 0.053418 | 0.031055 | 0.025587 | 0.138008 | 0.044898 | 0.1344 | 0.005502 | 0.013749 | 0.055461 | 0.023473 | 0.041068 | 0.018481 | 0.076049 | 0.010209 | 0.025182 | 0.036147 | 0.001911 | 0.165752 | 0.023254 | 0.019407 | 0.014806 | 0.035185 |
| 28.8055556 | 0.141241 | 0.00602 | 0.10273 | 0.035117 | 0.018601 | 0.091829 | 0.041981 | 0.130438 | 0.010105 | 0.019733 | 0.038042 | 0.020178 | 0.070603 | 0.009197 | 0.068154 | 0.003583 | 0.017648 | 0.026054 | 0.003464 | 0.194829 | 0.011305 | 0.038438 | 0.008793 | 0.063661 |
| 28.8194444 | 0.119717 | 0.004545 | 0.133907 | 0.064774 | 0.011497 | 0.045906 | 0.031809 | 0.112979 | 0.012853 | 0.024315 | 0.018256 | 0.014219 | 0.069614 | 0.005414 | 0.053473 | 0.004025 | 0.009178 | 0.017701 | 0.006391 | 0.194044 | 0.015758 | 0.054583 | 0.011638 | 0.073209 |
| 28.8333333 | 0.080201 | 0.003245 | 0.128428 | 0.084417 | 0.006456 | 0.036292 | 0.017533 | 0.084764 | 0.010862 | 0.023 | 0.006305 | 0.006967 | 0.040991 | 0.011683 | 0.043439 | 0.007329 | 0.008625 | 0.028926 | 0.009772 | 0.171827 | 0.027433 | 0.057053 | 0.013957 | 0.051308 |
| 28.8472222 | 0.048813 | 0.003683 | 0.086838 | 0.068774 | 0.007358 | 0.067371 | 0.012178 | 0.050338 | 0.011818 | 0.015434 | 0.009894 | 0.003806 | 0.018041 | 0.019364 | 0.047355 | 0.011078 | 0.009622 | 0.04562 | 0.010275 | 0.134091 | 0.036205 | 0.046626 | 0.010908 | 0.023568 |
| 28.8611111 | 0.067207 | 0.003799 | 0.04692 | 0.033811 | 0.011903 | 0.098655 | 0.018185 | 0.02405 | 0.022674 | 0.008315 | 0.030862 | 0.007126 | 0.016619 | 0.020305 | 0.068253 | 0.01346 | 0.006559 | 0.05901 | 0.007209 | 0.091492 | 0.040492 | 0.024966 | 0.007491 | 0.02338 |
| 28.875 | 0.138792 | 0.003662 | 0.057336 | 0.013072 | 0.014714 | 0.096403 | 0.024301 | 0.023649 | 0.037605 | 0.010215 | 0.058533 | 0.017258 | 0.0163 | 0.021483 | 0.097427 | 0.013228 | 0.005689 | 0.070393 | 0.005271 | 0.053704 | 0.03757 | 0.007122 | 0.007788 | 0.035742 |
| 28.8888889 | 0.205119 | 0.003735 | 0.100476 | 0.019719 | 0.019001 | 0.057471 | 0.025085 | 0.039444 | 0.05028 | 0.020874 | 0.078086 | 0.031337 | 0.017081 | 0.028386 | 0.123922 | 0.014156 | 0.009633 | 0.066634 | 0.008667 | 0.028389 | 0.029179 | 0.001975 | 0.005239 | 0.032715 |
| 28.9027778 | 0.224195 | 0.005019 | 0.131155 | 0.04061 | 0.027092 | 0.029142 | 0.022321 | 0.048684 | 0.057768 | 0.033936 | 0.082761 | 0.046631 | 0.038134 | 0.032734 | 0.136571 | 0.018426 | 0.013981 | 0.042332 | 0.013152 | 0.020327 | 0.021589 | 0.002593 | 0.002111 | 0.021031 |
| 28.9166667 | 0.200879 | 0.013513 | 0.134337 | 0.061143 | 0.036504 | 0.0539 | 0.019608 | 0.047359 | 0.059343 | 0.044351 | 0.071332 | 0.060214 | 0.064997 | 0.028554 | 0.128356 | 0.023834 | 0.015295 | 0.030074 | 0.016396 | 0.024395 | 0.018296 | 0.00215 | 0.002842 | 0.027214 |
| 28.9305556 | 0.161274 | 0.028639 | 0.11297 | 0.077447 | 0.04508 | 0.098665 | 0.020886 | 0.037414 | 0.053938 | 0.045672 | 0.048473 | 0.065172 | 0.073318 | 0.020922 | 0.104397 | 0.030588 | 0.014786 | 0.060933 | 0.020971 | 0.030374 | 0.01898 | 0.002748 | 0.009967 | 0.05835 |
| 28.9444444 | 0.127364 | 0.042866 | 0.082643 | 0.088797 | 0.047632 | 0.124826 | 0.027541 | 0.02437 | 0.040765 | 0.03484 | 0.025241 | 0.054656 | 0.06012 | 0.015513 | 0.074251 | 0.040864 | 0.015179 | 0.108931 | 0.023278 | 0.033499 | 0.022316 | 0.007999 | 0.02313 | 0.095377 |
| 28.9583333 | 0.100481 | 0.047501 | 0.056339 | 0.090155 | 0.04258 | 0.13412 | 0.036002 | 0.027804 | 0.026787 | 0.019938 | 0.010937 | 0.032394 | 0.035554 | 0.014053 | 0.045522 | 0.049702 | 0.015788 | 0.130727 | 0.019569 | 0.032992 | 0.024015 | 0.014091 | 0.034188 | 0.111568 |
| 28.9722222 | 0.067085 | 0.040359 | 0.034644 | 0.080242 | 0.034423 | 0.140682 | 0.043895 | 0.049865 | 0.020194 | 0.010559 | 0.007757 | 0.014 | 0.016147 | 0.015264 | 0.023557 | 0.051033 | 0.017029 | 0.116403 | 0.011858 | 0.028465 | 0.019886 | 0.012114 | 0.042614 | 0.092084 |
| 28.9861111 | 0.037936 | 0.028208 | 0.017491 | 0.065186 | 0.025044 | 0.144402 | 0.050281 | 0.069641 | 0.02032 | 0.00727 | 0.013688 | 0.009468 | 0.011961 | 0.01792 | 0.011014 | 0.047887 | 0.020888 | 0.093834 | 0.004672 | 0.020786 | 0.011678 | 0.006592 | 0.050131 | 0.050315 |
| 29 | 0.04409 | 0.015764 | 0.013894 | 0.049243 | 0.015788 | 0.126194 | 0.051391 | 0.077209 | 0.020575 | 0.005908 | 0.021743 | 0.011903 | 0.013444 | 0.020143 | 0.006545 | 0.043532 | 0.023009 | 0.088593 | 0.0011 | 0.012377 | 0.004704 | 0.006404 | 0.052314 | 0.0212 |
| 29.0138889 | 0.076068 | 0.005945 | 0.027322 | 0.036143 | 0.011848 | 0.086041 | 0.047899 | 0.077183 | 0.017473 | 0.004295 | 0.023624 | 0.009589 | 0.009793 | 0.017761 | 0.009994 | 0.035636 | 0.019211 | 0.090724 | 0.00054 | 0.010322 | 0.001343 | 0.007431 | 0.049641 | 0.020807 |
| 29.0277778 | 0.089908 | 0.004874 | 0.037444 | 0.033002 | 0.019855 | 0.055998 | 0.044967 | 0.079537 | 0.015006 | 0.004597 | 0.023612 | 0.010567 | 0.011713 | 0.01248 | 0.013137 | 0.02768 | 0.015236 | 0.085966 | 0.003144 | 0.012002 | 0.003298 | 0.012866 | 0.053804 | 0.024863 |
| 29.0416667 | 0.073998 | 0.022482 | 0.036673 | 0.050805 | 0.047443 | 0.070713 | 0.051141 | 0.101459 | 0.024023 | 0.012214 | 0.040833 | 0.039801 | 0.038937 | 0.018596 | 0.030555 | 0.035608 | 0.022326 | 0.087651 | 0.013251 | 0.016955 | 0.017104 | 0.034175 | 0.089718 | 0.044238 |
| 29.0555556 | 0.100652 | 0.057912 | 0.060045 | 0.087938 | 0.09896 | 0.14139 | 0.066539 | 0.149636 | 0.048097 | 0.029786 | 0.089711 | 0.105792 | 0.095196 | 0.045655 | 0.108729 | 0.065676 | 0.04345 | 0.111871 | 0.032016 | 0.044504 | 0.043869 | 0.06895 | 0.161181 | 0.110139 |
| 29.0694444 | 0.191173 | 0.081827 | 0.116092 | 0.115165 | 0.144932 | 0.22829 | 0.069758 | 0.176257 | 0.066326 | 0.049556 | 0.148109 | 0.172207 | 0.147826 | 0.073735 | 0.212345 | 0.091065 | 0.062193 | 0.130638 | 0.0488 | 0.088819 | 0.070518 | 0.085266 | 0.214188 | 0.172077 |
| 29.0833333 | 0.218015 | 0.069165 | 0.154837 | 0.108964 | 0.146237 | 0.25579 | 0.050193 | 0.138293 | 0.055831 | 0.056758 | 0.165463 | 0.185877 | 0.153205 | 0.073795 | 0.244959 | 0.083456 | 0.061097 | 0.107162 | 0.050822 | 0.109116 | 0.085631 | 0.063847 | 0.196163 | 0.166094 |
| 29.0972222 | 0.142881 | 0.034677 | 0.149624 | 0.079502 | 0.117471 | 0.212216 | 0.028594 | 0.070668 | 0.027103 | 0.046615 | 0.122297 | 0.152224 | 0.12032 | 0.049016 | 0.194936 | 0.05517 | 0.047307 | 0.063616 | 0.038482 | 0.08492 | 0.08864 | 0.03984 | 0.131468 | 0.109238 |
| 29.1111111 | 0.067142 | 0.011806 | 0.116467 | 0.045778 | 0.087505 | 0.147735 | 0.019036 | 0.027614 | 0.007808 | 0.027294 | 0.057042 | 0.115912 | 0.086577 | 0.022955 | 0.119514 | 0.031525 | 0.034555 | 0.036576 | 0.020966 | 0.041049 | 0.07921 | 0.035636 | 0.071386 | 0.049451 |
| 29.125 | 0.031422 | 0.010959 | 0.073279 | 0.019248 | 0.059819 | 0.081746 | 0.013554 | 0.024093 | 0.005574 | 0.013116 | 0.022527 | 0.088586 | 0.061381 | 0.009096 | 0.055795 | 0.017059 | 0.025049 | 0.026046 | 0.009207 | 0.021224 | 0.056717 | 0.029988 | 0.032128 | 0.019446 |
| 29.1388889 | 0.014144 | 0.013781 | 0.034508 | 0.008126 | 0.035281 | 0.0368 | 0.007802 | 0.03253 | 0.008688 | 0.015017 | 0.027374 | 0.061313 | 0.040032 | 0.007328 | 0.022426 | 0.008378 | 0.017207 | 0.021424 | 0.009164 | 0.036419 | 0.027918 | 0.016563 | 0.020517 | 0.027421 |
| 29.1527778 | 0.005412 | 0.011435 | 0.012309 | 0.009727 | 0.016399 | 0.041669 | 0.003935 | 0.034322 | 0.008861 | 0.026597 | 0.043731 | 0.035806 | 0.027301 | 0.007837 | 0.027673 | 0.00368 | 0.009384 | 0.016912 | 0.014347 | 0.055471 | 0.011515 | 0.013534 | 0.03157 | 0.049133 |
| 29.1666667 | 0.004097 | 0.010053 | 0.009371 | 0.013956 | 0.007796 | 0.073404 | 0.004437 | 0.038415 | 0.005546 | 0.036581 | 0.056599 | 0.016659 | 0.0402 | 0.005522 | 0.051558 | 0.003243 | 0.003893 | 0.011296 | 0.017426 | 0.062669 | 0.016298 | 0.024716 | 0.046245 | 0.069852 |
| 29.1805556 | 0.005534 | 0.009834 | 0.020813 | 0.01787 | 0.016379 | 0.098726 | 0.013373 | 0.047347 | 0.00573 | 0.044956 | 0.071794 | 0.009305 | 0.070225 | 0.004402 | 0.075835 | 0.010043 | 0.004171 | 0.005933 | 0.016735 | 0.066 | 0.026632 | 0.03794 | 0.060641 | 0.091107 |
| 29.1944444 | 0.006838 | 0.01175 | 0.045177 | 0.025975 | 0.053821 | 0.12541 | 0.033226 | 0.055083 | 0.019361 | 0.055163 | 0.084828 | 0.017127 | 0.064513 | 0.009837 | 0.098862 | 0.027472 | 0.006407 | 0.004416 | 0.014042 | 0.073231 | 0.031083 | 0.041634 | 0.068065 | 0.101445 |
| 29.2083333 | 0.012767 | 0.023846 | 0.069292 | 0.042198 | 0.116957 | 0.159498 | 0.045288 | 0.054008 | 0.041133 | 0.059374 | 0.080282 | 0.03492 | 0.023939 | 0.013949 | 0.090905 | 0.044239 | 0.012301 | 0.005662 | 0.018542 | 0.080233 | 0.026375 | 0.033372 | 0.054966 | 0.080092 |
| 29.2222222 | 0.023436 | 0.043199 | 0.070618 | 0.060678 | 0.145758 | 0.161827 | 0.034745 | 0.037108 | 0.053253 | 0.050086 | 0.063262 | 0.06533 | 0.029923 | 0.0452 | 0.085308 | 0.06021 | 0.017994 | 0.018067 | 0.021978 | 0.089176 | 0.041357 | 0.037864 | 0.036439 | 0.06553 |
| 29.2361111 | 0.026639 | 0.051491 | 0.051117 | 0.066042 | 0.111379 | 0.114863 | 0.026096 | 0.018555 | 0.046727 | 0.036358 | 0.042284 | 0.094665 | 0.091846 | 0.121564 | 0.099981 | 0.088324 | 0.025784 | 0.052268 | 0.027745 | 0.164145 | 0.075576 | 0.050571 | 0.028936 | 0.106084 |
| 29.25 | 0.016511 | 0.043554 | 0.028255 | 0.052629 | 0.072449 | 0.056698 | 0.029936 | 0.016763 | 0.029159 | 0.024786 | 0.02002 | 0.091138 | 0.139234 | 0.166997 | 0.096718 | 0.120387 | 0.054328 | 0.106124 | 0.063479 | 0.277873 | 0.099191 | 0.050075 | 0.046382 | 0.151069 |
| 29.2638889 | 0.005762 | 0.0346 | 0.010907 | 0.035114 | 0.058815 | 0.02141 | 0.032911 | 0.01558 | 0.014285 | 0.014914 | 0.007227 | 0.062528 | 0.145421 | 0.165016 | 0.144696 | 0.155473 | 0.086875 | 0.175646 | 0.110542 | 0.298195 | 0.142488 | 0.06615 | 0.121056 | 0.17794 |
| 29.2777778 | 0.004245 | 0.032198 | 0.002564 | 0.02502 | 0.054416 | 0.008315 | 0.031998 | 0.00896 | 0.007624 | 0.009399 | 0.004056 | 0.035661 | 0.13861 | 0.156444 | 0.220594 | 0.176851 | 0.105722 | 0.224505 | 0.143057 | 0.251389 | 0.185541 | 0.091144 | 0.205784 | 0.207168 |
| 29.2916667 | 0.005761 | 0.028494 | 0.001104 | 0.018578 | 0.0497 | 0.003542 | 0.027771 | 0.009194 | 0.00527 | 0.0073 | 0.006305 | 0.017759 | 0.137349 | 0.139282 | 0.233845 | 0.166191 | 0.119769 | 0.223997 | 0.156746 | 0.225076 | 0.182493 | 0.092074 | 0.224475 | 0.218192 |
| 29.3055556 | 0.004785 | 0.019848 | 0.003301 | 0.012432 | 0.039706 | 0.002737 | 0.020641 | 0.008758 | 0.005416 | 0.005966 | 0.010383 | 0.009474 | 0.121008 | 0.121483 | 0.190607 | 0.142229 | 0.128187 | 0.191441 | 0.150112 | 0.199316 | 0.151719 | 0.080851 | 0.201923 | 0.217921 |
| 29.3194444 | 0.00318 | 0.011351 | 0.011366 | 0.0077 | 0.027577 | 0.005671 | 0.011906 | 0.005327 | 0.010983 | 0.005971 | 0.010522 | 0.010372 | 0.094653 | 0.110254 | 0.123201 | 0.115055 | 0.122687 | 0.147848 | 0.133352 | 0.150471 | 0.118037 | 0.071375 | 0.175668 | 0.208218 |
| 29.3333333 | 0.004409 | 0.00664 | 0.023659 | 0.005533 | 0.020432 | 0.007126 | 0.004975 | 0.006851 | 0.021757 | 0.007604 | 0.01457 | 0.018047 | 0.079713 | 0.092396 | 0.068201 | 0.084473 | 0.100184 | 0.104929 | 0.107838 | 0.105217 | 0.090807 | 0.065767 | 0.140968 | 0.159717 |
| 29.3472222 | 0.008807 | 0.004682 | 0.03475 | 0.006261 | 0.015504 | 0.009348 | 0.002463 | 0.008677 | 0.032532 | 0.010327 | 0.027417 | 0.027171 | 0.069818 | 0.065679 | 0.039557 | 0.05721 | 0.069683 | 0.070909 | 0.07754 | 0.076296 | 0.070981 | 0.061131 | 0.097642 | 0.095671 |
| 29.3611111 | 0.012081 | 0.003229 | 0.039008 | 0.0083 | 0.009302 | 0.016116 | 0.001645 | 0.007536 | 0.036331 | 0.012687 | 0.038056 | 0.029585 | 0.061902 | 0.043542 | 0.029193 | 0.038936 | 0.045295 | 0.050655 | 0.055388 | 0.053873 | 0.05561 | 0.053862 | 0.061232 | 0.056917 |
| 29.375 | 0.011089 | 0.001556 | 0.033677 | 0.00911 | 0.008389 | 0.018546 | 0.00097 | 0.010643 | 0.030802 | 0.012998 | 0.038361 | 0.020566 | 0.057946 | 0.032433 | 0.026202 | 0.029647 | 0.032194 | 0.041575 | 0.044085 | 0.038123 | 0.042783 | 0.042949 | 0.040351 | 0.04027 |
| 29.3888889 | 0.007397 | 0.001599 | 0.024291 | 0.008735 | 0.01412 | 0.014336 | 0.000835 | 0.017078 | 0.020715 | 0.01264 | 0.02625 | 0.010967 | 0.052903 | 0.027861 | 0.02524 | 0.025418 | 0.026572 | 0.036489 | 0.041225 | 0.02962 | 0.033592 | 0.030865 | 0.028484 | 0.031407 |
| 29.4027778 | 0.00361 | 0.004598 | 0.017776 | 0.008594 | 0.018296 | 0.009302 | 0.00052 | 0.017345 | 0.011085 | 0.014134 | 0.013923 | 0.013715 | 0.044736 | 0.025145 | 0.024719 | 0.023185 | 0.024206 | 0.031801 | 0.04375 | 0.025099 | 0.028009 | 0.022671 | 0.020783 | 0.026592 |
| 29.4166667 | 0.001786 | 0.007627 | 0.01501 | 0.007995 | 0.018505 | 0.004879 | 0.000927 | 0.010398 | 0.005062 | 0.016499 | 0.01719 | 0.019066 | 0.038035 | 0.022306 | 0.023143 | 0.02179 | 0.022971 | 0.026926 | 0.043817 | 0.022014 | 0.025162 | 0.018602 | 0.016063 | 0.025135 |
| 29.4305556 | 0.002089 | 0.008873 | 0.014466 | 0.006776 | 0.015492 | 0.002089 | 0.001743 | 0.005971 | 0.007192 | 0.017972 | 0.021726 | 0.014601 | 0.033042 | 0.018962 | 0.019503 | 0.020784 | 0.021588 | 0.021241 | 0.037243 | 0.019139 | 0.023683 | 0.016357 | 0.013792 | 0.025315 |
| 29.4444444 | 0.002325 | 0.010004 | 0.016434 | 0.007359 | 0.009431 | 0.00454 | 0.003192 | 0.008478 | 0.019816 | 0.020097 | 0.017128 | 0.010747 | 0.0275 | 0.016103 | 0.015889 | 0.020894 | 0.019435 | 0.016256 | 0.028634 | 0.015523 | 0.02222 | 0.015271 | 0.013712 | 0.025211 |
| 29.4583333 | 0.003054 | 0.012169 | 0.019926 | 0.010957 | 0.007112 | 0.014353 | 0.007067 | 0.010917 | 0.037021 | 0.024196 | 0.020734 | 0.022981 | 0.022512 | 0.014366 | 0.014298 | 0.023628 | 0.016836 | 0.013612 | 0.02209 | 0.011135 | 0.02064 | 0.014602 | 0.014485 | 0.023207 |
| 29.4722222 | 0.005036 | 0.015503 | 0.02175 | 0.016085 | 0.011272 | 0.026492 | 0.01139 | 0.008546 | 0.0469 | 0.027848 | 0.035058 | 0.039917 | 0.019155 | 0.012479 | 0.013364 | 0.026928 | 0.013917 | 0.011903 | 0.017618 | 0.006559 | 0.019123 | 0.013044 | 0.014341 | 0.018483 |
| 29.4861111 | 0.007148 | 0.020039 | 0.021913 | 0.021631 | 0.016967 | 0.033505 | 0.01517 | 0.004681 | 0.043896 | 0.02836 | 0.041946 | 0.040331 | 0.017124 | 0.009924 | 0.011569 | 0.025479 | 0.010773 | 0.00978 | 0.014647 | 0.002828 | 0.017575 | 0.010429 | 0.012893 | 0.012836 |
| 29.5 | 0.009633 | 0.024906 | 0.023547 | 0.026847 | 0.02362 | 0.032329 | 0.019944 | 0.004026 | 0.033202 | 0.026273 | 0.032979 | 0.024168 | 0.015947 | 0.007643 | 0.00985 | 0.01855 | 0.00817 | 0.007925 | 0.01302 | 0.000892 | 0.01614 | 0.007844 | 0.010771 | 0.008594 |
| 29.5138889 | 0.0131 | 0.028107 | 0.026906 | 0.029303 | 0.029817 | 0.024211 | 0.024086 | 0.005752 | 0.019728 | 0.022873 | 0.019341 | 0.013394 | 0.014856 | 0.005546 | 0.009459 | 0.010861 | 0.006378 | 0.006511 | 0.011852 | 0.000805 | 0.014158 | 0.00579 | 0.008446 | 0.005673 |
| 29.5277778 | 0.025845 | 0.031458 | 0.03226 | 0.029982 | 0.033308 | 0.015563 | 0.024532 | 0.006002 | 0.009533 | 0.018826 | 0.017781 | 0.015245 | 0.013382 | 0.003593 | 0.010259 | 0.005829 | 0.004996 | 0.004674 | 0.009696 | 0.001843 | 0.010924 | 0.003734 | 0.006132 | 0.003293 |
| 29.5416667 | 0.087618 | 0.044134 | 0.048109 | 0.035658 | 0.042189 | 0.021764 | 0.023582 | 0.00394 | 0.016503 | 0.017063 | 0.021054 | 0.023885 | 0.011983 | 0.002734 | 0.010742 | 0.002912 | 0.003244 | 0.003023 | 0.00598 | 0.002842 | 0.006386 | 0.002653 | 0.003619 | 0.001967 |
| 29.5555556 | 0.212127 | 0.070113 | 0.078035 | 0.052185 | 0.062633 | 0.060794 | 0.027712 | 0.004824 | 0.042975 | 0.021022 | 0.021099 | 0.053781 | 0.010357 | 0.002745 | 0.008941 | 0.00149 | 0.002048 | 0.003326 | 0.002929 | 0.004894 | 0.003341 | 0.004749 | 0.002062 | 0.00251 |
| 29.5694444 | 0.327995 | 0.093814 | 0.100338 | 0.078504 | 0.083338 | 0.125833 | 0.034504 | 0.016092 | 0.069157 | 0.02688 | 0.042041 | 0.096243 | 0.007764 | 0.002652 | 0.005109 | 0.001818 | 0.003119 | 0.00417 | 0.002887 | 0.008068 | 0.004348 | 0.009125 | 0.001958 | 0.003481 |
| 29.5833333 | 0.373213 | 0.097774 | 0.099312 | 0.102512 | 0.090426 | 0.179915 | 0.035885 | 0.03417 | 0.079596 | 0.026825 | 0.093193 | 0.128137 | 0.005204 | 0.002247 | 0.001944 | 0.002385 | 0.00491 | 0.002907 | 0.004003 | 0.008672 | 0.005663 | 0.013652 | 0.00162 | 0.002891 |
| 29.5972222 | 0.346093 | 0.083161 | 0.088977 | 0.108018 | 0.084619 | 0.206408 | 0.033222 | 0.046533 | 0.076018 | 0.02203 | 0.139992 | 0.148344 | 0.004018 | 0.001697 | 0.000959 | 0.002457 | 0.005968 | 0.001143 | 0.004631 | 0.006396 | 0.00504 | 0.018217 | 0.001557 | 0.001377 |
| 29.6111111 | 0.268249 | 0.061348 | 0.074578 | 0.091685 | 0.070367 | 0.222153 | 0.031546 | 0.045812 | 0.062827 | 0.017538 | 0.130087 | 0.138935 | 0.003798 | 0.001346 | 0.001096 | 0.002487 | 0.00679 | 0.001069 | 0.004991 | 0.003862 | 0.004025 | 0.022204 | 0.002121 | 0.000438 |
| 29.625 | 0.176604 | 0.042865 | 0.046008 | 0.068852 | 0.046388 | 0.225253 | 0.027867 | 0.033367 | 0.039581 | 0.01321 | 0.075497 | 0.082226 | 0.003201 | 0.001173 | 0.00139 | 0.00259 | 0.007739 | 0.001619 | 0.005794 | 0.002158 | 0.003317 | 0.024017 | 0.002079 | 0.00026 |
| 29.6388889 | 0.099108 | 0.031156 | 0.025143 | 0.053213 | 0.02427 | 0.19739 | 0.018077 | 0.016405 | 0.020379 | 0.008038 | 0.068627 | 0.043357 | 0.001784 | 0.000752 | 0.002158 | 0.003067 | 0.008906 | 0.001489 | 0.008702 | 0.001552 | 0.003272 | 0.023571 | 0.001385 | 0.00044 |
| 29.6527778 | 0.04535 | 0.026286 | 0.033413 | 0.040779 | 0.02422 | 0.151467 | 0.013023 | 0.006766 | 0.023444 | 0.005939 | 0.121015 | 0.070912 | 0.000597 | 0.000249 | 0.003138 | 0.003587 | 0.009903 | 0.000982 | 0.012916 | 0.001541 | 0.003558 | 0.022288 | 0.000684 | 0.000848 |
| 29.6666667 | 0.020578 | 0.029187 | 0.037044 | 0.026137 | 0.037514 | 0.115634 | 0.019562 | 0.013933 | 0.032648 | 0.009408 | 0.145838 | 0.08738 | 0.000275 | 0.000227 | 0.003366 | 0.003568 | 0.010192 | 0.000842 | 0.015591 | 0.001291 | 0.003599 | 0.021252 | 0.00039 | 0.001175 |
| 29.6805556 | 0.016701 | 0.037171 | 0.020965 | 0.01508 | 0.038833 | 0.093149 | 0.022154 | 0.037627 | 0.025077 | 0.012132 | 0.109871 | 0.05325 | 0.000344 | 0.000637 | 0.002918 | 0.003257 | 0.009649 | 0.001176 | 0.016296 | 0.000745 | 0.003541 | 0.021342 | 0.000423 | 0.001976 |
| 29.6944444 | 0.030253 | 0.045339 | 0.011209 | 0.013217 | 0.026084 | 0.071298 | 0.013867 | 0.069739 | 0.01795 | 0.00904 | 0.058512 | 0.033264 | 0.00074 | 0.000684 | 0.002805 | 0.003456 | 0.00884 | 0.001361 | 0.017079 | 0.000508 | 0.004174 | 0.02448 | 0.001292 | 0.005308 |
| 29.7083333 | 0.070435 | 0.048596 | 0.010828 | 0.014225 | 0.012412 | 0.041459 | 0.008454 | 0.100784 | 0.032125 | 0.007227 | 0.051653 | 0.054973 | 0.000979 | 0.000626 | 0.003575 | 0.004666 | 0.008793 | 0.001033 | 0.019848 | 0.000979 | 0.005792 | 0.031957 | 0.003239 | 0.010864 |
| 29.7222222 | 0.116433 | 0.043504 | 0.010765 | 0.011622 | 0.006019 | 0.023467 | 0.012344 | 0.124747 | 0.050542 | 0.016406 | 0.092933 | 0.081844 | 0.000768 | 0.001117 | 0.004482 | 0.006306 | 0.009776 | 0.000658 | 0.024206 | 0.001648 | 0.007166 | 0.040018 | 0.004595 | 0.014073 |
| 29.7361111 | 0.143241 | 0.031357 | 0.019403 | 0.015906 | 0.006506 | 0.038024 | 0.017924 | 0.134604 | 0.053091 | 0.030887 | 0.126623 | 0.093193 | 0.00064 | 0.001206 | 0.004178 | 0.007066 | 0.010739 | 0.000736 | 0.025341 | 0.001476 | 0.007037 | 0.042635 | 0.004205 | 0.012257 |
| 29.75 | 0.16965 | 0.016378 | 0.035858 | 0.030474 | 0.007368 | 0.070575 | 0.021736 | 0.124451 | 0.037834 | 0.041063 | 0.116869 | 0.085073 | 0.000903 | 0.001046 | 0.00268 | 0.006669 | 0.011367 | 0.001381 | 0.023152 | 0.000769 | 0.005983 | 0.040499 | 0.002952 | 0.008291 |
| 29.7638889 | 0.196135 | 0.008072 | 0.039693 | 0.037438 | 0.007052 | 0.110456 | 0.0248 | 0.0946 | 0.018389 | 0.042496 | 0.076277 | 0.053181 | 0.00152 | 0.001448 | 0.0015 | 0.006411 | 0.012508 | 0.001985 | 0.031 | 0.000478 | 0.005373 | 0.038939 | 0.001968 | 0.005549 |
| 29.7777778 | 0.181187 | 0.010666 | 0.024539 | 0.032574 | 0.00701 | 0.153904 | 0.027218 | 0.050984 | 0.0132 | 0.032919 | 0.037002 | 0.026608 | 0.00148 | 0.001292 | 0.001773 | 0.007385 | 0.014531 | 0.00156 | 0.051558 | 0.000597 | 0.00598 | 0.03908 | 0.001678 | 0.004629 |
| 29.7916667 | 0.116927 | 0.01369 | 0.015032 | 0.023751 | 0.006059 | 0.186434 | 0.030303 | 0.028209 | 0.021936 | 0.017983 | 0.033282 | 0.038481 | 0.000987 | 0.000581 | 0.002945 | 0.008863 | 0.016888 | 0.000882 | 0.063042 | 0.001003 | 0.007215 | 0.038392 | 0.001818 | 0.004735 |
| 29.8055556 | 0.054387 | 0.011584 | 0.03502 | 0.014492 | 0.004575 | 0.191416 | 0.030954 | 0.051948 | 0.030332 | 0.008125 | 0.062878 | 0.063064 | 0.000795 | 0.000404 | 0.003359 | 0.009415 | 0.018517 | 0.000954 | 0.053828 | 0.001119 | 0.007386 | 0.035673 | 0.001547 | 0.004779 |
| 29.8194444 | 0.04233 | 0.008027 | 0.072088 | 0.013442 | 0.003806 | 0.164506 | 0.023029 | 0.087162 | 0.028731 | 0.010165 | 0.079488 | 0.067287 | 0.00056 | 0.000437 | 0.002742 | 0.009256 | 0.01976 | 0.000979 | 0.038546 | 0.000722 | 0.00616 | 0.033018 | 0.000839 | 0.004394 |
| 29.8333333 | 0.065653 | 0.00921 | 0.105542 | 0.020378 | 0.00571 | 0.116794 | 0.012518 | 0.103432 | 0.017857 | 0.018906 | 0.064907 | 0.060791 | 0.000785 | 0.000308 | 0.00234 | 0.009657 | 0.022478 | 0.000819 | 0.031371 | 0.000707 | 0.004924 | 0.033285 | 0.000671 | 0.00398 |
| 29.8472222 | 0.082592 | 0.015496 | 0.120505 | 0.019865 | 0.012696 | 0.059922 | 0.011193 | 0.105231 | 0.008278 | 0.024385 | 0.040152 | 0.051068 | 0.001517 | 0.000209 | 0.002472 | 0.010938 | 0.028399 | 0.000845 | 0.031757 | 0.001174 | 0.004434 | 0.036282 | 0.001154 | 0.003454 |
| 29.8611111 | 0.075739 | 0.016804 | 0.108763 | 0.013234 | 0.022263 | 0.034734 | 0.016386 | 0.099749 | 0.006228 | 0.022142 | 0.021809 | 0.034487 | 0.001969 | 0.000409 | 0.002605 | 0.012601 | 0.036887 | 0.000899 | 0.034947 | 0.001533 | 0.004416 | 0.039231 | 0.001706 | 0.002618 |
| 29.875 | 0.045737 | 0.010367 | 0.077458 | 0.018664 | 0.027499 | 0.07744 | 0.021037 | 0.089982 | 0.008663 | 0.01356 | 0.014692 | 0.018852 | 0.001918 | 0.000638 | 0.002422 | 0.013815 | 0.043091 | 0.000822 | 0.039941 | 0.00154 | 0.004522 | 0.038777 | 0.002002 | 0.001786 |
| 29.8888889 | 0.031605 | 0.006219 | 0.045072 | 0.035938 | 0.028318 | 0.137673 | 0.027247 | 0.074655 | 0.017983 | 0.009817 | 0.019288 | 0.016956 | 0.001354 | 0.000603 | 0.001742 | 0.013945 | 0.04386 | 0.000548 | 0.044935 | 0.001109 | 0.005451 | 0.035696 | 0.00186 | 0.001294 |
| 29.9027778 | 0.075225 | 0.011062 | 0.045568 | 0.051822 | 0.030812 | 0.159135 | 0.035364 | 0.057054 | 0.033567 | 0.016844 | 0.035833 | 0.029986 | 0.000783 | 0.000578 | 0.001038 | 0.013687 | 0.042232 | 0.000276 | 0.044585 | 0.000641 | 0.009171 | 0.036284 | 0.001665 | 0.001596 |
| 29.9166667 | 0.148018 | 0.018663 | 0.080969 | 0.062293 | 0.034419 | 0.127018 | 0.038859 | 0.049092 | 0.045638 | 0.022175 | 0.06401 | 0.051072 | 0.000917 | 0.000485 | 0.001051 | 0.013782 | 0.041715 | 0.000257 | 0.039563 | 0.000744 | 0.013655 | 0.040964 | 0.001885 | 0.00267 |
| 29.9305556 | 0.193857 | 0.021 | 0.113063 | 0.069873 | 0.038814 | 0.075565 | 0.034579 | 0.055991 | 0.050774 | 0.0168 | 0.093833 | 0.071094 | 0.00139 | 0.000324 | 0.00142 | 0.013749 | 0.042034 | 0.000505 | 0.040746 | 0.001083 | 0.013542 | 0.04185 | 0.002087 | 0.003171 |
| 29.9444444 | 0.190255 | 0.017439 | 0.126508 | 0.073202 | 0.047343 | 0.081988 | 0.027492 | 0.067196 | 0.052253 | 0.015086 | 0.106877 | 0.075503 | 0.001478 | 0.000307 | 0.001476 | 0.013277 | 0.041294 | 0.000683 | 0.055304 | 0.000995 | 0.009534 | 0.037288 | 0.001887 | 0.002301 |
| 29.9583333 | 0.153687 | 0.011414 | 0.125565 | 0.068118 | 0.052274 | 0.130174 | 0.023143 | 0.070258 | 0.051419 | 0.02867 | 0.100077 | 0.062519 | 0.001378 | 0.000411 | 0.001469 | 0.012514 | 0.038374 | 0.000843 | 0.076678 | 0.000775 | 0.006198 | 0.032984 | 0.00168 | 0.001184 |
| 29.9722222 | 0.112887 | 0.00631 | 0.110384 | 0.057165 | 0.047723 | 0.164042 | 0.022337 | 0.060116 | 0.0454 | 0.043851 | 0.083399 | 0.044942 | 0.00159 | 0.000593 | 0.00186 | 0.012304 | 0.034383 | 0.001197 | 0.089094 | 0.001074 | 0.004912 | 0.032242 | 0.001898 | 0.001054 |
| 29.9861111 | 0.077352 | 0.002935 | 0.089836 | 0.044981 | 0.040104 | 0.183115 | 0.021788 | 0.038605 | 0.030481 | 0.051351 | 0.056913 | 0.025569 | 0.002008 | 0.00063 | 0.002364 | 0.014209 | 0.033362 | 0.001365 | 0.082751 | 0.001776 | 0.00461 | 0.033103 | 0.002412 | 0.002125 |
| 30 | 0.044799 | 0.001079 | 0.072785 | 0.031736 | 0.030076 | 0.175648 | 0.015169 | 0.017371 | 0.01471 | 0.045258 | 0.02989 | 0.016554 | 0.002277 | 0.00038 | 0.002492 | 0.018544 | 0.038904 | 0.00112 | 0.066056 | 0.002479 | 0.004242 | 0.030987 | 0.002802 | 0.004094 |
| 30.0138889 | 0.018506 | 0.0023 | 0.056814 | 0.017522 | 0.015593 | 0.13331 | 0.007984 | 0.008826 | 0.011828 | 0.023271 | 0.025081 | 0.032663 | 0.002632 | 0.000228 | 0.002501 | 0.022848 | 0.050361 | 0.000751 | 0.052051 | 0.003289 | 0.003621 | 0.025101 | 0.00295 | 0.00599 |
| 30.0277778 | 0.006784 | 0.008874 | 0.044694 | 0.009041 | 0.006289 | 0.101497 | 0.008961 | 0.007495 | 0.01689 | 0.007681 | 0.032433 | 0.049314 | 0.003413 | 0.000368 | 0.002178 | 0.022161 | 0.055975 | 0.001159 | 0.041032 | 0.003457 | 0.002714 | 0.017654 | 0.002262 | 0.005728 |
| 30.0416667 | 0.020048 | 0.023718 | 0.048996 | 0.021918 | 0.01647 | 0.124003 | 0.013766 | 0.014212 | 0.018723 | 0.005664 | 0.035299 | 0.048728 | 0.003954 | 0.000513 | 0.002234 | 0.014411 | 0.039833 | 0.004163 | 0.025181 | 0.003387 | 0.004599 | 0.012545 | 0.002379 | 0.003745 |
| 30.0555556 | 0.058064 | 0.049086 | 0.076162 | 0.062348 | 0.047394 | 0.196529 | 0.023148 | 0.045783 | 0.030905 | 0.007904 | 0.072636 | 0.08179 | 0.003327 | 0.00067 | 0.004018 | 0.011804 | 0.024982 | 0.008994 | 0.020225 | 0.005697 | 0.013189 | 0.02511 | 0.005519 | 0.002753 |
| 30.0694444 | 0.090739 | 0.077491 | 0.111019 | 0.105254 | 0.079895 | 0.294484 | 0.033594 | 0.090809 | 0.060966 | 0.015288 | 0.157542 | 0.165383 | 0.00227 | 0.00098 | 0.005282 | 0.025473 | 0.047298 | 0.011454 | 0.047682 | 0.009145 | 0.025176 | 0.06727 | 0.008098 | 0.002382 |
| 30.0833333 | 0.086157 | 0.088716 | 0.128192 | 0.11791 | 0.092469 | 0.37035 | 0.035749 | 0.111262 | 0.074686 | 0.018857 | 0.204731 | 0.201535 | 0.002946 | 0.001708 | 0.004712 | 0.039821 | 0.081817 | 0.010552 | 0.078827 | 0.010795 | 0.03406 | 0.102863 | 0.007473 | 0.002698 |
| 30.0972222 | 0.057381 | 0.072463 | 0.119436 | 0.100583 | 0.082575 | 0.375531 | 0.030215 | 0.093112 | 0.052378 | 0.011685 | 0.160602 | 0.149837 | 0.00526 | 0.003 | 0.004652 | 0.039327 | 0.088944 | 0.008862 | 0.088422 | 0.010921 | 0.035595 | 0.095567 | 0.00616 | 0.005052 |

| | | | | | | | | | | | | | | | | | | | | | | | | |
|---|---|---|---|---|---|---|---|---|---|---|---|---|---|---|---|---|---|---|---|---|---|---|---|---|
| 30.1111111 | 0.035096 | 0.041546 | 0.094207 | 0.073059 | 0.060619 | 0.304642 | 0.019685 | 0.057678 | 0.02251 | 0.004688 | 0.077557 | 0.082373 | 0.006653 | 0.00416 | 0.005713 | 0.032805 | 0.073861 | 0.007543 | 0.092779 | 0.009923 | 0.029562 | 0.070751 | 0.005812 | 0.007756 |
| 30.125 | 0.021416 | 0.022073 | 0.062087 | 0.043245 | 0.035872 | 0.186195 | 0.009757 | 0.027522 | 0.011483 | 0.005158 | 0.034025 | 0.037251 | 0.006343 | 0.00455 | 0.006319 | 0.029917 | 0.058237 | 0.00638 | 0.088377 | 0.007891 | 0.020746 | 0.051939 | 0.005923 | 0.008747 |
| 30.1388889 | 0.009624 | 0.030225 | 0.030742 | 0.021138 | 0.015835 | 0.075822 | 0.004257 | 0.01627 | 0.017087 | 0.007957 | 0.049326 | 0.014291 | 0.005364 | 0.004328 | 0.005892 | 0.027217 | 0.048244 | 0.005354 | 0.065529 | 0.005881 | 0.013886 | 0.036401 | 0.005756 | 0.008097 |
| 30.1527778 | 0.004203 | 0.048343 | 0.010853 | 0.020537 | 0.008588 | 0.027944 | 0.003033 | 0.020944 | 0.026235 | 0.0102 | 0.073928 | 0.015324 | 0.004481 | 0.004111 | 0.005103 | 0.021137 | 0.037894 | 0.004622 | 0.042399 | 0.004615 | 0.010266 | 0.024802 | 0.005124 | 0.00683 |
| 30.1666667 | 0.006265 | 0.058419 | 0.006015 | 0.031849 | 0.013914 | 0.040939 | 0.00617 | 0.027547 | 0.036908 | 0.014532 | 0.084498 | 0.027872 | 0.003871 | 0.003527 | 0.004263 | 0.014654 | 0.026534 | 0.004123 | 0.031001 | 0.003894 | 0.008982 | 0.018469 | 0.004273 | 0.005713 |
| 30.1805556 | 0.009069 | 0.063371 | 0.008497 | 0.035201 | 0.021082 | 0.074462 | 0.009966 | 0.03747 | 0.053051 | 0.020876 | 0.098057 | 0.040118 | 0.002718 | 0.002136 | 0.003034 | 0.009909 | 0.017882 | 0.00352 | 0.02764 | 0.003559 | 0.009094 | 0.017031 | 0.003963 | 0.005627 |
| 30.1944444 | 0.007027 | 0.059583 | 0.011453 | 0.027 | 0.022326 | 0.107451 | 0.008765 | 0.054346 | 0.070834 | 0.024383 | 0.114753 | 0.047293 | 0.006114 | 0.001101 | 0.007367 | 0.005488 | 0.010825 | 0.002343 | 0.028849 | 0.005004 | 0.01227 | 0.02109 | 0.007791 | 0.0116 |
| 30.2083333 | 0.005531 | 0.042341 | 0.01511 | 0.014417 | 0.015388 | 0.140817 | 0.006822 | 0.066433 | 0.080027 | 0.019215 | 0.11863 | 0.044804 | 0.025214 | 0.00195 | 0.027553 | 0.004935 | 0.008823 | 0.002106 | 0.033437 | 0.011671 | 0.02196 | 0.034989 | 0.015251 | 0.030524 |
| 30.2222222 | 0.010519 | 0.020962 | 0.021697 | 0.005935 | 0.005201 | 0.173476 | 0.007985 | 0.06289 | 0.075874 | 0.01027 | 0.099049 | 0.036461 | 0.051308 | 0.00544 | 0.052739 | 0.01308 | 0.023328 | 0.003892 | 0.029619 | 0.02045 | 0.03168 | 0.048138 | 0.018469 | 0.048362 |
| 30.2361111 | 0.011425 | 0.00672 | 0.029025 | 0.007967 | 0.005381 | 0.194071 | 0.010946 | 0.047217 | 0.065923 | 0.011045 | 0.069277 | 0.036964 | 0.055307 | 0.007098 | 0.057016 | 0.021508 | 0.039054 | 0.004365 | 0.023142 | 0.019092 | 0.026395 | 0.037716 | 0.014075 | 0.040651 |
| 30.25 | 0.005823 | 0.00452 | 0.031575 | 0.017203 | 0.017329 | 0.187639 | 0.019574 | 0.025643 | 0.056303 | 0.02106 | 0.043949 | 0.049586 | 0.036433 | 0.004393 | 0.040427 | 0.023767 | 0.036333 | 0.003129 | 0.042927 | 0.009035 | 0.013336 | 0.021838 | 0.007551 | 0.028662 |
| 30.2638889 | 0.003897 | 0.008749 | 0.02898 | 0.027558 | 0.027208 | 0.156285 | 0.027133 | 0.013179 | 0.048317 | 0.025959 | 0.024914 | 0.059058 | 0.028059 | 0.004057 | 0.024068 | 0.027174 | 0.034423 | 0.0048 | 0.070753 | 0.006455 | 0.011839 | 0.029257 | 0.007113 | 0.041385 |
| 30.2777778 | 0.004788 | 0.008049 | 0.024924 | 0.03459 | 0.030237 | 0.119881 | 0.028804 | 0.022473 | 0.042296 | 0.021087 | 0.023518 | 0.059381 | 0.045166 | 0.009157 | 0.016205 | 0.03469 | 0.044364 | 0.010389 | 0.073375 | 0.014237 | 0.018529 | 0.045865 | 0.012482 | 0.059653 |
| 30.2916667 | 0.005738 | 0.00833 | 0.020364 | 0.03382 | 0.031875 | 0.087218 | 0.027943 | 0.037158 | 0.037322 | 0.012014 | 0.033808 | 0.056203 | 0.065666 | 0.017771 | 0.01383 | 0.041589 | 0.055015 | 0.017236 | 0.066938 | 0.021336 | 0.024168 | 0.060011 | 0.016847 | 0.065505 |
| 30.3055556 | 0.006838 | 0.017912 | 0.015088 | 0.026749 | 0.034981 | 0.056638 | 0.02324 | 0.037489 | 0.033059 | 0.00548 | 0.02848 | 0.051227 | 0.064743 | 0.029244 | 0.016365 | 0.043014 | 0.06222 | 0.024423 | 0.06982 | 0.022561 | 0.027432 | 0.073536 | 0.017973 | 0.060151 |
| 30.3194444 | 0.007288 | 0.023127 | 0.009159 | 0.017878 | 0.037337 | 0.030537 | 0.016163 | 0.023112 | 0.027277 | 0.005726 | 0.016867 | 0.040446 | 0.048633 | 0.040386 | 0.023152 | 0.03756 | 0.064442 | 0.030347 | 0.06959 | 0.024269 | 0.028761 | 0.080049 | 0.018141 | 0.04999 |
| 30.3333333 | 0.007557 | 0.015915 | 0.004261 | 0.010714 | 0.034114 | 0.014948 | 0.00894 | 0.013215 | 0.018244 | 0.008454 | 0.024634 | 0.022192 | 0.038147 | 0.046944 | 0.032808 | 0.034075 | 0.064594 | 0.031403 | 0.062864 | 0.032017 | 0.029568 | 0.075063 | 0.020534 | 0.044295 |
| 30.3472222 | 0.008011 | 0.011372 | 0.003291 | 0.007136 | 0.024311 | 0.017272 | 0.003239 | 0.020841 | 0.008551 | 0.008228 | 0.040823 | 0.008333 | 0.038599 | 0.046809 | 0.043252 | 0.040563 | 0.074105 | 0.028864 | 0.062152 | 0.042538 | 0.031728 | 0.062428 | 0.028712 | 0.0438 |
| 30.3611111 | 0.008392 | 0.016959 | 0.005547 | 0.005298 | 0.01265 | 0.032949 | 0.000693 | 0.029679 | 0.003396 | 0.00566 | 0.044939 | 0.007416 | 0.045732 | 0.048496 | 0.052878 | 0.049926 | 0.086088 | 0.027925 | 0.067868 | 0.049397 | 0.039214 | 0.050775 | 0.045433 | 0.046206 |
| 30.375 | 0.008165 | 0.021135 | 0.007482 | 0.003227 | 0.008346 | 0.046901 | 0.001661 | 0.027058 | 0.00242 | 0.003536 | 0.035363 | 0.00981 | 0.056015 | 0.068905 | 0.06577 | 0.050024 | 0.078575 | 0.030752 | 0.073192 | 0.051079 | 0.049825 | 0.044789 | 0.068032 | 0.052994 |
| 30.3888889 | 0.008082 | 0.016677 | 0.00738 | 0.002751 | 0.015026 | 0.04957 | 0.003825 | 0.017447 | 0.003753 | 0.004831 | 0.018713 | 0.008764 | 0.066999 | 0.090035 | 0.082651 | 0.041166 | 0.056217 | 0.036716 | 0.075986 | 0.053589 | 0.054077 | 0.041739 | 0.089145 | 0.062887 |
| 30.4027778 | 0.008811 | 0.008599 | 0.005593 | 0.004201 | 0.021851 | 0.041578 | 0.003374 | 0.00818 | 0.01044 | 0.00945 | 0.007781 | 0.005756 | 0.074436 | 0.079131 | 0.095487 | 0.034097 | 0.037291 | 0.048261 | 0.07341 | 0.061406 | 0.057298 | 0.037146 | 0.099165 | 0.0776 |
| 30.4166667 | 0.008845 | 0.004355 | 0.00302 | 0.004179 | 0.020779 | 0.028459 | 0.002352 | 0.004695 | 0.02002 | 0.014095 | 0.007683 | 0.002904 | 0.073494 | 0.055271 | 0.105439 | 0.030837 | 0.025949 | 0.064969 | 0.065664 | 0.069365 | 0.072879 | 0.030336 | 0.094585 | 0.096772 |
| 30.4305556 | 0.006823 | 0.004036 | 0.001553 | 0.003275 | 0.014019 | 0.015479 | 0.004706 | 0.004157 | 0.027774 | 0.017523 | 0.010374 | 0.002577 | 0.065068 | 0.048115 | 0.115558 | 0.02953 | 0.020321 | 0.075773 | 0.06374 | 0.070304 | 0.096969 | 0.024313 | 0.082652 | 0.101043 |
| 30.4444444 | 0.003825 | 0.006898 | 0.00263 | 0.005783 | 0.006796 | 0.010822 | 0.00901 | 0.004673 | 0.032228 | 0.020562 | 0.019967 | 0.00305 | 0.057135 | 0.0512 | 0.116027 | 0.03493 | 0.018538 | 0.074158 | 0.068504 | 0.06831 | 0.120673 | 0.021896 | 0.070634 | 0.086167 |
| 30.4583333 | 0.002283 | 0.016149 | 0.005006 | 0.011142 | 0.005062 | 0.017514 | 0.012423 | 0.00784 | 0.031297 | 0.023378 | 0.036586 | 0.002503 | 0.060314 | 0.05397 | 0.112205 | 0.046335 | 0.018516 | 0.067306 | 0.069411 | 0.074564 | 0.130552 | 0.023061 | 0.0668 | 0.076366 |
| 30.4722222 | 0.003965 | 0.028087 | 0.006798 | 0.015226 | 0.008567 | 0.026906 | 0.014677 | 0.008242 | 0.025537 | 0.02599 | 0.043692 | 0.002486 | 0.070442 | 0.061177 | 0.116596 | 0.057614 | 0.020038 | 0.062933 | 0.06472 | 0.094665 | 0.112459 | 0.025029 | 0.078783 | 0.088488 |
| 30.4861111 | 0.008582 | 0.03176 | 0.009159 | 0.016725 | 0.012568 | 0.034767 | 0.017612 | 0.004949 | 0.020788 | 0.02889 | 0.034776 | 0.002683 | 0.074358 | 0.071757 | 0.119683 | 0.068454 | 0.024673 | 0.060617 | 0.057399 | 0.119063 | 0.08328 | 0.024498 | 0.0992 | 0.116774 |
| 30.5 | 0.013025 | 0.024019 | 0.013575 | 0.016865 | 0.014811 | 0.043097 | 0.022624 | 0.005191 | 0.022473 | 0.031401 | 0.019204 | 0.004018 | 0.069131 | 0.074861 | 0.111077 | 0.077241 | 0.029589 | 0.060256 | 0.049762 | 0.136189 | 0.071168 | 0.022306 | 0.112851 | 0.144469 |
| 30.5138889 | 0.015591 | 0.013515 | 0.017981 | 0.016193 | 0.015942 | 0.050974 | 0.027507 | 0.011282 | 0.029598 | 0.031816 | 0.013141 | 0.009223 | 0.057494 | 0.072366 | 0.092601 | 0.078105 | 0.030092 | 0.065478 | 0.044384 | 0.152252 | 0.080968 | 0.021646 | 0.11663 | 0.156507 |
| 30.5277778 | 0.019701 | 0.007155 | 0.022719 | 0.017641 | 0.020735 | 0.055933 | 0.029535 | 0.018727 | 0.038227 | 0.030959 | 0.017838 | 0.019108 | 0.050327 | 0.073212 | 0.075445 | 0.078693 | 0.031089 | 0.075921 | 0.044651 | 0.166099 | 0.100625 | 0.023401 | 0.115049 | 0.144224 |
| 30.5416667 | 0.042842 | 0.0054 | 0.036603 | 0.029212 | 0.041577 | 0.063654 | 0.032542 | 0.021493 | 0.049544 | 0.033558 | 0.01795 | 0.036511 | 0.055544 | 0.079206 | 0.07944 | 0.097074 | 0.038822 | 0.089389 | 0.054227 | 0.169129 | 0.121907 | 0.03005 | 0.114574 | 0.129647 |
| 30.5555556 | 0.119185 | 0.014051 | 0.068727 | 0.056109 | 0.078774 | 0.081481 | 0.038683 | 0.016788 | 0.065724 | 0.040579 | 0.020584 | 0.066807 | 0.064846 | 0.09303 | 0.106071 | 0.134722 | 0.047321 | 0.1092 | 0.068098 | 0.170814 | 0.148999 | 0.043393 | 0.121835 | 0.140879 |
| 30.5694444 | 0.229245 | 0.03503 | 0.101419 | 0.086406 | 0.110458 | 0.102749 | 0.044657 | 0.012572 | 0.078846 | 0.045642 | 0.038438 | 0.100892 | 0.070432 | 0.110123 | 0.13262 | 0.16742 | 0.046876 | 0.13383 | 0.0734 | 0.185011 | 0.173782 | 0.056462 | 0.125829 | 0.162266 |
| 30.5833333 | 0.293372 | 0.052551 | 0.105249 | 0.103795 | 0.123374 | 0.116848 | 0.048404 | 0.025476 | 0.082822 | 0.046446 | 0.059192 | 0.115434 | 0.068545 | 0.112355 | 0.154454 | 0.177465 | 0.045094 | 0.146743 | 0.065539 | 0.207051 | 0.188333 | 0.064493 | 0.113938 | 0.165451 |
| 30.5972222 | 0.292303 | 0.056754 | 0.090414 | 0.106275 | 0.116237 | 0.122559 | 0.046402 | 0.053789 | 0.085465 | 0.046383 | 0.086899 | 0.10743 | 0.057014 | 0.096179 | 0.17432 | 0.170404 | 0.05017 | 0.136286 | 0.05241 | 0.210781 | 0.197276 | 0.074365 | 0.091795 | 0.146106 |
| 30.6111111 | 0.251719 | 0.052079 | 0.077902 | 0.090675 | 0.090102 | 0.123036 | 0.038532 | 0.072235 | 0.084913 | 0.044652 | 0.10723 | 0.089027 | 0.042929 | 0.07832 | 0.172939 | 0.149645 | 0.051997 | 0.109847 | 0.042806 | 0.181638 | 0.185587 | 0.084972 | 0.07036 | 0.11724 |
| 30.625 | 0.18222 | 0.04587 | 0.061508 | 0.056152 | 0.056434 | 0.117372 | 0.027478 | 0.063736 | 0.065501 | 0.037355 | 0.07639 | 0.061013 | 0.037468 | 0.073604 | 0.152091 | 0.120157 | 0.048481 | 0.084713 | 0.037117 | 0.139275 | 0.150587 | 0.084885 | 0.058424 | 0.093369 |
| 30.6388889 | 0.119127 | 0.042158 | 0.042462 | 0.023991 | 0.031867 | 0.110308 | 0.014809 | 0.036203 | 0.033814 | 0.023736 | 0.040444 | 0.031924 | 0.043479 | 0.077607 | 0.131108 | 0.095197 | 0.044812 | 0.070659 | 0.03146 | 0.107944 | 0.118262 | 0.075356 | 0.054737 | 0.07727 |
| 30.6527778 | 0.097498 | 0.03441 | 0.034369 | 0.014119 | 0.020797 | 0.109723 | 0.006256 | 0.020731 | 0.018101 | 0.011772 | 0.065739 | 0.022037 | 0.051696 | 0.072832 | 0.10799 | 0.077327 | 0.040283 | 0.060049 | 0.025481 | 0.090921 | 0.098275 | 0.064946 | 0.053888 | 0.064521 |
| 30.6666667 | 0.105809 | 0.022512 | 0.035018 | 0.013801 | 0.014242 | 0.108387 | 0.007487 | 0.032445 | 0.016708 | 0.011506 | 0.113936 | 0.026452 | 0.053126 | 0.057237 | 0.083402 | 0.060465 | 0.035257 | 0.046346 | 0.021319 | 0.081995 | 0.082717 | 0.055798 | 0.054329 | 0.055237 |
| 30.6805556 | 0.113954 | 0.018117 | 0.034166 | 0.011145 | 0.014874 | 0.094709 | 0.016506 | 0.046084 | 0.011108 | 0.01693 | 0.131776 | 0.021973 | 0.049468 | 0.042817 | 0.067554 | 0.047217 | 0.031492 | 0.034109 | 0.019973 | 0.073447 | 0.068319 | 0.047543 | 0.052081 | 0.050504 |
| 30.6944444 | 0.109259 | 0.019082 | 0.030611 | 0.016494 | 0.030467 | 0.071444 | 0.023285 | 0.044644 | 0.009458 | 0.015635 | 0.113935 | 0.01479 | 0.044936 | 0.034052 | 0.060602 | 0.044726 | 0.027974 | 0.026388 | 0.020191 | 0.05922 | 0.056782 | 0.041774 | 0.045606 | 0.048602 |
| 30.7083333 | 0.090365 | 0.017956 | 0.02808 | 0.026907 | 0.046301 | 0.049639 | 0.021852 | 0.031151 | 0.015305 | 0.008216 | 0.071772 | 0.022593 | 0.040652 | 0.031981 | 0.05688 | 0.050074 | 0.023023 | 0.022503 | 0.019881 | 0.041804 | 0.049935 | 0.042007 | 0.039222 | 0.048673 |
| 30.7222222 | 0.056656 | 0.018023 | 0.026072 | 0.028189 | 0.052242 | 0.033401 | 0.017087 | 0.018203 | 0.019158 | 0.004425 | 0.036062 | 0.0321 | 0.037279 | 0.035905 | 0.052464 | 0.055095 | 0.018816 | 0.021058 | 0.01808 | 0.026798 | 0.046809 | 0.048593 | 0.035364 | 0.049408 |
| 30.7361111 | 0.030196 | 0.019972 | 0.02268 | 0.019983 | 0.061185 | 0.020229 | 0.012032 | 0.026984 | 0.015135 | 0.010182 | 0.036305 | 0.029425 | 0.035251 | 0.03898 | 0.045757 | 0.054649 | 0.016626 | 0.019842 | 0.015615 | 0.017444 | 0.043596 | 0.055766 | 0.03233 | 0.048249 |
| 30.75 | 0.037546 | 0.02094 | 0.019483 | 0.014297 | 0.062695 | 0.010354 | 0.007406 | 0.059503 | 0.008034 | 0.021615 | 0.065956 | 0.01943 | 0.034246 | 0.038309 | 0.039345 | 0.047918 | 0.01455 | 0.017959 | 0.014252 | 0.015522 | 0.039487 | 0.057347 | 0.029031 | 0.044714 |
| 30.7638889 | 0.066266 | 0.024437 | 0.018078 | 0.01535 | 0.042452 | 0.011402 | 0.008344 | 0.096573 | 0.006909 | 0.032608 | 0.096065 | 0.00975 | 0.033289 | 0.035279 | 0.036463 | 0.03916 | 0.014016 | 0.016812 | 0.014814 | 0.019046 | 0.036132 | 0.052495 | 0.026193 | 0.03993 |
| 30.7777778 | 0.104434 | 0.030498 | 0.016555 | 0.015216 | 0.035022 | 0.030979 | 0.016646 | 0.119469 | 0.007761 | 0.03966 | 0.090267 | 0.005311 | 0.031064 | 0.030636 | 0.034552 | 0.032418 | 0.01749 | 0.017602 | 0.016599 | 0.022051 | 0.032381 | 0.043535 | 0.023529 | 0.034257 |
| 30.7916667 | 0.159833 | 0.033297 | 0.011364 | 0.011444 | 0.060625 | 0.057629 | 0.027685 | 0.12219 | 0.011693 | 0.042031 | 0.048825 | 0.005318 | 0.026809 | 0.025129 | 0.029194 | 0.027179 | 0.021873 | 0.019017 | 0.017775 | 0.022183 | 0.026825 | 0.032511 | 0.019911 | 0.028352 |
| 30.8055556 | 0.21053 | 0.032599 | 0.006669 | 0.008953 | 0.08059 | 0.075861 | 0.035379 | 0.10077 | 0.023368 | 0.040594 | 0.024861 | 0.00532 | 0.021409 | 0.019687 | 0.020911 | 0.021964 | 0.023365 | 0.01923 | 0.017669 | 0.020738 | 0.021227 | 0.023726 | 0.015739 | 0.023558 |
| 30.8194444 | 0.221502 | 0.02724 | 0.009465 | 0.009442 | 0.076531 | 0.087903 | 0.038045 | 0.05837 | 0.034772 | 0.035101 | 0.049558 | 0.004544 | 0.016577 | 0.015008 | 0.013664 | 0.01668 | 0.02168 | 0.018762 | 0.018162 | 0.0183 | 0.017922 | 0.022521 | 0.011436 | 0.020035 |
| 30.8333333 | 0.19533 | 0.017028 | 0.015583 | 0.010017 | 0.065362 | 0.095119 | 0.039122 | 0.039262 | 0.035153 | 0.025493 | 0.085747 | 0.005291 | 0.013481 | 0.01089 | 0.009654 | 0.01134 | 0.017742 | 0.017774 | 0.01937 | 0.014541 | 0.016857 | 0.028221 | 0.006243 | 0.017073 |
| 30.8472222 | 0.150963 | 0.012113 | 0.019717 | 0.00806 | 0.047118 | 0.095515 | 0.03507 | 0.06845 | 0.022351 | 0.013855 | 0.098151 | 0.005753 | 0.011898 | 0.00617 | 0.007975 | 0.005972 | 0.01185 | 0.015058 | 0.018148 | 0.010547 | 0.015661 | 0.032608 | 0.002471 | 0.014571 |
| 30.8611111 | 0.098983 | 0.013571 | 0.022182 | 0.006822 | 0.024635 | 0.086921 | 0.021497 | 0.106357 | 0.014053 | 0.010101 | 0.095847 | 0.009659 | 0.009574 | 0.004448 | 0.006589 | 0.003863 | 0.006339 | 0.012247 | 0.013614 | 0.013767 | 0.014698 | 0.029994 | 0.002972 | 0.013294 |
| 30.875 | 0.053537 | 0.010969 | 0.0226 | 0.011915 | 0.011192 | 0.07179 | 0.014591 | 0.129813 | 0.025235 | 0.021754 | 0.08749 | 0.023181 | 0.005718 | 0.010175 | 0.004653 | 0.007406 | 0.005588 | 0.011637 | 0.008465 | 0.024256 | 0.015754 | 0.023358 | 0.004834 | 0.013756 |
| 30.8888889 | 0.027551 | 0.00852 | 0.020854 | 0.018509 | 0.011917 | 0.062523 | 0.023961 | 0.140662 | 0.041549 | 0.039852 | 0.076161 | 0.04347 | 0.004382 | 0.019106 | 0.002681 | 0.012706 | 0.00802 | 0.011081 | 0.004413 | 0.026904 | 0.017107 | 0.015662 | 0.004631 | 0.014604 |
| 30.9027778 | 0.023232 | 0.0098 | 0.018402 | 0.020891 | 0.021283 | 0.070525 | 0.035087 | 0.132952 | 0.051039 | 0.054286 | 0.065532 | 0.06036 | 0.006969 | 0.025788 | 0.002982 | 0.016191 | 0.008806 | 0.007675 | 0.001873 | 0.016347 | 0.017057 | 0.008328 | 0.002888 | 0.013907 |
| 30.9166667 | 0.03248 | 0.008825 | 0.018254 | 0.024021 | 0.036096 | 0.095868 | 0.040363 | 0.108551 | 0.05708 | 0.064478 | 0.057424 | 0.064948 | 0.008006 | 0.026966 | 0.007957 | 0.016878 | 0.008533 | 0.003429 | 0.00167 | 0.008019 | 0.015617 | 0.006297 | 0.002035 | 0.010905 |
| 30.9305556 | 0.043827 | 0.010173 | 0.022333 | 0.031489 | 0.06404 | 0.118056 | 0.042268 | 0.079189 | 0.063171 | 0.068659 | 0.054261 | 0.05782 | 0.00554 | 0.021561 | 0.016103 | 0.016541 | 0.008688 | 0.00152 | 0.002672 | 0.009807 | 0.013108 | 0.010873 | 0.001989 | 0.006226 |
| 30.9444444 | 0.050194 | 0.022018 | 0.027652 | 0.037967 | 0.094703 | 0.120731 | 0.042205 | 0.056043 | 0.071048 | 0.060674 | 0.053757 | 0.04368 | 0.004688 | 0.013077 | 0.024617 | 0.017959 | 0.007923 | 0.002445 | 0.0025 | 0.009953 | 0.009057 | 0.015932 | 0.003627 | 0.004512 |
| 30.9583333 | 0.046405 | 0.04087 | 0.02675 | 0.038845 | 0.109348 | 0.101912 | 0.036307 | 0.043238 | 0.076737 | 0.043234 | 0.050061 | 0.026134 | 0.006516 | 0.005668 | 0.033291 | 0.018634 | 0.006119 | 0.005348 | 0.00122 | 0.010207 | 0.004664 | 0.018096 | 0.010399 | 0.009646 |
| 30.9722222 | 0.02905 | 0.046466 | 0.016404 | 0.034797 | 0.103065 | 0.060483 | 0.021973 | 0.037753 | 0.073825 | 0.025271 | 0.038137 | 0.012136 | 0.006658 | 0.001961 | 0.042518 | 0.016241 | 0.00551 | 0.008413 | 0.001365 | 0.020917 | 0.003806 | 0.019449 | 0.019441 | 0.018062 |
| 30.9861111 | 0.019579 | 0.033459 | 0.009178 | 0.025939 | 0.076336 | 0.032078 | 0.01052 | 0.031996 | 0.061276 | 0.013229 | 0.021029 | 0.011099 | 0.005008 | 0.001379 | 0.048294 | 0.013775 | 0.007147 | 0.010407 | 0.004482 | 0.038571 | 0.00751 | 0.021824 | 0.026154 | 0.024296 |
| 31 | 0.040328 | 0.030953 | 0.015148 | 0.014678 | 0.04439 | 0.05277 | 0.012541 | 0.021056 | 0.042258 | 0.013578 | 0.012831 | 0.021096 | 0.003191 | 0.001981 | 0.041979 | 0.01467 | 0.007201 | 0.009931 | 0.007608 | 0.054515 | 0.011817 | 0.02483 | 0.030314 | 0.025513 |
| 31.0138889 | 0.068296 | 0.049211 | 0.023583 | 0.005895 | 0.02071 | 0.088363 | 0.020954 | 0.011071 | 0.021336 | 0.02243 | 0.017809 | 0.029299 | 0.002626 | 0.003884 | 0.024535 | 0.016029 | 0.003672 | 0.006761 | 0.00806 | 0.055989 | 0.011923 | 0.027773 | 0.031278 | 0.021656 |
| 31.0277778 | 0.067821 | 0.062406 | 0.021707 | 0.002745 | 0.006052 | 0.095223 | 0.022112 | 0.009414 | 0.006144 | 0.027253 | 0.019665 | 0.025131 | 0.005824 | 0.006053 | 0.014809 | 0.01443 | 0.002628 | 0.005246 | 0.008071 | 0.039216 | 0.009118 | 0.031814 | 0.028099 | 0.016836 |
| 31.0416667 | 0.057426 | 0.046892 | 0.021785 | 0.013497 | 0.007275 | 0.073772 | 0.0182 | 0.013482 | 0.007776 | 0.021234 | 0.022661 | 0.024818 | 0.011608 | 0.009262 | 0.01607 | 0.018822 | 0.010856 | 0.008134 | 0.011289 | 0.019187 | 0.012415 | 0.041339 | 0.026167 | 0.019052 |
| 31.0555556 | 0.100461 | 0.037032 | 0.05253 | 0.046499 | 0.029863 | 0.094385 | 0.025773 | 0.032964 | 0.032591 | 0.012167 | 0.062449 | 0.064052 | 0.018595 | 0.020185 | 0.016085 | 0.04179 | 0.029734 | 0.014483 | 0.0223 | 0.009438 | 0.032108 | 0.065868 | 0.031902 | 0.031516 |
| 31.0694444 | 0.180084 | 0.07545 | 0.099423 | 0.081668 | 0.05579 | 0.181914 | 0.036119 | 0.072819 | 0.065211 | 0.011399 | 0.133177 | 0.123712 | 0.026616 | 0.03827 | 0.021178 | 0.073283 | 0.046995 | 0.020787 | 0.037878 | 0.009636 | 0.065029 | 0.097402 | 0.040611 | 0.042209 |
| 31.0833333 | 0.199817 | 0.109037 | 0.119239 | 0.086942 | 0.058362 | 0.225165 | 0.029481 | 0.104184 | 0.076926 | 0.012164 | 0.170743 | 0.140836 | 0.025263 | 0.040772 | 0.022597 | 0.076352 | 0.040244 | 0.018008 | 0.038559 | 0.02311 | 0.07554 | 0.091631 | 0.038945 | 0.033584 |
| 31.0972222 | 0.149688 | 0.092729 | 0.112462 | 0.065613 | 0.03826 | 0.206142 | 0.02183 | 0.110959 | 0.060797 | 0.007078 | 0.155661 | 0.111025 | 0.017827 | 0.028034 | 0.02786 | 0.046565 | 0.02541 | 0.01163 | 0.027173 | 0.047936 | 0.055208 | 0.053312 | 0.028992 | 0.020325 |
| 31.1111111 | 0.097758 | 0.055445 | 0.097468 | 0.036204 | 0.017574 | 0.196936 | 0.019222 | 0.10291 | 0.035537 | 0.003175 | 0.129814 | 0.078248 | 0.023071 | 0.028749 | 0.063069 | 0.028854 | 0.034439 | 0.01572 | 0.030368 | 0.058942 | 0.051475 | 0.041626 | 0.030851 | 0.025799 |
| 31.125 | 0.06443 | 0.026535 | 0.070736 | 0.017548 | 0.012811 | 0.17039 | 0.010046 | 0.085375 | 0.018887 | 0.005086 | 0.10082 | 0.056891 | 0.029785 | 0.031439 | 0.088967 | 0.036728 | 0.048934 | 0.021344 | 0.041613 | 0.044732 | 0.06304 | 0.051293 | 0.043317 | 0.031607 |
| 31.1388889 | 0.035993 | 0.020976 | 0.034966 | 0.021562 | 0.020223 | 0.100813 | 0.00326 | 0.062148 | 0.014425 | 0.008596 | 0.065491 | 0.037442 | 0.025072 | 0.020115 | 0.079515 | 0.047634 | 0.049576 | 0.018187 | 0.042519 | 0.024333 | 0.056382 | 0.05323 | 0.049219 | 0.025471 |
| 31.1527778 | 0.015517 | 0.036114 | 0.014594 | 0.029564 | 0.027616 | 0.038474 | 0.001944 | 0.040309 | 0.014432 | 0.009833 | 0.042172 | 0.023132 | 0.020276 | 0.009846 | 0.062529 | 0.05553 | 0.044681 | 0.011106 | 0.036421 | 0.017788 | 0.041133 | 0.051942 | 0.046559 | 0.022269 |
| 31.1666667 | 0.012358 | 0.056247 | 0.018279 | 0.031424 | 0.032649 | 0.019298 | 0.00162 | 0.022633 | 0.011711 | 0.00906 | 0.030506 | 0.034023 | 0.022453 | 0.008688 | 0.046595 | 0.060903 | 0.03844 | 0.005542 | 0.02878 | 0.027754 | 0.031656 | 0.048218 | 0.040806 | 0.026728 |
| 31.1805556 | 0.022861 | 0.072914 | 0.025103 | 0.034495 | 0.037155 | 0.028592 | 0.003149 | 0.013954 | 0.01428 | 0.007379 | 0.039147 | 0.068917 | 0.027774 | 0.012972 | 0.028223 | 0.060234 | 0.030999 | 0.004144 | 0.021347 | 0.043637 | 0.026344 | 0.042178 | 0.037323 | 0.03423 |
| 31.1944444 | 0.035515 | 0.080475 | 0.02473 | 0.037465 | 0.040162 | 0.04038 | 0.006181 | 0.02275 | 0.030226 | 0.006398 | 0.070105 | 0.095986 | 0.029444 | 0.018597 | 0.015291 | 0.056588 | 0.023003 | 0.005926 | 0.015336 | 0.05296 | 0.024096 | 0.038819 | 0.036522 | 0.042389 |
| 31.2083333 | 0.045371 | 0.075811 | 0.022236 | 0.037098 | 0.040712 | 0.047306 | 0.010643 | 0.042344 | 0.052464 | 0.00837 | 0.096741 | 0.103592 | 0.02903 | 0.024661 | 0.0158 | 0.053164 | 0.015632 | 0.00896 | 0.011851 | 0.052771 | 0.027802 | 0.037843 | 0.033537 | 0.050955 |
| 31.2222222 | 0.051679 | 0.072262 | 0.020136 | 0.035524 | 0.040412 | 0.052049 | 0.016888 | 0.058101 | 0.074855 | 0.014262 | 0.109043 | 0.100581 | 0.034777 | 0.032071 | 0.027072 | 0.051622 | 0.011256 | 0.014454 | 0.012233 | 0.048409 | 0.037306 | 0.03593 | 0.028128 | 0.057627 |
| 31.2361111 | 0.052879 | 0.082941 | 0.018219 | 0.03721 | 0.041856 | 0.05752 | 0.024998 | 0.06453 | 0.095262 | 0.021233 | 0.103455 | 0.092577 | 0.043449 | 0.037571 | 0.035355 | 0.052537 | 0.0099 | 0.020699 | 0.015691 | 0.037975 | 0.04724 | 0.033745 | 0.020482 | 0.055887 |
| 31.25 | 0.0456 | 0.088625 | 0.012846 | 0.041744 | 0.044122 | 0.064677 | 0.033389 | 0.063622 | 0.111196 | 0.022236 | 0.083258 | 0.082119 | 0.043176 | 0.035482 | 0.027754 | 0.049758 | 0.007214 | 0.021542 | 0.0188 | 0.021141 | 0.051746 | 0.034897 | 0.011872 | 0.041722 |
| 31.2638889 | 0.033651 | 0.069359 | 0.005726 | 0.043119 | 0.045116 | 0.068533 | 0.039836 | 0.061424 | 0.118117 | 0.019982 | 0.083094 | 0.068224 | 0.033617 | 0.027323 | 0.013775 | 0.040505 | 0.003238 | 0.016179 | 0.019282 | 0.012502 | 0.048773 | 0.037486 | 0.010731 | 0.020753 |
| 31.2777778 | 0.025432 | 0.044082 | 0.003003 | 0.037897 | 0.045457 | 0.064061 | 0.043066 | 0.063368 | 0.102482 | 0.025172 | 0.118156 | 0.051188 | 0.024018 | 0.017354 | 0.012871 | 0.028066 | 0.001298 | 0.010517 | 0.017278 | 0.015945 | 0.041534 | 0.036402 | 0.018767 | 0.00685 |
| 31.2916667 | 0.020555 | 0.029493 | 0.002907 | 0.029831 | 0.045727 | 0.055121 | 0.042342 | 0.066055 | 0.066953 | 0.031587 | 0.148324 | 0.032405 | 0.017848 | 0.008359 | 0.024137 | 0.015342 | 0.00118 | 0.007756 | 0.012569 | 0.017315 | 0.032067 | 0.031903 | 0.024419 | 0.004849 |
| 31.3055556 | 0.015941 | 0.017035 | 0.002217 | 0.022529 | 0.045683 | 0.050483 | 0.038173 | 0.061398 | 0.036899 | 0.029232 | 0.137761 | 0.015633 | 0.012996 | 0.004738 | 0.036588 | 0.009827 | 0.001356 | 0.005884 | 0.006431 | 0.011526 | 0.02314 | 0.027329 | 0.020584 | 0.006302 |
| 31.3194444 | 0.011307 | 0.011123 | 0.003839 | 0.015438 | 0.044887 | 0.052392 | 0.032618 | 0.047889 | 0.020478 | 0.021851 | 0.092428 | 0.009652 | 0.007791 | 0.005761 | 0.042524 | 0.016039 | 0.002213 | 0.004637 | 0.003146 | 0.005273 | 0.017052 | 0.024001 | 0.011861 | 0.010624 |
| 31.3333333 | 0.007113 | 0.023463 | 0.009781 | 0.008003 | 0.042063 | 0.054799 | 0.027642 | 0.028379 | 0.009997 | 0.013039 | 0.052033 | 0.015804 | 0.004018 | 0.006456 | 0.037563 | 0.025585 | 0.005449 | 0.00476 | 0.003033 | 0.00334 | 0.010113 | 0.020813 | 0.005408 | 0.018174 |
| 31.3472222 | 0.006558 | 0.038035 | 0.017149 | 0.00271 | 0.036481 | 0.050823 | 0.023063 | 0.01303 | 0.005411 | 0.006818 | 0.059263 | 0.025331 | 0.005014 | 0.004958 | 0.024162 | 0.030805 | 0.009321 | 0.004891 | 0.002682 | 0.003268 | 0.006233 | 0.016539 | 0.003684 | 0.020651 |
| 31.3611111 | 0.009272 | 0.035603 | 0.022829 | 0.001395 | 0.028815 | 0.039401 | 0.018665 | 0.011767 | 0.006416 | 0.00746 | 0.09543 | 0.030135 | 0.008725 | 0.00249 | 0.012116 | 0.032668 | 0.011295 | 0.004536 | 0.00274 | 0.003355 | 0.012444 | 0.011461 | 0.004663 | 0.013918 |
| 31.375 | 0.00842 | 0.021725 | 0.023345 | 0.003453 | 0.020857 | 0.024855 | 0.016115 | 0.018378 | 0.005797 | 0.009386 | 0.112118 | 0.029957 | 0.010977 | 0.001266 | 0.011675 | 0.03503 | 0.011419 | 0.003814 | 0.004293 | 0.004259 | 0.022331 | 0.00636 | 0.006542 | 0.005341 |
| 31.3888889 | 0.004643 | 0.00999 | 0.015821 | 0.008169 | 0.014529 | 0.011888 | 0.015003 | 0.021594 | 0.004348 | 0.008044 | 0.104166 | 0.030115 | 0.01001 | 0.002486 | 0.018664 | 0.03839 | 0.009874 | 0.00281 | 0.005056 | 0.005042 | 0.027311 | 0.002495 | 0.007133 | 0.003213 |
| 31.4027778 | 0.004777 | 0.006865 | 0.006819 | 0.013302 | 0.009411 | 0.004749 | 0.012813 | 0.018089 | 0.00677 | 0.005756 | 0.090386 | 0.028106 | 0.005843 | 0.007416 | 0.022476 | 0.039434 | 0.007782 | 0.001824 | 0.003582 | 0.011229 | 0.025707 | 0.001107 | 0.004858 | 0.003832 |
| 31.4166667 | 0.008609 | 0.010132 | 0.003841 | 0.015872 | 0.004494 | 0.005418 | 0.008775 | 0.010611 | 0.011103 | 0.003623 | 0.073645 | 0.01945 | 0.002502 | 0.016278 | 0.020182 | 0.035954 | 0.007043 | 0.001857 | 0.002031 | 0.02757 | 0.019783 | 0.001508 | 0.005745 | 0.002452 |
| 31.4305556 | 0.01132 | 0.012574 | 0.003152 | 0.015128 | 0.001624 | 0.010006 | 0.004594 | 0.004925 | 0.01707 | 0.002231 | 0.045457 | 0.019026 | 0.003052 | 0.023667 | 0.013984 | 0.02603 | 0.008889 | 0.002222 | 0.002527 | 0.048994 | 0.012798 | 0.002576 | 0.016424 | 0.003731 |
| 31.4444444 | 0.010334 | 0.012323 | 0.001708 | 0.013062 | 0.001566 | 0.013138 | 0.002543 | 0.004011 | 0.025464 | 0.002955 | 0.023197 | 0.039763 | 0.004607 | 0.022113 | 0.009347 | 0.015957 | 0.014562 | 0.003026 | 0.002604 | 0.066658 | 0.007535 | 0.003718 | 0.031151 | 0.015238 |
| 31.4583333 | 0.007637 | 0.013535 | 0.001483 | 0.012047 | 0.001891 | 0.012099 | 0.002821 | 0.004076 | 0.033059 | 0.005331 | 0.027852 | 0.061968 | 0.005722 | 0.013415 | 0.01129 | 0.021439 | 0.024457 | 0.007255 | 0.002971 | 0.071152 | 0.008245 | 0.003628 | 0.038669 | 0.03348 |
| 31.4722222 | 0.00823 | 0.018745 | 0.001976 | 0.013933 | 0.003144 | 0.007147 | 0.004213 | 0.002876 | 0.03676 | 0.007822 | 0.041534 | 0.066776 | 0.008305 | 0.012851 | 0.017813 | 0.042839 | 0.035148 | 0.01259 | 0.008025 | 0.05936 | 0.012835 | 0.002239 | 0.034882 | 0.044169 |
| 31.4861111 | 0.014364 | 0.028438 | 0.002878 | 0.020211 | 0.006853 | 0.003665 | 0.00714 | 0.002194 | 0.039467 | 0.009977 | 0.052303 | 0.059842 | 0.012524 | 0.029177 | 0.022197 | 0.06199 | 0.040387 | 0.014545 | 0.014939 | 0.036985 | 0.013055 | 0.00121 | 0.02413 | 0.038777 |
| 31.5 | 0.022979 | 0.040586 | 0.005336 | 0.028665 | 0.011474 | 0.005856 | 0.011271 | 0.002177 | 0.042792 | 0.011905 | 0.064475 | 0.04882 | 0.01659 | 0.048779 | 0.020448 | 0.065967 | 0.037228 | 0.012724 | 0.018237 | 0.017767 | 0.009525 | 0.001345 | 0.013032 | 0.025209 |
| 31.5138889 | 0.029668 | 0.050664 | 0.008283 | 0.035248 | 0.016488 | 0.011089 | 0.013669 | 0.002315 | 0.04271 | 0.013968 | 0.070884 | 0.034718 | 0.019563 | 0.05698 | 0.01379 | 0.058005 | 0.028389 | 0.008614 | 0.017904 | 0.015508 | 0.009109 | 0.004648 | 0.009861 | 0.016604 |
| 31.5277778 | 0.032908 | 0.05741 | 0.010363 | 0.040934 | 0.023309 | 0.016424 | 0.014443 | 0.003502 | 0.040083 | 0.01804 | 0.070305 | 0.022652 | 0.021951 | 0.05171 | 0.007129 | 0.047982 | 0.01888 | 0.005301 | 0.018507 | 0.02391 | 0.01685 | 0.015354 | 0.014058 | 0.016733 |
| 31.5416667 | 0.040474 | 0.065139 | 0.017197 | 0.050277 | 0.034285 | 0.020443 | 0.016603 | 0.006407 | 0.041083 | 0.02759 | 0.067069 | 0.025031 | 0.02601 | 0.038953 | 0.003819 | 0.043934 | 0.014872 | 0.008185 | 0.027175 | 0.026054 | 0.04298 | 0.042544 | 0.017483 | 0.027939 |
| 31.5555556 | 0.080283 | 0.074466 | 0.040839 | 0.061219 | 0.046469 | 0.021479 | 0.019991 | 0.012297 | 0.048835 | 0.041792 | 0.061485 | 0.048858 | 0.033978 | 0.032826 | 0.007193 | 0.049692 | 0.021632 | 0.019642 | 0.045563 | 0.016655 | 0.091921 | 0.087574 | 0.03617 | 0.046588 |
| 31.5694444 | 0.157978 | 0.077307 | 0.076915 | 0.06554 | 0.051897 | 0.020864 | 0.021934 | 0.02402 | 0.05977 | 0.053509 | 0.063178 | 0.083274 | 0.040965 | 0.040297 | 0.023127 | 0.058746 | 0.038514 | 0.03252 | 0.060223 | 0.011953 | 0.131574 | 0.114835 | 0.077229 | 0.061514 |
| 31.5833333 | 0.228151 | 0.071093 | 0.096571 | 0.060598 | 0.051673 | 0.026709 | 0.02227 | 0.043806 | 0.067063 | 0.057052 | 0.089815 | 0.109501 | 0.040274 | 0.052019 | 0.04706 | 0.062379 | 0.059484 | 0.037542 | 0.059588 | 0.024383 | 0.132458 | 0.097557 | 0.107629 | 0.069898 |
| 31.5972222 | 0.252251 | 0.060303 | 0.092931 | 0.050743 | 0.057942 | 0.04561 | 0.023123 | 0.068361 | 0.065992 | 0.049798 | 0.146349 | 0.122019 | 0.033731 | 0.05632 | 0.062139 | 0.061674 | 0.077815 | 0.034535 | 0.050833 | 0.034834 | 0.116685 | 0.0666 | 0.114531 | 0.072504 |
| 31.6111111 | 0.222975 | 0.047916 | 0.083653 | 0.041285 | 0.071224 | 0.067627 | 0.025634 | 0.081281 | 0.052717 | 0.033729 | 0.181767 | 0.115767 | 0.025415 | 0.045782 | 0.05819 | 0.056757 | 0.082049 | 0.030345 | 0.040653 | 0.030429 | 0.102962 | 0.049919 | 0.111146 | 0.066706 |

| | | | | | | | | | | | | | | | | | | | | | | | | |
|---|---|---|---|---|---|---|---|---|---|---|---|---|---|---|---|---|---|---|---|---|---|---|---|---|
| 31.625 | 0.163708 | 0.030738 | 0.069079 | 0.030223 | 0.072733 | 0.07411 | 0.030408 | 0.066594 | 0.028978 | 0.017606 | 0.137539 | 0.081898 | 0.015471 | 0.024527 | 0.037083 | 0.038246 | 0.057501 | 0.027731 | 0.027165 | 0.018452 | 0.080451 | 0.03973 | 0.095117 | 0.052451 |
| 31.6388889 | 0.118598 | 0.021411 | 0.045462 | 0.016844 | 0.059736 | 0.059068 | 0.037172 | 0.036712 | 0.01261 | 0.006958 | 0.070656 | 0.040296 | 0.007543 | 0.014826 | 0.021709 | 0.020921 | 0.030659 | 0.022143 | 0.013348 | 0.011885 | 0.04736 | 0.025393 | 0.065149 | 0.034144 |
| 31.6527778 | 0.111716 | 0.037728 | 0.022158 | 0.011428 | 0.048641 | 0.031848 | 0.041277 | 0.022249 | 0.015093 | 0.003105 | 0.076624 | 0.025432 | 0.006996 | 0.02524 | 0.036654 | 0.030448 | 0.041282 | 0.01222 | 0.008426 | 0.024189 | 0.021509 | 0.012666 | 0.034541 | 0.018632 |
| 31.6666667 | 0.135952 | 0.049957 | 0.008464 | 0.014358 | 0.042739 | 0.00995 | 0.037932 | 0.031385 | 0.017702 | 0.005981 | 0.146055 | 0.028114 | 0.010678 | 0.034645 | 0.067942 | 0.04911 | 0.061188 | 0.005439 | 0.013226 | 0.04608 | 0.018213 | 0.011763 | 0.021996 | 0.016807 |
| 31.6805556 | 0.163205 | 0.033507 | 0.007188 | 0.013785 | 0.037734 | 0.005241 | 0.031106 | 0.046016 | 0.012751 | 0.01024 | 0.194336 | 0.024138 | 0.013262 | 0.03117 | 0.089381 | 0.05855 | 0.057345 | 0.006702 | 0.017957 | 0.055465 | 0.026233 | 0.015911 | 0.030515 | 0.030766 |
| 31.6944444 | 0.171382 | 0.021618 | 0.01708 | 0.008924 | 0.034182 | 0.013295 | 0.026468 | 0.048862 | 0.016512 | 0.010941 | 0.174311 | 0.02873 | 0.013386 | 0.020898 | 0.090874 | 0.064783 | 0.041163 | 0.010503 | 0.018728 | 0.045796 | 0.029248 | 0.017139 | 0.039944 | 0.042343 |
| 31.7083333 | 0.165964 | 0.042821 | 0.030189 | 0.010809 | 0.03136 | 0.02619 | 0.022235 | 0.037526 | 0.029133 | 0.008422 | 0.118903 | 0.044698 | 0.012114 | 0.009804 | 0.071717 | 0.064408 | 0.022401 | 0.010642 | 0.015793 | 0.025594 | 0.029676 | 0.019672 | 0.041402 | 0.037454 |
| 31.7222222 | 0.156867 | 0.071616 | 0.035153 | 0.020234 | 0.026541 | 0.037855 | 0.014223 | 0.021552 | 0.038134 | 0.005317 | 0.068355 | 0.046142 | 0.012119 | 0.004941 | 0.03893 | 0.056568 | 0.014124 | 0.006733 | 0.008857 | 0.00962 | 0.034058 | 0.024272 | 0.036272 | 0.022119 |
| 31.7361111 | 0.139347 | 0.077701 | 0.027911 | 0.02414 | 0.016838 | 0.035478 | 0.009062 | 0.012527 | 0.033939 | 0.003299 | 0.038421 | 0.029207 | 0.013148 | 0.007254 | 0.019798 | 0.041621 | 0.025493 | 0.00529 | 0.004535 | 0.005058 | 0.041677 | 0.028218 | 0.022256 | 0.008816 |
| 31.75 | 0.105366 | 0.0607 | 0.014347 | 0.015699 | 0.012824 | 0.024095 | 0.011921 | 0.020181 | 0.019746 | 0.003042 | 0.041903 | 0.021482 | 0.012527 | 0.00827 | 0.031512 | 0.020754 | 0.041515 | 0.010303 | 0.008596 | 0.004799 | 0.050385 | 0.029755 | 0.010843 | 0.004228 |
| 31.7638889 | 0.05878 | 0.033992 | 0.006077 | 0.006055 | 0.02359 | 0.031806 | 0.012974 | 0.041038 | 0.014796 | 0.004709 | 0.065982 | 0.036573 | 0.009451 | 0.006687 | 0.051884 | 0.010709 | 0.048307 | 0.016191 | 0.015281 | 0.004463 | 0.058064 | 0.026157 | 0.012806 | 0.007717 |
| 31.7777778 | 0.026733 | 0.019048 | 0.006727 | 0.004896 | 0.02934 | 0.056416 | 0.008567 | 0.058268 | 0.021756 | 0.007824 | 0.087207 | 0.054445 | 0.005667 | 0.008185 | 0.061564 | 0.015526 | 0.04241 | 0.017222 | 0.017143 | 0.008049 | 0.058447 | 0.017089 | 0.017569 | 0.016426 |
| 31.7916667 | 0.03218 | 0.019255 | 0.007052 | 0.00529 | 0.01887 | 0.067484 | 0.005344 | 0.061126 | 0.021651 | 0.011781 | 0.092744 | 0.066997 | 0.005932 | 0.009983 | 0.056682 | 0.021229 | 0.025186 | 0.012527 | 0.014148 | 0.012589 | 0.046971 | 0.00988 | 0.017717 | 0.026755 |
| 31.8055556 | 0.060284 | 0.018621 | 0.005556 | 0.005106 | 0.007657 | 0.052173 | 0.00797 | 0.052386 | 0.014006 | 0.014785 | 0.067686 | 0.073373 | 0.012016 | 0.008218 | 0.040851 | 0.021032 | 0.014569 | 0.006792 | 0.009417 | 0.010071 | 0.026715 | 0.014238 | 0.017063 | 0.034882 |
| 31.8194444 | 0.089668 | 0.0189 | 0.010413 | 0.006649 | 0.005347 | 0.026562 | 0.01251 | 0.036856 | 0.014574 | 0.015424 | 0.042898 | 0.06557 | 0.018181 | 0.008907 | 0.021803 | 0.018425 | 0.022925 | 0.003219 | 0.00517 | 0.004255 | 0.011809 | 0.025363 | 0.013537 | 0.03503 |
| 31.8333333 | 0.112597 | 0.024573 | 0.021442 | 0.006896 | 0.005451 | 0.018577 | 0.013806 | 0.024817 | 0.021951 | 0.016664 | 0.070984 | 0.050465 | 0.019626 | 0.014699 | 0.012651 | 0.015184 | 0.032388 | 0.002448 | 0.004634 | 0.001955 | 0.011698 | 0.031092 | 0.009369 | 0.024409 |
| 31.8472222 | 0.130436 | 0.025059 | 0.031098 | 0.011246 | 0.007576 | 0.034308 | 0.012718 | 0.036708 | 0.022869 | 0.018702 | 0.127125 | 0.041017 | 0.016014 | 0.018872 | 0.026974 | 0.008683 | 0.030808 | 0.005249 | 0.00904 | 0.003557 | 0.015094 | 0.026809 | 0.016825 | 0.010888 |
| 31.8611111 | 0.157708 | 0.017098 | 0.034406 | 0.021601 | 0.015247 | 0.058644 | 0.012016 | 0.067083 | 0.015168 | 0.016339 | 0.158727 | 0.043772 | 0.008876 | 0.017447 | 0.057608 | 0.004415 | 0.022043 | 0.00992 | 0.014088 | 0.008707 | 0.013205 | 0.016361 | 0.034619 | 0.002844 |
| 31.875 | 0.195004 | 0.0106 | 0.03112 | 0.025705 | 0.01656 | 0.073912 | 0.01643 | 0.091283 | 0.012768 | 0.009326 | 0.162002 | 0.056253 | 0.00429 | 0.010806 | 0.088318 | 0.008164 | 0.010888 | 0.014985 | 0.017668 | 0.016193 | 0.010364 | 0.014161 | 0.050083 | 0.000727 |
| 31.8888889 | 0.225147 | 0.016755 | 0.026273 | 0.0226 | 0.009241 | 0.064505 | 0.026342 | 0.103993 | 0.022434 | 0.00615 | 0.153998 | 0.068726 | 0.006482 | 0.006838 | 0.105636 | 0.017591 | 0.004538 | 0.019794 | 0.020227 | 0.02612 | 0.017029 | 0.02767 | 0.060588 | 0.00134 |
| 31.9027778 | 0.237409 | 0.029155 | 0.026569 | 0.026412 | 0.005182 | 0.03571 | 0.035719 | 0.110086 | 0.034329 | 0.015481 | 0.148683 | 0.077289 | 0.010939 | 0.012179 | 0.106838 | 0.031648 | 0.002997 | 0.025359 | 0.02208 | 0.039306 | 0.036034 | 0.044963 | 0.069924 | 0.004791 |
| 31.9166667 | 0.222672 | 0.033963 | 0.034294 | 0.040021 | 0.004634 | 0.014259 | 0.038253 | 0.11148 | 0.043268 | 0.033408 | 0.149465 | 0.084113 | 0.014533 | 0.017788 | 0.100462 | 0.047767 | 0.004728 | 0.034353 | 0.024216 | 0.051654 | 0.050474 | 0.0569 | 0.081098 | 0.012855 |
| 31.9305556 | 0.183595 | 0.025873 | 0.04599 | 0.0474 | 0.003097 | 0.012066 | 0.031342 | 0.105923 | 0.048568 | 0.047938 | 0.143329 | 0.084054 | 0.018359 | 0.018687 | 0.093507 | 0.061791 | 0.014854 | 0.041888 | 0.026656 | 0.057414 | 0.04919 | 0.064296 | 0.097672 | 0.021547 |
| 31.9444444 | 0.140298 | 0.022057 | 0.051997 | 0.041806 | 0.002062 | 0.014851 | 0.018059 | 0.091872 | 0.047636 | 0.052206 | 0.12762 | 0.07059 | 0.022565 | 0.022634 | 0.088404 | 0.074179 | 0.031603 | 0.040269 | 0.030512 | 0.056606 | 0.040569 | 0.067758 | 0.120904 | 0.025941 |
| 31.9583333 | 0.102127 | 0.03653 | 0.043834 | 0.035173 | 0.002488 | 0.013931 | 0.011664 | 0.074459 | 0.041599 | 0.047608 | 0.108619 | 0.050374 | 0.026185 | 0.030883 | 0.082634 | 0.085321 | 0.048283 | 0.032402 | 0.035474 | 0.051007 | 0.034536 | 0.068519 | 0.136478 | 0.028326 |
| 31.9722222 | 0.065431 | 0.042172 | 0.024663 | 0.030937 | 0.002587 | 0.017593 | 0.021029 | 0.057732 | 0.034588 | 0.037126 | 0.081888 | 0.033023 | 0.027674 | 0.034019 | 0.074942 | 0.094367 | 0.062983 | 0.023119 | 0.038576 | 0.040302 | 0.033488 | 0.070025 | 0.13118 | 0.03311 |
| 31.9861111 | 0.033207 | 0.024932 | 0.008219 | 0.026163 | 0.002696 | 0.032586 | 0.03252 | 0.039714 | 0.027819 | 0.022483 | 0.04413 | 0.018857 | 0.026468 | 0.029774 | 0.067327 | 0.10129 | 0.071579 | 0.017792 | 0.039318 | 0.027249 | 0.035807 | 0.069656 | 0.114479 | 0.039015 |
| 32 | 0.025022 | 0.009022 | 0.002307 | 0.024032 | 0.002865 | 0.058583 | 0.035973 | 0.021347 | 0.018864 | 0.01008 | 0.021478 | 0.009794 | 0.023331 | 0.023449 | 0.05955 | 0.102575 | 0.066509 | 0.020838 | 0.037899 | 0.017374 | 0.038613 | 0.060823 | 0.096355 | 0.042805 |
| 32.0138889 | 0.04715 | 0.006884 | 0.001713 | 0.024116 | 0.003192 | 0.086105 | 0.035167 | 0.015819 | 0.008949 | 0.007772 | 0.037292 | 0.012038 | 0.01929 | 0.015081 | 0.049276 | 0.095142 | 0.049078 | 0.025594 | 0.032327 | 0.012746 | 0.039961 | 0.04657 | 0.080122 | 0.045745 |
| 32.0277778 | 0.063082 | 0.008202 | 0.005598 | 0.02395 | 0.008401 | 0.101775 | 0.033287 | 0.025135 | 0.003779 | 0.013019 | 0.057988 | 0.017576 | 0.016486 | 0.008597 | 0.039372 | 0.086015 | 0.033422 | 0.027125 | 0.024549 | 0.014773 | 0.044739 | 0.037093 | 0.07095 | 0.053458 |
| 32.0416667 | 0.056646 | 0.020848 | 0.028947 | 0.032611 | 0.021767 | 0.122151 | 0.044357 | 0.024543 | 0.010707 | 0.013037 | 0.051088 | 0.022237 | 0.020879 | 0.018924 | 0.040924 | 0.093845 | 0.035788 | 0.033091 | 0.024666 | 0.030122 | 0.067025 | 0.043878 | 0.073819 | 0.073738 |
| 32.0555556 | 0.081906 | 0.069958 | 0.086222 | 0.063512 | 0.039877 | 0.172973 | 0.066969 | 0.024394 | 0.032749 | 0.013818 | 0.063178 | 0.051086 | 0.040031 | 0.057716 | 0.056445 | 0.132983 | 0.06416 | 0.049802 | 0.040401 | 0.061071 | 0.112314 | 0.071558 | 0.091704 | 0.109392 |
| 32.0694444 | 0.146262 | 0.147757 | 0.152655 | 0.103404 | 0.05334 | 0.220368 | 0.070765 | 0.056791 | 0.056956 | 0.028466 | 0.134844 | 0.098267 | 0.062705 | 0.103311 | 0.067772 | 0.169788 | 0.098409 | 0.061205 | 0.060971 | 0.084546 | 0.14609 | 0.09988 | 0.107844 | 0.133096 |
| 32.0833333 | 0.170365 | 0.20552 | 0.178055 | 0.114951 | 0.054239 | 0.204378 | 0.050767 | 0.094743 | 0.062484 | 0.038182 | 0.209823 | 0.111983 | 0.060022 | 0.112195 | 0.05623 | 0.151968 | 0.097899 | 0.044399 | 0.062517 | 0.076133 | 0.124998 | 0.09587 | 0.096104 | 0.107764 |
| 32.0972222 | 0.153018 | 0.206372 | 0.149276 | 0.087599 | 0.0451 | 0.135774 | 0.026172 | 0.104039 | 0.045151 | 0.02761 | 0.24145 | 0.084432 | 0.032737 | 0.081597 | 0.027984 | 0.095527 | 0.058012 | 0.020908 | 0.040921 | 0.050902 | 0.068314 | 0.058964 | 0.060187 | 0.050995 |
| 32.1111111 | 0.134193 | 0.158302 | 0.090494 | 0.046396 | 0.03244 | 0.074896 | 0.009402 | 0.091466 | 0.023411 | 0.013174 | 0.234705 | 0.052869 | 0.013973 | 0.042987 | 0.007997 | 0.049828 | 0.021325 | 0.020667 | 0.016776 | 0.032278 | 0.026479 | 0.023637 | 0.025859 | 0.014968 |
| 32.125 | 0.099549 | 0.091047 | 0.042076 | 0.018791 | 0.018627 | 0.04266 | 0.002765 | 0.072234 | 0.016751 | 0.011421 | 0.200017 | 0.034946 | 0.014015 | 0.021613 | 0.005531 | 0.025403 | 0.014431 | 0.025859 | 0.007355 | 0.019502 | 0.01024 | 0.014332 | 0.007936 | 0.011297 |
| 32.1388889 | 0.062524 | 0.037267 | 0.029487 | 0.015684 | 0.008045 | 0.022502 | 0.001642 | 0.057794 | 0.019868 | 0.018143 | 0.154446 | 0.024744 | 0.0153 | 0.029827 | 0.008362 | 0.011288 | 0.021107 | 0.023371 | 0.01344 | 0.009502 | 0.004874 | 0.023995 | 0.002071 | 0.014052 |
| 32.1527778 | 0.041399 | 0.020587 | 0.030567 | 0.026747 | 0.00734 | 0.011235 | 0.003615 | 0.046825 | 0.017592 | 0.023647 | 0.101391 | 0.014932 | 0.009669 | 0.049284 | 0.009325 | 0.005809 | 0.02664 | 0.019355 | 0.023418 | 0.005885 | 0.003764 | 0.033954 | 0.002526 | 0.012593 |
| 32.1666667 | 0.025747 | 0.036216 | 0.023694 | 0.040911 | 0.016077 | 0.015251 | 0.007779 | 0.032717 | 0.016901 | 0.02493 | 0.047038 | 0.010957 | 0.004206 | 0.049853 | 0.008 | 0.009979 | 0.033078 | 0.013721 | 0.027949 | 0.00528 | 0.004682 | 0.035859 | 0.008093 | 0.010971 |
| 32.1805556 | 0.012371 | 0.0586 | 0.018046 | 0.056878 | 0.027154 | 0.022783 | 0.010369 | 0.021256 | 0.022209 | 0.021517 | 0.021759 | 0.016404 | 0.003596 | 0.029273 | 0.004543 | 0.017402 | 0.040213 | 0.008191 | 0.025043 | 0.00705 | 0.004376 | 0.030707 | 0.013961 | 0.010144 |
| 32.1944444 | 0.004435 | 0.072298 | 0.020308 | 0.070096 | 0.034129 | 0.023746 | 0.007994 | 0.014268 | 0.020219 | 0.015659 | 0.032135 | 0.020373 | 0.002962 | 0.012385 | 0.004126 | 0.022798 | 0.042061 | 0.005758 | 0.017128 | 0.015358 | 0.002637 | 0.019918 | 0.014766 | 0.010764 |
| 32.2083333 | 0.009372 | 0.078585 | 0.022827 | 0.074422 | 0.035798 | 0.025042 | 0.003662 | 0.007732 | 0.013943 | 0.011422 | 0.053605 | 0.020707 | 0.001056 | 0.013078 | 0.009909 | 0.025529 | 0.035113 | 0.008417 | 0.009874 | 0.022453 | 0.001244 | 0.01094 | 0.010692 | 0.011867 |
| 32.2222222 | 0.035089 | 0.082351 | 0.017538 | 0.069672 | 0.033904 | 0.031337 | 0.002228 | 0.005939 | 0.018261 | 0.012581 | 0.07599 | 0.025334 | 0.000659 | 0.013374 | 0.015082 | 0.025887 | 0.024627 | 0.012084 | 0.006775 | 0.018526 | 0.001062 | 0.014027 | 0.005588 | 0.011925 |
| 32.2361111 | 0.073214 | 0.092136 | 0.010008 | 0.05902 | 0.029313 | 0.037373 | 0.003846 | 0.012253 | 0.033926 | 0.019403 | 0.099163 | 0.036433 | 0.001454 | 0.012181 | 0.013253 | 0.025996 | 0.017448 | 0.009177 | 0.007298 | 0.011245 | 0.001109 | 0.020038 | 0.002102 | 0.011557 |
| 32.25 | 0.104846 | 0.115667 | 0.011059 | 0.04727 | 0.024123 | 0.0396 | 0.006946 | 0.023499 | 0.052663 | 0.028396 | 0.119823 | 0.056252 | 0.003045 | 0.027046 | 0.00707 | 0.027971 | 0.01517 | 0.006335 | 0.008847 | 0.014213 | 0.001589 | 0.015684 | 0.000922 | 0.01115 |
| 32.2638889 | 0.119306 | 0.146014 | 0.01554 | 0.039289 | 0.021843 | 0.042165 | 0.011102 | 0.038301 | 0.06925 | 0.035195 | 0.141865 | 0.091669 | 0.003806 | 0.045874 | 0.002279 | 0.029437 | 0.015226 | 0.01231 | 0.009444 | 0.02267 | 0.003699 | 0.009147 | 0.000791 | 0.009795 |
| 32.2777778 | 0.110364 | 0.159611 | 0.012717 | 0.036524 | 0.023412 | 0.048045 | 0.015326 | 0.05522 | 0.081087 | 0.037463 | 0.166359 | 0.133675 | 0.003033 | 0.049509 | 0.002949 | 0.026882 | 0.01514 | 0.018152 | 0.008541 | 0.025085 | 0.006091 | 0.012286 | 0.003092 | 0.007953 |
| 32.2916667 | 0.085914 | 0.143016 | 0.010975 | 0.037281 | 0.027029 | 0.055384 | 0.018052 | 0.069376 | 0.086702 | 0.037387 | 0.178131 | 0.157605 | 0.001906 | 0.041376 | 0.005392 | 0.021351 | 0.014145 | 0.016844 | 0.006856 | 0.022389 | 0.008182 | 0.019506 | 0.009861 | 0.006664 |
| 32.3055556 | 0.069435 | 0.114754 | 0.021138 | 0.039082 | 0.030852 | 0.060716 | 0.019046 | 0.077987 | 0.086776 | 0.03727 | 0.167221 | 0.157487 | 0.003122 | 0.032378 | 0.006936 | 0.016378 | 0.012427 | 0.012458 | 0.006813 | 0.021517 | 0.012179 | 0.019766 | 0.016323 | 0.006646 |
| 32.3194444 | 0.063188 | 0.100009 | 0.033188 | 0.039885 | 0.033676 | 0.062407 | 0.019044 | 0.082084 | 0.081819 | 0.036147 | 0.143304 | 0.142411 | 0.008115 | 0.0252 | 0.011453 | 0.013653 | 0.010501 | 0.008337 | 0.009535 | 0.02368 | 0.01833 | 0.01406 | 0.018634 | 0.00841 |
| 32.3333333 | 0.052894 | 0.089615 | 0.036435 | 0.037012 | 0.034163 | 0.060957 | 0.018681 | 0.082331 | 0.072178 | 0.032709 | 0.111204 | 0.118209 | 0.010425 | 0.017283 | 0.012336 | 0.010948 | 0.007897 | 0.004638 | 0.01263 | 0.023856 | 0.022333 | 0.009582 | 0.019803 | 0.011102 |
| 32.3472222 | 0.040055 | 0.063093 | 0.030038 | 0.029668 | 0.031889 | 0.056247 | 0.018712 | 0.076092 | 0.058971 | 0.027795 | 0.07972 | 0.087396 | 0.006739 | 0.009946 | 0.007419 | 0.006397 | 0.004527 | 0.001964 | 0.014045 | 0.020777 | 0.020937 | 0.009412 | 0.020445 | 0.013479 |
| 32.3611111 | 0.034547 | 0.032378 | 0.017163 | 0.019623 | 0.027911 | 0.048909 | 0.019344 | 0.063735 | 0.0427 | 0.023238 | 0.065915 | 0.048425 | 0.002845 | 0.007833 | 0.004715 | 0.005265 | 0.003144 | 0.003412 | 0.014954 | 0.020495 | 0.017935 | 0.013032 | 0.017857 | 0.015824 |
| 32.375 | 0.03579 | 0.017986 | 0.007012 | 0.010316 | 0.022719 | 0.042101 | 0.020078 | 0.049296 | 0.025121 | 0.019864 | 0.05512 | 0.023255 | 0.003164 | 0.012074 | 0.005863 | 0.015399 | 0.006796 | 0.010396 | 0.017583 | 0.028462 | 0.019389 | 0.018074 | 0.013023 | 0.019695 |
| 32.3888889 | 0.036543 | 0.028302 | 0.00651 | 0.008036 | 0.01683 | 0.037811 | 0.020781 | 0.034045 | 0.010744 | 0.018006 | 0.033823 | 0.036124 | 0.006996 | 0.014118 | 0.01043 | 0.032495 | 0.013288 | 0.01658 | 0.020471 | 0.039548 | 0.024376 | 0.021602 | 0.009723 | 0.024158 |
| 32.4027778 | 0.034179 | 0.053621 | 0.011814 | 0.011861 | 0.012444 | 0.033334 | 0.021352 | 0.01827 | 0.00475 | 0.01729 | 0.028703 | 0.0655 | 0.008957 | 0.012665 | 0.012373 | 0.042456 | 0.017718 | 0.014979 | 0.019879 | 0.042789 | 0.027332 | 0.023283 | 0.009575 | 0.025888 |
| 32.4166667 | 0.029051 | 0.076409 | 0.01373 | 0.014136 | 0.010985 | 0.025498 | 0.021179 | 0.008944 | 0.006261 | 0.016217 | 0.055275 | 0.090267 | 0.007087 | 0.017697 | 0.007658 | 0.04365 | 0.019743 | 0.009911 | 0.017071 | 0.03914 | 0.027717 | 0.024537 | 0.011681 | 0.025813 |
| 32.4305556 | 0.020152 | 0.091452 | 0.008856 | 0.016072 | 0.010427 | 0.01633 | 0.020094 | 0.012426 | 0.006909 | 0.014184 | 0.08743 | 0.108438 | 0.004667 | 0.02106 | 0.003543 | 0.045127 | 0.02132 | 0.007564 | 0.015288 | 0.036671 | 0.027719 | 0.024692 | 0.014596 | 0.026461 |
| 32.4444444 | 0.011666 | 0.096399 | 0.004142 | 0.021965 | 0.008112 | 0.010003 | 0.018542 | 0.021517 | 0.00569 | 0.011471 | 0.09441 | 0.101576 | 0.002734 | 0.01654 | 0.004096 | 0.0483 | 0.022967 | 0.008072 | 0.01393 | 0.035311 | 0.026679 | 0.021455 | 0.016346 | 0.026733 |
| 32.4583333 | 0.010769 | 0.089304 | 0.005058 | 0.026497 | 0.005411 | 0.007463 | 0.016539 | 0.027034 | 0.008702 | 0.007865 | 0.077427 | 0.065938 | 0.004062 | 0.023686 | 0.006362 | 0.053234 | 0.026955 | 0.012829 | 0.013468 | 0.036587 | 0.026362 | 0.016081 | 0.018276 | 0.028313 |
| 32.4722222 | 0.014207 | 0.070165 | 0.005714 | 0.021874 | 0.006051 | 0.006961 | 0.014156 | 0.027934 | 0.013294 | 0.0042 | 0.063086 | 0.034671 | 0.010849 | 0.047402 | 0.012578 | 0.062221 | 0.033288 | 0.021787 | 0.015637 | 0.044692 | 0.030188 | 0.014121 | 0.025483 | 0.035226 |
| 32.4861111 | 0.014314 | 0.043259 | 0.007029 | 0.012234 | 0.010587 | 0.006994 | 0.012777 | 0.025819 | 0.015011 | 0.002838 | 0.062067 | 0.024478 | 0.016046 | 0.062663 | 0.022255 | 0.066211 | 0.03618 | 0.027167 | 0.017646 | 0.051029 | 0.034894 | 0.018572 | 0.037996 | 0.043842 |
| 32.5 | 0.009124 | 0.019327 | 0.018145 | 0.009142 | 0.015777 | 0.007208 | 0.013708 | 0.021207 | 0.016662 | 0.004846 | 0.06197 | 0.022711 | 0.014799 | 0.056188 | 0.026714 | 0.056542 | 0.032135 | 0.024163 | 0.016597 | 0.046971 | 0.035574 | 0.025902 | 0.048356 | 0.047125 |
| 32.5138889 | 0.004956 | 0.01022 | 0.032691 | 0.013622 | 0.019085 | 0.007608 | 0.015926 | 0.014615 | 0.02031 | 0.008483 | 0.053096 | 0.017107 | 0.012673 | 0.042141 | 0.027456 | 0.041135 | 0.024258 | 0.017428 | 0.015203 | 0.038816 | 0.034902 | 0.031853 | 0.05063 | 0.046845 |
| 32.5277778 | 0.010336 | 0.019394 | 0.036566 | 0.021155 | 0.02202 | 0.008488 | 0.016816 | 0.009006 | 0.025587 | 0.011857 | 0.033429 | 0.015113 | 0.015914 | 0.035443 | 0.033355 | 0.031407 | 0.018763 | 0.011395 | 0.017697 | 0.035795 | 0.04171 | 0.038746 | 0.049461 | 0.052682 |
| 32.5416667 | 0.026732 | 0.041484 | 0.031238 | 0.035562 | 0.029632 | 0.010465 | 0.015746 | 0.011303 | 0.034599 | 0.016411 | 0.031314 | 0.03994 | 0.02877 | 0.04386 | 0.054665 | 0.034621 | 0.024157 | 0.00842 | 0.027925 | 0.043541 | 0.071521 | 0.056736 | 0.05942 | 0.078001 |
| 32.5555556 | 0.05554 | 0.076113 | 0.036918 | 0.058491 | 0.043574 | 0.011311 | 0.014767 | 0.020856 | 0.047116 | 0.026305 | 0.072147 | 0.088589 | 0.051328 | 0.071078 | 0.104869 | 0.059454 | 0.048346 | 0.013995 | 0.047877 | 0.060119 | 0.122239 | 0.085161 | 0.087664 | 0.123984 |
| 32.5694444 | 0.094404 | 0.114962 | 0.0708 | 0.079447 | 0.056541 | 0.008399 | 0.015218 | 0.026128 | 0.055799 | 0.041606 | 0.119947 | 0.125062 | 0.069478 | 0.10357 | 0.165732 | 0.102844 | 0.081172 | 0.031822 | 0.067661 | 0.073436 | 0.14706 | 0.101353 | 0.116517 | 0.151391 |
| 32.5833333 | 0.127891 | 0.138064 | 0.108942 | 0.089574 | 0.061765 | 0.010302 | 0.016469 | 0.027804 | 0.061351 | 0.054808 | 0.158898 | 0.136961 | 0.0683 | 0.124922 | 0.193731 | 0.139871 | 0.104546 | 0.053871 | 0.073576 | 0.079512 | 0.121069 | 0.096788 | 0.130291 | 0.131013 |
| 32.5972222 | 0.140466 | 0.141706 | 0.114722 | 0.089578 | 0.060266 | 0.024513 | 0.017214 | 0.035254 | 0.070418 | 0.058494 | 0.216051 | 0.158768 | 0.054349 | 0.124812 | 0.17109 | 0.14585 | 0.11452 | 0.05967 | 0.06949 | 0.085829 | 0.084752 | 0.087214 | 0.135779 | 0.09565 |
| 32.6111111 | 0.135145 | 0.138503 | 0.08833 | 0.077367 | 0.051422 | 0.038455 | 0.016048 | 0.044823 | 0.073609 | 0.052865 | 0.268317 | 0.190357 | 0.041504 | 0.091375 | 0.112708 | 0.126214 | 0.113263 | 0.043971 | 0.064207 | 0.083929 | 0.067771 | 0.079057 | 0.128784 | 0.073277 |
| 32.625 | 0.124685 | 0.125216 | 0.050617 | 0.047224 | 0.032819 | 0.043774 | 0.013131 | 0.050284 | 0.061798 | 0.042306 | 0.273351 | 0.194411 | 0.033105 | 0.043427 | 0.051808 | 0.104277 | 0.10104 | 0.025938 | 0.058632 | 0.05731 | 0.060281 | 0.068355 | 0.09361 | 0.053401 |
| 32.6388889 | 0.114214 | 0.088756 | 0.036443 | 0.020592 | 0.01673 | 0.046881 | 0.010903 | 0.053455 | 0.041547 | 0.029609 | 0.228329 | 0.170672 | 0.026804 | 0.023322 | 0.023409 | 0.074087 | 0.067604 | 0.01444 | 0.051399 | 0.02879 | 0.057373 | 0.057176 | 0.049524 | 0.028795 |
| 32.6527778 | 0.108569 | 0.047019 | 0.051015 | 0.026156 | 0.017531 | 0.046386 | 0.01241 | 0.052444 | 0.020784 | 0.017412 | 0.148607 | 0.129688 | 0.020531 | 0.03671 | 0.039586 | 0.037916 | 0.034166 | 0.014793 | 0.035231 | 0.034345 | 0.064697 | 0.050957 | 0.042125 | 0.015288 |
| 32.6666667 | 0.112806 | 0.022035 | 0.048633 | 0.049309 | 0.026131 | 0.037823 | 0.018278 | 0.042634 | 0.010713 | 0.010784 | 0.075256 | 0.079707 | 0.014023 | 0.051493 | 0.072495 | 0.026788 | 0.03649 | 0.036573 | 0.015101 | 0.057147 | 0.072562 | 0.050404 | 0.065888 | 0.019511 |
| 32.6805556 | 0.121617 | 0.011817 | 0.032696 | 0.062632 | 0.027506 | 0.025062 | 0.026311 | 0.025657 | 0.017077 | 0.013035 | 0.069167 | 0.040661 | 0.008798 | 0.041764 | 0.09275 | 0.038141 | 0.051796 | 0.053501 | 0.007998 | 0.063249 | 0.064673 | 0.048017 | 0.074231 | 0.019885 |
| 32.6944444 | 0.125513 | 0.015388 | 0.050178 | 0.056885 | 0.018962 | 0.015156 | 0.033816 | 0.011994 | 0.023987 | 0.019522 | 0.138575 | 0.035566 | 0.008332 | 0.030302 | 0.100674 | 0.053102 | 0.053957 | 0.035828 | 0.012051 | 0.05188 | 0.043405 | 0.039732 | 0.062492 | 0.011236 |
| 32.7083333 | 0.129212 | 0.029472 | 0.086414 | 0.036455 | 0.011259 | 0.014231 | 0.037422 | 0.010665 | 0.017915 | 0.022083 | 0.220816 | 0.06775 | 0.012312 | 0.053581 | 0.10553 | 0.057172 | 0.045271 | 0.013957 | 0.015532 | 0.032417 | 0.024008 | 0.035226 | 0.041525 | 0.011184 |
| 32.7222222 | 0.144203 | 0.033422 | 0.092518 | 0.016008 | 0.0165 | 0.022776 | 0.036749 | 0.017327 | 0.009851 | 0.017934 | 0.264199 | 0.108835 | 0.014517 | 0.081663 | 0.100243 | 0.037341 | 0.028004 | 0.021309 | 0.012125 | 0.018127 | 0.025328 | 0.036353 | 0.033014 | 0.023766 |
| 32.7361111 | 0.170026 | 0.020865 | 0.057763 | 0.011628 | 0.027758 | 0.032662 | 0.034075 | 0.026909 | 0.013886 | 0.010505 | 0.256163 | 0.122933 | 0.011484 | 0.074947 | 0.07641 | 0.019279 | 0.020297 | 0.043578 | 0.007027 | 0.025139 | 0.04858 | 0.032659 | 0.07191 | 0.037829 |
| 32.75 | 0.188975 | 0.017695 | 0.031217 | 0.024802 | 0.034803 | 0.034592 | 0.029124 | 0.036912 | 0.024236 | 0.004408 | 0.198928 | 0.094313 | 0.006014 | 0.043532 | 0.043299 | 0.03179 | 0.037012 | 0.059512 | 0.012357 | 0.045547 | 0.061677 | 0.020551 | 0.130224 | 0.049297 |
| 32.7638889 | 0.183579 | 0.035872 | 0.049392 | 0.037481 | 0.036396 | 0.026156 | 0.021371 | 0.040664 | 0.031861 | 0.00201 | 0.129045 | 0.053238 | 0.002849 | 0.024014 | 0.021371 | 0.046062 | 0.055749 | 0.058739 | 0.024554 | 0.057946 | 0.043633 | 0.011262 | 0.163471 | 0.063343 |
| 32.7777778 | 0.158241 | 0.050458 | 0.067371 | 0.036891 | 0.030769 | 0.0157 | 0.0129 | 0.033603 | 0.034161 | 0.00193 | 0.072273 | 0.036007 | 0.003263 | 0.040132 | 0.025095 | 0.034477 | 0.056264 | 0.039704 | 0.0332 | 0.05276 | 0.0174 | 0.015518 | 0.159048 | 0.078128 |
| 32.7916667 | 0.120704 | 0.042197 | 0.049213 | 0.025384 | 0.018227 | 0.019924 | 0.007532 | 0.020627 | 0.028944 | 0.00295 | 0.056204 | 0.031449 | 0.004342 | 0.064317 | 0.034978 | 0.014321 | 0.044359 | 0.022372 | 0.034891 | 0.031452 | 0.005612 | 0.025979 | 0.111961 | 0.079521 |
| 32.8055556 | 0.076617 | 0.043585 | 0.026911 | 0.01111 | 0.009849 | 0.039949 | 0.008702 | 0.017871 | 0.017106 | 0.0059 | 0.107705 | 0.0198 | 0.003688 | 0.06134 | 0.031537 | 0.00395 | 0.027323 | 0.027193 | 0.029714 | 0.016857 | 0.005469 | 0.032755 | 0.059265 | 0.056998 |
| 32.8194444 | 0.036722 | 0.080184 | 0.033263 | 0.002933 | 0.011606 | 0.057047 | 0.015808 | 0.02515 | 0.009493 | 0.00867 | 0.152439 | 0.011845 | 0.002182 | 0.038872 | 0.018851 | 0.006136 | 0.011707 | 0.037742 | 0.021986 | 0.028483 | 0.011271 | 0.031789 | 0.056784 | 0.02858 |
| 32.8333333 | 0.023932 | 0.112659 | 0.043828 | 0.002689 | 0.013084 | 0.06351 | 0.023239 | 0.022974 | 0.016194 | 0.009208 | 0.129391 | 0.016153 | 0.001622 | 0.033942 | 0.011072 | 0.018261 | 0.007616 | 0.032581 | 0.016774 | 0.048384 | 0.015198 | 0.023453 | 0.095018 | 0.021347 |
| 32.8472222 | 0.052292 | 0.114118 | 0.033443 | 0.004646 | 0.010665 | 0.066347 | 0.02572 | 0.022888 | 0.027177 | 0.007865 | 0.070124 | 0.0198 | 0.001837 | 0.059505 | 0.019709 | 0.027212 | 0.012896 | 0.018148 | 0.014325 | 0.06238 | 0.01304 | 0.014274 | 0.125643 | 0.031277 |
| 32.8611111 | 0.093516 | 0.100955 | 0.03416 | 0.003909 | 0.008038 | 0.071304 | 0.02272 | 0.04507 | 0.03021 | 0.011268 | 0.043372 | 0.017115 | 0.00184 | 0.07835 | 0.036445 | 0.023219 | 0.016988 | 0.008356 | 0.012283 | 0.071655 | 0.009007 | 0.009297 | 0.125774 | 0.040077 |
| 32.875 | 0.125234 | 0.077302 | 0.069271 | 0.001648 | 0.005487 | 0.076195 | 0.016842 | 0.073871 | 0.027602 | 0.02225 | 0.091096 | 0.025547 | 0.003737 | 0.061726 | 0.060612 | 0.017963 | 0.020975 | 0.008838 | 0.008757 | 0.071736 | 0.005139 | 0.008637 | 0.104891 | 0.040786 |
| 32.8888889 | 0.158001 | 0.04146 | 0.112149 | 0.002629 | 0.003195 | 0.070754 | 0.010783 | 0.089962 | 0.026487 | 0.02885 | 0.163869 | 0.055134 | 0.010434 | 0.034237 | 0.102834 | 0.022913 | 0.026988 | 0.012834 | 0.004433 | 0.062137 | 0.002152 | 0.010963 | 0.086835 | 0.035194 |
| 32.9027778 | 0.201368 | 0.021188 | 0.131972 | 0.008593 | 0.001411 | 0.051092 | 0.005923 | 0.092469 | 0.028005 | 0.027747 | 0.203249 | 0.089181 | 0.018675 | 0.036507 | 0.153673 | 0.036489 | 0.032645 | 0.011949 | 0.001905 | 0.049585 | 0.001252 | 0.015631 | 0.07948 | 0.030415 |
| 32.9166667 | 0.249474 | 0.037207 | 0.115754 | 0.01752 | 0.001123 | 0.026703 | 0.002661 | 0.087938 | 0.029749 | 0.027276 | 0.209229 | 0.118507 | 0.02122 | 0.071018 | 0.19293 | 0.052403 | 0.036282 | 0.007543 | 0.001039 | 0.040614 | 0.003534 | 0.022439 | 0.077052 | 0.030408 |
| 32.9305556 | 0.289383 | 0.073691 | 0.072208 | 0.026468 | 0.002275 | 0.010473 | 0.00173 | 0.082735 | 0.03243 | 0.027966 | 0.205308 | 0.143156 | 0.016996 | 0.095907 | 0.202843 | 0.066289 | 0.036438 | 0.006974 | 0.000665 | 0.036273 | 0.009103 | 0.03037 | 0.076506 | 0.033457 |
| 32.9444444 | 0.310946 | 0.109097 | 0.029409 | 0.02983 | 0.002463 | 0.007285 | 0.002656 | 0.079528 | 0.03505 | 0.024185 | 0.20891 | 0.151259 | 0.012782 | 0.082166 | 0.174592 | 0.077102 | 0.035789 | 0.012959 | 0.000788 | 0.0342 | 0.015218 | 0.036297 | 0.077758 | 0.036651 |
| 32.9583333 | 0.299317 | 0.123807 | 0.006483 | 0.022823 | 0.001157 | 0.009434 | 0.002614 | 0.074772 | 0.03449 | 0.014861 | 0.21823 | 0.133665 | 0.01245 | 0.051818 | 0.124587 | 0.086132 | 0.037638 | 0.02071 | 0.002113 | 0.031753 | 0.019244 | 0.037527 | 0.078945 | 0.037727 |
| 32.9722222 | 0.240916 | 0.117418 | 0.002338 | 0.011493 | 0.000626 | 0.008509 | 0.002234 | 0.065893 | 0.030569 | 0.006262 | 0.20932 | 0.101591 | 0.012721 | 0.032337 | 0.075687 | 0.092631 | 0.039793 | 0.024285 | 0.005009 | 0.026159 | 0.020394 | 0.03424 | 0.072474 | 0.03685 |
| 32.9861111 | 0.158319 | 0.111364 | 0.010464 | 0.007652 | 0.000946 | 0.005769 | 0.00394 | 0.056245 | 0.024864 | 0.00296 | 0.165444 | 0.072172 | 0.010155 | 0.022171 | 0.035034 | 0.088853 | 0.036945 | 0.019819 | 0.008377 | 0.016792 | 0.018283 | 0.027212 | 0.05771 | 0.034786 |
| 33 | 0.089317 | 0.103913 | 0.025742 | 0.01398 | 0.000885 | 0.006123 | 0.005602 | 0.04807 | 0.018829 | 0.00207 | 0.097512 | 0.045798 | 0.005441 | 0.01392 | 0.015807 | 0.068693 | 0.026275 | 0.010389 | 0.010832 | 0.00733 | 0.014644 | 0.017427 | 0.045073 | 0.031796 |
| 33.0138889 | 0.046213 | 0.082289 | 0.0386 | 0.022699 | 0.002024 | 0.007421 | 0.006572 | 0.038719 | 0.013856 | 0.001005 | 0.045035 | 0.020611 | 0.001756 | 0.010813 | 0.028025 | 0.038294 | 0.012269 | 0.004274 | 0.010535 | 0.001849 | 0.012929 | 0.008134 | 0.040002 | 0.029466 |
| 33.0277778 | 0.035935 | 0.052151 | 0.0387 | 0.023895 | 0.00347 | 0.007862 | 0.008239 | 0.02692 | 0.013157 | 0.001623 | 0.039291 | 0.006041 | 0.003972 | 0.011035 | 0.044738 | 0.017084 | 0.005035 | 0.005018 | 0.007547 | 0.00413 | 0.018405 | 0.004245 | 0.043769 | 0.034193 |
| 33.0416667 | 0.07209 | 0.038468 | 0.030075 | 0.018036 | 0.006363 | 0.017642 | 0.008355 | 0.022393 | 0.020583 | 0.007448 | 0.042122 | 0.014032 | 0.018248 | 0.02229 | 0.043278 | 0.030185 | 0.017186 | 0.01684 | 0.011305 | 0.021179 | 0.041176 | 0.010307 | 0.061315 | 0.05995 |
| 33.0555556 | 0.148082 | 0.076405 | 0.047037 | 0.021964 | 0.018183 | 0.041657 | 0.006119 | 0.038322 | 0.034337 | 0.020075 | 0.046749 | 0.047072 | 0.048501 | 0.061924 | 0.062302 | 0.091864 | 0.056559 | 0.045174 | 0.029871 | 0.057939 | 0.094499 | 0.031619 | 0.093155 | 0.118301 |
| 33.0694444 | 0.207042 | 0.157059 | 0.10282 | 0.039954 | 0.03441 | 0.064076 | 0.006311 | 0.066465 | 0.045556 | 0.032643 | 0.112065 | 0.088754 | 0.083396 | 0.113744 | 0.124767 | 0.169714 | 0.098376 | 0.07407 | 0.04746 | 0.101332 | 0.157204 | 0.060721 | 0.119495 | 0.172409 |
| 33.0833333 | 0.206253 | 0.211283 | 0.157128 | 0.050495 | 0.039365 | 0.068163 | 0.008923 | 0.081147 | 0.046599 | 0.035497 | 0.20162 | 0.113883 | 0.099438 | 0.139484 | 0.159018 | 0.199788 | 0.099583 | 0.074681 | 0.04182 | 0.119886 | 0.168117 | 0.076779 | 0.112856 | 0.171503 |
| 33.0972222 | 0.181407 | 0.198496 | 0.179279 | 0.042912 | 0.029788 | 0.056098 | 0.00712 | 0.073709 | 0.037066 | 0.025488 | 0.232922 | 0.109276 | 0.085266 | 0.110488 | 0.12432 | 0.165785 | 0.06334 | 0.043972 | 0.022344 | 0.100979 | 0.120139 | 0.07059 | 0.074729 | 0.13669 |
| 33.1111111 | 0.167006 | 0.143286 | 0.150295 | 0.025764 | 0.01544 | 0.037769 | 0.004092 | 0.055421 | 0.020705 | 0.014653 | 0.205387 | 0.081135 | 0.050913 | 0.058279 | 0.069981 | 0.102133 | 0.026405 | 0.019762 | 0.015126 | 0.066441 | 0.068018 | 0.044982 | 0.033073 | 0.103917 |
| 33.125 | 0.153574 | 0.083286 | 0.08846 | 0.01157 | 0.00815 | 0.019395 | 0.007127 | 0.035804 | 0.010631 | 0.020252 | 0.157884 | 0.046667 | 0.030001 | 0.052312 | 0.031134 | 0.044223 | 0.01019 | 0.02833 | 0.021084 | 0.035532 | 0.036116 | 0.021584 | 0.013623 | 0.070291 |

| | | | | | | | | | | | | | | | | | | | | | | | | |
|---|---|---|---|---|---|---|---|---|---|---|---|---|---|---|---|---|---|---|---|---|---|---|---|---|
| 33.1388889 | 0.129366 | 0.038042 | 0.072438 | 0.010542 | 0.012555 | 0.007439 | 0.0116 | 0.018814 | 0.014123 | 0.033913 | 0.112667 | 0.019179 | 0.037309 | 0.08776 | 0.011452 | 0.017793 | 0.014035 | 0.045693 | 0.026342 | 0.016568 | 0.016173 | 0.024786 | 0.016777 | 0.034238 |
| 33.1527778 | 0.092665 | 0.014978 | 0.111734 | 0.019055 | 0.019483 | 0.003357 | 0.012281 | 0.007217 | 0.016022 | 0.038609 | 0.074573 | 0.006179 | 0.04449 | 0.115125 | 0.009891 | 0.022642 | 0.021722 | 0.052525 | 0.028542 | 0.014721 | 0.008076 | 0.042661 | 0.023144 | 0.012737 |
| 33.1666667 | 0.054049 | 0.014853 | 0.136445 | 0.027499 | 0.023228 | 0.003203 | 0.011084 | 0.001908 | 0.009553 | 0.035755 | 0.04547 | 0.007118 | 0.033257 | 0.118323 | 0.015397 | 0.03241 | 0.025225 | 0.051097 | 0.029914 | 0.016707 | 0.011347 | 0.05609 | 0.024044 | 0.010413 |
| 33.1805556 | 0.027567 | 0.023322 | 0.127817 | 0.033852 | 0.028248 | 0.005978 | 0.010273 | 0.000864 | 0.00461 | 0.032786 | 0.024788 | 0.011058 | 0.014246 | 0.101903 | 0.015262 | 0.036153 | 0.027546 | 0.040079 | 0.028902 | 0.01094 | 0.01347 | 0.062685 | 0.021519 | 0.009163 |
| 33.1944444 | 0.015464 | 0.027515 | 0.109138 | 0.040243 | 0.034798 | 0.009445 | 0.009708 | 0.001024 | 0.005854 | 0.032276 | 0.010935 | 0.012497 | 0.005817 | 0.073908 | 0.008578 | 0.038975 | 0.029127 | 0.023223 | 0.023669 | 0.006618 | 0.008866 | 0.058438 | 0.016678 | 0.004204 |
| 33.2083333 | 0.013563 | 0.026757 | 0.074498 | 0.04597 | 0.038101 | 0.010916 | 0.008764 | 0.001066 | 0.010033 | 0.033406 | 0.007294 | 0.01345 | 0.01063 | 0.041521 | 0.003459 | 0.039245 | 0.029654 | 0.011291 | 0.015216 | 0.009995 | 0.006241 | 0.045424 | 0.009713 | 0.003764 |
| 33.2222222 | 0.018991 | 0.024161 | 0.043999 | 0.047713 | 0.037082 | 0.009597 | 0.008468 | 0.002142 | 0.013826 | 0.036126 | 0.016119 | 0.015606 | 0.016122 | 0.017209 | 0.003803 | 0.033637 | 0.028172 | 0.012588 | 0.007111 | 0.015128 | 0.009716 | 0.030915 | 0.004302 | 0.005789 |
| 33.2361111 | 0.029652 | 0.024823 | 0.065128 | 0.045799 | 0.034879 | 0.007085 | 0.011088 | 0.005811 | 0.017802 | 0.040885 | 0.034256 | 0.020237 | 0.015761 | 0.011015 | 0.005519 | 0.024957 | 0.024052 | 0.023147 | 0.003001 | 0.018029 | 0.011206 | 0.0168 | 0.004548 | 0.005398 |
| 33.25 | 0.048674 | 0.029289 | 0.112371 | 0.042767 | 0.033672 | 0.010144 | 0.017451 | 0.011744 | 0.022927 | 0.045592 | 0.060004 | 0.026857 | 0.011446 | 0.012478 | 0.005215 | 0.017844 | 0.019115 | 0.029023 | 0.003108 | 0.018595 | 0.006757 | 0.010759 | 0.005137 | 0.003075 |
| 33.2638889 | 0.093106 | 0.032133 | 0.127905 | 0.038101 | 0.032903 | 0.020107 | 0.024631 | 0.01779 | 0.027018 | 0.045861 | 0.090471 | 0.032214 | 0.00732 | 0.009336 | 0.003161 | 0.012475 | 0.015491 | 0.023564 | 0.006211 | 0.016847 | 0.001951 | 0.014886 | 0.005654 | 0.001125 |
| 33.2777778 | 0.153574 | 0.034796 | 0.096664 | 0.031497 | 0.031097 | 0.030029 | 0.028298 | 0.022187 | 0.027352 | 0.04036 | 0.117919 | 0.032896 | 0.00424 | 0.010132 | 0.004316 | 0.010031 | 0.013023 | 0.012755 | 0.010845 | 0.011497 | 0.001431 | 0.017428 | 0.014301 | 0.00114 |
| 33.2916667 | 0.174747 | 0.046 | 0.051203 | 0.027841 | 0.028306 | 0.037502 | 0.027191 | 0.025309 | 0.025915 | 0.033082 | 0.141206 | 0.032367 | 0.002719 | 0.02531 | 0.012607 | 0.013919 | 0.010096 | 0.008742 | 0.013276 | 0.007301 | 0.005068 | 0.012267 | 0.028191 | 0.003305 |
| 33.3055556 | 0.146862 | 0.064014 | 0.038312 | 0.029889 | 0.025605 | 0.0438 | 0.023599 | 0.027742 | 0.026451 | 0.027409 | 0.170221 | 0.040091 | 0.002941 | 0.041348 | 0.020804 | 0.019497 | 0.006913 | 0.013444 | 0.012231 | 0.009563 | 0.008354 | 0.006313 | 0.038133 | 0.006242 |
| 33.3194444 | 0.111834 | 0.077028 | 0.048896 | 0.033427 | 0.024507 | 0.046657 | 0.021092 | 0.029101 | 0.028214 | 0.023328 | 0.207116 | 0.055224 | 0.002713 | 0.043152 | 0.02201 | 0.020295 | 0.005334 | 0.015275 | 0.010517 | 0.010069 | 0.007613 | 0.008014 | 0.041876 | 0.006019 |
| 33.3333333 | 0.082665 | 0.079853 | 0.038711 | 0.033638 | 0.025072 | 0.043337 | 0.021684 | 0.029085 | 0.028561 | 0.018745 | 0.232077 | 0.066217 | 0.004356 | 0.033841 | 0.020231 | 0.020207 | 0.005392 | 0.012411 | 0.009544 | 0.007129 | 0.005494 | 0.013949 | 0.040272 | 0.004738 |
| 33.3472222 | 0.061724 | 0.075563 | 0.026188 | 0.027155 | 0.024952 | 0.03574 | 0.024722 | 0.028627 | 0.02769 | 0.011482 | 0.227741 | 0.072085 | 0.009074 | 0.020362 | 0.019266 | 0.022678 | 0.005675 | 0.011657 | 0.008169 | 0.007239 | 0.005399 | 0.016967 | 0.032859 | 0.008648 |
| 33.3611111 | 0.05559 | 0.062165 | 0.053883 | 0.015661 | 0.023143 | 0.029143 | 0.028615 | 0.028623 | 0.028418 | 0.004744 | 0.198846 | 0.080196 | 0.013424 | 0.009315 | 0.018688 | 0.024662 | 0.006357 | 0.014112 | 0.007005 | 0.010665 | 0.007643 | 0.017614 | 0.022227 | 0.015188 |
| 33.375 | 0.056373 | 0.042055 | 0.09572 | 0.006178 | 0.020931 | 0.024942 | 0.031991 | 0.02676 | 0.031313 | 0.00214 | 0.154565 | 0.0859 | 0.016288 | 0.00975 | 0.019877 | 0.025915 | 0.007848 | 0.018558 | 0.007475 | 0.016817 | 0.012655 | 0.018957 | 0.01418 | 0.01929 |
| 33.3888889 | 0.053602 | 0.022775 | 0.110455 | 0.00204 | 0.018942 | 0.020093 | 0.033922 | 0.021133 | 0.033066 | 0.002781 | 0.103323 | 0.07857 | 0.01652 | 0.024487 | 0.023035 | 0.026689 | 0.007902 | 0.023033 | 0.00782 | 0.024229 | 0.019123 | 0.020997 | 0.01172 | 0.019106 |
| 33.4027778 | 0.043484 | 0.008492 | 0.092649 | 0.002257 | 0.017413 | 0.014086 | 0.033564 | 0.013638 | 0.031908 | 0.006829 | 0.053641 | 0.057036 | 0.012204 | 0.042612 | 0.024798 | 0.022793 | 0.005097 | 0.024436 | 0.007033 | 0.028417 | 0.022819 | 0.021872 | 0.014906 | 0.014784 |
| 33.4166667 | 0.028197 | 0.004882 | 0.058951 | 0.006462 | 0.017457 | 0.00891 | 0.030455 | 0.006688 | 0.028213 | 0.012327 | 0.026157 | 0.031323 | 0.00582 | 0.049149 | 0.02341 | 0.01381 | 0.001853 | 0.022109 | 0.007405 | 0.027035 | 0.021399 | 0.020984 | 0.021946 | 0.009269 |
| 33.4305556 | 0.01367 | 0.007013 | 0.034917 | 0.013497 | 0.019393 | 0.004691 | 0.025823 | 0.004154 | 0.022535 | 0.01551 | 0.040232 | 0.02459 | 0.002118 | 0.038326 | 0.020918 | 0.005578 | 0.000391 | 0.01816 | 0.009677 | 0.022357 | 0.016695 | 0.018432 | 0.028598 | 0.004915 |
| 33.4444444 | 0.007754 | 0.009232 | 0.034445 | 0.019927 | 0.022287 | 0.002728 | 0.022369 | 0.007732 | 0.015964 | 0.015533 | 0.075922 | 0.045632 | 0.001248 | 0.021827 | 0.018396 | 0.002544 | 0.000179 | 0.013235 | 0.011912 | 0.016208 | 0.01037 | 0.013178 | 0.03043 | 0.002094 |
| 33.4583333 | 0.012469 | 0.012283 | 0.052588 | 0.023425 | 0.02546 | 0.005073 | 0.022371 | 0.012237 | 0.010079 | 0.012998 | 0.107359 | 0.07357 | 0.00138 | 0.019004 | 0.014412 | 0.004217 | 0.000629 | 0.007355 | 0.012854 | 0.00991 | 0.004358 | 0.006618 | 0.026216 | 0.000901 |
| 33.4722222 | 0.019364 | 0.013354 | 0.070184 | 0.024753 | 0.028481 | 0.008215 | 0.024863 | 0.014331 | 0.007063 | 0.010755 | 0.125186 | 0.088933 | 0.00241 | 0.027973 | 0.009833 | 0.006618 | 0.002117 | 0.003554 | 0.012818 | 0.007101 | 0.001808 | 0.003624 | 0.019776 | 0.00206 |
| 33.4861111 | 0.022552 | 0.010062 | 0.064303 | 0.026234 | 0.031115 | 0.006786 | 0.026933 | 0.014574 | 0.009781 | 0.012328 | 0.126833 | 0.087898 | 0.005437 | 0.028203 | 0.007731 | 0.008521 | 0.00423 | 0.00306 | 0.012989 | 0.010021 | 0.003178 | 0.003506 | 0.018783 | 0.006058 |
| 33.5 | 0.023633 | 0.006035 | 0.041976 | 0.029489 | 0.033604 | 0.004694 | 0.027599 | 0.012823 | 0.018548 | 0.017355 | 0.114909 | 0.072387 | 0.009442 | 0.018904 | 0.008204 | 0.009225 | 0.007005 | 0.003792 | 0.014186 | 0.017059 | 0.007438 | 0.003702 | 0.02568 | 0.01132 |
| 33.5138889 | 0.023094 | 0.008071 | 0.042796 | 0.032961 | 0.035196 | 0.008367 | 0.026912 | 0.00955 | 0.02863 | 0.022511 | 0.088472 | 0.044607 | 0.009644 | 0.019072 | 0.008335 | 0.009128 | 0.011016 | 0.005826 | 0.015394 | 0.025196 | 0.013526 | 0.00707 | 0.03458 | 0.015665 |
| 33.5277778 | 0.023386 | 0.017374 | 0.073453 | 0.033819 | 0.034952 | 0.014768 | 0.025294 | 0.005634 | 0.037695 | 0.026305 | 0.051465 | 0.02116 | 0.010109 | 0.029416 | 0.009437 | 0.018422 | 0.016117 | 0.010314 | 0.015229 | 0.03521 | 0.02445 | 0.016358 | 0.044801 | 0.022333 |
| 33.5416667 | 0.038249 | 0.035987 | 0.104112 | 0.034849 | 0.034636 | 0.023189 | 0.024975 | 0.00438 | 0.050832 | 0.031567 | 0.028977 | 0.02465 | 0.022484 | 0.038544 | 0.01885 | 0.042923 | 0.02351 | 0.014013 | 0.015362 | 0.053084 | 0.048224 | 0.036364 | 0.062944 | 0.041978 |
| 33.5555556 | 0.077079 | 0.063021 | 0.131771 | 0.043643 | 0.037531 | 0.035844 | 0.029013 | 0.009615 | 0.069132 | 0.044537 | 0.045744 | 0.053352 | 0.052252 | 0.047193 | 0.042901 | 0.072815 | 0.03448 | 0.013971 | 0.019133 | 0.079485 | 0.08607 | 0.068906 | 0.103932 | 0.081135 |
| 33.5694444 | 0.117781 | 0.081566 | 0.161554 | 0.061375 | 0.042843 | 0.047379 | 0.037775 | 0.017745 | 0.080686 | 0.067122 | 0.083294 | 0.080659 | 0.096826 | 0.061143 | 0.080274 | 0.095818 | 0.047492 | 0.020751 | 0.025923 | 0.102108 | 0.109466 | 0.095983 | 0.175806 | 0.120235 |
| 33.5833333 | 0.132262 | 0.082375 | 0.172599 | 0.081026 | 0.045984 | 0.05323 | 0.045397 | 0.024538 | 0.075573 | 0.081022 | 0.111232 | 0.086714 | 0.138495 | 0.085824 | 0.114802 | 0.113513 | 0.063304 | 0.044709 | 0.034206 | 0.114775 | 0.092178 | 0.099785 | 0.212762 | 0.13178 |
| 33.5972222 | 0.11948 | 0.08031 | 0.154709 | 0.095594 | 0.045288 | 0.059098 | 0.046383 | 0.031417 | 0.061625 | 0.071604 | 0.128887 | 0.080672 | 0.15207 | 0.115042 | 0.126623 | 0.135735 | 0.083651 | 0.069974 | 0.04455 | 0.129978 | 0.064473 | 0.088947 | 0.166012 | 0.108967 |
| 33.6111111 | 0.091099 | 0.088011 | 0.111401 | 0.091576 | 0.041908 | 0.065421 | 0.042295 | 0.037656 | 0.054009 | 0.052678 | 0.147515 | 0.077556 | 0.130002 | 0.125142 | 0.120466 | 0.149962 | 0.094681 | 0.07637 | 0.052752 | 0.142889 | 0.062517 | 0.077717 | 0.10739 | 0.073128 |
| 33.625 | 0.068968 | 0.105033 | 0.058312 | 0.059182 | 0.036941 | 0.061742 | 0.035186 | 0.04124 | 0.055251 | 0.034829 | 0.169051 | 0.080088 | 0.084737 | 0.106083 | 0.101055 | 0.127384 | 0.077555 | 0.064397 | 0.060225 | 0.12874 | 0.071414 | 0.068509 | 0.079432 | 0.065293 |
| 33.6388889 | 0.071482 | 0.124456 | 0.045625 | 0.024406 | 0.03061 | 0.047551 | 0.02515 | 0.044674 | 0.056017 | 0.018937 | 0.18063 | 0.087094 | 0.042226 | 0.07 | 0.059549 | 0.075457 | 0.053446 | 0.040735 | 0.069434 | 0.093823 | 0.061415 | 0.056846 | 0.058901 | 0.084287 |
| 33.6527778 | 0.085217 | 0.133564 | 0.085585 | 0.012823 | 0.021737 | 0.029254 | 0.014009 | 0.044897 | 0.052085 | 0.013596 | 0.171551 | 0.103242 | 0.0288 | 0.03756 | 0.029753 | 0.038506 | 0.046516 | 0.018545 | 0.067581 | 0.062798 | 0.038893 | 0.038577 | 0.038674 | 0.088533 |
| 33.6666667 | 0.096095 | 0.122212 | 0.130979 | 0.011442 | 0.012541 | 0.012451 | 0.005245 | 0.035496 | 0.045846 | 0.018546 | 0.150385 | 0.129313 | 0.039605 | 0.038788 | 0.050149 | 0.03434 | 0.035866 | 0.017212 | 0.047355 | 0.03525 | 0.025028 | 0.017912 | 0.02216 | 0.062689 |
| 33.6805556 | 0.115506 | 0.09429 | 0.144454 | 0.008678 | 0.008968 | 0.004846 | 0.001191 | 0.024815 | 0.038918 | 0.020914 | 0.130629 | 0.144286 | 0.041988 | 0.083049 | 0.089091 | 0.039612 | 0.020656 | 0.035098 | 0.028185 | 0.019521 | 0.016025 | 0.005024 | 0.023699 | 0.039686 |
| 33.6944444 | 0.151508 | 0.066941 | 0.107245 | 0.005535 | 0.011427 | 0.007858 | 0.00369 | 0.020393 | 0.031973 | 0.015016 | 0.10471 | 0.13461 | 0.026117 | 0.12927 | 0.098874 | 0.035418 | 0.023452 | 0.057951 | 0.032039 | 0.03642 | 0.007521 | 0.002481 | 0.042797 | 0.04788 |
| 33.7083333 | 0.195538 | 0.048595 | 0.055738 | 0.004598 | 0.014354 | 0.014108 | 0.013395 | 0.013851 | 0.025444 | 0.009167 | 0.059283 | 0.110746 | 0.021336 | 0.130748 | 0.079253 | 0.024678 | 0.037889 | 0.072617 | 0.042021 | 0.065518 | 0.005417 | 0.004266 | 0.061167 | 0.061261 |
| 33.7222222 | 0.247699 | 0.029589 | 0.05702 | 0.012589 | 0.016125 | 0.019118 | 0.025495 | 0.008403 | 0.02043 | 0.013269 | 0.031757 | 0.075734 | 0.039373 | 0.096445 | 0.047622 | 0.023492 | 0.053102 | 0.059299 | 0.034148 | 0.078067 | 0.006664 | 0.007889 | 0.076087 | 0.060896 |
| 33.7361111 | 0.320393 | 0.016512 | 0.108241 | 0.031266 | 0.019703 | 0.025242 | 0.034564 | 0.014191 | 0.018285 | 0.019915 | 0.062623 | 0.040797 | 0.048219 | 0.061828 | 0.02481 | 0.037538 | 0.065004 | 0.032732 | 0.01687 | 0.071256 | 0.007811 | 0.01462 | 0.088342 | 0.054273 |
| 33.75 | 0.383019 | 0.028552 | 0.15887 | 0.054654 | 0.024793 | 0.031608 | 0.040037 | 0.023744 | 0.017953 | 0.020582 | 0.122885 | 0.038286 | 0.032255 | 0.033583 | 0.035836 | 0.036568 | 0.053357 | 0.031715 | 0.011647 | 0.053296 | 0.008629 | 0.023498 | 0.086828 | 0.03559 |
| 33.7638889 | 0.375354 | 0.055618 | 0.175755 | 0.070457 | 0.028196 | 0.031539 | 0.042925 | 0.030125 | 0.018026 | 0.01813 | 0.171055 | 0.061879 | 0.018934 | 0.026542 | 0.069195 | 0.02617 | 0.033417 | 0.052064 | 0.023316 | 0.036628 | 0.010559 | 0.030778 | 0.061012 | 0.016944 |
| 33.7777778 | 0.302197 | 0.072711 | 0.153075 | 0.071384 | 0.02972 | 0.023614 | 0.041498 | 0.032138 | 0.020409 | 0.014756 | 0.182589 | 0.064283 | 0.029639 | 0.059536 | 0.092796 | 0.040435 | 0.050034 | 0.064589 | 0.038397 | 0.025382 | 0.015739 | 0.034609 | 0.030569 | 0.022777 |
| 33.7916667 | 0.223716 | 0.070553 | 0.109849 | 0.061818 | 0.030935 | 0.014393 | 0.036021 | 0.028305 | 0.025682 | 0.011968 | 0.146681 | 0.038488 | 0.052142 | 0.089225 | 0.11291 | 0.047763 | 0.083518 | 0.055986 | 0.051128 | 0.018317 | 0.01952 | 0.035505 | 0.031476 | 0.044137 |
| 33.8055556 | 0.162205 | 0.058619 | 0.067407 | 0.046418 | 0.03049 | 0.008275 | 0.031494 | 0.020201 | 0.031212 | 0.011841 | 0.084556 | 0.021798 | 0.068992 | 0.075652 | 0.130801 | 0.02902 | 0.092124 | 0.0321 | 0.055658 | 0.014763 | 0.01781 | 0.033036 | 0.062385 | 0.062428 |
| 33.8194444 | 0.09145 | 0.047938 | 0.034897 | 0.025421 | 0.024875 | 0.004459 | 0.029659 | 0.010945 | 0.035023 | 0.014176 | 0.035828 | 0.021741 | 0.077595 | 0.045555 | 0.107322 | 0.019635 | 0.073552 | 0.019452 | 0.044628 | 0.014204 | 0.012988 | 0.027202 | 0.092753 | 0.070174 |
| 33.8333333 | 0.046 | 0.039763 | 0.028324 | 0.012146 | 0.01464 | 0.005081 | 0.027832 | 0.004503 | 0.035973 | 0.0155 | 0.028285 | 0.018749 | 0.080419 | 0.03164 | 0.059955 | 0.037948 | 0.040064 | 0.039744 | 0.027049 | 0.016029 | 0.008695 | 0.019793 | 0.105632 | 0.065756 |
| 33.8472222 | 0.08208 | 0.035688 | 0.050362 | 0.019308 | 0.00587 | 0.014584 | 0.024215 | 0.003225 | 0.033266 | 0.013149 | 0.058742 | 0.010292 | 0.074177 | 0.030611 | 0.048529 | 0.072679 | 0.022307 | 0.076156 | 0.016082 | 0.017715 | 0.004938 | 0.01393 | 0.1056 | 0.05616 |
| 33.8611111 | 0.156792 | 0.030454 | 0.082188 | 0.033725 | 0.00193 | 0.032833 | 0.017722 | 0.007169 | 0.026554 | 0.007601 | 0.097986 | 0.005437 | 0.0548 | 0.025124 | 0.072654 | 0.098473 | 0.036753 | 0.101121 | 0.015375 | 0.016098 | 0.002193 | 0.011806 | 0.101633 | 0.046743 |
| 33.875 | 0.226298 | 0.016903 | 0.119935 | 0.045469 | 0.001907 | 0.057998 | 0.010167 | 0.01341 | 0.016512 | 0.004696 | 0.115914 | 0.006124 | 0.028396 | 0.013802 | 0.082895 | 0.097014 | 0.05082 | 0.106019 | 0.019541 | 0.009756 | 0.002136 | 0.012287 | 0.088148 | 0.031938 |
| 33.8888889 | 0.304362 | 0.007192 | 0.158914 | 0.055997 | 0.004438 | 0.082724 | 0.008507 | 0.02143 | 0.008201 | 0.008227 | 0.100805 | 0.005349 | 0.011857 | 0.007214 | 0.073018 | 0.078106 | 0.053616 | 0.095828 | 0.02097 | 0.003163 | 0.002981 | 0.012126 | 0.053704 | 0.020043 |
| 33.9027778 | 0.391117 | 0.007655 | 0.173456 | 0.061082 | 0.008764 | 0.095263 | 0.014684 | 0.035208 | 0.008648 | 0.012679 | 0.060587 | 0.003263 | 0.009882 | 0.012544 | 0.064011 | 0.060421 | 0.062508 | 0.079715 | 0.018762 | 0.003748 | 0.002805 | 0.009144 | 0.0298 | 0.031509 |
| 33.9166667 | 0.466581 | 0.009617 | 0.141734 | 0.056668 | 0.016225 | 0.089022 | 0.023195 | 0.051038 | 0.017049 | 0.016073 | 0.032887 | 0.005647 | 0.013474 | 0.027356 | 0.04713 | 0.049752 | 0.070967 | 0.063891 | 0.015899 | 0.013461 | 0.002138 | 0.004773 | 0.058618 | 0.056817 |
| 33.9305556 | 0.506804 | 0.018773 | 0.082972 | 0.042626 | 0.025665 | 0.068188 | 0.030999 | 0.061164 | 0.025749 | 0.019885 | 0.053737 | 0.010087 | 0.027901 | 0.041734 | 0.032513 | 0.043554 | 0.071857 | 0.051798 | 0.011159 | 0.028737 | 0.0032 | 0.002981 | 0.12129 | 0.076233 |
| 33.9444444 | 0.482858 | 0.040189 | 0.07086 | 0.021909 | 0.032453 | 0.046189 | 0.036632 | 0.066249 | 0.029202 | 0.021657 | 0.108208 | 0.012419 | 0.05103 | 0.049283 | 0.033969 | 0.034615 | 0.068832 | 0.042188 | 0.009644 | 0.045615 | 0.005436 | 0.005441 | 0.181701 | 0.087348 |
| 33.9583333 | 0.408386 | 0.059087 | 0.115953 | 0.008726 | 0.032794 | 0.031292 | 0.035905 | 0.067472 | 0.026472 | 0.018662 | 0.160317 | 0.013526 | 0.062306 | 0.050016 | 0.041126 | 0.022541 | 0.05752 | 0.032116 | 0.012871 | 0.063475 | 0.007013 | 0.010386 | 0.226067 | 0.09122 |
| 33.9722222 | 0.323145 | 0.065406 | 0.13249 | 0.011994 | 0.026196 | 0.025426 | 0.026413 | 0.062143 | 0.019802 | 0.011139 | 0.187242 | 0.014674 | 0.054142 | 0.044216 | 0.040612 | 0.01244 | 0.036384 | 0.021994 | 0.009694 | 0.081424 | 0.006319 | 0.01623 | 0.243876 | 0.086114 |
| 33.9861111 | 0.233386 | 0.06357 | 0.098151 | 0.020622 | 0.01599 | 0.024797 | 0.013714 | 0.052799 | 0.012253 | 0.006979 | 0.181121 | 0.014247 | 0.036219 | 0.032257 | 0.033168 | 0.007563 | 0.018762 | 0.012719 | 0.004529 | 0.091744 | 0.003777 | 0.021614 | 0.225001 | 0.069612 |
| 34 | 0.14082 | 0.055996 | 0.051504 | 0.026641 | 0.008068 | 0.020455 | 0.008974 | 0.043408 | 0.006152 | 0.010775 | 0.155605 | 0.010064 | 0.020058 | 0.017744 | 0.024964 | 0.010538 | 0.0194 | 0.005362 | 0.006042 | 0.08532 | 0.003756 | 0.0237 | 0.18052 | 0.045765 |
| 34.0138889 | 0.068859 | 0.04033 | 0.033143 | 0.027986 | 0.007032 | 0.0124 | 0.01368 | 0.034919 | 0.003196 | 0.014896 | 0.129539 | 0.005921 | 0.016922 | 0.007828 | 0.014439 | 0.01807 | 0.035126 | 0.003525 | 0.01035 | 0.065119 | 0.005508 | 0.020969 | 0.133314 | 0.027571 |
| 34.0277778 | 0.051541 | 0.023833 | 0.070664 | 0.020826 | 0.008313 | 0.010445 | 0.017341 | 0.030118 | 0.004559 | 0.013794 | 0.110805 | 0.01208 | 0.021709 | 0.005094 | 0.007251 | 0.018405 | 0.043975 | 0.005933 | 0.013099 | 0.043736 | 0.004813 | 0.016967 | 0.091006 | 0.02492 |
| 34.0416667 | 0.058558 | 0.024655 | 0.13411 | 0.014271 | 0.006828 | 0.023801 | 0.012662 | 0.034628 | 0.010016 | 0.010504 | 0.112586 | 0.037609 | 0.033496 | 0.011491 | 0.011943 | 0.02591 | 0.035899 | 0.012688 | 0.011113 | 0.034826 | 0.010406 | 0.020009 | 0.060735 | 0.044667 |
| 34.0555556 | 0.054015 | 0.052248 | 0.201167 | 0.024219 | 0.009761 | 0.050453 | 0.009256 | 0.051234 | 0.018678 | 0.014175 | 0.144917 | 0.078947 | 0.096019 | 0.041245 | 0.030263 | 0.085951 | 0.047122 | 0.0393 | 0.011814 | 0.054312 | 0.039818 | 0.034762 | 0.057581 | 0.096824 |
| 34.0694444 | 0.077193 | 0.087954 | 0.269401 | 0.041769 | 0.020988 | 0.070102 | 0.017575 | 0.06987 | 0.028256 | 0.02507 | 0.18307 | 0.116101 | 0.186538 | 0.084603 | 0.053705 | 0.178661 | 0.091908 | 0.080768 | 0.022776 | 0.094097 | 0.076665 | 0.049923 | 0.078824 | 0.159348 |
| 34.0833333 | 0.101453 | 0.102167 | 0.307907 | 0.044287 | 0.027986 | 0.063041 | 0.024388 | 0.073762 | 0.031429 | 0.029539 | 0.188051 | 0.130704 | 0.200226 | 0.099183 | 0.071975 | 0.218311 | 0.111911 | 0.098733 | 0.032473 | 0.112627 | 0.078621 | 0.047398 | 0.09502 | 0.168988 |
| 34.0972222 | 0.073666 | 0.089979 | 0.276548 | 0.027342 | 0.021335 | 0.034856 | 0.018815 | 0.059983 | 0.023079 | 0.023477 | 0.159624 | 0.125991 | 0.13651 | 0.0715 | 0.081775 | 0.181711 | 0.080559 | 0.078438 | 0.028531 | 0.092756 | 0.045459 | 0.028267 | 0.089514 | 0.115439 |
| 34.1111111 | 0.02883 | 0.065211 | 0.196863 | 0.011233 | 0.010572 | 0.012261 | 0.009364 | 0.039756 | 0.011697 | 0.013328 | 0.127615 | 0.109962 | 0.081289 | 0.033415 | 0.075979 | 0.125592 | 0.037951 | 0.040014 | 0.020071 | 0.060949 | 0.014736 | 0.010836 | 0.077091 | 0.058357 |
| 34.125 | 0.006374 | 0.039509 | 0.13693 | 0.012071 | 0.00975 | 0.009813 | 0.007111 | 0.021535 | 0.009505 | 0.006635 | 0.10141 | 0.081355 | 0.052758 | 0.019006 | 0.056212 | 0.087217 | 0.020031 | 0.014688 | 0.024829 | 0.038257 | 0.003457 | 0.005735 | 0.0688 | 0.028892 |
| 34.1388889 | 0.004586 | 0.018135 | 0.095665 | 0.017796 | 0.017373 | 0.013551 | 0.013298 | 0.008588 | 0.014237 | 0.008179 | 0.076854 | 0.04872 | 0.032003 | 0.029332 | 0.036632 | 0.057772 | 0.021985 | 0.017287 | 0.036388 | 0.024832 | 0.00309 | 0.008993 | 0.0582 | 0.016264 |
| 34.1527778 | 0.008755 | 0.007148 | 0.050715 | 0.018096 | 0.02244 | 0.012542 | 0.019212 | 0.002721 | 0.016486 | 0.012924 | 0.056442 | 0.023399 | 0.039228 | 0.044339 | 0.019446 | 0.029098 | 0.024235 | 0.029434 | 0.039607 | 0.016343 | 0.005625 | 0.013798 | 0.037652 | 0.01002 |
| 34.1666667 | 0.008281 | 0.010477 | 0.035066 | 0.016657 | 0.021712 | 0.010488 | 0.0198 | 0.001522 | 0.015618 | 0.0151 | 0.038341 | 0.009566 | 0.079679 | 0.051575 | 0.009913 | 0.01092 | 0.023638 | 0.036868 | 0.037599 | 0.009239 | 0.00707 | 0.017361 | 0.019241 | 0.010917 |
| 34.1805556 | 0.007581 | 0.020795 | 0.070796 | 0.017779 | 0.019764 | 0.011067 | 0.017399 | 0.001482 | 0.016481 | 0.015209 | 0.020309 | 0.007792 | 0.119464 | 0.051083 | 0.016571 | 0.009923 | 0.022038 | 0.041775 | 0.035589 | 0.003726 | 0.004937 | 0.01839 | 0.021559 | 0.017687 |
| 34.1944444 | 0.012587 | 0.029068 | 0.127038 | 0.021795 | 0.019999 | 0.014473 | 0.01516 | 0.00176 | 0.019727 | 0.014479 | 0.006963 | 0.011176 | 0.124959 | 0.047154 | 0.027868 | 0.013723 | 0.017883 | 0.045864 | 0.032282 | 0.001648 | 0.002471 | 0.015982 | 0.033454 | 0.022136 |
| 34.2083333 | 0.014553 | 0.030192 | 0.178201 | 0.023663 | 0.021472 | 0.018479 | 0.014072 | 0.001874 | 0.021771 | 0.013714 | 0.002006 | 0.01316 | 0.084242 | 0.039372 | 0.032483 | 0.012769 | 0.011445 | 0.046625 | 0.026576 | 0.001701 | 0.002227 | 0.011456 | 0.037507 | 0.022233 |
| 34.2222222 | 0.010108 | 0.025951 | 0.22601 | 0.022052 | 0.023229 | 0.021451 | 0.01451 | 0.00327 | 0.022566 | 0.015499 | 0.002103 | 0.015436 | 0.04135 | 0.027642 | 0.031709 | 0.008816 | 0.006213 | 0.040851 | 0.019398 | 0.002146 | 0.002352 | 0.00703 | 0.035879 | 0.022226 |
| 34.2361111 | 0.007309 | 0.02426 | 0.254695 | 0.020354 | 0.027048 | 0.023655 | 0.017886 | 0.006659 | 0.025282 | 0.022269 | 0.004394 | 0.020175 | 0.044024 | 0.016135 | 0.028515 | 0.007209 | 0.004699 | 0.030625 | 0.012947 | 0.003138 | 0.003658 | 0.003563 | 0.033722 | 0.025188 |
| 34.25 | 0.009307 | 0.028562 | 0.237648 | 0.021003 | 0.034812 | 0.026851 | 0.024637 | 0.007783 | 0.029151 | 0.032868 | 0.010202 | 0.025357 | 0.0597 | 0.007501 | 0.026381 | 0.01787 | 0.007022 | 0.020099 | 0.009831 | 0.005055 | 0.006713 | 0.001848 | 0.031132 | 0.030844 |
| 34.2638889 | 0.010337 | 0.037302 | 0.181272 | 0.028473 | 0.042783 | 0.031207 | 0.031307 | 0.005345 | 0.03165 | 0.042841 | 0.017802 | 0.029857 | 0.045977 | 0.00342 | 0.028175 | 0.0361 | 0.012537 | 0.01074 | 0.010881 | 0.006811 | 0.00995 | 0.003166 | 0.027616 | 0.036956 |
| 34.2777778 | 0.009157 | 0.049571 | 0.122506 | 0.05207 | 0.041145 | 0.034508 | 0.029952 | 0.003801 | 0.033491 | 0.050177 | 0.025537 | 0.034377 | 0.039777 | 0.005706 | 0.031352 | 0.037115 | 0.01913 | 0.005854 | 0.01309 | 0.005759 | 0.013399 | 0.00603 | 0.023538 | 0.041625 |
| 34.2916667 | 0.01268 | 0.06537 | 0.083839 | 0.083425 | 0.033342 | 0.035117 | 0.020576 | 0.005262 | 0.035353 | 0.056979 | 0.033216 | 0.038258 | 0.073424 | 0.013567 | 0.031468 | 0.021942 | 0.024192 | 0.007735 | 0.013827 | 0.004387 | 0.019147 | 0.008254 | 0.019945 | 0.045864 |
| 34.3055556 | 0.017236 | 0.081219 | 0.054146 | 0.088861 | 0.037326 | 0.031195 | 0.019471 | 0.005482 | 0.033818 | 0.059613 | 0.036919 | 0.035538 | 0.105499 | 0.022556 | 0.029397 | 0.01774 | 0.027546 | 0.007729 | 0.015194 | 0.005733 | 0.027005 | 0.010394 | 0.019289 | 0.049647 |
| 34.3194444 | 0.0162 | 0.093535 | 0.02723 | 0.055051 | 0.048442 | 0.021965 | 0.027435 | 0.006697 | 0.024552 | 0.04853 | 0.028587 | 0.028089 | 0.09835 | 0.02553 | 0.023085 | 0.023313 | 0.027332 | 0.004568 | 0.016661 | 0.00947 | 0.028799 | 0.00928 | 0.017461 | 0.044816 |
| 34.3333333 | 0.028698 | 0.111365 | 0.015692 | 0.015017 | 0.051548 | 0.018542 | 0.028995 | 0.011023 | 0.011522 | 0.02523 | 0.01609 | 0.040032 | 0.063092 | 0.020432 | 0.023932 | 0.020718 | 0.025205 | 0.015212 | 0.014023 | 0.026019 | 0.022195 | 0.011529 | 0.017165 | 0.028492 |
| 34.3472222 | 0.060276 | 0.14029 | 0.019993 | 0.006774 | 0.053878 | 0.036109 | 0.031692 | 0.014788 | 0.010387 | 0.011805 | 0.015886 | 0.06001 | 0.047962 | 0.017683 | 0.055725 | 0.016566 | 0.02068 | 0.043732 | 0.01249 | 0.05345 | 0.020644 | 0.025863 | 0.027415 | 0.017113 |
| 34.3611111 | 0.065351 | 0.15366 | 0.040517 | 0.017754 | 0.055699 | 0.062185 | 0.041501 | 0.022646 | 0.025585 | 0.022586 | 0.033848 | 0.049403 | 0.055244 | 0.024147 | 0.075628 | 0.026851 | 0.014451 | 0.063207 | 0.012261 | 0.057331 | 0.019949 | 0.032101 | 0.028366 | 0.013941 |
| 34.375 | 0.039023 | 0.126196 | 0.085721 | 0.015802 | 0.045153 | 0.066737 | 0.042749 | 0.027426 | 0.03726 | 0.038345 | 0.061606 | 0.031715 | 0.043615 | 0.03185 | 0.0437 | 0.040563 | 0.015631 | 0.052439 | 0.008979 | 0.032566 | 0.014014 | 0.019534 | 0.019675 | 0.010401 |
| 34.3888889 | 0.028892 | 0.078008 | 0.132356 | 0.006855 | 0.029352 | 0.049477 | 0.033826 | 0.019972 | 0.034189 | 0.041018 | 0.074888 | 0.034333 | 0.03462 | 0.036411 | 0.010725 | 0.039784 | 0.017245 | 0.027836 | 0.010938 | 0.016012 | 0.018009 | 0.006578 | 0.027442 | 0.011644 |
| 34.4027778 | 0.033504 | 0.039793 | 0.147215 | 0.008815 | 0.022305 | 0.032221 | 0.027895 | 0.010113 | 0.027577 | 0.03591 | 0.07494 | 0.041126 | 0.035872 | 0.041208 | 0.00523 | 0.031951 | 0.013775 | 0.0182 | 0.017117 | 0.01588 | 0.025608 | 0.00197 | 0.040093 | 0.014712 |
| 34.4166667 | 0.027529 | 0.018917 | 0.14305 | 0.016917 | 0.022385 | 0.020619 | 0.027142 | 0.006408 | 0.025168 | 0.027583 | 0.0833 | 0.041804 | 0.031566 | 0.044183 | 0.007618 | 0.0282 | 0.011065 | 0.024308 | 0.020815 | 0.017807 | 0.026223 | 0.001407 | 0.04307 | 0.016841 |
| 34.4305556 | 0.014465 | 0.016014 | 0.140904 | 0.021187 | 0.023376 | 0.01172 | 0.027258 | 0.010881 | 0.024952 | 0.016912 | 0.093107 | 0.034578 | 0.045593 | 0.041971 | 0.006759 | 0.028619 | 0.008852 | 0.025182 | 0.022603 | 0.021885 | 0.025794 | 0.001659 | 0.039742 | 0.022138 |
| 34.4444444 | 0.008362 | 0.023759 | 0.125424 | 0.023757 | 0.026571 | 0.009503 | 0.029662 | 0.018478 | 0.025401 | 0.010078 | 0.074126 | 0.021802 | 0.06274 | 0.035985 | 0.006042 | 0.030267 | 0.008064 | 0.01672 | 0.023105 | 0.030438 | 0.027274 | 0.003449 | 0.036065 | 0.028143 |
| 34.4583333 | 0.012896 | 0.027791 | 0.080814 | 0.028661 | 0.034555 | 0.013996 | 0.036323 | 0.019585 | 0.025575 | 0.010158 | 0.033447 | 0.010926 | 0.045808 | 0.028421 | 0.010875 | 0.028529 | 0.010058 | 0.010681 | 0.023428 | 0.037617 | 0.02997 | 0.007644 | 0.039983 | 0.030329 |
| 34.4722222 | 0.019314 | 0.023712 | 0.045523 | 0.035609 | 0.047115 | 0.021012 | 0.04529 | 0.014232 | 0.02534 | 0.013639 | 0.010229 | 0.004699 | 0.030033 | 0.022949 | 0.016051 | 0.021643 | 0.012209 | 0.013204 | 0.023373 | 0.036831 | 0.030965 | 0.010355 | 0.044944 | 0.026297 |
| 34.4861111 | 0.02174 | 0.01499 | 0.064035 | 0.043993 | 0.062073 | 0.032329 | 0.054209 | 0.009222 | 0.027357 | 0.017962 | 0.010695 | 0.002107 | 0.057492 | 0.023475 | 0.017962 | 0.014046 | 0.011972 | 0.0228 | 0.0224 | 0.030249 | 0.028764 | 0.00947 | 0.040306 | 0.018841 |
| 34.5 | 0.019218 | 0.007796 | 0.101706 | 0.054174 | 0.075381 | 0.047891 | 0.06091 | 0.006249 | 0.034628 | 0.023576 | 0.02164 | 0.001685 | 0.0834 | 0.026737 | 0.016251 | 0.010482 | 0.009267 | 0.032322 | 0.022018 | 0.024729 | 0.02508 | 0.008209 | 0.032625 | 0.012996 |
| 34.5138889 | 0.011852 | 0.007915 | 0.109262 | 0.063836 | 0.082309 | 0.058429 | 0.063933 | 0.003389 | 0.046223 | 0.030568 | 0.033944 | 0.002533 | 0.064038 | 0.024855 | 0.010056 | 0.011804 | 0.006347 | 0.032928 | 0.023626 | 0.023107 | 0.0219 | 0.009603 | 0.032246 | 0.012144 |
| 34.5277778 | 0.005144 | 0.015944 | 0.089681 | 0.065918 | 0.081353 | 0.058854 | 0.065751 | 0.002414 | 0.056893 | 0.040202 | 0.042998 | 0.006726 | 0.04698 | 0.015567 | 0.003962 | 0.019793 | 0.007905 | 0.02195 | 0.02732 | 0.027338 | 0.021206 | 0.015063 | 0.039275 | 0.018854 |
| 34.5416667 | 0.009155 | 0.032198 | 0.071772 | 0.059819 | 0.079456 | 0.057619 | 0.073327 | 0.007851 | 0.063664 | 0.056752 | 0.048119 | 0.01813 | 0.06933 | 0.005887 | 0.002891 | 0.04106 | 0.018666 | 0.012948 | 0.032572 | 0.04388 | 0.026758 | 0.025783 | 0.050889 | 0.036225 |
| 34.5555556 | 0.02839 | 0.057078 | 0.090981 | 0.061254 | 0.084084 | 0.062319 | 0.088774 | 0.019816 | 0.06683 | 0.078639 | 0.042728 | 0.036586 | 0.071105 | 0.006419 | 0.011994 | 0.073011 | 0.037603 | 0.015171 | 0.037618 | 0.07328 | 0.039788 | 0.038514 | 0.077313 | 0.067146 |
| 34.5694444 | 0.053459 | 0.078717 | 0.149723 | 0.082734 | 0.091489 | 0.072269 | 0.10211 | 0.031079 | 0.063531 | 0.093201 | 0.024194 | 0.052362 | 0.054119 | 0.022872 | 0.032623 | 0.095595 | 0.054412 | 0.023218 | 0.04054 | 0.104746 | 0.052943 | 0.04382 | 0.131461 | 0.099634 |
| 34.5833333 | 0.067377 | 0.081306 | 0.191818 | 0.112511 | 0.09408 | 0.075949 | 0.106564 | 0.033691 | 0.051683 | 0.091098 | 0.008254 | 0.055972 | 0.087321 | 0.052067 | 0.050641 | 0.10077 | 0.065317 | 0.055081 | 0.042572 | 0.125363 | 0.058842 | 0.040404 | 0.18847 | 0.117112 |
| 34.5972222 | 0.059915 | 0.067958 | 0.170507 | 0.133541 | 0.089632 | 0.058292 | 0.101846 | 0.027936 | 0.036196 | 0.078019 | 0.005143 | 0.050883 | 0.150237 | 0.079922 | 0.051178 | 0.102578 | 0.075999 | 0.11079 | 0.046172 | 0.129129 | 0.060339 | 0.039087 | 0.213898 | 0.12091 |
| 34.6111111 | 0.036032 | 0.052825 | 0.10896 | 0.127979 | 0.07559 | 0.032188 | 0.084234 | 0.019555 | 0.024944 | 0.065308 | 0.006028 | 0.044899 | 0.194952 | 0.092893 | 0.037307 | 0.104769 | 0.082292 | 0.140909 | 0.048886 | 0.11708 | 0.061857 | 0.043298 | 0.208059 | 0.112668 |
| 34.625 | 0.013891 | 0.042099 | 0.051343 | 0.081279 | 0.050823 | 0.01839 | 0.055725 | 0.011424 | 0.021922 | 0.056765 | 0.007242 | 0.039511 | 0.193594 | 0.088505 | 0.020535 | 0.099125 | 0.07882 | 0.125655 | 0.047805 | 0.091936 | 0.059597 | 0.047254 | 0.175422 | 0.094602 |
| 34.6388889 | 0.005697 | 0.036677 | 0.031455 | 0.049954 | 0.023895 | 0.013926 | 0.026246 | 0.005294 | 0.024875 | 0.049388 | 0.011139 | 0.033884 | 0.132745 | 0.060576 | 0.012163 | 0.087581 | 0.070663 | 0.083861 | 0.045008 | 0.064213 | 0.046083 | 0.047547 | 0.119179 | 0.078152 |

| | | | | | | | | | | | | | | | | | | | | | | | | |
|---|---|---|---|---|---|---|---|---|---|---|---|---|---|---|---|---|---|---|---|---|---|---|---|---|
| 34.6527778 | 0.009224 | 0.03418 | 0.060179 | 0.077305 | 0.010578 | 0.009966 | 0.010036 | 0.004374 | 0.02914 | 0.041471 | 0.016269 | 0.028543 | 0.067138 | 0.032389 | 0.015162 | 0.076068 | 0.063318 | 0.039614 | 0.040277 | 0.044407 | 0.023582 | 0.042849 | 0.064179 | 0.074711 |
| 34.6666667 | 0.017258 | 0.029333 | 0.115295 | 0.096299 | 0.013924 | 0.014604 | 0.008063 | 0.007136 | 0.030864 | 0.034179 | 0.020166 | 0.024107 | 0.071822 | 0.037099 | 0.016177 | 0.06243 | 0.055648 | 0.022165 | 0.031577 | 0.033661 | 0.006734 | 0.036249 | 0.036458 | 0.088848 |
| 34.6805556 | 0.02206 | 0.022723 | 0.171646 | 0.07824 | 0.015153 | 0.033149 | 0.009028 | 0.006976 | 0.029737 | 0.029964 | 0.023693 | 0.022613 | 0.127979 | 0.045474 | 0.010687 | 0.047051 | 0.044404 | 0.035903 | 0.021001 | 0.028211 | 0.004959 | 0.033394 | 0.027902 | 0.098572 |
| 34.6944444 | 0.027227 | 0.019351 | 0.199454 | 0.05115 | 0.011805 | 0.044787 | 0.016896 | 0.005856 | 0.029523 | 0.031217 | 0.031734 | 0.026329 | 0.161746 | 0.035672 | 0.005158 | 0.036496 | 0.031192 | 0.057885 | 0.011145 | 0.019201 | 0.012872 | 0.03263 | 0.023595 | 0.07171 |
| 34.7083333 | 0.038324 | 0.019495 | 0.187349 | 0.024915 | 0.019845 | 0.038577 | 0.032031 | 0.010454 | 0.031158 | 0.036339 | 0.042815 | 0.032843 | 0.142131 | 0.023994 | 0.005031 | 0.032306 | 0.020142 | 0.069177 | 0.003967 | 0.013207 | 0.022364 | 0.027589 | 0.042202 | 0.035767 |
| 34.7222222 | 0.053316 | 0.019775 | 0.139389 | 0.016387 | 0.036403 | 0.023075 | 0.0437 | 0.015545 | 0.031523 | 0.040937 | 0.047544 | 0.036413 | 0.082831 | 0.01437 | 0.013602 | 0.032951 | 0.013843 | 0.057912 | 0.005408 | 0.028202 | 0.029308 | 0.017692 | 0.088923 | 0.035202 |
| 34.7361111 | 0.078063 | 0.019328 | 0.082953 | 0.029002 | 0.050286 | 0.015725 | 0.048619 | 0.016044 | 0.029711 | 0.041932 | 0.040966 | 0.033313 | 0.041879 | 0.013177 | 0.022897 | 0.038024 | 0.011672 | 0.031571 | 0.017744 | 0.053984 | 0.030781 | 0.008032 | 0.126858 | 0.049513 |
| 34.75 | 0.115622 | 0.0202 | 0.081244 | 0.051655 | 0.058784 | 0.029788 | 0.048558 | 0.015197 | 0.027458 | 0.038925 | 0.026141 | 0.02373 | 0.064978 | 0.027157 | 0.025471 | 0.046343 | 0.010267 | 0.018412 | 0.030656 | 0.066728 | 0.02587 | 0.002897 | 0.135822 | 0.053623 |
| 34.7638889 | 0.149303 | 0.024156 | 0.130407 | 0.076099 | 0.062779 | 0.053427 | 0.046987 | 0.014986 | 0.026416 | 0.035064 | 0.012304 | 0.012673 | 0.111405 | 0.04416 | 0.027586 | 0.05547 | 0.00907 | 0.023593 | 0.035942 | 0.064854 | 0.015938 | 0.002851 | 0.134712 | 0.049175 |
| 34.7777778 | 0.159431 | 0.030396 | 0.161562 | 0.088813 | 0.064988 | 0.069675 | 0.04733 | 0.014609 | 0.027054 | 0.034432 | 0.010195 | 0.005094 | 0.125216 | 0.052891 | 0.032815 | 0.062647 | 0.011062 | 0.027854 | 0.036519 | 0.059516 | 0.007167 | 0.004609 | 0.138501 | 0.03388 |
| 34.7916667 | 0.141946 | 0.036009 | 0.15628 | 0.096865 | 0.067878 | 0.07458 | 0.050424 | 0.013195 | 0.028086 | 0.037899 | 0.020988 | 0.00329 | 0.099063 | 0.052365 | 0.028664 | 0.065468 | 0.016922 | 0.021269 | 0.036612 | 0.05294 | 0.004121 | 0.004869 | 0.14476 | 0.015032 |
| 34.8055556 | 0.108358 | 0.038635 | 0.146237 | 0.123536 | 0.071308 | 0.073939 | 0.055261 | 0.0113 | 0.02805 | 0.043867 | 0.035505 | 0.008002 | 0.053169 | 0.042999 | 0.014036 | 0.062162 | 0.024603 | 0.012683 | 0.036196 | 0.042643 | 0.003639 | 0.00386 | 0.142957 | 0.006166 |
| 34.8194444 | 0.074677 | 0.038638 | 0.137826 | 0.151368 | 0.074296 | 0.070336 | 0.060176 | 0.009185 | 0.027488 | 0.050566 | 0.043484 | 0.013465 | 0.033706 | 0.028524 | 0.004195 | 0.051692 | 0.033236 | 0.015869 | 0.034507 | 0.029433 | 0.004759 | 0.003135 | 0.119648 | 0.008852 |
| 34.8333333 | 0.045251 | 0.036162 | 0.108248 | 0.142513 | 0.075133 | 0.056322 | 0.062357 | 0.00657 | 0.027178 | 0.054958 | 0.04047 | 0.014658 | 0.071212 | 0.017234 | 0.003549 | 0.034001 | 0.040727 | 0.025926 | 0.033482 | 0.016109 | 0.008322 | 0.00258 | 0.073975 | 0.020681 |
| 34.8472222 | 0.024718 | 0.03001 | 0.057636 | 0.090916 | 0.07163 | 0.030485 | 0.059157 | 0.003924 | 0.026161 | 0.053212 | 0.030953 | 0.013645 | 0.124542 | 0.013062 | 0.012668 | 0.017087 | 0.043416 | 0.032546 | 0.03521 | 0.007123 | 0.012724 | 0.002013 | 0.06022 | 0.0398 |
| 34.8611111 | 0.030512 | 0.022101 | 0.032989 | 0.045246 | 0.062924 | 0.014619 | 0.050011 | 0.002334 | 0.024062 | 0.044486 | 0.021992 | 0.011649 | 0.145203 | 0.012698 | 0.027763 | 0.012347 | 0.041625 | 0.0347 | 0.038886 | 0.005863 | 0.016261 | 0.002363 | 0.12197 | 0.064387 |
| 34.875 | 0.065798 | 0.015926 | 0.066897 | 0.050842 | 0.049194 | 0.02745 | 0.03637 | 0.003259 | 0.021755 | 0.031713 | 0.014014 | 0.007334 | 0.137373 | 0.015708 | 0.039279 | 0.013538 | 0.039859 | 0.035974 | 0.040335 | 0.011488 | 0.017416 | 0.002749 | 0.185986 | 0.087235 |
| 34.8888889 | 0.112994 | 0.012197 | 0.130001 | 0.065921 | 0.032738 | 0.058111 | 0.020997 | 0.006385 | 0.019222 | 0.017981 | 0.009111 | 0.003345 | 0.12398 | 0.021511 | 0.047204 | 0.01194 | 0.041761 | 0.044734 | 0.034693 | 0.017772 | 0.014711 | 0.002175 | 0.192724 | 0.095967 |
| 34.9027778 | 0.159876 | 0.009993 | 0.186528 | 0.051841 | 0.017528 | 0.090887 | 0.008844 | 0.009648 | 0.01483 | 0.00905 | 0.014576 | 0.002681 | 0.109362 | 0.023663 | 0.050953 | 0.017108 | 0.044497 | 0.066256 | 0.022036 | 0.018839 | 0.009087 | 0.001462 | 0.16905 | 0.087669 |
| 34.9166667 | 0.203954 | 0.009679 | 0.219698 | 0.035785 | 0.007576 | 0.115599 | 0.005347 | 0.013609 | 0.00821 | 0.011639 | 0.026938 | 0.003152 | 0.091933 | 0.01723 | 0.044934 | 0.031507 | 0.041998 | 0.092265 | 0.009527 | 0.01244 | 0.006528 | 0.001698 | 0.121012 | 0.063822 |
| 34.9305556 | 0.240487 | 0.011798 | 0.234868 | 0.030059 | 0.006244 | 0.117689 | 0.009203 | 0.020406 | 0.004548 | 0.021768 | 0.034862 | 0.006033 | 0.07419 | 0.008748 | 0.029514 | 0.04428 | 0.034639 | 0.107543 | 0.003559 | 0.008704 | 0.008527 | 0.00331 | 0.07268 | 0.039283 |
| 34.9444444 | 0.26017 | 0.014302 | 0.230329 | 0.023254 | 0.01142 | 0.096911 | 0.013501 | 0.027614 | 0.007437 | 0.029995 | 0.033678 | 0.013617 | 0.057549 | 0.007502 | 0.013851 | 0.050631 | 0.028321 | 0.1066 | 0.002634 | 0.023228 | 0.008233 | 0.00501 | 0.092445 | 0.045159 |
| 34.9583333 | 0.249505 | 0.014649 | 0.17778 | 0.017226 | 0.016024 | 0.073346 | 0.013971 | 0.030254 | 0.012676 | 0.032344 | 0.024538 | 0.022271 | 0.046832 | 0.009616 | 0.009989 | 0.051045 | 0.023156 | 0.089892 | 0.005767 | 0.050601 | 0.005708 | 0.007219 | 0.169196 | 0.079207 |
| 34.9722222 | 0.203018 | 0.013399 | 0.099472 | 0.011501 | 0.015458 | 0.056726 | 0.009688 | 0.027517 | 0.016707 | 0.028958 | 0.012914 | 0.026451 | 0.042189 | 0.008986 | 0.020813 | 0.045214 | 0.01506 | 0.060613 | 0.015508 | 0.073482 | 0.008097 | 0.012074 | 0.235489 | 0.103501 |
| 34.9861111 | 0.13873 | 0.01337 | 0.081315 | 0.007585 | 0.010471 | 0.042082 | 0.005087 | 0.022284 | 0.018722 | 0.021653 | 0.00771 | 0.022216 | 0.036639 | 0.007375 | 0.035773 | 0.034825 | 0.013995 | 0.030451 | 0.02805 | 0.080698 | 0.014755 | 0.018664 | 0.251829 | 0.089991 |
| 35 | 0.078574 | 0.014248 | 0.148151 | 0.01353 | 0.005023 | 0.024244 | 0.006128 | 0.016441 | 0.018133 | 0.012653 | 0.0095 | 0.012079 | 0.023945 | 0.008433 | 0.042325 | 0.024848 | 0.027895 | 0.012923 | 0.036601 | 0.074425 | 0.019173 | 0.023514 | 0.216651 | 0.046225 |
| 35.0138889 | 0.035172 | 0.012465 | 0.216402 | 0.021735 | 0.002573 | 0.010049 | 0.010095 | 0.010096 | 0.01607 | 0.005085 | 0.010993 | 0.007123 | 0.011406 | 0.016432 | 0.035759 | 0.016917 | 0.042862 | 0.015268 | 0.040417 | 0.066108 | 0.018931 | 0.02397 | 0.172758 | 0.011815 |
| 35.0277778 | 0.015685 | 0.007496 | 0.224667 | 0.0225 | 0.002734 | 0.006627 | 0.010355 | 0.007047 | 0.015599 | 0.003442 | 0.015558 | 0.009632 | 0.00773 | 0.033871 | 0.020888 | 0.011341 | 0.051738 | 0.029743 | 0.042595 | 0.062344 | 0.01445 | 0.018806 | 0.14373 | 0.004336 |
| 35.0416667 | 0.022726 | 0.007109 | 0.163676 | 0.019172 | 0.007818 | 0.014455 | 0.008169 | 0.016271 | 0.020954 | 0.012898 | 0.041996 | 0.013331 | 0.015367 | 0.055521 | 0.022233 | 0.022326 | 0.065391 | 0.029787 | 0.042442 | 0.068684 | 0.010113 | 0.012487 | 0.133441 | 0.01712 |
| 35.0555556 | 0.073002 | 0.020235 | 0.102996 | 0.026326 | 0.022414 | 0.040085 | 0.01272 | 0.042218 | 0.0341 | 0.035729 | 0.104712 | 0.028646 | 0.050426 | 0.073319 | 0.06261 | 0.076529 | 0.094469 | 0.032748 | 0.032079 | 0.090675 | 0.015961 | 0.016449 | 0.149143 | 0.054891 |
| 35.0694444 | 0.148621 | 0.038699 | 0.148525 | 0.043921 | 0.036584 | 0.068272 | 0.024902 | 0.072481 | 0.049686 | 0.061333 | 0.18252 | 0.062166 | 0.116046 | 0.079341 | 0.121649 | 0.155676 | 0.122179 | 0.087527 | 0.023405 | 0.116582 | 0.030997 | 0.029526 | 0.165238 | 0.098449 |
| 35.0833333 | 0.178331 | 0.04379 | 0.256128 | 0.045088 | 0.036432 | 0.070162 | 0.032502 | 0.085214 | 0.055246 | 0.072165 | 0.219887 | 0.092597 | 0.174642 | 0.063696 | 0.151686 | 0.191689 | 0.11476 | 0.164496 | 0.035536 | 0.116329 | 0.037981 | 0.034518 | 0.149098 | 0.106341 |
| 35.0972222 | 0.144062 | 0.032193 | 0.327118 | 0.025729 | 0.023388 | 0.047395 | 0.027759 | 0.074233 | 0.044842 | 0.065126 | 0.191397 | 0.102634 | 0.192698 | 0.032554 | 0.140111 | 0.160621 | 0.074779 | 0.195493 | 0.047422 | 0.084686 | 0.02749 | 0.025595 | 0.108948 | 0.070406 |
| 35.1111111 | 0.092489 | 0.016269 | 0.329428 | 0.011459 | 0.011605 | 0.024747 | 0.015067 | 0.053063 | 0.027618 | 0.050496 | 0.137344 | 0.099708 | 0.156364 | 0.015381 | 0.110972 | 0.100697 | 0.033344 | 0.161979 | 0.038198 | 0.050051 | 0.012255 | 0.013174 | 0.074009 | 0.032398 |
| 35.125 | 0.059679 | 0.006366 | 0.270628 | 0.011528 | 0.013074 | 0.012668 | 0.006967 | 0.032899 | 0.013421 | 0.037259 | 0.098766 | 0.088253 | 0.087253 | 0.022175 | 0.073478 | 0.055114 | 0.010942 | 0.098148 | 0.019003 | 0.028253 | 0.006893 | 0.005028 | 0.049248 | 0.028296 |
| 35.1388889 | 0.047524 | 0.004374 | 0.19951 | 0.015448 | 0.02183 | 0.00628 | 0.009529 | 0.015735 | 0.006009 | 0.02785 | 0.077471 | 0.065342 | 0.0655 | 0.030858 | 0.038818 | 0.042631 | 0.0107 | 0.043744 | 0.008552 | 0.015309 | 0.007919 | 0.001581 | 0.028418 | 0.038624 |
| 35.1527778 | 0.040966 | 0.006794 | 0.155933 | 0.016993 | 0.026434 | 0.005562 | 0.014309 | 0.006511 | 0.007046 | 0.019883 | 0.063104 | 0.040074 | 0.098995 | 0.029296 | 0.021157 | 0.049723 | 0.021077 | 0.016826 | 0.013867 | 0.006927 | 0.007058 | 0.001817 | 0.012673 | 0.040093 |
| 35.1666667 | 0.030673 | 0.010631 | 0.131426 | 0.01742 | 0.026124 | 0.010506 | 0.016388 | 0.00532 | 0.012525 | 0.011864 | 0.049251 | 0.020082 | 0.116207 | 0.022935 | 0.014228 | 0.044439 | 0.021375 | 0.021944 | 0.026366 | 0.002902 | 0.005592 | 0.003244 | 0.006348 | 0.037961 |
| 35.1805556 | 0.016506 | 0.015011 | 0.10776 | 0.018162 | 0.02468 | 0.015154 | 0.017234 | 0.004846 | 0.016868 | 0.005556 | 0.037364 | 0.009781 | 0.112747 | 0.017093 | 0.010803 | 0.023807 | 0.013674 | 0.043079 | 0.04119 | 0.003362 | 0.006953 | 0.003059 | 0.008598 | 0.035064 |
| 35.1944444 | 0.006662 | 0.01798 | 0.082785 | 0.018709 | 0.024339 | 0.015822 | 0.017071 | 0.005656 | 0.018456 | 0.003303 | 0.030805 | 0.014268 | 0.105743 | 0.013781 | 0.007303 | 0.013717 | 0.017378 | 0.062125 | 0.059198 | 0.004802 | 0.010006 | 0.003224 | 0.010259 | 0.02838 |
| 35.2083333 | 0.00703 | 0.017971 | 0.057872 | 0.018735 | 0.026513 | 0.013474 | 0.017116 | 0.00788 | 0.0178 | 0.003998 | 0.028297 | 0.026461 | 0.084769 | 0.01424 | 0.004202 | 0.016532 | 0.024709 | 0.072336 | 0.077788 | 0.005233 | 0.012102 | 0.006329 | 0.008045 | 0.018953 |
| 35.2222222 | 0.011278 | 0.016598 | 0.034187 | 0.019512 | 0.031857 | 0.009689 | 0.019094 | 0.006067 | 0.016508 | 0.00584 | 0.026367 | 0.037687 | 0.049463 | 0.015855 | 0.00435 | 0.016575 | 0.022836 | 0.070905 | 0.090066 | 0.006774 | 0.012592 | 0.010226 | 0.004553 | 0.009801 |
| 35.2361111 | 0.014239 | 0.016575 | 0.016674 | 0.021214 | 0.0398 | 0.006436 | 0.02389 | 0.003107 | 0.015493 | 0.010017 | 0.022444 | 0.047407 | 0.025037 | 0.014263 | 0.004768 | 0.010227 | 0.014517 | 0.060276 | 0.089531 | 0.009857 | 0.011788 | 0.012235 | 0.002311 | 0.003488 |
| 35.25 | 0.017824 | 0.01864 | 0.014491 | 0.023061 | 0.048778 | 0.005056 | 0.030422 | 0.003596 | 0.014304 | 0.016291 | 0.016577 | 0.056216 | 0.033576 | 0.010798 | 0.002906 | 0.005736 | 0.007138 | 0.045835 | 0.078629 | 0.012104 | 0.009694 | 0.011673 | 0.004462 | 0.001701 |
| 35.2638889 | 0.023625 | 0.022024 | 0.027256 | 0.02449 | 0.056652 | 0.005443 | 0.036316 | 0.00426 | 0.013139 | 0.022363 | 0.011228 | 0.061486 | 0.04827 | 0.008877 | 0.001212 | 0.00874 | 0.006249 | 0.031145 | 0.065122 | 0.011947 | 0.007863 | 0.009665 | 0.012316 | 0.00626 |
| 35.2777778 | 0.030323 | 0.02577 | 0.044793 | 0.025009 | 0.061774 | 0.006875 | 0.039819 | 0.006112 | 0.012901 | 0.028377 | 0.009726 | 0.059017 | 0.040262 | 0.009805 | 0.003258 | 0.01221 | 0.007021 | 0.018105 | 0.053459 | 0.00924 | 0.008646 | 0.008306 | 0.023784 | 0.016964 |
| 35.2916667 | 0.034766 | 0.028751 | 0.066792 | 0.024713 | 0.063358 | 0.009373 | 0.040633 | 0.011268 | 0.013642 | 0.03594 | 0.012722 | 0.048567 | 0.03094 | 0.012774 | 0.011192 | 0.009555 | 0.004696 | 0.008014 | 0.04365 | 0.005225 | 0.011695 | 0.007299 | 0.033492 | 0.028619 |
| 35.3055556 | 0.03501 | 0.030317 | 0.105923 | 0.024057 | 0.062557 | 0.013844 | 0.040844 | 0.01492 | 0.014276 | 0.04401 | 0.017226 | 0.035984 | 0.055271 | 0.017012 | 0.023309 | 0.008127 | 0.004674 | 0.003151 | 0.034367 | 0.003217 | 0.01451 | 0.005035 | 0.037523 | 0.035953 |
| 35.3194444 | 0.032704 | 0.030264 | 0.154148 | 0.023457 | 0.061945 | 0.02099 | 0.043789 | 0.01476 | 0.014443 | 0.049975 | 0.020333 | 0.027896 | 0.103128 | 0.022638 | 0.034602 | 0.018509 | 0.013754 | 0.00323 | 0.025868 | 0.00338 | 0.015938 | 0.002264 | 0.036732 | 0.038843 |
| 35.3333333 | 0.032542 | 0.028903 | 0.195564 | 0.023455 | 0.061748 | 0.028051 | 0.049636 | 0.011462 | 0.015671 | 0.053368 | 0.021544 | 0.024702 | 0.140163 | 0.028309 | 0.041954 | 0.03686 | 0.028713 | 0.003112 | 0.020634 | 0.002803 | 0.016904 | 0.000911 | 0.033988 | 0.040024 |
| 35.3472222 | 0.036766 | 0.027725 | 0.237859 | 0.023641 | 0.060169 | 0.029456 | 0.054561 | 0.006126 | 0.018792 | 0.055319 | 0.021353 | 0.023044 | 0.151306 | 0.031938 | 0.046873 | 0.051144 | 0.039512 | 0.002844 | 0.020252 | 0.002507 | 0.018769 | 0.0018 | 0.031789 | 0.041205 |
| 35.3611111 | 0.046027 | 0.027393 | 0.284054 | 0.022698 | 0.056644 | 0.025409 | 0.054928 | 0.002106 | 0.022158 | 0.055153 | 0.020394 | 0.020611 | 0.139934 | 0.032907 | 0.050197 | 0.054399 | 0.039665 | 0.003744 | 0.02236 | 0.003362 | 0.021473 | 0.00423 | 0.031451 | 0.042034 |
| 35.375 | 0.060813 | 0.02688 | 0.332407 | 0.022833 | 0.051976 | 0.024523 | 0.051549 | 0.000924 | 0.024035 | 0.052149 | 0.019473 | 0.016716 | 0.119925 | 0.032773 | 0.050213 | 0.04906 | 0.032464 | 0.004053 | 0.023016 | 0.00347 | 0.023598 | 0.00624 | 0.033217 | 0.04113 |
| 35.3888889 | 0.072434 | 0.026131 | 0.353765 | 0.030734 | 0.046816 | 0.034894 | 0.048437 | 0.001474 | 0.025293 | 0.048764 | 0.01871 | 0.012448 | 0.103929 | 0.033449 | 0.047267 | 0.043832 | 0.026219 | 0.006044 | 0.021732 | 0.002394 | 0.024399 | 0.006531 | 0.036129 | 0.038689 |
| 35.4027778 | 0.07972 | 0.026204 | 0.309965 | 0.054925 | 0.042048 | 0.051066 | 0.047856 | 0.003379 | 0.027876 | 0.047197 | 0.017716 | 0.009151 | 0.095584 | 0.034092 | 0.041713 | 0.042214 | 0.023898 | 0.007761 | 0.02149 | 0.001826 | 0.023927 | 0.004969 | 0.036686 | 0.036484 |
| 35.4166667 | 0.079767 | 0.026804 | 0.213561 | 0.080592 | 0.039396 | 0.056567 | 0.049728 | 0.004767 | 0.031413 | 0.046209 | 0.016849 | 0.007203 | 0.086963 | 0.033354 | 0.032896 | 0.039565 | 0.021451 | 0.006072 | 0.023771 | 0.002913 | 0.022784 | 0.002619 | 0.032545 | 0.035436 |
| 35.4305556 | 0.052561 | 0.027025 | 0.104438 | 0.066465 | 0.039933 | 0.050153 | 0.05306 | 0.003479 | 0.033593 | 0.044224 | 0.016086 | 0.005186 | 0.070255 | 0.031944 | 0.0224 | 0.032236 | 0.016219 | 0.006932 | 0.027294 | 0.003079 | 0.022536 | 0.001423 | 0.026165 | 0.036399 |
| 35.4444444 | 0.028112 | 0.026991 | 0.047443 | 0.038055 | 0.043634 | 0.043893 | 0.05645 | 0.002359 | 0.033609 | 0.042146 | 0.015226 | 0.003626 | 0.042466 | 0.029347 | 0.011614 | 0.023886 | 0.012016 | 0.01402 | 0.031513 | 0.001934 | 0.023279 | 0.001225 | 0.020855 | 0.039296 |
| 35.4583333 | 0.045308 | 0.02652 | 0.091386 | 0.047978 | 0.049778 | 0.033054 | 0.059143 | 0.004389 | 0.033257 | 0.042156 | 0.015672 | 0.005186 | 0.030565 | 0.024383 | 0.005351 | 0.017547 | 0.010312 | 0.023294 | 0.0372 | 0.002244 | 0.023404 | 0.00202 | 0.017285 | 0.040123 |
| 35.4722222 | 0.058018 | 0.025571 | 0.199228 | 0.051999 | 0.057384 | 0.017905 | 0.061657 | 0.007025 | 0.035133 | 0.044357 | 0.017784 | 0.006343 | 0.064683 | 0.018766 | 0.006943 | 0.011892 | 0.008179 | 0.02958 | 0.043122 | 0.002913 | 0.022232 | 0.004921 | 0.015074 | 0.035616 |
| 35.4861111 | 0.039975 | 0.02586 | 0.311583 | 0.028027 | 0.065099 | 0.014582 | 0.064568 | 0.007103 | 0.040415 | 0.046479 | 0.021907 | 0.004856 | 0.109255 | 0.015707 | 0.008367 | 0.008703 | 0.008027 | 0.029955 | 0.046591 | 0.002181 | 0.020066 | 0.008393 | 0.01448 | 0.028358 |
| 35.5 | 0.031885 | 0.028542 | 0.409744 | 0.021553 | 0.071502 | 0.018293 | 0.067449 | 0.005662 | 0.047959 | 0.047225 | 0.030628 | 0.00635 | 0.122216 | 0.015783 | 0.005342 | 0.010777 | 0.014077 | 0.025739 | 0.046691 | 0.001066 | 0.017767 | 0.010813 | 0.014448 | 0.022694 |
| 35.5138889 | 0.037827 | 0.031591 | 0.47443 | 0.030254 | 0.075773 | 0.019341 | 0.069847 | 0.005964 | 0.053564 | 0.046273 | 0.044748 | 0.013375 | 0.100161 | 0.017318 | 0.002547 | 0.014858 | 0.022394 | 0.019945 | 0.044268 | 0.000652 | 0.014865 | 0.011288 | 0.013255 | 0.020658 |
| 35.5277778 | 0.031927 | 0.030416 | 0.464899 | 0.033095 | 0.078206 | 0.023445 | 0.072937 | 0.007266 | 0.052733 | 0.045482 | 0.058221 | 0.023453 | 0.054859 | 0.020453 | 0.003568 | 0.018249 | 0.026272 | 0.013599 | 0.040879 | 0.002658 | 0.01053 | 0.009784 | 0.013069 | 0.023979 |
| 35.5416667 | 0.023073 | 0.024532 | 0.369728 | 0.032953 | 0.082715 | 0.024162 | 0.080896 | 0.006904 | 0.047479 | 0.051908 | 0.060927 | 0.035274 | 0.030455 | 0.026043 | 0.01513 | 0.029164 | 0.029035 | 0.008506 | 0.038494 | 0.012664 | 0.00765 | 0.010385 | 0.019801 | 0.036559 |
| 35.5555556 | 0.0382 | 0.020803 | 0.211663 | 0.033561 | 0.091302 | 0.022067 | 0.092395 | 0.011574 | 0.043221 | 0.067476 | 0.048943 | 0.046726 | 0.059125 | 0.031106 | 0.040385 | 0.055938 | 0.040428 | 0.011713 | 0.035379 | 0.034262 | 0.010016 | 0.015195 | 0.036355 | 0.057269 |
| 35.5694444 | 0.071715 | 0.022278 | 0.127718 | 0.035162 | 0.094968 | 0.037111 | 0.09533 | 0.027652 | 0.041258 | 0.081071 | 0.027761 | 0.051641 | 0.116249 | 0.028827 | 0.066219 | 0.089361 | 0.05982 | 0.019083 | 0.030403 | 0.057987 | 0.017407 | 0.019943 | 0.061506 | 0.075061 |
| 35.5833333 | 0.102469 | 0.023943 | 0.225887 | 0.040702 | 0.090265 | 0.057223 | 0.088267 | 0.0475 | 0.041188 | 0.083727 | 0.015773 | 0.050072 | 0.179468 | 0.01705 | 0.080278 | 0.11032 | 0.075074 | 0.030859 | 0.036648 | 0.07317 | 0.028295 | 0.023504 | 0.094364 | 0.083961 |
| 35.5972222 | 0.118048 | 0.021485 | 0.333729 | 0.058894 | 0.085423 | 0.057814 | 0.081016 | 0.058522 | 0.042735 | 0.079184 | 0.020657 | 0.050559 | 0.220908 | 0.006334 | 0.081989 | 0.111026 | 0.078261 | 0.067251 | 0.063709 | 0.082954 | 0.041972 | 0.029876 | 0.130672 | 0.085812 |
| 35.6111111 | 0.107305 | 0.015594 | 0.322368 | 0.081493 | 0.076567 | 0.05618 | 0.07274 | 0.052096 | 0.043461 | 0.075966 | 0.024274 | 0.056834 | 0.2156 | 0.007591 | 0.07028 | 0.098228 | 0.071421 | 0.110357 | 0.092379 | 0.088645 | 0.054488 | 0.038886 | 0.160965 | 0.082388 |
| 35.625 | 0.085778 | 0.014782 | 0.263473 | 0.088945 | 0.058575 | 0.070657 | 0.061802 | 0.031551 | 0.041965 | 0.079143 | 0.021882 | 0.065629 | 0.180277 | 0.016745 | 0.045806 | 0.079112 | 0.057096 | 0.124603 | 0.103838 | 0.083707 | 0.05924 | 0.04632 | 0.177867 | 0.076442 |
| 35.6388889 | 0.081364 | 0.024117 | 0.209534 | 0.078725 | 0.039792 | 0.07643 | 0.052509 | 0.016736 | 0.041022 | 0.08743 | 0.022081 | 0.07231 | 0.120222 | 0.020416 | 0.02205 | 0.05568 | 0.034838 | 0.116332 | 0.103029 | 0.072433 | 0.054436 | 0.050104 | 0.178709 | 0.062378 |
| 35.6527778 | 0.089496 | 0.038658 | 0.168458 | 0.067664 | 0.02529 | 0.061639 | 0.044934 | 0.022248 | 0.041482 | 0.093759 | 0.022351 | 0.075483 | 0.064616 | 0.013004 | 0.017268 | 0.029791 | 0.016906 | 0.100659 | 0.092346 | 0.063115 | 0.041383 | 0.050706 | 0.172432 | 0.036108 |
| 35.6666667 | 0.088483 | 0.053223 | 0.134877 | 0.074964 | 0.014964 | 0.045704 | 0.037913 | 0.033946 | 0.039248 | 0.090355 | 0.017054 | 0.074622 | 0.066833 | 0.007089 | 0.032195 | 0.012642 | 0.01845 | 0.074266 | 0.068103 | 0.053887 | 0.025219 | 0.048442 | 0.168246 | 0.017249 |
| 35.6805556 | 0.0729 | 0.056548 | 0.091533 | 0.074208 | 0.010922 | 0.034583 | 0.033971 | 0.030359 | 0.033495 | 0.076844 | 0.008162 | 0.066179 | 0.122429 | 0.012569 | 0.046509 | 0.012888 | 0.027167 | 0.038196 | 0.037494 | 0.047623 | 0.018101 | 0.043448 | 0.156899 | 0.018691 |
| 35.6944444 | 0.048941 | 0.039579 | 0.050137 | 0.050291 | 0.014396 | 0.025164 | 0.034973 | 0.018965 | 0.027529 | 0.05965 | 0.004647 | 0.049864 | 0.179378 | 0.020954 | 0.045315 | 0.018616 | 0.02504 | 0.01203 | 0.014538 | 0.050078 | 0.018525 | 0.03593 | 0.130517 | 0.035885 |
| 35.7083333 | 0.024249 | 0.02052 | 0.057014 | 0.039931 | 0.02274 | 0.01613 | 0.038473 | 0.021187 | 0.022326 | 0.042055 | 0.008947 | 0.030662 | 0.184547 | 0.023496 | 0.032024 | 0.018241 | 0.014752 | 0.004459 | 0.006658 | 0.055734 | 0.025678 | 0.02586 | 0.109129 | 0.068643 |
| 35.7222222 | 0.010192 | 0.015771 | 0.118468 | 0.041614 | 0.032131 | 0.013074 | 0.041227 | 0.033365 | 0.017593 | 0.025391 | 0.014657 | 0.014145 | 0.128956 | 0.020089 | 0.016278 | 0.012895 | 0.011996 | 0.007841 | 0.009122 | 0.056132 | 0.044856 | 0.014024 | 0.110257 | 0.10536 |
| 35.7361111 | 0.011399 | 0.013441 | 0.199412 | 0.030886 | 0.040703 | 0.025302 | 0.043203 | 0.038285 | 0.014143 | 0.014733 | 0.017296 | 0.004181 | 0.063525 | 0.013763 | 0.006049 | 0.006876 | 0.025195 | 0.015924 | 0.0098 | 0.049993 | 0.054405 | 0.005695 | 0.11792 | 0.126333 |
| 35.75 | 0.018793 | 0.01025 | 0.28239 | 0.018697 | 0.048193 | 0.04106 | 0.046757 | 0.03195 | 0.012821 | 0.013533 | 0.015667 | 0.00182 | 0.046831 | 0.009499 | 0.003435 | 0.007108 | 0.043263 | 0.016104 | 0.012113 | 0.040248 | 0.043397 | 0.004214 | 0.11788 | 0.118886 |
| 35.7638889 | 0.028945 | 0.009502 | 0.365595 | 0.014836 | 0.054365 | 0.044233 | 0.052802 | 0.019505 | 0.014077 | 0.019224 | 0.011667 | 0.005044 | 0.073006 | 0.011616 | 0.007683 | 0.017775 | 0.054048 | 0.014043 | 0.032524 | 0.027596 | 0.029245 | 0.00444 | 0.105544 | 0.083918 |
| 35.7777778 | 0.037849 | 0.00741 | 0.404491 | 0.013837 | 0.058199 | 0.036073 | 0.058843 | 0.009478 | 0.017384 | 0.025833 | 0.009519 | 0.010098 | 0.102243 | 0.018918 | 0.017689 | 0.033819 | 0.056324 | 0.024728 | 0.061683 | 0.013354 | 0.021027 | 0.006263 | 0.076721 | 0.04271 |
| 35.7916667 | 0.030829 | 0.006851 | 0.336679 | 0.018606 | 0.05968 | 0.021 | 0.062317 | 0.007579 | 0.020935 | 0.03037 | 0.009996 | 0.014407 | 0.118428 | 0.027227 | 0.026494 | 0.04789 | 0.055006 | 0.03464 | 0.077973 | 0.00535 | 0.019831 | 0.012761 | 0.050273 | 0.02826 |
| 35.8055556 | 0.014984 | 0.009219 | 0.193829 | 0.028096 | 0.061534 | 0.009372 | 0.06427 | 0.008723 | 0.023371 | 0.034693 | 0.010502 | 0.016931 | 0.122991 | 0.032835 | 0.029253 | 0.056031 | 0.05555 | 0.031466 | 0.075353 | 0.008399 | 0.022867 | 0.019904 | 0.057829 | 0.044915 |
| 35.8194444 | 0.009443 | 0.012147 | 0.091897 | 0.037043 | 0.065528 | 0.012399 | 0.067036 | 0.007323 | 0.02489 | 0.040608 | 0.009243 | 0.017045 | 0.115801 | 0.031955 | 0.028951 | 0.057063 | 0.057795 | 0.019347 | 0.058934 | 0.016146 | 0.022929 | 0.022676 | 0.099179 | 0.061665 |
| 35.8333333 | 0.0126 | 0.01435 | 0.106572 | 0.045609 | 0.070575 | 0.026286 | 0.070991 | 0.008482 | 0.025662 | 0.046725 | 0.00571 | 0.015276 | 0.091225 | 0.024205 | 0.025394 | 0.052368 | 0.05723 | 0.013507 | 0.038113 | 0.023952 | 0.019065 | 0.019451 | 0.151747 | 0.061126 |
| 35.8472222 | 0.013953 | 0.014821 | 0.196467 | 0.051346 | 0.073798 | 0.041231 | 0.073331 | 0.016204 | 0.025107 | 0.049696 | 0.002813 | 0.012887 | 0.054359 | 0.014332 | 0.015505 | 0.043455 | 0.051251 | 0.02453 | 0.020147 | 0.033928 | 0.017064 | 0.013238 | 0.197258 | 0.042493 |
| 35.8611111 | 0.012353 | 0.013073 | 0.278145 | 0.052882 | 0.072984 | 0.052155 | 0.069789 | 0.025574 | 0.023594 | 0.04841 | 0.00426 | 0.010774 | 0.048956 | 0.006865 | 0.011263 | 0.03021 | 0.041887 | 0.037923 | 0.00873 | 0.044359 | 0.01758 | 0.00736 | 0.220323 | 0.026284 |
| 35.875 | 0.011726 | 0.00872 | 0.321341 | 0.058236 | 0.070185 | 0.059006 | 0.060798 | 0.030375 | 0.021958 | 0.04459 | 0.008029 | 0.008701 | 0.093877 | 0.004993 | 0.027004 | 0.015816 | 0.033045 | 0.04197 | 0.003709 | 0.048614 | 0.013932 | 0.003899 | 0.215709 | 0.033633 |
| 35.8888889 | 0.009811 | 0.005587 | 0.34009 | 0.077935 | 0.066258 | 0.070478 | 0.049009 | 0.027765 | 0.019975 | 0.03809 | 0.011541 | 0.005494 | 0.141035 | 0.009931 | 0.051829 | 0.009908 | 0.025101 | 0.040063 | 0.003902 | 0.046522 | 0.008747 | 0.005026 | 0.188218 | 0.04074 |
| 35.9027778 | 0.007835 | 0.009381 | 0.337384 | 0.10151 | 0.057353 | 0.080444 | 0.032649 | 0.023472 | 0.016704 | 0.026577 | 0.015449 | 0.003437 | 0.167725 | 0.015768 | 0.068574 | 0.015461 | 0.016281 | 0.034696 | 0.006862 | 0.04377 | 0.011256 | 0.008389 | 0.155177 | 0.028017 |
| 35.9166667 | 0.008522 | 0.01452 | 0.284424 | 0.094292 | 0.04119 | 0.066524 | 0.017222 | 0.029353 | 0.011078 | 0.014851 | 0.022005 | 0.00728 | 0.176496 | 0.015886 | 0.074618 | 0.024279 | 0.007589 | 0.025467 | 0.006404 | 0.041506 | 0.017005 | 0.010407 | 0.12726 | 0.017562 |
| 35.9305556 | 0.010465 | 0.015324 | 0.181053 | 0.05499 | 0.021653 | 0.042916 | 0.017338 | 0.048026 | 0.00638 | 0.015199 | 0.033869 | 0.016811 | 0.169067 | 0.012173 | 0.074053 | 0.02976 | 0.003068 | 0.015606 | 0.003888 | 0.03667 | 0.019046 | 0.010969 | 0.089222 | 0.023962 |
| 35.9444444 | 0.025411 | 0.0179 | 0.083132 | 0.03801 | 0.009148 | 0.042193 | 0.03407 | 0.067225 | 0.008842 | 0.029152 | 0.049764 | 0.027503 | 0.160596 | 0.009679 | 0.063567 | 0.030542 | 0.003635 | 0.01141 | 0.006401 | 0.026892 | 0.019521 | 0.011586 | 0.049071 | 0.028894 |
| 35.9583333 | 0.052487 | 0.023743 | 0.058266 | 0.064123 | 0.011196 | 0.062097 | 0.052602 | 0.074807 | 0.01735 | 0.045647 | 0.064118 | 0.034932 | 0.156316 | 0.008538 | 0.044725 | 0.026753 | 0.003969 | 0.015166 | 0.012074 | 0.013935 | 0.019182 | 0.01246 | 0.046308 | 0.019292 |
| 35.9722222 | 0.072255 | 0.024408 | 0.120588 | 0.091211 | 0.019625 | 0.078369 | 0.059832 | 0.071113 | 0.025729 | 0.05641 | 0.071611 | 0.037268 | 0.142277 | 0.006534 | 0.029906 | 0.017244 | 0.005111 | 0.025766 | 0.016035 | 0.006102 | 0.016181 | 0.010772 | 0.074182 | 0.00891 |
| 35.9861111 | 0.083011 | 0.015052 | 0.250391 | 0.08366 | 0.021156 | 0.082378 | 0.053762 | 0.061697 | 0.030606 | 0.059597 | 0.069779 | 0.036022 | 0.106092 | 0.004552 | 0.018765 | 0.010582 | 0.012602 | 0.039172 | 0.017612 | 0.005725 | 0.009773 | 0.006682 | 0.082742 | 0.010647 |
| 36 | 0.084761 | 0.006201 | 0.405461 | 0.046729 | 0.016582 | 0.081333 | 0.042074 | 0.049125 | 0.032318 | 0.057411 | 0.0583 | 0.032784 | 0.055421 | 0.004584 | 0.009756 | 0.019 | 0.023185 | 0.045081 | 0.020786 | 0.004705 | 0.006144 | 0.006311 | 0.05318 | 0.014793 |
| 36.0138889 | 0.071287 | 0.006864 | 0.502038 | 0.013656 | 0.013926 | 0.081842 | 0.034211 | 0.031574 | 0.032892 | 0.052963 | 0.040233 | 0.027086 | 0.02753 | 0.006183 | 0.009987 | 0.036003 | 0.032711 | 0.035287 | 0.031081 | 0.003149 | 0.012586 | 0.011647 | 0.03046 | 0.012285 |
| 36.0277778 | 0.048509 | 0.008532 | 0.496123 | 0.002391 | 0.017227 | 0.089986 | 0.034366 | 0.013928 | 0.034618 | 0.051612 | 0.027183 | 0.022321 | 0.034567 | 0.008793 | 0.015069 | 0.045973 | 0.034964 | 0.022181 | 0.040534 | 0.008741 | 0.021399 | 0.016134 | 0.060968 | 0.007366 |
| 36.0416667 | 0.040036 | 0.006323 | 0.417207 | 0.005544 | 0.030218 | 0.107801 | 0.043981 | 0.012301 | 0.043017 | 0.062373 | 0.038608 | 0.03343 | 0.043134 | 0.01616 | 0.020635 | 0.039047 | 0.034109 | 0.036189 | 0.037087 | 0.029864 | 0.01964 | 0.013971 | 0.130029 | 0.014747 |
| 36.0555556 | 0.066333 | 0.00759 | 0.338424 | 0.016397 | 0.053137 | 0.131582 | 0.062017 | 0.043048 | 0.06349 | 0.088394 | 0.091367 | 0.077013 | 0.073882 | 0.030399 | 0.050346 | 0.040047 | 0.052467 | 0.097569 | 0.052492 | 0.073134 | 0.022215 | 0.018811 | 0.210246 | 0.053541 |
| 36.0694444 | 0.106112 | 0.015718 | 0.298955 | 0.029206 | 0.073402 | 0.150307 | 0.077904 | 0.098952 | 0.09057 | 0.118317 | 0.164654 | 0.140047 | 0.157037 | 0.041705 | 0.108579 | 0.061817 | 0.069179 | 0.182623 | 0.109609 | 0.119888 | 0.049301 | 0.044921 | 0.274623 | 0.113753 |
| 36.0833333 | 0.117985 | 0.023301 | 0.283368 | 0.034541 | 0.075577 | 0.151296 | 0.079704 | 0.130682 | 0.112353 | 0.135018 | 0.194596 | 0.181696 | 0.21255 | 0.038559 | 0.142175 | 0.06422 | 0.049579 | 0.225834 | 0.149375 | 0.129959 | 0.07386 | 0.069468 | 0.274198 | 0.135174 |
| 36.0972222 | 0.09598 | 0.022291 | 0.276724 | 0.026021 | 0.057948 | 0.132682 | 0.066514 | 0.104829 | 0.12209 | 0.131356 | 0.161399 | 0.181881 | 0.16735 | 0.023465 | 0.124322 | 0.040313 | 0.021203 | 0.199654 | 0.132659 | 0.098156 | 0.06792 | 0.069648 | 0.209603 | 0.085206 |
| 36.1111111 | 0.067292 | 0.013023 | 0.254644 | 0.014262 | 0.032358 | 0.101433 | 0.047357 | 0.050257 | 0.117361 | 0.111752 | 0.112582 | 0.154114 | 0.0817 | 0.011479 | 0.089642 | 0.018842 | 0.017903 | 0.155922 | 0.101595 | 0.060074 | 0.048962 | 0.056537 | 0.145305 | 0.029802 |
| 36.125 | 0.048057 | 0.006315 | 0.204361 | 0.01518 | 0.01269 | 0.067418 | 0.031107 | 0.016326 | 0.099904 | 0.083944 | 0.082085 | 0.118983 | 0.027473 | 0.013324 | 0.062091 | 0.01647 | 0.037685 | 0.162069 | 0.11428 | 0.03605 | 0.0439 | 0.046589 | 0.107803 | 0.019213 |
| 36.1388889 | 0.035354 | 0.010095 | 0.148423 | 0.023511 | 0.007762 | 0.037373 | 0.017295 | 0.017878 | 0.074921 | 0.054062 | 0.067937 | 0.088824 | 0.006816 | 0.021107 | 0.03688 | 0.028112 | 0.061962 | 0.216028 | 0.155913 | 0.023386 | 0.054644 | 0.04237 | 0.085021 | 0.026135 |
| 36.1527778 | 0.026051 | 0.017231 | 0.108247 | 0.027816 | 0.01707 | 0.016021 | 0.010754 | 0.029298 | 0.047718 | 0.027211 | 0.060768 | 0.065053 | 0.004338 | 0.02308 | 0.015633 | 0.030935 | 0.07275 | 0.229147 | 0.151094 | 0.017531 | 0.060571 | 0.037211 | 0.069278 | 0.030212 |

| | | | | | | | | | | | | | | | | | | | | | | | | |
|---|---|---|---|---|---|---|---|---|---|---|---|---|---|---|---|---|---|---|---|---|---|---|---|---|
| 36.1666667 | 0.019734 | 0.020938 | 0.082677 | 0.027312 | 0.029903 | 0.008458 | 0.019469 | 0.030916 | 0.024316 | 0.010329 | 0.057501 | 0.045511 | 0.009718 | 0.019192 | 0.007644 | 0.018462 | 0.069568 | 0.164935 | 0.100971 | 0.014227 | 0.048384 | 0.027793 | 0.050871 | 0.036634 |
| 36.1805556 | 0.015687 | 0.021587 | 0.06382 | 0.024141 | 0.038612 | 0.010498 | 0.032685 | 0.024222 | 0.008947 | 0.006741 | 0.057303 | 0.03079 | 0.023908 | 0.014603 | 0.010292 | 0.007154 | 0.064118 | 0.092085 | 0.058413 | 0.011667 | 0.027119 | 0.018554 | 0.02784 | 0.05213 |
| 36.1944444 | 0.012324 | 0.020267 | 0.049868 | 0.019877 | 0.04304 | 0.01263 | 0.03951 | 0.014402 | 0.00261 | 0.00892 | 0.057919 | 0.022501 | 0.052214 | 0.0114 | 0.013926 | 0.004722 | 0.06175 | 0.051057 | 0.03291 | 0.010689 | 0.011593 | 0.014224 | 0.009541 | 0.074435 |
| 36.2083333 | 0.008404 | 0.018067 | 0.038802 | 0.016253 | 0.045489 | 0.012758 | 0.042541 | 0.006898 | 0.003471 | 0.010118 | 0.060399 | 0.019297 | 0.10122 | 0.008407 | 0.01297 | 0.005373 | 0.056213 | 0.031491 | 0.017055 | 0.01002 | 0.005973 | 0.015065 | 0.001988 | 0.085222 |
| 36.2222222 | 0.004289 | 0.016003 | 0.028453 | 0.014706 | 0.049785 | 0.013104 | 0.04644 | 0.007158 | 0.007994 | 0.012566 | 0.066163 | 0.015674 | 0.146123 | 0.005319 | 0.00785 | 0.007667 | 0.042237 | 0.025166 | 0.007912 | 0.007649 | 0.00439 | 0.017254 | 0.001148 | 0.073939 |
| 36.2361111 | 0.002327 | 0.014783 | 0.018981 | 0.015182 | 0.058055 | 0.014977 | 0.053249 | 0.015335 | 0.014229 | 0.019081 | 0.07181 | 0.009954 | 0.148087 | 0.003972 | 0.005844 | 0.010117 | 0.023694 | 0.027351 | 0.003421 | 0.004511 | 0.003914 | 0.01607 | 0.005234 | 0.057181 |
| 36.25 | 0.003092 | 0.014558 | 0.010724 | 0.01657 | 0.068268 | 0.018848 | 0.06179 | 0.024291 | 0.02086 | 0.028209 | 0.071749 | 0.009775 | 0.130483 | 0.006096 | 0.009777 | 0.010796 | 0.011317 | 0.033177 | 0.004112 | 0.002046 | 0.008402 | 0.01026 | 0.016256 | 0.051997 |
| 36.2638889 | 0.00408 | 0.015733 | 0.004523 | 0.018277 | 0.076244 | 0.023866 | 0.06951 | 0.026056 | 0.02689 | 0.036544 | 0.062954 | 0.019232 | 0.12256 | 0.010437 | 0.015966 | 0.012452 | 0.015373 | 0.036134 | 0.010441 | 0.00076 | 0.017398 | 0.00442 | 0.030514 | 0.054206 |
| 36.2777778 | 0.004715 | 0.019031 | 0.003098 | 0.020236 | 0.079642 | 0.027841 | 0.073952 | 0.017399 | 0.031672 | 0.042143 | 0.04674 | 0.031359 | 0.11185 | 0.013896 | 0.022257 | 0.016216 | 0.029188 | 0.030157 | 0.02167 | 0.000901 | 0.024344 | 0.003322 | 0.040246 | 0.053309 |
| 36.2916667 | 0.007501 | 0.023849 | 0.008656 | 0.022179 | 0.079516 | 0.028512 | 0.07401 | 0.010525 | 0.034089 | 0.045365 | 0.027697 | 0.040932 | 0.092374 | 0.015957 | 0.026571 | 0.016911 | 0.033646 | 0.016487 | 0.034946 | 0.002409 | 0.026533 | 0.005555 | 0.043976 | 0.04968 |
| 36.3055556 | 0.014181 | 0.028443 | 0.020364 | 0.023993 | 0.078873 | 0.025273 | 0.071272 | 0.019496 | 0.033734 | 0.047338 | 0.01266 | 0.048073 | 0.071936 | 0.017996 | 0.025768 | 0.011998 | 0.023602 | 0.00725 | 0.045865 | 0.006358 | 0.028942 | 0.008085 | 0.045967 | 0.051158 |
| 36.3194444 | 0.022892 | 0.031759 | 0.036724 | 0.025495 | 0.079558 | 0.018892 | 0.068091 | 0.034747 | 0.030956 | 0.048876 | 0.008783 | 0.053709 | 0.050812 | 0.019304 | 0.017278 | 0.012022 | 0.021098 | 0.009101 | 0.051611 | 0.013082 | 0.034533 | 0.00931 | 0.047762 | 0.060917 |
| 36.3333333 | 0.029116 | 0.033136 | 0.055064 | 0.026058 | 0.079652 | 0.010993 | 0.064608 | 0.042279 | 0.026414 | 0.049447 | 0.014729 | 0.057215 | 0.027013 | 0.018553 | 0.009076 | 0.024389 | 0.042807 | 0.012555 | 0.050968 | 0.018824 | 0.039754 | 0.00825 | 0.050873 | 0.077816 |
| 36.3472222 | 0.029921 | 0.032073 | 0.06936 | 0.025565 | 0.074801 | 0.004651 | 0.059489 | 0.039648 | 0.02102 | 0.047863 | 0.022047 | 0.054694 | 0.014641 | 0.016518 | 0.010301 | 0.039277 | 0.069765 | 0.011196 | 0.044576 | 0.020792 | 0.041126 | 0.00526 | 0.059038 | 0.102576 |
| 36.3611111 | 0.026561 | 0.029139 | 0.077686 | 0.024498 | 0.064783 | 0.002266 | 0.052772 | 0.027805 | 0.016413 | 0.044972 | 0.027471 | 0.045093 | 0.02249 | 0.014818 | 0.01267 | 0.047117 | 0.08047 | 0.006585 | 0.034707 | 0.020019 | 0.038915 | 0.002089 | 0.069582 | 0.127514 |
| 36.375 | 0.022634 | 0.025095 | 0.092802 | 0.022834 | 0.054404 | 0.002189 | 0.046261 | 0.014946 | 0.012895 | 0.041784 | 0.02994 | 0.03146 | 0.029356 | 0.01456 | 0.010707 | 0.046571 | 0.072902 | 0.003883 | 0.023282 | 0.017323 | 0.035097 | 0.0015 | 0.073786 | 0.135116 |
| 36.3888889 | 0.020357 | 0.020957 | 0.128086 | 0.020678 | 0.047065 | 0.002315 | 0.041162 | 0.013571 | 0.008503 | 0.037387 | 0.025034 | 0.016526 | 0.020702 | 0.015652 | 0.011769 | 0.039355 | 0.053639 | 0.008179 | 0.012445 | 0.011579 | 0.029499 | 0.003605 | 0.06813 | 0.119328 |
| 36.4027778 | 0.019122 | 0.018518 | 0.176074 | 0.019274 | 0.043852 | 0.004535 | 0.038532 | 0.022295 | 0.004335 | 0.031797 | 0.013911 | 0.007054 | 0.013438 | 0.016901 | 0.019531 | 0.029837 | 0.032824 | 0.016147 | 0.005042 | 0.00638 | 0.022034 | 0.004877 | 0.056444 | 0.088875 |
| 36.4166667 | 0.017434 | 0.017252 | 0.212866 | 0.019266 | 0.045312 | 0.010233 | 0.039435 | 0.030573 | 0.003345 | 0.027159 | 0.008112 | 0.007326 | 0.019422 | 0.018052 | 0.030316 | 0.022874 | 0.020696 | 0.022212 | 0.002031 | 0.00777 | 0.015133 | 0.004178 | 0.042332 | 0.05277 |
| 36.4305556 | 0.01447 | 0.014161 | 0.223031 | 0.019285 | 0.050132 | 0.011695 | 0.043569 | 0.034789 | 0.003264 | 0.024636 | 0.013133 | 0.01008 | 0.030698 | 0.019701 | 0.037348 | 0.020309 | 0.019218 | 0.023719 | 0.001593 | 0.011025 | 0.009954 | 0.003525 | 0.028067 | 0.022999 |
| 36.4444444 | 0.010221 | 0.009112 | 0.221305 | 0.017853 | 0.055896 | 0.006971 | 0.049598 | 0.036994 | 0.002916 | 0.023492 | 0.021722 | 0.010854 | 0.044892 | 0.021065 | 0.034449 | 0.019662 | 0.021495 | 0.021828 | 0.001726 | 0.009687 | 0.006701 | 0.003001 | 0.017042 | 0.008504 |
| 36.4583333 | 0.006323 | 0.005434 | 0.225948 | 0.015319 | 0.060963 | 0.006606 | 0.055599 | 0.040044 | 0.00573 | 0.022916 | 0.029297 | 0.011888 | 0.059697 | 0.020972 | 0.021441 | 0.017424 | 0.023503 | 0.020957 | 0.001743 | 0.00586 | 0.007293 | 0.001982 | 0.010887 | 0.010531 |
| 36.4722222 | 0.004568 | 0.005198 | 0.22652 | 0.013413 | 0.064621 | 0.01063 | 0.060243 | 0.043533 | 0.011446 | 0.02249 | 0.034667 | 0.013181 | 0.064216 | 0.020096 | 0.007815 | 0.013131 | 0.025398 | 0.021644 | 0.001501 | 0.002753 | 0.012362 | 0.002672 | 0.008957 | 0.032894 |
| 36.4861111 | 0.004647 | 0.008674 | 0.214838 | 0.01385 | 0.067281 | 0.014605 | 0.063234 | 0.044524 | 0.018222 | 0.022872 | 0.035649 | 0.012696 | 0.056609 | 0.019587 | 0.005432 | 0.007883 | 0.026288 | 0.01884 | 0.002562 | 0.001997 | 0.018725 | 0.007697 | 0.009096 | 0.067148 |
| 36.5 | 0.005806 | 0.016401 | 0.202488 | 0.017021 | 0.070079 | 0.018668 | 0.065325 | 0.04163 | 0.026396 | 0.026184 | 0.034325 | 0.00907 | 0.045337 | 0.019513 | 0.01461 | 0.003367 | 0.024666 | 0.010529 | 0.007887 | 0.004019 | 0.023017 | 0.013174 | 0.007912 | 0.089496 |
| 36.5138889 | 0.008014 | 0.024975 | 0.193322 | 0.02016 | 0.072339 | 0.021375 | 0.067407 | 0.034472 | 0.033536 | 0.032718 | 0.035548 | 0.004772 | 0.033567 | 0.019668 | 0.023079 | 0.001882 | 0.021903 | 0.005729 | 0.017117 | 0.007283 | 0.026407 | 0.011612 | 0.004974 | 0.081117 |
| 36.5277778 | 0.010834 | 0.025889 | 0.187334 | 0.018367 | 0.071873 | 0.023361 | 0.070235 | 0.021496 | 0.035968 | 0.040792 | 0.03841 | 0.005456 | 0.019614 | 0.020597 | 0.021796 | 0.008121 | 0.025864 | 0.015549 | 0.032188 | 0.01205 | 0.033409 | 0.010057 | 0.006977 | 0.048366 |
| 36.5416667 | 0.014083 | 0.018554 | 0.209111 | 0.011925 | 0.071126 | 0.029033 | 0.078771 | 0.019888 | 0.038556 | 0.051831 | 0.032329 | 0.016367 | 0.016659 | 0.024274 | 0.021295 | 0.029505 | 0.042973 | 0.044294 | 0.0623 | 0.024865 | 0.046321 | 0.0207 | 0.027081 | 0.018604 |
| 36.5555556 | 0.023922 | 0.01574 | 0.264674 | 0.013225 | 0.07341 | 0.040587 | 0.095109 | 0.05542 | 0.049972 | 0.067538 | 0.026007 | 0.038667 | 0.045528 | 0.031034 | 0.040478 | 0.064779 | 0.065945 | 0.089902 | 0.106991 | 0.051787 | 0.060724 | 0.037236 | 0.070085 | 0.013623 |
| 36.5694444 | 0.045349 | 0.024396 | 0.280196 | 0.026347 | 0.075757 | 0.052767 | 0.106753 | 0.108615 | 0.066802 | 0.082945 | 0.046397 | 0.067597 | 0.09493 | 0.033994 | 0.076959 | 0.097755 | 0.082858 | 0.140323 | 0.146118 | 0.08202 | 0.069644 | 0.047927 | 0.107464 | 0.032691 |
| 36.5833333 | 0.062924 | 0.033168 | 0.2121 | 0.033887 | 0.077031 | 0.060095 | 0.10431 | 0.139132 | 0.078481 | 0.091791 | 0.08138 | 0.092891 | 0.132014 | 0.028936 | 0.11202 | 0.109826 | 0.094915 | 0.187328 | 0.169385 | 0.098008 | 0.077416 | 0.052005 | 0.113083 | 0.061255 |
| 36.5972222 | 0.066584 | 0.031233 | 0.126817 | 0.033258 | 0.076452 | 0.062882 | 0.094669 | 0.135377 | 0.08262 | 0.092696 | 0.09741 | 0.107779 | 0.156461 | 0.025048 | 0.132521 | 0.100645 | 0.099069 | 0.226194 | 0.187563 | 0.09916 | 0.08972 | 0.052871 | 0.110596 | 0.088554 |
| 36.6111111 | 0.072277 | 0.023853 | 0.079967 | 0.033828 | 0.069357 | 0.062593 | 0.084215 | 0.105079 | 0.082082 | 0.088574 | 0.091448 | 0.114507 | 0.182194 | 0.031886 | 0.128424 | 0.083008 | 0.08709 | 0.244639 | 0.202935 | 0.094956 | 0.096137 | 0.052136 | 0.123613 | 0.103684 |
| 36.625 | 0.074542 | 0.020233 | 0.057643 | 0.030018 | 0.058239 | 0.061863 | 0.074961 | 0.060769 | 0.077306 | 0.086309 | 0.078498 | 0.116545 | 0.203247 | 0.038945 | 0.096763 | 0.067889 | 0.072869 | 0.245795 | 0.21383 | 0.086418 | 0.088204 | 0.053679 | 0.14325 | 0.10493 |
| 36.6388889 | 0.050346 | 0.021437 | 0.032632 | 0.019933 | 0.049039 | 0.065203 | 0.07115 | 0.030619 | 0.070429 | 0.089951 | 0.066168 | 0.11904 | 0.203987 | 0.029574 | 0.050906 | 0.061349 | 0.066633 | 0.242137 | 0.22062 | 0.069317 | 0.068316 | 0.060895 | 0.169509 | 0.095511 |
| 36.6527778 | 0.021785 | 0.023573 | 0.01237 | 0.010505 | 0.043395 | 0.071556 | 0.073936 | 0.040742 | 0.065033 | 0.095939 | 0.051903 | 0.119033 | 0.172752 | 0.019386 | 0.027986 | 0.059553 | 0.061545 | 0.229405 | 0.211686 | 0.047918 | 0.043573 | 0.06941 | 0.194301 | 0.065503 |
| 36.6666667 | 0.020184 | 0.023941 | 0.011317 | 0.006609 | 0.040364 | 0.074402 | 0.076714 | 0.068062 | 0.059128 | 0.097624 | 0.032115 | 0.105638 | 0.107797 | 0.022223 | 0.045012 | 0.055691 | 0.054031 | 0.198272 | 0.179334 | 0.031587 | 0.027008 | 0.07072 | 0.189094 | 0.035288 |
| 36.6805556 | 0.03122 | 0.023012 | 0.016392 | 0.007891 | 0.037035 | 0.070531 | 0.073659 | 0.077367 | 0.050189 | 0.093018 | 0.013656 | 0.082276 | 0.060349 | 0.020367 | 0.059118 | 0.046854 | 0.042227 | 0.150317 | 0.131479 | 0.024596 | 0.022293 | 0.061142 | 0.164766 | 0.047331 |
| 36.6944444 | 0.035037 | 0.022705 | 0.018345 | 0.011011 | 0.032643 | 0.06233 | 0.067163 | 0.064528 | 0.040941 | 0.086297 | 0.006967 | 0.061388 | 0.107926 | 0.015003 | 0.040134 | 0.031389 | 0.026231 | 0.098182 | 0.083028 | 0.028299 | 0.01925 | 0.046225 | 0.141397 | 0.081793 |
| 36.7083333 | 0.031362 | 0.02346 | 0.019066 | 0.015362 | 0.029024 | 0.051862 | 0.05974 | 0.037122 | 0.033554 | 0.07871 | 0.007792 | 0.046762 | 0.219303 | 0.019255 | 0.021454 | 0.016255 | 0.013766 | 0.0577 | 0.046164 | 0.042936 | 0.013499 | 0.032865 | 0.120885 | 0.074011 |
| 36.7222222 | 0.02609 | 0.023894 | 0.020349 | 0.020366 | 0.026801 | 0.039058 | 0.049736 | 0.017055 | 0.026969 | 0.066664 | 0.007234 | 0.033929 | 0.265908 | 0.026586 | 0.02926 | 0.013355 | 0.01265 | 0.035764 | 0.02449 | 0.060298 | 0.010212 | 0.020985 | 0.118646 | 0.04781 |
| 36.7361111 | 0.025591 | 0.023205 | 0.023717 | 0.024142 | 0.02567 | 0.026156 | 0.038534 | 0.0243 | 0.021115 | 0.050664 | 0.004502 | 0.021423 | 0.18347 | 0.030109 | 0.034254 | 0.017803 | 0.019732 | 0.025954 | 0.014328 | 0.069244 | 0.010201 | 0.010242 | 0.133575 | 0.08385 |
| 36.75 | 0.03171 | 0.021942 | 0.028017 | 0.026725 | 0.026416 | 0.017326 | 0.030992 | 0.048921 | 0.017418 | 0.036587 | 0.00187 | 0.012258 | 0.066709 | 0.029729 | 0.022714 | 0.015641 | 0.027352 | 0.01929 | 0.010572 | 0.066793 | 0.009628 | 0.003897 | 0.144544 | 0.151261 |
| 36.7638889 | 0.036731 | 0.02075 | 0.031794 | 0.029061 | 0.02965 | 0.014769 | 0.029896 | 0.070916 | 0.015785 | 0.028211 | 0.002427 | 0.007258 | 0.022308 | 0.027367 | 0.018295 | 0.009355 | 0.033688 | 0.012678 | 0.00991 | 0.054469 | 0.00993 | 0.002963 | 0.131019 | 0.17224 |
| 36.7777778 | 0.034143 | 0.020054 | 0.035202 | 0.031215 | 0.034354 | 0.017448 | 0.034508 | 0.075789 | 0.015436 | 0.024943 | 0.008359 | 0.005337 | 0.039286 | 0.027749 | 0.028648 | 0.010027 | 0.041962 | 0.007788 | 0.009596 | 0.031954 | 0.011596 | 0.004192 | 0.081518 | 0.137068 |
| 36.7916667 | 0.028274 | 0.020181 | 0.040541 | 0.03325 | 0.039685 | 0.022664 | 0.042015 | 0.05909 | 0.016049 | 0.025057 | 0.016948 | 0.006191 | 0.077259 | 0.036367 | 0.036573 | 0.017348 | 0.051837 | 0.009586 | 0.008304 | 0.014117 | 0.010311 | 0.003803 | 0.029653 | 0.076275 |
| 36.8055556 | 0.021714 | 0.020715 | 0.044241 | 0.035069 | 0.045653 | 0.02796 | 0.050019 | 0.030945 | 0.016816 | 0.027022 | 0.02339 | 0.009569 | 0.109366 | 0.045125 | 0.030365 | 0.026271 | 0.058566 | 0.01697 | 0.008143 | 0.015113 | 0.007611 | 0.004011 | 0.016593 | 0.043252 |
| 36.8194444 | 0.014881 | 0.020659 | 0.043994 | 0.035582 | 0.051669 | 0.031447 | 0.057166 | 0.017228 | 0.016665 | 0.02816 | 0.026771 | 0.013818 | 0.10526 | 0.034933 | 0.015989 | 0.035161 | 0.063255 | 0.019247 | 0.013915 | 0.017905 | 0.009701 | 0.010287 | 0.032918 | 0.063168 |
| 36.8333333 | 0.010303 | 0.019661 | 0.052035 | 0.03449 | 0.056993 | 0.032563 | 0.063161 | 0.03481 | 0.015678 | 0.025952 | 0.028709 | 0.016988 | 0.071303 | 0.016439 | 0.005956 | 0.042698 | 0.071231 | 0.012999 | 0.027649 | 0.012706 | 0.018453 | 0.019774 | 0.041203 | 0.103761 |
| 36.8472222 | 0.006555 | 0.018186 | 0.07085 | 0.033452 | 0.061311 | 0.032275 | 0.068508 | 0.064638 | 0.014738 | 0.021149 | 0.031293 | 0.018895 | 0.055409 | 0.008947 | 0.00531 | 0.048306 | 0.084888 | 0.011957 | 0.044677 | 0.009138 | 0.028923 | 0.025735 | 0.05408 | 0.145883 |
| 36.8611111 | 0.003437 | 0.016809 | 0.088793 | 0.032915 | 0.063472 | 0.031415 | 0.071111 | 0.082174 | 0.014441 | 0.015001 | 0.035183 | 0.019554 | 0.053424 | 0.004694 | 0.005386 | 0.051784 | 0.098368 | 0.024037 | 0.057033 | 0.011508 | 0.032287 | 0.025778 | 0.13081 | 0.165955 |
| 36.875 | 0.002764 | 0.016182 | 0.106718 | 0.030824 | 0.06243 | 0.029953 | 0.06839 | 0.078781 | 0.014977 | 0.009315 | 0.036997 | 0.018537 | 0.033803 | 0.003604 | 0.004699 | 0.051804 | 0.099827 | 0.035524 | 0.059734 | 0.018685 | 0.024542 | 0.021069 | 0.233297 | 0.121122 |
| 36.8888889 | 0.00391 | 0.016655 | 0.128489 | 0.026365 | 0.05839 | 0.02803 | 0.061801 | 0.058906 | 0.016091 | 0.007507 | 0.032711 | 0.016066 | 0.039978 | 0.011131 | 0.00555 | 0.046154 | 0.08639 | 0.034093 | 0.054038 | 0.028283 | 0.012013 | 0.015192 | 0.282008 | 0.055865 |

Supplementary Table S6. 1 hour integrated motion data in (a) lettuce subjected to 100 mM KCl

| IntervalMid | Control1 | Control2 | Control3 | Control4 | Control5 | Control6 | Control7 | Control8 | Control9 | Stress1 | Stress2 | Stress3 | Stress4 | Stress5 | Stress6 | Stress7 | Stress8 | Stress9 | Stress10 |
|---|---|---|---|---|---|---|---|---|---|---|---|---|---|---|---|---|---|---|---|
| 18.1111111 | 0.005512 | 0.025243 | 0.006489 | 0.008184 | 0.0143 | 0.011171 | 0.002568 | 0.028883 | 0.054189 | 0.007602 | 0.009236 | 0.006965 | 0.047401 | 0.005261 | 0.078547 | 0.019047 | 0.018691 | 0.016194 | 0.002523 |
| 18.125 | 0.004611 | 0.021333 | 0.003649 | 0.004569 | 0.012374 | 0.013106 | 0.000518 | 0.021173 | 0.050023 | 0.003374 | 0.00226 | 0.008701 | 0.032929 | 0.005552 | 0.070462 | 0.022338 | 0.029007 | 0.013626 | 0.001978 |
| 18.1388889 | 0.004644 | 0.01046 | 0.001063 | 0.001444 | 0.005346 | 0.008212 | 0.000844 | 0.010345 | 0.031553 | 0.002093 | 0.001433 | 0.007937 | 0.016742 | 0.003287 | 0.046237 | 0.015976 | 0.028401 | 0.008571 | 0.00226 |
| 18.1527778 | 0.003717 | 0.003601 | 0.0003 | 0.000303 | 0.001219 | 0.003884 | 0.002138 | 0.003818 | 0.016136 | 0.00341 | 0.001358 | 0.00457 | 0.007914 | 0.001442 | 0.0276 | 0.00952 | 0.019123 | 0.005183 | 0.001833 |
| 18.1666667 | 0.003314 | 0.001912 | 0.000547 | 0.000333 | 0.000182 | 0.002131 | 0.003194 | 0.001196 | 0.006951 | 0.006356 | 0.001106 | 0.001745 | 0.00389 | 0.00191 | 0.01692 | 0.005477 | 0.010155 | 0.003679 | 0.001548 |
| 18.1805556 | 0.004009 | 0.001607 | 0.001158 | 0.001361 | 0.000331 | 0.001382 | 0.001762 | 0.000634 | 0.00332 | 0.009951 | 0.001306 | 0.001301 | 0.002455 | 0.003213 | 0.009212 | 0.003474 | 0.005118 | 0.002913 | 0.001774 |
| 18.1944444 | 0.005574 | 0.001514 | 0.001583 | 0.001989 | 0.000971 | 0.001603 | 0.009363 | 0.001195 | 0.004506 | 0.015592 | 0.003256 | 0.003369 | 0.003681 | 0.004495 | 0.004304 | 0.004807 | 0.003275 | 0.004399 | 0.004063 |
| 18.2083333 | 0.006256 | 0.001931 | 0.00625 | 0.008202 | 0.00137 | 0.002499 | 0.010033 | 0.001863 | 0.004928 | 0.021158 | 0.004645 | 0.004938 | 0.005767 | 0.00812 | 0.003738 | 0.007191 | 0.003207 | 0.007419 | 0.007152 |
| 18.2222222 | 0.004295 | 0.003755 | 0.011542 | 0.018131 | 0.001165 | 0.002914 | 0.016832 | 0.001763 | 0.00278 | 0.023239 | 0.00311 | 0.003703 | 0.005759 | 0.010398 | 0.00399 | 0.005585 | 0.003681 | 0.006785 | 0.006618 |
| 18.2361111 | 0.002474 | 0.005578 | 0.007267 | 0.014876 | 0.001029 | 0.003111 | 0.007925 | 0.00186 | 0.003165 | 0.024239 | 0.003188 | 0.002612 | 0.004585 | 0.006514 | 0.004861 | 0.002656 | 0.006773 | 0.003357 | 0.003289 |
| 18.25 | 0.002923 | 0.005819 | 0.001846 | 0.00521 | 0.000808 | 0.003224 | 0.002209 | 0.002371 | 0.006977 | 0.026449 | 0.008372 | 0.003981 | 0.004971 | 0.00376 | 0.007401 | 0.002929 | 0.012471 | 0.001914 | 0.001543 |
| 18.2638889 | 0.002867 | 0.004061 | 0.001166 | 0.002109 | 0.000374 | 0.002573 | 0.001903 | 0.001877 | 0.008721 | 0.026703 | 0.013826 | 0.005589 | 0.005396 | 0.004997 | 0.007516 | 0.004031 | 0.014579 | 0.001897 | 0.001588 |
| 18.2777778 | 0.002278 | 0.002386 | 0.000711 | 0.001289 | 0.000643 | 0.001727 | 0.00152 | 0.000915 | 0.006301 | 0.024116 | 0.014571 | 0.005743 | 0.003629 | 0.005573 | 0.005041 | 0.003441 | 0.011188 | 0.001234 | 0.001382 |
| 18.2916667 | 0.002305 | 0.002294 | 0.00049 | 0.000665 | 0.00172 | 0.0017 | 0.001658 | 0.001071 | 0.004595 | 0.02153 | 0.012108 | 0.004866 | 0.001231 | 0.004684 | 0.004743 | 0.002202 | 0.006175 | 0.00051 | 0.000756 |
| 18.3055556 | 0.003282 | 0.002848 | 0.00074 | 0.000953 | 0.0023 | 0.002118 | 0.002 | 0.002511 | 0.008075 | 0.019373 | 0.00978 | 0.003391 | 0.000774 | 0.003228 | 0.004864 | 0.001634 | 0.002706 | 0.000358 | 0.000382 |
| 18.3194444 | 0.004288 | 0.004263 | 0.00072 | 0.001286 | 0.001696 | 0.001974 | 0.001693 | 0.003782 | 0.012963 | 0.016207 | 0.008187 | 0.002282 | 0.001277 | 0.002214 | 0.003706 | 0.001591 | 0.001714 | 0.000371 | 0.000517 |
| 18.3333333 | 0.003202 | 0.004181 | 0.000382 | 0.001018 | 0.001012 | 0.001365 | 0.001124 | 0.003 | 0.011832 | 0.013674 | 0.007274 | 0.002623 | 0.003189 | 0.001801 | 0.006545 | 0.001599 | 0.001818 | 0.001264 | 0.002002 |
| 18.3472222 | 0.001895 | 0.002394 | 0.00027 | 0.000519 | 0.001203 | 0.001053 | 0.001243 | 0.001246 | 0.00625 | 0.013801 | 0.007958 | 0.003189 | 0.009038 | 0.001278 | 0.013577 | 0.002365 | 0.002607 | 0.003314 | 0.00541 |
| 18.3611111 | 0.002626 | 0.002553 | 0.000187 | 0.000272 | 0.001451 | 0.000864 | 0.001507 | 0.000609 | 0.003114 | 0.012415 | 0.007017 | 0.005261 | 0.010888 | 0.002226 | 0.013845 | 0.005253 | 0.003768 | 0.003817 | 0.007775 |
| 18.375 | 0.003086 | 0.0034 | 0.000122 | 0.000185 | 0.000997 | 0.000678 | 0.001051 | 0.001213 | 0.002451 | 0.016241 | 0.012937 | 0.013622 | 0.020733 | 0.009091 | 0.020301 | 0.02075 | 0.00367 | 0.006371 | 0.017143 |
| 18.3888889 | 0.002742 | 0.00362 | 0.000244 | 0.000274 | 0.000471 | 0.000903 | 0.000639 | 0.002682 | 0.002617 | 0.033934 | 0.025433 | 0.02492 | 0.055107 | 0.018507 | 0.046995 | 0.047156 | 0.006916 | 0.013221 | 0.02909 |
| 18.4027778 | 0.003973 | 0.00475 | 0.000461 | 0.000773 | 0.000257 | 0.000922 | 0.001016 | 0.003752 | 0.003513 | 0.040516 | 0.02138 | 0.026423 | 0.071129 | 0.018455 | 0.052213 | 0.049399 | 0.01205 | 0.012908 | 0.021183 |
| 18.4166667 | 0.004009 | 0.004043 | 0.000499 | 0.001025 | 0.000266 | 0.000426 | 0.001404 | 0.002877 | 0.002945 | 0.022417 | 0.007869 | 0.015702 | 0.042651 | 0.009155 | 0.024134 | 0.02354 | 0.009045 | 0.005036 | 0.006615 |
| 18.4305556 | 0.002704 | 0.002476 | 0.000375 | 0.000835 | 0.000397 | 0.000264 | 0.001133 | 0.001165 | 0.001986 | 0.005448 | 0.0021 | 0.005929 | 0.011692 | 0.002209 | 0.00466 | 0.005223 | 0.002748 | 0.000894 | 0.001837 |
| 18.4444444 | 0.002545 | 0.002307 | 0.0006 | 0.00094 | 0.00068 | 0.000733 | 0.00062 | 0.000781 | 0.001692 | 0.001912 | 0.000999 | 0.002812 | 0.002841 | 0.00058 | 0.00294 | 0.001419 | 0.001265 | 0.000893 | 0.000719 |
| 18.4583333 | 0.003086 | 0.002815 | 0.001167 | 0.001226 | 0.001697 | 0.001763 | 0.000506 | 0.002508 | 0.002215 | 0.002394 | 0.000755 | 0.001871 | 0.003754 | 0.000845 | 0.006252 | 0.001387 | 0.002727 | 0.002123 | 0.001211 |
| 18.4722222 | 0.005479 | 0.005523 | 0.002381 | 0.002263 | 0.003577 | 0.003955 | 0.001369 | 0.00587 | 0.005763 | 0.001914 | 0.000932 | 0.002144 | 0.005463 | 0.001151 | 0.011134 | 0.003366 | 0.005039 | 0.004903 | 0.002241 |
| 18.4861111 | 0.007177 | 0.007946 | 0.004155 | 0.004486 | 0.005451 | 0.006985 | 0.003052 | 0.008695 | 0.009795 | 0.001733 | 0.002232 | 0.007897 | 0.006404 | 0.002212 | 0.014354 | 0.005378 | 0.007429 | 0.006864 | 0.002313 |
| 18.5 | 0.006554 | 0.007868 | 0.004932 | 0.005801 | 0.006412 | 0.008598 | 0.00424 | 0.009206 | 0.010723 | 0.003034 | 0.004524 | 0.007775 | 0.00609 | 0.005186 | 0.013379 | 0.005783 | 0.008743 | 0.006216 | 0.002396 |
| 18.5138889 | 0.008791 | 0.010495 | 0.004018 | 0.004683 | 0.006083 | 0.007603 | 0.003791 | 0.009127 | 0.012454 | 0.004665 | 0.008302 | 0.009341 | 0.00655 | 0.008179 | 0.012118 | 0.006293 | 0.009237 | 0.005611 | 0.003974 |
| 18.5277778 | 0.020039 | 0.0235 | 0.002264 | 0.002379 | 0.004796 | 0.005624 | 0.002913 | 0.012219 | 0.02078 | 0.00476 | 0.016741 | 0.011239 | 0.010632 | 0.011448 | 0.01578 | 0.008715 | 0.012193 | 0.007752 | 0.006699 |
| 18.5416667 | 0.050575 | 0.041805 | 0.001822 | 0.001172 | 0.00501 | 0.006639 | 0.002367 | 0.018968 | 0.038134 | 0.00397 | 0.042503 | 0.017897 | 0.018603 | 0.018244 | 0.02581 | 0.013859 | 0.022085 | 0.016004 | 0.010139 |
| 18.5555556 | 0.092245 | 0.056949 | 0.003672 | 0.003032 | 0.00842 | 0.012635 | 0.003248 | 0.022234 | 0.057188 | 0.010231 | 0.079562 | 0.030416 | 0.026744 | 0.026024 | 0.036296 | 0.021658 | 0.035504 | 0.029081 | 0.014192 |
| 18.5694444 | 0.101402 | 0.059885 | 0.004671 | 0.008823 | 0.013675 | 0.016759 | 0.006132 | 0.015645 | 0.055957 | 0.023038 | 0.086184 | 0.038298 | 0.030114 | 0.028056 | 0.036203 | 0.029869 | 0.037989 | 0.033899 | 0.017435 |
| 18.5833333 | 0.070484 | 0.04187 | 0.002937 | 0.013792 | 0.017638 | 0.012907 | 0.006236 | 0.006261 | 0.034294 | 0.027389 | 0.055934 | 0.031191 | 0.027459 | 0.024457 | 0.026936 | 0.03646 | 0.0267 | 0.02656 | 0.016587 |
| 18.5972222 | 0.039848 | 0.02034 | 0.000984 | 0.012704 | 0.01664 | 0.006641 | 0.00338 | 0.002515 | 0.017255 | 0.021142 | 0.026279 | 0.017009 | 0.020777 | 0.021789 | 0.018225 | 0.040375 | 0.015086 | 0.017598 | 0.011761 |
| 18.6111111 | 0.021997 | 0.008555 | 0.000518 | 0.009207 | 0.011863 | 0.003782 | 0.001634 | 0.001953 | 0.008869 | 0.013996 | 0.011932 | 0.007444 | 0.013617 | 0.022306 | 0.012361 | 0.042564 | 0.008937 | 0.011088 | 0.007555 |
| 18.625 | 0.011712 | 0.002345 | 0.000763 | 0.00673 | 0.007581 | 0.003264 | 0.000916 | 0.001279 | 0.004166 | 0.010687 | 0.007139 | 0.003559 | 0.009754 | 0.021902 | 0.008891 | 0.044013 | 0.005983 | 0.006178 | 0.005066 |
| 18.6388889 | 0.010665 | 0.001022 | 0.000772 | 0.004637 | 0.005781 | 0.003405 | 0.000457 | 0.001331 | 0.006458 | 0.010134 | 0.006218 | 0.002044 | 0.008873 | 0.01868 | 0.006879 | 0.040572 | 0.004891 | 0.002862 | 0.003047 |
| 18.6527778 | 0.012447 | 0.001397 | 0.000604 | 0.003763 | 0.005973 | 0.003582 | 0.000394 | 0.002128 | 0.012576 | 0.009317 | 0.004591 | 0.001421 | 0.007909 | 0.014767 | 0.005638 | 0.032186 | 0.00443 | 0.001124 | 0.00166 |
| 18.6666667 | 0.008942 | 0.001932 | 0.000695 | 0.00446 | 0.006029 | 0.002627 | 0.000546 | 0.001972 | 0.016431 | 0.007229 | 0.00256 | 0.002419 | 0.006013 | 0.011296 | 0.005834 | 0.025778 | 0.004008 | 0.001046 | 0.001284 |
| 18.6805556 | 0.005531 | 0.004209 | 0.000746 | 0.005334 | 0.004523 | 0.001483 | 0.000708 | 0.001693 | 0.015983 | 0.004141 | 0.002631 | 0.003987 | 0.004641 | 0.008373 | 0.006654 | 0.022226 | 0.00378 | 0.002512 | 0.001226 |
| 18.6944444 | 0.008995 | 0.006254 | 0.000685 | 0.005158 | 0.002205 | 0.001968 | 0.000848 | 0.001879 | 0.011096 | 0.001962 | 0.004606 | 0.004415 | 0.004989 | 0.006033 | 0.006745 | 0.017688 | 0.003829 | 0.00561 | 0.0008 |
| 18.7083333 | 0.010758 | 0.005329 | 0.001178 | 0.003878 | 0.000801 | 0.002432 | 0.000887 | 0.001907 | 0.008145 | 0.002278 | 0.00589 | 0.003993 | 0.006489 | 0.004103 | 0.005536 | 0.012148 | 0.003757 | 0.008644 | 0.000655 |
| 18.7222222 | 0.006726 | 0.002575 | 0.001851 | 0.002676 | 0.00067 | 0.001699 | 0.000711 | 0.002072 | 0.013952 | 0.003047 | 0.005266 | 0.003047 | 0.007217 | 0.002728 | 0.003332 | 0.006557 | 0.003317 | 0.010122 | 0.000731 |
| 18.7361111 | 0.003585 | 0.001272 | 0.001608 | 0.002679 | 0.000822 | 0.00088 | 0.000504 | 0.001923 | 0.018988 | 0.002552 | 0.003721 | 0.001772 | 0.005873 | 0.002108 | 0.001523 | 0.003498 | 0.002797 | 0.010563 | 0.000565 |
| 18.75 | 0.00236 | 0.002981 | 0.000834 | 0.002754 | 0.000695 | 0.000864 | 0.000614 | 0.001742 | 0.013973 | 0.001425 | 0.002451 | 0.000768 | 0.003328 | 0.002245 | 0.001596 | 0.005876 | 0.002459 | 0.010087 | 0.000471 |
| 18.7638889 | 0.001898 | 0.00664 | 0.000728 | 0.00191 | 0.000441 | 0.001665 | 0.000658 | 0.001509 | 0.005609 | 0.000536 | 0.001607 | 0.000311 | 0.001301 | 0.002969 | 0.00238 | 0.009817 | 0.002293 | 0.008038 | 0.000539 |
| 18.7777778 | 0.003721 | 0.009762 | 0.001253 | 0.001799 | 0.000567 | 0.002375 | 0.000412 | 0.001769 | 0.001949 | 0.000188 | 0.000982 | 0.000638 | 0.000437 | 0.003615 | 0.002643 | 0.010071 | 0.002056 | 0.004752 | 0.000473 |
| 18.7916667 | 0.00539 | 0.010544 | 0.001559 | 0.002402 | 0.001183 | 0.002444 | 0.000285 | 0.002543 | 0.004806 | 0.000204 | 0.000664 | 0.001829 | 0.000289 | 0.003509 | 0.004503 | 0.006528 | 0.001331 | 0.001834 | 0.000397 |
| 18.8055556 | 0.005036 | 0.008405 | 0.001125 | 0.002031 | 0.001542 | 0.001834 | 0.000569 | 0.002711 | 0.012367 | 0.000494 | 0.000667 | 0.003056 | 0.000458 | 0.002858 | 0.007935 | 0.002558 | 0.000684 | 0.000438 | 0.000472 |
| 18.8194444 | 0.006313 | 0.004701 | 0.000583 | 0.001963 | 0.001178 | 0.000976 | 0.001287 | 0.003514 | 0.018962 | 0.00078 | 0.000779 | 0.003456 | 0.00113 | 0.0024 | 0.009024 | 0.000642 | 0.000691 | 0.000416 | 0.000479 |
| 18.8333333 | 0.00875 | 0.002607 | 0.000635 | 0.002898 | 0.000531 | 0.000782 | 0.002181 | 0.004373 | 0.017709 | 0.000771 | 0.000782 | 0.002919 | 0.001647 | 0.002571 | 0.006647 | 0.000392 | 0.000583 | 0.001799 | 0.000302 |
| 18.8472222 | 0.007603 | 0.003729 | 0.00057 | 0.002805 | 0.000249 | 0.001379 | 0.002666 | 0.003242 | 0.010911 | 0.000672 | 0.00076 | 0.001886 | 0.002508 | 0.00308 | 0.003285 | 0.001111 | 0.000358 | 0.004828 | 0.000193 |
| 18.8611111 | 0.004241 | 0.007068 | 0.000465 | 0.001716 | 0.000274 | 0.002016 | 0.002171 | 0.002382 | 0.009095 | 0.000849 | 0.001026 | 0.001434 | 0.004351 | 0.003297 | 0.001616 | 0.002378 | 0.000736 | 0.00703 | 0.000475 |
| 18.875 | 0.003303 | 0.010182 | 0.001039 | 0.001433 | 0.000218 | 0.002099 | 0.001074 | 0.003621 | 0.011646 | 0.000891 | 0.001426 | 0.002042 | 0.004186 | 0.00305 | 0.002615 | 0.003697 | 0.001608 | 0.005488 | 0.000912 |
| 18.8888889 | 0.003235 | 0.011498 | 0.001417 | 0.001803 | 0.000325 | 0.001395 | 0.000418 | 0.003825 | 0.010003 | 0.00074 | 0.001513 | 0.002791 | 0.00378 | 0.002568 | 0.005706 | 0.005458 | 0.002348 | 0.004419 | 0.000827 |
| 18.9027778 | 0.00363 | 0.01131 | 0.001288 | 0.002097 | 0.001019 | 0.000712 | 0.000434 | 0.002436 | 0.009608 | 0.000707 | 0.00155 | 0.00314 | 0.007085 | 0.002059 | 0.010221 | 0.007838 | 0.002854 | 0.008842 | 0.000348 |
| 18.9166667 | 0.008313 | 0.01093 | 0.0018 | 0.002196 | 0.001846 | 0.000828 | 0.000618 | 0.003263 | 0.016037 | 0.000972 | 0.002001 | 0.003628 | 0.011052 | 0.00163 | 0.013403 | 0.009552 | 0.003778 | 0.014052 | 0.000353 |
| 18.9305556 | 0.013912 | 0.010741 | 0.002505 | 0.001907 | 0.002122 | 0.000809 | 0.000875 | 0.006009 | 0.020927 | 0.002225 | 0.002401 | 0.004154 | 0.013689 | 0.001461 | 0.012757 | 0.009702 | 0.005064 | 0.013113 | 0.001014 |
| 18.9444444 | 0.014649 | 0.012167 | 0.00213 | 0.002165 | 0.002031 | 0.000473 | 0.001008 | 0.006079 | 0.020537 | 0.00385 | 0.002186 | 0.003811 | 0.016736 | 0.001756 | 0.010604 | 0.008946 | 0.005697 | 0.007857 | 0.001599 |
| 18.9583333 | 0.013156 | 0.015452 | 0.001664 | 0.003362 | 0.002373 | 0.001126 | 0.001073 | 0.003341 | 0.020098 | 0.005236 | 0.001749 | 0.002754 | 0.019807 | 0.002172 | 0.010087 | 0.008166 | 0.005725 | 0.005268 | 0.001555 |
| 18.9722222 | 0.014789 | 0.016454 | 0.002916 | 0.004342 | 0.003838 | 0.003257 | 0.001748 | 0.002245 | 0.022334 | 0.00661 | 0.001601 | 0.002252 | 0.018896 | 0.002054 | 0.010438 | 0.007677 | 0.006334 | 0.004421 | 0.001004 |
| 18.9861111 | 0.016955 | 0.015766 | 0.005392 | 0.005144 | 0.005488 | 0.005248 | 0.002759 | 0.003092 | 0.022738 | 0.007744 | 0.00165 | 0.002488 | 0.01178 | 0.001473 | 0.009409 | 0.00716 | 0.007653 | 0.004115 | 0.000477 |
| 19 | 0.016311 | 0.017638 | 0.006885 | 0.005327 | 0.005492 | 0.005049 | 0.002827 | 0.003847 | 0.017375 | 0.008676 | 0.001779 | 0.003963 | 0.003863 | 0.00101 | 0.007385 | 0.006378 | 0.008711 | 0.008615 | 0.000244 |
| 19.0138889 | 0.018064 | 0.019112 | 0.006797 | 0.003592 | 0.004536 | 0.003236 | 0.002204 | 0.006679 | 0.012168 | 0.009492 | 0.002119 | 0.005206 | 0.00218 | 0.00067 | 0.007092 | 0.006194 | 0.009496 | 0.014747 | 0.000404 |
| 19.0277778 | 0.025499 | 0.015183 | 0.006498 | 0.002728 | 0.005163 | 0.002566 | 0.002257 | 0.013451 | 0.018685 | 0.011696 | 0.004518 | 0.007818 | 0.00878 | 0.001127 | 0.011832 | 0.00945 | 0.011664 | 0.019524 | 0.002441 |
| 19.0416667 | 0.038281 | 0.016351 | 0.007152 | 0.007254 | 0.008298 | 0.0063 | 0.002953 | 0.02079 | 0.041304 | 0.024932 | 0.016025 | 0.015508 | 0.025258 | 0.005595 | 0.028738 | 0.024271 | 0.019816 | 0.033581 | 0.010818 |
| 19.0555556 | 0.05144 | 0.027611 | 0.008483 | 0.016411 | 0.012955 | 0.013006 | 0.005401 | 0.022897 | 0.060824 | 0.048853 | 0.046965 | 0.028693 | 0.035253 | 0.012711 | 0.053214 | 0.049525 | 0.031032 | 0.051608 | 0.029136 |
| 19.0694444 | 0.047146 | 0.026164 | 0.00787 | 0.021439 | 0.016545 | 0.015095 | 0.008575 | 0.015687 | 0.051113 | 0.065611 | 0.09508 | 0.03924 | 0.038235 | 0.013251 | 0.050882 | 0.057386 | 0.028434 | 0.043478 | 0.051011 |
| 19.0833333 | 0.024122 | 0.013317 | 0.00459 | 0.014764 | 0.014503 | 0.009877 | 0.00774 | 0.008771 | 0.027235 | 0.064808 | 0.119483 | 0.039007 | 0.056138 | 0.012091 | 0.029815 | 0.034929 | 0.014353 | 0.024093 | 0.062513 |
| 19.0972222 | 0.006278 | 0.012861 | 0.002779 | 0.004784 | 0.008092 | 0.003795 | 0.004175 | 0.009211 | 0.019285 | 0.054972 | 0.082009 | 0.027159 | 0.059825 | 0.016626 | 0.027989 | 0.011179 | 0.007222 | 0.023373 | 0.054311 |
| 19.1111111 | 0.002009 | 0.018451 | 0.002577 | 0.001656 | 0.004539 | 0.001328 | 0.001902 | 0.006895 | 0.016091 | 0.038624 | 0.0265 | 0.011444 | 0.036754 | 0.013112 | 0.026323 | 0.004303 | 0.005059 | 0.023483 | 0.030113 |
| 19.125 | 0.003361 | 0.024718 | 0.002594 | 0.001577 | 0.004043 | 0.000883 | 0.0014 | 0.003674 | 0.010885 | 0.027081 | 0.011558 | 0.002357 | 0.023094 | 0.004604 | 0.013425 | 0.005569 | 0.0023 | 0.017316 | 0.011103 |
| 19.1388889 | 0.005155 | 0.03148 | 0.0037 | 0.001359 | 0.002975 | 0.001126 | 0.002017 | 0.005048 | 0.013758 | 0.027353 | 0.017083 | 0.000413 | 0.021372 | 0.001993 | 0.004481 | 0.007807 | 0.001005 | 0.013339 | 0.007883 |
| 19.1527778 | 0.004638 | 0.026937 | 0.003182 | 0.001543 | 0.001793 | 0.001272 | 0.001931 | 0.004792 | 0.011853 | 0.014542 | 0.012213 | 0.00044 | 0.016716 | 0.001202 | 0.001775 | 0.00722 | 0.000572 | 0.00975 | 0.006143 |
| 19.1666667 | 0.002686 | 0.016679 | 0.001215 | 0.001235 | 0.001513 | 0.001059 | 0.00122 | 0.001946 | 0.004021 | 0.001305 | 0.005373 | 0.000328 | 0.011528 | 0.000737 | 0.003527 | 0.00388 | 0.000441 | 0.008019 | 0.001716 |
| 19.1805556 | 0.001549 | 0.010139 | 0.000308 | 0.001 | 0.001385 | 0.000994 | 0.001101 | 0.000752 | 0.001751 | 0.001114 | 0.00282 | 0.000287 | 0.009186 | 0.000593 | 0.005274 | 0.00123 | 0.00042 | 0.008374 | 0.000486 |
| 19.1944444 | 0.001466 | 0.00675 | 0.000182 | 0.001058 | 0.000998 | 0.000876 | 0.001074 | 0.001252 | 0.002081 | 0.001463 | 0.001572 | 0.000741 | 0.007871 | 0.000301 | 0.004912 | 0.000358 | 0.000887 | 0.008377 | 0.000842 |
| 19.2083333 | 0.001593 | 0.007281 | 0.000222 | 0.000746 | 0.00089 | 0.00061 | 0.000859 | 0.001461 | 0.002022 | 0.002394 | 0.001024 | 0.001159 | 0.007406 | 0.000223 | 0.003847 | 0.000688 | 0.00169 | 0.007542 | 0.000732 |
| 19.2222222 | 0.001383 | 0.008862 | 0.0002 | 0.000431 | 0.00101 | 0.000297 | 0.000528 | 0.00102 | 0.001677 | 0.002641 | 0.000684 | 0.0011 | 0.00775 | 0.000261 | 0.003311 | 0.001146 | 0.001732 | 0.006362 | 0.000494 |
| 19.2361111 | 0.001765 | 0.006861 | 0.000323 | 0.000569 | 0.000944 | 8.77E-05 | 0.000284 | 0.000502 | 0.000833 | 0.001861 | 0.000422 | 0.001194 | 0.007697 | 0.000551 | 0.002834 | 0.001419 | 0.002407 | 0.004973 | 0.000386 |
| 19.25 | 0.003328 | 0.002875 | 0.000758 | 0.0008 | 0.000606 | 8.50E-05 | 0.000504 | 0.000361 | 0.000572 | 0.001853 | 0.000511 | 0.001644 | 0.006542 | 0.001095 | 0.001797 | 0.001945 | 0.004186 | 0.003272 | 0.000438 |
| 19.2638889 | 0.004816 | 0.000569 | 0.001204 | 0.000781 | 0.000474 | 0.000172 | 0.000916 | 0.000342 | 0.001362 | 0.002267 | 0.001057 | 0.001551 | 0.004917 | 0.001332 | 0.000861 | 0.002368 | 0.004306 | 0.00197 | 0.000779 |
| 19.2777778 | 0.004432 | 0.001888 | 0.001026 | 0.000791 | 0.000865 | 0.000257 | 0.000854 | 0.000372 | 0.001674 | 0.001757 | 0.00164 | 0.000833 | 0.003941 | 0.000998 | 0.000556 | 0.002107 | 0.002581 | 0.002384 | 0.001125 |
| 19.2916667 | 0.002513 | 0.007055 | 0.000407 | 0.00119 | 0.001332 | 0.000522 | 0.000704 | 0.000556 | 0.00152 | 0.001053 | 0.001938 | 0.000341 | 0.003847 | 0.000825 | 0.000478 | 0.001754 | 0.001129 | 0.003677 | 0.001206 |
| 19.3055556 | 0.001565 | 0.011864 | 0.000144 | 0.001332 | 0.001515 | 0.000937 | 0.001048 | 0.000709 | 0.002296 | 0.000628 | 0.002004 | 0.000346 | 0.003853 | 0.001303 | 0.000634 | 0.001901 | 0.000499 | 0.003877 | 0.001365 |

| | | | | | | | | | | | | | | | | | | | |
|---|---|---|---|---|---|---|---|---|---|---|---|---|---|---|---|---|---|---|---|
| 19.3194444 | 0.001393 | 0.013838 | 0.000505 | 0.000871 | 0.001184 | 0.001101 | 0.00123 | 0.000628 | 0.002987 | 0.000635 | 0.001971 | 0.000536 | 0.002866 | 0.001964 | 0.000725 | 0.002112 | 0.000295 | 0.003386 | 0.001781 |
| 19.3333333 | 0.001025 | 0.016175 | 0.001353 | 0.000464 | 0.00058 | 0.000829 | 0.00095 | 0.000689 | 0.003404 | 0.001276 | 0.001699 | 0.000763 | 0.001405 | 0.002839 | 0.000905 | 0.002001 | 0.000167 | 0.004334 | 0.001678 |
| 19.3472222 | 0.001825 | 0.020657 | 0.002151 | 0.000482 | 0.000396 | 0.000396 | 0.000838 | 0.00111 | 0.004903 | 0.00183 | 0.001225 | 0.000738 | 0.001022 | 0.0037 | 0.002143 | 0.001525 | 0.000422 | 0.006787 | 0.000952 |
| 19.3611111 | 0.004328 | 0.025802 | 0.00251 | 0.000934 | 0.000677 | 0.000258 | 0.001094 | 0.002054 | 0.007725 | 0.001829 | 0.001075 | 0.000743 | 0.001278 | 0.003003 | 0.004053 | 0.001066 | 0.001457 | 0.008258 | 0.000808 |
| 19.375 | 0.007224 | 0.028749 | 0.002691 | 0.001355 | 0.001122 | 0.000526 | 0.001362 | 0.003416 | 0.010683 | 0.001986 | 0.001278 | 0.001313 | 0.0014 | 0.001353 | 0.006359 | 0.001528 | 0.002565 | 0.008212 | 0.001529 |
| 19.3888889 | 0.007399 | 0.027602 | 0.002795 | 0.001251 | 0.00122 | 0.000713 | 0.001417 | 0.003766 | 0.011598 | 0.001936 | 0.001389 | 0.001675 | 0.002757 | 0.000842 | 0.009346 | 0.002764 | 0.002877 | 0.008552 | 0.002081 |
| 19.4027778 | 0.004317 | 0.025293 | 0.002581 | 0.000799 | 0.000933 | 0.000647 | 0.001336 | 0.00247 | 0.010483 | 0.00108 | 0.001402 | 0.001431 | 0.00634 | 0.000898 | 0.011925 | 0.003775 | 0.002507 | 0.009946 | 0.00172 |
| 19.4166667 | 0.001821 | 0.025959 | 0.002472 | 0.000928 | 0.00106 | 0.000923 | 0.001028 | 0.00127 | 0.010152 | 0.000517 | 0.001109 | 0.001015 | 0.01083 | 0.00077 | 0.01334 | 0.004252 | 0.002186 | 0.012293 | 0.001143 |
| 19.4305556 | 0.001616 | 0.027792 | 0.00308 | 0.002479 | 0.001576 | 0.001732 | 0.000972 | 0.00132 | 0.01153 | 0.000598 | 0.000782 | 0.000863 | 0.014066 | 0.000636 | 0.014286 | 0.004503 | 0.002486 | 0.015019 | 0.001343 |
| 19.4444444 | 0.00193 | 0.026602 | 0.003712 | 0.004172 | 0.001893 | 0.00236 | 0.001418 | 0.002111 | 0.012593 | 0.001076 | 0.000977 | 0.001544 | 0.014201 | 0.001034 | 0.01418 | 0.004659 | 0.003131 | 0.015962 | 0.00149 |
| 19.4583333 | 0.002427 | 0.024431 | 0.003857 | 0.004578 | 0.00256 | 0.002813 | 0.001621 | 0.003615 | 0.012968 | 0.002085 | 0.001129 | 0.002667 | 0.011617 | 0.002521 | 0.012392 | 0.004499 | 0.003524 | 0.014868 | 0.001011 |
| 19.4722222 | 0.004645 | 0.025252 | 0.004447 | 0.005371 | 0.004491 | 0.004471 | 0.002411 | 0.005872 | 0.013943 | 0.002661 | 0.001212 | 0.003038 | 0.009049 | 0.003256 | 0.010759 | 0.004012 | 0.003608 | 0.014236 | 0.000876 |
| 19.4861111 | 0.007506 | 0.02678 | 0.00556 | 0.007681 | 0.007048 | 0.007262 | 0.004378 | 0.007966 | 0.015214 | 0.002495 | 0.001523 | 0.002536 | 0.008471 | 0.002383 | 0.010328 | 0.003567 | 0.003534 | 0.014757 | 0.001507 |
| 19.5 | 0.008773 | 0.026273 | 0.005962 | 0.008976 | 0.00826 | 0.00872 | 0.005606 | 0.009041 | 0.016311 | 0.003091 | 0.001955 | 0.002525 | 0.00966 | 0.001962 | 0.010875 | 0.003917 | 0.00383 | 0.015269 | 0.002529 |
| 19.5138889 | 0.010514 | 0.027516 | 0.005286 | 0.006717 | 0.007106 | 0.007447 | 0.004381 | 0.009559 | 0.018683 | 0.005531 | 0.002765 | 0.004086 | 0.011465 | 0.003495 | 0.01259 | 0.005586 | 0.005291 | 0.015566 | 0.003666 |
| 19.5277778 | 0.018221 | 0.034267 | 0.005043 | 0.004084 | 0.005239 | 0.004942 | 0.002707 | 0.011039 | 0.02648 | 0.008931 | 0.00443 | 0.006541 | 0.014006 | 0.007277 | 0.014163 | 0.007723 | 0.008928 | 0.017233 | 0.005085 |
| 19.5416667 | 0.037576 | 0.042709 | 0.006582 | 0.003482 | 0.004903 | 0.00476 | 0.00229 | 0.013054 | 0.048004 | 0.014902 | 0.008907 | 0.010169 | 0.018971 | 0.012147 | 0.014011 | 0.009375 | 0.016768 | 0.025647 | 0.008056 |
| 19.5555556 | 0.058987 | 0.037327 | 0.00778 | 0.005422 | 0.008173 | 0.009588 | 0.003881 | 0.012772 | 0.068264 | 0.022963 | 0.021453 | 0.012829 | 0.025848 | 0.019777 | 0.012457 | 0.011446 | 0.026307 | 0.037664 | 0.014354 |
| 19.5694444 | 0.061582 | 0.019706 | 0.005488 | 0.010173 | 0.014864 | 0.016252 | 0.008549 | 0.009753 | 0.057873 | 0.0233 | 0.04149 | 0.012267 | 0.026337 | 0.026492 | 0.008441 | 0.014825 | 0.027481 | 0.035422 | 0.022344 |
| 19.5833333 | 0.045558 | 0.014204 | 0.002308 | 0.011809 | 0.017676 | 0.018084 | 0.011675 | 0.00699 | 0.029059 | 0.015736 | 0.053629 | 0.010228 | 0.016029 | 0.022952 | 0.003521 | 0.019572 | 0.018233 | 0.018012 | 0.027514 |
| 19.5972222 | 0.026327 | 0.025247 | 0.002392 | 0.009741 | 0.013306 | 0.013937 | 0.010587 | 0.006548 | 0.010962 | 0.01127 | 0.047609 | 0.006076 | 0.008812 | 0.015592 | 0.003688 | 0.024953 | 0.008514 | 0.006489 | 0.025515 |
| 19.6111111 | 0.014835 | 0.040742 | 0.003771 | 0.007047 | 0.00745 | 0.008469 | 0.007965 | 0.006717 | 0.005856 | 0.011476 | 0.03249 | 0.002351 | 0.012994 | 0.012716 | 0.008684 | 0.02627 | 0.003552 | 0.007223 | 0.016438 |
| 19.625 | 0.012858 | 0.048239 | 0.003632 | 0.003713 | 0.003313 | 0.005004 | 0.005356 | 0.005158 | 0.005087 | 0.012404 | 0.019399 | 0.002187 | 0.017442 | 0.010656 | 0.009396 | 0.022012 | 0.001991 | 0.009237 | 0.007421 |
| 19.6388889 | 0.012795 | 0.042294 | 0.002514 | 0.001532 | 0.001197 | 0.003493 | 0.003431 | 0.003635 | 0.005008 | 0.013381 | 0.011658 | 0.003277 | 0.013041 | 0.007139 | 0.007837 | 0.01807 | 0.002301 | 0.006664 | 0.002765 |
| 19.6527778 | 0.009226 | 0.027475 | 0.002581 | 0.000835 | 0.000657 | 0.002776 | 0.002186 | 0.005102 | 0.006111 | 0.015197 | 0.00879 | 0.003387 | 0.010522 | 0.004138 | 0.015153 | 0.017194 | 0.003115 | 0.004797 | 0.000975 |
| 19.6666667 | 0.004505 | 0.012385 | 0.00363 | 0.000758 | 0.001007 | 0.001832 | 0.001234 | 0.00823 | 0.013281 | 0.017232 | 0.008766 | 0.002342 | 0.021644 | 0.003069 | 0.019845 | 0.017175 | 0.004043 | 0.010019 | 0.000687 |
| 19.6805556 | 0.002553 | 0.005812 | 0.003077 | 0.001069 | 0.002538 | 0.00111 | 0.000502 | 0.009261 | 0.026832 | 0.017294 | 0.009149 | 0.002172 | 0.028993 | 0.003025 | 0.012401 | 0.016974 | 0.005255 | 0.017679 | 0.001014 |
| 19.6944444 | 0.00336 | 0.007988 | 0.002746 | 0.000912 | 0.004104 | 0.001362 | 0.000254 | 0.006363 | 0.039901 | 0.014718 | 0.008806 | 0.00356 | 0.020004 | 0.00251 | 0.007011 | 0.018334 | 0.005363 | 0.019467 | 0.001354 |
| 19.7083333 | 0.005933 | 0.011052 | 0.005046 | 0.000711 | 0.003362 | 0.001948 | 0.000434 | 0.003009 | 0.042144 | 0.012316 | 0.008356 | 0.004224 | 0.013334 | 0.001612 | 0.009488 | 0.020704 | 0.003656 | 0.014269 | 0.00152 |
| 19.7222222 | 0.009076 | 0.011145 | 0.006772 | 0.001387 | 0.002301 | 0.001998 | 0.00088 | 0.0028 | 0.026733 | 0.014232 | 0.008186 | 0.002847 | 0.019786 | 0.001794 | 0.01361 | 0.020338 | 0.002861 | 0.007169 | 0.001399 |
| 19.7361111 | 0.008316 | 0.014348 | 0.005665 | 0.00163 | 0.004271 | 0.001564 | 0.001602 | 0.002912 | 0.011142 | 0.019987 | 0.00781 | 0.001998 | 0.028489 | 0.002795 | 0.013394 | 0.016856 | 0.004023 | 0.004959 | 0.001174 |
| 19.75 | 0.00636 | 0.021755 | 0.003233 | 0.001602 | 0.006688 | 0.001531 | 0.001799 | 0.001798 | 0.012924 | 0.023279 | 0.006507 | 0.00408 | 0.031419 | 0.003149 | 0.011112 | 0.01338 | 0.00471 | 0.009423 | 0.001358 |
| 19.7638889 | 0.007193 | 0.022993 | 0.002101 | 0.00205 | 0.006665 | 0.001414 | 0.001225 | 0.00122 | 0.020303 | 0.019814 | 0.004034 | 0.005981 | 0.023382 | 0.002814 | 0.017533 | 0.011579 | 0.00429 | 0.015628 | 0.002083 |
| 19.7777778 | 0.006225 | 0.015212 | 0.003315 | 0.00155 | 0.004778 | 0.001216 | 0.001123 | 0.001338 | 0.0202 | 0.011846 | 0.001768 | 0.004309 | 0.019002 | 0.002605 | 0.030738 | 0.010649 | 0.004017 | 0.016335 | 0.002752 |
| 19.7916667 | 0.004884 | 0.010244 | 0.003407 | 0.000995 | 0.002839 | 0.001944 | 0.001664 | 0.001243 | 0.012566 | 0.0064 | 0.001712 | 0.001595 | 0.034411 | 0.002752 | 0.035189 | 0.008916 | 0.004046 | 0.012651 | 0.002549 |
| 19.8055556 | 0.005873 | 0.014678 | 0.002088 | 0.001297 | 0.002804 | 0.00261 | 0.00184 | 0.001114 | 0.005691 | 0.006452 | 0.003384 | 0.000748 | 0.040458 | 0.002961 | 0.025914 | 0.005742 | 0.002957 | 0.015304 | 0.001671 |
| 19.8194444 | 0.006216 | 0.0206 | 0.002539 | 0.001098 | 0.003854 | 0.002435 | 0.001272 | 0.002455 | 0.007136 | 0.006777 | 0.004154 | 0.000912 | 0.024422 | 0.00296 | 0.01347 | 0.002637 | 0.000994 | 0.02016 | 0.001395 |
| 19.8333333 | 0.005228 | 0.022891 | 0.003545 | 0.001475 | 0.003696 | 0.001572 | 0.000654 | 0.004183 | 0.012125 | 0.005326 | 0.003368 | 0.001402 | 0.016694 | 0.002535 | 0.009349 | 0.001362 | 0.000618 | 0.015229 | 0.002042 |
| 19.8472222 | 0.004694 | 0.023834 | 0.003934 | 0.00395 | 0.002245 | 0.000877 | 0.000626 | 0.003369 | 0.011133 | 0.003311 | 0.00194 | 0.001721 | 0.028177 | 0.001621 | 0.013933 | 0.001695 | 0.002202 | 0.011894 | 0.002529 |
| 19.8611111 | 0.006064 | 0.018695 | 0.004295 | 0.006288 | 0.00132 | 0.001144 | 0.000969 | 0.001823 | 0.013222 | 0.001891 | 0.001245 | 0.001205 | 0.041604 | 0.001065 | 0.018802 | 0.002453 | 0.004527 | 0.02151 | 0.002217 |
| 19.875 | 0.010885 | 0.011742 | 0.003657 | 0.005387 | 0.002837 | 0.001318 | 0.001199 | 0.003512 | 0.032719 | 0.002999 | 0.002412 | 0.00075 | 0.045622 | 0.001347 | 0.019359 | 0.00326 | 0.006162 | 0.031854 | 0.001545 |
| 19.8888889 | 0.015373 | 0.015367 | 0.002015 | 0.003738 | 0.006224 | 0.001535 | 0.001122 | 0.006465 | 0.048755 | 0.00492 | 0.003722 | 0.001027 | 0.037901 | 0.001359 | 0.014625 | 0.003607 | 0.006729 | 0.03081 | 0.001237 |
| 19.9027778 | 0.012739 | 0.023221 | 0.000946 | 0.006398 | 0.009159 | 0.003093 | 0.001561 | 0.007412 | 0.042663 | 0.005336 | 0.003626 | 0.001291 | 0.021344 | 0.001564 | 0.008357 | 0.002613 | 0.007425 | 0.019762 | 0.001372 |
| 19.9166667 | 0.008773 | 0.030018 | 0.000964 | 0.011075 | 0.010801 | 0.005446 | 0.002763 | 0.008659 | 0.02668 | 0.003957 | 0.002815 | 0.001641 | 0.00972 | 0.003173 | 0.008598 | 0.001792 | 0.007757 | 0.010518 | 0.001205 |
| 19.9305556 | 0.011227 | 0.035739 | 0.001955 | 0.01183 | 0.01113 | 0.006711 | 0.003098 | 0.012964 | 0.013592 | 0.001788 | 0.002338 | 0.002278 | 0.006612 | 0.005046 | 0.013022 | 0.002974 | 0.006664 | 0.011614 | 0.000568 |
| 19.9444444 | 0.016864 | 0.033247 | 0.003192 | 0.009074 | 0.010395 | 0.006082 | 0.002136 | 0.018531 | 0.00928 | 0.000648 | 0.0029 | 0.002093 | 0.005784 | 0.006038 | 0.013288 | 0.003948 | 0.005175 | 0.013205 | 0.000179 |
| 19.9583333 | 0.02221 | 0.024007 | 0.003642 | 0.007169 | 0.009588 | 0.00579 | 0.000697 | 0.021079 | 0.012382 | 0.000918 | 0.004133 | 0.001135 | 0.009978 | 0.006391 | 0.008714 | 0.002843 | 0.005227 | 0.008628 | 0.000185 |
| 19.9722222 | 0.02522 | 0.016306 | 0.003735 | 0.009029 | 0.009548 | 0.007892 | 0.001428 | 0.019102 | 0.015499 | 0.001836 | 0.004694 | 0.000621 | 0.017513 | 0.00683 | 0.004861 | 0.001687 | 0.007239 | 0.007185 | 0.000401 |
| 19.9861111 | 0.024441 | 0.013039 | 0.004001 | 0.013717 | 0.010249 | 0.010713 | 0.003922 | 0.015441 | 0.016024 | 0.002682 | 0.003837 | 0.000697 | 0.022098 | 0.008073 | 0.006905 | 0.002868 | 0.009525 | 0.012613 | 0.001057 |
| 20 | 0.022739 | 0.01399 | 0.003486 | 0.015481 | 0.011058 | 0.011289 | 0.005492 | 0.012641 | 0.01548 | 0.002484 | 0.002127 | 0.000753 | 0.023149 | 0.009866 | 0.014001 | 0.00637 | 0.010132 | 0.017953 | 0.001948 |
| 20.0138889 | 0.023499 | 0.019985 | 0.001896 | 0.011106 | 0.012094 | 0.010005 | 0.004386 | 0.012503 | 0.014553 | 0.002144 | 0.00102 | 0.000739 | 0.021779 | 0.011245 | 0.020823 | 0.010975 | 0.009112 | 0.018204 | 0.00196 |
| 20.0277778 | 0.026998 | 0.033737 | 0.000801 | 0.005137 | 0.013691 | 0.010218 | 0.002027 | 0.017081 | 0.017969 | 0.007233 | 0.003132 | 0.002242 | 0.018427 | 0.012169 | 0.025333 | 0.016034 | 0.008664 | 0.014986 | 0.002305 |
| 20.0416667 | 0.031134 | 0.055511 | 0.001248 | 0.004209 | 0.015552 | 0.013364 | 0.001368 | 0.024422 | 0.032715 | 0.025653 | 0.014894 | 0.009872 | 0.015214 | 0.014531 | 0.032883 | 0.023918 | 0.015514 | 0.016505 | 0.008161 |
| 20.0555556 | 0.036254 | 0.070222 | 0.002822 | 0.010333 | 0.018061 | 0.016413 | 0.003547 | 0.027693 | 0.050906 | 0.051685 | 0.04 | 0.022035 | 0.016206 | 0.01858 | 0.047531 | 0.035322 | 0.033016 | 0.02703 | 0.021378 |
| 20.0694444 | 0.03389 | 0.057422 | 0.004036 | 0.016496 | 0.018047 | 0.014852 | 0.005319 | 0.022421 | 0.053951 | 0.056226 | 0.056006 | 0.022445 | 0.020929 | 0.016687 | 0.060467 | 0.042101 | 0.037376 | 0.033329 | 0.028144 |
| 20.0833333 | 0.017643 | 0.028167 | 0.004121 | 0.0144 | 0.01118 | 0.008996 | 0.003952 | 0.013501 | 0.040968 | 0.034686 | 0.041958 | 0.013735 | 0.025239 | 0.009229 | 0.06133 | 0.037238 | 0.02347 | 0.023795 | 0.021768 |
| 20.0972222 | 0.004178 | 0.013618 | 0.004266 | 0.007183 | 0.00355 | 0.003393 | 0.001583 | 0.008414 | 0.025649 | 0.018894 | 0.023956 | 0.014998 | 0.023012 | 0.009118 | 0.0482 | 0.022456 | 0.02177 | 0.009355 | 0.017298 |
| 20.1111111 | 0.001604 | 0.015564 | 0.003681 | 0.002945 | 0.002198 | 0.001699 | 0.001287 | 0.00559 | 0.012975 | 0.013146 | 0.019638 | 0.017848 | 0.01633 | 0.01633 | 0.026853 | 0.010659 | 0.026795 | 0.003328 | 0.015059 |
| 20.125 | 0.001787 | 0.014011 | 0.00239 | 0.003174 | 0.003987 | 0.003726 | 0.003311 | 0.003155 | 0.004695 | 0.006834 | 0.014798 | 0.012567 | 0.014144 | 0.018929 | 0.011152 | 0.01045 | 0.019454 | 0.002839 | 0.009332 |
| 20.1388889 | 0.002005 | 0.007521 | 0.002253 | 0.003688 | 0.004437 | 0.005033 | 0.004534 | 0.0029 | 0.002359 | 0.004787 | 0.009205 | 0.006311 | 0.010934 | 0.01176 | 0.005567 | 0.010351 | 0.009651 | 0.002318 | 0.005887 |
| 20.1527778 | 0.002389 | 0.004396 | 0.001658 | 0.002347 | 0.002964 | 0.003463 | 0.003012 | 0.002187 | 0.001161 | 0.004739 | 0.005358 | 0.003331 | 0.005404 | 0.003703 | 0.003403 | 0.004999 | 0.007922 | 0.002541 | 0.003855 |
| 20.1666667 | 0.002958 | 0.004673 | 0.000803 | 0.001135 | 0.001776 | 0.00177 | 0.00123 | 0.001458 | 0.001154 | 0.002975 | 0.00185 | 0.001296 | 0.005037 | 0.001921 | 0.002382 | 0.001065 | 0.007989 | 0.002554 | 0.001589 |
| 20.1805556 | 0.003914 | 0.007643 | 0.001483 | 0.001044 | 0.001981 | 0.001316 | 0.000972 | 0.002551 | 0.003097 | 0.001982 | 0.000645 | 0.000983 | 0.005086 | 0.002173 | 0.003223 | 0.000466 | 0.004633 | 0.002147 | 0.00085 |
| 20.1944444 | 0.003761 | 0.008561 | 0.002209 | 0.001056 | 0.001831 | 0.000951 | 0.001086 | 0.002554 | 0.002878 | 0.001758 | 0.000367 | 0.000424 | 0.004526 | 0.002438 | 0.005028 | 0.001558 | 0.001576 | 0.003744 | 0.000855 |
| 20.2083333 | 0.003089 | 0.005302 | 0.00163 | 0.000755 | 0.001021 | 0.000602 | 0.000975 | 0.001235 | 0.001219 | 0.001405 | 0.000553 | 0.000541 | 0.006824 | 0.00344 | 0.004933 | 0.002104 | 0.000845 | 0.005263 | 0.00097 |
| 20.2222222 | 0.002718 | 0.002912 | 0.00079 | 0.000663 | 0.000594 | 0.000614 | 0.001092 | 0.000761 | 0.001138 | 0.001569 | 0.001582 | 0.000845 | 0.008526 | 0.004286 | 0.003131 | 0.001309 | 0.001674 | 0.004769 | 0.001482 |
| 20.2361111 | 0.002605 | 0.002639 | 0.000518 | 0.000848 | 0.000596 | 0.000514 | 0.001315 | 0.001444 | 0.001872 | 0.001879 | 0.002148 | 0.001016 | 0.008667 | 0.004161 | 0.003166 | 0.000576 | 0.002084 | 0.004387 | 0.002171 |
| 20.25 | 0.003392 | 0.002851 | 0.000514 | 0.000708 | 0.000593 | 0.000438 | 0.001653 | 0.002601 | 0.00197 | 0.00149 | 0.001708 | 0.000894 | 0.009024 | 0.003282 | 0.005784 | 0.00095 | 0.001356 | 0.005123 | 0.001914 |
| 20.2638889 | 0.003002 | 0.003716 | 0.000812 | 0.000337 | 0.0007 | 0.000787 | 0.001955 | 0.002634 | 0.002496 | 0.000957 | 0.00109 | 0.000769 | 0.00934 | 0.002549 | 0.009108 | 0.00189 | 0.000584 | 0.006506 | 0.00092 |
| 20.2777778 | 0.001612 | 0.006223 | 0.001194 | 0.000325 | 0.001388 | 0.001241 | 0.001613 | 0.002553 | 0.005028 | 0.00064 | 0.000771 | 0.000573 | 0.009244 | 0.00194 | 0.011401 | 0.002702 | 0.000678 | 0.007958 | 0.000382 |
| 20.2916667 | 0.0018 | 0.008349 | 0.001602 | 0.000888 | 0.002195 | 0.001635 | 0.001211 | 0.004705 | 0.008722 | 0.000467 | 0.000509 | 0.00034 | 0.009019 | 0.001064 | 0.012132 | 0.003063 | 0.001306 | 0.007967 | 0.00031 |
| 20.3055556 | 0.002585 | 0.007688 | 0.00213 | 0.001978 | 0.002442 | 0.002194 | 0.001657 | 0.006776 | 0.010829 | 0.000672 | 0.000486 | 0.000427 | 0.009291 | 0.001006 | 0.012359 | 0.003297 | 0.001535 | 0.005998 | 0.00107 |
| 20.3194444 | 0.002045 | 0.005255 | 0.001892 | 0.002332 | 0.002015 | 0.002072 | 0.002022 | 0.005474 | 0.010414 | 0.003522 | 0.004229 | 0.002198 | 0.013034 | 0.003309 | 0.015273 | 0.004866 | 0.001063 | 0.003521 | 0.005497 |
| 20.3333333 | 0.001568 | 0.003327 | 0.001763 | 0.001813 | 0.002107 | 0.001614 | 0.001736 | 0.003218 | 0.009367 | 0.016346 | 0.020966 | 0.00967 | 0.027904 | 0.007716 | 0.027646 | 0.01064 | 0.000425 | 0.003535 | 0.017277 |
| 20.3472222 | 0.002608 | 0.003447 | 0.002348 | 0.001551 | 0.002487 | 0.001846 | 0.001841 | 0.002719 | 0.0103 | 0.038159 | 0.048289 | 0.019076 | 0.052241 | 0.01396 | 0.050414 | 0.022309 | 0.000186 | 0.009947 | 0.03128 |
| 20.3611111 | 0.002802 | 0.006828 | 0.002813 | 0.002452 | 0.002615 | 0.003245 | 0.003283 | 0.003393 | 0.014663 | 0.041145 | 0.049649 | 0.016558 | 0.058903 | 0.017567 | 0.063163 | 0.031359 | 0.002185 | 0.019197 | 0.030686 |
| 20.375 | 0.002078 | 0.012561 | 0.004282 | 0.004871 | 0.004115 | 0.006015 | 0.005165 | 0.006119 | 0.020334 | 0.018649 | 0.019381 | 0.006071 | 0.038793 | 0.012784 | 0.050163 | 0.026778 | 0.005389 | 0.021252 | 0.014452 |
| 20.3888889 | 0.00325 | 0.016508 | 0.005662 | 0.0061 | 0.005485 | 0.007224 | 0.004712 | 0.008565 | 0.022925 | 0.003071 | 0.001323 | 0.002031 | 0.020422 | 0.005265 | 0.028155 | 0.014305 | 0.005375 | 0.015972 | 0.002661 |
| 20.4027778 | 0.005879 | 0.01738 | 0.005407 | 0.005764 | 0.005782 | 0.006775 | 0.002917 | 0.008613 | 0.022597 | 0.00173 | 0.00092 | 0.002249 | 0.01438 | 0.002589 | 0.016153 | 0.006277 | 0.003983 | 0.01131 | 0.001117 |
| 20.4166667 | 0.008764 | 0.019325 | 0.004814 | 0.006633 | 0.006417 | 0.007327 | 0.002605 | 0.0095 | 0.024549 | 0.003733 | 0.001983 | 0.002962 | 0.013329 | 0.004164 | 0.015528 | 0.004857 | 0.004802 | 0.010773 | 0.002277 |
| 20.4305556 | 0.011261 | 0.023968 | 0.004833 | 0.008597 | 0.007057 | 0.008341 | 0.003104 | 0.012973 | 0.030074 | 0.005832 | 0.003227 | 0.004477 | 0.01619 | 0.006176 | 0.020255 | 0.005708 | 0.006493 | 0.013067 | 0.003262 |
| 20.4444444 | 0.011383 | 0.025725 | 0.005223 | 0.010027 | 0.007437 | 0.00867 | 0.003241 | 0.015872 | 0.033251 | 0.005036 | 0.003664 | 0.005267 | 0.021741 | 0.005972 | 0.023029 | 0.005796 | 0.007832 | 0.015835 | 0.003768 |
| 20.4583333 | 0.009666 | 0.020747 | 0.005584 | 0.01088 | 0.007897 | 0.008302 | 0.00333 | 0.014722 | 0.029805 | 0.002962 | 0.004092 | 0.006114 | 0.028022 | 0.004872 | 0.021941 | 0.005605 | 0.00943 | 0.017603 | 0.004371 |
| 20.4722222 | 0.010189 | 0.014442 | 0.005909 | 0.011808 | 0.008492 | 0.007873 | 0.003797 | 0.01108 | 0.023695 | 0.002408 | 0.005073 | 0.007817 | 0.032362 | 0.005429 | 0.020685 | 0.006248 | 0.011569 | 0.018518 | 0.005405 |
| 20.4861111 | 0.013183 | 0.013114 | 0.006062 | 0.012228 | 0.009278 | 0.007886 | 0.004481 | 0.009018 | 0.019687 | 0.002534 | 0.00598 | 0.00894 | 0.031881 | 0.006972 | 0.020613 | 0.007 | 0.013312 | 0.018467 | 0.006111 |
| 20.5 | 0.014962 | 0.016321 | 0.00626 | 0.012085 | 0.010247 | 0.008389 | 0.005283 | 0.010218 | 0.01651 | 0.003288 | 0.006185 | 0.008704 | 0.025874 | 0.008108 | 0.019417 | 0.00723 | 0.013715 | 0.015991 | 0.006212 |
| 20.5138889 | 0.016189 | 0.021991 | 0.006975 | 0.011743 | 0.010498 | 0.009197 | 0.005829 | 0.012178 | 0.013957 | 0.007521 | 0.006263 | 0.009015 | 0.017168 | 0.009913 | 0.016952 | 0.008219 | 0.013685 | 0.011733 | 0.006743 |
| 20.5277778 | 0.02411 | 0.03123 | 0.00762 | 0.00978 | 0.009669 | 0.010314 | 0.005595 | 0.012642 | 0.018819 | 0.013959 | 0.008242 | 0.010684 | 0.010822 | 0.012611 | 0.017863 | 0.011968 | 0.016356 | 0.00995 | 0.008845 |
| 20.5416667 | 0.047871 | 0.048554 | 0.007643 | 0.009325 | 0.012879 | 0.013338 | 0.008159 | 0.014384 | 0.039049 | 0.021673 | 0.015732 | 0.013106 | 0.012739 | 0.016823 | 0.030777 | 0.022437 | 0.027432 | 0.014943 | 0.015824 |

| | | | | | | | | | | | | | | | | | | | |
|---|---|---|---|---|---|---|---|---|---|---|---|---|---|---|---|---|---|---|---|
| 20.5555556 | 0.078157 | 0.069301 | 0.007758 | 0.017681 | 0.024693 | 0.020869 | 0.018436 | 0.019303 | 0.075849 | 0.034951 | 0.02958 | 0.016601 | 0.028445 | 0.021093 | 0.061771 | 0.042174 | 0.044347 | 0.024207 | 0.024905 |
| 20.5694444 | 0.080915 | 0.071402 | 0.007468 | 0.027057 | 0.032908 | 0.026844 | 0.027416 | 0.022896 | 0.099866 | 0.050124 | 0.041247 | 0.018812 | 0.052626 | 0.018972 | 0.07989 | 0.052399 | 0.049819 | 0.027871 | 0.02488 |
| 20.5833333 | 0.049453 | 0.047665 | 0.004995 | 0.024517 | 0.027155 | 0.023045 | 0.024181 | 0.019202 | 0.076059 | 0.054738 | 0.040387 | 0.015822 | 0.063708 | 0.012759 | 0.053904 | 0.03727 | 0.038435 | 0.020704 | 0.016918 |
| 20.5972222 | 0.019577 | 0.020993 | 0.00402 | 0.015733 | 0.017155 | 0.014792 | 0.016359 | 0.01117 | 0.038737 | 0.044726 | 0.028705 | 0.008757 | 0.045809 | 0.009219 | 0.022069 | 0.015687 | 0.023196 | 0.010163 | 0.010492 |
| 20.6111111 | 0.012152 | 0.010898 | 0.008583 | 0.008983 | 0.009196 | 0.008388 | 0.011594 | 0.009301 | 0.038605 | 0.028802 | 0.016072 | 0.003008 | 0.022904 | 0.006967 | 0.022313 | 0.006322 | 0.011569 | 0.007235 | 0.007319 |
| 20.625 | 0.017005 | 0.01175 | 0.012382 | 0.004072 | 0.003173 | 0.004511 | 0.008336 | 0.011954 | 0.059876 | 0.017436 | 0.009124 | 0.00233 | 0.026614 | 0.004929 | 0.039195 | 0.008635 | 0.004582 | 0.011328 | 0.006321 |
| 20.6388889 | 0.019337 | 0.011094 | 0.009284 | 0.001585 | 0.000946 | 0.004239 | 0.005953 | 0.010233 | 0.065279 | 0.014242 | 0.008269 | 0.00583 | 0.039946 | 0.004391 | 0.047475 | 0.014264 | 0.002613 | 0.012906 | 0.006584 |
| 20.6527778 | 0.014845 | 0.015589 | 0.004231 | 0.002963 | 0.001326 | 0.005936 | 0.00378 | 0.01007 | 0.04504 | 0.015918 | 0.009307 | 0.008951 | 0.037575 | 0.004383 | 0.040372 | 0.01886 | 0.003846 | 0.008618 | 0.005784 |
| 20.6666667 | 0.016696 | 0.031115 | 0.003967 | 0.007642 | 0.002076 | 0.005413 | 0.002703 | 0.01687 | 0.023426 | 0.016447 | 0.007972 | 0.007819 | 0.023912 | 0.003246 | 0.026399 | 0.022772 | 0.00616 | 0.006393 | 0.003756 |
| 20.6805556 | 0.030234 | 0.040905 | 0.006399 | 0.012327 | 0.001808 | 0.00362 | 0.004716 | 0.021712 | 0.02814 | 0.011511 | 0.00474 | 0.003837 | 0.012243 | 0.001985 | 0.013632 | 0.024028 | 0.008069 | 0.011325 | 0.00367 |
| 20.6944444 | 0.031754 | 0.029217 | 0.006354 | 0.01268 | 0.001693 | 0.004471 | 0.006303 | 0.018261 | 0.043119 | 0.005997 | 0.003552 | 0.001252 | 0.015076 | 0.002286 | 0.010253 | 0.021297 | 0.007447 | 0.014561 | 0.005493 |
| 20.7083333 | 0.018112 | 0.014253 | 0.004415 | 0.009251 | 0.003605 | 0.005397 | 0.004399 | 0.010864 | 0.037868 | 0.006508 | 0.004506 | 0.001449 | 0.013387 | 0.002545 | 0.016031 | 0.014668 | 0.00639 | 0.011055 | 0.005801 |
| 20.7222222 | 0.011827 | 0.016486 | 0.004403 | 0.005479 | 0.005412 | 0.003637 | 0.002657 | 0.008708 | 0.020652 | 0.007037 | 0.003955 | 0.003907 | 0.038712 | 0.002765 | 0.014888 | 0.007499 | 0.009951 | 0.010578 | 0.004083 |
| 20.7361111 | 0.011756 | 0.017071 | 0.005619 | 0.002586 | 0.004548 | 0.002001 | 0.003219 | 0.011556 | 0.01909 | 0.005052 | 0.002255 | 0.007396 | 0.026508 | 0.003547 | 0.007012 | 0.003837 | 0.013056 | 0.01695 | 0.002145 |
| 20.75 | 0.009064 | 0.010958 | 0.004194 | 0.001452 | 0.002841 | 0.003223 | 0.003985 | 0.010402 | 0.03713 | 0.007912 | 0.001979 | 0.007959 | 0.017674 | 0.002668 | 0.006118 | 0.002278 | 0.009762 | 0.018566 | 0.000979 |
| 20.7638889 | 0.00622 | 0.014918 | 0.002145 | 0.001646 | 0.00342 | 0.005896 | 0.004667 | 0.008453 | 0.054129 | 0.014058 | 0.002412 | 0.004823 | 0.025987 | 0.001023 | 0.010777 | 0.001735 | 0.006993 | 0.012668 | 0.000871 |
| 20.7777778 | 0.003886 | 0.022687 | 0.00183 | 0.002597 | 0.005416 | 0.006172 | 0.004842 | 0.009164 | 0.050541 | 0.014997 | 0.002 | 0.003512 | 0.034266 | 0.001156 | 0.015715 | 0.003423 | 0.009129 | 0.012159 | 0.001776 |
| 20.7916667 | 0.001749 | 0.022425 | 0.002414 | 0.006559 | 0.006267 | 0.004041 | 0.003443 | 0.006644 | 0.028671 | 0.009165 | 0.001187 | 0.005757 | 0.03359 | 0.002859 | 0.019503 | 0.004793 | 0.00939 | 0.017546 | 0.0026 |
| 20.8055556 | 0.001031 | 0.013919 | 0.004811 | 0.012347 | 0.005448 | 0.003169 | 0.00245 | 0.005491 | 0.010989 | 0.004909 | 0.000987 | 0.00685 | 0.027308 | 0.0048 | 0.018953 | 0.003753 | 0.006209 | 0.016422 | 0.002231 |
| 20.8194444 | 0.001165 | 0.00866 | 0.005192 | 0.013735 | 0.003416 | 0.002969 | 0.002091 | 0.009006 | 0.008659 | 0.005515 | 0.001133 | 0.00507 | 0.015628 | 0.005918 | 0.013532 | 0.002632 | 0.00536 | 0.008579 | 0.002409 |
| 20.8333333 | 0.000912 | 0.014247 | 0.003226 | 0.009082 | 0.001393 | 0.00224 | 0.002209 | 0.009098 | 0.009418 | 0.005667 | 0.000895 | 0.004195 | 0.008975 | 0.006008 | 0.007313 | 0.004766 | 0.006327 | 0.003972 | 0.004413 |
| 20.8472222 | 0.000848 | 0.017641 | 0.004355 | 0.003456 | 0.000685 | 0.003028 | 0.004029 | 0.005469 | 0.005587 | 0.007629 | 0.00046 | 0.004709 | 0.016252 | 0.00536 | 0.003483 | 0.009812 | 0.005404 | 0.007575 | 0.00518 |
| 20.8611111 | 0.002243 | 0.012642 | 0.004875 | 0.002419 | 0.001443 | 0.004231 | 0.004423 | 0.004391 | 0.003564 | 0.012716 | 0.00084 | 0.003658 | 0.026037 | 0.004204 | 0.002861 | 0.014422 | 0.004231 | 0.016818 | 0.003654 |
| 20.875 | 0.003622 | 0.010523 | 0.004076 | 0.006755 | 0.003605 | 0.00354 | 0.002689 | 0.004944 | 0.010646 | 0.014025 | 0.002318 | 0.003326 | 0.024045 | 0.00297 | 0.004539 | 0.016719 | 0.004965 | 0.02522 | 0.003405 |
| 20.8888889 | 0.008446 | 0.010636 | 0.007627 | 0.011675 | 0.006791 | 0.002563 | 0.002244 | 0.008009 | 0.026929 | 0.008799 | 0.003447 | 0.005775 | 0.017579 | 0.002397 | 0.009519 | 0.016784 | 0.006865 | 0.026743 | 0.005448 |
| 20.9027778 | 0.022057 | 0.012933 | 0.012145 | 0.014005 | 0.009454 | 0.004161 | 0.003094 | 0.016674 | 0.04368 | 0.005243 | 0.003944 | 0.007188 | 0.025751 | 0.003351 | 0.018638 | 0.015518 | 0.009363 | 0.022409 | 0.007446 |
| 20.9166667 | 0.036141 | 0.029165 | 0.012766 | 0.014903 | 0.010872 | 0.007473 | 0.004115 | 0.024811 | 0.049963 | 0.010238 | 0.005423 | 0.005056 | 0.044592 | 0.00623 | 0.027351 | 0.014307 | 0.01131 | 0.017672 | 0.008246 |
| 20.9305556 | 0.040363 | 0.05009 | 0.010396 | 0.01486 | 0.011939 | 0.009245 | 0.004399 | 0.026897 | 0.043374 | 0.015496 | 0.008144 | 0.002382 | 0.055998 | 0.01033 | 0.035053 | 0.014992 | 0.011004 | 0.013601 | 0.008448 |
| 20.9444444 | 0.033848 | 0.057502 | 0.006133 | 0.013286 | 0.012749 | 0.007822 | 0.003083 | 0.023503 | 0.0316 | 0.013156 | 0.010521 | 0.002045 | 0.055046 | 0.014024 | 0.042924 | 0.017422 | 0.009207 | 0.008299 | 0.008177 |
| 20.9583333 | 0.021893 | 0.047675 | 0.002937 | 0.010783 | 0.012924 | 0.00421 | 0.00155 | 0.017544 | 0.02284 | 0.006877 | 0.011136 | 0.001892 | 0.043176 | 0.015876 | 0.044602 | 0.017945 | 0.008542 | 0.005708 | 0.005688 |
| 20.9722222 | 0.013043 | 0.028028 | 0.002396 | 0.008434 | 0.012719 | 0.001351 | 0.001431 | 0.012723 | 0.018299 | 0.002323 | 0.010302 | 0.001014 | 0.023488 | 0.015903 | 0.035288 | 0.014687 | 0.010355 | 0.009359 | 0.002373 |
| 20.9861111 | 0.014645 | 0.012551 | 0.003456 | 0.007925 | 0.012319 | 0.002088 | 0.001592 | 0.011908 | 0.016508 | 0.002155 | 0.009662 | 0.001259 | 0.010509 | 0.015292 | 0.020286 | 0.010623 | 0.013028 | 0.011996 | 0.0013 |
| 21 | 0.026208 | 0.009508 | 0.006836 | 0.011141 | 0.012242 | 0.005613 | 0.001199 | 0.015397 | 0.017958 | 0.006752 | 0.009825 | 0.0018 | 0.011803 | 0.014637 | 0.0098 | 0.009225 | 0.013749 | 0.008471 | 0.00237 |
| 21.0138889 | 0.03795 | 0.015407 | 0.010075 | 0.015242 | 0.011931 | 0.007673 | 0.002266 | 0.019324 | 0.021799 | 0.014822 | 0.009864 | 0.001316 | 0.014065 | 0.01333 | 0.009468 | 0.011311 | 0.012428 | 0.004183 | 0.005383 |
| 21.0277778 | 0.04315 | 0.025906 | 0.011509 | 0.015895 | 0.010959 | 0.008159 | 0.005373 | 0.02222 | 0.028774 | 0.023851 | 0.01025 | 0.001218 | 0.011085 | 0.010534 | 0.01993 | 0.016857 | 0.01246 | 0.005274 | 0.01011 |
| 21.0416667 | 0.045437 | 0.043062 | 0.0135 | 0.015326 | 0.012651 | 0.012932 | 0.007195 | 0.029411 | 0.044225 | 0.037474 | 0.015231 | 0.004113 | 0.009505 | 0.007186 | 0.045391 | 0.027911 | 0.018861 | 0.013346 | 0.0195 |
| 21.0555556 | 0.049157 | 0.063238 | 0.01674 | 0.016837 | 0.017958 | 0.020779 | 0.008933 | 0.042442 | 0.065249 | 0.051352 | 0.024264 | 0.008698 | 0.0184 | 0.0058 | 0.073336 | 0.0379 | 0.029724 | 0.026848 | 0.030192 |
| 21.0694444 | 0.046685 | 0.071392 | 0.018551 | 0.017203 | 0.020178 | 0.021626 | 0.012264 | 0.050687 | 0.070785 | 0.044984 | 0.025408 | 0.0099 | 0.03398 | 0.006139 | 0.070124 | 0.032677 | 0.03211 | 0.036916 | 0.027893 |
| 21.0833333 | 0.033157 | 0.059392 | 0.01501 | 0.013501 | 0.015388 | 0.013571 | 0.011065 | 0.043929 | 0.04878 | 0.021404 | 0.015316 | 0.006921 | 0.038376 | 0.004708 | 0.036635 | 0.016196 | 0.021239 | 0.035606 | 0.014695 |
| 21.0972222 | 0.020962 | 0.038278 | 0.008019 | 0.008751 | 0.008985 | 0.005545 | 0.005353 | 0.025409 | 0.020919 | 0.005978 | 0.007874 | 0.003453 | 0.029546 | 0.001893 | 0.011839 | 0.004692 | 0.009556 | 0.027339 | 0.007112 |
| 21.1111111 | 0.015884 | 0.020938 | 0.005132 | 0.005104 | 0.005813 | 0.003133 | 0.00179 | 0.012019 | 0.010573 | 0.003325 | 0.007701 | 0.001371 | 0.020015 | 0.000804 | 0.009208 | 0.001826 | 0.00712 | 0.01856 | 0.006361 |
| 21.125 | 0.010678 | 0.011057 | 0.007085 | 0.002555 | 0.004522 | 0.002965 | 0.002543 | 0.013221 | 0.01299 | 0.002355 | 0.00653 | 0.001086 | 0.015031 | 0.000979 | 0.00964 | 0.001547 | 0.009235 | 0.011814 | 0.004494 |
| 21.1388889 | 0.005302 | 0.007186 | 0.007542 | 0.001808 | 0.002995 | 0.002821 | 0.005883 | 0.015129 | 0.012542 | 0.000828 | 0.003186 | 0.003071 | 0.010864 | 0.001064 | 0.0054 | 0.001262 | 0.007886 | 0.008614 | 0.002003 |
| 21.1527778 | 0.004645 | 0.004518 | 0.005466 | 0.001842 | 0.001847 | 0.00393 | 0.008987 | 0.009155 | 0.007393 | 0.000573 | 0.001199 | 0.005275 | 0.005712 | 0.001014 | 0.001749 | 0.001382 | 0.003689 | 0.007432 | 0.001578 |
| 21.1666667 | 0.005727 | 0.0018 | 0.003083 | 0.001193 | 0.001815 | 0.005257 | 0.009674 | 0.004975 | 0.006695 | 0.000945 | 0.000702 | 0.004758 | 0.002505 | 0.001028 | 0.000508 | 0.001152 | 0.00185 | 0.006018 | 0.002159 |
| 21.1805556 | 0.00388 | 0.00122 | 0.001429 | 0.00065 | 0.001764 | 0.005745 | 0.008176 | 0.006767 | 0.01065 | 0.001823 | 0.001217 | 0.002326 | 0.002923 | 0.001225 | 0.001772 | 0.000538 | 0.003732 | 0.005368 | 0.002564 |
| 21.1944444 | 0.002055 | 0.00159 | 0.001056 | 0.00086 | 0.001206 | 0.005981 | 0.006723 | 0.007348 | 0.011561 | 0.002114 | 0.002445 | 0.00079 | 0.004985 | 0.002015 | 0.004671 | 0.000376 | 0.005293 | 0.006051 | 0.003928 |
| 21.2083333 | 0.003803 | 0.00182 | 0.001512 | 0.000857 | 0.001118 | 0.006062 | 0.006308 | 0.004976 | 0.007675 | 0.001303 | 0.002685 | 0.000568 | 0.004878 | 0.002272 | 0.005626 | 0.000616 | 0.003812 | 0.005066 | 0.005675 |
| 21.2222222 | 0.006653 | 0.003299 | 0.002548 | 0.00079 | 0.001494 | 0.0055 | 0.005764 | 0.005342 | 0.005505 | 0.000684 | 0.001564 | 0.001249 | 0.003895 | 0.001502 | 0.004542 | 0.00073 | 0.002087 | 0.003595 | 0.004607 |
| 21.2361111 | 0.007907 | 0.005302 | 0.002877 | 0.001306 | 0.001836 | 0.004526 | 0.004593 | 0.008278 | 0.007197 | 0.000645 | 0.000798 | 0.002552 | 0.006451 | 0.001596 | 0.005864 | 0.000585 | 0.002052 | 0.005668 | 0.002215 |
| 21.25 | 0.007918 | 0.00451 | 0.001826 | 0.001277 | 0.002191 | 0.003599 | 0.004152 | 0.009138 | 0.006041 | 0.000866 | 0.000997 | 0.002552 | 0.009657 | 0.002825 | 0.008029 | 0.000506 | 0.00226 | 0.008085 | 0.001533 |
| 21.2638889 | 0.007649 | 0.002286 | 0.000693 | 0.000512 | 0.002287 | 0.002873 | 0.004338 | 0.007044 | 0.002723 | 0.001409 | 0.001491 | 0.001824 | 0.008412 | 0.00398 | 0.008184 | 0.000483 | 0.002592 | 0.007738 | 0.00125 |
| 21.2777778 | 0.008101 | 0.001961 | 0.000168 | 0.0007 | 0.002096 | 0.002693 | 0.003241 | 0.003884 | 0.001963 | 0.002035 | 0.001394 | 0.00189 | 0.004729 | 0.003798 | 0.0078 | 0.000577 | 0.002904 | 0.006113 | 0.000992 |
| 21.2916667 | 0.008372 | 0.004023 | 0.0007 | 0.001649 | 0.002318 | 0.003776 | 0.001293 | 0.001535 | 0.002476 | 0.001932 | 0.000752 | 0.001525 | 0.003174 | 0.002148 | 0.007412 | 0.001065 | 0.002291 | 0.004096 | 0.001817 |
| 21.3055556 | 0.006469 | 0.006851 | 0.002212 | 0.001973 | 0.003155 | 0.005596 | 0.001278 | 0.001119 | 0.002862 | 0.001735 | 0.000627 | 0.001548 | 0.002739 | 0.000886 | 0.007468 | 0.001113 | 0.002122 | 0.002809 | 0.002104 |
| 21.3194444 | 0.005577 | 0.007792 | 0.002604 | 0.002816 | 0.003802 | 0.006068 | 0.00414 | 0.001638 | 0.00389 | 0.002237 | 0.001484 | 0.002769 | 0.002103 | 0.001147 | 0.010261 | 0.00093 | 0.00365 | 0.004175 | 0.00203 |
| 21.3333333 | 0.007554 | 0.0077 | 0.001543 | 0.004594 | 0.004164 | 0.005484 | 0.007717 | 0.002261 | 0.005761 | 0.002495 | 0.002718 | 0.003078 | 0.003635 | 0.001831 | 0.015123 | 0.001716 | 0.005351 | 0.008554 | 0.0047 |
| 21.3472222 | 0.007741 | 0.009011 | 0.001203 | 0.005416 | 0.00521 | 0.00716 | 0.010168 | 0.004468 | 0.005952 | 0.002058 | 0.003018 | 0.002313 | 0.0065 | 0.002077 | 0.017438 | 0.003013 | 0.005951 | 0.013335 | 0.009297 |
| 21.3611111 | 0.008155 | 0.013097 | 0.002493 | 0.0054 | 0.006746 | 0.011598 | 0.012041 | 0.008012 | 0.005629 | 0.001948 | 0.002165 | 0.002813 | 0.008608 | 0.002727 | 0.015758 | 0.004075 | 0.006551 | 0.014795 | 0.011448 |
| 21.375 | 0.011604 | 0.014882 | 0.004527 | 0.005013 | 0.007223 | 0.015018 | 0.013533 | 0.009354 | 0.005986 | 0.002824 | 0.001292 | 0.004701 | 0.009494 | 0.003953 | 0.013344 | 0.004246 | 0.007814 | 0.01238 | 0.009613 |
| 21.3888889 | 0.013291 | 0.011454 | 0.004914 | 0.003659 | 0.006137 | 0.015153 | 0.013401 | 0.007668 | 0.005007 | 0.003486 | 0.000865 | 0.005655 | 0.008747 | 0.004316 | 0.011351 | 0.003079 | 0.008614 | 0.00896 | 0.006605 |
| 21.4027778 | 0.013601 | 0.011841 | 0.003412 | 0.002894 | 0.005104 | 0.014192 | 0.012033 | 0.008503 | 0.007235 | 0.00311 | 0.000684 | 0.00561 | 0.007016 | 0.003795 | 0.008439 | 0.001837 | 0.008445 | 0.006459 | 0.005712 |
| 21.4166667 | 0.016171 | 0.021448 | 0.003084 | 0.005419 | 0.005944 | 0.014878 | 0.011943 | 0.016725 | 0.015686 | 0.002806 | 0.001148 | 0.006742 | 0.007118 | 0.004307 | 0.00676 | 0.001908 | 0.008211 | 0.005399 | 0.007657 |
| 21.4305556 | 0.020626 | 0.034702 | 0.005754 | 0.010577 | 0.008165 | 0.016693 | 0.013463 | 0.027187 | 0.023324 | 0.003426 | 0.002944 | 0.008871 | 0.01163 | 0.007016 | 0.010041 | 0.002806 | 0.008684 | 0.006846 | 0.010803 |
| 21.4444444 | 0.02563 | 0.043059 | 0.009456 | 0.013421 | 0.009282 | 0.017748 | 0.014397 | 0.029509 | 0.021761 | 0.004716 | 0.005364 | 0.009807 | 0.017902 | 0.010897 | 0.016839 | 0.003212 | 0.009718 | 0.010574 | 0.013044 |
| 21.4583333 | 0.028618 | 0.040721 | 0.01013 | 0.011278 | 0.008341 | 0.017558 | 0.013721 | 0.020451 | 0.014366 | 0.005761 | 0.006502 | 0.008661 | 0.019776 | 0.013186 | 0.020823 | 0.00272 | 0.01033 | 0.013849 | 0.013531 |
| 21.4722222 | 0.028714 | 0.03041 | 0.006742 | 0.007494 | 0.007252 | 0.016192 | 0.011781 | 0.00922 | 0.013318 | 0.005393 | 0.00564 | 0.0062 | 0.015583 | 0.012411 | 0.017522 | 0.002092 | 0.009632 | 0.014458 | 0.012387 |
| 21.4861111 | 0.02994 | 0.02235 | 0.002872 | 0.005869 | 0.007886 | 0.014376 | 0.010146 | 0.004536 | 0.014048 | 0.004036 | 0.004529 | 0.00416 | 0.010683 | 0.010558 | 0.01031 | 0.002532 | 0.008538 | 0.013077 | 0.010207 |
| 21.5 | 0.034952 | 0.023541 | 0.002525 | 0.007031 | 0.010116 | 0.01395 | 0.011063 | 0.008486 | 0.013765 | 0.003492 | 0.005653 | 0.003965 | 0.012306 | 0.010498 | 0.008312 | 0.00483 | 0.00933 | 0.013801 | 0.008656 |
| 21.5138889 | 0.042522 | 0.034136 | 0.006734 | 0.009297 | 0.012297 | 0.01532 | 0.014212 | 0.022792 | 0.028516 | 0.005742 | 0.008863 | 0.005731 | 0.0219 | 0.013158 | 0.014703 | 0.008411 | 0.012419 | 0.019098 | 0.008827 |
| 21.5277778 | 0.049892 | 0.048782 | 0.013922 | 0.010889 | 0.013642 | 0.016882 | 0.016997 | 0.035612 | 0.053972 | 0.01401 | 0.01242 | 0.008392 | 0.032033 | 0.017199 | 0.022484 | 0.011589 | 0.01512 | 0.025274 | 0.009223 |
| 21.5416667 | 0.062303 | 0.063757 | 0.019598 | 0.016535 | 0.018736 | 0.017677 | 0.018704 | 0.034171 | 0.082503 | 0.031423 | 0.017633 | 0.009708 | 0.037494 | 0.021613 | 0.029243 | 0.015391 | 0.020081 | 0.03212 | 0.009107 |
| 21.5555556 | 0.082485 | 0.078969 | 0.018443 | 0.034226 | 0.031938 | 0.01814 | 0.021443 | 0.026004 | 0.112374 | 0.057762 | 0.026349 | 0.00962 | 0.04738 | 0.026466 | 0.04612 | 0.02038 | 0.032485 | 0.034988 | 0.010979 |
| 21.5694444 | 0.086985 | 0.082707 | 0.012054 | 0.050367 | 0.040225 | 0.014809 | 0.020271 | 0.020379 | 0.117522 | 0.074031 | 0.032713 | 0.011366 | 0.063196 | 0.026959 | 0.064295 | 0.019969 | 0.039173 | 0.026124 | 0.014938 |
| 21.5833333 | 0.062008 | 0.062023 | 0.005697 | 0.043583 | 0.032485 | 0.009003 | 0.012981 | 0.015923 | 0.087572 | 0.059458 | 0.028565 | 0.013028 | 0.063562 | 0.018658 | 0.058725 | 0.012181 | 0.028108 | 0.017053 | 0.017024 |
| 21.5972222 | 0.032934 | 0.034609 | 0.003709 | 0.022356 | 0.020582 | 0.010005 | 0.011985 | 0.010323 | 0.044542 | 0.029997 | 0.016503 | 0.009116 | 0.038708 | 0.009559 | 0.033273 | 0.004686 | 0.012749 | 0.013439 | 0.014973 |
| 21.6111111 | 0.028441 | 0.032839 | 0.009098 | 0.008082 | 0.012993 | 0.014909 | 0.017435 | 0.008515 | 0.022067 | 0.009867 | 0.006054 | 0.00376 | 0.016528 | 0.009389 | 0.014689 | 0.002088 | 0.004879 | 0.007847 | 0.009941 |
| 21.625 | 0.043217 | 0.05137 | 0.015297 | 0.005859 | 0.007901 | 0.01505 | 0.016996 | 0.009459 | 0.037542 | 0.002769 | 0.002134 | 0.002313 | 0.019509 | 0.01381 | 0.01689 | 0.003488 | 0.002809 | 0.00288 | 0.006319 |
| 21.6388889 | 0.047564 | 0.054742 | 0.012689 | 0.007492 | 0.004297 | 0.009543 | 0.010132 | 0.008818 | 0.062426 | 0.004802 | 0.003728 | 0.003702 | 0.029249 | 0.012742 | 0.020283 | 0.006331 | 0.002821 | 0.001598 | 0.006721 |
| 21.6527778 | 0.03423 | 0.03501 | 0.008971 | 0.008931 | 0.003287 | 0.004793 | 0.00616 | 0.015661 | 0.067121 | 0.009926 | 0.007125 | 0.007839 | 0.02816 | 0.007554 | 0.013346 | 0.008374 | 0.003047 | 0.002292 | 0.007138 |
| 21.6666667 | 0.02859 | 0.028501 | 0.015147 | 0.009614 | 0.005935 | 0.006221 | 0.007286 | 0.02903 | 0.052254 | 0.011123 | 0.007377 | 0.01588 | 0.01909 | 0.007299 | 0.012743 | 0.008112 | 0.006209 | 0.003015 | 0.005226 |
| 21.6805556 | 0.038469 | 0.052649 | 0.020382 | 0.007724 | 0.008982 | 0.007811 | 0.0059 | 0.031589 | 0.028567 | 0.007545 | 0.00477 | 0.010777 | 0.011205 | 0.012472 | 0.01796 | 0.006326 | 0.012907 | 0.002339 | 0.005433 |
| 21.6944444 | 0.041258 | 0.070861 | 0.015824 | 0.005598 | 0.009036 | 0.005561 | 0.002531 | 0.018857 | 0.013505 | 0.006064 | 0.004312 | 0.006566 | 0.011082 | 0.015357 | 0.017246 | 0.006158 | 0.018033 | 0.00172 | 0.009594 |
| 21.7083333 | 0.026663 | 0.057149 | 0.00848 | 0.004149 | 0.005662 | 0.00339 | 0.003018 | 0.009193 | 0.018961 | 0.007904 | 0.00561 | 0.00442 | 0.012852 | 0.010868 | 0.023021 | 0.008075 | 0.019195 | 0.003723 | 0.013695 |
| 21.7222222 | 0.018307 | 0.033799 | 0.005098 | 0.002354 | 0.002198 | 0.003327 | 0.005303 | 0.014858 | 0.033865 | 0.006926 | 0.005219 | 0.005846 | 0.013762 | 0.00575 | 0.03883 | 0.008905 | 0.017841 | 0.007695 | 0.014542 |
| 21.7361111 | 0.035529 | 0.034294 | 0.007348 | 0.001217 | 0.001762 | 0.003907 | 0.005191 | 0.022227 | 0.042785 | 0.005766 | 0.003103 | 0.006148 | 0.02063 | 0.00826 | 0.040812 | 0.006523 | 0.013919 | 0.010044 | 0.01072 |
| 21.75 | 0.05539 | 0.05071 | 0.008402 | 0.001001 | 0.003406 | 0.003114 | 0.003272 | 0.019204 | 0.039403 | 0.008813 | 0.001683 | 0.007418 | 0.026496 | 0.011936 | 0.024687 | 0.004357 | 0.007627 | 0.008262 | 0.005794 |
| 21.7638889 | 0.05357 | 0.053076 | 0.006241 | 0.00245 | 0.00521 | 0.001757 | 0.003387 | 0.013191 | 0.024302 | 0.010222 | 0.002552 | 0.012678 | 0.022937 | 0.009614 | 0.014528 | 0.006955 | 0.002558 | 0.004339 | 0.005852 |
| 21.7777778 | 0.031902 | 0.035188 | 0.00653 | 0.004539 | 0.00474 | 0.001837 | 0.006004 | 0.017259 | 0.011088 | 0.006595 | 0.003852 | 0.016775 | 0.016418 | 0.00472 | 0.017092 | 0.011622 | 0.000845 | 0.002038 | 0.008478 |

| | | | | | | | | | | | | | | | | | | |
|---|---|---|---|---|---|---|---|---|---|---|---|---|---|---|---|---|---|---|
| 21.7916667 | 0.012815 | 0.017478 | 0.006981 | 0.004527 | 0.00239 | 0.00446 | 0.007872 | 0.026854 | 0.014038 | 0.003937 | 0.003226 | 0.015171 | 0.012049 | 0.002153 | 0.014664 | 0.012978 | 0.001619 | 0.002084 | 0.00698 |
| 21.8055556 | 0.008475 | 0.014746 | 0.007194 | 0.005714 | 0.001385 | 0.008944 | 0.007494 | 0.024059 | 0.031588 | 0.005436 | 0.002999 | 0.010593 | 0.017183 | 0.001418 | 0.012597 | 0.010751 | 0.004673 | 0.004395 | 0.005086 |
| 21.8194444 | 0.011442 | 0.016843 | 0.009471 | 0.012457 | 0.002239 | 0.011268 | 0.004947 | 0.014349 | 0.051428 | 0.005953 | 0.005795 | 0.010986 | 0.033617 | 0.001269 | 0.024832 | 0.007385 | 0.009107 | 0.008418 | 0.009933 |
| 21.8333333 | 0.027527 | 0.026321 | 0.010429 | 0.019971 | 0.003604 | 0.008438 | 0.002951 | 0.017054 | 0.064804 | 0.003931 | 0.008387 | 0.014192 | 0.046665 | 0.003327 | 0.035988 | 0.004088 | 0.012709 | 0.011066 | 0.015109 |
| 21.8472222 | 0.052353 | 0.046189 | 0.007334 | 0.021153 | 0.004245 | 0.003797 | 0.003817 | 0.02463 | 0.068319 | 0.002542 | 0.007854 | 0.010575 | 0.043304 | 0.00706 | 0.033021 | 0.003703 | 0.014543 | 0.010267 | 0.013568 |
| 21.8611111 | 0.065894 | 0.055925 | 0.003702 | 0.017007 | 0.00331 | 0.003186 | 0.00585 | 0.020484 | 0.057183 | 0.003526 | 0.005861 | 0.00638 | 0.026809 | 0.010218 | 0.023357 | 0.008361 | 0.015457 | 0.007049 | 0.008752 |
| 21.875 | 0.058816 | 0.046467 | 0.00377 | 0.011531 | 0.001551 | 0.007068 | 0.008474 | 0.011462 | 0.033211 | 0.007834 | 0.005357 | 0.010919 | 0.017639 | 0.012171 | 0.017278 | 0.014106 | 0.01606 | 0.006534 | 0.005166 |
| 21.8888889 | 0.04115 | 0.02973 | 0.004206 | 0.005902 | 0.000825 | 0.011725 | 0.010478 | 0.006978 | 0.013271 | 0.014803 | 0.007425 | 0.01553 | 0.020257 | 0.013716 | 0.01872 | 0.016728 | 0.016848 | 0.010247 | 0.005299 |
| 21.9027778 | 0.030674 | 0.017667 | 0.005815 | 0.003123 | 0.002304 | 0.014433 | 0.010039 | 0.008924 | 0.006886 | 0.02068 | 0.010755 | 0.011284 | 0.016946 | 0.015291 | 0.025546 | 0.016011 | 0.018232 | 0.009982 | 0.010078 |
| 21.9166667 | 0.033718 | 0.013865 | 0.011961 | 0.006571 | 0.006762 | 0.014416 | 0.008587 | 0.021787 | 0.007502 | 0.022973 | 0.013777 | 0.007149 | 0.015048 | 0.016346 | 0.03639 | 0.013216 | 0.01922 | 0.005558 | 0.017241 |
| 21.9305556 | 0.046292 | 0.016754 | 0.017745 | 0.013329 | 0.013839 | 0.012924 | 0.007352 | 0.036192 | 0.015825 | 0.027005 | 0.01525 | 0.012015 | 0.028963 | 0.016482 | 0.043914 | 0.009557 | 0.019106 | 0.004484 | 0.021153 |
| 21.9444444 | 0.061836 | 0.02601 | 0.019914 | 0.017195 | 0.020346 | 0.011873 | 0.006149 | 0.040957 | 0.030371 | 0.032908 | 0.013869 | 0.016568 | 0.043716 | 0.015781 | 0.040588 | 0.006977 | 0.017713 | 0.004589 | 0.018731 |
| 21.9583333 | 0.072349 | 0.039707 | 0.019386 | 0.015318 | 0.022447 | 0.011348 | 0.005044 | 0.03665 | 0.038629 | 0.030605 | 0.010197 | 0.013323 | 0.042855 | 0.014553 | 0.03331 | 0.006007 | 0.014359 | 0.002494 | 0.013429 |
| 21.9722222 | 0.070435 | 0.050449 | 0.016659 | 0.011082 | 0.019625 | 0.010168 | 0.004586 | 0.025331 | 0.033863 | 0.020271 | 0.006402 | 0.007245 | 0.027668 | 0.013215 | 0.026084 | 0.004203 | 0.009831 | 0.001069 | 0.009005 |
| 21.9861111 | 0.058935 | 0.054703 | 0.011763 | 0.0088 | 0.015571 | 0.008029 | 0.005011 | 0.01211 | 0.020616 | 0.010153 | 0.003276 | 0.003126 | 0.012514 | 0.012518 | 0.015989 | 0.002057 | 0.006266 | 0.001532 | 0.006767 |
| 22 | 0.046425 | 0.053194 | 0.00625 | 0.007821 | 0.012926 | 0.005372 | 0.005139 | 0.004409 | 0.009435 | 0.005457 | 0.001659 | 0.001915 | 0.007707 | 0.013139 | 0.006133 | 0.002008 | 0.004398 | 0.002353 | 0.007764 |
| 22.0138889 | 0.0374 | 0.044938 | 0.003231 | 0.006804 | 0.010814 | 0.00309 | 0.00371 | 0.007343 | 0.006411 | 0.008282 | 0.002423 | 0.002923 | 0.006745 | 0.014154 | 0.002688 | 0.002625 | 0.003552 | 0.003424 | 0.011714 |
| 22.0277778 | 0.037476 | 0.036471 | 0.003515 | 0.006969 | 0.009391 | 0.003715 | 0.001811 | 0.021847 | 0.006898 | 0.016364 | 0.004185 | 0.005683 | 0.008131 | 0.014236 | 0.009111 | 0.002393 | 0.004086 | 0.004357 | 0.016876 |
| 22.0416667 | 0.046877 | 0.041731 | 0.00877 | 0.00961 | 0.010266 | 0.008985 | 0.002575 | 0.044934 | 0.012805 | 0.026784 | 0.005216 | 0.010868 | 0.017849 | 0.01431 | 0.025669 | 0.004233 | 0.007595 | 0.003589 | 0.022122 |
| 22.0555556 | 0.049631 | 0.058153 | 0.019599 | 0.013444 | 0.012684 | 0.015659 | 0.006958 | 0.069763 | 0.034846 | 0.038008 | 0.004649 | 0.019018 | 0.032954 | 0.014972 | 0.046902 | 0.009534 | 0.01223 | 0.001714 | 0.026267 |
| 22.0694444 | 0.034225 | 0.068262 | 0.028582 | 0.014843 | 0.012783 | 0.017961 | 0.011279 | 0.082292 | 0.053268 | 0.052636 | 0.002967 | 0.026203 | 0.043857 | 0.01323 | 0.060342 | 0.012613 | 0.011827 | 0.000758 | 0.026547 |
| 22.0833333 | 0.015885 | 0.064044 | 0.02891 | 0.012736 | 0.008773 | 0.014042 | 0.012064 | 0.068585 | 0.040692 | 0.064277 | 0.002849 | 0.02577 | 0.039055 | 0.008208 | 0.052366 | 0.009787 | 0.006673 | 0.001439 | 0.022718 |
| 22.0972222 | 0.011061 | 0.047375 | 0.022289 | 0.009635 | 0.003762 | 0.007889 | 0.009467 | 0.038395 | 0.015883 | 0.056109 | 0.004377 | 0.018173 | 0.022398 | 0.003846 | 0.027631 | 0.004902 | 0.003843 | 0.002668 | 0.018061 |
| 22.1111111 | 0.010954 | 0.026923 | 0.014419 | 0.006591 | 0.001064 | 0.003981 | 0.005269 | 0.014726 | 0.004211 | 0.031772 | 0.004106 | 0.0101 | 0.014188 | 0.002109 | 0.016597 | 0.001834 | 0.003624 | 0.002876 | 0.013616 |
| 22.125 | 0.00981 | 0.012249 | 0.008344 | 0.003302 | 0.000543 | 0.00215 | 0.002864 | 0.006476 | 0.001517 | 0.011703 | 0.002068 | 0.004613 | 0.017545 | 0.001612 | 0.030886 | 0.001296 | 0.002351 | 0.001921 | 0.008681 |
| 22.1388889 | 0.009633 | 0.01006 | 0.004186 | 0.001674 | 0.001285 | 0.002066 | 0.003915 | 0.013189 | 0.004631 | 0.007326 | 0.000786 | 0.001701 | 0.019797 | 0.001036 | 0.045521 | 0.002437 | 0.00081 | 0.001544 | 0.004797 |
| 22.1527778 | 0.006521 | 0.020337 | 0.001828 | 0.002921 | 0.003066 | 0.004131 | 0.005483 | 0.028713 | 0.015492 | 0.014777 | 0.000619 | 0.001499 | 0.020105 | 0.000623 | 0.046835 | 0.003548 | 0.000731 | 0.004015 | 0.004816 |
| 22.1666667 | 0.005161 | 0.028565 | 0.002414 | 0.004682 | 0.004295 | 0.005016 | 0.00584 | 0.042078 | 0.027164 | 0.024526 | 0.000773 | 0.00294 | 0.021179 | 0.000774 | 0.041283 | 0.003891 | 0.001739 | 0.009021 | 0.007363 |
| 22.1805556 | 0.010479 | 0.030639 | 0.007205 | 0.004733 | 0.004349 | 0.003769 | 0.005067 | 0.049099 | 0.034614 | 0.030699 | 0.001027 | 0.004053 | 0.018316 | 0.000677 | 0.030856 | 0.00322 | 0.003799 | 0.01371 | 0.008153 |
| 22.1944444 | 0.02068 | 0.035827 | 0.01294 | 0.003592 | 0.004173 | 0.003106 | 0.003139 | 0.050799 | 0.038299 | 0.032505 | 0.001148 | 0.003553 | 0.010046 | 0.000243 | 0.0179 | 0.001675 | 0.007392 | 0.016609 | 0.00776 |
| 22.2083333 | 0.029696 | 0.042094 | 0.014855 | 0.002979 | 0.003496 | 0.003277 | 0.001553 | 0.04545 | 0.037186 | 0.031332 | 0.001164 | 0.002066 | 0.004152 | 0.000446 | 0.008255 | 0.000831 | 0.010404 | 0.018069 | 0.009215 |
| 22.2222222 | 0.030755 | 0.042802 | 0.012604 | 0.003472 | 0.002235 | 0.003347 | 0.0015 | 0.032705 | 0.030058 | 0.026992 | 0.001765 | 0.001255 | 0.005262 | 0.001704 | 0.003495 | 0.001563 | 0.010995 | 0.018304 | 0.011741 |
| 22.2361111 | 0.027972 | 0.041754 | 0.010537 | 0.004519 | 0.001906 | 0.004003 | 0.001911 | 0.01969 | 0.020115 | 0.020971 | 0.002096 | 0.002486 | 0.006038 | 0.003027 | 0.001611 | 0.002806 | 0.010701 | 0.016811 | 0.011565 |
| 22.25 | 0.02666 | 0.042239 | 0.011787 | 0.005609 | 0.003111 | 0.005901 | 0.001686 | 0.013594 | 0.012225 | 0.015901 | 0.001455 | 0.005743 | 0.004043 | 0.0033 | 0.001567 | 0.002912 | 0.010452 | 0.012684 | 0.007157 |
| 22.2638889 | 0.021724 | 0.033498 | 0.012808 | 0.005895 | 0.003763 | 0.00684 | 0.001099 | 0.012393 | 0.007389 | 0.010726 | 0.000691 | 0.00817 | 0.004717 | 0.00253 | 0.004779 | 0.001758 | 0.008732 | 0.006962 | 0.003059 |
| 22.2777778 | 0.012685 | 0.021341 | 0.009121 | 0.00511 | 0.002546 | 0.004777 | 0.000538 | 0.009501 | 0.006018 | 0.005916 | 0.000509 | 0.007193 | 0.007063 | 0.001373 | 0.010445 | 0.000785 | 0.00496 | 0.002675 | 0.002381 |
| 22.2916667 | 0.009094 | 0.026783 | 0.006127 | 0.004017 | 0.001358 | 0.003247 | 0.000256 | 0.006341 | 0.007306 | 0.005271 | 0.00062 | 0.004198 | 0.006227 | 0.000946 | 0.011955 | 0.001693 | 0.001699 | 0.000971 | 0.002906 |
| 22.3055556 | 0.009095 | 0.037114 | 0.009296 | 0.002489 | 0.00123 | 0.004813 | 0.00023 | 0.010407 | 0.008941 | 0.006288 | 0.000889 | 0.002122 | 0.003774 | 0.001496 | 0.008413 | 0.004845 | 0.001252 | 0.000727 | 0.005008 |
| 22.3194444 | 0.007144 | 0.028627 | 0.011398 | 0.001248 | 0.002015 | 0.005288 | 0.000364 | 0.01908 | 0.012149 | 0.004743 | 0.001374 | 0.002002 | 0.00311 | 0.001758 | 0.005965 | 0.007012 | 0.00361 | 0.001493 | 0.007751 |
| 22.3333333 | 0.00823 | 0.013675 | 0.00754 | 0.001068 | 0.003719 | 0.003111 | 0.000615 | 0.022422 | 0.017884 | 0.003826 | 0.002247 | 0.002662 | 0.003946 | 0.00144 | 0.006348 | 0.005415 | 0.008328 | 0.004281 | 0.006268 |
| 22.3472222 | 0.012353 | 0.011964 | 0.003402 | 0.001403 | 0.004493 | 0.00196 | 0.000877 | 0.01559 | 0.020364 | 0.00501 | 0.004327 | 0.002742 | 0.007858 | 0.001644 | 0.011102 | 0.003883 | 0.013196 | 0.008008 | 0.002897 |
| 22.3611111 | 0.0171 | 0.015668 | 0.003451 | 0.002489 | 0.003944 | 0.003177 | 0.000868 | 0.007269 | 0.015342 | 0.004424 | 0.005336 | 0.002436 | 0.012672 | 0.002245 | 0.016743 | 0.005272 | 0.015228 | 0.009287 | 0.002547 |
| 22.375 | 0.023419 | 0.015065 | 0.004986 | 0.003359 | 0.003917 | 0.004808 | 0.000412 | 0.00853 | 0.015701 | 0.002742 | 0.003758 | 0.00323 | 0.012006 | 0.002691 | 0.015068 | 0.005662 | 0.013816 | 0.007754 | 0.003905 |
| 22.3888889 | 0.028265 | 0.013099 | 0.00558 | 0.003656 | 0.004566 | 0.005728 | 0.000476 | 0.013737 | 0.029875 | 0.004566 | 0.002648 | 0.006487 | 0.006632 | 0.003787 | 0.008538 | 0.00362 | 0.011192 | 0.005687 | 0.00763 |
| 22.4027778 | 0.029368 | 0.014933 | 0.006525 | 0.004654 | 0.004924 | 0.006338 | 0.001577 | 0.014887 | 0.037964 | 0.01079 | 0.003192 | 0.011507 | 0.002961 | 0.005526 | 0.004552 | 0.002468 | 0.009859 | 0.00411 | 0.013664 |
| 22.4166667 | 0.026492 | 0.020153 | 0.007522 | 0.006762 | 0.003707 | 0.006228 | 0.002626 | 0.014174 | 0.025003 | 0.016832 | 0.002915 | 0.015609 | 0.00264 | 0.006312 | 0.005015 | 0.003151 | 0.009435 | 0.002445 | 0.016087 |
| 22.4305556 | 0.018705 | 0.021161 | 0.006853 | 0.009168 | 0.001897 | 0.005125 | 0.002835 | 0.016053 | 0.011725 | 0.016393 | 0.001394 | 0.016206 | 0.00522 | 0.005309 | 0.008251 | 0.003559 | 0.00789 | 0.001319 | 0.012568 |
| 22.4444444 | 0.011008 | 0.019896 | 0.00565 | 0.010201 | 0.001861 | 0.004986 | 0.002847 | 0.019122 | 0.013042 | 0.009814 | 0.0007 | 0.013646 | 0.011473 | 0.003535 | 0.010389 | 0.002466 | 0.004546 | 0.002042 | 0.006953 |
| 22.4583333 | 0.009565 | 0.024717 | 0.006759 | 0.009593 | 0.004017 | 0.006933 | 0.0038 | 0.020332 | 0.013658 | 0.004902 | 0.001036 | 0.01132 | 0.014794 | 0.002358 | 0.007942 | 0.00129 | 0.001373 | 0.00268 | 0.002685 |
| 22.4722222 | 0.011432 | 0.030296 | 0.009439 | 0.008437 | 0.006136 | 0.008555 | 0.005058 | 0.021946 | 0.017369 | 0.006051 | 0.001493 | 0.010483 | 0.01005 | 0.002187 | 0.005014 | 0.001594 | 0.001392 | 0.001751 | 0.001798 |
| 22.4861111 | 0.009249 | 0.024303 | 0.009576 | 0.007814 | 0.007649 | 0.007631 | 0.004845 | 0.023134 | 0.032356 | 0.010863 | 0.001842 | 0.009014 | 0.005294 | 0.002884 | 0.004642 | 0.002248 | 0.005691 | 0.002 | 0.006271 |
| 22.5 | 0.004182 | 0.010749 | 0.006339 | 0.007556 | 0.00983 | 0.006031 | 0.003594 | 0.016841 | 0.034613 | 0.015979 | 0.002428 | 0.005877 | 0.00587 | 0.004108 | 0.004712 | 0.001924 | 0.010927 | 0.004567 | 0.01297 |
| 22.5138889 | 0.007346 | 0.011291 | 0.004915 | 0.005498 | 0.009582 | 0.007042 | 0.002943 | 0.007943 | 0.019678 | 0.01844 | 0.003816 | 0.0033 | 0.006276 | 0.005972 | 0.008386 | 0.001145 | 0.012 | 0.006616 | 0.014314 |
| 22.5277778 | 0.021001 | 0.035576 | 0.009865 | 0.002729 | 0.005602 | 0.010182 | 0.003564 | 0.005261 | 0.017129 | 0.017979 | 0.007779 | 0.004144 | 0.009537 | 0.010505 | 0.016302 | 0.001977 | 0.009755 | 0.006772 | 0.010109 |
| 22.5416667 | 0.033292 | 0.070834 | 0.022042 | 0.004604 | 0.007161 | 0.013926 | 0.008677 | 0.010624 | 0.021926 | 0.020691 | 0.017211 | 0.011118 | 0.023283 | 0.022098 | 0.027397 | 0.005878 | 0.011216 | 0.007414 | 0.008567 |
| 22.5555556 | 0.042564 | 0.099577 | 0.038076 | 0.016348 | 0.019131 | 0.020358 | 0.020894 | 0.028233 | 0.033316 | 0.032475 | 0.02723 | 0.021726 | 0.043683 | 0.04151 | 0.041313 | 0.00815 | 0.019519 | 0.010449 | 0.01308 |
| 22.5694444 | 0.049021 | 0.099589 | 0.046187 | 0.033017 | 0.029832 | 0.025589 | 0.03006 | 0.042706 | 0.068739 | 0.045821 | 0.026631 | 0.026258 | 0.058738 | 0.051885 | 0.047844 | 0.005324 | 0.026301 | 0.014045 | 0.019599 |
| 22.5833333 | 0.04051 | 0.064004 | 0.037167 | 0.037343 | 0.028175 | 0.02099 | 0.024573 | 0.032058 | 0.083618 | 0.0442 | 0.015153 | 0.015609 | 0.05931 | 0.039057 | 0.044632 | 0.002912 | 0.022335 | 0.016734 | 0.022873 |
| 22.5972222 | 0.020702 | 0.029038 | 0.01986 | 0.023983 | 0.019922 | 0.010968 | 0.012623 | 0.015026 | 0.056298 | 0.026055 | 0.008053 | 0.011696 | 0.046157 | 0.018186 | 0.041333 | 0.005518 | 0.012408 | 0.01728 | 0.01794 |
| 22.6111111 | 0.010647 | 0.02978 | 0.013779 | 0.011186 | 0.014271 | 0.008674 | 0.010397 | 0.020382 | 0.026776 | 0.014017 | 0.012551 | 0.010881 | 0.025403 | 0.009669 | 0.037961 | 0.013032 | 0.009497 | 0.012987 | 0.012947 |
| 22.625 | 0.017643 | 0.050125 | 0.021874 | 0.011936 | 0.010921 | 0.015868 | 0.017713 | 0.037185 | 0.022227 | 0.023239 | 0.013198 | 0.021156 | 0.012553 | 0.014176 | 0.030367 | 0.022625 | 0.015783 | 0.006517 | 0.01679 |
| 22.6388889 | 0.026049 | 0.053536 | 0.026672 | 0.014521 | 0.007402 | 0.019483 | 0.021084 | 0.046876 | 0.044434 | 0.033709 | 0.010528 | 0.023029 | 0.011346 | 0.017513 | 0.022549 | 0.028049 | 0.017269 | 0.00343 | 0.015782 |
| 22.6527778 | 0.023037 | 0.033494 | 0.01887 | 0.011518 | 0.004014 | 0.013387 | 0.014305 | 0.038501 | 0.060972 | 0.027898 | 0.01722 | 0.017008 | 0.009288 | 0.012743 | 0.023847 | 0.022322 | 0.012302 | 0.004386 | 0.009397 |
| 22.6666667 | 0.014847 | 0.021822 | 0.010467 | 0.012825 | 0.00287 | 0.008852 | 0.008209 | 0.029847 | 0.044633 | 0.022796 | 0.019599 | 0.024209 | 0.008475 | 0.006239 | 0.030727 | 0.012623 | 0.016374 | 0.006628 | 0.012476 |
| 22.6805556 | 0.014611 | 0.036112 | 0.015175 | 0.017086 | 0.004698 | 0.009735 | 0.01089 | 0.051464 | 0.031656 | 0.025182 | 0.011208 | 0.029215 | 0.011917 | 0.003605 | 0.025621 | 0.013781 | 0.023604 | 0.005861 | 0.018948 |
| 22.6944444 | 0.019467 | 0.045521 | 0.022845 | 0.013394 | 0.006529 | 0.007596 | 0.011649 | 0.062508 | 0.038708 | 0.019911 | 0.006064 | 0.017809 | 0.019914 | 0.00353 | 0.019339 | 0.022806 | 0.018077 | 0.004077 | 0.016269 |
| 22.7083333 | 0.015321 | 0.029976 | 0.01845 | 0.008005 | 0.00629 | 0.006189 | 0.006867 | 0.042387 | 0.031722 | 0.009724 | 0.007995 | 0.008383 | 0.032593 | 0.002604 | 0.034656 | 0.030747 | 0.008388 | 0.006203 | 0.011329 |
| 22.7222222 | 0.007788 | 0.017163 | 0.011208 | 0.012708 | 0.005248 | 0.012688 | 0.006421 | 0.040567 | 0.025239 | 0.010628 | 0.009791 | 0.011293 | 0.040399 | 0.002282 | 0.058923 | 0.035853 | 0.008803 | 0.007562 | 0.016869 |
| 22.7361111 | 0.008501 | 0.0307 | 0.012616 | 0.021974 | 0.005272 | 0.019411 | 0.012279 | 0.05978 | 0.035569 | 0.024993 | 0.007123 | 0.02 | 0.037683 | 0.004629 | 0.077668 | 0.036518 | 0.014971 | 0.005409 | 0.025176 |
| 22.75 | 0.011284 | 0.050669 | 0.015962 | 0.022454 | 0.00675 | 0.017057 | 0.017623 | 0.060671 | 0.033454 | 0.029178 | 0.0051 | 0.022464 | 0.024318 | 0.006391 | 0.080961 | 0.03059 | 0.013533 | 0.00617 | 0.020507 |
| 22.7638889 | 0.010081 | 0.051051 | 0.012574 | 0.013078 | 0.008714 | 0.009138 | 0.017569 | 0.037403 | 0.02031 | 0.021078 | 0.00764 | 0.017686 | 0.012745 | 0.004978 | 0.055236 | 0.019435 | 0.010339 | 0.01383 | 0.012714 |
| 22.7777778 | 0.007214 | 0.031765 | 0.005848 | 0.006992 | 0.009645 | 0.004626 | 0.01241 | 0.024478 | 0.022076 | 0.023563 | 0.008547 | 0.020593 | 0.019112 | 0.002533 | 0.024851 | 0.00922 | 0.016475 | 0.022557 | 0.016204 |
| 22.7916667 | 0.00557 | 0.018319 | 0.002987 | 0.00956 | 0.007976 | 0.003766 | 0.006123 | 0.034373 | 0.031551 | 0.02204 | 0.006121 | 0.026635 | 0.031433 | 0.002027 | 0.015324 | 0.005002 | 0.017203 | 0.023997 | 0.015359 |
| 22.8055556 | 0.005973 | 0.019528 | 0.005179 | 0.011151 | 0.004252 | 0.004607 | 0.003313 | 0.031792 | 0.027799 | 0.013196 | 0.007315 | 0.02035 | 0.030051 | 0.003941 | 0.010409 | 0.005568 | 0.013087 | 0.017524 | 0.009048 |
| 22.8194444 | 0.00699 | 0.018336 | 0.014067 | 0.007911 | 0.001702 | 0.009095 | 0.002921 | 0.023143 | 0.028961 | 0.023535 | 0.012665 | 0.010577 | 0.017898 | 0.006575 | 0.005889 | 0.006455 | 0.02082 | 0.00871 | 0.015113 |
| 22.8333333 | 0.010084 | 0.025734 | 0.025968 | 0.007235 | 0.001368 | 0.013762 | 0.002092 | 0.045288 | 0.05665 | 0.046048 | 0.014865 | 0.012006 | 0.01398 | 0.009376 | 0.016226 | 0.006036 | 0.03046 | 0.004404 | 0.027767 |
| 22.8472222 | 0.01743 | 0.052413 | 0.032154 | 0.01314 | 0.001351 | 0.014497 | 0.002319 | 0.06869 | 0.08249 | 0.056791 | 0.010782 | 0.020307 | 0.025486 | 0.013087 | 0.040419 | 0.005768 | 0.029844 | 0.006676 | 0.029359 |
| 22.8611111 | 0.02485 | 0.077439 | 0.033464 | 0.018925 | 0.001587 | 0.012603 | 0.004648 | 0.057329 | 0.08058 | 0.046635 | 0.004814 | 0.021164 | 0.037064 | 0.016671 | 0.059607 | 0.004727 | 0.022545 | 0.010401 | 0.018121 |
| 22.875 | 0.026103 | 0.081917 | 0.032178 | 0.019101 | 0.002819 | 0.010482 | 0.00732 | 0.031654 | 0.063418 | 0.023325 | 0.002646 | 0.012894 | 0.037734 | 0.017201 | 0.05576 | 0.003055 | 0.015076 | 0.01166 | 0.011193 |
| 22.8888889 | 0.020184 | 0.068018 | 0.027121 | 0.015887 | 0.00503 | 0.010161 | 0.008974 | 0.019851 | 0.048643 | 0.006113 | 0.005096 | 0.006463 | 0.031514 | 0.014477 | 0.036137 | 0.004032 | 0.010881 | 0.010473 | 0.011901 |
| 22.9027778 | 0.012059 | 0.048635 | 0.020692 | 0.013984 | 0.008528 | 0.012061 | 0.008913 | 0.025118 | 0.038591 | 0.00391 | 0.009053 | 0.006001 | 0.024023 | 0.012589 | 0.017428 | 0.006032 | 0.010477 | 0.008747 | 0.009379 |
| 22.9166667 | 0.007468 | 0.028985 | 0.018469 | 0.01713 | 0.012375 | 0.014714 | 0.007903 | 0.038233 | 0.029648 | 0.012157 | 0.012605 | 0.006127 | 0.016472 | 0.013948 | 0.011665 | 0.005567 | 0.0122 | 0.008218 | 0.010096 |
| 22.9305556 | 0.006977 | 0.013856 | 0.021701 | 0.024151 | 0.015076 | 0.017806 | 0.00803 | 0.050948 | 0.023172 | 0.023859 | 0.014584 | 0.009212 | 0.01122 | 0.016167 | 0.022651 | 0.003354 | 0.0124 | 0.007137 | 0.023204 |
| 22.9444444 | 0.008685 | 0.009767 | 0.026173 | 0.029313 | 0.016443 | 0.020767 | 0.008883 | 0.054031 | 0.017888 | 0.033887 | 0.013986 | 0.017093 | 0.010445 | 0.016312 | 0.034332 | 0.001517 | 0.008899 | 0.004128 | 0.039604 |
| 22.9583333 | 0.01061 | 0.01012 | 0.026449 | 0.027553 | 0.016344 | 0.020017 | 0.008064 | 0.040816 | 0.011317 | 0.041415 | 0.011577 | 0.020284 | 0.01151 | 0.014366 | 0.029855 | 0.001745 | 0.004175 | 0.001854 | 0.045391 |
| 22.9722222 | 0.01009 | 0.009121 | 0.0202 | 0.019777 | 0.013893 | 0.014525 | 0.005 | 0.020904 | 0.009681 | 0.040785 | 0.008637 | 0.013834 | 0.011996 | 0.011629 | 0.015949 | 0.005044 | 0.002084 | 0.001878 | 0.037562 |
| 22.9861111 | 0.00651 | 0.008714 | 0.011214 | 0.010439 | 0.008947 | 0.007799 | 0.002095 | 0.012753 | 0.012789 | 0.025365 | 0.006102 | 0.00647 | 0.010197 | 0.008551 | 0.012885 | 0.010244 | 0.001947 | 0.001745 | 0.022572 |
| 23 | 0.004541 | 0.006735 | 0.004377 | 0.00472 | 0.00391 | 0.003516 | 0.001193 | 0.01502 | 0.011313 | 0.01248 | 0.004533 | 0.005794 | 0.007265 | 0.005093 | 0.027207 | 0.016806 | 0.001332 | 0.001332 | 0.010781 |
| 23.0138889 | 0.005465 | 0.008109 | 0.001829 | 0.005505 | 0.001176 | 0.004039 | 0.001021 | 0.012511 | 0.010872 | 0.016971 | 0.003611 | 0.005446 | 0.010069 | 0.002282 | 0.043816 | 0.023099 | 0.001354 | 0.0023 | 0.008699 |

| | | | | | | | | | | | | | | | | | | | |
|---|---|---|---|---|---|---|---|---|---|---|---|---|---|---|---|---|---|---|---|
| 23.0277778 | 0.007326 | 0.024075 | 0.002855 | 0.007009 | 0.000425 | 0.009255 | 0.001114 | 0.012482 | 0.018999 | 0.017814 | 0.003646 | 0.005308 | 0.017767 | 0.002388 | 0.047446 | 0.024051 | 0.004302 | 0.00356 | 0.008249 |
| 23.0416667 | 0.014159 | 0.050439 | 0.010044 | 0.00633 | 0.000888 | 0.016464 | 0.004871 | 0.028246 | 0.03049 | 0.015694 | 0.006657 | 0.014344 | 0.016767 | 0.007031 | 0.040589 | 0.020158 | 0.009542 | 0.004666 | 0.008331 |
| 23.0555556 | 0.022263 | 0.072961 | 0.025238 | 0.009862 | 0.002242 | 0.022552 | 0.014493 | 0.048366 | 0.040387 | 0.041314 | 0.013006 | 0.027076 | 0.015646 | 0.014229 | 0.032371 | 0.01799 | 0.01371 | 0.007016 | 0.018526 |
| 23.0694444 | 0.022011 | 0.078937 | 0.037384 | 0.016566 | 0.003988 | 0.024806 | 0.025401 | 0.055101 | 0.039716 | 0.086659 | 0.018491 | 0.030262 | 0.032844 | 0.017534 | 0.024764 | 0.019511 | 0.012252 | 0.009753 | 0.029268 |
| 23.0833333 | 0.016229 | 0.066206 | 0.036079 | 0.01987 | 0.005136 | 0.019405 | 0.029113 | 0.042843 | 0.024041 | 0.109827 | 0.018087 | 0.023697 | 0.049586 | 0.013518 | 0.016215 | 0.020433 | 0.007069 | 0.009865 | 0.028352 |
| 23.0972222 | 0.015959 | 0.049856 | 0.025622 | 0.020258 | 0.005657 | 0.009016 | 0.022861 | 0.024674 | 0.01006 | 0.099232 | 0.012498 | 0.015841 | 0.046585 | 0.006835 | 0.008265 | 0.01759 | 0.005216 | 0.00666 | 0.023773 |
| 23.1111111 | 0.022301 | 0.041101 | 0.015613 | 0.019235 | 0.006462 | 0.002775 | 0.013104 | 0.017424 | 0.007105 | 0.080016 | 0.008225 | 0.009326 | 0.029773 | 0.002873 | 0.005057 | 0.012397 | 0.005709 | 0.003366 | 0.023875 |
| 23.125 | 0.023558 | 0.035063 | 0.009054 | 0.014692 | 0.006552 | 0.00147 | 0.006138 | 0.018253 | 0.004732 | 0.058642 | 0.008436 | 0.003959 | 0.01299 | 0.002109 | 0.005599 | 0.007256 | 0.004424 | 0.003413 | 0.023504 |
| 23.1388889 | 0.014823 | 0.025049 | 0.00429 | 0.007418 | 0.004773 | 0.002712 | 0.002745 | 0.016075 | 0.001301 | 0.031507 | 0.008353 | 0.001161 | 0.009257 | 0.002163 | 0.005521 | 0.003995 | 0.002445 | 0.005627 | 0.017798 |
| 23.1527778 | 0.005338 | 0.012487 | 0.001914 | 0.003226 | 0.002908 | 0.007401 | 0.00137 | 0.009729 | 0.001237 | 0.013766 | 0.006007 | 0.000667 | 0.017216 | 0.001417 | 0.006307 | 0.00553 | 0.001391 | 0.005902 | 0.009955 |
| 23.1666667 | 0.003346 | 0.005291 | 0.002697 | 0.004522 | 0.004127 | 0.013919 | 0.001423 | 0.004897 | 0.00417 | 0.015988 | 0.00709 | 0.001239 | 0.026364 | 0.000865 | 0.011688 | 0.011799 | 0.001897 | 0.003998 | 0.008081 |
| 23.1805556 | 0.007037 | 0.006752 | 0.004455 | 0.00707 | 0.007439 | 0.017818 | 0.002764 | 0.006123 | 0.009106 | 0.029124 | 0.011989 | 0.001394 | 0.031555 | 0.001546 | 0.022798 | 0.019681 | 0.004618 | 0.003893 | 0.014455 |
| 23.1944444 | 0.010278 | 0.008036 | 0.004911 | 0.007679 | 0.008493 | 0.015436 | 0.00482 | 0.012882 | 0.015187 | 0.041471 | 0.014756 | 0.000925 | 0.032592 | 0.002321 | 0.037212 | 0.02707 | 0.008717 | 0.006341 | 0.0212 |
| 23.2083333 | 0.009569 | 0.004089 | 0.004575 | 0.007289 | 0.007115 | 0.00842 | 0.007072 | 0.022876 | 0.023047 | 0.051559 | 0.015269 | 0.000987 | 0.029468 | 0.002569 | 0.045904 | 0.030818 | 0.012575 | 0.008989 | 0.025463 |
| 23.2222222 | 0.008401 | 0.009266 | 0.006424 | 0.008567 | 0.006419 | 0.002613 | 0.008199 | 0.035624 | 0.031609 | 0.0576 | 0.015843 | 0.001296 | 0.024099 | 0.00333 | 0.042307 | 0.030768 | 0.015469 | 0.011827 | 0.028857 |
| 23.2361111 | 0.010337 | 0.030688 | 0.011278 | 0.011617 | 0.006109 | 0.001431 | 0.007522 | 0.045758 | 0.037313 | 0.053581 | 0.014269 | 0.001656 | 0.019061 | 0.004876 | 0.031396 | 0.030065 | 0.016951 | 0.014316 | 0.028158 |
| 23.25 | 0.012725 | 0.049258 | 0.015853 | 0.013531 | 0.004315 | 0.003395 | 0.00642 | 0.048033 | 0.038942 | 0.04069 | 0.009874 | 0.002882 | 0.014862 | 0.006411 | 0.020276 | 0.026664 | 0.017018 | 0.014558 | 0.023058 |
| 23.2638889 | 0.013436 | 0.054099 | 0.017371 | 0.013143 | 0.001998 | 0.005934 | 0.006686 | 0.045715 | 0.037309 | 0.026384 | 0.005179 | 0.004406 | 0.010445 | 0.007117 | 0.010884 | 0.017651 | 0.016326 | 0.012596 | 0.017322 |
| 23.2777778 | 0.012544 | 0.050501 | 0.01582 | 0.011228 | 0.001407 | 0.007771 | 0.007935 | 0.041064 | 0.032648 | 0.015624 | 0.008557 | 0.005949 | 0.007202 | 0.006159 | 0.006468 | 0.010669 | 0.011775 | 0.010032 | 0.01307 |
| 23.2916667 | 0.011007 | 0.041693 | 0.013195 | 0.009034 | 0.002505 | 0.007982 | 0.008223 | 0.031796 | 0.02687 | 0.018701 | 0.031337 | 0.014252 | 0.010222 | 0.008512 | 0.011523 | 0.017495 | 0.010747 | 0.014938 | 0.020879 |
| 23.3055556 | 0.009216 | 0.031434 | 0.011084 | 0.007491 | 0.002507 | 0.006098 | 0.007113 | 0.018717 | 0.021398 | 0.03258 | 0.059159 | 0.027957 | 0.016451 | 0.016186 | 0.020007 | 0.031634 | 0.02217 | 0.027765 | 0.040439 |
| 23.3194444 | 0.00566 | 0.019208 | 0.007987 | 0.005567 | 0.001846 | 0.00323 | 0.005812 | 0.010725 | 0.017154 | 0.031697 | 0.059105 | 0.02745 | 0.014651 | 0.017619 | 0.020277 | 0.033838 | 0.027922 | 0.029697 | 0.043764 |
| 23.3333333 | 0.001956 | 0.010067 | 0.004402 | 0.00272 | 0.002816 | 0.001594 | 0.004643 | 0.017751 | 0.021073 | 0.017637 | 0.03278 | 0.013614 | 0.008778 | 0.010501 | 0.01196 | 0.021259 | 0.019129 | 0.01679 | 0.025785 |
| 23.3472222 | 0.001273 | 0.013136 | 0.003752 | 0.000754 | 0.003426 | 0.001507 | 0.002837 | 0.033733 | 0.030218 | 0.012779 | 0.011666 | 0.006006 | 0.010081 | 0.00573 | 0.004515 | 0.007697 | 0.010055 | 0.005989 | 0.010969 |
| 23.3611111 | 0.003118 | 0.01826 | 0.004666 | 0.000766 | 0.002319 | 0.001556 | 0.001004 | 0.042403 | 0.028696 | 0.014016 | 0.006327 | 0.004955 | 0.011169 | 0.005881 | 0.004128 | 0.001927 | 0.005062 | 0.003549 | 0.006703 |
| 23.375 | 0.005497 | 0.013356 | 0.004101 | 0.002225 | 0.00147 | 0.002411 | 0.000414 | 0.037491 | 0.016757 | 0.011467 | 0.005169 | 0.00482 | 0.007251 | 0.005937 | 0.008604 | 0.002567 | 0.003655 | 0.002991 | 0.004505 |
| 23.3888889 | 0.007655 | 0.00933 | 0.002497 | 0.004682 | 0.001404 | 0.004555 | 0.000908 | 0.022301 | 0.011634 | 0.013941 | 0.004006 | 0.006005 | 0.005808 | 0.003772 | 0.012191 | 0.006269 | 0.009065 | 0.003527 | 0.002887 |
| 23.4027778 | 0.010594 | 0.020227 | 0.002353 | 0.007717 | 0.002266 | 0.006503 | 0.001892 | 0.01361 | 0.020275 | 0.025461 | 0.006922 | 0.00761 | 0.00565 | 0.002645 | 0.011521 | 0.010156 | 0.017929 | 0.007478 | 0.004883 |
| 23.4166667 | 0.014374 | 0.035453 | 0.005029 | 0.010474 | 0.004836 | 0.0076 | 0.003261 | 0.027258 | 0.031534 | 0.033587 | 0.011357 | 0.008619 | 0.004531 | 0.005457 | 0.007572 | 0.011475 | 0.021925 | 0.009967 | 0.008023 |
| 23.4305556 | 0.016472 | 0.03775 | 0.008355 | 0.011827 | 0.006628 | 0.007992 | 0.004971 | 0.048278 | 0.034425 | 0.032177 | 0.012439 | 0.008627 | 0.007576 | 0.009615 | 0.004288 | 0.009362 | 0.018148 | 0.009118 | 0.010222 |
| 23.4444444 | 0.014889 | 0.024339 | 0.009926 | 0.01103 | 0.005633 | 0.007674 | 0.006197 | 0.056815 | 0.02804 | 0.024414 | 0.010366 | 0.008316 | 0.01465 | 0.011996 | 0.00541 | 0.006204 | 0.010913 | 0.007188 | 0.013552 |
| 23.4583333 | 0.011798 | 0.009691 | 0.010254 | 0.00944 | 0.003571 | 0.006908 | 0.006517 | 0.05192 | 0.019174 | 0.014745 | 0.007877 | 0.009167 | 0.019074 | 0.012626 | 0.011407 | 0.00569 | 0.004933 | 0.00495 | 0.019768 |
| 23.4722222 | 0.012291 | 0.004657 | 0.011094 | 0.009437 | 0.002133 | 0.006813 | 0.007157 | 0.039545 | 0.014291 | 0.010979 | 0.008423 | 0.011683 | 0.018548 | 0.012914 | 0.019537 | 0.008239 | 0.001835 | 0.004103 | 0.024674 |
| 23.4861111 | 0.017303 | 0.004971 | 0.011937 | 0.010792 | 0.001274 | 0.007975 | 0.008602 | 0.024685 | 0.011385 | 0.016114 | 0.011113 | 0.01461 | 0.014546 | 0.013221 | 0.024285 | 0.010368 | 0.00106 | 0.004903 | 0.020097 |
| 23.5 | 0.020941 | 0.010073 | 0.010689 | 0.010989 | 0.000717 | 0.008822 | 0.009308 | 0.013671 | 0.007908 | 0.019689 | 0.010605 | 0.015104 | 0.009575 | 0.012077 | 0.01912 | 0.008714 | 0.000832 | 0.00535 | 0.011908 |
| 23.5138889 | 0.019302 | 0.020882 | 0.007167 | 0.010381 | 0.001606 | 0.00948 | 0.007735 | 0.012608 | 0.007711 | 0.014207 | 0.006672 | 0.011761 | 0.009245 | 0.009606 | 0.010561 | 0.00459 | 0.000629 | 0.004501 | 0.012634 |
| 23.5277778 | 0.017093 | 0.043628 | 0.00534 | 0.012232 | 0.005658 | 0.013292 | 0.00426 | 0.013846 | 0.01167 | 0.010928 | 0.005807 | 0.008012 | 0.010798 | 0.008337 | 0.009105 | 0.001831 | 0.001604 | 0.003745 | 0.013781 |
| 23.5416667 | 0.022457 | 0.084589 | 0.01371 | 0.016147 | 0.010774 | 0.019591 | 0.001616 | 0.023137 | 0.016536 | 0.024282 | 0.011648 | 0.010189 | 0.011811 | 0.011232 | 0.01087 | 0.001709 | 0.007948 | 0.006695 | 0.014056 |
| 23.5555556 | 0.036806 | 0.127882 | 0.034525 | 0.021959 | 0.011753 | 0.024056 | 0.005143 | 0.057673 | 0.03319 | 0.055284 | 0.020814 | 0.021564 | 0.025004 | 0.020369 | 0.020549 | 0.002922 | 0.023858 | 0.01554 | 0.027586 |
| 23.5694444 | 0.054013 | 0.140521 | 0.053707 | 0.030772 | 0.010991 | 0.024349 | 0.012664 | 0.090929 | 0.060811 | 0.085795 | 0.028958 | 0.041411 | 0.047664 | 0.032486 | 0.042039 | 0.002783 | 0.042286 | 0.022979 | 0.043564 |
| 23.5833333 | 0.061921 | 0.113137 | 0.055837 | 0.034972 | 0.015605 | 0.018736 | 0.014163 | 0.08852 | 0.071481 | 0.087836 | 0.032749 | 0.052818 | 0.057772 | 0.03651 | 0.052482 | 0.001808 | 0.048251 | 0.021847 | 0.041988 |
| 23.5972222 | 0.048617 | 0.067777 | 0.041776 | 0.028476 | 0.023573 | 0.010201 | 0.008579 | 0.057374 | 0.05895 | 0.055081 | 0.026333 | 0.037733 | 0.050573 | 0.024924 | 0.039428 | 0.003581 | 0.040223 | 0.016494 | 0.026995 |
| 23.6111111 | 0.025895 | 0.035075 | 0.02145 | 0.015523 | 0.027965 | 0.009188 | 0.005089 | 0.03096 | 0.034034 | 0.03128 | 0.015708 | 0.017591 | 0.035366 | 0.011715 | 0.021616 | 0.009568 | 0.029518 | 0.010218 | 0.023736 |
| 23.625 | 0.020435 | 0.038204 | 0.009421 | 0.008226 | 0.027356 | 0.01819 | 0.009437 | 0.038844 | 0.018105 | 0.051659 | 0.015662 | 0.018932 | 0.017586 | 0.011012 | 0.016945 | 0.017081 | 0.020987 | 0.00508 | 0.039055 |
| 23.6388889 | 0.032018 | 0.05982 | 0.01278 | 0.014205 | 0.024821 | 0.025094 | 0.015051 | 0.070048 | 0.030437 | 0.081005 | 0.019751 | 0.031923 | 0.005342 | 0.011027 | 0.02233 | 0.021637 | 0.01261 | 0.0046 | 0.05124 |
| 23.6527778 | 0.033526 | 0.067304 | 0.018842 | 0.021116 | 0.022721 | 0.019259 | 0.013722 | 0.096954 | 0.052058 | 0.074283 | 0.015074 | 0.03617 | 0.00357 | 0.006453 | 0.022926 | 0.020944 | 0.005357 | 0.004515 | 0.041723 |
| 23.6666667 | 0.023768 | 0.047547 | 0.016996 | 0.016583 | 0.020027 | 0.011947 | 0.007438 | 0.097621 | 0.053624 | 0.042699 | 0.011035 | 0.024545 | 0.004431 | 0.006228 | 0.023789 | 0.016804 | 0.002423 | 0.002938 | 0.027124 |
| 23.6805556 | 0.028056 | 0.030769 | 0.010322 | 0.010726 | 0.014208 | 0.014565 | 0.004557 | 0.067015 | 0.033115 | 0.028264 | 0.015927 | 0.01527 | 0.002676 | 0.009066 | 0.035969 | 0.01586 | 0.002659 | 0.003112 | 0.035338 |
| 23.6944444 | 0.04187 | 0.054113 | 0.010174 | 0.017827 | 0.007254 | 0.014445 | 0.007207 | 0.053096 | 0.021788 | 0.028419 | 0.01864 | 0.02876 | 0.006158 | 0.010785 | 0.040671 | 0.01833 | 0.004362 | 0.002967 | 0.045623 |
| 23.7083333 | 0.042232 | 0.080595 | 0.019945 | 0.02488 | 0.0059 | 0.007715 | 0.009759 | 0.071938 | 0.030983 | 0.021161 | 0.012343 | 0.039425 | 0.024722 | 0.008638 | 0.025585 | 0.016206 | 0.007227 | 0.002741 | 0.031864 |
| 23.7222222 | 0.027873 | 0.073549 | 0.029951 | 0.016875 | 0.011913 | 0.005547 | 0.00921 | 0.06842 | 0.030103 | 0.021424 | 0.007151 | 0.024685 | 0.05071 | 0.005909 | 0.013094 | 0.01022 | 0.009114 | 0.004954 | 0.0165 |
| 23.7361111 | 0.014185 | 0.044671 | 0.032636 | 0.008949 | 0.019133 | 0.010945 | 0.00599 | 0.042493 | 0.023179 | 0.039562 | 0.011563 | 0.011373 | 0.062809 | 0.011088 | 0.021846 | 0.009953 | 0.008507 | 0.007025 | 0.023829 |
| 23.75 | 0.012363 | 0.01983 | 0.024636 | 0.013017 | 0.021092 | 0.014946 | 0.002658 | 0.045024 | 0.037092 | 0.0556 | 0.015254 | 0.013623 | 0.056185 | 0.020275 | 0.038214 | 0.015594 | 0.006073 | 0.007983 | 0.041237 |
| 23.7638889 | 0.013662 | 0.019656 | 0.014339 | 0.01878 | 0.015954 | 0.010168 | 0.002222 | 0.076324 | 0.058539 | 0.05494 | 0.010958 | 0.017482 | 0.038744 | 0.024979 | 0.05008 | 0.020981 | 0.003256 | 0.009848 | 0.049304 |
| 23.7777778 | 0.009892 | 0.032964 | 0.015348 | 0.016376 | 0.007896 | 0.003808 | 0.003709 | 0.104308 | 0.070528 | 0.037098 | 0.007841 | 0.025665 | 0.017415 | 0.022358 | 0.052485 | 0.021709 | 0.001174 | 0.010256 | 0.036708 |
| 23.7916667 | 0.010215 | 0.045703 | 0.022228 | 0.010645 | 0.004027 | 0.002431 | 0.00477 | 0.103521 | 0.062002 | 0.015304 | 0.006927 | 0.03856 | 0.006386 | 0.012972 | 0.036473 | 0.013567 | 0.001046 | 0.006953 | 0.023087 |
| 23.8055556 | 0.014223 | 0.050484 | 0.021017 | 0.012422 | 0.004939 | 0.004499 | 0.006534 | 0.066573 | 0.034114 | 0.008236 | 0.005133 | 0.034668 | 0.010869 | 0.005258 | 0.015856 | 0.005645 | 0.003364 | 0.00615 | 0.028632 |
| 23.8194444 | 0.011626 | 0.035761 | 0.012335 | 0.016061 | 0.0052 | 0.013049 | 0.009717 | 0.041172 | 0.018639 | 0.008232 | 0.007966 | 0.019947 | 0.020634 | 0.005016 | 0.007168 | 0.004512 | 0.006592 | 0.009932 | 0.024098 |
| 23.8333333 | 0.007928 | 0.020571 | 0.008877 | 0.012262 | 0.003587 | 0.026547 | 0.01033 | 0.060491 | 0.023688 | 0.014819 | 0.014989 | 0.022703 | 0.028256 | 0.005517 | 0.005944 | 0.004771 | 0.006241 | 0.011022 | 0.013598 |
| 23.8472222 | 0.013441 | 0.025752 | 0.014752 | 0.010083 | 0.004424 | 0.032973 | 0.00638 | 0.07301 | 0.022085 | 0.032926 | 0.019703 | 0.034168 | 0.027838 | 0.004183 | 0.006503 | 0.006246 | 0.006342 | 0.007098 | 0.028543 |
| 23.8611111 | 0.02036 | 0.026197 | 0.017056 | 0.016017 | 0.009665 | 0.027454 | 0.002342 | 0.048677 | 0.015855 | 0.046997 | 0.018547 | 0.029532 | 0.018329 | 0.006565 | 0.007431 | 0.006861 | 0.015695 | 0.004394 | 0.051733 |
| 23.875 | 0.018312 | 0.016475 | 0.014668 | 0.023149 | 0.014924 | 0.016486 | 0.001302 | 0.034453 | 0.027598 | 0.055465 | 0.012191 | 0.015944 | 0.015359 | 0.012527 | 0.006974 | 0.005 | 0.027523 | 0.008359 | 0.056834 |
| 23.8888889 | 0.011069 | 0.02359 | 0.023161 | 0.026651 | 0.016411 | 0.008913 | 0.001303 | 0.062472 | 0.0486 | 0.056489 | 0.005423 | 0.013797 | 0.027846 | 0.017861 | 0.006876 | 0.005266 | 0.030988 | 0.014494 | 0.0458 |
| 23.9027778 | 0.005228 | 0.045467 | 0.038728 | 0.026027 | 0.015719 | 0.008661 | 0.002336 | 0.091701 | 0.055565 | 0.044762 | 0.002627 | 0.024784 | 0.040915 | 0.021518 | 0.009063 | 0.00983 | 0.024927 | 0.014293 | 0.033858 |
| 23.9166667 | 0.002709 | 0.065544 | 0.047236 | 0.021252 | 0.015105 | 0.01241 | 0.006 | 0.085919 | 0.042455 | 0.0353 | 0.004186 | 0.034983 | 0.043216 | 0.023352 | 0.012656 | 0.016288 | 0.01539 | 0.008025 | 0.02847 |
| 23.9305556 | 0.003096 | 0.075446 | 0.047746 | 0.013921 | 0.014313 | 0.01334 | 0.009708 | 0.059008 | 0.023459 | 0.034634 | 0.007693 | 0.034839 | 0.038082 | 0.021477 | 0.01605 | 0.020983 | 0.008762 | 0.003491 | 0.020834 |
| 23.9444444 | 0.003772 | 0.068189 | 0.042337 | 0.006504 | 0.012393 | 0.008334 | 0.010226 | 0.02963 | 0.010712 | 0.033337 | 0.010885 | 0.025396 | 0.031265 | 0.016227 | 0.016184 | 0.02 | 0.005385 | 0.003757 | 0.010404 |
| 23.9583333 | 0.003601 | 0.045947 | 0.033958 | 0.001954 | 0.009856 | 0.002554 | 0.009117 | 0.010992 | 0.00491 | 0.026888 | 0.012596 | 0.013074 | 0.02553 | 0.011505 | 0.012307 | 0.012166 | 0.003635 | 0.005228 | 0.008379 |
| 23.9722222 | 0.002882 | 0.022353 | 0.023396 | 0.00054 | 0.007931 | 0.002087 | 0.007566 | 0.008683 | 0.005672 | 0.018936 | 0.013493 | 0.007948 | 0.019251 | 0.00918 | 0.008241 | 0.00491 | 0.002492 | 0.004816 | 0.009485 |
| 23.9861111 | 0.001627 | 0.013304 | 0.012846 | 0.001923 | 0.006408 | 0.006869 | 0.004714 | 0.016201 | 0.010391 | 0.015299 | 0.015125 | 0.011088 | 0.012129 | 0.007173 | 0.006499 | 0.003273 | 0.001419 | 0.003035 | 0.008671 |
| 24 | 0.002134 | 0.020982 | 0.012881 | 0.006903 | 0.005041 | 0.013411 | 0.004095 | 0.025156 | 0.012733 | 0.015258 | 0.016535 | 0.013875 | 0.011419 | 0.004151 | 0.005972 | 0.004852 | 0.000936 | 0.002589 | 0.011933 |
| 24.0138889 | 0.00472 | 0.025594 | 0.023267 | 0.012153 | 0.005915 | 0.019689 | 0.008439 | 0.031413 | 0.011038 | 0.012802 | 0.014884 | 0.013529 | 0.018788 | 0.002203 | 0.004858 | 0.008302 | 0.001572 | 0.003548 | 0.01591 |
| 24.0277778 | 0.010002 | 0.023267 | 0.030043 | 0.012723 | 0.010118 | 0.028437 | 0.012024 | 0.028313 | 0.007667 | 0.008464 | 0.011767 | 0.010653 | 0.021587 | 0.002094 | 0.004873 | 0.010921 | 0.001836 | 0.003607 | 0.017223 |
| 24.0416667 | 0.022652 | 0.035049 | 0.02358 | 0.013333 | 0.012998 | 0.039828 | 0.009713 | 0.021743 | 0.009159 | 0.015082 | 0.013632 | 0.008413 | 0.016249 | 0.002107 | 0.011588 | 0.013777 | 0.002319 | 0.002722 | 0.020633 |
| 24.0555556 | 0.038052 | 0.057558 | 0.018605 | 0.023225 | 0.011725 | 0.047425 | 0.008456 | 0.027455 | 0.021167 | 0.039501 | 0.022028 | 0.015873 | 0.017431 | 0.003461 | 0.022928 | 0.020511 | 0.00555 | 0.002558 | 0.030168 |
| 24.0694444 | 0.042374 | 0.067018 | 0.029614 | 0.036639 | 0.0101 | 0.050373 | 0.012545 | 0.032921 | 0.033598 | 0.066739 | 0.030956 | 0.029575 | 0.025535 | 0.006679 | 0.026177 | 0.027019 | 0.008204 | 0.003022 | 0.041674 |
| 24.0833333 | 0.032139 | 0.061497 | 0.03949 | 0.039548 | 0.009727 | 0.050132 | 0.013942 | 0.022116 | 0.033185 | 0.083017 | 0.035632 | 0.032357 | 0.02318 | 0.008342 | 0.018091 | 0.02735 | 0.007407 | 0.003424 | 0.044147 |
| 24.0972222 | 0.017351 | 0.050943 | 0.037359 | 0.030485 | 0.008738 | 0.041885 | 0.010819 | 0.007785 | 0.022336 | 0.080653 | 0.03339 | 0.021976 | 0.011946 | 0.007141 | 0.010081 | 0.021267 | 0.005341 | 0.002896 | 0.038663 |
| 24.1111111 | 0.007429 | 0.037722 | 0.028775 | 0.017993 | 0.006221 | 0.025658 | 0.006824 | 0.002538 | 0.011311 | 0.053808 | 0.022553 | 0.01012 | 0.004788 | 0.004412 | 0.007917 | 0.012605 | 0.003427 | 0.002196 | 0.032414 |
| 24.125 | 0.004521 | 0.024318 | 0.016995 | 0.008448 | 0.003283 | 0.010781 | 0.003822 | 0.002213 | 0.004696 | 0.020762 | 0.009761 | 0.005532 | 0.002014 | 0.001827 | 0.008584 | 0.006502 | 0.002115 | 0.00462 | 0.023324 |
| 24.1388889 | 0.010086 | 0.012468 | 0.007589 | 0.006322 | 0.001508 | 0.006418 | 0.003746 | 0.004372 | 0.002089 | 0.008355 | 0.009857 | 0.006757 | 0.004972 | 0.001797 | 0.018379 | 0.010856 | 0.006864 | 0.01209 | 0.018216 |
| 24.1527778 | 0.025935 | 0.01229 | 0.005304 | 0.013558 | 0.003583 | 0.013961 | 0.004541 | 0.010336 | 0.00358 | 0.031176 | 0.036605 | 0.021275 | 0.028779 | 0.008521 | 0.060469 | 0.038432 | 0.031116 | 0.031404 | 0.034783 |
| 24.1666667 | 0.051514 | 0.039063 | 0.006002 | 0.027744 | 0.015941 | 0.031366 | 0.006302 | 0.013255 | 0.006344 | 0.070791 | 0.071391 | 0.057842 | 0.073562 | 0.02462 | 0.111037 | 0.073034 | 0.065704 | 0.055361 | 0.065442 |
| 24.1805556 | 0.073542 | 0.072729 | 0.005515 | 0.04276 | 0.035152 | 0.049926 | 0.010613 | 0.011064 | 0.009396 | 0.070225 | 0.066786 | 0.072415 | 0.093159 | 0.035266 | 0.104965 | 0.071208 | 0.069999 | 0.055692 | 0.066202 |
| 24.1944444 | 0.069236 | 0.073193 | 0.007749 | 0.048385 | 0.042042 | 0.058858 | 0.01333 | 0.013301 | 0.021735 | 0.037149 | 0.033059 | 0.039954 | 0.063152 | 0.024916 | 0.064876 | 0.043647 | 0.04289 | 0.034328 | 0.037815 |
| 24.2083333 | 0.042724 | 0.043321 | 0.012207 | 0.040612 | 0.030873 | 0.056701 | 0.012673 | 0.018516 | 0.038668 | 0.025751 | 0.016322 | 0.008399 | 0.032347 | 0.011085 | 0.051481 | 0.031282 | 0.024106 | 0.019182 | 0.025458 |
| 24.2222222 | 0.020477 | 0.018453 | 0.013411 | 0.025619 | 0.015817 | 0.044422 | 0.010845 | 0.016293 | 0.039641 | 0.021637 | 0.012841 | 0.005685 | 0.021884 | 0.007612 | 0.048135 | 0.027834 | 0.01838 | 0.015877 | 0.020808 |
| 24.2361111 | 0.010471 | 0.008798 | 0.009549 | 0.011856 | 0.006052 | 0.028433 | 0.007101 | 0.008207 | 0.024073 | 0.012717 | 0.011676 | 0.015018 | 0.013688 | 0.005324 | 0.043981 | 0.026423 | 0.014926 | 0.018905 | 0.011092 |
| 24.25 | 0.00528 | 0.004086 | 0.004792 | 0.006449 | 0.002422 | 0.017318 | 0.003295 | 0.002239 | 0.00915 | 0.013753 | 0.013737 | 0.018222 | 0.012435 | 0.003528 | 0.051015 | 0.031297 | 0.015017 | 0.023576 | 0.010672 |

| | | | | | | | | | | | | | | | | | | | |
|---|---|---|---|---|---|---|---|---|---|---|---|---|---|---|---|---|---|---|---|
| 24.2638889 | 0.004428 | 0.001699 | 0.003138 | 0.006895 | 0.003417 | 0.011993 | 0.002172 | 0.001157 | 0.002722 | 0.013797 | 0.011726 | 0.014933 | 0.012124 | 0.002933 | 0.040898 | 0.025685 | 0.01275 | 0.021053 | 0.012499 |
| 24.2777778 | 0.010419 | 0.002413 | 0.003153 | 0.006313 | 0.005323 | 0.008854 | 0.00243 | 0.002374 | 0.00295 | 0.008147 | 0.00673 | 0.012891 | 0.010758 | 0.002172 | 0.016238 | 0.01238 | 0.008623 | 0.01424 | 0.010454 |
| 24.2916667 | 0.017815 | 0.004745 | 0.004601 | 0.006782 | 0.005056 | 0.005868 | 0.003796 | 0.002455 | 0.006177 | 0.004563 | 0.00434 | 0.014001 | 0.015311 | 0.003053 | 0.004793 | 0.00741 | 0.008182 | 0.009257 | 0.009853 |
| 24.3055556 | 0.017672 | 0.005114 | 0.00511 | 0.0078 | 0.002794 | 0.006059 | 0.003824 | 0.001289 | 0.00801 | 0.004972 | 0.005248 | 0.01611 | 0.017197 | 0.004758 | 0.004331 | 0.009163 | 0.01007 | 0.006605 | 0.011596 |
| 24.3194444 | 0.011054 | 0.003169 | 0.003969 | 0.005972 | 0.000829 | 0.011277 | 0.002426 | 0.000575 | 0.007119 | 0.006087 | 0.006548 | 0.016132 | 0.014792 | 0.005219 | 0.008896 | 0.01077 | 0.011018 | 0.01033 | 0.013066 |
| 24.3333333 | 0.008025 | 0.001964 | 0.00566 | 0.005924 | 0.000884 | 0.016212 | 0.0037 | 0.000721 | 0.005715 | 0.004796 | 0.005907 | 0.012198 | 0.012116 | 0.003698 | 0.013055 | 0.009786 | 0.009527 | 0.017138 | 0.01281 |
| 24.3472222 | 0.009627 | 0.001358 | 0.007721 | 0.00892 | 0.003111 | 0.017002 | 0.006822 | 0.001311 | 0.005453 | 0.003388 | 0.004862 | 0.005891 | 0.008067 | 0.002421 | 0.014295 | 0.008402 | 0.007736 | 0.017397 | 0.012451 |
| 24.3611111 | 0.007976 | 0.000768 | 0.005624 | 0.009008 | 0.005245 | 0.015508 | 0.008394 | 0.001608 | 0.005914 | 0.005287 | 0.005398 | 0.001563 | 0.003781 | 0.003321 | 0.015474 | 0.008306 | 0.007837 | 0.011368 | 0.012484 |
| 24.375 | 0.005807 | 0.001204 | 0.003693 | 0.006536 | 0.004788 | 0.011528 | 0.006047 | 0.001897 | 0.006418 | 0.008064 | 0.006135 | 0.000829 | 0.002492 | 0.00465 | 0.016277 | 0.00831 | 0.008255 | 0.005908 | 0.011228 |
| 24.3888889 | 0.011962 | 0.003071 | 0.006416 | 0.009256 | 0.002542 | 0.007408 | 0.003451 | 0.002696 | 0.006565 | 0.007479 | 0.004983 | 0.003119 | 0.002322 | 0.003697 | 0.014069 | 0.006656 | 0.005968 | 0.005033 | 0.008241 |
| 24.4027778 | 0.02163 | 0.005345 | 0.008566 | 0.017228 | 0.001184 | 0.009103 | 0.005177 | 0.003131 | 0.005549 | 0.004828 | 0.002619 | 0.006574 | 0.002221 | 0.001534 | 0.008961 | 0.003731 | 0.002425 | 0.007818 | 0.00433 |
| 24.4166667 | 0.022187 | 0.005468 | 0.005785 | 0.020756 | 0.001708 | 0.013639 | 0.007319 | 0.00383 | 0.003219 | 0.003084 | 0.001261 | 0.008252 | 0.004668 | 0.000495 | 0.005016 | 0.002045 | 0.00146 | 0.011458 | 0.002392 |
| 24.4305556 | 0.015999 | 0.004095 | 0.002978 | 0.018569 | 0.002253 | 0.016111 | 0.00789 | 0.009059 | 0.003781 | 0.003617 | 0.002789 | 0.010861 | 0.010462 | 0.001484 | 0.008129 | 0.00427 | 0.003975 | 0.015726 | 0.005599 |
| 24.4444444 | 0.019286 | 0.004918 | 0.007097 | 0.017786 | 0.001561 | 0.015991 | 0.009574 | 0.02065 | 0.013334 | 0.008145 | 0.007097 | 0.017617 | 0.01806 | 0.004731 | 0.018335 | 0.0097 | 0.009014 | 0.021496 | 0.01166 |
| 24.4583333 | 0.031762 | 0.009645 | 0.016529 | 0.017912 | 0.000803 | 0.011417 | 0.012293 | 0.02932 | 0.026428 | 0.013478 | 0.010829 | 0.022537 | 0.021751 | 0.007686 | 0.025828 | 0.01356 | 0.013868 | 0.024793 | 0.014707 |
| 24.4722222 | 0.038322 | 0.014902 | 0.021341 | 0.017149 | 0.000638 | 0.006098 | 0.013623 | 0.025273 | 0.028781 | 0.014159 | 0.011414 | 0.020161 | 0.018192 | 0.007925 | 0.024301 | 0.013296 | 0.014906 | 0.021467 | 0.012896 |
| 24.4861111 | 0.031763 | 0.013352 | 0.016376 | 0.016978 | 0.000913 | 0.007653 | 0.01141 | 0.013481 | 0.018386 | 0.011698 | 0.010143 | 0.014761 | 0.012139 | 0.006597 | 0.018345 | 0.011443 | 0.012163 | 0.015359 | 0.009652 |
| 24.5 | 0.015937 | 0.006497 | 0.007635 | 0.013607 | 0.001611 | 0.013305 | 0.006672 | 0.0044 | 0.007312 | 0.009692 | 0.008861 | 0.011475 | 0.008105 | 0.005655 | 0.013661 | 0.01 | 0.009259 | 0.01031 | 0.007445 |
| 24.5138889 | 0.008895 | 0.009254 | 0.003248 | 0.008119 | 0.002431 | 0.013038 | 0.003302 | 0.001413 | 0.004674 | 0.008139 | 0.008093 | 0.009773 | 0.00615 | 0.005196 | 0.011218 | 0.008689 | 0.007787 | 0.006414 | 0.006047 |
| 24.5277778 | 0.025801 | 0.03103 | 0.002604 | 0.007209 | 0.002226 | 0.014165 | 0.00326 | 0.006349 | 0.013131 | 0.007517 | 0.009513 | 0.009115 | 0.00686 | 0.005376 | 0.013486 | 0.009696 | 0.009011 | 0.006331 | 0.006777 |
| 24.5416667 | 0.054919 | 0.058758 | 0.008478 | 0.008064 | 0.001414 | 0.024089 | 0.006898 | 0.027743 | 0.031518 | 0.010952 | 0.015222 | 0.013153 | 0.01421 | 0.008832 | 0.031362 | 0.019145 | 0.017355 | 0.015387 | 0.013085 |
| 24.5555556 | 0.072458 | 0.07296 | 0.031512 | 0.014921 | 0.00365 | 0.029191 | 0.016223 | 0.073262 | 0.053519 | 0.018186 | 0.022347 | 0.022899 | 0.028852 | 0.015192 | 0.071467 | 0.041106 | 0.03106 | 0.033634 | 0.024247 |
| 24.5694444 | 0.070319 | 0.070305 | 0.05828 | 0.033618 | 0.008766 | 0.022125 | 0.025393 | 0.128737 | 0.075195 | 0.022666 | 0.024056 | 0.032141 | 0.041513 | 0.018681 | 0.103542 | 0.061151 | 0.037736 | 0.044281 | 0.030992 |
| 24.5833333 | 0.057014 | 0.059139 | 0.061388 | 0.045454 | 0.012223 | 0.010271 | 0.024261 | 0.155294 | 0.086292 | 0.020449 | 0.018628 | 0.034131 | 0.040584 | 0.01666 | 0.090934 | 0.05766 | 0.030524 | 0.031508 | 0.026647 |
| 24.5972222 | 0.038918 | 0.046839 | 0.044656 | 0.036236 | 0.013053 | 0.005724 | 0.016558 | 0.1286 | 0.072518 | 0.014984 | 0.010946 | 0.027719 | 0.02788 | 0.012167 | 0.053172 | 0.037856 | 0.017834 | 0.010837 | 0.01654 |
| 24.6111111 | 0.024197 | 0.033216 | 0.026609 | 0.019023 | 0.012879 | 0.011245 | 0.009574 | 0.070586 | 0.041629 | 0.010293 | 0.005068 | 0.017998 | 0.015811 | 0.008048 | 0.026595 | 0.022079 | 0.009237 | 0.005913 | 0.007979 |
| 24.625 | 0.029497 | 0.017778 | 0.012725 | 0.014499 | 0.011917 | 0.020717 | 0.005454 | 0.030408 | 0.019728 | 0.007061 | 0.001942 | 0.010573 | 0.008854 | 0.004981 | 0.014639 | 0.013919 | 0.005653 | 0.015458 | 0.003102 |
| 24.6388889 | 0.04654 | 0.008702 | 0.00764 | 0.024862 | 0.010786 | 0.02733 | 0.006322 | 0.036504 | 0.022726 | 0.004828 | 0.001482 | 0.006697 | 0.004756 | 0.002992 | 0.00903 | 0.009649 | 0.004892 | 0.024173 | 0.001779 |
| 24.6527778 | 0.046729 | 0.01102 | 0.010978 | 0.035123 | 0.011959 | 0.022257 | 0.008637 | 0.076506 | 0.042024 | 0.00351 | 0.00303 | 0.005205 | 0.002546 | 0.002143 | 0.006414 | 0.007933 | 0.005459 | 0.024419 | 0.002856 |
| 24.6666667 | 0.027209 | 0.017573 | 0.011254 | 0.032614 | 0.017299 | 0.012928 | 0.00675 | 0.121013 | 0.059088 | 0.003389 | 0.00555 | 0.004601 | 0.00171 | 0.002174 | 0.005367 | 0.007843 | 0.006228 | 0.020166 | 0.004488 |
| 24.6805556 | 0.018938 | 0.02108 | 0.007619 | 0.020584 | 0.024684 | 0.013148 | 0.002642 | 0.139886 | 0.058525 | 0.004439 | 0.007631 | 0.003862 | 0.001803 | 0.002371 | 0.004872 | 0.008096 | 0.00662 | 0.015849 | 0.00535 |
| 24.6944444 | 0.037404 | 0.022447 | 0.009516 | 0.014293 | 0.027051 | 0.014264 | 0.002245 | 0.110848 | 0.037088 | 0.005717 | 0.007543 | 0.002793 | 0.002091 | 0.002166 | 0.004524 | 0.007847 | 0.006196 | 0.011757 | 0.004966 |
| 24.7083333 | 0.050988 | 0.024105 | 0.014926 | 0.011601 | 0.020154 | 0.009474 | 0.004152 | 0.06248 | 0.016945 | 0.006202 | 0.005 | 0.00161 | 0.002013 | 0.001508 | 0.004255 | 0.007061 | 0.004703 | 0.008145 | 0.003539 |
| 24.7222222 | 0.037987 | 0.023752 | 0.017519 | 0.007395 | 0.009449 | 0.007776 | 0.004044 | 0.0591 | 0.019283 | 0.005859 | 0.002166 | 0.001013 | 0.001697 | 0.000784 | 0.003884 | 0.006259 | 0.002839 | 0.006292 | 0.002233 |
| 24.7361111 | 0.019489 | 0.017559 | 0.020977 | 0.009784 | 0.003807 | 0.012232 | 0.003974 | 0.096382 | 0.035243 | 0.005275 | 0.001575 | 0.003755 | 0.001482 | 0.000473 | 0.002965 | 0.005805 | 0.001516 | 0.006638 | 0.00225 |
| 24.75 | 0.021807 | 0.008921 | 0.02366 | 0.015432 | 0.004393 | 0.020007 | 0.005544 | 0.118993 | 0.048992 | 0.004817 | 0.003577 | 0.015668 | 0.001384 | 0.0006 | 0.001513 | 0.005568 | 0.000994 | 0.007612 | 0.003419 |
| 24.7638889 | 0.043648 | 0.004494 | 0.018902 | 0.018484 | 0.005728 | 0.025843 | 0.006698 | 0.099436 | 0.047534 | 0.004474 | 0.00597 | 0.026851 | 0.001354 | 0.000921 | 0.000492 | 0.005185 | 0.001141 | 0.0073 | 0.004511 |
| 24.7777778 | 0.057075 | 0.006886 | 0.010742 | 0.020501 | 0.004471 | 0.019947 | 0.005694 | 0.057885 | 0.029098 | 0.004302 | 0.005885 | 0.021247 | 0.001472 | 0.001485 | 0.000341 | 0.004548 | 0.001772 | 0.005329 | 0.004745 |
| 24.7916667 | 0.046687 | 0.017554 | 0.005069 | 0.019552 | 0.002158 | 0.012134 | 0.00605 | 0.054402 | 0.02277 | 0.004596 | 0.003522 | 0.010899 | 0.001825 | 0.002287 | 0.000841 | 0.00397 | 0.002612 | 0.002672 | 0.004378 |
| 24.8055556 | 0.024974 | 0.038228 | 0.005221 | 0.013765 | 0.001619 | 0.016278 | 0.009631 | 0.09784 | 0.041018 | 0.005266 | 0.001961 | 0.008012 | 0.002247 | 0.002609 | 0.001838 | 0.003452 | 0.003086 | 0.001335 | 0.003833 |
| 24.8194444 | 0.007847 | 0.055786 | 0.010936 | 0.007575 | 0.001594 | 0.015678 | 0.009957 | 0.128972 | 0.053655 | 0.005484 | 0.001457 | 0.006801 | 0.002225 | 0.001801 | 0.002317 | 0.002451 | 0.002546 | 0.002404 | 0.003108 |
| 24.8333333 | 0.002003 | 0.043834 | 0.017402 | 0.004464 | 0.001578 | 0.011292 | 0.009571 | 0.115378 | 0.044538 | 0.004681 | 0.001508 | 0.004684 | 0.001561 | 0.001128 | 0.001721 | 0.001283 | 0.001541 | 0.004206 | 0.002296 |
| 24.8472222 | 0.002595 | 0.024745 | 0.020361 | 0.0043 | 0.003627 | 0.022037 | 0.015595 | 0.074534 | 0.028443 | 0.003382 | 0.003872 | 0.003006 | 0.000711 | 0.001846 | 0.000802 | 0.000918 | 0.001496 | 0.004607 | 0.001903 |
| 24.8611111 | 0.006379 | 0.037106 | 0.016281 | 0.004861 | 0.006757 | 0.038497 | 0.022662 | 0.035657 | 0.017558 | 0.002326 | 0.006661 | 0.002052 | 0.000247 | 0.002774 | 0.000383 | 0.000931 | 0.002318 | 0.00301 | 0.002234 |
| 24.875 | 0.018478 | 0.055714 | 0.009449 | 0.008358 | 0.009209 | 0.048082 | 0.023519 | 0.012566 | 0.010091 | 0.001751 | 0.005606 | 0.001346 | 0.000272 | 0.002884 | 0.000699 | 0.000786 | 0.002894 | 0.001314 | 0.002886 |
| 24.8888889 | 0.043705 | 0.054668 | 0.008203 | 0.024565 | 0.011628 | 0.051059 | 0.017673 | 0.003418 | 0.004473 | 0.001895 | 0.002929 | 0.000654 | 0.000469 | 0.002185 | 0.001325 | 0.001265 | 0.002927 | 0.001346 | 0.003187 |
| 24.9027778 | 0.077889 | 0.046645 | 0.009321 | 0.049545 | 0.018209 | 0.049206 | 0.00928 | 0.001624 | 0.002867 | 0.003078 | 0.0028 | 0.000229 | 0.000465 | 0.001334 | 0.001652 | 0.002373 | 0.002482 | 0.00216 | 0.003008 |
| 24.9166667 | 0.100195 | 0.04109 | 0.008555 | 0.065576 | 0.030383 | 0.041078 | 0.00338 | 0.001982 | 0.002364 | 0.004972 | 0.002937 | 0.000426 | 0.000603 | 0.001198 | 0.001203 | 0.002956 | 0.001633 | 0.002878 | 0.002716 |
| 24.9305556 | 0.096527 | 0.035664 | 0.01136 | 0.063095 | 0.041856 | 0.026123 | 0.002155 | 0.002679 | 0.002005 | 0.006504 | 0.003305 | 0.001 | 0.001316 | 0.001173 | 0.000694 | 0.002521 | 0.001007 | 0.003546 | 0.00242 |
| 24.9444444 | 0.075298 | 0.030065 | 0.017353 | 0.048537 | 0.046655 | 0.012285 | 0.00416 | 0.002652 | 0.003935 | 0.006485 | 0.006245 | 0.001102 | 0.001628 | 0.001438 | 0.001016 | 0.001653 | 0.00134 | 0.003873 | 0.002014 |
| 24.9583333 | 0.0482 | 0.025235 | 0.019317 | 0.03068 | 0.041016 | 0.010836 | 0.005481 | 0.003333 | 0.005989 | 0.004538 | 0.008031 | 0.000774 | 0.001336 | 0.002992 | 0.001157 | 0.001131 | 0.002613 | 0.003428 | 0.001812 |
| 24.9722222 | 0.023807 | 0.019782 | 0.017075 | 0.017113 | 0.025144 | 0.02152 | 0.003849 | 0.005741 | 0.005755 | 0.002033 | 0.006124 | 0.00087 | 0.002017 | 0.004174 | 0.000756 | 0.001096 | 0.004114 | 0.002549 | 0.002327 |
| 24.9861111 | 0.011934 | 0.014414 | 0.01514 | 0.017081 | 0.010721 | 0.030738 | 0.001984 | 0.006881 | 0.003565 | 0.001219 | 0.002965 | 0.001241 | 0.003697 | 0.003276 | 0.000789 | 0.001097 | 0.004645 | 0.002722 | 0.003441 |
| 25 | 0.012495 | 0.010002 | 0.01565 | 0.022546 | 0.006787 | 0.03151 | 0.002604 | 0.005844 | 0.002178 | 0.003083 | 0.001219 | 0.001186 | 0.004288 | 0.001515 | 0.000956 | 0.000878 | 0.003248 | 0.004564 | 0.004355 |
| 25.0138889 | 0.011448 | 0.007183 | 0.017468 | 0.02007 | 0.006443 | 0.028467 | 0.003716 | 0.004618 | 0.003773 | 0.006203 | 0.001613 | 0.00092 | 0.003071 | 0.00069 | 0.000781 | 0.000644 | 0.001687 | 0.006751 | 0.004166 |
| 25.0277778 | 0.013298 | 0.011209 | 0.01896 | 0.012776 | 0.009175 | 0.019809 | 0.003471 | 0.004191 | 0.004871 | 0.006604 | 0.003071 | 0.002167 | 0.001447 | 0.001236 | 0.00204 | 0.003366 | 0.00213 | 0.007824 | 0.004625 |
| 25.0416667 | 0.038531 | 0.024286 | 0.020817 | 0.019468 | 0.022634 | 0.01545 | 0.006159 | 0.006972 | 0.007974 | 0.007602 | 0.007175 | 0.010025 | 0.005328 | 0.005251 | 0.016055 | 0.01844 | 0.008353 | 0.009366 | 0.014257 |
| 25.0555556 | 0.073706 | 0.033391 | 0.022111 | 0.047474 | 0.036061 | 0.032516 | 0.016082 | 0.012292 | 0.019541 | 0.022664 | 0.019468 | 0.031953 | 0.023088 | 0.015814 | 0.057061 | 0.051917 | 0.024654 | 0.015205 | 0.037112 |
| 25.0694444 | 0.083425 | 0.028116 | 0.019974 | 0.064551 | 0.033646 | 0.055986 | 0.026709 | 0.012221 | 0.031429 | 0.04427 | 0.034537 | 0.053683 | 0.038864 | 0.025653 | 0.09226 | 0.077201 | 0.037956 | 0.01917 | 0.052382 |
| 25.0833333 | 0.064054 | 0.018473 | 0.015221 | 0.05105 | 0.02082 | 0.067591 | 0.029018 | 0.006759 | 0.033076 | 0.041703 | 0.031454 | 0.043287 | 0.030832 | 0.02131 | 0.069523 | 0.057682 | 0.028859 | 0.014229 | 0.03993 |
| 25.0972222 | 0.038555 | 0.014048 | 0.010205 | 0.029708 | 0.011359 | 0.061219 | 0.023203 | 0.004334 | 0.02515 | 0.020645 | 0.014636 | 0.017304 | 0.020224 | 0.010761 | 0.029974 | 0.024034 | 0.012671 | 0.010557 | 0.017385 |
| 25.1111111 | 0.019396 | 0.010127 | 0.005446 | 0.015892 | 0.006704 | 0.037112 | 0.014399 | 0.005555 | 0.013798 | 0.012908 | 0.007613 | 0.012355 | 0.025232 | 0.010445 | 0.02894 | 0.019783 | 0.011266 | 0.013396 | 0.008176 |
| 25.125 | 0.015624 | 0.007256 | 0.003924 | 0.0073 | 0.003437 | 0.01486 | 0.007409 | 0.005982 | 0.006274 | 0.016402 | 0.007652 | 0.018066 | 0.028299 | 0.013325 | 0.037964 | 0.025458 | 0.013268 | 0.014653 | 0.008756 |
| 25.1388889 | 0.030847 | 0.013803 | 0.008036 | 0.004144 | 0.002408 | 0.009965 | 0.006649 | 0.005339 | 0.006984 | 0.017236 | 0.008633 | 0.020206 | 0.022303 | 0.01072 | 0.038226 | 0.025838 | 0.011347 | 0.012104 | 0.009756 |
| 25.1527778 | 0.045002 | 0.024087 | 0.01324 | 0.007905 | 0.003845 | 0.014682 | 0.010814 | 0.005481 | 0.013492 | 0.015694 | 0.012424 | 0.020526 | 0.015341 | 0.007175 | 0.035818 | 0.025634 | 0.010865 | 0.009713 | 0.010724 |
| 25.1666667 | 0.043503 | 0.025057 | 0.015144 | 0.015824 | 0.005885 | 0.017235 | 0.013867 | 0.00653 | 0.019201 | 0.014419 | 0.014064 | 0.019647 | 0.0105 | 0.006188 | 0.027103 | 0.021364 | 0.011363 | 0.008271 | 0.010407 |
| 25.1805556 | 0.032662 | 0.016075 | 0.013728 | 0.023982 | 0.006523 | 0.015639 | 0.013683 | 0.006417 | 0.018353 | 0.011923 | 0.010245 | 0.018999 | 0.006168 | 0.006766 | 0.014279 | 0.012814 | 0.01145 | 0.006324 | 0.007997 |
| 25.1944444 | 0.019398 | 0.006023 | 0.009115 | 0.028354 | 0.005463 | 0.012886 | 0.01042 | 0.004617 | 0.010499 | 0.006446 | 0.006527 | 0.017332 | 0.003088 | 0.006218 | 0.005509 | 0.004621 | 0.009868 | 0.005333 | 0.007652 |
| 25.2083333 | 0.009767 | 0.001652 | 0.004624 | 0.027206 | 0.004436 | 0.013244 | 0.005436 | 0.003356 | 0.002748 | 0.00218 | 0.005754 | 0.012465 | 0.003253 | 0.003903 | 0.004369 | 0.002483 | 0.006259 | 0.006172 | 0.007233 |
| 25.2222222 | 0.011776 | 0.002431 | 0.004168 | 0.024207 | 0.00612 | 0.01913 | 0.002484 | 0.003433 | 0.003075 | 0.00296 | 0.006337 | 0.006246 | 0.006534 | 0.002316 | 0.007748 | 0.007038 | 0.002698 | 0.009785 | 0.00619 |
| 25.2361111 | 0.024856 | 0.004355 | 0.004266 | 0.022965 | 0.012136 | 0.027703 | 0.003948 | 0.00377 | 0.012298 | 0.006728 | 0.009232 | 0.002921 | 0.011266 | 0.002221 | 0.012529 | 0.01239 | 0.00124 | 0.011405 | 0.008948 |
| 25.25 | 0.039771 | 0.00624 | 0.00461 | 0.023645 | 0.020029 | 0.034817 | 0.008338 | 0.00406 | 0.023224 | 0.009402 | 0.008939 | 0.002971 | 0.012591 | 0.001532 | 0.012967 | 0.013274 | 0.000914 | 0.01227 | 0.010251 |
| 25.2638889 | 0.043977 | 0.008345 | 0.008015 | 0.023221 | 0.02296 | 0.037237 | 0.01159 | 0.004524 | 0.027091 | 0.010287 | 0.004432 | 0.004453 | 0.009732 | 0.000602 | 0.008552 | 0.010579 | 0.001199 | 0.006445 | 0.007313 |
| 25.2777778 | 0.029315 | 0.0091 | 0.009415 | 0.01716 | 0.015582 | 0.032777 | 0.0106 | 0.0048 | 0.021936 | 0.011033 | 0.003873 | 0.005878 | 0.00796 | 0.000504 | 0.005811 | 0.008571 | 0.002239 | 0.004833 | 0.005191 |
| 25.2916667 | 0.012845 | 0.008387 | 0.005827 | 0.01008 | 0.007728 | 0.024929 | 0.006015 | 0.004754 | 0.013112 | 0.012202 | 0.009742 | 0.004725 | 0.010084 | 0.001135 | 0.007305 | 0.009314 | 0.002686 | 0.009293 | 0.005914 |
| 25.3055556 | 0.01081 | 0.006712 | 0.002632 | 0.010367 | 0.009246 | 0.017552 | 0.00256 | 0.004536 | 0.00655 | 0.012706 | 0.014784 | 0.002287 | 0.012667 | 0.002733 | 0.008949 | 0.010607 | 0.003373 | 0.012838 | 0.006696 |
| 25.3194444 | 0.010064 | 0.004309 | 0.002699 | 0.010688 | 0.010177 | 0.010975 | 0.002013 | 0.004024 | 0.005147 | 0.01205 | 0.012735 | 0.002704 | 0.011215 | 0.003987 | 0.006904 | 0.010055 | 0.004715 | 0.010392 | 0.005218 |
| 25.3333333 | 0.009214 | 0.003341 | 0.003373 | 0.007951 | 0.005962 | 0.005879 | 0.002822 | 0.003377 | 0.00815 | 0.011314 | 0.008045 | 0.005182 | 0.006433 | 0.00327 | 0.003883 | 0.009141 | 0.004131 | 0.006676 | 0.003737 |
| 25.3472222 | 0.017236 | 0.003439 | 0.005963 | 0.011193 | 0.005134 | 0.002559 | 0.006439 | 0.003442 | 0.012382 | 0.010772 | 0.006128 | 0.006568 | 0.004352 | 0.002661 | 0.002871 | 0.009948 | 0.002701 | 0.005461 | 0.003445 |
| 25.3611111 | 0.024285 | 0.004311 | 0.010287 | 0.015731 | 0.008561 | 0.001665 | 0.010927 | 0.004547 | 0.015107 | 0.010696 | 0.005941 | 0.005334 | 0.008007 | 0.004412 | 0.002868 | 0.011137 | 0.004009 | 0.00438 | 0.002535 |
| 25.375 | 0.019664 | 0.006403 | 0.01013 | 0.012531 | 0.00869 | 0.004926 | 0.011549 | 0.005466 | 0.014317 | 0.011801 | 0.005174 | 0.006149 | 0.010431 | 0.006502 | 0.002735 | 0.009558 | 0.007226 | 0.003739 | 0.002507 |
| 25.3888889 | 0.009175 | 0.005959 | 0.005892 | 0.005411 | 0.004982 | 0.010066 | 0.008329 | 0.005158 | 0.010361 | 0.012912 | 0.007199 | 0.013014 | 0.007009 | 0.007534 | 0.004414 | 0.005336 | 0.009664 | 0.007312 | 0.00543 |
| 25.4027778 | 0.003721 | 0.003202 | 0.003559 | 0.002788 | 0.002658 | 0.011274 | 0.004698 | 0.004329 | 0.007608 | 0.013086 | 0.010917 | 0.016891 | 0.004754 | 0.007381 | 0.006503 | 0.0029 | 0.009241 | 0.009863 | 0.006441 |
| 25.4166667 | 0.005228 | 0.001872 | 0.003275 | 0.00643 | 0.002365 | 0.007294 | 0.003451 | 0.003916 | 0.010048 | 0.014204 | 0.010411 | 0.012152 | 0.007409 | 0.005431 | 0.006188 | 0.006472 | 0.005641 | 0.009197 | 0.006521 |
| 25.4305556 | 0.010404 | 0.001659 | 0.004087 | 0.011324 | 0.004401 | 0.002582 | 0.004619 | 0.003263 | 0.015296 | 0.015829 | 0.009245 | 0.007653 | 0.008385 | 0.00373 | 0.005703 | 0.012635 | 0.002199 | 0.011295 | 0.009578 |
| 25.4444444 | 0.012275 | 0.001972 | 0.007702 | 0.012518 | 0.008664 | 0.000721 | 0.006321 | 0.001984 | 0.016117 | 0.015845 | 0.009475 | 0.006827 | 0.007789 | 0.004447 | 0.004994 | 0.01433 | 0.001828 | 0.012082 | 0.009745 |
| 25.4583333 | 0.007816 | 0.002343 | 0.0109 | 0.00878 | 0.012321 | 0.001044 | 0.006785 | 0.000924 | 0.01029 | 0.017692 | 0.013076 | 0.005 | 0.007812 | 0.005916 | 0.006747 | 0.014593 | 0.003108 | 0.01351 | 0.009945 |
| 25.4722222 | 0.002636 | 0.003435 | 0.009377 | 0.004297 | 0.012647 | 0.003282 | 0.005669 | 0.001264 | 0.004151 | 0.022905 | 0.020681 | 0.00349 | 0.008376 | 0.006319 | 0.012451 | 0.016753 | 0.003812 | 0.019461 | 0.016198 |
| 25.4861111 | 0.001666 | 0.006554 | 0.005539 | 0.002322 | 0.009048 | 0.008145 | 0.003661 | 0.003177 | 0.002421 | 0.026377 | 0.022115 | 0.005852 | 0.01363 | 0.006561 | 0.014206 | 0.015118 | 0.005056 | 0.019911 | 0.021539 |

| | | | | | | | | | | | | | | | | | | | |
|---|---|---|---|---|---|---|---|---|---|---|---|---|---|---|---|---|---|---|---|
| 25.5 | 0.005212 | 0.00687 | 0.004059 | 0.002941 | 0.004278 | 0.012989 | 0.002223 | 0.004264 | 0.003158 | 0.025488 | 0.015719 | 0.010036 | 0.018553 | 0.007159 | 0.010693 | 0.009564 | 0.007218 | 0.01337 | 0.020955 |
| 25.5138889 | 0.011336 | 0.00349 | 0.00384 | 0.00744 | 0.001594 | 0.016315 | 0.003164 | 0.003754 | 0.007398 | 0.022035 | 0.011163 | 0.011018 | 0.016171 | 0.007069 | 0.009017 | 0.006353 | 0.008672 | 0.009669 | 0.018379 |
| 25.5277778 | 0.019221 | 0.004023 | 0.005491 | 0.012989 | 0.002648 | 0.020515 | 0.00712 | 0.003384 | 0.016585 | 0.021296 | 0.0137 | 0.009551 | 0.009405 | 0.006815 | 0.010993 | 0.007625 | 0.010312 | 0.014301 | 0.02093 |
| 25.5416667 | 0.029902 | 0.012324 | 0.016092 | 0.017202 | 0.010112 | 0.027729 | 0.014486 | 0.003827 | 0.024315 | 0.031577 | 0.022214 | 0.018822 | 0.007488 | 0.010337 | 0.017274 | 0.013957 | 0.016827 | 0.03129 | 0.035986 |
| 25.5555556 | 0.044443 | 0.02477 | 0.037616 | 0.02578 | 0.023389 | 0.0358 | 0.023544 | 0.004595 | 0.02606 | 0.048194 | 0.02852 | 0.045082 | 0.016901 | 0.01973 | 0.032228 | 0.025598 | 0.029116 | 0.050613 | 0.056434 |
| 25.5694444 | 0.059293 | 0.037788 | 0.060052 | 0.039946 | 0.030498 | 0.037829 | 0.029368 | 0.008312 | 0.02371 | 0.054161 | 0.024882 | 0.06417 | 0.031363 | 0.027323 | 0.047513 | 0.03343 | 0.038417 | 0.051972 | 0.06166 |
| 25.5833333 | 0.061406 | 0.049415 | 0.066896 | 0.047034 | 0.023083 | 0.031083 | 0.029535 | 0.019647 | 0.020581 | 0.047147 | 0.015366 | 0.058189 | 0.041029 | 0.025776 | 0.051889 | 0.030142 | 0.037477 | 0.042069 | 0.049677 |
| 25.5972222 | 0.039711 | 0.050974 | 0.052427 | 0.034778 | 0.011656 | 0.018141 | 0.022478 | 0.034184 | 0.016287 | 0.034076 | 0.006671 | 0.040774 | 0.039849 | 0.01966 | 0.050608 | 0.023331 | 0.027714 | 0.035774 | 0.034532 |
| 25.6111111 | 0.023673 | 0.038827 | 0.026886 | 0.021346 | 0.011446 | 0.010538 | 0.011979 | 0.042879 | 0.009851 | 0.018707 | 0.001994 | 0.023476 | 0.02701 | 0.01459 | 0.045244 | 0.018209 | 0.01515 | 0.029141 | 0.021788 |
| 25.625 | 0.040873 | 0.023875 | 0.012705 | 0.030263 | 0.021592 | 0.018748 | 0.008519 | 0.045717 | 0.007265 | 0.012158 | 0.002827 | 0.011166 | 0.012869 | 0.009888 | 0.032446 | 0.012815 | 0.008492 | 0.018868 | 0.011753 |
| 25.6388889 | 0.058215 | 0.012998 | 0.020532 | 0.044398 | 0.026563 | 0.028081 | 0.010802 | 0.046995 | 0.009226 | 0.017022 | 0.007346 | 0.00861 | 0.009708 | 0.005731 | 0.016833 | 0.006728 | 0.006831 | 0.01049 | 0.005011 |
| 25.6527778 | 0.053707 | 0.008557 | 0.027102 | 0.047638 | 0.020509 | 0.027298 | 0.009088 | 0.045134 | 0.008401 | 0.019354 | 0.01093 | 0.008322 | 0.017179 | 0.003315 | 0.008997 | 0.004496 | 0.007475 | 0.006594 | 0.0031 |
| 25.6666667 | 0.035531 | 0.01556 | 0.016894 | 0.039829 | 0.010795 | 0.024335 | 0.004029 | 0.03763 | 0.010616 | 0.015223 | 0.00976 | 0.0072 | 0.023326 | 0.002774 | 0.01623 | 0.010205 | 0.01472 | 0.004111 | 0.007684 |
| 25.6805556 | 0.022416 | 0.029411 | 0.005399 | 0.02454 | 0.006824 | 0.019407 | 0.001864 | 0.02522 | 0.022361 | 0.010574 | 0.009646 | 0.011879 | 0.019085 | 0.003129 | 0.028404 | 0.017961 | 0.017739 | 0.003436 | 0.014085 |
| 25.6944444 | 0.035881 | 0.04068 | 0.005508 | 0.021499 | 0.012111 | 0.010232 | 0.002096 | 0.021952 | 0.02972 | 0.008595 | 0.015308 | 0.017036 | 0.01046 | 0.003605 | 0.033427 | 0.022351 | 0.010644 | 0.006396 | 0.016526 |
| 25.7083333 | 0.049949 | 0.043351 | 0.008201 | 0.028177 | 0.018067 | 0.006115 | 0.001679 | 0.038576 | 0.020679 | 0.011774 | 0.018191 | 0.013351 | 0.008632 | 0.007989 | 0.030624 | 0.021919 | 0.008874 | 0.009223 | 0.016181 |
| 25.7222222 | 0.034909 | 0.033633 | 0.01061 | 0.022195 | 0.016442 | 0.007709 | 0.001543 | 0.056704 | 0.01073 | 0.01871 | 0.013061 | 0.007488 | 0.010877 | 0.015435 | 0.021073 | 0.014567 | 0.018988 | 0.008558 | 0.015434 |
| 25.7361111 | 0.01863 | 0.017579 | 0.020958 | 0.012166 | 0.009024 | 0.005103 | 0.001704 | 0.061752 | 0.016972 | 0.022557 | 0.006354 | 0.012951 | 0.008387 | 0.020656 | 0.011268 | 0.009359 | 0.028781 | 0.005617 | 0.012884 |
| 25.75 | 0.030971 | 0.012226 | 0.026193 | 0.01687 | 0.004503 | 0.002998 | 0.001384 | 0.052577 | 0.027876 | 0.016964 | 0.00451 | 0.023661 | 0.005838 | 0.01944 | 0.011062 | 0.014705 | 0.022958 | 0.002887 | 0.008825 |
| 25.7638889 | 0.055405 | 0.025931 | 0.016605 | 0.030827 | 0.010698 | 0.010288 | 0.001618 | 0.032492 | 0.02991 | 0.009735 | 0.005085 | 0.025651 | 0.008985 | 0.011916 | 0.016952 | 0.021283 | 0.012097 | 0.001229 | 0.009964 |
| 25.7777778 | 0.067874 | 0.043813 | 0.009231 | 0.042297 | 0.023854 | 0.020511 | 0.003368 | 0.014587 | 0.020794 | 0.014058 | 0.003949 | 0.017388 | 0.010373 | 0.008692 | 0.018541 | 0.021089 | 0.01721 | 0.000477 | 0.016714 |
| 25.7916667 | 0.05521 | 0.049327 | 0.013613 | 0.04245 | 0.028243 | 0.02926 | 0.006269 | 0.009641 | 0.011892 | 0.023489 | 0.002175 | 0.010199 | 0.00629 | 0.013047 | 0.013812 | 0.014674 | 0.028681 | 0.000633 | 0.018051 |
| 25.8055556 | 0.0292 | 0.040745 | 0.016482 | 0.030429 | 0.018012 | 0.039936 | 0.006951 | 0.014674 | 0.017731 | 0.026387 | 0.002376 | 0.014543 | 0.004179 | 0.014319 | 0.00738 | 0.007153 | 0.026746 | 0.000836 | 0.011243 |
| 25.8194444 | 0.022349 | 0.022767 | 0.012303 | 0.017236 | 0.009522 | 0.043458 | 0.004645 | 0.016522 | 0.026971 | 0.021754 | 0.00513 | 0.020929 | 0.006775 | 0.008769 | 0.005659 | 0.006189 | 0.014258 | 0.000541 | 0.004598 |
| 25.8333333 | 0.026622 | 0.011855 | 0.012397 | 0.010301 | 0.0137 | 0.03475 | 0.002782 | 0.010105 | 0.024922 | 0.012436 | 0.00689 | 0.019173 | 0.007742 | 0.005378 | 0.010716 | 0.011656 | 0.009206 | 0.000585 | 0.002944 |
| 25.8472222 | 0.019853 | 0.022234 | 0.018838 | 0.011588 | 0.020237 | 0.024884 | 0.005781 | 0.005412 | 0.015499 | 0.008182 | 0.005824 | 0.011456 | 0.005294 | 0.008775 | 0.018957 | 0.017681 | 0.014873 | 0.001482 | 0.004417 |
| 25.8611111 | 0.014448 | 0.041146 | 0.025449 | 0.017327 | 0.020386 | 0.023488 | 0.01692 | 0.009939 | 0.014744 | 0.016193 | 0.007273 | 0.009134 | 0.005172 | 0.011393 | 0.027513 | 0.023366 | 0.017365 | 0.002461 | 0.00644 |
| 25.875 | 0.01813 | 0.047717 | 0.030544 | 0.019994 | 0.015593 | 0.027335 | 0.027826 | 0.016236 | 0.028709 | 0.027872 | 0.013088 | 0.019584 | 0.008841 | 0.0091 | 0.034146 | 0.030319 | 0.010741 | 0.002397 | 0.009335 |
| 25.8888889 | 0.022324 | 0.036062 | 0.031288 | 0.018549 | 0.009076 | 0.026412 | 0.026374 | 0.015736 | 0.039447 | 0.035432 | 0.016802 | 0.028415 | 0.011421 | 0.006362 | 0.035925 | 0.034818 | 0.00353 | 0.001807 | 0.013096 |
| 25.9027778 | 0.027128 | 0.017882 | 0.029113 | 0.016061 | 0.010042 | 0.022117 | 0.017526 | 0.00912 | 0.034153 | 0.039179 | 0.014579 | 0.0256 | 0.010383 | 0.00837 | 0.03283 | 0.032026 | 0.003207 | 0.002829 | 0.015731 |
| 25.9166667 | 0.032495 | 0.006146 | 0.028023 | 0.015793 | 0.026281 | 0.020453 | 0.012209 | 0.003152 | 0.019204 | 0.039146 | 0.010204 | 0.018949 | 0.007619 | 0.015231 | 0.027605 | 0.022198 | 0.00722 | 0.004694 | 0.016518 |
| 25.9305556 | 0.03213 | 0.002792 | 0.023234 | 0.018832 | 0.041146 | 0.018639 | 0.01148 | 0.001684 | 0.008106 | 0.035277 | 0.009567 | 0.013239 | 0.004911 | 0.020615 | 0.022013 | 0.010562 | 0.010391 | 0.003976 | 0.017375 |
| 25.9444444 | 0.026055 | 0.004952 | 0.012552 | 0.02326 | 0.034288 | 0.012049 | 0.011884 | 0.00148 | 0.007394 | 0.027695 | 0.013706 | 0.008459 | 0.002803 | 0.020442 | 0.015865 | 0.003648 | 0.010407 | 0.002713 | 0.019012 |
| 25.9583333 | 0.020773 | 0.008462 | 0.005478 | 0.026642 | 0.017863 | 0.007736 | 0.010108 | 0.001118 | 0.00892 | 0.017167 | 0.018007 | 0.005065 | 0.001913 | 0.016719 | 0.008684 | 0.001993 | 0.008055 | 0.005066 | 0.019705 |
| 25.9722222 | 0.017562 | 0.010527 | 0.006869 | 0.024452 | 0.013222 | 0.015524 | 0.005797 | 0.002354 | 0.007024 | 0.00818 | 0.015769 | 0.002603 | 0.00213 | 0.012554 | 0.003372 | 0.002313 | 0.005339 | 0.007997 | 0.017724 |
| 25.9861111 | 0.011353 | 0.010854 | 0.008721 | 0.015047 | 0.016233 | 0.02843 | 0.003388 | 0.004694 | 0.005296 | 0.005868 | 0.009562 | 0.001689 | 0.00202 | 0.008984 | 0.002807 | 0.004952 | 0.003884 | 0.007571 | 0.01244 |
| 26 | 0.007515 | 0.010719 | 0.00641 | 0.007144 | 0.013933 | 0.035607 | 0.004752 | 0.00618 | 0.006365 | 0.007578 | 0.00881 | 0.002523 | 0.001171 | 0.005461 | 0.004198 | 0.00814 | 0.003253 | 0.004974 | 0.005922 |
| 26.0138889 | 0.014808 | 0.011058 | 0.004135 | 0.00832 | 0.008676 | 0.035095 | 0.006049 | 0.005568 | 0.009348 | 0.007825 | 0.010442 | 0.002331 | 0.000878 | 0.002419 | 0.004069 | 0.009115 | 0.002464 | 0.00293 | 0.002394 |
| 26.0277778 | 0.020585 | 0.009916 | 0.004255 | 0.010667 | 0.005751 | 0.026484 | 0.004456 | 0.00348 | 0.009384 | 0.005702 | 0.007854 | 0.002958 | 0.003615 | 0.002219 | 0.004766 | 0.011438 | 0.001778 | 0.00495 | 0.003263 |
| 26.0416667 | 0.021449 | 0.007976 | 0.005292 | 0.012535 | 0.008806 | 0.018098 | 0.005056 | 0.003185 | 0.010843 | 0.010133 | 0.008498 | 0.012409 | 0.016554 | 0.008852 | 0.014717 | 0.022388 | 0.005246 | 0.018918 | 0.012656 |
| 26.0555556 | 0.041313 | 0.011801 | 0.008831 | 0.027381 | 0.020684 | 0.030153 | 0.01446 | 0.007795 | 0.02847 | 0.028297 | 0.017859 | 0.03154 | 0.040653 | 0.021224 | 0.035781 | 0.041792 | 0.016875 | 0.045275 | 0.036852 |
| 26.0694444 | 0.073686 | 0.019249 | 0.013433 | 0.047391 | 0.037071 | 0.053687 | 0.024401 | 0.015307 | 0.056425 | 0.04687 | 0.0248 | 0.043244 | 0.053959 | 0.026868 | 0.043865 | 0.049396 | 0.025676 | 0.060964 | 0.054336 |
| 26.0833333 | 0.077492 | 0.02024 | 0.0149 | 0.052131 | 0.047997 | 0.055511 | 0.026241 | 0.018323 | 0.071951 | 0.041476 | 0.020961 | 0.03101 | 0.038631 | 0.018682 | 0.029359 | 0.031282 | 0.019624 | 0.050003 | 0.040326 |
| 26.0972222 | 0.048219 | 0.014438 | 0.018403 | 0.046562 | 0.042298 | 0.035891 | 0.030936 | 0.013238 | 0.064388 | 0.019565 | 0.011287 | 0.01201 | 0.018312 | 0.00763 | 0.020797 | 0.011061 | 0.010795 | 0.025896 | 0.016829 |
| 26.1111111 | 0.018356 | 0.007789 | 0.023862 | 0.035272 | 0.024799 | 0.015262 | 0.030657 | 0.007042 | 0.037438 | 0.012486 | 0.006574 | 0.007509 | 0.018664 | 0.003919 | 0.024317 | 0.007844 | 0.009917 | 0.012187 | 0.010925 |
| 26.125 | 0.006967 | 0.003981 | 0.018728 | 0.017007 | 0.018629 | 0.004681 | 0.013331 | 0.006886 | 0.017604 | 0.013045 | 0.0084 | 0.010412 | 0.025193 | 0.004266 | 0.024697 | 0.008996 | 0.008755 | 0.014283 | 0.013537 |
| 26.1388889 | 0.011589 | 0.003873 | 0.007788 | 0.006512 | 0.029289 | 0.004925 | 0.004248 | 0.009409 | 0.025647 | 0.009125 | 0.009328 | 0.013835 | 0.022385 | 0.004421 | 0.016519 | 0.006222 | 0.006739 | 0.014889 | 0.011684 |
| 26.1527778 | 0.021191 | 0.004069 | 0.006166 | 0.007219 | 0.036074 | 0.007775 | 0.01029 | 0.008978 | 0.039997 | 0.010275 | 0.010443 | 0.014895 | 0.01582 | 0.004149 | 0.00716 | 0.004216 | 0.007632 | 0.012677 | 0.010189 |
| 26.1666667 | 0.028554 | 0.003055 | 0.00863 | 0.012218 | 0.030594 | 0.008285 | 0.015095 | 0.006272 | 0.041673 | 0.011045 | 0.010433 | 0.009749 | 0.011991 | 0.002724 | 0.005201 | 0.003478 | 0.006461 | 0.01156 | 0.009098 |
| 26.1805556 | 0.030961 | 0.002835 | 0.007154 | 0.018024 | 0.022806 | 0.006709 | 0.015209 | 0.004317 | 0.034493 | 0.006801 | 0.008738 | 0.005132 | 0.009133 | 0.001147 | 0.006855 | 0.002645 | 0.004399 | 0.008624 | 0.006103 |
| 26.1944444 | 0.028716 | 0.002316 | 0.004258 | 0.02026 | 0.019677 | 0.004115 | 0.012096 | 0.003782 | 0.024638 | 0.003821 | 0.007234 | 0.005786 | 0.006668 | 0.000828 | 0.008104 | 0.002608 | 0.00605 | 0.007515 | 0.004916 |
| 26.2083333 | 0.025405 | 0.00141 | 0.003172 | 0.018589 | 0.018212 | 0.002534 | 0.008398 | 0.003447 | 0.017278 | 0.00339 | 0.003595 | 0.006707 | 0.007922 | 0.000823 | 0.005962 | 0.00181 | 0.007129 | 0.006793 | 0.003028 |
| 26.2222222 | 0.022766 | 0.002878 | 0.005744 | 0.015538 | 0.013015 | 0.003819 | 0.007459 | 0.003065 | 0.01463 | 0.003233 | 0.001579 | 0.005364 | 0.008664 | 0.000542 | 0.002234 | 0.000484 | 0.005035 | 0.005667 | 0.001772 |
| 26.2361111 | 0.021689 | 0.007327 | 0.011141 | 0.013525 | 0.006313 | 0.009202 | 0.011147 | 0.003436 | 0.016874 | 0.002538 | 0.001376 | 0.003756 | 0.005143 | 0.000633 | 0.001851 | 0.000266 | 0.004318 | 0.006 | 0.002657 |
| 26.25 | 0.022123 | 0.011704 | 0.015385 | 0.013293 | 0.003243 | 0.016533 | 0.016398 | 0.004435 | 0.022953 | 0.004111 | 0.001547 | 0.003176 | 0.002134 | 0.001406 | 0.002892 | 0.000895 | 0.00479 | 0.008171 | 0.002008 |
| 26.2638889 | 0.021516 | 0.011987 | 0.015147 | 0.014155 | 0.005169 | 0.019879 | 0.018005 | 0.005235 | 0.029574 | 0.008747 | 0.005884 | 0.003265 | 0.002636 | 0.001974 | 0.00418 | 0.002825 | 0.004334 | 0.008736 | 0.003537 |
| 26.2777778 | 0.018852 | 0.008918 | 0.01157 | 0.015439 | 0.008836 | 0.018022 | 0.015566 | 0.005613 | 0.032297 | 0.012914 | 0.012037 | 0.004322 | 0.003485 | 0.002118 | 0.007921 | 0.005038 | 0.004953 | 0.008207 | 0.009904 |
| 26.2916667 | 0.016158 | 0.007119 | 0.007818 | 0.016068 | 0.009014 | 0.016099 | 0.011662 | 0.005838 | 0.029423 | 0.012596 | 0.011665 | 0.006087 | 0.002656 | 0.002775 | 0.010196 | 0.005095 | 0.004304 | 0.008645 | 0.0136 |
| 26.3055556 | 0.014303 | 0.007595 | 0.005161 | 0.014832 | 0.005254 | 0.016303 | 0.007191 | 0.005582 | 0.0223 | 0.007187 | 0.006653 | 0.006168 | 0.002623 | 0.003446 | 0.007015 | 0.002901 | 0.001963 | 0.006361 | 0.010564 |
| 26.3194444 | 0.012755 | 0.006698 | 0.003923 | 0.012202 | 0.002671 | 0.015883 | 0.003393 | 0.004304 | 0.014333 | 0.002532 | 0.005342 | 0.003974 | 0.00419 | 0.002976 | 0.003047 | 0.000967 | 0.002075 | 0.005098 | 0.004988 |
| 26.3333333 | 0.011377 | 0.00415 | 0.004673 | 0.009755 | 0.003509 | 0.012733 | 0.002482 | 0.002526 | 0.010879 | 0.00464 | 0.005899 | 0.001745 | 0.004297 | 0.001662 | 0.002037 | 0.002154 | 0.004202 | 0.00675 | 0.002046 |
| 26.3472222 | 0.008054 | 0.003755 | 0.005492 | 0.007312 | 0.004354 | 0.008108 | 0.004029 | 0.001509 | 0.014673 | 0.011616 | 0.006594 | 0.002236 | 0.003552 | 0.001048 | 0.002284 | 0.006059 | 0.005353 | 0.005998 | 0.002939 |
| 26.3611111 | 0.005446 | 0.004137 | 0.004338 | 0.004665 | 0.003293 | 0.004516 | 0.004563 | 0.002147 | 0.019044 | 0.014241 | 0.007278 | 0.005045 | 0.003556 | 0.002129 | 0.002682 | 0.0078 | 0.006656 | 0.005637 | 0.003234 |
| 26.375 | 0.009441 | 0.003407 | 0.003628 | 0.00483 | 0.003871 | 0.002499 | 0.003058 | 0.004026 | 0.014575 | 0.009411 | 0.005562 | 0.005924 | 0.002213 | 0.003737 | 0.002685 | 0.005392 | 0.008772 | 0.007033 | 0.001665 |
| 26.3888889 | 0.013409 | 0.004984 | 0.00342 | 0.00763 | 0.008234 | 0.001729 | 0.001905 | 0.00555 | 0.008816 | 0.004429 | 0.005344 | 0.00398 | 0.000491 | 0.00416 | 0.001964 | 0.002466 | 0.007081 | 0.005851 | 0.001425 |
| 26.4027778 | 0.009852 | 0.008535 | 0.003893 | 0.007224 | 0.01106 | 0.002805 | 0.002205 | 0.005746 | 0.00863 | 0.005921 | 0.007525 | 0.002853 | 0.000313 | 0.003574 | 0.002053 | 0.00142 | 0.00286 | 0.005556 | 0.001727 |
| 26.4166667 | 0.007289 | 0.010059 | 0.009409 | 0.004785 | 0.008658 | 0.005519 | 0.004702 | 0.004663 | 0.009958 | 0.014549 | 0.009956 | 0.002457 | 0.002155 | 0.002732 | 0.002711 | 0.002835 | 0.000874 | 0.006212 | 0.001104 |
| 26.4305556 | 0.01316 | 0.007212 | 0.016283 | 0.007454 | 0.004281 | 0.008885 | 0.009555 | 0.002619 | 0.019556 | 0.023739 | 0.012729 | 0.002069 | 0.005747 | 0.002121 | 0.003733 | 0.006508 | 0.003116 | 0.005201 | 0.001515 |
| 26.4444444 | 0.018806 | 0.003667 | 0.017172 | 0.011946 | 0.002299 | 0.011615 | 0.014418 | 0.001145 | 0.031581 | 0.024637 | 0.013556 | 0.003768 | 0.008226 | 0.002888 | 0.00685 | 0.009954 | 0.010416 | 0.00839 | 0.003679 |
| 26.4583333 | 0.016612 | 0.003006 | 0.01262 | 0.011619 | 0.003717 | 0.010771 | 0.017636 | 0.000944 | 0.029374 | 0.018377 | 0.012594 | 0.007302 | 0.008042 | 0.005419 | 0.009756 | 0.011536 | 0.018084 | 0.014205 | 0.006307 |
| 26.4722222 | 0.011049 | 0.004176 | 0.008459 | 0.008267 | 0.006341 | 0.007758 | 0.018507 | 0.001902 | 0.01622 | 0.013878 | 0.01079 | 0.010707 | 0.006378 | 0.008582 | 0.008856 | 0.011668 | 0.021012 | 0.013389 | 0.00696 |
| 26.4861111 | 0.007312 | 0.007721 | 0.00878 | 0.004859 | 0.008084 | 0.008745 | 0.016015 | 0.004429 | 0.006852 | 0.015584 | 0.009795 | 0.011591 | 0.005704 | 0.010414 | 0.007832 | 0.012163 | 0.018912 | 0.008519 | 0.006603 |
| 26.5 | 0.00525 | 0.009313 | 0.011792 | 0.002403 | 0.00698 | 0.0121 | 0.010532 | 0.005805 | 0.004697 | 0.020233 | 0.013052 | 0.007703 | 0.007887 | 0.00909 | 0.011947 | 0.014253 | 0.013263 | 0.009825 | 0.009786 |
| 26.5138889 | 0.004363 | 0.005859 | 0.01252 | 0.002356 | 0.004184 | 0.012622 | 0.004711 | 0.004013 | 0.007242 | 0.022575 | 0.015327 | 0.002533 | 0.010532 | 0.006025 | 0.016099 | 0.015556 | 0.008546 | 0.014092 | 0.01376 |
| 26.5277778 | 0.007732 | 0.002778 | 0.009476 | 0.007238 | 0.002669 | 0.013128 | 0.003752 | 0.002224 | 0.020606 | 0.023033 | 0.012033 | 0.00275 | 0.013222 | 0.006196 | 0.01679 | 0.017194 | 0.010104 | 0.015254 | 0.016569 |
| 26.5416667 | 0.021285 | 0.003123 | 0.009024 | 0.020849 | 0.003553 | 0.021628 | 0.013637 | 0.005291 | 0.038124 | 0.03018 | 0.010874 | 0.012949 | 0.024136 | 0.01551 | 0.026012 | 0.029501 | 0.018847 | 0.020923 | 0.027002 |
| 26.5555556 | 0.046355 | 0.007653 | 0.021127 | 0.043767 | 0.007367 | 0.037489 | 0.033795 | 0.01354 | 0.045891 | 0.045371 | 0.01539 | 0.031202 | 0.04406 | 0.032116 | 0.045304 | 0.050302 | 0.030273 | 0.031404 | 0.042902 |
| 26.5694444 | 0.068793 | 0.015778 | 0.044649 | 0.066905 | 0.00804 | 0.04776 | 0.055599 | 0.019523 | 0.045638 | 0.053315 | 0.019122 | 0.043684 | 0.057425 | 0.044357 | 0.053948 | 0.058361 | 0.038159 | 0.033807 | 0.047904 |
| 26.5833333 | 0.075505 | 0.023111 | 0.063933 | 0.079096 | 0.007394 | 0.042857 | 0.068627 | 0.018766 | 0.040963 | 0.044938 | 0.016911 | 0.04321 | 0.056743 | 0.047564 | 0.04792 | 0.049592 | 0.040844 | 0.025855 | 0.038065 |
| 26.5972222 | 0.065814 | 0.024279 | 0.062875 | 0.073967 | 0.017006 | 0.026509 | 0.06192 | 0.014979 | 0.029318 | 0.02596 | 0.009954 | 0.036213 | 0.046447 | 0.04307 | 0.043349 | 0.038238 | 0.037247 | 0.016665 | 0.023187 |
| 26.6111111 | 0.043098 | 0.016507 | 0.03893 | 0.049268 | 0.031048 | 0.011427 | 0.035188 | 0.010132 | 0.01593 | 0.013674 | 0.006944 | 0.025563 | 0.027981 | 0.030017 | 0.041777 | 0.027772 | 0.024927 | 0.010179 | 0.010655 |
| 26.625 | 0.019808 | 0.006195 | 0.018747 | 0.023999 | 0.0378 | 0.008462 | 0.019114 | 0.005327 | 0.014715 | 0.022342 | 0.012264 | 0.013266 | 0.01388 | 0.015603 | 0.034906 | 0.015813 | 0.01361 | 0.005453 | 0.00501 |
| 26.6388889 | 0.009106 | 0.001028 | 0.025598 | 0.020287 | 0.037896 | 0.017199 | 0.034572 | 0.003369 | 0.027715 | 0.036194 | 0.016296 | 0.006106 | 0.016479 | 0.012533 | 0.021177 | 0.012036 | 0.014871 | 0.003258 | 0.00847 |
| 26.6527778 | 0.012219 | 0.000351 | 0.039075 | 0.025969 | 0.034104 | 0.027707 | 0.053967 | 0.002803 | 0.037003 | 0.035802 | 0.011343 | 0.006292 | 0.02459 | 0.02001 | 0.011649 | 0.022998 | 0.018789 | 0.003773 | 0.015348 |
| 26.6666667 | 0.019691 | 0.0012 | 0.039716 | 0.021981 | 0.0253 | 0.038506 | 0.049983 | 0.002111 | 0.026886 | 0.022012 | 0.006955 | 0.006541 | 0.024068 | 0.02365 | 0.018048 | 0.034092 | 0.01394 | 0.003985 | 0.020358 |
| 26.6805556 | 0.02125 | 0.00168 | 0.034304 | 0.011554 | 0.014182 | 0.049624 | 0.030444 | 0.002155 | 0.014459 | 0.006754 | 0.008645 | 0.003381 | 0.014007 | 0.017422 | 0.031146 | 0.03401 | 0.010397 | 0.004004 | 0.024257 |
| 26.6944444 | 0.014437 | 0.001336 | 0.022772 | 0.00707 | 0.006307 | 0.052994 | 0.022319 | 0.002424 | 0.020415 | 0.005152 | 0.008449 | 0.003623 | 0.004093 | 0.008929 | 0.037977 | 0.028116 | 0.012629 | 0.008238 | 0.026695 |
| 26.7083333 | 0.013762 | 0.001502 | 0.015913 | 0.013923 | 0.007185 | 0.044626 | 0.035996 | 0.003543 | 0.025424 | 0.017916 | 0.010439 | 0.006392 | 0.005262 | 0.006955 | 0.037521 | 0.022648 | 0.011895 | 0.014404 | 0.025355 |
| 26.7222222 | 0.030497 | 0.002374 | 0.027027 | 0.026407 | 0.017285 | 0.025757 | 0.04985 | 0.005354 | 0.015068 | 0.033208 | 0.019435 | 0.007048 | 0.016487 | 0.012124 | 0.031774 | 0.017489 | 0.014685 | 0.014628 | 0.020147 |

| | | | | | | | | | | | | | | | | | | | |
|---|---|---|---|---|---|---|---|---|---|---|---|---|---|---|---|---|---|---|---|
| 26.7361111 | 0.044211 | 0.002392 | 0.029262 | 0.029899 | 0.025052 | 0.015833 | 0.038224 | 0.006326 | 0.007166 | 0.039539 | 0.024154 | 0.007976 | 0.024801 | 0.017813 | 0.023552 | 0.010256 | 0.025496 | 0.009006 | 0.012375 |
| 26.75 | 0.034967 | 0.004967 | 0.013501 | 0.018149 | 0.023101 | 0.025934 | 0.019381 | 0.005522 | 0.011002 | 0.028279 | 0.016876 | 0.009499 | 0.019764 | 0.017899 | 0.015278 | 0.005264 | 0.030943 | 0.005324 | 0.006648 |
| 26.7638889 | 0.016334 | 0.013051 | 0.005375 | 0.006846 | 0.014812 | 0.035161 | 0.02655 | 0.002991 | 0.01885 | 0.019044 | 0.008743 | 0.009765 | 0.010109 | 0.011537 | 0.007707 | 0.008072 | 0.02252 | 0.007095 | 0.010607 |
| 26.7777778 | 0.00869 | 0.017473 | 0.009828 | 0.004368 | 0.006438 | 0.03448 | 0.04523 | 0.000887 | 0.023638 | 0.034353 | 0.010213 | 0.007773 | 0.008937 | 0.005478 | 0.003726 | 0.013288 | 0.012336 | 0.012111 | 0.023657 |
| 26.7916667 | 0.009393 | 0.011523 | 0.01526 | 0.002848 | 0.004526 | 0.028823 | 0.049995 | 0.000459 | 0.022409 | 0.048656 | 0.011064 | 0.007725 | 0.010299 | 0.005546 | 0.003952 | 0.012568 | 0.01461 | 0.01621 | 0.034786 |
| 26.8055556 | 0.01419 | 0.004351 | 0.014698 | 0.00529 | 0.011418 | 0.018582 | 0.036723 | 0.001362 | 0.016875 | 0.038299 | 0.008675 | 0.008668 | 0.00731 | 0.007173 | 0.003506 | 0.009115 | 0.017755 | 0.013203 | 0.034647 |
| 26.8194444 | 0.030295 | 0.002978 | 0.012132 | 0.0179 | 0.020298 | 0.016942 | 0.016295 | 0.00341 | 0.009335 | 0.022118 | 0.013516 | 0.006863 | 0.005403 | 0.005493 | 0.003882 | 0.013349 | 0.012618 | 0.005957 | 0.024002 |
| 26.8333333 | 0.045532 | 0.002968 | 0.014236 | 0.02733 | 0.020333 | 0.03236 | 0.003626 | 0.005207 | 0.004869 | 0.021524 | 0.018858 | 0.009456 | 0.007376 | 0.004218 | 0.010145 | 0.024783 | 0.012535 | 0.002663 | 0.012932 |
| 26.8472222 | 0.037894 | 0.004156 | 0.017095 | 0.020209 | 0.012673 | 0.049548 | 0.000721 | 0.005268 | 0.007847 | 0.025553 | 0.015029 | 0.019129 | 0.012904 | 0.006942 | 0.019579 | 0.033753 | 0.02263 | 0.002448 | 0.013389 |
| 26.8611111 | 0.020555 | 0.009306 | 0.012479 | 0.011799 | 0.00672 | 0.059735 | 0.001289 | 0.003614 | 0.010256 | 0.019139 | 0.009099 | 0.027724 | 0.02406 | 0.011549 | 0.026763 | 0.035105 | 0.029381 | 0.004625 | 0.024669 |
| 26.875 | 0.014707 | 0.013929 | 0.005497 | 0.018087 | 0.004018 | 0.065922 | 0.002917 | 0.001687 | 0.009236 | 0.013901 | 0.010427 | 0.028035 | 0.033693 | 0.015817 | 0.028005 | 0.030287 | 0.023985 | 0.008246 | 0.035276 |
| 26.8888889 | 0.015608 | 0.012266 | 0.004641 | 0.029092 | 0.006432 | 0.062632 | 0.008513 | 0.000886 | 0.013547 | 0.022224 | 0.018845 | 0.022281 | 0.031992 | 0.018724 | 0.022013 | 0.024736 | 0.013845 | 0.007727 | 0.038236 |
| 26.9027778 | 0.014712 | 0.006714 | 0.005305 | 0.032169 | 0.016648 | 0.045358 | 0.018692 | 0.001196 | 0.023195 | 0.037402 | 0.026427 | 0.016916 | 0.023812 | 0.019727 | 0.012639 | 0.0222 | 0.008545 | 0.004111 | 0.033148 |
| 26.9166667 | 0.012122 | 0.004478 | 0.007112 | 0.028299 | 0.02298 | 0.026862 | 0.029038 | 0.00293 | 0.029104 | 0.046896 | 0.025667 | 0.01437 | 0.017341 | 0.020507 | 0.005367 | 0.021986 | 0.011423 | 0.002898 | 0.024327 |
| 26.9305556 | 0.011707 | 0.004595 | 0.012726 | 0.024104 | 0.017444 | 0.017042 | 0.034072 | 0.005365 | 0.027556 | 0.043542 | 0.01779 | 0.012627 | 0.013846 | 0.02135 | 0.002039 | 0.021108 | 0.01862 | 0.005889 | 0.017677 |
| 26.9444444 | 0.015811 | 0.003819 | 0.015604 | 0.024719 | 0.008845 | 0.01053 | 0.030565 | 0.007315 | 0.022771 | 0.029592 | 0.011141 | 0.009444 | 0.012867 | 0.019824 | 0.001772 | 0.017155 | 0.023446 | 0.012545 | 0.015007 |
| 26.9583333 | 0.020838 | 0.004498 | 0.011814 | 0.030344 | 0.006753 | 0.005256 | 0.0201 | 0.009508 | 0.018865 | 0.016695 | 0.011276 | 0.00555 | 0.013231 | 0.015434 | 0.002957 | 0.011181 | 0.02307 | 0.018311 | 0.014756 |
| 26.9722222 | 0.019209 | 0.005228 | 0.006759 | 0.033033 | 0.012102 | 0.004323 | 0.010119 | 0.012275 | 0.014621 | 0.01269 | 0.015682 | 0.002462 | 0.011013 | 0.010245 | 0.004186 | 0.006427 | 0.017414 | 0.018667 | 0.014496 |
| 26.9861111 | 0.011397 | 0.003971 | 0.006766 | 0.025694 | 0.017931 | 0.005892 | 0.007161 | 0.013939 | 0.00891 | 0.015933 | 0.017864 | 0.001158 | 0.006271 | 0.006455 | 0.004419 | 0.004163 | 0.009053 | 0.013851 | 0.012903 |
| 27 | 0.008037 | 0.004543 | 0.008129 | 0.013034 | 0.019326 | 0.006479 | 0.009224 | 0.013461 | 0.007955 | 0.021472 | 0.014446 | 0.001155 | 0.004028 | 0.004794 | 0.003217 | 0.00275 | 0.004313 | 0.007129 | 0.011304 |
| 27.0138889 | 0.011795 | 0.007748 | 0.005589 | 0.006546 | 0.017952 | 0.006681 | 0.01028 | 0.011176 | 0.014862 | 0.023764 | 0.007918 | 0.000962 | 0.00341 | 0.004289 | 0.001733 | 0.001543 | 0.004603 | 0.004006 | 0.010954 |
| 27.0277778 | 0.013208 | 0.00863 | 0.003754 | 0.00719 | 0.016431 | 0.00682 | 0.008055 | 0.008596 | 0.018622 | 0.020684 | 0.00341 | 0.002132 | 0.007682 | 0.005178 | 0.005916 | 0.003984 | 0.006136 | 0.010178 | 0.014655 |
| 27.0416667 | 0.011426 | 0.007917 | 0.008855 | 0.010099 | 0.018649 | 0.006955 | 0.011465 | 0.010013 | 0.016815 | 0.020081 | 0.007774 | 0.009955 | 0.030506 | 0.010127 | 0.027693 | 0.019156 | 0.013101 | 0.027023 | 0.027815 |
| 27.0555556 | 0.018846 | 0.014774 | 0.021053 | 0.021685 | 0.022898 | 0.011462 | 0.027954 | 0.016467 | 0.027043 | 0.032414 | 0.021304 | 0.022418 | 0.065917 | 0.017123 | 0.055689 | 0.042447 | 0.02592 | 0.044033 | 0.043611 |
| 27.0694444 | 0.035351 | 0.028364 | 0.032507 | 0.040262 | 0.019989 | 0.018825 | 0.041389 | 0.020214 | 0.049103 | 0.043666 | 0.02797 | 0.024311 | 0.08158 | 0.017284 | 0.054013 | 0.044966 | 0.031189 | 0.038881 | 0.040619 |
| 27.0833333 | 0.043856 | 0.036013 | 0.034425 | 0.04699 | 0.010843 | 0.019891 | 0.036482 | 0.015098 | 0.062139 | 0.032782 | 0.019355 | 0.014085 | 0.060558 | 0.010637 | 0.028678 | 0.026864 | 0.021234 | 0.01632 | 0.01953 |
| 27.0972222 | 0.037928 | 0.033973 | 0.027457 | 0.035121 | 0.004199 | 0.014241 | 0.021689 | 0.007872 | 0.057028 | 0.026224 | 0.013876 | 0.009575 | 0.026816 | 0.007517 | 0.016616 | 0.020087 | 0.009335 | 0.005424 | 0.005395 |
| 27.1111111 | 0.024032 | 0.022935 | 0.016713 | 0.019054 | 0.003883 | 0.008219 | 0.010512 | 0.006057 | 0.043572 | 0.036243 | 0.019273 | 0.010717 | 0.013247 | 0.007075 | 0.020299 | 0.025445 | 0.00814 | 0.006313 | 0.004013 |
| 27.125 | 0.011783 | 0.015642 | 0.008997 | 0.008537 | 0.009632 | 0.008695 | 0.005669 | 0.005606 | 0.039944 | 0.027413 | 0.018123 | 0.007288 | 0.01944 | 0.004425 | 0.024745 | 0.025901 | 0.009943 | 0.012578 | 0.00836 |
| 27.1388889 | 0.012316 | 0.029595 | 0.009757 | 0.004124 | 0.020662 | 0.019845 | 0.00711 | 0.008206 | 0.039425 | 0.0131 | 0.014021 | 0.002998 | 0.02526 | 0.002916 | 0.025024 | 0.020815 | 0.008831 | 0.021439 | 0.012571 |
| 27.1527778 | 0.02572 | 0.05269 | 0.015342 | 0.005315 | 0.033268 | 0.026672 | 0.013993 | 0.021383 | 0.026953 | 0.016034 | 0.014435 | 0.001599 | 0.024415 | 0.002326 | 0.020973 | 0.014304 | 0.009057 | 0.018997 | 0.010308 |
| 27.1666667 | 0.028459 | 0.051162 | 0.030952 | 0.011678 | 0.035439 | 0.040368 | 0.017399 | 0.034244 | 0.011847 | 0.016339 | 0.009698 | 0.001514 | 0.021902 | 0.001682 | 0.015796 | 0.008328 | 0.015342 | 0.008992 | 0.008883 |
| 27.1805556 | 0.024388 | 0.038958 | 0.038743 | 0.021282 | 0.025178 | 0.066045 | 0.012953 | 0.042534 | 0.009172 | 0.009886 | 0.005665 | 0.002451 | 0.022629 | 0.003601 | 0.014786 | 0.006323 | 0.027206 | 0.004966 | 0.009236 |
| 27.1944444 | 0.03157 | 0.043294 | 0.017659 | 0.027114 | 0.015958 | 0.061009 | 0.009187 | 0.043235 | 0.026537 | 0.008123 | 0.010479 | 0.00576 | 0.026518 | 0.008374 | 0.018835 | 0.009544 | 0.036418 | 0.007818 | 0.008087 |
| 27.2083333 | 0.026267 | 0.031416 | 0.005349 | 0.029117 | 0.01721 | 0.045809 | 0.015149 | 0.027614 | 0.050222 | 0.01097 | 0.015057 | 0.008937 | 0.025445 | 0.011842 | 0.019336 | 0.012283 | 0.035603 | 0.008903 | 0.009991 |
| 27.2222222 | 0.012952 | 0.009681 | 0.010496 | 0.027782 | 0.02446 | 0.042069 | 0.023087 | 0.015125 | 0.059876 | 0.013804 | 0.014101 | 0.007041 | 0.017281 | 0.009901 | 0.012575 | 0.009015 | 0.023496 | 0.007945 | 0.0097 |
| 27.2361111 | 0.008754 | 0.004457 | 0.01195 | 0.019279 | 0.023906 | 0.025732 | 0.022224 | 0.014797 | 0.058109 | 0.017538 | 0.01636 | 0.002833 | 0.014603 | 0.004676 | 0.008487 | 0.005164 | 0.010225 | 0.011602 | 0.007899 |
| 27.25 | 0.008895 | 0.0094 | 0.019509 | 0.009457 | 0.016094 | 0.010788 | 0.019962 | 0.023166 | 0.066248 | 0.016028 | 0.016001 | 0.002414 | 0.018507 | 0.001982 | 0.008696 | 0.00476 | 0.005143 | 0.011714 | 0.007138 |
| 27.2638889 | 0.016027 | 0.02395 | 0.034745 | 0.004803 | 0.016851 | 0.009117 | 0.022652 | 0.03578 | 0.08712 | 0.016774 | 0.017585 | 0.00787 | 0.014844 | 0.005791 | 0.009305 | 0.006539 | 0.006247 | 0.011915 | 0.009071 |
| 27.2777778 | 0.017096 | 0.02862 | 0.043737 | 0.008985 | 0.018932 | 0.011124 | 0.025685 | 0.035475 | 0.088834 | 0.030715 | 0.03558 | 0.015891 | 0.00853 | 0.012506 | 0.015025 | 0.013264 | 0.012471 | 0.020354 | 0.019721 |
| 27.2916667 | 0.011086 | 0.017927 | 0.04213 | 0.023782 | 0.009523 | 0.010654 | 0.025671 | 0.022522 | 0.062469 | 0.032401 | 0.043122 | 0.017398 | 0.007965 | 0.013088 | 0.019998 | 0.018294 | 0.015761 | 0.019589 | 0.024786 |
| 27.3055556 | 0.015214 | 0.015154 | 0.035156 | 0.033372 | 0.003924 | 0.007075 | 0.02111 | 0.010083 | 0.033887 | 0.018833 | 0.025328 | 0.010845 | 0.007005 | 0.007457 | 0.014363 | 0.013157 | 0.011828 | 0.009834 | 0.014785 |
| 27.3194444 | 0.020284 | 0.018739 | 0.022537 | 0.023453 | 0.008197 | 0.003596 | 0.012887 | 0.003277 | 0.015275 | 0.015451 | 0.008633 | 0.004523 | 0.007991 | 0.004496 | 0.006346 | 0.006153 | 0.012725 | 0.007507 | 0.004586 |
| 27.3333333 | 0.014495 | 0.013685 | 0.009545 | 0.010713 | 0.014243 | 0.002821 | 0.009301 | 0.00125 | 0.008165 | 0.014441 | 0.004469 | 0.002254 | 0.012476 | 0.006271 | 0.005736 | 0.006532 | 0.019114 | 0.006929 | 0.001469 |
| 27.3472222 | 0.006485 | 0.006704 | 0.004737 | 0.010046 | 0.016476 | 0.003708 | 0.011183 | 0.00114 | 0.010206 | 0.011472 | 0.006715 | 0.002223 | 0.014389 | 0.006796 | 0.008271 | 0.008971 | 0.018796 | 0.003467 | 0.000731 |
| 27.3611111 | 0.002994 | 0.00698 | 0.005082 | 0.011002 | 0.013582 | 0.003047 | 0.0061 | 0.002093 | 0.009448 | 0.019943 | 0.014417 | 0.00266 | 0.012627 | 0.004151 | 0.008024 | 0.006343 | 0.011567 | 0.00239 | 0.000876 |
| 27.375 | 0.004054 | 0.007775 | 0.009696 | 0.014782 | 0.008705 | 0.003331 | 0.011206 | 0.005172 | 0.009067 | 0.027358 | 0.020113 | 0.002665 | 0.009558 | 0.002548 | 0.005286 | 0.003495 | 0.007711 | 0.002412 | 0.002214 |
| 27.3888889 | 0.010559 | 0.008711 | 0.017041 | 0.025406 | 0.005316 | 0.007686 | 0.018561 | 0.008363 | 0.018392 | 0.020304 | 0.017608 | 0.002313 | 0.005892 | 0.003683 | 0.002702 | 0.004651 | 0.00781 | 0.002151 | 0.002758 |
| 27.4027778 | 0.016427 | 0.012356 | 0.019321 | 0.029431 | 0.006941 | 0.010655 | 0.021561 | 0.008714 | 0.02748 | 0.00876 | 0.012501 | 0.002689 | 0.003571 | 0.004971 | 0.002514 | 0.005129 | 0.006076 | 0.002404 | 0.002128 |
| 27.4166667 | 0.014352 | 0.010984 | 0.01521 | 0.022073 | 0.010016 | 0.008228 | 0.016019 | 0.006361 | 0.026074 | 0.003169 | 0.010341 | 0.004418 | 0.006052 | 0.004894 | 0.0041 | 0.003487 | 0.004496 | 0.002736 | 0.002284 |
| 27.4305556 | 0.007445 | 0.005726 | 0.009769 | 0.011156 | 0.012597 | 0.003822 | 0.007942 | 0.003685 | 0.018117 | 0.007587 | 0.012026 | 0.007163 | 0.010724 | 0.004059 | 0.004808 | 0.002207 | 0.007242 | 0.004269 | 0.00239 |
| 27.4444444 | 0.002191 | 0.003707 | 0.006809 | 0.005454 | 0.016588 | 0.00178 | 0.003946 | 0.002123 | 0.010597 | 0.020921 | 0.014842 | 0.009593 | 0.013645 | 0.003664 | 0.004192 | 0.001785 | 0.013067 | 0.005276 | 0.00195 |
| 27.4583333 | 0.001788 | 0.004068 | 0.006691 | 0.005196 | 0.019842 | 0.002953 | 0.004308 | 0.001277 | 0.006095 | 0.029869 | 0.014251 | 0.009554 | 0.014068 | 0.004447 | 0.004442 | 0.00192 | 0.017053 | 0.00459 | 0.002911 |
| 27.4722222 | 0.005103 | 0.003068 | 0.008264 | 0.00549 | 0.020563 | 0.004409 | 0.003787 | 0.001027 | 0.006287 | 0.023438 | 0.009138 | 0.006897 | 0.012957 | 0.006347 | 0.006907 | 0.003137 | 0.017163 | 0.00632 | 0.003761 |
| 27.4861111 | 0.008553 | 0.001636 | 0.009899 | 0.008139 | 0.019287 | 0.003643 | 0.003489 | 0.00171 | 0.012052 | 0.010757 | 0.00464 | 0.004232 | 0.013407 | 0.008413 | 0.010553 | 0.005281 | 0.01615 | 0.007408 | 0.004196 |
| 27.5 | 0.009143 | 0.001553 | 0.010339 | 0.01214 | 0.016182 | 0.001827 | 0.006306 | 0.00273 | 0.020464 | 0.005503 | 0.004954 | 0.003687 | 0.016611 | 0.008422 | 0.012972 | 0.007127 | 0.015774 | 0.006378 | 0.007861 |
| 27.5138889 | 0.007786 | 0.001502 | 0.010797 | 0.012315 | 0.012585 | 0.001785 | 0.008687 | 0.003766 | 0.025623 | 0.005399 | 0.007337 | 0.005236 | 0.018666 | 0.005957 | 0.011819 | 0.007047 | 0.015468 | 0.008864 | 0.011872 |
| 27.5277778 | 0.008907 | 0.001124 | 0.013804 | 0.011143 | 0.011253 | 0.004958 | 0.010342 | 0.006708 | 0.024309 | 0.007055 | 0.00805 | 0.007611 | 0.018248 | 0.004313 | 0.008904 | 0.005855 | 0.015678 | 0.013411 | 0.011498 |
| 27.5416667 | 0.016247 | 0.001653 | 0.021571 | 0.018861 | 0.014113 | 0.015336 | 0.019855 | 0.014924 | 0.022993 | 0.018141 | 0.009625 | 0.012833 | 0.022818 | 0.007282 | 0.012483 | 0.009857 | 0.019351 | 0.023728 | 0.012938 |
| 27.5555556 | 0.030524 | 0.002547 | 0.035115 | 0.041903 | 0.020601 | 0.037622 | 0.043844 | 0.026641 | 0.029373 | 0.037491 | 0.016208 | 0.027197 | 0.035632 | 0.014769 | 0.024298 | 0.020316 | 0.02712 | 0.041186 | 0.023042 |
| 27.5694444 | 0.045686 | 0.004247 | 0.051436 | 0.072747 | 0.023437 | 0.063045 | 0.075786 | 0.031412 | 0.041127 | 0.053514 | 0.026794 | 0.038564 | 0.050045 | 0.021848 | 0.03225 | 0.02741 | 0.033582 | 0.048533 | 0.03404 |
| 27.5833333 | 0.052316 | 0.007833 | 0.06287 | 0.092051 | 0.018545 | 0.073158 | 0.09755 | 0.026088 | 0.048123 | 0.056384 | 0.03333 | 0.029193 | 0.061769 | 0.024337 | 0.028506 | 0.024913 | 0.035232 | 0.040604 | 0.034815 |
| 27.5972222 | 0.047692 | 0.010309 | 0.06134 | 0.081704 | 0.011631 | 0.062868 | 0.091472 | 0.017963 | 0.040639 | 0.042204 | 0.02537 | 0.012899 | 0.064531 | 0.022284 | 0.018694 | 0.018194 | 0.032055 | 0.029693 | 0.028115 |
| 27.6111111 | 0.034618 | 0.009965 | 0.045897 | 0.046437 | 0.006573 | 0.043297 | 0.059722 | 0.00974 | 0.021618 | 0.022526 | 0.016409 | 0.006367 | 0.049511 | 0.016959 | 0.009117 | 0.013049 | 0.024024 | 0.020217 | 0.020973 |
| 27.625 | 0.018503 | 0.00838 | 0.023666 | 0.020744 | 0.006307 | 0.025151 | 0.026772 | 0.00359 | 0.010345 | 0.02113 | 0.030426 | 0.007973 | 0.023927 | 0.009547 | 0.005234 | 0.010484 | 0.014154 | 0.011591 | 0.015512 |
| 27.6388889 | 0.007673 | 0.007886 | 0.010454 | 0.026634 | 0.010362 | 0.011612 | 0.01447 | 0.001718 | 0.01719 | 0.036511 | 0.04659 | 0.011143 | 0.007854 | 0.003428 | 0.009557 | 0.009559 | 0.006955 | 0.005289 | 0.012804 |
| 27.6527778 | 0.006145 | 0.008755 | 0.014449 | 0.04301 | 0.013125 | 0.005134 | 0.02076 | 0.001768 | 0.028256 | 0.035287 | 0.034432 | 0.0111 | 0.007089 | 0.00137 | 0.014029 | 0.007808 | 0.006797 | 0.002215 | 0.012781 |
| 27.6666667 | 0.009949 | 0.009585 | 0.019671 | 0.047042 | 0.01369 | 0.008306 | 0.03041 | 0.002791 | 0.029196 | 0.024359 | 0.02213 | 0.009805 | 0.009777 | 0.003077 | 0.01255 | 0.005765 | 0.011182 | 0.00158 | 0.014278 |
| 27.6805556 | 0.013048 | 0.010951 | 0.015516 | 0.037549 | 0.013952 | 0.019558 | 0.036783 | 0.004535 | 0.018464 | 0.038423 | 0.035735 | 0.009644 | 0.010222 | 0.008235 | 0.006999 | 0.006653 | 0.009872 | 0.001926 | 0.014869 |
| 27.6944444 | 0.011249 | 0.010373 | 0.00842 | 0.020286 | 0.012114 | 0.037145 | 0.039245 | 0.004964 | 0.010574 | 0.056184 | 0.041557 | 0.009077 | 0.010815 | 0.014313 | 0.002432 | 0.008724 | 0.007091 | 0.002338 | 0.012875 |
| 27.7083333 | 0.006054 | 0.008634 | 0.00372 | 0.00898 | 0.008664 | 0.056249 | 0.033149 | 0.003245 | 0.015744 | 0.044068 | 0.025474 | 0.006618 | 0.012379 | 0.018058 | 0.001328 | 0.006929 | 0.012068 | 0.001919 | 0.009898 |
| 27.7222222 | 0.003416 | 0.011302 | 0.003573 | 0.010765 | 0.011132 | 0.065318 | 0.019933 | 0.002157 | 0.018044 | 0.022641 | 0.018759 | 0.003368 | 0.013934 | 0.018546 | 0.001309 | 0.00409 | 0.015866 | 0.001668 | 0.008065 |
| 27.7361111 | 0.008357 | 0.01347 | 0.00757 | 0.014031 | 0.018144 | 0.056444 | 0.017157 | 0.004159 | 0.013487 | 0.024545 | 0.02811 | 0.00179 | 0.013243 | 0.01402 | 0.001179 | 0.006179 | 0.013078 | 0.002518 | 0.008024 |
| 27.75 | 0.020166 | 0.008986 | 0.012081 | 0.014497 | 0.022934 | 0.035117 | 0.032866 | 0.00715 | 0.015643 | 0.041119 | 0.031327 | 0.002135 | 0.008655 | 0.007835 | 0.001257 | 0.010436 | 0.008367 | 0.003253 | 0.009241 |
| 27.7638889 | 0.028035 | 0.003516 | 0.012043 | 0.012963 | 0.025429 | 0.016812 | 0.050274 | 0.007907 | 0.018974 | 0.044989 | 0.020559 | 0.00286 | 0.005279 | 0.008101 | 0.001787 | 0.012047 | 0.004887 | 0.00464 | 0.009305 |
| 27.7777778 | 0.020542 | 0.001943 | 0.007147 | 0.011842 | 0.019479 | 0.014978 | 0.054145 | 0.005635 | 0.013124 | 0.028779 | 0.009275 | 0.005049 | 0.009981 | 0.013724 | 0.002108 | 0.009694 | 0.007117 | 0.008176 | 0.006127 |
| 27.7916667 | 0.01061 | 0.002707 | 0.003495 | 0.021144 | 0.012293 | 0.028874 | 0.041802 | 0.00368 | 0.007045 | 0.013517 | 0.00745 | 0.009741 | 0.017753 | 0.017176 | 0.002025 | 0.005127 | 0.013968 | 0.011483 | 0.002356 |
| 27.8055556 | 0.014868 | 0.005605 | 0.003092 | 0.034004 | 0.01941 | 0.042943 | 0.021054 | 0.0046 | 0.008685 | 0.014256 | 0.007082 | 0.013487 | 0.019355 | 0.01604 | 0.001669 | 0.001722 | 0.017168 | 0.01192 | 0.001771 |
| 27.8194444 | 0.019824 | 0.009682 | 0.004492 | 0.036848 | 0.031934 | 0.045661 | 0.011673 | 0.005214 | 0.008745 | 0.013514 | 0.006721 | 0.012397 | 0.01369 | 0.012632 | 0.001452 | 0.000437 | 0.013374 | 0.010411 | 0.004451 |
| 27.8333333 | 0.013985 | 0.011119 | 0.005133 | 0.029075 | 0.036967 | 0.038054 | 0.024233 | 0.003368 | 0.005328 | 0.0121 | 0.016466 | 0.007004 | 0.012921 | 0.010473 | 0.002714 | 0.000457 | 0.00712 | 0.007627 | 0.009061 |
| 27.8472222 | 0.009848 | 0.007673 | 0.007159 | 0.016576 | 0.032596 | 0.022233 | 0.046558 | 0.001257 | 0.009047 | 0.027979 | 0.031226 | 0.002056 | 0.02805 | 0.012705 | 0.004668 | 0.001683 | 0.004684 | 0.003999 | 0.014882 |
| 27.8611111 | 0.013934 | 0.003535 | 0.016141 | 0.0062 | 0.022372 | 0.012284 | 0.057476 | 0.000626 | 0.017935 | 0.04623 | 0.037374 | 0.002247 | 0.047456 | 0.019385 | 0.006296 | 0.004632 | 0.009636 | 0.001543 | 0.021616 |
| 27.875 | 0.015136 | 0.002854 | 0.024558 | 0.00391 | 0.01246 | 0.025341 | 0.045919 | 0.001475 | 0.020781 | 0.044701 | 0.029749 | 0.00761 | 0.053678 | 0.025909 | 0.007456 | 0.007764 | 0.020047 | 0.00174 | 0.027514 |
| 27.8888889 | 0.017767 | 0.005629 | 0.031051 | 0.013822 | 0.011064 | 0.050329 | 0.02668 | 0.003257 | 0.015066 | 0.026091 | 0.014717 | 0.0125 | 0.043402 | 0.027091 | 0.006796 | 0.008228 | 0.02971 | 0.00506 | 0.02971 |
| 27.9027778 | 0.030049 | 0.014623 | 0.042289 | 0.030567 | 0.018208 | 0.06463 | 0.021631 | 0.003478 | 0.008367 | 0.009351 | 0.003972 | 0.011199 | 0.024334 | 0.023579 | 0.004154 | 0.006184 | 0.0343 | 0.01 | 0.027506 |
| 27.9166667 | 0.036768 | 0.027258 | 0.049973 | 0.041538 | 0.02371 | 0.058506 | 0.030575 | 0.003197 | 0.006613 | 0.005053 | 0.003394 | 0.00563 | 0.008594 | 0.019337 | 0.001633 | 0.004663 | 0.032545 | 0.014898 | 0.022487 |
| 27.9305556 | 0.033315 | 0.035277 | 0.043513 | 0.044753 | 0.02174 | 0.042266 | 0.040838 | 0.006304 | 0.008805 | 0.0111 | 0.010957 | 0.001934 | 0.002302 | 0.015909 | 0.000931 | 0.005435 | 0.024481 | 0.019443 | 0.01727 |
| 27.9444444 | 0.02997 | 0.034489 | 0.024829 | 0.046343 | 0.013261 | 0.03037 | 0.041774 | 0.011408 | 0.014545 | 0.026843 | 0.02001 | 0.002558 | 0.002983 | 0.012712 | 0.002415 | 0.007964 | 0.013485 | 0.022314 | 0.014241 |
| 27.9583333 | 0.024788 | 0.024408 | 0.011737 | 0.043625 | 0.006998 | 0.021872 | 0.030079 | 0.016038 | 0.019639 | 0.040667 | 0.022927 | 0.005349 | 0.006273 | 0.008242 | 0.005361 | 0.010804 | 0.005239 | 0.022878 | 0.01324 |

| | | | | | | | | | | | | | | | | | | | |
|---|---|---|---|---|---|---|---|---|---|---|---|---|---|---|---|---|---|---|---|
| 27.9722222 | 0.01436 | 0.013021 | 0.017711 | 0.031195 | 0.010111 | 0.013143 | 0.014857 | 0.017931 | 0.01579 | 0.036022 | 0.016461 | 0.006248 | 0.00823 | 0.004514 | 0.007844 | 0.011478 | 0.002499 | 0.022723 | 0.011814 |
| 27.9861111 | 0.008355 | 0.013044 | 0.027993 | 0.016842 | 0.017409 | 0.006038 | 0.010425 | 0.01436 | 0.008603 | 0.019878 | 0.007237 | 0.004191 | 0.007871 | 0.005047 | 0.008143 | 0.008199 | 0.003505 | 0.022775 | 0.008549 |
| 28 | 0.01242 | 0.021663 | 0.028037 | 0.015394 | 0.02206 | 0.003818 | 0.020077 | 0.007639 | 0.009704 | 0.011365 | 0.002945 | 0.00296 | 0.00724 | 0.005732 | 0.006607 | 0.003985 | 0.005335 | 0.022713 | 0.004693 |
| 28.0138889 | 0.017008 | 0.025558 | 0.022233 | 0.02516 | 0.021346 | 0.006597 | 0.03289 | 0.003658 | 0.019372 | 0.008673 | 0.003846 | 0.004398 | 0.007504 | 0.00297 | 0.005824 | 0.002208 | 0.005968 | 0.0215 | 0.00213 |
| 28.0277778 | 0.013823 | 0.018914 | 0.013716 | 0.026152 | 0.017972 | 0.008817 | 0.03648 | 0.005194 | 0.028573 | 0.007943 | 0.008242 | 0.004461 | 0.011113 | 0.002576 | 0.012634 | 0.007605 | 0.006088 | 0.019626 | 0.003463 |
| 28.0416667 | 0.013234 | 0.015012 | 0.01457 | 0.024199 | 0.020553 | 0.008241 | 0.028401 | 0.011933 | 0.026639 | 0.023018 | 0.015275 | 0.00922 | 0.027928 | 0.010642 | 0.037845 | 0.032073 | 0.010925 | 0.023712 | 0.012878 |
| 28.0555556 | 0.033907 | 0.036257 | 0.044471 | 0.04841 | 0.028655 | 0.011897 | 0.029038 | 0.020796 | 0.030728 | 0.056674 | 0.027462 | 0.028839 | 0.053274 | 0.024585 | 0.073278 | 0.067789 | 0.024404 | 0.033761 | 0.028761 |
| 28.0694444 | 0.06695 | 0.075331 | 0.091769 | 0.083988 | 0.028426 | 0.019968 | 0.044694 | 0.02593 | 0.054547 | 0.082335 | 0.042449 | 0.040748 | 0.056404 | 0.031077 | 0.079411 | 0.078064 | 0.038443 | 0.032768 | 0.033491 |
| 28.0833333 | 0.079768 | 0.100775 | 0.113365 | 0.087627 | 0.015713 | 0.020511 | 0.045018 | 0.022787 | 0.067171 | 0.06846 | 0.042374 | 0.027185 | 0.03398 | 0.021438 | 0.046908 | 0.049449 | 0.037239 | 0.017351 | 0.02218 |
| 28.0972222 | 0.061826 | 0.093013 | 0.086146 | 0.056952 | 0.006189 | 0.011458 | 0.025958 | 0.012435 | 0.060507 | 0.033434 | 0.025907 | 0.01495 | 0.020658 | 0.009204 | 0.02174 | 0.019298 | 0.020891 | 0.008035 | 0.016473 |
| 28.1111111 | 0.032127 | 0.057555 | 0.039389 | 0.025614 | 0.008057 | 0.003837 | 0.010207 | 0.005673 | 0.052783 | 0.019861 | 0.020782 | 0.016308 | 0.025257 | 0.007495 | 0.020919 | 0.015553 | 0.007935 | 0.010961 | 0.017825 |
| 28.125 | 0.012939 | 0.025965 | 0.017648 | 0.017513 | 0.01119 | 0.002286 | 0.003478 | 0.00689 | 0.041061 | 0.025462 | 0.030079 | 0.015642 | 0.028033 | 0.009965 | 0.020744 | 0.021058 | 0.00734 | 0.014619 | 0.012772 |
| 28.1388889 | 0.013219 | 0.024042 | 0.029271 | 0.028615 | 0.009273 | 0.002458 | 0.00183 | 0.006692 | 0.022719 | 0.028064 | 0.033117 | 0.009319 | 0.023922 | 0.009713 | 0.016173 | 0.020997 | 0.011223 | 0.013869 | 0.005475 |
| 28.1527778 | 0.024064 | 0.042905 | 0.049468 | 0.041374 | 0.006337 | 0.002906 | 0.002878 | 0.003364 | 0.009571 | 0.026401 | 0.029272 | 0.004236 | 0.016906 | 0.007216 | 0.012594 | 0.017915 | 0.012147 | 0.009601 | 0.002894 |
| 28.1666667 | 0.034149 | 0.064704 | 0.059782 | 0.048439 | 0.006538 | 0.003029 | 0.003244 | 0.001774 | 0.008147 | 0.027794 | 0.028414 | 0.003506 | 0.008983 | 0.004619 | 0.009335 | 0.015162 | 0.010276 | 0.005163 | 0.002768 |
| 28.1805556 | 0.040524 | 0.078754 | 0.056395 | 0.049734 | 0.008407 | 0.003387 | 0.004187 | 0.001779 | 0.009751 | 0.032143 | 0.030952 | 0.004457 | 0.004393 | 0.004032 | 0.006532 | 0.014008 | 0.009839 | 0.006159 | 0.003151 |
| 28.1944444 | 0.041907 | 0.077489 | 0.044629 | 0.043438 | 0.008365 | 0.007068 | 0.010455 | 0.001816 | 0.00926 | 0.034027 | 0.029887 | 0.007995 | 0.002753 | 0.005497 | 0.005019 | 0.013018 | 0.014227 | 0.01265 | 0.006595 |
| 28.2083333 | 0.03692 | 0.063071 | 0.034739 | 0.031658 | 0.006642 | 0.012199 | 0.018294 | 0.002181 | 0.014246 | 0.030978 | 0.023872 | 0.01367 | 0.002262 | 0.006635 | 0.005345 | 0.011962 | 0.020343 | 0.014084 | 0.010477 |
| 28.2222222 | 0.028238 | 0.045315 | 0.028953 | 0.020124 | 0.005144 | 0.014926 | 0.022239 | 0.004158 | 0.029362 | 0.024947 | 0.020006 | 0.012946 | 0.004808 | 0.005768 | 0.006278 | 0.011282 | 0.021256 | 0.010895 | 0.008949 |
| 28.2361111 | 0.021468 | 0.031757 | 0.024822 | 0.012463 | 0.003539 | 0.013937 | 0.021746 | 0.006445 | 0.044347 | 0.017338 | 0.016977 | 0.008169 | 0.006905 | 0.004625 | 0.005625 | 0.00895 | 0.016067 | 0.012849 | 0.006401 |
| 28.25 | 0.017956 | 0.021822 | 0.022667 | 0.008752 | 0.001693 | 0.011319 | 0.019512 | 0.006728 | 0.046348 | 0.00866 | 0.010577 | 0.008301 | 0.005928 | 0.005088 | 0.003483 | 0.005278 | 0.009035 | 0.011843 | 0.006375 |
| 28.2638889 | 0.015875 | 0.013472 | 0.024292 | 0.008208 | 0.001945 | 0.009988 | 0.019311 | 0.004946 | 0.03861 | 0.002926 | 0.007128 | 0.008403 | 0.007269 | 0.006034 | 0.002563 | 0.004125 | 0.004911 | 0.007636 | 0.005812 |
| 28.2777778 | 0.014846 | 0.008207 | 0.028024 | 0.009878 | 0.004522 | 0.0099 | 0.021051 | 0.002495 | 0.032347 | 0.00211 | 0.007912 | 0.008331 | 0.010393 | 0.00527 | 0.003223 | 0.004241 | 0.00698 | 0.010202 | 0.00945 |
| 28.2916667 | 0.015036 | 0.007018 | 0.030214 | 0.010817 | 0.006804 | 0.009641 | 0.021538 | 0.001056 | 0.030621 | 0.002078 | 0.00736 | 0.013145 | 0.009528 | 0.003566 | 0.003049 | 0.002864 | 0.010911 | 0.01117 | 0.015463 |
| 28.3055556 | 0.016084 | 0.009358 | 0.028725 | 0.008462 | 0.007139 | 0.008488 | 0.018968 | 0.001582 | 0.029945 | 0.00122 | 0.009151 | 0.013534 | 0.005498 | 0.004226 | 0.002646 | 0.002094 | 0.00955 | 0.009406 | 0.013849 |
| 28.3194444 | 0.017037 | 0.013027 | 0.02313 | 0.004232 | 0.005999 | 0.006625 | 0.014427 | 0.003969 | 0.024802 | 0.001566 | 0.014529 | 0.007692 | 0.002731 | 0.006194 | 0.003965 | 0.002309 | 0.007231 | 0.013717 | 0.010282 |
| 28.3333333 | 0.015514 | 0.013021 | 0.015324 | 0.001259 | 0.004982 | 0.004757 | 0.0103 | 0.005678 | 0.014888 | 0.003933 | 0.013042 | 0.005481 | 0.001918 | 0.005106 | 0.004382 | 0.002065 | 0.010137 | 0.015295 | 0.010541 |
| 28.3472222 | 0.00968 | 0.00906 | 0.008033 | 0.001177 | 0.004474 | 0.003142 | 0.007551 | 0.004276 | 0.011838 | 0.005771 | 0.00595 | 0.005178 | 0.001732 | 0.003273 | 0.002822 | 0.002651 | 0.012387 | 0.009153 | 0.007674 |
| 28.3611111 | 0.003721 | 0.008186 | 0.003302 | 0.004697 | 0.004011 | 0.001784 | 0.005315 | 0.003087 | 0.02011 | 0.005023 | 0.002409 | 0.004594 | 0.001391 | 0.003917 | 0.002688 | 0.003799 | 0.009389 | 0.004183 | 0.003416 |
| 28.375 | 0.002543 | 0.009035 | 0.001553 | 0.012084 | 0.00416 | 0.001241 | 0.003867 | 0.0052 | 0.025063 | 0.007238 | 0.004092 | 0.00892 | 0.001573 | 0.003632 | 0.003552 | 0.003139 | 0.004914 | 0.002095 | 0.003468 |
| 28.3888889 | 0.005771 | 0.007125 | 0.002675 | 0.021281 | 0.005302 | 0.00105 | 0.004968 | 0.00625 | 0.017546 | 0.016592 | 0.011904 | 0.013939 | 0.004069 | 0.002131 | 0.003386 | 0.00289 | 0.003146 | 0.00087 | 0.003884 |
| 28.4027778 | 0.0129 | 0.009366 | 0.009127 | 0.028013 | 0.006422 | 0.001212 | 0.009006 | 0.004715 | 0.011275 | 0.02509 | 0.020783 | 0.013048 | 0.008014 | 0.002309 | 0.004628 | 0.006183 | 0.003266 | 0.000842 | 0.002988 |
| 28.4166667 | 0.021494 | 0.018834 | 0.020269 | 0.02767 | 0.00641 | 0.002523 | 0.013266 | 0.004925 | 0.015499 | 0.024609 | 0.02202 | 0.008409 | 0.010735 | 0.003408 | 0.008106 | 0.009847 | 0.002902 | 0.000973 | 0.00422 |
| 28.4305556 | 0.025995 | 0.026888 | 0.029495 | 0.018901 | 0.004995 | 0.003451 | 0.013593 | 0.005163 | 0.014997 | 0.01735 | 0.014502 | 0.00458 | 0.0103 | 0.003586 | 0.009959 | 0.010481 | 0.003127 | 0.001298 | 0.00635 |
| 28.4444444 | 0.021287 | 0.023409 | 0.0292 | 0.009803 | 0.002723 | 0.002889 | 0.008692 | 0.004185 | 0.01079 | 0.009986 | 0.006098 | 0.0044 | 0.006988 | 0.002727 | 0.008624 | 0.008487 | 0.005159 | 0.001937 | 0.006832 |
| 28.4583333 | 0.012974 | 0.015055 | 0.017707 | 0.010511 | 0.001397 | 0.002049 | 0.004545 | 0.006664 | 0.019806 | 0.007006 | 0.002775 | 0.007575 | 0.003336 | 0.001573 | 0.006223 | 0.005387 | 0.008433 | 0.003383 | 0.00679 |
| 28.4722222 | 0.013776 | 0.018506 | 0.00954 | 0.015674 | 0.001812 | 0.001838 | 0.005396 | 0.010666 | 0.032491 | 0.009705 | 0.00493 | 0.011479 | 0.001759 | 0.000989 | 0.004732 | 0.002766 | 0.012518 | 0.006531 | 0.006807 |
| 28.4861111 | 0.018785 | 0.027337 | 0.014371 | 0.012828 | 0.002062 | 0.002118 | 0.005197 | 0.010974 | 0.031477 | 0.016031 | 0.014333 | 0.013174 | 0.00313 | 0.001536 | 0.004615 | 0.001853 | 0.015283 | 0.006863 | 0.005843 |
| 28.5 | 0.015283 | 0.026324 | 0.016332 | 0.008334 | 0.001585 | 0.002731 | 0.003069 | 0.009966 | 0.018354 | 0.020377 | 0.025321 | 0.010522 | 0.005814 | 0.002181 | 0.005183 | 0.002773 | 0.013414 | 0.004231 | 0.003862 |
| 28.5138889 | 0.008071 | 0.016 | 0.012165 | 0.014536 | 0.001715 | 0.003379 | 0.004065 | 0.003261 | 0.00782 | 0.017795 | 0.026442 | 0.005464 | 0.007338 | 0.001953 | 0.00625 | 0.004586 | 0.008424 | 0.004079 | 0.002964 |
| 28.5277778 | 0.008558 | 0.008096 | 0.01933 | 0.02609 | 0.003359 | 0.004481 | 0.008531 | 0.005584 | 0.007632 | 0.010526 | 0.016136 | 0.002777 | 0.008282 | 0.002897 | 0.009857 | 0.007897 | 0.006933 | 0.005687 | 0.006781 |
| 28.5416667 | 0.017504 | 0.011844 | 0.037741 | 0.040934 | 0.009933 | 0.008658 | 0.02097 | 0.014659 | 0.017369 | 0.00957 | 0.007019 | 0.009847 | 0.012313 | 0.009266 | 0.019203 | 0.015492 | 0.01758 | 0.008264 | 0.015535 |
| 28.5555556 | 0.027418 | 0.021127 | 0.055557 | 0.064719 | 0.024837 | 0.015958 | 0.042469 | 0.027472 | 0.038614 | 0.020765 | 0.007089 | 0.028529 | 0.021092 | 0.020778 | 0.03035 | 0.024726 | 0.034448 | 0.013556 | 0.023747 |
| 28.5694444 | 0.030611 | 0.0277 | 0.071229 | 0.0974 | 0.04087 | 0.021883 | 0.062158 | 0.035993 | 0.063446 | 0.036831 | 0.016379 | 0.040879 | 0.030195 | 0.028434 | 0.032227 | 0.026862 | 0.037711 | 0.017612 | 0.028987 |
| 28.5833333 | 0.03045 | 0.033433 | 0.085068 | 0.12868 | 0.049528 | 0.02525 | 0.072314 | 0.033878 | 0.07931 | 0.050804 | 0.033993 | 0.033304 | 0.035096 | 0.026886 | 0.024008 | 0.021955 | 0.028329 | 0.014326 | 0.03099 |
| 28.5972222 | 0.033112 | 0.037383 | 0.084664 | 0.127863 | 0.050623 | 0.026883 | 0.070544 | 0.024303 | 0.074639 | 0.054886 | 0.041231 | 0.017068 | 0.035124 | 0.022088 | 0.015156 | 0.017175 | 0.020299 | 0.0071 | 0.027113 |
| 28.6111111 | 0.033108 | 0.032101 | 0.064232 | 0.08591 | 0.041399 | 0.024407 | 0.057898 | 0.013085 | 0.046216 | 0.041982 | 0.028364 | 0.00689 | 0.029062 | 0.017439 | 0.009985 | 0.012669 | 0.013647 | 0.003869 | 0.018244 |
| 28.625 | 0.027951 | 0.018903 | 0.037668 | 0.037462 | 0.022846 | 0.019304 | 0.042128 | 0.007485 | 0.020048 | 0.02267 | 0.024689 | 0.004268 | 0.017766 | 0.010247 | 0.007135 | 0.007091 | 0.007665 | 0.00514 | 0.00946 |
| 28.6388889 | 0.024107 | 0.007993 | 0.019612 | 0.017573 | 0.008988 | 0.017497 | 0.028514 | 0.01025 | 0.01549 | 0.018005 | 0.046949 | 0.003941 | 0.008756 | 0.005119 | 0.004548 | 0.004209 | 0.007189 | 0.007351 | 0.005357 |
| 28.6527778 | 0.023471 | 0.006179 | 0.019566 | 0.028399 | 0.008592 | 0.020567 | 0.018797 | 0.011208 | 0.017243 | 0.029554 | 0.063608 | 0.005235 | 0.007514 | 0.006718 | 0.001939 | 0.003373 | 0.009473 | 0.008841 | 0.003859 |
| 28.6666667 | 0.022504 | 0.011583 | 0.029367 | 0.037992 | 0.014763 | 0.024227 | 0.015143 | 0.005788 | 0.016205 | 0.034505 | 0.047908 | 0.007486 | 0.00838 | 0.00833 | 0.001759 | 0.003081 | 0.008195 | 0.010132 | 0.005353 |
| 28.6805556 | 0.020981 | 0.016722 | 0.026976 | 0.029561 | 0.020773 | 0.023516 | 0.014432 | 0.001304 | 0.017215 | 0.022455 | 0.025866 | 0.006252 | 0.005405 | 0.005612 | 0.00458 | 0.006099 | 0.004287 | 0.01185 | 0.013125 |
| 28.6944444 | 0.019015 | 0.017254 | 0.014804 | 0.015578 | 0.025187 | 0.0165 | 0.011025 | 0.000321 | 0.019228 | 0.015331 | 0.031255 | 0.004098 | 0.00213 | 0.003335 | 0.007296 | 0.009511 | 0.002533 | 0.012872 | 0.017902 |
| 28.7083333 | 0.012708 | 0.014504 | 0.012516 | 0.007272 | 0.026019 | 0.010104 | 0.007924 | 0.000706 | 0.022206 | 0.028908 | 0.043584 | 0.005704 | 0.001681 | 0.004921 | 0.007833 | 0.009757 | 0.00297 | 0.013068 | 0.012446 |
| 28.7222222 | 0.00824 | 0.01117 | 0.018693 | 0.009786 | 0.02111 | 0.015463 | 0.015099 | 0.001994 | 0.027822 | 0.042358 | 0.031649 | 0.007445 | 0.002834 | 0.007066 | 0.006838 | 0.007445 | 0.002256 | 0.013847 | 0.005519 |
| 28.7361111 | 0.01794 | 0.008768 | 0.020391 | 0.022149 | 0.011602 | 0.029713 | 0.032696 | 0.003222 | 0.03091 | 0.036266 | 0.015555 | 0.006503 | 0.004897 | 0.007796 | 0.005808 | 0.00369 | 0.001797 | 0.013955 | 0.004234 |
| 28.75 | 0.032514 | 0.007359 | 0.014785 | 0.032178 | 0.006448 | 0.039151 | 0.046488 | 0.002825 | 0.025268 | 0.01954 | 0.020186 | 0.004414 | 0.005437 | 0.008153 | 0.005312 | 0.001127 | 0.004159 | 0.011931 | 0.004032 |
| 28.7638889 | 0.032984 | 0.006495 | 0.008907 | 0.024596 | 0.013916 | 0.034256 | 0.043374 | 0.002179 | 0.013344 | 0.012851 | 0.029819 | 0.00258 | 0.003502 | 0.006372 | 0.005432 | 0.001507 | 0.007468 | 0.009476 | 0.003776 |
| 28.7777778 | 0.020945 | 0.00635 | 0.009704 | 0.011904 | 0.029158 | 0.018058 | 0.024864 | 0.003277 | 0.004281 | 0.021224 | 0.029019 | 0.002896 | 0.001911 | 0.00447 | 0.006403 | 0.003306 | 0.008011 | 0.008135 | 0.006351 |
| 28.7916667 | 0.009333 | 0.007104 | 0.014932 | 0.014714 | 0.039344 | 0.006333 | 0.011599 | 0.004572 | 0.006999 | 0.029675 | 0.023706 | 0.005684 | 0.002883 | 0.005099 | 0.007046 | 0.003898 | 0.006717 | 0.008053 | 0.008378 |
| 28.8055556 | 0.005418 | 0.008623 | 0.026883 | 0.019617 | 0.038152 | 0.005812 | 0.018159 | 0.005339 | 0.017439 | 0.030092 | 0.021438 | 0.008159 | 0.005881 | 0.004094 | 0.006032 | 0.002601 | 0.009903 | 0.009862 | 0.006167 |
| 28.8194444 | 0.009369 | 0.010507 | 0.046331 | 0.012812 | 0.028535 | 0.006777 | 0.031176 | 0.007451 | 0.021941 | 0.025222 | 0.023196 | 0.008274 | 0.007941 | 0.001349 | 0.003828 | 0.001933 | 0.017131 | 0.014694 | 0.004931 |
| 28.8333333 | 0.014448 | 0.013472 | 0.048263 | 0.007487 | 0.016273 | 0.004558 | 0.036434 | 0.009604 | 0.017797 | 0.019832 | 0.025727 | 0.007255 | 0.007773 | 0.001962 | 0.003012 | 0.003862 | 0.021806 | 0.021131 | 0.010878 |
| 28.8472222 | 0.016663 | 0.018753 | 0.027962 | 0.013618 | 0.013547 | 0.003532 | 0.032948 | 0.007929 | 0.011442 | 0.018838 | 0.028218 | 0.007765 | 0.008244 | 0.006362 | 0.004903 | 0.006158 | 0.022781 | 0.024516 | 0.018969 |
| 28.8611111 | 0.016268 | 0.023756 | 0.019704 | 0.021159 | 0.025808 | 0.00528 | 0.022558 | 0.00467 | 0.009285 | 0.02378 | 0.032135 | 0.011838 | 0.012461 | 0.010746 | 0.006771 | 0.006653 | 0.022773 | 0.021949 | 0.024085 |
| 28.875 | 0.012019 | 0.024356 | 0.02846 | 0.022638 | 0.037353 | 0.007398 | 0.012331 | 0.005884 | 0.013819 | 0.03126 | 0.037566 | 0.019093 | 0.017854 | 0.013446 | 0.006591 | 0.0061 | 0.021571 | 0.016125 | 0.026526 |
| 28.8888889 | 0.01239 | 0.022397 | 0.034953 | 0.025758 | 0.038074 | 0.008934 | 0.016615 | 0.009718 | 0.024034 | 0.034544 | 0.041634 | 0.026232 | 0.018622 | 0.017407 | 0.005478 | 0.006242 | 0.018441 | 0.012464 | 0.025641 |
| 28.9027778 | 0.027576 | 0.023947 | 0.027893 | 0.037093 | 0.035939 | 0.010976 | 0.042974 | 0.010098 | 0.037734 | 0.029455 | 0.040472 | 0.027829 | 0.013488 | 0.023897 | 0.005431 | 0.007856 | 0.016736 | 0.013148 | 0.022428 |
| 28.9166667 | 0.047961 | 0.029425 | 0.02115 | 0.050889 | 0.033978 | 0.013505 | 0.075558 | 0.007029 | 0.05054 | 0.018585 | 0.035218 | 0.020229 | 0.006336 | 0.026962 | 0.006489 | 0.009665 | 0.018223 | 0.015788 | 0.019032 |
| 28.9305556 | 0.05554 | 0.033089 | 0.041427 | 0.056379 | 0.027305 | 0.014568 | 0.082908 | 0.009058 | 0.059566 | 0.008058 | 0.028864 | 0.009763 | 0.00184 | 0.022135 | 0.006187 | 0.009028 | 0.018625 | 0.016055 | 0.01535 |
| 28.9444444 | 0.045651 | 0.031323 | 0.069988 | 0.050053 | 0.018149 | 0.013877 | 0.058142 | 0.019103 | 0.061693 | 0.002682 | 0.020932 | 0.005506 | 0.001452 | 0.01246 | 0.003557 | 0.005229 | 0.015022 | 0.012153 | 0.010055 |
| 28.9583333 | 0.0263 | 0.02453 | 0.072185 | 0.038648 | 0.012331 | 0.01272 | 0.026814 | 0.027106 | 0.053654 | 0.003045 | 0.012184 | 0.004431 | 0.004433 | 0.004874 | 0.001351 | 0.001487 | 0.010862 | 0.007274 | 0.004853 |
| 28.9722222 | 0.01208 | 0.014682 | 0.046116 | 0.026485 | 0.009797 | 0.009866 | 0.019592 | 0.027756 | 0.03732 | 0.005758 | 0.006668 | 0.003643 | 0.010483 | 0.003608 | 0.001604 | 0.000593 | 0.009761 | 0.005629 | 0.003134 |
| 28.9861111 | 0.011999 | 0.006529 | 0.02242 | 0.01469 | 0.00676 | 0.00509 | 0.007327 | 0.022684 | 0.019642 | 0.007871 | 0.006421 | 0.00598 | 0.015811 | 0.004043 | 0.002564 | 0.001432 | 0.010028 | 0.00934 | 0.003537 |
| 29 | 0.016992 | 0.003831 | 0.030315 | 0.011504 | 0.006258 | 0.00273 | 0.011716 | 0.013753 | 0.014267 | 0.009385 | 0.008515 | 0.006258 | 0.014915 | 0.003526 | 0.002685 | 0.001765 | 0.008213 | 0.017108 | 0.003401 |
| 29.0138889 | 0.015887 | 0.003274 | 0.057494 | 0.017054 | 0.011936 | 0.005065 | 0.01538 | 0.006296 | 0.028443 | 0.011146 | 0.009223 | 0.004002 | 0.008879 | 0.004922 | 0.001987 | 0.001388 | 0.004648 | 0.024448 | 0.002976 |
| 29.0277778 | 0.011808 | 0.005123 | 0.07382 | 0.015828 | 0.013868 | 0.00674 | 0.015021 | 0.004483 | 0.043386 | 0.01512 | 0.010436 | 0.004572 | 0.004051 | 0.008916 | 0.004948 | 0.00465 | 0.002329 | 0.032298 | 0.003948 |
| 29.0416667 | 0.021291 | 0.021853 | 0.056794 | 0.019032 | 0.013405 | 0.006714 | 0.01287 | 0.014407 | 0.039438 | 0.03036 | 0.020013 | 0.014917 | 0.013082 | 0.017381 | 0.020918 | 0.01987 | 0.008164 | 0.057696 | 0.013851 |
| 29.0555556 | 0.054117 | 0.063546 | 0.042031 | 0.053529 | 0.027169 | 0.011565 | 0.019498 | 0.043564 | 0.044574 | 0.056399 | 0.038674 | 0.042316 | 0.038848 | 0.029956 | 0.044701 | 0.042991 | 0.026279 | 0.107688 | 0.034096 |
| 29.0694444 | 0.085684 | 0.113095 | 0.081708 | 0.092493 | 0.04518 | 0.018777 | 0.030601 | 0.068367 | 0.077965 | 0.064548 | 0.050057 | 0.064548 | 0.052307 | 0.036825 | 0.048843 | 0.048579 | 0.038546 | 0.136338 | 0.042104 |
| 29.0833333 | 0.087273 | 0.136149 | 0.125066 | 0.094969 | 0.043004 | 0.021826 | 0.030264 | 0.061893 | 0.096123 | 0.040025 | 0.040755 | 0.051785 | 0.036437 | 0.028085 | 0.028459 | 0.029535 | 0.02798 | 0.10429 | 0.025521 |
| 29.0972222 | 0.065865 | 0.12173 | 0.12252 | 0.068154 | 0.023564 | 0.01979 | 0.020647 | 0.040861 | 0.084836 | 0.013281 | 0.020521 | 0.022992 | 0.021897 | 0.013443 | 0.012911 | 0.013777 | 0.011343 | 0.04747 | 0.00877 |
| 29.1111111 | 0.036244 | 0.082293 | 0.100034 | 0.036652 | 0.009193 | 0.012226 | 0.011892 | 0.023979 | 0.070704 | 0.005312 | 0.00909 | 0.013127 | 0.02718 | 0.008645 | 0.010594 | 0.013107 | 0.005912 | 0.014361 | 0.00802 |
| 29.125 | 0.016724 | 0.039594 | 0.074675 | 0.015404 | 0.009804 | 0.005307 | 0.005872 | 0.011093 | 0.057298 | 0.003908 | 0.012391 | 0.018243 | 0.03317 | 0.010775 | 0.009691 | 0.014508 | 0.00557 | 0.010465 | 0.012965 |
| 29.1388889 | 0.022983 | 0.020607 | 0.046674 | 0.012107 | 0.015761 | 0.005155 | 0.004601 | 0.004544 | 0.0395 | 0.002368 | 0.018882 | 0.024181 | 0.032531 | 0.012643 | 0.007585 | 0.011542 | 0.004126 | 0.017012 | 0.015825 |
| 29.1527778 | 0.038997 | 0.027681 | 0.022624 | 0.02104 | 0.019399 | 0.005377 | 0.009002 | 0.008556 | 0.0216 | 0.001872 | 0.018027 | 0.027272 | 0.027965 | 0.013018 | 0.004673 | 0.006661 | 0.002322 | 0.0195 | 0.013869 |
| 29.1666667 | 0.05079 | 0.040297 | 0.015384 | 0.028552 | 0.022145 | 0.002512 | 0.012359 | 0.019542 | 0.008704 | 0.002041 | 0.012784 | 0.024128 | 0.018716 | 0.011867 | 0.002726 | 0.004526 | 0.001123 | 0.015127 | 0.007165 |
| 29.1805556 | 0.057676 | 0.04599 | 0.029076 | 0.027362 | 0.023388 | 0.001743 | 0.010154 | 0.03128 | 0.003354 | 0.003171 | 0.015211 | 0.017308 | 0.011144 | 0.010746 | 0.002766 | 0.004352 | 0.001441 | 0.010494 | 0.004794 |
| 29.1944444 | 0.056856 | 0.042939 | 0.056165 | 0.022857 | 0.020826 | 0.004273 | 0.008433 | 0.040561 | 0.00486 | 0.005539 | 0.031003 | 0.012431 | 0.009913 | 0.011234 | 0.004601 | 0.004763 | 0.003396 | 0.01208 | 0.012475 |

| | | | | | | | | | | | | | | | | | | | |
|---|---|---|---|---|---|---|---|---|---|---|---|---|---|---|---|---|---|---|---|
| 29.2083333 | 0.04931 | 0.038202 | 0.087694 | 0.022141 | 0.016089 | 0.007147 | 0.013095 | 0.042575 | 0.008131 | 0.010013 | 0.046593 | 0.010614 | 0.011482 | 0.012353 | 0.0077 | 0.006515 | 0.004976 | 0.01166 | 0.020204 |
| 29.2222222 | 0.037913 | 0.040843 | 0.109681 | 0.02094 | 0.011438 | 0.008396 | 0.021384 | 0.037639 | 0.011496 | 0.014881 | 0.046549 | 0.009821 | 0.01054 | 0.011502 | 0.007953 | 0.00563 | 0.003808 | 0.00955 | 0.016663 |
| 29.2361111 | 0.026555 | 0.054414 | 0.113301 | 0.015675 | 0.006606 | 0.008114 | 0.028075 | 0.029728 | 0.016019 | 0.016357 | 0.034248 | 0.008498 | 0.006572 | 0.00732 | 0.00497 | 0.002468 | 0.002123 | 0.016416 | 0.009058 |
| 29.25 | 0.018722 | 0.073485 | 0.10436 | 0.009417 | 0.002488 | 0.007808 | 0.026765 | 0.020658 | 0.02076 | 0.013692 | 0.0198 | 0.005411 | 0.002978 | 0.003639 | 0.002192 | 0.00093 | 0.002482 | 0.02654 | 0.01058 |
| 29.2638889 | 0.014709 | 0.081826 | 0.089746 | 0.0043 | 0.000809 | 0.00825 | 0.017196 | 0.012293 | 0.02423 | 0.010135 | 0.008761 | 0.003243 | 0.002891 | 0.004173 | 0.000973 | 0.000718 | 0.002988 | 0.032861 | 0.017724 |
| 29.2777778 | 0.012021 | 0.071016 | 0.066853 | 0.001786 | 0.001161 | 0.008703 | 0.008769 | 0.007965 | 0.025507 | 0.008142 | 0.004027 | 0.005381 | 0.005097 | 0.004793 | 0.000889 | 0.000813 | 0.003579 | 0.034327 | 0.018648 |
| 29.2916667 | 0.010708 | 0.05212 | 0.038939 | 0.001731 | 0.001369 | 0.009567 | 0.005104 | 0.007655 | 0.027799 | 0.008363 | 0.002984 | 0.006647 | 0.005538 | 0.0038 | 0.001686 | 0.001078 | 0.004158 | 0.028137 | 0.011764 |
| 29.3055556 | 0.013376 | 0.038743 | 0.019899 | 0.004566 | 0.001774 | 0.011767 | 0.003206 | 0.006917 | 0.034694 | 0.011297 | 0.002697 | 0.005824 | 0.003547 | 0.005082 | 0.003502 | 0.001769 | 0.003547 | 0.017041 | 0.007059 |
| 29.3194444 | 0.017829 | 0.031907 | 0.01222 | 0.012397 | 0.003716 | 0.013493 | 0.002884 | 0.006805 | 0.041943 | 0.014375 | 0.002956 | 0.008304 | 0.002269 | 0.007701 | 0.004596 | 0.002884 | 0.003488 | 0.008956 | 0.007259 |
| 29.3333333 | 0.019968 | 0.025646 | 0.007052 | 0.019877 | 0.004661 | 0.012491 | 0.004356 | 0.007066 | 0.04358 | 0.012621 | 0.005473 | 0.00947 | 0.002892 | 0.006983 | 0.003363 | 0.003382 | 0.004059 | 0.007393 | 0.007632 |
| 29.3472222 | 0.019283 | 0.017925 | 0.002522 | 0.020775 | 0.003613 | 0.009312 | 0.00581 | 0.004263 | 0.040061 | 0.007114 | 0.012101 | 0.006391 | 0.004051 | 0.003517 | 0.001362 | 0.002798 | 0.004318 | 0.008179 | 0.011682 |
| 29.3611111 | 0.017067 | 0.010859 | 0.001313 | 0.016047 | 0.002449 | 0.005955 | 0.006428 | 0.001678 | 0.034936 | 0.004052 | 0.015745 | 0.00613 | 0.004892 | 0.001168 | 0.000525 | 0.002113 | 0.004956 | 0.008254 | 0.016632 |
| 29.375 | 0.014997 | 0.006072 | 0.001234 | 0.009128 | 0.001779 | 0.00395 | 0.006039 | 0.001476 | 0.030705 | 0.00477 | 0.010857 | 0.008324 | 0.004737 | 0.001673 | 0.000936 | 0.002115 | 0.004971 | 0.008922 | 0.012877 |
| 29.3888889 | 0.01519 | 0.005646 | 0.000884 | 0.004583 | 0.001181 | 0.003998 | 0.005117 | 0.002176 | 0.029573 | 0.006119 | 0.00677 | 0.007267 | 0.003184 | 0.005036 | 0.002295 | 0.002399 | 0.003362 | 0.009226 | 0.006651 |
| 29.4027778 | 0.017959 | 0.01067 | 0.001333 | 0.005126 | 0.002042 | 0.00468 | 0.00437 | 0.004234 | 0.031032 | 0.0069 | 0.010805 | 0.007118 | 0.002532 | 0.008315 | 0.003299 | 0.002117 | 0.002268 | 0.007057 | 0.007694 |
| 29.4166667 | 0.019674 | 0.016887 | 0.002209 | 0.008223 | 0.00693 | 0.003892 | 0.003798 | 0.005707 | 0.031776 | 0.008802 | 0.016299 | 0.012264 | 0.003504 | 0.008014 | 0.002914 | 0.00159 | 0.002442 | 0.003756 | 0.010202 |
| 29.4305556 | 0.016085 | 0.018677 | 0.004235 | 0.010836 | 0.013494 | 0.002056 | 0.00458 | 0.004626 | 0.030773 | 0.013385 | 0.01855 | 0.017173 | 0.004503 | 0.004968 | 0.002219 | 0.001965 | 0.004572 | 0.001969 | 0.009079 |
| 29.4444444 | 0.00872 | 0.01554 | 0.009739 | 0.01336 | 0.016164 | 0.000967 | 0.0083 | 0.003024 | 0.028546 | 0.01886 | 0.01828 | 0.017849 | 0.008546 | 0.00253 | 0.002392 | 0.003068 | 0.010474 | 0.001574 | 0.007352 |
| 29.4583333 | 0.004616 | 0.010354 | 0.018002 | 0.01587 | 0.013334 | 0.000764 | 0.014191 | 0.001815 | 0.025697 | 0.021634 | 0.015535 | 0.01508 | 0.014893 | 0.002512 | 0.00322 | 0.003682 | 0.015862 | 0.000984 | 0.007372 |
| 29.4722222 | 0.004614 | 0.00575 | 0.023749 | 0.017335 | 0.008179 | 0.000642 | 0.018843 | 0.001368 | 0.022728 | 0.020813 | 0.012165 | 0.010299 | 0.016436 | 0.005157 | 0.003622 | 0.003202 | 0.016247 | 0.000817 | 0.009388 |
| 29.4861111 | 0.003359 | 0.002885 | 0.022526 | 0.017419 | 0.004605 | 0.000835 | 0.018682 | 0.00253 | 0.018224 | 0.018457 | 0.010631 | 0.005957 | 0.01149 | 0.008893 | 0.003062 | 0.00183 | 0.013356 | 0.002581 | 0.011894 |
| 29.5 | 0.002749 | 0.001349 | 0.01609 | 0.01582 | 0.003684 | 0.00096 | 0.013591 | 0.003369 | 0.01167 | 0.017183 | 0.012014 | 0.004875 | 0.004833 | 0.011066 | 0.002346 | 0.000733 | 0.011234 | 0.005707 | 0.012973 |
| 29.5138889 | 0.006951 | 0.000665 | 0.010885 | 0.014291 | 0.003102 | 0.001016 | 0.008105 | 0.002541 | 0.007108 | 0.018685 | 0.015376 | 0.008685 | 0.002099 | 0.011419 | 0.00271 | 0.001567 | 0.013116 | 0.007522 | 0.011678 |
| 29.5277778 | 0.014567 | 0.002189 | 0.012333 | 0.018003 | 0.005823 | 0.001579 | 0.00803 | 0.003074 | 0.010469 | 0.024256 | 0.019933 | 0.016216 | 0.006002 | 0.012466 | 0.004754 | 0.004807 | 0.01929 | 0.007814 | 0.010442 |
| 29.5416667 | 0.023707 | 0.008058 | 0.022581 | 0.032172 | 0.020185 | 0.003962 | 0.017303 | 0.008679 | 0.026672 | 0.035667 | 0.027384 | 0.023725 | 0.014556 | 0.01732 | 0.007879 | 0.009175 | 0.028914 | 0.012573 | 0.015162 |
| 29.5555556 | 0.03345 | 0.016161 | 0.041278 | 0.053211 | 0.043038 | 0.011343 | 0.03556 | 0.018097 | 0.050611 | 0.04991 | 0.037284 | 0.027939 | 0.022074 | 0.025499 | 0.009802 | 0.011887 | 0.03714 | 0.029139 | 0.024527 |
| 29.5694444 | 0.041425 | 0.021713 | 0.062512 | 0.069322 | 0.058 | 0.021053 | 0.055264 | 0.026081 | 0.065747 | 0.054646 | 0.044249 | 0.03018 | 0.025125 | 0.031903 | 0.008119 | 0.010679 | 0.03604 | 0.052063 | 0.02986 |
| 29.5833333 | 0.046355 | 0.023995 | 0.081404 | 0.074408 | 0.056342 | 0.025256 | 0.065681 | 0.028285 | 0.07078 | 0.046301 | 0.044882 | 0.034436 | 0.026307 | 0.033509 | 0.004005 | 0.007755 | 0.028084 | 0.060321 | 0.031982 |
| 29.5972222 | 0.045114 | 0.024466 | 0.089403 | 0.067789 | 0.04583 | 0.022625 | 0.060856 | 0.02673 | 0.07354 | 0.03686 | 0.039458 | 0.036283 | 0.02566 | 0.029387 | 0.001044 | 0.005338 | 0.019951 | 0.046235 | 0.032623 |
| 29.6111111 | 0.033234 | 0.023599 | 0.069703 | 0.051155 | 0.036848 | 0.017896 | 0.045245 | 0.025022 | 0.063844 | 0.029813 | 0.025105 | 0.024859 | 0.019723 | 0.018351 | 0.000806 | 0.003416 | 0.011458 | 0.028657 | 0.025853 |
| 29.625 | 0.016948 | 0.022809 | 0.032765 | 0.032294 | 0.034641 | 0.014123 | 0.030296 | 0.017976 | 0.038352 | 0.020169 | 0.017237 | 0.013695 | 0.010637 | 0.009374 | 0.002224 | 0.003981 | 0.006793 | 0.023299 | 0.014234 |
| 29.6388889 | 0.009036 | 0.02202 | 0.016934 | 0.016519 | 0.036678 | 0.0121 | 0.021132 | 0.009735 | 0.015875 | 0.009378 | 0.038394 | 0.021768 | 0.004869 | 0.012282 | 0.003789 | 0.006064 | 0.009535 | 0.027021 | 0.009974 |
| 29.6527778 | 0.012073 | 0.018649 | 0.034879 | 0.005682 | 0.035042 | 0.011923 | 0.017089 | 0.008587 | 0.007974 | 0.002924 | 0.06711 | 0.033513 | 0.005068 | 0.019631 | 0.00475 | 0.006384 | 0.012282 | 0.030262 | 0.01514 |
| 29.6666667 | 0.017628 | 0.01139 | 0.057434 | 0.001962 | 0.027131 | 0.013713 | 0.017711 | 0.007162 | 0.008127 | 0.001545 | 0.066376 | 0.033573 | 0.005662 | 0.021658 | 0.004048 | 0.004344 | 0.011858 | 0.027844 | 0.016579 |
| 29.6805556 | 0.021315 | 0.008877 | 0.062428 | 0.002787 | 0.015146 | 0.013669 | 0.019441 | 0.004682 | 0.010399 | 0.001463 | 0.037693 | 0.023623 | 0.003754 | 0.017487 | 0.002245 | 0.002047 | 0.01344 | 0.021264 | 0.010123 |
| 29.6944444 | 0.022032 | 0.019595 | 0.052698 | 0.005513 | 0.008234 | 0.013187 | 0.018085 | 0.005577 | 0.012981 | 0.003323 | 0.020213 | 0.01191 | 0.00293 | 0.009605 | 0.001164 | 0.001469 | 0.018762 | 0.014363 | 0.005425 |
| 29.7083333 | 0.015456 | 0.032475 | 0.03441 | 0.010174 | 0.014877 | 0.014588 | 0.012731 | 0.00694 | 0.01044 | 0.009057 | 0.030121 | 0.009783 | 0.00254 | 0.004757 | 0.001819 | 0.00262 | 0.022894 | 0.007899 | 0.004272 |
| 29.7222222 | 0.009956 | 0.036272 | 0.027384 | 0.014402 | 0.026926 | 0.010148 | 0.006695 | 0.007091 | 0.005199 | 0.014447 | 0.036289 | 0.014692 | 0.001971 | 0.006766 | 0.003959 | 0.005019 | 0.018561 | 0.003596 | 0.003868 |
| 29.7361111 | 0.014964 | 0.033606 | 0.04215 | 0.016236 | 0.036558 | 0.005147 | 0.005831 | 0.006345 | 0.005573 | 0.014856 | 0.023694 | 0.017445 | 0.003266 | 0.009579 | 0.004724 | 0.005372 | 0.011708 | 0.003462 | 0.005785 |
| 29.75 | 0.019585 | 0.029884 | 0.056092 | 0.0171 | 0.043849 | 0.009645 | 0.010867 | 0.004956 | 0.010769 | 0.011269 | 0.015564 | 0.016078 | 0.003507 | 0.009558 | 0.004157 | 0.00383 | 0.015544 | 0.003674 | 0.005857 |
| 29.7638889 | 0.016813 | 0.026847 | 0.056427 | 0.019135 | 0.043115 | 0.015464 | 0.017547 | 0.003048 | 0.016692 | 0.007339 | 0.025298 | 0.01195 | 0.002466 | 0.008308 | 0.005398 | 0.004811 | 0.023459 | 0.004699 | 0.004383 |
| 29.7777778 | 0.010896 | 0.022008 | 0.048726 | 0.022261 | 0.028376 | 0.011447 | 0.02432 | 0.001283 | 0.018378 | 0.0045 | 0.034814 | 0.006529 | 0.003941 | 0.005593 | 0.006574 | 0.006491 | 0.022645 | 0.00975 | 0.006454 |
| 29.7916667 | 0.006538 | 0.014946 | 0.034572 | 0.022289 | 0.014394 | 0.003188 | 0.029277 | 0.00042 | 0.013458 | 0.002633 | 0.033121 | 0.004307 | 0.005701 | 0.004736 | 0.004654 | 0.004918 | 0.014602 | 0.015085 | 0.006742 |
| 29.8055556 | 0.00937 | 0.010385 | 0.018491 | 0.016361 | 0.019971 | 0.000457 | 0.028957 | 0.001219 | 0.009815 | 0.003345 | 0.029152 | 0.008252 | 0.005026 | 0.008012 | 0.00293 | 0.003403 | 0.006521 | 0.017591 | 0.006105 |
| 29.8194444 | 0.019891 | 0.014261 | 0.01661 | 0.009504 | 0.033742 | 0.002037 | 0.024271 | 0.006344 | 0.013733 | 0.008743 | 0.031042 | 0.014079 | 0.003788 | 0.008684 | 0.003611 | 0.006007 | 0.003964 | 0.01704 | 0.013737 |
| 29.8333333 | 0.034451 | 0.02668 | 0.034543 | 0.009813 | 0.038232 | 0.007525 | 0.017066 | 0.019102 | 0.020673 | 0.018087 | 0.039997 | 0.016817 | 0.00624 | 0.00462 | 0.003249 | 0.008542 | 0.007604 | 0.012641 | 0.025419 |
| 29.8472222 | 0.047161 | 0.040283 | 0.056569 | 0.017338 | 0.033125 | 0.011852 | 0.008649 | 0.035899 | 0.023755 | 0.027307 | 0.050777 | 0.01551 | 0.014709 | 0.002076 | 0.001454 | 0.007605 | 0.014728 | 0.006511 | 0.029751 |
| 29.8611111 | 0.051076 | 0.047673 | 0.068572 | 0.024978 | 0.021127 | 0.008597 | 0.002911 | 0.044789 | 0.024457 | 0.031993 | 0.058007 | 0.012714 | 0.023971 | 0.004148 | 0.001095 | 0.005057 | 0.024071 | 0.004161 | 0.024738 |
| 29.875 | 0.044735 | 0.046743 | 0.064241 | 0.028327 | 0.012456 | 0.005533 | 0.001231 | 0.04081 | 0.026635 | 0.031049 | 0.060598 | 0.012519 | 0.027045 | 0.009603 | 0.002231 | 0.002705 | 0.033427 | 0.006567 | 0.016033 |
| 29.8888889 | 0.029912 | 0.039856 | 0.049665 | 0.027425 | 0.019599 | 0.004849 | 0.002211 | 0.031301 | 0.024388 | 0.027281 | 0.058661 | 0.016703 | 0.023674 | 0.017208 | 0.003884 | 0.002031 | 0.038937 | 0.009547 | 0.009582 |
| 29.9027778 | 0.01288 | 0.031661 | 0.033156 | 0.025718 | 0.034459 | 0.003523 | 0.006601 | 0.022514 | 0.014024 | 0.023507 | 0.052438 | 0.022038 | 0.016564 | 0.0254 | 0.005226 | 0.002879 | 0.037615 | 0.010902 | 0.006098 |
| 29.9166667 | 0.004214 | 0.02499 | 0.019455 | 0.026778 | 0.042216 | 0.003403 | 0.01415 | 0.015701 | 0.007283 | 0.019541 | 0.042726 | 0.022529 | 0.008525 | 0.030158 | 0.00422 | 0.002687 | 0.028938 | 0.010998 | 0.003674 |
| 29.9305556 | 0.005543 | 0.018512 | 0.014794 | 0.028987 | 0.044507 | 0.003192 | 0.020006 | 0.009052 | 0.010437 | 0.01424 | 0.030951 | 0.016356 | 0.004636 | 0.02791 | 0.003308 | 0.002061 | 0.017483 | 0.010333 | 0.003384 |
| 29.9444444 | 0.009975 | 0.012091 | 0.01161 | 0.02739 | 0.049752 | 0.002176 | 0.017221 | 0.006066 | 0.014744 | 0.007889 | 0.019516 | 0.008151 | 0.005844 | 0.019493 | 0.005879 | 0.004306 | 0.009389 | 0.00826 | 0.005024 |
| 29.9583333 | 0.013171 | 0.007195 | 0.010092 | 0.018828 | 0.048245 | 0.003117 | 0.01004 | 0.011311 | 0.017911 | 0.005278 | 0.009902 | 0.004396 | 0.007146 | 0.009412 | 0.008235 | 0.00806 | 0.005925 | 0.004529 | 0.005549 |
| 29.9722222 | 0.013074 | 0.003473 | 0.022634 | 0.008289 | 0.031693 | 0.006071 | 0.011193 | 0.016241 | 0.020373 | 0.009119 | 0.003517 | 0.004665 | 0.006208 | 0.00369 | 0.007342 | 0.009961 | 0.004549 | 0.001763 | 0.004083 |
| 29.9861111 | 0.008457 | 0.002208 | 0.040682 | 0.004874 | 0.014135 | 0.005888 | 0.018961 | 0.014932 | 0.018047 | 0.012447 | 0.000954 | 0.003884 | 0.004013 | 0.003128 | 0.004796 | 0.008828 | 0.003218 | 0.001182 | 0.002036 |
| 30 | 0.003771 | 0.00445 | 0.049818 | 0.007986 | 0.00829 | 0.00339 | 0.02456 | 0.010565 | 0.012549 | 0.010267 | 0.000384 | 0.003443 | 0.001897 | 0.00252 | 0.002585 | 0.005488 | 0.002989 | 0.001502 | 0.000857 |
| 30.0138889 | 0.003436 | 0.006731 | 0.047649 | 0.010509 | 0.011483 | 0.003724 | 0.026475 | 0.006557 | 0.008499 | 0.005374 | 0.000262 | 0.005303 | 0.001243 | 0.001188 | 0.001525 | 0.002487 | 0.005982 | 0.003072 | 0.000614 |
| 30.0277778 | 0.007858 | 0.008837 | 0.033685 | 0.010854 | 0.011455 | 0.004059 | 0.022649 | 0.010477 | 0.014696 | 0.004456 | 0.002076 | 0.009981 | 0.002978 | 0.002598 | 0.001336 | 0.003841 | 0.006995 | 0.005846 | 0.001949 |
| 30.0416667 | 0.024242 | 0.025491 | 0.025148 | 0.021454 | 0.015661 | 0.007842 | 0.017788 | 0.031483 | 0.047468 | 0.01564 | 0.011965 | 0.026224 | 0.009665 | 0.012594 | 0.001853 | 0.01478 | 0.013103 | 0.016586 | 0.009496 |
| 30.0555556 | 0.057091 | 0.06444 | 0.053114 | 0.048918 | 0.03865 | 0.020303 | 0.028278 | 0.057128 | 0.100485 | 0.03729 | 0.030038 | 0.056497 | 0.02369 | 0.034764 | 0.005179 | 0.033066 | 0.040195 | 0.037443 | 0.024096 |
| 30.0694444 | 0.086674 | 0.098355 | 0.109047 | 0.075688 | 0.056149 | 0.03092 | 0.056576 | 0.060016 | 0.132505 | 0.047408 | 0.040503 | 0.076678 | 0.035217 | 0.052675 | 0.008071 | 0.04233 | 0.061463 | 0.042227 | 0.030132 |
| 30.0833333 | 0.090248 | 0.098606 | 0.143563 | 0.081625 | 0.048954 | 0.03274 | 0.079758 | 0.03916 | 0.121265 | 0.031434 | 0.031719 | 0.062057 | 0.028937 | 0.044691 | 0.006214 | 0.031615 | 0.046701 | 0.027736 | 0.021395 |
| 30.0972222 | 0.074583 | 0.077296 | 0.131949 | 0.06072 | 0.03351 | 0.026764 | 0.073075 | 0.015632 | 0.088669 | 0.013579 | 0.014164 | 0.029674 | 0.014865 | 0.022077 | 0.006077 | 0.015662 | 0.019954 | 0.025065 | 0.01616 |
| 30.1111111 | 0.057187 | 0.060442 | 0.094759 | 0.030837 | 0.01985 | 0.016285 | 0.039983 | 0.003398 | 0.062128 | 0.011706 | 0.003961 | 0.015522 | 0.013855 | 0.012169 | 0.009243 | 0.01293 | 0.009804 | 0.02952 | 0.016724 |
| 30.125 | 0.038083 | 0.045609 | 0.052292 | 0.01485 | 0.008595 | 0.00769 | 0.020353 | 0.002164 | 0.047133 | 0.010732 | 0.003121 | 0.023372 | 0.021921 | 0.016781 | 0.011404 | 0.019451 | 0.012616 | 0.022852 | 0.012647 |
| 30.1388889 | 0.02057 | 0.025438 | 0.020075 | 0.017101 | 0.004346 | 0.005598 | 0.035259 | 0.004211 | 0.034999 | 0.005606 | 0.004466 | 0.034715 | 0.026121 | 0.021676 | 0.013575 | 0.023116 | 0.017003 | 0.012226 | 0.007008 |
| 30.1527778 | 0.022332 | 0.017589 | 0.010447 | 0.030546 | 0.008298 | 0.010153 | 0.055091 | 0.004024 | 0.019622 | 0.003203 | 0.004458 | 0.041868 | 0.023943 | 0.022317 | 0.015333 | 0.021526 | 0.019032 | 0.00571 | 0.004183 |
| 30.1666667 | 0.041537 | 0.030989 | 0.012022 | 0.043161 | 0.015248 | 0.015723 | 0.061347 | 0.002498 | 0.01055 | 0.003239 | 0.004725 | 0.048857 | 0.019746 | 0.022391 | 0.014655 | 0.017462 | 0.019081 | 0.003344 | 0.003328 |
| 30.1805556 | 0.055624 | 0.045972 | 0.014529 | 0.04691 | 0.01874 | 0.017811 | 0.060266 | 0.002253 | 0.016818 | 0.004948 | 0.007503 | 0.059664 | 0.017849 | 0.023807 | 0.012062 | 0.012434 | 0.018752 | 0.003112 | 0.002957 |
| 30.1944444 | 0.055949 | 0.049258 | 0.018509 | 0.040384 | 0.019407 | 0.016304 | 0.05233 | 0.00211 | 0.023577 | 0.006406 | 0.011799 | 0.068542 | 0.01757 | 0.024986 | 0.008001 | 0.007303 | 0.017605 | 0.003027 | 0.002926 |
| 30.2083333 | 0.056347 | 0.045323 | 0.023663 | 0.027237 | 0.022324 | 0.015714 | 0.037993 | 0.001905 | 0.022227 | 0.00517 | 0.014488 | 0.066515 | 0.016597 | 0.024601 | 0.004013 | 0.003937 | 0.014379 | 0.002725 | 0.004103 |
| 30.2222222 | 0.062057 | 0.033547 | 0.030538 | 0.016566 | 0.025776 | 0.020947 | 0.020817 | 0.005558 | 0.027059 | 0.004242 | 0.015133 | 0.053094 | 0.01375 | 0.022668 | 0.002484 | 0.004389 | 0.010664 | 0.003334 | 0.005871 |
| 30.2361111 | 0.062594 | 0.015355 | 0.040651 | 0.013469 | 0.026102 | 0.028073 | 0.007155 | 0.014416 | 0.040896 | 0.003912 | 0.016451 | 0.032938 | 0.009511 | 0.01883 | 0.00203 | 0.007445 | 0.007608 | 0.005337 | 0.005938 |
| 30.25 | 0.060814 | 0.006476 | 0.055358 | 0.013642 | 0.023147 | 0.029043 | 0.003126 | 0.023754 | 0.050393 | 0.004959 | 0.021445 | 0.014771 | 0.005603 | 0.012836 | 0.00148 | 0.008755 | 0.004461 | 0.00778 | 0.003796 |
| 30.2638889 | 0.060921 | 0.014653 | 0.072444 | 0.0184 | 0.018672 | 0.02481 | 0.006444 | 0.030812 | 0.050232 | 0.011608 | 0.029941 | 0.006049 | 0.003437 | 0.006123 | 0.001641 | 0.006598 | 0.002836 | 0.009578 | 0.001898 |
| 30.2777778 | 0.056937 | 0.030486 | 0.084712 | 0.029612 | 0.014885 | 0.023486 | 0.013811 | 0.0357 | 0.044175 | 0.018943 | 0.036655 | 0.004472 | 0.003643 | 0.001637 | 0.001865 | 0.003969 | 0.002724 | 0.009879 | 0.001294 |
| 30.2916667 | 0.048684 | 0.043732 | 0.086344 | 0.036306 | 0.011632 | 0.025166 | 0.023688 | 0.037082 | 0.038713 | 0.019947 | 0.035092 | 0.007313 | 0.005386 | 0.000742 | 0.002828 | 0.004193 | 0.002978 | 0.008041 | 0.001188 |
| 30.3055556 | 0.035452 | 0.05085 | 0.077539 | 0.032127 | 0.009242 | 0.02432 | 0.030345 | 0.034816 | 0.039876 | 0.016258 | 0.025362 | 0.014427 | 0.007192 | 0.002148 | 0.00513 | 0.007749 | 0.006549 | 0.005128 | 0.001755 |
| 30.3194444 | 0.018571 | 0.052397 | 0.064059 | 0.020913 | 0.008967 | 0.021607 | 0.029435 | 0.029623 | 0.051424 | 0.016267 | 0.017174 | 0.020151 | 0.009183 | 0.004402 | 0.006632 | 0.013206 | 0.014323 | 0.003737 | 0.002263 |
| 30.3333333 | 0.007797 | 0.048794 | 0.055162 | 0.010625 | 0.011402 | 0.019768 | 0.026002 | 0.023309 | 0.069799 | 0.025524 | 0.017553 | 0.018529 | 0.011081 | 0.0055 | 0.00732 | 0.018095 | 0.022446 | 0.004977 | 0.003695 |
| 30.3472222 | 0.007116 | 0.039839 | 0.052021 | 0.009699 | 0.016288 | 0.018703 | 0.028352 | 0.021185 | 0.079374 | 0.035703 | 0.02109 | 0.010568 | 0.011919 | 0.006728 | 0.00866 | 0.019392 | 0.024415 | 0.006984 | 0.007964 |
| 30.3611111 | 0.014288 | 0.030059 | 0.04795 | 0.016987 | 0.021832 | 0.01798 | 0.035317 | 0.026725 | 0.077074 | 0.036895 | 0.019298 | 0.005191 | 0.013306 | 0.011717 | 0.009436 | 0.016734 | 0.019158 | 0.007739 | 0.012507 |
| 30.375 | 0.023977 | 0.024915 | 0.038969 | 0.02308 | 0.026317 | 0.01707 | 0.03711 | 0.034054 | 0.071501 | 0.029437 | 0.013453 | 0.009226 | 0.01618 | 0.016791 | 0.007357 | 0.012077 | 0.013536 | 0.007211 | 0.013935 |
| 30.3888889 | 0.027134 | 0.023099 | 0.023693 | 0.022447 | 0.028054 | 0.013189 | 0.027142 | 0.031928 | 0.053013 | 0.017563 | 0.012795 | 0.018356 | 0.017712 | 0.016674 | 0.003781 | 0.006385 | 0.010403 | 0.005969 | 0.011113 |
| 30.4027778 | 0.021007 | 0.019333 | 0.01384 | 0.017241 | 0.025192 | 0.006523 | 0.012479 | 0.018979 | 0.026166 | 0.009894 | 0.015396 | 0.025528 | 0.014944 | 0.013057 | 0.001588 | 0.00187 | 0.007692 | 0.004309 | 0.006141 |
| 30.4166667 | 0.013218 | 0.011842 | 0.01549 | 0.012482 | 0.018354 | 0.002314 | 0.003846 | 0.009298 | 0.012759 | 0.012728 | 0.013455 | 0.025338 | 0.00874 | 0.010422 | 0.001521 | 0.000653 | 0.005068 | 0.002599 | 0.00288 |
| 30.4305556 | 0.014931 | 0.004962 | 0.013646 | 0.012831 | 0.010563 | 0.002734 | 0.003445 | 0.007657 | 0.010367 | 0.021206 | 0.016473 | 0.016551 | 0.003387 | 0.010245 | 0.003804 | 0.002066 | 0.004108 | 0.001373 | 0.002924 |

| | | | | | | | | | | | | | | | | | | | |
|---|---|---|---|---|---|---|---|---|---|---|---|---|---|---|---|---|---|---|---|
| 30.4444444 | 0.029174 | 0.003344 | 0.00673 | 0.01987 | 0.005364 | 0.005893 | 0.009315 | 0.005366 | 0.015224 | 0.025152 | 0.029558 | 0.00767 | 0.001704 | 0.010951 | 0.006641 | 0.005269 | 0.005824 | 0.001212 | 0.006839 |
| 30.4583333 | 0.047259 | 0.007251 | 0.004875 | 0.030578 | 0.004136 | 0.009173 | 0.016269 | 0.003099 | 0.028814 | 0.020399 | 0.038842 | 0.005206 | 0.003374 | 0.010998 | 0.007264 | 0.00804 | 0.008662 | 0.00191 | 0.014473 |
| 30.4722222 | 0.057651 | 0.012841 | 0.006678 | 0.038943 | 0.004659 | 0.00977 | 0.018701 | 0.002433 | 0.038433 | 0.012274 | 0.037369 | 0.00536 | 0.007523 | 0.010293 | 0.004861 | 0.007322 | 0.009729 | 0.002138 | 0.021919 |
| 30.4861111 | 0.055061 | 0.016126 | 0.006043 | 0.040091 | 0.003934 | 0.006808 | 0.016092 | 0.002974 | 0.030448 | 0.007207 | 0.029991 | 0.008738 | 0.01173 | 0.009975 | 0.001724 | 0.004028 | 0.007968 | 0.00133 | 0.023717 |
| 30.5 | 0.043702 | 0.01763 | 0.003614 | 0.035153 | 0.003181 | 0.003112 | 0.011518 | 0.006923 | 0.017548 | 0.008367 | 0.022066 | 0.01923 | 0.013064 | 0.011228 | 0.001487 | 0.00209 | 0.004886 | 0.000471 | 0.018291 |
| 30.5138889 | 0.032185 | 0.020942 | 0.003432 | 0.029644 | 0.00508 | 0.001468 | 0.00837 | 0.010197 | 0.021861 | 0.013364 | 0.016299 | 0.028675 | 0.010865 | 0.013505 | 0.004451 | 0.003382 | 0.0035 | 0.000276 | 0.009158 |
| 30.5277778 | 0.025959 | 0.0262 | 0.006764 | 0.027609 | 0.006223 | 0.00123 | 0.008616 | 0.008228 | 0.030189 | 0.016791 | 0.015303 | 0.028327 | 0.006276 | 0.014463 | 0.006863 | 0.006036 | 0.006213 | 0.000523 | 0.002868 |
| 30.5416667 | 0.028776 | 0.03051 | 0.015887 | 0.033024 | 0.012138 | 0.001068 | 0.013969 | 0.006212 | 0.024637 | 0.019221 | 0.023455 | 0.022014 | 0.002227 | 0.013378 | 0.005666 | 0.007696 | 0.012738 | 0.00166 | 0.002643 |
| 30.5555556 | 0.044475 | 0.033417 | 0.035349 | 0.049478 | 0.035331 | 0.002376 | 0.027118 | 0.013185 | 0.023518 | 0.02352 | 0.038234 | 0.01931 | 0.002308 | 0.013198 | 0.003829 | 0.009197 | 0.02195 | 0.005996 | 0.009383 |
| 30.5694444 | 0.067616 | 0.03682 | 0.067186 | 0.069933 | 0.062356 | 0.006974 | 0.047249 | 0.02832 | 0.039421 | 0.028346 | 0.04713 | 0.025252 | 0.008824 | 0.017318 | 0.006642 | 0.013691 | 0.031223 | 0.014704 | 0.024514 |
| 30.5833333 | 0.082043 | 0.044044 | 0.099919 | 0.079464 | 0.070419 | 0.010434 | 0.06292 | 0.046065 | 0.055234 | 0.032928 | 0.043597 | 0.036113 | 0.019819 | 0.024669 | 0.012945 | 0.021622 | 0.035801 | 0.021316 | 0.040937 |
| 30.5972222 | 0.079435 | 0.051385 | 0.109036 | 0.070047 | 0.063355 | 0.011553 | 0.061185 | 0.056623 | 0.059874 | 0.036128 | 0.031593 | 0.042213 | 0.0276 | 0.029283 | 0.016325 | 0.026923 | 0.033923 | 0.018521 | 0.045363 |
| 30.6111111 | 0.061215 | 0.049282 | 0.087824 | 0.048741 | 0.055385 | 0.014853 | 0.044438 | 0.052045 | 0.055052 | 0.033215 | 0.01671 | 0.035247 | 0.026076 | 0.026131 | 0.012297 | 0.021386 | 0.026341 | 0.010473 | 0.031594 |
| 30.625 | 0.03396 | 0.039738 | 0.054174 | 0.028111 | 0.051243 | 0.017685 | 0.023137 | 0.042393 | 0.047806 | 0.025204 | 0.010558 | 0.020094 | 0.017446 | 0.016062 | 0.007454 | 0.012969 | 0.014864 | 0.00486 | 0.017275 |
| 30.6388889 | 0.02043 | 0.030975 | 0.024318 | 0.012549 | 0.051013 | 0.017859 | 0.009984 | 0.034643 | 0.03936 | 0.017781 | 0.020359 | 0.014824 | 0.008299 | 0.006929 | 0.010206 | 0.01631 | 0.006913 | 0.003368 | 0.022519 |
| 30.6527778 | 0.025455 | 0.025217 | 0.011887 | 0.004074 | 0.050168 | 0.017016 | 0.010747 | 0.023998 | 0.025691 | 0.010159 | 0.037264 | 0.02572 | 0.002769 | 0.005411 | 0.01416 | 0.024226 | 0.007376 | 0.003085 | 0.036472 |
| 30.6666667 | 0.030226 | 0.020372 | 0.022221 | 0.002302 | 0.043808 | 0.014122 | 0.011846 | 0.012484 | 0.019733 | 0.003467 | 0.050696 | 0.034982 | 0.001268 | 0.007362 | 0.012383 | 0.02415 | 0.009255 | 0.002065 | 0.044431 |
| 30.6805556 | 0.032762 | 0.01315 | 0.04054 | 0.002911 | 0.031383 | 0.008182 | 0.006329 | 0.010935 | 0.026396 | 0.005223 | 0.053856 | 0.027596 | 0.002597 | 0.008393 | 0.006931 | 0.016021 | 0.007326 | 0.002764 | 0.040594 |
| 30.6944444 | 0.032998 | 0.010919 | 0.050515 | 0.005581 | 0.016799 | 0.005147 | 0.002032 | 0.021081 | 0.031794 | 0.014201 | 0.045683 | 0.014689 | 0.005217 | 0.010555 | 0.002727 | 0.006267 | 0.004523 | 0.006602 | 0.023792 |
| 30.7083333 | 0.027005 | 0.020219 | 0.054485 | 0.009157 | 0.012278 | 0.009972 | 0.001287 | 0.030158 | 0.034381 | 0.018856 | 0.030732 | 0.014753 | 0.007448 | 0.013035 | 0.001735 | 0.002383 | 0.003165 | 0.011421 | 0.011462 |
| 30.7222222 | 0.025511 | 0.028604 | 0.058758 | 0.01016 | 0.023028 | 0.015253 | 0.002158 | 0.032332 | 0.035024 | 0.01286 | 0.015132 | 0.023514 | 0.008279 | 0.012437 | 0.002531 | 0.005446 | 0.002723 | 0.014232 | 0.012684 |
| 30.7361111 | 0.031075 | 0.027129 | 0.045297 | 0.007142 | 0.035073 | 0.015904 | 0.004167 | 0.031138 | 0.025036 | 0.005724 | 0.007114 | 0.026399 | 0.007848 | 0.008183 | 0.004462 | 0.009465 | 0.002873 | 0.013648 | 0.011336 |
| 30.75 | 0.031591 | 0.018607 | 0.023399 | 0.005155 | 0.039457 | 0.016628 | 0.005328 | 0.030716 | 0.017057 | 0.006522 | 0.005771 | 0.02173 | 0.005671 | 0.004406 | 0.005428 | 0.009369 | 0.00427 | 0.010796 | 0.006221 |
| 30.7638889 | 0.027784 | 0.009136 | 0.027636 | 0.011007 | 0.043446 | 0.017376 | 0.003816 | 0.034972 | 0.027334 | 0.007311 | 0.004378 | 0.013896 | 0.00331 | 0.005353 | 0.004446 | 0.006341 | 0.006138 | 0.007358 | 0.007802 |
| 30.7777778 | 0.022012 | 0.006404 | 0.039211 | 0.020909 | 0.049357 | 0.011974 | 0.002509 | 0.040971 | 0.04405 | 0.006331 | 0.004171 | 0.008224 | 0.003905 | 0.008107 | 0.003397 | 0.003863 | 0.006834 | 0.004496 | 0.010168 |
| 30.7916667 | 0.012781 | 0.011467 | 0.030078 | 0.029858 | 0.04818 | 0.004098 | 0.004688 | 0.039633 | 0.048576 | 0.011167 | 0.004551 | 0.011396 | 0.005604 | 0.007957 | 0.003031 | 0.002454 | 0.005605 | 0.002956 | 0.008301 |
| 30.8055556 | 0.00736 | 0.018325 | 0.012444 | 0.035063 | 0.036082 | 0.001243 | 0.007496 | 0.03017 | 0.034332 | 0.017491 | 0.003713 | 0.020032 | 0.005027 | 0.004836 | 0.002639 | 0.001496 | 0.003088 | 0.002929 | 0.005298 |
| 30.8194444 | 0.007368 | 0.021963 | 0.007154 | 0.031747 | 0.019454 | 0.001429 | 0.007143 | 0.021536 | 0.015103 | 0.016394 | 0.006489 | 0.024854 | 0.003175 | 0.002857 | 0.002067 | 0.000989 | 0.00194 | 0.003798 | 0.004278 |
| 30.8333333 | 0.006783 | 0.021722 | 0.012647 | 0.019835 | 0.009158 | 0.001711 | 0.005344 | 0.016531 | 0.008669 | 0.010279 | 0.016309 | 0.025877 | 0.004103 | 0.005832 | 0.001933 | 0.001601 | 0.002451 | 0.004513 | 0.007013 |
| 30.8472222 | 0.005565 | 0.018597 | 0.016514 | 0.010927 | 0.011479 | 0.002992 | 0.007264 | 0.012019 | 0.018651 | 0.010211 | 0.031403 | 0.026895 | 0.009897 | 0.012352 | 0.002712 | 0.004969 | 0.002505 | 0.004761 | 0.010456 |
| 30.8611111 | 0.007327 | 0.014626 | 0.017984 | 0.015471 | 0.017886 | 0.008241 | 0.014452 | 0.007506 | 0.034803 | 0.015227 | 0.04557 | 0.026783 | 0.017393 | 0.018178 | 0.003696 | 0.009426 | 0.004069 | 0.00517 | 0.008976 |
| 30.875 | 0.013371 | 0.014733 | 0.026875 | 0.026271 | 0.017445 | 0.016042 | 0.020726 | 0.004792 | 0.042633 | 0.015682 | 0.052571 | 0.024175 | 0.022164 | 0.019637 | 0.004216 | 0.011477 | 0.008062 | 0.006245 | 0.005116 |
| 30.8888889 | 0.02002 | 0.023073 | 0.053621 | 0.035802 | 0.011665 | 0.017059 | 0.023882 | 0.003672 | 0.039011 | 0.013486 | 0.05352 | 0.022638 | 0.02258 | 0.016748 | 0.004907 | 0.011172 | 0.012135 | 0.007913 | 0.005042 |
| 30.9027778 | 0.023355 | 0.036993 | 0.087748 | 0.042432 | 0.012714 | 0.009223 | 0.02899 | 0.002779 | 0.031296 | 0.015092 | 0.052611 | 0.023579 | 0.019152 | 0.012638 | 0.005289 | 0.009304 | 0.016401 | 0.010044 | 0.006258 |
| 30.9166667 | 0.023467 | 0.046563 | 0.106071 | 0.044121 | 0.026674 | 0.002262 | 0.034759 | 0.003344 | 0.026868 | 0.021436 | 0.048017 | 0.022027 | 0.013181 | 0.009262 | 0.004147 | 0.00606 | 0.020357 | 0.011741 | 0.006021 |
| 30.9305556 | 0.018855 | 0.044218 | 0.1035 | 0.041932 | 0.042879 | 0.001867 | 0.036734 | 0.006681 | 0.027153 | 0.02997 | 0.038216 | 0.015876 | 0.006935 | 0.006692 | 0.004009 | 0.005582 | 0.022023 | 0.011473 | 0.006317 |
| 30.9444444 | 0.011882 | 0.032889 | 0.089464 | 0.037036 | 0.050747 | 0.006209 | 0.032629 | 0.013427 | 0.026789 | 0.034702 | 0.027828 | 0.009131 | 0.002555 | 0.004006 | 0.007579 | 0.009864 | 0.021624 | 0.009306 | 0.007835 |
| 30.9583333 | 0.01361 | 0.01876 | 0.068908 | 0.026141 | 0.048804 | 0.010893 | 0.019829 | 0.020776 | 0.020823 | 0.030512 | 0.019548 | 0.004569 | 0.000743 | 0.002423 | 0.011254 | 0.012927 | 0.019515 | 0.007171 | 0.008928 |
| 30.9722222 | 0.03026 | 0.008774 | 0.041517 | 0.014033 | 0.038444 | 0.011913 | 0.012334 | 0.022505 | 0.012364 | 0.019303 | 0.012669 | 0.001791 | 0.000656 | 0.00407 | 0.010988 | 0.010529 | 0.015369 | 0.006192 | 0.008063 |
| 30.9861111 | 0.053701 | 0.006206 | 0.02195 | 0.014224 | 0.023098 | 0.009999 | 0.026323 | 0.017745 | 0.011103 | 0.008013 | 0.007328 | 0.000959 | 0.000583 | 0.006658 | 0.007069 | 0.005924 | 0.01107 | 0.005737 | 0.005891 |
| 31 | 0.071142 | 0.007039 | 0.029252 | 0.021749 | 0.011225 | 0.007926 | 0.047445 | 0.01303 | 0.019494 | 0.00181 | 0.003503 | 0.002585 | 0.000418 | 0.006493 | 0.002943 | 0.004221 | 0.009212 | 0.005116 | 0.003783 |
| 31.0138889 | 0.07491 | 0.009758 | 0.051142 | 0.024316 | 0.010664 | 0.006631 | 0.05661 | 0.013006 | 0.02729 | 0.001514 | 0.001279 | 0.005889 | 0.001303 | 0.004228 | 0.001306 | 0.004138 | 0.008862 | 0.00411 | 0.002175 |
| 31.0277778 | 0.060596 | 0.010939 | 0.062447 | 0.022717 | 0.013022 | 0.008476 | 0.050485 | 0.015038 | 0.02575 | 0.005547 | 0.002843 | 0.010323 | 0.005177 | 0.006587 | 0.001683 | 0.005454 | 0.009542 | 0.004277 | 0.002397 |
| 31.0416667 | 0.042795 | 0.011092 | 0.046837 | 0.018795 | 0.015841 | 0.016949 | 0.032143 | 0.019862 | 0.019927 | 0.012135 | 0.013128 | 0.019553 | 0.016471 | 0.02103 | 0.005956 | 0.017575 | 0.016934 | 0.011459 | 0.008864 |
| 31.0555556 | 0.055513 | 0.018387 | 0.033006 | 0.023339 | 0.033125 | 0.026684 | 0.025185 | 0.026824 | 0.027197 | 0.019944 | 0.031684 | 0.034822 | 0.035672 | 0.040912 | 0.014619 | 0.040018 | 0.032217 | 0.023572 | 0.021405 |
| 31.0694444 | 0.08647 | 0.026386 | 0.059174 | 0.041948 | 0.053159 | 0.024745 | 0.050566 | 0.02341 | 0.045089 | 0.022503 | 0.041267 | 0.042366 | 0.046499 | 0.047285 | 0.018749 | 0.04958 | 0.041878 | 0.024362 | 0.025644 |
| 31.0833333 | 0.091028 | 0.027582 | 0.088771 | 0.052959 | 0.053501 | 0.012781 | 0.087438 | 0.013335 | 0.053747 | 0.015897 | 0.029339 | 0.031131 | 0.034031 | 0.032755 | 0.012043 | 0.032885 | 0.036017 | 0.015003 | 0.01779 |
| 31.0972222 | 0.073075 | 0.026699 | 0.088007 | 0.048469 | 0.038189 | 0.006612 | 0.10531 | 0.011239 | 0.053128 | 0.011955 | 0.010826 | 0.013811 | 0.015244 | 0.01373 | 0.003889 | 0.011545 | 0.022393 | 0.012829 | 0.014414 |
| 31.1111111 | 0.056391 | 0.026962 | 0.071253 | 0.03871 | 0.021872 | 0.008133 | 0.089953 | 0.012405 | 0.048886 | 0.018338 | 0.002057 | 0.007315 | 0.010221 | 0.007251 | 0.002467 | 0.004771 | 0.01031 | 0.014894 | 0.01557 |
| 31.125 | 0.04811 | 0.032449 | 0.052733 | 0.025851 | 0.011857 | 0.009401 | 0.049355 | 0.011371 | 0.038531 | 0.025852 | 0.001627 | 0.013463 | 0.013289 | 0.008907 | 0.003758 | 0.006696 | 0.00567 | 0.013315 | 0.011813 |
| 31.1388889 | 0.04421 | 0.036792 | 0.034514 | 0.012746 | 0.006157 | 0.011307 | 0.028255 | 0.010957 | 0.023771 | 0.027567 | 0.0056 | 0.025638 | 0.016751 | 0.011456 | 0.005735 | 0.012897 | 0.013069 | 0.014024 | 0.007924 |
| 31.1527778 | 0.029564 | 0.025625 | 0.021703 | 0.010339 | 0.003766 | 0.017561 | 0.05704 | 0.012749 | 0.012802 | 0.025865 | 0.01303 | 0.036622 | 0.019809 | 0.015669 | 0.009252 | 0.022492 | 0.026148 | 0.019306 | 0.008602 |
| 31.1666667 | 0.017251 | 0.014453 | 0.024682 | 0.023808 | 0.006635 | 0.025791 | 0.101528 | 0.01424 | 0.014596 | 0.023463 | 0.018457 | 0.039068 | 0.020178 | 0.020182 | 0.013134 | 0.029876 | 0.032869 | 0.02153 | 0.009566 |
| 31.1805556 | 0.028791 | 0.023869 | 0.033384 | 0.036983 | 0.009954 | 0.028404 | 0.120279 | 0.010695 | 0.024166 | 0.017823 | 0.016251 | 0.03097 | 0.018345 | 0.021811 | 0.016426 | 0.031596 | 0.029454 | 0.016676 | 0.007124 |
| 31.1944444 | 0.042027 | 0.036503 | 0.02918 | 0.035684 | 0.010071 | 0.02212 | 0.107926 | 0.004513 | 0.026175 | 0.010315 | 0.008742 | 0.01824 | 0.017131 | 0.020221 | 0.015957 | 0.024695 | 0.020593 | 0.009981 | 0.005993 |
| 31.2083333 | 0.038031 | 0.036495 | 0.015345 | 0.026284 | 0.008311 | 0.011456 | 0.076947 | 0.001247 | 0.016524 | 0.008436 | 0.003753 | 0.007278 | 0.015448 | 0.015538 | 0.010099 | 0.014438 | 0.011786 | 0.0055 | 0.008419 |
| 31.2222222 | 0.028257 | 0.034657 | 0.007178 | 0.018344 | 0.005036 | 0.004218 | 0.044648 | 0.001653 | 0.007305 | 0.009086 | 0.004006 | 0.003539 | 0.011206 | 0.008447 | 0.006416 | 0.01411 | 0.005975 | 0.002693 | 0.00856 |
| 31.2361111 | 0.02522 | 0.041087 | 0.005992 | 0.012708 | 0.001779 | 0.002566 | 0.022515 | 0.004571 | 0.007658 | 0.006709 | 0.004269 | 0.008892 | 0.006407 | 0.003268 | 0.007829 | 0.019793 | 0.002855 | 0.00153 | 0.004909 |
| 31.25 | 0.030427 | 0.049774 | 0.010241 | 0.013198 | 0.000958 | 0.002673 | 0.012581 | 0.009667 | 0.009052 | 0.005455 | 0.002617 | 0.017011 | 0.003675 | 0.002853 | 0.007021 | 0.019129 | 0.002185 | 0.00196 | 0.001637 |
| 31.2638889 | 0.040243 | 0.054329 | 0.022984 | 0.014708 | 0.002668 | 0.004183 | 0.01157 | 0.005595 | 0.006564 | 0.004711 | 0.001841 | 0.020369 | 0.002442 | 0.004276 | 0.003812 | 0.012512 | 0.003992 | 0.002764 | 0.000525 |
| 31.2777778 | 0.053926 | 0.055889 | 0.037136 | 0.016752 | 0.005828 | 0.006352 | 0.016044 | 0.002741 | 0.008818 | 0.002972 | 0.002409 | 0.016997 | 0.002824 | 0.003757 | 0.002623 | 0.005844 | 0.004862 | 0.003489 | 0.000234 |
| 31.2916667 | 0.067541 | 0.052025 | 0.043262 | 0.017908 | 0.00875 | 0.007472 | 0.022228 | 0.002416 | 0.017174 | 0.003753 | 0.003295 | 0.009594 | 0.005535 | 0.002393 | 0.002427 | 0.003256 | 0.003497 | 0.004088 | 9.87E-05 |
| 31.3055556 | 0.073477 | 0.041829 | 0.03914 | 0.016407 | 0.009439 | 0.007592 | 0.024081 | 0.004358 | 0.021525 | 0.005661 | 0.004778 | 0.005649 | 0.008403 | 0.002751 | 0.002596 | 0.006816 | 0.002893 | 0.004311 | 0.000165 |
| 31.3194444 | 0.06922 | 0.026813 | 0.02789 | 0.012459 | 0.008109 | 0.0068 | 0.018605 | 0.00545 | 0.016065 | 0.005296 | 0.006077 | 0.007301 | 0.009317 | 0.005933 | 0.005423 | 0.014681 | 0.004006 | 0.004148 | 0.000197 |
| 31.3333333 | 0.055396 | 0.013805 | 0.017218 | 0.00805 | 0.0065 | 0.005166 | 0.009583 | 0.004052 | 0.007277 | 0.002958 | 0.004866 | 0.006572 | 0.007804 | 0.00974 | 0.008762 | 0.019545 | 0.004447 | 0.003545 | 0.000648 |
| 31.3472222 | 0.033906 | 0.012073 | 0.012708 | 0.004123 | 0.005366 | 0.003411 | 0.003364 | 0.002297 | 0.00346 | 0.001461 | 0.003125 | 0.004106 | 0.004955 | 0.009333 | 0.008857 | 0.01723 | 0.003375 | 0.002524 | 0.001652 |
| 31.3611111 | 0.014597 | 0.013601 | 0.013708 | 0.001906 | 0.004017 | 0.004215 | 0.001839 | 0.002915 | 0.003388 | 0.001353 | 0.004946 | 0.005865 | 0.002534 | 0.005616 | 0.006117 | 0.01044 | 0.003612 | 0.001428 | 0.002315 |
| 31.375 | 0.007762 | 0.009712 | 0.018363 | 0.001776 | 0.002522 | 0.008402 | 0.002267 | 0.005396 | 0.002622 | 0.002332 | 0.008931 | 0.008963 | 0.001629 | 0.004396 | 0.003455 | 0.005581 | 0.006011 | 0.00136 | 0.002299 |
| 31.3888889 | 0.008582 | 0.009848 | 0.024194 | 0.001848 | 0.001777 | 0.013108 | 0.005209 | 0.007942 | 0.002533 | 0.005453 | 0.011374 | 0.009683 | 0.002478 | 0.006659 | 0.003456 | 0.007064 | 0.007329 | 0.002677 | 0.002076 |
| 31.4027778 | 0.008385 | 0.016664 | 0.028603 | 0.001541 | 0.001198 | 0.015483 | 0.009605 | 0.009106 | 0.00518 | 0.009026 | 0.011277 | 0.008023 | 0.002839 | 0.0084 | 0.004574 | 0.009947 | 0.006602 | 0.003605 | 0.001739 |
| 31.4166667 | 0.009422 | 0.01653 | 0.030592 | 0.001135 | 0.000902 | 0.013572 | 0.008953 | 0.00818 | 0.00775 | 0.010504 | 0.010227 | 0.005492 | 0.001846 | 0.007969 | 0.003636 | 0.008325 | 0.006123 | 0.003024 | 0.001481 |
| 31.4305556 | 0.015753 | 0.010608 | 0.02977 | 0.00207 | 0.002038 | 0.007864 | 0.007497 | 0.006449 | 0.00774 | 0.00949 | 0.01065 | 0.004588 | 0.001523 | 0.007071 | 0.002141 | 0.004716 | 0.006929 | 0.001718 | 0.001721 |
| 31.4444444 | 0.02555 | 0.012175 | 0.026841 | 0.005317 | 0.003736 | 0.002697 | 0.012608 | 0.006015 | 0.006277 | 0.00686 | 0.012759 | 0.006722 | 0.001532 | 0.007169 | 0.002689 | 0.003824 | 0.007947 | 0.000664 | 0.002289 |
| 31.4583333 | 0.03121 | 0.014468 | 0.022594 | 0.007884 | 0.004532 | 0.002045 | 0.015373 | 0.007493 | 0.004941 | 0.004623 | 0.01473 | 0.01065 | 0.001188 | 0.008652 | 0.003078 | 0.004202 | 0.008802 | 0.00036 | 0.002849 |
| 31.4722222 | 0.026961 | 0.008786 | 0.018234 | 0.006085 | 0.004016 | 0.005297 | 0.009575 | 0.009546 | 0.003926 | 0.005046 | 0.015413 | 0.013607 | 0.001964 | 0.011549 | 0.002005 | 0.003417 | 0.011018 | 0.000574 | 0.003415 |
| 31.4861111 | 0.019288 | 0.006653 | 0.016502 | 0.004061 | 0.003796 | 0.006287 | 0.003905 | 0.009887 | 0.002523 | 0.007914 | 0.014936 | 0.013234 | 0.003797 | 0.015047 | 0.001357 | 0.004214 | 0.014963 | 0.000802 | 0.00358 |
| 31.5 | 0.016462 | 0.015421 | 0.018022 | 0.007376 | 0.00648 | 0.004475 | 0.004786 | 0.007565 | 0.000925 | 0.010013 | 0.013845 | 0.00939 | 0.005085 | 0.018064 | 0.001219 | 0.007573 | 0.018264 | 0.001311 | 0.00261 |
| 31.5138889 | 0.012983 | 0.019425 | 0.019766 | 0.011527 | 0.011562 | 0.003619 | 0.006259 | 0.004565 | 0.000965 | 0.008821 | 0.012794 | 0.004805 | 0.005939 | 0.020783 | 0.000722 | 0.010293 | 0.018952 | 0.001865 | 0.001132 |
| 31.5277778 | 0.012705 | 0.016246 | 0.023006 | 0.014638 | 0.015486 | 0.00542 | 0.006743 | 0.003967 | 0.00515 | 0.007232 | 0.014612 | 0.004326 | 0.00897 | 0.024937 | 0.000857 | 0.012477 | 0.019156 | 0.001974 | 0.001197 |
| 31.5416667 | 0.028889 | 0.020781 | 0.039759 | 0.021652 | 0.015392 | 0.012552 | 0.014629 | 0.006979 | 0.015966 | 0.011943 | 0.026178 | 0.011394 | 0.01721 | 0.034039 | 0.00298 | 0.018231 | 0.023572 | 0.003716 | 0.004153 |
| 31.5555556 | 0.05177 | 0.023241 | 0.071432 | 0.034369 | 0.011304 | 0.019249 | 0.031655 | 0.010133 | 0.030438 | 0.022138 | 0.047751 | 0.024437 | 0.027549 | 0.047462 | 0.006202 | 0.027003 | 0.031415 | 0.007445 | 0.009341 |
| 31.5694444 | 0.07111 | 0.018373 | 0.093337 | 0.0523 | 0.01292 | 0.020983 | 0.057492 | 0.008027 | 0.038625 | 0.028722 | 0.059789 | 0.041215 | 0.031149 | 0.056367 | 0.008765 | 0.032355 | 0.035511 | 0.008105 | 0.015509 |
| 31.5833333 | 0.088225 | 0.019527 | 0.091071 | 0.069381 | 0.025975 | 0.018623 | 0.081585 | 0.004067 | 0.039405 | 0.029134 | 0.048144 | 0.05947 | 0.02698 | 0.054916 | 0.008894 | 0.032308 | 0.033772 | 0.005618 | 0.023096 |
| 31.5972222 | 0.097141 | 0.022233 | 0.075866 | 0.074179 | 0.03536 | 0.011188 | 0.079865 | 0.003368 | 0.036648 | 0.027772 | 0.025485 | 0.064791 | 0.022001 | 0.044651 | 0.00577 | 0.028696 | 0.028278 | 0.006517 | 0.02803 |
| 31.6111111 | 0.089863 | 0.015923 | 0.058218 | 0.062125 | 0.032919 | 0.004083 | 0.051291 | 0.003239 | 0.028882 | 0.022467 | 0.011749 | 0.04407 | 0.017675 | 0.027044 | 0.002445 | 0.019955 | 0.017937 | 0.01077 | 0.022298 |
| 31.625 | 0.068308 | 0.007424 | 0.042368 | 0.039764 | 0.030976 | 0.003459 | 0.022061 | 0.004856 | 0.02509 | 0.013676 | 0.015892 | 0.023469 | 0.01203 | 0.013273 | 0.002953 | 0.010685 | 0.009137 | 0.013436 | 0.012063 |
| 31.6388889 | 0.042511 | 0.006055 | 0.030914 | 0.018067 | 0.040271 | 0.00736 | 0.013817 | 0.010574 | 0.030367 | 0.011547 | 0.029166 | 0.029947 | 0.006816 | 0.016219 | 0.00719 | 0.010415 | 0.009642 | 0.012939 | 0.010779 |
| 31.6527778 | 0.021845 | 0.008212 | 0.020425 | 0.005457 | 0.052821 | 0.0102 | 0.025219 | 0.016777 | 0.030194 | 0.013641 | 0.039633 | 0.041368 | 0.003849 | 0.025409 | 0.011888 | 0.017749 | 0.012125 | 0.010029 | 0.016251 |
| 31.6666667 | 0.015988 | 0.008711 | 0.009099 | 0.002858 | 0.053018 | 0.008993 | 0.036798 | 0.019591 | 0.017578 | 0.00978 | 0.045948 | 0.037944 | 0.002248 | 0.026518 | 0.0143 | 0.024059 | 0.012173 | 0.006254 | 0.019688 |

| | | | | | | | | | | | | | | | | | | | |
|---|---|---|---|---|---|---|---|---|---|---|---|---|---|---|---|---|---|---|---|
| 31.6805556 | 0.026834 | 0.007439 | 0.003793 | 0.004647 | 0.044433 | 0.005917 | 0.037423 | 0.016397 | 0.01176 | 0.003939 | 0.049446 | 0.028521 | 0.001772 | 0.023094 | 0.012375 | 0.023339 | 0.014322 | 0.003051 | 0.020097 |
| 31.6944444 | 0.039259 | 0.007228 | 0.004519 | 0.008574 | 0.037918 | 0.003774 | 0.033934 | 0.010486 | 0.028835 | 0.002751 | 0.047848 | 0.020327 | 0.003855 | 0.022873 | 0.006966 | 0.015969 | 0.017499 | 0.001316 | 0.018929 |
| 31.7083333 | 0.041113 | 0.011904 | 0.006578 | 0.011769 | 0.030974 | 0.005001 | 0.033774 | 0.01136 | 0.049709 | 0.002886 | 0.039096 | 0.014398 | 0.007546 | 0.025175 | 0.003801 | 0.007644 | 0.018732 | 0.001029 | 0.016182 |
| 31.7222222 | 0.031694 | 0.017198 | 0.009104 | 0.0086 | 0.0194 | 0.010261 | 0.035623 | 0.019788 | 0.052758 | 0.001948 | 0.024573 | 0.010549 | 0.009876 | 0.02595 | 0.004072 | 0.003156 | 0.016171 | 0.00093 | 0.012058 |
| 31.7361111 | 0.016873 | 0.016737 | 0.009027 | 0.005156 | 0.011504 | 0.013114 | 0.033717 | 0.026535 | 0.041946 | 0.00243 | 0.013657 | 0.006598 | 0.009266 | 0.023235 | 0.004039 | 0.00342 | 0.01012 | 0.000735 | 0.007477 |
| 31.75 | 0.013626 | 0.014212 | 0.005151 | 0.009001 | 0.020352 | 0.011279 | 0.026145 | 0.025219 | 0.026383 | 0.004249 | 0.018362 | 0.004295 | 0.006431 | 0.015372 | 0.006411 | 0.0064 | 0.00771 | 0.000792 | 0.003974 |
| 31.7638889 | 0.027443 | 0.015082 | 0.00195 | 0.015782 | 0.041546 | 0.009954 | 0.015771 | 0.016638 | 0.016321 | 0.00645 | 0.032758 | 0.006695 | 0.003265 | 0.008658 | 0.011687 | 0.008528 | 0.011793 | 0.000818 | 0.003884 |
| 31.7777778 | 0.042419 | 0.018569 | 0.002462 | 0.019371 | 0.057588 | 0.011598 | 0.007907 | 0.007491 | 0.023271 | 0.01004 | 0.040301 | 0.009895 | 0.001578 | 0.012142 | 0.01313 | 0.006873 | 0.013674 | 0.001455 | 0.007296 |
| 31.7916667 | 0.051554 | 0.020969 | 0.005779 | 0.019041 | 0.060011 | 0.017708 | 0.007229 | 0.005623 | 0.043149 | 0.013906 | 0.035041 | 0.011021 | 0.002099 | 0.019574 | 0.009586 | 0.003562 | 0.009659 | 0.003076 | 0.010541 |
| 31.8055556 | 0.059355 | 0.018477 | 0.011085 | 0.014791 | 0.053103 | 0.02782 | 0.00993 | 0.014194 | 0.057746 | 0.014395 | 0.021036 | 0.010871 | 0.003598 | 0.021601 | 0.005359 | 0.002892 | 0.008505 | 0.004352 | 0.009915 |
| 31.8194444 | 0.057455 | 0.012348 | 0.016064 | 0.006826 | 0.046042 | 0.03393 | 0.012677 | 0.024445 | 0.056144 | 0.010653 | 0.009831 | 0.009251 | 0.005263 | 0.017399 | 0.002342 | 0.003569 | 0.014287 | 0.00441 | 0.007002 |
| 31.8333333 | 0.037172 | 0.0113 | 0.014223 | 0.00374 | 0.042797 | 0.033535 | 0.013933 | 0.026159 | 0.0406 | 0.005504 | 0.010864 | 0.006057 | 0.006876 | 0.010886 | 0.000923 | 0.002701 | 0.018501 | 0.003755 | 0.008908 |
| 31.8472222 | 0.01646 | 0.014612 | 0.007179 | 0.00995 | 0.042212 | 0.035965 | 0.011834 | 0.02478 | 0.022947 | 0.001938 | 0.018833 | 0.005855 | 0.007464 | 0.011388 | 0.001258 | 0.001479 | 0.015976 | 0.003131 | 0.015947 |
| 31.8611111 | 0.018791 | 0.014895 | 0.006615 | 0.019447 | 0.042602 | 0.045114 | 0.016216 | 0.025741 | 0.021188 | 0.002449 | 0.027421 | 0.012879 | 0.00582 | 0.021641 | 0.002063 | 0.001488 | 0.010773 | 0.002752 | 0.019146 |
| 31.875 | 0.038067 | 0.020334 | 0.01209 | 0.025512 | 0.040771 | 0.05346 | 0.02917 | 0.024253 | 0.033043 | 0.00821 | 0.035566 | 0.022054 | 0.003561 | 0.033175 | 0.002618 | 0.002227 | 0.008298 | 0.002287 | 0.014775 |
| 31.8888889 | 0.054495 | 0.035423 | 0.017479 | 0.025404 | 0.036635 | 0.052916 | 0.039415 | 0.019884 | 0.043929 | 0.015914 | 0.039934 | 0.025125 | 0.004418 | 0.038565 | 0.003118 | 0.002847 | 0.009988 | 0.0015 | 0.007532 |
| 31.9027778 | 0.062815 | 0.047633 | 0.025404 | 0.024208 | 0.033678 | 0.04627 | 0.046363 | 0.017727 | 0.052457 | 0.019851 | 0.037755 | 0.020278 | 0.007777 | 0.036012 | 0.003356 | 0.002652 | 0.01202 | 0.000782 | 0.003341 |
| 31.9166667 | 0.066312 | 0.051257 | 0.037762 | 0.025319 | 0.031725 | 0.043143 | 0.056916 | 0.019457 | 0.058821 | 0.017688 | 0.03026 | 0.013829 | 0.010771 | 0.028282 | 0.003552 | 0.001702 | 0.010849 | 0.000736 | 0.003805 |
| 31.9305556 | 0.065559 | 0.051071 | 0.051202 | 0.026633 | 0.028373 | 0.045929 | 0.068721 | 0.023809 | 0.059822 | 0.011091 | 0.021409 | 0.009854 | 0.01283 | 0.01883 | 0.004877 | 0.001492 | 0.007033 | 0.001324 | 0.004201 |
| 31.9444444 | 0.060921 | 0.055333 | 0.058155 | 0.026383 | 0.023123 | 0.047146 | 0.069643 | 0.029322 | 0.053527 | 0.005124 | 0.015338 | 0.00634 | 0.014207 | 0.009357 | 0.007353 | 0.00241 | 0.003717 | 0.002355 | 0.002307 |
| 31.9583333 | 0.050268 | 0.059377 | 0.051387 | 0.023438 | 0.01655 | 0.040212 | 0.056465 | 0.033222 | 0.044837 | 0.003527 | 0.013795 | 0.002846 | 0.014709 | 0.003119 | 0.009487 | 0.003224 | 0.004123 | 0.003903 | 0.001088 |
| 31.9722222 | 0.029674 | 0.047477 | 0.033985 | 0.015358 | 0.009486 | 0.028064 | 0.038647 | 0.030746 | 0.037268 | 0.00393 | 0.014268 | 0.000756 | 0.013832 | 0.001407 | 0.009995 | 0.002674 | 0.007822 | 0.005779 | 0.001215 |
| 31.9861111 | 0.013545 | 0.025879 | 0.016427 | 0.00895 | 0.004509 | 0.01547 | 0.021082 | 0.021599 | 0.023861 | 0.005605 | 0.013604 | 0.000169 | 0.010925 | 0.001268 | 0.00822 | 0.001118 | 0.011437 | 0.007549 | 0.001578 |
| 32 | 0.017157 | 0.013659 | 0.009847 | 0.014973 | 0.004536 | 0.006073 | 0.010808 | 0.012452 | 0.013066 | 0.008517 | 0.010734 | 0.000568 | 0.006419 | 0.003459 | 0.005052 | 0.000519 | 0.012874 | 0.008602 | 0.003337 |
| 32.0138889 | 0.027927 | 0.018153 | 0.018334 | 0.026505 | 0.007768 | 0.003139 | 0.016601 | 0.007522 | 0.021208 | 0.00811 | 0.007148 | 0.001228 | 0.002539 | 0.009174 | 0.002314 | 0.001188 | 0.011368 | 0.007735 | 0.006375 |
| 32.0277778 | 0.028731 | 0.026575 | 0.027601 | 0.028414 | 0.007759 | 0.004598 | 0.02682 | 0.007011 | 0.034753 | 0.005101 | 0.007357 | 0.003887 | 0.003872 | 0.010461 | 0.001575 | 0.004783 | 0.008848 | 0.007703 | 0.006164 |
| 32.0416667 | 0.023143 | 0.025144 | 0.028033 | 0.026992 | 0.008408 | 0.012258 | 0.024463 | 0.010457 | 0.032878 | 0.009798 | 0.01752 | 0.015688 | 0.016345 | 0.015758 | 0.005149 | 0.020078 | 0.015827 | 0.020559 | 0.007882 |
| 32.0555556 | 0.033635 | 0.033595 | 0.041762 | 0.049541 | 0.020727 | 0.027703 | 0.028337 | 0.015408 | 0.037743 | 0.028097 | 0.034919 | 0.032879 | 0.033155 | 0.044193 | 0.011884 | 0.042861 | 0.035979 | 0.045201 | 0.019189 |
| 32.0694444 | 0.063027 | 0.063821 | 0.070278 | 0.085319 | 0.034988 | 0.037936 | 0.054107 | 0.01887 | 0.066997 | 0.045579 | 0.039086 | 0.035196 | 0.03415 | 0.069228 | 0.013045 | 0.045662 | 0.047965 | 0.049978 | 0.024972 |
| 32.0833333 | 0.083847 | 0.087082 | 0.079348 | 0.095183 | 0.03713 | 0.039542 | 0.069834 | 0.019255 | 0.084641 | 0.042289 | 0.023785 | 0.019709 | 0.020147 | 0.056438 | 0.008556 | 0.027618 | 0.036656 | 0.029276 | 0.016811 |
| 32.0972222 | 0.082478 | 0.093206 | 0.073166 | 0.080481 | 0.032148 | 0.043809 | 0.06554 | 0.01587 | 0.076994 | 0.021913 | 0.010593 | 0.006298 | 0.012106 | 0.026069 | 0.007596 | 0.018434 | 0.016946 | 0.017674 | 0.011545 |
| 32.1111111 | 0.066115 | 0.093413 | 0.070731 | 0.058086 | 0.027314 | 0.046797 | 0.05792 | 0.010526 | 0.058036 | 0.008987 | 0.005917 | 0.002299 | 0.012605 | 0.013216 | 0.008303 | 0.020716 | 0.007964 | 0.019722 | 0.013976 |
| 32.125 | 0.04211 | 0.083499 | 0.060538 | 0.031306 | 0.021773 | 0.03754 | 0.041876 | 0.006019 | 0.029964 | 0.016328 | 0.002132 | 0.00315 | 0.012539 | 0.019058 | 0.007142 | 0.020828 | 0.010747 | 0.01845 | 0.01592 |
| 32.1388889 | 0.021422 | 0.05913 | 0.037932 | 0.012805 | 0.013043 | 0.022202 | 0.01902 | 0.003986 | 0.011896 | 0.030494 | 0.000556 | 0.009906 | 0.011337 | 0.026491 | 0.006203 | 0.01851 | 0.015783 | 0.014375 | 0.01703 |
| 32.1527778 | 0.018314 | 0.030086 | 0.016114 | 0.012952 | 0.007871 | 0.020419 | 0.007115 | 0.005852 | 0.0174 | 0.037111 | 0.001279 | 0.021044 | 0.009684 | 0.027544 | 0.005563 | 0.0149 | 0.01758 | 0.01161 | 0.017513 |
| 32.1666667 | 0.028782 | 0.010466 | 0.006633 | 0.021003 | 0.012089 | 0.034373 | 0.0106 | 0.009542 | 0.029306 | 0.037184 | 0.00336 | 0.02978 | 0.006597 | 0.023564 | 0.004762 | 0.009319 | 0.015288 | 0.009984 | 0.016643 |
| 32.1805556 | 0.037756 | 0.00394 | 0.009526 | 0.027346 | 0.018771 | 0.04556 | 0.01945 | 0.011561 | 0.035501 | 0.034835 | 0.007302 | 0.032959 | 0.003109 | 0.018821 | 0.004475 | 0.003901 | 0.011196 | 0.00859 | 0.015463 |
| 32.1944444 | 0.042776 | 0.005682 | 0.014875 | 0.03188 | 0.023005 | 0.048709 | 0.026468 | 0.012784 | 0.038635 | 0.029104 | 0.0101 | 0.0336 | 0.001771 | 0.015894 | 0.004443 | 0.001679 | 0.010494 | 0.00714 | 0.012772 |
| 32.2083333 | 0.048832 | 0.013184 | 0.016935 | 0.032854 | 0.024851 | 0.049435 | 0.028751 | 0.01578 | 0.04144 | 0.019321 | 0.008737 | 0.032859 | 0.003896 | 0.01554 | 0.003237 | 0.002445 | 0.01669 | 0.00603 | 0.007723 |
| 32.2222222 | 0.054619 | 0.02408 | 0.017768 | 0.030235 | 0.02386 | 0.046827 | 0.027222 | 0.019458 | 0.047447 | 0.009069 | 0.004693 | 0.026763 | 0.008342 | 0.019163 | 0.001824 | 0.00439 | 0.026152 | 0.004752 | 0.003033 |
| 32.2361111 | 0.055058 | 0.032114 | 0.019004 | 0.02801 | 0.020866 | 0.039402 | 0.02663 | 0.021932 | 0.05817 | 0.003687 | 0.002074 | 0.015283 | 0.012965 | 0.024986 | 0.001428 | 0.006375 | 0.029851 | 0.00283 | 0.001206 |
| 32.25 | 0.051655 | 0.036324 | 0.01979 | 0.02756 | 0.017463 | 0.030025 | 0.028986 | 0.023956 | 0.067395 | 0.003942 | 0.00183 | 0.006146 | 0.015936 | 0.027294 | 0.000999 | 0.006395 | 0.024004 | 0.001064 | 0.001196 |
| 32.2638889 | 0.047716 | 0.039336 | 0.019466 | 0.026509 | 0.014694 | 0.022659 | 0.035289 | 0.023468 | 0.070022 | 0.005884 | 0.001421 | 0.004051 | 0.016039 | 0.023105 | 0.000441 | 0.004205 | 0.013029 | 0.000357 | 0.001586 |
| 32.2777778 | 0.042191 | 0.040129 | 0.018446 | 0.023054 | 0.012402 | 0.018411 | 0.046457 | 0.018506 | 0.067879 | 0.008105 | 0.000938 | 0.004444 | 0.01276 | 0.013416 | 0.000543 | 0.002242 | 0.005474 | 0.000728 | 0.003173 |
| 32.2916667 | 0.036176 | 0.041271 | 0.016553 | 0.016323 | 0.011121 | 0.01619 | 0.057447 | 0.013466 | 0.063128 | 0.010233 | 0.001886 | 0.004308 | 0.007513 | 0.005977 | 0.000946 | 0.001832 | 0.005163 | 0.001109 | 0.00587 |
| 32.3055556 | 0.031152 | 0.04594 | 0.016114 | 0.010276 | 0.011456 | 0.014093 | 0.064641 | 0.011639 | 0.058024 | 0.009693 | 0.002655 | 0.00396 | 0.003881 | 0.006479 | 0.001023 | 0.001958 | 0.005061 | 0.001359 | 0.006012 |
| 32.3194444 | 0.027302 | 0.050765 | 0.021025 | 0.004062 | 0.012634 | 0.010368 | 0.069589 | 0.01122 | 0.052583 | 0.006188 | 0.00235 | 0.003044 | 0.003839 | 0.006986 | 0.001233 | 0.002934 | 0.004063 | 0.002671 | 0.00388 |
| 32.3333333 | 0.027269 | 0.051991 | 0.028578 | 0.003259 | 0.013473 | 0.00684 | 0.070868 | 0.010221 | 0.044628 | 0.004601 | 0.00291 | 0.003759 | 0.005436 | 0.007248 | 0.001747 | 0.005238 | 0.008435 | 0.00414 | 0.004336 |
| 32.3472222 | 0.030726 | 0.048858 | 0.032773 | 0.0048 | 0.013548 | 0.008123 | 0.068826 | 0.008351 | 0.03393 | 0.006974 | 0.004243 | 0.007332 | 0.005699 | 0.013469 | 0.001702 | 0.006745 | 0.014495 | 0.004134 | 0.007269 |
| 32.3611111 | 0.035391 | 0.045256 | 0.032657 | 0.007861 | 0.013971 | 0.015136 | 0.067526 | 0.007348 | 0.023417 | 0.009244 | 0.005078 | 0.011058 | 0.004073 | 0.019007 | 0.001251 | 0.006783 | 0.015553 | 0.00352 | 0.00887 |
| 32.375 | 0.040226 | 0.045992 | 0.030009 | 0.008958 | 0.015581 | 0.022041 | 0.065445 | 0.008611 | 0.017268 | 0.010595 | 0.006371 | 0.013889 | 0.0027 | 0.018462 | 0.001375 | 0.006947 | 0.011381 | 0.003983 | 0.007551 |
| 32.3888889 | 0.040574 | 0.048599 | 0.027306 | 0.006555 | 0.015823 | 0.022561 | 0.057941 | 0.00941 | 0.014327 | 0.012476 | 0.008765 | 0.017339 | 0.003422 | 0.014322 | 0.002156 | 0.008018 | 0.005653 | 0.005505 | 0.004252 |
| 32.4027778 | 0.034946 | 0.048222 | 0.026564 | 0.003644 | 0.012172 | 0.016947 | 0.044495 | 0.006816 | 0.009803 | 0.013467 | 0.010686 | 0.021983 | 0.005202 | 0.009805 | 0.002539 | 0.009017 | 0.00247 | 0.006349 | 0.001924 |
| 32.4166667 | 0.03257 | 0.045975 | 0.027083 | 0.004125 | 0.008258 | 0.011489 | 0.027776 | 0.003665 | 0.004304 | 0.01204 | 0.01104 | 0.025934 | 0.005843 | 0.007151 | 0.002123 | 0.008794 | 0.003662 | 0.005598 | 0.002051 |
| 32.4305556 | 0.041006 | 0.046503 | 0.027897 | 0.008985 | 0.00921 | 0.012153 | 0.012444 | 0.005158 | 0.003875 | 0.008226 | 0.011085 | 0.025507 | 0.005197 | 0.006676 | 0.002105 | 0.007828 | 0.007669 | 0.004196 | 0.001918 |
| 32.4444444 | 0.052543 | 0.048737 | 0.029635 | 0.014567 | 0.012972 | 0.016971 | 0.003597 | 0.009357 | 0.009518 | 0.003706 | 0.011881 | 0.019046 | 0.004674 | 0.00751 | 0.002781 | 0.006664 | 0.010788 | 0.002812 | 0.001362 |
| 32.4583333 | 0.053746 | 0.047105 | 0.030408 | 0.015749 | 0.013787 | 0.017699 | 0.002329 | 0.009264 | 0.014184 | 0.001292 | 0.013076 | 0.010003 | 0.004689 | 0.008766 | 0.00259 | 0.004476 | 0.010824 | 0.001391 | 0.002184 |
| 32.4722222 | 0.043671 | 0.040329 | 0.023526 | 0.010276 | 0.010987 | 0.011719 | 0.008574 | 0.004813 | 0.012892 | 0.002911 | 0.014012 | 0.005417 | 0.004596 | 0.009389 | 0.001321 | 0.002155 | 0.00822 | 0.00049 | 0.00293 |
| 32.4861111 | 0.030687 | 0.030482 | 0.012183 | 0.005401 | 0.00746 | 0.009405 | 0.021345 | 0.001886 | 0.007729 | 0.007911 | 0.013946 | 0.004971 | 0.004602 | 0.008416 | 0.00065 | 0.001165 | 0.004546 | 0.000405 | 0.002889 |
| 32.5 | 0.021444 | 0.021693 | 0.007845 | 0.00736 | 0.004573 | 0.017478 | 0.034185 | 0.002016 | 0.003414 | 0.013014 | 0.012333 | 0.00512 | 0.005157 | 0.00725 | 0.000923 | 0.001362 | 0.002107 | 0.000587 | 0.002734 |
| 32.5138889 | 0.020467 | 0.021235 | 0.008545 | 0.009079 | 0.00311 | 0.026578 | 0.040928 | 0.001978 | 0.003117 | 0.015611 | 0.009588 | 0.009497 | 0.00543 | 0.008789 | 0.001177 | 0.002349 | 0.002986 | 0.00055 | 0.001917 |
| 32.5277778 | 0.026984 | 0.030565 | 0.008973 | 0.006773 | 0.004084 | 0.027237 | 0.038536 | 0.001781 | 0.003819 | 0.01752 | 0.009399 | 0.017873 | 0.005828 | 0.015838 | 0.001423 | 0.004055 | 0.008292 | 0.00151 | 0.000951 |
| 32.5416667 | 0.037412 | 0.043409 | 0.021457 | 0.011074 | 0.008447 | 0.019067 | 0.026344 | 0.00417 | 0.007217 | 0.023952 | 0.018357 | 0.02988 | 0.009999 | 0.032131 | 0.001705 | 0.010337 | 0.019632 | 0.006865 | 0.001928 |
| 32.5555556 | 0.05346 | 0.052164 | 0.048897 | 0.029437 | 0.01583 | 0.017678 | 0.025166 | 0.008693 | 0.020669 | 0.03419 | 0.038493 | 0.045776 | 0.018325 | 0.050982 | 0.002418 | 0.020927 | 0.034256 | 0.014625 | 0.003389 |
| 32.5694444 | 0.07239 | 0.050772 | 0.066227 | 0.049991 | 0.020835 | 0.03279 | 0.055646 | 0.01453 | 0.037867 | 0.040567 | 0.05781 | 0.057252 | 0.023384 | 0.056892 | 0.004406 | 0.02641 | 0.042286 | 0.016044 | 0.003093 |
| 32.5833333 | 0.084184 | 0.042415 | 0.067211 | 0.068099 | 0.020403 | 0.046219 | 0.095203 | 0.022514 | 0.050269 | 0.042744 | 0.063696 | 0.063893 | 0.020297 | 0.050381 | 0.006337 | 0.024614 | 0.039673 | 0.00967 | 0.005177 |
| 32.5972222 | 0.084093 | 0.034118 | 0.068048 | 0.088514 | 0.020538 | 0.047378 | 0.114208 | 0.029969 | 0.056043 | 0.041712 | 0.057899 | 0.067563 | 0.012965 | 0.041137 | 0.00762 | 0.020914 | 0.032227 | 0.003584 | 0.01279 |
| 32.6111111 | 0.073924 | 0.027488 | 0.069012 | 0.09716 | 0.024659 | 0.041672 | 0.107995 | 0.031797 | 0.050854 | 0.029478 | 0.043703 | 0.054774 | 0.006336 | 0.028054 | 0.009021 | 0.014212 | 0.020335 | 0.00299 | 0.022158 |
| 32.625 | 0.057229 | 0.022126 | 0.064395 | 0.081496 | 0.030745 | 0.02928 | 0.086883 | 0.028573 | 0.037276 | 0.015453 | 0.024618 | 0.032564 | 0.003836 | 0.014452 | 0.007737 | 0.008044 | 0.012403 | 0.005849 | 0.029004 |
| 32.6388889 | 0.037818 | 0.016135 | 0.056521 | 0.051565 | 0.039225 | 0.013873 | 0.066458 | 0.023703 | 0.025093 | 0.016649 | 0.010285 | 0.03186 | 0.006764 | 0.011293 | 0.004308 | 0.009142 | 0.019539 | 0.007671 | 0.033009 |
| 32.6527778 | 0.022582 | 0.010033 | 0.047327 | 0.02404 | 0.047599 | 0.00724 | 0.052544 | 0.018348 | 0.01573 | 0.021664 | 0.007228 | 0.04805 | 0.009426 | 0.014992 | 0.00445 | 0.011095 | 0.02453 | 0.007366 | 0.032405 |
| 32.6666667 | 0.015808 | 0.006846 | 0.038259 | 0.010723 | 0.049695 | 0.006714 | 0.042264 | 0.014411 | 0.007525 | 0.019189 | 0.011741 | 0.054095 | 0.006649 | 0.016432 | 0.006311 | 0.009237 | 0.017072 | 0.006753 | 0.022509 |
| 32.6805556 | 0.012396 | 0.005166 | 0.030119 | 0.012974 | 0.044163 | 0.005839 | 0.033736 | 0.011386 | 0.006921 | 0.013688 | 0.017752 | 0.048439 | 0.003367 | 0.016869 | 0.00609 | 0.00756 | 0.010426 | 0.00528 | 0.012188 |
| 32.6944444 | 0.007531 | 0.003584 | 0.021824 | 0.018312 | 0.033857 | 0.007765 | 0.021489 | 0.007299 | 0.017243 | 0.007441 | 0.023887 | 0.039801 | 0.003195 | 0.020276 | 0.005272 | 0.007648 | 0.013072 | 0.003194 | 0.015725 |
| 32.7083333 | 0.004971 | 0.004208 | 0.014366 | 0.018943 | 0.023148 | 0.007253 | 0.016545 | 0.003739 | 0.031276 | 0.003577 | 0.029747 | 0.031266 | 0.003111 | 0.026937 | 0.00411 | 0.00761 | 0.022674 | 0.003188 | 0.028853 |
| 32.7222222 | 0.008712 | 0.006398 | 0.008143 | 0.017148 | 0.014492 | 0.008826 | 0.040476 | 0.001776 | 0.044358 | 0.002666 | 0.034858 | 0.025978 | 0.004475 | 0.032828 | 0.002213 | 0.005705 | 0.030393 | 0.003598 | 0.038501 |
| 32.7361111 | 0.014441 | 0.007621 | 0.004908 | 0.017337 | 0.007939 | 0.026133 | 0.077322 | 0.000998 | 0.055069 | 0.003004 | 0.036195 | 0.023548 | 0.008733 | 0.03306 | 0.001119 | 0.003854 | 0.030336 | 0.002532 | 0.039358 |
| 32.75 | 0.017335 | 0.007855 | 0.006807 | 0.019805 | 0.004779 | 0.052504 | 0.099148 | 0.002199 | 0.056315 | 0.006351 | 0.029871 | 0.018492 | 0.010808 | 0.027166 | 0.00243 | 0.005697 | 0.021885 | 0.001917 | 0.034228 |
| 32.7638889 | 0.016132 | 0.008117 | 0.011395 | 0.01774 | 0.005626 | 0.067369 | 0.093525 | 0.006656 | 0.040567 | 0.009914 | 0.018323 | 0.011343 | 0.007845 | 0.017869 | 0.005968 | 0.010165 | 0.01133 | 0.001889 | 0.025031 |
| 32.7777778 | 0.010524 | 0.009698 | 0.016002 | 0.012051 | 0.007974 | 0.059965 | 0.061777 | 0.012404 | 0.021482 | 0.010106 | 0.008049 | 0.010634 | 0.00564 | 0.00871 | 0.009136 | 0.012266 | 0.009647 | 0.001327 | 0.012904 |
| 32.7916667 | 0.005526 | 0.012505 | 0.018884 | 0.014596 | 0.008026 | 0.039618 | 0.026467 | 0.013722 | 0.022606 | 0.009736 | 0.003735 | 0.015983 | 0.007276 | 0.005073 | 0.009707 | 0.009761 | 0.015346 | 0.000869 | 0.004012 |
| 32.8055556 | 0.007131 | 0.01371 | 0.018617 | 0.028497 | 0.005608 | 0.019239 | 0.008259 | 0.008953 | 0.035657 | 0.012167 | 0.003743 | 0.018616 | 0.007486 | 0.008666 | 0.00781 | 0.005066 | 0.018135 | 0.001442 | 0.001791 |
| 32.8194444 | 0.010654 | 0.013089 | 0.016262 | 0.042046 | 0.007395 | 0.010018 | 0.00672 | 0.005625 | 0.04503 | 0.017586 | 0.00321 | 0.01417 | 0.005198 | 0.01461 | 0.00442 | 0.001641 | 0.013856 | 0.002616 | 0.002849 |
| 32.8333333 | 0.00978 | 0.013477 | 0.013469 | 0.045124 | 0.017898 | 0.014571 | 0.008903 | 0.008142 | 0.053226 | 0.025033 | 0.002601 | 0.009166 | 0.003174 | 0.018264 | 0.002399 | 0.000587 | 0.008732 | 0.003037 | 0.003209 |
| 32.8472222 | 0.011024 | 0.013639 | 0.010471 | 0.033878 | 0.031666 | 0.020039 | 0.009276 | 0.009469 | 0.055035 | 0.032561 | 0.005018 | 0.013435 | 0.001615 | 0.017953 | 0.002935 | 0.001542 | 0.011597 | 0.002227 | 0.002326 |
| 32.8611111 | 0.020261 | 0.011282 | 0.0063 | 0.022103 | 0.04313 | 0.019696 | 0.009285 | 0.005915 | 0.042372 | 0.035403 | 0.008518 | 0.020498 | 0.001114 | 0.013829 | 0.003247 | 0.004201 | 0.017456 | 0.00123 | 0.00148 |
| 32.875 | 0.031287 | 0.008713 | 0.004305 | 0.032215 | 0.052376 | 0.012933 | 0.008642 | 0.003272 | 0.022934 | 0.02894 | 0.009947 | 0.020273 | 0.00311 | 0.007736 | 0.004056 | 0.007255 | 0.017037 | 0.001454 | 0.00076 |
| 32.8888889 | 0.039521 | 0.00846 | 0.009368 | 0.053317 | 0.058739 | 0.007771 | 0.006275 | 0.005053 | 0.017344 | 0.015994 | 0.010825 | 0.015053 | 0.006524 | 0.005986 | 0.006939 | 0.008682 | 0.011705 | 0.003759 | 0.000492 |
| 32.9027778 | 0.047146 | 0.010787 | 0.018198 | 0.063899 | 0.059356 | 0.015257 | 0.00688 | 0.006572 | 0.0292 | 0.008967 | 0.012526 | 0.009067 | 0.007776 | 0.01322 | 0.008087 | 0.007637 | 0.008846 | 0.00676 | 0.001301 |

| | | | | | | | | | | | | | | | | | | | |
|---|---|---|---|---|---|---|---|---|---|---|---|---|---|---|---|---|---|---|---|
| 32.9166667 | 0.054508 | 0.015307 | 0.025085 | 0.063089 | 0.053125 | 0.027458 | 0.018549 | 0.00643 | 0.042809 | 0.012581 | 0.013554 | 0.004427 | 0.005579 | 0.02268 | 0.005933 | 0.005599 | 0.011646 | 0.008294 | 0.001812 |
| 32.9305556 | 0.058783 | 0.020998 | 0.027115 | 0.056192 | 0.040622 | 0.035844 | 0.03596 | 0.010966 | 0.058149 | 0.017339 | 0.012052 | 0.003704 | 0.002476 | 0.026102 | 0.004306 | 0.004905 | 0.0142 | 0.008445 | 0.001386 |
| 32.9444444 | 0.056843 | 0.025391 | 0.024214 | 0.045469 | 0.023435 | 0.043431 | 0.04828 | 0.018518 | 0.073264 | 0.017707 | 0.008193 | 0.007506 | 0.001227 | 0.022894 | 0.004865 | 0.005441 | 0.011123 | 0.007823 | 0.001323 |
| 32.9583333 | 0.046797 | 0.025194 | 0.019108 | 0.032255 | 0.01088 | 0.049268 | 0.049443 | 0.021655 | 0.07963 | 0.014075 | 0.00444 | 0.013488 | 0.001836 | 0.016201 | 0.00588 | 0.005296 | 0.005715 | 0.006401 | 0.001465 |
| 32.9722222 | 0.03091 | 0.020102 | 0.014555 | 0.019604 | 0.012771 | 0.046382 | 0.040251 | 0.021 | 0.080292 | 0.007425 | 0.002385 | 0.0196 | 0.00208 | 0.008836 | 0.006972 | 0.004674 | 0.004153 | 0.00519 | 0.001512 |
| 32.9861111 | 0.014804 | 0.013504 | 0.009795 | 0.011058 | 0.021933 | 0.031963 | 0.02833 | 0.0179 | 0.068091 | 0.002428 | 0.001542 | 0.023426 | 0.002699 | 0.003659 | 0.009624 | 0.005706 | 0.006489 | 0.005726 | 0.002495 |
| 33 | 0.007324 | 0.007453 | 0.007876 | 0.014347 | 0.028343 | 0.017142 | 0.015804 | 0.011382 | 0.041548 | 0.001605 | 0.001563 | 0.022682 | 0.005801 | 0.002727 | 0.014168 | 0.009036 | 0.008866 | 0.007595 | 0.004312 |
| 33.0138889 | 0.012397 | 0.003128 | 0.015062 | 0.032166 | 0.032777 | 0.017537 | 0.011073 | 0.00526 | 0.032809 | 0.00158 | 0.002049 | 0.017663 | 0.008101 | 0.003471 | 0.017803 | 0.012262 | 0.009415 | 0.008236 | 0.005401 |
| 33.0277778 | 0.019861 | 0.002725 | 0.02182 | 0.048901 | 0.034578 | 0.024829 | 0.019545 | 0.003273 | 0.051151 | 0.002572 | 0.003781 | 0.011979 | 0.007269 | 0.005116 | 0.016551 | 0.01055 | 0.009342 | 0.00761 | 0.004168 |
| 33.0416667 | 0.017972 | 0.009313 | 0.020254 | 0.04097 | 0.026447 | 0.02525 | 0.024418 | 0.003489 | 0.052538 | 0.012967 | 0.013751 | 0.019586 | 0.011429 | 0.017226 | 0.010825 | 0.009735 | 0.02277 | 0.015087 | 0.004405 |
| 33.0555556 | 0.022839 | 0.030995 | 0.026961 | 0.029951 | 0.017851 | 0.03816 | 0.03508 | 0.007217 | 0.040466 | 0.037529 | 0.034653 | 0.045696 | 0.026945 | 0.043789 | 0.00697 | 0.017866 | 0.058439 | 0.033675 | 0.009521 |
| 33.0694444 | 0.056555 | 0.070793 | 0.044937 | 0.055168 | 0.027597 | 0.069196 | 0.063218 | 0.017061 | 0.060324 | 0.058688 | 0.046289 | 0.059951 | 0.038187 | 0.060063 | 0.006273 | 0.020176 | 0.085589 | 0.039171 | 0.01069 |
| 33.0833333 | 0.10074 | 0.108545 | 0.052098 | 0.086799 | 0.046412 | 0.085957 | 0.074018 | 0.025438 | 0.090707 | 0.052734 | 0.032691 | 0.040802 | 0.027824 | 0.046075 | 0.005101 | 0.011493 | 0.075334 | 0.02365 | 0.007009 |
| 33.0972222 | 0.121296 | 0.123266 | 0.045929 | 0.087674 | 0.057744 | 0.082409 | 0.058519 | 0.026529 | 0.105092 | 0.028591 | 0.014012 | 0.016143 | 0.012616 | 0.020575 | 0.005247 | 0.006179 | 0.043568 | 0.01255 | 0.008634 |
| 33.1111111 | 0.101118 | 0.118747 | 0.037874 | 0.067554 | 0.055768 | 0.075257 | 0.042317 | 0.023556 | 0.110537 | 0.017337 | 0.009906 | 0.011545 | 0.013132 | 0.008432 | 0.007099 | 0.004861 | 0.019169 | 0.010535 | 0.012552 |
| 33.125 | 0.057405 | 0.101739 | 0.030798 | 0.042706 | 0.041197 | 0.070067 | 0.030617 | 0.020243 | 0.100015 | 0.025905 | 0.013308 | 0.015923 | 0.019128 | 0.012155 | 0.007714 | 0.004727 | 0.014399 | 0.008456 | 0.012629 |
| 33.1388889 | 0.023006 | 0.073689 | 0.024957 | 0.021228 | 0.022017 | 0.06711 | 0.019667 | 0.016446 | 0.076521 | 0.033367 | 0.014275 | 0.01835 | 0.019461 | 0.019188 | 0.006946 | 0.005652 | 0.022464 | 0.007903 | 0.010345 |
| 33.1527778 | 0.011348 | 0.03961 | 0.018978 | 0.010056 | 0.010059 | 0.06164 | 0.009718 | 0.009695 | 0.052953 | 0.029377 | 0.012874 | 0.021758 | 0.016538 | 0.022431 | 0.006651 | 0.004832 | 0.030111 | 0.007995 | 0.009053 |
| 33.1666667 | 0.016373 | 0.015017 | 0.011705 | 0.013527 | 0.015622 | 0.050504 | 0.004199 | 0.003153 | 0.034086 | 0.022221 | 0.01137 | 0.027788 | 0.012046 | 0.022475 | 0.007369 | 0.003708 | 0.033234 | 0.006138 | 0.01117 |
| 33.1805556 | 0.021992 | 0.011031 | 0.005476 | 0.024156 | 0.020811 | 0.035161 | 0.006819 | 0.000994 | 0.022394 | 0.020315 | 0.008867 | 0.032541 | 0.006053 | 0.019617 | 0.007152 | 0.005034 | 0.032952 | 0.003857 | 0.012823 |
| 33.1944444 | 0.024527 | 0.014749 | 0.005138 | 0.03379 | 0.026771 | 0.021024 | 0.014255 | 0.001388 | 0.014885 | 0.025609 | 0.006748 | 0.033391 | 0.001459 | 0.01651 | 0.005541 | 0.00664 | 0.030826 | 0.002839 | 0.009283 |
| 33.2083333 | 0.030958 | 0.018304 | 0.012525 | 0.043917 | 0.030514 | 0.021912 | 0.021753 | 0.004013 | 0.010729 | 0.031326 | 0.006004 | 0.033346 | 0.000844 | 0.015723 | 0.004474 | 0.005859 | 0.030129 | 0.002437 | 0.00417 |
| 33.2222222 | 0.041598 | 0.021677 | 0.022518 | 0.05111 | 0.032265 | 0.041667 | 0.026863 | 0.008781 | 0.018547 | 0.028011 | 0.004826 | 0.034917 | 0.002721 | 0.015147 | 0.003855 | 0.004188 | 0.031929 | 0.00319 | 0.001777 |
| 33.2361111 | 0.052429 | 0.02125 | 0.027976 | 0.047172 | 0.033839 | 0.062767 | 0.029309 | 0.013497 | 0.039686 | 0.017577 | 0.00537 | 0.034935 | 0.005116 | 0.011996 | 0.002567 | 0.003359 | 0.034507 | 0.004641 | 0.001915 |
| 33.25 | 0.059459 | 0.017715 | 0.027877 | 0.036486 | 0.034305 | 0.073393 | 0.030119 | 0.017577 | 0.064104 | 0.008625 | 0.007549 | 0.026404 | 0.007357 | 0.007328 | 0.001518 | 0.005069 | 0.034834 | 0.00419 | 0.003691 |
| 33.2638889 | 0.059045 | 0.010907 | 0.025514 | 0.027553 | 0.031001 | 0.074558 | 0.028676 | 0.020795 | 0.083886 | 0.00363 | 0.008452 | 0.015279 | 0.008154 | 0.004315 | 0.001129 | 0.008031 | 0.027804 | 0.002136 | 0.005431 |
| 33.2777778 | 0.051521 | 0.005318 | 0.022079 | 0.020822 | 0.023642 | 0.067769 | 0.024789 | 0.023288 | 0.093384 | 0.002571 | 0.008798 | 0.016012 | 0.006518 | 0.005872 | 0.001562 | 0.007815 | 0.015106 | 0.000833 | 0.005797 |
| 33.2916667 | 0.041613 | 0.005118 | 0.017359 | 0.013646 | 0.014946 | 0.057214 | 0.02014 | 0.026398 | 0.089026 | 0.00667 | 0.009074 | 0.023531 | 0.00416 | 0.011611 | 0.002374 | 0.004376 | 0.00856 | 0.001045 | 0.005745 |
| 33.3055556 | 0.030552 | 0.005173 | 0.011195 | 0.007157 | 0.007229 | 0.045333 | 0.014798 | 0.029418 | 0.078185 | 0.012473 | 0.008877 | 0.025628 | 0.003254 | 0.01736 | 0.001953 | 0.001818 | 0.011397 | 0.001927 | 0.005883 |
| 33.3194444 | 0.016629 | 0.006662 | 0.004836 | 0.005595 | 0.00273 | 0.028462 | 0.008043 | 0.028079 | 0.073195 | 0.014846 | 0.008716 | 0.02245 | 0.003729 | 0.01874 | 0.000872 | 0.001692 | 0.013488 | 0.003188 | 0.005508 |
| 33.3333333 | 0.005689 | 0.008988 | 0.001424 | 0.007323 | 0.002471 | 0.013066 | 0.003705 | 0.022686 | 0.074293 | 0.012327 | 0.007135 | 0.015855 | 0.004952 | 0.013437 | 0.000803 | 0.002817 | 0.01049 | 0.003888 | 0.005205 |
| 33.3472222 | 0.002956 | 0.011229 | 0.002726 | 0.006182 | 0.002818 | 0.008701 | 0.003763 | 0.018837 | 0.072333 | 0.007251 | 0.004856 | 0.007884 | 0.006743 | 0.007833 | 0.001766 | 0.005741 | 0.011474 | 0.003114 | 0.00561 |
| 33.3611111 | 0.004345 | 0.019719 | 0.006816 | 0.003187 | 0.003452 | 0.008778 | 0.003815 | 0.01628 | 0.059948 | 0.005541 | 0.006172 | 0.004005 | 0.008213 | 0.011594 | 0.002149 | 0.008852 | 0.020345 | 0.003014 | 0.005976 |
| 33.375 | 0.008238 | 0.026408 | 0.010189 | 0.003641 | 0.006809 | 0.00685 | 0.00336 | 0.012679 | 0.042017 | 0.008198 | 0.009946 | 0.00533 | 0.008832 | 0.020966 | 0.001608 | 0.009449 | 0.026864 | 0.004567 | 0.006147 |
| 33.3888889 | 0.015738 | 0.029761 | 0.011833 | 0.007209 | 0.012101 | 0.004843 | 0.005049 | 0.010341 | 0.032377 | 0.010123 | 0.012145 | 0.007669 | 0.008783 | 0.026562 | 0.002236 | 0.008479 | 0.023944 | 0.006881 | 0.006286 |
| 33.4027778 | 0.023737 | 0.038328 | 0.012151 | 0.010081 | 0.016841 | 0.006411 | 0.006204 | 0.009286 | 0.029483 | 0.009749 | 0.010354 | 0.009233 | 0.007675 | 0.025299 | 0.004407 | 0.006596 | 0.014197 | 0.009463 | 0.005644 |
| 33.4166667 | 0.026371 | 0.042916 | 0.012283 | 0.009813 | 0.017799 | 0.015093 | 0.004543 | 0.007395 | 0.021353 | 0.009747 | 0.005969 | 0.010685 | 0.005797 | 0.01884 | 0.005548 | 0.004209 | 0.006079 | 0.010946 | 0.004507 |
| 33.4305556 | 0.023912 | 0.037585 | 0.014859 | 0.008068 | 0.014743 | 0.027236 | 0.002852 | 0.005362 | 0.010855 | 0.012011 | 0.003778 | 0.012277 | 0.004442 | 0.011546 | 0.004475 | 0.00324 | 0.00486 | 0.010666 | 0.003905 |
| 33.4444444 | 0.022849 | 0.031848 | 0.019863 | 0.007868 | 0.011031 | 0.035319 | 0.002243 | 0.004021 | 0.008366 | 0.015154 | 0.002901 | 0.013511 | 0.003664 | 0.007205 | 0.003019 | 0.003735 | 0.004837 | 0.008933 | 0.003471 |
| 33.4583333 | 0.024436 | 0.029901 | 0.024285 | 0.008272 | 0.008167 | 0.03788 | 0.002865 | 0.002995 | 0.011004 | 0.016362 | 0.002325 | 0.01356 | 0.003345 | 0.007152 | 0.002274 | 0.004046 | 0.004531 | 0.006215 | 0.002929 |
| 33.4722222 | 0.026105 | 0.028428 | 0.026462 | 0.008144 | 0.00596 | 0.035426 | 0.00576 | 0.00251 | 0.011045 | 0.01491 | 0.004863 | 0.0125 | 0.00419 | 0.01059 | 0.001476 | 0.003974 | 0.009386 | 0.003946 | 0.002882 |
| 33.4861111 | 0.027471 | 0.026612 | 0.025934 | 0.007817 | 0.004671 | 0.031297 | 0.008051 | 0.002011 | 0.009078 | 0.011987 | 0.008514 | 0.010282 | 0.005597 | 0.0144 | 0.000707 | 0.003985 | 0.016296 | 0.002943 | 0.002741 |
| 33.5 | 0.026979 | 0.022927 | 0.022132 | 0.005967 | 0.003256 | 0.033759 | 0.006015 | 0.001518 | 0.008641 | 0.009183 | 0.010334 | 0.008202 | 0.00658 | 0.015486 | 0.000438 | 0.004184 | 0.018977 | 0.002654 | 0.001846 |
| 33.5138889 | 0.02215 | 0.015847 | 0.017239 | 0.002864 | 0.002253 | 0.042664 | 0.003091 | 0.002017 | 0.010818 | 0.008282 | 0.0115 | 0.008848 | 0.007325 | 0.014179 | 0.000794 | 0.005014 | 0.018159 | 0.002959 | 0.001446 |
| 33.5277778 | 0.015555 | 0.014538 | 0.016412 | 0.003772 | 0.002595 | 0.04727 | 0.002422 | 0.002519 | 0.011174 | 0.010363 | 0.015313 | 0.013134 | 0.007955 | 0.014871 | 0.002143 | 0.007309 | 0.020168 | 0.004609 | 0.002205 |
| 33.5416667 | 0.013595 | 0.032217 | 0.024321 | 0.014712 | 0.004142 | 0.03619 | 0.005816 | 0.005088 | 0.00991 | 0.016565 | 0.025881 | 0.021926 | 0.009506 | 0.024944 | 0.004601 | 0.012331 | 0.034994 | 0.009509 | 0.00382 |
| 33.5555556 | 0.021952 | 0.065091 | 0.040928 | 0.034992 | 0.009924 | 0.018857 | 0.019129 | 0.012294 | 0.018836 | 0.026657 | 0.040458 | 0.035048 | 0.014033 | 0.043829 | 0.007221 | 0.018503 | 0.058224 | 0.017661 | 0.006419 |
| 33.5694444 | 0.03993 | 0.085415 | 0.057007 | 0.056282 | 0.019286 | 0.018545 | 0.03805 | 0.019876 | 0.039018 | 0.03549 | 0.04663 | 0.047736 | 0.020423 | 0.057395 | 0.007698 | 0.020176 | 0.071086 | 0.022608 | 0.007779 |
| 33.5833333 | 0.059225 | 0.078646 | 0.062806 | 0.070523 | 0.025097 | 0.031439 | 0.048452 | 0.02342 | 0.060007 | 0.03777 | 0.041077 | 0.054833 | 0.023056 | 0.058434 | 0.005089 | 0.015967 | 0.073955 | 0.020067 | 0.00589 |
| 33.5972222 | 0.071894 | 0.067169 | 0.057896 | 0.072875 | 0.027209 | 0.039919 | 0.046136 | 0.023724 | 0.076035 | 0.031724 | 0.033774 | 0.054365 | 0.018989 | 0.053511 | 0.002738 | 0.009329 | 0.073783 | 0.014453 | 0.004091 |
| 33.6111111 | 0.075706 | 0.067156 | 0.047247 | 0.064609 | 0.030525 | 0.043103 | 0.037843 | 0.022797 | 0.079026 | 0.018166 | 0.029912 | 0.043554 | 0.01135 | 0.045796 | 0.00355 | 0.006231 | 0.063052 | 0.009061 | 0.003784 |
| 33.625 | 0.072651 | 0.070711 | 0.036636 | 0.053908 | 0.031951 | 0.044598 | 0.024669 | 0.020967 | 0.065874 | 0.00842 | 0.028532 | 0.023096 | 0.005323 | 0.034853 | 0.005708 | 0.011022 | 0.041609 | 0.00802 | 0.006354 |
| 33.6388889 | 0.063019 | 0.071288 | 0.028216 | 0.043796 | 0.030507 | 0.041204 | 0.01105 | 0.01896 | 0.046814 | 0.009734 | 0.028417 | 0.008648 | 0.005783 | 0.023287 | 0.006744 | 0.018375 | 0.020785 | 0.012782 | 0.014738 |
| 33.6527778 | 0.047227 | 0.068725 | 0.021603 | 0.032771 | 0.029864 | 0.035076 | 0.006447 | 0.018468 | 0.030232 | 0.009991 | 0.02751 | 0.009815 | 0.00914 | 0.013108 | 0.004996 | 0.019812 | 0.008342 | 0.015474 | 0.024172 |
| 33.6666667 | 0.028939 | 0.05905 | 0.01666 | 0.02286 | 0.027689 | 0.031594 | 0.005709 | 0.017848 | 0.015949 | 0.004861 | 0.024596 | 0.015305 | 0.007404 | 0.00689 | 0.00257 | 0.012908 | 0.006207 | 0.014725 | 0.028637 |
| 33.6805556 | 0.013734 | 0.037994 | 0.013132 | 0.014948 | 0.020314 | 0.028445 | 0.005159 | 0.012334 | 0.010459 | 0.001846 | 0.02027 | 0.018275 | 0.003115 | 0.008525 | 0.002722 | 0.004564 | 0.011234 | 0.013328 | 0.025208 |
| 33.6944444 | 0.011494 | 0.018071 | 0.011636 | 0.009062 | 0.012159 | 0.020655 | 0.009546 | 0.008454 | 0.02189 | 0.003474 | 0.015242 | 0.018674 | 0.001278 | 0.015548 | 0.002733 | 0.001038 | 0.016983 | 0.009763 | 0.015188 |
| 33.7083333 | 0.022366 | 0.017351 | 0.011274 | 0.009677 | 0.013584 | 0.0111 | 0.014251 | 0.016261 | 0.040276 | 0.008391 | 0.009992 | 0.016809 | 0.001581 | 0.021932 | 0.001707 | 0.000861 | 0.021094 | 0.004696 | 0.007628 |
| 33.7222222 | 0.031657 | 0.030908 | 0.010021 | 0.015631 | 0.02365 | 0.009783 | 0.01451 | 0.030678 | 0.05702 | 0.013186 | 0.004989 | 0.013004 | 0.004689 | 0.026827 | 0.003001 | 0.002367 | 0.02504 | 0.001172 | 0.010233 |
| 33.7361111 | 0.032179 | 0.042678 | 0.008195 | 0.018388 | 0.030424 | 0.016813 | 0.012241 | 0.044833 | 0.073645 | 0.013416 | 0.001904 | 0.009609 | 0.009397 | 0.030887 | 0.005437 | 0.004289 | 0.028264 | 0.000293 | 0.017987 |
| 33.75 | 0.028525 | 0.046682 | 0.007714 | 0.014315 | 0.029499 | 0.021873 | 0.009143 | 0.053484 | 0.084932 | 0.008816 | 0.001455 | 0.006496 | 0.010515 | 0.03076 | 0.004644 | 0.00396 | 0.028285 | 0.000431 | 0.020937 |
| 33.7638889 | 0.025 | 0.046966 | 0.009935 | 0.00848 | 0.023912 | 0.020645 | 0.008102 | 0.049598 | 0.083749 | 0.006793 | 0.001263 | 0.008802 | 0.007252 | 0.024303 | 0.003178 | 0.002277 | 0.023128 | 0.000371 | 0.017669 |
| 33.7777778 | 0.020112 | 0.046333 | 0.015584 | 0.0114 | 0.016147 | 0.017286 | 0.011104 | 0.033957 | 0.067771 | 0.010301 | 0.000887 | 0.016747 | 0.005223 | 0.015106 | 0.005399 | 0.002467 | 0.013226 | 0.000348 | 0.012203 |
| 33.7916667 | 0.014809 | 0.041457 | 0.023114 | 0.023686 | 0.008404 | 0.014121 | 0.015439 | 0.015985 | 0.03926 | 0.012715 | 0.001194 | 0.023424 | 0.006009 | 0.006948 | 0.008048 | 0.004151 | 0.00623 | 0.000808 | 0.006574 |
| 33.8055556 | 0.014134 | 0.032546 | 0.029036 | 0.033582 | 0.003376 | 0.009317 | 0.017875 | 0.004731 | 0.024221 | 0.011115 | 0.001126 | 0.024959 | 0.005758 | 0.00317 | 0.009211 | 0.006589 | 0.008215 | 0.002027 | 0.003381 |
| 33.8194444 | 0.01687 | 0.023921 | 0.031391 | 0.034539 | 0.003169 | 0.005725 | 0.017893 | 0.001445 | 0.04197 | 0.007932 | 0.000484 | 0.021805 | 0.00385 | 0.004378 | 0.0101 | 0.008991 | 0.011224 | 0.003048 | 0.003835 |
| 33.8333333 | 0.02007 | 0.016838 | 0.029158 | 0.028083 | 0.009071 | 0.009082 | 0.015231 | 0.002474 | 0.063854 | 0.004572 | 0.00103 | 0.015427 | 0.002041 | 0.004633 | 0.010139 | 0.008881 | 0.00791 | 0.002425 | 0.003728 |
| 33.8472222 | 0.027609 | 0.013072 | 0.021851 | 0.01669 | 0.021616 | 0.016839 | 0.009739 | 0.005072 | 0.066709 | 0.004143 | 0.003451 | 0.008123 | 0.001132 | 0.002589 | 0.009365 | 0.006202 | 0.004883 | 0.001914 | 0.003037 |
| 33.8611111 | 0.046643 | 0.022478 | 0.012739 | 0.012985 | 0.03962 | 0.022279 | 0.00747 | 0.007027 | 0.055518 | 0.009199 | 0.006263 | 0.005773 | 0.001638 | 0.001572 | 0.007324 | 0.003386 | 0.008858 | 0.003551 | 0.003838 |
| 33.875 | 0.074529 | 0.049534 | 0.011336 | 0.027494 | 0.059973 | 0.023199 | 0.012336 | 0.005522 | 0.033272 | 0.01708 | 0.009427 | 0.01267 | 0.002839 | 0.001846 | 0.003975 | 0.001747 | 0.015785 | 0.005494 | 0.003413 |
| 33.8888889 | 0.098661 | 0.078586 | 0.020852 | 0.046861 | 0.080862 | 0.021029 | 0.015342 | 0.004296 | 0.018445 | 0.024669 | 0.015335 | 0.023602 | 0.00278 | 0.002985 | 0.001393 | 0.001435 | 0.020444 | 0.006639 | 0.002096 |
| 33.9027778 | 0.107658 | 0.091876 | 0.030638 | 0.059897 | 0.1013 | 0.017531 | 0.011341 | 0.009203 | 0.034825 | 0.029504 | 0.023569 | 0.033446 | 0.002954 | 0.003003 | 0.000384 | 0.001884 | 0.020815 | 0.007254 | 0.002893 |
| 33.9166667 | 0.102899 | 0.096029 | 0.034802 | 0.068701 | 0.113811 | 0.012956 | 0.005462 | 0.016081 | 0.070514 | 0.028671 | 0.029105 | 0.037361 | 0.005502 | 0.003213 | 0.000884 | 0.002062 | 0.016067 | 0.005895 | 0.005153 |
| 33.9305556 | 0.094152 | 0.104896 | 0.036745 | 0.072075 | 0.110638 | 0.007159 | 0.002158 | 0.019429 | 0.103515 | 0.021829 | 0.028889 | 0.030058 | 0.007392 | 0.007505 | 0.001997 | 0.00202 | 0.008328 | 0.003745 | 0.006463 |
| 33.9444444 | 0.08011 | 0.113628 | 0.036214 | 0.067479 | 0.092846 | 0.003768 | 0.002 | 0.019015 | 0.123458 | 0.01222 | 0.025225 | 0.01674 | 0.006638 | 0.013784 | 0.003351 | 0.001567 | 0.003503 | 0.004276 | 0.005514 |
| 33.9583333 | 0.05942 | 0.110998 | 0.03019 | 0.055935 | 0.063906 | 0.00664 | 0.003341 | 0.016709 | 0.126441 | 0.005573 | 0.021164 | 0.008276 | 0.004554 | 0.01779 | 0.005677 | 0.000792 | 0.005075 | 0.006552 | 0.002984 |
| 33.9722222 | 0.038754 | 0.096847 | 0.019307 | 0.038842 | 0.033507 | 0.01034 | 0.004239 | 0.014407 | 0.120023 | 0.005834 | 0.017032 | 0.01145 | 0.003089 | 0.017464 | 0.005685 | 0.001112 | 0.008987 | 0.008084 | 0.001122 |
| 33.9861111 | 0.019699 | 0.080174 | 0.010198 | 0.02113 | 0.02679 | 0.007903 | 0.004619 | 0.011121 | 0.108258 | 0.010303 | 0.012992 | 0.021822 | 0.004212 | 0.013843 | 0.003318 | 0.002232 | 0.011736 | 0.008762 | 0.00181 |
| 34 | 0.010214 | 0.06424 | 0.011305 | 0.017724 | 0.049954 | 0.006913 | 0.008504 | 0.006416 | 0.086614 | 0.013894 | 0.009569 | 0.029448 | 0.006079 | 0.009922 | 0.003158 | 0.002281 | 0.012873 | 0.008839 | 0.004863 |
| 34.0138889 | 0.016144 | 0.04777 | 0.019862 | 0.032976 | 0.065307 | 0.017488 | 0.017359 | 0.004631 | 0.058549 | 0.014253 | 0.006839 | 0.03036 | 0.005309 | 0.007916 | 0.003408 | 0.003119 | 0.01119 | 0.007709 | 0.007654 |
| 34.0277778 | 0.019506 | 0.037434 | 0.022045 | 0.046916 | 0.049394 | 0.026171 | 0.023458 | 0.004938 | 0.039413 | 0.011162 | 0.008765 | 0.021484 | 0.003182 | 0.010731 | 0.004107 | 0.010327 | 0.010366 | 0.008038 | 0.00689 |
| 34.0416667 | 0.020951 | 0.048514 | 0.021221 | 0.039957 | 0.024919 | 0.027475 | 0.019455 | 0.007865 | 0.052352 | 0.018005 | 0.024081 | 0.01982 | 0.008067 | 0.024661 | 0.011654 | 0.028838 | 0.025236 | 0.022952 | 0.00944 |
| 34.0555556 | 0.050048 | 0.088466 | 0.035775 | 0.039301 | 0.024566 | 0.037644 | 0.017214 | 0.020425 | 0.094252 | 0.04733 | 0.054407 | 0.047556 | 0.0247 | 0.050305 | 0.023471 | 0.053226 | 0.058884 | 0.056172 | 0.022907 |
| 34.0694444 | 0.094497 | 0.136429 | 0.057344 | 0.069707 | 0.051109 | 0.049334 | 0.024719 | 0.033506 | 0.125974 | 0.073442 | 0.070241 | 0.070295 | 0.035005 | 0.064254 | 0.02593 | 0.058756 | 0.07674 | 0.073367 | 0.030709 |
| 34.0833333 | 0.113386 | 0.15253 | 0.067067 | 0.084285 | 0.072623 | 0.050742 | 0.027095 | 0.035386 | 0.134032 | 0.065101 | 0.047134 | 0.04965 | 0.02502 | 0.044241 | 0.016767 | 0.036223 | 0.053107 | 0.048908 | 0.021528 |
| 34.0972222 | 0.109139 | 0.129625 | 0.063872 | 0.064114 | 0.073351 | 0.047011 | 0.020857 | 0.031047 | 0.129938 | 0.033205 | 0.018397 | 0.020196 | 0.015218 | 0.017598 | 0.010276 | 0.016883 | 0.026403 | 0.022331 | 0.014112 |
| 34.1111111 | 0.099147 | 0.100048 | 0.051285 | 0.042187 | 0.061565 | 0.040121 | 0.012895 | 0.027843 | 0.114745 | 0.015179 | 0.014517 | 0.019808 | 0.018085 | 0.014905 | 0.011552 | 0.016516 | 0.029644 | 0.026027 | 0.017321 |
| 34.125 | 0.078695 | 0.079395 | 0.034974 | 0.026328 | 0.041458 | 0.032844 | 0.006022 | 0.025174 | 0.090941 | 0.025349 | 0.021003 | 0.031982 | 0.024103 | 0.020905 | 0.015289 | 0.018168 | 0.039058 | 0.036393 | 0.019847 |
| 34.1388889 | 0.048755 | 0.062997 | 0.020936 | 0.012165 | 0.01943 | 0.029621 | 0.005582 | 0.020254 | 0.064771 | 0.03837 | 0.022761 | 0.039062 | 0.03362 | 0.021454 | 0.019254 | 0.016131 | 0.039261 | 0.03896 | 0.017396 |

| | | | | | | | | | | | | | | | | | | | |
|---|---|---|---|---|---|---|---|---|---|---|---|---|---|---|---|---|---|---|---|
| 34.1527778 | 0.022888 | 0.047373 | 0.011292 | 0.004042 | 0.008004 | 0.029154 | 0.011681 | 0.013871 | 0.041665 | 0.037073 | 0.020085 | 0.038226 | 0.040507 | 0.020086 | 0.020488 | 0.014488 | 0.03558 | 0.03794 | 0.012486 |
| 34.1666667 | 0.008792 | 0.03277 | 0.006937 | 0.005313 | 0.01232 | 0.026844 | 0.018083 | 0.007908 | 0.024449 | 0.029401 | 0.018259 | 0.028752 | 0.033222 | 0.020297 | 0.018641 | 0.014019 | 0.031543 | 0.03354 | 0.008228 |
| 34.1805556 | 0.008019 | 0.020433 | 0.009447 | 0.012181 | 0.021952 | 0.02051 | 0.022094 | 0.003837 | 0.011543 | 0.02364 | 0.019745 | 0.017985 | 0.021721 | 0.023098 | 0.016877 | 0.014686 | 0.030671 | 0.028623 | 0.006544 |
| 34.1944444 | 0.020224 | 0.009924 | 0.016303 | 0.021186 | 0.023605 | 0.011172 | 0.022771 | 0.002209 | 0.004741 | 0.022924 | 0.020246 | 0.010111 | 0.015926 | 0.026466 | 0.014032 | 0.013962 | 0.031228 | 0.024722 | 0.005418 |
| 34.2083333 | 0.037448 | 0.003613 | 0.022549 | 0.031714 | 0.017926 | 0.006838 | 0.020722 | 0.001767 | 0.007281 | 0.027558 | 0.016623 | 0.00585 | 0.011427 | 0.029568 | 0.009355 | 0.010646 | 0.02916 | 0.019201 | 0.003512 |
| 34.2222222 | 0.048429 | 0.005176 | 0.028495 | 0.041328 | 0.017434 | 0.016157 | 0.021602 | 0.002547 | 0.01579 | 0.036465 | 0.012553 | 0.010195 | 0.005695 | 0.030806 | 0.005766 | 0.008004 | 0.025194 | 0.010881 | 0.003063 |
| 34.2361111 | 0.049387 | 0.013489 | 0.033805 | 0.048654 | 0.0254 | 0.034988 | 0.028282 | 0.006079 | 0.026577 | 0.045664 | 0.010484 | 0.021647 | 0.001792 | 0.027437 | 0.004084 | 0.007825 | 0.022363 | 0.005301 | 0.004845 |
| 34.25 | 0.043435 | 0.019731 | 0.03431 | 0.055286 | 0.033631 | 0.059223 | 0.03522 | 0.00901 | 0.035054 | 0.047015 | 0.007317 | 0.028624 | 0.000798 | 0.018842 | 0.002556 | 0.007195 | 0.019254 | 0.006921 | 0.005729 |
| 34.2638889 | 0.033442 | 0.016591 | 0.029167 | 0.059744 | 0.034903 | 0.08387 | 0.035822 | 0.007787 | 0.035027 | 0.036043 | 0.004279 | 0.024187 | 0.000585 | 0.009607 | 0.001032 | 0.00428 | 0.013896 | 0.012037 | 0.004553 |
| 34.2777778 | 0.02086 | 0.014262 | 0.022527 | 0.054966 | 0.025089 | 0.094612 | 0.030469 | 0.004692 | 0.03058 | 0.01853 | 0.004828 | 0.012416 | 0.00096 | 0.007345 | 0.000975 | 0.002297 | 0.007589 | 0.016892 | 0.002747 |
| 34.2916667 | 0.012804 | 0.036667 | 0.017767 | 0.040103 | 0.017104 | 0.084542 | 0.021908 | 0.003063 | 0.031768 | 0.006277 | 0.005798 | 0.004498 | 0.003092 | 0.009515 | 0.001921 | 0.002877 | 0.002906 | 0.018538 | 0.002268 |
| 34.3055556 | 0.015058 | 0.076872 | 0.013432 | 0.022968 | 0.028658 | 0.062635 | 0.01239 | 0.003967 | 0.04066 | 0.003554 | 0.005245 | 0.004704 | 0.006123 | 0.008479 | 0.002925 | 0.0029 | 0.001219 | 0.014846 | 0.004632 |
| 34.3194444 | 0.016239 | 0.090416 | 0.007981 | 0.010846 | 0.044432 | 0.043 | 0.006564 | 0.006495 | 0.050933 | 0.003622 | 0.00524 | 0.005851 | 0.007725 | 0.005602 | 0.003779 | 0.002491 | 0.00152 | 0.008652 | 0.007491 |
| 34.3333333 | 0.009638 | 0.068999 | 0.005203 | 0.006461 | 0.045294 | 0.028939 | 0.006235 | 0.00881 | 0.055234 | 0.00349 | 0.007008 | 0.003697 | 0.007648 | 0.003848 | 0.003667 | 0.002471 | 0.002398 | 0.003684 | 0.007751 |
| 34.3472222 | 0.005163 | 0.049232 | 0.00583 | 0.00516 | 0.0402 | 0.015769 | 0.007905 | 0.009682 | 0.053222 | 0.005392 | 0.009783 | 0.001485 | 0.008041 | 0.002699 | 0.003027 | 0.001673 | 0.002233 | 0.002209 | 0.006698 |
| 34.3611111 | 0.004406 | 0.044934 | 0.005808 | 0.00593 | 0.037523 | 0.01008 | 0.009168 | 0.009258 | 0.050284 | 0.008436 | 0.010956 | 0.00212 | 0.010555 | 0.003813 | 0.003633 | 0.00224 | 0.001978 | 0.004032 | 0.006642 |
| 34.375 | 0.002491 | 0.043922 | 0.005041 | 0.00898 | 0.034456 | 0.023392 | 0.009308 | 0.007853 | 0.04935 | 0.00997 | 0.009068 | 0.003347 | 0.014264 | 0.008182 | 0.005209 | 0.005145 | 0.004147 | 0.006441 | 0.00705 |
| 34.3888889 | 0.000881 | 0.040099 | 0.004531 | 0.008563 | 0.03067 | 0.047643 | 0.007468 | 0.005212 | 0.051552 | 0.007896 | 0.005282 | 0.004086 | 0.01656 | 0.012794 | 0.005212 | 0.007421 | 0.007204 | 0.008559 | 0.00672 |
| 34.4027778 | 0.003588 | 0.03832 | 0.003633 | 0.005899 | 0.028812 | 0.06513 | 0.004965 | 0.002564 | 0.056936 | 0.005598 | 0.004316 | 0.008799 | 0.017549 | 0.015376 | 0.003508 | 0.007831 | 0.009453 | 0.011577 | 0.006478 |
| 34.4166667 | 0.012797 | 0.047098 | 0.003253 | 0.008366 | 0.032924 | 0.067763 | 0.004497 | 0.002263 | 0.064781 | 0.009543 | 0.008681 | 0.017193 | 0.017736 | 0.015751 | 0.002633 | 0.007633 | 0.010663 | 0.01522 | 0.007511 |
| 34.4305556 | 0.02201 | 0.059925 | 0.005818 | 0.014941 | 0.040996 | 0.055136 | 0.00506 | 0.004033 | 0.070354 | 0.016548 | 0.012668 | 0.022289 | 0.014007 | 0.012752 | 0.002352 | 0.006632 | 0.00975 | 0.016971 | 0.008295 |
| 34.4444444 | 0.019604 | 0.056615 | 0.009766 | 0.014837 | 0.039667 | 0.033042 | 0.005631 | 0.004556 | 0.065355 | 0.018867 | 0.011947 | 0.020623 | 0.008499 | 0.007425 | 0.001517 | 0.004117 | 0.007019 | 0.014507 | 0.007081 |
| 34.4583333 | 0.008662 | 0.037506 | 0.010631 | 0.007249 | 0.029952 | 0.013878 | 0.009597 | 0.00259 | 0.054047 | 0.017347 | 0.009546 | 0.015736 | 0.007753 | 0.004973 | 0.00103 | 0.002301 | 0.004856 | 0.00933 | 0.005408 |
| 34.4722222 | 0.003886 | 0.022725 | 0.007332 | 0.002607 | 0.030765 | 0.007638 | 0.014831 | 0.001247 | 0.046699 | 0.016818 | 0.00901 | 0.012882 | 0.008351 | 0.005685 | 0.001341 | 0.00307 | 0.004362 | 0.004735 | 0.004607 |
| 34.4861111 | 0.008284 | 0.01766 | 0.004173 | 0.003861 | 0.042017 | 0.013053 | 0.015091 | 0.002999 | 0.039954 | 0.018048 | 0.009631 | 0.012207 | 0.005194 | 0.00476 | 0.002112 | 0.004578 | 0.004959 | 0.002263 | 0.003444 |
| 34.5 | 0.011073 | 0.015091 | 0.004584 | 0.006271 | 0.044125 | 0.020888 | 0.011586 | 0.006035 | 0.029376 | 0.020474 | 0.009443 | 0.01229 | 0.0025 | 0.003269 | 0.002852 | 0.005222 | 0.006899 | 0.001607 | 0.001612 |
| 34.5138889 | 0.007154 | 0.010857 | 0.005197 | 0.006977 | 0.030262 | 0.021945 | 0.010213 | 0.006801 | 0.020981 | 0.023978 | 0.00867 | 0.014685 | 0.003581 | 0.005781 | 0.00429 | 0.005858 | 0.010503 | 0.001678 | 0.00083 |
| 34.5277778 | 0.006536 | 0.008965 | 0.008397 | 0.009485 | 0.012874 | 0.014503 | 0.014125 | 0.004988 | 0.020019 | 0.027391 | 0.00966 | 0.019999 | 0.005109 | 0.012405 | 0.006386 | 0.007506 | 0.015057 | 0.003706 | 0.001127 |
| 34.5416667 | 0.017254 | 0.021295 | 0.022055 | 0.019257 | 0.007249 | 0.009713 | 0.022911 | 0.00285 | 0.029918 | 0.031611 | 0.017692 | 0.029384 | 0.004897 | 0.023618 | 0.007418 | 0.010871 | 0.022561 | 0.010463 | 0.002449 |
| 34.5555556 | 0.035384 | 0.048346 | 0.044783 | 0.03783 | 0.017382 | 0.023771 | 0.035366 | 0.003733 | 0.053529 | 0.037608 | 0.032195 | 0.041294 | 0.006138 | 0.041073 | 0.007865 | 0.015618 | 0.038907 | 0.021733 | 0.007087 |
| 34.5694444 | 0.056279 | 0.077542 | 0.069842 | 0.060764 | 0.036009 | 0.051245 | 0.049296 | 0.009828 | 0.081399 | 0.040928 | 0.04111 | 0.047056 | 0.011062 | 0.056708 | 0.009162 | 0.018519 | 0.055316 | 0.031604 | 0.012577 |
| 34.5833333 | 0.076387 | 0.10168 | 0.088334 | 0.080178 | 0.052456 | 0.070184 | 0.060134 | 0.015883 | 0.101277 | 0.038426 | 0.038706 | 0.044139 | 0.015905 | 0.06017 | 0.010819 | 0.016393 | 0.054554 | 0.033073 | 0.013419 |
| 34.5972222 | 0.087277 | 0.123964 | 0.088984 | 0.088833 | 0.059245 | 0.07497 | 0.062077 | 0.015764 | 0.112261 | 0.03224 | 0.033206 | 0.038769 | 0.015749 | 0.054531 | 0.011126 | 0.010911 | 0.04282 | 0.02691 | 0.009618 |
| 34.6111111 | 0.081611 | 0.145191 | 0.071116 | 0.079819 | 0.056908 | 0.07546 | 0.052742 | 0.013306 | 0.118565 | 0.023155 | 0.032883 | 0.033311 | 0.011235 | 0.046309 | 0.007776 | 0.005421 | 0.032336 | 0.017408 | 0.005204 |
| 34.625 | 0.062782 | 0.157149 | 0.045494 | 0.054027 | 0.048723 | 0.075922 | 0.036403 | 0.015011 | 0.122526 | 0.012396 | 0.036423 | 0.025337 | 0.005962 | 0.037499 | 0.004941 | 0.003047 | 0.021675 | 0.01113 | 0.004464 |
| 34.6388889 | 0.044215 | 0.154283 | 0.021091 | 0.024843 | 0.036241 | 0.071013 | 0.021348 | 0.018701 | 0.124414 | 0.007393 | 0.037549 | 0.014122 | 0.002775 | 0.031559 | 0.008654 | 0.006403 | 0.01168 | 0.018105 | 0.005292 |
| 34.6527778 | 0.035863 | 0.13682 | 0.006933 | 0.008974 | 0.025439 | 0.061342 | 0.015108 | 0.017709 | 0.120173 | 0.010131 | 0.034187 | 0.005792 | 0.00234 | 0.030069 | 0.012902 | 0.010219 | 0.008122 | 0.031507 | 0.004358 |
| 34.6666667 | 0.031272 | 0.108401 | 0.006309 | 0.008873 | 0.020465 | 0.050469 | 0.018967 | 0.010573 | 0.099559 | 0.012307 | 0.029223 | 0.00505 | 0.001905 | 0.030181 | 0.011763 | 0.00792 | 0.010296 | 0.035101 | 0.005331 |
| 34.6805556 | 0.01978 | 0.070543 | 0.010289 | 0.016208 | 0.014726 | 0.032379 | 0.026631 | 0.005336 | 0.062801 | 0.012512 | 0.025959 | 0.008089 | 0.000701 | 0.027255 | 0.007033 | 0.00376 | 0.011155 | 0.030221 | 0.00709 |
| 34.6944444 | 0.01208 | 0.036693 | 0.015419 | 0.028793 | 0.014925 | 0.025158 | 0.030816 | 0.008673 | 0.031602 | 0.014583 | 0.024271 | 0.012369 | 0.00106 | 0.018081 | 0.00309 | 0.002545 | 0.008147 | 0.025547 | 0.005674 |
| 34.7083333 | 0.022542 | 0.035573 | 0.022441 | 0.0358 | 0.032692 | 0.049831 | 0.032936 | 0.015761 | 0.032796 | 0.016816 | 0.022501 | 0.016537 | 0.003501 | 0.008471 | 0.001929 | 0.001878 | 0.004491 | 0.02298 | 0.004748 |
| 34.7222222 | 0.038992 | 0.062254 | 0.026717 | 0.027267 | 0.054324 | 0.078899 | 0.037798 | 0.021954 | 0.064954 | 0.014837 | 0.021531 | 0.01743 | 0.004712 | 0.006235 | 0.002773 | 0.001226 | 0.004189 | 0.020447 | 0.005955 |
| 34.7361111 | 0.045614 | 0.084835 | 0.025432 | 0.014422 | 0.068623 | 0.088093 | 0.04249 | 0.02635 | 0.098243 | 0.009279 | 0.022742 | 0.014738 | 0.003684 | 0.008744 | 0.007766 | 0.002584 | 0.007733 | 0.015755 | 0.005088 |
| 34.75 | 0.042176 | 0.089953 | 0.020242 | 0.006496 | 0.072686 | 0.078719 | 0.044562 | 0.02628 | 0.113575 | 0.008352 | 0.0235 | 0.010493 | 0.004433 | 0.009245 | 0.014648 | 0.005166 | 0.011028 | 0.008816 | 0.00361 |
| 34.7638889 | 0.03271 | 0.080504 | 0.013355 | 0.003909 | 0.061005 | 0.055765 | 0.046159 | 0.022469 | 0.116169 | 0.015839 | 0.020397 | 0.006033 | 0.006558 | 0.006136 | 0.016908 | 0.006086 | 0.0108 | 0.005886 | 0.003817 |
| 34.7777778 | 0.020979 | 0.06367 | 0.008538 | 0.006946 | 0.038357 | 0.027836 | 0.047336 | 0.019076 | 0.114299 | 0.025365 | 0.013958 | 0.003474 | 0.008094 | 0.004669 | 0.012688 | 0.004232 | 0.007482 | 0.012217 | 0.002865 |
| 34.7916667 | 0.011566 | 0.046016 | 0.013428 | 0.013942 | 0.016818 | 0.012042 | 0.047534 | 0.016266 | 0.10344 | 0.029774 | 0.006848 | 0.005828 | 0.010884 | 0.006967 | 0.006681 | 0.002118 | 0.004018 | 0.019508 | 0.001127 |
| 34.8055556 | 0.005803 | 0.030631 | 0.02742 | 0.021664 | 0.005438 | 0.017022 | 0.046549 | 0.011884 | 0.078878 | 0.027068 | 0.003864 | 0.011267 | 0.013683 | 0.007564 | 0.005498 | 0.001712 | 0.002808 | 0.017018 | 0.001079 |
| 34.8194444 | 0.002349 | 0.017799 | 0.038019 | 0.024712 | 0.00246 | 0.022212 | 0.043972 | 0.007957 | 0.049247 | 0.018 | 0.007903 | 0.014335 | 0.012964 | 0.004948 | 0.008182 | 0.00233 | 0.003407 | 0.010809 | 0.002159 |
| 34.8333333 | 0.000893 | 0.008191 | 0.037647 | 0.021759 | 0.002539 | 0.014519 | 0.04067 | 0.008639 | 0.024826 | 0.010338 | 0.014834 | 0.013414 | 0.009238 | 0.003575 | 0.008763 | 0.003137 | 0.004252 | 0.01408 | 0.002589 |
| 34.8472222 | 0.000864 | 0.00459 | 0.030123 | 0.016264 | 0.006122 | 0.007884 | 0.036344 | 0.012928 | 0.010611 | 0.013115 | 0.019728 | 0.010799 | 0.005772 | 0.004258 | 0.005832 | 0.003965 | 0.004697 | 0.021924 | 0.004516 |
| 34.8611111 | 0.001826 | 0.00614 | 0.02005 | 0.008975 | 0.013957 | 0.014289 | 0.029527 | 0.016159 | 0.009167 | 0.020171 | 0.02222 | 0.008515 | 0.004275 | 0.003292 | 0.002984 | 0.004421 | 0.004047 | 0.023157 | 0.010457 |
| 34.875 | 0.002106 | 0.00671 | 0.010618 | 0.003797 | 0.025073 | 0.027625 | 0.020588 | 0.017462 | 0.01404 | 0.024285 | 0.02501 | 0.005998 | 0.004233 | 0.001899 | 0.004174 | 0.003847 | 0.00228 | 0.018571 | 0.015645 |
| 34.8888889 | 0.002721 | 0.005483 | 0.010161 | 0.005653 | 0.035405 | 0.039589 | 0.010684 | 0.01913 | 0.01349 | 0.027806 | 0.030936 | 0.003803 | 0.004626 | 0.0036 | 0.008161 | 0.002718 | 0.000922 | 0.012629 | 0.014442 |
| 34.9027778 | 0.008702 | 0.009687 | 0.020661 | 0.010591 | 0.040529 | 0.049809 | 0.004258 | 0.021598 | 0.011365 | 0.030769 | 0.039948 | 0.004342 | 0.005447 | 0.006502 | 0.010083 | 0.002575 | 0.000674 | 0.006855 | 0.009177 |
| 34.9166667 | 0.018598 | 0.022219 | 0.031332 | 0.014189 | 0.046255 | 0.061237 | 0.004258 | 0.022792 | 0.022954 | 0.030475 | 0.048677 | 0.005617 | 0.006738 | 0.00832 | 0.007919 | 0.003877 | 0.000862 | 0.003971 | 0.005735 |
| 34.9305556 | 0.026958 | 0.034605 | 0.035695 | 0.015372 | 0.059099 | 0.074932 | 0.005139 | 0.021173 | 0.04506 | 0.026271 | 0.053409 | 0.004267 | 0.007967 | 0.009873 | 0.004093 | 0.005765 | 0.001311 | 0.00686 | 0.005997 |
| 34.9444444 | 0.033272 | 0.037502 | 0.038095 | 0.0147 | 0.072849 | 0.082788 | 0.004613 | 0.017642 | 0.064798 | 0.019979 | 0.052225 | 0.002687 | 0.009344 | 0.011713 | 0.002124 | 0.007072 | 0.002477 | 0.009435 | 0.007362 |
| 34.9583333 | 0.035082 | 0.029581 | 0.043318 | 0.015249 | 0.076714 | 0.079021 | 0.005684 | 0.013815 | 0.079513 | 0.014519 | 0.047249 | 0.003655 | 0.010753 | 0.012928 | 0.002009 | 0.007174 | 0.004101 | 0.006359 | 0.006983 |
| 34.9722222 | 0.026895 | 0.017121 | 0.043177 | 0.015258 | 0.061928 | 0.069018 | 0.006966 | 0.00905 | 0.087541 | 0.010807 | 0.041922 | 0.004527 | 0.01115 | 0.012764 | 0.001997 | 0.006617 | 0.00541 | 0.002378 | 0.006022 |
| 34.9861111 | 0.014637 | 0.008201 | 0.027627 | 0.010619 | 0.039576 | 0.055117 | 0.005642 | 0.00775 | 0.083029 | 0.007278 | 0.036917 | 0.004786 | 0.010532 | 0.010283 | 0.0019 | 0.006368 | 0.005639 | 0.003792 | 0.006493 |
| 35 | 0.013132 | 0.007916 | 0.012879 | 0.006296 | 0.046955 | 0.038145 | 0.005997 | 0.015415 | 0.065534 | 0.003729 | 0.031202 | 0.006704 | 0.008886 | 0.006735 | 0.002333 | 0.006475 | 0.004421 | 0.011238 | 0.007319 |
| 35.0138889 | 0.02216 | 0.018618 | 0.018602 | 0.009243 | 0.090235 | 0.023194 | 0.012646 | 0.025872 | 0.045249 | 0.001503 | 0.025005 | 0.00942 | 0.00544 | 0.008379 | 0.003956 | 0.00569 | 0.002308 | 0.018057 | 0.00698 |
| 35.0277778 | 0.026006 | 0.029364 | 0.033784 | 0.01392 | 0.122749 | 0.014814 | 0.020313 | 0.031254 | 0.032297 | 0.004436 | 0.024696 | 0.008398 | 0.003501 | 0.016748 | 0.005022 | 0.004792 | 0.002663 | 0.014983 | 0.004886 |
| 35.0416667 | 0.018245 | 0.027684 | 0.035777 | 0.012192 | 0.114439 | 0.018628 | 0.020028 | 0.026695 | 0.03581 | 0.020292 | 0.039262 | 0.011171 | 0.007337 | 0.033543 | 0.005814 | 0.011211 | 0.01365 | 0.017359 | 0.007445 |
| 35.0555556 | 0.014292 | 0.034077 | 0.028829 | 0.011596 | 0.082594 | 0.035745 | 0.012327 | 0.017454 | 0.060087 | 0.049375 | 0.071404 | 0.033422 | 0.017448 | 0.066444 | 0.013198 | 0.028557 | 0.032369 | 0.048556 | 0.021848 |
| 35.0694444 | 0.023931 | 0.062735 | 0.036161 | 0.017548 | 0.050684 | 0.050664 | 0.008089 | 0.019293 | 0.079564 | 0.07013 | 0.09311 | 0.054206 | 0.028454 | 0.091523 | 0.022056 | 0.039477 | 0.041814 | 0.078048 | 0.035749 |
| 35.0833333 | 0.032916 | 0.085729 | 0.051598 | 0.017614 | 0.025144 | 0.048552 | 0.008239 | 0.028021 | 0.074635 | 0.058315 | 0.067851 | 0.040684 | 0.026915 | 0.074016 | 0.017435 | 0.028225 | 0.032782 | 0.065417 | 0.030946 |
| 35.0972222 | 0.036393 | 0.092429 | 0.05411 | 0.010168 | 0.013029 | 0.038429 | 0.006427 | 0.027941 | 0.067551 | 0.026298 | 0.026316 | 0.017334 | 0.015386 | 0.033104 | 0.006373 | 0.013927 | 0.016267 | 0.034518 | 0.015314 |
| 35.1111111 | 0.038342 | 0.096821 | 0.04646 | 0.006497 | 0.020067 | 0.033675 | 0.005101 | 0.021078 | 0.067844 | 0.007409 | 0.012142 | 0.015627 | 0.014006 | 0.010147 | 0.002227 | 0.013997 | 0.011926 | 0.029725 | 0.006109 |
| 35.125 | 0.035859 | 0.098467 | 0.038613 | 0.009877 | 0.031415 | 0.035345 | 0.007187 | 0.013917 | 0.063588 | 0.010361 | 0.009499 | 0.021123 | 0.025036 | 0.011475 | 0.002023 | 0.015638 | 0.018378 | 0.046893 | 0.002518 |
| 35.1388889 | 0.031041 | 0.092357 | 0.030142 | 0.016166 | 0.033971 | 0.039187 | 0.010709 | 0.007476 | 0.057301 | 0.020812 | 0.004796 | 0.027156 | 0.035773 | 0.018082 | 0.00384 | 0.012442 | 0.022584 | 0.058805 | 0.001888 |
| 35.1527778 | 0.026867 | 0.080241 | 0.018907 | 0.017891 | 0.0282 | 0.040446 | 0.011264 | 0.004676 | 0.056944 | 0.02621 | 0.003636 | 0.037408 | 0.0389 | 0.019696 | 0.007805 | 0.009478 | 0.020925 | 0.059487 | 0.005036 |
| 35.1666667 | 0.021838 | 0.064778 | 0.008578 | 0.012489 | 0.015817 | 0.038221 | 0.00704 | 0.007418 | 0.061704 | 0.023731 | 0.003761 | 0.040442 | 0.034875 | 0.018202 | 0.009972 | 0.006439 | 0.014941 | 0.054601 | 0.00722 |
| 35.1805556 | 0.01511 | 0.04893 | 0.004586 | 0.005683 | 0.009796 | 0.034027 | 0.003586 | 0.011876 | 0.063275 | 0.016987 | 0.003797 | 0.031126 | 0.027485 | 0.015611 | 0.00854 | 0.003577 | 0.009047 | 0.044906 | 0.005205 |
| 35.1944444 | 0.008618 | 0.033977 | 0.007732 | 0.005356 | 0.02519 | 0.029082 | 0.007362 | 0.011988 | 0.05419 | 0.010488 | 0.003397 | 0.01817 | 0.019142 | 0.014369 | 0.006008 | 0.00245 | 0.005833 | 0.029038 | 0.002202 |
| 35.2083333 | 0.009393 | 0.018719 | 0.01526 | 0.015669 | 0.053708 | 0.022285 | 0.018665 | 0.008119 | 0.033898 | 0.007344 | 0.002825 | 0.010262 | 0.010442 | 0.019747 | 0.005664 | 0.002267 | 0.00436 | 0.013386 | 0.00191 |
| 35.2222222 | 0.020749 | 0.012217 | 0.025714 | 0.032114 | 0.076162 | 0.012502 | 0.032805 | 0.008109 | 0.017219 | 0.008346 | 0.004839 | 0.008866 | 0.004994 | 0.031205 | 0.007317 | 0.002846 | 0.004315 | 0.007763 | 0.004012 |
| 35.2361111 | 0.032836 | 0.021152 | 0.036501 | 0.043137 | 0.079326 | 0.007875 | 0.042818 | 0.014089 | 0.019406 | 0.011001 | 0.008367 | 0.010305 | 0.00657 | 0.038576 | 0.008219 | 0.003757 | 0.006193 | 0.015901 | 0.006318 |
| 35.25 | 0.038326 | 0.031874 | 0.04367 | 0.042058 | 0.065766 | 0.011264 | 0.044763 | 0.018676 | 0.028796 | 0.012038 | 0.009775 | 0.010716 | 0.010132 | 0.035386 | 0.006749 | 0.003299 | 0.008636 | 0.029839 | 0.006333 |
| 35.2638889 | 0.04032 | 0.033703 | 0.046878 | 0.035515 | 0.055244 | 0.011314 | 0.041064 | 0.019025 | 0.028155 | 0.010208 | 0.008074 | 0.009372 | 0.010102 | 0.022917 | 0.004496 | 0.001818 | 0.009423 | 0.042212 | 0.004911 |
| 35.2777778 | 0.042648 | 0.030646 | 0.046362 | 0.03403 | 0.056693 | 0.006415 | 0.036194 | 0.016596 | 0.016939 | 0.00734 | 0.004968 | 0.006196 | 0.006937 | 0.011227 | 0.004056 | 0.000917 | 0.007812 | 0.050097 | 0.004467 |
| 35.2916667 | 0.043961 | 0.027169 | 0.039792 | 0.039612 | 0.06158 | 0.005542 | 0.033541 | 0.011485 | 0.008833 | 0.00546 | 0.002365 | 0.003731 | 0.003354 | 0.011544 | 0.005278 | 0.001272 | 0.005356 | 0.048866 | 0.004553 |
| 35.3055556 | 0.039808 | 0.022033 | 0.028438 | 0.041922 | 0.059538 | 0.01074 | 0.029911 | 0.009412 | 0.015289 | 0.003851 | 0.001192 | 0.005301 | 0.00229 | 0.018577 | 0.006142 | 0.002276 | 0.003326 | 0.04066 | 0.004045 |
| 35.3194444 | 0.030064 | 0.01382 | 0.016387 | 0.0298 | 0.045584 | 0.01805 | 0.019987 | 0.012753 | 0.02727 | 0.001798 | 0.001618 | 0.007332 | 0.003536 | 0.02217 | 0.006605 | 0.003047 | 0.0018 | 0.032034 | 0.003148 |
| 35.3333333 | 0.020115 | 0.007187 | 0.007172 | 0.013301 | 0.024483 | 0.022341 | 0.009698 | 0.014546 | 0.032437 | 0.000937 | 0.002088 | 0.005632 | 0.004361 | 0.019921 | 0.008035 | 0.004099 | 0.001212 | 0.025468 | 0.002273 |
| 35.3472222 | 0.014983 | 0.006177 | 0.004383 | 0.006595 | 0.011997 | 0.021591 | 0.008505 | 0.013333 | 0.029773 | 0.002141 | 0.00146 | 0.004164 | 0.003282 | 0.013108 | 0.009628 | 0.004815 | 0.001214 | 0.019263 | 0.001531 |
| 35.3611111 | 0.014888 | 0.006688 | 0.005477 | 0.004936 | 0.017904 | 0.019085 | 0.013921 | 0.011937 | 0.024862 | 0.003472 | 0.000773 | 0.005596 | 0.001501 | 0.006107 | 0.008909 | 0.003434 | 0.001427 | 0.011967 | 0.001243 |
| 35.375 | 0.016049 | 0.0046 | 0.00526 | 0.005774 | 0.032507 | 0.017069 | 0.012959 | 0.00967 | 0.024825 | 0.002994 | 0.001482 | 0.006442 | 0.000707 | 0.002663 | 0.005554 | 0.001888 | 0.001679 | 0.005043 | 0.000999 |

| | | | | | | | | | | | | | | | | | | | |
|---|---|---|---|---|---|---|---|---|---|---|---|---|---|---|---|---|---|---|---|
| 35.3888889 | 0.015985 | 0.003407 | 0.006388 | 0.009535 | 0.044078 | 0.014649 | 0.005216 | 0.006485 | 0.03026 | 0.001677 | 0.003614 | 0.005818 | 0.001077 | 0.002113 | 0.002201 | 0.00206 | 0.001225 | 0.001216 | 0.000791 |
| 35.4027778 | 0.015862 | 0.005348 | 0.011505 | 0.010385 | 0.051035 | 0.01198 | 0.001573 | 0.004329 | 0.037382 | 0.00117 | 0.005483 | 0.005579 | 0.002705 | 0.001714 | 0.000682 | 0.002048 | 0.001105 | 0.00088 | 0.001349 |
| 35.4166667 | 0.016696 | 0.007345 | 0.015841 | 0.007734 | 0.054472 | 0.011176 | 0.00108 | 0.003861 | 0.042849 | 0.001332 | 0.005296 | 0.006466 | 0.004658 | 0.001686 | 0.000574 | 0.001228 | 0.003143 | 0.002847 | 0.001923 |
| 35.4305556 | 0.016997 | 0.00909 | 0.016672 | 0.004082 | 0.050032 | 0.013619 | 0.001375 | 0.003567 | 0.047108 | 0.002411 | 0.003477 | 0.008141 | 0.005256 | 0.003157 | 0.000919 | 0.000635 | 0.006169 | 0.005884 | 0.001796 |
| 35.4444444 | 0.015921 | 0.011227 | 0.016651 | 0.003203 | 0.036138 | 0.016382 | 0.002888 | 0.002203 | 0.055417 | 0.004538 | 0.002121 | 0.009463 | 0.00404 | 0.004576 | 0.000849 | 0.000426 | 0.007571 | 0.007794 | 0.001225 |
| 35.4583333 | 0.014987 | 0.011944 | 0.017406 | 0.008482 | 0.018766 | 0.014476 | 0.004394 | 0.00073 | 0.065268 | 0.006178 | 0.001741 | 0.00929 | 0.002191 | 0.005808 | 0.000413 | 0.000472 | 0.007015 | 0.007468 | 0.000761 |
| 35.4722222 | 0.01476 | 0.011604 | 0.016961 | 0.016697 | 0.007579 | 0.009673 | 0.008671 | 0.002103 | 0.074009 | 0.00632 | 0.00145 | 0.008079 | 0.001597 | 0.007441 | 0.000994 | 0.000671 | 0.006117 | 0.006165 | 0.000814 |
| 35.4861111 | 0.01358 | 0.012107 | 0.013023 | 0.022104 | 0.006087 | 0.008515 | 0.01477 | 0.006835 | 0.08409 | 0.006042 | 0.001123 | 0.007527 | 0.002361 | 0.007873 | 0.002962 | 0.001455 | 0.006197 | 0.005383 | 0.001055 |
| 35.5 | 0.010163 | 0.013527 | 0.00709 | 0.0226 | 0.006126 | 0.012613 | 0.017869 | 0.010866 | 0.087195 | 0.006582 | 0.001076 | 0.007989 | 0.003649 | 0.006089 | 0.003707 | 0.002619 | 0.007214 | 0.004854 | 0.001676 |
| 35.5138889 | 0.005616 | 0.014097 | 0.003404 | 0.020973 | 0.006387 | 0.018497 | 0.017578 | 0.01235 | 0.075756 | 0.007792 | 0.001136 | 0.009186 | 0.005014 | 0.003516 | 0.003291 | 0.00247 | 0.00865 | 0.003591 | 0.002596 |
| 35.5277778 | 0.0031 | 0.010357 | 0.004264 | 0.021398 | 0.009624 | 0.01899 | 0.018884 | 0.01254 | 0.054301 | 0.012082 | 0.004473 | 0.01591 | 0.006015 | 0.004573 | 0.005731 | 0.002787 | 0.013193 | 0.005347 | 0.002301 |
| 35.5416667 | 0.007686 | 0.009645 | 0.011729 | 0.028684 | 0.010777 | 0.018466 | 0.033757 | 0.010406 | 0.030947 | 0.027585 | 0.021977 | 0.037528 | 0.006948 | 0.015784 | 0.010636 | 0.008152 | 0.028656 | 0.018284 | 0.003425 |
| 35.5555556 | 0.023162 | 0.024491 | 0.025798 | 0.045845 | 0.016004 | 0.03314 | 0.065876 | 0.007073 | 0.025534 | 0.053277 | 0.04847 | 0.069666 | 0.008574 | 0.036233 | 0.012627 | 0.017177 | 0.048399 | 0.039409 | 0.010965 |
| 35.5694444 | 0.04302 | 0.043363 | 0.035949 | 0.06447 | 0.034803 | 0.053512 | 0.091551 | 0.007808 | 0.046198 | 0.065351 | 0.05537 | 0.082907 | 0.011487 | 0.053936 | 0.01113 | 0.022451 | 0.051686 | 0.052408 | 0.020697 |
| 35.5833333 | 0.057136 | 0.051981 | 0.040848 | 0.07496 | 0.057255 | 0.059121 | 0.091205 | 0.009352 | 0.069251 | 0.052056 | 0.038446 | 0.064574 | 0.015328 | 0.060202 | 0.010036 | 0.020212 | 0.040059 | 0.052231 | 0.024186 |
| 35.5972222 | 0.06125 | 0.05626 | 0.045949 | 0.076636 | 0.069459 | 0.056407 | 0.075785 | 0.007896 | 0.082807 | 0.033096 | 0.020667 | 0.040711 | 0.016379 | 0.053744 | 0.008332 | 0.013307 | 0.028501 | 0.045057 | 0.020127 |
| 35.6111111 | 0.052821 | 0.057414 | 0.044095 | 0.070218 | 0.066083 | 0.056338 | 0.060493 | 0.009517 | 0.090402 | 0.020961 | 0.01162 | 0.02827 | 0.01224 | 0.034117 | 0.004902 | 0.008171 | 0.019898 | 0.03073 | 0.012027 |
| 35.625 | 0.034904 | 0.047436 | 0.036603 | 0.058255 | 0.051289 | 0.05551 | 0.047889 | 0.013568 | 0.091249 | 0.012523 | 0.007829 | 0.022587 | 0.006237 | 0.018753 | 0.003745 | 0.011137 | 0.012946 | 0.01582 | 0.006299 |
| 35.6388889 | 0.015962 | 0.027698 | 0.03034 | 0.044519 | 0.033041 | 0.048801 | 0.035986 | 0.013343 | 0.086549 | 0.008251 | 0.007151 | 0.019824 | 0.002952 | 0.027242 | 0.005617 | 0.018203 | 0.010392 | 0.015723 | 0.007537 |
| 35.6527778 | 0.005615 | 0.010592 | 0.02496 | 0.033675 | 0.018055 | 0.037945 | 0.027391 | 0.008231 | 0.073845 | 0.009967 | 0.007515 | 0.018329 | 0.002776 | 0.039634 | 0.005814 | 0.019021 | 0.011398 | 0.027778 | 0.01144 |
| 35.6666667 | 0.009158 | 0.004019 | 0.017622 | 0.02773 | 0.009698 | 0.028776 | 0.025941 | 0.00513 | 0.055633 | 0.014076 | 0.006119 | 0.015176 | 0.002378 | 0.032856 | 0.003723 | 0.011824 | 0.012369 | 0.036314 | 0.011325 |
| 35.6805556 | 0.019309 | 0.006202 | 0.011836 | 0.021536 | 0.006064 | 0.022318 | 0.027748 | 0.00878 | 0.041181 | 0.015957 | 0.003119 | 0.009845 | 0.002172 | 0.016104 | 0.001874 | 0.004645 | 0.015309 | 0.038952 | 0.006988 |
| 35.6944444 | 0.026228 | 0.0143 | 0.013831 | 0.015504 | 0.008078 | 0.015025 | 0.027333 | 0.014227 | 0.028376 | 0.014778 | 0.000923 | 0.005442 | 0.003701 | 0.004414 | 0.000716 | 0.002391 | 0.01872 | 0.041523 | 0.004558 |
| 35.7083333 | 0.028537 | 0.025378 | 0.017033 | 0.018132 | 0.019711 | 0.007472 | 0.024584 | 0.016113 | 0.01635 | 0.010579 | 0.000463 | 0.002658 | 0.004368 | 0.001595 | 0.000291 | 0.002408 | 0.018013 | 0.046289 | 0.005408 |
| 35.7222222 | 0.029003 | 0.036252 | 0.014546 | 0.026388 | 0.039398 | 0.007326 | 0.021225 | 0.013391 | 0.008598 | 0.006095 | 0.001533 | 0.001638 | 0.003877 | 0.005329 | 0.000874 | 0.003857 | 0.01181 | 0.051822 | 0.004791 |
| 35.7361111 | 0.029521 | 0.046024 | 0.008745 | 0.02967 | 0.062322 | 0.017577 | 0.016962 | 0.01076 | 0.006766 | 0.006952 | 0.003601 | 0.003655 | 0.005756 | 0.011595 | 0.002872 | 0.006011 | 0.006566 | 0.05367 | 0.004828 |
| 35.75 | 0.029783 | 0.053468 | 0.003751 | 0.026258 | 0.082607 | 0.032853 | 0.012959 | 0.013779 | 0.013133 | 0.01211 | 0.005144 | 0.008127 | 0.009949 | 0.014987 | 0.005827 | 0.00591 | 0.00974 | 0.049742 | 0.008826 |
| 35.7638889 | 0.027631 | 0.056742 | 0.001426 | 0.01801 | 0.088156 | 0.045169 | 0.014351 | 0.016693 | 0.022742 | 0.015133 | 0.005238 | 0.014679 | 0.012626 | 0.012079 | 0.008663 | 0.00394 | 0.012817 | 0.039234 | 0.012507 |
| 35.7777778 | 0.021989 | 0.055588 | 0.00181 | 0.009258 | 0.073607 | 0.04672 | 0.020689 | 0.011488 | 0.03265 | 0.0137 | 0.004368 | 0.021682 | 0.011541 | 0.007626 | 0.011027 | 0.004284 | 0.008662 | 0.021814 | 0.012343 |
| 35.7916667 | 0.015183 | 0.050887 | 0.004145 | 0.007785 | 0.048877 | 0.040269 | 0.027085 | 0.005538 | 0.04306 | 0.00959 | 0.003179 | 0.024694 | 0.006893 | 0.009923 | 0.012747 | 0.007637 | 0.005074 | 0.010593 | 0.008822 |
| 35.8055556 | 0.010285 | 0.044497 | 0.007687 | 0.014552 | 0.023887 | 0.031996 | 0.031488 | 0.007827 | 0.050143 | 0.006061 | 0.001935 | 0.021803 | 0.003991 | 0.018662 | 0.013104 | 0.010051 | 0.005409 | 0.018165 | 0.004427 |
| 35.8194444 | 0.008081 | 0.038859 | 0.008152 | 0.023025 | 0.010278 | 0.0238 | 0.033337 | 0.01246 | 0.05191 | 0.005748 | 0.001167 | 0.016877 | 0.007107 | 0.028085 | 0.012485 | 0.009526 | 0.004456 | 0.032663 | 0.00195 |
| 35.8333333 | 0.008588 | 0.036594 | 0.008894 | 0.028285 | 0.013082 | 0.018094 | 0.032392 | 0.014255 | 0.050783 | 0.008336 | 0.001123 | 0.01361 | 0.012024 | 0.034278 | 0.01214 | 0.006623 | 0.00267 | 0.03945 | 0.001534 |
| 35.8472222 | 0.011649 | 0.038652 | 0.018242 | 0.028819 | 0.013869 | 0.018608 | 0.027327 | 0.016398 | 0.049403 | 0.011816 | 0.001374 | 0.011066 | 0.013891 | 0.034841 | 0.012385 | 0.003243 | 0.003183 | 0.036256 | 0.001749 |
| 35.8611111 | 0.01622 | 0.042263 | 0.024159 | 0.024807 | 0.012595 | 0.029769 | 0.016306 | 0.016128 | 0.04984 | 0.01406 | 0.001724 | 0.007972 | 0.011273 | 0.028668 | 0.012189 | 0.001553 | 0.005077 | 0.023127 | 0.001468 |
| 35.875 | 0.020897 | 0.044093 | 0.017421 | 0.017747 | 0.027863 | 0.049515 | 0.008202 | 0.013187 | 0.056774 | 0.013749 | 0.002127 | 0.004933 | 0.006188 | 0.017816 | 0.010268 | 0.001882 | 0.007976 | 0.012363 | 0.001004 |
| 35.8888889 | 0.024771 | 0.041643 | 0.016863 | 0.009632 | 0.051128 | 0.065942 | 0.012539 | 0.020595 | 0.072048 | 0.011477 | 0.002145 | 0.003029 | 0.00265 | 0.007988 | 0.00724 | 0.003191 | 0.011354 | 0.015659 | 0.001639 |
| 35.9027778 | 0.028453 | 0.034635 | 0.033698 | 0.003279 | 0.069726 | 0.070026 | 0.021298 | 0.036296 | 0.094176 | 0.00969 | 0.001802 | 0.002854 | 0.001767 | 0.003717 | 0.004245 | 0.004583 | 0.013216 | 0.0202 | 0.00321 |
| 35.9166667 | 0.032195 | 0.024778 | 0.050695 | 0.001589 | 0.08215 | 0.068158 | 0.027878 | 0.045958 | 0.118402 | 0.010763 | 0.002634 | 0.003853 | 0.002432 | 0.004071 | 0.001907 | 0.005111 | 0.013381 | 0.01653 | 0.00536 |
| 35.9305556 | 0.034639 | 0.013744 | 0.055216 | 0.004598 | 0.08493 | 0.066496 | 0.033123 | 0.047774 | 0.129457 | 0.014439 | 0.005532 | 0.005015 | 0.003514 | 0.005731 | 0.000969 | 0.003684 | 0.012827 | 0.009882 | 0.008666 |
| 35.9444444 | 0.03624 | 0.009701 | 0.053088 | 0.009432 | 0.084389 | 0.0616 | 0.035354 | 0.045803 | 0.1169 | 0.017999 | 0.009687 | 0.005936 | 0.004043 | 0.006361 | 0.001364 | 0.001836 | 0.011788 | 0.005562 | 0.0124 |
| 35.9583333 | 0.036819 | 0.017927 | 0.050919 | 0.011877 | 0.089789 | 0.057713 | 0.03285 | 0.043875 | 0.090783 | 0.019362 | 0.014362 | 0.006207 | 0.003473 | 0.005393 | 0.002051 | 0.001349 | 0.010734 | 0.008077 | 0.014135 |
| 35.9722222 | 0.031883 | 0.029501 | 0.046353 | 0.009099 | 0.091198 | 0.057501 | 0.026035 | 0.042078 | 0.065228 | 0.018462 | 0.018859 | 0.005495 | 0.002095 | 0.003356 | 0.001766 | 0.001834 | 0.009778 | 0.015892 | 0.012279 |
| 35.9861111 | 0.018966 | 0.03702 | 0.035985 | 0.004542 | 0.075738 | 0.050665 | 0.016792 | 0.034254 | 0.04615 | 0.015747 | 0.021656 | 0.004032 | 0.001145 | 0.002575 | 0.001426 | 0.004362 | 0.007997 | 0.0239 | 0.008577 |
| 36 | 0.010966 | 0.042386 | 0.020384 | 0.003735 | 0.04597 | 0.033649 | 0.008208 | 0.02052 | 0.033159 | 0.012489 | 0.021952 | 0.002511 | 0.002172 | 0.005335 | 0.0024 | 0.008025 | 0.005001 | 0.030766 | 0.006753 |
| 36.0138889 | 0.01991 | 0.048211 | 0.009792 | 0.00599 | 0.023483 | 0.016303 | 0.004735 | 0.014898 | 0.021972 | 0.011132 | 0.020779 | 0.002337 | 0.005386 | 0.008137 | 0.002666 | 0.009306 | 0.002409 | 0.03413 | 0.008374 |
| 36.0277778 | 0.026408 | 0.045565 | 0.011186 | 0.006387 | 0.023785 | 0.006644 | 0.004252 | 0.01646 | 0.018199 | 0.015273 | 0.021354 | 0.008894 | 0.006516 | 0.008503 | 0.003238 | 0.007879 | 0.005836 | 0.027429 | 0.008186 |
| 36.0416667 | 0.02892 | 0.037422 | 0.016471 | 0.011615 | 0.030078 | 0.01407 | 0.009308 | 0.020048 | 0.04422 | 0.033313 | 0.034409 | 0.037 | 0.013004 | 0.020642 | 0.011375 | 0.014601 | 0.026048 | 0.036052 | 0.011869 |
| 36.0555556 | 0.056141 | 0.051534 | 0.037192 | 0.027273 | 0.051144 | 0.047184 | 0.025923 | 0.041949 | 0.106948 | 0.066216 | 0.064539 | 0.086238 | 0.040073 | 0.05188 | 0.029343 | 0.038689 | 0.059878 | 0.081565 | 0.031244 |
| 36.0694444 | 0.084381 | 0.071699 | 0.066227 | 0.033576 | 0.091528 | 0.082866 | 0.037406 | 0.071956 | 0.157265 | 0.078176 | 0.0786 | 0.094586 | 0.06182 | 0.062904 | 0.039427 | 0.053366 | 0.06599 | 0.104878 | 0.044332 |
| 36.0833333 | 0.074108 | 0.060593 | 0.064675 | 0.02109 | 0.105127 | 0.088257 | 0.031192 | 0.08745 | 0.146603 | 0.053165 | 0.056977 | 0.05491 | 0.044703 | 0.041823 | 0.030094 | 0.040923 | 0.045298 | 0.06589 | 0.032283 |
| 36.0972222 | 0.044754 | 0.036228 | 0.043079 | 0.00972 | 0.089129 | 0.069875 | 0.017458 | 0.093076 | 0.105585 | 0.033238 | 0.039079 | 0.044025 | 0.023576 | 0.035123 | 0.024572 | 0.033323 | 0.047868 | 0.024414 | 0.021834 |
| 36.1111111 | 0.02354 | 0.024097 | 0.029879 | 0.009451 | 0.074365 | 0.050378 | 0.006749 | 0.083104 | 0.079757 | 0.027019 | 0.034079 | 0.048053 | 0.034946 | 0.043722 | 0.034437 | 0.038122 | 0.054192 | 0.022148 | 0.027781 |
| 36.125 | 0.010449 | 0.023371 | 0.023177 | 0.010449 | 0.064347 | 0.035472 | 0.00259 | 0.048485 | 0.060975 | 0.015862 | 0.019134 | 0.031828 | 0.051915 | 0.040548 | 0.041649 | 0.033304 | 0.035885 | 0.030482 | 0.031619 |
| 36.1388889 | 0.00365 | 0.027818 | 0.01712 | 0.008848 | 0.056834 | 0.026656 | 0.001768 | 0.023548 | 0.033612 | 0.008189 | 0.00646 | 0.018417 | 0.052506 | 0.029371 | 0.037974 | 0.02193 | 0.014499 | 0.032609 | 0.028677 |
| 36.1527778 | 0.003616 | 0.030845 | 0.009796 | 0.008738 | 0.050035 | 0.022806 | 0.00156 | 0.040054 | 0.014522 | 0.006471 | 0.004296 | 0.014919 | 0.044157 | 0.016695 | 0.029232 | 0.013573 | 0.003917 | 0.032786 | 0.023676 |
| 36.1666667 | 0.007523 | 0.031136 | 0.005271 | 0.010778 | 0.036876 | 0.017841 | 0.003343 | 0.074206 | 0.014273 | 0.004813 | 0.003504 | 0.012466 | 0.034009 | 0.005663 | 0.020298 | 0.007857 | 0.002147 | 0.032984 | 0.014985 |
| 36.1805556 | 0.013916 | 0.031074 | 0.009107 | 0.013331 | 0.020086 | 0.00987 | 0.007102 | 0.098415 | 0.019793 | 0.004202 | 0.002483 | 0.007172 | 0.025623 | 0.00097 | 0.01303 | 0.004072 | 0.003266 | 0.03637 | 0.006952 |
| 36.1944444 | 0.024326 | 0.02866 | 0.017534 | 0.015902 | 0.018066 | 0.004779 | 0.011317 | 0.108829 | 0.024971 | 0.004445 | 0.00207 | 0.003556 | 0.018784 | 0.000463 | 0.007345 | 0.002348 | 0.005545 | 0.041286 | 0.005877 |
| 36.2083333 | 0.037823 | 0.020876 | 0.026156 | 0.018954 | 0.035852 | 0.008892 | 0.014961 | 0.103676 | 0.026608 | 0.006305 | 0.003011 | 0.004003 | 0.013231 | 0.000909 | 0.003011 | 0.001847 | 0.008497 | 0.044974 | 0.008835 |
| 36.2222222 | 0.05036 | 0.010427 | 0.035748 | 0.022077 | 0.058136 | 0.021787 | 0.01658 | 0.086489 | 0.021944 | 0.009717 | 0.005725 | 0.003523 | 0.010175 | 0.00305 | 0.000904 | 0.001367 | 0.008839 | 0.045871 | 0.007574 |
| 36.2361111 | 0.059014 | 0.005184 | 0.046991 | 0.024802 | 0.07501 | 0.041272 | 0.015999 | 0.066803 | 0.014648 | 0.010912 | 0.007538 | 0.00355 | 0.009072 | 0.00646 | 0.001091 | 0.000953 | 0.006045 | 0.041463 | 0.004116 |
| 36.25 | 0.062167 | 0.008736 | 0.059079 | 0.027155 | 0.08431 | 0.051695 | 0.015704 | 0.050069 | 0.010672 | 0.010562 | 0.008406 | 0.007541 | 0.008267 | 0.008918 | 0.003059 | 0.001253 | 0.003324 | 0.029184 | 0.004499 |
| 36.2638889 | 0.061116 | 0.01692 | 0.067838 | 0.028994 | 0.084755 | 0.047092 | 0.017423 | 0.041278 | 0.014224 | 0.011497 | 0.010449 | 0.011054 | 0.006222 | 0.009606 | 0.005256 | 0.002297 | 0.003566 | 0.015646 | 0.004991 |
| 36.2777778 | 0.060912 | 0.026461 | 0.068679 | 0.030012 | 0.080108 | 0.040295 | 0.019931 | 0.039651 | 0.022549 | 0.012911 | 0.01264 | 0.011851 | 0.003977 | 0.009208 | 0.005103 | 0.003392 | 0.005921 | 0.014912 | 0.003441 |
| 36.2916667 | 0.062495 | 0.033743 | 0.061255 | 0.030217 | 0.081541 | 0.028073 | 0.021081 | 0.037363 | 0.031381 | 0.012973 | 0.013193 | 0.010436 | 0.005284 | 0.008601 | 0.00291 | 0.004132 | 0.007522 | 0.026633 | 0.003674 |
| 36.3055556 | 0.062507 | 0.033886 | 0.048806 | 0.029892 | 0.092375 | 0.014843 | 0.020612 | 0.035767 | 0.037027 | 0.012141 | 0.012316 | 0.006341 | 0.00996 | 0.009582 | 0.001269 | 0.005144 | 0.006872 | 0.036812 | 0.005257 |
| 36.3194444 | 0.060961 | 0.028043 | 0.03687 | 0.029932 | 0.103709 | 0.011819 | 0.019775 | 0.034934 | 0.034613 | 0.01199 | 0.011566 | 0.002803 | 0.013507 | 0.013528 | 0.00127 | 0.006751 | 0.004099 | 0.040126 | 0.004814 |
| 36.3333333 | 0.061082 | 0.023415 | 0.029162 | 0.0315 | 0.107598 | 0.015329 | 0.019971 | 0.031487 | 0.024749 | 0.012788 | 0.011796 | 0.002338 | 0.01268 | 0.018524 | 0.003231 | 0.008467 | 0.001433 | 0.040087 | 0.005046 |
| 36.3472222 | 0.062109 | 0.025313 | 0.025917 | 0.034574 | 0.096636 | 0.015966 | 0.021626 | 0.027989 | 0.01426 | 0.013745 | 0.012575 | 0.002573 | 0.007467 | 0.020741 | 0.006843 | 0.010114 | 0.001585 | 0.036487 | 0.008754 |
| 36.3611111 | 0.059308 | 0.031018 | 0.024695 | 0.037455 | 0.07 | 0.009613 | 0.023504 | 0.019936 | 0.006724 | 0.014223 | 0.012507 | 0.00323 | 0.003175 | 0.018652 | 0.009919 | 0.010884 | 0.004186 | 0.027782 | 0.011192 |
| 36.375 | 0.052573 | 0.034525 | 0.022593 | 0.038521 | 0.036909 | 0.005802 | 0.024488 | 0.013966 | 0.003241 | 0.013833 | 0.010457 | 0.004866 | 0.003417 | 0.013761 | 0.010117 | 0.009511 | 0.006851 | 0.016999 | 0.009117 |
| 36.3888889 | 0.044469 | 0.034958 | 0.02031 | 0.037948 | 0.015521 | 0.008846 | 0.025229 | 0.024215 | 0.00402 | 0.012453 | 0.007156 | 0.006492 | 0.004157 | 0.008674 | 0.007837 | 0.006797 | 0.008344 | 0.007813 | 0.004997 |
| 36.4027778 | 0.036609 | 0.035044 | 0.019727 | 0.036842 | 0.017369 | 0.012634 | 0.027357 | 0.041148 | 0.005788 | 0.011497 | 0.005126 | 0.008414 | 0.003562 | 0.006255 | 0.006137 | 0.00489 | 0.008761 | 0.004041 | 0.002836 |
| 36.4166667 | 0.030919 | 0.03541 | 0.020858 | 0.03568 | 0.028512 | 0.015618 | 0.030853 | 0.052948 | 0.006456 | 0.012659 | 0.005958 | 0.01047 | 0.003634 | 0.007257 | 0.006412 | 0.004419 | 0.008673 | 0.006825 | 0.002792 |
| 36.4305556 | 0.027963 | 0.033878 | 0.022352 | 0.03406 | 0.034426 | 0.019115 | 0.033952 | 0.056516 | 0.007007 | 0.015331 | 0.008536 | 0.011656 | 0.00488 | 0.009581 | 0.006824 | 0.00399 | 0.008538 | 0.012318 | 0.002267 |
| 36.4444444 | 0.026541 | 0.029087 | 0.02276 | 0.032081 | 0.035543 | 0.021008 | 0.035138 | 0.051622 | 0.008564 | 0.01734 | 0.010337 | 0.012239 | 0.00652 | 0.011524 | 0.00538 | 0.002499 | 0.008029 | 0.018072 | 0.002173 |
| 36.4583333 | 0.025937 | 0.023065 | 0.022316 | 0.030889 | 0.037117 | 0.021771 | 0.034012 | 0.044582 | 0.010803 | 0.017355 | 0.009873 | 0.012945 | 0.007774 | 0.01229 | 0.002661 | 0.000965 | 0.006976 | 0.023527 | 0.002996 |
| 36.4722222 | 0.027521 | 0.018549 | 0.022081 | 0.030953 | 0.044053 | 0.025559 | 0.032175 | 0.041436 | 0.013863 | 0.015901 | 0.008066 | 0.013543 | 0.00873 | 0.010808 | 0.000868 | 0.000438 | 0.006174 | 0.027457 | 0.002544 |
| 36.4861111 | 0.029756 | 0.015689 | 0.022001 | 0.031262 | 0.053441 | 0.032039 | 0.031241 | 0.043185 | 0.019477 | 0.013948 | 0.006517 | 0.013834 | 0.0088 | 0.007408 | 0.000548 | 0.000669 | 0.006221 | 0.028884 | 0.002062 |
| 36.5 | 0.026506 | 0.012842 | 0.021284 | 0.030833 | 0.058195 | 0.038222 | 0.030372 | 0.047374 | 0.026534 | 0.011724 | 0.005675 | 0.013628 | 0.006206 | 0.004018 | 0.001471 | 0.000929 | 0.006422 | 0.027568 | 0.003918 |
| 36.5138889 | 0.019818 | 0.010672 | 0.019809 | 0.030697 | 0.05725 | 0.041699 | 0.028917 | 0.051002 | 0.030538 | 0.011128 | 0.005707 | 0.013154 | 0.003393 | 0.002303 | 0.003762 | 0.000845 | 0.005987 | 0.025374 | 0.005613 |
| 36.5277778 | 0.021876 | 0.016947 | 0.020924 | 0.03335 | 0.047136 | 0.038606 | 0.029738 | 0.053619 | 0.027983 | 0.018985 | 0.010047 | 0.017109 | 0.003577 | 0.004236 | 0.005254 | 0.001482 | 0.008114 | 0.029996 | 0.004528 |
| 36.5416667 | 0.042607 | 0.044611 | 0.030412 | 0.040632 | 0.031913 | 0.026157 | 0.037619 | 0.052373 | 0.019734 | 0.045145 | 0.028078 | 0.033063 | 0.004302 | 0.013104 | 0.005367 | 0.005397 | 0.019769 | 0.050224 | 0.006517 |
| 36.5555556 | 0.076432 | 0.091902 | 0.044452 | 0.049001 | 0.034509 | 0.018524 | 0.050924 | 0.035952 | 0.020901 | 0.081287 | 0.051466 | 0.056726 | 0.008045 | 0.028518 | 0.009455 | 0.013237 | 0.039009 | 0.075985 | 0.018891 |
| 36.5694444 | 0.094506 | 0.121126 | 0.048539 | 0.048151 | 0.056188 | 0.025164 | 0.058803 | 0.019209 | 0.03922 | 0.09155 | 0.052893 | 0.065824 | 0.018459 | 0.042722 | 0.018342 | 0.01929 | 0.048831 | 0.080318 | 0.037291 |
| 36.5833333 | 0.078739 | 0.103409 | 0.038658 | 0.036031 | 0.079102 | 0.028149 | 0.055126 | 0.023988 | 0.049411 | 0.065773 | 0.034346 | 0.051052 | 0.029379 | 0.047639 | 0.02443 | 0.019406 | 0.040258 | 0.063318 | 0.048718 |
| 36.5972222 | 0.053167 | 0.06698 | 0.025591 | 0.024699 | 0.096367 | 0.023747 | 0.045802 | 0.030646 | 0.041458 | 0.035625 | 0.0211 | 0.0314 | 0.033717 | 0.042456 | 0.023994 | 0.016017 | 0.025588 | 0.049389 | 0.043826 |
| 36.6111111 | 0.04522 | 0.046864 | 0.017385 | 0.024362 | 0.103787 | 0.023036 | 0.03849 | 0.031355 | 0.037805 | 0.017902 | 0.018074 | 0.020646 | 0.028296 | 0.031666 | 0.018937 | 0.009947 | 0.016632 | 0.04034 | 0.025924 |

| | | | | | | | | | | | | | | | | | | | |
|---|---|---|---|---|---|---|---|---|---|---|---|---|---|---|---|---|---|---|---|
| 36.625 | 0.052033 | 0.044344 | 0.014517 | 0.036147 | 0.108391 | 0.027857 | 0.038489 | 0.03195 | 0.041974 | 0.007743 | 0.013599 | 0.015761 | 0.015397 | 0.018821 | 0.011759 | 0.006503 | 0.012931 | 0.02469 | 0.014185 |
| 36.6388889 | 0.060045 | 0.047388 | 0.015 | 0.053941 | 0.110212 | 0.03216 | 0.046405 | 0.021997 | 0.037176 | 0.002597 | 0.006018 | 0.012423 | 0.006701 | 0.008922 | 0.005706 | 0.011355 | 0.010811 | 0.014653 | 0.023208 |
| 36.6527778 | 0.062125 | 0.048057 | 0.017023 | 0.070943 | 0.100937 | 0.029689 | 0.053131 | 0.012645 | 0.020281 | 0.003103 | 0.002408 | 0.009572 | 0.006496 | 0.007117 | 0.003965 | 0.016363 | 0.008908 | 0.022663 | 0.039045 |
| 36.6666667 | 0.058036 | 0.04424 | 0.02084 | 0.085314 | 0.088403 | 0.026525 | 0.055797 | 0.020828 | 0.010646 | 0.008575 | 0.003035 | 0.008079 | 0.007537 | 0.010778 | 0.007311 | 0.014376 | 0.007679 | 0.03429 | 0.042939 |
| 36.6805556 | 0.050461 | 0.035281 | 0.027444 | 0.091867 | 0.072778 | 0.026593 | 0.059011 | 0.030372 | 0.019772 | 0.013953 | 0.004442 | 0.008781 | 0.01112 | 0.012976 | 0.013618 | 0.008068 | 0.007231 | 0.041824 | 0.032969 |
| 36.6944444 | 0.040963 | 0.022322 | 0.03464 | 0.078855 | 0.04963 | 0.021827 | 0.051506 | 0.027439 | 0.028287 | 0.013955 | 0.004542 | 0.01025 | 0.016851 | 0.012828 | 0.018321 | 0.003617 | 0.006726 | 0.050567 | 0.018787 |
| 36.7083333 | 0.028729 | 0.010971 | 0.035234 | 0.044601 | 0.025289 | 0.015031 | 0.03049 | 0.022784 | 0.020463 | 0.00853 | 0.003029 | 0.00978 | 0.014728 | 0.015947 | 0.017067 | 0.003069 | 0.005701 | 0.059961 | 0.014781 |
| 36.7222222 | 0.014706 | 0.009441 | 0.027026 | 0.022251 | 0.015828 | 0.020315 | 0.023056 | 0.029652 | 0.007703 | 0.005332 | 0.002759 | 0.006678 | 0.010019 | 0.022848 | 0.011301 | 0.00293 | 0.004526 | 0.067575 | 0.021977 |
| 36.7361111 | 0.006009 | 0.018232 | 0.018061 | 0.033045 | 0.032379 | 0.034757 | 0.035602 | 0.053018 | 0.002879 | 0.010386 | 0.004238 | 0.003413 | 0.014606 | 0.028821 | 0.00561 | 0.004282 | 0.003731 | 0.069335 | 0.028181 |
| 36.75 | 0.006885 | 0.029364 | 0.014181 | 0.048356 | 0.064822 | 0.041289 | 0.038856 | 0.085142 | 0.006467 | 0.018254 | 0.017379 | 0.002798 | 0.019585 | 0.031803 | 0.003223 | 0.008056 | 0.003078 | 0.059463 | 0.0264 |
| 36.7638889 | 0.011669 | 0.037294 | 0.015622 | 0.049561 | 0.093 | 0.038187 | 0.02383 | 0.104373 | 0.011525 | 0.022343 | 0.033413 | 0.003431 | 0.015265 | 0.030947 | 0.005167 | 0.009628 | 0.002875 | 0.044312 | 0.016622 |
| 36.7777778 | 0.011781 | 0.039859 | 0.021808 | 0.04071 | 0.107187 | 0.036261 | 0.010692 | 0.089173 | 0.009016 | 0.021985 | 0.031347 | 0.002515 | 0.009122 | 0.025189 | 0.009708 | 0.006538 | 0.003285 | 0.033863 | 0.008883 |
| 36.7916667 | 0.006744 | 0.036523 | 0.030288 | 0.027429 | 0.113219 | 0.039502 | 0.009312 | 0.053173 | 0.005437 | 0.018707 | 0.031599 | 0.001334 | 0.010555 | 0.016345 | 0.014442 | 0.003595 | 0.003072 | 0.024948 | 0.0109 |
| 36.8055556 | 0.003214 | 0.029452 | 0.035297 | 0.015229 | 0.115803 | 0.046879 | 0.013661 | 0.044144 | 0.007793 | 0.014564 | 0.031375 | 0.002479 | 0.016587 | 0.00843 | 0.015826 | 0.004527 | 0.003737 | 0.014209 | 0.012121 |
| 36.8194444 | 0.002504 | 0.022809 | 0.031993 | 0.008454 | 0.113118 | 0.050644 | 0.017086 | 0.073723 | 0.01069 | 0.011111 | 0.015951 | 0.005476 | 0.017032 | 0.004232 | 0.012023 | 0.004883 | 0.006542 | 0.011229 | 0.012625 |
| 36.8333333 | 0.002414 | 0.019467 | 0.023126 | 0.012924 | 0.095075 | 0.047602 | 0.016979 | 0.099037 | 0.011054 | 0.008787 | 0.006716 | 0.007914 | 0.011009 | 0.003597 | 0.006306 | 0.002966 | 0.008294 | 0.023068 | 0.018378 |
| 36.8472222 | 0.003743 | 0.018557 | 0.015213 | 0.026768 | 0.064225 | 0.042866 | 0.01294 | 0.097851 | 0.010136 | 0.008134 | 0.007445 | 0.00852 | 0.007875 | 0.007414 | 0.00539 | 0.001338 | 0.007471 | 0.039017 | 0.018929 |
| 36.8611111 | 0.004503 | 0.018342 | 0.008726 | 0.041707 | 0.040179 | 0.040415 | 0.007804 | 0.085419 | 0.013208 | 0.010686 | 0.009275 | 0.008049 | 0.008009 | 0.016117 | 0.012136 | 0.00111 | 0.005553 | 0.049446 | 0.013235 |
| 36.875 | 0.004902 | 0.019327 | 0.008698 | 0.052468 | 0.026416 | 0.040386 | 0.00777 | 0.068053 | 0.025368 | 0.016351 | 0.016379 | 0.007825 | 0.0076 | 0.024084 | 0.019693 | 0.001617 | 0.004085 | 0.052633 | 0.010185 |
| 36.8888889 | 0.006757 | 0.023295 | 0.018547 | 0.057134 | 0.014753 | 0.037763 | 0.014194 | 0.04071 | 0.055492 | 0.021824 | 0.033639 | 0.008047 | 0.013489 | 0.027007 | 0.021078 | 0.00328 | 0.003634 | 0.047606 | 0.008951 |
| 36.9027778 | 0.007045 | 0.029357 | 0.025496 | 0.055038 | 0.006773 | 0.032096 | 0.024367 | 0.027461 | 0.089516 | 0.023913 | 0.043697 | 0.008158 | 0.024676 | 0.02584 | 0.01629 | 0.008816 | 0.004067 | 0.036739 | 0.006408 |
| 36.9166667 | 0.010124 | 0.033609 | 0.027088 | 0.046698 | 0.009261 | 0.02846 | 0.036508 | 0.053487 | 0.096478 | 0.0225 | 0.039966 | 0.007925 | 0.031754 | 0.022519 | 0.009229 | 0.01617 | 0.005443 | 0.023095 | 0.004765 |
| 36.9305556 | 0.019379 | 0.032415 | 0.032191 | 0.035155 | 0.023524 | 0.033326 | 0.046482 | 0.090964 | 0.082504 | 0.019593 | 0.041019 | 0.007332 | 0.033024 | 0.017254 | 0.004336 | 0.020187 | 0.007089 | 0.010309 | 0.006678 |
| 36.9444444 | 0.02584 | 0.025383 | 0.042614 | 0.025898 | 0.045019 | 0.047342 | 0.051303 | 0.116821 | 0.067188 | 0.01692 | 0.035585 | 0.006292 | 0.030971 | 0.010569 | 0.003212 | 0.018486 | 0.008241 | 0.00412 | 0.008509 |
| 36.9583333 | 0.024615 | 0.01516 | 0.052211 | 0.02207 | 0.062198 | 0.056822 | 0.052175 | 0.141313 | 0.05351 | 0.014841 | 0.016989 | 0.004908 | 0.025457 | 0.004732 | 0.00429 | 0.012887 | 0.008537 | 0.005756 | 0.007612 |
| 36.9722222 | 0.017514 | 0.007875 | 0.051496 | 0.021305 | 0.065303 | 0.051288 | 0.048622 | 0.161261 | 0.041967 | 0.012734 | 0.006241 | 0.005276 | 0.016642 | 0.001462 | 0.005207 | 0.007536 | 0.008118 | 0.00977 | 0.012354 |
| 36.9861111 | 0.010492 | 0.010263 | 0.039367 | 0.020086 | 0.057526 | 0.037596 | 0.039738 | 0.157225 | 0.031589 | 0.010039 | 0.005224 | 0.010266 | 0.008779 | 0.000847 | 0.004614 | 0.005456 | 0.007711 | 0.013411 | 0.024763 |
| 37 | 0.013107 | 0.021078 | 0.020904 | 0.017745 | 0.046021 | 0.02436 | 0.027426 | 0.114603 | 0.02081 | 0.007106 | 0.004706 | 0.015852 | 0.005787 | 0.00226 | 0.00276 | 0.007551 | 0.006923 | 0.014848 | 0.03324 |
| 37.0138889 | 0.023576 | 0.032853 | 0.008014 | 0.014262 | 0.033619 | 0.014394 | 0.014642 | 0.062986 | 0.022479 | 0.004801 | 0.00347 | 0.014801 | 0.006575 | 0.005528 | 0.001665 | 0.011903 | 0.005427 | 0.01143 | 0.035937 |
| 37.0277778 | 0.026302 | 0.030126 | 0.005708 | 0.009375 | 0.026323 | 0.011147 | 0.006793 | 0.052877 | 0.038417 | 0.00588 | 0.003832 | 0.011972 | 0.00581 | 0.006642 | 0.003614 | 0.013531 | 0.008003 | 0.01167 | 0.033941 |
| 37.0416667 | 0.026415 | 0.032125 | 0.012403 | 0.005199 | 0.035575 | 0.021015 | 0.010262 | 0.052593 | 0.039963 | 0.016296 | 0.008119 | 0.022378 | 0.009424 | 0.016245 | 0.014791 | 0.013641 | 0.023378 | 0.033504 | 0.025123 |
| 37.0555556 | 0.047252 | 0.066559 | 0.03126 | 0.003105 | 0.060026 | 0.041369 | 0.022468 | 0.052898 | 0.038152 | 0.037522 | 0.010891 | 0.045964 | 0.032385 | 0.047025 | 0.040273 | 0.025639 | 0.049893 | 0.07579 | 0.023283 |
| 37.0694444 | 0.065835 | 0.080952 | 0.041559 | 0.006009 | 0.075026 | 0.052626 | 0.029693 | 0.098634 | 0.048074 | 0.04938 | 0.024285 | 0.051659 | 0.061417 | 0.061626 | 0.054139 | 0.041141 | 0.059413 | 0.096228 | 0.031476 |
| 37.0833333 | 0.049223 | 0.045474 | 0.029082 | 0.018756 | 0.069744 | 0.044237 | 0.026638 | 0.133947 | 0.03524 | 0.03659 | 0.062087 | 0.034134 | 0.060248 | 0.043502 | 0.040205 | 0.032623 | 0.043004 | 0.065981 | 0.028037 |
| 37.0972222 | 0.020106 | 0.010677 | 0.016028 | 0.034563 | 0.067869 | 0.027188 | 0.022564 | 0.117595 | 0.014198 | 0.022467 | 0.084891 | 0.027572 | 0.031593 | 0.03575 | 0.031155 | 0.015874 | 0.034163 | 0.031824 | 0.019519 |
| 37.1111111 | 0.006959 | 0.003325 | 0.016848 | 0.04009 | 0.078453 | 0.012585 | 0.022389 | 0.089858 | 0.019185 | 0.019458 | 0.06309 | 0.028403 | 0.015642 | 0.042988 | 0.033386 | 0.017052 | 0.034285 | 0.029855 | 0.024927 |
| 37.125 | 0.009953 | 0.010136 | 0.02313 | 0.034743 | 0.079346 | 0.004565 | 0.023042 | 0.069394 | 0.034059 | 0.014516 | 0.028002 | 0.019266 | 0.024105 | 0.04508 | 0.026713 | 0.023658 | 0.022847 | 0.033215 | 0.032679 |
| 37.1388889 | 0.018735 | 0.020106 | 0.026567 | 0.028323 | 0.067744 | 0.002475 | 0.022853 | 0.05211 | 0.042049 | 0.008213 | 0.012891 | 0.010292 | 0.039983 | 0.042521 | 0.016957 | 0.026136 | 0.008871 | 0.033077 | 0.037766 |
| 37.1527778 | 0.025593 | 0.029366 | 0.02785 | 0.024496 | 0.062819 | 0.002915 | 0.022894 | 0.036078 | 0.04382 | 0.004281 | 0.011638 | 0.006284 | 0.0489 | 0.033864 | 0.012482 | 0.025721 | 0.002462 | 0.036396 | 0.043658 |
| 37.1666667 | 0.027643 | 0.034778 | 0.029659 | 0.021625 | 0.065457 | 0.003984 | 0.022442 | 0.020181 | 0.039608 | 0.002261 | 0.008678 | 0.004292 | 0.044419 | 0.021543 | 0.009238 | 0.020276 | 0.001204 | 0.036764 | 0.042749 |
| 37.1805556 | 0.023347 | 0.034488 | 0.030234 | 0.018927 | 0.063587 | 0.004279 | 0.019425 | 0.010472 | 0.026839 | 0.002663 | 0.005032 | 0.004952 | 0.031422 | 0.011281 | 0.005618 | 0.011658 | 0.001747 | 0.032064 | 0.030224 |
| 37.1944444 | 0.014633 | 0.028069 | 0.024965 | 0.016618 | 0.051374 | 0.00306 | 0.014563 | 0.014399 | 0.016862 | 0.002511 | 0.005232 | 0.007977 | 0.018002 | 0.006629 | 0.00427 | 0.00457 | 0.00214 | 0.02624 | 0.014008 |
| 37.2083333 | 0.007486 | 0.018448 | 0.015144 | 0.016426 | 0.03144 | 0.001373 | 0.00919 | 0.023082 | 0.026598 | 0.001935 | 0.006548 | 0.011284 | 0.008078 | 0.008044 | 0.005138 | 0.001589 | 0.002179 | 0.021388 | 0.008603 |
| 37.2222222 | 0.007135 | 0.011567 | 0.006286 | 0.018324 | 0.013045 | 0.00043 | 0.004466 | 0.032234 | 0.048692 | 0.004023 | 0.005379 | 0.013121 | 0.002836 | 0.010084 | 0.006007 | 0.001277 | 0.001835 | 0.015353 | 0.016291 |
| 37.2361111 | 0.011824 | 0.009819 | 0.002563 | 0.020178 | 0.005553 | 0.000162 | 0.002324 | 0.044935 | 0.068916 | 0.007464 | 0.003594 | 0.012958 | 0.001386 | 0.008848 | 0.005777 | 0.00279 | 0.001704 | 0.009641 | 0.02591 |
| 37.25 | 0.01833 | 0.008494 | 0.004708 | 0.020715 | 0.009743 | 0.00051 | 0.0027 | 0.05749 | 0.084166 | 0.010164 | 0.003107 | 0.012333 | 0.002406 | 0.006058 | 0.004901 | 0.00525 | 0.003429 | 0.006581 | 0.028132 |
| 37.2638889 | 0.025133 | 0.00451 | 0.008852 | 0.019807 | 0.01734 | 0.001583 | 0.003406 | 0.064682 | 0.095541 | 0.012181 | 0.002639 | 0.012571 | 0.004809 | 0.008094 | 0.004001 | 0.006201 | 0.006392 | 0.005243 | 0.023146 |
| 37.2777778 | 0.028679 | 0.001628 | 0.011549 | 0.017085 | 0.022272 | 0.002619 | 0.00287 | 0.060811 | 0.102688 | 0.013269 | 0.00148 | 0.013131 | 0.007069 | 0.015907 | 0.003379 | 0.006203 | 0.008483 | 0.007442 | 0.015792 |
| 37.2916667 | 0.028011 | 0.001278 | 0.012057 | 0.01256 | 0.024179 | 0.002729 | 0.002051 | 0.05323 | 0.094535 | 0.012251 | 0.001199 | 0.012483 | 0.006416 | 0.022476 | 0.002405 | 0.006813 | 0.008402 | 0.014279 | 0.008869 |
| 37.3055556 | 0.024108 | 0.003165 | 0.009943 | 0.007805 | 0.023305 | 0.00175 | 0.001872 | 0.056031 | 0.07081 | 0.008178 | 0.001616 | 0.009592 | 0.005513 | 0.022484 | 0.001933 | 0.007331 | 0.005512 | 0.022169 | 0.00373 |
| 37.3194444 | 0.016306 | 0.00973 | 0.005817 | 0.006531 | 0.020269 | 0.000689 | 0.002731 | 0.069494 | 0.050066 | 0.003292 | 0.002556 | 0.005857 | 0.011299 | 0.016289 | 0.003859 | 0.00812 | 0.002805 | 0.028094 | 0.001561 |
| 37.3333333 | 0.007474 | 0.01691 | 0.002693 | 0.011347 | 0.017118 | 0.000374 | 0.005019 | 0.086064 | 0.034822 | 0.001848 | 0.003255 | 0.003918 | 0.020219 | 0.009203 | 0.005622 | 0.010273 | 0.002637 | 0.029195 | 0.001398 |
| 37.3472222 | 0.002503 | 0.017216 | 0.002958 | 0.019605 | 0.014875 | 0.000319 | 0.006593 | 0.093484 | 0.017296 | 0.005221 | 0.00316 | 0.004412 | 0.023749 | 0.007508 | 0.004837 | 0.011125 | 0.002833 | 0.022685 | 0.003291 |
| 37.3611111 | 0.003255 | 0.010844 | 0.005454 | 0.026826 | 0.013152 | 0.00075 | 0.006511 | 0.085276 | 0.005171 | 0.010877 | 0.004797 | 0.005188 | 0.020498 | 0.013012 | 0.004343 | 0.007338 | 0.003727 | 0.011506 | 0.007834 |
| 37.375 | 0.009657 | 0.00508 | 0.007721 | 0.031746 | 0.011443 | 0.00232 | 0.00603 | 0.066866 | 0.004379 | 0.014606 | 0.006872 | 0.004762 | 0.013871 | 0.022329 | 0.006564 | 0.003865 | 0.005401 | 0.005142 | 0.010441 |
| 37.3888889 | 0.018078 | 0.004971 | 0.009065 | 0.035445 | 0.009245 | 0.003945 | 0.00462 | 0.049399 | 0.01285 | 0.014371 | 0.005912 | 0.00435 | 0.007922 | 0.028899 | 0.0095 | 0.005481 | 0.005294 | 0.007669 | 0.007663 |
| 37.4027778 | 0.023408 | 0.00771 | 0.010129 | 0.038174 | 0.005652 | 0.003778 | 0.002621 | 0.042666 | 0.031806 | 0.011881 | 0.003156 | 0.00462 | 0.004175 | 0.028392 | 0.010089 | 0.006201 | 0.003539 | 0.011502 | 0.005483 |
| 37.41666667 | 0.02429 | 0.009975 | 0.011069 | 0.03875 | 0.004681 | 0.001945 | 0.002852 | 0.044543 | 0.054846 | 0.010166 | 0.001621 | 0.004806 | 0.001798 | 0.02335 | 0.008093 | 0.003937 | 0.001734 | 0.011067 | 0.007714 |
| 37.43055556 | 0.02299 | 0.01223 | 0.011911 | 0.037194 | 0.011453 | 0.000593 | 0.006112 | 0.046182 | 0.06856 | 0.010605 | 0.002044 | 0.004743 | 0.000553 | 0.017717 | 0.005477 | 0.003317 | 0.000713 | 0.008471 | 0.008186 |
| 37.44444444 | 0.02148 | 0.014034 | 0.013426 | 0.036343 | 0.020441 | 0.000615 | 0.011141 | 0.041055 | 0.065649 | 0.01192 | 0.003049 | 0.004877 | 0.00076 | 0.012996 | 0.004088 | 0.003075 | 0.000598 | 0.007969 | 0.006607 |
| 37.45833333 | 0.018876 | 0.012867 | 0.01567 | 0.03833 | 0.025765 | 0.000889 | 0.015879 | 0.025464 | 0.045108 | 0.011813 | 0.003178 | 0.005348 | 0.002922 | 0.010377 | 0.004136 | 0.002496 | 0.000929 | 0.011526 | 0.010062 |
| 37.47222222 | 0.014155 | 0.008737 | 0.016962 | 0.040906 | 0.029511 | 0.000641 | 0.016768 | 0.015666 | 0.024448 | 0.00986 | 0.002856 | 0.006315 | 0.006135 | 0.010609 | 0.004797 | 0.004829 | 0.001525 | 0.016237 | 0.018072 |
| 37.48611111 | 0.009101 | 0.005494 | 0.015958 | 0.039866 | 0.033855 | 0.00043 | 0.012449 | 0.030159 | 0.027276 | 0.007977 | 0.00263 | 0.007548 | 0.006801 | 0.012534 | 0.005519 | 0.008041 | 0.002218 | 0.018525 | 0.022543 |
| 37.5 | 0.00547 | 0.005298 | 0.013415 | 0.035018 | 0.03691 | 0.000992 | 0.006813 | 0.053564 | 0.046589 | 0.006983 | 0.002131 | 0.007398 | 0.004546 | 0.013032 | 0.006213 | 0.008944 | 0.002171 | 0.018446 | 0.017687 |
| 37.51388889 | 0.003325 | 0.006674 | 0.011464 | 0.030517 | 0.036439 | 0.001444 | 0.005696 | 0.071002 | 0.064361 | 0.006703 | 0.001983 | 0.005597 | 0.004085 | 0.011517 | 0.0066 | 0.008146 | 0.00178 | 0.019035 | 0.009582 |
| 37.52777778 | 0.006628 | 0.013178 | 0.014326 | 0.029445 | 0.029739 | 0.003378 | 0.014389 | 0.083301 | 0.076353 | 0.010747 | 0.003869 | 0.006135 | 0.008331 | 0.01333 | 0.006128 | 0.005413 | 0.005067 | 0.027079 | 0.007103 |
| 37.54166667 | 0.026232 | 0.037782 | 0.027397 | 0.033637 | 0.022411 | 0.012525 | 0.039959 | 0.079193 | 0.071099 | 0.026622 | 0.013912 | 0.014036 | 0.01817 | 0.025243 | 0.010037 | 0.00477 | 0.016723 | 0.049322 | 0.011197 |
| 37.55555556 | 0.063407 | 0.080751 | 0.047807 | 0.042235 | 0.036965 | 0.025997 | 0.077367 | 0.055045 | 0.050903 | 0.05076 | 0.030949 | 0.027612 | 0.030949 | 0.042872 | 0.020443 | 0.011517 | 0.032553 | 0.077269 | 0.033254 |
| 37.56944445 | 0.093087 | 0.112836 | 0.061798 | 0.046763 | 0.067652 | 0.030563 | 0.099588 | 0.047032 | 0.060125 | 0.062385 | 0.039302 | 0.035642 | 0.039284 | 0.050368 | 0.026365 | 0.020357 | 0.039179 | 0.084292 | 0.074315 |
| 37.58333334 | 0.095516 | 0.112289 | 0.062925 | 0.039769 | 0.091905 | 0.02267 | 0.093566 | 0.067911 | 0.107801 | 0.050457 | 0.038379 | 0.030527 | 0.040928 | 0.041506 | 0.023213 | 0.024269 | 0.030801 | 0.068039 | 0.098588 |

Supplementary Table S6. 1 hour integrated motion data in (a) lettuce subjected to Nutrient withdrawal

| IntervalMid | Control1 | Control2 | Control3 | Control4 | Control5 | Control6 | Control7 | Control8 | Control9 | Control10 | Control11 | Control12 | Stress1 | Stress2 | Stress3 | Stress4 | Stress5 | Stress6 | Stress7 | Stress8 | Stress9 | Stress10 | Stress11 | Stress12 |
|---|---|---|---|---|---|---|---|---|---|---|---|---|---|---|---|---|---|---|---|---|---|---|---|---|
| 18.1111111 | 0.027509 | 0.020401 | 0.016219 | 0.003804 | 0.014822 | 0.001633 | 0.016574 | 0.001892 | 0.047122 | 0.007611 | 0.006356 | 0.018381 | 0.035386 | 0.051357 | 0.000447 | 0.003621 | 0.003467 | 0.000178 | 0.002163 | 0.001763 | 0.056562 | 0.010662 | 0.06103 | 0.016547 |
| 18.125 | 0.013083 | 0.011243 | 0.010006 | 0.002767 | 0.006354 | 0.001563 | 0.005995 | 0.000962 | 0.034442 | 0.005058 | 0.004698 | 0.008786 | 0.026443 | 0.036486 | 0.000263 | 0.003325 | 0.00134 | 0.000249 | 0.001384 | 0.000811 | 0.045967 | 0.013556 | 0.035921 | 0.010827 |
| 18.1388889 | 0.005518 | 0.004912 | 0.008158 | 0.002457 | 0.002307 | 0.001823 | 0.001458 | 0.000744 | 0.019448 | 0.00263 | 0.004386 | 0.004156 | 0.015699 | 0.019752 | 0.000265 | 0.002447 | 0.000792 | 0.000223 | 0.001921 | 0.001008 | 0.033351 | 0.015258 | 0.018653 | 0.005612 |
| 18.1527778 | 0.002549 | 0.001898 | 0.00781 | 0.002686 | 0.000722 | 0.002619 | 0.001203 | 0.001098 | 0.011505 | 0.001029 | 0.004964 | 0.00212 | 0.007067 | 0.008762 | 0.000449 | 0.001508 | 0.000924 | 0.000185 | 0.002011 | 0.001276 | 0.025172 | 0.013874 | 0.007093 | 0.002974 |
| 18.1666667 | 0.00168 | 0.000881 | 0.00572 | 0.002929 | 0.000536 | 0.003628 | 0.001215 | 0.001138 | 0.007587 | 0.000342 | 0.005835 | 0.001573 | 0.003332 | 0.003637 | 0.000477 | 0.000965 | 0.00069 | 0.000189 | 0.001518 | 0.001024 | 0.023044 | 0.010868 | 0.001767 | 0.003036 |
| 18.1805556 | 0.001584 | 0.000858 | 0.00323 | 0.002746 | 0.000496 | 0.004031 | 0.000731 | 0.000753 | 0.006393 | 0.000272 | 0.006312 | 0.001846 | 0.004331 | 0.002309 | 0.000241 | 0.001162 | 0.000402 | 0.000219 | 0.000853 | 0.000608 | 0.023935 | 0.008349 | 0.002544 | 0.006409 |
| 18.1944444 | 0.001539 | 0.001271 | 0.002268 | 0.002269 | 0.000468 | 0.003506 | 0.000341 | 0.000646 | 0.006053 | 0.000546 | 0.006117 | 0.002003 | 0.007014 | 0.003209 | 0.00015 | 0.001617 | 0.000521 | 0.00026 | 0.000547 | 0.000725 | 0.024983 | 0.008531 | 0.006545 | 0.013544 |
| 18.2083333 | 0.001194 | 0.001779 | 0.001988 | 0.001937 | 0.000511 | 0.002643 | 0.000318 | 0.000741 | 0.00521 | 0.000899 | 0.005476 | 0.001496 | 0.008275 | 0.004729 | 0.00036 | 0.001732 | 0.000626 | 0.000229 | 0.000687 | 0.001352 | 0.024802 | 0.011073 | 0.007309 | 0.018408 |
| 18.2222222 | 0.000674 | 0.00212 | 0.001633 | 0.001894 | 0.000385 | 0.002029 | 0.000363 | 0.000566 | 0.003965 | 0.000969 | 0.004989 | 0.000681 | 0.007932 | 0.00556 | 0.000641 | 0.001664 | 0.000509 | 0.000266 | 0.000864 | 0.001729 | 0.023218 | 0.011605 | 0.006554 | 0.022072 |
| 18.2361111 | 0.0003 | 0.002026 | 0.001302 | 0.001931 | 0.000312 | 0.001736 | 0.000291 | 0.000254 | 0.003036 | 0.000736 | 0.005165 | 0.000206 | 0.00736 | 0.005445 | 0.000729 | 0.001658 | 0.000314 | 0.000471 | 0.000869 | 0.001633 | 0.020826 | 0.007458 | 0.011618 | 0.033977 |
| 18.25 | 0.000207 | 0.001395 | 0.000945 | 0.002008 | 0.000277 | 0.001742 | 0.00025 | 0.000118 | 0.002589 | 0.000447 | 0.005971 | 0.000159 | 0.006736 | 0.004881 | 0.000624 | 0.001418 | 0.000161 | 0.000672 | 0.000843 | 0.001519 | 0.018815 | 0.005354 | 0.018113 | 0.043121 |
| 18.2638889 | 0.000345 | 0.000801 | 0.000547 | 0.002153 | 0.000229 | 0.001928 | 0.000361 | 0.000149 | 0.002399 | 0.000434 | 0.007106 | 0.000239 | 0.005785 | 0.004178 | 0.000423 | 0.000911 | 0.000242 | 0.000759 | 0.000852 | 0.001722 | 0.018143 | 0.008188 | 0.019885 | 0.037786 |
| 18.2777778 | 0.00038 | 0.000848 | 0.000479 | 0.002041 | 0.000261 | 0.001989 | 0.000452 | 0.000181 | 0.002493 | 0.000586 | 0.008557 | 0.000356 | 0.005368 | 0.003368 | 0.000252 | 0.000702 | 0.000558 | 0.000739 | 0.000913 | 0.002204 | 0.018307 | 0.006981 | 0.018522 | 0.02672 |
| 18.2916667 | 0.000236 | 0.001326 | 0.000921 | 0.001652 | 0.000289 | 0.001897 | 0.000363 | 0.000236 | 0.002937 | 0.000525 | 0.010316 | 0.000458 | 0.005903 | 0.002678 | 0.000232 | 0.000632 | 0.000738 | 0.000718 | 0.000924 | 0.002535 | 0.017935 | 0.002622 | 0.016329 | 0.018581 |
| 18.3055556 | 0.000218 | 0.00186 | 0.001368 | 0.001602 | 0.000434 | 0.001953 | 0.000321 | 0.000287 | 0.003291 | 0.000597 | 0.012379 | 0.000361 | 0.006109 | 0.002443 | 0.000241 | 0.000513 | 0.00134 | 0.00087 | 0.000658 | 0.00228 | 0.016692 | 0.001168 | 0.014332 | 0.014432 |
| 18.3194444 | 0.000315 | 0.002421 | 0.001461 | 0.002063 | 0.000635 | 0.002113 | 0.000273 | 0.000251 | 0.003193 | 0.001208 | 0.015541 | 0.000145 | 0.00523 | 0.002523 | 0.000237 | 0.000724 | 0.005272 | 0.001049 | 0.000401 | 0.001688 | 0.015053 | 0.000979 | 0.013185 | 0.012263 |
| 18.3333333 | 0.000419 | 0.00268 | 0.001506 | 0.002441 | 0.000753 | 0.002027 | 0.000241 | 0.000183 | 0.002913 | 0.001879 | 0.019489 | 0.00012 | 0.004209 | 0.002446 | 0.000385 | 0.000966 | 0.012995 | 0.000915 | 0.000455 | 0.001222 | 0.013241 | 0.000818 | 0.012634 | 0.009765 |
| 18.3472222 | 0.000423 | 0.002122 | 0.001741 | 0.002435 | 0.000728 | 0.001768 | 0.000384 | 0.000135 | 0.002777 | 0.001926 | 0.021779 | 0.000348 | 0.0041 | 0.002241 | 0.000606 | 0.001346 | 0.016916 | 0.000678 | 0.000545 | 0.000793 | 0.011313 | 0.001216 | 0.012297 | 0.006728 |
| 18.3611111 | 0.000321 | 0.001288 | 0.002231 | 0.002285 | 0.000673 | 0.001571 | 0.000463 | 0.000143 | 0.003032 | 0.001465 | 0.02073 | 0.000644 | 0.005396 | 0.002339 | 0.000653 | 0.002229 | 0.012759 | 0.001007 | 0.000666 | 0.000741 | 0.00951 | 0.002419 | 0.012314 | 0.004343 |
| 18.375 | 0.000413 | 0.001337 | 0.002892 | 0.001984 | 0.00105 | 0.0013 | 0.000566 | 0.000316 | 0.003788 | 0.001237 | 0.016755 | 0.000874 | 0.007487 | 0.002651 | 0.000561 | 0.002507 | 0.00741 | 0.002117 | 0.000878 | 0.001493 | 0.008099 | 0.003293 | 0.012485 | 0.002963 |
| 18.3888889 | 0.000687 | 0.002206 | 0.003268 | 0.001605 | 0.00174 | 0.000957 | 0.000683 | 0.000662 | 0.004464 | 0.00141 | 0.012032 | 0.00096 | 0.008398 | 0.002691 | 0.000552 | 0.001844 | 0.005135 | 0.003252 | 0.000847 | 0.002149 | 0.007259 | 0.003511 | 0.012305 | 0.002026 |
| 18.4027778 | 0.000816 | 0.002899 | 0.003149 | 0.001537 | 0.002148 | 0.000854 | 0.000477 | 0.00084 | 0.004539 | 0.001635 | 0.008757 | 0.00094 | 0.007456 | 0.002512 | 0.000636 | 0.001764 | 0.004206 | 0.003331 | 0.000547 | 0.002105 | 0.007187 | 0.003967 | 0.011915 | 0.001263 |
| 18.4166667 | 0.000745 | 0.003051 | 0.002853 | 0.001808 | 0.002112 | 0.001043 | 0.00021 | 0.000712 | 0.004437 | 0.001956 | 0.007187 | 0.000805 | 0.00663 | 0.002467 | 0.000617 | 0.002526 | 0.0038 | 0.002463 | 0.000444 | 0.001925 | 0.007863 | 0.004409 | 0.012018 | 0.000904 |
| 18.4305556 | 0.000664 | 0.003092 | 0.002776 | 0.00203 | 0.001905 | 0.001264 | 0.000305 | 0.000567 | 0.004774 | 0.002581 | 0.006603 | 0.000545 | 0.007457 | 0.002587 | 0.000486 | 0.003175 | 0.003819 | 0.001733 | 0.000708 | 0.002252 | 0.008567 | 0.004815 | 0.012793 | 0.00079 |
| 18.4444444 | 0.00069 | 0.003075 | 0.002878 | 0.001965 | 0.001606 | 0.001456 | 0.000791 | 0.000556 | 0.005452 | 0.003175 | 0.006567 | 0.000692 | 0.008785 | 0.002486 | 0.000918 | 0.003133 | 0.003535 | 0.001279 | 0.00092 | 0.002869 | 0.008057 | 0.005547 | 0.013246 | 0.000911 |
| 18.4583333 | 0.000623 | 0.002276 | 0.002524 | 0.001995 | 0.001243 | 0.002 | 0.001271 | 0.000465 | 0.005777 | 0.002974 | 0.007111 | 0.001766 | 0.008568 | 0.001709 | 0.002811 | 0.002204 | 0.002476 | 0.001663 | 0.001679 | 0.002436 | 0.0054 | 0.006636 | 0.01195 | 0.002139 |
| 18.4722222 | 0.000329 | 0.001081 | 0.001511 | 0.002506 | 0.001037 | 0.002893 | 0.001681 | 0.000223 | 0.005734 | 0.00214 | 0.008053 | 0.003063 | 0.008033 | 0.000685 | 0.005162 | 0.001379 | 0.001413 | 0.003169 | 0.003251 | 0.001108 | 0.002143 | 0.008089 | 0.009816 | 0.0038 |
| 18.4861111 | 0.000917 | 0.002991 | 0.001305 | 0.002887 | 0.002706 | 0.003028 | 0.005025 | 0.00113 | 0.008361 | 0.003081 | 0.008078 | 0.002921 | 0.016345 | 0.002394 | 0.005246 | 0.002606 | 0.00277 | 0.00329 | 0.003269 | 0.000981 | 0.002047 | 0.010298 | 0.0112 | 0.003979 |
| 18.5 | 0.004121 | 0.012844 | 0.00435 | 0.002149 | 0.010639 | 0.002247 | 0.017661 | 0.008158 | 0.018499 | 0.008941 | 0.005836 | 0.002251 | 0.050471 | 0.009588 | 0.00282 | 0.006809 | 0.010053 | 0.003487 | 0.001762 | 0.00294 | 0.007505 | 0.020278 | 0.019887 | 0.007002 |
| 18.5138889 | 0.008602 | 0.027775 | 0.009099 | 0.000789 | 0.023844 | 0.002752 | 0.035863 | 0.019158 | 0.033815 | 0.020186 | 0.003004 | 0.003432 | 0.108777 | 0.015759 | 0.001382 | 0.011903 | 0.019222 | 0.007116 | 0.001755 | 0.004886 | 0.015171 | 0.046775 | 0.031724 | 0.016508 |
| 18.5277778 | 0.009987 | 0.034299 | 0.008565 | 0.000531 | 0.030896 | 0.005091 | 0.041448 | 0.019845 | 0.041909 | 0.03064 | 0.002623 | 0.00484 | 0.138761 | 0.013714 | 0.003473 | 0.0117 | 0.018773 | 0.009331 | 0.005103 | 0.004337 | 0.016003 | 0.072405 | 0.035503 | 0.019312 |
| 18.5416667 | 0.007575 | 0.026309 | 0.003364 | 0.001123 | 0.026361 | 0.006485 | 0.029974 | 0.010741 | 0.036777 | 0.032841 | 0.004137 | 0.004664 | 0.108834 | 0.008797 | 0.00617 | 0.006663 | 0.009042 | 0.006798 | 0.009451 | 0.002804 | 0.008771 | 0.070923 | 0.029129 | 0.010472 |
| 18.5555556 | 0.005622 | 0.0141 | 0.000738 | 0.001461 | 0.019363 | 0.00604 | 0.016778 | 0.005641 | 0.027076 | 0.028897 | 0.005314 | 0.003931 | 0.06556 | 0.008386 | 0.00527 | 0.003752 | 0.003509 | 0.00497 | 0.008249 | 0.002793 | 0.004641 | 0.050623 | 0.022925 | 0.004363 |
| 18.5694444 | 0.006343 | 0.00697 | 0.00047 | 0.002266 | 0.017004 | 0.005409 | 0.010224 | 0.005324 | 0.021172 | 0.025313 | 0.00578 | 0.002811 | 0.043897 | 0.012518 | 0.003321 | 0.003363 | 0.007137 | 0.006154 | 0.0039 | 0.004723 | 0.007712 | 0.031757 | 0.021963 | 0.003004 |
| 18.5833333 | 0.008314 | 0.00444 | 0.000523 | 0.003359 | 0.017521 | 0.005016 | 0.008195 | 0.004922 | 0.020109 | 0.023594 | 0.005918 | 0.001356 | 0.035854 | 0.016552 | 0.003591 | 0.003898 | 0.012459 | 0.006912 | 0.003567 | 0.007437 | 0.010138 | 0.016511 | 0.022347 | 0.001361 |
| 18.5972222 | 0.010371 | 0.004153 | 0.001222 | 0.003745 | 0.018363 | 0.004655 | 0.007836 | 0.003969 | 0.022517 | 0.022147 | 0.005821 | 0.000572 | 0.034342 | 0.017809 | 0.003818 | 0.004151 | 0.013957 | 0.006376 | 0.004983 | 0.008921 | 0.011757 | 0.008691 | 0.021246 | 0.000772 |
| 18.6111111 | 0.012067 | 0.005284 | 0.002331 | 0.003604 | 0.018482 | 0.00438 | 0.006977 | 0.003025 | 0.026445 | 0.020637 | 0.005809 | 0.000424 | 0.036476 | 0.016617 | 0.003075 | 0.003562 | 0.012528 | 0.006269 | 0.005613 | 0.009279 | 0.025494 | 0.011971 | 0.018941 | 0.001914 |
| 18.625 | 0.013018 | 0.007275 | 0.00373 | 0.003345 | 0.017448 | 0.003979 | 0.004911 | 0.002798 | 0.030444 | 0.019178 | 0.006603 | 0.000586 | 0.039196 | 0.013303 | 0.002469 | 0.003133 | 0.010977 | 0.007119 | 0.006255 | 0.0099 | 0.049817 | 0.012214 | 0.016637 | 0.00348 |
| 18.6388889 | 0.012829 | 0.009684 | 0.005034 | 0.003226 | 0.015579 | 0.003228 | 0.002435 | 0.004022 | 0.033958 | 0.017202 | 0.00817 | 0.00129 | 0.040284 | 0.009763 | 0.002158 | 0.002945 | 0.011441 | 0.008368 | 0.006659 | 0.010777 | 0.06356 | 0.007868 | 0.015898 | 0.0043 |
| 18.6527778 | 0.011287 | 0.011858 | 0.005728 | 0.003321 | 0.012992 | 0.002168 | 0.001219 | 0.006198 | 0.036188 | 0.01509 | 0.00923 | 0.001907 | 0.038991 | 0.008022 | 0.002259 | 0.003242 | 0.013274 | 0.009082 | 0.006565 | 0.010834 | 0.061241 | 0.010512 | 0.017271 | 0.004557 |
| 18.6666667 | 0.008969 | 0.012817 | 0.005402 | 0.003321 | 0.0097 | 0.001347 | 0.00167 | 0.007249 | 0.035946 | 0.013441 | 0.009096 | 0.001943 | 0.035407 | 0.007607 | 0.002447 | 0.003871 | 0.013818 | 0.008347 | 0.005571 | 0.009087 | 0.053484 | 0.014042 | 0.01907 | 0.004125 |
| 18.6805556 | 0.006544 | 0.01224 | 0.004168 | 0.002859 | 0.006128 | 0.001189 | 0.002078 | 0.006084 | 0.032696 | 0.011879 | 0.008455 | 0.001783 | 0.029753 | 0.007311 | 0.001928 | 0.004124 | 0.012004 | 0.006147 | 0.003439 | 0.005543 | 0.04482 | 0.009828 | 0.018847 | 0.002647 |
| 18.6944444 | 0.004358 | 0.010841 | 0.002738 | 0.002397 | 0.003193 | 0.001415 | 0.001626 | 0.004114 | 0.027626 | 0.009635 | 0.007916 | 0.001571 | 0.023378 | 0.007783 | 0.000968 | 0.004233 | 0.009332 | 0.003694 | 0.001241 | 0.002084 | 0.035044 | 0.004877 | 0.016439 | 0.001551 |
| 18.7083333 | 0.002383 | 0.009589 | 0.001623 | 0.002549 | 0.001405 | 0.001604 | 0.001059 | 0.002984 | 0.022519 | 0.006578 | 0.007545 | 0.001083 | 0.0179 | 0.008903 | 0.000523 | 0.004707 | 0.007484 | 0.002265 | 0.000381 | 0.000659 | 0.026129 | 0.006736 | 0.013444 | 0.001816 |
| 18.7222222 | 0.001088 | 0.008787 | 0.000883 | 0.002958 | 0.001192 | 0.001674 | 0.000793 | 0.002553 | 0.017883 | 0.003463 | 0.007322 | 0.000453 | 0.013668 | 0.009254 | 0.000496 | 0.004884 | 0.006093 | 0.001955 | 0.000742 | 0.000693 | 0.019666 | 0.010254 | 0.01063 | 0.002107 |
| 18.7361111 | 0.001158 | 0.008504 | 0.000511 | 0.003169 | 0.003139 | 0.001758 | 0.000533 | 0.00223 | 0.013648 | 0.001486 | 0.007256 | 0.000234 | 0.011154 | 0.008901 | 0.000427 | 0.004686 | 0.004449 | 0.002146 | 0.001613 | 0.001078 | 0.014224 | 0.010347 | 0.008357 | 0.001854 |
| 18.75 | 0.001785 | 0.00919 | 0.000413 | 0.002973 | 0.005826 | 0.001863 | 0.0003 | 0.002031 | 0.010034 | 0.001452 | 0.00728 | 0.000456 | 0.010669 | 0.007535 | 0.000441 | 0.005009 | 0.002632 | 0.00253 | 0.002339 | 0.002707 | 0.010154 | 0.007801 | 0.007051 | 0.001508 |
| 18.7638889 | 0.002081 | 0.010401 | 0.000433 | 0.002536 | 0.006978 | 0.002152 | 0.000371 | 0.002106 | 0.007522 | 0.002307 | 0.007154 | 0.000669 | 0.010963 | 0.004575 | 0.000607 | 0.005525 | 0.001061 | 0.003095 | 0.002241 | 0.005658 | 0.009346 | 0.004241 | 0.006164 | 0.001285 |
| 18.7777778 | 0.001899 | 0.010535 | 0.000493 | 0.0023 | 0.005857 | 0.003691 | 0.000395 | 0.002485 | 0.006534 | 0.002255 | 0.006705 | 0.000754 | 0.010629 | 0.001586 | 0.000689 | 0.005142 | 0.000363 | 0.003507 | 0.001464 | 0.006926 | 0.011282 | 0.003211 | 0.004526 | 0.001431 |
| 18.7916667 | 0.00125 | 0.008974 | 0.000551 | 0.00249 | 0.003651 | 0.006712 | 0.000248 | 0.002875 | 0.00649 | 0.002479 | 0.006233 | 0.000827 | 0.00973 | 0.001336 | 0.000571 | 0.004081 | 0.000442 | 0.003373 | 0.0008 | 0.004801 | 0.013826 | 0.006589 | 0.002477 | 0.001816 |
| 18.8055556 | 0.000543 | 0.006932 | 0.000633 | 0.003349 | 0.002069 | 0.008883 | 0.000394 | 0.002935 | 0.006269 | 0.00468 | 0.006464 | 0.000713 | 0.009115 | 0.004131 | 0.000335 | 0.002926 | 0.000654 | 0.002781 | 0.000546 | 0.002253 | 0.014769 | 0.010433 | 0.001849 | 0.002001 |
| 18.8194444 | 0.00036 | 0.005647 | 0.000643 | 0.00449 | 0.001685 | 0.008886 | 0.001101 | 0.002649 | 0.005557 | 0.006722 | 0.008296 | 0.000403 | 0.008919 | 0.006509 | 0.000143 | 0.001678 | 0.001649 | 0.002117 | 0.000351 | 0.002307 | 0.013045 | 0.009813 | 0.002881 | 0.00185 |
| 18.8333333 | 0.001039 | 0.005083 | 0.000419 | 0.004775 | 0.002462 | 0.007894 | 0.002269 | 0.002067 | 0.00449 | 0.006407 | 0.012181 | 0.000206 | 0.008601 | 0.006546 | 8.78E-05 | 0.000695 | 0.004851 | 0.001679 | 0.000297 | 0.003841 | 0.009607 | 0.006102 | 0.003853 | 0.001671 |
| 18.8472222 | 0.002292 | 0.005057 | 0.00019 | 0.004077 | 0.00446 | 0.006782 | 0.003705 | 0.001258 | 0.002856 | 0.003888 | 0.015757 | 0.000227 | 0.007708 | 0.006103 | 0.000119 | 0.000391 | 0.008844 | 0.001455 | 0.000707 | 0.005115 | 0.005967 | 0.00482 | 0.004014 | 0.001801 |
| 18.8611111 | 0.0036 | 0.005732 | 0.000118 | 0.003311 | 0.007557 | 0.005884 | 0.005368 | 0.000554 | 0.001637 | 0.002434 | 0.015377 | 0.000455 | 0.005899 | 0.005807 | 0.000223 | 0.000569 | 0.011893 | 0.001248 | 0.001362 | 0.005909 | 0.003176 | 0.005273 | 0.003654 | 0.002197 |
| 18.875 | 0.004822 | 0.006283 | 8.92E-05 | 0.002723 | 0.010796 | 0.005298 | 0.007598 | 0.00039 | 0.0023 | 0.004733 | 0.011756 | 0.000498 | 0.003569 | 0.004865 | 0.000329 | 0.000918 | 0.015201 | 0.000854 | 0.002194 | 0.006429 | 0.001889 | 0.004611 | 0.002716 | 0.002896 |
| 18.8888889 | 0.006199 | 0.005664 | 0.000116 | 0.002076 | 0.0127 | 0.004643 | 0.010484 | 0.00073 | 0.004036 | 0.00665 | 0.008427 | 0.000413 | 0.00338 | 0.003226 | 0.000344 | 0.001429 | 0.019405 | 0.000459 | 0.003439 | 0.00707 | 0.002062 | 0.0077 | 0.002598 | 0.004204 |
| 18.9027778 | 0.008237 | 0.004431 | 0.000356 | 0.001569 | 0.012817 | 0.003757 | 0.013278 | 0.00107 | 0.005877 | 0.004651 | 0.006865 | 0.000785 | 0.006704 | 0.002174 | 0.000522 | 0.001938 | 0.022499 | 0.000581 | 0.00456 | 0.007898 | 0.003345 | 0.017682 | 0.004485 | 0.006123 |
| 18.9166667 | 0.011003 | 0.00403 | 0.000851 | 0.001334 | 0.011787 | 0.002984 | 0.015634 | 0.001112 | 0.00788 | 0.001781 | 0.006596 | 0.001257 | 0.010352 | 0.003907 | 0.000977 | 0.002161 | 0.023082 | 0.00122 | 0.004681 | 0.009217 | 0.004496 | 0.029172 | 0.006969 | 0.00815 |
| 18.9305556 | 0.013441 | 0.005096 | 0.001219 | 0.001161 | 0.010273 | 0.002553 | 0.017723 | 0.000964 | 0.009403 | 0.000828 | 0.006853 | 0.001519 | 0.011659 | 0.007619 | 0.001393 | 0.002128 | 0.022228 | 0.001988 | 0.004122 | 0.011354 | 0.004056 | 0.038366 | 0.009141 | 0.009664 |
| 18.9444444 | 0.014617 | 0.006086 | 0.001232 | 0.00097 | 0.008694 | 0.002394 | 0.019483 | 0.000655 | 0.009713 | 0.001794 | 0.00716 | 0.00188 | 0.011248 | 0.009987 | 0.001618 | 0.002059 | 0.021146 | 0.002646 | 0.00391 | 0.013324 | 0.0048 | 0.046957 | 0.010884 | 0.010336 |
| 18.9583333 | 0.014309 | 0.0048 | 0.001282 | 0.000893 | 0.007119 | 0.002351 | 0.02023 | 0.000414 | 0.009186 | 0.005112 | 0.007484 | 0.002315 | 0.010893 | 0.008665 | 0.001686 | 0.001991 | 0.019518 | 0.002986 | 0.004524 | 0.014088 | 0.009096 | 0.056297 | 0.01177 | 0.010161 |
| 18.9722222 | 0.01227 | 0.002096 | 0.00158 | 0.000942 | 0.005648 | 0.002284 | 0.01902 | 0.000492 | 0.008827 | 0.009555 | 0.008075 | 0.002557 | 0.012041 | 0.005163 | 0.001467 | 0.001676 | 0.015949 | 0.00282 | 0.004843 | 0.013944 | 0.013221 | 0.066725 | 0.011435 | 0.009471 |
| 18.9861111 | 0.009454 | 0.001149 | 0.002286 | 0.00098 | 0.005035 | 0.001824 | 0.016853 | 0.000705 | 0.009948 | 0.013086 | 0.008767 | 0.00318 | 0.016638 | 0.00361 | 0.000911 | 0.001227 | 0.011469 | 0.002594 | 0.003575 | 0.013192 | 0.016287 | 0.077722 | 0.010454 | 0.009099 |
| 19 | 0.007945 | 0.005767 | 0.00428 | 0.0012 | 0.006472 | 0.001773 | 0.017528 | 0.001431 | 0.015711 | 0.018493 | 0.008124 | 0.006797 | 0.029666 | 0.00366 | 0.000858 | 0.002207 | 0.010007 | 0.003757 | 0.002018 | 0.011241 | 0.023701 | 0.084502 | 0.01138 | 0.0111 |
| 19.0138889 | 0.00851 | 0.01853 | 0.008534 | 0.002771 | 0.011062 | 0.005384 | 0.024898 | 0.002812 | 0.030583 | 0.031864 | 0.00646 | 0.016616 | 0.065389 | 0.006478 | 0.002677 | 0.004999 | 0.012439 | 0.007886 | 0.003077 | 0.007384 | 0.040981 | 0.083035 | 0.016633 | 0.017288 |
| 19.0277778 | 0.009403 | 0.031468 | 0.01442 | 0.00565 | 0.015641 | 0.013172 | 0.038025 | 0.009872 | 0.055654 | 0.049627 | 0.009929 | 0.031727 | 0.126927 | 0.01766 | 0.006499 | 0.005775 | 0.015214 | 0.012793 | 0.007871 | 0.003423 | 0.074708 | 0.084184 | 0.023293 | 0.023644 |
| 19.0416667 | 0.008901 | 0.028165 | 0.0152 | 0.007593 | 0.012974 | 0.017859 | 0.043914 | 0.018667 | 0.064282 | 0.05507 | 0.020982 | 0.03778 | 0.15593 | 0.026579 | 0.009249 | 0.004912 | 0.013849 | 0.012034 | 0.011846 | 0.002173 | 0.097808 | 0.088232 | 0.022099 | 0.0204 |
| 19.0555556 | 0.007418 | 0.015622 | 0.010546 | 0.007981 | 0.00739 | 0.014553 | 0.030278 | 0.017235 | 0.047496 | 0.037714 | 0.031666 | 0.026817 | 0.109116 | 0.020787 | 0.007001 | 0.010026 | 0.007949 | 0.00651 | 0.008991 | 0.004533 | 0.073009 | 0.068051 | 0.013493 | 0.011197 |
| 19.0694444 | 0.005817 | 0.012649 | 0.008272 | 0.007662 | 0.00633 | 0.009108 | 0.014852 | 0.013219 | 0.042283 | 0.01693 | 0.03646 | 0.016849 | 0.066855 | 0.010985 | 0.003152 | 0.022867 | 0.003203 | 0.004955 | 0.003836 | 0.008246 | 0.043735 | 0.035563 | 0.008841 | 0.008083 |
| 19.0833333 | 0.004379 | 0.012164 | 0.006137 | 0.006931 | 0.004684 | 0.006297 | 0.013518 | 0.010701 | 0.041769 | 0.013698 | 0.036319 | 0.014544 | 0.075839 | 0.009617 | 0.002482 | 0.030574 | 0.00296 | 0.006879 | 0.003126 | 0.008274 | 0.045239 | 0.021516 | 0.007897 | 0.007312 |
| 19.0972222 | 0.003128 | 0.007375 | 0.002789 | 0.005945 | 0.001899 | 0.004903 | 0.012441 | 0.004947 | 0.025306 | 0.015167 | 0.031943 | 0.009972 | 0.07397 | 0.012739 | 0.002322 | 0.023733 | 0.004987 | 0.006175 | 0.003299 | 0.004498 | 0.042555 | 0.013945 | 0.005166 | 0.004242 |
| 19.1111111 | 0.00197 | 0.003981 | 0.0011 | 0.004625 | 0.001229 | 0.003958 | 0.006968 | 0.001429 | 0.010403 | 0.010648 | 0.025032 | 0.004306 | 0.048854 | 0.010602 | 0.001217 | 0.012798 | 0.00678 | 0.003662 | 0.001952 | 0.001434 | 0.027141 | 0.005219 | 0.002788 | 0.001827 |
| 19.125 | 0.00099 | 0.00264 | 0.00036 | 0.002987 | 0.000928 | 0.00329 | 0.003156 | 0.001396 | 0.004277 | 0.005809 | 0.01899 | 0.001309 | 0.026675 | 0.005353 | 0.000597 | 0.006877 | 0.007502 | 0.001649 | 0.0009 | 0.000442 | 0.013173 | 0.002128 | 0.00221 | 0.000698 |
| 19.1388889 | 0.00038 | 0.002246 | 0.000121 | 0.001554 | 0.000432 | 0.002695 | 0.00113 | 0.001247 | 0.001908 | 0.002931 | 0.015279 | 0.000328 | 0.011432 | 0.003109 | 0.000455 | 0.003978 | 0.007339 | 0.000449 | 0.00034 | 0.000261 | 0.004661 | 0.003215 | 0.002112 | 0.000531 |
| 19.1527778 | 0.000196 | 0.002491 | 0.000182 | 0.000891 | 0.000188 | 0.002104 | 0.000367 | 0.000632 | 0.001052 | 0.001824 | 0.012674 | 0.000573 | 0.003805 | 0.005748 | 0.000507 | 0.002214 | 0.005739 | 0.000176 | 0.00022 | 0.000368 | 0.0026 | 0.004931 | 0.001546 | 0.001037 |
| 19.1666667 | 0.000215 | 0.002288 | 0.000687 | 0.000789 | 0.000366 | 0.001665 | 0.001427 | 0.000341 | 0.002322 | 0.001295 | 0.009984 | 0.001446 | 0.00707 | 0.035902 | 0.001397 | 0.005185 | 0.009052 | 0.000391 | 0.000759 | 0.001263 | 0.005116 | 0.009562 | 0.003171 | 0.003682 |
| 19.1805556 | 0.00042 | 0.001157 | 0.001633 | 0.000742 | 0.001756 | 0.001798 | 0.005362 | 0.000447 | 0.007649 | 0.001227 | 0.007808 | 0.001799 | 0.028196 | 0.094018 | 0.003815 | 0.015395 | 0.023177 | 0.00114 | 0.0012 | 0.002302 | 0.015606 | 0.022627 | 0.010517 | 0.009472 |
| 19.1944444 | 0.000751 | 0.001312 | 0.001974 | 0.001317 | 0.002872 | 0.002534 | 0.007676 | 0.000669 | 0.010098 | 0.001662 | 0.006581 | 0.00141 | 0.038666 | 0.09788 | 0.005807 | 0.018913 | 0.026652 | 0.003083 | 0.002607 | 0.003298 | 0.02106 | 0.030418 | 0.015922 | 0.013165 |
| 19.2083333 | 0.001664 | 0.004896 | 0.001494 | 0.002909 | 0.003109 | 0.002725 | 0.007042 | 0.000904 | 0.009336 | 0.003359 | 0.00494 | 0.001067 | 0.036112 | 0.069828 | 0.005074 | 0.015621 | 0.023915 | 0.003807 | 0.005927 | 0.005259 | 0.022259 | 0.025563 | 0.014034 | 0.010372 |
| 19.2222222 | 0.003683 | 0.009579 | 0.001414 | 0.003326 | 0.004457 | 0.001779 | 0.008919 | 0.00101 | 0.013551 | 0.007585 | 0.002454 | 0.001245 | 0.053465 | 0.099774 | 0.004565 | 0.019614 | 0.038805 | 0.003456 | 0.006032 | 0.005484 | 0.035842 | 0.025138 | 0.015104 | 0.007652 |
| 19.2361111 | 0.004897 | 0.011065 | 0.001608 | 0.001666 | 0.004734 | 0.000846 | 0.009838 | 0.001124 | 0.014415 | 0.00903 | 0.000791 | 0.002225 | 0.053677 | 0.104456 | 0.005906 | 0.018637 | 0.042031 | 0.004884 | 0.003162 | 0.004182 | 0.03502 | 0.023527 | 0.01836 | 0.007907 |
| 19.25 | 0.00404 | 0.008949 | 0.001443 | 0.000417 | 0.002987 | 0.000591 | 0.00572 | 0.000997 | 0.007661 | 0.005707 | 0.000371 | 0.002613 | 0.026447 | 0.047177 | 0.004863 | 0.008238 | 0.023339 | 0.004452 | 0.002206 | 0.004468 | 0.016013 | 0.012747 | 0.014072 | 0.00535 |
| 19.2638889 | 0.002482 | 0.005325 | 0.001313 | 0.000378 | 0.001307 | 0.000758 | 0.001778 | 0.000505 | 0.002073 | 0.00366 | 0.00032 | 0.00181 | 0.014257 | 0.009108 | 0.002061 | 0.001942 | 0.009004 | 0.001934 | 0.001631 | 0.004464 | 0.007201 | 0.007357 | 0.006828 | 0.002639 |
| 19.2777778 | 0.00161 | 0.002009 | 0.00114 | 0.00051 | 0.000624 | 0.000917 | 0.000734 | 0.000354 | 0.000602 | 0.003157 | 0.000479 | 0.001049 | 0.015738 | 0.00136 | 0.000856 | 0.000885 | 0.004372 | 0.00099 | 0.00059 | 0.004505 | 0.006973 | 0.007452 | 0.003261 | 0.002808 |
| 19.2916667 | 0.00189 | 0.001079 | 0.000828 | 0.000537 | 0.001169 | 0.00071 | 0.001042 | 0.000875 | 0.001672 | 0.002457 | 0.000519 | 0.001086 | 0.012308 | 0.001271 | 0.000785 | 0.000847 | 0.005253 | 0.001038 | 0.000801 | 0.006633 | 0.006858 | 0.005363 | 0.003361 | 0.002257 |
| 19.3055556 | 0.002904 | 0.003163 | 0.000911 | 0.000804 | 0.002114 | 0.000543 | 0.002526 | 0.001824 | 0.003532 | 0.003863 | 0.000437 | 0.001861 | 0.008873 | 0.003651 | 0.001297 | 0.000894 | 0.008774 | 0.001734 | 0.002425 | 0.009205 | 0.009949 | 0.00434 | 0.005826 | 0.001137 |
| 19.3194444 | 0.00373 | 0.006041 | 0.001096 | 0.001097 | 0.002553 | 0.000581 | 0.004164 | 0.00237 | 0.004892 | 0.006361 | 0.000557 | 0.002506 | 0.013991 | 0.007014 | 0.001943 | 0.00113 | 0.011397 | 0.002715 | 0.003691 | 0.009879 | 0.014994 | 0.006427 | 0.008639 | 0.001148 |
| 19.3333333 | 0.003849 | 0.007312 | 0.000746 | 0.000957 | 0.002438 | 0.000412 | 0.004616 | 0.002185 | 0.005337 | 0.007053 | 0.000587 | 0.002466 | 0.018719 | 0.008455 | 0.001843 | 0.000869 | 0.010905 | 0.002817 | 0.003378 | 0.008414 | 0.016218 | 0.007476 | 0.010011 | 0.001088 |
| 19.3472222 | 0.003574 | 0.006953 | 0.00025 | 0.00074 | 0.002338 | 0.000159 | 0.004485 | 0.001878 | 0.005304 | 0.006032 | 0.00061 | 0.001981 | 0.017906 | 0.007249 | 0.001253 | 0.000379 | 0.008849 | 0.002345 | 0.003015 | 0.007546 | 0.014725 | 0.006635 | 0.009982 | 0.000496 |
| 19.3611111 | 0.003385 | 0.006049 | 8.66E-05 | 0.00072 | 0.00245 | 9.46E-05 | 0.004515 | 0.001732 | 0.005237 | 0.004615 | 0.000851 | 0.001644 | 0.016017 | 0.005365 | 0.000884 | 0.000265 | 0.00765 | 0.002133 | 0.003698 | 0.010398 | 0.01416 | 0.006418 | 0.009682 | 0.000147 |
| 19.375 | 0.003532 | 0.005372 | 9.96E-05 | 0.000767 | 0.002663 | 0.000193 | 0.005016 | 0.001627 | 0.005566 | 0.003866 | 0.001179 | 0.001886 | 0.016653 | 0.004371 | 0.000846 | 0.000333 | 0.007814 | 0.002267 | 0.004342 | 0.01589 | 0.016454 | 0.007597 | 0.009712 | 0.0001 |
| 19.3888889 | 0.004061 | 0.005299 | 0.000161 | 0.000893 | 0.003017 | 0.00041 | 0.005884 | 0.001495 | 0.006641 | 0.004139 | 0.001543 | 0.002303 | 0.020044 | 0.004428 | 0.000943 | 0.000495 | 0.009267 | 0.00247 | 0.004023 | 0.018524 | 0.019382 | 0.009326 | 0.010033 | 0.000174 |
| 19.4027778 | 0.004816 | 0.005313 | 0.000254 | 0.001006 | 0.003359 | 0.000598 | 0.006498 | 0.001329 | 0.007932 | 0.004916 | 0.001909 | 0.002217 | 0.022189 | 0.004849 | 0.000945 | 0.000871 | 0.011572 | 0.002445 | 0.003333 | 0.014799 | 0.018556 | 0.009935 | 0.010489 | 0.000319 |
| 19.4166667 | 0.005773 | 0.004778 | 0.000273 | 0.000898 | 0.003457 | 0.000652 | 0.006564 | 0.001225 | 0.008603 | 0.005498 | 0.00226 | 0.001753 | 0.020288 | 0.004976 | 0.000889 | 0.001314 | 0.013152 | 0.00218 | 0.003359 | 0.007914 | 0.014668 | 0.009163 | 0.012078 | 0.000387 |
| 19.4305556 | 0.007112 | 0.004045 | 0.000306 | 0.000661 | 0.003406 | 0.00069 | 0.006668 | 0.001309 | 0.008858 | 0.005711 | 0.002585 | 0.00143 | 0.016834 | 0.004678 | 0.000823 | 0.00151 | 0.013512 | 0.001843 | 0.004292 | 0.00303 | 0.011895 | 0.008325 | 0.015413 | 0.000258 |
| 19.4444444 | 0.008526 | 0.003127 | 0.000402 | 0.000411 | 0.002954 | 0.000577 | 0.007152 | 0.001343 | 0.009584 | 0.00542 | 0.002659 | 0.00116 | 0.014675 | 0.003872 | 0.00096 | 0.001909 | 0.013012 | 0.001318 | 0.004589 | 0.001155 | 0.010989 | 0.007452 | 0.018231 | 0.000537 |
| 19.4583333 | 0.009184 | 0.002063 | 0.000359 | 0.000246 | 0.001784 | 0.000532 | 0.007656 | 0.001091 | 0.011116 | 0.00425 | 0.002272 | 0.001686 | 0.013045 | 0.002724 | 0.002181 | 0.003113 | 0.010867 | 0.001366 | 0.002968 | 0.001 | 0.009813 | 0.00587 | 0.018416 | 0.001806 |
| 19.4722222 | 0.008924 | 0.00144 | 0.000197 | 0.000311 | 0.000666 | 0.00092 | 0.008228 | 0.000696 | 0.013192 | 0.002631 | 0.001821 | 0.002993 | 0.011647 | 0.002563 | 0.003955 | 0.004268 | 0.008163 | 0.002273 | 0.000971 | 0.003091 | 0.008984 | 0.004677 | 0.017773 | 0.003195 |
| 19.4861111 | 0.009492 | 0.002975 | 0.000581 | 0.000367 | 0.002349 | 0.001295 | 0.01102 | 0.000479 | 0.018272 | 0.00316 | 0.00259 | 0.005633 | 0.016815 | 0.005804 | 0.004186 | 0.004043 | 0.009542 | 0.002579 | 0.001067 | 0.008073 | 0.01455 | 0.007273 | 0.019559 | 0.002799 |
| 19.5 | 0.014689 | 0.010541 | 0.002524 | 0.000243 | 0.009804 | 0.002853 | 0.021952 | 0.001024 | 0.0343 | 0.008907 | 0.006725 | 0.023269 | 0.040522 | 0.01416 | 0.002722 | 0.003041 | 0.018488 | 0.003829 | 0.003612 | 0.012981 | 0.032762 | 0.017217 | 0.024508 | 0.002053 |
| 19.5138889 | 0.024317 | 0.022378 | 0.005915 | 0.00049 | 0.020276 | 0.007062 | 0.035052 | 0.002161 | 0.062995 | 0.018801 | 0.017182 | 0.07972 | 0.079326 | 0.023446 | 0.001701 | 0.003807 | 0.029866 | 0.00684 | 0.006111 | 0.013054 | 0.053791 | 0.032172 | 0.029224 | 0.002797 |
| 19.5277778 | 0.030079 | 0.028708 | 0.008885 | 0.001563 | 0.026146 | 0.011995 | 0.036559 | 0.002898 | 0.084569 | 0.027907 | 0.03897 | 0.159135 | 0.095643 | 0.023729 | 0.003068 | 0.007525 | 0.031506 | 0.006654 | 0.005556 | 0.008343 | 0.056375 | 0.044609 | 0.026258 | 0.002983 |
| 19.5416667 | 0.027925 | 0.025543 | 0.010467 | 0.006019 | 0.025099 | 0.014715 | 0.028963 | 0.003285 | 0.080805 | 0.03272 | 0.064744 | 0.193585 | 0.071689 | 0.014695 | 0.005356 | 0.010736 | 0.021252 | 0.002915 | 0.003386 | 0.006172 | 0.041557 | 0.050905 | 0.015601 | 0.002519 |
| 19.5555556 | 0.024351 | 0.019738 | 0.012862 | 0.016866 | 0.022658 | 0.016529 | 0.01976 | 0.003815 | 0.063654 | 0.033422 | 0.078826 | 0.161824 | 0.042577 | 0.008138 | 0.004549 | 0.008572 | 0.010742 | 0.001373 | 0.002917 | 0.010692 | 0.028771 | 0.051024 | 0.007119 | 0.002374 |
| 19.5694444 | 0.022481 | 0.016762 | 0.017856 | 0.029727 | 0.022514 | 0.019702 | 0.013068 | 0.00403 | 0.050114 | 0.03102 | 0.103995 | 0.110904 | 0.029957 | 0.008273 | 0.002007 | 0.003877 | 0.006002 | 0.002869 | 0.003673 | 0.014067 | 0.023833 | 0.044711 | 0.003883 | 0.002773 |
| 19.5833333 | 0.018458 | 0.015856 | 0.023071 | 0.036272 | 0.022618 | 0.021899 | 0.009552 | 0.003335 | 0.041242 | 0.024871 | 0.135244 | 0.071345 | 0.025164 | 0.009963 | 0.001516 | 0.002186 | 0.003336 | 0.003995 | 0.003283 | 0.010022 | 0.020586 | 0.036307 | 0.00213 | 0.004196 |
| 19.5972222 | 0.01142 | 0.016458 | 0.026255 | 0.033382 | 0.020558 | 0.021895 | 0.006594 | 0.002116 | 0.035199 | 0.016546 | 0.116901 | 0.045848 | 0.02289 | 0.010661 | 0.001832 | 0.001684 | 0.001571 | 0.003573 | 0.001759 | 0.00642 | 0.016986 | 0.032917 | 0.000957 | 0.005738 |
| 19.6111111 | 0.005121 | 0.019315 | 0.026703 | 0.025816 | 0.01573 | 0.020299 | 0.00452 | 0.001273 | 0.030259 | 0.01075 | 0.076278 | 0.029879 | 0.022899 | 0.00949 | 0.001845 | 0.000638 | 0.001113 | 0.002846 | 0.000813 | 0.008109 | 0.015213 | 0.034387 | 0.002127 | 0.006858 |

| | | | | | | | | | | | | | | | | | | | | | | | | |
|---|---|---|---|---|---|---|---|---|---|---|---|---|---|---|---|---|---|---|---|---|---|---|---|---|
| 19.625 | 0.004543 | 0.023414 | 0.023699 | 0.019873 | 0.008684 | 0.018067 | 0.003945 | 0.001302 | 0.023622 | 0.009316 | 0.070273 | 0.017991 | 0.023977 | 0.006032 | 0.001925 | 0.000275 | 0.001565 | 0.002785 | 0.00129 | 0.008337 | 0.015961 | 0.033916 | 0.007442 | 0.007512 |
| 19.6388889 | 0.012102 | 0.026504 | 0.01919 | 0.015883 | 0.004177 | 0.017665 | 0.003905 | 0.001992 | 0.015437 | 0.010848 | 0.086697 | 0.009769 | 0.023696 | 0.004029 | 0.001603 | 0.000439 | 0.004161 | 0.003206 | 0.003475 | 0.005009 | 0.01748 | 0.027235 | 0.01532 | 0.007287 |
| 19.6527778 | 0.02298 | 0.027392 | 0.015552 | 0.01239 | 0.004889 | 0.018678 | 0.003565 | 0.0026 | 0.008746 | 0.013807 | 0.098049 | 0.004429 | 0.020557 | 0.004278 | 0.001024 | 0.000489 | 0.00773 | 0.00351 | 0.005192 | 0.002785 | 0.017873 | 0.016099 | 0.020651 | 0.005948 |
| 19.6666667 | 0.026785 | 0.026532 | 0.01304 | 0.010112 | 0.004743 | 0.018106 | 0.003395 | 0.002752 | 0.005653 | 0.016346 | 0.093303 | 0.0018 | 0.015089 | 0.004305 | 0.000799 | 0.000289 | 0.009264 | 0.00343 | 0.004756 | 0.004528 | 0.016986 | 0.006464 | 0.019799 | 0.004004 |
| 19.6805556 | 0.01901 | 0.025521 | 0.011004 | 0.008901 | 0.003365 | 0.015509 | 0.003789 | 0.002656 | 0.004843 | 0.015815 | 0.075021 | 0.001405 | 0.009648 | 0.006077 | 0.000718 | 0.000225 | 0.007967 | 0.003097 | 0.003647 | 0.008163 | 0.015957 | 0.002775 | 0.013997 | 0.002531 |
| 19.6944444 | 0.00918 | 0.025406 | 0.008742 | 0.007801 | 0.004712 | 0.013516 | 0.004115 | 0.002301 | 0.003811 | 0.010754 | 0.05404 | 0.001369 | 0.006767 | 0.009654 | 0.000422 | 0.000405 | 0.004691 | 0.002488 | 0.00228 | 0.009032 | 0.015426 | 0.002969 | 0.007043 | 0.001695 |
| 19.7083333 | 0.008313 | 0.025243 | 0.006212 | 0.006635 | 0.005871 | 0.013314 | 0.003741 | 0.00156 | 0.001999 | 0.005535 | 0.038206 | 0.002089 | 0.006559 | 0.010382 | 0.000249 | 0.000512 | 0.002303 | 0.001566 | 0.001363 | 0.006554 | 0.014473 | 0.002889 | 0.003124 | 0.00107 |
| 19.7222222 | 0.014452 | 0.023597 | 0.004046 | 0.005338 | 0.005647 | 0.013459 | 0.002846 | 0.000844 | 0.00083 | 0.005263 | 0.02846 | 0.003471 | 0.006931 | 0.006721 | 0.000173 | 0.000433 | 0.003197 | 0.00079 | 0.001648 | 0.007075 | 0.012645 | 0.002806 | 0.004499 | 0.000783 |
| 19.7361111 | 0.018142 | 0.020617 | 0.002849 | 0.003767 | 0.005554 | 0.012625 | 0.001913 | 0.000603 | 0.000741 | 0.007268 | 0.022363 | 0.005411 | 0.006634 | 0.003934 | 9.85E-05 | 0.000311 | 0.005325 | 0.000651 | 0.002044 | 0.013038 | 0.011097 | 0.006107 | 0.008421 | 0.000749 |
| 19.75 | 0.013052 | 0.017636 | 0.002691 | 0.002199 | 0.004898 | 0.010718 | 0.001322 | 0.00072 | 0.000659 | 0.006346 | 0.017581 | 0.007612 | 0.005821 | 0.004725 | 0.000214 | 0.00022 | 0.005775 | 0.000892 | 0.001911 | 0.017009 | 0.01055 | 0.012412 | 0.010632 | 0.000534 |
| 19.7638889 | 0.007733 | 0.015452 | 0.003273 | 0.001258 | 0.00317 | 0.008304 | 0.001492 | 0.000913 | 0.000403 | 0.004915 | 0.01347 | 0.008294 | 0.004912 | 0.00545 | 0.000379 | 0.000198 | 0.003869 | 0.001059 | 0.002132 | 0.014574 | 0.010525 | 0.017485 | 0.009237 | 0.000326 |
| 19.7777778 | 0.011976 | 0.013724 | 0.004038 | 0.001294 | 0.001445 | 0.005997 | 0.002426 | 0.001028 | 0.0004 | 0.008388 | 0.010824 | 0.007567 | 0.004243 | 0.0044 | 0.000319 | 0.000271 | 0.001667 | 0.001053 | 0.003205 | 0.008117 | 0.01039 | 0.019595 | 0.005431 | 0.000421 |
| 19.7916667 | 0.015627 | 0.011359 | 0.004538 | 0.001716 | 0.000926 | 0.00443 | 0.003331 | 0.001003 | 0.000683 | 0.013057 | 0.009617 | 0.006475 | 0.003764 | 0.002991 | 0.000181 | 0.000267 | 0.001301 | 0.000734 | 0.003324 | 0.003945 | 0.010027 | 0.018786 | 0.002061 | 0.000751 |
| 19.8055556 | 0.011085 | 0.0081 | 0.004575 | 0.001791 | 0.001263 | 0.003903 | 0.003552 | 0.000812 | 0.001851 | 0.013229 | 0.008633 | 0.004733 | 0.003134 | 0.00324 | 0.000271 | 0.000314 | 0.00146 | 0.000474 | 0.00255 | 0.006157 | 0.009397 | 0.013918 | 0.000736 | 0.001204 |
| 19.8194444 | 0.005758 | 0.005427 | 0.003972 | 0.001812 | 0.001359 | 0.003748 | 0.003275 | 0.000461 | 0.004672 | 0.009068 | 0.007396 | 0.002973 | 0.002549 | 0.004358 | 0.000432 | 0.000723 | 0.000976 | 0.00083 | 0.001782 | 0.009229 | 0.0083 | 0.006795 | 0.001608 | 0.001449 |
| 19.8333333 | 0.006559 | 0.003766 | 0.002928 | 0.001999 | 0.000935 | 0.002681 | 0.002646 | 0.000213 | 0.008341 | 0.005487 | 0.006513 | 0.004038 | 0.002631 | 0.003388 | 0.000437 | 0.001071 | 0.001108 | 0.00131 | 0.000913 | 0.00778 | 0.006663 | 0.002612 | 0.004557 | 0.001371 |
| 19.8472222 | 0.010921 | 0.002216 | 0.001965 | 0.001664 | 0.000724 | 0.00186 | 0.001601 | 0.000349 | 0.011483 | 0.0082 | 0.005972 | 0.009213 | 0.003164 | 0.001442 | 0.000336 | 0.000942 | 0.002759 | 0.001426 | 0.001031 | 0.004833 | 0.004537 | 0.001947 | 0.008898 | 0.001365 |
| 19.8611111 | 0.011267 | 0.001861 | 0.001429 | 0.000986 | 0.001387 | 0.003603 | 0.001273 | 0.000622 | 0.013709 | 0.015006 | 0.004864 | 0.017833 | 0.002953 | 0.001424 | 0.000241 | 0.00073 | 0.004749 | 0.001342 | 0.001632 | 0.006026 | 0.002217 | 0.001919 | 0.013562 | 0.001762 |
| 19.875 | 0.006489 | 0.003666 | 0.001393 | 0.000916 | 0.001909 | 0.006422 | 0.002988 | 0.000625 | 0.015358 | 0.019203 | 0.003153 | 0.028075 | 0.002168 | 0.003276 | 0.000361 | 0.000889 | 0.005658 | 0.001383 | 0.003197 | 0.011209 | 0.000732 | 0.002983 | 0.017826 | 0.002713 |
| 19.8888889 | 0.001899 | 0.006648 | 0.001232 | 0.001148 | 0.002036 | 0.00773 | 0.005598 | 0.000395 | 0.01716 | 0.017382 | 0.001489 | 0.035134 | 0.003411 | 0.006215 | 0.000721 | 0.001291 | 0.005652 | 0.001818 | 0.006447 | 0.01632 | 0.00063 | 0.005858 | 0.021541 | 0.004284 |
| 19.9027778 | 0.002593 | 0.011015 | 0.001776 | 0.001062 | 0.004279 | 0.007047 | 0.007027 | 0.000325 | 0.020056 | 0.011474 | 0.000761 | 0.03619 | 0.008111 | 0.00982 | 0.001078 | 0.001644 | 0.006972 | 0.002448 | 0.007798 | 0.019741 | 0.001361 | 0.01019 | 0.024601 | 0.006573 |
| 19.9166667 | 0.0104 | 0.016912 | 0.0051 | 0.000703 | 0.009405 | 0.005332 | 0.007433 | 0.000525 | 0.024093 | 0.006881 | 0.001383 | 0.035044 | 0.01467 | 0.011135 | 0.001487 | 0.002041 | 0.012032 | 0.00302 | 0.005178 | 0.021245 | 0.002912 | 0.018064 | 0.026527 | 0.009718 |
| 19.9305556 | 0.022688 | 0.022635 | 0.010334 | 0.00053 | 0.015287 | 0.004035 | 0.009016 | 0.000816 | 0.026665 | 0.007098 | 0.002104 | 0.036441 | 0.021088 | 0.008407 | 0.002033 | 0.002592 | 0.020652 | 0.003588 | 0.002507 | 0.019714 | 0.005419 | 0.033012 | 0.028716 | 0.013374 |
| 19.9444444 | 0.033148 | 0.026936 | 0.014613 | 0.000739 | 0.019579 | 0.003593 | 0.013301 | 0.001292 | 0.025182 | 0.011204 | 0.002187 | 0.041893 | 0.026113 | 0.005072 | 0.00221 | 0.002903 | 0.030037 | 0.004177 | 0.003247 | 0.015293 | 0.008019 | 0.054836 | 0.032117 | 0.0157 |
| 19.9583333 | 0.036111 | 0.029101 | 0.017219 | 0.000754 | 0.019021 | 0.003503 | 0.018733 | 0.001813 | 0.021266 | 0.016439 | 0.002348 | 0.049821 | 0.028882 | 0.003723 | 0.001697 | 0.002665 | 0.03704 | 0.004704 | 0.004859 | 0.010229 | 0.009776 | 0.075372 | 0.033602 | 0.015397 |
| 19.9722222 | 0.032355 | 0.028205 | 0.019685 | 0.000881 | 0.014189 | 0.003349 | 0.021845 | 0.001971 | 0.018219 | 0.021195 | 0.003247 | 0.054038 | 0.029403 | 0.004256 | 0.001169 | 0.002271 | 0.039395 | 0.005106 | 0.004054 | 0.006485 | 0.011014 | 0.083745 | 0.030802 | 0.013898 |
| 19.9861111 | 0.026921 | 0.024694 | 0.021418 | 0.001615 | 0.010304 | 0.003449 | 0.021394 | 0.00172 | 0.016802 | 0.024619 | 0.005471 | 0.049263 | 0.026957 | 0.005458 | 0.000944 | 0.00172 | 0.035618 | 0.005216 | 0.002245 | 0.004867 | 0.013091 | 0.077742 | 0.025217 | 0.012445 |
| 20 | 0.022396 | 0.020859 | 0.021174 | 0.002107 | 0.008373 | 0.005421 | 0.019806 | 0.001239 | 0.015799 | 0.029224 | 0.010075 | 0.042086 | 0.027988 | 0.005304 | 0.001233 | 0.001724 | 0.02671 | 0.004725 | 0.001875 | 0.004602 | 0.018825 | 0.064858 | 0.018722 | 0.01061 |
| 20.0138889 | 0.018426 | 0.018507 | 0.021351 | 0.001625 | 0.00621 | 0.013864 | 0.019594 | 0.001032 | 0.014672 | 0.042468 | 0.019636 | 0.044585 | 0.030291 | 0.003257 | 0.003469 | 0.004473 | 0.016397 | 0.003925 | 0.001831 | 0.00464 | 0.034542 | 0.051532 | 0.012808 | 0.008371 |
| 20.0277778 | 0.014919 | 0.016871 | 0.024277 | 0.000847 | 0.003337 | 0.028206 | 0.020821 | 0.000962 | 0.014114 | 0.058554 | 0.043089 | 0.062643 | 0.038298 | 0.001799 | 0.007772 | 0.009598 | 0.008248 | 0.004056 | 0.001995 | 0.004063 | 0.057069 | 0.040639 | 0.009234 | 0.008682 |
| 20.0416667 | 0.013258 | 0.014684 | 0.026054 | 0.003009 | 0.0012 | 0.033131 | 0.020925 | 0.002809 | 0.014382 | 0.055026 | 0.074329 | 0.077702 | 0.049192 | 0.001869 | 0.010502 | 0.012621 | 0.004474 | 0.004099 | 0.002943 | 0.005255 | 0.07608 | 0.034555 | 0.009806 | 0.018524 |
| 20.0555556 | 0.013199 | 0.011505 | 0.021991 | 0.007982 | 0.001614 | 0.023069 | 0.015647 | 0.006531 | 0.014228 | 0.031293 | 0.077623 | 0.057471 | 0.048527 | 0.002262 | 0.007762 | 0.00929 | 0.005295 | 0.002337 | 0.003222 | 0.011736 | 0.06804 | 0.031284 | 0.014756 | 0.032181 |
| 20.0694444 | 0.012131 | 0.007505 | 0.015026 | 0.009657 | 0.00343 | 0.012213 | 0.007442 | 0.007101 | 0.012085 | 0.016149 | 0.044715 | 0.026055 | 0.028962 | 0.003189 | 0.003673 | 0.004018 | 0.005855 | 0.001013 | 0.003267 | 0.015022 | 0.045231 | 0.027441 | 0.019544 | 0.031938 |
| 20.0833333 | 0.008219 | 0.004289 | 0.009664 | 0.007183 | 0.004059 | 0.006818 | 0.00395 | 0.004871 | 0.007756 | 0.016945 | 0.01384 | 0.02246 | 0.010914 | 0.002871 | 0.003437 | 0.003074 | 0.005351 | 0.001585 | 0.00288 | 0.011045 | 0.048534 | 0.022937 | 0.018748 | 0.019761 |
| 20.0972222 | 0.004651 | 0.002832 | 0.006483 | 0.004666 | 0.003529 | 0.004151 | 0.004318 | 0.003078 | 0.004629 | 0.014231 | 0.004712 | 0.025355 | 0.009556 | 0.001328 | 0.003785 | 0.00364 | 0.009426 | 0.001937 | 0.002619 | 0.010345 | 0.047817 | 0.01802 | 0.012764 | 0.009629 |
| 20.1111111 | 0.006283 | 0.002027 | 0.004574 | 0.003275 | 0.003257 | 0.00293 | 0.003353 | 0.002063 | 0.00703 | 0.006036 | 0.003455 | 0.019554 | 0.01117 | 0.000901 | 0.002824 | 0.003155 | 0.015982 | 0.001309 | 0.004048 | 0.012869 | 0.026187 | 0.013025 | 0.005828 | 0.004944 |
| 20.125 | 0.011168 | 0.001053 | 0.003337 | 0.002567 | 0.003421 | 0.002158 | 0.001725 | 0.00183 | 0.012533 | 0.002076 | 0.001685 | 0.013849 | 0.00808 | 0.001536 | 0.002055 | 0.002534 | 0.019858 | 0.001143 | 0.0047 | 0.012801 | 0.009769 | 0.009128 | 0.001831 | 0.003157 |
| 20.1388889 | 0.014587 | 0.000522 | 0.002535 | 0.002179 | 0.003189 | 0.001562 | 0.001574 | 0.001867 | 0.014841 | 0.002656 | 0.001203 | 0.009431 | 0.005021 | 0.002477 | 0.001543 | 0.00187 | 0.020255 | 0.001371 | 0.003346 | 0.011755 | 0.002326 | 0.006688 | 0.0015 | 0.002234 |
| 20.1527778 | 0.015368 | 0.000605 | 0.001844 | 0.001762 | 0.002084 | 0.001199 | 0.001472 | 0.001434 | 0.012755 | 0.004165 | 0.001223 | 0.007026 | 0.003134 | 0.002363 | 0.001225 | 0.001278 | 0.017169 | 0.001118 | 0.002118 | 0.011251 | 0.000736 | 0.004927 | 0.002558 | 0.001611 |
| 20.1666667 | 0.014589 | 0.000572 | 0.001111 | 0.001146 | 0.000978 | 0.001002 | 0.000841 | 0.00071 | 0.010375 | 0.005192 | 0.000887 | 0.006647 | 0.001876 | 0.005994 | 0.001294 | 0.001741 | 0.011736 | 0.000636 | 0.002171 | 0.010802 | 0.001337 | 0.00353 | 0.003822 | 0.001702 |
| 20.1805556 | 0.013298 | 0.000312 | 0.001149 | 0.00054 | 0.000732 | 0.00076 | 0.00191 | 0.000409 | 0.009721 | 0.006414 | 0.000947 | 0.005272 | 0.003268 | 0.046449 | 0.002313 | 0.009026 | 0.009162 | 0.000288 | 0.003085 | 0.010505 | 0.002911 | 0.00268 | 0.004224 | 0.00606 |
| 20.1944444 | 0.012054 | 0.000422 | 0.002562 | 0.000437 | 0.000695 | 0.000458 | 0.005586 | 0.00035 | 0.008609 | 0.007506 | 0.00167 | 0.003152 | 0.010317 | 0.128426 | 0.003068 | 0.025837 | 0.013655 | 0.000778 | 0.00375 | 0.010223 | 0.005132 | 0.004283 | 0.005309 | 0.01485 |
| 20.2083333 | 0.011175 | 0.001392 | 0.003795 | 0.001139 | 0.000692 | 0.00037 | 0.007262 | 0.000249 | 0.009989 | 0.005998 | 0.003151 | 0.005095 | 0.013352 | 0.138342 | 0.001923 | 0.029399 | 0.014688 | 0.001549 | 0.003156 | 0.007742 | 0.006015 | 0.005748 | 0.008637 | 0.015588 |
| 20.2222222 | 0.010408 | 0.003346 | 0.003502 | 0.002289 | 0.001408 | 0.000597 | 0.007555 | 0.000245 | 0.018303 | 0.004131 | 0.007161 | 0.012981 | 0.013068 | 0.102538 | 0.000662 | 0.023595 | 0.013501 | 0.001576 | 0.001826 | 0.003951 | 0.005681 | 0.004566 | 0.011112 | 0.011697 |
| 20.2361111 | 0.008979 | 0.005329 | 0.002707 | 0.002595 | 0.002334 | 0.000716 | 0.00973 | 0.000259 | 0.02531 | 0.005105 | 0.012514 | 0.021852 | 0.01944 | 0.139392 | 0.001312 | 0.031648 | 0.021425 | 0.001789 | 0.001462 | 0.001876 | 0.0075 | 0.003739 | 0.014439 | 0.015718 |
| 20.25 | 0.006233 | 0.005843 | 0.003 | 0.001899 | 0.002727 | 0.000492 | 0.007513 | 0.000496 | 0.022849 | 0.00418 | 0.013565 | 0.023373 | 0.020057 | 0.133136 | 0.002605 | 0.027813 | 0.023533 | 0.002203 | 0.001495 | 0.001207 | 0.010592 | 0.002508 | 0.017829 | 0.014651 |
| 20.2638889 | 0.003012 | 0.005044 | 0.00445 | 0.001566 | 0.00301 | 0.000354 | 0.002622 | 0.001426 | 0.017155 | 0.00135 | 0.010266 | 0.01917 | 0.013296 | 0.055334 | 0.002364 | 0.009803 | 0.015537 | 0.002216 | 0.000832 | 0.001143 | 0.010461 | 0.001872 | 0.01559 | 0.005628 |
| 20.2777778 | 0.00103 | 0.004402 | 0.005803 | 0.001673 | 0.003431 | 0.000463 | 0.001752 | 0.002506 | 0.015199 | 0.000892 | 0.006964 | 0.015179 | 0.009236 | 0.015631 | 0.001316 | 0.001858 | 0.011046 | 0.001957 | 0.000484 | 0.000994 | 0.007425 | 0.00249 | 0.011173 | 0.000847 |
| 20.2916667 | 0.001203 | 0.004676 | 0.006192 | 0.001497 | 0.003632 | 0.000482 | 0.003548 | 0.002663 | 0.01705 | 0.002343 | 0.005616 | 0.012602 | 0.007086 | 0.008902 | 0.000884 | 0.000783 | 0.012236 | 0.001546 | 0.001051 | 0.001266 | 0.004192 | 0.003367 | 0.00922 | 0.000159 |
| 20.3055556 | 0.003901 | 0.005479 | 0.005937 | 0.001118 | 0.00406 | 0.000386 | 0.00591 | 0.002168 | 0.021534 | 0.004706 | 0.006484 | 0.012879 | 0.00559 | 0.005033 | 0.00079 | 0.000474 | 0.016399 | 0.001075 | 0.002471 | 0.003036 | 0.002095 | 0.005281 | 0.008818 | 0.0003 |
| 20.3194444 | 0.008762 | 0.006061 | 0.005853 | 0.000906 | 0.005303 | 0.000413 | 0.008139 | 0.002023 | 0.02599 | 0.007314 | 0.010215 | 0.015748 | 0.004704 | 0.002148 | 0.000868 | 0.000239 | 0.020515 | 0.000672 | 0.003944 | 0.005173 | 0.001739 | 0.007377 | 0.009613 | 0.000426 |
| 20.3333333 | 0.013808 | 0.006103 | 0.006286 | 0.001015 | 0.006605 | 0.00062 | 0.010158 | 0.002408 | 0.027235 | 0.009214 | 0.019078 | 0.01863 | 0.004756 | 0.002956 | 0.001238 | 0.00024 | 0.023583 | 0.00071 | 0.004318 | 0.006266 | 0.003321 | 0.008673 | 0.012714 | 0.000321 |
| 20.3472222 | 0.016592 | 0.006026 | 0.007025 | 0.001283 | 0.006829 | 0.000936 | 0.012644 | 0.002781 | 0.025172 | 0.010056 | 0.030907 | 0.020316 | 0.005806 | 0.008263 | 0.001557 | 0.000541 | 0.027256 | 0.001198 | 0.003876 | 0.006173 | 0.006192 | 0.008497 | 0.017884 | 0.000182 |
| 20.3611111 | 0.016804 | 0.006554 | 0.007782 | 0.001326 | 0.006035 | 0.001319 | 0.015837 | 0.002827 | 0.021788 | 0.010151 | 0.03755 | 0.021495 | 0.007307 | 0.013192 | 0.001457 | 0.00104 | 0.031427 | 0.001666 | 0.003899 | 0.005679 | 0.008379 | 0.00734 | 0.021342 | 0.000227 |
| 20.375 | 0.016042 | 0.007948 | 0.008355 | 0.001093 | 0.005265 | 0.001815 | 0.018185 | 0.002733 | 0.020094 | 0.010245 | 0.036393 | 0.02304 | 0.008574 | 0.011508 | 0.001131 | 0.001255 | 0.033094 | 0.001831 | 0.005071 | 0.006066 | 0.00851 | 0.007426 | 0.019778 | 0.000369 |
| 20.3888889 | 0.015325 | 0.009655 | 0.008694 | 0.001184 | 0.005441 | 0.0025 | 0.018567 | 0.002741 | 0.022589 | 0.010754 | 0.032653 | 0.025237 | 0.009232 | 0.005726 | 0.000883 | 0.000963 | 0.029883 | 0.001829 | 0.006718 | 0.008387 | 0.007555 | 0.0105 | 0.016291 | 0.000327 |
| 20.4027778 | 0.01447 | 0.01056 | 0.008887 | 0.002129 | 0.006544 | 0.003118 | 0.017575 | 0.002774 | 0.028051 | 0.011637 | 0.029469 | 0.027631 | 0.009094 | 0.001838 | 0.00079 | 0.000598 | 0.022696 | 0.001755 | 0.007848 | 0.011543 | 0.007066 | 0.015369 | 0.017193 | 0.000159 |
| 20.4166667 | 0.012893 | 0.010257 | 0.009475 | 0.003781 | 0.007632 | 0.00334 | 0.016391 | 0.002608 | 0.032584 | 0.013427 | 0.026953 | 0.029936 | 0.008455 | 0.001881 | 0.000772 | 0.000608 | 0.015716 | 0.001591 | 0.008097 | 0.012939 | 0.007436 | 0.019699 | 0.024456 | 0.000179 |
| 20.4305556 | 0.010145 | 0.009856 | 0.010977 | 0.005506 | 0.008306 | 0.003592 | 0.016378 | 0.002458 | 0.034223 | 0.016629 | 0.026293 | 0.032514 | 0.008476 | 0.005066 | 0.000742 | 0.000958 | 0.013666 | 0.001501 | 0.007513 | 0.012438 | 0.008649 | 0.022588 | 0.032746 | 0.000543 |
| 20.4444444 | 0.006415 | 0.010422 | 0.012396 | 0.006822 | 0.008731 | 0.004693 | 0.01755 | 0.002683 | 0.033424 | 0.019865 | 0.02681 | 0.034916 | 0.009849 | 0.00828 | 0.000612 | 0.001173 | 0.015951 | 0.001411 | 0.006212 | 0.011856 | 0.009892 | 0.023771 | 0.034175 | 0.001019 |
| 20.4583333 | 0.00284 | 0.011125 | 0.012116 | 0.007081 | 0.008274 | 0.006761 | 0.017927 | 0.003054 | 0.030429 | 0.020583 | 0.025812 | 0.035748 | 0.011252 | 0.007736 | 0.001023 | 0.001063 | 0.016077 | 0.00116 | 0.004083 | 0.011019 | 0.00934 | 0.023185 | 0.026219 | 0.001015 |
| 20.4722222 | 0.000868 | 0.011486 | 0.010376 | 0.005699 | 0.007374 | 0.00877 | 0.015146 | 0.002929 | 0.025712 | 0.017397 | 0.024066 | 0.035247 | 0.01242 | 0.00454 | 0.002212 | 0.001045 | 0.011046 | 0.000841 | 0.00176 | 0.009179 | 0.007766 | 0.022932 | 0.014117 | 0.000927 |
| 20.4861111 | 0.002465 | 0.015755 | 0.010217 | 0.004328 | 0.010303 | 0.011112 | 0.009647 | 0.002493 | 0.021857 | 0.013183 | 0.025623 | 0.039536 | 0.019565 | 0.002418 | 0.00227 | 0.001006 | 0.006874 | 0.001539 | 0.0015 | 0.007276 | 0.010842 | 0.028176 | 0.00589 | 0.002523 |
| 20.5 | 0.00981 | 0.030358 | 0.015947 | 0.005459 | 0.022737 | 0.017398 | 0.005083 | 0.002874 | 0.022726 | 0.018708 | 0.033576 | 0.058617 | 0.044456 | 0.001452 | 0.001655 | 0.000854 | 0.010488 | 0.004466 | 0.004051 | 0.006177 | 0.023693 | 0.042956 | 0.006477 | 0.006529 |
| 20.5138889 | 0.024863 | 0.052386 | 0.027415 | 0.008611 | 0.038183 | 0.029249 | 0.004424 | 0.004137 | 0.027565 | 0.040824 | 0.046142 | 0.08599 | 0.07889 | 0.001324 | 0.00182 | 0.00123 | 0.019671 | 0.006954 | 0.006225 | 0.006271 | 0.041627 | 0.063061 | 0.013924 | 0.011041 |
| 20.5277778 | 0.041205 | 0.067138 | 0.036768 | 0.010951 | 0.040645 | 0.040187 | 0.006451 | 0.004922 | 0.029955 | 0.06604 | 0.054941 | 0.093517 | 0.084535 | 0.002844 | 0.002456 | 0.003442 | 0.02526 | 0.005337 | 0.005539 | 0.006849 | 0.047328 | 0.079475 | 0.017931 | 0.013722 |
| 20.5416667 | 0.042898 | 0.068819 | 0.038337 | 0.010779 | 0.031829 | 0.039585 | 0.006415 | 0.004912 | 0.027857 | 0.074041 | 0.05418 | 0.072375 | 0.05653 | 0.003867 | 0.003708 | 0.006311 | 0.023086 | 0.001954 | 0.003903 | 0.008417 | 0.03532 | 0.089839 | 0.013047 | 0.014774 |
| 20.5555556 | 0.029175 | 0.0691 | 0.035438 | 0.008952 | 0.028662 | 0.028453 | 0.003793 | 0.004838 | 0.026789 | 0.06164 | 0.047201 | 0.044795 | 0.035751 | 0.003314 | 0.003824 | 0.005678 | 0.018627 | 0.002216 | 0.003328 | 0.013094 | 0.023938 | 0.092617 | 0.008295 | 0.015227 |
| 20.5694444 | 0.013554 | 0.073984 | 0.031021 | 0.007072 | 0.043894 | 0.017898 | 0.00373 | 0.004331 | 0.027461 | 0.037445 | 0.037871 | 0.02428 | 0.035191 | 0.00299 | 0.002887 | 0.003342 | 0.014751 | 0.005285 | 0.003704 | 0.016472 | 0.021496 | 0.082165 | 0.009348 | 0.013465 |
| 20.5833333 | 0.00693 | 0.073713 | 0.023696 | 0.006513 | 0.06246 | 0.011306 | 0.006976 | 0.003019 | 0.022321 | 0.016447 | 0.027615 | 0.014142 | 0.038514 | 0.002453 | 0.002208 | 0.002604 | 0.00995 | 0.005897 | 0.00347 | 0.012847 | 0.02006 | 0.061738 | 0.009873 | 0.008488 |
| 20.5972222 | 0.013897 | 0.060862 | 0.014623 | 0.008458 | 0.053634 | 0.006461 | 0.008944 | 0.001826 | 0.011476 | 0.008477 | 0.018995 | 0.020955 | 0.038635 | 0.001285 | 0.001241 | 0.00155 | 0.010701 | 0.004036 | 0.002203 | 0.006252 | 0.017651 | 0.041652 | 0.00655 | 0.003856 |
| 20.6111111 | 0.023226 | 0.041517 | 0.007338 | 0.012079 | 0.0267 | 0.004343 | 0.006529 | 0.001261 | 0.005907 | 0.008477 | 0.012984 | 0.035864 | 0.036279 | 0.00115 | 0.001601 | 0.000585 | 0.020815 | 0.00374 | 0.003028 | 0.004108 | 0.017342 | 0.02718 | 0.004739 | 0.002859 |
| 20.625 | 0.019632 | 0.025222 | 0.003749 | 0.014817 | 0.009275 | 0.006998 | 0.004062 | 0.00102 | 0.009348 | 0.017231 | 0.008234 | 0.039619 | 0.030675 | 0.002204 | 0.003556 | 0.000984 | 0.030431 | 0.00574 | 0.005434 | 0.007065 | 0.018593 | 0.015354 | 0.006811 | 0.003852 |
| 20.6388889 | 0.013551 | 0.015139 | 0.004741 | 0.015466 | 0.006116 | 0.013875 | 0.006938 | 0.001153 | 0.014368 | 0.031935 | 0.004423 | 0.027489 | 0.019488 | 0.002874 | 0.00446 | 0.001519 | 0.02567 | 0.007327 | 0.005381 | 0.009574 | 0.018764 | 0.010995 | 0.009454 | 0.003992 |
| 20.6527778 | 0.025086 | 0.009446 | 0.009067 | 0.014279 | 0.00893 | 0.020604 | 0.009635 | 0.001625 | 0.016044 | 0.040189 | 0.003403 | 0.013352 | 0.010854 | 0.002616 | 0.003789 | 0.001934 | 0.017255 | 0.007306 | 0.004128 | 0.007535 | 0.01788 | 0.02431 | 0.011655 | 0.003845 |
| 20.6666667 | 0.041423 | 0.006029 | 0.012742 | 0.011356 | 0.008123 | 0.020307 | 0.006611 | 0.001914 | 0.013837 | 0.034513 | 0.005095 | 0.003892 | 0.015092 | 0.00211 | 0.003227 | 0.002821 | 0.027435 | 0.006757 | 0.004216 | 0.008388 | 0.017215 | 0.043718 | 0.011005 | 0.004742 |
| 20.6805556 | 0.042271 | 0.004086 | 0.013148 | 0.007787 | 0.006883 | 0.011657 | 0.003474 | 0.001645 | 0.008699 | 0.020396 | 0.005665 | 0.004688 | 0.022034 | 0.00183 | 0.003314 | 0.003703 | 0.044016 | 0.005912 | 0.003532 | 0.018863 | 0.016584 | 0.050716 | 0.013461 | 0.005459 |
| 20.6944444 | 0.030535 | 0.002666 | 0.011244 | 0.005407 | 0.015054 | 0.004545 | 0.005925 | 0.00107 | 0.007263 | 0.017708 | 0.00345 | 0.005022 | 0.018184 | 0.001494 | 0.003091 | 0.003784 | 0.048363 | 0.004048 | 0.006129 | 0.027815 | 0.015681 | 0.041817 | 0.02688 | 0.004209 |
| 20.7083333 | 0.016285 | 0.002163 | 0.008783 | 0.005089 | 0.027235 | 0.005155 | 0.010128 | 0.000537 | 0.012502 | 0.027099 | 0.002146 | 0.002464 | 0.010251 | 0.001146 | 0.001944 | 0.002873 | 0.038516 | 0.00178 | 0.000851 | 0.02588 | 0.014552 | 0.024107 | 0.036264 | 0.002766 |
| 20.7222222 | 0.011763 | 0.004492 | 0.006838 | 0.006537 | 0.030461 | 0.007522 | 0.010787 | 0.00023 | 0.016177 | 0.026624 | 0.005075 | 0.000663 | 0.010573 | 0.001798 | 0.0009 | 0.00147 | 0.021758 | 0.000927 | 0.001017 | 0.016856 | 0.012788 | 0.011435 | 0.02888 | 0.004206 |
| 20.7361111 | 0.020503 | 0.008954 | 0.005938 | 0.008298 | 0.022066 | 0.008315 | 0.008347 | 0.000219 | 0.012649 | 0.015034 | 0.011554 | 0.000651 | 0.016305 | 0.003102 | 0.00092 | 0.000821 | 0.017573 | 0.002009 | 0.001391 | 0.008304 | 0.010202 | 0.012905 | 0.01448 | 0.006749 |
| 20.75 | 0.03322 | 0.012814 | 0.004859 | 0.008821 | 0.011669 | 0.007656 | 0.004933 | 0.000221 | 0.007214 | 0.009627 | 0.018825 | 0.002899 | 0.018457 | 0.003134 | 0.001082 | 0.001488 | 0.035618 | 0.00321 | 0.001766 | 0.007413 | 0.007035 | 0.020786 | 0.008054 | 0.007825 |
| 20.7638889 | 0.03937 | 0.014058 | 0.002978 | 0.007883 | 0.009469 | 0.00563 | 0.003235 | 0.000162 | 0.007731 | 0.012523 | 0.021684 | 0.007329 | 0.013536 | 0.001843 | 0.000831 | 0.002484 | 0.053329 | 0.002811 | 0.001619 | 0.014017 | 0.004262 | 0.024125 | 0.012116 | 0.007571 |
| 20.7777778 | 0.028812 | 0.011974 | 0.002089 | 0.006512 | 0.012081 | 0.008269 | 0.006478 | 0.000331 | 0.012185 | 0.012695 | 0.01757 | 0.008886 | 0.007562 | 0.001015 | 0.001017 | 0.002191 | 0.049653 | 0.00145 | 0.000914 | 0.020562 | 0.003181 | 0.021296 | 0.018884 | 0.006277 |
| 20.7916667 | 0.015357 | 0.008045 | 0.002692 | 0.005684 | 0.011682 | 0.016842 | 0.010795 | 0.000724 | 0.012647 | 0.010081 | 0.009965 | 0.006952 | 0.0086 | 0.001174 | 0.001821 | 0.0015 | 0.029033 | 0.000689 | 0.000476 | 0.020549 | 0.003561 | 0.013942 | 0.025835 | 0.004654 |
| 20.8055556 | 0.023573 | 0.004558 | 0.003436 | 0.005177 | 0.009663 | 0.020975 | 0.009069 | 0.000854 | 0.007983 | 0.01687 | 0.003736 | 0.009547 | 0.011105 | 0.00129 | 0.002096 | 0.002337 | 0.019214 | 0.001032 | 0.000474 | 0.013322 | 0.003443 | 0.01214 | 0.03144 | 0.00342 |
| 20.8194444 | 0.039758 | 0.00232 | 0.003638 | 0.004241 | 0.009452 | 0.015831 | 0.004656 | 0.000556 | 0.004734 | 0.031629 | 0.002227 | 0.018169 | 0.008284 | 0.001762 | 0.00148 | 0.003107 | 0.032942 | 0.001618 | 0.000796 | 0.006461 | 0.002143 | 0.024759 | 0.030244 | 0.002514 |
| 20.8333333 | 0.039406 | 0.001214 | 0.003288 | 0.002944 | 0.009628 | 0.007589 | 0.004462 | 0.000544 | 0.005451 | 0.035473 | 0.005625 | 0.024578 | 0.005438 | 0.004354 | 0.001386 | 0.002277 | 0.044931 | 0.001346 | 0.00181 | 0.007267 | 0.001027 | 0.040153 | 0.021496 | 0.002074 |
| 20.8472222 | 0.024399 | 0.00132 | 0.002704 | 0.001995 | 0.008233 | 0.003528 | 0.006285 | 0.001198 | 0.004755 | 0.022347 | 0.010138 | 0.022407 | 0.006391 | 0.00882 | 0.002289 | 0.001711 | 0.037888 | 0.000527 | 0.002272 | 0.011286 | 0.000535 | 0.045711 | 0.011031 | 0.002715 |
| 20.8611111 | 0.011886 | 0.002631 | 0.002478 | 0.001432 | 0.005436 | 0.005463 | 0.006206 | 0.00146 | 0.003284 | 0.0089 | 0.01143 | 0.013577 | 0.007083 | 0.011982 | 0.002606 | 0.00263 | 0.020314 | 0.000407 | 0.001578 | 0.018838 | 0.000366 | 0.042818 | 0.004763 | 0.004547 |
| 20.875 | 0.014754 | 0.005188 | 0.003315 | 0.001301 | 0.003202 | 0.008918 | 0.005141 | 0.000897 | 0.004305 | 0.005094 | 0.009365 | 0.006883 | 0.00986 | 0.010636 | 0.001678 | 0.003339 | 0.011328 | 0.001333 | 0.001222 | 0.006184 | 0.000842 | 0.037491 | 0.003475 | 0.006925 |
| 20.8888889 | 0.029761 | 0.008759 | 0.005602 | 0.002662 | 0.003478 | 0.011268 | 0.004803 | 0.000637 | 0.004054 | 0.005677 | 0.006619 | 0.008892 | 0.016988 | 0.005799 | 0.000591 | 0.002984 | 0.022684 | 0.002674 | 0.001973 | 0.00396 | 0.002097 | 0.034551 | 0.005515 | 0.009104 |
| 20.9027778 | 0.046243 | 0.012106 | 0.009162 | 0.004898 | 0.005745 | 0.012101 | 0.005778 | 0.000751 | 0.002215 | 0.011597 | 0.005515 | 0.016999 | 0.022766 | 0.001949 | 0.000258 | 0.002506 | 0.040473 | 0.003629 | 0.003467 | 0.007738 | 0.003678 | 0.031658 | 0.009028 | 0.01079 |
| 20.9166667 | 0.05458 | 0.014081 | 0.0126 | 0.005449 | 0.007786 | 0.010051 | 0.007271 | 0.000739 | 0.001807 | 0.022582 | 0.006758 | 0.023796 | 0.026225 | 0.001134 | 0.000662 | 0.001912 | 0.047251 | 0.003548 | 0.005415 | 0.013059 | 0.004958 | 0.046277 | 0.010834 | 0.011516 |
| 20.9305556 | 0.05149 | 0.016017 | 0.014144 | 0.003808 | 0.008603 | 0.006228 | 0.008331 | 0.001474 | 0.004461 | 0.0307 | 0.009214 | 0.025796 | 0.029362 | 0.002089 | 0.001288 | 0.001213 | 0.04184 | 0.002434 | 0.007789 | 0.015901 | 0.005485 | 0.058955 | 0.009407 | 0.01103 |
| 20.9444444 | 0.040981 | 0.020735 | 0.014991 | 0.002086 | 0.008965 | 0.003177 | 0.009069 | 0.003789 | 0.012662 | 0.029599 | 0.012094 | 0.022764 | 0.0319 | 0.004247 | 0.001984 | 0.001237 | 0.031636 | 0.001133 | 0.010399 | 0.016618 | 0.005413 | 0.070493 | 0.007935 | 0.010415 |
| 20.9583333 | 0.02503 | 0.02767 | 0.017803 | 0.001792 | 0.00957 | 0.001796 | 0.009655 | 0.006708 | 0.0241 | 0.018467 | 0.01619 | 0.019438 | 0.034927 | 0.007411 | 0.003355 | 0.001321 | 0.023527 | 0.000885 | 0.012771 | 0.016171 | 0.005778 | 0.078443 | 0.011065 | 0.010608 |
| 20.9722222 | 0.011973 | 0.032598 | 0.022489 | 0.002096 | 0.010084 | 0.001627 | 0.009528 | 0.007737 | 0.03307 | 0.011613 | 0.021065 | 0.016972 | 0.03786 | 0.009911 | 0.005574 | 0.001133 | 0.01935 | 0.001982 | 0.014294 | 0.014167 | 0.007436 | 0.081181 | 0.019048 | 0.011338 |
| 20.9861111 | 0.011007 | 0.032258 | 0.02653 | 0.001839 | 0.009664 | 0.002179 | 0.008551 | 0.005433 | 0.035299 | 0.016933 | 0.024512 | 0.015662 | 0.036515 | 0.009084 | 0.007193 | 0.001512 | 0.018045 | 0.002925 | 0.013635 | 0.010077 | 0.010525 | 0.074289 | 0.027567 | 0.010876 |
| 21 | 0.011744 | 0.028266 | 0.028816 | 0.001862 | 0.008699 | 0.003434 | 0.009253 | 0.002197 | 0.032271 | 0.016904 | 0.025991 | 0.018762 | 0.029868 | 0.005918 | 0.006682 | 0.001667 | 0.017828 | 0.002365 | 0.01023 | 0.005908 | 0.014743 | 0.056286 | 0.032818 | 0.007263 |
| 21.0138889 | 0.010054 | 0.02539 | 0.030242 | 0.002832 | 0.007602 | 0.005903 | 0.014864 | 0.001098 | 0.031411 | 0.016784 | 0.02691 | 0.030882 | 0.022226 | 0.006609 | 0.004084 | 0.002549 | 0.017846 | 0.001277 | 0.005602 | 0.005052 | 0.020108 | 0.036852 | 0.033408 | 0.004253 |
| 21.0277778 | 0.018917 | 0.026603 | 0.030819 | 0.002817 | 0.005651 | 0.010161 | 0.023824 | 0.001602 | 0.034677 | 0.037632 | 0.02859 | 0.051627 | 0.017721 | 0.013207 | 0.002903 | 0.0062 | 0.019153 | 0.001147 | 0.00197 | 0.008585 | 0.027376 | 0.026545 | 0.032712 | 0.006631 |
| 21.0416667 | 0.037081 | 0.029663 | 0.031072 | 0.001554 | 0.004091 | 0.014355 | 0.031199 | 0.001812 | 0.035601 | 0.056805 | 0.03005 | 0.069592 | 0.019351 | 0.018911 | 0.005033 | 0.009446 | 0.021319 | 0.001934 | 0.002488 | 0.014432 | 0.034493 | 0.02599 | 0.028424 | 0.008782 |
| 21.0555556 | 0.047752 | 0.028647 | 0.030347 | 0.00185 | 0.005365 | 0.01436 | 0.032356 | 0.001526 | 0.030007 | 0.046463 | 0.027176 | 0.068002 | 0.027015 | 0.015986 | 0.005559 | 0.007703 | 0.021015 | 0.00396 | 0.007793 | 0.02108 | 0.034173 | 0.029143 | 0.022949 | 0.005852 |
| 21.0694444 | 0.039223 | 0.021648 | 0.025143 | 0.003529 | 0.00806 | 0.009848 | 0.025017 | 0.001817 | 0.02073 | 0.023434 | 0.018861 | 0.046347 | 0.032673 | 0.009022 | 0.002943 | 0.003443 | 0.017439 | 0.006323 | 0.010856 | 0.024289 | 0.023163 | 0.028866 | 0.014592 | 0.00304 |
| 21.0833333 | 0.019498 | 0.011832 | 0.015985 | 0.003411 | 0.008576 | 0.005013 | 0.013294 | 0.002952 | 0.010769 | 0.015401 | 0.010674 | 0.020881 | 0.029147 | 0.008572 | 0.001254 | 0.001546 | 0.011589 | 0.006251 | 0.008062 | 0.018352 | 0.010358 | 0.023062 | 0.006579 | 0.002781 |
| 21.0972222 | 0.008693 | 0.004259 | 0.00781 | 0.002927 | 0.006712 | 0.002418 | 0.004415 | 0.004133 | 0.004544 | 0.018354 | 0.006763 | 0.005354 | 0.019951 | 0.012143 | 0.001848 | 0.001539 | 0.005729 | 0.004413 | 0.005762 | 0.00809 | 0.005176 | 0.016078 | 0.005803 | 0.002881 |
| 21.1111111 | 0.011296 | 0.001796 | 0.003017 | 0.005952 | 0.005181 | 0.001378 | 0.001977 | 0.004838 | 0.00369 | 0.014929 | 0.005494 | 0.001282 | 0.011961 | 0.012541 | 0.003355 | 0.002033 | 0.003814 | 0.003185 | 0.005731 | 0.001991 | 0.005699 | 0.011974 | 0.008884 | 0.003461 |
| 21.125 | 0.014535 | 0.001471 | 0.00093 | 0.010378 | 0.005399 | 0.000967 | 0.004018 | 0.004534 | 0.00299 | 0.008316 | 0.004848 | 0.002195 | 0.006699 | 0.009151 | 0.00437 | 0.002886 | 0.005849 | 0.002484 | 0.004528 | 0.000665 | 0.005806 | 0.010811 | 0.009848 | 0.004429 |
| 21.1388889 | 0.012965 | 0.001313 | 0.000698 | 0.011936 | 0.005915 | 0.001028 | 0.005421 | 0.00276 | 0.002214 | 0.009311 | 0.005008 | 0.003995 | 0.0031 | 0.004984 | 0.004267 | 0.003344 | 0.009609 | 0.001661 | 0.002769 | 0.001008 | 0.004317 | 0.010734 | 0.007581 | 0.004519 |
| 21.1527778 | 0.010424 | 0.002188 | 0.001455 | 0.008993 | 0.004503 | 0.001204 | 0.004056 | 0.001375 | 0.003108 | 0.01513 | 0.005958 | 0.004177 | 0.00107 | 0.003834 | 0.003302 | 0.003021 | 0.013698 | 0.000897 | 0.001774 | 0.002395 | 0.002783 | 0.010251 | 0.003674 | 0.003811 |

| | | | | | | | | | | | | | | | | | | | | | | | | |
|---|---|---|---|---|---|---|---|---|---|---|---|---|---|---|---|---|---|---|---|---|---|---|---|---|
| 21.1666667 | 0.008845 | 0.003261 | 0.002241 | 0.00458 | 0.002226 | 0.001155 | 0.002293 | 0.001337 | 0.003136 | 0.018769 | 0.006762 | 0.003359 | 0.000613 | 0.004718 | 0.002211 | 0.002204 | 0.015557 | 0.00057 | 0.001944 | 0.00547 | 0.001578 | 0.009088 | 0.001094 | 0.002668 |
| 21.1805556 | 0.007768 | 0.004228 | 0.00303 | 0.002262 | 0.001184 | 0.001211 | 0.001924 | 0.001467 | 0.002313 | 0.017998 | 0.007361 | 0.004251 | 0.002043 | 0.00396 | 0.001887 | 0.001809 | 0.013809 | 0.000964 | 0.003831 | 0.008599 | 0.001804 | 0.00749 | 0.000946 | 0.001437 |
| 21.1944444 | 0.006716 | 0.005188 | 0.004217 | 0.002224 | 0.00095 | 0.001592 | 0.001676 | 0.002191 | 0.004624 | 0.011958 | 0.008211 | 0.003885 | 0.005784 | 0.003541 | 0.002765 | 0.002657 | 0.011712 | 0.002075 | 0.005966 | 0.010192 | 0.004585 | 0.005182 | 0.002905 | 0.001477 |
| 21.2083333 | 0.007401 | 0.004132 | 0.004723 | 0.00293 | 0.000992 | 0.001611 | 0.00183 | 0.002725 | 0.012445 | 0.007815 | 0.006998 | 0.003035 | 0.007856 | 0.004619 | 0.003797 | 0.004164 | 0.014119 | 0.002952 | 0.006823 | 0.01148 | 0.005485 | 0.003787 | 0.007188 | 0.002863 |
| 21.2222222 | 0.011516 | 0.002648 | 0.006218 | 0.0032 | 0.001128 | 0.001712 | 0.004068 | 0.002038 | 0.021849 | 0.012625 | 0.003985 | 0.006165 | 0.005801 | 0.004853 | 0.003298 | 0.004652 | 0.017915 | 0.002341 | 0.007232 | 0.013777 | 0.004315 | 0.005901 | 0.012327 | 0.003633 |
| 21.2361111 | 0.016282 | 0.003833 | 0.009141 | 0.004479 | 0.00178 | 0.002491 | 0.005547 | 0.002039 | 0.024497 | 0.015358 | 0.002266 | 0.009633 | 0.003711 | 0.006932 | 0.001723 | 0.00342 | 0.017382 | 0.00108 | 0.006944 | 0.014644 | 0.005106 | 0.008434 | 0.012995 | 0.002518 |
| 21.25 | 0.016598 | 0.004604 | 0.008749 | 0.005359 | 0.003223 | 0.002461 | 0.003775 | 0.002919 | 0.018592 | 0.010796 | 0.001918 | 0.007821 | 0.002399 | 0.009951 | 0.000921 | 0.001718 | 0.012747 | 0.000706 | 0.004388 | 0.011821 | 0.004075 | 0.007356 | 0.008187 | 0.001252 |
| 21.2638889 | 0.010963 | 0.002961 | 0.006131 | 0.005198 | 0.004632 | 0.001625 | 0.001406 | 0.002971 | 0.012621 | 0.01101 | 0.00304 | 0.003517 | 0.0008 | 0.009147 | 0.000892 | 0.001179 | 0.006415 | 0.000758 | 0.001465 | 0.007939 | 0.001169 | 0.004687 | 0.004707 | 0.001414 |
| 21.2777778 | 0.005786 | 0.001552 | 0.005192 | 0.005355 | 0.004967 | 0.001377 | 0.001555 | 0.002701 | 0.01456 | 0.014491 | 0.003515 | 0.003467 | 0.000728 | 0.006518 | 0.000812 | 0.001457 | 0.002447 | 0.000735 | 0.000985 | 0.006437 | 0.00041 | 0.003784 | 0.004139 | 0.001346 |
| 21.2916667 | 0.006543 | 0.001296 | 0.0058 | 0.005322 | 0.00425 | 0.001802 | 0.004574 | 0.002331 | 0.024503 | 0.013348 | 0.002304 | 0.010651 | 0.002544 | 0.005146 | 0.000758 | 0.001159 | 0.001613 | 0.000703 | 0.002564 | 0.007227 | 0.001194 | 0.005115 | 0.002634 | 0.000609 |
| 21.3055556 | 0.00768 | 0.001365 | 0.007189 | 0.005434 | 0.003615 | 0.002359 | 0.008316 | 0.00242 | 0.034947 | 0.016328 | 0.002034 | 0.02103 | 0.004164 | 0.00479 | 0.000952 | 0.000595 | 0.002516 | 0.000458 | 0.004586 | 0.007953 | 0.002253 | 0.007356 | 0.000861 | 0.000321 |
| 21.3194444 | 0.00516 | 0.002821 | 0.008996 | 0.005725 | 0.003536 | 0.002732 | 0.011009 | 0.004814 | 0.037352 | 0.023957 | 0.002754 | 0.024573 | 0.00453 | 0.004914 | 0.001473 | 0.000556 | 0.005275 | 0.000186 | 0.005455 | 0.007276 | 0.003212 | 0.010036 | 0.001457 | 0.000289 |
| 21.3333333 | 0.002584 | 0.00559 | 0.011511 | 0.005646 | 0.003967 | 0.003146 | 0.013001 | 0.007154 | 0.03237 | 0.021762 | 0.003218 | 0.019177 | 0.005241 | 0.006665 | 0.002275 | 0.00106 | 0.00846 | 0.000322 | 0.005485 | 0.006273 | 0.004188 | 0.013368 | 0.004439 | 0.000406 |
| 21.3472222 | 0.003917 | 0.007561 | 0.015093 | 0.00534 | 0.004567 | 0.004021 | 0.015276 | 0.005975 | 0.02711 | 0.011619 | 0.00367 | 0.01527 | 0.006004 | 0.011384 | 0.002792 | 0.001703 | 0.010611 | 0.000829 | 0.006555 | 0.006335 | 0.005248 | 0.015432 | 0.005881 | 0.000975 |
| 21.3611111 | 0.009276 | 0.008046 | 0.018901 | 0.005777 | 0.004938 | 0.005224 | 0.018343 | 0.002805 | 0.024684 | 0.005011 | 0.004638 | 0.021584 | 0.005572 | 0.016108 | 0.00243 | 0.001433 | 0.012036 | 0.001346 | 0.0088 | 0.007903 | 0.006123 | 0.014628 | 0.006119 | 0.001528 |
| 21.375 | 0.015393 | 0.007828 | 0.020958 | 0.007353 | 0.005191 | 0.006549 | 0.021602 | 0.001189 | 0.023798 | 0.010047 | 0.005795 | 0.035439 | 0.005969 | 0.015295 | 0.001734 | 0.000521 | 0.014665 | 0.00188 | 0.01072 | 0.010739 | 0.006621 | 0.013741 | 0.014015 | 0.001591 |
| 21.3888889 | 0.019662 | 0.008017 | 0.020354 | 0.009125 | 0.005318 | 0.008101 | 0.023425 | 0.002746 | 0.022846 | 0.030688 | 0.006768 | 0.045048 | 0.007983 | 0.009497 | 0.001721 | 0.000303 | 0.019311 | 0.002332 | 0.011078 | 0.013322 | 0.006841 | 0.015945 | 0.027775 | 0.001499 |
| 21.4027778 | 0.021646 | 0.009133 | 0.018374 | 0.009995 | 0.005155 | 0.009195 | 0.021799 | 0.005517 | 0.020586 | 0.049218 | 0.007731 | 0.042181 | 0.010209 | 0.004323 | 0.002823 | 0.00107 | 0.023111 | 0.002429 | 0.00995 | 0.014617 | 0.007403 | 0.020934 | 0.031871 | 0.001598 |
| 21.4166667 | 0.022552 | 0.011226 | 0.016032 | 0.009755 | 0.005121 | 0.009433 | 0.016833 | 0.006453 | 0.017112 | 0.046724 | 0.009062 | 0.032471 | 0.012485 | 0.001994 | 0.004761 | 0.002033 | 0.021763 | 0.002556 | 0.00835 | 0.015394 | 0.009173 | 0.027221 | 0.023688 | 0.002097 |
| 21.4305556 | 0.024124 | 0.01383 | 0.014005 | 0.008793 | 0.005705 | 0.009399 | 0.011678 | 0.005755 | 0.014548 | 0.029079 | 0.010902 | 0.02708 | 0.014741 | 0.001462 | 0.006086 | 0.002146 | 0.01491 | 0.003127 | 0.006758 | 0.015236 | 0.011092 | 0.032139 | 0.015644 | 0.00304 |
| 21.4444444 | 0.026903 | 0.015686 | 0.013702 | 0.006926 | 0.006462 | 0.008996 | 0.008579 | 0.005016 | 0.014226 | 0.011823 | 0.012671 | 0.030224 | 0.016186 | 0.001597 | 0.005338 | 0.001466 | 0.007086 | 0.003564 | 0.005096 | 0.012189 | 0.010632 | 0.033168 | 0.014429 | 0.003635 |
| 21.4583333 | 0.02897 | 0.015455 | 0.014245 | 0.004408 | 0.00647 | 0.007888 | 0.006822 | 0.004335 | 0.014752 | 0.00605 | 0.013259 | 0.03764 | 0.015943 | 0.001631 | 0.003303 | 0.000745 | 0.0027 | 0.003132 | 0.003695 | 0.006941 | 0.007599 | 0.032096 | 0.018636 | 0.003263 |
| 21.4722222 | 0.026586 | 0.014372 | 0.013459 | 0.002844 | 0.006074 | 0.007069 | 0.004514 | 0.003461 | 0.013913 | 0.014371 | 0.012681 | 0.039899 | 0.014465 | 0.001324 | 0.002152 | 0.000425 | 0.003405 | 0.002704 | 0.003388 | 0.003673 | 0.005416 | 0.033472 | 0.026982 | 0.003085 |
| 21.4861111 | 0.018335 | 0.017684 | 0.013564 | 0.004361 | 0.008398 | 0.008715 | 0.003534 | 0.003334 | 0.011163 | 0.030466 | 0.014614 | 0.036257 | 0.017005 | 0.000804 | 0.001721 | 0.00059 | 0.00919 | 0.004297 | 0.00478 | 0.006457 | 0.009395 | 0.043715 | 0.034566 | 0.004972 |
| 21.5 | 0.009142 | 0.031872 | 0.021383 | 0.011254 | 0.019099 | 0.015269 | 0.006051 | 0.004837 | 0.010235 | 0.046761 | 0.026769 | 0.044464 | 0.036119 | 0.000426 | 0.001875 | 0.000819 | 0.019356 | 0.008401 | 0.007133 | 0.016033 | 0.02318 | 0.074525 | 0.034331 | 0.009002 |
| 21.5138889 | 0.006142 | 0.055907 | 0.037408 | 0.020653 | 0.034044 | 0.024025 | 0.007029 | 0.006795 | 0.016398 | 0.065723 | 0.048635 | 0.072822 | 0.068792 | 0.001051 | 0.004111 | 0.001654 | 0.031064 | 0.013644 | 0.008578 | 0.027255 | 0.040292 | 0.121307 | 0.029331 | 0.012676 |
| 21.5277778 | 0.013346 | 0.071495 | 0.05011 | 0.023051 | 0.038589 | 0.025098 | 0.006548 | 0.007386 | 0.029163 | 0.084505 | 0.05925 | 0.092797 | 0.080291 | 0.001899 | 0.00614 | 0.004591 | 0.037745 | 0.016517 | 0.006862 | 0.030706 | 0.045041 | 0.140767 | 0.023688 | 0.013105 |
| 21.5416667 | 0.029411 | 0.066868 | 0.050607 | 0.016082 | 0.033485 | 0.016331 | 0.00719 | 0.006631 | 0.043018 | 0.084706 | 0.048034 | 0.081147 | 0.061009 | 0.001516 | 0.005904 | 0.008006 | 0.03931 | 0.015154 | 0.002904 | 0.023976 | 0.036263 | 0.110485 | 0.017063 | 0.011036 |
| 21.5555556 | 0.0434 | 0.0558 | 0.042068 | 0.00817 | 0.030223 | 0.006843 | 0.005996 | 0.005934 | 0.048925 | 0.061285 | 0.032408 | 0.054537 | 0.041462 | 0.000763 | 0.005508 | 0.007528 | 0.040741 | 0.013156 | 0.001581 | 0.015118 | 0.030149 | 0.072144 | 0.012312 | 0.009866 |
| 21.5694444 | 0.041849 | 0.047382 | 0.030595 | 0.004083 | 0.028548 | 0.003616 | 0.008457 | 0.005725 | 0.040915 | 0.033676 | 0.02154 | 0.030234 | 0.030954 | 0.000469 | 0.005294 | 0.003543 | 0.035505 | 0.011511 | 0.003235 | 0.008658 | 0.029048 | 0.058082 | 0.010137 | 0.008833 |
| 21.5833333 | 0.026945 | 0.038589 | 0.019303 | 0.001935 | 0.02414 | 0.007353 | 0.016888 | 0.005088 | 0.025963 | 0.032925 | 0.011153 | 0.017367 | 0.018461 | 0.001237 | 0.003638 | 0.000869 | 0.021471 | 0.008171 | 0.003159 | 0.005125 | 0.023168 | 0.053054 | 0.006802 | 0.005596 |
| 21.5972222 | 0.012153 | 0.028145 | 0.010026 | 0.000587 | 0.017811 | 0.013673 | 0.022327 | 0.003668 | 0.012755 | 0.060231 | 0.005199 | 0.024217 | 0.011264 | 0.00309 | 0.001792 | 0.000407 | 0.019971 | 0.00527 | 0.001439 | 0.007885 | 0.014505 | 0.03747 | 0.005128 | 0.003373 |
| 21.6111111 | 0.011328 | 0.015579 | 0.00462 | 0.000818 | 0.011378 | 0.016741 | 0.0181 | 0.002222 | 0.007766 | 0.065956 | 0.007721 | 0.033503 | 0.020609 | 0.004526 | 0.00212 | 0.00047 | 0.043473 | 0.005043 | 0.000777 | 0.015611 | 0.008137 | 0.022148 | 0.010664 | 0.003695 |
| 21.625 | 0.02082 | 0.008333 | 0.004479 | 0.00323 | 0.005807 | 0.013066 | 0.011905 | 0.002028 | 0.012556 | 0.042172 | 0.010347 | 0.025086 | 0.036186 | 0.0046 | 0.003706 | 0.000628 | 0.069076 | 0.005474 | 0.000845 | 0.021028 | 0.003505 | 0.026276 | 0.016749 | 0.00317 |
| 21.6388889 | 0.022591 | 0.01308 | 0.006824 | 0.006371 | 0.003881 | 0.007645 | 0.015616 | 0.002891 | 0.014835 | 0.038964 | 0.007862 | 0.012405 | 0.045425 | 0.003562 | 0.004184 | 0.000476 | 0.069124 | 0.003895 | 0.001731 | 0.01746 | 0.00147 | 0.035017 | 0.015493 | 0.003103 |
| 21.6527778 | 0.018446 | 0.021688 | 0.006501 | 0.007656 | 0.006614 | 0.009677 | 0.019916 | 0.002923 | 0.008986 | 0.053023 | 0.007891 | 0.008015 | 0.043599 | 0.002407 | 0.002807 | 0.000303 | 0.042104 | 0.001601 | 0.003734 | 0.010635 | 0.003128 | 0.031437 | 0.010137 | 0.005725 |
| 21.6666667 | 0.01566 | 0.025787 | 0.004292 | 0.006387 | 0.009776 | 0.01727 | 0.013237 | 0.001843 | 0.00342 | 0.045596 | 0.013978 | 0.007306 | 0.031987 | 0.001746 | 0.002003 | 0.000354 | 0.024582 | 0.000737 | 0.006146 | 0.013814 | 0.006299 | 0.024728 | 0.00558 | 0.007588 |
| 21.6805556 | 0.022791 | 0.024547 | 0.004703 | 0.003874 | 0.009681 | 0.021521 | 0.006976 | 0.000982 | 0.002451 | 0.023292 | 0.018198 | 0.014028 | 0.018332 | 0.001313 | 0.003628 | 0.000302 | 0.041587 | 0.000946 | 0.007765 | 0.025055 | 0.007404 | 0.024978 | 0.006161 | 0.006021 |
| 21.6944444 | 0.043721 | 0.019744 | 0.006411 | 0.001677 | 0.006165 | 0.018316 | 0.01334 | 0.000815 | 0.003679 | 0.017095 | 0.015856 | 0.025366 | 0.010058 | 0.000783 | 0.004531 | 0.00034 | 0.062756 | 0.002846 | 0.007516 | 0.032084 | 0.006772 | 0.032083 | 0.015656 | 0.003573 |
| 21.7083333 | 0.053206 | 0.013641 | 0.006345 | 0.000929 | 0.003055 | 0.010462 | 0.02325 | 0.001026 | 0.004542 | 0.034312 | 0.010382 | 0.029884 | 0.009618 | 0.000582 | 0.003045 | 0.000846 | 0.066288 | 0.005939 | 0.00563 | 0.028325 | 0.008168 | 0.038685 | 0.026289 | 0.003865 |
| 21.7222222 | 0.037024 | 0.007907 | 0.005411 | 0.001345 | 0.003732 | 0.004952 | 0.0228 | 0.001252 | 0.005692 | 0.052328 | 0.005273 | 0.021595 | 0.009147 | 0.001289 | 0.002532 | 0.001955 | 0.054986 | 0.006877 | 0.00335 | 0.015949 | 0.012568 | 0.03807 | 0.027129 | 0.004824 |
| 21.7361111 | 0.015035 | 0.004799 | 0.003825 | 0.001317 | 0.005871 | 0.006479 | 0.013399 | 0.001001 | 0.008556 | 0.046878 | 0.003459 | 0.012337 | 0.007534 | 0.002252 | 0.004123 | 0.002539 | 0.031561 | 0.004605 | 0.001624 | 0.00692 | 0.017552 | 0.030833 | 0.017862 | 0.003112 |
| 21.75 | 0.007716 | 0.004953 | 0.002474 | 0.001454 | 0.00689 | 0.012794 | 0.006391 | 0.000434 | 0.009374 | 0.027621 | 0.005695 | 0.019012 | 0.011337 | 0.002402 | 0.005079 | 0.001736 | 0.020947 | 0.002923 | 0.001204 | 0.009412 | 0.020041 | 0.019945 | 0.007556 | 0.001473 |
| 21.7638889 | 0.014309 | 0.007812 | 0.004224 | 0.002996 | 0.006393 | 0.017394 | 0.004083 | 0.000338 | 0.006425 | 0.025565 | 0.008918 | 0.029824 | 0.01786 | 0.001712 | 0.004353 | 0.000938 | 0.040113 | 0.003952 | 0.003202 | 0.01663 | 0.018308 | 0.009892 | 0.004633 | 0.003357 |
| 21.7777778 | 0.023792 | 0.013213 | 0.008765 | 0.004313 | 0.004035 | 0.015832 | 0.003536 | 0.000617 | 0.003246 | 0.035499 | 0.010143 | 0.026709 | 0.022468 | 0.00171 | 0.002841 | 0.001033 | 0.065391 | 0.004425 | 0.006649 | 0.022967 | 0.013918 | 0.005683 | 0.007301 | 0.005873 |
| 21.7916667 | 0.033029 | 0.019901 | 0.01333 | 0.004191 | 0.0022 | 0.010038 | 0.006829 | 0.000734 | 0.00259 | 0.029678 | 0.007684 | 0.016687 | 0.023177 | 0.003325 | 0.001602 | 0.001498 | 0.078584 | 0.003708 | 0.009794 | 0.02781 | 0.011411 | 0.006607 | 0.01336 | 0.004842 |
| 21.8055556 | 0.041428 | 0.027111 | 0.015704 | 0.00318 | 0.003721 | 0.005391 | 0.013732 | 0.001315 | 0.007885 | 0.017078 | 0.004859 | 0.018917 | 0.021748 | 0.004821 | 0.0016 | 0.002235 | 0.079359 | 0.004138 | 0.01159 | 0.027443 | 0.014243 | 0.014293 | 0.022577 | 0.00417 |
| 21.8194444 | 0.044589 | 0.03421 | 0.015434 | 0.001998 | 0.006916 | 0.003036 | 0.020339 | 0.003182 | 0.020153 | 0.024239 | 0.006259 | 0.030524 | 0.021573 | 0.004617 | 0.001877 | 0.003358 | 0.064691 | 0.005235 | 0.010296 | 0.020168 | 0.021005 | 0.029148 | 0.027572 | 0.008154 |
| 21.8333333 | 0.039565 | 0.036958 | 0.014352 | 0.001672 | 0.009792 | 0.002639 | 0.020909 | 0.006302 | 0.033789 | 0.047832 | 0.007499 | 0.0329 | 0.023251 | 0.003203 | 0.001587 | 0.005047 | 0.040885 | 0.005406 | 0.005994 | 0.009743 | 0.027985 | 0.040971 | 0.022934 | 0.011468 |
| 21.8472222 | 0.028009 | 0.033726 | 0.014374 | 0.002198 | 0.012394 | 0.003749 | 0.015917 | 0.010116 | 0.040362 | 0.063106 | 0.005571 | 0.02537 | 0.026752 | 0.002355 | 0.002247 | 0.006121 | 0.022103 | 0.004899 | 0.002623 | 0.002526 | 0.032494 | 0.044425 | 0.013954 | 0.010582 |
| 21.8611111 | 0.017101 | 0.031718 | 0.016328 | 0.002156 | 0.015019 | 0.004651 | 0.01429 | 0.01275 | 0.037238 | 0.057005 | 0.006741 | 0.020231 | 0.029615 | 0.004047 | 0.004243 | 0.004346 | 0.015959 | 0.005055 | 0.001672 | 0.002064 | 0.032726 | 0.044281 | 0.011962 | 0.009114 |
| 21.875 | 0.014723 | 0.036295 | 0.019987 | 0.001251 | 0.017738 | 0.008327 | 0.020196 | 0.012413 | 0.030769 | 0.03673 | 0.013872 | 0.022873 | 0.027694 | 0.008642 | 0.006291 | 0.00228 | 0.020246 | 0.006591 | 0.002083 | 0.007266 | 0.029276 | 0.045043 | 0.02004 | 0.00709 |
| 21.8888889 | 0.022084 | 0.043459 | 0.024404 | 0.000621 | 0.019987 | 0.014359 | 0.029706 | 0.009575 | 0.028379 | 0.018087 | 0.022082 | 0.029959 | 0.022244 | 0.013565 | 0.008093 | 0.002369 | 0.027801 | 0.008133 | 0.005961 | 0.014822 | 0.024892 | 0.050263 | 0.028196 | 0.003368 |
| 21.9027778 | 0.03267 | 0.047339 | 0.028239 | 0.001053 | 0.021098 | 0.016308 | 0.037053 | 0.00609 | 0.032114 | 0.010679 | 0.027872 | 0.03738 | 0.018305 | 0.015374 | 0.009757 | 0.003173 | 0.033026 | 0.008235 | 0.013387 | 0.0207 | 0.021018 | 0.061402 | 0.030834 | 0.000845 |
| 21.9166667 | 0.038403 | 0.047284 | 0.029512 | 0.002351 | 0.01948 | 0.013145 | 0.037455 | 0.004448 | 0.038207 | 0.010603 | 0.031447 | 0.043759 | 0.016618 | 0.013464 | 0.010384 | 0.006654 | 0.033557 | 0.007952 | 0.02115 | 0.023522 | 0.018463 | 0.071431 | 0.029764 | 0.002156 |
| 21.9305556 | 0.036284 | 0.043567 | 0.027044 | 0.003798 | 0.014567 | 0.008888 | 0.028605 | 0.006407 | 0.040616 | 0.019269 | 0.032113 | 0.04722 | 0.01559 | 0.009157 | 0.008658 | 0.012212 | 0.030377 | 0.007623 | 0.025891 | 0.023391 | 0.017396 | 0.069599 | 0.024864 | 0.007105 |
| 21.9444444 | 0.030528 | 0.035329 | 0.022134 | 0.004982 | 0.008803 | 0.005958 | 0.014921 | 0.010478 | 0.035788 | 0.042445 | 0.028477 | 0.043093 | 0.014654 | 0.004164 | 0.005056 | 0.01485 | 0.027248 | 0.006444 | 0.025762 | 0.019167 | 0.015783 | 0.055482 | 0.016109 | 0.011802 |
| 21.9583333 | 0.027186 | 0.025875 | 0.017427 | 0.005557 | 0.004868 | 0.004167 | 0.006455 | 0.01255 | 0.025635 | 0.067883 | 0.022021 | 0.03196 | 0.013913 | 0.001167 | 0.002154 | 0.013684 | 0.024384 | 0.0051 | 0.022709 | 0.013152 | 0.013672 | 0.038654 | 0.007074 | 0.013328 |
| 21.9722222 | 0.026862 | 0.019423 | 0.014056 | 0.004643 | 0.003588 | 0.003588 | 0.005041 | 0.010587 | 0.015788 | 0.075922 | 0.016062 | 0.021034 | 0.013906 | 0.000758 | 0.001753 | 0.010369 | 0.019129 | 0.004253 | 0.019528 | 0.009253 | 0.012952 | 0.022712 | 0.002028 | 0.012795 |
| 21.9861111 | 0.02517 | 0.016463 | 0.011996 | 0.002386 | 0.003617 | 0.004291 | 0.003789 | 0.006496 | 0.010034 | 0.060641 | 0.012425 | 0.01308 | 0.014501 | 0.001211 | 0.002756 | 0.005782 | 0.012659 | 0.002981 | 0.017102 | 0.007423 | 0.013101 | 0.012495 | 0.00091 | 0.011093 |
| 22 | 0.021756 | 0.015842 | 0.013103 | 0.001624 | 0.00357 | 0.005653 | 0.003035 | 0.00306 | 0.010266 | 0.036682 | 0.013694 | 0.008522 | 0.015527 | 0.002271 | 0.002706 | 0.002638 | 0.007803 | 0.00118 | 0.015212 | 0.006886 | 0.014064 | 0.015152 | 0.002687 | 0.007908 |
| 22.0138889 | 0.022445 | 0.016718 | 0.020095 | 0.006786 | 0.003166 | 0.008232 | 0.005972 | 0.001564 | 0.016236 | 0.025624 | 0.026073 | 0.01139 | 0.016838 | 0.004129 | 0.002411 | 0.003346 | 0.006288 | 0.001242 | 0.013659 | 0.008522 | 0.017065 | 0.017728 | 0.008946 | 0.00445 |
| 22.0277778 | 0.028963 | 0.018569 | 0.031297 | 0.016815 | 0.002055 | 0.012587 | 0.009933 | 0.002022 | 0.022308 | 0.036476 | 0.05031 | 0.024313 | 0.017285 | 0.00583 | 0.004874 | 0.005563 | 0.011129 | 0.004145 | 0.01177 | 0.013071 | 0.023113 | 0.013216 | 0.019764 | 0.002847 |
| 22.0416667 | 0.036297 | 0.023028 | 0.039511 | 0.021727 | 0.000733 | 0.015745 | 0.013426 | 0.005035 | 0.022142 | 0.052555 | 0.063988 | 0.040934 | 0.016435 | 0.005085 | 0.007918 | 0.005863 | 0.021909 | 0.006193 | 0.010456 | 0.020159 | 0.027402 | 0.018788 | 0.034515 | 0.003592 |
| 22.0555556 | 0.037924 | 0.02961 | 0.037195 | 0.014849 | 0.000633 | 0.014499 | 0.017246 | 0.008082 | 0.016843 | 0.047933 | 0.04435 | 0.049519 | 0.015283 | 0.004531 | 0.006658 | 0.004043 | 0.028181 | 0.004848 | 0.012354 | 0.027803 | 0.019975 | 0.036757 | 0.045435 | 0.004989 |
| 22.0694444 | 0.029618 | 0.030974 | 0.023208 | 0.007844 | 0.001636 | 0.010402 | 0.01947 | 0.007301 | 0.011903 | 0.02616 | 0.016494 | 0.043359 | 0.013668 | 0.00973 | 0.002608 | 0.002037 | 0.021392 | 0.004499 | 0.014309 | 0.03121 | 0.007795 | 0.046069 | 0.037545 | 0.004417 |
| 22.0833333 | 0.015331 | 0.023353 | 0.009816 | 0.011635 | 0.00207 | 0.006365 | 0.017163 | 0.008581 | 0.010077 | 0.018688 | 0.009646 | 0.025923 | 0.010849 | 0.015062 | 0.000973 | 0.002482 | 0.014085 | 0.007089 | 0.010771 | 0.025437 | 0.004966 | 0.036936 | 0.019791 | 0.003535 |
| 22.0972222 | 0.005582 | 0.011876 | 0.007125 | 0.015947 | 0.001443 | 0.003434 | 0.011572 | 0.015365 | 0.009697 | 0.038843 | 0.009673 | 0.015107 | 0.007969 | 0.01265 | 0.002598 | 0.006308 | 0.013636 | 0.00763 | 0.004799 | 0.013444 | 0.0067 | 0.021616 | 0.01579 | 0.004661 |
| 22.1111111 | 0.004217 | 0.004981 | 0.007495 | 0.013494 | 0.000874 | 0.001783 | 0.007771 | 0.019537 | 0.007715 | 0.065422 | 0.005667 | 0.018422 | 0.00649 | 0.008388 | 0.004856 | 0.009436 | 0.008248 | 0.005794 | 0.00208 | 0.006568 | 0.005927 | 0.011574 | 0.020398 | 0.004718 |
| 22.125 | 0.004489 | 0.004706 | 0.005091 | 0.00864 | 0.001107 | 0.001227 | 0.008933 | 0.017143 | 0.004743 | 0.077931 | 0.002446 | 0.019226 | 0.00589 | 0.011177 | 0.005731 | 0.008936 | 0.002233 | 0.005548 | 0.001963 | 0.009047 | 0.00499 | 0.007369 | 0.018453 | 0.002874 |
| 22.1388889 | 0.002824 | 0.004647 | 0.002645 | 0.00449 | 0.001401 | 0.001221 | 0.011084 | 0.010043 | 0.002918 | 0.072358 | 0.001109 | 0.011022 | 0.005236 | 0.018355 | 0.005033 | 0.005802 | 0.002239 | 0.00646 | 0.002507 | 0.013273 | 0.005447 | 0.006738 | 0.013204 | 0.001322 |
| 22.1527778 | 0.002337 | 0.003445 | 0.001453 | 0.002021 | 0.001053 | 0.00111 | 0.009152 | 0.004896 | 0.002458 | 0.054937 | 0.001008 | 0.004931 | 0.004136 | 0.021114 | 0.003588 | 0.00315 | 0.002415 | 0.006301 | 0.003671 | 0.015635 | 0.005858 | 0.007069 | 0.010514 | 0.000615 |
| 22.1666667 | 0.002417 | 0.004043 | 0.00101 | 0.000851 | 0.001081 | 0.000861 | 0.007366 | 0.007023 | 0.002791 | 0.033633 | 0.001294 | 0.005297 | 0.002454 | 0.016615 | 0.002115 | 0.004226 | 0.002416 | 0.005136 | 0.004865 | 0.018107 | 0.004861 | 0.006494 | 0.009547 | 0.000589 |
| 22.1805556 | 0.003358 | 0.005634 | 0.00061 | 0.000315 | 0.002158 | 0.001003 | 0.008335 | 0.009782 | 0.003344 | 0.017143 | 0.002303 | 0.006072 | 0.001121 | 0.009431 | 0.00108 | 0.006915 | 0.004578 | 0.003343 | 0.005229 | 0.019964 | 0.002813 | 0.00557 | 0.008858 | 0.00089 |
| 22.1944444 | 0.007735 | 0.006456 | 0.000279 | 0.000352 | 0.002767 | 0.001508 | 0.007152 | 0.007214 | 0.003658 | 0.016808 | 0.003845 | 0.00787 | 0.000816 | 0.004186 | 0.000826 | 0.007393 | 0.006186 | 0.001914 | 0.004848 | 0.020028 | 0.00117 | 0.00492 | 0.008432 | 0.001162 |
| 22.2083333 | 0.011767 | 0.005647 | 0.000149 | 0.000978 | 0.001921 | 0.001671 | 0.003626 | 0.002746 | 0.003586 | 0.027502 | 0.004639 | 0.011568 | 0.001155 | 0.002927 | 0.001124 | 0.004934 | 0.004788 | 0.002437 | 0.004885 | 0.019028 | 0.000584 | 0.003971 | 0.008156 | 0.001145 |
| 22.2222222 | 0.012025 | 0.003278 | 0.000192 | 0.001448 | 0.001031 | 0.00126 | 0.001993 | 0.001317 | 0.003623 | 0.029898 | 0.004245 | 0.009753 | 0.001739 | 0.004168 | 0.001343 | 0.002391 | 0.002643 | 0.003579 | 0.00622 | 0.017193 | 0.000508 | 0.002317 | 0.007347 | 0.000963 |
| 22.2361111 | 0.009018 | 0.001446 | 0.000294 | 0.00112 | 0.001009 | 0.000766 | 0.003404 | 0.003471 | 0.004145 | 0.019515 | 0.003333 | 0.006942 | 0.00241 | 0.004411 | 0.001358 | 0.002039 | 0.001832 | 0.003069 | 0.007924 | 0.014198 | 0.000857 | 0.00156 | 0.00564 | 0.00115 |
| 22.25 | 0.004868 | 0.001651 | 0.000232 | 0.000515 | 0.000853 | 0.000585 | 0.004486 | 0.006462 | 0.004573 | 0.010968 | 0.002717 | 0.011229 | 0.002832 | 0.003056 | 0.001043 | 0.004018 | 0.00318 | 0.001549 | 0.007938 | 0.01039 | 0.001649 | 0.002967 | 0.003524 | 0.001706 |
| 22.2638889 | 0.003368 | 0.002244 | 0.000207 | 0.000561 | 0.000582 | 0.000581 | 0.003729 | 0.006506 | 0.004138 | 0.008217 | 0.002579 | 0.014095 | 0.002698 | 0.002686 | 0.000763 | 0.008534 | 0.006537 | 0.000834 | 0.005769 | 0.006501 | 0.002105 | 0.004299 | 0.002557 | 0.001787 |
| 22.2777778 | 0.004394 | 0.002221 | 0.000704 | 0.001511 | 0.000942 | 0.000477 | 0.002422 | 0.004942 | 0.002844 | 0.007461 | 0.002476 | 0.009351 | 0.002265 | 0.004274 | 0.001385 | 0.010527 | 0.008777 | 0.000724 | 0.003232 | 0.003243 | 0.001932 | 0.003894 | 0.003737 | 0.001909 |
| 22.2916667 | 0.004491 | 0.002154 | 0.002064 | 0.002811 | 0.001275 | 0.00031 | 0.002221 | 0.006442 | 0.001383 | 0.011853 | 0.001757 | 0.005162 | 0.001755 | 0.004747 | 0.002511 | 0.008331 | 0.006913 | 0.0006 | 0.001847 | 0.001775 | 0.002014 | 0.003387 | 0.006027 | 0.002581 |
| 22.3055556 | 0.003619 | 0.001814 | 0.004214 | 0.003862 | 0.000966 | 0.000174 | 0.005168 | 0.007677 | 0.000972 | 0.013149 | 0.001611 | 0.006052 | 0.001161 | 0.002929 | 0.003377 | 0.009568 | 0.003167 | 0.000736 | 0.001518 | 0.0016 | 0.003269 | 0.00468 | 0.008006 | 0.002462 |
| 22.3194444 | 0.006149 | 0.001419 | 0.006498 | 0.00484 | 0.000757 | 0.000194 | 0.009671 | 0.005126 | 0.002518 | 0.011425 | 0.00397 | 0.006934 | 0.001778 | 0.00111 | 0.003926 | 0.013809 | 0.002238 | 0.000961 | 0.00203 | 0.002711 | 0.00537 | 0.007799 | 0.009082 | 0.002081 |
| 22.3333333 | 0.014186 | 0.002502 | 0.008586 | 0.006039 | 0.001414 | 0.000677 | 0.013117 | 0.00257 | 0.005752 | 0.022575 | 0.006922 | 0.008057 | 0.004658 | 0.001731 | 0.004154 | 0.013014 | 0.006733 | 0.002321 | 0.003912 | 0.00846 | 0.006875 | 0.011615 | 0.009461 | 0.002991 |
| 22.3472222 | 0.022876 | 0.005196 | 0.01046 | 0.007284 | 0.002364 | 0.00156 | 0.015018 | 0.003391 | 0.009879 | 0.039585 | 0.007458 | 0.013437 | 0.007762 | 0.005554 | 0.003914 | 0.007076 | 0.013837 | 0.005926 | 0.006828 | 0.017625 | 0.006602 | 0.015006 | 0.009644 | 0.004376 |
| 22.3611111 | 0.024959 | 0.006862 | 0.011595 | 0.008126 | 0.002552 | 0.002227 | 0.015001 | 0.008761 | 0.013623 | 0.040151 | 0.005695 | 0.021301 | 0.00887 | 0.009573 | 0.003366 | 0.002697 | 0.016701 | 0.010084 | 0.008785 | 0.024984 | 0.004914 | 0.017827 | 0.010056 | 0.00457 |
| 22.375 | 0.01933 | 0.006177 | 0.012287 | 0.008633 | 0.002293 | 0.002607 | 0.014013 | 0.015843 | 0.016026 | 0.024347 | 0.004116 | 0.027226 | 0.008704 | 0.010642 | 0.003773 | 0.001982 | 0.014422 | 0.012769 | 0.009118 | 0.028396 | 0.003711 | 0.02204 | 0.01229 | 0.004384 |
| 22.3888889 | 0.012638 | 0.004918 | 0.013729 | 0.009457 | 0.002972 | 0.003551 | 0.014441 | 0.017623 | 0.01738 | 0.010561 | 0.005168 | 0.027584 | 0.009134 | 0.009502 | 0.006227 | 0.002721 | 0.012249 | 0.013993 | 0.009681 | 0.028234 | 0.004497 | 0.029065 | 0.01727 | 0.006369 |
| 22.4027778 | 0.01067 | 0.004249 | 0.015345 | 0.010061 | 0.00418 | 0.005092 | 0.016384 | 0.011361 | 0.018167 | 0.012386 | 0.008657 | 0.023076 | 0.010891 | 0.00772 | 0.00903 | 0.004357 | 0.012728 | 0.014048 | 0.011655 | 0.02506 | 0.00645 | 0.037239 | 0.02089 | 0.011029 |
| 22.4166667 | 0.014309 | 0.004139 | 0.015679 | 0.00898 | 0.004098 | 0.006273 | 0.018204 | 0.0041 | 0.018999 | 0.032786 | 0.011856 | 0.018649 | 0.01308 | 0.005799 | 0.009254 | 0.004916 | 0.014896 | 0.012276 | 0.014489 | 0.020157 | 0.007897 | 0.044405 | 0.018232 | 0.015429 |
| 22.4305556 | 0.021905 | 0.004893 | 0.014704 | 0.006193 | 0.002586 | 0.006845 | 0.019543 | 0.003299 | 0.021127 | 0.053656 | 0.012388 | 0.018035 | 0.013574 | 0.004116 | 0.006755 | 0.003158 | 0.016222 | 0.008442 | 0.017583 | 0.016594 | 0.008411 | 0.049453 | 0.011584 | 0.016597 |
| 22.4444444 | 0.029707 | 0.0062 | 0.013791 | 0.003284 | 0.001067 | 0.00716 | 0.019724 | 0.00893 | 0.024632 | 0.051441 | 0.011044 | 0.020188 | 0.011536 | 0.003825 | 0.003561 | 0.001367 | 0.015182 | 0.003914 | 0.018985 | 0.01675 | 0.007907 | 0.052874 | 0.006012 | 0.014389 |
| 22.4583333 | 0.032826 | 0.006843 | 0.01453 | 0.002027 | 0.000668 | 0.007486 | 0.017677 | 0.016709 | 0.027597 | 0.028861 | 0.01039 | 0.02141 | 0.00891 | 0.005821 | 0.001399 | 0.001022 | 0.013571 | 0.000981 | 0.016003 | 0.019453 | 0.00598 | 0.057164 | 0.003914 | 0.010374 |
| 22.4722222 | 0.028766 | 0.006527 | 0.016591 | 0.002477 | 0.001202 | 0.007775 | 0.01477 | 0.018643 | 0.029679 | 0.013259 | 0.011852 | 0.019968 | 0.008627 | 0.008426 | 0.000939 | 0.001039 | 0.013911 | 0.000969 | 0.010288 | 0.020618 | 0.004002 | 0.065669 | 0.007379 | 0.008302 |
| 22.4861111 | 0.020755 | 0.007554 | 0.01931 | 0.004259 | 0.002034 | 0.008687 | 0.013104 | 0.012752 | 0.03747 | 0.012531 | 0.017062 | 0.018752 | 0.013325 | 0.007961 | 0.002849 | 0.001143 | 0.018945 | 0.003918 | 0.006745 | 0.018535 | 0.006231 | 0.082313 | 0.01503 | 0.010879 |
| 22.5 | 0.017323 | 0.013872 | 0.026519 | 0.010877 | 0.003663 | 0.012695 | 0.0139 | 0.007032 | 0.061975 | 0.025555 | 0.028864 | 0.024627 | 0.025566 | 0.005062 | 0.00721 | 0.00264 | 0.031684 | 0.010056 | 0.007682 | 0.017646 | 0.017412 | 0.113488 | 0.02035 | 0.015846 |
| 22.5138889 | 0.024794 | 0.028026 | 0.041306 | 0.023568 | 0.00579 | 0.019214 | 0.019146 | 0.009092 | 0.087738 | 0.071624 | 0.041345 | 0.040521 | 0.042992 | 0.005146 | 0.012071 | 0.004255 | 0.05048 | 0.019999 | 0.010959 | 0.021341 | 0.040025 | 0.155963 | 0.020149 | 0.018228 |
| 22.5277778 | 0.037621 | 0.04187 | 0.052625 | 0.032345 | 0.005455 | 0.022023 | 0.030875 | 0.017448 | 0.081234 | 0.119059 | 0.040993 | 0.058068 | 0.053799 | 0.007824 | 0.013783 | 0.004961 | 0.065196 | 0.028408 | 0.011861 | 0.0277 | 0.060217 | 0.16706 | 0.017383 | 0.015096 |
| 22.5416667 | 0.045411 | 0.042978 | 0.051061 | 0.029573 | 0.002494 | 0.020233 | 0.037893 | 0.020206 | 0.052883 | 0.11608 | 0.029142 | 0.063438 | 0.053015 | 0.00742 | 0.01116 | 0.005944 | 0.065907 | 0.028274 | 0.008673 | 0.033496 | 0.063647 | 0.117306 | 0.014212 | 0.008663 |
| 22.5555556 | 0.042018 | 0.03476 | 0.046213 | 0.021399 | 0.00185 | 0.018928 | 0.025779 | 0.013577 | 0.033236 | 0.076412 | 0.018003 | 0.049971 | 0.051046 | 0.004773 | 0.009321 | 0.005129 | 0.053918 | 0.023433 | 0.005565 | 0.033185 | 0.063379 | 0.050567 | 0.012301 | 0.00391 |
| 22.5694444 | 0.026286 | 0.028618 | 0.042697 | 0.013443 | 0.004688 | 0.01662 | 0.014563 | 0.007072 | 0.020833 | 0.035836 | 0.01092 | 0.026036 | 0.050482 | 0.004746 | 0.009675 | 0.004173 | 0.03398 | 0.017751 | 0.003942 | 0.026322 | 0.062445 | 0.019867 | 0.012877 | 0.003675 |
| 22.5833333 | 0.01414 | 0.027222 | 0.035134 | 0.006949 | 0.006473 | 0.010819 | 0.024636 | 0.003847 | 0.009362 | 0.01937 | 0.006312 | 0.012675 | 0.04379 | 0.00518 | 0.007534 | 0.007414 | 0.018145 | 0.010461 | 0.003587 | 0.018629 | 0.046306 | 0.028985 | 0.013108 | 0.00577 |
| 22.5972222 | 0.021683 | 0.027106 | 0.024349 | 0.004888 | 0.005931 | 0.005318 | 0.038495 | 0.00427 | 0.005157 | 0.037393 | 0.004266 | 0.016098 | 0.029318 | 0.003596 | 0.003954 | 0.00924 | 0.022376 | 0.007389 | 0.006226 | 0.014886 | 0.021889 | 0.049486 | 0.01082 | 0.006003 |
| 22.6111111 | 0.036312 | 0.028276 | 0.013672 | 0.005581 | 0.00604 | 0.003633 | 0.038932 | 0.011647 | 0.009502 | 0.06426 | 0.003703 | 0.015437 | 0.013316 | 0.003578 | 0.004001 | 0.006277 | 0.040689 | 0.009672 | 0.00675 | 0.023167 | 0.006915 | 0.060877 | 0.006568 | 0.006951 |
| 22.625 | 0.0392 | 0.032261 | 0.005845 | 0.004453 | 0.00761 | 0.004399 | 0.024629 | 0.016057 | 0.014105 | 0.056527 | 0.002854 | 0.011482 | 0.005892 | 0.005717 | 0.006833 | 0.002928 | 0.051868 | 0.008307 | 0.005518 | 0.038688 | 0.003948 | 0.059363 | 0.003908 | 0.011651 |
| 22.6388889 | 0.025956 | 0.037873 | 0.004004 | 0.004079 | 0.008727 | 0.003994 | 0.015359 | 0.011974 | 0.011381 | 0.037369 | 0.002169 | 0.02268 | 0.006717 | 0.006357 | 0.006866 | 0.001838 | 0.042579 | 0.006611 | 0.010388 | 0.040304 | 0.007082 | 0.049156 | 0.007073 | 0.014816 |
| 22.6527778 | 0.01546 | 0.042903 | 0.007612 | 0.007224 | 0.008285 | 0.002037 | 0.031583 | 0.014059 | 0.004988 | 0.057359 | 0.002338 | 0.038419 | 0.00887 | 0.004179 | 0.004754 | 0.003193 | 0.021106 | 0.013595 | 0.016772 | 0.025974 | 0.012104 | 0.037491 | 0.015265 | 0.012446 |
| 22.6666667 | 0.027978 | 0.044713 | 0.012559 | 0.010255 | 0.006297 | 0.000785 | 0.044637 | 0.020156 | 0.002771 | 0.077215 | 0.003704 | 0.04134 | 0.011901 | 0.002457 | 0.006929 | 0.005921 | 0.011749 | 0.020015 | 0.015577 | 0.020114 | 0.017083 | 0.02307 | 0.022967 | 0.007604 |
| 22.6805556 | 0.045099 | 0.042227 | 0.014673 | 0.011368 | 0.003325 | 0.000914 | 0.028255 | 0.015942 | 0.004418 | 0.058841 | 0.006741 | 0.027872 | 0.011849 | 0.003557 | 0.012099 | 0.008237 | 0.024938 | 0.016444 | 0.008914 | 0.030502 | 0.023256 | 0.014542 | 0.022953 | 0.006793 |
| 22.6944444 | 0.047451 | 0.036026 | 0.010762 | 0.010855 | 0.001144 | 0.00135 | 0.012967 | 0.006801 | 0.005011 | 0.027862 | 0.010332 | 0.013153 | 0.010623 | 0.004777 | 0.012818 | 0.008019 | 0.043224 | 0.008222 | 0.005712 | 0.037632 | 0.027908 | 0.028896 | 0.013947 | 0.011141 |

| | | | | | | | | | | | | | | | | | | | | | | | | |
|---|---|---|---|---|---|---|---|---|---|---|---|---|---|---|---|---|---|---|---|---|---|---|---|---|
| 22.7083333 | 0.040559 | 0.02652 | 0.00425 | 0.007755 | 0.000542 | 0.001389 | 0.025119 | 0.002251 | 0.005688 | 0.012679 | 0.012413 | 0.014187 | 0.018949 | 0.004401 | 0.007767 | 0.005473 | 0.049875 | 0.003783 | 0.007545 | 0.03433 | 0.025168 | 0.051677 | 0.006145 | 0.014814 |
| 22.7222222 | 0.027688 | 0.014658 | 0.001991 | 0.00432 | 0.000855 | 0.002154 | 0.048183 | 0.003187 | 0.010915 | 0.021072 | 0.011672 | 0.022284 | 0.028324 | 0.003011 | 0.003603 | 0.002757 | 0.046017 | 0.006529 | 0.009254 | 0.025019 | 0.016134 | 0.057263 | 0.007629 | 0.011965 |
| 22.7361111 | 0.016455 | 0.006337 | 0.003339 | 0.00463 | 0.001897 | 0.004421 | 0.054628 | 0.007793 | 0.016736 | 0.042934 | 0.008178 | 0.029851 | 0.02534 | 0.001362 | 0.005315 | 0.001882 | 0.035059 | 0.012219 | 0.008184 | 0.013721 | 0.007086 | 0.045367 | 0.013158 | 0.00793 |
| 22.75 | 0.021796 | 0.006596 | 0.003882 | 0.007085 | 0.002677 | 0.006129 | 0.034745 | 0.015078 | 0.017356 | 0.066524 | 0.0039 | 0.03488 | 0.014667 | 0.000867 | 0.009186 | 0.002215 | 0.020317 | 0.014947 | 0.005529 | 0.010754 | 0.003861 | 0.028707 | 0.015539 | 0.010599 |
| 22.7638889 | 0.037456 | 0.011948 | 0.002653 | 0.006517 | 0.002096 | 0.005752 | 0.017134 | 0.018899 | 0.013687 | 0.073838 | 0.002269 | 0.027028 | 0.005477 | 0.001842 | 0.010302 | 0.001766 | 0.014571 | 0.011868 | 0.00527 | 0.018372 | 0.007262 | 0.015343 | 0.013754 | 0.01359 |
| 22.7777778 | 0.045178 | 0.01712 | 0.002209 | 0.003844 | 0.001514 | 0.003733 | 0.0231 | 0.013091 | 0.009089 | 0.050681 | 0.004186 | 0.016372 | 0.001704 | 0.003474 | 0.008472 | 0.001667 | 0.024962 | 0.005435 | 0.005926 | 0.026874 | 0.010755 | 0.006946 | 0.010086 | 0.010077 |
| 22.7916667 | 0.041598 | 0.019597 | 0.003953 | 0.004223 | 0.002131 | 0.00269 | 0.024958 | 0.006829 | 0.005697 | 0.026836 | 0.006888 | 0.022756 | 0.003279 | 0.004556 | 0.005059 | 0.001782 | 0.040421 | 0.001482 | 0.005341 | 0.029237 | 0.012125 | 0.002762 | 0.005836 | 0.00692 |
| 22.8055556 | 0.028504 | 0.019308 | 0.006411 | 0.006757 | 0.002059 | 0.004541 | 0.015594 | 0.010793 | 0.003836 | 0.038643 | 0.007804 | 0.030554 | 0.008769 | 0.003868 | 0.002521 | 0.001226 | 0.05202 | 0.001968 | 0.008425 | 0.024238 | 0.013651 | 0.00464 | 0.007092 | 0.012418 |
| 22.8194444 | 0.012949 | 0.017921 | 0.009506 | 0.006852 | 0.001325 | 0.00702 | 0.020772 | 0.016956 | 0.003441 | 0.0571 | 0.006633 | 0.020126 | 0.016631 | 0.002014 | 0.002754 | 0.00174 | 0.059495 | 0.00556 | 0.015763 | 0.015702 | 0.012688 | 0.018712 | 0.017699 | 0.020456 |
| 22.8333333 | 0.00504 | 0.01768 | 0.013782 | 0.004787 | 0.001411 | 0.008836 | 0.042241 | 0.013555 | 0.004108 | 0.042664 | 0.00512 | 0.009958 | 0.024909 | 0.001412 | 0.003425 | 0.003837 | 0.059441 | 0.009767 | 0.021394 | 0.010844 | 0.008639 | 0.044198 | 0.026604 | 0.025707 |
| 22.8472222 | 0.004693 | 0.020635 | 0.017978 | 0.005276 | 0.001265 | 0.010009 | 0.05351 | 0.010422 | 0.007924 | 0.021207 | 0.005526 | 0.017652 | 0.02986 | 0.003416 | 0.002713 | 0.006159 | 0.047993 | 0.011752 | 0.02252 | 0.011504 | 0.006551 | 0.069729 | 0.025958 | 0.026444 |
| 22.8611111 | 0.010769 | 0.026732 | 0.021801 | 0.008623 | 0.001729 | 0.009728 | 0.039724 | 0.018621 | 0.017218 | 0.031852 | 0.009154 | 0.031623 | 0.030242 | 0.007708 | 0.003393 | 0.007119 | 0.034128 | 0.010621 | 0.018686 | 0.014237 | 0.009162 | 0.083753 | 0.018721 | 0.017682 |
| 22.875 | 0.022181 | 0.03355 | 0.026101 | 0.009603 | 0.004217 | 0.008601 | 0.017848 | 0.025645 | 0.029265 | 0.069445 | 0.014274 | 0.03779 | 0.029674 | 0.010892 | 0.007652 | 0.00635 | 0.028258 | 0.008486 | 0.010661 | 0.018745 | 0.014268 | 0.084736 | 0.01072 | 0.006739 |
| 22.8888889 | 0.039441 | 0.037381 | 0.030096 | 0.006464 | 0.006386 | 0.007897 | 0.005547 | 0.020609 | 0.03821 | 0.103013 | 0.016112 | 0.034697 | 0.029209 | 0.008354 | 0.01355 | 0.005851 | 0.02663 | 0.007868 | 0.004926 | 0.025671 | 0.0185 | 0.078593 | 0.009418 | 0.002935 |
| 22.9027778 | 0.052014 | 0.035922 | 0.032798 | 0.003039 | 0.006083 | 0.007686 | 0.003738 | 0.012443 | 0.03842 | 0.10386 | 0.013189 | 0.027272 | 0.026564 | 0.004362 | 0.017264 | 0.007747 | 0.024628 | 0.008704 | 0.003851 | 0.030797 | 0.019767 | 0.065468 | 0.017256 | 0.00296 |
| 22.9166667 | 0.044113 | 0.029605 | 0.034541 | 0.002409 | 0.004333 | 0.007732 | 0.007906 | 0.007484 | 0.032148 | 0.074376 | 0.009379 | 0.019912 | 0.020795 | 0.004943 | 0.017071 | 0.011815 | 0.024008 | 0.009788 | 0.003569 | 0.029498 | 0.018566 | 0.043406 | 0.023288 | 0.006033 |
| 22.9305556 | 0.02534 | 0.018394 | 0.034581 | 0.002633 | 0.002772 | 0.007966 | 0.016365 | 0.005256 | 0.02701 | 0.036497 | 0.007747 | 0.014281 | 0.013246 | 0.005734 | 0.015489 | 0.016086 | 0.023401 | 0.010693 | 0.006305 | 0.022559 | 0.016065 | 0.01974 | 0.018245 | 0.017547 |
| 22.9444444 | 0.011001 | 0.008207 | 0.031128 | 0.001728 | 0.002002 | 0.007922 | 0.024941 | 0.007067 | 0.025223 | 0.012826 | 0.008212 | 0.012149 | 0.006814 | 0.004987 | 0.015068 | 0.018648 | 0.019808 | 0.011291 | 0.013908 | 0.014965 | 0.011909 | 0.00637 | 0.008754 | 0.034606 |
| 22.9583333 | 0.004484 | 0.006621 | 0.024741 | 0.001106 | 0.00197 | 0.007517 | 0.025835 | 0.008242 | 0.024107 | 0.01066 | 0.010254 | 0.013876 | 0.003125 | 0.006889 | 0.014683 | 0.017 | 0.015793 | 0.011095 | 0.020745 | 0.008494 | 0.006622 | 0.003766 | 0.002925 | 0.044564 |
| 22.9722222 | 0.003141 | 0.007908 | 0.01765 | 0.001684 | 0.00229 | 0.007178 | 0.01663 | 0.006594 | 0.021316 | 0.013128 | 0.012739 | 0.016276 | 0.001551 | 0.011269 | 0.012919 | 0.010223 | 0.013925 | 0.009075 | 0.021866 | 0.004574 | 0.004087 | 0.002512 | 0.00111 | 0.039542 |
| 22.9861111 | 0.005184 | 0.005627 | 0.012294 | 0.00299 | 0.002725 | 0.006918 | 0.008627 | 0.004688 | 0.018404 | 0.022139 | 0.014137 | 0.016761 | 0.001296 | 0.014811 | 0.01081 | 0.00376 | 0.013491 | 0.005516 | 0.017225 | 0.004375 | 0.00463 | 0.001903 | 0.002349 | 0.025616 |
| 23 | 0.012288 | 0.004576 | 0.011762 | 0.005929 | 0.003474 | 0.006638 | 0.009523 | 0.00541 | 0.018607 | 0.046169 | 0.016126 | 0.016267 | 0.003699 | 0.015147 | 0.009047 | 0.001826 | 0.014768 | 0.002445 | 0.01165 | 0.006264 | 0.005145 | 0.004658 | 0.006881 | 0.013218 |
| 23.0138889 | 0.023295 | 0.009687 | 0.016832 | 0.009385 | 0.004435 | 0.007114 | 0.009408 | 0.009207 | 0.022652 | 0.073286 | 0.024473 | 0.020254 | 0.009092 | 0.013316 | 0.006464 | 0.00168 | 0.021545 | 0.002287 | 0.010482 | 0.015031 | 0.009311 | 0.005887 | 0.012985 | 0.006488 |
| 23.0277778 | 0.036554 | 0.01581 | 0.026303 | 0.00965 | 0.00484 | 0.009597 | 0.010575 | 0.013482 | 0.027014 | 0.099441 | 0.041528 | 0.03497 | 0.013005 | 0.011167 | 0.003099 | 0.001156 | 0.037615 | 0.005958 | 0.012202 | 0.030956 | 0.018573 | 0.006932 | 0.01996 | 0.004909 |
| 23.0416667 | 0.050477 | 0.018505 | 0.038074 | 0.007931 | 0.004627 | 0.012652 | 0.023302 | 0.023681 | 0.025997 | 0.12538 | 0.056949 | 0.057189 | 0.014154 | 0.007923 | 0.001801 | 0.002551 | 0.067705 | 0.01203 | 0.012989 | 0.0465 | 0.025414 | 0.016353 | 0.032052 | 0.007962 |
| 23.0555556 | 0.056895 | 0.021447 | 0.042799 | 0.007223 | 0.004264 | 0.012405 | 0.037991 | 0.047008 | 0.01964 | 0.131461 | 0.056757 | 0.07146 | 0.01572 | 0.007753 | 0.004707 | 0.007007 | 0.098713 | 0.01859 | 0.012269 | 0.060572 | 0.025018 | 0.032511 | 0.05125 | 0.014389 |
| 23.0694444 | 0.047071 | 0.024271 | 0.031708 | 0.005644 | 0.0032 | 0.008939 | 0.038734 | 0.069334 | 0.013651 | 0.099629 | 0.041124 | 0.064392 | 0.015212 | 0.014222 | 0.008648 | 0.008346 | 0.093122 | 0.018171 | 0.008393 | 0.066503 | 0.020001 | 0.044522 | 0.05974 | 0.017885 |
| 23.0833333 | 0.025097 | 0.021898 | 0.016186 | 0.002981 | 0.001811 | 0.006047 | 0.023588 | 0.062569 | 0.011053 | 0.048678 | 0.021395 | 0.040266 | 0.009106 | 0.01841 | 0.0088 | 0.008282 | 0.05155 | 0.012505 | 0.00453 | 0.052567 | 0.013112 | 0.043356 | 0.043508 | 0.012503 |
| 23.0972222 | 0.01008 | 0.01478 | 0.01329 | 0.002129 | 0.002371 | 0.004766 | 0.01079 | 0.034128 | 0.010897 | 0.023083 | 0.012117 | 0.018529 | 0.002731 | 0.01435 | 0.005114 | 0.014702 | 0.030253 | 0.015313 | 0.006268 | 0.027118 | 0.006368 | 0.031964 | 0.019798 | 0.006675 |
| 23.1111111 | 0.011162 | 0.007657 | 0.018365 | 0.002283 | 0.004798 | 0.003731 | 0.012963 | 0.022267 | 0.012198 | 0.036613 | 0.016775 | 0.013984 | 0.000742 | 0.007584 | 0.002354 | 0.019519 | 0.049961 | 0.023156 | 0.009548 | 0.010044 | 0.003239 | 0.01975 | 0.008619 | 0.009878 |
| 23.125 | 0.013999 | 0.003326 | 0.019025 | 0.002022 | 0.005695 | 0.002705 | 0.016481 | 0.032339 | 0.013621 | 0.057784 | 0.021514 | 0.020023 | 0.000757 | 0.005268 | 0.00285 | 0.016162 | 0.066908 | 0.025727 | 0.010162 | 0.007719 | 0.002483 | 0.011292 | 0.010089 | 0.014933 |
| 23.1388889 | 0.01194 | 0.001809 | 0.014842 | 0.00165 | 0.004011 | 0.002009 | 0.013602 | 0.041603 | 0.012466 | 0.067739 | 0.019846 | 0.022828 | 0.000809 | 0.00772 | 0.003039 | 0.010107 | 0.057926 | 0.0234 | 0.009639 | 0.012263 | 0.001701 | 0.006994 | 0.011532 | 0.014577 |
| 23.1527778 | 0.010255 | 0.001458 | 0.009535 | 0.00104 | 0.003261 | 0.001596 | 0.011457 | 0.04276 | 0.00846 | 0.070578 | 0.016482 | 0.018745 | 0.000776 | 0.010165 | 0.001748 | 0.004887 | 0.039795 | 0.01913 | 0.009412 | 0.015328 | 0.001803 | 0.004221 | 0.008564 | 0.009069 |
| 23.1666667 | 0.009724 | 0.001983 | 0.004559 | 0.000437 | 0.00403 | 0.001472 | 0.013694 | 0.039398 | 0.00467 | 0.067845 | 0.0144 | 0.011422 | 0.001139 | 0.01103 | 0.001232 | 0.001954 | 0.025857 | 0.014469 | 0.008957 | 0.016048 | 0.001917 | 0.00235 | 0.00552 | 0.005127 |
| 23.1805556 | 0.009844 | 0.003995 | 0.001317 | 0.000191 | 0.003571 | 0.001443 | 0.017918 | 0.034759 | 0.003056 | 0.062001 | 0.013309 | 0.005803 | 0.001513 | 0.010562 | 0.001272 | 0.001933 | 0.017902 | 0.009587 | 0.006815 | 0.016925 | 0.001558 | 0.003139 | 0.003923 | 0.009164 |
| 23.1944444 | 0.011138 | 0.005727 | 0.000309 | 0.000198 | 0.004214 | 0.001115 | 0.01821 | 0.031098 | 0.002585 | 0.055843 | 0.011749 | 0.003235 | 0.001338 | 0.008745 | 0.001013 | 0.002777 | 0.013522 | 0.004762 | 0.003511 | 0.018331 | 0.00248 | 0.005019 | 0.004099 | 0.015236 |
| 23.2083333 | 0.013853 | 0.005592 | 0.000324 | 0.000365 | 0.006773 | 0.000625 | 0.011606 | 0.028117 | 0.001762 | 0.047595 | 0.009215 | 0.001729 | 0.001241 | 0.006178 | 0.001278 | 0.002079 | 0.00985 | 0.001995 | 0.001421 | 0.019277 | 0.004166 | 0.005474 | 0.00485 | 0.016391 |
| 23.2222222 | 0.016048 | 0.004275 | 0.000288 | 0.000562 | 0.006328 | 0.000422 | 0.006799 | 0.025194 | 0.000805 | 0.035521 | 0.006424 | 0.000505 | 0.001302 | 0.003714 | 0.001773 | 0.000795 | 0.006491 | 0.002237 | 0.00108 | 0.019176 | 0.005269 | 0.004813 | 0.005045 | 0.011164 |
| 23.2361111 | 0.015616 | 0.002875 | 0.000202 | 0.00052 | 0.003255 | 0.000574 | 0.013221 | 0.02165 | 0.001133 | 0.02118 | 0.004044 | 0.000402 | 0.001392 | 0.002309 | 0.001364 | 0.002005 | 0.004679 | 0.002902 | 0.00094 | 0.018053 | 0.005745 | 0.004943 | 0.005083 | 0.00572 |
| 23.25 | 0.01273 | 0.001965 | 0.000737 | 0.000636 | 0.001679 | 0.00101 | 0.021892 | 0.01649 | 0.002576 | 0.009778 | 0.00196 | 0.001588 | 0.001854 | 0.002227 | 0.000867 | 0.007049 | 0.00424 | 0.002214 | 0.000451 | 0.015844 | 0.005418 | 0.006899 | 0.005063 | 0.006281 |
| 23.2638889 | 0.008677 | 0.001523 | 0.002154 | 0.001649 | 0.001249 | 0.00144 | 0.020804 | 0.00923 | 0.002557 | 0.007379 | 0.000839 | 0.00405 | 0.002443 | 0.002939 | 0.001156 | 0.013685 | 0.003676 | 0.001787 | 0.000217 | 0.012395 | 0.004396 | 0.009522 | 0.004739 | 0.006036 |
| 23.2777778 | 0.004722 | 0.001359 | 0.003299 | 0.002881 | 0.000754 | 0.001258 | 0.011596 | 0.003558 | 0.001049 | 0.011618 | 0.0011 | 0.008513 | 0.002477 | 0.003525 | 0.001639 | 0.017355 | 0.002676 | 0.001994 | 0.000316 | 0.008652 | 0.003393 | 0.011328 | 0.003904 | 0.003837 |
| 23.2916667 | 0.002095 | 0.001288 | 0.002769 | 0.003006 | 0.000513 | 0.001013 | 0.004314 | 0.002944 | 0.000337 | 0.015352 | 0.001822 | 0.014701 | 0.002077 | 0.003373 | 0.001853 | 0.014478 | 0.001913 | 0.001755 | 0.000442 | 0.005671 | 0.002922 | 0.011913 | 0.002357 | 0.004963 |
| 23.3055556 | 0.00139 | 0.001151 | 0.001449 | 0.001868 | 0.000433 | 0.001387 | 0.003419 | 0.004863 | 0.000805 | 0.014102 | 0.00202 | 0.018863 | 0.001576 | 0.002711 | 0.001626 | 0.008277 | 0.002228 | 0.002192 | 0.000686 | 0.003562 | 0.002325 | 0.011757 | 0.001214 | 0.00547 |
| 23.3194444 | 0.002249 | 0.001068 | 0.001127 | 0.000825 | 0.000471 | 0.001421 | 0.002965 | 0.00566 | 0.002276 | 0.009465 | 0.001649 | 0.018222 | 0.001162 | 0.002097 | 0.000962 | 0.006444 | 0.004242 | 0.002935 | 0.001335 | 0.001976 | 0.001924 | 0.012074 | 0.001276 | 0.006737 |
| 23.3333333 | 0.00421 | 0.001014 | 0.001631 | 0.000771 | 0.000752 | 0.000764 | 0.002021 | 0.005057 | 0.004418 | 0.009397 | 0.000951 | 0.015353 | 0.00122 | 0.002178 | 0.000619 | 0.007464 | 0.005388 | 0.002663 | 0.001636 | 0.001355 | 0.003759 | 0.013769 | 0.002553 | 0.013463 |
| 23.3472222 | 0.007248 | 0.000874 | 0.002533 | 0.000837 | 0.001437 | 0.000254 | 0.004398 | 0.007214 | 0.006198 | 0.014688 | 0.000558 | 0.013718 | 0.001687 | 0.003333 | 0.000987 | 0.005501 | 0.004657 | 0.00395 | 0.001823 | 0.003207 | 0.007251 | 0.016325 | 0.004972 | 0.018763 |
| 23.3611111 | 0.011052 | 0.00093 | 0.003998 | 0.000941 | 0.001599 | 0.000294 | 0.009333 | 0.012773 | 0.006911 | 0.018593 | 0.001098 | 0.014816 | 0.002194 | 0.005167 | 0.001299 | 0.003245 | 0.004018 | 0.008736 | 0.00284 | 0.008283 | 0.009644 | 0.018789 | 0.006795 | 0.016886 |
| 23.375 | 0.014842 | 0.001561 | 0.005598 | 0.001926 | 0.001729 | 0.000756 | 0.014365 | 0.015262 | 0.006982 | 0.020637 | 0.001825 | 0.018612 | 0.002428 | 0.006056 | 0.001172 | 0.002703 | 0.005493 | 0.013514 | 0.00373 | 0.015109 | 0.009445 | 0.020886 | 0.006849 | 0.011906 |
| 23.3888889 | 0.018318 | 0.002519 | 0.00706 | 0.003535 | 0.003924 | 0.002567 | 0.017638 | 0.012531 | 0.00731 | 0.025139 | 0.002116 | 0.023198 | 0.001837 | 0.004511 | 0.001205 | 0.003091 | 0.00942 | 0.014065 | 0.006381 | 0.021887 | 0.00718 | 0.022752 | 0.005536 | 0.009859 |
| 23.4027778 | 0.021235 | 0.003146 | 0.008354 | 0.004923 | 0.006147 | 0.00556 | 0.017462 | 0.00826 | 0.007769 | 0.031452 | 0.002675 | 0.025551 | 0.001096 | 0.001959 | 0.002756 | 0.005899 | 0.011479 | 0.010427 | 0.011379 | 0.027458 | 0.005272 | 0.024646 | 0.0039 | 0.014744 |
| 23.4166667 | 0.022355 | 0.003062 | 0.009094 | 0.005518 | 0.005124 | 0.005469 | 0.013972 | 0.00513 | 0.007746 | 0.033992 | 0.004129 | 0.023559 | 0.00119 | 0.000912 | 0.006494 | 0.008534 | 0.009834 | 0.005179 | 0.013733 | 0.028698 | 0.005467 | 0.026007 | 0.003459 | 0.023993 |
| 23.4305556 | 0.021761 | 0.002213 | 0.009736 | 0.005026 | 0.002994 | 0.003017 | 0.010616 | 0.003636 | 0.007857 | 0.032298 | 0.00633 | 0.01831 | 0.001492 | 0.001705 | 0.009711 | 0.007775 | 0.008502 | 0.001698 | 0.011736 | 0.024072 | 0.007391 | 0.026769 | 0.00495 | 0.02724 |
| 23.4444444 | 0.021902 | 0.001078 | 0.011566 | 0.00405 | 0.003189 | 0.003523 | 0.010662 | 0.00302 | 0.0098 | 0.032547 | 0.008313 | 0.013565 | 0.002046 | 0.003366 | 0.009512 | 0.004867 | 0.008195 | 0.001374 | 0.008785 | 0.01796 | 0.010033 | 0.028414 | 0.008106 | 0.020414 |
| 23.4583333 | 0.0239 | 0.000381 | 0.013899 | 0.003867 | 0.003505 | 0.005714 | 0.013121 | 0.004194 | 0.013803 | 0.037624 | 0.008756 | 0.011339 | 0.002792 | 0.005168 | 0.006734 | 0.002963 | 0.008019 | 0.003056 | 0.008223 | 0.014537 | 0.011997 | 0.030482 | 0.011856 | 0.012627 |
| 23.4722222 | 0.026152 | 0.001076 | 0.01399 | 0.004596 | 0.003199 | 0.006716 | 0.013957 | 0.007684 | 0.017944 | 0.043447 | 0.008146 | 0.011163 | 0.003054 | 0.006053 | 0.004055 | 0.002714 | 0.010173 | 0.006099 | 0.010706 | 0.014752 | 0.012989 | 0.032325 | 0.014348 | 0.014616 |
| 23.4861111 | 0.028615 | 0.005131 | 0.012247 | 0.005785 | 0.005743 | 0.007331 | 0.012978 | 0.010536 | 0.025458 | 0.046637 | 0.010274 | 0.01586 | 0.005547 | 0.00508 | 0.004573 | 0.004049 | 0.015959 | 0.008106 | 0.014983 | 0.020242 | 0.017957 | 0.041172 | 0.016117 | 0.027727 |
| 23.5 | 0.036722 | 0.0152 | 0.015183 | 0.009446 | 0.009202 | 0.010716 | 0.016227 | 0.013738 | 0.046193 | 0.055848 | 0.019937 | 0.03156 | 0.015284 | 0.00311 | 0.010013 | 0.007866 | 0.028247 | 0.007848 | 0.019519 | 0.034215 | 0.035954 | 0.071598 | 0.021306 | 0.041579 |
| 23.5138889 | 0.049458 | 0.030583 | 0.025394 | 0.015834 | 0.009575 | 0.017172 | 0.025709 | 0.021301 | 0.073154 | 0.087491 | 0.03356 | 0.054046 | 0.032452 | 0.001798 | 0.017245 | 0.010736 | 0.043417 | 0.008489 | 0.021238 | 0.046661 | 0.068116 | 0.121566 | 0.031929 | 0.042424 |
| 23.5277778 | 0.057255 | 0.04054 | 0.032536 | 0.019501 | 0.007026 | 0.020985 | 0.037088 | 0.029866 | 0.082343 | 0.136952 | 0.039197 | 0.065282 | 0.045286 | 0.003061 | 0.020163 | 0.016281 | 0.049144 | 0.011703 | 0.017443 | 0.040515 | 0.08514 | 0.144607 | 0.039117 | 0.027445 |
| 23.5416667 | 0.05861 | 0.035435 | 0.029116 | 0.018463 | 0.003409 | 0.020049 | 0.039618 | 0.032954 | 0.07331 | 0.159745 | 0.034217 | 0.057798 | 0.043205 | 0.005834 | 0.01777 | 0.027575 | 0.043039 | 0.017088 | 0.010937 | 0.023811 | 0.06969 | 0.111091 | 0.031877 | 0.012468 |
| 23.5555556 | 0.051418 | 0.02569 | 0.023289 | 0.016173 | 0.000991 | 0.016423 | 0.026243 | 0.028561 | 0.06352 | 0.125257 | 0.025215 | 0.042633 | 0.0344 | 0.00517 | 0.016002 | 0.028091 | 0.033778 | 0.023065 | 0.006128 | 0.014588 | 0.052621 | 0.064837 | 0.018271 | 0.012677 |
| 23.5694444 | 0.031989 | 0.023621 | 0.024252 | 0.013708 | 0.000328 | 0.011364 | 0.019743 | 0.017935 | 0.054879 | 0.065899 | 0.015558 | 0.027153 | 0.029474 | 0.002316 | 0.0184 | 0.017815 | 0.028442 | 0.023147 | 0.003941 | 0.013222 | 0.04665 | 0.043742 | 0.011575 | 0.020766 |
| 23.5833333 | 0.014102 | 0.022895 | 0.03079 | 0.009823 | 0.000938 | 0.011809 | 0.032911 | 0.007955 | 0.043304 | 0.031446 | 0.007715 | 0.012644 | 0.026162 | 0.001281 | 0.022074 | 0.022504 | 0.022144 | 0.013807 | 0.00537 | 0.012485 | 0.039493 | 0.039102 | 0.011037 | 0.020731 |
| 23.5972222 | 0.012815 | 0.019261 | 0.034552 | 0.005013 | 0.003342 | 0.017239 | 0.04003 | 0.00603 | 0.035972 | 0.045771 | 0.004283 | 0.005647 | 0.02062 | 0.001388 | 0.022826 | 0.039498 | 0.013536 | 0.006759 | 0.009317 | 0.009565 | 0.032098 | 0.035438 | 0.01236 | 0.017484 |
| 23.6111111 | 0.022841 | 0.018536 | 0.029104 | 0.0022 | 0.005973 | 0.015268 | 0.028208 | 0.008268 | 0.038257 | 0.076896 | 0.004859 | 0.010901 | 0.014845 | 0.001679 | 0.017981 | 0.039873 | 0.014531 | 0.010902 | 0.009707 | 0.007655 | 0.029526 | 0.025565 | 0.014036 | 0.02612 |
| 23.625 | 0.031701 | 0.020595 | 0.018357 | 0.002239 | 0.006126 | 0.00788 | 0.017296 | 0.007959 | 0.040178 | 0.072325 | 0.007069 | 0.024224 | 0.01278 | 0.003021 | 0.009471 | 0.020989 | 0.020943 | 0.016551 | 0.007904 | 0.008276 | 0.029463 | 0.013772 | 0.014751 | 0.037781 |
| 23.6388889 | 0.033679 | 0.021175 | 0.009871 | 0.003131 | 0.003779 | 0.005211 | 0.024507 | 0.005353 | 0.036306 | 0.043404 | 0.009221 | 0.037765 | 0.017852 | 0.006329 | 0.004596 | 0.006466 | 0.017378 | 0.016005 | 0.012307 | 0.00656 | 0.026655 | 0.011745 | 0.013514 | 0.036136 |
| 23.6527778 | 0.02614 | 0.020028 | 0.004535 | 0.003127 | 0.002019 | 0.006188 | 0.037224 | 0.004157 | 0.03249 | 0.039456 | 0.010987 | 0.038636 | 0.025686 | 0.008117 | 0.004972 | 0.008782 | 0.008818 | 0.010857 | 0.020447 | 0.003875 | 0.021662 | 0.022133 | 0.010151 | 0.0223 |
| 23.6666667 | 0.01363 | 0.020233 | 0.001496 | 0.002741 | 0.002553 | 0.005387 | 0.037424 | 0.007995 | 0.028453 | 0.06826 | 0.012134 | 0.021922 | 0.027922 | 0.006763 | 0.004338 | 0.022976 | 0.005149 | 0.007176 | 0.021792 | 0.00598 | 0.017807 | 0.036279 | 0.005514 | 0.016073 |
| 23.6805556 | 0.009942 | 0.022373 | 0.000704 | 0.004728 | 0.002836 | 0.002665 | 0.024394 | 0.014388 | 0.022236 | 0.086026 | 0.012275 | 0.008973 | 0.027357 | 0.005688 | 0.002062 | 0.035996 | 0.003228 | 0.01325 | 0.013453 | 0.010684 | 0.018678 | 0.049434 | 0.003627 | 0.030078 |
| 23.6944444 | 0.018691 | 0.02175 | 0.001987 | 0.008492 | 0.002593 | 0.001174 | 0.012827 | 0.017345 | 0.013907 | 0.063704 | 0.010994 | 0.012251 | 0.026304 | 0.005795 | 0.001829 | 0.033279 | 0.003512 | 0.024888 | 0.007656 | 0.014733 | 0.025902 | 0.061838 | 0.007856 | 0.050347 |
| 23.7083333 | 0.026622 | 0.01525 | 0.005614 | 0.010209 | 0.004594 | 0.001596 | 0.018633 | 0.015418 | 0.009357 | 0.033511 | 0.007822 | 0.018079 | 0.020193 | 0.005005 | 0.003363 | 0.018278 | 0.011838 | 0.028152 | 0.014189 | 0.017503 | 0.032878 | 0.066638 | 0.014089 | 0.056536 |
| 23.7222222 | 0.021556 | 0.006783 | 0.009719 | 0.008875 | 0.007766 | 0.002036 | 0.03442 | 0.009907 | 0.017235 | 0.043031 | 0.004189 | 0.021171 | 0.010749 | 0.002866 | 0.004523 | 0.012877 | 0.025225 | 0.019665 | 0.023522 | 0.016704 | 0.031033 | 0.052924 | 0.01577 | 0.042785 |
| 23.7361111 | 0.012157 | 0.002568 | 0.012486 | 0.006331 | 0.008066 | 0.003551 | 0.03774 | 0.005702 | 0.0286 | 0.082107 | 0.003151 | 0.021978 | 0.004968 | 0.001553 | 0.003851 | 0.02408 | 0.032153 | 0.01035 | 0.023949 | 0.012182 | 0.022801 | 0.028083 | 0.01046 | 0.034318 |
| 23.75 | 0.014593 | 0.005243 | 0.013102 | 0.003843 | 0.005512 | 0.008302 | 0.022841 | 0.00913 | 0.029865 | 0.111581 | 0.004964 | 0.014848 | 0.006312 | 0.002398 | 0.003855 | 0.034556 | 0.024049 | 0.009675 | 0.016235 | 0.006699 | 0.015839 | 0.015217 | 0.005354 | 0.043357 |
| 23.7638889 | 0.025884 | 0.014013 | 0.01166 | 0.001868 | 0.005042 | 0.013302 | 0.00955 | 0.017728 | 0.023656 | 0.116434 | 0.00702 | 0.009289 | 0.009799 | 0.003983 | 0.006937 | 0.029181 | 0.011832 | 0.01461 | 0.012497 | 0.002609 | 0.010272 | 0.01618 | 0.00821 | 0.043332 |
| 23.7777778 | 0.036599 | 0.024745 | 0.009548 | 0.000766 | 0.007893 | 0.014079 | 0.009127 | 0.02292 | 0.017739 | 0.092984 | 0.006876 | 0.011022 | 0.008991 | 0.004048 | 0.009714 | 0.014166 | 0.010915 | 0.017926 | 0.015395 | 0.000801 | 0.006903 | 0.012869 | 0.013444 | 0.026397 |
| 23.7916667 | 0.040764 | 0.030425 | 0.009174 | 0.000464 | 0.008813 | 0.009917 | 0.008454 | 0.020757 | 0.010768 | 0.049625 | 0.004394 | 0.009886 | 0.005016 | 0.004632 | 0.010071 | 0.006427 | 0.014843 | 0.018031 | 0.012889 | 0.000422 | 0.010856 | 0.008375 | 0.014466 | 0.019131 |
| 23.8055556 | 0.032729 | 0.027629 | 0.011753 | 0.001343 | 0.006089 | 0.005028 | 0.005766 | 0.013972 | 0.00481 | 0.020108 | 0.002742 | 0.0099 | 0.002038 | 0.008135 | 0.009368 | 0.007286 | 0.018288 | 0.016214 | 0.008173 | 0.001156 | 0.017903 | 0.016921 | 0.00997 | 0.036796 |
| 23.8194444 | 0.017164 | 0.021095 | 0.01503 | 0.003239 | 0.00556 | 0.004264 | 0.012526 | 0.006925 | 0.00346 | 0.013368 | 0.002771 | 0.016001 | 0.000939 | 0.011964 | 0.008839 | 0.006453 | 0.025911 | 0.012269 | 0.01403 | 0.003754 | 0.020709 | 0.03203 | 0.004732 | 0.059347 |
| 23.8333333 | 0.005514 | 0.01764 | 0.01758 | 0.00462 | 0.008257 | 0.007564 | 0.026428 | 0.00271 | 0.002677 | 0.009172 | 0.001988 | 0.028039 | 0.001703 | 0.015177 | 0.008954 | 0.005252 | 0.036207 | 0.007102 | 0.028083 | 0.008417 | 0.019087 | 0.041592 | 0.005097 | 0.068473 |
| 23.8472222 | 0.003435 | 0.018313 | 0.020374 | 0.004456 | 0.00721 | 0.011465 | 0.034509 | 0.0015 | 0.00388 | 0.012469 | 0.001459 | 0.043358 | 0.004919 | 0.01805 | 0.009475 | 0.009603 | 0.046973 | 0.004533 | 0.038196 | 0.01539 | 0.01582 | 0.04782 | 0.008254 | 0.062666 |
| 23.8611111 | 0.009556 | 0.020364 | 0.023082 | 0.002786 | 0.00506 | 0.013153 | 0.029838 | 0.00177 | 0.010322 | 0.033731 | 0.003839 | 0.049672 | 0.010804 | 0.020079 | 0.010531 | 0.014988 | 0.053492 | 0.00716 | 0.036384 | 0.024516 | 0.015084 | 0.065331 | 0.012434 | 0.045335 |
| 23.875 | 0.020938 | 0.023104 | 0.023964 | 0.001713 | 0.007674 | 0.012661 | 0.018196 | 0.003347 | 0.020727 | 0.06733 | 0.008876 | 0.041418 | 0.018889 | 0.021949 | 0.012134 | 0.017566 | 0.054326 | 0.014101 | 0.024936 | 0.032638 | 0.019611 | 0.094845 | 0.018607 | 0.027525 |
| 23.8888889 | 0.03322 | 0.027365 | 0.02239 | 0.003179 | 0.009914 | 0.013011 | 0.007774 | 0.005641 | 0.029712 | 0.104312 | 0.014416 | 0.023251 | 0.026507 | 0.022439 | 0.014053 | 0.018923 | 0.055696 | 0.021537 | 0.012189 | 0.035555 | 0.028676 | 0.109829 | 0.023199 | 0.016311 |
| 23.9027778 | 0.043713 | 0.030733 | 0.019537 | 0.004589 | 0.007415 | 0.015534 | 0.002133 | 0.010797 | 0.032586 | 0.137776 | 0.018102 | 0.007376 | 0.030425 | 0.019805 | 0.016967 | 0.018198 | 0.054216 | 0.023369 | 0.005324 | 0.032941 | 0.040079 | 0.093959 | 0.023497 | 0.010258 |
| 23.9166667 | 0.053222 | 0.028792 | 0.017021 | 0.003479 | 0.005597 | 0.017827 | 0.001482 | 0.016722 | 0.032675 | 0.142068 | 0.01907 | 0.002293 | 0.029592 | 0.015835 | 0.019975 | 0.014223 | 0.045278 | 0.016694 | 0.006199 | 0.02747 | 0.050887 | 0.061905 | 0.021034 | 0.007301 |
| 23.9305556 | 0.053952 | 0.021805 | 0.015642 | 0.002613 | 0.009974 | 0.017893 | 0.003901 | 0.013856 | 0.032412 | 0.099353 | 0.017133 | 0.004996 | 0.025342 | 0.013718 | 0.018983 | 0.008573 | 0.032369 | 0.007949 | 0.011151 | 0.019848 | 0.055267 | 0.035334 | 0.017549 | 0.008262 |
| 23.9444444 | 0.038092 | 0.013767 | 0.015604 | 0.003732 | 0.013256 | 0.014963 | 0.008092 | 0.006451 | 0.028846 | 0.046444 | 0.011728 | 0.012817 | 0.019729 | 0.013183 | 0.012522 | 0.004236 | 0.019428 | 0.005168 | 0.016602 | 0.010866 | 0.046621 | 0.021828 | 0.013334 | 0.015288 |
| 23.9583333 | 0.017708 | 0.006914 | 0.015548 | 0.003685 | 0.009233 | 0.009822 | 0.011014 | 0.005188 | 0.022634 | 0.027965 | 0.005959 | 0.021567 | 0.014281 | 0.011703 | 0.005059 | 0.00334 | 0.010272 | 0.00637 | 0.021853 | 0.005982 | 0.028661 | 0.0162 | 0.009262 | 0.025476 |
| 23.9722222 | 0.009396 | 0.002981 | 0.013044 | 0.002264 | 0.004972 | 0.005086 | 0.009738 | 0.00607 | 0.018416 | 0.040583 | 0.004723 | 0.029367 | 0.009833 | 0.00735 | 0.001454 | 0.005858 | 0.007174 | 0.005229 | 0.025255 | 0.006051 | 0.013083 | 0.01314 | 0.005918 | 0.031808 |
| 23.9861111 | 0.009512 | 0.00318 | 0.009422 | 0.002677 | 0.006572 | 0.00259 | 0.006505 | 0.005457 | 0.019401 | 0.047674 | 0.005831 | 0.039385 | 0.006001 | 0.003953 | 0.001583 | 0.009369 | 0.009341 | 0.002526 | 0.023557 | 0.004705 | 0.005683 | 0.009881 | 0.00341 | 0.03128 |
| 24 | 0.008967 | 0.004557 | 0.010101 | 0.004778 | 0.009911 | 0.003603 | 0.006415 | 0.009015 | 0.025057 | 0.035512 | 0.005822 | 0.048798 | 0.002997 | 0.006065 | 0.002837 | 0.013258 | 0.014976 | 0.001948 | 0.018129 | 0.003607 | 0.007578 | 0.005258 | 0.00328 | 0.025795 |
| 24.0138889 | 0.014241 | 0.003853 | 0.017057 | 0.006396 | 0.009226 | 0.009584 | 0.011951 | 0.015447 | 0.030373 | 0.035498 | 0.010396 | 0.052762 | 0.002173 | 0.008718 | 0.00477 | 0.019661 | 0.02542 | 0.002661 | 0.014265 | 0.009733 | 0.019425 | 0.00464 | 0.006096 | 0.019252 |
| 24.0277778 | 0.030446 | 0.004418 | 0.025834 | 0.006095 | 0.008292 | 0.019918 | 0.020678 | 0.016903 | 0.029189 | 0.068568 | 0.024114 | 0.057618 | 0.002098 | 0.009147 | 0.008882 | 0.028131 | 0.046778 | 0.003817 | 0.014244 | 0.022422 | 0.035523 | 0.012941 | 0.00979 | 0.016489 |
| 24.0416667 | 0.051776 | 0.009833 | 0.032509 | 0.00544 | 0.013696 | 0.028069 | 0.02974 | 0.017933 | 0.021423 | 0.09741 | 0.038705 | 0.073091 | 0.00142 | 0.009307 | 0.016985 | 0.038034 | 0.082801 | 0.007109 | 0.017712 | 0.038067 | 0.043967 | 0.024902 | 0.013649 | 0.020948 |
| 24.0555556 | 0.065411 | 0.01638 | 0.035556 | 0.007949 | 0.015626 | 0.02705 | 0.034761 | 0.03061 | 0.0144 | 0.082025 | 0.0461 | 0.085302 | 0.002713 | 0.007606 | 0.02421 | 0.046266 | 0.117732 | 0.012886 | 0.023243 | 0.054541 | 0.039578 | 0.03279 | 0.018741 | 0.031449 |
| 24.0694444 | 0.058956 | 0.018872 | 0.03283 | 0.011075 | 0.00958 | 0.019935 | 0.028202 | 0.044559 | 0.013185 | 0.040633 | 0.046272 | 0.078354 | 0.005412 | 0.005789 | 0.02231 | 0.037239 | 0.115676 | 0.01432 | 0.022877 | 0.063572 | 0.029422 | 0.034353 | 0.022423 | 0.036664 |
| 24.0833333 | 0.037368 | 0.015196 | 0.023627 | 0.008951 | 0.00868 | 0.01347 | 0.015742 | 0.040107 | 0.015387 | 0.011549 | 0.037023 | 0.058238 | 0.004916 | 0.006802 | 0.01411 | 0.025282 | 0.067928 | 0.008927 | 0.015157 | 0.055869 | 0.02053 | 0.029737 | 0.019166 | 0.025839 |
| 24.0972222 | 0.016506 | 0.007709 | 0.012371 | 0.004855 | 0.009696 | 0.008871 | 0.013841 | 0.021806 | 0.016755 | 0.002809 | 0.020096 | 0.031526 | 0.002499 | 0.006971 | 0.006411 | 0.038361 | 0.02932 | 0.006841 | 0.013368 | 0.036745 | 0.012664 | 0.02061 | 0.010578 | 0.019105 |
| 24.1111111 | 0.007171 | 0.002868 | 0.007065 | 0.005346 | 0.00706 | 0.005286 | 0.020294 | 0.011047 | 0.014961 | 0.002476 | 0.009047 | 0.011834 | 0.002653 | 0.004172 | 0.002519 | 0.048698 | 0.037819 | 0.011811 | 0.018501 | 0.018107 | 0.005861 | 0.011737 | 0.005734 | 0.031939 |
| 24.125 | 0.007128 | 0.002672 | 0.007617 | 0.007575 | 0.007511 | 0.003017 | 0.021857 | 0.017266 | 0.010335 | 0.002611 | 0.009405 | 0.010017 | 0.004563 | 0.001581 | 0.003283 | 0.035091 | 0.059133 | 0.018593 | 0.021031 | 0.006406 | 0.002132 | 0.007393 | 0.006361 | 0.038476 |
| 24.1388889 | 0.008178 | 0.002584 | 0.007397 | 0.006589 | 0.007621 | 0.002361 | 0.017727 | 0.024851 | 0.006377 | 0.001724 | 0.011167 | 0.015679 | 0.006174 | 0.000963 | 0.00496 | 0.017186 | 0.064916 | 0.020532 | 0.020717 | 0.002334 | 0.002253 | 0.007067 | 0.006814 | 0.027778 |
| 24.1527778 | 0.007836 | 0.001516 | 0.009703 | 0.003672 | 0.003981 | 0.002453 | 0.012617 | 0.023844 | 0.005332 | 0.001948 | 0.012024 | 0.019887 | 0.009103 | 0.012998 | 0.005386 | 0.011685 | 0.061376 | 0.017805 | 0.018514 | 0.001569 | 0.004491 | 0.007508 | 0.00976 | 0.019392 |
| 24.1666667 | 0.007245 | 0.002249 | 0.011858 | 0.002517 | 0.003053 | 0.003457 | 0.016549 | 0.027115 | 0.00665 | 0.014002 | 0.019415 | 0.023541 | 0.012075 | 0.051403 | 0.009458 | 0.022604 | 0.060415 | 0.018781 | 0.015457 | 0.001666 | 0.007117 | 0.006138 | 0.021236 | 0.028118 |
| 24.1805556 | 0.013877 | 0.005057 | 0.014294 | 0.007689 | 0.007254 | 0.0066 | 0.035077 | 0.039945 | 0.013055 | 0.046811 | 0.034292 | 0.03491 | 0.021276 | 0.072857 | 0.022931 | 0.028069 | 0.06149 | 0.017718 | 0.0261 | 0.014549 | 0.008607 | 0.006744 | 0.042355 | 0.036424 |
| 24.1944444 | 0.019258 | 0.006622 | 0.018699 | 0.012827 | 0.011366 | 0.008709 | 0.043547 | 0.04278 | 0.018799 | 0.07654 | 0.038414 | 0.053805 | 0.028004 | 0.051229 | 0.029775 | 0.014461 | 0.069664 | 0.014104 | 0.048399 | 0.033899 | 0.012759 | 0.014198 | 0.046652 | 0.027749 |
| 24.2083333 | 0.018737 | 0.005679 | 0.018679 | 0.010567 | 0.009206 | 0.007445 | 0.036286 | 0.032459 | 0.024683 | 0.062568 | 0.023578 | 0.056386 | 0.016005 | 0.030259 | 0.017382 | 0.009375 | 0.067008 | 0.016953 | 0.044811 | 0.02806 | 0.020231 | 0.021589 | 0.021803 | 0.032749 |
| 24.2222222 | 0.027056 | 0.005223 | 0.020213 | 0.00752 | 0.00495 | 0.005067 | 0.037667 | 0.029724 | 0.039082 | 0.022338 | 0.011227 | 0.039999 | 0.010883 | 0.051481 | 0.008991 | 0.038633 | 0.029962 | 0.019994 | 0.021891 | 0.008133 | 0.01783 | 0.018725 | 0.009316 | 0.05024 |
| 24.2361111 | 0.033394 | 0.005816 | 0.015244 | 0.004448 | 0.004055 | 0.003248 | 0.035575 | 0.025766 | 0.038673 | 0.010023 | 0.008437 | 0.034266 | 0.016328 | 0.097451 | 0.013195 | 0.07447 | 0.003514 | 0.020483 | 0.015499 | 0.001117 | 0.011214 | 0.011593 | 0.013956 | 0.052448 |

| | | | | | | | | | | | | | | | | | | | | | | | | |
|---|---|---|---|---|---|---|---|---|---|---|---|---|---|---|---|---|---|---|---|---|---|---|---|---|
| 24.25 | 0.033626 | 0.006039 | 0.005205 | 0.001545 | 0.003166 | 0.003172 | 0.021965 | 0.012034 | 0.018012 | 0.02004 | 0.007752 | 0.041242 | 0.012533 | 0.083808 | 0.013869 | 0.067248 | 0.003962 | 0.016842 | 0.014393 | 0.002596 | 0.011536 | 0.008811 | 0.01155 | 0.049653 |
| 24.2638889 | 0.03674 | 0.005367 | 0.003145 | 0.00205 | 0.001825 | 0.003552 | 0.016156 | 0.004868 | 0.00524 | 0.020285 | 0.006701 | 0.040824 | 0.008041 | 0.03049 | 0.008694 | 0.034637 | 0.005489 | 0.010382 | 0.00795 | 0.005028 | 0.010188 | 0.005434 | 0.006609 | 0.044883 |
| 24.2777778 | 0.037862 | 0.004089 | 0.003878 | 0.003948 | 0.001549 | 0.003377 | 0.015627 | 0.003048 | 0.003916 | 0.011751 | 0.005143 | 0.032011 | 0.007143 | 0.014672 | 0.004225 | 0.016591 | 0.004158 | 0.008648 | 0.006601 | 0.004851 | 0.005955 | 0.002117 | 0.004812 | 0.033684 |
| 24.2916667 | 0.036461 | 0.003566 | 0.005273 | 0.004898 | 0.002255 | 0.004749 | 0.010097 | 0.001361 | 0.003576 | 0.006609 | 0.005985 | 0.023184 | 0.006905 | 0.020995 | 0.001946 | 0.010089 | 0.001743 | 0.009549 | 0.009584 | 0.003359 | 0.005673 | 0.001693 | 0.004677 | 0.022115 |
| 24.3055556 | 0.036163 | 0.004453 | 0.00762 | 0.003504 | 0.004191 | 0.007676 | 0.005251 | 0.003141 | 0.002402 | 0.008616 | 0.009446 | 0.01707 | 0.022643 | 0.001616 | 0.005226 | 0.000752 | 0.008716 | 0.008446 | 0.004485 | 0.008534 | 0.002483 | 0.005994 | 0.022773 |
| 24.3194444 | 0.034646 | 0.005369 | 0.008425 | 0.002062 | 0.006532 | 0.00896 | 0.007383 | 0.005957 | 0.001579 | 0.012667 | 0.011984 | 0.011786 | 0.010729 | 0.018451 | 0.003742 | 0.002988 | 0.001159 | 0.005889 | 0.006779 | 0.00589 | 0.009633 | 0.004347 | 0.007269 | 0.035034 |
| 24.3333333 | 0.027973 | 0.004796 | 0.00683 | 0.002969 | 0.007444 | 0.007149 | 0.009988 | 0.005811 | 0.001178 | 0.012947 | 0.010874 | 0.006639 | 0.010332 | 0.013299 | 0.006913 | 0.005381 | 0.002804 | 0.002838 | 0.011635 | 0.006018 | 0.008487 | 0.00562 | 0.006015 | 0.035635 |
| 24.3472222 | 0.018845 | 0.003216 | 0.004349 | 0.003435 | 0.005674 | 0.004429 | 0.007276 | 0.003028 | 0.001058 | 0.008351 | 0.007186 | 0.005654 | 0.00774 | 0.009832 | 0.008949 | 0.008738 | 0.004433 | 0.001109 | 0.016535 | 0.00501 | 0.007814 | 0.005966 | 0.003165 | 0.020806 |
| 24.3611111 | 0.011289 | 0.001829 | 0.002614 | 0.002197 | 0.002877 | 0.003031 | 0.002768 | 0.001892 | 0.001097 | 0.005105 | 0.004352 | 0.008142 | 0.005267 | 0.008711 | 0.008362 | 0.008568 | 0.003884 | 0.000811 | 0.015605 | 0.003373 | 0.008916 | 0.006029 | 0.001167 | 0.007057 |
| 24.375 | 0.006549 | 0.000947 | 0.002555 | 0.001645 | 0.001726 | 0.003834 | 0.001864 | 0.005523 | 0.002612 | 0.00766 | 0.005471 | 0.007005 | 0.005667 | 0.010829 | 0.006571 | 0.005887 | 0.004234 | 0.000788 | 0.010952 | 0.005332 | 0.011946 | 0.005964 | 0.000892 | 0.01201 |
| 24.3888889 | 0.005497 | 0.000634 | 0.00441 | 0.002046 | 0.001699 | 0.006982 | 0.006623 | 0.014106 | 0.006831 | 0.008153 | 0.010533 | 0.004983 | 0.009191 | 0.013629 | 0.007266 | 0.00569 | 0.008853 | 0.001298 | 0.007153 | 0.013276 | 0.017945 | 0.007007 | 0.002569 | 0.043766 |
| 24.4027778 | 0.009056 | 0.000798 | 0.007408 | 0.00396 | 0.002504 | 0.011589 | 0.015324 | 0.024099 | 0.009854 | 0.00947 | 0.015922 | 0.009253 | 0.013165 | 0.014271 | 0.012167 | 0.00924 | 0.01478 | 0.00428 | 0.008869 | 0.024963 | 0.02644 | 0.011098 | 0.007567 | 0.073989 |
| 24.4166667 | 0.016389 | 0.001514 | 0.010369 | 0.007443 | 0.005148 | 0.015893 | 0.023026 | 0.028887 | 0.008803 | 0.022776 | 0.018279 | 0.014583 | 0.015547 | 0.013663 | 0.02007 | 0.01371 | 0.020012 | 0.008982 | 0.015918 | 0.036816 | 0.033607 | 0.018049 | 0.014795 | 0.069643 |
| 24.4305556 | 0.02446 | 0.002977 | 0.012301 | 0.009398 | 0.006969 | 0.017893 | 0.024536 | 0.025637 | 0.006205 | 0.04074 | 0.01752 | 0.016172 | 0.015817 | 0.012961 | 0.026135 | 0.016188 | 0.025349 | 0.011982 | 0.023364 | 0.045533 | 0.035326 | 0.024205 | 0.020065 | 0.040982 |
| 24.4444444 | 0.030036 | 0.004318 | 0.01277 | 0.008552 | 0.005322 | 0.016711 | 0.019984 | 0.017589 | 0.005282 | 0.052811 | 0.015033 | 0.016918 | 0.014486 | 0.012631 | 0.024757 | 0.01624 | 0.029426 | 0.010682 | 0.029104 | 0.049136 | 0.030826 | 0.026098 | 0.020228 | 0.017399 |
| 24.4583333 | 0.031372 | 0.004739 | 0.011931 | 0.006304 | 0.002982 | 0.01415 | 0.013332 | 0.008656 | 0.007468 | 0.057829 | 0.01196 | 0.018245 | 0.012652 | 0.012983 | 0.016621 | 0.015822 | 0.02915 | 0.005891 | 0.030152 | 0.046713 | 0.023876 | 0.023302 | 0.015961 | 0.011693 |
| 24.4722222 | 0.028277 | 0.00435 | 0.01044 | 0.004223 | 0.003025 | 0.012863 | 0.00824 | 0.002735 | 0.012261 | 0.054757 | 0.009628 | 0.019434 | 0.010827 | 0.01236 | 0.007705 | 0.01342 | 0.024873 | 0.002318 | 0.023436 | 0.038655 | 0.019481 | 0.018669 | 0.010699 | 0.01193 |
| 24.4861111 | 0.02418 | 0.004796 | 0.011403 | 0.004799 | 0.00293 | 0.015352 | 0.008535 | 0.001513 | 0.019305 | 0.0447 | 0.010312 | 0.020518 | 0.010754 | 0.008606 | 0.004396 | 0.008392 | 0.021978 | 0.001714 | 0.016646 | 0.029805 | 0.022213 | 0.018841 | 0.007569 | 0.011157 |
| 24.5 | 0.027047 | 0.009054 | 0.021774 | 0.009626 | 0.002865 | 0.025258 | 0.017606 | 0.00382 | 0.030718 | 0.03948 | 0.017034 | 0.025305 | 0.015363 | 0.004494 | 0.009878 | 0.00661 | 0.027342 | 0.002687 | 0.01902 | 0.028823 | 0.036689 | 0.030183 | 0.00794 | 0.020603 |
| 24.5138889 | 0.038913 | 0.016615 | 0.040842 | 0.015937 | 0.006164 | 0.040989 | 0.03267 | 0.010389 | 0.043502 | 0.05434 | 0.029821 | 0.036614 | 0.025106 | 0.005025 | 0.020967 | 0.011523 | 0.042368 | 0.00754 | 0.027327 | 0.038826 | 0.061974 | 0.049027 | 0.011044 | 0.037472 |
| 24.5277778 | 0.053453 | 0.020462 | 0.050983 | 0.018964 | 0.010098 | 0.049343 | 0.043926 | 0.018548 | 0.04705 | 0.08992 | 0.042361 | 0.04621 | 0.035374 | 0.009043 | 0.028919 | 0.020091 | 0.057255 | 0.015346 | 0.02978 | 0.052532 | 0.07789 | 0.056829 | 0.013428 | 0.044215 |
| 24.5416667 | 0.071393 | 0.015775 | 0.043884 | 0.018236 | 0.01126 | 0.043127 | 0.042631 | 0.022994 | 0.050386 | 0.123386 | 0.046496 | 0.044006 | 0.039142 | 0.010922 | 0.029712 | 0.028754 | 0.062946 | 0.021249 | 0.023321 | 0.05861 | 0.06752 | 0.045013 | 0.012409 | 0.034616 |
| 24.5555556 | 0.083523 | 0.009558 | 0.035012 | 0.01691 | 0.009493 | 0.032204 | 0.028436 | 0.024768 | 0.068676 | 0.130728 | 0.039517 | 0.033414 | 0.037058 | 0.007951 | 0.02802 | 0.03274 | 0.060753 | 0.023441 | 0.017232 | 0.052456 | 0.048415 | 0.029726 | 0.01087 | 0.019945 |
| 24.5694444 | 0.068465 | 0.009611 | 0.034737 | 0.015721 | 0.005459 | 0.023973 | 0.014738 | 0.024345 | 0.079283 | 0.108311 | 0.024591 | 0.022704 | 0.034877 | 0.004875 | 0.025511 | 0.027614 | 0.053431 | 0.020944 | 0.014328 | 0.040114 | 0.037996 | 0.021896 | 0.011686 | 0.010559 |
| 24.5833333 | 0.035124 | 0.01256 | 0.037801 | 0.013887 | 0.002413 | 0.017508 | 0.016654 | 0.018131 | 0.062329 | 0.0628 | 0.011434 | 0.013436 | 0.032355 | 0.004712 | 0.019316 | 0.016193 | 0.042068 | 0.014166 | 0.009313 | 0.02751 | 0.032576 | 0.017894 | 0.012874 | 0.011773 |
| 24.5972222 | 0.014211 | 0.013157 | 0.036603 | 0.012872 | 0.002193 | 0.012312 | 0.027865 | 0.009033 | 0.034238 | 0.030719 | 0.006202 | 0.006677 | 0.027382 | 0.003461 | 0.009907 | 0.007497 | 0.030222 | 0.006887 | 0.004329 | 0.015131 | 0.029585 | 0.016011 | 0.012712 | 0.02509 |
| 24.6111111 | 0.01735 | 0.011881 | 0.030526 | 0.016008 | 0.001797 | 0.008728 | 0.032764 | 0.003345 | 0.02109 | 0.042274 | 0.006552 | 0.005447 | 0.021315 | 0.001817 | 0.003786 | 0.008932 | 0.020691 | 0.004618 | 0.004412 | 0.0076 | 0.029788 | 0.015973 | 0.012371 | 0.041496 |
| 24.625 | 0.030453 | 0.0106 | 0.026006 | 0.021484 | 0.003147 | 0.0057 | 0.026264 | 0.001502 | 0.030546 | 0.071239 | 0.013703 | 0.007765 | 0.015795 | 0.002464 | 0.004976 | 0.018998 | 0.012322 | 0.008086 | 0.005654 | 0.011009 | 0.030573 | 0.014772 | 0.01262 | 0.049745 |
| 24.6388889 | 0.037477 | 0.009693 | 0.024084 | 0.022266 | 0.007669 | 0.002874 | 0.015895 | 0.001309 | 0.041356 | 0.092848 | 0.022492 | 0.007475 | 0.012524 | 0.002971 | 0.007873 | 0.027879 | 0.00568 | 0.010716 | 0.005515 | 0.020101 | 0.029322 | 0.01243 | 0.011976 | 0.03725 |
| 24.6527778 | 0.036784 | 0.009326 | 0.021049 | 0.019001 | 0.009318 | 0.001305 | 0.018979 | 0.003521 | 0.039965 | 0.096588 | 0.023861 | 0.008388 | 0.012106 | 0.001959 | 0.006296 | 0.024479 | 0.005053 | 0.010177 | 0.002246 | 0.026688 | 0.011076 | 0.009678 | 0.024533 |
| 24.6666667 | 0.031492 | 0.010179 | 0.017519 | 0.014915 | 0.006799 | 0.000635 | 0.034196 | 0.007237 | 0.032971 | 0.06757 | 0.020309 | 0.017903 | 0.012445 | 0.001241 | 0.004215 | 0.014357 | 0.010728 | 0.00527 | 0.02176 | 0.016549 | 0.024235 | 0.008955 | 0.006027 | 0.045887 |
| 24.6805556 | 0.02186 | 0.012904 | 0.013429 | 0.009114 | 0.003692 | 0.000919 | 0.038889 | 0.007419 | 0.023466 | 0.050786 | 0.014278 | 0.027101 | 0.011875 | 0.001514 | 0.007905 | 0.018208 | 0.017024 | 0.009757 | 0.033421 | 0.019062 | 0.02181 | 0.006158 | 0.003348 | 0.080192 |
| 24.6944444 | 0.018221 | 0.017665 | 0.007874 | 0.006264 | 0.003343 | 0.001409 | 0.0247 | 0.00777 | 0.011847 | 0.092068 | 0.009099 | 0.025591 | 0.010689 | 0.001412 | 0.012151 | 0.035719 | 0.023811 | 0.018573 | 0.036398 | 0.03643 | 0.017848 | 0.006574 | 0.004951 | 0.088537 |
| 24.7083333 | 0.029686 | 0.024749 | 0.005871 | 0.009218 | 0.005279 | 0.001149 | 0.013698 | 0.012962 | 0.004625 | 0.127929 | 0.008693 | 0.01496 | 0.009678 | 0.001489 | 0.011175 | 0.042286 | 0.027605 | 0.023807 | 0.027624 | 0.048654 | 0.012511 | 0.00979 | 0.008765 | 0.061751 |
| 24.7222222 | 0.044605 | 0.032538 | 0.009956 | 0.011957 | 0.004771 | 0.001501 | 0.02534 | 0.016328 | 0.013493 | 0.115674 | 0.007529 | 0.005076 | 0.009025 | 0.002421 | 0.007336 | 0.030724 | 0.0287 | 0.022035 | 0.013918 | 0.04221 | 0.008101 | 0.012422 | 0.011447 | 0.027925 |
| 24.7361111 | 0.052273 | 0.035177 | 0.014908 | 0.011158 | 0.003059 | 0.00245 | 0.042002 | 0.013466 | 0.029061 | 0.083403 | 0.006344 | 0.002202 | 0.007971 | 0.002414 | 0.003675 | 0.016011 | 0.02616 | 0.0157 | 0.006732 | 0.025177 | 0.00602 | 0.012862 | 0.011478 | 0.023689 |
| 24.75 | 0.047152 | 0.02916 | 0.017321 | 0.009659 | 0.004144 | 0.002792 | 0.046006 | 0.006937 | 0.024262 | 0.046305 | 0.009271 | 0.003408 | 0.005718 | 0.002068 | 0.00267 | 0.007727 | 0.018469 | 0.008624 | 0.008218 | 0.011797 | 0.0058 | 0.01129 | 0.008055 | 0.050177 |
| 24.7638889 | 0.027633 | 0.017391 | 0.018064 | 0.009378 | 0.004909 | 0.003337 | 0.039052 | 0.002665 | 0.013505 | 0.031796 | 0.010877 | 0.009636 | 0.003091 | 0.003778 | 0.004826 | 0.008863 | 0.008957 | 0.004251 | 0.010665 | 0.010358 | 0.005756 | 0.009104 | 0.003612 | 0.080098 |
| 24.7777778 | 0.015998 | 0.007561 | 0.017781 | 0.010149 | 0.003778 | 0.004959 | 0.022607 | 0.003866 | 0.025568 | 0.059366 | 0.008258 | 0.020962 | 0.002193 | 0.007162 | 0.008494 | 0.017743 | 0.005283 | 0.00588 | 0.01243 | 0.016724 | 0.004112 | 0.007125 | 0.001227 | 0.089283 |
| 24.7916667 | 0.027244 | 0.005913 | 0.016345 | 0.013085 | 0.005683 | 0.005323 | 0.009701 | 0.007291 | 0.04232 | 0.099504 | 0.007896 | 0.0305 | 0.003378 | 0.010756 | 0.013061 | 0.026777 | 0.008157 | 0.01208 | 0.017684 | 0.0236 | 0.004177 | 0.005408 | 0.00111 | 0.06998 |
| 24.8055556 | 0.040499 | 0.009229 | 0.014199 | 0.018365 | 0.009094 | 0.003239 | 0.009831 | 0.010082 | 0.040443 | 0.121962 | 0.01333 | 0.031016 | 0.005162 | 0.013186 | 0.017104 | 0.027982 | 0.009403 | 0.018214 | 0.025333 | 0.029913 | 0.010912 | 0.004486 | 0.001401 | 0.035269 |
| 24.8194444 | 0.039816 | 0.012385 | 0.01166 | 0.020406 | 0.007311 | 0.00193 | 0.009131 | 0.011906 | 0.028374 | 0.130293 | 0.015106 | 0.022378 | 0.006719 | 0.014878 | 0.016458 | 0.019293 | 0.006285 | 0.020957 | 0.029849 | 0.033695 | 0.021227 | 0.005718 | 0.001967 | 0.012598 |
| 24.8333333 | 0.033211 | 0.014336 | 0.009553 | 0.015017 | 0.004337 | 0.00347 | 0.006786 | 0.012397 | 0.020932 | 0.132391 | 0.008534 | 0.012269 | 0.008744 | 0.016288 | 0.016161 | 0.00931 | 0.005134 | 0.020001 | 0.028437 | 0.033153 | 0.029303 | 0.009283 | 0.003326 | 0.007473 |
| 24.8472222 | 0.031356 | 0.0148 | 0.008855 | 0.010528 | 0.005977 | 0.006055 | 0.014406 | 0.010412 | 0.018704 | 0.120701 | 0.002377 | 0.010603 | 0.011983 | 0.01811 | 0.01359 | 0.005359 | 0.009434 | 0.015758 | 0.022261 | 0.030025 | 0.033202 | 0.011685 | 0.004741 | 0.009099 |
| 24.8611111 | 0.038701 | 0.014588 | 0.010079 | 0.015965 | 0.006972 | 0.008061 | 0.026972 | 0.006112 | 0.016093 | 0.093045 | 0.001195 | 0.019101 | 0.016181 | 0.020572 | 0.013018 | 0.00664 | 0.015591 | 0.008954 | 0.01412 | 0.025374 | 0.035141 | 0.010677 | 0.00761 | 0.01621 |
| 24.875 | 0.051573 | 0.015417 | 0.014488 | 0.02765 | 0.004028 | 0.00914 | 0.034088 | 0.003042 | 0.013071 | 0.059485 | 0.003449 | 0.021585 | 0.020548 | 0.024051 | 0.015222 | 0.016753 | 0.020787 | 0.004093 | 0.010403 | 0.021766 | 0.039776 | 0.009422 | 0.012403 | 0.029595 |
| 24.8888889 | 0.061105 | 0.017752 | 0.02245 | 0.037336 | 0.001664 | 0.009687 | 0.032625 | 0.00512 | 0.013106 | 0.029944 | 0.008952 | 0.015916 | 0.024451 | 0.028946 | 0.019327 | 0.031099 | 0.025744 | 0.004152 | 0.014349 | 0.022076 | 0.049246 | 0.010955 | 0.016345 | 0.05044 |
| 24.9027778 | 0.065004 | 0.020057 | 0.029899 | 0.040157 | 0.001725 | 0.010604 | 0.025033 | 0.010561 | 0.017674 | 0.010587 | 0.016248 | 0.021814 | 0.028079 | 0.033924 | 0.024288 | 0.033922 | 0.030458 | 0.006535 | 0.021675 | 0.025608 | 0.062781 | 0.015111 | 0.017812 | 0.071436 |
| 24.9166667 | 0.064206 | 0.020852 | 0.03169 | 0.036446 | 0.001499 | 0.012125 | 0.014116 | 0.014959 | 0.023971 | 0.002358 | 0.023721 | 0.036635 | 0.030312 | 0.035195 | 0.027768 | 0.027444 | 0.032143 | 0.008002 | 0.024989 | 0.02753 | 0.07317 | 0.020943 | 0.017254 | 0.075792 |
| 24.9305556 | 0.054846 | 0.019389 | 0.028842 | 0.027266 | 0.000542 | 0.014181 | 0.005811 | 0.019807 | 0.02732 | 0.000798 | 0.029898 | 0.045332 | 0.029018 | 0.030096 | 0.027082 | 0.022292 | 0.028483 | 0.008433 | 0.021015 | 0.024313 | 0.071011 | 0.027884 | 0.015147 | 0.060784 |
| 24.9444444 | 0.03544 | 0.01463 | 0.024576 | 0.01422 | 0.000345 | 0.016222 | 0.004573 | 0.028901 | 0.026082 | 0.002659 | 0.029407 | 0.047983 | 0.025176 | 0.021346 | 0.023803 | 0.017583 | 0.022748 | 0.00901 | 0.015518 | 0.018971 | 0.059108 | 0.034577 | 0.010716 | 0.040355 |
| 24.9583333 | 0.016514 | 0.007971 | 0.019786 | 0.005574 | 0.000825 | 0.01747 | 0.004862 | 0.038496 | 0.021519 | 0.010436 | 0.019018 | 0.04643 | 0.022023 | 0.012987 | 0.020353 | 0.01041 | 0.019605 | 0.009587 | 0.015241 | 0.014541 | 0.045716 | 0.037616 | 0.004932 | 0.027857 |
| 24.9722222 | 0.011222 | 0.00343 | 0.016147 | 0.004539 | 0.001005 | 0.016386 | 0.003711 | 0.039869 | 0.015186 | 0.024391 | 0.009455 | 0.037284 | 0.020942 | 0.007575 | 0.015575 | 0.00421 | 0.018933 | 0.008907 | 0.019854 | 0.009944 | 0.034185 | 0.034391 | 0.001158 | 0.025855 |
| 24.9861111 | 0.013251 | 0.002541 | 0.015432 | 0.006251 | 0.000567 | 0.012178 | 0.005854 | 0.031442 | 0.008875 | 0.038865 | 0.01011 | 0.02118 | 0.020775 | 0.007639 | 0.009084 | 0.002351 | 0.018418 | 0.005869 | 0.024383 | 0.0048 | 0.026231 | 0.026288 | 0.001102 | 0.028594 |
| 25 | 0.011851 | 0.00251 | 0.017788 | 0.009525 | 0.001036 | 0.008613 | 0.012912 | 0.020683 | 0.005785 | 0.048492 | 0.010974 | 0.009597 | 0.020251 | 0.010097 | 0.006547 | 0.002884 | 0.016977 | 0.002198 | 0.024904 | 0.00211 | 0.023511 | 0.017504 | 0.004165 | 0.030941 |
| 25.0138889 | 0.016181 | 0.002587 | 0.022759 | 0.014535 | 0.003566 | 0.010649 | 0.022749 | 0.014789 | 0.009224 | 0.052904 | 0.013166 | 0.007414 | 0.018963 | 0.009739 | 0.006635 | 0.005756 | 0.014796 | 0.001044 | 0.022379 | 0.00633 | 0.026062 | 0.01291 | 0.009068 | 0.033281 |
| 25.0277778 | 0.034241 | 0.00587 | 0.02806 | 0.019001 | 0.005451 | 0.018541 | 0.033476 | 0.015137 | 0.020089 | 0.055015 | 0.029102 | 0.011979 | 0.015561 | 0.007375 | 0.016869 | 0.011374 | 0.015473 | 0.003129 | 0.020488 | 0.020401 | 0.032583 | 0.014673 | 0.014585 | 0.03993 |
| 25.0416667 | 0.058219 | 0.01217 | 0.030562 | 0.02312 | 0.003971 | 0.02818 | 0.042872 | 0.020367 | 0.02689 | 0.054659 | 0.049054 | 0.027349 | 0.009819 | 0.005671 | 0.032113 | 0.016028 | 0.024861 | 0.007753 | 0.018905 | 0.043456 | 0.041916 | 0.021776 | 0.021279 | 0.054618 |
| 25.0555556 | 0.074861 | 0.019375 | 0.030202 | 0.026299 | 0.001399 | 0.033685 | 0.045101 | 0.026004 | 0.020132 | 0.048206 | 0.057883 | 0.04407 | 0.005224 | 0.008739 | 0.041008 | 0.015908 | 0.03748 | 0.014039 | 0.013664 | 0.06382 | 0.048924 | 0.032231 | 0.028063 | 0.064584 |
| 25.0694444 | 0.072648 | 0.022905 | 0.027357 | 0.022693 | 0.000555 | 0.031409 | 0.033478 | 0.022727 | 0.018042 | 0.034029 | 0.052547 | 0.047826 | 0.003162 | 0.015413 | 0.034042 | 0.010365 | 0.036945 | 0.017791 | 0.0096 | 0.065515 | 0.043732 | 0.039556 | 0.026954 | 0.049008 |
| 25.0833333 | 0.051845 | 0.017565 | 0.020556 | 0.012322 | 0.001325 | 0.022583 | 0.016558 | 0.010914 | 0.029799 | 0.018016 | 0.037012 | 0.039266 | 0.003013 | 0.014179 | 0.017471 | 0.004429 | 0.021179 | 0.01408 | 0.01688 | 0.050476 | 0.025855 | 0.036534 | 0.016158 | 0.02619 |
| 25.0972222 | 0.025525 | 0.008537 | 0.011094 | 0.005767 | 0.003136 | 0.01146 | 0.011194 | 0.002465 | 0.036236 | 0.013209 | 0.020769 | 0.023994 | 0.005073 | 0.008279 | 0.008263 | 0.003505 | 0.009645 | 0.00779 | 0.029316 | 0.031642 | 0.009554 | 0.024825 | 0.008147 | 0.030551 |
| 25.1111111 | 0.012848 | 0.004724 | 0.004422 | 0.008211 | 0.005943 | 0.004159 | 0.015328 | 0.000692 | 0.02791 | 0.020113 | 0.016568 | 0.010399 | 0.007202 | 0.010415 | 0.013453 | 0.007964 | 0.01201 | 0.007511 | 0.034878 | 0.015466 | 0.004205 | 0.011649 | 0.008388 | 0.051506 |
| 25.125 | 0.016699 | 0.005063 | 0.003505 | 0.011666 | 0.008056 | 0.00325 | 0.016638 | 0.001079 | 0.013426 | 0.023549 | 0.023472 | 0.00577 | 0.007735 | 0.013825 | 0.019771 | 0.013011 | 0.014994 | 0.011211 | 0.033387 | 0.005732 | 0.004669 | 0.003651 | 0.009153 | 0.061341 |
| 25.1388889 | 0.019962 | 0.004355 | 0.00443 | 0.010245 | 0.007061 | 0.004166 | 0.013539 | 0.001559 | 0.005107 | 0.017186 | 0.025259 | 0.007376 | 0.007142 | 0.009915 | 0.017632 | 0.013866 | 0.011047 | 0.013054 | 0.027587 | 0.004085 | 0.004984 | 0.001708 | 0.006927 | 0.056178 |
| 25.1527778 | 0.015773 | 0.002627 | 0.003932 | 0.005795 | 0.00411 | 0.003212 | 0.010129 | 0.001462 | 0.006506 | 0.007529 | 0.018815 | 0.008495 | 0.005826 | 0.004739 | 0.011042 | 0.011421 | 0.005243 | 0.012417 | 0.021425 | 0.005339 | 0.003781 | 0.001815 | 0.003765 | 0.043133 |
| 25.1666667 | 0.008687 | 0.001617 | 0.003121 | 0.002732 | 0.003389 | 0.001687 | 0.00811 | 0.000947 | 0.009182 | 0.001566 | 0.013285 | 0.008428 | 0.004174 | 0.004321 | 0.00496 | 0.007697 | 0.002136 | 0.010635 | 0.018899 | 0.004953 | 0.001869 | 0.003138 | 0.002545 | 0.029168 |
| 25.1805556 | 0.003123 | 0.001459 | 0.002748 | 0.003415 | 0.005662 | 0.001365 | 0.006975 | 0.001216 | 0.006947 | 0.000694 | 0.01257 | 0.009829 | 0.002621 | 0.004401 | 0.002479 | 0.004025 | 0.001629 | 0.00885 | 0.018455 | 0.004135 | 0.000976 | 0.00475 | 0.003462 | 0.021305 |
| 25.1944444 | 0.00199 | 0.001685 | 0.002731 | 0.004152 | 0.00582 | 0.001409 | 0.005982 | 0.003192 | 0.004341 | 0.002548 | 0.015177 | 0.013585 | 0.001741 | 0.003873 | 0.003004 | 0.0025 | 0.001441 | 0.008068 | 0.01752 | 0.004821 | 0.001299 | 0.004228 | 0.004014 | 0.02196 |
| 25.2083333 | 0.005562 | 0.001779 | 0.002985 | 0.002566 | 0.003957 | 0.00146 | 0.004796 | 0.006355 | 0.005717 | 0.005034 | 0.017662 | 0.018912 | 0.001513 | 0.007548 | 0.002904 | 0.003791 | 0.001617 | 0.008016 | 0.016241 | 0.006941 | 0.003138 | 0.002708 | 0.006339 | 0.027358 |
| 25.2222222 | 0.012155 | 0.001625 | 0.00309 | 0.000841 | 0.004503 | 0.003173 | 0.002918 | 0.009345 | 0.006642 | 0.006694 | 0.018052 | 0.025083 | 0.001727 | 0.010851 | 0.001692 | 0.006326 | 0.001648 | 0.007888 | 0.015612 | 0.009397 | 0.006221 | 0.00299 | 0.011399 | 0.03016 |
| 25.2361111 | 0.017616 | 0.001413 | 0.00265 | 0.001166 | 0.004336 | 0.005924 | 0.001644 | 0.012047 | 0.004993 | 0.007121 | 0.01676 | 0.031132 | 0.002618 | 0.008906 | 0.001746 | 0.007839 | 0.000925 | 0.006878 | 0.015064 | 0.011674 | 0.007876 | 0.005057 | 0.014408 | 0.025812 |
| 25.25 | 0.018519 | 0.00116 | 0.001749 | 0.003892 | 0.002894 | 0.006709 | 0.003041 | 0.015318 | 0.003513 | 0.007064 | 0.014687 | 0.035506 | 0.00392 | 0.004713 | 0.00284 | 0.007189 | 0.000653 | 0.005031 | 0.012623 | 0.01383 | 0.00624 | 0.006039 | 0.013604 | 0.016887 |
| 25.2638889 | 0.014995 | 0.000711 | 0.000934 | 0.006656 | 0.003891 | 0.004518 | 0.005936 | 0.018108 | 0.003514 | 0.006637 | 0.012552 | 0.036616 | 0.004817 | 0.0015 | 0.003184 | 0.004911 | 0.001088 | 0.00261 | 0.007668 | 0.015692 | 0.003605 | 0.004663 | 0.011003 | 0.008205 |
| 25.2777778 | 0.009453 | 0.000478 | 0.000717 | 0.006769 | 0.004296 | 0.002045 | 0.007372 | 0.0184 | 0.00528 | 0.005568 | 0.010715 | 0.033169 | 0.005011 | 0.001201 | 0.002573 | 0.002593 | 0.001122 | 0.001193 | 0.003951 | 0.017366 | 0.003242 | 0.003066 | 0.008076 | 0.003881 |
| 25.2916667 | 0.004495 | 0.000914 | 0.000894 | 0.004459 | 0.003914 | 0.001165 | 0.005982 | 0.016215 | 0.00684 | 0.004528 | 0.009429 | 0.026546 | 0.004748 | 0.003754 | 0.001714 | 0.001343 | 0.000744 | 0.001605 | 0.005243 | 0.018761 | 0.003567 | 0.003102 | 0.006245 | 0.004227 |
| 25.3055556 | 0.0028 | 0.001563 | 0.001003 | 0.003329 | 0.005635 | 0.00127 | 0.003057 | 0.013855 | 0.00656 | 0.004348 | 0.008621 | 0.019818 | 0.004293 | 0.006607 | 0.001176 | 0.000989 | 0.000911 | 0.002332 | 0.007805 | 0.018856 | 0.003366 | 0.003941 | 0.00669 | 0.0038 |
| 25.3194444 | 0.00504 | 0.001859 | 0.001061 | 0.005475 | 0.006112 | 0.003026 | 0.001811 | 0.013621 | 0.007243 | 0.00512 | 0.007654 | 0.013836 | 0.004123 | 0.008207 | 0.001863 | 0.001419 | 0.00143 | 0.002124 | 0.007104 | 0.01793 | 0.005946 | 0.005326 | 0.009058 | 0.002773 |
| 25.3333333 | 0.007906 | 0.001916 | 0.001296 | 0.008291 | 0.004257 | 0.006515 | 0.003425 | 0.015197 | 0.010918 | 0.005957 | 0.006667 | 0.008007 | 0.004742 | 0.008675 | 0.004468 | 0.002481 | 0.001727 | 0.002422 | 0.00744 | 0.017324 | 0.010248 | 0.007843 | 0.011239 | 0.00599 |
| 25.3472222 | 0.008926 | 0.001874 | 0.001834 | 0.009622 | 0.003095 | 0.009833 | 0.005521 | 0.014823 | 0.013681 | 0.005895 | 0.006211 | 0.004404 | 0.006261 | 0.00804 | 0.00829 | 0.003847 | 0.001825 | 0.004401 | 0.010897 | 0.01696 | 0.011116 | 0.010181 | 0.011084 | 0.01271 |
| 25.3611111 | 0.008318 | 0.001637 | 0.002784 | 0.009644 | 0.003535 | 0.011523 | 0.006737 | 0.009648 | 0.011517 | 0.005164 | 0.006284 | 0.005728 | 0.008149 | 0.006601 | 0.011874 | 0.005829 | 0.002129 | 0.006052 | 0.010388 | 0.016289 | 0.00849 | 0.011466 | 0.008476 | 0.022321 |
| 25.375 | 0.007634 | 0.001602 | 0.004226 | 0.009039 | 0.004652 | 0.010752 | 0.008096 | 0.003712 | 0.006269 | 0.004488 | 0.006947 | 0.008549 | 0.009255 | 0.005132 | 0.01398 | 0.00786 | 0.002729 | 0.006146 | 0.005273 | 0.01595 | 0.005572 | 0.013139 | 0.005131 | 0.034023 |
| 25.3888889 | 0.007857 | 0.001988 | 0.005792 | 0.008415 | 0.005285 | 0.008 | 0.0102 | 0.001126 | 0.003106 | 0.004748 | 0.008539 | 0.007702 | 0.009015 | 0.004688 | 0.014145 | 0.007913 | 0.002954 | 0.005698 | 0.004506 | 0.01695 | 0.004418 | 0.016011 | 0.003449 | 0.042043 |
| 25.4027778 | 0.007274 | 0.002005 | 0.006937 | 0.008524 | 0.005319 | 0.006241 | 0.012728 | 0.001136 | 0.004767 | 0.007472 | 0.011531 | 0.004839 | 0.008445 | 0.006075 | 0.013176 | 0.005505 | 0.002479 | 0.005791 | 0.012945 | 0.019246 | 0.007913 | 0.019104 | 0.005478 | 0.045122 |
| 25.4166667 | 0.004954 | 0.001477 | 0.007568 | 0.010308 | 0.005237 | 0.007951 | 0.014644 | 0.004121 | 0.010137 | 0.01201 | 0.014753 | 0.005205 | 0.008657 | 0.007813 | 0.01248 | 0.003301 | 0.001962 | 0.006969 | 0.025622 | 0.021471 | 0.016892 | 0.02178 | 0.01051 | 0.048737 |
| 25.4305556 | 0.005412 | 0.001567 | 0.007922 | 0.013443 | 0.0049 | 0.012077 | 0.014414 | 0.009483 | 0.015469 | 0.014465 | 0.014945 | 0.008885 | 0.008674 | 0.00787 | 0.01278 | 0.003419 | 0.002172 | 0.008614 | 0.035469 | 0.022398 | 0.02506 | 0.02391 | 0.014728 | 0.050593 |
| 25.4444444 | 0.01196 | 0.002319 | 0.008266 | 0.016318 | 0.004239 | 0.015094 | 0.012401 | 0.013644 | 0.016419 | 0.013869 | 0.011322 | 0.011911 | 0.007085 | 0.00683 | 0.012772 | 0.005059 | 0.002705 | 0.009119 | 0.039573 | 0.02239 | 0.026456 | 0.024799 | 0.015483 | 0.041146 |
| 25.4583333 | 0.019726 | 0.002518 | 0.008399 | 0.017335 | 0.003817 | 0.016097 | 0.010658 | 0.015149 | 0.011897 | 0.013412 | 0.007472 | 0.013713 | 0.004834 | 0.005784 | 0.010777 | 0.006804 | 0.002454 | 0.007916 | 0.038186 | 0.022759 | 0.023671 | 0.024482 | 0.013885 | 0.02217 |
| 25.4722222 | 0.023777 | 0.002221 | 0.007985 | 0.0166 | 0.004007 | 0.017186 | 0.01059 | 0.014018 | 0.009232 | 0.01503 | 0.007413 | 0.016647 | 0.00471 | 0.005631 | 0.008637 | 0.00883 | 0.002498 | 0.00659 | 0.034307 | 0.024948 | 0.021591 | 0.025565 | 0.012493 | 0.007501 |
| 25.4861111 | 0.026106 | 0.002764 | 0.00795 | 0.017129 | 0.00526 | 0.017758 | 0.013671 | 0.010714 | 0.02187 | 0.021458 | 0.012359 | 0.021849 | 0.008349 | 0.007796 | 0.010315 | 0.012719 | 0.007027 | 0.006821 | 0.030272 | 0.030072 | 0.021939 | 0.03115 | 0.012908 | 0.008401 |
| 25.5 | 0.033686 | 0.005427 | 0.010373 | 0.023412 | 0.008181 | 0.021484 | 0.022805 | 0.006581 | 0.033471 | 0.039285 | 0.022815 | 0.031928 | 0.015293 | 0.010914 | 0.016465 | 0.019452 | 0.019558 | 0.009214 | 0.02672 | 0.039313 | 0.026072 | 0.045168 | 0.015514 | 0.028541 |
| 25.5138889 | 0.050926 | 0.009556 | 0.015404 | 0.0344 | 0.011595 | 0.035132 | 0.036526 | 0.004244 | 0.030012 | 0.065683 | 0.039341 | 0.047852 | 0.024628 | 0.011674 | 0.023636 | 0.030559 | 0.037622 | 0.013972 | 0.02445 | 0.054237 | 0.033222 | 0.071303 | 0.019042 | 0.05658 |
| 25.5277778 | 0.071675 | 0.011759 | 0.019864 | 0.039442 | 0.014061 | 0.047228 | 0.045241 | 0.003832 | 0.033489 | 0.082876 | 0.055965 | 0.059557 | 0.030787 | 0.008142 | 0.026379 | 0.044694 | 0.049088 | 0.020314 | 0.022483 | 0.068027 | 0.038614 | 0.092802 | 0.020272 | 0.064373 |
| 25.5416667 | 0.088641 | 0.010515 | 0.020104 | 0.033652 | 0.015055 | 0.043777 | 0.043904 | 0.00353 | 0.042301 | 0.08734 | 0.061217 | 0.056333 | 0.027905 | 0.003226 | 0.023126 | 0.054749 | 0.046661 | 0.025635 | 0.017086 | 0.067905 | 0.04076 | 0.088039 | 0.018335 | 0.043208 |
| 25.5555556 | 0.105625 | 0.008128 | 0.017757 | 0.025851 | 0.013233 | 0.037804 | 0.035384 | 0.006077 | 0.05544 | 0.092234 | 0.051149 | 0.043012 | 0.022321 | 0.001501 | 0.019238 | 0.052284 | 0.03882 | 0.026343 | 0.009436 | 0.056939 | 0.043059 | 0.070578 | 0.016466 | 0.018692 |
| 25.5694444 | 0.113877 | 0.006086 | 0.016066 | 0.020887 | 0.009078 | 0.035838 | 0.022309 | 0.009894 | 0.078942 | 0.097272 | 0.032499 | 0.031023 | 0.020477 | 0.001739 | 0.01685 | 0.036416 | 0.031741 | 0.021166 | 0.006026 | 0.045244 | 0.045014 | 0.058434 | 0.014506 | 0.01237 |
| 25.5833333 | 0.096174 | 0.004479 | 0.012725 | 0.01623 | 0.004621 | 0.026889 | 0.010254 | 0.009314 | 0.085739 | 0.089676 | 0.017459 | 0.02316 | 0.018256 | 0.001644 | 0.013344 | 0.017918 | 0.024376 | 0.013473 | 0.008167 | 0.032982 | 0.04279 | 0.04872 | 0.009751 | 0.020866 |
| 25.5972222 | 0.062496 | 0.003483 | 0.00857 | 0.011002 | 0.001861 | 0.0157 | 0.007486 | 0.006047 | 0.061268 | 0.061605 | 0.01852 | 0.015564 | 0.013681 | 0.000982 | 0.009869 | 0.007072 | 0.017768 | 0.00747 | 0.013675 | 0.021095 | 0.036769 | 0.040396 | 0.004853 | 0.028796 |
| 25.6111111 | 0.03088 | 0.002646 | 0.009441 | 0.008486 | 0.00125 | 0.010381 | 0.014596 | 0.005827 | 0.032193 | 0.031972 | 0.029092 | 0.010144 | 0.009847 | 0.000596 | 0.008713 | 0.007917 | 0.013763 | 0.003607 | 0.01971 | 0.011152 | 0.029674 | 0.033798 | 0.002732 | 0.025737 |
| 25.625 | 0.013918 | 0.001505 | 0.014838 | 0.012202 | 0.001258 | 0.008573 | 0.0223 | 0.008234 | 0.023301 | 0.023791 | 0.026036 | 0.010009 | 0.008845 | 0.001637 | 0.008408 | 0.021256 | 0.01203 | 0.006291 | 0.021443 | 0.006476 | 0.023451 | 0.026078 | 0.001721 | 0.017663 |
| 25.6388889 | 0.017939 | 0.000571 | 0.019665 | 0.016574 | 0.003139 | 0.005787 | 0.021286 | 0.009479 | 0.022784 | 0.030005 | 0.017854 | 0.019323 | 0.010925 | 0.003955 | 0.008513 | 0.041203 | 0.010792 | 0.007239 | 0.016673 | 0.009889 | 0.02133 | 0.017941 | 0.002435 | 0.021711 |
| 25.6527778 | 0.030434 | 0.001074 | 0.021749 | 0.013197 | 0.008142 | 0.003119 | 0.014385 | 0.011361 | 0.019171 | 0.04383 | 0.029226 | 0.026953 | 0.014802 | 0.006646 | 0.011378 | 0.047384 | 0.009333 | 0.01012 | 0.009062 | 0.015205 | 0.025424 | 0.012392 | 0.007253 | 0.040569 |
| 25.6666667 | 0.042107 | 0.002541 | 0.020971 | 0.006317 | 0.01314 | 0.004309 | 0.01695 | 0.016789 | 0.017069 | 0.055887 | 0.042754 | 0.022504 | 0.018024 | 0.008005 | 0.015289 | 0.029058 | 0.008194 | 0.008246 | 0.003992 | 0.013786 | 0.032908 | 0.008672 | 0.011054 | 0.061386 |
| 25.6805556 | 0.055233 | 0.002776 | 0.020005 | 0.005135 | 0.014396 | 0.007985 | 0.028448 | 0.017679 | 0.012836 | 0.045797 | 0.034418 | 0.013292 | 0.017603 | 0.006819 | 0.015434 | 0.013281 | 0.007497 | 0.011021 | 0.004867 | 0.008693 | 0.038134 | 0.007173 | 0.00844 | 0.067876 |
| 25.6944444 | 0.060176 | 0.001873 | 0.01952 | 0.008051 | 0.01125 | 0.011558 | 0.030433 | 0.01169 | 0.009019 | 0.023894 | 0.016121 | 0.013801 | 0.013014 | 0.003913 | 0.010126 | 0.020949 | 0.006621 | 0.012216 | 0.011244 | 0.009934 | 0.037328 | 0.014846 | 0.003969 | 0.047731 |
| 25.7083333 | 0.043487 | 0.001793 | 0.016096 | 0.010769 | 0.006009 | 0.011008 | 0.019702 | 0.010729 | 0.01356 | 0.022872 | 0.008601 | 0.020235 | 0.008344 | 0.002666 | 0.003766 | 0.034791 | 0.004343 | 0.027028 | 0.018672 | 0.02239 | 0.030911 | 0.029031 | 0.003309 | 0.024972 |
| 25.7222222 | 0.017409 | 0.00207 | 0.011231 | 0.01293 | 0.002432 | 0.006973 | 0.009949 | 0.013295 | 0.022619 | 0.04878 | 0.018786 | 0.02105 | 0.007096 | 0.004777 | 0.003067 | 0.034925 | 0.002735 | 0.02313 | 0.027387 | 0.036044 | 0.021794 | 0.036553 | 0.005296 | 0.035183 |
| 25.7361111 | 0.005903 | 0.001418 | 0.007445 | 0.013305 | 0.00201 | 0.006607 | 0.011337 | 0.016056 | 0.026557 | 0.076021 | 0.034789 | 0.015627 | 0.007779 | 0.007203 | 0.008145 | 0.020752 | 0.005285 | 0.015846 | 0.033015 | 0.039403 | 0.013357 | 0.035108 | 0.007441 | 0.066768 |
| 25.75 | 0.010579 | 0.001017 | 0.004705 | 0.009816 | 0.003605 | 0.00936 | 0.019006 | 0.018503 | 0.020894 | 0.082803 | 0.035304 | 0.008892 | 0.007374 | 0.007429 | 0.011972 | 0.007393 | 0.009156 | 0.009771 | 0.024342 | 0.03155 | 0.007369 | 0.032212 | 0.011195 | 0.088352 |
| 25.7638889 | 0.019064 | 0.001106 | 0.005006 | 0.006336 | 0.005686 | 0.009093 | 0.024818 | 0.012574 | 0.011514 | 0.06877 | 0.021042 | 0.006488 | 0.005184 | 0.00598 | 0.010151 | 0.007072 | 0.010389 | 0.005061 | 0.012623 | 0.017938 | 0.003391 | 0.03067 | 0.0167 | 0.077556 |
| 25.7777778 | 0.026467 | 0.00098 | 0.008512 | 0.010576 | 0.005032 | 0.005379 | 0.027751 | 0.005014 | 0.00564 | 0.040923 | 0.015061 | 0.00862 | 0.002812 | 0.004741 | 0.006273 | 0.013031 | 0.009554 | 0.002186 | 0.015873 | 0.007754 | 0.001245 | 0.028351 | 0.02114 | 0.045732 |

| | | | | | | | | | | | | | | | | | | | | | | | | |
|---|---|---|---|---|---|---|---|---|---|---|---|---|---|---|---|---|---|---|---|---|---|---|---|---|
| 25.7916667 | 0.029625 | 0.001898 | 0.01246 | 0.019892 | 0.00245 | 0.002726 | 0.027238 | 0.00486 | 0.003808 | 0.02583 | 0.021584 | 0.011598 | 0.002529 | 0.005202 | 0.003662 | 0.015717 | 0.008955 | 0.002578 | 0.028546 | 0.007383 | 0.000685 | 0.024563 | 0.018524 | 0.032719 |
| 25.8055556 | 0.02265 | 0.003847 | 0.015037 | 0.025807 | 0.00271 | 0.002504 | 0.022643 | 0.005415 | 0.002721 | 0.045986 | 0.02106 | 0.015956 | 0.004299 | 0.006936 | 0.002464 | 0.016051 | 0.009537 | 0.005253 | 0.03987 | 0.013831 | 0.000849 | 0.020639 | 0.009824 | 0.027833 |
| 25.8194444 | 0.015167 | 0.00504 | 0.016071 | 0.026405 | 0.00568 | 0.003316 | 0.016043 | 0.003008 | 0.001778 | 0.075401 | 0.013135 | 0.023614 | 0.005728 | 0.009592 | 0.002285 | 0.015404 | 0.010483 | 0.008088 | 0.046866 | 0.02074 | 0.002072 | 0.018144 | 0.00465 | 0.020129 |
| 25.8333333 | 0.025416 | 0.005201 | 0.015624 | 0.024701 | 0.007648 | 0.00473 | 0.009532 | 0.001264 | 0.002097 | 0.091119 | 0.014405 | 0.033278 | 0.006112 | 0.01416 | 0.002193 | 0.013325 | 0.010979 | 0.009999 | 0.048281 | 0.026361 | 0.004938 | 0.016863 | 0.002968 | 0.037405 |
| 25.8472222 | 0.046095 | 0.005787 | 0.012838 | 0.024321 | 0.005677 | 0.005876 | 0.004806 | 0.002146 | 0.003861 | 0.09036 | 0.023494 | 0.041792 | 0.006553 | 0.02014 | 0.004125 | 0.016025 | 0.01095 | 0.008146 | 0.045942 | 0.032443 | 0.009945 | 0.014522 | 0.003552 | 0.070139 |
| 25.8611111 | 0.05692 | 0.007874 | 0.009257 | 0.027314 | 0.003549 | 0.01058 | 0.002905 | 0.006769 | 0.010233 | 0.069614 | 0.028816 | 0.044842 | 0.007421 | 0.023996 | 0.013402 | 0.027994 | 0.011133 | 0.005894 | 0.04246 | 0.040601 | 0.017996 | 0.011312 | 0.008986 | 0.090679 |
| 25.875 | 0.054395 | 0.010227 | 0.008523 | 0.031089 | 0.005931 | 0.021062 | 0.002105 | 0.016076 | 0.02253 | 0.040971 | 0.025918 | 0.039577 | 0.007975 | 0.02425 | 0.028243 | 0.040513 | 0.012731 | 0.006241 | 0.038843 | 0.050078 | 0.028168 | 0.011081 | 0.01777 | 0.092729 |
| 25.8888889 | 0.046571 | 0.010359 | 0.01136 | 0.030381 | 0.009984 | 0.0292 | 0.001404 | 0.02769 | 0.036807 | 0.02219 | 0.017406 | 0.029732 | 0.00811 | 0.024038 | 0.034315 | 0.041446 | 0.016131 | 0.00519 | 0.039844 | 0.057778 | 0.038052 | 0.016108 | 0.023872 | 0.072547 |
| 25.9027778 | 0.039796 | 0.007673 | 0.01724 | 0.026349 | 0.012291 | 0.030354 | 0.001258 | 0.033035 | 0.04885 | 0.017766 | 0.010145 | 0.022209 | 0.010085 | 0.027296 | 0.027109 | 0.032452 | 0.021158 | 0.006639 | 0.049892 | 0.064687 | 0.046077 | 0.027013 | 0.021299 | 0.039906 |
| 25.9166667 | 0.037049 | 0.00388 | 0.02468 | 0.024337 | 0.011052 | 0.029563 | 0.002891 | 0.026655 | 0.053523 | 0.020855 | 0.010483 | 0.018313 | 0.014399 | 0.033551 | 0.021313 | 0.022569 | 0.027513 | 0.012549 | 0.062318 | 0.071868 | 0.05073 | 0.042502 | 0.016688 | 0.024988 |
| 25.9305556 | 0.034103 | 0.001575 | 0.026939 | 0.023923 | 0.00661 | 0.023742 | 0.008322 | 0.016407 | 0.045846 | 0.024615 | 0.017577 | 0.014438 | 0.017923 | 0.034409 | 0.021926 | 0.012892 | 0.032474 | 0.016711 | 0.060843 | 0.070742 | 0.046913 | 0.05308 | 0.018687 | 0.026269 |
| 25.9444444 | 0.023921 | 0.001592 | 0.020523 | 0.02238 | 0.002382 | 0.010776 | 0.015439 | 0.008947 | 0.028557 | 0.029035 | 0.024099 | 0.008216 | 0.019144 | 0.025205 | 0.020207 | 0.005267 | 0.032341 | 0.01545 | 0.042202 | 0.057827 | 0.034453 | 0.049518 | 0.020127 | 0.02021 |
| 25.9583333 | 0.012015 | 0.001921 | 0.011702 | 0.019313 | 0.000846 | 0.00516 | 0.019241 | 0.003863 | 0.014003 | 0.033424 | 0.024882 | 0.004557 | 0.018322 | 0.013465 | 0.015645 | 0.003289 | 0.02734 | 0.01078 | 0.020954 | 0.040966 | 0.023507 | 0.036815 | 0.016554 | 0.018722 |
| 25.9722222 | 0.005597 | 0.002188 | 0.005428 | 0.015141 | 0.001363 | 0.009855 | 0.016398 | 0.002093 | 0.010953 | 0.030762 | 0.019362 | 0.008657 | 0.015303 | 0.006842 | 0.012031 | 0.003303 | 0.021625 | 0.00653 | 0.00775 | 0.024552 | 0.019146 | 0.025446 | 0.01656 | 0.041355 |
| 25.9861111 | 0.004603 | 0.002598 | 0.002875 | 0.011427 | 0.002353 | 0.013906 | 0.008475 | 0.003964 | 0.017785 | 0.021765 | 0.011195 | 0.014663 | 0.011571 | 0.005514 | 0.010044 | 0.002119 | 0.018966 | 0.004141 | 0.004099 | 0.011175 | 0.019752 | 0.018928 | 0.022773 | 0.075435 |
| 26 | 0.010263 | 0.002502 | 0.005889 | 0.011026 | 0.005474 | 0.015991 | 0.002443 | 0.007442 | 0.029843 | 0.016884 | 0.007832 | 0.014828 | 0.009852 | 0.006443 | 0.010355 | 0.00184 | 0.019605 | 0.004529 | 0.007099 | 0.004553 | 0.024575 | 0.016802 | 0.030968 | 0.092621 |
| 26.0138889 | 0.02428 | 0.003335 | 0.012743 | 0.014627 | 0.012107 | 0.018178 | 0.002041 | 0.010321 | 0.042691 | 0.017987 | 0.013329 | 0.009489 | 0.00937 | 0.007421 | 0.011388 | 0.003394 | 0.018943 | 0.006326 | 0.011391 | 0.002994 | 0.030918 | 0.017978 | 0.038874 | 0.079932 |
| 26.0277778 | 0.045477 | 0.005202 | 0.017194 | 0.019546 | 0.018011 | 0.020819 | 0.005085 | 0.011652 | 0.054282 | 0.019199 | 0.021788 | 0.008737 | 0.006374 | 0.008109 | 0.010838 | 0.003322 | 0.013302 | 0.005622 | 0.014672 | 0.004888 | 0.03486 | 0.02146 | 0.045047 | 0.055696 |
| 26.0416667 | 0.069664 | 0.008412 | 0.015215 | 0.025538 | 0.019249 | 0.027086 | 0.011797 | 0.009879 | 0.061862 | 0.020043 | 0.026035 | 0.021099 | 0.002536 | 0.009443 | 0.013275 | 0.003324 | 0.009517 | 0.004367 | 0.020822 | 0.012697 | 0.036323 | 0.026593 | 0.043791 | 0.043552 |
| 26.0555556 | 0.08564 | 0.014888 | 0.010601 | 0.032549 | 0.015252 | 0.035621 | 0.021868 | 0.005995 | 0.05923 | 0.021911 | 0.030781 | 0.039212 | 0.003572 | 0.012369 | 0.023585 | 0.008747 | 0.015047 | 0.009031 | 0.029874 | 0.026873 | 0.037003 | 0.032132 | 0.031437 | 0.045066 |
| 26.0694444 | 0.0791 | 0.021172 | 0.009153 | 0.033659 | 0.009486 | 0.038315 | 0.026264 | 0.003211 | 0.04783 | 0.022625 | 0.037028 | 0.048818 | 0.007774 | 0.013707 | 0.031422 | 0.013197 | 0.022859 | 0.016676 | 0.029798 | 0.039016 | 0.034311 | 0.032594 | 0.01495 | 0.04299 |
| 26.0833333 | 0.051965 | 0.021505 | 0.00977 | 0.023744 | 0.009449 | 0.031425 | 0.01976 | 0.002001 | 0.035388 | 0.02287 | 0.029262 | 0.042349 | 0.008638 | 0.010328 | 0.02469 | 0.009474 | 0.022491 | 0.017983 | 0.01895 | 0.038998 | 0.025069 | 0.023046 | 0.004962 | 0.026944 |
| 26.0972222 | 0.022958 | 0.016101 | 0.008233 | 0.011195 | 0.014005 | 0.01799 | 0.009636 | 0.003164 | 0.021829 | 0.028741 | 0.018878 | 0.025316 | 0.00544 | 0.008937 | 0.012027 | 0.006188 | 0.015122 | 0.011983 | 0.015081 | 0.029515 | 0.012474 | 0.010004 | 0.002748 | 0.019813 |
| 26.1111111 | 0.009042 | 0.008784 | 0.004671 | 0.007284 | 0.014841 | 0.007427 | 0.003167 | 0.005869 | 0.01182 | 0.03534 | 0.034282 | 0.010283 | 0.002223 | 0.012501 | 0.007399 | 0.009211 | 0.007338 | 0.005101 | 0.021542 | 0.019148 | 0.00449 | 0.004131 | 0.001488 | 0.035764 |
| 26.125 | 0.010412 | 0.003802 | 0.003184 | 0.009096 | 0.011409 | 0.005135 | 0.001619 | 0.006981 | 0.014589 | 0.026858 | 0.056779 | 0.003023 | 0.00135 | 0.013835 | 0.008987 | 0.011086 | 0.002716 | 0.001876 | 0.024828 | 0.010971 | 0.004253 | 0.007012 | 0.002577 | 0.05074 |
| 26.1388889 | 0.013359 | 0.003512 | 0.004528 | 0.00878 | 0.007003 | 0.004945 | 0.003396 | 0.005873 | 0.024313 | 0.01164 | 0.058106 | 0.0019 | 0.001802 | 0.011929 | 0.009432 | 0.008917 | 0.001686 | 0.002828 | 0.022368 | 0.004845 | 0.006927 | 0.010376 | 0.009685 | 0.051525 |
| 26.1527778 | 0.011079 | 0.004798 | 0.004355 | 0.00626 | 0.003539 | 0.006861 | 0.005846 | 0.004046 | 0.030336 | 0.007574 | 0.042005 | 0.002551 | 0.001643 | 0.010448 | 0.007315 | 0.006403 | 0.001881 | 0.005818 | 0.019894 | 0.001582 | 0.008086 | 0.00905 | 0.017414 | 0.046953 |
| 26.1666667 | 0.006101 | 0.004542 | 0.002753 | 0.003394 | 0.003368 | 0.013509 | 0.006944 | 0.002731 | 0.026631 | 0.008237 | 0.020614 | 0.002191 | 0.001164 | 0.010726 | 0.004093 | 0.004742 | 0.001329 | 0.008385 | 0.019983 | 0.001252 | 0.006928 | 0.00538 | 0.018668 | 0.041642 |
| 26.1805556 | 0.002338 | 0.003532 | 0.0031 | 0.001196 | 0.005184 | 0.017477 | 0.006386 | 0.002425 | 0.014567 | 0.006834 | 0.00726 | 0.001395 | 0.001766 | 0.012353 | 0.001448 | 0.003237 | 0.000904 | 0.009948 | 0.022776 | 0.001676 | 0.003916 | 0.002441 | 0.012253 | 0.035931 |
| 26.1944444 | 0.001414 | 0.002829 | 0.00349 | 0.000962 | 0.00549 | 0.013519 | 0.005437 | 0.003792 | 0.005563 | 0.005393 | 0.004921 | 0.001449 | 0.002632 | 0.012691 | 0.000293 | 0.001989 | 0.001442 | 0.011381 | 0.026555 | 0.002104 | 0.001735 | 0.001824 | 0.006184 | 0.030933 |
| 26.2083333 | 0.001513 | 0.002648 | 0.002702 | 0.002677 | 0.003637 | 0.0067 | 0.005167 | 0.005059 | 0.005378 | 0.003507 | 0.006168 | 0.002491 | 0.002789 | 0.010111 | 0.000251 | 0.001768 | 0.002054 | 0.012971 | 0.028237 | 0.003418 | 0.002164 | 0.002023 | 0.00718 | 0.026616 |
| 26.2222222 | 0.003 | 0.0029 | 0.00346 | 0.005055 | 0.001753 | 0.003781 | 0.005101 | 0.003746 | 0.007341 | 0.001579 | 0.012492 | 0.003613 | 0.002554 | 0.00671 | 0.001087 | 0.002985 | 0.001753 | 0.013804 | 0.025377 | 0.005608 | 0.003092 | 0.001372 | 0.009524 | 0.021978 |
| 26.2361111 | 0.00517 | 0.003054 | 0.005881 | 0.006457 | 0.001693 | 0.004518 | 0.004209 | 0.002474 | 0.005587 | 0.000678 | 0.02025 | 0.004539 | 0.001853 | 0.00494 | 0.002756 | 0.004762 | 0.001174 | 0.013353 | 0.017871 | 0.007534 | 0.002784 | 0.000793 | 0.007905 | 0.016834 |
| 26.25 | 0.004799 | 0.002824 | 0.0078 | 0.006198 | 0.002342 | 0.004772 | 0.002402 | 0.002978 | 0.002498 | 0.000467 | 0.017106 | 0.005415 | 0.000851 | 0.003988 | 0.004003 | 0.005531 | 0.001654 | 0.012008 | 0.009028 | 0.008096 | 0.001837 | 0.001034 | 0.005444 | 0.012948 |
| 26.2638889 | 0.004654 | 0.002419 | 0.008313 | 0.004844 | 0.002419 | 0.004148 | 0.001331 | 0.00253 | 0.001295 | 0.000371 | 0.010021 | 0.005988 | 0.000267 | 0.002657 | 0.003455 | 0.004817 | 0.002625 | 0.010558 | 0.003214 | 0.007447 | 0.001412 | 0.001187 | 0.005382 | 0.013448 |
| 26.2777778 | 0.007717 | 0.001997 | 0.007897 | 0.003336 | 0.002171 | 0.004766 | 0.002121 | 0.003345 | 0.002114 | 0.000387 | 0.011834 | 0.006249 | 0.000262 | 0.002174 | 0.001856 | 0.002988 | 0.003031 | 0.009569 | 0.002653 | 0.006741 | 0.00175 | 0.000948 | 0.008078 | 0.018202 |
| 26.2916667 | 0.009397 | 0.001661 | 0.007436 | 0.00202 | 0.001707 | 0.006695 | 0.00316 | 0.007045 | 0.005762 | 0.000706 | 0.012331 | 0.006547 | 0.000312 | 0.002317 | 0.001086 | 0.001083 | 0.002954 | 0.008639 | 0.00789 | 0.006889 | 0.002017 | 0.001042 | 0.010566 | 0.022293 |
| 26.3055556 | 0.006874 | 0.001347 | 0.007197 | 0.001193 | 0.001006 | 0.008614 | 0.003335 | 0.00855 | 0.009529 | 0.001161 | 0.007962 | 0.006974 | 0.00028 | 0.001714 | 0.001009 | 0.000579 | 0.002593 | 0.00718 | 0.017068 | 0.007728 | 0.001498 | 0.001453 | 0.008826 | 0.021028 |
| 26.3194444 | 0.003123 | 0.000911 | 0.006631 | 0.001376 | 0.000828 | 0.009619 | 0.003132 | 0.005563 | 0.008209 | 0.001409 | 0.009351 | 0.00739 | 0.000619 | 0.001412 | 0.001188 | 0.001694 | 0.001954 | 0.005802 | 0.024709 | 0.00893 | 0.001356 | 0.001516 | 0.00512 | 0.015674 |
| 26.3333333 | 0.001912 | 0.000699 | 0.005492 | 0.001406 | 0.002159 | 0.010134 | 0.002702 | 0.003417 | 0.006564 | 0.002054 | 0.012661 | 0.007987 | 0.001364 | 0.001738 | 0.002959 | 0.003508 | 0.001198 | 0.005516 | 0.025737 | 0.010773 | 0.003503 | 0.001226 | 0.005072 | 0.012802 |
| 26.3472222 | 0.00382 | 0.001276 | 0.004129 | 0.001519 | 0.003881 | 0.011579 | 0.002257 | 0.00619 | 0.010704 | 0.004238 | 0.011428 | 0.009182 | 0.0022 | 0.001848 | 0.005922 | 0.005284 | 0.000936 | 0.00639 | 0.020587 | 0.013805 | 0.007524 | 0.001439 | 0.00801 | 0.017366 |
| 26.3611111 | 0.005835 | 0.002339 | 0.003041 | 0.003883 | 0.003448 | 0.014067 | 0.002183 | 0.009924 | 0.013925 | 0.008434 | 0.007811 | 0.01081 | 0.002634 | 0.0019 | 0.00796 | 0.00638 | 0.001329 | 0.007262 | 0.015028 | 0.01735 | 0.011547 | 0.002326 | 0.011021 | 0.02803 |
| 26.375 | 0.006448 | 0.003109 | 0.002802 | 0.007357 | 0.002146 | 0.015598 | 0.001811 | 0.010596 | 0.011542 | 0.012495 | 0.004622 | 0.01141 | 0.002506 | 0.00221 | 0.008522 | 0.007125 | 0.001996 | 0.007132 | 0.015136 | 0.019455 | 0.014606 | 0.003601 | 0.01496 | 0.038798 |
| 26.3888889 | 0.007627 | 0.003498 | 0.003818 | 0.010469 | 0.002989 | 0.014755 | 0.001702 | 0.007106 | 0.006427 | 0.013096 | 0.003855 | 0.010294 | 0.002737 | 0.003115 | 0.008631 | 0.008644 | 0.003025 | 0.006652 | 0.021217 | 0.019487 | 0.017184 | 0.005727 | 0.020485 | 0.043065 |
| 26.4027778 | 0.01189 | 0.003716 | 0.00527 | 0.014183 | 0.00441 | 0.012638 | 0.004 | 0.003464 | 0.003644 | 0.010962 | 0.007062 | 0.009374 | 0.003829 | 0.004539 | 0.008855 | 0.010434 | 0.004615 | 0.006573 | 0.02836 | 0.018569 | 0.019888 | 0.008686 | 0.025265 | 0.039004 |
| 26.4166667 | 0.019037 | 0.003659 | 0.005814 | 0.018542 | 0.005051 | 0.011731 | 0.007444 | 0.003512 | 0.006469 | 0.009971 | 0.012319 | 0.009706 | 0.004968 | 0.005558 | 0.008634 | 0.011629 | 0.006237 | 0.006475 | 0.032894 | 0.017778 | 0.022054 | 0.010925 | 0.025569 | 0.030306 |
| 26.4305556 | 0.025612 | 0.003248 | 0.005593 | 0.02148 | 0.005918 | 0.013535 | 0.009474 | 0.003663 | 0.010702 | 0.012209 | 0.016431 | 0.010053 | 0.005406 | 0.005639 | 0.007651 | 0.012548 | 0.006978 | 0.006119 | 0.033278 | 0.016961 | 0.022232 | 0.011327 | 0.019239 | 0.021649 |
| 26.4444444 | 0.027889 | 0.002523 | 0.005291 | 0.022033 | 0.006404 | 0.016944 | 0.010015 | 0.003056 | 0.015446 | 0.016928 | 0.019132 | 0.010018 | 0.004966 | 0.005092 | 0.006185 | 0.012602 | 0.006192 | 0.005875 | 0.028605 | 0.015781 | 0.019697 | 0.010778 | 0.010174 | 0.017949 |
| 26.4583333 | 0.025132 | 0.001569 | 0.005104 | 0.022151 | 0.005166 | 0.018973 | 0.009852 | 0.003851 | 0.021545 | 0.020479 | 0.018591 | 0.010403 | 0.003559 | 0.004289 | 0.004416 | 0.011078 | 0.003867 | 0.005747 | 0.021999 | 0.014707 | 0.015557 | 0.010324 | 0.005819 | 0.023166 |
| 26.4722222 | 0.020672 | 0.00096 | 0.005304 | 0.023358 | 0.00285 | 0.01816 | 0.009723 | 0.006279 | 0.019613 | 0.020038 | 0.011881 | 0.0113 | 0.001938 | 0.003663 | 0.003382 | 0.01 | 0.001751 | 0.005902 | 0.020633 | 0.014622 | 0.012796 | 0.010673 | 0.004836 | 0.036222 |
| 26.4861111 | 0.019199 | 0.001542 | 0.006694 | 0.026703 | 0.002173 | 0.016466 | 0.010548 | 0.00973 | 0.013712 | 0.01981 | 0.005302 | 0.01371 | 0.002188 | 0.003814 | 0.005133 | 0.012578 | 0.002846 | 0.007582 | 0.02712 | 0.017311 | 0.014811 | 0.014044 | 0.005462 | 0.050859 |
| 26.5 | 0.025824 | 0.003745 | 0.010088 | 0.036867 | 0.004546 | 0.017486 | 0.014458 | 0.013282 | 0.020488 | 0.028444 | 0.012888 | 0.021222 | 0.006019 | 0.004642 | 0.009707 | 0.020996 | 0.009067 | 0.0114 | 0.034569 | 0.024791 | 0.023664 | 0.023644 | 0.014506 | 0.061015 |
| 26.5138889 | 0.04503 | 0.007873 | 0.014633 | 0.053474 | 0.008174 | 0.022178 | 0.021727 | 0.020393 | 0.029454 | 0.050928 | 0.040024 | 0.033028 | 0.013166 | 0.005065 | 0.013615 | 0.036188 | 0.018993 | 0.016079 | 0.035126 | 0.036399 | 0.040085 | 0.03892 | 0.032755 | 0.065661 |
| 26.5277778 | 0.069782 | 0.010923 | 0.017137 | 0.061026 | 0.01061 | 0.025283 | 0.027285 | 0.036335 | 0.023155 | 0.082063 | 0.073983 | 0.0411 | 0.019682 | 0.004106 | 0.013387 | 0.052601 | 0.0255 | 0.018518 | 0.028825 | 0.045589 | 0.055739 | 0.049746 | 0.050188 | 0.061838 |
| 26.5416667 | 0.085465 | 0.0103 | 0.015488 | 0.051302 | 0.011923 | 0.022772 | 0.026999 | 0.051175 | 0.013095 | 0.113902 | 0.090069 | 0.03942 | 0.019373 | 0.002489 | 0.011298 | 0.058508 | 0.023828 | 0.017413 | 0.0229 | 0.042916 | 0.057129 | 0.047213 | 0.053044 | 0.045709 |
| 26.5555556 | 0.090202 | 0.009952 | 0.012482 | 0.038052 | 0.012195 | 0.01719 | 0.02321 | 0.049139 | 0.013545 | 0.13647 | 0.076212 | 0.029876 | 0.013609 | 0.002459 | 0.011716 | 0.050742 | 0.020145 | 0.016466 | 0.023268 | 0.030971 | 0.047386 | 0.03875 | 0.042968 | 0.024852 |
| 26.5694444 | 0.088202 | 0.011277 | 0.011788 | 0.030087 | 0.009666 | 0.009866 | 0.01896 | 0.035435 | 0.021262 | 0.138718 | 0.045328 | 0.019059 | 0.008952 | 0.003645 | 0.012627 | 0.036274 | 0.020266 | 0.017048 | 0.024934 | 0.021332 | 0.036588 | 0.033715 | 0.032628 | 0.012432 |
| 26.5833333 | 0.080029 | 0.01127 | 0.013141 | 0.021312 | 0.005233 | 0.005101 | 0.014063 | 0.020501 | 0.029129 | 0.122689 | 0.021822 | 0.010864 | 0.005349 | 0.003167 | 0.010614 | 0.020256 | 0.021697 | 0.015736 | 0.01977 | 0.016413 | 0.025384 | 0.030725 | 0.021525 | 0.015827 |
| 26.5972222 | 0.067765 | 0.009297 | 0.013765 | 0.01083 | 0.001883 | 0.007279 | 0.008209 | 0.010492 | 0.028855 | 0.090439 | 0.023313 | 0.005476 | 0.002461 | 0.001435 | 0.007378 | 0.008368 | 0.021691 | 0.012276 | 0.010181 | 0.014684 | 0.01494 | 0.027571 | 0.010876 | 0.029936 |
| 26.6111111 | 0.052196 | 0.006309 | 0.012376 | 0.005484 | 0.000445 | 0.00994 | 0.004171 | 0.010514 | 0.018908 | 0.047626 | 0.041365 | 0.003601 | 0.002148 | 0.000849 | 0.005836 | 0.00706 | 0.020514 | 0.009379 | 0.003977 | 0.015464 | 0.009447 | 0.024248 | 0.008542 | 0.038565 |
| 26.625 | 0.034152 | 0.003306 | 0.008587 | 0.005523 | 0.000223 | 0.007954 | 0.004761 | 0.015878 | 0.010484 | 0.028209 | 0.05012 | 0.007075 | 0.002745 | 0.002151 | 0.007124 | 0.012868 | 0.018002 | 0.007705 | 0.003821 | 0.017044 | 0.011514 | 0.021181 | 0.009462 | 0.030338 |
| 26.6388889 | 0.01816 | 0.001536 | 0.003316 | 0.006421 | 0.000978 | 0.004185 | 0.005251 | 0.024142 | 0.012196 | 0.047431 | 0.03504 | 0.015982 | 0.00491 | 0.005026 | 0.011972 | 0.016228 | 0.014946 | 0.006269 | 0.005613 | 0.018782 | 0.019961 | 0.020262 | 0.008285 | 0.026772 |
| 26.6527778 | 0.014367 | 0.001986 | 0.000446 | 0.007421 | 0.003008 | 0.002205 | 0.004804 | 0.028434 | 0.014372 | 0.075067 | 0.022846 | 0.026581 | 0.010328 | 0.007926 | 0.019357 | 0.012229 | 0.013624 | 0.004448 | 0.005637 | 0.019155 | 0.030398 | 0.021796 | 0.012254 | 0.050039 |
| 26.6666667 | 0.027482 | 0.003725 | 0.000769 | 0.007077 | 0.006708 | 0.0022 | 0.009983 | 0.018268 | 0.011627 | 0.095576 | 0.040715 | 0.033983 | 0.016399 | 0.009374 | 0.024884 | 0.007508 | 0.013571 | 0.002876 | 0.011268 | 0.016141 | 0.040711 | 0.02331 | 0.022055 | 0.069299 |
| 26.6805556 | 0.044635 | 0.005097 | 0.002019 | 0.009047 | 0.011688 | 0.002163 | 0.017098 | 0.00559 | 0.009376 | 0.101019 | 0.057825 | 0.03396 | 0.020555 | 0.009453 | 0.025574 | 0.011876 | 0.012615 | 0.003349 | 0.030708 | 0.009426 | 0.053302 | 0.023237 | 0.027025 | 0.056086 |
| 26.6944444 | 0.051899 | 0.005915 | 0.004439 | 0.018677 | 0.015016 | 0.005243 | 0.01758 | 0.003496 | 0.00867 | 0.07672 | 0.044012 | 0.024538 | 0.021485 | 0.008515 | 0.022163 | 0.02069 | 0.010785 | 0.006565 | 0.052365 | 0.004265 | 0.061971 | 0.0208 | 0.024837 | 0.026683 |
| 26.7083333 | 0.047785 | 0.006815 | 0.009165 | 0.028576 | 0.013476 | 0.015578 | 0.010606 | 0.005575 | 0.011586 | 0.039212 | 0.022853 | 0.012639 | 0.017664 | 0.006473 | 0.015704 | 0.02352 | 0.008946 | 0.010129 | 0.05769 | 0.00521 | 0.054082 | 0.016037 | 0.021694 | 0.010156 |
| 26.7222222 | 0.036284 | 0.00783 | 0.011274 | 0.030889 | 0.007844 | 0.024913 | 0.005815 | 0.005083 | 0.01012 | 0.013901 | 0.028652 | 0.010014 | 0.009713 | 0.00352 | 0.008459 | 0.017203 | 0.007065 | 0.010631 | 0.045888 | 0.007877 | 0.034652 | 0.010414 | 0.015957 | 0.015255 |
| 26.7361111 | 0.027155 | 0.008283 | 0.00881 | 0.025017 | 0.005494 | 0.025402 | 0.010778 | 0.003473 | 0.005631 | 0.003726 | 0.057073 | 0.017979 | 0.004075 | 0.001468 | 0.004983 | 0.009987 | 0.004982 | 0.007586 | 0.02864 | 0.007537 | 0.016342 | 0.006975 | 0.009023 | 0.026988 |
| 26.75 | 0.027312 | 0.007838 | 0.006192 | 0.014098 | 0.00987 | 0.020406 | 0.018862 | 0.003346 | 0.007772 | 0.000971 | 0.082081 | 0.026211 | 0.00457 | 0.001976 | 0.004855 | 0.013022 | 0.003057 | 0.003912 | 0.014025 | 0.004665 | 0.008648 | 0.009344 | 0.010777 | 0.03634 |
| 26.7638889 | 0.025357 | 0.006799 | 0.00339 | 0.008413 | 0.012994 | 0.012618 | 0.023457 | 0.003326 | 0.008486 | 0.000681 | 0.087562 | 0.026711 | 0.006098 | 0.003768 | 0.003338 | 0.023035 | 0.001903 | 0.001938 | 0.005544 | 0.002017 | 0.012679 | 0.013927 | 0.01846 | 0.046246 |
| 26.7777778 | 0.014156 | 0.005787 | 0.00111 | 0.014823 | 0.010812 | 0.005557 | 0.023487 | 0.003435 | 0.007547 | 0.001468 | 0.068967 | 0.020194 | 0.005158 | 0.004375 | 0.001271 | 0.028538 | 0.001432 | 0.001544 | 0.004522 | 0.001404 | 0.016133 | 0.01488 | 0.023232 | 0.057814 |
| 26.7916667 | 0.005526 | 0.004965 | 0.001043 | 0.024307 | 0.006137 | 0.002781 | 0.018457 | 0.00602 | 0.011692 | 0.00266 | 0.037073 | 0.010745 | 0.00299 | 0.003148 | 0.00125 | 0.023971 | 0.000973 | 0.002042 | 0.008033 | 0.001737 | 0.015018 | 0.012742 | 0.023213 | 0.064596 |
| 26.8055556 | 0.004127 | 0.003807 | 0.003642 | 0.025944 | 0.002577 | 0.002291 | 0.010919 | 0.008025 | 0.013677 | 0.00345 | 0.019526 | 0.005925 | 0.001795 | 0.001623 | 0.001763 | 0.014514 | 0.001036 | 0.00266 | 0.012501 | 0.001358 | 0.012014 | 0.010351 | 0.019458 | 0.064349 |
| 26.8194444 | 0.006168 | 0.002279 | 0.008573 | 0.019364 | 0.002904 | 0.005309 | 0.005041 | 0.008488 | 0.009666 | 0.004229 | 0.023407 | 0.010148 | 0.001504 | 0.000975 | 0.003134 | 0.008621 | 0.002613 | 0.002444 | 0.017255 | 0.001526 | 0.007252 | 0.010238 | 0.014419 | 0.064508 |
| 26.8333333 | 0.015307 | 0.001149 | 0.013452 | 0.011403 | 0.0048 | 0.014374 | 0.002622 | 0.007961 | 0.005752 | 0.005775 | 0.025671 | 0.018279 | 0.001919 | 0.001432 | 0.006767 | 0.008308 | 0.004991 | 0.00143 | 0.022679 | 0.004744 | 0.002958 | 0.012658 | 0.009039 | 0.06512 |
| 26.8472222 | 0.028665 | 0.001099 | 0.017744 | 0.010764 | 0.004294 | 0.024787 | 0.003797 | 0.010413 | 0.007492 | 0.007718 | 0.016208 | 0.024522 | 0.004944 | 0.003585 | 0.010034 | 0.011463 | 0.007283 | 0.000814 | 0.029679 | 0.010052 | 0.001993 | 0.015736 | 0.003994 | 0.062214 |
| 26.8611111 | 0.041533 | 0.001817 | 0.021844 | 0.020781 | 0.004117 | 0.028964 | 0.007969 | 0.022506 | 0.016835 | 0.010035 | 0.011938 | 0.026914 | 0.00938 | 0.009329 | 0.010842 | 0.016438 | 0.009857 | 0.001433 | 0.039127 | 0.014928 | 0.006653 | 0.018881 | 0.002704 | 0.06035 |
| 26.875 | 0.05375 | 0.002053 | 0.024325 | 0.036192 | 0.008856 | 0.024352 | 0.012746 | 0.034459 | 0.029304 | 0.013401 | 0.028987 | 0.02712 | 0.01058 | 0.016644 | 0.010671 | 0.021233 | 0.012985 | 0.00325 | 0.05027 | 0.018112 | 0.018144 | 0.021887 | 0.007688 | 0.063256 |
| 26.8888889 | 0.056989 | 0.001348 | 0.023062 | 0.047938 | 0.013776 | 0.017601 | 0.015058 | 0.036281 | 0.038602 | 0.017957 | 0.053545 | 0.027706 | 0.008168 | 0.020624 | 0.011313 | 0.023655 | 0.016592 | 0.005595 | 0.061571 | 0.019924 | 0.031822 | 0.023458 | 0.016653 | 0.063924 |
| 26.9027778 | 0.047623 | 0.000988 | 0.01858 | 0.051372 | 0.012691 | 0.016954 | 0.01383 | 0.029703 | 0.042107 | 0.021486 | 0.060998 | 0.027422 | 0.006294 | 0.021226 | 0.01275 | 0.024599 | 0.020935 | 0.007505 | 0.07024 | 0.021171 | 0.037942 | 0.023666 | 0.024507 | 0.053171 |
| 26.9166667 | 0.0333 | 0.001497 | 0.013448 | 0.047414 | 0.007831 | 0.021307 | 0.009923 | 0.018616 | 0.039574 | 0.019685 | 0.042608 | 0.021985 | 0.005304 | 0.017776 | 0.013712 | 0.024181 | 0.024949 | 0.008969 | 0.072313 | 0.021458 | 0.036226 | 0.023358 | 0.028985 | 0.034103 |
| 26.9305556 | 0.02254 | 0.001913 | 0.009025 | 0.037389 | 0.007259 | 0.023828 | 0.004805 | 0.008576 | 0.030002 | 0.012614 | 0.020464 | 0.011957 | 0.003651 | 0.009833 | 0.013381 | 0.020481 | 0.02773 | 0.010299 | 0.068408 | 0.021189 | 0.034359 | 0.022577 | 0.030851 | 0.018771 |
| 26.9444444 | 0.017685 | 0.00259 | 0.00547 | 0.023074 | 0.012339 | 0.0208 | 0.001243 | 0.003174 | 0.016072 | 0.006496 | 0.018353 | 0.004216 | 0.002071 | 0.002858 | 0.012552 | 0.014827 | 0.030717 | 0.011207 | 0.064164 | 0.022382 | 0.033544 | 0.023324 | 0.030916 | 0.012928 |
| 26.9583333 | 0.015197 | 0.003898 | 0.00362 | 0.009752 | 0.012962 | 0.013354 | 0.000856 | 0.001699 | 0.010081 | 0.005908 | 0.020459 | 0.00251 | 0.001957 | 0.000894 | 0.01204 | 0.009471 | 0.033359 | 0.011196 | 0.058461 | 0.024389 | 0.029746 | 0.025417 | 0.029112 | 0.013295 |
| 26.9722222 | 0.012355 | 0.003936 | 0.005563 | 0.00347 | 0.008183 | 0.006011 | 0.002019 | 0.00342 | 0.016218 | 0.009142 | 0.012654 | 0.003954 | 0.002977 | 0.001674 | 0.010648 | 0.00493 | 0.032371 | 0.009026 | 0.040629 | 0.023266 | 0.020529 | 0.023638 | 0.024702 | 0.015995 |
| 26.9861111 | 0.009833 | 0.002102 | 0.00898 | 0.0027 | 0.008189 | 0.002366 | 0.002146 | 0.00821 | 0.020199 | 0.012415 | 0.007802 | 0.005221 | 0.003335 | 0.002663 | 0.007421 | 0.002047 | 0.027316 | 0.004648 | 0.020033 | 0.0183 | 0.009865 | 0.016898 | 0.017936 | 0.021099 |
| 27 | 0.010276 | 0.000546 | 0.007683 | 0.004941 | 0.013358 | 0.005236 | 0.002363 | 0.009024 | 0.015368 | 0.013831 | 0.015465 | 0.004241 | 0.002817 | 0.003798 | 0.00404 | 0.001381 | 0.021609 | 0.001396 | 0.014284 | 0.014151 | 0.005908 | 0.010699 | 0.013642 | 0.028883 |
| 27.0138889 | 0.017635 | 0.000304 | 0.005946 | 0.012301 | 0.017721 | 0.016328 | 0.005007 | 0.007129 | 0.018427 | 0.012991 | 0.030987 | 0.003354 | 0.002784 | 0.004943 | 0.002296 | 0.002224 | 0.016742 | 0.00159 | 0.011563 | 0.014413 | 0.012141 | 0.00833 | 0.018259 | 0.036094 |
| 27.0277778 | 0.030349 | 0.000989 | 0.009632 | 0.020585 | 0.018895 | 0.027923 | 0.007472 | 0.00958 | 0.042131 | 0.011077 | 0.046868 | 0.006654 | 0.002574 | 0.007127 | 0.002311 | 0.005086 | 0.010745 | 0.002476 | 0.005958 | 0.016421 | 0.024155 | 0.007426 | 0.027199 | 0.040564 |
| 27.0416667 | 0.042183 | 0.002724 | 0.010789 | 0.027293 | 0.0176 | 0.031765 | 0.009273 | 0.011593 | 0.07718 | 0.011332 | 0.054493 | 0.010459 | 0.002121 | 0.011787 | 0.003221 | 0.007827 | 0.005608 | 0.001863 | 0.006713 | 0.018463 | 0.034874 | 0.006484 | 0.029432 | 0.047973 |
| 27.0555556 | 0.053225 | 0.008244 | 0.006623 | 0.03894 | 0.016834 | 0.030126 | 0.012336 | 0.011344 | 0.113516 | 0.014903 | 0.048997 | 0.011508 | 0.002376 | 0.01729 | 0.006203 | 0.006813 | 0.00668 | 0.00247 | 0.010554 | 0.024458 | 0.043205 | 0.008749 | 0.028498 | 0.060256 |
| 27.0694444 | 0.061624 | 0.016819 | 0.003097 | 0.052949 | 0.01513 | 0.025393 | 0.013502 | 0.00944 | 0.126376 | 0.017473 | 0.036178 | 0.010947 | 0.00178 | 0.019022 | 0.011534 | 0.005667 | 0.01194 | 0.00446 | 0.012973 | 0.031744 | 0.046867 | 0.013211 | 0.029864 | 0.064668 |
| 27.0833333 | 0.056842 | 0.020484 | 0.001921 | 0.054453 | 0.009649 | 0.016115 | 0.010281 | 0.006608 | 0.085002 | 0.015021 | 0.021856 | 0.008357 | 0.000718 | 0.01403 | 0.014013 | 0.006299 | 0.014488 | 0.004767 | 0.011978 | 0.032031 | 0.038598 | 0.013949 | 0.025853 | 0.053262 |
| 27.0972222 | 0.037104 | 0.017034 | 0.003094 | 0.04042 | 0.006335 | 0.007897 | 0.005017 | 0.008711 | 0.038138 | 0.009248 | 0.011207 | 0.004448 | 0.000774 | 0.006614 | 0.011696 | 0.005327 | 0.013265 | 0.00379 | 0.006687 | 0.025908 | 0.022426 | 0.011479 | 0.016382 | 0.034544 |
| 27.1111111 | 0.016592 | 0.01109 | 0.005846 | 0.023218 | 0.008292 | 0.008231 | 0.001851 | 0.015442 | 0.04761 | 0.004334 | 0.012387 | 0.002014 | 0.001699 | 0.003497 | 0.008159 | 0.00322 | 0.011207 | 0.002883 | 0.003629 | 0.018542 | 0.010285 | 0.009734 | 0.008247 | 0.018154 |
| 27.125 | 0.00865 | 0.005457 | 0.009056 | 0.012015 | 0.009608 | 0.012698 | 0.001913 | 0.017268 | 0.075934 | 0.00179 | 0.025426 | 0.001229 | 0.002659 | 0.004311 | 0.005319 | 0.001666 | 0.009013 | 0.001812 | 0.006942 | 0.011449 | 0.007354 | 0.00948 | 0.004675 | 0.007905 |
| 27.1388889 | 0.010819 | 0.002312 | 0.013079 | 0.007438 | 0.008759 | 0.013722 | 0.002069 | 0.011878 | 0.080636 | 0.000971 | 0.03949 | 0.001224 | 0.00237 | 0.005287 | 0.002817 | 0.001128 | 0.006455 | 0.001441 | 0.011449 | 0.005128 | 0.010399 | 0.010004 | 0.0075 | 0.005717 |
| 27.1527778 | 0.011164 | 0.002899 | 0.015962 | 0.006636 | 0.007956 | 0.010765 | 0.001098 | 0.006854 | 0.070557 | 0.000867 | 0.045575 | 0.002173 | 0.001374 | 0.006297 | 0.001244 | 0.001455 | 0.004264 | 0.002886 | 0.014097 | 0.001672 | 0.013323 | 0.010853 | 0.013307 | 0.00945 |
| 27.1666667 | 0.00708 | 0.005057 | 0.013291 | 0.006672 | 0.007291 | 0.006011 | 0.000305 | 0.007325 | 0.054658 | 0.001027 | 0.043895 | 0.00252 | 0.001415 | 0.008151 | 0.001407 | 0.001379 | 0.00286 | 0.004777 | 0.017155 | 0.001736 | 0.014778 | 0.011022 | 0.016217 | 0.013382 |
| 27.1805556 | 0.005031 | 0.006889 | 0.007242 | 0.006396 | 0.006342 | 0.00232 | 0.001092 | 0.009348 | 0.03931 | 0.002093 | 0.0375 | 0.002441 | 0.001666 | 0.009435 | 0.001812 | 0.001418 | 0.002459 | 0.005388 | 0.02106 | 0.002158 | 0.015703 | 0.009707 | 0.014274 | 0.013589 |
| 27.1944444 | 0.00556 | 0.009571 | 0.005644 | 0.008051 | 0.005989 | 0.002759 | 0.007839 | 0.014544 | 0.052924 | 0.012063 | 0.026721 | 0.008156 | 0.001674 | 0.007605 | 0.001236 | 0.003723 | 0.004046 | 0.004436 | 0.022474 | 0.00291 | 0.014121 | 0.009627 | 0.009312 | 0.009009 |
| 27.2083333 | 0.008098 | 0.010644 | 0.008605 | 0.013663 | 0.006089 | 0.00467 | 0.019076 | 0.030212 | 0.085629 | 0.028353 | 0.01384 | 0.018191 | 0.00382 | 0.008596 | 0.000796 | 0.009313 | 0.007996 | 0.002795 | 0.017099 | 0.008351 | 0.00863 | 0.014194 | 0.005372 | 0.004882 |
| 27.2222222 | 0.015533 | 0.008371 | 0.009588 | 0.019653 | 0.004537 | 0.006417 | 0.029615 | 0.039225 | 0.071016 | 0.033318 | 0.007147 | 0.02546 | 0.006934 | 0.015426 | 0.001396 | 0.013498 | 0.011409 | 0.001681 | 0.008572 | 0.014154 | 0.002965 | 0.018688 | 0.006658 | 0.004978 |
| 27.2361111 | 0.034315 | 0.011838 | 0.006641 | 0.034785 | 0.002583 | 0.010746 | 0.049633 | 0.048198 | 0.028351 | 0.044471 | 0.012122 | 0.043863 | 0.006089 | 0.018777 | 0.00171 | 0.012782 | 0.009799 | 0.001331 | 0.01048 | 0.013955 | 0.001953 | 0.014669 | 0.009079 | 0.006472 |
| 27.25 | 0.046434 | 0.01617 | 0.005278 | 0.051446 | 0.003821 | 0.014131 | 0.064983 | 0.066446 | 0.034362 | 0.068987 | 0.017055 | 0.061939 | 0.004013 | 0.024222 | 0.001733 | 0.015211 | 0.005779 | 0.00245 | 0.025621 | 0.015932 | 0.005711 | 0.009125 | 0.006584 | 0.011813 |
| 27.2638889 | 0.030743 | 0.010853 | 0.008732 | 0.040398 | 0.006439 | 0.011641 | 0.046406 | 0.056139 | 0.055107 | 0.060813 | 0.013625 | 0.046635 | 0.005021 | 0.032428 | 0.003054 | 0.020348 | 0.005004 | 0.005748 | 0.036335 | 0.021761 | 0.010277 | 0.010295 | 0.002907 | 0.018589 |
| 27.2777778 | 0.013574 | 0.006848 | 0.012119 | 0.016846 | 0.005101 | 0.007558 | 0.022395 | 0.026838 | 0.045232 | 0.030118 | 0.012019 | 0.025224 | 0.005358 | 0.028229 | 0.004047 | 0.017446 | 0.004999 | 0.009253 | 0.031166 | 0.020845 | 0.011537 | 0.010346 | 0.002312 | 0.017958 |
| 27.2916667 | 0.00774 | 0.007189 | 0.010912 | 0.00498 | 0.001648 | 0.007181 | 0.017023 | 0.01569 | 0.037895 | 0.016751 | 0.01152 | 0.020951 | 0.003867 | 0.016951 | 0.002748 | 0.010112 | 0.003185 | 0.010277 | 0.019653 | 0.013967 | 0.009639 | 0.006929 | 0.003486 | 0.011439 |
| 27.3055556 | 0.004605 | 0.005535 | 0.006276 | 0.001649 | 0.000777 | 0.006292 | 0.013413 | 0.0155 | 0.029386 | 0.014889 | 0.008565 | 0.015957 | 0.002522 | 0.010097 | 0.000918 | 0.006453 | 0.001476 | 0.008611 | 0.013253 | 0.007826 | 0.007433 | 0.003834 | 0.00386 | 0.006041 |
| 27.3194444 | 0.003951 | 0.00298 | 0.003904 | 0.002149 | 0.00189 | 0.002975 | 0.014278 | 0.011244 | 0.021431 | 0.025746 | 0.011621 | 0.0152 | 0.001427 | 0.007073 | 0.000836 | 0.005526 | 0.000665 | 0.006251 | 0.012703 | 0.004916 | 0.00594 | 0.001727 | 0.002795 | 0.004446 |

| | | | | | | | | | | | | | | | | | | | | | | | | |
|---|---|---|---|---|---|---|---|---|---|---|---|---|---|---|---|---|---|---|---|---|---|---|---|---|
| 27.3333333 | 0.003483 | 0.001487 | 0.003853 | 0.004599 | 0.003231 | 0.000938 | 0.02049 | 0.004414 | 0.032393 | 0.036239 | 0.019486 | 0.021317 | 0.000664 | 0.005849 | 0.002375 | 0.00474 | 0.000655 | 0.004926 | 0.014112 | 0.005114 | 0.004679 | 0.001446 | 0.001596 | 0.004563 |
| 27.3472222 | 0.004946 | 0.001488 | 0.005993 | 0.006408 | 0.004068 | 0.001194 | 0.017022 | 0.001048 | 0.032646 | 0.025853 | 0.021247 | 0.018876 | 0.000354 | 0.005853 | 0.003818 | 0.003819 | 0.000824 | 0.005025 | 0.015702 | 0.006495 | 0.003805 | 0.00167 | 0.001227 | 0.004446 |
| 27.3611111 | 0.007195 | 0.002339 | 0.015131 | 0.005429 | 0.003795 | 0.003021 | 0.010894 | 0.001162 | 0.018701 | 0.013701 | 0.014102 | 0.013854 | 0.000712 | 0.006989 | 0.003933 | 0.003543 | 0.000862 | 0.005983 | 0.017976 | 0.007677 | 0.004155 | 0.001046 | 0.002354 | 0.005494 |
| 27.375 | 0.008145 | 0.003126 | 0.023397 | 0.003733 | 0.003421 | 0.005011 | 0.01001 | 0.005074 | 0.014919 | 0.011421 | 0.009612 | 0.012161 | 0.001484 | 0.008294 | 0.002941 | 0.003982 | 0.001277 | 0.006585 | 0.019866 | 0.007663 | 0.005088 | 0.000611 | 0.005009 | 0.008793 |
| 27.3888889 | 0.009754 | 0.002436 | 0.020035 | 0.003396 | 0.004199 | 0.00664 | 0.009332 | 0.011857 | 0.01326 | 0.011575 | 0.014155 | 0.012223 | 0.001973 | 0.008382 | 0.001596 | 0.004108 | 0.001538 | 0.006316 | 0.019243 | 0.005921 | 0.005148 | 0.001473 | 0.006793 | 0.012498 |
| 27.4027778 | 0.011717 | 0.001226 | 0.011041 | 0.00376 | 0.004025 | 0.008828 | 0.008636 | 0.013968 | 0.005158 | 0.013985 | 0.016694 | 0.017614 | 0.001966 | 0.007423 | 0.000662 | 0.00357 | 0.001376 | 0.005663 | 0.016507 | 0.004046 | 0.004948 | 0.002413 | 0.006667 | 0.014393 |
| 27.4166667 | 0.009363 | 0.002537 | 0.006427 | 0.004129 | 0.002725 | 0.011984 | 0.011098 | 0.009842 | 0.005987 | 0.016102 | 0.01188 | 0.019839 | 0.001899 | 0.006471 | 0.000762 | 0.003397 | 0.001708 | 0.005132 | 0.01404 | 0.004608 | 0.005763 | 0.00219 | 0.006194 | 0.014562 |
| 27.4305556 | 0.004141 | 0.005185 | 0.006909 | 0.007733 | 0.00405 | 0.015078 | 0.021404 | 0.005999 | 0.015796 | 0.021409 | 0.009168 | 0.021272 | 0.001997 | 0.00598 | 0.002059 | 0.004132 | 0.002565 | 0.00486 | 0.013625 | 0.007894 | 0.007633 | 0.002205 | 0.006063 | 0.014536 |
| 27.4444444 | 0.005632 | 0.004911 | 0.008549 | 0.016082 | 0.007112 | 0.015907 | 0.02579 | 0.005903 | 0.018358 | 0.022998 | 0.01241 | 0.021488 | 0.002088 | 0.005925 | 0.003776 | 0.005301 | 0.002838 | 0.004474 | 0.014812 | 0.010852 | 0.009002 | 0.002866 | 0.006403 | 0.014427 |
| 27.4583333 | 0.014276 | 0.00244 | 0.008764 | 0.024094 | 0.007777 | 0.012882 | 0.014613 | 0.010203 | 0.01907 | 0.013264 | 0.01256 | 0.013831 | 0.001806 | 0.005972 | 0.003899 | 0.006161 | 0.001987 | 0.003183 | 0.015226 | 0.010192 | 0.007955 | 0.002693 | 0.007179 | 0.012135 |
| 27.4722222 | 0.018199 | 0.001237 | 0.008905 | 0.02459 | 0.006184 | 0.009569 | 0.007992 | 0.011089 | 0.022651 | 0.01439 | 0.007095 | 0.011841 | 0.001045 | 0.00573 | 0.00234 | 0.005853 | 0.000933 | 0.001496 | 0.014101 | 0.00659 | 0.00556 | 0.00177 | 0.008206 | 0.007595 |
| 27.4861111 | 0.011557 | 0.003003 | 0.010427 | 0.018626 | 0.005229 | 0.012593 | 0.021833 | 0.009638 | 0.035199 | 0.044677 | 0.003688 | 0.028131 | 0.001226 | 0.005685 | 0.001783 | 0.004932 | 0.001663 | 0.001171 | 0.014784 | 0.005212 | 0.005348 | 0.001624 | 0.010117 | 0.006704 |
| 27.5 | 0.008421 | 0.006092 | 0.012277 | 0.025381 | 0.006211 | 0.02233 | 0.043342 | 0.011769 | 0.065347 | 0.087695 | 0.008925 | 0.052322 | 0.003539 | 0.006954 | 0.005209 | 0.006053 | 0.005576 | 0.003048 | 0.021277 | 0.012287 | 0.009723 | 0.002951 | 0.013652 | 0.015826 |
| 27.5138889 | 0.02714 | 0.006598 | 0.014585 | 0.05248 | 0.008666 | 0.034392 | 0.047278 | 0.015732 | 0.075423 | 0.108778 | 0.025445 | 0.058081 | 0.006922 | 0.009821 | 0.012808 | 0.012402 | 0.010481 | 0.005458 | 0.03258 | 0.030102 | 0.015715 | 0.003869 | 0.017003 | 0.033096 |
| 27.5277778 | 0.057545 | 0.004757 | 0.018033 | 0.071167 | 0.009523 | 0.041444 | 0.033847 | 0.030204 | 0.045377 | 0.101588 | 0.045514 | 0.045643 | 0.01053 | 0.012264 | 0.018384 | 0.022809 | 0.012041 | 0.004946 | 0.041294 | 0.050058 | 0.018468 | 0.002726 | 0.017382 | 0.053253 |
| 27.5416667 | 0.079304 | 0.003858 | 0.021089 | 0.066174 | 0.006926 | 0.038617 | 0.023123 | 0.047023 | 0.013453 | 0.087714 | 0.055138 | 0.032262 | 0.01225 | 0.012211 | 0.016229 | 0.029326 | 0.010316 | 0.00241 | 0.041878 | 0.055411 | 0.019462 | 0.001257 | 0.015042 | 0.062609 |
| 27.5555556 | 0.089725 | 0.004852 | 0.021435 | 0.055223 | 0.004435 | 0.031064 | 0.02012 | 0.048854 | 0.005088 | 0.081949 | 0.046847 | 0.02083 | 0.010632 | 0.010989 | 0.010201 | 0.027825 | 0.009756 | 0.001681 | 0.037822 | 0.044584 | 0.021994 | 0.002708 | 0.012188 | 0.048907 |
| 27.5694444 | 0.083396 | 0.005992 | 0.018016 | 0.045614 | 0.003622 | 0.02191 | 0.018588 | 0.035775 | 0.010808 | 0.077965 | 0.026533 | 0.013888 | 0.008437 | 0.010355 | 0.006298 | 0.021767 | 0.010909 | 0.002603 | 0.03263 | 0.031346 | 0.02437 | 0.007671 | 0.011241 | 0.026716 |
| 27.5833333 | 0.061424 | 0.0062 | 0.012313 | 0.035489 | 0.003162 | 0.014555 | 0.017637 | 0.018941 | 0.018216 | 0.067388 | 0.015598 | 0.014615 | 0.006438 | 0.009887 | 0.003666 | 0.014547 | 0.009997 | 0.002853 | 0.025365 | 0.02033 | 0.023699 | 0.012661 | 0.01345 | 0.011813 |
| 27.5972222 | 0.037028 | 0.005811 | 0.006431 | 0.028523 | 0.004083 | 0.014155 | 0.018389 | 0.015761 | 0.022934 | 0.056186 | 0.024969 | 0.01854 | 0.004174 | 0.009266 | 0.001418 | 0.009164 | 0.006704 | 0.001692 | 0.018096 | 0.010931 | 0.020581 | 0.015462 | 0.017275 | 0.004816 |
| 27.6111111 | 0.017217 | 0.004359 | 0.002151 | 0.021371 | 0.005043 | 0.014458 | 0.018057 | 0.02604 | 0.025645 | 0.047371 | 0.038037 | 0.019183 | 0.002367 | 0.008822 | 0.001065 | 0.007263 | 0.00384 | 0.000559 | 0.013834 | 0.006306 | 0.018115 | 0.016336 | 0.020288 | 0.004226 |
| 27.625 | 0.00639 | 0.002153 | 0.002685 | 0.011254 | 0.003811 | 0.010637 | 0.01381 | 0.030636 | 0.029193 | 0.042738 | 0.038286 | 0.01702 | 0.00116 | 0.008588 | 0.002887 | 0.00866 | 0.00255 | 0.000393 | 0.011964 | 0.007799 | 0.018767 | 0.015139 | 0.021653 | 0.008481 |
| 27.6388889 | 0.003453 | 0.000761 | 0.006814 | 0.006687 | 0.003652 | 0.007316 | 0.007496 | 0.021858 | 0.031246 | 0.042617 | 0.023638 | 0.014432 | 0.000811 | 0.008598 | 0.007009 | 0.012923 | 0.002629 | 0.00049 | 0.010739 | 0.013426 | 0.022148 | 0.012755 | 0.022584 | 0.016676 |
| 27.6527778 | 0.003883 | 0.000595 | 0.007493 | 0.010775 | 0.005224 | 0.00493 | 0.004584 | 0.018639 | 0.025728 | 0.035702 | 0.013512 | 0.010054 | 0.00116 | 0.009193 | 0.011818 | 0.018028 | 0.004029 | 0.000413 | 0.009645 | 0.02008 | 0.026622 | 0.010945 | 0.024913 | 0.027338 |
| 27.6666667 | 0.005269 | 0.000861 | 0.006301 | 0.013597 | 0.004418 | 0.002035 | 0.006925 | 0.031161 | 0.014156 | 0.021706 | 0.027285 | 0.009216 | 0.002585 | 0.010574 | 0.015022 | 0.022039 | 0.006482 | 0.00024 | 0.008673 | 0.024784 | 0.03171 | 0.010243 | 0.027214 | 0.035782 |
| 27.6805556 | 0.008531 | 0.001045 | 0.007984 | 0.009734 | 0.004024 | 0.001498 | 0.008446 | 0.039049 | 0.005801 | 0.022393 | 0.044681 | 0.017833 | 0.0052 | 0.01189 | 0.015496 | 0.023717 | 0.008704 | 0.000272 | 0.007098 | 0.025119 | 0.035588 | 0.010018 | 0.025243 | 0.036031 |
| 27.6944444 | 0.016087 | 0.0017 | 0.007154 | 0.005765 | 0.008436 | 0.003239 | 0.006364 | 0.030752 | 0.007679 | 0.047928 | 0.039322 | 0.028731 | 0.007179 | 0.012142 | 0.013834 | 0.022632 | 0.009242 | 0.000548 | 0.004843 | 0.020744 | 0.035193 | 0.009813 | 0.018337 | 0.027754 |
| 27.7083333 | 0.018008 | 0.002555 | 0.004285 | 0.010622 | 0.01395 | 0.004722 | 0.007062 | 0.018743 | 0.020948 | 0.072093 | 0.021502 | 0.031537 | 0.00736 | 0.011374 | 0.011811 | 0.019601 | 0.008249 | 0.000786 | 0.003686 | 0.013377 | 0.029504 | 0.009136 | 0.010331 | 0.016342 |
| 27.7222222 | 0.011305 | 0.002716 | 0.005549 | 0.020245 | 0.016328 | 0.008258 | 0.013265 | 0.021391 | 0.034668 | 0.069993 | 0.018169 | 0.026135 | 0.006179 | 0.009628 | 0.009186 | 0.01535 | 0.006734 | 0.00133 | 0.004638 | 0.006657 | 0.019808 | 0.007434 | 0.004568 | 0.006991 |
| 27.7361111 | 0.011548 | 0.002781 | 0.008734 | 0.024928 | 0.013505 | 0.014685 | 0.018271 | 0.034676 | 0.038025 | 0.051067 | 0.032017 | 0.019058 | 0.004306 | 0.007103 | 0.005833 | 0.011021 | 0.004969 | 0.002459 | 0.006411 | 0.004638 | 0.009797 | 0.005301 | 0.003353 | 0.002555 |
| 27.75 | 0.017991 | 0.00326 | 0.010945 | 0.026969 | 0.007309 | 0.015879 | 0.017663 | 0.030956 | 0.028561 | 0.030629 | 0.046338 | 0.011864 | 0.00225 | 0.004672 | 0.004335 | 0.007925 | 0.003303 | 0.003467 | 0.007447 | 0.005898 | 0.005881 | 0.003574 | 0.007557 | 0.003551 |
| 27.7638889 | 0.020182 | 0.003728 | 0.014262 | 0.02616 | 0.004185 | 0.00929 | 0.013204 | 0.018265 | 0.015648 | 0.013815 | 0.053073 | 0.005928 | 0.00141 | 0.0026 | 0.005228 | 0.005908 | 0.002486 | 0.004065 | 0.006477 | 0.006016 | 0.009814 | 0.002351 | 0.014476 | 0.007669 |
| 27.7777778 | 0.015779 | 0.004233 | 0.017851 | 0.017617 | 0.005239 | 0.003194 | 0.007583 | 0.021557 | 0.015487 | 0.005164 | 0.049027 | 0.00333 | 0.003631 | 0.001953 | 0.00541 | 0.003521 | 0.002621 | 0.004676 | 0.003942 | 0.00497 | 0.015679 | 0.003837 | 0.020982 | 0.012482 |
| 27.7916667 | 0.008286 | 0.004702 | 0.017523 | 0.009389 | 0.00601 | 0.001157 | 0.006265 | 0.022341 | 0.019325 | 0.003765 | 0.03617 | 0.004375 | 0.00785 | 0.003974 | 0.003736 | 0.001457 | 0.003048 | 0.00527 | 0.003315 | 0.004303 | 0.020612 | 0.01086 | 0.023792 | 0.017261 |
| 27.8055556 | 0.00744 | 0.004452 | 0.015396 | 0.012303 | 0.005508 | 0.001054 | 0.013748 | 0.015637 | 0.014487 | 0.005248 | 0.023733 | 0.010829 | 0.012133 | 0.007532 | 0.00242 | 0.001848 | 0.002823 | 0.005814 | 0.006295 | 0.004553 | 0.024373 | 0.018564 | 0.021259 | 0.021431 |
| 27.8194444 | 0.01507 | 0.003735 | 0.01643 | 0.022967 | 0.007281 | 0.001694 | 0.023893 | 0.018369 | 0.00746 | 0.006911 | 0.015038 | 0.015408 | 0.016052 | 0.011516 | 0.004056 | 0.004093 | 0.001934 | 0.006369 | 0.010925 | 0.005897 | 0.025047 | 0.019515 | 0.017188 | 0.0252 |
| 27.8333333 | 0.024112 | 0.003387 | 0.019899 | 0.031804 | 0.011942 | 0.005113 | 0.029566 | 0.027694 | 0.006536 | 0.005852 | 0.008539 | 0.011142 | 0.018563 | 0.015836 | 0.00784 | 0.006693 | 0.002077 | 0.006607 | 0.01512 | 0.007215 | 0.023108 | 0.015519 | 0.015323 | 0.030116 |
| 27.8472222 | 0.031549 | 0.004016 | 0.02279 | 0.034295 | 0.014405 | 0.013024 | 0.03118 | 0.038161 | 0.014124 | 0.005625 | 0.003963 | 0.005267 | 0.018123 | 0.020262 | 0.010456 | 0.009022 | 0.003716 | 0.006685 | 0.017686 | 0.007746 | 0.019468 | 0.012989 | 0.016128 | 0.03599 |
| 27.8611111 | 0.039054 | 0.00574 | 0.02333 | 0.03095 | 0.01187 | 0.020498 | 0.028663 | 0.038367 | 0.02238 | 0.01123 | 0.002274 | 0.006655 | 0.015689 | 0.023741 | 0.010419 | 0.011367 | 0.005128 | 0.007483 | 0.019446 | 0.008594 | 0.014619 | 0.014055 | 0.018201 | 0.040812 |
| 27.875 | 0.048198 | 0.007348 | 0.021093 | 0.026728 | 0.008515 | 0.023542 | 0.02335 | 0.02403 | 0.025819 | 0.017932 | 0.003543 | 0.013891 | 0.014279 | 0.026157 | 0.008333 | 0.014526 | 0.005708 | 0.008782 | 0.022516 | 0.010678 | 0.010693 | 0.015974 | 0.019088 | 0.045425 |
| 27.8888889 | 0.057059 | 0.007647 | 0.017137 | 0.027336 | 0.008482 | 0.022528 | 0.019554 | 0.009801 | 0.025551 | 0.020096 | 0.004345 | 0.02048 | 0.013196 | 0.027122 | 0.005726 | 0.017854 | 0.005643 | 0.009323 | 0.027669 | 0.01336 | 0.009017 | 0.015032 | 0.018696 | 0.051796 |
| 27.9027778 | 0.061471 | 0.007024 | 0.013177 | 0.031582 | 0.010739 | 0.01947 | 0.017012 | 0.00535 | 0.021111 | 0.019588 | 0.003093 | 0.022671 | 0.009569 | 0.025263 | 0.003815 | 0.021206 | 0.00483 | 0.008847 | 0.03507 | 0.015537 | 0.008991 | 0.010851 | 0.018022 | 0.058843 |
| 27.9166667 | 0.060672 | 0.005973 | 0.009928 | 0.032155 | 0.012515 | 0.017212 | 0.01298 | 0.008982 | 0.015513 | 0.01893 | 0.001999 | 0.017814 | 0.004773 | 0.021862 | 0.003526 | 0.025559 | 0.004331 | 0.008532 | 0.044469 | 0.016962 | 0.009458 | 0.00607 | 0.015612 | 0.065007 |
| 27.9305556 | 0.054663 | 0.003972 | 0.006051 | 0.02577 | 0.011667 | 0.014229 | 0.007858 | 0.014815 | 0.013068 | 0.015549 | 0.001363 | 0.011287 | 0.001873 | 0.017984 | 0.004233 | 0.029278 | 0.005089 | 0.009112 | 0.053069 | 0.01767 | 0.009665 | 0.002657 | 0.009567 | 0.066554 |
| 27.9444444 | 0.042506 | 0.002763 | 0.002834 | 0.016187 | 0.007336 | 0.008094 | 0.005701 | 0.018137 | 0.01283 | 0.008607 | 0.002435 | 0.013546 | 0.000776 | 0.013775 | 0.004276 | 0.028749 | 0.006624 | 0.009046 | 0.056493 | 0.017101 | 0.009189 | 0.001054 | 0.003241 | 0.058436 |
| 27.9583333 | 0.02624 | 0.004129 | 0.001891 | 0.008605 | 0.00436 | 0.002245 | 0.005705 | 0.017736 | 0.009587 | 0.005899 | 0.007398 | 0.017972 | 0.000621 | 0.010375 | 0.003597 | 0.023786 | 0.008208 | 0.007689 | 0.053882 | 0.014934 | 0.008188 | 0.001944 | 0.00151 | 0.045607 |
| 27.9722222 | 0.015283 | 0.004376 | 0.002854 | 0.007274 | 0.007529 | 0.00267 | 0.009225 | 0.016259 | 0.009091 | 0.023918 | 0.013975 | 0.021632 | 0.001223 | 0.008152 | 0.002728 | 0.017852 | 0.00931 | 0.006446 | 0.048352 | 0.01101 | 0.007197 | 0.005177 | 0.002436 | 0.037685 |
| 27.9861111 | 0.018924 | 0.004296 | 0.008132 | 0.01086 | 0.010902 | 0.010131 | 0.027633 | 0.023725 | 0.022326 | 0.073494 | 0.018655 | 0.045282 | 0.001401 | 0.006101 | 0.001696 | 0.013127 | 0.009995 | 0.005393 | 0.04255 | 0.006354 | 0.006929 | 0.00867 | 0.002885 | 0.03454 |
| 28 | 0.02497 | 0.008828 | 0.015561 | 0.014268 | 0.007799 | 0.019498 | 0.050642 | 0.03686 | 0.030646 | 0.12354 | 0.015936 | 0.079213 | 0.000995 | 0.003665 | 0.000757 | 0.009086 | 0.01093 | 0.004335 | 0.036977 | 0.003725 | 0.008673 | 0.010544 | 0.002273 | 0.0314 |
| 28.0138889 | 0.02251 | 0.010688 | 0.01927 | 0.018225 | 0.004244 | 0.025005 | 0.048713 | 0.030831 | 0.022169 | 0.121702 | 0.012452 | 0.077071 | 0.001751 | 0.00274 | 0.000307 | 0.00552 | 0.011405 | 0.004584 | 0.030472 | 0.005927 | 0.01193 | 0.011027 | 0.00141 | 0.026757 |
| 28.0277778 | 0.025492 | 0.005327 | 0.019723 | 0.022961 | 0.005546 | 0.024829 | 0.023867 | 0.015418 | 0.016675 | 0.079669 | 0.019908 | 0.040471 | 0.004568 | 0.00357 | 0.00017 | 0.005725 | 0.00965 | 0.005834 | 0.023575 | 0.012184 | 0.014295 | 0.012479 | 0.002217 | 0.021812 |
| 28.0416667 | 0.03437 | 0.005595 | 0.019752 | 0.02935 | 0.00845 | 0.021493 | 0.008144 | 0.016708 | 0.013233 | 0.056028 | 0.025509 | 0.018988 | 0.007896 | 0.00451 | 0.000512 | 0.009022 | 0.00624 | 0.005257 | 0.021482 | 0.015962 | 0.015414 | 0.014604 | 0.00592 | 0.018354 |
| 28.0555556 | 0.035432 | 0.015888 | 0.018004 | 0.034709 | 0.012373 | 0.017684 | 0.011754 | 0.022015 | 0.006049 | 0.055268 | 0.019056 | 0.022643 | 0.0091 | 0.009386 | 0.002426 | 0.008023 | 0.004033 | 0.002972 | 0.029197 | 0.012711 | 0.01589 | 0.014666 | 0.01183 | 0.016452 |
| 28.0694444 | 0.028961 | 0.023617 | 0.01284 | 0.028088 | 0.014558 | 0.013784 | 0.01651 | 0.021368 | 0.002936 | 0.039865 | 0.012512 | 0.022116 | 0.0077 | 0.018635 | 0.005454 | 0.004005 | 0.00442 | 0.002163 | 0.043049 | 0.006887 | 0.015292 | 0.012459 | 0.015876 | 0.015695 |
| 28.0833333 | 0.017139 | 0.02132 | 0.00633 | 0.015783 | 0.011124 | 0.008548 | 0.012221 | 0.016533 | 0.002992 | 0.021959 | 0.013278 | 0.012186 | 0.00561 | 0.022256 | 0.006915 | 0.00319 | 0.005759 | 0.002705 | 0.051077 | 0.004464 | 0.015334 | 0.008822 | 0.015269 | 0.016012 |
| 28.0972222 | 0.006666 | 0.012135 | 0.00266 | 0.011903 | 0.005971 | 0.003822 | 0.00698 | 0.009679 | 0.00533 | 0.034174 | 0.014261 | 0.01009 | 0.004064 | 0.016644 | 0.005961 | 0.003177 | 0.006171 | 0.002357 | 0.047071 | 0.005033 | 0.015955 | 0.005034 | 0.011544 | 0.015806 |
| 28.1111111 | 0.005433 | 0.004658 | 0.005577 | 0.012664 | 0.00392 | 0.003349 | 0.008047 | 0.010206 | 0.011439 | 0.053515 | 0.009601 | 0.014426 | 0.002903 | 0.008553 | 0.004162 | 0.001671 | 0.005365 | 0.002123 | 0.036406 | 0.005568 | 0.013735 | 0.002242 | 0.007034 | 0.013222 |
| 28.125 | 0.008556 | 0.003369 | 0.013941 | 0.011334 | 0.003522 | 0.006186 | 0.008304 | 0.016986 | 0.017236 | 0.043224 | 0.006533 | 0.012726 | 0.001952 | 0.003322 | 0.00282 | 0.000744 | 0.004022 | 0.003075 | 0.025824 | 0.006017 | 0.009498 | 0.000777 | 0.00362 | 0.008753 |
| 28.1388889 | 0.011612 | 0.004096 | 0.02294 | 0.008428 | 0.00266 | 0.009498 | 0.004138 | 0.020093 | 0.018572 | 0.019174 | 0.010446 | 0.008408 | 0.001114 | 0.002377 | 0.002141 | 0.00106 | 0.003103 | 0.003349 | 0.018193 | 0.007718 | 0.006462 | 0.001017 | 0.001828 | 0.004754 |
| 28.1527778 | 0.012662 | 0.005065 | 0.02834 | 0.004907 | 0.002784 | 0.011474 | 0.004036 | 0.014048 | 0.01781 | 0.012988 | 0.013255 | 0.005949 | 0.001002 | 0.004202 | 0.00143 | 0.001239 | 0.002658 | 0.002715 | 0.0134 | 0.008938 | 0.004762 | 0.002593 | 0.00099 | 0.002481 |
| 28.1666667 | 0.010652 | 0.005876 | 0.02687 | 0.002647 | 0.003533 | 0.011603 | 0.006868 | 0.005828 | 0.016333 | 0.015867 | 0.011717 | 0.002776 | 0.002463 | 0.006489 | 0.000757 | 0.000834 | 0.002169 | 0.002191 | 0.009916 | 0.006642 | 0.003242 | 0.003906 | 0.000932 | 0.001208 |
| 28.1805556 | 0.00633 | 0.00501 | 0.019583 | 0.002891 | 0.003377 | 0.010366 | 0.007054 | 0.004873 | 0.013242 | 0.013749 | 0.009955 | 0.002562 | 0.004389 | 0.007891 | 0.000685 | 0.000898 | 0.001688 | 0.00138 | 0.006897 | 0.004942 | 0.001776 | 0.003353 | 0.001192 | 0.00051 |
| 28.1944444 | 0.001814 | 0.00325 | 0.012175 | 0.002949 | 0.002619 | 0.007913 | 0.005263 | 0.005674 | 0.009671 | 0.011598 | 0.00877 | 0.004443 | 0.00565 | 0.007926 | 0.000597 | 0.001347 | 0.001501 | 0.0006 | 0.004393 | 0.008864 | 0.001552 | 0.001728 | 0.001168 | 0.000352 |
| 28.2083333 | 0.001254 | 0.002504 | 0.008678 | 0.00243 | 0.002141 | 0.00563 | 0.004517 | 0.004087 | 0.007349 | 0.01615 | 0.008631 | 0.004001 | 0.006265 | 0.007417 | 0.000319 | 0.001711 | 0.001578 | 0.000482 | 0.002549 | 0.012851 | 0.003118 | 0.000826 | 0.001042 | 0.000232 |
| 28.2222222 | 0.003971 | 0.002832 | 0.010005 | 0.00411 | 0.001663 | 0.004674 | 0.006683 | 0.006319 | 0.006349 | 0.021758 | 0.009812 | 0.003996 | 0.006275 | 0.006951 | 0.000205 | 0.001977 | 0.001723 | 0.000953 | 0.001195 | 0.011478 | 0.004982 | 0.000652 | 0.001126 | 0.00011 |
| 28.2361111 | 0.005566 | 0.00272 | 0.01455 | 0.005998 | 0.001803 | 0.004317 | 0.006602 | 0.010695 | 0.006051 | 0.016671 | 0.010647 | 0.003794 | 0.005981 | 0.006522 | 0.000132 | 0.001937 | 0.00188 | 0.002082 | 0.000487 | 0.006599 | 0.005991 | 0.000832 | 0.00127 | 0.000217 |
| 28.25 | 0.005555 | 0.005438 | 0.017888 | 0.005716 | 0.004294 | 0.003616 | 0.007412 | 0.009527 | 0.008535 | 0.02029 | 0.007899 | 0.007899 | 0.005863 | 0.00627 | 0.00013 | 0.001491 | 0.002068 | 0.003457 | 0.00046 | 0.002154 | 0.00624 | 0.001843 | 0.001129 | 0.000334 |
| 28.2638889 | 0.007451 | 0.011918 | 0.015275 | 0.006891 | 0.007813 | 0.002652 | 0.016337 | 0.00601 | 0.015346 | 0.049921 | 0.004393 | 0.021095 | 0.006273 | 0.006262 | 0.000171 | 0.001063 | 0.002154 | 0.004684 | 0.00048 | 0.000922 | 0.00657 | 0.00448 | 0.000687 | 0.000283 |
| 28.2777778 | 0.008463 | 0.014509 | 0.009463 | 0.007762 | 0.007737 | 0.002198 | 0.01961 | 0.006024 | 0.018224 | 0.061315 | 0.003649 | 0.025107 | 0.00692 | 0.00652 | 0.000251 | 0.001261 | 0.002033 | 0.005937 | 0.000361 | 0.002586 | 0.00745 | 0.008468 | 0.000382 | 0.000329 |
| 28.2916667 | 0.007284 | 0.010103 | 0.009056 | 0.008011 | 0.005157 | 0.002487 | 0.015903 | 0.006883 | 0.014345 | 0.047159 | 0.00442 | 0.020222 | 0.00709 | 0.006546 | 0.000346 | 0.001842 | 0.001786 | 0.00703 | 0.000294 | 0.004913 | 0.007544 | 0.011215 | 0.000488 | 0.000669 |
| 28.3055556 | 0.009738 | 0.005242 | 0.01247 | 0.015324 | 0.004867 | 0.002618 | 0.021565 | 0.009719 | 0.01114 | 0.051808 | 0.007232 | 0.026446 | 0.006565 | 0.005857 | 0.000264 | 0.00211 | 0.00151 | 0.007615 | 0.00048 | 0.006174 | 0.006081 | 0.010432 | 0.000747 | 0.001264 |
| 28.3194444 | 0.010593 | 0.004487 | 0.010878 | 0.020135 | 0.005953 | 0.00232 | 0.021789 | 0.013053 | 0.00766 | 0.050321 | 0.007748 | 0.026705 | 0.005521 | 0.004928 | 0.000115 | 0.001979 | 0.001434 | 0.007794 | 0.001023 | 0.005739 | 0.004132 | 0.006978 | 0.000925 | 0.002072 |
| 28.3333333 | 0.009265 | 0.007323 | 0.00484 | 0.013565 | 0.009076 | 0.003197 | 0.013393 | 0.009816 | 0.004713 | 0.039892 | 0.007694 | 0.020062 | 0.004378 | 0.004435 | 0.000224 | 0.001761 | 0.001586 | 0.007812 | 0.001444 | 0.004209 | 0.002804 | 0.003816 | 0.000754 | 0.002765 |
| 28.3472222 | 0.010649 | 0.010597 | 0.002203 | 0.006945 | 0.012384 | 0.00389 | 0.014299 | 0.003504 | 0.004682 | 0.048927 | 0.009756 | 0.023647 | 0.00389 | 0.004523 | 0.000705 | 0.001717 | 0.001433 | 0.007296 | 0.001113 | 0.003342 | 0.002555 | 0.002749 | 0.000707 | 0.00308 |
| 28.3611111 | 0.00944 | 0.008593 | 0.004991 | 0.00623 | 0.009352 | 0.003052 | 0.014173 | 0.003094 | 0.004202 | 0.042533 | 0.008201 | 0.02116 | 0.004345 | 0.0049 | 0.001185 | 0.001819 | 0.000887 | 0.006155 | 0.000831 | 0.003876 | 0.003327 | 0.003649 | 0.001717 | 0.002881 |
| 28.375 | 0.008751 | 0.00326 | 0.008598 | 0.009472 | 0.003541 | 0.003746 | 0.009885 | 0.009123 | 0.002824 | 0.022285 | 0.0072 | 0.013734 | 0.005346 | 0.005408 | 0.00116 | 0.001888 | 0.000706 | 0.005274 | 0.001764 | 0.004713 | 0.004686 | 0.006064 | 0.003237 | 0.001923 |
| 28.3888889 | 0.009375 | 0.002102 | 0.010074 | 0.01239 | 0.002256 | 0.005798 | 0.008664 | 0.013816 | 0.00248 | 0.018716 | 0.007771 | 0.010917 | 0.00645 | 0.005982 | 0.000717 | 0.001755 | 0.000943 | 0.004822 | 0.00267 | 0.004636 | 0.005689 | 0.009001 | 0.004101 | 0.001037 |
| 28.4027778 | 0.006745 | 0.004915 | 0.00947 | 0.013325 | 0.003916 | 0.006904 | 0.007642 | 0.013315 | 0.003501 | 0.022066 | 0.004552 | 0.007731 | 0.007483 | 0.006233 | 0.000468 | 0.001318 | 0.000903 | 0.004204 | 0.002058 | 0.003624 | 0.005577 | 0.010591 | 0.003718 | 0.000744 |
| 28.4166667 | 0.006789 | 0.006702 | 0.008206 | 0.018251 | 0.004513 | 0.008014 | 0.006001 | 0.011928 | 0.005568 | 0.018338 | 0.00203 | 0.005187 | 0.008322 | 0.006187 | 0.000522 | 0.000949 | 0.00078 | 0.003702 | 0.000984 | 0.002689 | 0.004837 | 0.010243 | 0.00264 | 0.000466 |
| 28.4305556 | 0.009859 | 0.007075 | 0.007188 | 0.022937 | 0.004716 | 0.010292 | 0.006427 | 0.011589 | 0.008517 | 0.013403 | 0.002572 | 0.005151 | 0.008481 | 0.006211 | 0.000427 | 0.001111 | 0.000996 | 0.003882 | 0.001276 | 0.002542 | 0.004276 | 0.008521 | 0.002234 | 0.000331 |
| 28.4444444 | 0.009587 | 0.00788 | 0.006103 | 0.01722 | 0.006002 | 0.01277 | 0.012284 | 0.010531 | 0.011963 | 0.022042 | 0.004322 | 0.010443 | 0.007488 | 0.006352 | 0.000442 | 0.001614 | 0.00102 | 0.003915 | 0.002575 | 0.002645 | 0.003672 | 0.005787 | 0.002461 | 0.0003 |
| 28.4583333 | 0.007541 | 0.007335 | 0.006388 | 0.007004 | 0.005523 | 0.013722 | 0.016215 | 0.009784 | 0.012199 | 0.029544 | 0.005643 | 0.014165 | 0.005594 | 0.006567 | 0.001523 | 0.001951 | 0.000973 | 0.00272 | 0.002957 | 0.001892 | 0.002384 | 0.00296 | 0.001884 | 0.000658 |
| 28.4722222 | 0.006906 | 0.004947 | 0.007635 | 0.002453 | 0.00247 | 0.01274 | 0.013513 | 0.009994 | 0.009131 | 0.024181 | 0.003664 | 0.010243 | 0.003832 | 0.006642 | 0.003344 | 0.001734 | 0.001508 | 0.001333 | 0.002343 | 0.00064 | 0.001146 | 0.000987 | 0.000667 | 0.001631 |
| 28.4861111 | 0.006707 | 0.004832 | 0.006653 | 0.005228 | 0.0034 | 0.011208 | 0.016564 | 0.009047 | 0.012657 | 0.032692 | 0.002044 | 0.012072 | 0.004019 | 0.006368 | 0.003776 | 0.001003 | 0.002214 | 0.001298 | 0.003848 | 0.001529 | 0.001795 | 0.00114 | 0.001869 | 0.00352 |
| 28.5 | 0.00962 | 0.009553 | 0.006126 | 0.017722 | 0.01072 | 0.012622 | 0.030338 | 0.008922 | 0.02525 | 0.062735 | 0.003873 | 0.024237 | 0.007033 | 0.006389 | 0.005824 | 0.000786 | 0.004923 | 0.003353 | 0.011598 | 0.007007 | 0.00629 | 0.004269 | 0.008095 | 0.011846 |
| 28.5138889 | 0.024199 | 0.014062 | 0.013865 | 0.045763 | 0.017719 | 0.02158 | 0.039808 | 0.01694 | 0.02654 | 0.082203 | 0.012238 | 0.03031 | 0.0101 | 0.009028 | 0.013456 | 0.002004 | 0.010331 | 0.007207 | 0.025754 | 0.014338 | 0.013515 | 0.008025 | 0.01716 | 0.031489 |
| 28.5277778 | 0.050774 | 0.011438 | 0.030575 | 0.073636 | 0.019107 | 0.037721 | 0.033871 | 0.036892 | 0.025415 | 0.073615 | 0.024962 | 0.02246 | 0.009098 | 0.013696 | 0.018915 | 0.003485 | 0.011389 | 0.009604 | 0.038378 | 0.015789 | 0.017561 | 0.007555 | 0.022417 | 0.049611 |
| 28.5416667 | 0.071165 | 0.004802 | 0.042452 | 0.066914 | 0.016401 | 0.048448 | 0.023097 | 0.058747 | 0.046981 | 0.052835 | 0.025566 | 0.011217 | 0.005382 | 0.014962 | 0.014049 | 0.003478 | 0.007291 | 0.007432 | 0.039507 | 0.00988 | 0.013997 | 0.006067 | 0.020039 | 0.046357 |
| 28.5555556 | 0.070817 | 0.00227 | 0.035082 | 0.035201 | 0.014475 | 0.04305 | 0.021143 | 0.063329 | 0.055949 | 0.03435 | 0.013254 | 0.006046 | 0.002716 | 0.012021 | 0.005946 | 0.003325 | 0.005201 | 0.003928 | 0.031368 | 0.003601 | 0.006227 | 0.007725 | 0.013649 | 0.026677 |
| 28.5694444 | 0.057646 | 0.003784 | 0.017848 | 0.019614 | 0.014596 | 0.030201 | 0.024782 | 0.047914 | 0.034149 | 0.018512 | 0.004637 | 0.006653 | 0.002871 | 0.010616 | 0.004972 | 0.00552 | 0.004551 | 0.004828 | 0.02449 | 0.002251 | 0.002368 | 0.006915 | 0.009435 | 0.011727 |
| 28.5833333 | 0.042108 | 0.004592 | 0.011048 | 0.021439 | 0.014153 | 0.020658 | 0.025125 | 0.025726 | 0.020236 | 0.012814 | 0.003559 | 0.009347 | 0.006285 | 0.012371 | 0.008305 | 0.008413 | 0.003999 | 0.008189 | 0.021086 | 0.004431 | 0.003746 | 0.0049 | 0.00952 | 0.007294 |
| 28.5972222 | 0.026664 | 0.003883 | 0.01326 | 0.018839 | 0.012081 | 0.015365 | 0.020349 | 0.01299 | 0.024149 | 0.024235 | 0.004059 | 0.010685 | 0.008761 | 0.012825 | 0.008763 | 0.008571 | 0.004104 | 0.008141 | 0.017516 | 0.004667 | 0.00508 | 0.004966 | 0.010586 | 0.005676 |
| 28.6111111 | 0.01467 | 0.003208 | 0.012711 | 0.011824 | 0.008192 | 0.01211 | 0.013427 | 0.017692 | 0.020404 | 0.033731 | 0.005478 | 0.009818 | 0.007855 | 0.010361 | 0.006455 | 0.006327 | 0.003025 | 0.004503 | 0.013579 | 0.002276 | 0.004593 | 0.004428 | 0.010231 | 0.003848 |
| 28.625 | 0.007916 | 0.002975 | 0.008561 | 0.010849 | 0.003937 | 0.00843 | 0.007162 | 0.024591 | 0.011681 | 0.024759 | 0.009517 | 0.009005 | 0.006044 | 0.007507 | 0.004518 | 0.004547 | 0.001449 | 0.001481 | 0.011336 | 0.001344 | 0.003939 | 0.003867 | 0.010459 | 0.005361 |
| 28.6388889 | 0.005135 | 0.002649 | 0.004228 | 0.015227 | 0.002417 | 0.00542 | 0.005639 | 0.018808 | 0.014885 | 0.012782 | 0.013587 | 0.008866 | 0.005016 | 0.006222 | 0.003789 | 0.005117 | 0.000937 | 0.000848 | 0.01157 | 0.00255 | 0.004579 | 0.004509 | 0.014358 | 0.012309 |
| 28.6527778 | 0.005762 | 0.00199 | 0.001643 | 0.01998 | 0.002703 | 0.005316 | 0.011961 | 0.010298 | 0.019041 | 0.015025 | 0.011255 | 0.007776 | 0.005424 | 0.006838 | 0.004169 | 0.007857 | 0.00148 | 0.00145 | 0.013857 | 0.003876 | 0.007034 | 0.005968 | 0.023028 | 0.021943 |
| 28.6666667 | 0.008953 | 0.001417 | 0.001387 | 0.022207 | 0.002924 | 0.007459 | 0.019276 | 0.016722 | 0.013705 | 0.021039 | 0.007578 | 0.005048 | 0.006965 | 0.00843 | 0.005708 | 0.011218 | 0.002464 | 0.002909 | 0.017112 | 0.004796 | 0.009788 | 0.007199 | 0.03301 | 0.029148 |
| 28.6805556 | 0.011804 | 0.001341 | 0.003439 | 0.020008 | 0.003515 | 0.009914 | 0.020365 | 0.039058 | 0.008012 | 0.017091 | 0.011985 | 0.003167 | 0.00825 | 0.009864 | 0.007726 | 0.013219 | 0.003161 | 0.004295 | 0.019421 | 0.005485 | 0.011069 | 0.007347 | 0.037081 | 0.031381 |
| 28.6944444 | 0.010251 | 0.001487 | 0.007981 | 0.015463 | 0.003677 | 0.011856 | 0.016933 | 0.062642 | 0.010584 | 0.009907 | 0.015184 | 0.00536 | 0.008567 | 0.010479 | 0.008963 | 0.012295 | 0.003116 | 0.004466 | 0.019414 | 0.00512 | 0.010491 | 0.006408 | 0.032774 | 0.030049 |
| 28.7083333 | 0.006648 | 0.001215 | 0.009581 | 0.010549 | 0.002468 | 0.013062 | 0.010684 | 0.074289 | 0.014367 | 0.011522 | 0.009984 | 0.009992 | 0.008181 | 0.009854 | 0.007846 | 0.00865 | 0.002288 | 0.003246 | 0.016715 | 0.003047 | 0.00801 | 0.004723 | 0.024733 | 0.026292 |
| 28.7222222 | 0.009052 | 0.001052 | 0.008438 | 0.006949 | 0.001795 | 0.012736 | 0.007942 | 0.068231 | 0.012017 | 0.020332 | 0.003409 | 0.013082 | 0.007124 | 0.008308 | 0.004592 | 0.004159 | 0.001137 | 0.001568 | 0.012422 | 0.001267 | 0.004392 | 0.003072 | 0.016656 | 0.020349 |
| 28.7361111 | 0.013457 | 0.001427 | 0.012145 | 0.007635 | 0.003318 | 0.010557 | 0.014507 | 0.041822 | 0.006106 | 0.029196 | 0.000881 | 0.012166 | 0.005948 | 0.007159 | 0.001872 | 0.001713 | 0.000345 | 0.001017 | 0.008839 | 0.001694 | 0.00167 | 0.001901 | 0.009654 | 0.013549 |
| 28.75 | 0.010351 | 0.001238 | 0.016169 | 0.008417 | 0.005364 | 0.007633 | 0.021529 | 0.02167 | 0.001757 | 0.036234 | 0.00124 | 0.007558 | 0.005352 | 0.00716 | 0.001353 | 0.001704 | 9.79E-05 | 0.001653 | 0.00701 | 0.002984 | 0.000968 | 0.001484 | 0.004244 | 0.007636 |
| 28.7638889 | 0.007478 | 0.000578 | 0.017801 | 0.010243 | 0.007907 | 0.004558 | 0.023068 | 0.035953 | 0.003384 | 0.039443 | 0.002629 | 0.003082 | 0.004906 | 0.007453 | 0.001911 | 0.001765 | 0.000165 | 0.002718 | 0.006377 | 0.004035 | 0.001325 | 0.002495 | 0.002338 | 0.003758 |
| 28.7777778 | 0.013221 | 0.000446 | 0.018389 | 0.018391 | 0.011701 | 0.003346 | 0.021479 | 0.058147 | 0.010745 | 0.036084 | 0.005547 | 0.001736 | 0.003908 | 0.0067 | 0.002918 | 0.001596 | 0.000616 | 0.003985 | 0.006151 | 0.005352 | 0.001984 | 0.003729 | 0.005198 | 0.001685 |
| 28.7916667 | 0.01821 | 0.000818 | 0.014861 | 0.023034 | 0.013855 | 0.006752 | 0.016818 | 0.060339 | 0.016942 | 0.02793 | 0.006262 | 0.002289 | 0.002451 | 0.004525 | 0.004586 | 0.001258 | 0.001708 | 0.005516 | 0.005874 | 0.007254 | 0.003189 | 0.003085 | 0.010012 | 0.001157 |
| 28.8055556 | 0.013946 | 0.001573 | 0.010643 | 0.015887 | 0.011254 | 0.011253 | 0.009465 | 0.049974 | 0.015107 | 0.021949 | 0.00528 | 0.002221 | 0.002057 | 0.002283 | 0.00574 | 0.000963 | 0.002984 | 0.006852 | 0.005329 | 0.00904 | 0.004788 | 0.001825 | 0.014433 | 0.00239 |
| 28.8194444 | 0.009654 | 0.002633 | 0.009372 | 0.008895 | 0.006649 | 0.012373 | 0.003502 | 0.039761 | 0.008594 | 0.023353 | 0.009168 | 0.001657 | 0.003438 | 0.001639 | 0.005213 | 0.00169 | 0.00353 | 0.007249 | 0.004725 | 0.009702 | 0.005934 | 0.002287 | 0.016976 | 0.004352 |
| 28.8333333 | 0.017897 | 0.003774 | 0.008411 | 0.012484 | 0.006352 | 0.011297 | 0.002444 | 0.033287 | 0.006811 | 0.033264 | 0.01432 | 0.002761 | 0.004827 | 0.001982 | 0.003393 | 0.002195 | 0.003246 | 0.006908 | 0.004896 | 0.009495 | 0.006281 | 0.003184 | 0.017219 | 0.005562 |
| 28.8472222 | 0.032121 | 0.005125 | 0.006652 | 0.016889 | 0.011941 | 0.010697 | 0.005386 | 0.03051 | 0.010707 | 0.049322 | 0.014441 | 0.005946 | 0.00571 | 0.001818 | 0.001726 | 0.001653 | 0.002839 | 0.007254 | 0.00574 | 0.009847 | 0.005979 | 0.002387 | 0.017287 | 0.005938 |
| 28.8611111 | 0.04047 | 0.007269 | 0.006916 | 0.014056 | 0.018352 | 0.010901 | 0.009937 | 0.031799 | 0.011651 | 0.062342 | 0.010693 | 0.00841 | 0.007131 | 0.001643 | 0.001364 | 0.001558 | 0.00291 | 0.008554 | 0.005634 | 0.011729 | 0.005557 | 0.001035 | 0.017843 | 0.006701 |

| | | | | | | | | | | | | | | | | | | | | | | | | |
|---|---|---|---|---|---|---|---|---|---|---|---|---|---|---|---|---|---|---|---|---|---|---|---|---|
| 28.875 | 0.040122 | 0.009812 | 0.008737 | 0.010253 | 0.020003 | 0.010699 | 0.013767 | 0.034275 | 0.008954 | 0.062937 | 0.007199 | 0.007687 | 0.008681 | 0.002877 | 0.001952 | 0.002604 | 0.003142 | 0.00899 | 0.004082 | 0.014479 | 0.006028 | 0.000784 | 0.018251 | 0.008665 |
| 28.8888889 | 0.032736 | 0.012383 | 0.008242 | 0.012765 | 0.017466 | 0.009926 | 0.015313 | 0.034024 | 0.010212 | 0.050621 | 0.005178 | 0.00432 | 0.009698 | 0.005229 | 0.002343 | 0.003632 | 0.002477 | 0.008365 | 0.002178 | 0.016177 | 0.007421 | 0.001721 | 0.018197 | 0.01087 |
| 28.9027778 | 0.02572 | 0.013656 | 0.004977 | 0.019389 | 0.013496 | 0.009103 | 0.014638 | 0.029881 | 0.010368 | 0.03039 | 0.003643 | 0.001885 | 0.010024 | 0.007145 | 0.002312 | 0.003838 | 0.001143 | 0.009111 | 0.00149 | 0.016178 | 0.009206 | 0.002611 | 0.017847 | 0.011411 |
| 28.9166667 | 0.02442 | 0.009997 | 0.001818 | 0.024559 | 0.008652 | 0.007515 | 0.012791 | 0.022965 | 0.008737 | 0.013971 | 0.00508 | 0.002774 | 0.010511 | 0.008611 | 0.001747 | 0.003819 | 0.000769 | 0.012245 | 0.003059 | 0.015682 | 0.010678 | 0.002486 | 0.017896 | 0.009746 |
| 28.9305556 | 0.02495 | 0.004939 | 0.000728 | 0.025606 | 0.00422 | 0.004725 | 0.010574 | 0.014623 | 0.017104 | 0.012729 | 0.010413 | 0.005111 | 0.013089 | 0.011152 | 0.000918 | 0.004149 | 0.001852 | 0.014292 | 0.005292 | 0.014219 | 0.010848 | 0.002266 | 0.017851 | 0.007072 |
| 28.9444444 | 0.022238 | 0.004881 | 0.001029 | 0.022036 | 0.002708 | 0.001968 | 0.008455 | 0.006866 | 0.031094 | 0.020943 | 0.016184 | 0.007342 | 0.017621 | 0.014327 | 0.000895 | 0.004299 | 0.003084 | 0.012387 | 0.005832 | 0.010714 | 0.009516 | 0.002074 | 0.015729 | 0.004882 |
| 28.9583333 | 0.01558 | 0.007273 | 0.001148 | 0.016985 | 0.004394 | 0.000655 | 0.006781 | 0.002478 | 0.038384 | 0.024127 | 0.018368 | 0.00944 | 0.020468 | 0.015988 | 0.001276 | 0.00321 | 0.003279 | 0.008336 | 0.00466 | 0.006282 | 0.007094 | 0.002522 | 0.011897 | 0.003776 |
| 28.9722222 | 0.009314 | 0.008066 | 0.000714 | 0.016067 | 0.006696 | 0.000685 | 0.007183 | 0.001188 | 0.035372 | 0.018209 | 0.015404 | 0.010259 | 0.018468 | 0.016068 | 0.001133 | 0.001453 | 0.002901 | 0.004229 | 0.003487 | 0.002742 | 0.005132 | 0.004291 | 0.008168 | 0.004058 |
| 28.9861111 | 0.012289 | 0.006289 | 0.000691 | 0.022264 | 0.006901 | 0.002602 | 0.011233 | 0.002113 | 0.021924 | 0.022302 | 0.008847 | 0.007984 | 0.01261 | 0.015132 | 0.00071 | 0.000999 | 0.002779 | 0.001923 | 0.003573 | 0.001419 | 0.005731 | 0.004875 | 0.0048 | 0.006778 |
| 29 | 0.025283 | 0.004842 | 0.003042 | 0.030334 | 0.004242 | 0.0077 | 0.017656 | 0.007527 | 0.013652 | 0.056366 | 0.005358 | 0.00894 | 0.006129 | 0.012262 | 0.000684 | 0.002497 | 0.002712 | 0.002518 | 0.004636 | 0.002743 | 0.008132 | 0.003227 | 0.004149 | 0.012077 |
| 29.0138889 | 0.036646 | 0.005774 | 0.008451 | 0.037649 | 0.002651 | 0.012883 | 0.02315 | 0.014796 | 0.026881 | 0.105138 | 0.011602 | 0.021385 | 0.002208 | 0.00697 | 0.002384 | 0.006101 | 0.002557 | 0.004057 | 0.004635 | 0.005974 | 0.008762 | 0.002147 | 0.007287 | 0.016091 |
| 29.0277778 | 0.041112 | 0.005887 | 0.013101 | 0.043228 | 0.004322 | 0.013989 | 0.023808 | 0.021048 | 0.043775 | 0.132212 | 0.021574 | 0.039085 | 0.00129 | 0.002677 | 0.006535 | 0.011578 | 0.004259 | 0.005192 | 0.00365 | 0.008752 | 0.005776 | 0.003136 | 0.009848 | 0.014189 |
| 29.0416667 | 0.043589 | 0.006957 | 0.012659 | 0.03806 | 0.00451 | 0.014089 | 0.019599 | 0.028696 | 0.046474 | 0.109731 | 0.024692 | 0.045899 | 0.002507 | 0.002939 | 0.009611 | 0.014771 | 0.00714 | 0.005784 | 0.003019 | 0.009515 | 0.003218 | 0.006262 | 0.009474 | 0.009479 |
| 29.0555556 | 0.042895 | 0.00955 | 0.007678 | 0.022321 | 0.002682 | 0.016133 | 0.012999 | 0.033796 | 0.037571 | 0.062149 | 0.020631 | 0.031119 | 0.005451 | 0.008349 | 0.007151 | 0.010922 | 0.00622 | 0.00454 | 0.005386 | 0.009354 | 0.005588 | 0.011138 | 0.009223 | 0.008504 |
| 29.0694444 | 0.035078 | 0.007981 | 0.004689 | 0.010785 | 0.003249 | 0.014917 | 0.006226 | 0.027019 | 0.022355 | 0.05212 | 0.016179 | 0.015952 | 0.007819 | 0.017407 | 0.003048 | 0.004541 | 0.003156 | 0.002216 | 0.013948 | 0.007542 | 0.009178 | 0.012601 | 0.010082 | 0.010218 |
| 29.0833333 | 0.022544 | 0.003377 | 0.008248 | 0.012127 | 0.00619 | 0.008523 | 0.003284 | 0.016061 | 0.018308 | 0.079913 | 0.01414 | 0.024371 | 0.007846 | 0.02317 | 0.002484 | 0.002385 | 0.002201 | 0.001872 | 0.021415 | 0.004783 | 0.008051 | 0.008637 | 0.009349 | 0.010437 |
| 29.0972222 | 0.011046 | 0.001808 | 0.012234 | 0.0171 | 0.008211 | 0.00369 | 0.004422 | 0.015914 | 0.032226 | 0.097292 | 0.012215 | 0.038439 | 0.006161 | 0.021568 | 0.002788 | 0.002356 | 0.001631 | 0.003574 | 0.021433 | 0.005031 | 0.003771 | 0.007293 | 0.007199 | 0.009285 |
| 29.1111111 | 0.007053 | 0.004568 | 0.012042 | 0.017084 | 0.007609 | 0.004683 | 0.005756 | 0.02353 | 0.040794 | 0.093236 | 0.008933 | 0.040015 | 0.004528 | 0.016486 | 0.002109 | 0.001969 | 0.000804 | 0.004877 | 0.018077 | 0.007158 | 0.001156 | 0.011258 | 0.005613 | 0.008137 |
| 29.125 | 0.011303 | 0.008624 | 0.011865 | 0.015699 | 0.00518 | 0.007784 | 0.006314 | 0.030365 | 0.029302 | 0.082295 | 0.00521 | 0.033148 | 0.004465 | 0.011529 | 0.001426 | 0.001836 | 0.000623 | 0.00467 | 0.014536 | 0.007213 | 0.000929 | 0.013397 | 0.004733 | 0.007345 |
| 29.1388889 | 0.017749 | 0.011236 | 0.014621 | 0.015586 | 0.002653 | 0.009304 | 0.00673 | 0.030307 | 0.012387 | 0.067828 | 0.002391 | 0.02302 | 0.005476 | 0.007517 | 0.001075 | 0.001882 | 0.000729 | 0.003573 | 0.011911 | 0.004541 | 0.001678 | 0.011428 | 0.004138 | 0.006545 |
| 29.1527778 | 0.021043 | 0.012495 | 0.018307 | 0.01575 | 0.001383 | 0.009182 | 0.006776 | 0.02287 | 0.004823 | 0.048333 | 0.001412 | 0.012803 | 0.005994 | 0.004863 | 0.001291 | 0.002001 | 0.000862 | 0.002749 | 0.010811 | 0.001672 | 0.00253 | 0.008529 | 0.003503 | 0.0051 |
| 29.1666667 | 0.021492 | 0.012832 | 0.019481 | 0.016142 | 0.00134 | 0.008484 | 0.006666 | 0.013655 | 0.003699 | 0.026271 | 0.002609 | 0.005374 | 0.005294 | 0.003498 | 0.001272 | 0.001831 | 0.000807 | 0.002858 | 0.01052 | 0.000887 | 0.002581 | 0.006333 | 0.002234 | 0.003333 |
| 29.1805556 | 0.020501 | 0.01124 | 0.016976 | 0.017162 | 0.001793 | 0.008553 | 0.006995 | 0.007006 | 0.006762 | 0.011225 | 0.004757 | 0.002767 | 0.003202 | 0.0026 | 0.001053 | 0.001075 | 0.000537 | 0.003714 | 0.010315 | 0.002583 | 0.001928 | 0.004618 | 0.001409 | 0.002002 |
| 29.1944444 | 0.016502 | 0.008043 | 0.013867 | 0.018915 | 0.002285 | 0.009458 | 0.00778 | 0.00327 | 0.012555 | 0.01132 | 0.007152 | 0.003218 | 0.00121 | 0.002186 | 0.001712 | 0.000408 | 0.000288 | 0.004379 | 0.010165 | 0.006726 | 0.001689 | 0.002574 | 0.002321 | 0.001359 |
| 29.2083333 | 0.010889 | 0.004568 | 0.012528 | 0.020573 | 0.002216 | 0.010353 | 0.008279 | 0.001333 | 0.01716 | 0.017794 | 0.00976 | 0.002727 | 0.000909 | 0.002346 | 0.002474 | 0.000474 | 0.000371 | 0.004115 | 0.010042 | 0.011559 | 0.002076 | 0.000916 | 0.003178 | 0.001326 |
| 29.2222222 | 0.005688 | 0.002034 | 0.011456 | 0.02051 | 0.001401 | 0.010517 | 0.007649 | 0.002568 | 0.021601 | 0.018858 | 0.011746 | 0.001677 | 0.00103 | 0.002086 | 0.002364 | 0.000983 | 0.000637 | 0.003268 | 0.009877 | 0.014761 | 0.00269 | 0.000624 | 0.002687 | 0.001705 |
| 29.2361111 | 0.002759 | 0.001489 | 0.009874 | 0.017759 | 0.000608 | 0.010028 | 0.005697 | 0.006594 | 0.025853 | 0.013221 | 0.012015 | 0.002513 | 0.000663 | 0.001097 | 0.001541 | 0.001535 | 0.000546 | 0.002408 | 0.009923 | 0.015567 | 0.003372 | 0.001648 | 0.001583 | 0.002369 |
| 29.25 | 0.004508 | 0.002079 | 0.00842 | 0.012826 | 0.000308 | 0.009362 | 0.003002 | 0.009616 | 0.026933 | 0.006323 | 0.010484 | 0.004132 | 0.000451 | 0.000324 | 0.000715 | 0.0018 | 0.000317 | 0.001638 | 0.009993 | 0.014361 | 0.003549 | 0.004251 | 0.000953 | 0.003135 |
| 29.2638889 | 0.008411 | 0.001985 | 0.00688 | 0.007055 | 0.000775 | 0.008152 | 0.001487 | 0.008434 | 0.023708 | 0.004244 | 0.008388 | 0.004828 | 0.00059 | 0.00037 | 0.000581 | 0.0017 | 0.000375 | 0.001273 | 0.00964 | 0.012074 | 0.002744 | 0.007025 | 0.001114 | 0.00346 |
| 29.2777778 | 0.010704 | 0.001835 | 0.005035 | 0.00242 | 0.002441 | 0.005873 | 0.002155 | 0.005841 | 0.016741 | 0.008433 | 0.006569 | 0.004568 | 0.00071 | 0.001302 | 0.000855 | 0.001247 | 0.000605 | 0.00143 | 0.009073 | 0.009297 | 0.001825 | 0.007731 | 0.001205 | 0.003201 |
| 29.2916667 | 0.010783 | 0.002415 | 0.003734 | 0.001365 | 0.004132 | 0.003291 | 0.002333 | 0.006055 | 0.008417 | 0.014881 | 0.00535 | 0.004653 | 0.001014 | 0.002759 | 0.001487 | 0.000713 | 0.001125 | 0.001656 | 0.008822 | 0.006643 | 0.00221 | 0.006186 | 0.00115 | 0.002755 |
| 29.3055556 | 0.009941 | 0.002578 | 0.003274 | 0.004823 | 0.003948 | 0.00151 | 0.001942 | 0.00606 | 0.004187 | 0.017042 | 0.005279 | 0.006756 | 0.001632 | 0.003798 | 0.002075 | 0.000477 | 0.001516 | 0.002315 | 0.008458 | 0.005252 | 0.00299 | 0.004192 | 0.001126 | 0.001891 |
| 29.3194444 | 0.00923 | 0.002715 | 0.002905 | 0.011584 | 0.002138 | 0.000592 | 0.00373 | 0.006877 | 0.007044 | 0.011856 | 0.006452 | 0.010405 | 0.001471 | 0.003658 | 0.00159 | 0.000685 | 0.001186 | 0.004331 | 0.007185 | 0.005701 | 0.002492 | 0.003511 | 0.001528 | 0.001012 |
| 29.3333333 | 0.007721 | 0.003747 | 0.002894 | 0.017822 | 0.000621 | 0.000569 | 0.006142 | 0.014184 | 0.014375 | 0.007832 | 0.007973 | 0.013024 | 0.001258 | 0.002274 | 0.000874 | 0.001678 | 0.00096 | 0.006976 | 0.005557 | 0.006695 | 0.00223 | 0.003633 | 0.002396 | 0.000832 |
| 29.3472222 | 0.004593 | 0.004522 | 0.00399 | 0.019012 | 0.000198 | 0.002087 | 0.006958 | 0.021625 | 0.021241 | 0.0128 | 0.008696 | 0.012914 | 0.002411 | 0.001102 | 0.001098 | 0.002881 | 0.001282 | 0.008159 | 0.00472 | 0.006637 | 0.003195 | 0.003538 | 0.002282 | 0.000583 |
| 29.3611111 | 0.003692 | 0.003349 | 0.005746 | 0.015432 | 0.000558 | 0.00451 | 0.006455 | 0.021133 | 0.023396 | 0.017742 | 0.00845 | 0.010343 | 0.003543 | 0.001281 | 0.0013 | 0.003663 | 0.001176 | 0.007268 | 0.004746 | 0.005569 | 0.003497 | 0.003513 | 0.00127 | 0.00017 |
| 29.375 | 0.007461 | 0.002027 | 0.006966 | 0.011075 | 0.001311 | 0.005739 | 0.005211 | 0.015027 | 0.020065 | 0.014655 | 0.007889 | 0.00614 | 0.003465 | 0.00129 | 0.000815 | 0.004319 | 0.000605 | 0.005918 | 0.005027 | 0.004095 | 0.002585 | 0.003945 | 0.000539 | 0.000145 |
| 29.3888889 | 0.011407 | 0.002794 | 0.00707 | 0.008783 | 0.001554 | 0.005194 | 0.00417 | 0.009033 | 0.014552 | 0.007741 | 0.007432 | 0.002648 | 0.00315 | 0.000799 | 0.000347 | 0.005172 | 0.00029 | 0.005255 | 0.005201 | 0.002949 | 0.001772 | 0.004827 | 0.000171 | 0.000563 |
| 29.4027778 | 0.012875 | 0.00394 | 0.006517 | 0.010118 | 0.001161 | 0.004294 | 0.004623 | 0.006769 | 0.010127 | 0.003825 | 0.007228 | 0.00132 | 0.003553 | 0.00124 | 0.000221 | 0.006208 | 0.000229 | 0.004663 | 0.005139 | 0.002783 | 0.001744 | 0.005958 | 0.000132 | 0.001495 |
| 29.4166667 | 0.01396 | 0.004215 | 0.005292 | 0.014279 | 0.00125 | 0.004316 | 0.007288 | 0.009388 | 0.007335 | 0.003742 | 0.007534 | 0.001535 | 0.004017 | 0.002429 | 0.000289 | 0.00642 | 0.00028 | 0.00369 | 0.005114 | 0.003049 | 0.001899 | 0.006476 | 0.0003 | 0.002782 |
| 29.4305556 | 0.016216 | 0.003863 | 0.003522 | 0.018491 | 0.002572 | 0.005813 | 0.011761 | 0.015338 | 0.005766 | 0.003233 | 0.007985 | 0.003705 | 0.003508 | 0.0034 | 0.000663 | 0.004932 | 0.000734 | 0.002826 | 0.005437 | 0.002995 | 0.001625 | 0.005749 | 0.000806 | 0.003957 |
| 29.4444444 | 0.018715 | 0.002818 | 0.002007 | 0.020126 | 0.004862 | 0.0083 | 0.016296 | 0.021549 | 0.006155 | 0.002237 | 0.007658 | 0.006057 | 0.002222 | 0.003558 | 0.001594 | 0.00255 | 0.001814 | 0.002273 | 0.005983 | 0.002517 | 0.001064 | 0.004173 | 0.002068 | 0.00449 |
| 29.4583333 | 0.020133 | 0.001737 | 0.001331 | 0.017895 | 0.007077 | 0.011219 | 0.019178 | 0.026196 | 0.009306 | 0.003708 | 0.006745 | 0.005074 | 0.001172 | 0.002576 | 0.003329 | 0.001243 | 0.002813 | 0.001485 | 0.006507 | 0.001565 | 0.001137 | 0.002555 | 0.003241 | 0.004305 |
| 29.4722222 | 0.021741 | 0.00262 | 0.002198 | 0.016522 | 0.006893 | 0.016147 | 0.021221 | 0.031358 | 0.013807 | 0.008495 | 0.007043 | 0.002855 | 0.00077 | 0.001224 | 0.005098 | 0.002096 | 0.002727 | 0.000641 | 0.006425 | 0.000912 | 0.001838 | 0.001882 | 0.002806 | 0.003507 |
| 29.4861111 | 0.02658 | 0.006898 | 0.004761 | 0.024871 | 0.006169 | 0.026705 | 0.023794 | 0.041074 | 0.02114 | 0.020073 | 0.010252 | 0.002434 | 0.001404 | 0.00053 | 0.004545 | 0.003828 | 0.002032 | 0.000698 | 0.004749 | 0.001074 | 0.002184 | 0.003642 | 0.002797 | 0.002657 |
| 29.5 | 0.03679 | 0.011146 | 0.008604 | 0.040647 | 0.010686 | 0.037886 | 0.02369 | 0.056003 | 0.038026 | 0.043433 | 0.01511 | 0.003087 | 0.00477 | 0.000384 | 0.003863 | 0.003963 | 0.003236 | 0.002223 | 0.003324 | 0.002847 | 0.004028 | 0.008134 | 0.007647 | 0.004819 |
| 29.5138889 | 0.048521 | 0.010474 | 0.012866 | 0.047223 | 0.018778 | 0.03633 | 0.01636 | 0.071961 | 0.062103 | 0.069907 | 0.020204 | 0.005997 | 0.010266 | 0.000611 | 0.007454 | 0.003645 | 0.006391 | 0.004585 | 0.004256 | 0.006764 | 0.007813 | 0.011512 | 0.014187 | 0.010855 |
| 29.5277778 | 0.056244 | 0.007284 | 0.017124 | 0.035969 | 0.024232 | 0.023955 | 0.010532 | 0.084977 | 0.090369 | 0.084544 | 0.026011 | 0.009738 | 0.013455 | 0.001547 | 0.010403 | 0.005132 | 0.00657 | 0.005139 | 0.004359 | 0.007565 | 0.008342 | 0.00913 | 0.013753 | 0.014289 |
| 29.5416667 | 0.063586 | 0.006633 | 0.024081 | 0.021026 | 0.024635 | 0.015708 | 0.01271 | 0.092148 | 0.11147 | 0.081195 | 0.029154 | 0.010162 | 0.012759 | 0.003422 | 0.007678 | 0.004687 | 0.003514 | 0.003629 | 0.003129 | 0.004298 | 0.004367 | 0.005895 | 0.007876 | 0.011042 |
| 29.5555556 | 0.069058 | 0.007683 | 0.030016 | 0.018892 | 0.021598 | 0.016395 | 0.011298 | 0.084904 | 0.093682 | 0.060781 | 0.027083 | 0.008129 | 0.011349 | 0.005644 | 0.004366 | 0.001754 | 0.00179 | 0.003557 | 0.002443 | 0.002236 | 0.002241 | 0.007063 | 0.005533 | 0.006392 |
| 29.5694444 | 0.06401 | 0.007596 | 0.026541 | 0.024722 | 0.01735 | 0.019917 | 0.008213 | 0.063285 | 0.051399 | 0.037251 | 0.023077 | 0.006408 | 0.01005 | 0.006617 | 0.004064 | 0.000728 | 0.001625 | 0.005044 | 0.001468 | 0.001235 | 0.004191 | 0.007797 | 0.006822 | 0.004412 |
| 29.5833333 | 0.04965 | 0.006502 | 0.017678 | 0.027899 | 0.011538 | 0.020808 | 0.012581 | 0.039627 | 0.044638 | 0.021524 | 0.018679 | 0.004365 | 0.007741 | 0.005577 | 0.004023 | 0.001827 | 0.001654 | 0.005257 | 0.001305 | 0.000796 | 0.005904 | 0.006378 | 0.006578 | 0.002899 |
| 29.5972222 | 0.031181 | 0.004996 | 0.009323 | 0.029989 | 0.006786 | 0.017556 | 0.017349 | 0.020558 | 0.069512 | 0.013215 | 0.013908 | 0.004806 | 0.005609 | 0.003681 | 0.002645 | 0.004082 | 0.001366 | 0.004028 | 0.001808 | 0.00143 | 0.005553 | 0.004219 | 0.004469 | 0.002273 |
| 29.6111111 | 0.013777 | 0.002871 | 0.004326 | 0.03261 | 0.008832 | 0.011426 | 0.017728 | 0.016238 | 0.068179 | 0.011044 | 0.011337 | 0.011927 | 0.004156 | 0.002076 | 0.001536 | 0.006298 | 0.00152 | 0.002488 | 0.001662 | 0.00206 | 0.004136 | 0.002279 | 0.002228 | 0.004465 |
| 29.625 | 0.006389 | 0.00105 | 0.004377 | 0.032453 | 0.014606 | 0.005735 | 0.015281 | 0.032066 | 0.036384 | 0.011548 | 0.011756 | 0.01895 | 0.003347 | 0.000969 | 0.001881 | 0.005976 | 0.002128 | 0.001385 | 0.001266 | 0.002278 | 0.003117 | 0.003331 | 0.001308 | 0.005806 |
| 29.6388889 | 0.010573 | 0.000578 | 0.003756 | 0.02946 | 0.015516 | 0.002449 | 0.010117 | 0.052385 | 0.016729 | 0.01972 | 0.01206 | 0.018942 | 0.00423 | 0.001121 | 0.002082 | 0.003703 | 0.001758 | 0.001605 | 0.000771 | 0.003494 | 0.00441 | 0.007265 | 0.003353 | 0.004937 |
| 29.6527778 | 0.015508 | 0.001205 | 0.002047 | 0.025254 | 0.011328 | 0.001495 | 0.005431 | 0.065714 | 0.02393 | 0.034453 | 0.009549 | 0.013284 | 0.006341 | 0.002718 | 0.001294 | 0.002695 | 0.001191 | 0.002415 | 0.000353 | 0.005252 | 0.007759 | 0.010425 | 0.007766 | 0.006528 |
| 29.6666667 | 0.015513 | 0.002556 | 0.003147 | 0.020861 | 0.006949 | 0.00244 | 0.004948 | 0.0599 | 0.034705 | 0.039262 | 0.005271 | 0.007573 | 0.008281 | 0.00421 | 0.000585 | 0.00301 | 0.001054 | 0.002184 | 0.000352 | 0.004857 | 0.011208 | 0.009854 | 0.012061 | 0.010322 |
| 29.6805556 | 0.013123 | 0.003887 | 0.005193 | 0.017246 | 0.004152 | 0.003113 | 0.005501 | 0.037295 | 0.030364 | 0.02838 | 0.003267 | 0.008087 | 0.010273 | 0.00583 | 0.001004 | 0.00314 | 0.001499 | 0.001613 | 0.001105 | 0.003725 | 0.013725 | 0.007318 | 0.014799 | 0.012265 |
| 29.6944444 | 0.008356 | 0.004353 | 0.006765 | 0.013538 | 0.003147 | 0.002613 | 0.003636 | 0.042842 | 0.019313 | 0.016283 | 0.004626 | 0.012415 | 0.012612 | 0.008379 | 0.003019 | 0.003513 | 0.002918 | 0.002033 | 0.00323 | 0.00425 | 0.015195 | 0.006198 | 0.015547 | 0.011822 |
| 29.7083333 | 0.004308 | 0.005568 | 0.00812 | 0.009119 | 0.00425 | 0.002746 | 0.001818 | 0.085962 | 0.020656 | 0.018394 | 0.005233 | 0.012066 | 0.012879 | 0.009294 | 0.004003 | 0.003314 | 0.003204 | 0.002262 | 0.00514 | 0.004098 | 0.014021 | 0.005434 | 0.013242 | 0.008678 |
| 29.7222222 | 0.00557 | 0.008355 | 0.009024 | 0.005867 | 0.005461 | 0.004541 | 0.002659 | 0.12156 | 0.031088 | 0.027216 | 0.003908 | 0.008277 | 0.009452 | 0.006876 | 0.002893 | 0.003523 | 0.002337 | 0.002626 | 0.004139 | 0.003726 | 0.008936 | 0.003181 | 0.007647 | 0.004779 |
| 29.7361111 | 0.007193 | 0.009055 | 0.009168 | 0.005391 | 0.004747 | 0.008743 | 0.004651 | 0.129989 | 0.034733 | 0.028713 | 0.005541 | 0.011217 | 0.005703 | 0.004202 | 0.002229 | 0.005419 | 0.002661 | 0.00433 | 0.002674 | 0.006149 | 0.003682 | 0.001184 | 0.002548 | 0.003988 |
| 29.75 | 0.004763 | 0.005745 | 0.00788 | 0.006907 | 0.003563 | 0.013596 | 0.00476 | 0.116997 | 0.032954 | 0.025029 | 0.012205 | 0.022149 | 0.00475 | 0.003852 | 0.001842 | 0.005057 | 0.00251 | 0.00508 | 0.004565 | 0.008545 | 0.002046 | 0.000989 | 0.001354 | 0.004297 |
| 29.7638889 | 0.003752 | 0.003505 | 0.006094 | 0.008827 | 0.003085 | 0.015944 | 0.003277 | 0.084598 | 0.030195 | 0.021272 | 0.018806 | 0.031878 | 0.005333 | 0.004293 | 0.001143 | 0.002422 | 0.001163 | 0.00405 | 0.006294 | 0.009496 | 0.002815 | 0.002581 | 0.001941 | 0.002791 |
| 29.7777778 | 0.008664 | 0.005522 | 0.008181 | 0.010451 | 0.002499 | 0.014205 | 0.004025 | 0.043201 | 0.024082 | 0.018116 | 0.020779 | 0.033543 | 0.005884 | 0.003916 | 0.001359 | 0.001004 | 0.00048 | 0.002676 | 0.003862 | 0.009741 | 0.003368 | 0.004926 | 0.001724 | 0.000997 |
| 29.7916667 | 0.014109 | 0.007251 | 0.014009 | 0.010769 | 0.001437 | 0.00854 | 0.007828 | 0.018818 | 0.013438 | 0.017688 | 0.018981 | 0.02702 | 0.006153 | 0.003405 | 0.003114 | 0.001371 | 0.000609 | 0.001886 | 0.001113 | 0.009174 | 0.002872 | 0.006324 | 0.001013 | 0.000364 |
| 29.8055556 | 0.014524 | 0.006142 | 0.01593 | 0.008256 | 0.000552 | 0.003787 | 0.009906 | 0.031056 | 0.004441 | 0.022354 | 0.015537 | 0.018213 | 0.005682 | 0.003788 | 0.006167 | 0.003876 | 0.001554 | 0.002641 | 0.001262 | 0.009311 | 0.001811 | 0.005468 | 0.000542 | 0.000414 |
| 29.8194444 | 0.011078 | 0.004604 | 0.013185 | 0.004605 | 0.000361 | 0.003918 | 0.008747 | 0.063083 | 0.006611 | 0.033565 | 0.011166 | 0.015078 | 0.004441 | 0.004717 | 0.009131 | 0.006362 | 0.003492 | 0.004795 | 0.002479 | 0.01026 | 0.000982 | 0.00319 | 0.000803 | 0.001421 |
| 29.8333333 | 0.011582 | 0.004913 | 0.010829 | 0.004156 | 0.000377 | 0.007831 | 0.008286 | 0.082998 | 0.016242 | 0.052247 | 0.006659 | 0.019149 | 0.002942 | 0.005511 | 0.010431 | 0.005627 | 0.005205 | 0.007393 | 0.002899 | 0.010923 | 0.001288 | 0.002758 | 0.001756 | 0.004218 |
| 29.8472222 | 0.021151 | 0.006316 | 0.009798 | 0.007333 | 0.001346 | 0.013614 | 0.010165 | 0.083794 | 0.018366 | 0.070848 | 0.003086 | 0.024564 | 0.001568 | 0.006151 | 0.010069 | 0.003369 | 0.005932 | 0.01005 | 0.002588 | 0.011995 | 0.002874 | 0.005374 | 0.002871 | 0.007958 |
| 29.8611111 | 0.030111 | 0.005866 | 0.010531 | 0.01092 | 0.004746 | 0.019894 | 0.013625 | 0.076073 | 0.011516 | 0.077885 | 0.001841 | 0.02574 | 0.000712 | 0.006742 | 0.008483 | 0.002 | 0.006157 | 0.012399 | 0.002286 | 0.014132 | 0.004886 | 0.009029 | 0.003998 | 0.010175 |
| 29.875 | 0.032206 | 0.004313 | 0.01406 | 0.014648 | 0.010012 | 0.025151 | 0.014545 | 0.071154 | 0.008577 | 0.073072 | 0.004873 | 0.022408 | 0.000638 | 0.007156 | 0.005284 | 0.001775 | 0.00571 | 0.013466 | 0.002191 | 0.017303 | 0.005841 | 0.010789 | 0.005028 | 0.009716 |
| 29.8888889 | 0.030193 | 0.004289 | 0.019555 | 0.020913 | 0.014372 | 0.027493 | 0.011789 | 0.069066 | 0.011037 | 0.059778 | 0.012824 | 0.015084 | 0.002227 | 0.006187 | 0.001829 | 0.002102 | 0.004948 | 0.013204 | 0.003218 | 0.022644 | 0.0059 | 0.010493 | 0.006255 | 0.008985 |
| 29.9027778 | 0.024414 | 0.004693 | 0.021314 | 0.029535 | 0.014695 | 0.025584 | 0.013606 | 0.06217 | 0.011011 | 0.039446 | 0.023296 | 0.01097 | 0.006002 | 0.003764 | 0.00046 | 0.002104 | 0.004759 | 0.013523 | 0.006028 | 0.028919 | 0.006621 | 0.011431 | 0.00827 | 0.009838 |
| 29.9166667 | 0.017405 | 0.004299 | 0.013871 | 0.037234 | 0.010997 | 0.019095 | 0.022854 | 0.056181 | 0.009432 | 0.018608 | 0.031142 | 0.019181 | 0.009416 | 0.002281 | 0.000514 | 0.002524 | 0.004217 | 0.014219 | 0.00856 | 0.031531 | 0.006993 | 0.013024 | 0.009604 | 0.010105 |
| 29.9305556 | 0.011933 | 0.004622 | 0.007969 | 0.036004 | 0.006352 | 0.009872 | 0.031244 | 0.051433 | 0.007261 | 0.009272 | 0.027796 | 0.027651 | 0.010565 | 0.002011 | 0.000556 | 0.002952 | 0.002627 | 0.013114 | 0.009209 | 0.027464 | 0.005325 | 0.012096 | 0.008973 | 0.008286 |
| 29.9444444 | 0.007482 | 0.005135 | 0.014305 | 0.026839 | 0.005048 | 0.005063 | 0.02788 | 0.037799 | 0.004599 | 0.014616 | 0.013862 | 0.023841 | 0.010987 | 0.001302 | 0.000637 | 0.002679 | 0.001233 | 0.011529 | 0.009108 | 0.018792 | 0.002587 | 0.00946 | 0.007213 | 0.005792 |
| 29.9583333 | 0.00421 | 0.003817 | 0.023439 | 0.022004 | 0.007973 | 0.009114 | 0.017073 | 0.019735 | 0.006455 | 0.021582 | 0.004322 | 0.015219 | 0.010827 | 0.001466 | 0.000626 | 0.002288 | 0.001313 | 0.011051 | 0.008904 | 0.009578 | 0.001126 | 0.006596 | 0.004787 | 0.003231 |
| 29.9722222 | 0.003343 | 0.002335 | 0.028363 | 0.021568 | 0.010051 | 0.014874 | 0.012897 | 0.009494 | 0.012057 | 0.017576 | 0.004322 | 0.008893 | 0.008454 | 0.004011 | 0.000457 | 0.001735 | 0.002423 | 0.010257 | 0.008198 | 0.004552 | 0.00136 | 0.003617 | 0.002227 | 0.001605 |
| 29.9861111 | 0.005901 | 0.00251 | 0.028747 | 0.019584 | 0.007088 | 0.013881 | 0.011643 | 0.007938 | 0.013688 | 0.015857 | 0.005632 | 0.007199 | 0.004842 | 0.005978 | 0.000759 | 0.00099 | 0.002536 | 0.008232 | 0.007732 | 0.006163 | 0.001992 | 0.001653 | 0.00139 | 0.001895 |
| 30 | 0.014742 | 0.004291 | 0.019514 | 0.016321 | 0.004009 | 0.009406 | 0.009381 | 0.014782 | 0.012036 | 0.041444 | 0.00882 | 0.016179 | 0.002545 | 0.005057 | 0.001789 | 0.000999 | 0.001375 | 0.005532 | 0.008501 | 0.007986 | 0.003803 | 0.00149 | 0.00368 | 0.004339 |
| 30.0138889 | 0.028929 | 0.008657 | 0.010692 | 0.018278 | 0.006489 | 0.012052 | 0.009178 | 0.034937 | 0.018705 | 0.083982 | 0.018741 | 0.033261 | 0.001641 | 0.004447 | 0.002195 | 0.003436 | 0.001624 | 0.002928 | 0.008626 | 0.005115 | 0.005455 | 0.001962 | 0.006252 | 0.008113 |
| 30.0277778 | 0.040528 | 0.01521 | 0.014311 | 0.027758 | 0.011051 | 0.021815 | 0.009845 | 0.052331 | 0.028982 | 0.103929 | 0.027051 | 0.044535 | 0.002103 | 0.007947 | 0.003321 | 0.008168 | 0.005568 | 0.001806 | 0.006727 | 0.001586 | 0.004989 | 0.003169 | 0.005116 | 0.008841 |
| 30.0416667 | 0.047724 | 0.023012 | 0.01773 | 0.03769 | 0.014774 | 0.02964 | 0.013446 | 0.052118 | 0.028412 | 0.083435 | 0.026472 | 0.043194 | 0.003512 | 0.012752 | 0.00636 | 0.011514 | 0.01037 | 0.002639 | 0.006368 | 0.000744 | 0.004948 | 0.004423 | 0.002055 | 0.006421 |
| 30.0555556 | 0.05207 | 0.029175 | 0.012947 | 0.043023 | 0.01635 | 0.030386 | 0.014831 | 0.046708 | 0.020617 | 0.046826 | 0.02158 | 0.030591 | 0.003728 | 0.012568 | 0.00655 | 0.009568 | 0.010149 | 0.002988 | 0.006796 | 0.001353 | 0.004236 | 0.004398 | 0.000786 | 0.004685 |
| 30.0694444 | 0.048983 | 0.026194 | 0.006682 | 0.041158 | 0.013935 | 0.025379 | 0.008889 | 0.045145 | 0.013016 | 0.020946 | 0.015615 | 0.016647 | 0.003629 | 0.00749 | 0.00318 | 0.004282 | 0.005289 | 0.001776 | 0.004707 | 0.002164 | 0.001926 | 0.005248 | 0.001263 | 0.003893 |
| 30.0833333 | 0.035256 | 0.015409 | 0.004753 | 0.032707 | 0.009576 | 0.01962 | 0.00294 | 0.040596 | 0.006788 | 0.008244 | 0.008091 | 0.017097 | 0.003905 | 0.002948 | 0.000799 | 0.000953 | 0.001319 | 0.001376 | 0.003744 | 0.003449 | 0.000988 | 0.005845 | 0.001506 | 0.00312 |
| 30.0972222 | 0.017479 | 0.012112 | 0.007827 | 0.022235 | 0.005389 | 0.014505 | 0.001221 | 0.028426 | 0.005328 | 0.00732 | 0.003226 | 0.030058 | 0.003289 | 0.001659 | 0.00045 | 0.000518 | 0.000271 | 0.003056 | 0.005047 | 0.004658 | 0.00106 | 0.004067 | 0.000839 | 0.00203 |
| 30.1111111 | 0.010073 | 0.01652 | 0.011286 | 0.012458 | 0.003103 | 0.008858 | 0.000829 | 0.014679 | 0.009479 | 0.018889 | 0.004037 | 0.040111 | 0.002326 | 0.001825 | 0.000588 | 0.000672 | 0.00062 | 0.005143 | 0.005699 | 0.004811 | 0.001181 | 0.002073 | 0.00036 | 0.001489 |
| 30.125 | 0.016301 | 0.016737 | 0.013592 | 0.007266 | 0.00462 | 0.004113 | 0.001716 | 0.006846 | 0.013535 | 0.043437 | 0.0061 | 0.043193 | 0.001854 | 0.001265 | 0.000404 | 0.000479 | 0.000931 | 0.00603 | 0.005146 | 0.004181 | 0.00123 | 0.001024 | 0.000525 | 0.002311 |
| 30.1388889 | 0.023156 | 0.01559 | 0.015473 | 0.011568 | 0.007795 | 0.002637 | 0.006653 | 0.005054 | 0.015057 | 0.067185 | 0.006271 | 0.043364 | 0.001527 | 0.000492 | 0.000166 | 0.00046 | 0.000716 | 0.006003 | 0.003914 | 0.003434 | 0.00104 | 0.000959 | 0.000566 | 0.003243 |
| 30.1527778 | 0.022357 | 0.01437 | 0.016231 | 0.020683 | 0.00956 | 0.003523 | 0.013497 | 0.004114 | 0.016629 | 0.067453 | 0.006143 | 0.039885 | 0.001226 | 0.000341 | 0.000232 | 0.000862 | 0.000321 | 0.006141 | 0.002655 | 0.002998 | 0.00126 | 0.00203 | 0.000281 | 0.002586 |
| 30.1666667 | 0.016332 | 0.012626 | 0.016049 | 0.026272 | 0.009814 | 0.004101 | 0.015991 | 0.002996 | 0.019382 | 0.056211 | 0.008706 | 0.030113 | 0.001159 | 0.000468 | 0.000617 | 0.001186 | 0.000208 | 0.007086 | 0.001838 | 0.003727 | 0.002365 | 0.003336 | 0.000127 | 0.001898 |
| 30.1805556 | 0.009588 | 0.013734 | 0.014855 | 0.027901 | 0.009169 | 0.004059 | 0.013021 | 0.005062 | 0.023737 | 0.049389 | 0.014577 | 0.015515 | 0.001256 | 0.000747 | 0.001064 | 0.00107 | 0.000306 | 0.008583 | 0.001538 | 0.0062 | 0.003816 | 0.004124 | 0.000193 | 0.003316 |
| 30.1944444 | 0.00835 | 0.013599 | 0.012829 | 0.025491 | 0.007898 | 0.004136 | 0.012202 | 0.010826 | 0.028154 | 0.037131 | 0.019071 | 0.006944 | 0.001375 | 0.000796 | 0.001273 | 0.000655 | 0.000331 | 0.009751 | 0.001692 | 0.009473 | 0.00432 | 0.004541 | 0.000266 | 0.004791 |
| 30.2083333 | 0.016475 | 0.010536 | 0.013342 | 0.01587 | 0.006483 | 0.004362 | 0.014767 | 0.019441 | 0.029593 | 0.018582 | 0.019437 | 0.008915 | 0.001324 | 0.000695 | 0.001105 | 0.000285 | 0.000319 | 0.009123 | 0.00167 | 0.012067 | 0.003696 | 0.005078 | 0.00041 | 0.004345 |
| 30.2222222 | 0.027824 | 0.008218 | 0.018124 | 0.006359 | 0.005278 | 0.00551 | 0.012909 | 0.030705 | 0.029385 | 0.005247 | 0.02004 | 0.010734 | 0.000979 | 0.00106 | 0.000842 | 0.000331 | 0.000521 | 0.006478 | 0.001282 | 0.013257 | 0.00308 | 0.006121 | 0.000481 | 0.002979 |
| 30.2361111 | 0.033001 | 0.006719 | 0.024454 | 0.005208 | 0.004345 | 0.008787 | 0.007768 | 0.042864 | 0.030295 | 0.002616 | 0.020755 | 0.007483 | 0.000541 | 0.001156 | 0.001116 | 0.000842 | 0.000791 | 0.003812 | 0.001316 | 0.012624 | 0.003063 | 0.007285 | 0.000611 | 0.002602 |
| 30.25 | 0.028482 | 0.004534 | 0.028795 | 0.00597 | 0.003565 | 0.013067 | 0.004765 | 0.049796 | 0.030222 | 0.007103 | 0.018643 | 0.003836 | 0.000459 | 0.000685 | 0.001649 | 0.001502 | 0.000729 | 0.002424 | 0.001291 | 0.010327 | 0.003396 | 0.007681 | 0.001352 | 0.003872 |
| 30.2638889 | 0.018205 | 0.002824 | 0.028279 | 0.003573 | 0.002556 | 0.01515 | 0.003712 | 0.045841 | 0.027272 | 0.013359 | 0.016062 | 0.003152 | 0.00066 | 0.00043 | 0.001794 | 0.002112 | 0.00038 | 0.001839 | 0.000849 | 0.007748 | 0.003742 | 0.006882 | 0.002188 | 0.005561 |
| 30.2777778 | 0.008261 | 0.004421 | 0.022612 | 0.001802 | 0.001395 | 0.013313 | 0.00287 | 0.03457 | 0.023179 | 0.014968 | 0.013861 | 0.002913 | 0.000634 | 0.000717 | 0.001675 | 0.002442 | 0.000192 | 0.001332 | 0.00062 | 0.006069 | 0.003547 | 0.005222 | 0.002263 | 0.005909 |
| 30.2916667 | 0.003502 | 0.009489 | 0.014345 | 0.003176 | 0.000595 | 0.008101 | 0.001835 | 0.022611 | 0.017911 | 0.011476 | 0.010847 | 0.002757 | 0.000322 | 0.001672 | 0.001452 | 0.002222 | 0.000447 | 0.000963 | 0.000826 | 0.005318 | 0.00249 | 0.003322 | 0.001527 | 0.004057 |
| 30.3055556 | 0.003959 | 0.017305 | 0.006738 | 0.005609 | 0.000382 | 0.004521 | 0.002118 | 0.011393 | 0.011064 | 0.013332 | 0.007237 | 0.006186 | 0.000185 | 0.003104 | 0.00096 | 0.001763 | 0.000961 | 0.001098 | 0.002132 | 0.005061 | 0.001275 | 0.001672 | 0.000895 | 0.002257 |
| 30.3194444 | 0.003396 | 0.028156 | 0.003018 | 0.006773 | 0.000581 | 0.006498 | 0.004487 | 0.004542 | 0.005184 | 0.021609 | 0.004855 | 0.012148 | 0.000687 | 0.004421 | 0.000759 | 0.0014 | 0.00148 | 0.001382 | 0.004481 | 0.004586 | 0.000704 | 0.000634 | 0.001411 | 0.002898 |
| 30.3333333 | 0.001922 | 0.035199 | 0.002963 | 0.006756 | 0.000613 | 0.007803 | 0.005914 | 0.003811 | 0.003687 | 0.023545 | 0.005905 | 0.015937 | 0.001555 | 0.0051 | 0.001456 | 0.000923 | 0.001855 | 0.001276 | 0.006037 | 0.003825 | 0.000912 | 0.000341 | 0.002219 | 0.003585 |
| 30.3472222 | 0.004557 | 0.034272 | 0.002487 | 0.005926 | 0.000853 | 0.005729 | 0.004975 | 0.005314 | 0.007984 | 0.015332 | 0.009896 | 0.013391 | 0.001597 | 0.00467 | 0.00196 | 0.000373 | 0.001581 | 0.001009 | 0.005361 | 0.004115 | 0.002074 | 0.001091 | 0.002047 | 0.002426 |
| 30.3611111 | 0.010338 | 0.032056 | 0.001744 | 0.004493 | 0.002029 | 0.006291 | 0.005731 | 0.011411 | 0.013361 | 0.009895 | 0.013999 | 0.007052 | 0.001289 | 0.003094 | 0.001374 | 0.000234 | 0.001104 | 0.001439 | 0.002983 | 0.0063 | 0.004043 | 0.003305 | 0.002292 | 0.001987 |
| 30.375 | 0.0133 | 0.031626 | 0.002927 | 0.003153 | 0.003304 | 0.009784 | 0.009523 | 0.021623 | 0.016407 | 0.015158 | 0.015536 | 0.003547 | 0.002286 | 0.001338 | 0.000613 | 0.000597 | 0.00158 | 0.002758 | 0.001084 | 0.009071 | 0.005737 | 0.006155 | 0.003874 | 0.003582 |
| 30.3888889 | 0.010956 | 0.02892 | 0.004234 | 0.002395 | 0.004108 | 0.011818 | 0.012998 | 0.029212 | 0.021134 | 0.021963 | 0.014835 | 0.002823 | 0.003474 | 0.000835 | 0.000359 | 0.000976 | 0.002314 | 0.003948 | 0.000662 | 0.010324 | 0.006153 | 0.007853 | 0.004505 | 0.005154 |
| 30.4027778 | 0.007404 | 0.020463 | 0.004351 | 0.002462 | 0.004608 | 0.011752 | 0.013436 | 0.032845 | 0.027524 | 0.023385 | 0.014665 | 0.003665 | 0.003206 | 0.002071 | 0.000428 | 0.001008 | 0.002161 | 0.004231 | 0.000722 | 0.009424 | 0.005529 | 0.007664 | 0.003178 | 0.005181 |

| | | | | | | | | | | | | | | | | | | | | | | | | |
|---|---|---|---|---|---|---|---|---|---|---|---|---|---|---|---|---|---|---|---|---|---|---|---|---|
| 30.4166667 | 0.005361 | 0.009803 | 0.004164 | 0.003757 | 0.004451 | 0.011288 | 0.009692 | 0.035292 | 0.031239 | 0.017899 | 0.015369 | 0.007676 | 0.00213 | 0.003807 | 0.000763 | 0.000814 | 0.001274 | 0.003719 | 0.001075 | 0.007506 | 0.004527 | 0.006392 | 0.001429 | 0.00377 |
| 30.4305556 | 0.006065 | 0.004729 | 0.004482 | 0.006119 | 0.003312 | 0.01096 | 0.008794 | 0.036796 | 0.030898 | 0.011628 | 0.015161 | 0.010887 | 0.001196 | 0.004589 | 0.001042 | 0.000722 | 0.00059 | 0.002969 | 0.001288 | 0.00563 | 0.003284 | 0.004967 | 0.000518 | 0.002061 |
| 30.4444444 | 0.010212 | 0.006211 | 0.004865 | 0.008317 | 0.001964 | 0.010037 | 0.016463 | 0.036041 | 0.027678 | 0.014149 | 0.013962 | 0.008903 | 0.001022 | 0.004069 | 0.00154 | 0.000723 | 0.00066 | 0.0021 | 0.001107 | 0.003508 | 0.001957 | 0.00346 | 0.00061 | 0.0019 |
| 30.4583333 | 0.015591 | 0.009268 | 0.005293 | 0.008647 | 0.001769 | 0.008723 | 0.02091 | 0.03528 | 0.024795 | 0.015317 | 0.013638 | 0.004996 | 0.002458 | 0.00246 | 0.003316 | 0.000719 | 0.001011 | 0.001466 | 0.001742 | 0.001768 | 0.001847 | 0.001958 | 0.000773 | 0.003655 |
| 30.4722222 | 0.020906 | 0.013951 | 0.007187 | 0.007794 | 0.001778 | 0.010196 | 0.016788 | 0.038938 | 0.026621 | 0.013742 | 0.015508 | 0.004491 | 0.004344 | 0.00091 | 0.005663 | 0.000914 | 0.001118 | 0.001664 | 0.003668 | 0.001351 | 0.003026 | 0.000893 | 0.000579 | 0.005124 |
| 30.4861111 | 0.029123 | 0.024613 | 0.012767 | 0.011021 | 0.002224 | 0.020201 | 0.022669 | 0.05091 | 0.037005 | 0.028153 | 0.020478 | 0.00819 | 0.003724 | 0.001709 | 0.005789 | 0.00125 | 0.000796 | 0.001578 | 0.00446 | 0.00174 | 0.003003 | 0.001393 | 0.001493 | 0.004732 |
| 30.5 | 0.041483 | 0.044206 | 0.022747 | 0.024267 | 0.004928 | 0.041135 | 0.048618 | 0.064264 | 0.053517 | 0.051967 | 0.02612 | 0.017899 | 0.002329 | 0.006255 | 0.003523 | 0.001931 | 0.000725 | 0.001003 | 0.004026 | 0.004226 | 0.003016 | 0.004468 | 0.005426 | 0.007059 |
| 30.5138889 | 0.042888 | 0.070235 | 0.034001 | 0.037222 | 0.006565 | 0.056224 | 0.069807 | 0.068142 | 0.07694 | 0.064949 | 0.026801 | 0.036081 | 0.003111 | 0.011947 | 0.001609 | 0.003373 | 0.001392 | 0.000775 | 0.005686 | 0.008655 | 0.005105 | 0.007087 | 0.010657 | 0.015214 |
| 30.5277778 | 0.038993 | 0.087833 | 0.040006 | 0.031887 | 0.005997 | 0.05392 | 0.067654 | 0.067516 | 0.110715 | 0.062392 | 0.02143 | 0.051802 | 0.003218 | 0.013796 | 0.001955 | 0.004906 | 0.001957 | 0.001046 | 0.006095 | 0.009806 | 0.005081 | 0.006073 | 0.011646 | 0.020581 |
| 30.5416667 | 0.048568 | 0.08678 | 0.037414 | 0.020089 | 0.006504 | 0.048262 | 0.05516 | 0.073501 | 0.13723 | 0.049454 | 0.016487 | 0.049333 | 0.001964 | 0.010563 | 0.004161 | 0.004922 | 0.002026 | 0.001687 | 0.003666 | 0.006066 | 0.003328 | 0.004641 | 0.007678 | 0.016294 |
| 30.5555556 | 0.052962 | 0.074217 | 0.03201 | 0.021602 | 0.007482 | 0.041716 | 0.041202 | 0.083527 | 0.126634 | 0.037198 | 0.018527 | 0.031836 | 0.001832 | 0.005531 | 0.004259 | 0.004495 | 0.002142 | 0.003361 | 0.00231 | 0.003288 | 0.00325 | 0.004139 | 0.004755 | 0.008093 |
| 30.5694444 | 0.045957 | 0.057206 | 0.026179 | 0.029724 | 0.007033 | 0.027555 | 0.024745 | 0.085539 | 0.073872 | 0.027894 | 0.02369 | 0.013946 | 0.002747 | 0.002836 | 0.002088 | 0.004899 | 0.00287 | 0.006641 | 0.002396 | 0.004361 | 0.003151 | 0.003472 | 0.005068 | 0.003017 |
| 30.5833333 | 0.036798 | 0.040228 | 0.015773 | 0.034144 | 0.005918 | 0.01248 | 0.011237 | 0.08054 | 0.033567 | 0.018656 | 0.025061 | 0.006249 | 0.003461 | 0.002895 | 0.001117 | 0.003966 | 0.003516 | 0.007982 | 0.002896 | 0.005131 | 0.003759 | 0.004574 | 0.005074 | 0.002175 |
| 30.5972222 | 0.022838 | 0.026244 | 0.007304 | 0.03461 | 0.00573 | 0.005358 | 0.007515 | 0.074243 | 0.045035 | 0.011269 | 0.02095 | 0.011128 | 0.002913 | 0.002873 | 0.000972 | 0.003303 | 0.00307 | 0.005596 | 0.002555 | 0.003638 | 0.005014 | 0.006116 | 0.003753 | 0.00349 |
| 30.6111111 | 0.016095 | 0.013826 | 0.009415 | 0.032774 | 0.006149 | 0.00442 | 0.01105 | 0.065461 | 0.069131 | 0.005941 | 0.01315 | 0.019555 | 0.002016 | 0.002107 | 0.001048 | 0.00495 | 0.002527 | 0.00275 | 0.001789 | 0.002977 | 0.005348 | 0.006543 | 0.003047 | 0.003521 |
| 30.625 | 0.021442 | 0.005969 | 0.015151 | 0.029152 | 0.006084 | 0.005854 | 0.01563 | 0.053745 | 0.07372 | 0.003227 | 0.009692 | 0.022898 | 0.002119 | 0.001462 | 0.000945 | 0.005292 | 0.002797 | 0.001391 | 0.001807 | 0.004742 | 0.005934 | 0.006639 | 0.003152 | 0.002106 |
| 30.6388889 | 0.019468 | 0.007197 | 0.019234 | 0.024356 | 0.005779 | 0.007263 | 0.022475 | 0.035934 | 0.058852 | 0.002757 | 0.015339 | 0.021902 | 0.002771 | 0.001516 | 0.000761 | 0.003131 | 0.003226 | 0.001075 | 0.002024 | 0.007191 | 0.00664 | 0.006601 | 0.003353 | 0.001878 |
| 30.6527778 | 0.010615 | 0.013249 | 0.025056 | 0.020293 | 0.005566 | 0.004944 | 0.032284 | 0.018607 | 0.034451 | 0.002077 | 0.021974 | 0.018141 | 0.003341 | 0.002707 | 0.001642 | 0.001845 | 0.003608 | 0.001481 | 0.002207 | 0.008681 | 0.006587 | 0.006953 | 0.004026 | 0.002724 |
| 30.6666667 | 0.00967 | 0.016559 | 0.026138 | 0.017196 | 0.005488 | 0.00234 | 0.032207 | 0.01647 | 0.01687 | 0.001365 | 0.020559 | 0.010394 | 0.003806 | 0.004296 | 0.002981 | 0.002184 | 0.004257 | 0.002631 | 0.002682 | 0.008976 | 0.00647 | 0.007681 | 0.005189 | 0.003336 |
| 30.6805556 | 0.015102 | 0.016912 | 0.017019 | 0.014421 | 0.00622 | 0.00322 | 0.025541 | 0.022888 | 0.008811 | 0.002623 | 0.012104 | 0.005674 | 0.004108 | 0.005187 | 0.00351 | 0.001884 | 0.004733 | 0.004096 | 0.003128 | 0.008301 | 0.006324 | 0.008025 | 0.006006 | 0.002678 |
| 30.6944444 | 0.022712 | 0.01722 | 0.012429 | 0.012645 | 0.007851 | 0.006642 | 0.03188 | 0.02563 | 0.006075 | 0.005433 | 0.005548 | 0.01084 | 0.004142 | 0.005385 | 0.002885 | 0.000906 | 0.004575 | 0.005036 | 0.003144 | 0.006938 | 0.005594 | 0.007558 | 0.006462 | 0.001973 |
| 30.7083333 | 0.022501 | 0.016491 | 0.021604 | 0.012796 | 0.008901 | 0.009772 | 0.037191 | 0.029782 | 0.005303 | 0.008548 | 0.004933 | 0.018766 | 0.003849 | 0.005229 | 0.0018 | 0.001057 | 0.003785 | 0.004522 | 0.002693 | 0.005014 | 0.00417 | 0.006159 | 0.006764 | 0.003921 |
| 30.7222222 | 0.01618 | 0.013498 | 0.028854 | 0.013805 | 0.008094 | 0.008378 | 0.023448 | 0.031977 | 0.007655 | 0.012483 | 0.007387 | 0.022228 | 0.003094 | 0.004735 | 0.001722 | 0.002985 | 0.002325 | 0.002585 | 0.001825 | 0.002729 | 0.002526 | 0.004268 | 0.006176 | 0.00669 |
| 30.7361111 | 0.018125 | 0.009216 | 0.024508 | 0.013877 | 0.0059 | 0.008435 | 0.010033 | 0.024439 | 0.017428 | 0.016775 | 0.009567 | 0.019777 | 0.002145 | 0.004351 | 0.00306 | 0.005101 | 0.000853 | 0.000788 | 0.000859 | 0.001071 | 0.001999 | 0.003303 | 0.004981 | 0.006587 |
| 30.75 | 0.022148 | 0.006124 | 0.016004 | 0.013177 | 0.003691 | 0.01439 | 0.010438 | 0.024438 | 0.024564 | 0.01884 | 0.007755 | 0.012646 | 0.001489 | 0.004447 | 0.004425 | 0.005686 | 0.000227 | 0.000228 | 0.000276 | 0.000481 | 0.002576 | 0.003656 | 0.004376 | 0.004045 |
| 30.7638889 | 0.023987 | 0.004438 | 0.009505 | 0.011497 | 0.002199 | 0.013719 | 0.011622 | 0.041427 | 0.020693 | 0.017744 | 0.006 | 0.011641 | 0.001059 | 0.004225 | 0.005079 | 0.005273 | 0.000178 | 0.000439 | 0.000441 | 0.000707 | 0.002361 | 0.00405 | 0.004101 | 0.002519 |
| 30.7777778 | 0.031724 | 0.003112 | 0.011337 | 0.009112 | 0.001223 | 0.008319 | 0.009219 | 0.053291 | 0.012731 | 0.016665 | 0.01027 | 0.024381 | 0.000571 | 0.002778 | 0.005352 | 0.004881 | 0.000336 | 0.001014 | 0.001595 | 0.002002 | 0.001349 | 0.003681 | 0.003897 | 0.003076 |
| 30.7916667 | 0.043363 | 0.003441 | 0.020241 | 0.007704 | 0.000579 | 0.008979 | 0.012909 | 0.045494 | 0.00691 | 0.01708 | 0.015967 | 0.040327 | 0.000309 | 0.001069 | 0.005413 | 0.00478 | 0.000658 | 0.001695 | 0.003395 | 0.003849 | 0.001417 | 0.002748 | 0.004381 | 0.003639 |
| 30.8055556 | 0.044735 | 0.00381 | 0.024038 | 0.007634 | 0.001022 | 0.010708 | 0.030684 | 0.02622 | 0.005693 | 0.018007 | 0.016987 | 0.049913 | 0.000654 | 0.001006 | 0.005366 | 0.005034 | 0.000702 | 0.002441 | 0.004719 | 0.005162 | 0.002405 | 0.001538 | 0.005035 | 0.003486 |
| 30.8194444 | 0.029623 | 0.003102 | 0.017831 | 0.007325 | 0.002694 | 0.01052 | 0.052211 | 0.010771 | 0.009192 | 0.022048 | 0.012341 | 0.052392 | 0.001097 | 0.002801 | 0.005596 | 0.005487 | 0.000476 | 0.003496 | 0.005421 | 0.006475 | 0.003527 | 0.000864 | 0.004668 | 0.003708 |
| 30.8333333 | 0.015796 | 0.00423 | 0.007454 | 0.005421 | 0.004488 | 0.009743 | 0.05916 | 0.009802 | 0.015121 | 0.030802 | 0.007126 | 0.048403 | 0.000938 | 0.00495 | 0.006196 | 0.005743 | 0.00061 | 0.004482 | 0.006104 | 0.008246 | 0.004688 | 0.00144 | 0.003176 | 0.004586 |
| 30.8472222 | 0.021008 | 0.007011 | 0.001868 | 0.002737 | 0.006277 | 0.008826 | 0.055148 | 0.023354 | 0.022192 | 0.042739 | 0.008264 | 0.041661 | 0.00042 | 0.006257 | 0.006244 | 0.005295 | 0.00115 | 0.005036 | 0.006285 | 0.009562 | 0.005616 | 0.002473 | 0.001573 | 0.005473 |
| 30.8611111 | 0.040313 | 0.009825 | 0.001744 | 0.001838 | 0.008319 | 0.010232 | 0.049857 | 0.04486 | 0.028826 | 0.055131 | 0.012952 | 0.035735 | 0.000283 | 0.0066 | 0.004843 | 0.004103 | 0.001476 | 0.005051 | 0.006427 | 0.009748 | 0.006449 | 0.0033 | 0.00079 | 0.00622 |
| 30.875 | 0.062885 | 0.01529 | 0.002565 | 0.003871 | 0.00948 | 0.014067 | 0.040411 | 0.062298 | 0.030862 | 0.063837 | 0.014853 | 0.030391 | 0.001149 | 0.005781 | 0.002704 | 0.003072 | 0.001713 | 0.005266 | 0.008438 | 0.009358 | 0.00775 | 0.004828 | 0.001457 | 0.007405 |
| 30.8888889 | 0.074849 | 0.024399 | 0.002387 | 0.005953 | 0.007995 | 0.017174 | 0.025887 | 0.065523 | 0.027791 | 0.065506 | 0.01442 | 0.024285 | 0.002762 | 0.003529 | 0.001529 | 0.003287 | 0.002436 | 0.006697 | 0.011982 | 0.008563 | 0.009007 | 0.007124 | 0.003377 | 0.008589 |
| 30.9027778 | 0.065792 | 0.034483 | 0.001582 | 0.006216 | 0.004225 | 0.017408 | 0.015105 | 0.062922 | 0.02333 | 0.059695 | 0.013295 | 0.017104 | 0.004051 | 0.001433 | 0.001596 | 0.00435 | 0.003468 | 0.008681 | 0.014517 | 0.007284 | 0.009227 | 0.008877 | 0.005405 | 0.008289 |
| 30.9166667 | 0.046086 | 0.043126 | 0.002116 | 0.005682 | 0.001705 | 0.014671 | 0.01187 | 0.067282 | 0.017159 | 0.04661 | 0.012655 | 0.009805 | 0.004549 | 0.001391 | 0.001814 | 0.004819 | 0.004494 | 0.009552 | 0.014641 | 0.007071 | 0.008593 | 0.009419 | 0.006751 | 0.006609 |
| 30.9305556 | 0.027177 | 0.04708 | 0.003965 | 0.004976 | 0.002715 | 0.010212 | 0.011008 | 0.072974 | 0.010703 | 0.026164 | 0.013746 | 0.009 | 0.004474 | 0.002415 | 0.001617 | 0.004173 | 0.00515 | 0.008503 | 0.012797 | 0.008882 | 0.007009 | 0.008905 | 0.006834 | 0.004973 |
| 30.9444444 | 0.013356 | 0.04603 | 0.006381 | 0.004619 | 0.005401 | 0.006492 | 0.008089 | 0.068736 | 0.013338 | 0.012583 | 0.011774 | 0.018019 | 0.004373 | 0.002609 | 0.001399 | 0.003122 | 0.005123 | 0.006543 | 0.010131 | 0.011141 | 0.004364 | 0.008087 | 0.006092 | 0.004462 |
| 30.9583333 | 0.007183 | 0.040852 | 0.008928 | 0.004101 | 0.007553 | 0.004331 | 0.004389 | 0.053995 | 0.027325 | 0.019623 | 0.005373 | 0.028545 | 0.004696 | 0.001733 | 0.001579 | 0.002483 | 0.00488 | 0.005213 | 0.007831 | 0.012239 | 0.002231 | 0.008088 | 0.005695 | 0.00484 |
| 30.9722222 | 0.008521 | 0.030182 | 0.010337 | 0.002923 | 0.008139 | 0.002579 | 0.003571 | 0.036402 | 0.038845 | 0.033504 | 0.001689 | 0.032303 | 0.004912 | 0.0007 | 0.001375 | 0.002144 | 0.004728 | 0.004748 | 0.00632 | 0.011594 | 0.001941 | 0.009033 | 0.005162 | 0.004462 |
| 30.9861111 | 0.009787 | 0.017477 | 0.009031 | 0.003218 | 0.006237 | 0.002194 | 0.004087 | 0.026182 | 0.032612 | 0.034299 | 0.004123 | 0.024992 | 0.004609 | 0.00456 | 0.000997 | 0.002487 | 0.005121 | 0.004352 | 0.005682 | 0.009895 | 0.003216 | 0.009643 | 0.004249 | 0.003822 |
| 31 | 0.010124 | 0.010091 | 0.005403 | 0.006504 | 0.003215 | 0.006009 | 0.007822 | 0.031289 | 0.024829 | 0.02853 | 0.012467 | 0.016491 | 0.004488 | 0.020277 | 0.001926 | 0.00453 | 0.007474 | 0.003442 | 0.006716 | 0.009518 | 0.005342 | 0.009078 | 0.004632 | 0.006609 |
| 31.0138889 | 0.019976 | 0.007958 | 0.003192 | 0.010038 | 0.002355 | 0.01338 | 0.019913 | 0.050393 | 0.045727 | 0.042737 | 0.026742 | 0.027712 | 0.004129 | 0.038235 | 0.004171 | 0.005545 | 0.009856 | 0.002504 | 0.008596 | 0.009925 | 0.006919 | 0.007162 | 0.005962 | 0.01172 |
| 31.0277778 | 0.033905 | 0.007057 | 0.004581 | 0.011205 | 0.004564 | 0.021372 | 0.03251 | 0.070822 | 0.079851 | 0.070767 | 0.04628 | 0.050626 | 0.003618 | 0.035858 | 0.007184 | 0.005784 | 0.008174 | 0.003195 | 0.007388 | 0.007052 | 0.005656 | 0.004118 | 0.004851 | 0.011119 |
| 31.0416667 | 0.04239 | 0.010795 | 0.006461 | 0.011767 | 0.008557 | 0.029227 | 0.035202 | 0.080634 | 0.086952 | 0.076314 | 0.061667 | 0.056992 | 0.0041 | 0.017267 | 0.009007 | 0.008602 | 0.005542 | 0.004115 | 0.003744 | 0.002783 | 0.003299 | 0.002216 | 0.002021 | 0.005888 |
| 31.0555556 | 0.044107 | 0.023564 | 0.005296 | 0.013392 | 0.012707 | 0.031438 | 0.034773 | 0.077936 | 0.0619 | 0.05728 | 0.064804 | 0.046387 | 0.004482 | 0.005102 | 0.006699 | 0.008523 | 0.004964 | 0.003681 | 0.00229 | 0.001901 | 0.00262 | 0.003933 | 0.001586 | 0.003873 |
| 31.0694444 | 0.038318 | 0.042651 | 0.003223 | 0.0152 | 0.015256 | 0.024298 | 0.036497 | 0.066792 | 0.032949 | 0.039314 | 0.052232 | 0.036169 | 0.004918 | 0.001886 | 0.003445 | 0.003939 | 0.003341 | 0.004379 | 0.002131 | 0.002891 | 0.002387 | 0.007298 | 0.00284 | 0.00336 |
| 31.0833333 | 0.028404 | 0.057983 | 0.00354 | 0.016022 | 0.015879 | 0.016423 | 0.03054 | 0.05083 | 0.015046 | 0.030173 | 0.029122 | 0.028115 | 0.005518 | 0.001632 | 0.003141 | 0.00111 | 0.00156 | 0.00549 | 0.001277 | 0.002795 | 0.003067 | 0.008119 | 0.00276 | 0.002214 |
| 31.0972222 | 0.017725 | 0.060589 | 0.00423 | 0.015572 | 0.015072 | 0.012416 | 0.018308 | 0.03252 | 0.00836 | 0.022882 | 0.023547 | 0.016484 | 0.004529 | 0.002825 | 0.003212 | 0.000641 | 0.001522 | 0.005125 | 0.000991 | 0.001729 | 0.004283 | 0.006344 | 0.001504 | 0.002022 |
| 31.1111111 | 0.008684 | 0.051362 | 0.004282 | 0.014677 | 0.012851 | 0.009763 | 0.011935 | 0.017157 | 0.016507 | 0.014831 | 0.040478 | 0.007898 | 0.002839 | 0.003237 | 0.00272 | 0.000576 | 0.001922 | 0.004245 | 0.002002 | 0.001497 | 0.003511 | 0.004552 | 0.000779 | 0.001617 |
| 31.125 | 0.003415 | 0.037934 | 0.005256 | 0.013784 | 0.009666 | 0.008548 | 0.014011 | 0.017636 | 0.028558 | 0.007455 | 0.049747 | 0.010703 | 0.001694 | 0.001969 | 0.002615 | 0.000491 | 0.001746 | 0.003105 | 0.002839 | 0.002205 | 0.00203 | 0.003452 | 0.000576 | 0.001294 |
| 31.1388889 | 0.001283 | 0.027827 | 0.006832 | 0.012704 | 0.006475 | 0.008711 | 0.018253 | 0.003305 | 0.029417 | 0.002466 | 0.036369 | 0.018687 | 0.000825 | 0.00077 | 0.002408 | 0.000346 | 0.001183 | 0.001687 | 0.002284 | 0.002107 | 0.002359 | 0.002604 | 0.000297 | 0.002772 |
| 31.1527778 | 0.002827 | 0.02119 | 0.008336 | 0.010339 | 0.003507 | 0.008768 | 0.016784 | 0.004053 | 0.019752 | 0.000716 | 0.015658 | 0.025042 | 0.000401 | 0.001145 | 0.002157 | 0.00033 | 0.000998 | 0.000599 | 0.001268 | 0.001355 | 0.003632 | 0.001905 | 0.000216 | 0.00457 |
| 31.1666667 | 0.008839 | 0.013553 | 0.009352 | 0.006646 | 0.001892 | 0.007267 | 0.010758 | 0.007576 | 0.010156 | 0.002968 | 0.005387 | 0.029678 | 0.000401 | 0.001833 | 0.001912 | 0.000404 | 0.000991 | 0.000266 | 0.001316 | 0.001156 | 0.003356 | 0.001171 | 0.000256 | 0.004303 |
| 31.1805556 | 0.017617 | 0.011978 | 0.009631 | 0.005156 | 0.003271 | 0.004058 | 0.011946 | 0.010257 | 0.00873 | 0.012076 | 0.012038 | 0.032299 | 0.000471 | 0.001442 | 0.001258 | 0.000386 | 0.000692 | 0.000636 | 0.002686 | 0.001139 | 0.002509 | 0.001015 | 0.000603 | 0.002964 |
| 31.1944444 | 0.026122 | 0.026704 | 0.010064 | 0.006065 | 0.006085 | 0.002259 | 0.023381 | 0.009464 | 0.009 | 0.028372 | 0.029122 | 0.037128 | 0.000783 | 0.000707 | 0.000475 | 0.000422 | 0.00061 | 0.001217 | 0.003888 | 0.00124 | 0.003834 | 0.002368 | 0.00166 | 0.003429 |
| 31.2083333 | 0.032245 | 0.050272 | 0.012032 | 0.005903 | 0.007739 | 0.003814 | 0.033237 | 0.00541 | 0.00739 | 0.047397 | 0.032325 | 0.045325 | 0.001282 | 0.000723 | 0.000378 | 0.000422 | 0.000746 | 0.00146 | 0.003939 | 0.00196 | 0.005557 | 0.004197 | 0.002226 | 0.00519 |
| 31.2222222 | 0.035662 | 0.060346 | 0.014899 | 0.005581 | 0.00814 | 0.006273 | 0.035485 | 0.003267 | 0.016273 | 0.065005 | 0.022626 | 0.051557 | 0.001545 | 0.001502 | 0.00098 | 0.00036 | 0.000643 | 0.001281 | 0.003428 | 0.002671 | 0.005068 | 0.00542 | 0.001494 | 0.005249 |
| 31.2361111 | 0.03739 | 0.052726 | 0.01656 | 0.006363 | 0.008305 | 0.007956 | 0.034768 | 0.006658 | 0.043855 | 0.073235 | 0.032903 | 0.054379 | 0.001810 | 0.001655 | 0.001659 | 0.000445 | 0.000569 | 0.001192 | 0.003083 | 0.003482 | 0.00361 | 0.006331 | 0.000844 | 0.003337 |
| 31.25 | 0.039119 | 0.045333 | 0.016632 | 0.006835 | 0.008129 | 0.008566 | 0.035267 | 0.012394 | 0.07255 | 0.064646 | 0.056225 | 0.050242 | 0.001765 | 0.00114 | 0.001584 | 0.000801 | 0.001302 | 0.001782 | 0.003062 | 0.005168 | 0.002893 | 0.007285 | 0.00085 | 0.001712 |
| 31.2638889 | 0.039265 | 0.040853 | 0.015846 | 0.007221 | 0.006645 | 0.007671 | 0.035509 | 0.015959 | 0.071015 | 0.043062 | 0.057006 | 0.037485 | 0.001904 | 0.003886 | 0.000888 | 0.00109 | 0.002204 | 0.002554 | 0.00257 | 0.007123 | 0.002583 | 0.007947 | 0.000901 | 0.001083 |
| 31.2777778 | 0.036449 | 0.034582 | 0.014583 | 0.007963 | 0.003863 | 0.005409 | 0.02961 | 0.013893 | 0.041136 | 0.027123 | 0.035908 | 0.020926 | 0.001913 | 0.030276 | 0.000655 | 0.003545 | 0.002947 | 0.002127 | 0.002358 | 0.007432 | 0.002192 | 0.006935 | 0.002268 | 0.001612 |
| 31.2916667 | 0.032465 | 0.026281 | 0.013218 | 0.007284 | 0.001804 | 0.003044 | 0.016606 | 0.008082 | 0.024977 | 0.031194 | 0.015096 | 0.00927 | 0.002811 | 0.096118 | 0.002282 | 0.012254 | 0.004473 | 0.001172 | 0.005314 | 0.005418 | 0.003609 | 0.004497 | 0.005549 | 0.004442 |
| 31.3055556 | 0.027547 | 0.019063 | 0.011964 | 0.004153 | 0.00171 | 0.003675 | 0.005202 | 0.004723 | 0.037017 | 0.032948 | 0.00492 | 0.007954 | 0.004578 | 0.117418 | 0.006878 | 0.0189 | 0.008693 | 0.001781 | 0.007139 | 0.00452 | 0.00658 | 0.003913 | 0.011417 | 0.008656 |
| 31.3194444 | 0.022133 | 0.015419 | 0.011129 | 0.001457 | 0.002903 | 0.007853 | 0.00114 | 0.003204 | 0.043503 | 0.019173 | 0.003926 | 0.008288 | 0.009225 | 0.132375 | 0.011901 | 0.038803 | 0.025003 | 0.003908 | 0.011577 | 0.009954 | 0.012769 | 0.007613 | 0.025469 | 0.019723 |
| 31.3333333 | 0.018942 | 0.012408 | 0.01111 | 0.001568 | 0.004145 | 0.011488 | 0.002316 | 0.002142 | 0.028598 | 0.008199 | 0.008837 | 0.005298 | 0.012494 | 0.236101 | 0.012048 | 0.064588 | 0.041643 | 0.004349 | 0.022501 | 0.014489 | 0.023407 | 0.010941 | 0.034161 | 0.027853 |
| 31.3472222 | 0.019901 | 0.015106 | 0.012195 | 0.002814 | 0.003914 | 0.011657 | 0.006886 | 0.003678 | 0.014393 | 0.003622 | 0.014899 | 0.003076 | 0.008463 | 0.254594 | 0.013081 | 0.054255 | 0.046829 | 0.004756 | 0.025793 | 0.011325 | 0.029668 | 0.012385 | 0.03009 | 0.024104 |
| 31.3611111 | 0.022902 | 0.024164 | 0.013827 | 0.003487 | 0.002721 | 0.009093 | 0.010258 | 0.005582 | 0.018588 | 0.001583 | 0.016092 | 0.003988 | 0.004146 | 0.166712 | 0.016486 | 0.034579 | 0.048659 | 0.007648 | 0.019669 | 0.010084 | 0.024275 | 0.012796 | 0.030299 | 0.024367 |
| 31.375 | 0.023651 | 0.030195 | 0.014167 | 0.003471 | 0.00162 | 0.005536 | 0.00877 | 0.005124 | 0.024509 | 0.003921 | 0.011901 | 0.006587 | 0.002039 | 0.100171 | 0.011024 | 0.023319 | 0.033849 | 0.006784 | 0.010818 | 0.009348 | 0.011401 | 0.007817 | 0.024622 | 0.020119 |
| 31.3888889 | 0.020488 | 0.034377 | 0.012543 | 0.003214 | 0.000758 | 0.002518 | 0.006062 | 0.002972 | 0.01603 | 0.013614 | 0.00596 | 0.006135 | 0.000539 | 0.041285 | 0.002887 | 0.008656 | 0.011148 | 0.002861 | 0.003622 | 0.004122 | 0.002817 | 0.002464 | 0.009241 | 0.007067 |
| 31.4027778 | 0.014728 | 0.040025 | 0.009869 | 0.002836 | 0.000661 | 0.001819 | 0.005717 | 0.001426 | 0.008066 | 0.028958 | 0.004029 | 0.005168 | 0.000319 | 0.00765 | 0.000524 | 0.001401 | 0.002325 | 0.001546 | 0.00098 | 0.001752 | 0.000948 | 0.001411 | 0.00146 | |
| 31.4166667 | 0.009036 | 0.044331 | 0.00738 | 0.002104 | 0.001032 | 0.003827 | 0.007874 | 0.002006 | 0.01249 | 0.041788 | 0.007797 | 0.007614 | 0.000345 | 0.005025 | 0.000522 | 0.001103 | 0.00104 | 0.001497 | 0.000363 | 0.001749 | 0.002031 | 0.000832 | 0.000637 | 0.002194 |
| 31.4305556 | 0.00551 | 0.047923 | 0.006606 | 0.001609 | 0.001135 | 0.006315 | 0.010119 | 0.003467 | 0.018439 | 0.044065 | 0.012737 | 0.008207 | 0.000586 | 0.004502 | 0.00126 | 0.002084 | 0.001 | 0.000978 | 0.000292 | 0.002864 | 0.001319 | 0.001778 | 0.001618 | 0.002666 |
| 31.4444444 | 0.004838 | 0.052939 | 0.007942 | 0.001794 | 0.001006 | 0.007122 | 0.009512 | 0.004699 | 0.016441 | 0.033345 | 0.013358 | 0.005032 | 0.000888 | 0.003067 | 0.001412 | 0.002325 | 0.001064 | 0.000538 | 0.000623 | 0.00324 | 0.000969 | 0.0025 | 0.002709 | 0.001878 |
| 31.4583333 | 0.006302 | 0.057342 | 0.009273 | 0.002093 | 0.000837 | 0.006632 | 0.006267 | 0.004673 | 0.010184 | 0.01782 | 0.007657 | 0.003644 | 0.001679 | 0.001978 | 0.001365 | 0.001441 | 0.000675 | 0.001104 | 0.001887 | 0.002297 | 0.00201 | 0.001966 | 0.002672 | 0.001458 |
| 31.4722222 | 0.010999 | 0.05086 | 0.010028 | 0.002687 | 0.001286 | 0.008161 | 0.006422 | 0.005749 | 0.009088 | 0.010808 | 0.002447 | 0.00605 | 0.002833 | 0.002865 | 0.002275 | 0.000619 | 0.000519 | 0.001985 | 0.003026 | 0.001662 | 0.003012 | 0.001179 | 0.002227 | 0.001412 |
| 31.4861111 | 0.022904 | 0.037589 | 0.013055 | 0.008236 | 0.004588 | 0.016844 | 0.021485 | 0.01753 | 0.017533 | 0.02447 | 0.00428 | 0.011268 | 0.002424 | 0.003503 | 0.002157 | 0.000418 | 0.001246 | 0.001803 | 0.002542 | 0.002658 | 0.002633 | 0.002131 | 0.003184 | 0.002268 |
| 31.5 | 0.04161 | 0.046454 | 0.020265 | 0.022883 | 0.011134 | 0.031169 | 0.055305 | 0.043976 | 0.027749 | 0.056115 | 0.014833 | 0.020795 | 0.00157 | 0.003987 | 0.001416 | 0.000441 | 0.002595 | 0.001209 | 0.001386 | 0.004495 | 0.003114 | 0.004845 | 0.006666 | 0.007237 |
| 31.5138889 | 0.058772 | 0.07218 | 0.030424 | 0.036345 | 0.016314 | 0.038918 | 0.085036 | 0.068186 | 0.030385 | 0.079987 | 0.034003 | 0.0348 | 0.002388 | 0.007645 | 0.002184 | 0.001371 | 0.003533 | 0.001651 | 0.000776 | 0.005937 | 0.00559 | 0.006831 | 0.012195 | 0.015161 |
| 31.5277778 | 0.062719 | 0.081815 | 0.037689 | 0.033677 | 0.01715 | 0.032301 | 0.084989 | 0.071706 | 0.02776 | 0.081783 | 0.048494 | 0.05036 | 0.00301 | 0.011064 | 0.002571 | 0.003915 | 0.003007 | 0.00326 | 0.002121 | 0.005601 | 0.005667 | 0.005297 | 0.016489 | 0.018431 |
| 31.5416667 | 0.048674 | 0.077196 | 0.03633 | 0.020303 | 0.016822 | 0.02275 | 0.06872 | 0.065349 | 0.040023 | 0.076627 | 0.044309 | 0.059874 | 0.003178 | 0.010893 | 0.002151 | 0.005264 | 0.001922 | 0.004317 | 0.006091 | 0.003401 | 0.002923 | 0.00213 | 0.017129 | 0.013661 |
| 31.5555556 | 0.027094 | 0.074058 | 0.028435 | 0.011271 | 0.017014 | 0.025971 | 0.070434 | 0.066528 | 0.066232 | 0.068696 | 0.027956 | 0.053235 | 0.002863 | 0.007764 | 0.001838 | 0.005405 | 0.001944 | 0.003643 | 0.007319 | 0.002353 | 0.001887 | 0.001743 | 0.016102 | 0.006372 |
| 31.5694444 | 0.021589 | 0.079175 | 0.019015 | 0.008578 | 0.015545 | 0.034906 | 0.088894 | 0.071818 | 0.077241 | 0.051576 | 0.015211 | 0.03326 | 0.001654 | 0.003632 | 0.001354 | 0.007752 | 0.002905 | 0.003373 | 0.004715 | 0.003569 | 0.002098 | 0.00363 | 0.014167 | 0.002529 |
| 31.5833333 | 0.03527 | 0.086051 | 0.009632 | 0.008264 | 0.012059 | 0.037869 | 0.086051 | 0.073624 | 0.068338 | 0.033135 | 0.012953 | 0.014079 | 0.00119 | 0.001497 | 0.00134 | 0.008393 | 0.003201 | 0.004044 | 0.004112 | 0.004186 | 0.001623 | 0.005074 | 0.009633 | 0.003166 |
| 31.5972222 | 0.043386 | 0.079616 | 0.004645 | 0.009361 | 0.008189 | 0.034034 | 0.051983 | 0.068222 | 0.049607 | 0.018985 | 0.012249 | 0.007071 | 0.001005 | 0.001279 | 0.001354 | 0.005107 | 0.002181 | 0.003559 | 0.004827 | 0.002904 | 0.001226 | 0.005135 | 0.004358 | 0.005821 |
| 31.6111111 | 0.039871 | 0.061556 | 0.00805 | 0.011462 | 0.004847 | 0.02028 | 0.024581 | 0.054839 | 0.027529 | 0.009007 | 0.006678 | 0.013794 | 0.000945 | 0.001004 | 0.001142 | 0.003245 | 0.001255 | 0.002165 | 0.004298 | 0.001361 | 0.001048 | 0.004315 | 0.001309 | 0.007959 |
| 31.625 | 0.030889 | 0.04238 | 0.01597 | 0.013499 | 0.00253 | 0.009668 | 0.032495 | 0.040031 | 0.01891 | 0.006974 | 0.004103 | 0.021441 | 0.001575 | 0.000545 | 0.001143 | 0.003596 | 0.001266 | 0.001123 | 0.003412 | 0.001064 | 0.000584 | 0.003433 | 0.000529 | 0.008633 |
| 31.6388889 | 0.018439 | 0.024391 | 0.024056 | 0.013638 | 0.002324 | 0.012287 | 0.060941 | 0.029455 | 0.028013 | 0.01322 | 0.006784 | 0.023228 | 0.00231 | 0.000806 | 0.000848 | 0.002903 | 0.001266 | 0.000594 | 0.002558 | 0.001773 | 0.000332 | 0.002361 | 0.00092 | 0.007919 |
| 31.6527778 | 0.008572 | 0.012448 | 0.030543 | 0.01094 | 0.003393 | 0.013479 | 0.078373 | 0.022623 | 0.035006 | 0.021618 | 0.011726 | 0.02463 | 0.002689 | 0.001758 | 0.00054 | 0.001393 | 0.000798 | 0.000374 | 0.001729 | 0.001863 | 0.000642 | 0.001353 | 0.001088 | 0.006033 |
| 31.6666667 | 0.006463 | 0.015619 | 0.031414 | 0.006307 | 0.003649 | 0.008981 | 0.073441 | 0.017546 | 0.03027 | 0.02946 | 0.02087 | 0.025457 | 0.002345 | 0.002531 | 0.000723 | 0.000898 | 0.000435 | 0.000479 | 0.001212 | 0.001346 | 0.001286 | 0.001219 | 0.001005 | 0.003451 |
| 31.6805556 | 0.016969 | 0.029578 | 0.022176 | 0.003223 | 0.003122 | 0.004301 | 0.065665 | 0.014151 | 0.020941 | 0.036765 | 0.028181 | 0.02225 | 0.00151 | 0.002688 | 0.000718 | 0.001686 | 0.000284 | 0.001172 | 0.00085 | 0.001602 | 0.00216 | 0.00135 | 0.001998 | 0.002052 |
| 31.6944444 | 0.037962 | 0.041237 | 0.0097 | 0.003547 | 0.002091 | 0.001779 | 0.070069 | 0.013655 | 0.014007 | 0.042515 | 0.02946 | 0.013489 | 0.000735 | 0.002603 | 0.000346 | 0.002686 | 0.000281 | 0.002101 | 0.000745 | 0.001882 | 0.00294 | 0.000898 | 0.003595 | 0.001686 |
| 31.7083333 | 0.042441 | 0.045702 | 0.004071 | 0.004171 | 0.001331 | 0.002516 | 0.07483 | 0.017115 | 0.009772 | 0.047423 | 0.031609 | 0.005274 | 0.000316 | 0.002432 | 0.000182 | 0.002604 | 0.000844 | 0.002366 | 0.001515 | 0.001305 | 0.003031 | 0.000579 | 0.004313 | 0.001291 |
| 31.7222222 | 0.022448 | 0.052159 | 0.002546 | 0.005887 | 0.002076 | 0.004538 | 0.068961 | 0.021717 | 0.00773 | 0.052997 | 0.03318 | 0.003567 | 0.000341 | 0.00178 | 0.000347 | 0.001566 | 0.001885 | 0.001824 | 0.002295 | 0.000619 | 0.00239 | 0.001038 | 0.004019 | 0.001751 |
| 31.7361111 | 0.012153 | 0.06609 | 0.000886 | 0.010117 | 0.002484 | 0.005785 | 0.055424 | 0.022348 | 0.01135 | 0.058552 | 0.025337 | 0.003699 | 0.000504 | 0.000981 | 0.000636 | 0.001248 | 0.002705 | 0.000996 | 0.001979 | 0.000365 | 0.001441 | 0.001763 | 0.003275 | 0.002323 |
| 31.75 | 0.017175 | 0.078202 | 9.71E-05 | 0.011665 | 0.001525 | 0.005936 | 0.041154 | 0.019014 | 0.020147 | 0.063554 | 0.017219 | 0.003056 | 0.000493 | 0.000904 | 0.000682 | 0.002773 | 0.002705 | 0.000382 | 0.001027 | 0.00076 | 0.001099 | 0.001982 | 0.002151 | 0.002687 |
| 31.7638889 | 0.017927 | 0.081929 | 0.000706 | 0.007978 | 0.000963 | 0.006041 | 0.027969 | 0.013916 | 0.028327 | 0.066116 | 0.016751 | 0.002967 | 0.000546 | 0.001257 | 0.000755 | 0.005758 | 0.001839 | 0.000342 | 0.000552 | 0.00148 | 0.002494 | 0.001386 | 0.001179 | 0.003689 |
| 31.7777778 | 0.012004 | 0.080729 | 0.002283 | 0.004091 | 0.000784 | 0.006325 | 0.022445 | 0.008523 | 0.030916 | 0.06206 | 0.01155 | 0.005918 | 0.000706 | 0.0014 | 0.001353 | 0.007813 | 0.000796 | 0.000601 | 0.00103 | 0.001404 | 0.004849 | 0.00052 | 0.001281 | 0.005023 |
| 31.7916667 | 0.008885 | 0.078706 | 0.003682 | 0.00476 | 0.000459 | 0.005966 | 0.020432 | 0.004317 | 0.028069 | 0.056552 | 0.00355 | 0.015266 | 0.000554 | 0.001652 | 0.001871 | 0.006644 | 0.000291 | 0.000582 | 0.001713 | 0.000727 | 0.007349 | 9.21E-05 | 0.001925 | 0.005783 |
| 31.8055556 | 0.00788 | 0.071018 | 0.00436 | 0.006667 | 0.000468 | 0.004029 | 0.022424 | 0.002278 | 0.021333 | 0.058715 | 0.002481 | 0.028471 | 0.000337 | 0.002272 | 0.002076 | 0.004435 | 0.000235 | 0.000455 | 0.001812 | 0.00103 | 0.010134 | 0.000196 | 0.002078 | 0.005496 |
| 31.8194444 | 0.005202 | 0.054934 | 0.006835 | 0.004979 | 0.000817 | 0.001878 | 0.025903 | 0.002326 | 0.01252 | 0.067883 | 0.004136 | 0.042101 | 0.0006 | 0.002761 | 0.002606 | 0.00404 | 0.000424 | 0.000476 | 0.001242 | 0.002534 | 0.011556 | 0.000834 | 0.002648 | 0.005365 |
| 31.8333333 | 0.005222 | 0.037729 | 0.013552 | 0.002472 | 0.001413 | 0.002229 | 0.031587 | 0.002912 | 0.005862 | 0.074365 | 0.006445 | 0.057234 | 0.000883 | 0.003038 | 0.003306 | 0.004889 | 0.000866 | 0.00048 | 0.000565 | 0.003803 | 0.009587 | 0.001645 | 0.004248 | 0.006664 |
| 31.8472222 | 0.006897 | 0.026439 | 0.020036 | 0.00198 | 0.001578 | 0.004834 | 0.041201 | 0.004343 | 0.007152 | 0.07096 | 0.01123 | 0.068561 | 0.000631 | 0.003685 | 0.0034 | 0.005603 | 0.001234 | 0.000561 | 0.00043 | 0.003836 | 0.006089 | 0.002095 | 0.005298 | 0.0094 |
| 31.8611111 | 0.007588 | 0.02239 | 0.020697 | 0.002147 | 0.000908 | 0.008461 | 0.052322 | 0.00827 | 0.01267 | 0.060596 | 0.010996 | 0.067197 | 0.000295 | 0.004573 | 0.00294 | 0.005726 | 0.001403 | 0.000637 | 0.001197 | 0.003428 | 0.004595 | 0.002386 | 0.004796 | 0.012281 |
| 31.875 | 0.007672 | 0.023484 | 0.018364 | 0.003764 | 0.000297 | 0.011797 | 0.057601 | 0.014876 | 0.011231 | 0.045645 | 0.004535 | 0.057642 | 0.000351 | 0.005068 | 0.002445 | 0.005238 | 0.00146 | 0.001228 | 0.003017 | 0.003692 | 0.004694 | 0.002839 | 0.003327 | 0.01386 |
| 31.8888889 | 0.007919 | 0.026526 | 0.016973 | 0.006484 | 0.000557 | 0.012999 | 0.054434 | 0.022452 | 0.005345 | 0.027739 | 0.002311 | 0.047093 | 0.000597 | 0.00506 | 0.001674 | 0.004157 | 0.001449 | 0.003305 | 0.005303 | 0.004715 | 0.004755 | 0.003619 | 0.001719 | 0.01416 |
| 31.9027778 | 0.008009 | 0.02598 | 0.015539 | 0.007908 | 0.001712 | 0.011945 | 0.046841 | 0.027419 | 0.003915 | 0.01345 | 0.005214 | 0.035242 | 0.000714 | 0.005049 | 0.001022 | 0.002809 | 0.001605 | 0.006003 | 0.0071 | 0.005931 | 0.004601 | 0.004888 | 0.000692 | 0.014017 |
| 31.9166667 | 0.008739 | 0.020536 | 0.012215 | 0.007421 | 0.002938 | 0.009032 | 0.037755 | 0.026908 | 0.005645 | 0.005411 | 0.008365 | 0.023272 | 0.00055 | 0.005618 | 0.001564 | 0.001526 | 0.001669 | 0.007211 | 0.007132 | 0.00634 | 0.003751 | 0.005944 | 0.000849 | 0.013113 |
| 31.9305556 | 0.012602 | 0.013204 | 0.007256 | 0.005249 | 0.003666 | 0.005203 | 0.028998 | 0.021518 | 0.006906 | 0.002301 | 0.011206 | 0.012988 | 0.000252 | 0.00631 | 0.002407 | 0.000677 | 0.002222 | 0.006413 | 0.005405 | 0.005707 | 0.002236 | 0.00636 | 0.001245 | 0.01019 |
| 31.9444444 | 0.019176 | 0.009395 | 0.005502 | 0.0026 | 0.004116 | 0.004241 | 0.021502 | 0.013865 | 0.008081 | 0.003577 | 0.014035 | 0.00636 | 0.000419 | 0.006048 | 0.002002 | 0.000427 | 0.003941 | 0.005098 | 0.003845 | 0.004932 | 0.001428 | 0.006902 | 0.000842 | 0.005674 |

| | | | | | | | | | | | | | | | | | | | | | | | | |
|---|---|---|---|---|---|---|---|---|---|---|---|---|---|---|---|---|---|---|---|---|---|---|---|---|
| 31.9583333 | 0.02559 | 0.01613 | 0.009961 | 0.001113 | 0.004463 | 0.007572 | 0.015592 | 0.006422 | 0.01002 | 0.005696 | 0.016197 | 0.006852 | 0.001094 | 0.00507 | 0.000892 | 0.000316 | 0.004641 | 0.004279 | 0.003095 | 0.00462 | 0.001509 | 0.00726 | 0.00043 | 0.003435 |
| 31.9722222 | 0.030093 | 0.026913 | 0.014993 | 0.002267 | 0.00456 | 0.010714 | 0.012788 | 0.00272 | 0.010936 | 0.006671 | 0.013303 | 0.011085 | 0.001219 | 0.004356 | 0.000465 | 0.000272 | 0.003317 | 0.003614 | 0.002037 | 0.004808 | 0.001238 | 0.006706 | 0.000504 | 0.005008 |
| 31.9861111 | 0.028329 | 0.027039 | 0.013871 | 0.010914 | 0.00489 | 0.008951 | 0.016577 | 0.006682 | 0.008897 | 0.016302 | 0.009606 | 0.010267 | 0.000883 | 0.003347 | 0.000494 | 0.00053 | 0.001911 | 0.002564 | 0.000838 | 0.005491 | 0.001241 | 0.005901 | 0.00063 | 0.005005 |
| 32 | 0.017879 | 0.019001 | 0.009114 | 0.027524 | 0.005855 | 0.005784 | 0.025892 | 0.016693 | 0.011335 | 0.043729 | 0.01888 | 0.008921 | 0.001363 | 0.001651 | 0.000287 | 0.00129 | 0.001136 | 0.00164 | 0.000606 | 0.00648 | 0.002681 | 0.005269 | 0.001049 | 0.004232 |
| 32.0138889 | 0.010008 | 0.018967 | 0.009405 | 0.038889 | 0.005911 | 0.007514 | 0.033373 | 0.025277 | 0.023348 | 0.080638 | 0.037262 | 0.020663 | 0.002294 | 0.002824 | 0.001723 | 0.003817 | 0.001787 | 0.002916 | 0.001341 | 0.005582 | 0.00358 | 0.004117 | 0.001869 | 0.006055 |
| 32.0277778 | 0.013429 | 0.028599 | 0.014572 | 0.037525 | 0.004775 | 0.011177 | 0.034584 | 0.028413 | 0.036574 | 0.096982 | 0.046527 | 0.03878 | 0.004848 | 0.009635 | 0.006856 | 0.008608 | 0.005274 | 0.006702 | 0.003645 | 0.004623 | 0.005216 | 0.004148 | 0.003944 | 0.005847 |
| 32.0416667 | 0.021169 | 0.036853 | 0.017335 | 0.031996 | 0.005615 | 0.012497 | 0.031026 | 0.027542 | 0.042638 | 0.068056 | 0.041546 | 0.045777 | 0.00876 | 0.015712 | 0.011882 | 0.011737 | 0.008378 | 0.008128 | 0.007407 | 0.007132 | 0.009178 | 0.005215 | 0.006742 | 0.003945 |
| 32.0555556 | 0.022626 | 0.041122 | 0.015704 | 0.026097 | 0.009096 | 0.011056 | 0.027444 | 0.024417 | 0.038095 | 0.031672 | 0.03423 | 0.03829 | 0.007918 | 0.013253 | 0.010646 | 0.008814 | 0.006667 | 0.005988 | 0.009723 | 0.006958 | 0.008813 | 0.005043 | 0.005642 | 0.003188 |
| 32.0694444 | 0.015683 | 0.043516 | 0.011056 | 0.015609 | 0.011003 | 0.006768 | 0.023049 | 0.019359 | 0.026374 | 0.022765 | 0.035467 | 0.022438 | 0.00376 | 0.005751 | 0.004986 | 0.003564 | 0.002608 | 0.005842 | 0.008978 | 0.00335 | 0.004165 | 0.006251 | 0.00234 | 0.002092 |
| 32.0833333 | 0.01363 | 0.044484 | 0.006095 | 0.00843 | 0.009044 | 0.003493 | 0.014439 | 0.011591 | 0.014468 | 0.016166 | 0.042025 | 0.009465 | 0.002405 | 0.001068 | 0.001245 | 0.00142 | 0.000838 | 0.007519 | 0.006475 | 0.002159 | 0.001644 | 0.00821 | 0.001512 | 0.000938 |
| 32.0972222 | 0.021665 | 0.042912 | 0.005269 | 0.012002 | 0.005829 | 0.00519 | 0.006764 | 0.005926 | 0.007313 | 0.008071 | 0.037709 | 0.006821 | 0.002359 | 0.000172 | 0.000374 | 0.000888 | 0.000854 | 0.007413 | 0.003436 | 0.001826 | 0.000961 | 0.007853 | 0.001244 | 0.000712 |
| 32.1111111 | 0.028426 | 0.036876 | 0.007726 | 0.016385 | 0.004125 | 0.009195 | 0.005006 | 0.007827 | 0.007078 | 0.008217 | 0.020252 | 0.008693 | 0.002011 | 0.000321 | 0.000398 | 0.000389 | 0.000727 | 0.005836 | 0.001779 | 0.001374 | 0.000532 | 0.006182 | 0.000544 | 0.001105 |
| 32.125 | 0.030821 | 0.029043 | 0.009334 | 0.015665 | 0.00387 | 0.011529 | 0.005379 | 0.012088 | 0.00963 | 0.008145 | 0.008405 | 0.008654 | 0.001782 | 0.00036 | 0.000615 | 0.000353 | 0.000759 | 0.003893 | 0.002874 | 0.001291 | 0.000319 | 0.004355 | 0.000479 | 0.00184 |
| 32.1388889 | 0.033905 | 0.022832 | 0.011584 | 0.013164 | 0.003741 | 0.01064 | 0.003581 | 0.013291 | 0.010096 | 0.005744 | 0.007521 | 0.007993 | 0.001344 | 0.000233 | 0.000596 | 0.00047 | 0.00086 | 0.002376 | 0.004614 | 0.000841 | 0.000251 | 0.002825 | 0.001 | 0.001966 |
| 32.1527778 | 0.037261 | 0.016542 | 0.016149 | 0.011049 | 0.003026 | 0.006407 | 0.001309 | 0.011889 | 0.008122 | 0.003763 | 0.007037 | 0.008749 | 0.000721 | 8.70E-05 | 0.000371 | 0.000729 | 0.00059 | 0.001574 | 0.005228 | 0.000612 | 0.000237 | 0.00174 | 0.00152 | 0.001471 |
| 32.1666667 | 0.038332 | 0.009416 | 0.019499 | 0.01017 | 0.001807 | 0.002555 | 0.001957 | 0.008819 | 0.005044 | 0.005401 | 0.005057 | 0.010646 | 0.000344 | 7.99E-05 | 0.000405 | 0.001207 | 0.000304 | 0.001056 | 0.00502 | 0.000613 | 0.000188 | 0.000925 | 0.001196 | 0.002266 |
| 32.1805556 | 0.039643 | 0.008158 | 0.019532 | 0.0106 | 0.001255 | 0.002319 | 0.006349 | 0.005213 | 0.004354 | 0.009932 | 0.007538 | 0.013965 | 0.000495 | 0.000121 | 0.000548 | 0.00167 | 0.000153 | 0.00059 | 0.004591 | 0.000399 | 0.000203 | 0.000655 | 0.000806 | 0.004714 |
| 32.1944444 | 0.04296 | 0.015909 | 0.018343 | 0.012082 | 0.002673 | 0.003204 | 0.01272 | 0.002657 | 0.007456 | 0.013373 | 0.014 | 0.019734 | 0.000544 | 0.000285 | 0.000456 | 0.001847 | 9.85E-05 | 0.000562 | 0.004305 | 0.000251 | 0.000314 | 0.001096 | 0.001748 | 0.006925 |
| 32.2083333 | 0.043003 | 0.02502 | 0.017313 | 0.013797 | 0.005094 | 0.002392 | 0.01801 | 0.002333 | 0.011009 | 0.01404 | 0.01887 | 0.026337 | 0.000346 | 0.000713 | 0.000283 | 0.001904 | 0.000208 | 0.001305 | 0.004151 | 0.000324 | 0.000481 | 0.001496 | 0.003305 | 0.007692 |
| 32.2222222 | 0.035368 | 0.030272 | 0.016019 | 0.014607 | 0.006884 | 0.002063 | 0.020094 | 0.0043 | 0.013917 | 0.012613 | 0.023432 | 0.029742 | 0.000308 | 0.001049 | 0.00025 | 0.002282 | 0.000497 | 0.002384 | 0.003962 | 0.000488 | 0.000586 | 0.001436 | 0.004117 | 0.007309 |
| 32.2361111 | 0.025817 | 0.031739 | 0.016008 | 0.014039 | 0.007546 | 0.005136 | 0.020103 | 0.00799 | 0.01616 | 0.011469 | 0.030981 | 0.029421 | 0.000258 | 0.001152 | 0.000294 | 0.002911 | 0.000753 | 0.003198 | 0.00359 | 0.000711 | 0.000802 | 0.001263 | 0.004052 | 0.00599 |
| 32.25 | 0.017534 | 0.030231 | 0.018467 | 0.012632 | 0.006937 | 0.009097 | 0.020057 | 0.011992 | 0.015919 | 0.012286 | 0.034946 | 0.028434 | 0.000131 | 0.001277 | 0.000283 | 0.003251 | 0.000689 | 0.003299 | 0.002963 | 0.000821 | 0.000992 | 0.001159 | 0.003535 | 0.004098 |
| 32.2638889 | 0.009227 | 0.0286 | 0.020502 | 0.010957 | 0.005503 | 0.010264 | 0.019744 | 0.013731 | 0.012354 | 0.014675 | 0.02944 | 0.02701 | 6.62E-05 | 0.001318 | 0.000178 | 0.003077 | 0.00051 | 0.002614 | 0.002256 | 0.000655 | 0.000662 | 0.000876 | 0.002793 | 0.002676 |
| 32.2777778 | 0.00375 | 0.02751 | 0.018996 | 0.009437 | 0.004009 | 0.009102 | 0.018846 | 0.012739 | 0.008712 | 0.017764 | 0.016764 | 0.024509 | 9.26E-05 | 0.00105 | 0.000118 | 0.002564 | 0.000634 | 0.001803 | 0.002035 | 0.000428 | 0.00022 | 0.000526 | 0.001927 | 0.002538 |
| 32.2916667 | 0.003274 | 0.025356 | 0.014171 | 0.008314 | 0.00285 | 0.008386 | 0.017492 | 0.01022 | 0.008 | 0.020322 | 0.007707 | 0.021489 | 0.000116 | 0.000621 | 0.000165 | 0.002045 | 0.000894 | 0.001467 | 0.00236 | 0.000517 | 0.00019 | 0.000497 | 0.001201 | 0.003498 |
| 32.3055556 | 0.007365 | 0.022581 | 0.008713 | 0.007755 | 0.002154 | 0.010174 | 0.016801 | 0.006552 | 0.008412 | 0.021456 | 0.009364 | 0.017619 | 0.000251 | 0.000344 | 0.000491 | 0.001989 | 0.001125 | 0.001596 | 0.002458 | 0.001034 | 0.000593 | 0.0006 | 0.00091 | 0.004658 |
| 32.3194444 | 0.012584 | 0.019067 | 0.005591 | 0.007867 | 0.001612 | 0.014481 | 0.018365 | 0.006052 | 0.00825 | 0.021084 | 0.010246 | 0.012707 | 0.000809 | 0.000323 | 0.001409 | 0.002529 | 0.001619 | 0.001953 | 0.002023 | 0.001831 | 0.001438 | 0.001027 | 0.001068 | 0.005262 |
| 32.3333333 | 0.016326 | 0.014429 | 0.005978 | 0.008267 | 0.001088 | 0.020159 | 0.020589 | 0.010423 | 0.006899 | 0.019255 | 0.007606 | 0.007283 | 0.001501 | 0.000335 | 0.002386 | 0.00315 | 0.002184 | 0.00223 | 0.001549 | 0.002741 | 0.002273 | 0.002042 | 0.001386 | 0.005112 |
| 32.3472222 | 0.017699 | 0.009959 | 0.008204 | 0.008397 | 0.001018 | 0.025023 | 0.019695 | 0.012513 | 0.004638 | 0.016441 | 0.010011 | 0.003331 | 0.001791 | 0.000401 | 0.002573 | 0.00345 | 0.002247 | 0.002108 | 0.001296 | 0.003343 | 0.002535 | 0.002847 | 0.001551 | 0.00451 |
| 32.3611111 | 0.016816 | 0.006325 | 0.009264 | 0.007331 | 0.0012 | 0.026674 | 0.014798 | 0.010259 | 0.003065 | 0.013293 | 0.013872 | 0.00268 | 0.001595 | 0.000639 | 0.002029 | 0.003533 | 0.001922 | 0.001452 | 0.001097 | 0.002812 | 0.002431 | 0.003146 | 0.001534 | 0.004189 |
| 32.375 | 0.014115 | 0.003978 | 0.008078 | 0.004369 | 0.000933 | 0.024206 | 0.008319 | 0.008988 | 0.002686 | 0.009665 | 0.014686 | 0.003806 | 0.001227 | 0.000665 | 0.001502 | 0.003556 | 0.001661 | 0.000763 | 0.000876 | 0.001853 | 0.002689 | 0.003639 | 0.001745 | 0.004722 |
| 32.3888889 | 0.010076 | 0.003776 | 0.005924 | 0.001674 | 0.000429 | 0.018935 | 0.003981 | 0.010854 | 0.003453 | 0.005402 | 0.013462 | 0.003864 | 0.001103 | 0.000633 | 0.00139 | 0.003496 | 0.001592 | 0.000567 | 0.000683 | 0.00196 | 0.003413 | 0.00455 | 0.002176 | 0.005746 |
| 32.4027778 | 0.007854 | 0.007103 | 0.005423 | 0.001436 | 0.000312 | 0.014814 | 0.003325 | 0.010965 | 0.005851 | 0.002092 | 0.014574 | 0.002385 | 0.001367 | 0.001067 | 0.001415 | 0.003271 | 0.001622 | 0.000659 | 0.000743 | 0.001863 | 0.003827 | 0.005308 | 0.002172 | 0.006538 |
| 32.4166667 | 0.01027 | 0.014208 | 0.008482 | 0.001611 | 0.000322 | 0.01427 | 0.003018 | 0.008875 | 0.010307 | 0.000798 | 0.022433 | 0.001168 | 0.001613 | 0.001945 | 0.001355 | 0.00302 | 0.001654 | 0.000878 | 0.001275 | 0.001123 | 0.003513 | 0.005347 | 0.001684 | 0.006613 |
| 32.4305556 | 0.016566 | 0.021029 | 0.013393 | 0.001217 | 0.000521 | 0.015154 | 0.00305 | 0.011506 | 0.016411 | 0.000687 | 0.034125 | 0.001317 | 0.001356 | 0.002874 | 0.001331 | 0.002924 | 0.001589 | 0.000983 | 0.002132 | 0.001173 | 0.003006 | 0.004756 | 0.001358 | 0.00575 |
| 32.4444444 | 0.024167 | 0.023969 | 0.017926 | 0.00109 | 0.001883 | 0.014515 | 0.00596 | 0.016067 | 0.020066 | 0.001297 | 0.035334 | 0.001437 | 0.000863 | 0.00294 | 0.001307 | 0.002962 | 0.001457 | 0.00073 | 0.003152 | 0.001112 | 0.002489 | 0.003693 | 0.001193 | 0.004353 |
| 32.4583333 | 0.030814 | 0.023078 | 0.021745 | 0.000626 | 0.00402 | 0.011938 | 0.009563 | 0.015384 | 0.016601 | 0.002165 | 0.02237 | 0.001383 | 0.001238 | 0.001763 | 0.001153 | 0.002737 | 0.001324 | 0.000919 | 0.004182 | 0.00094 | 0.001743 | 0.002145 | 0.000953 | 0.002961 |
| 32.4722222 | 0.035882 | 0.018364 | 0.024318 | 0.001437 | 0.004837 | 0.010236 | 0.01454 | 0.010997 | 0.011562 | 0.005334 | 0.014808 | 0.002058 | 0.002619 | 0.000702 | 0.001299 | 0.001766 | 0.000989 | 0.001969 | 0.004905 | 0.001846 | 0.00135 | 0.000897 | 0.001065 | 0.002022 |
| 32.4861111 | 0.040544 | 0.012775 | 0.02602 | 0.007409 | 0.003706 | 0.012659 | 0.025716 | 0.010574 | 0.016251 | 0.01487 | 0.017569 | 0.003429 | 0.002918 | 0.001518 | 0.001369 | 0.00073 | 0.00064 | 0.002473 | 0.005146 | 0.002212 | 0.001631 | 0.001108 | 0.001428 | 0.002987 |
| 32.5 | 0.045707 | 0.016451 | 0.027116 | 0.019398 | 0.003539 | 0.017766 | 0.04751 | 0.021196 | 0.030314 | 0.026233 | 0.024303 | 0.010283 | 0.002115 | 0.004296 | 0.001561 | 0.000684 | 0.00101 | 0.001751 | 0.004969 | 0.00174 | 0.003555 | 0.003053 | 0.003061 | 0.007493 |
| 32.5138889 | 0.053012 | 0.032312 | 0.026689 | 0.030645 | 0.004455 | 0.019103 | 0.075327 | 0.040726 | 0.038093 | 0.031039 | 0.044381 | 0.027363 | 0.002134 | 0.007227 | 0.002812 | 0.000931 | 0.001813 | 0.000957 | 0.00427 | 0.002032 | 0.006653 | 0.004599 | 0.006295 | 0.013155 |
| 32.5277778 | 0.061559 | 0.049265 | 0.027992 | 0.033893 | 0.003289 | 0.013369 | 0.093319 | 0.060742 | 0.03073 | 0.033191 | 0.074142 | 0.051517 | 0.002471 | 0.008828 | 0.003033 | 0.001277 | 0.002075 | 0.001418 | 0.004056 | 0.002512 | 0.006254 | 0.004091 | 0.007071 | 0.013823 |
| 32.5416667 | 0.065187 | 0.068342 | 0.032381 | 0.03221 | 0.002052 | 0.005805 | 0.100595 | 0.074364 | 0.019381 | 0.042658 | 0.083894 | 0.068074 | 0.002857 | 0.008784 | 0.002447 | 0.001976 | 0.002169 | 0.002415 | 0.005611 | 0.002791 | 0.003015 | 0.003552 | 0.004504 | 0.008312 |
| 32.5555556 | 0.05939 | 0.099661 | 0.03216 | 0.029794 | 0.00293 | 0.003133 | 0.107716 | 0.076059 | 0.020164 | 0.063794 | 0.059147 | 0.065802 | 0.002826 | 0.006385 | 0.002514 | 0.002579 | 0.002337 | 0.002949 | 0.006124 | 0.002992 | 0.001959 | 0.003123 | 0.002966 | 0.003029 |
| 32.5694444 | 0.049525 | 0.117109 | 0.024033 | 0.024127 | 0.003586 | 0.004368 | 0.113527 | 0.067937 | 0.034042 | 0.08414 | 0.023669 | 0.055347 | 0.002381 | 0.002833 | 0.001933 | 0.004104 | 0.002721 | 0.004189 | 0.003562 | 0.003462 | 0.002549 | 0.002568 | 0.003523 | 0.00182 |
| 32.5833333 | 0.037306 | 0.099145 | 0.013442 | 0.01742 | 0.00356 | 0.005631 | 0.110768 | 0.054318 | 0.046843 | 0.087306 | 0.013729 | 0.044978 | 0.002806 | 0.000943 | 0.00161 | 0.005693 | 0.003599 | 0.005306 | 0.001225 | 0.00413 | 0.00242 | 0.003485 | 0.00358 | 0.002072 |
| 32.5972222 | 0.023427 | 0.078586 | 0.009435 | 0.016482 | 0.003645 | 0.007089 | 0.115547 | 0.037792 | 0.05198 | 0.081119 | 0.029426 | 0.033301 | 0.002947 | 0.000845 | 0.001724 | 0.005365 | 0.003823 | 0.004773 | 0.000921 | 0.003562 | 0.001303 | 0.004352 | 0.002403 | 0.001767 |
| 32.6111111 | 0.03129 | 0.072411 | 0.010535 | 0.019186 | 0.00261 | 0.011384 | 0.137616 | 0.032403 | 0.049411 | 0.072585 | 0.034218 | 0.022815 | 0.002456 | 0.000912 | 0.001334 | 0.004122 | 0.003243 | 0.003618 | 0.001283 | 0.002447 | 0.000426 | 0.004079 | 0.001364 | 0.001564 |
| 32.625 | 0.059571 | 0.065859 | 0.007029 | 0.020613 | 0.001715 | 0.017651 | 0.131628 | 0.043978 | 0.038314 | 0.053339 | 0.021209 | 0.01384 | 0.002451 | 0.00122 | 0.001232 | 0.003422 | 0.002834 | 0.002991 | 0.001651 | 0.002369 | 0.001067 | 0.003719 | 0.001385 | 0.002313 |
| 32.6388889 | 0.072601 | 0.056704 | 0.002676 | 0.021304 | 0.002591 | 0.019774 | 0.084726 | 0.051265 | 0.023869 | 0.030068 | 0.013739 | 0.006291 | 0.002736 | 0.002872 | 0.00173 | 0.003137 | 0.002685 | 0.00284 | 0.001936 | 0.002913 | 0.00322 | 0.003342 | 0.002434 | 0.003561 |
| 32.6527778 | 0.064865 | 0.047805 | 0.001802 | 0.019386 | 0.003226 | 0.017239 | 0.042234 | 0.039734 | 0.01127 | 0.027303 | 0.016071 | 0.003736 | 0.002413 | 0.004017 | 0.001815 | 0.002191 | 0.002156 | 0.002085 | 0.001601 | 0.002744 | 0.005223 | 0.002197 | 0.003546 | 0.003669 |
| 32.6666667 | 0.049909 | 0.039415 | 0.002508 | 0.012602 | 0.00249 | 0.013205 | 0.041458 | 0.021706 | 0.00391 | 0.048741 | 0.024122 | 0.008368 | 0.001424 | 0.003154 | 0.001056 | 0.000794 | 0.001406 | 0.000867 | 0.001009 | 0.00198 | 0.005756 | 0.000797 | 0.004082 | 0.002492 |
| 32.6805556 | 0.032782 | 0.032954 | 0.003617 | 0.006184 | 0.002383 | 0.008063 | 0.074767 | 0.010404 | 0.004209 | 0.066834 | 0.03944 | 0.017551 | 0.000863 | 0.002114 | 0.00061 | 0.000681 | 0.001544 | 0.000344 | 0.001585 | 0.002085 | 0.005668 | 0.000287 | 0.00465 | 0.001494 |
| 32.6944444 | 0.020026 | 0.026954 | 0.004505 | 0.006563 | 0.004355 | 0.003809 | 0.096701 | 0.007336 | 0.009783 | 0.066062 | 0.053831 | 0.0265 | 0.000754 | 0.001492 | 0.000729 | 0.001752 | 0.002055 | 0.00026 | 0.002777 | 0.002653 | 0.00505 | 0.000303 | 0.005103 | 0.000953 |
| 32.7083333 | 0.021699 | 0.016963 | 0.005268 | 0.010811 | 0.006559 | 0.002548 | 0.09409 | 0.010225 | 0.01509 | 0.059676 | 0.056207 | 0.030036 | 0.000607 | 0.001236 | 0.000567 | 0.002318 | 0.001594 | 0.00039 | 0.002762 | 0.002249 | 0.00301 | 0.000541 | 0.004655 | 0.000636 |
| 32.7222222 | 0.032306 | 0.007305 | 0.005423 | 0.011448 | 0.007166 | 0.003245 | 0.078357 | 0.016726 | 0.015849 | 0.056321 | 0.044762 | 0.026009 | 0.000943 | 0.002364 | 0.000303 | 0.002135 | 0.000765 | 0.000783 | 0.001599 | 0.001141 | 0.001421 | 0.000862 | 0.003375 | 0.001152 |
| 32.7361111 | 0.036136 | 0.007122 | 0.005025 | 0.007986 | 0.006745 | 0.002977 | 0.061484 | 0.023587 | 0.011233 | 0.051702 | 0.026007 | 0.019148 | 0.001898 | 0.004151 | 0.000696 | 0.00159 | 0.000919 | 0.001208 | 0.001171 | 0.000864 | 0.002355 | 0.001072 | 0.002177 | 0.002493 |
| 32.75 | 0.03462 | 0.015973 | 0.00466 | 0.006258 | 0.007085 | 0.001995 | 0.057619 | 0.026878 | 0.005177 | 0.048046 | 0.017119 | 0.015924 | 0.002861 | 0.005126 | 0.001613 | 0.000935 | 0.001896 | 0.001612 | 0.002562 | 0.002046 | 0.004393 | 0.001492 | 0.002488 | 0.003927 |
| 32.7638889 | 0.033014 | 0.031287 | 0.005246 | 0.008774 | 0.008929 | 0.001988 | 0.065531 | 0.026542 | 0.00292 | 0.048245 | 0.017025 | 0.018279 | 0.003187 | 0.004931 | 0.002197 | 0.000785 | 0.002604 | 0.001678 | 0.004057 | 0.00313 | 0.005774 | 0.001782 | 0.003821 | 0.004786 |
| 32.7777778 | 0.031917 | 0.048189 | 0.00618 | 0.012659 | 0.011287 | 0.002286 | 0.078017 | 0.024739 | 0.003819 | 0.051048 | 0.017447 | 0.023153 | 0.002669 | 0.004063 | 0.002176 | 0.001316 | 0.002526 | 0.001387 | 0.004241 | 0.003056 | 0.006568 | 0.001482 | 0.0041 | 0.004836 |
| 32.7916667 | 0.031292 | 0.057301 | 0.006325 | 0.015695 | 0.014176 | 0.003632 | 0.089911 | 0.020578 | 0.003945 | 0.054188 | 0.025706 | 0.025726 | 0.001691 | 0.003399 | 0.002009 | 0.002127 | 0.001851 | 0.001134 | 0.004068 | 0.002298 | 0.007438 | 0.000822 | 0.002812 | 0.004373 |
| 32.8055556 | 0.028428 | 0.058053 | 0.007107 | 0.018209 | 0.019063 | 0.005 | 0.102052 | 0.014556 | 0.002813 | 0.053123 | 0.034458 | 0.023511 | 0.00073 | 0.002937 | 0.001857 | 0.002567 | 0.000994 | 0.001053 | 0.004325 | 0.00127 | 0.008216 | 0.00043 | 0.002453 | 0.003879 |
| 32.8194444 | 0.02628 | 0.054431 | 0.010127 | 0.020697 | 0.024711 | 0.003565 | 0.121207 | 0.012279 | 0.002822 | 0.046759 | 0.035789 | 0.017724 | 0.000174 | 0.002573 | 0.001803 | 0.002571 | 0.000564 | 0.001095 | 0.004714 | 0.000555 | 0.008902 | 0.000487 | 0.007884 | 0.003946 |
| 32.8333333 | 0.027778 | 0.047938 | 0.016701 | 0.022189 | 0.025624 | 0.002077 | 0.138443 | 0.015858 | 0.003144 | 0.039961 | 0.03546 | 0.012597 | 8.01E-05 | 0.002542 | 0.001853 | 0.002503 | 0.000673 | 0.001059 | 0.004671 | 0.000454 | 0.009232 | 0.00041 | 0.015643 | 0.004704 |
| 32.8472222 | 0.030095 | 0.045568 | 0.026853 | 0.022297 | 0.020208 | 0.004031 | 0.143903 | 0.019294 | 0.003372 | 0.037186 | 0.038073 | 0.01299 | 0.000463 | 0.001996 | 0.001407 | 0.00215 | 0.000696 | 0.00074 | 0.003812 | 0.000614 | 0.007915 | 0.000577 | 0.013642 | 0.005659 |
| 32.8611111 | 0.029955 | 0.056019 | 0.036834 | 0.021692 | 0.013803 | 0.007747 | 0.150939 | 0.017395 | 0.008003 | 0.039458 | 0.042791 | 0.019141 | 0.001116 | 0.00085 | 0.000751 | 0.001299 | 0.000522 | 0.000456 | 0.002501 | 0.00067 | 0.005687 | 0.00136 | 0.005179 | 0.00653 |
| 32.875 | 0.023112 | 0.073649 | 0.043728 | 0.02139 | 0.010057 | 0.011287 | 0.1647 | 0.010961 | 0.015531 | 0.044817 | 0.043416 | 0.026487 | 0.001131 | 0.000452 | 0.00125 | 0.000783 | 0.000507 | 0.00092 | 0.001656 | 0.000997 | 0.004769 | 0.001643 | 0.002822 | 0.007903 |
| 32.8888889 | 0.012491 | 0.081124 | 0.04458 | 0.021547 | 0.008467 | 0.013771 | 0.177218 | 0.004472 | 0.016675 | 0.045573 | 0.038082 | 0.030891 | 0.000563 | 0.000638 | 0.002748 | 0.001358 | 0.000805 | 0.002197 | 0.001411 | 0.001868 | 0.00555 | 0.000979 | 0.004925 | 0.010089 |
| 32.9027778 | 0.006001 | 0.071321 | 0.038132 | 0.019884 | 0.007132 | 0.014045 | 0.181055 | 0.002446 | 0.009965 | 0.036549 | 0.026758 | 0.031423 | 0.000306 | 0.001167 | 0.00383 | 0.002389 | 0.001238 | 0.003096 | 0.001214 | 0.002137 | 0.006734 | 0.000358 | 0.00972 | 0.011294 |
| 32.9166667 | 0.004607 | 0.054823 | 0.02656 | 0.016153 | 0.004586 | 0.012511 | 0.156743 | 0.00451 | 0.004251 | 0.022895 | 0.014054 | 0.02857 | 0.000306 | 0.003611 | 0.003998 | 0.002772 | 0.001197 | 0.002731 | 0.000886 | 0.001415 | 0.007151 | 0.000378 | 0.019184 | 0.009923 |
| 32.9305556 | 0.007251 | 0.041424 | 0.013133 | 0.012928 | 0.002487 | 0.009821 | 0.106255 | 0.006546 | 0.00255 | 0.011346 | 0.016568 | 0.022805 | 0.000365 | 0.007586 | 0.003287 | 0.002076 | 0.001108 | 0.001689 | 0.000538 | 0.001096 | 0.006719 | 0.000666 | 0.023256 | 0.006971 |
| 32.9444444 | 0.012735 | 0.028992 | 0.005372 | 0.010392 | 0.002803 | 0.006312 | 0.064174 | 0.005152 | 0.002427 | 0.004256 | 0.03239 | 0.015205 | 0.000504 | 0.009922 | 0.002202 | 0.001226 | 0.001591 | 0.000883 | 0.000652 | 0.001705 | 0.0062 | 0.000692 | 0.014494 | 0.004131 |
| 32.9583333 | 0.017937 | 0.015239 | 0.003959 | 0.006675 | 0.003052 | 0.003829 | 0.038933 | 0.003602 | 0.006012 | 0.001607 | 0.044404 | 0.007495 | 0.000507 | 0.009703 | 0.001566 | 0.001233 | 0.001805 | 0.000517 | 0.00138 | 0.00229 | 0.005606 | 0.000495 | 0.00672 | 0.001969 |
| 32.9722222 | 0.021719 | 0.006091 | 0.004984 | 0.003046 | 0.003129 | 0.003971 | 0.018234 | 0.005376 | 0.011402 | 0.001893 | 0.046845 | 0.002786 | 0.000582 | 0.006845 | 0.001101 | 0.001563 | 0.001267 | 0.000493 | 0.002205 | 0.002302 | 0.004002 | 0.000727 | 0.012449 | 0.000693 |
| 32.9861111 | 0.019073 | 0.00488 | 0.012347 | 0.005147 | 0.00762 | 0.00661 | 0.004876 | 0.008979 | 0.010174 | 0.008532 | 0.03283 | 0.003806 | 0.000784 | 0.006511 | 0.00055 | 0.001384 | 0.001292 | 0.000528 | 0.003199 | 0.001728 | 0.002028 | 0.001038 | 0.021762 | 0.001187 |
| 33 | 0.011705 | 0.009538 | 0.02659 | 0.015157 | 0.014302 | 0.008753 | 0.003899 | 0.020727 | 0.009122 | 0.027648 | 0.021838 | 0.009512 | 0.002003 | 0.013373 | 0.000414 | 0.000794 | 0.002647 | 0.000501 | 0.004278 | 0.002406 | 0.002352 | 0.001044 | 0.019662 | 0.003949 |
| 33.0138889 | 0.009666 | 0.018136 | 0.040811 | 0.023567 | 0.017895 | 0.006193 | 0.014335 | 0.043517 | 0.023928 | 0.049113 | 0.030545 | 0.017407 | 0.003365 | 0.01477 | 0.001434 | 0.001894 | 0.003309 | 0.000681 | 0.005416 | 0.004686 | 0.004354 | 0.001826 | 0.008426 | 0.005946 |
| 33.0277778 | 0.011327 | 0.026038 | 0.051137 | 0.020952 | 0.019688 | 0.002487 | 0.023396 | 0.064168 | 0.045463 | 0.052603 | 0.099161 | 0.026941 | 0.005277 | 0.010372 | 0.005347 | 0.007065 | 0.004975 | 0.002414 | 0.006325 | 0.006158 | 0.005597 | 0.00517 | 0.003238 | 0.006066 |
| 33.0416667 | 0.015811 | 0.033197 | 0.058945 | 0.01143 | 0.020405 | 0.001961 | 0.017433 | 0.068107 | 0.05535 | 0.03425 | 0.194291 | 0.03312 | 0.010708 | 0.01349 | 0.010378 | 0.01256 | 0.009362 | 0.005685 | 0.004966 | 0.008801 | 0.007776 | 0.00943 | 0.006175 | 0.008259 |
| 33.0555556 | 0.022625 | 0.042865 | 0.05838 | 0.005475 | 0.01622 | 0.0026 | 0.006917 | 0.051592 | 0.049284 | 0.0134 | 0.17365 | 0.027895 | 0.01305 | 0.015773 | 0.010191 | 0.011625 | 0.009923 | 0.005749 | 0.002265 | 0.010818 | 0.008948 | 0.008569 | 0.007395 | 0.010236 |
| 33.0694444 | 0.025585 | 0.052348 | 0.042184 | 0.007076 | 0.010675 | 0.005708 | 0.004946 | 0.026624 | 0.030928 | 0.005356 | 0.086405 | 0.014517 | 0.007637 | 0.01041 | 0.005087 | 0.005816 | 0.005094 | 0.003219 | 0.001366 | 0.006966 | 0.00535 | 0.003901 | 0.010285 | 0.00788 |
| 33.0833333 | 0.022675 | 0.056133 | 0.022016 | 0.010131 | 0.012374 | 0.011233 | 0.007114 | 0.014327 | 0.012755 | 0.00345 | 0.036521 | 0.007408 | 0.002161 | 0.005267 | 0.001508 | 0.001789 | 0.001336 | 0.003227 | 0.00147 | 0.002293 | 0.001317 | 0.001663 | 0.019426 | 0.004295 |
| 33.0972222 | 0.015055 | 0.052639 | 0.016306 | 0.00993 | 0.01526 | 0.013675 | 0.007539 | 0.018485 | 0.005867 | 0.002602 | 0.014415 | 0.012529 | 0.001218 | 0.003209 | 0.001015 | 0.000688 | 0.000864 | 0.003901 | 0.001469 | 0.002116 | 0.000561 | 0.001457 | 0.023597 | 0.002251 |
| 33.1111111 | 0.006548 | 0.046187 | 0.021157 | 0.008224 | 0.010748 | 0.012099 | 0.006362 | 0.023371 | 0.006085 | 0.005377 | 0.01058 | 0.021582 | 0.002173 | 0.002266 | 0.001397 | 0.000448 | 0.001564 | 0.003457 | 0.002487 | 0.004 | 0.001218 | 0.00147 | 0.017983 | 0.001168 |
| 33.125 | 0.002469 | 0.04098 | 0.018145 | 0.007164 | 0.006518 | 0.010052 | 0.004495 | 0.021364 | 0.004745 | 0.009325 | 0.012427 | 0.028708 | 0.002562 | 0.001556 | 0.00133 | 0.00041 | 0.001733 | 0.002918 | 0.004442 | 0.00523 | 0.001424 | 0.001662 | 0.010237 | 0.000895 |
| 33.1388889 | 0.00322 | 0.037923 | 0.007515 | 0.007568 | 0.007915 | 0.009391 | 0.002596 | 0.014985 | 0.002102 | 0.011135 | 0.013415 | 0.029649 | 0.002238 | 0.000945 | 0.000713 | 0.000494 | 0.001303 | 0.002582 | 0.006212 | 0.005154 | 0.001035 | 0.001933 | 0.005295 | 0.001389 |
| 33.1527778 | 0.006557 | 0.035159 | 0.003924 | 0.009146 | 0.008363 | 0.010789 | 0.001315 | 0.007649 | 0.001174 | 0.010226 | 0.015933 | 0.022069 | 0.001807 | 0.000474 | 0.000378 | 0.000696 | 0.000999 | 0.002878 | 0.007373 | 0.004239 | 0.000715 | 0.002246 | 0.002881 | 0.001486 |
| 33.1666667 | 0.01061 | 0.030704 | 0.010765 | 0.010951 | 0.005599 | 0.014227 | 0.000883 | 0.005094 | 0.001568 | 0.007582 | 0.022882 | 0.01373 | 0.001242 | 0.000664 | 0.000824 | 0.001162 | 0.000737 | 0.003066 | 0.00741 | 0.00312 | 0.000527 | 0.001949 | 0.001497 | 0.001071 |
| 33.1805556 | 0.010071 | 0.025412 | 0.019691 | 0.012227 | 0.003254 | 0.017873 | 0.001651 | 0.008605 | 0.003007 | 0.00476 | 0.032359 | 0.009807 | 0.000752 | 0.001608 | 0.001839 | 0.002096 | 0.000434 | 0.002296 | 0.006016 | 0.002078 | 0.000453 | 0.001088 | 0.001262 | 0.001422 |
| 33.1944444 | 0.004974 | 0.020593 | 0.024694 | 0.012672 | 0.0043 | 0.019229 | 0.003552 | 0.010352 | 0.005386 | 0.002423 | 0.031014 | 0.008332 | 0.000782 | 0.002452 | 0.002926 | 0.003016 | 0.00049 | 0.001234 | 0.004633 | 0.001616 | 0.00066 | 0.000505 | 0.001966 | 0.002399 |
| 33.2083333 | 0.003295 | 0.015971 | 0.026567 | 0.012262 | 0.008924 | 0.016466 | 0.005754 | 0.007569 | 0.00749 | 0.001331 | 0.018933 | 0.00758 | 0.00089 | 0.002453 | 0.002666 | 0.003043 | 0.001043 | 0.001161 | 0.005012 | 0.002234 | 0.000708 | 0.000485 | 0.002014 | 0.002729 |
| 33.2222222 | 0.009109 | 0.011688 | 0.026335 | 0.011056 | 0.012584 | 0.009876 | 0.006488 | 0.007615 | 0.007939 | 0.001323 | 0.011277 | 0.007038 | 0.000871 | 0.002079 | 0.001132 | 0.002468 | 0.001657 | 0.001777 | 0.006376 | 0.003114 | 0.00049 | 0.000823 | 0.00153 | 0.002694 |
| 33.2361111 | 0.016453 | 0.008636 | 0.023072 | 0.0092 | 0.010956 | 0.005899 | 0.004852 | 0.013482 | 0.00631 | 0.000937 | 0.008248 | 0.006641 | 0.00101 | 0.002334 | 0.000627 | 0.002544 | 0.001311 | 0.001535 | 0.006346 | 0.002623 | 0.000578 | 0.000783 | 0.001996 | 0.00313 |
| 33.25 | 0.018709 | 0.007188 | 0.015385 | 0.007384 | 0.006864 | 0.006752 | 0.00403 | 0.019871 | 0.00443 | 0.000382 | 0.005956 | 0.006706 | 0.001368 | 0.002969 | 0.001298 | 0.003172 | 0.000468 | 0.000642 | 0.004878 | 0.001225 | 0.001031 | 0.000276 | 0.002868 | 0.003351 |
| 33.2638889 | 0.017205 | 0.007302 | 0.009883 | 0.00619 | 0.006924 | 0.005655 | 0.004193 | 0.022413 | 0.004621 | 0.000295 | 0.005048 | 0.007342 | 0.001429 | 0.002702 | 0.001395 | 0.00286 | 0.000672 | 0.000804 | 0.004148 | 0.001161 | 0.001015 | 0.000363 | 0.002554 | 0.002305 |
| 33.2777778 | 0.015748 | 0.008597 | 0.016001 | 0.005422 | 0.009637 | 0.004265 | 0.003775 | 0.02134 | 0.00712 | 0.001277 | 0.003507 | 0.008174 | 0.001394 | 0.001517 | 0.000978 | 0.001514 | 0.001572 | 0.001977 | 0.004478 | 0.002592 | 0.00045 | 0.001154 | 0.001359 | 0.001076 |
| 33.2916667 | 0.015043 | 0.009907 | 0.023685 | 0.004852 | 0.008435 | 0.00604 | 0.005971 | 0.01923 | 0.009399 | 0.003872 | 0.001615 | 0.008396 | 0.001541 | 0.000737 | 0.000755 | 0.000719 | 0.001514 | 0.002485 | 0.004429 | 0.003585 | 0.000596 | 0.00141 | 0.000761 | 0.000799 |
| 33.3055556 | 0.013627 | 0.010065 | 0.017712 | 0.004417 | 0.004403 | 0.00724 | 0.008723 | 0.017327 | 0.009649 | 0.005906 | 0.001084 | 0.007217 | 0.001187 | 0.000622 | 0.000571 | 0.00108 | 0.000728 | 0.002106 | 0.00448 | 0.003871 | 0.001765 | 0.000806 | 0.000916 | 0.000768 |
| 33.3194444 | 0.011772 | 0.009414 | 0.009363 | 0.003901 | 0.002295 | 0.007163 | 0.008155 | 0.015408 | 0.009239 | 0.005391 | 0.00099 | 0.004629 | 0.000785 | 0.000623 | 0.000611 | 0.001525 | 0.00042 | 0.002165 | 0.006046 | 0.004795 | 0.002734 | 0.000285 | 0.000873 | 0.00097 |
| 33.3333333 | 0.010035 | 0.00956 | 0.012835 | 0.003109 | 0.002145 | 0.007443 | 0.006536 | 0.014335 | 0.007963 | 0.003834 | 0.00068 | 0.002333 | 0.000845 | 0.000799 | 0.000721 | 0.001478 | 0.000428 | 0.002746 | 0.008258 | 0.006151 | 0.002803 | 0.000199 | 0.00039 | 0.001324 |
| 33.3472222 | 0.007379 | 0.011043 | 0.014097 | 0.001968 | 0.003871 | 0.007784 | 0.006031 | 0.014051 | 0.004745 | 0.003536 | 0.000541 | 0.00251 | 0.000872 | 0.00087 | 0.00104 | 0.001674 | 0.000573 | 0.002484 | 0.009051 | 0.006186 | 0.002625 | 0.000377 | 0.000317 | 0.0015 |
| 33.3611111 | 0.00443 | 0.012013 | 0.008309 | 0.001159 | 0.007728 | 0.007757 | 0.006219 | 0.013609 | 0.002249 | 0.00407 | 0.00055 | 0.003981 | 0.000936 | 0.000856 | 0.001681 | 0.002141 | 0.000826 | 0.001327 | 0.008322 | 0.004723 | 0.002556 | 0.000541 | 0.000577 | 0.00148 |
| 33.375 | 0.003825 | 0.011248 | 0.005588 | 0.001159 | 0.010146 | 0.008718 | 0.008323 | 0.013787 | 0.001618 | 0.003476 | 0.001347 | 0.003834 | 0.000923 | 0.00115 | 0.001477 | 0.0017 | 0.000944 | 0.000908 | 0.007765 | 0.003851 | 0.001825 | 0.000615 | 0.000959 | 0.001184 |
| 33.3888889 | 0.005885 | 0.010238 | 0.006015 | 0.002099 | 0.008598 | 0.011781 | 0.013421 | 0.015575 | 0.001489 | 0.003361 | 0.002911 | 0.00244 | 0.000585 | 0.001381 | 0.00051 | 0.000614 | 0.001059 | 0.001527 | 0.007322 | 0.004181 | 0.000796 | 0.000693 | 0.001602 | 0.000911 |
| 33.4027778 | 0.00774 | 0.010324 | 0.01166 | 0.004348 | 0.00586 | 0.015344 | 0.018401 | 0.018759 | 0.003268 | 0.005846 | 0.004817 | 0.002717 | 0.000418 | 0.001749 | 0.000214 | 0.000343 | 0.001123 | 0.001768 | 0.005687 | 0.004334 | 0.00071 | 0.000642 | 0.001399 | 0.000679 |
| 33.4166667 | 0.007367 | 0.010742 | 0.020085 | 0.006729 | 0.007453 | 0.017288 | 0.019602 | 0.023047 | 0.006772 | 0.008007 | 0.008097 | 0.005031 | 0.00036 | 0.002195 | 0.000494 | 0.000775 | 0.000846 | 0.001385 | 0.0037 | 0.004145 | 0.001086 | 0.000708 | 0.00076 | 0.000598 |
| 33.4305556 | 0.005723 | 0.011417 | 0.024056 | 0.00798 | 0.012488 | 0.017018 | 0.016117 | 0.027099 | 0.008828 | 0.007413 | 0.013027 | 0.006553 | 0.000369 | 0.002155 | 0.000426 | 0.001083 | 0.000453 | 0.001187 | 0.0028 | 0.003909 | 0.001163 | 0.000912 | 0.00088 | 0.000996 |
| 33.4444444 | 0.003977 | 0.014178 | 0.022958 | 0.007922 | 0.0127 | 0.014859 | 0.010622 | 0.028732 | 0.00763 | 0.004551 | 0.017798 | 0.005185 | 0.001088 | 0.002412 | 0.000376 | 0.000962 | 0.000411 | 0.001566 | 0.003166 | 0.003767 | 0.000896 | 0.0008 | 0.000901 | 0.001608 |
| 33.4583333 | 0.002282 | 0.01881 | 0.019583 | 0.006719 | 0.008174 | 0.012274 | 0.006751 | 0.027568 | 0.004486 | 0.001887 | 0.021611 | 0.002946 | 0.002819 | 0.003922 | 0.001208 | 0.000878 | 0.000946 | 0.002914 | 0.004828 | 0.004629 | 0.001022 | 0.001089 | 0.000849 | 0.001528 |
| 33.4722222 | 0.002697 | 0.020755 | 0.016667 | 0.005772 | 0.00737 | 0.011515 | 0.006743 | 0.024984 | 0.003282 | 0.002741 | 0.027913 | 0.001997 | 0.00461 | 0.005246 | 0.002406 | 0.001336 | 0.001575 | 0.004487 | 0.007283 | 0.005815 | 0.001825 | 0.002172 | 0.001423 | 0.00077 |
| 33.4861111 | 0.009864 | 0.016116 | 0.0151 | 0.00745 | 0.008152 | 0.013729 | 0.010759 | 0.021467 | 0.00662 | 0.008815 | 0.040863 | 0.003682 | 0.004411 | 0.004182 | 0.002183 | 0.001308 | 0.001312 | 0.004072 | 0.008915 | 0.004563 | 0.00271 | 0.002509 | 0.001955 | 0.000988 |

| | | | | | | | | | | | | | | | | | | | | | | | | |
|---|---|---|---|---|---|---|---|---|---|---|---|---|---|---|---|---|---|---|---|---|---|---|---|---|
| 33.5 | 0.024703 | 0.016454 | 0.016454 | 0.011624 | 0.007194 | 0.017032 | 0.015702 | 0.017461 | 0.0122 | 0.016782 | 0.055762 | 0.011184 | 0.002451 | 0.003139 | 0.001283 | 0.00068 | 0.000775 | 0.001968 | 0.008372 | 0.002843 | 0.00516 | 0.002623 | 0.003312 | 0.003097 |
| 33.5138889 | 0.040046 | 0.029833 | 0.024167 | 0.015259 | 0.00793 | 0.017356 | 0.017504 | 0.017702 | 0.015707 | 0.019798 | 0.066754 | 0.023194 | 0.001155 | 0.004997 | 0.001235 | 0.001037 | 0.000939 | 0.00138 | 0.005978 | 0.003404 | 0.007446 | 0.002799 | 0.005081 | 0.005059 |
| 33.5277778 | 0.048506 | 0.037449 | 0.040297 | 0.018331 | 0.007139 | 0.015118 | 0.015025 | 0.026477 | 0.018189 | 0.021059 | 0.075757 | 0.037199 | 0.002368 | 0.007399 | 0.002489 | 0.003067 | 0.002084 | 0.004519 | 0.003728 | 0.005132 | 0.006125 | 0.00352 | 0.004004 | 0.004031 |
| 33.5416667 | 0.050939 | 0.03207 | 0.059987 | 0.024366 | 0.005918 | 0.014921 | 0.0103 | 0.039128 | 0.020692 | 0.027119 | 0.090304 | 0.051517 | 0.004919 | 0.007299 | 0.004942 | 0.004766 | 0.00351 | 0.010913 | 0.002376 | 0.010136 | 0.004104 | 0.005698 | 0.00192 | 0.001335 |
| 33.5555556 | 0.049099 | 0.030236 | 0.070931 | 0.031769 | 0.006839 | 0.016764 | 0.006025 | 0.047769 | 0.018503 | 0.030201 | 0.131549 | 0.060506 | 0.00468 | 0.004932 | 0.004809 | 0.004404 | 0.003633 | 0.01429 | 0.00302 | 0.014567 | 0.003667 | 0.005329 | 0.003728 | 0.001315 |
| 33.5694444 | 0.039464 | 0.041409 | 0.06935 | 0.032195 | 0.007224 | 0.017738 | 0.005112 | 0.048265 | 0.012449 | 0.022896 | 0.192377 | 0.061681 | 0.002792 | 0.003436 | 0.002729 | 0.004139 | 0.003734 | 0.009791 | 0.005435 | 0.01096 | 0.003811 | 0.003225 | 0.007329 | 0.003744 |
| 33.5833333 | 0.025144 | 0.061206 | 0.063979 | 0.024043 | 0.008747 | 0.017223 | 0.013762 | 0.038332 | 0.009363 | 0.013693 | 0.211919 | 0.058913 | 0.002555 | 0.003211 | 0.002679 | 0.004623 | 0.004225 | 0.004067 | 0.004797 | 0.004082 | 0.003498 | 0.003118 | 0.00728 | 0.004716 |
| 33.5972222 | 0.015194 | 0.078312 | 0.058214 | 0.014126 | 0.009515 | 0.016755 | 0.031887 | 0.02143 | 0.009972 | 0.00989 | 0.161329 | 0.05359 | 0.002221 | 0.002028 | 0.002877 | 0.00432 | 0.003469 | 0.002341 | 0.002071 | 0.000845 | 0.002191 | 0.002957 | 0.004188 | 0.003702 |
| 33.6111111 | 0.012866 | 0.088735 | 0.049842 | 0.007959 | 0.008699 | 0.018064 | 0.045484 | 0.007984 | 0.011777 | 0.007992 | 0.080456 | 0.04291 | 0.001456 | 0.001269 | 0.001888 | 0.003147 | 0.00197 | 0.001836 | 0.002176 | 0.001149 | 0.002298 | 0.00182 | 0.001876 | 0.00253 |
| 33.625 | 0.015421 | 0.097946 | 0.039823 | 0.007301 | 0.007051 | 0.01979 | 0.046105 | 0.00359 | 0.015263 | 0.00585 | 0.03274 | 0.026596 | 0.002066 | 0.002864 | 0.001461 | 0.001773 | 0.001419 | 0.002006 | 0.004293 | 0.003416 | 0.004727 | 0.001473 | 0.001476 | 0.002848 |
| 33.6388889 | 0.01886 | 0.103786 | 0.030899 | 0.009498 | 0.003969 | 0.019519 | 0.041216 | 0.003444 | 0.019275 | 0.005477 | 0.075116 | 0.012505 | 0.003363 | 0.004934 | 0.00222 | 0.001235 | 0.002259 | 0.003174 | 0.005595 | 0.005469 | 0.006454 | 0.002226 | 0.001607 | 0.004268 |
| 33.6527778 | 0.022143 | 0.101656 | 0.02444 | 0.010411 | 0.002826 | 0.017691 | 0.040546 | 0.004373 | 0.020825 | 0.007507 | 0.160922 | 0.011384 | 0.003323 | 0.004873 | 0.002248 | 0.001095 | 0.002536 | 0.00338 | 0.005047 | 0.005222 | 0.005521 | 0.00238 | 0.00201 | 0.003965 |
| 33.6666667 | 0.025483 | 0.095233 | 0.018704 | 0.007447 | 0.004697 | 0.01542 | 0.044781 | 0.008184 | 0.017171 | 0.008805 | 0.170524 | 0.022074 | 0.001864 | 0.002895 | 0.001265 | 0.001132 | 0.001551 | 0.002293 | 0.003056 | 0.003057 | 0.002936 | 0.00143 | 0.003612 | 0.002315 |
| 33.6805556 | 0.027437 | 0.089218 | 0.012373 | 0.003469 | 0.009979 | 0.013286 | 0.050497 | 0.010594 | 0.011131 | 0.00664 | 0.109377 | 0.034004 | 0.000767 | 0.001512 | 0.000988 | 0.002326 | 0.000966 | 0.001034 | 0.001411 | 0.001421 | 0.001315 | 0.000759 | 0.005262 | 0.002036 |
| 33.6944444 | 0.025906 | 0.081087 | 0.008386 | 0.002935 | 0.017216 | 0.012371 | 0.054342 | 0.007311 | 0.014356 | 0.00489 | 0.060227 | 0.039625 | 0.000584 | 0.001586 | 0.001234 | 0.003799 | 0.001188 | 0.000336 | 0.000921 | 0.001737 | 0.001577 | 0.000737 | 0.005925 | 0.002532 |
| 33.7083333 | 0.01903 | 0.061495 | 0.007502 | 0.004007 | 0.023084 | 0.012181 | 0.051557 | 0.003663 | 0.02285 | 0.006634 | 0.033198 | 0.039194 | 0.000475 | 0.001418 | 0.001035 | 0.003945 | 0.001162 | 0.000246 | 0.000845 | 0.002203 | 0.001822 | 0.000573 | 0.005798 | 0.001835 |
| 33.7222222 | 0.009216 | 0.031491 | 0.005374 | 0.003801 | 0.026461 | 0.01143 | 0.041052 | 0.004072 | 0.023604 | 0.009037 | 0.019321 | 0.036477 | 0.000398 | 0.000749 | 0.000518 | 0.002743 | 0.000712 | 0.000313 | 0.001634 | 0.00162 | 0.001269 | 0.000343 | 0.004776 | 0.000764 |
| 33.7361111 | 0.00542 | 0.014027 | 0.003427 | 0.002451 | 0.027686 | 0.01039 | 0.028999 | 0.005867 | 0.020136 | 0.010917 | 0.037847 | 0.032137 | 0.00058 | 0.000669 | 0.000251 | 0.001368 | 0.000411 | 0.000648 | 0.002987 | 0.000801 | 0.000728 | 0.000316 | 0.003345 | 0.000687 |
| 33.75 | 0.011583 | 0.021014 | 0.006975 | 0.001717 | 0.029283 | 0.009424 | 0.021487 | 0.007388 | 0.016986 | 0.012668 | 0.075205 | 0.028422 | 0.000834 | 0.001024 | 0.000304 | 0.000546 | 0.000323 | 0.001462 | 0.003966 | 0.00051 | 0.00057 | 0.000239 | 0.002704 | 0.001065 |
| 33.7638889 | 0.020076 | 0.030909 | 0.014576 | 0.002789 | 0.033198 | 0.008169 | 0.016663 | 0.009384 | 0.016161 | 0.01406 | 0.10886 | 0.030537 | 0.001073 | 0.000912 | 0.00037 | 0.00031 | 0.00023 | 0.00238 | 0.00461 | 0.000866 | 0.001121 | 0.000139 | 0.003713 | 0.001557 |
| 33.7777778 | 0.024174 | 0.028302 | 0.025364 | 0.005366 | 0.038421 | 0.006663 | 0.010698 | 0.010879 | 0.016391 | 0.015458 | 0.13022 | 0.038266 | 0.001091 | 0.000512 | 0.000334 | 0.00038 | 0.000319 | 0.002983 | 0.004356 | 0.001311 | 0.001967 | 0.000164 | 0.005847 | 0.001892 |
| 33.7916667 | 0.027928 | 0.023256 | 0.041305 | 0.008514 | 0.045293 | 0.005196 | 0.005925 | 0.010059 | 0.014533 | 0.016949 | 0.175351 | 0.045939 | 0.000832 | 0.000735 | 0.000255 | 0.000378 | 0.000855 | 0.003074 | 0.00268 | 0.001409 | 0.002079 | 0.000315 | 0.007881 | 0.001798 |
| 33.8055556 | 0.041636 | 0.024172 | 0.059811 | 0.010153 | 0.055279 | 0.003169 | 0.007063 | 0.007108 | 0.013574 | 0.018427 | 0.238734 | 0.049507 | 0.000899 | 0.001601 | 0.000527 | 0.000484 | 0.001081 | 0.003106 | 0.000897 | 0.001745 | 0.002195 | 0.000682 | 0.009018 | 0.002333 |
| 33.8194444 | 0.066092 | 0.027762 | 0.075192 | 0.009508 | 0.065501 | 0.0027 | 0.013105 | 0.00471 | 0.012547 | 0.019139 | 0.216628 | 0.048424 | 0.001647 | 0.002649 | 0.001618 | 0.000804 | 0.000845 | 0.003758 | 0.000583 | 0.002929 | 0.00361 | 0.00187 | 0.008726 | 0.003894 |
| 33.8333333 | 0.084924 | 0.031189 | 0.0831 | 0.008432 | 0.06817 | 0.005763 | 0.020523 | 0.005586 | 0.007943 | 0.018326 | 0.125259 | 0.043071 | 0.002177 | 0.003348 | 0.002659 | 0.000815 | 0.00087 | 0.004497 | 0.001505 | 0.003859 | 0.005334 | 0.003525 | 0.007063 | 0.00519 |
| 33.8472222 | 0.086777 | 0.036279 | 0.08483 | 0.007507 | 0.058542 | 0.008582 | 0.027718 | 0.009333 | 0.005024 | 0.016695 | 0.064453 | 0.034767 | 0.001665 | 0.003157 | 0.002537 | 0.000705 | 0.000821 | 0.0043 | 0.002672 | 0.00332 | 0.006171 | 0.004278 | 0.005082 | 0.005488 |
| 33.8611111 | 0.074981 | 0.044379 | 0.08297 | 0.005887 | 0.046733 | 0.00925 | 0.032667 | 0.013328 | 0.009874 | 0.015229 | 0.089674 | 0.025688 | 0.000764 | 0.002342 | 0.001862 | 0.00111 | 0.001006 | 0.00318 | 0.003431 | 0.001911 | 0.006685 | 0.004028 | 0.003868 | 0.005195 |
| 33.875 | 0.061492 | 0.053751 | 0.076744 | 0.004543 | 0.044487 | 0.010304 | 0.034298 | 0.016009 | 0.016232 | 0.014074 | 0.172004 | 0.015895 | 0.000417 | 0.002901 | 0.001428 | 0.001482 | 0.001937 | 0.001696 | 0.003654 | 0.000831 | 0.007662 | 0.003212 | 0.003208 | 0.004452 |
| 33.8888889 | 0.055995 | 0.060469 | 0.069011 | 0.004453 | 0.043881 | 0.01288 | 0.038147 | 0.017387 | 0.018555 | 0.013173 | 0.200504 | 0.00821 | 0.000403 | 0.005175 | 0.001028 | 0.001118 | 0.002724 | 0.000582 | 0.003905 | 0.000593 | 0.00877 | 0.00189 | 0.00263 | 0.003599 |
| 33.9027778 | 0.052003 | 0.060081 | 0.06596 | 0.004587 | 0.031646 | 0.016232 | 0.050892 | 0.017715 | 0.020664 | 0.012723 | 0.149914 | 0.008276 | 0.000321 | 0.006408 | 0.000517 | 0.000907 | 0.002917 | 0.000291 | 0.004933 | 0.001007 | 0.009228 | 0.0009 | 0.002736 | 0.003395 |
| 33.9166667 | 0.046603 | 0.051273 | 0.070142 | 0.004218 | 0.015236 | 0.019828 | 0.067788 | 0.01696 | 0.026719 | 0.011653 | 0.096912 | 0.013603 | 0.000471 | 0.006832 | 0.000185 | 0.001443 | 0.003046 | 0.000438 | 0.006118 | 0.001706 | 0.009146 | 0.001151 | 0.003761 | 0.003682 |
| 33.9305556 | 0.040305 | 0.03552 | 0.076751 | 0.003509 | 0.008419 | 0.023124 | 0.077133 | 0.015466 | 0.03673 | 0.008442 | 0.073968 | 0.019085 | 0.000774 | 0.007871 | 0.000217 | 0.001746 | 0.003114 | 0.000467 | 0.005222 | 0.002076 | 0.009116 | 0.001978 | 0.004232 | 0.00431 |
| 33.9444444 | 0.030143 | 0.018878 | 0.075112 | 0.00241 | 0.00635 | 0.025787 | 0.075949 | 0.014031 | 0.045897 | 0.004288 | 0.052998 | 0.024101 | 0.000861 | 0.007884 | 0.000874 | 0.002132 | 0.002361 | 0.000603 | 0.004262 | 0.001582 | 0.009614 | 0.002151 | 0.002928 | 0.005186 |
| 33.9583333 | 0.021062 | 0.011728 | 0.059567 | 0.004008 | 0.003384 | 0.026339 | 0.065086 | 0.012838 | 0.051173 | 0.003837 | 0.027879 | 0.026686 | 0.000763 | 0.006294 | 0.001955 | 0.002716 | 0.001126 | 0.001033 | 0.007378 | 0.000714 | 0.010013 | 0.001703 | 0.00117 | 0.005693 |
| 33.9722222 | 0.01642 | 0.009989 | 0.038836 | 0.011945 | 0.003336 | 0.025347 | 0.044935 | 0.01068 | 0.049619 | 0.006415 | 0.017483 | 0.02444 | 0.001078 | 0.005276 | 0.002231 | 0.002328 | 0.000535 | 0.000968 | 0.010014 | 0.000599 | 0.008417 | 0.001612 | 0.001154 | 0.006265 |
| 33.9861111 | 0.017777 | 0.00816 | 0.029714 | 0.020103 | 0.004102 | 0.026333 | 0.031242 | 0.006518 | 0.033004 | 0.009785 | 0.027456 | 0.015996 | 0.00215 | 0.010524 | 0.001888 | 0.002424 | 0.001074 | 0.000647 | 0.006777 | 0.001364 | 0.005459 | 0.00153 | 0.005766 | 0.008096 |
| 34 | 0.02806 | 0.016543 | 0.035723 | 0.01794 | 0.009675 | 0.025634 | 0.039336 | 0.002907 | 0.018008 | 0.022124 | 0.045504 | 0.008802 | 0.002424 | 0.021272 | 0.002038 | 0.003864 | 0.002566 | 0.000837 | 0.00293 | 0.003719 | 0.004506 | 0.001902 | 0.013669 | 0.010492 |
| 34.0138889 | 0.040794 | 0.032753 | 0.044047 | 0.008475 | 0.027182 | 0.018995 | 0.058962 | 0.00201 | 0.029386 | 0.042304 | 0.05556 | 0.013401 | 0.001529 | 0.022513 | 0.002473 | 0.003941 | 0.003106 | 0.001393 | 0.0043 | 0.008988 | 0.006381 | 0.002886 | 0.014181 | 0.010087 |
| 34.0277778 | 0.045072 | 0.047739 | 0.047935 | 0.003008 | 0.042664 | 0.011161 | 0.068549 | 0.002754 | 0.049457 | 0.052346 | 0.049485 | 0.022262 | 0.00254 | 0.013498 | 0.00487 | 0.004076 | 0.003873 | 0.002759 | 0.008188 | 0.014864 | 0.007264 | 0.003555 | 0.010004 | 0.006837 |
| 34.0416667 | 0.04304 | 0.063206 | 0.052982 | 0.003242 | 0.036558 | 0.008934 | 0.06135 | 0.00575 | 0.057708 | 0.04426 | 0.031112 | 0.02333 | 0.006376 | 0.009331 | 0.0092 | 0.006156 | 0.00688 | 0.004096 | 0.00942 | 0.016213 | 0.005305 | 0.004982 | 0.012721 | 0.005955 |
| 34.0555556 | 0.043792 | 0.078416 | 0.061037 | 0.004859 | 0.024225 | 0.011519 | 0.047001 | 0.007637 | 0.057264 | 0.027739 | 0.014593 | 0.017336 | 0.008204 | 0.009034 | 0.009864 | 0.005817 | 0.007513 | 0.003136 | 0.006237 | 0.011972 | 0.004286 | 0.005173 | 0.012544 | 0.007476 |
| 34.0694444 | 0.045514 | 0.086309 | 0.063398 | 0.005222 | 0.035127 | 0.012097 | 0.034841 | 0.008118 | 0.048892 | 0.013673 | 0.009423 | 0.010198 | 0.005372 | 0.005985 | 0.006047 | 0.002739 | 0.004321 | 0.001403 | 0.002639 | 0.00784 | 0.004929 | 0.002989 | 0.005859 | 0.006945 |
| 34.0833333 | 0.044752 | 0.083241 | 0.053893 | 0.003841 | 0.05206 | 0.008626 | 0.025928 | 0.01385 | 0.033854 | 0.005622 | 0.008036 | 0.0051 | 0.001971 | 0.002827 | 0.0025 | 0.000759 | 0.001684 | 0.001711 | 0.002121 | 0.00825 | 0.004529 | 0.001445 | 0.005397 | 0.004746 |
| 34.0972222 | 0.04276 | 0.074636 | 0.03876 | 0.001858 | 0.048455 | 0.003767 | 0.01849 | 0.018803 | 0.019659 | 0.002167 | 0.005507 | 0.002504 | 0.00139 | 0.00132 | 0.001292 | 0.000664 | 0.001605 | 0.003585 | 0.002003 | 0.012507 | 0.003757 | 0.001019 | 0.012686 | 0.002487 |
| 34.1111111 | 0.037242 | 0.065979 | 0.026938 | 0.000819 | 0.033348 | 0.00116 | 0.012886 | 0.014516 | 0.009128 | 0.001827 | 0.004738 | 0.002922 | 0.002795 | 0.000508 | 0.001698 | 0.001784 | 0.002853 | 0.005652 | 0.001312 | 0.017852 | 0.003549 | 0.000876 | 0.019435 | 0.000945 |
| 34.125 | 0.030072 | 0.057259 | 0.02179 | 0.001371 | 0.021709 | 0.000933 | 0.009371 | 0.00664 | 0.003731 | 0.003675 | 0.00451 | 0.00531 | 0.003693 | 0.000766 | 0.001678 | 0.002641 | 0.003264 | 0.007426 | 0.002001 | 0.022338 | 0.00266 | 0.001434 | 0.0188 | 0.000715 |
| 34.1388889 | 0.02421 | 0.047569 | 0.024214 | 0.002649 | 0.015144 | 0.00077 | 0.008242 | 0.003327 | 0.004744 | 0.00527 | 0.008334 | 0.006921 | 0.003231 | 0.002361 | 0.00083 | 0.002332 | 0.002636 | 0.009889 | 0.003587 | 0.026479 | 0.001106 | 0.002423 | 0.012331 | 0.000704 |
| 34.1527778 | 0.016455 | 0.038968 | 0.028748 | 0.004073 | 0.010287 | 0.001145 | 0.008519 | 0.004659 | 0.005856 | 0.004712 | 0.019279 | 0.007206 | 0.002442 | 0.003951 | 0.000503 | 0.001644 | 0.002114 | 0.014506 | 0.004542 | 0.030311 | 0.000257 | 0.002705 | 0.005646 | 0.00061 |
| 34.1666667 | 0.007204 | 0.0339 | 0.024594 | 0.006508 | 0.01063 | 0.003113 | 0.006559 | 0.006627 | 0.004272 | 0.002647 | 0.034101 | 0.007423 | 0.001877 | 0.00496 | 0.000588 | 0.001127 | 0.001788 | 0.019907 | 0.004027 | 0.031153 | 0.000165 | 0.002098 | 0.002688 | 0.000587 |
| 34.1805556 | 0.003344 | 0.032253 | 0.012573 | 0.009994 | 0.014309 | 0.006038 | 0.003268 | 0.007606 | 0.004167 | 0.001091 | 0.047479 | 0.007769 | 0.001652 | 0.005841 | 0.000634 | 0.00062 | 0.001339 | 0.023429 | 0.002409 | 0.025522 | 0.000143 | 0.001109 | 0.003202 | 0.000815 |
| 34.1944444 | 0.005713 | 0.032456 | 0.005258 | 0.012742 | 0.012374 | 0.008517 | 0.002015 | 0.00757 | 0.007697 | 0.000723 | 0.054567 | 0.007805 | 0.002033 | 0.006018 | 0.000866 | 0.000388 | 0.000999 | 0.025119 | 0.001464 | 0.015045 | 0.000109 | 0.000494 | 0.003321 | 0.001897 |
| 34.2083333 | 0.007772 | 0.033033 | 0.006434 | 0.013843 | 0.011667 | 0.010673 | 0.002332 | 0.007343 | 0.012121 | 0.000739 | 0.056099 | 0.007877 | 0.003135 | 0.004842 | 0.001198 | 0.000754 | 0.000988 | 0.021821 | 0.002562 | 0.006042 | 0.000222 | 0.000558 | 0.002101 | 0.003397 |
| 34.2222222 | 0.008629 | 0.032415 | 0.008997 | 0.013883 | 0.020018 | 0.012775 | 0.003747 | 0.008338 | 0.01265 | 0.000826 | 0.0559 | 0.008461 | 0.004446 | 0.003189 | 0.001318 | 0.001062 | 0.001289 | 0.014711 | 0.005567 | 0.001911 | 0.001006 | 0.000533 | 0.000986 | 0.004814 |
| 34.2361111 | 0.009874 | 0.029364 | 0.008582 | 0.013961 | 0.028828 | 0.014003 | 0.004611 | 0.010127 | 0.007954 | 0.001363 | 0.055089 | 0.00878 | 0.004989 | 0.002391 | 0.001229 | 0.000864 | 0.001743 | 0.012873 | 0.009151 | 0.001202 | 0.002797 | 0.000216 | 0.000722 | 0.005775 |
| 34.25 | 0.010572 | 0.024571 | 0.005687 | 0.01391 | 0.031511 | 0.013658 | 0.003819 | 0.010511 | 0.003785 | 0.001837 | 0.053573 | 0.008003 | 0.004694 | 0.002172 | 0.000879 | 0.000522 | 0.002159 | 0.015505 | 0.011404 | 0.001224 | 0.004314 | 0.000182 | 0.001004 | 0.00509 |
| 34.2638889 | 0.010735 | 0.019726 | 0.002638 | 0.01194 | 0.025596 | 0.01221 | 0.003404 | 0.008754 | 0.003496 | 0.001963 | 0.052275 | 0.006757 | 0.004215 | 0.001499 | 0.000389 | 0.000467 | 0.002451 | 0.018099 | 0.011542 | 0.001462 | 0.004653 | 0.000495 | 0.001534 | 0.002746 |
| 34.2777778 | 0.011545 | 0.016753 | 0.001891 | 0.007872 | 0.016822 | 0.010877 | 0.003389 | 0.006741 | 0.002863 | 0.001975 | 0.050871 | 0.005664 | 0.003452 | 0.001027 | 0.000341 | 0.001051 | 0.002252 | 0.021304 | 0.010284 | 0.001466 | 0.004992 | 0.000735 | 0.002494 | 0.000875 |
| 34.2916667 | 0.014723 | 0.017365 | 0.00306 | 0.003737 | 0.021211 | 0.010118 | 0.002752 | 0.006509 | 0.002429 | 0.001418 | 0.048274 | 0.00393 | 0.002551 | 0.001459 | 0.000679 | 0.001909 | 0.00152 | 0.023984 | 0.008507 | 0.000732 | 0.005577 | 0.000884 | 0.002969 | 0.00074 |
| 34.3055556 | 0.01925 | 0.021525 | 0.003214 | 0.002071 | 0.036514 | 0.00991 | 0.001834 | 0.008244 | 0.004567 | 0.000959 | 0.043891 | 0.002307 | 0.002509 | 0.001634 | 0.000758 | 0.002081 | 0.001141 | 0.022328 | 0.007312 | 0.000308 | 0.005186 | 0.001294 | 0.001946 | 0.001342 |
| 34.3194444 | 0.0219 | 0.027615 | 0.001871 | 0.003304 | 0.045912 | 0.010079 | 0.000943 | 0.0106 | 0.006449 | 0.001866 | 0.037803 | 0.003132 | 0.00325 | 0.001102 | 0.000485 | 0.001426 | 0.001594 | 0.017157 | 0.00724 | 0.000569 | 0.003773 | 0.001763 | 0.000693 | 0.001567 |
| 34.3333333 | 0.022126 | 0.033755 | 0.001111 | 0.004975 | 0.043451 | 0.010526 | 0.000404 | 0.01219 | 0.005527 | 0.003432 | 0.031342 | 0.00579 | 0.003879 | 0.001295 | 0.000721 | 0.000784 | 0.002537 | 0.015146 | 0.008046 | 0.001918 | 0.002199 | 0.002138 | 0.000947 | 0.001178 |
| 34.3472222 | 0.019845 | 0.038201 | 0.001058 | 0.004871 | 0.031281 | 0.01135 | 0.000473 | 0.013128 | 0.00308 | 0.004556 | 0.026149 | 0.008378 | 0.004404 | 0.002676 | 0.002008 | 0.00088 | 0.003542 | 0.01943 | 0.009452 | 0.005292 | 0.001094 | 0.002779 | 0.002686 | 0.000598 |
| 34.3611111 | 0.014642 | 0.039994 | 0.001941 | 0.002827 | 0.015065 | 0.012 | 0.000778 | 0.014691 | 0.002893 | 0.004896 | 0.023662 | 0.010482 | 0.004554 | 0.003768 | 0.003148 | 0.001159 | 0.003861 | 0.025243 | 0.010204 | 0.008875 | 0.001347 | 0.003336 | 0.004173 | 0.000298 |
| 34.375 | 0.008904 | 0.040348 | 0.004455 | 0.000803 | 0.003794 | 0.012088 | 0.001177 | 0.017404 | 0.006003 | 0.004548 | 0.021377 | 0.012279 | 0.003607 | 0.003506 | 0.002879 | 0.00092 | 0.003186 | 0.025575 | 0.009326 | 0.008959 | 0.003103 | 0.003026 | 0.003553 | 0.000135 |
| 34.3888889 | 0.004987 | 0.040651 | 0.007032 | 0.000687 | 0.003808 | 0.011912 | 0.00214 | 0.020329 | 0.00951 | 0.004201 | 0.017448 | 0.013261 | 0.002693 | 0.002801 | 0.00223 | 0.000649 | 0.002822 | 0.020808 | 0.008634 | 0.006202 | 0.004382 | 0.002342 | 0.001649 | 0.000261 |
| 34.4027778 | 0.003492 | 0.040658 | 0.007987 | 0.002639 | 0.012603 | 0.012173 | 0.004349 | 0.02167 | 0.010192 | 0.003885 | 0.014802 | 0.012642 | 0.003026 | 0.0025 | 0.002487 | 0.000793 | 0.00358 | 0.016773 | 0.009358 | 0.004318 | 0.004116 | 0.002237 | 0.000685 | 0.000818 |
| 34.4166667 | 0.003345 | 0.040036 | 0.007563 | 0.004554 | 0.017455 | 0.013502 | 0.007908 | 0.021008 | 0.008713 | 0.003039 | 0.014574 | 0.011345 | 0.003277 | 0.001763 | 0.002484 | 0.000953 | 0.003922 | 0.014909 | 0.009276 | 0.003495 | 0.004076 | 0.002099 | 0.001137 | 0.001027 |
| 34.4305556 | 0.003241 | 0.038807 | 0.006944 | 0.003928 | 0.013107 | 0.014585 | 0.011952 | 0.02008 | 0.009753 | 0.001905 | 0.016556 | 0.011317 | 0.002662 | 0.00074 | 0.001322 | 0.001035 | 0.002896 | 0.014119 | 0.008306 | 0.002281 | 0.005238 | 0.001181 | 0.002066 | 0.000612 |
| 34.4444444 | 0.003911 | 0.037676 | 0.007448 | 0.001923 | 0.011952 | 0.013693 | 0.015755 | 0.01985 | 0.01501 | 0.001572 | 0.019896 | 0.0133 | 0.002925 | 0.000494 | 0.000972 | 0.001097 | 0.002351 | 0.013815 | 0.011264 | 0.001901 | 0.005628 | 0.000918 | 0.00234 | 0.000665 |
| 34.4583333 | 0.003828 | 0.037098 | 0.009948 | 0.000826 | 0.019254 | 0.011667 | 0.019002 | 0.019616 | 0.021196 | 0.001764 | 0.026758 | 0.015512 | 0.005034 | 0.001025 | 0.00281 | 0.000936 | 0.003723 | 0.012385 | 0.017567 | 0.003411 | 0.003792 | 0.002687 | 0.001856 | 0.001801 |
| 34.4722222 | 0.003104 | 0.035214 | 0.011721 | 0.000684 | 0.025727 | 0.01002 | 0.021206 | 0.01893 | 0.022946 | 0.002524 | 0.039177 | 0.015655 | 0.007002 | 0.002104 | 0.004571 | 0.000901 | 0.005232 | 0.009412 | 0.019898 | 0.00466 | 0.00179 | 0.004462 | 0.002062 | 0.002445 |
| 34.4861111 | 0.00874 | 0.028379 | 0.009129 | 0.001324 | 0.029084 | 0.009705 | 0.021184 | 0.018937 | 0.015944 | 0.007513 | 0.056102 | 0.012489 | 0.006562 | 0.002269 | 0.00363 | 0.00073 | 0.004365 | 0.005424 | 0.016624 | 0.003403 | 0.002961 | 0.003858 | 0.004193 | 0.00221 |
| 34.5 | 0.022074 | 0.015842 | 0.004644 | 0.004292 | 0.037588 | 0.01038 | 0.01672 | 0.019459 | 0.016129 | 0.017828 | 0.07164 | 0.008392 | 0.004147 | 0.001767 | 0.001807 | 0.000394 | 0.002398 | 0.002344 | 0.010063 | 0.001651 | 0.008343 | 0.003369 | 0.007477 | 0.003564 |
| 34.5138889 | 0.032531 | 0.007922 | 0.002693 | 0.010607 | 0.058513 | 0.010661 | 0.010957 | 0.019236 | 0.042457 | 0.026895 | 0.078812 | 0.01076 | 0.002481 | 0.00303 | 0.001227 | 0.000523 | 0.001407 | 0.001484 | 0.004515 | 0.001234 | 0.013804 | 0.004137 | 0.008789 | 0.005448 |
| 34.5277778 | 0.033073 | 0.013492 | 0.00579 | 0.01871 | 0.085144 | 0.0115 | 0.013592 | 0.023153 | 0.068662 | 0.027932 | 0.085405 | 0.021255 | 0.004545 | 0.005507 | 0.002198 | 0.001723 | 0.002187 | 0.002063 | 0.003188 | 0.002548 | 0.012161 | 0.003896 | 0.006594 | 0.00456 |
| 34.5416667 | 0.02927 | 0.026248 | 0.016332 | 0.02487 | 0.089209 | 0.01488 | 0.023173 | 0.03544 | 0.066517 | 0.021992 | 0.09315 | 0.03183 | 0.008464 | 0.006886 | 0.004721 | 0.002866 | 0.004521 | 0.003368 | 0.003049 | 0.005633 | 0.005534 | 0.003469 | 0.004228 | 0.002281 |
| 34.5555556 | 0.027825 | 0.038337 | 0.031146 | 0.024511 | 0.054669 | 0.019072 | 0.025993 | 0.048283 | 0.043707 | 0.015049 | 0.089623 | 0.037612 | 0.007522 | 0.00517 | 0.004974 | 0.003555 | 0.004836 | 0.003802 | 0.004985 | 0.005907 | 0.00307 | 0.003427 | 0.003389 | 0.002094 |
| 34.5694444 | 0.029543 | 0.045529 | 0.041882 | 0.017637 | 0.02605 | 0.020391 | 0.017585 | 0.054112 | 0.024272 | 0.011469 | 0.075832 | 0.039557 | 0.004018 | 0.00248 | 0.003296 | 0.005783 | 0.003894 | 0.003967 | 0.009605 | 0.003889 | 0.005048 | 0.003588 | 0.003564 | 0.003868 |
| 34.5833333 | 0.032629 | 0.046955 | 0.045753 | 0.009754 | 0.040353 | 0.019288 | 0.007458 | 0.054861 | 0.027422 | 0.013284 | 0.058965 | 0.03905 | 0.004322 | 0.001821 | 0.003163 | 0.00697 | 0.004214 | 0.004208 | 0.010367 | 0.003523 | 0.00593 | 0.003274 | 0.005084 | 0.004106 |
| 34.5972222 | 0.034468 | 0.04696 | 0.047828 | 0.005671 | 0.058904 | 0.017722 | 0.003433 | 0.054289 | 0.033871 | 0.018604 | 0.040722 | 0.03794 | 0.004344 | 0.001336 | 0.002411 | 0.004531 | 0.003175 | 0.002815 | 0.005947 | 0.002497 | 0.00387 | 0.002019 | 0.006268 | 0.002299 |
| 34.6111111 | 0.034568 | 0.048652 | 0.053395 | 0.006707 | 0.048343 | 0.016093 | 0.003166 | 0.05493 | 0.024305 | 0.021573 | 0.024343 | 0.036515 | 0.002367 | 0.000895 | 0.001364 | 0.00213 | 0.001601 | 0.001855 | 0.00308 | 0.001209 | 0.002182 | 0.001933 | 0.006577 | 0.001528 |
| 34.625 | 0.03335 | 0.050416 | 0.060225 | 0.010969 | 0.024406 | 0.014896 | 0.004958 | 0.055314 | 0.010146 | 0.019315 | 0.012213 | 0.032999 | 0.001779 | 0.001744 | 0.001919 | 0.00233 | 0.001945 | 0.002894 | 0.004101 | 0.001481 | 0.002484 | 0.002688 | 0.006877 | 0.002145 |
| 34.6388889 | 0.030988 | 0.048975 | 0.059894 | 0.013623 | 0.011274 | 0.014501 | 0.009217 | 0.050554 | 0.004118 | 0.014252 | 0.004928 | 0.026188 | 0.001538 | 0.002157 | 0.001922 | 0.002484 | 0.002097 | 0.00333 | 0.004145 | 0.001679 | 0.002385 | 0.002256 | 0.007034 | 0.001863 |
| 34.6527778 | 0.029529 | 0.046163 | 0.051079 | 0.011801 | 0.009938 | 0.014465 | 0.012415 | 0.039981 | 0.011129 | 0.010043 | 0.003579 | 0.017143 | 0.000998 | 0.00146 | 0.00113 | 0.001799 | 0.001337 | 0.002152 | 0.002411 | 0.002139 | 0.00263 | 0.001369 | 0.007197 | 0.001146 |
| 34.6666667 | 0.030004 | 0.046732 | 0.041214 | 0.00783 | 0.008889 | 0.013988 | 0.012623 | 0.026949 | 0.0273 | 0.007369 | 0.006446 | 0.008501 | 0.001299 | 0.00139 | 0.001379 | 0.002708 | 0.001484 | 0.000826 | 0.002337 | 0.003651 | 0.004374 | 0.001711 | 0.007223 | 0.001947 |
| 34.6805556 | 0.031535 | 0.048647 | 0.03436 | 0.004138 | 0.01609 | 0.013801 | 0.011989 | 0.013519 | 0.044483 | 0.005002 | 0.009283 | 0.005335 | 0.00164 | 0.002344 | 0.002016 | 0.004534 | 0.002131 | 0.000191 | 0.003695 | 0.004791 | 0.005792 | 0.002587 | 0.006734 | 0.003247 |
| 34.6944444 | 0.032635 | 0.046 | 0.029933 | 0.003898 | 0.043307 | 0.014219 | 0.012458 | 0.004777 | 0.055437 | 0.004354 | 0.009796 | 0.009303 | 0.00118 | 0.002868 | 0.001801 | 0.00511 | 0.002073 | 0.00033 | 0.004425 | 0.004721 | 0.005659 | 0.00224 | 0.006604 | 0.003374 |
| 34.7083333 | 0.033186 | 0.037678 | 0.026574 | 0.009685 | 0.077167 | 0.0143 | 0.01262 | 0.003909 | 0.054991 | 0.007317 | 0.007464 | 0.014908 | 0.00054 | 0.00242 | 0.001085 | 0.004212 | 0.001579 | 0.001109 | 0.003933 | 0.003818 | 0.004404 | 0.001108 | 0.007967 | 0.002355 |
| 34.7222222 | 0.034368 | 0.027139 | 0.023295 | 0.016244 | 0.092732 | 0.013846 | 0.009021 | 0.004136 | 0.046402 | 0.011669 | 0.007021 | 0.020096 | 0.000299 | 0.001708 | 0.000652 | 0.002642 | 0.001248 | 0.002306 | 0.00264 | 0.002651 | 0.003133 | 0.000622 | 0.008414 | 0.001121 |
| 34.7361111 | 0.036987 | 0.018592 | 0.018948 | 0.018162 | 0.082413 | 0.012672 | 0.006586 | 0.002552 | 0.039179 | 0.014371 | 0.019161 | 0.02736 | 0.000524 | 0.001327 | 0.000472 | 0.001407 | 0.000887 | 0.003662 | 0.001438 | 0.001557 | 0.002047 | 0.000762 | 0.005187 | 0.000869 |
| 34.75 | 0.041202 | 0.014165 | 0.012743 | 0.016826 | 0.057406 | 0.011116 | 0.013051 | 0.002773 | 0.037887 | 0.015411 | 0.038534 | 0.03486 | 0.001244 | 0.001192 | 0.000453 | 0.001366 | 0.000674 | 0.004981 | 0.001332 | 0.000739 | 0.001383 | 0.001034 | 0.002105 | 0.001985 |
| 34.7638889 | 0.045267 | 0.011842 | 0.007141 | 0.015444 | 0.033814 | 0.010461 | 0.021595 | 0.004716 | 0.03775 | 0.017203 | 0.044561 | 0.039219 | 0.002229 | 0.000922 | 0.000887 | 0.002224 | 0.001096 | 0.006226 | 0.002245 | 0.000585 | 0.00208 | 0.001285 | 0.003018 | 0.003737 |
| 34.7777778 | 0.046814 | 0.007654 | 0.005204 | 0.016034 | 0.023761 | 0.011512 | 0.024773 | 0.005483 | 0.032283 | 0.021068 | 0.035641 | 0.041221 | 0.003046 | 0.000549 | 0.001532 | 0.002855 | 0.001627 | 0.007364 | 0.003224 | 0.001076 | 0.00359 | 0.001595 | 0.005971 | 0.005402 |
| 34.7916667 | 0.044423 | 0.006206 | 0.005589 | 0.018183 | 0.030029 | 0.013013 | 0.02602 | 0.003972 | 0.023035 | 0.024441 | 0.030034 | 0.041186 | 0.003715 | 0.000527 | 0.002044 | 0.002935 | 0.002027 | 0.008168 | 0.004147 | 0.001555 | 0.005073 | 0.002347 | 0.008205 | 0.006465 |
| 34.8055556 | 0.037568 | 0.014491 | 0.005371 | 0.02028 | 0.044747 | 0.013486 | 0.02824 | 0.002651 | 0.01681 | 0.024834 | 0.034184 | 0.039269 | 0.004259 | 0.000798 | 0.00224 | 0.002629 | 0.002295 | 0.008337 | 0.004831 | 0.00167 | 0.006539 | 0.003475 | 0.008419 | 0.006557 |
| 34.8194444 | 0.028241 | 0.026969 | 0.005186 | 0.021833 | 0.052088 | 0.012765 | 0.029364 | 0.004779 | 0.012987 | 0.023076 | 0.038386 | 0.03721 | 0.004314 | 0.001378 | 0.001857 | 0.001878 | 0.001849 | 0.007681 | 0.004394 | 0.001351 | 0.007415 | 0.004097 | 0.006831 | 0.005857 |
| 34.8333333 | 0.019606 | 0.036649 | 0.006264 | 0.022804 | 0.048741 | 0.009878 | 0.027543 | 0.011287 | 0.007858 | 0.021783 | 0.037569 | 0.035059 | 0.00355 | 0.003261 | 0.00111 | 0.001656 | 0.001115 | 0.006295 | 0.002822 | 0.000895 | 0.007098 | 0.003608 | 0.004677 | 0.004503 |
| 34.8472222 | 0.011517 | 0.041986 | 0.007169 | 0.023595 | 0.049968 | 0.006067 | 0.024383 | 0.021234 | 0.005909 | 0.021922 | 0.031558 | 0.030922 | 0.002619 | 0.005395 | 0.000596 | 0.002391 | 0.000957 | 0.005255 | 0.001842 | 0.0008 | 0.006687 | 0.002868 | 0.003316 | 0.003505 |
| 34.8611111 | 0.006454 | 0.043565 | 0.006526 | 0.024373 | 0.062708 | 0.003998 | 0.022402 | 0.027842 | 0.010074 | 0.021958 | 0.021246 | 0.023449 | 0.002866 | 0.004573 | 0.000808 | 0.002187 | 0.001091 | 0.005955 | 0.003101 | 0.000839 | 0.007665 | 0.002965 | 0.00417 | 0.004624 |
| 34.875 | 0.009776 | 0.045353 | 0.005567 | 0.024105 | 0.08256 | 0.003344 | 0.02174 | 0.022931 | 0.017108 | 0.020687 | 0.010567 | 0.013353 | 0.004298 | 0.002165 | 0.002122 | 0.001637 | 0.001696 | 0.008432 | 0.006026 | 0.001146 | 0.009677 | 0.003675 | 0.007113 | 0.007483 |
| 34.8888889 | 0.018131 | 0.047174 | 0.00643 | 0.021594 | 0.099993 | 0.003246 | 0.019883 | 0.011207 | 0.025038 | 0.019142 | 0.006219 | 0.006259 | 0.005483 | 0.001454 | 0.00385 | 0.002468 | 0.002712 | 0.011019 | 0.009011 | 0.001914 | 0.011386 | 0.004137 | 0.009725 | 0.010227 |
| 34.9027778 | 0.023913 | 0.043926 | 0.00918 | 0.016382 | 0.103459 | 0.003937 | 0.014851 | 0.003438 | 0.032689 | 0.019288 | 0.011891 | 0.007469 | 0.005285 | 0.001421 | 0.004654 | 0.003059 | 0.002836 | 0.011841 | 0.011011 | 0.002329 | 0.012138 | 0.003632 | 0.010196 | 0.011662 |
| 34.9166667 | 0.024694 | 0.036441 | 0.012048 | 0.00945 | 0.092466 | 0.006079 | 0.007753 | 0.001906 | 0.038148 | 0.019225 | 0.017321 | 0.012022 | 0.003635 | 0.001743 | 0.004201 | 0.002432 | 0.001787 | 0.010638 | 0.011644 | 0.001671 | 0.012039 | 0.002074 | 0.008564 | 0.011141 |
| 34.9305556 | 0.021262 | 0.029208 | 0.012543 | 0.00372 | 0.075552 | 0.009908 | 0.004023 | 0.00355 | 0.042154 | 0.014019 | 0.015089 | 0.014534 | 0.001585 | 0.003405 | 0.003009 | 0.001366 | 0.001008 | 0.008426 | 0.010818 | 0.000798 | 0.011443 | 0.000794 | 0.005582 | 0.008951 |
| 34.9444444 | 0.016182 | 0.022163 | 0.00979 | 0.002241 | 0.058931 | 0.012887 | 0.006512 | 0.006993 | 0.044945 | 0.007396 | 0.012729 | 0.015123 | 0.000418 | 0.005051 | 0.001876 | 0.000543 | 0.001228 | 0.006196 | 0.008804 | 0.000778 | 0.010885 | 0.000564 | 0.002735 | 0.006568 |
| 34.9583333 | 0.012438 | 0.014624 | 0.007022 | 0.004926 | 0.043163 | 0.011345 | 0.01045 | 0.0103 | 0.045559 | 0.006053 | 0.018214 | 0.012928 | 0.000398 | 0.004561 | 0.001332 | 0.000248 | 0.001189 | 0.00449 | 0.006326 | 0.000762 | 0.010713 | 0.000606 | 0.001514 | 0.005404 |
| 34.9722222 | 0.013177 | 0.016451 | 0.007122 | 0.008241 | 0.031387 | 0.006903 | 0.012353 | 0.00966 | 0.043793 | 0.008128 | 0.037263 | 0.009654 | 0.000932 | 0.005785 | 0.001478 | 0.000326 | 0.000996 | 0.003966 | 0.003692 | 0.00063 | 0.010474 | 0.000798 | 0.00268 | 0.006077 |
| 34.9861111 | 0.019045 | 0.029827 | 0.007555 | 0.007354 | 0.03158 | 0.005935 | 0.010629 | 0.010565 | 0.034067 | 0.018543 | 0.066805 | 0.012204 | 0.001424 | 0.017827 | 0.002121 | 0.000612 | 0.002069 | 0.004571 | 0.002408 | 0.001311 | 0.008974 | 0.000874 | 0.006095 | 0.008471 |
| 35 | 0.023208 | 0.031277 | 0.0065 | 0.003919 | 0.047607 | 0.008827 | 0.00648 | 0.01832 | 0.019197 | 0.044846 | 0.085931 | 0.019401 | 0.002644 | 0.03558 | 0.002444 | 0.001039 | 0.003686 | 0.004365 | 0.002561 | 0.003228 | 0.006932 | 0.000706 | 0.010599 | 0.011337 |
| 35.0138889 | 0.019625 | 0.018758 | 0.006991 | 0.003148 | 0.067385 | 0.007904 | 0.006768 | 0.021865 | 0.013094 | 0.07378 | 0.078643 | 0.020369 | 0.004955 | 0.039357 | 0.001927 | 0.001715 | 0.003764 | 0.002526 | 0.005174 | 0.006467 | 0.006695 | 0.000993 | 0.014638 | 0.01253 |
| 35.0277778 | 0.010543 | 0.016584 | 0.008203 | 0.005817 | 0.071577 | 0.005238 | 0.012024 | 0.015876 | 0.014243 | 0.088978 | 0.052997 | 0.013075 | 0.005247 | 0.02375 | 0.002923 | 0.002593 | 0.003172 | 0.001777 | 0.01232 | 0.00731 | 0.006963 | 0.002779 | 0.014503 | 0.009338 |

| | | | | | | | | | | | | | | | | | | | | | | | | |
|---|---|---|---|---|---|---|---|---|---|---|---|---|---|---|---|---|---|---|---|---|---|---|---|---|
| 35.0416667 | 0.003036 | 0.025281 | 0.006157 | 0.010806 | 0.057795 | 0.008245 | 0.017425 | 0.008171 | 0.017106 | 0.09432 | 0.025733 | 0.00525 | 0.002806 | 0.008439 | 0.006693 | 0.002849 | 0.004732 | 0.003007 | 0.015667 | 0.007279 | 0.006019 | 0.006303 | 0.01026 | 0.006985 |
| 35.0555556 | 0.000751 | 0.030689 | 0.002395 | 0.013254 | 0.035614 | 0.012257 | 0.020455 | 0.0038 | 0.02244 | 0.093013 | 0.013093 | 0.001647 | 0.002051 | 0.00588 | 0.00813 | 0.002573 | 0.006109 | 0.00309 | 0.01046 | 0.009842 | 0.006382 | 0.007778 | 0.008539 | 0.010616 |
| 35.0694444 | 0.001145 | 0.02963 | 0.000635 | 0.011472 | 0.014915 | 0.013185 | 0.019635 | 0.00277 | 0.023904 | 0.078422 | 0.017688 | 0.001946 | 0.0042 | 0.005332 | 0.004799 | 0.003292 | 0.004205 | 0.002575 | 0.005066 | 0.008167 | 0.008689 | 0.004852 | 0.008913 | 0.01151 |
| 35.0833333 | 0.003362 | 0.02712 | 0.000997 | 0.010128 | 0.003827 | 0.013037 | 0.016567 | 0.002823 | 0.019099 | 0.053228 | 0.022984 | 0.004016 | 0.006535 | 0.003077 | 0.001814 | 0.004097 | 0.002357 | 0.003685 | 0.003624 | 0.003306 | 0.009855 | 0.001439 | 0.007207 | 0.006715 |
| 35.0972222 | 0.008307 | 0.027746 | 0.002899 | 0.010213 | 0.001113 | 0.013119 | 0.013645 | 0.002429 | 0.01328 | 0.027949 | 0.021795 | 0.004408 | 0.008252 | 0.00417 | 0.00179 | 0.002872 | 0.00284 | 0.004631 | 0.003757 | 0.002138 | 0.008752 | 0.00045 | 0.005983 | 0.002776 |
| 35.1111111 | 0.013209 | 0.031537 | 0.006041 | 0.009722 | 0.00286 | 0.013391 | 0.010326 | 0.003247 | 0.009708 | 0.010501 | 0.018272 | 0.00296 | 0.009931 | 0.007478 | 0.002093 | 0.001424 | 0.003142 | 0.004066 | 0.004083 | 0.002721 | 0.006645 | 0.000948 | 0.007063 | 0.001738 |
| 35.125 | 0.015473 | 0.034111 | 0.008295 | 0.008413 | 0.009717 | 0.012764 | 0.005623 | 0.00694 | 0.008427 | 0.004908 | 0.014654 | 0.002817 | 0.011846 | 0.010806 | 0.001572 | 0.001755 | 0.002381 | 0.002729 | 0.00463 | 0.002376 | 0.004644 | 0.001755 | 0.007484 | 0.002221 |
| 35.1388889 | 0.015202 | 0.031821 | 0.008369 | 0.007058 | 0.020585 | 0.011133 | 0.002283 | 0.011526 | 0.007714 | 0.010724 | 0.012982 | 0.004259 | 0.014071 | 0.011449 | 0.001812 | 0.001953 | 0.002314 | 0.001916 | 0.005807 | 0.002062 | 0.003741 | 0.001706 | 0.005102 | 0.002234 |
| 35.1527778 | 0.013706 | 0.027314 | 0.006957 | 0.006958 | 0.032048 | 0.009793 | 0.002757 | 0.014568 | 0.006266 | 0.023682 | 0.014758 | 0.005791 | 0.015186 | 0.007945 | 0.003053 | 0.001576 | 0.003396 | 0.002037 | 0.007343 | 0.002633 | 0.003893 | 0.000823 | 0.00201 | 0.001328 |
| 35.1666667 | 0.012583 | 0.025282 | 0.005473 | 0.008336 | 0.043151 | 0.00952 | 0.00378 | 0.015464 | 0.004816 | 0.036988 | 0.019259 | 0.006584 | 0.013527 | 0.003748 | 0.003262 | 0.001383 | 0.004139 | 0.001687 | 0.008368 | 0.002887 | 0.003308 | 0.000666 | 0.000902 | 0.001255 |
| 35.1805556 | 0.01256 | 0.025972 | 0.005454 | 0.010404 | 0.053702 | 0.009409 | 0.002712 | 0.014729 | 0.005113 | 0.043848 | 0.024239 | 0.006495 | 0.010312 | 0.001995 | 0.00187 | 0.001012 | 0.003862 | 0.000899 | 0.009468 | 0.002289 | 0.001697 | 0.001475 | 0.001435 | 0.00278 |
| 35.1944444 | 0.013624 | 0.026772 | 0.007454 | 0.013387 | 0.062125 | 0.009261 | 0.001685 | 0.012593 | 0.007804 | 0.04262 | 0.027294 | 0.005991 | 0.008325 | 0.002208 | 0.000666 | 0.001199 | 0.003439 | 0.000822 | 0.011624 | 0.001821 | 0.000645 | 0.002149 | 0.002196 | 0.004081 |
| 35.2083333 | 0.014632 | 0.026633 | 0.010593 | 0.016137 | 0.070052 | 0.010251 | 0.002466 | 0.009441 | 0.010458 | 0.036603 | 0.02757 | 0.005723 | 0.008259 | 0.003086 | 0.000313 | 0.001899 | 0.003335 | 0.000716 | 0.013983 | 0.001686 | 0.000391 | 0.002265 | 0.002609 | 0.004004 |
| 35.2222222 | 0.014033 | 0.024774 | 0.013537 | 0.018079 | 0.076662 | 0.012351 | 0.00308 | 0.007572 | 0.010185 | 0.02966 | 0.025769 | 0.005474 | 0.00868 | 0.003785 | 0.000219 | 0.002628 | 0.003143 | 0.00027 | 0.015151 | 0.00123 | 0.00079 | 0.0022 | 0.003139 | 0.003692 |
| 35.2361111 | 0.012098 | 0.019487 | 0.015577 | 0.020233 | 0.077995 | 0.014191 | 0.002151 | 0.008389 | 0.008763 | 0.024516 | 0.022874 | 0.00477 | 0.008817 | 0.00377 | 0.000213 | 0.003084 | 0.002807 | 7.53E-05 | 0.015038 | 0.000678 | 0.001859 | 0.00195 | 0.003741 | 0.003745 |
| 35.25 | 0.010589 | 0.011783 | 0.016304 | 0.021613 | 0.070863 | 0.014718 | 0.001255 | 0.009801 | 0.008971 | 0.022559 | 0.02046 | 0.003802 | 0.008935 | 0.002957 | 0.000516 | 0.002485 | 0.002872 | 0.000228 | 0.014452 | 0.001007 | 0.002494 | 0.001336 | 0.003438 | 0.003226 |
| 35.2638889 | 0.009592 | 0.005409 | 0.01474 | 0.021563 | 0.053194 | 0.013837 | 0.001755 | 0.00915 | 0.010199 | 0.02403 | 0.018842 | 0.002436 | 0.008805 | 0.001685 | 0.001243 | 0.001194 | 0.003488 | 0.000696 | 0.013774 | 0.002173 | 0.002066 | 0.001291 | 0.002377 | 0.001864 |
| 35.2777778 | 0.008082 | 0.002262 | 0.01071 | 0.021186 | 0.034042 | 0.012134 | 0.003625 | 0.00646 | 0.010327 | 0.029412 | 0.018673 | 0.001498 | 0.008183 | 0.001014 | 0.001673 | 0.000482 | 0.003981 | 0.000805 | 0.013711 | 0.002981 | 0.00173 | 0.002319 | 0.001402 | 0.001028 |
| 35.2916667 | 0.007091 | 0.001229 | 0.006288 | 0.020919 | 0.02319 | 0.010276 | 0.005843 | 0.003914 | 0.00963 | 0.038243 | 0.02167 | 0.001495 | 0.007514 | 0.001423 | 0.00129 | 0.000779 | 0.003804 | 0.000543 | 0.015716 | 0.002827 | 0.002753 | 0.003117 | 0.001056 | 0.001774 |
| 35.3055556 | 0.008085 | 0.001303 | 0.003327 | 0.018424 | 0.014687 | 0.008681 | 0.007032 | 0.002919 | 0.010308 | 0.046319 | 0.026818 | 0.001086 | 0.007457 | 0.001401 | 0.000728 | 0.001354 | 0.003276 | 0.000729 | 0.018791 | 0.002125 | 0.004776 | 0.002968 | 0.001616 | 0.002575 |
| 35.3194444 | 0.010026 | 0.002605 | 0.001832 | 0.012324 | 0.008347 | 0.007687 | 0.006811 | 0.003068 | 0.012206 | 0.048333 | 0.03142 | 0.000573 | 0.008456 | 0.000796 | 0.000611 | 0.001461 | 0.002956 | 0.001066 | 0.02 | 0.001536 | 0.00651 | 0.002238 | 0.002611 | 0.001865 |
| 35.3333333 | 0.011454 | 0.005199 | 0.001918 | 0.007006 | 0.009785 | 0.007596 | 0.005894 | 0.003812 | 0.013891 | 0.04333 | 0.033852 | 0.001172 | 0.010433 | 0.000616 | 0.000986 | 0.001009 | 0.003041 | 0.000857 | 0.019425 | 0.001491 | 0.007159 | 0.001356 | 0.0028 | 0.000672 |
| 35.3472222 | 0.012357 | 0.008156 | 0.003598 | 0.005016 | 0.01026 | 0.007887 | 0.005084 | 0.004534 | 0.015064 | 0.035457 | 0.033318 | 0.002399 | 0.013001 | 0.001031 | 0.00166 | 0.000704 | 0.003579 | 0.000759 | 0.019268 | 0.001976 | 0.006608 | 0.000945 | 0.001807 | 0.000515 |
| 35.3611111 | 0.012798 | 0.01061 | 0.005576 | 0.003273 | 0.009773 | 0.008044 | 0.004787 | 0.004444 | 0.016998 | 0.028737 | 0.031028 | 0.00328 | 0.01509 | 0.002196 | 0.002393 | 0.001191 | 0.004494 | 0.001519 | 0.019917 | 0.002698 | 0.005324 | 0.001375 | 0.001103 | 0.001381 |
| 35.375 | 0.01298 | 0.012855 | 0.006296 | 0.00138 | 0.013888 | 0.008763 | 0.004257 | 0.004477 | 0.020771 | 0.024943 | 0.029492 | 0.003647 | 0.015489 | 0.003094 | 0.002913 | 0.001889 | 0.005444 | 0.002164 | 0.020457 | 0.003221 | 0.004276 | 0.002218 | 0.001352 | 0.002046 |
| 35.3888889 | 0.013615 | 0.015207 | 0.005452 | 0.000957 | 0.014656 | 0.010372 | 0.002701 | 0.006873 | 0.023752 | 0.02434 | 0.029875 | 0.003868 | 0.014837 | 0.002994 | 0.003154 | 0.002186 | 0.006104 | 0.002207 | 0.020513 | 0.00312 | 0.003858 | 0.002745 | 0.001463 | 0.001831 |
| 35.4027778 | 0.015374 | 0.017817 | 0.004485 | 0.001007 | 0.008631 | 0.011695 | 0.001114 | 0.011877 | 0.023975 | 0.02495 | 0.032254 | 0.004026 | 0.014363 | 0.002409 | 0.003275 | 0.00238 | 0.006288 | 0.002284 | 0.020173 | 0.002557 | 0.003797 | 0.002739 | 0.001223 | 0.001338 |
| 35.4166667 | 0.018113 | 0.020604 | 0.004227 | 0.000881 | 0.004028 | 0.011788 | 0.000905 | 0.016452 | 0.021556 | 0.024362 | 0.036374 | 0.003518 | 0.013561 | 0.001481 | 0.002883 | 0.002194 | 0.005754 | 0.001969 | 0.018781 | 0.001762 | 0.004427 | 0.002064 | 0.000955 | 0.000958 |
| 35.4305556 | 0.020606 | 0.023323 | 0.003374 | 0.000915 | 0.007954 | 0.010133 | 0.001577 | 0.018386 | 0.014265 | 0.02208 | 0.043223 | 0.003788 | 0.012136 | 0.000695 | 0.00175 | 0.001275 | 0.004757 | 0.001045 | 0.016746 | 0.000828 | 0.005698 | 0.001024 | 0.000857 | 0.001068 |
| 35.4444444 | 0.022603 | 0.025184 | 0.0022 | 0.001459 | 0.021254 | 0.007356 | 0.001572 | 0.018687 | 0.006248 | 0.019959 | 0.051802 | 0.005774 | 0.012218 | 0.000492 | 0.001652 | 0.001365 | 0.004977 | 0.001233 | 0.017364 | 0.00113 | 0.00562 | 0.000877 | 0.001111 | 0.001509 |
| 35.4583333 | 0.025469 | 0.021847 | 0.002787 | 0.00538 | 0.039548 | 0.007177 | 0.001479 | 0.016584 | 0.009012 | 0.02003 | 0.051431 | 0.010323 | 0.015101 | 0.000342 | 0.004007 | 0.003653 | 0.007316 | 0.003636 | 0.021409 | 0.003494 | 0.003431 | 0.002393 | 0.001829 | 0.001356 |
| 35.4722222 | 0.031435 | 0.01178 | 0.006712 | 0.014476 | 0.057438 | 0.010249 | 0.002434 | 0.013822 | 0.020527 | 0.01972 | 0.041415 | 0.018835 | 0.01778 | 0.000525 | 0.006503 | 0.005825 | 0.009318 | 0.006049 | 0.024792 | 0.005393 | 0.001327 | 0.004323 | 0.002588 | 0.001063 |
| 35.4861111 | 0.040137 | 0.007811 | 0.017928 | 0.019751 | 0.080235 | 0.009774 | 0.002539 | 0.022639 | 0.030638 | 0.016681 | 0.050509 | 0.02288 | 0.016937 | 0.001414 | 0.006151 | 0.005821 | 0.007866 | 0.005668 | 0.023843 | 0.004261 | 0.002167 | 0.004339 | 0.002249 | 0.000864 |
| 35.5 | 0.044376 | 0.035468 | 0.040866 | 0.024056 | 0.120198 | 0.023246 | 0.001332 | 0.050822 | 0.077258 | 0.013976 | 0.112737 | 0.043172 | 0.013218 | 0.002982 | 0.003441 | 0.00458 | 0.004963 | 0.003141 | 0.01673 | 0.002181 | 0.007536 | 0.002564 | 0.002743 | 0.000622 |
| 35.5138889 | 0.039255 | 0.150805 | 0.080378 | 0.057291 | 0.170838 | 0.058178 | 0.008805 | 0.064776 | 0.154073 | 0.016471 | 0.216195 | 0.096382 | 0.008335 | 0.005025 | 0.001033 | 0.003348 | 0.004859 | 0.001335 | 0.009675 | 0.001547 | 0.014204 | 0.001398 | 0.005933 | 0.000978 |
| 35.5277778 | 0.035828 | 0.215106 | 0.092615 | 0.076463 | 0.164191 | 0.058628 | 0.032853 | 0.06078 | 0.145527 | 0.024564 | 0.242475 | 0.111631 | 0.003569 | 0.005101 | 0.001254 | 0.002276 | 0.004961 | 0.002019 | 0.010667 | 0.002488 | 0.015341 | 0.00107 | 0.008285 | 0.001736 |
| 35.5416667 | 0.036786 | 0.174449 | 0.047095 | 0.052421 | 0.110052 | 0.05741 | 0.052731 | 0.055779 | 0.112483 | 0.058993 | 0.174082 | 0.098894 | 0.001375 | 0.003445 | 0.004279 | 0.001554 | 0.003739 | 0.00411 | 0.012947 | 0.005846 | 0.009646 | 0.000942 | 0.006275 | 0.001794 |
| 35.5555556 | 0.031766 | 0.183224 | 0.013786 | 0.064869 | 0.082873 | 0.089557 | 0.050018 | 0.03606 | 0.167523 | 0.099997 | 0.128742 | 0.111968 | 0.003101 | 0.004136 | 0.007381 | 0.001321 | 0.004905 | 0.005036 | 0.010554 | 0.008822 | 0.003552 | 0.001664 | 0.003788 | 0.001523 |
| 35.5694444 | 0.028356 | 0.136758 | 0.020517 | 0.092444 | 0.063635 | 0.072766 | 0.039888 | 0.025318 | 0.169026 | 0.082456 | 0.094606 | 0.085036 | 0.006137 | 0.00617 | 0.007778 | 0.001166 | 0.006257 | 0.003794 | 0.008117 | 0.007401 | 0.001722 | 0.001888 | 0.004802 | 0.001512 |
| 35.5833333 | 0.032019 | 0.048212 | 0.033834 | 0.073086 | 0.054467 | 0.032825 | 0.033634 | 0.029439 | 0.074099 | 0.036319 | 0.065858 | 0.029739 | 0.006463 | 0.005052 | 0.005013 | 0.001324 | 0.004392 | 0.002242 | 0.007577 | 0.003681 | 0.003602 | 0.001468 | 0.00475 | 0.002016 |
| 35.5972222 | 0.035947 | 0.02799 | 0.042815 | 0.037769 | 0.047221 | 0.026163 | 0.028247 | 0.036124 | 0.013894 | 0.013629 | 0.068781 | 0.00956 | 0.00422 | 0.002495 | 0.001992 | 0.002131 | 0.001783 | 0.002326 | 0.006927 | 0.001796 | 0.006731 | 0.002117 | 0.002524 | 0.003453 |
| 35.6111111 | 0.037874 | 0.030395 | 0.041656 | 0.015347 | 0.040055 | 0.027923 | 0.023473 | 0.040603 | 0.005256 | 0.00803 | 0.062447 | 0.007288 | 0.002531 | 0.002276 | 0.001316 | 0.002663 | 0.000771 | 0.002694 | 0.005223 | 0.001814 | 0.00835 | 0.002713 | 0.001722 | 0.003515 |
| 35.625 | 0.037926 | 0.039043 | 0.03216 | 0.011265 | 0.068271 | 0.025094 | 0.01974 | 0.037057 | 0.008812 | 0.00774 | 0.040827 | 0.004829 | 0.002018 | 0.002452 | 0.001472 | 0.002614 | 0.000548 | 0.002096 | 0.002975 | 0.001666 | 0.007945 | 0.00252 | 0.001524 | 0.001883 |
| 35.6388889 | 0.036537 | 0.044165 | 0.023443 | 0.019568 | 0.104685 | 0.020447 | 0.015999 | 0.032019 | 0.013552 | 0.005518 | 0.024146 | 0.004149 | 0.001577 | 0.001422 | 0.001332 | 0.002462 | 0.0005 | 0.001463 | 0.001205 | 0.001327 | 0.007049 | 0.00232 | 0.001639 | 0.001472 |
| 35.6527778 | 0.034643 | 0.043205 | 0.019068 | 0.029 | 0.11759 | 0.01917 | 0.010933 | 0.027833 | 0.012781 | 0.005191 | 0.020416 | 0.004193 | 0.001007 | 0.000736 | 0.001477 | 0.002269 | 0.000599 | 0.001337 | 0.000575 | 0.001534 | 0.006691 | 0.002056 | 0.003116 | 0.002452 |
| 35.6666667 | 0.033131 | 0.045897 | 0.016564 | 0.034876 | 0.108187 | 0.021407 | 0.007186 | 0.027092 | 0.012186 | 0.012239 | 0.02629 | 0.003321 | 0.000602 | 0.000668 | 0.001951 | 0.002246 | 0.000627 | 0.001654 | 0.000559 | 0.002264 | 0.006815 | 0.001684 | 0.005066 | 0.003341 |
| 35.6805556 | 0.030823 | 0.067077 | 0.013714 | 0.036599 | 0.09372 | 0.023996 | 0.006888 | 0.025702 | 0.015155 | 0.022807 | 0.033053 | 0.004344 | 0.000586 | 0.001064 | 0.002468 | 0.002659 | 0.00111 | 0.002258 | 0.000774 | 0.003339 | 0.006936 | 0.00154 | 0.006302 | 0.003991 |
| 35.6944444 | 0.026967 | 0.102393 | 0.011152 | 0.033245 | 0.072052 | 0.025095 | 0.00657 | 0.021705 | 0.013788 | 0.031189 | 0.039644 | 0.005678 | 0.000651 | 0.002329 | 0.002574 | 0.003124 | 0.002133 | 0.002636 | 0.00121 | 0.004224 | 0.006627 | 0.001455 | 0.006341 | 0.004499 |
| 35.7083333 | 0.023704 | 0.123464 | 0.00996 | 0.021962 | 0.037749 | 0.025454 | 0.004883 | 0.019933 | 0.011018 | 0.034054 | 0.046565 | 0.004839 | 0.000481 | 0.003481 | 0.002089 | 0.00312 | 0.002874 | 0.002529 | 0.001574 | 0.004369 | 0.005917 | 0.001263 | 0.005444 | 0.004424 |
| 35.7222222 | 0.024289 | 0.116454 | 0.010062 | 0.011467 | 0.019067 | 0.025632 | 0.004117 | 0.020553 | 0.011394 | 0.030266 | 0.048135 | 0.00652 | 0.000559 | 0.00348 | 0.001245 | 0.002825 | 0.002832 | 0.002211 | 0.00201 | 0.003792 | 0.005182 | 0.001367 | 0.004312 | 0.003661 |
| 35.7361111 | 0.028651 | 0.098019 | 0.01147 | 0.014831 | 0.028914 | 0.026402 | 0.004439 | 0.021038 | 0.009988 | 0.02276 | 0.045146 | 0.012721 | 0.001017 | 0.002713 | 0.00064 | 0.002375 | 0.002156 | 0.001789 | 0.002367 | 0.002678 | 0.004447 | 0.001513 | 0.003444 | 0.002471 |
| 35.75 | 0.033237 | 0.083841 | 0.013097 | 0.025306 | 0.039348 | 0.026962 | 0.005963 | 0.017967 | 0.007754 | 0.014711 | 0.038177 | 0.021095 | 0.001462 | 0.001683 | 0.000652 | 0.00163 | 0.0014 | 0.001337 | 0.002185 | 0.001345 | 0.003641 | 0.00105 | 0.003112 | 0.001311 |
| 35.7638889 | 0.035995 | 0.073134 | 0.013473 | 0.029134 | 0.02935 | 0.025837 | 0.008283 | 0.010431 | 0.009669 | 0.008783 | 0.027001 | 0.027643 | 0.001945 | 0.000761 | 0.000731 | 0.000983 | 0.000839 | 0.000884 | 0.001725 | 0.000503 | 0.002879 | 0.000596 | 0.003255 | 0.00073 |
| 35.7777778 | 0.035409 | 0.056314 | 0.013574 | 0.019874 | 0.017457 | 0.026307 | 0.00824 | 0.005031 | 0.011749 | 0.00896 | 0.021728 | 0.027087 | 0.002723 | 0.000971 | 0.000465 | 0.000951 | 0.000458 | 0.000595 | 0.001123 | 0.000341 | 0.001919 | 0.00091 | 0.003423 | 0.001238 |
| 35.7916667 | 0.033449 | 0.036937 | 0.01415 | 0.010177 | 0.029155 | 0.028942 | 0.005764 | 0.004421 | 0.011907 | 0.015207 | 0.023912 | 0.022532 | 0.00366 | 0.002589 | 0.000427 | 0.00158 | 0.000288 | 0.000788 | 0.00074 | 0.000489 | 0.000842 | 0.00136 | 0.003329 | 0.002484 |
| 35.8055556 | 0.035837 | 0.028448 | 0.014201 | 0.014041 | 0.051166 | 0.027206 | 0.003281 | 0.008925 | 0.013782 | 0.022867 | 0.023257 | 0.019254 | 0.004486 | 0.004171 | 0.001206 | 0.002416 | 0.000171 | 0.001001 | 0.001433 | 0.000864 | 0.00023 | 0.001454 | 0.003016 | 0.002833 |
| 35.8194444 | 0.04261 | 0.031715 | 0.013785 | 0.01964 | 0.062975 | 0.018288 | 0.002548 | 0.017501 | 0.015631 | 0.027589 | 0.018528 | 0.016919 | 0.005621 | 0.004224 | 0.001876 | 0.002242 | 0.00064 | 0.001293 | 0.003546 | 0.001178 | 0.000682 | 0.002037 | 0.002264 | 0.001989 |
| 35.8333333 | 0.047384 | 0.038059 | 0.014695 | 0.019599 | 0.072021 | 0.008297 | 0.005363 | 0.022137 | 0.016237 | 0.027838 | 0.0155 | 0.015292 | 0.007365 | 0.003328 | 0.001391 | 0.001178 | 0.002043 | 0.002206 | 0.00694 | 0.001619 | 0.002281 | 0.003614 | 0.001757 | 0.001606 |
| 35.8472222 | 0.045154 | 0.037027 | 0.017475 | 0.021575 | 0.08598 | 0.003379 | 0.010842 | 0.021957 | 0.017881 | 0.025703 | 0.015118 | 0.013103 | 0.008697 | 0.004477 | 0.000638 | 0.000665 | 0.002922 | 0.002742 | 0.010005 | 0.002079 | 0.003381 | 0.004669 | 0.00205 | 0.001968 |
| 35.8611111 | 0.038172 | 0.024808 | 0.02013 | 0.026802 | 0.099426 | 0.002627 | 0.015978 | 0.020949 | 0.017815 | 0.023712 | 0.01506 | 0.008044 | 0.00915 | 0.007443 | 0.000615 | 0.00047 | 0.002658 | 0.002489 | 0.011576 | 0.002068 | 0.003244 | 0.004621 | 0.002292 | 0.002679 |
| 35.875 | 0.032208 | 0.010117 | 0.020976 | 0.027267 | 0.106361 | 0.002427 | 0.017482 | 0.020735 | 0.014488 | 0.023009 | 0.012934 | 0.003696 | 0.009507 | 0.007728 | 0.00125 | 0.000489 | 0.002521 | 0.002588 | 0.012493 | 0.002512 | 0.003445 | 0.004895 | 0.002229 | 0.003903 |
| 35.8888889 | 0.02912 | 0.002572 | 0.019465 | 0.021112 | 0.105482 | 0.002272 | 0.014232 | 0.021345 | 0.013511 | 0.022797 | 0.007454 | 0.003266 | 0.009507 | 0.005091 | 0.001646 | 0.000708 | 0.002392 | 0.002641 | 0.012123 | 0.00303 | 0.003958 | 0.004827 | 0.001605 | 0.004416 |
| 35.9027778 | 0.026717 | 0.001003 | 0.016375 | 0.017203 | 0.099293 | 0.002218 | 0.0094 | 0.020171 | 0.0134 | 0.021181 | 0.002378 | 0.003986 | 0.008832 | 0.003044 | 0.001089 | 0.000506 | 0.001685 | 0.001942 | 0.009928 | 0.002363 | 0.004005 | 0.003346 | 0.000686 | 0.003699 |
| 35.9166667 | 0.023222 | 0.000813 | 0.013652 | 0.016923 | 0.089634 | 0.002923 | 0.005086 | 0.01824 | 0.009423 | 0.0184 | 0.001306 | 0.004902 | 0.008768 | 0.002599 | 0.000695 | 0.00055 | 0.001662 | 0.002084 | 0.008533 | 0.002013 | 0.005049 | 0.002456 | 0.001229 | 0.003686 |
| 35.9305556 | 0.019995 | 0.000741 | 0.013179 | 0.013457 | 0.07193 | 0.006232 | 0.003614 | 0.017923 | 0.005984 | 0.015272 | 0.001622 | 0.006398 | 0.009385 | 0.003576 | 0.001318 | 0.001186 | 0.002667 | 0.003626 | 0.009027 | 0.003169 | 0.007509 | 0.002971 | 0.003547 | 0.004893 |
| 35.9444444 | 0.019056 | 0.00156 | 0.015656 | 0.007336 | 0.047852 | 0.011955 | 0.007886 | 0.017129 | 0.004413 | 0.013088 | 0.001083 | 0.007585 | 0.008728 | 0.004469 | 0.001628 | 0.001411 | 0.002885 | 0.004225 | 0.008859 | 0.003569 | 0.008915 | 0.002579 | 0.004824 | 0.004977 |
| 35.9583333 | 0.019548 | 0.005063 | 0.018613 | 0.004708 | 0.031044 | 0.017769 | 0.01409 | 0.014033 | 0.005845 | 0.014436 | 0.001547 | 0.006624 | 0.006241 | 0.003427 | 0.000959 | 0.000832 | 0.001709 | 0.002838 | 0.006521 | 0.00209 | 0.007846 | 0.001636 | 0.003664 | 0.003263 |
| 35.9722222 | 0.018084 | 0.009612 | 0.015862 | 0.006428 | 0.031581 | 0.01818 | 0.017622 | 0.013625 | 0.016502 | 0.022222 | 0.007362 | 0.013841 | 0.003362 | 0.003929 | 0.001231 | 0.001332 | 0.001021 | 0.001874 | 0.003967 | 0.001682 | 0.005602 | 0.001429 | 0.003131 | 0.003162 |
| 35.9861111 | 0.014307 | 0.009017 | 0.010242 | 0.006475 | 0.047483 | 0.012105 | 0.016893 | 0.021473 | 0.030526 | 0.040341 | 0.019678 | 0.038936 | 0.001514 | 0.012232 | 0.004321 | 0.005347 | 0.0029 | 0.003315 | 0.003432 | 0.005183 | 0.004042 | 0.001738 | 0.006073 | 0.007699 |
| 36 | 0.017639 | 0.013984 | 0.008028 | 0.010365 | 0.065901 | 0.008581 | 0.011659 | 0.02967 | 0.028321 | 0.061872 | 0.022322 | 0.045423 | 0.001317 | 0.02758 | 0.008388 | 0.010563 | 0.00655 | 0.004987 | 0.006504 | 0.011904 | 0.004302 | 0.003755 | 0.010778 | 0.014651 |
| 36.0138889 | 0.031269 | 0.040683 | 0.007724 | 0.018686 | 0.076918 | 0.009536 | 0.009161 | 0.028213 | 0.014858 | 0.07141 | 0.016823 | 0.033825 | 0.003788 | 0.038285 | 0.008367 | 0.010356 | 0.00782 | 0.004525 | 0.01692 | 0.019254 | 0.005183 | 0.005423 | 0.012621 | 0.017638 |
| 36.0277778 | 0.041005 | 0.062499 | 0.007228 | 0.018099 | 0.09132 | 0.010064 | 0.012095 | 0.026579 | 0.01117 | 0.081939 | 0.01863 | 0.036647 | 0.007264 | 0.031987 | 0.006348 | 0.006825 | 0.00605 | 0.004779 | 0.033886 | 0.021288 | 0.005613 | 0.004559 | 0.008781 | 0.013017 |
| 36.0416667 | 0.039328 | 0.050381 | 0.005533 | 0.016956 | 0.117062 | 0.015529 | 0.012218 | 0.033813 | 0.018721 | 0.113093 | 0.015476 | 0.047321 | 0.006498 | 0.015182 | 0.008423 | 0.006883 | 0.005351 | 0.006403 | 0.044342 | 0.016712 | 0.006781 | 0.004763 | 0.003632 | 0.00885 |
| 36.0555556 | 0.034799 | 0.032784 | 0.004526 | 0.020213 | 0.127255 | 0.019308 | 0.009655 | 0.034793 | 0.02431 | 0.138087 | 0.013559 | 0.05178 | 0.003112 | 0.004623 | 0.00955 | 0.006363 | 0.004744 | 0.005188 | 0.037197 | 0.012901 | 0.006539 | 0.006654 | 0.002112 | 0.010536 |
| 36.0694444 | 0.037458 | 0.04482 | 0.005758 | 0.019991 | 0.115631 | 0.011021 | 0.009498 | 0.022439 | 0.024951 | 0.124533 | 0.031612 | 0.047239 | 0.002788 | 0.002274 | 0.005485 | 0.003017 | 0.002795 | 0.002679 | 0.020269 | 0.009903 | 0.004777 | 0.00645 | 0.002422 | 0.009705 |
| 36.0833333 | 0.042108 | 0.062442 | 0.006892 | 0.018759 | 0.097682 | 0.005938 | 0.009386 | 0.012342 | 0.025605 | 0.089501 | 0.047782 | 0.049232 | 0.004467 | 0.001982 | 0.001876 | 0.001383 | 0.002395 | 0.00185 | 0.007231 | 0.005938 | 0.004485 | 0.00489 | 0.004417 | 0.005787 |
| 36.0972222 | 0.033716 | 0.046676 | 0.006453 | 0.019249 | 0.06439 | 0.019729 | 0.006208 | 0.009358 | 0.02095 | 0.068299 | 0.034709 | 0.051882 | 0.004751 | 0.002409 | 0.001306 | 0.001016 | 0.002447 | 0.001101 | 0.001769 | 0.003377 | 0.00378 | 0.004344 | 0.008173 | 0.004213 |
| 36.1111111 | 0.017191 | 0.017816 | 0.007133 | 0.020934 | 0.029967 | 0.040794 | 0.002701 | 0.006407 | 0.023407 | 0.063286 | 0.011605 | 0.056178 | 0.002936 | 0.004615 | 0.001321 | 0.001062 | 0.001821 | 0.000382 | 0.00176 | 0.001815 | 0.002419 | 0.003856 | 0.010287 | 0.003539 |
| 36.125 | 0.009527 | 0.007737 | 0.006283 | 0.021672 | 0.01343 | 0.042747 | 0.001817 | 0.004542 | 0.029913 | 0.058095 | 0.003011 | 0.053709 | 0.001407 | 0.007387 | 0.000795 | 0.001655 | 0.001621 | 0.000261 | 0.004234 | 0.001107 | 0.001902 | 0.002761 | 0.009364 | 0.002326 |
| 36.1388889 | 0.011568 | 0.012092 | 0.003803 | 0.019257 | 0.005802 | 0.026803 | 0.002077 | 0.009593 | 0.022291 | 0.052453 | 0.005909 | 0.030735 | 0.002553 | 0.008338 | 0.000467 | 0.00167 | 0.00159 | 0.00019 | 0.007187 | 0.001263 | 0.001719 | 0.001785 | 0.006815 | 0.001219 |
| 36.1527778 | 0.015538 | 0.042544 | 0.002847 | 0.016526 | 0.007709 | 0.016738 | 0.001934 | 0.023432 | 0.018996 | 0.045329 | 0.022955 | 0.034679 | 0.005266 | 0.006282 | 0.00104 | 0.001336 | 0.00195 | 0.00069 | 0.010758 | 0.002075 | 0.002218 | 0.000986 | 0.003929 | 0.000836 |
| 36.1666667 | 0.015506 | 0.092956 | 0.004212 | 0.01684 | 0.022513 | 0.012934 | 0.0046 | 0.043559 | 0.028747 | 0.030541 | 0.046473 | 0.084431 | 0.007527 | 0.00308 | 0.002013 | 0.001758 | 0.002922 | 0.001973 | 0.014715 | 0.003344 | 0.003584 | 0.000416 | 0.001468 | 0.00125 |
| 36.1805556 | 0.009703 | 0.106023 | 0.008999 | 0.01926 | 0.026968 | 0.011237 | 0.013362 | 0.056417 | 0.028141 | 0.02179 | 0.051349 | 0.103311 | 0.008106 | 0.001885 | 0.001977 | 0.00239 | 0.003521 | 0.002716 | 0.017105 | 0.003637 | 0.004578 | 0.000924 | 0.000709 | 0.001377 |
| 36.1944444 | 0.005673 | 0.070023 | 0.012416 | 0.021799 | 0.020026 | 0.023053 | 0.022041 | 0.051992 | 0.022123 | 0.028014 | 0.037154 | 0.060121 | 0.006937 | 0.001512 | 0.001137 | 0.001885 | 0.003045 | 0.002158 | 0.017042 | 0.002501 | 0.004254 | 0.002309 | 0.001702 | 0.000809 |
| 36.2083333 | 0.009107 | 0.03048 | 0.012997 | 0.022498 | 0.031169 | 0.03646 | 0.023374 | 0.038791 | 0.030138 | 0.033558 | 0.020348 | 0.027928 | 0.005375 | 0.001321 | 0.000928 | 0.000943 | 0.002129 | 0.00112 | 0.015766 | 0.001207 | 0.003208 | 0.003361 | 0.003039 | 0.000446 |
| 36.2222222 | 0.01816 | 0.02519 | 0.013894 | 0.019447 | 0.052432 | 0.037033 | 0.021007 | 0.023815 | 0.041267 | 0.024165 | 0.010296 | 0.049396 | 0.004825 | 0.002411 | 0.001055 | 0.00064 | 0.001709 | 0.000651 | 0.015001 | 0.000755 | 0.002422 | 0.003449 | 0.00338 | 0.000404 |
| 36.2361111 | 0.026923 | 0.045831 | 0.016025 | 0.014623 | 0.064312 | 0.036022 | 0.019634 | 0.012583 | 0.042373 | 0.01016 | 0.008909 | 0.082525 | 0.005774 | 0.003505 | 0.000744 | 0.000724 | 0.002117 | 0.001257 | 0.015586 | 0.001403 | 0.002546 | 0.002418 | 0.00242 | 0.000663 |
| 36.25 | 0.025952 | 0.048544 | 0.020075 | 0.015765 | 0.059867 | 0.033507 | 0.015891 | 0.019327 | 0.038184 | 0.004995 | 0.015903 | 0.075341 | 0.0072 | 0.003873 | 0.000453 | 0.000909 | 0.002585 | 0.002169 | 0.016453 | 0.00222 | 0.002904 | 0.001138 | 0.001529 | 0.000849 |
| 36.2638889 | 0.016887 | 0.040096 | 0.024349 | 0.021879 | 0.039558 | 0.020139 | 0.008278 | 0.040267 | 0.03982 | 0.004588 | 0.031229 | 0.057634 | 0.007147 | 0.004171 | 0.00105 | 0.001036 | 0.001916 | 0.001897 | 0.016122 | 0.001876 | 0.002391 | 0.001309 | 0.002403 | 0.001331 |
| 36.2777778 | 0.012064 | 0.036799 | 0.02288 | 0.023427 | 0.028794 | 0.011009 | 0.002351 | 0.047142 | 0.041173 | 0.008745 | 0.036043 | 0.051537 | 0.005533 | 0.004425 | 0.002371 | 0.0012 | 0.00073 | 0.001055 | 0.015409 | 0.000983 | 0.001919 | 0.002398 | 0.004215 | 0.002776 |
| 36.2916667 | 0.015821 | 0.038109 | 0.014175 | 0.020631 | 0.047649 | 0.015324 | 0.001125 | 0.030009 | 0.030887 | 0.017343 | 0.021302 | 0.049238 | 0.004736 | 0.003506 | 0.002542 | 0.00124 | 0.000637 | 0.001017 | 0.01639 | 0.001016 | 0.001749 | 0.0023 | 0.004284 | 0.002783 |
| 36.3055556 | 0.019316 | 0.052542 | 0.005851 | 0.019822 | 0.076857 | 0.022332 | 0.002171 | 0.012293 | 0.016371 | 0.028068 | 0.008829 | 0.067319 | 0.005516 | 0.001663 | 0.001385 | 0.001073 | 0.001509 | 0.00156 | 0.018594 | 0.002049 | 0.001488 | 0.001715 | 0.002436 | 0.001593 |
| 36.3194444 | 0.013413 | 0.04523 | 0.003009 | 0.020705 | 0.084823 | 0.026643 | 0.003816 | 0.012936 | 0.010582 | 0.044935 | 0.008513 | 0.063115 | 0.005797 | 0.000901 | 0.000673 | 0.000927 | 0.002002 | 0.001782 | 0.018848 | 0.002616 | 0.001354 | 0.001682 | 0.001115 | 0.001517 |
| 36.3333333 | 0.005838 | 0.016285 | 0.003092 | 0.019278 | 0.066362 | 0.021631 | 0.004947 | 0.022466 | 0.012083 | 0.064379 | 0.013283 | 0.027656 | 0.004643 | 0.001258 | 0.000781 | 0.000659 | 0.001488 | 0.00124 | 0.015917 | 0.001719 | 0.00103 | 0.001125 | 0.001402 | 0.001273 |
| 36.3472222 | 0.008338 | 0.009879 | 0.006032 | 0.014817 | 0.037584 | 0.010779 | 0.004891 | 0.030004 | 0.014222 | 0.079561 | 0.019196 | 0.014962 | 0.00365 | 0.001249 | 0.00107 | 0.000601 | 0.000888 | 0.00084 | 0.011951 | 0.000555 | 0.00102 | 0.000404 | 0.002226 | 0.000598 |
| 36.3611111 | 0.018661 | 0.036849 | 0.013957 | 0.012313 | 0.02546 | 0.00628 | 0.00477 | 0.041391 | 0.023539 | 0.09809 | 0.029976 | 0.045236 | 0.003904 | 0.000748 | 0.000964 | 0.000548 | 0.001074 | 0.000688 | 0.009776 | 0.000611 | 0.001021 | 0.000244 | 0.002157 | 0.000639 |
| 36.375 | 0.024601 | 0.066234 | 0.02033 | 0.015389 | 0.041921 | 0.010915 | 0.004078 | 0.05427 | 0.037499 | 0.112996 | 0.039768 | 0.085295 | 0.004793 | 0.000783 | 0.000998 | 0.00036 | 0.001683 | 0.000531 | 0.01126 | 0.00178 | 0.00051 | 0.000322 | 0.001111 | 0.001088 |
| 36.3888889 | 0.017115 | 0.056415 | 0.015643 | 0.0174 | 0.047055 | 0.019436 | 0.002673 | 0.048988 | 0.033635 | 0.095126 | 0.031202 | 0.080876 | 0.005618 | 0.001283 | 0.001522 | 0.000471 | 0.002076 | 0.000832 | 0.017891 | 0.002909 | 0.000125 | 0.000661 | 0.000267 | 0.001293 |
| 36.4027778 | 0.006594 | 0.032567 | 0.005778 | 0.019452 | 0.025865 | 0.032151 | 0.002954 | 0.026185 | 0.01993 | 0.053858 | 0.01553 | 0.065029 | 0.006275 | 0.001288 | 0.001496 | 0.000565 | 0.002099 | 0.001041 | 0.027811 | 0.003161 | 0.000111 | 0.001038 | 0.000285 | 0.000983 |
| 36.4166667 | 0.004031 | 0.023054 | 0.002655 | 0.022941 | 0.014122 | 0.041488 | 0.004143 | 0.013201 | 0.01574 | 0.028015 | 0.012097 | 0.06953 | 0.006816 | 0.00112 | 0.00084 | 0.000376 | 0.00199 | 0.000812 | 0.03372 | 0.002705 | 0.000457 | 0.000959 | 0.000955 | 0.000603 |
| 36.4305556 | 0.007074 | 0.01929 | 0.006664 | 0.017323 | 0.011925 | 0.034396 | 0.003874 | 0.018199 | 0.011764 | 0.026669 | 0.011541 | 0.050749 | 0.007309 | 0.001657 | 0.000321 | 0.00017 | 0.002052 | 0.000602 | 0.03251 | 0.002212 | 0.000861 | 0.000584 | 0.001553 | 0.000895 |
| 36.4444444 | 0.012968 | 0.033352 | 0.013083 | 0.007119 | 0.006808 | 0.021095 | 0.002955 | 0.02599 | 0.009932 | 0.031194 | 0.011643 | 0.019491 | 0.008563 | 0.002678 | 0.00065 | 0.000428 | 0.002867 | 0.001195 | 0.029111 | 0.002548 | 0.001137 | 0.0012 | 0.001378 | 0.001293 |
| 36.4583333 | 0.016134 | 0.055067 | 0.015918 | 0.00224 | 0.005167 | 0.014343 | 0.003573 | 0.026752 | 0.015438 | 0.027114 | 0.019201 | 0.015009 | 0.011382 | 0.003404 | 0.002364 | 0.001604 | 0.004845 | 0.0033 | 0.026894 | 0.004131 | 0.002142 | 0.003528 | 0.001504 | 0.001368 |
| 36.4722222 | 0.01671 | 0.072414 | 0.011379 | 0.002294 | 0.005849 | 0.010925 | 0.007581 | 0.022185 | 0.025624 | 0.020173 | 0.033996 | 0.034333 | 0.014142 | 0.002862 | 0.004234 | 0.00306 | 0.006659 | 0.005801 | 0.024498 | 0.005874 | 0.003593 | 0.005693 | 0.002586 | 0.002021 |
| 36.4861111 | 0.017725 | 0.089469 | 0.007896 | 0.004649 | 0.007714 | 0.006844 | 0.012663 | 0.017778 | 0.038459 | 0.014988 | 0.055398 | 0.059222 | 0.013846 | 0.001459 | 0.004089 | 0.003095 | 0.005398 | 0.006195 | 0.018843 | 0.005906 | 0.003408 | 0.00475 | 0.00258 | 0.00248 |
| 36.5 | 0.016946 | 0.092394 | 0.012765 | 0.004698 | 0.018584 | 0.005185 | 0.014915 | 0.017611 | 0.043532 | 0.021662 | 0.072257 | 0.060887 | 0.010268 | 0.0008 | 0.00215 | 0.001872 | 0.003551 | 0.004145 | 0.012393 | 0.003756 | 0.003716 | 0.003394 | 0.002491 | 0.001929 |
| 36.5138889 | 0.01614 | 0.082146 | 0.017019 | 0.002763 | 0.032287 | 0.007581 | 0.016925 | 0.025779 | 0.034895 | 0.078178 | 0.075467 | 0.033592 | 0.006084 | 0.000875 | 0.001468 | 0.001198 | 0.004586 | 0.002497 | 0.014904 | 0.001326 | 0.006467 | 0.004611 | 0.004114 | 0.001717 |
| 36.5277778 | 0.013629 | 0.064201 | 0.013825 | 0.006437 | 0.044202 | 0.015614 | 0.025303 | 0.043208 | 0.018012 | 0.194737 | 0.071172 | 0.012181 | 0.003182 | 0.000928 | 0.004702 | 0.002159 | 0.004862 | 0.003601 | 0.02436 | 0.001318 | 0.006189 | 0.004307 | 0.003918 | 0.001715 |
| 36.5416667 | 0.010417 | 0.034461 | 0.010031 | 0.02239 | 0.072991 | 0.02954 | 0.037261 | 0.059422 | 0.007674 | 0.272163 | 0.06663 | 0.014902 | 0.001688 | 0.001061 | 0.009471 | 0.003435 | 0.004059 | 0.005149 | 0.02785 | 0.003025 | 0.002262 | 0.002428 | 0.001514 | 0.001089 |
| 36.5555556 | 0.012734 | 0.010607 | 0.01168 | 0.041281 | 0.120396 | 0.040102 | 0.037888 | 0.060971 | 0.010722 | 0.220336 | 0.060111 | 0.021176 | 0.002012 | 0.001263 | 0.009616 | 0.003249 | 0.00327 | 0.004287 | 0.024149 | 0.003324 | 0.001654 | 0.001876 | 0.001378 | 0.00151 |
| 36.5694444 | 0.014668 | 0.010325 | 0.016076 | 0.04504 | 0.151314 | 0.036776 | 0.024081 | 0.051509 | 0.015326 | 0.109374 | 0.053525 | 0.015097 | 0.003271 | 0.001289 | 0.005008 | 0.003054 | 0.001389 | 0.003316 | 0.018558 | 0.002591 | 0.004688 | 0.001819 | 0.003331 | 0.00304 |

| | | | | | | | | | | | | | | | | | | | | | | | | |
|---|---|---|---|---|---|---|---|---|---|---|---|---|---|---|---|---|---|---|---|---|---|---|---|---|
| 36.5833333 | 0.015027 | 0.016935 | 0.02079 | 0.036608 | 0.137301 | 0.026331 | 0.011485 | 0.044542 | 0.014161 | 0.043899 | 0.05097 | 0.005615 | 0.002722 | 0.001277 | 0.001302 | 0.002973 | 0.000538 | 0.003489 | 0.012065 | 0.001894 | 0.0061 | 0.001436 | 0.004086 | 0.002759 |
| 36.5972222 | 0.023178 | 0.017532 | 0.026303 | 0.029379 | 0.092517 | 0.023526 | 0.007894 | 0.042393 | 0.010096 | 0.046128 | 0.042748 | 0.004436 | 0.001371 | 0.001646 | 0.000646 | 0.001938 | 0.001049 | 0.002819 | 0.006029 | 0.000808 | 0.005017 | 0.001102 | 0.003709 | 0.001586 |
| 36.6111111 | 0.035783 | 0.027468 | 0.029669 | 0.025185 | 0.043568 | 0.027742 | 0.007188 | 0.039752 | 0.005289 | 0.072252 | 0.028518 | 0.009508 | 0.001824 | 0.002825 | 0.001675 | 0.001172 | 0.001346 | 0.001474 | 0.002317 | 0.000144 | 0.00386 | 0.001291 | 0.0034 | 0.001976 |
| 36.625 | 0.040965 | 0.050073 | 0.027314 | 0.018973 | 0.015657 | 0.031218 | 0.005612 | 0.034588 | 0.003196 | 0.07682 | 0.019471 | 0.014296 | 0.003151 | 0.00381 | 0.00276 | 0.001104 | 0.001227 | 0.000567 | 0.001099 | 7.07E-05 | 0.003685 | 0.001345 | 0.002882 | 0.002638 |
| 36.6388889 | 0.036263 | 0.070069 | 0.021094 | 0.00992 | 0.01661 | 0.030932 | 0.004073 | 0.028203 | 0.00541 | 0.062223 | 0.017228 | 0.014835 | 0.003447 | 0.003502 | 0.003092 | 0.001354 | 0.000803 | 0.000314 | 0.001051 | 0.000333 | 0.004542 | 0.002036 | 0.0025 | 0.002503 |
| 36.6527778 | 0.02609 | 0.077057 | 0.015375 | 0.007585 | 0.02631 | 0.026101 | 0.003674 | 0.02349 | 0.008714 | 0.053754 | 0.019942 | 0.011026 | 0.002366 | 0.002224 | 0.003045 | 0.002074 | 0.000623 | 0.000731 | 0.001764 | 0.001316 | 0.006263 | 0.004228 | 0.003218 | 0.001943 |
| 36.6666667 | 0.014768 | 0.075317 | 0.011487 | 0.018347 | 0.042998 | 0.019458 | 0.003521 | 0.020099 | 0.011089 | 0.068832 | 0.022619 | 0.008453 | 0.001407 | 0.000949 | 0.003046 | 0.002919 | 0.00113 | 0.0013 | 0.003568 | 0.002459 | 0.00758 | 0.005761 | 0.004714 | 0.001964 |
| 36.6805556 | 0.008508 | 0.079179 | 0.009117 | 0.031325 | 0.073818 | 0.014905 | 0.002713 | 0.014132 | 0.012722 | 0.10698 | 0.019504 | 0.011909 | 0.001489 | 0.000963 | 0.0033 | 0.00335 | 0.001769 | 0.00137 | 0.004379 | 0.002969 | 0.007369 | 0.00566 | 0.006138 | 0.003513 |
| 36.6944444 | 0.01072 | 0.09479 | 0.007946 | 0.032964 | 0.103055 | 0.012922 | 0.004257 | 0.007408 | 0.013363 | 0.152699 | 0.012342 | 0.020782 | 0.002212 | 0.002408 | 0.004039 | 0.003776 | 0.002468 | 0.001534 | 0.004177 | 0.003479 | 0.006835 | 0.005375 | 0.007737 | 0.005771 |
| 36.7083333 | 0.017912 | 0.115313 | 0.006836 | 0.021484 | 0.110965 | 0.01185 | 0.008992 | 0.005651 | 0.012583 | 0.18509 | 0.006644 | 0.029553 | 0.003329 | 0.00404 | 0.004635 | 0.004353 | 0.003262 | 0.002278 | 0.00503 | 0.004379 | 0.007127 | 0.005152 | 0.009313 | 0.006973 |
| 36.7222222 | 0.02363 | 0.123951 | 0.006987 | 0.007769 | 0.097833 | 0.011638 | 0.012635 | 0.006723 | 0.01077 | 0.185522 | 0.007259 | 0.030674 | 0.003921 | 0.005244 | 0.004332 | 0.004552 | 0.003393 | 0.00291 | 0.006403 | 0.004843 | 0.00744 | 0.004335 | 0.009786 | 0.006373 |
| 36.7361111 | 0.02556 | 0.111661 | 0.009916 | 0.003174 | 0.070515 | 0.014053 | 0.011588 | 0.004811 | 0.009456 | 0.160325 | 0.013695 | 0.024187 | 0.003316 | 0.007086 | 0.003834 | 0.004742 | 0.002754 | 0.003183 | 0.00733 | 0.004858 | 0.007142 | 0.003319 | 0.009496 | 0.005 |
| 36.75 | 0.028002 | 0.087844 | 0.012172 | 0.005879 | 0.04265 | 0.018113 | 0.006807 | 0.002686 | 0.00981 | 0.128812 | 0.018169 | 0.018781 | 0.002037 | 0.009301 | 0.003698 | 0.005181 | 0.001936 | 0.003063 | 0.007725 | 0.00469 | 0.006175 | 0.002165 | 0.009009 | 0.003497 |
| 36.7638889 | 0.031001 | 0.063546 | 0.012615 | 0.009118 | 0.032061 | 0.022215 | 0.003288 | 0.005561 | 0.010725 | 0.104785 | 0.018957 | 0.016054 | 0.00101 | 0.009302 | 0.003323 | 0.005099 | 0.001189 | 0.002311 | 0.007401 | 0.003934 | 0.004384 | 0.001369 | 0.007809 | 0.002002 |
| 36.7777778 | 0.029427 | 0.041924 | 0.014297 | 0.009412 | 0.032097 | 0.023077 | 0.003501 | 0.01209 | 0.010978 | 0.107388 | 0.023198 | 0.010787 | 0.000956 | 0.006841 | 0.002579 | 0.004229 | 0.000598 | 0.001201 | 0.00613 | 0.002651 | 0.002151 | 0.002235 | 0.006159 | 0.001441 |
| 36.7916667 | 0.024511 | 0.027325 | 0.016329 | 0.00797 | 0.032386 | 0.019213 | 0.004025 | 0.01938 | 0.012201 | 0.114722 | 0.029968 | 0.0052 | 0.001614 | 0.003719 | 0.002056 | 0.003217 | 0.000292 | 0.000446 | 0.004 | 0.001333 | 0.000549 | 0.003653 | 0.004639 | 0.001759 |
| 36.8055556 | 0.021111 | 0.021545 | 0.016105 | 0.007165 | 0.030035 | 0.015654 | 0.003154 | 0.025726 | 0.015388 | 0.088817 | 0.029741 | 0.0048 | 0.002094 | 0.002054 | 0.002107 | 0.002865 | 0.000411 | 0.000275 | 0.001778 | 0.000422 | 0.000392 | 0.003734 | 0.003331 | 0.001976 |
| 36.8194444 | 0.019283 | 0.017743 | 0.015134 | 0.006526 | 0.023365 | 0.013583 | 0.004249 | 0.02937 | 0.018237 | 0.044609 | 0.020153 | 0.008239 | 0.002225 | 0.002551 | 0.002444 | 0.002936 | 0.000685 | 0.000251 | 0.00094 | 0.000192 | 0.000828 | 0.002999 | 0.002301 | 0.002002 |
| 36.8333333 | 0.017256 | 0.013388 | 0.01586 | 0.006833 | 0.016881 | 0.010507 | 0.008793 | 0.029481 | 0.020037 | 0.014613 | 0.008851 | 0.011394 | 0.003247 | 0.003113 | 0.002016 | 0.002457 | 0.001137 | 0.000937 | 0.002597 | 0.001064 | 0.001359 | 0.00323 | 0.002463 | 0.002727 |
| 36.8472222 | 0.014173 | 0.010431 | 0.016306 | 0.008713 | 0.013079 | 0.007692 | 0.012761 | 0.026422 | 0.021852 | 0.007427 | 0.004737 | 0.013266 | 0.005719 | 0.002451 | 0.0014 | 0.001825 | 0.002528 | 0.00256 | 0.00571 | 0.002451 | 0.003662 | 0.004182 | 0.004309 | 0.003821 |
| 36.8611111 | 0.01003 | 0.007101 | 0.01289 | 0.009888 | 0.012157 | 0.007278 | 0.012118 | 0.02252 | 0.022717 | 0.012482 | 0.008402 | 0.013724 | 0.00905 | 0.001087 | 0.005954 | 0.008567 | 0.003491 | 0.003215 | 0.007904 | 0.004013 | 0.013938 | 0.005474 | 0.00496 | 0.005568 |
| 36.875 | 0.010266 | 0.00531 | 0.009448 | 0.008706 | 0.017046 | 0.008196 | 0.008412 | 0.020848 | 0.022995 | 0.012816 | 0.012374 | 0.011597 | 0.010331 | 0.045691 | 0.016892 | 0.025714 | 0.006739 | 0.00526 | 0.008173 | 0.004355 | 0.042539 | 0.006033 | 0.014432 | 0.006976 |
| 36.8888889 | 0.017327 | 0.010552 | 0.012196 | 0.005412 | 0.027322 | 0.007621 | 0.005931 | 0.021556 | 0.024402 | 0.007026 | 0.015038 | 0.007402 | 0.007535 | 0.09734 | 0.020249 | 0.029536 | 0.009849 | 0.008014 | 0.00836 | 0.003596 | 0.057927 | 0.005044 | 0.026608 | 0.004772 |
| 36.9027778 | 0.023925 | 0.020967 | 0.018139 | 0.003216 | 0.035245 | 0.004801 | 0.006502 | 0.023718 | 0.026248 | 0.005783 | 0.021508 | 0.00352 | 0.00433 | 0.076005 | 0.012163 | 0.0152 | 0.005722 | 0.007524 | 0.008752 | 0.00426 | 0.041111 | 0.003718 | 0.023532 | 0.002636 |
| 36.9166667 | 0.024829 | 0.030782 | 0.022432 | 0.003853 | 0.03699 | 0.002792 | 0.007828 | 0.026931 | 0.027864 | 0.013155 | 0.030788 | 0.001456 | 0.003746 | 0.060843 | 0.008193 | 0.009881 | 0.001837 | 0.008143 | 0.007432 | 0.004377 | 0.035802 | 0.001951 | 0.022675 | 0.002927 |
| 36.9305556 | 0.021495 | 0.035999 | 0.024792 | 0.003593 | 0.036758 | 0.003659 | 0.006741 | 0.029009 | 0.030882 | 0.024284 | 0.035727 | 0.002315 | 0.007065 | 0.058855 | 0.007837 | 0.00996 | 0.001895 | 0.008309 | 0.007933 | 0.004916 | 0.033391 | 0.001028 | 0.022191 | 0.004392 |
| 36.9444444 | 0.017609 | 0.035645 | 0.025477 | 0.003668 | 0.034825 | 0.006688 | 0.004022 | 0.027853 | 0.034525 | 0.028009 | 0.033921 | 0.004811 | 0.010694 | 0.071969 | 0.011091 | 0.011436 | 0.002407 | 0.009736 | 0.011452 | 0.008699 | 0.035211 | 0.001242 | 0.024229 | 0.010425 |
| 36.9583333 | 0.015615 | 0.028782 | 0.023404 | 0.008745 | 0.024719 | 0.013369 | 0.002901 | 0.022595 | 0.034579 | 0.02134 | 0.031376 | 0.008787 | 0.007947 | 0.100076 | 0.020685 | 0.020803 | 0.002489 | 0.013865 | 0.011314 | 0.012532 | 0.053324 | 0.003909 | 0.03112 | 0.017458 |
| 36.9722222 | 0.01669 | 0.015425 | 0.017446 | 0.015292 | 0.010597 | 0.022765 | 0.003413 | 0.015012 | 0.029813 | 0.015904 | 0.032084 | 0.020529 | 0.003065 | 0.055291 | 0.019679 | 0.020239 | 0.001683 | 0.010988 | 0.007935 | 0.0118 | 0.039384 | 0.00761 | 0.018175 | 0.016487 |
| 36.9861111 | 0.021382 | 0.006909 | 0.010717 | 0.015818 | 0.009076 | 0.023045 | 0.003658 | 0.012479 | 0.025319 | 0.015659 | 0.028446 | 0.030524 | 0.003256 | 0.00694 | 0.011528 | 0.012513 | 0.001597 | 0.004919 | 0.005272 | 0.007699 | 0.020353 | 0.006959 | 0.004899 | 0.010505 |
| 37 | 0.027877 | 0.015143 | 0.008288 | 0.009694 | 0.029174 | 0.014333 | 0.002981 | 0.018842 | 0.02684 | 0.027922 | 0.015557 | 0.022186 | 0.00589 | 0.075892 | 0.014333 | 0.019926 | 0.002716 | 0.01246 | 0.007884 | 0.00599 | 0.06325 | 0.006062 | 0.032381 | 0.012072 |
| 37.0138889 | 0.034272 | 0.029199 | 0.009469 | 0.004583 | 0.057055 | 0.013291 | 0.004696 | 0.023944 | 0.029764 | 0.058162 | 0.006561 | 0.013287 | 0.028214 | 0.147198 | 0.025137 | 0.031894 | 0.006447 | 0.028707 | 0.063983 | 0.037758 | 0.093584 | 0.005834 | 0.058797 | 0.031516 |
| 37.0277778 | 0.040121 | 0.02565 | 0.009641 | 0.008804 | 0.073001 | 0.022186 | 0.013359 | 0.017871 | 0.024035 | 0.075387 | 0.012053 | 0.029327 | 0.067236 | 0.073252 | 0.050499 | 0.025945 | 0.010734 | 0.027506 | 0.17617 | 0.114141 | 0.041564 | 0.002381 | 0.028717 | 0.049147 |
| 37.0416667 | 0.042195 | 0.013331 | 0.011559 | 0.017699 | 0.098626 | 0.031158 | 0.020731 | 0.009604 | 0.014155 | 0.071118 | 0.015898 | 0.046935 | 0.069399 | 0.091266 | 0.071787 | 0.023553 | 0.011261 | 0.022392 | 0.210558 | 0.131529 | 0.04745 | 0.002176 | 0.024 | 0.043289 |
| 37.0555556 | 0.034065 | 0.035473 | 0.023022 | 0.019421 | 0.127875 | 0.037897 | 0.015931 | 0.021645 | 0.016171 | 0.08742 | 0.02923 | 0.05147 | 0.035477 | 0.183418 | 0.047668 | 0.028894 | 0.008151 | 0.02342 | 0.120962 | 0.057803 | 0.096236 | 0.005426 | 0.048687 | 0.027254 |
| 37.0694444 | 0.019265 | 0.077786 | 0.035826 | 0.015619 | 0.113447 | 0.051052 | 0.00663 | 0.044447 | 0.025347 | 0.118045 | 0.065504 | 0.059962 | 0.019935 | 0.189714 | 0.021681 | 0.023729 | 0.003921 | 0.021305 | 0.052987 | 0.011219 | 0.095664 | 0.004693 | 0.055705 | 0.017665 |
| 37.0833333 | 0.011874 | 0.07101 | 0.034664 | 0.017749 | 0.069357 | 0.049658 | 0.005808 | 0.044088 | 0.024047 | 0.095857 | 0.064341 | 0.060278 | 0.048439 | 0.194238 | 0.043131 | 0.01955 | 0.003666 | 0.026109 | 0.1326 | 0.025855 | 0.092419 | 0.001811 | 0.061348 | 0.027483 |
| 37.0972222 | 0.013214 | 0.031108 | 0.024326 | 0.027683 | 0.028507 | 0.02223 | 0.013642 | 0.032797 | 0.016343 | 0.043891 | 0.022588 | 0.055603 | 0.088333 | 0.096676 | 0.060447 | 0.011313 | 0.011098 | 0.023653 | 0.218115 | 0.036584 | 0.045542 | 0.000369 | 0.030331 | 0.032113 |
| 37.1111111 | 0.012584 | 0.018591 | 0.017746 | 0.029447 | 0.009264 | 0.01013 | 0.020805 | 0.026504 | 0.010554 | 0.023556 | 0.008074 | 0.04676 | 0.083191 | 0 | 0.041037 | 0.003643 | 0.014495 | 0.012572 | 0.147466 | 0.016821 | 0.019886 | 0.027679 | 0.037994 | 0.014945 |
| 37.125 | 0.00995 | 0.023922 | 0.014362 | 0.020167 | 0.009949 | 0.016194 | 0.022459 | 0.015416 | 0.007293 | 0.014011 | 0.013482 | 0.026413 | 0.04048 | 0.050645 | 0.021308 | 0.009016 | 0.006281 | 0.016736 | 0.059194 | 0.00343 | 0.072425 | 0.061039 | 0.108156 | 0.029973 |
| 37.1388889 | 0.010746 | 0.028143 | 0.009834 | 0.013568 | 0.009702 | 0.020071 | 0.020127 | 0.007559 | 0.006671 | 0.005161 | 0.007384 | 0.014341 | 0.0157 | 0.158353 | 0.017287 | 0.017617 | 0.002975 | 0.02994 | 0.035355 | 0.012549 | 0.115945 | 0.043704 | 0.113609 | 0.059234 |
| 37.1527778 | 0.01003 | 0.046321 | 0.011483 | 0.011215 | 0.013228 | 0.027531 | 0.01377 | 0.01437 | 0.009543 | 0.019933 | 0.014193 | 0.020784 | 0.015648 | 0.177462 | 0.02363 | 0.021974 | 0.006879 | 0.027821 | 0.020408 | 0.022836 | 0.104002 | 0.016386 | 0.055479 | 0.04395 |
| 37.1666667 | 0.005632 | 0.074652 | 0.021648 | 0.011687 | 0.023228 | 0.032067 | 0.008985 | 0.031728 | 0.019665 | 0.051368 | 0.045397 | 0.044755 | 0.011343 | 0.085525 | 0.022892 | 0.018353 | 0.006102 | 0.013068 | 0.011554 | 0.019405 | 0.050553 | 0.007833 | 0.013461 | 0.018695 |
| 37.1805556 | 0.00265 | 0.086831 | 0.028964 | 0.013218 | 0.025683 | 0.026639 | 0.011817 | 0.04357 | 0.030222 | 0.064892 | 0.069016 | 0.062311 | 0.003169 | 0.019949 | 0.009856 | 0.008382 | 0.002645 | 0.003741 | 0.008914 | 0.00774 | 0.010306 | 0.003361 | 0.002715 | 0.007578 |
| 37.1944444 | 0.002882 | 0.076567 | 0.027677 | 0.009911 | 0.018149 | 0.016328 | 0.017052 | 0.041863 | 0.031448 | 0.049718 | 0.068715 | 0.052957 | 0.000754 | 0.005988 | 0.002018 | 0.001934 | 0.000763 | 0.001504 | 0.008184 | 0.001641 | 0.000748 | 0.003828 | 0.002034 | 0.002215 |
| 37.2083333 | 0.008688 | 0.050222 | 0.019426 | 0.009182 | 0.021209 | 0.006074 | 0.016493 | 0.029009 | 0.023599 | 0.03339 | 0.045091 | 0.035923 | 0.001141 | 0.003152 | 0.000748 | 0.001351 | 0.000644 | 0.001204 | 0.008774 | 0.000773 | 0.001236 | 0.00413 | 0.001122 | 0.00191 |
| 37.2222222 | 0.015096 | 0.024734 | 0.011994 | 0.015961 | 0.039319 | 0.001985 | 0.011454 | 0.015716 | 0.014171 | 0.030369 | 0.017449 | 0.028215 | 0.002123 | 0.002575 | 0.000704 | 0.001022 | 0.001799 | 0.001032 | 0.010762 | 0.001339 | 0.002481 | 0.003324 | 0.00081 | 0.001269 |
| 37.2361111 | 0.013181 | 0.026544 | 0.016871 | 0.015785 | 0.036215 | 0.004411 | 0.011595 | 0.018428 | 0.01583 | 0.031818 | 0.025441 | 0.029361 | 0.002982 | 0.002215 | 0.000985 | 0.000576 | 0.002799 | 0.001456 | 0.012665 | 0.002527 | 0.003374 | 0.002961 | 0.000657 | 0.000834 |
| 37.25 | 0.007577 | 0.041427 | 0.027445 | 0.008209 | 0.014896 | 0.00818 | 0.015353 | 0.029508 | 0.024818 | 0.032913 | 0.054445 | 0.034657 | 0.00332 | 0.001468 | 0.00122 | 0.000675 | 0.002916 | 0.001854 | 0.013851 | 0.003577 | 0.003224 | 0.003292 | 0.000252 | 0.000667 |
| 37.2638889 | 0.004193 | 0.032712 | 0.027491 | 0.005585 | 0.017555 | 0.007943 | 0.01179 | 0.029752 | 0.021723 | 0.026614 | 0.055656 | 0.028888 | 0.003652 | 0.000863 | 0.001067 | 0.000867 | 0.002774 | 0.001983 | 0.014485 | 0.003789 | 0.002993 | 0.003763 | 9.77E-05 | 0.000665 |
| 37.2777778 | 0.003675 | 0.017324 | 0.023333 | 0.005637 | 0.034169 | 0.011994 | 0.005291 | 0.024508 | 0.013395 | 0.020685 | 0.047291 | 0.019266 | 0.003535 | 0.001418 | 0.000847 | 0.000856 | 0.002406 | 0.001809 | 0.013994 | 0.003036 | 0.002527 | 0.004338 | 0.000403 | 0.001575 |
| 37.2916667 | 0.005646 | 0.023303 | 0.025237 | 0.003218 | 0.033468 | 0.025451 | 0.003644 | 0.023876 | 0.015028 | 0.025659 | 0.059674 | 0.02154 | 0.002227 | 0.002933 | 0.001125 | 0.001126 | 0.001483 | 0.001231 | 0.012106 | 0.001557 | 0.001684 | 0.00532 | 0.001352 | 0.002591 |
| 37.3055556 | 0.007704 | 0.02501 | 0.0222 | 0.004923 | 0.030507 | 0.035437 | 0.006062 | 0.018937 | 0.015355 | 0.028445 | 0.060472 | 0.026177 | 0.0007 | 0.00384 | 0.001886 | 0.001844 | 0.000613 | 0.000992 | 0.009678 | 0.000578 | 0.001473 | 0.005863 | 0.002305 | 0.002498 |
| 37.3194444 | 0.010106 | 0.011249 | 0.011938 | 0.011259 | 0.033607 | 0.042726 | 0.014008 | 0.008079 | 0.010817 | 0.021928 | 0.041926 | 0.026079 | 0.000284 | 0.003144 | 0.001912 | 0.001909 | 0.00052 | 0.000777 | 0.007998 | 0.000994 | 0.00134 | 0.004981 | 0.001984 | 0.001727 |
| 37.3333333 | 0.010545 | 0.010823 | 0.008402 | 0.01184 | 0.025985 | 0.048578 | 0.032153 | 0.004258 | 0.007586 | 0.016895 | 0.040819 | 0.020079 | 0.000602 | 0.001946 | 0.001111 | 0.001151 | 0.001205 | 0.000308 | 0.006767 | 0.002157 | 0.000745 | 0.003723 | 0.000998 | 0.001084 |
| 37.3472222 | 0.008344 | 0.024679 | 0.011223 | 0.007088 | 0.018538 | 0.050163 | 0.063288 | 0.008703 | 0.003888 | 0.017267 | 0.052638 | 0.01025 | 0.001008 | 0.001515 | 0.000523 | 0.000554 | 0.002203 | 0.000331 | 0.004934 | 0.002991 | 0.000345 | 0.003315 | 0.000591 | 0.000716 |
| 37.3611111 | 0.008603 | 0.031262 | 0.01233 | 0.005217 | 0.019893 | 0.049566 | 0.081508 | 0.015149 | 0.002169 | 0.014214 | 0.054579 | 0.003656 | 0.001388 | 0.00156 | 0.000651 | 0.000635 | 0.003171 | 0.001073 | 0.003344 | 0.003794 | 0.000589 | 0.002773 | 0.000603 | 0.000934 |
| 37.375 | 0.008537 | 0.03431 | 0.01512 | 0.005414 | 0.027531 | 0.039067 | 0.061397 | 0.023764 | 0.005719 | 0.016233 | 0.059866 | 0.010211 | 0.001604 | 0.00164 | 0.001135 | 0.00114 | 0.003664 | 0.001929 | 0.00282 | 0.004486 | 0.001245 | 0.00138 | 0.000525 | 0.001537 |
| 37.3888889 | 0.006733 | 0.045088 | 0.024119 | 0.006449 | 0.046236 | 0.020673 | 0.029644 | 0.031803 | 0.015526 | 0.03439 | 0.080414 | 0.0351 | 0.001494 | 0.00187 | 0.001204 | 0.0012 | 0.00323 | 0.002002 | 0.002331 | 0.003916 | 0.001291 | 0.000492 | 0.000421 | 0.00132 |
| 37.4027778 | 0.00636 | 0.049034 | 0.030188 | 0.00846 | 0.059837 | 0.009603 | 0.012389 | 0.033032 | 0.022348 | 0.044744 | 0.091564 | 0.050973 | 0.0012 | 0.002252 | 0.000772 | 0.000709 | 0.002323 | 0.001232 | 0.001377 | 0.00262 | 0.000959 | 0.000959 | 0.000435 | 0.000606 |
| 37.4166667 | 0.0065 | 0.031964 | 0.021719 | 0.009638 | 0.043629 | 0.005673 | 0.005215 | 0.025519 | 0.015753 | 0.033176 | 0.067879 | 0.04171 | 0.000938 | 0.002289 | 0.000333 | 0.000332 | 0.001717 | 0.000432 | 0.000844 | 0.002067 | 0.000741 | 0.001779 | 0.00048 | 0.000516 |
| 37.4305556 | 0.010601 | 0.018482 | 0.010362 | 0.015794 | 0.024209 | 0.013349 | 0.006238 | 0.013151 | 0.011742 | 0.030088 | 0.034422 | 0.045658 | 0.000762 | 0.001797 | 0.001189 | 0.001042 | 0.001762 | 0.000188 | 0.000997 | 0.001946 | 0.001377 | 0.002351 | 0.000635 | 0.00084 |
| 37.4444444 | 0.014663 | 0.023728 | 0.008829 | 0.024322 | 0.032078 | 0.034669 | 0.009386 | 0.004636 | 0.020008 | 0.040373 | 0.02113 | 0.067877 | 0.001587 | 0.010022 | 0.009219 | 0.007657 | 0.002737 | 0.001895 | 0.00096 | 0.006384 | 0.012991 | 0.007614 | 0.002738 | 0.005129 |
| 37.4583333 | 0.013537 | 0.026373 | 0.012374 | 0.021148 | 0.034993 | 0.044559 | 0.006451 | 0.008749 | 0.021399 | 0.036246 | 0.029522 | 0.059244 | 0.00412 | 0.09556 | 0.02402 | 0.019394 | 0.006224 | 0.008625 | 0.005099 | 0.01603 | 0.052332 | 0.016707 | 0.010561 | 0.01517 |
| 37.4722222 | 0.021302 | 0.032356 | 0.023874 | 0.015046 | 0.029745 | 0.062614 | 0.00402 | 0.027454 | 0.021307 | 0.033452 | 0.069704 | 0.042696 | 0.006826 | 0.171511 | 0.027819 | 0.020509 | 0.010572 | 0.01298 | 0.00966 | 0.01867 | 0.07747 | 0.016324 | 0.021138 | 0.018453 |
| 37.4861111 | 0.042396 | 0.04451 | 0.036798 | 0.018304 | 0.027898 | 0.099497 | 0.004971 | 0.048855 | 0.036109 | 0.045789 | 0.112461 | 0.059124 | 0.008982 | 0.127255 | 0.024868 | 0.014421 | 0.010016 | 0.011702 | 0.008881 | 0.015496 | 0.066138 | 0.011457 | 0.02117 | 0.016667 |
| 37.5 | 0.059415 | 0.038232 | 0.035 | 0.017769 | 0.033626 | 0.089527 | 0.00456 | 0.058136 | 0.043799 | 0.041193 | 0.104103 | 0.060144 | 0.008789 | 0.07615 | 0.028441 | 0.016842 | 0.006921 | 0.010979 | 0.008927 | 0.014935 | 0.05902 | 0.011863 | 0.010234 | 0.018808 |
| 37.5138889 | 0.063318 | 0.022766 | 0.024867 | 0.01689 | 0.068735 | 0.039134 | 0.005556 | 0.053591 | 0.03221 | 0.018844 | 0.058008 | 0.030462 | 0.00498 | 0.034646 | 0.021166 | 0.01607 | 0.005311 | 0.006343 | 0.007408 | 0.011882 | 0.039965 | 0.009919 | 0.00408 | 0.013438 |
| 37.5277778 | 0.056959 | 0.019043 | 0.019137 | 0.025571 | 0.100885 | 0.013564 | 0.01135 | 0.04483 | 0.022665 | 0.005919 | 0.022503 | 0.008756 | 0.002824 | 0.001757 | 0.00746 | 0.007147 | 0.004189 | 0.002357 | 0.003784 | 0.005989 | 0.01269 | 0.00553 | 0.003493 | 0.003815 |
| 37.5416667 | 0.043477 | 0.022748 | 0.018521 | 0.032284 | 0.109832 | 0.018155 | 0.021645 | 0.04269 | 0.02564 | 0.004155 | 0.010677 | 0.002578 | 0.004055 | 0.002959 | 0.006118 | 0.002575 | 0.004747 | 0.00361 | 0.002896 | 0.003121 | 0.004571 | 0.002824 | 0.001662 | 0.000532 |
| 37.5555556 | 0.029537 | 0.021979 | 0.018935 | 0.030931 | 0.108326 | 0.028325 | 0.034526 | 0.04895 | 0.034386 | 0.004753 | 0.011904 | 0.001621 | 0.00395 | 0.004756 | 0.008096 | 0.002711 | 0.004537 | 0.004039 | 0.006828 | 0.002931 | 0.004109 | 0.002448 | 0.001408 | 0.001243 |
| 37.5694444 | 0.020822 | 0.015239 | 0.017103 | 0.026721 | 0.096525 | 0.028614 | 0.044109 | 0.056298 | 0.038863 | 0.004466 | 0.018299 | 0.002872 | 0.002624 | 0.007113 | 0.004911 | 0.003341 | 0.002837 | 0.004486 | 0.01424 | 0.002816 | 0.005638 | 0.003607 | 0.003512 | 0.003431 |
| 37.5833333 | 0.015707 | 0.013981 | 0.013692 | 0.025957 | 0.084132 | 0.023023 | 0.045642 | 0.057566 | 0.033608 | 0.004421 | 0.024966 | 0.007197 | 0.00286 | 0.006795 | 0.001343 | 0.00394 | 0.002435 | 0.005573 | 0.01798 | 0.002951 | 0.007321 | 0.003952 | 0.004532 | 0.004037 |
| 37.5972222 | 0.012753 | 0.021134 | 0.013295 | 0.025621 | 0.068676 | 0.021045 | 0.042663 | 0.053498 | 0.025469 | 0.005081 | 0.033153 | 0.009039 | 0.002901 | 0.003548 | 0.000449 | 0.003404 | 0.001747 | 0.004823 | 0.016039 | 0.002351 | 0.00662 | 0.002661 | 0.003575 | 0.002616 |
| 37.6111111 | 0.011903 | 0.022469 | 0.016901 | 0.020757 | 0.045911 | 0.024923 | 0.040066 | 0.049486 | 0.023055 | 0.003463 | 0.041698 | 0.005378 | 0.001949 | 0.001412 | 0.000359 | 0.002275 | 0.000814 | 0.002865 | 0.011959 | 0.001019 | 0.004874 | 0.000955 | 0.00234 | 0.001236 |
| 37.625 | 0.00877 | 0.016335 | 0.019374 | 0.018143 | 0.030816 | 0.028222 | 0.036272 | 0.04606 | 0.024084 | 0.001995 | 0.043463 | 0.002168 | 0.001199 | 0.001864 | 0.000157 | 0.001837 | 0.000843 | 0.001329 | 0.00829 | 0.000498 | 0.004018 | 0.000477 | 0.00139 | 0.000816 |
| 37.6388889 | 0.00443 | 0.010937 | 0.019 | 0.020528 | 0.027096 | 0.026327 | 0.03064 | 0.041718 | 0.023819 | 0.00284 | 0.040662 | 0.001821 | 0.000745 | 0.002812 | 0.000184 | 0.001511 | 0.001154 | 0.000486 | 0.005357 | 0.000477 | 0.003538 | 0.000905 | 0.000672 | 0.001532 |
| 37.6527778 | 0.003928 | 0.006501 | 0.020877 | 0.018102 | 0.022049 | 0.029612 | 0.026698 | 0.041224 | 0.025955 | 0.00426 | 0.044239 | 0.003277 | 0.000753 | 0.002672 | 0.000287 | 0.001144 | 0.001246 | 0.000314 | 0.003165 | 0.001139 | 0.003413 | 0.001376 | 0.000637 | 0.002282 |
| 37.6666667 | 0.005559 | 0.003597 | 0.025152 | 0.009609 | 0.02137 | 0.038007 | 0.026353 | 0.045541 | 0.030953 | 0.005554 | 0.0549 | 0.006176 | 0.000767 | 0.002069 | 0.000714 | 0.00147 | 0.001075 | 0.000707 | 0.002596 | 0.003118 | 0.004035 | 0.001354 | 0.001458 | 0.001815 |
| 37.6805556 | 0.006127 | 0.005294 | 0.025243 | 0.003825 | 0.031801 | 0.035139 | 0.026537 | 0.04548 | 0.0299 | 0.005474 | 0.061918 | 0.006161 | 0.000649 | 0.002933 | 0.00207 | 0.002461 | 0.001326 | 0.001633 | 0.004046 | 0.005534 | 0.004797 | 0.001893 | 0.003381 | 0.001496 |
| 37.6944444 | 0.004509 | 0.012109 | 0.021375 | 0.002344 | 0.042565 | 0.024424 | 0.024676 | 0.039278 | 0.022801 | 0.004318 | 0.061421 | 0.00356 | 0.000706 | 0.004802 | 0.003794 | 0.003138 | 0.001944 | 0.002351 | 0.005972 | 0.006475 | 0.004855 | 0.00357 | 0.005331 | 0.002645 |
| 37.7083333 | 0.002197 | 0.018419 | 0.019377 | 0.002192 | 0.052412 | 0.019804 | 0.022828 | 0.034696 | 0.018391 | 0.002592 | 0.059203 | 0.003359 | 0.000976 | 0.005274 | 0.004894 | 0.003404 | 0.002091 | 0.002572 | 0.007044 | 0.005868 | 0.004557 | 0.004543 | 0.005796 | 0.003662 |
| 37.7222222 | 0.001753 | 0.022157 | 0.01892 | 0.00321 | 0.061305 | 0.017043 | 0.022363 | 0.032284 | 0.016373 | 0.001185 | 0.056007 | 0.005894 | 0.00194 | 0.00452 | 0.005354 | 0.004228 | 0.002256 | 0.003241 | 0.007895 | 0.005631 | 0.00484 | 0.004288 | 0.005571 | 0.00365 |
| 37.7361111 | 0.003395 | 0.021659 | 0.018713 | 0.006378 | 0.066621 | 0.013751 | 0.021648 | 0.030917 | 0.015473 | 0.002513 | 0.051528 | 0.008073 | 0.003281 | 0.004096 | 0.005761 | 0.005583 | 0.002878 | 0.004436 | 0.008598 | 0.006269 | 0.005364 | 0.00381 | 0.005732 | 0.003407 |
| 37.75 | 0.004312 | 0.0137 | 0.020788 | 0.009138 | 0.071437 | 0.016845 | 0.019966 | 0.034935 | 0.019345 | 0.007169 | 0.049853 | 0.00704 | 0.003382 | 0.004003 | 0.005963 | 0.005944 | 0.002855 | 0.004652 | 0.007622 | 0.005957 | 0.004614 | 0.002963 | 0.005586 | 0.002666 |
| 37.7638889 | 0.003085 | 0.006581 | 0.023779 | 0.009797 | 0.069505 | 0.021513 | 0.018313 | 0.041777 | 0.023773 | 0.010871 | 0.048031 | 0.004738 | 0.002255 | 0.003363 | 0.004849 | 0.004354 | 0.001981 | 0.002975 | 0.004575 | 0.00367 | 0.002757 | 0.001924 | 0.003959 | 0.002094 |
| 37.7777778 | 0.00189 | 0.006735 | 0.025366 | 0.009903 | 0.052612 | 0.019425 | 0.016916 | 0.044558 | 0.022421 | 0.009086 | 0.041365 | 0.004651 | 0.002063 | 0.003272 | 0.002586 | 0.002206 | 0.001977 | 0.001541 | 0.002779 | 0.001758 | 0.002223 | 0.00203 | 0.002306 | 0.003741 |
| 37.7916667 | 0.002355 | 0.009197 | 0.025611 | 0.010174 | 0.027913 | 0.015345 | 0.016254 | 0.043858 | 0.018142 | 0.00421 | 0.033217 | 0.005931 | 0.002695 | 0.003835 | 0.001039 | 0.00108 | 0.002683 | 0.001745 | 0.003948 | 0.001754 | 0.002976 | 0.002917 | 0.002118 | 0.006105 |
| 37.8055556 | 0.003722 | 0.010024 | 0.024796 | 0.010927 | 0.010769 | 0.014989 | 0.016401 | 0.043111 | 0.015326 | 0.001658 | 0.027671 | 0.006633 | 0.002048 | 0.005238 | 0.002714 | 0.001928 | 0.002055 | 0.001603 | 0.004217 | 0.002368 | 0.002907 | 0.002775 | 0.002226 | 0.005551 |
| 37.8194444 | 0.004517 | 0.010959 | 0.023564 | 0.012083 | 0.006663 | 0.017105 | 0.016753 | 0.042741 | 0.014115 | 0.002249 | 0.023296 | 0.005899 | 0.002691 | 0.020677 | 0.013119 | 0.008003 | 0.002016 | 0.001086 | 0.002961 | 0.006728 | 0.006643 | 0.004324 | 0.002765 | 0.004109 |
| 37.8333333 | 0.004133 | 0.014751 | 0.022941 | 0.013391 | 0.005983 | 0.019437 | 0.018089 | 0.043265 | 0.014944 | 0.002945 | 0.018408 | 0.003653 | 0.006467 | 0.102721 | 0.030678 | 0.015624 | 0.008588 | 0.004773 | 0.003954 | 0.012485 | 0.026238 | 0.007745 | 0.014429 | 0.008535 |
| 37.8472222 | 0.002677 | 0.019835 | 0.023313 | 0.014568 | 0.005079 | 0.020477 | 0.02031 | 0.044607 | 0.017821 | 0.003127 | 0.012461 | 0.001424 | 0.007136 | 0.155851 | 0.032825 | 0.013502 | 0.013009 | 0.00847 | 0.006295 | 0.009797 | 0.03788 | 0.007359 | 0.027051 | 0.012042 |
| 37.8611111 | 0.001826 | 0.022508 | 0.023918 | 0.014684 | 0.009506 | 0.018791 | 0.020993 | 0.044332 | 0.020283 | 0.003647 | 0.006485 | 0.000498 | 0.004729 | 0.138445 | 0.022888 | 0.006944 | 0.014121 | 0.008402 | 0.006589 | 0.005248 | 0.037427 | 0.006684 | 0.03311 | 0.010267 |
| 37.875 | 0.002634 | 0.022309 | 0.023589 | 0.013642 | 0.016002 | 0.014304 | 0.018356 | 0.041354 | 0.019701 | 0.003872 | 0.00514 | 0.000383 | 0.005082 | 0.153284 | 0.023393 | 0.009316 | 0.01872 | 0.008348 | 0.005416 | 0.007679 | 0.045297 | 0.008001 | 0.037726 | 0.010676 |
| 37.8888889 | 0.004469 | 0.021079 | 0.02254 | 0.012662 | 0.019757 | 0.010823 | 0.013533 | 0.038322 | 0.01654 | 0.003395 | 0.007151 | 0.000623 | 0.005846 | 0.122264 | 0.02506 | 0.016326 | 0.014433 | 0.004981 | 0.004321 | 0.010114 | 0.035298 | 0.007126 | 0.023474 | 0.009165 |
| 37.9027778 | 0.006996 | 0.02005 | 0.021883 | 0.012187 | 0.022179 | 0.009372 | 0.008516 | 0.036377 | 0.014196 | 0.002601 | 0.008334 | 0.001446 | 0.004753 | 0.125306 | 0.02774 | 0.016641 | 0.015084 | 0.006285 | 0.00319 | 0.010208 | 0.039025 | 0.005979 | 0.019164 | 0.010148 |
| 37.9166667 | 0.009225 | 0.019242 | 0.021656 | 0.012156 | 0.026004 | 0.006627 | 0.004686 | 0.03322 | 0.013407 | 0.001826 | 0.010058 | 0.002428 | 0.002795 | 0.157439 | 0.02591 | 0.009063 | 0.020568 | 0.010487 | 0.003182 | 0.008647 | 0.053548 | 0.005659 | 0.029181 | 0.01367 |
| 37.9305556 | 0.010343 | 0.017339 | 0.021532 | 0.012866 | 0.030508 | 0.003048 | 0.003779 | 0.029194 | 0.013356 | 0.001456 | 0.012575 | 0.002403 | 0.001219 | 0.133338 | 0.017808 | 0.005213 | 0.016771 | 0.01015 | 0.007647 | 0.008048 | 0.047624 | 0.004223 | 0.028546 | 0.019034 |
| 37.9444444 | 0.01063 | 0.013766 | 0.021766 | 0.015131 | 0.028928 | 0.001732 | 0.006689 | 0.025625 | 0.016454 | 0.002035 | 0.014317 | 0.001346 | 0.001114 | 0.128911 | 0.015756 | 0.006296 | 0.01439 | 0.010284 | 0.010899 | 0.009173 | 0.044696 | 0.001603 | 0.026103 | 0.023419 |
| 37.9583333 | 0.011135 | 0.008919 | 0.022249 | 0.019223 | 0.017831 | 0.004798 | 0.012114 | 0.022317 | 0.022623 | 0.003288 | 0.015489 | 0.000489 | 0.00108 | 0.097433 | 0.015889 | 0.005358 | 0.015644 | 0.010949 | 0.007063 | 0.007995 | 0.046972 | 0.002213 | 0.023678 | 0.02105 |
| 37.9722222 | 0.012382 | 0.004823 | 0.022393 | 0.022898 | 0.008087 | 0.01152 | 0.017209 | 0.019057 | 0.026857 | 0.004291 | 0.017196 | 0.000305 | 0.001116 | 0.065421 | 0.01666 | 0.005145 | 0.017242 | 0.012537 | 0.003968 | 0.007882 | 0.051658 | 0.004429 | 0.027163 | 0.02265 |
