## Supplemental Table S6b for "Leaf movements as a quantitative metric for early stress detection"

**Supplementary Table S6.** 1 hour integrated motion data in **(b)** 100 mM NaCl treatment on different CEA crops and their respective controls

**Supplementary Table S6.** 1 hour integrated motion data in **(b)** 100 mM NaCl treatment on Mizuna

| IntervalMid | Stress1 | Stress10 | Stress11 | Stress12 | Stress2 | Stress3 | Stress4 | Stress5 | Stress6 | Stress7 | Stress8 | Stress9 | Control1 | Control10 | Control11 | Control12 | Control2 | Control3 | Control4 | Control5 | Control6 | Control7 | Control8 | Control9 |
|---|---|---|---|---|---|---|---|---|---|---|---|---|---|---|---|---|---|---|---|---|---|---|---|---|
| 18.1111111 | 0.01104 | 0.02848 | 0.003444 | 0.010391 | 0.016188 | 0.002079 | 0.004301 | 0.007573 | 0.003614 | 0.003509 | 0.007688 | 0.0012 | 0.007158 | 0.025405 | 0.003474 | 0.046765 | 0.043169 | 0.002846 | 0.008685 | 0.018651 | 0.008409 | 0.013989 | 0.012085 | 0.02373 |
| 18.125 | 0.015158 | 0.036827 | 0.001706 | 0.008536 | 0.004639 | 0.001379 | 0.002962 | 0.002755 | 0.004546 | 0.003572 | 0.010821 | 0.000544 | 0.00639 | 0.040852 | 0.004386 | 0.059874 | 0.01838 | 0.003715 | 0.01115 | 0.006926 | 0.010757 | 0.016594 | 0.017972 | 0.030622 |
| 18.1388889 | 0.020594 | 0.04087 | 0.0012 | 0.005398 | 0.002327 | 0.000966 | 0.002888 | 0.001898 | 0.00483 | 0.003146 | 0.011581 | 0.000547 | 0.009092 | 0.061977 | 0.003787 | 0.052416 | 0.017242 | 0.00389 | 0.01218 | 0.006478 | 0.010196 | 0.012732 | 0.018872 | 0.0279 |
| 18.1527778 | 0.016921 | 0.035717 | 0.001247 | 0.002352 | 0.001965 | 0.000526 | 0.002898 | 0.00238 | 0.004303 | 0.002657 | 0.009435 | 0.000308 | 0.008575 | 0.067048 | 0.001767 | 0.03262 | 0.020001 | 0.003539 | 0.009877 | 0.009496 | 0.007366 | 0.007004 | 0.014639 | 0.018699 |
| 18.1666667 | 0.010603 | 0.029009 | 0.00203 | 0.001857 | 0.000882 | 0.000317 | 0.002646 | 0.002434 | 0.003762 | 0.002257 | 0.007713 | 0.000309 | 0.007116 | 0.065637 | 0.002242 | 0.018104 | 0.014648 | 0.003162 | 0.007696 | 0.009076 | 0.004785 | 0.005343 | 0.009654 | 0.010059 |
| 18.1805556 | 0.006225 | 0.02341 | 0.002696 | 0.002491 | 0.001032 | 0.00054 | 0.001901 | 0.002592 | 0.003372 | 0.001694 | 0.00683 | 0.000662 | 0.00593 | 0.062916 | 0.003846 | 0.010151 | 0.010378 | 0.003146 | 0.006669 | 0.006324 | 0.00282 | 0.007686 | 0.006216 | 0.004557 |
| 18.1944444 | 0.004137 | 0.017237 | 0.002148 | 0.002452 | 0.001747 | 0.001149 | 0.001026 | 0.002771 | 0.002318 | 0.001202 | 0.005504 | 0.00075 | 0.004106 | 0.053611 | 0.004962 | 0.005364 | 0.00718 | 0.002731 | 0.005178 | 0.003392 | 0.00139 | 0.009437 | 0.003714 | 0.001987 |
| 18.2083333 | 0.002693 | 0.009375 | 0.001423 | 0.002799 | 0.001689 | 0.001535 | 0.000853 | 0.002264 | 0.001415 | 0.001485 | 0.003178 | 0.0008 | 0.002312 | 0.033593 | 0.006057 | 0.002839 | 0.004807 | 0.001716 | 0.003072 | 0.00294 | 0.00089 | 0.006765 | 0.001782 | 0.001921 |
| 18.2222222 | 0.001551 | 0.003367 | 0.002059 | 0.004631 | 0.002347 | 0.001241 | 0.001502 | 0.001348 | 0.002383 | 0.00245 | 0.001505 | 0.001357 | 0.002497 | 0.014002 | 0.006859 | 0.003596 | 0.007316 | 0.001314 | 0.002731 | 0.005746 | 0.001384 | 0.004126 | 0.001184 | 0.003801 |
| 18.2361111 | 0.001659 | 0.001727 | 0.002253 | 0.005324 | 0.00422 | 0.00069 | 0.002351 | 0.001238 | 0.003435 | 0.002978 | 0.001259 | 0.001649 | 0.003347 | 0.005568 | 0.006916 | 0.005381 | 0.010179 | 0.001451 | 0.003892 | 0.008337 | 0.001744 | 0.003656 | 0.001384 | 0.004954 |
| 18.25 | 0.002213 | 0.001677 | 0.001283 | 0.004161 | 0.005533 | 0.000545 | 0.002752 | 0.002005 | 0.00303 | 0.002891 | 0.001056 | 0.001381 | 0.002834 | 0.005633 | 0.00655 | 0.005504 | 0.007841 | 0.001144 | 0.004082 | 0.008375 | 0.00145 | 0.002939 | 0.001365 | 0.004021 |
| 18.2638889 | 0.001859 | 0.002626 | 0.000515 | 0.002529 | 0.005732 | 0.00068 | 0.002477 | 0.002515 | 0.001814 | 0.00262 | 0.00118 | 0.001067 | 0.001764 | 0.013658 | 0.006067 | 0.004015 | 0.004045 | 0.00061 | 0.002932 | 0.006583 | 0.000917 | 0.003834 | 0.001039 | 0.002144 |
| 18.2777778 | 0.000955 | 0.004333 | 0.000294 | 0.001069 | 0.005392 | 0.000561 | 0.001713 | 0.002257 | 0.000681 | 0.002114 | 0.002035 | 0.000656 | 0.001645 | 0.025162 | 0.005491 | 0.002341 | 0.003551 | 0.000386 | 0.001578 | 0.004041 | 0.000493 | 0.006067 | 0.000675 | 0.000883 |
| 18.2916667 | 0.00056 | 0.00428 | 0.000604 | 0.001408 | 0.005412 | 0.00035 | 0.000956 | 0.001634 | 0.000577 | 0.001649 | 0.00195 | 0.00025 | 0.001581 | 0.0224 | 0.005162 | 0.002768 | 0.004235 | 0.000473 | 0.001317 | 0.002075 | 0.000525 | 0.005256 | 0.00083 | 0.000895 |
| 18.3055556 | 0.001286 | 0.00499 | 0.001442 | 0.004103 | 0.006818 | 0.000392 | 0.000856 | 0.00156 | 0.001818 | 0.001849 | 0.001509 | 0.000204 | 0.001443 | 0.013792 | 0.005398 | 0.005957 | 0.003125 | 0.001009 | 0.002742 | 0.001702 | 0.001148 | 0.003367 | 0.001851 | 0.002223 |
| 18.3194444 | 0.002477 | 0.008128 | 0.002327 | 0.006661 | 0.009065 | 0.000582 | 0.001348 | 0.002276 | 0.0035 | 0.002504 | 0.002201 | 0.000408 | 0.002566 | 0.018849 | 0.005935 | 0.009819 | 0.003159 | 0.00183 | 0.004781 | 0.002645 | 0.001945 | 0.004659 | 0.003068 | 0.004701 |
| 18.3333333 | 0.00264 | 0.008781 | 0.002353 | 0.006477 | 0.010362 | 0.000629 | 0.001409 | 0.002704 | 0.003833 | 0.002495 | 0.002273 | 0.000439 | 0.003124 | 0.02183 | 0.005988 | 0.011535 | 0.00534 | 0.002056 | 0.005548 | 0.003483 | 0.002244 | 0.006034 | 0.003306 | 0.006245 |
| 18.3472222 | 0.001554 | 0.005258 | 0.001486 | 0.00386 | 0.00937 | 0.000457 | 0.000932 | 0.002245 | 0.002472 | 0.001498 | 0.001515 | 0.00044 | 0.001988 | 0.012607 | 0.004895 | 0.009686 | 0.004899 | 0.001356 | 0.004251 | 0.002783 | 0.001843 | 0.004766 | 0.002136 | 0.00511 |
| 18.3611111 | 0.00068 | 0.002115 | 0.000862 | 0.001817 | 0.006435 | 0.000442 | 0.000856 | 0.00155 | 0.001397 | 0.000662 | 0.00179 | 0.000814 | 0.001168 | 0.005693 | 0.003099 | 0.005835 | 0.002852 | 0.000692 | 0.002009 | 0.001503 | 0.0012 | 0.002438 | 0.000986 | 0.002464 |
| 18.375 | 0.000776 | 0.001857 | 0.00067 | 0.001562 | 0.004239 | 0.000535 | 0.001 | 0.001241 | 0.001327 | 0.000478 | 0.00231 | 0.001026 | 0.001392 | 0.007149 | 0.001947 | 0.00349 | 0.003258 | 0.000622 | 0.000574 | 0.00122 | 0.00087 | 0.000762 | 0.001059 | 0.0008 |
| 18.3888889 | 0.001011 | 0.001683 | 0.000682 | 0.001271 | 0.005205 | 0.000441 | 0.000728 | 0.00136 | 0.001002 | 0.000384 | 0.001849 | 0.000716 | 0.00141 | 0.008291 | 0.002164 | 0.00463 | 0.003207 | 0.000615 | 0.000821 | 0.001346 | 0.001091 | 0.001001 | 0.001148 | 0.001574 |
| 18.4027778 | 0.000797 | 0.001114 | 0.001305 | 0.001129 | 0.009276 | 0.000271 | 0.0004 | 0.001775 | 0.000706 | 0.000532 | 0.000996 | 0.000357 | 0.001011 | 0.005675 | 0.003347 | 0.008774 | 0.003068 | 0.000972 | 0.002458 | 0.002237 | 0.001869 | 0.003007 | 0.000695 | 0.004466 |
| 18.4166667 | 0.000588 | 0.002203 | 0.002067 | 0.001996 | 0.014754 | 0.000293 | 0.000538 | 0.002413 | 0.001288 | 0.001079 | 0.000769 | 0.000335 | 0.001212 | 0.005531 | 0.004674 | 0.014525 | 0.006441 | 0.002072 | 0.004475 | 0.003947 | 0.003036 | 0.004851 | 0.000483 | 0.008231 |
| 18.4305556 | 0.000911 | 0.003468 | 0.002233 | 0.002285 | 0.018766 | 0.000572 | 0.000847 | 0.003013 | 0.001832 | 0.001289 | 0.001138 | 0.000539 | 0.001823 | 0.007923 | 0.005359 | 0.019384 | 0.009588 | 0.002978 | 0.005756 | 0.004391 | 0.004077 | 0.004506 | 0.000539 | 0.010941 |
| 18.4444444 | 0.00124 | 0.002637 | 0.001673 | 0.001785 | 0.018207 | 0.000721 | 0.000688 | 0.003165 | 0.001394 | 0.001023 | 0.001104 | 0.000514 | 0.001709 | 0.007678 | 0.00494 | 0.019902 | 0.007659 | 0.002591 | 0.005429 | 0.003148 | 0.004406 | 0.002387 | 0.001016 | 0.010481 |
| 18.4583333 | 0.000907 | 0.001542 | 0.000851 | 0.002418 | 0.012631 | 0.000614 | 0.000365 | 0.002942 | 0.000969 | 0.001293 | 0.000677 | 0.000275 | 0.000991 | 0.00437 | 0.003525 | 0.01495 | 0.004689 | 0.00153 | 0.003326 | 0.00177 | 0.003838 | 0.000822 | 0.002153 | 0.006527 |
| 18.4722222 | 0.000735 | 0.002637 | 0.00055 | 0.003996 | 0.005603 | 0.000501 | 0.000527 | 0.00267 | 0.001759 | 0.002356 | 0.000815 | 0.000275 | 0.000755 | 0.002266 | 0.001756 | 0.007641 | 0.006586 | 0.001373 | 0.001558 | 0.001 | 0.002625 | 0.000848 | 0.003288 | 0.003534 |
| 18.4861111 | 0.001615 | 0.004448 | 0.000576 | 0.004658 | 0.001488 | 0.000487 | 0.001046 | 0.00223 | 0.002998 | 0.003395 | 0.001741 | 0.000635 | 0.000796 | 0.002572 | 0.000751 | 0.002533 | 0.009585 | 0.001538 | 0.001637 | 0.000724 | 0.001609 | 0.001015 | 0.003518 | 0.004344 |
| 18.5 | 0.002888 | 0.005809 | 0.00058 | 0.004029 | 0.000689 | 0.000737 | 0.001594 | 0.001485 | 0.003839 | 0.00416 | 0.002861 | 0.000956 | 0.00094 | 0.002417 | 0.001438 | 0.001082 | 0.008969 | 0.001343 | 0.001533 | 0.001203 | 0.001759 | 0.001086 | 0.002346 | 0.004783 |
| 18.5138889 | 0.003663 | 0.008803 | 0.000601 | 0.003496 | 0.002605 | 0.001156 | 0.001459 | 0.001343 | 0.00491 | 0.004823 | 0.00363 | 0.000855 | 0.001175 | 0.004669 | 0.002974 | 0.004143 | 0.00896 | 0.00188 | 0.000748 | 0.001829 | 0.002539 | 0.002178 | 0.001038 | 0.004863 |
| 18.5277778 | 0.003982 | 0.017868 | 0.001715 | 0.004871 | 0.011712 | 0.002582 | 0.000996 | 0.002399 | 0.007275 | 0.005752 | 0.004365 | 0.000979 | 0.003452 | 0.022034 | 0.002901 | 0.0184 | 0.019241 | 0.002819 | 0.002097 | 0.004781 | 0.002625 | 0.003118 | 0.001397 | 0.011023 |
| 18.5416667 | 0.004513 | 0.035526 | 0.005383 | 0.007793 | 0.030435 | 0.005106 | 0.00117 | 0.003209 | 0.01041 | 0.007808 | 0.006436 | 0.00121 | 0.013516 | 0.065972 | 0.00177 | 0.041414 | 0.036872 | 0.002757 | 0.007322 | 0.012844 | 0.003942 | 0.006779 | 0.00466 | 0.025406 |
| 18.5555556 | 0.005387 | 0.058315 | 0.008985 | 0.009712 | 0.046545 | 0.005254 | 0.002542 | 0.004823 | 0.013084 | 0.010887 | 0.013372 | 0.001456 | 0.028867 | 0.114914 | 0.002119 | 0.052112 | 0.043993 | 0.001716 | 0.014708 | 0.021055 | 0.008751 | 0.011309 | 0.011057 | 0.043808 |
| 18.5694444 | 0.005645 | 0.075364 | 0.00955 | 0.009487 | 0.044889 | 0.002918 | 0.005161 | 0.008365 | 0.014526 | 0.013051 | 0.024058 | 0.002793 | 0.037117 | 0.135262 | 0.003601 | 0.04255 | 0.035894 | 0.001501 | 0.020211 | 0.023236 | 0.013764 | 0.009368 | 0.017038 | 0.061157 |
| 18.5833333 | 0.004849 | 0.077102 | 0.008146 | 0.00931 | 0.02955 | 0.00188 | 0.006862 | 0.009697 | 0.014168 | 0.013342 | 0.030566 | 0.004357 | 0.034034 | 0.119538 | 0.004918 | 0.027061 | 0.023297 | 0.002905 | 0.020447 | 0.019742 | 0.014885 | 0.006414 | 0.019556 | 0.068683 |
| 18.5972222 | 0.003354 | 0.067074 | 0.006245 | 0.010211 | 0.015029 | 0.001991 | 0.00661 | 0.007187 | 0.011967 | 0.012419 | 0.029797 | 0.005124 | 0.028327 | 0.080442 | 0.005211 | 0.015312 | 0.013669 | 0.003948 | 0.016041 | 0.015725 | 0.012541 | 0.007308 | 0.018944 | 0.05927 |
| 18.6111111 | 0.001713 | 0.053843 | 0.004377 | 0.009532 | 0.007844 | 0.001525 | 0.005457 | 0.003476 | 0.008801 | 0.010489 | 0.024645 | 0.005041 | 0.026991 | 0.047972 | 0.004727 | 0.007958 | 0.007882 | 0.003699 | 0.010628 | 0.014243 | 0.009352 | 0.009206 | 0.016076 | 0.041596 |
| 18.625 | 0.000894 | 0.043045 | 0.002908 | 0.006953 | 0.005625 | 0.000993 | 0.00444 | 0.000845 | 0.006914 | 0.00862 | 0.01976 | 0.00441 | 0.029925 | 0.034355 | 0.004862 | 0.004289 | 0.005613 | 0.003181 | 0.006839 | 0.013216 | 0.007368 | 0.00933 | 0.013076 | 0.028876 |
| 18.6388889 | 0.000645 | 0.036401 | 0.002298 | 0.00522 | 0.004674 | 0.00105 | 0.00409 | 0.000149 | 0.007006 | 0.008005 | 0.017236 | 0.00405 | 0.033146 | 0.030597 | 0.006275 | 0.003283 | 0.007107 | 0.002807 | 0.005249 | 0.011045 | 0.007565 | 0.007018 | 0.012262 | 0.025149 |
| 18.6527778 | 0.000268 | 0.030032 | 0.002461 | 0.005522 | 0.003658 | 0.001262 | 0.004144 | 0.000377 | 0.007318 | 0.007565 | 0.014841 | 0.004034 | 0.031178 | 0.032549 | 0.008117 | 0.004306 | 0.01037 | 0.002324 | 0.004893 | 0.009184 | 0.009161 | 0.005237 | 0.012921 | 0.027257 |
| 18.6666667 | 0.00012 | 0.021717 | 0.002752 | 0.0067 | 0.002414 | 0.001338 | 0.004383 | 0.000726 | 0.006261 | 0.006291 | 0.011136 | 0.003806 | 0.02493 | 0.036453 | 0.008939 | 0.006405 | 0.012057 | 0.001764 | 0.004718 | 0.008057 | 0.009846 | 0.006254 | 0.012733 | 0.028974 |
| 18.6805556 | 0.000281 | 0.014747 | 0.003025 | 0.006983 | 0.00212 | 0.00152 | 0.004534 | 0.001392 | 0.004158 | 0.005027 | 0.007165 | 0.003071 | 0.019415 | 0.037451 | 0.008366 | 0.00783 | 0.011108 | 0.001393 | 0.004407 | 0.006549 | 0.008657 | 0.009142 | 0.011188 | 0.02631 |
| 18.6944444 | 0.00059 | 0.01068 | 0.003262 | 0.005644 | 0.00257 | 0.001545 | 0.003998 | 0.001936 | 0.002131 | 0.004109 | 0.00435 | 0.002079 | 0.016635 | 0.03298 | 0.006723 | 0.007496 | 0.008771 | 0.001082 | 0.003793 | 0.00481 | 0.00654 | 0.010525 | 0.009185 | 0.020563 |
| 18.7083333 | 0.001045 | 0.008033 | 0.003069 | 0.003479 | 0.001949 | 0.001 | 0.00304 | 0.001544 | 0.001228 | 0.00307 | 0.002787 | 0.001117 | 0.016338 | 0.024697 | 0.00458 | 0.006014 | 0.007347 | 0.000598 | 0.002807 | 0.003596 | 0.004079 | 0.008666 | 0.007498 | 0.015535 |
| 18.7222222 | 0.001077 | 0.00667 | 0.002301 | 0.002068 | 0.00099 | 0.000358 | 0.002823 | 0.000747 | 0.001653 | 0.002224 | 0.002235 | 0.000478 | 0.016401 | 0.016546 | 0.002848 | 0.00461 | 0.007546 | 0.000178 | 0.00191 | 0.003398 | 0.002029 | 0.005502 | 0.006606 | 0.013054 |
| 18.7361111 | 0.000895 | 0.006659 | 0.00142 | 0.002107 | 0.001073 | 0.00024 | 0.003633 | 0.000611 | 0.001804 | 0.002131 | 0.002362 | 0.000503 | 0.014461 | 0.010741 | 0.001868 | 0.003824 | 0.007779 | 0.000148 | 0.001743 | 0.004283 | 0.001288 | 0.003442 | 0.006476 | 0.013169 |
| 18.75 | 0.001615 | 0.00709 | 0.000915 | 0.002914 | 0.001388 | 0.000496 | 0.004385 | 0.001052 | 0.001279 | 0.00247 | 0.00258 | 0.000947 | 0.010965 | 0.006377 | 0.001579 | 0.003408 | 0.007501 | 0.000396 | 0.002242 | 0.005898 | 0.00134 | 0.003095 | 0.006719 | 0.014876 |
| 18.7638889 | 0.002663 | 0.008016 | 0.00082 | 0.003523 | 0.001723 | 0.000559 | 0.004008 | 0.001427 | 0.001119 | 0.002438 | 0.003058 | 0.001109 | 0.007436 | 0.003274 | 0.001714 | 0.003047 | 0.006712 | 0.000537 | 0.00269 | 0.007581 | 0.001207 | 0.003893 | 0.00709 | 0.015511 |
| 18.7777778 | 0.0029 | 0.009771 | 0.000936 | 0.003642 | 0.002861 | 0.000293 | 0.002905 | 0.001653 | 0.001144 | 0.002087 | 0.004047 | 0.00079 | 0.004781 | 0.001905 | 0.001576 | 0.002856 | 0.005551 | 0.000418 | 0.002643 | 0.008548 | 0.000747 | 0.004797 | 0.007873 | 0.013753 |
| 18.7916667 | 0.002607 | 0.010754 | 0.000967 | 0.00325 | 0.004406 | 0.000166 | 0.002057 | 0.002102 | 0.0008 | 0.001957 | 0.004652 | 0.000375 | 0.003768 | 0.001372 | 0.000923 | 0.003047 | 0.005145 | 0.000238 | 0.002418 | 0.008236 | 0.000398 | 0.004408 | 0.008658 | 0.011305 |
| 18.8055556 | 0.002477 | 0.009293 | 0.000754 | 0.002276 | 0.005233 | 0.000378 | 0.001317 | 0.00289 | 0.000666 | 0.001964 | 0.004016 | 0.00017 | 0.003441 | 0.001171 | 0.000485 | 0.00351 | 0.00542 | 0.000233 | 0.002257 | 0.006741 | 0.000389 | 0.002456 | 0.00804 | 0.00947 |
| 18.8194444 | 0.002441 | 0.006901 | 0.000787 | 0.001067 | 0.005077 | 0.000484 | 0.000839 | 0.003495 | 0.001228 | 0.001619 | 0.003004 | 0.000196 | 0.002763 | 0.001404 | 0.000854 | 0.00389 | 0.005078 | 0.000412 | 0.001855 | 0.00517 | 0.000706 | 0.000715 | 0.006063 | 0.007797 |
| 18.8333333 | 0.002184 | 0.006066 | 0.001305 | 0.000584 | 0.004777 | 0.00034 | 0.000975 | 0.003196 | 0.001875 | 0.001 | 0.002674 | 0.000272 | 0.003558 | 0.003366 | 0.001397 | 0.003859 | 0.004934 | 0.000591 | 0.001067 | 0.004831 | 0.001132 | 0.000349 | 0.003988 | 0.005517 |
| 18.8472222 | 0.001792 | 0.006081 | 0.0018 | 0.001425 | 0.004683 | 0.000281 | 0.000845 | 0.001901 | 0.00175 | 0.000626 | 0.002474 | 0.000396 | 0.005832 | 0.006968 | 0.001815 | 0.003414 | 0.005472 | 0.000609 | 0.000719 | 0.005277 | 0.001354 | 0.000513 | 0.002396 | 0.002755 |
| 18.8611111 | 0.00183 | 0.004345 | 0.002227 | 0.002799 | 0.004339 | 0.000392 | 0.000545 | 0.000895 | 0.000926 | 0.000563 | 0.001632 | 0.000747 | 0.005881 | 0.007192 | 0.002724 | 0.002814 | 0.004862 | 0.000423 | 0.001745 | 0.004748 | 0.001368 | 0.000986 | 0.001248 | 0.001013 |
| 18.875 | 0.006634 | 0.001794 | 0.002886 | 0.003808 | 0.003744 | 0.000765 | 0.000932 | 0.001306 | 0.000727 | 0.000445 | 0.001238 | 0.001114 | 0.003596 | 0.004005 | 0.003873 | 0.002336 | 0.003236 | 0.00035 | 0.002955 | 0.003152 | 0.001308 | 0.002576 | 0.000822 | 0.000838 |
| 18.8888889 | 0.018017 | 0.000901 | 0.003481 | 0.004602 | 0.003804 | 0.001425 | 0.001537 | 0.001962 | 0.001951 | 0.000661 | 0.001705 | 0.001127 | 0.003737 | 0.002235 | 0.004028 | 0.002253 | 0.002016 | 0.00061 | 0.002593 | 0.002099 | 0.001377 | 0.004041 | 0.000836 | 0.000966 |
| 18.9027778 | 0.023381 | 0.001197 | 0.003414 | 0.005875 | 0.004557 | 0.001646 | 0.001354 | 0.001693 | 0.003143 | 0.001465 | 0.001848 | 0.000748 | 0.008595 | 0.003167 | 0.00303 | 0.002601 | 0.002032 | 0.000706 | 0.001591 | 0.002719 | 0.001854 | 0.003777 | 0.000725 | 0.001143 |
| 18.9166667 | 0.017791 | 0.001617 | 0.002949 | 0.008225 | 0.005047 | 0.000995 | 0.000847 | 0.001597 | 0.00314 | 0.00175 | 0.002406 | 0.000421 | 0.014095 | 0.004657 | 0.002161 | 0.002917 | 0.003481 | 0.0004 | 0.001536 | 0.003837 | 0.002457 | 0.003052 | 0.00068 | 0.002216 |
| 18.9305556 | 0.013805 | 0.002184 | 0.00345 | 0.011597 | 0.00791 | 0.000427 | 0.000933 | 0.002143 | 0.003775 | 0.001856 | 0.002849 | 0.000433 | 0.014684 | 0.004458 | 0.002242 | 0.002379 | 0.004899 | 0.000227 | 0.001435 | 0.003374 | 0.002404 | 0.003364 | 0.001585 | 0.004785 |
| 18.9444444 | 0.014254 | 0.004088 | 0.005306 | 0.014772 | 0.012371 | 0.000655 | 0.001756 | 0.002073 | 0.00646 | 0.002961 | 0.002398 | 0.00095 | 0.011288 | 0.005759 | 0.003139 | 0.001476 | 0.004595 | 0.000593 | 0.000916 | 0.001641 | 0.001483 | 0.004181 | 0.004691 | 0.008834 |
| 18.9583333 | 0.014619 | 0.00643 | 0.007184 | 0.016351 | 0.011548 | 0.000913 | 0.003002 | 0.001439 | 0.008991 | 0.00442 | 0.002881 | 0.002158 | 0.01035 | 0.009898 | 0.004797 | 0.001918 | 0.004887 | 0.001238 | 0.001301 | 0.00078 | 0.000701 | 0.004894 | 0.008657 | 0.012564 |
| 18.9722222 | 0.014285 | 0.007874 | 0.00836 | 0.017022 | 0.006318 | 0.000642 | 0.003911 | 0.000969 | 0.009281 | 0.005538 | 0.003472 | 0.003272 | 0.015252 | 0.011862 | 0.00694 | 0.003644 | 0.008254 | 0.001676 | 0.002131 | 0.001848 | 0.000923 | 0.005763 | 0.011001 | 0.013895 |
| 18.9861111 | 0.013961 | 0.009544 | 0.009203 | 0.018241 | 0.002917 | 0.000317 | 0.004434 | 0.000612 | 0.008283 | 0.006427 | 0.00314 | 0.003799 | 0.021627 | 0.008797 | 0.008762 | 0.005436 | 0.012959 | 0.001709 | 0.00262 | 0.003971 | 0.001344 | 0.006218 | 0.011989 | 0.013699 |
| 19 | 0.014956 | 0.011599 | 0.009928 | 0.020101 | 0.004128 | 0.000198 | 0.004739 | 0.000487 | 0.007726 | 0.007209 | 0.003516 | 0.00484 | 0.022739 | 0.004243 | 0.009675 | 0.006668 | 0.019073 | 0.00146 | 0.002908 | 0.005502 | 0.000992 | 0.00562 | 0.012885 | 0.014187 |
| 19.0138889 | 0.015968 | 0.013011 | 0.010851 | 0.021765 | 0.008494 | 0.00067 | 0.004175 | 0.00092 | 0.007994 | 0.00765 | 0.004056 | 0.006255 | 0.01947 | 0.003492 | 0.009484 | 0.007195 | 0.026104 | 0.001072 | 0.002415 | 0.005597 | 0.000845 | 0.004827 | 0.012868 | 0.015479 |
| 19.0277778 | 0.014315 | 0.013534 | 0.012108 | 0.021604 | 0.012606 | 0.002678 | 0.002858 | 0.002416 | 0.007809 | 0.007391 | 0.003103 | 0.005281 | 0.018641 | 0.005096 | 0.008304 | 0.007295 | 0.027573 | 0.001607 | 0.001841 | 0.005591 | 0.002066 | 0.00403 | 0.010739 | 0.017386 |
| 19.0416667 | 0.01155 | 0.015816 | 0.013128 | 0.01908 | 0.017967 | 0.005244 | 0.001787 | 0.003297 | 0.007132 | 0.007538 | 0.001778 | 0.003924 | 0.023372 | 0.008122 | 0.006928 | 0.008371 | 0.022592 | 0.003233 | 0.00185 | 0.006792 | 0.003005 | 0.003598 | 0.008825 | 0.024898 |
| 19.0555556 | 0.012712 | 0.022122 | 0.014415 | 0.016877 | 0.028332 | 0.005219 | 0.002315 | 0.004853 | 0.007338 | 0.009921 | 0.001811 | 0.004925 | 0.031583 | 0.033384 | 0.007519 | 0.012489 | 0.022913 | 0.004043 | 0.005733 | 0.010786 | 0.003415 | 0.005858 | 0.010756 | 0.044919 |
| 19.0694444 | 0.017191 | 0.030974 | 0.015507 | 0.016661 | 0.036345 | 0.002969 | 0.005029 | 0.00904 | 0.007624 | 0.012495 | 0.004509 | 0.004342 | 0.037572 | 0.096457 | 0.010277 | 0.017931 | 0.032957 | 0.004567 | 0.01409 | 0.01923 | 0.006269 | 0.010741 | 0.013511 | 0.065622 |
| 19.0833333 | 0.017859 | 0.030918 | 0.013015 | 0.014275 | 0.030055 | 0.001889 | 0.007039 | 0.009853 | 0.006481 | 0.011217 | 0.007379 | 0.002528 | 0.03321 | 0.136167 | 0.010841 | 0.018204 | 0.039161 | 0.004623 | 0.015497 | 0.025902 | 0.008902 | 0.014244 | 0.011384 | 0.059702 |
| 19.0972222 | 0.011856 | 0.017104 | 0.00784 | 0.007966 | 0.014201 | 0.001985 | 0.006385 | 0.006578 | 0.004282 | 0.007246 | 0.005738 | 0.002322 | 0.018087 | 0.095388 | 0.006904 | 0.011225 | 0.032108 | 0.00277 | 0.008262 | 0.021987 | 0.006679 | 0.011313 | 0.006758 | 0.032124 |
| 19.1111111 | 0.004585 | 0.005624 | 0.005554 | 0.00387 | 0.004626 | 0.002057 | 0.005053 | 0.004554 | 0.002573 | 0.004906 | 0.00253 | 0.002924 | 0.005958 | 0.036045 | 0.002533 | 0.004134 | 0.019951 | 0.001375 | 0.003849 | 0.010787 | 0.003646 | 0.005667 | 0.003828 | 0.017509 |
| 19.125 | 0.001546 | 0.003512 | 0.006733 | 0.00475 | 0.005543 | 0.002118 | 0.00431 | 0.004147 | 0.001797 | 0.004748 | 0.001697 | 0.004083 | 0.004535 | 0.022161 | 0.001245 | 0.002395 | 0.012065 | 0.001473 | 0.0051 | 0.003963 | 0.00341 | 0.004934 | 0.002338 | 0.020887 |
| 19.1388889 | 0.002174 | 0.004554 | 0.007126 | 0.006076 | 0.007439 | 0.00174 | 0.00351 | 0.003414 | 0.001321 | 0.003979 | 0.001816 | 0.004594 | 0.005773 | 0.03951 | 0.001934 | 0.003424 | 0.008047 | 0.001925 | 0.008306 | 0.006055 | 0.003267 | 0.005606 | 0.001272 | 0.02311 |
| 19.1527778 | 0.003118 | 0.003855 | 0.005297 | 0.004889 | 0.005054 | 0.000977 | 0.002389 | 0.001872 | 0.001569 | 0.002132 | 0.002792 | 0.003397 | 0.00555 | 0.06682 | 0.003084 | 0.004018 | 0.005671 | 0.002687 | 0.012046 | 0.011277 | 0.00218 | 0.005 | 0.000454 | 0.020085 |
| 19.1666667 | 0.002417 | 0.002685 | 0.003375 | 0.002569 | 0.003355 | 0.000447 | 0.001525 | 0.000584 | 0.002434 | 0.001238 | 0.005373 | 0.001852 | 0.005729 | 0.095659 | 0.004616 | 0.004535 | 0.003757 | 0.003644 | 0.015291 | 0.013874 | 0.001115 | 0.00613 | 0.000121 | 0.017663 |
| 19.1805556 | 0.001274 | 0.00361 | 0.002851 | 0.001075 | 0.004371 | 0.000279 | 0.001294 | 0.000291 | 0.002519 | 0.001217 | 0.007317 | 0.001153 | 0.006328 | 0.111663 | 0.006561 | 0.006215 | 0.003685 | 0.004343 | 0.01638 | 0.013674 | 0.000438 | 0.008361 | 0.000205 | 0.017852 |
| 19.1944444 | 0.000945 | 0.005338 | 0.003985 | 0.001683 | 0.003901 | 0.000683 | 0.001953 | 0.000833 | 0.001661 | 0.001231 | 0.005988 | 0.001423 | 0.006573 | 0.110578 | 0.008575 | 0.008697 | 0.007422 | 0.003955 | 0.014495 | 0.010716 | 0.00039 | 0.007815 | 0.000282 | 0.017297 |
| 19.2083333 | 0.00147 | 0.011495 | 0.006719 | 0.004709 | 0.001667 | 0.002151 | 0.003741 | 0.002319 | 0.001435 | 0.002549 | 0.005845 | 0.00269 | 0.005389 | 0.093546 | 0.009558 | 0.010294 | 0.009669 | 0.002535 | 0.009808 | 0.00685 | 0.000829 | 0.005212 | 0.0005 | 0.011731 |
| 19.2222222 | 0.003344 | 0.020677 | 0.009012 | 0.007741 | 0.00148 | 0.003464 | 0.005216 | 0.003891 | 0.002306 | 0.004562 | 0.011103 | 0.003741 | 0.002914 | 0.059143 | 0.008839 | 0.010263 | 0.006866 | 0.001927 | 0.004411 | 0.007086 | 0.001308 | 0.006299 | 0.001502 | 0.005842 |
| 19.2361111 | 0.004615 | 0.020628 | 0.007981 | 0.006511 | 0.002418 | 0.002832 | 0.003973 | 0.003291 | 0.002557 | 0.00413 | 0.012478 | 0.002851 | 0.000832 | 0.033062 | 0.007985 | 0.008964 | 0.004157 | 0.002164 | 0.001582 | 0.00772 | 0.001906 | 0.008658 | 0.002761 | 0.005044 |
| 19.25 | 0.003397 | 0.011477 | 0.004289 | 0.003669 | 0.004262 | 0.001611 | 0.001938 | 0.001615 | 0.003214 | 0.003235 | 0.008118 | 0.001362 | 0.000768 | 0.033036 | 0.008527 | 0.00687 | 0.00609 | 0.001701 | 0.002628 | 0.005997 | 0.002967 | 0.006508 | 0.003291 | 0.009595 |
| 19.2638889 | 0.00232 | 0.005181 | 0.001868 | 0.004531 | 0.009374 | 0.001212 | 0.00221 | 0.000981 | 0.005757 | 0.005172 | 0.006843 | 0.001098 | 0.002337 | 0.038115 | 0.009475 | 0.004118 | 0.009532 | 0.001238 | 0.004863 | 0.007391 | 0.003951 | 0.004337 | 0.003276 | 0.018266 |
| 19.2777778 | 0.003191 | 0.003707 | 0.002409 | 0.005631 | 0.012457 | 0.000716 | 0.00288 | 0.000829 | 0.00712 | 0.006784 | 0.007061 | 0.001049 | 0.003266 | 0.031232 | 0.009553 | 0.002599 | 0.008375 | 0.00114 | 0.004958 | 0.008224 | 0.003991 | 0.005611 | 0.003443 | 0.020029 |
| 19.2916667 | 0.00326 | 0.012253 | 0.002885 | 0.004119 | 0.010647 | 0.000223 | 0.002423 | 0.000782 | 0.005973 | 0.006009 | 0.007169 | 0.000647 | 0.002553 | 0.019711 | 0.009047 | 0.003467 | 0.003892 | 0.000684 | 0.003243 | 0.004675 | 0.003474 | 0.00605 | 0.00394 | 0.010943 |
| 19.3055556 | 0.002115 | 0.021955 | 0.001677 | 0.002086 | 0.00845 | 0.000218 | 0.001667 | 0.001163 | 0.004713 | 0.004627 | 0.007615 | 0.000479 | 0.002289 | 0.013327 | 0.008062 | 0.003597 | 0.00148 | 0.000355 | 0.001907 | 0.00406 | 0.003422 | 0.005479 | 0.004284 | 0.002824 |
| 19.3194444 | 0.002186 | 0.016732 | 0.001137 | 0.001614 | 0.007253 | 0.000322 | 0.00088 | 0.002076 | 0.004016 | 0.003432 | 0.007072 | 0.000777 | 0.004762 | 0.011941 | 0.005642 | 0.002478 | 0.001489 | 0.000476 | 0.001664 | 0.008956 | 0.003836 | 0.006566 | 0.004271 | 0.002841 |
| 19.3333333 | 0.003667 | 0.015081 | 0.002051 | 0.002742 | 0.006283 | 0.000541 | 0.000381 | 0.002834 | 0.002577 | 0.002357 | 0.008289 | 0.001507 | 0.009123 | 0.013205 | 0.002949 | 0.001896 | 0.003488 | 0.00068 | 0.001916 | 0.014194 | 0.004003 | 0.008367 | 0.004135 | 0.00889 |
| 19.3472222 | 0.004004 | 0.025743 | 0.003125 | 0.00375 | 0.005792 | 0.000821 | 0.000352 | 0.003642 | 0.001212 | 0.001743 | 0.013737 | 0.00236 | 0.012044 | 0.016196 | 0.002568 | 0.001364 | 0.006235 | 0.001056 | 0.002164 | 0.019554 | 0.003278 | 0.009524 | 0.003506 | 0.01759 |
| 19.3611111 | 0.002255 | 0.033509 | 0.00365 | 0.004189 | 0.006113 | 0.000739 | 0.00061 | 0.004763 | 0.001504 | 0.001672 | 0.020592 | 0.002816 | 0.011463 | 0.019686 | 0.003656 | 0.000647 | 0.005881 | 0.001622 | 0.002467 | 0.021699 | 0.001936 | 0.010305 | 0.002572 | 0.025745 |
| 19.375 | 0.001094 | 0.031185 | 0.00353 | 0.003652 | 0.007133 | 0.000591 | 0.000987 | 0.004554 | 0.00189 | 0.001349 | 0.021644 | 0.002541 | 0.009202 | 0.020644 | 0.004568 | 0.000262 | 0.004134 | 0.001706 | 0.002743 | 0.017176 | 0.001452 | 0.010895 | 0.002585 | 0.026708 |
| 19.3888889 | 0.001995 | 0.021574 | 0.002951 | 0.00221 | 0.008513 | 0.000735 | 0.001133 | 0.002723 | 0.001417 | 0.000621 | 0.015463 | 0.002094 | 0.00868 | 0.017649 | 0.00532 | 0.000441 | 0.00341 | 0.001148 | 0.002505 | 0.010764 | 0.002123 | 0.010814 | 0.002704 | 0.018681 |
| 19.4027778 | 0.005298 | 0.012695 | 0.002099 | 0.001411 | 0.010369 | 0.000921 | 0.000985 | 0.00132 | 0.001868 | 0.000543 | 0.009591 | 0.002525 | 0.010869 | 0.012741 | 0.005627 | 0.000775 | 0.002481 | 0.000589 | 0.001692 | 0.007602 | 0.002639 | 0.009363 | 0.001801 | 0.009608 |
| 19.4166667 | 0.009975 | 0.009154 | 0.00119 | 0.001669 | 0.012695 | 0.001217 | 0.000689 | 0.001403 | 0.004046 | 0.002229 | 0.008479 | 0.003909 | 0.014095 | 0.009167 | 0.005773 | 0.000714 | 0.001442 | 0.000424 | 0.000774 | 0.012621 | 0.003017 | 0.006569 | 0.001224 | 0.00559 |
| 19.4305556 | 0.012771 | 0.010203 | 0.001023 | 0.002402 | 0.014324 | 0.001601 | 0.000342 | 0.00215 | 0.006787 | 0.005499 | 0.00954 | 0.005233 | 0.016491 | 0.008563 | 0.006277 | 0.000678 | 0.002076 | 0.000521 | 0.000197 | 0.015647 | 0.004317 | 0.0045 | 0.00109 | 0.005865 |
| 19.4444444 | 0.010873 | 0.012826 | 0.002246 | 0.003556 | 0.014415 | 0.00162 | 0.000327 | 0.003217 | 0.008862 | 0.008564 | 0.010232 | 0.005576 | 0.016679 | 0.010342 | 0.006594 | 0.001191 | 0.006505 | 0.000559 | 0.000183 | 0.012536 | 0.00667 | 0.005877 | 0.001157 | 0.008379 |
| 19.4583333 | 0.006447 | 0.013926 | 0.004416 | 0.004549 | 0.014765 | 0.001394 | 0.001044 | 0.005044 | 0.009941 | 0.010087 | 0.010703 | 0.005349 | 0.014743 | 0.013973 | 0.006213 | 0.002293 | 0.011001 | 0.000386 | 0.000695 | 0.017272 | 0.008878 | 0.009845 | 0.002846 | 0.011583 |
| 19.4722222 | 0.006542 | 0.01275 | 0.006549 | 0.004571 | 0.017566 | 0.001446 | 0.002582 | 0.006927 | 0.01023 | 0.01038 | 0.011319 | 0.005572 | 0.013242 | 0.019694 | 0.005895 | 0.003451 | 0.009739 | 0.000188 | 0.00138 | 0.025723 | 0.009663 | 0.012193 | 0.005765 | 0.013848 |
| 19.4861111 | 0.011656 | 0.010347 | 0.008158 | 0.003982 | 0.019947 | 0.001791 | 0.004402 | 0.007614 | 0.009813 | 0.010033 | 0.011617 | 0.005981 | 0.013771 | 0.025669 | 0.006361 | 0.003554 | 0.006901 | 0.000297 | 0.001567 | 0.026709 | 0.0094 | 0.011143 | 0.00835 | 0.014879 |
| 19.5 | 0.014908 | 0.007878 | 0.009387 | 0.003412 | 0.01765 | 0.002246 | 0.005564 | 0.007707 | 0.00837 | 0.009305 | 0.01124 | 0.006343 | 0.014595 | 0.02688 | 0.006473 | 0.00287 | 0.009321 | 0.000972 | 0.001063 | 0.022636 | 0.009171 | 0.009337 | 0.010204 | 0.01598 |
| 19.5138889 | 0.01273 | 0.006265 | 0.010894 | 0.002343 | 0.013049 | 0.002126 | 0.004888 | 0.007757 | 0.005495 | 0.007764 | 0.00971 | 0.006417 | 0.014109 | 0.025115 | 0.006097 | 0.002007 | 0.014128 | 0.002444 | 0.000693 | 0.019799 | 0.008544 | 0.008634 | 0.011806 | 0.01907 |
| 19.5277778 | 0.007995 | 0.004963 | 0.012673 | 0.001899 | 0.011138 | 0.003095 | 0.002953 | 0.006482 | 0.00312 | 0.004674 | 0.006866 | 0.004464 | 0.013078 | 0.027721 | 0.007644 | 0.001972 | 0.015285 | 0.004569 | 0.00073 | 0.02039 | 0.006165 | 0.007134 | 0.013322 | 0.025846 |
| 19.5416667 | 0.006009 | 0.004004 | 0.013151 | 0.003546 | 0.011674 | 0.006709 | 0.001531 | 0.005782 | 0.002872 | 0.002131 | 0.004758 | 0.002102 | 0.012287 | 0.030121 | 0.01114 | 0.004677 | 0.01415 | 0.005988 | 0.001233 | 0.018661 | 0.003063 | 0.005199 | 0.01457 | 0.034332 |
| 19.5555556 | 0.008066 | 0.007168 | 0.013549 | 0.003867 | 0.011607 | 0.00803 | 0.002238 | 0.007887 | 0.003103 | 0.004442 | 0.006103 | 0.001543 | 0.011669 | 0.028311 | 0.013292 | 0.009039 | 0.01523 | 0.004845 | 0.003751 | 0.012061 | 0.002525 | 0.005679 | 0.017559 | 0.039492 |
| 19.5694444 | 0.012278 | 0.01575 | 0.017009 | 0.003542 | 0.01172 | 0.004945 | 0.006149 | 0.009864 | 0.004442 | 0.011001 | 0.010352 | 0.002205 | 0.01208 | 0.033858 | 0.013804 | 0.013161 | 0.017864 | 0.002808 | 0.008256 | 0.007579 | 0.0057 | 0.007255 | 0.023651 | 0.042549 |
| 19.5833333 | 0.016726 | 0.023848 | 0.02164 | 0.007229 | 0.013655 | 0.002599 | 0.011038 | 0.01052 | 0.007025 | 0.015051 | 0.014574 | 0.005032 | 0.016019 | 0.051175 | 0.015446 | 0.017509 | 0.020656 | 0.003276 | 0.012401 | 0.008358 | 0.009834 | 0.007683 | 0.027583 | 0.047694 |

| | | | | | | | | | | | | | | | | | | | | | | | | |
|---|---|---|---|---|---|---|---|---|---|---|---|---|---|---|---|---|---|---|---|---|---|---|---|---|
| 19.5972222 | 0.019007 | 0.026726 | 0.022273 | 0.012453 | 0.014873 | 0.002746 | 0.014693 | 0.011224 | 0.007844 | 0.01363 | 0.0164 | 0.007935 | 0.02196 | 0.059185 | 0.018802 | 0.021253 | 0.024994 | 0.005806 | 0.015248 | 0.012842 | 0.011227 | 0.007881 | 0.024304 | 0.050858 |
| 19.6111111 | 0.017946 | 0.025858 | 0.017767 | 0.015792 | 0.01326 | 0.002666 | 0.016127 | 0.011029 | 0.00677 | 0.010693 | 0.015263 | 0.007888 | 0.023384 | 0.043475 | 0.021524 | 0.02201 | 0.029151 | 0.009179 | 0.017145 | 0.017087 | 0.00915 | 0.008852 | 0.017228 | 0.045957 |
| 19.625 | 0.015651 | 0.023963 | 0.012257 | 0.016742 | 0.010195 | 0.00182 | 0.015141 | 0.009658 | 0.005763 | 0.009147 | 0.012972 | 0.006258 | 0.019951 | 0.023971 | 0.021325 | 0.019278 | 0.029142 | 0.012286 | 0.017529 | 0.017616 | 0.006606 | 0.010075 | 0.011907 | 0.03399 |
| 19.6388889 | 0.014403 | 0.021069 | 0.009097 | 0.015045 | 0.007556 | 0.001464 | 0.014032 | 0.009049 | 0.005191 | 0.008546 | 0.011262 | 0.004863 | 0.015991 | 0.020888 | 0.018636 | 0.014715 | 0.024411 | 0.012928 | 0.01582 | 0.016086 | 0.006109 | 0.010027 | 0.009718 | 0.021509 |
| 19.6527778 | 0.013358 | 0.016888 | 0.008324 | 0.011736 | 0.006335 | 0.001673 | 0.013642 | 0.009197 | 0.004713 | 0.008184 | 0.010753 | 0.003951 | 0.011795 | 0.031012 | 0.01504 | 0.010543 | 0.020099 | 0.011148 | 0.012979 | 0.015055 | 0.007052 | 0.00846 | 0.00997 | 0.013334 |
| 19.6666667 | 0.012228 | 0.013989 | 0.009218 | 0.00887 | 0.006779 | 0.001843 | 0.012063 | 0.008342 | 0.004436 | 0.00809 | 0.011646 | 0.003621 | 0.007222 | 0.040639 | 0.012092 | 0.007656 | 0.021292 | 0.008819 | 0.010499 | 0.014852 | 0.008185 | 0.008224 | 0.012079 | 0.010049 |
| 19.6805556 | 0.011659 | 0.013688 | 0.010262 | 0.008331 | 0.008344 | 0.001755 | 0.009213 | 0.006718 | 0.004357 | 0.00845 | 0.012372 | 0.003945 | 0.004057 | 0.043805 | 0.010887 | 0.005923 | 0.022772 | 0.007309 | 0.008755 | 0.014148 | 0.007953 | 0.01046 | 0.014384 | 0.011525 |
| 19.6944444 | 0.011108 | 0.013235 | 0.009605 | 0.010292 | 0.010614 | 0.001593 | 0.007206 | 0.005253 | 0.004492 | 0.008923 | 0.011226 | 0.00413 | 0.002792 | 0.046111 | 0.010523 | 0.005432 | 0.019163 | 0.006883 | 0.007412 | 0.013129 | 0.005972 | 0.010248 | 0.015183 | 0.015079 |
| 19.7083333 | 0.010531 | 0.010666 | 0.007454 | 0.012789 | 0.013014 | 0.001459 | 0.006174 | 0.003555 | 0.004904 | 0.009121 | 0.008934 | 0.003818 | 0.002647 | 0.047598 | 0.009396 | 0.005647 | 0.016663 | 0.007168 | 0.006761 | 0.013691 | 0.005049 | 0.005724 | 0.015352 | 0.017322 |
| 19.7222222 | 0.010289 | 0.006452 | 0.005866 | 0.0142 | 0.014447 | 0.0012 | 0.004975 | 0.001608 | 0.005264 | 0.00885 | 0.006878 | 0.00348 | 0.003037 | 0.044071 | 0.00699 | 0.005363 | 0.019177 | 0.007944 | 0.007196 | 0.015619 | 0.006345 | 0.004295 | 0.016027 | 0.018083 |
| 19.7361111 | 0.00937 | 0.002862 | 0.005727 | 0.014762 | 0.015005 | 0.000738 | 0.003853 | 0.000446 | 0.004885 | 0.008003 | 0.005431 | 0.003464 | 0.003018 | 0.038146 | 0.003864 | 0.004042 | 0.02056 | 0.009276 | 0.008552 | 0.016771 | 0.007486 | 0.009711 | 0.016523 | 0.018091 |
| 19.75 | 0.006456 | 0.002387 | 0.005439 | 0.014947 | 0.015624 | 0.000292 | 0.003154 | 0.000317 | 0.003557 | 0.00685 | 0.005149 | 0.003156 | 0.002062 | 0.032581 | 0.001815 | 0.002155 | 0.016927 | 0.010607 | 0.01004 | 0.016091 | 0.006671 | 0.013451 | 0.016594 | 0.018026 |
| 19.7638889 | 0.002863 | 0.00638 | 0.004327 | 0.014701 | 0.015645 | 0.000119 | 0.002072 | 0.000692 | 0.001982 | 0.005954 | 0.006528 | 0.002366 | 0.001656 | 0.02889 | 0.002342 | 0.001116 | 0.012814 | 0.011044 | 0.010421 | 0.014148 | 0.005073 | 0.009847 | 0.0162 | 0.018337 |
| 19.7777778 | 0.001389 | 0.012255 | 0.003657 | 0.013231 | 0.014457 | 0.000252 | 0.001445 | 0.001751 | 0.000827 | 0.005777 | 0.00802 | 0.002533 | 0.002844 | 0.027005 | 0.004256 | 0.002159 | 0.01167 | 0.010487 | 0.00957 | 0.012407 | 0.004593 | 0.005833 | 0.014791 | 0.018595 |
| 19.7916667 | 0.002361 | 0.012529 | 0.003546 | 0.010111 | 0.013814 | 0.000445 | 0.003045 | 0.003697 | 0.000434 | 0.006041 | 0.007072 | 0.003946 | 0.004651 | 0.023703 | 0.005774 | 0.00434 | 0.010619 | 0.009158 | 0.008232 | 0.012222 | 0.004787 | 0.00446 | 0.013061 | 0.017912 |
| 19.8055556 | 0.002693 | 0.007648 | 0.003178 | 0.006768 | 0.014545 | 0.000393 | 0.005373 | 0.005761 | 0.000878 | 0.0058 | 0.004441 | 0.004685 | 0.006187 | 0.020785 | 0.006668 | 0.005993 | 0.006946 | 0.00714 | 0.006368 | 0.013222 | 0.004442 | 0.002789 | 0.011916 | 0.015032 |
| 19.8194444 | 0.0027 | 0.005678 | 0.002855 | 0.004505 | 0.014958 | 0.000379 | 0.006393 | 0.006896 | 0.001443 | 0.004668 | 0.003745 | 0.003765 | 0.007379 | 0.023406 | 0.007161 | 0.006706 | 0.003063 | 0.004905 | 0.004442 | 0.01349 | 0.004109 | 0.002887 | 0.010514 | 0.009936 |
| 19.8333333 | 0.005158 | 0.007937 | 0.003555 | 0.002978 | 0.013654 | 0.000774 | 0.006085 | 0.007289 | 0.001515 | 0.003283 | 0.005482 | 0.00264 | 0.008806 | 0.028407 | 0.007785 | 0.006985 | 0.002403 | 0.003285 | 0.003448 | 0.01189 | 0.00463 | 0.005281 | 0.00787 | 0.00541 |
| 19.8472222 | 0.008031 | 0.010091 | 0.004776 | 0.003048 | 0.010503 | 0.001204 | 0.005165 | 0.007886 | 0.001483 | 0.002057 | 0.007222 | 0.002603 | 0.011036 | 0.026576 | 0.008856 | 0.007519 | 0.003167 | 0.002282 | 0.003081 | 0.008589 | 0.005412 | 0.00547 | 0.004939 | 0.003408 |
| 19.8611111 | 0.010785 | 0.00803 | 0.004454 | 0.005996 | 0.006019 | 0.001548 | 0.004565 | 0.00832 | 0.001906 | 0.001225 | 0.006388 | 0.003153 | 0.013871 | 0.01633 | 0.009634 | 0.008529 | 0.004486 | 0.001281 | 0.002584 | 0.004532 | 0.005833 | 0.003118 | 0.003072 | 0.005659 |
| 19.875 | 0.013321 | 0.003105 | 0.002441 | 0.00943 | 0.00241 | 0.002269 | 0.004447 | 0.007842 | 0.002801 | 0.001486 | 0.003815 | 0.003644 | 0.016566 | 0.006458 | 0.009634 | 0.009563 | 0.007752 | 0.000748 | 0.001729 | 0.001713 | 0.005456 | 0.003007 | 0.002534 | 0.01252 |
| 19.8888889 | 0.013059 | 0.000843 | 0.000942 | 0.011475 | 0.001909 | 0.003235 | 0.004419 | 0.007463 | 0.004005 | 0.002726 | 0.003608 | 0.004178 | 0.018717 | 0.002548 | 0.010072 | 0.00955 | 0.008237 | 0.001333 | 0.000797 | 0.001068 | 0.004593 | 0.005945 | 0.002862 | 0.019719 |
| 19.9027778 | 0.012127 | 0.002282 | 0.000785 | 0.013614 | 0.004079 | 0.003709 | 0.004248 | 0.008042 | 0.005345 | 0.004277 | 0.005864 | 0.00466 | 0.019928 | 0.003965 | 0.011772 | 0.007959 | 0.004303 | 0.002245 | 0.000468 | 0.001523 | 0.004305 | 0.007728 | 0.003177 | 0.025959 |
| 19.9166667 | 0.014943 | 0.00556 | 0.001877 | 0.016627 | 0.007536 | 0.003741 | 0.003813 | 0.008662 | 0.006217 | 0.006116 | 0.008214 | 0.004573 | 0.020256 | 0.012338 | 0.013602 | 0.006147 | 0.005371 | 0.003187 | 0.000802 | 0.002519 | 0.003974 | 0.007805 | 0.002919 | 0.031735 |
| 19.9305556 | 0.020963 | 0.006838 | 0.004263 | 0.019726 | 0.010305 | 0.003688 | 0.003565 | 0.008832 | 0.006149 | 0.008203 | 0.009026 | 0.003468 | 0.021718 | 0.025133 | 0.013479 | 0.005911 | 0.014935 | 0.004754 | 0.001381 | 0.003764 | 0.003095 | 0.01001 | 0.002295 | 0.034804 |
| 19.9444444 | 0.025328 | 0.004277 | 0.006559 | 0.022161 | 0.011327 | 0.002962 | 0.00401 | 0.008432 | 0.005663 | 0.010305 | 0.008245 | 0.001929 | 0.025825 | 0.033627 | 0.011385 | 0.007561 | 0.022141 | 0.006425 | 0.002159 | 0.004481 | 0.00225 | 0.012789 | 0.00263 | 0.036753 |
| 19.9583333 | 0.026799 | 0.001967 | 0.007771 | 0.023818 | 0.01324 | 0.001551 | 0.004976 | 0.008129 | 0.005579 | 0.012262 | 0.007288 | 0.002017 | 0.029981 | 0.033563 | 0.010163 | 0.009934 | 0.02146 | 0.007534 | 0.003197 | 0.005012 | 0.001483 | 0.012008 | 0.005427 | 0.038907 |
| 19.9722222 | 0.028792 | 0.002313 | 0.00924 | 0.025081 | 0.017826 | 0.000798 | 0.006265 | 0.009378 | 0.006163 | 0.013873 | 0.005113 | 0.003727 | 0.031398 | 0.030979 | 0.011344 | 0.011915 | 0.021109 | 0.008354 | 0.004512 | 0.00758 | 0.001754 | 0.009739 | 0.010076 | 0.037789 |
| 19.9861111 | 0.033385 | 0.006926 | 0.012083 | 0.02619 | 0.022542 | 0.001321 | 0.007887 | 0.012312 | 0.00705 | 0.015087 | 0.004185 | 0.004196 | 0.033555 | 0.035307 | 0.013596 | 0.013456 | 0.027362 | 0.009534 | 0.006181 | 0.012957 | 0.003451 | 0.009245 | 0.01551 | 0.034818 |
| 20 | 0.040343 | 0.016504 | 0.015467 | 0.02665 | 0.025497 | 0.001963 | 0.009289 | 0.014279 | 0.007799 | 0.015805 | 0.00868 | 0.003125 | 0.039173 | 0.045366 | 0.015773 | 0.015107 | 0.03531 | 0.01122 | 0.009098 | 0.018323 | 0.005323 | 0.010277 | 0.022006 | 0.033479 |
| 20.0138889 | 0.046406 | 0.026171 | 0.01847 | 0.025733 | 0.027139 | 0.001876 | 0.009036 | 0.01186 | 0.007603 | 0.015819 | 0.013472 | 0.001922 | 0.043635 | 0.050676 | 0.017965 | 0.017063 | 0.03897 | 0.012678 | 0.014082 | 0.02084 | 0.007904 | 0.011506 | 0.029112 | 0.032714 |
| 20.0277778 | 0.047463 | 0.032268 | 0.019887 | 0.023692 | 0.027269 | 0.00292 | 0.005925 | 0.006269 | 0.005235 | 0.014915 | 0.013866 | 0.001627 | 0.043638 | 0.042509 | 0.02055 | 0.018336 | 0.042953 | 0.013059 | 0.019041 | 0.019129 | 0.011412 | 0.010984 | 0.034682 | 0.031132 |
| 20.0416667 | 0.045598 | 0.034883 | 0.01916 | 0.021625 | 0.029044 | 0.005098 | 0.002467 | 0.002617 | 0.002271 | 0.014352 | 0.013269 | 0.002131 | 0.042627 | 0.035805 | 0.021656 | 0.01843 | 0.054016 | 0.012603 | 0.022285 | 0.016505 | 0.013859 | 0.0093 | 0.036694 | 0.029283 |
| 20.0555556 | 0.04841 | 0.036869 | 0.021798 | 0.02105 | 0.040347 | 0.004868 | 0.004177 | 0.004399 | 0.002815 | 0.018321 | 0.018945 | 0.005003 | 0.044245 | 0.03365 | 0.019592 | 0.01828 | 0.069405 | 0.0141 | 0.026666 | 0.023468 | 0.016706 | 0.008629 | 0.037469 | 0.030525 |
| 20.0694444 | 0.0555 | 0.042567 | 0.026187 | 0.022142 | 0.054435 | 0.003588 | 0.010414 | 0.007361 | 0.007764 | 0.025111 | 0.03104 | 0.011705 | 0.045392 | 0.041226 | 0.016851 | 0.017571 | 0.071656 | 0.017403 | 0.03069 | 0.038518 | 0.020865 | 0.007529 | 0.038991 | 0.03465 |
| 20.0833333 | 0.049193 | 0.039389 | 0.021692 | 0.021174 | 0.046889 | 0.003432 | 0.013012 | 0.00595 | 0.010655 | 0.021897 | 0.030072 | 0.01291 | 0.034119 | 0.045 | 0.013167 | 0.022451 | 0.049439 | 0.016123 | 0.026458 | 0.039002 | 0.018409 | 0.004675 | 0.034664 | 0.030287 |
| 20.0972222 | 0.029881 | 0.025768 | 0.013481 | 0.013713 | 0.021763 | 0.002428 | 0.008247 | 0.002697 | 0.007497 | 0.014614 | 0.01884 | 0.008324 | 0.01913 | 0.031605 | 0.006793 | 0.023077 | 0.030102 | 0.009067 | 0.014244 | 0.02006 | 0.012687 | 0.002484 | 0.021427 | 0.016073 |
| 20.1111111 | 0.0197 | 0.025604 | 0.010072 | 0.004432 | 0.006798 | 0.001062 | 0.004161 | 0.001276 | 0.003362 | 0.014722 | 0.019456 | 0.006902 | 0.019718 | 0.01261 | 0.005953 | 0.022187 | 0.029801 | 0.003967 | 0.004794 | 0.005303 | 0.017424 | 0.002689 | 0.011201 | 0.005831 |
| 20.125 | 0.015283 | 0.027534 | 0.011187 | 0.00553 | 0.004516 | 0.000671 | 0.00634 | 0.001289 | 0.001896 | 0.010108 | 0.019942 | 0.006924 | 0.023437 | 0.00568 | 0.012289 | 0.021576 | 0.029474 | 0.003554 | 0.003466 | 0.007377 | 0.022885 | 0.002797 | 0.008967 | 0.002685 |
| 20.1388889 | 0.008049 | 0.024593 | 0.014906 | 0.015625 | 0.003917 | 0.000615 | 0.007994 | 0.00192 | 0.003732 | 0.002585 | 0.016156 | 0.006168 | 0.018367 | 0.009976 | 0.011884 | 0.01916 | 0.026747 | 0.003391 | 0.005064 | 0.016629 | 0.015881 | 0.002849 | 0.006987 | 0.003198 |
| 20.1527778 | 0.00437 | 0.033803 | 0.015144 | 0.018824 | 0.002031 | 0.000433 | 0.005201 | 0.003526 | 0.00587 | 0.001329 | 0.018734 | 0.003981 | 0.013296 | 0.022671 | 0.00727 | 0.013879 | 0.02299 | 0.005389 | 0.007908 | 0.016009 | 0.005598 | 0.00453 | 0.008239 | 0.004992 |
| 20.1666667 | 0.005865 | 0.033508 | 0.010629 | 0.011402 | 0.001102 | 0.0003 | 0.002343 | 0.004119 | 0.004796 | 0.001613 | 0.01913 | 0.00186 | 0.013006 | 0.036421 | 0.007347 | 0.007531 | 0.013109 | 0.011567 | 0.013256 | 0.008962 | 0.001324 | 0.004621 | 0.015878 | 0.005197 |
| 20.1805556 | 0.010208 | 0.020994 | 0.005462 | 0.004489 | 0.002107 | 0.000289 | 0.001892 | 0.002406 | 0.003196 | 0.001089 | 0.012571 | 0.002022 | 0.015435 | 0.040868 | 0.005969 | 0.004467 | 0.004473 | 0.015125 | 0.016303 | 0.008468 | 0.001062 | 0.002326 | 0.017536 | 0.00356 |
| 20.1944444 | 0.013678 | 0.022846 | 0.003009 | 0.002763 | 0.004352 | 0.000325 | 0.002009 | 0.001165 | 0.003544 | 0.000661 | 0.009118 | 0.003985 | 0.016884 | 0.031954 | 0.004343 | 0.005927 | 0.005629 | 0.013571 | 0.01491 | 0.009099 | 0.001858 | 0.001207 | 0.011863 | 0.002335 |
| 20.2083333 | 0.013449 | 0.028868 | 0.003799 | 0.003915 | 0.006913 | 0.000294 | 0.002479 | 0.002808 | 0.005564 | 0.000703 | 0.01323 | 0.005671 | 0.01708 | 0.021638 | 0.005734 | 0.006879 | 0.013247 | 0.010844 | 0.012039 | 0.009436 | 0.003659 | 0.002508 | 0.011172 | 0.002487 |
| 20.2222222 | 0.012186 | 0.023667 | 0.005207 | 0.007992 | 0.008887 | 0.00016 | 0.003353 | 0.007053 | 0.006168 | 0.001001 | 0.01485 | 0.005307 | 0.018186 | 0.020305 | 0.006694 | 0.005992 | 0.017837 | 0.008342 | 0.008975 | 0.011104 | 0.004561 | 0.004554 | 0.012175 | 0.002777 |
| 20.2361111 | 0.012539 | 0.014341 | 0.005987 | 0.011315 | 0.010853 | 0.000212 | 0.002791 | 0.010441 | 0.00345 | 0.001439 | 0.011652 | 0.004704 | 0.019154 | 0.024474 | 0.007304 | 0.005564 | 0.020844 | 0.006123 | 0.006259 | 0.009122 | 0.003316 | 0.005428 | 0.011211 | 0.003055 |
| 20.25 | 0.01135 | 0.007986 | 0.00667 | 0.009662 | 0.011909 | 0.000562 | 0.002083 | 0.010581 | 0.001025 | 0.00238 | 0.009473 | 0.005079 | 0.017813 | 0.027631 | 0.008347 | 0.005915 | 0.028137 | 0.004933 | 0.004847 | 0.004974 | 0.002155 | 0.004696 | 0.013322 | 0.002448 |
| 20.2638889 | 0.007016 | 0.007015 | 0.007818 | 0.006771 | 0.011228 | 0.000923 | 0.003904 | 0.009808 | 0.000717 | 0.004745 | 0.010322 | 0.005961 | 0.015437 | 0.018771 | 0.008614 | 0.006081 | 0.032907 | 0.004222 | 0.004323 | 0.005887 | 0.001577 | 0.003566 | 0.013448 | 0.001418 |
| 20.2777778 | 0.005302 | 0.01281 | 0.009088 | 0.006155 | 0.010936 | 0.000961 | 0.007304 | 0.010606 | 0.001719 | 0.008595 | 0.015526 | 0.007521 | 0.014836 | 0.006736 | 0.007926 | 0.005558 | 0.027259 | 0.003272 | 0.003872 | 0.009897 | 0.00154 | 0.002563 | 0.009365 | 0.001981 |
| 20.2916667 | 0.010325 | 0.020878 | 0.010272 | 0.005393 | 0.012614 | 0.000595 | 0.008442 | 0.012341 | 0.003862 | 0.011784 | 0.022283 | 0.009575 | 0.01655 | 0.009419 | 0.007594 | 0.005309 | 0.019103 | 0.003042 | 0.004075 | 0.010343 | 0.002813 | 0.002589 | 0.009535 | 0.005501 |
| 20.3055556 | 0.015917 | 0.022861 | 0.012012 | 0.002759 | 0.014408 | 0.000167 | 0.00581 | 0.013699 | 0.006058 | 0.01256 | 0.024788 | 0.011242 | 0.018425 | 0.01945 | 0.007927 | 0.006091 | 0.017776 | 0.003711 | 0.005087 | 0.013311 | 0.004059 | 0.004278 | 0.013815 | 0.010791 |
| 20.3194444 | 0.014198 | 0.018499 | 0.014485 | 0.001931 | 0.014982 | 6.20E-05 | 0.002418 | 0.01467 | 0.007339 | 0.013164 | 0.023522 | 0.012574 | 0.018396 | 0.01698 | 0.008052 | 0.007412 | 0.017928 | 0.00386 | 0.005556 | 0.021872 | 0.004987 | 0.005461 | 0.015792 | 0.013147 |
| 20.3333333 | 0.008551 | 0.015071 | 0.016912 | 0.005896 | 0.014803 | 0.000156 | 0.000772 | 0.015229 | 0.007546 | 0.014685 | 0.023089 | 0.013747 | 0.017191 | 0.006991 | 0.007811 | 0.008532 | 0.014955 | 0.003129 | 0.004946 | 0.025825 | 0.007343 | 0.006953 | 0.015286 | 0.011087 |
| 20.3472222 | 0.007234 | 0.015297 | 0.017705 | 0.010284 | 0.01369 | 0.000246 | 0.000781 | 0.014813 | 0.006947 | 0.013671 | 0.023958 | 0.01419 | 0.016946 | 0.003642 | 0.007648 | 0.009177 | 0.015215 | 0.001892 | 0.003795 | 0.02267 | 0.010763 | 0.009959 | 0.014363 | 0.008847 |
| 20.3611111 | 0.011256 | 0.0153 | 0.015978 | 0.008954 | 0.01202 | 0.000275 | 0.001551 | 0.014419 | 0.006068 | 0.010049 | 0.024629 | 0.013845 | 0.017458 | 0.003311 | 0.008618 | 0.009129 | 0.019241 | 0.000865 | 0.00249 | 0.021947 | 0.012533 | 0.010063 | 0.013417 | 0.008354 |
| 20.375 | 0.015598 | 0.011278 | 0.013475 | 0.003888 | 0.011426 | 0.000311 | 0.002319 | 0.016507 | 0.005167 | 0.008367 | 0.024366 | 0.013189 | 0.01783 | 0.008022 | 0.009679 | 0.009081 | 0.020737 | 0.00108 | 0.002135 | 0.024034 | 0.011996 | 0.007045 | 0.012126 | 0.007775 |
| 20.3888889 | 0.016666 | 0.004897 | 0.012482 | 0.002295 | 0.013367 | 0.001057 | 0.003889 | 0.01915 | 0.004527 | 0.010089 | 0.022445 | 0.012455 | 0.020775 | 0.021224 | 0.010925 | 0.010179 | 0.021427 | 0.002962 | 0.003612 | 0.019662 | 0.011223 | 0.007626 | 0.011902 | 0.008045 |
| 20.4027778 | 0.016671 | 0.001237 | 0.01275 | 0.006333 | 0.017813 | 0.002702 | 0.007779 | 0.019252 | 0.00453 | 0.012918 | 0.02017 | 0.011982 | 0.02815 | 0.031985 | 0.011725 | 0.012227 | 0.02664 | 0.005903 | 0.005328 | 0.011776 | 0.011125 | 0.008937 | 0.014504 | 0.010886 |
| 20.4166667 | 0.01975 | 0.000964 | 0.013578 | 0.011438 | 0.022032 | 0.004304 | 0.013355 | 0.018893 | 0.005341 | 0.016148 | 0.019218 | 0.012023 | 0.034662 | 0.035287 | 0.012733 | 0.013917 | 0.032225 | 0.008085 | 0.005912 | 0.011695 | 0.011586 | 0.007815 | 0.019541 | 0.013997 |
| 20.4305556 | 0.026088 | 0.001246 | 0.015997 | 0.012636 | 0.022807 | 0.006145 | 0.017824 | 0.018928 | 0.006923 | 0.020282 | 0.018754 | 0.012078 | 0.035126 | 0.042295 | 0.015207 | 0.014463 | 0.030536 | 0.009295 | 0.006152 | 0.018859 | 0.012613 | 0.01219 | 0.025682 | 0.014797 |
| 20.4444444 | 0.030433 | 0.002285 | 0.020544 | 0.011486 | 0.020752 | 0.007048 | 0.020365 | 0.018365 | 0.008995 | 0.023943 | 0.018811 | 0.011772 | 0.032408 | 0.057955 | 0.018043 | 0.014421 | 0.026774 | 0.0102 | 0.006508 | 0.025969 | 0.014111 | 0.019926 | 0.030983 | 0.015372 |
| 20.4583333 | 0.028647 | 0.005523 | 0.02498 | 0.014374 | 0.01937 | 0.005328 | 0.023099 | 0.017632 | 0.01093 | 0.026463 | 0.021415 | 0.011837 | 0.030348 | 0.068566 | 0.019772 | 0.015599 | 0.030913 | 0.010274 | 0.006852 | 0.033159 | 0.017007 | 0.023273 | 0.034956 | 0.01997 |
| 20.4722222 | 0.026101 | 0.010406 | 0.026611 | 0.021438 | 0.020137 | 0.003674 | 0.02589 | 0.016096 | 0.012352 | 0.028333 | 0.024847 | 0.012866 | 0.029202 | 0.067689 | 0.021449 | 0.018779 | 0.040121 | 0.008445 | 0.006293 | 0.040951 | 0.021164 | 0.024213 | 0.038076 | 0.028348 |
| 20.4861111 | 0.030504 | 0.017154 | 0.026158 | 0.024851 | 0.021204 | 0.003667 | 0.024947 | 0.012807 | 0.013417 | 0.02887 | 0.024852 | 0.013768 | 0.029839 | 0.069688 | 0.024227 | 0.021878 | 0.044988 | 0.004709 | 0.005743 | 0.042616 | 0.023428 | 0.023346 | 0.038984 | 0.035488 |
| 20.5 | 0.038413 | 0.027168 | 0.025446 | 0.022358 | 0.021447 | 0.004604 | 0.020326 | 0.009039 | 0.014201 | 0.02811 | 0.022667 | 0.014062 | 0.03203 | 0.081231 | 0.026298 | 0.02338 | 0.046226 | 0.002264 | 0.011504 | 0.037551 | 0.021516 | 0.018696 | 0.03768 | 0.039766 |
| 20.5138889 | 0.03927 | 0.037075 | 0.023281 | 0.020071 | 0.022215 | 0.005806 | 0.016038 | 0.006131 | 0.013275 | 0.024999 | 0.021743 | 0.013364 | 0.033099 | 0.086802 | 0.025795 | 0.023823 | 0.050064 | 0.003583 | 0.020808 | 0.03607 | 0.016834 | 0.012691 | 0.035653 | 0.043948 |
| 20.5277778 | 0.03122 | 0.038666 | 0.017943 | 0.019507 | 0.022565 | 0.004975 | 0.012525 | 0.003746 | 0.008709 | 0.017362 | 0.018766 | 0.009009 | 0.030511 | 0.075583 | 0.024953 | 0.023929 | 0.052093 | 0.006117 | 0.019643 | 0.038427 | 0.012576 | 0.007448 | 0.032681 | 0.043998 |
| 20.5416667 | 0.022731 | 0.030282 | 0.011908 | 0.01757 | 0.02026 | 0.003218 | 0.008929 | 0.002259 | 0.003933 | 0.009147 | 0.01166 | 0.003561 | 0.022926 | 0.058788 | 0.025165 | 0.02381 | 0.045853 | 0.006928 | 0.009581 | 0.03115 | 0.009652 | 0.004444 | 0.027893 | 0.034793 |
| 20.5555556 | 0.020345 | 0.021741 | 0.009959 | 0.016351 | 0.017381 | 0.002489 | 0.006944 | 0.00209 | 0.002863 | 0.008049 | 0.007013 | 0.00216 | 0.012404 | 0.050438 | 0.024235 | 0.022674 | 0.035959 | 0.00754 | 0.004089 | 0.014809 | 0.007712 | 0.003967 | 0.023764 | 0.026816 |
| 20.5694444 | 0.023046 | 0.022772 | 0.013594 | 0.019239 | 0.016738 | 0.002356 | 0.007935 | 0.003262 | 0.004566 | 0.013846 | 0.006978 | 0.003321 | 0.005852 | 0.058488 | 0.023878 | 0.022529 | 0.028906 | 0.010009 | 0.005569 | 0.010024 | 0.008359 | 0.005913 | 0.025289 | 0.033931 |
| 20.5833333 | 0.028521 | 0.0322 | 0.019316 | 0.023649 | 0.019584 | 0.004265 | 0.011088 | 0.005681 | 0.007573 | 0.018705 | 0.006914 | 0.003891 | 0.008953 | 0.077938 | 0.029422 | 0.027861 | 0.024651 | 0.014077 | 0.01035 | 0.028313 | 0.011939 | 0.010207 | 0.030651 | 0.05152 |
| 20.5972222 | 0.035873 | 0.037854 | 0.021975 | 0.024711 | 0.024576 | 0.006441 | 0.013817 | 0.007479 | 0.008952 | 0.018472 | 0.005635 | 0.00277 | 0.016892 | 0.08516 | 0.039339 | 0.036224 | 0.021711 | 0.018329 | 0.015505 | 0.051317 | 0.016152 | 0.01392 | 0.031936 | 0.062498 |
| 20.6111111 | 0.037863 | 0.031192 | 0.020527 | 0.021665 | 0.025136 | 0.00621 | 0.013053 | 0.006834 | 0.007633 | 0.01499 | 0.006294 | 0.001133 | 0.019372 | 0.070684 | 0.044629 | 0.040027 | 0.020214 | 0.020586 | 0.017616 | 0.055253 | 0.017967 | 0.015628 | 0.027343 | 0.056754 |
| 20.625 | 0.030662 | 0.021782 | 0.016728 | 0.017836 | 0.018365 | 0.004592 | 0.009036 | 0.004327 | 0.005492 | 0.010416 | 0.010642 | 0.00042 | 0.016329 | 0.047148 | 0.04144 | 0.037136 | 0.018341 | 0.019343 | 0.017814 | 0.044235 | 0.016421 | 0.014972 | 0.020271 | 0.037655 |
| 20.6388889 | 0.022741 | 0.018902 | 0.010504 | 0.01534 | 0.00984 | 0.002944 | 0.00489 | 0.002124 | 0.003554 | 0.005442 | 0.0148 | 0.000306 | 0.014751 | 0.028271 | 0.03453 | 0.030336 | 0.015246 | 0.01531 | 0.019372 | 0.033679 | 0.013646 | 0.012801 | 0.015533 | 0.020919 |
| 20.6527778 | 0.017834 | 0.017988 | 0.004163 | 0.013201 | 0.00507 | 0.002417 | 0.002218 | 0.001467 | 0.002368 | 0.00206 | 0.016354 | 0.000318 | 0.015055 | 0.021345 | 0.028881 | 0.022798 | 0.013544 | 0.011097 | 0.021509 | 0.027431 | 0.011057 | 0.011926 | 0.00846 | 0.019075 |
| 20.6666667 | 0.014102 | 0.016794 | 0.001388 | 0.010832 | 0.004209 | 0.003455 | 0.001854 | 0.00123 | 0.001974 | 0.001424 | 0.01859 | 0.000309 | 0.014947 | 0.024003 | 0.024864 | 0.017709 | 0.014987 | 0.008797 | 0.022135 | 0.021968 | 0.008375 | 0.011549 | 0.00455 | 0.027246 |
| 20.6805556 | 0.01245 | 0.016391 | 0.00113 | 0.008709 | 0.004892 | 0.004826 | 0.004197 | 0.001016 | 0.001766 | 0.001683 | 0.020613 | 0.000461 | 0.014148 | 0.028728 | 0.019348 | 0.015752 | 0.01708 | 0.008957 | 0.020713 | 0.013624 | 0.005821 | 0.009017 | 0.002632 | 0.035625 |
| 20.6944444 | 0.012155 | 0.014862 | 0.000914 | 0.006841 | 0.005335 | 0.004786 | 0.00676 | 0.001835 | 0.0013 | 0.001907 | 0.018721 | 0.001085 | 0.011764 | 0.032862 | 0.010857 | 0.014103 | 0.015413 | 0.010966 | 0.018669 | 0.005147 | 0.004399 | 0.004839 | 0.00252 | 0.041282 |
| 20.7083333 | 0.011712 | 0.012261 | 0.00154 | 0.005581 | 0.005076 | 0.003212 | 0.007168 | 0.003137 | 0.001165 | 0.001594 | 0.015861 | 0.001692 | 0.008054 | 0.037173 | 0.003858 | 0.011011 | 0.010986 | 0.013186 | 0.017814 | 0.002571 | 0.004269 | 0.00258 | 0.002351 | 0.044975 |
| 20.7222222 | 0.011975 | 0.009866 | 0.004675 | 0.005124 | 0.004872 | 0.001479 | 0.006179 | 0.004046 | 0.001373 | 0.000875 | 0.015069 | 0.001799 | 0.00625 | 0.040409 | 0.001948 | 0.007293 | 0.006685 | 0.0143 | 0.018489 | 0.007107 | 0.004481 | 0.003553 | 0.003225 | 0.045635 |
| 20.7361111 | 0.013125 | 0.00702 | 0.008615 | 0.004864 | 0.005243 | 0.001018 | 0.004586 | 0.00423 | 0.001253 | 0.000669 | 0.014068 | 0.001804 | 0.003929 | 0.040833 | 0.001627 | 0.003947 | 0.004183 | 0.01391 | 0.018935 | 0.015675 | 0.00472 | 0.004064 | 0.004528 | 0.042744 |
| 20.75 | 0.013093 | 0.00411 | 0.010869 | 0.004703 | 0.005991 | 0.001387 | 0.002344 | 0.003607 | 0.00107 | 0.000887 | 0.011549 | 0.001951 | 0.002041 | 0.036721 | 0.001071 | 0.001565 | 0.007185 | 0.012474 | 0.017265 | 0.023274 | 0.004951 | 0.003408 | 0.003626 | 0.036748 |
| 20.7638889 | 0.011834 | 0.003736 | 0.011183 | 0.004496 | 0.006629 | 0.001111 | 0.000603 | 0.002715 | 0.000919 | 0.001702 | 0.008814 | 0.001997 | 0.002613 | 0.027622 | 0.001553 | 0.000932 | 0.013879 | 0.011255 | 0.014637 | 0.026463 | 0.004562 | 0.004659 | 0.001723 | 0.028341 |
| 20.7777778 | 0.010813 | 0.003963 | 0.010377 | 0.003789 | 0.006863 | 0.000572 | 0.000675 | 0.002222 | 0.000533 | 0.00249 | 0.007369 | 0.001728 | 0.004855 | 0.018644 | 0.001979 | 0.002304 | 0.016893 | 0.010348 | 0.011926 | 0.024097 | 0.003909 | 0.007491 | 0.000827 | 0.021439 |
| 20.7916667 | 0.008971 | 0.003426 | 0.009038 | 0.00268 | 0.006848 | 0.000637 | 0.002153 | 0.002272 | 0.000313 | 0.002425 | 0.008251 | 0.001555 | 0.008435 | 0.015117 | 0.003303 | 0.004574 | 0.014909 | 0.008367 | 0.008726 | 0.017098 | 0.004197 | 0.008253 | 0.000665 | 0.018395 |
| 20.8055556 | 0.007115 | 0.006234 | 0.006911 | 0.001694 | 0.006753 | 0.000908 | 0.003093 | 0.002318 | 0.000475 | 0.001949 | 0.010343 | 0.002052 | 0.011256 | 0.016186 | 0.00667 | 0.006192 | 0.013625 | 0.004947 | 0.005041 | 0.010435 | 0.005246 | 0.006024 | 0.001492 | 0.017018 |
| 20.8194444 | 0.007886 | 0.010471 | 0.004299 | 0.001413 | 0.006456 | 0.000989 | 0.002915 | 0.002476 | 0.000502 | 0.001679 | 0.011362 | 0.002828 | 0.010614 | 0.017236 | 0.009072 | 0.006706 | 0.015462 | 0.003344 | 0.00326 | 0.007864 | 0.006002 | 0.004371 | 0.00323 | 0.014835 |
| 20.8333333 | 0.0088 | 0.012626 | 0.002617 | 0.001682 | 0.005383 | 0.000773 | 0.00348 | 0.003469 | 0.000313 | 0.001947 | 0.011031 | 0.002857 | 0.00912 | 0.014422 | 0.009143 | 0.007122 | 0.015183 | 0.005804 | 0.005947 | 0.006424 | 0.006161 | 0.005174 | 0.003847 | 0.01012 |
| 20.8472222 | 0.007202 | 0.012658 | 0.001949 | 0.001806 | 0.003156 | 0.000385 | 0.006027 | 0.005093 | 0.000421 | 0.001753 | 0.010001 | 0.002057 | 0.010447 | 0.008908 | 0.008697 | 0.008185 | 0.010221 | 0.009216 | 0.010856 | 0.004204 | 0.005271 | 0.006107 | 0.002971 | 0.007155 |
| 20.8611111 | 0.005233 | 0.008476 | 0.00163 | 0.001403 | 0.001751 | 0.000162 | 0.010347 | 0.007159 | 0.000554 | 0.002094 | 0.008215 | 0.00141 | 0.013751 | 0.008047 | 0.008704 | 0.009598 | 0.006441 | 0.011096 | 0.015812 | 0.003869 | 0.003062 | 0.004575 | 0.002561 | 0.012511 |
| 20.875 | 0.004104 | 0.004596 | 0.002444 | 0.001709 | 0.002606 | 0.000118 | 0.015116 | 0.010022 | 0.000937 | 0.00579 | 0.006082 | 0.001236 | 0.016401 | 0.013645 | 0.00892 | 0.011394 | 0.009208 | 0.012072 | 0.020325 | 0.004579 | 0.001357 | 0.00195 | 0.004012 | 0.021388 |
| 20.8888889 | 0.003001 | 0.007186 | 0.004054 | 0.004125 | 0.003298 | 0.000265 | 0.018681 | 0.013096 | 0.002763 | 0.010716 | 0.003748 | 0.001159 | 0.018275 | 0.01766 | 0.009517 | 0.013608 | 0.014869 | 0.013529 | 0.024766 | 0.005508 | 0.001203 | 0.000807 | 0.007866 | 0.026004 |
| 20.9027778 | 0.001925 | 0.01263 | 0.005161 | 0.007153 | 0.002492 | 0.000653 | 0.02101 | 0.015827 | 0.004945 | 0.014141 | 0.002811 | 0.001578 | 0.020613 | 0.01718 | 0.009839 | 0.014784 | 0.014862 | 0.015508 | 0.029429 | 0.008674 | 0.001013 | 0.001073 | 0.013423 | 0.026597 |
| 20.9166667 | 0.000991 | 0.018864 | 0.005933 | 0.009518 | 0.001678 | 0.000946 | 0.021419 | 0.018135 | 0.005412 | 0.017754 | 0.004692 | 0.002844 | 0.023498 | 0.016906 | 0.009243 | 0.013699 | 0.008483 | 0.016635 | 0.031876 | 0.012424 | 0.000532 | 0.002223 | 0.018428 | 0.027554 |
| 20.9305556 | 0.001239 | 0.025078 | 0.006655 | 0.012064 | 0.002786 | 0.001085 | 0.018349 | 0.018752 | 0.004337 | 0.023156 | 0.00667 | 0.003812 | 0.025834 | 0.020612 | 0.009037 | 0.011801 | 0.005417 | 0.016558 | 0.031869 | 0.012762 | 0.000583 | 0.00312 | 0.021255 | 0.032008 |
| 20.9444444 | 0.004406 | 0.026456 | 0.007388 | 0.015224 | 0.006633 | 0.001598 | 0.013521 | 0.017478 | 0.003166 | 0.027704 | 0.008022 | 0.003251 | 0.026952 | 0.027182 | 0.010091 | 0.011318 | 0.011976 | 0.016716 | 0.032619 | 0.011251 | 0.001999 | 0.002538 | 0.022294 | 0.0378 |
| 20.9583333 | 0.009151 | 0.023327 | 0.008412 | 0.018498 | 0.010739 | 0.002392 | 0.009998 | 0.016349 | 0.002793 | 0.028108 | 0.010163 | 0.001868 | 0.026734 | 0.031795 | 0.012015 | 0.012179 | 0.022841 | 0.018086 | 0.034644 | 0.013375 | 0.005265 | 0.001798 | 0.024169 | 0.040217 |
| 20.9722222 | 0.012332 | 0.021663 | 0.010253 | 0.020635 | 0.011866 | 0.002908 | 0.008534 | 0.015907 | 0.003467 | 0.026511 | 0.012663 | 0.000949 | 0.026516 | 0.034043 | 0.013757 | 0.012464 | 0.033254 | 0.019713 | 0.035055 | 0.019733 | 0.00968 | 0.002268 | 0.02892 | 0.040292 |
| 20.9861111 | 0.015386 | 0.024261 | 0.012755 | 0.020787 | 0.010498 | 0.003646 | 0.009198 | 0.015964 | 0.005675 | 0.026411 | 0.013928 | 0.000615 | 0.028852 | 0.039528 | 0.01528 | 0.011287 | 0.045203 | 0.020691 | 0.033333 | 0.02502 | 0.013321 | 0.003735 | 0.033925 | 0.040483 |
| 21 | 0.021887 | 0.027837 | 0.01436 | 0.019955 | 0.009346 | 0.00497 | 0.011028 | 0.016649 | 0.008725 | 0.027566 | 0.012438 | 0.00089 | 0.033952 | 0.048302 | 0.017725 | 0.010014 | 0.054615 | 0.021392 | 0.031733 | 0.026599 | 0.014945 | 0.005317 | 0.035429 | 0.040211 |
| 21.0138889 | 0.029841 | 0.029431 | 0.015655 | 0.019838 | 0.009001 | 0.006247 | 0.011228 | 0.0162 | 0.010581 | 0.027644 | 0.010029 | 0.002325 | 0.038898 | 0.051209 | 0.021251 | 0.010432 | 0.053207 | 0.022301 | 0.031431 | 0.028234 | 0.016303 | 0.005383 | 0.033538 | 0.038818 |
| 21.0277778 | 0.035175 | 0.030026 | 0.018463 | 0.021384 | 0.008846 | 0.005735 | 0.008879 | 0.012893 | 0.010773 | 0.026087 | 0.010331 | 0.004875 | 0.041645 | 0.04586 | 0.024884 | 0.012205 | 0.044197 | 0.022374 | 0.033377 | 0.035002 | 0.01746 | 0.003465 | 0.029207 | 0.0371 |
| 21.0416667 | 0.041568 | 0.030992 | 0.022987 | 0.025788 | 0.014167 | 0.003098 | 0.008662 | 0.009722 | 0.010883 | 0.026891 | 0.01451 | 0.005898 | 0.043289 | 0.04823 | 0.028298 | 0.014042 | 0.044332 | 0.021542 | 0.038685 | 0.052979 | 0.018386 | 0.001539 | 0.026116 | 0.038252 |
| 21.0555556 | 0.059669 | 0.034802 | 0.032717 | 0.035104 | 0.031516 | 0.002777 | 0.01538 | 0.011594 | 0.013418 | 0.035785 | 0.028468 | 0.006452 | 0.047235 | 0.071866 | 0.034639 | 0.017416 | 0.060969 | 0.022008 | 0.046169 | 0.080983 | 0.022273 | 0.001357 | 0.031674 | 0.049247 |
| 21.0694444 | 0.075026 | 0.045047 | 0.044105 | 0.046338 | 0.051646 | 0.006415 | 0.022137 | 0.015169 | 0.018646 | 0.047563 | 0.05497 | 0.012558 | 0.048086 | 0.101217 | 0.041236 | 0.023652 | 0.070669 | 0.024007 | 0.050607 | 0.102969 | 0.030025 | 0.002328 | 0.044152 | 0.063966 |
| 21.0833333 | 0.057244 | 0.044723 | 0.040444 | 0.047462 | 0.053486 | 0.008939 | 0.017663 | 0.012201 | 0.023316 | 0.041644 | 0.066381 | 0.019401 | 0.033724 | 0.104392 | 0.033567 | 0.027434 | 0.05079 | 0.022264 | 0.043168 | 0.093595 | 0.030669 | 0.003202 | 0.047958 | 0.060638 |
| 21.0972222 | 0.029835 | 0.034658 | 0.021551 | 0.030662 | 0.034 | 0.007027 | 0.009962 | 0.006473 | 0.019326 | 0.026671 | 0.044921 | 0.017182 | 0.015063 | 0.077311 | 0.017461 | 0.023903 | 0.024403 | 0.016557 | 0.026571 | 0.055609 | 0.023314 | 0.006353 | 0.031348 | 0.035334 |

| | | | | | | | | | | | | | | | | | | | | | | | | |
|---|---|---|---|---|---|---|---|---|---|---|---|---|---|---|---|---|---|---|---|---|---|---|---|---|
| 21.1111111 | 0.029852 | 0.043589 | 0.006494 | 0.013637 | 0.013388 | 0.003476 | 0.012696 | 0.006657 | 0.010813 | 0.032207 | 0.030024 | 0.009857 | 0.011276 | 0.050124 | 0.012451 | 0.018872 | 0.02498 | 0.015536 | 0.018745 | 0.030214 | 0.023853 | 0.013857 | 0.015999 | 0.01291 |
| 21.125 | 0.039075 | 0.043238 | 0.003575 | 0.012837 | 0.005259 | 0.002656 | 0.018637 | 0.009226 | 0.009374 | 0.045322 | 0.037671 | 0.006294 | 0.018753 | 0.041944 | 0.011999 | 0.019949 | 0.040135 | 0.015766 | 0.019814 | 0.022572 | 0.021459 | 0.02085 | 0.020368 | 0.010099 |
| 21.1388889 | 0.03695 | 0.027178 | 0.005959 | 0.013258 | 0.00527 | 0.004554 | 0.018521 | 0.008757 | 0.007403 | 0.046696 | 0.03735 | 0.005578 | 0.025762 | 0.043418 | 0.007298 | 0.028603 | 0.044044 | 0.010145 | 0.015862 | 0.012675 | 0.010285 | 0.024669 | 0.022364 | 0.016543 |
| 21.1527778 | 0.029188 | 0.032608 | 0.005271 | 0.009739 | 0.006488 | 0.006421 | 0.014256 | 0.006163 | 0.002901 | 0.04372 | 0.028298 | 0.003804 | 0.026442 | 0.037989 | 0.003334 | 0.040315 | 0.040056 | 0.004232 | 0.011253 | 0.00743 | 0.007513 | 0.028765 | 0.013088 | 0.024767 |
| 21.1666667 | 0.021647 | 0.039628 | 0.002525 | 0.010709 | 0.008995 | 0.006914 | 0.009087 | 0.005501 | 0.002628 | 0.040881 | 0.020159 | 0.001809 | 0.023029 | 0.024458 | 0.001742 | 0.043367 | 0.039119 | 0.002477 | 0.013638 | 0.008661 | 0.010793 | 0.031657 | 0.00464 | 0.03419 |
| 21.1805556 | 0.015896 | 0.035291 | 0.001838 | 0.010641 | 0.013074 | 0.005907 | 0.004619 | 0.008205 | 0.002791 | 0.032395 | 0.012461 | 0.000755 | 0.021846 | 0.015219 | 0.003221 | 0.034097 | 0.036265 | 0.004641 | 0.016622 | 0.010968 | 0.015929 | 0.028231 | 0.002187 | 0.038717 |
| 21.1944444 | 0.011468 | 0.033505 | 0.003264 | 0.008811 | 0.015676 | 0.00455 | 0.001786 | 0.010767 | 0.001503 | 0.017119 | 0.007054 | 0.002077 | 0.021957 | 0.0103 | 0.006727 | 0.020784 | 0.028632 | 0.007282 | 0.015376 | 0.012994 | 0.021021 | 0.020728 | 0.004148 | 0.034262 |
| 21.2083333 | 0.007117 | 0.028721 | 0.006659 | 0.010535 | 0.014611 | 0.004136 | 0.002057 | 0.01055 | 0.001 | 0.009371 | 0.004714 | 0.004798 | 0.017141 | 0.007292 | 0.008082 | 0.014967 | 0.028257 | 0.008613 | 0.02278 | 0.015121 | 0.019287 | 0.018741 | 0.004999 | 0.022222 |
| 21.2222222 | 0.00336 | 0.018757 | 0.009831 | 0.013864 | 0.009339 | 0.003914 | 0.00561 | 0.007155 | 0.002265 | 0.016119 | 0.008326 | 0.004745 | 0.013648 | 0.008905 | 0.011648 | 0.028766 | 0.043664 | 0.014912 | 0.050727 | 0.033035 | 0.017632 | 0.027845 | 0.004345 | 0.0165 |
| 21.2361111 | 0.001699 | 0.018273 | 0.010565 | 0.011961 | 0.003739 | 0.003653 | 0.008353 | 0.004406 | 0.003686 | 0.020785 | 0.010387 | 0.002659 | 0.024739 | 0.016776 | 0.014146 | 0.036701 | 0.047398 | 0.020236 | 0.066738 | 0.056191 | 0.025289 | 0.029105 | 0.013986 | 0.02285 |
| 21.25 | 0.00335 | 0.027672 | 0.009524 | 0.008162 | 0.003109 | 0.004486 | 0.006897 | 0.006235 | 0.004336 | 0.01322 | 0.007044 | 0.001838 | 0.030355 | 0.028529 | 0.012205 | 0.026535 | 0.026966 | 0.020674 | 0.051183 | 0.048725 | 0.028349 | 0.021629 | 0.028694 | 0.02087 |
| 21.2638889 | 0.009194 | 0.032548 | 0.006833 | 0.011706 | 0.009415 | 0.005052 | 0.005653 | 0.009013 | 0.006462 | 0.007637 | 0.008331 | 0.00241 | 0.017759 | 0.024381 | 0.016756 | 0.024912 | 0.016752 | 0.020982 | 0.032311 | 0.031607 | 0.017946 | 0.025393 | 0.024431 | 0.00845 |
| 21.2777778 | 0.018714 | 0.031998 | 0.00726 | 0.017539 | 0.01928 | 0.004294 | 0.007788 | 0.010596 | 0.008238 | 0.016083 | 0.019222 | 0.004055 | 0.012242 | 0.013876 | 0.023971 | 0.019961 | 0.026474 | 0.015017 | 0.020625 | 0.039773 | 0.010967 | 0.023 | 0.014332 | 0.00667 |
| 21.2916667 | 0.024574 | 0.033155 | 0.014344 | 0.020338 | 0.022846 | 0.004422 | 0.010504 | 0.014671 | 0.00843 | 0.028707 | 0.038001 | 0.00734 | 0.012304 | 0.020652 | 0.024357 | 0.009963 | 0.041124 | 0.009843 | 0.016015 | 0.057226 | 0.010755 | 0.010303 | 0.016468 | 0.016112 |
| 21.3055556 | 0.019087 | 0.036898 | 0.019295 | 0.022408 | 0.018083 | 0.005419 | 0.013026 | 0.017812 | 0.009996 | 0.033925 | 0.048528 | 0.010598 | 0.008542 | 0.026734 | 0.017673 | 0.00946 | 0.043123 | 0.009035 | 0.012505 | 0.05736 | 0.010954 | 0.005266 | 0.019721 | 0.02057 |
| 21.3194444 | 0.010252 | 0.034057 | 0.018645 | 0.022329 | 0.015243 | 0.00501 | 0.014704 | 0.015698 | 0.012841 | 0.034831 | 0.040274 | 0.012169 | 0.0099 | 0.020924 | 0.009295 | 0.010589 | 0.034849 | 0.005016 | 0.006759 | 0.046234 | 0.014204 | 0.006503 | 0.020255 | 0.017333 |
| 21.3333333 | 0.008846 | 0.024321 | 0.018767 | 0.01704 | 0.01958 | 0.003545 | 0.01433 | 0.014444 | 0.012589 | 0.035586 | 0.030859 | 0.013696 | 0.017003 | 0.018043 | 0.006541 | 0.009983 | 0.031854 | 0.001402 | 0.008988 | 0.047956 | 0.018537 | 0.007525 | 0.018582 | 0.017271 |
| 21.3472222 | 0.012384 | 0.019982 | 0.020524 | 0.009375 | 0.024038 | 0.002666 | 0.015196 | 0.01807 | 0.007871 | 0.033899 | 0.033165 | 0.01602 | 0.024565 | 0.023057 | 0.009298 | 0.008391 | 0.027699 | 0.00119 | 0.016426 | 0.059438 | 0.019124 | 0.007361 | 0.01375 | 0.0238 |
| 21.3611111 | 0.01578 | 0.02025 | 0.021455 | 0.005359 | 0.022586 | 0.002603 | 0.018718 | 0.020868 | 0.003994 | 0.029298 | 0.036733 | 0.017108 | 0.028519 | 0.027063 | 0.012131 | 0.007989 | 0.018633 | 0.002982 | 0.020583 | 0.061885 | 0.017948 | 0.00706 | 0.010476 | 0.029356 |
| 21.375 | 0.018791 | 0.016575 | 0.021341 | 0.006355 | 0.020562 | 0.002886 | 0.019889 | 0.018263 | 0.003516 | 0.024309 | 0.032554 | 0.016424 | 0.028496 | 0.027707 | 0.011585 | 0.00976 | 0.014857 | 0.005659 | 0.021218 | 0.05591 | 0.019497 | 0.007003 | 0.012574 | 0.028951 |
| 21.3888889 | 0.02204 | 0.011557 | 0.020049 | 0.007985 | 0.022194 | 0.003315 | 0.017535 | 0.0138 | 0.00498 | 0.01898 | 0.030837 | 0.015618 | 0.027326 | 0.032144 | 0.009026 | 0.01105 | 0.016714 | 0.008308 | 0.022496 | 0.049473 | 0.022001 | 0.00839 | 0.019046 | 0.027443 |
| 21.4027778 | 0.023286 | 0.01053 | 0.017544 | 0.006364 | 0.021227 | 0.003756 | 0.015845 | 0.010149 | 0.008688 | 0.012912 | 0.037143 | 0.014986 | 0.027616 | 0.042517 | 0.007864 | 0.010555 | 0.014291 | 0.010194 | 0.025171 | 0.042065 | 0.023011 | 0.0106 | 0.024566 | 0.030102 |
| 21.4166667 | 0.019181 | 0.010103 | 0.014705 | 0.004215 | 0.014454 | 0.003797 | 0.015422 | 0.00717 | 0.013201 | 0.008261 | 0.039713 | 0.013915 | 0.029402 | 0.05402 | 0.010305 | 0.010185 | 0.00921 | 0.012455 | 0.027567 | 0.037571 | 0.025609 | 0.011762 | 0.025423 | 0.034251 |
| 21.4305556 | 0.013685 | 0.007567 | 0.012793 | 0.006423 | 0.009986 | 0.003249 | 0.015406 | 0.007253 | 0.013279 | 0.006685 | 0.032919 | 0.012579 | 0.030138 | 0.059998 | 0.015038 | 0.012356 | 0.009882 | 0.016537 | 0.028596 | 0.038455 | 0.030053 | 0.011067 | 0.023952 | 0.034779 |
| 21.4444444 | 0.015782 | 0.007529 | 0.012932 | 0.012704 | 0.013228 | 0.002884 | 0.01697 | 0.010433 | 0.008766 | 0.008375 | 0.022866 | 0.01144 | 0.029033 | 0.059206 | 0.01856 | 0.016968 | 0.015495 | 0.020848 | 0.029219 | 0.034557 | 0.032232 | 0.00924 | 0.024729 | 0.031517 |
| 21.4583333 | 0.025175 | 0.010276 | 0.015461 | 0.018341 | 0.018677 | 0.003531 | 0.020135 | 0.012666 | 0.006909 | 0.013834 | 0.016664 | 0.010705 | 0.02838 | 0.060532 | 0.019481 | 0.020531 | 0.023233 | 0.023345 | 0.030847 | 0.023314 | 0.031432 | 0.007808 | 0.030923 | 0.027531 |
| 21.4722222 | 0.034017 | 0.013966 | 0.019273 | 0.019733 | 0.022522 | 0.00482 | 0.023195 | 0.011573 | 0.011738 | 0.023424 | 0.019186 | 0.010984 | 0.02828 | 0.071399 | 0.019302 | 0.020837 | 0.036408 | 0.024897 | 0.033884 | 0.017285 | 0.031612 | 0.006932 | 0.040246 | 0.029472 |
| 21.4861111 | 0.040075 | 0.018732 | 0.022456 | 0.018647 | 0.028541 | 0.006292 | 0.024514 | 0.009656 | 0.019449 | 0.033057 | 0.029504 | 0.013604 | 0.029346 | 0.085771 | 0.020591 | 0.020329 | 0.051273 | 0.027452 | 0.037755 | 0.02471 | 0.03623 | 0.005336 | 0.044335 | 0.045038 |
| 21.5 | 0.0446 | 0.022924 | 0.025249 | 0.019954 | 0.0366 | 0.008141 | 0.024757 | 0.010875 | 0.023675 | 0.0369 | 0.040698 | 0.018566 | 0.033771 | 0.089111 | 0.024923 | 0.021797 | 0.054894 | 0.031771 | 0.040737 | 0.038577 | 0.039223 | 0.003372 | 0.042345 | 0.068823 |
| 21.5138889 | 0.0467 | 0.024334 | 0.027449 | 0.022862 | 0.040282 | 0.008678 | 0.024583 | 0.013585 | 0.02116 | 0.033134 | 0.043549 | 0.021522 | 0.035952 | 0.076161 | 0.029717 | 0.024474 | 0.044893 | 0.035455 | 0.041829 | 0.041749 | 0.031963 | 0.001884 | 0.039193 | 0.0801 |
| 21.5277778 | 0.045248 | 0.019955 | 0.025864 | 0.021143 | 0.037599 | 0.005944 | 0.021632 | 0.011985 | 0.013912 | 0.024288 | 0.0341 | 0.017677 | 0.028368 | 0.057626 | 0.031144 | 0.024532 | 0.031839 | 0.035869 | 0.040458 | 0.029644 | 0.019266 | 0.001619 | 0.033164 | 0.070378 |
| 21.5416667 | 0.039548 | 0.011559 | 0.020615 | 0.014744 | 0.034617 | 0.002866 | 0.015758 | 0.006581 | 0.008033 | 0.018402 | 0.021135 | 0.010118 | 0.015869 | 0.047647 | 0.029661 | 0.021031 | 0.024106 | 0.0334 | 0.036947 | 0.022064 | 0.01208 | 0.002772 | 0.025757 | 0.057789 |
| 21.5555556 | 0.034286 | 0.006924 | 0.016696 | 0.010615 | 0.036193 | 0.002098 | 0.011449 | 0.003817 | 0.008755 | 0.020474 | 0.01521 | 0.006566 | 0.009027 | 0.048609 | 0.028195 | 0.017437 | 0.026176 | 0.028585 | 0.031162 | 0.033382 | 0.012278 | 0.003432 | 0.022502 | 0.052186 |
| 21.5694444 | 0.037156 | 0.009655 | 0.017 | 0.012422 | 0.039461 | 0.002758 | 0.011362 | 0.005315 | 0.014631 | 0.025543 | 0.016672 | 0.00902 | 0.010367 | 0.055218 | 0.02853 | 0.015842 | 0.037949 | 0.023894 | 0.02296 | 0.05062 | 0.014651 | 0.00563 | 0.022848 | 0.045572 |
| 21.5833333 | 0.047089 | 0.015765 | 0.019194 | 0.018258 | 0.040811 | 0.004267 | 0.014227 | 0.008132 | 0.020639 | 0.028437 | 0.020765 | 0.013542 | 0.015057 | 0.069274 | 0.031722 | 0.01761 | 0.048134 | 0.02452 | 0.017821 | 0.055334 | 0.01646 | 0.011627 | 0.02321 | 0.042961 |
| 21.5972222 | 0.052808 | 0.020708 | 0.020289 | 0.024372 | 0.040132 | 0.00506 | 0.018007 | 0.010685 | 0.02368 | 0.028742 | 0.025407 | 0.016144 | 0.018943 | 0.085523 | 0.037223 | 0.023701 | 0.047042 | 0.027704 | 0.017887 | 0.046255 | 0.018559 | 0.016535 | 0.022802 | 0.05323 |
| 21.6111111 | 0.046744 | 0.022555 | 0.019474 | 0.027445 | 0.037369 | 0.004281 | 0.019433 | 0.011299 | 0.021579 | 0.024643 | 0.028425 | 0.015035 | 0.019191 | 0.08523 | 0.038839 | 0.028666 | 0.038077 | 0.025419 | 0.017294 | 0.034358 | 0.019585 | 0.01656 | 0.024047 | 0.06052 |
| 21.625 | 0.037801 | 0.020701 | 0.016546 | 0.025812 | 0.032462 | 0.00295 | 0.017123 | 0.008994 | 0.015051 | 0.01616 | 0.025746 | 0.011448 | 0.01502 | 0.064869 | 0.0316 | 0.025733 | 0.025667 | 0.018335 | 0.014098 | 0.028062 | 0.018117 | 0.013712 | 0.02659 | 0.050615 |
| 21.6388889 | 0.033017 | 0.015591 | 0.012441 | 0.020376 | 0.025595 | 0.001758 | 0.01369 | 0.005634 | 0.007595 | 0.007772 | 0.018387 | 0.00778 | 0.012154 | 0.036967 | 0.020308 | 0.018065 | 0.013052 | 0.011668 | 0.011109 | 0.023265 | 0.015635 | 0.010678 | 0.0258 | 0.035418 |
| 21.6527778 | 0.022632 | 0.009578 | 0.007873 | 0.013851 | 0.016267 | 0.000981 | 0.010011 | 0.003003 | 0.004954 | 0.005849 | 0.009684 | 0.005161 | 0.015347 | 0.015306 | 0.011752 | 0.013136 | 0.008855 | 0.007108 | 0.009506 | 0.013945 | 0.011272 | 0.008526 | 0.018693 | 0.027117 |
| 21.6666667 | 0.013606 | 0.008042 | 0.003932 | 0.008545 | 0.007623 | 0.000497 | 0.005453 | 0.003109 | 0.009457 | 0.01142 | 0.00392 | 0.004 | 0.020283 | 0.010957 | 0.00754 | 0.012133 | 0.012654 | 0.006032 | 0.009806 | 0.005764 | 0.005907 | 0.006709 | 0.008428 | 0.027645 |
| 21.6805556 | 0.018808 | 0.012125 | 0.002876 | 0.005232 | 0.004509 | 0.000376 | 0.002335 | 0.004834 | 0.013457 | 0.015599 | 0.004512 | 0.003804 | 0.021168 | 0.019701 | 0.005915 | 0.011387 | 0.0123 | 0.010844 | 0.013491 | 0.003555 | 0.003817 | 0.003989 | 0.002378 | 0.03406 |
| 21.6944444 | 0.021645 | 0.011834 | 0.003589 | 0.003199 | 0.005793 | 0.00075 | 0.001612 | 0.004544 | 0.012793 | 0.014662 | 0.005307 | 0.003596 | 0.017702 | 0.02816 | 0.004349 | 0.008481 | 0.006914 | 0.01905 | 0.019794 | 0.00775 | 0.003786 | 0.002131 | 0.002425 | 0.04025 |
| 21.7083333 | 0.013614 | 0.00741 | 0.003019 | 0.001899 | 0.007419 | 0.00107 | 0.001659 | 0.003337 | 0.009762 | 0.011408 | 0.004972 | 0.003167 | 0.013584 | 0.031041 | 0.002188 | 0.00495 | 0.004017 | 0.023635 | 0.024168 | 0.017562 | 0.004236 | 0.004431 | 0.004055 | 0.04251 |
| 21.7222222 | 0.006304 | 0.009814 | 0.002469 | 0.001878 | 0.007328 | 0.001128 | 0.002391 | 0.002288 | 0.005453 | 0.006536 | 0.009822 | 0.002967 | 0.011629 | 0.030996 | 0.000948 | 0.003628 | 0.003189 | 0.022189 | 0.023732 | 0.023382 | 0.006884 | 0.007931 | 0.004505 | 0.041564 |
| 21.7361111 | 0.007007 | 0.015512 | 0.005012 | 0.003285 | 0.004985 | 0.001327 | 0.002444 | 0.001319 | 0.003273 | 0.003594 | 0.015914 | 0.003413 | 0.009898 | 0.031269 | 0.001122 | 0.004153 | 0.002543 | 0.018621 | 0.020021 | 0.019572 | 0.00988 | 0.006395 | 0.003894 | 0.038522 |
| 21.75 | 0.014912 | 0.016859 | 0.008571 | 0.00471 | 0.002573 | 0.001753 | 0.001744 | 0.00178 | 0.005254 | 0.005813 | 0.017565 | 0.003974 | 0.005922 | 0.028437 | 0.001994 | 0.003382 | 0.003204 | 0.015638 | 0.015714 | 0.011219 | 0.01162 | 0.003707 | 0.002963 | 0.032563 |
| 21.7638889 | 0.02334 | 0.012226 | 0.010323 | 0.004836 | 0.003111 | 0.002581 | 0.001675 | 0.003101 | 0.008553 | 0.010116 | 0.016413 | 0.004002 | 0.00229 | 0.018793 | 0.003437 | 0.002376 | 0.003633 | 0.013096 | 0.012719 | 0.004808 | 0.012737 | 0.006228 | 0.003736 | 0.024362 |
| 21.7777778 | 0.025367 | 0.005769 | 0.010521 | 0.003893 | 0.004802 | 0.003825 | 0.001686 | 0.003066 | 0.01115 | 0.013629 | 0.014327 | 0.004119 | 0.002455 | 0.012097 | 0.004483 | 0.004059 | 0.002534 | 0.010686 | 0.010551 | 0.002964 | 0.013137 | 0.009714 | 0.006423 | 0.016743 |
| 21.7916667 | 0.01866 | 0.005806 | 0.009076 | 0.002269 | 0.004018 | 0.004726 | 0.001939 | 0.001786 | 0.012329 | 0.015563 | 0.010155 | 0.004738 | 0.003955 | 0.015076 | 0.003754 | 0.005792 | 0.001379 | 0.007436 | 0.00683 | 0.003252 | 0.013278 | 0.009655 | 0.008177 | 0.011313 |
| 21.8055556 | 0.008615 | 0.014839 | 0.005617 | 0.001444 | 0.002379 | 0.004522 | 0.003569 | 0.001038 | 0.01223 | 0.015287 | 0.005113 | 0.004861 | 0.006147 | 0.018407 | 0.002154 | 0.005663 | 0.001734 | 0.003903 | 0.003243 | 0.005887 | 0.013817 | 0.007086 | 0.009348 | 0.007865 |
| 21.8194444 | 0.002896 | 0.02733 | 0.002487 | 0.002728 | 0.002451 | 0.003351 | 0.006947 | 0.000745 | 0.011855 | 0.013373 | 0.003649 | 0.003906 | 0.010475 | 0.016982 | 0.001875 | 0.005622 | 0.003451 | 0.0041 | 0.004302 | 0.010881 | 0.014975 | 0.003966 | 0.011555 | 0.004909 |
| 21.8333333 | 0.00319 | 0.035248 | 0.002589 | 0.004541 | 0.004051 | 0.001907 | 0.00918 | 0.001227 | 0.010576 | 0.010532 | 0.004941 | 0.002714 | 0.011443 | 0.013926 | 0.003099 | 0.007433 | 0.003436 | 0.010625 | 0.010517 | 0.013048 | 0.015285 | 0.001664 | 0.011584 | 0.004589 |
| 21.8472222 | 0.007133 | 0.036674 | 0.004867 | 0.006192 | 0.00592 | 0.000859 | 0.008392 | 0.003118 | 0.007095 | 0.006658 | 0.005522 | 0.001805 | 0.007824 | 0.010675 | 0.004206 | 0.011359 | 0.003286 | 0.021167 | 0.020328 | 0.008899 | 0.012257 | 0.000695 | 0.007164 | 0.011574 |
| 21.8611111 | 0.012883 | 0.034536 | 0.00683 | 0.008023 | 0.005069 | 0.000651 | 0.007068 | 0.004504 | 0.004514 | 0.004489 | 0.00572 | 0.001117 | 0.007095 | 0.010637 | 0.004842 | 0.015795 | 0.009826 | 0.030355 | 0.030037 | 0.005001 | 0.006641 | 0.000675 | 0.003644 | 0.022411 |
| 21.875 | 0.017494 | 0.029666 | 0.007783 | 0.009082 | 0.003639 | 0.001102 | 0.007968 | 0.004374 | 0.007953 | 0.008703 | 0.006574 | 0.000917 | 0.012318 | 0.018975 | 0.005698 | 0.018997 | 0.022198 | 0.034023 | 0.034946 | 0.008785 | 0.002924 | 0.000785 | 0.00504 | 0.032366 |
| 21.8888889 | 0.01919 | 0.021904 | 0.008356 | 0.009179 | 0.005258 | 0.001501 | 0.011445 | 0.005061 | 0.013822 | 0.016034 | 0.007243 | 0.001019 | 0.017702 | 0.033284 | 0.006432 | 0.020372 | 0.03056 | 0.033151 | 0.035452 | 0.014737 | 0.00373 | 0.000626 | 0.008425 | 0.041468 |
| 21.9027778 | 0.020804 | 0.012792 | 0.008841 | 0.009635 | 0.007394 | 0.001649 | 0.016037 | 0.008067 | 0.016224 | 0.019928 | 0.008097 | 0.000969 | 0.017548 | 0.043787 | 0.006752 | 0.020733 | 0.030339 | 0.031289 | 0.035002 | 0.017578 | 0.007338 | 0.000812 | 0.011173 | 0.048447 |
| 21.9166667 | 0.026311 | 0.006867 | 0.009118 | 0.011197 | 0.008726 | 0.002442 | 0.020675 | 0.01231 | 0.017951 | 0.023134 | 0.012311 | 0.001523 | 0.01659 | 0.047486 | 0.008088 | 0.020883 | 0.027338 | 0.030692 | 0.035553 | 0.019132 | 0.013467 | 0.00184 | 0.016091 | 0.052578 |
| 21.9305556 | 0.035593 | 0.007712 | 0.009426 | 0.01304 | 0.010837 | 0.003982 | 0.024654 | 0.016352 | 0.021821 | 0.030088 | 0.020171 | 0.003997 | 0.021626 | 0.051174 | 0.011339 | 0.020296 | 0.027988 | 0.032293 | 0.036648 | 0.020567 | 0.021375 | 0.002288 | 0.02386 | 0.054807 |
| 21.9444444 | 0.0439 | 0.012392 | 0.009995 | 0.015109 | 0.014094 | 0.004773 | 0.027467 | 0.019777 | 0.025015 | 0.038774 | 0.027771 | 0.007106 | 0.027847 | 0.054925 | 0.015131 | 0.019771 | 0.034362 | 0.034725 | 0.03747 | 0.023618 | 0.025595 | 0.00138 | 0.030702 | 0.054221 |
| 21.9583333 | 0.048414 | 0.015806 | 0.010835 | 0.01819 | 0.017191 | 0.004569 | 0.029107 | 0.021617 | 0.025203 | 0.046981 | 0.032553 | 0.008201 | 0.030185 | 0.053674 | 0.018005 | 0.021394 | 0.043711 | 0.035077 | 0.037771 | 0.030947 | 0.025263 | 0.001005 | 0.035275 | 0.051197 |
| 21.9722222 | 0.050653 | 0.017525 | 0.011403 | 0.021104 | 0.01953 | 0.00452 | 0.029081 | 0.021093 | 0.023555 | 0.051255 | 0.032886 | 0.007968 | 0.032882 | 0.048515 | 0.019588 | 0.023921 | 0.049004 | 0.03345 | 0.037947 | 0.040505 | 0.025138 | 0.002505 | 0.036649 | 0.049782 |
| 21.9861111 | 0.052449 | 0.022319 | 0.011478 | 0.023007 | 0.022079 | 0.005017 | 0.028072 | 0.021431 | 0.02192 | 0.049417 | 0.02963 | 0.008735 | 0.037223 | 0.0434 | 0.020235 | 0.024401 | 0.047588 | 0.032013 | 0.039507 | 0.048312 | 0.027452 | 0.0034 | 0.03654 | 0.050508 |
| 22 | 0.055277 | 0.031621 | 0.012161 | 0.026262 | 0.02595 | 0.005812 | 0.028246 | 0.024061 | 0.019826 | 0.046188 | 0.02922 | 0.010172 | 0.038517 | 0.041667 | 0.021266 | 0.023337 | 0.043855 | 0.031224 | 0.041186 | 0.054785 | 0.03198 | 0.002653 | 0.037441 | 0.050814 |
| 22.0138889 | 0.059883 | 0.037961 | 0.014115 | 0.031371 | 0.031005 | 0.005489 | 0.028938 | 0.026681 | 0.015903 | 0.04449 | 0.034289 | 0.009819 | 0.037113 | 0.042055 | 0.023704 | 0.023488 | 0.042862 | 0.030944 | 0.041256 | 0.061552 | 0.038467 | 0.002289 | 0.038022 | 0.053003 |
| 22.0277778 | 0.066228 | 0.037971 | 0.016928 | 0.035029 | 0.036465 | 0.003932 | 0.027716 | 0.028043 | 0.009534 | 0.040551 | 0.038989 | 0.006904 | 0.038054 | 0.041345 | 0.026444 | 0.025706 | 0.045814 | 0.03253 | 0.044419 | 0.06438 | 0.042212 | 0.001851 | 0.036783 | 0.058446 |
| 22.0416667 | 0.076109 | 0.036933 | 0.019669 | 0.036514 | 0.046269 | 0.003149 | 0.025006 | 0.026665 | 0.003968 | 0.032028 | 0.041299 | 0.004667 | 0.043971 | 0.041589 | 0.028101 | 0.028774 | 0.052055 | 0.036132 | 0.050741 | 0.065038 | 0.039018 | 0.001846 | 0.034433 | 0.06058 |
| 22.0555556 | 0.095724 | 0.041095 | 0.024473 | 0.038585 | 0.066116 | 0.004309 | 0.024048 | 0.024743 | 0.005789 | 0.028116 | 0.053069 | 0.008243 | 0.05763 | 0.045391 | 0.030611 | 0.032092 | 0.065324 | 0.04072 | 0.055609 | 0.073199 | 0.036424 | 0.002198 | 0.034781 | 0.057222 |
| 22.0694444 | 0.113452 | 0.053357 | 0.033285 | 0.045228 | 0.085642 | 0.008594 | 0.026115 | 0.024073 | 0.014619 | 0.037603 | 0.085855 | 0.021218 | 0.067508 | 0.044861 | 0.034262 | 0.033973 | 0.076464 | 0.045225 | 0.062848 | 0.083347 | 0.037239 | 0.005574 | 0.0417 | 0.057356 |
| 22.0833333 | 0.094396 | 0.060205 | 0.03706 | 0.047557 | 0.078578 | 0.013802 | 0.024775 | 0.018997 | 0.020225 | 0.042466 | 0.109552 | 0.032239 | 0.054593 | 0.03632 | 0.030752 | 0.031703 | 0.065081 | 0.043789 | 0.062937 | 0.08324 | 0.03022 | 0.015541 | 0.048891 | 0.055164 |
| 22.0972222 | 0.047792 | 0.043847 | 0.028869 | 0.030696 | 0.042607 | 0.014394 | 0.015149 | 0.010426 | 0.016464 | 0.029976 | 0.084273 | 0.02738 | 0.028555 | 0.036276 | 0.018133 | 0.026461 | 0.036478 | 0.030951 | 0.041346 | 0.068627 | 0.017484 | 0.018576 | 0.040197 | 0.034452 |
| 22.1111111 | 0.027411 | 0.019905 | 0.016722 | 0.014041 | 0.017085 | 0.009063 | 0.005716 | 0.007491 | 0.010635 | 0.026061 | 0.042004 | 0.016944 | 0.016484 | 0.048692 | 0.008459 | 0.022786 | 0.0215 | 0.013358 | 0.015922 | 0.04052 | 0.015711 | 0.01265 | 0.022047 | 0.014316 |
| 22.125 | 0.0428 | 0.012898 | 0.009971 | 0.014382 | 0.023645 | 0.004134 | 0.004222 | 0.009723 | 0.008512 | 0.036149 | 0.029227 | 0.010809 | 0.023885 | 0.055932 | 0.004953 | 0.022841 | 0.026252 | 0.003558 | 0.005701 | 0.019866 | 0.021344 | 0.01192 | 0.016629 | 0.015374 |
| 22.1388889 | 0.063022 | 0.016299 | 0.009063 | 0.011998 | 0.037683 | 0.004679 | 0.006345 | 0.011005 | 0.006319 | 0.03716 | 0.032113 | 0.006223 | 0.030521 | 0.053823 | 0.003568 | 0.02393 | 0.029785 | 0.002007 | 0.004109 | 0.018548 | 0.023577 | 0.00979 | 0.017553 | 0.021065 |
| 22.1527778 | 0.076152 | 0.020076 | 0.008462 | 0.00513 | 0.040348 | 0.006845 | 0.006916 | 0.011218 | 0.002862 | 0.032413 | 0.033464 | 0.004624 | 0.028565 | 0.050699 | 0.006472 | 0.021104 | 0.033196 | 0.002518 | 0.005241 | 0.013994 | 0.029858 | 0.004648 | 0.012996 | 0.022418 |
| 22.1666667 | 0.084988 | 0.023309 | 0.005204 | 0.004677 | 0.037214 | 0.007163 | 0.005566 | 0.01306 | 0.000948 | 0.033662 | 0.042956 | 0.006902 | 0.02639 | 0.047979 | 0.009721 | 0.013022 | 0.039717 | 0.003556 | 0.010707 | 0.0072 | 0.038647 | 0.007205 | 0.009471 | 0.023291 |
| 22.1805556 | 0.082624 | 0.019866 | 0.002055 | 0.006384 | 0.036547 | 0.006251 | 0.00332 | 0.015442 | 0.001038 | 0.034496 | 0.052057 | 0.009426 | 0.024068 | 0.042391 | 0.007797 | 0.004793 | 0.038962 | 0.003985 | 0.017263 | 0.011583 | 0.036721 | 0.010986 | 0.01055 | 0.020397 |
| 22.1944444 | 0.064983 | 0.012021 | 0.001264 | 0.008344 | 0.036205 | 0.0049 | 0.001418 | 0.014575 | 0.001562 | 0.029089 | 0.043561 | 0.009561 | 0.019151 | 0.030917 | 0.004436 | 0.001157 | 0.030997 | 0.003644 | 0.021088 | 0.013733 | 0.025119 | 0.007747 | 0.010741 | 0.01694 |
| 22.2083333 | 0.043286 | 0.010859 | 0.001172 | 0.008908 | 0.030854 | 0.003567 | 0.000815 | 0.013672 | 0.001838 | 0.02387 | 0.028208 | 0.007023 | 0.015295 | 0.015241 | 0.003399 | 0.002006 | 0.020491 | 0.004633 | 0.023399 | 0.007834 | 0.012069 | 0.00537 | 0.006516 | 0.01865 |
| 22.2222222 | 0.026985 | 0.01138 | 0.000606 | 0.00629 | 0.023395 | 0.002203 | 0.001698 | 0.016299 | 0.001682 | 0.022431 | 0.018769 | 0.003274 | 0.013729 | 0.006517 | 0.004366 | 0.004105 | 0.011309 | 0.006518 | 0.024422 | 0.003906 | 0.005535 | 0.008728 | 0.003519 | 0.016432 |
| 22.2361111 | 0.017597 | 0.006682 | 0.000956 | 0.00497 | 0.016889 | 0.001015 | 0.002841 | 0.018018 | 0.000916 | 0.018124 | 0.010869 | 0.001286 | 0.009616 | 0.00993 | 0.00529 | 0.003814 | 0.005397 | 0.006952 | 0.026703 | 0.005833 | 0.009419 | 0.010246 | 0.005382 | 0.010545 |
| 22.25 | 0.012845 | 0.005691 | 0.002651 | 0.007776 | 0.010396 | 0.000475 | 0.002693 | 0.015401 | 0.000562 | 0.009547 | 0.004725 | 0.001894 | 0.004671 | 0.017009 | 0.005279 | 0.002263 | 0.002605 | 0.008035 | 0.030719 | 0.010863 | 0.016158 | 0.007266 | 0.008048 | 0.013329 |
| 22.2638889 | 0.007658 | 0.01387 | 0.004131 | 0.012777 | 0.005408 | 0.000878 | 0.002218 | 0.010679 | 0.000773 | 0.003492 | 0.003264 | 0.003517 | 0.003468 | 0.022904 | 0.0059 | 0.003232 | 0.002855 | 0.01037 | 0.029669 | 0.012834 | 0.018835 | 0.004361 | 0.010669 | 0.017022 |
| 22.2777778 | 0.003294 | 0.026603 | 0.00463 | 0.017212 | 0.005799 | 0.001818 | 0.003804 | 0.008768 | 0.000537 | 0.001655 | 0.005835 | 0.006041 | 0.005364 | 0.028391 | 0.007424 | 0.006751 | 0.007292 | 0.0115 | 0.024594 | 0.009382 | 0.016377 | 0.002895 | 0.014162 | 0.013527 |
| 22.2916667 | 0.002083 | 0.03862 | 0.004709 | 0.019188 | 0.010923 | 0.002493 | 0.007422 | 0.010614 | 0.000208 | 0.001233 | 0.014712 | 0.009496 | 0.008258 | 0.03442 | 0.008536 | 0.009961 | 0.017569 | 0.013261 | 0.02245 | 0.009518 | 0.009653 | 0.002109 | 0.01583 | 0.007942 |
| 22.3055556 | 0.002304 | 0.048628 | 0.00405 | 0.01783 | 0.014312 | 0.002253 | 0.011525 | 0.014109 | 0.000654 | 0.000906 | 0.024945 | 0.01132 | 0.009161 | 0.042982 | 0.008531 | 0.011442 | 0.028621 | 0.01427 | 0.020441 | 0.015363 | 0.005756 | 0.003437 | 0.015626 | 0.007182 |
| 22.3194444 | 0.004799 | 0.051555 | 0.002251 | 0.013557 | 0.013528 | 0.0024 | 0.014801 | 0.016487 | 0.001279 | 0.001314 | 0.027828 | 0.010904 | 0.0081 | 0.051118 | 0.00657 | 0.011551 | 0.03325 | 0.010779 | 0.013539 | 0.018841 | 0.009282 | 0.006687 | 0.015516 | 0.013822 |
| 22.3333333 | 0.010383 | 0.040655 | 0.001055 | 0.007899 | 0.016238 | 0.004126 | 0.013847 | 0.012359 | 0.001106 | 0.004791 | 0.022185 | 0.011783 | 0.008988 | 0.053278 | 0.00331 | 0.010626 | 0.03151 | 0.006945 | 0.008036 | 0.018691 | 0.013527 | 0.006919 | 0.013662 | 0.023797 |
| 22.3472222 | 0.019754 | 0.023092 | 0.002694 | 0.003097 | 0.025427 | 0.005613 | 0.008843 | 0.006076 | 0.00194 | 0.014002 | 0.01709 | 0.015415 | 0.014501 | 0.050369 | 0.002013 | 0.009281 | 0.028995 | 0.008314 | 0.010898 | 0.018799 | 0.014841 | 0.003759 | 0.008019 | 0.034554 |
| 22.3611111 | 0.029299 | 0.015282 | 0.007148 | 0.003052 | 0.03124 | 0.006601 | 0.007806 | 0.005789 | 0.00661 | 0.026716 | 0.025281 | 0.019949 | 0.02142 | 0.044043 | 0.005083 | 0.009135 | 0.027958 | 0.012895 | 0.017151 | 0.021617 | 0.017658 | 0.004637 | 0.004199 | 0.041937 |
| 22.375 | 0.03206 | 0.018778 | 0.011114 | 0.008681 | 0.029639 | 0.007581 | 0.013317 | 0.009636 | 0.011418 | 0.029883 | 0.039012 | 0.022257 | 0.022509 | 0.032229 | 0.009583 | 0.010905 | 0.02651 | 0.014622 | 0.018372 | 0.026344 | 0.022612 | 0.011074 | 0.007517 | 0.040571 |
| 22.3888889 | 0.028784 | 0.019544 | 0.01172 | 0.013929 | 0.028012 | 0.007769 | 0.016015 | 0.011868 | 0.010484 | 0.018131 | 0.037528 | 0.020973 | 0.018714 | 0.016513 | 0.00925 | 0.011534 | 0.027426 | 0.014563 | 0.017043 | 0.03032 | 0.025011 | 0.015276 | 0.012027 | 0.031036 |
| 22.4027778 | 0.026653 | 0.01485 | 0.012033 | 0.014329 | 0.029397 | 0.00813 | 0.011593 | 0.007968 | 0.007592 | 0.007071 | 0.027245 | 0.020053 | 0.018604 | 0.007941 | 0.005031 | 0.008175 | 0.034611 | 0.019357 | 0.022883 | 0.033455 | 0.024995 | 0.016684 | 0.012332 | 0.022018 |
| 22.4166667 | 0.029769 | 0.010483 | 0.014595 | 0.013774 | 0.029102 | 0.009142 | 0.007942 | 0.004454 | 0.008989 | 0.005867 | 0.026725 | 0.022422 | 0.022554 | 0.014696 | 0.003749 | 0.003398 | 0.0418 | 0.026925 | 0.034946 | 0.032428 | 0.025649 | 0.020861 | 0.010444 | 0.022364 |
| 22.4305556 | 0.033728 | 0.007591 | 0.016311 | 0.01536 | 0.026069 | 0.009076 | 0.009177 | 0.008238 | 0.012431 | 0.007787 | 0.03222 | 0.026026 | 0.023865 | 0.0254 | 0.006391 | 0.000805 | 0.038637 | 0.030474 | 0.042636 | 0.025412 | 0.025556 | 0.021445 | 0.008794 | 0.028807 |
| 22.4444444 | 0.035157 | 0.011411 | 0.017701 | 0.016361 | 0.025569 | 0.00763 | 0.012482 | 0.01436 | 0.013799 | 0.009512 | 0.037881 | 0.029739 | 0.022014 | 0.028293 | 0.010073 | 0.000457 | 0.028768 | 0.03075 | 0.044059 | 0.02004 | 0.02471 | 0.014808 | 0.010359 | 0.035734 |
| 22.4583333 | 0.037686 | 0.023156 | 0.022856 | 0.015394 | 0.029164 | 0.00655 | 0.016462 | 0.017744 | 0.013913 | 0.013412 | 0.047921 | 0.034771 | 0.021681 | 0.027967 | 0.013473 | 0.00113 | 0.025479 | 0.031219 | 0.044283 | 0.019201 | 0.02594 | 0.011604 | 0.016366 | 0.042976 |
| 22.4722222 | 0.044019 | 0.032155 | 0.02912 | 0.013698 | 0.031211 | 0.006303 | 0.020839 | 0.018179 | 0.015491 | 0.020345 | 0.05962 | 0.040523 | 0.024769 | 0.029922 | 0.015316 | 0.001903 | 0.031612 | 0.032485 | 0.044105 | 0.016827 | 0.029059 | 0.015268 | 0.022569 | 0.048291 |
| 22.4861111 | 0.052964 | 0.033276 | 0.030438 | 0.012913 | 0.028957 | 0.006115 | 0.023449 | 0.017237 | 0.021467 | 0.029811 | 0.064752 | 0.044978 | 0.029361 | 0.033646 | 0.015405 | 0.002455 | 0.039895 | 0.035006 | 0.044971 | 0.012158 | 0.034001 | 0.017116 | 0.026623 | 0.050737 |
| 22.5 | 0.061293 | 0.03279 | 0.028618 | 0.013864 | 0.028627 | 0.005857 | 0.023777 | 0.016251 | 0.03118 | 0.038939 | 0.065253 | 0.04824 | 0.033622 | 0.03766 | 0.015832 | 0.002657 | 0.045756 | 0.037471 | 0.04783 | 0.011209 | 0.043415 | 0.013325 | 0.031234 | 0.054561 |
| 22.5138889 | 0.064569 | 0.033221 | 0.028848 | 0.015702 | 0.033232 | 0.004605 | 0.023403 | 0.015483 | 0.036699 | 0.040917 | 0.066229 | 0.050275 | 0.03828 | 0.040467 | 0.018971 | 0.002685 | 0.047662 | 0.038032 | 0.047617 | 0.01707 | 0.050276 | 0.0096 | 0.036891 | 0.059377 |
| 22.5277778 | 0.061868 | 0.032034 | 0.028826 | 0.015996 | 0.036615 | 0.003456 | 0.021538 | 0.013215 | 0.032489 | 0.033245 | 0.061248 | 0.047195 | 0.040812 | 0.039469 | 0.023761 | 0.003964 | 0.045061 | 0.037588 | 0.042284 | 0.026421 | 0.043101 | 0.009918 | 0.039069 | 0.058251 |
| 22.5416667 | 0.058433 | 0.029119 | 0.024852 | 0.013947 | 0.035184 | 0.004421 | 0.019052 | 0.010148 | 0.024127 | 0.02354 | 0.046898 | 0.03751 | 0.035989 | 0.032689 | 0.02538 | 0.006242 | 0.040289 | 0.035273 | 0.035653 | 0.030054 | 0.027998 | 0.009792 | 0.037463 | 0.048954 |
| 22.5555556 | 0.056281 | 0.025135 | 0.019629 | 0.012941 | 0.031619 | 0.004143 | 0.018049 | 0.010054 | 0.021303 | 0.022464 | 0.03336 | 0.027295 | 0.027059 | 0.022328 | 0.022982 | 0.005952 | 0.036261 | 0.02921 | 0.029578 | 0.022491 | 0.020268 | 0.005629 | 0.03684 | 0.036867 |
| 22.5694444 | 0.052099 | 0.022993 | 0.016945 | 0.016349 | 0.028237 | 0.002313 | 0.018592 | 0.012026 | 0.028165 | 0.031027 | 0.03115 | 0.024905 | 0.022011 | 0.014112 | 0.022192 | 0.003461 | 0.034929 | 0.023683 | 0.025777 | 0.01122 | 0.024921 | 0.002896 | 0.037974 | 0.028948 |
| 22.5833333 | 0.04953 | 0.022578 | 0.017611 | 0.022911 | 0.023951 | 0.002885 | 0.0232 | 0.012121 | 0.039078 | 0.039027 | 0.039306 | 0.030394 | 0.023272 | 0.012771 | 0.025188 | 0.002797 | 0.037478 | 0.027004 | 0.029011 | 0.008302 | 0.03456 | 0.005987 | 0.040377 | 0.029048 |
| 22.5972222 | 0.051385 | 0.020756 | 0.019236 | 0.027276 | 0.01908 | 0.005005 | 0.032084 | 0.010526 | 0.043127 | 0.03773 | 0.044954 | 0.034054 | 0.027856 | 0.017783 | 0.028497 | 0.005773 | 0.039921 | 0.039705 | 0.040324 | 0.018411 | 0.040021 | 0.010303 | 0.042829 | 0.034962 |
| 22.6111111 | 0.051201 | 0.020501 | 0.017678 | 0.026404 | 0.017281 | 0.005884 | 0.036207 | 0.009488 | 0.03617 | 0.027979 | 0.039563 | 0.030273 | 0.031395 | 0.024098 | 0.030316 | 0.011823 | 0.03691 | 0.04685 | 0.047576 | 0.033393 | 0.038956 | 0.009439 | 0.043378 | 0.038626 |

| | | | | | | | | | | | | | | | | | | | | | | | | |
|---|---|---|---|---|---|---|---|---|---|---|---|---|---|---|---|---|---|---|---|---|---|---|---|---|
| 22.625 | 0.04711 | 0.021662 | 0.013237 | 0.023068 | 0.019021 | 0.004821 | 0.028358 | 0.008839 | 0.023408 | 0.016784 | 0.028918 | 0.022281 | 0.032283 | 0.027647 | 0.031008 | 0.015829 | 0.031836 | 0.037644 | 0.039464 | 0.037111 | 0.031302 | 0.005327 | 0.042708 | 0.035935 |
| 22.6388889 | 0.041737 | 0.019735 | 0.00972 | 0.01983 | 0.01907 | 0.00296 | 0.015287 | 0.007641 | 0.012115 | 0.008686 | 0.021275 | 0.015655 | 0.031524 | 0.027924 | 0.030341 | 0.014794 | 0.029258 | 0.021834 | 0.023297 | 0.024819 | 0.020299 | 0.004256 | 0.040584 | 0.031365 |
| 22.6527778 | 0.034565 | 0.015382 | 0.008553 | 0.015986 | 0.013624 | 0.0015 | 0.006968 | 0.00544 | 0.005448 | 0.004275 | 0.016805 | 0.012299 | 0.02876 | 0.025816 | 0.027337 | 0.01024 | 0.026281 | 0.01015 | 0.010017 | 0.010804 | 0.011448 | 0.006858 | 0.033461 | 0.028404 |
| 22.6666667 | 0.024585 | 0.009732 | 0.008585 | 0.011501 | 0.006122 | 0.001174 | 0.003718 | 0.003236 | 0.004103 | 0.003917 | 0.011742 | 0.008984 | 0.024513 | 0.0217 | 0.021669 | 0.005524 | 0.018026 | 0.004139 | 0.004473 | 0.007505 | 0.006707 | 0.009952 | 0.019264 | 0.026126 |
| 22.6805556 | 0.012774 | 0.006123 | 0.007182 | 0.007398 | 0.002535 | 0.00191 | 0.003396 | 0.00342 | 0.007062 | 0.006044 | 0.006382 | 0.005559 | 0.020365 | 0.016642 | 0.015114 | 0.004657 | 0.010253 | 0.003107 | 0.004854 | 0.011213 | 0.007804 | 0.011188 | 0.010075 | 0.024041 |
| 22.6944444 | 0.006049 | 0.007443 | 0.003891 | 0.004447 | 0.003242 | 0.002414 | 0.005156 | 0.005249 | 0.010317 | 0.009143 | 0.004519 | 0.00619 | 0.016147 | 0.012866 | 0.0106 | 0.005404 | 0.01317 | 0.005575 | 0.005153 | 0.014656 | 0.012047 | 0.009893 | 0.017092 | 0.023381 |
| 22.7083333 | 0.008447 | 0.007566 | 0.001866 | 0.003505 | 0.003349 | 0.001864 | 0.006851 | 0.005531 | 0.01171 | 0.013741 | 0.005815 | 0.01016 | 0.011847 | 0.011394 | 0.008985 | 0.00514 | 0.019621 | 0.00894 | 0.004121 | 0.016474 | 0.01166 | 0.00656 | 0.027775 | 0.023493 |
| 22.7222222 | 0.010864 | 0.00534 | 0.001688 | 0.003934 | 0.002331 | 0.000724 | 0.007832 | 0.004721 | 0.010935 | 0.016446 | 0.005917 | 0.012654 | 0.007722 | 0.010511 | 0.007925 | 0.005996 | 0.017201 | 0.010429 | 0.00259 | 0.016334 | 0.007027 | 0.003401 | 0.028371 | 0.020921 |
| 22.7361111 | 0.007548 | 0.00429 | 0.002814 | 0.004002 | 0.002887 | 0.000179 | 0.008401 | 0.004969 | 0.007886 | 0.013783 | 0.004586 | 0.010238 | 0.005426 | 0.008284 | 0.005204 | 0.008392 | 0.010068 | 0.009423 | 0.002385 | 0.013197 | 0.00539 | 0.00373 | 0.020564 | 0.014095 |
| 22.75 | 0.003325 | 0.002682 | 0.005717 | 0.003031 | 0.003071 | 0.000167 | 0.008122 | 0.004178 | 0.004725 | 0.009312 | 0.005484 | 0.005474 | 0.006338 | 0.005235 | 0.002044 | 0.008636 | 0.007677 | 0.008275 | 0.004922 | 0.007379 | 0.006457 | 0.006103 | 0.011577 | 0.006582 |
| 22.7638889 | 0.001976 | 0.0022 | 0.007188 | 0.001729 | 0.002897 | 0.000227 | 0.007835 | 0.00223 | 0.002591 | 0.006169 | 0.008373 | 0.004275 | 0.007511 | 0.003287 | 0.001342 | 0.005863 | 0.00705 | 0.0086 | 0.007265 | 0.003092 | 0.006276 | 0.006072 | 0.005288 | 0.0032 |
| 22.7777778 | 0.002341 | 0.006545 | 0.005762 | 0.001803 | 0.00452 | 0.000525 | 0.008812 | 0.001534 | 0.001763 | 0.003602 | 0.009362 | 0.005209 | 0.005969 | 0.003528 | 0.002975 | 0.004338 | 0.005431 | 0.008646 | 0.007555 | 0.002908 | 0.005969 | 0.004016 | 0.001749 | 0.003252 |
| 22.7916667 | 0.002334 | 0.012855 | 0.004506 | 0.00371 | 0.006298 | 0.000918 | 0.010253 | 0.001506 | 0.002758 | 0.001928 | 0.008323 | 0.004235 | 0.005742 | 0.005156 | 0.003992 | 0.00376 | 0.005646 | 0.007246 | 0.007139 | 0.003334 | 0.005843 | 0.002941 | 0.001537 | 0.003359 |
| 22.8055556 | 0.001769 | 0.016785 | 0.003614 | 0.005595 | 0.007443 | 0.001122 | 0.009851 | 0.001475 | 0.003129 | 0.001426 | 0.006997 | 0.002103 | 0.010025 | 0.006199 | 0.003355 | 0.00288 | 0.004202 | 0.004945 | 0.00623 | 0.004764 | 0.004726 | 0.003398 | 0.004007 | 0.002812 |
| 22.8194444 | 0.003482 | 0.019372 | 0.002308 | 0.005247 | 0.008572 | 0.000929 | 0.006601 | 0.001385 | 0.001962 | 0.001142 | 0.004691 | 0.001241 | 0.012884 | 0.005248 | 0.002347 | 0.00329 | 0.00238 | 0.002639 | 0.003943 | 0.00949 | 0.003829 | 0.004239 | 0.00741 | 0.002171 |
| 22.8333333 | 0.010602 | 0.021769 | 0.001605 | 0.003273 | 0.00961 | 0.000814 | 0.003636 | 0.001246 | 0.001174 | 0.001162 | 0.00267 | 0.002526 | 0.010649 | 0.003099 | 0.001655 | 0.004118 | 0.007138 | 0.002385 | 0.002523 | 0.015397 | 0.004005 | 0.004726 | 0.011003 | 0.002809 |
| 22.8472222 | 0.019996 | 0.021873 | 0.001484 | 0.003086 | 0.010278 | 0.001125 | 0.00341 | 0.001044 | 0.00107 | 0.001667 | 0.003433 | 0.004135 | 0.008097 | 0.002061 | 0.002108 | 0.006031 | 0.015758 | 0.004878 | 0.004824 | 0.021553 | 0.00577 | 0.003639 | 0.014454 | 0.005997 |
| 22.8611111 | 0.026222 | 0.019802 | 0.001722 | 0.004934 | 0.010625 | 0.001291 | 0.004167 | 0.001336 | 0.000819 | 0.001578 | 0.005251 | 0.003987 | 0.010502 | 0.002696 | 0.003725 | 0.009526 | 0.021455 | 0.007964 | 0.008063 | 0.027231 | 0.009471 | 0.001931 | 0.016118 | 0.010536 |
| 22.875 | 0.028345 | 0.018979 | 0.002855 | 0.006371 | 0.011812 | 0.001334 | 0.005087 | 0.003204 | 0.001594 | 0.002409 | 0.008478 | 0.004281 | 0.017571 | 0.004347 | 0.004628 | 0.012211 | 0.025135 | 0.0105 | 0.009163 | 0.030747 | 0.011517 | 0.001926 | 0.014903 | 0.013824 |
| 22.8888889 | 0.029034 | 0.020925 | 0.006009 | 0.008484 | 0.014237 | 0.002295 | 0.007355 | 0.006284 | 0.006319 | 0.00763 | 0.017361 | 0.008171 | 0.028193 | 0.007655 | 0.005403 | 0.012991 | 0.030363 | 0.013653 | 0.010073 | 0.034467 | 0.008843 | 0.002563 | 0.013599 | 0.016147 |
| 22.9027778 | 0.031213 | 0.022612 | 0.010815 | 0.012947 | 0.015945 | 0.003912 | 0.010338 | 0.008881 | 0.013983 | 0.015691 | 0.028916 | 0.012337 | 0.038749 | 0.011984 | 0.008288 | 0.012849 | 0.037909 | 0.018238 | 0.01358 | 0.039668 | 0.009048 | 0.002952 | 0.013721 | 0.019354 |
| 22.9166667 | 0.034448 | 0.021484 | 0.015246 | 0.017674 | 0.014474 | 0.00435 | 0.012202 | 0.009999 | 0.01949 | 0.0231 | 0.034757 | 0.012652 | 0.040393 | 0.015181 | 0.012256 | 0.012897 | 0.044488 | 0.022617 | 0.020006 | 0.043473 | 0.019582 | 0.00327 | 0.012754 | 0.022127 |
| 22.9305556 | 0.037927 | 0.019891 | 0.018905 | 0.021144 | 0.011844 | 0.003284 | 0.013228 | 0.011027 | 0.021836 | 0.029166 | 0.033931 | 0.012976 | 0.034585 | 0.016174 | 0.014758 | 0.013027 | 0.045479 | 0.025336 | 0.028612 | 0.043997 | 0.035907 | 0.002821 | 0.011404 | 0.023515 |
| 22.9444444 | 0.04689 | 0.019928 | 0.022099 | 0.024575 | 0.014749 | 0.002083 | 0.01473 | 0.013882 | 0.025533 | 0.033778 | 0.034586 | 0.019367 | 0.030666 | 0.015079 | 0.01611 | 0.012027 | 0.041565 | 0.028721 | 0.03772 | 0.042087 | 0.052817 | 0.002509 | 0.012641 | 0.024044 |
| 22.9583333 | 0.061375 | 0.021443 | 0.022554 | 0.027555 | 0.025737 | 0.001541 | 0.01716 | 0.01633 | 0.033314 | 0.039145 | 0.043264 | 0.028584 | 0.033486 | 0.013368 | 0.019184 | 0.008421 | 0.037978 | 0.035779 | 0.044257 | 0.040632 | 0.065316 | 0.003928 | 0.01541 | 0.02347 |
| 22.9722222 | 0.070856 | 0.026751 | 0.018568 | 0.029808 | 0.038637 | 0.001891 | 0.020305 | 0.016021 | 0.04204 | 0.048405 | 0.054284 | 0.032669 | 0.040828 | 0.012281 | 0.023481 | 0.004764 | 0.038605 | 0.045519 | 0.048267 | 0.041454 | 0.069038 | 0.005922 | 0.017382 | 0.023923 |
| 22.9861111 | 0.074505 | 0.035159 | 0.01342 | 0.033552 | 0.045468 | 0.002633 | 0.023451 | 0.015446 | 0.045504 | 0.057429 | 0.059049 | 0.031761 | 0.045703 | 0.011661 | 0.023963 | 0.003498 | 0.042082 | 0.052591 | 0.05241 | 0.044836 | 0.068038 | 0.006322 | 0.018454 | 0.03013 |
| 23 | 0.082525 | 0.039246 | 0.012208 | 0.038357 | 0.046801 | 0.002533 | 0.025593 | 0.015931 | 0.040788 | 0.058719 | 0.060182 | 0.031174 | 0.046912 | 0.012222 | 0.021809 | 0.003224 | 0.044468 | 0.053843 | 0.056843 | 0.052125 | 0.065461 | 0.005411 | 0.019492 | 0.044261 |
| 23.0138889 | 0.096148 | 0.036568 | 0.016282 | 0.042666 | 0.05206 | 0.001441 | 0.024982 | 0.015911 | 0.032663 | 0.054447 | 0.062958 | 0.032332 | 0.046146 | 0.014833 | 0.023411 | 0.005328 | 0.042186 | 0.05246 | 0.059694 | 0.060431 | 0.058444 | 0.00465 | 0.021525 | 0.058637 |
| 23.0277778 | 0.10627 | 0.033156 | 0.022275 | 0.046875 | 0.065812 | 0.000405 | 0.022128 | 0.016497 | 0.02699 | 0.05207 | 0.070723 | 0.033052 | 0.044436 | 0.018342 | 0.029881 | 0.00796 | 0.037429 | 0.051592 | 0.059786 | 0.060222 | 0.048401 | 0.005396 | 0.025145 | 0.06482 |
| 23.0416667 | 0.107508 | 0.032008 | 0.026403 | 0.050658 | 0.085276 | 0.001019 | 0.021776 | 0.021371 | 0.024458 | 0.052072 | 0.081767 | 0.034038 | 0.047669 | 0.022524 | 0.037956 | 0.006588 | 0.040782 | 0.050224 | 0.058049 | 0.056051 | 0.040992 | 0.008244 | 0.030279 | 0.062826 |
| 23.0555556 | 0.107075 | 0.031334 | 0.029128 | 0.053577 | 0.109202 | 0.005163 | 0.027448 | 0.029205 | 0.02489 | 0.054094 | 0.091431 | 0.038519 | 0.061608 | 0.027611 | 0.044834 | 0.003631 | 0.054014 | 0.050452 | 0.059419 | 0.050458 | 0.037982 | 0.012634 | 0.038812 | 0.054416 |
| 23.0694444 | 0.110053 | 0.038576 | 0.034441 | 0.058286 | 0.1241 | 0.011663 | 0.035216 | 0.034027 | 0.026982 | 0.060763 | 0.104562 | 0.044859 | 0.075787 | 0.030926 | 0.049056 | 0.00281 | 0.061903 | 0.057995 | 0.066767 | 0.037158 | 0.036757 | 0.021202 | 0.04918 | 0.048865 |
| 23.0833333 | 0.093436 | 0.053967 | 0.03851 | 0.062534 | 0.105939 | 0.015745 | 0.032763 | 0.030952 | 0.025836 | 0.058552 | 0.106795 | 0.045445 | 0.063386 | 0.028855 | 0.041611 | 0.004025 | 0.0535 | 0.060218 | 0.065588 | 0.029096 | 0.029335 | 0.02936 | 0.055248 | 0.050689 |
| 23.0972222 | 0.049045 | 0.051257 | 0.030682 | 0.050955 | 0.058257 | 0.013606 | 0.01977 | 0.018795 | 0.017013 | 0.036163 | 0.07454 | 0.035409 | 0.032041 | 0.020842 | 0.02061 | 0.007327 | 0.032076 | 0.038736 | 0.041163 | 0.028799 | 0.018598 | 0.024316 | 0.049163 | 0.043355 |
| 23.1111111 | 0.020528 | 0.030689 | 0.015473 | 0.029189 | 0.02329 | 0.007526 | 0.01333 | 0.007292 | 0.007865 | 0.020439 | 0.034615 | 0.020382 | 0.018788 | 0.010996 | 0.007239 | 0.013975 | 0.015499 | 0.015669 | 0.017165 | 0.025112 | 0.017378 | 0.010954 | 0.031771 | 0.022635 |
| 23.125 | 0.026117 | 0.0222 | 0.00819 | 0.021317 | 0.026624 | 0.003329 | 0.014649 | 0.005935 | 0.007047 | 0.025861 | 0.024503 | 0.008465 | 0.022537 | 0.005646 | 0.010153 | 0.02242 | 0.018521 | 0.011925 | 0.013296 | 0.01681 | 0.019286 | 0.003858 | 0.020172 | 0.0066 |
| 23.1388889 | 0.037173 | 0.026001 | 0.007767 | 0.021832 | 0.042499 | 0.001893 | 0.013176 | 0.010136 | 0.007726 | 0.031457 | 0.031669 | 0.004258 | 0.026536 | 0.006956 | 0.011454 | 0.029193 | 0.029029 | 0.010638 | 0.011747 | 0.007613 | 0.016272 | 0.004907 | 0.018206 | 0.003319 |
| 23.1527778 | 0.042276 | 0.030177 | 0.005432 | 0.018531 | 0.042991 | 0.001852 | 0.010714 | 0.01432 | 0.007342 | 0.03304 | 0.034035 | 0.006581 | 0.029482 | 0.011275 | 0.005787 | 0.033107 | 0.034029 | 0.007051 | 0.008628 | 0.002568 | 0.017265 | 0.007456 | 0.016156 | 0.008673 |
| 23.1666667 | 0.044819 | 0.028357 | 0.00312 | 0.014124 | 0.029342 | 0.003019 | 0.009779 | 0.017516 | 0.010793 | 0.037449 | 0.032569 | 0.007016 | 0.029995 | 0.015549 | 0.003781 | 0.034698 | 0.033759 | 0.006078 | 0.008765 | 0.00187 | 0.025585 | 0.008172 | 0.01345 | 0.016248 |
| 23.1805556 | 0.04548 | 0.02094 | 0.002282 | 0.009113 | 0.017738 | 0.003943 | 0.007444 | 0.020686 | 0.017884 | 0.044686 | 0.030178 | 0.004289 | 0.027071 | 0.019578 | 0.00372 | 0.033798 | 0.030512 | 0.004803 | 0.007433 | 0.002813 | 0.031139 | 0.010502 | 0.012116 | 0.021993 |
| 23.1944444 | 0.048557 | 0.01839 | 0.002187 | 0.005499 | 0.01569 | 0.003611 | 0.004847 | 0.024717 | 0.023944 | 0.050314 | 0.026715 | 0.002918 | 0.022884 | 0.021928 | 0.005809 | 0.030237 | 0.029294 | 0.004609 | 0.008096 | 0.003136 | 0.030843 | 0.011496 | 0.012993 | 0.024309 |
| 23.2083333 | 0.052124 | 0.017686 | 0.004594 | 0.00544 | 0.014944 | 0.002864 | 0.005622 | 0.025733 | 0.021633 | 0.042927 | 0.019948 | 0.002273 | 0.019477 | 0.017844 | 0.011581 | 0.024814 | 0.028121 | 0.005287 | 0.012759 | 0.003978 | 0.030458 | 0.00652 | 0.014043 | 0.022207 |
| 23.2222222 | 0.050941 | 0.01304 | 0.009171 | 0.004431 | 0.011256 | 0.00232 | 0.008293 | 0.022536 | 0.012843 | 0.025565 | 0.009351 | 0.001429 | 0.01597 | 0.010386 | 0.009968 | 0.019785 | 0.019877 | 0.004377 | 0.013816 | 0.005904 | 0.028337 | 0.002276 | 0.011441 | 0.020067 |
| 23.2361111 | 0.046332 | 0.008855 | 0.013543 | 0.002287 | 0.00944 | 0.001786 | 0.009348 | 0.020586 | 0.006203 | 0.01452 | 0.003908 | 0.002451 | 0.01288 | 0.009434 | 0.003087 | 0.017828 | 0.008298 | 0.003843 | 0.008998 | 0.00532 | 0.020495 | 0.003019 | 0.008902 | 0.020752 |
| 23.25 | 0.039041 | 0.007019 | 0.013506 | 0.002334 | 0.010993 | 0.002194 | 0.008725 | 0.021447 | 0.002928 | 0.011837 | 0.006548 | 0.004803 | 0.010815 | 0.012364 | 0.002162 | 0.018388 | 0.001928 | 0.004265 | 0.005542 | 0.003759 | 0.009952 | 0.006598 | 0.011143 | 0.021047 |
| 23.2638889 | 0.028205 | 0.005514 | 0.009357 | 0.002952 | 0.013804 | 0.004196 | 0.008225 | 0.02085 | 0.001295 | 0.009843 | 0.008462 | 0.004666 | 0.007508 | 0.01111 | 0.008233 | 0.017592 | 0.002039 | 0.003367 | 0.00588 | 0.006012 | 0.004982 | 0.009544 | 0.011174 | 0.018008 |
| 23.2777778 | 0.019016 | 0.005468 | 0.008313 | 0.002837 | 0.018608 | 0.005049 | 0.009172 | 0.017321 | 0.001397 | 0.007854 | 0.008375 | 0.004826 | 0.004028 | 0.00725 | 0.015923 | 0.013501 | 0.008018 | 0.001881 | 0.005824 | 0.011797 | 0.007937 | 0.007453 | 0.00849 | 0.011805 |
| 23.2916667 | 0.015494 | 0.007328 | 0.010651 | 0.002139 | 0.022632 | 0.003187 | 0.011272 | 0.015189 | 0.002665 | 0.00864 | 0.012908 | 0.009747 | 0.003728 | 0.006085 | 0.015882 | 0.009335 | 0.018844 | 0.001993 | 0.005085 | 0.016545 | 0.009658 | 0.004523 | 0.013433 | 0.005609 |
| 23.3055556 | 0.011228 | 0.008588 | 0.01118 | 0.001892 | 0.019842 | 0.001593 | 0.012 | 0.016622 | 0.004621 | 0.010798 | 0.021321 | 0.015049 | 0.005825 | 0.008061 | 0.009695 | 0.008639 | 0.027796 | 0.006364 | 0.010531 | 0.01489 | 0.006988 | 0.006094 | 0.021574 | 0.002436 |
| 23.3194444 | 0.004983 | 0.007685 | 0.008697 | 0.004396 | 0.013414 | 0.001696 | 0.009942 | 0.016881 | 0.006298 | 0.012364 | 0.028236 | 0.016391 | 0.006528 | 0.010263 | 0.007383 | 0.009994 | 0.030203 | 0.014226 | 0.022315 | 0.007866 | 0.006354 | 0.008679 | 0.023084 | 0.001848 |
| 23.3333333 | 0.003246 | 0.010281 | 0.00815 | 0.012813 | 0.010833 | 0.001949 | 0.008606 | 0.014877 | 0.007026 | 0.013086 | 0.033072 | 0.01583 | 0.005712 | 0.010719 | 0.006488 | 0.009662 | 0.028252 | 0.019926 | 0.028085 | 0.002685 | 0.009384 | 0.012031 | 0.01964 | 0.003144 |
| 23.3472222 | 0.006101 | 0.02206 | 0.015041 | 0.023941 | 0.01027 | 0.001651 | 0.010674 | 0.013151 | 0.007173 | 0.012712 | 0.037171 | 0.015905 | 0.00865 | 0.007849 | 0.003975 | 0.006546 | 0.024584 | 0.021319 | 0.025258 | 0.001176 | 0.014501 | 0.01778 | 0.015974 | 0.00659 |
| 23.3611111 | 0.012337 | 0.032352 | 0.022269 | 0.026059 | 0.009966 | 0.001688 | 0.013142 | 0.009782 | 0.008283 | 0.012444 | 0.040427 | 0.016654 | 0.016099 | 0.004268 | 0.002343 | 0.00419 | 0.020864 | 0.021341 | 0.022722 | 0.002891 | 0.020065 | 0.021599 | 0.013059 | 0.010975 |
| 23.375 | 0.019697 | 0.031928 | 0.023566 | 0.018473 | 0.018011 | 0.003364 | 0.013628 | 0.005897 | 0.011522 | 0.014072 | 0.043632 | 0.017876 | 0.019848 | 0.003534 | 0.001726 | 0.007701 | 0.01792 | 0.021368 | 0.02235 | 0.009649 | 0.022506 | 0.021945 | 0.011063 | 0.010985 |
| 23.3888889 | 0.025288 | 0.024972 | 0.021927 | 0.012224 | 0.036591 | 0.005924 | 0.013214 | 0.006357 | 0.015945 | 0.017592 | 0.047337 | 0.02026 | 0.018254 | 0.003291 | 0.002489 | 0.014598 | 0.016019 | 0.021341 | 0.023749 | 0.01817 | 0.022528 | 0.020991 | 0.011909 | 0.007299 |
| 23.4027778 | 0.029938 | 0.019209 | 0.019735 | 0.011972 | 0.052005 | 0.006969 | 0.013617 | 0.010334 | 0.021145 | 0.022233 | 0.050852 | 0.024961 | 0.020561 | 0.005763 | 0.00481 | 0.016809 | 0.018087 | 0.021341 | 0.026834 | 0.023384 | 0.026396 | 0.018691 | 0.015208 | 0.010721 |
| 23.4166667 | 0.035336 | 0.017315 | 0.019921 | 0.014511 | 0.053819 | 0.006737 | 0.016732 | 0.014319 | 0.025621 | 0.026068 | 0.053391 | 0.0311 | 0.027983 | 0.018812 | 0.010892 | 0.01433 | 0.02535 | 0.020833 | 0.027945 | 0.025262 | 0.03545 | 0.015122 | 0.017691 | 0.022826 |
| 23.4305556 | 0.041467 | 0.016476 | 0.021294 | 0.015573 | 0.048627 | 0.007638 | 0.021133 | 0.016857 | 0.026271 | 0.025707 | 0.051832 | 0.034294 | 0.033355 | 0.045156 | 0.020531 | 0.012363 | 0.03243 | 0.020129 | 0.026367 | 0.026662 | 0.041863 | 0.012116 | 0.017935 | 0.036464 |
| 23.4444444 | 0.046427 | 0.015968 | 0.019783 | 0.014972 | 0.045097 | 0.01004 | 0.021765 | 0.016641 | 0.023785 | 0.022021 | 0.044785 | 0.033282 | 0.033962 | 0.066096 | 0.026465 | 0.012372 | 0.034233 | 0.019497 | 0.025775 | 0.030304 | 0.039952 | 0.011805 | 0.017162 | 0.046367 |
| 23.4583333 | 0.049383 | 0.016439 | 0.017983 | 0.01509 | 0.044335 | 0.012325 | 0.017876 | 0.015812 | 0.022135 | 0.020263 | 0.03897 | 0.031817 | 0.032255 | 0.057494 | 0.025842 | 0.014979 | 0.032805 | 0.019975 | 0.029476 | 0.034554 | 0.034959 | 0.013803 | 0.01736 | 0.049531 |
| 23.4722222 | 0.052088 | 0.017005 | 0.018845 | 0.01583 | 0.04563 | 0.013963 | 0.014456 | 0.01898 | 0.025176 | 0.02373 | 0.04184 | 0.031678 | 0.030921 | 0.034375 | 0.023857 | 0.019278 | 0.032013 | 0.023729 | 0.037134 | 0.03478 | 0.035073 | 0.01516 | 0.019972 | 0.048916 |
| 23.4861111 | 0.055978 | 0.014747 | 0.019638 | 0.015902 | 0.046898 | 0.015514 | 0.015233 | 0.022656 | 0.034144 | 0.031117 | 0.050367 | 0.033012 | 0.030124 | 0.022154 | 0.023746 | 0.022 | 0.031717 | 0.028664 | 0.044406 | 0.030714 | 0.04046 | 0.014704 | 0.024321 | 0.048772 |
| 23.5 | 0.058463 | 0.008654 | 0.018313 | 0.016065 | 0.045933 | 0.016597 | 0.018943 | 0.020691 | 0.043208 | 0.038764 | 0.055923 | 0.035124 | 0.03015 | 0.018789 | 0.024286 | 0.021822 | 0.03316 | 0.031072 | 0.047679 | 0.025947 | 0.044009 | 0.01441 | 0.028998 | 0.04835 |
| 23.5138889 | 0.055867 | 0.005421 | 0.016376 | 0.016216 | 0.043688 | 0.01576 | 0.021137 | 0.015264 | 0.045035 | 0.041628 | 0.05648 | 0.035423 | 0.032875 | 0.017329 | 0.024559 | 0.020927 | 0.03722 | 0.03389 | 0.048602 | 0.021521 | 0.041531 | 0.015268 | 0.032645 | 0.046247 |
| 23.5277778 | 0.050663 | 0.007196 | 0.014901 | 0.014113 | 0.039346 | 0.012101 | 0.01901 | 0.010157 | 0.038648 | 0.034988 | 0.051816 | 0.034525 | 0.037492 | 0.016336 | 0.024959 | 0.021844 | 0.040746 | 0.041262 | 0.050129 | 0.015604 | 0.033816 | 0.015075 | 0.035538 | 0.041966 |
| 23.5416667 | 0.047227 | 0.005956 | 0.012912 | 0.011245 | 0.033039 | 0.007394 | 0.015154 | 0.008421 | 0.027439 | 0.023327 | 0.042119 | 0.03353 | 0.038644 | 0.013646 | 0.024182 | 0.022807 | 0.043719 | 0.044966 | 0.047175 | 0.008688 | 0.025653 | 0.012615 | 0.038847 | 0.035005 |
| 23.5555556 | 0.043405 | 0.004302 | 0.011899 | 0.013265 | 0.030976 | 0.005742 | 0.013847 | 0.011635 | 0.019549 | 0.017735 | 0.034332 | 0.032363 | 0.035383 | 0.008726 | 0.020882 | 0.020093 | 0.046847 | 0.036304 | 0.03693 | 0.003551 | 0.024842 | 0.009634 | 0.040581 | 0.028007 |
| 23.5694444 | 0.038959 | 0.010088 | 0.017067 | 0.021049 | 0.033081 | 0.008416 | 0.01582 | 0.014935 | 0.022992 | 0.022036 | 0.034417 | 0.033682 | 0.034357 | 0.005306 | 0.017882 | 0.013911 | 0.049639 | 0.025257 | 0.030061 | 0.001327 | 0.032263 | 0.007797 | 0.042185 | 0.025979 |
| 23.5833333 | 0.039801 | 0.018061 | 0.026405 | 0.027352 | 0.031917 | 0.010409 | 0.020867 | 0.015546 | 0.035284 | 0.031324 | 0.041446 | 0.039988 | 0.041323 | 0.008727 | 0.019262 | 0.010282 | 0.05286 | 0.024295 | 0.032688 | 0.000606 | 0.041956 | 0.008849 | 0.046793 | 0.029592 |
| 23.5972222 | 0.046995 | 0.020425 | 0.031156 | 0.027903 | 0.026495 | 0.009423 | 0.027485 | 0.014538 | 0.044296 | 0.036075 | 0.047847 | 0.047475 | 0.053329 | 0.018873 | 0.02445 | 0.01338 | 0.049535 | 0.032721 | 0.037599 | 0.002348 | 0.048945 | 0.013414 | 0.051004 | 0.03405 |
| 23.6111111 | 0.052074 | 0.017951 | 0.028598 | 0.024977 | 0.022066 | 0.008905 | 0.030388 | 0.010724 | 0.039252 | 0.029388 | 0.045123 | 0.047419 | 0.057184 | 0.029109 | 0.028127 | 0.018279 | 0.034396 | 0.040481 | 0.035616 | 0.008861 | 0.046238 | 0.018048 | 0.050634 | 0.036412 |
| 23.625 | 0.049119 | 0.013484 | 0.021486 | 0.019857 | 0.021146 | 0.009182 | 0.027167 | 0.005893 | 0.025459 | 0.018458 | 0.034594 | 0.038752 | 0.045336 | 0.03331 | 0.027286 | 0.019606 | 0.019088 | 0.036895 | 0.025271 | 0.015668 | 0.034901 | 0.018864 | 0.046282 | 0.036243 |
| 23.6388889 | 0.04347 | 0.00807 | 0.013901 | 0.014671 | 0.021376 | 0.009049 | 0.02165 | 0.004414 | 0.016818 | 0.012907 | 0.023798 | 0.028361 | 0.027747 | 0.030863 | 0.023847 | 0.017919 | 0.014577 | 0.02236 | 0.014173 | 0.016359 | 0.026758 | 0.016676 | 0.042139 | 0.033389 |
| 23.6527778 | 0.039468 | 0.003976 | 0.009043 | 0.01162 | 0.019657 | 0.007725 | 0.01633 | 0.005515 | 0.012934 | 0.010015 | 0.016785 | 0.019781 | 0.017586 | 0.024946 | 0.020705 | 0.013652 | 0.017487 | 0.009034 | 0.009876 | 0.01086 | 0.020226 | 0.013893 | 0.037626 | 0.029225 |
| 23.6666667 | 0.031962 | 0.003788 | 0.005894 | 0.008851 | 0.015317 | 0.005192 | 0.009375 | 0.004541 | 0.008886 | 0.005659 | 0.011593 | 0.010941 | 0.015887 | 0.017577 | 0.01847 | 0.008541 | 0.018238 | 0.00478 | 0.0121 | 0.005129 | 0.010352 | 0.009895 | 0.027784 | 0.024693 |
| 23.6805556 | 0.018809 | 0.007417 | 0.003254 | 0.005051 | 0.009302 | 0.00538 | 0.004819 | 0.001585 | 0.00949 | 0.003228 | 0.00731 | 0.005655 | 0.015702 | 0.00987 | 0.014995 | 0.009157 | 0.012273 | 0.0046 | 0.016009 | 0.004615 | 0.00318 | 0.006623 | 0.016033 | 0.019854 |
| 23.6944444 | 0.007914 | 0.01042 | 0.002652 | 0.002198 | 0.004579 | 0.008948 | 0.008219 | 0.000393 | 0.016462 | 0.006052 | 0.008282 | 0.008502 | 0.015083 | 0.004314 | 0.009886 | 0.016885 | 0.006026 | 0.003796 | 0.017304 | 0.007331 | 0.001405 | 0.006762 | 0.012673 | 0.015809 |
| 23.7083333 | 0.00599 | 0.010727 | 0.003397 | 0.001861 | 0.004507 | 0.011293 | 0.015415 | 0.002078 | 0.02485 | 0.011083 | 0.012475 | 0.012398 | 0.017427 | 0.001739 | 0.005266 | 0.022411 | 0.005828 | 0.002457 | 0.014168 | 0.009511 | 0.003566 | 0.006724 | 0.017356 | 0.01305 |
| 23.7222222 | 0.010502 | 0.011498 | 0.003768 | 0.002702 | 0.007085 | 0.011801 | 0.020013 | 0.006468 | 0.030932 | 0.01563 | 0.01316 | 0.012948 | 0.021451 | 0.001468 | 0.002317 | 0.019525 | 0.010204 | 0.00184 | 0.008164 | 0.010791 | 0.010325 | 0.004272 | 0.019984 | 0.010765 |
| 23.7361111 | 0.015155 | 0.013458 | 0.005045 | 0.002864 | 0.008641 | 0.013759 | 0.019183 | 0.010189 | 0.032508 | 0.019934 | 0.010604 | 0.012174 | 0.019313 | 0.003112 | 0.002361 | 0.013662 | 0.015581 | 0.003734 | 0.004783 | 0.010313 | 0.016957 | 0.00246 | 0.017481 | 0.008461 |
| 23.75 | 0.016605 | 0.013383 | 0.006402 | 0.002395 | 0.008101 | 0.016647 | 0.015998 | 0.009241 | 0.027775 | 0.021794 | 0.007712 | 0.011033 | 0.01092 | 0.004597 | 0.004494 | 0.010124 | 0.017959 | 0.006647 | 0.005744 | 0.007208 | 0.01412 | 0.002147 | 0.013822 | 0.005589 |
| 23.7638889 | 0.012429 | 0.009662 | 0.006801 | 0.001855 | 0.005108 | 0.015167 | 0.014973 | 0.005133 | 0.017986 | 0.017996 | 0.004666 | 0.009155 | 0.007037 | 0.004572 | 0.004656 | 0.008101 | 0.016036 | 0.009135 | 0.005719 | 0.005783 | 0.008653 | 0.002514 | 0.012986 | 0.00461 |
| 23.7777778 | 0.005895 | 0.005358 | 0.009834 | 0.001408 | 0.001979 | 0.00946 | 0.014513 | 0.001833 | 0.010844 | 0.01092 | 0.002077 | 0.006995 | 0.009099 | 0.003824 | 0.002591 | 0.006476 | 0.010451 | 0.010183 | 0.004098 | 0.008683 | 0.008139 | 0.003279 | 0.013442 | 0.007422 |
| 23.7916667 | 0.002806 | 0.003564 | 0.012555 | 0.001835 | 0.003527 | 0.006293 | 0.013233 | 0.001963 | 0.011313 | 0.006894 | 0.001445 | 0.006675 | 0.009613 | 0.003746 | 0.001631 | 0.005455 | 0.004742 | 0.006983 | 0.004973 | 0.009378 | 0.007232 | 0.002577 | 0.0131 | 0.01079 |
| 23.8055556 | 0.00323 | 0.00308 | 0.009916 | 0.0029 | 0.009848 | 0.006358 | 0.011723 | 0.003504 | 0.013222 | 0.006681 | 0.002142 | 0.008008 | 0.008085 | 0.00417 | 0.002318 | 0.005275 | 0.003444 | 0.002525 | 0.008736 | 0.006307 | 0.007583 | 0.00113 | 0.014163 | 0.012121 |
| 23.8194444 | 0.007667 | 0.001872 | 0.006355 | 0.003305 | 0.015409 | 0.004697 | 0.008818 | 0.003093 | 0.012707 | 0.006409 | 0.002096 | 0.008539 | 0.008454 | 0.003004 | 0.002827 | 0.006624 | 0.006722 | 0.001023 | 0.010806 | 0.007446 | 0.0082 | 0.001768 | 0.015162 | 0.009072 |
| 23.8333333 | 0.013613 | 0.000657 | 0.00612 | 0.003589 | 0.017449 | 0.002361 | 0.004736 | 0.002081 | 0.013852 | 0.005964 | 0.001537 | 0.007419 | 0.011 | 0.000937 | 0.002621 | 0.010246 | 0.012351 | 0.000684 | 0.009243 | 0.016394 | 0.005217 | 0.002612 | 0.014819 | 0.003461 |
| 23.8472222 | 0.017571 | 0.000775 | 0.007312 | 0.005276 | 0.019124 | 0.002473 | 0.003178 | 0.003 | 0.01645 | 0.00634 | 0.00146 | 0.006601 | 0.013882 | 0.000415 | 0.002173 | 0.014321 | 0.015546 | 0.000833 | 0.007901 | 0.031747 | 0.003229 | 0.002233 | 0.015929 | 0.000725 |
| 23.8611111 | 0.020122 | 0.001469 | 0.007126 | 0.008136 | 0.0243 | 0.002411 | 0.006839 | 0.005181 | 0.015403 | 0.005491 | 0.002154 | 0.007587 | 0.017686 | 0.001524 | 0.001578 | 0.015323 | 0.016907 | 0.002964 | 0.009293 | 0.046443 | 0.006922 | 0.002799 | 0.018047 | 0.000615 |
| 23.875 | 0.022748 | 0.001297 | 0.005046 | 0.01014 | 0.031144 | 0.00128 | 0.013176 | 0.006637 | 0.011132 | 0.002964 | 0.005261 | 0.009942 | 0.023234 | 0.003872 | 0.001032 | 0.012914 | 0.021116 | 0.007709 | 0.013278 | 0.051603 | 0.014244 | 0.003526 | 0.01868 | 0.001032 |
| 23.8888889 | 0.027652 | 0.000687 | 0.003866 | 0.009882 | 0.035617 | 0.000838 | 0.017952 | 0.008126 | 0.008533 | 0.000959 | 0.011945 | 0.013575 | 0.030216 | 0.00818 | 0.001636 | 0.011149 | 0.026794 | 0.013376 | 0.017668 | 0.052905 | 0.020606 | 0.00399 | 0.018994 | 0.001485 |
| 23.9027778 | 0.035239 | 0.000736 | 0.008212 | 0.008637 | 0.037785 | 0.000924 | 0.019424 | 0.010915 | 0.00867 | 0.000438 | 0.021348 | 0.017885 | 0.03897 | 0.013955 | 0.003898 | 0.013644 | 0.029268 | 0.017479 | 0.020246 | 0.056859 | 0.026578 | 0.005387 | 0.021673 | 0.003315 |
| 23.9166667 | 0.043 | 0.000923 | 0.017507 | 0.009588 | 0.038301 | 0.00214 | 0.018426 | 0.014329 | 0.008697 | 0.001143 | 0.028874 | 0.020248 | 0.048526 | 0.019161 | 0.006931 | 0.018294 | 0.027637 | 0.0202 | 0.021697 | 0.057763 | 0.03319 | 0.006518 | 0.024982 | 0.008104 |
| 23.9305556 | 0.048304 | 0.000999 | 0.025202 | 0.013724 | 0.038216 | 0.005899 | 0.017929 | 0.019056 | 0.008223 | 0.00199 | 0.033808 | 0.021312 | 0.056611 | 0.023434 | 0.009652 | 0.02048 | 0.026814 | 0.023138 | 0.024259 | 0.054572 | 0.038093 | 0.006733 | 0.024129 | 0.015088 |
| 23.9444444 | 0.051173 | 0.003664 | 0.030079 | 0.018578 | 0.041715 | 0.012077 | 0.021119 | 0.023559 | 0.011194 | 0.005253 | 0.042119 | 0.025371 | 0.057906 | 0.028266 | 0.011958 | 0.020055 | 0.029084 | 0.028525 | 0.029851 | 0.051598 | 0.040917 | 0.007412 | 0.02033 | 0.023041 |
| 23.9583333 | 0.053867 | 0.009347 | 0.038204 | 0.022788 | 0.049098 | 0.018778 | 0.02689 | 0.024632 | 0.018548 | 0.014877 | 0.052444 | 0.030701 | 0.050838 | 0.033163 | 0.014325 | 0.019958 | 0.031024 | 0.035116 | 0.038095 | 0.049142 | 0.042833 | 0.008284 | 0.020125 | 0.028425 |
| 23.9722222 | 0.059356 | 0.013273 | 0.049592 | 0.026948 | 0.054907 | 0.021818 | 0.030602 | 0.023057 | 0.02697 | 0.025861 | 0.059242 | 0.031934 | 0.045645 | 0.036343 | 0.016841 | 0.021624 | 0.032139 | 0.038758 | 0.046785 | 0.048667 | 0.043847 | 0.007616 | 0.026254 | 0.029142 |
| 23.9861111 | 0.067809 | 0.014629 | 0.056241 | 0.031321 | 0.058535 | 0.020077 | 0.031775 | 0.021818 | 0.035101 | 0.036479 | 0.064244 | 0.030169 | 0.050437 | 0.038263 | 0.019427 | 0.024357 | 0.035197 | 0.041182 | 0.054611 | 0.050367 | 0.045231 | 0.006667 | 0.034762 | 0.030363 |
| 24 | 0.075706 | 0.016631 | 0.057573 | 0.034132 | 0.063103 | 0.017903 | 0.032936 | 0.025176 | 0.042248 | 0.050982 | 0.068574 | 0.029395 | 0.061201 | 0.039657 | 0.022709 | 0.02634 | 0.039926 | 0.045317 | 0.061434 | 0.047193 | 0.046871 | 0.007779 | 0.037063 | 0.035278 |
| 24.0138889 | 0.079681 | 0.018243 | 0.057038 | 0.033469 | 0.066845 | 0.0152 | 0.032444 | 0.029978 | 0.046159 | 0.064005 | 0.068392 | 0.029269 | 0.069863 | 0.039094 | 0.026784 | 0.026361 | 0.044614 | 0.049005 | 0.066567 | 0.038911 | 0.047646 | 0.009567 | 0.031539 | 0.040138 |
| 24.0277778 | 0.082205 | 0.016676 | 0.053174 | 0.034372 | 0.073616 | 0.009236 | 0.028997 | 0.027146 | 0.046993 | 0.071281 | 0.068784 | 0.027257 | 0.075233 | 0.03595 | 0.028461 | 0.026234 | 0.049928 | 0.049734 | 0.069124 | 0.038329 | 0.048675 | 0.010142 | 0.026799 | 0.042744 |
| 24.0416667 | 0.090624 | 0.012521 | 0.048681 | 0.044 | 0.096339 | 0.003965 | 0.027333 | 0.022352 | 0.045049 | 0.077746 | 0.077752 | 0.026857 | 0.080227 | 0.032873 | 0.028178 | 0.029422 | 0.057516 | 0.04749 | 0.067641 | 0.047165 | 0.050606 | 0.012951 | 0.026811 | 0.042887 |
| 24.0555556 | 0.103865 | 0.011514 | 0.047821 | 0.054244 | 0.131611 | 0.003286 | 0.032795 | 0.025624 | 0.04181 | 0.086512 | 0.093621 | 0.035957 | 0.084815 | 0.032537 | 0.032835 | 0.035969 | 0.070132 | 0.048317 | 0.064082 | 0.048944 | 0.050384 | 0.022039 | 0.031631 | 0.042681 |
| 24.0694444 | 0.107212 | 0.020467 | 0.052917 | 0.053462 | 0.145017 | 0.005557 | 0.041415 | 0.030532 | 0.043549 | 0.090209 | 0.108662 | 0.052005 | 0.08508 | 0.035193 | 0.041105 | 0.041898 | 0.078213 | 0.057212 | 0.062261 | 0.04115 | 0.045414 | 0.031019 | 0.039805 | 0.048599 |
| 24.0833333 | 0.085629 | 0.026963 | 0.053012 | 0.043647 | 0.115196 | 0.008087 | 0.040544 | 0.026656 | 0.044036 | 0.075048 | 0.100621 | 0.058274 | 0.069612 | 0.039775 | 0.0364 | 0.043036 | 0.067521 | 0.058271 | 0.055102 | 0.034384 | 0.034379 | 0.03025 | 0.041759 | 0.054596 |
| 24.0972222 | 0.045151 | 0.017274 | 0.034133 | 0.026533 | 0.066427 | 0.008509 | 0.028023 | 0.014515 | 0.031942 | 0.041545 | 0.059642 | 0.046044 | 0.038363 | 0.044462 | 0.021963 | 0.038944 | 0.045629 | 0.036117 | 0.034645 | 0.025773 | 0.025088 | 0.022022 | 0.028256 | 0.043887 |
| 24.1111111 | 0.017337 | 0.006033 | 0.011227 | 0.012509 | 0.029885 | 0.006086 | 0.01403 | 0.005299 | 0.015141 | 0.015382 | 0.021218 | 0.027222 | 0.020209 | 0.045567 | 0.019191 | 0.034284 | 0.02517 | 0.011269 | 0.01255 | 0.013828 | 0.029048 | 0.013967 | 0.012255 | 0.02188 |
| 24.125 | 0.017357 | 0.003802 | 0.001993 | 0.011801 | 0.020308 | 0.002698 | 0.004861 | 0.005075 | 0.007598 | 0.01473 | 0.014479 | 0.012934 | 0.027415 | 0.039378 | 0.023585 | 0.032816 | 0.015741 | 0.002862 | 0.004425 | 0.009266 | 0.032068 | 0.010927 | 0.011911 | 0.011829 |

| | | | | | | | | | | | | | | | | | | | | | | | | |
|---|---|---|---|---|---|---|---|---|---|---|---|---|---|---|---|---|---|---|---|---|---|---|---|---|
| 24.1388889 | 0.024347 | 0.008582 | 0.00249 | 0.014823 | 0.031123 | 0.000611 | 0.00172 | 0.00829 | 0.009924 | 0.025797 | 0.026191 | 0.008018 | 0.037993 | 0.029171 | 0.028842 | 0.033169 | 0.015335 | 0.003874 | 0.009015 | 0.011092 | 0.024287 | 0.010373 | 0.020574 | 0.016831 |
| 24.1527778 | 0.025232 | 0.020999 | 0.00584 | 0.018798 | 0.043111 | 9.60E-05 | 0.002672 | 0.011067 | 0.015299 | 0.034274 | 0.035192 | 0.013444 | 0.044964 | 0.022235 | 0.033433 | 0.033987 | 0.010811 | 0.010975 | 0.017118 | 0.009315 | 0.024305 | 0.006181 | 0.027198 | 0.022088 |
| 24.1666667 | 0.026135 | 0.030669 | 0.012484 | 0.026084 | 0.050661 | 7.94E-05 | 0.004515 | 0.013067 | 0.023785 | 0.042295 | 0.033905 | 0.01915 | 0.050068 | 0.019886 | 0.026379 | 0.03541 | 0.010021 | 0.019636 | 0.019582 | 0.006829 | 0.035972 | 0.00293 | 0.026825 | 0.02441 |
| 24.1805556 | 0.030347 | 0.026651 | 0.021973 | 0.030082 | 0.055967 | 0.000173 | 0.006558 | 0.017509 | 0.036259 | 0.052975 | 0.029031 | 0.020772 | 0.048815 | 0.018594 | 0.013671 | 0.031396 | 0.016843 | 0.021223 | 0.015823 | 0.00631 | 0.040093 | 0.005596 | 0.019654 | 0.029254 |
| 24.1944444 | 0.04346 | 0.018643 | 0.023043 | 0.027809 | 0.066955 | 0.001093 | 0.010052 | 0.028245 | 0.042338 | 0.062126 | 0.035841 | 0.023154 | 0.045964 | 0.016487 | 0.007285 | 0.018972 | 0.015208 | 0.017572 | 0.01461 | 0.004126 | 0.02933 | 0.007909 | 0.014052 | 0.033908 |
| 24.2083333 | 0.071441 | 0.022428 | 0.020527 | 0.026724 | 0.08709 | 0.010169 | 0.014168 | 0.038194 | 0.029853 | 0.058943 | 0.05434 | 0.02259 | 0.042129 | 0.014911 | 0.009334 | 0.009532 | 0.006027 | 0.01218 | 0.015911 | 0.003269 | 0.017677 | 0.005819 | 0.014435 | 0.031793 |
| 24.2222222 | 0.072085 | 0.027633 | 0.023778 | 0.023945 | 0.081286 | 0.025968 | 0.023679 | 0.040593 | 0.018047 | 0.061298 | 0.052285 | 0.025049 | 0.041121 | 0.013448 | 0.018018 | 0.006424 | 0.003165 | 0.005952 | 0.013946 | 0.004997 | 0.017439 | 0.004083 | 0.015394 | 0.025557 |
| 24.2361111 | 0.039611 | 0.021309 | 0.022584 | 0.027406 | 0.053185 | 0.049692 | 0.039191 | 0.030939 | 0.02538 | 0.069773 | 0.029194 | 0.039327 | 0.043581 | 0.00886 | 0.020693 | 0.00304 | 0.005469 | 0.002044 | 0.010706 | 0.004647 | 0.02407 | 0.008446 | 0.011899 | 0.022536 |
| 24.25 | 0.044277 | 0.024293 | 0.026026 | 0.033881 | 0.041634 | 0.069264 | 0.043776 | 0.017253 | 0.025382 | 0.044977 | 0.018267 | 0.041999 | 0.037241 | 0.004007 | 0.012874 | 0.001153 | 0.00614 | 0.001119 | 0.010015 | 0.002109 | 0.025546 | 0.014897 | 0.006778 | 0.019877 |
| 24.2638889 | 0.075736 | 0.042595 | 0.040566 | 0.029396 | 0.027805 | 0.049433 | 0.036468 | 0.020194 | 0.01048 | 0.018314 | 0.024533 | 0.026603 | 0.025199 | 0.004838 | 0.008619 | 0.002366 | 0.006792 | 0.001507 | 0.010486 | 0.001066 | 0.018296 | 0.014318 | 0.003916 | 0.014044 |
| 24.2777778 | 0.082953 | 0.056037 | 0.052977 | 0.040575 | 0.015927 | 0.020944 | 0.031284 | 0.030011 | 0.00449 | 0.028815 | 0.048789 | 0.023396 | 0.015138 | 0.009321 | 0.011451 | 0.005762 | 0.007595 | 0.00209 | 0.009436 | 0.001477 | 0.010295 | 0.011107 | 0.004122 | 0.009334 |
| 24.2916667 | 0.069105 | 0.062798 | 0.059349 | 0.057675 | 0.026119 | 0.022505 | 0.025483 | 0.031228 | 0.01219 | 0.048841 | 0.074536 | 0.037952 | 0.009415 | 0.011416 | 0.012738 | 0.010796 | 0.00675 | 0.003105 | 0.006934 | 0.003152 | 0.006914 | 0.00841 | 0.004368 | 0.00665 |
| 24.3055556 | 0.055601 | 0.067972 | 0.058573 | 0.059448 | 0.039987 | 0.032261 | 0.013589 | 0.029483 | 0.022885 | 0.057802 | 0.083217 | 0.052274 | 0.009822 | 0.007947 | 0.010969 | 0.012397 | 0.007007 | 0.006149 | 0.004384 | 0.006909 | 0.004453 | 0.004295 | 0.002568 | 0.006341 |
| 24.3194444 | 0.054812 | 0.070862 | 0.052451 | 0.060073 | 0.046726 | 0.034877 | 0.005017 | 0.028736 | 0.028063 | 0.057939 | 0.083013 | 0.059363 | 0.01074 | 0.002416 | 0.009608 | 0.010438 | 0.009288 | 0.00773 | 0.00225 | 0.00871 | 0.00328 | 0.003013 | 0.001666 | 0.010243 |
| 24.3333333 | 0.064092 | 0.072309 | 0.04489 | 0.062115 | 0.049321 | 0.02975 | 0.005969 | 0.028228 | 0.02779 | 0.055249 | 0.077516 | 0.05869 | 0.00791 | 0.003383 | 0.009479 | 0.013189 | 0.008155 | 0.005056 | 0.001083 | 0.005872 | 0.006103 | 0.004467 | 0.002266 | 0.014678 |
| 24.3472222 | 0.078193 | 0.074982 | 0.036395 | 0.059514 | 0.05495 | 0.020136 | 0.010534 | 0.027749 | 0.028334 | 0.055312 | 0.07109 | 0.051521 | 0.005492 | 0.011782 | 0.010256 | 0.020163 | 0.004676 | 0.003099 | 0.002975 | 0.005042 | 0.006558 | 0.010283 | 0.002833 | 0.013703 |
| 24.3611111 | 0.087505 | 0.081879 | 0.028782 | 0.051486 | 0.064207 | 0.013767 | 0.014749 | 0.026929 | 0.029644 | 0.059351 | 0.070989 | 0.042164 | 0.004505 | 0.018571 | 0.009813 | 0.02193 | 0.007317 | 0.006518 | 0.008682 | 0.012293 | 0.005383 | 0.018412 | 0.004487 | 0.008869 |
| 24.375 | 0.07981 | 0.088147 | 0.020956 | 0.0411 | 0.065676 | 0.012997 | 0.015808 | 0.021336 | 0.025323 | 0.057648 | 0.072263 | 0.034455 | 0.004222 | 0.018765 | 0.007882 | 0.018622 | 0.012142 | 0.012155 | 0.015529 | 0.019428 | 0.010979 | 0.019331 | 0.006471 | 0.007133 |
| 24.3888889 | 0.063492 | 0.08259 | 0.012066 | 0.030865 | 0.05508 | 0.014642 | 0.012546 | 0.011427 | 0.018146 | 0.04461 | 0.06851 | 0.030081 | 0.009572 | 0.01716 | 0.007275 | 0.018371 | 0.012156 | 0.015298 | 0.021669 | 0.019606 | 0.021585 | 0.016934 | 0.007147 | 0.006858 |
| 24.4027778 | 0.055207 | 0.067531 | 0.006102 | 0.024991 | 0.042238 | 0.015159 | 0.009294 | 0.005726 | 0.014285 | 0.030637 | 0.063146 | 0.026868 | 0.018642 | 0.017725 | 0.008611 | 0.022161 | 0.010611 | 0.018771 | 0.030658 | 0.018973 | 0.034541 | 0.02068 | 0.007893 | 0.005362 |
| 24.4166667 | 0.052802 | 0.057534 | 0.004113 | 0.024825 | 0.033753 | 0.013899 | 0.009521 | 0.007383 | 0.012535 | 0.024188 | 0.060179 | 0.023191 | 0.025607 | 0.017445 | 0.014972 | 0.025958 | 0.009661 | 0.025384 | 0.042957 | 0.019043 | 0.045971 | 0.025156 | 0.009952 | 0.007195 |
| 24.4305556 | 0.048601 | 0.053591 | 0.003834 | 0.024321 | 0.02856 | 0.012379 | 0.011702 | 0.011393 | 0.010986 | 0.023351 | 0.060412 | 0.017864 | 0.029904 | 0.014123 | 0.023009 | 0.028539 | 0.011507 | 0.030018 | 0.05013 | 0.017227 | 0.047259 | 0.023532 | 0.011674 | 0.013642 |
| 24.4444444 | 0.043144 | 0.049987 | 0.003976 | 0.019322 | 0.025213 | 0.011731 | 0.012365 | 0.013761 | 0.009293 | 0.024126 | 0.060809 | 0.012267 | 0.033489 | 0.011174 | 0.022517 | 0.02968 | 0.017264 | 0.030238 | 0.050116 | 0.015464 | 0.042331 | 0.021107 | 0.011654 | 0.021255 |
| 24.4583333 | 0.037647 | 0.046899 | 0.004138 | 0.013866 | 0.023126 | 0.011436 | 0.011519 | 0.01465 | 0.007385 | 0.025927 | 0.058225 | 0.010707 | 0.036876 | 0.012804 | 0.016047 | 0.02968 | 0.020487 | 0.031374 | 0.05075 | 0.013949 | 0.04435 | 0.022289 | 0.010679 | 0.026687 |
| 24.4722222 | 0.031613 | 0.045113 | 0.004423 | 0.011341 | 0.021541 | 0.010376 | 0.010838 | 0.014583 | 0.007168 | 0.030386 | 0.055337 | 0.011161 | 0.039434 | 0.018167 | 0.012012 | 0.030134 | 0.020674 | 0.03559 | 0.05482 | 0.011329 | 0.054436 | 0.024498 | 0.009855 | 0.027714 |
| 24.4861111 | 0.027222 | 0.045248 | 0.005442 | 0.011867 | 0.020955 | 0.0092 | 0.010691 | 0.013652 | 0.009838 | 0.035201 | 0.05447 | 0.010998 | 0.042634 | 0.021903 | 0.014643 | 0.031178 | 0.026087 | 0.037966 | 0.055851 | 0.008473 | 0.062097 | 0.026391 | 0.011082 | 0.025476 |
| 24.5 | 0.026554 | 0.047543 | 0.0071 | 0.015259 | 0.02227 | 0.009274 | 0.010726 | 0.012446 | 0.012356 | 0.035045 | 0.055747 | 0.011457 | 0.047713 | 0.020913 | 0.023198 | 0.031574 | 0.034923 | 0.035862 | 0.05215 | 0.00689 | 0.062928 | 0.030638 | 0.014841 | 0.024859 |
| 24.5138889 | 0.028794 | 0.051104 | 0.00864 | 0.020299 | 0.025708 | 0.009572 | 0.010419 | 0.010985 | 0.012902 | 0.030876 | 0.059213 | 0.012274 | 0.052477 | 0.017442 | 0.029934 | 0.032168 | 0.039811 | 0.034183 | 0.050334 | 0.006443 | 0.062291 | 0.038773 | 0.020502 | 0.029654 |
| 24.5277778 | 0.034322 | 0.053701 | 0.01079 | 0.02317 | 0.034078 | 0.008565 | 0.009913 | 0.010342 | 0.015163 | 0.029153 | 0.066366 | 0.013239 | 0.053744 | 0.015393 | 0.029399 | 0.033787 | 0.04398 | 0.03558 | 0.049807 | 0.006325 | 0.060231 | 0.046874 | 0.02687 | 0.041421 |
| 24.5416667 | 0.046281 | 0.055635 | 0.016484 | 0.024256 | 0.05217 | 0.008474 | 0.011563 | 0.014499 | 0.025567 | 0.035917 | 0.084424 | 0.016233 | 0.048574 | 0.014508 | 0.024341 | 0.032037 | 0.048143 | 0.034744 | 0.046145 | 0.006045 | 0.048685 | 0.048056 | 0.029189 | 0.054914 |
| 24.5555556 | 0.057214 | 0.056323 | 0.025615 | 0.027567 | 0.07155 | 0.011755 | 0.016981 | 0.023554 | 0.046496 | 0.051738 | 0.108922 | 0.023441 | 0.036788 | 0.011443 | 0.01956 | 0.022866 | 0.04708 | 0.028553 | 0.040365 | 0.005299 | 0.031896 | 0.039431 | 0.025346 | 0.056313 |
| 24.5694444 | 0.052948 | 0.051388 | 0.034331 | 0.031876 | 0.074273 | 0.015028 | 0.021775 | 0.030414 | 0.063621 | 0.063028 | 0.112249 | 0.031911 | 0.027656 | 0.010546 | 0.016696 | 0.01271 | 0.048524 | 0.023583 | 0.036715 | 0.005524 | 0.025933 | 0.028213 | 0.021612 | 0.045428 |
| 24.5833333 | 0.037339 | 0.039143 | 0.039054 | 0.034366 | 0.060696 | 0.013501 | 0.021526 | 0.03002 | 0.061352 | 0.057349 | 0.087422 | 0.034497 | 0.031096 | 0.023025 | 0.016431 | 0.013402 | 0.056725 | 0.025757 | 0.038745 | 0.007405 | 0.02837 | 0.025486 | 0.025662 | 0.040923 |
| 24.5972222 | 0.023474 | 0.024708 | 0.03834 | 0.033403 | 0.044994 | 0.008372 | 0.018241 | 0.025345 | 0.047826 | 0.042594 | 0.060665 | 0.029878 | 0.044769 | 0.04458 | 0.019312 | 0.026659 | 0.057734 | 0.032253 | 0.043977 | 0.009899 | 0.02544 | 0.029253 | 0.035085 | 0.050465 |
| 24.6111111 | 0.015083 | 0.014167 | 0.034608 | 0.02859 | 0.034614 | 0.00451 | 0.015752 | 0.021759 | 0.036924 | 0.030524 | 0.044209 | 0.022633 | 0.054089 | 0.057563 | 0.022871 | 0.038763 | 0.046261 | 0.036557 | 0.046679 | 0.011439 | 0.019928 | 0.02918 | 0.039724 | 0.064448 |
| 24.625 | 0.010424 | 0.007763 | 0.0297 | 0.021615 | 0.028787 | 0.00355 | 0.014849 | 0.019851 | 0.030426 | 0.022605 | 0.032245 | 0.01574 | 0.047644 | 0.06076 | 0.023136 | 0.036616 | 0.031912 | 0.032444 | 0.042349 | 0.011034 | 0.017535 | 0.022917 | 0.03525 | 0.070321 |
| 24.6388889 | 0.007796 | 0.004169 | 0.024286 | 0.015366 | 0.024685 | 0.003612 | 0.013473 | 0.017133 | 0.026255 | 0.01686 | 0.022904 | 0.010676 | 0.034397 | 0.060244 | 0.018384 | 0.024466 | 0.022216 | 0.020422 | 0.033966 | 0.010409 | 0.014954 | 0.013204 | 0.027193 | 0.062319 |
| 24.6527778 | 0.006649 | 0.003193 | 0.020178 | 0.011861 | 0.021269 | 0.003184 | 0.010587 | 0.012776 | 0.024405 | 0.013276 | 0.016825 | 0.008744 | 0.030296 | 0.048232 | 0.01143 | 0.015455 | 0.018133 | 0.008558 | 0.029798 | 0.009571 | 0.010502 | 0.005896 | 0.022761 | 0.046641 |
| 24.6666667 | 0.006349 | 0.004127 | 0.018363 | 0.01042 | 0.018234 | 0.002036 | 0.007954 | 0.009089 | 0.024325 | 0.012342 | 0.012607 | 0.008631 | 0.037911 | 0.026693 | 0.006453 | 0.014161 | 0.015375 | 0.003681 | 0.03021 | 0.006255 | 0.006045 | 0.003954 | 0.021906 | 0.029554 |
| 24.6805556 | 0.005735 | 0.006682 | 0.018069 | 0.009256 | 0.015196 | 0.00082 | 0.006696 | 0.007595 | 0.023245 | 0.012821 | 0.009156 | 0.007466 | 0.046686 | 0.013791 | 0.004291 | 0.014143 | 0.010102 | 0.005891 | 0.031125 | 0.003428 | 0.003368 | 0.005159 | 0.020121 | 0.016564 |
| 24.6944444 | 0.004862 | 0.009872 | 0.017394 | 0.007363 | 0.012386 | 0.000315 | 0.005706 | 0.006845 | 0.020081 | 0.013096 | 0.006817 | 0.005052 | 0.046363 | 0.011968 | 0.003567 | 0.00957 | 0.006432 | 0.009227 | 0.027381 | 0.005674 | 0.0027 | 0.00774 | 0.015495 | 0.014961 |
| 24.7083333 | 0.004487 | 0.010652 | 0.015525 | 0.005633 | 0.010527 | 0.000564 | 0.004322 | 0.006242 | 0.01648 | 0.012401 | 0.004826 | 0.003028 | 0.038261 | 0.015299 | 0.003274 | 0.003848 | 0.007602 | 0.00849 | 0.016455 | 0.010695 | 0.005215 | 0.007488 | 0.008637 | 0.020849 |
| 24.7222222 | 0.004536 | 0.008455 | 0.013072 | 0.00583 | 0.009908 | 0.001267 | 0.003307 | 0.006158 | 0.013422 | 0.010182 | 0.003215 | 0.0019 | 0.027736 | 0.020423 | 0.002928 | 0.001352 | 0.008358 | 0.004743 | 0.007498 | 0.014404 | 0.010985 | 0.005011 | 0.004747 | 0.024295 |
| 24.7361111 | 0.005113 | 0.006722 | 0.010781 | 0.007181 | 0.009947 | 0.001999 | 0.003297 | 0.006296 | 0.010812 | 0.006853 | 0.003693 | 0.001683 | 0.016367 | 0.019822 | 0.002386 | 0.001035 | 0.005394 | 0.001825 | 0.007951 | 0.015429 | 0.013186 | 0.004464 | 0.005644 | 0.023616 |
| 24.75 | 0.006908 | 0.006149 | 0.00983 | 0.006245 | 0.00958 | 0.002199 | 0.003383 | 0.006141 | 0.009169 | 0.004367 | 0.00516 | 0.002117 | 0.007673 | 0.019162 | 0.001648 | 0.001287 | 0.003656 | 0.001447 | 0.01113 | 0.013155 | 0.009057 | 0.005104 | 0.005075 | 0.02269 |
| 24.7638889 | 0.0096 | 0.005061 | 0.011817 | 0.003182 | 0.008313 | 0.001675 | 0.002347 | 0.005684 | 0.009148 | 0.00402 | 0.005058 | 0.002268 | 0.005967 | 0.026247 | 0.001091 | 0.001007 | 0.006797 | 0.002145 | 0.008617 | 0.008424 | 0.006003 | 0.004892 | 0.002555 | 0.022031 |
| 24.7777778 | 0.011404 | 0.004389 | 0.016055 | 0.00158 | 0.006774 | 0.000851 | 0.00102 | 0.005136 | 0.011214 | 0.005966 | 0.003723 | 0.001568 | 0.006788 | 0.031943 | 0.0009 | 0.000567 | 0.008959 | 0.001993 | 0.006112 | 0.004067 | 0.007047 | 0.004589 | 0.001341 | 0.01952 |
| 24.7916667 | 0.011106 | 0.005239 | 0.019003 | 0.002814 | 0.005645 | 0.000417 | 0.000638 | 0.004077 | 0.015228 | 0.00953 | 0.002946 | 0.000792 | 0.004474 | 0.028841 | 0.000752 | 0.000581 | 0.007046 | 0.002468 | 0.011586 | 0.002628 | 0.008916 | 0.003925 | 0.001701 | 0.016213 |
| 24.8055556 | 0.009863 | 0.006108 | 0.0182 | 0.005541 | 0.004998 | 0.000563 | 0.000608 | 0.002713 | 0.019024 | 0.012065 | 0.003401 | 0.000614 | 0.001477 | 0.02371 | 0.000791 | 0.001003 | 0.006092 | 0.003988 | 0.017967 | 0.003452 | 0.01024 | 0.003591 | 0.002501 | 0.014928 |
| 24.8194444 | 0.00926 | 0.005585 | 0.015079 | 0.005972 | 0.004808 | 0.000661 | 0.000316 | 0.002075 | 0.020649 | 0.011432 | 0.004603 | 0.001178 | 0.002263 | 0.023258 | 0.001624 | 0.00293 | 0.006579 | 0.003607 | 0.017215 | 0.003253 | 0.010937 | 0.007232 | 0.003356 | 0.015395 |
| 24.8333333 | 0.00934 | 0.004619 | 0.012867 | 0.003506 | 0.005142 | 0.000443 | 0.0004 | 0.002528 | 0.021405 | 0.009405 | 0.006152 | 0.003128 | 0.008415 | 0.023959 | 0.003437 | 0.005782 | 0.006269 | 0.003188 | 0.010795 | 0.005397 | 0.011582 | 0.010229 | 0.004847 | 0.015137 |
| 24.8472222 | 0.009725 | 0.004836 | 0.013025 | 0.001629 | 0.00603 | 0.000248 | 0.000961 | 0.003599 | 0.022858 | 0.008315 | 0.008213 | 0.005821 | 0.017318 | 0.021811 | 0.004979 | 0.006874 | 0.008237 | 0.006134 | 0.005244 | 0.018553 | 0.013355 | 0.006903 | 0.005487 | 0.013474 |
| 24.8611111 | 0.010257 | 0.006598 | 0.015027 | 0.002063 | 0.007329 | 0.000424 | 0.001218 | 0.004428 | 0.023971 | 0.007336 | 0.010506 | 0.007938 | 0.02648 | 0.01686 | 0.005205 | 0.005785 | 0.011084 | 0.009573 | 0.00543 | 0.035791 | 0.014648 | 0.002417 | 0.004017 | 0.011889 |
| 24.875 | 0.010867 | 0.008488 | 0.01794 | 0.003362 | 0.00884 | 0.001056 | 0.000938 | 0.004829 | 0.023564 | 0.005527 | 0.012301 | 0.009313 | 0.037767 | 0.009468 | 0.004262 | 0.005998 | 0.009551 | 0.01142 | 0.007831 | 0.042691 | 0.013158 | 0.001659 | 0.002043 | 0.010309 |
| 24.8888889 | 0.011501 | 0.009469 | 0.021315 | 0.00324 | 0.010495 | 0.001242 | 0.000914 | 0.005695 | 0.022184 | 0.003989 | 0.013758 | 0.010335 | 0.052394 | 0.003469 | 0.003642 | 0.009661 | 0.010537 | 0.012879 | 0.008136 | 0.041944 | 0.010577 | 0.002238 | 0.001263 | 0.007435 |
| 24.9027778 | 0.011829 | 0.010411 | 0.024243 | 0.002095 | 0.011919 | 0.001175 | 0.0019 | 0.007358 | 0.020462 | 0.003531 | 0.01505 | 0.010968 | 0.065261 | 0.001604 | 0.004749 | 0.013723 | 0.020293 | 0.01445 | 0.008074 | 0.042707 | 0.007134 | 0.001811 | 0.000924 | 0.0049 |
| 24.9166667 | 0.011451 | 0.010534 | 0.025204 | 0.002549 | 0.012601 | 0.001797 | 0.003438 | 0.008881 | 0.017463 | 0.003341 | 0.015121 | 0.010822 | 0.070153 | 0.002714 | 0.006179 | 0.015133 | 0.028602 | 0.014634 | 0.009886 | 0.03856 | 0.005342 | 0.00193 | 0.000447 | 0.007148 |
| 24.9305556 | 0.010213 | 0.008459 | 0.023505 | 0.005814 | 0.0127 | 0.001765 | 0.004363 | 0.009457 | 0.012669 | 0.002726 | 0.013621 | 0.010301 | 0.068469 | 0.008305 | 0.006991 | 0.015578 | 0.027446 | 0.011724 | 0.011839 | 0.028026 | 0.009206 | 0.003537 | 0.000249 | 0.015851 |
| 24.9444444 | 0.008415 | 0.005201 | 0.019883 | 0.007012 | 0.013015 | 0.001224 | 0.004333 | 0.009114 | 0.007377 | 0.001867 | 0.011678 | 0.010151 | 0.067003 | 0.017678 | 0.009024 | 0.018118 | 0.025706 | 0.011463 | 0.013878 | 0.033123 | 0.009707 | 0.006813 | 0.001257 | 0.026245 |
| 24.9583333 | 0.007231 | 0.003378 | 0.016446 | 0.00423 | 0.014566 | 0.002088 | 0.003605 | 0.008047 | 0.003519 | 0.002236 | 0.009968 | 0.010258 | 0.070391 | 0.028209 | 0.013022 | 0.02208 | 0.032551 | 0.019816 | 0.017909 | 0.054497 | 0.007401 | 0.013087 | 0.002542 | 0.031367 |
| 24.9722222 | 0.007515 | 0.004409 | 0.014735 | 0.002834 | 0.017652 | 0.003794 | 0.002964 | 0.005693 | 0.002896 | 0.005191 | 0.008979 | 0.009817 | 0.07635 | 0.043199 | 0.017162 | 0.024744 | 0.041461 | 0.028751 | 0.018622 | 0.067143 | 0.010636 | 0.017758 | 0.002998 | 0.030382 |
| 24.9861111 | 0.008065 | 0.005943 | 0.015307 | 0.003594 | 0.021161 | 0.00519 | 0.003906 | 0.004788 | 0.005469 | 0.010314 | 0.008807 | 0.008292 | 0.078593 | 0.065812 | 0.019757 | 0.026423 | 0.041154 | 0.030424 | 0.01452 | 0.070914 | 0.0125 | 0.016778 | 0.005563 | 0.028693 |
| 25 | 0.007668 | 0.006166 | 0.01848 | 0.005055 | 0.024029 | 0.005681 | 0.004684 | 0.007105 | 0.008405 | 0.015641 | 0.009123 | 0.006438 | 0.076622 | 0.086554 | 0.021431 | 0.028352 | 0.038512 | 0.029964 | 0.015658 | 0.079221 | 0.013527 | 0.014772 | 0.009163 | 0.027343 |
| 25.0138889 | 0.006725 | 0.004794 | 0.023481 | 0.009226 | 0.026327 | 0.004196 | 0.003226 | 0.006501 | 0.009263 | 0.017691 | 0.010911 | 0.005807 | 0.075574 | 0.086235 | 0.024199 | 0.029409 | 0.048576 | 0.032267 | 0.0274 | 0.088444 | 0.017008 | 0.012969 | 0.010371 | 0.022415 |
| 25.0277778 | 0.004587 | 0.003674 | 0.02638 | 0.015174 | 0.02441 | 0.002059 | 0.001474 | 0.002437 | 0.00794 | 0.016613 | 0.012465 | 0.005736 | 0.077178 | 0.063075 | 0.027541 | 0.028708 | 0.06352 | 0.036758 | 0.043884 | 0.095709 | 0.019756 | 0.009577 | 0.013359 | 0.014061 |
| 25.0416667 | 0.00592 | 0.00484 | 0.021499 | 0.017852 | 0.018854 | 0.001408 | 0.002842 | 0.003861 | 0.006451 | 0.017179 | 0.011697 | 0.005373 | 0.077766 | 0.045641 | 0.02868 | 0.029531 | 0.071676 | 0.042905 | 0.056472 | 0.097476 | 0.023038 | 0.006594 | 0.022609 | 0.009575 |
| 25.0555556 | 0.019522 | 0.005542 | 0.012465 | 0.017208 | 0.023625 | 0.002045 | 0.007872 | 0.01325 | 0.008843 | 0.022705 | 0.018221 | 0.008064 | 0.075502 | 0.057671 | 0.029225 | 0.036053 | 0.081175 | 0.052447 | 0.065395 | 0.079988 | 0.030148 | 0.004903 | 0.037515 | 0.017987 |
| 25.0694444 | 0.036784 | 0.003965 | 0.008791 | 0.017818 | 0.034536 | 0.003492 | 0.011136 | 0.020798 | 0.014682 | 0.028648 | 0.028746 | 0.012649 | 0.075554 | 0.080954 | 0.03305 | 0.044709 | 0.08807 | 0.064153 | 0.073402 | 0.050492 | 0.045702 | 0.007959 | 0.054408 | 0.033701 |
| 25.0833333 | 0.037615 | 0.002726 | 0.007185 | 0.019111 | 0.028891 | 0.004249 | 0.010828 | 0.017545 | 0.018272 | 0.026213 | 0.022071 | 0.013191 | 0.075484 | 0.068324 | 0.034841 | 0.045088 | 0.070511 | 0.061135 | 0.068899 | 0.032042 | 0.053247 | 0.014076 | 0.058514 | 0.036157 |
| 25.0972222 | 0.022253 | 0.002988 | 0.002957 | 0.01945 | 0.011984 | 0.003594 | 0.0106 | 0.008456 | 0.018103 | 0.016666 | 0.006519 | 0.012152 | 0.062909 | 0.029314 | 0.024412 | 0.036125 | 0.033822 | 0.039127 | 0.04417 | 0.024111 | 0.041931 | 0.013658 | 0.041925 | 0.020889 |
| 25.1111111 | 0.008577 | 0.00292 | 0.000688 | 0.020173 | 0.003638 | 0.00238 | 0.009979 | 0.003595 | 0.015942 | 0.007919 | 0.001774 | 0.012569 | 0.040866 | 0.013459 | 0.010194 | 0.029787 | 0.0122 | 0.024141 | 0.019227 | 0.013985 | 0.035264 | 0.009166 | 0.020481 | 0.006004 |
| 25.125 | 0.003603 | 0.002302 | 0.000372 | 0.019871 | 0.002769 | 0.001445 | 0.006572 | 0.003228 | 0.011174 | 0.003576 | 0.002562 | 0.010865 | 0.023581 | 0.019646 | 0.006474 | 0.030014 | 0.018933 | 0.019464 | 0.014468 | 0.006558 | 0.033805 | 0.005444 | 0.008309 | 0.001442 |
| 25.1388889 | 0.002109 | 0.005307 | 0.000336 | 0.017347 | 0.003876 | 0.00172 | 0.00248 | 0.002457 | 0.00537 | 0.004044 | 0.002211 | 0.007445 | 0.012604 | 0.027732 | 0.008708 | 0.030738 | 0.036163 | 0.01423 | 0.017327 | 0.007553 | 0.029649 | 0.002476 | 0.006689 | 0.001564 |
| 25.1527778 | 0.001264 | 0.016412 | 0.000727 | 0.014127 | 0.00934 | 0.002303 | 0.001137 | 0.001987 | 0.002744 | 0.007191 | 0.003308 | 0.005914 | 0.010355 | 0.022401 | 0.014628 | 0.030903 | 0.045845 | 0.019623 | 0.012442 | 0.010306 | 0.031311 | 0.003776 | 0.008362 | 0.00268 |
| 25.1666667 | 0.001002 | 0.027118 | 0.000675 | 0.010711 | 0.016696 | 0.002429 | 0.002083 | 0.004655 | 0.003468 | 0.010903 | 0.005503 | 0.005667 | 0.024172 | 0.008786 | 0.025148 | 0.031866 | 0.043098 | 0.030495 | 0.006708 | 0.011702 | 0.032407 | 0.007892 | 0.008381 | 0.005322 |
| 25.1805556 | 0.001229 | 0.024686 | 0.000619 | 0.007706 | 0.022237 | 0.00238 | 0.003034 | 0.009578 | 0.00594 | 0.014588 | 0.00779 | 0.00432 | 0.041679 | 0.007246 | 0.03341 | 0.030101 | 0.030817 | 0.027241 | 0.006945 | 0.012721 | 0.031101 | 0.00985 | 0.010702 | 0.007596 |
| 25.1944444 | 0.004236 | 0.012991 | 0.000702 | 0.007697 | 0.026493 | 0.001783 | 0.002669 | 0.014154 | 0.009278 | 0.01643 | 0.011165 | 0.002193 | 0.045323 | 0.017784 | 0.035264 | 0.023224 | 0.014987 | 0.016742 | 0.010469 | 0.013128 | 0.034164 | 0.010413 | 0.015707 | 0.00808 |
| 25.2083333 | 0.009938 | 0.004892 | 0.001496 | 0.009989 | 0.030706 | 0.001376 | 0.001637 | 0.016283 | 0.010342 | 0.014365 | 0.015009 | 0.001023 | 0.037717 | 0.027565 | 0.033106 | 0.016072 | 0.005638 | 0.013275 | 0.010237 | 0.013495 | 0.033427 | 0.01164 | 0.016882 | 0.007458 |
| 25.2222222 | 0.012723 | 0.005764 | 0.003962 | 0.010846 | 0.035688 | 0.001486 | 0.001483 | 0.01609 | 0.007801 | 0.008439 | 0.017087 | 0.000769 | 0.034137 | 0.024316 | 0.029666 | 0.012564 | 0.006732 | 0.012597 | 0.007949 | 0.011994 | 0.023516 | 0.011297 | 0.011544 | 0.006662 |
| 25.2361111 | 0.009842 | 0.009093 | 0.006513 | 0.009898 | 0.039618 | 0.001463 | 0.002199 | 0.015305 | 0.007603 | 0.00427 | 0.015909 | 0.000934 | 0.038595 | 0.012852 | 0.023994 | 0.00996 | 0.013366 | 0.007979 | 0.006867 | 0.008371 | 0.017702 | 0.008682 | 0.007531 | 0.008022 |
| 25.25 | 0.00475 | 0.008314 | 0.008029 | 0.008249 | 0.039445 | 0.001327 | 0.002126 | 0.01455 | 0.012783 | 0.005811 | 0.012054 | 0.001678 | 0.0472 | 0.008914 | 0.016897 | 0.005771 | 0.019362 | 0.002632 | 0.005253 | 0.010251 | 0.021513 | 0.007047 | 0.00888 | 0.012632 |
| 25.2638889 | 0.002017 | 0.00638 | 0.009347 | 0.005986 | 0.035043 | 0.001772 | 0.001814 | 0.013716 | 0.017533 | 0.007067 | 0.007054 | 0.002885 | 0.057795 | 0.015822 | 0.010563 | 0.002587 | 0.020239 | 0.000587 | 0.00543 | 0.015927 | 0.024423 | 0.006699 | 0.009491 | 0.017487 |
| 25.2777778 | 0.002273 | 0.006934 | 0.010457 | 0.003984 | 0.028804 | 0.002826 | 0.003214 | 0.012147 | 0.018883 | 0.004614 | 0.003521 | 0.004846 | 0.058799 | 0.02343 | 0.007754 | 0.003 | 0.021148 | 0.000853 | 0.007269 | 0.01321 | 0.025204 | 0.004672 | 0.006152 | 0.019652 |
| 25.2916667 | 0.003554 | 0.008027 | 0.010458 | 0.003074 | 0.022775 | 0.003313 | 0.005289 | 0.00964 | 0.019391 | 0.002718 | 0.004081 | 0.007204 | 0.048721 | 0.024006 | 0.006806 | 0.005547 | 0.025823 | 0.002661 | 0.011351 | 0.005824 | 0.027703 | 0.002071 | 0.004082 | 0.020253 |
| 25.3055556 | 0.005768 | 0.00602 | 0.008363 | 0.004035 | 0.017859 | 0.00409 | 0.006294 | 0.007621 | 0.021253 | 0.004103 | 0.007641 | 0.008089 | 0.041174 | 0.020387 | 0.00535 | 0.008347 | 0.023375 | 0.007922 | 0.017709 | 0.003235 | 0.024381 | 0.00179 | 0.00719 | 0.02088 |
| 25.3194444 | 0.009232 | 0.0022 | 0.004687 | 0.006088 | 0.013885 | 0.004867 | 0.005667 | 0.006458 | 0.023701 | 0.007097 | 0.010356 | 0.006726 | 0.038662 | 0.019465 | 0.005722 | 0.010535 | 0.012659 | 0.014848 | 0.020861 | 0.002325 | 0.012748 | 0.002025 | 0.010897 | 0.019567 |
| 25.3333333 | 0.011772 | 0.001347 | 0.001911 | 0.006014 | 0.010657 | 0.003882 | 0.004025 | 0.004962 | 0.02452 | 0.008736 | 0.009542 | 0.004075 | 0.034984 | 0.021284 | 0.005824 | 0.011326 | 0.006126 | 0.017181 | 0.019948 | 0.001701 | 0.003596 | 0.001116 | 0.011571 | 0.015591 |
| 25.3472222 | 0.010321 | 0.003153 | 0.001346 | 0.005198 | 0.009451 | 0.002685 | 0.003377 | 0.003625 | 0.02346 | 0.008479 | 0.00583 | 0.002148 | 0.029344 | 0.023738 | 0.005591 | 0.011133 | 0.005565 | 0.015041 | 0.019192 | 0.004172 | 0.003371 | 0.000597 | 0.010067 | 0.012926 |
| 25.3611111 | 0.005753 | 0.003343 | 0.001797 | 0.005366 | 0.011006 | 0.003405 | 0.004998 | 0.004854 | 0.021907 | 0.007885 | 0.003994 | 0.003169 | 0.025205 | 0.030519 | 0.007305 | 0.010545 | 0.010074 | 0.014098 | 0.019902 | 0.009866 | 0.006586 | 0.002569 | 0.00742 | 0.013779 |
| 25.375 | 0.003022 | 0.004126 | 0.002768 | 0.007922 | 0.013187 | 0.005323 | 0.007904 | 0.00907 | 0.020613 | 0.007563 | 0.006187 | 0.006289 | 0.025459 | 0.040595 | 0.00891 | 0.008612 | 0.016924 | 0.015617 | 0.02181 | 0.016744 | 0.006529 | 0.007296 | 0.005049 | 0.015565 |
| 25.3888889 | 0.003724 | 0.009691 | 0.003978 | 0.01325 | 0.013902 | 0.006469 | 0.010103 | 0.013415 | 0.01921 | 0.006899 | 0.009033 | 0.00885 | 0.030215 | 0.043856 | 0.011047 | 0.005757 | 0.015383 | 0.016486 | 0.024552 | 0.02206 | 0.005779 | 0.011997 | 0.005348 | 0.016308 |
| 25.4027778 | 0.005825 | 0.012951 | 0.003262 | 0.012072 | 0.013853 | 0.006404 | 0.011685 | 0.016646 | 0.01705 | 0.005178 | 0.01219 | 0.010572 | 0.035054 | 0.038329 | 0.01434 | 0.00509 | 0.00766 | 0.015067 | 0.02656 | 0.025413 | 0.010219 | 0.014763 | 0.009453 | 0.016625 |
| 25.4166667 | 0.008694 | 0.010508 | 0.002001 | 0.006605 | 0.014665 | 0.005831 | 0.013794 | 0.020267 | 0.0141 | 0.002932 | 0.017993 | 0.013356 | 0.037156 | 0.034982 | 0.014378 | 0.007416 | 0.004392 | 0.013715 | 0.026379 | 0.029073 | 0.018026 | 0.015786 | 0.014816 | 0.01742 |
| 25.4305556 | 0.011792 | 0.011227 | 0.003455 | 0.005174 | 0.016695 | 0.004725 | 0.015515 | 0.023106 | 0.011435 | 0.002497 | 0.025241 | 0.016816 | 0.040129 | 0.03951 | 0.009273 | 0.009743 | 0.008422 | 0.014695 | 0.025327 | 0.034674 | 0.024204 | 0.015629 | 0.018566 | 0.020029 |
| 25.4444444 | 0.01464 | 0.015675 | 0.00592 | 0.003224 | 0.019377 | 0.003175 | 0.016157 | 0.02261 | 0.008778 | 0.004999 | 0.031324 | 0.018886 | 0.045707 | 0.047918 | 0.004775 | 0.010252 | 0.020791 | 0.016764 | 0.026273 | 0.040161 | 0.029309 | 0.014872 | 0.020848 | 0.024979 |
| 25.4583333 | 0.018478 | 0.018534 | 0.007649 | 0.00422 | 0.022208 | 0.002498 | 0.016573 | 0.02067 | 0.004961 | 0.009382 | 0.03557 | 0.020346 | 0.049531 | 0.056301 | 0.006757 | 0.010416 | 0.034508 | 0.018362 | 0.02803 | 0.044705 | 0.035 | 0.014073 | 0.02164 | 0.030583 |
| 25.4722222 | 0.022864 | 0.02061 | 0.009802 | 0.014849 | 0.024848 | 0.003166 | 0.016265 | 0.020001 | 0.002356 | 0.01409 | 0.037894 | 0.021786 | 0.048935 | 0.063878 | 0.013605 | 0.011405 | 0.039367 | 0.020511 | 0.028095 | 0.047223 | 0.039972 | 0.014183 | 0.020994 | 0.034606 |
| 25.4861111 | 0.025072 | 0.021756 | 0.013304 | 0.026605 | 0.027254 | 0.004018 | 0.013833 | 0.019642 | 0.002725 | 0.017114 | 0.037818 | 0.021631 | 0.046767 | 0.072114 | 0.019874 | 0.012321 | 0.039342 | 0.024873 | 0.028053 | 0.047456 | 0.044783 | 0.016629 | 0.02212 | 0.038991 |
| 25.5 | 0.024355 | 0.019713 | 0.016848 | 0.028249 | 0.030236 | 0.004598 | 0.011262 | 0.019084 | 0.004315 | 0.017664 | 0.037246 | 0.020804 | 0.046449 | 0.078916 | 0.023056 | 0.012417 | 0.044804 | 0.030332 | 0.030403 | 0.05223 | 0.050003 | 0.022398 | 0.026493 | 0.048597 |
| 25.5138889 | 0.022653 | 0.015917 | 0.01831 | 0.023088 | 0.033066 | 0.004874 | 0.010917 | 0.018035 | 0.007472 | 0.01704 | 0.039154 | 0.02258 | 0.047926 | 0.07479 | 0.02271 | 0.012421 | 0.055193 | 0.033329 | 0.033572 | 0.061406 | 0.053976 | 0.029296 | 0.031267 | 0.056716 |
| 25.5277778 | 0.023997 | 0.01246 | 0.018394 | 0.021568 | 0.033241 | 0.003533 | 0.011184 | 0.014601 | 0.013261 | 0.015631 | 0.042854 | 0.026016 | 0.047465 | 0.058935 | 0.019506 | 0.013152 | 0.059916 | 0.032103 | 0.033256 | 0.069354 | 0.058038 | 0.03286 | 0.032874 | 0.054202 |
| 25.5416667 | 0.031784 | 0.010071 | 0.018776 | 0.023135 | 0.032536 | 0.001426 | 0.012191 | 0.010773 | 0.01974 | 0.014011 | 0.04685 | 0.027484 | 0.041352 | 0.04342 | 0.015597 | 0.013873 | 0.05603 | 0.027312 | 0.030716 | 0.071668 | 0.058821 | 0.030541 | 0.02991 | 0.046899 |
| 25.5555556 | 0.041542 | 0.01203 | 0.019768 | 0.021169 | 0.033767 | 0.001961 | 0.014022 | 0.009999 | 0.022723 | 0.013415 | 0.049994 | 0.027028 | 0.031224 | 0.032419 | 0.012078 | 0.012496 | 0.052524 | 0.021162 | 0.02935 | 0.065392 | 0.051257 | 0.024549 | 0.025533 | 0.042021 |
| 25.5694444 | 0.043257 | 0.020337 | 0.021762 | 0.017871 | 0.032313 | 0.004934 | 0.011801 | 0.010214 | 0.019414 | 0.012742 | 0.050704 | 0.027139 | 0.025642 | 0.024253 | 0.010791 | 0.00922 | 0.056897 | 0.017715 | 0.027198 | 0.059889 | 0.044005 | 0.020743 | 0.026317 | 0.039283 |
| 25.5833333 | 0.03669 | 0.031845 | 0.02556 | 0.019163 | 0.025064 | 0.008119 | 0.005553 | 0.009786 | 0.014976 | 0.014055 | 0.053173 | 0.029411 | 0.03381 | 0.022241 | 0.012701 | 0.009362 | 0.063396 | 0.020831 | 0.024981 | 0.065177 | 0.045033 | 0.021383 | 0.034623 | 0.038881 |
| 25.5972222 | 0.03023 | 0.039777 | 0.030285 | 0.01997 | 0.018012 | 0.011745 | 0.00324 | 0.01116 | 0.016436 | 0.020786 | 0.059009 | 0.032636 | 0.052095 | 0.029803 | 0.015262 | 0.017035 | 0.059197 | 0.02728 | 0.025529 | 0.064263 | 0.04909 | 0.021153 | 0.045497 | 0.042988 |
| 25.6111111 | 0.026234 | 0.038675 | 0.031569 | 0.015601 | 0.014893 | 0.015455 | 0.009632 | 0.013313 | 0.021239 | 0.02729 | 0.060453 | 0.034092 | 0.062922 | 0.046272 | 0.01605 | 0.026856 | 0.044137 | 0.02893 | 0.026443 | 0.041438 | 0.048454 | 0.01721 | 0.052726 | 0.050045 |
| 25.625 | 0.022388 | 0.030826 | 0.027248 | 0.012379 | 0.013556 | 0.017347 | 0.017998 | 0.012104 | 0.024597 | 0.028212 | 0.05689 | 0.035938 | 0.057918 | 0.064456 | 0.014872 | 0.029622 | 0.032362 | 0.023747 | 0.022888 | 0.02012 | 0.041306 | 0.014115 | 0.04999 | 0.055881 |
| 25.6388889 | 0.018113 | 0.022585 | 0.020925 | 0.011966 | 0.013204 | 0.01782 | 0.018653 | 0.007203 | 0.026323 | 0.027019 | 0.055897 | 0.042875 | 0.046222 | 0.073836 | 0.012453 | 0.023825 | 0.029754 | 0.014247 | 0.014034 | 0.013089 | 0.03074 | 0.013669 | 0.03645 | 0.058372 |

| | | | | | | | | | | | | | | | | | | | | | | | | |
|---|---|---|---|---|---|---|---|---|---|---|---|---|---|---|---|---|---|---|---|---|---|---|---|---|
| 25.6527778 | 0.015396 | 0.016437 | 0.015057 | 0.011441 | 0.014063 | 0.019075 | 0.013305 | 0.00381 | 0.023418 | 0.025306 | 0.057671 | 0.052688 | 0.037025 | 0.068258 | 0.008736 | 0.015557 | 0.026151 | 0.006477 | 0.007191 | 0.008761 | 0.022071 | 0.01444 | 0.021 | 0.055737 |
| 25.6666667 | 0.015846 | 0.010869 | 0.009676 | 0.009759 | 0.014135 | 0.019866 | 0.008235 | 0.004697 | 0.014784 | 0.019503 | 0.055384 | 0.057549 | 0.03324 | 0.050457 | 0.004884 | 0.010239 | 0.016034 | 0.005366 | 0.009245 | 0.004817 | 0.016113 | 0.015973 | 0.010312 | 0.046736 |
| 25.6805556 | 0.016365 | 0.005386 | 0.005404 | 0.008549 | 0.011475 | 0.016288 | 0.004706 | 0.007359 | 0.010364 | 0.010102 | 0.045675 | 0.051234 | 0.032037 | 0.030888 | 0.002873 | 0.007865 | 0.007356 | 0.006406 | 0.018179 | 0.002547 | 0.00869 | 0.017159 | 0.004661 | 0.034765 |
| 25.6944444 | 0.013949 | 0.002523 | 0.003253 | 0.008307 | 0.00701 | 0.009449 | 0.003918 | 0.012208 | 0.012489 | 0.00342 | 0.031235 | 0.036531 | 0.028165 | 0.015599 | 0.002308 | 0.005822 | 0.004682 | 0.005896 | 0.028045 | 0.002273 | 0.00524 | 0.015983 | 0.003792 | 0.023752 |
| 25.7083333 | 0.010479 | 0.003135 | 0.0039 | 0.00529 | 0.003275 | 0.00539 | 0.005333 | 0.016939 | 0.01059 | 0.005181 | 0.018673 | 0.024404 | 0.02131 | 0.008567 | 0.00235 | 0.003503 | 0.008214 | 0.005975 | 0.032646 | 0.002436 | 0.009693 | 0.011728 | 0.006588 | 0.013499 |
| 25.7222222 | 0.008055 | 0.003934 | 0.006418 | 0.002559 | 0.001552 | 0.006118 | 0.00546 | 0.016072 | 0.008015 | 0.013725 | 0.013514 | 0.020734 | 0.015551 | 0.014034 | 0.002058 | 0.002777 | 0.016859 | 0.005953 | 0.030415 | 0.001752 | 0.015055 | 0.007092 | 0.00786 | 0.005338 |
| 25.7361111 | 0.006632 | 0.00348 | 0.008568 | 0.00446 | 0.001879 | 0.00798 | 0.005134 | 0.012822 | 0.012074 | 0.019671 | 0.013419 | 0.02082 | 0.014205 | 0.021675 | 0.001176 | 0.004737 | 0.018634 | 0.003754 | 0.02442 | 0.001877 | 0.019437 | 0.004007 | 0.005236 | 0.002994 |
| 25.75 | 0.006154 | 0.003136 | 0.008981 | 0.006478 | 0.003954 | 0.007803 | 0.006094 | 0.013128 | 0.015405 | 0.017858 | 0.012279 | 0.017665 | 0.015899 | 0.02037 | 0.000852 | 0.006992 | 0.011018 | 0.001673 | 0.014881 | 0.002894 | 0.020655 | 0.001864 | 0.006009 | 0.006539 |
| 25.7638889 | 0.006688 | 0.002755 | 0.007194 | 0.005601 | 0.006727 | 0.006176 | 0.007308 | 0.016693 | 0.015146 | 0.013156 | 0.007658 | 0.010001 | 0.015947 | 0.014193 | 0.00086 | 0.00635 | 0.006879 | 0.002157 | 0.008242 | 0.007536 | 0.017879 | 0.001072 | 0.012967 | 0.011521 |
| 25.7777778 | 0.007483 | 0.003808 | 0.004108 | 0.00588 | 0.009552 | 0.005361 | 0.007277 | 0.01873 | 0.017339 | 0.012041 | 0.005013 | 0.005632 | 0.012852 | 0.008325 | 0.001283 | 0.003776 | 0.006904 | 0.003849 | 0.008584 | 0.019273 | 0.016795 | 0.001798 | 0.018032 | 0.014549 |
| 25.7916667 | 0.007565 | 0.007296 | 0.002276 | 0.006932 | 0.011555 | 0.005365 | 0.005079 | 0.015407 | 0.024265 | 0.014668 | 0.007758 | 0.008607 | 0.010048 | 0.004466 | 0.002701 | 0.002413 | 0.005301 | 0.005322 | 0.006838 | 0.034364 | 0.017704 | 0.003068 | 0.018618 | 0.016437 |
| 25.8055556 | 0.00667 | 0.009187 | 0.001669 | 0.005695 | 0.011131 | 0.004431 | 0.002612 | 0.008785 | 0.031078 | 0.016165 | 0.008561 | 0.010009 | 0.01001 | 0.004826 | 0.003454 | 0.002284 | 0.004275 | 0.006714 | 0.002621 | 0.045306 | 0.017264 | 0.004323 | 0.019352 | 0.017453 |
| 25.8194444 | 0.005858 | 0.009023 | 0.001687 | 0.004747 | 0.008008 | 0.003946 | 0.002784 | 0.003733 | 0.031651 | 0.013645 | 0.004897 | 0.006738 | 0.008616 | 0.007083 | 0.002638 | 0.001635 | 0.007585 | 0.007437 | 0.001715 | 0.045753 | 0.013875 | 0.005554 | 0.020665 | 0.016673 |
| 25.8333333 | 0.005985 | 0.00958 | 0.002279 | 0.004425 | 0.00392 | 0.005635 | 0.004458 | 0.00225 | 0.026347 | 0.008868 | 0.001711 | 0.004749 | 0.006572 | 0.008278 | 0.002808 | 0.000977 | 0.011512 | 0.006464 | 0.002617 | 0.038942 | 0.009567 | 0.006691 | 0.019067 | 0.015257 |
| 25.8472222 | 0.005886 | 0.008517 | 0.003207 | 0.005935 | 0.002153 | 0.007821 | 0.004591 | 0.002198 | 0.019472 | 0.004477 | 0.002709 | 0.006108 | 0.011028 | 0.01005 | 0.004901 | 0.001951 | 0.013436 | 0.004093 | 0.003199 | 0.034718 | 0.007471 | 0.006647 | 0.013682 | 0.013954 |
| 25.8611111 | 0.004581 | 0.005762 | 0.006503 | 0.010793 | 0.004672 | 0.011189 | 0.003375 | 0.002116 | 0.014449 | 0.002571 | 0.00735 | 0.0075 | 0.019601 | 0.01472 | 0.005604 | 0.003929 | 0.016518 | 0.003104 | 0.004754 | 0.037287 | 0.007687 | 0.00503 | 0.008857 | 0.013532 |
| 25.875 | 0.003275 | 0.005171 | 0.010668 | 0.013858 | 0.00951 | 0.017501 | 0.003485 | 0.002996 | 0.010845 | 0.004385 | 0.010942 | 0.00619 | 0.030238 | 0.020573 | 0.003954 | 0.005382 | 0.013505 | 0.004973 | 0.006808 | 0.043266 | 0.009422 | 0.003657 | 0.009101 | 0.013782 |
| 25.8888889 | 0.00332 | 0.00576 | 0.012531 | 0.010785 | 0.013729 | 0.02516 | 0.006034 | 0.003873 | 0.006977 | 0.00818 | 0.013507 | 0.005553 | 0.045283 | 0.021479 | 0.002667 | 0.007129 | 0.007389 | 0.007327 | 0.006234 | 0.047279 | 0.010978 | 0.002668 | 0.014833 | 0.012081 |
| 25.9027778 | 0.003935 | 0.007614 | 0.014209 | 0.005133 | 0.016578 | 0.030783 | 0.009192 | 0.004872 | 0.006197 | 0.011258 | 0.019273 | 0.00925 | 0.060667 | 0.014515 | 0.002162 | 0.010461 | 0.016882 | 0.008838 | 0.005735 | 0.048919 | 0.011071 | 0.002361 | 0.021971 | 0.00722 |
| 25.9166667 | 0.003445 | 0.013886 | 0.01815 | 0.007756 | 0.018873 | 0.030761 | 0.011439 | 0.008543 | 0.009907 | 0.011989 | 0.027892 | 0.013993 | 0.069985 | 0.0067 | 0.002241 | 0.013467 | 0.03904 | 0.011289 | 0.007794 | 0.050448 | 0.011346 | 0.005017 | 0.025317 | 0.004678 |
| 25.9305556 | 0.002643 | 0.021883 | 0.022961 | 0.020067 | 0.019605 | 0.025833 | 0.013873 | 0.014706 | 0.015354 | 0.01205 | 0.034804 | 0.016045 | 0.07398 | 0.006106 | 0.004051 | 0.014054 | 0.050538 | 0.016314 | 0.007815 | 0.049646 | 0.014262 | 0.008378 | 0.026435 | 0.007554 |
| 25.9444444 | 0.003049 | 0.026819 | 0.024291 | 0.028565 | 0.016227 | 0.021568 | 0.017981 | 0.019829 | 0.023083 | 0.015197 | 0.039445 | 0.019805 | 0.079773 | 0.011482 | 0.006947 | 0.013936 | 0.049008 | 0.02118 | 0.004535 | 0.04217 | 0.018542 | 0.009789 | 0.030307 | 0.010933 |
| 25.9583333 | 0.005058 | 0.030095 | 0.019807 | 0.023653 | 0.012636 | 0.021213 | 0.021927 | 0.022313 | 0.033179 | 0.022034 | 0.045617 | 0.027131 | 0.089511 | 0.018434 | 0.011276 | 0.0156 | 0.055581 | 0.022837 | 0.00228 | 0.035356 | 0.022555 | 0.011844 | 0.034598 | 0.014106 |
| 25.9722222 | 0.014345 | 0.035427 | 0.015636 | 0.012664 | 0.015762 | 0.023772 | 0.024379 | 0.025558 | 0.041911 | 0.030675 | 0.05386 | 0.031373 | 0.096294 | 0.024193 | 0.018455 | 0.019781 | 0.068472 | 0.022628 | 0.002879 | 0.043957 | 0.024901 | 0.016651 | 0.036148 | 0.019859 |
| 25.9861111 | 0.032518 | 0.039628 | 0.019422 | 0.005979 | 0.023415 | 0.025377 | 0.02744 | 0.030928 | 0.047219 | 0.037279 | 0.060371 | 0.032397 | 0.097422 | 0.026453 | 0.026135 | 0.025788 | 0.0699 | 0.023213 | 0.004206 | 0.066654 | 0.021864 | 0.022299 | 0.038596 | 0.026744 |
| 26 | 0.047361 | 0.038367 | 0.029343 | 0.006377 | 0.028858 | 0.022863 | 0.030321 | 0.036109 | 0.04973 | 0.037982 | 0.060817 | 0.033542 | 0.099932 | 0.026492 | 0.031559 | 0.03196 | 0.064607 | 0.026621 | 0.007816 | 0.083287 | 0.01676 | 0.027301 | 0.040913 | 0.032074 |
| 26.0138889 | 0.052169 | 0.034557 | 0.038393 | 0.015633 | 0.02993 | 0.01716 | 0.030472 | 0.03959 | 0.047887 | 0.033976 | 0.055535 | 0.032188 | 0.107085 | 0.027462 | 0.034645 | 0.036657 | 0.069456 | 0.031848 | 0.01681 | 0.079348 | 0.015873 | 0.035011 | 0.039873 | 0.03444 |
| 26.0277778 | 0.05013 | 0.031948 | 0.044243 | 0.027213 | 0.028713 | 0.011117 | 0.027663 | 0.039047 | 0.041322 | 0.030212 | 0.050566 | 0.028652 | 0.116567 | 0.027875 | 0.034216 | 0.038657 | 0.083793 | 0.036736 | 0.027163 | 0.061097 | 0.019965 | 0.046267 | 0.038231 | 0.034022 |
| 26.0416667 | 0.050024 | 0.033955 | 0.049027 | 0.033317 | 0.032148 | 0.007863 | 0.024941 | 0.03619 | 0.035195 | 0.030669 | 0.05354 | 0.027418 | 0.120514 | 0.028343 | 0.027039 | 0.037949 | 0.093896 | 0.042064 | 0.034657 | 0.037643 | 0.027083 | 0.057381 | 0.040312 | 0.035587 |
| 26.0555556 | 0.068396 | 0.043774 | 0.054712 | 0.036707 | 0.048341 | 0.010432 | 0.027906 | 0.038085 | 0.035522 | 0.035514 | 0.066083 | 0.030975 | 0.115204 | 0.041234 | 0.018126 | 0.036743 | 0.094633 | 0.050632 | 0.040365 | 0.018411 | 0.038035 | 0.068748 | 0.046249 | 0.048157 |
| 26.0694444 | 0.08674 | 0.058355 | 0.060785 | 0.040303 | 0.066227 | 0.01957 | 0.034023 | 0.044611 | 0.039134 | 0.038801 | 0.077554 | 0.035427 | 0.102657 | 0.066534 | 0.021 | 0.036581 | 0.08575 | 0.061702 | 0.044625 | 0.014478 | 0.051405 | 0.073358 | 0.052968 | 0.068088 |
| 26.0833333 | 0.066875 | 0.063848 | 0.059237 | 0.043815 | 0.061545 | 0.028891 | 0.033369 | 0.046008 | 0.037098 | 0.033902 | 0.071102 | 0.033788 | 0.084392 | 0.078827 | 0.031556 | 0.036736 | 0.066497 | 0.063774 | 0.048186 | 0.017231 | 0.053225 | 0.063454 | 0.056231 | 0.076862 |
| 26.0972222 | 0.027777 | 0.047345 | 0.044898 | 0.044487 | 0.034942 | 0.030208 | 0.023184 | 0.037219 | 0.029263 | 0.02295 | 0.045 | 0.026823 | 0.066034 | 0.068736 | 0.031735 | 0.036856 | 0.040643 | 0.043152 | 0.047996 | 0.013541 | 0.036178 | 0.048965 | 0.048625 | 0.066806 |
| 26.1111111 | 0.006199 | 0.022064 | 0.027712 | 0.03706 | 0.014659 | 0.024097 | 0.010676 | 0.023775 | 0.018723 | 0.01228 | 0.018946 | 0.021946 | 0.044115 | 0.046781 | 0.021006 | 0.036992 | 0.017795 | 0.018281 | 0.04005 | 0.0066 | 0.016069 | 0.03654 | 0.031665 | 0.043842 |
| 26.125 | 0.00162 | 0.012962 | 0.017089 | 0.021535 | 0.013612 | 0.016333 | 0.003351 | 0.011327 | 0.009093 | 0.005893 | 0.005857 | 0.02232 | 0.019954 | 0.021391 | 0.010375 | 0.036392 | 0.012103 | 0.01278 | 0.032114 | 0.002781 | 0.006175 | 0.023538 | 0.016765 | 0.019256 |
| 26.1388889 | 0.001706 | 0.020464 | 0.010277 | 0.012874 | 0.013417 | 0.010517 | 0.001366 | 0.003948 | 0.007236 | 0.006557 | 0.003663 | 0.024704 | 0.005705 | 0.005005 | 0.008331 | 0.034297 | 0.02601 | 0.012445 | 0.028315 | 0.002742 | 0.005686 | 0.0136 | 0.014078 | 0.006565 |
| 26.1527778 | 0.002689 | 0.029802 | 0.005442 | 0.020325 | 0.007015 | 0.007001 | 0.001338 | 0.003544 | 0.011553 | 0.01278 | 0.009154 | 0.023922 | 0.006126 | 0.003727 | 0.019421 | 0.032275 | 0.040296 | 0.011747 | 0.022752 | 0.004313 | 0.012707 | 0.008177 | 0.021784 | 0.008045 |
| 26.1666667 | 0.006005 | 0.033866 | 0.005785 | 0.025942 | 0.006469 | 0.0045 | 0.001498 | 0.007608 | 0.015009 | 0.018733 | 0.016751 | 0.016497 | 0.021517 | 0.015271 | 0.033341 | 0.030428 | 0.04065 | 0.017161 | 0.012967 | 0.005481 | 0.022399 | 0.00544 | 0.026975 | 0.019681 |
| 26.1805556 | 0.00893 | 0.0326 | 0.007936 | 0.018825 | 0.011558 | 0.002646 | 0.001777 | 0.014694 | 0.016549 | 0.021528 | 0.021441 | 0.007432 | 0.041945 | 0.036613 | 0.034319 | 0.024703 | 0.037925 | 0.027373 | 0.010894 | 0.008972 | 0.032129 | 0.012774 | 0.025268 | 0.037059 |
| 26.1944444 | 0.008872 | 0.028826 | 0.007894 | 0.008718 | 0.012665 | 0.002502 | 0.002042 | 0.0224 | 0.017492 | 0.022562 | 0.022975 | 0.003531 | 0.052368 | 0.05899 | 0.021646 | 0.013639 | 0.046775 | 0.046437 | 0.027423 | 0.025079 | 0.044342 | 0.035786 | 0.018583 | 0.053287 |
| 26.2083333 | 0.009136 | 0.024235 | 0.008004 | 0.002976 | 0.007848 | 0.003689 | 0.002674 | 0.027418 | 0.019481 | 0.02381 | 0.022043 | 0.002661 | 0.066328 | 0.053685 | 0.007313 | 0.011811 | 0.050858 | 0.049402 | 0.038423 | 0.038824 | 0.044505 | 0.049596 | 0.010931 | 0.053358 |
| 26.2222222 | 0.010747 | 0.01935 | 0.010092 | 0.003589 | 0.003484 | 0.003491 | 0.003082 | 0.027418 | 0.023238 | 0.025128 | 0.01873 | 0.002382 | 0.080835 | 0.038027 | 0.001199 | 0.027516 | 0.039054 | 0.038674 | 0.043156 | 0.029784 | 0.034078 | 0.039229 | 0.007834 | 0.041513 |
| 26.2361111 | 0.010669 | 0.017003 | 0.010613 | 0.010837 | 0.002127 | 0.002843 | 0.003405 | 0.02234 | 0.025344 | 0.02422 | 0.014075 | 0.002449 | 0.068158 | 0.059977 | 0.008517 | 0.035563 | 0.031488 | 0.046206 | 0.057086 | 0.015375 | 0.027432 | 0.026025 | 0.007855 | 0.043427 |
| 26.25 | 0.00927 | 0.02045 | 0.008279 | 0.021197 | 0.004299 | 0.004449 | 0.005621 | 0.016267 | 0.024798 | 0.022336 | 0.011348 | 0.001798 | 0.053166 | 0.077188 | 0.025192 | 0.024736 | 0.024683 | 0.03913 | 0.0501 | 0.008875 | 0.019994 | 0.0147 | 0.008135 | 0.04444 |
| 26.2638889 | 0.009003 | 0.027678 | 0.010335 | 0.029284 | 0.012465 | 0.005366 | 0.005623 | 0.012151 | 0.023812 | 0.022554 | 0.011794 | 0.002124 | 0.056598 | 0.053147 | 0.035136 | 0.013397 | 0.013959 | 0.014024 | 0.025527 | 0.00886 | 0.0219 | 0.006665 | 0.008592 | 0.025942 |
| 26.2777778 | 0.009194 | 0.032264 | 0.01612 | 0.030765 | 0.021961 | 0.004585 | 0.002551 | 0.009116 | 0.021025 | 0.022371 | 0.012005 | 0.005036 | 0.057722 | 0.022154 | 0.031895 | 0.00907 | 0.010773 | 0.005464 | 0.011777 | 0.010797 | 0.028448 | 0.006883 | 0.010116 | 0.009849 |
| 26.2916667 | 0.008365 | 0.034539 | 0.015987 | 0.022459 | 0.028596 | 0.005008 | 0.001763 | 0.005509 | 0.014492 | 0.018605 | 0.011401 | 0.008366 | 0.04717 | 0.010264 | 0.020926 | 0.006966 | 0.015669 | 0.00639 | 0.005688 | 0.010245 | 0.026704 | 0.011574 | 0.01222 | 0.008735 |
| 26.3055556 | 0.005841 | 0.03999 | 0.01112 | 0.010323 | 0.033712 | 0.006444 | 0.003356 | 0.002923 | 0.007035 | 0.013158 | 0.01437 | 0.008715 | 0.036143 | 0.016324 | 0.011866 | 0.004265 | 0.022166 | 0.008144 | 0.002036 | 0.009552 | 0.022899 | 0.017618 | 0.012857 | 0.011981 |
| 26.3194444 | 0.002468 | 0.045232 | 0.008997 | 0.00339 | 0.038038 | 0.007603 | 0.003889 | 0.003591 | 0.003599 | 0.009708 | 0.022192 | 0.00577 | 0.040807 | 0.023504 | 0.010031 | 0.003568 | 0.018924 | 0.010771 | 0.001108 | 0.009678 | 0.022993 | 0.016453 | 0.011291 | 0.012899 |
| 26.3333333 | 0.002314 | 0.043667 | 0.008828 | 0.002096 | 0.039616 | 0.008121 | 0.004051 | 0.007643 | 0.005077 | 0.009443 | 0.03079 | 0.004923 | 0.056207 | 0.021271 | 0.008842 | 0.006531 | 0.01188 | 0.012074 | 0.002756 | 0.009286 | 0.030015 | 0.009103 | 0.012471 | 0.011938 |
| 26.3472222 | 0.006061 | 0.036777 | 0.006449 | 0.004349 | 0.035067 | 0.008152 | 0.007897 | 0.01252 | 0.006959 | 0.009246 | 0.035059 | 0.009004 | 0.061364 | 0.014189 | 0.005859 | 0.008849 | 0.009711 | 0.013329 | 0.007972 | 0.006749 | 0.032997 | 0.009363 | 0.021196 | 0.01114 |
| 26.3611111 | 0.008356 | 0.029091 | 0.003776 | 0.00916 | 0.026629 | 0.007834 | 0.012433 | 0.011819 | 0.00518 | 0.006053 | 0.032945 | 0.012335 | 0.052121 | 0.008663 | 0.005532 | 0.007642 | 0.013591 | 0.015088 | 0.014621 | 0.003205 | 0.021512 | 0.018504 | 0.027341 | 0.011028 |
| 26.375 | 0.007047 | 0.02351 | 0.004903 | 0.016229 | 0.020294 | 0.006721 | 0.014094 | 0.005981 | 0.001846 | 0.002422 | 0.027224 | 0.01324 | 0.041949 | 0.005973 | 0.005408 | 0.005114 | 0.021629 | 0.014131 | 0.017015 | 0.000979 | 0.011311 | 0.0218 | 0.020216 | 0.008147 |
| 26.3888889 | 0.004686 | 0.020469 | 0.009151 | 0.022981 | 0.018014 | 0.005108 | 0.015956 | 0.002081 | 0.003789 | 0.003128 | 0.024695 | 0.015652 | 0.039701 | 0.006607 | 0.005942 | 0.003884 | 0.01892 | 0.011641 | 0.016114 | 0.000996 | 0.010765 | 0.014168 | 0.009373 | 0.003244 |
| 26.4027778 | 0.00369 | 0.018827 | 0.014297 | 0.024949 | 0.017485 | 0.003817 | 0.020345 | 0.003125 | 0.013083 | 0.008304 | 0.029878 | 0.018773 | 0.044149 | 0.006259 | 0.012472 | 0.00466 | 0.013759 | 0.013208 | 0.0178 | 0.002853 | 0.013212 | 0.005616 | 0.008345 | 0.002193 |
| 26.4166667 | 0.005011 | 0.01716 | 0.020265 | 0.023355 | 0.016195 | 0.002438 | 0.026121 | 0.008347 | 0.02251 | 0.013608 | 0.035271 | 0.02005 | 0.048899 | 0.004647 | 0.019357 | 0.006207 | 0.028599 | 0.018663 | 0.023101 | 0.004926 | 0.024476 | 0.00238 | 0.013922 | 0.006255 |
| 26.4305556 | 0.007776 | 0.013467 | 0.026994 | 0.023518 | 0.013779 | 0.00108 | 0.030413 | 0.013571 | 0.026607 | 0.016105 | 0.04031 | 0.021543 | 0.049699 | 0.007531 | 0.021623 | 0.006914 | 0.05348 | 0.022452 | 0.027744 | 0.006254 | 0.038549 | 0.003483 | 0.016871 | 0.01115 |
| 26.4444444 | 0.010539 | 0.00889 | 0.034041 | 0.027391 | 0.01297 | 0.000703 | 0.031983 | 0.015604 | 0.028114 | 0.016992 | 0.045008 | 0.025739 | 0.049664 | 0.013255 | 0.022401 | 0.006992 | 0.068957 | 0.022763 | 0.029396 | 0.007853 | 0.046855 | 0.007953 | 0.017601 | 0.014888 |
| 26.4583333 | 0.013142 | 0.007769 | 0.039202 | 0.030318 | 0.015581 | 0.001704 | 0.031787 | 0.017073 | 0.030821 | 0.018289 | 0.051249 | 0.031182 | 0.051463 | 0.021328 | 0.022271 | 0.007108 | 0.069754 | 0.022191 | 0.03074 | 0.009209 | 0.051304 | 0.012497 | 0.021287 | 0.019753 |
| 26.4722222 | 0.016125 | 0.011717 | 0.039495 | 0.029571 | 0.019312 | 0.004724 | 0.029656 | 0.020661 | 0.035052 | 0.02013 | 0.057763 | 0.035334 | 0.051475 | 0.030794 | 0.019607 | 0.007366 | 0.05825 | 0.022195 | 0.033154 | 0.008477 | 0.052793 | 0.015133 | 0.030381 | 0.026282 |
| 26.4861111 | 0.019305 | 0.017604 | 0.037813 | 0.027435 | 0.023122 | 0.00913 | 0.026088 | 0.026454 | 0.038815 | 0.02184 | 0.063489 | 0.036708 | 0.048685 | 0.038163 | 0.016973 | 0.008132 | 0.041683 | 0.022737 | 0.035489 | 0.006371 | 0.054318 | 0.01714 | 0.041556 | 0.03293 |
| 26.5 | 0.02285 | 0.020649 | 0.037746 | 0.024149 | 0.027906 | 0.012998 | 0.023948 | 0.032262 | 0.041081 | 0.023295 | 0.069048 | 0.035678 | 0.050491 | 0.046632 | 0.020297 | 0.009398 | 0.027903 | 0.024147 | 0.038273 | 0.00495 | 0.055798 | 0.018981 | 0.048716 | 0.039304 |
| 26.5138889 | 0.025886 | 0.017254 | 0.037413 | 0.021935 | 0.031131 | 0.014483 | 0.023357 | 0.032244 | 0.042058 | 0.023495 | 0.071877 | 0.033937 | 0.057812 | 0.054818 | 0.028383 | 0.009846 | 0.022043 | 0.025637 | 0.039105 | 0.005211 | 0.053761 | 0.022265 | 0.048976 | 0.043013 |
| 26.5277778 | 0.025981 | 0.010092 | 0.035693 | 0.020905 | 0.028833 | 0.012577 | 0.021335 | 0.025924 | 0.041553 | 0.020519 | 0.068812 | 0.032535 | 0.062446 | 0.056992 | 0.032695 | 0.008452 | 0.024955 | 0.024424 | 0.029519 | 0.008551 | 0.047412 | 0.028418 | 0.043538 | 0.042206 |
| 26.5416667 | 0.024114 | 0.005582 | 0.031631 | 0.016568 | 0.023362 | 0.011689 | 0.018547 | 0.020924 | 0.04112 | 0.018011 | 0.061861 | 0.031499 | 0.059421 | 0.057013 | 0.029563 | 0.00618 | 0.033768 | 0.019615 | 0.014584 | 0.017254 | 0.04107 | 0.034061 | 0.0368 | 0.040466 |
| 26.5555556 | 0.021906 | 0.006191 | 0.023651 | 0.011034 | 0.020488 | 0.018775 | 0.01869 | 0.020843 | 0.041681 | 0.022813 | 0.054144 | 0.031508 | 0.05123 | 0.056641 | 0.024002 | 0.004894 | 0.042882 | 0.015322 | 0.008314 | 0.026448 | 0.037235 | 0.03362 | 0.032344 | 0.038536 |
| 26.5694444 | 0.017851 | 0.012888 | 0.015835 | 0.010261 | 0.020193 | 0.030505 | 0.019818 | 0.020753 | 0.041132 | 0.031871 | 0.046918 | 0.033118 | 0.04552 | 0.050201 | 0.02243 | 0.005061 | 0.0516 | 0.015347 | 0.007205 | 0.028839 | 0.034083 | 0.026899 | 0.031022 | 0.032789 |
| 26.5833333 | 0.013901 | 0.02363 | 0.013727 | 0.014608 | 0.021663 | 0.036029 | 0.017626 | 0.016732 | 0.039449 | 0.035785 | 0.046806 | 0.036751 | 0.045477 | 0.042479 | 0.022288 | 0.007094 | 0.065068 | 0.019983 | 0.007895 | 0.029937 | 0.031013 | 0.022565 | 0.033444 | 0.027235 |
| 26.5972222 | 0.014218 | 0.032255 | 0.017652 | 0.017545 | 0.025324 | 0.033265 | 0.015009 | 0.013344 | 0.038849 | 0.031941 | 0.057245 | 0.044002 | 0.049299 | 0.043699 | 0.019103 | 0.011713 | 0.075246 | 0.027765 | 0.013281 | 0.029772 | 0.026955 | 0.029423 | 0.036282 | 0.029431 |
| 26.6111111 | 0.016715 | 0.032804 | 0.022942 | 0.012954 | 0.026843 | 0.027667 | 0.015818 | 0.015388 | 0.039004 | 0.022913 | 0.065513 | 0.050487 | 0.049307 | 0.048783 | 0.014568 | 0.01686 | 0.068115 | 0.034015 | 0.01843 | 0.021744 | 0.025336 | 0.043587 | 0.035511 | 0.036063 |
| 26.625 | 0.016096 | 0.024478 | 0.023407 | 0.006405 | 0.02368 | 0.021133 | 0.018414 | 0.018503 | 0.038042 | 0.013172 | 0.058045 | 0.047663 | 0.040895 | 0.047099 | 0.010471 | 0.01945 | 0.050886 | 0.033136 | 0.020618 | 0.021147 | 0.02843 | 0.050402 | 0.033723 | 0.03743 |
| 26.6388889 | 0.012118 | 0.012496 | 0.018084 | 0.004982 | 0.018081 | 0.014752 | 0.021952 | 0.019054 | 0.035926 | 0.010503 | 0.042345 | 0.038126 | 0.031507 | 0.041041 | 0.009037 | 0.018795 | 0.037384 | 0.024827 | 0.021773 | 0.039319 | 0.030907 | 0.043285 | 0.03298 | 0.033717 |
| 26.6527778 | 0.008545 | 0.00357 | 0.011034 | 0.004842 | 0.01204 | 0.012543 | 0.024963 | 0.018803 | 0.031063 | 0.017317 | 0.035764 | 0.03216 | 0.027143 | 0.039408 | 0.009785 | 0.015764 | 0.027491 | 0.015334 | 0.023727 | 0.051883 | 0.036178 | 0.032662 | 0.032607 | 0.031291 |
| 26.6666667 | 0.007336 | 0.001571 | 0.006694 | 0.005294 | 0.006873 | 0.0144 | 0.023197 | 0.016677 | 0.021765 | 0.024902 | 0.035967 | 0.030608 | 0.026847 | 0.040555 | 0.008814 | 0.010978 | 0.017635 | 0.009545 | 0.027001 | 0.045633 | 0.045005 | 0.026981 | 0.02854 | 0.027545 |
| 26.6805556 | 0.007352 | 0.003601 | 0.00788 | 0.008694 | 0.003607 | 0.014043 | 0.015938 | 0.012714 | 0.013451 | 0.02585 | 0.028885 | 0.026607 | 0.024892 | 0.03227 | 0.005299 | 0.005626 | 0.009201 | 0.005686 | 0.028447 | 0.036242 | 0.047661 | 0.024796 | 0.020058 | 0.019858 |
| 26.6944444 | 0.006869 | 0.005852 | 0.013194 | 0.009953 | 0.002182 | 0.008406 | 0.00779 | 0.008897 | 0.011168 | 0.022567 | 0.015354 | 0.016099 | 0.017313 | 0.01664 | 0.001682 | 0.002036 | 0.005236 | 0.002851 | 0.025668 | 0.025121 | 0.041662 | 0.022962 | 0.013359 | 0.012934 |
| 26.7083333 | 0.005964 | 0.006672 | 0.017606 | 0.006077 | 0.001457 | 0.002801 | 0.004313 | 0.005084 | 0.010518 | 0.018853 | 0.008496 | 0.006097 | 0.008239 | 0.008532 | 0.000625 | 0.001585 | 0.004452 | 0.00305 | 0.019932 | 0.012787 | 0.030399 | 0.018145 | 0.011109 | 0.008766 |
| 26.7222222 | 0.004658 | 0.005226 | 0.01537 | 0.003547 | 0.002271 | 0.001163 | 0.004199 | 0.002921 | 0.006764 | 0.014552 | 0.01041 | 0.005517 | 0.002817 | 0.008851 | 0.001389 | 0.002938 | 0.005613 | 0.003482 | 0.014644 | 0.007372 | 0.016237 | 0.010882 | 0.010336 | 0.006354 |
| 26.7361111 | 0.002799 | 0.004141 | 0.009691 | 0.007906 | 0.006516 | 0.001355 | 0.003028 | 0.005777 | 0.003968 | 0.009835 | 0.009782 | 0.008815 | 0.001461 | 0.011117 | 0.001609 | 0.003834 | 0.009979 | 0.002188 | 0.010482 | 0.010614 | 0.007768 | 0.004651 | 0.008537 | 0.004688 |
| 26.75 | 0.001339 | 0.004636 | 0.010769 | 0.012894 | 0.011332 | 0.00277 | 0.00217 | 0.012242 | 0.004786 | 0.006591 | 0.005001 | 0.009501 | 0.002661 | 0.011066 | 0.000891 | 0.003824 | 0.012743 | 0.002972 | 0.006611 | 0.02086 | 0.009317 | 0.001193 | 0.008955 | 0.003612 |
| 26.7638889 | 0.001637 | 0.007834 | 0.015287 | 0.010146 | 0.01141 | 0.005808 | 0.002165 | 0.015355 | 0.004238 | 0.005897 | 0.004534 | 0.006749 | 0.003035 | 0.00832 | 0.000836 | 0.004287 | 0.013163 | 0.004275 | 0.005039 | 0.030932 | 0.008648 | 0.000814 | 0.012617 | 0.004458 |
| 26.7777778 | 0.004375 | 0.014085 | 0.013559 | 0.004869 | 0.006684 | 0.008949 | 0.002038 | 0.010444 | 0.003237 | 0.006889 | 0.010019 | 0.004321 | 0.00167 | 0.009339 | 0.003123 | 0.005149 | 0.017527 | 0.002645 | 0.004571 | 0.036997 | 0.00371 | 0.001461 | 0.015889 | 0.007116 |
| 26.7916667 | 0.00658 | 0.015942 | 0.008733 | 0.004661 | 0.002433 | 0.011228 | 0.002847 | 0.003434 | 0.006312 | 0.008441 | 0.014253 | 0.007234 | 0.001451 | 0.013056 | 0.006589 | 0.004733 | 0.023167 | 0.00219 | 0.00464 | 0.04111 | 0.001123 | 0.001255 | 0.017005 | 0.009488 |
| 26.8055556 | 0.005512 | 0.009984 | 0.008813 | 0.006949 | 0.002702 | 0.011886 | 0.003392 | 0.002411 | 0.010241 | 0.010057 | 0.014199 | 0.010172 | 0.003572 | 0.014084 | 0.007334 | 0.003104 | 0.023045 | 0.008225 | 0.009481 | 0.04518 | 0.005934 | 0.001698 | 0.019056 | 0.010367 |
| 26.8194444 | 0.002743 | 0.003058 | 0.014983 | 0.008245 | 0.006781 | 0.012772 | 0.003395 | 0.00521 | 0.011915 | 0.010442 | 0.014314 | 0.009151 | 0.007911 | 0.012103 | 0.004435 | 0.002341 | 0.019724 | 0.018057 | 0.018542 | 0.047627 | 0.010816 | 0.003661 | 0.026999 | 0.009829 |
| 26.8333333 | 0.001439 | 0.001568 | 0.018166 | 0.006481 | 0.011113 | 0.00984 | 0.003277 | 0.005628 | 0.012229 | 0.009423 | 0.015286 | 0.006824 | 0.01188 | 0.008186 | 0.001722 | 0.002482 | 0.019436 | 0.0205 | 0.023056 | 0.047993 | 0.017714 | 0.00455 | 0.03988 | 0.008727 |
| 26.8472222 | 0.002324 | 0.004702 | 0.011398 | 0.004212 | 0.011906 | 0.006597 | 0.003644 | 0.005166 | 0.011541 | 0.008498 | 0.012329 | 0.004231 | 0.01427 | 0.005276 | 0.000941 | 0.002312 | 0.018965 | 0.014204 | 0.016573 | 0.05048 | 0.023028 | 0.003284 | 0.04926 | 0.008754 |
| 26.8611111 | 0.003675 | 0.006552 | 0.004936 | 0.006161 | 0.007602 | 0.00547 | 0.004484 | 0.006422 | 0.008762 | 0.007414 | 0.007784 | 0.003957 | 0.018252 | 0.003939 | 0.001589 | 0.002493 | 0.012756 | 0.014664 | 0.012314 | 0.053919 | 0.021532 | 0.004254 | 0.049391 | 0.008469 |
| 26.875 | 0.004924 | 0.005542 | 0.009447 | 0.011096 | 0.003875 | 0.005336 | 0.004498 | 0.005759 | 0.004867 | 0.004609 | 0.008894 | 0.007285 | 0.024498 | 0.004709 | 0.003207 | 0.002761 | 0.007915 | 0.026154 | 0.020837 | 0.052097 | 0.015048 | 0.008593 | 0.044834 | 0.005144 |
| 26.8888889 | 0.005934 | 0.006128 | 0.021225 | 0.015491 | 0.00411 | 0.004084 | 0.006071 | 0.004598 | 0.003584 | 0.001682 | 0.012176 | 0.009043 | 0.02714 | 0.00798 | 0.003301 | 0.004355 | 0.008797 | 0.033649 | 0.026395 | 0.044061 | 0.012223 | 0.010816 | 0.03779 | 0.001818 |
| 26.9027778 | 0.007267 | 0.00896 | 0.030461 | 0.015086 | 0.004921 | 0.004672 | 0.009491 | 0.005142 | 0.005188 | 0.000835 | 0.013839 | 0.008051 | 0.022947 | 0.008629 | 0.001924 | 0.008415 | 0.00986 | 0.030303 | 0.020776 | 0.036451 | 0.010212 | 0.010109 | 0.026216 | 0.002335 |
| 26.9166667 | 0.01004 | 0.010309 | 0.031626 | 0.01118 | 0.008489 | 0.010152 | 0.013166 | 0.006694 | 0.007759 | 0.00087 | 0.018771 | 0.009424 | 0.020319 | 0.007775 | 0.001184 | 0.012326 | 0.015806 | 0.021346 | 0.011639 | 0.034672 | 0.007377 | 0.01237 | 0.017798 | 0.003484 |
| 26.9305556 | 0.013375 | 0.008421 | 0.027676 | 0.011242 | 0.014852 | 0.015106 | 0.017724 | 0.011177 | 0.011532 | 0.001603 | 0.027002 | 0.013776 | 0.028429 | 0.012079 | 0.001355 | 0.014735 | 0.035751 | 0.011226 | 0.007289 | 0.032796 | 0.008872 | 0.01948 | 0.018254 | 0.003471 |
| 26.9444444 | 0.016093 | 0.00593 | 0.024902 | 0.016913 | 0.019489 | 0.015855 | 0.021551 | 0.016285 | 0.016707 | 0.002198 | 0.031603 | 0.016754 | 0.042539 | 0.020265 | 0.002951 | 0.016484 | 0.059939 | 0.006053 | 0.010527 | 0.024175 | 0.008873 | 0.026318 | 0.022151 | 0.005809 |
| 26.9583333 | 0.018291 | 0.008153 | 0.027585 | 0.020456 | 0.021973 | 0.015995 | 0.022985 | 0.018082 | 0.022288 | 0.001165 | 0.032091 | 0.016809 | 0.052873 | 0.0259 | 0.00522 | 0.017601 | 0.068839 | 0.008387 | 0.01402 | 0.012799 | 0.008601 | 0.02751 | 0.024836 | 0.008542 |
| 26.9722222 | 0.020676 | 0.016191 | 0.036949 | 0.016312 | 0.0228 | 0.018252 | 0.023232 | 0.018812 | 0.027555 | 0.001258 | 0.034396 | 0.016685 | 0.061471 | 0.023575 | 0.006539 | 0.01776 | 0.067081 | 0.014011 | 0.015215 | 0.006255 | 0.017902 | 0.023471 | 0.026631 | 0.007876 |
| 26.9861111 | 0.023741 | 0.025818 | 0.046626 | 0.010486 | 0.022968 | 0.022741 | 0.024161 | 0.022481 | 0.032692 | 0.00561 | 0.039555 | 0.017819 | 0.072666 | 0.013925 | 0.007662 | 0.018641 | 0.070585 | 0.021653 | 0.018208 | 0.00611 | 0.030583 | 0.021863 | 0.030407 | 0.005178 |
| 27 | 0.026522 | 0.03123 | 0.049403 | 0.01127 | 0.025389 | 0.027996 | 0.025916 | 0.026974 | 0.037236 | 0.012856 | 0.043321 | 0.019837 | 0.080323 | 0.00619 | 0.01013 | 0.02178 | 0.075017 | 0.030332 | 0.022241 | 0.00767 | 0.034872 | 0.027943 | 0.034395 | 0.003205 |
| 27.0138889 | 0.028289 | 0.029722 | 0.050033 | 0.017798 | 0.032739 | 0.030749 | 0.025924 | 0.027711 | 0.03914 | 0.018872 | 0.042556 | 0.022528 | 0.087279 | 0.004282 | 0.013605 | 0.02488 | 0.069561 | 0.03452 | 0.022364 | 0.006368 | 0.031335 | 0.03619 | 0.03226 | 0.002546 |
| 27.0277778 | 0.030153 | 0.026308 | 0.054406 | 0.023557 | 0.045337 | 0.029184 | 0.023751 | 0.026143 | 0.038965 | 0.022858 | 0.040844 | 0.02463 | 0.091363 | 0.006587 | 0.019478 | 0.024915 | 0.063784 | 0.037621 | 0.019239 | 0.002958 | 0.029217 | 0.041373 | 0.02513 | 0.002985 |
| 27.0416667 | 0.036477 | 0.028716 | 0.060322 | 0.026752 | 0.06638 | 0.02738 | 0.02653 | 0.028447 | 0.042584 | 0.029977 | 0.046269 | 0.026609 | 0.099528 | 0.019398 | 0.026933 | 0.023278 | 0.078598 | 0.043232 | 0.021306 | 0.002172 | 0.036469 | 0.043638 | 0.028044 | 0.009065 |
| 27.0555556 | 0.055539 | 0.042578 | 0.065659 | 0.030536 | 0.100556 | 0.031668 | 0.042232 | 0.041636 | 0.055023 | 0.045266 | 0.062312 | 0.028347 | 0.107274 | 0.042051 | 0.031317 | 0.024004 | 0.107747 | 0.042129 | 0.029356 | 0.003819 | 0.0546 | 0.042243 | 0.047341 | 0.018972 |
| 27.0694444 | 0.076078 | 0.064708 | 0.072492 | 0.036406 | 0.133645 | 0.041318 | 0.063879 | 0.055786 | 0.066387 | 0.059654 | 0.080354 | 0.027417 | 0.113704 | 0.062213 | 0.033404 | 0.027496 | 0.117334 | 0.037467 | 0.034103 | 0.003559 | 0.074054 | 0.033892 | 0.069897 | 0.026785 |
| 27.0833333 | 0.06986 | 0.077108 | 0.073112 | 0.044648 | 0.136059 | 0.048566 | 0.069411 | 0.050214 | 0.058172 | 0.057906 | 0.083489 | 0.022886 | 0.117382 | 0.074066 | 0.034605 | 0.028087 | 0.089867 | 0.035889 | 0.030958 | 0.002493 | 0.075355 | 0.018357 | 0.083247 | 0.035411 |
| 27.0972222 | 0.041302 | 0.062969 | 0.054936 | 0.049844 | 0.102099 | 0.045096 | 0.052099 | 0.030013 | 0.034999 | 0.043311 | 0.065087 | 0.01513 | 0.101347 | 0.074536 | 0.031329 | 0.022386 | 0.04677 | 0.028806 | 0.023052 | 0.003643 | 0.051984 | 0.006587 | 0.076168 | 0.040005 |
| 27.1111111 | 0.01904 | 0.032149 | 0.026783 | 0.041551 | 0.054486 | 0.031037 | 0.030014 | 0.014388 | 0.016 | 0.029064 | 0.03694 | 0.009184 | 0.065918 | 0.059937 | 0.025624 | 0.016483 | 0.016532 | 0.015861 | 0.015984 | 0.004108 | 0.022577 | 0.007171 | 0.052577 | 0.034243 |
| 27.125 | 0.01387 | 0.011423 | 0.008586 | 0.022574 | 0.019006 | 0.01673 | 0.015006 | 0.007338 | 0.006956 | 0.018122 | 0.015519 | 0.006763 | 0.029255 | 0.038447 | 0.023433 | 0.016772 | 0.00783 | 0.006095 | 0.010137 | 0.002379 | 0.007352 | 0.013576 | 0.033649 | 0.024062 |
| 27.1388889 | 0.0165 | 0.006136 | 0.006687 | 0.009416 | 0.009533 | 0.007947 | 0.006794 | 0.005842 | 0.007272 | 0.010615 | 0.009889 | 0.004445 | 0.007771 | 0.021007 | 0.018685 | 0.020479 | 0.013408 | 0.003168 | 0.003982 | 0.002015 | 0.004203 | 0.013306 | 0.028378 | 0.013755 |
| 27.1527778 | 0.013145 | 0.005909 | 0.012263 | 0.007584 | 0.02082 | 0.00486 | 0.00649 | 0.009094 | 0.013429 | 0.011722 | 0.016725 | 0.004352 | 0.003383 | 0.009112 | 0.012953 | 0.019626 | 0.021721 | 0.007955 | 0.002103 | 0.003952 | 0.00271 | 0.009415 | 0.029689 | 0.006382 |

| | | | | | | | | | | | | | | | | | | | | | | | | |
|---|---|---|---|---|---|---|---|---|---|---|---|---|---|---|---|---|---|---|---|---|---|---|---|---|
| 27.1666667 | 0.007843 | 0.00992 | 0.016825 | 0.011041 | 0.039281 | 0.008807 | 0.011653 | 0.011847 | 0.018787 | 0.019566 | 0.026956 | 0.008324 | 0.009114 | 0.003185 | 0.015562 | 0.013026 | 0.031642 | 0.015753 | 0.004571 | 0.004571 | 0.001946 | 0.008726 | 0.026378 | 0.005831 |
| 27.1805556 | 0.011862 | 0.018196 | 0.02001 | 0.018377 | 0.049959 | 0.014489 | 0.014882 | 0.014918 | 0.022731 | 0.024416 | 0.038596 | 0.01384 | 0.024591 | 0.006073 | 0.021637 | 0.010296 | 0.043125 | 0.019448 | 0.006596 | 0.004382 | 0.003927 | 0.00911 | 0.015485 | 0.011097 |
| 27.1944444 | 0.018469 | 0.024245 | 0.022051 | 0.02032 | 0.047895 | 0.014707 | 0.013468 | 0.023098 | 0.028813 | 0.022606 | 0.047915 | 0.017178 | 0.037459 | 0.015389 | 0.029127 | 0.015949 | 0.048115 | 0.021797 | 0.014915 | 0.007833 | 0.011448 | 0.012789 | 0.009737 | 0.019936 |
| 27.2083333 | 0.01967 | 0.023796 | 0.021149 | 0.01055 | 0.042387 | 0.010525 | 0.010401 | 0.029625 | 0.035814 | 0.019246 | 0.045521 | 0.016226 | 0.038243 | 0.017896 | 0.034362 | 0.020689 | 0.040049 | 0.027309 | 0.035207 | 0.011491 | 0.020908 | 0.01577 | 0.019535 | 0.02516 |
| 27.2222222 | 0.017022 | 0.018954 | 0.017844 | 0.003338 | 0.039262 | 0.005689 | 0.00747 | 0.029327 | 0.038301 | 0.018563 | 0.032198 | 0.01268 | 0.039014 | 0.013692 | 0.033146 | 0.017169 | 0.023171 | 0.030785 | 0.043929 | 0.011343 | 0.024618 | 0.016343 | 0.031833 | 0.01857 |
| 27.2361111 | 0.014017 | 0.013593 | 0.015093 | 0.006057 | 0.037029 | 0.002405 | 0.004141 | 0.027019 | 0.03521 | 0.017816 | 0.018652 | 0.009492 | 0.041631 | 0.018414 | 0.032248 | 0.011544 | 0.008878 | 0.024736 | 0.027225 | 0.010485 | 0.027245 | 0.017503 | 0.031624 | 0.007881 |
| 27.25 | 0.012853 | 0.009525 | 0.015546 | 0.011308 | 0.036488 | 0.001259 | 0.001385 | 0.027316 | 0.029946 | 0.013972 | 0.011373 | 0.007657 | 0.036072 | 0.027828 | 0.029623 | 0.011297 | 0.005945 | 0.014476 | 0.014961 | 0.010847 | 0.03327 | 0.014228 | 0.023347 | 0.003174 |
| 27.2638889 | 0.011419 | 0.007748 | 0.019732 | 0.01213 | 0.039768 | 0.001691 | 0.001138 | 0.029765 | 0.023712 | 0.008924 | 0.009895 | 0.006888 | 0.032059 | 0.029228 | 0.018871 | 0.013988 | 0.015685 | 0.012096 | 0.023405 | 0.011194 | 0.032918 | 0.006461 | 0.023552 | 0.002906 |
| 27.2777778 | 0.007899 | 0.009405 | 0.025062 | 0.008744 | 0.043129 | 0.003273 | 0.003111 | 0.032736 | 0.016 | 0.005378 | 0.011709 | 0.00705 | 0.039205 | 0.02079 | 0.006799 | 0.013543 | 0.025073 | 0.013948 | 0.028718 | 0.010576 | 0.023778 | 0.002862 | 0.035795 | 0.006196 |
| 27.2916667 | 0.004301 | 0.013802 | 0.027799 | 0.008382 | 0.04486 | 0.00411 | 0.004932 | 0.033566 | 0.008739 | 0.003162 | 0.014182 | 0.007686 | 0.046096 | 0.010419 | 0.001234 | 0.009578 | 0.019067 | 0.0132 | 0.020553 | 0.009402 | 0.017896 | 0.005523 | 0.039137 | 0.01114 |
| 27.3055556 | 0.003707 | 0.02013 | 0.028089 | 0.01233 | 0.047063 | 0.004208 | 0.005246 | 0.028815 | 0.004108 | 0.001445 | 0.016124 | 0.008432 | 0.044799 | 0.008952 | 0.001358 | 0.007953 | 0.011372 | 0.013964 | 0.018202 | 0.009611 | 0.017721 | 0.006632 | 0.03122 | 0.014218 |
| 27.3194444 | 0.007019 | 0.024741 | 0.03004 | 0.012657 | 0.048301 | 0.00628 | 0.005741 | 0.021205 | 0.002321 | 0.001075 | 0.017714 | 0.009801 | 0.04158 | 0.014948 | 0.002461 | 0.010221 | 0.015798 | 0.011883 | 0.018286 | 0.011512 | 0.015334 | 0.003958 | 0.036048 | 0.016308 |
| 27.3333333 | 0.011529 | 0.021317 | 0.033946 | 0.007517 | 0.046414 | 0.01031 | 0.008659 | 0.015741 | 0.003117 | 0.002712 | 0.01959 | 0.012642 | 0.041289 | 0.021988 | 0.005136 | 0.010636 | 0.017172 | 0.004451 | 0.009477 | 0.012136 | 0.009949 | 0.003741 | 0.052686 | 0.018984 |
| 27.3472222 | 0.014866 | 0.013762 | 0.037235 | 0.004669 | 0.043537 | 0.014318 | 0.013686 | 0.012404 | 0.005915 | 0.005195 | 0.023253 | 0.016488 | 0.03788 | 0.027766 | 0.013255 | 0.007295 | 0.016132 | 0.005357 | 0.007321 | 0.010722 | 0.008904 | 0.004647 | 0.060051 | 0.022851 |
| 27.3611111 | 0.017818 | 0.011164 | 0.039292 | 0.007597 | 0.042694 | 0.017478 | 0.017779 | 0.009773 | 0.009255 | 0.006455 | 0.028198 | 0.019517 | 0.031723 | 0.027733 | 0.01667 | 0.005246 | 0.021362 | 0.01644 | 0.017649 | 0.010536 | 0.015293 | 0.005779 | 0.053155 | 0.023878 |
| 27.375 | 0.020947 | 0.01286 | 0.04173 | 0.01201 | 0.042763 | 0.01927 | 0.018746 | 0.007836 | 0.011518 | 0.005638 | 0.031796 | 0.019946 | 0.033023 | 0.020366 | 0.010678 | 0.005477 | 0.020859 | 0.023217 | 0.027832 | 0.012796 | 0.022752 | 0.010577 | 0.042381 | 0.018234 |
| 27.3888889 | 0.023635 | 0.014328 | 0.046253 | 0.014989 | 0.040612 | 0.018498 | 0.017747 | 0.007268 | 0.01387 | 0.004929 | 0.034139 | 0.018034 | 0.042382 | 0.013006 | 0.007958 | 0.005063 | 0.011156 | 0.021263 | 0.029325 | 0.013304 | 0.022805 | 0.01096 | 0.04197 | 0.012384 |
| 27.4027778 | 0.026459 | 0.015193 | 0.049847 | 0.01795 | 0.035635 | 0.015159 | 0.017623 | 0.009379 | 0.017939 | 0.006576 | 0.036189 | 0.016731 | 0.053296 | 0.00985 | 0.012099 | 0.005504 | 0.006895 | 0.018963 | 0.026276 | 0.009668 | 0.017757 | 0.006101 | 0.04995 | 0.011011 |
| 27.4166667 | 0.029928 | 0.016148 | 0.048706 | 0.021053 | 0.030353 | 0.011223 | 0.019683 | 0.014077 | 0.022715 | 0.009917 | 0.037056 | 0.017974 | 0.064624 | 0.007196 | 0.018964 | 0.007066 | 0.00902 | 0.023338 | 0.027869 | 0.005208 | 0.017794 | 0.003208 | 0.05193 | 0.009271 |
| 27.4305556 | 0.031203 | 0.016918 | 0.044913 | 0.020835 | 0.025537 | 0.009329 | 0.022024 | 0.017181 | 0.028049 | 0.014281 | 0.03782 | 0.020449 | 0.071886 | 0.003962 | 0.026347 | 0.0069 | 0.010636 | 0.030781 | 0.031505 | 0.003133 | 0.02379 | 0.001556 | 0.050588 | 0.005889 |
| 27.4444444 | 0.029106 | 0.017282 | 0.042935 | 0.019311 | 0.019346 | 0.012123 | 0.023419 | 0.015586 | 0.033333 | 0.019034 | 0.040066 | 0.021946 | 0.064438 | 0.006472 | 0.032607 | 0.007373 | 0.021013 | 0.032584 | 0.026206 | 0.004173 | 0.030001 | 0.002468 | 0.052887 | 0.006159 |
| 27.4583333 | 0.02669 | 0.018404 | 0.04218 | 0.024247 | 0.013093 | 0.020058 | 0.025351 | 0.01369 | 0.036274 | 0.022322 | 0.042944 | 0.022497 | 0.047597 | 0.019145 | 0.039243 | 0.011498 | 0.036313 | 0.030857 | 0.016809 | 0.006211 | 0.035855 | 0.006479 | 0.055338 | 0.01232 |
| 27.4722222 | 0.025392 | 0.020315 | 0.039321 | 0.032787 | 0.010338 | 0.028645 | 0.029099 | 0.018825 | 0.036631 | 0.023932 | 0.044851 | 0.023595 | 0.039329 | 0.035762 | 0.046149 | 0.014261 | 0.043375 | 0.033282 | 0.015882 | 0.007443 | 0.043797 | 0.009903 | 0.055567 | 0.02002 |
| 27.4861111 | 0.026541 | 0.020677 | 0.035578 | 0.032747 | 0.012002 | 0.030926 | 0.032755 | 0.029704 | 0.03675 | 0.026276 | 0.045313 | 0.025594 | 0.042417 | 0.048492 | 0.050067 | 0.011797 | 0.042966 | 0.037796 | 0.019154 | 0.010031 | 0.050609 | 0.012364 | 0.053695 | 0.025943 |
| 27.5 | 0.02967 | 0.017818 | 0.034202 | 0.022246 | 0.016269 | 0.02621 | 0.03258 | 0.037829 | 0.037772 | 0.030135 | 0.045008 | 0.028483 | 0.048817 | 0.056418 | 0.050596 | 0.007698 | 0.035605 | 0.040769 | 0.021528 | 0.014563 | 0.050054 | 0.013726 | 0.053435 | 0.031809 |
| 27.5138889 | 0.030514 | 0.01341 | 0.035166 | 0.011939 | 0.021374 | 0.021084 | 0.02846 | 0.038661 | 0.03764 | 0.031325 | 0.04515 | 0.030995 | 0.056253 | 0.05777 | 0.050613 | 0.005545 | 0.02281 | 0.039341 | 0.023349 | 0.018685 | 0.044281 | 0.013697 | 0.061931 | 0.036014 |
| 27.5277778 | 0.02885 | 0.011867 | 0.034702 | 0.009912 | 0.025964 | 0.019675 | 0.023897 | 0.033744 | 0.034875 | 0.026718 | 0.047366 | 0.031065 | 0.058444 | 0.050215 | 0.050297 | 0.005537 | 0.012342 | 0.028867 | 0.020554 | 0.0224 | 0.041017 | 0.015361 | 0.073599 | 0.033856 |
| 27.5416667 | 0.029782 | 0.014297 | 0.029499 | 0.012946 | 0.028592 | 0.023924 | 0.022499 | 0.02884 | 0.031918 | 0.021201 | 0.053143 | 0.031375 | 0.050006 | 0.038847 | 0.043955 | 0.005742 | 0.00666 | 0.017341 | 0.014889 | 0.02488 | 0.040664 | 0.016992 | 0.070318 | 0.027297 |
| 27.5555556 | 0.03498 | 0.016158 | 0.020514 | 0.013407 | 0.028586 | 0.0355 | 0.023239 | 0.026526 | 0.033009 | 0.021743 | 0.05836 | 0.036317 | 0.039184 | 0.029395 | 0.031324 | 0.005108 | 0.005361 | 0.017866 | 0.011594 | 0.022962 | 0.040371 | 0.014783 | 0.056642 | 0.021235 |
| 27.5694444 | 0.039066 | 0.013093 | 0.012895 | 0.011404 | 0.027418 | 0.050112 | 0.021476 | 0.023856 | 0.037278 | 0.028239 | 0.055524 | 0.042593 | 0.03721 | 0.021713 | 0.021449 | 0.005194 | 0.00995 | 0.028706 | 0.010208 | 0.019556 | 0.041327 | 0.01233 | 0.048856 | 0.015357 |
| 27.5833333 | 0.040892 | 0.006688 | 0.013723 | 0.012074 | 0.02958 | 0.057859 | 0.018429 | 0.02073 | 0.039801 | 0.035085 | 0.049845 | 0.045545 | 0.049188 | 0.015079 | 0.01706 | 0.00839 | 0.02096 | 0.036987 | 0.008486 | 0.02313 | 0.040963 | 0.014703 | 0.044556 | 0.008591 |
| 27.5972222 | 0.042771 | 0.003869 | 0.023365 | 0.013651 | 0.034634 | 0.054585 | 0.019222 | 0.020117 | 0.039733 | 0.037464 | 0.048443 | 0.045837 | 0.06062 | 0.009015 | 0.015448 | 0.015008 | 0.041635 | 0.034288 | 0.008146 | 0.035782 | 0.035513 | 0.022852 | 0.042959 | 0.006327 |
| 27.6111111 | 0.040023 | 0.006884 | 0.031313 | 0.010658 | 0.03406 | 0.046884 | 0.020542 | 0.02255 | 0.038439 | 0.035644 | 0.046007 | 0.044756 | 0.050072 | 0.006711 | 0.015633 | 0.020548 | 0.06066 | 0.024598 | 0.011304 | 0.053359 | 0.0312 | 0.029951 | 0.040662 | 0.010439 |
| 27.625 | 0.031258 | 0.00888 | 0.028572 | 0.007682 | 0.024304 | 0.039834 | 0.015759 | 0.023136 | 0.03713 | 0.033684 | 0.037903 | 0.04189 | 0.031993 | 0.007894 | 0.014725 | 0.02081 | 0.058265 | 0.017629 | 0.014852 | 0.071054 | 0.036254 | 0.028107 | 0.031114 | 0.010992 |
| 27.6388889 | 0.02403 | 0.006968 | 0.023321 | 0.009073 | 0.014144 | 0.033928 | 0.009488 | 0.018717 | 0.035494 | 0.031586 | 0.028818 | 0.036574 | 0.029131 | 0.01258 | 0.013934 | 0.02072 | 0.040087 | 0.018327 | 0.016201 | 0.076488 | 0.045642 | 0.021368 | 0.024862 | 0.010128 |
| 27.6527778 | 0.023259 | 0.004955 | 0.027397 | 0.007622 | 0.011982 | 0.029246 | 0.007194 | 0.01362 | 0.029129 | 0.025925 | 0.026867 | 0.032589 | 0.029565 | 0.0273 | 0.015102 | 0.026878 | 0.025645 | 0.022248 | 0.0181 | 0.063284 | 0.048509 | 0.016288 | 0.027277 | 0.019163 |
| 27.6666667 | 0.025259 | 0.00478 | 0.031229 | 0.003299 | 0.014626 | 0.023663 | 0.005608 | 0.011323 | 0.017557 | 0.018506 | 0.028952 | 0.030164 | 0.021786 | 0.041026 | 0.012989 | 0.032325 | 0.02575 | 0.02345 | 0.023297 | 0.051428 | 0.042467 | 0.012281 | 0.029061 | 0.030047 |
| 27.6805556 | 0.024433 | 0.006026 | 0.021272 | 0.003483 | 0.014693 | 0.015881 | 0.003298 | 0.010568 | 0.007483 | 0.013237 | 0.025834 | 0.025036 | 0.012534 | 0.041178 | 0.006679 | 0.027539 | 0.033359 | 0.021956 | 0.029089 | 0.057081 | 0.033191 | 0.009438 | 0.028083 | 0.033028 |
| 27.6944444 | 0.020144 | 0.007686 | 0.0089 | 0.00752 | 0.010564 | 0.007918 | 0.00245 | 0.008019 | 0.002799 | 0.009958 | 0.018783 | 0.019023 | 0.006917 | 0.028314 | 0.002931 | 0.016845 | 0.032912 | 0.015723 | 0.030065 | 0.06524 | 0.024511 | 0.009316 | 0.028593 | 0.029996 |
| 27.7083333 | 0.013899 | 0.00906 | 0.005202 | 0.009215 | 0.006468 | 0.002574 | 0.003993 | 0.004525 | 0.001137 | 0.007388 | 0.011637 | 0.012076 | 0.004394 | 0.016652 | 0.003666 | 0.008202 | 0.02497 | 0.009333 | 0.026509 | 0.05787 | 0.017096 | 0.009503 | 0.027724 | 0.022355 |
| 27.7222222 | 0.007452 | 0.008161 | 0.004134 | 0.006043 | 0.005576 | 0.000716 | 0.004998 | 0.004302 | 0.001403 | 0.005175 | 0.00485 | 0.007598 | 0.004248 | 0.02412 | 0.003616 | 0.003916 | 0.02222 | 0.011138 | 0.021609 | 0.039254 | 0.012517 | 0.00656 | 0.023999 | 0.011447 |
| 27.7361111 | 0.00328 | 0.004198 | 0.003137 | 0.002389 | 0.006196 | 0.000545 | 0.003747 | 0.005362 | 0.004643 | 0.002945 | 0.002656 | 0.012878 | 0.004596 | 0.037112 | 0.002567 | 0.002506 | 0.028587 | 0.013155 | 0.015254 | 0.020364 | 0.009681 | 0.00402 | 0.021811 | 0.003687 |
| 27.75 | 0.001506 | 0.001275 | 0.003444 | 0.002224 | 0.005684 | 0.000503 | 0.002774 | 0.004891 | 0.008089 | 0.001016 | 0.005209 | 0.020194 | 0.006698 | 0.038489 | 0.003821 | 0.004036 | 0.035302 | 0.008716 | 0.009385 | 0.011418 | 0.010761 | 0.005506 | 0.017432 | 0.002544 |
| 27.7638889 | 0.000635 | 0.000939 | 0.006354 | 0.00356 | 0.003841 | 0.000719 | 0.002362 | 0.005364 | 0.006998 | 0.000392 | 0.00513 | 0.020123 | 0.008948 | 0.033886 | 0.004821 | 0.007086 | 0.035425 | 0.005503 | 0.0066 | 0.021839 | 0.015764 | 0.006614 | 0.009461 | 0.003901 |
| 27.7777778 | 0.000696 | 0.000817 | 0.008421 | 0.004051 | 0.001852 | 0.001596 | 0.002302 | 0.006952 | 0.004257 | 0.000708 | 0.003915 | 0.014613 | 0.005916 | 0.030735 | 0.003406 | 0.005913 | 0.028202 | 0.005958 | 0.006049 | 0.040401 | 0.017283 | 0.005917 | 0.005729 | 0.005972 |
| 27.7916667 | 0.002005 | 0.000637 | 0.006717 | 0.006846 | 0.000867 | 0.003175 | 0.002428 | 0.006283 | 0.005994 | 0.001344 | 0.007382 | 0.007529 | 0.003798 | 0.02928 | 0.001839 | 0.00241 | 0.018383 | 0.005912 | 0.006541 | 0.049552 | 0.016611 | 0.005357 | 0.00569 | 0.007939 |
| 27.8055556 | 0.002518 | 0.001116 | 0.005298 | 0.012064 | 0.00076 | 0.004737 | 0.002448 | 0.003609 | 0.009471 | 0.002907 | 0.010773 | 0.002102 | 0.003917 | 0.025373 | 0.001544 | 0.000905 | 0.016515 | 0.006907 | 0.007647 | 0.04665 | 0.017517 | 0.005385 | 0.004253 | 0.006912 |
| 27.8194444 | 0.003212 | 0.002556 | 0.007423 | 0.011986 | 0.000868 | 0.005566 | 0.002272 | 0.001753 | 0.010271 | 0.003789 | 0.010019 | 0.000415 | 0.002555 | 0.018119 | 0.001288 | 0.000568 | 0.023459 | 0.007516 | 0.009114 | 0.037456 | 0.019017 | 0.006438 | 0.001971 | 0.00328 |
| 27.8333333 | 0.00649 | 0.005067 | 0.014039 | 0.007131 | 0.001154 | 0.005312 | 0.00213 | 0.002859 | 0.008477 | 0.00262 | 0.007881 | 0.001325 | 0.002975 | 0.014575 | 0.000868 | 0.0008 | 0.026077 | 0.007121 | 0.010258 | 0.028606 | 0.021033 | 0.007214 | 0.001823 | 0.000923 |
| 27.8472222 | 0.006718 | 0.009461 | 0.021343 | 0.006485 | 0.000941 | 0.003488 | 0.003985 | 0.007051 | 0.005185 | 0.00163 | 0.005027 | 0.002625 | 0.005015 | 0.02389 | 0.00177 | 0.000843 | 0.018804 | 0.011155 | 0.008308 | 0.024058 | 0.021087 | 0.007148 | 0.002614 | 0.003092 |
| 27.8611111 | 0.005084 | 0.014761 | 0.024116 | 0.008311 | 0.000906 | 0.002176 | 0.006731 | 0.009897 | 0.002045 | 0.00131 | 0.002264 | 0.002757 | 0.004796 | 0.03814 | 0.002965 | 0.001127 | 0.01075 | 0.017455 | 0.004083 | 0.026587 | 0.017367 | 0.008346 | 0.004349 | 0.006625 |
| 27.875 | 0.009451 | 0.017008 | 0.021877 | 0.006148 | 0.002781 | 0.004271 | 0.007979 | 0.008328 | 0.000776 | 0.000948 | 0.0023 | 0.002369 | 0.004519 | 0.042434 | 0.002408 | 0.002353 | 0.009765 | 0.021114 | 0.002509 | 0.034938 | 0.011187 | 0.011074 | 0.010968 | 0.00578 |
| 27.8888889 | 0.012274 | 0.015284 | 0.01686 | 0.003985 | 0.005038 | 0.007592 | 0.007767 | 0.005141 | 0.00092 | 0.002117 | 0.004078 | 0.004398 | 0.012546 | 0.039861 | 0.002539 | 0.004041 | 0.013301 | 0.026983 | 0.004487 | 0.043124 | 0.006142 | 0.012533 | 0.018657 | 0.003477 |
| 27.9027778 | 0.007481 | 0.013349 | 0.013363 | 0.007981 | 0.004336 | 0.008728 | 0.00717 | 0.004515 | 0.001473 | 0.006617 | 0.005893 | 0.009378 | 0.028834 | 0.036962 | 0.005321 | 0.007338 | 0.013652 | 0.039325 | 0.010578 | 0.048231 | 0.007006 | 0.012926 | 0.022437 | 0.004218 |
| 27.9166667 | 0.004029 | 0.015432 | 0.015959 | 0.016046 | 0.00232 | 0.008187 | 0.006885 | 0.007442 | 0.00352 | 0.012903 | 0.008081 | 0.013255 | 0.041339 | 0.034272 | 0.008476 | 0.012062 | 0.010967 | 0.052591 | 0.021556 | 0.056864 | 0.014812 | 0.016256 | 0.024578 | 0.005396 |
| 27.9305556 | 0.00968 | 0.021245 | 0.025182 | 0.020575 | 0.003463 | 0.009096 | 0.007354 | 0.01167 | 0.007032 | 0.016597 | 0.011852 | 0.014018 | 0.04459 | 0.030881 | 0.011787 | 0.015931 | 0.009589 | 0.060234 | 0.03224 | 0.071684 | 0.024875 | 0.022897 | 0.026009 | 0.003869 |
| 27.9444444 | 0.022433 | 0.025453 | 0.038526 | 0.01794 | 0.005984 | 0.013676 | 0.009115 | 0.015813 | 0.011286 | 0.016959 | 0.018113 | 0.015295 | 0.044645 | 0.026501 | 0.014842 | 0.018302 | 0.007342 | 0.060302 | 0.035117 | 0.080176 | 0.029811 | 0.028391 | 0.025348 | 0.001266 |
| 27.9583333 | 0.031609 | 0.026049 | 0.050889 | 0.014985 | 0.009623 | 0.020401 | 0.011143 | 0.01996 | 0.015974 | 0.018332 | 0.026979 | 0.019923 | 0.048802 | 0.024942 | 0.016053 | 0.02043 | 0.008514 | 0.056989 | 0.032925 | 0.075666 | 0.030105 | 0.030695 | 0.025099 | 0.000422 |
| 27.9722222 | 0.033132 | 0.024916 | 0.056348 | 0.0202 | 0.020418 | 0.025808 | 0.012652 | 0.024851 | 0.019569 | 0.023572 | 0.038096 | 0.027512 | 0.06048 | 0.028414 | 0.017632 | 0.022408 | 0.022097 | 0.057316 | 0.036071 | 0.064552 | 0.033018 | 0.032043 | 0.026518 | 0.00093 |
| 27.9861111 | 0.032038 | 0.022318 | 0.056339 | 0.030941 | 0.03537 | 0.029251 | 0.01541 | 0.03249 | 0.022359 | 0.030258 | 0.046191 | 0.03438 | 0.077564 | 0.032103 | 0.019269 | 0.023016 | 0.03955 | 0.061141 | 0.044036 | 0.053883 | 0.038614 | 0.034394 | 0.031473 | 0.001203 |
| 28 | 0.031358 | 0.021396 | 0.060135 | 0.034788 | 0.046435 | 0.0326 | 0.018499 | 0.039554 | 0.026353 | 0.034731 | 0.047354 | 0.035743 | 0.090288 | 0.034269 | 0.018603 | 0.023394 | 0.048733 | 0.066793 | 0.050971 | 0.046444 | 0.04072 | 0.036337 | 0.03903 | 0.002342 |
| 28.0138889 | 0.032203 | 0.025817 | 0.07087 | 0.026338 | 0.058651 | 0.034476 | 0.018647 | 0.041376 | 0.028952 | 0.033777 | 0.04599 | 0.031899 | 0.085428 | 0.042604 | 0.019552 | 0.024718 | 0.051967 | 0.075694 | 0.056611 | 0.039923 | 0.037122 | 0.033783 | 0.037958 | 0.007796 |
| 28.0277778 | 0.033769 | 0.030453 | 0.077941 | 0.016749 | 0.072876 | 0.032459 | 0.014835 | 0.040303 | 0.025811 | 0.026428 | 0.046181 | 0.026875 | 0.071867 | 0.058746 | 0.024087 | 0.024198 | 0.057048 | 0.081449 | 0.060801 | 0.033805 | 0.028193 | 0.02696 | 0.027656 | 0.015553 |
| 28.0416667 | 0.038096 | 0.031711 | 0.076584 | 0.018566 | 0.077279 | 0.032609 | 0.010936 | 0.041195 | 0.022522 | 0.021231 | 0.050767 | 0.025979 | 0.078848 | 0.073193 | 0.030346 | 0.021017 | 0.070706 | 0.077593 | 0.066958 | 0.029656 | 0.01518 | 0.025825 | 0.024105 | 0.019567 |
| 28.0555556 | 0.050107 | 0.036727 | 0.080887 | 0.031254 | 0.077828 | 0.046228 | 0.015832 | 0.049483 | 0.029226 | 0.028427 | 0.061961 | 0.032647 | 0.115379 | 0.08489 | 0.038043 | 0.018168 | 0.090587 | 0.07197 | 0.077224 | 0.027658 | 0.004984 | 0.038743 | 0.03281 | 0.020277 |
| 28.0694444 | 0.06122 | 0.048772 | 0.094863 | 0.043706 | 0.084165 | 0.06745 | 0.026178 | 0.058873 | 0.040106 | 0.040023 | 0.074428 | 0.040888 | 0.158126 | 0.10789 | 0.041181 | 0.018245 | 0.101141 | 0.068935 | 0.081972 | 0.032634 | 0.003249 | 0.05885 | 0.040246 | 0.023232 |
| 28.0833333 | 0.057824 | 0.060116 | 0.09788 | 0.046504 | 0.087741 | 0.076524 | 0.028565 | 0.052113 | 0.041351 | 0.040961 | 0.077676 | 0.039796 | 0.160935 | 0.134149 | 0.03369 | 0.021407 | 0.08953 | 0.056872 | 0.068365 | 0.045331 | 0.008304 | 0.069315 | 0.037993 | 0.034323 |
| 28.0972222 | 0.045213 | 0.05891 | 0.077629 | 0.040664 | 0.077768 | 0.071272 | 0.021656 | 0.030142 | 0.034626 | 0.034908 | 0.072568 | 0.033109 | 0.107089 | 0.131504 | 0.020881 | 0.018716 | 0.062902 | 0.033145 | 0.039114 | 0.05436 | 0.013469 | 0.062929 | 0.028651 | 0.046027 |
| 28.1111111 | 0.033435 | 0.04107 | 0.050347 | 0.030666 | 0.052903 | 0.061579 | 0.013937 | 0.01283 | 0.027927 | 0.031397 | 0.067736 | 0.032639 | 0.044592 | 0.099385 | 0.011505 | 0.011127 | 0.035963 | 0.020279 | 0.023421 | 0.050768 | 0.012912 | 0.047178 | 0.017102 | 0.041028 |
| 28.125 | 0.024452 | 0.018032 | 0.034243 | 0.017583 | 0.028013 | 0.048684 | 0.009197 | 0.007713 | 0.021343 | 0.027989 | 0.060769 | 0.033874 | 0.016858 | 0.065576 | 0.00773 | 0.010759 | 0.015653 | 0.02564 | 0.028095 | 0.034553 | 0.010812 | 0.032197 | 0.010529 | 0.024181 |
| 28.1388889 | 0.019173 | 0.007057 | 0.025601 | 0.009458 | 0.014118 | 0.031427 | 0.007129 | 0.00737 | 0.012428 | 0.018747 | 0.048737 | 0.028176 | 0.020212 | 0.042177 | 0.006024 | 0.015089 | 0.006039 | 0.025277 | 0.024373 | 0.015052 | 0.011786 | 0.022664 | 0.012505 | 0.011051 |
| 28.1527778 | 0.015823 | 0.012509 | 0.014704 | 0.014336 | 0.010027 | 0.015323 | 0.011222 | 0.009699 | 0.005885 | 0.009171 | 0.032578 | 0.017354 | 0.030501 | 0.022616 | 0.00557 | 0.017964 | 0.007126 | 0.019929 | 0.016403 | 0.004445 | 0.012688 | 0.014187 | 0.01809 | 0.005768 |
| 28.1666667 | 0.010231 | 0.024066 | 0.006464 | 0.022259 | 0.018094 | 0.009497 | 0.018696 | 0.01341 | 0.007597 | 0.008324 | 0.017621 | 0.00794 | 0.033158 | 0.012103 | 0.010303 | 0.017068 | 0.017134 | 0.027289 | 0.025049 | 0.002913 | 0.010473 | 0.009543 | 0.01612 | 0.012942 |
| 28.1805556 | 0.007743 | 0.03272 | 0.008265 | 0.023141 | 0.03506 | 0.013337 | 0.023549 | 0.015756 | 0.013564 | 0.011879 | 0.015072 | 0.004465 | 0.025676 | 0.018387 | 0.019223 | 0.012894 | 0.028625 | 0.040102 | 0.042361 | 0.002275 | 0.005426 | 0.011394 | 0.007456 | 0.029987 |
| 28.1944444 | 0.016115 | 0.036359 | 0.014496 | 0.019334 | 0.049037 | 0.01703 | 0.026018 | 0.018026 | 0.019083 | 0.013954 | 0.022899 | 0.006066 | 0.012101 | 0.030973 | 0.027914 | 0.008271 | 0.029785 | 0.042562 | 0.049958 | 0.000893 | 0.002966 | 0.014062 | 0.004683 | 0.042288 |
| 28.2083333 | 0.025776 | 0.035194 | 0.020654 | 0.013913 | 0.052676 | 0.017831 | 0.022808 | 0.018686 | 0.021339 | 0.013316 | 0.030101 | 0.010158 | 0.003719 | 0.046682 | 0.029906 | 0.006724 | 0.024041 | 0.035335 | 0.043885 | 0.000673 | 0.00693 | 0.024864 | 0.010154 | 0.04843 |
| 28.2222222 | 0.025379 | 0.029951 | 0.026804 | 0.00876 | 0.048431 | 0.01702 | 0.013154 | 0.01885 | 0.01743 | 0.008335 | 0.033991 | 0.01434 | 0.004009 | 0.06122 | 0.023631 | 0.013756 | 0.021674 | 0.031558 | 0.039228 | 0.004031 | 0.009662 | 0.034375 | 0.018315 | 0.054342 |
| 28.2361111 | 0.017506 | 0.023702 | 0.028907 | 0.006993 | 0.041311 | 0.013152 | 0.005873 | 0.024418 | 0.0113 | 0.003024 | 0.035812 | 0.015972 | 0.006775 | 0.06329 | 0.018784 | 0.024087 | 0.02418 | 0.035357 | 0.047398 | 0.009567 | 0.006976 | 0.024372 | 0.024555 | 0.053319 |
| 28.25 | 0.009492 | 0.01919 | 0.024212 | 0.008897 | 0.033814 | 0.007461 | 0.005354 | 0.03346 | 0.007821 | 0.001437 | 0.033447 | 0.013098 | 0.010025 | 0.057938 | 0.01536 | 0.027056 | 0.029533 | 0.04154 | 0.059684 | 0.010757 | 0.006361 | 0.009347 | 0.026608 | 0.04451 |
| 28.2638889 | 0.004647 | 0.015569 | 0.015147 | 0.010425 | 0.034474 | 0.006754 | 0.005388 | 0.037466 | 0.005942 | 0.001255 | 0.023852 | 0.008326 | 0.018385 | 0.049418 | 0.009082 | 0.024199 | 0.03549 | 0.041837 | 0.059704 | 0.009368 | 0.006535 | 0.004258 | 0.024852 | 0.03425 |
| 28.2777778 | 0.002167 | 0.012921 | 0.007143 | 0.007802 | 0.051943 | 0.01305 | 0.003085 | 0.034713 | 0.005949 | 0.001459 | 0.011202 | 0.006538 | 0.028777 | 0.033007 | 0.005161 | 0.02178 | 0.037159 | 0.030489 | 0.043718 | 0.008888 | 0.004183 | 0.007326 | 0.021785 | 0.020737 |
| 28.2916667 | 0.003154 | 0.013341 | 0.005411 | 0.003342 | 0.081424 | 0.016046 | 0.003962 | 0.033298 | 0.005957 | 0.002444 | 0.007307 | 0.007762 | 0.029216 | 0.016573 | 0.006633 | 0.018274 | 0.026098 | 0.016921 | 0.02582 | 0.009069 | 0.00544 | 0.013685 | 0.015982 | 0.008076 |
| 28.3055556 | 0.008373 | 0.014184 | 0.010894 | 0.001933 | 0.100843 | 0.010692 | 0.010239 | 0.036059 | 0.004972 | 0.004567 | 0.015303 | 0.011991 | 0.020604 | 0.008983 | 0.008029 | 0.011001 | 0.010122 | 0.01125 | 0.015342 | 0.009863 | 0.010846 | 0.017412 | 0.010398 | 0.002591 |
| 28.3194444 | 0.013094 | 0.011881 | 0.016585 | 0.002118 | 0.097934 | 0.004756 | 0.016462 | 0.037192 | 0.005486 | 0.006909 | 0.022568 | 0.016185 | 0.015894 | 0.005569 | 0.007523 | 0.00568 | 0.004139 | 0.00917 | 0.01046 | 0.010756 | 0.012533 | 0.017511 | 0.009797 | 0.002083 |
| 28.3333333 | 0.015385 | 0.010759 | 0.017615 | 0.002719 | 0.094822 | 0.003199 | 0.017572 | 0.035593 | 0.005713 | 0.006437 | 0.021473 | 0.014794 | 0.014239 | 0.00392 | 0.008146 | 0.00704 | 0.003907 | 0.007388 | 0.012875 | 0.01151 | 0.007297 | 0.016507 | 0.008614 | 0.003436 |
| 28.3472222 | 0.019742 | 0.015855 | 0.024132 | 0.005416 | 0.10188 | 0.002837 | 0.016706 | 0.034339 | 0.00645 | 0.004312 | 0.019915 | 0.012181 | 0.016717 | 0.004626 | 0.006276 | 0.010884 | 0.005995 | 0.005803 | 0.018567 | 0.012577 | 0.00508 | 0.015808 | 0.010166 | 0.00401 |
| 28.3611111 | 0.024229 | 0.021674 | 0.040756 | 0.005915 | 0.102675 | 0.003185 | 0.018621 | 0.03441 | 0.006344 | 0.0038 | 0.022909 | 0.012424 | 0.026577 | 0.005528 | 0.00339 | 0.012102 | 0.007981 | 0.004798 | 0.020932 | 0.013938 | 0.011926 | 0.017626 | 0.012108 | 0.003519 |
| 28.375 | 0.022795 | 0.020604 | 0.05207 | 0.00514 | 0.093371 | 0.007661 | 0.02208 | 0.033156 | 0.005951 | 0.004998 | 0.025348 | 0.01321 | 0.026443 | 0.005733 | 0.002327 | 0.011399 | 0.007337 | 0.005393 | 0.016361 | 0.014168 | 0.018347 | 0.021606 | 0.009465 | 0.004011 |
| 28.3888889 | 0.018547 | 0.016014 | 0.052708 | 0.009511 | 0.07784 | 0.015191 | 0.023776 | 0.025969 | 0.007876 | 0.006288 | 0.024446 | 0.013618 | 0.017479 | 0.0042 | 0.004762 | 0.01345 | 0.00949 | 0.008709 | 0.009906 | 0.013008 | 0.018384 | 0.022505 | 0.008011 | 0.004596 |
| 28.4027778 | 0.016939 | 0.01438 | 0.054397 | 0.013889 | 0.056385 | 0.022204 | 0.021974 | 0.01402 | 0.007725 | 0.007773 | 0.023753 | 0.013672 | 0.021185 | 0.002137 | 0.012078 | 0.016153 | 0.010332 | 0.013927 | 0.008185 | 0.01135 | 0.016478 | 0.018255 | 0.011434 | 0.004767 |
| 28.4166667 | 0.017106 | 0.015928 | 0.062123 | 0.01273 | 0.036321 | 0.026054 | 0.018478 | 0.006315 | 0.005546 | 0.011629 | 0.025718 | 0.015701 | 0.040073 | 0.005788 | 0.017539 | 0.011917 | 0.008591 | 0.013247 | 0.005976 | 0.009242 | 0.016879 | 0.011207 | 0.024775 | 0.009767 |
| 28.4305556 | 0.017651 | 0.018159 | 0.070939 | 0.010915 | 0.028638 | 0.026556 | 0.01782 | 0.00939 | 0.00706 | 0.01837 | 0.028786 | 0.021429 | 0.061093 | 0.011853 | 0.022566 | 0.005876 | 0.013962 | 0.013716 | 0.00862 | 0.006081 | 0.022243 | 0.004642 | 0.037194 | 0.021295 |
| 28.4444444 | 0.019858 | 0.019361 | 0.070388 | 0.012913 | 0.034475 | 0.027372 | 0.021215 | 0.018721 | 0.014172 | 0.025559 | 0.031627 | 0.025979 | 0.073353 | 0.013096 | 0.034986 | 0.008559 | 0.023834 | 0.026905 | 0.020122 | 0.002652 | 0.031478 | 0.001599 | 0.040688 | 0.032547 |
| 28.4583333 | 0.023231 | 0.019979 | 0.056644 | 0.015999 | 0.038637 | 0.03148 | 0.026142 | 0.024512 | 0.022239 | 0.030771 | 0.034269 | 0.025952 | 0.077944 | 0.013083 | 0.04987 | 0.014383 | 0.032701 | 0.042012 | 0.028341 | 0.00075 | 0.036865 | 0.001499 | 0.04336 | 0.032391 |
| 28.4722222 | 0.025679 | 0.021407 | 0.041956 | 0.016415 | 0.02996 | 0.036519 | 0.030274 | 0.023907 | 0.026151 | 0.033296 | 0.037319 | 0.025009 | 0.081982 | 0.010484 | 0.059407 | 0.013839 | 0.039448 | 0.046947 | 0.025727 | 0.000312 | 0.034364 | 0.001792 | 0.04616 | 0.018778 |
| 28.4861111 | 0.027362 | 0.022065 | 0.036912 | 0.015596 | 0.019469 | 0.0375 | 0.031953 | 0.022818 | 0.025885 | 0.033501 | 0.039585 | 0.026093 | 0.084931 | 0.006033 | 0.067105 | 0.009613 | 0.039871 | 0.044963 | 0.020864 | 0.00094 | 0.029456 | 0.002652 | 0.048343 | 0.007995 |
| 28.5 | 0.028894 | 0.020144 | 0.040421 | 0.015912 | 0.017868 | 0.033674 | 0.031594 | 0.024497 | 0.025063 | 0.03215 | 0.039832 | 0.028641 | 0.085845 | 0.00561 | 0.075466 | 0.00719 | 0.034523 | 0.042482 | 0.021228 | 0.003243 | 0.027019 | 0.005125 | 0.050313 | 0.006829 |
| 28.5138889 | 0.029161 | 0.017402 | 0.045763 | 0.015818 | 0.020514 | 0.028716 | 0.030559 | 0.024653 | 0.026162 | 0.029643 | 0.041077 | 0.031766 | 0.092696 | 0.004577 | 0.080153 | 0.006549 | 0.027482 | 0.040998 | 0.021468 | 0.005997 | 0.026652 | 0.006992 | 0.048533 | 0.007062 |
| 28.5277778 | 0.028194 | 0.015421 | 0.045528 | 0.012955 | 0.018332 | 0.02488 | 0.026928 | 0.020218 | 0.027308 | 0.026053 | 0.044292 | 0.033776 | 0.107325 | 0.005224 | 0.077244 | 0.005687 | 0.020056 | 0.040237 | 0.017917 | 0.006933 | 0.030059 | 0.005965 | 0.042449 | 0.007165 |
| 28.5416667 | 0.028236 | 0.014161 | 0.041132 | 0.010052 | 0.010263 | 0.02388 | 0.019916 | 0.015178 | 0.025724 | 0.023171 | 0.047458 | 0.033793 | 0.117881 | 0.018844 | 0.066929 | 0.005731 | 0.011711 | 0.03862 | 0.012358 | 0.005565 | 0.037138 | 0.002993 | 0.041202 | 0.006998 |
| 28.5555556 | 0.028504 | 0.013416 | 0.038882 | 0.011568 | 0.00631 | 0.027023 | 0.014478 | 0.014053 | 0.022036 | 0.023305 | 0.049989 | 0.034724 | 0.110591 | 0.038419 | 0.056063 | 0.007319 | 0.006636 | 0.037561 | 0.006466 | 0.003734 | 0.040882 | 0.001925 | 0.052813 | 0.010884 |
| 28.5694444 | 0.026055 | 0.012698 | 0.036175 | 0.016283 | 0.013431 | 0.03128 | 0.014512 | 0.014658 | 0.019231 | 0.025566 | 0.050859 | 0.037571 | 0.089997 | 0.042651 | 0.051965 | 0.008432 | 0.008432 | 0.040209 | 0.002784 | 0.004126 | 0.042977 | 0.00432 | 0.062062 | 0.022112 |
| 28.5833333 | 0.023998 | 0.011161 | 0.031612 | 0.0198 | 0.025508 | 0.032496 | 0.018562 | 0.01641 | 0.020043 | 0.028696 | 0.050354 | 0.04081 | 0.081539 | 0.030827 | 0.0536 | 0.009467 | 0.011252 | 0.043795 | 0.004126 | 0.009332 | 0.05028 | 0.007176 | 0.054544 | 0.031286 |
| 28.5972222 | 0.025566 | 0.009348 | 0.030228 | 0.019497 | 0.032115 | 0.030334 | 0.022202 | 0.022173 | 0.024706 | 0.035195 | 0.048765 | 0.045962 | 0.090872 | 0.024923 | 0.053564 | 0.011528 | 0.010365 | 0.047301 | 0.008212 | 0.019956 | 0.058493 | 0.008867 | 0.052308 | 0.034165 |
| 28.6111111 | 0.026333 | 0.010231 | 0.026964 | 0.014112 | 0.0285 | 0.027355 | 0.021949 | 0.027004 | 0.030859 | 0.045866 | 0.046856 | 0.051714 | 0.087389 | 0.032739 | 0.044523 | 0.012366 | 0.012935 | 0.049486 | 0.010185 | 0.027083 | 0.063268 | 0.010138 | 0.060005 | 0.034811 |
| 28.625 | 0.022992 | 0.016832 | 0.016447 | 0.007503 | 0.019404 | 0.023473 | 0.018853 | 0.023688 | 0.034552 | 0.052647 | 0.043226 | 0.052376 | 0.056378 | 0.039739 | 0.031386 | 0.01044 | 0.024985 | 0.044474 | 0.008739 | 0.025061 | 0.065137 | 0.011794 | 0.054336 | 0.03554 |
| 28.6388889 | 0.019313 | 0.022357 | 0.007386 | 0.005056 | 0.014463 | 0.018443 | 0.016858 | 0.017338 | 0.033162 | 0.050902 | 0.032044 | 0.039967 | 0.024721 | 0.034458 | 0.027047 | 0.008 | 0.036475 | 0.034829 | 0.007607 | 0.022817 | 0.062041 | 0.013754 | 0.03936 | 0.038836 |
| 28.6527778 | 0.01939 | 0.01625 | 0.007033 | 0.003982 | 0.014504 | 0.015964 | 0.017755 | 0.016596 | 0.027203 | 0.044276 | 0.016152 | 0.0178 | 0.012232 | 0.022432 | 0.028704 | 0.006798 | 0.039717 | 0.02726 | 0.010623 | 0.023377 | 0.054236 | 0.014315 | 0.027395 | 0.045726 |
| 28.6666667 | 0.020653 | 0.006272 | 0.010462 | 0.002639 | 0.014003 | 0.01533 | 0.017287 | 0.016025 | 0.017327 | 0.032444 | 0.004651 | 0.004189 | 0.011174 | 0.017171 | 0.02391 | 0.006245 | 0.038681 | 0.021051 | 0.014801 | 0.020816 | 0.042643 | 0.012301 | 0.019569 | 0.053273 |

| | | | | | | | | | | | | | | | | | | | | | | | | |
|---|---|---|---|---|---|---|---|---|---|---|---|---|---|---|---|---|---|---|---|---|---|---|---|---|
| 28.6805556 | 0.017633 | 0.003644 | 0.011157 | 0.002878 | 0.011174 | 0.011864 | 0.011058 | 0.009503 | 0.007061 | 0.016578 | 0.002094 | 0.004322 | 0.015161 | 0.022867 | 0.011855 | 0.0055 | 0.031908 | 0.012179 | 0.014805 | 0.014454 | 0.026469 | 0.009294 | 0.015506 | 0.056084 |
| 28.6944444 | 0.009716 | 0.003884 | 0.014988 | 0.003048 | 0.007346 | 0.00641 | 0.004406 | 0.003128 | 0.001843 | 0.006383 | 0.003989 | 0.007851 | 0.02096 | 0.029438 | 0.002614 | 0.004066 | 0.020577 | 0.006107 | 0.012153 | 0.007343 | 0.011297 | 0.006654 | 0.013342 | 0.048771 |
| 28.7083333 | 0.004658 | 0.00496 | 0.022888 | 0.004621 | 0.008247 | 0.003887 | 0.003093 | 0.002661 | 0.000831 | 0.004689 | 0.006348 | 0.007399 | 0.024401 | 0.02543 | 0.001898 | 0.002312 | 0.014524 | 0.009275 | 0.01061 | 0.002599 | 0.006521 | 0.004457 | 0.010403 | 0.032889 |
| 28.7222222 | 0.006927 | 0.007451 | 0.022358 | 0.008962 | 0.014477 | 0.00484 | 0.003042 | 0.00731 | 0.000447 | 0.004866 | 0.01273 | 0.008564 | 0.025487 | 0.013277 | 0.004282 | 0.001194 | 0.011776 | 0.016296 | 0.010342 | 0.00126 | 0.009977 | 0.00483 | 0.006859 | 0.015667 |
| 28.7361111 | 0.008476 | 0.006583 | 0.012953 | 0.010299 | 0.015905 | 0.004253 | 0.002137 | 0.011342 | 0.000864 | 0.005518 | 0.021004 | 0.016827 | 0.023481 | 0.006056 | 0.005578 | 0.000786 | 0.009625 | 0.016929 | 0.009814 | 0.001552 | 0.01046 | 0.007018 | 0.00568 | 0.007936 |
| 28.75 | 0.005383 | 0.0066 | 0.005192 | 0.006126 | 0.011721 | 0.00186 | 0.002513 | 0.010634 | 0.002671 | 0.005296 | 0.023829 | 0.023446 | 0.016652 | 0.006983 | 0.008515 | 0.000638 | 0.013524 | 0.010778 | 0.007918 | 0.003815 | 0.012963 | 0.005897 | 0.005199 | 0.015894 |
| 28.7638889 | 0.002099 | 0.010872 | 0.00324 | 0.00171 | 0.011192 | 0.001339 | 0.0023 | 0.007253 | 0.003671 | 0.004186 | 0.018072 | 0.021697 | 0.008393 | 0.006448 | 0.008581 | 0.000558 | 0.018263 | 0.010909 | 0.005713 | 0.005696 | 0.022027 | 0.002646 | 0.003991 | 0.02655 |
| 28.7777778 | 0.000917 | 0.014125 | 0.005228 | 0.000712 | 0.015383 | 0.00271 | 0.001259 | 0.003785 | 0.003361 | 0.003525 | 0.00909 | 0.014802 | 0.003723 | 0.007827 | 0.005464 | 0.001011 | 0.022881 | 0.015633 | 0.004259 | 0.004856 | 0.020717 | 0.001303 | 0.003255 | 0.028362 |
| 28.7916667 | 0.002009 | 0.013269 | 0.010451 | 0.001258 | 0.017112 | 0.00457 | 0.003109 | 0.001321 | 0.00514 | 0.002297 | 0.003588 | 0.008919 | 0.001608 | 0.021565 | 0.004907 | 0.001729 | 0.034965 | 0.012015 | 0.003067 | 0.007113 | 0.013602 | 0.002022 | 0.00437 | 0.023775 |
| 28.8055556 | 0.004992 | 0.009182 | 0.016112 | 0.001507 | 0.012346 | 0.006289 | 0.007975 | 0.001104 | 0.009242 | 0.002405 | 0.00301 | 0.008233 | 0.000977 | 0.042138 | 0.003802 | 0.001487 | 0.046536 | 0.006737 | 0.001749 | 0.01289 | 0.013299 | 0.004368 | 0.008067 | 0.014557 |
| 28.8194444 | 0.008824 | 0.006738 | 0.016886 | 0.002314 | 0.007204 | 0.007835 | 0.010437 | 0.00246 | 0.01231 | 0.003231 | 0.004068 | 0.009709 | 0.002442 | 0.052559 | 0.002871 | 0.000861 | 0.044919 | 0.011375 | 0.0008 | 0.015582 | 0.012402 | 0.005631 | 0.010651 | 0.007309 |
| 28.8333333 | 0.011226 | 0.008582 | 0.012309 | 0.002618 | 0.010694 | 0.008723 | 0.00739 | 0.002552 | 0.012717 | 0.002446 | 0.004498 | 0.007327 | 0.003202 | 0.043685 | 0.005924 | 0.001164 | 0.035564 | 0.020397 | 0.000732 | 0.016204 | 0.015083 | 0.004703 | 0.010912 | 0.009717 |
| 28.8472222 | 0.010529 | 0.011608 | 0.006205 | 0.003041 | 0.017387 | 0.007334 | 0.003334 | 0.001337 | 0.010121 | 0.001745 | 0.007479 | 0.002776 | 0.003087 | 0.030725 | 0.01478 | 0.001942 | 0.031542 | 0.02954 | 0.000689 | 0.0184 | 0.018564 | 0.004492 | 0.009533 | 0.016888 |
| 28.8611111 | 0.0077 | 0.013221 | 0.002278 | 0.006156 | 0.015262 | 0.004253 | 0.00181 | 0.002334 | 0.006175 | 0.00342 | 0.009794 | 0.001308 | 0.013876 | 0.034157 | 0.024034 | 0.002851 | 0.030863 | 0.043795 | 0.000932 | 0.0207 | 0.015881 | 0.005546 | 0.008004 | 0.021343 |
| 28.875 | 0.00409 | 0.012752 | 0.002988 | 0.009381 | 0.010682 | 0.002978 | 0.002663 | 0.004078 | 0.003564 | 0.005074 | 0.007088 | 0.004569 | 0.033821 | 0.047344 | 0.025088 | 0.003277 | 0.023781 | 0.057248 | 0.002014 | 0.020679 | 0.014104 | 0.005866 | 0.013477 | 0.021359 |
| 28.8888889 | 0.00186 | 0.009609 | 0.00853 | 0.010729 | 0.017387 | 0.003966 | 0.008215 | 0.00521 | 0.002039 | 0.005094 | 0.005566 | 0.012417 | 0.040851 | 0.041507 | 0.019205 | 0.002415 | 0.014466 | 0.061652 | 0.003664 | 0.018207 | 0.011848 | 0.005253 | 0.024466 | 0.016228 |
| 28.9027778 | 0.002428 | 0.005765 | 0.015184 | 0.010682 | 0.035489 | 0.004496 | 0.01634 | 0.011103 | 0.002111 | 0.00755 | 0.009491 | 0.021044 | 0.032246 | 0.024875 | 0.01089 | 0.002106 | 0.011747 | 0.066086 | 0.008675 | 0.01544 | 0.009183 | 0.005696 | 0.031358 | 0.008436 |
| 28.9166667 | 0.003686 | 0.003054 | 0.020608 | 0.009575 | 0.055077 | 0.004114 | 0.020751 | 0.020139 | 0.006329 | 0.016362 | 0.01293 | 0.024091 | 0.019359 | 0.033706 | 0.006301 | 0.003584 | 0.011514 | 0.078991 | 0.01769 | 0.014224 | 0.011527 | 0.007099 | 0.036099 | 0.004352 |
| 28.9305556 | 0.004811 | 0.00395 | 0.026885 | 0.010194 | 0.070177 | 0.004501 | 0.02144 | 0.027455 | 0.012686 | 0.029506 | 0.014361 | 0.021591 | 0.015552 | 0.054883 | 0.00916 | 0.005909 | 0.008284 | 0.092805 | 0.027209 | 0.014216 | 0.013146 | 0.007722 | 0.045638 | 0.005618 |
| 28.9444444 | 0.006164 | 0.012605 | 0.035708 | 0.013391 | 0.083683 | 0.006599 | 0.021061 | 0.035036 | 0.017086 | 0.040214 | 0.020527 | 0.019766 | 0.033784 | 0.062561 | 0.013834 | 0.009379 | 0.004353 | 0.100563 | 0.035967 | 0.013482 | 0.014452 | 0.007235 | 0.054303 | 0.008227 |
| 28.9583333 | 0.007811 | 0.024519 | 0.047462 | 0.016785 | 0.092807 | 0.01035 | 0.022241 | 0.044259 | 0.020205 | 0.044722 | 0.035083 | 0.021455 | 0.062233 | 0.061124 | 0.017176 | 0.014222 | 0.001694 | 0.104844 | 0.045332 | 0.01124 | 0.023477 | 0.007027 | 0.054835 | 0.007558 |
| 28.9722222 | 0.011288 | 0.028858 | 0.05992 | 0.018882 | 0.099411 | 0.016301 | 0.027377 | 0.052454 | 0.025198 | 0.046576 | 0.053596 | 0.025295 | 0.078717 | 0.056631 | 0.020861 | 0.019567 | 0.001262 | 0.103941 | 0.052519 | 0.008781 | 0.031633 | 0.00864 | 0.057613 | 0.003512 |
| 28.9861111 | 0.017069 | 0.022187 | 0.065838 | 0.019135 | 0.116776 | 0.024433 | 0.032811 | 0.055562 | 0.031463 | 0.048986 | 0.065053 | 0.028993 | 0.081535 | 0.050524 | 0.023455 | 0.02401 | 0.002211 | 0.088182 | 0.051258 | 0.010262 | 0.030017 | 0.0106 | 0.062685 | 0.000635 |
| 29 | 0.026426 | 0.014665 | 0.063268 | 0.019426 | 0.135255 | 0.03252 | 0.03358 | 0.050686 | 0.035558 | 0.052473 | 0.06594 | 0.032164 | 0.078114 | 0.040299 | 0.024958 | 0.026879 | 0.005406 | 0.062466 | 0.048778 | 0.016339 | 0.023448 | 0.010794 | 0.058076 | 0.000443 |
| 29.0138889 | 0.040587 | 0.018486 | 0.059164 | 0.022042 | 0.137286 | 0.037391 | 0.029411 | 0.040438 | 0.036909 | 0.05928 | 0.064586 | 0.036657 | 0.07518 | 0.02578 | 0.026615 | 0.029066 | 0.011607 | 0.04419 | 0.059978 | 0.020648 | 0.013816 | 0.010064 | 0.052622 | 0.004451 |
| 29.0277778 | 0.054776 | 0.028208 | 0.058001 | 0.025516 | 0.126957 | 0.037965 | 0.02427 | 0.030231 | 0.037057 | 0.070269 | 0.065839 | 0.03949 | 0.084718 | 0.020797 | 0.02688 | 0.032467 | 0.017053 | 0.035731 | 0.075114 | 0.019132 | 0.00425 | 0.009567 | 0.051783 | 0.016304 |
| 29.0416667 | 0.062831 | 0.032542 | 0.058743 | 0.027305 | 0.116043 | 0.038964 | 0.024888 | 0.023478 | 0.038885 | 0.084466 | 0.071144 | 0.039752 | 0.107366 | 0.033376 | 0.027518 | 0.03889 | 0.02545 | 0.033458 | 0.081321 | 0.013149 | 0.003943 | 0.010171 | 0.056227 | 0.032019 |
| 29.0555556 | 0.068265 | 0.037738 | 0.062071 | 0.029452 | 0.114757 | 0.04629 | 0.032584 | 0.024021 | 0.04675 | 0.092981 | 0.082092 | 0.042786 | 0.134926 | 0.047107 | 0.035404 | 0.043725 | 0.04125 | 0.04449 | 0.09286 | 0.007716 | 0.014048 | 0.014183 | 0.072383 | 0.041353 |
| 29.0694444 | 0.079218 | 0.051868 | 0.068675 | 0.036304 | 0.113122 | 0.056077 | 0.0397 | 0.030378 | 0.055204 | 0.086875 | 0.088727 | 0.045022 | 0.159579 | 0.051715 | 0.050323 | 0.042799 | 0.055443 | 0.067086 | 0.114857 | 0.005864 | 0.025078 | 0.023286 | 0.085106 | 0.043586 |
| 29.0833333 | 0.081931 | 0.062664 | 0.069192 | 0.043153 | 0.092073 | 0.057997 | 0.037585 | 0.033081 | 0.05318 | 0.073232 | 0.078938 | 0.037461 | 0.16512 | 0.054862 | 0.053751 | 0.03878 | 0.055157 | 0.073097 | 0.12694 | 0.009613 | 0.026027 | 0.031713 | 0.074242 | 0.042966 |
| 29.0972222 | 0.059471 | 0.05611 | 0.057982 | 0.041428 | 0.058307 | 0.04984 | 0.026069 | 0.028783 | 0.042172 | 0.058889 | 0.058473 | 0.023353 | 0.141847 | 0.05521 | 0.03705 | 0.033799 | 0.043748 | 0.05165 | 0.109756 | 0.01963 | 0.017419 | 0.028652 | 0.048157 | 0.035128 |
| 29.1111111 | 0.028893 | 0.036026 | 0.045398 | 0.030158 | 0.027228 | 0.039041 | 0.014501 | 0.023804 | 0.030621 | 0.045016 | 0.040627 | 0.013546 | 0.10185 | 0.043572 | 0.021645 | 0.032634 | 0.037696 | 0.029855 | 0.06893 | 0.021357 | 0.00992 | 0.020222 | 0.02846 | 0.019911 |
| 29.125 | 0.010812 | 0.017498 | 0.038314 | 0.014424 | 0.009211 | 0.032163 | 0.011084 | 0.021879 | 0.023111 | 0.034445 | 0.029721 | 0.010345 | 0.067921 | 0.0256 | 0.017608 | 0.033842 | 0.040497 | 0.01984 | 0.035885 | 0.010205 | 0.006017 | 0.019584 | 0.023052 | 0.011315 |
| 29.1388889 | 0.005979 | 0.009881 | 0.032637 | 0.004459 | 0.008542 | 0.026091 | 0.012199 | 0.020052 | 0.018879 | 0.02322 | 0.023652 | 0.008886 | 0.045528 | 0.019529 | 0.012479 | 0.024733 | 0.039802 | 0.015346 | 0.03627 | 0.006255 | 0.002492 | 0.016966 | 0.022389 | 0.016934 |
| 29.1527778 | 0.013118 | 0.010454 | 0.024161 | 0.004644 | 0.017654 | 0.015393 | 0.009658 | 0.014564 | 0.014213 | 0.013018 | 0.016853 | 0.006494 | 0.02427 | 0.025515 | 0.007551 | 0.012771 | 0.029489 | 0.01263 | 0.045829 | 0.010315 | 0.003509 | 0.013632 | 0.01548 | 0.024278 |
| 29.1666667 | 0.02749 | 0.019003 | 0.012909 | 0.010077 | 0.030115 | 0.008096 | 0.007051 | 0.006831 | 0.008398 | 0.012899 | 0.010984 | 0.006408 | 0.010174 | 0.024338 | 0.010038 | 0.015418 | 0.01529 | 0.015618 | 0.041351 | 0.012248 | 0.009025 | 0.022876 | 0.007153 | 0.020343 |
| 29.1805556 | 0.038969 | 0.031431 | 0.005418 | 0.017011 | 0.042552 | 0.010293 | 0.009354 | 0.002402 | 0.003562 | 0.019087 | 0.014937 | 0.010011 | 0.012503 | 0.012704 | 0.016753 | 0.025274 | 0.007078 | 0.025769 | 0.049455 | 0.016559 | 0.011532 | 0.027484 | 0.006946 | 0.011415 |
| 29.1944444 | 0.046969 | 0.036994 | 0.006682 | 0.021455 | 0.05166 | 0.012785 | 0.010333 | 0.003307 | 0.001119 | 0.025473 | 0.022552 | 0.011928 | 0.01746 | 0.005096 | 0.029603 | 0.031732 | 0.009052 | 0.02739 | 0.061472 | 0.018459 | 0.009135 | 0.017964 | 0.012584 | 0.014361 |
| 29.2083333 | 0.053808 | 0.037856 | 0.012406 | 0.021748 | 0.058916 | 0.011607 | 0.007998 | 0.005795 | 0.001168 | 0.029499 | 0.022906 | 0.011515 | 0.014161 | 0.004371 | 0.042686 | 0.030082 | 0.01258 | 0.025194 | 0.056861 | 0.01355 | 0.005826 | 0.011378 | 0.017651 | 0.023342 |
| 29.2222222 | 0.056908 | 0.042174 | 0.018221 | 0.019661 | 0.06707 | 0.009912 | 0.006813 | 0.006536 | 0.002196 | 0.027674 | 0.019928 | 0.013342 | 0.008132 | 0.004073 | 0.04209 | 0.017355 | 0.013296 | 0.038126 | 0.065988 | 0.009091 | 0.004905 | 0.00954 | 0.020562 | 0.019049 |
| 29.2361111 | 0.053182 | 0.050114 | 0.021236 | 0.014969 | 0.074695 | 0.008467 | 0.006549 | 0.005473 | 0.002599 | 0.020834 | 0.021059 | 0.01718 | 0.006093 | 0.00684 | 0.031347 | 0.004944 | 0.020908 | 0.055401 | 0.08852 | 0.006347 | 0.007805 | 0.004866 | 0.028081 | 0.008652 |
| 29.25 | 0.04313 | 0.056979 | 0.020111 | 0.009323 | 0.078957 | 0.00883 | 0.005973 | 0.004541 | 0.004478 | 0.013648 | 0.025796 | 0.019248 | 0.007772 | 0.009736 | 0.021842 | 0.003339 | 0.038272 | 0.062392 | 0.095206 | 0.004421 | 0.012475 | 0.004579 | 0.045764 | 0.009944 |
| 29.2638889 | 0.035316 | 0.055963 | 0.020077 | 0.006877 | 0.081426 | 0.01259 | 0.007523 | 0.006818 | 0.009263 | 0.011298 | 0.02782 | 0.018065 | 0.009373 | 0.009589 | 0.014662 | 0.007005 | 0.049937 | 0.065592 | 0.097457 | 0.0035 | 0.014423 | 0.007686 | 0.059164 | 0.018989 |
| 29.2777778 | 0.032701 | 0.040647 | 0.023307 | 0.008681 | 0.083562 | 0.01599 | 0.010498 | 0.01209 | 0.012906 | 0.012394 | 0.020311 | 0.012731 | 0.012184 | 0.009791 | 0.008188 | 0.006409 | 0.047012 | 0.065824 | 0.107028 | 0.002756 | 0.015939 | 0.008268 | 0.051805 | 0.025134 |
| 29.2916667 | 0.028509 | 0.018063 | 0.024063 | 0.011681 | 0.084564 | 0.013685 | 0.010745 | 0.015954 | 0.012281 | 0.010029 | 0.009711 | 0.005839 | 0.013371 | 0.013676 | 0.00408 | 0.002566 | 0.036936 | 0.059153 | 0.108478 | 0.00368 | 0.018718 | 0.010958 | 0.036809 | 0.026491 |
| 29.3055556 | 0.021269 | 0.004099 | 0.02111 | 0.012653 | 0.0885 | 0.007939 | 0.008609 | 0.017057 | 0.009636 | 0.004629 | 0.006571 | 0.002384 | 0.00937 | 0.02013 | 0.003841 | 0.004018 | 0.026599 | 0.056335 | 0.095148 | 0.006462 | 0.015857 | 0.011499 | 0.034616 | 0.023771 |
| 29.3194444 | 0.015265 | 0.00335 | 0.01856 | 0.010943 | 0.098707 | 0.003986 | 0.007292 | 0.018286 | 0.008404 | 0.002224 | 0.007715 | 0.001593 | 0.006552 | 0.018105 | 0.006954 | 0.011126 | 0.017547 | 0.058745 | 0.077043 | 0.006167 | 0.007822 | 0.006667 | 0.042064 | 0.022768 |
| 29.3333333 | 0.013116 | 0.005801 | 0.01679 | 0.007539 | 0.111427 | 0.002969 | 0.008194 | 0.020763 | 0.009144 | 0.004188 | 0.007883 | 0.002154 | 0.005346 | 0.015007 | 0.009887 | 0.014235 | 0.012041 | 0.055944 | 0.067673 | 0.00312 | 0.00559 | 0.002896 | 0.047241 | 0.026136 |
| 29.3472222 | 0.01489 | 0.003679 | 0.015259 | 0.004954 | 0.120046 | 0.003897 | 0.012593 | 0.023206 | 0.010814 | 0.008965 | 0.006877 | 0.004496 | 0.003834 | 0.016965 | 0.015444 | 0.013432 | 0.012983 | 0.053347 | 0.072645 | 0.002407 | 0.013136 | 0.00167 | 0.047318 | 0.025185 |
| 29.3611111 | 0.018088 | 0.003424 | 0.015015 | 0.005104 | 0.11838 | 0.005866 | 0.021565 | 0.025263 | 0.012965 | 0.01487 | 0.009719 | 0.008379 | 0.005093 | 0.014519 | 0.026833 | 0.016286 | 0.018417 | 0.059572 | 0.082059 | 0.002501 | 0.021785 | 0.001257 | 0.044751 | 0.019113 |
| 29.375 | 0.019373 | 0.005004 | 0.016667 | 0.007118 | 0.11118 | 0.008018 | 0.030599 | 0.027683 | 0.015021 | 0.021216 | 0.01685 | 0.013445 | 0.005501 | 0.016079 | 0.038789 | 0.022964 | 0.023861 | 0.067504 | 0.07747 | 0.001606 | 0.024768 | 0.002065 | 0.042523 | 0.021729 |
| 29.3888889 | 0.018518 | 0.003544 | 0.021352 | 0.009737 | 0.109752 | 0.008979 | 0.034777 | 0.03275 | 0.016333 | 0.027755 | 0.022559 | 0.018393 | 0.004016 | 0.026149 | 0.047658 | 0.030901 | 0.02486 | 0.073634 | 0.066091 | 0.001796 | 0.030043 | 0.005385 | 0.043738 | 0.037482 |
| 29.4027778 | 0.018406 | 0.003823 | 0.030601 | 0.018476 | 0.12252 | 0.006739 | 0.038477 | 0.043937 | 0.01877 | 0.034022 | 0.030721 | 0.024385 | 0.005175 | 0.027215 | 0.056343 | 0.034487 | 0.022469 | 0.094471 | 0.086748 | 0.006608 | 0.042947 | 0.021172 | 0.042698 | 0.053142 |
| 29.4166667 | 0.02294 | 0.014398 | 0.038164 | 0.046046 | 0.166914 | 0.00427 | 0.049773 | 0.06025 | 0.024806 | 0.043699 | 0.054371 | 0.037223 | 0.009352 | 0.019774 | 0.071597 | 0.038261 | 0.022425 | 0.139849 | 0.151147 | 0.022969 | 0.051528 | 0.051805 | 0.034665 | 0.065621 |
| 29.4305556 | 0.031152 | 0.037113 | 0.053973 | 0.061623 | 0.18641 | 0.014559 | 0.058703 | 0.056955 | 0.032775 | 0.056031 | 0.076563 | 0.048013 | 0.026104 | 0.031709 | 0.089661 | 0.036927 | 0.022639 | 0.161156 | 0.176193 | 0.03461 | 0.048449 | 0.056194 | 0.030309 | 0.083468 |
| 29.4444444 | 0.028795 | 0.054452 | 0.088125 | 0.048979 | 0.117613 | 0.032217 | 0.042884 | 0.031404 | 0.028583 | 0.051378 | 0.060393 | 0.038084 | 0.045617 | 0.060951 | 0.076371 | 0.028082 | 0.016148 | 0.133387 | 0.132354 | 0.032506 | 0.035925 | 0.035008 | 0.028105 | 0.084141 |
| 29.4583333 | 0.013381 | 0.058873 | 0.102651 | 0.041096 | 0.038908 | 0.034863 | 0.015254 | 0.017824 | 0.011149 | 0.030298 | 0.026135 | 0.017391 | 0.040944 | 0.059334 | 0.031065 | 0.026812 | 0.010076 | 0.107794 | 0.102819 | 0.031497 | 0.024214 | 0.034059 | 0.017958 | 0.058562 |
| 29.4722222 | 0.007274 | 0.063778 | 0.085241 | 0.02815 | 0.016191 | 0.027695 | 0.008017 | 0.014753 | 0.006657 | 0.026416 | 0.021208 | 0.013001 | 0.029845 | 0.026681 | 0.015297 | 0.027277 | 0.014516 | 0.081759 | 0.077889 | 0.020955 | 0.025809 | 0.02988 | 0.01034 | 0.031787 |
| 29.4861111 | 0.016871 | 0.065829 | 0.069381 | 0.013763 | 0.026134 | 0.025938 | 0.019045 | 0.012261 | 0.017354 | 0.042604 | 0.039027 | 0.026543 | 0.031402 | 0.015387 | 0.041774 | 0.029152 | 0.021893 | 0.046985 | 0.037992 | 0.006159 | 0.032961 | 0.01004 | 0.015131 | 0.018219 |
| 29.5 | 0.030556 | 0.056018 | 0.064817 | 0.014148 | 0.036828 | 0.029627 | 0.031114 | 0.013553 | 0.024833 | 0.055266 | 0.048164 | 0.039243 | 0.036188 | 0.025093 | 0.068992 | 0.029617 | 0.025743 | 0.047354 | 0.027596 | 0.003268 | 0.039775 | 0.001856 | 0.023412 | 0.025658 |
| 29.5138889 | 0.042223 | 0.040666 | 0.064329 | 0.015563 | 0.036594 | 0.033553 | 0.036376 | 0.014389 | 0.026656 | 0.055134 | 0.044517 | 0.043576 | 0.039501 | 0.029062 | 0.07759 | 0.023759 | 0.027211 | 0.077392 | 0.041879 | 0.008519 | 0.048375 | 0.002376 | 0.03151 | 0.035379 |
| 29.5277778 | 0.046908 | 0.025213 | 0.063192 | 0.014048 | 0.028445 | 0.033572 | 0.032102 | 0.012722 | 0.027026 | 0.044273 | 0.036715 | 0.040825 | 0.041251 | 0.027405 | 0.070013 | 0.018433 | 0.02382 | 0.093725 | 0.050968 | 0.021146 | 0.055164 | 0.002219 | 0.047981 | 0.03606 |
| 29.5416667 | 0.043949 | 0.016537 | 0.061725 | 0.012007 | 0.018533 | 0.031157 | 0.022863 | 0.008856 | 0.027952 | 0.033311 | 0.02988 | 0.032971 | 0.039828 | 0.025456 | 0.055563 | 0.016534 | 0.015232 | 0.080091 | 0.043249 | 0.038303 | 0.055513 | 0.002372 | 0.060388 | 0.041676 |
| 29.5555556 | 0.036973 | 0.018691 | 0.059204 | 0.012093 | 0.014382 | 0.030306 | 0.01497 | 0.005653 | 0.028974 | 0.032559 | 0.025571 | 0.024088 | 0.033903 | 0.021522 | 0.042845 | 0.016511 | 0.007594 | 0.053769 | 0.024853 | 0.044947 | 0.045138 | 0.003701 | 0.056296 | 0.045527 |
| 29.5694444 | 0.027627 | 0.024298 | 0.048969 | 0.015054 | 0.012559 | 0.029385 | 0.011238 | 0.004772 | 0.027519 | 0.03781 | 0.022741 | 0.018174 | 0.028616 | 0.017065 | 0.037645 | 0.016432 | 0.009761 | 0.039965 | 0.011074 | 0.029429 | 0.034066 | 0.003842 | 0.049594 | 0.036879 |
| 29.5833333 | 0.023016 | 0.029568 | 0.035721 | 0.01621 | 0.012549 | 0.027194 | 0.013458 | 0.005744 | 0.025965 | 0.041778 | 0.025521 | 0.021002 | 0.032416 | 0.019591 | 0.041771 | 0.01897 | 0.0254 | 0.046971 | 0.007908 | 0.011369 | 0.036976 | 0.002769 | 0.051954 | 0.025812 |
| 29.5972222 | 0.030316 | 0.032806 | 0.031789 | 0.011789 | 0.017785 | 0.027554 | 0.020274 | 0.008601 | 0.030435 | 0.047565 | 0.035581 | 0.034248 | 0.041982 | 0.025385 | 0.04417 | 0.019874 | 0.044541 | 0.065797 | 0.008602 | 0.010626 | 0.053661 | 0.003353 | 0.063925 | 0.024708 |
| 29.6111111 | 0.0392 | 0.028035 | 0.035737 | 0.00704 | 0.021705 | 0.028222 | 0.024748 | 0.01305 | 0.038718 | 0.055518 | 0.043324 | 0.049576 | 0.042502 | 0.027837 | 0.035821 | 0.01546 | 0.051654 | 0.079642 | 0.009707 | 0.025398 | 0.065317 | 0.005493 | 0.070266 | 0.030739 |
| 29.625 | 0.033648 | 0.01606 | 0.036342 | 0.005523 | 0.018124 | 0.025353 | 0.02227 | 0.016822 | 0.043013 | 0.056529 | 0.040189 | 0.057207 | 0.033518 | 0.0278 | 0.022963 | 0.008635 | 0.046055 | 0.07463 | 0.011656 | 0.045571 | 0.055041 | 0.006069 | 0.066614 | 0.030948 |
| 29.6388889 | 0.018407 | 0.006271 | 0.031151 | 0.004764 | 0.011599 | 0.021844 | 0.018316 | 0.019017 | 0.041448 | 0.047721 | 0.028878 | 0.053349 | 0.030278 | 0.024625 | 0.016366 | 0.003033 | 0.036941 | 0.057348 | 0.011459 | 0.049159 | 0.039627 | 0.004811 | 0.060753 | 0.019827 |
| 29.6527778 | 0.011011 | 0.004239 | 0.028376 | 0.00547 | 0.008262 | 0.022124 | 0.018447 | 0.020125 | 0.038031 | 0.037172 | 0.01638 | 0.038946 | 0.033086 | 0.018631 | 0.019016 | 0.001577 | 0.024687 | 0.043115 | 0.008301 | 0.031714 | 0.035116 | 0.003552 | 0.047021 | 0.008179 |
| 29.6666667 | 0.014139 | 0.006866 | 0.029339 | 0.007487 | 0.008201 | 0.023233 | 0.018313 | 0.017932 | 0.034346 | 0.031244 | 0.009306 | 0.02235 | 0.026252 | 0.014517 | 0.017213 | 0.002698 | 0.023873 | 0.032459 | 0.006023 | 0.017066 | 0.029578 | 0.002491 | 0.029512 | 0.004439 |
| 29.6805556 | 0.016471 | 0.006609 | 0.022338 | 0.006657 | 0.008694 | 0.018202 | 0.012274 | 0.010944 | 0.027094 | 0.027109 | 0.009362 | 0.012438 | 0.012281 | 0.014167 | 0.011988 | 0.003498 | 0.042697 | 0.022844 | 0.006467 | 0.02097 | 0.018871 | 0.001228 | 0.015519 | 0.005031 |
| 29.6944444 | 0.011756 | 0.005445 | 0.01143 | 0.004413 | 0.006944 | 0.009019 | 0.006108 | 0.004777 | 0.016786 | 0.019302 | 0.009883 | 0.008025 | 0.005138 | 0.015207 | 0.018785 | 0.005191 | 0.051675 | 0.017345 | 0.008426 | 0.027623 | 0.009659 | 0.000476 | 0.006527 | 0.004309 |
| 29.7083333 | 0.004982 | 0.00739 | 0.011319 | 0.005653 | 0.00441 | 0.003759 | 0.006102 | 0.004076 | 0.008525 | 0.009263 | 0.006491 | 0.005196 | 0.003567 | 0.016296 | 0.026422 | 0.010195 | 0.045082 | 0.014722 | 0.009623 | 0.018319 | 0.00702 | 0.000386 | 0.00262 | 0.003613 |
| 29.7222222 | 0.002969 | 0.006578 | 0.012969 | 0.007568 | 0.007269 | 0.004013 | 0.005751 | 0.005252 | 0.003728 | 0.004076 | 0.004156 | 0.005017 | 0.003036 | 0.016968 | 0.024043 | 0.018339 | 0.041673 | 0.009913 | 0.008971 | 0.007259 | 0.008438 | 0.000476 | 0.00262 | 0.005305 |
| 29.7361111 | 0.005259 | 0.004077 | 0.008594 | 0.006712 | 0.015647 | 0.005521 | 0.002149 | 0.007194 | 0.001291 | 0.006145 | 0.006207 | 0.007549 | 0.00384 | 0.017482 | 0.017352 | 0.023827 | 0.03706 | 0.004852 | 0.007021 | 0.005345 | 0.009176 | 0.001218 | 0.00639 | 0.005198 |
| 29.75 | 0.007409 | 0.002782 | 0.005314 | 0.004998 | 0.021061 | 0.007071 | 0.002221 | 0.012129 | 0.001007 | 0.009346 | 0.00668 | 0.008972 | 0.004977 | 0.018003 | 0.009168 | 0.024597 | 0.025726 | 0.004122 | 0.004791 | 0.006308 | 0.013975 | 0.002468 | 0.009309 | 0.002589 |
| 29.7638889 | 0.006312 | 0.003967 | 0.004208 | 0.003736 | 0.022291 | 0.008892 | 0.006252 | 0.01502 | 0.00171 | 0.00829 | 0.003505 | 0.008817 | 0.004282 | 0.017548 | 0.004048 | 0.025146 | 0.021478 | 0.005706 | 0.003378 | 0.010825 | 0.017808 | 0.002763 | 0.008494 | 0.000889 |
| 29.7777778 | 0.003213 | 0.010177 | 0.003299 | 0.002791 | 0.026501 | 0.008648 | 0.007122 | 0.010607 | 0.001911 | 0.005402 | 0.001824 | 0.009399 | 0.002166 | 0.016883 | 0.003372 | 0.024722 | 0.024114 | 0.009067 | 0.002857 | 0.01959 | 0.012652 | 0.001797 | 0.008036 | 0.000476 |
| 29.7916667 | 0.00203 | 0.01553 | 0.003691 | 0.00257 | 0.031139 | 0.00542 | 0.007465 | 0.004178 | 0.002372 | 0.006971 | 0.003154 | 0.010716 | 0.001799 | 0.017772 | 0.003532 | 0.021332 | 0.020275 | 0.015349 | 0.002271 | 0.033876 | 0.007866 | 0.000916 | 0.009352 | 0.001609 |
| 29.8055556 | 0.004147 | 0.013725 | 0.007382 | 0.003246 | 0.031044 | 0.002845 | 0.011739 | 0.001315 | 0.003802 | 0.012765 | 0.004591 | 0.011108 | 0.003379 | 0.018944 | 0.004948 | 0.015273 | 0.012227 | 0.019206 | 0.001649 | 0.056822 | 0.0136 | 0.0009 | 0.009371 | 0.007639 |
| 29.8194444 | 0.006614 | 0.007924 | 0.012794 | 0.003124 | 0.028424 | 0.003216 | 0.013283 | 0.00113 | 0.004978 | 0.0162 | 0.00434 | 0.007862 | 0.003702 | 0.017692 | 0.004916 | 0.011719 | 0.008099 | 0.016533 | 0.001804 | 0.078895 | 0.020315 | 0.000925 | 0.010376 | 0.015997 |
| 29.8333333 | 0.00605 | 0.005858 | 0.01242 | 0.002594 | 0.026972 | 0.003446 | 0.010661 | 0.002248 | 0.00492 | 0.01546 | 0.003933 | 0.003425 | 0.004333 | 0.014206 | 0.002946 | 0.0162 | 0.012556 | 0.013657 | 0.002549 | 0.074149 | 0.02045 | 0.000943 | 0.014918 | 0.017037 |
| 29.8472222 | 0.003794 | 0.007649 | 0.008899 | 0.003106 | 0.028871 | 0.003104 | 0.006645 | 0.003448 | 0.004737 | 0.014367 | 0.003812 | 0.003497 | 0.010276 | 0.010217 | 0.002666 | 0.022899 | 0.019302 | 0.01691 | 0.002941 | 0.041845 | 0.020972 | 0.001151 | 0.019018 | 0.012431 |
| 29.8611111 | 0.003497 | 0.006869 | 0.014 | 0.006645 | 0.035416 | 0.005947 | 0.004082 | 0.005003 | 0.006663 | 0.016355 | 0.006109 | 0.010681 | 0.023293 | 0.006288 | 0.004715 | 0.023013 | 0.017688 | 0.024723 | 0.002944 | 0.019783 | 0.02468 | 0.00083 | 0.019314 | 0.012449 |
| 29.875 | 0.004917 | 0.003837 | 0.023729 | 0.014397 | 0.044377 | 0.01003 | 0.006659 | 0.006872 | 0.009748 | 0.021805 | 0.012385 | 0.023886 | 0.034648 | 0.003573 | 0.012296 | 0.015848 | 0.013094 | 0.033124 | 0.002927 | 0.017022 | 0.028892 | 0.000532 | 0.018213 | 0.017947 |
| 29.8888889 | 0.005765 | 0.004549 | 0.027187 | 0.019831 | 0.047739 | 0.010559 | 0.010134 | 0.006708 | 0.009648 | 0.026762 | 0.015485 | 0.031775 | 0.027462 | 0.003922 | 0.021629 | 0.009426 | 0.017526 | 0.040729 | 0.002271 | 0.025129 | 0.026677 | 0.000746 | 0.021699 | 0.019631 |
| 29.9027778 | 0.006733 | 0.006141 | 0.02493 | 0.01924 | 0.046582 | 0.009229 | 0.01227 | 0.006673 | 0.006472 | 0.028403 | 0.013848 | 0.028362 | 0.010594 | 0.005861 | 0.02134 | 0.008723 | 0.027489 | 0.04544 | 0.002419 | 0.043963 | 0.016785 | 0.000873 | 0.030509 | 0.013004 |
| 29.9166667 | 0.008803 | 0.005973 | 0.025506 | 0.016476 | 0.057834 | 0.010476 | 0.018992 | 0.011077 | 0.005766 | 0.029143 | 0.014679 | 0.023454 | 0.007179 | 0.007231 | 0.011758 | 0.010609 | 0.029828 | 0.047883 | 0.005539 | 0.04928 | 0.010146 | 0.000502 | 0.03802 | 0.005286 |
| 29.9305556 | 0.010451 | 0.009821 | 0.033548 | 0.014999 | 0.088974 | 0.015161 | 0.032112 | 0.017416 | 0.008972 | 0.029854 | 0.01902 | 0.02263 | 0.014579 | 0.008875 | 0.003971 | 0.0124 | 0.022382 | 0.057177 | 0.011315 | 0.034913 | 0.012879 | 0.000602 | 0.038265 | 0.003323 |
| 29.9444444 | 0.011097 | 0.017253 | 0.042134 | 0.016405 | 0.123908 | 0.020512 | 0.043524 | 0.021501 | 0.011706 | 0.028576 | 0.02325 | 0.024419 | 0.015975 | 0.010985 | 0.002752 | 0.013336 | 0.019954 | 0.074093 | 0.017308 | 0.027935 | 0.022899 | 0.001168 | 0.030403 | 0.005192 |
| 29.9583333 | 0.013129 | 0.026509 | 0.044845 | 0.020141 | 0.145182 | 0.024308 | 0.045721 | 0.024665 | 0.013543 | 0.025569 | 0.026516 | 0.02906 | 0.016138 | 0.011075 | 0.003707 | 0.014151 | 0.035256 | 0.083407 | 0.018295 | 0.043376 | 0.037145 | 0.001006 | 0.019631 | 0.007148 |
| 29.9722222 | 0.01734 | 0.040167 | 0.044603 | 0.024665 | 0.15343 | 0.026365 | 0.042825 | 0.028613 | 0.019049 | 0.025282 | 0.031573 | 0.037756 | 0.031389 | 0.007321 | 0.00833 | 0.016627 | 0.055767 | 0.079215 | 0.015917 | 0.066228 | 0.051176 | 0.001461 | 0.013429 | 0.013993 |
| 29.9861111 | 0.022091 | 0.056295 | 0.044157 | 0.029606 | 0.161823 | 0.026861 | 0.041681 | 0.032113 | 0.027525 | 0.033037 | 0.040046 | 0.048955 | 0.052848 | 0.004109 | 0.020937 | 0.022105 | 0.058141 | 0.071582 | 0.018092 | 0.069848 | 0.056656 | 0.003227 | 0.017394 | 0.030125 |
| 30 | 0.027063 | 0.064128 | 0.045662 | 0.034167 | 0.171448 | 0.026998 | 0.041821 | 0.033365 | 0.033423 | 0.048639 | 0.048478 | 0.057497 | 0.06999 | 0.005822 | 0.034374 | 0.027481 | 0.038407 | 0.069754 | 0.028861 | 0.047798 | 0.0514 | 0.003811 | 0.032494 | 0.045147 |
| 30.0138889 | 0.030423 | 0.055247 | 0.050461 | 0.035847 | 0.169527 | 0.027197 | 0.042803 | 0.032047 | 0.033703 | 0.059845 | 0.053579 | 0.060288 | 0.08405 | 0.009863 | 0.042318 | 0.029001 | 0.021214 | 0.076289 | 0.043256 | 0.020311 | 0.044539 | 0.003198 | 0.05131 | 0.049348 |
| 30.0277778 | 0.029065 | 0.038654 | 0.054483 | 0.033039 | 0.156559 | 0.026924 | 0.041487 | 0.029476 | 0.030128 | 0.055309 | 0.055165 | 0.059751 | 0.09639 | 0.009786 | 0.044604 | 0.026489 | 0.027847 | 0.090207 | 0.051974 | 0.010308 | 0.040172 | 0.002715 | 0.062362 | 0.048702 |
| 30.0416667 | 0.029602 | 0.033016 | 0.058207 | 0.028246 | 0.146735 | 0.028858 | 0.035558 | 0.028361 | 0.031536 | 0.046962 | 0.057502 | 0.061345 | 0.114191 | 0.005111 | 0.042582 | 0.022338 | 0.039155 | 0.100417 | 0.067309 | 0.012543 | 0.040655 | 0.002045 | 0.062572 | 0.049061 |
| 30.0555556 | 0.043617 | 0.047038 | 0.068236 | 0.030852 | 0.150638 | 0.040283 | 0.035074 | 0.032206 | 0.046029 | 0.055583 | 0.073343 | 0.069913 | 0.139803 | 0.012531 | 0.039604 | 0.021066 | 0.034323 | 0.090531 | 0.08946 | 0.010618 | 0.047709 | 0.004015 | 0.058851 | 0.048561 |
| 30.0694444 | 0.059747 | 0.067196 | 0.082766 | 0.04415 | 0.158748 | 0.055737 | 0.041188 | 0.039489 | 0.06228 | 0.077635 | 0.100059 | 0.078736 | 0.159804 | 0.033191 | 0.037257 | 0.020661 | 0.033512 | 0.06914 | 0.090201 | 0.007337 | 0.048886 | 0.005846 | 0.052379 | 0.042719 |
| 30.0833333 | 0.056032 | 0.069771 | 0.086659 | 0.060273 | 0.142089 | 0.055753 | 0.040028 | 0.042256 | 0.062656 | 0.088108 | 0.108852 | 0.073721 | 0.157392 | 0.049924 | 0.032537 | 0.027292 | 0.04963 | 0.053832 | 0.084193 | 0.010484 | 0.033885 | 0.003102 | 0.042306 | 0.034147 |
| 30.0972222 | 0.038578 | 0.047125 | 0.071635 | 0.063225 | 0.100987 | 0.041585 | 0.027597 | 0.03746 | 0.051567 | 0.080365 | 0.09066 | 0.058736 | 0.128419 | 0.050028 | 0.026205 | 0.037607 | 0.068862 | 0.043864 | 0.080018 | 0.015204 | 0.015156 | 0.001914 | 0.036943 | 0.024635 |
| 30.1111111 | 0.025015 | 0.022703 | 0.049033 | 0.041289 | 0.065251 | 0.029616 | 0.01242 | 0.031555 | 0.041549 | 0.069849 | 0.065742 | 0.046391 | 0.085551 | 0.027781 | 0.026979 | 0.034985 | 0.081234 | 0.029021 | 0.05475 | 0.012534 | 0.005877 | 0.003515 | 0.037573 | 0.012789 |
| 30.125 | 0.017636 | 0.020531 | 0.031013 | 0.015586 | 0.041311 | 0.021763 | 0.004433 | 0.028002 | 0.034019 | 0.060505 | 0.04599 | 0.036829 | 0.050317 | 0.011904 | 0.031581 | 0.029411 | 0.081723 | 0.011278 | 0.033046 | 0.008062 | 0.002364 | 0.006082 | 0.033979 | 0.008184 |
| 30.1388889 | 0.011928 | 0.027704 | 0.020662 | 0.005928 | 0.021754 | 0.012203 | 0.007079 | 0.022585 | 0.026225 | 0.041972 | 0.030394 | 0.027122 | 0.028248 | 0.022206 | 0.025382 | 0.021871 | 0.075216 | 0.005037 | 0.032253 | 0.014027 | 0.000728 | 0.011823 | 0.02304 | 0.011717 |
| 30.1527778 | 0.006341 | 0.029674 | 0.015406 | 0.003198 | 0.009221 | 0.005849 | 0.014065 | 0.01488 | 0.016934 | 0.022632 | 0.01735 | 0.017798 | 0.014375 | 0.037772 | 0.013899 | 0.011175 | 0.06369 | 0.006474 | 0.036968 | 0.021415 | 0.002493 | 0.018279 | 0.012964 | 0.017381 |
| 30.1666667 | 0.003907 | 0.033521 | 0.010139 | 0.004972 | 0.009001 | 0.008468 | 0.019113 | 0.008502 | 0.008961 | 0.023571 | 0.007534 | 0.009075 | 0.011598 | 0.038854 | 0.012235 | 0.01215 | 0.044142 | 0.009489 | 0.033014 | 0.01686 | 0.005831 | 0.020157 | 0.009729 | 0.020122 |
| 30.1805556 | 0.008354 | 0.039043 | 0.00629 | 0.014284 | 0.01916 | 0.012168 | 0.021249 | 0.007706 | 0.00829 | 0.036539 | 0.004264 | 0.004965 | 0.017625 | 0.02295 | 0.013183 | 0.018399 | 0.024436 | 0.022815 | 0.025619 | 0.008567 | 0.008171 | 0.015473 | 0.01219 | 0.018078 |

| | | | | | | | | | | | | | | | | | | | | | | | | |
|---|---|---|---|---|---|---|---|---|---|---|---|---|---|---|---|---|---|---|---|---|---|---|---|---|
| 30.1944444 | 0.016515 | 0.036619 | 0.008903 | 0.022983 | 0.030762 | 0.012728 | 0.020591 | 0.013334 | 0.013589 | 0.043251 | 0.008228 | 0.008993 | 0.026389 | 0.008071 | 0.013985 | 0.024082 | 0.012042 | 0.04094 | 0.040576 | 0.006537 | 0.013227 | 0.00936 | 0.014056 | 0.016494 |
| 30.2083333 | 0.021911 | 0.02791 | 0.015242 | 0.026108 | 0.038013 | 0.013085 | 0.018712 | 0.016612 | 0.016476 | 0.043262 | 0.013904 | 0.015022 | 0.033442 | 0.007991 | 0.023742 | 0.034436 | 0.013702 | 0.049339 | 0.061833 | 0.008023 | 0.022807 | 0.005239 | 0.010112 | 0.015731 |
| 30.2222222 | 0.021795 | 0.027143 | 0.021132 | 0.031178 | 0.04201 | 0.01311 | 0.017506 | 0.014628 | 0.014158 | 0.041252 | 0.017534 | 0.019026 | 0.034825 | 0.010779 | 0.036525 | 0.045492 | 0.028992 | 0.048468 | 0.066794 | 0.007625 | 0.027169 | 0.002264 | 0.008355 | 0.018648 |
| 30.2361111 | 0.018472 | 0.037628 | 0.027161 | 0.042748 | 0.044689 | 0.012856 | 0.015331 | 0.012779 | 0.010397 | 0.039008 | 0.019751 | 0.021481 | 0.032544 | 0.014811 | 0.040169 | 0.046196 | 0.038389 | 0.045905 | 0.068656 | 0.008577 | 0.018145 | 0.001935 | 0.018234 | 0.02975 |
| 30.25 | 0.016439 | 0.045185 | 0.031767 | 0.05017 | 0.045833 | 0.01297 | 0.011697 | 0.011717 | 0.007777 | 0.035715 | 0.01983 | 0.021061 | 0.032518 | 0.027484 | 0.035963 | 0.031954 | 0.034453 | 0.044551 | 0.074011 | 0.013415 | 0.008972 | 0.004668 | 0.032927 | 0.041595 |
| 30.2638889 | 0.01629 | 0.044172 | 0.031809 | 0.045958 | 0.044889 | 0.013401 | 0.009217 | 0.010149 | 0.005967 | 0.030729 | 0.016504 | 0.019163 | 0.035464 | 0.029441 | 0.031061 | 0.014992 | 0.030618 | 0.044212 | 0.078261 | 0.011462 | 0.009456 | 0.013367 | 0.042791 | 0.043067 |
| 30.2777778 | 0.015873 | 0.045522 | 0.027967 | 0.03951 | 0.043322 | 0.013061 | 0.008196 | 0.009493 | 0.004466 | 0.024811 | 0.010729 | 0.016678 | 0.030842 | 0.014024 | 0.021849 | 0.006132 | 0.028559 | 0.043956 | 0.077259 | 0.006268 | 0.011773 | 0.024229 | 0.045361 | 0.033774 |
| 30.2916667 | 0.015196 | 0.056539 | 0.024641 | 0.03969 | 0.042616 | 0.010443 | 0.00751 | 0.010166 | 0.002772 | 0.018514 | 0.00567 | 0.010956 | 0.018362 | 0.005137 | 0.012629 | 0.002845 | 0.019983 | 0.043041 | 0.069645 | 0.006224 | 0.01262 | 0.022765 | 0.042443 | 0.020709 |
| 30.3055556 | 0.0155 | 0.063778 | 0.023656 | 0.040548 | 0.040932 | 0.006224 | 0.006939 | 0.01014 | 0.001789 | 0.01271 | 0.004997 | 0.006665 | 0.014716 | 0.011185 | 0.012679 | 0.001872 | 0.014902 | 0.042548 | 0.061621 | 0.006145 | 0.014002 | 0.011335 | 0.041456 | 0.010376 |
| 30.3194444 | 0.016759 | 0.052397 | 0.022 | 0.033268 | 0.039316 | 0.003418 | 0.007095 | 0.010066 | 0.002835 | 0.009307 | 0.006193 | 0.006607 | 0.018909 | 0.02237 | 0.011337 | 0.002256 | 0.024179 | 0.043524 | 0.058432 | 0.006661 | 0.020733 | 0.007291 | 0.045584 | 0.008533 |
| 30.3333333 | 0.01793 | 0.037955 | 0.019366 | 0.021891 | 0.042833 | 0.003447 | 0.008742 | 0.012637 | 0.003735 | 0.008156 | 0.004372 | 0.005611 | 0.01904 | 0.029772 | 0.005227 | 0.005333 | 0.0314 | 0.04548 | 0.060217 | 0.005937 | 0.025364 | 0.015569 | 0.046578 | 0.014758 |
| 30.3472222 | 0.018631 | 0.036751 | 0.01931 | 0.016793 | 0.050843 | 0.004902 | 0.010963 | 0.016685 | 0.002656 | 0.006556 | 0.002428 | 0.007016 | 0.017312 | 0.030216 | 0.003284 | 0.014032 | 0.02272 | 0.04607 | 0.062999 | 0.00274 | 0.017963 | 0.02569 | 0.041904 | 0.018889 |
| 30.3611111 | 0.020217 | 0.034761 | 0.023739 | 0.018734 | 0.056544 | 0.00709 | 0.012396 | 0.018854 | 0.002564 | 0.005912 | 0.003215 | 0.012917 | 0.017408 | 0.03053 | 0.004073 | 0.026122 | 0.008731 | 0.042296 | 0.06032 | 0.001989 | 0.012049 | 0.024972 | 0.040757 | 0.015779 |
| 30.375 | 0.022967 | 0.021815 | 0.030605 | 0.020572 | 0.056768 | 0.010422 | 0.01369 | 0.019347 | 0.006657 | 0.010805 | 0.005401 | 0.020221 | 0.019946 | 0.035551 | 0.009583 | 0.0351 | 0.00781 | 0.039481 | 0.056018 | 0.002512 | 0.013999 | 0.014807 | 0.043449 | 0.014329 |
| 30.3888889 | 0.024129 | 0.008409 | 0.036786 | 0.020255 | 0.0547 | 0.013729 | 0.015565 | 0.0193 | 0.011869 | 0.01904 | 0.007918 | 0.025898 | 0.025148 | 0.041986 | 0.022688 | 0.038805 | 0.019331 | 0.046008 | 0.056433 | 0.002615 | 0.016313 | 0.008382 | 0.041016 | 0.018939 |
| 30.4027778 | 0.021934 | 0.002524 | 0.043673 | 0.020952 | 0.056904 | 0.015241 | 0.01663 | 0.017746 | 0.014218 | 0.024783 | 0.010667 | 0.027156 | 0.027608 | 0.047819 | 0.037211 | 0.040136 | 0.026053 | 0.057989 | 0.059628 | 0.004855 | 0.025173 | 0.008132 | 0.035121 | 0.021456 |
| 30.4166667 | 0.018024 | 0.002189 | 0.053459 | 0.023655 | 0.066609 | 0.015793 | 0.016133 | 0.015122 | 0.01524 | 0.029791 | 0.014834 | 0.027017 | 0.023345 | 0.05042 | 0.049079 | 0.04171 | 0.02177 | 0.064511 | 0.060582 | 0.007853 | 0.046138 | 0.007659 | 0.031099 | 0.016944 |
| 30.4305556 | 0.01481 | 0.002562 | 0.062501 | 0.024438 | 0.075914 | 0.01806 | 0.015502 | 0.013937 | 0.018831 | 0.038398 | 0.0208 | 0.029953 | 0.019457 | 0.048408 | 0.058198 | 0.042685 | 0.020219 | 0.064132 | 0.059457 | 0.010849 | 0.06163 | 0.00567 | 0.031991 | 0.009579 |
| 30.4444444 | 0.0134 | 0.001864 | 0.067856 | 0.020327 | 0.079618 | 0.022804 | 0.016244 | 0.015784 | 0.026235 | 0.047556 | 0.027667 | 0.036767 | 0.023334 | 0.045927 | 0.066171 | 0.04301 | 0.02592 | 0.0666 | 0.060292 | 0.019329 | 0.060126 | 0.005159 | 0.040787 | 0.007839 |
| 30.4583333 | 0.013619 | 0.006513 | 0.072781 | 0.015582 | 0.085912 | 0.027734 | 0.019356 | 0.021002 | 0.034893 | 0.05304 | 0.035081 | 0.045002 | 0.027867 | 0.048593 | 0.078129 | 0.043908 | 0.021307 | 0.071486 | 0.05634 | 0.037285 | 0.059692 | 0.00635 | 0.056497 | 0.015261 |
| 30.4722222 | 0.01453 | 0.023298 | 0.080419 | 0.015116 | 0.10315 | 0.030851 | 0.023873 | 0.026447 | 0.041584 | 0.055994 | 0.042302 | 0.051111 | 0.024788 | 0.054751 | 0.091391 | 0.04428 | 0.013298 | 0.070929 | 0.049024 | 0.065097 | 0.065413 | 0.005545 | 0.078086 | 0.030132 |
| 30.4861111 | 0.014977 | 0.045005 | 0.086624 | 0.017958 | 0.125822 | 0.031065 | 0.025922 | 0.027925 | 0.044069 | 0.057891 | 0.047398 | 0.053445 | 0.023645 | 0.058308 | 0.095805 | 0.041955 | 0.012978 | 0.068541 | 0.049346 | 0.097786 | 0.066695 | 0.003367 | 0.090976 | 0.047769 |
| 30.5 | 0.014734 | 0.058441 | 0.085033 | 0.020223 | 0.13853 | 0.027922 | 0.023715 | 0.027065 | 0.042348 | 0.057016 | 0.049589 | 0.053964 | 0.035579 | 0.059812 | 0.09404 | 0.035382 | 0.014329 | 0.070955 | 0.052869 | 0.126411 | 0.066308 | 0.00382 | 0.083825 | 0.061394 |
| 30.5138889 | 0.014125 | 0.058112 | 0.076085 | 0.020425 | 0.141304 | 0.023888 | 0.019828 | 0.026316 | 0.038096 | 0.051818 | 0.049987 | 0.055875 | 0.048907 | 0.061985 | 0.094878 | 0.027271 | 0.025739 | 0.072792 | 0.050874 | 0.13911 | 0.064437 | 0.007202 | 0.072418 | 0.072722 |
| 30.5277778 | 0.013618 | 0.04693 | 0.065224 | 0.019108 | 0.14514 | 0.021451 | 0.016503 | 0.024663 | 0.033001 | 0.043097 | 0.051457 | 0.061001 | 0.049002 | 0.070533 | 0.098647 | 0.02057 | 0.040514 | 0.061404 | 0.036773 | 0.124234 | 0.058972 | 0.011671 | 0.070609 | 0.082934 |
| 30.5416667 | 0.016461 | 0.032682 | 0.060078 | 0.01623 | 0.14796 | 0.022944 | 0.015128 | 0.023215 | 0.030783 | 0.035189 | 0.056346 | 0.06793 | 0.044337 | 0.08362 | 0.100947 | 0.01593 | 0.050287 | 0.046373 | 0.016388 | 0.080872 | 0.05489 | 0.015289 | 0.07524 | 0.086926 |
| 30.5555556 | 0.025171 | 0.0236 | 0.060072 | 0.01277 | 0.133112 | 0.026129 | 0.015983 | 0.024216 | 0.033282 | 0.03236 | 0.059292 | 0.068185 | 0.047068 | 0.088478 | 0.093569 | 0.014146 | 0.059401 | 0.046593 | 0.004703 | 0.040356 | 0.052679 | 0.017846 | 0.079625 | 0.081258 |
| 30.5694444 | 0.03834 | 0.023656 | 0.05438 | 0.010546 | 0.101417 | 0.023902 | 0.016033 | 0.025542 | 0.036212 | 0.032029 | 0.053415 | 0.057141 | 0.055389 | 0.083432 | 0.078232 | 0.015383 | 0.060872 | 0.057431 | 0.004831 | 0.028247 | 0.049064 | 0.016342 | 0.078155 | 0.062725 |
| 30.5833333 | 0.054792 | 0.031915 | 0.043315 | 0.009625 | 0.08477 | 0.018607 | 0.017578 | 0.024376 | 0.03626 | 0.031481 | 0.045249 | 0.046933 | 0.061337 | 0.074876 | 0.068082 | 0.018611 | 0.055188 | 0.071691 | 0.012895 | 0.029976 | 0.048925 | 0.009317 | 0.071426 | 0.037572 |
| 30.5972222 | 0.065795 | 0.040651 | 0.038607 | 0.008614 | 0.089009 | 0.016253 | 0.025292 | 0.020325 | 0.035522 | 0.033495 | 0.044763 | 0.048573 | 0.057157 | 0.068031 | 0.059792 | 0.022897 | 0.046155 | 0.084888 | 0.023194 | 0.041351 | 0.055128 | 0.0039 | 0.066809 | 0.018709 |
| 30.6111111 | 0.065361 | 0.039345 | 0.038225 | 0.007805 | 0.087784 | 0.0151 | 0.02815 | 0.014409 | 0.035994 | 0.038946 | 0.050453 | 0.056862 | 0.036859 | 0.063647 | 0.042398 | 0.025506 | 0.029917 | 0.08378 | 0.027116 | 0.048729 | 0.057051 | 0.002061 | 0.062541 | 0.009745 |
| 30.625 | 0.059323 | 0.029728 | 0.030632 | 0.006455 | 0.073132 | 0.012867 | 0.020795 | 0.010835 | 0.03617 | 0.044167 | 0.05489 | 0.061236 | 0.012688 | 0.061518 | 0.025734 | 0.022573 | 0.010812 | 0.073989 | 0.020443 | 0.039984 | 0.044807 | 0.002071 | 0.056333 | 0.006223 |
| 30.6388889 | 0.046351 | 0.019474 | 0.018197 | 0.003599 | 0.051779 | 0.011962 | 0.017618 | 0.011143 | 0.033942 | 0.0452 | 0.052028 | 0.05699 | 0.004347 | 0.060246 | 0.024624 | 0.015669 | 0.01081 | 0.067247 | 0.011184 | 0.033682 | 0.028885 | 0.00246 | 0.051131 | 0.005495 |
| 30.6527778 | 0.029403 | 0.011221 | 0.010897 | 0.002908 | 0.032247 | 0.013342 | 0.019874 | 0.01326 | 0.029903 | 0.040022 | 0.039539 | 0.046469 | 0.010505 | 0.047945 | 0.030057 | 0.008829 | 0.040894 | 0.054547 | 0.006135 | 0.030392 | 0.026312 | 0.002084 | 0.043379 | 0.006318 |
| 30.6666667 | 0.019865 | 0.007713 | 0.009575 | 0.004253 | 0.024211 | 0.013771 | 0.018255 | 0.017086 | 0.025622 | 0.030062 | 0.022831 | 0.035096 | 0.022013 | 0.025617 | 0.024095 | 0.004222 | 0.059116 | 0.036822 | 0.002859 | 0.024773 | 0.027606 | 0.004565 | 0.031466 | 0.009942 |
| 30.6805556 | 0.01615 | 0.00766 | 0.009095 | 0.00603 | 0.022349 | 0.012209 | 0.011265 | 0.016124 | 0.020007 | 0.018627 | 0.012206 | 0.025796 | 0.027328 | 0.013612 | 0.015878 | 0.002107 | 0.045936 | 0.023184 | 0.001079 | 0.019552 | 0.017898 | 0.00903 | 0.017994 | 0.018092 |
| 30.6944444 | 0.00994 | 0.005676 | 0.011687 | 0.011603 | 0.015872 | 0.009513 | 0.005225 | 0.012085 | 0.011449 | 0.009422 | 0.009119 | 0.017473 | 0.019559 | 0.015548 | 0.022282 | 0.002275 | 0.049654 | 0.013932 | 0.00145 | 0.011855 | 0.01164 | 0.011855 | 0.00891 | 0.023689 |
| 30.7083333 | 0.00331 | 0.004191 | 0.019996 | 0.017999 | 0.01138 | 0.006133 | 0.006598 | 0.017109 | 0.003882 | 0.006846 | 0.007812 | 0.011446 | 0.008083 | 0.015213 | 0.03316 | 0.00476 | 0.077283 | 0.008249 | 0.001594 | 0.005815 | 0.019265 | 0.01256 | 0.009359 | 0.022492 |
| 30.7222222 | 0.001231 | 0.004678 | 0.027023 | 0.01922 | 0.016605 | 0.005352 | 0.008603 | 0.021289 | 0.001685 | 0.008782 | 0.007853 | 0.011583 | 0.002807 | 0.009178 | 0.033707 | 0.007099 | 0.100555 | 0.004767 | 0.001153 | 0.008972 | 0.023106 | 0.00876 | 0.015992 | 0.016687 |
| 30.7361111 | 0.001618 | 0.004099 | 0.030695 | 0.018482 | 0.024129 | 0.008964 | 0.006348 | 0.013389 | 0.002443 | 0.010169 | 0.010467 | 0.017297 | 0.004496 | 0.005707 | 0.024884 | 0.007126 | 0.108543 | 0.003687 | 0.001249 | 0.022921 | 0.014437 | 0.006073 | 0.022064 | 0.009775 |
| 30.75 | 0.003171 | 0.00349 | 0.034187 | 0.018094 | 0.0302 | 0.011489 | 0.007706 | 0.006816 | 0.002883 | 0.012039 | 0.012112 | 0.021469 | 0.011692 | 0.003885 | 0.013748 | 0.005798 | 0.094652 | 0.005858 | 0.001479 | 0.045229 | 0.010386 | 0.01023 | 0.022811 | 0.008455 |
| 30.7638889 | 0.004773 | 0.004449 | 0.033574 | 0.013974 | 0.039694 | 0.008271 | 0.013152 | 0.011723 | 0.002023 | 0.012601 | 0.00982 | 0.017848 | 0.017034 | 0.002803 | 0.007162 | 0.004517 | 0.077923 | 0.008131 | 0.001957 | 0.063039 | 0.02268 | 0.011074 | 0.017452 | 0.011317 |
| 30.7777778 | 0.004129 | 0.00865 | 0.023701 | 0.007061 | 0.054267 | 0.004743 | 0.012913 | 0.015674 | 0.000876 | 0.01002 | 0.005294 | 0.009274 | 0.012113 | 0.006423 | 0.008748 | 0.003514 | 0.078378 | 0.006236 | 0.002688 | 0.063909 | 0.034388 | 0.007849 | 0.009577 | 0.012456 |
| 30.7916667 | 0.004231 | 0.01246 | 0.011953 | 0.002555 | 0.068365 | 0.005403 | 0.007531 | 0.011198 | 0.001051 | 0.011086 | 0.0016 | 0.00274 | 0.007065 | 0.010965 | 0.01141 | 0.002655 | 0.083003 | 0.003101 | 0.002289 | 0.053413 | 0.030949 | 0.008696 | 0.003826 | 0.012402 |
| 30.8055556 | 0.006295 | 0.013494 | 0.008569 | 0.001493 | 0.076018 | 0.005218 | 0.005746 | 0.008691 | 0.002032 | 0.015484 | 0.002022 | 0.002493 | 0.011536 | 0.009754 | 0.008236 | 0.00263 | 0.076258 | 0.004552 | 0.001043 | 0.043533 | 0.023183 | 0.010816 | 0.002186 | 0.011005 |
| 30.8194444 | 0.006384 | 0.014601 | 0.007223 | 0.00168 | 0.079979 | 0.003916 | 0.006614 | 0.007802 | 0.004426 | 0.015053 | 0.007903 | 0.009117 | 0.020192 | 0.007698 | 0.004961 | 0.00374 | 0.058297 | 0.008758 | 0.000668 | 0.033837 | 0.018674 | 0.010296 | 0.00482 | 0.008126 |
| 30.8333333 | 0.00418 | 0.011416 | 0.005027 | 0.003077 | 0.081614 | 0.004913 | 0.004551 | 0.004956 | 0.008591 | 0.010825 | 0.015883 | 0.018926 | 0.02693 | 0.008383 | 0.007744 | 0.004251 | 0.037733 | 0.011466 | 0.00142 | 0.021418 | 0.011938 | 0.007989 | 0.009643 | 0.007271 |
| 30.8472222 | 0.002112 | 0.006255 | 0.009484 | 0.007018 | 0.077313 | 0.00594 | 0.00183 | 0.004532 | 0.010439 | 0.007922 | 0.021121 | 0.025775 | 0.029558 | 0.006426 | 0.015271 | 0.003511 | 0.023752 | 0.01309 | 0.003137 | 0.010618 | 0.004258 | 0.009289 | 0.010705 | 0.010367 |
| 30.8611111 | 0.000832 | 0.007073 | 0.016793 | 0.011967 | 0.072152 | 0.006175 | 0.002606 | 0.004202 | 0.008059 | 0.00569 | 0.021369 | 0.024714 | 0.030108 | 0.003909 | 0.023505 | 0.003337 | 0.015076 | 0.017063 | 0.00498 | 0.003956 | 0.001281 | 0.012511 | 0.010159 | 0.015985 |
| 30.875 | 0.000672 | 0.009377 | 0.020579 | 0.016387 | 0.076789 | 0.005907 | 0.005076 | 0.003025 | 0.006296 | 0.003482 | 0.0169 | 0.015503 | 0.031362 | 0.003439 | 0.027518 | 0.00408 | 0.006581 | 0.022758 | 0.004612 | 0.001714 | 0.002394 | 0.009874 | 0.016939 | 0.021557 |
| 30.8888889 | 0.001515 | 0.014333 | 0.023366 | 0.021802 | 0.088373 | 0.003678 | 0.004815 | 0.002818 | 0.008586 | 0.003048 | 0.01087 | 0.010643 | 0.035273 | 0.004013 | 0.025699 | 0.004348 | 0.007885 | 0.026106 | 0.002956 | 0.001464 | 0.008165 | 0.006236 | 0.028214 | 0.025087 |
| 30.9027778 | 0.002273 | 0.026802 | 0.028098 | 0.027165 | 0.090128 | 0.001105 | 0.003523 | 0.004131 | 0.009813 | 0.003575 | 0.011656 | 0.012649 | 0.042219 | 0.00867 | 0.021456 | 0.003722 | 0.02284 | 0.030207 | 0.004046 | 0.001518 | 0.018944 | 0.007112 | 0.037693 | 0.026518 |
| 30.9166667 | 0.003053 | 0.033102 | 0.031352 | 0.029144 | 0.076814 | 0.000421 | 0.007054 | 0.00876 | 0.007107 | 0.006831 | 0.01813 | 0.008705 | 0.043927 | 0.013421 | 0.014042 | 0.003882 | 0.031605 | 0.036039 | 0.00804 | 0.002283 | 0.029647 | 0.008875 | 0.045332 | 0.024869 |
| 30.9305556 | 0.003025 | 0.023024 | 0.032218 | 0.028514 | 0.06674 | 0.002845 | 0.014652 | 0.01545 | 0.008072 | 0.01595 | 0.017079 | 0.003114 | 0.038373 | 0.011775 | 0.008914 | 0.006554 | 0.028931 | 0.037917 | 0.011275 | 0.002002 | 0.035055 | 0.009695 | 0.0518 | 0.023997 |
| 30.9444444 | 0.00294 | 0.015762 | 0.035447 | 0.028325 | 0.076977 | 0.009539 | 0.020437 | 0.022146 | 0.015088 | 0.027579 | 0.009333 | 0.002226 | 0.036882 | 0.011689 | 0.014589 | 0.009177 | 0.029845 | 0.037823 | 0.011011 | 0.001334 | 0.036053 | 0.009305 | 0.055375 | 0.028826 |
| 30.9583333 | 0.005771 | 0.024628 | 0.042039 | 0.03233 | 0.101425 | 0.018219 | 0.020436 | 0.024779 | 0.020338 | 0.035558 | 0.005931 | 0.004375 | 0.041638 | 0.018391 | 0.021916 | 0.008599 | 0.03793 | 0.03937 | 0.008653 | 0.002643 | 0.03769 | 0.006471 | 0.057972 | 0.030097 |
| 30.9722222 | 0.01103 | 0.038397 | 0.04979 | 0.045753 | 0.122373 | 0.025563 | 0.016161 | 0.020971 | 0.020907 | 0.038818 | 0.009123 | 0.010176 | 0.048449 | 0.02406 | 0.02624 | 0.007203 | 0.050245 | 0.038975 | 0.006778 | 0.005667 | 0.039241 | 0.003031 | 0.059158 | 0.025849 |
| 30.9861111 | 0.019762 | 0.054299 | 0.053419 | 0.071867 | 0.133728 | 0.031738 | 0.013619 | 0.018372 | 0.022827 | 0.046651 | 0.016071 | 0.019417 | 0.056507 | 0.029177 | 0.030986 | 0.009921 | 0.066657 | 0.03614 | 0.006701 | 0.008856 | 0.037872 | 0.001533 | 0.051503 | 0.028668 |
| 31 | 0.031467 | 0.069551 | 0.051213 | 0.100309 | 0.139053 | 0.03807 | 0.016725 | 0.023365 | 0.029547 | 0.065335 | 0.0242 | 0.031001 | 0.062972 | 0.038128 | 0.037205 | 0.018255 | 0.072813 | 0.038811 | 0.010719 | 0.012833 | 0.034126 | 0.001594 | 0.039588 | 0.034472 |
| 31.0138889 | 0.038233 | 0.076659 | 0.047457 | 0.115077 | 0.136603 | 0.041281 | 0.020703 | 0.02965 | 0.034637 | 0.079865 | 0.030074 | 0.042394 | 0.064402 | 0.045948 | 0.051983 | 0.031019 | 0.05177 | 0.051701 | 0.018824 | 0.016674 | 0.030409 | 0.002108 | 0.031694 | 0.032093 |
| 31.0277778 | 0.03771 | 0.075947 | 0.045349 | 0.118092 | 0.128023 | 0.038789 | 0.019419 | 0.0283 | 0.031547 | 0.077897 | 0.033042 | 0.050804 | 0.061967 | 0.053306 | 0.071979 | 0.049631 | 0.025148 | 0.07152 | 0.027828 | 0.019675 | 0.032924 | 0.00721 | 0.031697 | 0.029613 |
| 31.0416667 | 0.03985 | 0.073341 | 0.050381 | 0.121809 | 0.12425 | 0.035335 | 0.017015 | 0.020988 | 0.025834 | 0.074605 | 0.036042 | 0.055364 | 0.067366 | 0.058866 | 0.088483 | 0.066858 | 0.025 | 0.092114 | 0.036762 | 0.029772 | 0.041646 | 0.020156 | 0.046135 | 0.031899 |
| 31.0555556 | 0.051148 | 0.079668 | 0.06164 | 0.129696 | 0.131424 | 0.03872 | 0.023083 | 0.016088 | 0.027964 | 0.084086 | 0.044628 | 0.063476 | 0.085114 | 0.049834 | 0.099526 | 0.06826 | 0.04053 | 0.106921 | 0.042322 | 0.042379 | 0.043823 | 0.025713 | 0.060583 | 0.028788 |
| 31.0694444 | 0.058332 | 0.089982 | 0.06868 | 0.136133 | 0.147691 | 0.049859 | 0.033893 | 0.018024 | 0.035578 | 0.096634 | 0.060824 | 0.07123 | 0.099826 | 0.038602 | 0.093589 | 0.054072 | 0.044086 | 0.108846 | 0.043451 | 0.042403 | 0.035116 | 0.019408 | 0.05337 | 0.018748 |
| 31.0833333 | 0.05089 | 0.088031 | 0.062882 | 0.130925 | 0.155704 | 0.054672 | 0.035658 | 0.021582 | 0.036378 | 0.09251 | 0.076759 | 0.060322 | 0.095644 | 0.049048 | 0.079666 | 0.039923 | 0.03541 | 0.097147 | 0.047882 | 0.032443 | 0.025386 | 0.013208 | 0.040704 | 0.015598 |
| 31.0972222 | 0.043421 | 0.071465 | 0.043842 | 0.106637 | 0.13667 | 0.043134 | 0.029456 | 0.018014 | 0.027161 | 0.067734 | 0.084065 | 0.037559 | 0.062284 | 0.072967 | 0.07648 | 0.039396 | 0.030511 | 0.078202 | 0.051575 | 0.020143 | 0.015754 | 0.014602 | 0.038847 | 0.025879 |
| 31.1111111 | 0.039972 | 0.048207 | 0.023995 | 0.071312 | 0.105305 | 0.02591 | 0.023854 | 0.010307 | 0.016215 | 0.040289 | 0.078096 | 0.024504 | 0.021565 | 0.078531 | 0.066332 | 0.047451 | 0.031886 | 0.052115 | 0.044049 | 0.01396 | 0.008876 | 0.020762 | 0.038425 | 0.0318 |
| 31.125 | 0.030538 | 0.029365 | 0.014802 | 0.038884 | 0.081122 | 0.014572 | 0.020145 | 0.008837 | 0.010788 | 0.024114 | 0.063126 | 0.022783 | 0.004677 | 0.048456 | 0.042752 | 0.046533 | 0.034685 | 0.024008 | 0.027551 | 0.015927 | 0.010957 | 0.018929 | 0.038044 | 0.021095 |
| 31.1388889 | 0.019621 | 0.021013 | 0.013306 | 0.019467 | 0.059614 | 0.008442 | 0.016301 | 0.012007 | 0.008918 | 0.015958 | 0.050256 | 0.020597 | 0.002746 | 0.018637 | 0.022973 | 0.034034 | 0.034181 | 0.007793 | 0.014019 | 0.023071 | 0.013366 | 0.01408 | 0.038061 | 0.011185 |
| 31.1527778 | 0.011374 | 0.016614 | 0.010941 | 0.016007 | 0.035304 | 0.004738 | 0.009849 | 0.010304 | 0.006429 | 0.009493 | 0.043354 | 0.015992 | 0.00363 | 0.021319 | 0.020232 | 0.03737 | 0.030149 | 0.00654 | 0.011716 | 0.033143 | 0.01307 | 0.011055 | 0.033265 | 0.014251 |
| 31.1666667 | 0.007848 | 0.012551 | 0.006207 | 0.016052 | 0.015924 | 0.005351 | 0.004302 | 0.005362 | 0.005914 | 0.007088 | 0.034663 | 0.011877 | 0.006953 | 0.035617 | 0.030075 | 0.06562 | 0.026762 | 0.007576 | 0.013813 | 0.031526 | 0.012446 | 0.011147 | 0.021417 | 0.018833 |
| 31.1805556 | 0.010851 | 0.017597 | 0.004069 | 0.012251 | 0.009362 | 0.010884 | 0.004668 | 0.004974 | 0.008827 | 0.010948 | 0.021457 | 0.007724 | 0.011462 | 0.043393 | 0.035434 | 0.085736 | 0.024922 | 0.006843 | 0.011724 | 0.019354 | 0.011406 | 0.015972 | 0.011125 | 0.018834 |
| 31.1944444 | 0.013917 | 0.032153 | 0.004768 | 0.014955 | 0.010832 | 0.017038 | 0.007837 | 0.007786 | 0.011965 | 0.017908 | 0.017756 | 0.007833 | 0.020962 | 0.043193 | 0.037645 | 0.081267 | 0.028015 | 0.014361 | 0.011721 | 0.010198 | 0.014454 | 0.015255 | 0.017026 | 0.01605 |
| 31.2083333 | 0.013183 | 0.042927 | 0.005729 | 0.026512 | 0.009667 | 0.020908 | 0.009643 | 0.011134 | 0.014282 | 0.027121 | 0.025914 | 0.013287 | 0.042586 | 0.034229 | 0.041785 | 0.061347 | 0.029186 | 0.024696 | 0.01907 | 0.006143 | 0.020419 | 0.008743 | 0.036927 | 0.009706 |
| 31.2222222 | 0.010797 | 0.044792 | 0.008259 | 0.03745 | 0.006223 | 0.024093 | 0.01031 | 0.015633 | 0.015923 | 0.037048 | 0.03125 | 0.017666 | 0.064219 | 0.020607 | 0.044282 | 0.037392 | 0.018517 | 0.027941 | 0.024182 | 0.00522 | 0.026291 | 0.005855 | 0.044919 | 0.005992 |
| 31.2361111 | 0.009913 | 0.041674 | 0.011591 | 0.039682 | 0.01036 | 0.027113 | 0.010531 | 0.018472 | 0.016439 | 0.04377 | 0.032072 | 0.018593 | 0.060832 | 0.010032 | 0.046482 | 0.020994 | 0.011112 | 0.027907 | 0.026597 | 0.00402 | 0.031564 | 0.008026 | 0.03031 | 0.009373 |
| 31.25 | 0.011278 | 0.035317 | 0.012138 | 0.029319 | 0.021686 | 0.02736 | 0.010097 | 0.01854 | 0.015818 | 0.045439 | 0.036086 | 0.019366 | 0.040331 | 0.007183 | 0.05143 | 0.012532 | 0.021016 | 0.029081 | 0.031778 | 0.0035 | 0.033619 | 0.009054 | 0.017168 | 0.012233 |
| 31.2638889 | 0.012113 | 0.027862 | 0.010398 | 0.013183 | 0.031582 | 0.023947 | 0.008665 | 0.018644 | 0.014349 | 0.042096 | 0.042612 | 0.023134 | 0.02755 | 0.01011 | 0.055073 | 0.006915 | 0.032973 | 0.029832 | 0.033659 | 0.009284 | 0.035177 | 0.009184 | 0.01569 | 0.009911 |
| 31.2777778 | 0.009749 | 0.024051 | 0.009379 | 0.003915 | 0.035356 | 0.019088 | 0.005524 | 0.018182 | 0.012553 | 0.035241 | 0.045458 | 0.027767 | 0.033521 | 0.017992 | 0.051196 | 0.005817 | 0.035016 | 0.028324 | 0.029249 | 0.01884 | 0.040744 | 0.011997 | 0.01617 | 0.009045 |
| 31.2916667 | 0.005043 | 0.025272 | 0.009808 | 0.006145 | 0.033053 | 0.015009 | 0.002483 | 0.015365 | 0.011228 | 0.028075 | 0.044332 | 0.02881 | 0.054782 | 0.028702 | 0.041917 | 0.011609 | 0.044446 | 0.0281 | 0.026492 | 0.019183 | 0.037837 | 0.012684 | 0.013024 | 0.013141 |
| 31.3055556 | 0.00206 | 0.026824 | 0.010894 | 0.01269 | 0.03449 | 0.01228 | 0.001941 | 0.012639 | 0.011191 | 0.02318 | 0.045133 | 0.029205 | 0.070688 | 0.035167 | 0.033532 | 0.019823 | 0.060092 | 0.034208 | 0.033212 | 0.010334 | 0.021761 | 0.008727 | 0.006651 | 0.014915 |
| 31.3194444 | 0.001933 | 0.026751 | 0.011559 | 0.016806 | 0.048614 | 0.00984 | 0.001913 | 0.011206 | 0.011723 | 0.018209 | 0.046998 | 0.030901 | 0.06935 | 0.033231 | 0.026197 | 0.023392 | 0.047366 | 0.046446 | 0.047306 | 0.007052 | 0.022005 | 0.005579 | 0.004367 | 0.012557 |
| 31.3333333 | 0.001929 | 0.028242 | 0.010772 | 0.019241 | 0.064907 | 0.006753 | 0.001508 | 0.009674 | 0.01078 | 0.013157 | 0.043347 | 0.029433 | 0.060585 | 0.024374 | 0.01751 | 0.021838 | 0.021873 | 0.058549 | 0.058964 | 0.011543 | 0.03897 | 0.007693 | 0.008944 | 0.010492 |
| 31.3472222 | 0.001695 | 0.030424 | 0.008679 | 0.016837 | 0.071695 | 0.004628 | 0.001376 | 0.009394 | 0.007875 | 0.015093 | 0.033817 | 0.024379 | 0.056565 | 0.015279 | 0.01039 | 0.018584 | 0.013423 | 0.065952 | 0.062032 | 0.013944 | 0.043411 | 0.011316 | 0.017409 | 0.010634 |
| 31.3611111 | 0.001721 | 0.03208 | 0.006945 | 0.008591 | 0.074622 | 0.00608 | 0.000761 | 0.013799 | 0.00578 | 0.021435 | 0.024843 | 0.0201 | 0.060113 | 0.014319 | 0.009276 | 0.017368 | 0.015191 | 0.068604 | 0.058838 | 0.011955 | 0.035311 | 0.010123 | 0.027874 | 0.0138 |
| 31.375 | 0.002392 | 0.033782 | 0.00689 | 0.002767 | 0.082504 | 0.010827 | 0.001595 | 0.020982 | 0.006825 | 0.018853 | 0.019865 | 0.017952 | 0.064953 | 0.023381 | 0.012197 | 0.021029 | 0.024217 | 0.062464 | 0.049099 | 0.012098 | 0.02093 | 0.006118 | 0.037469 | 0.018923 |
| 31.3888889 | 0.003638 | 0.031466 | 0.007485 | 0.00386 | 0.091527 | 0.014802 | 0.0053 | 0.023124 | 0.009043 | 0.010261 | 0.013994 | 0.013746 | 0.06285 | 0.033004 | 0.012587 | 0.025926 | 0.028568 | 0.047347 | 0.035039 | 0.016767 | 0.008254 | 0.004721 | 0.045017 | 0.024397 |
| 31.4027778 | 0.003098 | 0.020816 | 0.008011 | 0.01224 | 0.096176 | 0.015573 | 0.009923 | 0.018134 | 0.009953 | 0.012188 | 0.007497 | 0.007847 | 0.053614 | 0.033282 | 0.011632 | 0.030131 | 0.023943 | 0.036108 | 0.027734 | 0.020397 | 0.01209 | 0.004031 | 0.050244 | 0.028532 |
| 31.4166667 | 0.001671 | 0.007298 | 0.009855 | 0.022854 | 0.099866 | 0.016211 | 0.013862 | 0.012657 | 0.010139 | 0.025817 | 0.006007 | 0.005784 | 0.037925 | 0.030447 | 0.018117 | 0.037587 | 0.02053 | 0.033974 | 0.028245 | 0.020829 | 0.030763 | 0.004801 | 0.047618 | 0.028373 |
| 31.4305556 | 0.002648 | 0.001229 | 0.013524 | 0.026343 | 0.10553 | 0.01986 | 0.016589 | 0.012778 | 0.011046 | 0.04035 | 0.00598 | 0.005737 | 0.017837 | 0.036485 | 0.034251 | 0.047541 | 0.029336 | 0.034078 | 0.028979 | 0.020336 | 0.051089 | 0.010579 | 0.042571 | 0.029824 |
| 31.4444444 | 0.006479 | 0.001445 | 0.018394 | 0.020047 | 0.112164 | 0.025314 | 0.018527 | 0.01684 | 0.013008 | 0.049908 | 0.004602 | 0.00464 | 0.010065 | 0.052262 | 0.058254 | 0.058129 | 0.039338 | 0.037778 | 0.033112 | 0.018413 | 0.064065 | 0.013232 | 0.051697 | 0.040267 |
| 31.4583333 | 0.011301 | 0.005041 | 0.023316 | 0.011397 | 0.118843 | 0.030047 | 0.021846 | 0.020369 | 0.01553 | 0.054052 | 0.00691 | 0.006544 | 0.023458 | 0.066542 | 0.085032 | 0.067161 | 0.033641 | 0.046757 | 0.043139 | 0.019076 | 0.068242 | 0.008634 | 0.069858 | 0.048547 |
| 31.4722222 | 0.013834 | 0.015124 | 0.02556 | 0.013045 | 0.1254 | 0.034012 | 0.026913 | 0.022113 | 0.018129 | 0.055079 | 0.014007 | 0.0131 | 0.046628 | 0.070703 | 0.1063 | 0.071002 | 0.042033 | 0.051385 | 0.047994 | 0.027137 | 0.068579 | 0.00606 | 0.07765 | 0.04658 |
| 31.4861111 | 0.012788 | 0.028969 | 0.02696 | 0.027325 | 0.131945 | 0.039594 | 0.031191 | 0.021508 | 0.020353 | 0.055401 | 0.021252 | 0.019205 | 0.065578 | 0.069195 | 0.120173 | 0.068053 | 0.07637 | 0.050223 | 0.044991 | 0.038299 | 0.066449 | 0.008895 | 0.07618 | 0.042035 |
| 31.5 | 0.011709 | 0.040812 | 0.032607 | 0.04619 | 0.138831 | 0.04665 | 0.034064 | 0.019888 | 0.021912 | 0.054498 | 0.027195 | 0.023998 | 0.07432 | 0.068385 | 0.120303 | 0.057209 | 0.097213 | 0.054032 | 0.043878 | 0.049428 | 0.061426 | 0.008772 | 0.082118 | 0.041654 |
| 31.5138889 | 0.013527 | 0.047277 | 0.039239 | 0.061438 | 0.144638 | 0.049192 | 0.035295 | 0.018009 | 0.02337 | 0.052124 | 0.033217 | 0.030971 | 0.076423 | 0.071236 | 0.10536 | 0.046771 | 0.095179 | 0.063228 | 0.04547 | 0.065045 | 0.05565 | 0.004928 | 0.093164 | 0.046345 |
| 31.5277778 | 0.016316 | 0.04463 | 0.039948 | 0.065768 | 0.145451 | 0.045761 | 0.035042 | 0.014781 | 0.025143 | 0.050074 | 0.038685 | 0.041105 | 0.07795 | 0.079566 | 0.095186 | 0.047628 | 0.092656 | 0.069639 | 0.045 | 0.089595 | 0.049065 | 0.003816 | 0.092739 | 0.055011 |
| 31.5416667 | 0.017401 | 0.033844 | 0.033195 | 0.058673 | 0.131566 | 0.043925 | 0.035375 | 0.012765 | 0.028265 | 0.051198 | 0.041534 | 0.052444 | 0.079969 | 0.091779 | 0.096804 | 0.055039 | 0.080154 | 0.07202 | 0.041952 | 0.112657 | 0.045237 | 0.004319 | 0.087214 | 0.063652 |
| 31.5555556 | 0.015213 | 0.021313 | 0.022861 | 0.043243 | 0.103293 | 0.044032 | 0.034401 | 0.013349 | 0.032373 | 0.053885 | 0.040182 | 0.057579 | 0.07938 | 0.098051 | 0.097395 | 0.056195 | 0.048651 | 0.072088 | 0.036732 | 0.108022 | 0.045034 | 0.005299 | 0.087858 | 0.066991 |
| 31.5694444 | 0.00963 | 0.011028 | 0.014837 | 0.023757 | 0.079676 | 0.037968 | 0.0283 | 0.013449 | 0.033721 | 0.051461 | 0.032991 | 0.048529 | 0.074838 | 0.093773 | 0.101518 | 0.052226 | 0.031975 | 0.073511 | 0.035411 | 0.072349 | 0.049459 | 0.011362 | 0.085744 | 0.067465 |
| 31.5833333 | 0.005782 | 0.01226 | 0.013615 | 0.012266 | 0.076325 | 0.025304 | 0.018379 | 0.012271 | 0.032768 | 0.046387 | 0.023411 | 0.027197 | 0.07325 | 0.086787 | 0.111386 | 0.047861 | 0.055958 | 0.078821 | 0.041833 | 0.038184 | 0.054948 | 0.015225 | 0.075477 | 0.069573 |
| 31.5972222 | 0.006371 | 0.030626 | 0.019571 | 0.016599 | 0.08463 | 0.014843 | 0.011114 | 0.010673 | 0.033817 | 0.047537 | 0.018958 | 0.014946 | 0.077192 | 0.07881 | 0.111979 | 0.03785 | 0.095942 | 0.07917 | 0.046545 | 0.031365 | 0.052311 | 0.015844 | 0.060559 | 0.068457 |
| 31.6111111 | 0.010706 | 0.051433 | 0.026727 | 0.030331 | 0.076311 | 0.011252 | 0.010279 | 0.008735 | 0.036132 | 0.051246 | 0.017984 | 0.017449 | 0.073616 | 0.068991 | 0.102668 | 0.023266 | 0.099179 | 0.071561 | 0.044352 | 0.046668 | 0.046681 | 0.020705 | 0.046151 | 0.060574 |
| 31.625 | 0.019405 | 0.074863 | 0.025644 | 0.039089 | 0.05163 | 0.010064 | 0.013959 | 0.007599 | 0.037489 | 0.05165 | 0.013184 | 0.014804 | 0.061466 | 0.059674 | 0.091315 | 0.012517 | 0.0796 | 0.066302 | 0.036876 | 0.056017 | 0.043693 | 0.017658 | 0.03911 | 0.050203 |
| 31.6388889 | 0.024297 | 0.102709 | 0.016327 | 0.036882 | 0.036925 | 0.008431 | 0.018912 | 0.00574 | 0.039734 | 0.054971 | 0.005951 | 0.009848 | 0.055095 | 0.05055 | 0.083862 | 0.008399 | 0.091462 | 0.06288 | 0.025937 | 0.050575 | 0.039341 | 0.009652 | 0.042445 | 0.040526 |
| 31.6527778 | 0.020525 | 0.111052 | 0.009956 | 0.035733 | 0.034033 | 0.009495 | 0.019116 | 0.002874 | 0.041039 | 0.058819 | 0.004254 | 0.007439 | 0.057412 | 0.042963 | 0.08112 | 0.005411 | 0.105857 | 0.052599 | 0.016502 | 0.043547 | 0.030837 | 0.008539 | 0.050348 | 0.033669 |
| 31.6666667 | 0.013581 | 0.098388 | 0.007195 | 0.046257 | 0.029414 | 0.012487 | 0.011596 | 0.002805 | 0.035835 | 0.05223 | 0.006573 | 0.002901 | 0.059342 | 0.035507 | 0.074185 | 0.002227 | 0.094288 | 0.038719 | 0.010077 | 0.036881 | 0.022191 | 0.009011 | 0.049533 | 0.029886 |
| 31.6805556 | 0.008809 | 0.073819 | 0.003683 | 0.059534 | 0.018452 | 0.01068 | 0.004143 | 0.003234 | 0.023792 | 0.03338 | 0.00543 | 0.001251 | 0.055152 | 0.021571 | 0.057743 | 0.00221 | 0.063348 | 0.028744 | 0.006102 | 0.029679 | 0.01445 | 0.00812 | 0.035453 | 0.02674 |
| 31.6944444 | 0.01114 | 0.043928 | 0.002885 | 0.063893 | 0.010104 | 0.006822 | 0.005047 | 0.003881 | 0.011627 | 0.016689 | 0.002947 | 0.003398 | 0.045057 | 0.009868 | 0.03844 | 0.004648 | 0.040071 | 0.021318 | 0.004241 | 0.021927 | 0.012165 | 0.005726 | 0.017708 | 0.022509 |

| | | | | | | | | | | | | | | | | | | | | | | | | |
|---|---|---|---|---|---|---|---|---|---|---|---|---|---|---|---|---|---|---|---|---|---|---|---|---|
| 31.7083333 | 0.017951 | 0.02822 | 0.004494 | 0.06241 | 0.015343 | 0.008998 | 0.012398 | 0.009902 | 0.007069 | 0.016316 | 0.003762 | 0.004773 | 0.029833 | 0.012531 | 0.021768 | 0.006862 | 0.06888 | 0.01534 | 0.002459 | 0.013752 | 0.029976 | 0.003455 | 0.009379 | 0.016893 |
| 31.7222222 | 0.023959 | 0.025991 | 0.00565 | 0.062473 | 0.029279 | 0.013563 | 0.019736 | 0.016461 | 0.007632 | 0.020399 | 0.004242 | 0.005089 | 0.015448 | 0.017552 | 0.00999 | 0.006741 | 0.117499 | 0.012962 | 0.000879 | 0.005702 | 0.054899 | 0.003734 | 0.008805 | 0.012289 |
| 31.7361111 | 0.028173 | 0.022064 | 0.006384 | 0.068065 | 0.041336 | 0.014659 | 0.022987 | 0.016869 | 0.006036 | 0.018681 | 0.004273 | 0.009338 | 0.010806 | 0.012894 | 0.006047 | 0.005222 | 0.138513 | 0.011712 | 0.000575 | 0.002922 | 0.056814 | 0.004435 | 0.006635 | 0.011584 |
| 31.75 | 0.030184 | 0.018168 | 0.006516 | 0.076394 | 0.051085 | 0.013025 | 0.021192 | 0.013928 | 0.002807 | 0.019983 | 0.008964 | 0.015351 | 0.013957 | 0.007903 | 0.008745 | 0.004233 | 0.127792 | 0.00823 | 0.002382 | 0.009894 | 0.035962 | 0.003472 | 0.005606 | 0.014204 |
| 31.7638889 | 0.030688 | 0.020938 | 0.008546 | 0.077894 | 0.067318 | 0.011642 | 0.016711 | 0.011988 | 0.00106 | 0.026309 | 0.015575 | 0.017749 | 0.011745 | 0.010334 | 0.021513 | 0.00421 | 0.093417 | 0.003503 | 0.006345 | 0.024725 | 0.01405 | 0.002919 | 0.007107 | 0.015174 |
| 31.7777778 | 0.028436 | 0.023783 | 0.012112 | 0.065498 | 0.091986 | 0.012734 | 0.013136 | 0.010648 | 0.001362 | 0.026748 | 0.020149 | 0.018545 | 0.006106 | 0.012871 | 0.032756 | 0.004645 | 0.055941 | 0.000906 | 0.007986 | 0.042972 | 0.004343 | 0.003833 | 0.011458 | 0.011266 |
| 31.7916667 | 0.021701 | 0.019184 | 0.010096 | 0.042704 | 0.11374 | 0.016091 | 0.010906 | 0.007005 | 0.003055 | 0.016527 | 0.022087 | 0.019724 | 0.006558 | 0.015274 | 0.023662 | 0.004106 | 0.033677 | 0.002785 | 0.00577 | 0.052069 | 0.004144 | 0.004067 | 0.017262 | 0.007645 |
| 31.8055556 | 0.014173 | 0.010944 | 0.004293 | 0.025077 | 0.122341 | 0.019811 | 0.007358 | 0.005233 | 0.0043 | 0.009252 | 0.020556 | 0.02159 | 0.007249 | 0.024354 | 0.010282 | 0.002665 | 0.024482 | 0.006618 | 0.004035 | 0.040413 | 0.007943 | 0.007602 | 0.018054 | 0.009088 |
| 31.8194444 | 0.010191 | 0.01147 | 0.001998 | 0.021682 | 0.121009 | 0.021686 | 0.004281 | 0.009593 | 0.005252 | 0.015557 | 0.017122 | 0.027209 | 0.005392 | 0.038786 | 0.009434 | 0.001665 | 0.02329 | 0.007579 | 0.004572 | 0.018044 | 0.014014 | 0.020609 | 0.017845 | 0.014704 |
| 31.8333333 | 0.00846 | 0.029932 | 0.00364 | 0.02595 | 0.119171 | 0.020323 | 0.006271 | 0.012352 | 0.007044 | 0.024423 | 0.014456 | 0.031996 | 0.003688 | 0.051321 | 0.009877 | 0.001643 | 0.027726 | 0.009274 | 0.006402 | 0.005836 | 0.018956 | 0.034791 | 0.025774 | 0.0221 |
| 31.8472222 | 0.006343 | 0.049432 | 0.004609 | 0.03572 | 0.119644 | 0.016594 | 0.010996 | 0.009698 | 0.008989 | 0.024421 | 0.016564 | 0.02385 | 0.009351 | 0.052968 | 0.006181 | 0.002693 | 0.025037 | 0.014539 | 0.007028 | 0.010346 | 0.016454 | 0.035187 | 0.031821 | 0.028471 |
| 31.8611111 | 0.005871 | 0.046069 | 0.004078 | 0.044058 | 0.121227 | 0.012904 | 0.01435 | 0.007348 | 0.010774 | 0.019516 | 0.019497 | 0.008991 | 0.028407 | 0.041787 | 0.003669 | 0.00281 | 0.0191 | 0.018561 | 0.004821 | 0.022383 | 0.010869 | 0.022452 | 0.027744 | 0.034194 |
| 31.875 | 0.008015 | 0.025208 | 0.005625 | 0.036716 | 0.118877 | 0.010624 | 0.016746 | 0.008613 | 0.012482 | 0.020182 | 0.017745 | 0.004423 | 0.045505 | 0.025553 | 0.002377 | 0.002361 | 0.023734 | 0.022808 | 0.002646 | 0.026584 | 0.011026 | 0.009142 | 0.030371 | 0.036732 |
| 31.8888889 | 0.007087 | 0.016088 | 0.006586 | 0.019156 | 0.10942 | 0.009956 | 0.016883 | 0.01056 | 0.013105 | 0.0263 | 0.025609 | 0.012461 | 0.045577 | 0.015934 | 0.002438 | 0.005823 | 0.033481 | 0.035083 | 0.005934 | 0.018964 | 0.010497 | 0.003317 | 0.043114 | 0.031316 |
| 31.9027778 | 0.003893 | 0.028598 | 0.006912 | 0.007928 | 0.09959 | 0.010878 | 0.012567 | 0.008995 | 0.012028 | 0.025641 | 0.043731 | 0.020916 | 0.036198 | 0.01591 | 0.009676 | 0.013321 | 0.035883 | 0.053662 | 0.01536 | 0.008869 | 0.005822 | 0.003321 | 0.048441 | 0.022001 |
| 31.9166667 | 0.003624 | 0.03953 | 0.01105 | 0.00905 | 0.098167 | 0.012561 | 0.006854 | 0.004887 | 0.010804 | 0.014931 | 0.041823 | 0.019425 | 0.034295 | 0.014647 | 0.02836 | 0.022133 | 0.030314 | 0.064147 | 0.022895 | 0.005138 | 0.007701 | 0.004 | 0.047487 | 0.012751 |
| 31.9305556 | 0.003122 | 0.041503 | 0.015994 | 0.015681 | 0.108789 | 0.012894 | 0.003259 | 0.002898 | 0.009895 | 0.009437 | 0.020642 | 0.019489 | 0.04125 | 0.007699 | 0.052332 | 0.029488 | 0.029515 | 0.061259 | 0.021931 | 0.008757 | 0.019381 | 0.005966 | 0.052156 | 0.006158 |
| 31.9444444 | 0.002025 | 0.044207 | 0.018414 | 0.021255 | 0.125664 | 0.011583 | 0.005353 | 0.006251 | 0.006938 | 0.020803 | 0.00938 | 0.02808 | 0.042077 | 0.003771 | 0.068572 | 0.031804 | 0.045229 | 0.050859 | 0.016803 | 0.013619 | 0.027493 | 0.009503 | 0.058894 | 0.006016 |
| 31.9583333 | 0.005234 | 0.048816 | 0.021397 | 0.0255 | 0.137698 | 0.012101 | 0.014634 | 0.014047 | 0.005724 | 0.037135 | 0.00943 | 0.035758 | 0.037721 | 0.005841 | 0.073828 | 0.032355 | 0.070429 | 0.04168 | 0.015299 | 0.01337 | 0.027738 | 0.010702 | 0.071013 | 0.007662 |
| 31.9722222 | 0.012741 | 0.049898 | 0.027424 | 0.028088 | 0.140686 | 0.01678 | 0.022819 | 0.024222 | 0.011436 | 0.044538 | 0.017973 | 0.042971 | 0.042971 | 0.009639 | 0.073172 | 0.036639 | 0.091508 | 0.041376 | 0.018294 | 0.009235 | 0.025174 | 0.00803 | 0.089236 | 0.005359 |
| 31.9861111 | 0.019267 | 0.042032 | 0.032092 | 0.024049 | 0.140334 | 0.022937 | 0.022737 | 0.03341 | 0.017872 | 0.046185 | 0.042245 | 0.054481 | 0.055514 | 0.014121 | 0.067427 | 0.041166 | 0.102334 | 0.044049 | 0.020564 | 0.010965 | 0.019537 | 0.005018 | 0.098723 | 0.002326 |
| 32 | 0.020746 | 0.029189 | 0.031251 | 0.01322 | 0.147156 | 0.027719 | 0.016485 | 0.035746 | 0.019256 | 0.056981 | 0.076434 | 0.068378 | 0.064161 | 0.019748 | 0.058877 | 0.04154 | 0.10557 | 0.044888 | 0.019158 | 0.023724 | 0.021495 | 0.013675 | 0.102873 | 0.001405 |
| 32.0138889 | 0.018934 | 0.018874 | 0.0292 | 0.00434 | 0.1629 | 0.031554 | 0.010575 | 0.032576 | 0.018013 | 0.079212 | 0.100105 | 0.072215 | 0.071156 | 0.025889 | 0.052326 | 0.039951 | 0.112179 | 0.04991 | 0.016452 | 0.04043 | 0.039626 | 0.02342 | 0.112139 | 0.00213 |
| 32.0277778 | 0.016026 | 0.012638 | 0.028563 | 0.00139 | 0.174467 | 0.03529 | 0.009464 | 0.032325 | 0.016601 | 0.094613 | 0.102395 | 0.064042 | 0.082608 | 0.029478 | 0.052073 | 0.040798 | 0.127384 | 0.058349 | 0.014135 | 0.054541 | 0.05112 | 0.013873 | 0.125512 | 0.007809 |
| 32.0416667 | 0.012443 | 0.010357 | 0.026281 | 0.002176 | 0.168036 | 0.043588 | 0.014852 | 0.038825 | 0.018126 | 0.102093 | 0.103125 | 0.059521 | 0.09654 | 0.032155 | 0.056375 | 0.043396 | 0.154564 | 0.062575 | 0.009737 | 0.069857 | 0.034997 | 0.003555 | 0.144262 | 0.021104 |
| 32.0555556 | 0.011701 | 0.018054 | 0.023952 | 0.007269 | 0.151759 | 0.061309 | 0.03007 | 0.050276 | 0.026288 | 0.116424 | 0.121099 | 0.06591 | 0.106543 | 0.040269 | 0.061099 | 0.043931 | 0.169919 | 0.059774 | 0.005314 | 0.086407 | 0.019164 | 0.011572 | 0.162787 | 0.034835 |
| 32.0694444 | 0.015543 | 0.036055 | 0.022669 | 0.013457 | 0.155045 | 0.078408 | 0.049287 | 0.058298 | 0.034938 | 0.137501 | 0.147417 | 0.074436 | 0.108954 | 0.049468 | 0.067683 | 0.045194 | 0.140286 | 0.057571 | 0.012027 | 0.084974 | 0.026214 | 0.02486 | 0.165655 | 0.033847 |
| 32.0833333 | 0.016798 | 0.042846 | 0.016174 | 0.014292 | 0.175049 | 0.076005 | 0.054415 | 0.055056 | 0.033062 | 0.142161 | 0.149939 | 0.068594 | 0.108018 | 0.048442 | 0.068765 | 0.049717 | 0.090434 | 0.060065 | 0.025692 | 0.060955 | 0.030031 | 0.026706 | 0.133952 | 0.018868 |
| 32.0972222 | 0.0106 | 0.025348 | 0.009878 | 0.012019 | 0.175103 | 0.053615 | 0.039478 | 0.043347 | 0.022414 | 0.117275 | 0.112909 | 0.048308 | 0.101796 | 0.035546 | 0.052545 | 0.051064 | 0.055338 | 0.053852 | 0.025821 | 0.040058 | 0.018899 | 0.028902 | 0.083624 | 0.007213 |
| 32.1111111 | 0.0054 | 0.0073 | 0.010884 | 0.014998 | 0.15372 | 0.030195 | 0.019547 | 0.029192 | 0.014153 | 0.086467 | 0.068109 | 0.030867 | 0.083075 | 0.022929 | 0.02648 | 0.045004 | 0.03837 | 0.03865 | 0.014706 | 0.026481 | 0.010483 | 0.045537 | 0.067926 | 0.006274 |
| 32.125 | 0.007475 | 0.003982 | 0.012462 | 0.023943 | 0.132874 | 0.016775 | 0.008017 | 0.017528 | 0.012637 | 0.066696 | 0.043396 | 0.023897 | 0.055696 | 0.014682 | 0.008745 | 0.032313 | 0.027412 | 0.031801 | 0.009857 | 0.014926 | 0.007715 | 0.058502 | 0.067709 | 0.008996 |
| 32.1388889 | 0.012876 | 0.003628 | 0.013882 | 0.02711 | 0.11407 | 0.011888 | 0.003487 | 0.010728 | 0.013028 | 0.055759 | 0.035281 | 0.023287 | 0.027557 | 0.008822 | 0.005303 | 0.019925 | 0.019381 | 0.033779 | 0.011596 | 0.013961 | 0.006425 | 0.053667 | 0.037413 | 0.009047 |
| 32.1527778 | 0.015707 | 0.002822 | 0.017858 | 0.024622 | 0.089265 | 0.011582 | 0.00253 | 0.00716 | 0.011094 | 0.048704 | 0.032172 | 0.024829 | 0.012528 | 0.009236 | 0.012917 | 0.02527 | 0.030041 | 0.030102 | 0.010029 | 0.017219 | 0.014738 | 0.041071 | 0.011487 | 0.012356 |
| 32.1666667 | 0.014172 | 0.005792 | 0.023355 | 0.027132 | 0.055576 | 0.010496 | 0.004668 | 0.00505 | 0.007199 | 0.038783 | 0.025278 | 0.025679 | 0.019078 | 0.009426 | 0.022961 | 0.045374 | 0.055136 | 0.01686 | 0.005257 | 0.014287 | 0.027869 | 0.027692 | 0.015139 | 0.018154 |
| 32.1805556 | 0.014273 | 0.012349 | 0.024551 | 0.032229 | 0.025843 | 0.006776 | 0.006588 | 0.006161 | 0.003724 | 0.025218 | 0.014956 | 0.022318 | 0.030596 | 0.005787 | 0.027016 | 0.060614 | 0.075823 | 0.005139 | 0.003664 | 0.010201 | 0.031035 | 0.020319 | 0.02782 | 0.017182 |
| 32.1944444 | 0.016814 | 0.019713 | 0.018266 | 0.031015 | 0.018896 | 0.005614 | 0.008169 | 0.01265 | 0.004045 | 0.012339 | 0.0149 | 0.013126 | 0.034121 | 0.004433 | 0.027633 | 0.064633 | 0.088081 | 0.001565 | 0.003722 | 0.011259 | 0.029313 | 0.018387 | 0.041806 | 0.0127 |
| 32.2083333 | 0.016182 | 0.024931 | 0.011158 | 0.024922 | 0.030489 | 0.009063 | 0.012196 | 0.020033 | 0.007488 | 0.007684 | 0.027388 | 0.005589 | 0.035538 | 0.009497 | 0.029463 | 0.060925 | 0.101395 | 0.004587 | 0.006753 | 0.010595 | 0.033049 | 0.018082 | 0.05687 | 0.010325 |
| 32.2222222 | 0.01397 | 0.026874 | 0.011939 | 0.022911 | 0.043797 | 0.012188 | 0.016127 | 0.024223 | 0.0095 | 0.013498 | 0.032484 | 0.006482 | 0.03874 | 0.018738 | 0.031553 | 0.053824 | 0.117897 | 0.011107 | 0.014126 | 0.007535 | 0.034904 | 0.024159 | 0.059041 | 0.011446 |
| 32.2361111 | 0.013453 | 0.02555 | 0.019543 | 0.025923 | 0.052716 | 0.013619 | 0.015749 | 0.027318 | 0.009238 | 0.021458 | 0.028358 | 0.009765 | 0.039421 | 0.0235 | 0.027268 | 0.037356 | 0.122707 | 0.015049 | 0.016274 | 0.008799 | 0.026206 | 0.02231 | 0.037698 | 0.012675 |
| 32.25 | 0.013717 | 0.024341 | 0.022427 | 0.028711 | 0.057614 | 0.014984 | 0.012272 | 0.031002 | 0.00843 | 0.025843 | 0.027245 | 0.010602 | 0.031154 | 0.023859 | 0.015898 | 0.019872 | 0.112868 | 0.012469 | 0.010278 | 0.011493 | 0.021943 | 0.015855 | 0.011753 | 0.014799 |
| 32.2638889 | 0.013676 | 0.025802 | 0.017714 | 0.029629 | 0.060082 | 0.015335 | 0.010075 | 0.03579 | 0.007414 | 0.026824 | 0.026907 | 0.010899 | 0.017187 | 0.026479 | 0.008072 | 0.016082 | 0.102093 | 0.006578 | 0.003905 | 0.016533 | 0.034149 | 0.01878 | 0.00389 | 0.022265 |
| 32.2777778 | 0.012895 | 0.02724 | 0.01348 | 0.030742 | 0.060209 | 0.01406 | 0.010719 | 0.04057 | 0.006084 | 0.02569 | 0.023301 | 0.010936 | 0.011669 | 0.028946 | 0.005843 | 0.01681 | 0.090001 | 0.004148 | 0.001561 | 0.02527 | 0.05166 | 0.018348 | 0.005576 | 0.02739 |
| 32.2916667 | 0.010825 | 0.027015 | 0.015787 | 0.032951 | 0.055466 | 0.011994 | 0.011335 | 0.040554 | 0.005258 | 0.023577 | 0.021097 | 0.00984 | 0.014762 | 0.025854 | 0.004189 | 0.016732 | 0.068957 | 0.006186 | 0.003007 | 0.026222 | 0.065494 | 0.020943 | 0.008664 | 0.029538 |
| 32.3055556 | 0.00805 | 0.026366 | 0.020198 | 0.033808 | 0.046639 | 0.008367 | 0.008177 | 0.033233 | 0.00518 | 0.018775 | 0.020689 | 0.007589 | 0.014094 | 0.018686 | 0.004792 | 0.021368 | 0.041205 | 0.005965 | 0.006512 | 0.01577 | 0.064418 | 0.027927 | 0.014975 | 0.027299 |
| 32.3194444 | 0.005978 | 0.024543 | 0.018237 | 0.031755 | 0.039184 | 0.003548 | 0.003557 | 0.023131 | 0.004364 | 0.012052 | 0.016615 | 0.004319 | 0.009585 | 0.011076 | 0.010723 | 0.028828 | 0.017997 | 0.005247 | 0.006951 | 0.007654 | 0.047737 | 0.026078 | 0.014508 | 0.018346 |
| 32.3333333 | 0.005964 | 0.02363 | 0.014505 | 0.033113 | 0.040792 | 0.00376 | 0.004617 | 0.020155 | 0.003964 | 0.01201 | 0.01498 | 0.001316 | 0.020872 | 0.006144 | 0.017243 | 0.029482 | 0.026638 | 0.016841 | 0.012037 | 0.010112 | 0.036138 | 0.036737 | 0.00882 | 0.016428 |
| 32.3472222 | 0.008737 | 0.030439 | 0.018623 | 0.047347 | 0.051493 | 0.011813 | 0.011232 | 0.025668 | 0.007649 | 0.021468 | 0.024747 | 0.000283 | 0.054088 | 0.006869 | 0.014191 | 0.019241 | 0.064608 | 0.036681 | 0.029129 | 0.016556 | 0.031474 | 0.060364 | 0.013022 | 0.021764 |
| 32.3611111 | 0.011775 | 0.040243 | 0.026288 | 0.064812 | 0.057614 | 0.017811 | 0.013264 | 0.026771 | 0.012817 | 0.027024 | 0.034111 | 0.000843 | 0.070863 | 0.01116 | 0.005063 | 0.008117 | 0.075158 | 0.041235 | 0.036049 | 0.017975 | 0.022273 | 0.059241 | 0.024835 | 0.023812 |
| 32.375 | 0.012007 | 0.041219 | 0.028403 | 0.064663 | 0.055672 | 0.013692 | 0.007458 | 0.019734 | 0.013469 | 0.021211 | 0.029506 | 0.001532 | 0.056998 | 0.013962 | 0.001653 | 0.007742 | 0.051517 | 0.029024 | 0.026841 | 0.013963 | 0.014702 | 0.036904 | 0.032352 | 0.020928 |
| 32.3888889 | 0.010031 | 0.03494 | 0.024438 | 0.050845 | 0.061019 | 0.006466 | 0.002439 | 0.01586 | 0.010486 | 0.01582 | 0.018551 | 0.002125 | 0.046035 | 0.015163 | 0.008998 | 0.022718 | 0.048524 | 0.021568 | 0.024057 | 0.013493 | 0.026575 | 0.024758 | 0.038017 | 0.016163 |
| 32.4027778 | 0.008402 | 0.02911 | 0.020274 | 0.041309 | 0.077027 | 0.004213 | 0.002847 | 0.018778 | 0.008368 | 0.018312 | 0.012078 | 0.002784 | 0.045042 | 0.017564 | 0.026487 | 0.046945 | 0.066704 | 0.02194 | 0.029075 | 0.020457 | 0.046782 | 0.028734 | 0.044786 | 0.016053 |
| 32.4166667 | 0.009465 | 0.025952 | 0.020296 | 0.038306 | 0.086796 | 0.005583 | 0.005939 | 0.021393 | 0.00909 | 0.022451 | 0.013365 | 0.002152 | 0.035049 | 0.021015 | 0.04389 | 0.057357 | 0.063012 | 0.017726 | 0.027433 | 0.027844 | 0.050027 | 0.029884 | 0.048934 | 0.022225 |
| 32.4305556 | 0.012436 | 0.025937 | 0.023409 | 0.036417 | 0.083322 | 0.007657 | 0.008916 | 0.021434 | 0.012312 | 0.026142 | 0.023507 | 0.001615 | 0.017286 | 0.023715 | 0.05276 | 0.050127 | 0.033829 | 0.009621 | 0.017934 | 0.030441 | 0.047784 | 0.017089 | 0.052986 | 0.030971 |
| 32.4444444 | 0.014503 | 0.03044 | 0.027622 | 0.037667 | 0.082379 | 0.01059 | 0.012103 | 0.024101 | 0.016625 | 0.035567 | 0.042009 | 0.003647 | 0.006625 | 0.025088 | 0.056332 | 0.045728 | 0.011338 | 0.004652 | 0.01158 | 0.028858 | 0.043969 | 0.005848 | 0.056375 | 0.03505 |
| 32.4583333 | 0.015082 | 0.039933 | 0.032618 | 0.049181 | 0.096323 | 0.014769 | 0.017874 | 0.032103 | 0.021036 | 0.052787 | 0.0654 | 0.006318 | 0.008733 | 0.025585 | 0.05659 | 0.041346 | 0.013363 | 0.005561 | 0.015385 | 0.025726 | 0.030818 | 0.012338 | 0.049873 | 0.029571 |
| 32.4722222 | 0.015 | 0.051875 | 0.037045 | 0.071094 | 0.116653 | 0.01913 | 0.024839 | 0.04031 | 0.025203 | 0.069427 | 0.086448 | 0.008899 | 0.022939 | 0.027833 | 0.053464 | 0.031967 | 0.033636 | 0.016139 | 0.025086 | 0.027069 | 0.023864 | 0.04026 | 0.038986 | 0.023193 |
| 32.4861111 | 0.015585 | 0.062072 | 0.039609 | 0.089246 | 0.131577 | 0.024889 | 0.029398 | 0.045405 | 0.029779 | 0.077443 | 0.099409 | 0.015048 | 0.045 | 0.034117 | 0.053473 | 0.030087 | 0.064896 | 0.035543 | 0.036907 | 0.042986 | 0.02351 | 0.076785 | 0.045702 | 0.027936 |
| 32.5 | 0.017963 | 0.0705 | 0.041653 | 0.096995 | 0.142924 | 0.037074 | 0.034867 | 0.05033 | 0.035154 | 0.079599 | 0.108433 | 0.025827 | 0.071053 | 0.044386 | 0.062026 | 0.036688 | 0.103822 | 0.057299 | 0.050979 | 0.060973 | 0.025036 | 0.097484 | 0.072635 | 0.044592 |
| 32.5138889 | 0.021128 | 0.079092 | 0.043791 | 0.111547 | 0.156794 | 0.052973 | 0.043831 | 0.05186 | 0.038934 | 0.08216 | 0.116016 | 0.033879 | 0.098575 | 0.061014 | 0.077871 | 0.043516 | 0.145037 | 0.072181 | 0.061918 | 0.063255 | 0.037509 | 0.096156 | 0.096489 | 0.061766 |
| 32.5277778 | 0.021945 | 0.082395 | 0.043097 | 0.123184 | 0.176745 | 0.060194 | 0.049156 | 0.046201 | 0.037979 | 0.083783 | 0.115034 | 0.034894 | 0.121667 | 0.083754 | 0.094484 | 0.051887 | 0.189482 | 0.079167 | 0.066666 | 0.060002 | 0.060687 | 0.083572 | 0.103743 | 0.071985 |
| 32.5416667 | 0.019665 | 0.076922 | 0.039141 | 0.105419 | 0.192059 | 0.056242 | 0.046963 | 0.040759 | 0.033071 | 0.081015 | 0.105583 | 0.036129 | 0.132341 | 0.104602 | 0.104148 | 0.063499 | 0.232765 | 0.084515 | 0.066374 | 0.059664 | 0.080015 | 0.071799 | 0.106064 | 0.08318 |
| 32.5555556 | 0.018023 | 0.065604 | 0.036096 | 0.079348 | 0.182854 | 0.048939 | 0.040868 | 0.043564 | 0.027409 | 0.076135 | 0.095121 | 0.043829 | 0.124626 | 0.116341 | 0.107166 | 0.067066 | 0.23804 | 0.083266 | 0.058309 | 0.055367 | 0.086682 | 0.072692 | 0.104599 | 0.099171 |
| 32.5694444 | 0.019148 | 0.048931 | 0.036406 | 0.064681 | 0.164394 | 0.042008 | 0.03298 | 0.05192 | 0.022085 | 0.070475 | 0.087316 | 0.054157 | 0.112609 | 0.11256 | 0.107937 | 0.058368 | 0.197169 | 0.074444 | 0.048279 | 0.039955 | 0.089564 | 0.065493 | 0.088405 | 0.103583 |
| 32.5833333 | 0.022647 | 0.031484 | 0.042645 | 0.051691 | 0.165211 | 0.038684 | 0.026989 | 0.056034 | 0.017393 | 0.064909 | 0.079712 | 0.058171 | 0.116533 | 0.097777 | 0.10619 | 0.046793 | 0.163608 | 0.071516 | 0.04708 | 0.029249 | 0.088597 | 0.035654 | 0.067469 | 0.0876 |
| 32.5972222 | 0.028569 | 0.020012 | 0.052709 | 0.036387 | 0.174494 | 0.042157 | 0.026537 | 0.050919 | 0.015773 | 0.063285 | 0.069114 | 0.053865 | 0.123356 | 0.089801 | 0.100658 | 0.03726 | 0.158763 | 0.0776 | 0.050483 | 0.035207 | 0.079733 | 0.017951 | 0.059286 | 0.067055 |
| 32.6111111 | 0.034729 | 0.016985 | 0.053625 | 0.025291 | 0.169045 | 0.046342 | 0.025983 | 0.040246 | 0.019534 | 0.065183 | 0.060103 | 0.050456 | 0.111675 | 0.091588 | 0.094245 | 0.028381 | 0.160849 | 0.076558 | 0.045961 | 0.038135 | 0.068308 | 0.024529 | 0.072915 | 0.061837 |
| 32.625 | 0.038196 | 0.01919 | 0.041241 | 0.025788 | 0.147586 | 0.044156 | 0.020459 | 0.027745 | 0.025775 | 0.065417 | 0.059455 | 0.055021 | 0.083423 | 0.09274 | 0.092408 | 0.018052 | 0.164233 | 0.062189 | 0.033366 | 0.02981 | 0.056754 | 0.04319 | 0.101006 | 0.075522 |
| 32.6388889 | 0.038694 | 0.019291 | 0.02697 | 0.031259 | 0.116709 | 0.043765 | 0.013033 | 0.017099 | 0.030706 | 0.065146 | 0.059626 | 0.060903 | 0.052638 | 0.084106 | 0.096507 | 0.008547 | 0.181744 | 0.05155 | 0.020662 | 0.023346 | 0.046031 | 0.065523 | 0.120372 | 0.090858 |
| 32.6527778 | 0.03663 | 0.013419 | 0.017823 | 0.029385 | 0.081536 | 0.05157 | 0.006232 | 0.013012 | 0.032444 | 0.066777 | 0.051758 | 0.061447 | 0.031119 | 0.066311 | 0.104275 | 0.004729 | 0.204624 | 0.054423 | 0.01033 | 0.02237 | 0.036915 | 0.070438 | 0.126518 | 0.091597 |
| 32.6666667 | 0.033541 | 0.005663 | 0.01297 | 0.020316 | 0.048565 | 0.050589 | 0.002932 | 0.010764 | 0.028625 | 0.063005 | 0.044651 | 0.057102 | 0.018433 | 0.047187 | 0.103522 | 0.005999 | 0.18929 | 0.059693 | 0.005484 | 0.021903 | 0.028206 | 0.057051 | 0.12093 | 0.071191 |
| 32.6805556 | 0.028786 | 0.004082 | 0.009299 | 0.016513 | 0.026079 | 0.033213 | 0.007004 | 0.006936 | 0.020509 | 0.049534 | 0.038219 | 0.045646 | 0.012471 | 0.027861 | 0.075075 | 0.008701 | 0.120421 | 0.054637 | 0.00484 | 0.018966 | 0.02044 | 0.039567 | 0.094645 | 0.035686 |
| 32.6944444 | 0.020127 | 0.011687 | 0.006651 | 0.019897 | 0.017216 | 0.022649 | 0.019175 | 0.00761 | 0.011648 | 0.032201 | 0.022605 | 0.026336 | 0.016029 | 0.01128 | 0.036801 | 0.011032 | 0.056906 | 0.042973 | 0.003656 | 0.014025 | 0.012611 | 0.022128 | 0.063557 | 0.009684 |
| 32.7083333 | 0.011923 | 0.020412 | 0.009271 | 0.01788 | 0.013552 | 0.022951 | 0.034064 | 0.013181 | 0.006123 | 0.019016 | 0.013279 | 0.013386 | 0.021353 | 0.006007 | 0.025335 | 0.012477 | 0.052255 | 0.029344 | 0.004348 | 0.013788 | 0.009646 | 0.013821 | 0.040367 | 0.005273 |
| 32.7222222 | 0.007647 | 0.01619 | 0.014527 | 0.012373 | 0.008448 | 0.015727 | 0.043109 | 0.019848 | 0.007809 | 0.011977 | 0.026403 | 0.020407 | 0.016458 | 0.016067 | 0.033084 | 0.011858 | 0.06873 | 0.01592 | 0.007543 | 0.015025 | 0.019632 | 0.01065 | 0.02798 | 0.013157 |
| 32.7361111 | 0.005294 | 0.006307 | 0.015313 | 0.013979 | 0.003845 | 0.00834 | 0.046505 | 0.028826 | 0.013062 | 0.00661 | 0.045828 | 0.033174 | 0.010162 | 0.029134 | 0.033054 | 0.009422 | 0.058623 | 0.014806 | 0.009735 | 0.009954 | 0.02904 | 0.006283 | 0.050912 | 0.024315 |
| 32.75 | 0.004037 | 0.004812 | 0.010583 | 0.018729 | 0.001792 | 0.009162 | 0.045314 | 0.041909 | 0.014934 | 0.008267 | 0.051088 | 0.036293 | 0.011142 | 0.03 | 0.022306 | 0.007854 | 0.036792 | 0.022565 | 0.0105 | 0.008293 | 0.029807 | 0.00646 | 0.107029 | 0.031674 |
| 32.7638889 | 0.00303 | 0.00812 | 0.004608 | 0.021986 | 0.001034 | 0.010251 | 0.034353 | 0.053292 | 0.012151 | 0.02346 | 0.041351 | 0.03585 | 0.009576 | 0.021467 | 0.010505 | 0.007132 | 0.031913 | 0.022879 | 0.011432 | 0.017195 | 0.034351 | 0.011611 | 0.149064 | 0.029853 |
| 32.7777778 | 0.002788 | 0.010945 | 0.001298 | 0.026431 | 0.003674 | 0.010362 | 0.019241 | 0.056048 | 0.007235 | 0.035808 | 0.027171 | 0.037556 | 0.003878 | 0.0118 | 0.003163 | 0.005133 | 0.047492 | 0.0138 | 0.012983 | 0.028368 | 0.043849 | 0.021301 | 0.14157 | 0.020966 |
| 32.7916667 | 0.004487 | 0.008592 | 0.002009 | 0.022177 | 0.014693 | 0.01046 | 0.011592 | 0.052978 | 0.00538 | 0.031402 | 0.019459 | 0.03795 | 0.001366 | 0.006809 | 0.002223 | 0.002051 | 0.067048 | 0.006085 | 0.013705 | 0.033002 | 0.045515 | 0.028734 | 0.103153 | 0.012976 |
| 32.8055556 | 0.004814 | 0.003216 | 0.004629 | 0.014603 | 0.027812 | 0.011218 | 0.01161 | 0.046638 | 0.007663 | 0.022749 | 0.01683 | 0.029449 | 0.003823 | 0.010594 | 0.004004 | 0.000613 | 0.07445 | 0.005724 | 0.012292 | 0.03014 | 0.035097 | 0.027826 | 0.075581 | 0.011698 |
| 32.8194444 | 0.002441 | 0.004537 | 0.006051 | 0.022042 | 0.031198 | 0.013605 | 0.012112 | 0.035676 | 0.008407 | 0.021654 | 0.015162 | 0.016319 | 0.005866 | 0.020708 | 0.004138 | 0.002116 | 0.074139 | 0.010392 | 0.010053 | 0.024467 | 0.019389 | 0.024015 | 0.06646 | 0.015333 |
| 32.8333333 | 0.000905 | 0.012853 | 0.008641 | 0.028967 | 0.030049 | 0.014786 | 0.00949 | 0.023692 | 0.00657 | 0.024026 | 0.017632 | 0.011708 | 0.007868 | 0.03086 | 0.005094 | 0.006556 | 0.077356 | 0.014247 | 0.009293 | 0.020866 | 0.007383 | 0.021144 | 0.061468 | 0.014945 |
| 32.8472222 | 0.001609 | 0.018157 | 0.010742 | 0.022452 | 0.036783 | 0.012318 | 0.005183 | 0.015188 | 0.005137 | 0.025953 | 0.018541 | 0.011198 | 0.017393 | 0.034581 | 0.006188 | 0.01236 | 0.084368 | 0.01136 | 0.010543 | 0.024414 | 0.009217 | 0.019184 | 0.059711 | 0.012365 |
| 32.8611111 | 0.003024 | 0.015576 | 0.008865 | 0.014211 | 0.048396 | 0.008423 | 0.002921 | 0.010078 | 0.004245 | 0.028487 | 0.01697 | 0.009867 | 0.027742 | 0.02713 | 0.005245 | 0.016162 | 0.087192 | 0.008528 | 0.011705 | 0.036515 | 0.020423 | 0.018585 | 0.068166 | 0.012842 |
| 32.875 | 0.00446 | 0.009981 | 0.008814 | 0.012692 | 0.061375 | 0.006964 | 0.005163 | 0.006642 | 0.002999 | 0.030252 | 0.019519 | 0.012667 | 0.032943 | 0.016424 | 0.003692 | 0.018906 | 0.077867 | 0.009612 | 0.011868 | 0.04899 | 0.029931 | 0.017356 | 0.081594 | 0.010014 |
| 32.8888889 | 0.005753 | 0.006569 | 0.009964 | 0.012214 | 0.083776 | 0.00881 | 0.008911 | 0.005497 | 0.003444 | 0.02746 | 0.025915 | 0.017966 | 0.038333 | 0.012376 | 0.003266 | 0.022264 | 0.064507 | 0.015491 | 0.01639 | 0.04892 | 0.033181 | 0.015701 | 0.086536 | 0.008416 |
| 32.9027778 | 0.005674 | 0.00693 | 0.007233 | 0.010426 | 0.110329 | 0.011904 | 0.010154 | 0.006339 | 0.00642 | 0.019864 | 0.0341 | 0.024195 | 0.042123 | 0.009863 | 0.00706 | 0.025306 | 0.055923 | 0.032548 | 0.02649 | 0.036352 | 0.026319 | 0.017711 | 0.081895 | 0.016367 |
| 32.9166667 | 0.004303 | 0.007026 | 0.005144 | 0.016217 | 0.126208 | 0.013802 | 0.0077 | 0.005993 | 0.008259 | 0.012089 | 0.034013 | 0.026132 | 0.043199 | 0.006558 | 0.012605 | 0.029166 | 0.049841 | 0.049423 | 0.03461 | 0.029192 | 0.018284 | 0.02152 | 0.070653 | 0.030043 |
| 32.9305556 | 0.002495 | 0.005712 | 0.008458 | 0.026482 | 0.133718 | 0.013956 | 0.004768 | 0.005007 | 0.007616 | 0.009582 | 0.025503 | 0.019423 | 0.052305 | 0.006387 | 0.014619 | 0.033137 | 0.051573 | 0.058087 | 0.041777 | 0.030651 | 0.024 | 0.022178 | 0.061787 | 0.040233 |
| 32.9444444 | 0.002556 | 0.009309 | 0.014624 | 0.032119 | 0.142643 | 0.01466 | 0.006387 | 0.008506 | 0.00694 | 0.01349 | 0.02605 | 0.013875 | 0.074594 | 0.006086 | 0.014203 | 0.034611 | 0.066752 | 0.06488 | 0.053751 | 0.034134 | 0.041637 | 0.022973 | 0.06425 | 0.041535 |
| 32.9583333 | 0.00746 | 0.017857 | 0.020526 | 0.034596 | 0.155085 | 0.018311 | 0.012049 | 0.017181 | 0.008052 | 0.022598 | 0.036549 | 0.016312 | 0.097242 | 0.002915 | 0.016001 | 0.038033 | 0.086472 | 0.073993 | 0.067262 | 0.047631 | 0.064809 | 0.026552 | 0.066446 | 0.038393 |
| 32.9722222 | 0.01383 | 0.027369 | 0.025824 | 0.040533 | 0.16642 | 0.025013 | 0.018546 | 0.025958 | 0.010793 | 0.038498 | 0.049299 | 0.023731 | 0.109746 | 0.007466 | 0.020133 | 0.050836 | 0.10713 | 0.088337 | 0.075702 | 0.067839 | 0.088494 | 0.026655 | 0.068467 | 0.038238 |
| 32.9861111 | 0.016309 | 0.03541 | 0.029276 | 0.047694 | 0.167392 | 0.034096 | 0.022846 | 0.032603 | 0.013837 | 0.059026 | 0.060737 | 0.029925 | 0.117475 | 0.024975 | 0.025515 | 0.067275 | 0.133195 | 0.102711 | 0.076074 | 0.082405 | 0.103579 | 0.021651 | 0.081297 | 0.04211 |
| 33 | 0.015559 | 0.042859 | 0.031188 | 0.052754 | 0.159597 | 0.042453 | 0.023819 | 0.037535 | 0.015741 | 0.072326 | 0.06699 | 0.030516 | 0.127143 | 0.041702 | 0.035636 | 0.078033 | 0.152679 | 0.106014 | 0.072242 | 0.091766 | 0.106568 | 0.019395 | 0.093768 | 0.042987 |
| 33.0138889 | 0.014793 | 0.050893 | 0.033693 | 0.059332 | 0.159599 | 0.046982 | 0.024615 | 0.045154 | 0.01599 | 0.072201 | 0.077952 | 0.032279 | 0.145777 | 0.049805 | 0.045726 | 0.088532 | 0.15781 | 0.097612 | 0.068715 | 0.095573 | 0.090339 | 0.02339 | 0.094663 | 0.045079 |
| 33.0277778 | 0.015214 | 0.052296 | 0.03511 | 0.065924 | 0.168911 | 0.049652 | 0.028058 | 0.060592 | 0.015406 | 0.066518 | 0.09628 | 0.038347 | 0.170978 | 0.053253 | 0.045384 | 0.101446 | 0.1642 | 0.088486 | 0.066491 | 0.084976 | 0.06245 | 0.025159 | 0.094625 | 0.057571 |
| 33.0416667 | 0.017722 | 0.053273 | 0.033119 | 0.073719 | 0.174304 | 0.056081 | 0.033841 | 0.077029 | 0.018732 | 0.066555 | 0.109508 | 0.043467 | 0.185212 | 0.051542 | 0.038648 | 0.112613 | 0.167865 | 0.086888 | 0.068144 | 0.070889 | 0.045679 | 0.019166 | 0.08541 | 0.065789 |
| 33.0555556 | 0.024061 | 0.067664 | 0.035278 | 0.082928 | 0.182743 | 0.071044 | 0.039383 | 0.083546 | 0.029592 | 0.079317 | 0.127375 | 0.055442 | 0.179701 | 0.047852 | 0.036653 | 0.102171 | 0.142905 | 0.094685 | 0.073055 | 0.074823 | 0.03769 | 0.01515 | 0.061025 | 0.07604 |
| 33.0694444 | 0.032503 | 0.083515 | 0.046799 | 0.074688 | 0.211885 | 0.085058 | 0.03877 | 0.085297 | 0.041049 | 0.093758 | 0.15358 | 0.070966 | 0.161688 | 0.047171 | 0.039549 | 0.063099 | 0.091594 | 0.109151 | 0.077345 | 0.087599 | 0.027161 | 0.023427 | 0.059965 | 0.103296 |
| 33.0833333 | 0.033197 | 0.076202 | 0.053078 | 0.042699 | 0.231235 | 0.081115 | 0.027335 | 0.086639 | 0.041294 | 0.088238 | 0.152773 | 0.070528 | 0.153142 | 0.045251 | 0.047112 | 0.045335 | 0.040603 | 0.112692 | 0.072595 | 0.091469 | 0.020675 | 0.03946 | 0.078348 | 0.100779 |
| 33.0972222 | 0.020537 | 0.043233 | 0.042791 | 0.021424 | 0.20111 | 0.060511 | 0.012479 | 0.080168 | 0.029177 | 0.064487 | 0.108244 | 0.053195 | 0.144323 | 0.03657 | 0.058602 | 0.065636 | 0.01974 | 0.083145 | 0.051182 | 0.079751 | 0.01964 | 0.052905 | 0.077131 | 0.066893 |
| 33.1111111 | 0.00643 | 0.019192 | 0.025344 | 0.030347 | 0.150497 | 0.038714 | 0.005468 | 0.071396 | 0.016225 | 0.04296 | 0.04947 | 0.031797 | 0.104413 | 0.025465 | 0.066112 | 0.08587 | 0.037727 | 0.035923 | 0.024694 | 0.054904 | 0.01716 | 0.068625 | 0.061763 | 0.061038 |
| 33.125 | 0.002066 | 0.025901 | 0.011754 | 0.053012 | 0.116112 | 0.025887 | 0.006654 | 0.063914 | 0.009668 | 0.03141 | 0.01759 | 0.018695 | 0.061238 | 0.023276 | 0.061841 | 0.087227 | 0.058768 | 0.011206 | 0.012407 | 0.030329 | 0.020373 | 0.084949 | 0.047724 | 0.071353 |
| 33.1388889 | 0.002746 | 0.040349 | 0.004093 | 0.07491 | 0.097069 | 0.020219 | 0.009589 | 0.049846 | 0.007342 | 0.023905 | 0.02268 | 0.015055 | 0.050548 | 0.019551 | 0.040594 | 0.060076 | 0.051216 | 0.019447 | 0.014381 | 0.017929 | 0.021205 | 0.076291 | 0.034031 | 0.065407 |
| 33.1527778 | 0.004128 | 0.04593 | 0.002163 | 0.083864 | 0.080385 | 0.014431 | 0.009145 | 0.028317 | 0.005093 | 0.016508 | 0.039759 | 0.012267 | 0.046803 | 0.010927 | 0.023945 | 0.035758 | 0.026991 | 0.028225 | 0.017091 | 0.024434 | 0.014408 | 0.05264 | 0.018231 | 0.047353 |
| 33.1666667 | 0.009823 | 0.045793 | 0.006033 | 0.082513 | 0.053616 | 0.00746 | 0.005525 | 0.010542 | 0.003196 | 0.009936 | 0.052855 | 0.006863 | 0.042472 | 0.013887 | 0.02822 | 0.055599 | 0.024243 | 0.023575 | 0.022274 | 0.035854 | 0.015693 | 0.059358 | 0.005935 | 0.02855 |
| 33.1805556 | 0.017972 | 0.042494 | 0.01218 | 0.08969 | 0.027407 | 0.004752 | 0.003851 | 0.007421 | 0.004765 | 0.004934 | 0.061677 | 0.002483 | 0.067228 | 0.022233 | 0.034754 | 0.083422 | 0.058028 | 0.030133 | 0.032127 | 0.038423 | 0.018185 | 0.089098 | 0.002295 | 0.014902 |
| 33.1944444 | 0.021486 | 0.039966 | 0.015312 | 0.104072 | 0.02174 | 0.007839 | 0.006346 | 0.017056 | 0.007616 | 0.002847 | 0.067916 | 0.001562 | 0.092744 | 0.022809 | 0.038999 | 0.086156 | 0.094728 | 0.038467 | 0.03265 | 0.051131 | 0.012787 | 0.099216 | 0.004886 | 0.00608 |
| 33.2083333 | 0.020365 | 0.039327 | 0.014061 | 0.109128 | 0.028338 | 0.012454 | 0.009277 | 0.035179 | 0.00996 | 0.004715 | 0.068375 | 0.001671 | 0.085451 | 0.018136 | 0.045283 | 0.078057 | 0.106807 | 0.030724 | 0.023891 | 0.081893 | 0.009316 | 0.090094 | 0.012065 | 0.003543 |

| | | | | | | | | | | | | | | | | | | | | | | | | |
|---|---|---|---|---|---|---|---|---|---|---|---|---|---|---|---|---|---|---|---|---|---|---|---|---|
| 33.2222222 | 0.019704 | 0.039143 | 0.011248 | 0.103469 | 0.034829 | 0.016313 | 0.010681 | 0.053772 | 0.012936 | 0.007871 | 0.060411 | 0.002325 | 0.067121 | 0.016502 | 0.047001 | 0.072427 | 0.100111 | 0.021183 | 0.017675 | 0.098468 | 0.016695 | 0.080873 | 0.019988 | 0.00438 |
| 33.2361111 | 0.019614 | 0.040925 | 0.00977 | 0.095076 | 0.04628 | 0.018932 | 0.010674 | 0.061254 | 0.01592 | 0.010066 | 0.052717 | 0.004117 | 0.055128 | 0.017983 | 0.040676 | 0.066027 | 0.079151 | 0.013359 | 0.013439 | 0.081502 | 0.031094 | 0.075198 | 0.022592 | 0.006776 |
| 33.25 | 0.017844 | 0.044808 | 0.008679 | 0.080097 | 0.060078 | 0.018223 | 0.008081 | 0.058272 | 0.017318 | 0.010574 | 0.050382 | 0.005191 | 0.041788 | 0.018673 | 0.030608 | 0.049867 | 0.043959 | 0.009363 | 0.011932 | 0.055002 | 0.041023 | 0.064671 | 0.015877 | 0.012668 |
| 33.2638889 | 0.015003 | 0.046309 | 0.006016 | 0.052459 | 0.064106 | 0.013459 | 0.004433 | 0.05408 | 0.016123 | 0.008877 | 0.047035 | 0.005929 | 0.022834 | 0.018124 | 0.021579 | 0.028012 | 0.016592 | 0.015728 | 0.015662 | 0.045986 | 0.040321 | 0.048792 | 0.007951 | 0.018553 |
| 33.2777778 | 0.012959 | 0.041801 | 0.003442 | 0.024845 | 0.058016 | 0.007664 | 0.002587 | 0.053313 | 0.013556 | 0.006577 | 0.037096 | 0.007081 | 0.013363 | 0.014811 | 0.012447 | 0.010645 | 0.009089 | 0.01682 | 0.013297 | 0.056289 | 0.034709 | 0.043324 | 0.007127 | 0.016198 |
| 33.2916667 | 0.012325 | 0.034896 | 0.00333 | 0.014423 | 0.053977 | 0.003926 | 0.001978 | 0.051165 | 0.011479 | 0.005179 | 0.0267 | 0.007019 | 0.019836 | 0.008719 | 0.005397 | 0.004372 | 0.011836 | 0.009041 | 0.005659 | 0.067356 | 0.035162 | 0.045057 | 0.006029 | 0.008385 |
| 33.3055556 | 0.011771 | 0.030937 | 0.004606 | 0.018533 | 0.058152 | 0.00204 | 0.001122 | 0.04312 | 0.009729 | 0.003907 | 0.024249 | 0.006192 | 0.028685 | 0.005383 | 0.004933 | 0.009947 | 0.023526 | 0.005746 | 0.00586 | 0.062153 | 0.041724 | 0.037483 | 0.00544 | 0.006377 |
| 33.3194444 | 0.010751 | 0.029133 | 0.004909 | 0.022008 | 0.063568 | 0.000975 | 0.000439 | 0.036123 | 0.007509 | 0.00203 | 0.026501 | 0.006483 | 0.033958 | 0.006708 | 0.00763 | 0.016715 | 0.025201 | 0.005771 | 0.009071 | 0.04791 | 0.046394 | 0.027705 | 0.00978 | 0.008783 |
| 33.3333333 | 0.009952 | 0.027633 | 0.004375 | 0.021929 | 0.06227 | 0.000969 | 0.000305 | 0.034094 | 0.005684 | 0.0009 | 0.027237 | 0.006971 | 0.038525 | 0.006364 | 0.008412 | 0.013416 | 0.019586 | 0.009685 | 0.010736 | 0.044928 | 0.048366 | 0.03416 | 0.013477 | 0.007806 |
| 33.3472222 | 0.009555 | 0.028642 | 0.004811 | 0.03111 | 0.056602 | 0.001702 | 0.000563 | 0.031235 | 0.005882 | 0.000898 | 0.029554 | 0.005301 | 0.03793 | 0.005943 | 0.014295 | 0.008765 | 0.02586 | 0.023363 | 0.019768 | 0.059623 | 0.051017 | 0.047419 | 0.01069 | 0.010876 |
| 33.3611111 | 0.009724 | 0.034217 | 0.006733 | 0.055444 | 0.05539 | 0.002196 | 0.001979 | 0.028709 | 0.007969 | 0.001524 | 0.040105 | 0.003173 | 0.025812 | 0.013785 | 0.03137 | 0.014156 | 0.025117 | 0.032978 | 0.028704 | 0.072221 | 0.052824 | 0.050264 | 0.007445 | 0.028624 |
| 33.375 | 0.01156 | 0.042151 | 0.009019 | 0.079403 | 0.063284 | 0.003931 | 0.004924 | 0.032699 | 0.010507 | 0.003728 | 0.058258 | 0.004472 | 0.010801 | 0.025459 | 0.046324 | 0.025185 | 0.023604 | 0.021799 | 0.023552 | 0.064255 | 0.051891 | 0.046355 | 0.013058 | 0.045299 |
| 33.3888889 | 0.015883 | 0.047251 | 0.010486 | 0.085012 | 0.076259 | 0.007833 | 0.007881 | 0.040384 | 0.013192 | 0.006174 | 0.072145 | 0.008671 | 0.006227 | 0.030498 | 0.048633 | 0.034136 | 0.038098 | 0.007866 | 0.018815 | 0.053369 | 0.054167 | 0.043453 | 0.018515 | 0.043726 |
| 33.4027778 | 0.020917 | 0.048656 | 0.010865 | 0.078515 | 0.088181 | 0.011585 | 0.009928 | 0.046284 | 0.016472 | 0.007949 | 0.073375 | 0.014328 | 0.01463 | 0.028861 | 0.047735 | 0.040075 | 0.056241 | 0.013022 | 0.033812 | 0.062554 | 0.06305 | 0.043502 | 0.018725 | 0.033422 |
| 33.4166667 | 0.023374 | 0.052984 | 0.010877 | 0.074614 | 0.097773 | 0.014702 | 0.012309 | 0.053135 | 0.019439 | 0.01003 | 0.074464 | 0.023187 | 0.036096 | 0.029649 | 0.05425 | 0.048058 | 0.067516 | 0.028176 | 0.06078 | 0.085588 | 0.069079 | 0.03438 | 0.0225 | 0.032636 |
| 33.4305556 | 0.022363 | 0.064079 | 0.011488 | 0.076284 | 0.103809 | 0.018283 | 0.015459 | 0.063912 | 0.0207 | 0.011385 | 0.088912 | 0.035162 | 0.061226 | 0.036237 | 0.064839 | 0.057171 | 0.062913 | 0.034461 | 0.07874 | 0.099024 | 0.077214 | 0.01735 | 0.030371 | 0.044184 |
| 33.4444444 | 0.020303 | 0.076588 | 0.013356 | 0.084564 | 0.103497 | 0.020854 | 0.018034 | 0.071022 | 0.021176 | 0.011352 | 0.107765 | 0.045517 | 0.071164 | 0.041367 | 0.06978 | 0.063791 | 0.040057 | 0.031372 | 0.081194 | 0.095675 | 0.096613 | 0.012514 | 0.035567 | 0.05321 |
| 33.4583333 | 0.019816 | 0.082322 | 0.017474 | 0.093446 | 0.0998 | 0.020708 | 0.020407 | 0.07104 | 0.02251 | 0.010774 | 0.114301 | 0.049773 | 0.065081 | 0.04246 | 0.069423 | 0.071524 | 0.016077 | 0.033939 | 0.077521 | 0.096255 | 0.115858 | 0.02158 | 0.038607 | 0.053718 |
| 33.4722222 | 0.020647 | 0.079616 | 0.023591 | 0.093819 | 0.100728 | 0.01943 | 0.024506 | 0.071854 | 0.024239 | 0.012454 | 0.107902 | 0.053631 | 0.062822 | 0.046554 | 0.073187 | 0.083948 | 0.013675 | 0.046037 | 0.073482 | 0.107468 | 0.124048 | 0.032318 | 0.044961 | 0.056629 |
| 33.4861111 | 0.021487 | 0.073913 | 0.028 | 0.092692 | 0.110139 | 0.020365 | 0.030961 | 0.072654 | 0.025829 | 0.020537 | 0.101669 | 0.06372 | 0.071934 | 0.056153 | 0.082298 | 0.096906 | 0.036694 | 0.0559 | 0.071598 | 0.107782 | 0.118835 | 0.040693 | 0.051588 | 0.064565 |
| 33.5 | 0.022976 | 0.068864 | 0.028533 | 0.096321 | 0.122337 | 0.02393 | 0.035853 | 0.068576 | 0.026468 | 0.033332 | 0.103682 | 0.073527 | 0.083075 | 0.065469 | 0.08896 | 0.101159 | 0.073252 | 0.053128 | 0.07218 | 0.096519 | 0.107546 | 0.053052 | 0.054185 | 0.070963 |
| 33.5138889 | 0.025353 | 0.065296 | 0.02867 | 0.097903 | 0.130812 | 0.027799 | 0.033296 | 0.060498 | 0.023637 | 0.041716 | 0.107542 | 0.076495 | 0.093803 | 0.073776 | 0.090311 | 0.093738 | 0.114099 | 0.047219 | 0.072182 | 0.101027 | 0.096 | 0.055573 | 0.056804 | 0.078897 |
| 33.5277778 | 0.02661 | 0.066088 | 0.031396 | 0.091923 | 0.136694 | 0.032571 | 0.024652 | 0.05088 | 0.018767 | 0.043382 | 0.103839 | 0.075251 | 0.110105 | 0.088527 | 0.090287 | 0.075881 | 0.148788 | 0.057461 | 0.075584 | 0.121358 | 0.078384 | 0.039298 | 0.063053 | 0.093571 |
| 33.5416667 | 0.026421 | 0.072553 | 0.035346 | 0.080013 | 0.146439 | 0.040416 | 0.019476 | 0.050309 | 0.016776 | 0.046165 | 0.098781 | 0.07877 | 0.127001 | 0.106476 | 0.088186 | 0.055075 | 0.161987 | 0.075647 | 0.083035 | 0.131126 | 0.061498 | 0.024773 | 0.069409 | 0.106886 |
| 33.5555556 | 0.027053 | 0.075348 | 0.038099 | 0.065897 | 0.153157 | 0.049743 | 0.024363 | 0.056073 | 0.018467 | 0.051717 | 0.09179 | 0.083286 | 0.124629 | 0.102929 | 0.082186 | 0.050624 | 0.144307 | 0.079541 | 0.08704 | 0.120325 | 0.064494 | 0.027001 | 0.070494 | 0.104754 |
| 33.5694444 | 0.029718 | 0.061078 | 0.038851 | 0.050257 | 0.146024 | 0.056768 | 0.034034 | 0.050124 | 0.021606 | 0.057804 | 0.070372 | 0.073446 | 0.103907 | 0.070866 | 0.082548 | 0.058302 | 0.112084 | 0.07297 | 0.088142 | 0.099563 | 0.081934 | 0.034932 | 0.070129 | 0.088529 |
| 33.5833333 | 0.03335 | 0.03614 | 0.038414 | 0.035508 | 0.149539 | 0.060289 | 0.04022 | 0.038918 | 0.024777 | 0.06785 | 0.04764 | 0.058489 | 0.08965 | 0.044948 | 0.094931 | 0.058415 | 0.096817 | 0.072207 | 0.093896 | 0.086554 | 0.083666 | 0.035859 | 0.073598 | 0.076132 |
| 33.5972222 | 0.032553 | 0.021846 | 0.037698 | 0.026761 | 0.186646 | 0.062117 | 0.039112 | 0.033803 | 0.026817 | 0.079321 | 0.046166 | 0.05958 | 0.085589 | 0.046508 | 0.10929 | 0.058204 | 0.093392 | 0.081023 | 0.099214 | 0.084064 | 0.059261 | 0.03228 | 0.073357 | 0.076921 |
| 33.6111111 | 0.025617 | 0.0235 | 0.035637 | 0.025093 | 0.220872 | 0.065453 | 0.032646 | 0.03109 | 0.026537 | 0.083137 | 0.061592 | 0.071208 | 0.0807 | 0.063915 | 0.11507 | 0.058006 | 0.072918 | 0.094823 | 0.086568 | 0.072964 | 0.044094 | 0.022668 | 0.06688 | 0.076585 |
| 33.625 | 0.018192 | 0.030143 | 0.031349 | 0.029111 | 0.208373 | 0.073315 | 0.027069 | 0.032897 | 0.025354 | 0.081236 | 0.071502 | 0.074622 | 0.084184 | 0.085966 | 0.114073 | 0.04943 | 0.043564 | 0.101696 | 0.066469 | 0.051219 | 0.06075 | 0.016919 | 0.063326 | 0.066992 |
| 33.6388889 | 0.01193 | 0.043182 | 0.027082 | 0.038191 | 0.163725 | 0.081258 | 0.025538 | 0.03793 | 0.026056 | 0.082222 | 0.066091 | 0.068366 | 0.101823 | 0.100356 | 0.113686 | 0.040436 | 0.035856 | 0.095915 | 0.064132 | 0.041708 | 0.081886 | 0.018056 | 0.062085 | 0.066323 |
| 33.6527778 | 0.007967 | 0.068547 | 0.023887 | 0.049654 | 0.124978 | 0.076989 | 0.021441 | 0.035192 | 0.027674 | 0.082335 | 0.054597 | 0.061489 | 0.113828 | 0.083853 | 0.120008 | 0.031716 | 0.048197 | 0.085736 | 0.064789 | 0.040782 | 0.084166 | 0.013064 | 0.057038 | 0.069232 |
| 33.6666667 | 0.008293 | 0.080599 | 0.020024 | 0.053899 | 0.107484 | 0.060285 | 0.013597 | 0.023787 | 0.027168 | 0.072637 | 0.04694 | 0.058564 | 0.111423 | 0.048382 | 0.132449 | 0.024622 | 0.052056 | 0.068007 | 0.048767 | 0.027312 | 0.076247 | 0.005499 | 0.048466 | 0.053944 |
| 33.6805556 | 0.008389 | 0.055936 | 0.015071 | 0.042031 | 0.102696 | 0.04181 | 0.010375 | 0.0118 | 0.023361 | 0.055227 | 0.043036 | 0.060041 | 0.101484 | 0.031505 | 0.13623 | 0.024187 | 0.034442 | 0.040471 | 0.030046 | 0.009771 | 0.068152 | 0.005512 | 0.036603 | 0.040459 |
| 33.6944444 | 0.005076 | 0.023474 | 0.009585 | 0.023851 | 0.099376 | 0.02288 | 0.007158 | 0.004348 | 0.016819 | 0.035508 | 0.035251 | 0.059078 | 0.07658 | 0.022737 | 0.118715 | 0.020894 | 0.016081 | 0.03282 | 0.030767 | 0.004689 | 0.045811 | 0.01215 | 0.020917 | 0.036456 |
| 33.7083333 | 0.003107 | 0.012328 | 0.005009 | 0.020198 | 0.088493 | 0.011027 | 0.002069 | 0.002877 | 0.01049 | 0.017038 | 0.020408 | 0.045562 | 0.043299 | 0.010928 | 0.075065 | 0.01048 | 0.019415 | 0.066264 | 0.046685 | 0.01104 | 0.019021 | 0.018651 | 0.010627 | 0.021572 |
| 33.7222222 | 0.004593 | 0.010731 | 0.004914 | 0.026397 | 0.056511 | 0.013137 | 0.004234 | 0.004613 | 0.007163 | 0.008588 | 0.011866 | 0.025851 | 0.034035 | 0.011341 | 0.030647 | 0.004038 | 0.034316 | 0.110755 | 0.067349 | 0.020008 | 0.014609 | 0.024504 | 0.014465 | 0.007191 |
| 33.7361111 | 0.004695 | 0.011361 | 0.008565 | 0.020306 | 0.033148 | 0.015152 | 0.014517 | 0.007428 | 0.00528 | 0.011371 | 0.019911 | 0.022758 | 0.045377 | 0.022243 | 0.014077 | 0.004044 | 0.039034 | 0.123141 | 0.088924 | 0.02434 | 0.025604 | 0.028069 | 0.023875 | 0.003667 |
| 33.75 | 0.002528 | 0.015441 | 0.010285 | 0.008346 | 0.051354 | 0.01025 | 0.017095 | 0.010492 | 0.003578 | 0.014376 | 0.030983 | 0.04062 | 0.043413 | 0.031035 | 0.016159 | 0.004829 | 0.030686 | 0.098399 | 0.096983 | 0.023198 | 0.048444 | 0.025079 | 0.030993 | 0.005439 |
| 33.7638889 | 0.001408 | 0.013019 | 0.009704 | 0.008802 | 0.077122 | 0.006274 | 0.014073 | 0.012836 | 0.004287 | 0.0162 | 0.03452 | 0.056379 | 0.02685 | 0.02417 | 0.026884 | 0.00438 | 0.024188 | 0.063978 | 0.090066 | 0.016158 | 0.065597 | 0.020153 | 0.037246 | 0.011626 |
| 33.7777778 | 0.001382 | 0.008953 | 0.008771 | 0.023115 | 0.087321 | 0.006607 | 0.019295 | 0.014041 | 0.007226 | 0.022545 | 0.031091 | 0.050708 | 0.012154 | 0.021042 | 0.033127 | 0.003755 | 0.025535 | 0.03617 | 0.072446 | 0.007841 | 0.055191 | 0.020953 | 0.039052 | 0.016311 |
| 33.7916667 | 0.001323 | 0.010716 | 0.006002 | 0.036864 | 0.092801 | 0.009076 | 0.017381 | 0.013089 | 0.00995 | 0.029064 | 0.020445 | 0.030536 | 0.003496 | 0.03844 | 0.024327 | 0.007672 | 0.023051 | 0.019436 | 0.050235 | 0.008222 | 0.041408 | 0.027222 | 0.033953 | 0.01543 |
| 33.8055556 | 0.00226 | 0.009909 | 0.00426 | 0.031447 | 0.095879 | 0.009294 | 0.013865 | 0.010865 | 0.011635 | 0.027055 | 0.012198 | 0.01928 | 0.001123 | 0.047957 | 0.017281 | 0.017159 | 0.01937 | 0.018778 | 0.039069 | 0.012508 | 0.035036 | 0.034168 | 0.031399 | 0.012121 |
| 33.8194444 | 0.004121 | 0.007292 | 0.005879 | 0.015082 | 0.096001 | 0.006184 | 0.02385 | 0.010155 | 0.012351 | 0.021138 | 0.009751 | 0.015406 | 0.002888 | 0.048902 | 0.0314 | 0.031542 | 0.019611 | 0.026664 | 0.044856 | 0.011223 | 0.026469 | 0.042275 | 0.039518 | 0.014702 |
| 33.8333333 | 0.005571 | 0.005808 | 0.006347 | 0.009061 | 0.098869 | 0.005725 | 0.031262 | 0.010982 | 0.012417 | 0.020781 | 0.007699 | 0.009963 | 0.009978 | 0.057248 | 0.057313 | 0.049362 | 0.017765 | 0.032098 | 0.05498 | 0.008338 | 0.017451 | 0.049335 | 0.051225 | 0.028482 |
| 33.8472222 | 0.008648 | 0.003934 | 0.004101 | 0.023564 | 0.102698 | 0.0093 | 0.027191 | 0.009719 | 0.011282 | 0.022633 | 0.006564 | 0.008568 | 0.02989 | 0.071368 | 0.077149 | 0.062188 | 0.024244 | 0.035351 | 0.053739 | 0.007353 | 0.012452 | 0.044461 | 0.056924 | 0.044474 |
| 33.8611111 | 0.013601 | 0.002545 | 0.001842 | 0.047927 | 0.100637 | 0.012098 | 0.015862 | 0.005685 | 0.009162 | 0.024998 | 0.007959 | 0.011458 | 0.062547 | 0.089289 | 0.08714 | 0.062673 | 0.03132 | 0.040522 | 0.04342 | 0.00524 | 0.016188 | 0.026348 | 0.05801 | 0.055713 |
| 33.875 | 0.015223 | 0.002158 | 0.000764 | 0.0547 | 0.099854 | 0.010275 | 0.005851 | 0.003033 | 0.007495 | 0.027 | 0.0169 | 0.022893 | 0.092522 | 0.107623 | 0.089711 | 0.052521 | 0.023213 | 0.042629 | 0.036288 | 0.004669 | 0.017784 | 0.010221 | 0.0575 | 0.070499 |
| 33.8888889 | 0.011606 | 0.005085 | 0.001137 | 0.042447 | 0.106611 | 0.00537 | 0.007292 | 0.00347 | 0.00652 | 0.020223 | 0.023834 | 0.029939 | 0.101241 | 0.123061 | 0.085988 | 0.03873 | 0.015952 | 0.035567 | 0.034451 | 0.014019 | 0.012326 | 0.006598 | 0.055592 | 0.087792 |
| 33.9027778 | 0.008464 | 0.008694 | 0.002165 | 0.03499 | 0.115609 | 0.003699 | 0.018039 | 0.003848 | 0.006368 | 0.009922 | 0.016542 | 0.022476 | 0.091191 | 0.119922 | 0.075215 | 0.027468 | 0.020309 | 0.024343 | 0.035326 | 0.028067 | 0.013724 | 0.010391 | 0.05276 | 0.089981 |
| 33.9166667 | 0.009792 | 0.012415 | 0.001993 | 0.043875 | 0.120865 | 0.004106 | 0.029044 | 0.002737 | 0.006488 | 0.006255 | 0.010318 | 0.025143 | 0.083572 | 0.095365 | 0.071313 | 0.024778 | 0.027665 | 0.019402 | 0.041637 | 0.031101 | 0.017187 | 0.017015 | 0.050658 | 0.074982 |
| 33.9305556 | 0.010971 | 0.022024 | 0.002008 | 0.059619 | 0.123676 | 0.005104 | 0.03309 | 0.003486 | 0.004807 | 0.006229 | 0.016878 | 0.044395 | 0.090717 | 0.07418 | 0.075745 | 0.024849 | 0.03279 | 0.025344 | 0.050677 | 0.034188 | 0.014888 | 0.024419 | 0.054725 | 0.061574 |
| 33.9444444 | 0.007451 | 0.033488 | 0.004913 | 0.068635 | 0.126552 | 0.010593 | 0.025563 | 0.007604 | 0.003326 | 0.00812 | 0.025376 | 0.062064 | 0.10451 | 0.068511 | 0.076861 | 0.019975 | 0.038464 | 0.039412 | 0.055228 | 0.053268 | 0.012015 | 0.025119 | 0.06403 | 0.062288 |
| 33.9583333 | 0.003695 | 0.038842 | 0.009899 | 0.069047 | 0.125666 | 0.01677 | 0.013658 | 0.012498 | 0.004299 | 0.010194 | 0.031095 | 0.070511 | 0.110146 | 0.074839 | 0.083504 | 0.020183 | 0.045533 | 0.052047 | 0.053579 | 0.073045 | 0.010248 | 0.016704 | 0.068659 | 0.076532 |
| 33.9722222 | 0.005225 | 0.039209 | 0.015481 | 0.067317 | 0.119876 | 0.019169 | 0.008675 | 0.014926 | 0.005195 | 0.013817 | 0.036477 | 0.072057 | 0.104354 | 0.088943 | 0.095357 | 0.031023 | 0.056543 | 0.048442 | 0.050572 | 0.08063 | 0.010176 | 0.008295 | 0.069043 | 0.096783 |
| 33.9861111 | 0.010783 | 0.041195 | 0.021905 | 0.072237 | 0.118487 | 0.021014 | 0.009179 | 0.015012 | 0.00702 | 0.023629 | 0.04181 | 0.07139 | 0.0936 | 0.108777 | 0.103513 | 0.04511 | 0.074726 | 0.03619 | 0.054637 | 0.085503 | 0.01606 | 0.005894 | 0.073281 | 0.117844 |
| 34 | 0.018158 | 0.050479 | 0.02895 | 0.086964 | 0.126821 | 0.025654 | 0.012951 | 0.020433 | 0.011632 | 0.036572 | 0.049878 | 0.071639 | 0.087173 | 0.12645 | 0.10837 | 0.053969 | 0.092066 | 0.040275 | 0.068251 | 0.102995 | 0.031478 | 0.008999 | 0.075864 | 0.136531 |
| 34.0138889 | 0.023826 | 0.06404 | 0.035553 | 0.102631 | 0.147002 | 0.031573 | 0.022961 | 0.034883 | 0.016635 | 0.05261 | 0.062301 | 0.076727 | 0.083247 | 0.138048 | 0.107787 | 0.056406 | 0.098234 | 0.062889 | 0.082977 | 0.141352 | 0.049074 | 0.020902 | 0.069614 | 0.14782 |
| 34.0277778 | 0.022886 | 0.074755 | 0.041291 | 0.116945 | 0.178277 | 0.036879 | 0.035614 | 0.053579 | 0.020063 | 0.069081 | 0.076331 | 0.090013 | 0.078888 | 0.148343 | 0.104122 | 0.057301 | 0.105533 | 0.09841 | 0.098316 | 0.176774 | 0.073775 | 0.03709 | 0.063999 | 0.151955 |
| 34.0416667 | 0.01826 | 0.083113 | 0.044094 | 0.130691 | 0.206095 | 0.044369 | 0.043701 | 0.068364 | 0.023241 | 0.083959 | 0.089185 | 0.108379 | 0.082167 | 0.142856 | 0.10118 | 0.057876 | 0.11334 | 0.133883 | 0.12052 | 0.17064 | 0.135861 | 0.04586 | 0.071453 | 0.172318 |
| 34.0555556 | 0.019483 | 0.089165 | 0.044136 | 0.135465 | 0.232765 | 0.057396 | 0.04761 | 0.082095 | 0.030871 | 0.097956 | 0.104519 | 0.125743 | 0.098099 | 0.141985 | 0.096219 | 0.053087 | 0.105723 | 0.147748 | 0.145166 | 0.117244 | 0.171017 | 0.051065 | 0.095743 | 0.156014 |
| 34.0694444 | 0.025188 | 0.084726 | 0.045429 | 0.13332 | 0.2563 | 0.072099 | 0.049238 | 0.098675 | 0.040149 | 0.103169 | 0.116632 | 0.134932 | 0.119778 | 0.158616 | 0.085493 | 0.044369 | 0.10058 | 0.12672 | 0.149655 | 0.065612 | 0.119234 | 0.048707 | 0.121336 | 0.100239 |
| 34.0833333 | 0.023609 | 0.065058 | 0.044103 | 0.118741 | 0.251683 | 0.076423 | 0.040601 | 0.105755 | 0.039483 | 0.09539 | 0.110399 | 0.127398 | 0.11749 | 0.158794 | 0.073561 | 0.041656 | 0.095683 | 0.08392 | 0.122407 | 0.063739 | 0.082752 | 0.033615 | 0.126846 | 0.138059 |
| 34.0972222 | 0.01486 | 0.037697 | 0.035381 | 0.079081 | 0.219628 | 0.065323 | 0.022142 | 0.103421 | 0.027898 | 0.079303 | 0.08668 | 0.102233 | 0.090382 | 0.138662 | 0.064656 | 0.037976 | 0.07014 | 0.04916 | 0.089821 | 0.086541 | 0.091832 | 0.034308 | 0.11508 | 0.19976 |
| 34.1111111 | 0.0077 | 0.015804 | 0.02433 | 0.033444 | 0.175175 | 0.0492 | 0.009112 | 0.094025 | 0.015617 | 0.06397 | 0.061007 | 0.073067 | 0.06878 | 0.112539 | 0.060165 | 0.033997 | 0.03664 | 0.024818 | 0.062592 | 0.10447 | 0.086074 | 0.047551 | 0.097486 | 0.182678 |
| 34.125 | 0.004537 | 0.006528 | 0.016062 | 0.011605 | 0.132517 | 0.037948 | 0.005498 | 0.073936 | 0.008205 | 0.050971 | 0.041487 | 0.050811 | 0.05276 | 0.101101 | 0.06745 | 0.041875 | 0.015388 | 0.015285 | 0.045564 | 0.114082 | 0.067537 | 0.048813 | 0.079723 | 0.163913 |
| 34.1388889 | 0.002918 | 0.007992 | 0.011148 | 0.011047 | 0.095574 | 0.031198 | 0.004117 | 0.049762 | 0.004192 | 0.036082 | 0.025584 | 0.034204 | 0.039001 | 0.110382 | 0.072958 | 0.04751 | 0.012213 | 0.029856 | 0.051806 | 0.095745 | 0.058659 | 0.051408 | 0.062475 | 0.170279 |
| 34.1527778 | 0.001855 | 0.015892 | 0.008737 | 0.011396 | 0.052481 | 0.023643 | 0.00378 | 0.025969 | 0.002945 | 0.02192 | 0.012078 | 0.020215 | 0.027984 | 0.123651 | 0.062478 | 0.043159 | 0.016037 | 0.042443 | 0.050282 | 0.061015 | 0.063064 | 0.048869 | 0.051345 | 0.162904 |
| 34.1666667 | 0.002283 | 0.023565 | 0.006481 | 0.012388 | 0.024381 | 0.013716 | 0.006342 | 0.012158 | 0.004505 | 0.012355 | 0.005938 | 0.011365 | 0.020864 | 0.127539 | 0.052216 | 0.037065 | 0.026469 | 0.036598 | 0.040131 | 0.062247 | 0.054954 | 0.02861 | 0.053851 | 0.143377 |
| 34.1805556 | 0.003791 | 0.024902 | 0.004564 | 0.025212 | 0.038663 | 0.006103 | 0.010291 | 0.020018 | 0.00748 | 0.006759 | 0.012448 | 0.015263 | 0.036888 | 0.129901 | 0.048534 | 0.029967 | 0.042185 | 0.043203 | 0.063609 | 0.089672 | 0.027623 | 0.015294 | 0.059553 | 0.136608 |
| 34.1944444 | 0.006205 | 0.022662 | 0.006418 | 0.051272 | 0.069573 | 0.00534 | 0.014969 | 0.041008 | 0.011498 | 0.005754 | 0.025687 | 0.030113 | 0.072063 | 0.131879 | 0.037871 | 0.019132 | 0.04524 | 0.066442 | 0.093653 | 0.100683 | 0.013482 | 0.018153 | 0.056924 | 0.130373 |
| 34.2083333 | 0.008906 | 0.021029 | 0.009998 | 0.079619 | 0.088167 | 0.011595 | 0.020557 | 0.058462 | 0.01516 | 0.009337 | 0.035218 | 0.042754 | 0.093901 | 0.11883 | 0.019363 | 0.009056 | 0.040557 | 0.081362 | 0.096649 | 0.117845 | 0.023296 | 0.026647 | 0.047199 | 0.106281 |
| 34.2222222 | 0.010453 | 0.018555 | 0.010866 | 0.091385 | 0.092654 | 0.017896 | 0.024123 | 0.063701 | 0.016794 | 0.011513 | 0.036776 | 0.047028 | 0.086023 | 0.097755 | 0.006074 | 0.007378 | 0.031739 | 0.088573 | 0.089619 | 0.152686 | 0.049347 | 0.039277 | 0.037255 | 0.077359 |
| 34.2361111 | 0.012334 | 0.016073 | 0.01028 | 0.083761 | 0.089598 | 0.018536 | 0.024376 | 0.060735 | 0.017263 | 0.011976 | 0.03209 | 0.044752 | 0.061298 | 0.079531 | 0.002787 | 0.009443 | 0.016364 | 0.084442 | 0.072073 | 0.153189 | 0.078991 | 0.045418 | 0.031969 | 0.055772 |
| 34.25 | 0.018413 | 0.019565 | 0.008985 | 0.079807 | 0.087176 | 0.015123 | 0.03005 | 0.069503 | 0.018145 | 0.027356 | 0.028698 | 0.035444 | 0.037059 | 0.069429 | 0.007448 | 0.009211 | 0.010144 | 0.055043 | 0.042251 | 0.093855 | 0.084708 | 0.031484 | 0.032127 | 0.043441 |
| 34.2638889 | 0.028428 | 0.034476 | 0.006449 | 0.094106 | 0.079345 | 0.012978 | 0.060259 | 0.122384 | 0.020122 | 0.064203 | 0.030716 | 0.019659 | 0.023124 | 0.079128 | 0.020215 | 0.013478 | 0.016001 | 0.028174 | 0.031803 | 0.036625 | 0.059931 | 0.02606 | 0.038009 | 0.050065 |
| 34.2777778 | 0.030365 | 0.043601 | 0.004941 | 0.102964 | 0.094718 | 0.015431 | 0.078116 | 0.160578 | 0.019378 | 0.075087 | 0.02525 | 0.007126 | 0.020738 | 0.099399 | 0.031129 | 0.022184 | 0.02507 | 0.028494 | 0.034676 | 0.03474 | 0.037366 | 0.045349 | 0.044165 | 0.068939 |
| 34.2916667 | 0.018771 | 0.036099 | 0.009051 | 0.080476 | 0.125739 | 0.016485 | 0.053021 | 0.123868 | 0.014812 | 0.057283 | 0.010389 | 0.010754 | 0.020268 | 0.096998 | 0.025116 | 0.018802 | 0.02837 | 0.03803 | 0.02979 | 0.083542 | 0.043122 | 0.043987 | 0.042158 | 0.075003 |
| 34.3055556 | 0.006914 | 0.030748 | 0.019955 | 0.05821 | 0.093766 | 0.013667 | 0.031328 | 0.076832 | 0.009578 | 0.057788 | 0.00383 | 0.019568 | 0.019886 | 0.073119 | 0.01021 | 0.006783 | 0.027698 | 0.042003 | 0.029207 | 0.121432 | 0.055728 | 0.022824 | 0.03496 | 0.063362 |
| 34.3194444 | 0.00326 | 0.035189 | 0.027115 | 0.072517 | 0.042696 | 0.010157 | 0.019419 | 0.044509 | 0.006717 | 0.04858 | 0.003754 | 0.016026 | 0.016193 | 0.050683 | 0.003889 | 0.004725 | 0.029399 | 0.041353 | 0.032509 | 0.087396 | 0.049062 | 0.013181 | 0.029326 | 0.045762 |
| 34.3333333 | 0.005557 | 0.046236 | 0.024639 | 0.098898 | 0.047803 | 0.005541 | 0.011181 | 0.030235 | 0.009294 | 0.020686 | 0.002605 | 0.007995 | 0.007236 | 0.031041 | 0.003354 | 0.007566 | 0.027214 | 0.045834 | 0.039997 | 0.038911 | 0.036455 | 0.007268 | 0.021558 | 0.028423 |
| 34.3472222 | 0.009503 | 0.053029 | 0.019925 | 0.10972 | 0.073666 | 0.002216 | 0.015697 | 0.04054 | 0.012276 | 0.006461 | 0.003388 | 0.00618 | 0.002797 | 0.016671 | 0.002317 | 0.005341 | 0.026331 | 0.043197 | 0.039587 | 0.040845 | 0.048677 | 0.012916 | 0.010371 | 0.016412 |
| 34.3611111 | 0.012108 | 0.051207 | 0.017599 | 0.108927 | 0.082275 | 0.002998 | 0.021682 | 0.046901 | 0.013531 | 0.003488 | 0.008331 | 0.01264 | 0.006215 | 0.009123 | 0.002783 | 0.002539 | 0.027893 | 0.039854 | 0.038502 | 0.081894 | 0.084398 | 0.038491 | 0.003029 | 0.011041 |
| 34.375 | 0.013441 | 0.045479 | 0.015311 | 0.102333 | 0.076382 | 0.006656 | 0.023005 | 0.042242 | 0.014966 | 0.003123 | 0.013726 | 0.01978 | 0.010369 | 0.003765 | 0.003371 | 0.002569 | 0.02135 | 0.037024 | 0.043547 | 0.10345 | 0.109932 | 0.059904 | 0.001311 | 0.006894 |
| 34.3888889 | 0.01568 | 0.036468 | 0.011266 | 0.092278 | 0.076011 | 0.008347 | 0.019993 | 0.037432 | 0.014743 | 0.004846 | 0.012445 | 0.016542 | 0.007905 | 0.003319 | 0.003115 | 0.004835 | 0.015316 | 0.022623 | 0.028295 | 0.075438 | 0.106237 | 0.060648 | 0.001076 | 0.003433 |
| 34.4027778 | 0.020185 | 0.029497 | 0.0076 | 0.08339 | 0.086528 | 0.005771 | 0.018347 | 0.042578 | 0.013778 | 0.011487 | 0.007046 | 0.007012 | 0.004122 | 0.004546 | 0.004043 | 0.006036 | 0.023805 | 0.017552 | 0.013822 | 0.050214 | 0.085372 | 0.054482 | 0.001568 | 0.003556 |
| 34.4166667 | 0.023589 | 0.0275 | 0.00616 | 0.077498 | 0.091915 | 0.00351 | 0.021269 | 0.053853 | 0.014175 | 0.019273 | 0.00746 | 0.005669 | 0.00595 | 0.013366 | 0.011132 | 0.010993 | 0.031606 | 0.031164 | 0.030994 | 0.068045 | 0.068366 | 0.055774 | 0.004368 | 0.012254 |
| 34.4305556 | 0.023277 | 0.028984 | 0.007072 | 0.074891 | 0.086073 | 0.006581 | 0.025575 | 0.061174 | 0.01619 | 0.021268 | 0.020793 | 0.024788 | 0.016613 | 0.044635 | 0.027396 | 0.028465 | 0.027076 | 0.054892 | 0.058103 | 0.118553 | 0.058636 | 0.059272 | 0.00928 | 0.033251 |
| 34.4444444 | 0.023243 | 0.033743 | 0.010749 | 0.077983 | 0.082059 | 0.015765 | 0.026862 | 0.063577 | 0.018511 | 0.019554 | 0.044006 | 0.058196 | 0.036682 | 0.083083 | 0.043314 | 0.048189 | 0.037564 | 0.064697 | 0.069108 | 0.171618 | 0.056324 | 0.067636 | 0.014296 | 0.056106 |
| 34.4583333 | 0.027519 | 0.042044 | 0.016316 | 0.087735 | 0.090812 | 0.026722 | 0.02468 | 0.063284 | 0.019632 | 0.019332 | 0.062624 | 0.083967 | 0.052317 | 0.10244 | 0.0435 | 0.052559 | 0.067121 | 0.066265 | 0.073779 | 0.201351 | 0.067903 | 0.086969 | 0.016006 | 0.065444 |
| 34.4722222 | 0.033747 | 0.051836 | 0.021765 | 0.097898 | 0.109416 | 0.034567 | 0.023276 | 0.060407 | 0.020719 | 0.023655 | 0.070321 | 0.096468 | 0.057181 | 0.09812 | 0.02566 | 0.038543 | 0.083888 | 0.085215 | 0.085207 | 0.204035 | 0.087557 | 0.100501 | 0.013336 | 0.061293 |
| 34.4861111 | 0.03705 | 0.059055 | 0.027275 | 0.101937 | 0.129626 | 0.039559 | 0.026963 | 0.061693 | 0.02409 | 0.030968 | 0.077628 | 0.104208 | 0.062728 | 0.086983 | 0.012046 | 0.02198 | 0.084211 | 0.102404 | 0.094401 | 0.205446 | 0.111048 | 0.103282 | 0.011736 | 0.059687 |
| 34.5 | 0.037935 | 0.059806 | 0.033613 | 0.102582 | 0.143896 | 0.043336 | 0.033297 | 0.068246 | 0.028655 | 0.036811 | 0.087117 | 0.108675 | 0.070762 | 0.087448 | 0.030962 | 0.024252 | 0.083276 | 0.095535 | 0.096054 | 0.223815 | 0.119459 | 0.103422 | 0.017245 | 0.070014 |
| 34.5138889 | 0.043402 | 0.055688 | 0.038313 | 0.098873 | 0.151903 | 0.045368 | 0.038196 | 0.073973 | 0.031647 | 0.040167 | 0.089858 | 0.108095 | 0.073077 | 0.1017 | 0.07362 | 0.045412 | 0.082266 | 0.077311 | 0.095694 | 0.228804 | 0.09012 | 0.094463 | 0.028793 | 0.08472 |
| 34.5277778 | 0.054349 | 0.05505 | 0.037379 | 0.09113 | 0.159373 | 0.048 | 0.041162 | 0.073934 | 0.031269 | 0.040564 | 0.08441 | 0.1064 | 0.075567 | 0.134228 | 0.110327 | 0.06746 | 0.084557 | 0.066726 | 0.101402 | 0.186342 | 0.053997 | 0.075008 | 0.0424 | 0.106088 |
| 34.5416667 | 0.061436 | 0.061631 | 0.036049 | 0.089321 | 0.166123 | 0.056273 | 0.042339 | 0.066875 | 0.028156 | 0.04173 | 0.07689 | 0.102257 | 0.084759 | 0.181634 | 0.135582 | 0.079774 | 0.097448 | 0.071635 | 0.113269 | 0.131167 | 0.084758 | 0.048986 | 0.053633 | 0.141091 |
| 34.5555556 | 0.059226 | 0.067149 | 0.043414 | 0.087222 | 0.162342 | 0.066107 | 0.042029 | 0.059783 | 0.025582 | 0.049522 | 0.069304 | 0.088452 | 0.085797 | 0.209249 | 0.144142 | 0.080976 | 0.1139 | 0.080473 | 0.116019 | 0.117693 | 0.156197 | 0.028693 | 0.057517 | 0.167869 |
| 34.5694444 | 0.050953 | 0.062107 | 0.050034 | 0.073754 | 0.141063 | 0.065124 | 0.0417 | 0.05657 | 0.024494 | 0.057582 | 0.057799 | 0.070425 | 0.072218 | 0.203761 | 0.139583 | 0.08069 | 0.120639 | 0.080954 | 0.108897 | 0.123439 | 0.163374 | 0.032476 | 0.057966 | 0.168092 |
| 34.5833333 | 0.043666 | 0.048042 | 0.04392 | 0.055436 | 0.116625 | 0.052971 | 0.041158 | 0.056556 | 0.023831 | 0.058975 | 0.045427 | 0.055901 | 0.06064 | 0.195025 | 0.14786 | 0.089132 | 0.118811 | 0.079714 | 0.096412 | 0.111031 | 0.135309 | 0.051018 | 0.062256 | 0.159287 |
| 34.5972222 | 0.043982 | 0.039593 | 0.033401 | 0.045129 | 0.109659 | 0.043853 | 0.036787 | 0.06038 | 0.024044 | 0.057633 | 0.043345 | 0.047606 | 0.064129 | 0.198442 | 0.168984 | 0.100657 | 0.112294 | 0.079304 | 0.080824 | 0.095468 | 0.130895 | 0.063115 | 0.069045 | 0.162211 |
| 34.6111111 | 0.051672 | 0.042807 | 0.028665 | 0.04464 | 0.118529 | 0.045407 | 0.028016 | 0.064096 | 0.023167 | 0.05419 | 0.051967 | 0.04656 | 0.077904 | 0.205581 | 0.172765 | 0.108247 | 0.098906 | 0.080352 | 0.071809 | 0.094686 | 0.119332 | 0.070499 | 0.071926 | 0.172063 |
| 34.625 | 0.05947 | 0.04266 | 0.029444 | 0.042081 | 0.125436 | 0.049698 | 0.021537 | 0.059868 | 0.020499 | 0.050553 | 0.05433 | 0.041128 | 0.084913 | 0.205261 | 0.147169 | 0.107134 | 0.084617 | 0.079905 | 0.06111 | 0.101912 | 0.077095 | 0.082716 | 0.066852 | 0.171773 |
| 34.6388889 | 0.058523 | 0.030911 | 0.030883 | 0.030264 | 0.119453 | 0.047794 | 0.019262 | 0.048754 | 0.017871 | 0.051644 | 0.045346 | 0.025691 | 0.078326 | 0.187101 | 0.114074 | 0.094193 | 0.076218 | 0.068659 | 0.035652 | 0.102342 | 0.031963 | 0.088595 | 0.059483 | 0.160478 |
| 34.6527778 | 0.047693 | 0.015236 | 0.028878 | 0.016615 | 0.101983 | 0.040117 | 0.012863 | 0.039199 | 0.014862 | 0.055223 | 0.036714 | 0.013328 | 0.067644 | 0.14518 | 0.092795 | 0.080183 | 0.069208 | 0.051627 | 0.012857 | 0.093107 | 0.012029 | 0.071963 | 0.055242 | 0.134637 |
| 34.6666667 | 0.032839 | 0.005476 | 0.02306 | 0.009929 | 0.07787 | 0.030076 | 0.005786 | 0.033046 | 0.014628 | 0.056498 | 0.029694 | 0.007776 | 0.059651 | 0.093376 | 0.082374 | 0.070416 | 0.050461 | 0.040265 | 0.00667 | 0.080132 | 0.006899 | 0.051595 | 0.049401 | 0.093696 |
| 34.6805556 | 0.016843 | 0.007189 | 0.015014 | 0.010311 | 0.050088 | 0.019069 | 0.005157 | 0.024945 | 0.018585 | 0.049325 | 0.022658 | 0.007172 | 0.052906 | 0.058821 | 0.073594 | 0.059357 | 0.022829 | 0.045896 | 0.009375 | 0.061747 | 0.004481 | 0.049461 | 0.038725 | 0.063238 |
| 34.6944444 | 0.005882 | 0.017188 | 0.007093 | 0.017274 | 0.023223 | 0.00892 | 0.004953 | 0.01315 | 0.020958 | 0.033976 | 0.016963 | 0.008512 | 0.042936 | 0.046488 | 0.05748 | 0.042017 | 0.005446 | 0.063916 | 0.021688 | 0.034579 | 0.005289 | 0.046827 | 0.024899 | 0.051198 |
| 34.7083333 | 0.004552 | 0.0283 | 0.002433 | 0.020282 | 0.008731 | 0.005382 | 0.003577 | 0.007035 | 0.018019 | 0.017688 | 0.010313 | 0.006766 | 0.027319 | 0.03903 | 0.034375 | 0.028063 | 0.001565 | 0.075708 | 0.036935 | 0.016988 | 0.01019 | 0.031216 | 0.013961 | 0.04013 |
| 34.7222222 | 0.006281 | 0.032695 | 0.001222 | 0.020318 | 0.016869 | 0.010483 | 0.002432 | 0.009979 | 0.011533 | 0.007837 | 0.005596 | 0.005512 | 0.022444 | 0.023673 | 0.027711 | 0.037427 | 0.013249 | 0.077684 | 0.046965 | 0.032126 | 0.01502 | 0.028999 | 0.014409 | 0.022676 |

| | | | | | | | | | | | | | | | | | | | | | | | | |
|---|---|---|---|---|---|---|---|---|---|---|---|---|---|---|---|---|---|---|---|---|---|---|---|---|
| 34.7361111 | 0.008115 | 0.024212 | 0.001249 | 0.029296 | 0.044332 | 0.014689 | 0.001476 | 0.013544 | 0.00541 | 0.009594 | 0.007465 | 0.004488 | 0.037437 | 0.01106 | 0.03804 | 0.048642 | 0.046747 | 0.061168 | 0.052164 | 0.066173 | 0.013541 | 0.040965 | 0.022795 | 0.00946 |
| 34.75 | 0.011242 | 0.015068 | 0.003586 | 0.032715 | 0.073713 | 0.010954 | 0.004101 | 0.016014 | 0.001871 | 0.017751 | 0.011648 | 0.002061 | 0.047933 | 0.013277 | 0.032307 | 0.03431 | 0.077956 | 0.036465 | 0.049811 | 0.0851 | 0.008579 | 0.04276 | 0.027575 | 0.006527 |
| 34.7638889 | 0.012104 | 0.024663 | 0.008726 | 0.021426 | 0.088936 | 0.004627 | 0.009981 | 0.019721 | 0.0008 | 0.021954 | 0.013482 | 0.001887 | 0.040828 | 0.022441 | 0.022193 | 0.013102 | 0.09267 | 0.033501 | 0.034725 | 0.080131 | 0.014831 | 0.039553 | 0.027878 | 0.008699 |
| 34.7777778 | 0.009458 | 0.040667 | 0.013146 | 0.011023 | 0.090391 | 0.002443 | 0.01478 | 0.023424 | 0.001772 | 0.022027 | 0.0134 | 0.004691 | 0.025182 | 0.027628 | 0.024633 | 0.004263 | 0.122605 | 0.040351 | 0.015173 | 0.06855 | 0.021711 | 0.041671 | 0.029215 | 0.011351 |
| 34.7916667 | 0.008305 | 0.035378 | 0.01422 | 0.01399 | 0.085904 | 0.005411 | 0.015195 | 0.025035 | 0.00287 | 0.029034 | 0.013102 | 0.007155 | 0.013632 | 0.025583 | 0.023099 | 0.004436 | 0.165128 | 0.031572 | 0.006936 | 0.061604 | 0.013169 | 0.048426 | 0.030466 | 0.0105 |
| 34.8055556 | 0.013092 | 0.018093 | 0.012704 | 0.029869 | 0.080929 | 0.010783 | 0.010321 | 0.023123 | 0.002032 | 0.041275 | 0.014194 | 0.007915 | 0.013983 | 0.02012 | 0.022786 | 0.008507 | 0.170457 | 0.014021 | 0.00872 | 0.056396 | 0.01058 | 0.052467 | 0.026922 | 0.006137 |
| 34.8194444 | 0.021694 | 0.01236 | 0.010016 | 0.047485 | 0.080243 | 0.01272 | 0.004195 | 0.019179 | 0.000924 | 0.045246 | 0.014437 | 0.009319 | 0.020024 | 0.012469 | 0.040108 | 0.021628 | 0.121478 | 0.00553 | 0.020819 | 0.051119 | 0.023584 | 0.048888 | 0.019184 | 0.005033 |
| 34.8333333 | 0.026686 | 0.011859 | 0.007115 | 0.045576 | 0.085794 | 0.010044 | 0.001956 | 0.017701 | 0.000702 | 0.041018 | 0.014864 | 0.014507 | 0.020303 | 0.012718 | 0.061401 | 0.037741 | 0.055188 | 0.009162 | 0.037175 | 0.049424 | 0.026571 | 0.043999 | 0.016642 | 0.016642 |
| 34.8472222 | 0.022617 | 0.013081 | 0.005712 | 0.025683 | 0.090221 | 0.008111 | 0.004732 | 0.019934 | 0.000848 | 0.035713 | 0.021182 | 0.024532 | 0.01235 | 0.026573 | 0.055618 | 0.036719 | 0.020513 | 0.016441 | 0.047072 | 0.052944 | 0.014265 | 0.042224 | 0.023054 | 0.034827 |
| 34.8611111 | 0.012282 | 0.013796 | 0.006503 | 0.01057 | 0.081907 | 0.007914 | 0.009716 | 0.021113 | 0.001195 | 0.032067 | 0.028082 | 0.032629 | 0.004292 | 0.026865 | 0.028701 | 0.022329 | 0.01659 | 0.023435 | 0.051513 | 0.054743 | 0.007639 | 0.041972 | 0.025498 | 0.037482 |
| 34.875 | 0.003868 | 0.015088 | 0.007831 | 0.014706 | 0.061156 | 0.006748 | 0.011028 | 0.018721 | 0.001627 | 0.031193 | 0.023874 | 0.027904 | 0.002213 | 0.011404 | 0.014424 | 0.013235 | 0.013186 | 0.029154 | 0.050071 | 0.04643 | 0.011302 | 0.039555 | 0.016969 | 0.022432 |
| 34.8888889 | 0.002181 | 0.02741 | 0.007809 | 0.037315 | 0.042041 | 0.004607 | 0.008477 | 0.019018 | 0.003284 | 0.033015 | 0.013868 | 0.017021 | 0.00256 | 0.00698 | 0.017526 | 0.01382 | 0.007872 | 0.02985 | 0.043196 | 0.033739 | 0.021522 | 0.030669 | 0.007014 | 0.013026 |
| 34.9027778 | 0.006511 | 0.031377 | 0.006544 | 0.055135 | 0.034494 | 0.002987 | 0.008151 | 0.026669 | 0.005966 | 0.030295 | 0.010573 | 0.013458 | 0.005745 | 0.01595 | 0.017974 | 0.01481 | 0.015786 | 0.029726 | 0.03837 | 0.031351 | 0.02607 | 0.023176 | 0.005101 | 0.020941 |
| 34.9166667 | 0.014202 | 0.020932 | 0.006333 | 0.057554 | 0.033416 | 0.002848 | 0.011159 | 0.035584 | 0.007308 | 0.022022 | 0.010519 | 0.012041 | 0.022148 | 0.032982 | 0.017811 | 0.017135 | 0.042701 | 0.034976 | 0.04368 | 0.045352 | 0.016832 | 0.029669 | 0.013095 | 0.039243 |
| 34.9305556 | 0.018698 | 0.019152 | 0.008883 | 0.058282 | 0.035096 | 0.004506 | 0.013479 | 0.040723 | 0.006046 | 0.021352 | 0.01185 | 0.012487 | 0.045213 | 0.058263 | 0.034637 | 0.03237 | 0.075583 | 0.041258 | 0.056778 | 0.066441 | 0.015835 | 0.038563 | 0.024239 | 0.062427 |
| 34.9444444 | 0.013982 | 0.030583 | 0.012395 | 0.065375 | 0.03954 | 0.007259 | 0.013846 | 0.042849 | 0.003217 | 0.026133 | 0.016717 | 0.016172 | 0.058009 | 0.085206 | 0.052149 | 0.044503 | 0.100151 | 0.04403 | 0.064959 | 0.077973 | 0.031783 | 0.03482 | 0.030289 | 0.085364 |
| 34.9583333 | 0.010245 | 0.041376 | 0.014248 | 0.077567 | 0.043378 | 0.009985 | 0.014221 | 0.044615 | 0.001033 | 0.025019 | 0.016977 | 0.012799 | 0.062026 | 0.108889 | 0.053446 | 0.039515 | 0.109417 | 0.046114 | 0.060712 | 0.077573 | 0.038427 | 0.030578 | 0.031428 | 0.102663 |
| 34.9722222 | 0.017977 | 0.044037 | 0.015151 | 0.088896 | 0.047926 | 0.012107 | 0.016069 | 0.049542 | 0.000764 | 0.021827 | 0.013556 | 0.005112 | 0.059862 | 0.14757 | 0.062528 | 0.036295 | 0.099305 | 0.049173 | 0.051943 | 0.083687 | 0.033991 | 0.035214 | 0.031772 | 0.124293 |
| 34.9861111 | 0.026998 | 0.046145 | 0.01774 | 0.094373 | 0.055621 | 0.012811 | 0.020514 | 0.057138 | 0.003432 | 0.02449 | 0.016677 | 0.007275 | 0.051044 | 0.182662 | 0.09216 | 0.041232 | 0.077969 | 0.05604 | 0.049529 | 0.11204 | 0.040403 | 0.039991 | 0.033655 | 0.14794 |
| 35 | 0.031503 | 0.051922 | 0.021994 | 0.096482 | 0.067823 | 0.01106 | 0.026183 | 0.065579 | 0.010099 | 0.035429 | 0.026918 | 0.024634 | 0.044643 | 0.17524 | 0.117283 | 0.046713 | 0.061874 | 0.068625 | 0.055792 | 0.150737 | 0.055425 | 0.043954 | 0.033534 | 0.150356 |
| 35.0138889 | 0.037653 | 0.055345 | 0.025312 | 0.09923 | 0.083565 | 0.010235 | 0.029361 | 0.07196 | 0.017166 | 0.048803 | 0.037065 | 0.043262 | 0.050451 | 0.164443 | 0.127624 | 0.053664 | 0.058785 | 0.085422 | 0.065752 | 0.176 | 0.074401 | 0.055408 | 0.028694 | 0.140658 |
| 35.0277778 | 0.043633 | 0.056124 | 0.027413 | 0.104049 | 0.095181 | 0.017311 | 0.029097 | 0.073977 | 0.019181 | 0.054448 | 0.039885 | 0.05039 | 0.074996 | 0.181576 | 0.134174 | 0.058419 | 0.066916 | 0.102201 | 0.07685 | 0.16376 | 0.094372 | 0.078859 | 0.027403 | 0.142919 |
| 35.0416667 | 0.044352 | 0.059596 | 0.031402 | 0.113876 | 0.10465 | 0.035924 | 0.030388 | 0.080029 | 0.018471 | 0.05428 | 0.039015 | 0.05884 | 0.123532 | 0.219023 | 0.136355 | 0.05976 | 0.092506 | 0.101658 | 0.09396 | 0.106142 | 0.095624 | 0.099307 | 0.043038 | 0.155021 |
| 35.0555556 | 0.041065 | 0.067573 | 0.035778 | 0.129414 | 0.118745 | 0.063609 | 0.036332 | 0.097667 | 0.023807 | 0.064578 | 0.049267 | 0.082255 | 0.181774 | 0.2673 | 0.1331 | 0.061034 | 0.157149 | 0.081029 | 0.121708 | 0.052798 | 0.093741 | 0.092301 | 0.083745 | 0.141451 |
| 35.0694444 | 0.034444 | 0.071982 | 0.03368 | 0.139889 | 0.134907 | 0.080504 | 0.038466 | 0.114898 | 0.035655 | 0.089662 | 0.064906 | 0.102923 | 0.203964 | 0.258025 | 0.122315 | 0.062096 | 0.213495 | 0.069027 | 0.141416 | 0.063978 | 0.101608 | 0.055762 | 0.117236 | 0.107562 |
| 35.0833333 | 0.022379 | 0.061685 | 0.025968 | 0.125019 | 0.135985 | 0.066122 | 0.028523 | 0.116153 | 0.041596 | 0.112169 | 0.065425 | 0.092657 | 0.174163 | 0.180978 | 0.106245 | 0.071209 | 0.182123 | 0.066159 | 0.12562 | 0.100309 | 0.112164 | 0.018913 | 0.106726 | 0.104291 |
| 35.0972222 | 0.010276 | 0.040426 | 0.018672 | 0.085979 | 0.108691 | 0.03572 | 0.013561 | 0.110891 | 0.036981 | 0.119684 | 0.055455 | 0.062287 | 0.122547 | 0.131003 | 0.088057 | 0.081775 | 0.108205 | 0.046303 | 0.081664 | 0.088619 | 0.1192 | 0.012924 | 0.077017 | 0.113608 |
| 35.1111111 | 0.005803 | 0.019065 | 0.01264 | 0.048233 | 0.076896 | 0.013498 | 0.004977 | 0.11358 | 0.028874 | 0.111843 | 0.044018 | 0.041565 | 0.06969 | 0.123978 | 0.069589 | 0.075631 | 0.05548 | 0.022085 | 0.039597 | 0.061203 | 0.135628 | 0.026152 | 0.059242 | 0.109474 |
| 35.125 | 0.006407 | 0.006648 | 0.007525 | 0.021746 | 0.060091 | 0.006925 | 0.003253 | 0.117154 | 0.01831 | 0.09039 | 0.029516 | 0.031198 | 0.029525 | 0.121855 | 0.060901 | 0.061484 | 0.021973 | 0.02354 | 0.028845 | 0.060089 | 0.150823 | 0.030834 | 0.053791 | 0.104741 |
| 35.1388889 | 0.007514 | 0.005164 | 0.003832 | 0.007984 | 0.047463 | 0.013451 | 0.003643 | 0.111043 | 0.007792 | 0.062085 | 0.017283 | 0.019424 | 0.010229 | 0.121917 | 0.054018 | 0.046648 | 0.008098 | 0.043728 | 0.046677 | 0.056467 | 0.123652 | 0.025545 | 0.052399 | 0.10214 |
| 35.1527778 | 0.008104 | 0.004729 | 0.001778 | 0.007919 | 0.028729 | 0.017835 | 0.003367 | 0.090896 | 0.004763 | 0.036678 | 0.00986 | 0.019232 | 0.008536 | 0.122433 | 0.041353 | 0.026314 | 0.010952 | 0.053455 | 0.058782 | 0.034386 | 0.064149 | 0.018981 | 0.048719 | 0.099375 |
| 35.1666667 | 0.008837 | 0.002105 | 0.001888 | 0.01174 | 0.017892 | 0.013509 | 0.002688 | 0.062975 | 0.007637 | 0.019084 | 0.006764 | 0.013069 | 0.023028 | 0.113322 | 0.029722 | 0.012504 | 0.020555 | 0.053475 | 0.057564 | 0.02196 | 0.021986 | 0.012352 | 0.043288 | 0.088369 |
| 35.1805556 | 0.012097 | 0.00213 | 0.003543 | 0.018074 | 0.028161 | 0.01047 | 0.004741 | 0.039768 | 0.011455 | 0.008901 | 0.011413 | 0.013453 | 0.048222 | 0.085368 | 0.023295 | 0.011224 | 0.034053 | 0.071737 | 0.063782 | 0.034549 | 0.012889 | 0.010982 | 0.037563 | 0.06337 |
| 35.1944444 | 0.01724 | 0.005975 | 0.005976 | 0.031136 | 0.04241 | 0.013018 | 0.010977 | 0.02177 | 0.015833 | 0.007335 | 0.021005 | 0.026401 | 0.072594 | 0.052981 | 0.021401 | 0.010782 | 0.049274 | 0.114171 | 0.079381 | 0.049533 | 0.021161 | 0.020124 | 0.033112 | 0.038418 |
| 35.2083333 | 0.019732 | 0.010993 | 0.008773 | 0.048263 | 0.051107 | 0.016709 | 0.017817 | 0.016303 | 0.020282 | 0.013225 | 0.027511 | 0.041129 | 0.085044 | 0.045512 | 0.016827 | 0.011301 | 0.054465 | 0.148079 | 0.083411 | 0.051979 | 0.027608 | 0.029613 | 0.033716 | 0.033037 |
| 35.2222222 | 0.017896 | 0.013045 | 0.010184 | 0.064554 | 0.059638 | 0.017872 | 0.022029 | 0.031365 | 0.022544 | 0.017881 | 0.028239 | 0.048848 | 0.08499 | 0.065869 | 0.01848 | 0.027828 | 0.041781 | 0.14213 | 0.066863 | 0.048113 | 0.020605 | 0.028392 | 0.039463 | 0.046856 |
| 35.2361111 | 0.016006 | 0.012128 | 0.009815 | 0.07134 | 0.066332 | 0.016455 | 0.02433 | 0.049604 | 0.021726 | 0.017507 | 0.02683 | 0.049885 | 0.076116 | 0.0914 | 0.037137 | 0.048991 | 0.023503 | 0.101339 | 0.037843 | 0.041903 | 0.009663 | 0.016534 | 0.043261 | 0.063147 |
| 35.25 | 0.017036 | 0.009803 | 0.009671 | 0.063968 | 0.067301 | 0.014103 | 0.025118 | 0.059975 | 0.018938 | 0.014965 | 0.025842 | 0.04524 | 0.05814 | 0.102928 | 0.053982 | 0.063562 | 0.014731 | 0.051681 | 0.019785 | 0.027203 | 0.006381 | 0.005756 | 0.03925 | 0.06777 |
| 35.2638889 | 0.018257 | 0.006401 | 0.010298 | 0.052382 | 0.062203 | 0.011662 | 0.023201 | 0.067337 | 0.015603 | 0.010198 | 0.023638 | 0.036655 | 0.036178 | 0.096633 | 0.055339 | 0.062403 | 0.012583 | 0.020428 | 0.022365 | 0.014245 | 0.008634 | 0.003817 | 0.03067 | 0.060931 |
| 35.2777778 | 0.017186 | 0.004634 | 0.011055 | 0.049297 | 0.054511 | 0.008854 | 0.018509 | 0.072449 | 0.012462 | 0.005989 | 0.01899 | 0.025752 | 0.017472 | 0.081044 | 0.048761 | 0.049259 | 0.010625 | 0.022626 | 0.020068 | 0.013374 | 0.016624 | 0.004774 | 0.024775 | 0.051532 |
| 35.2916667 | 0.015404 | 0.0074 | 0.01215 | 0.053753 | 0.047887 | 0.005799 | 0.012652 | 0.07497 | 0.009123 | 0.007855 | 0.012609 | 0.013627 | 0.006807 | 0.064052 | 0.04073 | 0.033201 | 0.010431 | 0.043947 | 0.010339 | 0.013918 | 0.019687 | 0.006612 | 0.02282 | 0.043725 |
| 35.3055556 | 0.014769 | 0.012669 | 0.013128 | 0.053112 | 0.04387 | 0.003927 | 0.008065 | 0.074032 | 0.00575 | 0.012915 | 0.006449 | 0.006179 | 0.002999 | 0.045947 | 0.032505 | 0.018254 | 0.010754 | 0.047565 | 0.008047 | 0.019327 | 0.011837 | 0.007594 | 0.019615 | 0.034961 |
| 35.3194444 | 0.015039 | 0.014938 | 0.012695 | 0.040816 | 0.04148 | 0.004331 | 0.007347 | 0.068607 | 0.003686 | 0.015942 | 0.002531 | 0.007608 | 0.001263 | 0.027386 | 0.025546 | 0.008238 | 0.008044 | 0.03077 | 0.015004 | 0.025728 | 0.004774 | 0.004865 | 0.012754 | 0.023514 |
| 35.3333333 | 0.015033 | 0.012458 | 0.010729 | 0.022656 | 0.038376 | 0.005274 | 0.010212 | 0.062594 | 0.003019 | 0.015291 | 0.001309 | 0.011544 | 0.001367 | 0.014928 | 0.022083 | 0.00551 | 0.006477 | 0.021833 | 0.028098 | 0.019818 | 0.002046 | 0.003009 | 0.005893 | 0.013251 |
| 35.3472222 | 0.013912 | 0.01153 | 0.008997 | 0.008434 | 0.034497 | 0.004872 | 0.01291 | 0.062745 | 0.002765 | 0.011312 | 0.002203 | 0.013554 | 0.003427 | 0.011931 | 0.021348 | 0.007695 | 0.007089 | 0.021596 | 0.028337 | 0.009813 | 0.002081 | 0.004155 | 0.002801 | 0.008164 |
| 35.3611111 | 0.012418 | 0.018766 | 0.009533 | 0.00609 | 0.0332 | 0.005135 | 0.013118 | 0.070593 | 0.002969 | 0.005993 | 0.003406 | 0.01308 | 0.011901 | 0.010254 | 0.023299 | 0.01366 | 0.00799 | 0.035428 | 0.017709 | 0.012596 | 0.009662 | 0.013291 | 0.002229 | 0.00531 |
| 35.375 | 0.012544 | 0.031038 | 0.012193 | 0.019415 | 0.037774 | 0.008837 | 0.012237 | 0.077467 | 0.005026 | 0.003053 | 0.00359 | 0.009174 | 0.031147 | 0.011947 | 0.02909 | 0.023669 | 0.014999 | 0.065971 | 0.021768 | 0.035869 | 0.027946 | 0.035397 | 0.00175 | 0.004448 |
| 35.3888889 | 0.015069 | 0.040257 | 0.015217 | 0.043471 | 0.04772 | 0.015492 | 0.012306 | 0.0747 | 0.009508 | 0.005393 | 0.005009 | 0.008241 | 0.049327 | 0.026203 | 0.035305 | 0.032459 | 0.024191 | 0.079188 | 0.036711 | 0.071964 | 0.049519 | 0.058522 | 0.001848 | 0.008644 |
| 35.4027778 | 0.018273 | 0.044276 | 0.016786 | 0.062802 | 0.059187 | 0.023757 | 0.013523 | 0.06809 | 0.014344 | 0.010299 | 0.00829 | 0.014087 | 0.053775 | 0.038576 | 0.036431 | 0.032564 | 0.02666 | 0.067507 | 0.04754 | 0.101544 | 0.068416 | 0.068827 | 0.002437 | 0.012642 |
| 35.4166667 | 0.020245 | 0.046478 | 0.016404 | 0.070122 | 0.068255 | 0.031734 | 0.015032 | 0.068478 | 0.017434 | 0.015323 | 0.010386 | 0.019344 | 0.047062 | 0.036756 | 0.033444 | 0.026467 | 0.022164 | 0.060006 | 0.058334 | 0.108413 | 0.084571 | 0.067416 | 0.003676 | 0.014041 |
| 35.4305556 | 0.021549 | 0.049561 | 0.015656 | 0.076665 | 0.074229 | 0.034793 | 0.0163 | 0.075321 | 0.019262 | 0.020959 | 0.012271 | 0.022422 | 0.038308 | 0.032237 | 0.03486 | 0.025066 | 0.013764 | 0.071444 | 0.07371 | 0.109828 | 0.094177 | 0.064437 | 0.006994 | 0.017196 |
| 35.4444444 | 0.024341 | 0.055554 | 0.016059 | 0.093335 | 0.078918 | 0.032552 | 0.017555 | 0.082459 | 0.021352 | 0.026628 | 0.017635 | 0.030357 | 0.036371 | 0.035425 | 0.045407 | 0.033915 | 0.00968 | 0.09559 | 0.09178 | 0.133227 | 0.096681 | 0.069628 | 0.012098 | 0.022863 |
| 35.4583333 | 0.029675 | 0.06634 | 0.01806 | 0.115686 | 0.084691 | 0.031743 | 0.018784 | 0.088845 | 0.024234 | 0.030438 | 0.025958 | 0.044375 | 0.042954 | 0.042901 | 0.058839 | 0.047276 | 0.015499 | 0.120397 | 0.115978 | 0.16211 | 0.102564 | 0.085928 | 0.016705 | 0.028027 |
| 35.4722222 | 0.036467 | 0.078198 | 0.02109 | 0.130078 | 0.093362 | 0.034933 | 0.019531 | 0.095709 | 0.02632 | 0.032583 | 0.033982 | 0.058785 | 0.045905 | 0.046924 | 0.066636 | 0.057037 | 0.026494 | 0.13247 | 0.139814 | 0.164783 | 0.120561 | 0.110099 | 0.018712 | 0.029421 |
| 35.4861111 | 0.042533 | 0.086114 | 0.023514 | 0.132127 | 0.101129 | 0.038132 | 0.021074 | 0.098998 | 0.025306 | 0.035331 | 0.040786 | 0.069366 | 0.036942 | 0.04596 | 0.068424 | 0.061485 | 0.034023 | 0.127686 | 0.145395 | 0.141014 | 0.143208 | 0.12945 | 0.019298 | 0.028023 |
| 35.5 | 0.046061 | 0.08997 | 0.024844 | 0.132117 | 0.103715 | 0.038871 | 0.022853 | 0.097615 | 0.023481 | 0.036787 | 0.046678 | 0.075378 | 0.028626 | 0.045026 | 0.070697 | 0.064892 | 0.03662 | 0.122539 | 0.135803 | 0.124863 | 0.152835 | 0.133481 | 0.023301 | 0.029282 |
| 35.5138889 | 0.047932 | 0.090921 | 0.02701 | 0.13606 | 0.106698 | 0.037886 | 0.021516 | 0.094271 | 0.025432 | 0.034811 | 0.052131 | 0.079438 | 0.036204 | 0.051342 | 0.078061 | 0.072178 | 0.04236 | 0.129756 | 0.125087 | 0.13338 | 0.142718 | 0.128985 | 0.033843 | 0.037798 |
| 35.5277778 | 0.051005 | 0.088584 | 0.031911 | 0.142926 | 0.114292 | 0.038684 | 0.017191 | 0.085274 | 0.029713 | 0.032798 | 0.058043 | 0.084326 | 0.060429 | 0.07026 | 0.089724 | 0.082615 | 0.05949 | 0.147053 | 0.115032 | 0.151178 | 0.134089 | 0.122985 | 0.048309 | 0.055502 |
| 35.5416667 | 0.055518 | 0.087285 | 0.03983 | 0.149691 | 0.121997 | 0.046379 | 0.014428 | 0.067438 | 0.033027 | 0.032957 | 0.059969 | 0.084029 | 0.082166 | 0.095847 | 0.100368 | 0.091965 | 0.082328 | 0.167026 | 0.107121 | 0.194198 | 0.146704 | 0.11998 | 0.059876 | 0.078007 |
| 35.5555556 | 0.058463 | 0.093156 | 0.047324 | 0.145404 | 0.122903 | 0.057487 | 0.014366 | 0.04941 | 0.03595 | 0.033827 | 0.055488 | 0.075977 | 0.08197 | 0.110847 | 0.109833 | 0.095219 | 0.095562 | 0.173985 | 0.105815 | 0.242413 | 0.165507 | 0.13163 | 0.063487 | 0.094005 |
| 35.5694444 | 0.057211 | 0.093985 | 0.048464 | 0.12338 | 0.111859 | 0.060581 | 0.015995 | 0.04709 | 0.038761 | 0.033571 | 0.042265 | 0.06618 | 0.067998 | 0.110911 | 0.132022 | 0.088926 | 0.094103 | 0.164035 | 0.106089 | 0.231705 | 0.163425 | 0.153688 | 0.063356 | 0.101033 |
| 35.5833333 | 0.054993 | 0.080826 | 0.041904 | 0.093971 | 0.097625 | 0.052789 | 0.02162 | 0.05333 | 0.039845 | 0.030706 | 0.036385 | 0.056755 | 0.054511 | 0.109615 | 0.167256 | 0.076201 | 0.086075 | 0.1472 | 0.096756 | 0.183228 | 0.143755 | 0.166792 | 0.067954 | 0.108683 |
| 35.5972222 | 0.057487 | 0.067589 | 0.034543 | 0.073139 | 0.09345 | 0.04221 | 0.027784 | 0.050138 | 0.038511 | 0.026979 | 0.045702 | 0.052396 | 0.042731 | 0.115231 | 0.192087 | 0.066571 | 0.080628 | 0.130052 | 0.087159 | 0.157883 | 0.123501 | 0.17407 | 0.077975 | 0.118032 |
| 35.6111111 | 0.065185 | 0.063866 | 0.035866 | 0.068595 | 0.095713 | 0.03767 | 0.02888 | 0.042484 | 0.035312 | 0.025722 | 0.060225 | 0.047623 | 0.033284 | 0.123362 | 0.19473 | 0.07019 | 0.074628 | 0.115937 | 0.091016 | 0.161006 | 0.089647 | 0.181845 | 0.08784 | 0.118488 |
| 35.625 | 0.071949 | 0.062255 | 0.045487 | 0.074169 | 0.092662 | 0.041463 | 0.028578 | 0.039538 | 0.030501 | 0.02787 | 0.061435 | 0.032505 | 0.029381 | 0.125869 | 0.173424 | 0.07989 | 0.063724 | 0.106027 | 0.089192 | 0.15842 | 0.052747 | 0.175153 | 0.090421 | 0.106948 |
| 35.6388889 | 0.068688 | 0.050517 | 0.052231 | 0.068067 | 0.084653 | 0.052783 | 0.034033 | 0.038117 | 0.025741 | 0.031467 | 0.054069 | 0.02327 | 0.030233 | 0.105644 | 0.137033 | 0.076457 | 0.056347 | 0.096371 | 0.073533 | 0.132601 | 0.04015 | 0.14153 | 0.082101 | 0.08969 |
| 35.6527778 | 0.048404 | 0.032024 | 0.0477 | 0.043462 | 0.075189 | 0.069067 | 0.041967 | 0.036555 | 0.02328 | 0.034919 | 0.048726 | 0.030234 | 0.03191 | 0.08591 | 0.112167 | 0.064073 | 0.06069 | 0.078516 | 0.063182 | 0.104713 | 0.045592 | 0.091317 | 0.068923 | 0.072852 |
| 35.6666667 | 0.026907 | 0.02141 | 0.035235 | 0.019255 | 0.056997 | 0.079353 | 0.044726 | 0.037248 | 0.02289 | 0.039416 | 0.041281 | 0.036128 | 0.033624 | 0.076458 | 0.101487 | 0.054875 | 0.067705 | 0.055957 | 0.061032 | 0.089726 | 0.060357 | 0.054544 | 0.054555 | 0.063636 |
| 35.6805556 | 0.026129 | 0.015827 | 0.02358 | 0.006774 | 0.032196 | 0.075497 | 0.041326 | 0.034806 | 0.022377 | 0.042571 | 0.026799 | 0.02913 | 0.033186 | 0.080743 | 0.087295 | 0.037566 | 0.065777 | 0.03777 | 0.057553 | 0.07562 | 0.073934 | 0.040463 | 0.044072 | 0.066316 |
| 35.6944444 | 0.030547 | 0.011256 | 0.016068 | 0.0043 | 0.014667 | 0.067194 | 0.034569 | 0.025707 | 0.018602 | 0.037709 | 0.010922 | 0.015838 | 0.027912 | 0.096083 | 0.06357 | 0.015814 | 0.055728 | 0.022272 | 0.045302 | 0.056421 | 0.069104 | 0.036103 | 0.045745 | 0.076398 |
| 35.7083333 | 0.028099 | 0.018406 | 0.009748 | 0.011201 | 0.008718 | 0.060808 | 0.027487 | 0.014776 | 0.011599 | 0.022449 | 0.002342 | 0.007415 | 0.01987 | 0.109076 | 0.035572 | 0.008422 | 0.039472 | 0.014572 | 0.0285 | 0.038013 | 0.044884 | 0.034245 | 0.049109 | 0.083874 |
| 35.7222222 | 0.032181 | 0.031051 | 0.005155 | 0.025583 | 0.009488 | 0.041885 | 0.018267 | 0.006215 | 0.006433 | 0.014903 | 0.00097 | 0.005413 | 0.010929 | 0.10291 | 0.015884 | 0.008634 | 0.022052 | 0.024896 | 0.029729 | 0.020787 | 0.023133 | 0.029456 | 0.042725 | 0.078392 |
| 35.7361111 | 0.043252 | 0.030322 | 0.00609 | 0.039914 | 0.01772 | 0.022476 | 0.007826 | 0.0027 | 0.005634 | 0.015078 | 0.003886 | 0.005304 | 0.004542 | 0.082254 | 0.011313 | 0.008752 | 0.01172 | 0.040351 | 0.054495 | 0.012821 | 0.01181 | 0.020801 | 0.033459 | 0.061775 |
| 35.75 | 0.046791 | 0.018866 | 0.009626 | 0.050078 | 0.033963 | 0.025809 | 0.002991 | 0.002732 | 0.006489 | 0.013373 | 0.009679 | 0.008116 | 0.004051 | 0.064644 | 0.01599 | 0.008209 | 0.008217 | 0.040446 | 0.072899 | 0.028275 | 0.00951 | 0.014591 | 0.029296 | 0.046279 |
| 35.7638889 | 0.036371 | 0.011816 | 0.011622 | 0.059011 | 0.048205 | 0.033605 | 0.004327 | 0.00191 | 0.006827 | 0.006982 | 0.014643 | 0.01446 | 0.006276 | 0.048932 | 0.02069 | 0.006471 | 0.007341 | 0.026373 | 0.075258 | 0.054439 | 0.012498 | 0.014898 | 0.026591 | 0.031425 |
| 35.7777778 | 0.021337 | 0.012761 | 0.013332 | 0.066051 | 0.055223 | 0.034868 | 0.008163 | 0.001136 | 0.006363 | 0.005327 | 0.014306 | 0.019682 | 0.009979 | 0.030225 | 0.020508 | 0.008604 | 0.014493 | 0.022449 | 0.086396 | 0.070736 | 0.011741 | 0.016147 | 0.017043 | 0.019409 |
| 35.7916667 | 0.012784 | 0.013382 | 0.016527 | 0.062193 | 0.059945 | 0.035038 | 0.014142 | 0.003927 | 0.004779 | 0.010268 | 0.00879 | 0.019244 | 0.011904 | 0.026154 | 0.013976 | 0.009756 | 0.029794 | 0.039083 | 0.107171 | 0.076154 | 0.016875 | 0.01221 | 0.008114 | 0.025232 |
| 35.8055556 | 0.012754 | 0.01022 | 0.019801 | 0.046361 | 0.063864 | 0.027603 | 0.02074 | 0.009475 | 0.003112 | 0.017837 | 0.004936 | 0.011979 | 0.007662 | 0.041878 | 0.010864 | 0.006194 | 0.030979 | 0.056963 | 0.117722 | 0.069716 | 0.030186 | 0.010612 | 0.010795 | 0.041365 |
| 35.8194444 | 0.015951 | 0.009672 | 0.020729 | 0.032725 | 0.063865 | 0.014107 | 0.024945 | 0.013944 | 0.004506 | 0.022548 | 0.004147 | 0.004714 | 0.003492 | 0.052508 | 0.019124 | 0.005499 | 0.019268 | 0.062176 | 0.081199 | 0.052703 | 0.04441 | 0.014607 | 0.015657 | 0.047316 |
| 35.8333333 | 0.017971 | 0.011582 | 0.018811 | 0.030802 | 0.062836 | 0.004896 | 0.025693 | 0.017534 | 0.008616 | 0.019417 | 0.004754 | 0.003866 | 0.003831 | 0.04598 | 0.022446 | 0.007776 | 0.0219 | 0.05449 | 0.05476 | 0.049899 | 0.05815 | 0.015948 | 0.014789 | 0.03941 |
| 35.8472222 | 0.018944 | 0.018658 | 0.015339 | 0.033333 | 0.064112 | 0.001463 | 0.027622 | 0.021794 | 0.010259 | 0.011203 | 0.009008 | 0.007534 | 0.011839 | 0.034046 | 0.012728 | 0.0071 | 0.029937 | 0.050476 | 0.047284 | 0.07628 | 0.062394 | 0.018858 | 0.014651 | 0.027456 |
| 35.8611111 | 0.018593 | 0.028961 | 0.012043 | 0.028586 | 0.062194 | 0.000733 | 0.031986 | 0.021981 | 0.008061 | 0.006614 | 0.013632 | 0.011701 | 0.022274 | 0.028624 | 0.003487 | 0.00515 | 0.023452 | 0.062836 | 0.057002 | 0.111263 | 0.052191 | 0.029886 | 0.018854 | 0.021967 |
| 35.875 | 0.014566 | 0.030334 | 0.0103 | 0.019815 | 0.053904 | 0.001317 | 0.030468 | 0.015243 | 0.004651 | 0.010382 | 0.014453 | 0.011098 | 0.019389 | 0.030109 | 0.00199 | 0.005447 | 0.013414 | 0.064418 | 0.062855 | 0.115788 | 0.035204 | 0.044511 | 0.023132 | 0.023662 |
| 35.8888889 | 0.009453 | 0.027031 | 0.011285 | 0.018421 | 0.044736 | 0.001746 | 0.022883 | 0.010513 | 0.003332 | 0.023102 | 0.012075 | 0.006257 | 0.011191 | 0.03488 | 0.00612 | 0.008396 | 0.014215 | 0.037083 | 0.049427 | 0.079248 | 0.017595 | 0.06074 | 0.024417 | 0.028735 |
| 35.9027778 | 0.009356 | 0.026777 | 0.014791 | 0.024895 | 0.041993 | 0.001713 | 0.015244 | 0.014163 | 0.006078 | 0.035993 | 0.00843 | 0.00389 | 0.016118 | 0.03927 | 0.012989 | 0.013716 | 0.021403 | 0.01557 | 0.030537 | 0.056171 | 0.029892 | 0.074992 | 0.025028 | 0.034145 |
| 35.9166667 | 0.013723 | 0.025295 | 0.018063 | 0.03214 | 0.045318 | 0.001351 | 0.008396 | 0.020104 | 0.008351 | 0.03293 | 0.004889 | 0.004871 | 0.027627 | 0.034333 | 0.01757 | 0.016306 | 0.026136 | 0.013293 | 0.021063 | 0.050364 | 0.082415 | 0.07629 | 0.026387 | 0.034411 |
| 35.9305556 | 0.017552 | 0.024513 | 0.01928 | 0.038019 | 0.048368 | 0.001728 | 0.003281 | 0.023624 | 0.006333 | 0.020081 | 0.00343 | 0.00528 | 0.031055 | 0.01933 | 0.017819 | 0.011105 | 0.032285 | 0.010798 | 0.023271 | 0.042824 | 0.16986 | 0.066471 | 0.026064 | 0.02707 |
| 35.9444444 | 0.018145 | 0.032053 | 0.020107 | 0.047011 | 0.04888 | 0.002825 | 0.001706 | 0.027965 | 0.003646 | 0.017379 | 0.005814 | 0.004362 | 0.027481 | 0.008837 | 0.021292 | 0.0075 | 0.048236 | 0.008552 | 0.02985 | 0.069271 | 0.216775 | 0.058749 | 0.025324 | 0.017029 |
| 35.9583333 | 0.016622 | 0.045171 | 0.022402 | 0.060088 | 0.04876 | 0.002568 | 0.002828 | 0.036702 | 0.004811 | 0.023679 | 0.009031 | 0.003464 | 0.024495 | 0.00794 | 0.031242 | 0.014139 | 0.069406 | 0.008667 | 0.036289 | 0.109065 | 0.136848 | 0.067097 | 0.027698 | 0.009868 |
| 35.9722222 | 0.018359 | 0.056268 | 0.026465 | 0.073499 | 0.052242 | 0.00168 | 0.006092 | 0.044991 | 0.009044 | 0.031038 | 0.01142 | 0.00545 | 0.024996 | 0.008705 | 0.035942 | 0.020133 | 0.08005 | 0.010968 | 0.043479 | 0.138129 | 0.050862 | 0.091885 | 0.032467 | 0.005857 |
| 35.9861111 | 0.026142 | 0.060646 | 0.030948 | 0.08592 | 0.062412 | 0.002468 | 0.010985 | 0.051138 | 0.012844 | 0.040517 | 0.013907 | 0.009346 | 0.03158 | 0.014317 | 0.024924 | 0.01594 | 0.071444 | 0.022007 | 0.053806 | 0.164175 | 0.035913 | 0.112503 | 0.03925 | 0.00817 |
| 36 | 0.035174 | 0.062862 | 0.033943 | 0.102286 | 0.074801 | 0.004234 | 0.013276 | 0.060231 | 0.014785 | 0.049653 | 0.015734 | 0.010297 | 0.04257 | 0.027678 | 0.011971 | 0.006932 | 0.057035 | 0.046469 | 0.073175 | 0.17347 | 0.039 | 0.113079 | 0.046958 | 0.017797 |
| 36.0138889 | 0.040354 | 0.074468 | 0.035259 | 0.127066 | 0.081764 | 0.004966 | 0.011746 | 0.071988 | 0.014094 | 0.051378 | 0.018738 | 0.01341 | 0.051584 | 0.032089 | 0.009985 | 0.002421 | 0.061352 | 0.075129 | 0.092923 | 0.172586 | 0.047032 | 0.100248 | 0.052967 | 0.021203 |
| 36.0277778 | 0.040443 | 0.097801 | 0.037987 | 0.152521 | 0.083649 | 0.006235 | 0.009717 | 0.083095 | 0.01157 | 0.045456 | 0.023695 | 0.022614 | 0.060192 | 0.027819 | 0.012722 | 0.011407 | 0.095113 | 0.101729 | 0.09576 | 0.179216 | 0.052659 | 0.097183 | 0.061885 | 0.019614 |
| 36.0416667 | 0.038869 | 0.120206 | 0.050755 | 0.165117 | 0.085483 | 0.015783 | 0.006293 | 0.08765 | 0.013331 | 0.047442 | 0.030364 | 0.040402 | 0.076275 | 0.045529 | 0.02938 | 0.038305 | 0.14214 | 0.125278 | 0.09704 | 0.177439 | 0.051185 | 0.106475 | 0.075662 | 0.034703 |
| 36.0555556 | 0.03818 | 0.123417 | 0.072913 | 0.168362 | 0.097421 | 0.038296 | 0.003814 | 0.083854 | 0.023619 | 0.070754 | 0.044436 | 0.071321 | 0.099355 | 0.085755 | 0.070202 | 0.074989 | 0.162963 | 0.142684 | 0.12091 | 0.156361 | 0.060044 | 0.126213 | 0.088348 | 0.06436 |
| 36.0694444 | 0.033558 | 0.111544 | 0.081768 | 0.162935 | 0.115374 | 0.058315 | 0.006436 | 0.073219 | 0.033281 | 0.098623 | 0.060918 | 0.095083 | 0.117379 | 0.10932 | 0.10659 | 0.097739 | 0.139024 | 0.138033 | 0.136035 | 0.145506 | 0.098305 | 0.17725 | 0.095132 | 0.081981 |
| 36.0833333 | 0.022452 | 0.098425 | 0.065815 | 0.133254 | 0.112652 | 0.055498 | 0.008879 | 0.06223 | 0.030735 | 0.10953 | 0.062261 | 0.086188 | 0.125763 | 0.096929 | 0.104516 | 0.089337 | 0.098893 | 0.109942 | 0.114793 | 0.167116 | 0.131488 | 0.204405 | 0.093511 | 0.07552 |
| 36.0972222 | 0.010005 | 0.077769 | 0.042328 | 0.090214 | 0.087837 | 0.035593 | 0.006677 | 0.067955 | 0.020514 | 0.104883 | 0.049279 | 0.05747 | 0.124815 | 0.066691 | 0.078259 | 0.067676 | 0.075007 | 0.106498 | 0.090974 | 0.190997 | 0.122775 | 0.158292 | 0.08354 | 0.056159 |
| 36.1111111 | 0.003946 | 0.049372 | 0.02606 | 0.057581 | 0.0612 | 0.01767 | 0.004105 | 0.084225 | 0.012893 | 0.095671 | 0.036126 | 0.036445 | 0.121162 | 0.042309 | 0.059498 | 0.053853 | 0.062026 | 0.12195 | 0.08692 | 0.17711 | 0.105698 | 0.113326 | 0.063059 | 0.03762 |
| 36.125 | 0.004347 | 0.026243 | 0.017255 | 0.038829 | 0.041057 | 0.010185 | 0.006318 | 0.09059 | 0.009721 | 0.088593 | 0.028367 | 0.028062 | 0.118649 | 0.038639 | 0.053061 | 0.050741 | 0.038864 | 0.092805 | 0.078648 | 0.131655 | 0.099642 | 0.097096 | 0.033982 | 0.027611 |
| 36.1388889 | 0.004715 | 0.0143 | 0.011305 | 0.025319 | 0.025991 | 0.010295 | 0.011795 | 0.086098 | 0.007687 | 0.076168 | 0.025221 | 0.026518 | 0.108069 | 0.061262 | 0.052103 | 0.052526 | 0.015841 | 0.050355 | 0.051046 | 0.076439 | 0.085618 | 0.074658 | 0.013706 | 0.033303 |
| 36.1527778 | 0.004886 | 0.008045 | 0.005548 | 0.013826 | 0.012929 | 0.011726 | 0.014941 | 0.069378 | 0.004475 | 0.052356 | 0.021411 | 0.027194 | 0.089639 | 0.093073 | 0.051199 | 0.046364 | 0.010264 | 0.054416 | 0.030268 | 0.032512 | 0.064171 | 0.048729 | 0.009576 | 0.049628 |
| 36.1666667 | 0.006813 | 0.007875 | 0.002028 | 0.012843 | 0.009446 | 0.011903 | 0.013883 | 0.039836 | 0.002036 | 0.025299 | 0.012769 | 0.023367 | 0.07488 | 0.10365 | 0.046918 | 0.032653 | 0.011197 | 0.0613 | 0.02954 | 0.022377 | 0.046496 | 0.045664 | 0.009004 | 0.054818 |
| 36.1805556 | 0.00937 | 0.020067 | 0.001254 | 0.020955 | 0.02418 | 0.010943 | 0.010637 | 0.014497 | 0.00323 | 0.009299 | 0.005342 | 0.013306 | 0.065915 | 0.090994 | 0.040151 | 0.020158 | 0.009571 | 0.044165 | 0.029601 | 0.041535 | 0.036272 | 0.066525 | 0.012561 | 0.043483 |
| 36.1944444 | 0.011629 | 0.033075 | 0.000978 | 0.025978 | 0.044072 | 0.008132 | 0.006108 | 0.008194 | 0.007343 | 0.010579 | 0.005094 | 0.00473 | 0.058808 | 0.07865 | 0.033143 | 0.009817 | 0.012698 | 0.039705 | 0.032385 | 0.066333 | 0.032002 | 0.081116 | 0.022083 | 0.034111 |
| 36.2083333 | 0.012684 | 0.036414 | 0.000727 | 0.027698 | 0.053079 | 0.00442 | 0.003795 | 0.018708 | 0.011182 | 0.018436 | 0.008075 | 0.004116 | 0.050425 | 0.074477 | 0.028007 | 0.003574 | 0.025312 | 0.061247 | 0.043613 | 0.073508 | 0.029399 | 0.08415 | 0.030814 | 0.031589 |
| 36.2222222 | 0.012127 | 0.037336 | 0.001519 | 0.033177 | 0.055529 | 0.003398 | 0.00734 | 0.039372 | 0.012923 | 0.024658 | 0.011512 | 0.010151 | 0.038447 | 0.070403 | 0.025805 | 0.003005 | 0.040261 | 0.086704 | 0.052438 | 0.05858 | 0.022449 | 0.084383 | 0.035536 | 0.024594 |
| 36.2361111 | 0.010093 | 0.041449 | 0.004571 | 0.039837 | 0.060596 | 0.005792 | 0.013149 | 0.059577 | 0.014006 | 0.028964 | 0.016721 | 0.020104 | 0.026685 | 0.061881 | 0.025456 | 0.003514 | 0.047849 | 0.090585 | 0.046696 | 0.033175 | 0.016037 | 0.085819 | 0.031395 | 0.014774 |

| | | | | | | | | | | | | | | | | | | | | | | | | |
|---|---|---|---|---|---|---|---|---|---|---|---|---|---|---|---|---|---|---|---|---|---|---|---|---|
| 36.25 | 0.008052 | 0.046131 | 0.008509 | 0.041316 | 0.066135 | 0.008805 | 0.017287 | 0.068132 | 0.015418 | 0.029681 | 0.02258 | 0.030518 | 0.024686 | 0.054476 | 0.026726 | 0.003859 | 0.039685 | 0.068076 | 0.033076 | 0.021277 | 0.016476 | 0.089106 | 0.023219 | 0.013444 |
| 36.2638889 | 0.009106 | 0.055386 | 0.010949 | 0.04091 | 0.0677 | 0.010711 | 0.019248 | 0.069364 | 0.015486 | 0.02687 | 0.026974 | 0.036575 | 0.03311 | 0.054892 | 0.028701 | 0.005141 | 0.019643 | 0.033894 | 0.040588 | 0.040684 | 0.01532 | 0.083854 | 0.019751 | 0.022604 |
| 36.2777778 | 0.013051 | 0.073634 | 0.011593 | 0.044211 | 0.064705 | 0.011148 | 0.019519 | 0.072041 | 0.013855 | 0.023324 | 0.028883 | 0.036903 | 0.039215 | 0.054246 | 0.02877 | 0.007807 | 0.005544 | 0.016025 | 0.058298 | 0.062061 | 0.007651 | 0.070171 | 0.014339 | 0.0318 |
| 36.2916667 | 0.015779 | 0.083853 | 0.011262 | 0.044146 | 0.055859 | 0.009929 | 0.019194 | 0.074895 | 0.011758 | 0.020513 | 0.028479 | 0.034005 | 0.035999 | 0.041257 | 0.025562 | 0.01271 | 0.001554 | 0.021991 | 0.054115 | 0.058083 | 0.004608 | 0.064192 | 0.007039 | 0.032116 |
| 36.3055556 | 0.015354 | 0.07207 | 0.011402 | 0.030752 | 0.042904 | 0.007467 | 0.018567 | 0.073339 | 0.009761 | 0.016646 | 0.026334 | 0.030018 | 0.029396 | 0.025315 | 0.020952 | 0.018571 | 0.003089 | 0.026368 | 0.035748 | 0.038852 | 0.013862 | 0.073481 | 0.011168 | 0.02856 |
| 36.3194444 | 0.013949 | 0.053769 | 0.012738 | 0.014689 | 0.031343 | 0.004354 | 0.016749 | 0.068464 | 0.007726 | 0.011092 | 0.022581 | 0.025351 | 0.020943 | 0.018958 | 0.017582 | 0.025238 | 0.006485 | 0.016988 | 0.021721 | 0.020442 | 0.02736 | 0.087565 | 0.022965 | 0.028035 |
| 36.3333333 | 0.013515 | 0.049782 | 0.01535 | 0.00964 | 0.025049 | 0.001802 | 0.013407 | 0.066677 | 0.005814 | 0.007206 | 0.017714 | 0.021183 | 0.016713 | 0.016287 | 0.014394 | 0.027165 | 0.012533 | 0.010696 | 0.012083 | 0.011418 | 0.027282 | 0.0816 | 0.024745 | 0.026555 |
| 36.3472222 | 0.013229 | 0.059971 | 0.018846 | 0.017992 | 0.02355 | 0.000765 | 0.01072 | 0.071548 | 0.005824 | 0.007671 | 0.014007 | 0.019824 | 0.027883 | 0.009241 | 0.008809 | 0.018904 | 0.029344 | 0.022185 | 0.005078 | 0.013591 | 0.023443 | 0.049236 | 0.014355 | 0.020171 |
| 36.3611111 | 0.0123 | 0.065502 | 0.020412 | 0.037529 | 0.024106 | 0.0011 | 0.011838 | 0.080235 | 0.008278 | 0.010913 | 0.013461 | 0.02137 | 0.036234 | 0.004552 | 0.003778 | 0.011364 | 0.047399 | 0.04131 | 0.007402 | 0.016496 | 0.032299 | 0.023303 | 0.005333 | 0.016215 |
| 36.375 | 0.01187 | 0.063558 | 0.018023 | 0.052758 | 0.025488 | 0.002163 | 0.016052 | 0.086811 | 0.0102 | 0.012928 | 0.014301 | 0.021933 | 0.023726 | 0.010207 | 0.002701 | 0.011664 | 0.05145 | 0.042886 | 0.017752 | 0.01081 | 0.036707 | 0.019492 | 0.002296 | 0.020824 |
| 36.3888889 | 0.013033 | 0.077943 | 0.015249 | 0.061293 | 0.027761 | 0.002487 | 0.019488 | 0.090291 | 0.009683 | 0.012463 | 0.013854 | 0.018786 | 0.009285 | 0.018376 | 0.003609 | 0.013502 | 0.051023 | 0.028403 | 0.019148 | 0.005419 | 0.041805 | 0.02657 | 0.00157 | 0.027968 |
| 36.4027778 | 0.015494 | 0.104991 | 0.01619 | 0.075487 | 0.033806 | 0.002955 | 0.021871 | 0.096253 | 0.008806 | 0.012449 | 0.012837 | 0.015221 | 0.015223 | 0.016471 | 0.005483 | 0.0178 | 0.057941 | 0.024314 | 0.015802 | 0.01713 | 0.055467 | 0.043224 | 0.002347 | 0.03029 |
| 36.4166667 | 0.018577 | 0.112441 | 0.022323 | 0.097689 | 0.061739 | 0.012382 | 0.029969 | 0.117928 | 0.012623 | 0.015377 | 0.012659 | 0.020738 | 0.049038 | 0.016454 | 0.008327 | 0.033807 | 0.059894 | 0.042816 | 0.017065 | 0.078071 | 0.047916 | 0.072571 | 0.022747 | 0.024022 |
| 36.4305556 | 0.030519 | 0.117577 | 0.039567 | 0.144848 | 0.164046 | 0.055131 | 0.065281 | 0.196099 | 0.024226 | 0.022477 | 0.02656 | 0.056921 | 0.136707 | 0.066235 | 0.011278 | 0.073083 | 0.075598 | 0.058415 | 0.024699 | 0.249442 | 0.032894 | 0.181237 | 0.140387 | 0.032761 |

**Supplementary Table S6.** 1 hour integrated motion data in **(b)** 100 mM NaCl treatment on Radish

| IntervalMid | Stress1 | Stress10 | Stress11 | Stress12 | Stress2 | Stress3 | Stress4 | Stress5 | Stress6 | Stress7 | Stress8 | Stress9 | Control1 | Control10 | Control11 | Control12 | Control2 | Control3 | Control4 | Control5 | Control6 | Control7 | Control8 | Control9 |
|---|---|---|---|---|---|---|---|---|---|---|---|---|---|---|---|---|---|---|---|---|---|---|---|---|
| 18.1944444 | 0.02231 | 0.015595 | 0.004569 | 0.033411 | 0.030353 | 0.00033 | 0.044666 | 0.004306 | 0.001225 | 0.030497 | 0.000348 | 0.004338 | 0.011232 | 0.00687 | 0.016445 | 0.007246 | 0.003624 | 0.003714 | 0.001388 | 0.011849 | 0.002383 | 0.002669 | 0.005994 | 0.013593 |
| 18.2083333 | 0.024576 | 0.023723 | 0.009997 | 0.041291 | 0.029783 | 0.000225 | 0.053213 | 0.005756 | 0.001003 | 0.033088 | 0.000108 | 0.003628 | 0.013275 | 0.004675 | 0.022998 | 0.011585 | 0.002891 | 0.001474 | 0.001218 | 0.008681 | 0.00288 | 0.002569 | 0.003922 | 0.011613 |
| 18.2222222 | 0.022462 | 0.029992 | 0.017183 | 0.04844 | 0.034765 | 0.000504 | 0.058171 | 0.005423 | 0.00141 | 0.024942 | 0.000174 | 0.002712 | 0.014751 | 0.002691 | 0.023168 | 0.012976 | 0.002108 | 0.00161 | 0.001756 | 0.005038 | 0.003054 | 0.002472 | 0.001982 | 0.010157 |
| 18.2361111 | 0.019183 | 0.03355 | 0.022178 | 0.053139 | 0.050382 | 0.001739 | 0.059411 | 0.005108 | 0.003704 | 0.016419 | 0.000434 | 0.003542 | 0.013971 | 0.001789 | 0.022574 | 0.014503 | 0.001121 | 0.003167 | 0.003253 | 0.004155 | 0.003161 | 0.004927 | 0.002559 | 0.006652 |
| 18.25 | 0.016542 | 0.039816 | 0.022974 | 0.056389 | 0.060202 | 0.003458 | 0.057354 | 0.004531 | 0.00774 | 0.014208 | 0.001012 | 0.005734 | 0.011973 | 0.002791 | 0.024205 | 0.016776 | 0.000376 | 0.004306 | 0.007112 | 0.007902 | 0.003578 | 0.010047 | 0.003377 | 0.002845 |
| 18.2638889 | 0.01392 | 0.049561 | 0.021319 | 0.059812 | 0.049172 | 0.004603 | 0.049264 | 0.004012 | 0.011752 | 0.014826 | 0.001945 | 0.007252 | 0.011942 | 0.00458 | 0.023319 | 0.014921 | 0.000313 | 0.003978 | 0.008436 | 0.009635 | 0.004265 | 0.011472 | 0.002884 | 0.003038 |
| 18.2777778 | 0.010661 | 0.056814 | 0.018998 | 0.063244 | 0.028891 | 0.004442 | 0.034543 | 0.005504 | 0.01319 | 0.014836 | 0.00337 | 0.006686 | 0.013558 | 0.004669 | 0.018045 | 0.008307 | 0.001089 | 0.002374 | 0.0053 | 0.006247 | 0.005016 | 0.007555 | 0.001996 | 0.006446 |
| 18.2916667 | 0.007846 | 0.052761 | 0.016411 | 0.068476 | 0.016494 | 0.003069 | 0.018956 | 0.009554 | 0.010567 | 0.016848 | 0.004883 | 0.004722 | 0.013615 | 0.002624 | 0.01278 | 0.002934 | 0.002804 | 0.000919 | 0.002906 | 0.003519 | 0.00553 | 0.002919 | 0.001289 | 0.009469 |
| 18.3055556 | 0.008278 | 0.041935 | 0.015087 | 0.076818 | 0.018213 | 0.001841 | 0.008354 | 0.017942 | 0.005909 | 0.020957 | 0.005571 | 0.003935 | 0.011768 | 0.000882 | 0.011217 | 0.001603 | 0.004275 | 0.001043 | 0.002606 | 0.002975 | 0.006079 | 0.002105 | 0.001739 | 0.009987 |
| 18.3194444 | 0.012883 | 0.038728 | 0.017141 | 0.080667 | 0.031009 | 0.002167 | 0.006286 | 0.032822 | 0.004186 | 0.022493 | 0.00541 | 0.006446 | 0.010334 | 0.001151 | 0.012041 | 0.001802 | 0.004467 | 0.002676 | 0.002916 | 0.002787 | 0.006861 | 0.004704 | 0.003312 | 0.008053 |
| 18.3333333 | 0.018259 | 0.042756 | 0.020499 | 0.070813 | 0.045764 | 0.003805 | 0.013803 | 0.048367 | 0.007778 | 0.019107 | 0.005728 | 0.011493 | 0.009975 | 0.002704 | 0.011423 | 0.002672 | 0.004054 | 0.003966 | 0.003821 | 0.003904 | 0.006749 | 0.006657 | 0.004063 | 0.005282 |
| 18.3472222 | 0.021805 | 0.044558 | 0.02079 | 0.056025 | 0.057356 | 0.006097 | 0.027339 | 0.053216 | 0.014452 | 0.012453 | 0.007705 | 0.014968 | 0.009797 | 0.003841 | 0.007938 | 0.002638 | 0.004207 | 0.003315 | 0.003267 | 0.003723 | 0.004718 | 0.005141 | 0.00306 | 0.002846 |
| 18.3611111 | 0.024734 | 0.047428 | 0.01834 | 0.050022 | 0.066998 | 0.00973 | 0.039219 | 0.042189 | 0.020948 | 0.00658 | 0.010556 | 0.015722 | 0.00938 | 0.004371 | 0.003704 | 0.001469 | 0.005075 | 0.001562 | 0.001779 | 0.001768 | 0.002144 | 0.003135 | 0.001508 | 0.001706 |
| 18.375 | 0.02684 | 0.06351 | 0.015261 | 0.053559 | 0.072503 | 0.014502 | 0.043189 | 0.02246 | 0.025939 | 0.005798 | 0.012511 | 0.020061 | 0.00878 | 0.005725 | 0.001911 | 0.001717 | 0.00508 | 0.000822 | 0.001984 | 0.001586 | 0.000851 | 0.002797 | 0.000905 | 0.003429 |
| 18.3888889 | 0.024605 | 0.089656 | 0.012317 | 0.069273 | 0.065097 | 0.017681 | 0.038022 | 0.007666 | 0.029152 | 0.010385 | 0.013258 | 0.025818 | 0.008113 | 0.007879 | 0.002777 | 0.003711 | 0.003745 | 0.001647 | 0.003496 | 0.002971 | 0.000565 | 0.002754 | 0.001253 | 0.006698 |
| 18.4027778 | 0.016823 | 0.093002 | 0.010084 | 0.084385 | 0.041273 | 0.016772 | 0.027058 | 0.002634 | 0.029981 | 0.01511 | 0.012467 | 0.023264 | 0.007438 | 0.009214 | 0.003779 | 0.006072 | 0.002687 | 0.003002 | 0.005659 | 0.004944 | 0.000878 | 0.004163 | 0.00163 | 0.008064 |
| 18.4166667 | 0.008222 | 0.05871 | 0.006967 | 0.068183 | 0.025192 | 0.012647 | 0.015743 | 0.002687 | 0.030755 | 0.015185 | 0.009998 | 0.014731 | 0.006776 | 0.008894 | 0.003587 | 0.007424 | 0.002857 | 0.003902 | 0.007213 | 0.006378 | 0.00147 | 0.006017 | 0.001707 | 0.007805 |
| 18.4305556 | 0.003438 | 0.037358 | 0.004628 | 0.030455 | 0.033539 | 0.008135 | 0.008244 | 0.00556 | 0.033148 | 0.010823 | 0.007242 | 0.006814 | 0.005892 | 0.006702 | 0.003348 | 0.006068 | 0.004342 | 0.003496 | 0.005819 | 0.005244 | 0.001427 | 0.004992 | 0.001307 | 0.007783 |
| 18.4444444 | 0.0037 | 0.04499 | 0.006389 | 0.006956 | 0.034329 | 0.004957 | 0.006383 | 0.010669 | 0.032711 | 0.005064 | 0.005094 | 0.00216 | 0.004895 | 0.004147 | 0.005493 | 0.002767 | 0.006079 | 0.001916 | 0.002404 | 0.002483 | 0.000902 | 0.00228 | 0.000582 | 0.008407 |
| 18.4583333 | 0.008421 | 0.04717 | 0.008232 | 0.005586 | 0.029989 | 0.00374 | 0.008997 | 0.012332 | 0.027414 | 0.001162 | 0.004512 | 0.003579 | 0.004716 | 0.004727 | 0.009205 | 0.001313 | 0.005924 | 0.000951 | 0.001698 | 0.002657 | 0.001196 | 0.002204 | 0.000294 | 0.010663 |
| 18.4722222 | 0.011687 | 0.062756 | 0.00609 | 0.016162 | 0.04285 | 0.003921 | 0.012739 | 0.010343 | 0.020416 | 0.00084 | 0.004898 | 0.01105 | 0.0047 | 0.007451 | 0.012547 | 0.00139 | 0.003618 | 0.000919 | 0.003266 | 0.005152 | 0.002081 | 0.003649 | 0.000779 | 0.01432 |
| 18.4861111 | 0.009573 | 0.07402 | 0.002643 | 0.020583 | 0.047017 | 0.004342 | 0.015073 | 0.012743 | 0.014343 | 0.002905 | 0.005401 | 0.016787 | 0.003583 | 0.006565 | 0.017411 | 0.00282 | 0.001629 | 0.001194 | 0.004167 | 0.005462 | 0.001918 | 0.004711 | 0.001683 | 0.01662 |
| 18.5 | 0.007391 | 0.056236 | 0.001387 | 0.018868 | 0.035864 | 0.004991 | 0.016152 | 0.013849 | 0.010713 | 0.00472 | 0.007155 | 0.014559 | 0.003069 | 0.002916 | 0.023311 | 0.007267 | 0.001713 | 0.002361 | 0.005695 | 0.005264 | 0.001275 | 0.007626 | 0.002344 | 0.017032 |
| 18.5138889 | 0.010277 | 0.05785 | 0.002952 | 0.023053 | 0.024625 | 0.006007 | 0.015562 | 0.00969 | 0.007858 | 0.005811 | 0.009573 | 0.013306 | 0.003459 | 0.002081 | 0.02665 | 0.011236 | 0.002703 | 0.003955 | 0.005825 | 0.005039 | 0.001071 | 0.009182 | 0.002334 | 0.01697 |
| 18.5277778 | 0.013769 | 0.058439 | 0.006186 | 0.022789 | 0.018509 | 0.006291 | 0.011635 | 0.008025 | 0.004597 | 0.007373 | 0.00958 | 0.014823 | 0.002767 | 0.006603 | 0.029652 | 0.013659 | 0.002398 | 0.00604 | 0.003152 | 0.002892 | 0.001192 | 0.006163 | 0.00193 | 0.016455 |
| 18.5416667 | 0.012357 | 0.042773 | 0.007903 | 0.027796 | 0.019464 | 0.004697 | 0.006829 | 0.008317 | 0.003963 | 0.009315 | 0.006587 | 0.00956 | 0.002034 | 0.017758 | 0.034458 | 0.015781 | 0.003025 | 0.009761 | 0.002279 | 0.001295 | 0.00205 | 0.002795 | 0.002971 | 0.016462 |
| 18.5555556 | 0.009183 | 0.047014 | 0.006056 | 0.044617 | 0.020053 | 0.00212 | 0.006758 | 0.007035 | 0.005436 | 0.01118 | 0.004853 | 0.005787 | 0.002356 | 0.029188 | 0.035298 | 0.015612 | 0.007959 | 0.014024 | 0.005009 | 0.000937 | 0.002221 | 0.001746 | 0.004596 | 0.019806 |
| 18.5694444 | 0.006768 | 0.042568 | 0.002733 | 0.041407 | 0.013554 | 0.001285 | 0.011744 | 0.009691 | 0.004751 | 0.012739 | 0.005568 | 0.007318 | 0.004364 | 0.030757 | 0.030347 | 0.013848 | 0.013438 | 0.016114 | 0.008486 | 0.001013 | 0.0021 | 0.002273 | 0.004648 | 0.026895 |
| 18.5833333 | 0.005064 | 0.028647 | 0.000823 | 0.022193 | 0.007968 | 0.003167 | 0.015144 | 0.021054 | 0.003015 | 0.013002 | 0.006717 | 0.006312 | 0.009205 | 0.021728 | 0.025631 | 0.013197 | 0.01545 | 0.015079 | 0.01079 | 0.002813 | 0.005311 | 0.00557 | 0.006215 | 0.033249 |
| 18.5972222 | 0.006258 | 0.024492 | 0.000346 | 0.016849 | 0.008222 | 0.005504 | 0.013145 | 0.02824 | 0.004302 | 0.011605 | 0.007614 | 0.004498 | 0.014892 | 0.011108 | 0.024366 | 0.013935 | 0.015184 | 0.012325 | 0.011906 | 0.005663 | 0.011166 | 0.011693 | 0.009977 | 0.035767 |
| 18.6111111 | 0.011216 | 0.027043 | 0.002963 | 0.015277 | 0.00658 | 0.005839 | 0.008576 | 0.021067 | 0.006903 | 0.009579 | 0.007742 | 0.005606 | 0.020156 | 0.007003 | 0.023237 | 0.016435 | 0.016245 | 0.009219 | 0.011011 | 0.006338 | 0.015218 | 0.016264 | 0.013788 | 0.035577 |
| 18.625 | 0.015184 | 0.04394 | 0.008154 | 0.009024 | 0.003627 | 0.004213 | 0.005063 | 0.008772 | 0.006339 | 0.007509 | 0.007969 | 0.010914 | 0.026847 | 0.007616 | 0.019417 | 0.022224 | 0.020758 | 0.007693 | 0.008185 | 0.005428 | 0.015153 | 0.015632 | 0.019339 | 0.035459 |
| 18.6388889 | 0.012497 | 0.049049 | 0.008976 | 0.009094 | 0.002371 | 0.002698 | 0.003735 | 0.004705 | 0.003298 | 0.005705 | 0.009125 | 0.012366 | 0.035538 | 0.010248 | 0.01607 | 0.030896 | 0.024932 | 0.01009 | 0.006166 | 0.00707 | 0.012513 | 0.012623 | 0.026519 | 0.038332 |
| 18.6527778 | 0.006569 | 0.034317 | 0.005033 | 0.008037 | 0.001173 | 0.00223 | 0.003131 | 0.00988 | 0.001829 | 0.004707 | 0.008915 | 0.006559 | 0.041294 | 0.01367 | 0.012901 | 0.035994 | 0.024279 | 0.014299 | 0.006041 | 0.010219 | 0.009679 | 0.009791 | 0.028659 | 0.039943 |
| 18.6666667 | 0.004188 | 0.023956 | 0.003175 | 0.002913 | 0.002217 | 0.001654 | 0.002182 | 0.012669 | 0.002783 | 0.004714 | 0.005755 | 0.00452 | 0.037803 | 0.013837 | 0.008975 | 0.030026 | 0.019616 | 0.01519 | 0.006386 | 0.010225 | 0.00661 | 0.005656 | 0.019889 | 0.031248 |
| 18.6805556 | 0.004489 | 0.020369 | 0.007349 | 0.004833 | 0.005913 | 0.001009 | 0.001775 | 0.008326 | 0.003271 | 0.0056 | 0.002855 | 0.009741 | 0.027634 | 0.009557 | 0.012594 | 0.016298 | 0.014601 | 0.012234 | 0.006128 | 0.006761 | 0.00317 | 0.003015 | 0.008389 | 0.016862 |
| 18.6944444 | 0.003734 | 0.014361 | 0.01031 | 0.012085 | 0.008898 | 0.000967 | 0.001897 | 0.00377 | 0.003071 | 0.006387 | 0.00168 | 0.014075 | 0.017836 | 0.004206 | 0.021537 | 0.006349 | 0.011774 | 0.008515 | 0.004692 | 0.003038 | 0.000886 | 0.003438 | 0.004995 | 0.013107 |
| 18.7083333 | 0.002246 | 0.010736 | 0.006259 | 0.012074 | 0.010151 | 0.001178 | 0.001966 | 0.004713 | 0.005661 | 0.006154 | 0.001126 | 0.011516 | 0.011312 | 0.002381 | 0.020495 | 0.005928 | 0.010446 | 0.005833 | 0.00264 | 0.00186 | 0.001351 | 0.004149 | 0.004793 | 0.019192 |
| 18.7222222 | 0.00161 | 0.01946 | 0.003712 | 0.00723 | 0.007926 | 0.001926 | 0.003441 | 0.007597 | 0.010058 | 0.005803 | 0.001416 | 0.00659 | 0.008991 | 0.005571 | 0.011774 | 0.010399 | 0.008991 | 0.003906 | 0.001261 | 0.002611 | 0.003836 | 0.005053 | 0.003889 | 0.016694 |
| 18.7361111 | 0.001421 | 0.02561 | 0.00476 | 0.006862 | 0.005722 | 0.002929 | 0.007453 | 0.009355 | 0.012152 | 0.006506 | 0.001641 | 0.005134 | 0.00974 | 0.01017 | 0.009188 | 0.013743 | 0.007679 | 0.002247 | 0.00078 | 0.002278 | 0.004455 | 0.005613 | 0.005739 | 0.012074 |
| 18.75 | 0.001002 | 0.01661 | 0.003808 | 0.008581 | 0.009974 | 0.003205 | 0.011054 | 0.014946 | 0.013206 | 0.007485 | 0.000961 | 0.009101 | 0.00985 | 0.010114 | 0.010597 | 0.011285 | 0.006371 | 0.001382 | 0.001195 | 0.001043 | 0.004772 | 0.007526 | 0.008336 | 0.022206 |
| 18.7638889 | 0.000773 | 0.005935 | 0.001615 | 0.017447 | 0.014663 | 0.00275 | 0.010598 | 0.018297 | 0.017719 | 0.00753 | 0.00062 | 0.020053 | 0.007377 | 0.005552 | 0.008176 | 0.008239 | 0.004483 | 0.001009 | 0.001953 | 0.001197 | 0.007326 | 0.010675 | 0.008916 | 0.028977 |
| 18.7777778 | 0.002192 | 0.005291 | 0.005848 | 0.039091 | 0.016383 | 0.002745 | 0.007637 | 0.011793 | 0.023412 | 0.006347 | 0.001342 | 0.031765 | 0.003615 | 0.001979 | 0.007676 | 0.013202 | 0.003005 | 0.001055 | 0.001889 | 0.002889 | 0.007327 | 0.010448 | 0.007796 | 0.019787 |
| 18.7916667 | 0.01048 | 0.021304 | 0.020847 | 0.071671 | 0.023687 | 0.005277 | 0.008785 | 0.005743 | 0.024276 | 0.004941 | 0.005808 | 0.037651 | 0.001094 | 0.004434 | 0.012469 | 0.014925 | 0.00244 | 0.00182 | 0.001491 | 0.004625 | 0.009319 | 0.014022 | 0.00694 | 0.011141 |
| 18.8055556 | 0.023047 | 0.042535 | 0.029371 | 0.102765 | 0.038943 | 0.010742 | 0.015001 | 0.005621 | 0.022353 | 0.007357 | 0.015727 | 0.038371 | 0.001205 | 0.013841 | 0.013216 | 0.009792 | 0.002014 | 0.002474 | 0.001987 | 0.004908 | 0.015912 | 0.020911 | 0.008086 | 0.009649 |
| 18.8194444 | 0.022543 | 0.039885 | 0.023657 | 0.092053 | 0.042254 | 0.014814 | 0.01677 | 0.011568 | 0.020878 | 0.011248 | 0.021414 | 0.031063 | 0.002011 | 0.017756 | 0.009918 | 0.009122 | 0.00173 | 0.002118 | 0.002867 | 0.003429 | 0.015079 | 0.017574 | 0.009006 | 0.012738 |
| 18.8333333 | 0.011744 | 0.024794 | 0.027016 | 0.048624 | 0.023801 | 0.012778 | 0.01495 | 0.024463 | 0.017134 | 0.008664 | 0.016477 | 0.015994 | 0.002545 | 0.011992 | 0.009015 | 0.008334 | 0.002207 | 0.001532 | 0.002486 | 0.003082 | 0.011579 | 0.014161 | 0.007543 | 0.019062 |
| 18.8472222 | 0.007618 | 0.020581 | 0.03954 | 0.033173 | 0.012125 | 0.006653 | 0.01989 | 0.028241 | 0.010057 | 0.003861 | 0.008237 | 0.005307 | 0.004234 | 0.010501 | 0.007332 | 0.006161 | 0.003389 | 0.002182 | 0.001825 | 0.00552 | 0.013041 | 0.017307 | 0.005345 | 0.020577 |
| 18.8611111 | 0.006056 | 0.03659 | 0.033864 | 0.033056 | 0.023611 | 0.002317 | 0.027822 | 0.038212 | 0.003695 | 0.002554 | 0.004634 | 0.003307 | 0.004786 | 0.010453 | 0.006432 | 0.008258 | 0.004193 | 0.003153 | 0.002638 | 0.007024 | 0.011358 | 0.0163 | 0.003461 | 0.015085 |
| 18.875 | 0.003025 | 0.079872 | 0.017385 | 0.026287 | 0.036878 | 0.003319 | 0.030751 | 0.077138 | 0.001767 | 0.002153 | 0.005928 | 0.002205 | 0.004489 | 0.00768 | 0.013618 | 0.007401 | 0.003389 | 0.003198 | 0.003233 | 0.00569 | 0.009612 | 0.01492 | 0.002828 | 0.00906 |
| 18.8888889 | 0.003824 | 0.101279 | 0.018386 | 0.043645 | 0.035455 | 0.008303 | 0.029406 | 0.087393 | 0.00484 | 0.002732 | 0.006468 | 0.001605 | 0.004385 | 0.007515 | 0.018335 | 0.003156 | 0.001666 | 0.002593 | 0.002283 | 0.003574 | 0.008723 | 0.012266 | 0.005338 | 0.005178 |
| 18.9027778 | 0.007231 | 0.072534 | 0.031257 | 0.086164 | 0.032156 | 0.011629 | 0.026993 | 0.055819 | 0.011903 | 0.004372 | 0.005355 | 0.002562 | 0.004207 | 0.007334 | 0.012985 | 0.002708 | 0.000819 | 0.001482 | 0.001036 | 0.002382 | 0.006703 | 0.008801 | 0.00736 | 0.003399 |
| 18.9166667 | 0.008463 | 0.05573 | 0.032536 | 0.125835 | 0.041357 | 0.010404 | 0.026811 | 0.035677 | 0.018243 | 0.006209 | 0.005704 | 0.004559 | 0.006674 | 0.011101 | 0.010256 | 0.006484 | 0.001867 | 0.000451 | 0.000624 | 0.002564 | 0.008192 | 0.012078 | 0.006809 | 0.005543 |
| 18.9305556 | 0.0067 | 0.084129 | 0.018102 | 0.13804 | 0.050684 | 0.008412 | 0.029798 | 0.036586 | 0.020369 | 0.011592 | 0.006806 | 0.01042 | 0.007185 | 0.0147 | 0.010885 | 0.010831 | 0.004721 | 0.000297 | 0.001208 | 0.002625 | 0.007885 | 0.011818 | 0.009427 | 0.011058 |
| 18.9444444 | 0.008515 | 0.115046 | 0.005582 | 0.121432 | 0.041581 | 0.009488 | 0.030399 | 0.056846 | 0.020176 | 0.018557 | 0.007572 | 0.012865 | 0.003969 | 0.010989 | 0.007174 | 0.009527 | 0.007068 | 0.000686 | 0.002407 | 0.002041 | 0.005225 | 0.006244 | 0.013225 | 0.014511 |
| 18.9583333 | 0.019437 | 0.100164 | 0.003234 | 0.098908 | 0.022165 | 0.01108 | 0.024215 | 0.059782 | 0.02059 | 0.020425 | 0.008696 | 0.008419 | 0.002367 | 0.006408 | 0.004313 | 0.006594 | 0.006533 | 0.000865 | 0.002769 | 0.00302 | 0.004042 | 0.00382 | 0.010345 | 0.011517 |
| 18.9722222 | 0.023767 | 0.051849 | 0.003093 | 0.075069 | 0.013427 | 0.009597 | 0.014413 | 0.041788 | 0.021305 | 0.017286 | 0.00915 | 0.00719 | 0.003588 | 0.004078 | 0.007897 | 0.00835 | 0.003765 | 0.001336 | 0.001731 | 0.0052 | 0.004118 | 0.004082 | 0.004223 | 0.006142 |
| 18.9861111 | 0.014326 | 0.021442 | 0.008691 | 0.048965 | 0.013483 | 0.007094 | 0.008444 | 0.05006 | 0.015612 | 0.011029 | 0.007898 | 0.01282 | 0.006252 | 0.004849 | 0.011911 | 0.009228 | 0.001463 | 0.002302 | 0.000627 | 0.005058 | 0.004988 | 0.005562 | 0.001648 | 0.003565 |
| 19 | 0.007512 | 0.018484 | 0.025687 | 0.048937 | 0.008696 | 0.00607 | 0.012931 | 0.052458 | 0.00977 | 0.006183 | 0.004854 | 0.01534 | 0.006389 | 0.006378 | 0.008761 | 0.00833 | 0.001283 | 0.002019 | 0.000642 | 0.003274 | 0.003805 | 0.004642 | 0.001871 | 0.003578 |
| 19.0138889 | 0.006552 | 0.022442 | 0.036491 | 0.071926 | 0.006808 | 0.005837 | 0.024203 | 0.027084 | 0.015439 | 0.006646 | 0.002502 | 0.013059 | 0.003309 | 0.004953 | 0.003122 | 0.011084 | 0.001973 | 0.001584 | 0.002117 | 0.00461 | 0.001724 | 0.003058 | 0.00194 | 0.006453 |
| 19.0277778 | 0.007118 | 0.026189 | 0.027025 | 0.075427 | 0.022458 | 0.005581 | 0.033199 | 0.01022 | 0.020199 | 0.010994 | 0.003828 | 0.01662 | 0.000911 | 0.006734 | 0.00535 | 0.018747 | 0.002506 | 0.003806 | 0.004045 | 0.010169 | 0.002133 | 0.006209 | 0.002429 | 0.007475 |
| 19.0416667 | 0.006987 | 0.027755 | 0.019438 | 0.067541 | 0.047891 | 0.003981 | 0.036868 | 0.016983 | 0.015866 | 0.017051 | 0.006827 | 0.016226 | 0.002667 | 0.015553 | 0.016424 | 0.02944 | 0.00574 | 0.007964 | 0.005742 | 0.018033 | 0.007895 | 0.012399 | 0.007378 | 0.007858 |
| 19.0555556 | 0.006911 | 0.024632 | 0.025685 | 0.081304 | 0.060773 | 0.002107 | 0.033566 | 0.03122 | 0.013576 | 0.015908 | 0.009489 | 0.006679 | 0.010545 | 0.025967 | 0.030984 | 0.042537 | 0.014428 | 0.013169 | 0.009932 | 0.02782 | 0.017407 | 0.015404 | 0.017472 | 0.018207 |
| 19.0694444 | 0.00702 | 0.018384 | 0.023541 | 0.103285 | 0.049209 | 0.002227 | 0.024088 | 0.029253 | 0.019721 | 0.007649 | 0.011414 | 0.002964 | 0.02088 | 0.032555 | 0.042136 | 0.053371 | 0.023131 | 0.017246 | 0.017724 | 0.035868 | 0.021426 | 0.010545 | 0.026816 | 0.033799 |
| 19.0833333 | 0.005376 | 0.012599 | 0.01334 | 0.101917 | 0.02523 | 0.004711 | 0.016358 | 0.029475 | 0.031719 | 0.003881 | 0.012483 | 0.007111 | 0.025322 | 0.030248 | 0.040058 | 0.047624 | 0.022438 | 0.01406 | 0.022411 | 0.031255 | 0.019607 | 0.003512 | 0.028302 | 0.039791 |
| 19.0972222 | 0.005116 | 0.011107 | 0.01552 | 0.078478 | 0.008677 | 0.009143 | 0.015153 | 0.040099 | 0.04081 | 0.003266 | 0.013197 | 0.009579 | 0.020608 | 0.021085 | 0.02636 | 0.025002 | 0.013325 | 0.006237 | 0.019332 | 0.014846 | 0.018951 | 0.002405 | 0.022032 | 0.034362 |
| 19.1111111 | 0.004948 | 0.018235 | 0.022711 | 0.057017 | 0.011822 | 0.010275 | 0.016256 | 0.03212 | 0.034425 | 0.002544 | 0.012785 | 0.006745 | 0.011747 | 0.015197 | 0.015253 | 0.006644 | 0.006282 | 0.002243 | 0.014022 | 0.00457 | 0.01531 | 0.006375 | 0.013764 | 0.025333 |
| 19.125 | 0.004694 | 0.033564 | 0.017347 | 0.044036 | 0.035914 | 0.006551 | 0.016826 | 0.018497 | 0.019402 | 0.003172 | 0.010666 | 0.004461 | 0.004958 | 0.015576 | 0.011355 | 0.001245 | 0.00508 | 0.001279 | 0.010871 | 0.004992 | 0.007645 | 0.011894 | 0.006485 | 0.016331 |
| 19.1388889 | 0.005965 | 0.04656 | 0.007922 | 0.029495 | 0.057639 | 0.004211 | 0.019684 | 0.014228 | 0.017322 | 0.002578 | 0.007321 | 0.00565 | 0.003254 | 0.016577 | 0.009553 | 0.000791 | 0.004527 | 0.000775 | 0.007104 | 0.004985 | 0.003272 | 0.015346 | 0.002575 | 0.0091 |
| 19.1527778 | 0.004888 | 0.046609 | 0.00592 | 0.027223 | 0.052244 | 0.00445 | 0.024889 | 0.008448 | 0.022615 | 0.001104 | 0.003507 | 0.005884 | 0.005805 | 0.015508 | 0.008893 | 0.00103 | 0.00256 | 0.000535 | 0.003146 | 0.002862 | 0.001795 | 0.014448 | 0.002514 | 0.006284 |
| 19.1666667 | 0.002313 | 0.035035 | 0.006829 | 0.051685 | 0.026864 | 0.003685 | 0.028074 | 0.003712 | 0.016933 | 0.000819 | 0.001305 | 0.004175 | 0.009055 | 0.013682 | 0.009619 | 0.001634 | 0.001209 | 0.000214 | 0.002805 | 0.003691 | 0.001268 | 0.01018 | 0.002271 | 0.007834 |
| 19.1805556 | 0.002835 | 0.018618 | 0.011199 | 0.071847 | 0.006708 | 0.002707 | 0.02638 | 0.003161 | 0.008356 | 0.001277 | 0.000907 | 0.004845 | 0.010361 | 0.01148 | 0.008402 | 0.002098 | 0.001451 | 9.24E-05 | 0.004688 | 0.006212 | 0.002279 | 0.004905 | 0.001462 | 0.010851 |
| 19.1944444 | 0.005789 | 0.010865 | 0.017305 | 0.057529 | 0.005098 | 0.004185 | 0.021523 | 0.004847 | 0.009192 | 0.00269 | 0.000888 | 0.00761 | 0.009932 | 0.009816 | 0.004464 | 0.001836 | 0.002033 | 0.000725 | 0.004529 | 0.005614 | 0.002446 | 0.00206 | 0.001855 | 0.011104 |
| 19.2083333 | 0.00776 | 0.022503 | 0.0203 | 0.032699 | 0.017714 | 0.006986 | 0.019223 | 0.006564 | 0.011406 | 0.004571 | 0.001398 | 0.007796 | 0.009259 | 0.009304 | 0.001223 | 0.001852 | 0.002383 | 0.002876 | 0.003389 | 0.004176 | 0.002487 | 0.00353 | 0.002402 | 0.006891 |
| 19.2222222 | 0.008465 | 0.036205 | 0.021847 | 0.036844 | 0.034429 | 0.008461 | 0.023747 | 0.010412 | 0.00933 | 0.005607 | 0.001751 | 0.00477 | 0.009058 | 0.007749 | 0.0008 | 0.003068 | 0.004239 | 0.005446 | 0.005162 | 0.006945 | 0.005099 | 0.007 | 0.002664 | 0.002771 |
| 19.2361111 | 0.010987 | 0.028998 | 0.019599 | 0.057345 | 0.047661 | 0.008031 | 0.033597 | 0.021754 | 0.007469 | 0.005694 | 0.00161 | 0.00188 | 0.009746 | 0.004581 | 0.001978 | 0.003956 | 0.005816 | 0.006075 | 0.006809 | 0.007784 | 0.006659 | 0.008021 | 0.003064 | 0.002274 |
| 19.25 | 0.018054 | 0.02068 | 0.012268 | 0.062456 | 0.052032 | 0.007615 | 0.043847 | 0.037625 | 0.007024 | 0.005162 | 0.00205 | 0.003486 | 0.011253 | 0.003513 | 0.002186 | 0.007378 | 0.004886 | 0.004087 | 0.005452 | 0.006528 | 0.004451 | 0.006857 | 0.003307 | 0.002111 |
| 19.2638889 | 0.025916 | 0.040015 | 0.005899 | 0.052614 | 0.042219 | 0.00851 | 0.046824 | 0.046455 | 0.005107 | 0.004485 | 0.003553 | 0.010627 | 0.011391 | 0.005199 | 0.001272 | 0.014592 | 0.004914 | 0.001522 | 0.004835 | 0.0104 | 0.002483 | 0.007573 | 0.002646 | 0.000937 |
| 19.2777778 | 0.024755 | 0.062854 | 0.00487 | 0.038572 | 0.022389 | 0.008522 | 0.035346 | 0.040845 | 0.004673 | 0.003926 | 0.004932 | 0.015539 | 0.008603 | 0.004881 | 0.003187 | 0.017524 | 0.00682 | 0.000691 | 0.004868 | 0.012221 | 0.003762 | 0.009955 | 0.001263 | 0.000521 |
| 19.2916667 | 0.013657 | 0.062675 | 0.009761 | 0.025778 | 0.008919 | 0.005589 | 0.0177 | 0.025778 | 0.011964 | 0.004226 | 0.004143 | 0.011972 | 0.004956 | 0.002426 | 0.009425 | 0.013466 | 0.006088 | 0.001503 | 0.00317 | 0.007219 | 0.005734 | 0.010382 | 0.000403 | 0.001173 |
| 19.3055556 | 0.005108 | 0.05337 | 0.019608 | 0.022959 | 0.00763 | 0.002335 | 0.008432 | 0.010914 | 0.020413 | 0.006024 | 0.002011 | 0.006494 | 0.003156 | 0.002066 | 0.01403 | 0.008076 | 0.003462 | 0.002612 | 0.002819 | 0.003872 | 0.005662 | 0.008773 | 0.000351 | 0.002562 |
| 19.3194444 | 0.009569 | 0.05011 | 0.03214 | 0.040121 | 0.006847 | 0.00323 | 0.009085 | 0.004029 | 0.017737 | 0.007514 | 0.003381 | 0.00909 | 0.003408 | 0.003838 | 0.011067 | 0.004243 | 0.002553 | 0.003676 | 0.005363 | 0.005663 | 0.004222 | 0.008058 | 0.001172 | 0.002613 |
| 19.3333333 | 0.026852 | 0.052162 | 0.044095 | 0.078899 | 0.008308 | 0.010747 | 0.020909 | 0.009074 | 0.015655 | 0.007025 | 0.013069 | 0.020424 | 0.004053 | 0.004995 | 0.008798 | 0.002534 | 0.002826 | 0.004015 | 0.006887 | 0.006668 | 0.002446 | 0.007679 | 0.003415 | 0.001082 |
| 19.3472222 | 0.04715 | 0.053862 | 0.050992 | 0.120009 | 0.026819 | 0.022444 | 0.040415 | 0.023931 | 0.032103 | 0.006583 | 0.028719 | 0.0325 | 0.003457 | 0.004247 | 0.01368 | 0.003018 | 0.004066 | 0.002665 | 0.004984 | 0.004674 | 0.001959 | 0.004896 | 0.005761 | 0.000411 |
| 19.3611111 | 0.056339 | 0.046435 | 0.051281 | 0.126978 | 0.056801 | 0.031473 | 0.05443 | 0.039465 | 0.049901 | 0.008635 | 0.042965 | 0.042127 | 0.002089 | 0.002634 | 0.013597 | 0.00292 | 0.005212 | 0.001416 | 0.003016 | 0.004075 | 0.004089 | 0.002046 | 0.00654 | 0.001447 |
| 19.375 | 0.053166 | 0.028357 | 0.044906 | 0.094847 | 0.069478 | 0.033571 | 0.062226 | 0.045804 | 0.051921 | 0.0122 | 0.053333 | 0.049385 | 0.002227 | 0.0016 | 0.007744 | 0.002256 | 0.004046 | 0.001324 | 0.00247 | 0.004052 | 0.00529 | 0.002637 | 0.00664 | 0.006878 |
| 19.3888889 | 0.046185 | 0.014894 | 0.032488 | 0.058437 | 0.050518 | 0.028722 | 0.072523 | 0.041903 | 0.047074 | 0.014509 | 0.055705 | 0.048942 | 0.004567 | 0.001129 | 0.006411 | 0.003465 | 0.001847 | 0.001463 | 0.002192 | 0.003053 | 0.003843 | 0.005324 | 0.007209 | 0.017224 |
| 19.4027778 | 0.035856 | 0.013111 | 0.018185 | 0.040127 | 0.0237 | 0.019073 | 0.084232 | 0.033607 | 0.040596 | 0.013913 | 0.048617 | 0.043795 | 0.00733 | 0.001189 | 0.006781 | 0.004036 | 0.001351 | 0.001792 | 0.002407 | 0.002585 | 0.003254 | 0.009303 | 0.007166 | 0.023502 |
| 19.4166667 | 0.023546 | 0.019796 | 0.00953 | 0.039542 | 0.016159 | 0.008963 | 0.087805 | 0.025337 | 0.032361 | 0.010915 | 0.039687 | 0.039124 | 0.009286 | 0.002087 | 0.009203 | 0.004268 | 0.002976 | 0.001533 | 0.001854 | 0.001871 | 0.003345 | 0.015041 | 0.005608 | 0.019834 |
| 19.4305556 | 0.014858 | 0.030358 | 0.014846 | 0.05142 | 0.022334 | 0.002758 | 0.075142 | 0.017361 | 0.026332 | 0.007081 | 0.030823 | 0.030691 | 0.011095 | 0.002498 | 0.017968 | 0.006344 | 0.003892 | 0.000696 | 0.001341 | 0.002551 | 0.002121 | 0.019337 | 0.00412 | 0.012638 |
| 19.4444444 | 0.010137 | 0.030801 | 0.029605 | 0.064291 | 0.023885 | 0.00159 | 0.046477 | 0.00897 | 0.022534 | 0.004306 | 0.02029 | 0.0188 | 0.013713 | 0.002116 | 0.023955 | 0.007339 | 0.002642 | 0.000288 | 0.002701 | 0.006371 | 0.001344 | 0.018259 | 0.004958 | 0.009613 |
| 19.4583333 | 0.006882 | 0.022095 | 0.032011 | 0.064583 | 0.018575 | 0.003376 | 0.018329 | 0.003056 | 0.020953 | 0.004052 | 0.011383 | 0.010304 | 0.016511 | 0.003344 | 0.020903 | 0.007621 | 0.001156 | 0.0008 | 0.004688 | 0.010233 | 0.002417 | 0.013382 | 0.008709 | 0.012102 |
| 19.4722222 | 0.004537 | 0.019351 | 0.022661 | 0.054639 | 0.011569 | 0.005242 | 0.007205 | 0.004717 | 0.026823 | 0.004986 | 0.008895 | 0.010377 | 0.017344 | 0.006526 | 0.015264 | 0.010523 | 0.002022 | 0.002529 | 0.004518 | 0.008872 | 0.00469 | 0.010225 | 0.013176 | 0.015742 |
| 19.4861111 | 0.003316 | 0.024913 | 0.028974 | 0.042746 | 0.009821 | 0.004639 | 0.007805 | 0.00936 | 0.040548 | 0.004613 | 0.010463 | 0.019442 | 0.016035 | 0.009124 | 0.011974 | 0.016146 | 0.006715 | 0.005041 | 0.004342 | 0.007339 | 0.008069 | 0.012269 | 0.014953 | 0.015127 |
| 19.5 | 0.003024 | 0.024265 | 0.042924 | 0.025188 | 0.021851 | 0.002418 | 0.00789 | 0.007405 | 0.053167 | 0.002635 | 0.015401 | 0.028614 | 0.014963 | 0.009936 | 0.009717 | 0.021069 | 0.011735 | 0.007294 | 0.006495 | 0.011384 | 0.012565 | 0.017658 | 0.01363 | 0.012296 |
| 19.5138889 | 0.004609 | 0.01264 | 0.040728 | 0.010453 | 0.039248 | 0.002414 | 0.009112 | 0.004048 | 0.053205 | 0.002032 | 0.030918 | 0.027538 | 0.015157 | 0.010828 | 0.006826 | 0.021269 | 0.012049 | 0.008087 | 0.00634 | 0.013294 | 0.015956 | 0.020459 | 0.011053 | 0.014006 |
| 19.5277778 | 0.00657 | 0.006274 | 0.025887 | 0.011519 | 0.045309 | 0.006729 | 0.021987 | 0.01078 | 0.037714 | 0.005351 | 0.052724 | 0.018705 | 0.014985 | 0.014825 | 0.006284 | 0.018208 | 0.009479 | 0.008113 | 0.003612 | 0.013389 | 0.01528 | 0.016613 | 0.008325 | 0.020053 |
| 19.5416667 | 0.006496 | 0.007036 | 0.015853 | 0.02121 | 0.037309 | 0.012466 | 0.044809 | 0.019588 | 0.01689 | 0.009129 | 0.060958 | 0.009094 | 0.013506 | 0.019775 | 0.012708 | 0.015775 | 0.010957 | 0.009877 | 0.003415 | 0.019948 | 0.011129 | 0.008834 | 0.006376 | 0.023299 |
| 19.5555556 | 0.009275 | 0.008986 | 0.031489 | 0.032625 | 0.022204 | 0.013322 | 0.051316 | 0.017067 | 0.004663 | 0.009583 | 0.047712 | 0.003547 | 0.01385 | 0.020243 | 0.026425 | 0.014394 | 0.017836 | 0.01447 | 0.006711 | 0.028845 | 0.007082 | 0.003457 | 0.006836 | 0.019777 |
| 19.5694444 | 0.01548 | 0.00961 | 0.057101 | 0.045699 | 0.014719 | 0.009975 | 0.030759 | 0.008977 | 0.004843 | 0.008408 | 0.027827 | 0.003657 | 0.018985 | 0.019412 | 0.043839 | 0.015197 | 0.024324 | 0.019974 | 0.010434 | 0.03118 | 0.005909 | 0.002445 | 0.010443 | 0.013453 |
| 19.5833333 | 0.015322 | 0.008991 | 0.057705 | 0.04809 | 0.025475 | 0.008918 | 0.017527 | 0.00788 | 0.015491 | 0.007092 | 0.013167 | 0.008007 | 0.026853 | 0.026858 | 0.056118 | 0.020038 | 0.026735 | 0.02257 | 0.012025 | 0.02907 | 0.00763 | 0.005141 | 0.0146 | 0.011838 |
| 19.5972222 | 0.008806 | 0.012508 | 0.034854 | 0.028604 | 0.045237 | 0.010143 | 0.027846 | 0.018712 | 0.029588 | 0.005061 | 0.009629 | 0.015459 | 0.03418 | 0.04593 | 0.050835 | 0.026697 | 0.02623 | 0.021416 | 0.011825 | 0.029818 | 0.0117 | 0.012805 | 0.017675 | 0.020574 |
| 19.6111111 | 0.004133 | 0.013824 | 0.016574 | 0.006901 | 0.059501 | 0.008917 | 0.036322 | 0.030698 | 0.031513 | 0.003408 | 0.023073 | 0.02236 | 0.038945 | 0.069362 | 0.029668 | 0.031901 | 0.02518 | 0.019876 | 0.011547 | 0.033092 | 0.016722 | 0.023113 | 0.022563 | 0.039791 |
| 19.625 | 0.004647 | 0.009811 | 0.018446 | 0.000797 | 0.060732 | 0.005239 | 0.029553 | 0.029174 | 0.019312 | 0.003721 | 0.040114 | 0.025079 | 0.039233 | 0.088235 | 0.013656 | 0.034015 | 0.024794 | 0.019426 | 0.011634 | 0.032644 | 0.018359 | 0.02645 | 0.03238 | 0.05909 |
| 19.6388889 | 0.006299 | 0.006368 | 0.026676 | 0.003727 | 0.045537 | 0.003568 | 0.017087 | 0.017563 | 0.008085 | 0.004818 | 0.036745 | 0.027341 | 0.035569 | 0.097079 | 0.01105 | 0.032699 | 0.024213 | 0.01819 | 0.011311 | 0.028382 | 0.015133 | 0.018539 | 0.037372 | 0.060882 |
| 19.6527778 | 0.007418 | 0.006125 | 0.030331 | 0.013634 | 0.021515 | 0.005777 | 0.009153 | 0.011866 | 0.007783 | 0.004464 | 0.022626 | 0.027302 | 0.03049 | 0.088526 | 0.014252 | 0.029651 | 0.021343 | 0.015317 | 0.009958 | 0.027991 | 0.008977 | 0.009445 | 0.024676 | 0.042135 |
| 19.6666667 | 0.011902 | 0.009262 | 0.028736 | 0.032499 | 0.011298 | 0.008314 | 0.005375 | 0.012258 | 0.013105 | 0.002726 | 0.021857 | 0.016562 | 0.023899 | 0.060864 | 0.015901 | 0.025797 | 0.01663 | 0.011292 | 0.007634 | 0.031082 | 0.004886 | 0.008109 | 0.011996 | 0.0196 |

| | | | | | | | | | | | | | | | | | | | | | | | | |
|---|---|---|---|---|---|---|---|---|---|---|---|---|---|---|---|---|---|---|---|---|---|---|---|---|
| 19.6805556 | 0.01362 | 0.013625 | 0.018287 | 0.050227 | 0.018055 | 0.007544 | 0.004306 | 0.009492 | 0.011817 | 0.002001 | 0.023087 | 0.00563 | 0.014923 | 0.028841 | 0.010836 | 0.019586 | 0.013011 | 0.006814 | 0.005474 | 0.028988 | 0.004578 | 0.009234 | 0.014745 | 0.006083 |
| 19.6944444 | 0.007978 | 0.016648 | 0.01678 | 0.06402 | 0.020212 | 0.004387 | 0.009602 | 0.007763 | 0.007173 | 0.00314 | 0.014242 | 0.002698 | 0.006487 | 0.017295 | 0.005386 | 0.011138 | 0.011998 | 0.003608 | 0.005303 | 0.021396 | 0.005156 | 0.007167 | 0.01561 | 0.003161 |
| 19.7083333 | 0.00487 | 0.016052 | 0.04212 | 0.076128 | 0.012842 | 0.00173 | 0.017249 | 0.008112 | 0.010821 | 0.00356 | 0.008107 | 0.004728 | 0.003877 | 0.030917 | 0.00468 | 0.005106 | 0.013489 | 0.002145 | 0.007155 | 0.01484 | 0.009066 | 0.006478 | 0.016704 | 0.009319 |
| 19.7222222 | 0.009981 | 0.013212 | 0.071206 | 0.071708 | 0.006219 | 0.001166 | 0.017158 | 0.011763 | 0.018913 | 0.00261 | 0.009995 | 0.013061 | 0.009265 | 0.042668 | 0.010614 | 0.003071 | 0.015294 | 0.001329 | 0.00761 | 0.012959 | 0.014743 | 0.007504 | 0.030845 | 0.020201 |
| 19.7361111 | 0.017688 | 0.011735 | 0.080564 | 0.050298 | 0.007033 | 0.002379 | 0.017923 | 0.02537 | 0.025831 | 0.002139 | 0.012657 | 0.018175 | 0.017394 | 0.032053 | 0.020035 | 0.003681 | 0.014824 | 0.000972 | 0.004538 | 0.014766 | 0.01516 | 0.005948 | 0.038537 | 0.026456 |
| 19.75 | 0.023479 | 0.018629 | 0.07168 | 0.032955 | 0.018408 | 0.004269 | 0.030162 | 0.03874 | 0.036366 | 0.001893 | 0.014152 | 0.01168 | 0.020873 | 0.022486 | 0.02021 | 0.009495 | 0.013049 | 0.00169 | 0.00145 | 0.014929 | 0.008931 | 0.003819 | 0.025838 | 0.021566 |
| 19.7638889 | 0.025414 | 0.030846 | 0.055468 | 0.029793 | 0.037686 | 0.006618 | 0.035821 | 0.037367 | 0.048807 | 0.00179 | 0.015023 | 0.008785 | 0.018774 | 0.035512 | 0.010942 | 0.016642 | 0.011939 | 0.00299 | 0.001888 | 0.00971 | 0.003988 | 0.003039 | 0.010409 | 0.012718 |
| 19.7777778 | 0.025135 | 0.033183 | 0.04686 | 0.039697 | 0.059994 | 0.010906 | 0.028078 | 0.033072 | 0.052733 | 0.00274 | 0.014935 | 0.016482 | 0.015318 | 0.039385 | 0.004717 | 0.018187 | 0.011417 | 0.003206 | 0.00413 | 0.004566 | 0.004743 | 0.003492 | 0.005263 | 0.012416 |
| 19.7916667 | 0.034216 | 0.03395 | 0.050139 | 0.074421 | 0.092948 | 0.018642 | 0.024152 | 0.058724 | 0.050536 | 0.00387 | 0.025811 | 0.023522 | 0.013652 | 0.022687 | 0.0032 | 0.016812 | 0.010962 | 0.002369 | 0.004267 | 0.004499 | 0.00748 | 0.005746 | 0.010691 | 0.021805 |
| 19.8055556 | 0.051204 | 0.057399 | 0.054261 | 0.135192 | 0.130527 | 0.026771 | 0.028231 | 0.102762 | 0.047057 | 0.008868 | 0.050099 | 0.037295 | 0.013256 | 0.01358 | 0.003346 | 0.01721 | 0.010824 | 0.001657 | 0.002325 | 0.004637 | 0.013258 | 0.009232 | 0.023206 | 0.030925 |
| 19.8194444 | 0.050207 | 0.085992 | 0.039863 | 0.167673 | 0.141381 | 0.02655 | 0.031478 | 0.092846 | 0.034512 | 0.01697 | 0.05985 | 0.054311 | 0.011542 | 0.013195 | 0.007187 | 0.01802 | 0.011449 | 0.001144 | 0.001112 | 0.00324 | 0.018178 | 0.011192 | 0.027357 | 0.02979 |
| 19.8333333 | 0.026708 | 0.078025 | 0.022229 | 0.138963 | 0.109277 | 0.016914 | 0.031029 | 0.05317 | 0.016746 | 0.017475 | 0.045248 | 0.052088 | 0.007559 | 0.01048 | 0.010935 | 0.015 | 0.012817 | 0.001812 | 0.000929 | 0.003947 | 0.01525 | 0.008991 | 0.018195 | 0.021548 |
| 19.8472222 | 0.008033 | 0.041605 | 0.031902 | 0.093099 | 0.06244 | 0.01033 | 0.030127 | 0.054281 | 0.006669 | 0.015518 | 0.028562 | 0.03782 | 0.003307 | 0.009306 | 0.00971 | 0.009191 | 0.01404 | 0.005512 | 0.000922 | 0.004783 | 0.010093 | 0.00671 | 0.010532 | 0.013246 |
| 19.8611111 | 0.004988 | 0.013721 | 0.048677 | 0.053475 | 0.050815 | 0.013156 | 0.028334 | 0.0518 | 0.003572 | 0.017753 | 0.020253 | 0.025769 | 0.000938 | 0.011675 | 0.006444 | 0.007114 | 0.013897 | 0.010708 | 0.000716 | 0.003439 | 0.008915 | 0.007477 | 0.008975 | 0.006068 |
| 19.875 | 0.004872 | 0.005537 | 0.048589 | 0.034201 | 0.066563 | 0.0178 | 0.0245 | 0.025998 | 0.00254 | 0.014732 | 0.015424 | 0.0143 | 0.000451 | 0.010672 | 0.007806 | 0.010352 | 0.012374 | 0.013257 | 0.000491 | 0.003608 | 0.010554 | 0.008261 | 0.013815 | 0.003126 |
| 19.8888889 | 0.003736 | 0.006703 | 0.033998 | 0.064252 | 0.075013 | 0.016647 | 0.021386 | 0.0167 | 0.008752 | 0.006973 | 0.010435 | 0.005311 | 0.000448 | 0.005447 | 0.013399 | 0.014048 | 0.011122 | 0.009886 | 0.000974 | 0.007415 | 0.011864 | 0.007388 | 0.030012 | 0.005989 |
| 19.9027778 | 0.005877 | 0.006841 | 0.031516 | 0.097988 | 0.083194 | 0.011517 | 0.020847 | 0.025446 | 0.028025 | 0.002802 | 0.005839 | 0.004814 | 0.000583 | 0.004181 | 0.01803 | 0.013584 | 0.01134 | 0.006434 | 0.002258 | 0.012199 | 0.009565 | 0.004716 | 0.039061 | 0.014082 |
| 19.9166667 | 0.006937 | 0.013704 | 0.051035 | 0.076115 | 0.087407 | 0.008928 | 0.022672 | 0.036296 | 0.045332 | 0.004941 | 0.004016 | 0.01176 | 0.001497 | 0.008035 | 0.017659 | 0.008639 | 0.01227 | 0.010624 | 0.003495 | 0.015591 | 0.005432 | 0.003063 | 0.029948 | 0.018998 |
| 19.9305556 | 0.007349 | 0.0448 | 0.050095 | 0.044612 | 0.062474 | 0.008241 | 0.027611 | 0.048256 | 0.042854 | 0.012654 | 0.005998 | 0.020616 | 0.002783 | 0.011258 | 0.012332 | 0.004817 | 0.012716 | 0.015711 | 0.003588 | 0.015628 | 0.005377 | 0.005928 | 0.018242 | 0.013868 |
| 19.9444444 | 0.011894 | 0.092736 | 0.025225 | 0.072643 | 0.033594 | 0.005161 | 0.03337 | 0.059761 | 0.030005 | 0.02545 | 0.010457 | 0.027955 | 0.003499 | 0.010707 | 0.009901 | 0.003563 | 0.012756 | 0.014209 | 0.002211 | 0.011086 | 0.010206 | 0.011445 | 0.009969 | 0.006144 |
| 19.9583333 | 0.012067 | 0.119233 | 0.011339 | 0.12837 | 0.044533 | 0.002917 | 0.03256 | 0.055353 | 0.021147 | 0.039796 | 0.012997 | 0.028675 | 0.002874 | 0.01228 | 0.011936 | 0.001845 | 0.012941 | 0.008253 | 0.000721 | 0.005473 | 0.014946 | 0.016129 | 0.006026 | 0.004279 |
| 19.9722222 | 0.010224 | 0.090591 | 0.009133 | 0.148877 | 0.072511 | 0.004247 | 0.023047 | 0.031956 | 0.019024 | 0.045432 | 0.010957 | 0.019211 | 0.001808 | 0.019218 | 0.01163 | 0.003778 | 0.01264 | 0.002851 | 0.000587 | 0.003692 | 0.015585 | 0.016993 | 0.008875 | 0.010057 |
| 19.9861111 | 0.014437 | 0.043371 | 0.011424 | 0.120238 | 0.077292 | 0.00466 | 0.012221 | 0.010984 | 0.02377 | 0.040417 | 0.007376 | 0.007893 | 0.003755 | 0.019685 | 0.007139 | 0.011631 | 0.011109 | 0.001666 | 0.001283 | 0.003593 | 0.013327 | 0.01395 | 0.012167 | 0.015102 |
| 20 | 0.013983 | 0.031334 | 0.016367 | 0.067187 | 0.055268 | 0.003765 | 0.009013 | 0.006079 | 0.031578 | 0.029903 | 0.004855 | 0.0037 | 0.008985 | 0.012093 | 0.005398 | 0.019395 | 0.009065 | 0.005805 | 0.001722 | 0.004708 | 0.013046 | 0.011288 | 0.013498 | 0.011369 |
| 20.0138889 | 0.009312 | 0.032723 | 0.016556 | 0.025793 | 0.028811 | 0.005687 | 0.013221 | 0.009896 | 0.031248 | 0.019707 | 0.003046 | 0.003715 | 0.012214 | 0.008855 | 0.012629 | 0.021844 | 0.00844 | 0.015556 | 0.001817 | 0.010352 | 0.015304 | 0.011346 | 0.016894 | 0.004705 |
| 20.0277778 | 0.011413 | 0.027402 | 0.020961 | 0.012571 | 0.025177 | 0.006308 | 0.01627 | 0.020518 | 0.020013 | 0.020412 | 0.001996 | 0.005374 | 0.011276 | 0.016874 | 0.020003 | 0.016233 | 0.010385 | 0.027271 | 0.001798 | 0.015819 | 0.0169 | 0.013785 | 0.018513 | 0.001706 |
| 20.0416667 | 0.011826 | 0.027409 | 0.029163 | 0.018576 | 0.037695 | 0.00368 | 0.015209 | 0.033539 | 0.015142 | 0.027689 | 0.002978 | 0.007806 | 0.012741 | 0.03757 | 0.0195 | 0.013186 | 0.012007 | 0.038209 | 0.002912 | 0.020589 | 0.020562 | 0.020437 | 0.015583 | 0.005198 |
| 20.0555556 | 0.008752 | 0.026669 | 0.02242 | 0.020816 | 0.040317 | 0.003499 | 0.012393 | 0.034895 | 0.021784 | 0.025248 | 0.005738 | 0.007529 | 0.020853 | 0.067974 | 0.021022 | 0.024097 | 0.009347 | 0.052919 | 0.00776 | 0.031098 | 0.029613 | 0.035516 | 0.011545 | 0.012811 |
| 20.0694444 | 0.01022 | 0.016409 | 0.014168 | 0.012114 | 0.028824 | 0.005716 | 0.00819 | 0.025775 | 0.02588 | 0.015159 | 0.008052 | 0.004476 | 0.026623 | 0.090206 | 0.028906 | 0.03941 | 0.004469 | 0.062821 | 0.013966 | 0.042721 | 0.034545 | 0.049873 | 0.017482 | 0.012463 |
| 20.0833333 | 0.010792 | 0.007924 | 0.022233 | 0.006977 | 0.015995 | 0.007614 | 0.004268 | 0.01874 | 0.02681 | 0.010419 | 0.009766 | 0.001671 | 0.020945 | 0.081505 | 0.028129 | 0.04435 | 0.002396 | 0.051774 | 0.015533 | 0.042835 | 0.024172 | 0.042349 | 0.03413 | 0.008082 |
| 20.0972222 | 0.013265 | 0.009588 | 0.024793 | 0.014721 | 0.009269 | 0.007845 | 0.004854 | 0.01547 | 0.030466 | 0.01435 | 0.011645 | 0.001083 | 0.011116 | 0.046366 | 0.017514 | 0.032256 | 0.003273 | 0.026473 | 0.012087 | 0.029375 | 0.013537 | 0.02415 | 0.034213 | 0.010844 |
| 20.1111111 | 0.01873 | 0.009477 | 0.017274 | 0.028066 | 0.005419 | 0.006175 | 0.009658 | 0.011377 | 0.033813 | 0.021326 | 0.012484 | 0.002432 | 0.00696 | 0.020538 | 0.013864 | 0.022695 | 0.003556 | 0.011992 | 0.006882 | 0.015108 | 0.019091 | 0.021834 | 0.024277 | 0.014353 |
| 20.125 | 0.016717 | 0.005342 | 0.022938 | 0.038631 | 0.002951 | 0.004406 | 0.011681 | 0.00756 | 0.03451 | 0.02675 | 0.01131 | 0.00456 | 0.007154 | 0.025862 | 0.01641 | 0.021154 | 0.002003 | 0.01962 | 0.003627 | 0.012185 | 0.023932 | 0.021757 | 0.040965 | 0.017267 |
| 20.1388889 | 0.0084 | 0.00618 | 0.035508 | 0.048477 | 0.002465 | 0.004749 | 0.008238 | 0.005423 | 0.031957 | 0.027115 | 0.00921 | 0.007389 | 0.013021 | 0.039943 | 0.019778 | 0.016706 | 0.00064 | 0.032162 | 0.002696 | 0.015051 | 0.016071 | 0.012407 | 0.069559 | 0.029978 |
| 20.1527778 | 0.002396 | 0.009901 | 0.037151 | 0.060068 | 0.003268 | 0.005277 | 0.006069 | 0.003587 | 0.024709 | 0.018198 | 0.007355 | 0.010785 | 0.02407 | 0.038005 | 0.021228 | 0.022676 | 0.000476 | 0.033469 | 0.003147 | 0.01109 | 0.007351 | 0.0076 | 0.066743 | 0.046433 |
| 20.1666667 | 0.002617 | 0.009912 | 0.029137 | 0.073939 | 0.00699 | 0.00367 | 0.008855 | 0.002539 | 0.015436 | 0.009438 | 0.005677 | 0.012138 | 0.029225 | 0.023667 | 0.013494 | 0.028561 | 0.000688 | 0.023811 | 0.006409 | 0.00606 | 0.005683 | 0.005728 | 0.04037 | 0.041212 |
| 20.1805556 | 0.008091 | 0.005767 | 0.019826 | 0.088238 | 0.0108 | 0.003615 | 0.012395 | 0.002934 | 0.008042 | 0.011583 | 0.004625 | 0.010134 | 0.025028 | 0.010331 | 0.005206 | 0.019595 | 0.000988 | 0.011058 | 0.008229 | 0.01012 | 0.005983 | 0.002149 | 0.037683 | 0.024272 |
| 20.1944444 | 0.015264 | 0.002575 | 0.014115 | 0.089077 | 0.01178 | 0.006395 | 0.014624 | 0.005799 | 0.003386 | 0.015986 | 0.005052 | 0.0076 | 0.019398 | 0.005951 | 0.004873 | 0.009338 | 0.001423 | 0.005536 | 0.005228 | 0.016895 | 0.005496 | 0.003633 | 0.051818 | 0.023355 |
| 20.2083333 | 0.020376 | 0.001848 | 0.01281 | 0.067831 | 0.01207 | 0.008203 | 0.016535 | 0.011037 | 0.001397 | 0.013623 | 0.006219 | 0.005916 | 0.017604 | 0.009569 | 0.008567 | 0.006206 | 0.002011 | 0.010543 | 0.002405 | 0.020173 | 0.008688 | 0.011425 | 0.041106 | 0.020947 |
| 20.2222222 | 0.022687 | 0.002106 | 0.016722 | 0.039612 | 0.015983 | 0.008342 | 0.018667 | 0.017368 | 0.001669 | 0.008338 | 0.007072 | 0.004472 | 0.016691 | 0.014756 | 0.01201 | 0.004937 | 0.002647 | 0.016554 | 0.00332 | 0.018236 | 0.012534 | 0.016658 | 0.024406 | 0.009864 |
| 20.2361111 | 0.024543 | 0.005922 | 0.026811 | 0.02104 | 0.027542 | 0.009282 | 0.021089 | 0.027101 | 0.003309 | 0.008802 | 0.007858 | 0.004742 | 0.012178 | 0.016243 | 0.01117 | 0.00275 | 0.002879 | 0.013445 | 0.003828 | 0.012891 | 0.012504 | 0.012567 | 0.032333 | 0.00886 |
| 20.25 | 0.027372 | 0.016127 | 0.039681 | 0.014087 | 0.04274 | 0.011193 | 0.023432 | 0.037121 | 0.005906 | 0.01427 | 0.008679 | 0.007036 | 0.00558 | 0.011839 | 0.011423 | 0.003553 | 0.002863 | 0.006807 | 0.001906 | 0.012624 | 0.010147 | 0.007522 | 0.040925 | 0.010248 |
| 20.2638889 | 0.028302 | 0.032931 | 0.049861 | 0.019101 | 0.052193 | 0.012487 | 0.025782 | 0.039427 | 0.009089 | 0.015516 | 0.009415 | 0.009073 | 0.001529 | 0.006522 | 0.016429 | 0.007506 | 0.003404 | 0.006404 | 0.001301 | 0.015829 | 0.006057 | 0.008484 | 0.031494 | 0.009288 |
| 20.2777778 | 0.023472 | 0.050395 | 0.052247 | 0.033533 | 0.051895 | 0.011574 | 0.026864 | 0.034829 | 0.010037 | 0.011935 | 0.010415 | 0.010481 | 0.001514 | 0.007662 | 0.015227 | 0.01474 | 0.004135 | 0.009486 | 0.001993 | 0.016683 | 0.003741 | 0.009333 | 0.031391 | 0.018343 |
| 20.2916667 | 0.014309 | 0.053959 | 0.042717 | 0.040569 | 0.044215 | 0.007365 | 0.02307 | 0.025246 | 0.00692 | 0.00839 | 0.011287 | 0.01135 | 0.00442 | 0.012471 | 0.010694 | 0.023646 | 0.004216 | 0.010727 | 0.002198 | 0.01655 | 0.00786 | 0.00971 | 0.054886 | 0.031434 |
| 20.3055556 | 0.006416 | 0.036756 | 0.025925 | 0.027516 | 0.029308 | 0.003666 | 0.014466 | 0.017679 | 0.002873 | 0.005617 | 0.010618 | 0.009178 | 0.009146 | 0.014837 | 0.012437 | 0.026963 | 0.004386 | 0.009054 | 0.00254 | 0.013825 | 0.013973 | 0.013362 | 0.061147 | 0.028652 |
| 20.3194444 | 0.003395 | 0.014354 | 0.012509 | 0.009629 | 0.016822 | 0.003209 | 0.00861 | 0.026175 | 0.004642 | 0.004348 | 0.00907 | 0.005269 | 0.01295 | 0.014213 | 0.01411 | 0.021864 | 0.005381 | 0.005314 | 0.004165 | 0.007813 | 0.015196 | 0.016095 | 0.042843 | 0.017999 |
| 20.3333333 | 0.00489 | 0.005689 | 0.007613 | 0.003299 | 0.020049 | 0.006245 | 0.014277 | 0.038648 | 0.012722 | 0.006737 | 0.009826 | 0.00614 | 0.013056 | 0.012022 | 0.013032 | 0.014298 | 0.006404 | 0.003395 | 0.006432 | 0.003535 | 0.01216 | 0.014064 | 0.04072 | 0.020932 |
| 20.3472222 | 0.010152 | 0.014616 | 0.009614 | 0.009107 | 0.02291 | 0.015123 | 0.034071 | 0.030496 | 0.020301 | 0.010914 | 0.013848 | 0.013946 | 0.010981 | 0.008798 | 0.012389 | 0.01206 | 0.006226 | 0.00344 | 0.005778 | 0.002601 | 0.011146 | 0.012224 | 0.04936 | 0.029862 |
| 20.3611111 | 0.019448 | 0.036093 | 0.014259 | 0.033255 | 0.021969 | 0.025286 | 0.060811 | 0.019067 | 0.026705 | 0.013169 | 0.017878 | 0.024883 | 0.010075 | 0.005743 | 0.012112 | 0.015461 | 0.005304 | 0.003118 | 0.003125 | 0.002572 | 0.014064 | 0.013261 | 0.041165 | 0.029088 |
| 20.375 | 0.034263 | 0.062002 | 0.02213 | 0.071274 | 0.037287 | 0.031009 | 0.081615 | 0.036927 | 0.034543 | 0.012798 | 0.020783 | 0.0354 | 0.010946 | 0.005672 | 0.011513 | 0.018667 | 0.005099 | 0.005358 | 0.003593 | 0.003368 | 0.015611 | 0.011954 | 0.024173 | 0.023076 |
| 20.3888889 | 0.055663 | 0.083549 | 0.03975 | 0.096948 | 0.05654 | 0.030407 | 0.088315 | 0.078182 | 0.043506 | 0.011146 | 0.023044 | 0.042619 | 0.012218 | 0.008911 | 0.008861 | 0.019006 | 0.004923 | 0.011687 | 0.006844 | 0.007791 | 0.013738 | 0.008465 | 0.022847 | 0.032343 |
| 20.4027778 | 0.072611 | 0.092584 | 0.058252 | 0.095073 | 0.058635 | 0.020704 | 0.080375 | 0.106795 | 0.051381 | 0.008204 | 0.029642 | 0.04484 | 0.013158 | 0.011679 | 0.005438 | 0.015936 | 0.004723 | 0.017293 | 0.009214 | 0.014192 | 0.011396 | 0.007608 | 0.03407 | 0.05081 |
| 20.4166667 | 0.068772 | 0.082078 | 0.05824 | 0.068959 | 0.044272 | 0.009026 | 0.066202 | 0.085714 | 0.059792 | 0.005144 | 0.045359 | 0.043468 | 0.015118 | 0.011366 | 0.004123 | 0.009468 | 0.006284 | 0.018365 | 0.009017 | 0.016806 | 0.010409 | 0.009073 | 0.037416 | 0.050043 |
| 20.4305556 | 0.048346 | 0.055778 | 0.045507 | 0.035718 | 0.023512 | 0.006377 | 0.053796 | 0.035772 | 0.072861 | 0.003401 | 0.049732 | 0.039694 | 0.02027 | 0.01106 | 0.005532 | 0.004625 | 0.008219 | 0.017684 | 0.008329 | 0.017232 | 0.012375 | 0.010663 | 0.031061 | 0.02996 |
| 20.4444444 | 0.027287 | 0.028718 | 0.031016 | 0.015206 | 0.007545 | 0.007559 | 0.038657 | 0.012363 | 0.081617 | 0.002457 | 0.029291 | 0.034431 | 0.027408 | 0.015251 | 0.012487 | 0.006879 | 0.008019 | 0.018387 | 0.008547 | 0.019511 | 0.018949 | 0.0144 | 0.025362 | 0.013386 |
| 20.4583333 | 0.012321 | 0.013072 | 0.018211 | 0.016532 | 0.001739 | 0.006327 | 0.020576 | 0.015759 | 0.07204 | 0.004135 | 0.014505 | 0.028903 | 0.031337 | 0.022224 | 0.023385 | 0.014947 | 0.00677 | 0.01883 | 0.007609 | 0.020573 | 0.027471 | 0.020172 | 0.018366 | 0.012377 |
| 20.4722222 | 0.004678 | 0.013052 | 0.018553 | 0.04961 | 0.003478 | 0.007218 | 0.010314 | 0.019878 | 0.044385 | 0.007849 | 0.028165 | 0.024049 | 0.029707 | 0.026192 | 0.031755 | 0.023632 | 0.006432 | 0.018196 | 0.006669 | 0.019984 | 0.032412 | 0.022434 | 0.007391 | 0.012797 |
| 20.4861111 | 0.003894 | 0.017513 | 0.022835 | 0.098398 | 0.007388 | 0.013019 | 0.012922 | 0.022602 | 0.019751 | 0.010129 | 0.048464 | 0.023715 | 0.026335 | 0.025586 | 0.035829 | 0.027191 | 0.006551 | 0.017209 | 0.008908 | 0.021587 | 0.030799 | 0.018748 | 0.009863 | 0.014787 |
| 20.5 | 0.004449 | 0.014462 | 0.016224 | 0.105825 | 0.007841 | 0.020437 | 0.018714 | 0.026832 | 0.019378 | 0.008393 | 0.051636 | 0.031112 | 0.024209 | 0.024839 | 0.037039 | 0.022677 | 0.006209 | 0.015271 | 0.012096 | 0.02285 | 0.02335 | 0.01085 | 0.033258 | 0.034512 |
| 20.5138889 | 0.005945 | 0.014614 | 0.008484 | 0.059126 | 0.00584 | 0.025257 | 0.014435 | 0.025286 | 0.03386 | 0.003801 | 0.038685 | 0.042687 | 0.021985 | 0.029431 | 0.036773 | 0.016444 | 0.005249 | 0.013562 | 0.012054 | 0.021966 | 0.01288 | 0.004972 | 0.053052 | 0.052353 |
| 20.5277778 | 0.012524 | 0.021727 | 0.008034 | 0.024207 | 0.011431 | 0.026857 | 0.006568 | 0.015119 | 0.040801 | 0.001862 | 0.020783 | 0.047427 | 0.018359 | 0.038664 | 0.034145 | 0.017127 | 0.003178 | 0.014548 | 0.009475 | 0.022802 | 0.006496 | 0.006589 | 0.047215 | 0.047332 |
| 20.5416667 | 0.01614 | 0.020088 | 0.013534 | 0.032265 | 0.018361 | 0.019918 | 0.00609 | 0.014176 | 0.03205 | 0.005534 | 0.01434 | 0.039417 | 0.014482 | 0.044172 | 0.028467 | 0.020982 | 0.002266 | 0.016966 | 0.006396 | 0.023135 | 0.008318 | 0.010296 | 0.02891 | 0.031461 |
| 20.5555556 | 0.011368 | 0.010075 | 0.022149 | 0.047358 | 0.015206 | 0.009962 | 0.011929 | 0.032984 | 0.015936 | 0.0113 | 0.023393 | 0.025692 | 0.013539 | 0.041722 | 0.024052 | 0.020243 | 0.005771 | 0.019233 | 0.004447 | 0.019426 | 0.009418 | 0.008556 | 0.016536 | 0.018651 |
| 20.5694444 | 0.005997 | 0.005044 | 0.023764 | 0.042273 | 0.013494 | 0.012303 | 0.0202 | 0.040479 | 0.004973 | 0.011876 | 0.033695 | 0.012758 | 0.018488 | 0.039187 | 0.02647 | 0.020423 | 0.011035 | 0.023547 | 0.006589 | 0.017745 | 0.007584 | 0.006535 | 0.00851 | 0.010574 |
| 20.5833333 | 0.006169 | 0.008042 | 0.019365 | 0.023895 | 0.019311 | 0.020526 | 0.019345 | 0.026964 | 0.004595 | 0.006284 | 0.039965 | 0.004567 | 0.028995 | 0.046935 | 0.037353 | 0.031478 | 0.013942 | 0.028105 | 0.013792 | 0.024432 | 0.012719 | 0.012597 | 0.003608 | 0.006061 |
| 20.5972222 | 0.009563 | 0.013543 | 0.023415 | 0.01447 | 0.024788 | 0.024503 | 0.015211 | 0.038933 | 0.011504 | 0.001947 | 0.036974 | 0.002584 | 0.040381 | 0.06634 | 0.048414 | 0.050477 | 0.014227 | 0.026523 | 0.02506 | 0.03504 | 0.022991 | 0.020819 | 0.002758 | 0.010262 |
| 20.6111111 | 0.009073 | 0.0158 | 0.035187 | 0.016004 | 0.023613 | 0.023273 | 0.03351 | 0.067846 | 0.013077 | 0.002588 | 0.023166 | 0.003246 | 0.046076 | 0.085423 | 0.046302 | 0.057167 | 0.013205 | 0.018569 | 0.037265 | 0.040691 | 0.027344 | 0.022758 | 0.006433 | 0.030183 |
| 20.625 | 0.006288 | 0.013132 | 0.045588 | 0.026655 | 0.014621 | 0.01617 | 0.050596 | 0.056851 | 0.006939 | 0.005432 | 0.016903 | 0.005309 | 0.044715 | 0.085308 | 0.032186 | 0.042463 | 0.011955 | 0.009912 | 0.046857 | 0.037511 | 0.02453 | 0.017945 | 0.022336 | 0.055103 |
| 20.6388889 | 0.008386 | 0.008428 | 0.051553 | 0.049723 | 0.005451 | 0.008236 | 0.033617 | 0.022932 | 0.003819 | 0.008394 | 0.021974 | 0.006193 | 0.04124 | 0.059632 | 0.017024 | 0.025123 | 0.010471 | 0.005782 | 0.053166 | 0.02783 | 0.023866 | 0.010399 | 0.048524 | 0.060147 |
| 20.6527778 | 0.010334 | 0.004415 | 0.051042 | 0.065966 | 0.003658 | 0.008181 | 0.01461 | 0.008334 | 0.007516 | 0.011698 | 0.031992 | 0.006934 | 0.037347 | 0.030394 | 0.006217 | 0.013511 | 0.008866 | 0.006397 | 0.04749 | 0.017738 | 0.029171 | 0.005894 | 0.060167 | 0.040803 |
| 20.6666667 | 0.006498 | 0.004028 | 0.041415 | 0.080216 | 0.008946 | 0.018812 | 0.012413 | 0.012106 | 0.011404 | 0.01634 | 0.046121 | 0.011868 | 0.029118 | 0.028085 | 0.004033 | 0.006919 | 0.008206 | 0.006569 | 0.026093 | 0.009676 | 0.033281 | 0.005635 | 0.044636 | 0.018739 |
| 20.6805556 | 0.003368 | 0.008337 | 0.026987 | 0.144745 | 0.013611 | 0.032464 | 0.01233 | 0.018017 | 0.009833 | 0.020602 | 0.054003 | 0.014013 | 0.016311 | 0.050049 | 0.007505 | 0.009268 | 0.008328 | 0.00784 | 0.011836 | 0.00353 | 0.027752 | 0.00615 | 0.02128 | 0.01298 |
| 20.6944444 | 0.004595 | 0.009447 | 0.019574 | 0.226726 | 0.014301 | 0.039113 | 0.019742 | 0.014824 | 0.008766 | 0.01828 | 0.053316 | 0.00921 | 0.005486 | 0.071008 | 0.009088 | 0.012061 | 0.007681 | 0.012209 | 0.017378 | 0.000951 | 0.015934 | 0.00495 | 0.006724 | 0.018438 |
| 20.7083333 | 0.006207 | 0.005895 | 0.027176 | 0.220825 | 0.017152 | 0.039646 | 0.035957 | 0.011385 | 0.01191 | 0.011933 | 0.04757 | 0.004426 | 0.002888 | 0.076803 | 0.008403 | 0.008242 | 0.00641 | 0.013295 | 0.027465 | 0.003572 | 0.012551 | 0.002985 | 0.003817 | 0.017337 |
| 20.7222222 | 0.00555 | 0.003653 | 0.030865 | 0.147827 | 0.01975 | 0.031614 | 0.039146 | 0.014128 | 0.01721 | 0.014141 | 0.031005 | 0.005042 | 0.007683 | 0.058405 | 0.00948 | 0.005305 | 0.006777 | 0.016471 | 0.029584 | 0.009422 | 0.018828 | 0.00369 | 0.010916 | 0.010808 |
| 20.7361111 | 0.003541 | 0.00272 | 0.023401 | 0.083538 | 0.015934 | 0.01616 | 0.023134 | 0.011255 | 0.02735 | 0.027254 | 0.011635 | 0.00868 | 0.013164 | 0.036779 | 0.013007 | 0.011885 | 0.009123 | 0.023423 | 0.022223 | 0.013517 | 0.018263 | 0.008054 | 0.018994 | 0.014521 |
| 20.75 | 0.004453 | 0.0056 | 0.031967 | 0.038013 | 0.012015 | 0.013256 | 0.013249 | 0.007411 | 0.029768 | 0.053878 | 0.006042 | 0.01133 | 0.015153 | 0.047261 | 0.016296 | 0.025061 | 0.010656 | 0.021065 | 0.011225 | 0.012412 | 0.011302 | 0.011094 | 0.021779 | 0.028144 |
| 20.7638889 | 0.01025 | 0.015014 | 0.056175 | 0.013507 | 0.011631 | 0.02487 | 0.022801 | 0.015479 | 0.018429 | 0.080906 | 0.010428 | 0.015252 | 0.015473 | 0.06341 | 0.020626 | 0.037702 | 0.00965 | 0.012641 | 0.006904 | 0.009022 | 0.011661 | 0.008547 | 0.021606 | 0.03639 |
| 20.7777778 | 0.021284 | 0.027616 | 0.080174 | 0.016685 | 0.012735 | 0.034611 | 0.040417 | 0.032168 | 0.023825 | 0.08191 | 0.013914 | 0.02365 | 0.016157 | 0.054141 | 0.021174 | 0.043561 | 0.007313 | 0.009205 | 0.013354 | 0.012497 | 0.016358 | 0.004636 | 0.019457 | 0.032456 |
| 20.7916667 | 0.041251 | 0.043677 | 0.098279 | 0.053799 | 0.019106 | 0.029591 | 0.057114 | 0.057028 | 0.058752 | 0.064937 | 0.019611 | 0.04011 | 0.015809 | 0.041281 | 0.014749 | 0.03727 | 0.005757 | 0.009236 | 0.022502 | 0.019733 | 0.019407 | 0.004319 | 0.018004 | 0.024016 |
| 20.8055556 | 0.068761 | 0.065586 | 0.105992 | 0.126632 | 0.031863 | 0.013186 | 0.057216 | 0.085752 | 0.098354 | 0.062648 | 0.024876 | 0.067651 | 0.012778 | 0.040983 | 0.007096 | 0.024098 | 0.005832 | 0.006987 | 0.02515 | 0.020194 | 0.020713 | 0.004213 | 0.019379 | 0.021674 |
| 20.8194444 | 0.084793 | 0.081359 | 0.093676 | 0.165928 | 0.034413 | 0.002475 | 0.037753 | 0.085864 | 0.117305 | 0.072578 | 0.017544 | 0.079599 | 0.008265 | 0.040889 | 0.004215 | 0.012523 | 0.006048 | 0.004108 | 0.020513 | 0.015796 | 0.018569 | 0.002603 | 0.01855 | 0.023285 |
| 20.8333333 | 0.073265 | 0.084628 | 0.063607 | 0.111333 | 0.020289 | 0.001245 | 0.030736 | 0.049929 | 0.107919 | 0.054013 | 0.005141 | 0.054863 | 0.004295 | 0.027112 | 0.011879 | 0.004889 | 0.004837 | 0.006655 | 0.013764 | 0.013289 | 0.012641 | 0.002083 | 0.012989 | 0.01818 |
| 20.8472222 | 0.04481 | 0.084291 | 0.04061 | 0.038465 | 0.007839 | 0.002817 | 0.041763 | 0.022736 | 0.081751 | 0.035011 | 0.002455 | 0.026528 | 0.002508 | 0.01457 | 0.026339 | 0.002575 | 0.003166 | 0.02022 | 0.009447 | 0.013181 | 0.007898 | 0.00213 | 0.008598 | 0.008458 |
| 20.8611111 | 0.019763 | 0.072381 | 0.024046 | 0.014277 | 0.005342 | 0.0086 | 0.044878 | 0.018156 | 0.057263 | 0.061245 | 0.00715 | 0.015401 | 0.004488 | 0.020532 | 0.031482 | 0.00527 | 0.002679 | 0.043553 | 0.006521 | 0.010773 | 0.0124 | 0.003224 | 0.013108 | 0.002321 |
| 20.875 | 0.009262 | 0.042276 | 0.011605 | 0.015125 | 0.006287 | 0.020981 | 0.033769 | 0.015907 | 0.039867 | 0.094444 | 0.010898 | 0.010586 | 0.008302 | 0.026881 | 0.021947 | 0.009214 | 0.003336 | 0.057434 | 0.005213 | 0.006246 | 0.025806 | 0.007143 | 0.021833 | 0.001863 |
| 20.8888889 | 0.008074 | 0.025656 | 0.011235 | 0.025826 | 0.005851 | 0.029471 | 0.0209 | 0.016451 | 0.023156 | 0.102761 | 0.010075 | 0.0097 | 0.011149 | 0.022358 | 0.017698 | 0.010534 | 0.004295 | 0.04853 | 0.011118 | 0.006083 | 0.038408 | 0.01324 | 0.025456 | 0.003378 |
| 20.9027778 | 0.012115 | 0.042796 | 0.011885 | 0.040971 | 0.009533 | 0.024994 | 0.015365 | 0.01989 | 0.010862 | 0.100551 | 0.005588 | 0.011826 | 0.013388 | 0.017795 | 0.025638 | 0.008921 | 0.005147 | 0.029113 | 0.023716 | 0.013192 | 0.041879 | 0.019639 | 0.021457 | 0.003782 |
| 20.9166667 | 0.019608 | 0.054347 | 0.013326 | 0.039683 | 0.019006 | 0.020146 | 0.015278 | 0.018172 | 0.01409 | 0.0762 | 0.002125 | 0.012096 | 0.014329 | 0.017047 | 0.029584 | 0.006104 | 0.005296 | 0.013806 | 0.033516 | 0.024042 | 0.034843 | 0.021739 | 0.011706 | 0.006674 |
| 20.9305556 | 0.021081 | 0.039392 | 0.016256 | 0.024566 | 0.022911 | 0.03846 | 0.017176 | 0.011947 | 0.020864 | 0.036541 | 0.001536 | 0.009345 | 0.011787 | 0.019037 | 0.027501 | 0.0061 | 0.004182 | 0.013375 | 0.033634 | 0.029579 | 0.02638 | 0.019762 | 0.005092 | 0.008294 |
| 20.9444444 | 0.015262 | 0.036943 | 0.01247 | 0.025018 | 0.014519 | 0.067636 | 0.020801 | 0.011469 | 0.018692 | 0.028767 | 0.002372 | 0.005548 | 0.006987 | 0.022488 | 0.023841 | 0.014566 | 0.002244 | 0.034059 | 0.026914 | 0.024213 | 0.024801 | 0.019006 | 0.004352 | 0.004544 |
| 20.9583333 | 0.009497 | 0.053605 | 0.007051 | 0.047491 | 0.005213 | 0.067928 | 0.024596 | 0.021994 | 0.022482 | 0.041456 | 0.005378 | 0.004588 | 0.004072 | 0.023619 | 0.0156 | 0.024739 | 0.001007 | 0.054554 | 0.020876 | 0.014076 | 0.020837 | 0.016216 | 0.00431 | 0.002114 |
| 20.9722222 | 0.010493 | 0.054206 | 0.007659 | 0.067586 | 0.006452 | 0.046111 | 0.02872 | 0.033917 | 0.036789 | 0.068541 | 0.00849 | 0.009326 | 0.004852 | 0.021058 | 0.013622 | 0.025661 | 0.000974 | 0.052866 | 0.018546 | 0.007672 | 0.012849 | 0.009666 | 0.009464 | 0.003398 |
| 20.9861111 | 0.018021 | 0.040307 | 0.00986 | 0.064326 | 0.013478 | 0.034065 | 0.037556 | 0.037597 | 0.047142 | 0.105984 | 0.008143 | 0.015839 | 0.006564 | 0.017382 | 0.030188 | 0.022055 | 0.000837 | 0.036706 | 0.018639 | 0.008406 | 0.013823 | 0.00573 | 0.02018 | 0.005389 |
| 21 | 0.022528 | 0.028697 | 0.008865 | 0.040084 | 0.018077 | 0.028011 | 0.041708 | 0.028657 | 0.042403 | 0.103277 | 0.005129 | 0.016114 | 0.005148 | 0.019231 | 0.054689 | 0.022316 | 0.000431 | 0.02106 | 0.019195 | 0.012912 | 0.017288 | 0.004134 | 0.028605 | 0.00988 |
| 21.0138889 | 0.016985 | 0.018019 | 0.010102 | 0.015429 | 0.01552 | 0.017896 | 0.029348 | 0.017124 | 0.025778 | 0.06701 | 0.002566 | 0.010107 | 0.002566 | 0.028604 | 0.072308 | 0.025784 | 0.00068 | 0.011084 | 0.018408 | 0.014367 | 0.012036 | 0.001906 | 0.037355 | 0.018476 |
| 21.0277778 | 0.008312 | 0.008526 | 0.019436 | 0.006811 | 0.008357 | 0.012333 | 0.024578 | 0.019999 | 0.014388 | 0.034891 | 0.002817 | 0.007376 | 0.003167 | 0.036601 | 0.080881 | 0.028846 | 0.002197 | 0.006862 | 0.016248 | 0.01297 | 0.009939 | 0.002724 | 0.051088 | 0.025968 |
| 21.0416667 | 0.006039 | 0.006108 | 0.032234 | 0.012249 | 0.00392 | 0.019507 | 0.033643 | 0.030626 | 0.02053 | 0.033907 | 0.005387 | 0.009595 | 0.01005 | 0.038986 | 0.085772 | 0.032852 | 0.002858 | 0.012647 | 0.018305 | 0.019294 | 0.026177 | 0.012331 | 0.069367 | 0.03204 |
| 21.0555556 | 0.006908 | 0.011255 | 0.037412 | 0.016592 | 0.003938 | 0.029556 | 0.026211 | 0.036576 | 0.030414 | 0.058785 | 0.007582 | 0.010865 | 0.02553 | 0.039261 | 0.08495 | 0.042589 | 0.003439 | 0.024462 | 0.028396 | 0.050418 | 0.047706 | 0.028146 | 0.08686 | 0.045888 |
| 21.0694444 | 0.005509 | 0.018979 | 0.03187 | 0.012441 | 0.004263 | 0.025933 | 0.011527 | 0.040395 | 0.027604 | 0.066408 | 0.008445 | 0.010258 | 0.038864 | 0.037991 | 0.07465 | 0.054155 | 0.007204 | 0.026437 | 0.034847 | 0.083602 | 0.046208 | 0.037318 | 0.083475 | 0.064205 |
| 21.0833333 | 0.003954 | 0.025397 | 0.023881 | 0.010261 | 0.005867 | 0.013446 | 0.009086 | 0.041433 | 0.020213 | 0.057977 | 0.009155 | 0.008964 | 0.036944 | 0.035114 | 0.05372 | 0.057136 | 0.009808 | 0.015511 | 0.028162 | 0.073666 | 0.024876 | 0.033447 | 0.055518 | 0.065979 |
| 21.0972222 | 0.004476 | 0.028245 | 0.022057 | 0.015502 | 0.011734 | 0.008559 | 0.009009 | 0.034673 | 0.01444 | 0.047733 | 0.010366 | 0.007251 | 0.024309 | 0.031064 | 0.028208 | 0.052616 | 0.009416 | 0.004701 | 0.01873 | 0.040035 | 0.010908 | 0.021571 | 0.029124 | 0.045781 |
| 21.1111111 | 0.008738 | 0.028079 | 0.024519 | 0.020346 | 0.015829 | 0.010105 | 0.006303 | 0.022502 | 0.008342 | 0.031801 | 0.010835 | 0.005466 | 0.01125 | 0.02176 | 0.013228 | 0.0487 | 0.009506 | 0.002987 | 0.014275 | 0.018013 | 0.009617 | 0.009477 | 0.013922 | 0.021575 |
| 21.125 | 0.014137 | 0.023936 | 0.022837 | 0.022863 | 0.014 | 0.008169 | 0.004032 | 0.010519 | 0.004388 | 0.018333 | 0.009387 | 0.003697 | 0.002855 | 0.010092 | 0.008659 | 0.044339 | 0.009046 | 0.008413 | 0.010949 | 0.006727 | 0.01054 | 0.003833 | 0.008263 | 0.01016 |
| 21.1388889 | 0.014408 | 0.015247 | 0.015983 | 0.026126 | 0.01013 | 0.004004 | 0.002691 | 0.004985 | 0.005714 | 0.017225 | 0.007569 | 0.002422 | 0.00198 | 0.004282 | 0.012307 | 0.034673 | 0.006033 | 0.014993 | 0.007644 | 0.002682 | 0.012058 | 0.004904 | 0.013287 | 0.009629 |
| 21.1527778 | 0.009683 | 0.011826 | 0.009797 | 0.027191 | 0.008017 | 0.001941 | 0.003557 | 0.006843 | 0.014497 | 0.017747 | 0.006713 | 0.002115 | 0.006737 | 0.006259 | 0.024248 | 0.020912 | 0.002319 | 0.019229 | 0.005766 | 0.004372 | 0.018406 | 0.0083 | 0.017607 | 0.007814 |
| 21.1666667 | 0.004459 | 0.018516 | 0.006878 | 0.0208 | 0.008527 | 0.001576 | 0.007021 | 0.008025 | 0.029392 | 0.011627 | 0.006156 | 0.002333 | 0.010808 | 0.016319 | 0.032319 | 0.00851 | 0.000552 | 0.019295 | 0.004564 | 0.007183 | 0.026743 | 0.011572 | 0.013477 | 0.009864 |
| 21.1805556 | 0.00189 | 0.020368 | 0.007303 | 0.010907 | 0.010101 | 0.00175 | 0.010116 | 0.010261 | 0.04744 | 0.008058 | 0.00526 | 0.002366 | 0.011566 | 0.027654 | 0.034485 | 0.003357 | 0.000435 | 0.014047 | 0.003805 | 0.007459 | 0.027423 | 0.013076 | 0.012568 | 0.027952 |

| | | | | | | | | | | | | | | | | | | | | | | | | |
|---|---|---|---|---|---|---|---|---|---|---|---|---|---|---|---|---|---|---|---|---|---|---|---|---|
| 21.1944444 | 0.00268 | 0.015244 | 0.012814 | 0.007158 | 0.010743 | 0.002095 | 0.012256 | 0.018385 | 0.062949 | 0.00994 | 0.004308 | 0.002105 | 0.011302 | 0.029017 | 0.037496 | 0.007128 | 0.000409 | 0.007088 | 0.004554 | 0.005374 | 0.018069 | 0.011279 | 0.024164 | 0.052448 |
| 21.2083333 | 0.005593 | 0.021417 | 0.023828 | 0.007161 | 0.011221 | 0.004146 | 0.017288 | 0.024403 | 0.066802 | 0.030393 | 0.004076 | 0.001879 | 0.011826 | 0.01885 | 0.034504 | 0.014387 | 0.00019 | 0.004455 | 0.005173 | 0.005992 | 0.00981 | 0.006912 | 0.033929 | 0.062505 |
| 21.2222222 | 0.009173 | 0.041064 | 0.034583 | 0.011348 | 0.013975 | 0.009826 | 0.02504 | 0.025426 | 0.057439 | 0.063537 | 0.004924 | 0.001837 | 0.013841 | 0.007705 | 0.020674 | 0.020019 | 0.000756 | 0.006685 | 0.003732 | 0.01142 | 0.014052 | 0.005981 | 0.031332 | 0.051179 |
| 21.2361111 | 0.012702 | 0.059627 | 0.038815 | 0.027894 | 0.018497 | 0.015748 | 0.030494 | 0.026046 | 0.038866 | 0.072555 | 0.006335 | 0.001828 | 0.017972 | 0.006291 | 0.01002 | 0.022556 | 0.002064 | 0.009809 | 0.00254 | 0.014096 | 0.024979 | 0.010729 | 0.02375 | 0.028558 |
| 21.25 | 0.016616 | 0.069057 | 0.038846 | 0.050638 | 0.021977 | 0.019883 | 0.034557 | 0.02602 | 0.021889 | 0.058896 | 0.008001 | 0.001905 | 0.021244 | 0.013536 | 0.009149 | 0.019965 | 0.002963 | 0.011834 | 0.003576 | 0.009838 | 0.030332 | 0.012077 | 0.015422 | 0.012641 |
| 21.2638889 | 0.018801 | 0.071357 | 0.038661 | 0.071196 | 0.022758 | 0.025069 | 0.038385 | 0.025543 | 0.015738 | 0.04078 | 0.009657 | 0.002495 | 0.018056 | 0.021012 | 0.010109 | 0.01293 | 0.003461 | 0.009545 | 0.004438 | 0.007311 | 0.02472 | 0.00765 | 0.009617 | 0.015013 |
| 21.2777778 | 0.015475 | 0.071524 | 0.034605 | 0.082469 | 0.019562 | 0.023232 | 0.035559 | 0.026109 | 0.012436 | 0.022049 | 0.009329 | 0.003247 | 0.009701 | 0.022395 | 0.011141 | 0.006592 | 0.003028 | 0.004274 | 0.003502 | 0.012873 | 0.013972 | 0.005553 | 0.012539 | 0.027481 |
| 21.2916667 | 0.006081 | 0.035839 | 0.015322 | 0.042076 | 0.008362 | 0.009106 | 0.015703 | 0.013296 | 0.004795 | 0.006835 | 0.004143 | 0.001758 | 0.003471 | 0.016614 | 0.012522 | 0.006322 | 0.001926 | 0.003636 | 0.004276 | 0.019605 | 0.010909 | 0.00771 | 0.019002 | 0.03427 |
| 21.3055556 | 0.004851 | 0.040365 | 0.013164 | 0.04015 | 0.010029 | 0.002967 | 0.015675 | 0.017892 | 0.009669 | 0.021628 | 0.009763 | 0.010349 | 0.001329 | 0.009717 | 0.012103 | 0.012118 | 0.002255 | 0.005885 | 0.00698 | 0.019939 | 0.014038 | 0.009412 | 0.01863 | 0.028648 |
| 21.3194444 | 0.001047 | 0.006097 | 0.000731 | 0.000841 | 0.003121 | 0.0007 | 0.00513 | 0.005475 | 0.004177 | 0.009117 | 0.005889 | 0.0062 | 0.00132 | 0.009242 | 0.008987 | 0.01721 | 0.003424 | 0.006158 | 0.006251 | 0.01442 | 0.014088 | 0.007944 | 0.012787 | 0.015267 |
| 21.3333333 | 0.003794 | 0.017276 | 0.001743 | 0.003578 | 0.008301 | 0.001932 | 0.009911 | 0.012851 | 0.006775 | 0.013715 | 0.010318 | 0.01095 | 0.001984 | 0.010606 | 0.005663 | 0.017308 | 0.003491 | 0.006119 | 0.003434 | 0.008463 | 0.011962 | 0.004181 | 0.007299 | 0.008236 |
| 21.3472222 | 0.007856 | 0.022007 | 0.003769 | 0.007499 | 0.014226 | 0.004307 | 0.00764 | 0.014833 | 0.006788 | 0.008793 | 0.007634 | 0.011698 | 0.002005 | 0.007147 | 0.008863 | 0.015185 | 0.002805 | 0.005777 | 0.002227 | 0.006974 | 0.010788 | 0.001077 | 0.00359 | 0.012467 |
| 21.3611111 | 0.010985 | 0.015596 | 0.006992 | 0.010124 | 0.018281 | 0.00913 | 0.008097 | 0.011008 | 0.01098 | 0.010583 | 0.004614 | 0.015487 | 0.001966 | 0.003805 | 0.019369 | 0.015359 | 0.003368 | 0.003774 | 0.001119 | 0.009623 | 0.010476 | 0.001372 | 0.002485 | 0.015222 |
| 21.375 | 0.01015 | 0.007249 | 0.009747 | 0.010902 | 0.01621 | 0.012803 | 0.013558 | 0.004635 | 0.014337 | 0.015034 | 0.003365 | 0.018259 | 0.002622 | 0.003858 | 0.026174 | 0.018145 | 0.005477 | 0.002366 | 0.000973 | 0.01179 | 0.009971 | 0.004041 | 0.003082 | 0.012223 |
| 21.3888889 | 0.006395 | 0.006515 | 0.012727 | 0.011021 | 0.011532 | 0.012249 | 0.017088 | 0.003092 | 0.013872 | 0.019317 | 0.006477 | 0.022791 | 0.004878 | 0.004038 | 0.02196 | 0.01955 | 0.005702 | 0.002281 | 0.002319 | 0.010257 | 0.008427 | 0.004599 | 0.003831 | 0.015472 |
| 21.4027778 | 0.003955 | 0.012608 | 0.01578 | 0.010954 | 0.008299 | 0.009693 | 0.017359 | 0.008303 | 0.011873 | 0.023471 | 0.010045 | 0.030897 | 0.009134 | 0.00558 | 0.015465 | 0.020015 | 0.002999 | 0.002639 | 0.00277 | 0.007388 | 0.0071 | 0.002506 | 0.007125 | 0.027024 |
| 21.4166667 | 0.004727 | 0.020664 | 0.016925 | 0.01257 | 0.009288 | 0.008165 | 0.018362 | 0.019678 | 0.012129 | 0.027994 | 0.010256 | 0.036759 | 0.013089 | 0.011175 | 0.016413 | 0.024202 | 0.00095 | 0.004353 | 0.002922 | 0.007452 | 0.007195 | 0.001176 | 0.013054 | 0.037048 |
| 21.4305556 | 0.006539 | 0.022599 | 0.015807 | 0.017761 | 0.015175 | 0.008953 | 0.021175 | 0.032227 | 0.015589 | 0.03289 | 0.006867 | 0.035983 | 0.014534 | 0.01757 | 0.021786 | 0.029614 | 0.000778 | 0.004851 | 0.002894 | 0.01038 | 0.008198 | 0.001162 | 0.018494 | 0.042872 |
| 21.4444444 | 0.007797 | 0.015993 | 0.01542 | 0.024947 | 0.023404 | 0.011474 | 0.023014 | 0.040186 | 0.020399 | 0.036933 | 0.003761 | 0.034181 | 0.01354 | 0.020353 | 0.024871 | 0.029848 | 0.00116 | 0.004223 | 0.001687 | 0.013135 | 0.009726 | 0.002061 | 0.022521 | 0.044527 |
| 21.4583333 | 0.009441 | 0.00751 | 0.018431 | 0.031815 | 0.031582 | 0.014077 | 0.022186 | 0.041678 | 0.023477 | 0.039071 | 0.00548 | 0.035529 | 0.011751 | 0.018685 | 0.024221 | 0.024763 | 0.001405 | 0.006543 | 0.001777 | 0.015001 | 0.011559 | 0.003918 | 0.024517 | 0.039504 |
| 21.4722222 | 0.01313 | 0.004765 | 0.024531 | 0.04092 | 0.042067 | 0.01542 | 0.020909 | 0.034427 | 0.023739 | 0.042822 | 0.009658 | 0.036591 | 0.01066 | 0.014801 | 0.024004 | 0.019594 | 0.001223 | 0.009545 | 0.005374 | 0.018563 | 0.013016 | 0.005436 | 0.023435 | 0.029458 |
| 21.4861111 | 0.019432 | 0.012634 | 0.032365 | 0.054369 | 0.055322 | 0.016045 | 0.022256 | 0.019337 | 0.024056 | 0.056063 | 0.013136 | 0.034598 | 0.011272 | 0.013626 | 0.030494 | 0.020314 | 0.001121 | 0.01123 | 0.013413 | 0.026054 | 0.014288 | 0.004499 | 0.019951 | 0.019328 |
| 21.5 | 0.026075 | 0.028617 | 0.040142 | 0.066646 | 0.063694 | 0.017854 | 0.024317 | 0.006744 | 0.026839 | 0.077754 | 0.016219 | 0.031286 | 0.013473 | 0.020173 | 0.044999 | 0.026734 | 0.001777 | 0.015922 | 0.02409 | 0.033114 | 0.015841 | 0.003639 | 0.015068 | 0.011686 |
| 21.5138889 | 0.028291 | 0.036663 | 0.045053 | 0.070143 | 0.059714 | 0.019813 | 0.021732 | 0.00265 | 0.028075 | 0.087395 | 0.017284 | 0.029451 | 0.014737 | 0.031154 | 0.057665 | 0.031053 | 0.003008 | 0.022829 | 0.02859 | 0.030347 | 0.016347 | 0.006531 | 0.008582 | 0.008418 |
| 21.5277778 | 0.02361 | 0.027277 | 0.045421 | 0.064024 | 0.045249 | 0.017398 | 0.014334 | 0.005166 | 0.022594 | 0.068635 | 0.012837 | 0.02806 | 0.012262 | 0.036018 | 0.05837 | 0.027893 | 0.004552 | 0.021884 | 0.02087 | 0.016932 | 0.013968 | 0.009089 | 0.004475 | 0.01141 |
| 21.5416667 | 0.015815 | 0.012334 | 0.042694 | 0.055249 | 0.030833 | 0.010104 | 0.007795 | 0.017655 | 0.013575 | 0.035141 | 0.005466 | 0.025762 | 0.006999 | 0.031826 | 0.051785 | 0.02174 | 0.006303 | 0.013416 | 0.008851 | 0.004944 | 0.009914 | 0.008638 | 0.003866 | 0.012783 |
| 21.5555556 | 0.010826 | 0.00568 | 0.039762 | 0.049794 | 0.024912 | 0.003458 | 0.006666 | 0.039528 | 0.008016 | 0.010784 | 0.001298 | 0.024929 | 0.003614 | 0.026096 | 0.047689 | 0.022581 | 0.009526 | 0.009277 | 0.002938 | 0.002782 | 0.007242 | 0.00838 | 0.005567 | 0.009671 |
| 21.5694444 | 0.011388 | 0.007571 | 0.038118 | 0.049745 | 0.026681 | 0.000776 | 0.010721 | 0.061004 | 0.009223 | 0.005489 | 0.000681 | 0.025484 | 0.005936 | 0.025884 | 0.053373 | 0.035675 | 0.013251 | 0.013211 | 0.004989 | 0.008748 | 0.005235 | 0.009924 | 0.015652 | 0.015575 |
| 21.5833333 | 0.017873 | 0.013822 | 0.040467 | 0.059652 | 0.03161 | 0.002753 | 0.018696 | 0.074451 | 0.018095 | 0.017674 | 0.000952 | 0.026346 | 0.013383 | 0.034806 | 0.069593 | 0.05495 | 0.011013 | 0.018675 | 0.014821 | 0.019379 | 0.004701 | 0.012166 | 0.031283 | 0.034799 |
| 21.5972222 | 0.028779 | 0.024996 | 0.051878 | 0.081622 | 0.036562 | 0.010236 | 0.026556 | 0.074771 | 0.033702 | 0.039227 | 0.001743 | 0.030309 | 0.021888 | 0.057022 | 0.088423 | 0.067455 | 0.00643 | 0.019323 | 0.031469 | 0.02982 | 0.008346 | 0.014566 | 0.041658 | 0.061187 |
| 21.6111111 | 0.040628 | 0.040145 | 0.070235 | 0.103621 | 0.040078 | 0.01753 | 0.025642 | 0.059935 | 0.049111 | 0.066033 | 0.004691 | 0.039916 | 0.028852 | 0.080993 | 0.098723 | 0.068698 | 0.009502 | 0.016567 | 0.051229 | 0.036118 | 0.010339 | 0.017713 | 0.043422 | 0.087298 |
| 21.625 | 0.049386 | 0.05452 | 0.086538 | 0.111469 | 0.040837 | 0.01693 | 0.01479 | 0.038682 | 0.055202 | 0.096793 | 0.010726 | 0.056205 | 0.035121 | 0.081226 | 0.086648 | 0.062922 | 0.017253 | 0.014564 | 0.067364 | 0.037424 | 0.013563 | 0.020859 | 0.040068 | 0.09786 |
| 21.6388889 | 0.05134 | 0.064674 | 0.095059 | 0.102005 | 0.038595 | 0.010045 | 0.006472 | 0.019621 | 0.049146 | 0.109095 | 0.016159 | 0.06784 | 0.041263 | 0.058931 | 0.054497 | 0.051207 | 0.021643 | 0.013587 | 0.07131 | 0.033164 | 0.029013 | 0.022137 | 0.034583 | 0.084566 |
| 21.6527778 | 0.04602 | 0.067836 | 0.091667 | 0.076553 | 0.035045 | 0.004181 | 0.008821 | 0.00965 | 0.032936 | 0.082464 | 0.013464 | 0.056923 | 0.0443 | 0.032098 | 0.024426 | 0.034593 | 0.018504 | 0.010709 | 0.05668 | 0.023135 | 0.047404 | 0.02146 | 0.02576 | 0.055963 |
| 21.6666667 | 0.035868 | 0.061863 | 0.074861 | 0.042814 | 0.029866 | 0.005012 | 0.014866 | 0.013688 | 0.016459 | 0.037681 | 0.005166 | 0.030432 | 0.041805 | 0.017886 | 0.015476 | 0.017704 | 0.011075 | 0.005977 | 0.029544 | 0.011457 | 0.049813 | 0.019283 | 0.014118 | 0.028036 |
| 21.6805556 | 0.024926 | 0.050504 | 0.051004 | 0.018014 | 0.022742 | 0.00906 | 0.018322 | 0.020698 | 0.009617 | 0.012496 | 0.002604 | 0.01001 | 0.037939 | 0.030875 | 0.024263 | 0.006623 | 0.005813 | 0.003525 | 0.011606 | 0.005151 | 0.032533 | 0.013994 | 0.008681 | 0.020183 |
| 21.6944444 | 0.017687 | 0.040844 | 0.029743 | 0.012719 | 0.017865 | 0.008868 | 0.01532 | 0.018868 | 0.00817 | 0.01391 | 0.006724 | 0.002295 | 0.03634 | 0.05694 | 0.0319 | 0.004241 | 0.003224 | 0.003516 | 0.011899 | 0.00507 | 0.014613 | 0.007059 | 0.014605 | 0.033509 |
| 21.7083333 | 0.015349 | 0.0357 | 0.016151 | 0.016358 | 0.019283 | 0.011533 | 0.010992 | 0.012353 | 0.012828 | 0.016291 | 0.011439 | 0.003522 | 0.035107 | 0.07032 | 0.030624 | 0.005945 | 0.001736 | 0.005644 | 0.013773 | 0.004787 | 0.010657 | 0.004178 | 0.022053 | 0.044095 |
| 21.7222222 | 0.017147 | 0.033631 | 0.007908 | 0.016107 | 0.027269 | 0.022497 | 0.016734 | 0.013985 | 0.026311 | 0.01132 | 0.013066 | 0.009926 | 0.033889 | 0.058914 | 0.020652 | 0.006327 | 0.00109 | 0.010211 | 0.008479 | 0.003197 | 0.012446 | 0.006144 | 0.023511 | 0.036894 |
| 21.7361111 | 0.021329 | 0.034813 | 0.005162 | 0.014585 | 0.038406 | 0.029041 | 0.02774 | 0.02572 | 0.033676 | 0.013333 | 0.011216 | 0.014698 | 0.033043 | 0.032233 | 0.01436 | 0.005908 | 0.001758 | 0.012134 | 0.007428 | 0.005788 | 0.012632 | 0.008091 | 0.017608 | 0.021212 |
| 21.75 | 0.025048 | 0.040487 | 0.010431 | 0.016286 | 0.045655 | 0.02735 | 0.033267 | 0.027701 | 0.029297 | 0.020844 | 0.008514 | 0.013881 | 0.031379 | 0.020276 | 0.024249 | 0.012295 | 0.004004 | 0.009993 | 0.015412 | 0.01022 | 0.010439 | 0.00739 | 0.01473 | 0.019606 |
| 21.7638889 | 0.025849 | 0.046794 | 0.017485 | 0.016086 | 0.04341 | 0.026602 | 0.028079 | 0.015292 | 0.023993 | 0.020461 | 0.006553 | 0.011998 | 0.02993 | 0.028874 | 0.033363 | 0.029967 | 0.006393 | 0.009495 | 0.024445 | 0.010655 | 0.006065 | 0.005006 | 0.028211 | 0.037852 |
| 21.7777778 | 0.021814 | 0.048356 | 0.021112 | 0.010352 | 0.032866 | 0.030073 | 0.016466 | 0.007957 | 0.020569 | 0.01205 | 0.005739 | 0.015354 | 0.029691 | 0.032881 | 0.025825 | 0.049176 | 0.007119 | 0.013749 | 0.028669 | 0.009186 | 0.005115 | 0.005468 | 0.045307 | 0.056673 |
| 21.7916667 | 0.012728 | 0.043761 | 0.023446 | 0.006128 | 0.018948 | 0.034406 | 0.007389 | 0.009806 | 0.01645 | 0.006715 | 0.006722 | 0.023813 | 0.027257 | 0.027345 | 0.022123 | 0.057708 | 0.006113 | 0.017856 | 0.028534 | 0.0099 | 0.012017 | 0.010869 | 0.05274 | 0.064141 |
| 21.8055556 | 0.006364 | 0.033057 | 0.025029 | 0.011113 | 0.008873 | 0.032516 | 0.003626 | 0.011802 | 0.011097 | 0.005343 | 0.008832 | 0.029242 | 0.018059 | 0.026906 | 0.045423 | 0.05525 | 0.005186 | 0.013688 | 0.026638 | 0.012339 | 0.019515 | 0.014976 | 0.058209 | 0.065701 |
| 21.8194444 | 0.008812 | 0.019207 | 0.022949 | 0.021077 | 0.008096 | 0.021362 | 0.005504 | 0.010518 | 0.00569 | 0.005961 | 0.00925 | 0.025153 | 0.007942 | 0.033369 | 0.088671 | 0.050321 | 0.00534 | 0.007038 | 0.032131 | 0.015715 | 0.021765 | 0.013989 | 0.07324 | 0.067117 |
| 21.8333333 | 0.010447 | 0.012945 | 0.017423 | 0.025702 | 0.012636 | 0.011774 | 0.011375 | 0.011443 | 0.003949 | 0.011921 | 0.006102 | 0.014226 | 0.004614 | 0.035685 | 0.126349 | 0.046768 | 0.004786 | 0.006248 | 0.039633 | 0.018276 | 0.020455 | 0.01077 | 0.087446 | 0.058654 |
| 21.8472222 | 0.009735 | 0.016638 | 0.01398 | 0.025333 | 0.017116 | 0.015434 | 0.01899 | 0.013883 | 0.006622 | 0.01657 | 0.003474 | 0.008618 | 0.003301 | 0.024331 | 0.127728 | 0.038805 | 0.002761 | 0.005058 | 0.03643 | 0.015035 | 0.014492 | 0.006754 | 0.073971 | 0.03277 |
| 21.8611111 | 0.017877 | 0.02032 | 0.015923 | 0.027501 | 0.020256 | 0.025367 | 0.026412 | 0.010843 | 0.00972 | 0.011896 | 0.004379 | 0.014222 | 0.002395 | 0.014693 | 0.093289 | 0.023468 | 0.001317 | 0.002557 | 0.046362 | 0.00799 | 0.011332 | 0.004012 | 0.041369 | 0.014886 |
| 21.875 | 0.027683 | 0.0238 | 0.021567 | 0.03196 | 0.020922 | 0.02917 | 0.027784 | 0.004954 | 0.010777 | 0.006192 | 0.004795 | 0.020458 | 0.006687 | 0.024214 | 0.051392 | 0.012534 | 0.000919 | 0.003026 | 0.074602 | 0.008217 | 0.022211 | 0.006494 | 0.031866 | 0.025715 |
| 21.8888889 | 0.021854 | 0.027653 | 0.026249 | 0.03271 | 0.0184 | 0.024841 | 0.018683 | 0.005309 | 0.009606 | 0.007153 | 0.003587 | 0.019936 | 0.016066 | 0.033227 | 0.023377 | 0.017215 | 0.000492 | 0.008719 | 0.080855 | 0.014953 | 0.039089 | 0.012223 | 0.055056 | 0.043903 |
| 21.9027778 | 0.010768 | 0.02704 | 0.025505 | 0.028907 | 0.014869 | 0.016804 | 0.007537 | 0.013294 | 0.008555 | 0.00692 | 0.003754 | 0.016175 | 0.02307 | 0.026483 | 0.022791 | 0.030792 | 0.000279 | 0.017871 | 0.056665 | 0.015538 | 0.0467 | 0.017868 | 0.075618 | 0.04727 |
| 21.9166667 | 0.012752 | 0.026949 | 0.021049 | 0.02498 | 0.013812 | 0.010767 | 0.004358 | 0.015835 | 0.01165 | 0.005346 | 0.005 | 0.014158 | 0.01996 | 0.015414 | 0.043011 | 0.04555 | 0.00047 | 0.023532 | 0.029868 | 0.009705 | 0.042426 | 0.025517 | 0.06784 | 0.035391 |
| 21.9305556 | 0.019856 | 0.030771 | 0.016321 | 0.023343 | 0.015771 | 0.008874 | 0.006247 | 0.013894 | 0.015447 | 0.013691 | 0.00415 | 0.01548 | 0.010151 | 0.007265 | 0.063606 | 0.063413 | 0.000605 | 0.020959 | 0.013037 | 0.004739 | 0.031551 | 0.034321 | 0.042361 | 0.019072 |
| 21.9444444 | 0.024946 | 0.026576 | 0.012281 | 0.023872 | 0.019871 | 0.008948 | 0.009171 | 0.019371 | 0.014113 | 0.035457 | 0.002229 | 0.017317 | 0.003583 | 0.0021 | 0.06716 | 0.08098 | 0.000477 | 0.014863 | 0.005065 | 0.003386 | 0.01924 | 0.036486 | 0.020512 | 0.012069 |
| 21.9583333 | 0.029907 | 0.016235 | 0.011157 | 0.027088 | 0.025618 | 0.00936 | 0.010787 | 0.018351 | 0.011694 | 0.060348 | 0.001868 | 0.014554 | 0.002794 | 0.003094 | 0.043765 | 0.08521 | 0.000361 | 0.011865 | 0.003666 | 0.004849 | 0.010549 | 0.028572 | 0.014306 | 0.015146 |
| 21.9722222 | 0.031581 | 0.01291 | 0.013967 | 0.030674 | 0.03168 | 0.012169 | 0.007458 | 0.011075 | 0.015321 | 0.071771 | 0.001538 | 0.00731 | 0.00273 | 0.011645 | 0.015139 | 0.071709 | 0.000433 | 0.008861 | 0.004422 | 0.005633 | 0.00617 | 0.015246 | 0.025205 | 0.024597 |
| 21.9861111 | 0.029281 | 0.016784 | 0.019194 | 0.030682 | 0.035163 | 0.018691 | 0.004722 | 0.016637 | 0.024854 | 0.069671 | 0.001788 | 0.004841 | 0.003322 | 0.021549 | 0.01102 | 0.047248 | 0.000495 | 0.007783 | 0.00506 | 0.003929 | 0.003771 | 0.005861 | 0.040916 | 0.039334 |
| 22 | 0.021414 | 0.033549 | 0.02568 | 0.029066 | 0.033655 | 0.026071 | 0.008799 | 0.025765 | 0.035563 | 0.069232 | 0.004221 | 0.014855 | 0.007086 | 0.025155 | 0.023342 | 0.023323 | 0.000664 | 0.008383 | 0.004565 | 0.001646 | 0.004176 | 0.004817 | 0.043551 | 0.049135 |
| 22.0138889 | 0.010179 | 0.053983 | 0.031357 | 0.030165 | 0.029113 | 0.030667 | 0.013188 | 0.030585 | 0.043838 | 0.071633 | 0.006834 | 0.030941 | 0.010649 | 0.02162 | 0.032581 | 0.015948 | 0.000909 | 0.005747 | 0.004985 | 0.001782 | 0.008403 | 0.004947 | 0.027771 | 0.046788 |
| 22.0277778 | 0.003987 | 0.058791 | 0.0363 | 0.037547 | 0.029187 | 0.032423 | 0.012697 | 0.041233 | 0.047575 | 0.067617 | 0.008173 | 0.039231 | 0.012195 | 0.017611 | 0.029605 | 0.030558 | 0.001472 | 0.003337 | 0.009497 | 0.005895 | 0.016467 | 0.006236 | 0.013038 | 0.038111 |
| 22.0416667 | 0.0052 | 0.059415 | 0.04877 | 0.061703 | 0.044261 | 0.040149 | 0.014976 | 0.065504 | 0.052169 | 0.066905 | 0.010142 | 0.04567 | 0.020459 | 0.020671 | 0.022792 | 0.053792 | 0.00173 | 0.009157 | 0.019982 | 0.017412 | 0.031045 | 0.015993 | 0.019463 | 0.032262 |
| 22.0555556 | 0.015585 | 0.070378 | 0.066987 | 0.098139 | 0.076618 | 0.05932 | 0.031933 | 0.097822 | 0.066958 | 0.085142 | 0.014006 | 0.065662 | 0.041441 | 0.030564 | 0.031566 | 0.053864 | 0.002312 | 0.02239 | 0.031885 | 0.036743 | 0.047356 | 0.031931 | 0.042719 | 0.039744 |
| 22.0694444 | 0.029454 | 0.077188 | 0.071314 | 0.108514 | 0.108976 | 0.072556 | 0.054751 | 0.107563 | 0.079431 | 0.103171 | 0.01651 | 0.08449 | 0.062649 | 0.038208 | 0.056375 | 0.035987 | 0.005862 | 0.023697 | 0.032119 | 0.049821 | 0.048355 | 0.045884 | 0.066116 | 0.062447 |
| 22.0833333 | 0.035667 | 0.062885 | 0.057109 | 0.078626 | 0.116633 | 0.064228 | 0.062211 | 0.078798 | 0.070079 | 0.096644 | 0.014452 | 0.076637 | 0.07214 | 0.0412 | 0.079125 | 0.044835 | 0.012159 | 0.015054 | 0.018526 | 0.045641 | 0.032467 | 0.052941 | 0.076687 | 0.086867 |
| 22.0972222 | 0.034483 | 0.038757 | 0.03679 | 0.042857 | 0.096486 | 0.046212 | 0.055157 | 0.038278 | 0.047081 | 0.077062 | 0.008979 | 0.046697 | 0.068568 | 0.04394 | 0.079207 | 0.069329 | 0.018495 | 0.016739 | 0.00646 | 0.033853 | 0.016272 | 0.048913 | 0.071027 | 0.091773 |
| 22.1111111 | 0.032049 | 0.028887 | 0.017991 | 0.026391 | 0.071753 | 0.032087 | 0.044693 | 0.013623 | 0.0266 | 0.059958 | 0.00485 | 0.020615 | 0.050893 | 0.043968 | 0.057401 | 0.073809 | 0.019139 | 0.017837 | 0.003716 | 0.025259 | 0.007176 | 0.031423 | 0.052209 | 0.066919 |
| 22.125 | 0.030227 | 0.044963 | 0.005887 | 0.025702 | 0.054988 | 0.020399 | 0.031883 | 0.007282 | 0.012285 | 0.047056 | 0.006123 | 0.015487 | 0.027361 | 0.039372 | 0.028907 | 0.064763 | 0.011464 | 0.010736 | 0.005386 | 0.01959 | 0.003132 | 0.015051 | 0.029687 | 0.030804 |
| 22.1388889 | 0.026049 | 0.06993 | 0.003022 | 0.028261 | 0.032936 | 0.009438 | 0.01808 | 0.008142 | 0.004825 | 0.0351 | 0.010209 | 0.028041 | 0.010291 | 0.033582 | 0.012253 | 0.060773 | 0.005054 | 0.009828 | 0.010125 | 0.015073 | 0.004167 | 0.015185 | 0.023602 | 0.017278 |
| 22.1527778 | 0.018727 | 0.075532 | 0.006366 | 0.027158 | 0.011384 | 0.003867 | 0.01494 | 0.009188 | 0.003675 | 0.021759 | 0.012109 | 0.044086 | 0.002395 | 0.027672 | 0.015877 | 0.061488 | 0.006652 | 0.015691 | 0.014425 | 0.01154 | 0.008577 | 0.024672 | 0.036251 | 0.033241 |
| 22.1666667 | 0.009931 | 0.052828 | 0.010705 | 0.025005 | 0.005311 | 0.006208 | 0.02075 | 0.019378 | 0.003401 | 0.010932 | 0.01125 | 0.048203 | 0.000627 | 0.019103 | 0.023287 | 0.052752 | 0.008938 | 0.02003 | 0.014144 | 0.009236 | 0.010249 | 0.032366 | 0.040975 | 0.047978 |
| 22.1805556 | 0.00505 | 0.022759 | 0.013304 | 0.024514 | 0.007148 | 0.010295 | 0.017476 | 0.04095 | 0.00247 | 0.005227 | 0.009196 | 0.036069 | 0.001033 | 0.009794 | 0.025247 | 0.031829 | 0.007016 | 0.020231 | 0.011176 | 0.008594 | 0.007008 | 0.033438 | 0.028899 | 0.039578 |
| 22.1944444 | 0.009635 | 0.009434 | 0.01518 | 0.024913 | 0.010805 | 0.010196 | 0.010023 | 0.039998 | 0.004649 | 0.003025 | 0.006114 | 0.020052 | 0.002106 | 0.006494 | 0.023717 | 0.014124 | 0.003918 | 0.016768 | 0.006659 | 0.009142 | 0.003645 | 0.025607 | 0.014711 | 0.019935 |
| 22.2083333 | 0.017256 | 0.007893 | 0.017597 | 0.024637 | 0.011301 | 0.006638 | 0.012107 | 0.024867 | 0.008789 | 0.002514 | 0.004001 | 0.016319 | 0.003773 | 0.007124 | 0.018943 | 0.011058 | 0.002762 | 0.011765 | 0.002789 | 0.010693 | 0.004615 | 0.012987 | 0.011942 | 0.012901 |
| 22.2222222 | 0.018701 | 0.0107 | 0.020701 | 0.021898 | 0.006794 | 0.003813 | 0.015779 | 0.034214 | 0.011133 | 0.00247 | 0.005794 | 0.027172 | 0.005763 | 0.006182 | 0.013805 | 0.019882 | 0.003578 | 0.00788 | 0.001462 | 0.013403 | 0.008484 | 0.005554 | 0.02302 | 0.023634 |
| 22.2361111 | 0.013494 | 0.012754 | 0.022602 | 0.016776 | 0.004125 | 0.005139 | 0.014678 | 0.039386 | 0.010985 | 0.003213 | 0.008133 | 0.037618 | 0.007807 | 0.007444 | 0.011361 | 0.030102 | 0.004844 | 0.005057 | 0.001069 | 0.014674 | 0.011997 | 0.007991 | 0.036231 | 0.036899 |
| 22.25 | 0.007101 | 0.008335 | 0.02084 | 0.012158 | 0.007423 | 0.009101 | 0.012247 | 0.022978 | 0.009747 | 0.003856 | 0.006515 | 0.037633 | 0.008816 | 0.011698 | 0.012138 | 0.031058 | 0.006357 | 0.00379 | 0.001418 | 0.011018 | 0.014513 | 0.012623 | 0.041053 | 0.039492 |
| 22.2638889 | 0.006173 | 0.004489 | 0.015751 | 0.009733 | 0.014932 | 0.012431 | 0.008128 | 0.017475 | 0.00869 | 0.002644 | 0.003417 | 0.029888 | 0.007715 | 0.012502 | 0.014396 | 0.021371 | 0.007334 | 0.006272 | 0.002789 | 0.005718 | 0.016458 | 0.01312 | 0.037026 | 0.026945 |
| 22.2777778 | 0.00905 | 0.008255 | 0.01043 | 0.008301 | 0.018877 | 0.013679 | 0.004809 | 0.025483 | 0.008144 | 0.001269 | 0.00304 | 0.01992 | 0.005061 | 0.008708 | 0.016706 | 0.009987 | 0.007011 | 0.00842 | 0.003075 | 0.00444 | 0.017339 | 0.010241 | 0.027739 | 0.010119 |
| 22.2916667 | 0.010098 | 0.0159 | 0.00791 | 0.0072 | 0.015769 | 0.012827 | 0.005397 | 0.025944 | 0.007877 | 0.00153 | 0.004594 | 0.012032 | 0.002647 | 0.008682 | 0.01778 | 0.005685 | 0.006342 | 0.005835 | 0.003504 | 0.004912 | 0.016034 | 0.005619 | 0.015103 | 0.004619 |
| 22.3055556 | 0.010125 | 0.01643 | 0.007708 | 0.007602 | 0.010571 | 0.011042 | 0.00495 | 0.023821 | 0.00771 | 0.002961 | 0.005816 | 0.007455 | 0.00131 | 0.016045 | 0.015455 | 0.006117 | 0.006839 | 0.003133 | 0.006356 | 0.004035 | 0.012813 | 0.002848 | 0.007614 | 0.00855 |
| 22.3194444 | 0.009933 | 0.014684 | 0.006518 | 0.009378 | 0.009289 | 0.009537 | 0.002626 | 0.02025 | 0.007117 | 0.004605 | 0.006312 | 0.004928 | 0.001254 | 0.020222 | 0.010224 | 0.005389 | 0.007769 | 0.005715 | 0.006399 | 0.003679 | 0.009863 | 0.004151 | 0.01337 | 0.018684 |
| 22.3333333 | 0.007265 | 0.016868 | 0.00376 | 0.010353 | 0.013982 | 0.008446 | 0.00164 | 0.014169 | 0.004615 | 0.004838 | 0.005681 | 0.002921 | 0.00287 | 0.015252 | 0.006707 | 0.006602 | 0.007396 | 0.010321 | 0.004197 | 0.004583 | 0.008623 | 0.005184 | 0.024807 | 0.037078 |
| 22.3472222 | 0.005763 | 0.016151 | 0.002608 | 0.009574 | 0.020129 | 0.007024 | 0.001698 | 0.01481 | 0.002224 | 0.00298 | 0.003949 | 0.002018 | 0.004614 | 0.007534 | 0.007854 | 0.013851 | 0.008567 | 0.010456 | 0.005326 | 0.005541 | 0.008387 | 0.003176 | 0.034593 | 0.056434 |
| 22.3611111 | 0.007531 | 0.018278 | 0.003447 | 0.00817 | 0.018576 | 0.004664 | 0.003129 | 0.018188 | 0.002264 | 0.001145 | 0.002279 | 0.002096 | 0.004151 | 0.003462 | 0.01101 | 0.020729 | 0.010143 | 0.006083 | 0.005692 | 0.006073 | 0.007819 | 0.001181 | 0.039987 | 0.060258 |
| 22.375 | 0.007054 | 0.021299 | 0.003212 | 0.006835 | 0.010354 | 0.002167 | 0.00416 | 0.023832 | 0.002192 | 0.000589 | 0.001178 | 0.001808 | 0.002031 | 0.003306 | 0.011969 | 0.018274 | 0.006819 | 0.001928 | 0.00426 | 0.005999 | 0.006227 | 0.001669 | 0.036525 | 0.042242 |
| 22.3888889 | 0.005919 | 0.020697 | 0.002429 | 0.004812 | 0.004754 | 0.001477 | 0.003938 | 0.025161 | 0.001292 | 0.001323 | 0.00076 | 0.00237 | 0.001328 | 0.004513 | 0.010999 | 0.009334 | 0.002532 | 0.001336 | 0.005809 | 0.005475 | 0.004128 | 0.004207 | 0.025644 | 0.018434 |
| 22.4027778 | 0.005502 | 0.024566 | 0.004488 | 0.00416 | 0.005101 | 0.00305 | 0.005859 | 0.021333 | 0.002287 | 0.005351 | 0.000852 | 0.005696 | 0.003858 | 0.006 | 0.011159 | 0.003875 | 0.002545 | 0.003909 | 0.009364 | 0.00578 | 0.003799 | 0.006899 | 0.014301 | 0.004627 |
| 22.4166667 | 0.005642 | 0.028362 | 0.008725 | 0.006845 | 0.009639 | 0.004915 | 0.008688 | 0.022718 | 0.006063 | 0.012429 | 0.001301 | 0.010839 | 0.00836 | 0.005239 | 0.013485 | 0.006415 | 0.005213 | 0.006827 | 0.010466 | 0.007542 | 0.005291 | 0.007654 | 0.007265 | 0.001205 |
| 22.4305556 | 0.00996 | 0.023867 | 0.011707 | 0.010655 | 0.013358 | 0.006387 | 0.009275 | 0.025692 | 0.009742 | 0.018789 | 0.003408 | 0.016326 | 0.012609 | 0.00215 | 0.018511 | 0.017122 | 0.006681 | 0.008882 | 0.009347 | 0.010038 | 0.005423 | 0.006032 | 0.005909 | 0.002018 |
| 22.4444444 | 0.01305 | 0.015885 | 0.013539 | 0.016278 | 0.017096 | 0.009534 | 0.010014 | 0.023438 | 0.011686 | 0.022968 | 0.006808 | 0.022969 | 0.015254 | 0.004989 | 0.02723 | 0.03452 | 0.00569 | 0.012216 | 0.01035 | 0.012663 | 0.004713 | 0.004488 | 0.007397 | 0.002762 |
| 22.4583333 | 0.010925 | 0.012586 | 0.016164 | 0.02405 | 0.024543 | 0.015037 | 0.012967 | 0.017281 | 0.013266 | 0.025699 | 0.009668 | 0.031917 | 0.015781 | 0.011462 | 0.035935 | 0.048894 | 0.003376 | 0.018607 | 0.014246 | 0.014591 | 0.005124 | 0.004586 | 0.006734 | 0.003532 |
| 22.4722222 | 0.008461 | 0.013849 | 0.018833 | 0.030688 | 0.03128 | 0.020361 | 0.016686 | 0.016571 | 0.014992 | 0.02701 | 0.011762 | 0.042536 | 0.015193 | 0.009479 | 0.038479 | 0.04959 | 0.001412 | 0.02583 | 0.017558 | 0.01472 | 0.005343 | 0.00474 | 0.005212 | 0.008985 |
| 22.4861111 | 0.009826 | 0.01667 | 0.020545 | 0.035987 | 0.033669 | 0.023042 | 0.020064 | 0.027346 | 0.01677 | 0.027508 | 0.013465 | 0.051812 | 0.015853 | 0.005451 | 0.034881 | 0.041225 | 0.000742 | 0.0299 | 0.018372 | 0.013657 | 0.004312 | 0.004189 | 0.0112 | 0.026244 |
| 22.5 | 0.01549 | 0.020909 | 0.023045 | 0.04196 | 0.035135 | 0.024393 | 0.023065 | 0.043306 | 0.01925 | 0.029445 | 0.01449 | 0.05627 | 0.017964 | 0.013525 | 0.030409 | 0.034625 | 0.000531 | 0.030164 | 0.017882 | 0.013117 | 0.00316 | 0.003997 | 0.032117 | 0.054254 |
| 22.5138889 | 0.021359 | 0.023825 | 0.027052 | 0.047774 | 0.037071 | 0.026077 | 0.023783 | 0.052319 | 0.021633 | 0.03421 | 0.013823 | 0.054217 | 0.017905 | 0.026885 | 0.028107 | 0.031545 | 0.002095 | 0.029704 | 0.017069 | 0.012928 | 0.002314 | 0.004404 | 0.058414 | 0.074543 |
| 22.5277778 | 0.020015 | 0.019383 | 0.028585 | 0.048013 | 0.034353 | 0.025552 | 0.019873 | 0.04541 | 0.020336 | 0.037932 | 0.010524 | 0.044814 | 0.01238 | 0.027123 | 0.02522 | 0.025804 | 0.009794 | 0.028617 | 0.015754 | 0.011082 | 0.003191 | 0.004508 | 0.067509 | 0.069346 |
| 22.5416667 | 0.011403 | 0.009252 | 0.024552 | 0.039505 | 0.025188 | 0.021036 | 0.013768 | 0.026719 | 0.013511 | 0.033846 | 0.005896 | 0.031075 | 0.005579 | 0.014779 | 0.017867 | 0.014262 | 0.028662 | 0.023596 | 0.013671 | 0.00681 | 0.007854 | 0.003795 | 0.056324 | 0.044687 |
| 22.5555556 | 0.005438 | 0.00366 | 0.01873 | 0.029729 | 0.015653 | 0.014959 | 0.010048 | 0.018326 | 0.005668 | 0.02218 | 0.003213 | 0.021404 | 0.003535 | 0.008136 | 0.008344 | 0.005781 | 0.053185 | 0.01585 | 0.011778 | 0.003047 | 0.010302 | 0.003801 | 0.039438 | 0.019897 |
| 22.5694444 | 0.005109 | 0.00783 | 0.018909 | 0.033053 | 0.013685 | 0.012129 | 0.010153 | 0.024193 | 0.003164 | 0.013708 | 0.003658 | 0.023183 | 0.0073 | 0.019058 | 0.006357 | 0.011836 | 0.060442 | 0.010161 | 0.012401 | 0.004468 | 0.006847 | 0.007318 | 0.022716 | 0.010704 |
| 22.5833333 | 0.008989 | 0.019211 | 0.029929 | 0.054703 | 0.022898 | 0.016993 | 0.01552 | 0.02301 | 0.008646 | 0.021196 | 0.005702 | 0.037057 | 0.015616 | 0.043354 | 0.021152 | 0.031412 | 0.040348 | 0.008436 | 0.016862 | 0.012281 | 0.006021 | 0.015998 | 0.01131 | 0.015651 |
| 22.5972222 | 0.018562 | 0.032595 | 0.049217 | 0.081071 | 0.038201 | 0.029053 | 0.027825 | 0.027535 | 0.02116 | 0.045805 | 0.008194 | 0.049475 | 0.02287 | 0.071681 | 0.044596 | 0.053375 | 0.019507 | 0.009532 | 0.025004 | 0.021858 | 0.011848 | 0.027254 | 0.012984 | 0.016409 |
| 22.6111111 | 0.030508 | 0.041609 | 0.067571 | 0.094793 | 0.050025 | 0.041236 | 0.041166 | 0.055479 | 0.035539 | 0.073428 | 0.010466 | 0.04503 | 0.026857 | 0.099092 | 0.055616 | 0.068876 | 0.025066 | 0.012584 | 0.033902 | 0.0282 | 0.018724 | 0.034786 | 0.021217 | 0.014551 |
| 22.625 | 0.041576 | 0.04484 | 0.077674 | 0.095788 | 0.056652 | 0.045046 | 0.046352 | 0.081591 | 0.046408 | 0.090512 | 0.012399 | 0.032449 | 0.032693 | 0.114035 | 0.052513 | 0.077004 | 0.049734 | 0.017807 | 0.039586 | 0.032656 | 0.024273 | 0.036256 | 0.022408 | 0.028821 |
| 22.6388889 | 0.047073 | 0.046521 | 0.078523 | 0.09512 | 0.060426 | 0.039584 | 0.043364 | 0.084451 | 0.050851 | 0.096467 | 0.016133 | 0.030776 | 0.04063 | 0.100398 | 0.050495 | 0.075529 | 0.068825 | 0.024194 | 0.041155 | 0.038484 | 0.030176 | 0.035391 | 0.013856 | 0.050092 |
| 22.6527778 | 0.041064 | 0.048329 | 0.070935 | 0.092616 | 0.056938 | 0.032808 | 0.037479 | 0.065972 | 0.048762 | 0.090829 | 0.021844 | 0.041798 | 0.043568 | 0.063156 | 0.053 | 0.060359 | 0.066603 | 0.027778 | 0.03803 | 0.040414 | 0.034709 | 0.034073 | 0.009399 | 0.057145 |
| 22.6666667 | 0.027732 | 0.050968 | 0.058603 | 0.078381 | 0.04606 | 0.030287 | 0.030733 | 0.035868 | 0.042787 | 0.070053 | 0.02519 | 0.050799 | 0.03789 | 0.028444 | 0.054691 | 0.039221 | 0.047758 | 0.024618 | 0.02968 | 0.03303 | 0.034842 | 0.030216 | 0.017664 | 0.048688 |
| 22.6805556 | 0.016739 | 0.055095 | 0.045819 | 0.054035 | 0.037321 | 0.029426 | 0.023086 | 0.022581 | 0.034878 | 0.039127 | 0.024492 | 0.048434 | 0.02856 | 0.017461 | 0.058409 | 0.0226 | 0.02718 | 0.016787 | 0.01934 | 0.02318 | 0.030872 | 0.02405 | 0.024177 | 0.030194 |
| 22.6944444 | 0.014139 | 0.058515 | 0.034305 | 0.034054 | 0.037239 | 0.026656 | 0.014369 | 0.033677 | 0.02674 | 0.01704 | 0.022733 | 0.040275 | 0.022191 | 0.025501 | 0.066769 | 0.012243 | 0.012873 | 0.010871 | 0.012034 | 0.017041 | 0.025468 | 0.017668 | 0.02425 | 0.013303 |

| | | | | | | | | | | | | | | | | | | | | | | | | |
|---|---|---|---|---|---|---|---|---|---|---|---|---|---|---|---|---|---|---|---|---|---|---|---|---|
| 22.7083333 | 0.019455 | 0.059619 | 0.025321 | 0.026273 | 0.045333 | 0.022007 | 0.006182 | 0.030285 | 0.020443 | 0.015616 | 0.022359 | 0.033407 | 0.021075 | 0.032576 | 0.072354 | 0.009775 | 0.005955 | 0.011658 | 0.009021 | 0.013773 | 0.020997 | 0.011251 | 0.033578 | 0.011634 |
| 22.7222222 | 0.026292 | 0.058239 | 0.019593 | 0.02375 | 0.057827 | 0.01821 | 0.001719 | 0.016744 | 0.018726 | 0.016723 | 0.024918 | 0.031443 | 0.023776 | 0.03621 | 0.064731 | 0.020625 | 0.004226 | 0.019841 | 0.009761 | 0.012562 | 0.018685 | 0.005929 | 0.052447 | 0.018239 |
| 22.7361111 | 0.028199 | 0.056202 | 0.018183 | 0.01635 | 0.069205 | 0.01775 | 0.002686 | 0.01489 | 0.021591 | 0.009908 | 0.029068 | 0.032866 | 0.028064 | 0.041914 | 0.038766 | 0.043376 | 0.00425 | 0.028915 | 0.016798 | 0.013453 | 0.018065 | 0.004468 | 0.066136 | 0.025135 |
| 22.75 | 0.024655 | 0.055368 | 0.020476 | 0.006938 | 0.072721 | 0.020204 | 0.0078 | 0.010055 | 0.02437 | 0.007553 | 0.028841 | 0.031736 | 0.031566 | 0.039163 | 0.019317 | 0.065667 | 0.003989 | 0.029943 | 0.029401 | 0.016938 | 0.017831 | 0.006412 | 0.063543 | 0.031845 |
| 22.7638889 | 0.020681 | 0.052782 | 0.020944 | 0.002877 | 0.065638 | 0.023543 | 0.014222 | 0.004079 | 0.024166 | 0.011794 | 0.02208 | 0.02687 | 0.030327 | 0.025493 | 0.035008 | 0.068902 | 0.003098 | 0.022152 | 0.037087 | 0.018736 | 0.017796 | 0.007642 | 0.041393 | 0.029762 |
| 22.7777778 | 0.021098 | 0.047853 | 0.015226 | 0.00278 | 0.051061 | 0.025699 | 0.021101 | 0.00525 | 0.021852 | 0.014655 | 0.012631 | 0.020738 | 0.023708 | 0.023266 | 0.060244 | 0.050221 | 0.002451 | 0.013617 | 0.031561 | 0.014653 | 0.017588 | 0.005534 | 0.022285 | 0.01715 |
| 22.7916667 | 0.026928 | 0.039662 | 0.007695 | 0.003095 | 0.034959 | 0.026295 | 0.026857 | 0.00924 | 0.019333 | 0.011857 | 0.005538 | 0.015063 | 0.017977 | 0.035098 | 0.060695 | 0.03239 | 0.002834 | 0.011083 | 0.019703 | 0.008749 | 0.016126 | 0.002965 | 0.025167 | 0.008043 |
| 22.8055556 | 0.031993 | 0.023299 | 0.005866 | 0.00312 | 0.024187 | 0.028016 | 0.027523 | 0.00847 | 0.018653 | 0.006159 | 0.00388 | 0.010686 | 0.016639 | 0.034416 | 0.048561 | 0.02465 | 0.003675 | 0.012176 | 0.011241 | 0.004713 | 0.011547 | 0.003494 | 0.029256 | 0.010419 |
| 22.8194444 | 0.026492 | 0.010436 | 0.009426 | 0.00509 | 0.019538 | 0.032541 | 0.022465 | 0.011448 | 0.021063 | 0.002691 | 0.006074 | 0.006125 | 0.015606 | 0.018096 | 0.0446 | 0.018623 | 0.003775 | 0.009728 | 0.006611 | 0.003777 | 0.00618 | 0.005683 | 0.02208 | 0.014319 |
| 22.8333333 | 0.01424 | 0.013164 | 0.009374 | 0.00764 | 0.015314 | 0.036131 | 0.014689 | 0.037029 | 0.023443 | 0.004955 | 0.009342 | 0.003934 | 0.010876 | 0.007052 | 0.040244 | 0.012086 | 0.002792 | 0.006149 | 0.006094 | 0.006727 | 0.007485 | 0.007344 | 0.012961 | 0.016299 |
| 22.8472222 | 0.010544 | 0.02009 | 0.00547 | 0.00816 | 0.008494 | 0.032034 | 0.00691 | 0.05612 | 0.020627 | 0.015611 | 0.011489 | 0.00744 | 0.005001 | 0.011104 | 0.024717 | 0.006827 | 0.001442 | 0.009916 | 0.011593 | 0.010247 | 0.01653 | 0.008221 | 0.005741 | 0.022262 |
| 22.8611111 | 0.020105 | 0.021099 | 0.006275 | 0.008616 | 0.002559 | 0.021378 | 0.001924 | 0.040513 | 0.013836 | 0.026064 | 0.011518 | 0.011731 | 0.002209 | 0.021739 | 0.009075 | 0.006587 | 0.000658 | 0.014665 | 0.020397 | 0.011264 | 0.025291 | 0.00862 | 0.004132 | 0.025566 |
| 22.875 | 0.030906 | 0.018983 | 0.009525 | 0.011203 | 0.002332 | 0.014032 | 0.001791 | 0.023371 | 0.0091 | 0.025247 | 0.010721 | 0.015079 | 0.001407 | 0.022467 | 0.007236 | 0.012982 | 0.000822 | 0.012128 | 0.027928 | 0.010703 | 0.025792 | 0.008242 | 0.006709 | 0.018087 |
| 22.8888889 | 0.027604 | 0.019105 | 0.010647 | 0.015465 | 0.005994 | 0.013518 | 0.003787 | 0.019092 | 0.009339 | 0.018431 | 0.009681 | 0.020724 | 0.001764 | 0.016678 | 0.024369 | 0.01952 | 0.001288 | 0.00676 | 0.031351 | 0.008139 | 0.017775 | 0.006324 | 0.011565 | 0.007189 |
| 22.9027778 | 0.012665 | 0.023789 | 0.012089 | 0.01966 | 0.006518 | 0.014661 | 0.003792 | 0.018924 | 0.011358 | 0.015014 | 0.006495 | 0.026316 | 0.004811 | 0.02381 | 0.047368 | 0.023544 | 0.001414 | 0.005042 | 0.030893 | 0.004419 | 0.007939 | 0.003312 | 0.032623 | 0.003789 |
| 22.9166667 | 0.005595 | 0.026578 | 0.015897 | 0.023613 | 0.00412 | 0.011746 | 0.003071 | 0.020535 | 0.010027 | 0.015488 | 0.003294 | 0.02753 | 0.008444 | 0.037593 | 0.061258 | 0.033249 | 0.001444 | 0.00688 | 0.025883 | 0.003864 | 0.002972 | 0.001192 | 0.075943 | 0.006791 |
| 22.9305556 | 0.013881 | 0.019694 | 0.020336 | 0.027061 | 0.006399 | 0.006152 | 0.003498 | 0.021163 | 0.005597 | 0.014838 | 0.003415 | 0.025554 | 0.008053 | 0.039402 | 0.074028 | 0.044815 | 0.001444 | 0.007611 | 0.014838 | 0.00672 | 0.00334 | 0.001464 | 0.108527 | 0.008839 |
| 22.9444444 | 0.018933 | 0.008551 | 0.022048 | 0.028037 | 0.014333 | 0.003182 | 0.006258 | 0.037484 | 0.002451 | 0.009395 | 0.003122 | 0.023811 | 0.004178 | 0.031599 | 0.075691 | 0.04908 | 0.001105 | 0.006063 | 0.005787 | 0.010685 | 0.005669 | 0.003408 | 0.093575 | 0.012534 |
| 22.9583333 | 0.013863 | 0.006508 | 0.020766 | 0.027326 | 0.020494 | 0.002032 | 0.01196 | 0.05405 | 0.001663 | 0.00463 | 0.001274 | 0.024063 | 0.0022 | 0.026727 | 0.057537 | 0.047894 | 0.000558 | 0.004103 | 0.004532 | 0.01265 | 0.009186 | 0.006347 | 0.047781 | 0.023968 |
| 22.9722222 | 0.014805 | 0.017688 | 0.021535 | 0.029762 | 0.020266 | 0.002383 | 0.014224 | 0.045886 | 0.003 | 0.008018 | 0.00098 | 0.02641 | 0.004176 | 0.029054 | 0.036168 | 0.034978 | 0.000329 | 0.002457 | 0.005014 | 0.009455 | 0.013204 | 0.009065 | 0.021407 | 0.037037 |
| 22.9861111 | 0.018772 | 0.033043 | 0.025723 | 0.036592 | 0.016576 | 0.007927 | 0.009018 | 0.04692 | 0.008959 | 0.018065 | 0.004044 | 0.029621 | 0.007466 | 0.0324 | 0.018272 | 0.022152 | 0.000653 | 0.004232 | 0.00718 | 0.004875 | 0.015228 | 0.010037 | 0.0282 | 0.040103 |
| 23 | 0.020475 | 0.045835 | 0.028196 | 0.041878 | 0.015584 | 0.017923 | 0.004431 | 0.05973 | 0.017493 | 0.031733 | 0.009446 | 0.029469 | 0.009204 | 0.029089 | 0.013887 | 0.030984 | 0.000912 | 0.016074 | 0.014372 | 0.004127 | 0.01384 | 0.009232 | 0.0423 | 0.029929 |
| 23.0138889 | 0.028487 | 0.05805 | 0.02988 | 0.041775 | 0.021531 | 0.026075 | 0.005702 | 0.049247 | 0.022778 | 0.041542 | 0.011979 | 0.019848 | 0.009351 | 0.017572 | 0.016583 | 0.044641 | 0.000831 | 0.037732 | 0.025753 | 0.004557 | 0.010213 | 0.006846 | 0.051071 | 0.024335 |
| 23.0277778 | 0.040865 | 0.067898 | 0.034547 | 0.044442 | 0.03315 | 0.028322 | 0.007825 | 0.034796 | 0.027041 | 0.041727 | 0.010696 | 0.00727 | 0.009401 | 0.009709 | 0.012823 | 0.048192 | 0.001359 | 0.053579 | 0.03962 | 0.00434 | 0.007929 | 0.003733 | 0.053133 | 0.040306 |
| 23.0416667 | 0.055113 | 0.07243 | 0.042843 | 0.062553 | 0.05117 | 0.030571 | 0.013118 | 0.051659 | 0.040128 | 0.047034 | 0.012703 | 0.010442 | 0.011891 | 0.013458 | 0.014522 | 0.040101 | 0.001828 | 0.049951 | 0.053098 | 0.011824 | 0.012995 | 0.003142 | 0.051307 | 0.066731 |
| 23.0555556 | 0.071662 | 0.079124 | 0.05546 | 0.092946 | 0.077346 | 0.042834 | 0.027588 | 0.093157 | 0.061432 | 0.066031 | 0.022283 | 0.040531 | 0.019009 | 0.014213 | 0.027435 | 0.034374 | 0.004036 | 0.03909 | 0.063542 | 0.037732 | 0.024714 | 0.007354 | 0.054893 | 0.079325 |
| 23.0694444 | 0.077369 | 0.090132 | 0.065679 | 0.113914 | 0.093126 | 0.060069 | 0.045853 | 0.108758 | 0.072746 | 0.07857 | 0.031478 | 0.080389 | 0.029042 | 0.009565 | 0.043276 | 0.060063 | 0.009875 | 0.037364 | 0.068779 | 0.068329 | 0.032243 | 0.011741 | 0.057076 | 0.070903 |
| 23.0833333 | 0.064061 | 0.088148 | 0.060332 | 0.110166 | 0.085181 | 0.063081 | 0.058275 | 0.07274 | 0.064706 | 0.073577 | 0.031502 | 0.096604 | 0.037448 | 0.015153 | 0.053994 | 0.096056 | 0.01295 | 0.035868 | 0.063273 | 0.074958 | 0.030368 | 0.012313 | 0.04618 | 0.060309 |
| 23.0972222 | 0.046345 | 0.059132 | 0.042298 | 0.08696 | 0.074169 | 0.049838 | 0.063067 | 0.027272 | 0.04994 | 0.066657 | 0.024063 | 0.087249 | 0.04028 | 0.029618 | 0.054694 | 0.10558 | 0.011621 | 0.021973 | 0.045767 | 0.059216 | 0.023853 | 0.011849 | 0.027945 | 0.05384 |
| 23.1111111 | 0.037282 | 0.026343 | 0.027613 | 0.060526 | 0.067394 | 0.040303 | 0.065629 | 0.010443 | 0.041233 | 0.065466 | 0.01418 | 0.069247 | 0.036191 | 0.039472 | 0.046535 | 0.098067 | 0.009923 | 0.006992 | 0.028426 | 0.039782 | 0.018923 | 0.01102 | 0.019406 | 0.035684 |
| 23.125 | 0.034209 | 0.012718 | 0.020958 | 0.043046 | 0.086125 | 0.040232 | 0.063984 | 0.019898 | 0.037197 | 0.061481 | 0.00726 | 0.044923 | 0.026949 | 0.041753 | 0.033611 | 0.077625 | 0.006971 | 0.002146 | 0.018663 | 0.022716 | 0.01534 | 0.007439 | 0.03447 | 0.023541 |
| 23.1388889 | 0.028767 | 0.016711 | 0.017227 | 0.033858 | 0.075233 | 0.035667 | 0.050343 | 0.034757 | 0.029213 | 0.048832 | 0.006147 | 0.022374 | 0.016427 | 0.045093 | 0.019819 | 0.042635 | 0.00317 | 0.002887 | 0.014627 | 0.011417 | 0.010178 | 0.004694 | 0.061235 | 0.05143 |
| 23.1527778 | 0.019345 | 0.037901 | 0.012679 | 0.028197 | 0.0415 | 0.022801 | 0.026928 | 0.03555 | 0.018201 | 0.032088 | 0.00622 | 0.019442 | 0.008276 | 0.051178 | 0.010844 | 0.017054 | 0.001282 | 0.007228 | 0.014308 | 0.011003 | 0.006114 | 0.004779 | 0.070023 | 0.088356 |
| 23.1666667 | 0.013687 | 0.049033 | 0.00741 | 0.025545 | 0.014865 | 0.011779 | 0.010306 | 0.022974 | 0.014975 | 0.019976 | 0.00491 | 0.031437 | 0.003962 | 0.050086 | 0.012223 | 0.010927 | 0.001164 | 0.012184 | 0.01427 | 0.015821 | 0.006205 | 0.003268 | 0.054225 | 0.084097 |
| 23.1805556 | 0.015319 | 0.031024 | 0.004117 | 0.024728 | 0.021482 | 0.009175 | 0.012157 | 0.017128 | 0.020742 | 0.012146 | 0.004312 | 0.03479 | 0.002095 | 0.037296 | 0.025045 | 0.018818 | 0.001682 | 0.013145 | 0.011822 | 0.015228 | 0.007102 | 0.001394 | 0.028868 | 0.047716 |
| 23.1944444 | 0.019688 | 0.011415 | 0.003578 | 0.021846 | 0.063511 | 0.013071 | 0.020561 | 0.023168 | 0.025236 | 0.005228 | 0.00342 | 0.024403 | 0.001666 | 0.018871 | 0.03733 | 0.028511 | 0.003093 | 0.011006 | 0.007394 | 0.009324 | 0.005189 | 0.00133 | 0.009854 | 0.022638 |
| 23.2083333 | 0.022272 | 0.006427 | 0.002911 | 0.016181 | 0.097468 | 0.018095 | 0.022595 | 0.024975 | 0.022474 | 0.001462 | 0.002013 | 0.012371 | 0.002203 | 0.00824 | 0.042846 | 0.027083 | 0.004136 | 0.00821 | 0.004397 | 0.004149 | 0.004288 | 0.001046 | 0.004958 | 0.025325 |
| 23.2222222 | 0.019931 | 0.012493 | 0.002317 | 0.010689 | 0.078479 | 0.018419 | 0.016368 | 0.016886 | 0.015383 | 0.001078 | 0.001555 | 0.010797 | 0.002393 | 0.009872 | 0.043921 | 0.020045 | 0.003644 | 0.007081 | 0.005634 | 0.002906 | 0.007264 | 0.000535 | 0.009055 | 0.040654 |
| 23.2361111 | 0.012667 | 0.032145 | 0.004466 | 0.007116 | 0.032421 | 0.013916 | 0.01265 | 0.008233 | 0.009315 | 0.001492 | 0.002359 | 0.021054 | 0.003031 | 0.00953 | 0.037961 | 0.014088 | 0.002863 | 0.007638 | 0.008748 | 0.004676 | 0.010915 | 0.000496 | 0.015389 | 0.062404 |
| 23.25 | 0.006391 | 0.048742 | 0.007729 | 0.005089 | 0.01486 | 0.010371 | 0.023836 | 0.007144 | 0.0058 | 0.001806 | 0.002999 | 0.031226 | 0.004129 | 0.005977 | 0.03103 | 0.00812 | 0.002344 | 0.006766 | 0.0081 | 0.005922 | 0.01354 | 0.001406 | 0.019996 | 0.07779 |
| 23.2638889 | 0.006621 | 0.041418 | 0.009624 | 0.00312 | 0.029641 | 0.00828 | 0.03362 | 0.007662 | 0.003321 | 0.002823 | 0.002225 | 0.031907 | 0.004749 | 0.008216 | 0.020824 | 0.005673 | 0.00183 | 0.00397 | 0.005048 | 0.004607 | 0.016071 | 0.003029 | 0.019239 | 0.06906 |
| 23.2777778 | 0.010367 | 0.0211 | 0.008874 | 0.001741 | 0.035646 | 0.005507 | 0.026208 | 0.004157 | 0.00263 | 0.004168 | 0.001669 | 0.024674 | 0.005517 | 0.010748 | 0.013844 | 0.013342 | 0.001221 | 0.001991 | 0.004361 | 0.002627 | 0.017651 | 0.003083 | 0.014895 | 0.044153 |
| 23.2916667 | 0.009507 | 0.006923 | 0.005119 | 0.002499 | 0.027518 | 0.005583 | 0.011952 | 0.004043 | 0.005192 | 0.003914 | 0.001472 | 0.016655 | 0.005222 | 0.009471 | 0.024396 | 0.026143 | 0.000646 | 0.001508 | 0.004346 | 0.002197 | 0.016784 | 0.00262 | 0.010346 | 0.023366 |
| 23.3055556 | 0.004927 | 0.003112 | 0.001961 | 0.00394 | 0.027876 | 0.008714 | 0.004047 | 0.008302 | 0.009043 | 0.003932 | 0.00101 | 0.011398 | 0.003185 | 0.006184 | 0.027852 | 0.031364 | 0.000561 | 0.00165 | 0.003902 | 0.001978 | 0.014523 | 0.004586 | 0.006077 | 0.022142 |
| 23.3194444 | 0.00263 | 0.003193 | 0.002615 | 0.004223 | 0.02746 | 0.009534 | 0.004494 | 0.009253 | 0.010659 | 0.005258 | 0.001302 | 0.00774 | 0.001791 | 0.003135 | 0.013264 | 0.026607 | 0.000619 | 0.002675 | 0.003809 | 0.001492 | 0.012602 | 0.007652 | 0.005832 | 0.038239 |
| 23.3333333 | 0.002089 | 0.002086 | 0.005205 | 0.004023 | 0.027945 | 0.008327 | 0.00964 | 0.011662 | 0.009058 | 0.004315 | 0.002153 | 0.00452 | 0.001962 | 0.002534 | 0.005718 | 0.017995 | 0.000596 | 0.003795 | 0.004256 | 0.00252 | 0.010834 | 0.009482 | 0.012368 | 0.055526 |
| 23.3472222 | 0.002958 | 0.000739 | 0.007366 | 0.004294 | 0.027754 | 0.00803 | 0.016486 | 0.019865 | 0.006169 | 0.001585 | 0.002116 | 0.002686 | 0.002967 | 0.00213 | 0.008993 | 0.009899 | 0.001652 | 0.002972 | 0.004937 | 0.004153 | 0.008197 | 0.008779 | 0.020507 | 0.06156 |
| 23.3611111 | 0.006198 | 0.000543 | 0.008473 | 0.004294 | 0.025137 | 0.007653 | 0.019175 | 0.021052 | 0.003798 | 0.000848 | 0.001501 | 0.003242 | 0.004838 | 0.001009 | 0.012209 | 0.005458 | 0.004071 | 0.002584 | 0.004782 | 0.003998 | 0.005189 | 0.005531 | 0.025767 | 0.053376 |
| 23.375 | 0.009868 | 0.000568 | 0.00841 | 0.004047 | 0.038569 | 0.006596 | 0.014451 | 0.011604 | 0.002195 | 0.001743 | 0.002252 | 0.003482 | 0.006982 | 0.001175 | 0.012276 | 0.006763 | 0.005359 | 0.004171 | 0.005598 | 0.003257 | 0.002899 | 0.002312 | 0.026635 | 0.039355 |
| 23.3888889 | 0.01094 | 0.001737 | 0.008144 | 0.004774 | 0.03876 | 0.006891 | 0.007655 | 0.004148 | 0.002513 | 0.002342 | 0.004986 | 0.003653 | 0.008577 | 0.002458 | 0.008399 | 0.01267 | 0.003954 | 0.004102 | 0.008179 | 0.004435 | 0.002669 | 0.001968 | 0.022742 | 0.027636 |
| 23.4027778 | 0.009258 | 0.007413 | 0.009205 | 0.006551 | 0.021206 | 0.009422 | 0.00448 | 0.006316 | 0.006251 | 0.003999 | 0.008259 | 0.008186 | 0.010198 | 0.002665 | 0.006161 | 0.019751 | 0.002872 | 0.004022 | 0.011416 | 0.00656 | 0.006199 | 0.004219 | 0.016679 | 0.018541 |
| 23.4166667 | 0.006949 | 0.017164 | 0.011301 | 0.009159 | 0.018662 | 0.012516 | 0.006471 | 0.016491 | 0.011205 | 0.007356 | 0.009578 | 0.014988 | 0.012222 | 0.002095 | 0.011737 | 0.024068 | 0.00411 | 0.00737 | 0.015175 | 0.008152 | 0.011679 | 0.00608 | 0.010568 | 0.011493 |
| 23.4305556 | 0.006849 | 0.02349 | 0.012515 | 0.013462 | 0.018174 | 0.01483 | 0.012896 | 0.031064 | 0.014411 | 0.010903 | 0.007503 | 0.020059 | 0.013384 | 0.003018 | 0.022006 | 0.024239 | 0.005348 | 0.010736 | 0.017202 | 0.008985 | 0.015442 | 0.006106 | 0.004995 | 0.007115 |
| 23.4444444 | 0.010894 | 0.022228 | 0.012638 | 0.019518 | 0.017293 | 0.016319 | 0.021873 | 0.044884 | 0.01644 | 0.014273 | 0.003632 | 0.024209 | 0.012914 | 0.004302 | 0.030978 | 0.024468 | 0.005511 | 0.01192 | 0.016579 | 0.009049 | 0.016996 | 0.005317 | 0.001713 | 0.005243 |
| 23.4583333 | 0.016314 | 0.01887 | 0.013227 | 0.025058 | 0.03189 | 0.017897 | 0.030946 | 0.054126 | 0.019291 | 0.017291 | 0.001465 | 0.029358 | 0.012073 | 0.00518 | 0.036427 | 0.029646 | 0.005584 | 0.013637 | 0.016307 | 0.009179 | 0.018007 | 0.004939 | 0.001405 | 0.005936 |
| 23.4722222 | 0.019186 | 0.017869 | 0.015477 | 0.028392 | 0.043998 | 0.02086 | 0.038814 | 0.059282 | 0.023435 | 0.019482 | 0.003149 | 0.03522 | 0.012433 | 0.00626 | 0.039626 | 0.0362 | 0.005951 | 0.017763 | 0.018593 | 0.01013 | 0.019307 | 0.005328 | 0.004041 | 0.01205 |
| 23.4861111 | 0.019681 | 0.017091 | 0.019209 | 0.030703 | 0.040967 | 0.025328 | 0.043417 | 0.060067 | 0.027366 | 0.021866 | 0.008111 | 0.039752 | 0.01367 | 0.008545 | 0.041594 | 0.038032 | 0.006107 | 0.022518 | 0.022979 | 0.011184 | 0.021069 | 0.005704 | 0.010775 | 0.025359 |
| 23.5 | 0.021181 | 0.01412 | 0.023201 | 0.033449 | 0.037087 | 0.030151 | 0.041521 | 0.056956 | 0.029148 | 0.026207 | 0.013806 | 0.042916 | 0.013908 | 0.016141 | 0.041433 | 0.037369 | 0.005025 | 0.024974 | 0.028125 | 0.011786 | 0.023383 | 0.005396 | 0.018964 | 0.040946 |
| 23.5138889 | 0.024359 | 0.009551 | 0.025618 | 0.036519 | 0.039848 | 0.033781 | 0.034582 | 0.049924 | 0.02809 | 0.031256 | 0.016833 | 0.045174 | 0.011883 | 0.026225 | 0.038996 | 0.036669 | 0.002812 | 0.024536 | 0.031933 | 0.012181 | 0.024564 | 0.004717 | 0.024722 | 0.052827 |
| 23.5277778 | 0.025502 | 0.005469 | 0.023719 | 0.039117 | 0.042919 | 0.033353 | 0.02474 | 0.036391 | 0.023288 | 0.032462 | 0.015294 | 0.044755 | 0.009642 | 0.029946 | 0.034725 | 0.032851 | 0.005277 | 0.021594 | 0.031317 | 0.011851 | 0.021656 | 0.003647 | 0.026778 | 0.056942 |
| 23.5416667 | 0.024568 | 0.005474 | 0.017422 | 0.040122 | 0.041316 | 0.025859 | 0.012808 | 0.019094 | 0.014083 | 0.028146 | 0.011173 | 0.041008 | 0.016644 | 0.027683 | 0.027459 | 0.024513 | 0.030563 | 0.01679 | 0.026166 | 0.008411 | 0.013829 | 0.002256 | 0.024717 | 0.046203 |
| 23.5555556 | 0.026573 | 0.009052 | 0.011864 | 0.040917 | 0.039465 | 0.015279 | 0.004023 | 0.006358 | 0.005369 | 0.022452 | 0.009715 | 0.036641 | 0.031259 | 0.022434 | 0.01697 | 0.01649 | 0.091632 | 0.013244 | 0.0208 | 0.007313 | 0.006214 | 0.002455 | 0.019096 | 0.024323 |
| 23.5694444 | 0.031327 | 0.009118 | 0.014461 | 0.046061 | 0.043642 | 0.010768 | 0.003592 | 0.007266 | 0.002372 | 0.021169 | 0.013501 | 0.034753 | 0.03198 | 0.015751 | 0.009625 | 0.020557 | 0.145475 | 0.014357 | 0.02021 | 0.010945 | 0.002947 | 0.005607 | 0.012928 | 0.01173 |
| 23.5833333 | 0.031466 | 0.00667 | 0.026169 | 0.059075 | 0.054762 | 0.016708 | 0.012758 | 0.023932 | 0.007204 | 0.02966 | 0.01965 | 0.0358 | 0.018323 | 0.01695 | 0.013883 | 0.044789 | 0.127549 | 0.02038 | 0.02655 | 0.018993 | 0.002871 | 0.009715 | 0.01158 | 0.010683 |
| 23.5972222 | 0.025411 | 0.009653 | 0.03901 | 0.074738 | 0.064106 | 0.032696 | 0.030985 | 0.047986 | 0.022947 | 0.04806 | 0.02551 | 0.038859 | 0.01358 | 0.0343 | 0.028246 | 0.076234 | 0.065207 | 0.030078 | 0.039046 | 0.029586 | 0.005322 | 0.012779 | 0.017535 | 0.013269 |
| 23.6111111 | 0.024825 | 0.019687 | 0.047704 | 0.081334 | 0.065605 | 0.05274 | 0.055907 | 0.066139 | 0.045042 | 0.067837 | 0.030435 | 0.041217 | 0.022047 | 0.061572 | 0.042875 | 0.091609 | 0.046727 | 0.039366 | 0.053387 | 0.038804 | 0.007266 | 0.015368 | 0.028291 | 0.03181 |
| 23.625 | 0.035021 | 0.034365 | 0.051357 | 0.076757 | 0.063034 | 0.06332 | 0.077646 | 0.067701 | 0.060004 | 0.083081 | 0.030251 | 0.038852 | 0.040002 | 0.079658 | 0.050998 | 0.089829 | 0.088079 | 0.042512 | 0.061695 | 0.042657 | 0.007327 | 0.017637 | 0.03861 | 0.063718 |
| 23.6388889 | 0.045805 | 0.044791 | 0.047981 | 0.064925 | 0.055166 | 0.059007 | 0.078595 | 0.049777 | 0.059937 | 0.091606 | 0.02128 | 0.031508 | 0.0514 | 0.074015 | 0.051368 | 0.083715 | 0.125694 | 0.038355 | 0.059392 | 0.038973 | 0.007159 | 0.018098 | 0.044753 | 0.080336 |
| 23.6527778 | 0.039876 | 0.042946 | 0.037931 | 0.046333 | 0.038461 | 0.051573 | 0.058851 | 0.025187 | 0.049293 | 0.087722 | 0.009367 | 0.024067 | 0.052009 | 0.050787 | 0.046041 | 0.070402 | 0.117231 | 0.02861 | 0.048705 | 0.028346 | 0.007113 | 0.015981 | 0.04664 | 0.06615 |
| 23.6666667 | 0.023214 | 0.037282 | 0.025917 | 0.024827 | 0.02578 | 0.050987 | 0.041823 | 0.015794 | 0.036782 | 0.071532 | 0.003623 | 0.0223 | 0.043899 | 0.026314 | 0.038697 | 0.04899 | 0.079343 | 0.015401 | 0.034685 | 0.016151 | 0.009952 | 0.012646 | 0.044347 | 0.037374 |
| 23.6805556 | 0.026621 | 0.038438 | 0.016901 | 0.009794 | 0.035198 | 0.051089 | 0.035609 | 0.01974 | 0.026967 | 0.049848 | 0.00654 | 0.027744 | 0.032185 | 0.009713 | 0.030302 | 0.032433 | 0.040479 | 0.007645 | 0.022078 | 0.008394 | 0.017988 | 0.010443 | 0.03778 | 0.01615 |
| 23.6944444 | 0.038553 | 0.042794 | 0.012845 | 0.0041 | 0.051879 | 0.044439 | 0.026836 | 0.020597 | 0.020378 | 0.02727 | 0.014056 | 0.038006 | 0.022146 | 0.002863 | 0.021668 | 0.021679 | 0.013926 | 0.010673 | 0.012614 | 0.007224 | 0.027967 | 0.00916 | 0.028481 | 0.006388 |
| 23.7083333 | 0.028722 | 0.044991 | 0.011914 | 0.005572 | 0.051883 | 0.033748 | 0.013575 | 0.016781 | 0.015882 | 0.011834 | 0.020909 | 0.045664 | 0.017747 | 0.003577 | 0.017789 | 0.011146 | 0.003131 | 0.011918 | 0.007079 | 0.00987 | 0.033293 | 0.006574 | 0.018459 | 0.003696 |
| 23.7222222 | 0.012131 | 0.048429 | 0.012328 | 0.016232 | 0.042236 | 0.024055 | 0.008613 | 0.01586 | 0.011956 | 0.009239 | 0.025331 | 0.044803 | 0.017887 | 0.005548 | 0.020482 | 0.006605 | 0.002417 | 0.013996 | 0.00754 | 0.012466 | 0.031458 | 0.003337 | 0.010449 | 0.00744 |
| 23.7361111 | 0.014915 | 0.054727 | 0.014834 | 0.030619 | 0.038815 | 0.020576 | 0.018688 | 0.022639 | 0.008639 | 0.008618 | 0.030526 | 0.043109 | 0.020492 | 0.00566 | 0.02485 | 0.014237 | 0.003704 | 0.027952 | 0.011148 | 0.015175 | 0.026592 | 0.001444 | 0.00481 | 0.022004 |
| 23.75 | 0.02735 | 0.053758 | 0.020536 | 0.040169 | 0.046225 | 0.023524 | 0.035496 | 0.03485 | 0.00803 | 0.006548 | 0.03908 | 0.048339 | 0.024312 | 0.009589 | 0.028335 | 0.02883 | 0.004364 | 0.037333 | 0.012156 | 0.019797 | 0.022292 | 0.000898 | 0.003419 | 0.046623 |
| 23.7638889 | 0.023353 | 0.037964 | 0.029448 | 0.046358 | 0.057162 | 0.026149 | 0.041868 | 0.037828 | 0.011437 | 0.012196 | 0.047172 | 0.054939 | 0.027478 | 0.02064 | 0.03399 | 0.041138 | 0.003457 | 0.033847 | 0.011269 | 0.023139 | 0.018272 | 0.001978 | 0.008623 | 0.063415 |
| 23.7777778 | 0.012909 | 0.018717 | 0.037441 | 0.05343 | 0.059428 | 0.025685 | 0.030148 | 0.022991 | 0.017006 | 0.018944 | 0.046699 | 0.052126 | 0.027805 | 0.032615 | 0.037437 | 0.045377 | 0.002421 | 0.028387 | 0.009423 | 0.019638 | 0.015509 | 0.004492 | 0.018161 | 0.058784 |
| 23.7916667 | 0.012796 | 0.010466 | 0.037951 | 0.058255 | 0.04819 | 0.026995 | 0.012703 | 0.012751 | 0.020956 | 0.016506 | 0.034014 | 0.036453 | 0.02396 | 0.039589 | 0.02872 | 0.040043 | 0.002888 | 0.023926 | 0.007571 | 0.011275 | 0.016332 | 0.005098 | 0.024323 | 0.044876 |
| 23.8055556 | 0.015378 | 0.009138 | 0.030547 | 0.054473 | 0.029935 | 0.035032 | 0.00391 | 0.020273 | 0.02154 | 0.0087 | 0.016667 | 0.017157 | 0.015799 | 0.042274 | 0.01507 | 0.031622 | 0.00264 | 0.015679 | 0.01128 | 0.004984 | 0.018775 | 0.003432 | 0.022692 | 0.033623 |
| 23.8194444 | 0.024761 | 0.009387 | 0.019436 | 0.042423 | 0.019849 | 0.043038 | 0.003447 | 0.025929 | 0.019785 | 0.004307 | 0.005687 | 0.005418 | 0.00661 | 0.041733 | 0.012711 | 0.025424 | 0.002267 | 0.010312 | 0.017434 | 0.004944 | 0.016249 | 0.001994 | 0.018195 | 0.024981 |
| 23.8333333 | 0.038991 | 0.018238 | 0.008936 | 0.0267 | 0.023996 | 0.039686 | 0.005643 | 0.021697 | 0.017258 | 0.008412 | 0.004409 | 0.003671 | 0.002052 | 0.033674 | 0.019768 | 0.020198 | 0.002924 | 0.018695 | 0.019783 | 0.008726 | 0.012421 | 0.002201 | 0.018617 | 0.0179 |
| 23.8472222 | 0.043016 | 0.026783 | 0.005077 | 0.013286 | 0.028501 | 0.025169 | 0.007354 | 0.017695 | 0.015887 | 0.019545 | 0.006862 | 0.007014 | 0.001945 | 0.017976 | 0.022924 | 0.013989 | 0.003109 | 0.026973 | 0.021301 | 0.009635 | 0.021572 | 0.005075 | 0.021824 | 0.012629 |
| 23.8611111 | 0.040234 | 0.02141 | 0.006817 | 0.011195 | 0.022679 | 0.009935 | 0.00678 | 0.019722 | 0.015754 | 0.028398 | 0.006866 | 0.009484 | 0.003745 | 0.007592 | 0.020267 | 0.006926 | 0.003639 | 0.021897 | 0.024631 | 0.006881 | 0.036012 | 0.006194 | 0.017755 | 0.009271 |
| 23.875 | 0.041014 | 0.010933 | 0.007044 | 0.02094 | 0.013057 | 0.002967 | 0.004129 | 0.023183 | 0.013273 | 0.026769 | 0.004919 | 0.007927 | 0.007858 | 0.010668 | 0.014911 | 0.003899 | 0.005757 | 0.011574 | 0.027126 | 0.004986 | 0.034607 | 0.00352 | 0.010352 | 0.007062 |
| 23.8888889 | 0.046175 | 0.011016 | 0.005589 | 0.031444 | 0.009139 | 0.002462 | 0.002591 | 0.017278 | 0.007353 | 0.019174 | 0.00293 | 0.00447 | 0.012747 | 0.018469 | 0.010267 | 0.007529 | 0.00776 | 0.004136 | 0.024076 | 0.005819 | 0.020225 | 0.001992 | 0.013036 | 0.00604 |
| 23.9027778 | 0.048356 | 0.014122 | 0.006494 | 0.037371 | 0.009112 | 0.003126 | 0.003028 | 0.007358 | 0.003248 | 0.015574 | 0.001283 | 0.004123 | 0.014728 | 0.024484 | 0.013135 | 0.012566 | 0.007754 | 0.001087 | 0.013895 | 0.006724 | 0.008054 | 0.002991 | 0.023195 | 0.004789 |
| 23.9166667 | 0.03701 | 0.023649 | 0.011362 | 0.041355 | 0.00603 | 0.005084 | 0.002656 | 0.003702 | 0.005389 | 0.019072 | 0.001066 | 0.005862 | 0.012708 | 0.029936 | 0.019318 | 0.014371 | 0.005975 | 0.002072 | 0.005456 | 0.006516 | 0.003268 | 0.003739 | 0.030756 | 0.003645 |
| 23.9305556 | 0.017808 | 0.037927 | 0.017473 | 0.046436 | 0.002293 | 0.009335 | 0.004144 | 0.002356 | 0.010631 | 0.023661 | 0.002997 | 0.004774 | 0.009252 | 0.03294 | 0.02236 | 0.014955 | 0.003814 | 0.006072 | 0.002203 | 0.00885 | 0.005685 | 0.002774 | 0.032061 | 0.005921 |
| 23.9444444 | 0.009685 | 0.034322 | 0.018695 | 0.051223 | 0.001598 | 0.012417 | 0.010897 | 0.00508 | 0.012903 | 0.019954 | 0.006787 | 0.003236 | 0.008126 | 0.030718 | 0.020385 | 0.027132 | 0.002915 | 0.008298 | 0.002444 | 0.011139 | 0.014247 | 0.001475 | 0.031563 | 0.010718 |
| 23.9583333 | 0.014765 | 0.015578 | 0.014388 | 0.05131 | 0.001505 | 0.010553 | 0.017514 | 0.015772 | 0.010518 | 0.011053 | 0.011394 | 0.004818 | 0.010963 | 0.026641 | 0.014272 | 0.052113 | 0.00413 | 0.006678 | 0.002208 | 0.007594 | 0.024064 | 0.000845 | 0.035556 | 0.019231 |
| 23.9722222 | 0.021458 | 0.003086 | 0.011157 | 0.049178 | 0.001841 | 0.005899 | 0.015384 | 0.026241 | 0.005818 | 0.007871 | 0.014453 | 0.005126 | 0.015758 | 0.023933 | 0.017062 | 0.068674 | 0.006624 | 0.007988 | 0.001706 | 0.002471 | 0.029604 | 0.001377 | 0.035891 | 0.027218 |
| 23.9861111 | 0.027625 | 0.00132 | 0.012055 | 0.049602 | 0.003538 | 0.004968 | 0.008805 | 0.024653 | 0.002196 | 0.007292 | 0.013603 | 0.00357 | 0.019914 | 0.017538 | 0.034239 | 0.067011 | 0.00937 | 0.012027 | 0.001352 | 0.000644 | 0.029096 | 0.003616 | 0.023254 | 0.024689 |
| 24 | 0.036957 | 0.002661 | 0.013828 | 0.048762 | 0.011325 | 0.010136 | 0.006803 | 0.016036 | 0.001892 | 0.004399 | 0.009098 | 0.006264 | 0.02382 | 0.010324 | 0.049078 | 0.056753 | 0.012079 | 0.010262 | 0.003296 | 0.001478 | 0.023514 | 0.006314 | 0.008729 | 0.021013 |
| 24.0138889 | 0.04532 | 0.008952 | 0.012523 | 0.042697 | 0.030527 | 0.015355 | 0.006157 | 0.019641 | 0.005186 | 0.004752 | 0.004705 | 0.015173 | 0.028322 | 0.011277 | 0.049594 | 0.050647 | 0.014932 | 0.007873 | 0.00892 | 0.004681 | 0.014038 | 0.006697 | 0.002983 | 0.02285 |
| 24.0277778 | 0.043815 | 0.027789 | 0.008011 | 0.039909 | 0.044585 | 0.013868 | 0.006849 | 0.038728 | 0.012307 | 0.013824 | 0.005676 | 0.02818 | 0.033712 | 0.012462 | 0.044769 | 0.049056 | 0.022474 | 0.016453 | 0.017545 | 0.009709 | 0.007437 | 0.004765 | 0.002576 | 0.0232 |
| 24.0416667 | 0.036534 | 0.051306 | 0.005454 | 0.052752 | 0.041544 | 0.013913 | 0.019891 | 0.060566 | 0.026213 | 0.039224 | 0.016199 | 0.044332 | 0.04416 | 0.010896 | 0.043868 | 0.04864 | 0.050099 | 0.04333 | 0.033962 | 0.017278 | 0.012469 | 0.00334 | 0.007082 | 0.030608 |
| 24.0555556 | 0.045737 | 0.067003 | 0.009959 | 0.084043 | 0.046967 | 0.028638 | 0.045506 | 0.081364 | 0.046378 | 0.082893 | 0.035727 | 0.06257 | 0.064495 | 0.01735 | 0.042862 | 0.056496 | 0.107663 | 0.076051 | 0.056652 | 0.031945 | 0.02581 | 0.002606 | 0.019804 | 0.053199 |
| 24.0694444 | 0.064785 | 0.074216 | 0.019159 | 0.10821 | 0.072878 | 0.04256 | 0.060959 | 0.095091 | 0.061446 | 0.112424 | 0.046582 | 0.078864 | 0.085057 | 0.028525 | 0.038098 | 0.06479 | 0.168972 | 0.077047 | 0.063741 | 0.05123 | 0.035581 | 0.002805 | 0.034123 | 0.069534 |
| 24.0833333 | 0.06014 | 0.075902 | 0.025853 | 0.097118 | 0.09885 | 0.036579 | 0.05272 | 0.086265 | 0.060515 | 0.100603 | 0.037295 | 0.083336 | 0.084534 | 0.032937 | 0.030757 | 0.052743 | 0.206394 | 0.045878 | 0.049076 | 0.057035 | 0.036595 | 0.005405 | 0.038111 | 0.063257 |
| 24.0972222 | 0.035826 | 0.066936 | 0.028915 | 0.068952 | 0.111362 | 0.021341 | 0.040373 | 0.059633 | 0.045656 | 0.072195 | 0.025899 | 0.069319 | 0.061168 | 0.030204 | 0.021256 | 0.029694 | 0.205859 | 0.020131 | 0.032347 | 0.041688 | 0.033989 | 0.006458 | 0.034565 | 0.048157 |
| 24.1111111 | 0.020506 | 0.043828 | 0.033497 | 0.049628 | 0.10123 | 0.011404 | 0.042993 | 0.029849 | 0.028038 | 0.051027 | 0.022636 | 0.046781 | 0.036249 | 0.02757 | 0.012186 | 0.019737 | 0.150545 | 0.011681 | 0.022905 | 0.024031 | 0.030894 | 0.004652 | 0.032914 | 0.034222 |
| 24.125 | 0.02019 | 0.027915 | 0.03956 | 0.037369 | 0.072623 | 0.009641 | 0.051467 | 0.012171 | 0.016167 | 0.035899 | 0.0223 | 0.025893 | 0.023886 | 0.027367 | 0.005808 | 0.020587 | 0.072039 | 0.009107 | 0.016531 | 0.017663 | 0.025385 | 0.002998 | 0.029921 | 0.022823 |
| 24.1388889 | 0.018467 | 0.029235 | 0.039031 | 0.026501 | 0.050072 | 0.013387 | 0.054914 | 0.01832 | 0.011378 | 0.025168 | 0.019409 | 0.013629 | 0.018973 | 0.029088 | 0.002869 | 0.01845 | 0.027651 | 0.005624 | 0.010042 | 0.015207 | 0.016113 | 0.00289 | 0.02072 | 0.016134 |
| 24.1527778 | 0.013469 | 0.025508 | 0.02872 | 0.020904 | 0.03904 | 0.017367 | 0.057003 | 0.035687 | 0.012535 | 0.017795 | 0.011641 | 0.020013 | 0.012568 | 0.033287 | 0.004179 | 0.011245 | 0.028637 | 0.002956 | 0.004168 | 0.010074 | 0.012009 | 0.002009 | 0.009765 | 0.010207 |
| 24.1666667 | 0.01743 | 0.013515 | 0.015929 | 0.021257 | 0.024562 | 0.016698 | 0.058608 | 0.04267 | 0.018945 | 0.013303 | 0.006312 | 0.03469 | 0.00577 | 0.03488 | 0.006192 | 0.005532 | 0.041989 | 0.004277 | 0.001361 | 0.00639 | 0.002998 | 0.000842 | 0.002801 | 0.005906 |
| 24.1805556 | 0.019542 | 0.005245 | 0.006847 | 0.018558 | 0.009463 | 0.011383 | 0.052651 | 0.031853 | 0.026852 | 0.010179 | 0.004948 | 0.035884 | 0.004233 | 0.027762 | 0.005278 | 0.007087 | 0.039763 | 0.008586 | 0.002519 | 0.007544 | 0.002122 | 0.001181 | 0.00146 | 0.006442 |
| 24.1944444 | 0.010884 | 0.021982 | 0.004006 | 0.011395 | 0.005159 | 0.00591 | 0.033277 | 0.022597 | 0.025802 | 0.009816 | 0.002967 | 0.026606 | 0.007355 | 0.015736 | 0.003135 | 0.014558 | 0.024227 | 0.012711 | 0.005808 | 0.010167 | 0.00163 | 0.00189 | 0.002446 | 0.006073 |
| 24.2083333 | 0.014885 | 0.050767 | 0.015127 | 0.017744 | 0.019834 | 0.010444 | 0.01114 | 0.031305 | 0.018567 | 0.012777 | 0.010616 | 0.023758 | 0.010117 | 0.007938 | 0.002221 | 0.021775 | 0.020797 | 0.015446 | 0.009266 | 0.011555 | 0.002177 | 0.001419 | 0.002589 | 0.006405 |

| | | | | | | | | | | | | | | | | | | | | | | | | |
|---|---|---|---|---|---|---|---|---|---|---|---|---|---|---|---|---|---|---|---|---|---|---|---|---|
| 24.2222222 | 0.064691 | 0.088522 | 0.057742 | 0.065777 | 0.070685 | 0.017693 | 0.011104 | 0.032756 | 0.024845 | 0.043273 | 0.02529 | 0.029869 | 0.010189 | 0.006697 | 0.002905 | 0.023024 | 0.04124 | 0.016643 | 0.012414 | 0.014667 | 0.002246 | 0.00053 | 0.002746 | 0.014293 |
| 24.2361111 | 0.10484 | 0.081093 | 0.096469 | 0.096033 | 0.130894 | 0.020842 | 0.029199 | 0.033699 | 0.034648 | 0.083526 | 0.038905 | 0.031431 | 0.008274 | 0.007779 | 0.005681 | 0.016364 | 0.049617 | 0.014837 | 0.013461 | 0.019047 | 0.001878 | 0.000577 | 0.003855 | 0.026117 |
| 24.25 | 0.073218 | 0.049768 | 0.07673 | 0.06653 | 0.116093 | 0.023028 | 0.029779 | 0.041486 | 0.031122 | 0.068135 | 0.04301 | 0.029732 | 0.005636 | 0.008034 | 0.00999 | 0.008085 | 0.027705 | 0.009361 | 0.011033 | 0.016708 | 0.002912 | 0.00168 | 0.006506 | 0.036325 |
| 24.2638889 | 0.055595 | 0.040547 | 0.040243 | 0.062738 | 0.065529 | 0.019555 | 0.016172 | 0.037696 | 0.033522 | 0.042271 | 0.030401 | 0.050716 | 0.00285 | 0.006558 | 0.01197 | 0.007444 | 0.006905 | 0.003816 | 0.007108 | 0.008549 | 0.003423 | 0.002853 | 0.007915 | 0.04089 |
| 24.2777778 | 0.096266 | 0.057532 | 0.027071 | 0.078561 | 0.098209 | 0.022474 | 0.017317 | 0.052951 | 0.042006 | 0.052837 | 0.033789 | 0.085857 | 0.001101 | 0.003762 | 0.009709 | 0.015454 | 0.002348 | 0.001535 | 0.004211 | 0.003319 | 0.002993 | 0.003559 | 0.005791 | 0.036109 |
| 24.2916667 | 0.12755 | 0.075334 | 0.01883 | 0.060065 | 0.160891 | 0.025391 | 0.029031 | 0.070297 | 0.039448 | 0.061162 | 0.051817 | 0.095117 | 0.002493 | 0.002073 | 0.006375 | 0.026948 | 0.002888 | 0.00094 | 0.00367 | 0.003164 | 0.004032 | 0.004172 | 0.003487 | 0.024285 |
| 24.3055556 | 0.113762 | 0.0716 | 0.010236 | 0.030159 | 0.160818 | 0.018328 | 0.038986 | 0.057373 | 0.033964 | 0.055736 | 0.051046 | 0.075666 | 0.006172 | 0.004989 | 0.005551 | 0.036148 | 0.00331 | 0.00065 | 0.005256 | 0.006722 | 0.00619 | 0.004544 | 0.00236 | 0.011867 |
| 24.3194444 | 0.076281 | 0.051169 | 0.005467 | 0.013051 | 0.120687 | 0.011865 | 0.043884 | 0.037292 | 0.032025 | 0.047368 | 0.035894 | 0.051429 | 0.007542 | 0.012025 | 0.007835 | 0.035502 | 0.005546 | 0.001124 | 0.00715 | 0.010694 | 0.006383 | 0.00479 | 0.002561 | 0.006971 |
| 24.3333333 | 0.047903 | 0.030359 | 0.005158 | 0.010236 | 0.08406 | 0.009247 | 0.042369 | 0.024958 | 0.030439 | 0.039351 | 0.022658 | 0.032807 | 0.004678 | 0.01647 | 0.011142 | 0.024323 | 0.00915 | 0.002397 | 0.007448 | 0.012661 | 0.004386 | 0.005825 | 0.007186 | 0.013548 |
| 24.3472222 | 0.031899 | 0.016162 | 0.008894 | 0.011686 | 0.061693 | 0.011062 | 0.03842 | 0.021391 | 0.030081 | 0.032993 | 0.016517 | 0.023554 | 0.002068 | 0.013902 | 0.013798 | 0.01502 | 0.012798 | 0.004516 | 0.005028 | 0.012293 | 0.003983 | 0.007404 | 0.015403 | 0.025073 |
| 24.3611111 | 0.02129 | 0.011017 | 0.013405 | 0.013411 | 0.048593 | 0.015297 | 0.035494 | 0.022425 | 0.031011 | 0.028502 | 0.014974 | 0.023732 | 0.003388 | 0.008553 | 0.015802 | 0.016434 | 0.016868 | 0.00689 | 0.002389 | 0.008776 | 0.005256 | 0.008196 | 0.022192 | 0.031817 |
| 24.375 | 0.015124 | 0.010453 | 0.014013 | 0.015388 | 0.038583 | 0.01821 | 0.031514 | 0.022707 | 0.02909 | 0.0236 | 0.014299 | 0.024386 | 0.006439 | 0.005741 | 0.01583 | 0.022803 | 0.018316 | 0.007535 | 0.002309 | 0.004863 | 0.005278 | 0.007559 | 0.024496 | 0.029617 |
| 24.3888889 | 0.011463 | 0.009293 | 0.010503 | 0.016318 | 0.028391 | 0.017884 | 0.025715 | 0.02033 | 0.02275 | 0.017594 | 0.012399 | 0.020579 | 0.008287 | 0.003945 | 0.013136 | 0.026731 | 0.014087 | 0.005427 | 0.00254 | 0.005278 | 0.004774 | 0.005939 | 0.023099 | 0.024325 |
| 24.4027778 | 0.009064 | 0.008106 | 0.00661 | 0.014825 | 0.020752 | 0.015713 | 0.021023 | 0.017146 | 0.016401 | 0.013139 | 0.010307 | 0.015437 | 0.008697 | 0.002616 | 0.010165 | 0.02824 | 0.007187 | 0.002576 | 0.001949 | 0.009666 | 0.005752 | 0.004314 | 0.019691 | 0.022486 |
| 24.4166667 | 0.007721 | 0.006987 | 0.004555 | 0.010966 | 0.01621 | 0.013642 | 0.017966 | 0.014064 | 0.013192 | 0.01097 | 0.009029 | 0.011809 | 0.009118 | 0.003887 | 0.0088 | 0.029924 | 0.003286 | 0.002008 | 0.002308 | 0.014694 | 0.007766 | 0.003046 | 0.015478 | 0.022388 |
| 24.4305556 | 0.006747 | 0.005582 | 0.004227 | 0.0073 | 0.012979 | 0.012489 | 0.015884 | 0.011032 | 0.012192 | 0.009868 | 0.008232 | 0.009574 | 0.011184 | 0.004955 | 0.008098 | 0.033763 | 0.006162 | 0.004784 | 0.004334 | 0.018468 | 0.009323 | 0.00174 | 0.012235 | 0.021686 |
| 24.4444444 | 0.005813 | 0.004499 | 0.004798 | 0.006048 | 0.010742 | 0.012705 | 0.01515 | 0.008754 | 0.012122 | 0.009433 | 0.007639 | 0.008311 | 0.014713 | 0.004493 | 0.006009 | 0.038839 | 0.013982 | 0.008947 | 0.007358 | 0.019675 | 0.009989 | 0.000957 | 0.01021 | 0.022074 |
| 24.4583333 | 0.005163 | 0.004025 | 0.005292 | 0.00614 | 0.009525 | 0.013919 | 0.015126 | 0.007326 | 0.012255 | 0.009513 | 0.007168 | 0.007882 | 0.017165 | 0.004813 | 0.003143 | 0.040817 | 0.020094 | 0.011733 | 0.009863 | 0.018884 | 0.01019 | 0.00162 | 0.008831 | 0.023875 |
| 24.4722222 | 0.004787 | 0.003755 | 0.005388 | 0.006464 | 0.008497 | 0.016458 | 0.014234 | 0.006232 | 0.011195 | 0.009856 | 0.006496 | 0.00783 | 0.016833 | 0.006882 | 0.001954 | 0.038035 | 0.020502 | 0.013077 | 0.011156 | 0.017607 | 0.010227 | 0.003014 | 0.008671 | 0.024828 |
| 24.4861111 | 0.00483 | 0.003865 | 0.006338 | 0.007624 | 0.007515 | 0.024583 | 0.012136 | 0.00587 | 0.008396 | 0.010884 | 0.005967 | 0.008389 | 0.014823 | 0.009785 | 0.001775 | 0.03345 | 0.015618 | 0.015102 | 0.012971 | 0.016597 | 0.010746 | 0.003662 | 0.010697 | 0.024122 |
| 24.5 | 0.005562 | 0.005161 | 0.009302 | 0.009723 | 0.007313 | 0.035665 | 0.01017 | 0.006994 | 0.006859 | 0.013067 | 0.006511 | 0.010136 | 0.012547 | 0.011502 | 0.001285 | 0.030379 | 0.008078 | 0.019032 | 0.018556 | 0.016601 | 0.012888 | 0.002854 | 0.014801 | 0.022763 |
| 24.5138889 | 0.005779 | 0.006668 | 0.012823 | 0.011223 | 0.007342 | 0.037861 | 0.009414 | 0.008099 | 0.009945 | 0.015145 | 0.007415 | 0.012272 | 0.010986 | 0.011212 | 0.001782 | 0.031371 | 0.002643 | 0.022893 | 0.026415 | 0.016599 | 0.015058 | 0.001603 | 0.019281 | 0.022381 |
| 24.5277778 | 0.004114 | 0.005735 | 0.015174 | 0.011589 | 0.007912 | 0.031276 | 0.00958 | 0.006934 | 0.016489 | 0.016119 | 0.006771 | 0.013911 | 0.012381 | 0.00951 | 0.003991 | 0.033288 | 0.004563 | 0.023421 | 0.030462 | 0.014968 | 0.013124 | 0.00187 | 0.019513 | 0.021528 |
| 24.5416667 | 0.002677 | 0.003073 | 0.018554 | 0.013219 | 0.014161 | 0.026082 | 0.011743 | 0.005641 | 0.025138 | 0.018796 | 0.005704 | 0.01698 | 0.018276 | 0.007846 | 0.007218 | 0.026564 | 0.012889 | 0.020001 | 0.028832 | 0.013809 | 0.007213 | 0.003651 | 0.013374 | 0.017346 |
| 24.5555556 | 0.004399 | 0.002391 | 0.024928 | 0.01715 | 0.026879 | 0.025453 | 0.016582 | 0.007268 | 0.034932 | 0.024591 | 0.007805 | 0.022602 | 0.026378 | 0.007715 | 0.01036 | 0.013223 | 0.014896 | 0.014783 | 0.024821 | 0.016617 | 0.002659 | 0.004357 | 0.006967 | 0.011842 |
| 24.5694444 | 0.007232 | 0.003671 | 0.030148 | 0.0205 | 0.034303 | 0.025149 | 0.019837 | 0.008891 | 0.039622 | 0.028008 | 0.013087 | 0.026796 | 0.032084 | 0.010813 | 0.00916 | 0.012902 | 0.011996 | 0.012834 | 0.022654 | 0.022905 | 0.002551 | 0.002915 | 0.006998 | 0.012455 |
| 24.5833333 | 0.007834 | 0.003831 | 0.028833 | 0.019993 | 0.028163 | 0.021683 | 0.018391 | 0.006949 | 0.035511 | 0.024852 | 0.017244 | 0.025705 | 0.032673 | 0.023223 | 0.010904 | 0.040974 | 0.014002 | 0.019155 | 0.025616 | 0.030392 | 0.007847 | 0.002064 | 0.018137 | 0.028898 |
| 24.5972222 | 0.006752 | 0.002216 | 0.021637 | 0.015758 | 0.01581 | 0.016162 | 0.014051 | 0.003189 | 0.02824 | 0.018573 | 0.017249 | 0.020146 | 0.029375 | 0.05113 | 0.029493 | 0.080522 | 0.012857 | 0.031968 | 0.032772 | 0.038246 | 0.019103 | 0.002889 | 0.038706 | 0.061465 |
| 24.6111111 | 0.005285 | 0.001095 | 0.014051 | 0.010922 | 0.006732 | 0.010062 | 0.007908 | 0.00122 | 0.023471 | 0.013734 | 0.013746 | 0.013271 | 0.026702 | 0.081506 | 0.053067 | 0.094503 | 0.017563 | 0.043376 | 0.038196 | 0.044448 | 0.030839 | 0.003551 | 0.054226 | 0.090789 |
| 24.625 | 0.003845 | 0.001778 | 0.010028 | 0.007583 | 0.00267 | 0.004588 | 0.002796 | 0.001664 | 0.020528 | 0.01126 | 0.008968 | 0.007845 | 0.029927 | 0.090961 | 0.064694 | 0.084652 | 0.046302 | 0.046907 | 0.038351 | 0.04665 | 0.03644 | 0.005181 | 0.060666 | 0.10005 |
| 24.6388889 | 0.002657 | 0.002704 | 0.00985 | 0.005909 | 0.002849 | 0.003409 | 0.002867 | 0.002141 | 0.018635 | 0.010139 | 0.005128 | 0.005304 | 0.03932 | 0.079964 | 0.060928 | 0.07945 | 0.085312 | 0.041682 | 0.035394 | 0.044159 | 0.036418 | 0.008216 | 0.063808 | 0.091298 |
| 24.6527778 | 0.001588 | 0.002232 | 0.011587 | 0.005023 | 0.005963 | 0.00724 | 0.007582 | 0.001612 | 0.019523 | 0.009536 | 0.003051 | 0.005865 | 0.047285 | 0.062861 | 0.040201 | 0.077912 | 0.107662 | 0.031849 | 0.031706 | 0.036453 | 0.035738 | 0.009802 | 0.056165 | 0.076773 |
| 24.6666667 | 0.000742 | 0.00315 | 0.012237 | 0.003815 | 0.008361 | 0.010907 | 0.01347 | 0.001403 | 0.021229 | 0.009316 | 0.002187 | 0.010306 | 0.047157 | 0.044073 | 0.016939 | 0.069963 | 0.107122 | 0.023966 | 0.026925 | 0.024631 | 0.035378 | 0.009698 | 0.036641 | 0.064851 |
| 24.6805556 | 0.000569 | 0.004907 | 0.01046 | 0.002367 | 0.008125 | 0.011351 | 0.016315 | 0.001971 | 0.020616 | 0.00874 | 0.001844 | 0.014477 | 0.041058 | 0.025058 | 0.014183 | 0.059847 | 0.095637 | 0.020179 | 0.021156 | 0.012579 | 0.030602 | 0.008957 | 0.016823 | 0.049035 |
| 24.6944444 | 0.001016 | 0.003341 | 0.007362 | 0.001198 | 0.007024 | 0.009616 | 0.01475 | 0.00214 | 0.01787 | 0.007059 | 0.00179 | 0.012295 | 0.03422 | 0.011952 | 0.033899 | 0.051289 | 0.082889 | 0.016743 | 0.015732 | 0.004468 | 0.020012 | 0.007955 | 0.00864 | 0.029237 |
| 24.7083333 | 0.001031 | 0.00155 | 0.004283 | 0.000462 | 0.005524 | 0.007301 | 0.010935 | 0.001585 | 0.014177 | 0.004956 | 0.00215 | 0.0073 | 0.029301 | 0.010736 | 0.058047 | 0.041495 | 0.071576 | 0.012935 | 0.011346 | 0.002035 | 0.009295 | 0.007026 | 0.016269 | 0.017524 |
| 24.7222222 | 0.000901 | 0.001228 | 0.00198 | 0.000176 | 0.003016 | 0.005636 | 0.007006 | 0.00087 | 0.010913 | 0.003401 | 0.002717 | 0.004015 | 0.027025 | 0.016286 | 0.068971 | 0.032392 | 0.061225 | 0.013104 | 0.009221 | 0.005224 | 0.00458 | 0.005705 | 0.030483 | 0.01523 |
| 24.7361111 | 0.002065 | 0.001253 | 0.000809 | 0.000144 | 0.001775 | 0.004836 | 0.00397 | 0.000533 | 0.009033 | 0.002735 | 0.003096 | 0.002295 | 0.026563 | 0.017334 | 0.066699 | 0.031653 | 0.050012 | 0.019968 | 0.010463 | 0.012135 | 0.007444 | 0.004048 | 0.042266 | 0.01211 |
| 24.75 | 0.003893 | 0.002125 | 0.000546 | 0.000292 | 0.003531 | 0.004594 | 0.001771 | 0.000855 | 0.008331 | 0.002763 | 0.003221 | 0.001377 | 0.026606 | 0.010996 | 0.053498 | 0.037541 | 0.035478 | 0.029682 | 0.013421 | 0.018491 | 0.015202 | 0.003156 | 0.047824 | 0.011382 |
| 24.7638889 | 0.004836 | 0.002918 | 0.000642 | 0.000453 | 0.006047 | 0.004908 | 0.000486 | 0.001156 | 0.008536 | 0.003213 | 0.003193 | 0.000734 | 0.02507 | 0.008178 | 0.031066 | 0.045546 | 0.018718 | 0.034111 | 0.014918 | 0.02181 | 0.021923 | 0.003233 | 0.040264 | 0.017497 |
| 24.7777778 | 0.004146 | 0.002748 | 0.000815 | 0.001018 | 0.007621 | 0.006017 | 0.000356 | 0.000827 | 0.009694 | 0.003971 | 0.003153 | 0.000801 | 0.019154 | 0.019597 | 0.025991 | 0.055525 | 0.007551 | 0.029068 | 0.013385 | 0.023026 | 0.024375 | 0.003619 | 0.020751 | 0.015571 |
| 24.7916667 | 0.002525 | 0.001678 | 0.001048 | 0.002817 | 0.009074 | 0.007471 | 0.001127 | 0.000578 | 0.01103 | 0.004622 | 0.003267 | 0.001614 | 0.010366 | 0.038338 | 0.049961 | 0.055074 | 0.005762 | 0.018241 | 0.009302 | 0.023137 | 0.025298 | 0.004043 | 0.010756 | 0.012686 |
| 24.8055556 | 0.001897 | 0.000686 | 0.001454 | 0.004811 | 0.009586 | 0.008327 | 0.002428 | 0.001445 | 0.011238 | 0.00487 | 0.003631 | 0.002604 | 0.005287 | 0.053251 | 0.06769 | 0.038955 | 0.005465 | 0.008309 | 0.004883 | 0.021182 | 0.026362 | 0.004357 | 0.01856 | 0.022813 |
| 24.8194444 | 0.001753 | 0.000558 | 0.002427 | 0.005737 | 0.006738 | 0.008405 | 0.003807 | 0.003245 | 0.010362 | 0.005054 | 0.004266 | 0.003599 | 0.009936 | 0.05637 | 0.05863 | 0.024632 | 0.004181 | 0.003944 | 0.002616 | 0.015273 | 0.024696 | 0.004731 | 0.026324 | 0.029936 |
| 24.8333333 | 0.001343 | 0.001007 | 0.004087 | 0.005985 | 0.004288 | 0.008075 | 0.004863 | 0.005221 | 0.009407 | 0.005425 | 0.005015 | 0.004598 | 0.027554 | 0.042684 | 0.035844 | 0.019571 | 0.007537 | 0.008558 | 0.001919 | 0.007958 | 0.018241 | 0.004901 | 0.027961 | 0.026643 |
| 24.8472222 | 0.002166 | 0.001255 | 0.00567 | 0.006092 | 0.00721 | 0.007638 | 0.005451 | 0.006681 | 0.008627 | 0.005592 | 0.005603 | 0.005115 | 0.053397 | 0.020775 | 0.020343 | 0.01896 | 0.013292 | 0.019771 | 0.001801 | 0.003879 | 0.009576 | 0.004108 | 0.035379 | 0.025631 |
| 24.8611111 | 0.003757 | 0.000987 | 0.006425 | 0.0059 | 0.011683 | 0.007178 | 0.005529 | 0.006962 | 0.007768 | 0.004953 | 0.005782 | 0.004834 | 0.070653 | 0.00685 | 0.027801 | 0.018332 | 0.015052 | 0.027016 | 0.004104 | 0.002234 | 0.003726 | 0.002509 | 0.046524 | 0.030956 |
| 24.875 | 0.005357 | 0.000573 | 0.006393 | 0.005279 | 0.013616 | 0.006848 | 0.005457 | 0.00613 | 0.006809 | 0.003683 | 0.005586 | 0.00428 | 0.065263 | 0.00393 | 0.040675 | 0.015877 | 0.011801 | 0.026088 | 0.008182 | 0.002138 | 0.002345 | 0.001238 | 0.047385 | 0.033851 |
| 24.8888889 | 0.008056 | 0.000853 | 0.006035 | 0.004856 | 0.01329 | 0.007323 | 0.00635 | 0.005238 | 0.006545 | 0.00301 | 0.005355 | 0.004352 | 0.043221 | 0.003494 | 0.046145 | 0.014307 | 0.007234 | 0.023576 | 0.010338 | 0.005248 | 0.001961 | 0.000969 | 0.037103 | 0.032538 |
| 24.9027778 | 0.01108 | 0.002652 | 0.005996 | 0.005125 | 0.01223 | 0.008849 | 0.009484 | 0.005149 | 0.007622 | 0.003871 | 0.00504 | 0.005253 | 0.020752 | 0.007116 | 0.050195 | 0.013635 | 0.003718 | 0.022143 | 0.008572 | 0.009849 | 0.001537 | 0.000938 | 0.022086 | 0.028218 |
| 24.9166667 | 0.011654 | 0.005418 | 0.006494 | 0.005453 | 0.011714 | 0.010207 | 0.01457 | 0.005574 | 0.009006 | 0.005529 | 0.004058 | 0.006203 | 0.009379 | 0.019813 | 0.046999 | 0.010545 | 0.002467 | 0.018016 | 0.005505 | 0.010192 | 0.002516 | 0.000996 | 0.008884 | 0.01903 |
| 24.9305556 | 0.009567 | 0.007431 | 0.006906 | 0.005364 | 0.012068 | 0.010059 | 0.019211 | 0.005877 | 0.009313 | 0.006331 | 0.002528 | 0.006435 | 0.013826 | 0.035853 | 0.037046 | 0.006935 | 0.005706 | 0.011593 | 0.005013 | 0.006569 | 0.005329 | 0.002018 | 0.003852 | 0.012633 |
| 24.9444444 | 0.007525 | 0.007783 | 0.006585 | 0.005441 | 0.014573 | 0.008529 | 0.021333 | 0.005717 | 0.008625 | 0.005612 | 0.001722 | 0.006156 | 0.028323 | 0.04672 | 0.028856 | 0.00929 | 0.012131 | 0.010886 | 0.005316 | 0.005415 | 0.010924 | 0.004148 | 0.005566 | 0.012678 |
| 24.9583333 | 0.006643 | 0.006969 | 0.00551 | 0.005937 | 0.021164 | 0.006479 | 0.020816 | 0.004773 | 0.007318 | 0.004098 | 0.002227 | 0.006043 | 0.045843 | 0.049817 | 0.022672 | 0.017966 | 0.01882 | 0.015111 | 0.004263 | 0.006282 | 0.015795 | 0.007031 | 0.00971 | 0.010737 |
| 24.9722222 | 0.005649 | 0.00609 | 0.003939 | 0.006261 | 0.0267 | 0.004143 | 0.018326 | 0.003223 | 0.004983 | 0.00272 | 0.002814 | 0.006153 | 0.063967 | 0.045561 | 0.014145 | 0.025791 | 0.026569 | 0.013052 | 0.003977 | 0.005448 | 0.017249 | 0.009574 | 0.016783 | 0.005398 |
| 24.9861111 | 0.003726 | 0.005844 | 0.002294 | 0.006359 | 0.023749 | 0.002347 | 0.014644 | 0.002175 | 0.002246 | 0.002001 | 0.002221 | 0.006075 | 0.077994 | 0.033594 | 0.010767 | 0.031655 | 0.043139 | 0.0079 | 0.004604 | 0.003839 | 0.019139 | 0.010119 | 0.023555 | 0.002313 |
| 25 | 0.001847 | 0.006295 | 0.001541 | 0.00714 | 0.016865 | 0.002407 | 0.01092 | 0.002649 | 0.001379 | 0.002656 | 0.001058 | 0.005837 | 0.078756 | 0.017567 | 0.024664 | 0.043459 | 0.071909 | 0.007413 | 0.003636 | 0.002369 | 0.017677 | 0.008365 | 0.02535 | 0.003205 |
| 25.0138889 | 0.002524 | 0.007258 | 0.002355 | 0.008858 | 0.013121 | 0.002781 | 0.009032 | 0.004997 | 0.003996 | 0.005652 | 0.001067 | 0.006295 | 0.061279 | 0.00604 | 0.046455 | 0.056799 | 0.105258 | 0.008012 | 0.002916 | 0.002634 | 0.010729 | 0.006326 | 0.022721 | 0.013656 |
| 25.0277778 | 0.006593 | 0.008236 | 0.00349 | 0.010034 | 0.012623 | 0.002237 | 0.009516 | 0.008995 | 0.008245 | 0.010331 | 0.002971 | 0.00772 | 0.038823 | 0.002244 | 0.052687 | 0.062139 | 0.120642 | 0.010779 | 0.006489 | 0.007446 | 0.008164 | 0.006596 | 0.015923 | 0.034678 |
| 25.0416667 | 0.010864 | 0.006716 | 0.003435 | 0.008225 | 0.011062 | 0.001549 | 0.008016 | 0.011843 | 0.008552 | 0.012036 | 0.004936 | 0.007293 | 0.028265 | 0.004669 | 0.038636 | 0.060592 | 0.102105 | 0.026753 | 0.01896 | 0.018875 | 0.014654 | 0.009688 | 0.010833 | 0.055711 |
| 25.0555556 | 0.009545 | 0.006007 | 0.004993 | 0.004849 | 0.008858 | 0.001973 | 0.005993 | 0.009511 | 0.007325 | 0.009498 | 0.004343 | 0.005197 | 0.028424 | 0.01292 | 0.024607 | 0.060939 | 0.068486 | 0.060665 | 0.042279 | 0.03846 | 0.027275 | 0.01331 | 0.015023 | 0.067018 |
| 25.0694444 | 0.004534 | 0.011337 | 0.009582 | 0.003658 | 0.010123 | 0.003453 | 0.008083 | 0.003934 | 0.012969 | 0.009385 | 0.004184 | 0.005203 | 0.024408 | 0.023305 | 0.026613 | 0.072754 | 0.040217 | 0.080481 | 0.057471 | 0.05947 | 0.033716 | 0.012426 | 0.021484 | 0.06403 |
| 25.0833333 | 0.002474 | 0.013967 | 0.011613 | 0.003164 | 0.009736 | 0.003723 | 0.007444 | 0.001883 | 0.019875 | 0.011172 | 0.008761 | 0.005053 | 0.019007 | 0.030403 | 0.038445 | 0.084173 | 0.045459 | 0.059781 | 0.049838 | 0.063227 | 0.025435 | 0.007188 | 0.017183 | 0.050475 |
| 25.0972222 | 0.002565 | 0.009652 | 0.008116 | 0.00168 | 0.006406 | 0.004896 | 0.00272 | 0.004542 | 0.018906 | 0.00849 | 0.014246 | 0.002916 | 0.025687 | 0.031009 | 0.046549 | 0.076169 | 0.099764 | 0.030227 | 0.034747 | 0.039794 | 0.014588 | 0.003629 | 0.007682 | 0.030038 |
| 25.1111111 | 0.004424 | 0.008707 | 0.004096 | 0.001617 | 0.00429 | 0.010188 | 0.000422 | 0.006974 | 0.012993 | 0.003807 | 0.015968 | 0.001346 | 0.031622 | 0.025355 | 0.040834 | 0.053669 | 0.166919 | 0.017816 | 0.02532 | 0.011929 | 0.009809 | 0.001937 | 0.00498 | 0.017174 |
| 25.125 | 0.007629 | 0.013739 | 0.004748 | 0.004122 | 0.002336 | 0.016928 | 0.001275 | 0.007336 | 0.007556 | 0.001255 | 0.014442 | 0.000847 | 0.037658 | 0.018729 | 0.026008 | 0.029861 | 0.196018 | 0.014021 | 0.022081 | 0.01071 | 0.007229 | 0.000878 | 0.005243 | 0.024429 |
| 25.1388889 | 0.01027 | 0.01839 | 0.008146 | 0.007746 | 0.000946 | 0.019191 | 0.005051 | 0.007721 | 0.003639 | 0.000483 | 0.011445 | 0.001245 | 0.054006 | 0.017867 | 0.021711 | 0.020036 | 0.159421 | 0.010713 | 0.020804 | 0.036053 | 0.008651 | 0.001061 | 0.007811 | 0.033015 |
| 25.1527778 | 0.012852 | 0.021264 | 0.010783 | 0.011424 | 0.001242 | 0.014871 | 0.010365 | 0.008418 | 0.002125 | 0.000576 | 0.007688 | 0.002567 | 0.050175 | 0.024317 | 0.024242 | 0.028596 | 0.087071 | 0.00652 | 0.01579 | 0.059186 | 0.010937 | 0.001765 | 0.016636 | 0.030815 |
| 25.1666667 | 0.015054 | 0.023952 | 0.013088 | 0.01488 | 0.003047 | 0.008404 | 0.01158 | 0.007989 | 0.003834 | 0.00115 | 0.00404 | 0.004205 | 0.03658 | 0.031174 | 0.016329 | 0.029989 | 0.03902 | 0.003002 | 0.009385 | 0.056064 | 0.008237 | 0.00396 | 0.02372 | 0.02206 |
| 25.1805556 | 0.015514 | 0.024785 | 0.014583 | 0.016503 | 0.004842 | 0.006949 | 0.008144 | 0.005932 | 0.006006 | 0.002014 | 0.003289 | 0.005552 | 0.054774 | 0.027208 | 0.006854 | 0.016452 | 0.047669 | 0.003393 | 0.007265 | 0.032511 | 0.0084 | 0.005618 | 0.01938 | 0.012321 |
| 25.1944444 | 0.013407 | 0.022359 | 0.011882 | 0.014568 | 0.004082 | 0.011685 | 0.007151 | 0.003068 | 0.005951 | 0.002373 | 0.00587 | 0.005989 | 0.074512 | 0.015212 | 0.00719 | 0.007292 | 0.098838 | 0.008321 | 0.008706 | 0.012917 | 0.009622 | 0.005861 | 0.008255 | 0.013712 |
| 25.2083333 | 0.009587 | 0.018524 | 0.007084 | 0.009601 | 0.002903 | 0.015857 | 0.00924 | 0.001071 | 0.003793 | 0.001723 | 0.008477 | 0.00518 | 0.072614 | 0.011027 | 0.010824 | 0.005809 | 0.154055 | 0.013244 | 0.009022 | 0.006702 | 0.007977 | 0.009308 | 0.003272 | 0.023677 |
| 25.2222222 | 0.005718 | 0.014599 | 0.005525 | 0.004465 | 0.004939 | 0.015528 | 0.009822 | 0.000696 | 0.002765 | 0.000946 | 0.009282 | 0.003453 | 0.065524 | 0.016164 | 0.010411 | 0.004057 | 0.181243 | 0.014646 | 0.007577 | 0.005479 | 0.012118 | 0.011643 | 0.004997 | 0.021713 |
| 25.2361111 | 0.002855 | 0.010802 | 0.005486 | 0.002566 | 0.007924 | 0.010071 | 0.008392 | 0.000808 | 0.005269 | 0.00086 | 0.008799 | 0.0021 | 0.049007 | 0.020188 | 0.007543 | 0.003739 | 0.159806 | 0.012274 | 0.006132 | 0.004467 | 0.018083 | 0.007436 | 0.005798 | 0.013578 |
| 25.25 | 0.001293 | 0.008282 | 0.006521 | 0.004639 | 0.009292 | 0.00666 | 0.005946 | 0.00054 | 0.008778 | 0.000863 | 0.007863 | 0.00212 | 0.02459 | 0.017671 | 0.006357 | 0.007809 | 0.101352 | 0.007515 | 0.004412 | 0.006248 | 0.014881 | 0.003275 | 0.003986 | 0.021981 |
| 25.2638889 | 0.001329 | 0.007133 | 0.00887 | 0.008201 | 0.009808 | 0.014841 | 0.003568 | 0.0003 | 0.011179 | 0.000616 | 0.006605 | 0.002725 | 0.012815 | 0.011996 | 0.005568 | 0.009856 | 0.04703 | 0.005388 | 0.003586 | 0.011096 | 0.00626 | 0.003275 | 0.001792 | 0.037017 |
| 25.2777778 | 0.002535 | 0.00678 | 0.008297 | 0.010312 | 0.009123 | 0.025022 | 0.00222 | 0.000579 | 0.012636 | 0.000927 | 0.00483 | 0.002829 | 0.020269 | 0.013619 | 0.003702 | 0.00935 | 0.036004 | 0.007086 | 0.00436 | 0.014501 | 0.00218 | 0.005906 | 0.002038 | 0.044374 |
| 25.2916667 | 0.003546 | 0.0074 | 0.006696 | 0.010472 | 0.00609 | 0.024191 | 0.001342 | 0.001081 | 0.0128 | 0.002108 | 0.002998 | 0.002408 | 0.035328 | 0.020911 | 0.002941 | 0.014143 | 0.072332 | 0.007369 | 0.004408 | 0.0237 | 0.003526 | 0.010942 | 0.004957 | 0.048211 |
| 25.3055556 | 0.00405 | 0.007848 | 0.009521 | 0.01034 | 0.003364 | 0.016721 | 0.00096 | 0.001266 | 0.011613 | 0.003233 | 0.00161 | 0.001929 | 0.04723 | 0.019949 | 0.002629 | 0.020598 | 0.11727 | 0.006564 | 0.004457 | 0.037512 | 0.00589 | 0.01529 | 0.01005 | 0.046337 |
| 25.3194444 | 0.004661 | 0.00654 | 0.014899 | 0.010973 | 0.002529 | 0.011209 | 0.002004 | 0.001442 | 0.009623 | 0.003142 | 0.000742 | 0.001553 | 0.051175 | 0.011338 | 0.002682 | 0.020986 | 0.131249 | 0.007659 | 0.005779 | 0.038218 | 0.006368 | 0.014584 | 0.015188 | 0.034483 |
| 25.3333333 | 0.005129 | 0.004876 | 0.016364 | 0.011775 | 0.001561 | 0.008625 | 0.003685 | 0.001936 | 0.007134 | 0.001953 | 0.000277 | 0.001029 | 0.046798 | 0.007688 | 0.005395 | 0.017613 | 0.1057 | 0.00761 | 0.006354 | 0.030785 | 0.004775 | 0.01012 | 0.013926 | 0.020219 |
| 25.3472222 | 0.004513 | 0.005565 | 0.01239 | 0.011647 | 0.001124 | 0.006641 | 0.0047 | 0.002373 | 0.004711 | 0.000952 | 0.00021 | 0.000472 | 0.033963 | 0.01298 | 0.008237 | 0.014868 | 0.061917 | 0.004357 | 0.006043 | 0.027489 | 0.002947 | 0.006529 | 0.00818 | 0.012033 |
| 25.3611111 | 0.00282 | 0.008311 | 0.007486 | 0.01103 | 0.002092 | 0.004522 | 0.004736 | 0.002541 | 0.003036 | 0.001046 | 0.000476 | 0.000274 | 0.016641 | 0.019925 | 0.00927 | 0.013053 | 0.0265 | 0.001955 | 0.007198 | 0.022279 | 0.002194 | 0.005603 | 0.007249 | 0.007302 |
| 25.375 | 0.001377 | 0.010253 | 0.005587 | 0.0112 | 0.003553 | 0.003264 | 0.004136 | 0.002547 | 0.001984 | 0.002128 | 0.000621 | 0.00074 | 0.006757 | 0.020098 | 0.010156 | 0.010867 | 0.014085 | 0.00164 | 0.010418 | 0.012871 | 0.001857 | 0.004339 | 0.012168 | 0.007344 |
| 25.3888889 | 0.001152 | 0.010077 | 0.006507 | 0.011768 | 0.003291 | 0.004013 | 0.003334 | 0.002754 | 0.00104 | 0.003563 | 0.000873 | 0.00206 | 0.008326 | 0.012914 | 0.011339 | 0.009956 | 0.020153 | 0.001856 | 0.014206 | 0.009683 | 0.00162 | 0.002764 | 0.018516 | 0.018283 |
| 25.4027778 | 0.002055 | 0.009383 | 0.00834 | 0.011779 | 0.001861 | 0.006721 | 0.002441 | 0.003412 | 0.000614 | 0.004872 | 0.001699 | 0.003785 | 0.013027 | 0.005028 | 0.011377 | 0.011549 | 0.027054 | 0.003468 | 0.016896 | 0.014657 | 0.002883 | 0.003366 | 0.023461 | 0.03012 |
| 25.4166667 | 0.003837 | 0.009933 | 0.010771 | 0.011481 | 0.003678 | 0.009823 | 0.001627 | 0.004013 | 0.001432 | 0.006025 | 0.00291 | 0.005537 | 0.015015 | 0.001103 | 0.008817 | 0.012792 | 0.026078 | 0.005785 | 0.017854 | 0.019491 | 0.005364 | 0.002913 | 0.024303 | 0.033142 |
| 25.4305556 | 0.005785 | 0.011025 | 0.013879 | 0.011357 | 0.009281 | 0.011366 | 0.001212 | 0.004175 | 0.002911 | 0.006899 | 0.004809 | 0.007376 | 0.01246 | 0.000221 | 0.006382 | 0.013343 | 0.018968 | 0.007815 | 0.017717 | 0.020125 | 0.00797 | 0.001446 | 0.01982 | 0.032249 |
| 25.4444444 | 0.006778 | 0.010975 | 0.015928 | 0.011243 | 0.01356 | 0.010604 | 0.001488 | 0.004101 | 0.004355 | 0.00725 | 0.007396 | 0.009032 | 0.007713 | 0.000266 | 0.008595 | 0.01716 | 0.009615 | 0.00967 | 0.017915 | 0.017239 | 0.010473 | 0.002151 | 0.015121 | 0.029245 |
| 25.4583333 | 0.006532 | 0.00977 | 0.015083 | 0.010642 | 0.013221 | 0.008185 | 0.002494 | 0.003972 | 0.00606 | 0.00726 | 0.009744 | 0.010011 | 0.00742 | 0.000795 | 0.014649 | 0.024584 | 0.005383 | 0.011693 | 0.019599 | 0.014875 | 0.012164 | 0.0032 | 0.015715 | 0.023472 |
| 25.4722222 | 0.005565 | 0.00836 | 0.012479 | 0.009482 | 0.011284 | 0.00559 | 0.003878 | 0.003908 | 0.008085 | 0.007319 | 0.011044 | 0.009953 | 0.011471 | 0.002088 | 0.020427 | 0.032613 | 0.010729 | 0.014698 | 0.022084 | 0.018208 | 0.013012 | 0.00263 | 0.020048 | 0.019441 |
| 25.4861111 | 0.004938 | 0.007484 | 0.011147 | 0.008762 | 0.012034 | 0.003921 | 0.005377 | 0.003705 | 0.010103 | 0.007796 | 0.010948 | 0.009254 | 0.014775 | 0.003627 | 0.024079 | 0.038838 | 0.018615 | 0.018434 | 0.024295 | 0.027737 | 0.013992 | 0.001629 | 0.023138 | 0.01847 |
| 25.5 | 0.005599 | 0.007529 | 0.011837 | 0.009067 | 0.016278 | 0.003178 | 0.007258 | 0.002922 | 0.01231 | 0.008816 | 0.009903 | 0.009014 | 0.015445 | 0.005291 | 0.024943 | 0.040055 | 0.022378 | 0.021496 | 0.025545 | 0.038754 | 0.015015 | 0.002308 | 0.022299 | 0.018447 |
| 25.5138889 | 0.006165 | 0.007068 | 0.012022 | 0.008802 | 0.019137 | 0.002506 | 0.008552 | 0.002845 | 0.014501 | 0.0091 | 0.007877 | 0.00899 | 0.013332 | 0.006878 | 0.021915 | 0.035517 | 0.020707 | 0.022393 | 0.024224 | 0.044919 | 0.014816 | 0.00427 | 0.020244 | 0.019788 |
| 25.5277778 | 0.004347 | 0.004446 | 0.009336 | 0.007232 | 0.014991 | 0.002812 | 0.007167 | 0.00529 | 0.015429 | 0.007383 | 0.005497 | 0.007299 | 0.008772 | 0.007187 | 0.016694 | 0.027229 | 0.012823 | 0.019994 | 0.018948 | 0.042818 | 0.012275 | 0.004299 | 0.016344 | 0.022301 |
| 25.5416667 | 0.00212 | 0.001519 | 0.005934 | 0.007255 | 0.007766 | 0.004103 | 0.004555 | 0.00735 | 0.014232 | 0.006283 | 0.005726 | 0.004577 | 0.004221 | 0.005321 | 0.013379 | 0.015797 | 0.007135 | 0.014657 | 0.012104 | 0.034847 | 0.007281 | 0.004472 | 0.008702 | 0.023 |
| 25.5555556 | 0.003562 | 0.000724 | 0.006148 | 0.01071 | 0.00644 | 0.003565 | 0.004528 | 0.006074 | 0.011265 | 0.009004 | 0.007446 | 0.003949 | 0.003537 | 0.002733 | 0.016002 | 0.008441 | 0.012562 | 0.009895 | 0.010261 | 0.02929 | 0.00325 | 0.007118 | 0.00253 | 0.020121 |
| 25.5694444 | 0.009574 | 0.001006 | 0.009893 | 0.014183 | 0.014469 | 0.002357 | 0.007151 | 0.006763 | 0.008502 | 0.012897 | 0.006199 | 0.005139 | 0.007991 | 0.003646 | 0.025699 | 0.018922 | 0.017886 | 0.010943 | 0.018224 | 0.034846 | 0.002497 | 0.0074 | 0.001084 | 0.017261 |
| 25.5833333 | 0.018492 | 0.001155 | 0.013983 | 0.013431 | 0.030808 | 0.002611 | 0.010638 | 0.013369 | 0.009506 | 0.014817 | 0.003877 | 0.005857 | 0.014603 | 0.014109 | 0.033805 | 0.049039 | 0.013829 | 0.023105 | 0.035348 | 0.049943 | 0.004986 | 0.010022 | 0.005955 | 0.022131 |
| 25.5972222 | 0.028085 | 0.002198 | 0.017079 | 0.009168 | 0.051302 | 0.003732 | 0.015197 | 0.020787 | 0.0157 | 0.016024 | 0.005149 | 0.00617 | 0.023453 | 0.041315 | 0.028151 | 0.08012 | 0.013168 | 0.048454 | 0.056635 | 0.064792 | 0.011164 | 0.020778 | 0.017575 | 0.042697 |
| 25.6111111 | 0.034846 | 0.003831 | 0.019041 | 0.005581 | 0.067628 | 0.005906 | 0.019805 | 0.023894 | 0.022934 | 0.016252 | 0.008272 | 0.006396 | 0.037693 | 0.07998 | 0.012953 | 0.09579 | 0.024213 | 0.073087 | 0.067747 | 0.06888 | 0.014899 | 0.026797 | 0.032677 | 0.071371 |
| 25.625 | 0.035523 | 0.004439 | 0.019154 | 0.005215 | 0.073463 | 0.009086 | 0.022079 | 0.022666 | 0.026883 | 0.013903 | 0.008504 | 0.006301 | 0.054065 | 0.105114 | 0.003963 | 0.10144 | 0.039516 | 0.075651 | 0.057713 | 0.05966 | 0.010718 | 0.019859 | 0.04759 | 0.078894 |
| 25.6388889 | 0.031035 | 0.004096 | 0.017334 | 0.007162 | 0.067309 | 0.012555 | 0.022623 | 0.019278 | 0.028759 | 0.010259 | 0.006352 | 0.006447 | 0.064273 | 0.098108 | 0.006017 | 0.107149 | 0.052232 | 0.056628 | 0.035552 | 0.045789 | 0.003964 | 0.010847 | 0.053809 | 0.055543 |
| 25.6527778 | 0.022839 | 0.003766 | 0.015992 | 0.008908 | 0.049172 | 0.015739 | 0.022702 | 0.015413 | 0.029785 | 0.006864 | 0.005757 | 0.007291 | 0.064492 | 0.069298 | 0.01698 | 0.107642 | 0.056473 | 0.031717 | 0.015814 | 0.033844 | 0.003853 | 0.008261 | 0.046814 | 0.028799 |
| 25.6666667 | 0.012571 | 0.004282 | 0.018592 | 0.008895 | 0.02611 | 0.017219 | 0.020802 | 0.012108 | 0.027952 | 0.003879 | 0.007412 | 0.008523 | 0.05652 | 0.041935 | 0.034013 | 0.09114 | 0.05082 | 0.013373 | 0.008091 | 0.025511 | 0.010331 | 0.010034 | 0.032285 | 0.021122 |
| 25.6805556 | 0.00824 | 0.005939 | 0.024097 | 0.007235 | 0.016803 | 0.015693 | 0.016275 | 0.009453 | 0.022661 | 0.001676 | 0.010492 | 0.009503 | 0.044465 | 0.025367 | 0.044842 | 0.065206 | 0.043244 | 0.008426 | 0.011585 | 0.020208 | 0.014185 | 0.011672 | 0.016541 | 0.027307 |
| 25.6944444 | 0.012202 | 0.007416 | 0.026058 | 0.004985 | 0.027008 | 0.012192 | 0.011919 | 0.006807 | 0.015735 | 0.000721 | 0.016302 | 0.009908 | 0.03666 | 0.014755 | 0.033822 | 0.047731 | 0.039484 | 0.014712 | 0.015027 | 0.015114 | 0.012216 | 0.009943 | 0.006634 | 0.02972 |
| 25.7083333 | 0.015449 | 0.007064 | 0.020358 | 0.003238 | 0.037288 | 0.008915 | 0.010897 | 0.004514 | 0.008973 | 0.000967 | 0.024474 | 0.009778 | 0.037767 | 0.008031 | 0.02341 | 0.039937 | 0.039711 | 0.022947 | 0.01162 | 0.009649 | 0.007156 | 0.005724 | 0.005332 | 0.01923 |
| 25.7222222 | 0.012901 | 0.004811 | 0.010781 | 0.002016 | 0.035792 | 0.006572 | 0.014637 | 0.003135 | 0.003889 | 0.002408 | 0.030275 | 0.009351 | 0.043642 | 0.007714 | 0.041318 | 0.035008 | 0.039875 | 0.020684 | 0.006819 | 0.006632 | 0.003871 | 0.002216 | 0.006211 | 0.006164 |

| | | | | | | | | | | | | | | | | | | | | | | | | |
|---|---|---|---|---|---|---|---|---|---|---|---|---|---|---|---|---|---|---|---|---|---|---|---|---|
| 25.7361111 | 0.007009 | 0.002335 | 0.005773 | 0.000957 | 0.025797 | 0.005036 | 0.023445 | 0.001966 | 0.001612 | 0.005258 | 0.029887 | 0.008431 | 0.048668 | 0.007493 | 0.055117 | 0.034576 | 0.035308 | 0.019531 | 0.008212 | 0.008917 | 0.007682 | 0.001102 | 0.005593 | 0.007755 |
| 25.75 | 0.003986 | 0.001845 | 0.006128 | 0.000573 | 0.016403 | 0.004433 | 0.032682 | 0.00125 | 0.001811 | 0.008969 | 0.02477 | 0.006248 | 0.050676 | 0.008602 | 0.037631 | 0.041435 | 0.027207 | 0.039398 | 0.014133 | 0.015077 | 0.015942 | 0.002661 | 0.006835 | 0.026531 |
| 25.7638889 | 0.009002 | 0.003115 | 0.006186 | 0.00136 | 0.026436 | 0.004393 | 0.034495 | 0.001928 | 0.004615 | 0.011806 | 0.017546 | 0.003361 | 0.049907 | 0.024013 | 0.023763 | 0.049962 | 0.019298 | 0.058373 | 0.017512 | 0.019722 | 0.025037 | 0.004868 | 0.011768 | 0.046286 |
| 25.7777778 | 0.019596 | 0.003822 | 0.008997 | 0.002649 | 0.056183 | 0.004558 | 0.028916 | 0.002244 | 0.009204 | 0.012716 | 0.00903 | 0.001709 | 0.044743 | 0.046549 | 0.043587 | 0.051269 | 0.013488 | 0.054963 | 0.015718 | 0.018297 | 0.03114 | 0.005217 | 0.017895 | 0.049408 |
| 25.7916667 | 0.025712 | 0.003173 | 0.016591 | 0.003145 | 0.077903 | 0.005059 | 0.021259 | 0.00179 | 0.011902 | 0.012301 | 0.003907 | 0.002173 | 0.034886 | 0.058607 | 0.060156 | 0.045012 | 0.00931 | 0.040189 | 0.013645 | 0.011186 | 0.027381 | 0.004038 | 0.023684 | 0.0354 |
| 25.8055556 | 0.02483 | 0.003621 | 0.021388 | 0.00273 | 0.082129 | 0.005746 | 0.013583 | 0.002782 | 0.011439 | 0.011202 | 0.005443 | 0.003792 | 0.022794 | 0.060234 | 0.052873 | 0.040646 | 0.005315 | 0.026301 | 0.013964 | 0.007049 | 0.01713 | 0.002441 | 0.031092 | 0.019821 |
| 25.8194444 | 0.025027 | 0.006617 | 0.020218 | 0.002273 | 0.075817 | 0.006296 | 0.006763 | 0.004716 | 0.010215 | 0.009701 | 0.006902 | 0.005781 | 0.010739 | 0.061764 | 0.047476 | 0.040117 | 0.002281 | 0.014915 | 0.015294 | 0.010416 | 0.008912 | 0.001494 | 0.039853 | 0.015333 |
| 25.8333333 | 0.026256 | 0.008763 | 0.016969 | 0.001879 | 0.055916 | 0.006475 | 0.004363 | 0.005683 | 0.00964 | 0.007775 | 0.004386 | 0.007856 | 0.00454 | 0.060087 | 0.045605 | 0.037274 | 0.001541 | 0.014783 | 0.01753 | 0.011662 | 0.006165 | 0.001905 | 0.043325 | 0.01743 |
| 25.8472222 | 0.020855 | 0.007761 | 0.013912 | 0.001235 | 0.029159 | 0.006259 | 0.007544 | 0.005751 | 0.009104 | 0.004891 | 0.002744 | 0.00895 | 0.007213 | 0.046785 | 0.039703 | 0.031318 | 0.001637 | 0.034055 | 0.017919 | 0.007042 | 0.00949 | 0.002806 | 0.034866 | 0.01396 |
| 25.8611111 | 0.011962 | 0.005295 | 0.010725 | 0.000628 | 0.012731 | 0.006214 | 0.01143 | 0.006422 | 0.008734 | 0.00237 | 0.00442 | 0.008877 | 0.010321 | 0.025716 | 0.029169 | 0.023409 | 0.001455 | 0.043814 | 0.016555 | 0.003616 | 0.014978 | 0.00634 | 0.018489 | 0.009309 |
| 25.875 | 0.012152 | 0.003121 | 0.00713 | 0.000759 | 0.019265 | 0.006917 | 0.012245 | 0.008015 | 0.00897 | 0.002031 | 0.008028 | 0.008627 | 0.009312 | 0.011689 | 0.02739 | 0.014044 | 0.001945 | 0.030708 | 0.017965 | 0.003497 | 0.017439 | 0.009313 | 0.009583 | 0.011252 |
| 25.8888889 | 0.020113 | 0.001508 | 0.00395 | 0.002063 | 0.04626 | 0.008118 | 0.008976 | 0.008307 | 0.008152 | 0.002983 | 0.014018 | 0.008281 | 0.011209 | 0.012044 | 0.041715 | 0.007932 | 0.002098 | 0.024791 | 0.018907 | 0.002971 | 0.017205 | 0.009097 | 0.010584 | 0.011605 |
| 25.9027778 | 0.027295 | 0.000578 | 0.003774 | 0.004225 | 0.075033 | 0.009253 | 0.005498 | 0.00597 | 0.005233 | 0.003311 | 0.021485 | 0.007521 | 0.019639 | 0.015822 | 0.053179 | 0.009445 | 0.001953 | 0.021981 | 0.014823 | 0.002521 | 0.017769 | 0.016482 | 0.007886 | 0.011717 |
| 25.9166667 | 0.033909 | 0.000394 | 0.006515 | 0.006491 | 0.094235 | 0.010203 | 0.008283 | 0.003248 | 0.002744 | 0.002829 | 0.024585 | 0.006791 | 0.027733 | 0.021728 | 0.045148 | 0.015713 | 0.003955 | 0.017466 | 0.010356 | 0.007636 | 0.020144 | 0.027599 | 0.002709 | 0.01291 |
| 25.9305556 | 0.038836 | 0.000533 | 0.007736 | 0.008084 | 0.104064 | 0.010581 | 0.014615 | 0.002124 | 0.00489 | 0.00376 | 0.020483 | 0.006645 | 0.028131 | 0.037619 | 0.024498 | 0.020887 | 0.00723 | 0.024711 | 0.009917 | 0.015948 | 0.021552 | 0.02726 | 0.003012 | 0.011721 |
| 25.9444444 | 0.037779 | 0.001636 | 0.005444 | 0.008721 | 0.09556 | 0.010004 | 0.018411 | 0.001605 | 0.0139 | 0.005863 | 0.013805 | 0.007071 | 0.0224 | 0.057408 | 0.011313 | 0.021931 | 0.01016 | 0.024609 | 0.01028 | 0.016847 | 0.019476 | 0.018683 | 0.009199 | 0.013989 |
| 25.9583333 | 0.030249 | 0.00385 | 0.003445 | 0.008752 | 0.067187 | 0.008759 | 0.019499 | 0.001444 | 0.025864 | 0.006043 | 0.010031 | 0.007455 | 0.01696 | 0.064741 | 0.006879 | 0.026079 | 0.012999 | 0.02568 | 0.007318 | 0.009067 | 0.015772 | 0.012426 | 0.021085 | 0.014864 |
| 25.9722222 | 0.0206 | 0.004821 | 0.003424 | 0.008474 | 0.039874 | 0.006515 | 0.0192 | 0.003543 | 0.031204 | 0.004538 | 0.010358 | 0.006635 | 0.016074 | 0.05763 | 0.011051 | 0.039517 | 0.016276 | 0.038594 | 0.005634 | 0.002789 | 0.013821 | 0.009089 | 0.032003 | 0.01066 |
| 25.9861111 | 0.011362 | 0.00375 | 0.002739 | 0.008029 | 0.021654 | 0.003631 | 0.016636 | 0.007543 | 0.025934 | 0.003926 | 0.011337 | 0.004221 | 0.019293 | 0.051676 | 0.021502 | 0.049891 | 0.020366 | 0.036564 | 0.011125 | 0.004765 | 0.013929 | 0.006765 | 0.027639 | 0.014104 |
| 26 | 0.009095 | 0.003749 | 0.001414 | 0.007419 | 0.019847 | 0.001807 | 0.010571 | 0.00968 | 0.017796 | 0.005423 | 0.009462 | 0.002123 | 0.021328 | 0.050418 | 0.028338 | 0.050524 | 0.023294 | 0.040697 | 0.022429 | 0.016265 | 0.014924 | 0.0074 | 0.020755 | 0.030524 |
| 26.0138889 | 0.020082 | 0.004624 | 0.001537 | 0.006448 | 0.045436 | 0.001691 | 0.005177 | 0.007024 | 0.01271 | 0.008106 | 0.005697 | 0.003077 | 0.016813 | 0.043987 | 0.032916 | 0.054819 | 0.02255 | 0.065555 | 0.036018 | 0.033579 | 0.016643 | 0.012104 | 0.031083 | 0.046274 |
| 26.0277778 | 0.030955 | 0.003305 | 0.002272 | 0.004387 | 0.070046 | 0.003655 | 0.00637 | 0.00438 | 0.008107 | 0.009082 | 0.002505 | 0.007578 | 0.009811 | 0.030178 | 0.033051 | 0.063106 | 0.021547 | 0.070739 | 0.045252 | 0.046068 | 0.019493 | 0.018537 | 0.040676 | 0.048886 |
| 26.0416667 | 0.027616 | 0.003348 | 0.006115 | 0.005409 | 0.058511 | 0.005767 | 0.015049 | 0.005527 | 0.004091 | 0.00796 | 0.001992 | 0.011679 | 0.014412 | 0.019236 | 0.023205 | 0.05921 | 0.029084 | 0.062592 | 0.054056 | 0.059134 | 0.018395 | 0.02907 | 0.044846 | 0.045267 |
| 26.0555556 | 0.017739 | 0.010894 | 0.020653 | 0.016234 | 0.038127 | 0.004713 | 0.031242 | 0.006659 | 0.004832 | 0.012755 | 0.005293 | 0.009526 | 0.032903 | 0.016958 | 0.014671 | 0.050497 | 0.045156 | 0.07189 | 0.071482 | 0.08759 | 0.014137 | 0.039624 | 0.049224 | 0.046787 |
| 26.0694444 | 0.018203 | 0.023076 | 0.038805 | 0.029824 | 0.053333 | 0.002174 | 0.049187 | 0.010624 | 0.006822 | 0.026569 | 0.012438 | 0.007464 | 0.047297 | 0.019169 | 0.017426 | 0.051872 | 0.050693 | 0.082787 | 0.083732 | 0.110591 | 0.01597 | 0.033632 | 0.0428 | 0.04485 |
| 26.0833333 | 0.029171 | 0.02985 | 0.048062 | 0.03117 | 0.084942 | 0.001844 | 0.055067 | 0.018357 | 0.00448 | 0.036328 | 0.019882 | 0.014496 | 0.044473 | 0.022639 | 0.020422 | 0.056054 | 0.033128 | 0.070868 | 0.076304 | 0.096841 | 0.020036 | 0.017668 | 0.029226 | 0.028312 |
| 26.0972222 | 0.036228 | 0.027856 | 0.050165 | 0.018866 | 0.098347 | 0.002898 | 0.044577 | 0.021839 | 0.004062 | 0.034699 | 0.019826 | 0.022468 | 0.029259 | 0.022465 | 0.022851 | 0.055888 | 0.012047 | 0.043233 | 0.056802 | 0.054577 | 0.019787 | 0.014157 | 0.015502 | 0.016442 |
| 26.1111111 | 0.035961 | 0.022306 | 0.045856 | 0.006488 | 0.096752 | 0.00343 | 0.027811 | 0.018746 | 0.009359 | 0.027262 | 0.011798 | 0.02394 | 0.015164 | 0.016502 | 0.030951 | 0.054128 | 0.005653 | 0.02177 | 0.038437 | 0.017854 | 0.016274 | 0.027397 | 0.005576 | 0.030968 |
| 26.125 | 0.032555 | 0.015555 | 0.034974 | 0.002423 | 0.096385 | 0.0056 | 0.013002 | 0.01259 | 0.011894 | 0.018669 | 0.005848 | 0.020144 | 0.009646 | 0.013038 | 0.03766 | 0.050588 | 0.006925 | 0.012449 | 0.028447 | 0.004109 | 0.011915 | 0.0369 | 0.006044 | 0.047328 |
| 26.1388889 | 0.026242 | 0.008407 | 0.024022 | 0.002764 | 0.094249 | 0.008304 | 0.006164 | 0.006141 | 0.007682 | 0.012229 | 0.009026 | 0.012068 | 0.00839 | 0.01854 | 0.029489 | 0.034562 | 0.00895 | 0.008887 | 0.024899 | 0.003678 | 0.008662 | 0.026748 | 0.010577 | 0.037293 |
| 26.1527778 | 0.018038 | 0.006381 | 0.015557 | 0.005739 | 0.079778 | 0.007252 | 0.012759 | 0.002615 | 0.003182 | 0.008008 | 0.016957 | 0.006803 | 0.006191 | 0.029575 | 0.016309 | 0.018031 | 0.011463 | 0.00685 | 0.019533 | 0.005381 | 0.006025 | 0.015742 | 0.014998 | 0.018553 |
| 26.1666667 | 0.01074 | 0.010031 | 0.009476 | 0.012168 | 0.061483 | 0.004538 | 0.027107 | 0.004003 | 0.001897 | 0.005016 | 0.022145 | 0.010931 | 0.004589 | 0.037213 | 0.014135 | 0.017238 | 0.013875 | 0.005182 | 0.011252 | 0.008114 | 0.005013 | 0.026789 | 0.027179 | 0.015311 |
| 26.1805556 | 0.006168 | 0.012779 | 0.006438 | 0.019332 | 0.050199 | 0.004173 | 0.036853 | 0.007497 | 0.002493 | 0.006049 | 0.019237 | 0.017301 | 0.004104 | 0.03435 | 0.011726 | 0.015526 | 0.01578 | 0.0032 | 0.005541 | 0.02006 | 0.007912 | 0.043493 | 0.040867 | 0.016462 |
| 26.1944444 | 0.00445 | 0.013239 | 0.005955 | 0.023009 | 0.042132 | 0.005232 | 0.033022 | 0.009783 | 0.007921 | 0.009972 | 0.011204 | 0.018819 | 0.005483 | 0.021248 | 0.01227 | 0.012882 | 0.016689 | 0.002158 | 0.003782 | 0.039066 | 0.009386 | 0.03499 | 0.037085 | 0.011843 |
| 26.2083333 | 0.003747 | 0.014401 | 0.008886 | 0.020441 | 0.028213 | 0.00481 | 0.018624 | 0.009134 | 0.01944 | 0.01143 | 0.010257 | 0.016763 | 0.011082 | 0.012122 | 0.021887 | 0.014878 | 0.016165 | 0.002346 | 0.004951 | 0.045818 | 0.005735 | 0.022538 | 0.018538 | 0.017645 |
| 26.2222222 | 0.003053 | 0.01602 | 0.014492 | 0.013401 | 0.012604 | 0.003289 | 0.011097 | 0.005821 | 0.030292 | 0.0091 | 0.016924 | 0.011829 | 0.013462 | 0.015633 | 0.023507 | 0.011582 | 0.013226 | 0.004548 | 0.007678 | 0.034168 | 0.005212 | 0.026878 | 0.005469 | 0.029876 |
| 26.2361111 | 0.004102 | 0.015891 | 0.01729 | 0.006436 | 0.005563 | 0.002812 | 0.017126 | 0.005007 | 0.032897 | 0.006346 | 0.020817 | 0.005454 | 0.00832 | 0.014715 | 0.016851 | 0.007141 | 0.008856 | 0.009856 | 0.007074 | 0.019315 | 0.007818 | 0.025009 | 0.002329 | 0.026021 |
| 26.25 | 0.005575 | 0.013523 | 0.014276 | 0.002238 | 0.00749 | 0.002859 | 0.022532 | 0.011483 | 0.027826 | 0.00589 | 0.019481 | 0.001959 | 0.003425 | 0.008236 | 0.018347 | 0.005557 | 0.005207 | 0.010042 | 0.005225 | 0.020333 | 0.00573 | 0.023987 | 0.001901 | 0.012292 |
| 26.2638889 | 0.004141 | 0.010511 | 0.009958 | 0.000566 | 0.014009 | 0.001958 | 0.021876 | 0.024679 | 0.019942 | 0.006733 | 0.017356 | 0.001993 | 0.003173 | 0.011343 | 0.021337 | 0.004359 | 0.003201 | 0.005449 | 0.007653 | 0.032172 | 0.001721 | 0.02865 | 0.004244 | 0.008939 |
| 26.2777778 | 0.002709 | 0.008324 | 0.008466 | 0.000952 | 0.015521 | 0.001422 | 0.020463 | 0.038939 | 0.010847 | 0.007026 | 0.015029 | 0.001987 | 0.00521 | 0.019696 | 0.015756 | 0.005967 | 0.004083 | 0.004422 | 0.00775 | 0.028971 | 0.000722 | 0.019752 | 0.012711 | 0.011464 |
| 26.2916667 | 0.004698 | 0.007054 | 0.008872 | 0.003635 | 0.009028 | 0.00411 | 0.017845 | 0.043807 | 0.006388 | 0.006953 | 0.010342 | 0.001258 | 0.006801 | 0.02763 | 0.009169 | 0.007232 | 0.006741 | 0.00376 | 0.003999 | 0.018461 | 0.003005 | 0.006958 | 0.020733 | 0.013443 |
| 26.3055556 | 0.006896 | 0.006957 | 0.008377 | 0.007366 | 0.005151 | 0.00891 | 0.01208 | 0.036572 | 0.012074 | 0.007042 | 0.01004 | 0.002653 | 0.006369 | 0.034106 | 0.007126 | 0.004523 | 0.006674 | 0.001869 | 0.003214 | 0.02534 | 0.008592 | 0.004969 | 0.020279 | 0.017514 |
| 26.3194444 | 0.005829 | 0.008624 | 0.00641 | 0.009731 | 0.011208 | 0.012155 | 0.006241 | 0.026021 | 0.020971 | 0.006787 | 0.019667 | 0.00756 | 0.004135 | 0.034951 | 0.010238 | 0.001945 | 0.00429 | 0.001094 | 0.00346 | 0.035237 | 0.015349 | 0.009643 | 0.01707 | 0.019298 |
| 26.3333333 | 0.004044 | 0.011327 | 0.007756 | 0.010461 | 0.025808 | 0.013123 | 0.005733 | 0.018459 | 0.025135 | 0.006354 | 0.029501 | 0.013343 | 0.001843 | 0.030223 | 0.014513 | 0.003036 | 0.004521 | 0.002548 | 0.002885 | 0.032219 | 0.019694 | 0.012057 | 0.017555 | 0.014368 |
| 26.3472222 | 0.006005 | 0.01265 | 0.014409 | 0.011714 | 0.04347 | 0.01222 | 0.010637 | 0.014108 | 0.023798 | 0.006163 | 0.032133 | 0.016307 | 0.001825 | 0.024403 | 0.013316 | 0.007815 | 0.007282 | 0.007198 | 0.003819 | 0.025015 | 0.021226 | 0.009391 | 0.015375 | 0.00708 |
| 26.3611111 | 0.009948 | 0.01037 | 0.021735 | 0.014595 | 0.05458 | 0.009177 | 0.016792 | 0.009885 | 0.020961 | 0.006225 | 0.031804 | 0.016054 | 0.004251 | 0.020377 | 0.007376 | 0.011406 | 0.009092 | 0.011626 | 0.003544 | 0.01862 | 0.021951 | 0.005437 | 0.007931 | 0.004858 |
| 26.375 | 0.012222 | 0.005701 | 0.028012 | 0.017835 | 0.052974 | 0.00547 | 0.022418 | 0.006098 | 0.021564 | 0.007266 | 0.031548 | 0.015522 | 0.004574 | 0.017506 | 0.002953 | 0.01169 | 0.009467 | 0.011437 | 0.003272 | 0.01239 | 0.021323 | 0.004361 | 0.002553 | 0.004481 |
| 26.3888889 | 0.013094 | 0.003778 | 0.032776 | 0.020302 | 0.044607 | 0.003323 | 0.027433 | 0.007314 | 0.024269 | 0.008652 | 0.028931 | 0.016932 | 0.003241 | 0.015653 | 0.002026 | 0.014232 | 0.00998 | 0.006793 | 0.008193 | 0.012516 | 0.017402 | 0.007766 | 0.003007 | 0.002953 |
| 26.4027778 | 0.013706 | 0.008307 | 0.034707 | 0.021997 | 0.038768 | 0.002967 | 0.030224 | 0.012091 | 0.023605 | 0.00835 | 0.022 | 0.019515 | 0.005391 | 0.015403 | 0.003092 | 0.022079 | 0.010773 | 0.003727 | 0.015548 | 0.019719 | 0.012191 | 0.013732 | 0.007477 | 0.004586 |
| 26.4166667 | 0.014817 | 0.015191 | 0.03488 | 0.022736 | 0.038719 | 0.003676 | 0.030673 | 0.017045 | 0.01889 | 0.007289 | 0.012426 | 0.020992 | 0.009908 | 0.014384 | 0.00559 | 0.031436 | 0.010876 | 0.006365 | 0.020214 | 0.022642 | 0.008567 | 0.016549 | 0.012345 | 0.007097 |
| 26.4305556 | 0.016567 | 0.016994 | 0.033785 | 0.022469 | 0.041256 | 0.004685 | 0.030412 | 0.024338 | 0.014779 | 0.008611 | 0.00452 | 0.01985 | 0.014166 | 0.009423 | 0.00851 | 0.038664 | 0.010416 | 0.010344 | 0.021247 | 0.018152 | 0.006363 | 0.012576 | 0.015812 | 0.006749 |
| 26.4444444 | 0.017744 | 0.012925 | 0.032852 | 0.021482 | 0.041734 | 0.005185 | 0.028991 | 0.032449 | 0.014746 | 0.01208 | 0.002995 | 0.016668 | 0.018081 | 0.004715 | 0.009443 | 0.042306 | 0.010164 | 0.012338 | 0.020083 | 0.012715 | 0.00462 | 0.00617 | 0.018736 | 0.009156 |
| 26.4583333 | 0.017479 | 0.008911 | 0.034324 | 0.019943 | 0.039435 | 0.005059 | 0.026599 | 0.034833 | 0.018416 | 0.014963 | 0.008873 | 0.013761 | 0.021247 | 0.004802 | 0.00718 | 0.040004 | 0.009928 | 0.013778 | 0.018032 | 0.011623 | 0.002629 | 0.002997 | 0.02243 | 0.014149 |
| 26.4722222 | 0.017012 | 0.010248 | 0.036607 | 0.018274 | 0.037683 | 0.004822 | 0.025731 | 0.031967 | 0.023066 | 0.017086 | 0.018648 | 0.012945 | 0.0213 | 0.004429 | 0.004126 | 0.03236 | 0.009245 | 0.015951 | 0.016933 | 0.0162 | 0.001341 | 0.002012 | 0.024217 | 0.012842 |
| 26.4861111 | 0.018107 | 0.01539 | 0.035808 | 0.016712 | 0.040003 | 0.004836 | 0.027392 | 0.028479 | 0.025443 | 0.018429 | 0.028112 | 0.013276 | 0.017319 | 0.00241 | 0.00492 | 0.025089 | 0.008715 | 0.018596 | 0.019307 | 0.02296 | 0.00219 | 0.001398 | 0.021854 | 0.007821 |
| 26.5 | 0.02074 | 0.018428 | 0.030899 | 0.014432 | 0.046192 | 0.004889 | 0.029195 | 0.025206 | 0.025262 | 0.017237 | 0.034602 | 0.012745 | 0.012918 | 0.002081 | 0.01069 | 0.023612 | 0.008098 | 0.020588 | 0.025228 | 0.027795 | 0.003589 | 0.004781 | 0.018534 | 0.011255 |
| 26.5138889 | 0.022492 | 0.01668 | 0.024386 | 0.0107 | 0.050371 | 0.003897 | 0.027268 | 0.021271 | 0.024585 | 0.012887 | 0.03559 | 0.010757 | 0.009655 | 0.00267 | 0.01622 | 0.026226 | 0.005698 | 0.021385 | 0.029382 | 0.029671 | 0.003734 | 0.016405 | 0.014925 | 0.024596 |
| 26.5277778 | 0.0207 | 0.012857 | 0.019547 | 0.006846 | 0.045768 | 0.002608 | 0.021534 | 0.015939 | 0.023628 | 0.007008 | 0.029836 | 0.008303 | 0.006144 | 0.003312 | 0.01631 | 0.023834 | 0.00242 | 0.020341 | 0.026075 | 0.024861 | 0.00268 | 0.023984 | 0.011288 | 0.033429 |
| 26.5416667 | 0.017039 | 0.012474 | 0.02042 | 0.005569 | 0.03475 | 0.00316 | 0.016157 | 0.010527 | 0.023559 | 0.003045 | 0.021351 | 0.008633 | 0.00392 | 0.002537 | 0.013914 | 0.015417 | 0.000739 | 0.018144 | 0.017905 | 0.01532 | 0.003495 | 0.018838 | 0.010444 | 0.028532 |
| 26.5555556 | 0.017017 | 0.018229 | 0.027783 | 0.007246 | 0.026937 | 0.003781 | 0.014557 | 0.0084 | 0.023707 | 0.002813 | 0.015366 | 0.012321 | 0.006564 | 0.000943 | 0.016975 | 0.008396 | 0.000416 | 0.018071 | 0.013232 | 0.009322 | 0.00836 | 0.019112 | 0.010033 | 0.019874 |
| 26.5694444 | 0.025105 | 0.025261 | 0.033751 | 0.009222 | 0.031323 | 0.002835 | 0.016144 | 0.011532 | 0.019472 | 0.003343 | 0.011449 | 0.01528 | 0.015361 | 0.00298 | 0.023892 | 0.014217 | 0.001687 | 0.023049 | 0.018089 | 0.014798 | 0.0107 | 0.023596 | 0.012507 | 0.018601 |
| 26.5833333 | 0.042515 | 0.026206 | 0.030438 | 0.011619 | 0.052196 | 0.002813 | 0.018923 | 0.021589 | 0.012846 | 0.002172 | 0.00704 | 0.016112 | 0.027945 | 0.016248 | 0.023554 | 0.047462 | 0.005907 | 0.034301 | 0.034618 | 0.046107 | 0.010058 | 0.020894 | 0.023494 | 0.02601 |
| 26.5972222 | 0.059865 | 0.022202 | 0.019857 | 0.015778 | 0.074971 | 0.007691 | 0.023067 | 0.036187 | 0.010767 | 0.000947 | 0.003299 | 0.015242 | 0.039305 | 0.047601 | 0.013522 | 0.098694 | 0.011483 | 0.049908 | 0.059408 | 0.093258 | 0.017909 | 0.033661 | 0.03653 | 0.039108 |
| 26.6111111 | 0.060303 | 0.018757 | 0.010201 | 0.018388 | 0.071947 | 0.016953 | 0.026467 | 0.043041 | 0.014559 | 0.000628 | 0.002779 | 0.012728 | 0.047607 | 0.083142 | 0.003626 | 0.135178 | 0.017347 | 0.06024 | 0.079235 | 0.122442 | 0.032351 | 0.059864 | 0.047753 | 0.051947 |
| 26.625 | 0.041642 | 0.014485 | 0.010897 | 0.017075 | 0.041153 | 0.025011 | 0.026333 | 0.03663 | 0.021477 | 0.000832 | 0.003064 | 0.01134 | 0.057537 | 0.096139 | 0.003408 | 0.143747 | 0.025499 | 0.053938 | 0.083218 | 0.13019 | 0.042029 | 0.064923 | 0.056043 | 0.05383 |
| 26.6388889 | 0.018872 | 0.008712 | 0.022025 | 0.014177 | 0.018395 | 0.028694 | 0.023754 | 0.027133 | 0.028429 | 0.001187 | 0.003928 | 0.013607 | 0.069897 | 0.079183 | 0.011253 | 0.132321 | 0.035531 | 0.037303 | 0.074282 | 0.126053 | 0.046525 | 0.048562 | 0.053534 | 0.03647 |
| 26.6527778 | 0.008724 | 0.006991 | 0.02866 | 0.012239 | 0.019411 | 0.026481 | 0.019461 | 0.020743 | 0.030644 | 0.00139 | 0.009772 | 0.017579 | 0.077004 | 0.046056 | 0.015883 | 0.107247 | 0.043274 | 0.02419 | 0.055394 | 0.100568 | 0.046994 | 0.02818 | 0.041375 | 0.015418 |
| 26.6666667 | 0.010724 | 0.011447 | 0.020521 | 0.011773 | 0.017472 | 0.01931 | 0.013892 | 0.013445 | 0.026128 | 0.00154 | 0.015747 | 0.019042 | 0.074 | 0.018206 | 0.013238 | 0.078996 | 0.045407 | 0.013819 | 0.028807 | 0.060081 | 0.040806 | 0.013413 | 0.031064 | 0.007567 |
| 26.6805556 | 0.010532 | 0.017832 | 0.009418 | 0.013147 | 0.008837 | 0.012323 | 0.010524 | 0.006903 | 0.0191 | 0.00181 | 0.016578 | 0.017211 | 0.064612 | 0.006223 | 0.011132 | 0.054731 | 0.041524 | 0.005932 | 0.00942 | 0.026973 | 0.031795 | 0.008075 | 0.027215 | 0.010487 |
| 26.6944444 | 0.011298 | 0.020414 | 0.010226 | 0.016729 | 0.011478 | 0.009232 | 0.013203 | 0.008237 | 0.01368 | 0.001857 | 0.014381 | 0.015196 | 0.054886 | 0.005375 | 0.014429 | 0.033884 | 0.034471 | 0.005027 | 0.004575 | 0.011734 | 0.025717 | 0.009791 | 0.027099 | 0.024251 |
| 26.7083333 | 0.01934 | 0.0168 | 0.012345 | 0.019412 | 0.026787 | 0.009337 | 0.02245 | 0.015852 | 0.012391 | 0.00149 | 0.01078 | 0.015579 | 0.048478 | 0.009417 | 0.022599 | 0.016452 | 0.027738 | 0.006475 | 0.006101 | 0.010774 | 0.020972 | 0.013452 | 0.027612 | 0.036016 |
| 26.7222222 | 0.020301 | 0.00922 | 0.009089 | 0.017251 | 0.037748 | 0.010328 | 0.037849 | 0.024409 | 0.016254 | 0.002107 | 0.009439 | 0.018721 | 0.048705 | 0.014833 | 0.031815 | 0.007988 | 0.022264 | 0.007955 | 0.013489 | 0.021206 | 0.015641 | 0.012035 | 0.024667 | 0.032738 |
| 26.7361111 | 0.013475 | 0.004387 | 0.012869 | 0.012017 | 0.035204 | 0.011873 | 0.051599 | 0.03164 | 0.020595 | 0.003593 | 0.011754 | 0.022678 | 0.056819 | 0.018775 | 0.038716 | 0.016875 | 0.018279 | 0.018371 | 0.024143 | 0.038812 | 0.011952 | 0.006765 | 0.01748 | 0.025737 |
| 26.75 | 0.017208 | 0.005386 | 0.020382 | 0.008449 | 0.047242 | 0.014894 | 0.050172 | 0.034517 | 0.020751 | 0.003407 | 0.013053 | 0.025297 | 0.069109 | 0.021018 | 0.047471 | 0.039667 | 0.015902 | 0.036657 | 0.031192 | 0.044527 | 0.011279 | 0.004534 | 0.008887 | 0.02456 |
| 26.7638889 | 0.030645 | 0.005155 | 0.024615 | 0.008837 | 0.079246 | 0.01972 | 0.036756 | 0.030389 | 0.018154 | 0.004171 | 0.012319 | 0.027399 | 0.080679 | 0.026678 | 0.053977 | 0.061444 | 0.014123 | 0.04544 | 0.03431 | 0.034409 | 0.013448 | 0.005681 | 0.00574 | 0.030746 |
| 26.7777778 | 0.036831 | 0.00351 | 0.029291 | 0.011866 | 0.089289 | 0.026061 | 0.024637 | 0.022041 | 0.015728 | 0.010037 | 0.011914 | 0.029416 | 0.086426 | 0.040197 | 0.046983 | 0.076382 | 0.012235 | 0.037388 | 0.032908 | 0.021765 | 0.017343 | 0.006927 | 0.0107 | 0.038138 |
| 26.7916667 | 0.029049 | 0.006096 | 0.032711 | 0.013736 | 0.057313 | 0.030417 | 0.016264 | 0.014486 | 0.01472 | 0.016503 | 0.010333 | 0.028537 | 0.085492 | 0.057469 | 0.032108 | 0.083165 | 0.010911 | 0.025242 | 0.029449 | 0.014816 | 0.021487 | 0.006499 | 0.011811 | 0.036016 |
| 26.8055556 | 0.015434 | 0.008698 | 0.03188 | 0.012851 | 0.023165 | 0.028166 | 0.009149 | 0.00835 | 0.01497 | 0.01836 | 0.007429 | 0.023695 | 0.079231 | 0.06806 | 0.027832 | 0.075305 | 0.009972 | 0.026169 | 0.024862 | 0.02338 | 0.02293 | 0.005256 | 0.010752 | 0.026468 |
| 26.8194444 | 0.009122 | 0.007517 | 0.028591 | 0.011069 | 0.014847 | 0.020182 | 0.008516 | 0.003435 | 0.014907 | 0.016337 | 0.005075 | 0.017856 | 0.0638 | 0.065657 | 0.039275 | 0.060068 | 0.008464 | 0.047689 | 0.018245 | 0.053204 | 0.019949 | 0.005419 | 0.016588 | 0.021596 |
| 26.8333333 | 0.017048 | 0.005037 | 0.023236 | 0.009413 | 0.024196 | 0.010714 | 0.017797 | 0.001233 | 0.012491 | 0.01325 | 0.004341 | 0.013591 | 0.040967 | 0.057293 | 0.045981 | 0.049746 | 0.007086 | 0.068129 | 0.011685 | 0.091126 | 0.017556 | 0.007082 | 0.017681 | 0.029335 |
| 26.8472222 | 0.030246 | 0.003509 | 0.015974 | 0.007659 | 0.059918 | 0.004172 | 0.026603 | 0.002174 | 0.007124 | 0.010464 | 0.007768 | 0.011628 | 0.020887 | 0.054237 | 0.030973 | 0.045276 | 0.006658 | 0.056532 | 0.005994 | 0.108939 | 0.018644 | 0.008457 | 0.010454 | 0.040142 |
| 26.8611111 | 0.036002 | 0.003246 | 0.008438 | 0.006183 | 0.09038 | 0.003261 | 0.025753 | 0.006094 | 0.003772 | 0.006659 | 0.014111 | 0.011451 | 0.010597 | 0.051985 | 0.022687 | 0.04269 | 0.006571 | 0.026605 | 0.004415 | 0.091455 | 0.01548 | 0.015016 | 0.005861 | 0.036948 |
| 26.875 | 0.032452 | 0.003966 | 0.003678 | 0.005281 | 0.074591 | 0.006684 | 0.01691 | 0.013186 | 0.007238 | 0.004284 | 0.020395 | 0.011541 | 0.006371 | 0.039685 | 0.041346 | 0.034891 | 0.005826 | 0.010633 | 0.007005 | 0.058572 | 0.006847 | 0.027498 | 0.009857 | 0.022631 |
| 26.8888889 | 0.020551 | 0.00532 | 0.003764 | 0.004515 | 0.035884 | 0.010221 | 0.008696 | 0.022529 | 0.012838 | 0.005134 | 0.024276 | 0.011477 | 0.007078 | 0.020147 | 0.04951 | 0.019251 | 0.004766 | 0.013883 | 0.007589 | 0.038051 | 0.004065 | 0.027218 | 0.014417 | 0.012727 |
| 26.9027778 | 0.013621 | 0.005736 | 0.005776 | 0.003041 | 0.012012 | 0.009713 | 0.009272 | 0.031004 | 0.01454 | 0.004358 | 0.020927 | 0.011527 | 0.017481 | 0.007841 | 0.036451 | 0.008803 | 0.004917 | 0.021997 | 0.006614 | 0.025312 | 0.00944 | 0.017171 | 0.011386 | 0.012868 |
| 26.9166667 | 0.02605 | 0.006068 | 0.006625 | 0.001804 | 0.012581 | 0.006285 | 0.015434 | 0.035582 | 0.012786 | 0.002332 | 0.012557 | 0.011958 | 0.029833 | 0.010855 | 0.032615 | 0.012805 | 0.006294 | 0.024769 | 0.008946 | 0.020314 | 0.018151 | 0.024932 | 0.010909 | 0.016775 |
| 26.9305556 | 0.046729 | 0.008534 | 0.006307 | 0.002503 | 0.029863 | 0.005948 | 0.021187 | 0.036648 | 0.011295 | 0.00172 | 0.009573 | 0.013101 | 0.036069 | 0.021813 | 0.037521 | 0.021529 | 0.007121 | 0.023442 | 0.016353 | 0.051192 | 0.027085 | 0.045017 | 0.018888 | 0.014712 |
| 26.9444444 | 0.055068 | 0.010498 | 0.004668 | 0.004277 | 0.058511 | 0.01031 | 0.027189 | 0.034717 | 0.011521 | 0.000967 | 0.012956 | 0.014255 | 0.035953 | 0.036202 | 0.03262 | 0.023542 | 0.006506 | 0.015627 | 0.027137 | 0.10685 | 0.032926 | 0.050245 | 0.029727 | 0.01296 |
| 26.9583333 | 0.046215 | 0.010461 | 0.002822 | 0.005716 | 0.091535 | 0.014954 | 0.033569 | 0.028032 | 0.010645 | 0.001388 | 0.014301 | 0.014528 | 0.030138 | 0.056521 | 0.01872 | 0.027048 | 0.005139 | 0.005112 | 0.035911 | 0.142154 | 0.033925 | 0.039362 | 0.04438 | 0.023415 |
| 26.9722222 | 0.036657 | 0.011597 | 0.003055 | 0.006319 | 0.117018 | 0.01628 | 0.035531 | 0.017058 | 0.006709 | 0.004221 | 0.011551 | 0.013922 | 0.022706 | 0.073419 | 0.015475 | 0.040967 | 0.003977 | 0.003031 | 0.036857 | 0.130975 | 0.02941 | 0.027501 | 0.056751 | 0.027901 |
| 26.9861111 | 0.034007 | 0.012768 | 0.003687 | 0.006895 | 0.119761 | 0.014095 | 0.02942 | 0.008008 | 0.004185 | 0.006158 | 0.008248 | 0.013141 | 0.019025 | 0.069633 | 0.037442 | 0.049204 | 0.003312 | 0.010796 | 0.024867 | 0.076436 | 0.022963 | 0.023208 | 0.04537 | 0.018423 |
| 27 | 0.030144 | 0.01371 | 0.002899 | 0.008682 | 0.099292 | 0.010747 | 0.018608 | 0.010092 | 0.006786 | 0.00484 | 0.007007 | 0.012462 | 0.023414 | 0.051078 | 0.061949 | 0.051901 | 0.002395 | 0.022944 | 0.014755 | 0.030975 | 0.020621 | 0.023654 | 0.01754 | 0.011436 |
| 27.0138889 | 0.02587 | 0.019082 | 0.002427 | 0.010408 | 0.07825 | 0.008778 | 0.009073 | 0.021113 | 0.010213 | 0.003481 | 0.007822 | 0.011729 | 0.03526 | 0.043841 | 0.059109 | 0.063789 | 0.002239 | 0.031246 | 0.024667 | 0.033215 | 0.018725 | 0.019323 | 0.004028 | 0.010664 |
| 27.0277778 | 0.0257 | 0.024572 | 0.003042 | 0.011922 | 0.074972 | 0.007039 | 0.008866 | 0.027825 | 0.008603 | 0.006441 | 0.007754 | 0.008909 | 0.045957 | 0.053269 | 0.037275 | 0.073488 | 0.007714 | 0.03433 | 0.038318 | 0.061809 | 0.012894 | 0.01057 | 0.010766 | 0.017288 |
| 27.0416667 | 0.028727 | 0.019548 | 0.004863 | 0.018103 | 0.073895 | 0.007705 | 0.025676 | 0.029656 | 0.012274 | 0.017072 | 0.012504 | 0.011682 | 0.052617 | 0.064898 | 0.029812 | 0.074316 | 0.028078 | 0.04357 | 0.048004 | 0.102282 | 0.013733 | 0.011151 | 0.026804 | 0.019357 |
| 27.0555556 | 0.031998 | 0.01523 | 0.011151 | 0.030069 | 0.057393 | 0.017668 | 0.053791 | 0.045012 | 0.033408 | 0.036499 | 0.026128 | 0.031111 | 0.058759 | 0.072051 | 0.047501 | 0.081515 | 0.054678 | 0.057454 | 0.064707 | 0.144475 | 0.027869 | 0.029637 | 0.037935 | 0.020995 |
| 27.0694444 | 0.031462 | 0.027433 | 0.019942 | 0.038155 | 0.040429 | 0.030167 | 0.076625 | 0.067129 | 0.047136 | 0.057445 | 0.031885 | 0.050323 | 0.054638 | 0.073529 | 0.068326 | 0.09511 | 0.054763 | 0.05738 | 0.07018 | 0.159032 | 0.036745 | 0.058845 | 0.036923 | 0.037457 |
| 27.0833333 | 0.026892 | 0.034946 | 0.024349 | 0.033822 | 0.038943 | 0.029767 | 0.077446 | 0.06373 | 0.034507 | 0.07084 | 0.022323 | 0.049985 | 0.035806 | 0.072566 | 0.072975 | 0.106868 | 0.028652 | 0.040765 | 0.047569 | 0.127607 | 0.029301 | 0.088471 | 0.02772 | 0.048916 |
| 27.0972222 | 0.023257 | 0.026105 | 0.025287 | 0.021612 | 0.049917 | 0.017065 | 0.055579 | 0.037616 | 0.017719 | 0.069751 | 0.009968 | 0.039215 | 0.019672 | 0.070822 | 0.061618 | 0.114228 | 0.008456 | 0.023332 | 0.01988 | 0.066432 | 0.01934 | 0.087919 | 0.019815 | 0.042492 |
| 27.1111111 | 0.019673 | 0.020942 | 0.025854 | 0.011423 | 0.055058 | 0.005618 | 0.031739 | 0.016351 | 0.012256 | 0.052528 | 0.00357 | 0.03011 | 0.015168 | 0.062196 | 0.053469 | 0.102769 | 0.002547 | 0.017814 | 0.010772 | 0.022299 | 0.014239 | 0.056032 | 0.018318 | 0.027889 |
| 27.125 | 0.011627 | 0.023771 | 0.02633 | 0.006682 | 0.03917 | 0.001762 | 0.01983 | 0.012964 | 0.01274 | 0.030354 | 0.004377 | 0.021988 | 0.013325 | 0.054031 | 0.050516 | 0.073155 | 0.001511 | 0.02175 | 0.01746 | 0.016916 | 0.012884 | 0.051854 | 0.013068 | 0.022637 |
| 27.1388889 | 0.00483 | 0.02697 | 0.028039 | 0.005654 | 0.017939 | 0.002929 | 0.017625 | 0.020132 | 0.013199 | 0.013579 | 0.011618 | 0.013458 | 0.010272 | 0.050612 | 0.034536 | 0.039722 | 0.002267 | 0.019787 | 0.025609 | 0.02188 | 0.009713 | 0.062288 | 0.004274 | 0.021376 |
| 27.1527778 | 0.00471 | 0.026203 | 0.029933 | 0.004799 | 0.009175 | 0.00498 | 0.019013 | 0.026408 | 0.01276 | 0.007332 | 0.02353 | 0.006579 | 0.014076 | 0.035922 | 0.023148 | 0.017687 | 0.00583 | 0.00969 | 0.024161 | 0.018007 | 0.006194 | 0.040315 | 0.00191 | 0.014931 |
| 27.1666667 | 0.005432 | 0.018478 | 0.027481 | 0.004741 | 0.014398 | 0.004956 | 0.016323 | 0.025673 | 0.01195 | 0.012123 | 0.037893 | 0.004898 | 0.021505 | 0.015327 | 0.035357 | 0.014013 | 0.008785 | 0.003861 | 0.014277 | 0.022105 | 0.009134 | 0.016125 | 0.005889 | 0.017039 |
| 27.1805556 | 0.007505 | 0.009825 | 0.018732 | 0.009034 | 0.041226 | 0.003562 | 0.010621 | 0.015704 | 0.009771 | 0.017572 | 0.049958 | 0.008571 | 0.017319 | 0.008751 | 0.044545 | 0.012326 | 0.007211 | 0.009417 | 0.005204 | 0.041264 | 0.012495 | 0.010847 | 0.014016 | 0.017149 |
| 27.1944444 | 0.011874 | 0.0085 | 0.008573 | 0.013862 | 0.069652 | 0.003824 | 0.011704 | 0.006287 | 0.006966 | 0.015117 | 0.049435 | 0.011795 | 0.008097 | 0.012358 | 0.031381 | 0.007556 | 0.00339 | 0.018943 | 0.001734 | 0.05517 | 0.011951 | 0.011467 | 0.018297 | 0.01012 |
| 27.2083333 | 0.010271 | 0.01184 | 0.002807 | 0.013814 | 0.061672 | 0.005861 | 0.016488 | 0.008547 | 0.00516 | 0.011274 | 0.034139 | 0.011993 | 0.009675 | 0.019394 | 0.0134 | 0.00588 | 0.001432 | 0.017361 | 0.001397 | 0.053263 | 0.012659 | 0.024185 | 0.014568 | 0.012642 |
| 27.2222222 | 0.004937 | 0.010421 | 0.001993 | 0.009222 | 0.030285 | 0.006306 | 0.014065 | 0.019981 | 0.004087 | 0.018505 | 0.01751 | 0.009041 | 0.018199 | 0.022519 | 0.006449 | 0.008214 | 0.002409 | 0.009374 | 0.001436 | 0.041596 | 0.014284 | 0.039107 | 0.00963 | 0.01865 |
| 27.2361111 | 0.004018 | 0.005796 | 0.004563 | 0.003925 | 0.010839 | 0.003948 | 0.007202 | 0.03052 | 0.002817 | 0.03322 | 0.01563 | 0.005284 | 0.022245 | 0.016078 | 0.006403 | 0.012893 | 0.006548 | 0.005616 | 0.001502 | 0.028756 | 0.011116 | 0.030968 | 0.0063 | 0.014566 |

| | | | | | | | | | | | | | | | | | | | | | | | | |
|---|---|---|---|---|---|---|---|---|---|---|---|---|---|---|---|---|---|---|---|---|---|---|---|---|
| 27.25 | 0.006036 | 0.005671 | 0.009932 | 0.001849 | 0.008488 | 0.002474 | 0.004714 | 0.033188 | 0.001258 | 0.041954 | 0.028239 | 0.005967 | 0.018277 | 0.011883 | 0.010549 | 0.013394 | 0.013897 | 0.004992 | 0.002498 | 0.018499 | 0.008583 | 0.013211 | 0.005958 | 0.008621 |
| 27.2638889 | 0.007001 | 0.005421 | 0.017192 | 0.003819 | 0.01266 | 0.004419 | 0.006666 | 0.028448 | 0.000366 | 0.040884 | 0.04176 | 0.010228 | 0.01134 | 0.012023 | 0.01986 | 0.008213 | 0.021951 | 0.004968 | 0.004649 | 0.012476 | 0.011631 | 0.0085 | 0.006479 | 0.015345 |
| 27.2777778 | 0.006487 | 0.005047 | 0.02223 | 0.00698 | 0.015193 | 0.006684 | 0.009759 | 0.021404 | 0.000168 | 0.035024 | 0.049177 | 0.013933 | 0.006543 | 0.011637 | 0.030587 | 0.003066 | 0.027903 | 0.003276 | 0.00465 | 0.016324 | 0.015216 | 0.010057 | 0.004718 | 0.025754 |
| 27.2916667 | 0.005585 | 0.011581 | 0.021299 | 0.009391 | 0.02539 | 0.00703 | 0.013398 | 0.016549 | 0.000675 | 0.029276 | 0.049208 | 0.015073 | 0.004006 | 0.015494 | 0.035854 | 0.000974 | 0.031498 | 0.00151 | 0.002278 | 0.018756 | 0.015994 | 0.010645 | 0.003479 | 0.021248 |
| 27.3055556 | 0.005217 | 0.020163 | 0.016602 | 0.012466 | 0.04525 | 0.006631 | 0.016804 | 0.014692 | 0.002202 | 0.022164 | 0.040779 | 0.011883 | 0.002748 | 0.021762 | 0.032207 | 0.000337 | 0.032954 | 0.002257 | 0.001275 | 0.016619 | 0.013881 | 0.016544 | 0.003636 | 0.014139 |
| 27.3194444 | 0.004774 | 0.024742 | 0.011563 | 0.017152 | 0.045426 | 0.005854 | 0.017933 | 0.011084 | 0.003195 | 0.011886 | 0.025942 | 0.006746 | 0.003869 | 0.024443 | 0.024073 | 0.000896 | 0.032169 | 0.004305 | 0.001802 | 0.027698 | 0.0102 | 0.03967 | 0.002744 | 0.024134 |
| 27.3333333 | 0.003497 | 0.02572 | 0.007121 | 0.023016 | 0.025973 | 0.004081 | 0.015082 | 0.009483 | 0.002441 | 0.004414 | 0.011746 | 0.006105 | 0.008642 | 0.02393 | 0.017618 | 0.00332 | 0.031097 | 0.006693 | 0.002459 | 0.044387 | 0.006357 | 0.071684 | 0.00086 | 0.035663 |
| 27.3472222 | 0.00275 | 0.02448 | 0.004312 | 0.028764 | 0.01802 | 0.002065 | 0.008883 | 0.016725 | 0.002146 | 0.003191 | 0.007425 | 0.011265 | 0.015931 | 0.021024 | 0.014889 | 0.004733 | 0.032719 | 0.010534 | 0.003295 | 0.049138 | 0.004443 | 0.076662 | 0.000682 | 0.031012 |
| 27.3611111 | 0.004348 | 0.023155 | 0.005904 | 0.032734 | 0.018647 | 0.001649 | 0.004297 | 0.023464 | 0.005543 | 0.003669 | 0.01573 | 0.019633 | 0.021642 | 0.015493 | 0.01478 | 0.004211 | 0.036984 | 0.015266 | 0.004469 | 0.041627 | 0.006073 | 0.048533 | 0.002665 | 0.016552 |
| 27.375 | 0.007133 | 0.022925 | 0.012419 | 0.034863 | 0.024912 | 0.003527 | 0.00687 | 0.024418 | 0.012802 | 0.006143 | 0.02757 | 0.029409 | 0.021335 | 0.010547 | 0.017277 | 0.006564 | 0.039234 | 0.017858 | 0.005659 | 0.025141 | 0.008834 | 0.018979 | 0.005145 | 0.005839 |
| 27.3888889 | 0.009422 | 0.022288 | 0.020083 | 0.034592 | 0.047314 | 0.006823 | 0.015101 | 0.026489 | 0.021409 | 0.013087 | 0.034234 | 0.037328 | 0.016685 | 0.009992 | 0.023068 | 0.013108 | 0.036604 | 0.016072 | 0.0069 | 0.013645 | 0.008041 | 0.006166 | 0.006843 | 0.003273 |
| 27.4027778 | 0.01151 | 0.019207 | 0.026308 | 0.031336 | 0.062359 | 0.010266 | 0.023223 | 0.03066 | 0.028392 | 0.019977 | 0.037169 | 0.041443 | 0.013756 | 0.011389 | 0.02892 | 0.019634 | 0.033417 | 0.011234 | 0.008259 | 0.021027 | 0.005241 | 0.003875 | 0.008698 | 0.004827 |
| 27.4166667 | 0.012237 | 0.014451 | 0.030907 | 0.02729 | 0.053875 | 0.013194 | 0.027635 | 0.032843 | 0.032153 | 0.021116 | 0.038408 | 0.044148 | 0.012939 | 0.010074 | 0.029088 | 0.024795 | 0.032479 | 0.007699 | 0.009064 | 0.033658 | 0.004355 | 0.004586 | 0.012163 | 0.008054 |
| 27.4305556 | 0.011288 | 0.011428 | 0.034581 | 0.024806 | 0.038305 | 0.015344 | 0.02828 | 0.032746 | 0.034288 | 0.017375 | 0.036558 | 0.045415 | 0.009325 | 0.006352 | 0.023902 | 0.032757 | 0.02945 | 0.008743 | 0.009039 | 0.035766 | 0.003994 | 0.004302 | 0.015917 | 0.008424 |
| 27.4444444 | 0.010794 | 0.012955 | 0.037966 | 0.024526 | 0.030655 | 0.016881 | 0.027902 | 0.031526 | 0.036859 | 0.014727 | 0.032573 | 0.042224 | 0.005596 | 0.003708 | 0.02038 | 0.043356 | 0.02182 | 0.013008 | 0.009156 | 0.031834 | 0.005499 | 0.003886 | 0.018166 | 0.00552 |
| 27.4583333 | 0.01097 | 0.018116 | 0.041168 | 0.025115 | 0.033434 | 0.018792 | 0.02823 | 0.029447 | 0.037695 | 0.017243 | 0.029221 | 0.035807 | 0.008002 | 0.004604 | 0.0222 | 0.047287 | 0.013021 | 0.016244 | 0.010007 | 0.03032 | 0.009374 | 0.008447 | 0.018509 | 0.006222 |
| 27.4722222 | 0.011428 | 0.022337 | 0.044056 | 0.024651 | 0.04112 | 0.021415 | 0.028582 | 0.027062 | 0.03425 | 0.023876 | 0.028291 | 0.029901 | 0.01247 | 0.00675 | 0.024799 | 0.04307 | 0.007072 | 0.016762 | 0.010736 | 0.031854 | 0.012455 | 0.017037 | 0.017654 | 0.011762 |
| 27.4861111 | 0.013064 | 0.022339 | 0.045389 | 0.0232 | 0.048069 | 0.024136 | 0.0292 | 0.027564 | 0.028848 | 0.031948 | 0.029266 | 0.026673 | 0.01298 | 0.006939 | 0.025178 | 0.040169 | 0.004634 | 0.01815 | 0.011041 | 0.032446 | 0.012874 | 0.023973 | 0.017162 | 0.016877 |
| 27.5 | 0.015912 | 0.020412 | 0.043707 | 0.02262 | 0.050612 | 0.026052 | 0.031188 | 0.038849 | 0.025828 | 0.03992 | 0.029558 | 0.025494 | 0.009958 | 0.006843 | 0.026724 | 0.043048 | 0.004422 | 0.022367 | 0.01195 | 0.030929 | 0.010537 | 0.021897 | 0.0182 | 0.017954 |
| 27.5138889 | 0.018442 | 0.020354 | 0.039121 | 0.023361 | 0.047582 | 0.02535 | 0.033146 | 0.060714 | 0.026032 | 0.045055 | 0.02528 | 0.02276 | 0.006409 | 0.00762 | 0.030189 | 0.048675 | 0.005762 | 0.025194 | 0.013172 | 0.028612 | 0.008653 | 0.011796 | 0.02121 | 0.014841 |
| 27.5277778 | 0.018829 | 0.022108 | 0.033359 | 0.023864 | 0.042208 | 0.020628 | 0.033413 | 0.076486 | 0.025665 | 0.044745 | 0.015744 | 0.016172 | 0.003911 | 0.008415 | 0.031918 | 0.049625 | 0.009302 | 0.022807 | 0.01302 | 0.026115 | 0.011818 | 0.006766 | 0.024053 | 0.009529 |
| 27.5416667 | 0.01764 | 0.025381 | 0.029188 | 0.025429 | 0.039808 | 0.01305 | 0.032534 | 0.073706 | 0.020722 | 0.043753 | 0.007989 | 0.008983 | 0.001893 | 0.009886 | 0.029928 | 0.041296 | 0.018006 | 0.015434 | 0.011069 | 0.021893 | 0.020409 | 0.013302 | 0.02221 | 0.004999 |
| 27.5555556 | 0.01789 | 0.029679 | 0.028273 | 0.029864 | 0.039497 | 0.005779 | 0.029676 | 0.058626 | 0.012732 | 0.045899 | 0.005976 | 0.004755 | 0.000737 | 0.013086 | 0.025025 | 0.029452 | 0.037006 | 0.007997 | 0.009446 | 0.016078 | 0.029213 | 0.028819 | 0.015512 | 0.003452 |
| 27.5694444 | 0.021254 | 0.030029 | 0.028953 | 0.035835 | 0.038736 | 0.002008 | 0.023584 | 0.044391 | 0.009139 | 0.044921 | 0.006539 | 0.002769 | 0.003245 | 0.019212 | 0.021715 | 0.024196 | 0.058691 | 0.006454 | 0.011711 | 0.019705 | 0.028077 | 0.0387 | 0.011218 | 0.006993 |
| 27.5833333 | 0.029453 | 0.02267 | 0.028881 | 0.044272 | 0.043945 | 0.001692 | 0.017239 | 0.036571 | 0.008353 | 0.039524 | 0.007502 | 0.003943 | 0.014265 | 0.0336 | 0.025246 | 0.04107 | 0.05614 | 0.010412 | 0.02002 | 0.04625 | 0.017727 | 0.0307 | 0.0136 | 0.017842 |
| 27.5972222 | 0.040923 | 0.011113 | 0.026517 | 0.051772 | 0.05354 | 0.004251 | 0.013994 | 0.030962 | 0.006743 | 0.03429 | 0.008146 | 0.009129 | 0.030258 | 0.062126 | 0.031873 | 0.076716 | 0.030472 | 0.016514 | 0.033315 | 0.088328 | 0.016904 | 0.032333 | 0.022886 | 0.034994 |
| 27.6111111 | 0.045783 | 0.003352 | 0.020926 | 0.04832 | 0.054131 | 0.012529 | 0.014082 | 0.024654 | 0.012948 | 0.029929 | 0.010316 | 0.01083 | 0.03904 | 0.095364 | 0.03381 | 0.104147 | 0.020178 | 0.023999 | 0.048074 | 0.120105 | 0.028489 | 0.0617 | 0.042751 | 0.04945 |
| 27.625 | 0.040253 | 0.002881 | 0.01507 | 0.035478 | 0.035462 | 0.02638 | 0.018048 | 0.023737 | 0.027725 | 0.026768 | 0.014935 | 0.013075 | 0.043172 | 0.107364 | 0.030349 | 0.118206 | 0.039913 | 0.03343 | 0.064952 | 0.126985 | 0.033524 | 0.082602 | 0.067429 | 0.05082 |
| 27.6388889 | 0.034031 | 0.009465 | 0.013512 | 0.024279 | 0.022666 | 0.04157 | 0.027755 | 0.03432 | 0.04249 | 0.025419 | 0.016559 | 0.025405 | 0.052577 | 0.085173 | 0.026879 | 0.122462 | 0.070137 | 0.047572 | 0.080882 | 0.118887 | 0.025383 | 0.079948 | 0.07984 | 0.042445 |
| 27.6527778 | 0.035118 | 0.0226 | 0.01246 | 0.017386 | 0.021472 | 0.052059 | 0.039485 | 0.04729 | 0.049816 | 0.023988 | 0.01997 | 0.038546 | 0.060786 | 0.047413 | 0.030478 | 0.115567 | 0.090273 | 0.063309 | 0.080884 | 0.102734 | 0.022221 | 0.069287 | 0.068329 | 0.034974 |
| 27.6666667 | 0.040128 | 0.035537 | 0.009341 | 0.012031 | 0.028674 | 0.050739 | 0.04496 | 0.038669 | 0.04708 | 0.018169 | 0.026978 | 0.046066 | 0.057952 | 0.017923 | 0.041677 | 0.100755 | 0.091561 | 0.065847 | 0.05537 | 0.07267 | 0.03004 | 0.045132 | 0.037632 | 0.029156 |
| 27.6805556 | 0.039129 | 0.040217 | 0.010189 | 0.008662 | 0.039332 | 0.037506 | 0.041208 | 0.018889 | 0.039224 | 0.009963 | 0.02457 | 0.042934 | 0.045536 | 0.006095 | 0.051504 | 0.079374 | 0.079774 | 0.054202 | 0.026383 | 0.035353 | 0.031735 | 0.017056 | 0.018024 | 0.026051 |
| 27.6944444 | 0.027077 | 0.036006 | 0.012218 | 0.009195 | 0.0443 | 0.01984 | 0.031325 | 0.013429 | 0.030643 | 0.006414 | 0.016015 | 0.027468 | 0.031244 | 0.010256 | 0.049616 | 0.055939 | 0.065798 | 0.039177 | 0.013228 | 0.00988 | 0.02356 | 0.004717 | 0.023001 | 0.029175 |
| 27.7083333 | 0.013441 | 0.028963 | 0.008942 | 0.013387 | 0.039787 | 0.007565 | 0.023882 | 0.015891 | 0.023295 | 0.006078 | 0.009456 | 0.013579 | 0.02184 | 0.022458 | 0.031833 | 0.031409 | 0.056394 | 0.022048 | 0.011956 | 0.007675 | 0.013476 | 0.003123 | 0.023257 | 0.031711 |
| 27.7222222 | 0.008662 | 0.023558 | 0.007157 | 0.016402 | 0.028439 | 0.004185 | 0.023795 | 0.03095 | 0.017219 | 0.009383 | 0.006128 | 0.008432 | 0.01921 | 0.034376 | 0.016917 | 0.011025 | 0.052677 | 0.010079 | 0.016308 | 0.028084 | 0.00566 | 0.003572 | 0.014119 | 0.030237 |
| 27.7361111 | 0.008051 | 0.019029 | 0.012941 | 0.014449 | 0.013971 | 0.005052 | 0.025916 | 0.052914 | 0.011663 | 0.019837 | 0.0065 | 0.00807 | 0.021821 | 0.037122 | 0.014265 | 0.009239 | 0.050433 | 0.012156 | 0.022516 | 0.058292 | 0.003534 | 0.012642 | 0.017582 | 0.028083 |
| 27.75 | 0.013452 | 0.015145 | 0.017331 | 0.009392 | 0.00834 | 0.010657 | 0.026214 | 0.060267 | 0.006818 | 0.028122 | 0.005243 | 0.007847 | 0.028605 | 0.027107 | 0.01897 | 0.029728 | 0.047529 | 0.022177 | 0.025234 | 0.063475 | 0.006044 | 0.024339 | 0.026313 | 0.026664 |
| 27.7638889 | 0.03347 | 0.013227 | 0.014473 | 0.005726 | 0.022285 | 0.020524 | 0.026429 | 0.06027 | 0.00287 | 0.027436 | 0.003896 | 0.006058 | 0.037536 | 0.013117 | 0.050203 | 0.057775 | 0.047645 | 0.028176 | 0.022928 | 0.038944 | 0.00986 | 0.020863 | 0.027263 | 0.024054 |
| 27.7777778 | 0.054136 | 0.016522 | 0.012673 | 0.006382 | 0.046302 | 0.026394 | 0.029876 | 0.067743 | 0.001141 | 0.023508 | 0.007317 | 0.00871 | 0.043833 | 0.009465 | 0.101501 | 0.066671 | 0.052432 | 0.026138 | 0.021003 | 0.029867 | 0.014444 | 0.012985 | 0.023909 | 0.017679 |
| 27.7916667 | 0.05655 | 0.026442 | 0.01841 | 0.009368 | 0.063094 | 0.025287 | 0.036695 | 0.076518 | 0.001243 | 0.02095 | 0.01259 | 0.022222 | 0.045228 | 0.022267 | 0.139207 | 0.059512 | 0.05588 | 0.024654 | 0.022685 | 0.039416 | 0.016084 | 0.009128 | 0.016434 | 0.017662 |
| 27.8055556 | 0.047214 | 0.032719 | 0.029266 | 0.010259 | 0.069016 | 0.023418 | 0.039948 | 0.061903 | 0.00293 | 0.020622 | 0.018212 | 0.042224 | 0.042282 | 0.045911 | 0.137161 | 0.057035 | 0.051757 | 0.029646 | 0.022944 | 0.04026 | 0.015437 | 0.004548 | 0.010569 | 0.030334 |
| 27.8194444 | 0.044062 | 0.028307 | 0.038425 | 0.007126 | 0.068009 | 0.0265 | 0.032201 | 0.032741 | 0.008219 | 0.021806 | 0.025535 | 0.0608 | 0.036043 | 0.066952 | 0.087258 | 0.054662 | 0.041966 | 0.030978 | 0.01599 | 0.045903 | 0.0135 | 0.002329 | 0.013646 | 0.036296 |
| 27.8333333 | 0.045124 | 0.020703 | 0.0394 | 0.003156 | 0.051645 | 0.031517 | 0.017833 | 0.024527 | 0.014144 | 0.02115 | 0.034701 | 0.069389 | 0.028785 | 0.073031 | 0.041588 | 0.052525 | 0.03146 | 0.021108 | 0.007391 | 0.071556 | 0.008487 | 0.002846 | 0.02068 | 0.027992 |
| 27.8472222 | 0.041095 | 0.012581 | 0.031171 | 0.002062 | 0.025205 | 0.031374 | 0.007025 | 0.022858 | 0.016104 | 0.016624 | 0.043122 | 0.061067 | 0.023729 | 0.065179 | 0.043767 | 0.049717 | 0.022914 | 0.010155 | 0.006656 | 0.096084 | 0.004719 | 0.005558 | 0.029162 | 0.023892 |
| 27.8611111 | 0.033621 | 0.008183 | 0.017254 | 0.003417 | 0.011646 | 0.0253 | 0.002258 | 0.024479 | 0.014204 | 0.010783 | 0.047937 | 0.044547 | 0.022283 | 0.050819 | 0.053449 | 0.037888 | 0.016853 | 0.005247 | 0.008374 | 0.0993 | 0.0081 | 0.015081 | 0.035103 | 0.025385 |
| 27.875 | 0.024349 | 0.011468 | 0.00685 | 0.005433 | 0.016029 | 0.018129 | 0.000897 | 0.046102 | 0.012928 | 0.007609 | 0.051225 | 0.031993 | 0.021538 | 0.032823 | 0.039628 | 0.024793 | 0.011827 | 0.011962 | 0.007585 | 0.087754 | 0.014388 | 0.021422 | 0.030992 | 0.02299 |
| 27.8888889 | 0.016719 | 0.014716 | 0.00475 | 0.006697 | 0.029873 | 0.012334 | 0.001297 | 0.055548 | 0.018022 | 0.008959 | 0.051856 | 0.02302 | 0.017681 | 0.015453 | 0.021778 | 0.015389 | 0.006998 | 0.02657 | 0.006297 | 0.071079 | 0.011408 | 0.014016 | 0.02124 | 0.019026 |
| 27.9027778 | 0.011036 | 0.012121 | 0.005037 | 0.005927 | 0.029908 | 0.006843 | 0.005599 | 0.041232 | 0.028935 | 0.012866 | 0.042712 | 0.015777 | 0.010111 | 0.005584 | 0.017386 | 0.012064 | 0.006452 | 0.025156 | 0.004248 | 0.048655 | 0.005584 | 0.012193 | 0.016698 | 0.014866 |
| 27.9166667 | 0.00957 | 0.00784 | 0.005153 | 0.003505 | 0.018626 | 0.004759 | 0.015378 | 0.023686 | 0.037994 | 0.015226 | 0.025419 | 0.010661 | 0.004209 | 0.008857 | 0.025053 | 0.023016 | 0.013701 | 0.011183 | 0.004662 | 0.027974 | 0.008071 | 0.025571 | 0.018442 | 0.017913 |
| 27.9305556 | 0.015117 | 0.0112 | 0.007215 | 0.002942 | 0.018406 | 0.011047 | 0.023938 | 0.011886 | 0.03216 | 0.014171 | 0.016083 | 0.010453 | 0.003989 | 0.026433 | 0.037037 | 0.041203 | 0.023808 | 0.003979 | 0.008298 | 0.030823 | 0.012323 | 0.036073 | 0.01604 | 0.025221 |
| 27.9444444 | 0.017962 | 0.018033 | 0.015225 | 0.008006 | 0.019281 | 0.020493 | 0.02392 | 0.012764 | 0.019334 | 0.010696 | 0.020814 | 0.018666 | 0.004512 | 0.047082 | 0.051246 | 0.054439 | 0.02964 | 0.00343 | 0.019051 | 0.066847 | 0.009664 | 0.028896 | 0.01216 | 0.019593 |
| 27.9583333 | 0.015629 | 0.021029 | 0.027465 | 0.015946 | 0.021239 | 0.022605 | 0.016572 | 0.026101 | 0.028766 | 0.006241 | 0.031089 | 0.031558 | 0.002974 | 0.060734 | 0.063943 | 0.063412 | 0.02848 | 0.00974 | 0.039335 | 0.108412 | 0.006761 | 0.01628 | 0.020476 | 0.013515 |
| 27.9722222 | 0.016737 | 0.021021 | 0.030669 | 0.018158 | 0.026329 | 0.015127 | 0.008256 | 0.037407 | 0.062631 | 0.002366 | 0.041421 | 0.037351 | 0.001574 | 0.066918 | 0.054356 | 0.070719 | 0.023304 | 0.024724 | 0.053635 | 0.124225 | 0.013422 | 0.018407 | 0.036386 | 0.016552 |
| 27.9861111 | 0.023458 | 0.018024 | 0.020999 | 0.014371 | 0.025894 | 0.006267 | 0.003756 | 0.040876 | 0.090478 | 0.000663 | 0.040018 | 0.027272 | 0.00108 | 0.071615 | 0.030609 | 0.072605 | 0.018922 | 0.036259 | 0.049588 | 0.113544 | 0.023643 | 0.026803 | 0.049658 | 0.015075 |
| 28 | 0.033716 | 0.012722 | 0.010079 | 0.012774 | 0.041321 | 0.003847 | 0.004454 | 0.039788 | 0.090833 | 0.001104 | 0.025185 | 0.011577 | 0.002017 | 0.082143 | 0.036421 | 0.070365 | 0.017015 | 0.033532 | 0.034865 | 0.082331 | 0.030916 | 0.02697 | 0.056435 | 0.011203 |
| 28.0138889 | 0.04618 | 0.00891 | 0.005226 | 0.01461 | 0.069046 | 0.008526 | 0.008744 | 0.03319 | 0.079899 | 0.003604 | 0.011636 | 0.003426 | 0.002922 | 0.080074 | 0.060541 | 0.06534 | 0.017576 | 0.022732 | 0.019244 | 0.040349 | 0.033301 | 0.02422 | 0.054996 | 0.012821 |
| 28.0277778 | 0.053085 | 0.008396 | 0.00677 | 0.018692 | 0.066122 | 0.016086 | 0.012365 | 0.027113 | 0.076405 | 0.010142 | 0.005769 | 0.002087 | 0.003609 | 0.053971 | 0.075541 | 0.051417 | 0.02046 | 0.01386 | 0.011768 | 0.01838 | 0.031997 | 0.030034 | 0.046537 | 0.020219 |
| 28.0416667 | 0.050579 | 0.00957 | 0.014746 | 0.035261 | 0.034036 | 0.016188 | 0.012178 | 0.038612 | 0.07923 | 0.024648 | 0.008595 | 0.002884 | 0.011145 | 0.031854 | 0.077914 | 0.040254 | 0.028825 | 0.021497 | 0.026936 | 0.042544 | 0.024497 | 0.030162 | 0.031401 | 0.024834 |
| 28.0555556 | 0.048147 | 0.016191 | 0.021814 | 0.070329 | 0.021688 | 0.015326 | 0.017395 | 0.083811 | 0.084594 | 0.04825 | 0.014561 | 0.003911 | 0.02871 | 0.030875 | 0.065554 | 0.049616 | 0.038135 | 0.047551 | 0.065989 | 0.110335 | 0.016505 | 0.019978 | 0.017216 | 0.019165 |
| 28.0694444 | 0.044759 | 0.027464 | 0.017932 | 0.091671 | 0.050812 | 0.029957 | 0.032443 | 0.118893 | 0.080438 | 0.075572 | 0.01625 | 0.005197 | 0.038086 | 0.041294 | 0.052312 | 0.070186 | 0.031376 | 0.061799 | 0.100906 | 0.175403 | 0.0231 | 0.031111 | 0.013415 | 0.017685 |
| 28.0833333 | 0.033167 | 0.037054 | 0.012014 | 0.069932 | 0.092312 | 0.0463 | 0.040716 | 0.082753 | 0.059438 | 0.090053 | 0.013268 | 0.008782 | 0.026774 | 0.051094 | 0.040981 | 0.080853 | 0.014426 | 0.054728 | 0.104809 | 0.185081 | 0.031794 | 0.062839 | 0.023057 | 0.036557 |
| 28.0972222 | 0.022338 | 0.042621 | 0.013741 | 0.035127 | 0.105341 | 0.049299 | 0.035002 | 0.025134 | 0.034836 | 0.07256 | 0.007281 | 0.014284 | 0.011309 | 0.049769 | 0.023715 | 0.079426 | 0.008503 | 0.046119 | 0.080394 | 0.142024 | 0.026021 | 0.07568 | 0.038326 | 0.05466 |
| 28.1111111 | 0.017233 | 0.042347 | 0.01295 | 0.023613 | 0.076686 | 0.041519 | 0.025125 | 0.010051 | 0.018663 | 0.034597 | 0.006333 | 0.015382 | 0.004734 | 0.034262 | 0.011017 | 0.07449 | 0.010754 | 0.034692 | 0.045256 | 0.077232 | 0.012412 | 0.051012 | 0.039435 | 0.041691 |
| 28.125 | 0.011466 | 0.039269 | 0.007625 | 0.030972 | 0.034376 | 0.025307 | 0.020317 | 0.01424 | 0.016421 | 0.009289 | 0.014553 | 0.010766 | 0.00667 | 0.01908 | 0.017156 | 0.066892 | 0.011429 | 0.018196 | 0.02024 | 0.030354 | 0.00651 | 0.026644 | 0.027085 | 0.021859 |
| 28.1388889 | 0.005428 | 0.03943 | 0.007308 | 0.037345 | 0.011497 | 0.010392 | 0.023136 | 0.012043 | 0.027001 | 0.005327 | 0.022621 | 0.00811 | 0.013535 | 0.022019 | 0.026376 | 0.058407 | 0.011392 | 0.007439 | 0.009337 | 0.017331 | 0.00988 | 0.023427 | 0.022444 | 0.016971 |
| 28.1527778 | 0.003855 | 0.043263 | 0.011064 | 0.031959 | 0.006806 | 0.008434 | 0.03172 | 0.006144 | 0.037981 | 0.011686 | 0.024313 | 0.01746 | 0.020015 | 0.042098 | 0.018637 | 0.047787 | 0.01243 | 0.006472 | 0.004401 | 0.017361 | 0.011809 | 0.02304 | 0.036528 | 0.016528 |
| 28.1666667 | 0.004628 | 0.037107 | 0.012646 | 0.017545 | 0.004796 | 0.016659 | 0.043166 | 0.009246 | 0.042088 | 0.025967 | 0.019927 | 0.043884 | 0.020045 | 0.056172 | 0.010139 | 0.02846 | 0.013243 | 0.011241 | 0.004185 | 0.029166 | 0.010743 | 0.027834 | 0.048724 | 0.021747 |
| 28.1805556 | 0.003993 | 0.022623 | 0.012031 | 0.005993 | 0.004711 | 0.022551 | 0.051187 | 0.035408 | 0.041584 | 0.045087 | 0.011643 | 0.066315 | 0.011886 | 0.046671 | 0.011972 | 0.010851 | 0.011771 | 0.012839 | 0.008607 | 0.055052 | 0.008755 | 0.027236 | 0.038753 | 0.018896 |
| 28.1944444 | 0.002596 | 0.019187 | 0.012265 | 0.003063 | 0.008533 | 0.020083 | 0.047191 | 0.056321 | 0.03717 | 0.050791 | 0.00537 | 0.058052 | 0.005498 | 0.025009 | 0.012278 | 0.00634 | 0.008842 | 0.014947 | 0.017653 | 0.071493 | 0.006684 | 0.015811 | 0.039861 | 0.010013 |
| 28.2083333 | 0.001397 | 0.018381 | 0.013632 | 0.002881 | 0.01445 | 0.012448 | 0.030033 | 0.043569 | 0.02583 | 0.036901 | 0.005872 | 0.034718 | 0.006626 | 0.017448 | 0.010603 | 0.005624 | 0.006046 | 0.024288 | 0.027762 | 0.066265 | 0.007925 | 0.014048 | 0.073091 | 0.018055 |
| 28.2222222 | 0.000633 | 0.010581 | 0.014254 | 0.002802 | 0.022498 | 0.005768 | 0.012601 | 0.027406 | 0.011971 | 0.035781 | 0.010425 | 0.021706 | 0.007616 | 0.024873 | 0.00952 | 0.00546 | 0.003583 | 0.023772 | 0.029575 | 0.051259 | 0.010701 | 0.016116 | 0.095738 | 0.03234 |
| 28.2361111 | 0.000509 | 0.004794 | 0.01347 | 0.002973 | 0.02913 | 0.004002 | 0.004537 | 0.020115 | 0.004977 | 0.05668 | 0.013287 | 0.017861 | 0.004661 | 0.024713 | 0.00626 | 0.010772 | 0.004359 | 0.011819 | 0.017834 | 0.040108 | 0.00989 | 0.01627 | 0.081874 | 0.027959 |
| 28.25 | 0.000949 | 0.003315 | 0.011212 | 0.008761 | 0.0289 | 0.006136 | 0.001772 | 0.018916 | 0.004014 | 0.054749 | 0.010277 | 0.012462 | 0.008048 | 0.013375 | 0.005451 | 0.012699 | 0.015443 | 0.006924 | 0.009455 | 0.034326 | 0.010573 | 0.028899 | 0.052153 | 0.018648 |
| 28.2638889 | 0.017261 | 0.007325 | 0.009585 | 0.038299 | 0.054954 | 0.016099 | 0.006288 | 0.060495 | 0.009116 | 0.055396 | 0.009586 | 0.004855 | 0.02114 | 0.007567 | 0.007269 | 0.014252 | 0.043047 | 0.010632 | 0.012528 | 0.033385 | 0.016296 | 0.040156 | 0.03038 | 0.031381 |
| 28.2777778 | 0.065397 | 0.020097 | 0.024243 | 0.079738 | 0.103343 | 0.037693 | 0.011708 | 0.144016 | 0.015947 | 0.075563 | 0.039911 | 0.007801 | 0.029591 | 0.014635 | 0.007677 | 0.025649 | 0.061984 | 0.026471 | 0.015894 | 0.049098 | 0.022066 | 0.031433 | 0.020417 | 0.039114 |
| 28.2916667 | 0.10754 | 0.044766 | 0.072032 | 0.110353 | 0.134319 | 0.052228 | 0.007555 | 0.178528 | 0.014043 | 0.055969 | 0.094947 | 0.015889 | 0.035229 | 0.025482 | 0.015073 | 0.02772 | 0.054006 | 0.052944 | 0.01527 | 0.066952 | 0.021265 | 0.014903 | 0.022785 | 0.02429 |
| 28.3055556 | 0.089521 | 0.052712 | 0.093732 | 0.119012 | 0.159483 | 0.05047 | 0.006501 | 0.14462 | 0.014558 | 0.065802 | 0.102195 | 0.011505 | 0.02926 | 0.02777 | 0.023116 | 0.014534 | 0.042976 | 0.055516 | 0.013579 | 0.059464 | 0.014653 | 0.007248 | 0.039564 | 0.012683 |
| 28.3194444 | 0.035297 | 0.026901 | 0.049269 | 0.082509 | 0.138108 | 0.040962 | 0.008741 | 0.102174 | 0.016705 | 0.109855 | 0.050758 | 0.002852 | 0.016946 | 0.022037 | 0.016522 | 0.006267 | 0.027869 | 0.030281 | 0.015429 | 0.031695 | 0.013019 | 0.010414 | 0.049402 | 0.014267 |
| 28.3333333 | 0.004943 | 0.009866 | 0.011661 | 0.031773 | 0.065444 | 0.022721 | 0.00583 | 0.052183 | 0.009916 | 0.077883 | 0.012884 | 0.000723 | 0.012375 | 0.017896 | 0.013292 | 0.00547 | 0.00897 | 0.016229 | 0.012342 | 0.014809 | 0.016273 | 0.018514 | 0.044124 | 0.022262 |
| 28.3472222 | 0.001894 | 0.00699 | 0.004953 | 0.013382 | 0.018116 | 0.008369 | 0.003099 | 0.01367 | 0.004714 | 0.030401 | 0.004061 | 0.001655 | 0.013903 | 0.014546 | 0.022406 | 0.004889 | 0.003475 | 0.011999 | 0.005389 | 0.007282 | 0.020057 | 0.025101 | 0.03992 | 0.028081 |
| 28.3611111 | 0.004832 | 0.003452 | 0.003836 | 0.017964 | 0.005707 | 0.005427 | 0.002329 | 0.004097 | 0.003974 | 0.031216 | 0.002374 | 0.002502 | 0.009283 | 0.00855 | 0.0229 | 0.00339 | 0.004366 | 0.008933 | 0.002715 | 0.004098 | 0.022505 | 0.023307 | 0.040306 | 0.029274 |
| 28.375 | 0.008735 | 0.003418 | 0.005766 | 0.029412 | 0.004941 | 0.003828 | 0.003221 | 0.003346 | 0.003413 | 0.042911 | 0.004291 | 0.003264 | 0.004526 | 0.004704 | 0.012705 | 0.004556 | 0.005145 | 0.010713 | 0.002655 | 0.005323 | 0.018746 | 0.013511 | 0.037017 | 0.025481 |
| 28.3888889 | 0.011416 | 0.005843 | 0.00744 | 0.04223 | 0.00933 | 0.002438 | 0.005354 | 0.005478 | 0.004539 | 0.048689 | 0.007322 | 0.007458 | 0.006323 | 0.004609 | 0.013879 | 0.011067 | 0.006536 | 0.012432 | 0.006874 | 0.00618 | 0.011107 | 0.006082 | 0.02338 | 0.01502 |
| 28.4027778 | 0.010525 | 0.013772 | 0.009341 | 0.050534 | 0.012159 | 0.003581 | 0.006895 | 0.007842 | 0.007367 | 0.047949 | 0.01099 | 0.016038 | 0.01039 | 0.005778 | 0.034143 | 0.020424 | 0.006122 | 0.013537 | 0.015125 | 0.008372 | 0.006082 | 0.007039 | 0.011581 | 0.006457 |
| 28.4166667 | 0.006549 | 0.021864 | 0.012143 | 0.052571 | 0.008475 | 0.006439 | 0.008411 | 0.010802 | 0.009886 | 0.039854 | 0.015552 | 0.02574 | 0.011526 | 0.007219 | 0.047315 | 0.028981 | 0.003828 | 0.011642 | 0.020588 | 0.015307 | 0.004114 | 0.011977 | 0.016522 | 0.005864 |
| 28.4305556 | 0.003173 | 0.024202 | 0.015751 | 0.053193 | 0.002837 | 0.010578 | 0.01091 | 0.013799 | 0.011818 | 0.026022 | 0.020209 | 0.034286 | 0.009447 | 0.007852 | 0.032031 | 0.033044 | 0.00351 | 0.00756 | 0.018346 | 0.02253 | 0.003935 | 0.016662 | 0.030043 | 0.006435 |
| 28.4444444 | 0.004425 | 0.023583 | 0.019658 | 0.055858 | 0.003013 | 0.015456 | 0.01382 | 0.0135 | 0.014681 | 0.011509 | 0.024767 | 0.041629 | 0.005685 | 0.005676 | 0.012437 | 0.031715 | 0.005938 | 0.005963 | 0.012217 | 0.027977 | 0.004921 | 0.021391 | 0.043994 | 0.007107 |
| 28.4583333 | 0.009846 | 0.024102 | 0.023301 | 0.056114 | 0.010002 | 0.01902 | 0.015338 | 0.014238 | 0.020038 | 0.004237 | 0.029154 | 0.047553 | 0.005236 | 0.003868 | 0.009194 | 0.029248 | 0.00803 | 0.009803 | 0.008372 | 0.033464 | 0.004955 | 0.026117 | 0.057687 | 0.012247 |
| 28.4722222 | 0.015071 | 0.027673 | 0.026221 | 0.051666 | 0.019273 | 0.021328 | 0.015014 | 0.022067 | 0.026831 | 0.010497 | 0.032641 | 0.051035 | 0.010678 | 0.008465 | 0.012222 | 0.028937 | 0.008573 | 0.017426 | 0.008968 | 0.039608 | 0.006689 | 0.026931 | 0.064502 | 0.018566 |
| 28.4861111 | 0.017889 | 0.033231 | 0.028665 | 0.046095 | 0.027131 | 0.024252 | 0.015091 | 0.032611 | 0.032001 | 0.029659 | 0.035682 | 0.052448 | 0.017962 | 0.015792 | 0.021598 | 0.030085 | 0.008186 | 0.024341 | 0.011253 | 0.046822 | 0.009964 | 0.020938 | 0.059955 | 0.019802 |
| 28.5 | 0.019064 | 0.037222 | 0.031133 | 0.040414 | 0.033111 | 0.028259 | 0.016884 | 0.039387 | 0.033823 | 0.052349 | 0.038522 | 0.053682 | 0.023138 | 0.019721 | 0.04037 | 0.032602 | 0.007771 | 0.02611 | 0.018108 | 0.055951 | 0.010624 | 0.01172 | 0.04913 | 0.015297 |
| 28.5138889 | 0.017958 | 0.036764 | 0.033106 | 0.03544 | 0.036865 | 0.030706 | 0.018297 | 0.04207 | 0.030958 | 0.065615 | 0.038596 | 0.055087 | 0.02549 | 0.021907 | 0.057785 | 0.035639 | 0.008123 | 0.021546 | 0.023128 | 0.062358 | 0.007379 | 0.005295 | 0.040467 | 0.009967 |
| 28.5277778 | 0.014723 | 0.030773 | 0.032792 | 0.031426 | 0.0367 | 0.028827 | 0.01723 | 0.04013 | 0.023187 | 0.064735 | 0.033895 | 0.052084 | 0.02559 | 0.025682 | 0.063533 | 0.034485 | 0.006905 | 0.021532 | 0.026682 | 0.053756 | 0.008104 | 0.007879 | 0.036249 | 0.006001 |
| 28.5416667 | 0.013363 | 0.021613 | 0.030342 | 0.028685 | 0.033979 | 0.024976 | 0.016638 | 0.036327 | 0.013437 | 0.059027 | 0.026471 | 0.042611 | 0.022667 | 0.028213 | 0.056385 | 0.026979 | 0.006219 | 0.006377 | 0.02829 | 0.030874 | 0.021405 | 0.026786 | 0.033015 | 0.003291 |
| 28.5555556 | 0.014598 | 0.014589 | 0.02689 | 0.029027 | 0.030192 | 0.022079 | 0.01894 | 0.034169 | 0.00599 | 0.055784 | 0.019518 | 0.031691 | 0.015791 | 0.022501 | 0.040976 | 0.021192 | 0.009914 | 0.00725 | 0.028353 | 0.011229 | 0.036109 | 0.061176 | 0.027607 | 0.003509 |
| 28.5694444 | 0.014921 | 0.010895 | 0.02208 | 0.032287 | 0.02431 | 0.018759 | 0.01951 | 0.029978 | 0.004353 | 0.050642 | 0.013511 | 0.020811 | 0.009652 | 0.013289 | 0.026105 | 0.027613 | 0.011833 | 0.006812 | 0.029354 | 0.006066 | 0.037321 | 0.086528 | 0.022409 | 0.005302 |
| 28.5833333 | 0.014967 | 0.008241 | 0.021101 | 0.04091 | 0.020053 | 0.017021 | 0.016686 | 0.025082 | 0.006889 | 0.042633 | 0.007617 | 0.010259 | 0.009831 | 0.014033 | 0.015617 | 0.044 | 0.009037 | 0.006587 | 0.037296 | 0.019162 | 0.026457 | 0.070562 | 0.022621 | 0.007039 |
| 28.5972222 | 0.018032 | 0.007439 | 0.027043 | 0.053169 | 0.022209 | 0.021404 | 0.015752 | 0.025372 | 0.006354 | 0.0367 | 0.002824 | 0.006775 | 0.013825 | 0.031884 | 0.007466 | 0.054132 | 0.009972 | 0.01697 | 0.055814 | 0.050112 | 0.02417 | 0.040993 | 0.027526 | 0.016461 |
| 28.6111111 | 0.02247 | 0.011574 | 0.03183 | 0.056495 | 0.027515 | 0.029402 | 0.016978 | 0.034851 | 0.006141 | 0.033002 | 0.001235 | 0.010527 | 0.017019 | 0.059189 | 0.012472 | 0.055697 | 0.012255 | 0.037263 | 0.074577 | 0.087651 | 0.036675 | 0.057757 | 0.034832 | 0.039907 |
| 28.625 | 0.02469 | 0.020876 | 0.032707 | 0.048193 | 0.031219 | 0.036128 | 0.016905 | 0.049916 | 0.01552 | 0.028002 | 0.004087 | 0.009976 | 0.022372 | 0.072853 | 0.038153 | 0.0654 | 0.03835 | 0.059983 | 0.077139 | 0.112586 | 0.037268 | 0.08568 | 0.045619 | 0.063876 |
| 28.6388889 | 0.025737 | 0.031028 | 0.03467 | 0.038392 | 0.034407 | 0.040395 | 0.016925 | 0.05452 | 0.029138 | 0.021289 | 0.010992 | 0.00977 | 0.032486 | 0.062976 | 0.063132 | 0.076889 | 0.05503 | 0.071695 | 0.063424 | 0.113876 | 0.026163 | 0.085007 | 0.052076 | 0.071574 |
| 28.6527778 | 0.028563 | 0.04235 | 0.034441 | 0.029749 | 0.040398 | 0.041637 | 0.018587 | 0.044681 | 0.041539 | 0.015256 | 0.019817 | 0.023134 | 0.042222 | 0.040557 | 0.066077 | 0.071454 | 0.065369 | 0.064491 | 0.047985 | 0.099606 | 0.031024 | 0.088454 | 0.041757 | 0.061426 |
| 28.6666667 | 0.033811 | 0.055599 | 0.02351 | 0.021315 | 0.048645 | 0.040285 | 0.020021 | 0.039399 | 0.051541 | 0.010411 | 0.029046 | 0.037486 | 0.046729 | 0.01892 | 0.050706 | 0.065377 | 0.064256 | 0.044912 | 0.034323 | 0.07672 | 0.0468 | 0.080888 | 0.020348 | 0.039334 |
| 28.6805556 | 0.038933 | 0.064884 | 0.008492 | 0.016126 | 0.052922 | 0.036936 | 0.019722 | 0.042571 | 0.054205 | 0.006252 | 0.034958 | 0.039215 | 0.046188 | 0.011714 | 0.02968 | 0.063791 | 0.056753 | 0.024653 | 0.017645 | 0.045767 | 0.045349 | 0.045781 | 0.006006 | 0.016647 |
| 28.6944444 | 0.040793 | 0.064803 | 0.002841 | 0.01538 | 0.048888 | 0.030166 | 0.017226 | 0.042183 | 0.050462 | 0.003887 | 0.031378 | 0.034161 | 0.042788 | 0.016776 | 0.013072 | 0.042341 | 0.047305 | 0.013088 | 0.010695 | 0.022221 | 0.023973 | 0.016789 | 0.006327 | 0.004306 |
| 28.7083333 | 0.040798 | 0.050009 | 0.007317 | 0.018182 | 0.040571 | 0.020794 | 0.01297 | 0.036204 | 0.044663 | 0.005865 | 0.021009 | 0.028994 | 0.038019 | 0.01834 | 0.006715 | 0.014927 | 0.034412 | 0.010219 | 0.019607 | 0.022492 | 0.006515 | 0.00725 | 0.012638 | 0.003036 |
| 28.7222222 | 0.043064 | 0.026231 | 0.020479 | 0.021848 | 0.035015 | 0.011939 | 0.009572 | 0.026271 | 0.040613 | 0.014934 | 0.01329 | 0.028797 | 0.033831 | 0.015248 | 0.006142 | 0.005678 | 0.021719 | 0.016871 | 0.031576 | 0.040708 | 0.003898 | 0.005764 | 0.014649 | 0.008294 |
| 28.7361111 | 0.048615 | 0.010209 | 0.036226 | 0.022086 | 0.034719 | 0.005174 | 0.009201 | 0.032363 | 0.039545 | 0.028713 | 0.01273 | 0.034883 | 0.032443 | 0.010879 | 0.00761 | 0.009529 | 0.014173 | 0.03224 | 0.044325 | 0.069343 | 0.009631 | 0.010768 | 0.019138 | 0.017675 |
| 28.75 | 0.054612 | 0.008178 | 0.043044 | 0.017605 | 0.036377 | 0.001489 | 0.011356 | 0.060511 | 0.036337 | 0.036257 | 0.017428 | 0.038155 | 0.033848 | 0.01227 | 0.013615 | 0.018196 | 0.01014 | 0.042857 | 0.056104 | 0.108091 | 0.020347 | 0.035997 | 0.034214 | 0.028754 |

| | | | | | | | | | | | | | | | | | | | | | | | | |
|---|---|---|---|---|---|---|---|---|---|---|---|---|---|---|---|---|---|---|---|---|---|---|---|---|
| 28.7638889 | 0.054716 | 0.013293 | 0.040781 | 0.012126 | 0.036142 | 0.003724 | 0.015557 | 0.078954 | 0.029281 | 0.034112 | 0.023516 | 0.033592 | 0.037472 | 0.030002 | 0.020841 | 0.036569 | 0.007083 | 0.036502 | 0.059674 | 0.128694 | 0.029497 | 0.068319 | 0.054081 | 0.04052 |
| 28.7777778 | 0.045564 | 0.020661 | 0.035504 | 0.010602 | 0.031698 | 0.013114 | 0.022238 | 0.080858 | 0.023426 | 0.032579 | 0.028377 | 0.025786 | 0.043137 | 0.051055 | 0.024112 | 0.059421 | 0.005883 | 0.018774 | 0.051873 | 0.10036 | 0.028727 | 0.068497 | 0.067226 | 0.051321 |
| 28.7916667 | 0.03388 | 0.02387 | 0.031526 | 0.012199 | 0.022079 | 0.023715 | 0.029687 | 0.083943 | 0.023462 | 0.035303 | 0.029604 | 0.019921 | 0.047996 | 0.056975 | 0.021154 | 0.070469 | 0.006592 | 0.005859 | 0.0345 | 0.048763 | 0.02104 | 0.039552 | 0.060565 | 0.052253 |
| 28.8055556 | 0.026465 | 0.019498 | 0.028614 | 0.01194 | 0.011183 | 0.027834 | 0.0352 | 0.088989 | 0.025695 | 0.037731 | 0.028079 | 0.019018 | 0.048599 | 0.049747 | 0.013665 | 0.06949 | 0.006767 | 0.009065 | 0.017561 | 0.029039 | 0.011462 | 0.015511 | 0.037416 | 0.038662 |
| 28.8194444 | 0.02223 | 0.016455 | 0.023024 | 0.008296 | 0.005599 | 0.026115 | 0.038794 | 0.084534 | 0.023123 | 0.037436 | 0.028408 | 0.021324 | 0.045681 | 0.041552 | 0.007435 | 0.056796 | 0.00502 | 0.028665 | 0.014935 | 0.055927 | 0.007172 | 0.010531 | 0.016884 | 0.019271 |
| 28.8333333 | 0.018256 | 0.014136 | 0.018109 | 0.004202 | 0.00809 | 0.022731 | 0.041 | 0.069649 | 0.017667 | 0.033805 | 0.031365 | 0.022259 | 0.041626 | 0.036056 | 0.007172 | 0.040236 | 0.003176 | 0.039202 | 0.025434 | 0.09441 | 0.013416 | 0.018872 | 0.010044 | 0.009248 |
| 28.8472222 | 0.013658 | 0.018353 | 0.019675 | 0.002426 | 0.01472 | 0.019671 | 0.040635 | 0.054755 | 0.011832 | 0.027197 | 0.036265 | 0.022788 | 0.037534 | 0.028985 | 0.01376 | 0.038861 | 0.004072 | 0.026895 | 0.033879 | 0.115334 | 0.023779 | 0.030645 | 0.010481 | 0.011714 |
| 28.8611111 | 0.010815 | 0.031921 | 0.023584 | 0.001679 | 0.018185 | 0.018757 | 0.03913 | 0.035804 | 0.007742 | 0.020718 | 0.042038 | 0.028264 | 0.032939 | 0.018025 | 0.020585 | 0.050766 | 0.00462 | 0.018405 | 0.034265 | 0.123302 | 0.031803 | 0.05313 | 0.01254 | 0.016252 |
| 28.875 | 0.012504 | 0.034264 | 0.019991 | 0.000799 | 0.015372 | 0.018358 | 0.038299 | 0.019158 | 0.008441 | 0.014368 | 0.041845 | 0.037132 | 0.027252 | 0.009459 | 0.017761 | 0.054121 | 0.004127 | 0.018909 | 0.029698 | 0.12768 | 0.033879 | 0.075539 | 0.019741 | 0.018348 |
| 28.8888889 | 0.016351 | 0.034745 | 0.011377 | 0.001215 | 0.009659 | 0.017649 | 0.032688 | 0.027245 | 0.00874 | 0.012716 | 0.031131 | 0.042288 | 0.024055 | 0.009032 | 0.014535 | 0.04245 | 0.005575 | 0.01449 | 0.031212 | 0.125377 | 0.028001 | 0.076736 | 0.029652 | 0.01674 |
| 28.9027778 | 0.019243 | 0.038456 | 0.010948 | 0.002291 | 0.005802 | 0.018268 | 0.019542 | 0.04055 | 0.005358 | 0.019893 | 0.016673 | 0.04189 | 0.025969 | 0.013137 | 0.020051 | 0.025754 | 0.004716 | 0.01211 | 0.039604 | 0.122107 | 0.018614 | 0.071894 | 0.037236 | 0.011943 |
| 28.9166667 | 0.021276 | 0.032505 | 0.018256 | 0.002779 | 0.003911 | 0.019326 | 0.008123 | 0.0325 | 0.003818 | 0.021455 | 0.010589 | 0.040362 | 0.028623 | 0.023792 | 0.021412 | 0.012217 | 0.002456 | 0.016986 | 0.038945 | 0.11605 | 0.011469 | 0.068962 | 0.036506 | 0.008603 |
| 28.9305556 | 0.022008 | 0.034918 | 0.024906 | 0.005887 | 0.005113 | 0.019045 | 0.004431 | 0.016405 | 0.004932 | 0.015531 | 0.014422 | 0.038791 | 0.028112 | 0.042498 | 0.016351 | 0.013382 | 0.002604 | 0.019999 | 0.026234 | 0.082896 | 0.007071 | 0.064878 | 0.024173 | 0.007413 |
| 28.9444444 | 0.019125 | 0.043919 | 0.028182 | 0.014148 | 0.013178 | 0.01647 | 0.00451 | 0.013151 | 0.004686 | 0.01808 | 0.016726 | 0.033569 | 0.02492 | 0.064216 | 0.019339 | 0.036514 | 0.002372 | 0.015899 | 0.020246 | 0.044411 | 0.004856 | 0.054624 | 0.013089 | 0.005325 |
| 28.9583333 | 0.012977 | 0.042561 | 0.028885 | 0.022605 | 0.023615 | 0.011112 | 0.005592 | 0.02574 | 0.012587 | 0.028893 | 0.012161 | 0.025177 | 0.019784 | 0.076192 | 0.031104 | 0.066539 | 0.001728 | 0.019131 | 0.032608 | 0.04468 | 0.004406 | 0.030741 | 0.020033 | 0.006387 |
| 28.9722222 | 0.010651 | 0.032932 | 0.02865 | 0.027989 | 0.027116 | 0.007009 | 0.006826 | 0.042308 | 0.028708 | 0.030883 | 0.011269 | 0.016082 | 0.013753 | 0.069563 | 0.035467 | 0.081022 | 0.002237 | 0.040576 | 0.048428 | 0.061965 | 0.005257 | 0.017579 | 0.041201 | 0.017827 |
| 28.9861111 | 0.016173 | 0.018658 | 0.025494 | 0.032907 | 0.021052 | 0.010041 | 0.006801 | 0.063094 | 0.032349 | 0.021704 | 0.021535 | 0.008449 | 0.009662 | 0.05698 | 0.027652 | 0.080742 | 0.002001 | 0.06478 | 0.056756 | 0.075711 | 0.004896 | 0.030329 | 0.058032 | 0.029645 |
| 29 | 0.022383 | 0.007336 | 0.019206 | 0.036303 | 0.011792 | 0.019098 | 0.005805 | 0.094505 | 0.025308 | 0.01613 | 0.036665 | 0.010082 | 0.007894 | 0.054674 | 0.028606 | 0.078752 | 0.00093 | 0.067395 | 0.055672 | 0.095556 | 0.004081 | 0.036371 | 0.058343 | 0.025699 |
| 29.0138889 | 0.021628 | 0.005488 | 0.01085 | 0.036814 | 0.009812 | 0.030436 | 0.006182 | 0.111706 | 0.02085 | 0.012893 | 0.04959 | 0.024303 | 0.008227 | 0.063333 | 0.047177 | 0.076294 | 0.004423 | 0.051703 | 0.044584 | 0.105722 | 0.008229 | 0.021031 | 0.04137 | 0.011306 |
| 29.0277778 | 0.013356 | 0.009443 | 0.003554 | 0.040348 | 0.016521 | 0.038519 | 0.014124 | 0.089188 | 0.02297 | 0.005797 | 0.052281 | 0.037464 | 0.018526 | 0.069481 | 0.05542 | 0.073945 | 0.017553 | 0.040499 | 0.029358 | 0.079096 | 0.014288 | 0.006118 | 0.022566 | 0.009329 |
| 29.0416667 | 0.007947 | 0.01317 | 0.004903 | 0.058967 | 0.017033 | 0.040125 | 0.021321 | 0.057178 | 0.037293 | 0.004601 | 0.041884 | 0.037602 | 0.037275 | 0.056867 | 0.037441 | 0.075847 | 0.02992 | 0.028969 | 0.019988 | 0.056236 | 0.013778 | 0.004306 | 0.020565 | 0.023057 |
| 29.0555556 | 0.01257 | 0.023748 | 0.015497 | 0.094337 | 0.018325 | 0.039865 | 0.027156 | 0.073866 | 0.057726 | 0.018971 | 0.030298 | 0.027056 | 0.045389 | 0.034246 | 0.017705 | 0.077164 | 0.025818 | 0.017047 | 0.036497 | 0.100562 | 0.007664 | 0.015465 | 0.035881 | 0.026929 |
| 29.0694444 | 0.024561 | 0.046797 | 0.019923 | 0.110662 | 0.037099 | 0.039356 | 0.048145 | 0.119076 | 0.068779 | 0.047509 | 0.020787 | 0.014343 | 0.035664 | 0.031908 | 0.01818 | 0.067696 | 0.012041 | 0.021673 | 0.07349 | 0.165543 | 0.007456 | 0.036005 | 0.057596 | 0.022597 |
| 29.0833333 | 0.046418 | 0.069704 | 0.014918 | 0.076439 | 0.05293 | 0.03561 | 0.066763 | 0.10457 | 0.067601 | 0.08058 | 0.010771 | 0.009395 | 0.021182 | 0.047263 | 0.02186 | 0.060254 | 0.003963 | 0.029654 | 0.085643 | 0.191549 | 0.016008 | 0.06431 | 0.073477 | 0.03599 |
| 29.0972222 | 0.065977 | 0.068338 | 0.014915 | 0.028155 | 0.050195 | 0.029022 | 0.060545 | 0.066934 | 0.058866 | 0.098339 | 0.008453 | 0.011616 | 0.014301 | 0.054193 | 0.02042 | 0.064753 | 0.002062 | 0.023304 | 0.054194 | 0.187413 | 0.018759 | 0.08944 | 0.091023 | 0.054607 |
| 29.1111111 | 0.065224 | 0.047533 | 0.013842 | 0.018575 | 0.03615 | 0.021673 | 0.038304 | 0.070917 | 0.046609 | 0.092895 | 0.011778 | 0.010632 | 0.014439 | 0.044981 | 0.022181 | 0.061934 | 0.000904 | 0.009739 | 0.020513 | 0.153329 | 0.013909 | 0.077356 | 0.111106 | 0.048123 |
| 29.125 | 0.053663 | 0.046602 | 0.009475 | 0.043848 | 0.022426 | 0.015659 | 0.017675 | 0.055692 | 0.037084 | 0.084828 | 0.011044 | 0.008373 | 0.015563 | 0.027068 | 0.021854 | 0.053925 | 0.001189 | 0.003468 | 0.013986 | 0.084881 | 0.011582 | 0.044592 | 0.096807 | 0.027173 |
| 29.1388889 | 0.045019 | 0.062145 | 0.011151 | 0.065752 | 0.018213 | 0.010394 | 0.006379 | 0.018019 | 0.0319 | 0.082475 | 0.007717 | 0.015398 | 0.012923 | 0.012925 | 0.016656 | 0.050558 | 0.00389 | 0.005283 | 0.024811 | 0.02984 | 0.01108 | 0.035011 | 0.053933 | 0.021055 |
| 29.1527778 | 0.034568 | 0.068846 | 0.012475 | 0.0679 | 0.018737 | 0.00751 | 0.003116 | 0.003043 | 0.026153 | 0.066644 | 0.009008 | 0.025951 | 0.007809 | 0.008461 | 0.021498 | 0.048978 | 0.005408 | 0.005746 | 0.047147 | 0.014845 | 0.017383 | 0.026679 | 0.040563 | 0.019766 |
| 29.1666667 | 0.018591 | 0.061529 | 0.009961 | 0.059381 | 0.015595 | 0.009906 | 0.003828 | 0.015416 | 0.015964 | 0.044039 | 0.012171 | 0.028664 | 0.006121 | 0.009353 | 0.040496 | 0.047005 | 0.00363 | 0.003127 | 0.05946 | 0.017702 | 0.026506 | 0.014196 | 0.066595 | 0.010564 |
| 29.1805556 | 0.006782 | 0.03703 | 0.007654 | 0.04834 | 0.011143 | 0.009641 | 0.005697 | 0.048201 | 0.008197 | 0.054114 | 0.008909 | 0.0241 | 0.006604 | 0.008887 | 0.044171 | 0.033902 | 0.001851 | 0.003194 | 0.046299 | 0.035822 | 0.02768 | 0.013905 | 0.090156 | 0.006984 |
| 29.1944444 | 0.006907 | 0.015015 | 0.008002 | 0.038624 | 0.012885 | 0.009376 | 0.008335 | 0.066498 | 0.010264 | 0.080524 | 0.00415 | 0.015914 | 0.005136 | 0.005696 | 0.029252 | 0.012493 | 0.002437 | 0.007746 | 0.030599 | 0.046183 | 0.024562 | 0.01429 | 0.087544 | 0.0105 |
| 29.2083333 | 0.013794 | 0.01222 | 0.012447 | 0.031995 | 0.018193 | 0.018115 | 0.010502 | 0.050342 | 0.01481 | 0.081896 | 0.003768 | 0.008221 | 0.002836 | 0.006908 | 0.021922 | 0.002911 | 0.003062 | 0.012807 | 0.02674 | 0.02955 | 0.019678 | 0.014828 | 0.065282 | 0.011783 |
| 29.2222222 | 0.018387 | 0.010708 | 0.019509 | 0.027845 | 0.019084 | 0.024791 | 0.013396 | 0.025017 | 0.014835 | 0.071272 | 0.003945 | 0.005856 | 0.002975 | 0.012874 | 0.022406 | 0.003894 | 0.003002 | 0.015348 | 0.026117 | 0.012199 | 0.013579 | 0.022228 | 0.035339 | 0.011029 |
| 29.2361111 | 0.019192 | 0.003856 | 0.027017 | 0.024914 | 0.015032 | 0.021103 | 0.017528 | 0.014797 | 0.011754 | 0.057333 | 0.006066 | 0.010923 | 0.006546 | 0.01218 | 0.018791 | 0.006013 | 0.00272 | 0.017038 | 0.018585 | 0.011013 | 0.008065 | 0.036786 | 0.013709 | 0.01607 |
| 29.25 | 0.019402 | 0.000876 | 0.032893 | 0.022605 | 0.010669 | 0.016568 | 0.019805 | 0.014557 | 0.007557 | 0.03399 | 0.013273 | 0.021325 | 0.010565 | 0.00536 | 0.013607 | 0.009184 | 0.001759 | 0.01839 | 0.007518 | 0.011769 | 0.008065 | 0.052364 | 0.011154 | 0.027474 |
| 29.2638889 | 0.018943 | 0.000925 | 0.035397 | 0.021622 | 0.008332 | 0.018521 | 0.021596 | 0.016844 | 0.004095 | 0.024556 | 0.019991 | 0.028774 | 0.012279 | 0.00497 | 0.009226 | 0.008325 | 0.001535 | 0.013997 | 0.005072 | 0.0129 | 0.015344 | 0.049888 | 0.018865 | 0.031913 |
| 29.2777778 | 0.014487 | 0.003131 | 0.034959 | 0.022188 | 0.007387 | 0.020386 | 0.024316 | 0.01628 | 0.002043 | 0.047192 | 0.020761 | 0.026103 | 0.0109 | 0.012785 | 0.008499 | 0.004021 | 0.002852 | 0.006489 | 0.012398 | 0.028269 | 0.019926 | 0.028147 | 0.024331 | 0.021287 |
| 29.2916667 | 0.007911 | 0.009812 | 0.032603 | 0.02314 | 0.007636 | 0.01903 | 0.024215 | 0.01087 | 0.002109 | 0.077046 | 0.017632 | 0.017613 | 0.009041 | 0.018332 | 0.024059 | 0.004107 | 0.004055 | 0.003721 | 0.01951 | 0.046125 | 0.014931 | 0.021271 | 0.025343 | 0.011682 |
| 29.3055556 | 0.005263 | 0.021094 | 0.028987 | 0.022713 | 0.009669 | 0.019811 | 0.021848 | 0.010271 | 0.005432 | 0.088683 | 0.014955 | 0.012677 | 0.010316 | 0.014009 | 0.051966 | 0.007875 | 0.004738 | 0.003928 | 0.019415 | 0.050239 | 0.007268 | 0.030748 | 0.033054 | 0.012143 |
| 29.3194444 | 0.004703 | 0.029735 | 0.025553 | 0.01907 | 0.013811 | 0.020365 | 0.019009 | 0.012813 | 0.010157 | 0.077582 | 0.013055 | 0.013274 | 0.011585 | 0.006171 | 0.058632 | 0.011987 | 0.00612 | 0.00408 | 0.017243 | 0.042231 | 0.005915 | 0.025128 | 0.05491 | 0.010039 |
| 29.3333333 | 0.005006 | 0.025498 | 0.02137 | 0.011257 | 0.018517 | 0.017112 | 0.015294 | 0.014646 | 0.014205 | 0.04645 | 0.010686 | 0.015746 | 0.008581 | 0.00331 | 0.042373 | 0.017305 | 0.007276 | 0.005281 | 0.018053 | 0.033488 | 0.009598 | 0.012117 | 0.073291 | 0.005458 |
| 29.3472222 | 0.007846 | 0.015607 | 0.01276 | 0.006146 | 0.020124 | 0.01183 | 0.010596 | 0.015075 | 0.017875 | 0.022639 | 0.008508 | 0.018056 | 0.004045 | 0.003631 | 0.042858 | 0.018135 | 0.00597 | 0.007386 | 0.017216 | 0.030757 | 0.012052 | 0.010316 | 0.06967 | 0.00558 |
| 29.3611111 | 0.008633 | 0.015803 | 0.005713 | 0.011059 | 0.016186 | 0.006132 | 0.005395 | 0.011516 | 0.021545 | 0.031712 | 0.007049 | 0.018375 | 0.003346 | 0.004195 | 0.059369 | 0.011153 | 0.003294 | 0.00845 | 0.011158 | 0.032905 | 0.010013 | 0.01474 | 0.047407 | 0.006381 |
| 29.375 | 0.010988 | 0.020102 | 0.007771 | 0.020194 | 0.008494 | 0.002482 | 0.002739 | 0.011164 | 0.022391 | 0.061785 | 0.005992 | 0.014355 | 0.006666 | 0.005349 | 0.053001 | 0.004127 | 0.00237 | 0.007382 | 0.004482 | 0.039877 | 0.005255 | 0.024429 | 0.024023 | 0.008991 |
| 29.3888889 | 0.019852 | 0.023276 | 0.014245 | 0.030477 | 0.007478 | 0.001911 | 0.002942 | 0.013848 | 0.020008 | 0.090728 | 0.008412 | 0.00851 | 0.011635 | 0.009067 | 0.024763 | 0.002512 | 0.004657 | 0.004973 | 0.002836 | 0.047093 | 0.00274 | 0.040247 | 0.014308 | 0.019908 |
| 29.4027778 | 0.023537 | 0.036712 | 0.023403 | 0.050223 | 0.025688 | 0.005698 | 0.004234 | 0.012377 | 0.02793 | 0.109339 | 0.012739 | 0.004516 | 0.018325 | 0.014117 | 0.006828 | 0.00541 | 0.009977 | 0.00314 | 0.00417 | 0.054833 | 0.002511 | 0.051417 | 0.01823 | 0.03359 |
| 29.4166667 | 0.017623 | 0.048608 | 0.029666 | 0.073498 | 0.037072 | 0.012324 | 0.00955 | 0.021778 | 0.033194 | 0.105052 | 0.013383 | 0.009419 | 0.023107 | 0.018403 | 0.009391 | 0.016833 | 0.014737 | 0.003241 | 0.005165 | 0.060565 | 0.001697 | 0.041846 | 0.019872 | 0.035976 |
| 29.4305556 | 0.014941 | 0.040025 | 0.021257 | 0.063656 | 0.037654 | 0.012227 | 0.018397 | 0.039236 | 0.03098 | 0.075072 | 0.019732 | 0.020722 | 0.020761 | 0.027417 | 0.023692 | 0.034335 | 0.012942 | 0.003088 | 0.004458 | 0.045727 | 0.000819 | 0.021068 | 0.021404 | 0.023179 |
| 29.4444444 | 0.016468 | 0.022074 | 0.008298 | 0.025944 | 0.04655 | 0.005517 | 0.022349 | 0.033609 | 0.040582 | 0.046933 | 0.027541 | 0.021025 | 0.014394 | 0.038181 | 0.027714 | 0.039996 | 0.006351 | 0.002632 | 0.00325 | 0.019688 | 0.001346 | 0.014489 | 0.035496 | 0.015483 |
| 29.4583333 | 0.014083 | 0.014914 | 0.008873 | 0.019825 | 0.032241 | 0.003915 | 0.01909 | 0.017805 | 0.035063 | 0.031811 | 0.020268 | 0.014903 | 0.010281 | 0.036373 | 0.023717 | 0.029808 | 0.002561 | 0.006438 | 0.003525 | 0.007582 | 0.002787 | 0.014111 | 0.043663 | 0.021656 |
| 29.4722222 | 0.012655 | 0.022644 | 0.020992 | 0.046324 | 0.007112 | 0.008844 | 0.015536 | 0.018031 | 0.016908 | 0.02279 | 0.011758 | 0.015243 | 0.010016 | 0.026954 | 0.040114 | 0.019412 | 0.002172 | 0.014042 | 0.004816 | 0.012124 | 0.002881 | 0.009472 | 0.038184 | 0.025889 |
| 29.4861111 | 0.015444 | 0.0295 | 0.034018 | 0.065131 | 0.002293 | 0.012314 | 0.013492 | 0.023621 | 0.013033 | 0.019421 | 0.014319 | 0.016777 | 0.011566 | 0.026051 | 0.066147 | 0.022164 | 0.002172 | 0.019137 | 0.010055 | 0.035831 | 0.003587 | 0.011222 | 0.030981 | 0.024289 |
| 29.5 | 0.016676 | 0.029619 | 0.04334 | 0.068725 | 0.003886 | 0.013872 | 0.011463 | 0.02625 | 0.015702 | 0.021225 | 0.017892 | 0.016788 | 0.012322 | 0.034336 | 0.07851 | 0.039064 | 0.001854 | 0.018176 | 0.017313 | 0.073753 | 0.007653 | 0.021974 | 0.026599 | 0.020127 |
| 29.5138889 | 0.016508 | 0.031393 | 0.048131 | 0.068512 | 0.006279 | 0.017367 | 0.010367 | 0.027313 | 0.020703 | 0.034374 | 0.016196 | 0.018122 | 0.01149 | 0.038513 | 0.081865 | 0.061044 | 0.001291 | 0.013832 | 0.019593 | 0.102484 | 0.009393 | 0.028741 | 0.027621 | 0.013757 |
| 29.5277778 | 0.017733 | 0.036792 | 0.047075 | 0.069185 | 0.012788 | 0.02052 | 0.00883 | 0.024932 | 0.024975 | 0.055089 | 0.012182 | 0.015774 | 0.010037 | 0.028608 | 0.084216 | 0.075937 | 0.004172 | 0.009721 | 0.015723 | 0.100063 | 0.006762 | 0.022429 | 0.03054 | 0.007361 |
| 29.5416667 | 0.021517 | 0.042532 | 0.038211 | 0.068497 | 0.020203 | 0.020646 | 0.00686 | 0.020777 | 0.024272 | 0.069815 | 0.009331 | 0.009474 | 0.009094 | 0.01301 | 0.078939 | 0.071038 | 0.012829 | 0.007071 | 0.009912 | 0.071508 | 0.006338 | 0.021729 | 0.027255 | 0.006601 |
| 29.5555556 | 0.027198 | 0.042736 | 0.024787 | 0.072136 | 0.023301 | 0.016964 | 0.007528 | 0.020177 | 0.019707 | 0.071123 | 0.008616 | 0.010206 | 0.010076 | 0.003224 | 0.062531 | 0.056809 | 0.02102 | 0.007094 | 0.0075 | 0.045087 | 0.007685 | 0.041252 | 0.01855 | 0.011675 |
| 29.5694444 | 0.030519 | 0.034047 | 0.014105 | 0.084577 | 0.021145 | 0.009435 | 0.009718 | 0.025466 | 0.012874 | 0.056421 | 0.008772 | 0.017101 | 0.014563 | 0.00607 | 0.047558 | 0.064319 | 0.024182 | 0.011664 | 0.0112 | 0.037702 | 0.00692 | 0.050143 | 0.015281 | 0.016347 |
| 29.5833333 | 0.032286 | 0.019986 | 0.009855 | 0.099929 | 0.018866 | 0.003194 | 0.010636 | 0.03121 | 0.009003 | 0.039621 | 0.008811 | 0.020265 | 0.02112 | 0.024243 | 0.045443 | 0.084942 | 0.027725 | 0.021254 | 0.0232 | 0.051811 | 0.007407 | 0.032488 | 0.022705 | 0.023328 |
| 29.5972222 | 0.036328 | 0.008333 | 0.01557 | 0.111116 | 0.022187 | 0.003984 | 0.0111 | 0.031341 | 0.015395 | 0.038536 | 0.011186 | 0.02143 | 0.02561 | 0.041916 | 0.060419 | 0.082877 | 0.033783 | 0.02952 | 0.042195 | 0.079597 | 0.01328 | 0.03255 | 0.041729 | 0.032692 |
| 29.6111111 | 0.039276 | 0.006952 | 0.028051 | 0.108347 | 0.028415 | 0.011581 | 0.011964 | 0.027482 | 0.027078 | 0.046242 | 0.017248 | 0.027298 | 0.027237 | 0.045842 | 0.083981 | 0.063201 | 0.028593 | 0.032362 | 0.064388 | 0.102031 | 0.018955 | 0.066141 | 0.072528 | 0.048669 |
| 29.625 | 0.037226 | 0.017621 | 0.036941 | 0.086626 | 0.031573 | 0.021652 | 0.013108 | 0.022972 | 0.032902 | 0.047675 | 0.026265 | 0.037748 | 0.028989 | 0.043112 | 0.106357 | 0.048961 | 0.011492 | 0.033714 | 0.081266 | 0.104728 | 0.01827 | 0.082132 | 0.102679 | 0.077309 |
| 29.6388889 | 0.031911 | 0.031669 | 0.044832 | 0.056355 | 0.031043 | 0.031475 | 0.014834 | 0.016229 | 0.032893 | 0.041594 | 0.038217 | 0.048311 | 0.031347 | 0.036516 | 0.125118 | 0.044586 | 0.00305 | 0.028438 | 0.07782 | 0.090235 | 0.0161 | 0.075122 | 0.111321 | 0.0929 |
| 29.6527778 | 0.02703 | 0.050459 | 0.054367 | 0.028449 | 0.027979 | 0.040344 | 0.01726 | 0.007712 | 0.033583 | 0.033704 | 0.052278 | 0.053201 | 0.030462 | 0.023467 | 0.120425 | 0.047718 | 0.004774 | 0.015335 | 0.058723 | 0.077522 | 0.015955 | 0.073876 | 0.087554 | 0.075864 |
| 29.6666667 | 0.024498 | 0.07552 | 0.051993 | 0.011925 | 0.018345 | 0.045829 | 0.018831 | 0.005298 | 0.036672 | 0.025198 | 0.061102 | 0.048581 | 0.024478 | 0.011464 | 0.0821 | 0.048157 | 0.010759 | 0.007363 | 0.042331 | 0.070198 | 0.015669 | 0.05424 | 0.04442 | 0.039756 |
| 29.6805556 | 0.022788 | 0.085998 | 0.032637 | 0.012596 | 0.008335 | 0.046498 | 0.019232 | 0.012979 | 0.038175 | 0.015442 | 0.057467 | 0.040271 | 0.016538 | 0.01852 | 0.037114 | 0.038199 | 0.019285 | 0.006724 | 0.029712 | 0.051379 | 0.013436 | 0.024941 | 0.018328 | 0.018091 |
| 29.6944444 | 0.02177 | 0.077035 | 0.017786 | 0.025876 | 0.008123 | 0.043331 | 0.020261 | 0.022428 | 0.034631 | 0.007204 | 0.047302 | 0.037358 | 0.009711 | 0.046135 | 0.018928 | 0.025742 | 0.021466 | 0.010687 | 0.016968 | 0.025517 | 0.009327 | 0.023667 | 0.026153 | 0.026974 |
| 29.7083333 | 0.024404 | 0.060408 | 0.015957 | 0.03038 | 0.009348 | 0.037116 | 0.021058 | 0.029184 | 0.026875 | 0.003879 | 0.040056 | 0.038613 | 0.005059 | 0.06141 | 0.042432 | 0.018996 | 0.020257 | 0.018228 | 0.011692 | 0.009041 | 0.005271 | 0.036777 | 0.041802 | 0.038722 |
| 29.7222222 | 0.030855 | 0.036331 | 0.020357 | 0.030164 | 0.012009 | 0.029827 | 0.018957 | 0.040596 | 0.019375 | 0.004958 | 0.035548 | 0.039219 | 0.00308 | 0.046217 | 0.076993 | 0.019841 | 0.017156 | 0.023039 | 0.022022 | 0.008563 | 0.002658 | 0.035734 | 0.046236 | 0.036534 |
| 29.7361111 | 0.036488 | 0.017622 | 0.030035 | 0.046734 | 0.026315 | 0.025378 | 0.015435 | 0.056709 | 0.017243 | 0.012531 | 0.026774 | 0.036689 | 0.003655 | 0.02109 | 0.068134 | 0.029292 | 0.012871 | 0.026803 | 0.04401 | 0.019699 | 0.001982 | 0.022517 | 0.042781 | 0.024084 |
| 29.75 | 0.03735 | 0.023619 | 0.030975 | 0.052782 | 0.044412 | 0.024657 | 0.015756 | 0.061004 | 0.020438 | 0.027769 | 0.015538 | 0.028306 | 0.006048 | 0.013354 | 0.049296 | 0.045202 | 0.016249 | 0.032376 | 0.060408 | 0.029324 | 0.006038 | 0.02755 | 0.028318 | 0.018434 |
| 29.7638889 | 0.03455 | 0.035678 | 0.021636 | 0.044053 | 0.051703 | 0.024383 | 0.020765 | 0.046784 | 0.024902 | 0.036911 | 0.012578 | 0.015505 | 0.009866 | 0.032161 | 0.066452 | 0.066642 | 0.019552 | 0.034746 | 0.058356 | 0.03889 | 0.021587 | 0.061501 | 0.014989 | 0.042047 |
| 29.7777778 | 0.033321 | 0.029866 | 0.012978 | 0.046414 | 0.041755 | 0.022611 | 0.024107 | 0.029387 | 0.028558 | 0.033807 | 0.016498 | 0.006307 | 0.015454 | 0.0536 | 0.057764 | 0.088025 | 0.017067 | 0.033542 | 0.046771 | 0.051038 | 0.03471 | 0.079677 | 0.027122 | 0.068607 |
| 29.7916667 | 0.036156 | 0.027102 | 0.008757 | 0.058045 | 0.026656 | 0.020367 | 0.022575 | 0.021643 | 0.032183 | 0.029194 | 0.018795 | 0.006109 | 0.021929 | 0.051799 | 0.021288 | 0.089214 | 0.01803 | 0.032719 | 0.040931 | 0.054129 | 0.033204 | 0.053452 | 0.049467 | 0.056497 |
| 29.8055556 | 0.040068 | 0.040321 | 0.0103 | 0.064039 | 0.026481 | 0.018283 | 0.022246 | 0.022819 | 0.038761 | 0.026918 | 0.015891 | 0.008565 | 0.026279 | 0.034975 | 0.014444 | 0.067463 | 0.022371 | 0.032697 | 0.04097 | 0.046976 | 0.03588 | 0.029021 | 0.055588 | 0.025536 |
| 29.8194444 | 0.041965 | 0.047329 | 0.015443 | 0.050965 | 0.040147 | 0.015532 | 0.030474 | 0.019208 | 0.047553 | 0.02421 | 0.008394 | 0.007595 | 0.026033 | 0.023869 | 0.032635 | 0.050377 | 0.017831 | 0.030129 | 0.037326 | 0.0417 | 0.036213 | 0.030777 | 0.052901 | 0.014646 |
| 29.8333333 | 0.041954 | 0.04385 | 0.022665 | 0.027477 | 0.054158 | 0.013104 | 0.042429 | 0.007387 | 0.050391 | 0.024223 | 0.003764 | 0.00531 | 0.021823 | 0.021525 | 0.059506 | 0.052562 | 0.01063 | 0.021029 | 0.027298 | 0.037985 | 0.022844 | 0.027332 | 0.050716 | 0.016433 |
| 29.8472222 | 0.039041 | 0.044162 | 0.029469 | 0.014815 | 0.067123 | 0.013653 | 0.04627 | 0.009694 | 0.043368 | 0.028599 | 0.009624 | 0.005081 | 0.016532 | 0.022899 | 0.064382 | 0.063196 | 0.014775 | 0.010832 | 0.016114 | 0.025911 | 0.013097 | 0.013813 | 0.042507 | 0.016033 |
| 29.8611111 | 0.027754 | 0.060039 | 0.033196 | 0.015877 | 0.07599 | 0.016674 | 0.047131 | 0.027537 | 0.030481 | 0.031856 | 0.022895 | 0.007243 | 0.012482 | 0.024204 | 0.053793 | 0.068741 | 0.018344 | 0.010498 | 0.010565 | 0.018457 | 0.011448 | 0.015414 | 0.028487 | 0.023132 |
| 29.875 | 0.013366 | 0.085349 | 0.034597 | 0.020205 | 0.058324 | 0.018257 | 0.058071 | 0.031818 | 0.016763 | 0.030994 | 0.031996 | 0.008515 | 0.010324 | 0.018528 | 0.048861 | 0.060992 | 0.013377 | 0.018971 | 0.017197 | 0.032126 | 0.015186 | 0.047366 | 0.015357 | 0.042146 |
| 29.8888889 | 0.010542 | 0.082019 | 0.033847 | 0.026026 | 0.022479 | 0.015752 | 0.074579 | 0.017603 | 0.00905 | 0.027625 | 0.031973 | 0.009329 | 0.009513 | 0.012088 | 0.027473 | 0.042337 | 0.008977 | 0.027337 | 0.027568 | 0.055498 | 0.021373 | 0.094366 | 0.005997 | 0.055837 |
| 29.9027778 | 0.01978 | 0.050154 | 0.026846 | 0.036573 | 0.011472 | 0.011268 | 0.070244 | 0.008842 | 0.006751 | 0.02196 | 0.030099 | 0.012941 | 0.009246 | 0.016558 | 0.01578 | 0.021352 | 0.011682 | 0.025629 | 0.022672 | 0.072042 | 0.023457 | 0.118907 | 0.00282 | 0.049842 |
| 29.9166667 | 0.030323 | 0.038999 | 0.015143 | 0.046782 | 0.035468 | 0.008057 | 0.040146 | 0.010379 | 0.003236 | 0.015404 | 0.033774 | 0.017464 | 0.009584 | 0.026349 | 0.02155 | 0.016234 | 0.019863 | 0.013857 | 0.011408 | 0.076391 | 0.023186 | 0.109728 | 0.006153 | 0.032533 |
| 29.9305556 | 0.033662 | 0.053375 | 0.009343 | 0.053348 | 0.066722 | 0.006882 | 0.02494 | 0.007762 | 0.003503 | 0.011027 | 0.042037 | 0.019803 | 0.010838 | 0.030229 | 0.026057 | 0.042949 | 0.025093 | 0.005312 | 0.013839 | 0.070998 | 0.020864 | 0.087746 | 0.014059 | 0.01829 |
| 29.9444444 | 0.028738 | 0.05417 | 0.012339 | 0.06956 | 0.083018 | 0.007413 | 0.032441 | 0.006545 | 0.011314 | 0.012556 | 0.046071 | 0.020878 | 0.012315 | 0.029529 | 0.041019 | 0.082976 | 0.020386 | 0.007838 | 0.024798 | 0.054823 | 0.012547 | 0.075342 | 0.025302 | 0.0133 |
| 29.9583333 | 0.021104 | 0.040192 | 0.017406 | 0.093168 | 0.086115 | 0.007149 | 0.028201 | 0.023088 | 0.019861 | 0.026205 | 0.042559 | 0.023334 | 0.012398 | 0.031902 | 0.04126 | 0.110944 | 0.009659 | 0.020103 | 0.034868 | 0.030716 | 0.005419 | 0.086195 | 0.039295 | 0.016086 |
| 29.9722222 | 0.014667 | 0.039459 | 0.024103 | 0.09543 | 0.084572 | 0.004223 | 0.017312 | 0.050554 | 0.023432 | 0.043561 | 0.038611 | 0.026017 | 0.010173 | 0.037804 | 0.017695 | 0.124312 | 0.003042 | 0.036852 | 0.037801 | 0.017737 | 0.007076 | 0.101622 | 0.048613 | 0.016828 |
| 29.9861111 | 0.016888 | 0.043368 | 0.033836 | 0.06623 | 0.074316 | 0.001479 | 0.020515 | 0.062312 | 0.020075 | 0.042547 | 0.037614 | 0.027343 | 0.007105 | 0.041276 | 0.005972 | 0.123642 | 0.00156 | 0.048862 | 0.028019 | 0.01785 | 0.014156 | 0.081856 | 0.047752 | 0.012571 |
| 30 | 0.028794 | 0.039683 | 0.040387 | 0.034395 | 0.053191 | 0.0011 | 0.022484 | 0.050725 | 0.011993 | 0.026839 | 0.040171 | 0.035463 | 0.005724 | 0.039275 | 0.022302 | 0.114109 | 0.001743 | 0.050579 | 0.013919 | 0.013224 | 0.023356 | 0.043637 | 0.044257 | 0.01396 |
| 30.0138889 | 0.033511 | 0.034965 | 0.037371 | 0.034392 | 0.030203 | 0.001572 | 0.017673 | 0.040068 | 0.004961 | 0.023418 | 0.038002 | 0.054183 | 0.005843 | 0.041316 | 0.061561 | 0.109482 | 0.0033 | 0.041731 | 0.012572 | 0.010509 | 0.027591 | 0.028594 | 0.041621 | 0.019339 |
| 30.0277778 | 0.023274 | 0.031785 | 0.029246 | 0.050776 | 0.012458 | 0.001975 | 0.018397 | 0.045979 | 0.006623 | 0.025849 | 0.022261 | 0.061112 | 0.007041 | 0.052096 | 0.100304 | 0.105344 | 0.004471 | 0.026435 | 0.040944 | 0.031493 | 0.018923 | 0.02449 | 0.043741 | 0.015401 |
| 30.0416667 | 0.017645 | 0.038282 | 0.030846 | 0.049657 | 0.015353 | 0.007238 | 0.022288 | 0.041485 | 0.02427 | 0.024068 | 0.012766 | 0.039771 | 0.014527 | 0.058599 | 0.119945 | 0.103606 | 0.003531 | 0.02389 | 0.087434 | 0.062385 | 0.01102 | 0.027641 | 0.049674 | 0.012909 |
| 30.0555556 | 0.039809 | 0.05418 | 0.047173 | 0.052415 | 0.053169 | 0.022151 | 0.027304 | 0.027927 | 0.049537 | 0.041748 | 0.023589 | 0.020947 | 0.023947 | 0.059452 | 0.111531 | 0.090432 | 0.003697 | 0.038187 | 0.111402 | 0.074981 | 0.011025 | 0.049405 | 0.047103 | 0.028605 |
| 30.0694444 | 0.073126 | 0.057448 | 0.057528 | 0.057851 | 0.094018 | 0.034958 | 0.026219 | 0.048891 | 0.060584 | 0.077652 | 0.030646 | 0.025258 | 0.02083 | 0.06487 | 0.074278 | 0.044396 | 0.008191 | 0.043642 | 0.097653 | 0.086463 | 0.016258 | 0.070598 | 0.031391 | 0.047405 |
| 30.0833333 | 0.083317 | 0.040197 | 0.044315 | 0.038961 | 0.087321 | 0.032795 | 0.019027 | 0.070974 | 0.054166 | 0.098407 | 0.02029 | 0.028593 | 0.010135 | 0.063497 | 0.040661 | 0.007487 | 0.014238 | 0.029738 | 0.058848 | 0.094606 | 0.032952 | 0.074063 | 0.021863 | 0.049501 |
| 30.0972222 | 0.07167 | 0.023712 | 0.022361 | 0.023713 | 0.052036 | 0.023548 | 0.027752 | 0.056991 | 0.043732 | 0.094877 | 0.010251 | 0.025231 | 0.005585 | 0.040782 | 0.028357 | 0.003328 | 0.015141 | 0.013136 | 0.021127 | 0.063701 | 0.035942 | 0.049878 | 0.027139 | 0.033084 |
| 30.1111111 | 0.05766 | 0.022864 | 0.011361 | 0.021485 | 0.03282 | 0.016112 | 0.049477 | 0.034375 | 0.032797 | 0.088555 | 0.013854 | 0.030605 | 0.005078 | 0.019835 | 0.032296 | 0.015874 | 0.008934 | 0.012221 | 0.006708 | 0.030245 | 0.016866 | 0.029305 | 0.023103 | 0.017508 |
| 30.125 | 0.048097 | 0.031222 | 0.011426 | 0.014817 | 0.031388 | 0.008906 | 0.059796 | 0.015875 | 0.020863 | 0.07756 | 0.026482 | 0.042023 | 0.002963 | 0.015434 | 0.036669 | 0.03426 | 0.002751 | 0.018966 | 0.014444 | 0.021681 | 0.008018 | 0.0488 | 0.015394 | 0.01886 |
| 30.1388889 | 0.040847 | 0.029637 | 0.013564 | 0.010343 | 0.030329 | 0.004957 | 0.050964 | 0.020398 | 0.013195 | 0.056523 | 0.038067 | 0.046711 | 0.000872 | 0.009503 | 0.03548 | 0.041241 | 0.000803 | 0.015503 | 0.032362 | 0.014071 | 0.012764 | 0.06682 | 0.017564 | 0.018753 |
| 30.1527778 | 0.031849 | 0.024075 | 0.010269 | 0.01709 | 0.025678 | 0.008929 | 0.03195 | 0.038244 | 0.008123 | 0.046176 | 0.041127 | 0.041735 | 0.000501 | 0.004592 | 0.048403 | 0.03522 | 0.000911 | 0.007961 | 0.035287 | 0.009897 | 0.016002 | 0.054807 | 0.026097 | 0.008853 |
| 30.1666667 | 0.01989 | 0.036678 | 0.003688 | 0.032669 | 0.025441 | 0.014555 | 0.016147 | 0.04572 | 0.005122 | 0.042975 | 0.032935 | 0.030285 | 0.000704 | 0.012045 | 0.049286 | 0.02614 | 0.009047 | 0.004858 | 0.024045 | 0.009465 | 0.016837 | 0.048611 | 0.037313 | 0.005749 |
| 30.1805556 | 0.010502 | 0.058132 | 0.001543 | 0.05517 | 0.029155 | 0.013756 | 0.008868 | 0.056401 | 0.007241 | 0.028511 | 0.020325 | 0.019355 | 0.001397 | 0.021671 | 0.024981 | 0.017077 | 0.019502 | 0.004761 | 0.025225 | 0.013879 | 0.013931 | 0.064478 | 0.042296 | 0.01375 |
| 30.1944444 | 0.009018 | 0.072831 | 0.004025 | 0.079156 | 0.033599 | 0.008535 | 0.008442 | 0.048322 | 0.007638 | 0.011959 | 0.010386 | 0.015756 | 0.002654 | 0.022662 | 0.017857 | 0.012385 | 0.014472 | 0.005035 | 0.026352 | 0.031364 | 0.007708 | 0.069203 | 0.0446 | 0.020321 |
| 30.2083333 | 0.012774 | 0.07025 | 0.007411 | 0.095661 | 0.033106 | 0.005747 | 0.011328 | 0.025058 | 0.005268 | 0.011346 | 0.003854 | 0.015478 | 0.002879 | 0.019255 | 0.034817 | 0.010949 | 0.004537 | 0.005686 | 0.020158 | 0.057347 | 0.008023 | 0.047401 | 0.044667 | 0.014268 |
| 30.2222222 | 0.016145 | 0.047587 | 0.012214 | 0.099742 | 0.032149 | 0.005198 | 0.01322 | 0.025085 | 0.007616 | 0.019907 | 0.001285 | 0.012491 | 0.002511 | 0.019127 | 0.048038 | 0.009563 | 0.003637 | 0.010949 | 0.027095 | 0.078801 | 0.012743 | 0.023451 | 0.040052 | 0.004592 |
| 30.2361111 | 0.01736 | 0.026352 | 0.019384 | 0.097677 | 0.039061 | 0.00347 | 0.010978 | 0.028828 | 0.01307 | 0.020022 | 0.001697 | 0.008468 | 0.002872 | 0.026955 | 0.041181 | 0.011338 | 0.00451 | 0.017394 | 0.034497 | 0.083049 | 0.016389 | 0.020314 | 0.032484 | 0.003606 |
| 30.25 | 0.017515 | 0.021886 | 0.026662 | 0.095756 | 0.042314 | 0.003637 | 0.01262 | 0.024883 | 0.015149 | 0.022415 | 0.00334 | 0.007492 | 0.004164 | 0.035651 | 0.036006 | 0.012811 | 0.002456 | 0.017937 | 0.02447 | 0.077335 | 0.021201 | 0.035427 | 0.019961 | 0.00631 |
| 30.2638889 | 0.017974 | 0.022469 | 0.033281 | 0.0859 | 0.03492 | 0.006975 | 0.022165 | 0.045188 | 0.014548 | 0.042409 | 0.005489 | 0.008151 | 0.00745 | 0.030639 | 0.043297 | 0.016107 | 0.0025 | 0.011543 | 0.009684 | 0.072446 | 0.024563 | 0.040433 | 0.015865 | 0.005642 |

| | | | | | | | | | | | | | | | | | | | | | | | | |
|---|---|---|---|---|---|---|---|---|---|---|---|---|---|---|---|---|---|---|---|---|---|---|---|---|
| 30.2777778 | 0.018049 | 0.018135 | 0.04089 | 0.068682 | 0.02638 | 0.009087 | 0.023976 | 0.073175 | 0.013244 | 0.060749 | 0.00655 | 0.00785 | 0.011764 | 0.018238 | 0.034653 | 0.0229 | 0.005183 | 0.005051 | 0.003579 | 0.054285 | 0.020405 | 0.027588 | 0.030225 | 0.003735 |
| 30.2916667 | 0.017471 | 0.013008 | 0.047586 | 0.053159 | 0.019797 | 0.007545 | 0.015502 | 0.063611 | 0.010736 | 0.059416 | 0.005037 | 0.006526 | 0.015651 | 0.011209 | 0.0187 | 0.026006 | 0.005935 | 0.004684 | 0.008007 | 0.031498 | 0.011567 | 0.016272 | 0.045533 | 0.002437 |
| 30.3055556 | 0.017494 | 0.009015 | 0.047774 | 0.04136 | 0.012361 | 0.00383 | 0.01755 | 0.031315 | 0.008001 | 0.046269 | 0.003319 | 0.003994 | 0.018678 | 0.008234 | 0.018522 | 0.029053 | 0.007826 | 0.009516 | 0.020069 | 0.039884 | 0.00985 | 0.012656 | 0.049305 | 0.001329 |
| 30.3194444 | 0.018388 | 0.006224 | 0.040813 | 0.03044 | 0.005377 | 0.00186 | 0.03266 | 0.016728 | 0.005051 | 0.032452 | 0.004928 | 0.001718 | 0.019269 | 0.007749 | 0.017229 | 0.029744 | 0.012456 | 0.016965 | 0.030604 | 0.061352 | 0.014644 | 0.011976 | 0.042708 | 0.001619 |
| 30.3333333 | 0.018403 | 0.005116 | 0.031508 | 0.017935 | 0.001763 | 0.003254 | 0.037094 | 0.020976 | 0.00302 | 0.019496 | 0.008057 | 0.000792 | 0.015059 | 0.009182 | 0.011887 | 0.02283 | 0.014263 | 0.019621 | 0.024886 | 0.059397 | 0.016064 | 0.013246 | 0.026713 | 0.003325 |
| 30.3472222 | 0.014217 | 0.004934 | 0.021517 | 0.014219 | 0.002281 | 0.004458 | 0.024798 | 0.021295 | 0.00256 | 0.018361 | 0.009678 | 0.001244 | 0.009577 | 0.007832 | 0.022283 | 0.020704 | 0.010824 | 0.012785 | 0.013236 | 0.041507 | 0.014134 | 0.014694 | 0.013916 | 0.006103 |
| 30.3611111 | 0.007334 | 0.007319 | 0.011345 | 0.026393 | 0.009298 | 0.003477 | 0.011385 | 0.02313 | 0.001563 | 0.037835 | 0.009227 | 0.003611 | 0.010246 | 0.004043 | 0.042919 | 0.02937 | 0.00567 | 0.005448 | 0.016745 | 0.021912 | 0.010856 | 0.012186 | 0.019179 | 0.007196 |
| 30.375 | 0.00427 | 0.01086 | 0.00702 | 0.034237 | 0.022228 | 0.0024 | 0.005548 | 0.023483 | 0.002089 | 0.054742 | 0.006263 | 0.0056 | 0.0142 | 0.003618 | 0.044036 | 0.029791 | 0.003576 | 0.005372 | 0.021981 | 0.011039 | 0.005841 | 0.006158 | 0.036011 | 0.005079 |
| 30.3888889 | 0.006108 | 0.012391 | 0.011774 | 0.026388 | 0.035694 | 0.001801 | 0.004165 | 0.021381 | 0.005038 | 0.057885 | 0.00273 | 0.00595 | 0.014875 | 0.004307 | 0.033793 | 0.021421 | 0.004721 | 0.007092 | 0.017753 | 0.016281 | 0.001995 | 0.001588 | 0.047557 | 0.006535 |
| 30.4027778 | 0.009154 | 0.013771 | 0.019517 | 0.01413 | 0.049023 | 0.001956 | 0.003503 | 0.039787 | 0.005265 | 0.061423 | 0.001669 | 0.00733 | 0.013298 | 0.005326 | 0.047563 | 0.029032 | 0.005427 | 0.006708 | 0.014071 | 0.024443 | 0.001137 | 0.001356 | 0.047198 | 0.013822 |
| 30.4166667 | 0.009887 | 0.01411 | 0.025831 | 0.008632 | 0.051632 | 0.003083 | 0.008241 | 0.045668 | 0.002722 | 0.06195 | 0.001572 | 0.009245 | 0.011969 | 0.01322 | 0.059511 | 0.039594 | 0.003765 | 0.006424 | 0.014861 | 0.03128 | 0.001891 | 0.004053 | 0.039591 | 0.023121 |
| 30.4305556 | 0.006615 | 0.013771 | 0.030778 | 0.007971 | 0.035715 | 0.003622 | 0.020287 | 0.038003 | 0.001835 | 0.048384 | 0.001127 | 0.009568 | 0.010324 | 0.026412 | 0.038891 | 0.034588 | 0.001569 | 0.005544 | 0.017614 | 0.043235 | 0.003753 | 0.006083 | 0.035065 | 0.034124 |
| 30.4444444 | 0.0028 | 0.018447 | 0.034468 | 0.010592 | 0.0181 | 0.004574 | 0.029867 | 0.05905 | 0.001843 | 0.031118 | 0.002815 | 0.010436 | 0.007966 | 0.038841 | 0.013342 | 0.029812 | 0.001129 | 0.003321 | 0.019466 | 0.052125 | 0.005512 | 0.005074 | 0.035459 | 0.048176 |
| 30.4583333 | 0.002204 | 0.026688 | 0.037055 | 0.024332 | 0.01228 | 0.00691 | 0.023926 | 0.068889 | 0.001642 | 0.023939 | 0.006634 | 0.012875 | 0.00608 | 0.047475 | 0.00503 | 0.034493 | 0.00181 | 0.001567 | 0.019673 | 0.050659 | 0.004561 | 0.003202 | 0.02853 | 0.058581 |
| 30.4722222 | 0.00462 | 0.033017 | 0.040062 | 0.039659 | 0.013298 | 0.008809 | 0.011793 | 0.043054 | 0.00184 | 0.027578 | 0.009178 | 0.014066 | 0.005199 | 0.051907 | 0.007588 | 0.043493 | 0.002322 | 0.002145 | 0.020961 | 0.049285 | 0.002415 | 0.004506 | 0.017284 | 0.055862 |
| 30.4861111 | 0.007142 | 0.038037 | 0.045307 | 0.049534 | 0.013342 | 0.009425 | 0.013904 | 0.016859 | 0.002502 | 0.039885 | 0.009046 | 0.012894 | 0.004756 | 0.05387 | 0.009177 | 0.051336 | 0.002501 | 0.006164 | 0.02511 | 0.057691 | 0.002425 | 0.007761 | 0.021134 | 0.055172 |
| 30.5 | 0.008153 | 0.045466 | 0.052618 | 0.066721 | 0.009063 | 0.009217 | 0.023986 | 0.006106 | 0.005515 | 0.050585 | 0.007954 | 0.010595 | 0.003641 | 0.0515 | 0.008714 | 0.049862 | 0.004959 | 0.011724 | 0.02714 | 0.065324 | 0.003743 | 0.007955 | 0.03736 | 0.064994 |
| 30.5138889 | 0.008253 | 0.053781 | 0.060414 | 0.097913 | 0.003993 | 0.008783 | 0.031551 | 0.005218 | 0.011513 | 0.052942 | 0.007303 | 0.008385 | 0.002012 | 0.043525 | 0.016431 | 0.038487 | 0.01338 | 0.015361 | 0.022695 | 0.070257 | 0.004314 | 0.005615 | 0.05325 | 0.064085 |
| 30.5277778 | 0.007874 | 0.062918 | 0.062232 | 0.125899 | 0.009073 | 0.009156 | 0.037067 | 0.007065 | 0.017473 | 0.063542 | 0.008172 | 0.009504 | 0.001421 | 0.033338 | 0.02287 | 0.036771 | 0.026778 | 0.016292 | 0.014819 | 0.081493 | 0.003574 | 0.006843 | 0.069693 | 0.054351 |
| 30.5416667 | 0.008742 | 0.071291 | 0.053254 | 0.130875 | 0.024195 | 0.009441 | 0.040886 | 0.012761 | 0.01926 | 0.082123 | 0.009605 | 0.014 | 0.001409 | 0.02107 | 0.01727 | 0.05476 | 0.040707 | 0.01489 | 0.008015 | 0.089783 | 0.002552 | 0.012529 | 0.07845 | 0.04534 |
| 30.5555556 | 0.011328 | 0.065734 | 0.035927 | 0.120664 | 0.033331 | 0.006805 | 0.043245 | 0.020219 | 0.015852 | 0.078469 | 0.008467 | 0.014096 | 0.00202 | 0.009113 | 0.018931 | 0.073481 | 0.048088 | 0.011674 | 0.0081 | 0.092231 | 0.002659 | 0.017006 | 0.073893 | 0.028307 |
| 30.5694444 | 0.013683 | 0.047129 | 0.016751 | 0.107942 | 0.029832 | 0.003843 | 0.039029 | 0.019031 | 0.010221 | 0.051369 | 0.008047 | 0.007652 | 0.006956 | 0.002693 | 0.043085 | 0.082711 | 0.053328 | 0.009483 | 0.021598 | 0.100562 | 0.002588 | 0.014 | 0.066135 | 0.016447 |
| 30.5833333 | 0.019582 | 0.025713 | 0.008724 | 0.097205 | 0.024668 | 0.003336 | 0.026917 | 0.01091 | 0.008192 | 0.028416 | 0.009844 | 0.002267 | 0.015649 | 0.003922 | 0.086888 | 0.08052 | 0.060712 | 0.012343 | 0.046342 | 0.108831 | 0.003761 | 0.022389 | 0.057766 | 0.026673 |
| 30.5972222 | 0.030382 | 0.010834 | 0.014859 | 0.085355 | 0.027716 | 0.005741 | 0.015856 | 0.008129 | 0.01242 | 0.020681 | 0.007942 | 0.003053 | 0.022309 | 0.014031 | 0.133066 | 0.058524 | 0.054969 | 0.021157 | 0.072217 | 0.097667 | 0.00953 | 0.061662 | 0.047803 | 0.048647 |
| 30.6111111 | 0.035995 | 0.006962 | 0.023565 | 0.064154 | 0.035663 | 0.015127 | 0.011273 | 0.007782 | 0.018482 | 0.023875 | 0.007609 | 0.009134 | 0.025803 | 0.027254 | 0.158104 | 0.035079 | 0.035757 | 0.032956 | 0.091905 | 0.065526 | 0.01741 | 0.099979 | 0.040725 | 0.060683 |
| 30.625 | 0.03197 | 0.015115 | 0.038322 | 0.047991 | 0.039206 | 0.027745 | 0.014104 | 0.007595 | 0.021124 | 0.034613 | 0.015353 | 0.01688 | 0.027804 | 0.031661 | 0.156985 | 0.032229 | 0.020973 | 0.041025 | 0.101311 | 0.044306 | 0.021421 | 0.11018 | 0.04033 | 0.055774 |
| 30.6388889 | 0.022118 | 0.04826 | 0.060426 | 0.039045 | 0.0311 | 0.034979 | 0.023798 | 0.014284 | 0.01882 | 0.049044 | 0.023318 | 0.023653 | 0.026987 | 0.024062 | 0.124598 | 0.032319 | 0.019962 | 0.034539 | 0.09585 | 0.056432 | 0.021025 | 0.093037 | 0.043787 | 0.049132 |
| 30.6527778 | 0.011734 | 0.081173 | 0.066117 | 0.025779 | 0.016877 | 0.037215 | 0.034352 | 0.028337 | 0.016424 | 0.069581 | 0.028413 | 0.03091 | 0.022567 | 0.015272 | 0.066631 | 0.019515 | 0.021993 | 0.017956 | 0.078424 | 0.075091 | 0.023654 | 0.054326 | 0.037553 | 0.044683 |
| 30.6666667 | 0.008873 | 0.084555 | 0.043782 | 0.026225 | 0.008135 | 0.039485 | 0.035245 | 0.045761 | 0.019332 | 0.098357 | 0.035879 | 0.042566 | 0.016527 | 0.015219 | 0.026517 | 0.008756 | 0.014984 | 0.010679 | 0.058862 | 0.059819 | 0.025677 | 0.031485 | 0.026546 | 0.029469 |
| 30.6805556 | 0.010218 | 0.078818 | 0.023208 | 0.039933 | 0.00571 | 0.04128 | 0.023915 | 0.051689 | 0.023735 | 0.118355 | 0.040232 | 0.057093 | 0.009843 | 0.014854 | 0.030449 | 0.005952 | 0.015117 | 0.011056 | 0.039167 | 0.035899 | 0.016696 | 0.045453 | 0.035763 | 0.018132 |
| 30.6944444 | 0.007275 | 0.071767 | 0.029747 | 0.04582 | 0.011265 | 0.038731 | 0.011868 | 0.039569 | 0.024266 | 0.11138 | 0.035277 | 0.061145 | 0.003792 | 0.008703 | 0.054017 | 0.010592 | 0.031019 | 0.007179 | 0.023651 | 0.044165 | 0.007513 | 0.061061 | 0.057211 | 0.030738 |
| 30.7083333 | 0.004078 | 0.056438 | 0.041671 | 0.040162 | 0.025962 | 0.033552 | 0.009877 | 0.021596 | 0.02029 | 0.071361 | 0.031043 | 0.049125 | 0.002281 | 0.007944 | 0.061991 | 0.032736 | 0.042752 | 0.004674 | 0.021654 | 0.046614 | 0.010112 | 0.060914 | 0.068726 | 0.049444 |
| 30.7222222 | 0.002921 | 0.037586 | 0.034389 | 0.029855 | 0.040135 | 0.029244 | 0.016904 | 0.015243 | 0.014274 | 0.02659 | 0.034473 | 0.034629 | 0.007326 | 0.014636 | 0.045516 | 0.061647 | 0.037799 | 0.010071 | 0.019458 | 0.025319 | 0.014338 | 0.057521 | 0.066376 | 0.057057 |
| 30.7361111 | 0.002129 | 0.02208 | 0.019121 | 0.032496 | 0.039876 | 0.025309 | 0.019514 | 0.018427 | 0.01256 | 0.013734 | 0.038632 | 0.026842 | 0.017741 | 0.022228 | 0.051957 | 0.063496 | 0.020887 | 0.019781 | 0.016721 | 0.012014 | 0.011179 | 0.045051 | 0.052602 | 0.048721 |
| 30.75 | 0.003118 | 0.020724 | 0.021613 | 0.055216 | 0.027424 | 0.021668 | 0.015486 | 0.016249 | 0.016419 | 0.022915 | 0.038455 | 0.024396 | 0.030131 | 0.024582 | 0.079296 | 0.040226 | 0.011053 | 0.031259 | 0.02884 | 0.0168 | 0.012124 | 0.028207 | 0.032622 | 0.031163 |
| 30.7638889 | 0.00666 | 0.036636 | 0.038514 | 0.079489 | 0.022197 | 0.020081 | 0.02468 | 0.016512 | 0.022168 | 0.041566 | 0.035882 | 0.023933 | 0.038684 | 0.029434 | 0.070136 | 0.02113 | 0.017739 | 0.039713 | 0.041472 | 0.027006 | 0.028845 | 0.036168 | 0.024332 | 0.035006 |
| 30.7777778 | 0.012225 | 0.050248 | 0.047041 | 0.086231 | 0.022838 | 0.019504 | 0.051357 | 0.020347 | 0.027163 | 0.071845 | 0.032038 | 0.023187 | 0.041075 | 0.041778 | 0.055669 | 0.009866 | 0.022821 | 0.040793 | 0.03991 | 0.026391 | 0.051533 | 0.067628 | 0.053485 | 0.061663 |
| 30.7916667 | 0.018643 | 0.04755 | 0.044614 | 0.072682 | 0.021527 | 0.017457 | 0.070962 | 0.032563 | 0.025345 | 0.092278 | 0.027938 | 0.023301 | 0.040164 | 0.047294 | 0.048673 | 0.006037 | 0.015749 | 0.040299 | 0.025777 | 0.016449 | 0.070865 | 0.090759 | 0.097153 | 0.071047 |
| 30.8055556 | 0.02762 | 0.035139 | 0.047084 | 0.057754 | 0.024286 | 0.013835 | 0.060891 | 0.051194 | 0.015413 | 0.078524 | 0.02558 | 0.025218 | 0.03758 | 0.039123 | 0.03329 | 0.012384 | 0.006158 | 0.044527 | 0.015602 | 0.007361 | 0.080233 | 0.077165 | 0.104414 | 0.054783 |
| 30.8194444 | 0.035881 | 0.023453 | 0.05815 | 0.049497 | 0.024045 | 0.010407 | 0.032539 | 0.046714 | 0.006577 | 0.048916 | 0.025731 | 0.027558 | 0.030976 | 0.021495 | 0.033453 | 0.021625 | 0.003167 | 0.048253 | 0.015869 | 0.004291 | 0.066183 | 0.040019 | 0.078688 | 0.034476 |
| 30.8333333 | 0.037007 | 0.016016 | 0.063183 | 0.033983 | 0.020014 | 0.00948 | 0.014872 | 0.027907 | 0.005511 | 0.032751 | 0.030792 | 0.029325 | 0.018556 | 0.008393 | 0.039633 | 0.028462 | 0.00754 | 0.036507 | 0.011843 | 0.004976 | 0.034554 | 0.023959 | 0.051076 | 0.021877 |
| 30.8472222 | 0.034714 | 0.022661 | 0.057018 | 0.016492 | 0.027264 | 0.012847 | 0.009772 | 0.016062 | 0.007076 | 0.035503 | 0.03896 | 0.030939 | 0.011324 | 0.007048 | 0.039631 | 0.041485 | 0.018229 | 0.020593 | 0.007666 | 0.007262 | 0.017675 | 0.024305 | 0.033162 | 0.022154 |
| 30.8611111 | 0.027889 | 0.061773 | 0.044503 | 0.008164 | 0.052101 | 0.018156 | 0.005632 | 0.008308 | 0.005155 | 0.063757 | 0.043545 | 0.031688 | 0.017943 | 0.011344 | 0.034406 | 0.063132 | 0.02908 | 0.025014 | 0.009253 | 0.012874 | 0.030389 | 0.017287 | 0.024108 | 0.024173 |
| 30.875 | 0.015879 | 0.114931 | 0.029821 | 0.008899 | 0.087213 | 0.020426 | 0.002592 | 0.015349 | 0.002831 | 0.09883 | 0.043737 | 0.032348 | 0.024935 | 0.021314 | 0.018472 | 0.079317 | 0.028894 | 0.036729 | 0.010768 | 0.012613 | 0.048314 | 0.013159 | 0.024519 | 0.021755 |
| 30.8888889 | 0.010087 | 0.149956 | 0.019692 | 0.010914 | 0.102662 | 0.019685 | 0.009689 | 0.035071 | 0.003165 | 0.101201 | 0.041664 | 0.036246 | 0.024042 | 0.027056 | 0.011524 | 0.085409 | 0.020597 | 0.03623 | 0.019795 | 0.00549 | 0.052114 | 0.026256 | 0.030092 | 0.024666 |
| 30.9027778 | 0.009315 | 0.149483 | 0.017177 | 0.009051 | 0.081516 | 0.019701 | 0.02553 | 0.045622 | 0.007207 | 0.070419 | 0.036301 | 0.040618 | 0.019965 | 0.021768 | 0.022403 | 0.0791 | 0.018199 | 0.026695 | 0.032554 | 0.008086 | 0.045583 | 0.055579 | 0.024477 | 0.03383 |
| 30.9166667 | 0.006316 | 0.106614 | 0.016989 | 0.012902 | 0.050454 | 0.021057 | 0.029417 | 0.059308 | 0.009455 | 0.051551 | 0.031122 | 0.034392 | 0.012636 | 0.027489 | 0.060359 | 0.062604 | 0.024272 | 0.015477 | 0.031278 | 0.021165 | 0.040599 | 0.085101 | 0.02039 | 0.048202 |
| 30.9305556 | 0.003897 | 0.049801 | 0.013002 | 0.022991 | 0.025795 | 0.020698 | 0.018525 | 0.074568 | 0.006749 | 0.09021 | 0.030878 | 0.01926 | 0.008414 | 0.049801 | 0.11687 | 0.048613 | 0.026992 | 0.007434 | 0.017208 | 0.031016 | 0.039283 | 0.103749 | 0.044286 | 0.062855 |
| 30.9444444 | 0.006796 | 0.015206 | 0.009307 | 0.032506 | 0.015256 | 0.018215 | 0.009032 | 0.062445 | 0.006719 | 0.152535 | 0.028428 | 0.01391 | 0.017175 | 0.064117 | 0.108875 | 0.039756 | 0.018105 | 0.005207 | 0.005693 | 0.037674 | 0.035413 | 0.115544 | 0.078251 | 0.066397 |
| 30.9583333 | 0.014373 | 0.011935 | 0.010679 | 0.050378 | 0.034019 | 0.018635 | 0.00557 | 0.035319 | 0.011412 | 0.187869 | 0.017862 | 0.018119 | 0.028906 | 0.064774 | 0.056887 | 0.036648 | 0.006107 | 0.007928 | 0.003679 | 0.052732 | 0.028037 | 0.113842 | 0.094865 | 0.056681 |
| 30.9722222 | 0.020134 | 0.032967 | 0.009873 | 0.059322 | 0.069799 | 0.021541 | 0.004234 | 0.0175 | 0.017768 | 0.192329 | 0.008525 | 0.016 | 0.033499 | 0.060322 | 0.037707 | 0.042415 | 0.000971 | 0.015109 | 0.004901 | 0.076317 | 0.017662 | 0.096332 | 0.091588 | 0.038088 |
| 30.9861111 | 0.02129 | 0.061527 | 0.009254 | 0.041199 | 0.101339 | 0.021098 | 0.006101 | 0.009857 | 0.020741 | 0.162446 | 0.007973 | 0.007896 | 0.03648 | 0.053465 | 0.039826 | 0.056313 | 0.00082 | 0.021884 | 0.005253 | 0.092859 | 0.009556 | 0.086834 | 0.078533 | 0.025102 |
| 31 | 0.020016 | 0.088901 | 0.022278 | 0.064921 | 0.115006 | 0.017651 | 0.013732 | 0.013545 | 0.019124 | 0.132072 | 0.010189 | 0.002804 | 0.037205 | 0.051673 | 0.067418 | 0.072652 | 0.002453 | 0.02158 | 0.007103 | 0.093002 | 0.009822 | 0.084765 | 0.072678 | 0.028172 |
| 31.0138889 | 0.016969 | 0.112444 | 0.041812 | 0.159624 | 0.108026 | 0.013088 | 0.021269 | 0.026689 | 0.015076 | 0.123956 | 0.007655 | 0.002272 | 0.030544 | 0.052191 | 0.096 | 0.086713 | 0.004339 | 0.014052 | 0.016674 | 0.092792 | 0.011264 | 0.069898 | 0.081308 | 0.03289 |
| 31.0277778 | 0.01086 | 0.115962 | 0.054329 | 0.204252 | 0.090141 | 0.007681 | 0.020474 | 0.035257 | 0.008492 | 0.098082 | 0.003918 | 0.002603 | 0.021238 | 0.05026 | 0.092492 | 0.10223 | 0.005203 | 0.013443 | 0.040961 | 0.116862 | 0.012134 | 0.052115 | 0.096837 | 0.030172 |
| 31.0416667 | 0.009056 | 0.104801 | 0.059025 | 0.163099 | 0.064422 | 0.009074 | 0.011948 | 0.029528 | 0.005762 | 0.048037 | 0.008264 | 0.001912 | 0.022564 | 0.052266 | 0.100048 | 0.11366 | 0.012951 | 0.037533 | 0.079101 | 0.163482 | 0.019253 | 0.048083 | 0.108489 | 0.031344 |
| 31.0555556 | 0.011221 | 0.090894 | 0.057376 | 0.122746 | 0.034953 | 0.02847 | 0.010424 | 0.017026 | 0.012195 | 0.013021 | 0.020808 | 0.001501 | 0.033035 | 0.05805 | 0.123116 | 0.099845 | 0.036609 | 0.070532 | 0.124105 | 0.200501 | 0.025137 | 0.051055 | 0.10244 | 0.039731 |
| 31.0694444 | 0.008125 | 0.061655 | 0.051253 | 0.087841 | 0.020033 | 0.060026 | 0.026488 | 0.01718 | 0.018811 | 0.02329 | 0.031641 | 0.004113 | 0.038625 | 0.062438 | 0.102733 | 0.075774 | 0.06351 | 0.074747 | 0.1419 | 0.192247 | 0.021225 | 0.0462 | 0.073151 | 0.039191 |
| 31.0833333 | 0.007828 | 0.031572 | 0.036808 | 0.046926 | 0.022297 | 0.076463 | 0.045041 | 0.028162 | 0.019821 | 0.08537 | 0.031766 | 0.008459 | 0.031397 | 0.058578 | 0.054639 | 0.069929 | 0.066726 | 0.048996 | 0.105429 | 0.140559 | 0.010648 | 0.037098 | 0.035894 | 0.023319 |
| 31.0972222 | 0.009617 | 0.019146 | 0.017946 | 0.020077 | 0.034389 | 0.059511 | 0.042567 | 0.039927 | 0.016355 | 0.15619 | 0.0252 | 0.00916 | 0.01864 | 0.04502 | 0.03533 | 0.052478 | 0.045588 | 0.03275 | 0.050664 | 0.082141 | 0.003489 | 0.031322 | 0.020853 | 0.008954 |
| 31.1111111 | 0.006118 | 0.011626 | 0.007377 | 0.020248 | 0.041807 | 0.029091 | 0.030521 | 0.06687 | 0.01177 | 0.189415 | 0.020866 | 0.006833 | 0.017483 | 0.02698 | 0.043184 | 0.017795 | 0.03162 | 0.036176 | 0.019331 | 0.04018 | 0.002836 | 0.023074 | 0.043277 | 0.008442 |
| 31.125 | 0.005124 | 0.00912 | 0.005475 | 0.04048 | 0.027766 | 0.013371 | 0.038055 | 0.081449 | 0.012867 | 0.169874 | 0.015902 | 0.004775 | 0.018942 | 0.013299 | 0.07868 | 0.005608 | 0.035491 | 0.0372 | 0.018805 | 0.029386 | 0.003554 | 0.028316 | 0.07882 | 0.015497 |
| 31.1388889 | 0.011584 | 0.011261 | 0.004443 | 0.057454 | 0.016059 | 0.011763 | 0.057029 | 0.05878 | 0.018525 | 0.118118 | 0.007688 | 0.00327 | 0.012736 | 0.013685 | 0.110503 | 0.011714 | 0.039361 | 0.033102 | 0.039568 | 0.047785 | 0.002993 | 0.052027 | 0.084208 | 0.017696 |
| 31.1527778 | 0.017438 | 0.011564 | 0.002542 | 0.077204 | 0.013428 | 0.014328 | 0.060433 | 0.049614 | 0.025148 | 0.085442 | 0.002148 | 0.005807 | 0.012214 | 0.01895 | 0.094558 | 0.01563 | 0.031412 | 0.027393 | 0.053721 | 0.055408 | 0.002176 | 0.056432 | 0.052221 | 0.011063 |
| 31.1666667 | 0.013569 | 0.024421 | 0.002793 | 0.096422 | 0.008199 | 0.01769 | 0.043477 | 0.052143 | 0.023034 | 0.063357 | 0.002668 | 0.011641 | 0.012965 | 0.019835 | 0.062927 | 0.017123 | 0.023533 | 0.017574 | 0.04157 | 0.033707 | 0.005483 | 0.039947 | 0.02426 | 0.004076 |
| 31.1805556 | 0.00999 | 0.056074 | 0.004415 | 0.081074 | 0.006221 | 0.017919 | 0.023214 | 0.035152 | 0.017498 | 0.043866 | 0.007694 | 0.011592 | 0.009009 | 0.021433 | 0.048489 | 0.020818 | 0.035868 | 0.009379 | 0.03477 | 0.012496 | 0.013727 | 0.029164 | 0.025024 | 0.003904 |
| 31.1944444 | 0.017427 | 0.067905 | 0.005075 | 0.045886 | 0.007514 | 0.011726 | 0.014757 | 0.017358 | 0.027371 | 0.063044 | 0.013488 | 0.006814 | 0.009782 | 0.02775 | 0.042518 | 0.023947 | 0.054376 | 0.009936 | 0.047425 | 0.007916 | 0.020174 | 0.027217 | 0.038629 | 0.006667 |
| 31.2083333 | 0.022294 | 0.040595 | 0.003754 | 0.049521 | 0.015575 | 0.006929 | 0.014686 | 0.010718 | 0.037666 | 0.10647 | 0.016199 | 0.004754 | 0.009342 | 0.036114 | 0.036187 | 0.016444 | 0.060685 | 0.013462 | 0.045684 | 0.011254 | 0.016451 | 0.026764 | 0.049966 | 0.007994 |
| 31.2222222 | 0.015914 | 0.021285 | 0.002213 | 0.094262 | 0.028276 | 0.012365 | 0.013092 | 0.009842 | 0.032464 | 0.130078 | 0.014314 | 0.004222 | 0.004273 | 0.031475 | 0.052666 | 0.008015 | 0.055267 | 0.012758 | 0.02589 | 0.0226 | 0.008878 | 0.022374 | 0.062375 | 0.006236 |
| 31.2361111 | 0.009636 | 0.035328 | 0.002186 | 0.124791 | 0.037035 | 0.018833 | 0.008388 | 0.007502 | 0.023116 | 0.115066 | 0.008186 | 0.008752 | 0.00193 | 0.021263 | 0.073751 | 0.008083 | 0.036407 | 0.012923 | 0.011019 | 0.03591 | 0.009844 | 0.025746 | 0.075621 | 0.002314 |
| 31.25 | 0.006688 | 0.052222 | 0.003423 | 0.113489 | 0.040851 | 0.015633 | 0.003827 | 0.005329 | 0.015647 | 0.092315 | 0.004218 | 0.017746 | 0.002004 | 0.020246 | 0.055613 | 0.006973 | 0.014777 | 0.014145 | 0.012077 | 0.042266 | 0.010057 | 0.038622 | 0.080823 | 0.00045 |
| 31.2638889 | 0.004696 | 0.047362 | 0.006878 | 0.078529 | 0.051227 | 0.008205 | 0.00297 | 0.003815 | 0.009582 | 0.107139 | 0.007905 | 0.01812 | 0.002599 | 0.024816 | 0.043024 | 0.004408 | 0.006915 | 0.010105 | 0.026773 | 0.043737 | 0.005038 | 0.043894 | 0.073714 | 0.002645 |
| 31.2777778 | 0.005943 | 0.031866 | 0.014641 | 0.049197 | 0.063604 | 0.006006 | 0.003437 | 0.001576 | 0.008503 | 0.141724 | 0.013463 | 0.01282 | 0.00453 | 0.028915 | 0.066922 | 0.004721 | 0.017839 | 0.004627 | 0.046573 | 0.04338 | 0.005121 | 0.042313 | 0.058088 | 0.009167 |
| 31.2916667 | 0.006771 | 0.025873 | 0.023683 | 0.035829 | 0.056146 | 0.008533 | 0.006859 | 0.011313 | 0.011848 | 0.148072 | 0.016353 | 0.012414 | 0.006532 | 0.020872 | 0.066651 | 0.006261 | 0.033928 | 0.002761 | 0.058235 | 0.039332 | 0.008414 | 0.034463 | 0.039678 | 0.014599 |
| 31.3055556 | 0.005102 | 0.03333 | 0.026071 | 0.028681 | 0.039534 | 0.011109 | 0.015645 | 0.035951 | 0.015729 | 0.116225 | 0.017797 | 0.012394 | 0.005309 | 0.026056 | 0.038674 | 0.01188 | 0.030441 | 0.002365 | 0.052277 | 0.032622 | 0.009356 | 0.029451 | 0.023433 | 0.013715 |
| 31.3194444 | 0.003142 | 0.045568 | 0.024897 | 0.022311 | 0.028209 | 0.011844 | 0.017527 | 0.048485 | 0.017083 | 0.068828 | 0.016732 | 0.009363 | 0.003937 | 0.060272 | 0.037717 | 0.020232 | 0.018061 | 0.002164 | 0.033467 | 0.029381 | 0.008795 | 0.033574 | 0.014219 | 0.012036 |
| 31.3333333 | 0.001833 | 0.053045 | 0.027943 | 0.015773 | 0.018321 | 0.009611 | 0.009918 | 0.034928 | 0.014763 | 0.036686 | 0.012024 | 0.007229 | 0.007471 | 0.073683 | 0.04964 | 0.018688 | 0.024115 | 0.001781 | 0.015745 | 0.031202 | 0.010713 | 0.030658 | 0.010142 | 0.01056 |
| 31.3472222 | 0.001418 | 0.050861 | 0.029662 | 0.008642 | 0.012583 | 0.006917 | 0.006248 | 0.014175 | 0.012197 | 0.043234 | 0.005857 | 0.005343 | 0.011642 | 0.04941 | 0.048801 | 0.010048 | 0.037737 | 0.000943 | 0.009599 | 0.036036 | 0.01363 | 0.024432 | 0.006426 | 0.005529 |
| 31.3611111 | 0.00137 | 0.041197 | 0.023665 | 0.005108 | 0.009104 | 0.007061 | 0.013539 | 0.00535 | 0.012273 | 0.072967 | 0.002314 | 0.002822 | 0.011673 | 0.033032 | 0.050734 | 0.00543 | 0.041256 | 0.001844 | 0.009976 | 0.038957 | 0.013562 | 0.023727 | 0.003298 | 0.005839 |
| 31.375 | 0.00197 | 0.029588 | 0.015434 | 0.006205 | 0.007639 | 0.007842 | 0.019235 | 0.01144 | 0.015549 | 0.092708 | 0.002823 | 0.002282 | 0.008441 | 0.043963 | 0.061803 | 0.003843 | 0.034151 | 0.005408 | 0.00894 | 0.03978 | 0.010204 | 0.032417 | 0.002748 | 0.017323 |
| 31.3888889 | 0.003653 | 0.016539 | 0.010586 | 0.010816 | 0.011667 | 0.006322 | 0.012989 | 0.019027 | 0.020423 | 0.077277 | 0.004771 | 0.002914 | 0.004448 | 0.057195 | 0.064349 | 0.005921 | 0.019098 | 0.007977 | 0.011816 | 0.049911 | 0.00671 | 0.042332 | 0.003701 | 0.032221 |
| 31.4027778 | 0.005023 | 0.012647 | 0.006738 | 0.015814 | 0.019681 | 0.005161 | 0.007586 | 0.01682 | 0.023035 | 0.043795 | 0.006969 | 0.004673 | 0.003304 | 0.053808 | 0.04337 | 0.008648 | 0.007965 | 0.006278 | 0.018719 | 0.05808 | 0.007607 | 0.044477 | 0.005783 | 0.040891 |
| 31.4166667 | 0.005868 | 0.028307 | 0.005174 | 0.012859 | 0.028751 | 0.007736 | 0.011881 | 0.014815 | 0.022375 | 0.042249 | 0.009093 | 0.00942 | 0.005376 | 0.041088 | 0.023865 | 0.006271 | 0.010473 | 0.004721 | 0.021765 | 0.047391 | 0.007818 | 0.053054 | 0.00978 | 0.03954 |
| 31.4305556 | 0.00718 | 0.04661 | 0.009515 | 0.007278 | 0.035347 | 0.011213 | 0.014136 | 0.01938 | 0.020078 | 0.058315 | 0.008509 | 0.013413 | 0.008004 | 0.036327 | 0.035793 | 0.003753 | 0.017786 | 0.006317 | 0.021902 | 0.03943 | 0.004154 | 0.069972 | 0.015721 | 0.034331 |
| 31.4444444 | 0.006356 | 0.052065 | 0.014863 | 0.00632 | 0.046724 | 0.010774 | 0.011713 | 0.026357 | 0.018626 | 0.066951 | 0.006465 | 0.012146 | 0.010373 | 0.042326 | 0.062646 | 0.004462 | 0.021099 | 0.007175 | 0.023176 | 0.045298 | 0.001766 | 0.080341 | 0.023134 | 0.030515 |
| 31.4583333 | 0.003037 | 0.048285 | 0.016582 | 0.00759 | 0.061204 | 0.006295 | 0.010175 | 0.034682 | 0.018407 | 0.07118 | 0.008204 | 0.008616 | 0.012784 | 0.047746 | 0.085596 | 0.011266 | 0.022444 | 0.005885 | 0.025716 | 0.053968 | 0.004001 | 0.084476 | 0.028799 | 0.027795 |
| 31.4722222 | 0.001576 | 0.048545 | 0.017663 | 0.011547 | 0.067655 | 0.002084 | 0.005071 | 0.035447 | 0.017522 | 0.066342 | 0.014342 | 0.008514 | 0.014385 | 0.045302 | 0.100334 | 0.020815 | 0.026192 | 0.003839 | 0.02892 | 0.062852 | 0.008139 | 0.10114 | 0.029383 | 0.028156 |
| 31.4861111 | 0.003018 | 0.067534 | 0.023685 | 0.022333 | 0.06974 | 0.003535 | 0.001935 | 0.022149 | 0.015754 | 0.063497 | 0.018614 | 0.012314 | 0.014302 | 0.040276 | 0.103914 | 0.020294 | 0.031442 | 0.00231 | 0.03208 | 0.073716 | 0.008255 | 0.127153 | 0.026218 | 0.038173 |
| 31.5 | 0.003384 | 0.102506 | 0.032653 | 0.04567 | 0.062746 | 0.011352 | 0.00345 | 0.011885 | 0.013867 | 0.07568 | 0.017191 | 0.017317 | 0.013642 | 0.043111 | 0.096624 | 0.017915 | 0.0328 | 0.002008 | 0.034527 | 0.089055 | 0.006785 | 0.138269 | 0.024077 | 0.06186 |
| 31.5138889 | 0.002585 | 0.135008 | 0.038781 | 0.073596 | 0.0413 | 0.018464 | 0.00562 | 0.018359 | 0.011347 | 0.085125 | 0.013307 | 0.020158 | 0.01427 | 0.056676 | 0.083746 | 0.027359 | 0.030963 | 0.002954 | 0.036079 | 0.111669 | 0.005772 | 0.138554 | 0.024464 | 0.088733 |
| 31.5277778 | 0.003692 | 0.148223 | 0.044318 | 0.0859 | 0.021549 | 0.018854 | 0.012496 | 0.023891 | 0.007314 | 0.071584 | 0.008457 | 0.017813 | 0.017341 | 0.06144 | 0.070048 | 0.038303 | 0.030931 | 0.004845 | 0.040411 | 0.1235 | 0.003691 | 0.128311 | 0.02383 | 0.08939 |
| 31.5416667 | 0.004326 | 0.134259 | 0.051841 | 0.08112 | 0.011984 | 0.014913 | 0.025242 | 0.019936 | 0.003416 | 0.044683 | 0.003688 | 0.011914 | 0.020442 | 0.038911 | 0.049959 | 0.045215 | 0.028106 | 0.006712 | 0.050393 | 0.09772 | 0.003725 | 0.091518 | 0.020845 | 0.055735 |
| 31.5555556 | 0.002649 | 0.096105 | 0.054255 | 0.071463 | 0.008063 | 0.012088 | 0.040677 | 0.018605 | 0.001627 | 0.026468 | 0.001227 | 0.007154 | 0.019494 | 0.013113 | 0.026585 | 0.058868 | 0.019556 | 0.008065 | 0.065447 | 0.051282 | 0.004819 | 0.054634 | 0.027031 | 0.028123 |
| 31.5694444 | 0.001746 | 0.05049 | 0.045547 | 0.06133 | 0.0073 | 0.014589 | 0.053637 | 0.020295 | 0.002794 | 0.030378 | 0.000683 | 0.005303 | 0.016073 | 0.005638 | 0.011581 | 0.07523 | 0.014239 | 0.009997 | 0.087925 | 0.022144 | 0.006622 | 0.057997 | 0.043267 | 0.023183 |
| 31.5833333 | 0.00223 | 0.023575 | 0.035091 | 0.059138 | 0.009349 | 0.021465 | 0.055867 | 0.019469 | 0.005525 | 0.042232 | 0.001143 | 0.004057 | 0.014758 | 0.003926 | 0.009711 | 0.07377 | 0.026509 | 0.0138 | 0.104574 | 0.026924 | 0.009481 | 0.098956 | 0.044239 | 0.020368 |
| 31.5972222 | 0.001735 | 0.032192 | 0.035558 | 0.07204 | 0.015052 | 0.024442 | 0.043284 | 0.017885 | 0.005164 | 0.040659 | 0.002985 | 0.002844 | 0.015048 | 0.002327 | 0.012437 | 0.058242 | 0.046671 | 0.020405 | 0.10181 | 0.047637 | 0.008539 | 0.129265 | 0.028109 | 0.015474 |
| 31.6111111 | 0.001838 | 0.057222 | 0.045285 | 0.089163 | 0.023261 | 0.021978 | 0.024763 | 0.018737 | 0.004194 | 0.025462 | 0.006785 | 0.004702 | 0.014203 | 0.007784 | 0.025448 | 0.045992 | 0.04805 | 0.030712 | 0.096449 | 0.081126 | 0.006562 | 0.116317 | 0.015555 | 0.012335 |
| 31.625 | 0.007207 | 0.057166 | 0.051361 | 0.105934 | 0.03378 | 0.024708 | 0.015602 | 0.021303 | 0.008347 | 0.019606 | 0.011776 | 0.009757 | 0.012971 | 0.023874 | 0.06224 | 0.052465 | 0.041079 | 0.040134 | 0.103299 | 0.128702 | 0.009875 | 0.083914 | 0.009211 | 0.015976 |
| 31.6388889 | 0.018064 | 0.056041 | 0.04835 | 0.127199 | 0.048703 | 0.039754 | 0.024175 | 0.02191 | 0.014584 | 0.052813 | 0.018166 | 0.015523 | 0.013404 | 0.048148 | 0.100146 | 0.095032 | 0.037054 | 0.040825 | 0.108647 | 0.147914 | 0.017571 | 0.052145 | 0.005329 | 0.017576 |
| 31.6527778 | 0.029852 | 0.091675 | 0.043032 | 0.126799 | 0.070443 | 0.057432 | 0.039591 | 0.018096 | 0.021044 | 0.111727 | 0.026583 | 0.020015 | 0.014576 | 0.061735 | 0.129837 | 0.139328 | 0.028967 | 0.030522 | 0.088091 | 0.136757 | 0.023504 | 0.024255 | 0.013131 | 0.014005 |
| 31.6666667 | 0.035764 | 0.119202 | 0.033782 | 0.085739 | 0.092174 | 0.06208 | 0.045553 | 0.020329 | 0.031255 | 0.147914 | 0.036896 | 0.021505 | 0.011054 | 0.050441 | 0.138054 | 0.165411 | 0.019335 | 0.016259 | 0.049415 | 0.115099 | 0.022354 | 0.016508 | 0.02702 | 0.015532 |
| 31.6805556 | 0.032144 | 0.112768 | 0.017817 | 0.046312 | 0.089047 | 0.053837 | 0.037635 | 0.031749 | 0.044782 | 0.145083 | 0.046873 | 0.020398 | 0.005813 | 0.027652 | 0.089294 | 0.170961 | 0.019952 | 0.010393 | 0.031722 | 0.085068 | 0.016144 | 0.037539 | 0.030225 | 0.029436 |
| 31.6944444 | 0.022368 | 0.09221 | 0.005397 | 0.047818 | 0.055891 | 0.040926 | 0.019918 | 0.033422 | 0.053006 | 0.128335 | 0.050443 | 0.019388 | 0.006459 | 0.018047 | 0.035655 | 0.111713 | 0.038996 | 0.012342 | 0.040371 | 0.055981 | 0.010768 | 0.080092 | 0.027776 | 0.070128 |
| 31.7083333 | 0.011667 | 0.072481 | 0.002982 | 0.066488 | 0.023631 | 0.02795 | 0.006715 | 0.022329 | 0.049932 | 0.120463 | 0.045622 | 0.019867 | 0.008614 | 0.024994 | 0.034914 | 0.066014 | 0.064138 | 0.0107 | 0.0409 | 0.036815 | 0.008982 | 0.112855 | 0.023085 | 0.099099 |
| 31.7222222 | 0.004829 | 0.065006 | 0.006506 | 0.062014 | 0.008311 | 0.019764 | 0.011565 | 0.011165 | 0.041537 | 0.12596 | 0.039733 | 0.020915 | 0.011982 | 0.021827 | 0.051096 | 0.127845 | 0.069861 | 0.008747 | 0.024599 | 0.038542 | 0.010231 | 0.099841 | 0.01823 | 0.07589 |
| 31.7361111 | 0.003268 | 0.077819 | 0.010127 | 0.035467 | 0.008319 | 0.01844 | 0.031585 | 0.005377 | 0.036876 | 0.132062 | 0.041067 | 0.022179 | 0.020287 | 0.014452 | 0.052896 | 0.174239 | 0.045091 | 0.017466 | 0.018126 | 0.054107 | 0.01164 | 0.054411 | 0.01536 | 0.035555 |
| 31.75 | 0.009072 | 0.110456 | 0.013274 | 0.022402 | 0.034568 | 0.017081 | 0.039877 | 0.005573 | 0.037964 | 0.117729 | 0.044076 | 0.022506 | 0.027538 | 0.026564 | 0.042208 | 0.112117 | 0.017197 | 0.032345 | 0.034195 | 0.064151 | 0.011088 | 0.020838 | 0.01025 | 0.018619 |
| 31.7638889 | 0.015319 | 0.14211 | 0.017111 | 0.041542 | 0.085592 | 0.010827 | 0.024019 | 0.006445 | 0.039862 | 0.086487 | 0.037522 | 0.020556 | 0.032577 | 0.040829 | 0.045103 | 0.037159 | 0.008723 | 0.043601 | 0.062073 | 0.063365 | 0.00932 | 0.010532 | 0.009266 | 0.025902 |
| 31.7777778 | 0.013067 | 0.147728 | 0.020373 | 0.070514 | 0.107513 | 0.005597 | 0.00903 | 0.004549 | 0.041353 | 0.064385 | 0.026214 | 0.019971 | 0.036439 | 0.040169 | 0.069473 | 0.019301 | 0.007624 | 0.046898 | 0.085264 | 0.059774 | 0.007738 | 0.006737 | 0.01752 | 0.034054 |

| | | | | | | | | | | | | | | | | | | | | | | | | |
|---|---|---|---|---|---|---|---|---|---|---|---|---|---|---|---|---|---|---|---|---|---|---|---|---|
| 31.7916667 | 0.013101 | 0.131148 | 0.023453 | 0.094851 | 0.080406 | 0.005481 | 0.009827 | 0.004786 | 0.043401 | 0.064908 | 0.017963 | 0.024216 | 0.035314 | 0.033606 | 0.088431 | 0.042734 | 0.006815 | 0.039873 | 0.089562 | 0.056734 | 0.008738 | 0.013607 | 0.030804 | 0.031945 |
| 31.8055556 | 0.018643 | 0.107246 | 0.029389 | 0.11377 | 0.047273 | 0.007662 | 0.015897 | 0.010682 | 0.04417 | 0.078403 | 0.012375 | 0.030384 | 0.031561 | 0.03055 | 0.089816 | 0.080854 | 0.016434 | 0.02881 | 0.074787 | 0.046566 | 0.013069 | 0.032402 | 0.043028 | 0.025635 |
| 31.8194444 | 0.020975 | 0.065278 | 0.037103 | 0.114071 | 0.042176 | 0.007641 | 0.012303 | 0.022891 | 0.049259 | 0.07739 | 0.009505 | 0.033641 | 0.027485 | 0.031414 | 0.063814 | 0.099694 | 0.031949 | 0.021123 | 0.047681 | 0.027994 | 0.017845 | 0.04029 | 0.047311 | 0.020313 |
| 31.8333333 | 0.016012 | 0.025949 | 0.043545 | 0.10554 | 0.080584 | 0.006206 | 0.004702 | 0.041942 | 0.062791 | 0.047516 | 0.009618 | 0.031526 | 0.017914 | 0.030103 | 0.028967 | 0.091175 | 0.032559 | 0.015297 | 0.02528 | 0.014006 | 0.023186 | 0.032361 | 0.043742 | 0.018241 |
| 31.8472222 | 0.006882 | 0.028773 | 0.049544 | 0.104784 | 0.119653 | 0.007066 | 0.007423 | 0.057389 | 0.071629 | 0.01987 | 0.012156 | 0.026518 | 0.009687 | 0.022644 | 0.011174 | 0.077578 | 0.017573 | 0.010181 | 0.031261 | 0.008223 | 0.032498 | 0.02587 | 0.038335 | 0.014247 |
| 31.8611111 | 0.002141 | 0.043578 | 0.052279 | 0.09908 | 0.136543 | 0.008067 | 0.01309 | 0.055533 | 0.063253 | 0.017553 | 0.015831 | 0.025121 | 0.01605 | 0.020147 | 0.007656 | 0.062809 | 0.008519 | 0.009525 | 0.060334 | 0.00389 | 0.051164 | 0.03772 | 0.031931 | 0.009685 |
| 31.875 | 0.006075 | 0.043738 | 0.053823 | 0.078196 | 0.140609 | 0.006836 | 0.025045 | 0.049578 | 0.044568 | 0.018247 | 0.01932 | 0.030669 | 0.031777 | 0.024038 | 0.01731 | 0.047037 | 0.017332 | 0.011294 | 0.09214 | 0.005416 | 0.067827 | 0.069767 | 0.025343 | 0.01363 |
| 31.8888889 | 0.017647 | 0.047542 | 0.052944 | 0.045758 | 0.112672 | 0.007568 | 0.04448 | 0.05797 | 0.031561 | 0.01197 | 0.021833 | 0.039274 | 0.039155 | 0.021761 | 0.030253 | 0.036507 | 0.032138 | 0.010036 | 0.11097 | 0.015149 | 0.055286 | 0.102227 | 0.020376 | 0.020641 |
| 31.9027778 | 0.024575 | 0.051203 | 0.036157 | 0.035 | 0.072209 | 0.009525 | 0.044387 | 0.066255 | 0.033239 | 0.017474 | 0.025181 | 0.043674 | 0.030487 | 0.01842 | 0.035454 | 0.028232 | 0.039348 | 0.011246 | 0.108996 | 0.02711 | 0.025961 | 0.124007 | 0.01406 | 0.024478 |
| 31.9166667 | 0.018652 | 0.050805 | 0.01544 | 0.08451 | 0.046669 | 0.007323 | 0.041961 | 0.055541 | 0.040714 | 0.046707 | 0.03163 | 0.040553 | 0.019404 | 0.017647 | 0.032642 | 0.022983 | 0.043908 | 0.015203 | 0.095261 | 0.036969 | 0.02049 | 0.144873 | 0.016071 | 0.031253 |
| 31.9305556 | 0.008435 | 0.048644 | 0.007489 | 0.141046 | 0.032109 | 0.003617 | 0.055774 | 0.031849 | 0.037412 | 0.08677 | 0.037319 | 0.032248 | 0.013698 | 0.015536 | 0.022131 | 0.021364 | 0.055519 | 0.012667 | 0.080638 | 0.042121 | 0.040305 | 0.179895 | 0.029984 | 0.042605 |
| 31.9444444 | 0.002526 | 0.047444 | 0.003672 | 0.147517 | 0.018626 | 0.004464 | 0.052696 | 0.023664 | 0.02403 | 0.115776 | 0.037139 | 0.022372 | 0.00905 | 0.013418 | 0.023831 | 0.016969 | 0.064211 | 0.009174 | 0.070516 | 0.049362 | 0.051693 | 0.208075 | 0.031683 | 0.048739 |
| 31.9583333 | 0.009567 | 0.062718 | 0.00276 | 0.130662 | 0.009897 | 0.0061 | 0.036083 | 0.037928 | 0.018103 | 0.128292 | 0.031639 | 0.014204 | 0.010975 | 0.012469 | 0.036385 | 0.013929 | 0.052959 | 0.010937 | 0.061467 | 0.064571 | 0.042806 | 0.201042 | 0.018614 | 0.046066 |
| 31.9722222 | 0.027557 | 0.077397 | 0.009561 | 0.115112 | 0.016771 | 0.003626 | 0.024489 | 0.041523 | 0.02468 | 0.125431 | 0.024065 | 0.009764 | 0.020794 | 0.012856 | 0.041739 | 0.018927 | 0.030459 | 0.018755 | 0.041977 | 0.077559 | 0.030567 | 0.186939 | 0.016055 | 0.038706 |
| 31.9861111 | 0.034449 | 0.059509 | 0.021084 | 0.108449 | 0.022217 | 0.003748 | 0.012796 | 0.033139 | 0.031948 | 0.111903 | 0.01706 | 0.007365 | 0.028501 | 0.009661 | 0.048371 | 0.021961 | 0.014101 | 0.031328 | 0.017605 | 0.077092 | 0.018746 | 0.182424 | 0.028995 | 0.024997 |
| 32 | 0.025525 | 0.06079 | 0.032914 | 0.116381 | 0.021859 | 0.008937 | 0.007055 | 0.042434 | 0.041292 | 0.10267 | 0.011434 | 0.005146 | 0.032369 | 0.004297 | 0.041604 | 0.019738 | 0.006294 | 0.032495 | 0.006454 | 0.064017 | 0.010616 | 0.171835 | 0.040597 | 0.01095 |
| 32.0138889 | 0.015959 | 0.104181 | 0.040907 | 0.127899 | 0.047676 | 0.013078 | 0.005397 | 0.06738 | 0.048272 | 0.109271 | 0.006826 | 0.005603 | 0.035314 | 0.003802 | 0.027643 | 0.022148 | 0.004029 | 0.019766 | 0.017101 | 0.050716 | 0.017455 | 0.147481 | 0.043053 | 0.006426 |
| 32.0277778 | 0.015659 | 0.113744 | 0.04433 | 0.119348 | 0.101561 | 0.011699 | 0.002744 | 0.077859 | 0.036702 | 0.139999 | 0.004114 | 0.010097 | 0.029716 | 0.00965 | 0.052504 | 0.034202 | 0.011454 | 0.011914 | 0.044173 | 0.049446 | 0.029409 | 0.11622 | 0.033877 | 0.006997 |
| 32.0416667 | 0.034218 | 0.111928 | 0.056356 | 0.093735 | 0.150735 | 0.008505 | 0.00279 | 0.066645 | 0.016702 | 0.188366 | 0.005455 | 0.016521 | 0.020423 | 0.01667 | 0.101718 | 0.059372 | 0.035079 | 0.027214 | 0.080416 | 0.056306 | 0.033587 | 0.100158 | 0.025075 | 0.011625 |
| 32.0555556 | 0.058441 | 0.145019 | 0.077795 | 0.076682 | 0.164801 | 0.012822 | 0.002982 | 0.056294 | 0.00814 | 0.208239 | 0.014196 | 0.021755 | 0.024606 | 0.022364 | 0.115694 | 0.09292 | 0.062998 | 0.052787 | 0.121763 | 0.058769 | 0.027223 | 0.099907 | 0.037965 | 0.020496 |
| 32.0694444 | 0.062463 | 0.138941 | 0.080276 | 0.072681 | 0.158697 | 0.020452 | 0.003691 | 0.045214 | 0.012537 | 0.193978 | 0.027853 | 0.020061 | 0.031594 | 0.028013 | 0.083776 | 0.110223 | 0.079551 | 0.057189 | 0.136648 | 0.052423 | 0.013491 | 0.090033 | 0.048852 | 0.022774 |
| 32.0833333 | 0.050562 | 0.069637 | 0.058778 | 0.070387 | 0.175601 | 0.018271 | 0.008377 | 0.028255 | 0.015444 | 0.15357 | 0.034102 | 0.011166 | 0.026398 | 0.029561 | 0.05422 | 0.093818 | 0.079453 | 0.040555 | 0.104318 | 0.044852 | 0.004555 | 0.067155 | 0.037664 | 0.017467 |
| 32.0972222 | 0.032446 | 0.020686 | 0.037001 | 0.05455 | 0.166485 | 0.012088 | 0.011298 | 0.029032 | 0.012692 | 0.075221 | 0.026054 | 0.003765 | 0.016821 | 0.024886 | 0.054683 | 0.054955 | 0.054543 | 0.021479 | 0.056915 | 0.045296 | 0.005464 | 0.048268 | 0.031817 | 0.011983 |
| 32.1111111 | 0.012827 | 0.019417 | 0.025413 | 0.025374 | 0.087562 | 0.009637 | 0.014269 | 0.04859 | 0.011509 | 0.034448 | 0.014587 | 0.002754 | 0.017911 | 0.017549 | 0.048771 | 0.019811 | 0.030511 | 0.015764 | 0.02843 | 0.046545 | 0.00748 | 0.034221 | 0.042312 | 0.007573 |
| 32.125 | 0.00454 | 0.033495 | 0.017627 | 0.010131 | 0.020473 | 0.00606 | 0.023125 | 0.063881 | 0.009494 | 0.067981 | 0.012463 | 0.004342 | 0.031307 | 0.010248 | 0.034151 | 0.00722 | 0.039356 | 0.014947 | 0.022833 | 0.038671 | 0.005517 | 0.023019 | 0.053287 | 0.005091 |
| 32.1388889 | 0.00555 | 0.041061 | 0.008698 | 0.014026 | 0.008181 | 0.003908 | 0.021526 | 0.064849 | 0.009999 | 0.112844 | 0.016946 | 0.003964 | 0.041122 | 0.006437 | 0.03519 | 0.010663 | 0.053468 | 0.014891 | 0.021098 | 0.026139 | 0.002992 | 0.032237 | 0.0636 | 0.008829 |
| 32.1527778 | 0.004093 | 0.032303 | 0.002599 | 0.014569 | 0.01632 | 0.004565 | 0.011772 | 0.045167 | 0.015229 | 0.141664 | 0.018335 | 0.003474 | 0.0385 | 0.007606 | 0.028095 | 0.013954 | 0.052633 | 0.027765 | 0.017522 | 0.017495 | 0.002252 | 0.065735 | 0.080176 | 0.016258 |
| 32.1666667 | 0.001715 | 0.037145 | 0.001579 | 0.012713 | 0.025386 | 0.003916 | 0.018966 | 0.028589 | 0.018442 | 0.170476 | 0.013348 | 0.00859 | 0.025909 | 0.007283 | 0.012106 | 0.014967 | 0.047605 | 0.033565 | 0.019233 | 0.017 | 0.003637 | 0.09112 | 0.090129 | 0.015425 |
| 32.1805556 | 0.00135 | 0.069315 | 0.002207 | 0.014163 | 0.032217 | 0.002142 | 0.033032 | 0.042451 | 0.018732 | 0.194003 | 0.009976 | 0.016956 | 0.011805 | 0.004878 | 0.010152 | 0.014726 | 0.041766 | 0.021618 | 0.016651 | 0.017856 | 0.004843 | 0.097519 | 0.083729 | 0.008389 |
| 32.1944444 | 0.002564 | 0.083634 | 0.003167 | 0.013405 | 0.030456 | 0.002354 | 0.027986 | 0.056699 | 0.019839 | 0.194707 | 0.013068 | 0.020852 | 0.004725 | 0.006073 | 0.017822 | 0.009896 | 0.03844 | 0.013389 | 0.01457 | 0.012648 | 0.005227 | 0.119345 | 0.069177 | 0.005537 |
| 32.2083333 | 0.007296 | 0.066582 | 0.005581 | 0.014687 | 0.023466 | 0.006066 | 0.010726 | 0.046217 | 0.028902 | 0.176272 | 0.015187 | 0.020663 | 0.003476 | 0.009288 | 0.019396 | 0.006898 | 0.04004 | 0.014481 | 0.024288 | 0.009789 | 0.009369 | 0.176217 | 0.050047 | 0.005857 |
| 32.2222222 | 0.017185 | 0.05013 | 0.010158 | 0.017017 | 0.035526 | 0.010036 | 0.00317 | 0.02424 | 0.039873 | 0.141398 | 0.012206 | 0.01939 | 0.00257 | 0.013848 | 0.024566 | 0.012059 | 0.039613 | 0.014612 | 0.031515 | 0.015074 | 0.011723 | 0.217248 | 0.028442 | 0.008462 |
| 32.2361111 | 0.034084 | 0.050465 | 0.013945 | 0.01279 | 0.061889 | 0.011169 | 0.00463 | 0.017669 | 0.032976 | 0.090381 | 0.006778 | 0.017128 | 0.001588 | 0.020397 | 0.049146 | 0.021823 | 0.026908 | 0.012986 | 0.024416 | 0.020573 | 0.007071 | 0.194092 | 0.012477 | 0.010641 |
| 32.25 | 0.049585 | 0.065759 | 0.013481 | 0.00549 | 0.077257 | 0.010029 | 0.004299 | 0.026911 | 0.020217 | 0.042682 | 0.00249 | 0.015631 | 0.002876 | 0.030716 | 0.062545 | 0.037494 | 0.013847 | 0.011925 | 0.016418 | 0.023238 | 0.003904 | 0.152253 | 0.005075 | 0.010844 |
| 32.2638889 | 0.044614 | 0.077131 | 0.011177 | 0.0025 | 0.075785 | 0.008665 | 0.004237 | 0.02362 | 0.027144 | 0.025705 | 0.001426 | 0.015646 | 0.003888 | 0.038058 | 0.049048 | 0.051488 | 0.016796 | 0.009615 | 0.013025 | 0.024905 | 0.003441 | 0.124482 | 0.004383 | 0.012714 |
| 32.2777778 | 0.024098 | 0.058776 | 0.009422 | 0.004756 | 0.071369 | 0.008706 | 0.007901 | 0.00977 | 0.034892 | 0.040978 | 0.002944 | 0.013976 | 0.00365 | 0.027164 | 0.061013 | 0.050403 | 0.025231 | 0.005352 | 0.009901 | 0.02147 | 0.003888 | 0.081104 | 0.010725 | 0.013404 |
| 32.2916667 | 0.014658 | 0.0323 | 0.006991 | 0.00713 | 0.081577 | 0.01009 | 0.01143 | 0.007157 | 0.027911 | 0.082298 | 0.006214 | 0.008376 | 0.004757 | 0.016468 | 0.098287 | 0.040627 | 0.032966 | 0.00183 | 0.011178 | 0.015798 | 0.007374 | 0.04494 | 0.019903 | 0.011419 |
| 32.3055556 | 0.016007 | 0.042144 | 0.004455 | 0.005921 | 0.087092 | 0.008831 | 0.015097 | 0.014023 | 0.022815 | 0.147371 | 0.011354 | 0.005747 | 0.004456 | 0.020378 | 0.088124 | 0.033976 | 0.045895 | 0.001044 | 0.010621 | 0.017707 | 0.008174 | 0.059771 | 0.026422 | 0.00913 |
| 32.3194444 | 0.017133 | 0.082115 | 0.006653 | 0.006153 | 0.06633 | 0.0065 | 0.023423 | 0.021381 | 0.020189 | 0.182526 | 0.017905 | 0.014384 | 0.004216 | 0.017142 | 0.038744 | 0.0272 | 0.051453 | 0.001529 | 0.011398 | 0.020392 | 0.005936 | 0.083154 | 0.030427 | 0.007901 |
| 32.3333333 | 0.020519 | 0.127332 | 0.013226 | 0.010702 | 0.03568 | 0.008235 | 0.029124 | 0.024844 | 0.015295 | 0.185788 | 0.022942 | 0.028075 | 0.009102 | 0.014042 | 0.010231 | 0.023381 | 0.036759 | 0.003195 | 0.020634 | 0.021674 | 0.00561 | 0.088514 | 0.032867 | 0.014572 |
| 32.3472222 | 0.020061 | 0.152613 | 0.017076 | 0.012447 | 0.013561 | 0.008315 | 0.020665 | 0.019211 | 0.015012 | 0.20735 | 0.021268 | 0.031018 | 0.012458 | 0.018068 | 0.011051 | 0.033752 | 0.019447 | 0.004946 | 0.025928 | 0.023905 | 0.003551 | 0.084866 | 0.031641 | 0.024837 |
| 32.3611111 | 0.013325 | 0.136629 | 0.016508 | 0.008503 | 0.008702 | 0.006493 | 0.012107 | 0.022927 | 0.01571 | 0.202475 | 0.012188 | 0.021758 | 0.008019 | 0.010404 | 0.015092 | 0.039887 | 0.015777 | 0.003928 | 0.023676 | 0.015825 | 0.001365 | 0.050622 | 0.022957 | 0.022335 |
| 32.375 | 0.009919 | 0.098028 | 0.014883 | 0.004375 | 0.018607 | 0.006748 | 0.020997 | 0.035805 | 0.01018 | 0.142098 | 0.004667 | 0.013559 | 0.002658 | 0.003113 | 0.017055 | 0.033001 | 0.013293 | 0.002421 | 0.022494 | 0.006992 | 0.001988 | 0.01493 | 0.011179 | 0.010373 |
| 32.3888889 | 0.013609 | 0.062928 | 0.013711 | 0.003786 | 0.027183 | 0.005007 | 0.025171 | 0.028419 | 0.005726 | 0.075655 | 0.005001 | 0.009091 | 0.00241 | 0.005489 | 0.037868 | 0.034676 | 0.009485 | 0.003329 | 0.021243 | 0.012662 | 0.002923 | 0.018987 | 0.008484 | 0.003753 |
| 32.4027778 | 0.022919 | 0.036704 | 0.012732 | 0.004717 | 0.036248 | 0.00262 | 0.013215 | 0.013814 | 0.01022 | 0.065436 | 0.009395 | 0.004667 | 0.004434 | 0.006442 | 0.06726 | 0.044906 | 0.013887 | 0.004111 | 0.015771 | 0.025591 | 0.004965 | 0.052859 | 0.019125 | 0.00552 |
| 32.4166667 | 0.033085 | 0.017974 | 0.010791 | 0.003959 | 0.056931 | 0.002894 | 0.006049 | 0.013507 | 0.019755 | 0.103324 | 0.015279 | 0.002284 | 0.004988 | 0.012518 | 0.072547 | 0.04392 | 0.018255 | 0.003926 | 0.009669 | 0.034227 | 0.005613 | 0.089643 | 0.032475 | 0.012217 |
| 32.4305556 | 0.036487 | 0.009391 | 0.006817 | 0.004241 | 0.081088 | 0.003624 | 0.004599 | 0.015591 | 0.028311 | 0.126908 | 0.021698 | 0.003833 | 0.003494 | 0.032622 | 0.04674 | 0.033982 | 0.020366 | 0.00337 | 0.007912 | 0.035768 | 0.00364 | 0.107714 | 0.039021 | 0.02001 |
| 32.4444444 | 0.0313 | 0.011754 | 0.003363 | 0.00968 | 0.099561 | 0.00303 | 0.003277 | 0.014256 | 0.035137 | 0.124425 | 0.027089 | 0.006656 | 0.001532 | 0.048138 | 0.026308 | 0.030267 | 0.024669 | 0.001959 | 0.009865 | 0.034528 | 0.003905 | 0.10562 | 0.04007 | 0.021668 |
| 32.4583333 | 0.021082 | 0.016795 | 0.004147 | 0.0175 | 0.109162 | 0.002797 | 0.004114 | 0.01308 | 0.039654 | 0.108373 | 0.033024 | 0.008238 | 0.000966 | 0.035822 | 0.033964 | 0.031129 | 0.027054 | 0.001091 | 0.01219 | 0.034491 | 0.005459 | 0.098224 | 0.037595 | 0.015922 |
| 32.4722222 | 0.011062 | 0.027513 | 0.008865 | 0.025094 | 0.109211 | 0.004584 | 0.009575 | 0.009451 | 0.040668 | 0.107236 | 0.038162 | 0.01033 | 0.003713 | 0.013254 | 0.042803 | 0.035122 | 0.037304 | 0.002933 | 0.015066 | 0.037068 | 0.004714 | 0.107098 | 0.031931 | 0.013939 |
| 32.4861111 | 0.005256 | 0.053332 | 0.014579 | 0.032826 | 0.100473 | 0.007117 | 0.015392 | 0.006542 | 0.041054 | 0.134422 | 0.037418 | 0.014245 | 0.01098 | 0.016155 | 0.044385 | 0.04941 | 0.064283 | 0.0081 | 0.019083 | 0.042161 | 0.003742 | 0.133514 | 0.023952 | 0.02445 |
| 32.5 | 0.003001 | 0.082236 | 0.018 | 0.037916 | 0.079436 | 0.008627 | 0.013839 | 0.011916 | 0.041547 | 0.153744 | 0.02882 | 0.017004 | 0.020902 | 0.033667 | 0.05446 | 0.074267 | 0.086631 | 0.014125 | 0.021947 | 0.049247 | 0.005399 | 0.157717 | 0.014438 | 0.039299 |
| 32.5138889 | 0.00353 | 0.085541 | 0.017907 | 0.037955 | 0.056911 | 0.007827 | 0.013303 | 0.01909 | 0.041649 | 0.149459 | 0.016117 | 0.013422 | 0.030433 | 0.034149 | 0.070977 | 0.099137 | 0.092886 | 0.017543 | 0.022082 | 0.056263 | 0.006225 | 0.171576 | 0.007502 | 0.041067 |
| 32.5277778 | 0.005788 | 0.057489 | 0.016197 | 0.034822 | 0.050118 | 0.005569 | 0.0208 | 0.015536 | 0.043219 | 0.123922 | 0.007165 | 0.007254 | 0.037683 | 0.020956 | 0.082726 | 0.103396 | 0.093992 | 0.018069 | 0.021809 | 0.058655 | 0.00793 | 0.174631 | 0.010084 | 0.030718 |
| 32.5416667 | 0.005359 | 0.026885 | 0.014186 | 0.030601 | 0.048732 | 0.004835 | 0.028711 | 0.009472 | 0.045067 | 0.089117 | 0.006381 | 0.007026 | 0.040656 | 0.008613 | 0.085132 | 0.075729 | 0.083607 | 0.018994 | 0.027289 | 0.050817 | 0.013352 | 0.158483 | 0.023108 | 0.025285 |
| 32.5555556 | 0.004425 | 0.017845 | 0.012726 | 0.028328 | 0.05132 | 0.00894 | 0.033657 | 0.011942 | 0.041148 | 0.087225 | 0.010982 | 0.009549 | 0.04064 | 0.006396 | 0.075984 | 0.041406 | 0.063204 | 0.025053 | 0.047433 | 0.03727 | 0.012985 | 0.130207 | 0.0357 | 0.030033 |
| 32.5694444 | 0.008994 | 0.030194 | 0.016995 | 0.034556 | 0.066056 | 0.022044 | 0.042204 | 0.019605 | 0.029761 | 0.1147 | 0.016137 | 0.00863 | 0.041468 | 0.024912 | 0.062293 | 0.038874 | 0.055582 | 0.039071 | 0.083616 | 0.035532 | 0.010804 | 0.131315 | 0.037072 | 0.042041 |
| 32.5833333 | 0.015181 | 0.04601 | 0.033897 | 0.056388 | 0.062792 | 0.034956 | 0.03821 | 0.027166 | 0.019514 | 0.111446 | 0.017445 | 0.004971 | 0.039741 | 0.045684 | 0.056348 | 0.076936 | 0.069989 | 0.056181 | 0.105464 | 0.056419 | 0.016589 | 0.149195 | 0.033349 | 0.049829 |
| 32.5972222 | 0.017651 | 0.056938 | 0.058297 | 0.090362 | 0.03414 | 0.029805 | 0.015423 | 0.029699 | 0.016754 | 0.068228 | 0.01228 | 0.003429 | 0.030983 | 0.039205 | 0.055822 | 0.118583 | 0.084293 | 0.065194 | 0.094551 | 0.090804 | 0.022802 | 0.132437 | 0.036126 | 0.03784 |
| 32.6111111 | 0.022539 | 0.081602 | 0.07276 | 0.119487 | 0.018226 | 0.016939 | 0.004142 | 0.027661 | 0.018307 | 0.02856 | 0.007782 | 0.010386 | 0.021103 | 0.030589 | 0.065773 | 0.131689 | 0.072703 | 0.064858 | 0.082007 | 0.118393 | 0.025516 | 0.085057 | 0.046968 | 0.020149 |
| 32.625 | 0.029691 | 0.118379 | 0.072797 | 0.136327 | 0.044752 | 0.017412 | 0.006606 | 0.020986 | 0.020449 | 0.010903 | 0.014073 | 0.024566 | 0.017056 | 0.042149 | 0.087102 | 0.11871 | 0.050285 | 0.06366 | 0.086687 | 0.126972 | 0.030589 | 0.039958 | 0.057767 | 0.017398 |
| 32.6388889 | 0.023632 | 0.131879 | 0.074287 | 0.141673 | 0.111672 | 0.020454 | 0.009106 | 0.015015 | 0.024108 | 0.010137 | 0.026793 | 0.0367 | 0.013444 | 0.049927 | 0.086515 | 0.089511 | 0.04339 | 0.061941 | 0.098145 | 0.121848 | 0.039372 | 0.029845 | 0.065454 | 0.020072 |
| 32.6527778 | 0.014635 | 0.106927 | 0.081954 | 0.138768 | 0.166437 | 0.01225 | 0.014573 | 0.016297 | 0.029837 | 0.017921 | 0.035286 | 0.039349 | 0.009658 | 0.046314 | 0.052436 | 0.079414 | 0.054713 | 0.054867 | 0.099535 | 0.112022 | 0.042048 | 0.059047 | 0.087454 | 0.015122 |
| 32.6666667 | 0.014776 | 0.062885 | 0.078971 | 0.133365 | 0.190546 | 0.00411 | 0.023273 | 0.017366 | 0.036058 | 0.019212 | 0.036805 | 0.033441 | 0.015372 | 0.034948 | 0.023776 | 0.086367 | 0.057686 | 0.039051 | 0.083378 | 0.097852 | 0.031831 | 0.086666 | 0.115713 | 0.012454 |
| 32.6805556 | 0.021526 | 0.032424 | 0.06051 | 0.116773 | 0.222114 | 0.004507 | 0.031828 | 0.014423 | 0.039053 | 0.013347 | 0.034461 | 0.026491 | 0.0266 | 0.026681 | 0.023371 | 0.082241 | 0.039645 | 0.022767 | 0.058421 | 0.082036 | 0.016486 | 0.066471 | 0.116164 | 0.019625 |
| 32.6944444 | 0.037545 | 0.029736 | 0.042884 | 0.088802 | 0.222666 | 0.0076 | 0.035171 | 0.011817 | 0.032889 | 0.020272 | 0.030335 | 0.024177 | 0.030906 | 0.029765 | 0.032764 | 0.080757 | 0.040054 | 0.023224 | 0.032281 | 0.070532 | 0.006785 | 0.026539 | 0.091729 | 0.021799 |
| 32.7083333 | 0.04472 | 0.040815 | 0.034873 | 0.063818 | 0.14307 | 0.009448 | 0.031992 | 0.008118 | 0.019663 | 0.044905 | 0.025899 | 0.027832 | 0.021827 | 0.022115 | 0.039821 | 0.076338 | 0.064188 | 0.033329 | 0.012231 | 0.063538 | 0.004946 | 0.013675 | 0.06081 | 0.015809 |
| 32.7222222 | 0.035597 | 0.048504 | 0.029254 | 0.045559 | 0.050268 | 0.010866 | 0.025938 | 0.007408 | 0.014802 | 0.068213 | 0.023633 | 0.03432 | 0.012488 | 0.010896 | 0.047167 | 0.066018 | 0.065682 | 0.030751 | 0.010668 | 0.061097 | 0.005639 | 0.01361 | 0.033241 | 0.020195 |
| 32.7361111 | 0.020711 | 0.03639 | 0.019317 | 0.031617 | 0.011319 | 0.012571 | 0.019096 | 0.015431 | 0.024115 | 0.068729 | 0.025146 | 0.03666 | 0.022967 | 0.010618 | 0.05229 | 0.056593 | 0.038839 | 0.028319 | 0.022777 | 0.06427 | 0.011495 | 0.022634 | 0.018166 | 0.031993 |
| 32.75 | 0.019693 | 0.019027 | 0.010994 | 0.022505 | 0.041468 | 0.014452 | 0.012163 | 0.024621 | 0.03262 | 0.054189 | 0.030292 | 0.032775 | 0.041312 | 0.012901 | 0.050788 | 0.045964 | 0.023183 | 0.038855 | 0.023003 | 0.066703 | 0.027977 | 0.036669 | 0.020103 | 0.029273 |
| 32.7638889 | 0.041349 | 0.014905 | 0.013154 | 0.015598 | 0.123015 | 0.015731 | 0.005748 | 0.027268 | 0.032881 | 0.058148 | 0.036166 | 0.030279 | 0.043465 | 0.021176 | 0.040556 | 0.031689 | 0.026565 | 0.036093 | 0.015048 | 0.061269 | 0.051305 | 0.034944 | 0.034992 | 0.016843 |
| 32.7777778 | 0.051048 | 0.02192 | 0.018746 | 0.008898 | 0.20856 | 0.017144 | 0.007811 | 0.019259 | 0.029839 | 0.090399 | 0.039502 | 0.037478 | 0.031074 | 0.026581 | 0.02684 | 0.018561 | 0.025558 | 0.019758 | 0.021596 | 0.048931 | 0.076142 | 0.026454 | 0.047895 | 0.010955 |
| 32.7916667 | 0.039064 | 0.053398 | 0.017234 | 0.013863 | 0.26087 | 0.019305 | 0.013213 | 0.02871 | 0.020143 | 0.12301 | 0.038936 | 0.053575 | 0.016719 | 0.020612 | 0.016734 | 0.029773 | 0.020649 | 0.014648 | 0.028967 | 0.036492 | 0.095499 | 0.015261 | 0.047878 | 0.008844 |
| 32.8055556 | 0.028375 | 0.098386 | 0.010573 | 0.031177 | 0.219776 | 0.020885 | 0.009754 | 0.056046 | 0.017118 | 0.133156 | 0.035533 | 0.072282 | 0.006865 | 0.015531 | 0.010742 | 0.057575 | 0.019647 | 0.023107 | 0.020302 | 0.032867 | 0.097067 | 0.00717 | 0.038539 | 0.013031 |
| 32.8194444 | 0.027827 | 0.140387 | 0.007872 | 0.03977 | 0.114217 | 0.018072 | 0.012351 | 0.056061 | 0.045156 | 0.129679 | 0.030651 | 0.079588 | 0.00546 | 0.027319 | 0.014182 | 0.070011 | 0.017574 | 0.032873 | 0.011804 | 0.038306 | 0.078606 | 0.012694 | 0.033033 | 0.034081 |
| 32.8333333 | 0.031503 | 0.146103 | 0.015193 | 0.037296 | 0.051955 | 0.009386 | 0.02599 | 0.054491 | 0.093059 | 0.121901 | 0.027032 | 0.066011 | 0.010996 | 0.054044 | 0.018282 | 0.055922 | 0.013549 | 0.033609 | 0.025096 | 0.038226 | 0.054693 | 0.018048 | 0.031022 | 0.059509 |
| 32.8472222 | 0.022238 | 0.103024 | 0.028928 | 0.042823 | 0.039074 | 0.006122 | 0.045376 | 0.087927 | 0.135119 | 0.105213 | 0.02878 | 0.047866 | 0.01636 | 0.06424 | 0.021261 | 0.032621 | 0.009464 | 0.026328 | 0.053481 | 0.02499 | 0.035385 | 0.01634 | 0.022061 | 0.059048 |
| 32.8611111 | 0.007942 | 0.074232 | 0.040067 | 0.041856 | 0.036718 | 0.012492 | 0.073056 | 0.115226 | 0.151798 | 0.077418 | 0.030472 | 0.040403 | 0.017105 | 0.040855 | 0.041437 | 0.01787 | 0.009484 | 0.015644 | 0.071966 | 0.014439 | 0.022994 | 0.02703 | 0.010345 | 0.038235 |
| 32.875 | 0.003558 | 0.07527 | 0.044013 | 0.026003 | 0.042864 | 0.012826 | 0.082435 | 0.094648 | 0.135262 | 0.050641 | 0.026734 | 0.04181 | 0.020416 | 0.018431 | 0.06169 | 0.010407 | 0.018887 | 0.006174 | 0.076945 | 0.018011 | 0.01573 | 0.051685 | 0.005643 | 0.034958 |
| 32.8888889 | 0.00231 | 0.059421 | 0.046803 | 0.024413 | 0.047384 | 0.004646 | 0.05356 | 0.065614 | 0.098809 | 0.043527 | 0.023564 | 0.046096 | 0.033122 | 0.027295 | 0.06584 | 0.009765 | 0.024812 | 0.010167 | 0.081945 | 0.025402 | 0.015263 | 0.068937 | 0.007442 | 0.047007 |
| 32.9027778 | 0.004215 | 0.066874 | 0.052095 | 0.030824 | 0.034144 | 0.000319 | 0.020537 | 0.056207 | 0.062259 | 0.055612 | 0.022907 | 0.044248 | 0.048497 | 0.051439 | 0.06001 | 0.020431 | 0.02578 | 0.029584 | 0.088792 | 0.025128 | 0.030228 | 0.088895 | 0.008392 | 0.051733 |
| 32.9166667 | 0.011637 | 0.148324 | 0.053384 | 0.034137 | 0.022566 | 0.00027 | 0.009419 | 0.035694 | 0.040626 | 0.05987 | 0.022579 | 0.030165 | 0.053271 | 0.066621 | 0.045392 | 0.037437 | 0.040632 | 0.04798 | 0.090516 | 0.016947 | 0.048173 | 0.124287 | 0.006323 | 0.052731 |
| 32.9305556 | 0.017202 | 0.203901 | 0.047882 | 0.043931 | 0.023785 | 0.000474 | 0.00742 | 0.017184 | 0.035398 | 0.039747 | 0.019775 | 0.015387 | 0.038426 | 0.063351 | 0.035976 | 0.045196 | 0.058893 | 0.05702 | 0.087368 | 0.008463 | 0.052616 | 0.162064 | 0.0075 | 0.057885 |
| 32.9444444 | 0.016038 | 0.185157 | 0.040537 | 0.051346 | 0.025989 | 0.001117 | 0.006174 | 0.017061 | 0.035594 | 0.019907 | 0.014376 | 0.019556 | 0.019061 | 0.046831 | 0.042081 | 0.038935 | 0.062647 | 0.058203 | 0.080106 | 0.008284 | 0.052039 | 0.182956 | 0.014457 | 0.058383 |
| 32.9583333 | 0.011873 | 0.155279 | 0.033702 | 0.044793 | 0.033341 | 0.001931 | 0.010061 | 0.014934 | 0.02713 | 0.02876 | 0.012406 | 0.037308 | 0.017867 | 0.032337 | 0.053078 | 0.029393 | 0.05756 | 0.054914 | 0.071993 | 0.020347 | 0.053256 | 0.188146 | 0.014635 | 0.048964 |
| 32.9722222 | 0.008186 | 0.132042 | 0.024142 | 0.030739 | 0.055578 | 0.002455 | 0.018893 | 0.011326 | 0.015875 | 0.0601 | 0.014238 | 0.042306 | 0.03058 | 0.029874 | 0.049391 | 0.024146 | 0.053487 | 0.058344 | 0.067929 | 0.045728 | 0.053029 | 0.208736 | 0.018248 | 0.04168 |
| 32.9861111 | 0.005174 | 0.098668 | 0.013347 | 0.019977 | 0.086734 | 0.002995 | 0.022099 | 0.015406 | 0.017485 | 0.096408 | 0.016494 | 0.030341 | 0.044448 | 0.033795 | 0.027379 | 0.024848 | 0.054269 | 0.069002 | 0.06249 | 0.076447 | 0.051488 | 0.243641 | 0.049589 | 0.039324 |
| 33 | 0.004548 | 0.095013 | 0.011673 | 0.014284 | 0.092959 | 0.00508 | 0.016729 | 0.02009 | 0.027466 | 0.126052 | 0.015403 | 0.021515 | 0.057573 | 0.030864 | 0.012091 | 0.035064 | 0.056885 | 0.059442 | 0.045317 | 0.102525 | 0.047572 | 0.246306 | 0.080802 | 0.031144 |
| 33.0138889 | 0.006628 | 0.151104 | 0.02345 | 0.020844 | 0.089048 | 0.009142 | 0.009976 | 0.026656 | 0.030222 | 0.121705 | 0.011212 | 0.018071 | 0.064977 | 0.022393 | 0.016906 | 0.060673 | 0.061121 | 0.028028 | 0.026783 | 0.121696 | 0.041798 | 0.209411 | 0.085685 | 0.015911 |
| 33.0277778 | 0.009464 | 0.181422 | 0.042886 | 0.046028 | 0.087311 | 0.011712 | 0.005803 | 0.034371 | 0.020821 | 0.090588 | 0.009613 | 0.01939 | 0.050827 | 0.015762 | 0.020791 | 0.103032 | 0.077325 | 0.03164 | 0.019489 | 0.133123 | 0.039387 | 0.184022 | 0.078684 | 0.004797 |
| 33.0416667 | 0.013289 | 0.164075 | 0.05521 | 0.076197 | 0.053727 | 0.009676 | 0.005746 | 0.024756 | 0.023447 | 0.096053 | 0.015764 | 0.031932 | 0.029201 | 0.013346 | 0.017371 | 0.139672 | 0.108807 | 0.085935 | 0.019955 | 0.136552 | 0.037891 | 0.192188 | 0.053412 | 0.002202 |
| 33.0555556 | 0.016487 | 0.15241 | 0.042991 | 0.09384 | 0.053355 | 0.008657 | 0.007701 | 0.026309 | 0.035729 | 0.150779 | 0.026088 | 0.03774 | 0.029475 | 0.01193 | 0.027038 | 0.146765 | 0.138488 | 0.127401 | 0.03905 | 0.12948 | 0.031728 | 0.194414 | 0.023955 | 0.004352 |
| 33.0694444 | 0.016368 | 0.11946 | 0.024447 | 0.087002 | 0.112084 | 0.01245 | 0.011782 | 0.06432 | 0.026768 | 0.179757 | 0.027076 | 0.025712 | 0.03607 | 0.006624 | 0.043561 | 0.126182 | 0.139479 | 0.122659 | 0.057804 | 0.103137 | 0.023288 | 0.156195 | 0.023817 | 0.007379 |
| 33.0833333 | 0.013855 | 0.056809 | 0.031351 | 0.059443 | 0.157401 | 0.013403 | 0.020339 | 0.085375 | 0.010754 | 0.155172 | 0.017397 | 0.011802 | 0.034602 | 0.004404 | 0.04924 | 0.080203 | 0.107763 | 0.085436 | 0.043578 | 0.056059 | 0.018677 | 0.085229 | 0.03196 | 0.00605 |
| 33.0972222 | 0.011308 | 0.018399 | 0.04752 | 0.037639 | 0.152651 | 0.009186 | 0.023383 | 0.065078 | 0.007983 | 0.109943 | 0.010247 | 0.006886 | 0.028757 | 0.010485 | 0.045935 | 0.034547 | 0.06388 | 0.035637 | 0.017517 | 0.019813 | 0.017357 | 0.034513 | 0.03157 | 0.002142 |
| 33.1111111 | 0.00971 | 0.017068 | 0.045375 | 0.028795 | 0.114343 | 0.008198 | 0.016503 | 0.039539 | 0.009328 | 0.064025 | 0.011871 | 0.007746 | 0.020001 | 0.022385 | 0.03733 | 0.027094 | 0.032472 | 0.016812 | 0.008688 | 0.013465 | 0.011678 | 0.038249 | 0.03124 | 0.00253 |
| 33.125 | 0.007537 | 0.019926 | 0.031576 | 0.02019 | 0.102315 | 0.012077 | 0.015202 | 0.028012 | 0.008175 | 0.069555 | 0.015323 | 0.007111 | 0.025097 | 0.037409 | 0.025041 | 0.0406 | 0.030229 | 0.033119 | 0.009912 | 0.013178 | 0.007679 | 0.065602 | 0.040572 | 0.008261 |
| 33.1388889 | 0.004383 | 0.018306 | 0.021841 | 0.013891 | 0.129642 | 0.013606 | 0.022098 | 0.021672 | 0.008351 | 0.128344 | 0.015405 | 0.005734 | 0.047568 | 0.058085 | 0.011793 | 0.043035 | 0.039062 | 0.036533 | 0.006978 | 0.007678 | 0.013604 | 0.113675 | 0.07716 | 0.017904 |
| 33.1527778 | 0.004161 | 0.036206 | 0.020014 | 0.018067 | 0.116074 | 0.012133 | 0.026517 | 0.011962 | 0.015487 | 0.191514 | 0.013333 | 0.008576 | 0.063904 | 0.067771 | 0.014618 | 0.03843 | 0.039934 | 0.020802 | 0.00559 | 0.006669 | 0.026154 | 0.180584 | 0.116941 | 0.029127 |
| 33.1666667 | 0.006892 | 0.073583 | 0.02016 | 0.027631 | 0.062639 | 0.010242 | 0.028418 | 0.003147 | 0.032864 | 0.253216 | 0.010621 | 0.010385 | 0.063252 | 0.049483 | 0.041097 | 0.040783 | 0.040045 | 0.008595 | 0.011218 | 0.013724 | 0.03786 | 0.218775 | 0.108987 | 0.036322 |
| 33.1805556 | 0.008203 | 0.102996 | 0.016778 | 0.026981 | 0.055984 | 0.007398 | 0.029684 | 0.000363 | 0.053681 | 0.329523 | 0.007328 | 0.007458 | 0.054479 | 0.030042 | 0.059939 | 0.05066 | 0.041958 | 0.008352 | 0.014506 | 0.022936 | 0.037381 | 0.218127 | 0.0587 | 0.036591 |
| 33.1944444 | 0.006592 | 0.113862 | 0.012102 | 0.014917 | 0.063212 | 0.004656 | 0.030001 | 0.001786 | 0.063472 | 0.407795 | 0.00479 | 0.006084 | 0.044331 | 0.028599 | 0.047794 | 0.053107 | 0.041682 | 0.013296 | 0.009418 | 0.027727 | 0.030924 | 0.212422 | 0.015491 | 0.037229 |
| 33.2083333 | 0.003297 | 0.118112 | 0.009901 | 0.007099 | 0.04081 | 0.003561 | 0.03076 | 0.006236 | 0.06444 | 0.383629 | 0.007729 | 0.008878 | 0.037006 | 0.035019 | 0.021249 | 0.042024 | 0.040471 | 0.012766 | 0.006367 | 0.027938 | 0.02893 | 0.226766 | 0.002199 | 0.042334 |
| 33.2222222 | 0.000951 | 0.100891 | 0.008087 | 0.008887 | 0.01828 | 0.005194 | 0.032902 | 0.015781 | 0.066795 | 0.251137 | 0.014491 | 0.01237 | 0.032885 | 0.033118 | 0.008149 | 0.037267 | 0.040829 | 0.015009 | 0.01223 | 0.025771 | 0.021082 | 0.23833 | 0.004433 | 0.043764 |
| 33.2361111 | 0.000874 | 0.067709 | 0.005325 | 0.011999 | 0.012732 | 0.006763 | 0.035943 | 0.031288 | 0.066335 | 0.170346 | 0.017999 | 0.014543 | 0.026332 | 0.023081 | 0.010466 | 0.051217 | 0.04158 | 0.021285 | 0.014072 | 0.022752 | 0.00957 | 0.205555 | 0.005695 | 0.040144 |
| 33.25 | 0.002547 | 0.068709 | 0.00519 | 0.009112 | 0.017397 | 0.004563 | 0.037345 | 0.043165 | 0.055834 | 0.157606 | 0.016246 | 0.014499 | 0.01705 | 0.022975 | 0.012613 | 0.066707 | 0.039921 | 0.019286 | 0.013167 | 0.02112 | 0.009952 | 0.135724 | 0.005614 | 0.036299 |
| 33.2638889 | 0.004743 | 0.07696 | 0.005957 | 0.005946 | 0.017882 | 0.002482 | 0.034848 | 0.040835 | 0.032372 | 0.110262 | 0.011733 | 0.012582 | 0.011066 | 0.030395 | 0.01555 | 0.064355 | 0.033503 | 0.010294 | 0.020851 | 0.019965 | 0.016492 | 0.075614 | 0.011366 | 0.031202 |
| 33.2777778 | 0.005361 | 0.050148 | 0.004447 | 0.009696 | 0.018827 | 0.003104 | 0.028933 | 0.029073 | 0.01613 | 0.057049 | 0.006556 | 0.009384 | 0.008966 | 0.036498 | 0.022373 | 0.043175 | 0.022928 | 0.005775 | 0.022132 | 0.01667 | 0.02046 | 0.043812 | 0.018952 | 0.023295 |
| 33.2916667 | 0.004903 | 0.023086 | 0.003754 | 0.013866 | 0.017806 | 0.003059 | 0.025205 | 0.016834 | 0.027785 | 0.073911 | 0.003065 | 0.004854 | 0.007208 | 0.036207 | 0.020852 | 0.024436 | 0.012615 | 0.013983 | 0.01277 | 0.014217 | 0.023824 | 0.022721 | 0.018694 | 0.015021 |

| | | | | | | | | | | | | | | | | | | | | | | | | |
|---|---|---|---|---|---|---|---|---|---|---|---|---|---|---|---|---|---|---|---|---|---|---|---|---|
| 33.3055556 | 0.005709 | 0.028593 | 0.004215 | 0.012337 | 0.016774 | 0.002898 | 0.031257 | 0.009163 | 0.048416 | 0.117698 | 0.003434 | 0.003103 | 0.009248 | 0.022711 | 0.017553 | 0.030158 | 0.007156 | 0.021762 | 0.006222 | 0.015889 | 0.02389 | 0.007085 | 0.01038 | 0.010179 |
| 33.3194444 | 0.007016 | 0.058043 | 0.003281 | 0.007426 | 0.019735 | 0.007071 | 0.048134 | 0.009254 | 0.062591 | 0.13169 | 0.005469 | 0.005508 | 0.016222 | 0.01434 | 0.025292 | 0.04357 | 0.005151 | 0.016034 | 0.004317 | 0.017935 | 0.016726 | 0.003804 | 0.004395 | 0.010153 |
| 33.3333333 | 0.011319 | 0.089254 | 0.001941 | 0.007368 | 0.014331 | 0.012329 | 0.067341 | 0.013791 | 0.073044 | 0.148335 | 0.005545 | 0.008324 | 0.019877 | 0.029429 | 0.030778 | 0.040004 | 0.00321 | 0.012661 | 0.006559 | 0.016558 | 0.008053 | 0.01565 | 0.008784 | 0.016097 |
| 33.3472222 | 0.01708 | 0.120111 | 0.001822 | 0.013897 | 0.011435 | 0.014169 | 0.075114 | 0.014351 | 0.074663 | 0.193079 | 0.003399 | 0.011352 | 0.015557 | 0.046816 | 0.027758 | 0.022948 | 0.003176 | 0.018279 | 0.010523 | 0.01509 | 0.005943 | 0.042078 | 0.018626 | 0.027517 |
| 33.3611111 | 0.014664 | 0.143444 | 0.002959 | 0.018173 | 0.013998 | 0.0154 | 0.068267 | 0.008527 | 0.065151 | 0.231897 | 0.002274 | 0.01514 | 0.008743 | 0.050475 | 0.021591 | 0.010836 | 0.007236 | 0.017715 | 0.011952 | 0.018536 | 0.006644 | 0.0668 | 0.026568 | 0.034476 |
| 33.375 | 0.007986 | 0.128072 | 0.004679 | 0.016952 | 0.032119 | 0.016572 | 0.059426 | 0.002352 | 0.052225 | 0.22756 | 0.003668 | 0.017355 | 0.008178 | 0.056528 | 0.012051 | 0.017293 | 0.012646 | 0.011395 | 0.013281 | 0.026556 | 0.004879 | 0.073609 | 0.02768 | 0.03 |
| 33.3888889 | 0.005388 | 0.073768 | 0.006367 | 0.011673 | 0.077477 | 0.016993 | 0.054703 | 0.000151 | 0.040327 | 0.185426 | 0.004828 | 0.015548 | 0.012594 | 0.066 | 0.005348 | 0.033928 | 0.015407 | 0.01233 | 0.017349 | 0.032926 | 0.004026 | 0.074605 | 0.02025 | 0.024212 |
| 33.4027778 | 0.004682 | 0.038244 | 0.007379 | 0.00613 | 0.091951 | 0.01602 | 0.048119 | 6.86E-05 | 0.027036 | 0.134363 | 0.004704 | 0.010882 | 0.015534 | 0.068298 | 0.005647 | 0.051115 | 0.015866 | 0.022699 | 0.021659 | 0.032756 | 0.006535 | 0.088379 | 0.012616 | 0.026468 |
| 33.4166667 | 0.004036 | 0.061052 | 0.007018 | 0.00449 | 0.050565 | 0.013994 | 0.038613 | 7.88E-05 | 0.013445 | 0.086709 | 0.004303 | 0.007534 | 0.016481 | 0.064709 | 0.009658 | 0.065304 | 0.016401 | 0.033447 | 0.024562 | 0.030892 | 0.008582 | 0.111958 | 0.013651 | 0.03317 |
| 33.4305556 | 0.003573 | 0.102807 | 0.005377 | 0.003812 | 0.019031 | 0.013683 | 0.032767 | 0.000344 | 0.006016 | 0.046567 | 0.003952 | 0.010182 | 0.017552 | 0.051775 | 0.0141 | 0.069117 | 0.017531 | 0.03467 | 0.025941 | 0.034018 | 0.007147 | 0.1293 | 0.021958 | 0.038259 |
| 33.4444444 | 0.002899 | 0.111782 | 0.003232 | 0.004909 | 0.030786 | 0.016121 | 0.034009 | 0.001247 | 0.008553 | 0.041545 | 0.003916 | 0.017596 | 0.020125 | 0.03311 | 0.015492 | 0.064298 | 0.019172 | 0.025822 | 0.02359 | 0.040171 | 0.004946 | 0.136234 | 0.031187 | 0.042951 |
| 33.4583333 | 0.002658 | 0.077866 | 0.001861 | 0.010251 | 0.066616 | 0.020124 | 0.037074 | 0.002685 | 0.016765 | 0.078782 | 0.004328 | 0.021012 | 0.024556 | 0.020805 | 0.021198 | 0.058988 | 0.023814 | 0.016196 | 0.018832 | 0.047733 | 0.004508 | 0.142368 | 0.027576 | 0.050224 |
| 33.4722222 | 0.004138 | 0.043979 | 0.002242 | 0.01563 | 0.117436 | 0.022413 | 0.035564 | 0.002837 | 0.025736 | 0.124392 | 0.005465 | 0.017536 | 0.029672 | 0.017106 | 0.036913 | 0.052511 | 0.035468 | 0.012913 | 0.017626 | 0.059653 | 0.005006 | 0.147549 | 0.010946 | 0.054092 |
| 33.4861111 | 0.007473 | 0.040237 | 0.003372 | 0.018777 | 0.18468 | 0.021227 | 0.028858 | 0.001507 | 0.03418 | 0.152423 | 0.007931 | 0.010666 | 0.034325 | 0.017732 | 0.0554 | 0.041393 | 0.051533 | 0.014876 | 0.021427 | 0.0718 | 0.005679 | 0.149013 | 0.003136 | 0.049083 |
| 33.5 | 0.010429 | 0.044027 | 0.003695 | 0.01808 | 0.249435 | 0.018462 | 0.019232 | 0.000741 | 0.040873 | 0.153015 | 0.01116 | 0.006299 | 0.037879 | 0.014959 | 0.070617 | 0.030953 | 0.066107 | 0.01794 | 0.025264 | 0.078261 | 0.006053 | 0.155716 | 0.003949 | 0.043431 |
| 33.5138889 | 0.010169 | 0.056519 | 0.003297 | 0.011324 | 0.287276 | 0.015483 | 0.009188 | 0.002469 | 0.046999 | 0.138629 | 0.01263 | 0.007652 | 0.03997 | 0.020869 | 0.077528 | 0.022284 | 0.078513 | 0.020001 | 0.025225 | 0.085744 | 0.00497 | 0.161654 | 0.00839 | 0.047801 |
| 33.5277778 | 0.008086 | 0.073359 | 0.006367 | 0.003772 | 0.299616 | 0.012771 | 0.002923 | 0.009579 | 0.057692 | 0.130669 | 0.011152 | 0.007722 | 0.039197 | 0.043262 | 0.069822 | 0.015477 | 0.084448 | 0.021246 | 0.021694 | 0.09275 | 0.003988 | 0.147821 | 0.023689 | 0.058319 |
| 33.5416667 | 0.010276 | 0.059471 | 0.01636 | 0.001329 | 0.263513 | 0.011516 | 0.004583 | 0.021609 | 0.069633 | 0.127938 | 0.008012 | 0.004941 | 0.034152 | 0.062918 | 0.058059 | 0.012928 | 0.075059 | 0.023389 | 0.016759 | 0.08181 | 0.004905 | 0.123289 | 0.042834 | 0.063543 |
| 33.5555556 | 0.021569 | 0.034 | 0.02839 | 0.004581 | 0.186498 | 0.013039 | 0.011896 | 0.02805 | 0.069263 | 0.120772 | 0.004483 | 0.004372 | 0.027004 | 0.076897 | 0.054597 | 0.010793 | 0.05524 | 0.031618 | 0.018462 | 0.061025 | 0.005516 | 0.102172 | 0.054761 | 0.063549 |
| 33.5694444 | 0.041953 | 0.044776 | 0.027555 | 0.022415 | 0.121146 | 0.018693 | 0.01612 | 0.019305 | 0.057794 | 0.111068 | 0.002223 | 0.004166 | 0.01817 | 0.088637 | 0.053219 | 0.010334 | 0.037643 | 0.052701 | 0.041062 | 0.058148 | 0.01036 | 0.092885 | 0.057344 | 0.068983 |
| 33.5833333 | 0.053448 | 0.070361 | 0.024246 | 0.056483 | 0.063717 | 0.024534 | 0.013153 | 0.007185 | 0.048609 | 0.107475 | 0.001889 | 0.00363 | 0.022356 | 0.090876 | 0.053201 | 0.018524 | 0.027102 | 0.083065 | 0.081199 | 0.075755 | 0.022965 | 0.116459 | 0.059967 | 0.078922 |
| 33.5972222 | 0.043806 | 0.075608 | 0.047414 | 0.085149 | 0.019809 | 0.022054 | 0.007437 | 0.003474 | 0.044649 | 0.107243 | 0.004154 | 0.005616 | 0.067853 | 0.081914 | 0.051816 | 0.03169 | 0.023892 | 0.101723 | 0.103537 | 0.096318 | 0.035952 | 0.141474 | 0.068232 | 0.081688 |
| 33.6111111 | 0.03318 | 0.072347 | 0.083438 | 0.097894 | 0.008343 | 0.013457 | 0.005685 | 0.004564 | 0.040961 | 0.110432 | 0.010643 | 0.012147 | 0.118762 | 0.07448 | 0.051611 | 0.040425 | 0.025928 | 0.095159 | 0.08795 | 0.109761 | 0.0492 | 0.127801 | 0.082097 | 0.081174 |
| 33.625 | 0.026033 | 0.101765 | 0.110724 | 0.094662 | 0.017024 | 0.0102 | 0.010207 | 0.006168 | 0.034354 | 0.124235 | 0.019156 | 0.022409 | 0.115911 | 0.072029 | 0.067963 | 0.045365 | 0.030867 | 0.075119 | 0.077624 | 0.116635 | 0.062469 | 0.113188 | 0.087007 | 0.08477 |
| 33.6388889 | 0.019627 | 0.162146 | 0.127495 | 0.076539 | 0.057249 | 0.01156 | 0.017318 | 0.005714 | 0.024813 | 0.137359 | 0.025922 | 0.032846 | 0.07737 | 0.058709 | 0.088048 | 0.045924 | 0.032079 | 0.055487 | 0.099132 | 0.117444 | 0.067188 | 0.11758 | 0.059344 | 0.080811 |
| 33.6527778 | 0.023345 | 0.172042 | 0.124004 | 0.059572 | 0.128661 | 0.008804 | 0.023223 | 0.00761 | 0.017767 | 0.14167 | 0.031409 | 0.045486 | 0.039093 | 0.034508 | 0.080544 | 0.038385 | 0.022945 | 0.035317 | 0.113254 | 0.111881 | 0.053324 | 0.11183 | 0.022478 | 0.063248 |
| 33.6666667 | 0.037913 | 0.111517 | 0.097149 | 0.047835 | 0.193639 | 0.005986 | 0.027155 | 0.013011 | 0.018765 | 0.143953 | 0.035767 | 0.057344 | 0.014916 | 0.015057 | 0.045528 | 0.024667 | 0.015562 | 0.016088 | 0.099226 | 0.102192 | 0.024573 | 0.095685 | 0.008573 | 0.041934 |
| 33.6805556 | 0.045814 | 0.08725 | 0.063501 | 0.029363 | 0.207059 | 0.006864 | 0.028753 | 0.017686 | 0.02315 | 0.140553 | 0.03611 | 0.063401 | 0.014837 | 0.006771 | 0.018149 | 0.011395 | 0.01559 | 0.007558 | 0.064162 | 0.090044 | 0.007156 | 0.081802 | 0.015312 | 0.02448 |
| 33.6944444 | 0.043793 | 0.154412 | 0.040776 | 0.009373 | 0.147866 | 0.006705 | 0.02934 | 0.022182 | 0.024015 | 0.123153 | 0.032255 | 0.063553 | 0.048048 | 0.007975 | 0.01938 | 0.003774 | 0.009902 | 0.009842 | 0.03132 | 0.077246 | 0.012956 | 0.059676 | 0.048095 | 0.014667 |
| 33.7083333 | 0.063458 | 0.174229 | 0.040736 | 0.001686 | 0.121281 | 0.008432 | 0.030323 | 0.021857 | 0.018952 | 0.095667 | 0.027113 | 0.050472 | 0.121489 | 0.020776 | 0.029243 | 0.004456 | 0.004136 | 0.01697 | 0.027312 | 0.068823 | 0.029741 | 0.032571 | 0.106023 | 0.015474 |
| 33.7222222 | 0.095771 | 0.156503 | 0.062291 | 0.003797 | 0.206936 | 0.013396 | 0.027755 | 0.015426 | 0.011712 | 0.068419 | 0.023538 | 0.027501 | 0.197306 | 0.033259 | 0.023129 | 0.009081 | 0.007454 | 0.020457 | 0.036746 | 0.068228 | 0.044333 | 0.027578 | 0.136515 | 0.022909 |
| 33.7361111 | 0.105025 | 0.219852 | 0.080712 | 0.009284 | 0.29042 | 0.015812 | 0.021885 | 0.015796 | 0.011375 | 0.048957 | 0.022058 | 0.011802 | 0.21639 | 0.026758 | 0.012733 | 0.020151 | 0.022294 | 0.014544 | 0.040978 | 0.075045 | 0.051626 | 0.041915 | 0.10394 | 0.021778 |
| 33.75 | 0.098707 | 0.200058 | 0.075956 | 0.01448 | 0.26657 | 0.011832 | 0.017053 | 0.026125 | 0.012638 | 0.04248 | 0.022099 | 0.011126 | 0.166153 | 0.015478 | 0.019505 | 0.046549 | 0.049741 | 0.009137 | 0.04703 | 0.083802 | 0.058044 | 0.041495 | 0.056534 | 0.009793 |
| 33.7638889 | 0.094222 | 0.080856 | 0.056584 | 0.01638 | 0.152328 | 0.008852 | 0.01658 | 0.036847 | 0.011016 | 0.054098 | 0.022981 | 0.01653 | 0.082226 | 0.0127 | 0.029147 | 0.066804 | 0.053433 | 0.020001 | 0.05811 | 0.08498 | 0.068064 | 0.024453 | 0.047956 | 0.006925 |
| 33.7777778 | 0.071266 | 0.029338 | 0.053046 | 0.016496 | 0.045385 | 0.010468 | 0.023589 | 0.042598 | 0.018495 | 0.088888 | 0.022975 | 0.024762 | 0.032691 | 0.011186 | 0.023906 | 0.056206 | 0.036711 | 0.044943 | 0.062337 | 0.076423 | 0.07885 | 0.020238 | 0.064861 | 0.020332 |
| 33.7916667 | 0.03629 | 0.023359 | 0.073651 | 0.015767 | 0.02013 | 0.008468 | 0.025549 | 0.043027 | 0.03445 | 0.138111 | 0.021649 | 0.042154 | 0.033629 | 0.015667 | 0.015659 | 0.028677 | 0.049302 | 0.059656 | 0.060092 | 0.061534 | 0.089457 | 0.027603 | 0.06583 | 0.04111 |
| 33.8055556 | 0.035996 | 0.032378 | 0.07834 | 0.013637 | 0.036773 | 0.007491 | 0.013132 | 0.041473 | 0.054256 | 0.175547 | 0.020202 | 0.059588 | 0.039459 | 0.033136 | 0.019651 | 0.017803 | 0.072354 | 0.053114 | 0.06784 | 0.042362 | 0.093411 | 0.030406 | 0.055247 | 0.063133 |
| 33.8194444 | 0.078884 | 0.057229 | 0.049923 | 0.023258 | 0.068787 | 0.010708 | 0.006406 | 0.038207 | 0.078048 | 0.186681 | 0.019662 | 0.064385 | 0.043582 | 0.052178 | 0.023782 | 0.025733 | 0.082723 | 0.039753 | 0.081302 | 0.024292 | 0.084757 | 0.037862 | 0.050662 | 0.076776 |
| 33.8333333 | 0.140838 | 0.075135 | 0.024264 | 0.055874 | 0.108448 | 0.01466 | 0.014154 | 0.02964 | 0.096657 | 0.175027 | 0.021703 | 0.057655 | 0.058617 | 0.053039 | 0.01415 | 0.029009 | 0.095365 | 0.028966 | 0.08534 | 0.013399 | 0.074739 | 0.053829 | 0.035955 | 0.073469 |
| 33.8472222 | 0.188815 | 0.095857 | 0.027847 | 0.083643 | 0.108252 | 0.020023 | 0.022667 | 0.017776 | 0.115154 | 0.150155 | 0.026193 | 0.047697 | 0.063191 | 0.041875 | 0.006729 | 0.024619 | 0.107665 | 0.022088 | 0.073872 | 0.010272 | 0.079052 | 0.063123 | 0.019775 | 0.062982 |
| 33.8611111 | 0.207931 | 0.105467 | 0.048048 | 0.068515 | 0.072759 | 0.022927 | 0.024657 | 0.009265 | 0.141155 | 0.126059 | 0.029859 | 0.038913 | 0.042325 | 0.033702 | 0.013422 | 0.019949 | 0.100308 | 0.023503 | 0.054595 | 0.010028 | 0.099668 | 0.053608 | 0.024063 | 0.060397 |
| 33.875 | 0.210821 | 0.115275 | 0.054807 | 0.028651 | 0.064915 | 0.020695 | 0.018845 | 0.007104 | 0.14851 | 0.108128 | 0.031487 | 0.025021 | 0.031341 | 0.024832 | 0.024603 | 0.015594 | 0.065187 | 0.027036 | 0.049778 | 0.007663 | 0.11543 | 0.030101 | 0.029585 | 0.05763 |
| 33.8888889 | 0.196657 | 0.064923 | 0.042488 | 0.008228 | 0.127946 | 0.014623 | 0.009786 | 0.011345 | 0.136488 | 0.091116 | 0.031801 | 0.008302 | 0.051632 | 0.013841 | 0.036902 | 0.010199 | 0.0348 | 0.021262 | 0.057286 | 0.004622 | 0.092527 | 0.018962 | 0.029045 | 0.038506 |
| 33.9027778 | 0.181718 | 0.112075 | 0.025111 | 0.00602 | 0.211277 | 0.008061 | 0.004776 | 0.02136 | 0.131115 | 0.071475 | 0.031029 | 0.00497 | 0.078543 | 0.007931 | 0.052409 | 0.004432 | 0.033675 | 0.010392 | 0.055095 | 0.005006 | 0.041705 | 0.031333 | 0.031968 | 0.018171 |
| 33.9166667 | 0.162419 | 0.032879 | 0.011298 | 0.012898 | 0.267771 | 0.005693 | 0.008939 | 0.032901 | 0.12263 | 0.048596 | 0.029871 | 0.009854 | 0.090912 | 0.014605 | 0.059736 | 0.005577 | 0.036888 | 0.006527 | 0.04704 | 0.010412 | 0.012599 | 0.048738 | 0.032978 | 0.023833 |
| 33.9305556 | 0.105967 | 0.147641 | 0.006383 | 0.029512 | 0.307425 | 0.012414 | 0.021976 | 0.046808 | 0.108725 | 0.024718 | 0.027258 | 0.017153 | 0.081565 | 0.027948 | 0.052882 | 0.014601 | 0.034445 | 0.01443 | 0.041249 | 0.018705 | 0.01012 | 0.064396 | 0.034736 | 0.042164 |
| 33.9444444 | 0.049175 | 0.302564 | 0.006574 | 0.043629 | 0.263452 | 0.022484 | 0.029703 | 0.060309 | 0.089537 | 0.007661 | 0.021492 | 0.032969 | 0.067303 | 0.027108 | 0.041949 | 0.01754 | 0.038947 | 0.025874 | 0.030129 | 0.018235 | 0.010563 | 0.080903 | 0.048678 | 0.040093 |
| 33.9583333 | 0.031138 | 0.349344 | 0.014559 | 0.04288 | 0.172921 | 0.027596 | 0.033871 | 0.069115 | 0.054371 | 0.001881 | 0.012931 | 0.040771 | 0.066885 | 0.012702 | 0.033885 | 0.018523 | 0.049714 | 0.025855 | 0.015769 | 0.011499 | 0.013751 | 0.096784 | 0.072275 | 0.021071 |
| 33.9722222 | 0.030968 | 0.227509 | 0.023908 | 0.027189 | 0.17252 | 0.028463 | 0.040419 | 0.071048 | 0.038745 | 0.006365 | 0.007437 | 0.037922 | 0.074556 | 0.009711 | 0.023971 | 0.031556 | 0.054952 | 0.015306 | 0.006487 | 0.009408 | 0.02708 | 0.122701 | 0.085357 | 0.015068 |
| 33.9861111 | 0.02258 | 0.069136 | 0.064477 | 0.01491 | 0.186535 | 0.02419 | 0.029394 | 0.052728 | 0.063363 | 0.019776 | 0.00957 | 0.04211 | 0.078004 | 0.024368 | 0.01708 | 0.045562 | 0.049847 | 0.00678 | 0.005607 | 0.012016 | 0.046883 | 0.154287 | 0.074593 | 0.031973 |
| 34 | 0.021607 | 0.011351 | 0.146114 | 0.025254 | 0.122894 | 0.027539 | 0.012869 | 0.026411 | 0.088283 | 0.033794 | 0.010362 | 0.052158 | 0.069398 | 0.043574 | 0.016838 | 0.048135 | 0.052778 | 0.003439 | 0.007656 | 0.026056 | 0.055367 | 0.186309 | 0.042351 | 0.053489 |
| 34.0138889 | 0.030688 | 0.043275 | 0.18117 | 0.046906 | 0.045702 | 0.03649 | 0.009272 | 0.020453 | 0.074195 | 0.047427 | 0.007163 | 0.058373 | 0.04936 | 0.05373 | 0.028241 | 0.035905 | 0.081439 | 0.005329 | 0.013268 | 0.044203 | 0.052228 | 0.223177 | 0.014881 | 0.066309 |
| 34.0277778 | 0.026767 | 0.093213 | 0.172923 | 0.046251 | 0.031006 | 0.02782 | 0.010082 | 0.03748 | 0.03817 | 0.096303 | 0.013487 | 0.058252 | 0.029351 | 0.049363 | 0.055828 | 0.020119 | 0.128252 | 0.008159 | 0.030858 | 0.056728 | 0.081498 | 0.245178 | 0.010515 | 0.074225 |
| 34.0416667 | 0.036652 | 0.14107 | 0.183458 | 0.02925 | 0.057159 | 0.013272 | 0.014491 | 0.059228 | 0.028977 | 0.195786 | 0.034131 | 0.056097 | 0.022247 | 0.048262 | 0.067296 | 0.021348 | 0.14523 | 0.013463 | 0.042136 | 0.067582 | 0.16132 | 0.238853 | 0.021865 | 0.07611 |
| 34.0555556 | 0.082302 | 0.177043 | 0.190393 | 0.044186 | 0.066497 | 0.009089 | 0.016013 | 0.060766 | 0.042543 | 0.280514 | 0.047158 | 0.051211 | 0.026152 | 0.057853 | 0.051985 | 0.030955 | 0.109287 | 0.029481 | 0.031431 | 0.068896 | 0.203145 | 0.218462 | 0.046979 | 0.068993 |
| 34.0694444 | 0.113086 | 0.156624 | 0.15917 | 0.082262 | 0.069234 | 0.006294 | 0.012238 | 0.042706 | 0.052816 | 0.304848 | 0.034173 | 0.038576 | 0.02386 | 0.059752 | 0.038339 | 0.029657 | 0.073547 | 0.050374 | 0.016662 | 0.05331 | 0.153085 | 0.202986 | 0.075716 | 0.060626 |
| 34.0833333 | 0.088393 | 0.090455 | 0.114735 | 0.089896 | 0.063451 | 0.0077 | 0.008495 | 0.022564 | 0.048292 | 0.283091 | 0.01373 | 0.022692 | 0.013064 | 0.05247 | 0.044148 | 0.020717 | 0.056873 | 0.055929 | 0.017339 | 0.030457 | 0.084253 | 0.167998 | 0.087024 | 0.055244 |
| 34.0972222 | 0.046299 | 0.037092 | 0.096294 | 0.069405 | 0.040238 | 0.023107 | 0.007605 | 0.010235 | 0.030567 | 0.229634 | 0.006427 | 0.011382 | 0.010131 | 0.044505 | 0.059661 | 0.024864 | 0.037706 | 0.035479 | 0.028268 | 0.016637 | 0.041209 | 0.095497 | 0.08051 | 0.049203 |
| 34.1111111 | 0.025724 | 0.018266 | 0.090829 | 0.048219 | 0.037668 | 0.037658 | 0.011979 | 0.011715 | 0.016099 | 0.148128 | 0.007374 | 0.006195 | 0.014044 | 0.035035 | 0.073089 | 0.047378 | 0.038878 | 0.011978 | 0.034901 | 0.011798 | 0.023273 | 0.046316 | 0.070044 | 0.034495 |
| 34.125 | 0.02101 | 0.021589 | 0.069141 | 0.034416 | 0.059633 | 0.032456 | 0.016537 | 0.01583 | 0.008344 | 0.073739 | 0.007338 | 0.005011 | 0.015555 | 0.023495 | 0.085441 | 0.063804 | 0.059904 | 0.004539 | 0.036062 | 0.005936 | 0.02662 | 0.052508 | 0.050111 | 0.021647 |
| 34.1388889 | 0.014758 | 0.021058 | 0.038324 | 0.028243 | 0.069451 | 0.018047 | 0.013373 | 0.013248 | 0.003755 | 0.062373 | 0.006135 | 0.004282 | 0.016545 | 0.022444 | 0.076069 | 0.060309 | 0.061555 | 0.012949 | 0.031707 | 0.006217 | 0.027963 | 0.072077 | 0.039262 | 0.033926 |
| 34.1527778 | 0.00574 | 0.01402 | 0.034821 | 0.029125 | 0.054472 | 0.01288 | 0.005543 | 0.010703 | 0.003754 | 0.093574 | 0.005465 | 0.003343 | 0.022513 | 0.03867 | 0.040504 | 0.048915 | 0.047209 | 0.022673 | 0.025297 | 0.015642 | 0.020363 | 0.090516 | 0.076304 | 0.060971 |
| 34.1666667 | 0.005418 | 0.013214 | 0.06713 | 0.029242 | 0.035601 | 0.019452 | 0.002182 | 0.010651 | 0.011265 | 0.135061 | 0.005308 | 0.005385 | 0.02932 | 0.057258 | 0.02282 | 0.041758 | 0.033959 | 0.016903 | 0.021857 | 0.026011 | 0.016073 | 0.106994 | 0.127067 | 0.082093 |
| 34.1805556 | 0.01885 | 0.021073 | 0.10084 | 0.020895 | 0.031806 | 0.029994 | 0.009493 | 0.013037 | 0.027426 | 0.190987 | 0.005341 | 0.009623 | 0.026692 | 0.064568 | 0.041244 | 0.03605 | 0.024332 | 0.007234 | 0.016346 | 0.039005 | 0.024148 | 0.113531 | 0.111529 | 0.091265 |
| 34.1944444 | 0.055661 | 0.035975 | 0.114631 | 0.013063 | 0.038211 | 0.032615 | 0.017927 | 0.012434 | 0.049385 | 0.222962 | 0.004436 | 0.012333 | 0.01712 | 0.062418 | 0.064505 | 0.026305 | 0.019542 | 0.00663 | 0.011821 | 0.050018 | 0.035326 | 0.127911 | 0.045982 | 0.085776 |
| 34.2083333 | 0.101543 | 0.056008 | 0.109662 | 0.013531 | 0.162738 | 0.022183 | 0.012742 | 0.010395 | 0.067523 | 0.223244 | 0.002688 | 0.013653 | 0.011796 | 0.055817 | 0.071351 | 0.016046 | 0.013991 | 0.009673 | 0.012395 | 0.047264 | 0.030199 | 0.15089 | 0.013955 | 0.074919 |
| 34.2222222 | 0.114756 | 0.070388 | 0.080035 | 0.020236 | 0.333782 | 0.01104 | 0.006721 | 0.011027 | 0.07479 | 0.206824 | 0.001596 | 0.015869 | 0.011754 | 0.044796 | 0.062699 | 0.011898 | 0.010781 | 0.007594 | 0.017355 | 0.029763 | 0.016564 | 0.161541 | 0.01118 | 0.078109 |
| 34.2361111 | 0.081858 | 0.070251 | 0.044594 | 0.031751 | 0.258154 | 0.009745 | 0.008798 | 0.0061 | 0.071358 | 0.135842 | 0.002531 | 0.018347 | 0.020087 | 0.031482 | 0.044584 | 0.021676 | 0.02177 | 0.009767 | 0.046664 | 0.022389 | 0.010771 | 0.143003 | 0.014634 | 0.085285 |
| 34.25 | 0.042189 | 0.058801 | 0.045563 | 0.032314 | 0.09758 | 0.011641 | 0.00964 | 0.002026 | 0.058142 | 0.051236 | 0.00595 | 0.018666 | 0.042691 | 0.025784 | 0.026776 | 0.035899 | 0.037196 | 0.036654 | 0.119935 | 0.044157 | 0.009736 | 0.098347 | 0.0383 | 0.064046 |
| 34.2638889 | 0.043306 | 0.04419 | 0.078479 | 0.026701 | 0.156625 | 0.010396 | 0.006969 | 0.00533 | 0.041062 | 0.031291 | 0.009674 | 0.01504 | 0.050981 | 0.029003 | 0.027288 | 0.034157 | 0.047665 | 0.061015 | 0.148183 | 0.052139 | 0.018432 | 0.051125 | 0.047192 | 0.033121 |
| 34.2777778 | 0.04759 | 0.031497 | 0.102667 | 0.036743 | 0.260072 | 0.007516 | 0.006535 | 0.013529 | 0.026684 | 0.060974 | 0.01014 | 0.012308 | 0.037614 | 0.028843 | 0.046134 | 0.026633 | 0.047657 | 0.061261 | 0.108154 | 0.039418 | 0.037469 | 0.04139 | 0.022183 | 0.038184 |
| 34.2916667 | 0.022409 | 0.022459 | 0.097532 | 0.047357 | 0.196407 | 0.012335 | 0.012482 | 0.025026 | 0.016711 | 0.078004 | 0.007662 | 0.014594 | 0.031584 | 0.02667 | 0.05655 | 0.020533 | 0.031395 | 0.060075 | 0.091582 | 0.053995 | 0.04549 | 0.085558 | 0.019091 | 0.058972 |
| 34.3055556 | 0.014142 | 0.026171 | 0.067551 | 0.046024 | 0.083324 | 0.020639 | 0.019606 | 0.028574 | 0.021421 | 0.051281 | 0.006237 | 0.01346 | 0.025923 | 0.023669 | 0.048959 | 0.013182 | 0.018927 | 0.044551 | 0.063533 | 0.065042 | 0.032324 | 0.113967 | 0.040385 | 0.054041 |
| 34.3194444 | 0.027708 | 0.031021 | 0.0309 | 0.03009 | 0.044413 | 0.038687 | 0.02268 | 0.018919 | 0.036107 | 0.016755 | 0.006918 | 0.007438 | 0.013992 | 0.018605 | 0.033387 | 0.017148 | 0.013674 | 0.01897 | 0.022901 | 0.045176 | 0.01424 | 0.081937 | 0.047478 | 0.049177 |
| 34.3333333 | 0.03985 | 0.022791 | 0.015708 | 0.0178 | 0.048536 | 0.072251 | 0.02414 | 0.007803 | 0.036673 | 0.009002 | 0.008 | 0.003636 | 0.00798 | 0.015113 | 0.018759 | 0.02625 | 0.011068 | 0.011614 | 0.013349 | 0.027603 | 0.007187 | 0.034353 | 0.052284 | 0.060314 |
| 34.3472222 | 0.047692 | 0.032068 | 0.037439 | 0.030141 | 0.041855 | 0.085553 | 0.024999 | 0.005639 | 0.030019 | 0.018607 | 0.009972 | 0.002278 | 0.00741 | 0.013055 | 0.011094 | 0.028303 | 0.01802 | 0.013821 | 0.011774 | 0.020537 | 0.010712 | 0.022045 | 0.044757 | 0.052734 |
| 34.3611111 | 0.046692 | 0.056352 | 0.084817 | 0.041908 | 0.042243 | 0.075088 | 0.022837 | 0.010471 | 0.032424 | 0.026204 | 0.012553 | 0.001666 | 0.007636 | 0.014387 | 0.014026 | 0.024907 | 0.029897 | 0.014797 | 0.009867 | 0.021279 | 0.013666 | 0.044331 | 0.024598 | 0.026516 |
| 34.375 | 0.03704 | 0.055499 | 0.124495 | 0.034106 | 0.073003 | 0.061016 | 0.017881 | 0.016399 | 0.038126 | 0.025379 | 0.012011 | 0.003048 | 0.00637 | 0.015218 | 0.022316 | 0.020031 | 0.032011 | 0.014079 | 0.007483 | 0.024365 | 0.008875 | 0.057156 | 0.016746 | 0.012878 |
| 34.3888889 | 0.022683 | 0.039521 | 0.131348 | 0.021934 | 0.104576 | 0.0469 | 0.011467 | 0.020361 | 0.039827 | 0.017184 | 0.007529 | 0.010513 | 0.004801 | 0.01078 | 0.030096 | 0.015377 | 0.024333 | 0.012233 | 0.004806 | 0.02097 | 0.002683 | 0.042327 | 0.015411 | 0.017104 |
| 34.4027778 | 0.011302 | 0.032375 | 0.110116 | 0.018072 | 0.094175 | 0.037188 | 0.007026 | 0.023473 | 0.03782 | 0.007123 | 0.003014 | 0.022042 | 0.008322 | 0.010182 | 0.037307 | 0.010767 | 0.024677 | 0.009562 | 0.003699 | 0.01769 | 0.002045 | 0.025763 | 0.015481 | 0.017178 |
| 34.4166667 | 0.01569 | 0.024516 | 0.073105 | 0.019338 | 0.042081 | 0.030389 | 0.007891 | 0.02768 | 0.035502 | 0.002684 | 0.001906 | 0.030794 | 0.018549 | 0.02176 | 0.038554 | 0.009303 | 0.034641 | 0.006352 | 0.003894 | 0.024137 | 0.005388 | 0.032868 | 0.027509 | 0.016961 |
| 34.4305556 | 0.042325 | 0.015351 | 0.042359 | 0.02283 | 0.030639 | 0.020642 | 0.011728 | 0.030799 | 0.032362 | 0.002407 | 0.004073 | 0.037794 | 0.02714 | 0.039477 | 0.02741 | 0.014548 | 0.038289 | 0.003391 | 0.003676 | 0.039213 | 0.007062 | 0.061173 | 0.063836 | 0.040466 |
| 34.4444444 | 0.08034 | 0.015387 | 0.026386 | 0.02849 | 0.127778 | 0.010041 | 0.015912 | 0.028865 | 0.02655 | 0.00424 | 0.006194 | 0.041685 | 0.028254 | 0.054087 | 0.014215 | 0.023062 | 0.033959 | 0.001765 | 0.002954 | 0.052336 | 0.005695 | 0.092218 | 0.104739 | 0.068495 |
| 34.4583333 | 0.128444 | 0.021356 | 0.013795 | 0.032727 | 0.245826 | 0.004659 | 0.017783 | 0.021353 | 0.018255 | 0.013772 | 0.005658 | 0.031213 | 0.031941 | 0.06632 | 0.013097 | 0.027079 | 0.043992 | 0.004718 | 0.004023 | 0.063032 | 0.006947 | 0.118313 | 0.124724 | 0.076149 |
| 34.4722222 | 0.169762 | 0.03616 | 0.008889 | 0.030858 | 0.221156 | 0.004184 | 0.015015 | 0.013574 | 0.009371 | 0.033977 | 0.003233 | 0.015384 | 0.048208 | 0.087163 | 0.01581 | 0.025611 | 0.080188 | 0.012146 | 0.009263 | 0.077621 | 0.013246 | 0.1542 | 0.111264 | 0.082255 |
| 34.4861111 | 0.147789 | 0.059073 | 0.015211 | 0.022833 | 0.108055 | 0.011448 | 0.009285 | 0.010892 | 0.005062 | 0.057442 | 0.002239 | 0.01663 | 0.076173 | 0.11374 | 0.014367 | 0.023062 | 0.118347 | 0.0187 | 0.015616 | 0.087096 | 0.019915 | 0.221955 | 0.073287 | 0.099294 |
| 34.5 | 0.082354 | 0.072059 | 0.026053 | 0.014288 | 0.068469 | 0.020689 | 0.004089 | 0.009507 | 0.007001 | 0.072538 | 0.003779 | 0.02957 | 0.113017 | 0.131482 | 0.014751 | 0.024385 | 0.129199 | 0.022025 | 0.021865 | 0.085531 | 0.023945 | 0.281759 | 0.056907 | 0.10873 |
| 34.5138889 | 0.052735 | 0.062421 | 0.037144 | 0.007897 | 0.099961 | 0.019492 | 0.003416 | 0.007585 | 0.010724 | 0.073356 | 0.00505 | 0.035085 | 0.157883 | 0.143097 | 0.021815 | 0.032599 | 0.122713 | 0.022745 | 0.031028 | 0.084026 | 0.026968 | 0.280849 | 0.072153 | 0.110626 |
| 34.5277778 | 0.051205 | 0.043634 | 0.041222 | 0.003335 | 0.117238 | 0.011953 | 0.007377 | 0.011133 | 0.017844 | 0.067124 | 0.004739 | 0.027732 | 0.196133 | 0.143357 | 0.036009 | 0.043376 | 0.11695 | 0.021949 | 0.044006 | 0.081264 | 0.032027 | 0.243084 | 0.07798 | 0.106521 |
| 34.5416667 | 0.033655 | 0.037872 | 0.030971 | 0.004556 | 0.080794 | 0.005595 | 0.012817 | 0.01813 | 0.031453 | 0.059003 | 0.004411 | 0.016951 | 0.205141 | 0.11721 | 0.047088 | 0.048199 | 0.099938 | 0.022935 | 0.055586 | 0.066625 | 0.03669 | 0.184424 | 0.069471 | 0.080983 |
| 34.5555556 | 0.013638 | 0.044626 | 0.018124 | 0.015224 | 0.046281 | 0.002608 | 0.015384 | 0.0207 | 0.041776 | 0.045901 | 0.004605 | 0.010851 | 0.185098 | 0.08283 | 0.042549 | 0.045068 | 0.067225 | 0.02794 | 0.061972 | 0.049184 | 0.034857 | 0.12451 | 0.065303 | 0.049911 |
| 34.5694444 | 0.010252 | 0.057467 | 0.024649 | 0.031771 | 0.073752 | 0.004376 | 0.011024 | 0.016891 | 0.040107 | 0.034566 | 0.004481 | 0.008427 | 0.139399 | 0.071267 | 0.029427 | 0.041261 | 0.042523 | 0.032393 | 0.076178 | 0.047144 | 0.028101 | 0.108931 | 0.060621 | 0.039394 |
| 34.5833333 | 0.01582 | 0.07628 | 0.045742 | 0.042244 | 0.109578 | 0.010497 | 0.00607 | 0.010679 | 0.032463 | 0.032191 | 0.004868 | 0.006456 | 0.103781 | 0.084896 | 0.020933 | 0.04156 | 0.038088 | 0.033629 | 0.096746 | 0.064306 | 0.025932 | 0.135705 | 0.06293 | 0.049969 |
| 34.5972222 | 0.021239 | 0.091843 | 0.071026 | 0.043138 | 0.085454 | 0.020231 | 0.010114 | 0.014542 | 0.02609 | 0.035795 | 0.007577 | 0.004458 | 0.110061 | 0.109941 | 0.016726 | 0.046035 | 0.044471 | 0.040069 | 0.106759 | 0.086975 | 0.037345 | 0.164039 | 0.068575 | 0.066916 |
| 34.6111111 | 0.028831 | 0.097461 | 0.120186 | 0.041542 | 0.058043 | 0.028886 | 0.018868 | 0.03507 | 0.022863 | 0.043514 | 0.01141 | 0.006453 | 0.109214 | 0.134144 | 0.021819 | 0.055278 | 0.05051 | 0.054697 | 0.11235 | 0.098987 | 0.054187 | 0.158415 | 0.064149 | 0.076476 |
| 34.625 | 0.040715 | 0.106823 | 0.177317 | 0.040389 | 0.08105 | 0.031898 | 0.027115 | 0.053499 | 0.022261 | 0.056144 | 0.012786 | 0.012351 | 0.078214 | 0.131164 | 0.045091 | 0.063901 | 0.046488 | 0.063627 | 0.115415 | 0.097393 | 0.058011 | 0.114152 | 0.062394 | 0.072744 |
| 34.6388889 | 0.040291 | 0.119289 | 0.185907 | 0.040209 | 0.112064 | 0.027105 | 0.031146 | 0.052442 | 0.024187 | 0.071917 | 0.011256 | 0.01655 | 0.049149 | 0.096717 | 0.064419 | 0.063147 | 0.038692 | 0.054658 | 0.110209 | 0.087925 | 0.046448 | 0.060385 | 0.057797 | 0.058883 |
| 34.6527778 | 0.02208 | 0.114643 | 0.147406 | 0.048943 | 0.124766 | 0.022369 | 0.029513 | 0.038067 | 0.033003 | 0.087465 | 0.010731 | 0.02017 | 0.033897 | 0.065658 | 0.057443 | 0.052448 | 0.036929 | 0.037615 | 0.104834 | 0.078286 | 0.026638 | 0.027217 | 0.035907 | 0.045823 |
| 34.6666667 | 0.018007 | 0.09399 | 0.104257 | 0.066532 | 0.106099 | 0.031465 | 0.032182 | 0.024967 | 0.048536 | 0.092431 | 0.013053 | 0.026434 | 0.02609 | 0.043634 | 0.037051 | 0.038512 | 0.031138 | 0.027059 | 0.095483 | 0.07433 | 0.011863 | 0.013118 | 0.013246 | 0.030485 |
| 34.6805556 | 0.039867 | 0.062501 | 0.072975 | 0.068383 | 0.085368 | 0.046918 | 0.039169 | 0.022075 | 0.06053 | 0.076899 | 0.015896 | 0.030299 | 0.018073 | 0.021158 | 0.017723 | 0.026718 | 0.019664 | 0.018365 | 0.070543 | 0.074153 | 0.011704 | 0.009142 | 0.005314 | 0.015814 |
| 34.6944444 | 0.054832 | 0.028966 | 0.047261 | 0.044944 | 0.097526 | 0.050847 | 0.042791 | 0.026093 | 0.062604 | 0.051814 | 0.01804 | 0.02959 | 0.02007 | 0.013463 | 0.007963 | 0.019898 | 0.020162 | 0.008264 | 0.039925 | 0.077627 | 0.018304 | 0.015734 | 0.010151 | 0.016339 |
| 34.7083333 | 0.049189 | 0.016695 | 0.027968 | 0.029189 | 0.109072 | 0.042693 | 0.04116 | 0.029091 | 0.062174 | 0.04666 | 0.019211 | 0.027059 | 0.065115 | 0.01753 | 0.010735 | 0.018744 | 0.026322 | 0.005085 | 0.017654 | 0.085591 | 0.023372 | 0.031015 | 0.030644 | 0.020461 |
| 34.7222222 | 0.033318 | 0.037775 | 0.04421 | 0.034202 | 0.112166 | 0.032738 | 0.033839 | 0.027809 | 0.068466 | 0.068066 | 0.019253 | 0.030266 | 0.120464 | 0.019084 | 0.01258 | 0.020913 | 0.024009 | 0.009316 | 0.005736 | 0.092514 | 0.026812 | 0.053119 | 0.069808 | 0.01869 |
| 34.7361111 | 0.035747 | 0.081555 | 0.098114 | 0.041286 | 0.115024 | 0.026605 | 0.023609 | 0.023346 | 0.080602 | 0.085503 | 0.018958 | 0.037002 | 0.128079 | 0.025736 | 0.007501 | 0.021604 | 0.028913 | 0.013197 | 0.004797 | 0.099938 | 0.026414 | 0.071016 | 0.11015 | 0.02017 |
| 34.75 | 0.061638 | 0.085371 | 0.122956 | 0.049225 | 0.086973 | 0.025533 | 0.016422 | 0.019209 | 0.092174 | 0.086366 | 0.018505 | 0.037773 | 0.101306 | 0.038674 | 0.005282 | 0.018038 | 0.036999 | 0.01177 | 0.0176 | 0.104767 | 0.018445 | 0.0576 | 0.127324 | 0.022106 |
| 34.7638889 | 0.072116 | 0.046799 | 0.082539 | 0.043177 | 0.06264 | 0.024793 | 0.015281 | 0.0185 | 0.095241 | 0.085693 | 0.016835 | 0.028867 | 0.062865 | 0.043618 | 0.009114 | 0.011979 | 0.032905 | 0.013786 | 0.043172 | 0.090546 | 0.008481 | 0.025944 | 0.114378 | 0.016701 |
| 34.7777778 | 0.055644 | 0.058614 | 0.040317 | 0.031269 | 0.082044 | 0.017401 | 0.014869 | 0.021296 | 0.085303 | 0.090598 | 0.014055 | 0.019086 | 0.050399 | 0.032413 | 0.012958 | 0.006917 | 0.036862 | 0.025282 | 0.070002 | 0.06014 | 0.004492 | 0.018256 | 0.086337 | 0.015347 |
| 34.7916667 | 0.032658 | 0.111693 | 0.056807 | 0.037253 | 0.154777 | 0.010239 | 0.013087 | 0.02659 | 0.067512 | 0.104645 | 0.011819 | 0.028926 | 0.081151 | 0.018246 | 0.018184 | 0.006187 | 0.055169 | 0.031419 | 0.086461 | 0.033811 | 0.011008 | 0.028612 | 0.069263 | 0.023349 |
| 34.8055556 | 0.05255 | 0.143475 | 0.120219 | 0.035039 | 0.309721 | 0.020227 | 0.010365 | 0.032076 | 0.047763 | 0.127166 | 0.010579 | 0.043185 | 0.116276 | 0.018773 | 0.029373 | 0.008771 | 0.066644 | 0.025683 | 0.095621 | 0.017106 | 0.027164 | 0.035569 | 0.068382 | 0.025658 |

| | | | | | | | | | | | | | | | | | | | | | | | | |
|---|---|---|---|---|---|---|---|---|---|---|---|---|---|---|---|---|---|---|---|---|---|---|---|---|
| 34.8194444 | 0.097886 | 0.158143 | 0.183346 | 0.017015 | 0.371596 | 0.040941 | 0.004797 | 0.037835 | 0.02584 | 0.142062 | 0.009546 | 0.035779 | 0.128192 | 0.024853 | 0.040134 | 0.010337 | 0.053155 | 0.018379 | 0.108328 | 0.008179 | 0.043423 | 0.031932 | 0.065594 | 0.02144 |
| 34.8333333 | 0.085473 | 0.158588 | 0.229615 | 0.007068 | 0.251237 | 0.049527 | 0.005951 | 0.048396 | 0.01276 | 0.139626 | 0.00683 | 0.02107 | 0.117724 | 0.02242 | 0.040291 | 0.008212 | 0.021318 | 0.017101 | 0.11095 | 0.009974 | 0.050655 | 0.022854 | 0.052472 | 0.015261 |
| 34.8472222 | 0.038991 | 0.13103 | 0.30632 | 0.009199 | 0.163659 | 0.038402 | 0.026767 | 0.065168 | 0.020287 | 0.126273 | 0.003341 | 0.027519 | 0.096792 | 0.020497 | 0.028482 | 0.003979 | 0.003461 | 0.022161 | 0.083277 | 0.017289 | 0.054482 | 0.021679 | 0.041289 | 0.010839 |
| 34.8611111 | 0.015618 | 0.101993 | 0.353878 | 0.008645 | 0.124316 | 0.01839 | 0.05532 | 0.071024 | 0.036534 | 0.10894 | 0.001642 | 0.048911 | 0.069038 | 0.029608 | 0.014168 | 0.001313 | 0.002743 | 0.028381 | 0.044896 | 0.016225 | 0.060278 | 0.025345 | 0.039453 | 0.01633 |
| 34.875 | 0.007497 | 0.094822 | 0.288132 | 0.010375 | 0.079099 | 0.007099 | 0.061964 | 0.048112 | 0.050493 | 0.095897 | 0.000999 | 0.054914 | 0.036417 | 0.038733 | 0.007207 | 0.002737 | 0.004192 | 0.03427 | 0.018319 | 0.006442 | 0.057618 | 0.023089 | 0.038901 | 0.020877 |
| 34.8888889 | 0.005301 | 0.087066 | 0.196858 | 0.017517 | 0.116298 | 0.006695 | 0.049214 | 0.021836 | 0.060351 | 0.085093 | 0.001704 | 0.046531 | 0.023753 | 0.033311 | 0.005873 | 0.009439 | 0.007273 | 0.042996 | 0.005783 | 0.001983 | 0.045529 | 0.013884 | 0.039056 | 0.013835 |
| 34.9027778 | 0.007145 | 0.061735 | 0.125013 | 0.033709 | 0.179044 | 0.013421 | 0.035507 | 0.01814 | 0.057729 | 0.063866 | 0.006008 | 0.04909 | 0.0534 | 0.017691 | 0.006622 | 0.020188 | 0.012227 | 0.048693 | 0.004187 | 0.003846 | 0.035853 | 0.004967 | 0.044916 | 0.006927 |
| 34.9166667 | 0.007662 | 0.040548 | 0.056919 | 0.084804 | 0.168103 | 0.025583 | 0.023178 | 0.024204 | 0.040321 | 0.044768 | 0.013501 | 0.071214 | 0.094736 | 0.006921 | 0.012282 | 0.026936 | 0.018276 | 0.040574 | 0.004722 | 0.010804 | 0.028786 | 0.006157 | 0.049744 | 0.006937 |
| 34.9305556 | 0.01186 | 0.052045 | 0.026847 | 0.150403 | 0.115745 | 0.033844 | 0.018993 | 0.02704 | 0.019218 | 0.042352 | 0.014562 | 0.083731 | 0.11774 | 0.005309 | 0.021065 | 0.022099 | 0.021779 | 0.0211 | 0.004292 | 0.027206 | 0.018053 | 0.019746 | 0.048749 | 0.006273 |
| 34.9444444 | 0.025132 | 0.072629 | 0.039401 | 0.187114 | 0.091138 | 0.031502 | 0.028878 | 0.027863 | 0.006236 | 0.041417 | 0.006412 | 0.066483 | 0.129656 | 0.007991 | 0.024576 | 0.012304 | 0.01987 | 0.006636 | 0.00358 | 0.039465 | 0.008159 | 0.038143 | 0.047981 | 0.003713 |
| 34.9583333 | 0.049579 | 0.067803 | 0.068387 | 0.166133 | 0.096449 | 0.022987 | 0.051013 | 0.029313 | 0.00763 | 0.030612 | 0.002565 | 0.041538 | 0.137228 | 0.014243 | 0.020279 | 0.007531 | 0.022231 | 0.011079 | 0.002541 | 0.036769 | 0.003471 | 0.05914 | 0.052921 | 0.005115 |
| 34.9722222 | 0.094311 | 0.043411 | 0.089065 | 0.127535 | 0.110006 | 0.023928 | 0.074693 | 0.030261 | 0.020284 | 0.01684 | 0.003947 | 0.021733 | 0.145358 | 0.026299 | 0.014396 | 0.007475 | 0.032491 | 0.028782 | 0.004411 | 0.036363 | 0.002527 | 0.077498 | 0.059696 | 0.015909 |
| 34.9861111 | 0.125531 | 0.038353 | 0.084726 | 0.124421 | 0.111937 | 0.040582 | 0.080511 | 0.02507 | 0.032409 | 0.012672 | 0.003937 | 0.010557 | 0.136913 | 0.039225 | 0.014235 | 0.009183 | 0.035552 | 0.03745 | 0.010972 | 0.048623 | 0.005608 | 0.09277 | 0.056561 | 0.029367 |
| 35 | 0.102233 | 0.072284 | 0.072527 | 0.111436 | 0.074813 | 0.060173 | 0.063639 | 0.020451 | 0.034087 | 0.017812 | 0.00558 | 0.018199 | 0.107158 | 0.04421 | 0.024673 | 0.016342 | 0.027301 | 0.026711 | 0.017664 | 0.059904 | 0.009599 | 0.11092 | 0.043163 | 0.036592 |
| 35.0138889 | 0.067406 | 0.104095 | 0.063357 | 0.096228 | 0.042432 | 0.059834 | 0.042056 | 0.019495 | 0.025993 | 0.020128 | 0.007163 | 0.040142 | 0.097002 | 0.043824 | 0.049779 | 0.03444 | 0.021571 | 0.011472 | 0.019261 | 0.049845 | 0.013991 | 0.115595 | 0.02813 | 0.04275 |
| 35.0277778 | 0.063477 | 0.106533 | 0.046624 | 0.143318 | 0.058069 | 0.041426 | 0.0262 | 0.021737 | 0.018734 | 0.027778 | 0.008581 | 0.048365 | 0.104545 | 0.041267 | 0.083126 | 0.058217 | 0.032123 | 0.004605 | 0.014356 | 0.021943 | 0.026417 | 0.109108 | 0.021557 | 0.051688 |
| 35.0416667 | 0.069065 | 0.0892 | 0.059782 | 0.205936 | 0.086211 | 0.047276 | 0.029292 | 0.046031 | 0.025163 | 0.072505 | 0.015506 | 0.037046 | 0.102381 | 0.033864 | 0.105913 | 0.066273 | 0.055652 | 0.012159 | 0.012155 | 0.006593 | 0.058628 | 0.130067 | 0.031548 | 0.056355 |
| 35.0555556 | 0.0776 | 0.059067 | 0.129741 | 0.168057 | 0.100188 | 0.076249 | 0.059869 | 0.080328 | 0.035945 | 0.162342 | 0.020375 | 0.023643 | 0.090447 | 0.03544 | 0.117209 | 0.050766 | 0.074218 | 0.032219 | 0.024424 | 0.009932 | 0.096634 | 0.157667 | 0.099294 | 0.049549 |
| 35.0694444 | 0.085207 | 0.026303 | 0.200562 | 0.070025 | 0.104104 | 0.082664 | 0.077403 | 0.091636 | 0.032697 | 0.202173 | 0.014108 | 0.011926 | 0.062926 | 0.057078 | 0.117909 | 0.026907 | 0.071371 | 0.044854 | 0.041051 | 0.019339 | 0.118472 | 0.154538 | 0.216217 | 0.054253 |
| 35.0833333 | 0.058042 | 0.009618 | 0.231434 | 0.033107 | 0.103939 | 0.064063 | 0.059197 | 0.07321 | 0.023636 | 0.130469 | 0.005387 | 0.00641 | 0.030375 | 0.065282 | 0.097487 | 0.008322 | 0.043641 | 0.040411 | 0.044663 | 0.030142 | 0.133627 | 0.135812 | 0.230503 | 0.070189 |
| 35.0972222 | 0.02087 | 0.012329 | 0.243126 | 0.027201 | 0.096735 | 0.050427 | 0.038712 | 0.038581 | 0.01634 | 0.04861 | 0.00413 | 0.007613 | 0.012588 | 0.04086 | 0.071971 | 0.001552 | 0.021776 | 0.024194 | 0.035371 | 0.03905 | 0.121618 | 0.093471 | 0.139662 | 0.059364 |
| 35.1111111 | 0.011253 | 0.017322 | 0.256193 | 0.016714 | 0.076517 | 0.056447 | 0.039171 | 0.013129 | 0.00992 | 0.017405 | 0.006936 | 0.009831 | 0.006138 | 0.018849 | 0.054603 | 0.004436 | 0.029347 | 0.011328 | 0.022958 | 0.044221 | 0.071845 | 0.038353 | 0.079665 | 0.039391 |
| 35.125 | 0.012942 | 0.021455 | 0.244418 | 0.030266 | 0.055149 | 0.075205 | 0.050527 | 0.006339 | 0.00905 | 0.009612 | 0.0105 | 0.009467 | 0.005978 | 0.014439 | 0.035941 | 0.017943 | 0.041566 | 0.01026 | 0.013984 | 0.04689 | 0.043093 | 0.010081 | 0.052118 | 0.033006 |
| 35.1388889 | 0.01535 | 0.035983 | 0.181726 | 0.044876 | 0.034858 | 0.085331 | 0.054929 | 0.004784 | 0.012882 | 0.010978 | 0.015051 | 0.010457 | 0.009741 | 0.015487 | 0.029588 | 0.039929 | 0.038376 | 0.008264 | 0.013038 | 0.046526 | 0.044394 | 0.014117 | 0.051007 | 0.025504 |
| 35.1527778 | 0.021577 | 0.050677 | 0.126618 | 0.034152 | 0.031399 | 0.071368 | 0.045027 | 0.004658 | 0.018011 | 0.011204 | 0.017217 | 0.018264 | 0.009375 | 0.025846 | 0.045189 | 0.060495 | 0.024924 | 0.005896 | 0.021964 | 0.040382 | 0.031252 | 0.038851 | 0.063866 | 0.023645 |
| 35.1666667 | 0.027661 | 0.0472 | 0.107094 | 0.021187 | 0.068971 | 0.041333 | 0.027328 | 0.006823 | 0.022417 | 0.014942 | 0.01402 | 0.037463 | 0.004934 | 0.045986 | 0.055829 | 0.072906 | 0.012084 | 0.013992 | 0.033585 | 0.029298 | 0.013518 | 0.075415 | 0.056649 | 0.037084 |
| 35.1805556 | 0.026154 | 0.03227 | 0.07772 | 0.02169 | 0.091465 | 0.017224 | 0.012919 | 0.006899 | 0.023282 | 0.027781 | 0.007099 | 0.058563 | 0.002515 | 0.060977 | 0.052363 | 0.073239 | 0.006672 | 0.020424 | 0.035373 | 0.017401 | 0.008343 | 0.115772 | 0.048471 | 0.053162 |
| 35.1944444 | 0.017174 | 0.017725 | 0.043823 | 0.022185 | 0.057038 | 0.012117 | 0.00623 | 0.007965 | 0.020542 | 0.040937 | 0.002409 | 0.06084 | 0.002027 | 0.06375 | 0.051656 | 0.068321 | 0.007284 | 0.016457 | 0.023553 | 0.008652 | 0.012188 | 0.137445 | 0.060066 | 0.059779 |
| 35.2083333 | 0.007601 | 0.012254 | 0.02415 | 0.015541 | 0.030573 | 0.016179 | 0.004887 | 0.012782 | 0.017536 | 0.055063 | 0.002474 | 0.050606 | 0.004261 | 0.059306 | 0.058638 | 0.063734 | 0.012016 | 0.01769 | 0.009638 | 0.004312 | 0.037044 | 0.131331 | 0.086218 | 0.057859 |
| 35.2222222 | 0.004993 | 0.02011 | 0.014154 | 0.013098 | 0.049294 | 0.016878 | 0.003776 | 0.017479 | 0.01571 | 0.068392 | 0.005295 | 0.050664 | 0.009922 | 0.053938 | 0.068415 | 0.05745 | 0.016346 | 0.026732 | 0.002958 | 0.003692 | 0.070571 | 0.115442 | 0.083417 | 0.054994 |
| 35.2361111 | 0.012083 | 0.025517 | 0.015868 | 0.014514 | 0.058302 | 0.018372 | 0.006723 | 0.018816 | 0.012777 | 0.072907 | 0.00969 | 0.058255 | 0.012498 | 0.05016 | 0.075815 | 0.050384 | 0.012792 | 0.034299 | 0.003187 | 0.004028 | 0.080532 | 0.102239 | 0.053938 | 0.056541 |
| 35.25 | 0.034724 | 0.018308 | 0.019887 | 0.01798 | 0.04108 | 0.021208 | 0.016793 | 0.018317 | 0.007529 | 0.071429 | 0.010928 | 0.051199 | 0.009951 | 0.047656 | 0.074262 | 0.04475 | 0.008827 | 0.034657 | 0.009052 | 0.003204 | 0.062415 | 0.097244 | 0.037701 | 0.060408 |
| 35.2638889 | 0.052177 | 0.011365 | 0.020365 | 0.033283 | 0.027367 | 0.017346 | 0.022992 | 0.016407 | 0.003088 | 0.069127 | 0.007949 | 0.029902 | 0.01134 | 0.046771 | 0.063148 | 0.043498 | 0.017153 | 0.023706 | 0.016124 | 0.00325 | 0.032423 | 0.091503 | 0.033149 | 0.05882 |
| 35.2777778 | 0.037709 | 0.019825 | 0.016964 | 0.047924 | 0.02581 | 0.009911 | 0.022332 | 0.012039 | 0.002628 | 0.060652 | 0.009161 | 0.021757 | 0.018201 | 0.045459 | 0.049729 | 0.04424 | 0.029677 | 0.012603 | 0.020481 | 0.003822 | 0.011101 | 0.069227 | 0.030252 | 0.050327 |
| 35.2916667 | 0.035205 | 0.038964 | 0.012316 | 0.050819 | 0.054235 | 0.006235 | 0.019623 | 0.006871 | 0.005666 | 0.039135 | 0.015963 | 0.037584 | 0.021323 | 0.041498 | 0.039011 | 0.040696 | 0.036368 | 0.013762 | 0.022395 | 0.004796 | 0.012437 | 0.039319 | 0.027 | 0.041367 |
| 35.3055556 | 0.056382 | 0.049899 | 0.020859 | 0.053223 | 0.079194 | 0.006175 | 0.015151 | 0.004651 | 0.010169 | 0.018138 | 0.019955 | 0.054972 | 0.018585 | 0.039133 | 0.032475 | 0.035753 | 0.03677 | 0.020955 | 0.021485 | 0.012361 | 0.025467 | 0.022083 | 0.023589 | 0.034951 |
| 35.3194444 | 0.048261 | 0.052054 | 0.028306 | 0.05474 | 0.061426 | 0.004999 | 0.010635 | 0.00916 | 0.014566 | 0.009495 | 0.015936 | 0.057254 | 0.014477 | 0.042567 | 0.028266 | 0.03429 | 0.031653 | 0.024019 | 0.01757 | 0.025971 | 0.032963 | 0.024204 | 0.020965 | 0.031608 |
| 35.3333333 | 0.019064 | 0.05552 | 0.030648 | 0.043987 | 0.039237 | 0.002671 | 0.00858 | 0.01285 | 0.017439 | 0.008399 | 0.00904 | 0.049094 | 0.010928 | 0.046787 | 0.021343 | 0.034487 | 0.022943 | 0.022252 | 0.012872 | 0.032652 | 0.031659 | 0.032999 | 0.023446 | 0.030617 |
| 35.3472222 | 0.004249 | 0.060212 | 0.060229 | 0.025013 | 0.043889 | 0.002636 | 0.007946 | 0.008425 | 0.018488 | 0.010535 | 0.007426 | 0.039817 | 0.006865 | 0.045163 | 0.011851 | 0.034958 | 0.012863 | 0.016706 | 0.008951 | 0.025248 | 0.020957 | 0.041953 | 0.035763 | 0.029541 |
| 35.3611111 | 0.002712 | 0.064126 | 0.092847 | 0.010983 | 0.051874 | 0.003249 | 0.006819 | 0.004112 | 0.018 | 0.010112 | 0.009262 | 0.03194 | 0.005423 | 0.042275 | 0.008266 | 0.036901 | 0.009751 | 0.010271 | 0.005501 | 0.01372 | 0.012846 | 0.0628 | 0.052858 | 0.031853 |
| 35.375 | 0.004599 | 0.065926 | 0.087069 | 0.013598 | 0.041651 | 0.002207 | 0.004939 | 0.002831 | 0.016509 | 0.011608 | 0.008458 | 0.021783 | 0.009086 | 0.042312 | 0.012589 | 0.04049 | 0.017166 | 0.01042 | 0.004747 | 0.009699 | 0.018352 | 0.092267 | 0.080857 | 0.038657 |
| 35.3888889 | 0.005201 | 0.058957 | 0.060204 | 0.028881 | 0.031525 | 0.001141 | 0.002517 | 0.002497 | 0.015155 | 0.018468 | 0.005163 | 0.011446 | 0.010291 | 0.041715 | 0.017442 | 0.043138 | 0.022174 | 0.014892 | 0.007134 | 0.011461 | 0.020666 | 0.10621 | 0.124747 | 0.043451 |
| 35.4027778 | 0.004095 | 0.041601 | 0.035348 | 0.034961 | 0.022991 | 0.001939 | 0.000956 | 0.006147 | 0.01358 | 0.021529 | 0.002533 | 0.009831 | 0.006415 | 0.040279 | 0.021465 | 0.042053 | 0.017939 | 0.014494 | 0.007581 | 0.01179 | 0.012598 | 0.097765 | 0.147285 | 0.042305 |
| 35.4166667 | 0.003156 | 0.024298 | 0.018445 | 0.027064 | 0.014664 | 0.005215 | 0.001955 | 0.013656 | 0.010607 | 0.018472 | 0.001295 | 0.018664 | 0.00435 | 0.039481 | 0.025833 | 0.039113 | 0.013288 | 0.009253 | 0.005392 | 0.0098 | 0.005982 | 0.088166 | 0.140281 | 0.038571 |
| 35.4305556 | 0.003761 | 0.014913 | 0.010537 | 0.020806 | 0.024267 | 0.012106 | 0.00591 | 0.023259 | 0.006919 | 0.013879 | 0.000559 | 0.030952 | 0.007166 | 0.037864 | 0.026408 | 0.038762 | 0.018619 | 0.005 | 0.005319 | 0.009776 | 0.00504 | 0.094408 | 0.149012 | 0.038542 |
| 35.4444444 | 0.007394 | 0.012701 | 0.009144 | 0.023674 | 0.03468 | 0.022866 | 0.012169 | 0.032536 | 0.004429 | 0.012152 | 0.000502 | 0.038563 | 0.014146 | 0.036384 | 0.021158 | 0.040685 | 0.036774 | 0.004258 | 0.009 | 0.01434 | 0.012191 | 0.11305 | 0.171547 | 0.042427 |
| 35.4583333 | 0.013485 | 0.012813 | 0.022083 | 0.031166 | 0.024169 | 0.034067 | 0.018326 | 0.036944 | 0.004322 | 0.01395 | 0.000914 | 0.037207 | 0.02639 | 0.040216 | 0.012493 | 0.039526 | 0.055049 | 0.005023 | 0.013562 | 0.021374 | 0.027409 | 0.131966 | 0.184317 | 0.048029 |
| 35.4722222 | 0.019625 | 0.010029 | 0.057678 | 0.038446 | 0.017183 | 0.039839 | 0.020493 | 0.033955 | 0.006095 | 0.017007 | 0.001008 | 0.032382 | 0.040415 | 0.048906 | 0.005644 | 0.034316 | 0.05649 | 0.004917 | 0.016333 | 0.026564 | 0.039883 | 0.142951 | 0.188626 | 0.052511 |
| 35.4861111 | 0.025367 | 0.006522 | 0.105244 | 0.041405 | 0.024806 | 0.040008 | 0.01752 | 0.025545 | 0.008164 | 0.020214 | 0.000918 | 0.029856 | 0.048444 | 0.056431 | 0.00501 | 0.031136 | 0.048714 | 0.00491 | 0.018059 | 0.028819 | 0.042683 | 0.147693 | 0.184425 | 0.054006 |
| 35.5 | 0.033256 | 0.010526 | 0.136725 | 0.03824 | 0.032908 | 0.042089 | 0.01277 | 0.018194 | 0.009626 | 0.025347 | 0.000913 | 0.029428 | 0.053456 | 0.064372 | 0.006293 | 0.035234 | 0.049902 | 0.007536 | 0.020413 | 0.032063 | 0.04184 | 0.147528 | 0.186287 | 0.057529 |
| 35.5138889 | 0.044791 | 0.021721 | 0.138912 | 0.030661 | 0.046423 | 0.052886 | 0.010158 | 0.017869 | 0.01061 | 0.037015 | 0.001784 | 0.02967 | 0.067024 | 0.075101 | 0.007258 | 0.04536 | 0.057777 | 0.012538 | 0.023349 | 0.041347 | 0.046699 | 0.144252 | 0.198825 | 0.067621 |
| 35.5277778 | 0.058569 | 0.030786 | 0.122552 | 0.018294 | 0.069837 | 0.076357 | 0.011011 | 0.023479 | 0.012438 | 0.052915 | 0.004442 | 0.025027 | 0.086681 | 0.074696 | 0.013409 | 0.052553 | 0.064954 | 0.016533 | 0.025976 | 0.054766 | 0.0628 | 0.141044 | 0.203745 | 0.071259 |
| 35.5416667 | 0.071229 | 0.032789 | 0.089406 | 0.008943 | 0.106243 | 0.10461 | 0.013693 | 0.028862 | 0.01661 | 0.054736 | 0.008003 | 0.015596 | 0.090562 | 0.061724 | 0.023606 | 0.040589 | 0.069418 | 0.017613 | 0.02909 | 0.063172 | 0.078121 | 0.124701 | 0.189769 | 0.061494 |
| 35.5555556 | 0.073871 | 0.028352 | 0.058194 | 0.009894 | 0.117478 | 0.116457 | 0.011618 | 0.029475 | 0.019561 | 0.036465 | 0.009399 | 0.011332 | 0.070031 | 0.063458 | 0.03127 | 0.015182 | 0.072704 | 0.015528 | 0.032637 | 0.06243 | 0.077619 | 0.100735 | 0.158868 | 0.061127 |
| 35.5694444 | 0.058169 | 0.017306 | 0.057343 | 0.009606 | 0.094759 | 0.112653 | 0.00445 | 0.026118 | 0.017543 | 0.029018 | 0.006431 | 0.009734 | 0.05646 | 0.081986 | 0.030431 | 0.007944 | 0.083136 | 0.011849 | 0.036126 | 0.059759 | 0.071709 | 0.092859 | 0.128072 | 0.079362 |
| 35.5833333 | 0.033492 | 0.01053 | 0.074547 | 0.006351 | 0.133901 | 0.114141 | 0.002738 | 0.027772 | 0.014972 | 0.041784 | 0.00283 | 0.004837 | 0.07335 | 0.093455 | 0.018565 | 0.026508 | 0.09747 | 0.013588 | 0.045893 | 0.062413 | 0.074594 | 0.100211 | 0.117724 | 0.104909 |
| 35.5972222 | 0.019578 | 0.018654 | 0.086327 | 0.011964 | 0.177812 | 0.12113 | 0.008 | 0.03698 | 0.014756 | 0.05 | 0.002161 | 0.003629 | 0.089957 | 0.093157 | 0.006052 | 0.049084 | 0.107625 | 0.027852 | 0.06224 | 0.069409 | 0.093914 | 0.107304 | 0.130088 | 0.125253 |
| 35.6111111 | 0.019803 | 0.030541 | 0.093679 | 0.024906 | 0.111794 | 0.119043 | 0.012898 | 0.043982 | 0.014341 | 0.050842 | 0.001673 | 0.007048 | 0.085912 | 0.090306 | 0.005854 | 0.061498 | 0.10657 | 0.052947 | 0.072124 | 0.074063 | 0.135492 | 0.097034 | 0.137691 | 0.127234 |
| 35.625 | 0.027288 | 0.039163 | 0.104866 | 0.038186 | 0.034262 | 0.106905 | 0.015172 | 0.046143 | 0.013308 | 0.041392 | 0.000575 | 0.007001 | 0.092118 | 0.089906 | 0.016391 | 0.070034 | 0.086518 | 0.072374 | 0.070561 | 0.068502 | 0.1652 | 0.073915 | 0.124705 | 0.115324 |
| 35.6388889 | 0.039134 | 0.051169 | 0.112372 | 0.046948 | 0.011496 | 0.095553 | 0.017186 | 0.044766 | 0.012607 | 0.024524 | 0.00167 | 0.006005 | 0.116659 | 0.087464 | 0.028256 | 0.072817 | 0.060831 | 0.065593 | 0.065335 | 0.057015 | 0.144293 | 0.054406 | 0.109246 | 0.102035 |
| 35.6527778 | 0.050726 | 0.065747 | 0.101775 | 0.04602 | 0.025746 | 0.089878 | 0.019674 | 0.043378 | 0.011927 | 0.025994 | 0.004053 | 0.009108 | 0.123415 | 0.074668 | 0.036837 | 0.0613 | 0.045251 | 0.037661 | 0.061956 | 0.048134 | 0.09375 | 0.042435 | 0.095468 | 0.088801 |
| 35.6666667 | 0.05757 | 0.070976 | 0.06512 | 0.033604 | 0.075745 | 0.084712 | 0.020524 | 0.03987 | 0.012385 | 0.045316 | 0.007103 | 0.013296 | 0.103412 | 0.0537 | 0.041771 | 0.049126 | 0.041228 | 0.02241 | 0.058593 | 0.043418 | 0.04739 | 0.03603 | 0.074234 | 0.076595 |
| 35.6805556 | 0.051662 | 0.05809 | 0.032984 | 0.018714 | 0.120398 | 0.076147 | 0.017885 | 0.024378 | 0.01664 | 0.064468 | 0.013868 | 0.023292 | 0.079496 | 0.037348 | 0.039499 | 0.050806 | 0.049069 | 0.024623 | 0.053608 | 0.042554 | 0.022792 | 0.031269 | 0.051287 | 0.067326 |
| 35.6944444 | 0.033351 | 0.034392 | 0.048766 | 0.011621 | 0.122274 | 0.062961 | 0.013069 | 0.011489 | 0.023814 | 0.075103 | 0.02113 | 0.035307 | 0.055811 | 0.032115 | 0.026861 | 0.062432 | 0.060311 | 0.021757 | 0.049023 | 0.042303 | 0.016759 | 0.02619 | 0.031009 | 0.059771 |
| 35.7083333 | 0.017853 | 0.017004 | 0.080919 | 0.009606 | 0.07924 | 0.043544 | 0.012677 | 0.020357 | 0.029287 | 0.075353 | 0.024653 | 0.034946 | 0.029561 | 0.031396 | 0.011783 | 0.076377 | 0.048454 | 0.024186 | 0.045566 | 0.041347 | 0.010315 | 0.019664 | 0.017048 | 0.052077 |
| 35.7222222 | 0.016979 | 0.013879 | 0.066881 | 0.015438 | 0.039266 | 0.024748 | 0.020297 | 0.040981 | 0.031955 | 0.068932 | 0.025354 | 0.026471 | 0.024205 | 0.02787 | 0.005321 | 0.081512 | 0.031963 | 0.036837 | 0.041722 | 0.036657 | 0.006259 | 0.011367 | 0.020256 | 0.042395 |
| 35.7361111 | 0.02638 | 0.022877 | 0.033702 | 0.034322 | 0.028671 | 0.014787 | 0.028488 | 0.058446 | 0.034821 | 0.057854 | 0.02368 | 0.023369 | 0.044415 | 0.023217 | 0.004394 | 0.071863 | 0.049069 | 0.041537 | 0.036355 | 0.026905 | 0.018987 | 0.009281 | 0.032426 | 0.031362 |
| 35.75 | 0.024899 | 0.039338 | 0.043596 | 0.060072 | 0.031806 | 0.010636 | 0.029481 | 0.070513 | 0.03912 | 0.046893 | 0.022902 | 0.028994 | 0.059329 | 0.016697 | 0.003883 | 0.060413 | 0.07355 | 0.035576 | 0.029887 | 0.026283 | 0.04695 | 0.017462 | 0.044609 | 0.019998 |
| 35.7638889 | 0.013516 | 0.054386 | 0.066959 | 0.085081 | 0.054676 | 0.009324 | 0.024389 | 0.078506 | 0.044895 | 0.036674 | 0.02335 | 0.038964 | 0.055564 | 0.01113 | 0.00751 | 0.061705 | 0.082724 | 0.029755 | 0.026733 | 0.038238 | 0.07033 | 0.022428 | 0.062498 | 0.010195 |
| 35.7777778 | 0.007824 | 0.062789 | 0.077752 | 0.097646 | 0.068261 | 0.011703 | 0.017807 | 0.086565 | 0.053207 | 0.023226 | 0.021496 | 0.049313 | 0.038317 | 0.009858 | 0.013709 | 0.07138 | 0.073359 | 0.032882 | 0.030487 | 0.04479 | 0.076806 | 0.018555 | 0.073666 | 0.006081 |
| 35.7916667 | 0.011495 | 0.070571 | 0.110797 | 0.083807 | 0.058456 | 0.014131 | 0.012999 | 0.100501 | 0.065179 | 0.017387 | 0.01916 | 0.057116 | 0.01963 | 0.006894 | 0.017876 | 0.073455 | 0.041621 | 0.036566 | 0.036066 | 0.042845 | 0.067329 | 0.01653 | 0.067365 | 0.010159 |
| 35.8055556 | 0.018117 | 0.078451 | 0.157545 | 0.049079 | 0.064706 | 0.016751 | 0.010946 | 0.110655 | 0.077703 | 0.026281 | 0.017655 | 0.060254 | 0.015474 | 0.005899 | 0.021334 | 0.06579 | 0.013741 | 0.030015 | 0.037186 | 0.04028 | 0.04564 | 0.021621 | 0.05524 | 0.019074 |
| 35.8194444 | 0.041504 | 0.075541 | 0.176148 | 0.027952 | 0.084659 | 0.022585 | 0.009982 | 0.105617 | 0.082275 | 0.040853 | 0.01648 | 0.06058 | 0.026209 | 0.00883 | 0.026408 | 0.055805 | 0.007941 | 0.016191 | 0.034917 | 0.036126 | 0.021978 | 0.027329 | 0.051155 | 0.026084 |
| 35.8333333 | 0.091331 | 0.06228 | 0.166588 | 0.033814 | 0.078749 | 0.026323 | 0.006678 | 0.09463 | 0.072667 | 0.055278 | 0.01796 | 0.060739 | 0.036229 | 0.00936 | 0.029217 | 0.04621 | 0.010209 | 0.006125 | 0.033225 | 0.030458 | 0.008469 | 0.030411 | 0.056846 | 0.028423 |
| 35.8472222 | 0.157767 | 0.049073 | 0.161409 | 0.041038 | 0.082983 | 0.023631 | 0.004528 | 0.084523 | 0.053015 | 0.068941 | 0.025891 | 0.057558 | 0.035791 | 0.007491 | 0.030251 | 0.037102 | 0.011687 | 0.005103 | 0.035172 | 0.025564 | 0.006197 | 0.035609 | 0.067755 | 0.025602 |
| 35.8611111 | 0.229397 | 0.041482 | 0.165969 | 0.038006 | 0.180443 | 0.017632 | 0.009743 | 0.074044 | 0.034313 | 0.080229 | 0.038312 | 0.044441 | 0.02579 | 0.010966 | 0.03319 | 0.029957 | 0.012483 | 0.006428 | 0.043056 | 0.021996 | 0.007646 | 0.037728 | 0.074815 | 0.018856 |
| 35.875 | 0.215977 | 0.040521 | 0.155931 | 0.026626 | 0.414294 | 0.015801 | 0.018682 | 0.064319 | 0.021795 | 0.081711 | 0.043632 | 0.025615 | 0.013233 | 0.018772 | 0.031002 | 0.025792 | 0.012101 | 0.011616 | 0.054406 | 0.017887 | 0.013065 | 0.02854 | 0.074596 | 0.012241 |
| 35.8888889 | 0.11604 | 0.04803 | 0.119014 | 0.015316 | 0.508469 | 0.025312 | 0.027864 | 0.055515 | 0.013524 | 0.074228 | 0.038916 | 0.0365 | 0.011341 | 0.022478 | 0.026277 | 0.021011 | 0.016534 | 0.02622 | 0.062158 | 0.015553 | 0.017875 | 0.015315 | 0.082133 | 0.007059 |
| 35.9027778 | 0.043432 | 0.053725 | 0.07509 | 0.021251 | 0.26019 | 0.041959 | 0.040459 | 0.051948 | 0.017063 | 0.072056 | 0.032171 | 0.084066 | 0.032387 | 0.02572 | 0.02731 | 0.015774 | 0.027369 | 0.040179 | 0.06276 | 0.021481 | 0.015159 | 0.007559 | 0.099242 | 0.005153 |
| 35.9166667 | 0.014277 | 0.073415 | 0.048131 | 0.051133 | 0.074984 | 0.058123 | 0.055099 | 0.057474 | 0.032575 | 0.083945 | 0.029977 | 0.136064 | 0.078095 | 0.029678 | 0.027112 | 0.014959 | 0.037061 | 0.043836 | 0.058688 | 0.026719 | 0.017794 | 0.006406 | 0.113316 | 0.009577 |
| 35.9305556 | 0.004523 | 0.110635 | 0.031225 | 0.096908 | 0.108406 | 0.065476 | 0.063388 | 0.064112 | 0.035825 | 0.101264 | 0.028234 | 0.150037 | 0.135624 | 0.029225 | 0.019356 | 0.015993 | 0.039583 | 0.043495 | 0.052193 | 0.018574 | 0.033135 | 0.012571 | 0.124324 | 0.017556 |
| 35.9444444 | 0.007347 | 0.105951 | 0.013736 | 0.124022 | 0.124268 | 0.057057 | 0.064951 | 0.058698 | 0.039163 | 0.102284 | 0.018673 | 0.096258 | 0.174804 | 0.021515 | 0.016877 | 0.01241 | 0.038184 | 0.045135 | 0.04445 | 0.008536 | 0.047773 | 0.022298 | 0.133801 | 0.024922 |
| 35.9583333 | 0.01798 | 0.058262 | 0.028408 | 0.114694 | 0.089535 | 0.044574 | 0.064746 | 0.037192 | 0.06774 | 0.078567 | 0.017168 | 0.040059 | 0.164955 | 0.010483 | 0.028777 | 0.007585 | 0.036066 | 0.044795 | 0.037823 | 0.006331 | 0.057567 | 0.027934 | 0.133419 | 0.028131 |
| 35.9722222 | 0.032975 | 0.024046 | 0.088153 | 0.087848 | 0.069244 | 0.045474 | 0.060131 | 0.020725 | 0.092624 | 0.044007 | 0.028158 | 0.045406 | 0.12416 | 0.008721 | 0.041136 | 0.007404 | 0.030688 | 0.040624 | 0.033653 | 0.007585 | 0.069325 | 0.028807 | 0.117372 | 0.024482 |
| 35.9861111 | 0.044464 | 0.02381 | 0.101798 | 0.054152 | 0.052427 | 0.064735 | 0.052404 | 0.026421 | 0.088258 | 0.030571 | 0.041336 | 0.095718 | 0.0974 | 0.015571 | 0.045371 | 0.013203 | 0.019566 | 0.039007 | 0.031665 | 0.018837 | 0.082204 | 0.025719 | 0.093744 | 0.019247 |
| 36 | 0.050181 | 0.032341 | 0.081713 | 0.026552 | 0.033124 | 0.078808 | 0.043797 | 0.035972 | 0.060375 | 0.046145 | 0.048185 | 0.124296 | 0.084804 | 0.021047 | 0.045432 | 0.027195 | 0.013532 | 0.042816 | 0.031806 | 0.035064 | 0.088779 | 0.01967 | 0.075666 | 0.01677 |
| 36.0138889 | 0.056607 | 0.033763 | 0.092705 | 0.026402 | 0.044193 | 0.058096 | 0.033733 | 0.040875 | 0.044944 | 0.056959 | 0.03607 | 0.100194 | 0.066997 | 0.025725 | 0.045829 | 0.050954 | 0.022218 | 0.04086 | 0.031334 | 0.039778 | 0.087252 | 0.028613 | 0.071019 | 0.013536 |
| 36.0277778 | 0.068497 | 0.029489 | 0.104505 | 0.058686 | 0.065635 | 0.036104 | 0.02366 | 0.055479 | 0.06303 | 0.04991 | 0.039278 | 0.076312 | 0.058004 | 0.033612 | 0.051537 | 0.084422 | 0.030992 | 0.029013 | 0.02281 | 0.037989 | 0.090794 | 0.066609 | 0.083836 | 0.018176 |
| 36.0416667 | 0.07624 | 0.030324 | 0.108524 | 0.087367 | 0.086908 | 0.059062 | 0.017054 | 0.079549 | 0.071729 | 0.063666 | 0.086415 | 0.061203 | 0.08476 | 0.040219 | 0.066787 | 0.122423 | 0.038874 | 0.025317 | 0.018405 | 0.041896 | 0.105185 | 0.109392 | 0.105794 | 0.033703 |
| 36.0555556 | 0.075782 | 0.050547 | 0.075375 | 0.070996 | 0.110668 | 0.106723 | 0.019288 | 0.091509 | 0.052604 | 0.100797 | 0.115363 | 0.041705 | 0.137361 | 0.038652 | 0.09098 | 0.132727 | 0.051496 | 0.033002 | 0.028705 | 0.048239 | 0.112562 | 0.120941 | 0.117253 | 0.037368 |
| 36.0694444 | 0.070307 | 0.061286 | 0.042257 | 0.053214 | 0.114579 | 0.134107 | 0.021558 | 0.077393 | 0.036792 | 0.090992 | 0.092242 | 0.029534 | 0.163115 | 0.030343 | 0.107453 | 0.099085 | 0.060512 | 0.032357 | 0.030142 | 0.041326 | 0.09996 | 0.102785 | 0.098223 | 0.026235 |
| 36.0833333 | 0.054884 | 0.044556 | 0.038695 | 0.086029 | 0.082833 | 0.131975 | 0.019618 | 0.044927 | 0.028161 | 0.048426 | 0.064606 | 0.018628 | 0.146735 | 0.022317 | 0.097884 | 0.058405 | 0.066426 | 0.01939 | 0.015867 | 0.021539 | 0.07714 | 0.073763 | 0.053817 | 0.017298 |
| 36.0972222 | 0.034447 | 0.020762 | 0.032195 | 0.123688 | 0.050056 | 0.130064 | 0.024862 | 0.017826 | 0.014524 | 0.028491 | 0.043393 | 0.014492 | 0.126335 | 0.019373 | 0.070799 | 0.032571 | 0.070967 | 0.006914 | 0.004438 | 0.00742 | 0.051889 | 0.038072 | 0.020437 | 0.014521 |
| 36.1111111 | 0.016778 | 0.006525 | 0.032284 | 0.121532 | 0.064469 | 0.142438 | 0.038674 | 0.011627 | 0.006056 | 0.013564 | 0.025039 | 0.019527 | 0.110763 | 0.018005 | 0.045378 | 0.017434 | 0.070152 | 0.002379 | 0.003483 | 0.004761 | 0.028823 | 0.015998 | 0.013739 | 0.014963 |
| 36.125 | 0.00541 | 0.006386 | 0.043695 | 0.088527 | 0.098395 | 0.143397 | 0.052338 | 0.012442 | 0.007176 | 0.002629 | 0.01535 | 0.020743 | 0.086653 | 0.013756 | 0.025706 | 0.021176 | 0.058814 | 0.003277 | 0.005718 | 0.009818 | 0.023724 | 0.018201 | 0.017987 | 0.012231 |
| 36.1388889 | 0.005357 | 0.013426 | 0.070157 | 0.046421 | 0.103408 | 0.102365 | 0.058665 | 0.010069 | 0.011748 | 0.003188 | 0.009961 | 0.016242 | 0.059679 | 0.015852 | 0.015305 | 0.053856 | 0.043787 | 0.005567 | 0.004458 | 0.016002 | 0.033142 | 0.02493 | 0.021515 | 0.00853 |
| 36.1527778 | 0.014882 | 0.01733 | 0.101332 | 0.015954 | 0.08328 | 0.041495 | 0.054117 | 0.007322 | 0.018572 | 0.006782 | 0.0131 | 0.012518 | 0.039887 | 0.02874 | 0.024801 | 0.087715 | 0.032365 | 0.009436 | 0.005329 | 0.01646 | 0.036851 | 0.028225 | 0.024239 | 0.009845 |
| 36.1666667 | 0.0316 | 0.016322 | 0.122445 | 0.009599 | 0.060875 | 0.014551 | 0.038651 | 0.009782 | 0.025709 | 0.014911 | 0.019264 | 0.013758 | 0.028749 | 0.035849 | 0.043877 | 0.086694 | 0.018167 | 0.013814 | 0.010659 | 0.01274 | 0.031495 | 0.028495 | 0.022375 | 0.011101 |
| 36.1805556 | 0.048026 | 0.026661 | 0.125087 | 0.020558 | 0.042662 | 0.021556 | 0.021139 | 0.023802 | 0.0225 | 0.027489 | 0.019645 | 0.01701 | 0.023012 | 0.029919 | 0.052891 | 0.065863 | 0.007787 | 0.017251 | 0.017566 | 0.010466 | 0.025134 | 0.020815 | 0.013161 | 0.015376 |
| 36.1944444 | 0.049685 | 0.05562 | 0.104439 | 0.031537 | 0.024426 | 0.023354 | 0.014189 | 0.041374 | 0.014543 | 0.034425 | 0.018463 | 0.020351 | 0.016913 | 0.023784 | 0.053251 | 0.054251 | 0.014116 | 0.018415 | 0.024298 | 0.012341 | 0.026641 | 0.011484 | 0.008174 | 0.020603 |
| 36.2083333 | 0.03757 | 0.081 | 0.085636 | 0.037177 | 0.014715 | 0.02293 | 0.016613 | 0.049709 | 0.013696 | 0.030375 | 0.025954 | 0.028635 | 0.008977 | 0.030905 | 0.052502 | 0.054739 | 0.033117 | 0.013216 | 0.027674 | 0.014339 | 0.037015 | 0.009414 | 0.015281 | 0.019728 |
| 36.2222222 | 0.024063 | 0.087232 | 0.091752 | 0.039625 | 0.020874 | 0.044378 | 0.016057 | 0.045248 | 0.011188 | 0.023405 | 0.04571 | 0.042225 | 0.005722 | 0.039493 | 0.052547 | 0.058892 | 0.052969 | 0.009805 | 0.026755 | 0.01298 | 0.046615 | 0.007951 | 0.026811 | 0.020287 |
| 36.2361111 | 0.01294 | 0.080309 | 0.107671 | 0.037566 | 0.038489 | 0.061906 | 0.013804 | 0.03055 | 0.007379 | 0.019572 | 0.067833 | 0.049642 | 0.014344 | 0.031688 | 0.053786 | 0.059326 | 0.057779 | 0.011586 | 0.023778 | 0.008621 | 0.048477 | 0.008563 | 0.03005 | 0.022235 |
| 36.25 | 0.010454 | 0.070398 | 0.108183 | 0.03504 | 0.066176 | 0.051334 | 0.017804 | 0.01449 | 0.009988 | 0.018553 | 0.085636 | 0.046397 | 0.034785 | 0.020229 | 0.054757 | 0.055185 | 0.039187 | 0.010578 | 0.024396 | 0.003591 | 0.040816 | 0.017877 | 0.021805 | 0.023794 |
| 36.2638889 | 0.024242 | 0.055154 | 0.092523 | 0.036316 | 0.080971 | 0.036813 | 0.020551 | 0.007386 | 0.019349 | 0.013558 | 0.093186 | 0.03273 | 0.053508 | 0.018292 | 0.051853 | 0.050102 | 0.017124 | 0.013013 | 0.028825 | 0.000861 | 0.030524 | 0.026063 | 0.010814 | 0.026949 |
| 36.2777778 | 0.043514 | 0.033408 | 0.081728 | 0.07023 | 0.060836 | 0.039821 | 0.016138 | 0.00793 | 0.023748 | 0.007865 | 0.072717 | 0.012896 | 0.054715 | 0.022628 | 0.042366 | 0.041918 | 0.013427 | 0.016905 | 0.030021 | 0.001122 | 0.027952 | 0.023497 | 0.007698 | 0.027021 |
| 36.2916667 | 0.05304 | 0.015537 | 0.074614 | 0.10534 | 0.029182 | 0.051187 | 0.009119 | 0.009493 | 0.02055 | 0.006457 | 0.03727 | 0.002522 | 0.042268 | 0.028452 | 0.02873 | 0.029641 | 0.021071 | 0.013325 | 0.027134 | 0.003273 | 0.034283 | 0.014276 | 0.014219 | 0.024108 |
| 36.3055556 | 0.049893 | 0.005665 | 0.055475 | 0.069222 | 0.015855 | 0.054209 | 0.00656 | 0.011703 | 0.029642 | 0.010758 | 0.026995 | 0.001139 | 0.033389 | 0.035579 | 0.018586 | 0.020483 | 0.026319 | 0.009208 | 0.025208 | 0.004069 | 0.050086 | 0.008951 | 0.018032 | 0.024626 |
| 36.3194444 | 0.040048 | 0.005356 | 0.031622 | 0.048325 | 0.017181 | 0.04787 | 0.010251 | 0.015876 | 0.045526 | 0.02742 | 0.054021 | 0.000942 | 0.027361 | 0.041804 | 0.015943 | 0.022204 | 0.025547 | 0.011544 | 0.027061 | 0.004769 | 0.070287 | 0.008948 | 0.011686 | 0.031413 |

| | | | | | | | | | | | | | | | | | | | | | | | | |
|---|---|---|---|---|---|---|---|---|---|---|---|---|---|---|---|---|---|---|---|---|---|---|---|---|
| 36.3333333 | 0.030149 | 0.015827 | 0.015608 | 0.0924 | 0.021078 | 0.040187 | 0.011255 | 0.020063 | 0.03807 | 0.044752 | 0.093194 | 0.001619 | 0.016449 | 0.038608 | 0.015153 | 0.031628 | 0.018508 | 0.018204 | 0.030606 | 0.010035 | 0.090447 | 0.013342 | 0.004864 | 0.036516 |
| 36.3472222 | 0.019334 | 0.023632 | 0.007954 | 0.10303 | 0.019086 | 0.028042 | 0.008913 | 0.023428 | 0.018616 | 0.045343 | 0.111014 | 0.007514 | 0.009635 | 0.025688 | 0.010003 | 0.035223 | 0.010732 | 0.022002 | 0.031012 | 0.015462 | 0.105706 | 0.02294 | 0.006524 | 0.029714 |
| 36.3611111 | 0.009501 | 0.016499 | 0.007212 | 0.06479 | 0.011312 | 0.016932 | 0.008036 | 0.026005 | 0.020442 | 0.030871 | 0.10072 | 0.016653 | 0.010422 | 0.01244 | 0.005215 | 0.027991 | 0.0111 | 0.020033 | 0.027414 | 0.015392 | 0.09654 | 0.026694 | 0.010102 | 0.015133 |
| 36.375 | 0.009015 | 0.007539 | 0.013801 | 0.037368 | 0.009555 | 0.017159 | 0.006718 | 0.023631 | 0.033046 | 0.023649 | 0.073299 | 0.017013 | 0.009564 | 0.004657 | 0.005966 | 0.019753 | 0.018137 | 0.016568 | 0.026337 | 0.012653 | 0.063685 | 0.021703 | 0.00782 | 0.005952 |

**Supplementary Table S6.** 1 hour integrated motion data in **(b)** 100 mM NaCl treatment on Amaranth

| IntervalMid | Control1 | Control2 | Control3 | Control4 | Control5 | Control6 | Control7 | Control8 | Control9 | Control10 | Control11 | Control12 | Stress1 | Stress2 | Stress4 | Stress5 | Stress7 | Stress8 | Stress9 | Stress10 | Stress11 | Stress12 |
|---|---|---|---|---|---|---|---|---|---|---|---|---|---|---|---|---|---|---|---|---|---|---|
| 18.1111111 | 0.004178 | 0.000537 | 0.001834 | 0.000947 | 0.001224 | 0.0014 | 0.001489 | 0.001139 | 0.002503 | 0.001219 | 0.001265 | 0.007736 | 0.014277 | 0.012243 | 0.014708 | 0.013575 | 0.007003 | 0.013172 | 0.017341 | 0.008397 | 0.015159 | 0.012894 |
| 18.125 | 0.004113 | 0.000417 | 0.00072 | 0.001116 | 0.001422 | 0.001213 | 0.001468 | 0.000944 | 0.000949 | 0.001123 | 0.002538 | 0.006986 | 0.013652 | 0.012544 | 0.01406 | 0.013939 | 0.010933 | 0.014182 | 0.016652 | 0.00978 | 0.014389 | 0.01123 |
| 18.1388889 | 0.004189 | 0.000626 | 0.000478 | 0.00239 | 0.002439 | 0.00172 | 0.002372 | 0.001391 | 0.00106 | 0.001597 | 0.004233 | 0.005349 | 0.010571 | 0.009697 | 0.010959 | 0.011308 | 0.011996 | 0.01359 | 0.013571 | 0.008535 | 0.010972 | 0.006891 |
| 18.1527778 | 0.002958 | 0.000887 | 0.001205 | 0.004419 | 0.003096 | 0.002022 | 0.003011 | 0.001788 | 0.002103 | 0.001931 | 0.005521 | 0.003776 | 0.005766 | 0.005312 | 0.006338 | 0.006907 | 0.009751 | 0.010365 | 0.009 | 0.005187 | 0.005829 | 0.003844 |
| 18.1666667 | 0.002309 | 0.000842 | 0.002031 | 0.006181 | 0.003211 | 0.001875 | 0.003436 | 0.002036 | 0.002939 | 0.001855 | 0.006248 | 0.003004 | 0.001889 | 0.001635 | 0.00246 | 0.002619 | 0.00562 | 0.006224 | 0.00482 | 0.001813 | 0.002259 | 0.005803 |
| 18.1805556 | 0.001864 | 0.000912 | 0.002114 | 0.00715 | 0.003212 | 0.001625 | 0.004004 | 0.002223 | 0.003391 | 0.001813 | 0.006468 | 0.003019 | 0.001305 | 0.001099 | 0.001837 | 0.001338 | 0.002637 | 0.004593 | 0.003617 | 0.001221 | 0.001639 | 0.008296 |
| 18.1944444 | 0.001291 | 0.00151 | 0.001386 | 0.006983 | 0.003344 | 0.001578 | 0.004694 | 0.002496 | 0.003707 | 0.002101 | 0.006506 | 0.003404 | 0.002688 | 0.00331 | 0.004043 | 0.003346 | 0.002403 | 0.00608 | 0.005561 | 0.003351 | 0.002301 | 0.007662 |
| 18.2083333 | 0.001908 | 0.001982 | 0.000696 | 0.005756 | 0.003247 | 0.00164 | 0.004878 | 0.002758 | 0.003906 | 0.002246 | 0.006257 | 0.003447 | 0.003777 | 0.004999 | 0.005442 | 0.004951 | 0.002604 | 0.0068 | 0.006864 | 0.00485 | 0.003025 | 0.007275 |
| 18.2222222 | 0.002597 | 0.001763 | 0.000556 | 0.004312 | 0.002727 | 0.001741 | 0.004515 | 0.002903 | 0.003851 | 0.001919 | 0.005414 | 0.002922 | 0.003299 | 0.003951 | 0.004186 | 0.003958 | 0.00238 | 0.005258 | 0.005248 | 0.003957 | 0.00333 | 0.010322 |
| 18.2361111 | 0.002065 | 0.001272 | 0.001094 | 0.003606 | 0.002306 | 0.002096 | 0.004268 | 0.003067 | 0.003623 | 0.001702 | 0.004634 | 0.002511 | 0.003078 | 0.002715 | 0.003269 | 0.003104 | 0.003635 | 0.005341 | 0.004059 | 0.003291 | 0.004656 | 0.014308 |
| 18.25 | 0.001109 | 0.00134 | 0.002251 | 0.003711 | 0.002533 | 0.002648 | 0.004382 | 0.003182 | 0.003467 | 0.002143 | 0.004899 | 0.002766 | 0.003791 | 0.003195 | 0.003807 | 0.003379 | 0.004685 | 0.006662 | 0.004431 | 0.003297 | 0.005384 | 0.014214 |
| 18.2638889 | 0.001098 | 0.002191 | 0.003092 | 0.003913 | 0.003135 | 0.002923 | 0.004628 | 0.003074 | 0.003476 | 0.002869 | 0.005463 | 0.003292 | 0.003002 | 0.002665 | 0.002843 | 0.002282 | 0.003412 | 0.005083 | 0.003201 | 0.002113 | 0.00329 | 0.01038 |
| 18.2777778 | 0.002092 | 0.002903 | 0.003092 | 0.003734 | 0.003348 | 0.002796 | 0.004628 | 0.00296 | 0.003434 | 0.003134 | 0.004908 | 0.003451 | 0.001757 | 0.001247 | 0.001174 | 0.000869 | 0.002065 | 0.002741 | 0.001758 | 0.000915 | 0.001454 | 0.007372 |
| 18.2916667 | 0.002147 | 0.002524 | 0.002823 | 0.002995 | 0.002753 | 0.002488 | 0.003885 | 0.00276 | 0.003015 | 0.002659 | 0.003265 | 0.002937 | 0.002347 | 0.001527 | 0.001558 | 0.001542 | 0.002428 | 0.003222 | 0.002813 | 0.001137 | 0.002052 | 0.006665 |
| 18.3055556 | 0.001143 | 0.001386 | 0.002489 | 0.001808 | 0.001707 | 0.001984 | 0.002536 | 0.001991 | 0.002229 | 0.001802 | 0.001533 | 0.001909 | 0.003937 | 0.003207 | 0.003465 | 0.003521 | 0.003437 | 0.005231 | 0.00498 | 0.002749 | 0.003748 | 0.006704 |
| 18.3194444 | 0.000888 | 0.000805 | 0.002145 | 0.001041 | 0.001207 | 0.001474 | 0.001637 | 0.001123 | 0.001683 | 0.001432 | 0.000872 | 0.001306 | 0.005236 | 0.005004 | 0.005356 | 0.005311 | 0.004136 | 0.006793 | 0.006659 | 0.004674 | 0.005198 | 0.006548 |
| 18.3333333 | 0.001662 | 0.001491 | 0.002074 | 0.000781 | 0.001676 | 0.001367 | 0.001759 | 0.001035 | 0.001784 | 0.001949 | 0.001585 | 0.001721 | 0.005608 | 0.005941 | 0.005969 | 0.005877 | 0.003863 | 0.006897 | 0.007144 | 0.005347 | 0.005509 | 0.005443 |
| 18.3472222 | 0.002259 | 0.002136 | 0.001968 | 0.000701 | 0.002001 | 0.001449 | 0.001959 | 0.001316 | 0.001878 | 0.002308 | 0.002129 | 0.002027 | 0.003993 | 0.004537 | 0.004085 | 0.004082 | 0.00296 | 0.004609 | 0.005264 | 0.003804 | 0.003908 | 0.003653 |
| 18.3611111 | 0.001666 | 0.001503 | 0.001545 | 0.001218 | 0.001292 | 0.001243 | 0.00133 | 0.001102 | 0.001321 | 0.001608 | 0.001416 | 0.001283 | 0.002355 | 0.002371 | 0.001992 | 0.001977 | 0.003689 | 0.002471 | 0.002634 | 0.001459 | 0.002682 | 0.003703 |
| 18.375 | 0.000817 | 0.000523 | 0.001131 | 0.00189 | 0.000378 | 0.000909 | 0.000583 | 0.000675 | 0.000666 | 0.000713 | 0.000579 | 0.000461 | 0.003211 | 0.002504 | 0.002617 | 0.002435 | 0.005862 | 0.003194 | 0.002367 | 0.000611 | 0.003969 | 0.005833 |
| 18.3888889 | 0.000535 | 0.000357 | 0.000941 | 0.002119 | 0.000233 | 0.000882 | 0.000661 | 0.000681 | 0.000673 | 0.000648 | 0.000577 | 0.00045 | 0.004409 | 0.003541 | 0.00391 | 0.003682 | 0.007152 | 0.004597 | 0.003468 | 0.001109 | 0.005852 | 0.007379 |
| 18.4027778 | 0.000569 | 0.000944 | 0.000917 | 0.002154 | 0.000755 | 0.001157 | 0.001294 | 0.001049 | 0.001179 | 0.00118 | 0.000883 | 0.000897 | 0.004192 | 0.003421 | 0.003777 | 0.003654 | 0.006908 | 0.004693 | 0.003549 | 0.001772 | 0.006517 | 0.007114 |
| 18.4166667 | 0.000815 | 0.001529 | 0.00102 | 0.002325 | 0.001291 | 0.001474 | 0.001755 | 0.001402 | 0.001576 | 0.001663 | 0.000898 | 0.001281 | 0.003123 | 0.002262 | 0.002509 | 0.002507 | 0.005634 | 0.003531 | 0.002182 | 0.001699 | 0.005706 | 0.00548 |
| 18.4305556 | 0.000883 | 0.001698 | 0.001198 | 0.002655 | 0.001512 | 0.001733 | 0.001823 | 0.001741 | 0.001646 | 0.001896 | 0.000817 | 0.001406 | 0.001888 | 0.00093 | 0.001061 | 0.00108 | 0.004282 | 0.001766 | 0.000949 | 0.000939 | 0.00402 | 0.003579 |
| 18.4444444 | 0.001026 | 0.002219 | 0.001819 | 0.003785 | 0.00222 | 0.002601 | 0.002557 | 0.003723 | 0.002107 | 0.002703 | 0.001732 | 0.002114 | 0.001584 | 0.001081 | 0.001137 | 0.001068 | 0.003 | 0.001604 | 0.002537 | 0.00112 | 0.00309 | 0.002744 |
| 18.4583333 | 0.001205 | 0.004524 | 0.002985 | 0.006619 | 0.004861 | 0.005354 | 0.005915 | 0.009901 | 0.003947 | 0.005254 | 0.005561 | 0.005079 | 0.003607 | 0.003956 | 0.003726 | 0.003427 | 0.003123 | 0.005058 | 0.007921 | 0.003621 | 0.005188 | 0.004799 |
| 18.4722222 | 0.000781 | 0.008649 | 0.005803 | 0.007134 | 0.009096 | 0.008068 | 0.012663 | 0.016781 | 0.004712 | 0.008269 | 0.012141 | 0.008658 | 0.005995 | 0.008074 | 0.007399 | 0.005885 | 0.005716 | 0.010316 | 0.013494 | 0.00656 | 0.008415 | 0.008287 |
| 18.4861111 | 0.005136 | 0.014694 | 0.016936 | 0.009255 | 0.012957 | 0.007259 | 0.022487 | 0.016104 | 0.004277 | 0.009304 | 0.015338 | 0.00797 | 0.006282 | 0.010716 | 0.01205 | 0.00568 | 0.007517 | 0.011056 | 0.014806 | 0.006986 | 0.007784 | 0.008157 |
| 18.5 | 0.016787 | 0.022152 | 0.0336 | 0.019319 | 0.015232 | 0.005192 | 0.031366 | 0.009456 | 0.004776 | 0.012065 | 0.010962 | 0.003922 | 0.00482 | 0.012337 | 0.019416 | 0.004801 | 0.006987 | 0.006465 | 0.014168 | 0.006202 | 0.007909 | 0.007386 |
| 18.5138889 | 0.021931 | 0.022716 | 0.03142 | 0.019844 | 0.012937 | 0.004105 | 0.026852 | 0.007263 | 0.006992 | 0.017194 | 0.007328 | 0.00195 | 0.004484 | 0.013168 | 0.021133 | 0.005494 | 0.006104 | 0.003227 | 0.016569 | 0.005775 | 0.008746 | 0.01156 |
| 18.5277778 | 0.017488 | 0.018369 | 0.022994 | 0.015122 | 0.008739 | 0.00534 | 0.018091 | 0.010795 | 0.00971 | 0.016231 | 0.007181 | 0.002112 | 0.013574 | 0.013592 | 0.011863 | 0.010604 | 0.006368 | 0.005428 | 0.020466 | 0.009671 | 0.012721 | 0.015294 |
| 18.5416667 | 0.015574 | 0.021357 | 0.037728 | 0.025028 | 0.009506 | 0.011012 | 0.024058 | 0.014914 | 0.006822 | 0.012185 | 0.005091 | 0.001992 | 0.038456 | 0.026028 | 0.013156 | 0.031261 | 0.019163 | 0.021487 | 0.031048 | 0.028579 | 0.038208 | 0.022142 |
| 18.5555556 | 0.013905 | 0.022441 | 0.045257 | 0.029093 | 0.009482 | 0.014456 | 0.026174 | 0.016221 | 0.001794 | 0.010015 | 0.003724 | 0.00269 | 0.067667 | 0.053777 | 0.038155 | 0.062486 | 0.048206 | 0.052738 | 0.053017 | 0.05541 | 0.073633 | 0.039504 |
| 18.5694444 | 0.00773 | 0.012869 | 0.027743 | 0.016027 | 0.004828 | 0.010333 | 0.013807 | 0.010684 | 0.001214 | 0.005724 | 0.002774 | 0.00558 | 0.073883 | 0.066535 | 0.055381 | 0.071529 | 0.061575 | 0.068659 | 0.061149 | 0.061286 | 0.080501 | 0.046977 |
| 18.5833333 | 0.002856 | 0.004315 | 0.009757 | 0.004949 | 0.00156 | 0.004378 | 0.004644 | 0.003867 | 0.002821 | 0.001895 | 0.0017 | 0.0086 | 0.051009 | 0.047975 | 0.040875 | 0.048661 | 0.041156 | 0.049416 | 0.040191 | 0.039041 | 0.052431 | 0.031019 |
| 18.5972222 | 0.002706 | 0.003457 | 0.002298 | 0.003412 | 0.001068 | 0.001814 | 0.004227 | 0.002662 | 0.003651 | 0.000825 | 0.003216 | 0.008863 | 0.022673 | 0.021646 | 0.016403 | 0.02081 | 0.017095 | 0.020031 | 0.015421 | 0.014608 | 0.02017 | 0.011431 |
| 18.6111111 | 0.004307 | 0.005164 | 0.001455 | 0.003896 | 0.001569 | 0.001291 | 0.006351 | 0.002771 | 0.002707 | 0.000649 | 0.004974 | 0.006999 | 0.006956 | 0.006916 | 0.004792 | 0.006351 | 0.012063 | 0.005428 | 0.007143 | 0.006327 | 0.006239 | 0.004151 |
| 18.625 | 0.004951 | 0.005334 | 0.002105 | 0.00273 | 0.002096 | 0.000849 | 0.007297 | 0.002102 | 0.001526 | 0.000681 | 0.005163 | 0.006175 | 0.004305 | 0.003908 | 0.005383 | 0.003935 | 0.014253 | 0.005457 | 0.01031 | 0.007983 | 0.007679 | 0.006321 |
| 18.6388889 | 0.004765 | 0.004393 | 0.001949 | 0.001709 | 0.002592 | 0.000754 | 0.006747 | 0.002295 | 0.001494 | 0.000762 | 0.00469 | 0.00728 | 0.008305 | 0.007194 | 0.009743 | 0.00755 | 0.014286 | 0.01106 | 0.016009 | 0.011093 | 0.013425 | 0.011567 |
| 18.6527778 | 0.003695 | 0.002877 | 0.00114 | 0.000961 | 0.002916 | 0.000689 | 0.004911 | 0.002557 | 0.001857 | 0.000701 | 0.003895 | 0.008224 | 0.013212 | 0.011577 | 0.013608 | 0.012148 | 0.014287 | 0.016714 | 0.021075 | 0.014428 | 0.018138 | 0.016912 |
| 18.6666667 | 0.002023 | 0.001396 | 0.000795 | 0.000559 | 0.002432 | 0.000561 | 0.002404 | 0.002016 | 0.001664 | 0.000554 | 0.002523 | 0.007264 | 0.016324 | 0.014759 | 0.016243 | 0.015329 | 0.014129 | 0.020747 | 0.023889 | 0.01748 | 0.021296 | 0.020866 |
| 18.6805556 | 0.001101 | 0.000765 | 0.000773 | 0.000755 | 0.001622 | 0.000889 | 0.000574 | 0.0012 | 0.001233 | 0.000764 | 0.001165 | 0.005076 | 0.017718 | 0.016627 | 0.01736 | 0.016948 | 0.013848 | 0.022557 | 0.023851 | 0.019559 | 0.02276 | 0.022811 |
| 18.6944444 | 0.000905 | 0.000507 | 0.000589 | 0.000703 | 0.001103 | 0.001053 | 9.19E-05 | 0.000761 | 0.001044 | 0.000744 | 0.001171 | 0.003191 | 0.018977 | 0.01822 | 0.018523 | 0.01833 | 0.015204 | 0.023971 | 0.022928 | 0.021425 | 0.024205 | 0.024288 |
| 18.7083333 | 0.000592 | 0.000258 | 0.000815 | 0.000464 | 0.00065 | 0.000655 | 0.000199 | 0.000653 | 0.000916 | 0.000405 | 0.00246 | 0.001886 | 0.021643 | 0.020818 | 0.021468 | 0.020957 | 0.019123 | 0.027165 | 0.023937 | 0.024166 | 0.027618 | 0.027226 |
| 18.7222222 | 0.000359 | 0.000408 | 0.001526 | 0.000833 | 0.000334 | 0.00025 | 0.000374 | 0.000694 | 0.000808 | 0.000336 | 0.002786 | 0.00125 | 0.024974 | 0.024044 | 0.025132 | 0.02415 | 0.02348 | 0.030845 | 0.026563 | 0.026761 | 0.032144 | 0.030647 |
| 18.7361111 | 0.000376 | 0.000731 | 0.001777 | 0.001218 | 0.000292 | 0.000259 | 0.000484 | 0.000638 | 0.000641 | 0.000394 | 0.001784 | 0.001041 | 0.025419 | 0.024575 | 0.025549 | 0.02446 | 0.024178 | 0.030477 | 0.026758 | 0.025814 | 0.033268 | 0.030713 |
| 18.75 | 0.000782 | 0.001128 | 0.001137 | 0.000924 | 0.000627 | 0.000764 | 0.000632 | 0.000426 | 0.000597 | 0.000661 | 0.00174 | 0.000911 | 0.020433 | 0.019461 | 0.020303 | 0.019367 | 0.019081 | 0.023908 | 0.021452 | 0.019348 | 0.027334 | 0.024643 |
| 18.7638889 | 0.001467 | 0.002057 | 0.000364 | 0.00042 | 0.001384 | 0.001339 | 0.001261 | 0.000346 | 0.001076 | 0.001395 | 0.003087 | 0.001054 | 0.012417 | 0.011102 | 0.01201 | 0.011241 | 0.010731 | 0.014475 | 0.012792 | 0.010498 | 0.016857 | 0.014743 |
| 18.7777778 | 0.001372 | 0.002497 | 0.00013 | 0.000248 | 0.001528 | 0.001427 | 0.002037 | 0.00047 | 0.001451 | 0.001592 | 0.004075 | 0.000898 | 0.005867 | 0.004557 | 0.005139 | 0.004639 | 0.004089 | 0.006812 | 0.005465 | 0.004179 | 0.007704 | 0.006375 |
| 18.7916667 | 0.001074 | 0.002014 | 0.000554 | 0.000304 | 0.001123 | 0.001452 | 0.002924 | 0.000945 | 0.001544 | 0.001261 | 0.004467 | 0.000661 | 0.002482 | 0.001847 | 0.001953 | 0.001793 | 0.001957 | 0.002891 | 0.002122 | 0.001498 | 0.003149 | 0.002231 |
| 18.8055556 | 0.001889 | 0.002146 | 0.001544 | 0.000682 | 0.001501 | 0.001939 | 0.004351 | 0.001973 | 0.001965 | 0.001765 | 0.005078 | 0.001465 | 0.001602 | 0.001862 | 0.001686 | 0.001835 | 0.002549 | 0.002122 | 0.002045 | 0.000693 | 0.002655 | 0.001268 |
| 18.8194444 | 0.002737 | 0.002921 | 0.002088 | 0.00111 | 0.002225 | 0.002132 | 0.005256 | 0.002761 | 0.002259 | 0.002612 | 0.005227 | 0.002542 | 0.001438 | 0.002081 | 0.001565 | 0.001871 | 0.002219 | 0.001906 | 0.001891 | 0.000638 | 0.002756 | 0.001096 |
| 18.8333333 | 0.002462 | 0.003145 | 0.001736 | 0.00102 | 0.002133 | 0.001668 | 0.004873 | 0.002619 | 0.001922 | 0.002526 | 0.004071 | 0.002433 | 0.000838 | 0.001505 | 0.000764 | 0.001096 | 0.00127 | 0.000951 | 0.000772 | 0.000685 | 0.001847 | 0.000624 |
| 18.8472222 | 0.001889 | 0.002797 | 0.001616 | 0.000848 | 0.00153 | 0.001419 | 0.004085 | 0.002176 | 0.001617 | 0.001817 | 0.002293 | 0.001629 | 0.00143 | 0.001926 | 0.001396 | 0.001802 | 0.002811 | 0.001315 | 0.001575 | 0.001391 | 0.002485 | 0.00102 |
| 18.8611111 | 0.00196 | 0.002486 | 0.00204 | 0.00095 | 0.001189 | 0.001549 | 0.003351 | 0.001808 | 0.001766 | 0.001361 | 0.001009 | 0.001396 | 0.004049 | 0.004052 | 0.004167 | 0.004526 | 0.006868 | 0.003824 | 0.004928 | 0.004067 | 0.005686 | 0.003181 |
| 18.875 | 0.002485 | 0.00254 | 0.002027 | 0.000943 | 0.001201 | 0.001349 | 0.002622 | 0.001109 | 0.001892 | 0.001315 | 0.000593 | 0.001934 | 0.006312 | 0.005866 | 0.006337 | 0.006756 | 0.010145 | 0.005438 | 0.007579 | 0.00674 | 0.008614 | 0.005429 |
| 18.8888889 | 0.002946 | 0.002976 | 0.00134 | 0.000789 | 0.001345 | 0.000853 | 0.002301 | 0.000379 | 0.001759 | 0.00143 | 0.000732 | 0.002694 | 0.0062 | 0.005605 | 0.005922 | 0.006619 | 0.010279 | 0.004009 | 0.007576 | 0.007044 | 0.008933 | 0.005836 |
| 18.9027778 | 0.003218 | 0.003507 | 0.000668 | 0.000674 | 0.001314 | 0.000569 | 0.002434 | 0.000238 | 0.001582 | 0.001354 | 0.00085 | 0.003102 | 0.00427 | 0.003679 | 0.004078 | 0.004742 | 0.007707 | 0.00161 | 0.006178 | 0.005556 | 0.007031 | 0.004764 |
| 18.9166667 | 0.003458 | 0.003764 | 0.000476 | 0.000485 | 0.000912 | 0.000514 | 0.002339 | 0.000442 | 0.001452 | 0.000855 | 0.000617 | 0.003002 | 0.002256 | 0.001576 | 0.002497 | 0.002752 | 0.00445 | 0.000932 | 0.004876 | 0.003584 | 0.00448 | 0.003532 |
| 18.9305556 | 0.003811 | 0.003603 | 0.000784 | 0.000333 | 0.000457 | 0.000664 | 0.001822 | 0.000865 | 0.001481 | 0.00029 | 0.000305 | 0.002724 | 0.0017 | 0.000572 | 0.002225 | 0.002001 | 0.002816 | 0.003129 | 0.004625 | 0.002485 | 0.003187 | 0.003288 |
| 18.9444444 | 0.00437 | 0.003321 | 0.001198 | 0.000502 | 0.000516 | 0.001006 | 0.001511 | 0.001706 | 0.001872 | 0.000211 | 0.000473 | 0.002851 | 0.002842 | 0.001092 | 0.0033 | 0.002785 | 0.00355 | 0.007697 | 0.005703 | 0.003146 | 0.004064 | 0.004554 |
| 18.9583333 | 0.005161 | 0.003305 | 0.001087 | 0.000746 | 0.001061 | 0.001032 | 0.001736 | 0.002335 | 0.002263 | 0.000566 | 0.001083 | 0.003606 | 0.003877 | 0.00198 | 0.003975 | 0.003542 | 0.004708 | 0.011237 | 0.006481 | 0.004377 | 0.005322 | 0.005912 |
| 18.9722222 | 0.006099 | 0.003296 | 0.000567 | 0.000686 | 0.001501 | 0.000575 | 0.002139 | 0.002164 | 0.002159 | 0.000826 | 0.001333 | 0.004359 | 0.002906 | 0.001774 | 0.002847 | 0.002616 | 0.004157 | 0.010808 | 0.005216 | 0.004024 | 0.004534 | 0.005201 |
| 18.9861111 | 0.00673 | 0.00277 | 0.000305 | 0.000385 | 0.001513 | 0.000241 | 0.002146 | 0.00164 | 0.001712 | 0.0006 | 0.00086 | 0.004407 | 0.001664 | 0.001613 | 0.001992 | 0.0017 | 0.002203 | 0.007342 | 0.002872 | 0.00232 | 0.002538 | 0.002924 |
| 19 | 0.005899 | 0.001846 | 0.000955 | 0.001016 | 0.001071 | 0.000925 | 0.001436 | 0.001933 | 0.001135 | 0.000556 | 0.001473 | 0.00308 | 0.003188 | 0.003128 | 0.003464 | 0.003232 | 0.001376 | 0.003849 | 0.002615 | 0.002105 | 0.002903 | 0.001987 |
| 19.0138889 | 0.004743 | 0.002312 | 0.005293 | 0.006284 | 0.00051 | 0.004377 | 0.001638 | 0.006612 | 0.00094 | 0.000952 | 0.005872 | 0.003181 | 0.006524 | 0.0043 | 0.004422 | 0.006554 | 0.001684 | 0.003317 | 0.004864 | 0.005027 | 0.00743 | 0.002352 |
| 19.0277778 | 0.009265 | 0.006952 | 0.017006 | 0.019048 | 0.001194 | 0.011552 | 0.006414 | 0.017367 | 0.001348 | 0.001736 | 0.013211 | 0.008625 | 0.009855 | 0.004439 | 0.005678 | 0.00896 | 0.006774 | 0.004423 | 0.005761 | 0.00877 | 0.012605 | 0.005832 |
| 19.0416667 | 0.014252 | 0.011874 | 0.027773 | 0.029588 | 0.002655 | 0.017185 | 0.011637 | 0.025423 | 0.002958 | 0.002362 | 0.016221 | 0.014526 | 0.009814 | 0.006386 | 0.012298 | 0.008593 | 0.021781 | 0.00664 | 0.005957 | 0.009777 | 0.010926 | 0.01562 |
| 19.0555556 | 0.012548 | 0.012745 | 0.02581 | 0.025947 | 0.004449 | 0.01352 | 0.014592 | 0.020265 | 0.006346 | 0.004307 | 0.010525 | 0.012334 | 0.007682 | 0.006966 | 0.015309 | 0.007843 | 0.026462 | 0.00901 | 0.006936 | 0.009775 | 0.00459 | 0.019406 |
| 19.0694444 | 0.013247 | 0.01516 | 0.022617 | 0.020523 | 0.008104 | 0.007072 | 0.022216 | 0.009984 | 0.007754 | 0.009246 | 0.004611 | 0.006545 | 0.008133 | 0.005932 | 0.012786 | 0.006527 | 0.015658 | 0.008837 | 0.008377 | 0.009014 | 0.002332 | 0.015185 |
| 19.0833333 | 0.013611 | 0.016446 | 0.02357 | 0.021393 | 0.009655 | 0.00599 | 0.026195 | 0.006243 | 0.005209 | 0.010047 | 0.003307 | 0.005345 | 0.007599 | 0.008119 | 0.014228 | 0.004182 | 0.009873 | 0.009797 | 0.010016 | 0.005233 | 0.003412 | 0.015092 |
| 19.0972222 | 0.006847 | 0.009547 | 0.015554 | 0.015897 | 0.005815 | 0.004119 | 0.01701 | 0.004039 | 0.002764 | 0.004963 | 0.00182 | 0.004478 | 0.006847 | 0.009531 | 0.013254 | 0.005225 | 0.005665 | 0.010953 | 0.007076 | 0.003259 | 0.005636 | 0.013741 |
| 19.1111111 | 0.001682 | 0.002336 | 0.004728 | 0.006936 | 0.001717 | 0.000927 | 0.00611 | 0.001337 | 0.00199 | 0.001705 | 0.000949 | 0.002333 | 0.008096 | 0.00869 | 0.009268 | 0.007385 | 0.002752 | 0.009981 | 0.002097 | 0.003492 | 0.007687 | 0.010173 |
| 19.125 | 0.001565 | 0.000705 | 0.001038 | 0.003453 | 0.000729 | 0.000396 | 0.00193 | 0.001455 | 0.001851 | 0.001209 | 0.002693 | 0.002054 | 0.007167 | 0.007717 | 0.007216 | 0.006068 | 0.005632 | 0.007886 | 0.001671 | 0.002979 | 0.006581 | 0.008812 |
| 19.1388889 | 0.00207 | 0.000921 | 0.00096 | 0.003077 | 0.00088 | 0.00108 | 0.001016 | 0.002707 | 0.002832 | 0.001088 | 0.005239 | 0.002619 | 0.003809 | 0.004921 | 0.00431 | 0.002992 | 0.006879 | 0.004398 | 0.005156 | 0.004494 | 0.003663 | 0.005867 |
| 19.1527778 | 0.001475 | 0.001009 | 0.001537 | 0.00294 | 0.000869 | 0.001671 | 0.000725 | 0.003312 | 0.004719 | 0.001065 | 0.007098 | 0.00293 | 0.001458 | 0.002213 | 0.001721 | 0.00113 | 0.004399 | 0.0021 | 0.00859 | 0.007516 | 0.002099 | 0.002528 |
| 19.1666667 | 0.00065 | 0.000936 | 0.002017 | 0.002347 | 0.000794 | 0.001734 | 0.001325 | 0.002973 | 0.006121 | 0.001027 | 0.007266 | 0.00291 | 0.00096 | 0.001729 | 0.001031 | 0.000693 | 0.002252 | 0.002156 | 0.008186 | 0.008248 | 0.001654 | 0.001542 |
| 19.1805556 | 0.000395 | 0.000658 | 0.002011 | 0.001643 | 0.000738 | 0.001445 | 0.002324 | 0.002232 | 0.006215 | 0.000974 | 0.006254 | 0.002888 | 0.001059 | 0.001179 | 0.000723 | 0.000753 | 0.002987 | 0.003021 | 0.004405 | 0.006354 | 0.001336 | 0.001661 |
| 19.1944444 | 0.000666 | 0.000545 | 0.001584 | 0.00127 | 0.000854 | 0.001235 | 0.003249 | 0.001648 | 0.005272 | 0.001012 | 0.005238 | 0.003101 | 0.001566 | 0.000952 | 0.000826 | 0.001362 | 0.003677 | 0.00466 | 0.001536 | 0.004026 | 0.002339 | 0.001864 |
| 19.2083333 | 0.000751 | 0.000695 | 0.001025 | 0.001222 | 0.001101 | 0.001212 | 0.004121 | 0.001287 | 0.004131 | 0.001121 | 0.004324 | 0.003518 | 0.002106 | 0.00274 | 0.00185 | 0.002059 | 0.002179 | 0.00682 | 0.001657 | 0.002446 | 0.004525 | 0.002534 |
| 19.2222222 | 0.000487 | 0.00086 | 0.000526 | 0.001218 | 0.001189 | 0.001245 | 0.004653 | 0.000997 | 0.003117 | 0.001123 | 0.003537 | 0.00403 | 0.002313 | 0.004532 | 0.002737 | 0.002463 | 0.000864 | 0.008034 | 0.002624 | 0.001792 | 0.006397 | 0.003763 |
| 19.2361111 | 0.000306 | 0.000918 | 0.000296 | 0.001138 | 0.000924 | 0.001214 | 0.004296 | 0.000767 | 0.002083 | 0.000939 | 0.002964 | 0.004446 | 0.002798 | 0.005022 | 0.003031 | 0.003133 | 0.000841 | 0.008258 | 0.002864 | 0.002108 | 0.007936 | 0.003973 |
| 19.25 | 0.00035 | 0.000807 | 0.000445 | 0.001011 | 0.000456 | 0.001085 | 0.002852 | 0.000642 | 0.001194 | 0.000669 | 0.002343 | 0.004525 | 0.003286 | 0.005119 | 0.003054 | 0.003733 | 0.001541 | 0.007597 | 0.002728 | 0.002563 | 0.009111 | 0.002592 |
| 19.2638889 | 0.000416 | 0.000514 | 0.000955 | 0.001049 | 0.000145 | 0.001129 | 0.001407 | 0.00074 | 0.00085 | 0.000521 | 0.001601 | 0.004565 | 0.002593 | 0.005832 | 0.002499 | 0.003195 | 0.002805 | 0.005434 | 0.00195 | 0.002033 | 0.008733 | 0.001527 |
| 19.2777778 | 0.000597 | 0.000492 | 0.001503 | 0.00145 | 0.000314 | 0.001553 | 0.001228 | 0.00121 | 0.001112 | 0.000776 | 0.001379 | 0.005232 | 0.001521 | 0.005221 | 0.001703 | 0.002298 | 0.004418 | 0.003361 | 0.001099 | 0.001263 | 0.007528 | 0.001312 |
| 19.2916667 | 0.001166 | 0.000933 | 0.001456 | 0.001872 | 0.000868 | 0.001929 | 0.001871 | 0.001812 | 0.001424 | 0.001342 | 0.002092 | 0.006321 | 0.002331 | 0.003205 | 0.00249 | 0.003063 | 0.007921 | 0.004363 | 0.002288 | 0.002067 | 0.00814 | 0.001847 |
| 19.3055556 | 0.001525 | 0.001059 | 0.000833 | 0.001871 | 0.001298 | 0.001871 | 0.001834 | 0.002164 | 0.001238 | 0.001591 | 0.00268 | 0.006718 | 0.005344 | 0.004451 | 0.005338 | 0.005733 | 0.012976 | 0.008454 | 0.005692 | 0.004391 | 0.010894 | 0.004707 |
| 19.3194444 | 0.001478 | 0.000691 | 0.00034 | 0.001585 | 0.001514 | 0.001626 | 0.001037 | 0.00224 | 0.000685 | 0.001374 | 0.002347 | 0.006032 | 0.009229 | 0.008689 | 0.008911 | 0.00899 | 0.016245 | 0.012529 | 0.00931 | 0.006931 | 0.013958 | 0.009587 |
| 19.3333333 | 0.001298 | 0.000445 | 0.000263 | 0.001147 | 0.001664 | 0.001297 | 0.000781 | 0.001972 | 0.000433 | 0.001062 | 0.001645 | 0.004669 | 0.012398 | 0.012185 | 0.011563 | 0.011335 | 0.015417 | 0.013496 | 0.011529 | 0.008725 | 0.015766 | 0.013876 |
| 19.3472222 | 0.000764 | 0.000478 | 0.000292 | 0.000589 | 0.001475 | 0.000818 | 0.001809 | 0.001241 | 0.000889 | 0.000688 | 0.000902 | 0.002875 | 0.012839 | 0.012495 | 0.01159 | 0.010904 | 0.010579 | 0.010437 | 0.011157 | 0.008091 | 0.014309 | 0.014755 |
| 19.3611111 | 0.000206 | 0.000911 | 0.000433 | 0.000215 | 0.000866 | 0.000471 | 0.003435 | 0.000642 | 0.001618 | 0.000294 | 0.00045 | 0.001358 | 0.009615 | 0.008709 | 0.008206 | 0.006774 | 0.005453 | 0.005361 | 0.007435 | 0.00509 | 0.008542 | 0.010701 |
| 19.375 | 5.84E-05 | 0.001694 | 0.00072 | 0.000102 | 0.000517 | 0.000348 | 0.003378 | 0.000917 | 0.002081 | 0.000196 | 0.000857 | 0.001317 | 0.004665 | 0.004137 | 0.004023 | 0.003238 | 0.004736 | 0.002445 | 0.003901 | 0.003785 | 0.004157 | 0.005151 |
| 19.3888889 | 7.98E-05 | 0.002101 | 0.000651 | 0.000245 | 0.000845 | 0.00033 | 0.002761 | 0.001456 | 0.00206 | 0.000258 | 0.001407 | 0.002677 | 0.001336 | 0.002306 | 0.002362 | 0.004133 | 0.006941 | 0.002647 | 0.004728 | 0.005041 | 0.005763 | 0.002803 |
| 19.4027778 | 0.000134 | 0.001582 | 0.000368 | 0.000681 | 0.001166 | 0.000299 | 0.004584 | 0.001138 | 0.001562 | 0.000336 | 0.001107 | 0.004002 | 0.00067 | 0.001783 | 0.001904 | 0.00548 | 0.007289 | 0.002484 | 0.007299 | 0.004973 | 0.007878 | 0.002528 |
| 19.4166667 | 0.000274 | 0.000813 | 0.000261 | 0.000996 | 0.001159 | 0.000237 | 0.006488 | 0.000405 | 0.001022 | 0.000623 | 0.000494 | 0.004831 | 0.001217 | 0.001388 | 0.001307 | 0.004469 | 0.005295 | 0.001324 | 0.008228 | 0.002941 | 0.00725 | 0.001591 |
| 19.4305556 | 0.00028 | 0.000765 | 0.000364 | 0.000785 | 0.001148 | 0.00029 | 0.007019 | 0.000366 | 0.000775 | 0.000842 | 0.000603 | 0.005726 | 0.00172 | 0.002502 | 0.001812 | 0.003288 | 0.003171 | 0.000678 | 0.007895 | 0.001288 | 0.006472 | 0.001029 |
| 19.4444444 | 0.00026 | 0.000858 | 0.00063 | 0.001133 | 0.001319 | 0.000919 | 0.007053 | 0.002141 | 0.000745 | 0.001402 | 0.000868 | 0.006104 | 0.001417 | 0.003856 | 0.002481 | 0.003425 | 0.002241 | 0.000998 | 0.006207 | 0.00102 | 0.006526 | 0.002176 |
| 19.4583333 | 0.000664 | 0.001174 | 0.001214 | 0.003933 | 0.003105 | 0.003552 | 0.005041 | 0.007749 | 0.001781 | 0.003642 | 0.002137 | 0.004278 | 0.000782 | 0.004054 | 0.002475 | 0.004266 | 0.001813 | 0.001929 | 0.003852 | 0.001724 | 0.005839 | 0.004659 |
| 19.4722222 | 0.0026 | 0.002252 | 0.002854 | 0.007746 | 0.006563 | 0.007784 | 0.002585 | 0.015068 | 0.004079 | 0.006952 | 0.006676 | 0.002462 | 0.001116 | 0.004988 | 0.003474 | 0.004131 | 0.001022 | 0.003074 | 0.004285 | 0.001819 | 0.004367 | 0.007695 |

| | | | | | | | | | | | | | | | | | | | | | | |
|---|---|---|---|---|---|---|---|---|---|---|---|---|---|---|---|---|---|---|---|---|---|---|
| 19.4861111 | 0.004815 | 0.002468 | 0.004647 | 0.008317 | 0.007789 | 0.009247 | 0.001938 | 0.016414 | 0.005079 | 0.008084 | 0.0095 | 0.002365 | 0.003689 | 0.009485 | 0.007014 | 0.003737 | 0.001207 | 0.005867 | 0.008748 | 0.002646 | 0.005408 | 0.011897 |
| 19.5 | 0.003874 | 0.001397 | 0.004649 | 0.005197 | 0.00519 | 0.006091 | 0.001871 | 0.010086 | 0.003486 | 0.005695 | 0.006463 | 0.002342 | 0.009097 | 0.016789 | 0.011834 | 0.006181 | 0.002816 | 0.009738 | 0.014405 | 0.006996 | 0.00861 | 0.018011 |
| 19.5138889 | 0.001928 | 0.000755 | 0.003314 | 0.001986 | 0.002105 | 0.002282 | 0.002842 | 0.003479 | 0.001536 | 0.002419 | 0.003796 | 0.003916 | 0.013384 | 0.020779 | 0.013809 | 0.009419 | 0.003633 | 0.011487 | 0.016773 | 0.01034 | 0.009467 | 0.022001 |
| 19.5277778 | 0.002456 | 0.001669 | 0.002599 | 0.000827 | 0.001205 | 0.000658 | 0.004497 | 0.000965 | 0.001013 | 0.001028 | 0.005229 | 0.006341 | 0.012711 | 0.017154 | 0.01143 | 0.009051 | 0.004713 | 0.01053 | 0.013773 | 0.00846 | 0.006663 | 0.019526 |
| 19.5416667 | 0.004098 | 0.003477 | 0.003739 | 0.000805 | 0.001508 | 0.001155 | 0.005097 | 0.00103 | 0.001095 | 0.001321 | 0.007219 | 0.00725 | 0.012283 | 0.011784 | 0.010495 | 0.009031 | 0.005427 | 0.011942 | 0.010786 | 0.007347 | 0.005577 | 0.015618 |
| 19.5555556 | 0.0052 | 0.003932 | 0.005436 | 0.00052 | 0.001427 | 0.002795 | 0.004159 | 0.001783 | 0.001198 | 0.001309 | 0.006371 | 0.007018 | 0.017826 | 0.012629 | 0.015284 | 0.014131 | 0.004237 | 0.017915 | 0.012924 | 0.012177 | 0.010853 | 0.017069 |
| 19.5694444 | 0.005553 | 0.002482 | 0.004463 | 0.000602 | 0.001358 | 0.002921 | 0.002083 | 0.002212 | 0.001425 | 0.000812 | 0.003144 | 0.006245 | 0.022735 | 0.014889 | 0.017753 | 0.017187 | 0.004243 | 0.018875 | 0.01402 | 0.01556 | 0.015283 | 0.018713 |
| 19.5833333 | 0.005536 | 0.000982 | 0.002213 | 0.001883 | 0.001222 | 0.002055 | 0.00061 | 0.003045 | 0.002154 | 0.000768 | 0.001034 | 0.005634 | 0.017616 | 0.01128 | 0.012364 | 0.012376 | 0.007964 | 0.012677 | 0.009806 | 0.011633 | 0.011872 | 0.013175 |
| 19.5972222 | 0.00534 | 0.00051 | 0.002133 | 0.003181 | 0.00087 | 0.002415 | 0.000551 | 0.00407 | 0.003575 | 0.001057 | 0.001282 | 0.005203 | 0.008538 | 0.008581 | 0.0089 | 0.008943 | 0.015996 | 0.011385 | 0.007537 | 0.009207 | 0.009179 | 0.008077 |
| 19.6111111 | 0.004645 | 0.000646 | 0.002767 | 0.003479 | 0.001026 | 0.002388 | 0.000747 | 0.003734 | 0.004208 | 0.001135 | 0.001688 | 0.004024 | 0.006453 | 0.011602 | 0.011747 | 0.01179 | 0.020921 | 0.015861 | 0.008551 | 0.012489 | 0.012849 | 0.010336 |
| 19.625 | 0.003609 | 0.000559 | 0.003114 | 0.003476 | 0.0013 | 0.001416 | 0.000551 | 0.002701 | 0.003721 | 0.000719 | 0.0012 | 0.00208 | 0.008787 | 0.013394 | 0.013396 | 0.013429 | 0.018092 | 0.017796 | 0.007269 | 0.014048 | 0.015519 | 0.012687 |
| 19.6388889 | 0.002788 | 0.000345 | 0.003671 | 0.00393 | 0.001483 | 0.000823 | 0.000862 | 0.002282 | 0.003121 | 0.000302 | 0.000708 | 0.000871 | 0.009846 | 0.011495 | 0.011665 | 0.01137 | 0.011994 | 0.016279 | 0.004399 | 0.011843 | 0.014197 | 0.011511 |
| 19.6527778 | 0.002401 | 0.000364 | 0.003169 | 0.004533 | 0.001677 | 0.000635 | 0.001639 | 0.002545 | 0.002578 | 0.000252 | 0.000562 | 0.000775 | 0.00944 | 0.008839 | 0.009299 | 0.008518 | 0.005752 | 0.013668 | 0.002583 | 0.009022 | 0.011781 | 0.009576 |
| 19.6666667 | 0.002128 | 0.000317 | 0.00153 | 0.004692 | 0.001489 | 0.000515 | 0.001505 | 0.002988 | 0.001941 | 0.000338 | 0.000521 | 0.000767 | 0.008737 | 0.006979 | 0.007664 | 0.006512 | 0.001585 | 0.011314 | 0.0022 | 0.007089 | 0.010171 | 0.008209 |
| 19.6805556 | 0.001575 | 0.000234 | 0.000432 | 0.004231 | 0.000936 | 0.000468 | 0.001057 | 0.00316 | 0.001232 | 0.000309 | 0.00046 | 0.000603 | 0.008627 | 0.006272 | 0.007128 | 0.005876 | 0.00095 | 0.010213 | 0.003581 | 0.006444 | 0.009888 | 0.008029 |
| 19.6944444 | 0.000864 | 0.000326 | 0.000492 | 0.003151 | 0.000453 | 0.000557 | 0.001585 | 0.00283 | 0.000796 | 0.000222 | 0.000323 | 0.000501 | 0.008464 | 0.005587 | 0.006936 | 0.005722 | 0.002662 | 0.009956 | 0.005687 | 0.006399 | 0.009498 | 0.008401 |
| 19.7083333 | 0.000444 | 0.000342 | 0.000939 | 0.001718 | 0.000395 | 0.000559 | 0.002056 | 0.001787 | 0.000874 | 0.000348 | 0.000201 | 0.000387 | 0.007139 | 0.00408 | 0.006142 | 0.005056 | 0.004006 | 0.009041 | 0.006366 | 0.005994 | 0.007901 | 0.007896 |
| 19.7222222 | 0.00042 | 0.000271 | 0.001173 | 0.000905 | 0.000573 | 0.000329 | 0.001572 | 0.000694 | 0.001189 | 0.000418 | 0.0002 | 0.000325 | 0.005333 | 0.002713 | 0.004907 | 0.004152 | 0.003962 | 0.007409 | 0.005424 | 0.005318 | 0.006056 | 0.00626 |
| 19.7361111 | 0.00048 | 0.000491 | 0.001253 | 0.000938 | 0.00061 | 0.000341 | 0.000884 | 0.000712 | 0.001438 | 0.000439 | 0.000546 | 0.000324 | 0.004734 | 0.002734 | 0.004327 | 0.003951 | 0.003949 | 0.006693 | 0.004868 | 0.00526 | 0.005551 | 0.004939 |
| 19.75 | 0.000357 | 0.00062 | 0.001068 | 0.001057 | 0.000715 | 0.00051 | 0.000608 | 0.001553 | 0.001205 | 0.000522 | 0.000899 | 0.000759 | 0.00504 | 0.003356 | 0.004323 | 0.004155 | 0.004353 | 0.006661 | 0.005212 | 0.005617 | 0.005992 | 0.004538 |
| 19.7638889 | 0.000269 | 0.000647 | 0.000973 | 0.001282 | 0.0012 | 0.000548 | 0.000823 | 0.002078 | 0.001043 | 0.000715 | 0.000869 | 0.002093 | 0.004208 | 0.002571 | 0.00334 | 0.003215 | 0.003591 | 0.005091 | 0.004493 | 0.005022 | 0.005076 | 0.00356 |
| 19.7777778 | 0.00039 | 0.000859 | 0.001415 | 0.00131 | 0.001256 | 0.000723 | 0.001282 | 0.001587 | 0.001425 | 0.00111 | 0.000694 | 0.003233 | 0.002171 | 0.001297 | 0.001652 | 0.001661 | 0.001814 | 0.002414 | 0.002325 | 0.003086 | 0.002557 | 0.001851 |
| 19.7916667 | 0.000656 | 0.000654 | 0.001652 | 0.000758 | 0.000718 | 0.000692 | 0.001396 | 0.001056 | 0.001352 | 0.000972 | 0.000388 | 0.003068 | 0.000925 | 0.001503 | 0.000843 | 0.001303 | 0.00073 | 0.000987 | 0.000695 | 0.001378 | 0.000706 | 0.00088 |
| 19.8055556 | 0.001269 | 0.000391 | 0.001535 | 0.000354 | 0.00049 | 0.000378 | 0.001636 | 0.001847 | 0.001167 | 0.000602 | 0.000304 | 0.002417 | 0.001504 | 0.001694 | 0.00099 | 0.0012 | 0.000884 | 0.000899 | 0.001146 | 0.000884 | 0.000865 | 0.00104 |
| 19.8194444 | 0.001644 | 0.000477 | 0.001298 | 0.000268 | 0.000378 | 0.00018 | 0.002573 | 0.003009 | 0.001229 | 0.000594 | 0.000374 | 0.002095 | 0.003192 | 0.001734 | 0.002392 | 0.001283 | 0.002522 | 0.001923 | 0.003993 | 0.001114 | 0.002625 | 0.002842 |
| 19.8333333 | 0.001204 | 0.000387 | 0.001054 | 0.00043 | 0.000214 | 0.000202 | 0.004108 | 0.002939 | 0.00087 | 0.000466 | 0.000317 | 0.002146 | 0.003781 | 0.003607 | 0.004876 | 0.002945 | 0.005213 | 0.004159 | 0.007889 | 0.002218 | 0.004116 | 0.005932 |
| 19.8472222 | 0.000911 | 0.000394 | 0.001201 | 0.001335 | 0.00026 | 0.000488 | 0.005849 | 0.001845 | 0.000709 | 0.000219 | 0.000305 | 0.003276 | 0.003272 | 0.005599 | 0.00622 | 0.004159 | 0.006734 | 0.005388 | 0.0104 | 0.003053 | 0.003733 | 0.007698 |
| 19.8611111 | 0.001537 | 0.00127 | 0.001401 | 0.002642 | 0.000634 | 0.000849 | 0.005563 | 0.000956 | 0.001534 | 0.000301 | 0.000426 | 0.006378 | 0.004596 | 0.00483 | 0.005187 | 0.002883 | 0.005792 | 0.003766 | 0.010318 | 0.002403 | 0.004337 | 0.006356 |
| 19.875 | 0.002491 | 0.002762 | 0.001116 | 0.00322 | 0.001263 | 0.001056 | 0.003773 | 0.001017 | 0.002505 | 0.00064 | 0.000934 | 0.011314 | 0.006956 | 0.002321 | 0.002731 | 0.001437 | 0.003325 | 0.00213 | 0.008487 | 0.00208 | 0.008548 | 0.003234 |
| 19.8888889 | 0.002945 | 0.003707 | 0.000571 | 0.002532 | 0.001424 | 0.001207 | 0.003802 | 0.002536 | 0.002367 | 0.000631 | 0.001417 | 0.015996 | 0.007575 | 0.000712 | 0.000772 | 0.00195 | 0.001168 | 0.002855 | 0.00671 | 0.003159 | 0.012656 | 0.001074 |
| 19.9027778 | 0.00318 | 0.003785 | 0.000233 | 0.001178 | 0.00103 | 0.001361 | 0.004956 | 0.005099 | 0.001428 | 0.00033 | 0.001425 | 0.017519 | 0.007049 | 0.000546 | 0.000198 | 0.002915 | 0.000468 | 0.003484 | 0.005565 | 0.004189 | 0.013341 | 0.000625 |
| 19.9166667 | 0.004223 | 0.003715 | 0.000207 | 0.000473 | 0.000989 | 0.001403 | 0.006552 | 0.006609 | 0.001129 | 0.000313 | 0.001547 | 0.015712 | 0.007232 | 0.000668 | 0.000572 | 0.004073 | 0.000533 | 0.004457 | 0.004248 | 0.004476 | 0.011514 | 0.001319 |
| 19.9305556 | 0.005981 | 0.003726 | 0.000462 | 0.001125 | 0.001417 | 0.001346 | 0.009598 | 0.006186 | 0.001754 | 0.000593 | 0.002145 | 0.012597 | 0.008483 | 0.000741 | 0.001811 | 0.006285 | 0.000512 | 0.008076 | 0.005257 | 0.004621 | 0.009142 | 0.003 |
| 19.9444444 | 0.007132 | 0.003233 | 0.001041 | 0.002354 | 0.0014 | 0.001226 | 0.012761 | 0.005543 | 0.002432 | 0.000622 | 0.002453 | 0.009875 | 0.009775 | 0.001236 | 0.003157 | 0.007796 | 0.001023 | 0.012638 | 0.01077 | 0.005271 | 0.007403 | 0.0044 |
| 19.9583333 | 0.007117 | 0.002175 | 0.00159 | 0.002972 | 0.000822 | 0.001111 | 0.014041 | 0.005562 | 0.002753 | 0.000519 | 0.001707 | 0.009191 | 0.01035 | 0.001502 | 0.003643 | 0.007323 | 0.00266 | 0.015849 | 0.015598 | 0.007923 | 0.006282 | 0.004247 |
| 19.9722222 | 0.006781 | 0.001383 | 0.001557 | 0.002968 | 0.00045 | 0.001285 | 0.012886 | 0.005492 | 0.002825 | 0.000615 | 0.000749 | 0.01089 | 0.0105 | 0.001556 | 0.003697 | 0.006538 | 0.004351 | 0.01741 | 0.01577 | 0.012303 | 0.005757 | 0.003382 |
| 19.9861111 | 0.00664 | 0.001469 | 0.001136 | 0.00306 | 0.000676 | 0.001682 | 0.010286 | 0.005066 | 0.002717 | 0.000579 | 0.000836 | 0.012534 | 0.010473 | 0.002145 | 0.003608 | 0.006889 | 0.004288 | 0.017307 | 0.013254 | 0.01463 | 0.006063 | 0.00287 |
| 20 | 0.005789 | 0.001628 | 0.0012 | 0.002891 | 0.000982 | 0.001613 | 0.006989 | 0.003765 | 0.001945 | 0.000781 | 0.001698 | 0.010529 | 0.009376 | 0.002031 | 0.002504 | 0.00714 | 0.003074 | 0.014426 | 0.009299 | 0.0124 | 0.005809 | 0.002058 |
| 20.0138889 | 0.003794 | 0.001696 | 0.002313 | 0.002558 | 0.002098 | 0.002576 | 0.004627 | 0.004934 | 0.001421 | 0.002621 | 0.005091 | 0.007117 | 0.006228 | 0.001891 | 0.001763 | 0.005416 | 0.002917 | 0.008339 | 0.004997 | 0.007053 | 0.004133 | 0.001692 |
| 20.0277778 | 0.00262 | 0.003385 | 0.004033 | 0.003841 | 0.005212 | 0.006165 | 0.006329 | 0.012366 | 0.002833 | 0.006238 | 0.011528 | 0.00862 | 0.004137 | 0.002833 | 0.002682 | 0.003839 | 0.003673 | 0.003612 | 0.0031 | 0.003284 | 0.003454 | 0.002889 |
| 20.0416667 | 0.002577 | 0.00533 | 0.004688 | 0.005445 | 0.007203 | 0.008719 | 0.009724 | 0.018695 | 0.004399 | 0.008171 | 0.013358 | 0.01142 | 0.009133 | 0.007591 | 0.007737 | 0.007969 | 0.008875 | 0.009147 | 0.008667 | 0.008209 | 0.009535 | 0.008784 |
| 20.0555556 | 0.001589 | 0.00412 | 0.003549 | 0.004273 | 0.005237 | 0.007341 | 0.008879 | 0.016142 | 0.003826 | 0.005943 | 0.007892 | 0.008449 | 0.017916 | 0.017245 | 0.017278 | 0.015188 | 0.019493 | 0.021426 | 0.019544 | 0.020357 | 0.020964 | 0.019601 |
| 20.0694444 | 0.000456 | 0.001825 | 0.002442 | 0.001665 | 0.002104 | 0.004421 | 0.00518 | 0.008422 | 0.002071 | 0.002344 | 0.003603 | 0.003674 | 0.017871 | 0.018997 | 0.018526 | 0.015035 | 0.022207 | 0.023435 | 0.020288 | 0.024369 | 0.022452 | 0.021427 |
| 20.0833333 | 0.000357 | 0.001669 | 0.002477 | 0.000376 | 0.000835 | 0.00209 | 0.004762 | 0.003488 | 0.000742 | 0.000482 | 0.003706 | 0.002803 | 0.009155 | 0.010429 | 0.009774 | 0.008132 | 0.012877 | 0.014005 | 0.009588 | 0.015405 | 0.011428 | 0.011812 |
| 20.0972222 | 0.000392 | 0.001925 | 0.002892 | 0.000342 | 0.000707 | 0.000748 | 0.006336 | 0.002379 | 0.000223 | 0.000146 | 0.003861 | 0.003459 | 0.003232 | 0.005058 | 0.004373 | 0.00298 | 0.007131 | 0.006632 | 0.003528 | 0.006109 | 0.002798 | 0.005016 |
| 20.1111111 | 0.000327 | 0.001247 | 0.003237 | 0.000419 | 0.00054 | 0.000585 | 0.005164 | 0.001922 | 0.000205 | 0.000221 | 0.002821 | 0.002528 | 0.001549 | 0.004594 | 0.003582 | 0.002648 | 0.009962 | 0.004371 | 0.004618 | 0.002499 | 0.000988 | 0.003202 |
| 20.125 | 0.000319 | 0.000471 | 0.003182 | 0.000385 | 0.000556 | 0.001002 | 0.00339 | 0.002053 | 0.00036 | 0.000258 | 0.002211 | 0.001679 | 0.000679 | 0.0044 | 0.003434 | 0.005679 | 0.011818 | 0.004164 | 0.004559 | 0.001171 | 0.001061 | 0.002094 |
| 20.1388889 | 0.000204 | 0.000218 | 0.002235 | 0.000614 | 0.000892 | 0.00095 | 0.002519 | 0.002119 | 0.000705 | 0.000278 | 0.00237 | 0.002057 | 0.000728 | 0.003762 | 0.003334 | 0.006258 | 0.013067 | 0.003958 | 0.004016 | 0.000786 | 0.002238 | 0.001298 |
| 20.1527778 | 0.000314 | 0.000491 | 0.000907 | 0.000869 | 0.001226 | 0.000511 | 0.001081 | 0.001898 | 0.001205 | 0.000377 | 0.002423 | 0.001774 | 0.001328 | 0.002877 | 0.002702 | 0.003008 | 0.015375 | 0.00202 | 0.005443 | 0.00156 | 0.005074 | 0.000726 |
| 20.1666667 | 0.00092 | 0.000911 | 0.000197 | 0.000801 | 0.001394 | 0.000524 | 0.000556 | 0.001567 | 0.001529 | 0.000394 | 0.001993 | 0.001523 | 0.001501 | 0.002398 | 0.002097 | 0.000947 | 0.01253 | 0.000906 | 0.004677 | 0.002517 | 0.00756 | 0.000858 |
| 20.1805556 | 0.001426 | 0.001494 | 0.00018 | 0.000745 | 0.001586 | 0.000856 | 0.001611 | 0.000906 | 0.001669 | 0.000528 | 0.001717 | 0.003788 | 0.001479 | 0.002411 | 0.001917 | 0.00098 | 0.005975 | 0.001382 | 0.002568 | 0.002467 | 0.008636 | 0.001839 |
| 20.1944444 | 0.001243 | 0.002172 | 0.000274 | 0.000951 | 0.001678 | 0.001028 | 0.00338 | 0.000539 | 0.001667 | 0.000824 | 0.001912 | 0.007378 | 0.001892 | 0.002269 | 0.001639 | 0.001298 | 0.002214 | 0.00106 | 0.002735 | 0.001549 | 0.009052 | 0.003062 |
| 20.2083333 | 0.000654 | 0.002309 | 0.000223 | 0.001096 | 0.001297 | 0.000956 | 0.005169 | 0.000441 | 0.001474 | 0.000705 | 0.002073 | 0.009432 | 0.00242 | 0.001623 | 0.001071 | 0.001505 | 0.002223 | 0.000635 | 0.002509 | 0.00118 | 0.009069 | 0.004186 |
| 20.2222222 | 0.000694 | 0.001655 | 0.000408 | 0.001023 | 0.000851 | 0.00101 | 0.004931 | 0.000731 | 0.001646 | 0.00038 | 0.002019 | 0.008458 | 0.002501 | 0.000801 | 0.000805 | 0.001322 | 0.003531 | 0.001177 | 0.001059 | 0.00189 | 0.008591 | 0.004987 |
| 20.2361111 | 0.001865 | 0.000763 | 0.000968 | 0.00081 | 0.000965 | 0.001302 | 0.002504 | 0.001174 | 0.002325 | 0.000592 | 0.002105 | 0.004945 | 0.002375 | 0.001076 | 0.001451 | 0.001916 | 0.003267 | 0.002303 | 0.001201 | 0.003417 | 0.006077 | 0.006086 |
| 20.25 | 0.002882 | 0.000472 | 0.001268 | 0.00057 | 0.001174 | 0.001095 | 0.000632 | 0.000982 | 0.002383 | 0.000964 | 0.002162 | 0.001942 | 0.003729 | 0.003338 | 0.003519 | 0.004648 | 0.002457 | 0.00377 | 0.003348 | 0.005741 | 0.003807 | 0.007959 |
| 20.2638889 | 0.002931 | 0.000853 | 0.001251 | 0.000489 | 0.000989 | 0.00061 | 0.000643 | 0.000344 | 0.001488 | 0.000973 | 0.0021 | 0.001138 | 0.005142 | 0.004904 | 0.004926 | 0.005572 | 0.002715 | 0.004008 | 0.00402 | 0.006193 | 0.0045 | 0.007635 |
| 20.2777778 | 0.002953 | 0.000976 | 0.001717 | 0.000372 | 0.000873 | 0.000889 | 0.001591 | 0.0002 | 0.000859 | 0.001065 | 0.002415 | 0.00118 | 0.005059 | 0.004501 | 0.004801 | 0.005064 | 0.003449 | 0.005068 | 0.004569 | 0.004959 | 0.005236 | 0.005216 |
| 20.2916667 | 0.003347 | 0.00092 | 0.00234 | 0.000185 | 0.000994 | 0.001537 | 0.001634 | 0.000578 | 0.001117 | 0.001283 | 0.002854 | 0.001882 | 0.006724 | 0.005476 | 0.006666 | 0.008698 | 0.006183 | 0.009557 | 0.009145 | 0.006323 | 0.007448 | 0.005269 |
| 20.3055556 | 0.003393 | 0.000894 | 0.002098 | 0.00023 | 0.000836 | 0.001482 | 0.001181 | 0.000671 | 0.00163 | 0.001008 | 0.002702 | 0.002787 | 0.007807 | 0.006551 | 0.008204 | 0.011708 | 0.007867 | 0.010929 | 0.011478 | 0.006874 | 0.00997 | 0.005774 |
| 20.3194444 | 0.002643 | 0.001059 | 0.00124 | 0.000544 | 0.00049 | 0.000818 | 0.002304 | 0.000704 | 0.001562 | 0.000477 | 0.001751 | 0.002674 | 0.006013 | 0.004921 | 0.006066 | 0.008398 | 0.005816 | 0.007838 | 0.008534 | 0.005825 | 0.008671 | 0.004301 |
| 20.3333333 | 0.001425 | 0.002157 | 0.000893 | 0.000953 | 0.000481 | 0.000456 | 0.00457 | 0.001486 | 0.001008 | 0.000339 | 0.001061 | 0.001629 | 0.006705 | 0.004387 | 0.005235 | 0.005087 | 0.00511 | 0.008134 | 0.008369 | 0.009273 | 0.009323 | 0.004229 |
| 20.3472222 | 0.00055 | 0.002912 | 0.001023 | 0.001176 | 0.000587 | 0.000653 | 0.007198 | 0.002178 | 0.000839 | 0.000387 | 0.001596 | 0.000779 | 0.008528 | 0.005813 | 0.006684 | 0.00566 | 0.006225 | 0.009987 | 0.010978 | 0.012653 | 0.013193 | 0.004452 |
| 20.3611111 | 0.000406 | 0.002023 | 0.000927 | 0.000861 | 0.000503 | 0.000648 | 0.009677 | 0.001806 | 0.000883 | 0.000375 | 0.001965 | 0.001021 | 0.006485 | 0.005153 | 0.00579 | 0.005043 | 0.00532 | 0.007289 | 0.008243 | 0.00928 | 0.012655 | 0.003599 |
| 20.375 | 0.000566 | 0.000946 | 0.000939 | 0.000346 | 0.000547 | 0.000407 | 0.010526 | 0.001464 | 0.000566 | 0.000533 | 0.001661 | 0.001712 | 0.003854 | 0.004139 | 0.004665 | 0.003367 | 0.004647 | 0.003648 | 0.002642 | 0.003891 | 0.009403 | 0.00464 |
| 20.3888889 | 0.000532 | 0.000793 | 0.000907 | 0.00028 | 0.000574 | 0.000386 | 0.010375 | 0.002455 | 0.00031 | 0.000586 | 0.002565 | 0.002431 | 0.003415 | 0.004439 | 0.005008 | 0.00251 | 0.005045 | 0.002411 | 0.001692 | 0.002838 | 0.00988 | 0.005966 |
| 20.4027778 | 0.000483 | 0.000945 | 0.000492 | 0.000484 | 0.000364 | 0.000431 | 0.010745 | 0.003854 | 0.000436 | 0.000392 | 0.004973 | 0.003191 | 0.005151 | 0.004126 | 0.00454 | 0.00135 | 0.004681 | 0.001522 | 0.004383 | 0.004874 | 0.012017 | 0.006028 |
| 20.4166667 | 0.001056 | 0.001581 | 0.000259 | 0.000492 | 0.000227 | 0.000474 | 0.011369 | 0.003901 | 0.000547 | 0.000334 | 0.006455 | 0.003768 | 0.008957 | 0.005375 | 0.005537 | 0.001709 | 0.006417 | 0.00225 | 0.005409 | 0.004723 | 0.010802 | 0.007792 |
| 20.4305556 | 0.00208 | 0.002428 | 0.000373 | 0.000313 | 0.000342 | 0.000548 | 0.01299 | 0.002748 | 0.000464 | 0.000695 | 0.005188 | 0.004328 | 0.010745 | 0.010053 | 0.009698 | 0.004797 | 0.011728 | 0.006461 | 0.003647 | 0.003085 | 0.006151 | 0.012568 |
| 20.4444444 | 0.003273 | 0.002176 | 0.000555 | 0.000739 | 0.001112 | 0.00102 | 0.016722 | 0.002233 | 0.000758 | 0.002189 | 0.002871 | 0.004624 | 0.009005 | 0.014511 | 0.013047 | 0.00773 | 0.016667 | 0.009821 | 0.002929 | 0.003229 | 0.002008 | 0.017813 |
| 20.4583333 | 0.004034 | 0.00183 | 0.000823 | 0.002518 | 0.002872 | 0.003017 | 0.020215 | 0.005053 | 0.001815 | 0.005032 | 0.003389 | 0.00354 | 0.006121 | 0.015061 | 0.012258 | 0.006947 | 0.01776 | 0.008505 | 0.00443 | 0.002877 | 0.000827 | 0.020119 |
| 20.4722222 | 0.003028 | 0.002256 | 0.001702 | 0.004004 | 0.004493 | 0.00623 | 0.02054 | 0.010135 | 0.002845 | 0.007491 | 0.007212 | 0.002541 | 0.004398 | 0.0115 | 0.007719 | 0.004559 | 0.014453 | 0.00511 | 0.004651 | 0.002688 | 0.002051 | 0.016652 |
| 20.4861111 | 0.001678 | 0.002015 | 0.002836 | 0.003403 | 0.003783 | 0.00699 | 0.019084 | 0.010424 | 0.002403 | 0.006884 | 0.008504 | 0.002406 | 0.007291 | 0.007909 | 0.005374 | 0.006529 | 0.009233 | 0.007435 | 0.005725 | 0.006352 | 0.007749 | 0.010293 |
| 20.5 | 0.002463 | 0.001546 | 0.003068 | 0.002885 | 0.001796 | 0.00398 | 0.018103 | 0.006011 | 0.001276 | 0.00351 | 0.004874 | 0.001681 | 0.013467 | 0.009838 | 0.010514 | 0.012734 | 0.008077 | 0.017942 | 0.012625 | 0.012159 | 0.018442 | 0.009108 |
| 20.5138889 | 0.003809 | 0.001644 | 0.002676 | 0.003798 | 0.001158 | 0.001055 | 0.016552 | 0.003899 | 0.001053 | 0.000816 | 0.001704 | 0.001471 | 0.014869 | 0.01313 | 0.015054 | 0.015357 | 0.009422 | 0.024589 | 0.018076 | 0.013933 | 0.025115 | 0.010251 |
| 20.5277778 | 0.004126 | 0.001802 | 0.002968 | 0.004061 | 0.001132 | 0.000762 | 0.013186 | 0.003243 | 0.000922 | 0.000133 | 0.001159 | 0.001516 | 0.011218 | 0.010828 | 0.012379 | 0.011951 | 0.00926 | 0.017462 | 0.013456 | 0.011058 | 0.019296 | 0.008941 |
| 20.5416667 | 0.005074 | 0.00256 | 0.004559 | 0.003244 | 0.001035 | 0.002464 | 0.009105 | 0.001961 | 0.000724 | 0.000289 | 0.001683 | 0.001639 | 0.014336 | 0.011502 | 0.014371 | 0.014108 | 0.01573 | 0.010996 | 0.009783 | 0.014527 | 0.011698 | 0.012879 |
| 20.5555556 | 0.006969 | 0.003357 | 0.006031 | 0.002562 | 0.001141 | 0.003921 | 0.006023 | 0.001442 | 0.000729 | 0.000464 | 0.002249 | 0.003259 | 0.022751 | 0.018913 | 0.023554 | 0.023159 | 0.025563 | 0.016123 | 0.014743 | 0.023279 | 0.013688 | 0.019545 |
| 20.5694444 | 0.008341 | 0.002611 | 0.004912 | 0.002601 | 0.000999 | 0.003267 | 0.004883 | 0.00221 | 0.000962 | 0.00036 | 0.001689 | 0.005731 | 0.019953 | 0.017428 | 0.021241 | 0.020769 | 0.022071 | 0.018075 | 0.016377 | 0.020589 | 0.015066 | 0.017183 |
| 20.5833333 | 0.009125 | 0.001175 | 0.002076 | 0.003185 | 0.000666 | 0.002144 | 0.004869 | 0.004305 | 0.001903 | 0.000221 | 0.000843 | 0.007659 | 0.013175 | 0.010548 | 0.012711 | 0.012134 | 0.014383 | 0.017929 | 0.0173 | 0.013774 | 0.015537 | 0.013744 |
| 20.5972222 | 0.009824 | 0.000464 | 0.000705 | 0.003434 | 0.000574 | 0.001701 | 0.003896 | 0.004946 | 0.002163 | 0.000345 | 0.000968 | 0.00822 | 0.015236 | 0.010825 | 0.012279 | 0.011932 | 0.014811 | 0.021177 | 0.022115 | 0.015091 | 0.019696 | 0.015363 |
| 20.6111111 | 0.00965 | 0.000746 | 0.001065 | 0.002872 | 0.001188 | 0.001073 | 0.002443 | 0.002991 | 0.001243 | 0.000981 | 0.001005 | 0.007404 | 0.013361 | 0.00909 | 0.009857 | 0.009898 | 0.011156 | 0.015948 | 0.01937 | 0.011932 | 0.016837 | 0.011437 |
| 20.625 | 0.007548 | 0.002191 | 0.001559 | 0.001859 | 0.002407 | 0.000964 | 0.001966 | 0.001212 | 0.000764 | 0.002265 | 0.000712 | 0.006025 | 0.006689 | 0.005303 | 0.005694 | 0.005943 | 0.004514 | 0.006631 | 0.009533 | 0.005294 | 0.00816 | 0.005759 |
| 20.6388889 | 0.004235 | 0.003614 | 0.001803 | 0.001049 | 0.003264 | 0.001452 | 0.001552 | 0.000967 | 0.001008 | 0.003251 | 0.001054 | 0.005082 | 0.006421 | 0.006434 | 0.00652 | 0.007828 | 0.004415 | 0.004119 | 0.003342 | 0.005594 | 0.004276 | 0.006663 |
| 20.6527778 | 0.001646 | 0.003733 | 0.002055 | 0.001022 | 0.003278 | 0.00166 | 0.000742 | 0.001153 | 0.001134 | 0.003191 | 0.001315 | 0.005143 | 0.008552 | 0.007095 | 0.006888 | 0.009483 | 0.005144 | 0.004793 | 0.004024 | 0.006563 | 0.005058 | 0.008315 |
| 20.6666667 | 0.000661 | 0.002966 | 0.002167 | 0.001376 | 0.002765 | 0.001424 | 0.000397 | 0.000852 | 0.00085 | 0.002441 | 0.000911 | 0.005775 | 0.008018 | 0.005067 | 0.005086 | 0.007916 | 0.003427 | 0.004907 | 0.006785 | 0.004625 | 0.005526 | 0.007835 |
| 20.6805556 | 0.000449 | 0.002087 | 0.00194 | 0.001189 | 0.002105 | 0.000925 | 0.000799 | 0.000688 | 0.000436 | 0.001627 | 0.000388 | 0.005583 | 0.006464 | 0.003708 | 0.004144 | 0.006347 | 0.002326 | 0.005797 | 0.008903 | 0.003598 | 0.005192 | 0.007631 |
| 20.6944444 | 0.000199 | 0.001234 | 0.001445 | 0.000663 | 0.001483 | 0.000485 | 0.001501 | 0.000881 | 0.000276 | 0.001022 | 0.000151 | 0.00425 | 0.005496 | 0.004135 | 0.004901 | 0.006194 | 0.002842 | 0.007301 | 0.010738 | 0.004331 | 0.005618 | 0.007074 |
| 20.7083333 | 0.000217 | 0.000462 | 0.000947 | 0.00059 | 0.001012 | 0.000284 | 0.001883 | 0.000878 | 0.000266 | 0.000684 | 0.000143 | 0.002501 | 0.005052 | 0.004924 | 0.005834 | 0.006368 | 0.003648 | 0.007738 | 0.010981 | 0.005245 | 0.006034 | 0.004966 |
| 20.7222222 | 0.000663 | 0.000197 | 0.000659 | 0.000801 | 0.000918 | 0.000455 | 0.001501 | 0.001181 | 0.000351 | 0.000823 | 0.000313 | 0.001227 | 0.004832 | 0.004474 | 0.005181 | 0.005386 | 0.003062 | 0.006094 | 0.008763 | 0.004838 | 0.00496 | 0.002548 |
| 20.7361111 | 0.00109 | 0.000485 | 0.000426 | 0.000669 | 0.001149 | 0.000857 | 0.000738 | 0.002072 | 0.000844 | 0.00129 | 0.000825 | 0.001239 | 0.004827 | 0.002667 | 0.003023 | 0.003303 | 0.002043 | 0.003307 | 0.005556 | 0.002812 | 0.003085 | 0.001419 |
| 20.75 | 0.001104 | 0.000776 | 0.000196 | 0.000559 | 0.001194 | 0.000992 | 0.000342 | 0.002272 | 0.001298 | 0.00143 | 0.001208 | 0.001979 | 0.004584 | 0.00114 | 0.001079 | 0.001358 | 0.002845 | 0.00186 | 0.003103 | 0.001169 | 0.002677 | 0.001312 |
| 20.7638889 | 0.000677 | 0.000623 | 0.000218 | 0.00088 | 0.000758 | 0.000638 | 0.000415 | 0.00131 | 0.001148 | 0.000934 | 0.000886 | 0.002442 | 0.003906 | 0.001346 | 0.000504 | 0.000354 | 0.00464 | 0.002384 | 0.001593 | 0.001037 | 0.004011 | 0.001211 |
| 20.7777778 | 0.000312 | 0.000307 | 0.000583 | 0.001108 | 0.000373 | 0.000296 | 0.000609 | 0.000384 | 0.000641 | 0.000388 | 0.000523 | 0.002562 | 0.003545 | 0.002699 | 0.001132 | 0.000447 | 0.005943 | 0.003419 | 0.001047 | 0.001056 | 0.004053 | 0.001136 |
| 20.7916667 | 0.000263 | 0.000218 | 0.000909 | 0.000945 | 0.000503 | 0.000437 | 0.000759 | 0.0007 | 0.000424 | 0.00036 | 0.000796 | 0.002652 | 0.004094 | 0.004113 | 0.00277 | 0.001598 | 0.007277 | 0.004615 | 0.001953 | 0.001772 | 0.002983 | 0.002234 |
| 20.8055556 | 0.000397 | 0.000465 | 0.000861 | 0.000913 | 0.000715 | 0.00069 | 0.001735 | 0.002751 | 0.000637 | 0.000515 | 0.000858 | 0.003251 | 0.005194 | 0.004779 | 0.004637 | 0.002527 | 0.008464 | 0.005216 | 0.003328 | 0.003352 | 0.003879 | 0.004138 |
| 20.8194444 | 0.000879 | 0.00152 | 0.001276 | 0.001472 | 0.000611 | 0.000578 | 0.003963 | 0.005812 | 0.0006 | 0.000405 | 0.000579 | 0.005033 | 0.005632 | 0.004336 | 0.005262 | 0.002132 | 0.008306 | 0.003762 | 0.002988 | 0.003876 | 0.004408 | 0.004799 |
| 20.8333333 | 0.001862 | 0.00338 | 0.002432 | 0.002336 | 0.000396 | 0.000406 | 0.005965 | 0.007709 | 0.000494 | 0.000347 | 0.000975 | 0.00809 | 0.004766 | 0.00307 | 0.004112 | 0.002213 | 0.006584 | 0.003264 | 0.002503 | 0.003032 | 0.003118 | 0.003504 |
| 20.8472222 | 0.003652 | 0.005131 | 0.002628 | 0.002849 | 0.000426 | 0.000456 | 0.00777 | 0.007195 | 0.001077 | 0.000712 | 0.001924 | 0.011823 | 0.003982 | 0.001982 | 0.002534 | 0.003909 | 0.004113 | 0.007947 | 0.006592 | 0.002138 | 0.004431 | 0.00226 |
| 20.8611111 | 0.006123 | 0.00608 | 0.002168 | 0.003194 | 0.000665 | 0.000549 | 0.016431 | 0.005606 | 0.001984 | 0.001074 | 0.002667 | 0.015088 | 0.00517 | 0.002704 | 0.002896 | 0.005461 | 0.00211 | 0.015346 | 0.016201 | 0.004073 | 0.010237 | 0.003486 |
| 20.875 | 0.008483 | 0.006159 | 0.00353 | 0.004925 | 0.000932 | 0.000849 | 0.025543 | 0.004871 | 0.002649 | 0.001026 | 0.00334 | 0.016669 | 0.008761 | 0.004983 | 0.005128 | 0.006129 | 0.002035 | 0.020128 | 0.025148 | 0.010097 | 0.016241 | 0.006238 |

| | | | | | | | | | | | | | | | | | | | | | | |
|---|---|---|---|---|---|---|---|---|---|---|---|---|---|---|---|---|---|---|---|---|---|---|
| 20.8888889 | 0.010175 | 0.005438 | 0.005658 | 0.007567 | 0.001196 | 0.001359 | 0.017942 | 0.005089 | 0.002939 | 0.00069 | 0.004748 | 0.016245 | 0.013898 | 0.007244 | 0.007188 | 0.007416 | 0.004407 | 0.020421 | 0.026717 | 0.016158 | 0.019161 | 0.008527 |
| 20.9027778 | 0.011333 | 0.004364 | 0.00554 | 0.008121 | 0.001502 | 0.001709 | 0.005502 | 0.005117 | 0.003248 | 0.000536 | 0.006963 | 0.014806 | 0.019341 | 0.009736 | 0.00902 | 0.0112 | 0.007232 | 0.018443 | 0.024472 | 0.018774 | 0.02084 | 0.01072 |
| 20.9166667 | 0.012418 | 0.003655 | 0.00334 | 0.0063 | 0.001956 | 0.001857 | 0.002891 | 0.00485 | 0.003749 | 0.000918 | 0.008948 | 0.013565 | 0.024131 | 0.012967 | 0.011192 | 0.017091 | 0.006556 | 0.016897 | 0.025398 | 0.019188 | 0.022984 | 0.013296 |
| 20.9305556 | 0.012934 | 0.002996 | 0.002051 | 0.004556 | 0.002205 | 0.001813 | 0.004435 | 0.004454 | 0.003562 | 0.001191 | 0.009058 | 0.012167 | 0.028247 | 0.015394 | 0.012524 | 0.021424 | 0.003742 | 0.01607 | 0.028388 | 0.019513 | 0.024602 | 0.01494 |
| 20.9444444 | 0.012451 | 0.00214 | 0.002814 | 0.004463 | 0.002049 | 0.001624 | 0.00843 | 0.003597 | 0.002661 | 0.000865 | 0.006738 | 0.009662 | 0.032065 | 0.015622 | 0.012007 | 0.021122 | 0.003795 | 0.015783 | 0.028962 | 0.020982 | 0.023948 | 0.014405 |
| 20.9583333 | 0.011527 | 0.001983 | 0.003626 | 0.005479 | 0.001945 | 0.001379 | 0.016447 | 0.002582 | 0.002039 | 0.000463 | 0.003819 | 0.006344 | 0.033395 | 0.014014 | 0.010076 | 0.017118 | 0.004984 | 0.015889 | 0.025894 | 0.025395 | 0.020046 | 0.012097 |
| 20.9722222 | 0.010387 | 0.002593 | 0.003819 | 0.005791 | 0.001746 | 0.001035 | 0.025577 | 0.001952 | 0.002043 | 0.000385 | 0.002317 | 0.003082 | 0.028492 | 0.011056 | 0.007922 | 0.012004 | 0.004119 | 0.015734 | 0.021158 | 0.028285 | 0.013797 | 0.009004 |
| 20.9861111 | 0.009033 | 0.003176 | 0.003791 | 0.004968 | 0.001245 | 0.000758 | 0.028521 | 0.001916 | 0.002678 | 0.000546 | 0.001982 | 0.0011 | 0.018618 | 0.007497 | 0.006454 | 0.00746 | 0.002416 | 0.015677 | 0.016899 | 0.021939 | 0.008483 | 0.006235 |
| 21 | 0.006976 | 0.003186 | 0.003689 | 0.003536 | 0.000881 | 0.000961 | 0.022311 | 0.001782 | 0.003334 | 0.001069 | 0.002378 | 0.001296 | 0.009364 | 0.00437 | 0.005404 | 0.004316 | 0.0021 | 0.015924 | 0.012791 | 0.010428 | 0.006472 | 0.004536 |
| 21.0138889 | 0.004365 | 0.002573 | 0.004168 | 0.002744 | 0.001935 | 0.003105 | 0.012194 | 0.003769 | 0.003025 | 0.003171 | 0.005342 | 0.004136 | 0.004205 | 0.002775 | 0.004003 | 0.002318 | 0.003846 | 0.012997 | 0.007917 | 0.00274 | 0.005156 | 0.002952 |
| 21.0277778 | 0.003259 | 0.00292 | 0.005154 | 0.004403 | 0.004974 | 0.007633 | 0.004856 | 0.01068 | 0.003366 | 0.006745 | 0.008001 | 0.008967 | 0.003341 | 0.003082 | 0.002482 | 0.001325 | 0.006088 | 0.007847 | 0.006185 | 0.002386 | 0.003004 | 0.001598 |
| 21.0416667 | 0.003254 | 0.003968 | 0.005756 | 0.006427 | 0.006937 | 0.010553 | 0.002352 | 0.015722 | 0.005167 | 0.00835 | 0.006894 | 0.011385 | 0.003394 | 0.003396 | 0.002966 | 0.001322 | 0.006084 | 0.006815 | 0.008522 | 0.007997 | 0.003641 | 0.001849 |
| 21.0555556 | 0.00206 | 0.003204 | 0.005453 | 0.005527 | 0.005558 | 0.008991 | 0.00285 | 0.011402 | 0.005239 | 0.006233 | 0.006563 | 0.008282 | 0.001825 | 0.002942 | 0.005567 | 0.001438 | 0.003818 | 0.008953 | 0.008607 | 0.014265 | 0.006247 | 0.003528 |
| 21.0694444 | 0.00071 | 0.002167 | 0.005129 | 0.002923 | 0.00288 | 0.005318 | 0.008417 | 0.006399 | 0.002925 | 0.00305 | 0.009599 | 0.004066 | 0.001085 | 0.002534 | 0.006307 | 0.001601 | 0.00332 | 0.008827 | 0.006394 | 0.014999 | 0.00579 | 0.004453 |
| 21.0833333 | 0.000601 | 0.002704 | 0.004671 | 0.001078 | 0.001122 | 0.002345 | 0.019887 | 0.008141 | 0.001253 | 0.001234 | 0.013443 | 0.00404 | 0.001906 | 0.002434 | 0.004144 | 0.002148 | 0.005706 | 0.008005 | 0.004299 | 0.00886 | 0.004016 | 0.003655 |
| 21.0972222 | 0.000848 | 0.002797 | 0.002788 | 0.00035 | 0.000462 | 0.000857 | 0.028553 | 0.007493 | 0.00095 | 0.000671 | 0.011573 | 0.006289 | 0.002308 | 0.002673 | 0.001862 | 0.00216 | 0.007312 | 0.011298 | 0.001999 | 0.003482 | 0.004277 | 0.004165 |
| 21.1111111 | 0.000861 | 0.002021 | 0.000979 | 0.000274 | 0.000277 | 0.000448 | 0.024663 | 0.004245 | 0.000696 | 0.000474 | 0.00854 | 0.006734 | 0.002407 | 0.003336 | 0.00103 | 0.001589 | 0.006568 | 0.013475 | 0.00196 | 0.003666 | 0.00472 | 0.006183 |
| 21.125 | 0.000824 | 0.002735 | 0.00055 | 0.000535 | 0.00017 | 0.000425 | 0.013072 | 0.004333 | 0.000515 | 0.000231 | 0.015458 | 0.005174 | 0.002713 | 0.004265 | 0.001962 | 0.001318 | 0.005052 | 0.010447 | 0.004558 | 0.00413 | 0.00644 | 0.006601 |
| 21.1388889 | 0.000959 | 0.005131 | 0.000904 | 0.00122 | 0.000165 | 0.000474 | 0.008069 | 0.003868 | 0.001044 | 0.000102 | 0.021541 | 0.002971 | 0.002254 | 0.003801 | 0.003336 | 0.001235 | 0.003449 | 0.008176 | 0.00518 | 0.002519 | 0.009938 | 0.004754 |
| 21.1527778 | 0.000889 | 0.00577 | 0.001204 | 0.001937 | 0.000268 | 0.000916 | 0.010709 | 0.002404 | 0.002274 | 0.000206 | 0.015335 | 0.001148 | 0.001429 | 0.002148 | 0.002911 | 0.001636 | 0.001803 | 0.007967 | 0.004022 | 0.002031 | 0.009223 | 0.002289 |
| 21.1666667 | 0.000533 | 0.003875 | 0.000865 | 0.001634 | 0.000473 | 0.001949 | 0.011071 | 0.002828 | 0.002863 | 0.000519 | 0.008363 | 0.000442 | 0.002508 | 0.001398 | 0.001652 | 0.00273 | 0.001013 | 0.005625 | 0.005 | 0.003164 | 0.007087 | 0.001288 |
| 21.1805556 | 0.000639 | 0.002427 | 0.000699 | 0.001053 | 0.000678 | 0.002829 | 0.009338 | 0.004446 | 0.001922 | 0.000778 | 0.010588 | 0.000948 | 0.005324 | 0.001036 | 0.00175 | 0.004082 | 0.001495 | 0.002314 | 0.005378 | 0.00295 | 0.011578 | 0.001937 |
| 21.1944444 | 0.001698 | 0.002194 | 0.000588 | 0.001444 | 0.000532 | 0.002291 | 0.01259 | 0.004706 | 0.000771 | 0.000603 | 0.010653 | 0.002453 | 0.006705 | 0.000734 | 0.002069 | 0.005789 | 0.002731 | 0.001386 | 0.004826 | 0.001979 | 0.016617 | 0.002939 |
| 21.2083333 | 0.003612 | 0.002989 | 0.000222 | 0.002197 | 0.000428 | 0.001522 | 0.016798 | 0.00268 | 0.000531 | 0.000445 | 0.006445 | 0.004613 | 0.005965 | 0.001457 | 0.001367 | 0.007085 | 0.003747 | 0.001282 | 0.005719 | 0.003173 | 0.01435 | 0.003202 |
| 21.2222222 | 0.005157 | 0.00403 | 7.08E-05 | 0.002881 | 0.000809 | 0.002029 | 0.013831 | 0.001242 | 0.000648 | 0.000682 | 0.006567 | 0.006365 | 0.005507 | 0.002151 | 0.000723 | 0.006376 | 0.003552 | 0.001809 | 0.004773 | 0.004956 | 0.007709 | 0.00217 |
| 21.2361111 | 0.005561 | 0.003244 | 0.000309 | 0.003381 | 0.001054 | 0.001877 | 0.00915 | 0.002124 | 0.000478 | 0.000733 | 0.006092 | 0.00634 | 0.004905 | 0.001536 | 0.000596 | 0.003713 | 0.002719 | 0.003145 | 0.003129 | 0.005773 | 0.003063 | 0.000758 |
| 21.25 | 0.005556 | 0.001524 | 0.000877 | 0.003299 | 0.000857 | 0.001079 | 0.007396 | 0.003304 | 0.000244 | 0.000431 | 0.004105 | 0.004905 | 0.002835 | 0.000595 | 0.00064 | 0.002165 | 0.002305 | 0.002895 | 0.004412 | 0.006091 | 0.003515 | 0.00024 |
| 21.2638889 | 0.005838 | 0.000871 | 0.000994 | 0.003241 | 0.000763 | 0.000979 | 0.00814 | 0.004312 | 0.000358 | 0.000336 | 0.002792 | 0.003933 | 0.001177 | 0.000656 | 0.001256 | 0.00422 | 0.002075 | 0.002048 | 0.004427 | 0.005629 | 0.00587 | 0.000514 |
| 21.2777778 | 0.006025 | 0.00073 | 0.000687 | 0.003841 | 0.000961 | 0.000795 | 0.012976 | 0.006148 | 0.000803 | 0.000569 | 0.001826 | 0.003944 | 0.00075 | 0.001934 | 0.002045 | 0.007472 | 0.001898 | 0.001768 | 0.001938 | 0.004787 | 0.005957 | 0.00099 |
| 21.2916667 | 0.004991 | 0.000421 | 0.000654 | 0.004194 | 0.001079 | 0.000494 | 0.013446 | 0.00694 | 0.001087 | 0.000724 | 0.003432 | 0.003868 | 0.000736 | 0.003103 | 0.002142 | 0.008896 | 0.002155 | 0.001208 | 0.001356 | 0.004509 | 0.003656 | 0.001121 |
| 21.3055556 | 0.002899 | 0.00036 | 0.000883 | 0.00348 | 0.000846 | 0.000604 | 0.007883 | 0.00495 | 0.000825 | 0.000655 | 0.005108 | 0.003034 | 0.001255 | 0.002449 | 0.001828 | 0.00797 | 0.002945 | 0.001205 | 0.001968 | 0.003461 | 0.002315 | 0.001205 |
| 21.3194444 | 0.00156 | 0.000661 | 0.000909 | 0.002641 | 0.000546 | 0.000725 | 0.005685 | 0.002202 | 0.000413 | 0.000552 | 0.004091 | 0.002071 | 0.002164 | 0.001108 | 0.00137 | 0.005695 | 0.003452 | 0.002626 | 0.001399 | 0.001333 | 0.002544 | 0.001556 |
| 21.3333333 | 0.001213 | 0.001547 | 0.000931 | 0.002566 | 0.00058 | 0.001302 | 0.005318 | 0.001684 | 0.000812 | 0.000689 | 0.003976 | 0.001954 | 0.003457 | 0.000831 | 0.001474 | 0.004466 | 0.002676 | 0.005245 | 0.002029 | 0.002065 | 0.003271 | 0.001765 |
| 21.3472222 | 0.00108 | 0.002456 | 0.001485 | 0.002526 | 0.000787 | 0.002633 | 0.006369 | 0.003288 | 0.002013 | 0.001068 | 0.006141 | 0.002208 | 0.005137 | 0.000859 | 0.002759 | 0.004711 | 0.001777 | 0.00864 | 0.005731 | 0.007149 | 0.006111 | 0.002073 |
| 21.3611111 | 0.0015 | 0.002491 | 0.001755 | 0.001965 | 0.000786 | 0.002652 | 0.014037 | 0.004532 | 0.003066 | 0.001138 | 0.006529 | 0.002161 | 0.006479 | 0.000739 | 0.003256 | 0.004497 | 0.001398 | 0.010753 | 0.009173 | 0.011916 | 0.008149 | 0.001878 |
| 21.375 | 0.002319 | 0.002921 | 0.001347 | 0.001637 | 0.000666 | 0.001243 | 0.023304 | 0.004974 | 0.00371 | 0.000933 | 0.005533 | 0.002956 | 0.008013 | 0.000975 | 0.002158 | 0.004017 | 0.001287 | 0.009709 | 0.009651 | 0.012899 | 0.006385 | 0.000804 |
| 21.3888889 | 0.002974 | 0.004528 | 0.001104 | 0.001847 | 0.000649 | 0.000676 | 0.025927 | 0.006251 | 0.004095 | 0.000857 | 0.007546 | 0.005209 | 0.010822 | 0.001306 | 0.001692 | 0.005027 | 0.002204 | 0.007442 | 0.010118 | 0.012989 | 0.004446 | 0.000541 |
| 21.4027778 | 0.003557 | 0.006021 | 0.001299 | 0.002252 | 0.000696 | 0.001395 | 0.02124 | 0.008468 | 0.0042 | 0.00076 | 0.012757 | 0.008113 | 0.014607 | 0.002037 | 0.00245 | 0.007187 | 0.003861 | 0.007013 | 0.013952 | 0.015476 | 0.005929 | 0.001414 |
| 21.4166667 | 0.00508 | 0.006782 | 0.001404 | 0.003137 | 0.001175 | 0.002627 | 0.014225 | 0.010317 | 0.004488 | 0.000932 | 0.017483 | 0.011733 | 0.01797 | 0.004089 | 0.002882 | 0.008915 | 0.004242 | 0.008175 | 0.020371 | 0.018859 | 0.010315 | 0.001651 |
| 21.4305556 | 0.007918 | 0.006417 | 0.001403 | 0.004443 | 0.002002 | 0.003527 | 0.009859 | 0.010958 | 0.005264 | 0.0013 | 0.018096 | 0.016016 | 0.018739 | 0.006175 | 0.002435 | 0.009465 | 0.002575 | 0.008674 | 0.025763 | 0.019761 | 0.01477 | 0.001319 |
| 21.4444444 | 0.010992 | 0.004165 | 0.001315 | 0.004407 | 0.001864 | 0.00291 | 0.00963 | 0.009115 | 0.005356 | 0.001377 | 0.012764 | 0.018514 | 0.01708 | 0.006752 | 0.002929 | 0.009735 | 0.001338 | 0.008976 | 0.025474 | 0.018749 | 0.017537 | 0.001476 |
| 21.4583333 | 0.013105 | 0.001836 | 0.001717 | 0.003277 | 0.001414 | 0.002431 | 0.012982 | 0.006359 | 0.003565 | 0.002298 | 0.006996 | 0.016497 | 0.016577 | 0.007536 | 0.00635 | 0.011563 | 0.002823 | 0.012366 | 0.018685 | 0.019447 | 0.019744 | 0.00212 |
| 21.4722222 | 0.01185 | 0.001038 | 0.003807 | 0.003434 | 0.002152 | 0.004669 | 0.020257 | 0.006569 | 0.001323 | 0.004142 | 0.007038 | 0.010698 | 0.01882 | 0.009271 | 0.011144 | 0.014482 | 0.007578 | 0.019485 | 0.010667 | 0.021855 | 0.021172 | 0.004771 |
| 21.4861111 | 0.007871 | 0.000969 | 0.006634 | 0.003295 | 0.002279 | 0.006097 | 0.032071 | 0.006338 | 0.001346 | 0.004311 | 0.00768 | 0.006264 | 0.021404 | 0.009965 | 0.013414 | 0.015564 | 0.013026 | 0.026568 | 0.007668 | 0.021587 | 0.020608 | 0.008373 |
| 21.5 | 0.005893 | 0.001997 | 0.007778 | 0.002318 | 0.001088 | 0.003855 | 0.041695 | 0.004831 | 0.003961 | 0.002619 | 0.005023 | 0.007361 | 0.021225 | 0.009587 | 0.012029 | 0.013754 | 0.015652 | 0.02867 | 0.010978 | 0.016727 | 0.020152 | 0.008949 |
| 21.5138889 | 0.006192 | 0.004664 | 0.005871 | 0.003114 | 0.000683 | 0.001821 | 0.037846 | 0.006251 | 0.006282 | 0.001604 | 0.003256 | 0.011389 | 0.016874 | 0.00856 | 0.008576 | 0.010801 | 0.015691 | 0.024392 | 0.015288 | 0.011099 | 0.019172 | 0.007001 |
| 21.5277778 | 0.006586 | 0.005734 | 0.002662 | 0.004269 | 0.000851 | 0.001727 | 0.022659 | 0.006631 | 0.005991 | 0.001512 | 0.003912 | 0.013949 | 0.010671 | 0.006067 | 0.006173 | 0.007726 | 0.012308 | 0.015718 | 0.013202 | 0.007189 | 0.01315 | 0.005084 |
| 21.5416667 | 0.006445 | 0.00405 | 0.001372 | 0.003803 | 0.000423 | 0.001913 | 0.009019 | 0.00414 | 0.004499 | 0.001535 | 0.00488 | 0.014122 | 0.011785 | 0.008204 | 0.011781 | 0.010612 | 0.010128 | 0.010314 | 0.01095 | 0.010087 | 0.010572 | 0.008186 |
| 21.5555556 | 0.004981 | 0.003048 | 0.00237 | 0.00284 | 0.000532 | 0.002481 | 0.002323 | 0.002397 | 0.00361 | 0.001343 | 0.004614 | 0.012174 | 0.022572 | 0.019784 | 0.025522 | 0.02305 | 0.017312 | 0.015322 | 0.020746 | 0.022362 | 0.021454 | 0.018559 |
| 21.5694444 | 0.003048 | 0.002149 | 0.003031 | 0.002202 | 0.001375 | 0.002432 | 0.00192 | 0.002759 | 0.003522 | 0.000599 | 0.002947 | 0.009369 | 0.028236 | 0.029127 | 0.033361 | 0.031175 | 0.023227 | 0.020553 | 0.032135 | 0.030256 | 0.03112 | 0.024095 |
| 21.5833333 | 0.002812 | 0.000823 | 0.002475 | 0.001668 | 0.001304 | 0.001803 | 0.00661 | 0.005137 | 0.004328 | 0.000392 | 0.001334 | 0.008516 | 0.020879 | 0.025898 | 0.027771 | 0.025821 | 0.016264 | 0.015297 | 0.030309 | 0.025799 | 0.026915 | 0.016874 |
| 21.5972222 | 0.004298 | 0.000419 | 0.002567 | 0.001067 | 0.000677 | 0.002355 | 0.011465 | 0.007314 | 0.005055 | 0.001119 | 0.00117 | 0.010488 | 0.009734 | 0.015311 | 0.016589 | 0.014748 | 0.006395 | 0.007422 | 0.019211 | 0.016649 | 0.015552 | 0.006766 |
| 21.6111111 | 0.005564 | 0.000402 | 0.003278 | 0.000627 | 0.000691 | 0.002954 | 0.012827 | 0.006969 | 0.004601 | 0.001797 | 0.001753 | 0.012908 | 0.004399 | 0.00596 | 0.00756 | 0.005917 | 0.003632 | 0.006336 | 0.00839 | 0.008734 | 0.006186 | 0.002311 |
| 21.625 | 0.005085 | 0.00038 | 0.003036 | 0.000945 | 0.00076 | 0.002686 | 0.013377 | 0.004643 | 0.003415 | 0.001909 | 0.001969 | 0.012693 | 0.004054 | 0.001615 | 0.002886 | 0.001868 | 0.003338 | 0.007927 | 0.003635 | 0.003932 | 0.002592 | 0.002344 |
| 21.6388889 | 0.002981 | 0.000286 | 0.001779 | 0.002146 | 0.000504 | 0.002041 | 0.013935 | 0.002189 | 0.002351 | 0.001584 | 0.001734 | 0.009259 | 0.002904 | 0.001438 | 0.002214 | 0.001804 | 0.002359 | 0.005531 | 0.004618 | 0.002256 | 0.001773 | 0.005422 |
| 21.6527778 | 0.001547 | 0.000191 | 0.00114 | 0.003526 | 0.000352 | 0.001415 | 0.011665 | 0.001168 | 0.002241 | 0.001227 | 0.001384 | 0.004943 | 0.001077 | 0.002565 | 0.003451 | 0.003269 | 0.003524 | 0.00199 | 0.006222 | 0.001786 | 0.001012 | 0.009773 |
| 21.6666667 | 0.001716 | 0.000135 | 0.001847 | 0.004084 | 0.000252 | 0.001066 | 0.007517 | 0.001689 | 0.003343 | 0.000891 | 0.000907 | 0.002075 | 0.000686 | 0.003656 | 0.004982 | 0.00489 | 0.004444 | 0.00165 | 0.004686 | 0.002361 | 0.00144 | 0.012012 |
| 21.6805556 | 0.001559 | 0.000352 | 0.001899 | 0.003181 | 0.000227 | 0.000705 | 0.004041 | 0.002412 | 0.004465 | 0.000509 | 0.000511 | 0.001242 | 0.002158 | 0.00532 | 0.006927 | 0.007069 | 0.002944 | 0.004097 | 0.003346 | 0.005487 | 0.003953 | 0.013119 |
| 21.6944444 | 0.001501 | 0.000768 | 0.001168 | 0.001505 | 0.000428 | 0.000287 | 0.001791 | 0.00238 | 0.004048 | 0.000324 | 0.000427 | 0.001096 | 0.005652 | 0.007689 | 0.009312 | 0.009711 | 0.001195 | 0.008009 | 0.00705 | 0.010616 | 0.007616 | 0.014878 |
| 21.7083333 | 0.002994 | 0.00082 | 0.00143 | 0.0006 | 0.000569 | 0.000281 | 0.000859 | 0.001599 | 0.002222 | 0.000339 | 0.00062 | 0.001274 | 0.00817 | 0.009703 | 0.010922 | 0.011191 | 0.001345 | 0.011445 | 0.0125 | 0.014652 | 0.010086 | 0.016179 |
| 21.7222222 | 0.004303 | 0.000566 | 0.002267 | 0.000482 | 0.000711 | 0.000691 | 0.000621 | 0.00107 | 0.001416 | 0.000224 | 0.001267 | 0.003075 | 0.007608 | 0.010374 | 0.010655 | 0.010377 | 0.004262 | 0.012343 | 0.015297 | 0.015021 | 0.009526 | 0.015561 |
| 21.7361111 | 0.00426 | 0.000372 | 0.002236 | 0.000423 | 0.001141 | 0.001032 | 0.000367 | 0.001411 | 0.002432 | 0.000381 | 0.001933 | 0.005091 | 0.006015 | 0.009278 | 0.008469 | 0.00775 | 0.007855 | 0.010491 | 0.014101 | 0.010683 | 0.006003 | 0.012194 |
| 21.75 | 0.004395 | 0.000476 | 0.002188 | 0.000515 | 0.0018 | 0.001308 | 0.000534 | 0.001877 | 0.003298 | 0.001505 | 0.002385 | 0.005584 | 0.004094 | 0.006251 | 0.005656 | 0.005051 | 0.007831 | 0.007268 | 0.010134 | 0.006438 | 0.004247 | 0.007634 |
| 21.7638889 | 0.005447 | 0.000774 | 0.003647 | 0.000789 | 0.002286 | 0.00147 | 0.001338 | 0.001735 | 0.00431 | 0.002961 | 0.002626 | 0.004909 | 0.003618 | 0.006525 | 0.007464 | 0.0071 | 0.005634 | 0.009325 | 0.01286 | 0.010497 | 0.010276 | 0.008923 |
| 21.7777778 | 0.006249 | 0.000906 | 0.004698 | 0.001204 | 0.002036 | 0.001136 | 0.002027 | 0.001119 | 0.005635 | 0.003347 | 0.00211 | 0.0031 | 0.008953 | 0.015246 | 0.017009 | 0.016172 | 0.006826 | 0.021023 | 0.026257 | 0.019699 | 0.021789 | 0.01983 |
| 21.7916667 | 0.006338 | 0.001011 | 0.003987 | 0.001668 | 0.00148 | 0.000658 | 0.002235 | 0.000988 | 0.004984 | 0.002574 | 0.001527 | 0.0014 | 0.017836 | 0.025646 | 0.027813 | 0.025739 | 0.00859 | 0.033459 | 0.037473 | 0.024845 | 0.031251 | 0.031699 |
| 21.8055556 | 0.005862 | 0.001007 | 0.002624 | 0.002091 | 0.001495 | 0.00064 | 0.001708 | 0.001609 | 0.002783 | 0.001409 | 0.002288 | 0.001665 | 0.024117 | 0.030574 | 0.033312 | 0.029796 | 0.008698 | 0.037588 | 0.037561 | 0.024504 | 0.03426 | 0.036626 |
| 21.8194444 | 0.003845 | 0.000885 | 0.001331 | 0.002045 | 0.001655 | 0.000737 | 0.001815 | 0.002547 | 0.002534 | 0.000992 | 0.004313 | 0.003243 | 0.025389 | 0.030654 | 0.032901 | 0.028391 | 0.013124 | 0.033318 | 0.03042 | 0.022456 | 0.031673 | 0.0352 |
| 21.8333333 | 0.002471 | 0.00077 | 0.000584 | 0.001467 | 0.001165 | 0.000629 | 0.004539 | 0.003769 | 0.005123 | 0.001898 | 0.006172 | 0.005258 | 0.019852 | 0.026263 | 0.026866 | 0.022054 | 0.020482 | 0.022626 | 0.019264 | 0.020581 | 0.024813 | 0.028242 |
| 21.8472222 | 0.005227 | 0.000839 | 0.000618 | 0.001661 | 0.00075 | 0.000868 | 0.012319 | 0.00462 | 0.006692 | 0.002556 | 0.005521 | 0.008451 | 0.011333 | 0.017127 | 0.015922 | 0.012218 | 0.020556 | 0.011804 | 0.01054 | 0.015846 | 0.01358 | 0.016027 |
| 21.8611111 | 0.009643 | 0.001191 | 0.000743 | 0.003117 | 0.001295 | 0.001754 | 0.0281 | 0.004592 | 0.005154 | 0.002131 | 0.002751 | 0.013321 | 0.010087 | 0.008645 | 0.007543 | 0.007206 | 0.014241 | 0.010363 | 0.015229 | 0.009957 | 0.005459 | 0.009025 |
| 21.875 | 0.012164 | 0.001305 | 0.000852 | 0.004757 | 0.002167 | 0.002868 | 0.032708 | 0.004979 | 0.002886 | 0.00158 | 0.00148 | 0.016539 | 0.015925 | 0.008638 | 0.009137 | 0.012429 | 0.015799 | 0.016978 | 0.028982 | 0.012151 | 0.006893 | 0.016836 |
| 21.8888889 | 0.012529 | 0.000894 | 0.001697 | 0.005829 | 0.002718 | 0.003443 | 0.021579 | 0.007229 | 0.002471 | 0.001638 | 0.002091 | 0.012682 | 0.022294 | 0.016576 | 0.016621 | 0.021714 | 0.027469 | 0.023712 | 0.037306 | 0.022406 | 0.012919 | 0.031706 |
| 21.9027778 | 0.012448 | 0.000535 | 0.00257 | 0.006358 | 0.003071 | 0.003121 | 0.01932 | 0.009502 | 0.004103 | 0.0025 | 0.003231 | 0.005389 | 0.028658 | 0.026666 | 0.024416 | 0.031079 | 0.038458 | 0.029896 | 0.039314 | 0.032177 | 0.021222 | 0.04506 |
| 21.9166667 | 0.013847 | 0.000883 | 0.002312 | 0.006436 | 0.003301 | 0.002309 | 0.018564 | 0.008228 | 0.005522 | 0.003663 | 0.005278 | 0.002943 | 0.034058 | 0.034767 | 0.030353 | 0.038664 | 0.044171 | 0.036683 | 0.04164 | 0.036611 | 0.03101 | 0.053317 |
| 21.9305556 | 0.015668 | 0.001365 | 0.001298 | 0.006205 | 0.002846 | 0.001368 | 0.010677 | 0.004619 | 0.004977 | 0.003575 | 0.007815 | 0.007195 | 0.035692 | 0.036574 | 0.0311 | 0.038833 | 0.045 | 0.040368 | 0.042048 | 0.035314 | 0.036049 | 0.052056 |
| 21.9444444 | 0.015147 | 0.001391 | 0.001193 | 0.006222 | 0.00154 | 0.00056 | 0.00503 | 0.003545 | 0.003606 | 0.00197 | 0.008883 | 0.016351 | 0.033533 | 0.031378 | 0.026134 | 0.032024 | 0.040772 | 0.03921 | 0.037073 | 0.03123 | 0.034454 | 0.042062 |
| 21.9583333 | 0.011606 | 0.00184 | 0.002998 | 0.006518 | 0.000726 | 0.000229 | 0.004725 | 0.0058 | 0.00387 | 0.000928 | 0.008488 | 0.023711 | 0.030083 | 0.023961 | 0.018966 | 0.024411 | 0.033129 | 0.036285 | 0.030858 | 0.027857 | 0.031304 | 0.029673 |
| 21.9722222 | 0.006955 | 0.002669 | 0.004949 | 0.006481 | 0.001318 | 0.000283 | 0.010645 | 0.007709 | 0.005408 | 0.001253 | 0.011817 | 0.023612 | 0.025577 | 0.01765 | 0.012348 | 0.018335 | 0.027585 | 0.033241 | 0.02795 | 0.025536 | 0.029574 | 0.019721 |
| 21.9861111 | 0.003653 | 0.002389 | 0.004712 | 0.006067 | 0.002377 | 0.000376 | 0.017154 | 0.007045 | 0.004507 | 0.001916 | 0.019733 | 0.018905 | 0.018752 | 0.013474 | 0.007028 | 0.012365 | 0.028769 | 0.028926 | 0.026662 | 0.022456 | 0.027955 | 0.013062 |
| 22 | 0.001853 | 0.001722 | 0.00345 | 0.004959 | 0.003164 | 0.001119 | 0.01892 | 0.004343 | 0.002313 | 0.002702 | 0.023176 | 0.013263 | 0.011671 | 0.011981 | 0.004337 | 0.006546 | 0.033475 | 0.022913 | 0.022289 | 0.018409 | 0.026445 | 0.009992 |
| 22.0138889 | 0.001494 | 0.002682 | 0.003788 | 0.003208 | 0.004614 | 0.003828 | 0.015821 | 0.003059 | 0.002994 | 0.00474 | 0.015912 | 0.016127 | 0.007024 | 0.01154 | 0.004742 | 0.002616 | 0.034199 | 0.015716 | 0.014429 | 0.014232 | 0.024829 | 0.010611 |
| 22.0277778 | 0.003092 | 0.003877 | 0.005666 | 0.002957 | 0.006692 | 0.007865 | 0.010426 | 0.006129 | 0.004323 | 0.007642 | 0.007052 | 0.030094 | 0.006178 | 0.011257 | 0.006909 | 0.001717 | 0.028997 | 0.009659 | 0.008126 | 0.010922 | 0.022786 | 0.013204 |
| 22.0416667 | 0.004411 | 0.003715 | 0.007112 | 0.004127 | 0.006753 | 0.010009 | 0.009755 | 0.007122 | 0.004136 | 0.008243 | 0.004418 | 0.032333 | 0.005865 | 0.010737 | 0.008216 | 0.002438 | 0.022061 | 0.008434 | 0.00497 | 0.008409 | 0.018737 | 0.012543 |
| 22.0555556 | 0.005145 | 0.005054 | 0.007963 | 0.003669 | 0.003883 | 0.008531 | 0.019585 | 0.006962 | 0.003768 | 0.005338 | 0.004353 | 0.018326 | 0.009387 | 0.008868 | 0.007271 | 0.006064 | 0.016497 | 0.008377 | 0.00695 | 0.006738 | 0.015213 | 0.011165 |
| 22.0694444 | 0.006476 | 0.009536 | 0.009364 | 0.002694 | 0.001467 | 0.005304 | 0.038338 | 0.014167 | 0.002903 | 0.002452 | 0.011054 | 0.012908 | 0.020034 | 0.010044 | 0.007654 | 0.013303 | 0.01226 | 0.005557 | 0.01757 | 0.007836 | 0.022852 | 0.018823 |
| 22.0833333 | 0.005606 | 0.013318 | 0.009721 | 0.00327 | 0.001163 | 0.002806 | 0.055814 | 0.019161 | 0.002105 | 0.001734 | 0.026359 | 0.017636 | 0.019778 | 0.014875 | 0.009567 | 0.016337 | 0.00769 | 0.003682 | 0.025135 | 0.007944 | 0.030476 | 0.027286 |
| 22.0972222 | 0.003412 | 0.010742 | 0.007265 | 0.003158 | 0.001161 | 0.001994 | 0.054772 | 0.013193 | 0.00187 | 0.00254 | 0.034962 | 0.015248 | 0.009585 | 0.016181 | 0.010663 | 0.014937 | 0.00484 | 0.00527 | 0.020707 | 0.006319 | 0.020978 | 0.02427 |
| 22.1111111 | 0.003776 | 0.006251 | 0.003949 | 0.001755 | 0.001257 | 0.00171 | 0.033566 | 0.007619 | 0.00176 | 0.004395 | 0.02473 | 0.010939 | 0.006308 | 0.014564 | 0.01328 | 0.016863 | 0.00854 | 0.01258 | 0.014281 | 0.01049 | 0.00983 | 0.01782 |
| 22.125 | 0.00694 | 0.007393 | 0.002126 | 0.001117 | 0.001655 | 0.001409 | 0.013854 | 0.009637 | 0.002654 | 0.004436 | 0.012894 | 0.011721 | 0.004429 | 0.014876 | 0.017855 | 0.019968 | 0.015088 | 0.022937 | 0.012662 | 0.019539 | 0.006733 | 0.017965 |
| 22.1388889 | 0.009304 | 0.00742 | 0.001484 | 0.001407 | 0.001924 | 0.002149 | 0.00937 | 0.010461 | 0.002664 | 0.002423 | 0.01401 | 0.012659 | 0.001221 | 0.017539 | 0.022834 | 0.019598 | 0.018299 | 0.02656 | 0.011982 | 0.02688 | 0.006651 | 0.021586 |
| 22.1527778 | 0.007491 | 0.004949 | 0.000902 | 0.001676 | 0.002186 | 0.002622 | 0.012562 | 0.00694 | 0.001918 | 0.001264 | 0.01303 | 0.01724 | 0.002957 | 0.021202 | 0.026519 | 0.019722 | 0.022031 | 0.02342 | 0.009815 | 0.030618 | 0.014409 | 0.023016 |
| 22.1666667 | 0.00511 | 0.005556 | 0.000434 | 0.001953 | 0.001907 | 0.001574 | 0.012426 | 0.004214 | 0.002029 | 0.001495 | 0.00594 | 0.017003 | 0.01165 | 0.023571 | 0.026732 | 0.022791 | 0.035244 | 0.023606 | 0.009149 | 0.030528 | 0.025817 | 0.022957 |
| 22.1805556 | 0.00402 | 0.005137 | 0.000582 | 0.002899 | 0.000959 | 0.000834 | 0.008285 | 0.003094 | 0.002722 | 0.002748 | 0.003266 | 0.011411 | 0.022847 | 0.022955 | 0.023818 | 0.022791 | 0.05182 | 0.027962 | 0.011252 | 0.025582 | 0.028354 | 0.023132 |
| 22.1944444 | 0.003172 | 0.003134 | 0.000786 | 0.004771 | 0.000382 | 0.001502 | 0.005628 | 0.002184 | 0.004694 | 0.00324 | 0.007025 | 0.013679 | 0.028562 | 0.020918 | 0.02038 | 0.018756 | 0.053211 | 0.028363 | 0.012811 | 0.015599 | 0.020917 | 0.022425 |
| 22.2083333 | 0.003119 | 0.004625 | 0.000607 | 0.006299 | 0.000466 | 0.002012 | 0.007921 | 0.001612 | 0.004544 | 0.002127 | 0.01165 | 0.013995 | 0.024869 | 0.020191 | 0.018081 | 0.017031 | 0.041311 | 0.024059 | 0.012663 | 0.006775 | 0.012312 | 0.020031 |
| 22.2222222 | 0.003568 | 0.008243 | 0.001003 | 0.006287 | 0.000547 | 0.00143 | 0.015434 | 0.002559 | 0.001759 | 0.001121 | 0.012635 | 0.007644 | 0.013933 | 0.019592 | 0.015269 | 0.015509 | 0.033396 | 0.019276 | 0.014841 | 0.010554 | 0.008918 | 0.015881 |
| 22.2361111 | 0.004868 | 0.007309 | 0.002666 | 0.005564 | 0.000537 | 0.001374 | 0.023101 | 0.005662 | 0.002219 | 0.001189 | 0.013034 | 0.003332 | 0.004446 | 0.016674 | 0.010746 | 0.009494 | 0.032446 | 0.013897 | 0.021684 | 0.027223 | 0.010818 | 0.010314 |
| 22.25 | 0.005653 | 0.003299 | 0.004539 | 0.004928 | 0.000515 | 0.002599 | 0.021466 | 0.010353 | 0.007344 | 0.001465 | 0.014171 | 0.002809 | 0.004651 | 0.013846 | 0.007541 | 0.005425 | 0.030875 | 0.008976 | 0.025822 | 0.035364 | 0.01523 | 0.005539 |
| 22.2638889 | 0.004895 | 0.002988 | 0.004907 | 0.004344 | 0.000393 | 0.003497 | 0.010807 | 0.013932 | 0.011588 | 0.001033 | 0.011376 | 0.004461 | 0.010492 | 0.013267 | 0.007663 | 0.008267 | 0.026873 | 0.008573 | 0.018703 | 0.022279 | 0.017034 | 0.003038 |
| 22.2777778 | 0.004923 | 0.005953 | 0.003304 | 0.004195 | 0.000529 | 0.002989 | 0.00405 | 0.013966 | 0.00928 | 0.000653 | 0.006517 | 0.007718 | 0.011875 | 0.012179 | 0.008459 | 0.011846 | 0.023692 | 0.011537 | 0.007492 | 0.010136 | 0.014206 | 0.001965 |

| | | | | | | | | | | | | | | | | | | | | | | |
|---|---|---|---|---|---|---|---|---|---|---|---|---|---|---|---|---|---|---|---|---|---|---|
| 22.2916667 | 0.006305 | 0.007543 | 0.001357 | 0.004156 | 0.00071 | 0.001626 | 0.007422 | 0.011503 | 0.004394 | 0.000989 | 0.005551 | 0.010033 | 0.0121 | 0.009843 | 0.0083 | 0.011623 | 0.022933 | 0.01402 | 0.007826 | 0.010407 | 0.011054 | 0.002587 |
| 22.3055556 | 0.006178 | 0.005672 | 0.001154 | 0.003057 | 0.000802 | 0.00086 | 0.013926 | 0.008701 | 0.002872 | 0.001042 | 0.008112 | 0.007209 | 0.017423 | 0.009508 | 0.008551 | 0.009949 | 0.021784 | 0.015427 | 0.020825 | 0.015335 | 0.010109 | 0.00505 |
| 22.3194444 | 0.003817 | 0.004951 | 0.001939 | 0.001352 | 0.001461 | 0.001218 | 0.016539 | 0.00643 | 0.003028 | 0.000757 | 0.008246 | 0.002725 | 0.01906 | 0.01183 | 0.009175 | 0.008166 | 0.01589 | 0.015315 | 0.030452 | 0.02355 | 0.008678 | 0.006797 |
| 22.3333333 | 0.001958 | 0.006722 | 0.001731 | 0.000534 | 0.002166 | 0.001285 | 0.016127 | 0.004766 | 0.004126 | 0.00095 | 0.004486 | 0.002629 | 0.011758 | 0.01341 | 0.008362 | 0.006155 | 0.007593 | 0.012717 | 0.02491 | 0.023585 | 0.005558 | 0.006593 |
| 22.3472222 | 0.002022 | 0.006006 | 0.001316 | 0.000409 | 0.002009 | 0.000749 | 0.014972 | 0.003039 | 0.004527 | 0.000893 | 0.002414 | 0.008932 | 0.00427 | 0.010965 | 0.005212 | 0.004491 | 0.003537 | 0.008423 | 0.012064 | 0.01316 | 0.002448 | 0.005527 |
| 22.3611111 | 0.002474 | 0.00532 | 0.002266 | 0.000573 | 0.001465 | 0.000562 | 0.012068 | 0.001316 | 0.003156 | 0.000391 | 0.003351 | 0.01774 | 0.003451 | 0.006052 | 0.003196 | 0.002878 | 0.004714 | 0.004237 | 0.005391 | 0.005186 | 0.000728 | 0.004573 |
| 22.375 | 0.001864 | 0.007275 | 0.003318 | 0.001351 | 0.001335 | 0.001019 | 0.008798 | 0.001711 | 0.003002 | 0.000399 | 0.005786 | 0.017709 | 0.005422 | 0.003736 | 0.005014 | 0.001531 | 0.007248 | 0.00227 | 0.004661 | 0.003773 | 0.000722 | 0.00549 |
| 22.3888889 | 0.001082 | 0.006582 | 0.002805 | 0.00157 | 0.00155 | 0.001695 | 0.008844 | 0.00588 | 0.002968 | 0.000899 | 0.006001 | 0.009617 | 0.005658 | 0.003626 | 0.006145 | 0.001205 | 0.009298 | 0.003586 | 0.008105 | 0.007156 | 0.001953 | 0.008735 |
| 22.4027778 | 0.001849 | 0.003026 | 0.00233 | 0.001024 | 0.001444 | 0.001508 | 0.011542 | 0.011116 | 0.003202 | 0.000928 | 0.005969 | 0.004758 | 0.004824 | 0.003539 | 0.004036 | 0.001551 | 0.012505 | 0.005995 | 0.016566 | 0.011062 | 0.003581 | 0.01025 |
| 22.4166667 | 0.004247 | 0.001193 | 0.002884 | 0.000961 | 0.001014 | 0.001067 | 0.01413 | 0.011881 | 0.005679 | 0.000761 | 0.013458 | 0.005143 | 0.007334 | 0.005848 | 0.001985 | 0.00341 | 0.020089 | 0.006866 | 0.019771 | 0.012241 | 0.005346 | 0.00745 |
| 22.4305556 | 0.006878 | 0.002066 | 0.002391 | 0.000893 | 0.000992 | 0.001276 | 0.017765 | 0.008704 | 0.007502 | 0.001015 | 0.023936 | 0.013204 | 0.017943 | 0.006891 | 0.002382 | 0.006355 | 0.033678 | 0.007501 | 0.012039 | 0.01287 | 0.008164 | 0.004796 |
| 22.4444444 | 0.008576 | 0.003308 | 0.001744 | 0.000728 | 0.001274 | 0.001687 | 0.02204 | 0.006752 | 0.006926 | 0.001504 | 0.028815 | 0.029962 | 0.026152 | 0.006348 | 0.004425 | 0.006452 | 0.04353 | 0.009685 | 0.004223 | 0.014797 | 0.011326 | 0.005243 |
| 22.4583333 | 0.00881 | 0.003931 | 0.003729 | 0.001693 | 0.002386 | 0.003928 | 0.020507 | 0.005353 | 0.00467 | 0.003575 | 0.023978 | 0.038722 | 0.021011 | 0.00935 | 0.006461 | 0.005581 | 0.0405 | 0.011082 | 0.003119 | 0.016262 | 0.011745 | 0.005995 |
| 22.4722222 | 0.007526 | 0.00651 | 0.007336 | 0.003534 | 0.00467 | 0.008083 | 0.012205 | 0.003142 | 0.001916 | 0.006693 | 0.012001 | 0.02657 | 0.010995 | 0.009848 | 0.007914 | 0.007668 | 0.033739 | 0.008865 | 0.002801 | 0.015244 | 0.009097 | 0.005398 |
| 22.4861111 | 0.007172 | 0.008905 | 0.010359 | 0.003906 | 0.005405 | 0.009498 | 0.006563 | 0.00245 | 0.00076 | 0.006708 | 0.00481 | 0.009941 | 0.011072 | 0.008846 | 0.009506 | 0.009524 | 0.029692 | 0.004899 | 0.002187 | 0.01048 | 0.005727 | 0.004595 |
| 22.5 | 0.00911 | 0.00661 | 0.011 | 0.003038 | 0.003332 | 0.006734 | 0.010223 | 0.003243 | 0.00124 | 0.003621 | 0.006739 | 0.004791 | 0.022732 | 0.0086 | 0.012581 | 0.010869 | 0.023503 | 0.003011 | 0.004354 | 0.007557 | 0.003041 | 0.005053 |
| 22.5138889 | 0.010154 | 0.002879 | 0.008584 | 0.003599 | 0.001223 | 0.00351 | 0.020769 | 0.004059 | 0.002098 | 0.001778 | 0.007964 | 0.006144 | 0.033209 | 0.007303 | 0.016505 | 0.015764 | 0.013892 | 0.004149 | 0.005225 | 0.011727 | 0.002501 | 0.006248 |
| 22.5277778 | 0.008854 | 0.002283 | 0.00654 | 0.004374 | 0.00087 | 0.002248 | 0.032104 | 0.003969 | 0.002939 | 0.002009 | 0.004116 | 0.009948 | 0.029261 | 0.01318 | 0.020988 | 0.025598 | 0.008753 | 0.00501 | 0.005964 | 0.017425 | 0.005904 | 0.010121 |
| 22.5416667 | 0.006489 | 0.003546 | 0.007767 | 0.00341 | 0.000853 | 0.002905 | 0.038631 | 0.003046 | 0.004363 | 0.002352 | 0.001092 | 0.019264 | 0.022688 | 0.027015 | 0.031524 | 0.040211 | 0.019101 | 0.01003 | 0.016766 | 0.025598 | 0.016829 | 0.022951 |
| 22.5555556 | 0.00368 | 0.003278 | 0.009903 | 0.001769 | 0.000783 | 0.003997 | 0.03696 | 0.002136 | 0.005998 | 0.001955 | 0.000993 | 0.025722 | 0.034465 | 0.040527 | 0.045915 | 0.052576 | 0.038194 | 0.025971 | 0.033836 | 0.038436 | 0.031826 | 0.039586 |
| 22.5694444 | 0.001358 | 0.002246 | 0.009048 | 0.000727 | 0.001529 | 0.003318 | 0.027249 | 0.001911 | 0.006666 | 0.00103 | 0.001379 | 0.017635 | 0.045048 | 0.041722 | 0.046193 | 0.048802 | 0.043117 | 0.035433 | 0.037304 | 0.041748 | 0.033335 | 0.042401 |
| 22.5833333 | 0.000475 | 0.002947 | 0.005645 | 0.000426 | 0.002027 | 0.001997 | 0.015614 | 0.002154 | 0.006568 | 0.000412 | 0.00166 | 0.005883 | 0.03475 | 0.028517 | 0.02855 | 0.029549 | 0.028749 | 0.024849 | 0.023148 | 0.029369 | 0.01893 | 0.028722 |
| 22.5972222 | 0.000397 | 0.003767 | 0.004233 | 0.000744 | 0.001738 | 0.001849 | 0.008997 | 0.001857 | 0.005184 | 0.000271 | 0.00226 | 0.002584 | 0.018553 | 0.015467 | 0.010573 | 0.012478 | 0.012926 | 0.010361 | 0.008114 | 0.015829 | 0.008492 | 0.01449 |
| 22.6111111 | 0.000786 | 0.003416 | 0.00531 | 0.001633 | 0.001548 | 0.001294 | 0.007583 | 0.001113 | 0.002764 | 0.000314 | 0.002366 | 0.003756 | 0.009 | 0.0107 | 0.002342 | 0.006861 | 0.006268 | 0.004732 | 0.001812 | 0.010423 | 0.007357 | 0.009307 |
| 22.625 | 0.002209 | 0.002546 | 0.0051 | 0.002068 | 0.001259 | 0.000323 | 0.007795 | 0.000897 | 0.001688 | 0.000477 | 0.00151 | 0.003331 | 0.00407 | 0.009792 | 0.001222 | 0.007711 | 0.004451 | 0.003151 | 0.000572 | 0.009021 | 0.006926 | 0.010051 |
| 22.6388889 | 0.004393 | 0.001598 | 0.002842 | 0.001429 | 0.000587 | 0.000172 | 0.006987 | 0.001377 | 0.002163 | 0.000578 | 0.000621 | 0.001654 | 0.001449 | 0.008019 | 0.00247 | 0.008813 | 0.002721 | 0.002062 | 0.000405 | 0.007688 | 0.005236 | 0.011488 |
| 22.6527778 | 0.006822 | 0.001328 | 0.001171 | 0.000588 | 0.000221 | 0.000499 | 0.004247 | 0.001463 | 0.002287 | 0.000855 | 0.000533 | 0.00152 | 0.002068 | 0.005926 | 0.003145 | 0.008102 | 0.001347 | 0.003193 | 0.000642 | 0.006204 | 0.00324 | 0.010558 |
| 22.6666667 | 0.00866 | 0.001392 | 0.001131 | 0.000382 | 0.000222 | 0.000677 | 0.00227 | 0.000927 | 0.001936 | 0.001042 | 0.000705 | 0.002067 | 0.002949 | 0.0037 | 0.002361 | 0.005318 | 0.00228 | 0.005131 | 0.001763 | 0.005013 | 0.002089 | 0.006342 |
| 22.6805556 | 0.008462 | 0.001009 | 0.001097 | 0.000699 | 0.00034 | 0.000542 | 0.004243 | 0.0012 | 0.003212 | 0.000715 | 0.000664 | 0.002529 | 0.002287 | 0.002062 | 0.001156 | 0.002516 | 0.005519 | 0.00545 | 0.003366 | 0.003934 | 0.002363 | 0.002762 |
| 22.6944444 | 0.005421 | 0.000476 | 0.000914 | 0.000775 | 0.000445 | 0.000485 | 0.009003 | 0.002898 | 0.005755 | 0.000422 | 0.000634 | 0.002574 | 0.001998 | 0.002636 | 0.000738 | 0.002037 | 0.0088 | 0.003709 | 0.003676 | 0.003127 | 0.002172 | 0.003289 |
| 22.7083333 | 0.004168 | 0.000488 | 0.001577 | 0.000862 | 0.000437 | 0.000529 | 0.014324 | 0.00437 | 0.005928 | 0.000611 | 0.00073 | 0.001607 | 0.002358 | 0.002885 | 0.000603 | 0.00225 | 0.010078 | 0.001589 | 0.002587 | 0.003751 | 0.001415 | 0.005132 |
| 22.7222222 | 0.009706 | 0.001075 | 0.002112 | 0.001572 | 0.000626 | 0.000369 | 0.016385 | 0.003931 | 0.003625 | 0.001064 | 0.001133 | 0.000775 | 0.003754 | 0.002056 | 0.000423 | 0.002008 | 0.009526 | 0.00071 | 0.002039 | 0.005197 | 0.00178 | 0.006165 |
| 22.7361111 | 0.016539 | 0.001347 | 0.001788 | 0.002113 | 0.000734 | 0.000135 | 0.011704 | 0.002228 | 0.002824 | 0.001234 | 0.001097 | 0.001423 | 0.004013 | 0.002209 | 0.000323 | 0.002048 | 0.007024 | 0.000965 | 0.002815 | 0.005122 | 0.002989 | 0.006132 |
| 22.75 | 0.017536 | 0.000998 | 0.002319 | 0.001619 | 0.000572 | 0.000225 | 0.00615 | 0.001335 | 0.004884 | 0.001229 | 0.000721 | 0.002884 | 0.002567 | 0.00293 | 0.000312 | 0.001938 | 0.005454 | 0.002596 | 0.003936 | 0.00308 | 0.00461 | 0.004311 |
| 22.7638889 | 0.01373 | 0.000537 | 0.004322 | 0.00077 | 0.000501 | 0.000787 | 0.007959 | 0.001619 | 0.007208 | 0.001507 | 0.001294 | 0.00347 | 0.002614 | 0.00297 | 0.000477 | 0.001218 | 0.01095 | 0.004855 | 0.004482 | 0.001111 | 0.005877 | 0.002291 |
| 22.7777778 | 0.009203 | 0.000274 | 0.005649 | 0.000418 | 0.000487 | 0.00154 | 0.013186 | 0.001617 | 0.007828 | 0.001851 | 0.003146 | 0.002728 | 0.003004 | 0.002377 | 0.000879 | 0.000504 | 0.01714 | 0.006444 | 0.004229 | 0.000513 | 0.004787 | 0.002418 |
| 22.7916667 | 0.00526 | 0.000245 | 0.005332 | 0.000472 | 0.000319 | 0.001859 | 0.012089 | 0.001051 | 0.006303 | 0.001754 | 0.00516 | 0.00154 | 0.002404 | 0.001808 | 0.000976 | 0.0006 | 0.013676 | 0.006951 | 0.002927 | 0.000476 | 0.004562 | 0.004621 |
| 22.8055556 | 0.00309 | 0.000496 | 0.003852 | 0.001248 | 0.000125 | 0.001636 | 0.007284 | 0.00122 | 0.003838 | 0.001183 | 0.005551 | 0.000686 | 0.001881 | 0.001431 | 0.000781 | 0.001428 | 0.007448 | 0.006211 | 0.00314 | 0.000763 | 0.009098 | 0.007411 |
| 22.8194444 | 0.005556 | 0.001102 | 0.002469 | 0.002636 | 0.000123 | 0.00138 | 0.008717 | 0.002761 | 0.003545 | 0.000583 | 0.004499 | 0.000514 | 0.001622 | 0.000948 | 0.001305 | 0.002458 | 0.010085 | 0.004364 | 0.006874 | 0.002055 | 0.012323 | 0.009312 |
| 22.8333333 | 0.012092 | 0.001509 | 0.002933 | 0.003558 | 0.000176 | 0.001033 | 0.016157 | 0.004239 | 0.00478 | 0.000226 | 0.00369 | 0.001564 | 0.001616 | 0.000588 | 0.002121 | 0.003493 | 0.016586 | 0.002154 | 0.009322 | 0.003591 | 0.008519 | 0.008166 |
| 22.8472222 | 0.018085 | 0.001219 | 0.003452 | 0.003513 | 0.000176 | 0.00051 | 0.023944 | 0.003935 | 0.003968 | 0.000283 | 0.003343 | 0.004196 | 0.001869 | 0.001286 | 0.001906 | 0.003723 | 0.01412 | 0.001037 | 0.008516 | 0.00491 | 0.005176 | 0.006153 |
| 22.8611111 | 0.019319 | 0.000848 | 0.00314 | 0.003373 | 0.000335 | 0.000338 | 0.027469 | 0.002117 | 0.002888 | 0.001021 | 0.003235 | 0.008229 | 0.003348 | 0.00338 | 0.002001 | 0.00326 | 0.010766 | 0.00605 | 0.008707 | 0.006787 | 0.009564 | 0.009189 |
| 22.875 | 0.016598 | 0.001355 | 0.004924 | 0.003987 | 0.000804 | 0.000797 | 0.022631 | 0.000643 | 0.00531 | 0.002451 | 0.004572 | 0.011765 | 0.007088 | 0.006147 | 0.004257 | 0.00509 | 0.016009 | 0.014969 | 0.011219 | 0.009105 | 0.015796 | 0.014645 |
| 22.8888889 | 0.013182 | 0.002985 | 0.007046 | 0.004768 | 0.001474 | 0.001541 | 0.01257 | 0.001146 | 0.009019 | 0.00431 | 0.005807 | 0.011181 | 0.013432 | 0.008422 | 0.006713 | 0.009272 | 0.018091 | 0.016893 | 0.014588 | 0.011485 | 0.018286 | 0.015586 |
| 22.9027778 | 0.009142 | 0.004764 | 0.005203 | 0.005056 | 0.00223 | 0.002149 | 0.005987 | 0.003885 | 0.010807 | 0.006196 | 0.004241 | 0.006344 | 0.019984 | 0.009609 | 0.007107 | 0.011672 | 0.017559 | 0.014033 | 0.018134 | 0.013929 | 0.017956 | 0.012978 |
| 22.9166667 | 0.005194 | 0.005067 | 0.002261 | 0.00488 | 0.003019 | 0.002396 | 0.008646 | 0.008397 | 0.011319 | 0.007379 | 0.00327 | 0.002018 | 0.022898 | 0.009785 | 0.005777 | 0.011301 | 0.02097 | 0.012686 | 0.021646 | 0.015653 | 0.016405 | 0.011733 |
| 22.9305556 | 0.004546 | 0.003377 | 0.00217 | 0.004645 | 0.003503 | 0.001801 | 0.018349 | 0.014759 | 0.010578 | 0.006717 | 0.007764 | 0.001893 | 0.021836 | 0.009016 | 0.003723 | 0.010087 | 0.020377 | 0.011547 | 0.02472 | 0.014741 | 0.014681 | 0.013161 |
| 22.9444444 | 0.0069 | 0.002012 | 0.002782 | 0.005175 | 0.003222 | 0.000902 | 0.026555 | 0.021794 | 0.00818 | 0.003968 | 0.018251 | 0.006034 | 0.019517 | 0.007849 | 0.001735 | 0.008923 | 0.015334 | 0.011394 | 0.026245 | 0.011939 | 0.012946 | 0.015091 |
| 22.9583333 | 0.008665 | 0.002274 | 0.002404 | 0.006592 | 0.002586 | 0.000598 | 0.027447 | 0.026366 | 0.006542 | 0.00117 | 0.027851 | 0.012991 | 0.020046 | 0.00783 | 0.001324 | 0.008474 | 0.013775 | 0.013786 | 0.024737 | 0.011745 | 0.012485 | 0.015943 |
| 22.9722222 | 0.008649 | 0.002265 | 0.00242 | 0.007719 | 0.002539 | 0.000614 | 0.02258 | 0.028629 | 0.006827 | 0.000625 | 0.028746 | 0.019949 | 0.026226 | 0.010421 | 0.002545 | 0.009892 | 0.018275 | 0.016995 | 0.02097 | 0.014099 | 0.014963 | 0.01605 |
| 22.9861111 | 0.007869 | 0.002403 | 0.00324 | 0.007487 | 0.002792 | 0.000962 | 0.017094 | 0.027667 | 0.007393 | 0.00172 | 0.023396 | 0.022702 | 0.035143 | 0.014862 | 0.003381 | 0.011978 | 0.025869 | 0.018643 | 0.017898 | 0.014521 | 0.018237 | 0.015004 |
| 23 | 0.005608 | 0.002876 | 0.003131 | 0.005502 | 0.002251 | 0.001311 | 0.013493 | 0.019163 | 0.006735 | 0.001963 | 0.015287 | 0.019426 | 0.037804 | 0.017609 | 0.003315 | 0.010925 | 0.034456 | 0.015223 | 0.01552 | 0.012156 | 0.017567 | 0.011326 |
| 23.0138889 | 0.003966 | 0.002305 | 0.002529 | 0.003323 | 0.002556 | 0.002716 | 0.01039 | 0.009941 | 0.004218 | 0.002441 | 0.008683 | 0.011888 | 0.027184 | 0.015384 | 0.00274 | 0.0066 | 0.041066 | 0.009979 | 0.010019 | 0.007533 | 0.011353 | 0.006849 |
| 23.0277778 | 0.005628 | 0.001627 | 0.003886 | 0.003532 | 0.004989 | 0.006395 | 0.008234 | 0.009396 | 0.001842 | 0.005162 | 0.008205 | 0.004666 | 0.013656 | 0.009485 | 0.002644 | 0.004302 | 0.034519 | 0.010918 | 0.005183 | 0.004184 | 0.005223 | 0.007409 |
| 23.0416667 | 0.005565 | 0.002647 | 0.005768 | 0.004425 | 0.005597 | 0.008416 | 0.010087 | 0.009677 | 0.002012 | 0.006171 | 0.007502 | 0.002454 | 0.012485 | 0.004523 | 0.003672 | 0.005474 | 0.017127 | 0.012459 | 0.006535 | 0.003864 | 0.003984 | 0.01374 |
| 23.0555556 | 0.002409 | 0.007313 | 0.005864 | 0.003101 | 0.003171 | 0.005933 | 0.016508 | 0.011111 | 0.006502 | 0.004601 | 0.003638 | 0.004914 | 0.019721 | 0.006342 | 0.005008 | 0.006334 | 0.014255 | 0.008486 | 0.00937 | 0.006495 | 0.006541 | 0.021573 |
| 23.0694444 | 0.003502 | 0.011575 | 0.004831 | 0.001185 | 0.001484 | 0.003518 | 0.025419 | 0.022525 | 0.013609 | 0.004402 | 0.005504 | 0.007137 | 0.029595 | 0.016866 | 0.0069 | 0.007695 | 0.031932 | 0.00715 | 0.016718 | 0.012775 | 0.009331 | 0.031498 |
| 23.0833333 | 0.011613 | 0.011203 | 0.003226 | 0.000784 | 0.002795 | 0.009269 | 0.033575 | 0.027538 | 0.014352 | 0.003951 | 0.016535 | 0.005541 | 0.047111 | 0.024617 | 0.010797 | 0.010544 | 0.049435 | 0.013262 | 0.046416 | 0.013411 | 0.030243 | 0.049109 |
| 23.0972222 | 0.015862 | 0.010441 | 0.00219 | 0.001346 | 0.004118 | 0.019621 | 0.036615 | 0.016849 | 0.007599 | 0.003182 | 0.021056 | 0.003301 | 0.059953 | 0.020364 | 0.014038 | 0.011057 | 0.055119 | 0.017178 | 0.076272 | 0.00682 | 0.067377 | 0.066626 |
| 23.1111111 | 0.011273 | 0.008426 | 0.002099 | 0.001359 | 0.002274 | 0.019528 | 0.029932 | 0.009072 | 0.003288 | 0.004684 | 0.012578 | 0.002724 | 0.043008 | 0.012092 | 0.011856 | 0.007799 | 0.040604 | 0.011659 | 0.05703 | 0.002366 | 0.063744 | 0.057113 |
| 23.125 | 0.009419 | 0.004298 | 0.001476 | 0.000943 | 0.001094 | 0.009935 | 0.01632 | 0.011353 | 0.004508 | 0.005535 | 0.007328 | 0.002445 | 0.017821 | 0.009282 | 0.010932 | 0.004877 | 0.02982 | 0.004402 | 0.017112 | 0.001821 | 0.0303 | 0.028024 |
| 23.1388889 | 0.011861 | 0.002619 | 0.002275 | 0.001424 | 0.001631 | 0.00331 | 0.009258 | 0.011491 | 0.005744 | 0.003568 | 0.009853 | 0.002473 | 0.015694 | 0.013499 | 0.016236 | 0.0046 | 0.058864 | 0.00248 | 0.008983 | 0.002699 | 0.015724 | 0.015902 |
| 23.1527778 | 0.011259 | 0.004647 | 0.005657 | 0.00234 | 0.003123 | 0.001358 | 0.012355 | 0.006237 | 0.004278 | 0.002071 | 0.017093 | 0.00483 | 0.02099 | 0.022261 | 0.016582 | 0.004325 | 0.095873 | 0.004375 | 0.018926 | 0.003359 | 0.008242 | 0.016432 |
| 23.1666667 | 0.00729 | 0.010371 | 0.006435 | 0.002121 | 0.0074 | 0.000722 | 0.01412 | 0.002348 | 0.002166 | 0.002907 | 0.026232 | 0.011104 | 0.021502 | 0.032467 | 0.009109 | 0.004663 | 0.101127 | 0.009297 | 0.020521 | 0.005322 | 0.005928 | 0.011914 |
| 23.1805556 | 0.006352 | 0.012398 | 0.005023 | 0.001883 | 0.010727 | 0.000918 | 0.010465 | 0.004092 | 0.001174 | 0.002995 | 0.026684 | 0.015469 | 0.016187 | 0.03805 | 0.00389 | 0.005303 | 0.072769 | 0.010714 | 0.013783 | 0.012141 | 0.011672 | 0.010256 |
| 23.1944444 | 0.011748 | 0.008422 | 0.006852 | 0.003458 | 0.008746 | 0.002157 | 0.007068 | 0.012904 | 0.001549 | 0.002245 | 0.017348 | 0.012399 | 0.014184 | 0.029723 | 0.007551 | 0.006236 | 0.034953 | 0.005745 | 0.007492 | 0.019626 | 0.013188 | 0.014522 |
| 23.2083333 | 0.015722 | 0.008593 | 0.008502 | 0.004922 | 0.004749 | 0.00396 | 0.008987 | 0.0225 | 0.00367 | 0.002204 | 0.01574 | 0.007021 | 0.021937 | 0.016055 | 0.015061 | 0.006813 | 0.018631 | 0.003272 | 0.004706 | 0.016234 | 0.017102 | 0.012964 |
| 23.2222222 | 0.012735 | 0.011674 | 0.007523 | 0.004724 | 0.002709 | 0.005494 | 0.018481 | 0.023757 | 0.00781 | 0.001489 | 0.029298 | 0.006312 | 0.023369 | 0.009436 | 0.015984 | 0.007772 | 0.014635 | 0.005968 | 0.00762 | 0.012259 | 0.028452 | 0.013546 |
| 23.2361111 | 0.006236 | 0.008927 | 0.007553 | 0.003334 | 0.002149 | 0.005651 | 0.034684 | 0.01665 | 0.011661 | 0.000651 | 0.040098 | 0.007362 | 0.021158 | 0.012706 | 0.018428 | 0.012972 | 0.014164 | 0.008666 | 0.016613 | 0.019477 | 0.03477 | 0.038421 |
| 23.25 | 0.002048 | 0.003592 | 0.008109 | 0.001543 | 0.00293 | 0.004186 | 0.048329 | 0.009026 | 0.011418 | 0.000676 | 0.035936 | 0.006193 | 0.021424 | 0.025278 | 0.031719 | 0.013452 | 0.035317 | 0.007042 | 0.02339 | 0.019201 | 0.025658 | 0.072465 |
| 23.2638889 | 0.003521 | 0.001594 | 0.006187 | 0.001154 | 0.003015 | 0.002375 | 0.046074 | 0.006126 | 0.006611 | 0.000854 | 0.02177 | 0.005639 | 0.025236 | 0.025815 | 0.040534 | 0.009921 | 0.058822 | 0.004554 | 0.027159 | 0.011821 | 0.011485 | 0.078561 |
| 23.2777778 | 0.007089 | 0.000861 | 0.002693 | 0.00243 | 0.001391 | 0.001153 | 0.030396 | 0.006283 | 0.002258 | 0.000839 | 0.010741 | 0.00661 | 0.028471 | 0.013374 | 0.032852 | 0.010134 | 0.045549 | 0.004965 | 0.035619 | 0.009617 | 0.007609 | 0.055665 |
| 23.2916667 | 0.006481 | 0.001069 | 0.000845 | 0.002896 | 0.00122 | 0.001274 | 0.015021 | 0.006084 | 0.001162 | 0.000798 | 0.010851 | 0.00581 | 0.01821 | 0.008551 | 0.01925 | 0.008015 | 0.020518 | 0.006641 | 0.043782 | 0.005921 | 0.009408 | 0.028241 |
| 23.3055556 | 0.002628 | 0.002851 | 0.001056 | 0.001716 | 0.003138 | 0.003485 | 0.011397 | 0.004883 | 0.001162 | 0.00143 | 0.01313 | 0.0051 | 0.009103 | 0.010154 | 0.014892 | 0.008695 | 0.024322 | 0.008994 | 0.044538 | 0.004321 | 0.01013 | 0.009405 |
| 23.3194444 | 0.001021 | 0.003319 | 0.002015 | 0.001487 | 0.003785 | 0.005286 | 0.017041 | 0.005189 | 0.001752 | 0.003418 | 0.010602 | 0.008215 | 0.016478 | 0.01655 | 0.025296 | 0.021767 | 0.038185 | 0.021154 | 0.049628 | 0.017071 | 0.019725 | 0.010658 |
| 23.3333333 | 0.002497 | 0.002218 | 0.003072 | 0.003862 | 0.002874 | 0.00509 | 0.018037 | 0.009695 | 0.003221 | 0.003992 | 0.007412 | 0.008486 | 0.029603 | 0.02378 | 0.038513 | 0.034552 | 0.033714 | 0.040067 | 0.061938 | 0.037979 | 0.03353 | 0.027458 |
| 23.3472222 | 0.004133 | 0.001319 | 0.003304 | 0.006165 | 0.002856 | 0.006881 | 0.022552 | 0.015945 | 0.005152 | 0.003348 | 0.009076 | 0.006252 | 0.03235 | 0.019255 | 0.035126 | 0.028019 | 0.02353 | 0.036367 | 0.057428 | 0.036669 | 0.030425 | 0.027637 |
| 23.3611111 | 0.004063 | 0.001951 | 0.002685 | 0.00491 | 0.002785 | 0.010288 | 0.042197 | 0.018585 | 0.006004 | 0.004586 | 0.013557 | 0.008563 | 0.020351 | 0.009902 | 0.01793 | 0.011117 | 0.026252 | 0.016431 | 0.031052 | 0.021192 | 0.022446 | 0.011545 |
| 23.375 | 0.002519 | 0.003538 | 0.003098 | 0.001961 | 0.002646 | 0.009531 | 0.059531 | 0.017741 | 0.00554 | 0.004615 | 0.013428 | 0.009788 | 0.01236 | 0.006779 | 0.005854 | 0.003273 | 0.025788 | 0.010618 | 0.010036 | 0.025246 | 0.024511 | 0.005434 |
| 23.3888889 | 0.001624 | 0.003487 | 0.006655 | 0.001286 | 0.002534 | 0.005561 | 0.055775 | 0.015513 | 0.006057 | 0.002817 | 0.009472 | 0.00823 | 0.020983 | 0.004994 | 0.004239 | 0.002623 | 0.015151 | 0.013107 | 0.006514 | 0.031157 | 0.023489 | 0.006651 |
| 23.4027778 | 0.00344 | 0.002805 | 0.010935 | 0.002345 | 0.001537 | 0.00309 | 0.036945 | 0.011248 | 0.006326 | 0.002041 | 0.005566 | 0.009178 | 0.030683 | 0.003991 | 0.004387 | 0.003874 | 0.008286 | 0.01359 | 0.006708 | 0.021699 | 0.016266 | 0.007074 |
| 23.4166667 | 0.007344 | 0.002682 | 0.010456 | 0.002526 | 0.000794 | 0.002834 | 0.020045 | 0.007669 | 0.004338 | 0.002064 | 0.002496 | 0.01254 | 0.031413 | 0.006556 | 0.0024 | 0.006228 | 0.008694 | 0.014628 | 0.004279 | 0.010028 | 0.010011 | 0.009291 |
| 23.4305556 | 0.010729 | 0.002107 | 0.00605 | 0.00154 | 0.000556 | 0.002866 | 0.010425 | 0.008428 | 0.002135 | 0.001972 | 0.001415 | 0.015337 | 0.026998 | 0.009681 | 0.001672 | 0.007176 | 0.016822 | 0.015742 | 0.003325 | 0.007379 | 0.007766 | 0.010934 |
| 23.4444444 | 0.012108 | 0.002035 | 0.003836 | 0.000781 | 0.00044 | 0.002168 | 0.005173 | 0.010534 | 0.001467 | 0.001686 | 0.002513 | 0.015059 | 0.017272 | 0.010728 | 0.002358 | 0.006562 | 0.033 | 0.013796 | 0.002791 | 0.01533 | 0.009726 | 0.009888 |
| 23.4583333 | 0.0124 | 0.00219 | 0.00596 | 0.001359 | 0.000445 | 0.0029 | 0.00271 | 0.008388 | 0.00153 | 0.00283 | 0.005537 | 0.011054 | 0.006151 | 0.010175 | 0.002781 | 0.004661 | 0.053421 | 0.008374 | 0.001842 | 0.025217 | 0.011642 | 0.009403 |
| 23.4722222 | 0.012719 | 0.002715 | 0.007773 | 0.001989 | 0.000596 | 0.005629 | 0.002402 | 0.003761 | 0.002465 | 0.005255 | 0.009977 | 0.006362 | 0.010119 | 0.009658 | 0.004352 | 0.002886 | 0.064574 | 0.004184 | 0.001583 | 0.026343 | 0.009705 | 0.010894 |
| 23.4861111 | 0.013266 | 0.004099 | 0.005938 | 0.002603 | 0.001782 | 0.005757 | 0.004957 | 0.001965 | 0.005568 | 0.00506 | 0.011265 | 0.00897 | 0.035229 | 0.010749 | 0.006574 | 0.002403 | 0.058703 | 0.004507 | 0.001784 | 0.01927 | 0.007671 | 0.012456 |
| 23.5 | 0.0145 | 0.003513 | 0.003659 | 0.005267 | 0.004327 | 0.003207 | 0.01155 | 0.003434 | 0.009254 | 0.003217 | 0.007066 | 0.021935 | 0.061765 | 0.01272 | 0.007531 | 0.001784 | 0.046774 | 0.00977 | 0.001668 | 0.015331 | 0.010334 | 0.012047 |
| 23.5138889 | 0.016176 | 0.002141 | 0.005362 | 0.007664 | 0.006058 | 0.002524 | 0.020935 | 0.007004 | 0.010169 | 0.004087 | 0.003988 | 0.035034 | 0.062665 | 0.012318 | 0.008172 | 0.000969 | 0.035807 | 0.015514 | 0.001955 | 0.016751 | 0.014083 | 0.009586 |
| 23.5277778 | 0.017258 | 0.004442 | 0.01069 | 0.007449 | 0.005468 | 0.002789 | 0.031013 | 0.009461 | 0.008752 | 0.006099 | 0.005857 | 0.03741 | 0.040157 | 0.00917 | 0.010549 | 0.002933 | 0.023693 | 0.013459 | 0.003841 | 0.017432 | 0.012748 | 0.007949 |
| 23.5416667 | 0.016956 | 0.00803 | 0.015158 | 0.005876 | 0.003506 | 0.001865 | 0.038606 | 0.007694 | 0.008156 | 0.006818 | 0.009492 | 0.031077 | 0.020231 | 0.006775 | 0.018125 | 0.012084 | 0.016596 | 0.010914 | 0.006439 | 0.013142 | 0.008446 | 0.007057 |
| 23.5555556 | 0.014445 | 0.009539 | 0.014842 | 0.004083 | 0.002028 | 0.001275 | 0.038863 | 0.003877 | 0.008561 | 0.006163 | 0.013029 | 0.023025 | 0.026361 | 0.009126 | 0.029249 | 0.023974 | 0.02653 | 0.019352 | 0.013784 | 0.011333 | 0.008235 | 0.007423 |
| 23.5694444 | 0.00984 | 0.008545 | 0.009778 | 0.003357 | 0.002179 | 0.003449 | 0.03246 | 0.001423 | 0.008568 | 0.004355 | 0.013855 | 0.015249 | 0.048512 | 0.014465 | 0.031319 | 0.028138 | 0.039493 | 0.027336 | 0.023991 | 0.019252 | 0.008526 | 0.012782 |
| 23.5833333 | 0.005047 | 0.005069 | 0.007118 | 0.00368 | 0.002068 | 0.008109 | 0.024925 | 0.000627 | 0.007348 | 0.002822 | 0.009208 | 0.009237 | 0.040514 | 0.014941 | 0.020374 | 0.023509 | 0.032975 | 0.022803 | 0.023247 | 0.023523 | 0.009623 | 0.017766 |
| 23.5972222 | 0.002089 | 0.003031 | 0.010231 | 0.003321 | 0.001267 | 0.01142 | 0.018958 | 0.000851 | 0.004559 | 0.00257 | 0.003545 | 0.00745 | 0.018522 | 0.008686 | 0.012833 | 0.015091 | 0.018569 | 0.011367 | 0.012197 | 0.016063 | 0.017619 | 0.014845 |
| 23.6111111 | 0.002235 | 0.003508 | 0.011106 | 0.002002 | 0.001395 | 0.010898 | 0.013987 | 0.00147 | 0.002092 | 0.002554 | 0.002015 | 0.008762 | 0.020172 | 0.002967 | 0.013004 | 0.007858 | 0.012382 | 0.002978 | 0.004467 | 0.007731 | 0.021659 | 0.008983 |
| 23.625 | 0.005807 | 0.002987 | 0.006432 | 0.001448 | 0.001259 | 0.007779 | 0.008709 | 0.002855 | 0.001051 | 0.001632 | 0.003648 | 0.009889 | 0.02026 | 0.002224 | 0.009081 | 0.004693 | 0.014204 | 0.001191 | 0.00277 | 0.00715 | 0.016919 | 0.006807 |
| 23.6388889 | 0.00965 | 0.002391 | 0.002531 | 0.002419 | 0.000513 | 0.004435 | 0.005033 | 0.00637 | 0.001176 | 0.000604 | 0.005352 | 0.009542 | 0.010701 | 0.002746 | 0.003833 | 0.003508 | 0.014445 | 0.00276 | 0.002693 | 0.008185 | 0.011993 | 0.006225 |
| 23.6527778 | 0.009173 | 0.003875 | 0.002834 | 0.002653 | 0.000466 | 0.002452 | 0.006291 | 0.009072 | 0.003016 | 0.000492 | 0.004851 | 0.008189 | 0.00707 | 0.002113 | 0.002381 | 0.001855 | 0.008983 | 0.003471 | 0.003434 | 0.005229 | 0.009598 | 0.004067 |
| 23.6666667 | 0.005831 | 0.003969 | 0.004369 | 0.001362 | 0.000797 | 0.001949 | 0.009171 | 0.008019 | 0.005222 | 0.000712 | 0.002745 | 0.005898 | 0.005783 | 0.001831 | 0.002333 | 0.001226 | 0.007717 | 0.002783 | 0.003343 | 0.002408 | 0.00702 | 0.002293 |
| 23.6805556 | 0.006028 | 0.002932 | 0.004742 | 0.000762 | 0.000697 | 0.002087 | 0.008109 | 0.004664 | 0.005525 | 0.000531 | 0.001107 | 0.002947 | 0.004475 | 0.001818 | 0.003883 | 0.002799 | 0.021369 | 0.002452 | 0.002196 | 0.001807 | 0.00391 | 0.002302 |

| | | | | | | | | | | | | | | | | | | | | | | |
|---|---|---|---|---|---|---|---|---|---|---|---|---|---|---|---|---|---|---|---|---|---|---|
| 23.6944444 | 0.01071 | 0.005169 | 0.003774 | 0.001733 | 0.000382 | 0.002366 | 0.004119 | 0.00202 | 0.004717 | 0.000265 | 0.000586 | 0.000826 | 0.007304 | 0.002212 | 0.005942 | 0.004541 | 0.042867 | 0.001959 | 0.002054 | 0.002786 | 0.003104 | 0.002865 |
| 23.7083333 | 0.014729 | 0.007865 | 0.001907 | 0.003631 | 0.000406 | 0.002723 | 0.00137 | 0.002148 | 0.003971 | 0.000403 | 0.001387 | 0.000381 | 0.011395 | 0.002768 | 0.0059 | 0.004345 | 0.047589 | 0.001547 | 0.002649 | 0.00344 | 0.003913 | 0.003887 |
| 23.7222222 | 0.014765 | 0.007247 | 0.000513 | 0.005093 | 0.00058 | 0.002967 | 0.001777 | 0.003398 | 0.002463 | 0.000872 | 0.003465 | 0.001134 | 0.010126 | 0.00281 | 0.004283 | 0.002436 | 0.031869 | 0.002361 | 0.005056 | 0.003496 | 0.005185 | 0.004625 |
| 23.7361111 | 0.011258 | 0.004229 | 0.000242 | 0.004803 | 0.00055 | 0.002906 | 0.004798 | 0.003338 | 0.001262 | 0.001205 | 0.004892 | 0.002963 | 0.008464 | 0.003453 | 0.004047 | 0.000802 | 0.017156 | 0.002298 | 0.00842 | 0.005302 | 0.00846 | 0.003576 |
| 23.75 | 0.006596 | 0.002587 | 0.000392 | 0.003178 | 0.000457 | 0.002557 | 0.00864 | 0.001892 | 0.001862 | 0.00114 | 0.003973 | 0.005247 | 0.01081 | 0.004759 | 0.006137 | 0.000668 | 0.021998 | 0.001064 | 0.009811 | 0.007566 | 0.013368 | 0.002165 |
| 23.7638889 | 0.002905 | 0.004625 | 0.000679 | 0.001482 | 0.000601 | 0.00205 | 0.011798 | 0.000846 | 0.002915 | 0.00085 | 0.00198 | 0.0069 | 0.012734 | 0.005051 | 0.007873 | 0.001263 | 0.039756 | 0.00145 | 0.009497 | 0.006308 | 0.016465 | 0.004351 |
| 23.7777778 | 0.001774 | 0.00659 | 0.001819 | 0.000468 | 0.001049 | 0.001417 | 0.012925 | 0.001639 | 0.003425 | 0.000599 | 0.000702 | 0.007526 | 0.011817 | 0.003784 | 0.006938 | 0.002059 | 0.045232 | 0.004318 | 0.008837 | 0.004344 | 0.015967 | 0.010744 |
| 23.7916667 | 0.004127 | 0.006031 | 0.002999 | 0.000205 | 0.001539 | 0.000965 | 0.011498 | 0.00326 | 0.00393 | 0.000413 | 0.000468 | 0.006788 | 0.008226 | 0.002301 | 0.004243 | 0.003554 | 0.031261 | 0.008112 | 0.007605 | 0.005871 | 0.013184 | 0.016024 |
| 23.8055556 | 0.008351 | 0.004769 | 0.002685 | 0.000372 | 0.001697 | 0.001398 | 0.009831 | 0.003964 | 0.00438 | 0.000213 | 0.001312 | 0.00448 | 0.008846 | 0.001374 | 0.002357 | 0.005685 | 0.013779 | 0.010551 | 0.006343 | 0.007333 | 0.008747 | 0.01645 |
| 23.8194444 | 0.012254 | 0.003909 | 0.002835 | 0.000636 | 0.001447 | 0.00243 | 0.011784 | 0.003175 | 0.003507 | 0.000118 | 0.002366 | 0.002383 | 0.019275 | 0.001784 | 0.003365 | 0.008149 | 0.004687 | 0.011221 | 0.005631 | 0.006768 | 0.004083 | 0.014827 |
| 23.8333333 | 0.015364 | 0.002781 | 0.005303 | 0.000712 | 0.00101 | 0.002798 | 0.017593 | 0.00204 | 0.002167 | 0.000216 | 0.002605 | 0.002116 | 0.026287 | 0.004298 | 0.006294 | 0.010472 | 0.004115 | 0.012272 | 0.003679 | 0.005649 | 0.00173 | 0.014407 |
| 23.8472222 | 0.014899 | 0.001975 | 0.007501 | 0.001236 | 0.000525 | 0.002261 | 0.022034 | 0.002438 | 0.002423 | 0.000512 | 0.002675 | 0.002007 | 0.020194 | 0.00627 | 0.007612 | 0.011321 | 0.009549 | 0.013105 | 0.001238 | 0.003996 | 0.001333 | 0.014356 |
| 23.8611111 | 0.010379 | 0.003245 | 0.007799 | 0.002756 | 0.000141 | 0.001498 | 0.021094 | 0.004171 | 0.002917 | 0.001029 | 0.002877 | 0.001515 | 0.019378 | 0.004923 | 0.006248 | 0.010619 | 0.014706 | 0.009874 | 0.001144 | 0.001964 | 0.002505 | 0.013054 |
| 23.875 | 0.008998 | 0.004945 | 0.007334 | 0.00404 | 0.000169 | 0.00076 | 0.016223 | 0.0052 | 0.002241 | 0.001634 | 0.002878 | 0.001818 | 0.030737 | 0.00465 | 0.004513 | 0.009441 | 0.01469 | 0.004279 | 0.003224 | 0.000907 | 0.006199 | 0.01342 |
| 23.8888889 | 0.00926 | 0.004559 | 0.006989 | 0.004302 | 0.000437 | 0.000485 | 0.011348 | 0.005273 | 0.002791 | 0.002394 | 0.002369 | 0.004268 | 0.031875 | 0.010347 | 0.004178 | 0.007538 | 0.013449 | 0.002877 | 0.004214 | 0.00129 | 0.010509 | 0.014991 |
| 23.9027778 | 0.007551 | 0.00315 | 0.007127 | 0.004223 | 0.000696 | 0.001683 | 0.010399 | 0.006445 | 0.006826 | 0.00357 | 0.002196 | 0.00947 | 0.019971 | 0.017619 | 0.004763 | 0.00493 | 0.014379 | 0.004912 | 0.004688 | 0.002105 | 0.012793 | 0.014606 |
| 23.9166667 | 0.012832 | 0.003838 | 0.00744 | 0.004544 | 0.001607 | 0.004568 | 0.01564 | 0.009671 | 0.011738 | 0.004815 | 0.002982 | 0.013966 | 0.018447 | 0.020825 | 0.005833 | 0.003902 | 0.016963 | 0.004549 | 0.007698 | 0.00356 | 0.013984 | 0.021897 |
| 23.9305556 | 0.026393 | 0.00634 | 0.008321 | 0.005819 | 0.003396 | 0.007896 | 0.023484 | 0.013994 | 0.014253 | 0.005272 | 0.004492 | 0.014136 | 0.035245 | 0.019646 | 0.007313 | 0.00545 | 0.021638 | 0.003699 | 0.010429 | 0.005945 | 0.015731 | 0.036303 |
| 23.9444444 | 0.036627 | 0.007304 | 0.010091 | 0.008057 | 0.005052 | 0.009787 | 0.025006 | 0.018493 | 0.014043 | 0.004535 | 0.008929 | 0.010588 | 0.059436 | 0.017058 | 0.008034 | 0.006504 | 0.028826 | 0.004879 | 0.011405 | 0.008735 | 0.01828 | 0.033067 |
| 23.9583333 | 0.035586 | 0.006024 | 0.009929 | 0.010745 | 0.005852 | 0.009023 | 0.018067 | 0.021831 | 0.011974 | 0.003147 | 0.013757 | 0.008311 | 0.078486 | 0.016005 | 0.007147 | 0.005242 | 0.041528 | 0.004165 | 0.012414 | 0.011573 | 0.02299 | 0.017325 |
| 23.9722222 | 0.026193 | 0.004757 | 0.006397 | 0.012841 | 0.005953 | 0.006497 | 0.012545 | 0.024343 | 0.009715 | 0.002269 | 0.015071 | 0.011884 | 0.080829 | 0.016475 | 0.004478 | 0.00357 | 0.05639 | 0.001594 | 0.012786 | 0.013521 | 0.030301 | 0.021666 |
| 23.9861111 | 0.015247 | 0.004139 | 0.004526 | 0.012253 | 0.005733 | 0.004242 | 0.014735 | 0.028437 | 0.008173 | 0.002467 | 0.015949 | 0.017959 | 0.063625 | 0.01699 | 0.002858 | 0.00307 | 0.060388 | 0.000428 | 0.012847 | 0.014954 | 0.03707 | 0.036475 |
| 24 | 0.007288 | 0.003188 | 0.007859 | 0.007787 | 0.004963 | 0.002432 | 0.017888 | 0.030161 | 0.006396 | 0.002693 | 0.015604 | 0.018408 | 0.03657 | 0.016068 | 0.004172 | 0.003939 | 0.049709 | 0.002003 | 0.013317 | 0.01603 | 0.037119 | 0.034766 |
| 24.0138889 | 0.00586 | 0.003871 | 0.010647 | 0.003241 | 0.003351 | 0.001912 | 0.014289 | 0.022092 | 0.004042 | 0.002461 | 0.010511 | 0.011159 | 0.01625 | 0.012298 | 0.004118 | 0.006131 | 0.029175 | 0.008111 | 0.011447 | 0.014231 | 0.026289 | 0.020692 |
| 24.0277778 | 0.008634 | 0.0069 | 0.007697 | 0.003218 | 0.002372 | 0.004019 | 0.007114 | 0.009369 | 0.002604 | 0.003071 | 0.006355 | 0.00343 | 0.012032 | 0.00738 | 0.00177 | 0.007973 | 0.012929 | 0.01536 | 0.007305 | 0.010458 | 0.013916 | 0.008164 |
| 24.0416667 | 0.007734 | 0.007139 | 0.002857 | 0.004314 | 0.00238 | 0.004656 | 0.004446 | 0.007754 | 0.003393 | 0.003566 | 0.007647 | 0.004742 | 0.011264 | 0.007314 | 0.002537 | 0.007121 | 0.011817 | 0.014789 | 0.007248 | 0.012212 | 0.014431 | 0.002041 |
| 24.0555556 | 0.00551 | 0.004721 | 0.000896 | 0.003684 | 0.003211 | 0.007275 | 0.011344 | 0.023221 | 0.009645 | 0.005721 | 0.011009 | 0.019427 | 0.006555 | 0.011891 | 0.007383 | 0.007179 | 0.013023 | 0.015748 | 0.011495 | 0.026688 | 0.024092 | 0.002893 |
| 24.0694444 | 0.009117 | 0.005902 | 0.001297 | 0.004901 | 0.005887 | 0.018942 | 0.033588 | 0.042919 | 0.020143 | 0.011775 | 0.01388 | 0.041393 | 0.007142 | 0.013138 | 0.010002 | 0.009532 | 0.010985 | 0.032022 | 0.012318 | 0.047797 | 0.03156 | 0.009309 |
| 24.0833333 | 0.012156 | 0.010284 | 0.003328 | 0.007578 | 0.008458 | 0.025499 | 0.064635 | 0.048429 | 0.024372 | 0.014396 | 0.018441 | 0.054043 | 0.014773 | 0.012333 | 0.009612 | 0.009319 | 0.015284 | 0.055394 | 0.009877 | 0.055922 | 0.040029 | 0.012783 |
| 24.0972222 | 0.008491 | 0.012909 | 0.006046 | 0.008116 | 0.009 | 0.016031 | 0.075508 | 0.035498 | 0.019465 | 0.009951 | 0.028713 | 0.046552 | 0.02599 | 0.017538 | 0.011074 | 0.008925 | 0.016827 | 0.071772 | 0.010093 | 0.0459 | 0.05029 | 0.0095 |
| 24.1111111 | 0.006343 | 0.010048 | 0.006046 | 0.005978 | 0.007262 | 0.00691 | 0.052963 | 0.017585 | 0.011258 | 0.005197 | 0.031857 | 0.02651 | 0.03148 | 0.020292 | 0.012062 | 0.009338 | 0.010535 | 0.064404 | 0.011686 | 0.031326 | 0.04667 | 0.005109 |
| 24.125 | 0.012504 | 0.005685 | 0.006588 | 0.003092 | 0.004122 | 0.009577 | 0.023705 | 0.012016 | 0.005928 | 0.005679 | 0.018916 | 0.011221 | 0.024736 | 0.012935 | 0.008996 | 0.006313 | 0.009593 | 0.038685 | 0.011378 | 0.018244 | 0.026029 | 0.002294 |
| 24.1388889 | 0.019004 | 0.006385 | 0.010056 | 0.001128 | 0.003072 | 0.013864 | 0.015981 | 0.019282 | 0.008558 | 0.008693 | 0.009825 | 0.009003 | 0.015159 | 0.006999 | 0.004798 | 0.002313 | 0.009487 | 0.020524 | 0.010611 | 0.007325 | 0.008977 | 0.001323 |
| 24.1527778 | 0.015754 | 0.007463 | 0.008951 | 0.00063 | 0.004548 | 0.012596 | 0.025687 | 0.023594 | 0.014519 | 0.008317 | 0.014533 | 0.011826 | 0.010173 | 0.004908 | 0.00217 | 0.00143 | 0.006682 | 0.012008 | 0.012037 | 0.001723 | 0.006232 | 0.001552 |
| 24.1666667 | 0.009203 | 0.005599 | 0.005392 | 0.001868 | 0.004496 | 0.008372 | 0.031884 | 0.020071 | 0.016907 | 0.004623 | 0.022424 | 0.012895 | 0.007741 | 0.003183 | 0.000762 | 0.003043 | 0.008151 | 0.006383 | 0.016027 | 0.00051 | 0.009005 | 0.002547 |
| 24.1805556 | 0.009377 | 0.005771 | 0.0053 | 0.003763 | 0.002677 | 0.003807 | 0.024051 | 0.016723 | 0.013823 | 0.001595 | 0.025879 | 0.013526 | 0.005615 | 0.005079 | 0.000241 | 0.004188 | 0.013956 | 0.003928 | 0.019376 | 0.000319 | 0.012017 | 0.003242 |
| 24.1944444 | 0.008674 | 0.006587 | 0.006717 | 0.003923 | 0.002148 | 0.000955 | 0.012969 | 0.017954 | 0.007148 | 0.000722 | 0.019089 | 0.01287 | 0.004149 | 0.00961 | 0.000389 | 0.003118 | 0.020454 | 0.002978 | 0.017868 | 0.001208 | 0.017182 | 0.002456 |
| 24.2083333 | 0.004427 | 0.006285 | 0.006853 | 0.002172 | 0.001993 | 0.001042 | 0.0095 | 0.021789 | 0.004188 | 0.00145 | 0.006228 | 0.008455 | 0.006518 | 0.014897 | 0.000596 | 0.001246 | 0.022255 | 0.003964 | 0.010675 | 0.005497 | 0.024548 | 0.001818 |
| 24.2222222 | 0.004146 | 0.009655 | 0.005833 | 0.000697 | 0.00133 | 0.003189 | 0.010517 | 0.023242 | 0.007076 | 0.002854 | 0.00195 | 0.004089 | 0.015324 | 0.021145 | 0.000928 | 0.000468 | 0.018303 | 0.007588 | 0.006473 | 0.013786 | 0.02896 | 0.001734 |
| 24.2361111 | 0.005062 | 0.011401 | 0.007229 | 0.000729 | 0.001901 | 0.005428 | 0.020293 | 0.02034 | 0.007928 | 0.003027 | 0.004032 | 0.002968 | 0.027123 | 0.025995 | 0.003412 | 0.000739 | 0.011856 | 0.011764 | 0.014937 | 0.024449 | 0.023344 | 0.002474 |
| 24.25 | 0.004171 | 0.007501 | 0.006877 | 0.002279 | 0.003385 | 0.006523 | 0.031593 | 0.015906 | 0.005556 | 0.001864 | 0.005122 | 0.004602 | 0.036671 | 0.023336 | 0.009077 | 0.002488 | 0.005777 | 0.016855 | 0.030614 | 0.034054 | 0.010764 | 0.005557 |
| 24.2638889 | 0.004099 | 0.004907 | 0.003565 | 0.004704 | 0.004331 | 0.005921 | 0.024912 | 0.01208 | 0.005669 | 0.001285 | 0.008652 | 0.011573 | 0.040691 | 0.014187 | 0.014263 | 0.006623 | 0.002853 | 0.02039 | 0.042534 | 0.039424 | 0.003951 | 0.010695 |
| 24.2777778 | 0.005438 | 0.005065 | 0.00235 | 0.006651 | 0.003965 | 0.003655 | 0.010825 | 0.010023 | 0.005214 | 0.00168 | 0.011321 | 0.017926 | 0.037615 | 0.012876 | 0.013947 | 0.011446 | 0.003956 | 0.014874 | 0.045058 | 0.039599 | 0.006967 | 0.015901 |
| 24.2916667 | 0.005752 | 0.006207 | 0.002509 | 0.007274 | 0.002628 | 0.002026 | 0.010188 | 0.011606 | 0.003379 | 0.001612 | 0.008582 | 0.015645 | 0.029121 | 0.022216 | 0.008658 | 0.013579 | 0.005921 | 0.006939 | 0.038675 | 0.035593 | 0.0098 | 0.018339 |
| 24.3055556 | 0.004402 | 0.005981 | 0.003102 | 0.00728 | 0.002006 | 0.002024 | 0.018459 | 0.015691 | 0.005277 | 0.001071 | 0.004366 | 0.010162 | 0.01836 | 0.025555 | 0.004354 | 0.011369 | 0.007753 | 0.007544 | 0.028594 | 0.031215 | 0.01072 | 0.016826 |
| 24.3194444 | 0.004255 | 0.004722 | 0.00533 | 0.006766 | 0.002795 | 0.002099 | 0.022296 | 0.019099 | 0.009362 | 0.000991 | 0.002189 | 0.008662 | 0.009264 | 0.016956 | 0.003832 | 0.00618 | 0.009801 | 0.009517 | 0.017533 | 0.028366 | 0.019744 | 0.011628 |
| 24.3333333 | 0.006579 | 0.003975 | 0.008606 | 0.005355 | 0.003567 | 0.003117 | 0.022619 | 0.019143 | 0.009792 | 0.001436 | 0.001341 | 0.009382 | 0.00529 | 0.007882 | 0.005183 | 0.001827 | 0.011162 | 0.006191 | 0.007612 | 0.025872 | 0.032182 | 0.005892 |
| 24.3472222 | 0.008638 | 0.003691 | 0.009868 | 0.00361 | 0.003004 | 0.004696 | 0.028298 | 0.014655 | 0.005857 | 0.002303 | 0.001861 | 0.007514 | 0.007503 | 0.007097 | 0.006933 | 0.000957 | 0.012069 | 0.003462 | 0.004266 | 0.024746 | 0.035086 | 0.003775 |
| 24.3611111 | 0.007337 | 0.004273 | 0.009239 | 0.002021 | 0.001879 | 0.00486 | 0.040405 | 0.009638 | 0.003103 | 0.002374 | 0.005025 | 0.003827 | 0.013249 | 0.01055 | 0.007941 | 0.00289 | 0.012029 | 0.0039 | 0.008845 | 0.023879 | 0.024652 | 0.00396 |
| 24.375 | 0.004498 | 0.00374 | 0.009466 | 0.0011 | 0.001405 | 0.004233 | 0.047046 | 0.009857 | 0.003466 | 0.001716 | 0.007568 | 0.001837 | 0.016842 | 0.011201 | 0.007188 | 0.00623 | 0.010021 | 0.005083 | 0.016118 | 0.020155 | 0.012436 | 0.007057 |
| 24.3888889 | 0.003705 | 0.003717 | 0.010708 | 0.000833 | 0.00181 | 0.004097 | 0.041236 | 0.013373 | 0.004182 | 0.001792 | 0.005998 | 0.002584 | 0.016695 | 0.008518 | 0.005586 | 0.009254 | 0.00819 | 0.006291 | 0.02238 | 0.013372 | 0.010561 | 0.013514 |
| 24.4027778 | 0.004682 | 0.006871 | 0.010471 | 0.001446 | 0.003091 | 0.004775 | 0.032001 | 0.01417 | 0.005104 | 0.002644 | 0.002769 | 0.005293 | 0.016245 | 0.00498 | 0.004971 | 0.009726 | 0.0099 | 0.00978 | 0.026289 | 0.006009 | 0.010942 | 0.016921 |
| 24.4166667 | 0.009104 | 0.009585 | 0.010041 | 0.003433 | 0.004384 | 0.005599 | 0.026056 | 0.011934 | 0.008233 | 0.003574 | 0.001105 | 0.007784 | 0.016503 | 0.005057 | 0.005327 | 0.007289 | 0.014352 | 0.021316 | 0.025798 | 0.001787 | 0.01066 | 0.015259 |
| 24.4305556 | 0.017762 | 0.007273 | 0.011859 | 0.005399 | 0.004074 | 0.005672 | 0.021046 | 0.01046 | 0.01204 | 0.00377 | 0.001075 | 0.00924 | 0.016319 | 0.010233 | 0.005094 | 0.004034 | 0.018462 | 0.029893 | 0.020096 | 0.001228 | 0.011682 | 0.012472 |
| 24.4444444 | 0.023937 | 0.003864 | 0.010112 | 0.00596 | 0.002403 | 0.00467 | 0.017118 | 0.011186 | 0.012468 | 0.002785 | 0.003162 | 0.01015 | 0.017843 | 0.013569 | 0.00352 | 0.002104 | 0.021181 | 0.020256 | 0.011597 | 0.001825 | 0.008059 | 0.010206 |
| 24.4583333 | 0.02166 | 0.004352 | 0.005212 | 0.004992 | 0.00221 | 0.002978 | 0.018566 | 0.010418 | 0.009556 | 0.002645 | 0.00749 | 0.009751 | 0.022606 | 0.010311 | 0.002026 | 0.002232 | 0.02267 | 0.009819 | 0.004993 | 0.00393 | 0.00312 | 0.008262 |
| 24.4722222 | 0.012656 | 0.004686 | 0.004678 | 0.003134 | 0.004035 | 0.001886 | 0.027427 | 0.006415 | 0.007361 | 0.004136 | 0.012199 | 0.008294 | 0.023811 | 0.005238 | 0.001724 | 0.004326 | 0.022874 | 0.01466 | 0.00278 | 0.008316 | 0.001847 | 0.005912 |
| 24.4861111 | 0.008004 | 0.00507 | 0.006576 | 0.002966 | 0.004448 | 0.002067 | 0.038586 | 0.004532 | 0.008286 | 0.003762 | 0.019099 | 0.009216 | 0.017715 | 0.004758 | 0.001894 | 0.007343 | 0.023447 | 0.017959 | 0.004846 | 0.013216 | 0.005016 | 0.004824 |
| 24.5 | 0.017495 | 0.008144 | 0.007397 | 0.005334 | 0.005142 | 0.00392 | 0.041276 | 0.008353 | 0.010397 | 0.002131 | 0.023613 | 0.014794 | 0.012375 | 0.007714 | 0.003078 | 0.009028 | 0.025915 | 0.011639 | 0.009384 | 0.017432 | 0.012753 | 0.008875 |
| 24.5138889 | 0.036599 | 0.00845 | 0.006044 | 0.007267 | 0.009164 | 0.007393 | 0.032561 | 0.014112 | 0.010854 | 0.002087 | 0.016652 | 0.021062 | 0.010478 | 0.008248 | 0.004439 | 0.007721 | 0.028202 | 0.009879 | 0.011282 | 0.019223 | 0.017881 | 0.013946 |
| 24.5277778 | 0.050555 | 0.005989 | 0.006096 | 0.007489 | 0.011758 | 0.009983 | 0.020576 | 0.016587 | 0.009901 | 0.002568 | 0.005891 | 0.024392 | 0.007078 | 0.005691 | 0.00426 | 0.005433 | 0.026415 | 0.014932 | 0.007969 | 0.016636 | 0.016514 | 0.012472 |
| 24.5416667 | 0.048933 | 0.006989 | 0.016754 | 0.007114 | 0.010092 | 0.009939 | 0.012585 | 0.01428 | 0.008956 | 0.00314 | 0.001743 | 0.026229 | 0.004762 | 0.006382 | 0.007094 | 0.008468 | 0.018416 | 0.016476 | 0.007257 | 0.011331 | 0.011872 | 0.011151 |
| 24.5555556 | 0.034945 | 0.009828 | 0.031949 | 0.005613 | 0.006971 | 0.008487 | 0.013817 | 0.008987 | 0.007946 | 0.003366 | 0.001855 | 0.026553 | 0.006029 | 0.011944 | 0.014298 | 0.016856 | 0.009804 | 0.011183 | 0.012986 | 0.007064 | 0.006034 | 0.015833 |
| 24.5694444 | 0.017421 | 0.010642 | 0.033659 | 0.003535 | 0.004492 | 0.007955 | 0.025657 | 0.004544 | 0.007507 | 0.002277 | 0.003625 | 0.023967 | 0.006722 | 0.013326 | 0.01495 | 0.019222 | 0.007771 | 0.005505 | 0.012599 | 0.005308 | 0.004642 | 0.015637 |
| 24.5833333 | 0.005701 | 0.010226 | 0.020953 | 0.003167 | 0.003541 | 0.008597 | 0.036393 | 0.002396 | 0.008842 | 0.001226 | 0.009686 | 0.020727 | 0.008947 | 0.008516 | 0.010799 | 0.01157 | 0.010246 | 0.004572 | 0.007557 | 0.006032 | 0.012211 | 0.00853 |
| 24.5972222 | 0.003533 | 0.009468 | 0.009113 | 0.002971 | 0.003869 | 0.008689 | 0.034237 | 0.002665 | 0.009918 | 0.001202 | 0.015569 | 0.018581 | 0.013136 | 0.004205 | 0.013131 | 0.00391 | 0.011078 | 0.006108 | 0.009142 | 0.007148 | 0.02102 | 0.003583 |
| 24.6111111 | 0.003648 | 0.008662 | 0.007243 | 0.002044 | 0.003198 | 0.00776 | 0.024196 | 0.004669 | 0.007852 | 0.001099 | 0.013022 | 0.015737 | 0.012908 | 0.005374 | 0.014134 | 0.002222 | 0.007198 | 0.006379 | 0.008844 | 0.005544 | 0.021112 | 0.002958 |
| 24.625 | 0.002167 | 0.006881 | 0.009992 | 0.002462 | 0.001743 | 0.006701 | 0.014704 | 0.005358 | 0.004184 | 0.000694 | 0.006001 | 0.011611 | 0.008562 | 0.013054 | 0.008705 | 0.003652 | 0.002666 | 0.004264 | 0.003711 | 0.003853 | 0.016308 | 0.004962 |
| 24.6388889 | 0.001064 | 0.003579 | 0.011132 | 0.00324 | 0.001958 | 0.006004 | 0.007889 | 0.007624 | 0.002032 | 0.000547 | 0.00283 | 0.008381 | 0.00959 | 0.0176 | 0.005084 | 0.004528 | 0.001404 | 0.002242 | 0.001148 | 0.004204 | 0.015172 | 0.006153 |
| 24.6527778 | 0.001044 | 0.001419 | 0.00938 | 0.002816 | 0.002279 | 0.00508 | 0.006934 | 0.012266 | 0.002033 | 0.000431 | 0.003873 | 0.007501 | 0.017201 | 0.013247 | 0.004826 | 0.003322 | 0.00107 | 0.001231 | 0.000803 | 0.003894 | 0.016489 | 0.00447 |
| 24.6666667 | 0.002137 | 0.001608 | 0.005543 | 0.002007 | 0.001274 | 0.003643 | 0.011225 | 0.011078 | 0.002863 | 0.000389 | 0.005822 | 0.007345 | 0.016335 | 0.006749 | 0.003765 | 0.001375 | 0.000708 | 0.000407 | 0.000731 | 0.002941 | 0.014934 | 0.00206 |
| 24.6805556 | 0.002646 | 0.001697 | 0.004524 | 0.002615 | 0.000475 | 0.00241 | 0.013755 | 0.00674 | 0.002286 | 0.000327 | 0.007634 | 0.005155 | 0.008628 | 0.002737 | 0.001692 | 0.000781 | 0.000883 | 0.00076 | 0.001093 | 0.002455 | 0.011173 | 0.001108 |
| 24.6944444 | 0.003007 | 0.00176 | 0.007307 | 0.004225 | 0.000274 | 0.001798 | 0.011396 | 0.008337 | 0.001288 | 0.000112 | 0.008107 | 0.002658 | 0.008197 | 0.001838 | 0.000691 | 0.002163 | 0.001741 | 0.002572 | 0.002693 | 0.003413 | 0.009183 | 0.002053 |
| 24.7083333 | 0.004811 | 0.003294 | 0.008458 | 0.004301 | 0.000357 | 0.001472 | 0.007005 | 0.010849 | 0.001666 | 0.000133 | 0.006857 | 0.002651 | 0.010829 | 0.00372 | 0.000749 | 0.004241 | 0.003185 | 0.004486 | 0.004912 | 0.006305 | 0.009446 | 0.003426 |
| 24.7222222 | 0.005584 | 0.004264 | 0.005753 | 0.00285 | 0.000684 | 0.00124 | 0.005743 | 0.0082 | 0.001823 | 0.000497 | 0.005849 | 0.003077 | 0.009949 | 0.005712 | 0.001583 | 0.004967 | 0.00424 | 0.004872 | 0.006047 | 0.008317 | 0.011498 | 0.003837 |
| 24.7361111 | 0.004238 | 0.003233 | 0.002809 | 0.001349 | 0.000835 | 0.001408 | 0.00743 | 0.004022 | 0.001081 | 0.000817 | 0.005447 | 0.004482 | 0.00746 | 0.006063 | 0.002554 | 0.003821 | 0.004484 | 0.003641 | 0.005822 | 0.008128 | 0.014472 | 0.003958 |
| 24.75 | 0.002561 | 0.001792 | 0.002608 | 0.00089 | 0.00072 | 0.002051 | 0.005858 | 0.002586 | 0.00093 | 0.000777 | 0.005009 | 0.008537 | 0.005252 | 0.005364 | 0.002619 | 0.001879 | 0.004059 | 0.001871 | 0.00478 | 0.006884 | 0.020401 | 0.004198 |
| 24.7638889 | 0.001703 | 0.002089 | 0.003673 | 0.001861 | 0.000457 | 0.00247 | 0.003912 | 0.003959 | 0.001973 | 0.000481 | 0.004093 | 0.011536 | 0.003456 | 0.00412 | 0.001745 | 0.000689 | 0.003152 | 0.000719 | 0.003274 | 0.005133 | 0.029518 | 0.003665 |
| 24.7777778 | 0.002973 | 0.00449 | 0.003115 | 0.003375 | 0.000269 | 0.00199 | 0.009264 | 0.00558 | 0.002445 | 0.000293 | 0.002862 | 0.010956 | 0.001819 | 0.002667 | 0.00072 | 0.000592 | 0.001945 | 0.000569 | 0.001871 | 0.003271 | 0.031021 | 0.00229 |
| 24.7916667 | 0.007271 | 0.007847 | 0.002373 | 0.00445 | 0.000528 | 0.002691 | 0.019601 | 0.005134 | 0.002745 | 0.000578 | 0.001721 | 0.008071 | 0.001575 | 0.001505 | 0.000319 | 0.000823 | 0.001078 | 0.000719 | 0.002015 | 0.001679 | 0.02304 | 0.001214 |
| 24.8055556 | 0.013634 | 0.011619 | 0.004264 | 0.004538 | 0.000919 | 0.006583 | 0.028817 | 0.003178 | 0.004905 | 0.001315 | 0.00126 | 0.004442 | 0.003681 | 0.000849 | 0.000362 | 0.000824 | 0.001443 | 0.000594 | 0.004363 | 0.000778 | 0.015798 | 0.001786 |
| 24.8194444 | 0.022809 | 0.01328 | 0.005822 | 0.00365 | 0.000819 | 0.009677 | 0.032431 | 0.002534 | 0.007011 | 0.002004 | 0.002024 | 0.002831 | 0.006299 | 0.000844 | 0.000572 | 0.000629 | 0.002693 | 0.000531 | 0.006942 | 0.000736 | 0.011632 | 0.003788 |
| 24.8333333 | 0.033745 | 0.010621 | 0.004359 | 0.002257 | 0.00073 | 0.008493 | 0.028183 | 0.003135 | 0.007625 | 0.002154 | 0.002568 | 0.00705 | 0.00736 | 0.001444 | 0.001015 | 0.001007 | 0.003799 | 0.000519 | 0.008103 | 0.000743 | 0.006982 | 0.005075 |
| 24.8472222 | 0.034071 | 0.006173 | 0.00469 | 0.001241 | 0.001054 | 0.004973 | 0.018553 | 0.002656 | 0.00743 | 0.001838 | 0.004181 | 0.016733 | 0.007147 | 0.002102 | 0.001277 | 0.001899 | 0.004947 | 0.000731 | 0.008559 | 0.00038 | 0.003065 | 0.004988 |
| 24.8611111 | 0.01949 | 0.003733 | 0.011053 | 0.001221 | 0.001045 | 0.002303 | 0.008925 | 0.001378 | 0.006666 | 0.001387 | 0.009709 | 0.024757 | 0.006972 | 0.002935 | 0.002337 | 0.00226 | 0.007176 | 0.001821 | 0.009391 | 0.000246 | 0.00248 | 0.004256 |
| 24.875 | 0.008713 | 0.003734 | 0.016939 | 0.001641 | 0.000998 | 0.001408 | 0.004648 | 0.001863 | 0.005764 | 0.001 | 0.01756 | 0.02177 | 0.007485 | 0.004388 | 0.004718 | 0.001662 | 0.010738 | 0.003082 | 0.010591 | 0.000822 | 0.003261 | 0.003511 |
| 24.8888889 | 0.008805 | 0.003321 | 0.014616 | 0.002039 | 0.001402 | 0.002084 | 0.007377 | 0.005318 | 0.005026 | 0.000824 | 0.023778 | 0.014375 | 0.008437 | 0.006263 | 0.00514 | 0.000888 | 0.01528 | 0.0038 | 0.011264 | 0.001935 | 0.002776 | 0.003996 |
| 24.9027778 | 0.009738 | 0.004106 | 0.00768 | 0.002882 | 0.00195 | 0.005931 | 0.014984 | 0.010607 | 0.00504 | 0.00105 | 0.024497 | 0.018627 | 0.009891 | 0.008321 | 0.003097 | 0.001152 | 0.021006 | 0.004947 | 0.010806 | 0.003371 | 0.002114 | 0.00681 |
| 24.9166667 | 0.013623 | 0.008071 | 0.003754 | 0.004111 | 0.002723 | 0.012969 | 0.024918 | 0.015858 | 0.00632 | 0.001702 | 0.02017 | 0.025933 | 0.012431 | 0.010918 | 0.002234 | 0.002381 | 0.02846 | 0.007123 | 0.010158 | 0.005558 | 0.003804 | 0.012635 |
| 24.9305556 | 0.021638 | 0.011729 | 0.004326 | 0.004831 | 0.003969 | 0.019597 | 0.033626 | 0.019911 | 0.007866 | 0.002698 | 0.014575 | 0.019206 | 0.016036 | 0.014637 | 0.002395 | 0.004129 | 0.03737 | 0.009616 | 0.010724 | 0.008652 | 0.006533 | 0.019952 |
| 24.9444444 | 0.029723 | 0.012116 | 0.006023 | 0.005184 | 0.005893 | 0.02035 | 0.036914 | 0.022406 | 0.007677 | 0.004371 | 0.009788 | 0.010955 | 0.019742 | 0.018779 | 0.00316 | 0.006472 | 0.045576 | 0.011662 | 0.012087 | 0.012342 | 0.008022 | 0.023912 |
| 24.9583333 | 0.04178 | 0.00868 | 0.007439 | 0.006453 | 0.008114 | 0.015067 | 0.034408 | 0.023582 | 0.006455 | 0.006741 | 0.008134 | 0.021641 | 0.022158 | 0.021236 | 0.005942 | 0.00872 | 0.049673 | 0.012708 | 0.012471 | 0.015611 | 0.008812 | 0.02108 |
| 24.9722222 | 0.054707 | 0.003855 | 0.009896 | 0.008178 | 0.009568 | 0.009749 | 0.027036 | 0.026337 | 0.008041 | 0.008611 | 0.009937 | 0.047284 | 0.022729 | 0.020714 | 0.008187 | 0.010569 | 0.04739 | 0.012564 | 0.011165 | 0.016988 | 0.00971 | 0.013621 |
| 24.9861111 | 0.05526 | 0.001 | 0.013681 | 0.008313 | 0.009728 | 0.007326 | 0.015659 | 0.031332 | 0.012195 | 0.008839 | 0.011385 | 0.069528 | 0.021798 | 0.016969 | 0.007725 | 0.012005 | 0.040214 | 0.010702 | 0.008563 | 0.017145 | 0.010682 | 0.00645 |
| 25 | 0.039459 | 0.000367 | 0.017339 | 0.006771 | 0.008635 | 0.006042 | 0.005905 | 0.032618 | 0.014083 | 0.007014 | 0.008993 | 0.071274 | 0.018711 | 0.009892 | 0.005159 | 0.010772 | 0.030578 | 0.00694 | 0.005676 | 0.016846 | 0.011004 | 0.00389 |
| 25.0138889 | 0.018787 | 0.000893 | 0.017879 | 0.004156 | 0.00553 | 0.004233 | 0.001652 | 0.026097 | 0.011357 | 0.004497 | 0.006513 | 0.050981 | 0.012256 | 0.006436 | 0.003707 | 0.00624 | 0.018474 | 0.006596 | 0.00676 | 0.014143 | 0.008153 | 0.007092 |
| 25.0277778 | 0.006878 | 0.002222 | 0.014018 | 0.002349 | 0.002454 | 0.002722 | 0.002751 | 0.017179 | 0.006437 | 0.003865 | 0.00956 | 0.026281 | 0.005597 | 0.010795 | 0.005038 | 0.0024 | 0.007948 | 0.010976 | 0.010394 | 0.011219 | 0.003356 | 0.009774 |
| 25.0416667 | 0.005196 | 0.002915 | 0.009051 | 0.002161 | 0.002141 | 0.003887 | 0.010549 | 0.01946 | 0.005184 | 0.004475 | 0.012031 | 0.014991 | 0.005803 | 0.011472 | 0.00635 | 0.004437 | 0.007443 | 0.012281 | 0.011182 | 0.015275 | 0.007852 | 0.009932 |
| 25.0555556 | 0.008885 | 0.00391 | 0.005237 | 0.00365 | 0.003711 | 0.009881 | 0.022597 | 0.037151 | 0.01214 | 0.008245 | 0.007773 | 0.022556 | 0.012388 | 0.006178 | 0.007818 | 0.011317 | 0.014617 | 0.012002 | 0.013125 | 0.023734 | 0.024253 | 0.013967 |
| 25.0694444 | 0.014505 | 0.006857 | 0.002654 | 0.00849 | 0.005733 | 0.016297 | 0.035037 | 0.055166 | 0.025013 | 0.0173 | 0.004503 | 0.043763 | 0.017525 | 0.005143 | 0.007027 | 0.014286 | 0.016031 | 0.011833 | 0.013904 | 0.024617 | 0.039898 | 0.017429 |
| 25.0833333 | 0.017147 | 0.00901 | 0.001601 | 0.01268 | 0.005807 | 0.015388 | 0.049868 | 0.055372 | 0.03294 | 0.02197 | 0.010247 | 0.057673 | 0.019715 | 0.009376 | 0.00285 | 0.010861 | 0.009658 | 0.008611 | 0.009957 | 0.01744 | 0.049527 | 0.014069 |

| | | | | | | | | | | | | | | | | | | | | | | |
|---|---|---|---|---|---|---|---|---|---|---|---|---|---|---|---|---|---|---|---|---|---|---|
| 25.0972222 | 0.013319 | 0.009466 | 0.002284 | 0.012414 | 0.004436 | 0.008349 | 0.06297 | 0.036341 | 0.027192 | 0.015398 | 0.019565 | 0.047567 | 0.020878 | 0.012312 | 0.002643 | 0.006975 | 0.006397 | 0.00976 | 0.009571 | 0.012364 | 0.048146 | 0.008056 |
| 25.1111111 | 0.00934 | 0.008361 | 0.006113 | 0.008329 | 0.003855 | 0.004478 | 0.061074 | 0.016021 | 0.014597 | 0.006997 | 0.022407 | 0.02595 | 0.01906 | 0.009315 | 0.007205 | 0.004726 | 0.008987 | 0.017131 | 0.013287 | 0.012414 | 0.031613 | 0.004994 |
| 25.125 | 0.015235 | 0.006198 | 0.01048 | 0.004027 | 0.002731 | 0.008509 | 0.042296 | 0.010907 | 0.009442 | 0.00614 | 0.015192 | 0.012184 | 0.014287 | 0.004386 | 0.009487 | 0.002744 | 0.009726 | 0.019665 | 0.015249 | 0.014087 | 0.014157 | 0.007674 |
| 25.1388889 | 0.02216 | 0.007984 | 0.009136 | 0.003972 | 0.001821 | 0.014146 | 0.02154 | 0.017938 | 0.014014 | 0.009119 | 0.008154 | 0.015279 | 0.008894 | 0.002834 | 0.006945 | 0.001808 | 0.007465 | 0.012639 | 0.014117 | 0.014277 | 0.005472 | 0.011299 |
| 25.1527778 | 0.01777 | 0.014224 | 0.00598 | 0.006227 | 0.003029 | 0.014136 | 0.015551 | 0.022734 | 0.018325 | 0.0095 | 0.015867 | 0.029582 | 0.004058 | 0.002993 | 0.003476 | 0.001923 | 0.008366 | 0.004181 | 0.010803 | 0.012536 | 0.002441 | 0.010624 |
| 25.1666667 | 0.009357 | 0.014396 | 0.008109 | 0.006852 | 0.003577 | 0.008494 | 0.021697 | 0.017404 | 0.017472 | 0.007136 | 0.031705 | 0.038858 | 0.001257 | 0.003142 | 0.002778 | 0.003079 | 0.010364 | 0.004494 | 0.006777 | 0.010297 | 0.003211 | 0.006248 |
| 25.1805556 | 0.005112 | 0.010602 | 0.008761 | 0.0047 | 0.001911 | 0.002678 | 0.021263 | 0.007039 | 0.014943 | 0.004842 | 0.031421 | 0.033254 | 0.000798 | 0.004091 | 0.005285 | 0.006196 | 0.008397 | 0.018149 | 0.00487 | 0.006732 | 0.005362 | 0.002695 |
| 25.1944444 | 0.005635 | 0.010934 | 0.006248 | 0.001644 | 0.001327 | 0.001288 | 0.011521 | 0.004942 | 0.016761 | 0.004937 | 0.016335 | 0.019111 | 0.001666 | 0.005843 | 0.00935 | 0.008084 | 0.004785 | 0.043539 | 0.008939 | 0.003187 | 0.00541 | 0.003637 |
| 25.2083333 | 0.010838 | 0.01083 | 0.009292 | 0.001422 | 0.003635 | 0.00348 | 0.004679 | 0.014693 | 0.021206 | 0.007021 | 0.009042 | 0.012208 | 0.002109 | 0.006908 | 0.01124 | 0.005679 | 0.00377 | 0.059512 | 0.015472 | 0.002218 | 0.003085 | 0.005516 |
| 25.2222222 | 0.013974 | 0.01182 | 0.014591 | 0.003648 | 0.006206 | 0.005836 | 0.00882 | 0.022958 | 0.016585 | 0.00803 | 0.007072 | 0.015286 | 0.003385 | 0.005367 | 0.008642 | 0.002762 | 0.005337 | 0.044954 | 0.014708 | 0.002162 | 0.001968 | 0.004334 |
| 25.2361111 | 0.009615 | 0.010699 | 0.013128 | 0.004034 | 0.006689 | 0.006872 | 0.024828 | 0.022314 | 0.007864 | 0.006354 | 0.005225 | 0.01601 | 0.009144 | 0.002774 | 0.003944 | 0.002444 | 0.006328 | 0.02245 | 0.014781 | 0.002094 | 0.004166 | 0.004198 |
| 25.25 | 0.003835 | 0.004692 | 0.008439 | 0.002603 | 0.006419 | 0.007695 | 0.043128 | 0.02057 | 0.005546 | 0.003974 | 0.009591 | 0.018126 | 0.014421 | 0.002006 | 0.001568 | 0.002585 | 0.005771 | 0.02503 | 0.02369 | 0.002538 | 0.007821 | 0.011314 |
| 25.2638889 | 0.002622 | 0.00219 | 0.007354 | 0.002951 | 0.006077 | 0.008134 | 0.05109 | 0.022651 | 0.008935 | 0.003021 | 0.013579 | 0.035127 | 0.01311 | 0.002446 | 0.002754 | 0.002946 | 0.004185 | 0.037059 | 0.023342 | 0.005436 | 0.009115 | 0.019954 |
| 25.2777778 | 0.005097 | 0.004239 | 0.006327 | 0.003847 | 0.004077 | 0.006241 | 0.044836 | 0.023756 | 0.018213 | 0.003402 | 0.009845 | 0.046162 | 0.007499 | 0.004627 | 0.004264 | 0.002842 | 0.002295 | 0.02732 | 0.012396 | 0.011372 | 0.007392 | 0.018034 |
| 25.2916667 | 0.005574 | 0.005051 | 0.005139 | 0.002501 | 0.001778 | 0.003079 | 0.030903 | 0.019636 | 0.020413 | 0.003938 | 0.005536 | 0.031619 | 0.003202 | 0.00925 | 0.003334 | 0.002061 | 0.001288 | 0.010909 | 0.012525 | 0.015616 | 0.004974 | 0.01017 |
| 25.3055556 | 0.004082 | 0.003149 | 0.006752 | 0.001233 | 0.00104 | 0.001392 | 0.019867 | 0.013169 | 0.011391 | 0.003272 | 0.004247 | 0.01756 | 0.003118 | 0.010486 | 0.001401 | 0.001655 | 0.000997 | 0.009279 | 0.024877 | 0.014993 | 0.00285 | 0.009362 |
| 25.3194444 | 0.006018 | 0.002096 | 0.009575 | 0.002344 | 0.000647 | 0.002264 | 0.018273 | 0.011564 | 0.005208 | 0.001672 | 0.003652 | 0.022219 | 0.008085 | 0.007529 | 0.000749 | 0.000813 | 0.002253 | 0.009362 | 0.032473 | 0.012553 | 0.002111 | 0.011439 |
| 25.3333333 | 0.009003 | 0.001968 | 0.0148 | 0.003819 | 0.001004 | 0.004713 | 0.025744 | 0.01676 | 0.004704 | 0.000936 | 0.005935 | 0.022843 | 0.015808 | 0.007985 | 0.001377 | 0.000693 | 0.006436 | 0.005748 | 0.030039 | 0.012931 | 0.002353 | 0.008202 |
| 25.3472222 | 0.008395 | 0.002465 | 0.019943 | 0.00419 | 0.0018 | 0.005219 | 0.033309 | 0.019634 | 0.007218 | 0.001485 | 0.008685 | 0.014036 | 0.017111 | 0.009937 | 0.003488 | 0.001275 | 0.01047 | 0.006565 | 0.023916 | 0.016199 | 0.002501 | 0.003174 |
| 25.3611111 | 0.005481 | 0.004676 | 0.017809 | 0.004604 | 0.002502 | 0.003552 | 0.03205 | 0.016877 | 0.008155 | 0.001887 | 0.009437 | 0.014541 | 0.011895 | 0.007624 | 0.007066 | 0.0014 | 0.010385 | 0.008324 | 0.018334 | 0.019158 | 0.003909 | 0.000849 |
| 25.375 | 0.003802 | 0.005913 | 0.009381 | 0.005252 | 0.004946 | 0.003199 | 0.026022 | 0.015875 | 0.004578 | 0.00115 | 0.010429 | 0.017427 | 0.010132 | 0.003918 | 0.009084 | 0.001256 | 0.007932 | 0.008061 | 0.013066 | 0.020461 | 0.006575 | 0.000915 |
| 25.3888889 | 0.004576 | 0.005429 | 0.005188 | 0.005546 | 0.006469 | 0.004417 | 0.025561 | 0.017613 | 0.001874 | 0.000799 | 0.013119 | 0.014293 | 0.013718 | 0.003061 | 0.00664 | 0.002393 | 0.007478 | 0.009868 | 0.008501 | 0.020443 | 0.00913 | 0.001856 |
| 25.4027778 | 0.009752 | 0.006117 | 0.011022 | 0.006185 | 0.004162 | 0.005577 | 0.032284 | 0.018053 | 0.004657 | 0.001977 | 0.017797 | 0.010947 | 0.019036 | 0.004178 | 0.003527 | 0.007121 | 0.009977 | 0.014038 | 0.008788 | 0.019157 | 0.012588 | 0.002724 |
| 25.4166667 | 0.018808 | 0.008594 | 0.019582 | 0.008196 | 0.001503 | 0.007112 | 0.039293 | 0.015208 | 0.010409 | 0.003473 | 0.023745 | 0.009881 | 0.025866 | 0.004787 | 0.004993 | 0.012795 | 0.015324 | 0.017638 | 0.013662 | 0.016439 | 0.017648 | 0.005084 |
| 25.4305556 | 0.023219 | 0.010841 | 0.021247 | 0.010948 | 0.001945 | 0.009564 | 0.040811 | 0.014495 | 0.012805 | 0.004298 | 0.028363 | 0.011822 | 0.033774 | 0.0052 | 0.007614 | 0.014778 | 0.024557 | 0.018554 | 0.016371 | 0.012569 | 0.020166 | 0.009927 |
| 25.4444444 | 0.01903 | 0.010011 | 0.01629 | 0.011236 | 0.003739 | 0.011388 | 0.036733 | 0.018794 | 0.011479 | 0.00372 | 0.030391 | 0.014425 | 0.037218 | 0.004257 | 0.006543 | 0.012201 | 0.035 | 0.015027 | 0.012417 | 0.007621 | 0.014744 | 0.011064 |
| 25.4583333 | 0.012613 | 0.006211 | 0.012391 | 0.00713 | 0.003985 | 0.010364 | 0.030867 | 0.017744 | 0.01089 | 0.002554 | 0.035691 | 0.013534 | 0.032734 | 0.003236 | 0.005131 | 0.006892 | 0.041643 | 0.008236 | 0.007305 | 0.003702 | 0.008396 | 0.007482 |
| 25.4722222 | 0.011689 | 0.003235 | 0.011746 | 0.002927 | 0.003905 | 0.006684 | 0.029124 | 0.009445 | 0.013517 | 0.002523 | 0.037531 | 0.010241 | 0.022986 | 0.005769 | 0.007529 | 0.002545 | 0.043469 | 0.003016 | 0.01049 | 0.002809 | 0.009857 | 0.010336 |
| 25.4861111 | 0.021534 | 0.002775 | 0.011439 | 0.001676 | 0.003481 | 0.004666 | 0.035146 | 0.003465 | 0.019796 | 0.00257 | 0.030037 | 0.011824 | 0.013214 | 0.007439 | 0.008711 | 0.001807 | 0.041583 | 0.002387 | 0.016315 | 0.003163 | 0.01096 | 0.017965 |
| 25.5 | 0.038809 | 0.00302 | 0.015979 | 0.002098 | 0.003093 | 0.005825 | 0.043392 | 0.002937 | 0.024626 | 0.00236 | 0.028691 | 0.021031 | 0.008314 | 0.005455 | 0.005576 | 0.003507 | 0.037184 | 0.004614 | 0.013 | 0.003753 | 0.007407 | 0.019458 |
| 25.5138889 | 0.05086 | 0.002922 | 0.025708 | 0.004642 | 0.006364 | 0.005817 | 0.046558 | 0.00838 | 0.022705 | 0.002618 | 0.031756 | 0.032652 | 0.006554 | 0.00374 | 0.003657 | 0.005392 | 0.033072 | 0.005356 | 0.005213 | 0.00308 | 0.004934 | 0.015718 |
| 25.5277778 | 0.053163 | 0.003962 | 0.026354 | 0.007234 | 0.011917 | 0.003967 | 0.044703 | 0.019856 | 0.016055 | 0.002851 | 0.030792 | 0.040962 | 0.004333 | 0.00409 | 0.003411 | 0.007853 | 0.029064 | 0.003696 | 0.001888 | 0.002892 | 0.005052 | 0.010259 |
| 25.5416667 | 0.051061 | 0.006887 | 0.015003 | 0.007489 | 0.015583 | 0.003382 | 0.042175 | 0.02975 | 0.009477 | 0.002882 | 0.028118 | 0.04249 | 0.005342 | 0.007838 | 0.004317 | 0.013813 | 0.019269 | 0.004807 | 0.00639 | 0.0087 | 0.008495 | 0.005381 |
| 25.5555556 | 0.041933 | 0.009387 | 0.005517 | 0.006973 | 0.014008 | 0.003959 | 0.042764 | 0.027965 | 0.004959 | 0.00224 | 0.024345 | 0.035463 | 0.010526 | 0.014993 | 0.007869 | 0.021443 | 0.00884 | 0.010627 | 0.016258 | 0.017782 | 0.014539 | 0.00589 |
| 25.5694444 | 0.024867 | 0.008698 | 0.003617 | 0.006819 | 0.007934 | 0.004567 | 0.044457 | 0.015413 | 0.002011 | 0.001404 | 0.020726 | 0.022938 | 0.011344 | 0.019487 | 0.007889 | 0.022104 | 0.0071 | 0.013047 | 0.017909 | 0.021373 | 0.016539 | 0.010244 |
| 25.5833333 | 0.011997 | 0.005583 | 0.007097 | 0.007467 | 0.002938 | 0.005106 | 0.041083 | 0.008704 | 0.000712 | 0.001015 | 0.016793 | 0.012915 | 0.005958 | 0.017791 | 0.005431 | 0.013774 | 0.00632 | 0.007922 | 0.010802 | 0.016098 | 0.010692 | 0.008631 |
| 25.5972222 | 0.008263 | 0.003279 | 0.011249 | 0.00773 | 0.001476 | 0.004822 | 0.030732 | 0.011697 | 0.001629 | 0.000935 | 0.011381 | 0.010369 | 0.001654 | 0.013551 | 0.007318 | 0.005212 | 0.002591 | 0.00299 | 0.009321 | 0.007981 | 0.004634 | 0.003533 |
| 25.6111111 | 0.007443 | 0.002994 | 0.009008 | 0.005702 | 0.001375 | 0.003668 | 0.01861 | 0.010149 | 0.003355 | 0.001561 | 0.006872 | 0.013894 | 0.001005 | 0.008364 | 0.009083 | 0.002375 | 0.001644 | 0.002805 | 0.011506 | 0.004668 | 0.005112 | 0.003152 |
| 25.625 | 0.005118 | 0.003003 | 0.007037 | 0.002792 | 0.001369 | 0.003056 | 0.009999 | 0.008568 | 0.003447 | 0.001809 | 0.008502 | 0.020861 | 0.00132 | 0.003285 | 0.007023 | 0.002406 | 0.003682 | 0.003082 | 0.009585 | 0.004612 | 0.007097 | 0.00343 |
| 25.6388889 | 0.004719 | 0.002318 | 0.01436 | 0.001469 | 0.002122 | 0.003746 | 0.004652 | 0.015444 | 0.003977 | 0.001162 | 0.012976 | 0.029006 | 0.002218 | 0.001322 | 0.003755 | 0.001689 | 0.005798 | 0.001945 | 0.005553 | 0.002883 | 0.005427 | 0.002398 |
| 25.6527778 | 0.009956 | 0.002592 | 0.020245 | 0.001479 | 0.002833 | 0.004157 | 0.002259 | 0.018354 | 0.005159 | 0.001007 | 0.012294 | 0.032609 | 0.006453 | 0.001943 | 0.002507 | 0.001174 | 0.006962 | 0.001925 | 0.004499 | 0.000919 | 0.002753 | 0.003709 |
| 25.6666667 | 0.014835 | 0.003489 | 0.014791 | 0.001903 | 0.002884 | 0.002651 | 0.004626 | 0.012005 | 0.00346 | 0.001352 | 0.008908 | 0.02575 | 0.012155 | 0.002995 | 0.004593 | 0.002155 | 0.007323 | 0.003238 | 0.007045 | 0.001427 | 0.002554 | 0.00681 |
| 25.6805556 | 0.013671 | 0.003077 | 0.006089 | 0.003799 | 0.003583 | 0.000751 | 0.010293 | 0.011984 | 0.002731 | 0.00142 | 0.007436 | 0.013407 | 0.013387 | 0.003119 | 0.007456 | 0.003227 | 0.006131 | 0.003383 | 0.007827 | 0.003684 | 0.003431 | 0.008019 |
| 25.6944444 | 0.008414 | 0.003092 | 0.002128 | 0.005388 | 0.003902 | 0.000365 | 0.015113 | 0.017865 | 0.004787 | 0.00109 | 0.0054 | 0.009974 | 0.008462 | 0.002278 | 0.008714 | 0.003169 | 0.003581 | 0.001961 | 0.005087 | 0.004471 | 0.002973 | 0.005508 |
| 25.7083333 | 0.003665 | 0.004827 | 0.002137 | 0.004301 | 0.004098 | 0.000611 | 0.016836 | 0.013818 | 0.005063 | 0.00056 | 0.00525 | 0.022779 | 0.002602 | 0.001565 | 0.008123 | 0.002197 | 0.001833 | 0.000855 | 0.002183 | 0.00262 | 0.001867 | 0.003423 |
| 25.7222222 | 0.001542 | 0.005801 | 0.00444 | 0.003346 | 0.005794 | 0.000462 | 0.016187 | 0.005081 | 0.003443 | 0.000169 | 0.011331 | 0.040762 | 0.000324 | 0.001042 | 0.006488 | 0.001016 | 0.001817 | 0.00078 | 0.001552 | 0.00167 | 0.001485 | 0.00542 |
| 25.7361111 | 0.000884 | 0.004762 | 0.005613 | 0.002904 | 0.005313 | 0.000456 | 0.014715 | 0.002556 | 0.001716 | 0.000229 | 0.015775 | 0.042386 | 0.000662 | 0.000679 | 0.004832 | 0.000392 | 0.003194 | 0.001657 | 0.002718 | 0.004525 | 0.002048 | 0.006989 |
| 25.75 | 0.000439 | 0.002678 | 0.004113 | 0.001833 | 0.002683 | 0.000778 | 0.013084 | 0.004916 | 0.001335 | 0.000548 | 0.011851 | 0.027144 | 0.001254 | 0.001 | 0.003168 | 0.000587 | 0.005909 | 0.002897 | 0.003669 | 0.008262 | 0.003207 | 0.004694 |
| 25.7638889 | 0.000436 | 0.00135 | 0.00333 | 0.001431 | 0.002221 | 0.001714 | 0.011425 | 0.005265 | 0.002352 | 0.000556 | 0.006808 | 0.023113 | 0.002168 | 0.001431 | 0.001517 | 0.001293 | 0.008313 | 0.003976 | 0.004466 | 0.007211 | 0.004892 | 0.002523 |
| 25.7777778 | 0.001754 | 0.002426 | 0.005041 | 0.001569 | 0.002752 | 0.003431 | 0.010395 | 0.003058 | 0.002195 | 0.000324 | 0.010596 | 0.03848 | 0.003637 | 0.001301 | 0.000626 | 0.002162 | 0.009951 | 0.005128 | 0.006382 | 0.003332 | 0.007368 | 0.004063 |
| 25.7916667 | 0.005643 | 0.00554 | 0.00863 | 0.002253 | 0.002426 | 0.004841 | 0.012277 | 0.003379 | 0.001413 | 0.000384 | 0.021003 | 0.050302 | 0.002899 | 0.001799 | 0.000467 | 0.002743 | 0.012165 | 0.006266 | 0.008758 | 0.002136 | 0.010337 | 0.006869 |
| 25.8055556 | 0.012061 | 0.009573 | 0.012622 | 0.003161 | 0.003079 | 0.005242 | 0.019189 | 0.004199 | 0.001835 | 0.000607 | 0.026963 | 0.044429 | 0.001848 | 0.004029 | 0.001486 | 0.002973 | 0.016051 | 0.007969 | 0.010742 | 0.002664 | 0.013186 | 0.00909 |
| 25.8194444 | 0.016278 | 0.012281 | 0.014206 | 0.003771 | 0.004002 | 0.004591 | 0.029449 | 0.003645 | 0.002465 | 0.000697 | 0.022465 | 0.024765 | 0.006027 | 0.007598 | 0.005379 | 0.004162 | 0.02225 | 0.012305 | 0.0128 | 0.003373 | 0.016679 | 0.013625 |
| 25.8333333 | 0.013582 | 0.010957 | 0.011822 | 0.004328 | 0.003352 | 0.003282 | 0.037018 | 0.003314 | 0.003084 | 0.001048 | 0.011545 | 0.01001 | 0.015575 | 0.011473 | 0.011697 | 0.006979 | 0.029503 | 0.019199 | 0.015721 | 0.006606 | 0.021116 | 0.022475 |
| 25.8472222 | 0.006627 | 0.006306 | 0.007013 | 0.004338 | 0.001867 | 0.002229 | 0.034863 | 0.003599 | 0.004542 | 0.001641 | 0.007089 | 0.017899 | 0.027228 | 0.014444 | 0.01759 | 0.009993 | 0.035364 | 0.024639 | 0.019043 | 0.011149 | 0.023946 | 0.032541 |
| 25.8611111 | 0.002082 | 0.003315 | 0.004612 | 0.003919 | 0.000985 | 0.002858 | 0.02321 | 0.004915 | 0.007153 | 0.001708 | 0.012791 | 0.041843 | 0.038826 | 0.016524 | 0.021511 | 0.012493 | 0.038432 | 0.02512 | 0.021788 | 0.012203 | 0.02377 | 0.036267 |
| 25.875 | 0.003334 | 0.003577 | 0.006521 | 0.0037 | 0.001127 | 0.006254 | 0.013802 | 0.007234 | 0.009744 | 0.001245 | 0.018521 | 0.061341 | 0.04824 | 0.019017 | 0.02355 | 0.015474 | 0.038595 | 0.021891 | 0.024059 | 0.011248 | 0.02275 | 0.030273 |
| 25.8888889 | 0.008708 | 0.003609 | 0.008108 | 0.003385 | 0.001177 | 0.01082 | 0.01845 | 0.009611 | 0.010219 | 0.001104 | 0.02816 | 0.058353 | 0.053911 | 0.022422 | 0.022575 | 0.019204 | 0.035926 | 0.01881 | 0.026056 | 0.013224 | 0.02271 | 0.020227 |
| 25.9027778 | 0.016251 | 0.004398 | 0.008965 | 0.002934 | 0.001112 | 0.012778 | 0.035971 | 0.01141 | 0.009336 | 0.00218 | 0.041302 | 0.032926 | 0.055733 | 0.025451 | 0.018996 | 0.023085 | 0.031771 | 0.016448 | 0.027007 | 0.018319 | 0.023373 | 0.011289 |
| 25.9166667 | 0.027258 | 0.006616 | 0.012506 | 0.003926 | 0.002168 | 0.010898 | 0.049811 | 0.014006 | 0.009768 | 0.004916 | 0.037484 | 0.012879 | 0.052915 | 0.02661 | 0.016076 | 0.026346 | 0.028985 | 0.012314 | 0.025986 | 0.024126 | 0.023499 | 0.00526 |
| 25.9305556 | 0.038654 | 0.007158 | 0.017912 | 0.006152 | 0.005166 | 0.007137 | 0.045831 | 0.01629 | 0.011967 | 0.008332 | 0.018757 | 0.011827 | 0.045867 | 0.025282 | 0.015043 | 0.026652 | 0.029677 | 0.006693 | 0.022554 | 0.028265 | 0.021138 | 0.003948 |
| 25.9444444 | 0.043485 | 0.006719 | 0.021705 | 0.006534 | 0.008207 | 0.003204 | 0.032639 | 0.016238 | 0.014886 | 0.010539 | 0.005258 | 0.010463 | 0.037087 | 0.021162 | 0.01367 | 0.021834 | 0.032563 | 0.004153 | 0.016998 | 0.027299 | 0.014409 | 0.006599 |
| 25.9583333 | 0.041551 | 0.006927 | 0.021615 | 0.00515 | 0.008499 | 0.001551 | 0.027337 | 0.018177 | 0.018649 | 0.010566 | 0.002889 | 0.006388 | 0.030212 | 0.014744 | 0.01003 | 0.013709 | 0.031706 | 0.005718 | 0.01166 | 0.021926 | 0.007647 | 0.009707 |
| 25.9722222 | 0.035273 | 0.006855 | 0.01862 | 0.004851 | 0.006766 | 0.003792 | 0.032812 | 0.022593 | 0.022451 | 0.009123 | 0.007537 | 0.010105 | 0.028332 | 0.008139 | 0.006673 | 0.007059 | 0.024401 | 0.006125 | 0.010387 | 0.018538 | 0.007524 | 0.01034 |
| 25.9861111 | 0.025692 | 0.006997 | 0.013833 | 0.005779 | 0.004718 | 0.007367 | 0.040227 | 0.023262 | 0.023456 | 0.007435 | 0.013538 | 0.014954 | 0.028298 | 0.003589 | 0.006943 | 0.004663 | 0.015605 | 0.003453 | 0.013878 | 0.019106 | 0.010457 | 0.007999 |
| 26 | 0.015835 | 0.00821 | 0.00779 | 0.00603 | 0.00271 | 0.007662 | 0.041028 | 0.018372 | 0.019589 | 0.005262 | 0.019066 | 0.01675 | 0.023178 | 0.002158 | 0.00858 | 0.004442 | 0.008502 | 0.001613 | 0.015397 | 0.018893 | 0.011882 | 0.004887 |
| 26.0138889 | 0.008575 | 0.007983 | 0.004991 | 0.004172 | 0.002364 | 0.007308 | 0.03185 | 0.010229 | 0.01123 | 0.003454 | 0.027165 | 0.018347 | 0.012692 | 0.004094 | 0.006548 | 0.003187 | 0.004107 | 0.004242 | 0.009583 | 0.015144 | 0.014529 | 0.004117 |
| 26.0277778 | 0.006432 | 0.004758 | 0.008928 | 0.003416 | 0.003747 | 0.010246 | 0.018396 | 0.009285 | 0.003792 | 0.005325 | 0.03877 | 0.018451 | 0.00441 | 0.007871 | 0.003706 | 0.002526 | 0.004461 | 0.012278 | 0.005208 | 0.009268 | 0.018988 | 0.006394 |
| 26.0416667 | 0.013759 | 0.001614 | 0.014299 | 0.006801 | 0.004599 | 0.01412 | 0.012209 | 0.035573 | 0.004512 | 0.009343 | 0.047137 | 0.020146 | 0.004558 | 0.012504 | 0.006616 | 0.006148 | 0.006483 | 0.023657 | 0.015373 | 0.004752 | 0.021764 | 0.010953 |
| 26.0555556 | 0.031117 | 0.001928 | 0.015024 | 0.008087 | 0.008399 | 0.021771 | 0.017767 | 0.054054 | 0.011311 | 0.013051 | 0.040778 | 0.033137 | 0.011194 | 0.017431 | 0.011429 | 0.010316 | 0.009822 | 0.030785 | 0.025107 | 0.005489 | 0.019826 | 0.014672 |
| 26.0694444 | 0.044624 | 0.004966 | 0.0135 | 0.007941 | 0.013746 | 0.027152 | 0.030671 | 0.053651 | 0.014586 | 0.022004 | 0.021333 | 0.061161 | 0.017133 | 0.018822 | 0.009572 | 0.008025 | 0.019824 | 0.023448 | 0.01915 | 0.007447 | 0.012631 | 0.011955 |
| 26.0833333 | 0.037984 | 0.007008 | 0.013115 | 0.013067 | 0.011603 | 0.019587 | 0.048479 | 0.065224 | 0.011537 | 0.028398 | 0.007788 | 0.079399 | 0.021061 | 0.01571 | 0.00427 | 0.003174 | 0.026458 | 0.012077 | 0.011731 | 0.01064 | 0.0094 | 0.008506 |
| 26.0972222 | 0.018936 | 0.006398 | 0.009969 | 0.016961 | 0.004988 | 0.01079 | 0.062823 | 0.048074 | 0.007813 | 0.018719 | 0.008983 | 0.056702 | 0.029249 | 0.011895 | 0.003027 | 0.003614 | 0.017975 | 0.01212 | 0.008755 | 0.017208 | 0.013868 | 0.010945 |
| 26.1111111 | 0.010504 | 0.005486 | 0.006946 | 0.012394 | 0.002259 | 0.012494 | 0.052008 | 0.016679 | 0.005615 | 0.008306 | 0.014847 | 0.02541 | 0.041312 | 0.00964 | 0.00273 | 0.007713 | 0.00929 | 0.014228 | 0.010415 | 0.015065 | 0.013239 | 0.011443 |
| 26.125 | 0.018847 | 0.010438 | 0.010481 | 0.007034 | 0.004135 | 0.020598 | 0.027104 | 0.014923 | 0.005786 | 0.010255 | 0.014613 | 0.030081 | 0.048447 | 0.010736 | 0.002796 | 0.011508 | 0.012321 | 0.01252 | 0.013425 | 0.006566 | 0.00644 | 0.007585 |
| 26.1388889 | 0.02862 | 0.021823 | 0.011134 | 0.010002 | 0.007383 | 0.027407 | 0.020549 | 0.023513 | 0.012879 | 0.013661 | 0.008938 | 0.052089 | 0.045133 | 0.01426 | 0.007272 | 0.014884 | 0.016528 | 0.010506 | 0.011035 | 0.004889 | 0.002264 | 0.003929 |
| 26.1527778 | 0.029373 | 0.02608 | 0.008254 | 0.015359 | 0.008749 | 0.02424 | 0.022018 | 0.022156 | 0.025296 | 0.011739 | 0.006127 | 0.062159 | 0.031922 | 0.014778 | 0.014135 | 0.017837 | 0.016329 | 0.008149 | 0.008176 | 0.008638 | 0.002177 | 0.003718 |
| 26.1666667 | 0.026209 | 0.0161 | 0.011164 | 0.015276 | 0.006702 | 0.014367 | 0.013573 | 0.015778 | 0.032499 | 0.007937 | 0.011791 | 0.049719 | 0.015237 | 0.010846 | 0.019978 | 0.020115 | 0.015482 | 0.00542 | 0.00703 | 0.015656 | 0.003463 | 0.00971 |
| 26.1805556 | 0.022038 | 0.00476 | 0.012482 | 0.01112 | 0.002935 | 0.011449 | 0.005931 | 0.017647 | 0.025588 | 0.006605 | 0.024047 | 0.027317 | 0.006807 | 0.006841 | 0.023083 | 0.021782 | 0.014555 | 0.002897 | 0.005064 | 0.025589 | 0.003358 | 0.021206 |
| 26.1944444 | 0.015199 | 0.002555 | 0.010664 | 0.006923 | 0.001108 | 0.011746 | 0.005105 | 0.019611 | 0.011063 | 0.008282 | 0.025566 | 0.029343 | 0.013977 | 0.007088 | 0.022863 | 0.022307 | 0.012659 | 0.002246 | 0.002849 | 0.031836 | 0.002314 | 0.0327 |
| 26.2083333 | 0.013363 | 0.004427 | 0.009059 | 0.004349 | 0.002007 | 0.011032 | 0.008891 | 0.023395 | 0.004779 | 0.009633 | 0.015319 | 0.045745 | 0.029941 | 0.012625 | 0.020076 | 0.022299 | 0.011049 | 0.002523 | 0.002399 | 0.026338 | 0.001761 | 0.035921 |
| 26.2222222 | 0.015591 | 0.003588 | 0.005379 | 0.002583 | 0.003483 | 0.017341 | 0.02284 | 0.028685 | 0.015335 | 0.009431 | 0.009422 | 0.041487 | 0.038306 | 0.019649 | 0.017173 | 0.022807 | 0.011605 | 0.002231 | 0.006121 | 0.014512 | 0.001826 | 0.02759 |
| 26.2361111 | 0.012971 | 0.003422 | 0.00602 | 0.001935 | 0.004546 | 0.023457 | 0.046646 | 0.023624 | 0.028947 | 0.009883 | 0.011361 | 0.044898 | 0.032489 | 0.021876 | 0.017376 | 0.023659 | 0.014458 | 0.003625 | 0.012724 | 0.007006 | 0.003391 | 0.015263 |
| 26.25 | 0.013921 | 0.009221 | 0.014676 | 0.003034 | 0.005699 | 0.021326 | 0.064909 | 0.018123 | 0.02302 | 0.011015 | 0.021142 | 0.069196 | 0.021042 | 0.017298 | 0.020269 | 0.02533 | 0.017229 | 0.008579 | 0.014302 | 0.009545 | 0.008843 | 0.008716 |
| 26.2638889 | 0.019947 | 0.015478 | 0.020856 | 0.003208 | 0.00734 | 0.015034 | 0.059074 | 0.020342 | 0.008363 | 0.010526 | 0.022987 | 0.059927 | 0.015671 | 0.010499 | 0.020786 | 0.027509 | 0.017633 | 0.014587 | 0.008766 | 0.026016 | 0.017111 | 0.008339 |
| 26.2777778 | 0.017235 | 0.014092 | 0.01547 | 0.002484 | 0.010268 | 0.008496 | 0.035855 | 0.019509 | 0.007813 | 0.009205 | 0.012773 | 0.03417 | 0.022151 | 0.007052 | 0.016298 | 0.027408 | 0.017076 | 0.0169 | 0.003396 | 0.041933 | 0.020486 | 0.013075 |
| 26.2916667 | 0.013175 | 0.007335 | 0.008398 | 0.00542 | 0.012569 | 0.007771 | 0.019633 | 0.015352 | 0.018414 | 0.009028 | 0.009869 | 0.034897 | 0.032212 | 0.007808 | 0.010952 | 0.023576 | 0.018352 | 0.012241 | 0.007929 | 0.044129 | 0.015358 | 0.018819 |
| 26.3055556 | 0.019722 | 0.003094 | 0.006186 | 0.011769 | 0.010215 | 0.014682 | 0.014439 | 0.016892 | 0.021389 | 0.007414 | 0.012986 | 0.030579 | 0.033505 | 0.009997 | 0.009552 | 0.0188 | 0.021085 | 0.004813 | 0.026754 | 0.040498 | 0.008716 | 0.019114 |
| 26.3194444 | 0.019815 | 0.003362 | 0.005648 | 0.016785 | 0.007675 | 0.020978 | 0.011014 | 0.020217 | 0.012534 | 0.003446 | 0.013451 | 0.024615 | 0.026163 | 0.01134 | 0.011637 | 0.016872 | 0.023384 | 0.003697 | 0.044213 | 0.039152 | 0.004657 | 0.018635 |
| 26.3333333 | 0.010407 | 0.003607 | 0.005436 | 0.016558 | 0.00748 | 0.020911 | 0.007714 | 0.021524 | 0.005366 | 0.002349 | 0.015463 | 0.036374 | 0.018839 | 0.012105 | 0.014157 | 0.018654 | 0.026165 | 0.0111 | 0.041104 | 0.04297 | 0.002965 | 0.023058 |
| 26.3472222 | 0.007526 | 0.002668 | 0.003246 | 0.012096 | 0.006524 | 0.017223 | 0.008887 | 0.023173 | 0.004084 | 0.005458 | 0.014729 | 0.041415 | 0.018729 | 0.013999 | 0.014939 | 0.020973 | 0.031152 | 0.019364 | 0.024801 | 0.046373 | 0.004799 | 0.027477 |
| 26.3611111 | 0.008664 | 0.00368 | 0.005786 | 0.008824 | 0.007767 | 0.014031 | 0.015384 | 0.024621 | 0.007823 | 0.007567 | 0.008009 | 0.034867 | 0.026974 | 0.016736 | 0.013054 | 0.019001 | 0.035558 | 0.020827 | 0.013993 | 0.041316 | 0.006268 | 0.024212 |
| 26.375 | 0.009268 | 0.006583 | 0.018438 | 0.009012 | 0.008714 | 0.012663 | 0.021924 | 0.025276 | 0.013718 | 0.006402 | 0.007541 | 0.032213 | 0.038513 | 0.017298 | 0.009964 | 0.012391 | 0.033235 | 0.017422 | 0.012742 | 0.032166 | 0.004224 | 0.013736 |
| 26.3888889 | 0.012117 | 0.009348 | 0.027444 | 0.011077 | 0.006656 | 0.011794 | 0.024808 | 0.02597 | 0.015991 | 0.005195 | 0.019016 | 0.033051 | 0.044036 | 0.014947 | 0.008784 | 0.006508 | 0.02348 | 0.014775 | 0.01446 | 0.025968 | 0.002433 | 0.004423 |
| 26.4027778 | 0.017848 | 0.012002 | 0.019501 | 0.012635 | 0.006483 | 0.010154 | 0.024589 | 0.025295 | 0.013446 | 0.00621 | 0.034921 | 0.026551 | 0.040556 | 0.013345 | 0.010386 | 0.005681 | 0.016044 | 0.013552 | 0.014649 | 0.02358 | 0.002441 | 0.001723 |
| 26.4166667 | 0.024279 | 0.014287 | 0.006703 | 0.012506 | 0.009917 | 0.009886 | 0.023358 | 0.022924 | 0.010444 | 0.007811 | 0.047293 | 0.015113 | 0.034986 | 0.014846 | 0.012639 | 0.005711 | 0.020193 | 0.011966 | 0.015621 | 0.023483 | 0.002038 | 0.002378 |
| 26.4305556 | 0.026057 | 0.015399 | 0.00611 | 0.012238 | 0.014398 | 0.014481 | 0.021774 | 0.024575 | 0.012449 | 0.008099 | 0.057921 | 0.010733 | 0.033888 | 0.016609 | 0.013867 | 0.002929 | 0.033201 | 0.010139 | 0.022126 | 0.024768 | 0.001641 | 0.00215 |
| 26.4444444 | 0.022128 | 0.015783 | 0.019008 | 0.012703 | 0.014996 | 0.023193 | 0.018166 | 0.029435 | 0.016845 | 0.006796 | 0.073906 | 0.013599 | 0.032086 | 0.014364 | 0.014207 | 0.001108 | 0.045555 | 0.00887 | 0.033555 | 0.025194 | 0.003484 | 0.003213 |
| 26.4583333 | 0.020327 | 0.013568 | 0.034517 | 0.012225 | 0.010442 | 0.028916 | 0.012057 | 0.034082 | 0.01736 | 0.004257 | 0.082418 | 0.017284 | 0.021116 | 0.008295 | 0.013147 | 0.001949 | 0.050287 | 0.006608 | 0.040417 | 0.01982 | 0.007956 | 0.008646 |
| 26.4722222 | 0.024946 | 0.008193 | 0.0362 | 0.009851 | 0.006048 | 0.025224 | 0.005398 | 0.036886 | 0.016151 | 0.00222 | 0.068234 | 0.016972 | 0.011047 | 0.003517 | 0.009129 | 0.00378 | 0.049464 | 0.003529 | 0.033854 | 0.010039 | 0.011623 | 0.013665 |
| 26.4861111 | 0.031797 | 0.006143 | 0.024169 | 0.008861 | 0.007077 | 0.018101 | 0.003044 | 0.034018 | 0.01936 | 0.001875 | 0.054569 | 0.011892 | 0.015943 | 0.002594 | 0.004626 | 0.004146 | 0.044016 | 0.003827 | 0.021815 | 0.004717 | 0.01071 | 0.012433 |

| | | | | | | | | | | | | | | | | | | | | | | |
|---|---|---|---|---|---|---|---|---|---|---|---|---|---|---|---|---|---|---|---|---|---|---|
| 26.5 | 0.042741 | 0.010073 | 0.013221 | 0.01149 | 0.014535 | 0.020153 | 0.008026 | 0.03109 | 0.026267 | 0.003429 | 0.055918 | 0.006622 | 0.025856 | 0.002758 | 0.00386 | 0.004261 | 0.033452 | 0.00768 | 0.015847 | 0.007083 | 0.006891 | 0.011061 |
| 26.5138889 | 0.058581 | 0.011461 | 0.011372 | 0.01301 | 0.023408 | 0.033842 | 0.016354 | 0.030425 | 0.028469 | 0.008819 | 0.057624 | 0.019033 | 0.027854 | 0.00315 | 0.005228 | 0.005064 | 0.026376 | 0.010377 | 0.010845 | 0.01094 | 0.004086 | 0.022973 |
| 26.5277778 | 0.062824 | 0.008622 | 0.014305 | 0.009652 | 0.024426 | 0.042435 | 0.020924 | 0.023411 | 0.02149 | 0.01536 | 0.058207 | 0.059219 | 0.029114 | 0.005606 | 0.004613 | 0.005705 | 0.023081 | 0.008357 | 0.005559 | 0.010295 | 0.004888 | 0.041731 |
| 26.5416667 | 0.051266 | 0.008807 | 0.011535 | 0.004676 | 0.016899 | 0.030763 | 0.017699 | 0.011284 | 0.014891 | 0.01766 | 0.056259 | 0.090169 | 0.034736 | 0.012595 | 0.00431 | 0.0131 | 0.015639 | 0.008514 | 0.00932 | 0.013882 | 0.012817 | 0.041744 |
| 26.5555556 | 0.040579 | 0.009078 | 0.005698 | 0.001874 | 0.012702 | 0.013286 | 0.014851 | 0.004017 | 0.01323 | 0.015193 | 0.045059 | 0.082543 | 0.045087 | 0.025605 | 0.010227 | 0.029087 | 0.012459 | 0.021576 | 0.016276 | 0.0298 | 0.029325 | 0.027368 |
| 26.5694444 | 0.034209 | 0.005498 | 0.006792 | 0.001711 | 0.014107 | 0.006318 | 0.02369 | 0.002422 | 0.011608 | 0.009198 | 0.028385 | 0.052244 | 0.049856 | 0.037495 | 0.016464 | 0.039539 | 0.021467 | 0.03625 | 0.014922 | 0.041215 | 0.044602 | 0.021306 |
| 26.5833333 | 0.026114 | 0.001847 | 0.009573 | 0.003563 | 0.013423 | 0.005393 | 0.036087 | 0.001936 | 0.011065 | 0.004292 | 0.013976 | 0.022087 | 0.032762 | 0.035967 | 0.012468 | 0.033327 | 0.025607 | 0.034891 | 0.00864 | 0.033911 | 0.043807 | 0.017293 |
| 26.5972222 | 0.017779 | 0.000742 | 0.008433 | 0.005779 | 0.01194 | 0.004446 | 0.036338 | 0.004621 | 0.011935 | 0.0054 | 0.006516 | 0.009189 | 0.011396 | 0.020425 | 0.004675 | 0.016897 | 0.016588 | 0.026928 | 0.003942 | 0.016596 | 0.027834 | 0.010369 |
| 26.6111111 | 0.010396 | 0.002169 | 0.007596 | 0.006154 | 0.012385 | 0.005201 | 0.024016 | 0.013247 | 0.01044 | 0.008664 | 0.003863 | 0.015223 | 0.011895 | 0.007139 | 0.002588 | 0.005099 | 0.006384 | 0.023165 | 0.002363 | 0.004345 | 0.011761 | 0.011171 |
| 26.625 | 0.005974 | 0.004782 | 0.007533 | 0.004721 | 0.010632 | 0.011313 | 0.010312 | 0.01696 | 0.006879 | 0.007349 | 0.004722 | 0.023494 | 0.022784 | 0.006961 | 0.003669 | 0.002602 | 0.001759 | 0.014738 | 0.0037 | 0.001507 | 0.003769 | 0.017569 |
| 26.6388889 | 0.01057 | 0.006889 | 0.006394 | 0.003526 | 0.008568 | 0.017293 | 0.003069 | 0.011849 | 0.003951 | 0.003405 | 0.012768 | 0.021169 | 0.023822 | 0.012157 | 0.004156 | 0.002315 | 0.001603 | 0.006546 | 0.005261 | 0.002636 | 0.003091 | 0.024202 |
| 26.6527778 | 0.022199 | 0.007684 | 0.003777 | 0.003429 | 0.011039 | 0.013619 | 0.005513 | 0.013211 | 0.002405 | 0.002903 | 0.027902 | 0.015495 | 0.029708 | 0.014933 | 0.003456 | 0.003211 | 0.001968 | 0.008751 | 0.005295 | 0.007716 | 0.007232 | 0.029689 |
| 26.6666667 | 0.031291 | 0.006969 | 0.001887 | 0.003299 | 0.012347 | 0.006572 | 0.02454 | 0.019027 | 0.002678 | 0.003647 | 0.042713 | 0.022012 | 0.051971 | 0.014147 | 0.003488 | 0.008484 | 0.00291 | 0.012341 | 0.004891 | 0.013704 | 0.010563 | 0.029843 |
| 26.6805556 | 0.030032 | 0.00458 | 0.003464 | 0.002356 | 0.008275 | 0.007996 | 0.050499 | 0.01521 | 0.004454 | 0.004328 | 0.044267 | 0.034581 | 0.051576 | 0.010963 | 0.005234 | 0.010935 | 0.005008 | 0.011519 | 0.005865 | 0.013171 | 0.010204 | 0.024608 |
| 26.6944444 | 0.019364 | 0.001993 | 0.007435 | 0.001341 | 0.007231 | 0.012599 | 0.05561 | 0.009445 | 0.004767 | 0.007746 | 0.028036 | 0.036902 | 0.030525 | 0.008781 | 0.008383 | 0.009341 | 0.005125 | 0.01039 | 0.00893 | 0.010767 | 0.009466 | 0.01928 |
| 26.7083333 | 0.013562 | 0.00118 | 0.011357 | 0.00113 | 0.012584 | 0.012175 | 0.038033 | 0.012642 | 0.003029 | 0.008511 | 0.015751 | 0.028109 | 0.035774 | 0.008879 | 0.013851 | 0.013213 | 0.006758 | 0.010569 | 0.012386 | 0.01062 | 0.01027 | 0.016648 |
| 26.7222222 | 0.017946 | 0.001439 | 0.011858 | 0.001544 | 0.012274 | 0.007531 | 0.017493 | 0.016871 | 0.001522 | 0.004873 | 0.018434 | 0.014597 | 0.049814 | 0.009783 | 0.021237 | 0.019224 | 0.012913 | 0.010163 | 0.014149 | 0.009347 | 0.010888 | 0.015435 |
| 26.7361111 | 0.020104 | 0.001317 | 0.008201 | 0.001768 | 0.006634 | 0.004144 | 0.006241 | 0.013948 | 0.000967 | 0.002053 | 0.020596 | 0.006681 | 0.055975 | 0.01064 | 0.026897 | 0.020975 | 0.019507 | 0.007953 | 0.014009 | 0.011157 | 0.010312 | 0.014407 |
| 26.75 | 0.015193 | 0.000932 | 0.007579 | 0.001546 | 0.005355 | 0.005346 | 0.006307 | 0.011206 | 0.001466 | 0.003002 | 0.021669 | 0.011202 | 0.05596 | 0.011509 | 0.025526 | 0.021162 | 0.022787 | 0.004799 | 0.013096 | 0.017006 | 0.009506 | 0.013441 |
| 26.7638889 | 0.007864 | 0.001453 | 0.01386 | 0.00144 | 0.00536 | 0.005183 | 0.011831 | 0.016337 | 0.001874 | 0.007547 | 0.019809 | 0.022464 | 0.040576 | 0.011165 | 0.01708 | 0.020631 | 0.019608 | 0.004142 | 0.012053 | 0.015626 | 0.009042 | 0.012013 |
| 26.7777778 | 0.00422 | 0.003854 | 0.017993 | 0.001443 | 0.004181 | 0.003651 | 0.020349 | 0.021661 | 0.001222 | 0.010768 | 0.010625 | 0.0341 | 0.039284 | 0.00827 | 0.008506 | 0.017393 | 0.01222 | 0.008459 | 0.01059 | 0.00823 | 0.008399 | 0.009399 |
| 26.7916667 | 0.009577 | 0.007743 | 0.012345 | 0.001265 | 0.003435 | 0.007293 | 0.032233 | 0.017986 | 0.000856 | 0.008749 | 0.005353 | 0.042458 | 0.057862 | 0.004831 | 0.007873 | 0.011463 | 0.005888 | 0.014669 | 0.008381 | 0.005267 | 0.006979 | 0.005691 |
| 26.8055556 | 0.023466 | 0.010953 | 0.006808 | 0.00161 | 0.002826 | 0.013177 | 0.044283 | 0.009211 | 0.001486 | 0.003996 | 0.010944 | 0.045686 | 0.054975 | 0.005382 | 0.013232 | 0.005299 | 0.003641 | 0.018597 | 0.005854 | 0.005436 | 0.004942 | 0.002772 |
| 26.8194444 | 0.038264 | 0.010758 | 0.011223 | 0.003104 | 0.004065 | 0.015135 | 0.052314 | 0.003504 | 0.002005 | 0.001041 | 0.025032 | 0.042448 | 0.029476 | 0.010949 | 0.013758 | 0.002341 | 0.00606 | 0.018007 | 0.00392 | 0.005241 | 0.003052 | 0.003099 |
| 26.8333333 | 0.040971 | 0.00705 | 0.015125 | 0.004565 | 0.008391 | 0.012083 | 0.053157 | 0.002785 | 0.002714 | 0.000433 | 0.030602 | 0.029062 | 0.018359 | 0.016916 | 0.008683 | 0.003758 | 0.009351 | 0.013046 | 0.003111 | 0.004661 | 0.001937 | 0.005675 |
| 26.8472222 | 0.028841 | 0.002816 | 0.010656 | 0.004895 | 0.011288 | 0.007678 | 0.044824 | 0.004738 | 0.004509 | 0.000641 | 0.022583 | 0.019915 | 0.028338 | 0.017994 | 0.006035 | 0.007601 | 0.011888 | 0.006687 | 0.003397 | 0.003212 | 0.001863 | 0.007231 |
| 26.8611111 | 0.013063 | 0.001049 | 0.005188 | 0.004755 | 0.009286 | 0.006593 | 0.028136 | 0.005954 | 0.006062 | 0.001778 | 0.018351 | 0.027557 | 0.030018 | 0.014049 | 0.007475 | 0.011146 | 0.015033 | 0.002572 | 0.004353 | 0.002784 | 0.002634 | 0.007336 |
| 26.875 | 0.003275 | 0.002134 | 0.005667 | 0.00478 | 0.005904 | 0.010013 | 0.011296 | 0.004054 | 0.006445 | 0.004207 | 0.018299 | 0.030929 | 0.016888 | 0.009718 | 0.009774 | 0.01107 | 0.018028 | 0.002196 | 0.005428 | 0.005522 | 0.003559 | 0.007536 |
| 26.8888889 | 0.002895 | 0.003952 | 0.011002 | 0.005678 | 0.005081 | 0.015952 | 0.002739 | 0.001539 | 0.006496 | 0.007021 | 0.029015 | 0.031935 | 0.009165 | 0.008123 | 0.011521 | 0.007941 | 0.019515 | 0.003689 | 0.006516 | 0.010435 | 0.004237 | 0.008 |
| 26.9027778 | 0.010868 | 0.00443 | 0.015617 | 0.007094 | 0.006428 | 0.021354 | 0.003333 | 0.001791 | 0.007333 | 0.008898 | 0.053884 | 0.055324 | 0.022596 | 0.009114 | 0.012304 | 0.006176 | 0.018991 | 0.005352 | 0.007892 | 0.016272 | 0.004943 | 0.008245 |
| 26.9166667 | 0.023383 | 0.004906 | 0.017735 | 0.007575 | 0.008737 | 0.02475 | 0.010252 | 0.004301 | 0.009828 | 0.010095 | 0.066794 | 0.085181 | 0.058191 | 0.012047 | 0.012274 | 0.006943 | 0.017851 | 0.007114 | 0.009364 | 0.023265 | 0.005798 | 0.007884 |
| 26.9305556 | 0.036693 | 0.0079 | 0.019803 | 0.007156 | 0.011441 | 0.026582 | 0.01977 | 0.006845 | 0.014599 | 0.011471 | 0.058773 | 0.089663 | 0.083478 | 0.015173 | 0.012845 | 0.008859 | 0.016742 | 0.008381 | 0.01004 | 0.029646 | 0.006117 | 0.006721 |
| 26.9444444 | 0.048363 | 0.013371 | 0.023895 | 0.007071 | 0.013106 | 0.027283 | 0.028415 | 0.008734 | 0.020817 | 0.012374 | 0.04536 | 0.071156 | 0.07768 | 0.016028 | 0.014878 | 0.010587 | 0.014304 | 0.008148 | 0.009095 | 0.031221 | 0.005284 | 0.005425 |
| 26.9583333 | 0.055767 | 0.01956 | 0.02828 | 0.007808 | 0.012922 | 0.026159 | 0.031967 | 0.011949 | 0.026092 | 0.013027 | 0.040751 | 0.058269 | 0.056502 | 0.014189 | 0.017706 | 0.010834 | 0.010818 | 0.006513 | 0.006669 | 0.026867 | 0.003658 | 0.005136 |
| 26.9722222 | 0.055897 | 0.023212 | 0.030174 | 0.008518 | 0.012142 | 0.022877 | 0.028543 | 0.019665 | 0.031103 | 0.014359 | 0.041393 | 0.049188 | 0.033214 | 0.010402 | 0.020267 | 0.009257 | 0.007181 | 0.004634 | 0.003845 | 0.020464 | 0.002116 | 0.006046 |
| 26.9861111 | 0.049805 | 0.02161 | 0.030757 | 0.009103 | 0.010887 | 0.020041 | 0.021189 | 0.030552 | 0.035056 | 0.014917 | 0.031671 | 0.034712 | 0.014133 | 0.006063 | 0.021608 | 0.006964 | 0.003566 | 0.00359 | 0.001786 | 0.015923 | 0.001406 | 0.007277 |
| 27 | 0.040613 | 0.014625 | 0.028447 | 0.009308 | 0.007931 | 0.018951 | 0.014939 | 0.035264 | 0.032292 | 0.012743 | 0.015741 | 0.046219 | 0.006465 | 0.002718 | 0.019467 | 0.004894 | 0.001937 | 0.004304 | 0.001471 | 0.013118 | 0.002373 | 0.007667 |
| 27.0138889 | 0.028463 | 0.008073 | 0.018858 | 0.006847 | 0.004187 | 0.016592 | 0.012556 | 0.027727 | 0.022788 | 0.008393 | 0.011323 | 0.078883 | 0.010148 | 0.001758 | 0.012932 | 0.002806 | 0.003752 | 0.0074 | 0.003415 | 0.010066 | 0.005736 | 0.005772 |
| 27.0277778 | 0.016055 | 0.008578 | 0.008086 | 0.002797 | 0.001961 | 0.01301 | 0.009694 | 0.016517 | 0.012549 | 0.004861 | 0.015646 | 0.079927 | 0.013727 | 0.00274 | 0.006732 | 0.001459 | 0.005057 | 0.010247 | 0.004755 | 0.007901 | 0.009135 | 0.003224 |
| 27.0416667 | 0.010842 | 0.010702 | 0.003283 | 0.002825 | 0.004179 | 0.015214 | 0.004376 | 0.020467 | 0.008995 | 0.007627 | 0.014155 | 0.043543 | 0.009607 | 0.003556 | 0.007056 | 0.003383 | 0.005609 | 0.008295 | 0.005071 | 0.010203 | 0.007875 | 0.005527 |
| 27.0555556 | 0.015466 | 0.007755 | 0.00406 | 0.010161 | 0.012571 | 0.02942 | 0.001973 | 0.047244 | 0.016141 | 0.016553 | 0.012446 | 0.019673 | 0.004666 | 0.003857 | 0.015309 | 0.008427 | 0.011601 | 0.004155 | 0.006763 | 0.015802 | 0.004021 | 0.012233 |
| 27.0694444 | 0.023019 | 0.006068 | 0.01176 | 0.022376 | 0.021164 | 0.050709 | 0.008261 | 0.07993 | 0.026757 | 0.022726 | 0.019624 | 0.040593 | 0.007115 | 0.006053 | 0.025696 | 0.012524 | 0.022237 | 0.003312 | 0.008395 | 0.02236 | 0.003832 | 0.016182 |
| 27.0833333 | 0.025109 | 0.009941 | 0.018845 | 0.031643 | 0.019854 | 0.057352 | 0.026805 | 0.090361 | 0.029285 | 0.020834 | 0.026066 | 0.084628 | 0.012688 | 0.013625 | 0.026381 | 0.012599 | 0.030352 | 0.005451 | 0.013131 | 0.030095 | 0.00785 | 0.014697 |
| 27.0972222 | 0.019441 | 0.009947 | 0.014716 | 0.029549 | 0.011046 | 0.036774 | 0.04655 | 0.066092 | 0.021851 | 0.012895 | 0.029242 | 0.108994 | 0.013077 | 0.021554 | 0.01494 | 0.008744 | 0.0306 | 0.007875 | 0.019291 | 0.03152 | 0.012425 | 0.011011 |
| 27.1111111 | 0.011746 | 0.006384 | 0.010226 | 0.017408 | 0.007671 | 0.0151 | 0.048692 | 0.031075 | 0.011585 | 0.00576 | 0.031418 | 0.095544 | 0.014046 | 0.023069 | 0.005408 | 0.007105 | 0.023488 | 0.007691 | 0.01761 | 0.021055 | 0.013732 | 0.008575 |
| 27.125 | 0.01238 | 0.008555 | 0.015799 | 0.007454 | 0.011967 | 0.016303 | 0.033409 | 0.018762 | 0.005136 | 0.005268 | 0.02757 | 0.055363 | 0.02389 | 0.021799 | 0.006371 | 0.012019 | 0.014976 | 0.004868 | 0.010821 | 0.009194 | 0.011706 | 0.006355 |
| 27.1388889 | 0.019621 | 0.010347 | 0.02036 | 0.00763 | 0.013752 | 0.030374 | 0.016928 | 0.029546 | 0.005644 | 0.008602 | 0.018445 | 0.025769 | 0.033935 | 0.021116 | 0.012908 | 0.016807 | 0.00912 | 0.002315 | 0.005596 | 0.005211 | 0.009128 | 0.004643 |
| 27.1527778 | 0.023472 | 0.006372 | 0.015127 | 0.012651 | 0.008591 | 0.040995 | 0.013906 | 0.040791 | 0.013882 | 0.009365 | 0.010773 | 0.03845 | 0.037809 | 0.020278 | 0.020565 | 0.016764 | 0.00484 | 0.002151 | 0.002637 | 0.004326 | 0.00729 | 0.007247 |
| 27.1666667 | 0.029577 | 0.003235 | 0.0138 | 0.013267 | 0.002805 | 0.034673 | 0.022637 | 0.037827 | 0.024616 | 0.006806 | 0.006096 | 0.070302 | 0.036051 | 0.019926 | 0.026205 | 0.014364 | 0.002121 | 0.004291 | 0.002083 | 0.002721 | 0.005951 | 0.01095 |
| 27.1805556 | 0.04314 | 0.005535 | 0.029576 | 0.00802 | 0.005795 | 0.019134 | 0.028671 | 0.022635 | 0.024759 | 0.006261 | 0.005273 | 0.090791 | 0.029069 | 0.02029 | 0.02619 | 0.013 | 0.001467 | 0.006412 | 0.002799 | 0.004307 | 0.00492 | 0.010741 |
| 27.1944444 | 0.045691 | 0.009567 | 0.040515 | 0.003772 | 0.017649 | 0.013923 | 0.025166 | 0.009322 | 0.014374 | 0.0107 | 0.013564 | 0.091447 | 0.023599 | 0.017988 | 0.021713 | 0.015107 | 0.001503 | 0.006817 | 0.002291 | 0.01707 | 0.004605 | 0.008694 |
| 27.2083333 | 0.027953 | 0.009633 | 0.029881 | 0.004587 | 0.021919 | 0.013416 | 0.016415 | 0.007858 | 0.007395 | 0.013732 | 0.031511 | 0.064778 | 0.024861 | 0.010567 | 0.018105 | 0.019629 | 0.001762 | 0.005804 | 0.001994 | 0.046345 | 0.005157 | 0.007683 |
| 27.2222222 | 0.011953 | 0.007083 | 0.024932 | 0.00663 | 0.013867 | 0.011749 | 0.009295 | 0.012276 | 0.006835 | 0.009911 | 0.057198 | 0.032138 | 0.028841 | 0.003506 | 0.019903 | 0.023677 | 0.002155 | 0.00422 | 0.004367 | 0.075064 | 0.005919 | 0.006171 |
| 27.2361111 | 0.007139 | 0.0085 | 0.03609 | 0.005705 | 0.007426 | 0.009489 | 0.005086 | 0.012057 | 0.012231 | 0.005373 | 0.063148 | 0.033946 | 0.026304 | 0.005185 | 0.026187 | 0.024653 | 0.003354 | 0.003589 | 0.005673 | 0.079105 | 0.004697 | 0.003138 |
| 27.25 | 0.00515 | 0.012097 | 0.033953 | 0.003212 | 0.004561 | 0.006003 | 0.003414 | 0.008011 | 0.016114 | 0.005731 | 0.045933 | 0.054912 | 0.016718 | 0.0143 | 0.029432 | 0.021221 | 0.004103 | 0.005337 | 0.004655 | 0.058138 | 0.003381 | 0.001721 |
| 27.2638889 | 0.009168 | 0.009851 | 0.018522 | 0.003156 | 0.001706 | 0.007258 | 0.004039 | 0.00911 | 0.009291 | 0.007642 | 0.045978 | 0.04648 | 0.011293 | 0.019402 | 0.025443 | 0.017866 | 0.004347 | 0.00642 | 0.004971 | 0.030377 | 0.004364 | 0.001955 |
| 27.2777778 | 0.014053 | 0.00439 | 0.015165 | 0.005975 | 0.002463 | 0.010764 | 0.004701 | 0.017182 | 0.00416 | 0.007382 | 0.045491 | 0.03263 | 0.016819 | 0.015157 | 0.020594 | 0.020719 | 0.007149 | 0.0051 | 0.004716 | 0.014511 | 0.004395 | 0.005171 |
| 27.2916667 | 0.010251 | 0.003639 | 0.018052 | 0.0084 | 0.005547 | 0.008172 | 0.008498 | 0.026822 | 0.008773 | 0.005863 | 0.023178 | 0.049472 | 0.02897 | 0.007931 | 0.020778 | 0.025239 | 0.011297 | 0.004987 | 0.002575 | 0.014701 | 0.003851 | 0.016369 |
| 27.3055556 | 0.005403 | 0.006566 | 0.027213 | 0.009786 | 0.006732 | 0.003075 | 0.016274 | 0.032984 | 0.013858 | 0.00578 | 0.008173 | 0.056688 | 0.038996 | 0.003543 | 0.024382 | 0.021407 | 0.013483 | 0.00541 | 0.001276 | 0.022312 | 0.003827 | 0.02956 |
| 27.3194444 | 0.004759 | 0.006607 | 0.048312 | 0.013599 | 0.008276 | 0.00464 | 0.024965 | 0.034698 | 0.014339 | 0.007454 | 0.006231 | 0.033704 | 0.038946 | 0.002473 | 0.025808 | 0.011236 | 0.013682 | 0.003895 | 0.001352 | 0.030608 | 0.003337 | 0.035025 |
| 27.3333333 | 0.003822 | 0.004165 | 0.050643 | 0.017483 | 0.009653 | 0.012752 | 0.033988 | 0.029658 | 0.01036 | 0.007203 | 0.005167 | 0.012997 | 0.029536 | 0.004281 | 0.021453 | 0.005613 | 0.013062 | 0.001925 | 0.002854 | 0.037568 | 0.002883 | 0.033008 |
| 27.3472222 | 0.002845 | 0.004803 | 0.030792 | 0.014972 | 0.007152 | 0.020296 | 0.038383 | 0.021065 | 0.006129 | 0.004117 | 0.004855 | 0.010023 | 0.021486 | 0.007993 | 0.015174 | 0.008534 | 0.011932 | 0.001879 | 0.004525 | 0.039422 | 0.002313 | 0.029215 |
| 27.3611111 | 0.004096 | 0.008308 | 0.021733 | 0.008524 | 0.003238 | 0.02303 | 0.033149 | 0.0245 | 0.009142 | 0.003341 | 0.005433 | 0.019039 | 0.020053 | 0.010697 | 0.011398 | 0.014586 | 0.010847 | 0.004947 | 0.003885 | 0.035706 | 0.003129 | 0.026316 |
| 27.375 | 0.010051 | 0.010991 | 0.025242 | 0.0049 | 0.003818 | 0.020541 | 0.024854 | 0.041355 | 0.014306 | 0.007768 | 0.00864 | 0.030579 | 0.020824 | 0.011907 | 0.01 | 0.016585 | 0.011799 | 0.008729 | 0.003043 | 0.032578 | 0.005903 | 0.022471 |
| 27.3888889 | 0.022848 | 0.012068 | 0.022603 | 0.006851 | 0.010201 | 0.013068 | 0.024558 | 0.053268 | 0.014059 | 0.013747 | 0.017338 | 0.032102 | 0.022316 | 0.014547 | 0.010286 | 0.013011 | 0.017275 | 0.009633 | 0.004278 | 0.029327 | 0.007087 | 0.017891 |
| 27.4027778 | 0.034248 | 0.013075 | 0.021623 | 0.011742 | 0.016788 | 0.005294 | 0.032023 | 0.048011 | 0.013415 | 0.013998 | 0.030189 | 0.02106 | 0.029056 | 0.018354 | 0.011373 | 0.007464 | 0.026273 | 0.008347 | 0.004994 | 0.02227 | 0.005787 | 0.014794 |
| 27.4166667 | 0.035312 | 0.015436 | 0.034728 | 0.013833 | 0.018191 | 0.002661 | 0.040197 | 0.032399 | 0.01418 | 0.00755 | 0.046716 | 0.011088 | 0.040898 | 0.019844 | 0.012264 | 0.003816 | 0.035303 | 0.007954 | 0.003708 | 0.017459 | 0.004849 | 0.012701 |
| 27.4305556 | 0.028949 | 0.018473 | 0.044463 | 0.011781 | 0.013383 | 0.006113 | 0.047662 | 0.022067 | 0.015003 | 0.002673 | 0.063679 | 0.013845 | 0.049295 | 0.01919 | 0.013425 | 0.004987 | 0.042151 | 0.008921 | 0.002502 | 0.021641 | 0.004943 | 0.011686 |
| 27.4444444 | 0.022356 | 0.020322 | 0.035474 | 0.008447 | 0.0077 | 0.014716 | 0.055227 | 0.0224 | 0.01983 | 0.003606 | 0.074596 | 0.026521 | 0.046312 | 0.019317 | 0.014024 | 0.011596 | 0.045642 | 0.010953 | 0.003341 | 0.03284 | 0.005531 | 0.013188 |
| 27.4583333 | 0.019662 | 0.021704 | 0.021281 | 0.00515 | 0.006899 | 0.022439 | 0.057127 | 0.028459 | 0.024716 | 0.007415 | 0.078537 | 0.035797 | 0.033442 | 0.01761 | 0.012965 | 0.018141 | 0.046384 | 0.015356 | 0.004331 | 0.040306 | 0.007718 | 0.015432 |
| 27.4722222 | 0.021464 | 0.024997 | 0.018221 | 0.003564 | 0.010604 | 0.022634 | 0.049666 | 0.04192 | 0.023928 | 0.014676 | 0.079043 | 0.034596 | 0.019493 | 0.011146 | 0.012842 | 0.017057 | 0.046504 | 0.020032 | 0.003558 | 0.036569 | 0.010389 | 0.015597 |
| 27.4861111 | 0.028479 | 0.028423 | 0.015365 | 0.005592 | 0.016855 | 0.017509 | 0.041836 | 0.05053 | 0.022304 | 0.024403 | 0.079852 | 0.031054 | 0.012448 | 0.00563 | 0.015199 | 0.010387 | 0.047858 | 0.019227 | 0.003342 | 0.028008 | 0.009398 | 0.012837 |
| 27.5 | 0.045208 | 0.027944 | 0.009729 | 0.008274 | 0.021427 | 0.011825 | 0.043288 | 0.036416 | 0.019659 | 0.026303 | 0.082536 | 0.033953 | 0.01602 | 0.003651 | 0.017385 | 0.005961 | 0.047904 | 0.012474 | 0.004697 | 0.022789 | 0.004832 | 0.00768 |
| 27.5138889 | 0.067101 | 0.022571 | 0.015041 | 0.008454 | 0.021279 | 0.006564 | 0.054385 | 0.020489 | 0.014051 | 0.018281 | 0.072317 | 0.040979 | 0.025614 | 0.002867 | 0.015912 | 0.00511 | 0.042011 | 0.005274 | 0.0071 | 0.020583 | 0.001283 | 0.003498 |
| 27.5277778 | 0.073784 | 0.014847 | 0.02179 | 0.007624 | 0.020222 | 0.003899 | 0.067318 | 0.019897 | 0.011851 | 0.010023 | 0.050148 | 0.0473 | 0.032243 | 0.004456 | 0.010394 | 0.005952 | 0.030519 | 0.004049 | 0.013646 | 0.015717 | 0.003439 | 0.004278 |
| 27.5416667 | 0.059542 | 0.009518 | 0.018245 | 0.006599 | 0.020473 | 0.003604 | 0.071916 | 0.020549 | 0.017943 | 0.005343 | 0.043265 | 0.053272 | 0.028427 | 0.014439 | 0.011484 | 0.015467 | 0.019961 | 0.016783 | 0.029421 | 0.012457 | 0.017295 | 0.011769 |
| 27.5555556 | 0.040675 | 0.006877 | 0.012273 | 0.004079 | 0.020092 | 0.003016 | 0.064402 | 0.016674 | 0.02658 | 0.00792 | 0.047087 | 0.055088 | 0.024908 | 0.033611 | 0.023347 | 0.03473 | 0.020365 | 0.039223 | 0.048527 | 0.018322 | 0.038121 | 0.023129 |
| 27.5694444 | 0.025491 | 0.003914 | 0.009828 | 0.001587 | 0.016313 | 0.006898 | 0.052217 | 0.011151 | 0.024667 | 0.013995 | 0.042035 | 0.045013 | 0.026827 | 0.042606 | 0.027752 | 0.040787 | 0.022833 | 0.047571 | 0.049749 | 0.021048 | 0.043588 | 0.025747 |
| 27.5833333 | 0.014726 | 0.001453 | 0.009118 | 0.001562 | 0.008698 | 0.01359 | 0.048044 | 0.006898 | 0.015181 | 0.012312 | 0.032814 | 0.02913 | 0.022161 | 0.028504 | 0.019855 | 0.025171 | 0.015684 | 0.03345 | 0.029181 | 0.015888 | 0.026026 | 0.016479 |
| 27.5972222 | 0.009618 | 0.000478 | 0.00754 | 0.003061 | 0.002981 | 0.012485 | 0.042782 | 0.006707 | 0.011404 | 0.006908 | 0.027485 | 0.019598 | 0.019755 | 0.012503 | 0.014434 | 0.011778 | 0.012379 | 0.014857 | 0.014447 | 0.017197 | 0.009142 | 0.009269 |
| 27.6111111 | 0.006614 | 0.000175 | 0.005266 | 0.003639 | 0.002901 | 0.007752 | 0.023279 | 0.005745 | 0.009565 | 0.003093 | 0.02125 | 0.016752 | 0.017252 | 0.013844 | 0.012584 | 0.011256 | 0.01577 | 0.00526 | 0.0176 | 0.019938 | 0.007494 | 0.010236 |
| 27.625 | 0.004028 | 0.000248 | 0.004784 | 0.002466 | 0.004976 | 0.006491 | 0.005794 | 0.004541 | 0.006317 | 0.001125 | 0.01306 | 0.017921 | 0.008119 | 0.019392 | 0.007305 | 0.010838 | 0.014939 | 0.004358 | 0.019599 | 0.014726 | 0.008504 | 0.015057 |
| 27.6388889 | 0.004399 | 0.000968 | 0.007328 | 0.001689 | 0.005388 | 0.004981 | 0.002115 | 0.004889 | 0.007185 | 0.000767 | 0.018013 | 0.022052 | 0.009351 | 0.016138 | 0.003868 | 0.006116 | 0.009586 | 0.0065 | 0.011544 | 0.012727 | 0.005172 | 0.014665 |
| 27.6527778 | 0.006802 | 0.002386 | 0.01434 | 0.002192 | 0.003928 | 0.004279 | 0.007204 | 0.005738 | 0.007653 | 0.001307 | 0.04254 | 0.024329 | 0.016191 | 0.00903 | 0.008495 | 0.003594 | 0.004401 | 0.007925 | 0.005395 | 0.012533 | 0.002022 | 0.009235 |
| 27.6666667 | 0.008103 | 0.0036 | 0.015752 | 0.00237 | 0.002503 | 0.008514 | 0.016063 | 0.005912 | 0.00652 | 0.002065 | 0.067187 | 0.020322 | 0.012328 | 0.003803 | 0.015228 | 0.003821 | 0.001877 | 0.008704 | 0.008847 | 0.010408 | 0.001017 | 0.01377 |
| 27.6805556 | 0.007323 | 0.003562 | 0.009592 | 0.002589 | 0.001566 | 0.014667 | 0.024993 | 0.00426 | 0.009583 | 0.001863 | 0.063883 | 0.012359 | 0.008845 | 0.001988 | 0.016456 | 0.003061 | 0.001035 | 0.009052 | 0.01628 | 0.016552 | 0.001423 | 0.02776 |
| 27.6944444 | 0.004777 | 0.002228 | 0.009381 | 0.00239 | 0.002532 | 0.017094 | 0.03232 | 0.006531 | 0.0106 | 0.003961 | 0.041351 | 0.011405 | 0.015975 | 0.003599 | 0.0117 | 0.001568 | 0.000875 | 0.008561 | 0.020742 | 0.021942 | 0.001898 | 0.033672 |
| 27.7083333 | 0.002169 | 0.001181 | 0.010176 | 0.001436 | 0.006539 | 0.014339 | 0.032181 | 0.010006 | 0.005548 | 0.009435 | 0.043742 | 0.023088 | 0.020367 | 0.006273 | 0.006239 | 0.000936 | 0.001437 | 0.007206 | 0.018473 | 0.01665 | 0.001833 | 0.025925 |
| 27.7222222 | 0.001475 | 0.001362 | 0.005457 | 0.001339 | 0.0084 | 0.00938 | 0.019803 | 0.007315 | 0.003253 | 0.010397 | 0.069193 | 0.035614 | 0.02334 | 0.006671 | 0.005469 | 0.000873 | 0.002115 | 0.005254 | 0.011448 | 0.012945 | 0.001184 | 0.014159 |
| 27.7361111 | 0.001719 | 0.001849 | 0.004522 | 0.001327 | 0.005339 | 0.005089 | 0.01127 | 0.00302 | 0.006167 | 0.0072 | 0.07128 | 0.036174 | 0.031656 | 0.005127 | 0.008445 | 0.001209 | 0.002856 | 0.002938 | 0.006032 | 0.015228 | 0.000667 | 0.006283 |
| 27.75 | 0.002057 | 0.001931 | 0.010832 | 0.000736 | 0.001948 | 0.002198 | 0.022306 | 0.002865 | 0.008338 | 0.010823 | 0.04437 | 0.023626 | 0.026518 | 0.004157 | 0.010441 | 0.00221 | 0.003997 | 0.001479 | 0.003951 | 0.013586 | 0.001032 | 0.002764 |
| 27.7638889 | 0.005054 | 0.001419 | 0.016314 | 0.000549 | 0.001423 | 0.000808 | 0.04029 | 0.003904 | 0.008496 | 0.012428 | 0.018491 | 0.013621 | 0.010044 | 0.004019 | 0.009621 | 0.003948 | 0.005046 | 0.002435 | 0.00327 | 0.008286 | 0.001712 | 0.00341 |
| 27.7777778 | 0.012508 | 0.001142 | 0.015294 | 0.00096 | 0.001694 | 0.000819 | 0.052334 | 0.004144 | 0.010084 | 0.005135 | 0.008878 | 0.024061 | 0.002898 | 0.003834 | 0.007164 | 0.00621 | 0.00538 | 0.005145 | 0.00261 | 0.007119 | 0.002032 | 0.007937 |
| 27.7916667 | 0.023864 | 0.001595 | 0.011744 | 0.001328 | 0.002232 | 0.000997 | 0.055545 | 0.006179 | 0.009759 | 0.000794 | 0.015523 | 0.049049 | 0.005305 | 0.004542 | 0.004603 | 0.008436 | 0.005121 | 0.006953 | 0.001656 | 0.006672 | 0.001887 | 0.013359 |
| 27.8055556 | 0.033584 | 0.001996 | 0.011407 | 0.00135 | 0.003827 | 0.001369 | 0.046783 | 0.00705 | 0.006195 | 0.0018 | 0.029671 | 0.069012 | 0.01003 | 0.007854 | 0.002427 | 0.010127 | 0.005128 | 0.006316 | 0.000699 | 0.007802 | 0.001482 | 0.016501 |
| 27.8194444 | 0.032382 | 0.002216 | 0.01126 | 0.001362 | 0.004713 | 0.004154 | 0.027805 | 0.007158 | 0.005375 | 0.006214 | 0.041652 | 0.062537 | 0.014591 | 0.012727 | 0.001366 | 0.010679 | 0.005628 | 0.00405 | 0.000387 | 0.018555 | 0.000978 | 0.015559 |
| 27.8333333 | 0.019225 | 0.002518 | 0.01054 | 0.001419 | 0.006735 | 0.009321 | 0.01269 | 0.011788 | 0.005023 | 0.011088 | 0.055416 | 0.037347 | 0.019066 | 0.015002 | 0.001621 | 0.009937 | 0.006184 | 0.002335 | 0.001118 | 0.029609 | 0.000937 | 0.011841 |
| 27.8472222 | 0.007453 | 0.002816 | 0.011454 | 0.001263 | 0.012047 | 0.014543 | 0.010576 | 0.016206 | 0.00459 | 0.010018 | 0.063739 | 0.037488 | 0.022832 | 0.012915 | 0.001351 | 0.009516 | 0.007491 | 0.00286 | 0.002251 | 0.02637 | 0.001781 | 0.009554 |
| 27.8611111 | 0.004331 | 0.002545 | 0.008883 | 0.001116 | 0.018062 | 0.017117 | 0.012527 | 0.016257 | 0.007747 | 0.006144 | 0.044132 | 0.067728 | 0.026 | 0.009542 | 0.001827 | 0.011398 | 0.010665 | 0.004172 | 0.002602 | 0.014506 | 0.002176 | 0.010628 |
| 27.875 | 0.005888 | 0.001539 | 0.003845 | 0.001301 | 0.022251 | 0.016237 | 0.022648 | 0.015108 | 0.010267 | 0.006374 | 0.023554 | 0.093091 | 0.029631 | 0.00819 | 0.005859 | 0.015 | 0.014992 | 0.004033 | 0.001777 | 0.00636 | 0.001372 | 0.01262 |
| 27.8888889 | 0.016407 | 0.000625 | 0.002355 | 0.001526 | 0.025492 | 0.015634 | 0.043166 | 0.014366 | 0.010644 | 0.008843 | 0.034446 | 0.098227 | 0.034197 | 0.009847 | 0.011772 | 0.017365 | 0.018647 | 0.002959 | 0.000737 | 0.007141 | 0.000462 | 0.012972 |

| | | | | | | | | | | | | | | | | | | | | | | |
|---|---|---|---|---|---|---|---|---|---|---|---|---|---|---|---|---|---|---|---|---|---|---|
| 27.9027778 | 0.036808 | 0.001159 | 0.008285 | 0.001584 | 0.028084 | 0.018424 | 0.056766 | 0.012875 | 0.012628 | 0.010173 | 0.039206 | 0.074364 | 0.039205 | 0.014157 | 0.016901 | 0.017 | 0.020521 | 0.002291 | 0.000368 | 0.019977 | 0.000177 | 0.012251 |
| 27.9166667 | 0.057843 | 0.004259 | 0.017443 | 0.002893 | 0.02702 | 0.023854 | 0.056156 | 0.010973 | 0.017441 | 0.011287 | 0.024047 | 0.035093 | 0.043532 | 0.02057 | 0.0204 | 0.016306 | 0.020496 | 0.001971 | 0.000379 | 0.037734 | 0.000184 | 0.01224 |
| 27.9305556 | 0.070022 | 0.009586 | 0.015904 | 0.007083 | 0.021916 | 0.032446 | 0.053699 | 0.012352 | 0.024169 | 0.013761 | 0.02924 | 0.016077 | 0.044911 | 0.027077 | 0.022338 | 0.018421 | 0.019237 | 0.001388 | 0.000598 | 0.049728 | 0.000133 | 0.013399 |
| 27.9444444 | 0.069963 | 0.014161 | 0.007719 | 0.012392 | 0.015188 | 0.040316 | 0.052585 | 0.020291 | 0.030357 | 0.018295 | 0.059857 | 0.012498 | 0.041654 | 0.028794 | 0.023585 | 0.021644 | 0.017508 | 0.000755 | 0.001059 | 0.051329 | 6.10E-05 | 0.015392 |
| 27.9583333 | 0.062132 | 0.015271 | 0.010168 | 0.016003 | 0.00854 | 0.039479 | 0.040116 | 0.03105 | 0.035439 | 0.023746 | 0.08901 | 0.00746 | 0.035427 | 0.023529 | 0.024392 | 0.021134 | 0.015608 | 0.000414 | 0.001629 | 0.042753 | 0.000171 | 0.018233 |
| 27.9722222 | 0.049273 | 0.015391 | 0.023885 | 0.017375 | 0.004011 | 0.032485 | 0.028159 | 0.039108 | 0.040473 | 0.027447 | 0.097074 | 0.007045 | 0.030244 | 0.014623 | 0.024494 | 0.016567 | 0.014024 | 0.000427 | 0.002334 | 0.029972 | 0.000663 | 0.021318 |
| 27.9861111 | 0.033382 | 0.017268 | 0.037326 | 0.01695 | 0.003884 | 0.026249 | 0.033195 | 0.043993 | 0.043295 | 0.02733 | 0.079976 | 0.013956 | 0.028188 | 0.006697 | 0.023749 | 0.012098 | 0.013124 | 0.000621 | 0.002745 | 0.018317 | 0.001154 | 0.022596 |
| 28 | 0.02039 | 0.019488 | 0.041795 | 0.014024 | 0.00734 | 0.02055 | 0.042693 | 0.042507 | 0.039103 | 0.024052 | 0.051226 | 0.026266 | 0.026309 | 0.004028 | 0.020745 | 0.009474 | 0.011687 | 0.001487 | 0.002534 | 0.00905 | 0.001748 | 0.019691 |
| 28.0138889 | 0.012084 | 0.019107 | 0.03604 | 0.008134 | 0.011177 | 0.01267 | 0.042301 | 0.032089 | 0.026084 | 0.018612 | 0.025637 | 0.042628 | 0.021388 | 0.007042 | 0.014664 | 0.006865 | 0.007929 | 0.005066 | 0.003984 | 0.003843 | 0.004487 | 0.012369 |
| 28.0277778 | 0.0063 | 0.015512 | 0.021693 | 0.003623 | 0.013121 | 0.004888 | 0.033969 | 0.019466 | 0.011682 | 0.012032 | 0.009809 | 0.055097 | 0.01581 | 0.009297 | 0.010144 | 0.004956 | 0.003817 | 0.009298 | 0.006584 | 0.00282 | 0.00795 | 0.005141 |
| 28.0416667 | 0.007506 | 0.014915 | 0.008372 | 0.00359 | 0.01031 | 0.006543 | 0.020812 | 0.018611 | 0.005955 | 0.009761 | 0.00475 | 0.053475 | 0.015416 | 0.008042 | 0.014518 | 0.008209 | 0.005314 | 0.008713 | 0.007193 | 0.004762 | 0.007967 | 0.00439 |
| 28.0555556 | 0.023319 | 0.027976 | 0.004306 | 0.009304 | 0.011725 | 0.025576 | 0.00771 | 0.038742 | 0.011902 | 0.016269 | 0.008987 | 0.039131 | 0.022469 | 0.009 | 0.028997 | 0.016966 | 0.01336 | 0.006027 | 0.008043 | 0.011653 | 0.00719 | 0.009088 |
| 28.0694444 | 0.048617 | 0.046988 | 0.006856 | 0.022985 | 0.022032 | 0.053627 | 0.007219 | 0.069913 | 0.024182 | 0.02485 | 0.019207 | 0.020707 | 0.03266 | 0.012631 | 0.049782 | 0.025633 | 0.023142 | 0.004938 | 0.007769 | 0.02091 | 0.006215 | 0.011555 |
| 28.0833333 | 0.062795 | 0.045104 | 0.01386 | 0.038288 | 0.021874 | 0.064487 | 0.03162 | 0.086477 | 0.030032 | 0.023556 | 0.028235 | 0.01479 | 0.037135 | 0.013603 | 0.068018 | 0.027741 | 0.030586 | 0.004569 | 0.004238 | 0.028064 | 0.004113 | 0.010619 |
| 28.0972222 | 0.052058 | 0.022635 | 0.015411 | 0.043895 | 0.009691 | 0.044222 | 0.078749 | 0.071906 | 0.024484 | 0.0125 | 0.033409 | 0.024192 | 0.029435 | 0.011401 | 0.067994 | 0.019571 | 0.031875 | 0.00567 | 0.001715 | 0.030655 | 0.004777 | 0.009069 |
| 28.1111111 | 0.02702 | 0.006506 | 0.009667 | 0.035068 | 0.002781 | 0.019218 | 0.103721 | 0.037996 | 0.015731 | 0.00421 | 0.033366 | 0.023076 | 0.014794 | 0.008909 | 0.046988 | 0.009313 | 0.027399 | 0.005233 | 0.001524 | 0.026234 | 0.005137 | 0.006776 |
| 28.125 | 0.013684 | 0.004262 | 0.008815 | 0.018303 | 0.006191 | 0.016323 | 0.07766 | 0.016723 | 0.007976 | 0.004113 | 0.025358 | 0.017371 | 0.004757 | 0.006993 | 0.02219 | 0.007342 | 0.02097 | 0.002823 | 0.003192 | 0.018083 | 0.002686 | 0.005029 |
| 28.1388889 | 0.022185 | 0.005087 | 0.012301 | 0.0077 | 0.012367 | 0.025678 | 0.037868 | 0.022253 | 0.004343 | 0.006391 | 0.013423 | 0.029984 | 0.002453 | 0.004936 | 0.008538 | 0.00851 | 0.013205 | 0.002482 | 0.005759 | 0.012305 | 0.000732 | 0.00349 |
| 28.1527778 | 0.03557 | 0.005158 | 0.013138 | 0.009636 | 0.011561 | 0.03394 | 0.014767 | 0.037179 | 0.009738 | 0.007507 | 0.006629 | 0.046361 | 0.003687 | 0.005939 | 0.008573 | 0.006564 | 0.005457 | 0.0034 | 0.007072 | 0.011459 | 0.000164 | 0.002945 |
| 28.1666667 | 0.040327 | 0.005588 | 0.01072 | 0.014708 | 0.006996 | 0.036395 | 0.009835 | 0.04447 | 0.019241 | 0.007673 | 0.00851 | 0.050693 | 0.008816 | 0.013059 | 0.01502 | 0.003519 | 0.001817 | 0.00328 | 0.007102 | 0.013639 | 8.57E-05 | 0.00608 |
| 28.1805556 | 0.039647 | 0.004543 | 0.007165 | 0.01362 | 0.004526 | 0.025693 | 0.019062 | 0.035857 | 0.022994 | 0.007876 | 0.011393 | 0.046718 | 0.01924 | 0.022092 | 0.019159 | 0.001513 | 0.002892 | 0.002078 | 0.006856 | 0.012593 | 0.000226 | 0.009302 |
| 28.1944444 | 0.042608 | 0.00241 | 0.005087 | 0.007504 | 0.006996 | 0.013494 | 0.031763 | 0.018534 | 0.01733 | 0.009636 | 0.013149 | 0.038483 | 0.029962 | 0.025684 | 0.01986 | 0.002983 | 0.004749 | 0.00121 | 0.006378 | 0.009986 | 0.000997 | 0.007841 |
| 28.2083333 | 0.051196 | 0.001857 | 0.007096 | 0.002347 | 0.014658 | 0.01478 | 0.039352 | 0.011036 | 0.008443 | 0.011467 | 0.018558 | 0.025574 | 0.035186 | 0.020406 | 0.019161 | 0.009722 | 0.004895 | 0.001611 | 0.005538 | 0.016727 | 0.002029 | 0.004536 |
| 28.2222222 | 0.05133 | 0.00438 | 0.009791 | 0.0019 | 0.016447 | 0.014478 | 0.038298 | 0.01149 | 0.004488 | 0.010138 | 0.031296 | 0.015957 | 0.031795 | 0.010216 | 0.017886 | 0.016646 | 0.003985 | 0.002074 | 0.005014 | 0.030713 | 0.002437 | 0.004483 |
| 28.2361111 | 0.036398 | 0.008286 | 0.007502 | 0.003778 | 0.010013 | 0.008498 | 0.028676 | 0.008557 | 0.005757 | 0.005692 | 0.050451 | 0.022152 | 0.02008 | 0.003033 | 0.017282 | 0.017598 | 0.002637 | 0.001472 | 0.005272 | 0.039418 | 0.002 | 0.005855 |
| 28.25 | 0.021797 | 0.008598 | 0.003414 | 0.004603 | 0.007185 | 0.00858 | 0.016388 | 0.004146 | 0.007044 | 0.006771 | 0.054621 | 0.034858 | 0.008743 | 0.004841 | 0.018356 | 0.015238 | 0.002442 | 0.000908 | 0.00617 | 0.03845 | 0.001278 | 0.004986 |
| 28.2638889 | 0.018721 | 0.007332 | 0.003122 | 0.004782 | 0.008306 | 0.009811 | 0.013258 | 0.005155 | 0.005458 | 0.016205 | 0.038571 | 0.034865 | 0.006925 | 0.01472 | 0.021388 | 0.015588 | 0.004765 | 0.001784 | 0.007636 | 0.034016 | 0.001416 | 0.003785 |
| 28.2777778 | 0.022797 | 0.010517 | 0.010068 | 0.006751 | 0.007304 | 0.010514 | 0.019043 | 0.012459 | 0.003054 | 0.018274 | 0.03636 | 0.020669 | 0.016946 | 0.024288 | 0.026673 | 0.019621 | 0.007184 | 0.003656 | 0.009439 | 0.034258 | 0.002793 | 0.00467 |
| 28.2916667 | 0.020351 | 0.012787 | 0.026323 | 0.009412 | 0.009231 | 0.012776 | 0.016678 | 0.019592 | 0.005249 | 0.011617 | 0.044085 | 0.009833 | 0.034864 | 0.026805 | 0.032607 | 0.024959 | 0.008322 | 0.005167 | 0.010685 | 0.039614 | 0.004084 | 0.006759 |
| 28.3055556 | 0.010805 | 0.009348 | 0.042444 | 0.008745 | 0.013042 | 0.012767 | 0.0119 | 0.020902 | 0.014709 | 0.009788 | 0.029488 | 0.009091 | 0.046493 | 0.023412 | 0.033676 | 0.029422 | 0.010065 | 0.005626 | 0.01071 | 0.048017 | 0.00434 | 0.007295 |
| 28.3194444 | 0.011138 | 0.007693 | 0.040783 | 0.00628 | 0.010596 | 0.009661 | 0.023816 | 0.020518 | 0.024791 | 0.013185 | 0.011007 | 0.011627 | 0.045292 | 0.019013 | 0.02776 | 0.029508 | 0.013133 | 0.005003 | 0.009698 | 0.05655 | 0.003726 | 0.00791 |
| 28.3333333 | 0.023256 | 0.008194 | 0.027722 | 0.008539 | 0.006472 | 0.006111 | 0.042866 | 0.024118 | 0.027708 | 0.024005 | 0.010497 | 0.022686 | 0.03963 | 0.016848 | 0.021976 | 0.024212 | 0.016165 | 0.003438 | 0.008062 | 0.057612 | 0.00239 | 0.016271 |
| 28.3472222 | 0.028516 | 0.006638 | 0.030027 | 0.015614 | 0.007306 | 0.006084 | 0.052477 | 0.030393 | 0.022305 | 0.039005 | 0.012311 | 0.033865 | 0.034255 | 0.01672 | 0.022103 | 0.019576 | 0.017349 | 0.0017 | 0.006264 | 0.051959 | 0.001243 | 0.02968 |
| 28.3611111 | 0.021358 | 0.007735 | 0.039434 | 0.018123 | 0.009431 | 0.012043 | 0.048748 | 0.03327 | 0.01468 | 0.036399 | 0.007926 | 0.029555 | 0.028497 | 0.017362 | 0.024899 | 0.021139 | 0.0161 | 0.000803 | 0.005281 | 0.046564 | 0.001156 | 0.038205 |
| 28.375 | 0.014939 | 0.010698 | 0.031239 | 0.011879 | 0.00962 | 0.02132 | 0.037029 | 0.029709 | 0.015267 | 0.018282 | 0.004046 | 0.017719 | 0.025107 | 0.018438 | 0.0252 | 0.026135 | 0.014358 | 0.000531 | 0.005701 | 0.042188 | 0.001126 | 0.040221 |
| 28.3888889 | 0.01728 | 0.01068 | 0.023804 | 0.006277 | 0.010076 | 0.026671 | 0.031463 | 0.027495 | 0.025857 | 0.00736 | 0.006178 | 0.013197 | 0.029051 | 0.020496 | 0.023085 | 0.026181 | 0.015263 | 0.000538 | 0.006201 | 0.038797 | 0.000996 | 0.037808 |
| 28.4027778 | 0.032849 | 0.01038 | 0.034481 | 0.009897 | 0.012473 | 0.025234 | 0.039607 | 0.03221 | 0.036989 | 0.007146 | 0.016271 | 0.018744 | 0.041129 | 0.024808 | 0.022287 | 0.018628 | 0.020188 | 0.001921 | 0.004718 | 0.037432 | 0.002571 | 0.030663 |
| 28.4166667 | 0.04466 | 0.013038 | 0.036663 | 0.01951 | 0.018191 | 0.023362 | 0.055022 | 0.040389 | 0.039476 | 0.01536 | 0.031374 | 0.026416 | 0.056734 | 0.031511 | 0.025921 | 0.010019 | 0.027368 | 0.005691 | 0.003018 | 0.03798 | 0.006663 | 0.024431 |
| 28.4305556 | 0.036296 | 0.016607 | 0.01961 | 0.024383 | 0.024666 | 0.025505 | 0.069445 | 0.049648 | 0.033964 | 0.025518 | 0.045571 | 0.029259 | 0.067465 | 0.037415 | 0.033199 | 0.007989 | 0.033249 | 0.009965 | 0.004517 | 0.038955 | 0.011153 | 0.032468 |
| 28.4444444 | 0.02676 | 0.017686 | 0.009445 | 0.018986 | 0.025589 | 0.027416 | 0.079282 | 0.053929 | 0.029157 | 0.025636 | 0.055318 | 0.027158 | 0.068682 | 0.037831 | 0.038369 | 0.01455 | 0.035332 | 0.010452 | 0.006807 | 0.036514 | 0.01143 | 0.049168 |
| 28.4583333 | 0.030723 | 0.016066 | 0.015179 | 0.009411 | 0.020541 | 0.025058 | 0.081925 | 0.044863 | 0.028565 | 0.015539 | 0.063407 | 0.024747 | 0.065037 | 0.032095 | 0.037367 | 0.023635 | 0.032303 | 0.007817 | 0.005707 | 0.02879 | 0.007917 | 0.052358 |
| 28.4722222 | 0.044397 | 0.014661 | 0.019469 | 0.003847 | 0.013941 | 0.021734 | 0.074565 | 0.028855 | 0.027604 | 0.008091 | 0.072604 | 0.026679 | 0.063917 | 0.026043 | 0.030691 | 0.028667 | 0.024639 | 0.007291 | 0.00389 | 0.021196 | 0.006178 | 0.042762 |
| 28.4861111 | 0.060737 | 0.017554 | 0.020228 | 0.005294 | 0.011149 | 0.02444 | 0.062047 | 0.019738 | 0.025778 | 0.007687 | 0.081693 | 0.033606 | 0.064885 | 0.026025 | 0.022964 | 0.027586 | 0.019035 | 0.006682 | 0.004245 | 0.022747 | 0.004879 | 0.035849 |
| 28.5 | 0.068601 | 0.026226 | 0.02818 | 0.008514 | 0.015317 | 0.035752 | 0.053834 | 0.022201 | 0.028869 | 0.010424 | 0.0912 | 0.040749 | 0.058341 | 0.02851 | 0.018136 | 0.020382 | 0.020326 | 0.003211 | 0.004347 | 0.03365 | 0.002252 | 0.030977 |
| 28.5138889 | 0.065088 | 0.036143 | 0.034472 | 0.006918 | 0.023409 | 0.047829 | 0.050523 | 0.032695 | 0.036774 | 0.021843 | 0.097042 | 0.04438 | 0.043125 | 0.024538 | 0.015074 | 0.010457 | 0.022661 | 0.001589 | 0.004978 | 0.037651 | 0.001469 | 0.021914 |
| 28.5277778 | 0.058258 | 0.035025 | 0.028257 | 0.0036 | 0.030145 | 0.048157 | 0.046979 | 0.037809 | 0.04026 | 0.037657 | 0.085817 | 0.046186 | 0.026195 | 0.013662 | 0.014147 | 0.005877 | 0.018793 | 0.005513 | 0.009434 | 0.024773 | 0.004336 | 0.024179 |
| 28.5416667 | 0.050538 | 0.020801 | 0.015776 | 0.002561 | 0.031039 | 0.034539 | 0.041817 | 0.028582 | 0.033763 | 0.044091 | 0.058126 | 0.05034 | 0.010632 | 0.007874 | 0.011189 | 0.011611 | 0.013037 | 0.016761 | 0.019734 | 0.015214 | 0.013671 | 0.051417 |
| 28.5555556 | 0.04358 | 0.009253 | 0.007385 | 0.001983 | 0.025381 | 0.018285 | 0.042568 | 0.013671 | 0.023745 | 0.034925 | 0.03498 | 0.055945 | 0.002992 | 0.013526 | 0.007485 | 0.019707 | 0.014925 | 0.030629 | 0.032151 | 0.017502 | 0.026039 | 0.065554 |
| 28.5694444 | 0.03804 | 0.004886 | 0.004787 | 0.000949 | 0.01586 | 0.007784 | 0.047082 | 0.003932 | 0.016972 | 0.01744 | 0.029529 | 0.053096 | 0.00343 | 0.017002 | 0.010541 | 0.018256 | 0.018805 | 0.034556 | 0.034889 | 0.017516 | 0.030172 | 0.040486 |
| 28.5833333 | 0.03102 | 0.002604 | 0.002911 | 0.002709 | 0.007343 | 0.003386 | 0.042562 | 0.004253 | 0.010496 | 0.008042 | 0.035197 | 0.042852 | 0.003396 | 0.010427 | 0.014699 | 0.009844 | 0.014888 | 0.024995 | 0.024544 | 0.020649 | 0.022581 | 0.014814 |
| 28.5972222 | 0.022774 | 0.002303 | 0.001802 | 0.006637 | 0.005372 | 0.003715 | 0.031315 | 0.007143 | 0.003993 | 0.007528 | 0.037127 | 0.037288 | 0.001677 | 0.004128 | 0.016755 | 0.00576 | 0.006893 | 0.011894 | 0.011494 | 0.024481 | 0.011027 | 0.019631 |
| 28.6111111 | 0.01344 | 0.002105 | 0.006358 | 0.007271 | 0.007939 | 0.003718 | 0.020252 | 0.00432 | 0.004081 | 0.005577 | 0.028954 | 0.03379 | 0.002973 | 0.002649 | 0.015335 | 0.005043 | 0.001854 | 0.00409 | 0.004228 | 0.016706 | 0.003464 | 0.035402 |
| 28.625 | 0.005508 | 0.001344 | 0.015548 | 0.005561 | 0.011351 | 0.003349 | 0.012988 | 0.00429 | 0.00724 | 0.005313 | 0.0178 | 0.024261 | 0.007802 | 0.002724 | 0.00817 | 0.005485 | 0.000843 | 0.002312 | 0.00273 | 0.014292 | 0.001545 | 0.029222 |
| 28.6388889 | 0.003654 | 0.001498 | 0.01613 | 0.004637 | 0.013587 | 0.007622 | 0.012333 | 0.007454 | 0.006886 | 0.00875 | 0.019833 | 0.012439 | 0.008577 | 0.004149 | 0.002606 | 0.011043 | 0.00162 | 0.003232 | 0.003588 | 0.025149 | 0.002625 | 0.012731 |
| 28.6527778 | 0.006835 | 0.001407 | 0.012643 | 0.003514 | 0.016738 | 0.015387 | 0.012944 | 0.007045 | 0.006043 | 0.007276 | 0.037819 | 0.005267 | 0.004192 | 0.005734 | 0.001562 | 0.01698 | 0.002987 | 0.004199 | 0.004395 | 0.024052 | 0.003895 | 0.012726 |
| 28.6666667 | 0.009225 | 0.000586 | 0.021287 | 0.001856 | 0.024571 | 0.02053 | 0.009558 | 0.009234 | 0.007824 | 0.002314 | 0.047042 | 0.002997 | 0.002197 | 0.007622 | 0.002112 | 0.01668 | 0.003216 | 0.004431 | 0.004421 | 0.017467 | 0.004225 | 0.02379 |
| 28.6805556 | 0.007927 | 0.000269 | 0.027506 | 0.002438 | 0.023801 | 0.017555 | 0.008072 | 0.012973 | 0.011416 | 0.00347 | 0.04016 | 0.002074 | 0.007968 | 0.008278 | 0.003016 | 0.00995 | 0.002684 | 0.003329 | 0.003218 | 0.025156 | 0.003028 | 0.034165 |
| 28.6944444 | 0.005511 | 0.000317 | 0.018876 | 0.00543 | 0.013193 | 0.008963 | 0.009171 | 0.009293 | 0.010425 | 0.010707 | 0.027716 | 0.002188 | 0.023194 | 0.005966 | 0.002947 | 0.00433 | 0.004305 | 0.002085 | 0.001883 | 0.025093 | 0.00188 | 0.03486 |
| 28.7083333 | 0.004617 | 0.00037 | 0.010451 | 0.006651 | 0.008331 | 0.003114 | 0.010335 | 0.003391 | 0.004216 | 0.015267 | 0.017925 | 0.004273 | 0.028816 | 0.005315 | 0.002301 | 0.005879 | 0.007115 | 0.002898 | 0.002346 | 0.01536 | 0.00302 | 0.021533 |
| 28.7222222 | 0.003547 | 0.000768 | 0.011686 | 0.00647 | 0.008852 | 0.002729 | 0.018866 | 0.005185 | 0.000924 | 0.012023 | 0.025796 | 0.006229 | 0.022975 | 0.006702 | 0.002701 | 0.00913 | 0.008391 | 0.004232 | 0.003407 | 0.019118 | 0.004519 | 0.012929 |
| 28.7361111 | 0.003346 | 0.001042 | 0.015628 | 0.006404 | 0.008365 | 0.003844 | 0.027883 | 0.008271 | 0.00135 | 0.005545 | 0.045542 | 0.006432 | 0.026852 | 0.006215 | 0.004183 | 0.009611 | 0.007486 | 0.003669 | 0.00284 | 0.032178 | 0.00389 | 0.021002 |
| 28.75 | 0.006694 | 0.000742 | 0.018355 | 0.004263 | 0.005946 | 0.00379 | 0.024145 | 0.006085 | 0.002804 | 0.002417 | 0.057449 | 0.004569 | 0.025349 | 0.00367 | 0.005186 | 0.007378 | 0.005275 | 0.001911 | 0.001493 | 0.031953 | 0.002058 | 0.029806 |
| 28.7638889 | 0.007829 | 0.00046 | 0.019087 | 0.001918 | 0.006104 | 0.002939 | 0.015488 | 0.006844 | 0.003245 | 0.006025 | 0.049453 | 0.002618 | 0.012498 | 0.004264 | 0.004218 | 0.004917 | 0.002983 | 0.001076 | 0.001359 | 0.020907 | 0.001236 | 0.032271 |
| 28.7777778 | 0.005982 | 0.000521 | 0.016823 | 0.00259 | 0.00707 | 0.0025 | 0.020001 | 0.011929 | 0.003529 | 0.014381 | 0.025016 | 0.004395 | 0.007897 | 0.011833 | 0.002026 | 0.005796 | 0.001425 | 0.001706 | 0.002239 | 0.022545 | 0.001813 | 0.029109 |
| 28.7916667 | 0.008914 | 0.000509 | 0.011907 | 0.004238 | 0.006141 | 0.003534 | 0.038468 | 0.015433 | 0.00738 | 0.018322 | 0.011218 | 0.009159 | 0.01969 | 0.019915 | 0.000576 | 0.0069 | 0.001231 | 0.002328 | 0.002872 | 0.028837 | 0.002388 | 0.016953 |
| 28.8055556 | 0.014689 | 0.000701 | 0.008465 | 0.004022 | 0.00646 | 0.006428 | 0.051462 | 0.015391 | 0.012166 | 0.013294 | 0.018712 | 0.014004 | 0.04543 | 0.021986 | 0.000799 | 0.005502 | 0.002337 | 0.002394 | 0.003077 | 0.020535 | 0.002508 | 0.007431 |
| 28.8194444 | 0.018245 | 0.002055 | 0.015409 | 0.00265 | 0.006931 | 0.011011 | 0.048806 | 0.012806 | 0.013229 | 0.005854 | 0.028548 | 0.017508 | 0.064212 | 0.019041 | 0.002902 | 0.004665 | 0.004308 | 0.002661 | 0.003338 | 0.012254 | 0.002726 | 0.010583 |
| 28.8333333 | 0.021819 | 0.003859 | 0.027361 | 0.001918 | 0.005315 | 0.016444 | 0.035202 | 0.009836 | 0.010018 | 0.002942 | 0.033249 | 0.017463 | 0.058026 | 0.014395 | 0.006712 | 0.007695 | 0.007115 | 0.003204 | 0.003621 | 0.020409 | 0.003118 | 0.017904 |
| 28.8472222 | 0.026141 | 0.004814 | 0.03031 | 0.005456 | 0.003505 | 0.019838 | 0.02244 | 0.005741 | 0.00788 | 0.002838 | 0.041654 | 0.012047 | 0.03864 | 0.012064 | 0.011765 | 0.014767 | 0.010585 | 0.003244 | 0.003388 | 0.036425 | 0.003169 | 0.021864 |
| 28.8611111 | 0.029273 | 0.005677 | 0.026175 | 0.012427 | 0.005021 | 0.017763 | 0.020494 | 0.002309 | 0.006945 | 0.005108 | 0.050766 | 0.010586 | 0.02699 | 0.013475 | 0.01751 | 0.023312 | 0.014191 | 0.002349 | 0.002592 | 0.04766 | 0.002527 | 0.022498 |
| 28.875 | 0.030578 | 0.00836 | 0.021011 | 0.012755 | 0.009753 | 0.012321 | 0.018935 | 0.002874 | 0.005647 | 0.009914 | 0.040723 | 0.023173 | 0.027606 | 0.016899 | 0.020884 | 0.031806 | 0.016676 | 0.001185 | 0.00167 | 0.042102 | 0.001399 | 0.023497 |
| 28.8888889 | 0.029291 | 0.012492 | 0.014671 | 0.007799 | 0.010508 | 0.009274 | 0.018314 | 0.005557 | 0.012144 | 0.014427 | 0.024481 | 0.04266 | 0.034339 | 0.020649 | 0.020167 | 0.040171 | 0.017117 | 0.00104 | 0.000871 | 0.022465 | 0.000591 | 0.028136 |
| 28.9027778 | 0.027776 | 0.015525 | 0.012738 | 0.007149 | 0.007658 | 0.011945 | 0.031469 | 0.011411 | 0.025351 | 0.016762 | 0.031042 | 0.062508 | 0.04096 | 0.024623 | 0.018752 | 0.045938 | 0.0162 | 0.001785 | 0.000428 | 0.006802 | 0.000715 | 0.033166 |
| 28.9166667 | 0.031744 | 0.016874 | 0.013376 | 0.007015 | 0.013145 | 0.020606 | 0.045016 | 0.024957 | 0.034136 | 0.01754 | 0.038958 | 0.080381 | 0.042834 | 0.03051 | 0.017382 | 0.046085 | 0.014254 | 0.00222 | 0.000621 | 0.003994 | 0.001402 | 0.033753 |
| 28.9305556 | 0.042312 | 0.01674 | 0.010547 | 0.003903 | 0.022161 | 0.030645 | 0.044136 | 0.042912 | 0.034782 | 0.020547 | 0.037814 | 0.084766 | 0.040238 | 0.038832 | 0.0151 | 0.039839 | 0.011341 | 0.002535 | 0.001299 | 0.012925 | 0.00222 | 0.030172 |
| 28.9444444 | 0.053507 | 0.014714 | 0.013485 | 0.002192 | 0.022659 | 0.036331 | 0.040111 | 0.054697 | 0.031832 | 0.026158 | 0.049397 | 0.069903 | 0.037573 | 0.043739 | 0.01265 | 0.028472 | 0.008404 | 0.003304 | 0.002223 | 0.026129 | 0.003193 | 0.025126 |
| 28.9583333 | 0.059066 | 0.012626 | 0.030562 | 0.003473 | 0.01706 | 0.03771 | 0.048828 | 0.055001 | 0.030529 | 0.031317 | 0.074547 | 0.044481 | 0.036764 | 0.038593 | 0.009752 | 0.016271 | 0.005916 | 0.004383 | 0.003229 | 0.035096 | 0.004338 | 0.019605 |
| 28.9722222 | 0.058886 | 0.012814 | 0.051373 | 0.00799 | 0.010517 | 0.035079 | 0.061871 | 0.052697 | 0.034594 | 0.036253 | 0.09465 | 0.021293 | 0.035049 | 0.025915 | 0.008485 | 0.007574 | 0.003449 | 0.004932 | 0.003663 | 0.037552 | 0.005034 | 0.014719 |
| 28.9861111 | 0.053984 | 0.015701 | 0.056924 | 0.014056 | 0.004658 | 0.02846 | 0.057688 | 0.056835 | 0.043145 | 0.04231 | 0.089465 | 0.01339 | 0.029943 | 0.013461 | 0.009446 | 0.003567 | 0.001561 | 0.004192 | 0.002982 | 0.035214 | 0.004474 | 0.011623 |
| 29 | 0.042531 | 0.019579 | 0.043076 | 0.017339 | 0.00132 | 0.021354 | 0.034107 | 0.061893 | 0.046996 | 0.043401 | 0.060913 | 0.017518 | 0.022238 | 0.010829 | 0.0078 | 0.001972 | 0.00199 | 0.003877 | 0.002547 | 0.028599 | 0.00402 | 0.00896 |
| 29.0138889 | 0.025714 | 0.021315 | 0.022343 | 0.014885 | 0.001753 | 0.014993 | 0.012559 | 0.054777 | 0.038976 | 0.035741 | 0.031021 | 0.019972 | 0.013462 | 0.018731 | 0.006019 | 0.002028 | 0.005243 | 0.007136 | 0.004914 | 0.01806 | 0.00672 | 0.005801 |
| 29.0277778 | 0.010886 | 0.020595 | 0.011223 | 0.009493 | 0.003474 | 0.00985 | 0.011263 | 0.036657 | 0.024917 | 0.026068 | 0.021226 | 0.020529 | 0.006537 | 0.022397 | 0.006607 | 0.004186 | 0.008152 | 0.012388 | 0.008717 | 0.008107 | 0.011414 | 0.004363 |
| 29.0416667 | 0.010185 | 0.021902 | 0.010244 | 0.008972 | 0.006629 | 0.012598 | 0.030643 | 0.029185 | 0.018264 | 0.022224 | 0.021793 | 0.020743 | 0.006167 | 0.016029 | 0.010265 | 0.005532 | 0.007562 | 0.011941 | 0.008027 | 0.005926 | 0.011098 | 0.004605 |
| 29.0555556 | 0.034399 | 0.033777 | 0.010037 | 0.018943 | 0.017233 | 0.030856 | 0.050502 | 0.045252 | 0.029167 | 0.026519 | 0.022767 | 0.01975 | 0.012637 | 0.0143 | 0.024841 | 0.00855 | 0.00769 | 0.005926 | 0.00382 | 0.013423 | 0.005884 | 0.005964 |
| 29.0694444 | 0.078156 | 0.054386 | 0.01747 | 0.034574 | 0.032431 | 0.057206 | 0.049362 | 0.074149 | 0.050317 | 0.032297 | 0.042047 | 0.035302 | 0.022628 | 0.025389 | 0.041976 | 0.018918 | 0.012158 | 0.002393 | 0.00195 | 0.026888 | 0.002592 | 0.009787 |
| 29.0833333 | 0.102299 | 0.063214 | 0.031904 | 0.045517 | 0.038073 | 0.072266 | 0.042348 | 0.094275 | 0.061211 | 0.029364 | 0.061336 | 0.063944 | 0.032589 | 0.035689 | 0.048198 | 0.030896 | 0.018515 | 0.002152 | 0.001896 | 0.041041 | 0.001894 | 0.014598 |
| 29.0972222 | 0.078151 | 0.053927 | 0.034791 | 0.048548 | 0.028511 | 0.069486 | 0.061014 | 0.087546 | 0.056023 | 0.016649 | 0.05483 | 0.073006 | 0.035017 | 0.029195 | 0.047243 | 0.029807 | 0.025808 | 0.002353 | 0.001993 | 0.045771 | 0.002062 | 0.019093 |
| 29.1111111 | 0.035221 | 0.040742 | 0.02092 | 0.040982 | 0.013448 | 0.052849 | 0.094683 | 0.056217 | 0.042836 | 0.007014 | 0.033192 | 0.0634 | 0.024706 | 0.014881 | 0.044081 | 0.015668 | 0.03179 | 0.001921 | 0.001714 | 0.034271 | 0.001854 | 0.022809 |
| 29.125 | 0.013916 | 0.02928 | 0.010436 | 0.025364 | 0.005043 | 0.028141 | 0.102723 | 0.025042 | 0.027301 | 0.008985 | 0.016859 | 0.042804 | 0.011349 | 0.013293 | 0.037896 | 0.006509 | 0.033123 | 0.000959 | 0.000989 | 0.016216 | 0.000976 | 0.025839 |
| 29.1388889 | 0.021184 | 0.018722 | 0.016653 | 0.011668 | 0.004532 | 0.016149 | 0.080022 | 0.014756 | 0.013199 | 0.01355 | 0.020386 | 0.018686 | 0.007071 | 0.019238 | 0.028789 | 0.006643 | 0.029871 | 0.000435 | 0.000992 | 0.007161 | 0.000432 | 0.026539 |
| 29.1527778 | 0.043872 | 0.010816 | 0.023988 | 0.010135 | 0.004511 | 0.02682 | 0.047125 | 0.021148 | 0.009591 | 0.014499 | 0.035789 | 0.010801 | 0.007675 | 0.01926 | 0.017548 | 0.006695 | 0.023795 | 0.000373 | 0.001601 | 0.007481 | 0.000224 | 0.020426 |
| 29.1666667 | 0.069373 | 0.009809 | 0.020169 | 0.028459 | 0.002625 | 0.04018 | 0.024392 | 0.02937 | 0.023204 | 0.014013 | 0.048412 | 0.016548 | 0.005573 | 0.01549 | 0.009408 | 0.005557 | 0.018809 | 0.000634 | 0.00151 | 0.00635 | 0.000447 | 0.01127 |
| 29.1805556 | 0.076416 | 0.025262 | 0.011305 | 0.047488 | 0.001065 | 0.039905 | 0.031186 | 0.033951 | 0.04237 | 0.016851 | 0.048462 | 0.020877 | 0.004458 | 0.015431 | 0.011033 | 0.007443 | 0.018119 | 0.001843 | 0.001198 | 0.003699 | 0.001649 | 0.011046 |
| 29.1944444 | 0.057482 | 0.057423 | 0.009382 | 0.039329 | 0.007019 | 0.025843 | 0.058904 | 0.035525 | 0.046012 | 0.021415 | 0.030719 | 0.018468 | 0.012293 | 0.020501 | 0.013904 | 0.014877 | 0.020409 | 0.003815 | 0.002278 | 0.00889 | 0.003574 | 0.020094 |
| 29.2083333 | 0.034773 | 0.066064 | 0.02055 | 0.019523 | 0.025442 | 0.012774 | 0.081419 | 0.034531 | 0.028753 | 0.018147 | 0.009983 | 0.011262 | 0.03113 | 0.022456 | 0.009219 | 0.025284 | 0.022382 | 0.005349 | 0.003894 | 0.023858 | 0.005155 | 0.027794 |
| 29.2222222 | 0.026952 | 0.036676 | 0.032119 | 0.01264 | 0.038218 | 0.013351 | 0.078096 | 0.031692 | 0.010307 | 0.009792 | 0.003922 | 0.005473 | 0.051664 | 0.016888 | 0.003986 | 0.030126 | 0.021295 | 0.006437 | 0.005032 | 0.038397 | 0.006281 | 0.033799 |
| 29.2361111 | 0.041801 | 0.021423 | 0.026998 | 0.016336 | 0.03044 | 0.016188 | 0.047857 | 0.025744 | 0.006792 | 0.0052 | 0.009467 | 0.005245 | 0.056702 | 0.01162 | 0.005621 | 0.026115 | 0.015864 | 0.007679 | 0.006112 | 0.04341 | 0.007514 | 0.041185 |
| 29.25 | 0.049767 | 0.031085 | 0.013257 | 0.030081 | 0.01635 | 0.011546 | 0.023202 | 0.015364 | 0.012679 | 0.006765 | 0.017973 | 0.006926 | 0.04005 | 0.013297 | 0.012157 | 0.020826 | 0.008662 | 0.007918 | 0.006356 | 0.04308 | 0.007831 | 0.043826 |
| 29.2638889 | 0.030535 | 0.034741 | 0.009056 | 0.037836 | 0.007643 | 0.005175 | 0.026847 | 0.006827 | 0.014165 | 0.015952 | 0.023954 | 0.014727 | 0.018914 | 0.019313 | 0.015772 | 0.020728 | 0.004332 | 0.005379 | 0.004283 | 0.04437 | 0.005429 | 0.036347 |
| 29.2777778 | 0.018805 | 0.025656 | 0.014411 | 0.023789 | 0.005774 | 0.003815 | 0.036549 | 0.004589 | 0.009617 | 0.0363 | 0.0188 | 0.031714 | 0.01146 | 0.024696 | 0.012368 | 0.025075 | 0.00423 | 0.002488 | 0.001985 | 0.047904 | 0.002629 | 0.026827 |
| 29.2916667 | 0.021508 | 0.013612 | 0.023691 | 0.013019 | 0.009925 | 0.009776 | 0.027516 | 0.007503 | 0.007096 | 0.050187 | 0.012164 | 0.043675 | 0.009913 | 0.026505 | 0.007793 | 0.030187 | 0.005665 | 0.002217 | 0.002034 | 0.04951 | 0.002527 | 0.019107 |

| | | | | | | | | | | | | | | | | | | | | | | |
|---|---|---|---|---|---|---|---|---|---|---|---|---|---|---|---|---|---|---|---|---|---|---|
| 29.3055556 | 0.030564 | 0.011293 | 0.032995 | 0.016049 | 0.010359 | 0.014614 | 0.019429 | 0.016526 | 0.012542 | 0.034545 | 0.01654 | 0.044484 | 0.012529 | 0.024378 | 0.008543 | 0.034351 | 0.005152 | 0.002083 | 0.002096 | 0.044314 | 0.002617 | 0.011435 |
| 29.3194444 | 0.044595 | 0.018363 | 0.033524 | 0.022786 | 0.0091 | 0.010219 | 0.032511 | 0.019899 | 0.03253 | 0.018549 | 0.01603 | 0.043082 | 0.033625 | 0.020826 | 0.012888 | 0.037857 | 0.004266 | 0.001589 | 0.001603 | 0.03469 | 0.00188 | 0.012285 |
| 29.3333333 | 0.044138 | 0.024202 | 0.022652 | 0.028816 | 0.013895 | 0.004038 | 0.040451 | 0.012504 | 0.047059 | 0.017885 | 0.007434 | 0.040581 | 0.057832 | 0.018272 | 0.014421 | 0.039342 | 0.00778 | 0.002775 | 0.002508 | 0.028462 | 0.002418 | 0.015345 |
| 29.3472222 | 0.031208 | 0.023926 | 0.010208 | 0.020003 | 0.019792 | 0.009055 | 0.026784 | 0.013414 | 0.033636 | 0.013594 | 0.005732 | 0.036415 | 0.063464 | 0.017581 | 0.010859 | 0.036849 | 0.013149 | 0.003662 | 0.003022 | 0.03125 | 0.002897 | 0.011179 |
| 29.3611111 | 0.023021 | 0.018675 | 0.007023 | 0.007372 | 0.025859 | 0.025253 | 0.010631 | 0.030333 | 0.014558 | 0.019267 | 0.012326 | 0.036959 | 0.054515 | 0.021651 | 0.007536 | 0.030248 | 0.014754 | 0.003019 | 0.001852 | 0.040611 | 0.002081 | 0.008345 |
| 29.375 | 0.022002 | 0.010688 | 0.012767 | 0.006004 | 0.034421 | 0.035228 | 0.004256 | 0.053183 | 0.009427 | 0.032769 | 0.019059 | 0.039701 | 0.041608 | 0.032408 | 0.007884 | 0.022281 | 0.013205 | 0.001974 | 0.000752 | 0.047133 | 0.001031 | 0.014816 |
| 29.3888889 | 0.017378 | 0.005408 | 0.01467 | 0.006936 | 0.039663 | 0.028751 | 0.004233 | 0.060074 | 0.015577 | 0.031541 | 0.015979 | 0.036856 | 0.029734 | 0.044445 | 0.008649 | 0.016361 | 0.012033 | 0.000999 | 0.00069 | 0.046225 | 0.000643 | 0.030238 |
| 29.4027778 | 0.008057 | 0.007689 | 0.023496 | 0.01015 | 0.035705 | 0.018185 | 0.009993 | 0.040691 | 0.03156 | 0.017583 | 0.006949 | 0.029187 | 0.020713 | 0.049972 | 0.009173 | 0.014317 | 0.013826 | 0.00073 | 0.00086 | 0.04192 | 0.000764 | 0.045721 |
| 29.4166667 | 0.009854 | 0.013262 | 0.050153 | 0.014063 | 0.025278 | 0.019056 | 0.022207 | 0.02654 | 0.040553 | 0.005906 | 0.003547 | 0.0234 | 0.021155 | 0.047438 | 0.013741 | 0.016716 | 0.019571 | 0.001615 | 0.000925 | 0.039477 | 0.001034 | 0.052997 |
| 29.4305556 | 0.028993 | 0.016789 | 0.056658 | 0.012904 | 0.013427 | 0.030924 | 0.036061 | 0.041736 | 0.033341 | 0.004293 | 0.009418 | 0.024557 | 0.036166 | 0.040176 | 0.022891 | 0.022015 | 0.026459 | 0.002125 | 0.001445 | 0.039159 | 0.001475 | 0.053807 |
| 29.4444444 | 0.054546 | 0.017439 | 0.043867 | 0.008964 | 0.004737 | 0.040539 | 0.049953 | 0.063136 | 0.026569 | 0.009464 | 0.02509 | 0.029824 | 0.062166 | 0.034455 | 0.031716 | 0.026703 | 0.02756 | 0.003779 | 0.004199 | 0.036086 | 0.004139 | 0.053301 |
| 29.4583333 | 0.066085 | 0.014127 | 0.050587 | 0.005305 | 0.003427 | 0.040262 | 0.064438 | 0.068746 | 0.031398 | 0.017854 | 0.046751 | 0.032289 | 0.080227 | 0.033577 | 0.036866 | 0.027477 | 0.021191 | 0.011089 | 0.010713 | 0.026787 | 0.01197 | 0.052275 |
| 29.4722222 | 0.051594 | 0.007973 | 0.050195 | 0.002612 | 0.009637 | 0.037287 | 0.075476 | 0.065429 | 0.042969 | 0.027747 | 0.066791 | 0.031073 | 0.069175 | 0.031478 | 0.036323 | 0.02278 | 0.015758 | 0.018791 | 0.016258 | 0.014591 | 0.018675 | 0.044104 |
| 29.4861111 | 0.029349 | 0.003526 | 0.023969 | 0.003839 | 0.024241 | 0.040539 | 0.082353 | 0.057295 | 0.050521 | 0.033846 | 0.082293 | 0.031982 | 0.045139 | 0.026957 | 0.030839 | 0.015403 | 0.018468 | 0.016974 | 0.014413 | 0.008969 | 0.015523 | 0.03094 |
| 29.5 | 0.025881 | 0.002815 | 0.015438 | 0.009895 | 0.046271 | 0.047595 | 0.087258 | 0.04263 | 0.046479 | 0.035971 | 0.090031 | 0.037488 | 0.036756 | 0.02626 | 0.027525 | 0.010799 | 0.026104 | 0.008505 | 0.007356 | 0.016417 | 0.0074 | 0.023811 |
| 29.5138889 | 0.041926 | 0.004591 | 0.031095 | 0.014997 | 0.067549 | 0.04922 | 0.088403 | 0.027146 | 0.031467 | 0.039861 | 0.081886 | 0.043558 | 0.041321 | 0.027866 | 0.028887 | 0.009764 | 0.031129 | 0.003877 | 0.00291 | 0.030871 | 0.00386 | 0.027177 |
| 29.5277778 | 0.062605 | 0.009604 | 0.0361 | 0.016366 | 0.073993 | 0.043019 | 0.083578 | 0.014974 | 0.018232 | 0.040896 | 0.067822 | 0.048428 | 0.043439 | 0.024507 | 0.029322 | 0.008099 | 0.029982 | 0.003759 | 0.003422 | 0.038432 | 0.003586 | 0.033729 |
| 29.5416667 | 0.068662 | 0.018939 | 0.022473 | 0.013967 | 0.060556 | 0.037339 | 0.071444 | 0.008437 | 0.019759 | 0.032415 | 0.067555 | 0.054675 | 0.033874 | 0.015087 | 0.022515 | 0.007506 | 0.021206 | 0.00844 | 0.010596 | 0.029897 | 0.00678 | 0.025013 |
| 29.5555556 | 0.051556 | 0.027379 | 0.008256 | 0.008021 | 0.041497 | 0.034829 | 0.052692 | 0.009067 | 0.026598 | 0.022351 | 0.072428 | 0.057661 | 0.016082 | 0.008948 | 0.010577 | 0.012805 | 0.010355 | 0.021146 | 0.024917 | 0.013373 | 0.017907 | 0.015001 |
| 29.5694444 | 0.028113 | 0.026744 | 0.004333 | 0.002597 | 0.030204 | 0.026344 | 0.033791 | 0.011223 | 0.027699 | 0.019717 | 0.061836 | 0.049692 | 0.008456 | 0.007915 | 0.002797 | 0.014375 | 0.005242 | 0.027995 | 0.031186 | 0.007283 | 0.025094 | 0.026829 |
| 29.5833333 | 0.011908 | 0.01871 | 0.007365 | 0.000617 | 0.026988 | 0.011993 | 0.021149 | 0.009536 | 0.025288 | 0.018738 | 0.040041 | 0.039512 | 0.017843 | 0.00856 | 0.001138 | 0.010731 | 0.004234 | 0.01873 | 0.020009 | 0.019819 | 0.01735 | 0.031627 |
| 29.5972222 | 0.006411 | 0.0098 | 0.014982 | 0.000792 | 0.026887 | 0.004967 | 0.013486 | 0.007742 | 0.017844 | 0.016859 | 0.020479 | 0.038282 | 0.026984 | 0.015747 | 0.002508 | 0.013963 | 0.006387 | 0.009307 | 0.00855 | 0.036784 | 0.00852 | 0.018409 |
| 29.6111111 | 0.006728 | 0.006416 | 0.01718 | 0.001438 | 0.021498 | 0.007742 | 0.009622 | 0.008773 | 0.010689 | 0.015851 | 0.011518 | 0.041411 | 0.024377 | 0.02004 | 0.005617 | 0.01691 | 0.011938 | 0.010084 | 0.00841 | 0.036898 | 0.00982 | 0.01652 |
| 29.625 | 0.005568 | 0.014405 | 0.011378 | 0.002038 | 0.010296 | 0.010888 | 0.009796 | 0.010445 | 0.012403 | 0.011569 | 0.0244 | 0.042489 | 0.014046 | 0.013562 | 0.009141 | 0.010669 | 0.015993 | 0.010048 | 0.008525 | 0.020985 | 0.010538 | 0.022476 |
| 29.6388889 | 0.009737 | 0.026714 | 0.009102 | 0.002849 | 0.005791 | 0.018753 | 0.011601 | 0.01468 | 0.014088 | 0.006737 | 0.049687 | 0.040614 | 0.007726 | 0.009212 | 0.010968 | 0.005637 | 0.014619 | 0.005637 | 0.004772 | 0.016107 | 0.006152 | 0.020764 |
| 29.6527778 | 0.019027 | 0.032705 | 0.007269 | 0.002985 | 0.007643 | 0.028546 | 0.012306 | 0.017196 | 0.011784 | 0.011145 | 0.057473 | 0.034471 | 0.007965 | 0.010786 | 0.00939 | 0.009508 | 0.008961 | 0.003945 | 0.003616 | 0.026057 | 0.004107 | 0.015407 |
| 29.6666667 | 0.021573 | 0.029573 | 0.003137 | 0.003101 | 0.013531 | 0.022819 | 0.009504 | 0.011885 | 0.012933 | 0.020249 | 0.036689 | 0.021973 | 0.0045 | 0.009996 | 0.005304 | 0.014699 | 0.003647 | 0.00497 | 0.004638 | 0.023955 | 0.004996 | 0.010962 |
| 29.6805556 | 0.013828 | 0.019282 | 0.005561 | 0.005142 | 0.031566 | 0.009733 | 0.00557 | 0.005023 | 0.010192 | 0.017934 | 0.019716 | 0.012328 | 0.003749 | 0.005693 | 0.003103 | 0.015242 | 0.002198 | 0.00464 | 0.004231 | 0.023115 | 0.004687 | 0.008956 |
| 29.6944444 | 0.006713 | 0.010194 | 0.009559 | 0.005472 | 0.039831 | 0.003852 | 0.005649 | 0.002789 | 0.005288 | 0.011516 | 0.032753 | 0.020059 | 0.010796 | 0.002515 | 0.00567 | 0.013343 | 0.003733 | 0.002935 | 0.002639 | 0.031548 | 0.002935 | 0.009301 |
| 29.7083333 | 0.007111 | 0.01238 | 0.007026 | 0.00295 | 0.026167 | 0.004381 | 0.007948 | 0.003631 | 0.005043 | 0.012279 | 0.049092 | 0.039979 | 0.019189 | 0.002919 | 0.010531 | 0.012641 | 0.004674 | 0.002606 | 0.002387 | 0.023437 | 0.002365 | 0.009283 |
| 29.7222222 | 0.00988 | 0.023635 | 0.006932 | 0.001019 | 0.01882 | 0.006749 | 0.007494 | 0.004658 | 0.003571 | 0.012829 | 0.048458 | 0.055285 | 0.032239 | 0.006615 | 0.013627 | 0.014418 | 0.003399 | 0.004584 | 0.004331 | 0.012456 | 0.004075 | 0.00828 |
| 29.7361111 | 0.014452 | 0.032785 | 0.012951 | 0.00033 | 0.020731 | 0.006494 | 0.004916 | 0.005063 | 0.00549 | 0.014755 | 0.03831 | 0.058426 | 0.047184 | 0.012519 | 0.011344 | 0.015255 | 0.001617 | 0.007396 | 0.006968 | 0.023398 | 0.006911 | 0.008511 |
| 29.75 | 0.022183 | 0.037047 | 0.018566 | 0.00079 | 0.020218 | 0.003698 | 0.003937 | 0.004946 | 0.010597 | 0.014349 | 0.02184 | 0.048204 | 0.045574 | 0.018378 | 0.006253 | 0.011249 | 0.001503 | 0.009161 | 0.008364 | 0.043219 | 0.008846 | 0.009736 |
| 29.7638889 | 0.027092 | 0.039878 | 0.019886 | 0.002715 | 0.016175 | 0.002456 | 0.004275 | 0.004953 | 0.010322 | 0.012241 | 0.011498 | 0.0268 | 0.02563 | 0.020974 | 0.004122 | 0.006488 | 0.003373 | 0.009087 | 0.007877 | 0.048466 | 0.008802 | 0.011821 |
| 29.7777778 | 0.022059 | 0.039521 | 0.018785 | 0.004829 | 0.010203 | 0.004587 | 0.007008 | 0.005745 | 0.010523 | 0.015799 | 0.020249 | 0.013062 | 0.010309 | 0.017932 | 0.003139 | 0.007175 | 0.005577 | 0.007965 | 0.006525 | 0.034689 | 0.007681 | 0.015902 |
| 29.7916667 | 0.013278 | 0.033698 | 0.016974 | 0.004449 | 0.009668 | 0.006909 | 0.012815 | 0.009155 | 0.016189 | 0.013811 | 0.037445 | 0.025625 | 0.005987 | 0.010841 | 0.002347 | 0.008867 | 0.005905 | 0.007248 | 0.005772 | 0.014893 | 0.007042 | 0.020267 |
| 29.8055556 | 0.017008 | 0.02614 | 0.012353 | 0.002655 | 0.014922 | 0.007317 | 0.012616 | 0.012807 | 0.018125 | 0.011797 | 0.050902 | 0.051116 | 0.003921 | 0.007698 | 0.00433 | 0.006415 | 0.004469 | 0.007011 | 0.005512 | 0.004399 | 0.006766 | 0.021517 |
| 29.8194444 | 0.032831 | 0.01994 | 0.00847 | 0.002032 | 0.01551 | 0.006712 | 0.013139 | 0.00995 | 0.012726 | 0.012917 | 0.054113 | 0.061112 | 0.006069 | 0.012188 | 0.007821 | 0.004784 | 0.002972 | 0.006536 | 0.005005 | 0.003725 | 0.006098 | 0.017883 |
| 29.8333333 | 0.046189 | 0.015083 | 0.007224 | 0.002607 | 0.010067 | 0.006101 | 0.028832 | 0.00543 | 0.01039 | 0.007722 | 0.038891 | 0.043975 | 0.017501 | 0.019908 | 0.013457 | 0.010598 | 0.00198 | 0.006298 | 0.004993 | 0.005082 | 0.005715 | 0.012543 |
| 29.8472222 | 0.052789 | 0.0112 | 0.004492 | 0.004264 | 0.006644 | 0.006065 | 0.04517 | 0.008621 | 0.016762 | 0.006025 | 0.022733 | 0.025639 | 0.034846 | 0.027704 | 0.023312 | 0.021566 | 0.002669 | 0.006853 | 0.006147 | 0.012419 | 0.006223 | 0.011738 |
| 29.8611111 | 0.052185 | 0.007573 | 0.002154 | 0.004104 | 0.006047 | 0.006712 | 0.046812 | 0.011084 | 0.024917 | 0.009202 | 0.027593 | 0.040592 | 0.0438 | 0.033577 | 0.029655 | 0.031798 | 0.005911 | 0.0071 | 0.007147 | 0.029396 | 0.006575 | 0.015225 |
| 29.875 | 0.043578 | 0.004141 | 0.005061 | 0.001781 | 0.011453 | 0.007335 | 0.032139 | 0.010072 | 0.026615 | 0.010892 | 0.030018 | 0.071708 | 0.04362 | 0.037604 | 0.026233 | 0.040266 | 0.009746 | 0.006204 | 0.006756 | 0.04633 | 0.005861 | 0.017745 |
| 29.8888889 | 0.03536 | 0.001683 | 0.01662 | 0.000854 | 0.024189 | 0.008116 | 0.011591 | 0.017367 | 0.025393 | 0.014762 | 0.022422 | 0.084806 | 0.051412 | 0.044154 | 0.021166 | 0.047512 | 0.012362 | 0.005156 | 0.005656 | 0.057615 | 0.004731 | 0.018836 |
| 29.9027778 | 0.035296 | 0.001404 | 0.034236 | 0.001574 | 0.038946 | 0.011152 | 0.005367 | 0.031282 | 0.026839 | 0.02315 | 0.036432 | 0.081699 | 0.069498 | 0.054443 | 0.021147 | 0.050068 | 0.014051 | 0.004669 | 0.004817 | 0.061338 | 0.004083 | 0.021499 |
| 29.9166667 | 0.038885 | 0.004345 | 0.044152 | 0.003465 | 0.046057 | 0.018598 | 0.007097 | 0.043496 | 0.029429 | 0.032528 | 0.076661 | 0.072451 | 0.088062 | 0.057875 | 0.024638 | 0.046593 | 0.015868 | 0.004351 | 0.004298 | 0.051512 | 0.003894 | 0.028121 |
| 29.9305556 | 0.040398 | 0.010243 | 0.038168 | 0.004689 | 0.04102 | 0.031116 | 0.005344 | 0.051304 | 0.034228 | 0.036605 | 0.119538 | 0.056405 | 0.095726 | 0.044615 | 0.030389 | 0.041948 | 0.018394 | 0.003765 | 0.003794 | 0.03265 | 0.003591 | 0.03807 |
| 29.9444444 | 0.045602 | 0.018786 | 0.025254 | 0.004875 | 0.032828 | 0.044038 | 0.014076 | 0.059222 | 0.038912 | 0.036038 | 0.131383 | 0.037618 | 0.08478 | 0.024716 | 0.042127 | 0.03767 | 0.020656 | 0.002612 | 0.002813 | 0.016273 | 0.002624 | 0.046475 |
| 29.9583333 | 0.05696 | 0.031026 | 0.02111 | 0.007107 | 0.027679 | 0.048974 | 0.041744 | 0.073963 | 0.036214 | 0.036365 | 0.098559 | 0.025017 | 0.064403 | 0.013047 | 0.055911 | 0.032512 | 0.021469 | 0.001443 | 0.001772 | 0.008101 | 0.001435 | 0.049027 |
| 29.9722222 | 0.068962 | 0.045899 | 0.034873 | 0.008533 | 0.026161 | 0.044957 | 0.064635 | 0.091079 | 0.028517 | 0.037509 | 0.049852 | 0.018839 | 0.045339 | 0.011617 | 0.057064 | 0.024583 | 0.020518 | 0.001632 | 0.002168 | 0.008599 | 0.001435 | 0.044959 |
| 29.9861111 | 0.075692 | 0.054239 | 0.053339 | 0.005724 | 0.023581 | 0.038642 | 0.062034 | 0.096232 | 0.025236 | 0.038435 | 0.016897 | 0.012156 | 0.028084 | 0.01375 | 0.042337 | 0.013454 | 0.017152 | 0.003512 | 0.004027 | 0.012747 | 0.003131 | 0.035728 |
| 30 | 0.068183 | 0.048984 | 0.057288 | 0.003319 | 0.019459 | 0.033246 | 0.043456 | 0.0824 | 0.028437 | 0.036899 | 0.004723 | 0.009782 | 0.013139 | 0.013005 | 0.021668 | 0.007588 | 0.01165 | 0.006781 | 0.006581 | 0.013381 | 0.006204 | 0.02353 |
| 30.0138889 | 0.04674 | 0.036936 | 0.051768 | 0.005322 | 0.018266 | 0.025689 | 0.023084 | 0.060366 | 0.029406 | 0.028731 | 0.006914 | 0.026443 | 0.006056 | 0.008562 | 0.010644 | 0.011249 | 0.005778 | 0.012082 | 0.010493 | 0.008367 | 0.011062 | 0.012092 |
| 30.0277778 | 0.025773 | 0.026415 | 0.037615 | 0.007428 | 0.0171 | 0.015975 | 0.019561 | 0.045195 | 0.024071 | 0.017778 | 0.01521 | 0.04897 | 0.00902 | 0.004301 | 0.014103 | 0.017239 | 0.003485 | 0.020039 | 0.016662 | 0.005282 | 0.018373 | 0.009067 |
| 30.0416667 | 0.020492 | 0.017998 | 0.015827 | 0.009637 | 0.018197 | 0.015447 | 0.029572 | 0.049242 | 0.025334 | 0.015071 | 0.016224 | 0.041844 | 0.014087 | 0.004235 | 0.018985 | 0.020891 | 0.007226 | 0.028381 | 0.024136 | 0.010488 | 0.026535 | 0.016183 |
| 30.0555556 | 0.038208 | 0.009972 | 0.007982 | 0.01938 | 0.028776 | 0.035725 | 0.03914 | 0.074127 | 0.045184 | 0.027981 | 0.018839 | 0.022582 | 0.012681 | 0.004715 | 0.018224 | 0.018673 | 0.010287 | 0.032119 | 0.029184 | 0.015765 | 0.030825 | 0.021391 |
| 30.0694444 | 0.065536 | 0.003717 | 0.02262 | 0.037083 | 0.045574 | 0.06339 | 0.054517 | 0.099267 | 0.073548 | 0.04414 | 0.041158 | 0.019037 | 0.009662 | 0.007194 | 0.013371 | 0.014938 | 0.009383 | 0.029215 | 0.028309 | 0.013103 | 0.028411 | 0.015469 |
| 30.0833333 | 0.072242 | 0.002296 | 0.033263 | 0.052812 | 0.051998 | 0.074414 | 0.063965 | 0.105103 | 0.093289 | 0.045345 | 0.062819 | 0.025035 | 0.015726 | 0.01831 | 0.01055 | 0.021334 | 0.011916 | 0.023237 | 0.023179 | 0.013128 | 0.022707 | 0.007285 |
| 30.0972222 | 0.051569 | 0.006228 | 0.025233 | 0.051015 | 0.038956 | 0.070963 | 0.073873 | 0.09052 | 0.094401 | 0.033994 | 0.06265 | 0.044381 | 0.023361 | 0.026073 | 0.016058 | 0.025798 | 0.018832 | 0.018065 | 0.017972 | 0.023349 | 0.018077 | 0.006605 |
| 30.1111111 | 0.025233 | 0.01243 | 0.023567 | 0.031732 | 0.019545 | 0.062459 | 0.097807 | 0.063117 | 0.069467 | 0.018099 | 0.048416 | 0.046169 | 0.020159 | 0.019029 | 0.021241 | 0.016433 | 0.024859 | 0.013578 | 0.013423 | 0.028304 | 0.014076 | 0.008637 |
| 30.125 | 0.011555 | 0.015995 | 0.025906 | 0.013455 | 0.008852 | 0.045032 | 0.102456 | 0.03275 | 0.031422 | 0.008166 | 0.028406 | 0.02271 | 0.013391 | 0.00947 | 0.015483 | 0.010952 | 0.027012 | 0.009107 | 0.008973 | 0.018321 | 0.009592 | 0.007944 |
| 30.1388889 | 0.018248 | 0.014547 | 0.016468 | 0.005652 | 0.010229 | 0.024543 | 0.069533 | 0.017692 | 0.009945 | 0.012301 | 0.014305 | 0.014753 | 0.016361 | 0.011289 | 0.009796 | 0.016339 | 0.022502 | 0.005862 | 0.005751 | 0.009673 | 0.006079 | 0.004954 |
| 30.1527778 | 0.041902 | 0.008825 | 0.00762 | 0.009637 | 0.018952 | 0.022693 | 0.035749 | 0.02469 | 0.015255 | 0.019966 | 0.020637 | 0.017483 | 0.024769 | 0.020771 | 0.015373 | 0.019342 | 0.012216 | 0.004138 | 0.004173 | 0.01554 | 0.004253 | 0.003063 |
| 30.1666667 | 0.073811 | 0.005221 | 0.005187 | 0.023229 | 0.027676 | 0.039059 | 0.039472 | 0.032882 | 0.034142 | 0.024568 | 0.043918 | 0.013243 | 0.025163 | 0.031593 | 0.021315 | 0.017108 | 0.00456 | 0.003107 | 0.003342 | 0.024056 | 0.003282 | 0.003785 |
| 30.1805556 | 0.088958 | 0.009426 | 0.009883 | 0.028179 | 0.028942 | 0.044848 | 0.066195 | 0.030695 | 0.051797 | 0.033057 | 0.064514 | 0.007404 | 0.018009 | 0.04379 | 0.020193 | 0.017861 | 0.003247 | 0.002601 | 0.002897 | 0.025025 | 0.002773 | 0.005702 |
| 30.1944444 | 0.063324 | 0.016273 | 0.029103 | 0.017782 | 0.019709 | 0.030344 | 0.079524 | 0.024817 | 0.055394 | 0.036885 | 0.064454 | 0.009058 | 0.014687 | 0.053036 | 0.016021 | 0.025667 | 0.004181 | 0.002839 | 0.003135 | 0.023243 | 0.00293 | 0.009378 |
| 30.2083333 | 0.027674 | 0.020441 | 0.046659 | 0.010487 | 0.008635 | 0.013236 | 0.062772 | 0.019783 | 0.038974 | 0.022842 | 0.041216 | 0.018304 | 0.021508 | 0.051199 | 0.011234 | 0.033698 | 0.007895 | 0.003677 | 0.003985 | 0.024463 | 0.003696 | 0.016203 |
| 30.2222222 | 0.018075 | 0.020897 | 0.036146 | 0.009762 | 0.004984 | 0.007002 | 0.031829 | 0.015293 | 0.017602 | 0.009562 | 0.015559 | 0.025316 | 0.031965 | 0.036658 | 0.006543 | 0.031968 | 0.013471 | 0.003915 | 0.004232 | 0.030425 | 0.003845 | 0.025167 |
| 30.2361111 | 0.031404 | 0.016037 | 0.016931 | 0.0167 | 0.006499 | 0.006955 | 0.019213 | 0.009413 | 0.007357 | 0.008224 | 0.0112 | 0.020164 | 0.035236 | 0.018108 | 0.004645 | 0.022093 | 0.017612 | 0.002996 | 0.00305 | 0.035932 | 0.002994 | 0.030252 |
| 30.25 | 0.043213 | 0.007931 | 0.015737 | 0.035909 | 0.009557 | 0.007193 | 0.026475 | 0.004596 | 0.00676 | 0.008351 | 0.021049 | 0.014612 | 0.029959 | 0.006669 | 0.006556 | 0.017058 | 0.020457 | 0.002143 | 0.001593 | 0.034883 | 0.002339 | 0.028346 |
| 30.2638889 | 0.033294 | 0.00328 | 0.018194 | 0.040356 | 0.013835 | 0.007562 | 0.033213 | 0.006057 | 0.010568 | 0.01252 | 0.024585 | 0.028006 | 0.024739 | 0.005411 | 0.008302 | 0.024366 | 0.019565 | 0.002412 | 0.001922 | 0.031637 | 0.002705 | 0.026306 |
| 30.2777778 | 0.02166 | 0.003664 | 0.017137 | 0.023368 | 0.015769 | 0.008119 | 0.028428 | 0.013534 | 0.010641 | 0.023154 | 0.016403 | 0.042567 | 0.027107 | 0.012812 | 0.008779 | 0.034849 | 0.013276 | 0.004381 | 0.004639 | 0.032565 | 0.004837 | 0.027657 |
| 30.2916667 | 0.0188 | 0.005033 | 0.023695 | 0.015713 | 0.014408 | 0.01371 | 0.016822 | 0.020855 | 0.007021 | 0.034181 | 0.012135 | 0.034318 | 0.031606 | 0.027639 | 0.008833 | 0.036841 | 0.006387 | 0.006307 | 0.006805 | 0.034393 | 0.006813 | 0.025201 |
| 30.3055556 | 0.022521 | 0.007668 | 0.038075 | 0.016046 | 0.016199 | 0.021446 | 0.015089 | 0.020932 | 0.011205 | 0.03578 | 0.024828 | 0.014908 | 0.026403 | 0.041094 | 0.0079 | 0.032336 | 0.003276 | 0.005291 | 0.005609 | 0.030361 | 0.00563 | 0.014746 |
| 30.3194444 | 0.03589 | 0.010361 | 0.045978 | 0.016203 | 0.018902 | 0.021712 | 0.027587 | 0.013876 | 0.022819 | 0.022503 | 0.036837 | 0.004829 | 0.01315 | 0.04203 | 0.00685 | 0.029547 | 0.00261 | 0.002384 | 0.002718 | 0.020217 | 0.002652 | 0.008168 |
| 30.3333333 | 0.034035 | 0.008514 | 0.031934 | 0.022163 | 0.017006 | 0.016935 | 0.038673 | 0.011656 | 0.033046 | 0.010485 | 0.030172 | 0.007272 | 0.003476 | 0.033307 | 0.007932 | 0.031948 | 0.003822 | 0.000633 | 0.001238 | 0.012171 | 0.000966 | 0.013736 |
| 30.3472222 | 0.016645 | 0.004254 | 0.018646 | 0.018389 | 0.019271 | 0.016215 | 0.040461 | 0.024318 | 0.03703 | 0.010794 | 0.014102 | 0.016788 | 0.005313 | 0.026156 | 0.01165 | 0.034091 | 0.007262 | 0.000369 | 0.001231 | 0.012221 | 0.000791 | 0.017574 |
| 30.3611111 | 0.010497 | 0.001743 | 0.025413 | 0.006781 | 0.025485 | 0.021936 | 0.033708 | 0.045589 | 0.034078 | 0.012535 | 0.005059 | 0.027533 | 0.019299 | 0.023665 | 0.015907 | 0.030061 | 0.009544 | 0.000792 | 0.001275 | 0.017625 | 0.001044 | 0.012316 |
| 30.375 | 0.024194 | 0.0016 | 0.022852 | 0.004479 | 0.024806 | 0.028644 | 0.020279 | 0.059346 | 0.026252 | 0.016326 | 0.011099 | 0.035026 | 0.034553 | 0.022635 | 0.019838 | 0.023533 | 0.009759 | 0.002162 | 0.001777 | 0.024168 | 0.002119 | 0.012283 |
| 30.3888889 | 0.047061 | 0.004785 | 0.017021 | 0.005869 | 0.018291 | 0.03223 | 0.012385 | 0.055346 | 0.020863 | 0.026525 | 0.030355 | 0.04053 | 0.039805 | 0.022304 | 0.024118 | 0.020615 | 0.011137 | 0.003437 | 0.002721 | 0.030308 | 0.003446 | 0.026237 |
| 30.4027778 | 0.054805 | 0.010933 | 0.03705 | 0.00487 | 0.016659 | 0.033488 | 0.017831 | 0.040551 | 0.024749 | 0.03763 | 0.046355 | 0.047237 | 0.038369 | 0.022997 | 0.027875 | 0.022578 | 0.01574 | 0.003236 | 0.002916 | 0.035095 | 0.003378 | 0.044079 |
| 30.4166667 | 0.045046 | 0.015844 | 0.059691 | 0.004385 | 0.024253 | 0.033009 | 0.029018 | 0.032261 | 0.034997 | 0.043692 | 0.046005 | 0.054254 | 0.041429 | 0.022669 | 0.028467 | 0.026199 | 0.022597 | 0.003062 | 0.003186 | 0.037859 | 0.003147 | 0.055091 |
| 30.4305556 | 0.03769 | 0.017127 | 0.059849 | 0.004647 | 0.028264 | 0.03141 | 0.035148 | 0.037787 | 0.04363 | 0.040246 | 0.037504 | 0.057844 | 0.054649 | 0.023039 | 0.025725 | 0.027947 | 0.029055 | 0.004472 | 0.004548 | 0.040777 | 0.004631 | 0.055607 |
| 30.4444444 | 0.040206 | 0.016819 | 0.044818 | 0.005782 | 0.020447 | 0.030493 | 0.032429 | 0.053062 | 0.044353 | 0.031626 | 0.035512 | 0.055426 | 0.073309 | 0.031869 | 0.022751 | 0.026967 | 0.033099 | 0.006692 | 0.006243 | 0.047253 | 0.007288 | 0.048663 |
| 30.4583333 | 0.049942 | 0.016935 | 0.025839 | 0.014597 | 0.011584 | 0.032593 | 0.025986 | 0.078794 | 0.034014 | 0.026034 | 0.045607 | 0.049461 | 0.084165 | 0.047344 | 0.021211 | 0.022692 | 0.033968 | 0.010196 | 0.008469 | 0.05171 | 0.010829 | 0.041177 |
| 30.4722222 | 0.062018 | 0.018889 | 0.01173 | 0.026867 | 0.014806 | 0.040545 | 0.024251 | 0.109375 | 0.018272 | 0.024194 | 0.06231 | 0.045728 | 0.077316 | 0.05591 | 0.0217 | 0.016331 | 0.032854 | 0.014378 | 0.010936 | 0.045138 | 0.013752 | 0.034905 |
| 30.4861111 | 0.066684 | 0.024126 | 0.011619 | 0.025706 | 0.029734 | 0.055439 | 0.031638 | 0.132712 | 0.011444 | 0.026952 | 0.077132 | 0.046009 | 0.061333 | 0.049303 | 0.025853 | 0.011891 | 0.032228 | 0.014883 | 0.011239 | 0.032026 | 0.013167 | 0.027202 |
| 30.5 | 0.059135 | 0.030779 | 0.023682 | 0.014658 | 0.03697 | 0.070669 | 0.045428 | 0.133362 | 0.0219 | 0.039329 | 0.084386 | 0.049178 | 0.050391 | 0.035147 | 0.031615 | 0.011548 | 0.032513 | 0.011381 | 0.009138 | 0.022826 | 0.010149 | 0.018868 |
| 30.5138889 | 0.049088 | 0.034991 | 0.032705 | 0.011319 | 0.029986 | 0.072159 | 0.058538 | 0.100781 | 0.041198 | 0.054275 | 0.078227 | 0.05491 | 0.043747 | 0.029683 | 0.033909 | 0.013176 | 0.030924 | 0.008022 | 0.007088 | 0.020923 | 0.007595 | 0.015365 |
| 30.5277778 | 0.043297 | 0.034222 | 0.027335 | 0.012072 | 0.033841 | 0.059326 | 0.063817 | 0.058994 | 0.050107 | 0.059698 | 0.06407 | 0.061707 | 0.039397 | 0.034339 | 0.031622 | 0.011724 | 0.024919 | 0.008306 | 0.008391 | 0.022763 | 0.007737 | 0.016929 |
| 30.5416667 | 0.035291 | 0.029268 | 0.021881 | 0.008418 | 0.049301 | 0.046218 | 0.057163 | 0.031452 | 0.041634 | 0.056147 | 0.058743 | 0.06519 | 0.038205 | 0.028861 | 0.021802 | 0.011174 | 0.015847 | 0.016087 | 0.017252 | 0.021129 | 0.014254 | 0.023266 |
| 30.5555556 | 0.022091 | 0.023805 | 0.025755 | 0.004641 | 0.049853 | 0.036967 | 0.040596 | 0.01668 | 0.029092 | 0.049105 | 0.060537 | 0.056906 | 0.032195 | 0.012754 | 0.013438 | 0.022184 | 0.013566 | 0.030613 | 0.032142 | 0.012853 | 0.027188 | 0.023462 |
| 30.5694444 | 0.0178 | 0.019046 | 0.020845 | 0.005913 | 0.04076 | 0.02772 | 0.022645 | 0.00906 | 0.023197 | 0.043083 | 0.05379 | 0.035741 | 0.022004 | 0.010439 | 0.018598 | 0.033425 | 0.020232 | 0.040157 | 0.040676 | 0.004238 | 0.036742 | 0.013436 |
| 30.5833333 | 0.029953 | 0.017398 | 0.015473 | 0.012939 | 0.039618 | 0.017133 | 0.01213 | 0.007947 | 0.024005 | 0.042965 | 0.043013 | 0.0188 | 0.019228 | 0.021306 | 0.022888 | 0.027394 | 0.021304 | 0.034583 | 0.03388 | 0.006152 | 0.033046 | 0.008271 |
| 30.5972222 | 0.039052 | 0.021157 | 0.02542 | 0.013356 | 0.036079 | 0.007953 | 0.009113 | 0.010982 | 0.022137 | 0.040102 | 0.033898 | 0.017423 | 0.020974 | 0.024466 | 0.019855 | 0.014231 | 0.013743 | 0.019502 | 0.018654 | 0.017236 | 0.019505 | 0.007342 |
| 30.6111111 | 0.032033 | 0.023967 | 0.031044 | 0.006641 | 0.024183 | 0.004804 | 0.008908 | 0.009574 | 0.012098 | 0.023864 | 0.019251 | 0.022021 | 0.025674 | 0.015777 | 0.017627 | 0.007496 | 0.006222 | 0.007117 | 0.00663 | 0.021861 | 0.007368 | 0.009135 |
| 30.625 | 0.021429 | 0.020463 | 0.021975 | 0.006381 | 0.01981 | 0.005192 | 0.008782 | 0.005811 | 0.008872 | 0.009972 | 0.013668 | 0.024758 | 0.034267 | 0.013642 | 0.015802 | 0.004541 | 0.002637 | 0.001806 | 0.001618 | 0.0159 | 0.001855 | 0.010064 |
| 30.6388889 | 0.016326 | 0.014157 | 0.012876 | 0.007039 | 0.01971 | 0.011797 | 0.007569 | 0.005346 | 0.013293 | 0.010117 | 0.03281 | 0.024896 | 0.032895 | 0.023386 | 0.014126 | 0.002288 | 0.001762 | 0.001266 | 0.00134 | 0.01836 | 0.001172 | 0.006146 |
| 30.6527778 | 0.017858 | 0.009957 | 0.012314 | 0.003986 | 0.01113 | 0.023766 | 0.006623 | 0.004741 | 0.013301 | 0.014744 | 0.050761 | 0.01888 | 0.022089 | 0.024385 | 0.012868 | 0.003559 | 0.001089 | 0.002613 | 0.002439 | 0.035217 | 0.002474 | 0.005232 |
| 30.6666667 | 0.034383 | 0.008405 | 0.013143 | 0.002773 | 0.003214 | 0.02463 | 0.007762 | 0.003142 | 0.015991 | 0.026039 | 0.039357 | 0.009566 | 0.0157 | 0.011751 | 0.010625 | 0.010197 | 0.000788 | 0.002868 | 0.0023 | 0.034905 | 0.00275 | 0.006525 |
| 30.6805556 | 0.056231 | 0.00638 | 0.009611 | 0.002263 | 0.00776 | 0.015508 | 0.010358 | 0.002205 | 0.019707 | 0.035505 | 0.023142 | 0.00444 | 0.035428 | 0.004037 | 0.00934 | 0.01878 | 0.000662 | 0.001529 | 0.001078 | 0.025459 | 0.00147 | 0.008943 |
| 30.6944444 | 0.059441 | 0.003086 | 0.007115 | 0.002331 | 0.015035 | 0.007326 | 0.010921 | 0.002464 | 0.01333 | 0.027361 | 0.034665 | 0.002683 | 0.063897 | 0.006249 | 0.014205 | 0.023645 | 0.000405 | 0.000467 | 0.000358 | 0.039659 | 0.000468 | 0.020328 |

| | | | | | | | | | | | | | | | | | | | | | | |
|---|---|---|---|---|---|---|---|---|---|---|---|---|---|---|---|---|---|---|---|---|---|---|
| 30.7083333 | 0.037638 | 0.00093 | 0.013557 | 0.006249 | 0.018163 | 0.003239 | 0.007532 | 0.003789 | 0.004955 | 0.016534 | 0.051405 | 0.005539 | 0.05205 | 0.009904 | 0.025008 | 0.020328 | 0.000411 | 0.000219 | 0.000224 | 0.04508 | 0.000145 | 0.030983 |
| 30.7222222 | 0.019865 | 0.000531 | 0.019498 | 0.01368 | 0.032544 | 0.004034 | 0.003735 | 0.008047 | 0.007466 | 0.017688 | 0.043547 | 0.010277 | 0.020416 | 0.010547 | 0.031661 | 0.012321 | 0.000515 | 0.000386 | 0.000362 | 0.02469 | 0.000242 | 0.022124 |
| 30.7361111 | 0.032324 | 0.00078 | 0.013298 | 0.020003 | 0.048055 | 0.00464 | 0.002573 | 0.015344 | 0.014581 | 0.019554 | 0.023278 | 0.007733 | 0.01849 | 0.007828 | 0.026823 | 0.01143 | 0.00084 | 0.000848 | 0.000858 | 0.015457 | 0.000791 | 0.008907 |
| 30.75 | 0.056289 | 0.000772 | 0.008783 | 0.021032 | 0.037912 | 0.002773 | 0.004134 | 0.015569 | 0.011551 | 0.013042 | 0.009946 | 0.003286 | 0.037616 | 0.004747 | 0.016646 | 0.019685 | 0.001778 | 0.001568 | 0.001487 | 0.036417 | 0.001503 | 0.009667 |
| 30.7638889 | 0.064378 | 0.00093 | 0.01888 | 0.017064 | 0.014659 | 0.001758 | 0.007269 | 0.010403 | 0.007192 | 0.010778 | 0.010287 | 0.005559 | 0.032953 | 0.004355 | 0.01007 | 0.025666 | 0.002992 | 0.002196 | 0.002068 | 0.067639 | 0.002161 | 0.013834 |
| 30.7777778 | 0.046114 | 0.002459 | 0.031428 | 0.010736 | 0.00706 | 0.002417 | 0.008737 | 0.011709 | 0.01264 | 0.018562 | 0.023577 | 0.015884 | 0.011256 | 0.009076 | 0.008396 | 0.021369 | 0.004104 | 0.002672 | 0.002754 | 0.072368 | 0.002879 | 0.011745 |
| 30.7916667 | 0.02264 | 0.005243 | 0.025614 | 0.006094 | 0.016338 | 0.002291 | 0.006577 | 0.014302 | 0.020718 | 0.025257 | 0.039995 | 0.029356 | 0.014285 | 0.014743 | 0.009289 | 0.012182 | 0.004813 | 0.002966 | 0.00323 | 0.044675 | 0.003371 | 0.00711 |
| 30.8055556 | 0.026431 | 0.00829 | 0.012986 | 0.004727 | 0.02948 | 0.001251 | 0.00611 | 0.012324 | 0.019835 | 0.019508 | 0.052254 | 0.03788 | 0.028925 | 0.017813 | 0.010942 | 0.011105 | 0.004622 | 0.002684 | 0.002933 | 0.028673 | 0.003076 | 0.00825 |
| 30.8194444 | 0.045737 | 0.010589 | 0.019306 | 0.00387 | 0.040412 | 0.001505 | 0.008844 | 0.008914 | 0.012324 | 0.009672 | 0.055153 | 0.03847 | 0.024011 | 0.018034 | 0.011929 | 0.019229 | 0.004039 | 0.002086 | 0.002253 | 0.047126 | 0.002324 | 0.01713 |
| 30.8333333 | 0.046405 | 0.011191 | 0.046021 | 0.002681 | 0.041491 | 0.004429 | 0.009519 | 0.007653 | 0.012815 | 0.010546 | 0.044555 | 0.03266 | 0.012407 | 0.013138 | 0.011185 | 0.024479 | 0.004199 | 0.002152 | 0.00225 | 0.060994 | 0.002302 | 0.027274 |
| 30.8472222 | 0.028799 | 0.010777 | 0.071813 | 0.001757 | 0.027647 | 0.010205 | 0.010486 | 0.008106 | 0.015575 | 0.01642 | 0.025986 | 0.02298 | 0.010439 | 0.006687 | 0.007786 | 0.022712 | 0.00531 | 0.003332 | 0.003282 | 0.04356 | 0.003554 | 0.034105 |
| 30.8611111 | 0.011995 | 0.011195 | 0.068279 | 0.001566 | 0.020243 | 0.018507 | 0.010763 | 0.005966 | 0.016828 | 0.017434 | 0.009872 | 0.014089 | 0.01352 | 0.004167 | 0.003473 | 0.020003 | 0.006251 | 0.005103 | 0.004789 | 0.022972 | 0.005455 | 0.036023 |
| 30.875 | 0.004017 | 0.012853 | 0.045073 | 0.001873 | 0.031627 | 0.02774 | 0.010934 | 0.004164 | 0.019682 | 0.01466 | 0.005512 | 0.015665 | 0.019702 | 0.010217 | 0.002514 | 0.020564 | 0.005888 | 0.006774 | 0.006232 | 0.02863 | 0.007212 | 0.034493 |
| 30.8888889 | 0.006139 | 0.01573 | 0.029386 | 0.002145 | 0.041105 | 0.033991 | 0.011826 | 0.00779 | 0.015533 | 0.013809 | 0.009812 | 0.022652 | 0.02927 | 0.025565 | 0.00449 | 0.023851 | 0.004423 | 0.008073 | 0.00745 | 0.051133 | 0.008543 | 0.032729 |
| 30.9027778 | 0.014455 | 0.020517 | 0.025067 | 0.002301 | 0.038875 | 0.033612 | 0.008538 | 0.017279 | 0.00614 | 0.016902 | 0.01507 | 0.019262 | 0.040615 | 0.043692 | 0.004613 | 0.029308 | 0.002758 | 0.008791 | 0.008227 | 0.061004 | 0.009229 | 0.030858 |
| 30.9166667 | 0.02811 | 0.025579 | 0.024959 | 0.001901 | 0.030137 | 0.029198 | 0.00993 | 0.028929 | 0.003539 | 0.021812 | 0.028646 | 0.015021 | 0.050904 | 0.056771 | 0.004812 | 0.037048 | 0.001446 | 0.008677 | 0.008363 | 0.040273 | 0.009064 | 0.028889 |
| 30.9305556 | 0.051567 | 0.028515 | 0.032001 | 0.001784 | 0.02035 | 0.028239 | 0.016467 | 0.041842 | 0.009335 | 0.026675 | 0.047579 | 0.024696 | 0.05431 | 0.057436 | 0.009555 | 0.042918 | 0.001362 | 0.007948 | 0.007952 | 0.01784 | 0.008378 | 0.027623 |
| 30.9444444 | 0.083379 | 0.030267 | 0.052845 | 0.002575 | 0.012066 | 0.033466 | 0.014137 | 0.063732 | 0.014471 | 0.035542 | 0.05804 | 0.046033 | 0.051043 | 0.046263 | 0.014794 | 0.041672 | 0.003187 | 0.007048 | 0.007155 | 0.019794 | 0.007567 | 0.025157 |
| 30.9583333 | 0.109122 | 0.032248 | 0.069128 | 0.003238 | 0.00737 | 0.040971 | 0.009906 | 0.092996 | 0.020034 | 0.049118 | 0.0708 | 0.076795 | 0.047362 | 0.031936 | 0.017481 | 0.03547 | 0.006441 | 0.006017 | 0.006023 | 0.020633 | 0.006466 | 0.020302 |
| 30.9722222 | 0.111846 | 0.0338 | 0.058688 | 0.002479 | 0.01026 | 0.04434 | 0.02089 | 0.102346 | 0.029815 | 0.059111 | 0.089234 | 0.103899 | 0.045144 | 0.019583 | 0.019782 | 0.028843 | 0.010387 | 0.004643 | 0.004596 | 0.011572 | 0.004878 | 0.015399 |
| 30.9861111 | 0.093667 | 0.033754 | 0.037594 | 0.001161 | 0.024479 | 0.041786 | 0.040692 | 0.07852 | 0.036392 | 0.0624 | 0.090505 | 0.109731 | 0.043242 | 0.010603 | 0.02186 | 0.020515 | 0.01402 | 0.0031 | 0.003078 | 0.010311 | 0.00319 | 0.012691 |
| 31 | 0.066854 | 0.032044 | 0.03259 | 0.000662 | 0.039304 | 0.036483 | 0.052443 | 0.05198 | 0.037304 | 0.05831 | 0.068974 | 0.099418 | 0.039402 | 0.005297 | 0.020656 | 0.010326 | 0.015862 | 0.002749 | 0.002511 | 0.016295 | 0.002684 | 0.011649 |
| 31.0138889 | 0.037629 | 0.028402 | 0.029621 | 0.002324 | 0.037585 | 0.028621 | 0.050844 | 0.046299 | 0.034483 | 0.04344 | 0.037827 | 0.082816 | 0.032064 | 0.00246 | 0.014817 | 0.003744 | 0.014457 | 0.005504 | 0.004484 | 0.019521 | 0.004985 | 0.009179 |
| 31.0277778 | 0.016481 | 0.023833 | 0.022868 | 0.006277 | 0.029615 | 0.019785 | 0.049407 | 0.05106 | 0.03032 | 0.025173 | 0.013169 | 0.061098 | 0.024734 | 0.001201 | 0.009948 | 0.002026 | 0.011004 | 0.010219 | 0.007898 | 0.018683 | 0.009088 | 0.004924 |
| 31.0416667 | 0.018215 | 0.021826 | 0.031737 | 0.007097 | 0.040951 | 0.019995 | 0.055789 | 0.059243 | 0.034196 | 0.019824 | 0.006831 | 0.035206 | 0.022579 | 0.003693 | 0.011886 | 0.003458 | 0.009993 | 0.011947 | 0.008896 | 0.019047 | 0.010603 | 0.001853 |
| 31.0555556 | 0.04198 | 0.023468 | 0.039649 | 0.006901 | 0.063917 | 0.042196 | 0.059313 | 0.084979 | 0.052296 | 0.034897 | 0.018786 | 0.012998 | 0.026952 | 0.011642 | 0.018759 | 0.010035 | 0.013512 | 0.009485 | 0.007423 | 0.025356 | 0.008317 | 0.003186 |
| 31.0694444 | 0.06329 | 0.029338 | 0.027588 | 0.019735 | 0.06779 | 0.08611 | 0.055769 | 0.130236 | 0.077472 | 0.057569 | 0.042347 | 0.006542 | 0.045248 | 0.020227 | 0.026833 | 0.019918 | 0.021867 | 0.007614 | 0.007253 | 0.042626 | 0.006565 | 0.011265 |
| 31.0833333 | 0.05659 | 0.043087 | 0.016554 | 0.03985 | 0.055182 | 0.113808 | 0.039164 | 0.162323 | 0.096244 | 0.066103 | 0.058004 | 0.012458 | 0.06566 | 0.023585 | 0.033529 | 0.026864 | 0.036623 | 0.007344 | 0.007935 | 0.063045 | 0.006538 | 0.022448 |
| 31.0972222 | 0.030123 | 0.059406 | 0.027782 | 0.044827 | 0.041194 | 0.095426 | 0.019033 | 0.147698 | 0.095636 | 0.053911 | 0.049951 | 0.017392 | 0.052328 | 0.019753 | 0.032286 | 0.025689 | 0.049411 | 0.005047 | 0.005739 | 0.063383 | 0.004778 | 0.028099 |
| 31.1111111 | 0.012937 | 0.064033 | 0.048992 | 0.032468 | 0.028354 | 0.056741 | 0.020754 | 0.097698 | 0.07151 | 0.030359 | 0.027619 | 0.018782 | 0.02738 | 0.011342 | 0.020947 | 0.016748 | 0.047099 | 0.002498 | 0.002791 | 0.040747 | 0.002714 | 0.024022 |
| 31.125 | 0.008759 | 0.056124 | 0.053262 | 0.016728 | 0.017781 | 0.02629 | 0.036546 | 0.050385 | 0.038438 | 0.012737 | 0.012462 | 0.018358 | 0.032951 | 0.004601 | 0.009083 | 0.00854 | 0.03077 | 0.002651 | 0.002659 | 0.018425 | 0.002986 | 0.01448 |
| 31.1388889 | 0.006443 | 0.045281 | 0.034764 | 0.006627 | 0.011158 | 0.017672 | 0.038401 | 0.020746 | 0.014771 | 0.011861 | 0.013631 | 0.014368 | 0.045693 | 0.0017 | 0.006451 | 0.008203 | 0.015916 | 0.00236 | 0.002425 | 0.012573 | 0.00255 | 0.006478 |
| 31.1527778 | 0.009594 | 0.035727 | 0.013932 | 0.00708 | 0.014056 | 0.03206 | 0.025304 | 0.010599 | 0.009331 | 0.01935 | 0.018998 | 0.009729 | 0.041136 | 0.001709 | 0.012622 | 0.01181 | 0.016319 | 0.001165 | 0.001225 | 0.017477 | 0.001263 | 0.002711 |
| 31.1666667 | 0.023925 | 0.027226 | 0.007636 | 0.022298 | 0.022304 | 0.050661 | 0.023501 | 0.020616 | 0.020846 | 0.024217 | 0.024047 | 0.009677 | 0.02892 | 0.006682 | 0.02002 | 0.013767 | 0.02374 | 0.001941 | 0.00191 | 0.023358 | 0.002105 | 0.004396 |
| 31.1805556 | 0.042337 | 0.018969 | 0.011742 | 0.040773 | 0.029736 | 0.059743 | 0.043341 | 0.037303 | 0.037738 | 0.02507 | 0.04188 | 0.015868 | 0.018738 | 0.018236 | 0.023316 | 0.016979 | 0.023043 | 0.004995 | 0.004908 | 0.028781 | 0.005344 | 0.014699 |
| 31.1944444 | 0.04904 | 0.010192 | 0.014606 | 0.03843 | 0.041571 | 0.054192 | 0.070416 | 0.048483 | 0.047677 | 0.026057 | 0.066003 | 0.023722 | 0.011153 | 0.031544 | 0.021589 | 0.025121 | 0.014333 | 0.009394 | 0.009136 | 0.032446 | 0.009875 | 0.029742 |
| 31.2083333 | 0.036658 | 0.007657 | 0.01109 | 0.022727 | 0.048258 | 0.036453 | 0.075634 | 0.048392 | 0.043686 | 0.027336 | 0.075019 | 0.024738 | 0.006613 | 0.036617 | 0.015091 | 0.034624 | 0.006959 | 0.012618 | 0.012134 | 0.030724 | 0.012971 | 0.041515 |
| 31.2222222 | 0.020113 | 0.020412 | 0.008023 | 0.021513 | 0.038358 | 0.017091 | 0.044541 | 0.038964 | 0.026672 | 0.023047 | 0.060126 | 0.015879 | 0.008431 | 0.029017 | 0.008486 | 0.039356 | 0.006822 | 0.012229 | 0.01161 | 0.024375 | 0.012237 | 0.048496 |
| 31.2361111 | 0.020268 | 0.041644 | 0.010275 | 0.02187 | 0.026888 | 0.006262 | 0.015834 | 0.032603 | 0.008907 | 0.012821 | 0.034519 | 0.006667 | 0.011015 | 0.015958 | 0.009452 | 0.038524 | 0.009157 | 0.008588 | 0.007944 | 0.020424 | 0.008258 | 0.047946 |
| 31.25 | 0.035065 | 0.060858 | 0.027198 | 0.020249 | 0.023837 | 0.00565 | 0.017525 | 0.034639 | 0.003287 | 0.003648 | 0.02318 | 0.005254 | 0.007637 | 0.009393 | 0.0168 | 0.034791 | 0.009481 | 0.004368 | 0.003901 | 0.02274 | 0.00417 | 0.03207 |
| 31.2638889 | 0.039365 | 0.077317 | 0.062164 | 0.029975 | 0.026859 | 0.006299 | 0.030233 | 0.038843 | 0.009444 | 0.003401 | 0.022509 | 0.005389 | 0.005316 | 0.008301 | 0.021498 | 0.030958 | 0.009784 | 0.002949 | 0.002505 | 0.02745 | 0.003392 | 0.010031 |
| 31.2777778 | 0.026338 | 0.084766 | 0.08272 | 0.026006 | 0.031528 | 0.010268 | 0.030696 | 0.035131 | 0.018963 | 0.01175 | 0.019401 | 0.003363 | 0.01209 | 0.009847 | 0.019174 | 0.029387 | 0.010786 | 0.003994 | 0.003092 | 0.030179 | 0.004832 | 0.011953 |
| 31.2916667 | 0.015917 | 0.075076 | 0.06566 | 0.010665 | 0.033352 | 0.02525 | 0.019838 | 0.021493 | 0.02438 | 0.02167 | 0.023252 | 0.003706 | 0.02404 | 0.02112 | 0.011313 | 0.031282 | 0.009466 | 0.003791 | 0.002974 | 0.032917 | 0.004403 | 0.047832 |
| 31.3055556 | 0.011269 | 0.057256 | 0.038271 | 0.006949 | 0.029567 | 0.035683 | 0.018478 | 0.011435 | 0.02367 | 0.02865 | 0.037894 | 0.012862 | 0.033406 | 0.035799 | 0.00655 | 0.036931 | 0.007025 | 0.004344 | 0.004185 | 0.039811 | 0.004595 | 0.080272 |
| 31.3194444 | 0.007205 | 0.041994 | 0.034406 | 0.010376 | 0.01971 | 0.028454 | 0.024536 | 0.00899 | 0.023231 | 0.028567 | 0.04045 | 0.032641 | 0.035072 | 0.045364 | 0.012163 | 0.043051 | 0.009806 | 0.009082 | 0.008847 | 0.046561 | 0.009344 | 0.069636 |
| 31.3333333 | 0.00924 | 0.028476 | 0.036528 | 0.014768 | 0.02136 | 0.015837 | 0.020522 | 0.013125 | 0.026319 | 0.02385 | 0.022415 | 0.0518 | 0.029454 | 0.050478 | 0.019223 | 0.0444 | 0.016261 | 0.012986 | 0.012403 | 0.046214 | 0.013396 | 0.039233 |
| 31.3472222 | 0.01711 | 0.016457 | 0.02897 | 0.01428 | 0.037237 | 0.010711 | 0.009664 | 0.036432 | 0.026029 | 0.027714 | 0.007394 | 0.059915 | 0.024658 | 0.05122 | 0.018611 | 0.040842 | 0.019772 | 0.011776 | 0.011315 | 0.039314 | 0.012231 | 0.033326 |
| 31.3611111 | 0.026895 | 0.008828 | 0.030156 | 0.008067 | 0.043809 | 0.016319 | 0.002698 | 0.062761 | 0.01979 | 0.044878 | 0.00367 | 0.060271 | 0.029165 | 0.04808 | 0.012846 | 0.036825 | 0.019361 | 0.007466 | 0.00735 | 0.03012 | 0.008116 | 0.039563 |
| 31.375 | 0.034803 | 0.004839 | 0.030873 | 0.00666 | 0.033325 | 0.029228 | 0.00426 | 0.063817 | 0.014718 | 0.051859 | 0.006421 | 0.055897 | 0.040792 | 0.039586 | 0.007385 | 0.034375 | 0.016386 | 0.004073 | 0.00392 | 0.021706 | 0.004391 | 0.027468 |
| 31.3888889 | 0.040663 | 0.00249 | 0.024041 | 0.015367 | 0.020937 | 0.03968 | 0.015389 | 0.04907 | 0.016089 | 0.03456 | 0.013917 | 0.045213 | 0.047085 | 0.026883 | 0.005236 | 0.031427 | 0.011814 | 0.005327 | 0.004418 | 0.01632 | 0.004653 | 0.0141 |
| 31.4027778 | 0.049686 | 0.002308 | 0.021527 | 0.028506 | 0.015203 | 0.041341 | 0.027975 | 0.03581 | 0.020581 | 0.01599 | 0.018331 | 0.032318 | 0.043593 | 0.017179 | 0.00593 | 0.026502 | 0.007492 | 0.010384 | 0.008839 | 0.016192 | 0.009627 | 0.021773 |
| 31.4166667 | 0.060057 | 0.00338 | 0.015404 | 0.030013 | 0.011828 | 0.037243 | 0.032387 | 0.031965 | 0.018371 | 0.010544 | 0.019324 | 0.023952 | 0.038718 | 0.017806 | 0.009584 | 0.021601 | 0.005402 | 0.014636 | 0.013221 | 0.021344 | 0.014491 | 0.033733 |
| 31.4305556 | 0.065517 | 0.003458 | 0.007793 | 0.0241 | 0.011114 | 0.033764 | 0.031621 | 0.048818 | 0.015879 | 0.015377 | 0.022642 | 0.020804 | 0.035496 | 0.03049 | 0.018018 | 0.01963 | 0.007234 | 0.015545 | 0.014549 | 0.027991 | 0.01545 | 0.035889 |
| 31.4444444 | 0.068191 | 0.002558 | 0.018491 | 0.027146 | 0.013342 | 0.03435 | 0.030403 | 0.080417 | 0.029663 | 0.023889 | 0.029284 | 0.018076 | 0.030046 | 0.047114 | 0.028971 | 0.020626 | 0.013542 | 0.014333 | 0.013198 | 0.031644 | 0.014058 | 0.03334 |
| 31.4583333 | 0.073407 | 0.00198 | 0.04844 | 0.022757 | 0.017531 | 0.040037 | 0.025616 | 0.08999 | 0.047449 | 0.026467 | 0.041325 | 0.013422 | 0.026079 | 0.055815 | 0.036993 | 0.022019 | 0.024037 | 0.013575 | 0.011235 | 0.032965 | 0.013171 | 0.031784 |
| 31.4722222 | 0.080374 | 0.00227 | 0.074939 | 0.010526 | 0.017461 | 0.050893 | 0.018455 | 0.069207 | 0.055174 | 0.022164 | 0.060665 | 0.011341 | 0.036699 | 0.053833 | 0.039695 | 0.022735 | 0.03721 | 0.012213 | 0.00876 | 0.035578 | 0.011117 | 0.033545 |
| 31.4861111 | 0.08522 | 0.006551 | 0.083361 | 0.007453 | 0.014221 | 0.065845 | 0.016384 | 0.057611 | 0.059523 | 0.018198 | 0.081203 | 0.017993 | 0.067159 | 0.047354 | 0.037461 | 0.024879 | 0.047981 | 0.007339 | 0.004856 | 0.041188 | 0.006249 | 0.032516 |
| 31.5 | 0.084997 | 0.015783 | 0.074623 | 0.006399 | 0.025831 | 0.076736 | 0.020387 | 0.068335 | 0.054621 | 0.022674 | 0.094286 | 0.030561 | 0.100446 | 0.045531 | 0.033475 | 0.029475 | 0.052611 | 0.002688 | 0.002289 | 0.049322 | 0.002968 | 0.022518 |
| 31.5138889 | 0.080877 | 0.023636 | 0.054851 | 0.003059 | 0.051516 | 0.070926 | 0.025453 | 0.085581 | 0.036261 | 0.039057 | 0.090309 | 0.039194 | 0.110042 | 0.052491 | 0.032004 | 0.032279 | 0.052089 | 0.002496 | 0.003077 | 0.057858 | 0.003378 | 0.013112 |
| 31.5277778 | 0.076089 | 0.027878 | 0.03753 | 0.001674 | 0.072111 | 0.050928 | 0.029708 | 0.101607 | 0.028766 | 0.060642 | 0.067396 | 0.041407 | 0.087341 | 0.060249 | 0.031377 | 0.027228 | 0.047709 | 0.003363 | 0.00394 | 0.063926 | 0.003333 | 0.011383 |
| 31.5416667 | 0.073782 | 0.027929 | 0.027901 | 0.001399 | 0.074717 | 0.03316 | 0.033237 | 0.09784 | 0.043232 | 0.075508 | 0.046194 | 0.044543 | 0.05009 | 0.056185 | 0.023137 | 0.014991 | 0.037781 | 0.007279 | 0.007953 | 0.061341 | 0.005065 | 0.017397 |
| 31.5555556 | 0.077123 | 0.020124 | 0.019802 | 0.002369 | 0.055759 | 0.025035 | 0.031854 | 0.06068 | 0.046242 | 0.075953 | 0.040432 | 0.050259 | 0.023207 | 0.03852 | 0.012453 | 0.004677 | 0.021452 | 0.018007 | 0.018932 | 0.047125 | 0.013735 | 0.026631 |
| 31.5694444 | 0.077195 | 0.011563 | 0.010233 | 0.006234 | 0.038499 | 0.02106 | 0.02495 | 0.027006 | 0.029591 | 0.064928 | 0.045688 | 0.055322 | 0.017396 | 0.020836 | 0.010357 | 0.00558 | 0.007545 | 0.024286 | 0.024678 | 0.029282 | 0.020403 | 0.024764 |
| 31.5833333 | 0.060721 | 0.008434 | 0.008332 | 0.014853 | 0.041294 | 0.013698 | 0.017032 | 0.022958 | 0.024068 | 0.056123 | 0.060687 | 0.063165 | 0.028867 | 0.017178 | 0.008231 | 0.014267 | 0.00342 | 0.018432 | 0.017855 | 0.019042 | 0.016744 | 0.016595 |
| 31.5972222 | 0.032797 | 0.005866 | 0.019708 | 0.023385 | 0.042069 | 0.011524 | 0.00936 | 0.022722 | 0.021122 | 0.047714 | 0.069026 | 0.067819 | 0.038769 | 0.022647 | 0.00274 | 0.016904 | 0.004092 | 0.00968 | 0.008762 | 0.01747 | 0.009393 | 0.013879 |
| 31.6111111 | 0.012284 | 0.003008 | 0.028237 | 0.02064 | 0.03285 | 0.027841 | 0.004877 | 0.019572 | 0.017592 | 0.037158 | 0.047498 | 0.058653 | 0.028137 | 0.018783 | 0.001282 | 0.011158 | 0.004897 | 0.006226 | 0.005365 | 0.013738 | 0.006008 | 0.012601 |
| 31.625 | 0.009609 | 0.002339 | 0.021081 | 0.01575 | 0.022163 | 0.038872 | 0.005392 | 0.022533 | 0.024605 | 0.034591 | 0.025575 | 0.040079 | 0.017367 | 0.011331 | 0.003197 | 0.010621 | 0.005061 | 0.006456 | 0.005528 | 0.008304 | 0.0061 | 0.009897 |
| 31.6388889 | 0.011397 | 0.004977 | 0.016271 | 0.015909 | 0.010484 | 0.030366 | 0.00624 | 0.022693 | 0.023499 | 0.033269 | 0.040397 | 0.020737 | 0.035223 | 0.015359 | 0.006173 | 0.01796 | 0.004383 | 0.006534 | 0.005588 | 0.013344 | 0.006178 | 0.00727 |
| 31.6527778 | 0.020074 | 0.00888 | 0.028523 | 0.010527 | 0.008509 | 0.030385 | 0.005101 | 0.022336 | 0.021623 | 0.02463 | 0.077397 | 0.007365 | 0.051309 | 0.016058 | 0.01059 | 0.020125 | 0.00381 | 0.00592 | 0.005248 | 0.023523 | 0.005649 | 0.004838 |
| 31.6666667 | 0.032906 | 0.012197 | 0.045478 | 0.003816 | 0.010065 | 0.038062 | 0.003597 | 0.023876 | 0.023931 | 0.023841 | 0.098221 | 0.002305 | 0.039483 | 0.009752 | 0.016979 | 0.018899 | 0.003404 | 0.005388 | 0.004993 | 0.02334 | 0.005444 | 0.002456 |
| 31.6805556 | 0.030843 | 0.012135 | 0.058286 | 0.004983 | 0.011928 | 0.031841 | 0.002406 | 0.014722 | 0.014587 | 0.02661 | 0.075354 | 0.001893 | 0.032067 | 0.007282 | 0.023675 | 0.024777 | 0.002436 | 0.004766 | 0.004472 | 0.013022 | 0.005226 | 0.001469 |
| 31.6944444 | 0.02334 | 0.009533 | 0.062077 | 0.009496 | 0.028255 | 0.017447 | 0.002295 | 0.004487 | 0.005084 | 0.019682 | 0.040398 | 0.004366 | 0.033245 | 0.007584 | 0.027584 | 0.031479 | 0.001116 | 0.003957 | 0.003858 | 0.009648 | 0.004529 | 0.002211 |
| 31.7083333 | 0.025007 | 0.008738 | 0.052575 | 0.012086 | 0.043506 | 0.007204 | 0.004333 | 0.004598 | 0.005765 | 0.018509 | 0.049523 | 0.0066 | 0.022455 | 0.015842 | 0.028149 | 0.03068 | 0.000435 | 0.003602 | 0.003785 | 0.016109 | 0.004031 | 0.003895 |
| 31.7222222 | 0.043256 | 0.006323 | 0.034863 | 0.025104 | 0.045106 | 0.002415 | 0.00604 | 0.005582 | 0.006587 | 0.014664 | 0.087483 | 0.005789 | 0.017506 | 0.028336 | 0.026744 | 0.024531 | 0.001192 | 0.003228 | 0.003427 | 0.015313 | 0.003551 | 0.005412 |
| 31.7361111 | 0.048388 | 0.002981 | 0.017054 | 0.036542 | 0.037122 | 0.000981 | 0.005085 | 0.005674 | 0.005027 | 0.004853 | 0.095457 | 0.005688 | 0.036195 | 0.032326 | 0.022451 | 0.016367 | 0.003712 | 0.002758 | 0.002805 | 0.008234 | 0.002897 | 0.004906 |
| 31.75 | 0.028043 | 0.004291 | 0.008891 | 0.026155 | 0.023368 | 0.001718 | 0.00366 | 0.008962 | 0.005181 | 0.005179 | 0.064774 | 0.006928 | 0.058633 | 0.028541 | 0.015474 | 0.013381 | 0.006799 | 0.003903 | 0.003862 | 0.007881 | 0.003701 | 0.002714 |
| 31.7638889 | 0.02097 | 0.011422 | 0.011781 | 0.019434 | 0.017679 | 0.002937 | 0.002321 | 0.013932 | 0.006899 | 0.011963 | 0.030971 | 0.006577 | 0.058087 | 0.021366 | 0.010443 | 0.02068 | 0.007835 | 0.005483 | 0.005336 | 0.015844 | 0.005166 | 0.001546 |
| 31.7777778 | 0.033338 | 0.018516 | 0.012676 | 0.017339 | 0.015825 | 0.003396 | 0.002758 | 0.016514 | 0.007948 | 0.016057 | 0.025899 | 0.009022 | 0.034543 | 0.01246 | 0.009952 | 0.029631 | 0.006147 | 0.005072 | 0.00488 | 0.024457 | 0.004902 | 0.002112 |
| 31.7916667 | 0.039541 | 0.018941 | 0.008164 | 0.007059 | 0.015785 | 0.003556 | 0.005311 | 0.016627 | 0.005303 | 0.014986 | 0.048947 | 0.018678 | 0.018319 | 0.005693 | 0.011463 | 0.028797 | 0.003979 | 0.003828 | 0.003671 | 0.021295 | 0.003751 | 0.003015 |
| 31.8055556 | 0.032124 | 0.012811 | 0.005136 | 0.00273 | 0.026774 | 0.00414 | 0.004709 | 0.014007 | 0.003759 | 0.016706 | 0.069012 | 0.031206 | 0.022841 | 0.004643 | 0.012271 | 0.022185 | 0.002856 | 0.00322 | 0.003031 | 0.012518 | 0.003079 | 0.004807 |
| 31.8194444 | 0.021933 | 0.00755 | 0.007011 | 0.004726 | 0.031503 | 0.005291 | 0.002546 | 0.008263 | 0.005783 | 0.020943 | 0.071469 | 0.04215 | 0.022158 | 0.009619 | 0.011946 | 0.025821 | 0.001834 | 0.002556 | 0.002336 | 0.013599 | 0.002365 | 0.009455 |
| 31.8333333 | 0.023952 | 0.011763 | 0.016997 | 0.006132 | 0.024494 | 0.008174 | 0.00549 | 0.005233 | 0.012796 | 0.019385 | 0.062014 | 0.048722 | 0.012791 | 0.017799 | 0.01069 | 0.034535 | 0.000949 | 0.001465 | 0.001357 | 0.014335 | 0.001424 | 0.015804 |
| 31.8472222 | 0.043396 | 0.027257 | 0.031285 | 0.006145 | 0.026617 | 0.012932 | 0.012739 | 0.007696 | 0.020945 | 0.016605 | 0.042413 | 0.049831 | 0.014193 | 0.0254 | 0.009255 | 0.028945 | 0.001046 | 0.00105 | 0.001136 | 0.011204 | 0.001227 | 0.020292 |
| 31.8611111 | 0.063063 | 0.038417 | 0.040546 | 0.005959 | 0.02768 | 0.017796 | 0.01635 | 0.009358 | 0.016115 | 0.018705 | 0.018992 | 0.048444 | 0.026284 | 0.03162 | 0.008318 | 0.023026 | 0.000924 | 0.002104 | 0.002148 | 0.023885 | 0.002332 | 0.022757 |
| 31.875 | 0.068667 | 0.028276 | 0.040956 | 0.006745 | 0.014476 | 0.022979 | 0.011266 | 0.009441 | 0.008421 | 0.021043 | 0.006468 | 0.04495 | 0.041143 | 0.037697 | 0.007795 | 0.039358 | 0.000298 | 0.004045 | 0.003734 | 0.051935 | 0.004125 | 0.026399 |
| 31.8888889 | 0.060177 | 0.011217 | 0.038692 | 0.007633 | 0.009335 | 0.029985 | 0.004961 | 0.021918 | 0.010855 | 0.026703 | 0.011204 | 0.038171 | 0.051972 | 0.045761 | 0.007607 | 0.058 | 0.000384 | 0.005844 | 0.005234 | 0.075784 | 0.005701 | 0.033147 |
| 31.9027778 | 0.04606 | 0.006925 | 0.035971 | 0.007222 | 0.023991 | 0.036623 | 0.003856 | 0.042408 | 0.016243 | 0.038404 | 0.023754 | 0.032618 | 0.053821 | 0.059306 | 0.007487 | 0.057406 | 0.001781 | 0.006827 | 0.006177 | 0.081128 | 0.006545 | 0.044852 |
| 31.9166667 | 0.037272 | 0.017222 | 0.034575 | 0.005186 | 0.040135 | 0.039146 | 0.003919 | 0.050776 | 0.022935 | 0.042327 | 0.034818 | 0.03543 | 0.052723 | 0.071667 | 0.007413 | 0.050843 | 0.004622 | 0.006669 | 0.006149 | 0.060737 | 0.006389 | 0.059667 |
| 31.9305556 | 0.038667 | 0.034384 | 0.040513 | 0.003086 | 0.040612 | 0.038849 | 0.004517 | 0.046899 | 0.035217 | 0.03649 | 0.045531 | 0.046904 | 0.054983 | 0.073899 | 0.008036 | 0.050396 | 0.00879 | 0.00512 | 0.004887 | 0.033244 | 0.004903 | 0.069878 |
| 31.9444444 | 0.052815 | 0.044337 | 0.043184 | 0.001807 | 0.03661 | 0.039528 | 0.005373 | 0.042511 | 0.053449 | 0.032387 | 0.057367 | 0.058095 | 0.058454 | 0.072759 | 0.008785 | 0.051302 | 0.013775 | 0.002583 | 0.0027 | 0.027123 | 0.002465 | 0.071651 |
| 31.9583333 | 0.079277 | 0.040129 | 0.035072 | 0.001764 | 0.034799 | 0.044452 | 0.006178 | 0.041753 | 0.068068 | 0.03193 | 0.069119 | 0.056584 | 0.060099 | 0.070471 | 0.006969 | 0.048128 | 0.017711 | 0.001104 | 0.001309 | 0.03403 | 0.001103 | 0.065378 |
| 31.9722222 | 0.101588 | 0.027689 | 0.027647 | 0.002811 | 0.027882 | 0.052117 | 0.011283 | 0.038577 | 0.071909 | 0.034978 | 0.07441 | 0.03915 | 0.059538 | 0.060655 | 0.003379 | 0.04071 | 0.019725 | 0.0019 | 0.001891 | 0.030992 | 0.002052 | 0.055207 |
| 31.9861111 | 0.101319 | 0.017493 | 0.035625 | 0.003941 | 0.019211 | 0.054045 | 0.021966 | 0.024981 | 0.071807 | 0.041562 | 0.065607 | 0.01831 | 0.05678 | 0.045201 | 0.002179 | 0.031793 | 0.021605 | 0.002748 | 0.002622 | 0.017893 | 0.0031 | 0.048733 |
| 32 | 0.083538 | 0.015329 | 0.058652 | 0.004652 | 0.017137 | 0.045588 | 0.032077 | 0.009294 | 0.073125 | 0.042565 | 0.047228 | 0.008252 | 0.051213 | 0.028342 | 0.004723 | 0.023944 | 0.022901 | 0.002727 | 0.002648 | 0.006441 | 0.00295 | 0.045155 |
| 32.0138889 | 0.06203 | 0.016715 | 0.074427 | 0.004094 | 0.024099 | 0.029892 | 0.036067 | 0.012136 | 0.068334 | 0.03064 | 0.02653 | 0.009016 | 0.042108 | 0.01554 | 0.008432 | 0.015441 | 0.019699 | 0.005583 | 0.005126 | 0.002124 | 0.005153 | 0.036194 |
| 32.0277778 | 0.045352 | 0.014926 | 0.066782 | 0.004915 | 0.029327 | 0.015055 | 0.044126 | 0.028642 | 0.053527 | 0.016354 | 0.010206 | 0.012422 | 0.032348 | 0.014895 | 0.009721 | 0.006953 | 0.012597 | 0.01089 | 0.00925 | 0.00174 | 0.009784 | 0.021436 |
| 32.0416667 | 0.04634 | 0.010809 | 0.042372 | 0.007374 | 0.029954 | 0.013906 | 0.055086 | 0.045567 | 0.042561 | 0.016973 | 0.009133 | 0.015283 | 0.030172 | 0.018428 | 0.012359 | 0.004975 | 0.010081 | 0.010833 | 0.00915 | 0.005039 | 0.010026 | 0.01048 |
| 32.0555556 | 0.072145 | 0.008227 | 0.021807 | 0.006792 | 0.04752 | 0.036044 | 0.051403 | 0.068794 | 0.047994 | 0.041295 | 0.028462 | 0.019872 | 0.038833 | 0.013019 | 0.020831 | 0.011839 | 0.015401 | 0.007912 | 0.007601 | 0.01723 | 0.00828 | 0.00926 |
| 32.0694444 | 0.093157 | 0.006944 | 0.021626 | 0.007242 | 0.077865 | 0.073954 | 0.045809 | 0.099174 | 0.063755 | 0.077346 | 0.059204 | 0.025354 | 0.049226 | 0.004859 | 0.029255 | 0.021597 | 0.021317 | 0.008471 | 0.0082 | 0.035043 | 0.009492 | 0.012105 |
| 32.0833333 | 0.076659 | 0.007235 | 0.028357 | 0.013602 | 0.078893 | 0.100998 | 0.044006 | 0.124174 | 0.070585 | 0.092329 | 0.073068 | 0.028082 | 0.047711 | 0.003249 | 0.031788 | 0.023123 | 0.025646 | 0.006494 | 0.005729 | 0.047528 | 0.007404 | 0.011054 |
| 32.0972222 | 0.044059 | 0.010531 | 0.020201 | 0.021421 | 0.052254 | 0.093617 | 0.028812 | 0.125399 | 0.059542 | 0.073691 | 0.053124 | 0.032034 | 0.029564 | 0.006985 | 0.028095 | 0.013818 | 0.031566 | 0.002432 | 0.001995 | 0.045585 | 0.002931 | 0.008995 |

| | | | | | | | | | | | | | | | | | | | | | | |
|---|---|---|---|---|---|---|---|---|---|---|---|---|---|---|---|---|---|---|---|---|---|---|
| 32.1111111 | 0.025619 | 0.014944 | 0.009821 | 0.02225 | 0.0336 | 0.056302 | 0.012165 | 0.100428 | 0.039647 | 0.041419 | 0.023367 | 0.033695 | 0.014513 | 0.011285 | 0.018381 | 0.007331 | 0.033533 | 0.002414 | 0.002207 | 0.030181 | 0.002564 | 0.01395 |
| 32.125 | 0.016359 | 0.018142 | 0.010885 | 0.014183 | 0.022127 | 0.023588 | 0.010844 | 0.069498 | 0.019782 | 0.016839 | 0.012576 | 0.023052 | 0.022133 | 0.015081 | 0.007934 | 0.013529 | 0.025809 | 0.004921 | 0.004528 | 0.013714 | 0.005029 | 0.021786 |
| 32.1388889 | 0.00999 | 0.017903 | 0.010154 | 0.006382 | 0.012229 | 0.020713 | 0.016379 | 0.042073 | 0.008553 | 0.010325 | 0.018446 | 0.011804 | 0.03743 | 0.018254 | 0.003141 | 0.02417 | 0.014155 | 0.006835 | 0.006296 | 0.007616 | 0.006825 | 0.018689 |
| 32.1527778 | 0.014532 | 0.012892 | 0.005013 | 0.003359 | 0.013851 | 0.035051 | 0.0139 | 0.023543 | 0.011864 | 0.018653 | 0.022627 | 0.015043 | 0.040487 | 0.018566 | 0.001775 | 0.029426 | 0.006412 | 0.006647 | 0.006314 | 0.009675 | 0.006376 | 0.010659 |
| 32.1666667 | 0.031583 | 0.005908 | 0.002827 | 0.005977 | 0.026495 | 0.04744 | 0.009029 | 0.030341 | 0.024239 | 0.03027 | 0.019041 | 0.030777 | 0.03067 | 0.015879 | 0.000964 | 0.027697 | 0.005409 | 0.004659 | 0.004472 | 0.012486 | 0.004335 | 0.013446 |
| 32.1805556 | 0.054263 | 0.002138 | 0.004128 | 0.014215 | 0.048204 | 0.050108 | 0.015107 | 0.056341 | 0.036471 | 0.040967 | 0.016773 | 0.05064 | 0.016759 | 0.012091 | 0.000927 | 0.021484 | 0.009792 | 0.002241 | 0.002029 | 0.015016 | 0.002136 | 0.015238 |
| 32.1944444 | 0.068672 | 0.004051 | 0.008443 | 0.018068 | 0.067721 | 0.040553 | 0.023066 | 0.075948 | 0.040885 | 0.047687 | 0.022445 | 0.062042 | 0.006965 | 0.007725 | 0.001685 | 0.017669 | 0.014894 | 0.000588 | 0.000515 | 0.014815 | 0.000741 | 0.016601 |
| 32.2083333 | 0.058291 | 0.009479 | 0.008626 | 0.012328 | 0.063263 | 0.023785 | 0.021614 | 0.075627 | 0.036876 | 0.043237 | 0.031833 | 0.055158 | 0.003357 | 0.00444 | 0.004366 | 0.020789 | 0.016269 | 8.55E-05 | 0.000185 | 0.009662 | 0.000322 | 0.034322 |
| 32.2222222 | 0.030014 | 0.013331 | 0.008053 | 0.004803 | 0.036514 | 0.010347 | 0.013213 | 0.058593 | 0.03173 | 0.031355 | 0.037329 | 0.035992 | 0.004225 | 0.005514 | 0.009094 | 0.025646 | 0.013681 | 0.000306 | 0.000434 | 0.009513 | 0.000467 | 0.048619 |
| 32.2361111 | 0.013074 | 0.011548 | 0.020827 | 0.001347 | 0.017147 | 0.008587 | 0.005884 | 0.043479 | 0.030709 | 0.020489 | 0.029623 | 0.016907 | 0.006428 | 0.010312 | 0.013379 | 0.026364 | 0.010617 | 0.000778 | 0.000906 | 0.018886 | 0.000815 | 0.043364 |
| 32.25 | 0.016662 | 0.006779 | 0.041935 | 0.00654 | 0.024756 | 0.018069 | 0.006878 | 0.048868 | 0.03178 | 0.014338 | 0.015317 | 0.006024 | 0.00788 | 0.013905 | 0.014295 | 0.025491 | 0.009195 | 0.001073 | 0.001131 | 0.022107 | 0.001124 | 0.034598 |
| 32.2638889 | 0.016474 | 0.005857 | 0.052491 | 0.0163 | 0.043851 | 0.029981 | 0.012626 | 0.07029 | 0.02951 | 0.019005 | 0.013058 | 0.004576 | 0.007076 | 0.01182 | 0.011445 | 0.029968 | 0.00928 | 0.001823 | 0.001714 | 0.013609 | 0.002095 | 0.032012 |
| 32.2777778 | 0.012645 | 0.009689 | 0.043795 | 0.015713 | 0.058763 | 0.033573 | 0.012548 | 0.078425 | 0.023574 | 0.034521 | 0.016519 | 0.006249 | 0.003797 | 0.013369 | 0.007782 | 0.04213 | 0.01103 | 0.003236 | 0.003077 | 0.009772 | 0.003772 | 0.02921 |
| 32.2916667 | 0.027476 | 0.015162 | 0.026132 | 0.006949 | 0.067467 | 0.027058 | 0.008636 | 0.051204 | 0.019899 | 0.044234 | 0.023827 | 0.007069 | 0.003286 | 0.029334 | 0.006394 | 0.051213 | 0.014444 | 0.003077 | 0.00302 | 0.018915 | 0.003545 | 0.022423 |
| 32.3055556 | 0.046595 | 0.020937 | 0.014245 | 0.001598 | 0.063034 | 0.018497 | 0.01419 | 0.02116 | 0.024827 | 0.034805 | 0.038634 | 0.007055 | 0.010134 | 0.048784 | 0.007362 | 0.04758 | 0.018436 | 0.002616 | 0.002352 | 0.030328 | 0.002573 | 0.018056 |
| 32.3194444 | 0.046273 | 0.025916 | 0.016855 | 0.000675 | 0.044683 | 0.014633 | 0.025367 | 0.019116 | 0.034566 | 0.018737 | 0.039412 | 0.008287 | 0.021453 | 0.05625 | 0.009177 | 0.036488 | 0.021084 | 0.004928 | 0.004384 | 0.034153 | 0.004493 | 0.018566 |
| 32.3333333 | 0.029742 | 0.029752 | 0.03468 | 0.001315 | 0.023314 | 0.018079 | 0.025533 | 0.019553 | 0.036347 | 0.009333 | 0.023259 | 0.015091 | 0.03032 | 0.048064 | 0.011512 | 0.025347 | 0.021312 | 0.007758 | 0.007431 | 0.031077 | 0.007063 | 0.016011 |
| 32.3472222 | 0.012457 | 0.032929 | 0.05259 | 0.001958 | 0.012086 | 0.029766 | 0.017675 | 0.02082 | 0.033582 | 0.015569 | 0.015082 | 0.029501 | 0.03286 | 0.030189 | 0.013808 | 0.018693 | 0.018866 | 0.008262 | 0.008259 | 0.027124 | 0.00723 | 0.008461 |
| 32.3611111 | 0.003668 | 0.036004 | 0.048957 | 0.003741 | 0.012609 | 0.043991 | 0.014539 | 0.040756 | 0.035656 | 0.044196 | 0.027454 | 0.04659 | 0.031483 | 0.012859 | 0.01407 | 0.019176 | 0.013133 | 0.005749 | 0.005898 | 0.02718 | 0.004791 | 0.003756 |
| 32.375 | 0.0086 | 0.039403 | 0.026371 | 0.00699 | 0.023356 | 0.050963 | 0.015919 | 0.063934 | 0.034412 | 0.068341 | 0.048637 | 0.058346 | 0.030289 | 0.00413 | 0.012052 | 0.023385 | 0.006342 | 0.002993 | 0.003013 | 0.030147 | 0.002533 | 0.005087 |
| 32.3888889 | 0.031864 | 0.040716 | 0.008548 | 0.007451 | 0.039586 | 0.048521 | 0.017207 | 0.07323 | 0.033205 | 0.063114 | 0.059283 | 0.062656 | 0.031379 | 0.008281 | 0.009372 | 0.027044 | 0.003369 | 0.003569 | 0.00305 | 0.032477 | 0.003395 | 0.005967 |
| 32.4027778 | 0.064786 | 0.036997 | 0.007145 | 0.004303 | 0.038611 | 0.043335 | 0.018158 | 0.069675 | 0.037142 | 0.03718 | 0.053893 | 0.06013 | 0.035469 | 0.024032 | 0.006459 | 0.031529 | 0.005184 | 0.005496 | 0.004473 | 0.033359 | 0.005211 | 0.008316 |
| 32.4166667 | 0.087563 | 0.030057 | 0.01984 | 0.004362 | 0.020138 | 0.041605 | 0.019598 | 0.069577 | 0.030901 | 0.015637 | 0.041702 | 0.052248 | 0.043336 | 0.042705 | 0.003286 | 0.037916 | 0.009251 | 0.006865 | 0.005566 | 0.034525 | 0.006461 | 0.024954 |
| 32.4305556 | 0.086405 | 0.022817 | 0.039288 | 0.009536 | 0.007355 | 0.046022 | 0.021958 | 0.081364 | 0.025332 | 0.01042 | 0.034574 | 0.044772 | 0.05469 | 0.05519 | 0.001892 | 0.044315 | 0.014166 | 0.00825 | 0.006927 | 0.038523 | 0.007888 | 0.052002 |
| 32.4444444 | 0.068631 | 0.015869 | 0.048963 | 0.009946 | 0.005103 | 0.055897 | 0.023088 | 0.088915 | 0.040095 | 0.012585 | 0.037319 | 0.041238 | 0.066073 | 0.060597 | 0.003963 | 0.05139 | 0.018862 | 0.008777 | 0.008238 | 0.047437 | 0.008191 | 0.069382 |
| 32.4583333 | 0.054191 | 0.010774 | 0.035676 | 0.006958 | 0.013288 | 0.062627 | 0.01969 | 0.082201 | 0.050148 | 0.020998 | 0.04534 | 0.041123 | 0.074689 | 0.06388 | 0.006377 | 0.057377 | 0.021092 | 0.0065 | 0.007971 | 0.059396 | 0.005643 | 0.067922 |
| 32.4722222 | 0.054702 | 0.014546 | 0.022705 | 0.010646 | 0.031628 | 0.061876 | 0.015983 | 0.077247 | 0.037774 | 0.030202 | 0.052898 | 0.046807 | 0.07821 | 0.067498 | 0.005914 | 0.054888 | 0.01978 | 0.002746 | 0.006059 | 0.067963 | 0.002564 | 0.056494 |
| 32.4861111 | 0.069186 | 0.033664 | 0.023096 | 0.012094 | 0.054312 | 0.061788 | 0.019068 | 0.074493 | 0.029285 | 0.03432 | 0.062861 | 0.058901 | 0.077958 | 0.070871 | 0.004915 | 0.044868 | 0.017881 | 0.002365 | 0.005137 | 0.069247 | 0.002984 | 0.060112 |
| 32.5 | 0.087762 | 0.05142 | 0.023948 | 0.006659 | 0.073626 | 0.061776 | 0.027894 | 0.063929 | 0.03519 | 0.039816 | 0.07673 | 0.071151 | 0.080067 | 0.073827 | 0.00734 | 0.038464 | 0.018002 | 0.004742 | 0.005325 | 0.063419 | 0.005025 | 0.082259 |
| 32.5138889 | 0.094375 | 0.046287 | 0.035427 | 0.005399 | 0.074764 | 0.052114 | 0.035245 | 0.055131 | 0.03807 | 0.046573 | 0.083944 | 0.081072 | 0.080423 | 0.070769 | 0.015442 | 0.036425 | 0.01803 | 0.004973 | 0.00466 | 0.050046 | 0.004824 | 0.074678 |
| 32.5277778 | 0.080942 | 0.030864 | 0.047938 | 0.009768 | 0.050857 | 0.03591 | 0.03632 | 0.039416 | 0.04392 | 0.048523 | 0.076556 | 0.090356 | 0.071902 | 0.056355 | 0.026136 | 0.02885 | 0.015806 | 0.006901 | 0.007028 | 0.032651 | 0.006531 | 0.040323 |
| 32.5416667 | 0.059274 | 0.025889 | 0.041242 | 0.013172 | 0.019458 | 0.023984 | 0.03063 | 0.015803 | 0.052118 | 0.052206 | 0.054516 | 0.093561 | 0.054185 | 0.037219 | 0.034566 | 0.015934 | 0.011148 | 0.017353 | 0.017122 | 0.019647 | 0.015785 | 0.026159 |
| 32.5555556 | 0.043959 | 0.028398 | 0.023156 | 0.011477 | 0.016731 | 0.019668 | 0.018764 | 0.0066 | 0.037265 | 0.069096 | 0.030919 | 0.080944 | 0.029572 | 0.021808 | 0.040014 | 0.009427 | 0.006813 | 0.030519 | 0.029524 | 0.014471 | 0.027546 | 0.031079 |
| 32.5694444 | 0.041197 | 0.024568 | 0.015459 | 0.006492 | 0.055201 | 0.018657 | 0.009393 | 0.00911 | 0.015598 | 0.088189 | 0.031291 | 0.055312 | 0.009394 | 0.013369 | 0.039373 | 0.010452 | 0.006571 | 0.035191 | 0.033932 | 0.01431 | 0.032614 | 0.038817 |
| 32.5833333 | 0.045689 | 0.014377 | 0.023096 | 0.003994 | 0.091218 | 0.013894 | 0.009932 | 0.008976 | 0.009816 | 0.087684 | 0.056714 | 0.035646 | 0.007453 | 0.012589 | 0.032332 | 0.009263 | 0.00805 | 0.030499 | 0.029185 | 0.016489 | 0.029841 | 0.037432 |
| 32.5972222 | 0.038148 | 0.006678 | 0.026349 | 0.005305 | 0.081647 | 0.009755 | 0.012941 | 0.006762 | 0.013768 | 0.065303 | 0.069867 | 0.031547 | 0.015591 | 0.012212 | 0.02538 | 0.007379 | 0.007363 | 0.024127 | 0.022971 | 0.016277 | 0.024895 | 0.029725 |
| 32.6111111 | 0.01718 | 0.004326 | 0.018393 | 0.006815 | 0.047246 | 0.01751 | 0.012251 | 0.009226 | 0.024669 | 0.059949 | 0.049872 | 0.034168 | 0.01948 | 0.006515 | 0.020919 | 0.010378 | 0.00448 | 0.019609 | 0.018832 | 0.010182 | 0.02074 | 0.019632 |
| 32.625 | 0.011249 | 0.007532 | 0.007612 | 0.010751 | 0.023634 | 0.023747 | 0.008465 | 0.010588 | 0.031418 | 0.092794 | 0.022864 | 0.03644 | 0.036172 | 0.00131 | 0.017042 | 0.015806 | 0.001957 | 0.015121 | 0.014844 | 0.009759 | 0.016095 | 0.008928 |
| 32.6388889 | 0.034492 | 0.010762 | 0.009917 | 0.014351 | 0.015285 | 0.02185 | 0.004709 | 0.008085 | 0.025394 | 0.107312 | 0.021507 | 0.036034 | 0.069425 | 0.004647 | 0.013608 | 0.018927 | 0.002034 | 0.010558 | 0.010706 | 0.02223 | 0.011163 | 0.009865 |
| 32.6527778 | 0.050062 | 0.011213 | 0.028093 | 0.012852 | 0.017672 | 0.028603 | 0.004717 | 0.01545 | 0.014463 | 0.077552 | 0.042496 | 0.028523 | 0.072078 | 0.014032 | 0.011321 | 0.015228 | 0.003838 | 0.006866 | 0.007147 | 0.03535 | 0.007197 | 0.02485 |
| 32.6666667 | 0.033151 | 0.011382 | 0.036588 | 0.013677 | 0.039158 | 0.032302 | 0.007648 | 0.025724 | 0.010298 | 0.044538 | 0.061615 | 0.015801 | 0.048059 | 0.016087 | 0.010267 | 0.00977 | 0.005817 | 0.004049 | 0.004195 | 0.036121 | 0.004181 | 0.05242 |
| 32.6805556 | 0.014273 | 0.009925 | 0.02451 | 0.014932 | 0.06926 | 0.019845 | 0.007615 | 0.019998 | 0.014017 | 0.033979 | 0.067556 | 0.007281 | 0.057079 | 0.01444 | 0.011192 | 0.008237 | 0.006406 | 0.003128 | 0.002976 | 0.028496 | 0.002889 | 0.085718 |
| 32.6944444 | 0.018353 | 0.005954 | 0.013426 | 0.010711 | 0.06615 | 0.006975 | 0.003796 | 0.006922 | 0.015103 | 0.030155 | 0.054273 | 0.005549 | 0.084858 | 0.023216 | 0.014287 | 0.008413 | 0.005344 | 0.004547 | 0.004048 | 0.032329 | 0.003965 | 0.096322 |
| 32.7083333 | 0.026508 | 0.00285 | 0.018111 | 0.006467 | 0.052027 | 0.002886 | 0.003708 | 0.004624 | 0.014904 | 0.02454 | 0.030986 | 0.007128 | 0.086878 | 0.032857 | 0.018245 | 0.014687 | 0.004658 | 0.006022 | 0.005449 | 0.044928 | 0.00545 | 0.068466 |
| 32.7222222 | 0.0203 | 0.003551 | 0.03045 | 0.008491 | 0.056025 | 0.003099 | 0.005012 | 0.008865 | 0.01883 | 0.044499 | 0.028382 | 0.008655 | 0.062515 | 0.029744 | 0.020623 | 0.022426 | 0.00536 | 0.005613 | 0.005418 | 0.041445 | 0.005268 | 0.034592 |
| 32.7361111 | 0.026705 | 0.004123 | 0.041638 | 0.015761 | 0.045447 | 0.004959 | 0.009397 | 0.013244 | 0.0165 | 0.061518 | 0.061222 | 0.007863 | 0.043307 | 0.020312 | 0.01894 | 0.018334 | 0.006974 | 0.003544 | 0.003914 | 0.032233 | 0.003295 | 0.033265 |
| 32.75 | 0.035107 | 0.006012 | 0.045214 | 0.017812 | 0.034688 | 0.009587 | 0.022274 | 0.019584 | 0.006759 | 0.037117 | 0.090121 | 0.006156 | 0.045721 | 0.017516 | 0.014 | 0.01036 | 0.008445 | 0.001355 | 0.001957 | 0.046971 | 0.001206 | 0.056614 |
| 32.7638889 | 0.033228 | 0.008474 | 0.031723 | 0.009372 | 0.039202 | 0.013688 | 0.027607 | 0.017808 | 0.008177 | 0.014961 | 0.072789 | 0.005252 | 0.039795 | 0.024243 | 0.008949 | 0.012189 | 0.008792 | 0.000442 | 0.00077 | 0.068486 | 0.000516 | 0.069069 |
| 32.7777778 | 0.053812 | 0.007024 | 0.019485 | 0.00217 | 0.02777 | 0.011697 | 0.018727 | 0.011953 | 0.021072 | 0.016113 | 0.034399 | 0.009045 | 0.035537 | 0.030455 | 0.004835 | 0.015406 | 0.008736 | 0.000915 | 0.001029 | 0.063342 | 0.001037 | 0.058316 |
| 32.7916667 | 0.065727 | 0.010527 | 0.036169 | 0.001287 | 0.009959 | 0.00562 | 0.009755 | 0.011683 | 0.024685 | 0.014354 | 0.023724 | 0.020926 | 0.058358 | 0.030839 | 0.001977 | 0.011107 | 0.009223 | 0.002238 | 0.002418 | 0.038279 | 0.00228 | 0.036005 |
| 32.8055556 | 0.042264 | 0.016806 | 0.058127 | 0.004095 | 0.008258 | 0.00274 | 0.006346 | 0.00868 | 0.017627 | 0.011371 | 0.037265 | 0.037594 | 0.08178 | 0.034256 | 0.000984 | 0.006974 | 0.009318 | 0.003196 | 0.003374 | 0.031523 | 0.003212 | 0.017047 |
| 32.8194444 | 0.027534 | 0.01574 | 0.049461 | 0.010582 | 0.017293 | 0.004363 | 0.005929 | 0.004516 | 0.014995 | 0.016041 | 0.038486 | 0.051337 | 0.090984 | 0.043143 | 0.000697 | 0.011467 | 0.007907 | 0.002846 | 0.00298 | 0.046951 | 0.00288 | 0.012565 |
| 32.8333333 | 0.041302 | 0.015225 | 0.025575 | 0.012709 | 0.023507 | 0.004607 | 0.01002 | 0.005253 | 0.022215 | 0.01986 | 0.022849 | 0.05195 | 0.082825 | 0.049979 | 0.000483 | 0.022196 | 0.005465 | 0.001784 | 0.001996 | 0.044647 | 0.001876 | 0.020451 |
| 32.8472222 | 0.062237 | 0.01709 | 0.017733 | 0.007685 | 0.019276 | 0.005321 | 0.017962 | 0.007224 | 0.02767 | 0.029593 | 0.01637 | 0.033961 | 0.052582 | 0.058252 | 0.000648 | 0.030771 | 0.003311 | 0.001021 | 0.001201 | 0.025756 | 0.001176 | 0.029888 |
| 32.8611111 | 0.081723 | 0.019111 | 0.021013 | 0.005952 | 0.011535 | 0.013863 | 0.025504 | 0.008175 | 0.019627 | 0.046398 | 0.018633 | 0.011724 | 0.021983 | 0.064187 | 0.000621 | 0.036372 | 0.001978 | 0.000972 | 0.000807 | 0.024073 | 0.001221 | 0.035804 |
| 32.875 | 0.101696 | 0.02527 | 0.017325 | 0.012269 | 0.008113 | 0.026675 | 0.027125 | 0.007965 | 0.010823 | 0.047603 | 0.021572 | 0.005172 | 0.012214 | 0.045046 | 0.000268 | 0.04493 | 0.001144 | 0.001342 | 0.000817 | 0.041155 | 0.001669 | 0.039982 |
| 32.8888889 | 0.112303 | 0.025466 | 0.011601 | 0.017203 | 0.010417 | 0.034385 | 0.018237 | 0.00845 | 0.015427 | 0.036625 | 0.031297 | 0.011599 | 0.024707 | 0.019798 | 0.000304 | 0.056695 | 0.001138 | 0.001793 | 0.001229 | 0.053175 | 0.002011 | 0.04307 |
| 32.9027778 | 0.103356 | 0.029005 | 0.010026 | 0.012756 | 0.014961 | 0.033526 | 0.007941 | 0.010171 | 0.019992 | 0.022705 | 0.027707 | 0.019954 | 0.047696 | 0.016001 | 0.001183 | 0.065299 | 0.002555 | 0.002635 | 0.002179 | 0.05473 | 0.002593 | 0.043938 |
| 32.9166667 | 0.081097 | 0.049158 | 0.007938 | 0.007312 | 0.023644 | 0.033265 | 0.005827 | 0.01478 | 0.013963 | 0.018926 | 0.014056 | 0.028402 | 0.061804 | 0.020161 | 0.002307 | 0.069929 | 0.004904 | 0.003621 | 0.003197 | 0.048954 | 0.003428 | 0.044225 |
| 32.9305556 | 0.060834 | 0.065382 | 0.009557 | 0.009515 | 0.025178 | 0.04394 | 0.007059 | 0.027447 | 0.007506 | 0.035003 | 0.009775 | 0.03609 | 0.056643 | 0.028548 | 0.00244 | 0.074329 | 0.007855 | 0.003465 | 0.003181 | 0.030575 | 0.003325 | 0.04746 |
| 32.9444444 | 0.052428 | 0.068647 | 0.015378 | 0.012661 | 0.019792 | 0.058349 | 0.006615 | 0.04326 | 0.010114 | 0.040169 | 0.009384 | 0.044176 | 0.039604 | 0.041088 | 0.001415 | 0.075473 | 0.011189 | 0.001933 | 0.001993 | 0.015161 | 0.001941 | 0.053082 |
| 32.9583333 | 0.054303 | 0.067732 | 0.018689 | 0.012582 | 0.030314 | 0.062933 | 0.007413 | 0.058712 | 0.022436 | 0.024166 | 0.00851 | 0.063349 | 0.023403 | 0.054984 | 0.000609 | 0.06696 | 0.013957 | 0.000666 | 0.000786 | 0.016265 | 0.000702 | 0.058167 |
| 32.9722222 | 0.062048 | 0.059134 | 0.022052 | 0.011228 | 0.035677 | 0.056448 | 0.013353 | 0.080513 | 0.025081 | 0.013476 | 0.014221 | 0.094573 | 0.014186 | 0.066657 | 0.000809 | 0.050963 | 0.016756 | 0.000552 | 0.000347 | 0.017741 | 0.000471 | 0.061912 |
| 32.9861111 | 0.069054 | 0.039334 | 0.028162 | 0.01145 | 0.021169 | 0.045851 | 0.021294 | 0.094099 | 0.026748 | 0.021834 | 0.036562 | 0.117268 | 0.013616 | 0.066748 | 0.001158 | 0.036077 | 0.019345 | 0.000606 | 0.000315 | 0.010804 | 0.000475 | 0.061698 |
| 33 | 0.059306 | 0.024366 | 0.032491 | 0.022953 | 0.009845 | 0.03559 | 0.026669 | 0.083651 | 0.058872 | 0.034904 | 0.065832 | 0.105416 | 0.019542 | 0.055585 | 0.001214 | 0.024747 | 0.017949 | 0.001436 | 0.001058 | 0.005001 | 0.001169 | 0.05327 |
| 33.0138889 | 0.03228 | 0.036283 | 0.02611 | 0.048812 | 0.014221 | 0.025532 | 0.028668 | 0.065001 | 0.089296 | 0.031172 | 0.078279 | 0.061228 | 0.026232 | 0.037966 | 0.002726 | 0.013232 | 0.011071 | 0.005604 | 0.004432 | 0.003423 | 0.004833 | 0.03652 |
| 33.0277778 | 0.011431 | 0.056478 | 0.014533 | 0.067019 | 0.031255 | 0.015679 | 0.025578 | 0.046928 | 0.091317 | 0.020924 | 0.062442 | 0.032072 | 0.029813 | 0.019598 | 0.007187 | 0.005926 | 0.005086 | 0.013469 | 0.010788 | 0.002428 | 0.012124 | 0.017668 |
| 33.0416667 | 0.01699 | 0.047866 | 0.01149 | 0.058924 | 0.038507 | 0.013735 | 0.019072 | 0.033704 | 0.089575 | 0.022212 | 0.040421 | 0.030153 | 0.031498 | 0.007203 | 0.012176 | 0.007003 | 0.006252 | 0.021071 | 0.017372 | 0.00173 | 0.019881 | 0.005203 |
| 33.0555556 | 0.052398 | 0.039032 | 0.010933 | 0.033845 | 0.02524 | 0.030019 | 0.026047 | 0.041357 | 0.100685 | 0.034624 | 0.049391 | 0.020338 | 0.036146 | 0.002338 | 0.01372 | 0.008766 | 0.010707 | 0.02348 | 0.020718 | 0.004504 | 0.023009 | 0.001349 |
| 33.0694444 | 0.094301 | 0.03955 | 0.00951 | 0.010908 | 0.010205 | 0.063789 | 0.045126 | 0.072186 | 0.11587 | 0.062391 | 0.081489 | 0.007239 | 0.044569 | 0.00641 | 0.011527 | 0.014739 | 0.011815 | 0.020809 | 0.019911 | 0.015052 | 0.020458 | 0.002757 |
| 33.0833333 | 0.109746 | 0.019616 | 0.007846 | 0.006281 | 0.009759 | 0.100336 | 0.039868 | 0.114337 | 0.110223 | 0.08903 | 0.090526 | 0.004926 | 0.045253 | 0.023805 | 0.009369 | 0.022082 | 0.007699 | 0.017153 | 0.017155 | 0.02721 | 0.016565 | 0.009751 |
| 33.0972222 | 0.092639 | 0.009171 | 0.004105 | 0.02248 | 0.022494 | 0.111884 | 0.022861 | 0.144401 | 0.08104 | 0.088065 | 0.065085 | 0.010317 | 0.029339 | 0.039285 | 0.009269 | 0.016459 | 0.004804 | 0.014536 | 0.014488 | 0.02698 | 0.014195 | 0.018402 |
| 33.1111111 | 0.060082 | 0.020072 | 0.002313 | 0.041814 | 0.033831 | 0.082157 | 0.026291 | 0.13861 | 0.047122 | 0.056898 | 0.031876 | 0.025245 | 0.017646 | 0.031411 | 0.008819 | 0.010675 | 0.010233 | 0.011821 | 0.011437 | 0.015425 | 0.011758 | 0.019059 |
| 33.125 | 0.032629 | 0.029521 | 0.002773 | 0.042162 | 0.02827 | 0.038086 | 0.024613 | 0.106629 | 0.019775 | 0.021565 | 0.019099 | 0.038531 | 0.028136 | 0.01691 | 0.006212 | 0.016055 | 0.018923 | 0.008011 | 0.007469 | 0.006034 | 0.007861 | 0.01434 |
| 33.1388889 | 0.016927 | 0.034087 | 0.005829 | 0.024373 | 0.013586 | 0.020428 | 0.013513 | 0.072631 | 0.008876 | 0.009985 | 0.028402 | 0.041513 | 0.04107 | 0.02228 | 0.002941 | 0.023056 | 0.023693 | 0.003938 | 0.0037 | 0.00528 | 0.00374 | 0.015152 |
| 33.1527778 | 0.01141 | 0.03514 | 0.010722 | 0.012804 | 0.010454 | 0.034158 | 0.009278 | 0.039489 | 0.01689 | 0.017471 | 0.034237 | 0.042845 | 0.04056 | 0.039113 | 0.001123 | 0.025183 | 0.023424 | 0.001277 | 0.001498 | 0.009403 | 0.001271 | 0.024159 |
| 33.1666667 | 0.023101 | 0.033692 | 0.015212 | 0.021844 | 0.02004 | 0.054877 | 0.00953 | 0.017949 | 0.032551 | 0.032145 | 0.028978 | 0.04705 | 0.030646 | 0.052055 | 0.001014 | 0.021918 | 0.020481 | 0.001469 | 0.001716 | 0.014369 | 0.00158 | 0.029449 |
| 33.1805556 | 0.052833 | 0.028709 | 0.02163 | 0.035005 | 0.034095 | 0.06171 | 0.01817 | 0.022044 | 0.04354 | 0.049899 | 0.022335 | 0.046534 | 0.016533 | 0.056776 | 0.002361 | 0.017508 | 0.018332 | 0.004443 | 0.004129 | 0.01873 | 0.004342 | 0.026794 |
| 33.1944444 | 0.074927 | 0.019539 | 0.026904 | 0.036441 | 0.045858 | 0.048626 | 0.027052 | 0.03457 | 0.040144 | 0.063615 | 0.017399 | 0.037463 | 0.00763 | 0.051085 | 0.005384 | 0.018703 | 0.017841 | 0.00847 | 0.007501 | 0.021326 | 0.007999 | 0.027093 |
| 33.2083333 | 0.06419 | 0.009196 | 0.022443 | 0.026116 | 0.049204 | 0.024486 | 0.026146 | 0.042385 | 0.02415 | 0.063117 | 0.016575 | 0.026427 | 0.014676 | 0.039917 | 0.009148 | 0.028374 | 0.016799 | 0.012165 | 0.010961 | 0.022804 | 0.01152 | 0.033645 |
| 33.2222222 | 0.036114 | 0.003097 | 0.011789 | 0.012495 | 0.040926 | 0.007101 | 0.017901 | 0.053497 | 0.009315 | 0.045417 | 0.027012 | 0.018075 | 0.035047 | 0.029555 | 0.011976 | 0.043964 | 0.013052 | 0.014628 | 0.01361 | 0.023362 | 0.014148 | 0.055798 |
| 33.2361111 | 0.020233 | 0.001614 | 0.008044 | 0.01145 | 0.027922 | 0.008149 | 0.011632 | 0.065936 | 0.003845 | 0.022104 | 0.04222 | 0.010216 | 0.050488 | 0.020372 | 0.012136 | 0.057124 | 0.008671 | 0.014211 | 0.013583 | 0.019618 | 0.014097 | 0.084538 |
| 33.25 | 0.025691 | 0.001525 | 0.012275 | 0.017647 | 0.022144 | 0.021081 | 0.018064 | 0.065083 | 0.009517 | 0.009552 | 0.038548 | 0.00664 | 0.045129 | 0.011763 | 0.008807 | 0.056591 | 0.007719 | 0.010072 | 0.009933 | 0.010773 | 0.010248 | 0.085745 |
| 33.2638889 | 0.044744 | 0.003892 | 0.011216 | 0.017128 | 0.022659 | 0.03134 | 0.028845 | 0.048024 | 0.025519 | 0.01674 | 0.018106 | 0.013374 | 0.024601 | 0.005629 | 0.00473 | 0.040398 | 0.010584 | 0.005363 | 0.005301 | 0.003868 | 0.005553 | 0.059867 |
| 33.2777778 | 0.055797 | 0.010106 | 0.008657 | 0.012471 | 0.022483 | 0.032292 | 0.030224 | 0.027985 | 0.041312 | 0.042215 | 0.007964 | 0.022162 | 0.011611 | 0.004437 | 0.004074 | 0.021007 | 0.014803 | 0.004749 | 0.004108 | 0.002191 | 0.004793 | 0.029012 |
| 33.2916667 | 0.039403 | 0.013938 | 0.017859 | 0.01152 | 0.017567 | 0.024958 | 0.021419 | 0.017281 | 0.043967 | 0.065802 | 0.015265 | 0.022533 | 0.009734 | 0.008955 | 0.005509 | 0.013841 | 0.017495 | 0.006466 | 0.005502 | 0.004814 | 0.006373 | 0.012659 |
| 33.3055556 | 0.017443 | 0.009592 | 0.029604 | 0.017437 | 0.011967 | 0.016144 | 0.011078 | 0.020221 | 0.036867 | 0.065435 | 0.029682 | 0.01732 | 0.011554 | 0.01698 | 0.00475 | 0.023841 | 0.015984 | 0.005408 | 0.004826 | 0.013729 | 0.00517 | 0.016292 |
| 33.3194444 | 0.012954 | 0.005501 | 0.031468 | 0.021144 | 0.011188 | 0.012738 | 0.009922 | 0.02549 | 0.033115 | 0.045862 | 0.034878 | 0.016526 | 0.024799 | 0.022959 | 0.002954 | 0.042084 | 0.011039 | 0.002981 | 0.002892 | 0.023883 | 0.002715 | 0.021739 |
| 33.3333333 | 0.012684 | 0.013103 | 0.024721 | 0.015419 | 0.011899 | 0.014816 | 0.0139 | 0.021567 | 0.033491 | 0.022893 | 0.035003 | 0.021291 | 0.04165 | 0.022437 | 0.003433 | 0.054399 | 0.007086 | 0.00329 | 0.00322 | 0.028602 | 0.003147 | 0.015837 |
| 33.3472222 | 0.023236 | 0.034193 | 0.014654 | 0.00745 | 0.01797 | 0.018428 | 0.013911 | 0.020904 | 0.035739 | 0.011917 | 0.063037 | 0.026301 | 0.045187 | 0.015013 | 0.005529 | 0.052868 | 0.006626 | 0.00574 | 0.005135 | 0.027562 | 0.005602 | 0.007515 |
| 33.3611111 | 0.045008 | 0.050806 | 0.006447 | 0.003629 | 0.030579 | 0.021515 | 0.010431 | 0.022411 | 0.040621 | 0.021643 | 0.102789 | 0.032256 | 0.036583 | 0.009942 | 0.006946 | 0.04096 | 0.007899 | 0.007523 | 0.006343 | 0.021704 | 0.007226 | 0.00716 |
| 33.375 | 0.052235 | 0.041902 | 0.004703 | 0.002745 | 0.037886 | 0.024455 | 0.006257 | 0.02429 | 0.042999 | 0.042203 | 0.110831 | 0.039209 | 0.025325 | 0.017535 | 0.006934 | 0.027851 | 0.008201 | 0.00744 | 0.00626 | 0.014956 | 0.007073 | 0.007653 |
| 33.3888889 | 0.04322 | 0.020248 | 0.010027 | 0.002705 | 0.038524 | 0.028476 | 0.004054 | 0.051441 | 0.043402 | 0.055361 | 0.099172 | 0.042967 | 0.01979 | 0.027321 | 0.006094 | 0.018832 | 0.007143 | 0.006108 | 0.005573 | 0.013732 | 0.005927 | 0.005467 |
| 33.4027778 | 0.031888 | 0.008215 | 0.020235 | 0.00323 | 0.035993 | 0.036355 | 0.005734 | 0.088724 | 0.041161 | 0.051705 | 0.082048 | 0.045197 | 0.025021 | 0.02707 | 0.005327 | 0.012693 | 0.005847 | 0.005015 | 0.005188 | 0.019715 | 0.005176 | 0.005005 |
| 33.4166667 | 0.024778 | 0.015009 | 0.032895 | 0.005068 | 0.031932 | 0.049626 | 0.011898 | 0.102551 | 0.02941 | 0.039954 | 0.058662 | 0.052006 | 0.039035 | 0.024539 | 0.005408 | 0.008969 | 0.005231 | 0.005366 | 0.005798 | 0.027573 | 0.005701 | 0.006912 |
| 33.4305556 | 0.02725 | 0.031134 | 0.045589 | 0.006759 | 0.031057 | 0.063826 | 0.020275 | 0.105323 | 0.016248 | 0.030454 | 0.047854 | 0.061727 | 0.054987 | 0.030086 | 0.007667 | 0.014082 | 0.00667 | 0.008186 | 0.008341 | 0.030884 | 0.008409 | 0.013803 |
| 33.4444444 | 0.042325 | 0.033026 | 0.054322 | 0.005897 | 0.034059 | 0.069489 | 0.025131 | 0.121489 | 0.017401 | 0.024094 | 0.045158 | 0.067623 | 0.068553 | 0.043703 | 0.012932 | 0.018886 | 0.010179 | 0.013733 | 0.01309 | 0.027319 | 0.014085 | 0.032862 |
| 33.4583333 | 0.061466 | 0.020336 | 0.051064 | 0.005094 | 0.038408 | 0.062557 | 0.023795 | 0.14702 | 0.032659 | 0.022042 | 0.039289 | 0.072061 | 0.07932 | 0.044382 | 0.018686 | 0.025797 | 0.01266 | 0.020003 | 0.018103 | 0.019397 | 0.020721 | 0.0648 |
| 33.4722222 | 0.077064 | 0.009563 | 0.031419 | 0.007699 | 0.038891 | 0.054139 | 0.023683 | 0.158289 | 0.06039 | 0.027436 | 0.043424 | 0.078894 | 0.084754 | 0.026604 | 0.019993 | 0.044821 | 0.011799 | 0.022562 | 0.019743 | 0.013837 | 0.022593 | 0.087097 |
| 33.4861111 | 0.088683 | 0.008076 | 0.010318 | 0.008053 | 0.039268 | 0.056089 | 0.03177 | 0.139775 | 0.091042 | 0.039273 | 0.062952 | 0.085669 | 0.079824 | 0.01769 | 0.01448 | 0.043848 | 0.008107 | 0.017315 | 0.015107 | 0.01654 | 0.016106 | 0.082858 |
| 33.5 | 0.092248 | 0.016076 | 0.013514 | 0.006749 | 0.053107 | 0.059243 | 0.042662 | 0.106615 | 0.094403 | 0.053771 | 0.090284 | 0.094614 | 0.070817 | 0.020628 | 0.007514 | 0.030845 | 0.007715 | 0.00905 | 0.008163 | 0.026695 | 0.008142 | 0.065821 |

| | | | | | | | | | | | | | | | | | | | | | | |
|---|---|---|---|---|---|---|---|---|---|---|---|---|---|---|---|---|---|---|---|---|---|---|
| 33.5138889 | 0.083231 | 0.02403 | 0.03799 | 0.011884 | 0.073064 | 0.051882 | 0.049023 | 0.080664 | 0.061612 | 0.065976 | 0.110678 | 0.106938 | 0.066013 | 0.019724 | 0.006297 | 0.035052 | 0.01719 | 0.006874 | 0.0064 | 0.039296 | 0.00723 | 0.045425 |
| 33.5277778 | 0.069603 | 0.021417 | 0.055327 | 0.019595 | 0.080118 | 0.039321 | 0.049897 | 0.056688 | 0.03061 | 0.073689 | 0.109204 | 0.119271 | 0.060806 | 0.021699 | 0.007447 | 0.035805 | 0.029763 | 0.007623 | 0.006955 | 0.048686 | 0.008865 | 0.028606 |
| 33.5416667 | 0.071584 | 0.020218 | 0.053747 | 0.025295 | 0.075768 | 0.027499 | 0.052323 | 0.031009 | 0.021593 | 0.078545 | 0.085126 | 0.12699 | 0.049822 | 0.041674 | 0.006853 | 0.030861 | 0.032313 | 0.006739 | 0.006373 | 0.052422 | 0.007139 | 0.032119 |
| 33.5555556 | 0.086882 | 0.034491 | 0.042275 | 0.027097 | 0.068632 | 0.017789 | 0.056913 | 0.012125 | 0.015834 | 0.073453 | 0.051113 | 0.109791 | 0.036523 | 0.0657 | 0.00935 | 0.034029 | 0.023777 | 0.009435 | 0.009757 | 0.048921 | 0.00742 | 0.03578 |
| 33.5694444 | 0.088585 | 0.060508 | 0.028519 | 0.0205 | 0.061845 | 0.013067 | 0.05041 | 0.003399 | 0.019828 | 0.058886 | 0.026336 | 0.066827 | 0.029042 | 0.074176 | 0.013655 | 0.031909 | 0.013806 | 0.014189 | 0.014531 | 0.03483 | 0.01174 | 0.019653 |
| 33.5833333 | 0.069686 | 0.093101 | 0.015202 | 0.009673 | 0.055243 | 0.013538 | 0.0326 | 0.0029 | 0.024344 | 0.05105 | 0.023479 | 0.038762 | 0.024115 | 0.070302 | 0.012853 | 0.025336 | 0.009299 | 0.013796 | 0.013783 | 0.021825 | 0.012844 | 0.012807 |
| 33.5972222 | 0.039002 | 0.111797 | 0.012987 | 0.010526 | 0.042501 | 0.015342 | 0.015442 | 0.004245 | 0.011726 | 0.050631 | 0.039434 | 0.046472 | 0.017107 | 0.056826 | 0.007674 | 0.036399 | 0.012717 | 0.008614 | 0.00866 | 0.026455 | 0.008421 | 0.014802 |
| 33.6111111 | 0.019972 | 0.098378 | 0.024366 | 0.030209 | 0.028592 | 0.016423 | 0.006172 | 0.00345 | 0.007204 | 0.042614 | 0.042394 | 0.057905 | 0.011013 | 0.038262 | 0.003565 | 0.047478 | 0.023284 | 0.003987 | 0.004025 | 0.039424 | 0.003886 | 0.00997 |
| 33.625 | 0.025912 | 0.070127 | 0.033351 | 0.044585 | 0.023804 | 0.016589 | 0.003605 | 0.004453 | 0.01982 | 0.033844 | 0.025197 | 0.041109 | 0.014954 | 0.030758 | 0.003367 | 0.032155 | 0.032234 | 0.003455 | 0.003056 | 0.038574 | 0.003382 | 0.008053 |
| 33.6388889 | 0.038807 | 0.039675 | 0.030156 | 0.03107 | 0.02419 | 0.016753 | 0.00436 | 0.008345 | 0.024405 | 0.034959 | 0.016464 | 0.020156 | 0.048847 | 0.034103 | 0.005131 | 0.013223 | 0.029626 | 0.005549 | 0.004768 | 0.02241 | 0.005276 | 0.005904 |
| 33.6527778 | 0.060105 | 0.017993 | 0.017213 | 0.015043 | 0.021599 | 0.01666 | 0.011408 | 0.012535 | 0.012126 | 0.028433 | 0.009682 | 0.01572 | 0.096565 | 0.034465 | 0.006078 | 0.009776 | 0.019899 | 0.006926 | 0.006128 | 0.018603 | 0.00654 | 0.006946 |
| 33.6666667 | 0.088908 | 0.012327 | 0.007014 | 0.021077 | 0.021633 | 0.01281 | 0.017622 | 0.018056 | 0.002732 | 0.013421 | 0.013638 | 0.012951 | 0.093808 | 0.024997 | 0.005759 | 0.007894 | 0.013514 | 0.006379 | 0.005831 | 0.030867 | 0.006249 | 0.017194 |
| 33.6805556 | 0.102736 | 0.015205 | 0.013246 | 0.026532 | 0.024728 | 0.006926 | 0.012578 | 0.019887 | 0.001669 | 0.011448 | 0.040491 | 0.00727 | 0.057806 | 0.012246 | 0.004777 | 0.00432 | 0.011543 | 0.004732 | 0.004444 | 0.031796 | 0.004754 | 0.028792 |
| 33.6944444 | 0.109043 | 0.031652 | 0.039705 | 0.022174 | 0.018021 | 0.003523 | 0.00871 | 0.013848 | 0.001254 | 0.01833 | 0.070072 | 0.005577 | 0.056163 | 0.01535 | 0.003863 | 0.003248 | 0.011643 | 0.00333 | 0.003029 | 0.033693 | 0.003225 | 0.032988 |
| 33.7083333 | 0.12776 | 0.042098 | 0.065849 | 0.025311 | 0.01104 | 0.002267 | 0.015177 | 0.008173 | 0.008115 | 0.015864 | 0.081058 | 0.011696 | 0.060652 | 0.029866 | 0.002962 | 0.004418 | 0.012608 | 0.002084 | 0.001775 | 0.05356 | 0.001897 | 0.023171 |
| 33.7222222 | 0.136123 | 0.027625 | 0.056672 | 0.033097 | 0.015131 | 0.001603 | 0.019865 | 0.00588 | 0.025505 | 0.014027 | 0.055356 | 0.024346 | 0.039288 | 0.033072 | 0.002157 | 0.007258 | 0.013459 | 0.00088 | 0.000694 | 0.064459 | 0.000775 | 0.01345 |
| 33.7361111 | 0.101214 | 0.009876 | 0.029625 | 0.034038 | 0.022594 | 0.002142 | 0.018364 | 0.004896 | 0.032792 | 0.018938 | 0.040275 | 0.034005 | 0.029263 | 0.029811 | 0.002743 | 0.007845 | 0.014851 | 0.001438 | 0.00103 | 0.049212 | 0.001419 | 0.022399 |
| 33.75 | 0.04516 | 0.006123 | 0.021984 | 0.023186 | 0.020242 | 0.004429 | 0.01617 | 0.016741 | 0.023302 | 0.024682 | 0.078453 | 0.025883 | 0.052823 | 0.039887 | 0.004536 | 0.005748 | 0.016293 | 0.003755 | 0.00298 | 0.039002 | 0.003842 | 0.039485 |
| 33.7638889 | 0.016477 | 0.014608 | 0.024614 | 0.011242 | 0.016571 | 0.006313 | 0.011945 | 0.02871 | 0.022057 | 0.035094 | 0.116813 | 0.01272 | 0.066458 | 0.051736 | 0.005008 | 0.003304 | 0.014855 | 0.004834 | 0.004101 | 0.048702 | 0.005146 | 0.050123 |
| 33.7777778 | 0.033609 | 0.02327 | 0.022772 | 0.008835 | 0.018892 | 0.005185 | 0.006596 | 0.024477 | 0.027587 | 0.034884 | 0.112282 | 0.012928 | 0.042748 | 0.046678 | 0.003539 | 0.001999 | 0.010476 | 0.003582 | 0.003258 | 0.045698 | 0.00406 | 0.054475 |
| 33.7916667 | 0.074942 | 0.024794 | 0.016902 | 0.008393 | 0.01582 | 0.002402 | 0.009325 | 0.022802 | 0.019989 | 0.043035 | 0.076053 | 0.011755 | 0.031092 | 0.038899 | 0.00229 | 0.002555 | 0.006855 | 0.002158 | 0.002251 | 0.030666 | 0.002511 | 0.052515 |
| 33.8055556 | 0.083165 | 0.02862 | 0.011343 | 0.007962 | 0.018285 | 0.002227 | 0.013197 | 0.020346 | 0.008948 | 0.067337 | 0.03536 | 0.011925 | 0.068098 | 0.040377 | 0.002233 | 0.003739 | 0.005882 | 0.001973 | 0.002234 | 0.03421 | 0.00208 | 0.044576 |
| 33.8194444 | 0.056118 | 0.033807 | 0.008722 | 0.006887 | 0.034111 | 0.005987 | 0.008958 | 0.009908 | 0.013376 | 0.062886 | 0.016077 | 0.020011 | 0.114106 | 0.044431 | 0.002293 | 0.003286 | 0.005912 | 0.002148 | 0.002331 | 0.04868 | 0.002235 | 0.030438 |
| 33.8333333 | 0.06036 | 0.024947 | 0.013442 | 0.008937 | 0.052446 | 0.012272 | 0.009152 | 0.005476 | 0.029213 | 0.044967 | 0.025834 | 0.016282 | 0.129456 | 0.048278 | 0.00157 | 0.003794 | 0.005027 | 0.001569 | 0.001602 | 0.045046 | 0.001767 | 0.018012 |
| 33.8472222 | 0.101859 | 0.009549 | 0.028441 | 0.015205 | 0.055133 | 0.019591 | 0.01913 | 0.008778 | 0.036489 | 0.036001 | 0.048249 | 0.010314 | 0.107692 | 0.051823 | 0.001046 | 0.008623 | 0.003331 | 0.001051 | 0.000965 | 0.030057 | 0.001079 | 0.016612 |
| 33.8611111 | 0.133807 | 0.006592 | 0.040069 | 0.016833 | 0.034537 | 0.024778 | 0.025908 | 0.009182 | 0.026799 | 0.029327 | 0.067871 | 0.016512 | 0.062531 | 0.044183 | 0.001681 | 0.015629 | 0.00197 | 0.001607 | 0.001431 | 0.035816 | 0.001459 | 0.020451 |
| 33.875 | 0.134763 | 0.012226 | 0.043876 | 0.019368 | 0.014551 | 0.026571 | 0.02536 | 0.004218 | 0.015695 | 0.027187 | 0.088948 | 0.019684 | 0.042002 | 0.032417 | 0.002655 | 0.0235 | 0.001368 | 0.002633 | 0.002487 | 0.062762 | 0.002603 | 0.025151 |
| 33.8888889 | 0.113665 | 0.023964 | 0.044039 | 0.025645 | 0.014579 | 0.027891 | 0.028909 | 0.001836 | 0.012486 | 0.02662 | 0.108676 | 0.012648 | 0.051096 | 0.044411 | 0.004129 | 0.033832 | 0.002884 | 0.004507 | 0.004503 | 0.08028 | 0.004611 | 0.028577 |
| 33.9027778 | 0.086856 | 0.045897 | 0.031433 | 0.02232 | 0.020308 | 0.030883 | 0.038243 | 0.010117 | 0.012314 | 0.022635 | 0.109132 | 0.014496 | 0.044502 | 0.070277 | 0.007116 | 0.053257 | 0.008248 | 0.007967 | 0.007983 | 0.074204 | 0.008183 | 0.031386 |
| 33.9166667 | 0.063122 | 0.058074 | 0.015169 | 0.012 | 0.017919 | 0.03447 | 0.047483 | 0.028511 | 0.019529 | 0.021172 | 0.089499 | 0.04124 | 0.04257 | 0.087583 | 0.010654 | 0.078858 | 0.015241 | 0.011465 | 0.011224 | 0.056574 | 0.011648 | 0.036443 |
| 33.9305556 | 0.043799 | 0.045626 | 0.013661 | 0.011119 | 0.017917 | 0.038059 | 0.054562 | 0.042802 | 0.028174 | 0.02661 | 0.064418 | 0.071646 | 0.086305 | 0.096134 | 0.01194 | 0.088946 | 0.019613 | 0.012519 | 0.011929 | 0.041993 | 0.012587 | 0.040278 |
| 33.9444444 | 0.029575 | 0.027245 | 0.015719 | 0.015759 | 0.029559 | 0.044371 | 0.051617 | 0.050316 | 0.038926 | 0.02717 | 0.045676 | 0.082889 | 0.138152 | 0.095127 | 0.010049 | 0.084517 | 0.021097 | 0.010812 | 0.010064 | 0.031203 | 0.010867 | 0.037197 |
| 33.9583333 | 0.023663 | 0.027211 | 0.013601 | 0.015575 | 0.035835 | 0.054642 | 0.035745 | 0.065674 | 0.057394 | 0.020782 | 0.041621 | 0.080869 | 0.149778 | 0.088437 | 0.007531 | 0.087384 | 0.02333 | 0.008712 | 0.007989 | 0.021157 | 0.008615 | 0.024597 |
| 33.9722222 | 0.026276 | 0.051757 | 0.013033 | 0.012681 | 0.022531 | 0.060581 | 0.01964 | 0.086444 | 0.072888 | 0.018718 | 0.04642 | 0.06743 | 0.122555 | 0.088125 | 0.006656 | 0.095891 | 0.02781 | 0.00816 | 0.007516 | 0.015534 | 0.007627 | 0.011181 |
| 33.9861111 | 0.033035 | 0.085986 | 0.009729 | 0.009898 | 0.006748 | 0.053295 | 0.013797 | 0.090747 | 0.084679 | 0.027043 | 0.052196 | 0.041967 | 0.084287 | 0.096656 | 0.007509 | 0.094892 | 0.031841 | 0.008741 | 0.008224 | 0.01516 | 0.007793 | 0.008396 |
| 34 | 0.037137 | 0.104653 | 0.006384 | 0.007159 | 0.007626 | 0.037942 | 0.014741 | 0.076478 | 0.085782 | 0.032016 | 0.055312 | 0.022096 | 0.054928 | 0.102957 | 0.008351 | 0.081569 | 0.032082 | 0.008178 | 0.007994 | 0.015611 | 0.007424 | 0.016353 |
| 34.0138889 | 0.036563 | 0.091759 | 0.005809 | 0.006288 | 0.021097 | 0.022395 | 0.015441 | 0.053129 | 0.059211 | 0.027315 | 0.050166 | 0.023361 | 0.03614 | 0.092722 | 0.0062 | 0.060818 | 0.027078 | 0.006209 | 0.005695 | 0.012039 | 0.00557 | 0.025344 |
| 34.0277778 | 0.036374 | 0.053125 | 0.004453 | 0.006763 | 0.026022 | 0.010022 | 0.01439 | 0.026307 | 0.023457 | 0.018043 | 0.038749 | 0.045962 | 0.024243 | 0.064211 | 0.004391 | 0.038844 | 0.017129 | 0.007728 | 0.005292 | 0.005903 | 0.0062 | 0.021232 |
| 34.0416667 | 0.048144 | 0.018678 | 0.009266 | 0.010592 | 0.018929 | 0.008197 | 0.011333 | 0.014494 | 0.014064 | 0.018792 | 0.03316 | 0.074092 | 0.018002 | 0.036548 | 0.008084 | 0.022546 | 0.007695 | 0.013169 | 0.009286 | 0.002043 | 0.011151 | 0.009952 |
| 34.0555556 | 0.083946 | 0.016351 | 0.0235 | 0.023309 | 0.01451 | 0.024945 | 0.012384 | 0.033958 | 0.040295 | 0.048124 | 0.048178 | 0.089745 | 0.017753 | 0.020164 | 0.011988 | 0.016655 | 0.004885 | 0.01564 | 0.012789 | 0.00316 | 0.014212 | 0.006127 |
| 34.0694444 | 0.133573 | 0.035998 | 0.032387 | 0.035243 | 0.024342 | 0.058037 | 0.017211 | 0.079895 | 0.093896 | 0.095431 | 0.086454 | 0.089974 | 0.022096 | 0.009522 | 0.00977 | 0.021544 | 0.005599 | 0.013235 | 0.012498 | 0.01172 | 0.012335 | 0.014849 |
| 34.0833333 | 0.165618 | 0.040822 | 0.026873 | 0.028486 | 0.052275 | 0.093175 | 0.016932 | 0.131944 | 0.139939 | 0.124927 | 0.122323 | 0.07328 | 0.023786 | 0.007504 | 0.004062 | 0.026632 | 0.004876 | 0.008914 | 0.00937 | 0.021779 | 0.008281 | 0.029304 |
| 34.0972222 | 0.157199 | 0.024264 | 0.016987 | 0.011048 | 0.073118 | 0.109173 | 0.01375 | 0.162259 | 0.141285 | 0.107178 | 0.121107 | 0.052196 | 0.015782 | 0.013806 | 0.001785 | 0.020707 | 0.003855 | 0.004787 | 0.005342 | 0.019037 | 0.004352 | 0.029128 |
| 34.1111111 | 0.11398 | 0.013082 | 0.010565 | 0.003793 | 0.064543 | 0.091948 | 0.014198 | 0.150808 | 0.10242 | 0.057629 | 0.082804 | 0.0545 | 0.013383 | 0.018614 | 0.005114 | 0.012586 | 0.003976 | 0.003766 | 0.004118 | 0.010632 | 0.003467 | 0.018651 |
| 34.125 | 0.060366 | 0.0125 | 0.007191 | 0.007413 | 0.04611 | 0.051598 | 0.020064 | 0.114473 | 0.050632 | 0.020149 | 0.040451 | 0.069303 | 0.032579 | 0.019503 | 0.010223 | 0.017746 | 0.003331 | 0.007338 | 0.007309 | 0.013074 | 0.007318 | 0.020377 |
| 34.1388889 | 0.027992 | 0.009317 | 0.00539 | 0.015831 | 0.035835 | 0.028744 | 0.022469 | 0.082833 | 0.022136 | 0.012718 | 0.02969 | 0.070931 | 0.054564 | 0.021724 | 0.012033 | 0.03398 | 0.001826 | 0.0109 | 0.010018 | 0.015972 | 0.011203 | 0.026845 |
| 34.1527778 | 0.037636 | 0.00886 | 0.003342 | 0.021882 | 0.030224 | 0.043069 | 0.022116 | 0.053897 | 0.035198 | 0.022399 | 0.056044 | 0.060399 | 0.062493 | 0.019815 | 0.008922 | 0.048303 | 0.001357 | 0.009355 | 0.007688 | 0.010714 | 0.009749 | 0.018833 |
| 34.1666667 | 0.068319 | 0.023603 | 0.00466 | 0.021965 | 0.037745 | 0.057543 | 0.026194 | 0.028693 | 0.060187 | 0.037147 | 0.087528 | 0.037326 | 0.057699 | 0.012983 | 0.004245 | 0.051556 | 0.001843 | 0.00472 | 0.003506 | 0.006153 | 0.004893 | 0.006562 |
| 34.1805556 | 0.077408 | 0.038828 | 0.015429 | 0.02251 | 0.05749 | 0.048104 | 0.028748 | 0.028447 | 0.068545 | 0.051406 | 0.093416 | 0.024005 | 0.038337 | 0.01399 | 0.001414 | 0.042307 | 0.00213 | 0.001973 | 0.001933 | 0.009241 | 0.001877 | 0.004371 |
| 34.1944444 | 0.056865 | 0.032512 | 0.02995 | 0.021396 | 0.062918 | 0.030088 | 0.026672 | 0.045552 | 0.058724 | 0.058108 | 0.073712 | 0.031308 | 0.024864 | 0.026193 | 0.0007 | 0.025756 | 0.001857 | 0.001654 | 0.001915 | 0.01943 | 0.001566 | 0.009658 |
| 34.2083333 | 0.043338 | 0.014978 | 0.032309 | 0.013312 | 0.046494 | 0.020101 | 0.023644 | 0.053809 | 0.039125 | 0.052394 | 0.052863 | 0.031098 | 0.035699 | 0.043084 | 0.00048 | 0.010344 | 0.002683 | 0.001473 | 0.001493 | 0.026555 | 0.001428 | 0.017541 |
| 34.2222222 | 0.058082 | 0.006102 | 0.020121 | 0.007914 | 0.023469 | 0.021358 | 0.01896 | 0.052315 | 0.02045 | 0.038488 | 0.044577 | 0.040995 | 0.036338 | 0.048939 | 0.000212 | 0.002577 | 0.005411 | 0.001396 | 0.00132 | 0.022037 | 0.001108 | 0.023426 |
| 34.2361111 | 0.075574 | 0.012723 | 0.012788 | 0.007693 | 0.008423 | 0.025712 | 0.011701 | 0.052639 | 0.00842 | 0.026359 | 0.034845 | 0.062392 | 0.022637 | 0.041509 | 0.000717 | 0.00461 | 0.006612 | 0.001635 | 0.001453 | 0.011827 | 0.000889 | 0.019676 |
| 34.25 | 0.069005 | 0.047648 | 0.014409 | 0.008044 | 0.006583 | 0.026804 | 0.011007 | 0.053552 | 0.007616 | 0.020888 | 0.020804 | 0.050541 | 0.02161 | 0.039722 | 0.002276 | 0.016686 | 0.004423 | 0.001372 | 0.00122 | 0.006884 | 0.000849 | 0.010685 |
| 34.2638889 | 0.042557 | 0.087137 | 0.011645 | 0.011376 | 0.013446 | 0.024557 | 0.021099 | 0.048411 | 0.020676 | 0.017801 | 0.015584 | 0.039313 | 0.020361 | 0.040821 | 0.004163 | 0.031446 | 0.007028 | 0.001922 | 0.001887 | 0.00686 | 0.002327 | 0.009526 |
| 34.2777778 | 0.02035 | 0.0738 | 0.010017 | 0.011651 | 0.025767 | 0.019108 | 0.031111 | 0.041394 | 0.041578 | 0.015167 | 0.01623 | 0.063168 | 0.019496 | 0.028824 | 0.006495 | 0.039231 | 0.018613 | 0.005098 | 0.00484 | 0.01154 | 0.005708 | 0.021534 |
| 34.2916667 | 0.018456 | 0.051805 | 0.01392 | 0.007858 | 0.040779 | 0.017387 | 0.032986 | 0.039679 | 0.056588 | 0.017504 | 0.022528 | 0.072858 | 0.026329 | 0.023864 | 0.008788 | 0.039616 | 0.024453 | 0.00861 | 0.007931 | 0.026234 | 0.008987 | 0.042735 |
| 34.3055556 | 0.022162 | 0.055147 | 0.014182 | 0.006651 | 0.051986 | 0.028058 | 0.02694 | 0.036374 | 0.053848 | 0.023852 | 0.030935 | 0.059926 | 0.031953 | 0.037037 | 0.008031 | 0.03476 | 0.015538 | 0.008226 | 0.007404 | 0.043525 | 0.008673 | 0.058315 |
| 34.3194444 | 0.018976 | 0.041273 | 0.010648 | 0.007151 | 0.053974 | 0.042621 | 0.016112 | 0.023315 | 0.036657 | 0.031215 | 0.028266 | 0.057929 | 0.034674 | 0.040632 | 0.004391 | 0.027202 | 0.007775 | 0.004691 | 0.003831 | 0.052525 | 0.005057 | 0.059487 |
| 34.3333333 | 0.022998 | 0.056394 | 0.010021 | 0.008735 | 0.049066 | 0.045783 | 0.013551 | 0.012477 | 0.029412 | 0.038087 | 0.025409 | 0.069391 | 0.033933 | 0.024918 | 0.002221 | 0.023174 | 0.012366 | 0.002624 | 0.001364 | 0.051151 | 0.002733 | 0.0498 |
| 34.3472222 | 0.02743 | 0.104368 | 0.015351 | 0.01053 | 0.043843 | 0.037507 | 0.025054 | 0.020647 | 0.031388 | 0.040443 | 0.039761 | 0.084088 | 0.029333 | 0.01335 | 0.002326 | 0.02681 | 0.018113 | 0.002751 | 0.001052 | 0.041568 | 0.002816 | 0.035713 |
| 34.3611111 | 0.022122 | 0.109545 | 0.024772 | 0.007556 | 0.036388 | 0.030316 | 0.033845 | 0.046788 | 0.031057 | 0.038614 | 0.064545 | 0.094367 | 0.021024 | 0.024901 | 0.001924 | 0.035666 | 0.017834 | 0.002319 | 0.000927 | 0.02869 | 0.002517 | 0.021631 |
| 34.375 | 0.028041 | 0.072795 | 0.034551 | 0.002288 | 0.028561 | 0.032792 | 0.030043 | 0.06913 | 0.052812 | 0.038292 | 0.077076 | 0.092082 | 0.015252 | 0.034596 | 0.000863 | 0.044572 | 0.015877 | 0.001093 | 0.000753 | 0.017284 | 0.001179 | 0.009971 |
| 34.3888889 | 0.045684 | 0.032387 | 0.039507 | 0.000532 | 0.031953 | 0.042254 | 0.021239 | 0.068262 | 0.088256 | 0.041336 | 0.070313 | 0.073483 | 0.019409 | 0.024841 | 0.000665 | 0.05165 | 0.015965 | 0.000688 | 0.001331 | 0.0119 | 0.000411 | 0.005248 |
| 34.4027778 | 0.053552 | 0.014635 | 0.036934 | 0.001657 | 0.044199 | 0.048672 | 0.013484 | 0.050891 | 0.0955 | 0.046014 | 0.059559 | 0.049095 | 0.033037 | 0.018767 | 0.001111 | 0.05814 | 0.018709 | 0.001103 | 0.002088 | 0.014709 | 0.000683 | 0.011528 |
| 34.4166667 | 0.047876 | 0.029976 | 0.031847 | 0.003227 | 0.049674 | 0.048091 | 0.007863 | 0.03813 | 0.078434 | 0.053014 | 0.054614 | 0.037651 | 0.060393 | 0.028335 | 0.001385 | 0.064444 | 0.022598 | 0.001242 | 0.002139 | 0.022352 | 0.000902 | 0.020228 |
| 34.4305556 | 0.040455 | 0.047292 | 0.02875 | 0.003462 | 0.045155 | 0.04701 | 0.005584 | 0.04384 | 0.061583 | 0.064471 | 0.053098 | 0.047108 | 0.095149 | 0.047249 | 0.001295 | 0.070191 | 0.027364 | 0.000777 | 0.001461 | 0.029948 | 0.000498 | 0.043008 |
| 34.4444444 | 0.04292 | 0.034175 | 0.025249 | 0.002905 | 0.039657 | 0.048977 | 0.00627 | 0.066 | 0.044778 | 0.074486 | 0.051367 | 0.071858 | 0.114709 | 0.062121 | 0.00138 | 0.073374 | 0.033288 | 0.001036 | 0.001279 | 0.03378 | 0.001098 | 0.092023 |
| 34.4583333 | 0.05829 | 0.018136 | 0.020942 | 0.004154 | 0.035477 | 0.050537 | 0.006697 | 0.096423 | 0.033536 | 0.074521 | 0.05164 | 0.100532 | 0.101581 | 0.065467 | 0.003016 | 0.072833 | 0.03847 | 0.004503 | 0.003728 | 0.031772 | 0.004971 | 0.119573 |
| 34.4722222 | 0.082552 | 0.029392 | 0.024164 | 0.007212 | 0.03419 | 0.051343 | 0.007659 | 0.120358 | 0.036599 | 0.066723 | 0.064427 | 0.117337 | 0.062333 | 0.059681 | 0.006565 | 0.070039 | 0.040199 | 0.010625 | 0.008519 | 0.025727 | 0.010491 | 0.10217 |
| 34.4861111 | 0.108004 | 0.046233 | 0.039746 | 0.006987 | 0.040835 | 0.053156 | 0.01206 | 0.11067 | 0.047813 | 0.061143 | 0.095698 | 0.119946 | 0.037458 | 0.049144 | 0.009364 | 0.063584 | 0.038288 | 0.013998 | 0.011537 | 0.02092 | 0.012766 | 0.072259 |
| 34.5 | 0.120897 | 0.049223 | 0.054789 | 0.006776 | 0.053411 | 0.049919 | 0.016794 | 0.080014 | 0.069564 | 0.066211 | 0.127926 | 0.124789 | 0.057312 | 0.041051 | 0.008325 | 0.053771 | 0.035303 | 0.011193 | 0.009776 | 0.021635 | 0.009831 | 0.054589 |
| 34.5138889 | 0.10778 | 0.046805 | 0.065886 | 0.012238 | 0.06404 | 0.037322 | 0.018092 | 0.068692 | 0.089405 | 0.074775 | 0.128799 | 0.13512 | 0.107322 | 0.037966 | 0.004708 | 0.045506 | 0.033055 | 0.006009 | 0.005331 | 0.02616 | 0.005448 | 0.050339 |
| 34.5277778 | 0.079936 | 0.049105 | 0.076391 | 0.016078 | 0.066749 | 0.022164 | 0.017353 | 0.061906 | 0.086456 | 0.073624 | 0.098644 | 0.134599 | 0.147831 | 0.040066 | 0.002264 | 0.039208 | 0.031461 | 0.003557 | 0.002272 | 0.029392 | 0.004279 | 0.048638 |
| 34.5416667 | 0.069115 | 0.057901 | 0.077583 | 0.014427 | 0.066414 | 0.010533 | 0.017837 | 0.036657 | 0.071371 | 0.060892 | 0.076756 | 0.118569 | 0.14298 | 0.047637 | 0.003201 | 0.029721 | 0.028167 | 0.004125 | 0.003123 | 0.028651 | 0.004279 | 0.046834 |
| 34.5555556 | 0.088014 | 0.068252 | 0.069394 | 0.010225 | 0.067486 | 0.003578 | 0.018621 | 0.015377 | 0.067919 | 0.044216 | 0.081103 | 0.096808 | 0.092556 | 0.049836 | 0.008469 | 0.016385 | 0.018856 | 0.008352 | 0.008916 | 0.023545 | 0.005486 | 0.042162 |
| 34.5694444 | 0.109778 | 0.069133 | 0.054136 | 0.006054 | 0.072852 | 0.001009 | 0.014956 | 0.015801 | 0.084778 | 0.038367 | 0.091912 | 0.072769 | 0.042472 | 0.036005 | 0.013833 | 0.007786 | 0.008965 | 0.014522 | 0.015141 | 0.017517 | 0.010631 | 0.028176 |
| 34.5833333 | 0.105623 | 0.055127 | 0.039767 | 0.003826 | 0.083832 | 0.003696 | 0.009847 | 0.023944 | 0.092941 | 0.052511 | 0.08639 | 0.065077 | 0.040855 | 0.020457 | 0.012333 | 0.007775 | 0.008022 | 0.014317 | 0.014085 | 0.020704 | 0.012318 | 0.011851 |
| 34.5972222 | 0.080554 | 0.047358 | 0.027322 | 0.004275 | 0.083022 | 0.009652 | 0.009983 | 0.020625 | 0.06673 | 0.068092 | 0.067492 | 0.079167 | 0.062839 | 0.015585 | 0.006259 | 0.007428 | 0.013459 | 0.007873 | 0.007458 | 0.038674 | 0.007366 | 0.005831 |
| 34.6111111 | 0.044588 | 0.054337 | 0.020721 | 0.005783 | 0.066225 | 0.010255 | 0.012682 | 0.011412 | 0.031777 | 0.058651 | 0.045994 | 0.068647 | 0.056873 | 0.012402 | 0.003077 | 0.006356 | 0.018399 | 0.004422 | 0.00382 | 0.056374 | 0.00395 | 0.011015 |
| 34.625 | 0.017151 | 0.048915 | 0.024825 | 0.005697 | 0.047002 | 0.008902 | 0.013512 | 0.011275 | 0.023579 | 0.037253 | 0.027359 | 0.02928 | 0.056026 | 0.007799 | 0.003551 | 0.008109 | 0.016106 | 0.005877 | 0.004964 | 0.050189 | 0.00518 | 0.024379 |
| 34.6388889 | 0.01365 | 0.04146 | 0.027459 | 0.005018 | 0.030495 | 0.014658 | 0.015596 | 0.015087 | 0.028155 | 0.023909 | 0.024902 | 0.010561 | 0.070109 | 0.007252 | 0.004887 | 0.009703 | 0.008946 | 0.008104 | 0.00709 | 0.032076 | 0.007502 | 0.039195 |
| 34.6527778 | 0.021784 | 0.056286 | 0.031685 | 0.004252 | 0.031521 | 0.02011 | 0.015722 | 0.019014 | 0.022867 | 0.015626 | 0.028581 | 0.012055 | 0.048388 | 0.01022 | 0.006597 | 0.012255 | 0.007901 | 0.009679 | 0.008767 | 0.042463 | 0.009296 | 0.044815 |
| 34.6666667 | 0.024166 | 0.062202 | 0.027398 | 0.002786 | 0.040011 | 0.018815 | 0.00809 | 0.025614 | 0.018316 | 0.016928 | 0.019441 | 0.010122 | 0.020637 | 0.026095 | 0.008561 | 0.011411 | 0.012208 | 0.010951 | 0.010276 | 0.064536 | 0.010906 | 0.034408 |
| 34.6805556 | 0.028256 | 0.045669 | 0.016133 | 0.00153 | 0.031388 | 0.015941 | 0.008579 | 0.023913 | 0.010689 | 0.026339 | 0.007138 | 0.011425 | 0.037311 | 0.06026 | 0.010157 | 0.010478 | 0.015556 | 0.011908 | 0.011399 | 0.048694 | 0.012248 | 0.018053 |
| 34.6944444 | 0.058597 | 0.050158 | 0.033665 | 0.001886 | 0.017838 | 0.017246 | 0.03316 | 0.017741 | 0.00633 | 0.035104 | 0.002997 | 0.014642 | 0.053456 | 0.076093 | 0.010911 | 0.017334 | 0.016595 | 0.012129 | 0.011585 | 0.02632 | 0.012695 | 0.018938 |
| 34.7083333 | 0.09575 | 0.077332 | 0.066853 | 0.004935 | 0.01648 | 0.022351 | 0.05946 | 0.024348 | 0.023245 | 0.03733 | 0.004513 | 0.011269 | 0.054127 | 0.059191 | 0.010638 | 0.023415 | 0.015583 | 0.011331 | 0.010725 | 0.047155 | 0.011712 | 0.030279 |
| 34.7222222 | 0.11097 | 0.075622 | 0.0677 | 0.007454 | 0.028302 | 0.023387 | 0.061878 | 0.045994 | 0.038253 | 0.035849 | 0.018687 | 0.007471 | 0.075306 | 0.043044 | 0.009897 | 0.022314 | 0.013512 | 0.010238 | 0.009611 | 0.075774 | 0.010154 | 0.030607 |
| 34.7361111 | 0.092086 | 0.049158 | 0.036431 | 0.006846 | 0.032609 | 0.014915 | 0.051223 | 0.049816 | 0.033948 | 0.048306 | 0.044821 | 0.011475 | 0.067765 | 0.037945 | 0.008775 | 0.016454 | 0.010884 | 0.009048 | 0.008457 | 0.063832 | 0.008803 | 0.021534 |
| 34.75 | 0.052057 | 0.050609 | 0.013966 | 0.004852 | 0.020612 | 0.006546 | 0.037009 | 0.037366 | 0.031373 | 0.053385 | 0.053471 | 0.021501 | 0.028268 | 0.033488 | 0.006845 | 0.008678 | 0.007893 | 0.00719 | 0.006845 | 0.032053 | 0.007227 | 0.012066 |
| 34.7638889 | 0.043959 | 0.075339 | 0.026304 | 0.002424 | 0.014276 | 0.006442 | 0.028909 | 0.046587 | 0.031715 | 0.031098 | 0.034382 | 0.030651 | 0.01808 | 0.024177 | 0.004768 | 0.006289 | 0.005391 | 0.005275 | 0.005217 | 0.033237 | 0.005499 | 0.00618 |
| 34.7777778 | 0.060132 | 0.070647 | 0.050953 | 0.002773 | 0.017995 | 0.008611 | 0.036386 | 0.045362 | 0.022673 | 0.017113 | 0.022198 | 0.033426 | 0.030667 | 0.011088 | 0.003056 | 0.015977 | 0.003198 | 0.003795 | 0.003738 | 0.061529 | 0.003923 | 0.006872 |
| 34.7916667 | 0.04838 | 0.037407 | 0.055189 | 0.006682 | 0.015275 | 0.011019 | 0.039887 | 0.021288 | 0.011862 | 0.027752 | 0.038392 | 0.031242 | 0.047034 | 0.012607 | 0.001751 | 0.027102 | 0.001963 | 0.002412 | 0.002306 | 0.06977 | 0.002359 | 0.011145 |
| 34.8055556 | 0.031838 | 0.012361 | 0.044506 | 0.008819 | 0.011128 | 0.014507 | 0.029506 | 0.008177 | 0.007303 | 0.04091 | 0.057559 | 0.024426 | 0.063403 | 0.035404 | 0.001418 | 0.028893 | 0.003191 | 0.001714 | 0.001682 | 0.04986 | 0.001479 | 0.012848 |
| 34.8194444 | 0.043444 | 0.007985 | 0.028059 | 0.007292 | 0.016867 | 0.016349 | 0.024766 | 0.013228 | 0.010061 | 0.038791 | 0.048362 | 0.014644 | 0.063693 | 0.050422 | 0.001272 | 0.026556 | 0.005053 | 0.001435 | 0.001455 | 0.026679 | 0.001167 | 0.013174 |
| 34.8333333 | 0.05447 | 0.024552 | 0.014977 | 0.012902 | 0.023037 | 0.014109 | 0.02531 | 0.025563 | 0.020229 | 0.029835 | 0.02905 | 0.021776 | 0.040666 | 0.045077 | 0.000845 | 0.025517 | 0.006282 | 0.001179 | 0.001328 | 0.024055 | 0.000992 | 0.015897 |
| 34.8472222 | 0.043539 | 0.0439 | 0.015943 | 0.023019 | 0.027235 | 0.008432 | 0.021227 | 0.03466 | 0.030381 | 0.032022 | 0.043597 | 0.041319 | 0.032804 | 0.037275 | 0.000866 | 0.031375 | 0.008551 | 0.001543 | 0.001765 | 0.032271 | 0.001487 | 0.021507 |
| 34.8611111 | 0.024509 | 0.038835 | 0.017116 | 0.017953 | 0.040484 | 0.007106 | 0.028981 | 0.036975 | 0.026189 | 0.03563 | 0.086612 | 0.050474 | 0.073185 | 0.0332 | 0.000713 | 0.041629 | 0.012789 | 0.00156 | 0.001656 | 0.026295 | 0.001822 | 0.02879 |
| 34.875 | 0.020145 | 0.026307 | 0.009865 | 0.009322 | 0.051077 | 0.017753 | 0.036093 | 0.027173 | 0.020721 | 0.025016 | 0.125771 | 0.051255 | 0.117766 | 0.031409 | 0.000393 | 0.044911 | 0.019308 | 0.000904 | 0.000913 | 0.024698 | 0.001173 | 0.035862 |
| 34.8888889 | 0.047747 | 0.031639 | 0.003272 | 0.009829 | 0.043878 | 0.035658 | 0.028839 | 0.015782 | 0.032085 | 0.028628 | 0.117026 | 0.054815 | 0.118425 | 0.034944 | 0.00069 | 0.045161 | 0.027569 | 0.000633 | 0.000671 | 0.052501 | 0.000426 | 0.04132 |
| 34.9027778 | 0.087844 | 0.029454 | 0.001852 | 0.007955 | 0.040206 | 0.049362 | 0.031211 | 0.017639 | 0.043117 | 0.05699 | 0.059677 | 0.062785 | 0.086553 | 0.045669 | 0.001151 | 0.049118 | 0.035766 | 0.00074 | 0.000662 | 0.088114 | 0.000329 | 0.043878 |

| | | | | | | | | | | | | | | | | | | | | | | |
|---|---|---|---|---|---|---|---|---|---|---|---|---|---|---|---|---|---|---|---|---|---|---|
| 34.9166667 | 0.0971 | 0.016838 | 0.003795 | 0.00412 | 0.043733 | 0.055389 | 0.030585 | 0.020919 | 0.035106 | 0.067272 | 0.034143 | 0.060951 | 0.057249 | 0.061424 | 0.001177 | 0.059167 | 0.042769 | 0.001086 | 0.000752 | 0.103975 | 0.000809 | 0.041603 |
| 34.9305556 | 0.066283 | 0.027746 | 0.009194 | 0.007917 | 0.040377 | 0.060439 | 0.023573 | 0.017199 | 0.020494 | 0.050227 | 0.04291 | 0.050043 | 0.059242 | 0.073019 | 0.001332 | 0.078764 | 0.04762 | 0.001859 | 0.001481 | 0.085816 | 0.001794 | 0.035898 |
| 34.9444444 | 0.032003 | 0.057814 | 0.016435 | 0.018097 | 0.04278 | 0.067067 | 0.043472 | 0.026894 | 0.034183 | 0.045607 | 0.03062 | 0.04369 | 0.092103 | 0.074592 | 0.002089 | 0.098886 | 0.050905 | 0.00272 | 0.002613 | 0.04481 | 0.002848 | 0.03157 |
| 34.9583333 | 0.019702 | 0.077675 | 0.021504 | 0.021805 | 0.05732 | 0.065738 | 0.076983 | 0.060948 | 0.071663 | 0.061822 | 0.023076 | 0.039331 | 0.118705 | 0.07143 | 0.003332 | 0.107255 | 0.053264 | 0.003614 | 0.003813 | 0.020904 | 0.003699 | 0.0309 |
| 34.9722222 | 0.021573 | 0.063922 | 0.024616 | 0.014445 | 0.067154 | 0.053115 | 0.103216 | 0.107494 | 0.087551 | 0.072292 | 0.043746 | 0.056149 | 0.114218 | 0.068277 | 0.005138 | 0.102432 | 0.052741 | 0.005157 | 0.005346 | 0.021029 | 0.005022 | 0.031824 |
| 34.9861111 | 0.049027 | 0.035692 | 0.025262 | 0.005176 | 0.052817 | 0.040306 | 0.125071 | 0.149908 | 0.070839 | 0.07068 | 0.065727 | 0.099301 | 0.093513 | 0.062252 | 0.006852 | 0.091043 | 0.047981 | 0.006728 | 0.00663 | 0.017614 | 0.006479 | 0.031933 |
| 35 | 0.091738 | 0.040672 | 0.019246 | 0.00299 | 0.023816 | 0.03755 | 0.14298 | 0.156866 | 0.049835 | 0.052249 | 0.068007 | 0.131177 | 0.071866 | 0.051808 | 0.006565 | 0.077381 | 0.039266 | 0.005708 | 0.005651 | 0.007796 | 0.005672 | 0.029336 |
| 35.0138889 | 0.111881 | 0.061955 | 0.015504 | 0.008437 | 0.00981 | 0.03922 | 0.14011 | 0.106264 | 0.0577 | 0.02738 | 0.053644 | 0.141371 | 0.0526 | 0.040822 | 0.0046 | 0.058237 | 0.028772 | 0.004383 | 0.004023 | 0.004863 | 0.004234 | 0.022713 |
| 35.0277778 | 0.104516 | 0.052163 | 0.023998 | 0.013228 | 0.010101 | 0.033443 | 0.093664 | 0.043895 | 0.071433 | 0.022107 | 0.040695 | 0.135714 | 0.038702 | 0.032767 | 0.003863 | 0.034655 | 0.020139 | 0.0065 | 0.004761 | 0.007095 | 0.0056 | 0.015204 |
| 35.0416667 | 0.084956 | 0.041694 | 0.026792 | 0.010817 | 0.017046 | 0.026443 | 0.039759 | 0.027436 | 0.049362 | 0.026556 | 0.040269 | 0.122655 | 0.032007 | 0.028762 | 0.00409 | 0.020564 | 0.018057 | 0.007219 | 0.00529 | 0.014477 | 0.006405 | 0.01419 |
| 35.0555556 | 0.075235 | 0.077957 | 0.017269 | 0.007066 | 0.05272 | 0.035966 | 0.021437 | 0.053 | 0.036795 | 0.041217 | 0.060568 | 0.116922 | 0.028777 | 0.02674 | 0.005321 | 0.025554 | 0.021256 | 0.006007 | 0.0051 | 0.028129 | 0.006405 | 0.020541 |
| 35.0694444 | 0.084983 | 0.121932 | 0.019608 | 0.006547 | 0.11068 | 0.064886 | 0.017145 | 0.086979 | 0.077523 | 0.086719 | 0.104948 | 0.112488 | 0.021357 | 0.021667 | 0.007012 | 0.036649 | 0.018095 | 0.005655 | 0.004761 | 0.038888 | 0.006985 | 0.025426 |
| 35.0833333 | 0.101247 | 0.136611 | 0.033649 | 0.007302 | 0.144397 | 0.092376 | 0.016968 | 0.105885 | 0.127551 | 0.128729 | 0.149581 | 0.080221 | 0.011058 | 0.013238 | 0.005412 | 0.033495 | 0.012351 | 0.003641 | 0.002833 | 0.040219 | 0.004582 | 0.02239 |
| 35.0972222 | 0.099539 | 0.126114 | 0.032755 | 0.014254 | 0.138993 | 0.095978 | 0.029508 | 0.112326 | 0.136216 | 0.123378 | 0.149152 | 0.027863 | 0.005736 | 0.00944 | 0.003225 | 0.019456 | 0.016237 | 0.00154 | 0.001411 | 0.031674 | 0.001369 | 0.013331 |
| 35.1111111 | 0.071774 | 0.088952 | 0.016067 | 0.023802 | 0.107813 | 0.072655 | 0.035674 | 0.112299 | 0.1033 | 0.081366 | 0.097151 | 0.010496 | 0.008672 | 0.014563 | 0.003246 | 0.014153 | 0.018665 | 0.000984 | 0.001106 | 0.017521 | 0.000605 | 0.005702 |
| 35.125 | 0.037774 | 0.04424 | 0.004021 | 0.023824 | 0.061936 | 0.036416 | 0.030758 | 0.10259 | 0.060256 | 0.036612 | 0.040069 | 0.030151 | 0.021116 | 0.017059 | 0.003262 | 0.017466 | 0.011859 | 0.001261 | 0.001406 | 0.006786 | 0.001311 | 0.003004 |
| 35.1388889 | 0.021555 | 0.022697 | 0.002508 | 0.017986 | 0.026589 | 0.015263 | 0.023125 | 0.082281 | 0.03 | 0.018931 | 0.018322 | 0.050073 | 0.036232 | 0.015195 | 0.005368 | 0.017016 | 0.008349 | 0.0026 | 0.002665 | 0.002156 | 0.002829 | 0.005389 |
| 35.1527778 | 0.041634 | 0.044248 | 0.003764 | 0.012376 | 0.020579 | 0.018765 | 0.014961 | 0.053625 | 0.014107 | 0.033571 | 0.033924 | 0.047933 | 0.040318 | 0.014293 | 0.011351 | 0.011613 | 0.013187 | 0.003671 | 0.003605 | 0.001018 | 0.003901 | 0.016088 |
| 35.1666667 | 0.087132 | 0.090018 | 0.006079 | 0.006943 | 0.033196 | 0.02438 | 0.01482 | 0.02752 | 0.013926 | 0.053191 | 0.056903 | 0.041291 | 0.035168 | 0.009803 | 0.017894 | 0.006591 | 0.015843 | 0.003318 | 0.003439 | 0.002983 | 0.003443 | 0.029874 |
| 35.1805556 | 0.114493 | 0.103119 | 0.017659 | 0.007957 | 0.052769 | 0.021127 | 0.022345 | 0.023605 | 0.026039 | 0.062287 | 0.063018 | 0.06393 | 0.031234 | 0.010617 | 0.020497 | 0.005827 | 0.012093 | 0.002043 | 0.002551 | 0.009758 | 0.002029 | 0.034119 |
| 35.1944444 | 0.100216 | 0.063852 | 0.032895 | 0.013669 | 0.072587 | 0.014498 | 0.02455 | 0.044121 | 0.034971 | 0.064301 | 0.056152 | 0.086225 | 0.029296 | 0.019699 | 0.014842 | 0.005972 | 0.007207 | 0.000918 | 0.001377 | 0.020709 | 0.000876 | 0.026903 |
| 35.2083333 | 0.062935 | 0.030411 | 0.033118 | 0.013423 | 0.073998 | 0.007827 | 0.02055 | 0.065755 | 0.032163 | 0.057795 | 0.052845 | 0.070386 | 0.02951 | 0.034636 | 0.009283 | 0.009515 | 0.003905 | 0.000788 | 0.000827 | 0.029203 | 0.000791 | 0.019754 |
| 35.2222222 | 0.03151 | 0.042486 | 0.019923 | 0.009386 | 0.049987 | 0.003382 | 0.017976 | 0.079411 | 0.019912 | 0.049482 | 0.056577 | 0.045515 | 0.027869 | 0.047411 | 0.014524 | 0.028606 | 0.004693 | 0.001716 | 0.001615 | 0.029938 | 0.001634 | 0.018541 |
| 35.2361111 | 0.023627 | 0.065271 | 0.014808 | 0.011293 | 0.027482 | 0.001346 | 0.017065 | 0.082218 | 0.012879 | 0.045435 | 0.055954 | 0.060237 | 0.022574 | 0.04264 | 0.020501 | 0.059134 | 0.010942 | 0.002923 | 0.002808 | 0.022826 | 0.002743 | 0.013958 |
| 35.25 | 0.036376 | 0.061257 | 0.024298 | 0.013492 | 0.028335 | 0.003966 | 0.017119 | 0.060054 | 0.016405 | 0.039918 | 0.044371 | 0.079863 | 0.026087 | 0.028032 | 0.018996 | 0.079611 | 0.018719 | 0.003767 | 0.003549 | 0.011537 | 0.003631 | 0.008546 |
| 35.2638889 | 0.044612 | 0.051349 | 0.039763 | 0.008147 | 0.028534 | 0.017995 | 0.023978 | 0.026011 | 0.017258 | 0.025533 | 0.025491 | 0.054669 | 0.031643 | 0.013336 | 0.013547 | 0.073542 | 0.022439 | 0.003986 | 0.003726 | 0.004156 | 0.003856 | 0.012647 |
| 35.2777778 | 0.034616 | 0.083041 | 0.069094 | 0.00878 | 0.017746 | 0.042744 | 0.033744 | 0.01159 | 0.023919 | 0.009053 | 0.01342 | 0.042757 | 0.023674 | 0.011479 | 0.007238 | 0.048446 | 0.019775 | 0.003393 | 0.003169 | 0.006786 | 0.003139 | 0.016402 |
| 35.2916667 | 0.022904 | 0.116636 | 0.095505 | 0.016555 | 0.024002 | 0.063374 | 0.02817 | 0.022986 | 0.047615 | 0.006369 | 0.023155 | 0.083159 | 0.0219 | 0.025037 | 0.002887 | 0.026535 | 0.012526 | 0.002221 | 0.002022 | 0.018449 | 0.001995 | 0.011803 |
| 35.3055556 | 0.022532 | 0.104372 | 0.077101 | 0.014912 | 0.045574 | 0.065335 | 0.014983 | 0.045336 | 0.074548 | 0.016054 | 0.043239 | 0.110512 | 0.035416 | 0.038233 | 0.001197 | 0.019129 | 0.005396 | 0.001117 | 0.001092 | 0.033625 | 0.001072 | 0.005165 |
| 35.3194444 | 0.026215 | 0.075942 | 0.048705 | 0.007394 | 0.063425 | 0.05456 | 0.014358 | 0.063245 | 0.088194 | 0.027225 | 0.041259 | 0.087133 | 0.040053 | 0.038239 | 0.001234 | 0.021802 | 0.00243 | 0.000652 | 0.001077 | 0.044992 | 0.000468 | 0.002195 |
| 35.3333333 | 0.028194 | 0.05031 | 0.053725 | 0.007683 | 0.073482 | 0.051078 | 0.015875 | 0.071872 | 0.085853 | 0.037898 | 0.020845 | 0.045722 | 0.029474 | 0.028943 | 0.001361 | 0.025951 | 0.002398 | 0.000615 | 0.00112 | 0.047764 | 0.000232 | 0.007233 |
| 35.3472222 | 0.03405 | 0.027711 | 0.04913 | 0.010593 | 0.075915 | 0.059604 | 0.012133 | 0.078926 | 0.077455 | 0.05005 | 0.010972 | 0.028818 | 0.01487 | 0.022288 | 0.000999 | 0.029564 | 0.002168 | 0.000793 | 0.000814 | 0.042042 | 0.000472 | 0.015137 |
| 35.3611111 | 0.049954 | 0.017224 | 0.035636 | 0.007684 | 0.069774 | 0.070342 | 0.015251 | 0.089468 | 0.069714 | 0.061765 | 0.014479 | 0.054362 | 0.008222 | 0.019453 | 0.00098 | 0.035359 | 0.002868 | 0.001305 | 0.000914 | 0.030905 | 0.000957 | 0.017437 |
| 35.375 | 0.063394 | 0.027835 | 0.04741 | 0.004941 | 0.05696 | 0.075426 | 0.021753 | 0.093679 | 0.061713 | 0.067356 | 0.032737 | 0.100657 | 0.018795 | 0.034718 | 0.000844 | 0.043719 | 0.005737 | 0.001269 | 0.000905 | 0.01981 | 0.000969 | 0.027281 |
| 35.3888889 | 0.053006 | 0.050913 | 0.050352 | 0.006553 | 0.043376 | 0.064843 | 0.022453 | 0.080358 | 0.052349 | 0.064521 | 0.057832 | 0.133213 | 0.042677 | 0.057863 | 0.000411 | 0.050968 | 0.009053 | 0.000694 | 0.000694 | 0.013453 | 0.000596 | 0.046544 |
| 35.4027778 | 0.025989 | 0.059955 | 0.03391 | 0.009481 | 0.033719 | 0.040294 | 0.020297 | 0.054802 | 0.044245 | 0.05801 | 0.064996 | 0.129345 | 0.066828 | 0.067267 | 0.000318 | 0.055052 | 0.012491 | 0.000529 | 0.000866 | 0.013161 | 0.00046 | 0.057269 |
| 35.4166667 | 0.008141 | 0.049796 | 0.020496 | 0.009805 | 0.033358 | 0.023803 | 0.018143 | 0.03317 | 0.042646 | 0.051828 | 0.054777 | 0.097538 | 0.079755 | 0.063241 | 0.000312 | 0.059872 | 0.01631 | 0.000654 | 0.001084 | 0.018208 | 0.000385 | 0.062283 |
| 35.4305556 | 0.009021 | 0.035073 | 0.014903 | 0.008508 | 0.042272 | 0.028876 | 0.01635 | 0.027554 | 0.047937 | 0.0438 | 0.044119 | 0.066 | 0.080145 | 0.054822 | 0.000317 | 0.06711 | 0.01896 | 0.000959 | 0.001272 | 0.024964 | 0.000547 | 0.066251 |
| 35.4444444 | 0.031956 | 0.026282 | 0.020171 | 0.007055 | 0.051792 | 0.049 | 0.014719 | 0.041999 | 0.055702 | 0.036053 | 0.043201 | 0.045655 | 0.086166 | 0.062399 | 0.000683 | 0.0723 | 0.019047 | 0.001321 | 0.001501 | 0.027961 | 0.001422 | 0.062463 |
| 35.4583333 | 0.076258 | 0.041256 | 0.030141 | 0.007908 | 0.055819 | 0.068961 | 0.011821 | 0.069703 | 0.069503 | 0.034709 | 0.051701 | 0.03832 | 0.115463 | 0.084363 | 0.001457 | 0.074352 | 0.016343 | 0.002148 | 0.002059 | 0.026395 | 0.003021 | 0.052466 |
| 35.4722222 | 0.116632 | 0.070114 | 0.037075 | 0.009155 | 0.055391 | 0.074042 | 0.007483 | 0.098366 | 0.087797 | 0.041103 | 0.06309 | 0.049187 | 0.151967 | 0.098255 | 0.002035 | 0.076449 | 0.011823 | 0.003221 | 0.002428 | 0.025111 | 0.003606 | 0.046653 |
| 35.4861111 | 0.125806 | 0.067149 | 0.042526 | 0.005354 | 0.055442 | 0.067058 | 0.005859 | 0.132307 | 0.096024 | 0.051849 | 0.075909 | 0.08062 | 0.161599 | 0.090176 | 0.001888 | 0.078678 | 0.008202 | 0.003121 | 0.002221 | 0.02679 | 0.002766 | 0.05113 |
| 35.5 | 0.108108 | 0.046037 | 0.046174 | 0.003243 | 0.062804 | 0.066244 | 0.01823 | 0.161925 | 0.093277 | 0.060407 | 0.091531 | 0.113715 | 0.132852 | 0.068616 | 0.002009 | 0.075907 | 0.007594 | 0.003027 | 0.002932 | 0.029225 | 0.003151 | 0.057201 |
| 35.5138889 | 0.089944 | 0.05891 | 0.048804 | 0.006636 | 0.073301 | 0.068818 | 0.04635 | 0.155291 | 0.081941 | 0.067428 | 0.108822 | 0.120347 | 0.084916 | 0.053971 | 0.002324 | 0.063475 | 0.008277 | 0.003526 | 0.003655 | 0.031802 | 0.003972 | 0.05578 |
| 35.5277778 | 0.088145 | 0.084182 | 0.058555 | 0.008083 | 0.071799 | 0.058553 | 0.062793 | 0.120494 | 0.064564 | 0.069724 | 0.125918 | 0.114076 | 0.042241 | 0.045127 | 0.003251 | 0.043885 | 0.006359 | 0.003974 | 0.003925 | 0.036039 | 0.003667 | 0.050859 |
| 35.5416667 | 0.098846 | 0.078894 | 0.068478 | 0.005449 | 0.059865 | 0.040707 | 0.044668 | 0.088211 | 0.059594 | 0.060614 | 0.132469 | 0.119291 | 0.021719 | 0.02914 | 0.00947 | 0.022686 | 0.006885 | 0.009209 | 0.009192 | 0.040397 | 0.007049 | 0.044353 |
| 35.5555556 | 0.097972 | 0.051563 | 0.061815 | 0.004314 | 0.051013 | 0.027448 | 0.01873 | 0.062314 | 0.073928 | 0.060024 | 0.132631 | 0.113719 | 0.029397 | 0.013635 | 0.020975 | 0.010167 | 0.016981 | 0.021242 | 0.020705 | 0.036576 | 0.017276 | 0.033727 |
| 35.5694444 | 0.070521 | 0.024882 | 0.053722 | 0.004805 | 0.048489 | 0.019342 | 0.020491 | 0.038384 | 0.081957 | 0.084433 | 0.129769 | 0.077642 | 0.052827 | 0.010312 | 0.028067 | 0.011333 | 0.026554 | 0.029783 | 0.028493 | 0.021874 | 0.025954 | 0.023311 |
| 35.5833333 | 0.042828 | 0.013289 | 0.069777 | 0.003777 | 0.042961 | 0.013611 | 0.034616 | 0.023684 | 0.064434 | 0.098421 | 0.089532 | 0.049401 | 0.073497 | 0.022918 | 0.024411 | 0.01241 | 0.025659 | 0.026343 | 0.025071 | 0.013437 | 0.024256 | 0.020651 |
| 35.5972222 | 0.039926 | 0.012883 | 0.085878 | 0.004447 | 0.030679 | 0.010905 | 0.028861 | 0.015252 | 0.034276 | 0.066668 | 0.049111 | 0.063164 | 0.072834 | 0.036463 | 0.015381 | 0.007533 | 0.019806 | 0.016342 | 0.015578 | 0.024473 | 0.015761 | 0.029552 |
| 35.6111111 | 0.039549 | 0.009003 | 0.065702 | 0.010033 | 0.027068 | 0.01328 | 0.01305 | 0.006898 | 0.018905 | 0.0298 | 0.049447 | 0.071216 | 0.052705 | 0.031611 | 0.008522 | 0.005932 | 0.016593 | 0.008613 | 0.008155 | 0.047822 | 0.008554 | 0.041689 |
| 35.625 | 0.034056 | 0.008133 | 0.029361 | 0.014381 | 0.033803 | 0.019748 | 0.010287 | 0.010632 | 0.028475 | 0.020002 | 0.043447 | 0.040765 | 0.036091 | 0.028357 | 0.004951 | 0.009479 | 0.017337 | 0.004692 | 0.004377 | 0.061431 | 0.004679 | 0.036621 |
| 35.6388889 | 0.046717 | 0.01048 | 0.017897 | 0.010495 | 0.028766 | 0.022638 | 0.010168 | 0.017977 | 0.030168 | 0.023135 | 0.037366 | 0.023884 | 0.03291 | 0.036517 | 0.00301 | 0.012933 | 0.018355 | 0.002653 | 0.002564 | 0.045906 | 0.002651 | 0.01936 |
| 35.6527778 | 0.062705 | 0.010006 | 0.037409 | 0.004581 | 0.016565 | 0.019144 | 0.008411 | 0.01467 | 0.020638 | 0.035846 | 0.039283 | 0.027032 | 0.053674 | 0.036461 | 0.002171 | 0.013022 | 0.016013 | 0.001856 | 0.001931 | 0.038782 | 0.001997 | 0.010258 |
| 35.6666667 | 0.089779 | 0.026207 | 0.065097 | 0.003484 | 0.017642 | 0.013204 | 0.012134 | 0.022264 | 0.020647 | 0.039426 | 0.039039 | 0.016591 | 0.08836 | 0.028003 | 0.001654 | 0.008738 | 0.010758 | 0.001362 | 0.001436 | 0.06454 | 0.001674 | 0.005097 |
| 35.6805556 | 0.115205 | 0.053014 | 0.075297 | 0.004094 | 0.019869 | 0.008163 | 0.016469 | 0.034365 | 0.014476 | 0.03291 | 0.058605 | 0.012863 | 0.098491 | 0.021569 | 0.000985 | 0.00411 | 0.00558 | 0.000895 | 0.000822 | 0.065627 | 0.001162 | 0.005584 |
| 35.6944444 | 0.085833 | 0.059927 | 0.051443 | 0.008995 | 0.021331 | 0.010553 | 0.01785 | 0.023058 | 0.007213 | 0.047472 | 0.082006 | 0.0207 | 0.085405 | 0.016375 | 0.00071 | 0.004265 | 0.002398 | 0.001177 | 0.000808 | 0.034939 | 0.001268 | 0.009858 |
| 35.7083333 | 0.036043 | 0.047558 | 0.030819 | 0.018081 | 0.033542 | 0.023047 | 0.015805 | 0.012906 | 0.010733 | 0.084437 | 0.079801 | 0.022429 | 0.064918 | 0.011809 | 0.000921 | 0.006039 | 0.002863 | 0.001701 | 0.001185 | 0.032461 | 0.001667 | 0.008033 |
| 35.7222222 | 0.01596 | 0.035129 | 0.044965 | 0.017121 | 0.03548 | 0.029635 | 0.015901 | 0.025577 | 0.012092 | 0.106446 | 0.049725 | 0.033875 | 0.035351 | 0.014739 | 0.00189 | 0.005586 | 0.008493 | 0.002686 | 0.002226 | 0.059178 | 0.002445 | 0.0101 |
| 35.7361111 | 0.02098 | 0.024731 | 0.056169 | 0.008065 | 0.024236 | 0.022364 | 0.014795 | 0.042244 | 0.007249 | 0.084427 | 0.032109 | 0.047398 | 0.01372 | 0.019398 | 0.003493 | 0.004078 | 0.017016 | 0.004279 | 0.003873 | 0.062993 | 0.003796 | 0.01834 |
| 35.75 | 0.021956 | 0.012833 | 0.049694 | 0.004011 | 0.018925 | 0.013966 | 0.012226 | 0.041231 | 0.006331 | 0.051342 | 0.055051 | 0.036052 | 0.034285 | 0.017388 | 0.004377 | 0.003338 | 0.024231 | 0.005025 | 0.004626 | 0.034488 | 0.004448 | 0.028644 |
| 35.7638889 | 0.012083 | 0.00918 | 0.031003 | 0.003782 | 0.015307 | 0.013269 | 0.013797 | 0.030634 | 0.012298 | 0.060097 | 0.071379 | 0.020435 | 0.075046 | 0.018306 | 0.003985 | 0.004036 | 0.026319 | 0.004356 | 0.003981 | 0.015135 | 0.003816 | 0.041268 |
| 35.7777778 | 0.01075 | 0.018019 | 0.01709 | 0.005595 | 0.017984 | 0.02183 | 0.018509 | 0.027554 | 0.0143 | 0.07956 | 0.057578 | 0.020404 | 0.070111 | 0.018881 | 0.003117 | 0.00564 | 0.021695 | 0.00318 | 0.00298 | 0.028806 | 0.002754 | 0.052277 |
| 35.7916667 | 0.013906 | 0.039848 | 0.027401 | 0.007504 | 0.022326 | 0.027003 | 0.036679 | 0.022192 | 0.009593 | 0.072175 | 0.033673 | 0.024136 | 0.035664 | 0.009616 | 0.002544 | 0.006538 | 0.014271 | 0.00244 | 0.00251 | 0.053048 | 0.00222 | 0.048769 |
| 35.8055556 | 0.010947 | 0.051547 | 0.041644 | 0.006236 | 0.022106 | 0.022552 | 0.066275 | 0.015465 | 0.010901 | 0.059284 | 0.01537 | 0.024312 | 0.013601 | 0.012602 | 0.002246 | 0.008676 | 0.008325 | 0.002212 | 0.002338 | 0.056059 | 0.002237 | 0.028662 |
| 35.8194444 | 0.008042 | 0.034065 | 0.056387 | 0.006209 | 0.022004 | 0.01477 | 0.087208 | 0.012353 | 0.021997 | 0.038839 | 0.012766 | 0.023631 | 0.010504 | 0.037784 | 0.001895 | 0.019626 | 0.004403 | 0.002044 | 0.001991 | 0.035419 | 0.002184 | 0.018131 |
| 35.8333333 | 0.007994 | 0.01642 | 0.07139 | 0.009873 | 0.01271 | 0.009902 | 0.082215 | 0.008141 | 0.032363 | 0.017518 | 0.015723 | 0.025211 | 0.033862 | 0.061563 | 0.001605 | 0.033617 | 0.001897 | 0.001878 | 0.001726 | 0.016977 | 0.001948 | 0.024095 |
| 35.8472222 | 0.019062 | 0.014507 | 0.062364 | 0.011663 | 0.005355 | 0.016987 | 0.049738 | 0.005589 | 0.039038 | 0.010716 | 0.028008 | 0.021678 | 0.076197 | 0.066177 | 0.001798 | 0.033978 | 0.001043 | 0.002086 | 0.002002 | 0.018325 | 0.00211 | 0.02208 |
| 35.8611111 | 0.058325 | 0.016594 | 0.03796 | 0.010143 | 0.009073 | 0.036182 | 0.017261 | 0.007737 | 0.03396 | 0.00803 | 0.064393 | 0.011914 | 0.122928 | 0.055229 | 0.002273 | 0.026356 | 0.002221 | 0.002452 | 0.002349 | 0.021287 | 0.002617 | 0.013173 |
| 35.875 | 0.103009 | 0.020113 | 0.024067 | 0.009441 | 0.018748 | 0.057274 | 0.022207 | 0.010291 | 0.023896 | 0.006707 | 0.107201 | 0.005732 | 0.156843 | 0.042165 | 0.002368 | 0.026945 | 0.004612 | 0.002355 | 0.001998 | 0.019479 | 0.002608 | 0.018238 |
| 35.8888889 | 0.096222 | 0.02873 | 0.022684 | 0.008793 | 0.034124 | 0.069154 | 0.052853 | 0.008225 | 0.027306 | 0.004226 | 0.122194 | 0.010444 | 0.162321 | 0.029902 | 0.002381 | 0.034825 | 0.007011 | 0.002297 | 0.001683 | 0.034183 | 0.002297 | 0.02903 |
| 35.9027778 | 0.044818 | 0.036794 | 0.031731 | 0.005917 | 0.059306 | 0.066044 | 0.05677 | 0.008315 | 0.027261 | 0.00791 | 0.096051 | 0.025043 | 0.145888 | 0.019092 | 0.003206 | 0.043656 | 0.008631 | 0.003316 | 0.002647 | 0.064208 | 0.00291 | 0.033302 |
| 35.9166667 | 0.017792 | 0.028589 | 0.050463 | 0.002534 | 0.078787 | 0.06134 | 0.032827 | 0.013202 | 0.018044 | 0.026015 | 0.050647 | 0.041549 | 0.123876 | 0.01225 | 0.004565 | 0.05332 | 0.009538 | 0.005004 | 0.004328 | 0.087452 | 0.004422 | 0.035201 |
| 35.9305556 | 0.027385 | 0.010288 | 0.067054 | 0.001201 | 0.063756 | 0.066755 | 0.020382 | 0.012804 | 0.017879 | 0.031774 | 0.026899 | 0.054718 | 0.102492 | 0.006498 | 0.005508 | 0.064016 | 0.010285 | 0.006162 | 0.005361 | 0.091708 | 0.005721 | 0.039103 |
| 35.9444444 | 0.050974 | 0.007793 | 0.061715 | 0.001052 | 0.032262 | 0.062269 | 0.024695 | 0.010574 | 0.02831 | 0.014654 | 0.035497 | 0.054281 | 0.077829 | 0.00485 | 0.00588 | 0.073789 | 0.010789 | 0.00659 | 0.005598 | 0.077891 | 0.006411 | 0.045202 |
| 35.9583333 | 0.085489 | 0.032342 | 0.034003 | 0.001965 | 0.014854 | 0.042147 | 0.023566 | 0.01778 | 0.037335 | 0.0023 | 0.053131 | 0.035628 | 0.045501 | 0.013367 | 0.005857 | 0.08211 | 0.011324 | 0.006705 | 0.005577 | 0.05153 | 0.006663 | 0.040877 |
| 35.9722222 | 0.121407 | 0.080991 | 0.015637 | 0.007018 | 0.011302 | 0.025471 | 0.0143 | 0.036283 | 0.042513 | 0.005665 | 0.072531 | 0.029323 | 0.021057 | 0.028496 | 0.005431 | 0.089783 | 0.012402 | 0.006642 | 0.005526 | 0.025853 | 0.006551 | 0.026663 |
| 35.9861111 | 0.136965 | 0.124377 | 0.020514 | 0.014048 | 0.017394 | 0.020763 | 0.014686 | 0.066255 | 0.05019 | 0.011404 | 0.096864 | 0.060448 | 0.023363 | 0.04188 | 0.004242 | 0.094987 | 0.013489 | 0.006558 | 0.005561 | 0.01 | 0.006393 | 0.021656 |
| 36 | 0.105745 | 0.116246 | 0.02746 | 0.015201 | 0.034713 | 0.032609 | 0.032809 | 0.090546 | 0.048829 | 0.014032 | 0.10794 | 0.106653 | 0.040462 | 0.049808 | 0.002508 | 0.092108 | 0.013435 | 0.007234 | 0.006146 | 0.002401 | 0.006991 | 0.02446 |
| 36.0138889 | 0.068553 | 0.062147 | 0.0235 | 0.011213 | 0.058493 | 0.05927 | 0.054743 | 0.090041 | 0.038127 | 0.017616 | 0.090377 | 0.134798 | 0.051529 | 0.049748 | 0.001227 | 0.077842 | 0.011081 | 0.010045 | 0.008189 | 0.000869 | 0.009433 | 0.025287 |
| 36.0277778 | 0.085513 | 0.023034 | 0.013282 | 0.01223 | 0.074658 | 0.086967 | 0.061956 | 0.060579 | 0.030984 | 0.021393 | 0.058565 | 0.147267 | 0.048194 | 0.043169 | 0.000651 | 0.057892 | 0.007444 | 0.013555 | 0.010517 | 0.003506 | 0.012445 | 0.027058 |
| 36.0416667 | 0.110215 | 0.046865 | 0.006828 | 0.015467 | 0.081478 | 0.104356 | 0.044352 | 0.025123 | 0.019477 | 0.030064 | 0.03859 | 0.168391 | 0.030959 | 0.037892 | 0.002018 | 0.045254 | 0.0058 | 0.012105 | 0.009088 | 0.011127 | 0.010911 | 0.036194 |
| 36.0555556 | 0.100606 | 0.113035 | 0.008082 | 0.01323 | 0.105446 | 0.110932 | 0.016904 | 0.038396 | 0.023459 | 0.059102 | 0.047356 | 0.18672 | 0.014383 | 0.036363 | 0.00611 | 0.051235 | 0.004893 | 0.006363 | 0.005093 | 0.02105 | 0.005136 | 0.053473 |
| 36.0694444 | 0.07704 | 0.149345 | 0.013315 | 0.015412 | 0.147224 | 0.103697 | 0.005739 | 0.108021 | 0.076218 | 0.108756 | 0.087662 | 0.178015 | 0.014572 | 0.033109 | 0.009817 | 0.063522 | 0.006169 | 0.005687 | 0.005854 | 0.021955 | 0.00418 | 0.074536 |
| 36.0833333 | 0.074492 | 0.133205 | 0.034903 | 0.02253 | 0.160713 | 0.074335 | 0.006793 | 0.175884 | 0.138528 | 0.142792 | 0.139522 | 0.136795 | 0.028161 | 0.02342 | 0.010841 | 0.055796 | 0.013719 | 0.01132 | 0.011428 | 0.014267 | 0.009927 | 0.08268 |
| 36.0972222 | 0.104663 | 0.098644 | 0.066553 | 0.020172 | 0.134587 | 0.037063 | 0.006158 | 0.194771 | 0.148265 | 0.13124 | 0.153698 | 0.086131 | 0.039412 | 0.016019 | 0.010859 | 0.029762 | 0.019294 | 0.014769 | 0.01432 | 0.011293 | 0.013889 | 0.063437 |
| 36.1111111 | 0.111068 | 0.067708 | 0.06862 | 0.010443 | 0.096172 | 0.016581 | 0.006906 | 0.170988 | 0.111731 | 0.086626 | 0.109629 | 0.064136 | 0.043517 | 0.020142 | 0.010097 | 0.009274 | 0.018136 | 0.012719 | 0.012029 | 0.01188 | 0.012177 | 0.033241 |
| 36.125 | 0.068994 | 0.049161 | 0.04168 | 0.003466 | 0.062396 | 0.016529 | 0.012172 | 0.1309 | 0.073724 | 0.042517 | 0.049588 | 0.063401 | 0.045019 | 0.024592 | 0.0069 | 0.005166 | 0.016433 | 0.008358 | 0.007774 | 0.009392 | 0.007941 | 0.01401 |
| 36.1388889 | 0.032443 | 0.038792 | 0.017759 | 0.001708 | 0.038151 | 0.022334 | 0.013666 | 0.092722 | 0.052947 | 0.017602 | 0.02165 | 0.053643 | 0.041184 | 0.017335 | 0.002945 | 0.009947 | 0.01503 | 0.004397 | 0.004127 | 0.006432 | 0.004105 | 0.01448 |
| 36.1527778 | 0.03969 | 0.030645 | 0.010475 | 0.003205 | 0.020488 | 0.01951 | 0.01012 | 0.064981 | 0.039226 | 0.014036 | 0.036994 | 0.03424 | 0.027501 | 0.005757 | 0.00118 | 0.017809 | 0.010391 | 0.001778 | 0.001692 | 0.005076 | 0.001618 | 0.01908 |
| 36.1666667 | 0.068465 | 0.048461 | 0.016497 | 0.005376 | 0.017613 | 0.013077 | 0.009258 | 0.041435 | 0.02241 | 0.027452 | 0.066316 | 0.034306 | 0.026712 | 0.006347 | 0.001142 | 0.026435 | 0.005095 | 0.000645 | 0.000705 | 0.003662 | 0.000642 | 0.012065 |
| 36.1805556 | 0.08275 | 0.08767 | 0.026896 | 0.005709 | 0.044316 | 0.017519 | 0.011214 | 0.022473 | 0.010125 | 0.048693 | 0.07679 | 0.068879 | 0.059919 | 0.015873 | 0.001197 | 0.029437 | 0.004626 | 0.000722 | 0.001169 | 0.00183 | 0.000958 | 0.006727 |
| 36.1944444 | 0.069892 | 0.098942 | 0.035188 | 0.005635 | 0.087795 | 0.030706 | 0.008913 | 0.010866 | 0.012328 | 0.07126 | 0.060768 | 0.107504 | 0.092352 | 0.020991 | 0.001685 | 0.025358 | 0.007732 | 0.00142 | 0.002027 | 0.003714 | 0.001742 | 0.010941 |
| 36.2083333 | 0.039044 | 0.067136 | 0.035439 | 0.009594 | 0.113534 | 0.034529 | 0.003416 | 0.010244 | 0.019995 | 0.082114 | 0.031442 | 0.110487 | 0.086258 | 0.025168 | 0.003218 | 0.023139 | 0.01052 | 0.001914 | 0.002278 | 0.014148 | 0.002016 | 0.01979 |
| 36.2222222 | 0.021702 | 0.031887 | 0.026043 | 0.015764 | 0.098562 | 0.021555 | 0.00471 | 0.018252 | 0.022615 | 0.071036 | 0.010233 | 0.069991 | 0.052211 | 0.027717 | 0.005359 | 0.030847 | 0.012094 | 0.001835 | 0.001939 | 0.029115 | 0.001639 | 0.022417 |
| 36.2361111 | 0.023527 | 0.032655 | 0.014064 | 0.015787 | 0.054649 | 0.00753 | 0.010981 | 0.018694 | 0.020401 | 0.043633 | 0.004203 | 0.036488 | 0.02121 | 0.022381 | 0.006183 | 0.044581 | 0.012642 | 0.001435 | 0.001455 | 0.038318 | 0.001103 | 0.015712 |
| 36.25 | 0.031435 | 0.055485 | 0.005801 | 0.010812 | 0.022864 | 0.005803 | 0.009827 | 0.013601 | 0.016002 | 0.01797 | 0.00701 | 0.045669 | 0.007385 | 0.014524 | 0.00444 | 0.051912 | 0.01213 | 0.001097 | 0.001156 | 0.038458 | 0.000828 | 0.01014 |
| 36.2638889 | 0.055636 | 0.056564 | 0.001915 | 0.01163 | 0.014611 | 0.012796 | 0.012119 | 0.01102 | 0.013874 | 0.006385 | 0.015303 | 0.048439 | 0.010409 | 0.010245 | 0.002387 | 0.04313 | 0.010527 | 0.001057 | 0.001146 | 0.033478 | 0.00093 | 0.007064 |
| 36.2777778 | 0.081907 | 0.041019 | 0.001134 | 0.017684 | 0.010176 | 0.030429 | 0.027439 | 0.009888 | 0.019524 | 0.007263 | 0.025331 | 0.029105 | 0.026064 | 0.01113 | 0.002044 | 0.023562 | 0.008386 | 0.00122 | 0.001248 | 0.02803 | 0.001141 | 0.004647 |
| 36.2916667 | 0.086673 | 0.060124 | 0.002243 | 0.019225 | 0.013963 | 0.057898 | 0.030378 | 0.011518 | 0.036706 | 0.020921 | 0.031152 | 0.032699 | 0.030591 | 0.033414 | 0.002579 | 0.007736 | 0.007876 | 0.001278 | 0.001185 | 0.024576 | 0.001178 | 0.007093 |
| 36.3055556 | 0.065008 | 0.095952 | 0.005742 | 0.014456 | 0.027423 | 0.070771 | 0.017053 | 0.018777 | 0.06132 | 0.04051 | 0.031891 | 0.063634 | 0.02538 | 0.070963 | 0.003157 | 0.005467 | 0.010568 | 0.001339 | 0.001156 | 0.023571 | 0.001332 | 0.013389 |

| | | | | | | | | | | | | | | | | | | | | | | |
|---|---|---|---|---|---|---|---|---|---|---|---|---|---|---|---|---|---|---|---|---|---|---|
| 36.3194444 | 0.030899 | 0.094333 | 0.013798 | 0.013116 | 0.041936 | 0.066015 | 0.014664 | 0.03405 | 0.08404 | 0.051771 | 0.034132 | 0.093757 | 0.040359 | 0.067318 | 0.003565 | 0.020887 | 0.014186 | 0.001777 | 0.00146 | 0.024854 | 0.001839 | 0.019938 |
| 36.3333333 | 0.014342 | 0.06512 | 0.02656 | 0.017819 | 0.057177 | 0.074604 | 0.029111 | 0.050533 | 0.094201 | 0.051808 | 0.042923 | 0.109111 | 0.065086 | 0.039003 | 0.003242 | 0.044479 | 0.01526 | 0.002337 | 0.001825 | 0.02723 | 0.002228 | 0.025691 |
| 36.3472222 | 0.017409 | 0.044205 | 0.03644 | 0.022195 | 0.067056 | 0.093439 | 0.047811 | 0.055449 | 0.08335 | 0.046455 | 0.052978 | 0.106437 | 0.071751 | 0.046975 | 0.002688 | 0.058053 | 0.013556 | 0.002592 | 0.001987 | 0.028271 | 0.00223 | 0.029226 |
| 36.3611111 | 0.020027 | 0.038301 | 0.035528 | 0.021507 | 0.067104 | 0.088337 | 0.057491 | 0.050883 | 0.062954 | 0.043653 | 0.056258 | 0.083925 | 0.059643 | 0.062677 | 0.003195 | 0.059118 | 0.010895 | 0.002656 | 0.002062 | 0.025572 | 0.002208 | 0.030193 |
| 36.375 | 0.014806 | 0.033255 | 0.028006 | 0.016133 | 0.059665 | 0.063653 | 0.049173 | 0.046304 | 0.051221 | 0.048961 | 0.052994 | 0.049957 | 0.040131 | 0.053066 | 0.004092 | 0.050524 | 0.007188 | 0.002595 | 0.001902 | 0.020074 | 0.002126 | 0.030155 |
| 36.3888889 | 0.015331 | 0.025529 | 0.021629 | 0.01227 | 0.052769 | 0.04772 | 0.033255 | 0.043941 | 0.0507 | 0.06272 | 0.050872 | 0.027724 | 0.022796 | 0.04415 | 0.003436 | 0.03525 | 0.004843 | 0.002181 | 0.001234 | 0.01582 | 0.001444 | 0.030851 |
| 36.4027778 | 0.028005 | 0.017946 | 0.018405 | 0.014258 | 0.055839 | 0.043864 | 0.023475 | 0.043751 | 0.054626 | 0.077056 | 0.055582 | 0.034878 | 0.020627 | 0.054632 | 0.001895 | 0.028412 | 0.008949 | 0.001365 | 0.00076 | 0.016022 | 0.000868 | 0.032458 |
| 36.4166667 | 0.030926 | 0.011377 | 0.018016 | 0.020212 | 0.062331 | 0.047614 | 0.014708 | 0.042762 | 0.057196 | 0.078647 | 0.068734 | 0.066527 | 0.040189 | 0.061301 | 0.001655 | 0.04209 | 0.017971 | 0.000686 | 0.001041 | 0.020886 | 0.0013 | 0.034288 |
| 36.4305556 | 0.020482 | 0.008055 | 0.020006 | 0.023502 | 0.058414 | 0.062881 | 0.007035 | 0.04107 | 0.058545 | 0.067407 | 0.086808 | 0.111595 | 0.071494 | 0.051233 | 0.002122 | 0.06669 | 0.026835 | 0.000461 | 0.001056 | 0.02716 | 0.001477 | 0.036543 |
| 36.4444444 | 0.01983 | 0.006665 | 0.020835 | 0.018055 | 0.049437 | 0.08387 | 0.004851 | 0.04438 | 0.0607 | 0.055309 | 0.096313 | 0.149364 | 0.101685 | 0.03281 | 0.002304 | 0.078911 | 0.03145 | 0.001644 | 0.00161 | 0.030972 | 0.002217 | 0.040756 |
| 36.4583333 | 0.02607 | 0.00728 | 0.014246 | 0.010842 | 0.044618 | 0.092029 | 0.005671 | 0.054585 | 0.062605 | 0.048881 | 0.085872 | 0.151368 | 0.125128 | 0.01631 | 0.004703 | 0.07515 | 0.029056 | 0.006721 | 0.005438 | 0.029992 | 0.006884 | 0.047525 |
| 36.4722222 | 0.022604 | 0.009321 | 0.004529 | 0.008755 | 0.043528 | 0.076899 | 0.01504 | 0.070115 | 0.063293 | 0.050863 | 0.063902 | 0.122141 | 0.145782 | 0.020379 | 0.009446 | 0.073821 | 0.021844 | 0.01317 | 0.010334 | 0.023445 | 0.012526 | 0.055288 |
| 36.4861111 | 0.012224 | 0.009928 | 0.0013 | 0.010996 | 0.052629 | 0.049101 | 0.034887 | 0.092108 | 0.066171 | 0.060554 | 0.060926 | 0.092676 | 0.171317 | 0.052027 | 0.011314 | 0.075254 | 0.01554 | 0.013892 | 0.010928 | 0.015572 | 0.012588 | 0.062803 |
| 36.5 | 0.011136 | 0.016404 | 0.002256 | 0.022163 | 0.074191 | 0.027395 | 0.048987 | 0.126697 | 0.074872 | 0.070397 | 0.098257 | 0.078412 | 0.195995 | 0.097859 | 0.008952 | 0.061983 | 0.013535 | 0.00905 | 0.007598 | 0.012408 | 0.008191 | 0.06767 |
| 36.5138889 | 0.031631 | 0.029944 | 0.002562 | 0.033722 | 0.096658 | 0.027636 | 0.041153 | 0.160977 | 0.092196 | 0.072753 | 0.151135 | 0.087253 | 0.197946 | 0.125848 | 0.006731 | 0.0456 | 0.013467 | 0.005307 | 0.005186 | 0.015856 | 0.005091 | 0.065154 |
| 36.5277778 | 0.069103 | 0.036843 | 0.0019 | 0.035353 | 0.113118 | 0.06047 | 0.021212 | 0.162686 | 0.113877 | 0.073535 | 0.179267 | 0.120405 | 0.160366 | 0.11485 | 0.008795 | 0.042094 | 0.009799 | 0.007001 | 0.007595 | 0.025423 | 0.006594 | 0.054223 |
| 36.5416667 | 0.111472 | 0.032331 | 0.001466 | 0.030274 | 0.115779 | 0.102905 | 0.00944 | 0.108898 | 0.123822 | 0.084838 | 0.174974 | 0.142997 | 0.095961 | 0.072267 | 0.018055 | 0.031602 | 0.008759 | 0.016113 | 0.017455 | 0.034554 | 0.014614 | 0.036752 |
| 36.5555556 | 0.126707 | 0.024955 | 0.007599 | 0.024671 | 0.092555 | 0.114003 | 0.013612 | 0.033517 | 0.119755 | 0.091097 | 0.142752 | 0.126847 | 0.038515 | 0.030436 | 0.031822 | 0.021383 | 0.019871 | 0.030459 | 0.032131 | 0.031183 | 0.027626 | 0.017622 |
| 36.5694444 | 0.088971 | 0.016644 | 0.025872 | 0.02471 | 0.05668 | 0.081124 | 0.031421 | 0.003678 | 0.104271 | 0.083193 | 0.090907 | 0.077805 | 0.012446 | 0.015635 | 0.038956 | 0.030648 | 0.029897 | 0.039746 | 0.040259 | 0.017253 | 0.036859 | 0.00601 |
| 36.5833333 | 0.061468 | 0.009265 | 0.05137 | 0.032139 | 0.031434 | 0.031547 | 0.0392 | 0.019843 | 0.080993 | 0.078031 | 0.053369 | 0.025162 | 0.01971 | 0.015065 | 0.032799 | 0.031978 | 0.025839 | 0.035436 | 0.034171 | 0.012924 | 0.034095 | 0.008402 |
| 36.5972222 | 0.072314 | 0.006543 | 0.068989 | 0.043889 | 0.02361 | 0.012441 | 0.031544 | 0.061646 | 0.069591 | 0.066867 | 0.044784 | 0.014715 | 0.040215 | 0.019755 | 0.019937 | 0.01773 | 0.014969 | 0.022223 | 0.020607 | 0.027779 | 0.022374 | 0.021945 |
| 36.6111111 | 0.07347 | 0.00604 | 0.06816 | 0.050108 | 0.035226 | 0.029892 | 0.039918 | 0.098368 | 0.058687 | 0.048143 | 0.052235 | 0.048412 | 0.051369 | 0.027588 | 0.009257 | 0.012495 | 0.006504 | 0.010137 | 0.009323 | 0.054092 | 0.010719 | 0.03236 |
| 36.625 | 0.098743 | 0.015421 | 0.058908 | 0.044943 | 0.053625 | 0.046503 | 0.07448 | 0.08716 | 0.032624 | 0.031019 | 0.049582 | 0.068043 | 0.045466 | 0.031705 | 0.00344 | 0.01934 | 0.002288 | 0.003739 | 0.003482 | 0.071302 | 0.004161 | 0.027067 |
| 36.6388889 | 0.135594 | 0.032089 | 0.049838 | 0.03978 | 0.054654 | 0.039739 | 0.083732 | 0.047793 | 0.018168 | 0.012472 | 0.039915 | 0.048125 | 0.029979 | 0.045799 | 0.001834 | 0.022296 | 0.001007 | 0.002713 | 0.002512 | 0.053184 | 0.002957 | 0.012008 |
| 36.6527778 | 0.115816 | 0.046732 | 0.037924 | 0.039616 | 0.033879 | 0.024878 | 0.054744 | 0.0359 | 0.024862 | 0.008986 | 0.057025 | 0.034055 | 0.033306 | 0.061572 | 0.002722 | 0.016829 | 0.001559 | 0.00434 | 0.004105 | 0.025386 | 0.004363 | 0.003973 |
| 36.6666667 | 0.068356 | 0.058913 | 0.027358 | 0.041052 | 0.013737 | 0.014398 | 0.067617 | 0.042952 | 0.025837 | 0.016655 | 0.068397 | 0.036889 | 0.067665 | 0.047388 | 0.004766 | 0.010691 | 0.00315 | 0.006572 | 0.006558 | 0.026714 | 0.006228 | 0.00981 |
| 36.6805556 | 0.030185 | 0.047134 | 0.020616 | 0.042872 | 0.007781 | 0.006748 | 0.099231 | 0.044027 | 0.023036 | 0.01787 | 0.042183 | 0.02748 | 0.095228 | 0.024898 | 0.00746 | 0.010416 | 0.004887 | 0.008763 | 0.00912 | 0.027896 | 0.00824 | 0.023071 |
| 36.6944444 | 0.041587 | 0.038354 | 0.015093 | 0.034346 | 0.014979 | 0.005381 | 0.078995 | 0.058495 | 0.033071 | 0.026707 | 0.02354 | 0.023438 | 0.077338 | 0.029623 | 0.009202 | 0.011927 | 0.005218 | 0.009705 | 0.01014 | 0.01996 | 0.009254 | 0.031908 |
| 36.7083333 | 0.109495 | 0.060094 | 0.012759 | 0.020771 | 0.035275 | 0.011389 | 0.06308 | 0.05227 | 0.034429 | 0.036867 | 0.037164 | 0.048144 | 0.053502 | 0.046008 | 0.008801 | 0.008679 | 0.003541 | 0.008943 | 0.008936 | 0.031914 | 0.008491 | 0.032036 |
| 36.7222222 | 0.134507 | 0.053458 | 0.012581 | 0.015005 | 0.058973 | 0.020945 | 0.072598 | 0.022167 | 0.020707 | 0.030455 | 0.058879 | 0.059669 | 0.082647 | 0.050151 | 0.007384 | 0.009233 | 0.001683 | 0.007748 | 0.007038 | 0.050494 | 0.007133 | 0.025406 |
| 36.7361111 | 0.089009 | 0.036391 | 0.008643 | 0.010102 | 0.059834 | 0.024081 | 0.054724 | 0.008659 | 0.01332 | 0.031324 | 0.07075 | 0.035157 | 0.125058 | 0.033588 | 0.006569 | 0.010539 | 0.001382 | 0.00743 | 0.006015 | 0.051132 | 0.006603 | 0.027422 |
| 36.75 | 0.067226 | 0.049932 | 0.013641 | 0.003924 | 0.03294 | 0.022567 | 0.032174 | 0.006177 | 0.014082 | 0.042431 | 0.05422 | 0.020444 | 0.102922 | 0.017516 | 0.006708 | 0.007355 | 0.001491 | 0.008037 | 0.005989 | 0.03162 | 0.007108 | 0.052392 |
| 36.7638889 | 0.075385 | 0.05631 | 0.029861 | 0.003256 | 0.013644 | 0.017911 | 0.041379 | 0.007102 | 0.014204 | 0.033605 | 0.027638 | 0.021838 | 0.074768 | 0.017974 | 0.007072 | 0.010082 | 0.001009 | 0.008483 | 0.006111 | 0.020673 | 0.00766 | 0.075624 |
| 36.7777778 | 0.054282 | 0.054168 | 0.028186 | 0.006808 | 0.011477 | 0.00844 | 0.049361 | 0.008367 | 0.010561 | 0.019015 | 0.030147 | 0.021813 | 0.114324 | 0.013715 | 0.006845 | 0.01596 | 0.000963 | 0.007975 | 0.005999 | 0.034396 | 0.007376 | 0.073356 |
| 36.7916667 | 0.038452 | 0.055742 | 0.015375 | 0.00884 | 0.0142 | 0.004843 | 0.032352 | 0.007428 | 0.00588 | 0.020333 | 0.050669 | 0.031357 | 0.135163 | 0.004602 | 0.006004 | 0.015251 | 0.00159 | 0.007045 | 0.006209 | 0.040732 | 0.00666 | 0.05068 |
| 36.8055556 | 0.04034 | 0.054391 | 0.010921 | 0.009237 | 0.019222 | 0.006142 | 0.021947 | 0.010337 | 0.006633 | 0.030213 | 0.070817 | 0.038247 | 0.119815 | 0.00387 | 0.005238 | 0.011244 | 0.002345 | 0.006486 | 0.006812 | 0.027379 | 0.006252 | 0.022905 |
| 36.8194444 | 0.040416 | 0.047154 | 0.010534 | 0.007915 | 0.02581 | 0.007089 | 0.046977 | 0.019516 | 0.010342 | 0.032091 | 0.081515 | 0.033192 | 0.128566 | 0.009463 | 0.005071 | 0.013093 | 0.002896 | 0.006466 | 0.007216 | 0.02386 | 0.006245 | 0.005865 |
| 36.8333333 | 0.076929 | 0.032657 | 0.016976 | 0.004106 | 0.032664 | 0.011846 | 0.100744 | 0.023971 | 0.010108 | 0.017047 | 0.06074 | 0.033728 | 0.13101 | 0.011867 | 0.005441 | 0.023704 | 0.002962 | 0.006754 | 0.007182 | 0.050329 | 0.006459 | 0.004176 |
| 36.8472222 | 0.101201 | 0.018443 | 0.030316 | 0.002904 | 0.030587 | 0.020041 | 0.159232 | 0.019835 | 0.009717 | 0.01094 | 0.023263 | 0.039576 | 0.08562 | 0.012854 | 0.005878 | 0.034743 | 0.003063 | 0.006746 | 0.006557 | 0.077044 | 0.006541 | 0.010211 |
| 36.8611111 | 0.053338 | 0.012024 | 0.048362 | 0.006056 | 0.021404 | 0.028995 | 0.188484 | 0.013737 | 0.014362 | 0.020279 | 0.012067 | 0.040624 | 0.044445 | 0.024807 | 0.005944 | 0.035694 | 0.003819 | 0.006057 | 0.005365 | 0.064962 | 0.00616 | 0.017932 |
| 36.875 | 0.014035 | 0.021873 | 0.063029 | 0.008582 | 0.013946 | 0.037992 | 0.162947 | 0.014038 | 0.023901 | 0.021227 | 0.036493 | 0.028516 | 0.060187 | 0.042683 | 0.005638 | 0.032992 | 0.00481 | 0.005184 | 0.004196 | 0.037237 | 0.005465 | 0.027277 |
| 36.8888889 | 0.020365 | 0.037574 | 0.062359 | 0.010781 | 0.014052 | 0.038967 | 0.09322 | 0.022775 | 0.030948 | 0.022179 | 0.084262 | 0.011708 | 0.072922 | 0.053892 | 0.005364 | 0.043401 | 0.005542 | 0.004887 | 0.00365 | 0.047249 | 0.005006 | 0.034779 |
| 36.9027778 | 0.034131 | 0.04276 | 0.049388 | 0.017539 | 0.027491 | 0.026981 | 0.037949 | 0.022433 | 0.025526 | 0.026795 | 0.128745 | 0.009962 | 0.061705 | 0.053034 | 0.005339 | 0.068273 | 0.00592 | 0.005055 | 0.003696 | 0.090094 | 0.00501 | 0.036533 |
| 36.9166667 | 0.04556 | 0.034805 | 0.034218 | 0.024651 | 0.032896 | 0.013369 | 0.054263 | 0.017898 | 0.018078 | 0.024253 | 0.163895 | 0.021604 | 0.07559 | 0.042546 | 0.004989 | 0.093868 | 0.00639 | 0.004608 | 0.003565 | 0.117127 | 0.004818 | 0.034568 |
| 36.9305556 | 0.068812 | 0.023492 | 0.025137 | 0.030311 | 0.02274 | 0.011839 | 0.107477 | 0.030336 | 0.021482 | 0.021864 | 0.187722 | 0.052174 | 0.112038 | 0.030957 | 0.003584 | 0.112938 | 0.007557 | 0.002978 | 0.002608 | 0.112473 | 0.003686 | 0.032833 |
| 36.9444444 | 0.114748 | 0.026207 | 0.026466 | 0.034591 | 0.027284 | 0.032317 | 0.119986 | 0.046202 | 0.028388 | 0.016304 | 0.163996 | 0.094284 | 0.143437 | 0.023555 | 0.001728 | 0.126303 | 0.009163 | 0.00142 | 0.001419 | 0.087644 | 0.001974 | 0.030451 |
| 36.9583333 | 0.136111 | 0.023498 | 0.036445 | 0.031741 | 0.046106 | 0.076368 | 0.085397 | 0.054626 | 0.025768 | 0.015008 | 0.095489 | 0.095082 | 0.154086 | 0.02079 | 0.001054 | 0.131148 | 0.010707 | 0.001392 | 0.001289 | 0.056353 | 0.001189 | 0.02547 |
| 36.9722222 | 0.105567 | 0.014155 | 0.048922 | 0.024545 | 0.058001 | 0.124971 | 0.047194 | 0.066047 | 0.019065 | 0.025791 | 0.039439 | 0.049182 | 0.140211 | 0.022541 | 0.001939 | 0.12357 | 0.012256 | 0.002298 | 0.002173 | 0.033788 | 0.001892 | 0.020701 |
| 36.9861111 | 0.058458 | 0.036683 | 0.051618 | 0.018395 | 0.045502 | 0.149543 | 0.02943 | 0.079647 | 0.029846 | 0.045278 | 0.023516 | 0.027291 | 0.110291 | 0.028091 | 0.003192 | 0.105088 | 0.013839 | 0.002862 | 0.002852 | 0.023048 | 0.00275 | 0.018038 |
| 37 | 0.032534 | 0.088605 | 0.046137 | 0.024099 | 0.029655 | 0.146796 | 0.023915 | 0.083117 | 0.048191 | 0.076575 | 0.017675 | 0.048476 | 0.076539 | 0.035038 | 0.003289 | 0.079331 | 0.014122 | 0.002336 | 0.002345 | 0.018537 | 0.002286 | 0.014893 |
| 37.0138889 | 0.060766 | 0.135837 | 0.04892 | 0.041062 | 0.047952 | 0.130468 | 0.031723 | 0.071709 | 0.065669 | 0.096226 | 0.011506 | 0.056037 | 0.050646 | 0.038331 | 0.002783 | 0.049886 | 0.01105 | 0.002966 | 0.002195 | 0.016541 | 0.002447 | 0.00916 |
| 37.0277778 | 0.120041 | 0.149484 | 0.054729 | 0.042988 | 0.073692 | 0.1049 | 0.046139 | 0.053945 | 0.088871 | 0.087687 | 0.014389 | 0.040185 | 0.040257 | 0.036911 | 0.003273 | 0.02445 | 0.006697 | 0.006135 | 0.003776 | 0.016908 | 0.005095 | 0.004211 |
| 37.0416667 | 0.158927 | 0.114879 | 0.052796 | 0.024999 | 0.093726 | 0.081346 | 0.038287 | 0.044877 | 0.098949 | 0.074085 | 0.037436 | 0.047749 | 0.046295 | 0.037168 | 0.00423 | 0.012492 | 0.007481 | 0.007253 | 0.004635 | 0.023578 | 0.006375 | 0.007933 |
| 37.0555556 | 0.17443 | 0.069598 | 0.051185 | 0.012609 | 0.122353 | 0.074676 | 0.042535 | 0.055725 | 0.10442 | 0.08143 | 0.097587 | 0.099553 | 0.057192 | 0.040584 | 0.005767 | 0.015534 | 0.013715 | 0.007181 | 0.005305 | 0.036466 | 0.007068 | 0.019961 |
| 37.0694444 | 0.177274 | 0.059539 | 0.05544 | 0.025436 | 0.152634 | 0.086292 | 0.090602 | 0.08878 | 0.126367 | 0.109016 | 0.161329 | 0.150411 | 0.057985 | 0.040011 | 0.006168 | 0.021353 | 0.017851 | 0.007966 | 0.005857 | 0.046072 | 0.008593 | 0.029433 |
| 37.0833333 | 0.176054 | 0.073296 | 0.054112 | 0.059233 | 0.174199 | 0.097451 | 0.133025 | 0.126281 | 0.145729 | 0.124237 | 0.177864 | 0.142154 | 0.046865 | 0.036719 | 0.004106 | 0.019375 | 0.01639 | 0.005567 | 0.004152 | 0.040542 | 0.006196 | 0.030645 |
| 37.0972222 | 0.169222 | 0.074017 | 0.042495 | 0.087283 | 0.1656 | 0.087431 | 0.128066 | 0.132416 | 0.13899 | 0.119544 | 0.141184 | 0.102637 | 0.03258 | 0.044473 | 0.00319 | 0.013978 | 0.012754 | 0.002437 | 0.002962 | 0.021553 | 0.002623 | 0.025649 |
| 37.1111111 | 0.129475 | 0.049377 | 0.030586 | 0.083076 | 0.12524 | 0.059562 | 0.081181 | 0.10903 | 0.111494 | 0.114858 | 0.081945 | 0.079247 | 0.01835 | 0.058855 | 0.003479 | 0.009499 | 0.009701 | 0.00192 | 0.003029 | 0.00587 | 0.001952 | 0.021966 |
| 37.125 | 0.064782 | 0.022503 | 0.02057 | 0.051121 | 0.080394 | 0.032771 | 0.033446 | 0.093628 | 0.07993 | 0.098145 | 0.035152 | 0.06397 | 0.011594 | 0.058244 | 0.00271 | 0.007168 | 0.006803 | 0.00195 | 0.00266 | 0.001424 | 0.002005 | 0.022456 |
| 37.1388889 | 0.041371 | 0.016946 | 0.011762 | 0.020861 | 0.047848 | 0.01748 | 0.020345 | 0.088357 | 0.05246 | 0.062218 | 0.022528 | 0.043839 | 0.022829 | 0.04121 | 0.001392 | 0.012806 | 0.003941 | 0.001743 | 0.002264 | 0.001313 | 0.001802 | 0.017682 |
| 37.1527778 | 0.088019 | 0.036774 | 0.009774 | 0.012436 | 0.023796 | 0.017381 | 0.0431 | 0.073718 | 0.032886 | 0.028824 | 0.041057 | 0.025457 | 0.041424 | 0.02127 | 0.000682 | 0.025381 | 0.0023 | 0.001358 | 0.001743 | 0.001293 | 0.001412 | 0.009062 |
| 37.1666667 | 0.157687 | 0.060174 | 0.013364 | 0.022168 | 0.013434 | 0.034245 | 0.074869 | 0.044597 | 0.019338 | 0.019449 | 0.063812 | 0.026403 | 0.0509 | 0.009504 | 0.00088 | 0.036308 | 0.002186 | 0.000706 | 0.001002 | 0.002234 | 0.000805 | 0.007133 |
| 37.1805556 | 0.188568 | 0.057081 | 0.014137 | 0.034425 | 0.033002 | 0.05245 | 0.080479 | 0.022744 | 0.008816 | 0.030318 | 0.06461 | 0.047793 | 0.044282 | 0.010968 | 0.000941 | 0.038701 | 0.002674 | 0.000325 | 0.000784 | 0.005118 | 0.000631 | 0.007659 |
| 37.1944444 | 0.148847 | 0.030319 | 0.0128 | 0.038942 | 0.07708 | 0.054282 | 0.0599 | 0.031621 | 0.012411 | 0.043998 | 0.039395 | 0.068217 | 0.024432 | 0.020201 | 0.000607 | 0.032507 | 0.002849 | 0.000386 | 0.001065 | 0.011465 | 0.00085 | 0.006166 |
| 37.2083333 | 0.086852 | 0.009342 | 0.015513 | 0.036271 | 0.117578 | 0.038832 | 0.037514 | 0.045233 | 0.040471 | 0.047204 | 0.020531 | 0.069198 | 0.008524 | 0.030903 | 0.000357 | 0.0272 | 0.002375 | 0.000411 | 0.000916 | 0.023027 | 0.000732 | 0.008005 |
| 37.2222222 | 0.079105 | 0.008453 | 0.020975 | 0.028359 | 0.119673 | 0.021531 | 0.019305 | 0.039211 | 0.075019 | 0.03395 | 0.030004 | 0.048036 | 0.004858 | 0.040157 | 0.000265 | 0.032527 | 0.001374 | 0.000553 | 0.000601 | 0.036523 | 0.000643 | 0.011985 |
| 37.2361111 | 0.101282 | 0.023361 | 0.022779 | 0.017516 | 0.07886 | 0.017593 | 0.008129 | 0.024316 | 0.088802 | 0.031809 | 0.036304 | 0.03749 | 0.006134 | 0.042504 | 0.000509 | 0.04823 | 0.000673 | 0.001256 | 0.001142 | 0.045832 | 0.001342 | 0.014605 |
| 37.25 | 0.083804 | 0.031842 | 0.017636 | 0.011124 | 0.037219 | 0.028887 | 0.011075 | 0.01086 | 0.078247 | 0.042649 | 0.020503 | 0.052408 | 0.009548 | 0.035706 | 0.000974 | 0.063335 | 0.000874 | 0.002249 | 0.002274 | 0.050233 | 0.002461 | 0.017179 |
| 37.2638889 | 0.037813 | 0.037042 | 0.009218 | 0.008969 | 0.027928 | 0.051223 | 0.017704 | 0.003716 | 0.057133 | 0.030568 | 0.008879 | 0.045726 | 0.009335 | 0.026667 | 0.001429 | 0.065074 | 0.001324 | 0.003117 | 0.003157 | 0.050738 | 0.003392 | 0.019055 |
| 37.2777778 | 0.009269 | 0.056044 | 0.00379 | 0.010542 | 0.02675 | 0.07716 | 0.016049 | 0.002564 | 0.03803 | 0.015411 | 0.016315 | 0.024251 | 0.008881 | 0.020268 | 0.001774 | 0.050891 | 0.001706 | 0.003744 | 0.003623 | 0.047172 | 0.003954 | 0.016223 |
| 37.2916667 | 0.0127 | 0.050209 | 0.004738 | 0.014709 | 0.022131 | 0.097694 | 0.014427 | 0.004704 | 0.031336 | 0.024614 | 0.024357 | 0.032088 | 0.019435 | 0.014385 | 0.001971 | 0.030358 | 0.002154 | 0.004147 | 0.003914 | 0.040357 | 0.004226 | 0.009118 |
| 37.3055556 | 0.040468 | 0.030278 | 0.010621 | 0.011384 | 0.039323 | 0.100778 | 0.02344 | 0.00865 | 0.039119 | 0.041235 | 0.031398 | 0.056107 | 0.029278 | 0.008125 | 0.00217 | 0.01529 | 0.002303 | 0.004265 | 0.004085 | 0.03239 | 0.004304 | 0.006283 |
| 37.3194444 | 0.069745 | 0.032355 | 0.01684 | 0.009465 | 0.063427 | 0.083099 | 0.032178 | 0.015181 | 0.048874 | 0.05174 | 0.067597 | 0.086315 | 0.023229 | 0.00355 | 0.002031 | 0.011642 | 0.00165 | 0.003828 | 0.003824 | 0.025523 | 0.003912 | 0.010351 |
| 37.3333333 | 0.073869 | 0.026266 | 0.020191 | 0.0245 | 0.071213 | 0.062686 | 0.030144 | 0.028072 | 0.054509 | 0.052762 | 0.117391 | 0.128052 | 0.016027 | 0.003356 | 0.001202 | 0.019023 | 0.001174 | 0.002834 | 0.003001 | 0.020777 | 0.002921 | 0.014944 |
| 37.3472222 | 0.057813 | 0.014596 | 0.020187 | 0.048011 | 0.066722 | 0.052482 | 0.019814 | 0.041745 | 0.061837 | 0.045809 | 0.134133 | 0.155594 | 0.031511 | 0.006282 | 0.000504 | 0.03178 | 0.002046 | 0.002021 | 0.002246 | 0.018494 | 0.002063 | 0.017029 |
| 37.3611111 | 0.068973 | 0.02327 | 0.017817 | 0.058993 | 0.058573 | 0.047785 | 0.007711 | 0.045109 | 0.067113 | 0.040591 | 0.110442 | 0.146379 | 0.060722 | 0.006069 | 0.000796 | 0.042089 | 0.002851 | 0.001904 | 0.002147 | 0.018493 | 0.001994 | 0.018647 |
| 37.375 | 0.097998 | 0.041864 | 0.014589 | 0.049617 | 0.054197 | 0.048452 | 0.002937 | 0.037748 | 0.064218 | 0.042877 | 0.064745 | 0.12097 | 0.080005 | 0.006213 | 0.001535 | 0.044109 | 0.002554 | 0.002056 | 0.002367 | 0.019486 | 0.002304 | 0.021378 |
| 37.3888889 | 0.082499 | 0.056836 | 0.01222 | 0.034805 | 0.056202 | 0.057517 | 0.008337 | 0.028303 | 0.05688 | 0.049533 | 0.030355 | 0.099013 | 0.077803 | 0.015763 | 0.001806 | 0.038271 | 0.001994 | 0.002112 | 0.002154 | 0.019112 | 0.002395 | 0.024092 |
| 37.4027778 | 0.056284 | 0.063377 | 0.012596 | 0.027922 | 0.060483 | 0.064303 | 0.018391 | 0.025701 | 0.04665 | 0.055525 | 0.033816 | 0.082247 | 0.05517 | 0.030578 | 0.001452 | 0.033071 | 0.002021 | 0.00318 | 0.001694 | 0.015395 | 0.002436 | 0.025208 |
| 37.4166667 | 0.076521 | 0.056824 | 0.015745 | 0.027846 | 0.059427 | 0.056852 | 0.027584 | 0.030848 | 0.036554 | 0.057057 | 0.064915 | 0.0831 | 0.026522 | 0.044252 | 0.000829 | 0.035921 | 0.002715 | 0.006334 | 0.002613 | 0.009525 | 0.002357 | 0.024566 |
| 37.4305556 | 0.088651 | 0.036605 | 0.019759 | 0.03154 | 0.051402 | 0.035656 | 0.033301 | 0.033791 | 0.035976 | 0.056777 | 0.102929 | 0.110626 | 0.008289 | 0.052018 | 0.000336 | 0.044732 | 0.002677 | 0.006949 | 0.003346 | 0.007958 | 0.001512 | 0.025201 |
| 37.4444444 | 0.069231 | 0.021359 | 0.019901 | 0.035905 | 0.041495 | 0.018273 | 0.029327 | 0.030693 | 0.048792 | 0.063049 | 0.132615 | 0.136242 | 0.011265 | 0.047878 | 0.000576 | 0.052669 | 0.001307 | 0.006051 | 0.00319 | 0.01293 | 0.001391 | 0.031439 |
| 37.4583333 | 0.092175 | 0.023186 | 0.013358 | 0.033802 | 0.034711 | 0.019162 | 0.019479 | 0.036406 | 0.070561 | 0.074233 | 0.135165 | 0.12768 | 0.022607 | 0.038697 | 0.002361 | 0.058655 | 0.001487 | 0.010012 | 0.003715 | 0.015255 | 0.004478 | 0.042541 |
| 37.4722222 | 0.161386 | 0.024262 | 0.007165 | 0.024755 | 0.033065 | 0.020996 | 0.021311 | 0.057248 | 0.091405 | 0.083894 | 0.106573 | 0.112069 | 0.022446 | 0.03723 | 0.004639 | 0.069924 | 0.003922 | 0.011979 | 0.003132 | 0.012407 | 0.007148 | 0.057182 |
| 37.4861111 | 0.202854 | 0.023728 | 0.004644 | 0.018551 | 0.039548 | 0.032871 | 0.026088 | 0.075008 | 0.104841 | 0.099526 | 0.072869 | 0.119799 | 0.017048 | 0.043417 | 0.004663 | 0.088584 | 0.004806 | 0.007488 | 0.001934 | 0.014376 | 0.004795 | 0.075581 |
| 37.5 | 0.184663 | 0.041803 | 0.002182 | 0.022254 | 0.052134 | 0.081855 | 0.027409 | 0.086167 | 0.113529 | 0.12579 | 0.0631 | 0.137419 | 0.031817 | 0.053176 | 0.002531 | 0.103365 | 0.003667 | 0.002645 | 0.001085 | 0.023215 | 0.001284 | 0.091381 |
| 37.5138889 | 0.129365 | 0.070994 | 0.001916 | 0.034444 | 0.062091 | 0.129305 | 0.066681 | 0.111158 | 0.120108 | 0.147175 | 0.080332 | 0.150213 | 0.074152 | 0.061945 | 0.001195 | 0.106465 | 0.00332 | 0.000763 | 0.000638 | 0.030999 | 0.000305 | 0.095926 |
| 37.5277778 | 0.08341 | 0.083 | 0.004122 | 0.048644 | 0.065484 | 0.140189 | 0.138274 | 0.132128 | 0.12419 | 0.151089 | 0.115638 | 0.159037 | 0.087933 | 0.062603 | 0.004164 | 0.100821 | 0.006539 | 0.003382 | 0.003537 | 0.032424 | 0.002458 | 0.089133 |
| 37.5416667 | 0.076359 | 0.074752 | 0.006659 | 0.060629 | 0.067086 | 0.1474 | 0.161992 | 0.096375 | 0.12846 | 0.137057 | 0.163257 | 0.170561 | 0.051077 | 0.050776 | 0.015757 | 0.084107 | 0.018023 | 0.014513 | 0.014808 | 0.026108 | 0.012029 | 0.076227 |
| 37.5555556 | 0.085848 | 0.057841 | 0.010158 | 0.065698 | 0.071041 | 0.158174 | 0.111572 | 0.038276 | 0.123936 | 0.110059 | 0.197839 | 0.16202 | 0.024208 | 0.028433 | 0.032352 | 0.054142 | 0.035276 | 0.031566 | 0.032169 | 0.014436 | 0.027918 | 0.056658 |
| 37.5694444 | 0.068346 | 0.03927 | 0.010225 | 0.06426 | 0.073272 | 0.147742 | 0.055442 | 0.020002 | 0.101916 | 0.088495 | 0.169001 | 0.10744 | 0.02193 | 0.013509 | 0.038971 | 0.026113 | 0.042754 | 0.039747 | 0.040258 | 0.009373 | 0.036996 | 0.03569 |
| 37.5833333 | 0.035997 | 0.022798 | 0.008221 | 0.069543 | 0.073994 | 0.137021 | 0.048008 | 0.026541 | 0.07584 | 0.086533 | 0.08907 | 0.041359 | 0.01723 | 0.012474 | 0.029914 | 0.018821 | 0.033341 | 0.031767 | 0.031451 | 0.022483 | 0.031245 | 0.029865 |
| 37.5972222 | 0.02432 | 0.008766 | 0.012817 | 0.07568 | 0.076896 | 0.135953 | 0.049883 | 0.044162 | 0.053836 | 0.089585 | 0.034269 | 0.00821 | 0.024059 | 0.011957 | 0.016389 | 0.02695 | 0.018653 | 0.017728 | 0.016893 | 0.050134 | 0.018518 | 0.04109 |
| 37.6111111 | 0.035274 | 0.005486 | 0.015104 | 0.056486 | 0.077514 | 0.109101 | 0.052045 | 0.043755 | 0.030374 | 0.074602 | 0.031912 | 0.007269 | 0.060075 | 0.008736 | 0.007503 | 0.030675 | 0.009092 | 0.007797 | 0.007115 | 0.083821 | 0.008572 | 0.052954 |
| 37.625 | 0.053969 | 0.023503 | 0.010187 | 0.028971 | 0.059807 | 0.05437 | 0.099813 | 0.022499 | 0.012792 | 0.046661 | 0.059598 | 0.011276 | 0.079875 | 0.007609 | 0.003257 | 0.025633 | 0.004717 | 0.002927 | 0.002402 | 0.096626 | 0.003307 | 0.050741 |
| 37.6388889 | 0.048191 | 0.057265 | 0.008491 | 0.023752 | 0.033883 | 0.019182 | 0.138191 | 0.014501 | 0.007807 | 0.027247 | 0.062847 | 0.033149 | 0.050375 | 0.00915 | 0.001697 | 0.018264 | 0.002982 | 0.001003 | 0.000584 | 0.063428 | 0.001115 | 0.032102 |
| 37.6527778 | 0.039182 | 0.087121 | 0.006625 | 0.028047 | 0.024969 | 0.013713 | 0.143179 | 0.016779 | 0.010341 | 0.020123 | 0.040777 | 0.08728 | 0.02895 | 0.00936 | 0.001654 | 0.013801 | 0.002712 | 0.000776 | 0.00028 | 0.034264 | 0.000758 | 0.016243 |
| 37.6666667 | 0.051301 | 0.088615 | 0.007062 | 0.034279 | 0.027589 | 0.011713 | 0.134921 | 0.031561 | 0.016801 | 0.029843 | 0.044673 | 0.112008 | 0.035483 | 0.007435 | 0.002119 | 0.018052 | 0.003277 | 0.001293 | 0.00038 | 0.049638 | 0.001266 | 0.023917 |
| 37.6805556 | 0.045227 | 0.065795 | 0.02274 | 0.03727 | 0.045915 | 0.008432 | 0.091951 | 0.035358 | 0.017536 | 0.041717 | 0.04232 | 0.074169 | 0.036647 | 0.00735 | 0.002052 | 0.024855 | 0.003693 | 0.00152 | 0.000537 | 0.054166 | 0.001581 | 0.043154 |
| 37.6944444 | 0.0305 | 0.07595 | 0.050873 | 0.026672 | 0.064625 | 0.010965 | 0.071477 | 0.014708 | 0.012626 | 0.031809 | 0.028794 | 0.028675 | 0.045245 | 0.0102 | 0.001249 | 0.020043 | 0.003203 | 0.001061 | 0.00057 | 0.032758 | 0.001193 | 0.049263 |
| 37.7083333 | 0.029746 | 0.107467 | 0.086976 | 0.014341 | 0.051759 | 0.027666 | 0.098123 | 0.003506 | 0.00845 | 0.017705 | 0.042859 | 0.032159 | 0.059271 | 0.012839 | 0.001283 | 0.011694 | 0.002078 | 0.001053 | 0.000883 | 0.032642 | 0.00116 | 0.036903 |

| | | | | | | | | | | | | | | | | | | | | | | |
|---|---|---|---|---|---|---|---|---|---|---|---|---|---|---|---|---|---|---|---|---|---|---|
| 37.7222222 | 0.029305 | 0.081125 | 0.114034 | 0.009112 | 0.023461 | 0.041186 | 0.102995 | 0.007624 | 0.00581 | 0.013509 | 0.078328 | 0.057886 | 0.05349 | 0.01435 | 0.003817 | 0.014072 | 0.002354 | 0.003303 | 0.002837 | 0.050716 | 0.003384 | 0.01784 |
| 37.7361111 | 0.035572 | 0.042972 | 0.100456 | 0.007944 | 0.01237 | 0.031139 | 0.064682 | 0.018923 | 0.011217 | 0.027236 | 0.095507 | 0.055422 | 0.031161 | 0.015909 | 0.008186 | 0.020073 | 0.005462 | 0.007664 | 0.006586 | 0.054423 | 0.007791 | 0.006195 |
| 37.75 | 0.042779 | 0.057474 | 0.056382 | 0.008714 | 0.023949 | 0.013707 | 0.068407 | 0.034448 | 0.022563 | 0.044885 | 0.068795 | 0.035268 | 0.013622 | 0.017432 | 0.012436 | 0.021073 | 0.009574 | 0.012245 | 0.010645 | 0.037735 | 0.012399 | 0.005332 |
| 37.7638889 | 0.034986 | 0.059336 | 0.019469 | 0.007866 | 0.032581 | 0.007391 | 0.10019 | 0.041229 | 0.023646 | 0.050755 | 0.05432 | 0.020333 | 0.013796 | 0.01794 | 0.015154 | 0.015909 | 0.012503 | 0.015487 | 0.013736 | 0.028573 | 0.015526 | 0.007223 |
| 37.7777778 | 0.027063 | 0.030367 | 0.007154 | 0.005721 | 0.022865 | 0.008337 | 0.06326 | 0.032532 | 0.013113 | 0.054653 | 0.077288 | 0.014663 | 0.021189 | 0.014838 | 0.015781 | 0.009359 | 0.013579 | 0.016677 | 0.015397 | 0.038797 | 0.016525 | 0.006001 |
| 37.7916667 | 0.021358 | 0.027781 | 0.014177 | 0.010309 | 0.008693 | 0.007052 | 0.021323 | 0.016443 | 0.010943 | 0.042261 | 0.099044 | 0.01874 | 0.021414 | 0.009381 | 0.014565 | 0.015046 | 0.012889 | 0.015837 | 0.015423 | 0.040749 | 0.015679 | 0.005038 |
| 37.8055556 | 0.010645 | 0.0503 | 0.024899 | 0.019387 | 0.007434 | 0.008998 | 0.01989 | 0.007923 | 0.020365 | 0.029722 | 0.102833 | 0.023804 | 0.019102 | 0.008439 | 0.01228 | 0.039497 | 0.011091 | 0.013628 | 0.013842 | 0.023913 | 0.013705 | 0.009998 |
| 37.8194444 | 0.016594 | 0.060687 | 0.031142 | 0.024539 | 0.011681 | 0.018845 | 0.031563 | 0.009574 | 0.02614 | 0.036172 | 0.085374 | 0.023565 | 0.038306 | 0.008922 | 0.00962 | 0.069663 | 0.008809 | 0.010873 | 0.011165 | 0.019772 | 0.011138 | 0.021441 |
| 37.8333333 | 0.06084 | 0.04833 | 0.03432 | 0.019815 | 0.006636 | 0.029387 | 0.057087 | 0.010091 | 0.024841 | 0.034312 | 0.052979 | 0.025964 | 0.084068 | 0.008707 | 0.006989 | 0.085378 | 0.006276 | 0.007992 | 0.007978 | 0.046345 | 0.008275 | 0.031304 |
| 37.8472222 | 0.104532 | 0.031162 | 0.038459 | 0.010544 | 0.002026 | 0.034275 | 0.069177 | 0.01093 | 0.0202 | 0.02306 | 0.026279 | 0.031883 | 0.125065 | 0.018365 | 0.004625 | 0.076755 | 0.003601 | 0.005101 | 0.004656 | 0.0668 | 0.005437 | 0.032461 |
| 37.8611111 | 0.105462 | 0.022779 | 0.043761 | 0.012518 | 0.003146 | 0.028338 | 0.054025 | 0.015639 | 0.013362 | 0.022191 | 0.023079 | 0.037309 | 0.135861 | 0.034927 | 0.002751 | 0.055243 | 0.001501 | 0.002565 | 0.001894 | 0.047623 | 0.003082 | 0.029744 |
| 37.875 | 0.087445 | 0.023366 | 0.047142 | 0.024807 | 0.006454 | 0.015703 | 0.02939 | 0.018039 | 0.008684 | 0.026586 | 0.041429 | 0.057655 | 0.116922 | 0.043922 | 0.001509 | 0.044041 | 0.000875 | 0.000934 | 0.000615 | 0.030897 | 0.001495 | 0.030897 |
| 37.8888889 | 0.077493 | 0.02299 | 0.047992 | 0.033287 | 0.013964 | 0.013087 | 0.013529 | 0.019245 | 0.01358 | 0.030815 | 0.062111 | 0.079841 | 0.080677 | 0.040502 | 0.000786 | 0.05558 | 0.001415 | 0.000436 | 0.000769 | 0.066679 | 0.000651 | 0.0379 |
| 37.9027778 | 0.102505 | 0.015316 | 0.045298 | 0.026482 | 0.024792 | 0.03356 | 0.023976 | 0.020957 | 0.022901 | 0.042936 | 0.078098 | 0.067355 | 0.044408 | 0.03145 | 0.000453 | 0.085958 | 0.002021 | 0.000538 | 0.0012 | 0.118987 | 0.000328 | 0.048551 |

**Supplementary Table S6.** 1 hour integrated motion data in **(b)** 100 mM NaCl treatment on Tomato

| IntervalMid | Control1 | Control2 | Control3 | Control4 | Control5 | Control6 | Control7 | Control8 | Control9 | Control10 | Control11 | Control12 | Stress1 | Stress2 | Stress3 | Stress4 | Stress5 | Stress6 | Stress7 | Stress8 | Stress9 | Stress10 |
|---|---|---|---|---|---|---|---|---|---|---|---|---|---|---|---|---|---|---|---|---|---|---|
| 18.11111111 | 0.026317 | 0.006916 | 0.006197 | 0.001845 | 0.002906 | 0.010435 | 0.016882 | 0.018099 | 0.018578 | 0.00312 | 0.009722 | 0.033248 | 0.011067 | 0.012811 | 0.023687 | 0.005988 | 0.005565 | 0.012234 | 0.001851 | 0.002404 | 0.012047 | 0.018367 |
| 18.125 | 0.026227 | 0.001702 | 0.010517 | 0.0034 | 0.005059 | 0.012409 | 0.016269 | 0.03129 | 0.021279 | 0.002762 | 0.013989 | 0.014097 | 0.012216 | 0.016096 | 0.02293 | 0.009005 | 0.008696 | 0.017905 | 0.002945 | 0.003097 | 0.014466 | 0.014372 |
| 18.13888889 | 0.022644 | 0.000355 | 0.013034 | 0.0062 | 0.006249 | 0.014111 | 0.016833 | 0.036054 | 0.020687 | 0.002025 | 0.015353 | 0.003076 | 0.01165 | 0.017536 | 0.018274 | 0.00981 | 0.009538 | 0.019562 | 0.003231 | 0.003479 | 0.01664 | 0.013238 |
| 18.15277778 | 0.016653 | 0.000277 | 0.012756 | 0.00827 | 0.006315 | 0.014224 | 0.015193 | 0.032414 | 0.017074 | 0.00108 | 0.012599 | 0.000535 | 0.009522 | 0.01721 | 0.013396 | 0.009414 | 0.009722 | 0.019362 | 0.00366 | 0.003928 | 0.016599 | 0.011694 |
| 18.16666667 | 0.011483 | 0.001018 | 0.011202 | 0.008468 | 0.005067 | 0.012716 | 0.011918 | 0.024685 | 0.012436 | 0.0004 | 0.008596 | 0.000702 | 0.007054 | 0.015187 | 0.009575 | 0.008622 | 0.00976 | 0.01833 | 0.004417 | 0.004747 | 0.014742 | 0.009389 |
| 18.18055556 | 0.00761 | 0.002188 | 0.008678 | 0.00657 | 0.002914 | 0.010335 | 0.008621 | 0.017034 | 0.008646 | 0.000174 | 0.006463 | 0.001464 | 0.005019 | 0.011991 | 0.006658 | 0.007355 | 0.009098 | 0.016298 | 0.004744 | 0.00513 | 0.01182 | 0.006561 |
| 18.19444444 | 0.004426 | 0.003243 | 0.0058 | 0.003799 | 0.001082 | 0.008042 | 0.005898 | 0.010943 | 0.006096 | 8.10E-05 | 0.006077 | 0.001838 | 0.003753 | 0.009023 | 0.004562 | 0.005895 | 0.007875 | 0.013847 | 0.004513 | 0.004687 | 0.008819 | 0.004301 |
| 18.20833333 | 0.003243 | 0.004135 | 0.0045 | 0.001984 | 0.000296 | 0.007057 | 0.0045 | 0.0073 | 0.004372 | 0.000112 | 0.005675 | 0.001462 | 0.003054 | 0.00701 | 0.003236 | 0.004603 | 0.006611 | 0.01166 | 0.004046 | 0.003722 | 0.006455 | 0.003263 |
| 18.22222222 | 0.005505 | 0.005425 | 0.005296 | 0.001766 | 0.000246 | 0.007743 | 0.00498 | 0.006533 | 0.003401 | 0.000393 | 0.004635 | 0.001121 | 0.002417 | 0.005624 | 0.00266 | 0.003847 | 0.005715 | 0.010088 | 0.003644 | 0.002786 | 0.00461 | 0.002576 |
| 18.23611111 | 0.010019 | 0.006879 | 0.00702 | 0.002556 | 0.000509 | 0.009143 | 0.006581 | 0.007081 | 0.003213 | 0.000558 | 0.003808 | 0.001367 | 0.001931 | 0.004804 | 0.00286 | 0.003906 | 0.005409 | 0.009376 | 0.003546 | 0.002385 | 0.003389 | 0.001805 |
| 18.25 | 0.014113 | 0.006961 | 0.007951 | 0.00311 | 0.000625 | 0.009618 | 0.007634 | 0.006343 | 0.003081 | 0.000513 | 0.003062 | 0.001929 | 0.002052 | 0.004794 | 0.003325 | 0.004195 | 0.005282 | 0.008868 | 0.003509 | 0.002531 | 0.003093 | 0.001539 |
| 18.26388889 | 0.017074 | 0.005721 | 0.007692 | 0.002958 | 0.000501 | 0.00888 | 0.007699 | 0.004128 | 0.002275 | 0.000523 | 0.001705 | 0.002423 | 0.002432 | 0.005107 | 0.003483 | 0.004066 | 0.004834 | 0.007818 | 0.003061 | 0.002672 | 0.003234 | 0.001699 |
| 18.27777778 | 0.020027 | 0.005134 | 0.008229 | 0.003253 | 0.000587 | 0.008891 | 0.008622 | 0.002786 | 0.001376 | 0.000598 | 0.000491 | 0.002134 | 0.002237 | 0.004862 | 0.003579 | 0.003805 | 0.004326 | 0.006707 | 0.002322 | 0.002392 | 0.003007 | 0.001516 |
| 18.29166667 | 0.022241 | 0.005601 | 0.009747 | 0.004314 | 0.001022 | 0.010607 | 0.011553 | 0.003205 | 0.00124 | 0.000903 | 0.000248 | 0.001732 | 0.001445 | 0.003957 | 0.003837 | 0.003635 | 0.004024 | 0.006127 | 0.001672 | 0.001814 | 0.002332 | 0.000881 |
| 18.30555556 | 0.021933 | 0.005852 | 0.009884 | 0.004721 | 0.001453 | 0.012501 | 0.014647 | 0.004179 | 0.001814 | 0.001321 | 0.000893 | 0.001806 | 0.000785 | 0.003087 | 0.004055 | 0.003357 | 0.003756 | 0.005751 | 0.001297 | 0.00131 | 0.001746 | 0.000328 |
| 18.31944444 | 0.019815 | 0.005423 | 0.008386 | 0.003957 | 0.001435 | 0.012962 | 0.01511 | 0.00479 | 0.002705 | 0.001482 | 0.002052 | 0.001776 | 0.000945 | 0.002964 | 0.004376 | 0.003391 | 0.003608 | 0.005483 | 0.001424 | 0.001344 | 0.001804 | 0.000391 |
| 18.33333333 | 0.017446 | 0.004575 | 0.006401 | 0.002649 | 0.001009 | 0.011509 | 0.012034 | 0.004635 | 0.003342 | 0.001477 | 0.002734 | 0.001318 | 0.001733 | 0.003367 | 0.004739 | 0.003903 | 0.003573 | 0.005401 | 0.001834 | 0.001885 | 0.002425 | 0.001126 |
| 18.34722222 | 0.014892 | 0.003339 | 0.004336 | 0.001312 | 0.000589 | 0.008686 | 0.007177 | 0.003671 | 0.003075 | 0.001241 | 0.002184 | 0.000774 | 0.001974 | 0.003202 | 0.004512 | 0.003881 | 0.00317 | 0.004904 | 0.001813 | 0.002025 | 0.002662 | 0.001686 |
| 18.36111111 | 0.012135 | 0.001953 | 0.003177 | 0.000576 | 0.000428 | 0.005839 | 0.003122 | 0.002771 | 0.002135 | 0.000774 | 0.0013 | 0.00059 | 0.001258 | 0.002296 | 0.003768 | 0.002952 | 0.002348 | 0.003624 | 0.001342 | 0.001393 | 0.002183 | 0.001486 |
| 18.375 | 0.009993 | 0.000918 | 0.003774 | 0.000537 | 0.000419 | 0.004059 | 0.001373 | 0.00264 | 0.001472 | 0.000604 | 0.001341 | 0.000871 | 0.00058 | 0.001629 | 0.003219 | 0.00204 | 0.001649 | 0.002162 | 0.001005 | 0.000742 | 0.001763 | 0.001133 |
| 18.38888889 | 0.008771 | 0.000502 | 0.005266 | 0.001115 | 0.000735 | 0.003542 | 0.002102 | 0.002907 | 0.00152 | 0.00098 | 0.002126 | 0.001773 | 0.000593 | 0.00156 | 0.003052 | 0.001723 | 0.001435 | 0.00141 | 0.001047 | 0.000634 | 0.001681 | 0.001116 |
| 18.40277778 | 0.007688 | 0.000464 | 0.005519 | 0.002025 | 0.00121 | 0.003405 | 0.003666 | 0.002827 | 0.001741 | 0.001278 | 0.002522 | 0.002023 | 0.000955 | 0.001519 | 0.002924 | 0.001717 | 0.001503 | 0.001427 | 0.001262 | 0.000877 | 0.001558 | 0.001088 |
| 18.41666667 | 0.006151 | 0.000496 | 0.003458 | 0.001905 | 0.001129 | 0.002358 | 0.003638 | 0.001888 | 0.001302 | 0.001017 | 0.001986 | 0.001479 | 0.001089 | 0.001052 | 0.002469 | 0.001509 | 0.001263 | 0.001222 | 0.001259 | 0.000985 | 0.001253 | 0.00071 |
| 18.43055556 | 0.004967 | 0.000747 | 0.00104 | 0.001331 | 0.000974 | 0.000979 | 0.001984 | 0.000903 | 0.000749 | 0.000787 | 0.001292 | 0.00154 | 0.000877 | 0.000712 | 0.001751 | 0.001031 | 0.000746 | 0.000775 | 0.00099 | 0.000769 | 0.000954 | 0.000273 |
| 18.44444444 | 0.005587 | 0.001145 | 0.000341 | 0.001527 | 0.000999 | 0.000465 | 0.000781 | 0.000742 | 0.000784 | 0.000746 | 0.001361 | 0.001558 | 0.000688 | 0.000945 | 0.001211 | 0.000647 | 0.000619 | 0.000934 | 0.000818 | 0.000529 | 0.000919 | 0.000205 |
| 18.45833333 | 0.00853 | 0.001247 | 0.00115 | 0.001268 | 0.000972 | 0.000648 | 0.000678 | 0.00129 | 0.00118 | 0.000876 | 0.002337 | 0.002126 | 0.000873 | 0.001169 | 0.000966 | 0.000528 | 0.00083 | 0.000996 | 0.000937 | 0.000742 | 0.001099 | 0.000384 |
| 18.47222222 | 0.013076 | 0.003969 | 0.003996 | 0.000697 | 0.001025 | 0.00093 | 0.001322 | 0.004274 | 0.002785 | 0.001914 | 0.0028 | 0.003822 | 0.001137 | 0.001985 | 0.00076 | 0.001127 | 0.001147 | 0.001546 | 0.001431 | 0.002222 | 0.001064 | 0.000962 |
| 18.48611111 | 0.017285 | 0.014762 | 0.009582 | 0.002139 | 0.002665 | 0.001799 | 0.002449 | 0.010953 | 0.005043 | 0.00257 | 0.00532 | 0.008697 | 0.001528 | 0.004813 | 0.000776 | 0.003788 | 0.002903 | 0.005083 | 0.002331 | 0.005602 | 0.001543 | 0.006028 |
| 18.5 | 0.018565 | 0.032858 | 0.014211 | 0.005042 | 0.004695 | 0.005098 | 0.005189 | 0.015658 | 0.004962 | 0.003111 | 0.015211 | 0.028744 | 0.001989 | 0.006115 | 0.001765 | 0.008244 | 0.00581 | 0.009855 | 0.003081 | 0.00889 | 0.00332 | 0.020004 |
| 18.51388889 | 0.015635 | 0.036261 | 0.011617 | 0.005602 | 0.005409 | 0.007004 | 0.008389 | 0.012002 | 0.003306 | 0.003734 | 0.018625 | 0.053194 | 0.001873 | 0.005396 | 0.003168 | 0.009814 | 0.005571 | 0.009714 | 0.002927 | 0.00799 | 0.003697 | 0.030039 |
| 18.52777778 | 0.013882 | 0.023628 | 0.007125 | 0.010099 | 0.011299 | 0.012915 | 0.011708 | 0.006688 | 0.00554 | 0.007439 | 0.018679 | 0.060662 | 0.002613 | 0.006597 | 0.004693 | 0.007171 | 0.002803 | 0.005346 | 0.002975 | 0.004729 | 0.003573 | 0.026911 |
| 18.54166667 | 0.023525 | 0.018741 | 0.019953 | 0.029243 | 0.02597 | 0.037468 | 0.026778 | 0.014645 | 0.018506 | 0.02257 | 0.039374 | 0.075355 | 0.005213 | 0.006294 | 0.008735 | 0.005067 | 0.003461 | 0.003546 | 0.005388 | 0.003777 | 0.005985 | 0.026329 |
| 18.55555556 | 0.040974 | 0.023359 | 0.049146 | 0.051651 | 0.040273 | 0.067212 | 0.049766 | 0.037697 | 0.039006 | 0.041286 | 0.057226 | 0.091184 | 0.007453 | 0.005147 | 0.015391 | 0.004089 | 0.009064 | 0.007053 | 0.008515 | 0.003667 | 0.00696 | 0.026021 |
| 18.56944444 | 0.046316 | 0.033755 | 0.063578 | 0.05659 | 0.042186 | 0.076898 | 0.058112 | 0.054402 | 0.047959 | 0.047602 | 0.048508 | 0.068334 | 0.008199 | 0.007674 | 0.019285 | 0.003245 | 0.014372 | 0.014352 | 0.007911 | 0.003806 | 0.005534 | 0.014404 |
| 18.58333333 | 0.031091 | 0.034038 | 0.04962 | 0.043438 | 0.030748 | 0.065878 | 0.050954 | 0.045411 | 0.036295 | 0.039098 | 0.02963 | 0.03022 | 0.008347 | 0.01005 | 0.016409 | 0.002969 | 0.013766 | 0.01994 | 0.003985 | 0.004321 | 0.004618 | 0.003826 |
| 18.59722222 | 0.014734 | 0.019351 | 0.027308 | 0.026154 | 0.0151 | 0.047819 | 0.040032 | 0.021616 | 0.017162 | 0.023116 | 0.013723 | 0.009954 | 0.007685 | 0.008176 | 0.010269 | 0.002342 | 0.009946 | 0.020255 | 0.001129 | 0.003133 | 0.003683 | 0.001085 |
| 18.61111111 | 0.017971 | 0.006933 | 0.013024 | 0.013442 | 0.004869 | 0.030711 | 0.02843 | 0.005413 | 0.004901 | 0.009825 | 0.004273 | 0.00471 | 0.006354 | 0.004179 | 0.005946 | 0.001894 | 0.00709 | 0.016433 | 0.000913 | 0.001503 | 0.002615 | 0.000802 |
| 18.625 | 0.031592 | 0.002153 | 0.005539 | 0.005657 | 0.00252 | 0.016828 | 0.017644 | 0.001287 | 0.001207 | 0.003662 | 0.001428 | 0.005301 | 0.005015 | 0.001494 | 0.003531 | 0.001889 | 0.004939 | 0.010736 | 0.001919 | 0.000954 | 0.002021 | 0.000917 |
| 18.63888889 | 0.03636 | 0.003013 | 0.002381 | 0.001912 | 0.003707 | 0.008532 | 0.012036 | 0.002243 | 0.001161 | 0.00159 | 0.002253 | 0.006107 | 0.003375 | 0.000888 | 0.002063 | 0.001542 | 0.003457 | 0.006032 | 0.002392 | 0.00068 | 0.001507 | 0.000931 |
| 18.65277778 | 0.031158 | 0.005071 | 0.002848 | 0.001144 | 0.005156 | 0.00645 | 0.012781 | 0.004699 | 0.003046 | 0.00145 | 0.004177 | 0.007052 | 0.00189 | 0.001427 | 0.00283 | 0.001236 | 0.002747 | 0.003636 | 0.001964 | 0.000369 | 0.001125 | 0.000846 |
| 18.66666667 | 0.023745 | 0.005684 | 0.003265 | 0.001019 | 0.006212 | 0.007375 | 0.01493 | 0.006626 | 0.007124 | 0.003872 | 0.006084 | 0.00899 | 0.001327 | 0.00187 | 0.00635 | 0.001278 | 0.002135 | 0.002497 | 0.001187 | 0.000341 | 0.001149 | 0.000914 |
| 18.68055556 | 0.018128 | 0.005718 | 0.002449 | 0.000602 | 0.006986 | 0.006338 | 0.012218 | 0.006722 | 0.012142 | 0.007844 | 0.007387 | 0.011161 | 0.00106 | 0.001953 | 0.010798 | 0.001022 | 0.00126 | 0.001529 | 0.000676 | 0.001078 | 0.000956 | 0.000893 |
| 18.69444444 | 0.01525 | 0.007024 | 0.002631 | 0.001095 | 0.007732 | 0.004915 | 0.009062 | 0.006357 | 0.01713 | 0.011638 | 0.00801 | 0.012539 | 0.000573 | 0.001589 | 0.013817 | 0.00044 | 0.000886 | 0.00078 | 0.000983 | 0.002516 | 0.000513 | 0.00094 |
| 18.70833333 | 0.014098 | 0.009962 | 0.005335 | 0.003672 | 0.009164 | 0.009668 | 0.018679 | 0.007957 | 0.022164 | 0.014753 | 0.009026 | 0.013277 | 0.000648 | 0.001729 | 0.014353 | 0.000473 | 0.00216 | 0.001858 | 0.001726 | 0.002958 | 0.000404 | 0.001177 |
| 18.72222222 | 0.012337 | 0.012937 | 0.009444 | 0.007728 | 0.011649 | 0.02036 | 0.045348 | 0.011768 | 0.026164 | 0.016967 | 0.011104 | 0.01424 | 0.00133 | 0.003439 | 0.012952 | 0.00106 | 0.005216 | 0.007084 | 0.002284 | 0.001993 | 0.000507 | 0.001323 |
| 18.73611111 | 0.008815 | 0.014082 | 0.01173 | 0.010643 | 0.01351 | 0.029288 | 0.070239 | 0.014416 | 0.026542 | 0.017239 | 0.012687 | 0.014891 | 0.001927 | 0.005249 | 0.01098 | 0.001295 | 0.009028 | 0.015703 | 0.00246 | 0.001988 | 0.000482 | 0.001061 |
| 18.75 | 0.004554 | 0.012805 | 0.011338 | 0.010823 | 0.012971 | 0.029325 | 0.067114 | 0.013702 | 0.022394 | 0.014177 | 0.012022 | 0.013762 | 0.002761 | 0.006925 | 0.009652 | 0.002351 | 0.012364 | 0.024756 | 0.002409 | 0.003291 | 0.000734 | 0.000947 |
| 18.76388889 | 0.002527 | 0.010289 | 0.009999 | 0.009495 | 0.01076 | 0.021456 | 0.042949 | 0.01075 | 0.015343 | 0.008107 | 0.009681 | 0.010943 | 0.004031 | 0.008885 | 0.009244 | 0.005266 | 0.014499 | 0.03093 | 0.002061 | 0.003702 | 0.001958 | 0.001699 |
| 18.77777778 | 0.005319 | 0.007865 | 0.008051 | 0.007784 | 0.008263 | 0.011254 | 0.019437 | 0.006473 | 0.007779 | 0.004485 | 0.007007 | 0.00777 | 0.005476 | 0.009485 | 0.008217 | 0.006732 | 0.014752 | 0.030409 | 0.001425 | 0.002905 | 0.003567 | 0.002125 |
| 18.79166667 | 0.011137 | 0.005628 | 0.00509 | 0.005445 | 0.005921 | 0.004842 | 0.00831 | 0.003053 | 0.003255 | 0.006244 | 0.004737 | 0.005447 | 0.007144 | 0.008387 | 0.005042 | 0.004366 | 0.012539 | 0.023066 | 0.001235 | 0.002034 | 0.005454 | 0.001975 |
| 18.80555556 | 0.015877 | 0.003408 | 0.002971 | 0.003142 | 0.004316 | 0.005669 | 0.015205 | 0.002805 | 0.002068 | 0.00691 | 0.003509 | 0.004671 | 0.008232 | 0.007142 | 0.002144 | 0.002385 | 0.009213 | 0.014042 | 0.001262 | 0.001508 | 0.008647 | 0.002543 |
| 18.81944444 | 0.018083 | 0.001544 | 0.003486 | 0.002158 | 0.003818 | 0.008855 | 0.031978 | 0.003347 | 0.002124 | 0.004281 | 0.003059 | 0.005036 | 0.008087 | 0.006276 | 0.002257 | 0.00439 | 0.007402 | 0.007684 | 0.000822 | 0.001173 | 0.012303 | 0.003649 |
| 18.83333333 | 0.01981 | 0.000978 | 0.004659 | 0.002289 | 0.003558 | 0.008906 | 0.041413 | 0.003074 | 0.004191 | 0.003582 | 0.002123 | 0.005721 | 0.007338 | 0.005172 | 0.003372 | 0.007121 | 0.007597 | 0.004208 | 0.000433 | 0.001104 | 0.013832 | 0.004187 |
| 18.84722222 | 0.022924 | 0.002211 | 0.003498 | 0.002286 | 0.002451 | 0.006141 | 0.032448 | 0.003221 | 0.006418 | 0.005946 | 0.001258 | 0.006593 | 0.006302 | 0.003635 | 0.002813 | 0.007057 | 0.007861 | 0.002826 | 0.000758 | 0.001954 | 0.01227 | 0.003797 |
| 18.86111111 | 0.024048 | 0.003866 | 0.00124 | 0.001693 | 0.001338 | 0.003678 | 0.015524 | 0.003024 | 0.006492 | 0.008264 | 0.001958 | 0.007245 | 0.004883 | 0.002184 | 0.001766 | 0.00468 | 0.007114 | 0.004691 | 0.001562 | 0.003213 | 0.009255 | 0.003054 |
| 18.875 | 0.019043 | 0.004267 | 0.001113 | 0.001343 | 0.00154 | 0.002621 | 0.006855 | 0.00228 | 0.005642 | 0.008734 | 0.002999 | 0.00631 | 0.003524 | 0.001452 | 0.002542 | 0.002015 | 0.005419 | 0.009075 | 0.002039 | 0.003309 | 0.006768 | 0.002698 |
| 18.88888889 | 0.012201 | 0.003247 | 0.003554 | 0.002364 | 0.002008 | 0.004248 | 0.012762 | 0.0034 | 0.004353 | 0.00677 | 0.002732 | 0.003468 | 0.002577 | 0.001005 | 0.004512 | 0.000909 | 0.003273 | 0.012294 | 0.00201 | 0.00242 | 0.00506 | 0.002756 |
| 18.90277778 | 0.011881 | 0.001656 | 0.007248 | 0.004949 | 0.001686 | 0.010226 | 0.031273 | 0.00637 | 0.002345 | 0.00345 | 0.001519 | 0.001654 | 0.001574 | 0.001029 | 0.00631 | 0.001916 | 0.002817 | 0.01393 | 0.00194 | 0.001844 | 0.002959 | 0.002526 |
| 18.91666667 | 0.0164 | 0.001109 | 0.010059 | 0.008212 | 0.001224 | 0.016378 | 0.045154 | 0.008544 | 0.001203 | 0.001924 | 0.000634 | 0.002803 | 0.001138 | 0.002827 | 0.007797 | 0.003929 | 0.004446 | 0.015863 | 0.002043 | 0.001966 | 0.001641 | 0.001667 |
| 18.93055556 | 0.014274 | 0.002677 | 0.010038 | 0.010204 | 0.001224 | 0.01882 | 0.04215 | 0.008727 | 0.002477 | 0.002849 | 0.000468 | 0.004454 | 0.002389 | 0.005512 | 0.008599 | 0.005475 | 0.005857 | 0.018557 | 0.002028 | 0.002264 | 0.003034 | 0.000749 |
| 18.94444444 | 0.008616 | 0.005519 | 0.007668 | 0.009388 | 0.001428 | 0.019396 | 0.032026 | 0.008104 | 0.004466 | 0.003943 | 0.000641 | 0.005076 | 0.004414 | 0.00703 | 0.008321 | 0.005894 | 0.007008 | 0.02185 | 0.001454 | 0.001913 | 0.005365 | 0.000585 |
| 18.95833333 | 0.007404 | 0.008889 | 0.005372 | 0.007151 | 0.001108 | 0.020351 | 0.024792 | 0.009742 | 0.004634 | 0.003998 | 0.001164 | 0.005238 | 0.006473 | 0.007035 | 0.007343 | 0.005491 | 0.009218 | 0.025195 | 0.000605 | 0.001085 | 0.006768 | 0.001151 |
| 18.97222222 | 0.011288 | 0.012073 | 0.00516 | 0.005945 | 0.00104 | 0.020932 | 0.020881 | 0.014226 | 0.002808 | 0.003736 | 0.001266 | 0.004902 | 0.008954 | 0.006936 | 0.006428 | 0.004878 | 0.011619 | 0.0275 | 0.000232 | 0.000747 | 0.00706 | 0.001547 |
| 18.98611111 | 0.025405 | 0.013359 | 0.006818 | 0.006108 | 0.002357 | 0.019651 | 0.017648 | 0.017164 | 0.001341 | 0.003904 | 0.000676 | 0.004523 | 0.01135 | 0.007656 | 0.005677 | 0.004063 | 0.012406 | 0.027028 | 0.000269 | 0.000893 | 0.005973 | 0.001428 |
| 19 | 0.034402 | 0.012154 | 0.008425 | 0.006251 | 0.003539 | 0.018325 | 0.013212 | 0.015769 | 0.001379 | 0.003839 | 0.000483 | 0.004096 | 0.012879 | 0.008092 | 0.004636 | 0.002768 | 0.011009 | 0.023364 | 0.000326 | 0.001193 | 0.004012 | 0.001013 |
| 19.01388889 | 0.024083 | 0.008689 | 0.007336 | 0.005622 | 0.003032 | 0.018389 | 0.010113 | 0.011295 | 0.001456 | 0.004103 | 0.001944 | 0.007601 | 0.013396 | 0.00728 | 0.002943 | 0.001822 | 0.008012 | 0.017818 | 0.000679 | 0.002214 | 0.003176 | 0.002682 |
| 19.02777778 | 0.017084 | 0.009352 | 0.004342 | 0.004824 | 0.003429 | 0.017519 | 0.011414 | 0.005577 | 0.001033 | 0.005904 | 0.006766 | 0.022446 | 0.013173 | 0.005868 | 0.00142 | 0.003126 | 0.004195 | 0.01093 | 0.001371 | 0.004668 | 0.00542 | 0.011016 |
| 19.04166667 | 0.024878 | 0.013044 | 0.003113 | 0.003612 | 0.005363 | 0.013223 | 0.011989 | 0.001833 | 0.002624 | 0.007007 | 0.010577 | 0.03083 | 0.011726 | 0.005148 | 0.001 | 0.005728 | 0.001376 | 0.004493 | 0.001554 | 0.007104 | 0.007932 | 0.018503 |
| 19.05555556 | 0.024837 | 0.018338 | 0.005245 | 0.002441 | 0.006553 | 0.006445 | 0.013905 | 0.001466 | 0.007969 | 0.007385 | 0.012094 | 0.036686 | 0.008454 | 0.006004 | 0.001371 | 0.005235 | 0.000805 | 0.00145 | 0.001075 | 0.005958 | 0.008097 | 0.020672 |
| 19.06944444 | 0.012147 | 0.029281 | 0.008782 | 0.003456 | 0.006041 | 0.002372 | 0.021637 | 0.002344 | 0.012372 | 0.00787 | 0.017121 | 0.054341 | 0.007321 | 0.006334 | 0.002411 | 0.002544 | 0.001024 | 0.001569 | 0.00099 | 0.00352 | 0.010107 | 0.029143 |
| 19.08333333 | 0.006121 | 0.027809 | 0.008283 | 0.004921 | 0.004609 | 0.002298 | 0.021726 | 0.005915 | 0.012035 | 0.005405 | 0.01852 | 0.043549 | 0.009226 | 0.004373 | 0.005767 | 0.001855 | 0.00103 | 0.002063 | 0.001198 | 0.00362 | 0.012543 | 0.034087 |
| 19.09722222 | 0.008348 | 0.012827 | 0.003961 | 0.00475 | 0.006922 | 0.003526 | 0.015335 | 0.012691 | 0.010425 | 0.001876 | 0.013061 | 0.012383 | 0.008753 | 0.003281 | 0.011088 | 0.002536 | 0.000829 | 0.002156 | 0.000893 | 0.002882 | 0.010023 | 0.022686 |
| 19.11111111 | 0.009531 | 0.003973 | 0.001765 | 0.005174 | 0.01118 | 0.006843 | 0.01691 | 0.017489 | 0.010398 | 0.000819 | 0.011525 | 0.004762 | 0.006478 | 0.004861 | 0.013402 | 0.003265 | 0.000764 | 0.002412 | 0.000797 | 0.001162 | 0.006073 | 0.010112 |
| 19.125 | 0.008995 | 0.001741 | 0.002628 | 0.006817 | 0.012451 | 0.012047 | 0.024353 | 0.01832 | 0.010329 | 0.000716 | 0.014062 | 0.010823 | 0.005655 | 0.00787 | 0.01198 | 0.003992 | 0.000885 | 0.002224 | 0.002016 | 0.001671 | 0.004494 | 0.005918 |
| 19.13888889 | 0.008739 | 0.001493 | 0.003876 | 0.008113 | 0.011915 | 0.016034 | 0.02971 | 0.017044 | 0.008901 | 0.000949 | 0.012727 | 0.013413 | 0.005525 | 0.01111 | 0.010183 | 0.004951 | 0.000872 | 0.001967 | 0.004094 | 0.003509 | 0.004283 | 0.004892 |
| 19.15277778 | 0.009091 | 0.003165 | 0.004703 | 0.008607 | 0.010189 | 0.017787 | 0.031109 | 0.015447 | 0.007487 | 0.002062 | 0.011065 | 0.014893 | 0.004898 | 0.013347 | 0.008372 | 0.005561 | 0.001205 | 0.00195 | 0.005283 | 0.004748 | 0.004153 | 0.003827 |
| 19.16666667 | 0.010056 | 0.006987 | 0.005917 | 0.008454 | 0.007733 | 0.018018 | 0.029532 | 0.014757 | 0.007161 | 0.003256 | 0.012997 | 0.014004 | 0.003808 | 0.013515 | 0.005951 | 0.004558 | 0.001311 | 0.001496 | 0.004696 | 0.004629 | 0.003461 | 0.002702 |
| 19.18055556 | 0.010831 | 0.011145 | 0.007207 | 0.00743 | 0.005776 | 0.016247 | 0.024955 | 0.014232 | 0.006865 | 0.003378 | 0.011403 | 0.009098 | 0.002973 | 0.011195 | 0.00366 | 0.002685 | 0.001014 | 0.000958 | 0.003307 | 0.003588 | 0.002438 | 0.002029 |
| 19.19444444 | 0.010293 | 0.012513 | 0.007075 | 0.005762 | 0.004154 | 0.012837 | 0.01881 | 0.012323 | 0.006046 | 0.002883 | 0.00741 | 0.005567 | 0.002729 | 0.007947 | 0.002664 | 0.001579 | 0.001144 | 0.001044 | 0.002351 | 0.002525 | 0.001696 | 0.001972 |
| 19.20833333 | 0.00918 | 0.011265 | 0.00572 | 0.004647 | 0.002942 | 0.010175 | 0.014016 | 0.009576 | 0.005997 | 0.003397 | 0.007866 | 0.006178 | 0.002789 | 0.005767 | 0.002929 | 0.001571 | 0.001627 | 0.001712 | 0.002074 | 0.001934 | 0.001551 | 0.002197 |
| 19.22222222 | 0.009149 | 0.01024 | 0.005168 | 0.004632 | 0.002828 | 0.009168 | 0.011233 | 0.008412 | 0.007096 | 0.004626 | 0.010583 | 0.008947 | 0.002805 | 0.004843 | 0.003334 | 0.001716 | 0.00195 | 0.002303 | 0.002058 | 0.001686 | 0.001834 | 0.00246 |
| 19.23611111 | 0.010058 | 0.010293 | 0.006004 | 0.00506 | 0.003465 | 0.008636 | 0.008843 | 0.009173 | 0.007869 | 0.005011 | 0.010813 | 0.007703 | 0.002562 | 0.003699 | 0.003149 | 0.001393 | 0.001816 | 0.002292 | 0.001843 | 0.00136 | 0.002067 | 0.00261 |
| 19.25 | 0.010566 | 0.010507 | 0.006982 | 0.005354 | 0.003845 | 0.00808 | 0.00669 | 0.009696 | 0.007562 | 0.004675 | 0.009069 | 0.003882 | 0.002019 | 0.003407 | 0.002486 | 0.001005 | 0.001488 | 0.001857 | 0.001492 | 0.000879 | 0.001974 | 0.002455 |
| 19.26388889 | 0.010132 | 0.010173 | 0.007368 | 0.005348 | 0.003429 | 0.007784 | 0.005535 | 0.008892 | 0.006766 | 0.004452 | 0.008041 | 0.002197 | 0.001536 | 0.007086 | 0.001629 | 0.001072 | 0.001429 | 0.001534 | 0.001439 | 0.000509 | 0.001703 | 0.002112 |
| 19.27777778 | 0.008867 | 0.009111 | 0.006928 | 0.004637 | 0.002153 | 0.007154 | 0.004538 | 0.007259 | 0.005705 | 0.003919 | 0.007569 | 0.001996 | 0.001541 | 0.01106 | 0.001007 | 0.001479 | 0.001587 | 0.001456 | 0.001649 | 0.000433 | 0.001506 | 0.001854 |
| 19.29166667 | 0.007671 | 0.008259 | 0.006027 | 0.003547 | 0.000792 | 0.006221 | 0.003379 | 0.005737 | 0.004656 | 0.003208 | 0.006835 | 0.001875 | 0.001862 | 0.009497 | 0.001053 | 0.001902 | 0.001549 | 0.001398 | 0.001675 | 0.000571 | 0.001361 | 0.001616 |
| 19.30555556 | 0.008211 | 0.009131 | 0.006507 | 0.003704 | 0.000898 | 0.006731 | 0.003644 | 0.005656 | 0.004861 | 0.003723 | 0.006763 | 0.00236 | 0.001863 | 0.004564 | 0.001528 | 0.002118 | 0.001301 | 0.001296 | 0.001422 | 0.000579 | 0.001021 | 0.001193 |
| 19.31944444 | 0.010392 | 0.011473 | 0.00864 | 0.005507 | 0.002487 | 0.009131 | 0.005995 | 0.007091 | 0.006547 | 0.00584 | 0.008287 | 0.005343 | 0.001479 | 0.001369 | 0.001986 | 0.002089 | 0.001189 | 0.001266 | 0.001171 | 0.000398 | 0.000612 | 0.000702 |
| 19.33333333 | 0.011914 | 0.012987 | 0.009803 | 0.007073 | 0.003758 | 0.011286 | 0.008627 | 0.008071 | 0.00795 | 0.007962 | 0.010234 | 0.009472 | 0.001159 | 0.000721 | 0.002203 | 0.001944 | 0.001142 | 0.001234 | 0.000994 | 0.000285 | 0.00046 | 0.000341 |
| 19.34722222 | 0.011154 | 0.011844 | 0.008824 | 0.006643 | 0.003468 | 0.011058 | 0.00872 | 0.007317 | 0.007362 | 0.007688 | 0.009681 | 0.008763 | 0.000846 | 0.000636 | 0.001912 | 0.001485 | 0.000799 | 0.000847 | 0.000642 | 0.000355 | 0.000351 | 0.00019 |
| 19.36111111 | 0.008295 | 0.007943 | 0.00662 | 0.004332 | 0.002182 | 0.008387 | 0.005598 | 0.005187 | 0.004906 | 0.00456 | 0.006109 | 0.00446 | 0.000412 | 0.000284 | 0.001108 | 0.000723 | 0.000428 | 0.000329 | 0.00027 | 0.000665 | 0.00021 | 0.00025 |
| 19.375 | 0.004567 | 0.003714 | 0.003635 | 0.001808 | 0.001661 | 0.005113 | 0.002445 | 0.002631 | 0.002256 | 0.001426 | 0.003083 | 0.003463 | 0.000159 | 9.52E-05 | 0.000347 | 0.000259 | 0.000331 | 0.000222 | 0.00015 | 0.000938 | 0.000305 | 0.000316 |
| 19.38888889 | 0.002018 | 0.001606 | 0.001375 | 0.000465 | 0.001882 | 0.002654 | 0.001629 | 0.001147 | 0.000734 | 0.000347 | 0.002116 | 0.00328 | 0.000426 | 0.000346 | 0.00021 | 0.000437 | 0.000479 | 0.000519 | 0.000267 | 0.000745 | 0.000454 | 0.000382 |
| 19.40277778 | 0.001959 | 0.001849 | 0.001681 | 0.000335 | 0.001415 | 0.00111 | 0.001653 | 0.00188 | 0.00041 | 0.000413 | 0.001424 | 0.004584 | 0.001218 | 0.001046 | 0.00071 | 0.001113 | 0.000974 | 0.001072 | 0.000736 | 0.000574 | 0.000755 | 0.000682 |
| 19.41666667 | 0.002901 | 0.003337 | 0.002915 | 0.000693 | 0.000601 | 0.000503 | 0.001295 | 0.003602 | 0.000771 | 0.00062 | 0.000675 | 0.008809 | 0.001741 | 0.001516 | 0.001203 | 0.001621 | 0.001281 | 0.001322 | 0.001114 | 0.00081 | 0.001205 | 0.000914 |
| 19.43055556 | 0.00263 | 0.003108 | 0.002402 | 0.000698 | 0.000595 | 0.00066 | 0.001052 | 0.003677 | 0.000974 | 0.000572 | 0.000404 | 0.008922 | 0.001293 | 0.001102 | 0.001035 | 0.001287 | 0.000894 | 0.000927 | 0.000841 | 0.000761 | 0.000993 | 0.000717 |
| 19.44444444 | 0.001307 | 0.001659 | 0.001146 | 0.000453 | 0.001444 | 0.000907 | 0.001359 | 0.002111 | 0.000674 | 0.000483 | 0.000972 | 0.005374 | 0.00061 | 0.000749 | 0.000859 | 0.000944 | 0.000458 | 0.000621 | 0.000513 | 0.000607 | 0.000429 | 0.000697 |
| 19.45833333 | 0.000371 | 0.001998 | 0.000888 | 0.000363 | 0.001877 | 0.000864 | 0.002378 | 0.000929 | 0.000447 | 0.001008 | 0.002408 | 0.004889 | 0.000475 | 0.001275 | 0.001248 | 0.001379 | 0.000385 | 0.000656 | 0.000578 | 0.00077 | 0.000278 | 0.001045 |
| 19.47222222 | 0.000657 | 0.003343 | 0.001234 | 0.000476 | 0.001218 | 0.000905 | 0.003182 | 0.001174 | 0.000454 | 0.00162 | 0.002922 | 0.005419 | 0.000457 | 0.001832 | 0.001226 | 0.001437 | 0.000324 | 0.000466 | 0.000479 | 0.000662 | 0.000249 | 0.00107 |

| | | | | | | | | | | | | | | | | | | | | | | |
|---|---|---|---|---|---|---|---|---|---|---|---|---|---|---|---|---|---|---|---|---|---|---|
| 19.48611111 | 0.002337 | 0.004041 | 0.002631 | 0.001209 | 0.000483 | 0.001192 | 0.002637 | 0.002728 | 0.000386 | 0.0016 | 0.001836 | 0.005174 | 0.00027 | 0.002244 | 0.000749 | 0.000773 | 0.000249 | 0.00025 | 0.000221 | 0.000288 | 0.000201 | 0.000941 |
| 19.5 | 0.004665 | 0.003598 | 0.004773 | 0.002487 | 0.000323 | 0.00123 | 0.001395 | 0.004573 | 0.000684 | 0.002025 | 0.001072 | 0.006918 | 0.000219 | 0.003162 | 0.000757 | 0.000408 | 0.000444 | 0.000546 | 0.000295 | 0.000223 | 0.00031 | 0.001184 |
| 19.51388889 | 0.005678 | 0.00285 | 0.005131 | 0.002971 | 0.000812 | 0.001394 | 0.001519 | 0.005062 | 0.001268 | 0.003072 | 0.000848 | 0.006031 | 0.000445 | 0.00401 | 0.000934 | 0.000372 | 0.000849 | 0.001001 | 0.00048 | 0.000406 | 0.000618 | 0.001701 |
| 19.52777778 | 0.004834 | 0.002857 | 0.005247 | 0.002667 | 0.00184 | 0.001514 | 0.00341 | 0.004498 | 0.001254 | 0.00304 | 0.001314 | 0.002581 | 0.000937 | 0.004244 | 0.001755 | 0.000715 | 0.001258 | 0.00095 | 0.00087 | 0.000527 | 0.001513 | 0.003038 |
| 19.54166667 | 0.005911 | 0.003791 | 0.012307 | 0.005895 | 0.003191 | 0.004426 | 0.005021 | 0.007225 | 0.002628 | 0.003801 | 0.002857 | 0.003922 | 0.001426 | 0.004683 | 0.006181 | 0.002563 | 0.002854 | 0.001196 | 0.002895 | 0.000525 | 0.003433 | 0.006027 |
| 19.55555556 | 0.011425 | 0.007281 | 0.025006 | 0.013967 | 0.005825 | 0.012781 | 0.00804 | 0.014591 | 0.006881 | 0.008776 | 0.003683 | 0.011486 | 0.001424 | 0.005372 | 0.013682 | 0.005437 | 0.005677 | 0.002487 | 0.006474 | 0.00127 | 0.004774 | 0.00863 |
| 19.56944444 | 0.015904 | 0.009656 | 0.027563 | 0.018926 | 0.00863 | 0.015993 | 0.011932 | 0.018903 | 0.00876 | 0.01432 | 0.004456 | 0.017828 | 0.001588 | 0.005503 | 0.017344 | 0.006361 | 0.005563 | 0.00273 | 0.007467 | 0.00322 | 0.003459 | 0.007535 |
| 19.58333333 | 0.013331 | 0.008625 | 0.016008 | 0.014565 | 0.007429 | 0.010183 | 0.009748 | 0.013858 | 0.005991 | 0.013436 | 0.007634 | 0.014349 | 0.002118 | 0.005624 | 0.013459 | 0.004049 | 0.00322 | 0.001765 | 0.004464 | 0.004107 | 0.001324 | 0.003913 |
| 19.59722222 | 0.006447 | 0.010513 | 0.006862 | 0.006843 | 0.004153 | 0.006926 | 0.004531 | 0.00643 | 0.004587 | 0.007219 | 0.012006 | 0.008084 | 0.001928 | 0.005664 | 0.0067 | 0.001343 | 0.003408 | 0.001629 | 0.002101 | 0.002742 | 0.000999 | 0.001424 |
| 19.61111111 | 0.00245 | 0.013081 | 0.006798 | 0.004143 | 0.004389 | 0.008031 | 0.00409 | 0.005059 | 0.005995 | 0.003328 | 0.014476 | 0.010773 | 0.001485 | 0.004272 | 0.002897 | 0.001071 | 0.003948 | 0.001468 | 0.001896 | 0.001232 | 0.001327 | 0.000643 |
| 19.625 | 0.003392 | 0.011693 | 0.008344 | 0.005362 | 0.006343 | 0.008685 | 0.00657 | 0.006311 | 0.006682 | 0.004057 | 0.014565 | 0.01857 | 0.00133 | 0.002022 | 0.004505 | 0.002409 | 0.003132 | 0.001645 | 0.001557 | 0.000773 | 0.000811 | 0.000506 |
| 19.63888889 | 0.005349 | 0.009075 | 0.008518 | 0.006401 | 0.006959 | 0.008317 | 0.008562 | 0.00607 | 0.006281 | 0.005552 | 0.01344 | 0.024923 | 0.002154 | 0.00064 | 0.01155 | 0.003201 | 0.003632 | 0.005398 | 0.001069 | 0.000581 | 0.000393 | 0.000837 |
| 19.65277778 | 0.005743 | 0.006699 | 0.008144 | 0.006669 | 0.00683 | 0.006894 | 0.008947 | 0.005697 | 0.005828 | 0.005843 | 0.011567 | 0.029207 | 0.005176 | 0.000778 | 0.021894 | 0.002394 | 0.006725 | 0.014621 | 0.001428 | 0.00065 | 0.001123 | 0.001885 |
| 19.66666667 | 0.005032 | 0.004341 | 0.007431 | 0.006634 | 0.006508 | 0.004482 | 0.006506 | 0.006457 | 0.005598 | 0.005441 | 0.009166 | 0.030693 | 0.009424 | 0.002206 | 0.033782 | 0.00214 | 0.011998 | 0.028914 | 0.002214 | 0.00154 | 0.003112 | 0.00367 |
| 19.68055556 | 0.003998 | 0.002135 | 0.006827 | 0.006515 | 0.006114 | 0.001972 | 0.004304 | 0.008721 | 0.005398 | 0.004691 | 0.006658 | 0.028823 | 0.014482 | 0.004421 | 0.045835 | 0.004962 | 0.018357 | 0.046453 | 0.002349 | 0.002414 | 0.006545 | 0.005877 |
| 19.69444444 | 0.003327 | 0.000705 | 0.006652 | 0.006306 | 0.005638 | 0.000856 | 0.006988 | 0.012378 | 0.005087 | 0.003659 | 0.004352 | 0.024993 | 0.021491 | 0.00669 | 0.053572 | 0.008117 | 0.023527 | 0.061533 | 0.001723 | 0.002073 | 0.011911 | 0.008704 |
| 19.70833333 | 0.003608 | 0.000585 | 0.007097 | 0.006208 | 0.005446 | 0.002665 | 0.011144 | 0.016606 | 0.005073 | 0.002725 | 0.002911 | 0.021219 | 0.030014 | 0.007823 | 0.052902 | 0.009908 | 0.023754 | 0.062986 | 0.001012 | 0.001246 | 0.01901 | 0.013084 |
| 19.72222222 | 0.004958 | 0.001697 | 0.008398 | 0.006913 | 0.006151 | 0.008802 | 0.009677 | 0.020031 | 0.006099 | 0.002555 | 0.003269 | 0.017814 | 0.036473 | 0.007566 | 0.045703 | 0.012905 | 0.018547 | 0.048444 | 0.001464 | 0.001856 | 0.025097 | 0.018506 |
| 19.73611111 | 0.006867 | 0.003917 | 0.010661 | 0.00892 | 0.007869 | 0.019041 | 0.010456 | 0.02117 | 0.008451 | 0.003317 | 0.005101 | 0.014211 | 0.036697 | 0.006943 | 0.037244 | 0.018047 | 0.011837 | 0.029496 | 0.00343 | 0.003662 | 0.026235 | 0.021605 |
| 19.75 | 0.008428 | 0.007354 | 0.013783 | 0.011139 | 0.009818 | 0.033664 | 0.037335 | 0.019484 | 0.011012 | 0.004282 | 0.006929 | 0.010284 | 0.030435 | 0.00632 | 0.029962 | 0.023626 | 0.00608 | 0.013863 | 0.004627 | 0.004494 | 0.022078 | 0.019781 |
| 19.76388889 | 0.00874 | 0.010994 | 0.018084 | 0.011831 | 0.010597 | 0.047267 | 0.078773 | 0.016137 | 0.012247 | 0.00436 | 0.007895 | 0.00677 | 0.020725 | 0.005859 | 0.023108 | 0.026025 | 0.00231 | 0.005833 | 0.004006 | 0.003601 | 0.015904 | 0.014695 |
| 19.77777778 | 0.007697 | 0.012385 | 0.019536 | 0.01092 | 0.009474 | 0.04696 | 0.089628 | 0.012438 | 0.011604 | 0.003433 | 0.007972 | 0.00434 | 0.011028 | 0.00642 | 0.016166 | 0.02241 | 0.000927 | 0.006603 | 0.003243 | 0.002284 | 0.010785 | 0.009853 |
| 19.79166667 | 0.00591 | 0.010048 | 0.014224 | 0.009335 | 0.007123 | 0.033105 | 0.065454 | 0.008928 | 0.009266 | 0.002143 | 0.007067 | 0.002933 | 0.004685 | 0.007633 | 0.009856 | 0.014197 | 0.001185 | 0.006706 | 0.002892 | 0.001452 | 0.007106 | 0.006421 |
| 19.80555556 | 0.004154 | 0.005559 | 0.008371 | 0.008166 | 0.005232 | 0.017105 | 0.033129 | 0.00618 | 0.006166 | 0.000961 | 0.005545 | 0.002096 | 0.00303 | 0.008105 | 0.004754 | 0.006628 | 0.003457 | 0.005689 | 0.002017 | 0.000852 | 0.003771 | 0.003457 |
| 19.81944444 | 0.003485 | 0.002526 | 0.007194 | 0.008091 | 0.005086 | 0.00679 | 0.013863 | 0.00464 | 0.003716 | 0.000327 | 0.004401 | 0.0015 | 0.002838 | 0.008021 | 0.002796 | 0.004537 | 0.00676 | 0.011502 | 0.000802 | 0.000521 | 0.001572 | 0.001169 |
| 19.83333333 | 0.004285 | 0.001902 | 0.007864 | 0.008077 | 0.005778 | 0.005876 | 0.020615 | 0.003986 | 0.002338 | 0.000383 | 0.003916 | 0.000957 | 0.00266 | 0.008467 | 0.005866 | 0.006647 | 0.007804 | 0.016261 | 0.000217 | 0.00089 | 0.001013 | 0.000477 |
| 19.84722222 | 0.005679 | 0.002636 | 0.00729 | 0.006921 | 0.005643 | 0.011974 | 0.050109 | 0.003824 | 0.001644 | 0.000697 | 0.003631 | 0.000398 | 0.003229 | 0.008883 | 0.010378 | 0.008075 | 0.005331 | 0.011511 | 0.000342 | 0.00168 | 0.001125 | 0.000535 |
| 19.86111111 | 0.006785 | 0.00419 | 0.005848 | 0.005186 | 0.004674 | 0.016308 | 0.07549 | 0.004327 | 0.001702 | 0.000937 | 0.003541 | 0.000197 | 0.002607 | 0.007842 | 0.010513 | 0.006324 | 0.003216 | 0.006806 | 0.000673 | 0.001599 | 0.001699 | 0.00065 |
| 19.875 | 0.00712 | 0.004478 | 0.004819 | 0.003953 | 0.003724 | 0.012113 | 0.060731 | 0.005053 | 0.002403 | 0.001305 | 0.003983 | 0.00048 | 0.001431 | 0.005936 | 0.007233 | 0.003665 | 0.003809 | 0.00806 | 0.000752 | 0.00065 | 0.002063 | 0.00101 |
| 19.88888889 | 0.006523 | 0.002833 | 0.003628 | 0.003123 | 0.002928 | 0.00723 | 0.031687 | 0.004854 | 0.002794 | 0.001734 | 0.004754 | 0.000671 | 0.001006 | 0.005257 | 0.006963 | 0.003195 | 0.00388 | 0.008252 | 0.000476 | 0.000263 | 0.001829 | 0.001037 |
| 19.90277778 | 0.005456 | 0.001099 | 0.001901 | 0.002039 | 0.001905 | 0.009148 | 0.03432 | 0.00358 | 0.002256 | 0.001574 | 0.005116 | 0.000725 | 0.000536 | 0.006292 | 0.008226 | 0.002972 | 0.002551 | 0.005602 | 0.000251 | 0.000482 | 0.001395 | 0.000762 |
| 19.91666667 | 0.004591 | 0.00038 | 0.000675 | 0.000896 | 0.000927 | 0.010301 | 0.040239 | 0.002291 | 0.001412 | 0.000807 | 0.00473 | 0.001327 | 0.000945 | 0.007286 | 0.006784 | 0.002934 | 0.002215 | 0.005275 | 0.000464 | 0.000597 | 0.001926 | 0.000971 |
| 19.93055556 | 0.00413 | 0.001008 | 0.000613 | 0.000263 | 0.000574 | 0.006869 | 0.02515 | 0.001698 | 0.001141 | 0.000238 | 0.004248 | 0.00242 | 0.002797 | 0.00717 | 0.007962 | 0.005824 | 0.002672 | 0.006734 | 0.000867 | 0.000745 | 0.004217 | 0.002312 |
| 19.94444444 | 0.003666 | 0.003518 | 0.000868 | 0.000111 | 0.000572 | 0.003493 | 0.009467 | 0.001276 | 0.001234 | 0.000264 | 0.004133 | 0.003792 | 0.004869 | 0.006647 | 0.014045 | 0.009896 | 0.002272 | 0.006217 | 0.001048 | 0.000879 | 0.00628 | 0.004694 |
| 19.95833333 | 0.003007 | 0.007388 | 0.000986 | 0.000261 | 0.00071 | 0.003645 | 0.005261 | 0.00078 | 0.000938 | 0.000488 | 0.00371 | 0.006119 | 0.007704 | 0.005723 | 0.01907 | 0.012684 | 0.002706 | 0.007938 | 0.001161 | 0.000731 | 0.006812 | 0.007445 |
| 19.97222222 | 0.003077 | 0.009128 | 0.001747 | 0.000879 | 0.001456 | 0.006986 | 0.008131 | 0.001181 | 0.000365 | 0.000579 | 0.002524 | 0.009914 | 0.013468 | 0.003588 | 0.020852 | 0.014028 | 0.005722 | 0.015245 | 0.001178 | 0.000731 | 0.007655 | 0.010934 |
| 19.98611111 | 0.004713 | 0.006402 | 0.002715 | 0.001875 | 0.002684 | 0.010127 | 0.011519 | 0.004719 | 0.000603 | 0.000821 | 0.001132 | 0.014498 | 0.021248 | 0.001779 | 0.022237 | 0.013879 | 0.008882 | 0.022132 | 0.000872 | 0.00136 | 0.011322 | 0.015474 |
| 20 | 0.006783 | 0.002406 | 0.00318 | 0.00275 | 0.0036 | 0.011614 | 0.014249 | 0.009873 | 0.00165 | 0.00112 | 0.000539 | 0.018815 | 0.025913 | 0.001744 | 0.025535 | 0.013048 | 0.009821 | 0.024278 | 0.001175 | 0.002561 | 0.018647 | 0.019937 |
| 20.01388889 | 0.00689 | 0.002237 | 0.004308 | 0.004104 | 0.004412 | 0.013087 | 0.017388 | 0.009988 | 0.002695 | 0.00116 | 0.001262 | 0.023161 | 0.024221 | 0.001839 | 0.030992 | 0.013795 | 0.009415 | 0.024318 | 0.002588 | 0.003963 | 0.027351 | 0.02236 |
| 20.02777778 | 0.004428 | 0.006672 | 0.0073 | 0.007402 | 0.006644 | 0.016237 | 0.021769 | 0.009744 | 0.005936 | 0.00205 | 0.002544 | 0.026545 | 0.019325 | 0.001625 | 0.036454 | 0.016724 | 0.0087 | 0.024578 | 0.003723 | 0.004951 | 0.030843 | 0.021425 |
| 20.04166667 | 0.003128 | 0.011026 | 0.010804 | 0.011941 | 0.010294 | 0.020373 | 0.026482 | 0.017931 | 0.013222 | 0.004852 | 0.003431 | 0.025206 | 0.016229 | 0.001973 | 0.037582 | 0.018307 | 0.007328 | 0.022795 | 0.002997 | 0.00479 | 0.02445 | 0.018175 |
| 20.05555556 | 0.004875 | 0.010841 | 0.011624 | 0.013994 | 0.012211 | 0.022037 | 0.027902 | 0.027213 | 0.020517 | 0.007379 | 0.003854 | 0.01929 | 0.016548 | 0.003175 | 0.033243 | 0.014576 | 0.006255 | 0.019275 | 0.001329 | 0.003527 | 0.013604 | 0.014661 |
| 20.06944444 | 0.005245 | 0.006637 | 0.008776 | 0.011141 | 0.009672 | 0.018454 | 0.023243 | 0.028994 | 0.02244 | 0.006239 | 0.005065 | 0.014287 | 0.018013 | 0.005089 | 0.028528 | 0.007848 | 0.007943 | 0.019703 | 0.000566 | 0.002049 | 0.006613 | 0.011916 |
| 20.08333333 | 0.0028 | 0.002305 | 0.005312 | 0.006024 | 0.004615 | 0.012106 | 0.015074 | 0.024864 | 0.019162 | 0.003022 | 0.008969 | 0.013316 | 0.017408 | 0.006478 | 0.026261 | 0.003075 | 0.011975 | 0.024802 | 0.000419 | 0.001139 | 0.004661 | 0.009908 |
| 20.09722222 | 0.00088 | 0.000535 | 0.004414 | 0.002882 | 0.001308 | 0.008129 | 0.008733 | 0.01915 | 0.015354 | 0.001615 | 0.014086 | 0.013997 | 0.013838 | 0.006264 | 0.023063 | 0.001018 | 0.014399 | 0.027611 | 0.000391 | 0.000951 | 0.004516 | 0.0082 |
| 20.11111111 | 0.000469 | 0.000793 | 0.005711 | 0.002467 | 0.00081 | 0.008021 | 0.006043 | 0.014666 | 0.013435 | 0.001552 | 0.015891 | 0.013057 | 0.008894 | 0.004702 | 0.017899 | 0.000268 | 0.013161 | 0.02429 | 0.000408 | 0.001035 | 0.004265 | 0.006623 |
| 20.125 | 0.000811 | 0.001636 | 0.007226 | 0.002931 | 0.001309 | 0.009722 | 0.005179 | 0.012137 | 0.012252 | 0.001331 | 0.013684 | 0.010268 | 0.004546 | 0.002791 | 0.014282 | 0.000476 | 0.009624 | 0.017002 | 0.000449 | 0.001103 | 0.003864 | 0.005187 |
| 20.13888889 | 0.001727 | 0.002188 | 0.007608 | 0.003065 | 0.001609 | 0.010751 | 0.004497 | 0.010891 | 0.010672 | 0.00117 | 0.010597 | 0.007023 | 0.001881 | 0.001211 | 0.013219 | 0.001893 | 0.006186 | 0.009467 | 0.000904 | 0.001111 | 0.003227 | 0.003664 |
| 20.15277778 | 0.003025 | 0.002072 | 0.00634 | 0.002783 | 0.00153 | 0.009851 | 0.003726 | 0.00988 | 0.008392 | 0.001153 | 0.008076 | 0.003768 | 0.001284 | 0.000335 | 0.011999 | 0.003493 | 0.004524 | 0.005729 | 0.001681 | 0.000961 | 0.002171 | 0.002132 |
| 20.16666667 | 0.004593 | 0.002058 | 0.004495 | 0.002563 | 0.001257 | 0.007938 | 0.003169 | 0.008921 | 0.006153 | 0.00091 | 0.006082 | 0.00145 | 0.001652 | 0.000196 | 0.008381 | 0.00314 | 0.004346 | 0.006151 | 0.001839 | 0.000628 | 0.001213 | 0.001085 |
| 20.18055556 | 0.006179 | 0.003003 | 0.003699 | 0.002972 | 0.001308 | 0.007122 | 0.003062 | 0.008174 | 0.005077 | 0.000622 | 0.005171 | 0.000704 | 0.00138 | 0.00044 | 0.004032 | 0.00225 | 0.004172 | 0.006826 | 0.001111 | 0.000259 | 0.000996 | 0.000511 |
| 20.19444444 | 0.00704 | 0.004145 | 0.003932 | 0.003685 | 0.001708 | 0.007902 | 0.003515 | 0.007213 | 0.004778 | 0.000734 | 0.005221 | 0.000659 | 0.000934 | 0.000606 | 0.002144 | 0.003829 | 0.003372 | 0.005522 | 0.000447 | 7.54E-05 | 0.001463 | 0.000347 |
| 20.20833333 | 0.006712 | 0.004192 | 0.004201 | 0.003791 | 0.001795 | 0.008915 | 0.004039 | 0.005807 | 0.004053 | 0.000676 | 0.00523 | 0.001125 | 0.001042 | 0.000474 | 0.001679 | 0.005663 | 0.002516 | 0.003664 | 0.000557 | 0.0001 | 0.001998 | 0.000638 |
| 20.22222222 | 0.005848 | 0.003466 | 0.004401 | 0.003318 | 0.001591 | 0.00926 | 0.004051 | 0.004845 | 0.002973 | 0.000267 | 0.005153 | 0.001765 | 0.001164 | 0.000443 | 0.001025 | 0.005126 | 0.002465 | 0.003316 | 0.001504 | 0.000295 | 0.002571 | 0.000894 |
| 20.23611111 | 0.005522 | 0.003182 | 0.004464 | 0.002933 | 0.001722 | 0.008822 | 0.003931 | 0.00544 | 0.002813 | 0.000309 | 0.00568 | 0.002332 | 0.001482 | 0.001041 | 0.000889 | 0.003173 | 0.003456 | 0.004763 | 0.002756 | 0.000779 | 0.003768 | 0.001397 |
| 20.25 | 0.005714 | 0.003556 | 0.003833 | 0.002778 | 0.002172 | 0.007815 | 0.004133 | 0.007602 | 0.004149 | 0.000969 | 0.006738 | 0.002743 | 0.001749 | 0.00174 | 0.001049 | 0.001538 | 0.004541 | 0.006538 | 0.00342 | 0.001257 | 0.005118 | 0.002468 |
| 20.26388889 | 0.006034 | 0.00432 | 0.003265 | 0.002825 | 0.002523 | 0.006712 | 0.004482 | 0.010192 | 0.005923 | 0.001395 | 0.007203 | 0.002069 | 0.00118 | 0.001464 | 0.001885 | 0.001136 | 0.004825 | 0.006927 | 0.003358 | 0.001137 | 0.005676 | 0.003067 |
| 20.27777778 | 0.006837 | 0.005606 | 0.00397 | 0.003246 | 0.002717 | 0.006071 | 0.004725 | 0.011487 | 0.00658 | 0.001239 | 0.006571 | 0.001168 | 0.000542 | 0.000638 | 0.004182 | 0.001525 | 0.00436 | 0.005377 | 0.003191 | 0.000815 | 0.005688 | 0.002921 |
| 20.29166667 | 0.008254 | 0.007145 | 0.005432 | 0.004034 | 0.002953 | 0.006297 | 0.005091 | 0.010623 | 0.005813 | 0.001372 | 0.006152 | 0.001218 | 0.000642 | 0.000289 | 0.007079 | 0.001873 | 0.003427 | 0.00282 | 0.003006 | 0.00087 | 0.005497 | 0.002453 |
| 20.30555556 | 0.009406 | 0.007836 | 0.006187 | 0.004585 | 0.002937 | 0.006811 | 0.005583 | 0.008331 | 0.004351 | 0.002031 | 0.006891 | 0.001017 | 0.001144 | 0.000364 | 0.009381 | 0.001866 | 0.002524 | 0.001034 | 0.002452 | 0.001013 | 0.004567 | 0.001667 |
| 20.31944444 | 0.009417 | 0.007083 | 0.005515 | 0.004097 | 0.002112 | 0.006521 | 0.00553 | 0.005695 | 0.002804 | 0.002274 | 0.007985 | 0.00057 | 0.001592 | 0.000447 | 0.010596 | 0.001686 | 0.002139 | 0.000455 | 0.001547 | 0.000895 | 0.002721 | 0.000926 |
| 20.33333333 | 0.008883 | 0.006168 | 0.004442 | 0.003285 | 0.001299 | 0.006026 | 0.005352 | 0.004139 | 0.002229 | 0.002451 | 0.008761 | 0.000574 | 0.001861 | 0.000394 | 0.011015 | 0.001763 | 0.001861 | 0.000434 | 0.000674 | 0.00063 | 0.001215 | 0.000929 |
| 20.34722222 | 0.009021 | 0.006876 | 0.004609 | 0.003838 | 0.002052 | 0.006853 | 0.006436 | 0.004878 | 0.003502 | 0.003911 | 0.008964 | 0.000795 | 0.002106 | 0.000338 | 0.010992 | 0.002206 | 0.001188 | 0.001277 | 0.000222 | 0.000535 | 0.001343 | 0.001644 |
| 20.36111111 | 0.009449 | 0.008667 | 0.005706 | 0.005394 | 0.003891 | 0.008362 | 0.00841 | 0.006581 | 0.005218 | 0.005906 | 0.007587 | 0.001416 | 0.00252 | 0.000538 | 0.010636 | 0.002535 | 0.000832 | 0.002907 | 0.000257 | 0.000626 | 0.002859 | 0.002581 |
| 20.375 | 0.008851 | 0.009187 | 0.005962 | 0.005856 | 0.004727 | 0.008461 | 0.009289 | 0.006542 | 0.005223 | 0.006534 | 0.004636 | 0.001773 | 0.003031 | 0.000936 | 0.01051 | 0.002503 | 0.001407 | 0.004492 | 0.000629 | 0.000555 | 0.005036 | 0.003803 |
| 20.38888889 | 0.007181 | 0.007437 | 0.004716 | 0.004518 | 0.003823 | 0.006605 | 0.008065 | 0.003964 | 0.00351 | 0.005768 | 0.002031 | 0.001315 | 0.003557 | 0.001248 | 0.010613 | 0.002048 | 0.002323 | 0.005931 | 0.001101 | 0.000251 | 0.00694 | 0.005002 |
| 20.40277778 | 0.005601 | 0.004907 | 0.003242 | 0.00287 | 0.002577 | 0.004438 | 0.005727 | 0.001654 | 0.001806 | 0.005039 | 0.000993 | 0.000739 | 0.004481 | 0.001327 | 0.010126 | 0.001136 | 0.003178 | 0.007625 | 0.001357 | 6.71E-05 | 0.007837 | 0.005765 |
| 20.41666667 | 0.005368 | 0.004001 | 0.003622 | 0.003004 | 0.003014 | 0.004157 | 0.004502 | 0.001264 | 0.001723 | 0.005664 | 0.001374 | 0.000911 | 0.006373 | 0.00095 | 0.010259 | 0.00053 | 0.004254 | 0.009729 | 0.001493 | 0.0002 | 0.008639 | 0.007138 |
| 20.43055556 | 0.007064 | 0.005728 | 0.006386 | 0.005564 | 0.005809 | 0.006147 | 0.005415 | 0.001477 | 0.003407 | 0.008259 | 0.002942 | 0.002603 | 0.008373 | 0.000355 | 0.01368 | 0.001467 | 0.005752 | 0.01204 | 0.001713 | 0.000551 | 0.010179 | 0.009676 |
| 20.44444444 | 0.008702 | 0.007747 | 0.008527 | 0.007886 | 0.008446 | 0.007506 | 0.005833 | 0.002065 | 0.004191 | 0.010874 | 0.003859 | 0.004696 | 0.008727 | 0.000264 | 0.020187 | 0.004629 | 0.007699 | 0.014847 | 0.001952 | 0.000793 | 0.011977 | 0.012043 |
| 20.45833333 | 0.006496 | 0.006067 | 0.006334 | 0.00612 | 0.007155 | 0.005467 | 0.004482 | 0.005245 | 0.003983 | 0.010064 | 0.004043 | 0.004251 | 0.008381 | 0.000597 | 0.027508 | 0.009636 | 0.01018 | 0.018791 | 0.002447 | 0.001006 | 0.014347 | 0.01395 |
| 20.47222222 | 0.003807 | 0.003578 | 0.003472 | 0.00346 | 0.004148 | 0.003583 | 0.004757 | 0.013055 | 0.006973 | 0.006206 | 0.007485 | 0.002287 | 0.010251 | 0.001395 | 0.035358 | 0.015592 | 0.012946 | 0.022792 | 0.003572 | 0.001883 | 0.017893 | 0.016728 |
| 20.48611111 | 0.006744 | 0.005992 | 0.00521 | 0.005513 | 0.004846 | 0.005919 | 0.007643 | 0.021941 | 0.012439 | 0.002825 | 0.013559 | 0.003231 | 0.014638 | 0.003078 | 0.043553 | 0.021269 | 0.015112 | 0.024792 | 0.005146 | 0.003676 | 0.021617 | 0.020571 |
| 20.5 | 0.012315 | 0.010648 | 0.008034 | 0.009123 | 0.007826 | 0.009983 | 0.012186 | 0.02874 | 0.017134 | 0.001121 | 0.017416 | 0.009672 | 0.019225 | 0.004956 | 0.049311 | 0.024517 | 0.016158 | 0.025637 | 0.006543 | 0.005595 | 0.025191 | 0.024373 |
| 20.51388889 | 0.016408 | 0.013125 | 0.007705 | 0.009682 | 0.008739 | 0.012758 | 0.018329 | 0.034886 | 0.021342 | 0.000607 | 0.017164 | 0.019882 | 0.02286 | 0.006272 | 0.051162 | 0.024657 | 0.016599 | 0.028128 | 0.006872 | 0.0068 | 0.03016 | 0.028102 |
| 20.52777778 | 0.0174 | 0.01029 | 0.007145 | 0.00734 | 0.006421 | 0.010644 | 0.020952 | 0.038394 | 0.023959 | 0.003341 | 0.012204 | 0.023463 | 0.027326 | 0.00714 | 0.050098 | 0.022975 | 0.016389 | 0.032717 | 0.005435 | 0.007459 | 0.036234 | 0.033363 |
| 20.54166667 | 0.012214 | 0.007992 | 0.015771 | 0.009112 | 0.006617 | 0.009808 | 0.016515 | 0.032405 | 0.019997 | 0.012356 | 0.008492 | 0.016917 | 0.034123 | 0.007406 | 0.046982 | 0.019349 | 0.014388 | 0.035707 | 0.003366 | 0.007363 | 0.039814 | 0.039953 |
| 20.55555556 | 0.004236 | 0.012657 | 0.030453 | 0.017201 | 0.012398 | 0.017645 | 0.014017 | 0.020056 | 0.010248 | 0.023311 | 0.011101 | 0.011438 | 0.039782 | 0.007525 | 0.045138 | 0.014928 | 0.011333 | 0.03369 | 0.002241 | 0.005756 | 0.03736 | 0.042511 |
| 20.56944444 | 0.007377 | 0.014877 | 0.028259 | 0.016761 | 0.012188 | 0.019513 | 0.014804 | 0.017719 | 0.006069 | 0.024878 | 0.012839 | 0.010181 | 0.041427 | 0.009112 | 0.049055 | 0.012848 | 0.010869 | 0.029601 | 0.002202 | 0.003544 | 0.031051 | 0.037171 |
| 20.58333333 | 0.026765 | 0.017443 | 0.013855 | 0.009974 | 0.006609 | 0.019133 | 0.010366 | 0.029382 | 0.013368 | 0.015066 | 0.016781 | 0.011547 | 0.040672 | 0.01198 | 0.057366 | 0.012584 | 0.014087 | 0.028654 | 0.003316 | 0.002719 | 0.02966 | 0.031444 |
| 20.59722222 | 0.045362 | 0.025651 | 0.010296 | 0.011824 | 0.007291 | 0.031075 | 0.007681 | 0.040844 | 0.025179 | 0.005151 | 0.02798 | 0.015271 | 0.036476 | 0.014423 | 0.060071 | 0.011827 | 0.013969 | 0.022849 | 0.004416 | 0.003156 | 0.036591 | 0.033904 |
| 20.61111111 | 0.046549 | 0.02776 | 0.011316 | 0.015192 | 0.009616 | 0.039678 | 0.01529 | 0.042825 | 0.035022 | 0.00278 | 0.03357 | 0.01881 | 0.028054 | 0.015259 | 0.047709 | 0.009037 | 0.007334 | 0.009901 | 0.003839 | 0.003753 | 0.040505 | 0.038073 |
| 20.625 | 0.032476 | 0.021939 | 0.010934 | 0.014257 | 0.009464 | 0.036231 | 0.029473 | 0.036564 | 0.038224 | 0.004778 | 0.02577 | 0.022531 | 0.019076 | 0.013309 | 0.026667 | 0.004348 | 0.002201 | 0.004056 | 0.003333 | 0.003966 | 0.029474 | 0.029503 |
| 20.63888889 | 0.016977 | 0.014074 | 0.014832 | 0.012126 | 0.009657 | 0.029693 | 0.044462 | 0.025547 | 0.027721 | 0.005724 | 0.012766 | 0.021389 | 0.013824 | 0.011328 | 0.011451 | 0.001683 | 0.00248 | 0.004965 | 0.004217 | 0.002832 | 0.014405 | 0.011949 |
| 20.65277778 | 0.011651 | 0.007244 | 0.02391 | 0.008285 | 0.008269 | 0.023911 | 0.051446 | 0.014656 | 0.012079 | 0.004595 | 0.004666 | 0.012384 | 0.013605 | 0.013224 | 0.01074 | 0.004266 | 0.005728 | 0.01066 | 0.004477 | 0.001557 | 0.009192 | 0.003942 |
| 20.66666667 | 0.012691 | 0.005896 | 0.032295 | 0.004735 | 0.005023 | 0.016982 | 0.039651 | 0.008607 | 0.005698 | 0.003271 | 0.005057 | 0.004501 | 0.014396 | 0.015788 | 0.022403 | 0.00903 | 0.007562 | 0.015684 | 0.003416 | 0.001689 | 0.009682 | 0.007585 |
| 20.68055556 | 0.010765 | 0.010662 | 0.030854 | 0.003758 | 0.003571 | 0.009686 | 0.017703 | 0.009813 | 0.007539 | 0.002837 | 0.012107 | 0.004055 | 0.012162 | 0.015097 | 0.030265 | 0.009826 | 0.005788 | 0.010734 | 0.002538 | 0.002155 | 0.00945 | 0.01103 |
| 20.69444444 | 0.007642 | 0.014678 | 0.022443 | 0.004104 | 0.003423 | 0.006421 | 0.005724 | 0.01437 | 0.012362 | 0.003067 | 0.016976 | 0.010029 | 0.008813 | 0.014036 | 0.025845 | 0.006421 | 0.003223 | 0.004714 | 0.002962 | 0.002067 | 0.006661 | 0.007852 |
| 20.70833333 | 0.00871 | 0.011667 | 0.016206 | 0.004272 | 0.00244 | 0.008083 | 0.003498 | 0.014061 | 0.01249 | 0.003285 | 0.012572 | 0.018114 | 0.009003 | 0.01815 | 0.020485 | 0.004666 | 0.002578 | 0.006294 | 0.003436 | 0.002006 | 0.009319 | 0.00335 |
| 20.72222222 | 0.011028 | 0.005514 | 0.015122 | 0.004028 | 0.001321 | 0.00937 | 0.004166 | 0.012628 | 0.008247 | 0.002741 | 0.005235 | 0.022057 | 0.009317 | 0.021951 | 0.020801 | 0.006783 | 0.003567 | 0.009662 | 0.002346 | 0.002037 | 0.015864 | 0.002836 |
| 20.73611111 | 0.008957 | 0.004785 | 0.014787 | 0.003043 | 0.00147 | 0.005805 | 0.009161 | 0.017358 | 0.008033 | 0.001633 | 0.002798 | 0.018873 | 0.006763 | 0.014909 | 0.016814 | 0.007274 | 0.004182 | 0.00884 | 0.001185 | 0.001493 | 0.013778 | 0.003806 |
| 20.75 | 0.005082 | 0.009146 | 0.010626 | 0.00148 | 0.002314 | 0.002407 | 0.014254 | 0.019614 | 0.007979 | 0.000928 | 0.004302 | 0.010522 | 0.005989 | 0.004275 | 0.013299 | 0.005832 | 0.004387 | 0.008098 | 0.000957 | 0.000881 | 0.009147 | 0.005827 |
| 20.76388889 | 0.007176 | 0.010201 | 0.007182 | 0.000738 | 0.002091 | 0.003191 | 0.012019 | 0.012162 | 0.00628 | 0.0006 | 0.006183 | 0.004737 | 0.00561 | 0.003722 | 0.021239 | 0.006181 | 0.003245 | 0.006625 | 0.000685 | 0.000539 | 0.007173 | 0.007738 |
| 20.77777778 | 0.021019 | 0.005795 | 0.010588 | 0.002523 | 0.001272 | 0.006531 | 0.006991 | 0.004446 | 0.007005 | 0.000348 | 0.004967 | 0.004812 | 0.005721 | 0.01189 | 0.021508 | 0.007193 | 0.001633 | 0.004181 | 0.001015 | 0.000552 | 0.003463 | 0.006748 |
| 20.79166667 | 0.031592 | 0.002724 | 0.015764 | 0.006239 | 0.001391 | 0.01073 | 0.007353 | 0.00346 | 0.008733 | 0.000579 | 0.002578 | 0.006524 | 0.00628 | 0.016387 | 0.012051 | 0.009437 | 0.002735 | 0.005756 | 0.001799 | 0.000775 | 0.004126 | 0.00697 |
| 20.80555556 | 0.028926 | 0.003953 | 0.013841 | 0.008431 | 0.002592 | 0.011103 | 0.008707 | 0.005471 | 0.013168 | 0.001397 | 0.002942 | 0.007734 | 0.005747 | 0.011956 | 0.008515 | 0.008632 | 0.006444 | 0.012655 | 0.00162 | 0.000834 | 0.010488 | 0.011329 |
| 20.81944444 | 0.031763 | 0.005091 | 0.006608 | 0.007488 | 0.002884 | 0.007133 | 0.006887 | 0.005791 | 0.014253 | 0.001942 | 0.003797 | 0.005459 | 0.007661 | 0.00624 | 0.008342 | 0.003837 | 0.007117 | 0.014614 | 0.001155 | 0.001379 | 0.011555 | 0.012539 |
| 20.83333333 | 0.030647 | 0.004496 | 0.001496 | 0.005085 | 0.001646 | 0.004444 | 0.004512 | 0.005484 | 0.011188 | 0.002117 | 0.003469 | 0.002567 | 0.008013 | 0.007004 | 0.012797 | 0.002538 | 0.00661 | 0.014324 | 0.002177 | 0.001962 | 0.008431 | 0.008717 |
| 20.84722222 | 0.027037 | 0.003729 | 0.000422 | 0.00259 | 0.000827 | 0.004182 | 0.002405 | 0.008885 | 0.013575 | 0.002224 | 0.002478 | 0.003444 | 0.006512 | 0.014541 | 0.014769 | 0.003221 | 0.010241 | 0.023243 | 0.004443 | 0.002435 | 0.007386 | 0.006601 |
| 20.86111111 | 0.037416 | 0.003138 | 0.001086 | 0.0014 | 0.002088 | 0.004073 | 0.00243 | 0.011238 | 0.015706 | 0.001733 | 0.001711 | 0.004507 | 0.00586 | 0.017546 | 0.012343 | 0.003693 | 0.009137 | 0.021705 | 0.005886 | 0.003709 | 0.00595 | 0.004977 |
| 20.875 | 0.03474 | 0.002729 | 0.001342 | 0.002749 | 0.004678 | 0.002677 | 0.003703 | 0.016517 | 0.022783 | 0.001289 | 0.002369 | 0.005384 | 0.005248 | 0.012286 | 0.015205 | 0.005093 | 0.004468 | 0.011903 | 0.00548 | 0.004645 | 0.004485 | 0.002464 |

| | | | | | | | | | | | | | | | | | | | | | | |
|---|---|---|---|---|---|---|---|---|---|---|---|---|---|---|---|---|---|---|---|---|---|---|
| 20.88888889 | 0.029561 | 0.005195 | 0.002495 | 0.005401 | 0.00597 | 0.000794 | 0.003507 | 0.026143 | 0.034967 | 0.001719 | 0.002911 | 0.006258 | 0.007308 | 0.008138 | 0.015211 | 0.004643 | 0.004588 | 0.012008 | 0.00427 | 0.003958 | 0.004229 | 0.002799 |
| 20.90277778 | 0.036934 | 0.014127 | 0.006701 | 0.00773 | 0.006059 | 0.000113 | 0.004278 | 0.023836 | 0.029369 | 0.003343 | 0.004248 | 0.008691 | 0.007317 | 0.007751 | 0.015527 | 0.004333 | 0.008056 | 0.019211 | 0.004368 | 0.002918 | 0.005446 | 0.005104 |
| 20.91666667 | 0.031146 | 0.017453 | 0.010803 | 0.00896 | 0.006438 | 0.001969 | 0.005158 | 0.017974 | 0.017613 | 0.005665 | 0.00858 | 0.014361 | 0.004015 | 0.010082 | 0.022722 | 0.007031 | 0.009116 | 0.02291 | 0.004664 | 0.002714 | 0.008633 | 0.005718 |
| 20.93055556 | 0.01941 | 0.008522 | 0.01138 | 0.009527 | 0.007511 | 0.008281 | 0.007087 | 0.01887 | 0.017563 | 0.007915 | 0.010443 | 0.014537 | 0.005537 | 0.011535 | 0.024765 | 0.007822 | 0.005374 | 0.015042 | 0.003096 | 0.002591 | 0.008995 | 0.007606 |
| 20.94444444 | 0.016603 | 0.002648 | 0.009448 | 0.011328 | 0.009664 | 0.015276 | 0.014577 | 0.015269 | 0.013576 | 0.009102 | 0.006829 | 0.009125 | 0.01109 | 0.012149 | 0.021308 | 0.006242 | 0.003757 | 0.010262 | 0.001552 | 0.00207 | 0.008016 | 0.015696 |
| 20.95833333 | 0.024194 | 0.006796 | 0.007879 | 0.014099 | 0.010822 | 0.015783 | 0.020333 | 0.012717 | 0.007087 | 0.007927 | 0.004677 | 0.009271 | 0.013428 | 0.01252 | 0.024747 | 0.007746 | 0.006474 | 0.016915 | 0.001315 | 0.002427 | 0.014707 | 0.021577 |
| 20.97222222 | 0.033395 | 0.016637 | 0.009483 | 0.015146 | 0.009236 | 0.011436 | 0.020872 | 0.022139 | 0.015018 | 0.004752 | 0.009906 | 0.014103 | 0.014004 | 0.010683 | 0.043066 | 0.0119 | 0.010213 | 0.026935 | 0.001531 | 0.004056 | 0.02337 | 0.018034 |
| 20.98611111 | 0.026595 | 0.019843 | 0.013949 | 0.014386 | 0.00745 | 0.0105 | 0.023185 | 0.03482 | 0.032981 | 0.002346 | 0.018594 | 0.015115 | 0.019058 | 0.011722 | 0.067637 | 0.015315 | 0.010853 | 0.029157 | 0.002015 | 0.005251 | 0.022124 | 0.01544 |
| 21 | 0.014073 | 0.011111 | 0.017616 | 0.014679 | 0.007757 | 0.016763 | 0.028688 | 0.038022 | 0.042947 | 0.003728 | 0.02252 | 0.013488 | 0.027276 | 0.018132 | 0.073765 | 0.017124 | 0.009585 | 0.025819 | 0.002722 | 0.005261 | 0.016985 | 0.023677 |
| 21.01388889 | 0.018529 | 0.004652 | 0.019222 | 0.016855 | 0.00891 | 0.024142 | 0.034656 | 0.032822 | 0.03877 | 0.00534 | 0.021301 | 0.014883 | 0.032166 | 0.025334 | 0.053252 | 0.018247 | 0.012143 | 0.030505 | 0.003339 | 0.005509 | 0.021867 | 0.035789 |
| 21.02777778 | 0.03589 | 0.008682 | 0.019172 | 0.017953 | 0.00859 | 0.024728 | 0.038626 | 0.031092 | 0.032022 | 0.003504 | 0.020304 | 0.0196 | 0.029909 | 0.027452 | 0.035506 | 0.020535 | 0.018949 | 0.045188 | 0.003163 | 0.006976 | 0.033509 | 0.03841 |
| 21.04166667 | 0.044963 | 0.01536 | 0.015936 | 0.016001 | 0.007499 | 0.019622 | 0.038573 | 0.035462 | 0.0326 | 0.001903 | 0.021806 | 0.022879 | 0.023833 | 0.02498 | 0.044556 | 0.022351 | 0.023097 | 0.054387 | 0.002399 | 0.007752 | 0.035692 | 0.031436 |
| 21.05555556 | 0.039801 | 0.013711 | 0.011097 | 0.012775 | 0.007687 | 0.014147 | 0.033302 | 0.036749 | 0.036061 | 0.001964 | 0.02544 | 0.021872 | 0.021413 | 0.026566 | 0.05827 | 0.018019 | 0.018504 | 0.043124 | 0.002278 | 0.005888 | 0.025276 | 0.025031 |
| 21.06944444 | 0.032407 | 0.00877 | 0.010434 | 0.010648 | 0.008313 | 0.011192 | 0.023052 | 0.033229 | 0.036672 | 0.001765 | 0.029417 | 0.021638 | 0.023812 | 0.034334 | 0.050499 | 0.008983 | 0.012352 | 0.026822 | 0.002514 | 0.00317 | 0.013071 | 0.022751 |
| 21.08333333 | 0.031542 | 0.00857 | 0.016186 | 0.010325 | 0.007587 | 0.012346 | 0.012291 | 0.031081 | 0.035803 | 0.001582 | 0.030033 | 0.024877 | 0.023927 | 0.036768 | 0.029737 | 0.003705 | 0.011829 | 0.022398 | 0.003259 | 0.00275 | 0.005981 | 0.020272 |
| 21.09722222 | 0.034721 | 0.011732 | 0.023266 | 0.011059 | 0.006304 | 0.01755 | 0.008635 | 0.030838 | 0.032601 | 0.001529 | 0.025426 | 0.027653 | 0.018122 | 0.026685 | 0.01627 | 0.004621 | 0.013834 | 0.024768 | 0.004692 | 0.004173 | 0.00428 | 0.015533 |
| 21.11111111 | 0.035246 | 0.019618 | 0.024251 | 0.011948 | 0.006603 | 0.022584 | 0.013002 | 0.029826 | 0.026662 | 0.001763 | 0.01874 | 0.025932 | 0.012123 | 0.014551 | 0.017151 | 0.008502 | 0.014949 | 0.027762 | 0.005892 | 0.005021 | 0.007763 | 0.012281 |
| 21.125 | 0.029459 | 0.022758 | 0.017514 | 0.012007 | 0.008176 | 0.023122 | 0.019103 | 0.027794 | 0.02147 | 0.002557 | 0.014656 | 0.019631 | 0.009263 | 0.009982 | 0.023398 | 0.012611 | 0.015677 | 0.031875 | 0.007247 | 0.005138 | 0.013535 | 0.012681 |
| 21.13888889 | 0.019664 | 0.014795 | 0.009369 | 0.01076 | 0.008848 | 0.020853 | 0.023456 | 0.024055 | 0.018395 | 0.002657 | 0.013872 | 0.012456 | 0.00619 | 0.009348 | 0.024915 | 0.015007 | 0.016079 | 0.034879 | 0.008664 | 0.004731 | 0.015262 | 0.013456 |
| 21.15277778 | 0.012475 | 0.007995 | 0.006275 | 0.00907 | 0.008005 | 0.019163 | 0.025781 | 0.019103 | 0.016311 | 0.001949 | 0.013593 | 0.007382 | 0.002935 | 0.008323 | 0.024486 | 0.016174 | 0.014709 | 0.031422 | 0.008625 | 0.003554 | 0.011885 | 0.011835 |
| 21.16666667 | 0.011208 | 0.008606 | 0.007655 | 0.007912 | 0.006954 | 0.017935 | 0.024615 | 0.015314 | 0.014364 | 0.001561 | 0.012672 | 0.005037 | 0.002593 | 0.008104 | 0.026777 | 0.017846 | 0.012072 | 0.023537 | 0.006415 | 0.002467 | 0.008917 | 0.009273 |
| 21.18055556 | 0.012441 | 0.010329 | 0.009606 | 0.007505 | 0.006772 | 0.016114 | 0.019759 | 0.013549 | 0.012381 | 0.001785 | 0.012485 | 0.00493 | 0.003401 | 0.008689 | 0.026607 | 0.018859 | 0.010244 | 0.018128 | 0.003151 | 0.001929 | 0.008195 | 0.007806 |
| 21.19444444 | 0.011854 | 0.007495 | 0.01058 | 0.007406 | 0.007035 | 0.014078 | 0.014081 | 0.012786 | 0.010708 | 0.00216 | 0.01287 | 0.006171 | 0.002583 | 0.007495 | 0.019219 | 0.016714 | 0.009567 | 0.016741 | 0.001027 | 0.001406 | 0.006462 | 0.006939 |
| 21.20833333 | 0.009088 | 0.002917 | 0.010203 | 0.006713 | 0.006404 | 0.012247 | 0.010221 | 0.011723 | 0.009198 | 0.00205 | 0.011885 | 0.007905 | 0.001525 | 0.004921 | 0.009158 | 0.011435 | 0.008833 | 0.015705 | 0.00075 | 0.000974 | 0.003343 | 0.005689 |
| 21.22222222 | 0.00626 | 0.001917 | 0.008199 | 0.004689 | 0.003936 | 0.010233 | 0.007933 | 0.009233 | 0.006558 | 0.001307 | 0.008651 | 0.008717 | 0.001222 | 0.003221 | 0.003036 | 0.006088 | 0.007764 | 0.012901 | 0.000927 | 0.000906 | 0.001188 | 0.003642 |
| 21.23611111 | 0.004372 | 0.003624 | 0.005365 | 0.002437 | 0.001957 | 0.007309 | 0.005457 | 0.004988 | 0.003412 | 0.001038 | 0.004363 | 0.006586 | 0.00106 | 0.002607 | 0.001761 | 0.005933 | 0.006553 | 0.0097 | 0.000872 | 0.000816 | 0.000844 | 0.002067 |
| 21.25 | 0.002628 | 0.004082 | 0.002631 | 0.002015 | 0.002831 | 0.003782 | 0.002951 | 0.001954 | 0.002446 | 0.002034 | 0.00216 | 0.003227 | 0.001547 | 0.001862 | 0.002038 | 0.011905 | 0.004677 | 0.006311 | 0.000742 | 0.000505 | 0.00217 | 0.002685 |
| 21.26388889 | 0.001712 | 0.005046 | 0.001221 | 0.002791 | 0.003995 | 0.001764 | 0.001931 | 0.002465 | 0.003577 | 0.002964 | 0.003026 | 0.002158 | 0.00365 | 0.001811 | 0.002877 | 0.014344 | 0.002471 | 0.003584 | 0.000627 | 0.000452 | 0.004125 | 0.003607 |
| 21.27777778 | 0.002414 | 0.008125 | 0.001361 | 0.002734 | 0.003683 | 0.001508 | 0.002097 | 0.00436 | 0.004809 | 0.002612 | 0.003946 | 0.002337 | 0.005857 | 0.003937 | 0.005994 | 0.008269 | 0.001588 | 0.003876 | 0.001189 | 0.001004 | 0.005182 | 0.00316 |
| 21.29166667 | 0.002676 | 0.00847 | 0.001174 | 0.002057 | 0.00337 | 0.001535 | 0.001895 | 0.006103 | 0.005519 | 0.001838 | 0.003803 | 0.002473 | 0.006564 | 0.006378 | 0.011553 | 0.005225 | 0.002154 | 0.005004 | 0.002693 | 0.001809 | 0.00523 | 0.002732 |
| 21.30555556 | 0.002053 | 0.005673 | 0.000665 | 0.0019 | 0.003691 | 0.002662 | 0.002316 | 0.007392 | 0.005591 | 0.001729 | 0.006089 | 0.003845 | 0.006963 | 0.00744 | 0.017219 | 0.011712 | 0.002891 | 0.004938 | 0.003995 | 0.002307 | 0.005425 | 0.003422 |
| 21.31944444 | 0.002721 | 0.0052 | 0.001315 | 0.002221 | 0.00398 | 0.005173 | 0.005055 | 0.007459 | 0.005528 | 0.001987 | 0.011894 | 0.006192 | 0.00802 | 0.008034 | 0.019484 | 0.017654 | 0.003609 | 0.00553 | 0.004205 | 0.002614 | 0.006518 | 0.004872 |
| 21.33333333 | 0.005903 | 0.00844 | 0.003064 | 0.002961 | 0.004274 | 0.007154 | 0.008903 | 0.007992 | 0.00697 | 0.00246 | 0.015107 | 0.007439 | 0.008882 | 0.009172 | 0.019089 | 0.017537 | 0.004202 | 0.006643 | 0.00405 | 0.003452 | 0.007625 | 0.006013 |
| 21.34722222 | 0.010115 | 0.011328 | 0.005068 | 0.004338 | 0.005084 | 0.008052 | 0.012685 | 0.011121 | 0.010299 | 0.00338 | 0.013796 | 0.007368 | 0.008604 | 0.011188 | 0.01972 | 0.016477 | 0.004222 | 0.006371 | 0.004047 | 0.004484 | 0.007456 | 0.005831 |
| 21.36111111 | 0.012441 | 0.011232 | 0.006365 | 0.005604 | 0.00557 | 0.009065 | 0.016068 | 0.014761 | 0.013651 | 0.003923 | 0.01414 | 0.007486 | 0.007924 | 0.013521 | 0.020587 | 0.016313 | 0.004134 | 0.005826 | 0.003895 | 0.004527 | 0.00694 | 0.005177 |
| 21.375 | 0.012321 | 0.010602 | 0.006805 | 0.006135 | 0.005179 | 0.010673 | 0.018613 | 0.015736 | 0.015277 | 0.003381 | 0.0166 | 0.007701 | 0.008658 | 0.014717 | 0.020211 | 0.015469 | 0.004712 | 0.00755 | 0.003767 | 0.003692 | 0.00792 | 0.006114 |
| 21.38888889 | 0.011381 | 0.010701 | 0.00736 | 0.006297 | 0.004978 | 0.012087 | 0.019982 | 0.014865 | 0.014789 | 0.002245 | 0.018224 | 0.00787 | 0.010922 | 0.014434 | 0.020886 | 0.014629 | 0.00588 | 0.010701 | 0.003753 | 0.002954 | 0.009649 | 0.007995 |
| 21.40277778 | 0.010971 | 0.009532 | 0.008402 | 0.006466 | 0.005715 | 0.012902 | 0.019832 | 0.014387 | 0.01342 | 0.001363 | 0.022146 | 0.00869 | 0.012563 | 0.014307 | 0.023914 | 0.015853 | 0.006799 | 0.012239 | 0.003657 | 0.002955 | 0.010511 | 0.008956 |
| 21.41666667 | 0.012017 | 0.00807 | 0.009215 | 0.007024 | 0.007068 | 0.01376 | 0.018317 | 0.014288 | 0.013607 | 0.001566 | 0.027288 | 0.00961 | 0.012636 | 0.015337 | 0.027509 | 0.01915 | 0.007004 | 0.011842 | 0.003961 | 0.0039 | 0.011035 | 0.009661 |
| 21.43055556 | 0.014225 | 0.008623 | 0.009365 | 0.008127 | 0.008373 | 0.0148 | 0.015779 | 0.014812 | 0.016634 | 0.003019 | 0.02792 | 0.010741 | 0.012459 | 0.01679 | 0.030095 | 0.020394 | 0.007227 | 0.011796 | 0.004883 | 0.005105 | 0.011718 | 0.01096 |
| 21.44444444 | 0.016822 | 0.009689 | 0.010289 | 0.00981 | 0.009637 | 0.015876 | 0.014237 | 0.018395 | 0.022068 | 0.004694 | 0.028727 | 0.013414 | 0.013481 | 0.018912 | 0.031673 | 0.017483 | 0.008498 | 0.013045 | 0.005951 | 0.006019 | 0.012345 | 0.012388 |
| 21.45833333 | 0.020617 | 0.010452 | 0.014619 | 0.01324 | 0.012292 | 0.018797 | 0.017701 | 0.025175 | 0.028023 | 0.00589 | 0.037584 | 0.017699 | 0.016867 | 0.0232 | 0.032975 | 0.015056 | 0.010561 | 0.014951 | 0.006973 | 0.007058 | 0.01367 | 0.014107 |
| 21.47222222 | 0.025711 | 0.012885 | 0.021401 | 0.018537 | 0.016541 | 0.024276 | 0.025952 | 0.031481 | 0.032212 | 0.006872 | 0.049406 | 0.023673 | 0.022317 | 0.028765 | 0.035878 | 0.017402 | 0.012567 | 0.018171 | 0.007821 | 0.00826 | 0.015832 | 0.016412 |
| 21.48611111 | 0.03058 | 0.016035 | 0.025883 | 0.022772 | 0.019766 | 0.028703 | 0.032169 | 0.0368 | 0.036197 | 0.007412 | 0.05278 | 0.031225 | 0.028246 | 0.033626 | 0.043275 | 0.022777 | 0.015062 | 0.024802 | 0.00883 | 0.009629 | 0.018919 | 0.019972 |
| 21.5 | 0.035349 | 0.016967 | 0.027638 | 0.024657 | 0.020923 | 0.030502 | 0.034128 | 0.045273 | 0.043692 | 0.007334 | 0.046619 | 0.038694 | 0.034388 | 0.038416 | 0.056164 | 0.026287 | 0.018809 | 0.034442 | 0.011041 | 0.011837 | 0.024103 | 0.026072 |
| 21.51388889 | 0.041322 | 0.017084 | 0.030896 | 0.026735 | 0.022284 | 0.033324 | 0.037631 | 0.055515 | 0.053237 | 0.007696 | 0.041545 | 0.046277 | 0.040518 | 0.043565 | 0.070214 | 0.025894 | 0.022763 | 0.04335 | 0.014053 | 0.014508 | 0.030846 | 0.034057 |
| 21.52777778 | 0.045632 | 0.01943 | 0.033707 | 0.028745 | 0.024284 | 0.036576 | 0.040611 | 0.059084 | 0.059467 | 0.008511 | 0.044226 | 0.055189 | 0.044835 | 0.047103 | 0.073748 | 0.024617 | 0.024993 | 0.049561 | 0.015563 | 0.01614 | 0.037381 | 0.0429 |
| 21.54166667 | 0.04251 | 0.021608 | 0.026982 | 0.024322 | 0.021213 | 0.032097 | 0.032824 | 0.052372 | 0.057738 | 0.006872 | 0.045333 | 0.058671 | 0.045777 | 0.048013 | 0.059903 | 0.023899 | 0.023469 | 0.05105 | 0.013462 | 0.015227 | 0.042665 | 0.051166 |
| 21.55555556 | 0.034781 | 0.020233 | 0.014318 | 0.012574 | 0.012474 | 0.020033 | 0.019601 | 0.044121 | 0.049383 | 0.004484 | 0.035878 | 0.049781 | 0.043169 | 0.050664 | 0.044699 | 0.023722 | 0.020642 | 0.047347 | 0.009184 | 0.011599 | 0.0463 | 0.054667 |
| 21.56944444 | 0.036189 | 0.020546 | 0.015686 | 0.007673 | 0.00683 | 0.016441 | 0.022485 | 0.046667 | 0.044957 | 0.0055 | 0.030042 | 0.039206 | 0.038516 | 0.060856 | 0.047177 | 0.025609 | 0.023256 | 0.046784 | 0.007801 | 0.009189 | 0.048669 | 0.050794 |
| 21.58333333 | 0.051137 | 0.02906 | 0.031881 | 0.016698 | 0.004489 | 0.029441 | 0.035837 | 0.051544 | 0.048067 | 0.006193 | 0.032143 | 0.035209 | 0.029142 | 0.064447 | 0.058037 | 0.02889 | 0.028261 | 0.049038 | 0.010274 | 0.011103 | 0.047548 | 0.0424 |
| 21.59722222 | 0.058169 | 0.035348 | 0.036013 | 0.022935 | 0.007159 | 0.038939 | 0.033934 | 0.042205 | 0.048519 | 0.003706 | 0.024725 | 0.030142 | 0.01557 | 0.04743 | 0.06166 | 0.031507 | 0.027297 | 0.044324 | 0.010998 | 0.012197 | 0.04032 | 0.033706 |
| 21.61111111 | 0.043015 | 0.027795 | 0.022032 | 0.018762 | 0.017516 | 0.027859 | 0.019569 | 0.026887 | 0.039064 | 0.001062 | 0.01183 | 0.021686 | 0.009003 | 0.028811 | 0.059446 | 0.028666 | 0.020069 | 0.032073 | 0.007674 | 0.008316 | 0.029966 | 0.027139 |
| 21.625 | 0.023725 | 0.015178 | 0.01108 | 0.015592 | 0.021499 | 0.012821 | 0.010749 | 0.017925 | 0.023344 | 0.000286 | 0.005193 | 0.013014 | 0.01077 | 0.021181 | 0.054672 | 0.02071 | 0.010611 | 0.017166 | 0.004386 | 0.004017 | 0.020214 | 0.021916 |
| 21.63888889 | 0.014246 | 0.007072 | 0.008694 | 0.014296 | 0.014284 | 0.009897 | 0.013797 | 0.013025 | 0.011516 | 0.000402 | 0.002953 | 0.007403 | 0.013031 | 0.016092 | 0.047526 | 0.013948 | 0.004796 | 0.007982 | 0.00409 | 0.002916 | 0.013057 | 0.014576 |
| 21.65277778 | 0.009313 | 0.004321 | 0.008481 | 0.010099 | 0.006929 | 0.01118 | 0.017937 | 0.010896 | 0.009825 | 0.000624 | 0.002501 | 0.008368 | 0.013839 | 0.011115 | 0.035595 | 0.00785 | 0.00559 | 0.010374 | 0.005543 | 0.003319 | 0.009759 | 0.007055 |
| 21.66666667 | 0.007358 | 0.007479 | 0.00901 | 0.005596 | 0.00442 | 0.008017 | 0.013098 | 0.012501 | 0.013626 | 0.001661 | 0.003076 | 0.011715 | 0.01369 | 0.00789 | 0.018732 | 0.002711 | 0.009076 | 0.01753 | 0.005337 | 0.002784 | 0.008365 | 0.004039 |
| 21.68055556 | 0.011179 | 0.012238 | 0.009184 | 0.003898 | 0.003435 | 0.003612 | 0.006098 | 0.012517 | 0.013322 | 0.003094 | 0.003663 | 0.010336 | 0.011531 | 0.00653 | 0.005797 | 0.001153 | 0.009506 | 0.017304 | 0.003288 | 0.001449 | 0.006719 | 0.005253 |
| 21.69444444 | 0.015355 | 0.010552 | 0.008433 | 0.004734 | 0.002329 | 0.004602 | 0.00709 | 0.007812 | 0.00715 | 0.003484 | 0.003778 | 0.005289 | 0.007509 | 0.005657 | 0.002479 | 0.00086 | 0.006126 | 0.008995 | 0.00253 | 0.000828 | 0.004649 | 0.0066 |
| 21.70833333 | 0.01451 | 0.004198 | 0.006636 | 0.006095 | 0.003097 | 0.009261 | 0.015365 | 0.003729 | 0.002073 | 0.002522 | 0.005765 | 0.002298 | 0.006605 | 0.003872 | 0.00463 | 0.001578 | 0.002741 | 0.002735 | 0.002378 | 0.000695 | 0.002368 | 0.004879 |
| 21.72222222 | 0.013303 | 0.003737 | 0.003776 | 0.006111 | 0.00433 | 0.009968 | 0.019682 | 0.00242 | 0.001202 | 0.001233 | 0.009672 | 0.002756 | 0.008905 | 0.002962 | 0.0047 | 0.005145 | 0.001538 | 0.001597 | 0.001755 | 0.000846 | 0.000841 | 0.003115 |
| 21.73611111 | 0.013964 | 0.008823 | 0.001498 | 0.003787 | 0.00353 | 0.005569 | 0.013928 | 0.00194 | 0.002649 | 0.000765 | 0.009856 | 0.003855 | 0.007301 | 0.005207 | 0.005971 | 0.006963 | 0.001673 | 0.001085 | 0.002016 | 0.001279 | 0.000775 | 0.004231 |
| 21.75 | 0.011101 | 0.009676 | 0.002374 | 0.001237 | 0.001956 | 0.002022 | 0.006033 | 0.002271 | 0.003737 | 0.000888 | 0.005761 | 0.00317 | 0.002812 | 0.007528 | 0.014154 | 0.005376 | 0.001484 | 0.000796 | 0.001833 | 0.001202 | 0.001383 | 0.005064 |
| 21.76388889 | 0.010569 | 0.009324 | 0.006096 | 0.001691 | 0.00157 | 0.004562 | 0.005428 | 0.003513 | 0.004053 | 0.00225 | 0.002913 | 0.002676 | 0.001298 | 0.006225 | 0.016843 | 0.00514 | 0.001091 | 0.001424 | 0.00111 | 0.000958 | 0.001457 | 0.003509 |
| 21.77777778 | 0.021764 | 0.015905 | 0.00938 | 0.004903 | 0.004028 | 0.011435 | 0.011829 | 0.007049 | 0.005706 | 0.006768 | 0.004158 | 0.005098 | 0.003382 | 0.005286 | 0.015131 | 0.005255 | 0.00152 | 0.002182 | 0.001497 | 0.001143 | 0.001535 | 0.001527 |
| 21.79166667 | 0.029366 | 0.02147 | 0.011966 | 0.010568 | 0.010614 | 0.01581 | 0.018163 | 0.011266 | 0.005456 | 0.011992 | 0.009876 | 0.007647 | 0.006265 | 0.005192 | 0.022252 | 0.003561 | 0.001941 | 0.002392 | 0.001536 | 0.001561 | 0.003158 | 0.003327 |
| 21.80555556 | 0.019928 | 0.015883 | 0.015758 | 0.01816 | 0.017406 | 0.016451 | 0.020084 | 0.011798 | 0.002834 | 0.014391 | 0.016295 | 0.008343 | 0.00669 | 0.004162 | 0.023618 | 0.002277 | 0.001822 | 0.002026 | 0.000917 | 0.002062 | 0.005688 | 0.010101 |
| 21.81944444 | 0.010603 | 0.008653 | 0.018849 | 0.022784 | 0.020075 | 0.017521 | 0.020974 | 0.010522 | 0.006413 | 0.014633 | 0.018252 | 0.008908 | 0.005273 | 0.005819 | 0.014905 | 0.002776 | 0.001812 | 0.002564 | 0.000841 | 0.002249 | 0.007505 | 0.01462 |
| 21.83333333 | 0.020268 | 0.014995 | 0.016532 | 0.019503 | 0.015542 | 0.019093 | 0.021054 | 0.010463 | 0.015854 | 0.013978 | 0.013236 | 0.009422 | 0.004421 | 0.006661 | 0.010577 | 0.004616 | 0.002407 | 0.003827 | 0.001588 | 0.001847 | 0.006798 | 0.010663 |
| 21.84722222 | 0.035263 | 0.026017 | 0.008687 | 0.009956 | 0.006434 | 0.016406 | 0.0167 | 0.008967 | 0.016726 | 0.011566 | 0.006029 | 0.008869 | 0.003637 | 0.005279 | 0.010966 | 0.008126 | 0.00258 | 0.004274 | 0.003584 | 0.001959 | 0.004609 | 0.005628 |
| 21.86111111 | 0.033463 | 0.023145 | 0.002329 | 0.002746 | 0.00157 | 0.010418 | 0.010821 | 0.005962 | 0.011993 | 0.007151 | 0.003994 | 0.008503 | 0.002332 | 0.004858 | 0.016283 | 0.009086 | 0.001368 | 0.00267 | 0.00483 | 0.002824 | 0.004058 | 0.004716 |
| 21.875 | 0.026038 | 0.014759 | 0.001119 | 0.001878 | 0.001498 | 0.009113 | 0.012131 | 0.005626 | 0.012799 | 0.003086 | 0.006101 | 0.007476 | 0.001512 | 0.003289 | 0.020178 | 0.006996 | 0.000665 | 0.001652 | 0.0044 | 0.003177 | 0.004649 | 0.006069 |
| 21.88888889 | 0.021756 | 0.01106 | 0.002843 | 0.003259 | 0.002854 | 0.009913 | 0.019856 | 0.005924 | 0.010301 | 0.001016 | 0.006347 | 0.009494 | 0.002565 | 0.001379 | 0.017441 | 0.007176 | 0.001712 | 0.005819 | 0.005064 | 0.003728 | 0.008225 | 0.012162 |
| 21.90277778 | 0.016781 | 0.006192 | 0.00717 | 0.006417 | 0.006768 | 0.006324 | 0.018888 | 0.005033 | 0.006121 | 0.002122 | 0.005419 | 0.017737 | 0.006754 | 0.002934 | 0.018658 | 0.009943 | 0.004195 | 0.012244 | 0.006784 | 0.005521 | 0.017141 | 0.017338 |
| 21.91666667 | 0.01716 | 0.002406 | 0.011613 | 0.011141 | 0.010914 | 0.004754 | 0.012069 | 0.009258 | 0.014147 | 0.008324 | 0.007566 | 0.017899 | 0.012378 | 0.010981 | 0.026005 | 0.013491 | 0.005481 | 0.013082 | 0.006325 | 0.007379 | 0.023702 | 0.016513 |
| 21.93055556 | 0.02431 | 0.005069 | 0.010344 | 0.011863 | 0.010544 | 0.010992 | 0.010318 | 0.019449 | 0.028077 | 0.013834 | 0.007314 | 0.007506 | 0.015491 | 0.019619 | 0.031845 | 0.011295 | 0.005833 | 0.012345 | 0.004151 | 0.007813 | 0.02004 | 0.014089 |
| 21.94444444 | 0.027992 | 0.013073 | 0.006886 | 0.009851 | 0.008918 | 0.019946 | 0.020152 | 0.02656 | 0.028952 | 0.010805 | 0.005178 | 0.006631 | 0.015589 | 0.017601 | 0.030653 | 0.004364 | 0.011387 | 0.022703 | 0.00538 | 0.007853 | 0.014601 | 0.014262 |
| 21.95833333 | 0.019062 | 0.014457 | 0.009404 | 0.011134 | 0.011495 | 0.023612 | 0.043736 | 0.026935 | 0.021183 | 0.004311 | 0.010294 | 0.01944 | 0.019191 | 0.012885 | 0.027302 | 0.006986 | 0.020574 | 0.041367 | 0.01078 | 0.010008 | 0.019978 | 0.019868 |
| 21.97222222 | 0.010394 | 0.00589 | 0.015628 | 0.014484 | 0.012916 | 0.021853 | 0.049495 | 0.02794 | 0.02499 | 0.003182 | 0.018978 | 0.029782 | 0.029134 | 0.020806 | 0.037638 | 0.021216 | 0.02384 | 0.050131 | 0.013841 | 0.013261 | 0.030797 | 0.027996 |
| 21.98611111 | 0.02259 | 0.008226 | 0.019324 | 0.016189 | 0.008861 | 0.020716 | 0.026222 | 0.035277 | 0.040635 | 0.006736 | 0.02081 | 0.035019 | 0.036158 | 0.034753 | 0.065531 | 0.032122 | 0.02116 | 0.04481 | 0.013438 | 0.014958 | 0.03397 | 0.031256 |
| 22 | 0.048003 | 0.027452 | 0.020402 | 0.017585 | 0.009072 | 0.021083 | 0.012522 | 0.043166 | 0.049865 | 0.009127 | 0.020614 | 0.04202 | 0.035552 | 0.040475 | 0.086205 | 0.029501 | 0.022477 | 0.042523 | 0.01605 | 0.016926 | 0.031313 | 0.028941 |
| 22.01388889 | 0.052542 | 0.036935 | 0.02276 | 0.021087 | 0.016496 | 0.021435 | 0.028958 | 0.045163 | 0.046548 | 0.008796 | 0.03113 | 0.047163 | 0.03652 | 0.04341 | 0.078262 | 0.021413 | 0.030001 | 0.055453 | 0.021517 | 0.020771 | 0.031752 | 0.027323 |
| 22.02777778 | 0.032425 | 0.025355 | 0.025966 | 0.023856 | 0.018735 | 0.023441 | 0.054084 | 0.043999 | 0.046541 | 0.007744 | 0.046685 | 0.046904 | 0.0435 | 0.052438 | 0.059295 | 0.020973 | 0.035974 | 0.074253 | 0.023287 | 0.023206 | 0.034491 | 0.029908 |
| 22.04166667 | 0.019691 | 0.016259 | 0.025888 | 0.021812 | 0.011012 | 0.025893 | 0.060949 | 0.04178 | 0.053611 | 0.006062 | 0.048897 | 0.042347 | 0.045803 | 0.056397 | 0.051299 | 0.02539 | 0.033115 | 0.07482 | 0.019474 | 0.021507 | 0.034517 | 0.035083 |
| 22.05555556 | 0.033643 | 0.020761 | 0.026349 | 0.017768 | 0.004505 | 0.025512 | 0.046351 | 0.036671 | 0.052954 | 0.003336 | 0.03813 | 0.036652 | 0.036506 | 0.046979 | 0.046854 | 0.024908 | 0.021247 | 0.050212 | 0.0132 | 0.016583 | 0.030129 | 0.038354 |
| 22.06944444 | 0.059275 | 0.023521 | 0.034439 | 0.016776 | 0.002702 | 0.0246 | 0.028232 | 0.030694 | 0.043312 | 0.001148 | 0.03031 | 0.030834 | 0.022388 | 0.033155 | 0.030456 | 0.018689 | 0.008669 | 0.022274 | 0.00711 | 0.011327 | 0.021576 | 0.035556 |
| 22.08333333 | 0.059925 | 0.015396 | 0.038459 | 0.016779 | 0.002164 | 0.026197 | 0.023392 | 0.026166 | 0.035668 | 0.000845 | 0.02718 | 0.02132 | 0.010244 | 0.021752 | 0.013657 | 0.011842 | 0.002434 | 0.009128 | 0.003262 | 0.007649 | 0.011341 | 0.026866 |
| 22.09722222 | 0.033077 | 0.005588 | 0.027193 | 0.014439 | 0.005498 | 0.027755 | 0.029089 | 0.02181 | 0.032625 | 0.001913 | 0.025342 | 0.010483 | 0.00345 | 0.012614 | 0.009524 | 0.009737 | 0.002985 | 0.010376 | 0.004688 | 0.006441 | 0.006937 | 0.018848 |
| 22.11111111 | 0.010741 | 0.003294 | 0.016459 | 0.012452 | 0.010895 | 0.02599 | 0.032867 | 0.018979 | 0.030989 | 0.002478 | 0.025495 | 0.007237 | 0.005752 | 0.00937 | 0.013693 | 0.015258 | 0.00977 | 0.021479 | 0.011936 | 0.008145 | 0.013635 | 0.018482 |
| 22.125 | 0.006538 | 0.003996 | 0.015322 | 0.011392 | 0.012055 | 0.020083 | 0.029618 | 0.018181 | 0.028302 | 0.002547 | 0.020607 | 0.009231 | 0.012834 | 0.013871 | 0.031907 | 0.024624 | 0.018837 | 0.035953 | 0.01837 | 0.009804 | 0.021845 | 0.022314 |
| 22.13888889 | 0.010314 | 0.002865 | 0.01323 | 0.008055 | 0.007397 | 0.011686 | 0.021064 | 0.014638 | 0.021902 | 0.004811 | 0.011848 | 0.008315 | 0.013477 | 0.016688 | 0.048395 | 0.02849 | 0.021198 | 0.040482 | 0.016757 | 0.008082 | 0.019526 | 0.020318 |
| 22.15277778 | 0.009638 | 0.00331 | 0.007083 | 0.003882 | 0.003536 | 0.004449 | 0.010633 | 0.007512 | 0.011897 | 0.008179 | 0.007395 | 0.005668 | 0.006777 | 0.012486 | 0.041855 | 0.023042 | 0.015272 | 0.029838 | 0.009394 | 0.004315 | 0.01062 | 0.011751 |
| 22.16666667 | 0.00656 | 0.006026 | 0.002494 | 0.002495 | 0.004546 | 0.000859 | 0.003114 | 0.002406 | 0.00376 | 0.009112 | 0.007789 | 0.003241 | 0.001519 | 0.008095 | 0.025128 | 0.014025 | 0.008654 | 0.015456 | 0.003414 | 0.001532 | 0.004301 | 0.004404 |
| 22.18055556 | 0.005176 | 0.007726 | 0.002253 | 0.003444 | 0.006262 | 0.000149 | 0.002253 | 0.001075 | 0.001436 | 0.007489 | 0.008661 | 0.002633 | 0.000858 | 0.006355 | 0.013549 | 0.007707 | 0.00542 | 0.008585 | 0.001779 | 0.000602 | 0.0031 | 0.001905 |
| 22.19444444 | 0.00338 | 0.005702 | 0.004431 | 0.004596 | 0.006874 | 0.000214 | 0.005993 | 0.000821 | 0.002427 | 0.005424 | 0.007272 | 0.004684 | 0.002972 | 0.004833 | 0.007451 | 0.004758 | 0.003826 | 0.006382 | 0.002441 | 0.000949 | 0.004509 | 0.002193 |
| 22.20833333 | 0.00263 | 0.00261 | 0.004465 | 0.005286 | 0.007786 | 0.000641 | 0.007447 | 0.000735 | 0.002779 | 0.004303 | 0.006576 | 0.004858 | 0.005258 | 0.004033 | 0.004778 | 0.002754 | 0.002526 | 0.003936 | 0.002792 | 0.001376 | 0.00427 | 0.002984 |
| 22.22222222 | 0.002814 | 0.004387 | 0.002282 | 0.005746 | 0.009055 | 0.002137 | 0.005492 | 0.001069 | 0.002083 | 0.004299 | 0.005113 | 0.002084 | 0.005259 | 0.003342 | 0.00746 | 0.00152 | 0.001759 | 0.002917 | 0.003193 | 0.001213 | 0.003003 | 0.004065 |
| 22.23611111 | 0.004491 | 0.009803 | 0.002919 | 0.006763 | 0.010234 | 0.004592 | 0.00594 | 0.003125 | 0.003065 | 0.005181 | 0.004551 | 0.002286 | 0.00527 | 0.003605 | 0.011572 | 0.001542 | 0.001244 | 0.002641 | 0.004975 | 0.001226 | 0.003227 | 0.005088 |
| 22.25 | 0.008421 | 0.014305 | 0.006924 | 0.008072 | 0.010291 | 0.007592 | 0.008806 | 0.007296 | 0.006929 | 0.006189 | 0.009411 | 0.006556 | 0.005282 | 0.005947 | 0.010548 | 0.001258 | 0.000664 | 0.001928 | 0.007051 | 0.001454 | 0.003642 | 0.005367 |
| 22.26388889 | 0.011811 | 0.017182 | 0.009933 | 0.008704 | 0.009187 | 0.010704 | 0.009176 | 0.011448 | 0.012645 | 0.00663 | 0.014836 | 0.008401 | 0.00306 | 0.00546 | 0.007168 | 0.001431 | 0.000419 | 0.001667 | 0.0076 | 0.00153 | 0.004168 | 0.005877 |
| 22.27777778 | 0.012901 | 0.017586 | 0.010988 | 0.009033 | 0.008781 | 0.012757 | 0.007704 | 0.014343 | 0.017731 | 0.006839 | 0.014461 | 0.006467 | 0.002177 | 0.002159 | 0.007064 | 0.003206 | 0.000932 | 0.001741 | 0.006994 | 0.001964 | 0.006641 | 0.007373 |

| | | | | | | | | | | | | | | | | | | | | | | |
|---|---|---|---|---|---|---|---|---|---|---|---|---|---|---|---|---|---|---|---|---|---|---|
| 22.29166667 | 0.012091 | 0.014351 | 0.011916 | 0.009494 | 0.009637 | 0.012949 | 0.007698 | 0.016474 | 0.019328 | 0.007105 | 0.013288 | 0.005767 | 0.004206 | 0.001961 | 0.008536 | 0.004925 | 0.001832 | 0.001647 | 0.005706 | 0.002241 | 0.008222 | 0.00751 |
| 22.30555556 | 0.010385 | 0.011032 | 0.01157 | 0.009544 | 0.010224 | 0.01206 | 0.008675 | 0.017938 | 0.017948 | 0.006647 | 0.016121 | 0.008144 | 0.006067 | 0.005566 | 0.008343 | 0.005898 | 0.002275 | 0.001485 | 0.003313 | 0.001586 | 0.005819 | 0.004696 |
| 22.31944444 | 0.007931 | 0.010361 | 0.009378 | 0.009263 | 0.009864 | 0.011246 | 0.008673 | 0.018335 | 0.016184 | 0.004956 | 0.014933 | 0.011335 | 0.006599 | 0.008436 | 0.007997 | 0.007177 | 0.002339 | 0.002017 | 0.001035 | 0.000725 | 0.002807 | 0.001996 |
| 22.33333333 | 0.005078 | 0.009149 | 0.008456 | 0.008958 | 0.008891 | 0.010651 | 0.007801 | 0.01784 | 0.014926 | 0.00318 | 0.008008 | 0.013014 | 0.007508 | 0.008395 | 0.008425 | 0.008275 | 0.002609 | 0.003479 | 0.000465 | 0.000666 | 0.003406 | 0.002959 |
| 22.34722222 | 0.003123 | 0.005913 | 0.008925 | 0.007566 | 0.006909 | 0.009381 | 0.007054 | 0.016175 | 0.013296 | 0.002243 | 0.005893 | 0.012332 | 0.007944 | 0.0078 | 0.008139 | 0.007525 | 0.003152 | 0.005186 | 0.000939 | 0.00112 | 0.00607 | 0.006276 |
| 22.36111111 | 0.002331 | 0.003264 | 0.006726 | 0.004338 | 0.003804 | 0.006119 | 0.006366 | 0.012673 | 0.009907 | 0.002247 | 0.012007 | 0.009749 | 0.005803 | 0.007738 | 0.007827 | 0.005562 | 0.003991 | 0.006298 | 0.001051 | 0.001069 | 0.00614 | 0.006945 |
| 22.375 | 0.003483 | 0.002332 | 0.003874 | 0.001713 | 0.001677 | 0.0027 | 0.00531 | 0.008769 | 0.004979 | 0.004345 | 0.016577 | 0.00569 | 0.004354 | 0.007875 | 0.00931 | 0.00481 | 0.00487 | 0.00657 | 0.000841 | 0.000833 | 0.004682 | 0.004295 |
| 22.38888889 | 0.006417 | 0.003418 | 0.003838 | 0.001655 | 0.001398 | 0.001573 | 0.003463 | 0.006337 | 0.001346 | 0.007583 | 0.012114 | 0.001917 | 0.007041 | 0.009556 | 0.012523 | 0.006767 | 0.005551 | 0.007337 | 0.001517 | 0.001053 | 0.005198 | 0.002524 |
| 22.40277778 | 0.007193 | 0.005403 | 0.003601 | 0.002464 | 0.001224 | 0.001022 | 0.00212 | 0.004061 | 0.000637 | 0.008831 | 0.005446 | 0.000956 | 0.010933 | 0.013273 | 0.016626 | 0.01049 | 0.006607 | 0.009883 | 0.003771 | 0.002281 | 0.007069 | 0.003954 |
| 22.41666667 | 0.005119 | 0.005141 | 0.00359 | 0.002962 | 0.001002 | 0.000903 | 0.003139 | 0.001974 | 0.000938 | 0.008622 | 0.004985 | 0.001718 | 0.011638 | 0.015954 | 0.020938 | 0.01293 | 0.008082 | 0.012491 | 0.00647 | 0.00461 | 0.009248 | 0.007323 |
| 22.43055556 | 0.004247 | 0.003279 | 0.005884 | 0.003279 | 0.001388 | 0.002436 | 0.004459 | 0.001713 | 0.000892 | 0.009753 | 0.007 | 0.001734 | 0.009536 | 0.015009 | 0.025185 | 0.013142 | 0.009204 | 0.013064 | 0.008296 | 0.006418 | 0.011888 | 0.010107 |
| 22.44444444 | 0.005629 | 0.003053 | 0.007206 | 0.003189 | 0.001486 | 0.003718 | 0.003505 | 0.001407 | 0.000994 | 0.012594 | 0.006527 | 0.00097 | 0.009094 | 0.014209 | 0.029279 | 0.014107 | 0.010213 | 0.013371 | 0.010024 | 0.007607 | 0.014362 | 0.011404 |
| 22.45833333 | 0.005678 | 0.004051 | 0.004898 | 0.002437 | 0.001042 | 0.002986 | 0.002525 | 0.000559 | 0.002242 | 0.015442 | 0.004156 | 0.002004 | 0.013543 | 0.018741 | 0.03211 | 0.017669 | 0.01197 | 0.016322 | 0.012701 | 0.009709 | 0.016783 | 0.012977 |
| 22.47222222 | 0.003747 | 0.004023 | 0.001697 | 0.001279 | 0.000953 | 0.001334 | 0.004293 | 0.000506 | 0.005343 | 0.01708 | 0.003045 | 0.0063 | 0.020093 | 0.025826 | 0.03416 | 0.02102 | 0.013913 | 0.021142 | 0.015448 | 0.012595 | 0.021249 | 0.016435 |
| 22.48611111 | 0.003978 | 0.002535 | 0.000937 | 0.000736 | 0.001811 | 0.000412 | 0.006431 | 0.00187 | 0.008683 | 0.017644 | 0.006011 | 0.009631 | 0.024322 | 0.030262 | 0.039483 | 0.020514 | 0.015671 | 0.025795 | 0.017541 | 0.015608 | 0.027907 | 0.020515 |
| 22.5 | 0.007847 | 0.002988 | 0.003391 | 0.001893 | 0.002802 | 0.000986 | 0.007351 | 0.007302 | 0.013049 | 0.017269 | 0.013395 | 0.009775 | 0.027681 | 0.034459 | 0.049344 | 0.017414 | 0.019179 | 0.032144 | 0.01965 | 0.019179 | 0.034174 | 0.024058 |
| 22.51388889 | 0.013779 | 0.008702 | 0.009034 | 0.003927 | 0.002537 | 0.002523 | 0.008138 | 0.016425 | 0.021154 | 0.016579 | 0.022804 | 0.012556 | 0.033892 | 0.042842 | 0.056254 | 0.0165 | 0.024788 | 0.044264 | 0.021858 | 0.022442 | 0.039236 | 0.028827 |
| 22.52777778 | 0.017984 | 0.011728 | 0.011135 | 0.005433 | 0.005035 | 0.005348 | 0.008311 | 0.020343 | 0.028167 | 0.019267 | 0.027838 | 0.020154 | 0.040875 | 0.050958 | 0.053611 | 0.018735 | 0.028244 | 0.058282 | 0.021841 | 0.023009 | 0.044857 | 0.036066 |
| 22.54166667 | 0.016211 | 0.010203 | 0.01055 | 0.011567 | 0.016934 | 0.014372 | 0.012632 | 0.013455 | 0.02672 | 0.028945 | 0.022986 | 0.019347 | 0.043837 | 0.052603 | 0.04507 | 0.022029 | 0.026956 | 0.064155 | 0.017224 | 0.020277 | 0.051047 | 0.044976 |
| 22.55555556 | 0.01756 | 0.010059 | 0.012256 | 0.022203 | 0.032255 | 0.023226 | 0.016883 | 0.008509 | 0.024702 | 0.038575 | 0.017042 | 0.008491 | 0.043369 | 0.05321 | 0.040187 | 0.027289 | 0.027016 | 0.06547 | 0.010757 | 0.016461 | 0.055658 | 0.054226 |
| 22.56944444 | 0.036628 | 0.011454 | 0.016946 | 0.020913 | 0.032338 | 0.022915 | 0.0232 | 0.02301 | 0.036608 | 0.033974 | 0.029597 | 0.007452 | 0.042915 | 0.063687 | 0.047826 | 0.033216 | 0.033683 | 0.071197 | 0.009103 | 0.015857 | 0.057332 | 0.059376 |
| 22.58333333 | 0.062299 | 0.023493 | 0.03052 | 0.013144 | 0.01677 | 0.026512 | 0.051656 | 0.04614 | 0.051975 | 0.017733 | 0.046715 | 0.018129 | 0.038722 | 0.073673 | 0.061887 | 0.03328 | 0.039176 | 0.072629 | 0.013281 | 0.019485 | 0.056376 | 0.055743 |
| 22.59722222 | 0.068685 | 0.037653 | 0.038708 | 0.016871 | 0.007063 | 0.03425 | 0.075701 | 0.050901 | 0.051554 | 0.005913 | 0.039805 | 0.026982 | 0.026591 | 0.061427 | 0.057826 | 0.025737 | 0.035015 | 0.062349 | 0.018165 | 0.022013 | 0.053213 | 0.045343 |
| 22.61111111 | 0.05077 | 0.034863 | 0.03016 | 0.021506 | 0.009954 | 0.029 | 0.060351 | 0.038564 | 0.042174 | 0.002564 | 0.025239 | 0.029531 | 0.016493 | 0.039159 | 0.037695 | 0.01723 | 0.024952 | 0.048427 | 0.018696 | 0.019557 | 0.045888 | 0.033546 |
| 22.625 | 0.028768 | 0.028461 | 0.020238 | 0.020368 | 0.015827 | 0.017763 | 0.033673 | 0.027612 | 0.038763 | 0.001933 | 0.021795 | 0.024614 | 0.018564 | 0.029277 | 0.029804 | 0.011977 | 0.015137 | 0.034795 | 0.012362 | 0.012524 | 0.035124 | 0.025982 |
| 22.63888889 | 0.020996 | 0.028742 | 0.016026 | 0.017629 | 0.017779 | 0.014464 | 0.027921 | 0.022221 | 0.035803 | 0.001879 | 0.020247 | 0.014569 | 0.028186 | 0.027974 | 0.03285 | 0.008332 | 0.009005 | 0.023989 | 0.00503 | 0.005047 | 0.026045 | 0.025474 |
| 22.65277778 | 0.021938 | 0.024294 | 0.012448 | 0.01195 | 0.01356 | 0.01584 | 0.033559 | 0.0188 | 0.023589 | 0.002811 | 0.014339 | 0.011263 | 0.03425 | 0.025457 | 0.034334 | 0.006585 | 0.009063 | 0.021357 | 0.002025 | 0.001625 | 0.020207 | 0.025178 |
| 22.66666667 | 0.018496 | 0.012781 | 0.009109 | 0.006242 | 0.006126 | 0.014258 | 0.028802 | 0.016654 | 0.012203 | 0.002755 | 0.007786 | 0.017132 | 0.031436 | 0.023511 | 0.037987 | 0.007425 | 0.012633 | 0.024099 | 0.001673 | 0.001243 | 0.016727 | 0.020614 |
| 22.68055556 | 0.010112 | 0.003419 | 0.008458 | 0.00442 | 0.001546 | 0.008049 | 0.015507 | 0.014789 | 0.011109 | 0.001415 | 0.006147 | 0.019411 | 0.023514 | 0.025415 | 0.039202 | 0.009524 | 0.015051 | 0.026849 | 0.003476 | 0.002134 | 0.015524 | 0.016206 |
| 22.69444444 | 0.006131 | 0.003647 | 0.008062 | 0.004469 | 0.000871 | 0.003128 | 0.007408 | 0.010482 | 0.01361 | 0.001416 | 0.008563 | 0.012556 | 0.013333 | 0.022175 | 0.026694 | 0.008788 | 0.014182 | 0.024715 | 0.004855 | 0.003291 | 0.011855 | 0.011118 |
| 22.70833333 | 0.010845 | 0.010168 | 0.006682 | 0.003358 | 0.001007 | 0.005036 | 0.010507 | 0.004912 | 0.011805 | 0.003172 | 0.009645 | 0.00473 | 0.006096 | 0.0117 | 0.014515 | 0.007902 | 0.009836 | 0.015309 | 0.003921 | 0.00288 | 0.010544 | 0.007437 |
| 22.72222222 | 0.017804 | 0.015152 | 0.007744 | 0.004477 | 0.001673 | 0.010092 | 0.022399 | 0.003446 | 0.006799 | 0.004782 | 0.008517 | 0.003205 | 0.006819 | 0.006198 | 0.013061 | 0.009929 | 0.004827 | 0.007653 | 0.003337 | 0.001514 | 0.015548 | 0.011559 |
| 22.73611111 | 0.017472 | 0.017749 | 0.01162 | 0.010066 | 0.004737 | 0.011299 | 0.025048 | 0.006064 | 0.003442 | 0.004815 | 0.006322 | 0.009482 | 0.007223 | 0.006576 | 0.013341 | 0.010611 | 0.002251 | 0.007763 | 0.002933 | 0.000864 | 0.013469 | 0.014253 |
| 22.75 | 0.009828 | 0.018592 | 0.01462 | 0.015104 | 0.009374 | 0.007632 | 0.014547 | 0.006531 | 0.002375 | 0.003275 | 0.003091 | 0.018514 | 0.004269 | 0.006196 | 0.012264 | 0.010685 | 0.001482 | 0.007805 | 0.003009 | 0.001071 | 0.007118 | 0.009469 |
| 22.76388889 | 0.006113 | 0.015965 | 0.013981 | 0.013865 | 0.00998 | 0.003808 | 0.006867 | 0.004204 | 0.004012 | 0.001504 | 0.00186 | 0.018804 | 0.002656 | 0.005242 | 0.011403 | 0.008777 | 0.002041 | 0.007457 | 0.004808 | 0.002386 | 0.004605 | 0.007014 |
| 22.77777778 | 0.009774 | 0.009179 | 0.010607 | 0.00731 | 0.006626 | 0.004337 | 0.006051 | 0.004511 | 0.008074 | 0.000997 | 0.003879 | 0.010395 | 0.003167 | 0.003696 | 0.012367 | 0.008407 | 0.003795 | 0.009302 | 0.005043 | 0.002905 | 0.004 | 0.006335 |
| 22.79166667 | 0.014205 | 0.002809 | 0.007218 | 0.002281 | 0.006402 | 0.005605 | 0.009565 | 0.006345 | 0.008238 | 0.002183 | 0.00457 | 0.004053 | 0.008229 | 0.004439 | 0.008358 | 0.011717 | 0.003811 | 0.00747 | 0.004416 | 0.001339 | 0.00426 | 0.004707 |
| 22.80555556 | 0.012976 | 0.002083 | 0.004928 | 0.001373 | 0.007454 | 0.003193 | 0.012168 | 0.005765 | 0.007383 | 0.004373 | 0.005508 | 0.005472 | 0.015217 | 0.012731 | 0.003833 | 0.00982 | 0.001924 | 0.003926 | 0.004246 | 0.001197 | 0.009835 | 0.006644 |
| 22.81944444 | 0.007582 | 0.004674 | 0.003775 | 0.00252 | 0.006025 | 0.002704 | 0.011093 | 0.004225 | 0.010756 | 0.005359 | 0.00975 | 0.007952 | 0.015187 | 0.019986 | 0.004937 | 0.00735 | 0.001702 | 0.004594 | 0.003675 | 0.002682 | 0.019494 | 0.007616 |
| 22.83333333 | 0.004023 | 0.006033 | 0.003331 | 0.004247 | 0.005967 | 0.007648 | 0.011071 | 0.004798 | 0.010836 | 0.003894 | 0.009627 | 0.005597 | 0.008461 | 0.020161 | 0.010313 | 0.010885 | 0.003709 | 0.00789 | 0.003641 | 0.003479 | 0.017719 | 0.004489 |
| 22.84722222 | 0.006471 | 0.008212 | 0.002974 | 0.004571 | 0.005783 | 0.011462 | 0.01065 | 0.007768 | 0.009573 | 0.001737 | 0.008307 | 0.006035 | 0.00537 | 0.016742 | 0.014489 | 0.012497 | 0.004271 | 0.007855 | 0.005028 | 0.003843 | 0.008902 | 0.004011 |
| 22.86111111 | 0.014565 | 0.01309 | 0.002047 | 0.003195 | 0.004417 | 0.009893 | 0.00729 | 0.007777 | 0.008679 | 0.000928 | 0.010326 | 0.007041 | 0.008598 | 0.01161 | 0.013668 | 0.014044 | 0.003471 | 0.007342 | 0.008324 | 0.003115 | 0.008356 | 0.007849 |
| 22.875 | 0.014655 | 0.012675 | 0.002182 | 0.003701 | 0.005611 | 0.009009 | 0.00507 | 0.006339 | 0.006885 | 0.001411 | 0.015693 | 0.007131 | 0.009189 | 0.009974 | 0.013944 | 0.014064 | 0.004837 | 0.012131 | 0.00765 | 0.001626 | 0.011728 | 0.008674 |
| 22.88888889 | 0.011362 | 0.009741 | 0.005072 | 0.006815 | 0.005701 | 0.009328 | 0.009131 | 0.007948 | 0.00917 | 0.001463 | 0.024292 | 0.009965 | 0.005535 | 0.00976 | 0.013774 | 0.010403 | 0.00667 | 0.014805 | 0.003328 | 0.00149 | 0.012169 | 0.006379 |
| 22.90277778 | 0.01914 | 0.014228 | 0.00742 | 0.007295 | 0.003532 | 0.006212 | 0.012347 | 0.006839 | 0.008629 | 0.001628 | 0.020641 | 0.007563 | 0.008298 | 0.011528 | 0.012181 | 0.011313 | 0.005735 | 0.010931 | 0.00294 | 0.003687 | 0.017042 | 0.007402 |
| 22.91666667 | 0.020062 | 0.015387 | 0.006836 | 0.004433 | 0.002655 | 0.004132 | 0.011159 | 0.003522 | 0.005105 | 0.003451 | 0.009924 | 0.006917 | 0.017266 | 0.022897 | 0.021378 | 0.01715 | 0.004757 | 0.008101 | 0.007102 | 0.008238 | 0.025316 | 0.015735 |
| 22.93055556 | 0.012163 | 0.00978 | 0.006469 | 0.004523 | 0.004392 | 0.007086 | 0.020317 | 0.004434 | 0.014806 | 0.003982 | 0.008136 | 0.015133 | 0.019513 | 0.032707 | 0.042657 | 0.023497 | 0.00969 | 0.017501 | 0.011408 | 0.012164 | 0.021348 | 0.02576 |
| 22.94444444 | 0.009318 | 0.007962 | 0.010247 | 0.010322 | 0.009139 | 0.01107 | 0.02539 | 0.011218 | 0.033243 | 0.002216 | 0.012139 | 0.020942 | 0.013658 | 0.027319 | 0.050508 | 0.020074 | 0.015834 | 0.031028 | 0.010093 | 0.010492 | 0.008284 | 0.024957 |
| 22.95833333 | 0.015983 | 0.010655 | 0.016027 | 0.015949 | 0.011922 | 0.012051 | 0.013811 | 0.018879 | 0.035987 | 0.001059 | 0.015911 | 0.031369 | 0.013789 | 0.023413 | 0.04509 | 0.017397 | 0.01404 | 0.026888 | 0.00647 | 0.00753 | 0.009935 | 0.01666 |
| 22.97222222 | 0.036741 | 0.020302 | 0.018839 | 0.016129 | 0.011431 | 0.012981 | 0.008991 | 0.020439 | 0.028999 | 0.001791 | 0.017026 | 0.050901 | 0.027484 | 0.039166 | 0.04719 | 0.031482 | 0.01078 | 0.015438 | 0.009457 | 0.013805 | 0.024663 | 0.016052 |
| 22.98611111 | 0.04862 | 0.030326 | 0.019026 | 0.015223 | 0.014174 | 0.016422 | 0.024937 | 0.024587 | 0.038135 | 0.005361 | 0.023573 | 0.048054 | 0.04348 | 0.059119 | 0.055415 | 0.044547 | 0.020148 | 0.029502 | 0.017854 | 0.025154 | 0.029943 | 0.028396 |
| 23 | 0.041945 | 0.026745 | 0.021877 | 0.019569 | 0.019931 | 0.023091 | 0.058955 | 0.039438 | 0.057942 | 0.010475 | 0.036832 | 0.024528 | 0.047666 | 0.060135 | 0.064078 | 0.042349 | 0.038433 | 0.068147 | 0.026883 | 0.032426 | 0.027533 | 0.041164 |
| 23.01388889 | 0.042587 | 0.021082 | 0.030568 | 0.02898 | 0.022174 | 0.030763 | 0.075758 | 0.049595 | 0.065125 | 0.013264 | 0.040107 | 0.026411 | 0.046254 | 0.052847 | 0.074526 | 0.039562 | 0.045296 | 0.087763 | 0.032226 | 0.035004 | 0.032431 | 0.044404 |
| 23.02777778 | 0.055439 | 0.030019 | 0.042435 | 0.036059 | 0.02029 | 0.035847 | 0.054441 | 0.043177 | 0.058535 | 0.010728 | 0.027273 | 0.050479 | 0.051678 | 0.057222 | 0.081602 | 0.045131 | 0.037634 | 0.068888 | 0.028918 | 0.031921 | 0.043724 | 0.044024 |
| 23.04166667 | 0.062222 | 0.044141 | 0.051122 | 0.034688 | 0.017975 | 0.037495 | 0.031561 | 0.034375 | 0.054597 | 0.005184 | 0.021602 | 0.05687 | 0.055835 | 0.063276 | 0.076787 | 0.046448 | 0.028199 | 0.04132 | 0.020854 | 0.024569 | 0.047893 | 0.043676 |
| 23.05555556 | 0.062381 | 0.045762 | 0.053747 | 0.031634 | 0.017297 | 0.034022 | 0.032447 | 0.036576 | 0.055865 | 0.002564 | 0.032235 | 0.044226 | 0.044744 | 0.053184 | 0.061862 | 0.037081 | 0.02194 | 0.029073 | 0.016661 | 0.017369 | 0.040179 | 0.041669 |
| 23.06944444 | 0.068162 | 0.03091 | 0.053858 | 0.035006 | 0.018649 | 0.030227 | 0.045719 | 0.04268 | 0.051763 | 0.002237 | 0.043147 | 0.028404 | 0.023263 | 0.029382 | 0.044241 | 0.024638 | 0.018015 | 0.027863 | 0.015855 | 0.013501 | 0.02812 | 0.040216 |
| 23.08333333 | 0.067043 | 0.011792 | 0.045548 | 0.03414 | 0.015829 | 0.030052 | 0.054387 | 0.04086 | 0.040764 | 0.001109 | 0.041837 | 0.013246 | 0.009715 | 0.009765 | 0.025676 | 0.014069 | 0.014126 | 0.026053 | 0.011505 | 0.011534 | 0.017211 | 0.037303 |
| 23.09722222 | 0.044946 | 0.014151 | 0.025638 | 0.019781 | 0.007089 | 0.028951 | 0.051462 | 0.029678 | 0.030726 | 0.001624 | 0.022893 | 0.005554 | 0.008275 | 0.004828 | 0.008945 | 0.006828 | 0.010667 | 0.020941 | 0.00545 | 0.008839 | 0.009882 | 0.027212 |
| 23.11111111 | 0.025007 | 0.04167 | 0.015314 | 0.008906 | 0.007615 | 0.025853 | 0.042421 | 0.020802 | 0.028608 | 0.006614 | 0.019683 | 0.010845 | 0.014315 | 0.011977 | 0.014041 | 0.012108 | 0.012221 | 0.020649 | 0.009558 | 0.010191 | 0.01316 | 0.018711 |
| 23.125 | 0.025456 | 0.049129 | 0.023097 | 0.012426 | 0.019342 | 0.024277 | 0.035324 | 0.023432 | 0.032738 | 0.01292 | 0.048181 | 0.020153 | 0.025211 | 0.023073 | 0.045028 | 0.029066 | 0.019128 | 0.031123 | 0.026051 | 0.017629 | 0.028469 | 0.022849 |
| 23.13888889 | 0.029921 | 0.030593 | 0.02628 | 0.016531 | 0.022552 | 0.023607 | 0.028546 | 0.029015 | 0.032929 | 0.013867 | 0.056416 | 0.019245 | 0.028699 | 0.027665 | 0.06214 | 0.034749 | 0.024814 | 0.041443 | 0.035072 | 0.02189 | 0.03844 | 0.028687 |
| 23.15277778 | 0.025506 | 0.02174 | 0.015138 | 0.01225 | 0.013152 | 0.021105 | 0.019755 | 0.026738 | 0.024205 | 0.008672 | 0.035927 | 0.011761 | 0.020697 | 0.022459 | 0.047957 | 0.024584 | 0.024404 | 0.038283 | 0.02498 | 0.017009 | 0.029397 | 0.022401 |
| 23.16666667 | 0.017181 | 0.015601 | 0.005066 | 0.005203 | 0.004506 | 0.018043 | 0.013922 | 0.019583 | 0.012736 | 0.003239 | 0.021664 | 0.009799 | 0.011341 | 0.015418 | 0.032961 | 0.015523 | 0.019571 | 0.027962 | 0.010092 | 0.008629 | 0.013334 | 0.010858 |
| 23.18055556 | 0.010402 | 0.008603 | 0.002478 | 0.001538 | 0.001807 | 0.015425 | 0.014025 | 0.012557 | 0.004757 | 0.001294 | 0.018066 | 0.011545 | 0.005674 | 0.010901 | 0.026913 | 0.010548 | 0.013394 | 0.020417 | 0.004134 | 0.003766 | 0.005548 | 0.004527 |
| 23.19444444 | 0.008165 | 0.009326 | 0.00267 | 0.001843 | 0.00331 | 0.011333 | 0.015707 | 0.005961 | 0.0012 | 0.001484 | 0.024416 | 0.010615 | 0.004304 | 0.006513 | 0.016122 | 0.005251 | 0.007924 | 0.014439 | 0.006374 | 0.003796 | 0.004505 | 0.002496 |
| 23.20833333 | 0.008129 | 0.009699 | 0.00395 | 0.003546 | 0.007124 | 0.006598 | 0.014105 | 0.002809 | 0.000494 | 0.001752 | 0.025785 | 0.007455 | 0.005143 | 0.003527 | 0.005527 | 0.001727 | 0.004943 | 0.009619 | 0.00825 | 0.00483 | 0.003412 | 0.002041 |
| 23.22222222 | 0.005275 | 0.006546 | 0.004385 | 0.003803 | 0.008515 | 0.004574 | 0.009371 | 0.005137 | 0.000506 | 0.002013 | 0.023565 | 0.004604 | 0.003844 | 0.002465 | 0.006195 | 0.002324 | 0.004251 | 0.008497 | 0.005984 | 0.003308 | 0.002029 | 0.002014 |
| 23.23611111 | 0.002629 | 0.005216 | 0.004558 | 0.002377 | 0.006656 | 0.004461 | 0.004559 | 0.007597 | 0.001251 | 0.003132 | 0.028905 | 0.002491 | 0.001528 | 0.001227 | 0.010903 | 0.004173 | 0.004095 | 0.008403 | 0.003508 | 0.00101 | 0.001601 | 0.001523 |
| 23.25 | 0.002609 | 0.005146 | 0.004161 | 0.001452 | 0.005676 | 0.003546 | 0.001706 | 0.00626 | 0.003073 | 0.00482 | 0.022316 | 0.001078 | 0.001359 | 0.000361 | 0.009928 | 0.003921 | 0.00287 | 0.005872 | 0.003715 | 0.000897 | 0.002184 | 0.001341 |
| 23.26388889 | 0.003292 | 0.004709 | 0.002495 | 0.001457 | 0.005948 | 0.00326 | 0.001497 | 0.004962 | 0.004729 | 0.005306 | 0.007676 | 0.000951 | 0.002805 | 0.00047 | 0.009927 | 0.003182 | 0.00167 | 0.00368 | 0.005132 | 0.002338 | 0.00322 | 0.000981 |
| 23.27777778 | 0.00259 | 0.006124 | 0.001815 | 0.001263 | 0.004313 | 0.004042 | 0.003993 | 0.00429 | 0.005316 | 0.00368 | 0.002289 | 0.001195 | 0.002773 | 0.001264 | 0.014483 | 0.002965 | 0.001748 | 0.00295 | 0.005252 | 0.002793 | 0.002793 | 0.000522 |
| 23.29166667 | 0.001404 | 0.006531 | 0.001765 | 0.001341 | 0.002078 | 0.00301 | 0.00617 | 0.002505 | 0.004374 | 0.001612 | 0.001962 | 0.001822 | 0.002287 | 0.00299 | 0.014623 | 0.00258 | 0.002134 | 0.002294 | 0.003541 | 0.001969 | 0.002461 | 0.00153 |
| 23.30555556 | 0.00291 | 0.005464 | 0.002429 | 0.002384 | 0.002769 | 0.001567 | 0.005478 | 0.001385 | 0.002238 | 0.001166 | 0.003558 | 0.003478 | 0.004952 | 0.005446 | 0.009057 | 0.004319 | 0.003377 | 0.003222 | 0.0024 | 0.001791 | 0.004536 | 0.003529 |
| 23.31944444 | 0.005815 | 0.004504 | 0.002866 | 0.003468 | 0.00446 | 0.001816 | 0.005203 | 0.002303 | 0.001083 | 0.001934 | 0.007812 | 0.003512 | 0.007031 | 0.006629 | 0.004194 | 0.006442 | 0.003897 | 0.003575 | 0.002897 | 0.001984 | 0.00509 | 0.003837 |
| 23.33333333 | 0.006718 | 0.007779 | 0.002693 | 0.003776 | 0.00436 | 0.003957 | 0.005526 | 0.007296 | 0.003261 | 0.001886 | 0.008938 | 0.002695 | 0.007211 | 0.006053 | 0.005856 | 0.006975 | 0.003801 | 0.003907 | 0.006189 | 0.003003 | 0.005039 | 0.004294 |
| 23.34722222 | 0.008367 | 0.016411 | 0.005166 | 0.005313 | 0.005189 | 0.010525 | 0.005834 | 0.013578 | 0.007234 | 0.002284 | 0.008787 | 0.00406 | 0.01098 | 0.007532 | 0.014111 | 0.00985 | 0.006773 | 0.008032 | 0.012488 | 0.005924 | 0.008604 | 0.007342 |
| 23.36111111 | 0.011926 | 0.017867 | 0.009905 | 0.008123 | 0.008366 | 0.017292 | 0.007892 | 0.014705 | 0.008415 | 0.006407 | 0.010511 | 0.007027 | 0.015 | 0.010724 | 0.016896 | 0.012054 | 0.00938 | 0.010342 | 0.013784 | 0.006867 | 0.009699 | 0.007552 |
| 23.375 | 0.012608 | 0.011407 | 0.012178 | 0.009201 | 0.010004 | 0.016613 | 0.007404 | 0.01248 | 0.007637 | 0.008347 | 0.010915 | 0.011432 | 0.014153 | 0.012546 | 0.01207 | 0.009368 | 0.008144 | 0.008407 | 0.009632 | 0.005456 | 0.008209 | 0.004906 |
| 23.38888889 | 0.011989 | 0.010927 | 0.010055 | 0.009659 | 0.010035 | 0.012055 | 0.00403 | 0.012816 | 0.01046 | 0.006286 | 0.013817 | 0.015922 | 0.012947 | 0.014539 | 0.013738 | 0.008159 | 0.007441 | 0.009668 | 0.009933 | 0.006329 | 0.010775 | 0.006358 |
| 23.40277778 | 0.012283 | 0.014809 | 0.006941 | 0.010553 | 0.010381 | 0.010918 | 0.00452 | 0.014162 | 0.015369 | 0.007837 | 0.017499 | 0.016922 | 0.013619 | 0.016525 | 0.023646 | 0.011581 | 0.010602 | 0.015537 | 0.014233 | 0.009067 | 0.014723 | 0.012014 |
| 23.41666667 | 0.012025 | 0.014397 | 0.007373 | 0.010542 | 0.010683 | 0.013076 | 0.01147 | 0.012642 | 0.016158 | 0.010174 | 0.01359 | 0.014432 | 0.013799 | 0.017088 | 0.030313 | 0.01487 | 0.014571 | 0.019137 | 0.014626 | 0.010302 | 0.016134 | 0.017065 |
| 23.43055556 | 0.013906 | 0.014024 | 0.011075 | 0.011681 | 0.012199 | 0.014767 | 0.017416 | 0.011609 | 0.014641 | 0.009882 | 0.009539 | 0.012944 | 0.014664 | 0.01899 | 0.030892 | 0.015795 | 0.016042 | 0.01864 | 0.011973 | 0.010496 | 0.017597 | 0.020107 |
| 23.44444444 | 0.020212 | 0.018633 | 0.014816 | 0.016244 | 0.016017 | 0.01506 | 0.01629 | 0.015616 | 0.017247 | 0.009688 | 0.017798 | 0.016342 | 0.017825 | 0.023447 | 0.032053 | 0.016541 | 0.017272 | 0.020154 | 0.013199 | 0.012321 | 0.02142 | 0.022659 |
| 23.45833333 | 0.027645 | 0.022517 | 0.019966 | 0.022079 | 0.02093 | 0.015975 | 0.014471 | 0.023481 | 0.024102 | 0.010785 | 0.031309 | 0.023247 | 0.021161 | 0.027877 | 0.038699 | 0.019825 | 0.021376 | 0.026566 | 0.018908 | 0.016134 | 0.026556 | 0.025999 |
| 23.47222222 | 0.032158 | 0.02237 | 0.028901 | 0.026187 | 0.024763 | 0.019077 | 0.020612 | 0.031435 | 0.031225 | 0.013264 | 0.035629 | 0.029469 | 0.024026 | 0.032367 | 0.051657 | 0.026091 | 0.026721 | 0.034448 | 0.025071 | 0.020626 | 0.033383 | 0.032098 |
| 23.48611111 | 0.034189 | 0.023813 | 0.036163 | 0.028651 | 0.026244 | 0.024005 | 0.033252 | 0.037142 | 0.038411 | 0.016146 | 0.034759 | 0.035359 | 0.03014 | 0.041044 | 0.068919 | 0.033586 | 0.030842 | 0.042389 | 0.030439 | 0.025558 | 0.042072 | 0.041303 |
| 23.5 | 0.037243 | 0.03044 | 0.036531 | 0.030898 | 0.026234 | 0.029317 | 0.04136 | 0.040824 | 0.04805 | 0.017828 | 0.041863 | 0.045475 | 0.040928 | 0.054922 | 0.086085 | 0.04153 | 0.035339 | 0.052295 | 0.035726 | 0.030992 | 0.048001 | 0.049986 |
| 23.51388889 | 0.042821 | 0.038028 | 0.036632 | 0.032196 | 0.026249 | 0.03352 | 0.039824 | 0.043731 | 0.057729 | 0.016491 | 0.054444 | 0.0595 | 0.051453 | 0.068445 | 0.097891 | 0.049336 | 0.041635 | 0.063218 | 0.039985 | 0.035695 | 0.048907 | 0.055393 |
| 23.52777778 | 0.048493 | 0.042418 | 0.042786 | 0.030379 | 0.024452 | 0.034522 | 0.038483 | 0.044901 | 0.062387 | 0.0115 | 0.059077 | 0.069296 | 0.057137 | 0.077409 | 0.103784 | 0.056476 | 0.045369 | 0.070484 | 0.040144 | 0.037577 | 0.05091 | 0.059807 |
| 23.54166667 | 0.04899 | 0.04221 | 0.044338 | 0.024436 | 0.016799 | 0.029328 | 0.045549 | 0.041773 | 0.061465 | 0.00659 | 0.05511 | 0.069605 | 0.056106 | 0.077558 | 0.103148 | 0.064998 | 0.041876 | 0.069665 | 0.034365 | 0.035427 | 0.057533 | 0.06544 |
| 23.55555556 | 0.044009 | 0.036425 | 0.033281 | 0.018122 | 0.00744 | 0.019765 | 0.05343 | 0.038568 | 0.060016 | 0.004401 | 0.051188 | 0.063497 | 0.048065 | 0.064211 | 0.084788 | 0.069126 | 0.034677 | 0.063414 | 0.025022 | 0.029751 | 0.059382 | 0.066541 |
| 23.56944444 | 0.043084 | 0.028316 | 0.024651 | 0.01714 | 0.005661 | 0.01427 | 0.053849 | 0.043775 | 0.06385 | 0.006332 | 0.043615 | 0.05511 | 0.041361 | 0.048328 | 0.052722 | 0.060331 | 0.033078 | 0.063263 | 0.016188 | 0.022797 | 0.049225 | 0.056934 |
| 23.58333333 | 0.051169 | 0.026634 | 0.02837 | 0.021351 | 0.011638 | 0.019237 | 0.052115 | 0.050131 | 0.067271 | 0.013896 | 0.029649 | 0.048564 | 0.043229 | 0.05256 | 0.036141 | 0.046072 | 0.038688 | 0.071602 | 0.011661 | 0.020075 | 0.040741 | 0.046971 |
| 23.59722222 | 0.068211 | 0.047627 | 0.035511 | 0.028278 | 0.01815 | 0.031389 | 0.054736 | 0.044645 | 0.061421 | 0.017585 | 0.026762 | 0.044078 | 0.044788 | 0.070473 | 0.039058 | 0.033195 | 0.040963 | 0.072506 | 0.014603 | 0.027692 | 0.049397 | 0.049624 |
| 23.61111111 | 0.081363 | 0.077565 | 0.039129 | 0.035917 | 0.019525 | 0.036858 | 0.054536 | 0.036406 | 0.050444 | 0.011825 | 0.037886 | 0.046646 | 0.038129 | 0.069281 | 0.040941 | 0.018855 | 0.034605 | 0.061958 | 0.020359 | 0.035586 | 0.05942 | 0.051663 |
| 23.625 | 0.064799 | 0.070162 | 0.033796 | 0.036412 | 0.016208 | 0.030663 | 0.03983 | 0.040699 | 0.044627 | 0.007595 | 0.035059 | 0.050814 | 0.030689 | 0.050648 | 0.039732 | 0.011426 | 0.023734 | 0.047914 | 0.020622 | 0.027083 | 0.045895 | 0.036864 |
| 23.63888889 | 0.032523 | 0.035164 | 0.020617 | 0.02551 | 0.0133 | 0.02402 | 0.025077 | 0.046907 | 0.043839 | 0.011034 | 0.015824 | 0.03477 | 0.029475 | 0.039548 | 0.037058 | 0.013367 | 0.013287 | 0.029688 | 0.016397 | 0.011094 | 0.019418 | 0.016365 |
| 23.65277778 | 0.018909 | 0.019662 | 0.012098 | 0.012449 | 0.01304 | 0.022432 | 0.031154 | 0.034751 | 0.036267 | 0.017841 | 0.009814 | 0.012957 | 0.032112 | 0.042656 | 0.034616 | 0.01414 | 0.011205 | 0.02125 | 0.011663 | 0.003954 | 0.005427 | 0.010382 |
| 23.66666667 | 0.019847 | 0.021785 | 0.01239 | 0.006003 | 0.010206 | 0.021495 | 0.041566 | 0.014989 | 0.019999 | 0.022855 | 0.013519 | 0.008583 | 0.032897 | 0.049912 | 0.043995 | 0.015278 | 0.015985 | 0.028498 | 0.008936 | 0.006024 | 0.010587 | 0.023914 |
| 23.68055556 | 0.016733 | 0.018404 | 0.014225 | 0.006303 | 0.004527 | 0.016682 | 0.035782 | 0.009433 | 0.006215 | 0.01799 | 0.010078 | 0.009365 | 0.023769 | 0.040688 | 0.055053 | 0.016846 | 0.015362 | 0.030118 | 0.008228 | 0.010663 | 0.023497 | 0.036499 |

| | | | | | | | | | | | | | | | | | | | | | | |
|---|---|---|---|---|---|---|---|---|---|---|---|---|---|---|---|---|---|---|---|---|---|---|
| 23.69444444 | 0.008596 | 0.009882 | 0.011057 | 0.007389 | 0.002754 | 0.011003 | 0.023936 | 0.018516 | 0.002898 | 0.00898 | 0.009843 | 0.004885 | 0.010018 | 0.01946 | 0.046822 | 0.014983 | 0.010187 | 0.0247 | 0.005883 | 0.011029 | 0.023788 | 0.027682 |
| 23.70833333 | 0.009877 | 0.014461 | 0.005745 | 0.006608 | 0.004167 | 0.008874 | 0.017548 | 0.026659 | 0.004303 | 0.006265 | 0.010121 | 0.003567 | 0.005282 | 0.007931 | 0.025431 | 0.010598 | 0.008298 | 0.021381 | 0.004513 | 0.007687 | 0.017469 | 0.015318 |
| 23.72222222 | 0.021795 | 0.02732 | 0.002557 | 0.005677 | 0.004645 | 0.007131 | 0.01409 | 0.01865 | 0.004183 | 0.005689 | 0.006465 | 0.005956 | 0.004504 | 0.003486 | 0.011815 | 0.006667 | 0.008 | 0.014667 | 0.006032 | 0.004764 | 0.02236 | 0.017938 |
| 23.73611111 | 0.022892 | 0.022997 | 0.002963 | 0.004095 | 0.005231 | 0.004079 | 0.012592 | 0.005899 | 0.006271 | 0.005028 | 0.005876 | 0.009799 | 0.004196 | 0.002319 | 0.012019 | 0.006639 | 0.005506 | 0.008531 | 0.006997 | 0.002733 | 0.022631 | 0.016772 |
| 23.75 | 0.01473 | 0.010502 | 0.005831 | 0.004485 | 0.005613 | 0.002506 | 0.010879 | 0.003852 | 0.009412 | 0.00345 | 0.00619 | 0.025998 | 0.005735 | 0.005333 | 0.012634 | 0.009641 | 0.002402 | 0.009299 | 0.007839 | 0.001915 | 0.017312 | 0.009079 |
| 23.76388889 | 0.01027 | 0.018046 | 0.007316 | 0.007696 | 0.004552 | 0.004167 | 0.007972 | 0.004983 | 0.011448 | 0.004393 | 0.009901 | 0.039948 | 0.004679 | 0.008402 | 0.007842 | 0.01568 | 0.002213 | 0.010547 | 0.009364 | 0.002705 | 0.015581 | 0.009127 |
| 23.77777778 | 0.006738 | 0.029646 | 0.01069 | 0.010833 | 0.005746 | 0.006237 | 0.00547 | 0.009457 | 0.017193 | 0.013664 | 0.019906 | 0.033461 | 0.004144 | 0.014499 | 0.008041 | 0.019923 | 0.005136 | 0.01389 | 0.009558 | 0.004894 | 0.012035 | 0.010207 |
| 23.79166667 | 0.007951 | 0.025149 | 0.014228 | 0.012779 | 0.007527 | 0.008165 | 0.008606 | 0.024264 | 0.020237 | 0.019379 | 0.022963 | 0.023476 | 0.004607 | 0.018333 | 0.011123 | 0.015769 | 0.007343 | 0.016037 | 0.009726 | 0.005135 | 0.010158 | 0.012332 |
| 23.80555556 | 0.014527 | 0.020311 | 0.010412 | 0.010565 | 0.006782 | 0.01274 | 0.018534 | 0.029849 | 0.026204 | 0.016633 | 0.017105 | 0.040298 | 0.003975 | 0.016635 | 0.007918 | 0.01293 | 0.006008 | 0.013558 | 0.009231 | 0.003169 | 0.008052 | 0.015605 |
| 23.81944444 | 0.01712 | 0.020972 | 0.003618 | 0.004869 | 0.004782 | 0.012433 | 0.018611 | 0.023287 | 0.034832 | 0.014768 | 0.022598 | 0.059045 | 0.006085 | 0.015722 | 0.007676 | 0.016382 | 0.003877 | 0.014964 | 0.010836 | 0.002968 | 0.005873 | 0.012995 |
| 23.83333333 | 0.011418 | 0.025004 | 0.002149 | 0.001794 | 0.002753 | 0.005669 | 0.009711 | 0.020568 | 0.029057 | 0.017929 | 0.025397 | 0.039428 | 0.01212 | 0.01967 | 0.015767 | 0.015003 | 0.002948 | 0.01538 | 0.019129 | 0.004078 | 0.007394 | 0.009668 |
| 23.84722222 | 0.005597 | 0.024288 | 0.005913 | 0.002912 | 0.003813 | 0.00185 | 0.006511 | 0.024123 | 0.024744 | 0.02782 | 0.022906 | 0.029994 | 0.016873 | 0.02688 | 0.016562 | 0.010585 | 0.002008 | 0.010294 | 0.020592 | 0.004532 | 0.009183 | 0.010861 |
| 23.86111111 | 0.008561 | 0.021921 | 0.008288 | 0.003915 | 0.006694 | 0.00495 | 0.011041 | 0.032879 | 0.01949 | 0.030144 | 0.030186 | 0.03694 | 0.013277 | 0.024216 | 0.010338 | 0.010787 | 0.002757 | 0.007474 | 0.011792 | 0.005692 | 0.010264 | 0.01513 |
| 23.875 | 0.019808 | 0.029599 | 0.007494 | 0.002283 | 0.006269 | 0.00893 | 0.016736 | 0.036329 | 0.014236 | 0.023906 | 0.026552 | 0.030384 | 0.006748 | 0.015383 | 0.012795 | 0.017971 | 0.005088 | 0.008379 | 0.008082 | 0.007074 | 0.01187 | 0.014005 |
| 23.88888889 | 0.022405 | 0.031241 | 0.00747 | 0.001679 | 0.005062 | 0.00607 | 0.014616 | 0.034628 | 0.017134 | 0.017092 | 0.024584 | 0.025833 | 0.00555 | 0.012806 | 0.0218 | 0.024822 | 0.004667 | 0.005559 | 0.01158 | 0.00705 | 0.013707 | 0.011009 |
| 23.90277778 | 0.014276 | 0.020385 | 0.007622 | 0.003164 | 0.0062 | 0.003408 | 0.00977 | 0.031912 | 0.016513 | 0.007484 | 0.028653 | 0.018124 | 0.007057 | 0.014025 | 0.028627 | 0.018313 | 0.003608 | 0.005598 | 0.010654 | 0.006832 | 0.021119 | 0.013257 |
| 23.91666667 | 0.013887 | 0.014345 | 0.006003 | 0.002935 | 0.007016 | 0.005725 | 0.005641 | 0.032078 | 0.020733 | 0.006568 | 0.022091 | 0.014257 | 0.008786 | 0.010435 | 0.023406 | 0.00638 | 0.007984 | 0.02086 | 0.005558 | 0.010081 | 0.021862 | 0.018641 |
| 23.93055556 | 0.014459 | 0.014362 | 0.007519 | 0.002583 | 0.007734 | 0.006431 | 0.002201 | 0.028948 | 0.018867 | 0.014688 | 0.021677 | 0.030471 | 0.011297 | 0.014715 | 0.015515 | 0.007562 | 0.014223 | 0.037803 | 0.004866 | 0.019085 | 0.00914 | 0.02848 |
| 23.94444444 | 0.013974 | 0.019095 | 0.014239 | 0.005317 | 0.009177 | 0.009421 | 0.011042 | 0.027855 | 0.018714 | 0.015671 | 0.028376 | 0.046002 | 0.019197 | 0.037707 | 0.035178 | 0.027305 | 0.016723 | 0.038525 | 0.007625 | 0.021909 | 0.008368 | 0.029465 |
| 23.95833333 | 0.024403 | 0.03469 | 0.016336 | 0.008589 | 0.011112 | 0.017603 | 0.028759 | 0.036084 | 0.037701 | 0.01183 | 0.041077 | 0.05212 | 0.025688 | 0.047997 | 0.064573 | 0.047902 | 0.018173 | 0.032385 | 0.015995 | 0.014534 | 0.025889 | 0.018779 |
| 23.97222222 | 0.042007 | 0.053413 | 0.0153 | 0.012607 | 0.012103 | 0.018577 | 0.035634 | 0.028295 | 0.047433 | 0.017563 | 0.051038 | 0.062396 | 0.02373 | 0.037576 | 0.064209 | 0.051573 | 0.022868 | 0.036529 | 0.024838 | 0.013247 | 0.037098 | 0.015991 |
| 23.98611111 | 0.054131 | 0.052583 | 0.024739 | 0.019779 | 0.014762 | 0.013627 | 0.038321 | 0.019585 | 0.04294 | 0.027033 | 0.04395 | 0.058641 | 0.024918 | 0.037413 | 0.053214 | 0.050701 | 0.030792 | 0.051186 | 0.024342 | 0.021187 | 0.030468 | 0.027883 |
| 24 | 0.055475 | 0.038579 | 0.038051 | 0.02681 | 0.021929 | 0.017124 | 0.054206 | 0.035608 | 0.04298 | 0.031495 | 0.040726 | 0.033901 | 0.03413 | 0.051989 | 0.065056 | 0.057321 | 0.037378 | 0.062776 | 0.019812 | 0.026682 | 0.023621 | 0.04278 |
| 24.01388889 | 0.056744 | 0.039666 | 0.038406 | 0.030289 | 0.028647 | 0.033418 | 0.069625 | 0.058302 | 0.052105 | 0.036685 | 0.049692 | 0.02383 | 0.044704 | 0.065974 | 0.092939 | 0.062403 | 0.039107 | 0.064156 | 0.019023 | 0.026283 | 0.029775 | 0.046595 |
| 24.02777778 | 0.065896 | 0.049528 | 0.031735 | 0.031893 | 0.032095 | 0.050601 | 0.064105 | 0.067436 | 0.068154 | 0.042964 | 0.068624 | 0.04775 | 0.051352 | 0.071006 | 0.107145 | 0.064743 | 0.040029 | 0.06286 | 0.0242 | 0.027279 | 0.041176 | 0.040982 |
| 24.04166667 | 0.070651 | 0.048897 | 0.036479 | 0.035432 | 0.030299 | 0.052788 | 0.049134 | 0.060168 | 0.078601 | 0.0424 | 0.092122 | 0.075692 | 0.052474 | 0.066573 | 0.092545 | 0.067486 | 0.042222 | 0.062408 | 0.029006 | 0.030355 | 0.044859 | 0.037569 |
| 24.05555556 | 0.063272 | 0.042184 | 0.055483 | 0.041931 | 0.025679 | 0.04385 | 0.049641 | 0.05104 | 0.072572 | 0.03412 | 0.082469 | 0.074478 | 0.048274 | 0.057903 | 0.065563 | 0.059265 | 0.037891 | 0.053109 | 0.025989 | 0.026959 | 0.03825 | 0.040382 |
| 24.06944444 | 0.058925 | 0.042668 | 0.077678 | 0.050731 | 0.024192 | 0.037309 | 0.065071 | 0.051385 | 0.055862 | 0.023688 | 0.038377 | 0.050668 | 0.041776 | 0.047611 | 0.044755 | 0.038434 | 0.027324 | 0.035739 | 0.016221 | 0.019049 | 0.028133 | 0.04355 |
| 24.08333333 | 0.061534 | 0.043115 | 0.083717 | 0.053536 | 0.020456 | 0.036204 | 0.076425 | 0.052277 | 0.03841 | 0.014283 | 0.011585 | 0.027092 | 0.03661 | 0.045055 | 0.039679 | 0.024935 | 0.022315 | 0.029118 | 0.011044 | 0.013515 | 0.016603 | 0.036679 |
| 24.09722222 | 0.055759 | 0.032831 | 0.062304 | 0.040437 | 0.011322 | 0.036354 | 0.074154 | 0.044402 | 0.027759 | 0.010916 | 0.021145 | 0.015437 | 0.043256 | 0.068847 | 0.06711 | 0.036945 | 0.027725 | 0.039382 | 0.01513 | 0.016163 | 0.006559 | 0.019462 |
| 24.11111111 | 0.046082 | 0.024857 | 0.036921 | 0.02454 | 0.008916 | 0.036286 | 0.064685 | 0.034773 | 0.031379 | 0.019674 | 0.045374 | 0.017034 | 0.054254 | 0.087312 | 0.091824 | 0.051812 | 0.031902 | 0.050155 | 0.016395 | 0.019594 | 0.003883 | 0.010206 |
| 24.125 | 0.047438 | 0.030937 | 0.032282 | 0.023393 | 0.021324 | 0.039455 | 0.057622 | 0.034765 | 0.045841 | 0.035276 | 0.060438 | 0.027365 | 0.044948 | 0.066972 | 0.076684 | 0.039373 | 0.026604 | 0.049215 | 0.014047 | 0.012314 | 0.014878 | 0.019259 |
| 24.13888889 | 0.054097 | 0.038688 | 0.035697 | 0.029382 | 0.036576 | 0.044049 | 0.054255 | 0.042119 | 0.05372 | 0.040496 | 0.05507 | 0.035097 | 0.036237 | 0.057906 | 0.070685 | 0.025372 | 0.027083 | 0.05441 | 0.012842 | 0.005943 | 0.041641 | 0.036641 |
| 24.15277778 | 0.050603 | 0.035313 | 0.029249 | 0.026708 | 0.035138 | 0.039722 | 0.048077 | 0.043235 | 0.044478 | 0.028333 | 0.03444 | 0.032373 | 0.045653 | 0.071098 | 0.080038 | 0.027344 | 0.035728 | 0.070424 | 0.0105 | 0.005513 | 0.060348 | 0.046233 |
| 24.16666667 | 0.033449 | 0.023241 | 0.02114 | 0.016147 | 0.01808 | 0.0255 | 0.036286 | 0.032538 | 0.027558 | 0.011715 | 0.013186 | 0.023051 | 0.051217 | 0.067716 | 0.071044 | 0.028906 | 0.034844 | 0.07064 | 0.011975 | 0.00783 | 0.056268 | 0.044038 |
| 24.18055556 | 0.014615 | 0.011309 | 0.020179 | 0.008519 | 0.004837 | 0.014154 | 0.024199 | 0.019031 | 0.016904 | 0.003801 | 0.004732 | 0.015254 | 0.050529 | 0.056035 | 0.054732 | 0.028046 | 0.026462 | 0.057959 | 0.021311 | 0.014878 | 0.047941 | 0.04382 |
| 24.19444444 | 0.005528 | 0.006233 | 0.016265 | 0.006804 | 0.002893 | 0.0134 | 0.016808 | 0.011614 | 0.01443 | 0.004094 | 0.007784 | 0.012522 | 0.051573 | 0.050283 | 0.041522 | 0.027678 | 0.021549 | 0.049918 | 0.031684 | 0.023173 | 0.046126 | 0.050483 |
| 24.20833333 | 0.005546 | 0.007287 | 0.007004 | 0.00712 | 0.003958 | 0.016859 | 0.013712 | 0.009915 | 0.012995 | 0.005023 | 0.009046 | 0.012111 | 0.051299 | 0.046074 | 0.033027 | 0.026091 | 0.019721 | 0.044255 | 0.036046 | 0.027459 | 0.04643 | 0.055559 |
| 24.22222222 | 0.006736 | 0.008022 | 0.005339 | 0.007006 | 0.002806 | 0.015382 | 0.011869 | 0.009734 | 0.009022 | 0.003566 | 0.004895 | 0.010508 | 0.047667 | 0.039541 | 0.034937 | 0.02308 | 0.018105 | 0.036987 | 0.034248 | 0.026398 | 0.046394 | 0.055602 |
| 24.23611111 | 0.005336 | 0.00732 | 0.012072 | 0.006435 | 0.001288 | 0.009868 | 0.009851 | 0.009322 | 0.006033 | 0.002717 | 0.003622 | 0.008784 | 0.04165 | 0.032865 | 0.045358 | 0.020174 | 0.016491 | 0.030367 | 0.030638 | 0.023481 | 0.046522 | 0.053494 |
| 24.25 | 0.003651 | 0.008728 | 0.01414 | 0.005739 | 0.001031 | 0.007185 | 0.008032 | 0.009341 | 0.006499 | 0.003813 | 0.006082 | 0.00946 | 0.035483 | 0.03029 | 0.052903 | 0.018347 | 0.015114 | 0.026064 | 0.028677 | 0.021157 | 0.046082 | 0.051328 |
| 24.26388889 | 0.002284 | 0.009988 | 0.00839 | 0.0048 | 0.001459 | 0.008507 | 0.006529 | 0.009411 | 0.007341 | 0.003904 | 0.007609 | 0.011655 | 0.03071 | 0.029744 | 0.05238 | 0.016805 | 0.013235 | 0.022146 | 0.026703 | 0.018261 | 0.041743 | 0.047082 |
| 24.27777778 | 0.001058 | 0.007357 | 0.003506 | 0.003509 | 0.001132 | 0.00905 | 0.005055 | 0.00779 | 0.005908 | 0.002639 | 0.010339 | 0.012277 | 0.027256 | 0.027264 | 0.046853 | 0.014638 | 0.010607 | 0.017523 | 0.021594 | 0.014216 | 0.032907 | 0.039862 |
| 24.29166667 | 0.001242 | 0.003974 | 0.002977 | 0.001906 | 0.000551 | 0.005962 | 0.003681 | 0.004321 | 0.00383 | 0.002594 | 0.012759 | 0.010027 | 0.024595 | 0.023508 | 0.038932 | 0.011966 | 0.007953 | 0.012702 | 0.013553 | 0.010162 | 0.023509 | 0.032133 |
| 24.30555556 | 0.003303 | 0.003753 | 0.003263 | 0.001065 | 0.000556 | 0.002301 | 0.002371 | 0.002052 | 0.00285 | 0.002404 | 0.008809 | 0.006882 | 0.021833 | 0.019812 | 0.030304 | 0.009085 | 0.005591 | 0.00814 | 0.006267 | 0.006549 | 0.01598 | 0.025166 |
| 24.31944444 | 0.005895 | 0.004104 | 0.00408 | 0.001873 | 0.000769 | 0.001593 | 0.001945 | 0.003075 | 0.002143 | 0.001326 | 0.003577 | 0.005301 | 0.01899 | 0.016259 | 0.021975 | 0.00572 | 0.003203 | 0.004025 | 0.003407 | 0.003252 | 0.010016 | 0.018132 |
| 24.33333333 | 0.005376 | 0.003508 | 0.00531 | 0.003057 | 0.001352 | 0.003673 | 0.004613 | 0.005859 | 0.001699 | 0.001086 | 0.005429 | 0.004317 | 0.016686 | 0.013401 | 0.014245 | 0.002717 | 0.001518 | 0.00194 | 0.003785 | 0.001056 | 0.005128 | 0.011191 |
| 24.34722222 | 0.00328 | 0.004691 | 0.004642 | 0.003479 | 0.002165 | 0.007659 | 0.009346 | 0.009123 | 0.003233 | 0.001866 | 0.010881 | 0.002691 | 0.014853 | 0.011676 | 0.007999 | 0.001789 | 0.001736 | 0.002905 | 0.003732 | 0.00058 | 0.002217 | 0.005318 |
| 24.36111111 | 0.004196 | 0.006484 | 0.004761 | 0.003405 | 0.003104 | 0.011252 | 0.011842 | 0.011647 | 0.006362 | 0.003541 | 0.011181 | 0.001746 | 0.013164 | 0.010117 | 0.00379 | 0.002452 | 0.003128 | 0.004636 | 0.003477 | 0.001243 | 0.003101 | 0.002442 |
| 24.375 | 0.006422 | 0.008321 | 0.007464 | 0.003692 | 0.004871 | 0.01253 | 0.012336 | 0.012125 | 0.009247 | 0.00489 | 0.008238 | 0.002004 | 0.011607 | 0.007861 | 0.002774 | 0.00279 | 0.004307 | 0.005262 | 0.004845 | 0.002485 | 0.00748 | 0.004112 |
| 24.38888889 | 0.009449 | 0.012946 | 0.008713 | 0.00504 | 0.007255 | 0.013629 | 0.016096 | 0.011707 | 0.011111 | 0.005705 | 0.011311 | 0.004104 | 0.00991 | 0.005466 | 0.00483 | 0.002738 | 0.004688 | 0.004998 | 0.006767 | 0.003533 | 0.011777 | 0.007029 |
| 24.40277778 | 0.014553 | 0.018063 | 0.008314 | 0.007038 | 0.008803 | 0.015955 | 0.022779 | 0.011916 | 0.012131 | 0.00727 | 0.019092 | 0.008038 | 0.007547 | 0.003279 | 0.00582 | 0.003136 | 0.004012 | 0.004373 | 0.006565 | 0.00321 | 0.011687 | 0.006862 |
| 24.41666667 | 0.019028 | 0.0199 | 0.010659 | 0.009072 | 0.009435 | 0.016938 | 0.024634 | 0.012409 | 0.013176 | 0.00937 | 0.021554 | 0.010833 | 0.005341 | 0.001742 | 0.00361 | 0.003495 | 0.002415 | 0.003441 | 0.003749 | 0.001844 | 0.007332 | 0.003796 |
| 24.43055556 | 0.0213 | 0.021849 | 0.01509 | 0.012141 | 0.011396 | 0.01558 | 0.020194 | 0.013507 | 0.01465 | 0.01097 | 0.018311 | 0.011989 | 0.00428 | 0.001617 | 0.001711 | 0.002598 | 0.001137 | 0.001968 | 0.001613 | 0.001266 | 0.0032 | 0.001205 |
| 24.44444444 | 0.024132 | 0.028931 | 0.018131 | 0.018345 | 0.016429 | 0.016096 | 0.018768 | 0.018385 | 0.016896 | 0.011956 | 0.020641 | 0.015417 | 0.003881 | 0.002624 | 0.002989 | 0.001535 | 0.001546 | 0.001093 | 0.002687 | 0.002015 | 0.001261 | 0.001027 |
| 24.45833333 | 0.030267 | 0.038595 | 0.023329 | 0.028422 | 0.023987 | 0.022052 | 0.027054 | 0.028747 | 0.022245 | 0.013578 | 0.033123 | 0.023522 | 0.004048 | 0.004612 | 0.005922 | 0.002452 | 0.003439 | 0.002397 | 0.005632 | 0.003271 | 0.001571 | 0.002818 |
| 24.47222222 | 0.039179 | 0.044206 | 0.036439 | 0.040263 | 0.032399 | 0.03065 | 0.039396 | 0.04072 | 0.032456 | 0.017066 | 0.047296 | 0.03385 | 0.005618 | 0.008181 | 0.009152 | 0.004697 | 0.006199 | 0.005462 | 0.009111 | 0.004996 | 0.005148 | 0.006377 |
| 24.48611111 | 0.049982 | 0.046877 | 0.052712 | 0.050575 | 0.04054 | 0.037937 | 0.048126 | 0.05023 | 0.045788 | 0.022638 | 0.057552 | 0.043442 | 0.008226 | 0.012645 | 0.012667 | 0.007233 | 0.009201 | 0.009052 | 0.012912 | 0.007408 | 0.010283 | 0.010615 |
| 24.5 | 0.064917 | 0.055511 | 0.062309 | 0.057385 | 0.04797 | 0.047063 | 0.059637 | 0.059417 | 0.061215 | 0.031202 | 0.069345 | 0.053599 | 0.010261 | 0.015872 | 0.015909 | 0.009975 | 0.011478 | 0.012248 | 0.016591 | 0.00987 | 0.013684 | 0.013181 |
| 24.51388889 | 0.085161 | 0.071528 | 0.067136 | 0.06202 | 0.053846 | 0.062514 | 0.088351 | 0.071002 | 0.079988 | 0.04207 | 0.085953 | 0.067888 | 0.011208 | 0.016886 | 0.018718 | 0.012123 | 0.012794 | 0.014656 | 0.019988 | 0.011743 | 0.014953 | 0.014001 |
| 24.52777778 | 0.102493 | 0.083971 | 0.075592 | 0.066269 | 0.056336 | 0.07613 | 0.125485 | 0.08061 | 0.100561 | 0.050733 | 0.099485 | 0.087043 | 0.012563 | 0.016758 | 0.023173 | 0.012641 | 0.01393 | 0.017109 | 0.023383 | 0.013891 | 0.017637 | 0.017065 |
| 24.54166667 | 0.103012 | 0.083809 | 0.083783 | 0.064979 | 0.050804 | 0.070491 | 0.125841 | 0.08035 | 0.115548 | 0.051206 | 0.099437 | 0.106163 | 0.016965 | 0.016368 | 0.034355 | 0.011311 | 0.016531 | 0.021688 | 0.02853 | 0.018647 | 0.026918 | 0.027378 |
| 24.55555556 | 0.085223 | 0.074262 | 0.079901 | 0.053697 | 0.035051 | 0.046847 | 0.075225 | 0.071245 | 0.110263 | 0.040269 | 0.082115 | 0.104959 | 0.025099 | 0.016483 | 0.050681 | 0.010464 | 0.022057 | 0.030926 | 0.036913 | 0.026466 | 0.040313 | 0.043094 |
| 24.56944444 | 0.066728 | 0.064235 | 0.067065 | 0.041384 | 0.019557 | 0.030926 | 0.029271 | 0.065715 | 0.08487 | 0.027649 | 0.051997 | 0.069833 | 0.032386 | 0.020134 | 0.06021 | 0.014071 | 0.029784 | 0.045013 | 0.04282 | 0.033081 | 0.048321 | 0.054466 |
| 24.58333333 | 0.062578 | 0.060206 | 0.060692 | 0.040008 | 0.017404 | 0.036215 | 0.031606 | 0.071458 | 0.069654 | 0.02867 | 0.021942 | 0.029168 | 0.034778 | 0.028018 | 0.05906 | 0.020722 | 0.035533 | 0.056692 | 0.040896 | 0.033807 | 0.048772 | 0.055848 |
| 24.59722222 | 0.071568 | 0.076495 | 0.06066 | 0.046201 | 0.023332 | 0.052619 | 0.071344 | 0.078302 | 0.080004 | 0.039312 | 0.016232 | 0.02301 | 0.033603 | 0.035101 | 0.055349 | 0.023659 | 0.033831 | 0.055527 | 0.033452 | 0.029403 | 0.046096 | 0.050914 |
| 24.61111111 | 0.080021 | 0.106867 | 0.052667 | 0.04445 | 0.024172 | 0.059069 | 0.103952 | 0.067978 | 0.089005 | 0.044169 | 0.028926 | 0.057197 | 0.029851 | 0.037563 | 0.058455 | 0.020814 | 0.025665 | 0.044097 | 0.024332 | 0.023833 | 0.044221 | 0.047635 |
| 24.625 | 0.074977 | 0.117249 | 0.034973 | 0.030435 | 0.020623 | 0.038958 | 0.082338 | 0.044361 | 0.076517 | 0.032848 | 0.030558 | 0.092447 | 0.02514 | 0.039777 | 0.075844 | 0.016301 | 0.019078 | 0.035987 | 0.018814 | 0.020194 | 0.045533 | 0.050237 |
| 24.63888889 | 0.057844 | 0.095492 | 0.019815 | 0.015018 | 0.014691 | 0.011381 | 0.034819 | 0.029046 | 0.056783 | 0.01195 | 0.023781 | 0.075668 | 0.024255 | 0.049772 | 0.09402 | 0.013645 | 0.018073 | 0.037012 | 0.018283 | 0.019326 | 0.045913 | 0.051633 |
| 24.65277778 | 0.036338 | 0.053913 | 0.01248 | 0.005413 | 0.006379 | 0.004286 | 0.015092 | 0.02691 | 0.039767 | 0.008307 | 0.014792 | 0.027988 | 0.025696 | 0.060671 | 0.087536 | 0.0135 | 0.019004 | 0.039752 | 0.018303 | 0.018971 | 0.039872 | 0.044463 |
| 24.66666667 | 0.024029 | 0.031312 | 0.009738 | 0.003292 | 0.002827 | 0.012849 | 0.017484 | 0.030982 | 0.029356 | 0.023652 | 0.015052 | 0.00741 | 0.024814 | 0.060969 | 0.062315 | 0.015076 | 0.01752 | 0.035699 | 0.015691 | 0.016738 | 0.030885 | 0.03346 |
| 24.68055556 | 0.023604 | 0.039111 | 0.011162 | 0.00733 | 0.003077 | 0.01659 | 0.025605 | 0.031448 | 0.0309 | 0.033789 | 0.026075 | 0.005635 | 0.02436 | 0.057497 | 0.041258 | 0.017904 | 0.013777 | 0.02708 | 0.01131 | 0.013017 | 0.024781 | 0.025636 |
| 24.69444444 | 0.018347 | 0.038419 | 0.018119 | 0.016104 | 0.002771 | 0.01102 | 0.023541 | 0.021885 | 0.030157 | 0.026729 | 0.022698 | 0.00788 | 0.026454 | 0.05663 | 0.030746 | 0.022694 | 0.010956 | 0.02066 | 0.007931 | 0.009992 | 0.022006 | 0.021553 |
| 24.70833333 | 0.007307 | 0.024069 | 0.022827 | 0.019839 | 0.003299 | 0.007772 | 0.014156 | 0.011021 | 0.019157 | 0.011645 | 0.00999 | 0.010678 | 0.028004 | 0.054038 | 0.027122 | 0.02825 | 0.010384 | 0.018403 | 0.006692 | 0.00888 | 0.020379 | 0.018803 |
| 24.72222222 | 0.003428 | 0.015127 | 0.016581 | 0.012188 | 0.00474 | 0.010791 | 0.01083 | 0.007228 | 0.012753 | 0.006184 | 0.006297 | 0.007875 | 0.02631 | 0.04831 | 0.027096 | 0.031426 | 0.010931 | 0.018569 | 0.007072 | 0.009148 | 0.018701 | 0.016421 |
| 24.73611111 | 0.008 | 0.011757 | 0.012845 | 0.011867 | 0.005692 | 0.01548 | 0.007735 | 0.006585 | 0.016661 | 0.010362 | 0.010965 | 0.007374 | 0.021605 | 0.042661 | 0.029909 | 0.031388 | 0.011391 | 0.019149 | 0.007932 | 0.010052 | 0.017655 | 0.015639 |
| 24.75 | 0.010157 | 0.010197 | 0.015716 | 0.015775 | 0.005757 | 0.013881 | 0.008821 | 0.004579 | 0.016979 | 0.015572 | 0.015583 | 0.009642 | 0.016266 | 0.040354 | 0.034552 | 0.028521 | 0.011359 | 0.01947 | 0.00831 | 0.010771 | 0.017947 | 0.017237 |
| 24.76388889 | 0.008338 | 0.010019 | 0.010959 | 0.011348 | 0.009018 | 0.006462 | 0.014383 | 0.004268 | 0.014966 | 0.015966 | 0.014568 | 0.01205 | 0.012421 | 0.040696 | 0.036688 | 0.022939 | 0.010965 | 0.019613 | 0.007725 | 0.010378 | 0.01883 | 0.019573 |
| 24.77777778 | 0.009407 | 0.011983 | 0.006723 | 0.010697 | 0.014472 | 0.002768 | 0.014751 | 0.004105 | 0.022286 | 0.014208 | 0.015926 | 0.01605 | 0.010497 | 0.038276 | 0.032589 | 0.016381 | 0.010399 | 0.019435 | 0.005918 | 0.008618 | 0.018073 | 0.019548 |
| 24.79166667 | 0.017021 | 0.026289 | 0.011486 | 0.010421 | 0.015343 | 0.003775 | 0.012777 | 0.004194 | 0.02774 | 0.01318 | 0.027438 | 0.016734 | 0.009721 | 0.030592 | 0.022842 | 0.011374 | 0.009909 | 0.018749 | 0.003672 | 0.006654 | 0.013495 | 0.015209 |
| 24.80555556 | 0.02535 | 0.036156 | 0.013076 | 0.005228 | 0.010215 | 0.005866 | 0.013757 | 0.009659 | 0.037636 | 0.013813 | 0.031409 | 0.017118 | 0.009615 | 0.020735 | 0.011096 | 0.009208 | 0.009657 | 0.017573 | 0.003144 | 0.006273 | 0.007139 | 0.008108 |
| 24.81944444 | 0.023338 | 0.031208 | 0.008093 | 0.004358 | 0.006737 | 0.011872 | 0.015932 | 0.014967 | 0.052893 | 0.0292 | 0.020273 | 0.029861 | 0.009923 | 0.011739 | 0.003357 | 0.009926 | 0.009138 | 0.015855 | 0.005231 | 0.007338 | 0.003685 | 0.002939 |
| 24.83333333 | 0.022369 | 0.037199 | 0.005684 | 0.009493 | 0.007835 | 0.018154 | 0.020006 | 0.019954 | 0.045874 | 0.039902 | 0.020462 | 0.035141 | 0.01042 | 0.006557 | 0.002151 | 0.012631 | 0.007639 | 0.013138 | 0.008608 | 0.00789 | 0.003613 | 0.002625 |
| 24.84722222 | 0.021992 | 0.041638 | 0.006143 | 0.011834 | 0.006267 | 0.018568 | 0.023891 | 0.02398 | 0.030844 | 0.023697 | 0.039249 | 0.023521 | 0.011223 | 0.008177 | 0.005666 | 0.015737 | 0.00553 | 0.009635 | 0.01152 | 0.007629 | 0.005853 | 0.005928 |
| 24.86111111 | 0.016632 | 0.033637 | 0.005273 | 0.009675 | 0.007532 | 0.014823 | 0.019171 | 0.025435 | 0.026233 | 0.008104 | 0.056603 | 0.017052 | 0.011955 | 0.015486 | 0.012895 | 0.017619 | 0.003952 | 0.006988 | 0.011315 | 0.007056 | 0.007957 | 0.008051 |
| 24.875 | 0.015153 | 0.027682 | 0.005376 | 0.012723 | 0.016049 | 0.008418 | 0.009409 | 0.027211 | 0.030546 | 0.007428 | 0.05126 | 0.021597 | 0.011363 | 0.022493 | 0.0183 | 0.016953 | 0.003529 | 0.006738 | 0.007438 | 0.005269 | 0.006472 | 0.006375 |
| 24.88888889 | 0.018205 | 0.033453 | 0.006342 | 0.013307 | 0.021405 | 0.005038 | 0.0094 | 0.02529 | 0.032492 | 0.008634 | 0.036054 | 0.026163 | 0.009103 | 0.021416 | 0.014766 | 0.013202 | 0.003583 | 0.007731 | 0.00605 | 0.003058 | 0.004244 | 0.004161 |
| 24.90277778 | 0.028042 | 0.039303 | 0.004478 | 0.005394 | 0.019891 | 0.011451 | 0.017137 | 0.023489 | 0.023836 | 0.012297 | 0.027723 | 0.028054 | 0.005915 | 0.014111 | 0.00898 | 0.007697 | 0.002683 | 0.006677 | 0.008255 | 0.003203 | 0.003377 | 0.004344 |
| 24.91666667 | 0.026814 | 0.028379 | 0.002514 | 0.003247 | 0.016325 | 0.020378 | 0.018068 | 0.021596 | 0.018709 | 0.020019 | 0.012611 | 0.030092 | 0.003361 | 0.014495 | 0.012189 | 0.003165 | 0.001455 | 0.004296 | 0.007368 | 0.003812 | 0.003108 | 0.008023 |
| 24.93055556 | 0.014327 | 0.019446 | 0.009019 | 0.012897 | 0.010709 | 0.021068 | 0.015572 | 0.027995 | 0.023347 | 0.018286 | 0.010441 | 0.031354 | 0.003841 | 0.024066 | 0.018084 | 0.001519 | 0.002267 | 0.006444 | 0.00549 | 0.002458 | 0.004594 | 0.009313 |
| 24.94444444 | 0.023293 | 0.031866 | 0.020642 | 0.02298 | 0.004071 | 0.022951 | 0.013876 | 0.044424 | 0.039292 | 0.007216 | 0.026787 | 0.03114 | 0.006764 | 0.030508 | 0.018427 | 0.002585 | 0.005614 | 0.014438 | 0.005453 | 0.002683 | 0.00957 | 0.008494 |
| 24.95833333 | 0.047063 | 0.051116 | 0.021159 | 0.018456 | 0.004429 | 0.026618 | 0.009848 | 0.040729 | 0.043531 | 0.008495 | 0.027579 | 0.027877 | 0.008664 | 0.03224 | 0.021107 | 0.007771 | 0.010127 | 0.022703 | 0.006851 | 0.006515 | 0.016652 | 0.01493 |
| 24.97222222 | 0.053081 | 0.041914 | 0.010936 | 0.006687 | 0.012163 | 0.02237 | 0.021922 | 0.020267 | 0.024512 | 0.025927 | 0.02804 | 0.027234 | 0.009528 | 0.036457 | 0.03343 | 0.018148 | 0.015273 | 0.028757 | 0.014777 | 0.014548 | 0.020226 | 0.02277 |
| 24.98611111 | 0.048797 | 0.022587 | 0.004628 | 0.002856 | 0.018657 | 0.021369 | 0.060604 | 0.022515 | 0.013025 | 0.037569 | 0.043444 | 0.027942 | 0.012868 | 0.046299 | 0.049055 | 0.026757 | 0.021202 | 0.035294 | 0.024848 | 0.02378 | 0.020302 | 0.025147 |
| 25 | 0.057172 | 0.034959 | 0.013293 | 0.011348 | 0.021807 | 0.033109 | 0.091816 | 0.054396 | 0.024733 | 0.037818 | 0.041657 | 0.040333 | 0.019702 | 0.056551 | 0.056855 | 0.028344 | 0.026348 | 0.043453 | 0.02592 | 0.026458 | 0.020454 | 0.024773 |
| 25.01388889 | 0.062651 | 0.058335 | 0.041547 | 0.032665 | 0.030394 | 0.047384 | 0.081421 | 0.085631 | 0.04565 | 0.044009 | 0.024964 | 0.058435 | 0.026153 | 0.059566 | 0.053785 | 0.028228 | 0.027889 | 0.04825 | 0.017516 | 0.022227 | 0.022236 | 0.024562 |
| 25.02777778 | 0.052317 | 0.060449 | 0.073328 | 0.053297 | 0.043888 | 0.047926 | 0.045635 | 0.093101 | 0.062078 | 0.060283 | 0.03755 | 0.051554 | 0.026614 | 0.053664 | 0.04692 | 0.030306 | 0.023287 | 0.041024 | 0.007867 | 0.01571 | 0.026208 | 0.026262 |
| 25.04166667 | 0.047555 | 0.050144 | 0.080051 | 0.062002 | 0.052762 | 0.036711 | 0.02465 | 0.080687 | 0.073218 | 0.073484 | 0.081855 | 0.035627 | 0.019366 | 0.039496 | 0.039926 | 0.028229 | 0.013787 | 0.024337 | 0.005475 | 0.009154 | 0.028137 | 0.026802 |
| 25.05555556 | 0.064211 | 0.051482 | 0.059881 | 0.060771 | 0.054662 | 0.033012 | 0.038357 | 0.064825 | 0.075947 | 0.071464 | 0.094353 | 0.032157 | 0.010747 | 0.023601 | 0.033491 | 0.018073 | 0.005455 | 0.012648 | 0.008276 | 0.005119 | 0.025378 | 0.025181 |
| 25.06944444 | 0.088638 | 0.058526 | 0.037231 | 0.051972 | 0.047247 | 0.041259 | 0.073503 | 0.053972 | 0.064411 | 0.053611 | 0.053721 | 0.032652 | 0.009109 | 0.019119 | 0.031269 | 0.01024 | 0.005161 | 0.011393 | 0.008807 | 0.005374 | 0.02293 | 0.02549 |
| 25.08333333 | 0.086896 | 0.051217 | 0.028505 | 0.040405 | 0.032394 | 0.049803 | 0.102313 | 0.044891 | 0.044975 | 0.031442 | 0.015081 | 0.030985 | 0.01229 | 0.025478 | 0.029982 | 0.010444 | 0.011568 | 0.016777 | 0.010803 | 0.010286 | 0.022334 | 0.025895 |

| | | | | | | | | | | | | | | | | | | | | | | |
|---|---|---|---|---|---|---|---|---|---|---|---|---|---|---|---|---|---|---|---|---|---|---|
| 25.09722222 | 0.056581 | 0.037134 | 0.031388 | 0.033199 | 0.019511 | 0.053726 | 0.11373 | 0.036286 | 0.034259 | 0.018946 | 0.016085 | 0.031212 | 0.012605 | 0.028629 | 0.025245 | 0.009779 | 0.014059 | 0.022461 | 0.017304 | 0.017379 | 0.018765 | 0.021226 |
| 25.11111111 | 0.042193 | 0.040019 | 0.037418 | 0.036556 | 0.018185 | 0.059184 | 0.116598 | 0.035165 | 0.040968 | 0.02759 | 0.047032 | 0.037667 | 0.010564 | 0.026004 | 0.019362 | 0.00519 | 0.010745 | 0.022452 | 0.021488 | 0.020808 | 0.012837 | 0.01397 |
| 25.125 | 0.059058 | 0.058614 | 0.044813 | 0.048189 | 0.030491 | 0.067725 | 0.11416 | 0.04467 | 0.057825 | 0.052274 | 0.076031 | 0.045792 | 0.011051 | 0.025839 | 0.016796 | 0.002722 | 0.009245 | 0.019836 | 0.021899 | 0.019877 | 0.010231 | 0.010633 |
| 25.13888889 | 0.077633 | 0.066891 | 0.054231 | 0.054363 | 0.043312 | 0.066993 | 0.096438 | 0.055312 | 0.066851 | 0.066576 | 0.078917 | 0.047079 | 0.01415 | 0.029503 | 0.018714 | 0.004943 | 0.011942 | 0.01926 | 0.021613 | 0.018379 | 0.012495 | 0.01256 |
| 25.15277778 | 0.073451 | 0.056758 | 0.061106 | 0.047642 | 0.042938 | 0.052 | 0.066154 | 0.056113 | 0.058847 | 0.055935 | 0.061935 | 0.040277 | 0.016708 | 0.031905 | 0.021054 | 0.008948 | 0.015755 | 0.019852 | 0.021327 | 0.017702 | 0.016438 | 0.016159 |
| 25.16666667 | 0.049815 | 0.041268 | 0.058584 | 0.035927 | 0.032908 | 0.035968 | 0.039863 | 0.047753 | 0.043976 | 0.035184 | 0.04672 | 0.031907 | 0.017634 | 0.031223 | 0.020494 | 0.011 | 0.017435 | 0.019779 | 0.020617 | 0.017034 | 0.019464 | 0.018713 |
| 25.18055556 | 0.027602 | 0.026841 | 0.047231 | 0.028648 | 0.025135 | 0.029267 | 0.02641 | 0.036761 | 0.034008 | 0.022257 | 0.04051 | 0.026506 | 0.018058 | 0.030204 | 0.018351 | 0.010533 | 0.016099 | 0.018765 | 0.018315 | 0.015372 | 0.021379 | 0.019847 |
| 25.19444444 | 0.021312 | 0.014656 | 0.036261 | 0.026463 | 0.023204 | 0.030007 | 0.024794 | 0.027671 | 0.029004 | 0.018327 | 0.039045 | 0.023656 | 0.019034 | 0.03131 | 0.018172 | 0.010135 | 0.013324 | 0.017883 | 0.015138 | 0.01305 | 0.02243 | 0.020357 |
| 25.20833333 | 0.028924 | 0.011851 | 0.030505 | 0.02443 | 0.022014 | 0.031974 | 0.028866 | 0.023215 | 0.024436 | 0.016483 | 0.037775 | 0.021622 | 0.0209 | 0.034174 | 0.021571 | 0.012458 | 0.011192 | 0.017868 | 0.013391 | 0.01143 | 0.022687 | 0.021135 |
| 25.22222222 | 0.03555 | 0.019055 | 0.026244 | 0.019619 | 0.017915 | 0.031108 | 0.031465 | 0.023044 | 0.020093 | 0.013692 | 0.03374 | 0.019931 | 0.023599 | 0.037166 | 0.026224 | 0.017313 | 0.010784 | 0.018805 | 0.013699 | 0.011165 | 0.022646 | 0.021942 |
| 25.23611111 | 0.030891 | 0.024333 | 0.020136 | 0.014674 | 0.013987 | 0.028241 | 0.030218 | 0.023515 | 0.017891 | 0.0114 | 0.027912 | 0.019094 | 0.026565 | 0.038465 | 0.028291 | 0.02134 | 0.011847 | 0.020373 | 0.014913 | 0.011923 | 0.022699 | 0.022134 |
| 25.25 | 0.020055 | 0.020186 | 0.014741 | 0.013646 | 0.013486 | 0.025321 | 0.025836 | 0.021699 | 0.016239 | 0.010419 | 0.023617 | 0.018771 | 0.027755 | 0.036376 | 0.026132 | 0.021648 | 0.012901 | 0.021184 | 0.015234 | 0.012651 | 0.021924 | 0.021153 |
| 25.26388889 | 0.012949 | 0.013695 | 0.014991 | 0.015651 | 0.013507 | 0.022554 | 0.019896 | 0.018417 | 0.013295 | 0.010169 | 0.022406 | 0.018131 | 0.025482 | 0.030818 | 0.021467 | 0.019186 | 0.012564 | 0.019795 | 0.013469 | 0.011933 | 0.019182 | 0.018613 |
| 25.27777778 | 0.01255 | 0.013297 | 0.020034 | 0.015942 | 0.010858 | 0.020284 | 0.015522 | 0.016706 | 0.011006 | 0.010408 | 0.023116 | 0.017469 | 0.021154 | 0.023318 | 0.01726 | 0.016718 | 0.011 | 0.01645 | 0.010359 | 0.009508 | 0.014854 | 0.015346 |
| 25.29166667 | 0.01373 | 0.015019 | 0.022872 | 0.013566 | 0.008279 | 0.018973 | 0.014405 | 0.017007 | 0.011189 | 0.010639 | 0.022833 | 0.017087 | 0.017638 | 0.016481 | 0.014363 | 0.014785 | 0.009308 | 0.012724 | 0.007477 | 0.006709 | 0.010226 | 0.012017 |
| 25.30555556 | 0.011503 | 0.012442 | 0.019774 | 0.011917 | 0.008517 | 0.017064 | 0.013925 | 0.015711 | 0.012318 | 0.010089 | 0.01858 | 0.015699 | 0.015703 | 0.011972 | 0.011285 | 0.012367 | 0.008246 | 0.010209 | 0.005369 | 0.005097 | 0.006258 | 0.008514 |
| 25.31944444 | 0.007397 | 0.007934 | 0.014111 | 0.011042 | 0.008401 | 0.012379 | 0.009961 | 0.010717 | 0.011863 | 0.008803 | 0.011254 | 0.012175 | 0.013458 | 0.008616 | 0.007153 | 0.010028 | 0.007776 | 0.00914 | 0.003447 | 0.004862 | 0.003545 | 0.005862 |
| 25.33333333 | 0.004943 | 0.004556 | 0.009637 | 0.007287 | 0.005361 | 0.005998 | 0.005044 | 0.004976 | 0.009111 | 0.006946 | 0.00523 | 0.007784 | 0.00954 | 0.005203 | 0.003462 | 0.008275 | 0.00675 | 0.0077 | 0.001912 | 0.004315 | 0.002224 | 0.005167 |
| 25.34722222 | 0.004339 | 0.002963 | 0.005996 | 0.002944 | 0.002363 | 0.001783 | 0.003637 | 0.001653 | 0.005861 | 0.004588 | 0.002768 | 0.004484 | 0.005396 | 0.00385 | 0.00218 | 0.00565 | 0.004467 | 0.004443 | 0.001538 | 0.002564 | 0.001455 | 0.005488 |
| 25.36111111 | 0.004842 | 0.002802 | 0.002954 | 0.001868 | 0.001801 | 0.001286 | 0.003393 | 0.00059 | 0.00407 | 0.002807 | 0.00386 | 0.002636 | 0.002841 | 0.005135 | 0.002594 | 0.002771 | 0.002034 | 0.0018 | 0.001498 | 0.001072 | 0.001258 | 0.004691 |
| 25.375 | 0.005898 | 0.002546 | 0.001424 | 0.001817 | 0.002244 | 0.001784 | 0.002738 | 0.00101 | 0.003243 | 0.002516 | 0.006865 | 0.001803 | 0.002621 | 0.006639 | 0.003675 | 0.002 | 0.001096 | 0.002509 | 0.001891 | 0.000921 | 0.002948 | 0.003 |
| 25.38888889 | 0.005574 | 0.002458 | 0.000873 | 0.002116 | 0.002376 | 0.002148 | 0.00338 | 0.001783 | 0.002065 | 0.00245 | 0.008861 | 0.001409 | 0.006565 | 0.009723 | 0.006949 | 0.003016 | 0.00252 | 0.005507 | 0.00496 | 0.002938 | 0.005987 | 0.002897 |
| 25.40277778 | 0.006412 | 0.003542 | 0.00176 | 0.003868 | 0.002422 | 0.004385 | 0.007323 | 0.001838 | 0.001428 | 0.001682 | 0.009784 | 0.00173 | 0.014147 | 0.016275 | 0.012621 | 0.005773 | 0.005798 | 0.009532 | 0.010636 | 0.007518 | 0.008816 | 0.004183 |
| 25.41666667 | 0.010781 | 0.00677 | 0.006372 | 0.00629 | 0.001963 | 0.008119 | 0.014338 | 0.002732 | 0.002931 | 0.001873 | 0.011324 | 0.003337 | 0.020241 | 0.023579 | 0.018197 | 0.01128 | 0.010029 | 0.014685 | 0.015649 | 0.012076 | 0.012031 | 0.006392 |
| 25.43055556 | 0.013663 | 0.011063 | 0.013894 | 0.009153 | 0.002529 | 0.012429 | 0.022268 | 0.00512 | 0.006452 | 0.004089 | 0.013655 | 0.00564 | 0.022447 | 0.028389 | 0.022996 | 0.017969 | 0.015052 | 0.020979 | 0.018452 | 0.014988 | 0.017323 | 0.012569 |
| 25.44444444 | 0.015028 | 0.017415 | 0.018743 | 0.014237 | 0.00733 | 0.017538 | 0.031068 | 0.007631 | 0.010769 | 0.007276 | 0.016003 | 0.008434 | 0.025074 | 0.033181 | 0.030723 | 0.023472 | 0.020476 | 0.027738 | 0.021989 | 0.019149 | 0.024705 | 0.023822 |
| 25.45833333 | 0.021072 | 0.027342 | 0.020856 | 0.021774 | 0.014394 | 0.022835 | 0.040963 | 0.011803 | 0.015412 | 0.011055 | 0.019416 | 0.012122 | 0.033446 | 0.043559 | 0.044344 | 0.029038 | 0.025264 | 0.034942 | 0.028623 | 0.027612 | 0.032652 | 0.036026 |
| 25.47222222 | 0.033485 | 0.037261 | 0.028327 | 0.030275 | 0.019723 | 0.02877 | 0.051255 | 0.020758 | 0.023235 | 0.01724 | 0.026839 | 0.018513 | 0.045502 | 0.059337 | 0.060355 | 0.039949 | 0.030321 | 0.045017 | 0.03567 | 0.037688 | 0.041693 | 0.045953 |
| 25.48611111 | 0.049383 | 0.045855 | 0.043065 | 0.040254 | 0.026245 | 0.036778 | 0.061255 | 0.033812 | 0.038291 | 0.028224 | 0.039666 | 0.029812 | 0.054555 | 0.074695 | 0.07171 | 0.056465 | 0.038847 | 0.060213 | 0.040446 | 0.043413 | 0.053846 | 0.056332 |
| 25.5 | 0.067081 | 0.057974 | 0.05662 | 0.052285 | 0.039245 | 0.046783 | 0.071792 | 0.048367 | 0.05962 | 0.044379 | 0.056783 | 0.044537 | 0.057379 | 0.084415 | 0.0765 | 0.066825 | 0.049604 | 0.076995 | 0.043712 | 0.044924 | 0.064803 | 0.068541 |
| 25.51388889 | 0.087035 | 0.075536 | 0.065694 | 0.064953 | 0.054305 | 0.057907 | 0.087577 | 0.065253 | 0.082061 | 0.064201 | 0.075248 | 0.059057 | 0.056372 | 0.097046 | 0.081144 | 0.064406 | 0.055588 | 0.087493 | 0.047103 | 0.048114 | 0.066408 | 0.076407 |
| 25.52777778 | 0.104982 | 0.09128 | 0.07801 | 0.07519 | 0.060713 | 0.069388 | 0.111293 | 0.085404 | 0.101522 | 0.08167 | 0.090883 | 0.068478 | 0.055791 | 0.115744 | 0.089612 | 0.061989 | 0.052837 | 0.086316 | 0.047768 | 0.051135 | 0.058677 | 0.074105 |
| 25.54166667 | 0.110627 | 0.096336 | 0.093137 | 0.077188 | 0.054746 | 0.074155 | 0.124399 | 0.099051 | 0.113984 | 0.084147 | 0.098486 | 0.070387 | 0.051664 | 0.10933 | 0.086643 | 0.062514 | 0.043186 | 0.071256 | 0.03583 | 0.040824 | 0.046948 | 0.062813 |
| 25.55555556 | 0.100508 | 0.089357 | 0.097125 | 0.067423 | 0.043646 | 0.064596 | 0.102816 | 0.095891 | 0.115768 | 0.067999 | 0.098178 | 0.072462 | 0.040024 | 0.067749 | 0.055021 | 0.047891 | 0.033012 | 0.048144 | 0.017503 | 0.022261 | 0.032018 | 0.046396 |
| 25.56944444 | 0.087508 | 0.079898 | 0.086494 | 0.055462 | 0.038342 | 0.050936 | 0.061882 | 0.087643 | 0.10922 | 0.050894 | 0.100626 | 0.080561 | 0.031854 | 0.034437 | 0.022926 | 0.02299 | 0.029393 | 0.034065 | 0.010054 | 0.01541 | 0.021027 | 0.033732 |
| 25.58333333 | 0.080749 | 0.075075 | 0.073009 | 0.05358 | 0.040072 | 0.047571 | 0.036949 | 0.087905 | 0.100055 | 0.056379 | 0.113548 | 0.084033 | 0.037019 | 0.034729 | 0.015113 | 0.011249 | 0.031847 | 0.036069 | 0.008492 | 0.012796 | 0.027286 | 0.036082 |
| 25.59722222 | 0.071478 | 0.07509 | 0.066206 | 0.058465 | 0.04093 | 0.051628 | 0.04069 | 0.089444 | 0.092263 | 0.080468 | 0.125835 | 0.074288 | 0.046412 | 0.049014 | 0.022745 | 0.009979 | 0.034142 | 0.042755 | 0.005792 | 0.010395 | 0.045101 | 0.047741 |
| 25.61111111 | 0.064602 | 0.091738 | 0.069897 | 0.06459 | 0.03718 | 0.057792 | 0.065074 | 0.079791 | 0.085476 | 0.09546 | 0.113609 | 0.063003 | 0.043836 | 0.056172 | 0.042326 | 0.017871 | 0.03152 | 0.045127 | 0.00385 | 0.021151 | 0.057165 | 0.054453 |
| 25.625 | 0.072483 | 0.125825 | 0.074371 | 0.067048 | 0.032132 | 0.066393 | 0.089737 | 0.054793 | 0.072926 | 0.079407 | 0.074145 | 0.058363 | 0.02678 | 0.046478 | 0.057448 | 0.048059 | 0.025572 | 0.046248 | 0.001463 | 0.031067 | 0.053033 | 0.054363 |
| 25.63888889 | 0.078045 | 0.138953 | 0.06061 | 0.050416 | 0.02692 | 0.058919 | 0.080481 | 0.031708 | 0.061569 | 0.039666 | 0.033339 | 0.042461 | 0.012389 | 0.029762 | 0.055597 | 0.074761 | 0.01763 | 0.037645 | 0.001837 | 0.022901 | 0.03374 | 0.048321 |
| 25.65277778 | 0.06101 | 0.097654 | 0.033513 | 0.021131 | 0.018606 | 0.031163 | 0.042312 | 0.025223 | 0.051601 | 0.010548 | 0.010208 | 0.026782 | 0.0095 | 0.01818 | 0.039177 | 0.057502 | 0.008499 | 0.018902 | 0.005034 | 0.00976 | 0.014357 | 0.034839 |
| 25.66666667 | 0.031596 | 0.037597 | 0.015809 | 0.007409 | 0.009876 | 0.011762 | 0.016873 | 0.022554 | 0.027866 | 0.007462 | 0.006572 | 0.045158 | 0.015503 | 0.020383 | 0.02153 | 0.0299 | 0.004239 | 0.009697 | 0.007808 | 0.006973 | 0.006666 | 0.021514 |
| 25.68055556 | 0.014282 | 0.016655 | 0.015617 | 0.01697 | 0.006845 | 0.008286 | 0.015201 | 0.017499 | 0.011925 | 0.021961 | 0.01613 | 0.057884 | 0.028271 | 0.043388 | 0.01983 | 0.031419 | 0.004016 | 0.006602 | 0.011498 | 0.007118 | 0.00549 | 0.013614 |
| 25.69444444 | 0.012957 | 0.028559 | 0.023187 | 0.031958 | 0.007529 | 0.018481 | 0.036309 | 0.01215 | 0.016518 | 0.030862 | 0.019805 | 0.040202 | 0.029764 | 0.048351 | 0.029338 | 0.029893 | 0.003462 | 0.003991 | 0.011968 | 0.004355 | 0.006039 | 0.01036 |
| 25.70833333 | 0.012693 | 0.029722 | 0.02376 | 0.029392 | 0.007244 | 0.026494 | 0.044708 | 0.011702 | 0.0191 | 0.0278 | 0.014148 | 0.026186 | 0.01839 | 0.026684 | 0.030385 | 0.015572 | 0.004281 | 0.006491 | 0.010622 | 0.003869 | 0.007305 | 0.011835 |
| 25.72222222 | 0.00879 | 0.016682 | 0.013856 | 0.014029 | 0.005491 | 0.023107 | 0.034292 | 0.016671 | 0.018174 | 0.030916 | 0.015209 | 0.029095 | 0.012652 | 0.020958 | 0.018955 | 0.008827 | 0.006896 | 0.011226 | 0.01002 | 0.00482 | 0.008 | 0.009964 |
| 25.73611111 | 0.006335 | 0.019225 | 0.00637 | 0.007943 | 0.003843 | 0.026923 | 0.043048 | 0.016699 | 0.02931 | 0.028121 | 0.01454 | 0.052227 | 0.010163 | 0.02101 | 0.012887 | 0.010732 | 0.007003 | 0.009976 | 0.00885 | 0.003703 | 0.013498 | 0.003764 |
| 25.75 | 0.006037 | 0.023856 | 0.006138 | 0.008403 | 0.004728 | 0.025093 | 0.038475 | 0.01771 | 0.042442 | 0.021888 | 0.00935 | 0.064031 | 0.010701 | 0.019507 | 0.014742 | 0.018381 | 0.005994 | 0.005337 | 0.014696 | 0.001284 | 0.016947 | 0.001731 |
| 25.76388889 | 0.005452 | 0.018976 | 0.004287 | 0.007025 | 0.004882 | 0.017524 | 0.018496 | 0.019448 | 0.032174 | 0.030157 | 0.012274 | 0.045592 | 0.020895 | 0.027399 | 0.020643 | 0.026301 | 0.007534 | 0.005723 | 0.016854 | 0.000197 | 0.013041 | 0.002439 |
| 25.77777778 | 0.00637 | 0.017512 | 0.002341 | 0.007425 | 0.002264 | 0.019081 | 0.019961 | 0.019102 | 0.021442 | 0.034181 | 0.024519 | 0.03128 | 0.025806 | 0.023677 | 0.026777 | 0.026069 | 0.007173 | 0.005924 | 0.012174 | 0.003255 | 0.013752 | 0.001885 |
| 25.79166667 | 0.009231 | 0.011129 | 0.004665 | 0.006398 | 0.00106 | 0.019157 | 0.0282 | 0.025907 | 0.033889 | 0.029403 | 0.03668 | 0.040935 | 0.016501 | 0.0114 | 0.02104 | 0.019428 | 0.003787 | 0.003881 | 0.013901 | 0.008898 | 0.019988 | 0.004556 |
| 25.80555556 | 0.012718 | 0.010051 | 0.007118 | 0.007774 | 0.002959 | 0.017881 | 0.030287 | 0.026388 | 0.034646 | 0.02187 | 0.03514 | 0.053547 | 0.009744 | 0.004116 | 0.014646 | 0.011712 | 0.003888 | 0.006383 | 0.012023 | 0.008698 | 0.021011 | 0.014797 |
| 25.81944444 | 0.017829 | 0.024481 | 0.00724 | 0.01509 | 0.009079 | 0.017063 | 0.025875 | 0.018731 | 0.025012 | 0.017783 | 0.026904 | 0.043601 | 0.010494 | 0.004006 | 0.025097 | 0.005766 | 0.007962 | 0.009943 | 0.009208 | 0.007418 | 0.014093 | 0.022552 |
| 25.83333333 | 0.027082 | 0.042471 | 0.007835 | 0.015163 | 0.016307 | 0.017396 | 0.022393 | 0.018957 | 0.032778 | 0.028825 | 0.031922 | 0.023618 | 0.009129 | 0.007711 | 0.031869 | 0.004787 | 0.009509 | 0.011552 | 0.013473 | 0.012095 | 0.008577 | 0.017262 |
| 25.84722222 | 0.030321 | 0.042024 | 0.008206 | 0.007135 | 0.015975 | 0.015651 | 0.032188 | 0.019829 | 0.03016 | 0.035215 | 0.037319 | 0.00956 | 0.010168 | 0.008195 | 0.017004 | 0.004789 | 0.008971 | 0.012253 | 0.011517 | 0.012968 | 0.007728 | 0.010967 |
| 25.86111111 | 0.024273 | 0.035604 | 0.007986 | 0.003582 | 0.00887 | 0.009593 | 0.032567 | 0.01664 | 0.013377 | 0.027282 | 0.022022 | 0.002601 | 0.013506 | 0.009223 | 0.005882 | 0.005063 | 0.008198 | 0.013228 | 0.00443 | 0.007684 | 0.007698 | 0.010084 |
| 25.875 | 0.024118 | 0.045427 | 0.01234 | 0.009413 | 0.003237 | 0.007077 | 0.020237 | 0.016779 | 0.0073 | 0.024572 | 0.010245 | 0.007485 | 0.010863 | 0.017025 | 0.01501 | 0.007495 | 0.005788 | 0.01739 | 0.004887 | 0.006846 | 0.011499 | 0.011257 |
| 25.88888889 | 0.024914 | 0.050695 | 0.017236 | 0.018366 | 0.006314 | 0.0064 | 0.024049 | 0.022663 | 0.004501 | 0.024198 | 0.011806 | 0.023108 | 0.005519 | 0.026213 | 0.020518 | 0.009306 | 0.003185 | 0.012993 | 0.012491 | 0.012502 | 0.019274 | 0.027198 |
| 25.90277778 | 0.02715 | 0.059466 | 0.015009 | 0.018391 | 0.018019 | 0.008447 | 0.029227 | 0.021609 | 0.001383 | 0.021461 | 0.023719 | 0.028956 | 0.006794 | 0.030308 | 0.015577 | 0.015722 | 0.005251 | 0.01176 | 0.018938 | 0.012895 | 0.022582 | 0.04432 |
| 25.91666667 | 0.02715 | 0.055819 | 0.012012 | 0.012756 | 0.022819 | 0.011855 | 0.017692 | 0.012981 | 0.002556 | 0.029286 | 0.045243 | 0.019021 | 0.017152 | 0.027266 | 0.021087 | 0.025088 | 0.013142 | 0.031877 | 0.017414 | 0.009517 | 0.014722 | 0.034038 |
| 25.93055556 | 0.014807 | 0.029649 | 0.012143 | 0.009902 | 0.014908 | 0.009601 | 0.007143 | 0.009018 | 0.007558 | 0.033681 | 0.040678 | 0.012652 | 0.026472 | 0.025948 | 0.026237 | 0.026669 | 0.02057 | 0.05382 | 0.009909 | 0.008677 | 0.004327 | 0.013615 |
| 25.94444444 | 0.009199 | 0.021543 | 0.012789 | 0.008278 | 0.008659 | 0.005515 | 0.010895 | 0.013464 | 0.012107 | 0.021637 | 0.021242 | 0.014232 | 0.030306 | 0.053981 | 0.031209 | 0.026477 | 0.023287 | 0.057152 | 0.00538 | 0.008194 | 0.005819 | 0.008774 |
| 25.95833333 | 0.015174 | 0.031759 | 0.018848 | 0.00813 | 0.013798 | 0.011647 | 0.028705 | 0.028755 | 0.01316 | 0.01348 | 0.024306 | 0.015692 | 0.034151 | 0.088784 | 0.059241 | 0.035614 | 0.018482 | 0.04351 | 0.008068 | 0.012834 | 0.022873 | 0.023071 |
| 25.97222222 | 0.017965 | 0.041288 | 0.030298 | 0.005686 | 0.017376 | 0.031462 | 0.057359 | 0.041704 | 0.024805 | 0.036636 | 0.055102 | 0.010846 | 0.038416 | 0.095568 | 0.102482 | 0.053462 | 0.010957 | 0.030138 | 0.019556 | 0.022086 | 0.044221 | 0.048587 |
| 25.98611111 | 0.027756 | 0.040022 | 0.033539 | 0.005767 | 0.017113 | 0.044204 | 0.08055 | 0.045163 | 0.055927 | 0.080937 | 0.079897 | 0.008151 | 0.045887 | 0.08933 | 0.130926 | 0.067798 | 0.016031 | 0.035979 | 0.035393 | 0.031516 | 0.055125 | 0.065981 |
| 26 | 0.056706 | 0.0441 | 0.021286 | 0.017072 | 0.030548 | 0.038323 | 0.082694 | 0.049694 | 0.083925 | 0.096639 | 0.078582 | 0.021277 | 0.057794 | 0.083307 | 0.115443 | 0.068458 | 0.03644 | 0.068154 | 0.04472 | 0.038576 | 0.055891 | 0.070275 |
| 26.01388889 | 0.073683 | 0.058677 | 0.022009 | 0.03662 | 0.04311 | 0.035006 | 0.064399 | 0.063939 | 0.082403 | 0.078243 | 0.101534 | 0.044634 | 0.063731 | 0.079918 | 0.06954 | 0.058098 | 0.055613 | 0.102537 | 0.04461 | 0.04003 | 0.054317 | 0.07251 |
| 26.02777778 | 0.056683 | 0.061397 | 0.059729 | 0.058329 | 0.038721 | 0.045203 | 0.046105 | 0.082641 | 0.057591 | 0.065591 | 0.119068 | 0.059167 | 0.065844 | 0.086003 | 0.040599 | 0.045008 | 0.059598 | 0.108114 | 0.040636 | 0.038132 | 0.05478 | 0.080258 |
| 26.04166667 | 0.031876 | 0.045907 | 0.089782 | 0.061865 | 0.024943 | 0.046997 | 0.054274 | 0.080177 | 0.036038 | 0.06751 | 0.080904 | 0.053445 | 0.065904 | 0.091856 | 0.04524 | 0.035303 | 0.04977 | 0.08603 | 0.034817 | 0.035188 | 0.050613 | 0.083754 |
| 26.05555556 | 0.016889 | 0.023548 | 0.063762 | 0.036614 | 0.012121 | 0.029149 | 0.071638 | 0.046169 | 0.026682 | 0.052681 | 0.032559 | 0.033131 | 0.05307 | 0.080083 | 0.056553 | 0.02785 | 0.035839 | 0.051609 | 0.024717 | 0.0268 | 0.035586 | 0.065594 |
| 26.06944444 | 0.007289 | 0.016204 | 0.026272 | 0.013229 | 0.009624 | 0.012682 | 0.067378 | 0.015356 | 0.020566 | 0.025453 | 0.007488 | 0.016708 | 0.037621 | 0.056938 | 0.054254 | 0.02143 | 0.022881 | 0.021702 | 0.013826 | 0.013913 | 0.018819 | 0.032582 |
| 26.08333333 | 0.00318 | 0.024158 | 0.021502 | 0.008551 | 0.012988 | 0.008172 | 0.049777 | 0.01004 | 0.0134 | 0.015996 | 0.001954 | 0.015287 | 0.028628 | 0.035165 | 0.043902 | 0.01683 | 0.011311 | 0.007922 | 0.007787 | 0.006564 | 0.007582 | 0.009316 |
| 26.09722222 | 0.01369 | 0.025504 | 0.021298 | 0.011118 | 0.011283 | 0.01188 | 0.040949 | 0.011612 | 0.011372 | 0.016691 | 0.008317 | 0.014291 | 0.02116 | 0.0221 | 0.032074 | 0.012113 | 0.00414 | 0.005334 | 0.005993 | 0.007549 | 0.001725 | 0.003415 |
| 26.11111111 | 0.042718 | 0.032365 | 0.022544 | 0.023511 | 0.010616 | 0.024793 | 0.050877 | 0.018933 | 0.021108 | 0.026924 | 0.022458 | 0.012938 | 0.017264 | 0.018998 | 0.020683 | 0.006755 | 0.002006 | 0.006004 | 0.005913 | 0.010328 | 0.001528 | 0.003446 |
| 26.125 | 0.067394 | 0.055702 | 0.041311 | 0.040831 | 0.022333 | 0.038254 | 0.068653 | 0.038383 | 0.037743 | 0.050324 | 0.037239 | 0.023495 | 0.015164 | 0.018073 | 0.012475 | 0.00301 | 0.002327 | 0.008093 | 0.007065 | 0.010867 | 0.005419 | 0.005889 |
| 26.13888889 | 0.06529 | 0.060639 | 0.061509 | 0.048564 | 0.034561 | 0.043528 | 0.073032 | 0.052843 | 0.050004 | 0.064234 | 0.044885 | 0.032641 | 0.011812 | 0.015492 | 0.010643 | 0.002432 | 0.004754 | 0.010766 | 0.008936 | 0.010471 | 0.0096 | 0.011044 |
| 26.15277778 | 0.047232 | 0.041687 | 0.067507 | 0.047444 | 0.03399 | 0.040367 | 0.059389 | 0.048995 | 0.051984 | 0.057151 | 0.044208 | 0.031469 | 0.009511 | 0.013612 | 0.013808 | 0.004491 | 0.009844 | 0.012502 | 0.010884 | 0.010443 | 0.012529 | 0.015173 |
| 26.16666667 | 0.032706 | 0.026941 | 0.0607 | 0.042915 | 0.026439 | 0.030793 | 0.039863 | 0.03287 | 0.044817 | 0.04042 | 0.037909 | 0.026639 | 0.009833 | 0.014015 | 0.016411 | 0.006852 | 0.014533 | 0.013068 | 0.01252 | 0.010536 | 0.015405 | 0.018055 |
| 26.18055556 | 0.023611 | 0.02025 | 0.048004 | 0.03404 | 0.019766 | 0.020942 | 0.025744 | 0.016852 | 0.033219 | 0.026224 | 0.031016 | 0.019748 | 0.011039 | 0.014904 | 0.01492 | 0.007336 | 0.015189 | 0.013404 | 0.013071 | 0.010178 | 0.017953 | 0.019976 |
| 26.19444444 | 0.014009 | 0.0148 | 0.034175 | 0.019894 | 0.01256 | 0.015035 | 0.018984 | 0.006849 | 0.022655 | 0.016186 | 0.026007 | 0.010995 | 0.01041 | 0.013383 | 0.010706 | 0.006417 | 0.013066 | 0.013908 | 0.012069 | 0.009111 | 0.017646 | 0.018803 |
| 26.20833333 | 0.005756 | 0.01471 | 0.020526 | 0.00721 | 0.006771 | 0.010329 | 0.01439 | 0.002352 | 0.015422 | 0.008061 | 0.019621 | 0.004134 | 0.008305 | 0.010393 | 0.007507 | 0.005632 | 0.011021 | 0.013897 | 0.009814 | 0.007375 | 0.013726 | 0.014201 |
| 26.22222222 | 0.003415 | 0.01666 | 0.010725 | 0.00249 | 0.008585 | 0.006027 | 0.009297 | 0.002831 | 0.009421 | 0.005218 | 0.01121 | 0.002911 | 0.006621 | 0.009064 | 0.006489 | 0.005108 | 0.00973 | 0.012262 | 0.006783 | 0.00481 | 0.008404 | 0.009021 |
| 26.23611111 | 0.003742 | 0.012257 | 0.010091 | 0.004756 | 0.012743 | 0.005605 | 0.008656 | 0.007742 | 0.007138 | 0.009041 | 0.007207 | 0.007023 | 0.005196 | 0.009166 | 0.004898 | 0.003581 | 0.008481 | 0.008961 | 0.003825 | 0.002744 | 0.004187 | 0.004764 |
| 26.25 | 0.005078 | 0.00943 | 0.013168 | 0.010512 | 0.013122 | 0.006681 | 0.01235 | 0.010779 | 0.010564 | 0.01202 | 0.008279 | 0.011094 | 0.003537 | 0.007368 | 0.0046 | 0.003049 | 0.006527 | 0.005408 | 0.003396 | 0.003409 | 0.003633 | 0.003218 |
| 26.26388889 | 0.008119 | 0.013128 | 0.012974 | 0.015786 | 0.011428 | 0.005871 | 0.012797 | 0.007606 | 0.011738 | 0.010163 | 0.007037 | 0.010415 | 0.00374 | 0.003944 | 0.009585 | 0.007347 | 0.0037 | 0.00272 | 0.005824 | 0.005712 | 0.006733 | 0.006689 |
| 26.27777778 | 0.007247 | 0.012183 | 0.012193 | 0.016643 | 0.009253 | 0.004656 | 0.009724 | 0.00273 | 0.009131 | 0.007403 | 0.003616 | 0.00642 | 0.006141 | 0.002367 | 0.014554 | 0.01346 | 0.002572 | 0.002181 | 0.00736 | 0.007052 | 0.009167 | 0.01106 |
| 26.29166667 | 0.00277 | 0.006117 | 0.012854 | 0.013725 | 0.006959 | 0.004644 | 0.009243 | 0.001287 | 0.008444 | 0.006473 | 0.002009 | 0.002702 | 0.007819 | 0.002813 | 0.015343 | 0.016565 | 0.005316 | 0.003899 | 0.006541 | 0.007004 | 0.008321 | 0.01119 |
| 26.30555556 | 0.004231 | 0.005883 | 0.014145 | 0.012666 | 0.006737 | 0.007892 | 0.014239 | 0.004402 | 0.010345 | 0.006758 | 0.002206 | 0.000896 | 0.007261 | 0.003424 | 0.015186 | 0.016519 | 0.010087 | 0.004678 | 0.004864 | 0.005967 | 0.005856 | 0.008161 |
| 26.31944444 | 0.012961 | 0.009977 | 0.01754 | 0.017001 | 0.010121 | 0.015009 | 0.019453 | 0.010477 | 0.013772 | 0.008679 | 0.004102 | 0.000737 | 0.005477 | 0.00415 | 0.015367 | 0.014491 | 0.012782 | 0.003507 | 0.003592 | 0.00423 | 0.00481 | 0.005387 |
| 26.33333333 | 0.01969 | 0.011454 | 0.024129 | 0.023295 | 0.014262 | 0.021065 | 0.018419 | 0.017013 | 0.018531 | 0.013002 | 0.007427 | 0.001727 | 0.004895 | 0.006173 | 0.013683 | 0.010994 | 0.009121 | 0.003177 | 0.003647 | 0.002908 | 0.006192 | 0.004448 |
| 26.34722222 | 0.021883 | 0.012422 | 0.030272 | 0.026487 | 0.016994 | 0.021838 | 0.016316 | 0.022407 | 0.021858 | 0.016985 | 0.012732 | 0.003088 | 0.006819 | 0.00923 | 0.009426 | 0.006813 | 0.004048 | 0.004368 | 0.00474 | 0.003107 | 0.00764 | 0.004347 |
| 26.36111111 | 0.023758 | 0.018256 | 0.029724 | 0.02525 | 0.018443 | 0.01993 | 0.019863 | 0.024477 | 0.019195 | 0.01804 | 0.020398 | 0.007213 | 0.009028 | 0.010448 | 0.005142 | 0.004234 | 0.004168 | 0.004984 | 0.005668 | 0.004379 | 0.007332 | 0.004156 |
| 26.375 | 0.023513 | 0.021899 | 0.021643 | 0.020054 | 0.016421 | 0.018376 | 0.02225 | 0.021173 | 0.012232 | 0.016503 | 0.025499 | 0.013491 | 0.009033 | 0.00843 | 0.003737 | 0.00498 | 0.00466 | 0.004804 | 0.006282 | 0.00552 | 0.006976 | 0.005265 |
| 26.38888889 | 0.019524 | 0.017662 | 0.014354 | 0.01323 | 0.010851 | 0.016395 | 0.018188 | 0.01552 | 0.008326 | 0.013292 | 0.024852 | 0.014317 | 0.008574 | 0.007075 | 0.005872 | 0.008264 | 0.00348 | 0.006287 | 0.008053 | 0.006482 | 0.009021 | 0.009327 |
| 26.40277778 | 0.017237 | 0.017072 | 0.016129 | 0.010892 | 0.008449 | 0.015268 | 0.017759 | 0.013702 | 0.01074 | 0.011258 | 0.023428 | 0.013626 | 0.012281 | 0.01176 | 0.01119 | 0.012946 | 0.005154 | 0.010616 | 0.012446 | 0.00908 | 0.013908 | 0.016264 |
| 26.41666667 | 0.019581 | 0.024506 | 0.023547 | 0.016439 | 0.013085 | 0.018004 | 0.027232 | 0.016483 | 0.017048 | 0.014215 | 0.024377 | 0.018258 | 0.02185 | 0.022153 | 0.020152 | 0.019436 | 0.009431 | 0.016373 | 0.018916 | 0.014662 | 0.020089 | 0.024508 |
| 26.43055556 | 0.023305 | 0.029516 | 0.026002 | 0.023921 | 0.019422 | 0.022311 | 0.036255 | 0.019149 | 0.023173 | 0.020401 | 0.024833 | 0.022974 | 0.034418 | 0.033841 | 0.032072 | 0.027977 | 0.014108 | 0.022706 | 0.025106 | 0.021631 | 0.025359 | 0.031541 |
| 26.44444444 | 0.027716 | 0.029998 | 0.022837 | 0.027254 | 0.023317 | 0.024975 | 0.039233 | 0.021073 | 0.027026 | 0.025629 | 0.025215 | 0.024788 | 0.046409 | 0.046277 | 0.044316 | 0.03693 | 0.020313 | 0.030793 | 0.030227 | 0.027836 | 0.030417 | 0.037709 |
| 26.45833333 | 0.035166 | 0.035017 | 0.025011 | 0.030677 | 0.028039 | 0.030058 | 0.048317 | 0.027614 | 0.031892 | 0.032881 | 0.031667 | 0.029151 | 0.057099 | 0.062839 | 0.055503 | 0.04511 | 0.028804 | 0.041597 | 0.035556 | 0.034093 | 0.03931 | 0.048355 |
| 26.47222222 | 0.044701 | 0.046594 | 0.037163 | 0.040407 | 0.03606 | 0.041731 | 0.071207 | 0.040549 | 0.042338 | 0.046614 | 0.047467 | 0.041554 | 0.067702 | 0.084368 | 0.066265 | 0.051507 | 0.037587 | 0.054732 | 0.041799 | 0.041184 | 0.051581 | 0.064606 |
| 26.48611111 | 0.055613 | 0.059827 | 0.052864 | 0.056308 | 0.046609 | 0.056515 | 0.096238 | 0.057093 | 0.059462 | 0.064632 | 0.06997 | 0.062063 | 0.07671 | 0.108454 | 0.078299 | 0.056687 | 0.044937 | 0.06981 | 0.048115 | 0.047588 | 0.058874 | 0.0757 |

| | | | | | | | | | | | | | | | | | | | | | | |
|---|---|---|---|---|---|---|---|---|---|---|---|---|---|---|---|---|---|---|---|---|---|---|
| 26.5 | 0.074361 | 0.078583 | 0.069877 | 0.076691 | 0.060436 | 0.069614 | 0.111203 | 0.079115 | 0.083819 | 0.085002 | 0.095314 | 0.087087 | 0.082089 | 0.13267 | 0.09433 | 0.065647 | 0.050948 | 0.086383 | 0.053106 | 0.052862 | 0.056977 | 0.075795 |
| 26.51388889 | 0.10466 | 0.110377 | 0.093147 | 0.100132 | 0.076904 | 0.080242 | 0.119682 | 0.109329 | 0.113399 | 0.108795 | 0.123682 | 0.109941 | 0.080964 | 0.150537 | 0.112297 | 0.076618 | 0.053359 | 0.099339 | 0.05607 | 0.05691 | 0.053005 | 0.076133 |
| 26.52777778 | 0.127276 | 0.14035 | 0.119656 | 0.121094 | 0.087838 | 0.087308 | 0.125922 | 0.140127 | 0.137101 | 0.133112 | 0.150327 | 0.121137 | 0.070245 | 0.147142 | 0.11761 | 0.07591 | 0.047574 | 0.095594 | 0.053741 | 0.057456 | 0.052982 | 0.080491 |
| 26.54166667 | 0.11781 | 0.139285 | 0.135862 | 0.128315 | 0.082368 | 0.086098 | 0.120647 | 0.147029 | 0.13946 | 0.14339 | 0.151494 | 0.115133 | 0.051551 | 0.110152 | 0.091453 | 0.055633 | 0.033939 | 0.069941 | 0.041281 | 0.051389 | 0.04937 | 0.075184 |
| 26.55555556 | 0.082134 | 0.102154 | 0.116698 | 0.105445 | 0.061508 | 0.071157 | 0.09261 | 0.104307 | 0.103744 | 0.114711 | 0.109705 | 0.0891 | 0.028409 | 0.056114 | 0.048859 | 0.026784 | 0.018537 | 0.037368 | 0.02155 | 0.037673 | 0.034343 | 0.05407 |
| 26.56944444 | 0.042515 | 0.050523 | 0.062811 | 0.055156 | 0.034277 | 0.042545 | 0.048145 | 0.051296 | 0.051739 | 0.054683 | 0.052416 | 0.048231 | 0.017284 | 0.023372 | 0.031783 | 0.011265 | 0.010685 | 0.017466 | 0.007201 | 0.020136 | 0.018863 | 0.027829 |
| 26.58333333 | 0.019501 | 0.016885 | 0.023324 | 0.018231 | 0.01433 | 0.023149 | 0.028283 | 0.029728 | 0.029896 | 0.022205 | 0.024727 | 0.022982 | 0.019522 | 0.015681 | 0.030213 | 0.022277 | 0.01564 | 0.019749 | 0.005648 | 0.010221 | 0.021128 | 0.017209 |
| 26.59722222 | 0.028836 | 0.039571 | 0.031446 | 0.024699 | 0.009058 | 0.037932 | 0.063491 | 0.032737 | 0.048857 | 0.059014 | 0.033763 | 0.029961 | 0.019938 | 0.024488 | 0.032249 | 0.05015 | 0.024944 | 0.03577 | 0.011659 | 0.013121 | 0.036621 | 0.028486 |
| 26.61111111 | 0.052989 | 0.111328 | 0.071205 | 0.054717 | 0.017495 | 0.062536 | 0.104374 | 0.067713 | 0.093801 | 0.122208 | 0.046627 | 0.052894 | 0.031146 | 0.058595 | 0.064213 | 0.074074 | 0.024887 | 0.042912 | 0.019669 | 0.017558 | 0.040559 | 0.040637 |
| 26.625 | 0.060838 | 0.158359 | 0.094615 | 0.07388 | 0.04324 | 0.056601 | 0.087574 | 0.093205 | 0.117779 | 0.130601 | 0.050238 | 0.071198 | 0.049766 | 0.097569 | 0.091768 | 0.081422 | 0.016523 | 0.035665 | 0.020624 | 0.016517 | 0.028875 | 0.040021 |
| 26.63888889 | 0.051121 | 0.140973 | 0.075481 | 0.061168 | 0.057759 | 0.027147 | 0.041313 | 0.066197 | 0.085544 | 0.080406 | 0.045749 | 0.062485 | 0.051106 | 0.105844 | 0.078104 | 0.065732 | 0.009275 | 0.025788 | 0.010511 | 0.010894 | 0.016601 | 0.029313 |
| 26.65277778 | 0.031638 | 0.084504 | 0.039776 | 0.027652 | 0.036036 | 0.010578 | 0.028458 | 0.025296 | 0.034122 | 0.037459 | 0.029113 | 0.032703 | 0.033279 | 0.071485 | 0.044679 | 0.041897 | 0.004658 | 0.015899 | 0.002271 | 0.004341 | 0.009478 | 0.014465 |
| 26.66666667 | 0.012559 | 0.029448 | 0.015239 | 0.007494 | 0.012513 | 0.013112 | 0.035651 | 0.018702 | 0.018338 | 0.029733 | 0.016507 | 0.010612 | 0.015547 | 0.031663 | 0.022348 | 0.035731 | 0.00355 | 0.012228 | 0.000696 | 0.003395 | 0.008546 | 0.004809 |
| 26.68055556 | 0.010667 | 0.017411 | 0.015382 | 0.019645 | 0.016871 | 0.021405 | 0.040321 | 0.049941 | 0.047464 | 0.050075 | 0.009145 | 0.006142 | 0.014348 | 0.023347 | 0.022582 | 0.033818 | 0.005373 | 0.01461 | 0.000652 | 0.006254 | 0.011558 | 0.005546 |
| 26.69444444 | 0.021885 | 0.042749 | 0.03448 | 0.048615 | 0.030393 | 0.03078 | 0.050331 | 0.076226 | 0.077648 | 0.078304 | 0.017017 | 0.010091 | 0.016376 | 0.023571 | 0.022689 | 0.022671 | 0.005003 | 0.012425 | 0.002206 | 0.007711 | 0.011118 | 0.009431 |
| 26.70833333 | 0.026819 | 0.056036 | 0.035271 | 0.050595 | 0.024847 | 0.026033 | 0.045345 | 0.064053 | 0.063727 | 0.067577 | 0.041318 | 0.009192 | 0.009636 | 0.015461 | 0.019645 | 0.017211 | 0.004412 | 0.011579 | 0.003373 | 0.007288 | 0.009748 | 0.011372 |
| 26.72222222 | 0.018502 | 0.038486 | 0.013925 | 0.029363 | 0.01343 | 0.014105 | 0.040545 | 0.046566 | 0.043677 | 0.044021 | 0.04978 | 0.004876 | 0.011791 | 0.026166 | 0.037815 | 0.020063 | 0.00812 | 0.017911 | 0.003767 | 0.005824 | 0.015072 | 0.016263 |
| 26.73611111 | 0.008368 | 0.021816 | 0.004264 | 0.022422 | 0.016404 | 0.012018 | 0.049912 | 0.042327 | 0.053813 | 0.044434 | 0.044796 | 0.009835 | 0.025826 | 0.051466 | 0.053319 | 0.017911 | 0.010093 | 0.020469 | 0.00593 | 0.006121 | 0.015765 | 0.01763 |
| 26.75 | 0.007041 | 0.030077 | 0.005917 | 0.019813 | 0.01618 | 0.01153 | 0.037064 | 0.030085 | 0.049069 | 0.038515 | 0.032401 | 0.019997 | 0.026815 | 0.047099 | 0.033789 | 0.007371 | 0.009007 | 0.015492 | 0.005347 | 0.007114 | 0.009574 | 0.015729 |
| 26.76388889 | 0.013912 | 0.039706 | 0.008843 | 0.016034 | 0.009202 | 0.008753 | 0.010916 | 0.026769 | 0.028005 | 0.032828 | 0.027278 | 0.020805 | 0.016015 | 0.019956 | 0.01371 | 0.002865 | 0.012804 | 0.016052 | 0.004161 | 0.005254 | 0.008451 | 0.018306 |
| 26.77777778 | 0.019769 | 0.034139 | 0.020105 | 0.02777 | 0.012888 | 0.005621 | 0.006316 | 0.029547 | 0.019514 | 0.029892 | 0.043051 | 0.018746 | 0.01516 | 0.005958 | 0.018318 | 0.008117 | 0.018701 | 0.021245 | 0.008564 | 0.00413 | 0.012002 | 0.019858 |
| 26.79166667 | 0.013349 | 0.025895 | 0.026196 | 0.033137 | 0.024203 | 0.013091 | 0.025963 | 0.022265 | 0.016522 | 0.019161 | 0.037998 | 0.016948 | 0.01971 | 0.012036 | 0.034445 | 0.016227 | 0.015425 | 0.018287 | 0.013012 | 0.006073 | 0.020897 | 0.023415 |
| 26.80555556 | 0.004455 | 0.019217 | 0.018879 | 0.023289 | 0.026239 | 0.022198 | 0.043549 | 0.018488 | 0.014423 | 0.010136 | 0.024854 | 0.011178 | 0.016582 | 0.033243 | 0.052549 | 0.022353 | 0.009529 | 0.012281 | 0.010492 | 0.006565 | 0.02836 | 0.031112 |
| 26.81944444 | 0.005329 | 0.01826 | 0.011086 | 0.01652 | 0.016705 | 0.018764 | 0.045315 | 0.016543 | 0.015331 | 0.007819 | 0.027966 | 0.009602 | 0.013026 | 0.061083 | 0.062238 | 0.024374 | 0.01295 | 0.012226 | 0.004412 | 0.004262 | 0.025969 | 0.027834 |
| 26.83333333 | 0.008 | 0.020577 | 0.006753 | 0.010062 | 0.007571 | 0.020026 | 0.041468 | 0.013413 | 0.022228 | 0.014424 | 0.029608 | 0.009759 | 0.011748 | 0.068372 | 0.056134 | 0.025792 | 0.012977 | 0.011839 | 0.00298 | 0.004787 | 0.020979 | 0.019446 |
| 26.84722222 | 0.006253 | 0.019408 | 0.005211 | 0.009443 | 0.006535 | 0.01965 | 0.035439 | 0.021718 | 0.025135 | 0.019793 | 0.03383 | 0.014516 | 0.007721 | 0.042854 | 0.038562 | 0.028034 | 0.007596 | 0.011035 | 0.004722 | 0.006381 | 0.019749 | 0.026023 |
| 26.86111111 | 0.006026 | 0.021683 | 0.004635 | 0.014891 | 0.010074 | 0.021296 | 0.046262 | 0.027488 | 0.025601 | 0.019609 | 0.035012 | 0.023034 | 0.008966 | 0.01832 | 0.029458 | 0.03403 | 0.010181 | 0.017498 | 0.005775 | 0.00413 | 0.020179 | 0.029088 |
| 26.875 | 0.01269 | 0.029701 | 0.00483 | 0.018281 | 0.011927 | 0.033443 | 0.057836 | 0.034347 | 0.030136 | 0.021817 | 0.038744 | 0.019514 | 0.010873 | 0.016861 | 0.031461 | 0.040321 | 0.012431 | 0.023388 | 0.008386 | 0.001176 | 0.028688 | 0.030912 |
| 26.88888889 | 0.019499 | 0.038536 | 0.00784 | 0.021756 | 0.013021 | 0.036781 | 0.055468 | 0.056551 | 0.037766 | 0.031404 | 0.035355 | 0.014659 | 0.00635 | 0.027713 | 0.027623 | 0.031459 | 0.008962 | 0.017564 | 0.01001 | 0.001277 | 0.041058 | 0.050218 |
| 26.90277778 | 0.017701 | 0.035962 | 0.010395 | 0.020372 | 0.015834 | 0.034391 | 0.06163 | 0.052663 | 0.046766 | 0.03713 | 0.03126 | 0.026701 | 0.005378 | 0.040491 | 0.032571 | 0.020753 | 0.010592 | 0.010143 | 0.007257 | 0.002813 | 0.032024 | 0.052008 |
| 26.91666667 | 0.014695 | 0.034205 | 0.012144 | 0.021365 | 0.020888 | 0.025231 | 0.050715 | 0.03006 | 0.037647 | 0.024606 | 0.038648 | 0.03273 | 0.009565 | 0.040985 | 0.051522 | 0.032214 | 0.012528 | 0.010197 | 0.003556 | 0.003308 | 0.015023 | 0.028174 |
| 26.93055556 | 0.016486 | 0.039593 | 0.016488 | 0.026858 | 0.022908 | 0.011128 | 0.023308 | 0.031812 | 0.024074 | 0.009539 | 0.034401 | 0.036891 | 0.011011 | 0.030009 | 0.047815 | 0.037342 | 0.010177 | 0.012606 | 0.001682 | 0.005638 | 0.011038 | 0.010583 |
| 26.94444444 | 0.023297 | 0.037818 | 0.023381 | 0.022137 | 0.019789 | 0.006875 | 0.009384 | 0.064881 | 0.037208 | 0.01389 | 0.027115 | 0.062518 | 0.020778 | 0.042982 | 0.040019 | 0.029591 | 0.016737 | 0.02377 | 0.005861 | 0.012304 | 0.017769 | 0.013596 |
| 26.95833333 | 0.040006 | 0.048094 | 0.037145 | 0.022479 | 0.015936 | 0.014809 | 0.018214 | 0.097065 | 0.057577 | 0.042003 | 0.023745 | 0.060768 | 0.047586 | 0.080799 | 0.066405 | 0.048752 | 0.03122 | 0.043685 | 0.020043 | 0.0177 | 0.040022 | 0.033932 |
| 26.97222222 | 0.049112 | 0.063175 | 0.044641 | 0.037698 | 0.018118 | 0.0328 | 0.053422 | 0.081229 | 0.0564 | 0.078822 | 0.029438 | 0.030203 | 0.068125 | 0.11538 | 0.100016 | 0.085264 | 0.037677 | 0.053771 | 0.037556 | 0.018209 | 0.059915 | 0.05249 |
| 26.98611111 | 0.038709 | 0.058006 | 0.03429 | 0.043795 | 0.030392 | 0.045803 | 0.079948 | 0.05672 | 0.056052 | 0.090757 | 0.043255 | 0.020625 | 0.076224 | 0.139486 | 0.122711 | 0.104855 | 0.034814 | 0.054629 | 0.048884 | 0.024619 | 0.058764 | 0.0557 |
| 27 | 0.035285 | 0.054606 | 0.030798 | 0.043356 | 0.044167 | 0.041228 | 0.069228 | 0.082297 | 0.081333 | 0.073716 | 0.073622 | 0.044314 | 0.083548 | 0.144348 | 0.133829 | 0.096617 | 0.035118 | 0.065887 | 0.052165 | 0.038584 | 0.044401 | 0.047924 |
| 27.01388889 | 0.055251 | 0.071376 | 0.062765 | 0.060112 | 0.052755 | 0.036038 | 0.054218 | 0.121968 | 0.106967 | 0.060354 | 0.121064 | 0.07868 | 0.083478 | 0.127634 | 0.127361 | 0.071639 | 0.042925 | 0.085118 | 0.049629 | 0.045972 | 0.03499 | 0.047208 |
| 27.02777778 | 0.079892 | 0.093311 | 0.112497 | 0.085508 | 0.06234 | 0.047533 | 0.069335 | 0.117972 | 0.106686 | 0.066411 | 0.117803 | 0.071331 | 0.073617 | 0.106528 | 0.114072 | 0.050886 | 0.049642 | 0.092579 | 0.044845 | 0.043353 | 0.040369 | 0.063454 |
| 27.04166667 | 0.0809 | 0.093496 | 0.123487 | 0.09684 | 0.0731 | 0.060697 | 0.092926 | 0.084959 | 0.088152 | 0.080582 | 0.068942 | 0.039592 | 0.065263 | 0.088671 | 0.097813 | 0.038214 | 0.044088 | 0.079195 | 0.039678 | 0.036201 | 0.04793 | 0.07945 |
| 27.05555556 | 0.060217 | 0.070319 | 0.083879 | 0.08845 | 0.07728 | 0.057014 | 0.088274 | 0.065448 | 0.069414 | 0.082876 | 0.047701 | 0.03295 | 0.057875 | 0.066439 | 0.071096 | 0.027806 | 0.027872 | 0.051377 | 0.031328 | 0.027032 | 0.043024 | 0.07246 |
| 27.06944444 | 0.043945 | 0.043771 | 0.036901 | 0.065507 | 0.067675 | 0.040791 | 0.056563 | 0.061703 | 0.056737 | 0.066518 | 0.053668 | 0.037256 | 0.041772 | 0.044376 | 0.041409 | 0.02059 | 0.015447 | 0.028118 | 0.02045 | 0.018052 | 0.027071 | 0.045657 |
| 27.08333333 | 0.039451 | 0.02064 | 0.014098 | 0.039264 | 0.041634 | 0.026108 | 0.025546 | 0.053692 | 0.043086 | 0.042375 | 0.054265 | 0.028334 | 0.020397 | 0.028403 | 0.020196 | 0.016626 | 0.012113 | 0.018604 | 0.011455 | 0.012203 | 0.012569 | 0.019487 |
| 27.09722222 | 0.045949 | 0.014133 | 0.013065 | 0.026521 | 0.016208 | 0.023748 | 0.017355 | 0.036664 | 0.028207 | 0.023331 | 0.044807 | 0.015202 | 0.007685 | 0.014318 | 0.009754 | 0.010511 | 0.011097 | 0.014362 | 0.005608 | 0.009874 | 0.006812 | 0.005789 |
| 27.11111111 | 0.062757 | 0.036236 | 0.029084 | 0.036065 | 0.013847 | 0.035618 | 0.037509 | 0.026204 | 0.025556 | 0.025893 | 0.043273 | 0.017357 | 0.007223 | 0.005337 | 0.009179 | 0.005364 | 0.008818 | 0.009801 | 0.003679 | 0.010214 | 0.005831 | 0.003465 |
| 27.125 | 0.075144 | 0.066896 | 0.052899 | 0.055474 | 0.032086 | 0.052458 | 0.070102 | 0.033496 | 0.039935 | 0.052553 | 0.056936 | 0.035041 | 0.007572 | 0.007402 | 0.008932 | 0.005793 | 0.007568 | 0.008165 | 0.006147 | 0.012777 | 0.008223 | 0.006901 |
| 27.13888889 | 0.074414 | 0.077852 | 0.066717 | 0.064227 | 0.047031 | 0.059784 | 0.086869 | 0.04892 | 0.055327 | 0.07564 | 0.067245 | 0.049226 | 0.007694 | 0.016317 | 0.006057 | 0.006502 | 0.008992 | 0.011762 | 0.010578 | 0.015875 | 0.015147 | 0.014939 |
| 27.15277778 | 0.065552 | 0.065901 | 0.06786 | 0.054738 | 0.043493 | 0.051927 | 0.076408 | 0.054764 | 0.055943 | 0.070617 | 0.061513 | 0.050474 | 0.012917 | 0.024346 | 0.0078 | 0.007727 | 0.01171 | 0.016889 | 0.014539 | 0.017605 | 0.020234 | 0.022571 |
| 27.16666667 | 0.054517 | 0.048232 | 0.060932 | 0.037831 | 0.031206 | 0.039085 | 0.053077 | 0.045981 | 0.045985 | 0.048022 | 0.048527 | 0.047783 | 0.018473 | 0.028485 | 0.011642 | 0.010987 | 0.013611 | 0.019197 | 0.01771 | 0.017738 | 0.020512 | 0.025273 |
| 27.18055556 | 0.044351 | 0.035538 | 0.050145 | 0.028571 | 0.023404 | 0.031326 | 0.035584 | 0.031133 | 0.036266 | 0.031717 | 0.037914 | 0.042606 | 0.021157 | 0.029734 | 0.014 | 0.01235 | 0.014182 | 0.018686 | 0.01995 | 0.017501 | 0.020418 | 0.026447 |
| 27.19444444 | 0.034403 | 0.023244 | 0.042376 | 0.028444 | 0.018805 | 0.025757 | 0.027683 | 0.01962 | 0.029038 | 0.024127 | 0.030183 | 0.034041 | 0.023491 | 0.030944 | 0.017228 | 0.011544 | 0.014224 | 0.018355 | 0.020692 | 0.018253 | 0.022999 | 0.02865 |
| 27.20833333 | 0.024754 | 0.011103 | 0.040685 | 0.028733 | 0.014584 | 0.0182 | 0.023531 | 0.014671 | 0.022188 | 0.016425 | 0.024947 | 0.024028 | 0.028332 | 0.034358 | 0.022251 | 0.010813 | 0.014216 | 0.019995 | 0.020773 | 0.020066 | 0.026889 | 0.030984 |
| 27.22222222 | 0.016519 | 0.005784 | 0.038816 | 0.024136 | 0.012878 | 0.012849 | 0.020197 | 0.013239 | 0.015555 | 0.010085 | 0.021029 | 0.013691 | 0.03465 | 0.038195 | 0.027422 | 0.012011 | 0.014204 | 0.022616 | 0.021133 | 0.021477 | 0.028606 | 0.031523 |
| 27.23611111 | 0.009861 | 0.004786 | 0.029654 | 0.016595 | 0.012274 | 0.012337 | 0.018033 | 0.011142 | 0.010446 | 0.008534 | 0.015842 | 0.00727 | 0.036444 | 0.037467 | 0.029767 | 0.015237 | 0.014618 | 0.024163 | 0.020711 | 0.020936 | 0.026374 | 0.029211 |
| 27.25 | 0.005365 | 0.003381 | 0.014975 | 0.010312 | 0.008706 | 0.010818 | 0.016331 | 0.007575 | 0.006894 | 0.006846 | 0.010683 | 0.004409 | 0.029958 | 0.030434 | 0.027025 | 0.017925 | 0.015453 | 0.023302 | 0.018091 | 0.017786 | 0.021113 | 0.023715 |
| 27.26388889 | 0.002952 | 0.003422 | 0.006144 | 0.006264 | 0.003781 | 0.005735 | 0.012432 | 0.003767 | 0.003973 | 0.004468 | 0.008399 | 0.003161 | 0.017167 | 0.018734 | 0.018751 | 0.016165 | 0.014738 | 0.019512 | 0.013512 | 0.012244 | 0.014551 | 0.015532 |
| 27.27777778 | 0.002613 | 0.005266 | 0.008281 | 0.003633 | 0.001082 | 0.001774 | 0.006589 | 0.001306 | 0.002304 | 0.004395 | 0.007514 | 0.00364 | 0.009137 | 0.009663 | 0.009001 | 0.009174 | 0.010908 | 0.013292 | 0.008747 | 0.00589 | 0.008906 | 0.007478 |
| 27.29166667 | 0.003561 | 0.006192 | 0.011548 | 0.003675 | 0.001226 | 0.001403 | 0.003776 | 0.001092 | 0.002378 | 0.003594 | 0.004995 | 0.003855 | 0.01223 | 0.00934 | 0.003386 | 0.002559 | 0.006404 | 0.007421 | 0.005712 | 0.001855 | 0.006608 | 0.004182 |
| 27.30555556 | 0.003016 | 0.004454 | 0.010313 | 0.006506 | 0.004189 | 0.004471 | 0.005606 | 0.001664 | 0.003026 | 0.002624 | 0.0023 | 0.003818 | 0.013388 | 0.009141 | 0.002428 | 0.001876 | 0.004639 | 0.004498 | 0.004976 | 0.001122 | 0.007683 | 0.006322 |
| 27.31944444 | 0.00171 | 0.001878 | 0.008049 | 0.008643 | 0.006589 | 0.007994 | 0.008349 | 0.001482 | 0.003019 | 0.003025 | 0.002668 | 0.005241 | 0.007071 | 0.004898 | 0.004073 | 0.005305 | 0.005114 | 0.003656 | 0.005673 | 0.001993 | 0.009521 | 0.009339 |
| 27.33333333 | 0.001636 | 0.001301 | 0.006253 | 0.007366 | 0.004729 | 0.006655 | 0.009009 | 0.000836 | 0.00176 | 0.003484 | 0.004357 | 0.005014 | 0.001726 | 0.002268 | 0.005427 | 0.007435 | 0.00526 | 0.003019 | 0.006165 | 0.003056 | 0.009706 | 0.009872 |
| 27.34722222 | 0.003165 | 0.003291 | 0.004884 | 0.005642 | 0.002996 | 0.003298 | 0.007054 | 0.001127 | 0.001602 | 0.003177 | 0.003693 | 0.002451 | 0.000581 | 0.00214 | 0.004416 | 0.005907 | 0.004201 | 0.002555 | 0.004966 | 0.002501 | 0.008063 | 0.008888 |
| 27.36111111 | 0.00648 | 0.007515 | 0.005052 | 0.008968 | 0.007513 | 0.006057 | 0.006045 | 0.003936 | 0.005065 | 0.004119 | 0.002396 | 0.002581 | 0.000987 | 0.002588 | 0.002665 | 0.003022 | 0.002743 | 0.002396 | 0.00282 | 0.000926 | 0.006158 | 0.008676 |
| 27.375 | 0.010801 | 0.012295 | 0.007657 | 0.015976 | 0.014562 | 0.013276 | 0.010542 | 0.008686 | 0.010439 | 0.009207 | 0.004338 | 0.006521 | 0.000906 | 0.002936 | 0.001699 | 0.002652 | 0.001447 | 0.001897 | 0.002684 | 0.000573 | 0.004769 | 0.008489 |
| 27.38888889 | 0.015801 | 0.017077 | 0.012394 | 0.020102 | 0.016942 | 0.016546 | 0.017895 | 0.013276 | 0.015172 | 0.014745 | 0.009879 | 0.011333 | 0.002048 | 0.00381 | 0.001982 | 0.00591 | 0.000946 | 0.002086 | 0.00557 | 0.001769 | 0.003149 | 0.00586 |
| 27.40277778 | 0.021191 | 0.022974 | 0.018505 | 0.020633 | 0.017472 | 0.016238 | 0.021544 | 0.015895 | 0.018565 | 0.018916 | 0.01626 | 0.016825 | 0.006545 | 0.005235 | 0.004798 | 0.011307 | 0.002345 | 0.004544 | 0.009466 | 0.00484 | 0.003029 | 0.004043 |
| 27.41666667 | 0.027039 | 0.029382 | 0.025612 | 0.022889 | 0.022948 | 0.018728 | 0.022231 | 0.016987 | 0.0218 | 0.024902 | 0.020025 | 0.023239 | 0.013403 | 0.010884 | 0.009366 | 0.016823 | 0.005637 | 0.009027 | 0.012935 | 0.009359 | 0.007081 | 0.007208 |
| 27.43055556 | 0.033684 | 0.03481 | 0.034288 | 0.029359 | 0.031752 | 0.023974 | 0.026848 | 0.020589 | 0.02732 | 0.032486 | 0.023603 | 0.028655 | 0.021118 | 0.0214 | 0.014747 | 0.021904 | 0.010136 | 0.015101 | 0.016968 | 0.014323 | 0.013466 | 0.012854 |
| 27.44444444 | 0.039366 | 0.039493 | 0.043099 | 0.037584 | 0.038963 | 0.029012 | 0.037418 | 0.029091 | 0.035811 | 0.04111 | 0.033369 | 0.033984 | 0.03161 | 0.031959 | 0.024703 | 0.029017 | 0.015205 | 0.022246 | 0.023904 | 0.021167 | 0.020465 | 0.020373 |
| 27.45833333 | 0.043472 | 0.046024 | 0.047473 | 0.044968 | 0.045022 | 0.035483 | 0.051191 | 0.040196 | 0.046083 | 0.054069 | 0.047992 | 0.043491 | 0.048786 | 0.043366 | 0.04361 | 0.041259 | 0.020621 | 0.030343 | 0.035031 | 0.032025 | 0.029902 | 0.032218 |
| 27.47222222 | 0.049698 | 0.0574 | 0.047592 | 0.051784 | 0.052348 | 0.045276 | 0.06653 | 0.051631 | 0.057792 | 0.070281 | 0.060884 | 0.057497 | 0.072117 | 0.061827 | 0.068978 | 0.05687 | 0.027024 | 0.041099 | 0.048485 | 0.045479 | 0.042132 | 0.047368 |
| 27.48611111 | 0.061709 | 0.073674 | 0.054113 | 0.061773 | 0.062307 | 0.05552 | 0.082234 | 0.064727 | 0.072575 | 0.083412 | 0.072526 | 0.071775 | 0.090018 | 0.089277 | 0.091649 | 0.068205 | 0.035404 | 0.057868 | 0.061449 | 0.057886 | 0.05284 | 0.062736 |
| 27.5 | 0.080453 | 0.092291 | 0.082079 | 0.081476 | 0.077336 | 0.064854 | 0.098085 | 0.084098 | 0.094103 | 0.098351 | 0.090787 | 0.082936 | 0.093916 | 0.120985 | 0.108106 | 0.075844 | 0.045957 | 0.080407 | 0.073305 | 0.066908 | 0.057933 | 0.077395 |
| 27.51388889 | 0.103236 | 0.112763 | 0.131553 | 0.111598 | 0.096302 | 0.077761 | 0.115591 | 0.112072 | 0.123416 | 0.128349 | 0.117796 | 0.092442 | 0.093463 | 0.151521 | 0.120873 | 0.089174 | 0.054386 | 0.097505 | 0.082328 | 0.072784 | 0.059614 | 0.091312 |
| 27.52777778 | 0.121732 | 0.134653 | 0.165777 | 0.135351 | 0.108408 | 0.092758 | 0.131916 | 0.140144 | 0.151655 | 0.166028 | 0.139884 | 0.100576 | 0.092785 | 0.166657 | 0.127145 | 0.104385 | 0.052891 | 0.097606 | 0.084771 | 0.075036 | 0.061445 | 0.100155 |
| 27.54166667 | 0.11676 | 0.138323 | 0.140828 | 0.123893 | 0.095523 | 0.09176 | 0.129496 | 0.14356 | 0.155438 | 0.167215 | 0.133515 | 0.092879 | 0.080024 | 0.142681 | 0.117001 | 0.10356 | 0.039774 | 0.082324 | 0.076397 | 0.06827 | 0.062096 | 0.098263 |
| 27.55555556 | 0.074012 | 0.095688 | 0.079953 | 0.07191 | 0.056014 | 0.06085 | 0.089868 | 0.097764 | 0.110934 | 0.103133 | 0.092266 | 0.063744 | 0.05049 | 0.091091 | 0.079509 | 0.075916 | 0.021942 | 0.057878 | 0.053481 | 0.04769 | 0.056218 | 0.082954 |
| 27.56944444 | 0.025801 | 0.039211 | 0.043581 | 0.020581 | 0.019671 | 0.024636 | 0.037179 | 0.042534 | 0.054082 | 0.037319 | 0.042225 | 0.032483 | 0.027363 | 0.061819 | 0.031542 | 0.036689 | 0.010296 | 0.032214 | 0.026359 | 0.021684 | 0.041678 | 0.056322 |
| 27.58333333 | 0.012347 | 0.01747 | 0.035015 | 0.007781 | 0.009864 | 0.019307 | 0.023896 | 0.029886 | 0.035951 | 0.032147 | 0.016983 | 0.022509 | 0.02988 | 0.050805 | 0.01108 | 0.018137 | 0.014198 | 0.021838 | 0.009978 | 0.008241 | 0.028447 | 0.032763 |
| 27.59722222 | 0.036135 | 0.02953 | 0.05079 | 0.032292 | 0.023326 | 0.049263 | 0.062899 | 0.055719 | 0.060304 | 0.084343 | 0.033028 | 0.041418 | 0.030035 | 0.043676 | 0.026018 | 0.029931 | 0.031206 | 0.03636 | 0.01019 | 0.015391 | 0.030603 | 0.032616 |
| 27.61111111 | 0.067706 | 0.056765 | 0.096886 | 0.07771 | 0.047155 | 0.081412 | 0.103287 | 0.111747 | 0.115063 | 0.151943 | 0.071286 | 0.074906 | 0.021858 | 0.060421 | 0.049945 | 0.056324 | 0.044352 | 0.060962 | 0.024369 | 0.029144 | 0.044556 | 0.052723 |
| 27.625 | 0.071858 | 0.087078 | 0.117722 | 0.106043 | 0.068955 | 0.075996 | 0.098025 | 0.145383 | 0.144653 | 0.154379 | 0.093606 | 0.102624 | 0.027487 | 0.081765 | 0.059668 | 0.078191 | 0.045008 | 0.075532 | 0.035017 | 0.027611 | 0.050068 | 0.067773 |
| 27.63888889 | 0.0544 | 0.105303 | 0.079868 | 0.078696 | 0.06804 | 0.048552 | 0.06051 | 0.103471 | 0.107027 | 0.089739 | 0.080313 | 0.098328 | 0.03513 | 0.093595 | 0.063267 | 0.068489 | 0.036483 | 0.070864 | 0.031516 | 0.015006 | 0.04104 | 0.065547 |
| 27.65277778 | 0.029377 | 0.080176 | 0.032943 | 0.027266 | 0.038082 | 0.027809 | 0.035785 | 0.049409 | 0.05414 | 0.036663 | 0.047593 | 0.058173 | 0.032104 | 0.082604 | 0.066388 | 0.035152 | 0.019875 | 0.049949 | 0.019857 | 0.00808 | 0.028149 | 0.048107 |
| 27.66666667 | 0.009506 | 0.036417 | 0.017198 | 0.007365 | 0.013032 | 0.017052 | 0.038469 | 0.032033 | 0.030878 | 0.026711 | 0.033388 | 0.032053 | 0.019836 | 0.042136 | 0.046935 | 0.018495 | 0.007465 | 0.027012 | 0.008507 | 0.005207 | 0.015291 | 0.023712 |
| 27.68055556 | 0.00479 | 0.022548 | 0.018537 | 0.010999 | 0.0064 | 0.016419 | 0.039124 | 0.032636 | 0.033506 | 0.039324 | 0.030157 | 0.032144 | 0.010337 | 0.016078 | 0.015006 | 0.016003 | 0.00784 | 0.01563 | 0.004039 | 0.004759 | 0.005139 | 0.008731 |
| 27.69444444 | 0.008692 | 0.022065 | 0.025137 | 0.020211 | 0.008331 | 0.014327 | 0.034782 | 0.052846 | 0.055991 | 0.055223 | 0.022544 | 0.028421 | 0.016582 | 0.028718 | 0.005857 | 0.012807 | 0.009289 | 0.013715 | 0.006133 | 0.007337 | 0.003725 | 0.005 |
| 27.70833333 | 0.019832 | 0.024629 | 0.031378 | 0.029565 | 0.016234 | 0.013118 | 0.028366 | 0.056671 | 0.059187 | 0.047599 | 0.035955 | 0.032007 | 0.025795 | 0.039944 | 0.016083 | 0.017786 | 0.006965 | 0.012133 | 0.007629 | 0.006493 | 0.0073 | 0.0071 |
| 27.72222222 | 0.025573 | 0.038217 | 0.025274 | 0.031613 | 0.019953 | 0.014684 | 0.014423 | 0.040073 | 0.039305 | 0.026123 | 0.046892 | 0.031731 | 0.020955 | 0.031849 | 0.028351 | 0.025619 | 0.008317 | 0.018141 | 0.006039 | 0.002976 | 0.00819 | 0.011813 |
| 27.73611111 | 0.027897 | 0.041753 | 0.018318 | 0.032843 | 0.020857 | 0.011046 | 0.013125 | 0.048967 | 0.038787 | 0.020841 | 0.053355 | 0.033257 | 0.02242 | 0.047755 | 0.038998 | 0.021939 | 0.012426 | 0.026093 | 0.007854 | 0.001411 | 0.007239 | 0.011194 |
| 27.75 | 0.03416 | 0.045719 | 0.013778 | 0.024262 | 0.022816 | 0.010515 | 0.023214 | 0.054848 | 0.042236 | 0.020159 | 0.067805 | 0.049332 | 0.040898 | 0.065294 | 0.036487 | 0.017515 | 0.011495 | 0.022447 | 0.013796 | 0.005665 | 0.009078 | 0.011938 |
| 27.76388889 | 0.025885 | 0.057648 | 0.009868 | 0.01126 | 0.018644 | 0.011618 | 0.022251 | 0.035481 | 0.027957 | 0.020723 | 0.053656 | 0.043722 | 0.044653 | 0.04039 | 0.01881 | 0.016615 | 0.006884 | 0.015258 | 0.014539 | 0.012943 | 0.010968 | 0.015521 |
| 27.77777778 | 0.017272 | 0.059472 | 0.013237 | 0.011969 | 0.010643 | 0.007833 | 0.019633 | 0.036064 | 0.029133 | 0.01534 | 0.05917 | 0.029196 | 0.027427 | 0.010623 | 0.010138 | 0.01395 | 0.006123 | 0.009521 | 0.009141 | 0.012308 | 0.007619 | 0.013769 |
| 27.79166667 | 0.021909 | 0.067295 | 0.012432 | 0.012127 | 0.009023 | 0.010478 | 0.030891 | 0.056808 | 0.051193 | 0.006698 | 0.077116 | 0.021951 | 0.012554 | 0.004611 | 0.014956 | 0.015108 | 0.008571 | 0.00721 | 0.006952 | 0.005709 | 0.007197 | 0.01114 |
| 27.80555556 | 0.018602 | 0.067024 | 0.005461 | 0.007805 | 0.010507 | 0.018916 | 0.034913 | 0.055933 | 0.050861 | 0.007976 | 0.051751 | 0.012777 | 0.011439 | 0.008078 | 0.017115 | 0.012907 | 0.009027 | 0.011113 | 0.007502 | 0.002154 | 0.018334 | 0.016902 |
| 27.81944444 | 0.010246 | 0.043084 | 0.003906 | 0.01214 | 0.016803 | 0.018161 | 0.024092 | 0.054172 | 0.044781 | 0.009213 | 0.026633 | 0.00593 | 0.022606 | 0.020702 | 0.015293 | 0.015243 | 0.007171 | 0.015119 | 0.005082 | 0.002697 | 0.027833 | 0.032191 |
| 27.83333333 | 0.008287 | 0.022271 | 0.007984 | 0.016269 | 0.028446 | 0.013644 | 0.019039 | 0.056223 | 0.054234 | 0.006588 | 0.019695 | 0.01075 | 0.039946 | 0.046726 | 0.012897 | 0.025608 | 0.004841 | 0.015187 | 0.00551 | 0.005855 | 0.022487 | 0.037517 |
| 27.84722222 | 0.009653 | 0.010795 | 0.0112 | 0.013253 | 0.023136 | 0.011798 | 0.018433 | 0.041413 | 0.05011 | 0.010111 | 0.018664 | 0.025471 | 0.051796 | 0.068123 | 0.011534 | 0.021783 | 0.002845 | 0.01195 | 0.009937 | 0.009355 | 0.016142 | 0.023798 |
| 27.86111111 | 0.014178 | 0.01172 | 0.010415 | 0.00989 | 0.010975 | 0.007769 | 0.016967 | 0.035124 | 0.045911 | 0.016902 | 0.025765 | 0.030542 | 0.041579 | 0.058242 | 0.012171 | 0.00917 | 0.001309 | 0.006862 | 0.010181 | 0.008924 | 0.017684 | 0.012671 |
| 27.875 | 0.015774 | 0.027917 | 0.01266 | 0.009949 | 0.00852 | 0.007474 | 0.023109 | 0.040767 | 0.045573 | 0.014526 | 0.041473 | 0.028635 | 0.021878 | 0.035496 | 0.010631 | 0.005882 | 0.000672 | 0.004136 | 0.008252 | 0.005105 | 0.014789 | 0.01556 |
| 27.88888889 | 0.011479 | 0.035725 | 0.019787 | 0.019828 | 0.008721 | 0.014069 | 0.026069 | 0.051693 | 0.046447 | 0.01254 | 0.047696 | 0.045011 | 0.018226 | 0.034374 | 0.007311 | 0.01112 | 0.000894 | 0.005991 | 0.006872 | 0.003917 | 0.014788 | 0.019594 |

| | | | | | | | | | | | | | | | | | | | | | | |
|---|---|---|---|---|---|---|---|---|---|---|---|---|---|---|---|---|---|---|---|---|---|---|
| 27.90277778 | 0.009047 | 0.0352 | 0.021101 | 0.025078 | 0.012671 | 0.018168 | 0.020523 | 0.060801 | 0.063374 | 0.019266 | 0.035715 | 0.063333 | 0.0223 | 0.037349 | 0.012693 | 0.019349 | 0.002706 | 0.009158 | 0.009144 | 0.004643 | 0.030116 | 0.030788 |
| 27.91666667 | 0.014422 | 0.03462 | 0.015296 | 0.015821 | 0.012984 | 0.017525 | 0.01919 | 0.04918 | 0.059455 | 0.022383 | 0.023607 | 0.05312 | 0.034478 | 0.048283 | 0.025436 | 0.019794 | 0.008017 | 0.017981 | 0.021705 | 0.005677 | 0.042306 | 0.057045 |
| 27.93055556 | 0.027231 | 0.042342 | 0.007864 | 0.00939 | 0.019203 | 0.024697 | 0.023928 | 0.027096 | 0.027613 | 0.016308 | 0.015832 | 0.033943 | 0.053913 | 0.071071 | 0.025049 | 0.010853 | 0.016458 | 0.040581 | 0.034633 | 0.011744 | 0.03406 | 0.069129 |
| 27.94444444 | 0.02936 | 0.065129 | 0.007927 | 0.009568 | 0.029028 | 0.029436 | 0.02103 | 0.026145 | 0.02116 | 0.01852 | 0.02009 | 0.025353 | 0.053376 | 0.06616 | 0.014758 | 0.016355 | 0.02409 | 0.061057 | 0.034183 | 0.018065 | 0.018226 | 0.056757 |
| 27.95833333 | 0.023476 | 0.05566 | 0.017266 | 0.014906 | 0.017112 | 0.019275 | 0.008752 | 0.063267 | 0.059764 | 0.043342 | 0.052912 | 0.031648 | 0.035903 | 0.045895 | 0.019594 | 0.039402 | 0.025372 | 0.056797 | 0.0236 | 0.01579 | 0.013496 | 0.041535 |
| 27.97222222 | 0.040675 | 0.027928 | 0.03747 | 0.036629 | 0.012172 | 0.012713 | 0.014515 | 0.113525 | 0.106404 | 0.07965 | 0.071032 | 0.04989 | 0.02854 | 0.059487 | 0.050805 | 0.064474 | 0.02186 | 0.037636 | 0.019542 | 0.013676 | 0.024822 | 0.038742 |
| 27.98611111 | 0.069308 | 0.038376 | 0.067075 | 0.06657 | 0.029217 | 0.025212 | 0.045092 | 0.131027 | 0.122061 | 0.102371 | 0.061016 | 0.064338 | 0.041792 | 0.105666 | 0.102982 | 0.093752 | 0.022391 | 0.033822 | 0.035096 | 0.022723 | 0.04866 | 0.05743 |
| 28 | 0.068953 | 0.068492 | 0.069965 | 0.066827 | 0.044325 | 0.05228 | 0.06979 | 0.098434 | 0.1038 | 0.099188 | 0.082008 | 0.064163 | 0.069406 | 0.148413 | 0.139012 | 0.114769 | 0.030734 | 0.052103 | 0.062017 | 0.037505 | 0.071001 | 0.087772 |
| 28.01388889 | 0.049195 | 0.081309 | 0.040193 | 0.037349 | 0.052558 | 0.06627 | 0.075681 | 0.064783 | 0.090894 | 0.087472 | 0.10561 | 0.05082 | 0.091432 | 0.156234 | 0.127235 | 0.107202 | 0.042009 | 0.074661 | 0.079494 | 0.050682 | 0.077195 | 0.101037 |
| 28.02777778 | 0.044776 | 0.072084 | 0.021699 | 0.022847 | 0.056749 | 0.05373 | 0.06877 | 0.082943 | 0.118597 | 0.099293 | 0.081419 | 0.042736 | 0.094561 | 0.126048 | 0.092184 | 0.084743 | 0.048668 | 0.086898 | 0.076038 | 0.058631 | 0.070049 | 0.092132 |
| 28.04166667 | 0.050871 | 0.05822 | 0.040183 | 0.04982 | 0.064379 | 0.04107 | 0.057866 | 0.117637 | 0.150278 | 0.126994 | 0.046489 | 0.056486 | 0.089918 | 0.094541 | 0.069501 | 0.067223 | 0.044681 | 0.079955 | 0.056918 | 0.054097 | 0.058857 | 0.077488 |
| 28.05555556 | 0.049556 | 0.052338 | 0.077327 | 0.091606 | 0.078642 | 0.04779 | 0.051767 | 0.111378 | 0.13373 | 0.122341 | 0.035966 | 0.078364 | 0.084541 | 0.077951 | 0.060571 | 0.052611 | 0.030355 | 0.056355 | 0.035556 | 0.037243 | 0.046037 | 0.060445 |
| 28.06944444 | 0.047475 | 0.054425 | 0.099586 | 0.105904 | 0.0877 | 0.056057 | 0.04217 | 0.070824 | 0.08132 | 0.090211 | 0.033144 | 0.077645 | 0.070047 | 0.06172 | 0.052881 | 0.037679 | 0.016148 | 0.034424 | 0.022574 | 0.021922 | 0.032765 | 0.040968 |
| 28.08333333 | 0.045685 | 0.045102 | 0.099232 | 0.086156 | 0.065177 | 0.045231 | 0.027453 | 0.03097 | 0.033023 | 0.06392 | 0.022005 | 0.048252 | 0.043269 | 0.040861 | 0.042471 | 0.025301 | 0.010325 | 0.024786 | 0.018803 | 0.015169 | 0.020917 | 0.024333 |
| 28.09722222 | 0.039434 | 0.02273 | 0.083815 | 0.05202 | 0.029056 | 0.024195 | 0.017685 | 0.010818 | 0.008457 | 0.044916 | 0.015035 | 0.016761 | 0.018248 | 0.019916 | 0.029122 | 0.013958 | 0.009426 | 0.019854 | 0.016691 | 0.011253 | 0.011185 | 0.013359 |
| 28.11111111 | 0.039985 | 0.020956 | 0.069813 | 0.037752 | 0.018685 | 0.015016 | 0.020721 | 0.010589 | 0.009041 | 0.036874 | 0.02815 | 0.014101 | 0.008988 | 0.007773 | 0.014496 | 0.006252 | 0.007806 | 0.011944 | 0.011092 | 0.006327 | 0.005073 | 0.008168 |
| 28.125 | 0.050786 | 0.049989 | 0.075275 | 0.057785 | 0.032793 | 0.026152 | 0.038566 | 0.023808 | 0.028162 | 0.04352 | 0.057838 | 0.040477 | 0.007071 | 0.004951 | 0.004404 | 0.005751 | 0.005486 | 0.004392 | 0.00536 | 0.003356 | 0.003676 | 0.007686 |
| 28.13888889 | 0.062235 | 0.079314 | 0.092603 | 0.083107 | 0.063576 | 0.045177 | 0.058899 | 0.03855 | 0.048681 | 0.054745 | 0.07861 | 0.061243 | 0.004193 | 0.005579 | 0.00287 | 0.006984 | 0.005632 | 0.003396 | 0.005162 | 0.005983 | 0.005009 | 0.010875 |
| 28.15277778 | 0.062791 | 0.080351 | 0.099807 | 0.082899 | 0.079867 | 0.054597 | 0.066789 | 0.041315 | 0.051954 | 0.057841 | 0.072159 | 0.053592 | 0.005195 | 0.009883 | 0.008666 | 0.010732 | 0.009312 | 0.00878 | 0.010467 | 0.012432 | 0.007227 | 0.016184 |
| 28.16666667 | 0.050001 | 0.058849 | 0.088938 | 0.06063 | 0.0611 | 0.051111 | 0.059742 | 0.036538 | 0.041602 | 0.047594 | 0.048197 | 0.032157 | 0.010217 | 0.016438 | 0.016831 | 0.017545 | 0.013847 | 0.01505 | 0.016791 | 0.017738 | 0.010522 | 0.021621 |
| 28.18055556 | 0.035439 | 0.039196 | 0.071496 | 0.04138 | 0.037315 | 0.039886 | 0.043476 | 0.033918 | 0.033029 | 0.031954 | 0.026879 | 0.01889 | 0.015805 | 0.022463 | 0.022096 | 0.020317 | 0.016008 | 0.017212 | 0.020554 | 0.018884 | 0.015151 | 0.025736 |
| 28.19444444 | 0.026727 | 0.03018 | 0.059093 | 0.037735 | 0.032962 | 0.027214 | 0.026945 | 0.031275 | 0.032705 | 0.025487 | 0.01726 | 0.019346 | 0.021499 | 0.027856 | 0.024981 | 0.019808 | 0.015347 | 0.015793 | 0.021917 | 0.017948 | 0.020785 | 0.0283 |
| 28.20833333 | 0.021302 | 0.024556 | 0.051941 | 0.03864 | 0.03605 | 0.02101 | 0.019913 | 0.026031 | 0.035776 | 0.029309 | 0.015378 | 0.025124 | 0.028172 | 0.033557 | 0.030386 | 0.022747 | 0.015338 | 0.016524 | 0.023854 | 0.020103 | 0.026055 | 0.029912 |
| 28.22222222 | 0.01785 | 0.017937 | 0.045569 | 0.032487 | 0.027217 | 0.022085 | 0.02393 | 0.022486 | 0.033357 | 0.028152 | 0.014857 | 0.024909 | 0.036621 | 0.039979 | 0.03953 | 0.030203 | 0.017039 | 0.02265 | 0.027112 | 0.026128 | 0.029133 | 0.030552 |
| 28.23611111 | 0.017123 | 0.012415 | 0.036974 | 0.024635 | 0.013893 | 0.022571 | 0.029172 | 0.022528 | 0.024942 | 0.017801 | 0.014404 | 0.017076 | 0.043704 | 0.043648 | 0.046612 | 0.035822 | 0.019228 | 0.029192 | 0.028282 | 0.029982 | 0.028675 | 0.028896 |
| 28.25 | 0.016622 | 0.009058 | 0.025987 | 0.022314 | 0.010599 | 0.017665 | 0.025979 | 0.021411 | 0.017751 | 0.010852 | 0.014262 | 0.010326 | 0.042576 | 0.040161 | 0.044371 | 0.033335 | 0.018589 | 0.028883 | 0.024164 | 0.026028 | 0.023753 | 0.023346 |
| 28.26388889 | 0.014013 | 0.006533 | 0.015032 | 0.021004 | 0.013213 | 0.01181 | 0.017654 | 0.016488 | 0.015175 | 0.011208 | 0.013904 | 0.009442 | 0.031279 | 0.029326 | 0.033711 | 0.024433 | 0.015161 | 0.022319 | 0.016268 | 0.016469 | 0.015399 | 0.014842 |
| 28.27777778 | 0.011765 | 0.005263 | 0.007629 | 0.015775 | 0.011556 | 0.009503 | 0.012865 | 0.011269 | 0.01406 | 0.010139 | 0.014648 | 0.009805 | 0.016571 | 0.016674 | 0.022722 | 0.016882 | 0.011928 | 0.016029 | 0.009411 | 0.008299 | 0.007718 | 0.007428 |
| 28.29166667 | 0.01332 | 0.007033 | 0.00437 | 0.009412 | 0.006103 | 0.008212 | 0.011137 | 0.008414 | 0.011862 | 0.005564 | 0.017027 | 0.007749 | 0.00753 | 0.009164 | 0.016918 | 0.014565 | 0.009695 | 0.012167 | 0.00634 | 0.004874 | 0.005082 | 0.004228 |
| 28.30555556 | 0.017375 | 0.010308 | 0.002556 | 0.005098 | 0.002502 | 0.005267 | 0.007207 | 0.005695 | 0.008733 | 0.003298 | 0.018185 | 0.005464 | 0.00742 | 0.008497 | 0.014604 | 0.01381 | 0.007132 | 0.008735 | 0.006026 | 0.004266 | 0.007183 | 0.004257 |
| 28.31944444 | 0.01882 | 0.01052 | 0.001548 | 0.002882 | 0.001407 | 0.002993 | 0.00262 | 0.002735 | 0.004994 | 0.002891 | 0.016696 | 0.003803 | 0.010504 | 0.009703 | 0.011584 | 0.01074 | 0.004593 | 0.006217 | 0.006572 | 0.003727 | 0.00925 | 0.004751 |
| 28.33333333 | 0.01545 | 0.006392 | 0.002044 | 0.003123 | 0.002031 | 0.002186 | 0.001189 | 0.001327 | 0.003293 | 0.003002 | 0.013951 | 0.002189 | 0.008828 | 0.007663 | 0.007548 | 0.007018 | 0.003338 | 0.005647 | 0.007653 | 0.0032 | 0.008585 | 0.004954 |
| 28.34722222 | 0.010192 | 0.002498 | 0.002974 | 0.005379 | 0.004452 | 0.002797 | 0.002394 | 0.001175 | 0.005998 | 0.006289 | 0.009536 | 0.001785 | 0.005463 | 0.003738 | 0.004498 | 0.005357 | 0.002385 | 0.00429 | 0.009022 | 0.003491 | 0.00701 | 0.005306 |
| 28.36111111 | 0.005904 | 0.00351 | 0.005731 | 0.008913 | 0.006423 | 0.006467 | 0.005904 | 0.004213 | 0.011001 | 0.010917 | 0.006207 | 0.003874 | 0.00778 | 0.001938 | 0.002523 | 0.004467 | 0.001319 | 0.002393 | 0.008816 | 0.003339 | 0.006677 | 0.005379 |
| 28.375 | 0.006308 | 0.010029 | 0.012713 | 0.01439 | 0.008717 | 0.010761 | 0.010367 | 0.010916 | 0.017714 | 0.01567 | 0.008467 | 0.010179 | 0.011852 | 0.002284 | 0.002254 | 0.003106 | 0.001142 | 0.002609 | 0.005779 | 0.0024 | 0.006703 | 0.004601 |
| 28.38888889 | 0.013938 | 0.019811 | 0.02169 | 0.021721 | 0.016012 | 0.014632 | 0.01393 | 0.016859 | 0.0268 | 0.021318 | 0.01148 | 0.017845 | 0.013034 | 0.003835 | 0.004787 | 0.003465 | 0.002035 | 0.00353 | 0.003832 | 0.002892 | 0.005114 | 0.003245 |
| 28.40277778 | 0.024346 | 0.029577 | 0.028003 | 0.028913 | 0.026619 | 0.020678 | 0.019462 | 0.021202 | 0.035343 | 0.027498 | 0.011956 | 0.02242 | 0.014568 | 0.00787 | 0.008441 | 0.006626 | 0.00583 | 0.007328 | 0.007039 | 0.005462 | 0.005571 | 0.004579 |
| 28.41666667 | 0.033366 | 0.037838 | 0.030276 | 0.033638 | 0.033691 | 0.02695 | 0.027363 | 0.026209 | 0.041 | 0.034186 | 0.01448 | 0.02629 | 0.019739 | 0.014759 | 0.012813 | 0.011797 | 0.013041 | 0.016289 | 0.012902 | 0.009464 | 0.012175 | 0.011565 |
| 28.43055556 | 0.039525 | 0.044918 | 0.031017 | 0.036921 | 0.038661 | 0.02975 | 0.032567 | 0.030009 | 0.044826 | 0.03839 | 0.01957 | 0.032581 | 0.027758 | 0.023031 | 0.019692 | 0.019386 | 0.019925 | 0.024585 | 0.019019 | 0.014802 | 0.020342 | 0.020328 |
| 28.44444444 | 0.043136 | 0.051987 | 0.035292 | 0.044305 | 0.049692 | 0.03184 | 0.036129 | 0.031747 | 0.049716 | 0.041405 | 0.025186 | 0.04107 | 0.03718 | 0.033009 | 0.030157 | 0.029856 | 0.024913 | 0.029199 | 0.025601 | 0.021271 | 0.027308 | 0.027647 |
| 28.45833333 | 0.049633 | 0.062171 | 0.050049 | 0.060071 | 0.066637 | 0.039538 | 0.046458 | 0.037071 | 0.05863 | 0.05417 | 0.034036 | 0.053206 | 0.051273 | 0.049203 | 0.045828 | 0.043342 | 0.032231 | 0.037426 | 0.033891 | 0.029918 | 0.037893 | 0.037846 |
| 28.47222222 | 0.063462 | 0.07738 | 0.075184 | 0.08047 | 0.08233 | 0.053256 | 0.064537 | 0.048868 | 0.071289 | 0.079619 | 0.049625 | 0.070224 | 0.074314 | 0.075513 | 0.067277 | 0.059527 | 0.045079 | 0.055089 | 0.046556 | 0.042788 | 0.053333 | 0.055479 |
| 28.48611111 | 0.082045 | 0.096128 | 0.099784 | 0.099018 | 0.097178 | 0.067725 | 0.082907 | 0.063913 | 0.087158 | 0.10587 | 0.071391 | 0.087878 | 0.101453 | 0.10904 | 0.090735 | 0.075974 | 0.059227 | 0.075605 | 0.063141 | 0.05932 | 0.067439 | 0.0765 |
| 28.5 | 0.10001 | 0.116082 | 0.11703 | 0.117912 | 0.116489 | 0.079969 | 0.100553 | 0.082227 | 0.107877 | 0.127079 | 0.095608 | 0.100679 | 0.121476 | 0.142446 | 0.113352 | 0.091975 | 0.067177 | 0.089116 | 0.079269 | 0.075199 | 0.076467 | 0.092423 |
| 28.51388889 | 0.112949 | 0.134737 | 0.130444 | 0.138735 | 0.135042 | 0.089478 | 0.118618 | 0.105921 | 0.130214 | 0.145743 | 0.117129 | 0.108142 | 0.123831 | 0.169053 | 0.136835 | 0.109505 | 0.064805 | 0.093928 | 0.08848 | 0.082961 | 0.080141 | 0.09972 |
| 28.52777778 | 0.116203 | 0.14854 | 0.147199 | 0.150002 | 0.136218 | 0.092153 | 0.128171 | 0.131118 | 0.149123 | 0.162273 | 0.132103 | 0.113515 | 0.109455 | 0.190312 | 0.157133 | 0.122384 | 0.053374 | 0.091261 | 0.086475 | 0.078847 | 0.077513 | 0.099299 |
| 28.54166667 | 0.104661 | 0.147142 | 0.161739 | 0.14507 | 0.113315 | 0.080778 | 0.115844 | 0.140143 | 0.152446 | 0.164472 | 0.139934 | 0.114323 | 0.094144 | 0.201084 | 0.16234 | 0.124491 | 0.038371 | 0.080755 | 0.080945 | 0.068793 | 0.073694 | 0.10086 |
| 28.55555556 | 0.076735 | 0.114884 | 0.133424 | 0.113154 | 0.075275 | 0.052479 | 0.075241 | 0.106189 | 0.11285 | 0.124912 | 0.127942 | 0.092752 | 0.068398 | 0.163932 | 0.13723 | 0.112068 | 0.02488 | 0.065918 | 0.082256 | 0.055489 | 0.069863 | 0.105725 |
| 28.56944444 | 0.044197 | 0.061418 | 0.072475 | 0.058146 | 0.034855 | 0.023646 | 0.032053 | 0.048482 | 0.046629 | 0.059356 | 0.083467 | 0.046753 | 0.042241 | 0.09535 | 0.078188 | 0.079553 | 0.015918 | 0.048525 | 0.075861 | 0.033434 | 0.057982 | 0.092034 |
| 28.58333333 | 0.033554 | 0.035034 | 0.034554 | 0.021641 | 0.016708 | 0.012623 | 0.018665 | 0.028661 | 0.034912 | 0.040789 | 0.038929 | 0.013655 | 0.05173 | 0.061745 | 0.023908 | 0.050412 | 0.013628 | 0.033299 | 0.048534 | 0.011597 | 0.043741 | 0.060699 |
| 28.59722222 | 0.059109 | 0.062581 | 0.033739 | 0.023834 | 0.028202 | 0.020726 | 0.033951 | 0.06958 | 0.1002 | 0.093161 | 0.040057 | 0.019516 | 0.055406 | 0.047551 | 0.018943 | 0.049454 | 0.013745 | 0.030078 | 0.0245 | 0.005526 | 0.041729 | 0.040671 |
| 28.61111111 | 0.09301 | 0.108856 | 0.075797 | 0.057696 | 0.046545 | 0.039442 | 0.06295 | 0.132997 | 0.157817 | 0.144276 | 0.082699 | 0.050133 | 0.036181 | 0.035384 | 0.04902 | 0.066594 | 0.00974 | 0.033101 | 0.019885 | 0.008478 | 0.045545 | 0.036393 |
| 28.625 | 0.097832 | 0.140494 | 0.1135 | 0.091241 | 0.060775 | 0.052811 | 0.082033 | 0.162681 | 0.150888 | 0.130254 | 0.127762 | 0.089867 | 0.039859 | 0.050704 | 0.067478 | 0.079377 | 0.006498 | 0.033522 | 0.021701 | 0.010709 | 0.038168 | 0.035013 |
| 28.63888889 | 0.079429 | 0.150131 | 0.08919 | 0.077043 | 0.058088 | 0.057458 | 0.077084 | 0.137289 | 0.102918 | 0.071176 | 0.141019 | 0.121126 | 0.053933 | 0.084599 | 0.059965 | 0.070064 | 0.010107 | 0.035094 | 0.020584 | 0.009563 | 0.021505 | 0.034865 |
| 28.65277778 | 0.058419 | 0.122522 | 0.035445 | 0.037536 | 0.034055 | 0.044988 | 0.04936 | 0.084654 | 0.059582 | 0.026979 | 0.099221 | 0.097468 | 0.052962 | 0.111031 | 0.047693 | 0.040862 | 0.011643 | 0.03153 | 0.015495 | 0.004561 | 0.015107 | 0.030439 |
| 28.66666667 | 0.036366 | 0.068926 | 0.009735 | 0.027879 | 0.017962 | 0.02558 | 0.024392 | 0.050637 | 0.039683 | 0.028401 | 0.052124 | 0.048642 | 0.044198 | 0.100526 | 0.042274 | 0.014899 | 0.006411 | 0.018398 | 0.008357 | 0.003032 | 0.023019 | 0.018755 |
| 28.68055556 | 0.015874 | 0.027389 | 0.011834 | 0.028222 | 0.019148 | 0.024408 | 0.033899 | 0.039804 | 0.035362 | 0.052543 | 0.053062 | 0.040157 | 0.031532 | 0.060115 | 0.03063 | 0.010733 | 0.003163 | 0.011293 | 0.003049 | 0.006793 | 0.021386 | 0.015081 |
| 28.69444444 | 0.005946 | 0.010765 | 0.023404 | 0.024589 | 0.019093 | 0.027338 | 0.058153 | 0.046911 | 0.05848 | 0.06422 | 0.044505 | 0.03437 | 0.019083 | 0.039125 | 0.014311 | 0.015763 | 0.003987 | 0.009613 | 0.005231 | 0.011711 | 0.008027 | 0.016246 |
| 28.70833333 | 0.00896 | 0.012832 | 0.044994 | 0.041815 | 0.015834 | 0.026072 | 0.063589 | 0.063375 | 0.073484 | 0.062456 | 0.024611 | 0.020548 | 0.017164 | 0.034042 | 0.007615 | 0.015437 | 0.007581 | 0.009707 | 0.014794 | 0.0131 | 0.006422 | 0.00986 |
| 28.72222222 | 0.013354 | 0.020652 | 0.046427 | 0.035386 | 0.017476 | 0.018368 | 0.044688 | 0.048013 | 0.052653 | 0.071942 | 0.04419 | 0.022588 | 0.018355 | 0.029943 | 0.010769 | 0.010869 | 0.01221 | 0.019165 | 0.019788 | 0.009922 | 0.015247 | 0.004573 |
| 28.73611111 | 0.010605 | 0.020692 | 0.030245 | 0.04381 | 0.018613 | 0.014944 | 0.033247 | 0.029286 | 0.048395 | 0.075093 | 0.056209 | 0.021736 | 0.011339 | 0.047599 | 0.026578 | 0.005232 | 0.011215 | 0.024018 | 0.013949 | 0.008774 | 0.014993 | 0.011801 |
| 28.75 | 0.006349 | 0.020535 | 0.034646 | 0.035301 | 0.012234 | 0.027338 | 0.044977 | 0.046582 | 0.045809 | 0.053771 | 0.062366 | 0.01678 | 0.010773 | 0.058513 | 0.041783 | 0.003261 | 0.005622 | 0.01493 | 0.009348 | 0.014046 | 0.009312 | 0.031247 |
| 28.76388889 | 0.003714 | 0.033765 | 0.033539 | 0.025376 | 0.008391 | 0.02652 | 0.038925 | 0.048944 | 0.026431 | 0.047097 | 0.086191 | 0.014819 | 0.022267 | 0.042609 | 0.053447 | 0.004698 | 0.00211 | 0.007272 | 0.014456 | 0.018751 | 0.01005 | 0.03915 |
| 28.77777778 | 0.005872 | 0.041981 | 0.02049 | 0.017097 | 0.012349 | 0.012248 | 0.021001 | 0.034422 | 0.033775 | 0.058452 | 0.065731 | 0.015621 | 0.023278 | 0.04373 | 0.029873 | 0.007742 | 0.002698 | 0.006704 | 0.018501 | 0.015963 | 0.010815 | 0.024256 |
| 28.79166667 | 0.009153 | 0.040645 | 0.024346 | 0.028532 | 0.017658 | 0.01136 | 0.026932 | 0.038324 | 0.039901 | 0.04842 | 0.04472 | 0.019547 | 0.017776 | 0.056231 | 0.035803 | 0.008995 | 0.005004 | 0.004669 | 0.013126 | 0.009639 | 0.010911 | 0.008909 |
| 28.80555556 | 0.007112 | 0.040498 | 0.030611 | 0.040783 | 0.019283 | 0.018798 | 0.029253 | 0.033535 | 0.033945 | 0.034965 | 0.063703 | 0.029188 | 0.021844 | 0.041925 | 0.026885 | 0.008183 | 0.005372 | 0.00241 | 0.006689 | 0.00603 | 0.01316 | 0.007656 |
| 28.81944444 | 0.007288 | 0.031567 | 0.024562 | 0.03005 | 0.013393 | 0.019589 | 0.020712 | 0.02856 | 0.054592 | 0.04933 | 0.058454 | 0.033558 | 0.022509 | 0.023931 | 0.013455 | 0.006047 | 0.004121 | 0.005216 | 0.006691 | 0.00713 | 0.014056 | 0.009886 |
| 28.83333333 | 0.009259 | 0.025743 | 0.012025 | 0.017597 | 0.010239 | 0.021091 | 0.029925 | 0.039378 | 0.054656 | 0.047189 | 0.04713 | 0.038116 | 0.016167 | 0.034872 | 0.016798 | 0.00391 | 0.004879 | 0.008537 | 0.009094 | 0.007341 | 0.010916 | 0.005662 |
| 28.84722222 | 0.010611 | 0.035751 | 0.007978 | 0.019034 | 0.022061 | 0.025954 | 0.036792 | 0.045999 | 0.041173 | 0.028567 | 0.050895 | 0.063137 | 0.009214 | 0.041362 | 0.021832 | 0.011806 | 0.007097 | 0.00681 | 0.010678 | 0.004737 | 0.006943 | 0.00239 |
| 28.86111111 | 0.011078 | 0.038781 | 0.010317 | 0.015329 | 0.035622 | 0.021831 | 0.025531 | 0.045713 | 0.057285 | 0.036933 | 0.045764 | 0.075147 | 0.007936 | 0.031112 | 0.021707 | 0.024517 | 0.007506 | 0.002968 | 0.008714 | 0.004924 | 0.006271 | 0.003094 |
| 28.875 | 0.007258 | 0.02736 | 0.005507 | 0.015837 | 0.039037 | 0.02112 | 0.027477 | 0.036641 | 0.056546 | 0.044916 | 0.056302 | 0.072264 | 0.013078 | 0.040597 | 0.025956 | 0.022602 | 0.006388 | 0.003061 | 0.006989 | 0.007488 | 0.009781 | 0.009552 |
| 28.88888889 | 0.010051 | 0.034995 | 0.00149 | 0.024923 | 0.036684 | 0.02294 | 0.033236 | 0.030289 | 0.046449 | 0.047394 | 0.05912 | 0.062387 | 0.024742 | 0.07221 | 0.028303 | 0.012346 | 0.005395 | 0.00672 | 0.008292 | 0.006224 | 0.022931 | 0.026318 |
| 28.90277778 | 0.023366 | 0.047521 | 0.005277 | 0.032247 | 0.029892 | 0.021071 | 0.034338 | 0.031015 | 0.047415 | 0.060977 | 0.029359 | 0.046999 | 0.046069 | 0.081535 | 0.023913 | 0.007427 | 0.006272 | 0.010779 | 0.009413 | 0.00642 | 0.038313 | 0.041485 |
| 28.91666667 | 0.030993 | 0.03384 | 0.016499 | 0.04916 | 0.026012 | 0.02969 | 0.036512 | 0.031229 | 0.039578 | 0.051234 | 0.017059 | 0.052675 | 0.055496 | 0.04764 | 0.018897 | 0.003924 | 0.008516 | 0.014095 | 0.016314 | 0.015662 | 0.03854 | 0.041584 |
| 28.93055556 | 0.02191 | 0.015951 | 0.031908 | 0.05361 | 0.019826 | 0.029709 | 0.059164 | 0.033074 | 0.031584 | 0.026204 | 0.01864 | 0.040821 | 0.048241 | 0.015695 | 0.017344 | 0.003455 | 0.009573 | 0.015056 | 0.026684 | 0.02992 | 0.02328 | 0.028196 |
| 28.94444444 | 0.007502 | 0.00855 | 0.036184 | 0.034851 | 0.007807 | 0.016198 | 0.031437 | 0.036007 | 0.038982 | 0.026981 | 0.052945 | 0.016914 | 0.041983 | 0.030512 | 0.016764 | 0.006557 | 0.010922 | 0.017227 | 0.035755 | 0.040073 | 0.008242 | 0.015861 |
| 28.95833333 | 0.010165 | 0.014858 | 0.030123 | 0.016639 | 0.009625 | 0.010843 | 0.017501 | 0.042407 | 0.063327 | 0.056911 | 0.141321 | 0.02464 | 0.046338 | 0.087864 | 0.019562 | 0.012654 | 0.014414 | 0.024301 | 0.043178 | 0.037646 | 0.010466 | 0.017361 |
| 28.97222222 | 0.04672 | 0.042451 | 0.046336 | 0.024644 | 0.031853 | 0.01953 | 0.020243 | 0.052529 | 0.094326 | 0.097722 | 0.1807 | 0.040498 | 0.068946 | 0.151833 | 0.038711 | 0.037986 | 0.021414 | 0.036653 | 0.043492 | 0.029446 | 0.035401 | 0.03529 |
| 28.98611111 | 0.09771 | 0.077792 | 0.092788 | 0.070896 | 0.062389 | 0.041471 | 0.036046 | 0.072318 | 0.124532 | 0.106846 | 0.128999 | 0.039441 | 0.093398 | 0.187675 | 0.078978 | 0.077713 | 0.037043 | 0.058766 | 0.041324 | 0.025419 | 0.066437 | 0.06537 |
| 29 | 0.104766 | 0.092827 | 0.125316 | 0.11552 | 0.087746 | 0.050183 | 0.059225 | 0.097095 | 0.137133 | 0.073896 | 0.06222 | 0.049498 | 0.091079 | 0.17765 | 0.12108 | 0.102186 | 0.059469 | 0.083611 | 0.043591 | 0.029183 | 0.079576 | 0.084124 |
| 29.01388889 | 0.068515 | 0.080274 | 0.111948 | 0.120966 | 0.100714 | 0.039211 | 0.075944 | 0.101337 | 0.119877 | 0.054268 | 0.031701 | 0.087346 | 0.068234 | 0.130066 | 0.133856 | 0.106488 | 0.074886 | 0.091173 | 0.049864 | 0.044596 | 0.071621 | 0.076474 |
| 29.02777778 | 0.051021 | 0.065259 | 0.08507 | 0.099961 | 0.098985 | 0.042538 | 0.085818 | 0.093414 | 0.104444 | 0.087774 | 0.024232 | 0.111294 | 0.05371 | 0.06828 | 0.116154 | 0.101674 | 0.07199 | 0.081835 | 0.061985 | 0.064926 | 0.056981 | 0.059627 |
| 29.04166667 | 0.065012 | 0.069363 | 0.088391 | 0.087427 | 0.0958 | 0.065204 | 0.088853 | 0.096046 | 0.11487 | 0.133118 | 0.013289 | 0.108862 | 0.060278 | 0.074431 | 0.089652 | 0.090576 | 0.054412 | 0.071227 | 0.071481 | 0.072536 | 0.04635 | 0.049083 |
| 29.05555556 | 0.075724 | 0.0764 | 0.105891 | 0.098393 | 0.104611 | 0.07459 | 0.07283 | 0.09717 | 0.12629 | 0.131117 | 0.008565 | 0.087934 | 0.078237 | 0.080427 | 0.066996 | 0.072913 | 0.035419 | 0.05984 | 0.065008 | 0.061302 | 0.041463 | 0.045619 |
| 29.06944444 | 0.069562 | 0.068846 | 0.09905 | 0.110722 | 0.106479 | 0.059105 | 0.04113 | 0.086817 | 0.112518 | 0.085906 | 0.018379 | 0.048301 | 0.078546 | 0.077297 | 0.05095 | 0.05251 | 0.023473 | 0.04441 | 0.047471 | 0.042508 | 0.03882 | 0.043939 |
| 29.08333333 | 0.056486 | 0.04884 | 0.067023 | 0.09413 | 0.073196 | 0.035037 | 0.01795 | 0.064287 | 0.073685 | 0.037155 | 0.032751 | 0.019475 | 0.04977 | 0.055134 | 0.037408 | 0.034776 | 0.017523 | 0.029862 | 0.02888 | 0.024443 | 0.034478 | 0.039255 |
| 29.09722222 | 0.046798 | 0.035039 | 0.036243 | 0.049678 | 0.036268 | 0.01512 | 0.011775 | 0.036994 | 0.030694 | 0.013163 | 0.031512 | 0.014269 | 0.020669 | 0.028231 | 0.024255 | 0.019912 | 0.011056 | 0.017234 | 0.013462 | 0.01055 | 0.027071 | 0.031336 |
| 29.11111111 | 0.050536 | 0.049253 | 0.033201 | 0.018042 | 0.028704 | 0.008021 | 0.014316 | 0.019308 | 0.013496 | 0.019366 | 0.024196 | 0.013764 | 0.009612 | 0.012615 | 0.014021 | 0.009322 | 0.005351 | 0.007385 | 0.004365 | 0.004195 | 0.017763 | 0.021803 |
| 29.125 | 0.063813 | 0.081691 | 0.065969 | 0.03124 | 0.033544 | 0.017556 | 0.028422 | 0.018391 | 0.02697 | 0.041618 | 0.041964 | 0.019022 | 0.011127 | 0.011316 | 0.008623 | 0.004976 | 0.004282 | 0.003755 | 0.002848 | 0.003188 | 0.010136 | 0.014024 |
| 29.13888889 | 0.074323 | 0.100828 | 0.110795 | 0.070848 | 0.051326 | 0.034662 | 0.050975 | 0.041406 | 0.046322 | 0.055853 | 0.061614 | 0.03142 | 0.019925 | 0.021565 | 0.009533 | 0.005309 | 0.004969 | 0.005932 | 0.007143 | 0.005033 | 0.007126 | 0.011793 |
| 29.15277778 | 0.077747 | 0.092864 | 0.127533 | 0.088741 | 0.062863 | 0.042969 | 0.059849 | 0.062566 | 0.051973 | 0.05384 | 0.055307 | 0.038125 | 0.030202 | 0.035801 | 0.015464 | 0.011026 | 0.00868 | 0.012144 | 0.012443 | 0.009205 | 0.009052 | 0.015101 |
| 29.16666667 | 0.069716 | 0.069719 | 0.103765 | 0.066256 | 0.049925 | 0.039305 | 0.047515 | 0.055523 | 0.045284 | 0.046578 | 0.036413 | 0.033795 | 0.035608 | 0.043922 | 0.021336 | 0.01841 | 0.013072 | 0.016253 | 0.01512 | 0.012837 | 0.01342 | 0.019343 |
| 29.18055556 | 0.053914 | 0.048991 | 0.068249 | 0.032421 | 0.034312 | 0.032701 | 0.030662 | 0.036538 | 0.037223 | 0.040868 | 0.021596 | 0.024378 | 0.035866 | 0.044483 | 0.023007 | 0.020284 | 0.013787 | 0.015046 | 0.017041 | 0.015653 | 0.017499 | 0.021189 |
| 29.19444444 | 0.041441 | 0.037243 | 0.052108 | 0.018729 | 0.028717 | 0.027245 | 0.026212 | 0.0244 | 0.034087 | 0.0346 | 0.014463 | 0.024544 | 0.036796 | 0.044583 | 0.022207 | 0.018179 | 0.014529 | 0.014769 | 0.0224 | 0.020341 | 0.020797 | 0.022575 |
| 29.20833333 | 0.034289 | 0.029872 | 0.053628 | 0.024728 | 0.029114 | 0.023684 | 0.03202 | 0.020808 | 0.033158 | 0.025766 | 0.013142 | 0.035362 | 0.042898 | 0.049475 | 0.024465 | 0.019694 | 0.018586 | 0.019961 | 0.03022 | 0.026662 | 0.023494 | 0.026583 |
| 29.22222222 | 0.028015 | 0.020685 | 0.050585 | 0.029381 | 0.025748 | 0.023691 | 0.033573 | 0.0208 | 0.030206 | 0.020379 | 0.015553 | 0.038282 | 0.053807 | 0.058356 | 0.031835 | 0.026583 | 0.023702 | 0.026926 | 0.035254 | 0.031578 | 0.025758 | 0.031705 |
| 29.23611111 | 0.020981 | 0.010754 | 0.034766 | 0.021834 | 0.017073 | 0.025047 | 0.026712 | 0.019652 | 0.026219 | 0.022715 | 0.018119 | 0.025628 | 0.064913 | 0.066628 | 0.039281 | 0.031264 | 0.025306 | 0.029184 | 0.034587 | 0.032711 | 0.02713 | 0.032604 |
| 29.25 | 0.015772 | 0.005836 | 0.016751 | 0.011173 | 0.010368 | 0.022338 | 0.020478 | 0.01691 | 0.022665 | 0.023649 | 0.017834 | 0.012149 | 0.070257 | 0.068633 | 0.039885 | 0.027673 | 0.022986 | 0.026843 | 0.029672 | 0.029655 | 0.026233 | 0.026751 |
| 29.26388889 | 0.012839 | 0.006173 | 0.006279 | 0.008038 | 0.009632 | 0.015497 | 0.018971 | 0.016489 | 0.018436 | 0.015166 | 0.015265 | 0.007739 | 0.066093 | 0.061089 | 0.032943 | 0.018112 | 0.019666 | 0.023911 | 0.023685 | 0.023025 | 0.022705 | 0.017969 |
| 29.27777778 | 0.009372 | 0.006208 | 0.002971 | 0.008732 | 0.010002 | 0.010311 | 0.017642 | 0.017206 | 0.013592 | 0.005271 | 0.013265 | 0.006904 | 0.0535 | 0.047254 | 0.023945 | 0.009477 | 0.015825 | 0.020115 | 0.017729 | 0.014613 | 0.01739 | 0.011321 |
| 29.29166667 | 0.006426 | 0.003929 | 0.004472 | 0.006555 | 0.006437 | 0.009197 | 0.013453 | 0.013897 | 0.01027 | 0.002014 | 0.013451 | 0.005292 | 0.038237 | 0.034169 | 0.017463 | 0.00639 | 0.010563 | 0.013716 | 0.011772 | 0.007704 | 0.011661 | 0.00763 |

| | | | | | | | | | | | | | | | | | | | | | | |
|---|---|---|---|---|---|---|---|---|---|---|---|---|---|---|---|---|---|---|---|---|---|---|
| 29.30555556 | 0.007296 | 0.001998 | 0.00872 | 0.004022 | 0.002072 | 0.007756 | 0.007717 | 0.008842 | 0.008951 | 0.003723 | 0.014102 | 0.004268 | 0.026422 | 0.024897 | 0.013746 | 0.00749 | 0.006466 | 0.008307 | 0.007887 | 0.004562 | 0.007623 | 0.005578 |
| 29.31944444 | 0.009674 | 0.001341 | 0.01091 | 0.003403 | 0.001725 | 0.004495 | 0.003027 | 0.007637 | 0.006369 | 0.008211 | 0.012744 | 0.006594 | 0.02016 | 0.018589 | 0.01136 | 0.008651 | 0.00666 | 0.008258 | 0.007877 | 0.004458 | 0.007014 | 0.005306 |
| 29.33333333 | 0.007798 | 0.003299 | 0.010663 | 0.004728 | 0.006019 | 0.003273 | 0.001286 | 0.007085 | 0.004652 | 0.011367 | 0.00972 | 0.009881 | 0.01761 | 0.014892 | 0.009868 | 0.008243 | 0.008586 | 0.010403 | 0.010156 | 0.005305 | 0.008627 | 0.006466 |
| 29.34722222 | 0.007204 | 0.010928 | 0.012109 | 0.009491 | 0.012993 | 0.004695 | 0.002376 | 0.005476 | 0.009269 | 0.012161 | 0.005815 | 0.008118 | 0.013931 | 0.012197 | 0.008446 | 0.007838 | 0.006782 | 0.008254 | 0.011752 | 0.005071 | 0.008919 | 0.00662 |
| 29.36111111 | 0.017138 | 0.021256 | 0.015561 | 0.015195 | 0.017927 | 0.007391 | 0.004354 | 0.006601 | 0.01563 | 0.016206 | 0.003526 | 0.004419 | 0.009431 | 0.00792 | 0.00534 | 0.007837 | 0.002847 | 0.003674 | 0.010593 | 0.00345 | 0.00597 | 0.004748 |
| 29.375 | 0.028738 | 0.02862 | 0.01836 | 0.018488 | 0.019582 | 0.01036 | 0.006125 | 0.007319 | 0.018563 | 0.021314 | 0.004633 | 0.008339 | 0.011842 | 0.005409 | 0.003416 | 0.006122 | 0.001356 | 0.001706 | 0.006425 | 0.002595 | 0.0025 | 0.002469 |
| 29.38888889 | 0.031617 | 0.032748 | 0.022138 | 0.021632 | 0.023076 | 0.013469 | 0.009727 | 0.007347 | 0.020765 | 0.022403 | 0.006234 | 0.021992 | 0.018736 | 0.006281 | 0.006168 | 0.003416 | 0.000915 | 0.001106 | 0.002361 | 0.002621 | 0.001642 | 0.001281 |
| 29.40277778 | 0.030544 | 0.037003 | 0.031862 | 0.029257 | 0.032181 | 0.020036 | 0.017666 | 0.013499 | 0.026344 | 0.025105 | 0.009931 | 0.036938 | 0.021161 | 0.006147 | 0.010341 | 0.003685 | 0.003092 | 0.004287 | 0.00146 | 0.00259 | 0.002778 | 0.002233 |
| 29.41666667 | 0.033123 | 0.043056 | 0.045606 | 0.041594 | 0.045289 | 0.030569 | 0.027909 | 0.023379 | 0.034843 | 0.034069 | 0.017751 | 0.047961 | 0.022479 | 0.008276 | 0.014459 | 0.008662 | 0.011184 | 0.014818 | 0.00422 | 0.005351 | 0.005822 | 0.005959 |
| 29.43055556 | 0.039435 | 0.049653 | 0.055027 | 0.052753 | 0.057332 | 0.038374 | 0.035685 | 0.029191 | 0.042157 | 0.044253 | 0.024597 | 0.05659 | 0.030427 | 0.018023 | 0.021345 | 0.017718 | 0.021202 | 0.026696 | 0.012279 | 0.013123 | 0.011834 | 0.011805 |
| 29.44444444 | 0.046081 | 0.05739 | 0.058797 | 0.059254 | 0.065887 | 0.04004 | 0.040909 | 0.032748 | 0.048549 | 0.052527 | 0.02866 | 0.065446 | 0.044518 | 0.03282 | 0.031307 | 0.029395 | 0.029028 | 0.03445 | 0.025668 | 0.024262 | 0.02028 | 0.01945 |
| 29.45833333 | 0.054512 | 0.069923 | 0.066317 | 0.068437 | 0.076405 | 0.043209 | 0.048979 | 0.042334 | 0.059512 | 0.064191 | 0.036982 | 0.079165 | 0.061802 | 0.051054 | 0.04305 | 0.043262 | 0.037083 | 0.042636 | 0.040563 | 0.037333 | 0.030165 | 0.029569 |
| 29.47222222 | 0.067205 | 0.087735 | 0.084997 | 0.090362 | 0.095679 | 0.054993 | 0.062367 | 0.06064 | 0.077142 | 0.081178 | 0.05657 | 0.098804 | 0.084307 | 0.077526 | 0.057619 | 0.062552 | 0.049681 | 0.057809 | 0.055283 | 0.052899 | 0.04164 | 0.043016 |
| 29.48611111 | 0.082001 | 0.107076 | 0.111485 | 0.122375 | 0.120839 | 0.07174 | 0.078658 | 0.080865 | 0.097552 | 0.096357 | 0.085557 | 0.118335 | 0.114619 | 0.116833 | 0.077119 | 0.089195 | 0.064087 | 0.077132 | 0.072602 | 0.070385 | 0.055314 | 0.060429 |
| 29.5 | 0.097419 | 0.129154 | 0.141972 | 0.151374 | 0.143092 | 0.086969 | 0.096913 | 0.098835 | 0.122652 | 0.111329 | 0.117681 | 0.13063 | 0.142883 | 0.163535 | 0.102057 | 0.118299 | 0.071753 | 0.09116 | 0.091622 | 0.085102 | 0.069245 | 0.080496 |
| 29.51388889 | 0.114791 | 0.156506 | 0.171952 | 0.172579 | 0.155719 | 0.09899 | 0.116691 | 0.12337 | 0.155927 | 0.137997 | 0.147451 | 0.137101 | 0.151668 | 0.205121 | 0.127007 | 0.140575 | 0.070935 | 0.096588 | 0.10337 | 0.091878 | 0.079503 | 0.098697 |
| 29.52777778 | 0.127472 | 0.174438 | 0.194578 | 0.192982 | 0.151967 | 0.106346 | 0.12947 | 0.156578 | 0.186148 | 0.165368 | 0.165741 | 0.137303 | 0.140526 | 0.232785 | 0.137083 | 0.148976 | 0.064582 | 0.096126 | 0.100621 | 0.091878 | 0.085668 | 0.110639 |
| 29.54166667 | 0.117447 | 0.155686 | 0.192426 | 0.195179 | 0.1248 | 0.102689 | 0.120156 | 0.16913 | 0.186813 | 0.15698 | 0.162529 | 0.119274 | 0.116781 | 0.229454 | 0.129672 | 0.137722 | 0.052402 | 0.090698 | 0.086035 | 0.088634 | 0.090146 | 0.111383 |
| 29.55555556 | 0.072741 | 0.094932 | 0.128732 | 0.133909 | 0.071713 | 0.076376 | 0.082221 | 0.124428 | 0.1301 | 0.096241 | 0.127868 | 0.079772 | 0.075919 | 0.162743 | 0.106255 | 0.102558 | 0.035467 | 0.076714 | 0.071876 | 0.076565 | 0.086746 | 0.094868 |
| 29.56944444 | 0.033157 | 0.040736 | 0.063605 | 0.05706 | 0.022985 | 0.036217 | 0.036996 | 0.050998 | 0.056097 | 0.037904 | 0.071354 | 0.0352 | 0.042037 | 0.08304 | 0.060234 | 0.053047 | 0.018443 | 0.054388 | 0.057784 | 0.051082 | 0.064887 | 0.061034 |
| 29.58333333 | 0.027101 | 0.027351 | 0.062393 | 0.035854 | 0.015831 | 0.01761 | 0.025589 | 0.026146 | 0.043622 | 0.033115 | 0.027736 | 0.013386 | 0.048532 | 0.06837 | 0.019038 | 0.018926 | 0.006712 | 0.033868 | 0.037038 | 0.025113 | 0.034727 | 0.028682 |
| 29.59722222 | 0.043521 | 0.04693 | 0.06501 | 0.037872 | 0.035005 | 0.039643 | 0.062587 | 0.068534 | 0.103957 | 0.075195 | 0.036856 | 0.041937 | 0.052699 | 0.060141 | 0.012084 | 0.023036 | 0.003632 | 0.029274 | 0.022482 | 0.015813 | 0.020609 | 0.023884 |
| 29.61111111 | 0.093388 | 0.099375 | 0.075064 | 0.058167 | 0.049204 | 0.075479 | 0.100962 | 0.139815 | 0.180973 | 0.137995 | 0.104362 | 0.112405 | 0.035599 | 0.037302 | 0.02517 | 0.048214 | 0.006839 | 0.042866 | 0.019886 | 0.023372 | 0.031052 | 0.047053 |
| 29.625 | 0.122205 | 0.138079 | 0.104296 | 0.08394 | 0.040316 | 0.083741 | 0.099992 | 0.182585 | 0.192983 | 0.151187 | 0.162278 | 0.158532 | 0.045826 | 0.05252 | 0.039504 | 0.066873 | 0.009872 | 0.053661 | 0.019476 | 0.031949 | 0.055184 | 0.075744 |
| 29.63888889 | 0.088128 | 0.115877 | 0.091746 | 0.066023 | 0.019085 | 0.068669 | 0.074135 | 0.149246 | 0.120654 | 0.094653 | 0.159236 | 0.135218 | 0.069916 | 0.073425 | 0.043665 | 0.059451 | 0.010886 | 0.04855 | 0.016718 | 0.026813 | 0.068475 | 0.083263 |
| 29.65277778 | 0.048017 | 0.066181 | 0.046387 | 0.033846 | 0.01038 | 0.045134 | 0.044588 | 0.07293 | 0.045393 | 0.050884 | 0.102813 | 0.071005 | 0.064914 | 0.071374 | 0.035601 | 0.031631 | 0.012249 | 0.039415 | 0.01449 | 0.013452 | 0.051535 | 0.061898 |
| 29.66666667 | 0.024241 | 0.032645 | 0.02152 | 0.021963 | 0.013905 | 0.023469 | 0.027025 | 0.02349 | 0.020519 | 0.045968 | 0.054463 | 0.023162 | 0.039017 | 0.062871 | 0.031337 | 0.021789 | 0.011583 | 0.032479 | 0.013813 | 0.00845 | 0.024626 | 0.033574 |
| 29.68055556 | 0.011184 | 0.02514 | 0.016742 | 0.014228 | 0.01738 | 0.017204 | 0.021181 | 0.019448 | 0.02738 | 0.04919 | 0.059714 | 0.011632 | 0.020819 | 0.045688 | 0.035295 | 0.028145 | 0.007004 | 0.022781 | 0.012696 | 0.008096 | 0.015585 | 0.019748 |
| 29.69444444 | 0.011387 | 0.023914 | 0.015236 | 0.016076 | 0.015782 | 0.01491 | 0.026473 | 0.035842 | 0.058018 | 0.074842 | 0.054723 | 0.022753 | 0.013854 | 0.027056 | 0.029766 | 0.022972 | 0.002999 | 0.014403 | 0.014066 | 0.003899 | 0.013194 | 0.014295 |
| 29.70833333 | 0.009799 | 0.015181 | 0.022156 | 0.032568 | 0.013467 | 0.013364 | 0.037208 | 0.035454 | 0.072019 | 0.09297 | 0.042347 | 0.046852 | 0.014567 | 0.026886 | 0.016572 | 0.01855 | 0.002179 | 0.010687 | 0.015211 | 0.003334 | 0.009144 | 0.014072 |
| 29.72222222 | 0.003421 | 0.01213 | 0.022298 | 0.03368 | 0.009978 | 0.016303 | 0.028485 | 0.028438 | 0.059027 | 0.084105 | 0.061505 | 0.051125 | 0.018311 | 0.023771 | 0.01524 | 0.029502 | 0.004903 | 0.012847 | 0.009955 | 0.007098 | 0.015349 | 0.029097 |
| 29.73611111 | 0.001205 | 0.013679 | 0.014291 | 0.0304 | 0.013529 | 0.018865 | 0.018808 | 0.03802 | 0.064434 | 0.087833 | 0.060979 | 0.053446 | 0.01415 | 0.024876 | 0.017531 | 0.034536 | 0.006448 | 0.019976 | 0.003919 | 0.008389 | 0.022743 | 0.038537 |
| 29.75 | 0.001616 | 0.014467 | 0.015405 | 0.031814 | 0.021369 | 0.027337 | 0.040705 | 0.034679 | 0.068439 | 0.073424 | 0.059303 | 0.086033 | 0.013571 | 0.045778 | 0.019821 | 0.021648 | 0.004048 | 0.020112 | 0.0031 | 0.009686 | 0.017111 | 0.028715 |
| 29.76388889 | 0.001568 | 0.013063 | 0.014372 | 0.020516 | 0.024341 | 0.039771 | 0.061747 | 0.015884 | 0.053481 | 0.050843 | 0.082002 | 0.082432 | 0.022859 | 0.043232 | 0.024278 | 0.010139 | 0.004241 | 0.017408 | 0.006537 | 0.016169 | 0.010358 | 0.025277 |
| 29.77777778 | 0.001248 | 0.007985 | 0.01076 | 0.012143 | 0.022252 | 0.032299 | 0.051149 | 0.00877 | 0.058172 | 0.072546 | 0.069188 | 0.045365 | 0.024022 | 0.029373 | 0.017229 | 0.007535 | 0.006589 | 0.01981 | 0.008336 | 0.017593 | 0.012081 | 0.035284 |
| 29.79166667 | 0.003536 | 0.007044 | 0.011893 | 0.007137 | 0.013315 | 0.012618 | 0.036276 | 0.015233 | 0.079992 | 0.079098 | 0.043137 | 0.041969 | 0.022396 | 0.043799 | 0.007184 | 0.007432 | 0.007705 | 0.01805 | 0.005475 | 0.009654 | 0.0113 | 0.032493 |
| 29.80555556 | 0.008209 | 0.011318 | 0.014553 | 0.00894 | 0.007666 | 0.010934 | 0.033796 | 0.018245 | 0.073332 | 0.062845 | 0.049714 | 0.044285 | 0.029956 | 0.052862 | 0.004231 | 0.013192 | 0.009267 | 0.014052 | 0.003088 | 0.005854 | 0.009512 | 0.023332 |
| 29.81944444 | 0.012029 | 0.017892 | 0.023473 | 0.025805 | 0.007864 | 0.025218 | 0.035984 | 0.017621 | 0.06934 | 0.080485 | 0.042962 | 0.039071 | 0.030194 | 0.039849 | 0.00354 | 0.01616 | 0.00869 | 0.014464 | 0.002247 | 0.007506 | 0.012687 | 0.032321 |
| 29.83333333 | 0.013272 | 0.02118 | 0.028649 | 0.029823 | 0.008196 | 0.029396 | 0.039535 | 0.017549 | 0.084014 | 0.078277 | 0.022914 | 0.04707 | 0.01722 | 0.040064 | 0.0062 | 0.013257 | 0.004609 | 0.013673 | 0.002643 | 0.007781 | 0.014323 | 0.035358 |
| 29.84722222 | 0.009816 | 0.025338 | 0.023407 | 0.019145 | 0.006859 | 0.029115 | 0.045984 | 0.011043 | 0.0674 | 0.047928 | 0.035687 | 0.043259 | 0.007998 | 0.039321 | 0.008054 | 0.024158 | 0.003965 | 0.013474 | 0.003741 | 0.008043 | 0.017419 | 0.027541 |
| 29.86111111 | 0.007134 | 0.027751 | 0.01675 | 0.013762 | 0.007239 | 0.040484 | 0.049105 | 0.010841 | 0.050578 | 0.04647 | 0.056504 | 0.030827 | 0.010428 | 0.029943 | 0.004508 | 0.046498 | 0.007755 | 0.01363 | 0.00424 | 0.010584 | 0.016052 | 0.029274 |
| 29.875 | 0.01132 | 0.019018 | 0.01672 | 0.017137 | 0.010587 | 0.034717 | 0.040416 | 0.016883 | 0.055195 | 0.043041 | 0.048297 | 0.03744 | 0.024029 | 0.051745 | 0.001698 | 0.050852 | 0.008104 | 0.010483 | 0.00973 | 0.012384 | 0.010534 | 0.025594 |
| 29.88888889 | 0.016379 | 0.023284 | 0.018998 | 0.025663 | 0.009787 | 0.015286 | 0.029456 | 0.030311 | 0.058594 | 0.031897 | 0.026495 | 0.040799 | 0.053111 | 0.096539 | 0.006451 | 0.029059 | 0.005466 | 0.01649 | 0.017842 | 0.015894 | 0.01493 | 0.01784 |
| 29.90277778 | 0.02579 | 0.03961 | 0.016889 | 0.03107 | 0.010848 | 0.00979 | 0.018497 | 0.050387 | 0.076429 | 0.043504 | 0.01328 | 0.028238 | 0.07442 | 0.110897 | 0.01913 | 0.010115 | 0.006645 | 0.026168 | 0.017502 | 0.019668 | 0.01642 | 0.016893 |
| 29.91666667 | 0.046555 | 0.040034 | 0.024449 | 0.047805 | 0.026302 | 0.011682 | 0.013134 | 0.0432 | 0.083547 | 0.046432 | 0.021444 | 0.021612 | 0.063313 | 0.06698 | 0.033704 | 0.010088 | 0.010504 | 0.022512 | 0.009684 | 0.016033 | 0.014056 | 0.015987 |
| 29.93055556 | 0.052248 | 0.035711 | 0.03456 | 0.063622 | 0.044109 | 0.018806 | 0.026453 | 0.016615 | 0.051935 | 0.027508 | 0.057653 | 0.045538 | 0.037832 | 0.018636 | 0.050733 | 0.019854 | 0.013957 | 0.0161 | 0.003515 | 0.008583 | 0.027742 | 0.026253 |
| 29.94444444 | 0.034972 | 0.035394 | 0.029926 | 0.04559 | 0.051716 | 0.028287 | 0.044252 | 0.01757 | 0.015987 | 0.013528 | 0.091888 | 0.103053 | 0.028754 | 0.030911 | 0.069603 | 0.034357 | 0.021776 | 0.023204 | 0.005572 | 0.008353 | 0.045327 | 0.049304 |
| 29.95833333 | 0.027855 | 0.026833 | 0.019651 | 0.021951 | 0.056758 | 0.03636 | 0.058093 | 0.056448 | 0.026641 | 0.032307 | 0.088179 | 0.12475 | 0.049939 | 0.089202 | 0.067288 | 0.049675 | 0.036592 | 0.043477 | 0.018991 | 0.025286 | 0.045303 | 0.060868 |
| 29.97222222 | 0.030626 | 0.028995 | 0.033204 | 0.03833 | 0.062935 | 0.052116 | 0.076953 | 0.100092 | 0.092626 | 0.082035 | 0.06608 | 0.082556 | 0.07803 | 0.133095 | 0.038109 | 0.05753 | 0.049373 | 0.06547 | 0.036773 | 0.05016 | 0.036108 | 0.057439 |
| 29.98611111 | 0.039301 | 0.050951 | 0.070149 | 0.082631 | 0.06676 | 0.075811 | 0.088451 | 0.114732 | 0.156694 | 0.108165 | 0.063426 | 0.044698 | 0.082686 | 0.135671 | 0.018578 | 0.061354 | 0.052353 | 0.078269 | 0.043181 | 0.057935 | 0.039477 | 0.060823 |
| 30 | 0.06392 | 0.065955 | 0.090526 | 0.105042 | 0.069741 | 0.081926 | 0.075506 | 0.112241 | 0.165088 | 0.090425 | 0.086756 | 0.051117 | 0.067188 | 0.118836 | 0.03119 | 0.071074 | 0.051212 | 0.080577 | 0.038055 | 0.050054 | 0.059288 | 0.077884 |
| 30.01388889 | 0.07728 | 0.07389 | 0.102853 | 0.105695 | 0.078701 | 0.056059 | 0.060573 | 0.113749 | 0.130714 | 0.083702 | 0.128257 | 0.084998 | 0.047003 | 0.105509 | 0.062577 | 0.083895 | 0.052934 | 0.081076 | 0.040314 | 0.046646 | 0.077029 | 0.091815 |
| 30.02777778 | 0.067503 | 0.077944 | 0.130282 | 0.113001 | 0.088226 | 0.033812 | 0.073923 | 0.124207 | 0.103309 | 0.108684 | 0.163895 | 0.11295 | 0.038631 | 0.103071 | 0.095277 | 0.091694 | 0.057938 | 0.085704 | 0.055742 | 0.051201 | 0.075533 | 0.088868 |
| 30.04166667 | 0.058886 | 0.076354 | 0.140476 | 0.116624 | 0.085014 | 0.045677 | 0.102426 | 0.131749 | 0.102975 | 0.124943 | 0.164026 | 0.113764 | 0.057466 | 0.111256 | 0.115067 | 0.084624 | 0.059798 | 0.083726 | 0.071372 | 0.0571 | 0.060429 | 0.074509 |
| 30.05555556 | 0.060774 | 0.074937 | 0.112211 | 0.104178 | 0.072617 | 0.070813 | 0.101927 | 0.099713 | 0.111001 | 0.117173 | 0.133498 | 0.093504 | 0.091035 | 0.119564 | 0.109592 | 0.064191 | 0.050932 | 0.064988 | 0.073218 | 0.057527 | 0.048011 | 0.061681 |
| 30.06944444 | 0.065723 | 0.063472 | 0.075319 | 0.08889 | 0.055439 | 0.07387 | 0.060145 | 0.050339 | 0.10482 | 0.091238 | 0.0907 | 0.061953 | 0.097588 | 0.10427 | 0.084936 | 0.047823 | 0.036897 | 0.04137 | 0.057783 | 0.049748 | 0.039505 | 0.05082 |
| 30.08333333 | 0.067385 | 0.042923 | 0.051196 | 0.068681 | 0.035232 | 0.052526 | 0.025893 | 0.028689 | 0.078808 | 0.049492 | 0.045151 | 0.029637 | 0.061238 | 0.060147 | 0.054042 | 0.038349 | 0.025861 | 0.025919 | 0.034864 | 0.036068 | 0.028879 | 0.03562 |
| 30.09722222 | 0.073411 | 0.032666 | 0.040349 | 0.035232 | 0.025615 | 0.029917 | 0.019883 | 0.021359 | 0.047879 | 0.016421 | 0.013794 | 0.01416 | 0.021778 | 0.019779 | 0.025308 | 0.026846 | 0.017115 | 0.016935 | 0.015134 | 0.02217 | 0.018229 | 0.019193 |
| 30.11111111 | 0.094626 | 0.04729 | 0.045825 | 0.013025 | 0.021409 | 0.026128 | 0.020784 | 0.018491 | 0.031297 | 0.008607 | 0.015798 | 0.020451 | 0.010935 | 0.005254 | 0.008117 | 0.013194 | 0.00932 | 0.010728 | 0.004326 | 0.014429 | 0.010774 | 0.007722 |
| 30.125 | 0.11614 | 0.07526 | 0.065333 | 0.023176 | 0.016572 | 0.041219 | 0.033504 | 0.018182 | 0.030229 | 0.014459 | 0.047144 | 0.040388 | 0.018381 | 0.007524 | 0.002592 | 0.003949 | 0.003555 | 0.007015 | 0.001967 | 0.013523 | 0.006993 | 0.002589 |
| 30.13888889 | 0.11375 | 0.085189 | 0.089531 | 0.048265 | 0.021006 | 0.050996 | 0.052923 | 0.021999 | 0.034125 | 0.026005 | 0.077742 | 0.052565 | 0.026362 | 0.015812 | 0.003537 | 0.001581 | 0.000919 | 0.005819 | 0.004055 | 0.015218 | 0.006953 | 0.00343 |
| 30.15277778 | 0.089994 | 0.06978 | 0.101322 | 0.061084 | 0.024491 | 0.039205 | 0.051121 | 0.037742 | 0.036509 | 0.031248 | 0.080191 | 0.04521 | 0.028828 | 0.024024 | 0.007347 | 0.002495 | 0.001491 | 0.006467 | 0.007961 | 0.015339 | 0.008065 | 0.007141 |
| 30.16666667 | 0.062025 | 0.046697 | 0.087646 | 0.055789 | 0.025253 | 0.019217 | 0.030869 | 0.052747 | 0.036958 | 0.027604 | 0.056134 | 0.032686 | 0.027428 | 0.026735 | 0.010184 | 0.006212 | 0.003534 | 0.007536 | 0.009423 | 0.012592 | 0.007495 | 0.00888 |
| 30.18055556 | 0.038673 | 0.029439 | 0.060764 | 0.04146 | 0.028789 | 0.009431 | 0.015656 | 0.054204 | 0.036409 | 0.027318 | 0.028125 | 0.025001 | 0.02502 | 0.024417 | 0.01202 | 0.011779 | 0.004547 | 0.006947 | 0.007895 | 0.009302 | 0.006337 | 0.00808 |
| 30.19444444 | 0.022404 | 0.017645 | 0.04231 | 0.025885 | 0.023387 | 0.00941 | 0.012325 | 0.042519 | 0.034528 | 0.027872 | 0.010224 | 0.01775 | 0.025186 | 0.025881 | 0.016631 | 0.016244 | 0.005158 | 0.005692 | 0.009076 | 0.009615 | 0.008053 | 0.009318 |
| 30.20833333 | 0.014225 | 0.007784 | 0.039449 | 0.016111 | 0.012206 | 0.009432 | 0.012396 | 0.031162 | 0.0338 | 0.023007 | 0.002332 | 0.01178 | 0.031143 | 0.036071 | 0.023526 | 0.019614 | 0.00904 | 0.008802 | 0.015999 | 0.015611 | 0.013501 | 0.015214 |
| 30.22222222 | 0.01121 | 0.002975 | 0.045325 | 0.014658 | 0.008346 | 0.008417 | 0.009125 | 0.027883 | 0.036608 | 0.018562 | 0.001178 | 0.010689 | 0.043583 | 0.050089 | 0.027864 | 0.022605 | 0.017062 | 0.017864 | 0.024631 | 0.025079 | 0.019842 | 0.021723 |
| 30.23611111 | 0.007699 | 0.002809 | 0.044951 | 0.014259 | 0.009443 | 0.010678 | 0.006507 | 0.020028 | 0.035954 | 0.019592 | 0.002951 | 0.008651 | 0.059242 | 0.06094 | 0.029061 | 0.023613 | 0.023888 | 0.025911 | 0.029026 | 0.030988 | 0.024484 | 0.023734 |
| 30.25 | 0.007185 | 0.006031 | 0.030574 | 0.00898 | 0.007327 | 0.011699 | 0.007468 | 0.012438 | 0.024678 | 0.020512 | 0.004027 | 0.005414 | 0.069903 | 0.062678 | 0.028794 | 0.019571 | 0.023134 | 0.025308 | 0.028052 | 0.027706 | 0.025834 | 0.020762 |
| 30.26388889 | 0.014599 | 0.012969 | 0.012672 | 0.003261 | 0.003042 | 0.008692 | 0.007046 | 0.015023 | 0.009859 | 0.015569 | 0.004495 | 0.005759 | 0.067189 | 0.052575 | 0.025273 | 0.011131 | 0.015994 | 0.018388 | 0.023992 | 0.018996 | 0.022191 | 0.015314 |
| 30.27777778 | 0.020135 | 0.014481 | 0.003371 | 0.002387 | 0.001405 | 0.006665 | 0.005316 | 0.013142 | 0.00285 | 0.007322 | 0.004572 | 0.006018 | 0.05285 | 0.035484 | 0.018117 | 0.004141 | 0.009536 | 0.013069 | 0.018111 | 0.011083 | 0.014879 | 0.009942 |
| 30.29166667 | 0.014726 | 0.008039 | 0.001301 | 0.005098 | 0.002771 | 0.006395 | 0.004865 | 0.007897 | 0.003051 | 0.002548 | 0.005584 | 0.004768 | 0.036947 | 0.021148 | 0.012136 | 0.00344 | 0.006806 | 0.011308 | 0.011037 | 0.005395 | 0.008798 | 0.006626 |
| 30.30555556 | 0.007713 | 0.003709 | 0.001755 | 0.007691 | 0.004792 | 0.006724 | 0.005102 | 0.007348 | 0.003693 | 0.002429 | 0.010358 | 0.003392 | 0.025269 | 0.0142 | 0.01062 | 0.006782 | 0.005399 | 0.009833 | 0.005538 | 0.002979 | 0.00678 | 0.006404 |
| 30.31944444 | 0.00946 | 0.004535 | 0.005916 | 0.006394 | 0.005028 | 0.006306 | 0.005894 | 0.008251 | 0.00435 | 0.003948 | 0.015997 | 0.002923 | 0.016448 | 0.010409 | 0.010204 | 0.008327 | 0.003812 | 0.007624 | 0.004348 | 0.003621 | 0.007258 | 0.007971 |
| 30.33333333 | 0.012219 | 0.005659 | 0.014397 | 0.004221 | 0.003857 | 0.005772 | 0.004549 | 0.007683 | 0.00669 | 0.005246 | 0.019954 | 0.003176 | 0.00912 | 0.007087 | 0.006936 | 0.006419 | 0.002918 | 0.006114 | 0.005146 | 0.004523 | 0.007836 | 0.008319 |
| 30.34722222 | 0.009917 | 0.006343 | 0.022963 | 0.00535 | 0.005133 | 0.006044 | 0.002441 | 0.004242 | 0.009737 | 0.007398 | 0.020016 | 0.002911 | 0.008013 | 0.008102 | 0.002608 | 0.004512 | 0.002811 | 0.005036 | 0.003909 | 0.005359 | 0.007782 | 0.006633 |
| 30.36111111 | 0.011801 | 0.01221 | 0.027877 | 0.009164 | 0.011383 | 0.007837 | 0.004122 | 0.003097 | 0.013655 | 0.011614 | 0.012907 | 0.004323 | 0.015875 | 0.01251 | 0.000751 | 0.005236 | 0.00297 | 0.003263 | 0.001653 | 0.006216 | 0.007005 | 0.004932 |
| 30.375 | 0.022062 | 0.023441 | 0.030571 | 0.016382 | 0.019169 | 0.015154 | 0.009897 | 0.009267 | 0.019442 | 0.014847 | 0.005159 | 0.007596 | 0.026771 | 0.014784 | 0.000987 | 0.006648 | 0.003493 | 0.001405 | 0.001624 | 0.005512 | 0.005118 | 0.004227 |
| 30.38888889 | 0.033951 | 0.035457 | 0.035822 | 0.024957 | 0.025228 | 0.024617 | 0.01733 | 0.021819 | 0.028462 | 0.016863 | 0.005823 | 0.014694 | 0.036672 | 0.016437 | 0.00334 | 0.005404 | 0.003381 | 0.00196 | 0.002727 | 0.004341 | 0.002816 | 0.002977 |
| 30.40277778 | 0.04239 | 0.046546 | 0.045278 | 0.031846 | 0.031947 | 0.027652 | 0.022455 | 0.031047 | 0.039435 | 0.022927 | 0.014128 | 0.027555 | 0.044269 | 0.020774 | 0.009136 | 0.004873 | 0.00484 | 0.007558 | 0.003189 | 0.006037 | 0.002157 | 0.002191 |
| 30.41666667 | 0.042862 | 0.052311 | 0.053723 | 0.039655 | 0.042144 | 0.02685 | 0.02613 | 0.035427 | 0.046127 | 0.033245 | 0.024975 | 0.040801 | 0.045913 | 0.02464 | 0.01438 | 0.009707 | 0.01049 | 0.015408 | 0.005276 | 0.010731 | 0.00331 | 0.004398 |
| 30.43055556 | 0.035381 | 0.050687 | 0.057684 | 0.051363 | 0.055297 | 0.032332 | 0.03458 | 0.044615 | 0.047729 | 0.042416 | 0.034016 | 0.049426 | 0.041012 | 0.025202 | 0.015986 | 0.016021 | 0.015268 | 0.019701 | 0.009691 | 0.014993 | 0.005329 | 0.007984 |
| 30.44444444 | 0.032229 | 0.051394 | 0.065193 | 0.065396 | 0.06835 | 0.04476 | 0.050144 | 0.056946 | 0.053246 | 0.050974 | 0.042194 | 0.056774 | 0.038945 | 0.028656 | 0.019127 | 0.02251 | 0.018208 | 0.022311 | 0.016475 | 0.018309 | 0.010613 | 0.013854 |
| 30.45833333 | 0.044762 | 0.065316 | 0.085416 | 0.081535 | 0.078519 | 0.055548 | 0.069475 | 0.064538 | 0.06818 | 0.064064 | 0.054419 | 0.071029 | 0.053622 | 0.045827 | 0.030142 | 0.03479 | 0.026512 | 0.031108 | 0.028875 | 0.02471 | 0.022213 | 0.025969 |
| 30.47222222 | 0.072019 | 0.092505 | 0.11558 | 0.105041 | 0.089466 | 0.062015 | 0.086699 | 0.070821 | 0.089618 | 0.083059 | 0.071309 | 0.092623 | 0.088479 | 0.080136 | 0.048684 | 0.054585 | 0.041639 | 0.047199 | 0.046176 | 0.036997 | 0.038697 | 0.042504 |
| 30.48611111 | 0.103312 | 0.126359 | 0.141463 | 0.136724 | 0.110726 | 0.070025 | 0.097238 | 0.084581 | 0.114407 | 0.108301 | 0.092387 | 0.114193 | 0.132417 | 0.12403 | 0.068533 | 0.078083 | 0.056553 | 0.064237 | 0.062798 | 0.053235 | 0.055245 | 0.059087 |
| 30.5 | 0.126695 | 0.158811 | 0.151849 | 0.168588 | 0.140574 | 0.085254 | 0.10693 | 0.108002 | 0.141521 | 0.137432 | 0.119587 | 0.132704 | 0.166291 | 0.171114 | 0.08424 | 0.101617 | 0.065886 | 0.080049 | 0.07921 | 0.071067 | 0.066961 | 0.076095 |
| 30.51388889 | 0.139038 | 0.182248 | 0.154048 | 0.18763 | 0.159933 | 0.105278 | 0.124037 | 0.130394 | 0.16807 | 0.161898 | 0.150987 | 0.151125 | 0.178341 | 0.222771 | 0.095892 | 0.122241 | 0.070687 | 0.096084 | 0.097583 | 0.088557 | 0.07404 | 0.095862 |
| 30.52777778 | 0.139248 | 0.188517 | 0.160162 | 0.185217 | 0.159631 | 0.114387 | 0.132509 | 0.135201 | 0.186797 | 0.1751 | 0.176805 | 0.165321 | 0.170994 | 0.268807 | 0.10755 | 0.135871 | 0.070608 | 0.108219 | 0.110193 | 0.102809 | 0.081008 | 0.11428 |
| 30.54166667 | 0.117581 | 0.157938 | 0.155723 | 0.16191 | 0.139823 | 0.095665 | 0.104627 | 0.114989 | 0.175067 | 0.158493 | 0.177173 | 0.151493 | 0.142452 | 0.26676 | 0.115244 | 0.128031 | 0.06304 | 0.108806 | 0.111157 | 0.109551 | 0.090198 | 0.122531 |
| 30.55555556 | 0.075723 | 0.095583 | 0.107855 | 0.10506 | 0.090978 | 0.056367 | 0.051616 | 0.070672 | 0.111194 | 0.10027 | 0.128171 | 0.095902 | 0.091074 | 0.182203 | 0.095114 | 0.088719 | 0.049347 | 0.094347 | 0.098849 | 0.093213 | 0.094053 | 0.110719 |
| 30.56944444 | 0.054808 | 0.054569 | 0.057935 | 0.04694 | 0.042462 | 0.025506 | 0.022504 | 0.030949 | 0.053521 | 0.053138 | 0.061682 | 0.035509 | 0.065205 | 0.114679 | 0.053974 | 0.041363 | 0.03228 | 0.067835 | 0.066867 | 0.050839 | 0.073715 | 0.073462 |
| 30.58333333 | 0.054317 | 0.047883 | 0.048268 | 0.02902 | 0.035292 | 0.02256 | 0.040144 | 0.058473 | 0.058287 | 0.04504 | 0.034795 | 0.016202 | 0.094119 | 0.133776 | 0.031666 | 0.015891 | 0.017447 | 0.040894 | 0.028565 | 0.01612 | 0.037374 | 0.031835 |
| 30.59722222 | 0.051916 | 0.073506 | 0.043534 | 0.034682 | 0.06351 | 0.053831 | 0.088115 | 0.143368 | 0.116556 | 0.057528 | 0.067533 | 0.054227 | 0.095622 | 0.118075 | 0.031859 | 0.023224 | 0.01612 | 0.033879 | 0.014034 | 0.011638 | 0.016114 | 0.016109 |
| 30.61111111 | 0.081517 | 0.134648 | 0.061914 | 0.074738 | 0.100285 | 0.097355 | 0.133294 | 0.190844 | 0.206437 | 0.097908 | 0.104144 | 0.124066 | 0.064746 | 0.074182 | 0.050729 | 0.053009 | 0.024062 | 0.04876 | 0.024604 | 0.026187 | 0.013825 | 0.030259 |
| 30.625 | 0.106766 | 0.15931 | 0.099358 | 0.124174 | 0.107178 | 0.112139 | 0.135447 | 0.15476 | 0.237086 | 0.123927 | 0.080856 | 0.166017 | 0.080525 | 0.087519 | 0.067908 | 0.068215 | 0.023951 | 0.061267 | 0.036872 | 0.042254 | 0.025123 | 0.058971 |
| 30.63888889 | 0.08406 | 0.117397 | 0.094867 | 0.110529 | 0.076301 | 0.097107 | 0.095396 | 0.072817 | 0.167423 | 0.089189 | 0.064928 | 0.154541 | 0.095719 | 0.092516 | 0.050512 | 0.051476 | 0.017891 | 0.060852 | 0.038463 | 0.043363 | 0.045592 | 0.080372 |
| 30.65277778 | 0.050215 | 0.065399 | 0.053852 | 0.053299 | 0.048306 | 0.066794 | 0.053574 | 0.026613 | 0.075559 | 0.052871 | 0.083601 | 0.095478 | 0.060992 | 0.067825 | 0.021289 | 0.025135 | 0.011626 | 0.047436 | 0.033087 | 0.026043 | 0.054273 | 0.074486 |
| 30.66666667 | 0.02515 | 0.03696 | 0.020383 | 0.022243 | 0.035831 | 0.030196 | 0.032491 | 0.035857 | 0.022621 | 0.04902 | 0.092873 | 0.030539 | 0.020667 | 0.0441 | 0.006769 | 0.008313 | 0.006304 | 0.025029 | 0.028294 | 0.009676 | 0.037885 | 0.048982 |
| 30.68055556 | 0.013855 | 0.029582 | 0.012191 | 0.017189 | 0.037232 | 0.007243 | 0.024686 | 0.047839 | 0.021851 | 0.051155 | 0.126166 | 0.013339 | 0.008154 | 0.028063 | 0.011859 | 0.004224 | 0.005921 | 0.010043 | 0.024003 | 0.007528 | 0.019974 | 0.043031 |
| 30.69444444 | 0.011169 | 0.023963 | 0.012823 | 0.0177 | 0.065138 | 0.00189 | 0.017098 | 0.045033 | 0.050186 | 0.077349 | 0.161499 | 0.018946 | 0.009927 | 0.030013 | 0.030765 | 0.005413 | 0.005175 | 0.008014 | 0.016994 | 0.01196 | 0.018478 | 0.049132 |

| | | | | | | | | | | | | | | | | | | | | | | |
|---|---|---|---|---|---|---|---|---|---|---|---|---|---|---|---|---|---|---|---|---|---|---|
| 30.70833333 | 0.008226 | 0.015181 | 0.016356 | 0.026141 | 0.074132 | 0.003875 | 0.010414 | 0.03001 | 0.061153 | 0.09914 | 0.119218 | 0.016999 | 0.00835 | 0.024005 | 0.037189 | 0.005673 | 0.002662 | 0.009448 | 0.015142 | 0.015193 | 0.014696 | 0.034444 |
| 30.72222222 | 0.010103 | 0.017602 | 0.027341 | 0.029827 | 0.041966 | 0.007114 | 0.00667 | 0.020377 | 0.044081 | 0.07197 | 0.056602 | 0.031926 | 0.008914 | 0.0213 | 0.030227 | 0.006374 | 0.00267 | 0.010106 | 0.016245 | 0.012752 | 0.01007 | 0.027152 |
| 30.73611111 | 0.010622 | 0.019493 | 0.031181 | 0.023243 | 0.025177 | 0.015539 | 0.013567 | 0.036183 | 0.03942 | 0.046255 | 0.048097 | 0.061351 | 0.014064 | 0.053827 | 0.022406 | 0.007727 | 0.003248 | 0.017495 | 0.010423 | 0.009377 | 0.01407 | 0.034775 |
| 30.75 | 0.007782 | 0.017818 | 0.01911 | 0.015948 | 0.034606 | 0.024526 | 0.030534 | 0.044794 | 0.058717 | 0.049453 | 0.043681 | 0.059305 | 0.019008 | 0.073134 | 0.014436 | 0.011505 | 0.00273 | 0.026077 | 0.004973 | 0.009644 | 0.012213 | 0.026167 |
| 30.76388889 | 0.007835 | 0.027515 | 0.008625 | 0.01344 | 0.036745 | 0.023356 | 0.033806 | 0.02933 | 0.060666 | 0.038009 | 0.037574 | 0.052688 | 0.017789 | 0.048027 | 0.015645 | 0.013718 | 0.004687 | 0.027441 | 0.004095 | 0.007319 | 0.007569 | 0.02068 |
| 30.77777778 | 0.005873 | 0.032405 | 0.01066 | 0.011467 | 0.027625 | 0.028497 | 0.038066 | 0.023191 | 0.039361 | 0.030457 | 0.052529 | 0.085442 | 0.02013 | 0.042762 | 0.025078 | 0.014211 | 0.009485 | 0.025609 | 0.004198 | 0.006676 | 0.010257 | 0.037949 |
| 30.79166667 | 0.004678 | 0.030361 | 0.010954 | 0.011336 | 0.018158 | 0.036727 | 0.049573 | 0.033331 | 0.028748 | 0.051025 | 0.064565 | 0.087854 | 0.035013 | 0.078931 | 0.029714 | 0.021269 | 0.0107 | 0.020473 | 0.007347 | 0.008906 | 0.011797 | 0.038569 |
| 30.80555556 | 0.006344 | 0.02617 | 0.010595 | 0.014623 | 0.012418 | 0.026824 | 0.040103 | 0.034088 | 0.026982 | 0.049891 | 0.083251 | 0.065781 | 0.038929 | 0.081604 | 0.024614 | 0.020858 | 0.00909 | 0.019502 | 0.011321 | 0.01325 | 0.013671 | 0.021293 |
| 30.81944444 | 0.005157 | 0.026265 | 0.020118 | 0.015426 | 0.010638 | 0.023293 | 0.042855 | 0.042101 | 0.038589 | 0.042955 | 0.083038 | 0.068153 | 0.02423 | 0.054405 | 0.01821 | 0.010885 | 0.011809 | 0.02838 | 0.009923 | 0.027806 | 0.020176 | 0.024319 |
| 30.83333333 | 0.00289 | 0.045392 | 0.021304 | 0.015037 | 0.012825 | 0.03957 | 0.069697 | 0.060911 | 0.063559 | 0.054555 | 0.06078 | 0.049689 | 0.014957 | 0.056426 | 0.014717 | 0.009776 | 0.012573 | 0.03084 | 0.008629 | 0.037241 | 0.021561 | 0.028292 |
| 30.84722222 | 0.00328 | 0.053802 | 0.016508 | 0.015177 | 0.01454 | 0.038148 | 0.062443 | 0.048185 | 0.058951 | 0.043023 | 0.057222 | 0.031535 | 0.01314 | 0.053644 | 0.015125 | 0.015141 | 0.009512 | 0.020699 | 0.014618 | 0.025475 | 0.022982 | 0.016564 |
| 30.86111111 | 0.004735 | 0.03649 | 0.025143 | 0.023285 | 0.012004 | 0.024616 | 0.031088 | 0.031052 | 0.048328 | 0.035643 | 0.055933 | 0.038044 | 0.012655 | 0.040989 | 0.023204 | 0.021125 | 0.012717 | 0.015395 | 0.024143 | 0.01818 | 0.027763 | 0.007378 |
| 30.875 | 0.005297 | 0.028769 | 0.032191 | 0.034212 | 0.008579 | 0.028331 | 0.018826 | 0.043798 | 0.050286 | 0.051049 | 0.082747 | 0.028007 | 0.030022 | 0.063721 | 0.021483 | 0.020817 | 0.01665 | 0.017292 | 0.026626 | 0.027539 | 0.021152 | 0.005074 |
| 30.88888889 | 0.008386 | 0.03821 | 0.031895 | 0.032749 | 0.01254 | 0.026666 | 0.01472 | 0.043574 | 0.046276 | 0.053778 | 0.139327 | 0.014342 | 0.056673 | 0.095344 | 0.008037 | 0.014125 | 0.013199 | 0.016011 | 0.01666 | 0.029062 | 0.010272 | 0.008255 |
| 30.90277778 | 0.009954 | 0.05568 | 0.028053 | 0.027749 | 0.031082 | 0.016904 | 0.012339 | 0.027548 | 0.053804 | 0.056055 | 0.128802 | 0.015568 | 0.078834 | 0.119813 | 0.01133 | 0.010713 | 0.012247 | 0.018605 | 0.005423 | 0.020883 | 0.008704 | 0.015243 |
| 30.91666667 | 0.007491 | 0.07841 | 0.023381 | 0.03246 | 0.045859 | 0.013446 | 0.017622 | 0.037321 | 0.046956 | 0.049836 | 0.075008 | 0.015896 | 0.098069 | 0.140416 | 0.038597 | 0.013345 | 0.020488 | 0.028377 | 0.004266 | 0.016517 | 0.007397 | 0.018663 |
| 30.93055556 | 0.008379 | 0.069554 | 0.020688 | 0.03709 | 0.038711 | 0.011048 | 0.02377 | 0.049179 | 0.01885 | 0.023919 | 0.086722 | 0.017637 | 0.099187 | 0.149331 | 0.080023 | 0.030625 | 0.02754 | 0.032079 | 0.006952 | 0.01272 | 0.008018 | 0.02398 |
| 30.94444444 | 0.014471 | 0.031625 | 0.019536 | 0.031072 | 0.020666 | 0.017898 | 0.029308 | 0.032465 | 0.012843 | 0.008241 | 0.117805 | 0.033827 | 0.088044 | 0.160069 | 0.124805 | 0.059245 | 0.027367 | 0.023289 | 0.008594 | 0.013718 | 0.021993 | 0.044336 |
| 30.95833333 | 0.023181 | 0.018771 | 0.019942 | 0.024091 | 0.016469 | 0.03542 | 0.044204 | 0.014634 | 0.039124 | 0.012787 | 0.08401 | 0.064641 | 0.086688 | 0.161069 | 0.137993 | 0.074133 | 0.02411 | 0.016767 | 0.024483 | 0.037282 | 0.038328 | 0.061602 |
| 30.97222222 | 0.029192 | 0.043217 | 0.033147 | 0.035596 | 0.040062 | 0.055345 | 0.062161 | 0.034832 | 0.080692 | 0.041799 | 0.042383 | 0.084317 | 0.096432 | 0.150154 | 0.09729 | 0.0635 | 0.029649 | 0.031632 | 0.051498 | 0.069506 | 0.041823 | 0.054218 |
| 30.98611111 | 0.036428 | 0.078032 | 0.07624 | 0.080841 | 0.077361 | 0.083097 | 0.080831 | 0.092444 | 0.117231 | 0.090109 | 0.041547 | 0.085022 | 0.107322 | 0.148558 | 0.039795 | 0.045041 | 0.052038 | 0.06372 | 0.060164 | 0.080099 | 0.036084 | 0.036226 |
| 31 | 0.05322 | 0.097948 | 0.109373 | 0.124898 | 0.108642 | 0.106744 | 0.104574 | 0.127156 | 0.133479 | 0.125083 | 0.055541 | 0.08387 | 0.105044 | 0.147964 | 0.013902 | 0.041286 | 0.073864 | 0.087593 | 0.05009 | 0.072979 | 0.035542 | 0.032083 |
| 31.01388889 | 0.070069 | 0.108974 | 0.103266 | 0.128334 | 0.119677 | 0.098012 | 0.112961 | 0.111355 | 0.124719 | 0.134748 | 0.103692 | 0.087944 | 0.079177 | 0.131972 | 0.029372 | 0.055265 | 0.076419 | 0.088363 | 0.045079 | 0.066366 | 0.047965 | 0.050674 |
| 31.02777778 | 0.074935 | 0.119107 | 0.089403 | 0.112809 | 0.110229 | 0.075779 | 0.104938 | 0.112926 | 0.118462 | 0.13438 | 0.166542 | 0.095029 | 0.044842 | 0.100878 | 0.068766 | 0.076543 | 0.064726 | 0.080436 | 0.055893 | 0.068399 | 0.063339 | 0.076731 |
| 31.04166667 | 0.073988 | 0.111293 | 0.089774 | 0.104734 | 0.096752 | 0.076183 | 0.096265 | 0.161516 | 0.134568 | 0.130398 | 0.184662 | 0.092923 | 0.038421 | 0.083763 | 0.10392 | 0.091117 | 0.051609 | 0.078666 | 0.0737 | 0.075588 | 0.067968 | 0.088295 |
| 31.05555556 | 0.090474 | 0.099205 | 0.100742 | 0.1072 | 0.08575 | 0.0911 | 0.084875 | 0.170647 | 0.14208 | 0.116593 | 0.158376 | 0.079366 | 0.074384 | 0.10385 | 0.110078 | 0.088096 | 0.04211 | 0.076424 | 0.078617 | 0.076947 | 0.060319 | 0.08064 |
| 31.06944444 | 0.121724 | 0.101648 | 0.114639 | 0.113599 | 0.068879 | 0.095453 | 0.067863 | 0.097603 | 0.120412 | 0.09395 | 0.112783 | 0.060406 | 0.1011 | 0.119505 | 0.090508 | 0.070038 | 0.035112 | 0.066348 | 0.063499 | 0.065553 | 0.045388 | 0.061493 |
| 31.08333333 | 0.122389 | 0.084618 | 0.111461 | 0.101763 | 0.041571 | 0.076715 | 0.045264 | 0.025224 | 0.083824 | 0.059047 | 0.063883 | 0.041873 | 0.071989 | 0.087422 | 0.059251 | 0.045905 | 0.027923 | 0.050712 | 0.039972 | 0.044097 | 0.027834 | 0.038343 |
| 31.09722222 | 0.084336 | 0.039955 | 0.080164 | 0.065403 | 0.017706 | 0.042095 | 0.020635 | 0.005944 | 0.05167 | 0.028398 | 0.025505 | 0.034719 | 0.028123 | 0.037967 | 0.026859 | 0.021468 | 0.018174 | 0.030821 | 0.020287 | 0.022939 | 0.012225 | 0.017431 |
| 31.11111111 | 0.051342 | 0.018217 | 0.054102 | 0.039849 | 0.014864 | 0.018914 | 0.008799 | 0.011404 | 0.03738 | 0.020747 | 0.017227 | 0.042356 | 0.012743 | 0.012309 | 0.009529 | 0.006293 | 0.008066 | 0.01312 | 0.009547 | 0.010284 | 0.00326 | 0.004619 |
| 31.125 | 0.046004 | 0.040096 | 0.060876 | 0.048874 | 0.032908 | 0.025723 | 0.021291 | 0.021568 | 0.041312 | 0.021538 | 0.034577 | 0.053521 | 0.015925 | 0.009033 | 0.008261 | 0.002592 | 0.002356 | 0.005485 | 0.007585 | 0.006978 | 0.002259 | 0.002208 |
| 31.13888889 | 0.055074 | 0.071862 | 0.087652 | 0.073367 | 0.055592 | 0.050728 | 0.049429 | 0.033748 | 0.054474 | 0.03544 | 0.049714 | 0.056203 | 0.022609 | 0.014887 | 0.010628 | 0.002302 | 0.002072 | 0.004471 | 0.01077 | 0.009731 | 0.006214 | 0.006685 |
| 31.15277778 | 0.061056 | 0.082974 | 0.105975 | 0.087058 | 0.064665 | 0.068637 | 0.068545 | 0.048103 | 0.066398 | 0.053734 | 0.052101 | 0.052748 | 0.028612 | 0.021944 | 0.011348 | 0.003973 | 0.005381 | 0.007153 | 0.014624 | 0.014211 | 0.011357 | 0.012124 |
| 31.16666667 | 0.06352 | 0.075077 | 0.100145 | 0.082637 | 0.056436 | 0.065 | 0.065354 | 0.065779 | 0.069078 | 0.053368 | 0.04905 | 0.051707 | 0.032499 | 0.025502 | 0.010229 | 0.007893 | 0.008955 | 0.01183 | 0.016628 | 0.017174 | 0.013864 | 0.014875 |
| 31.18055556 | 0.064201 | 0.060862 | 0.080483 | 0.067536 | 0.042077 | 0.046136 | 0.049512 | 0.082679 | 0.064673 | 0.043642 | 0.040476 | 0.053785 | 0.035416 | 0.026957 | 0.00906 | 0.011659 | 0.010535 | 0.013779 | 0.017532 | 0.018059 | 0.013337 | 0.015852 |
| 31.19444444 | 0.054572 | 0.046268 | 0.064501 | 0.051649 | 0.033575 | 0.028683 | 0.033781 | 0.088182 | 0.061266 | 0.036473 | 0.030389 | 0.052661 | 0.041375 | 0.032285 | 0.011614 | 0.015092 | 0.011033 | 0.013104 | 0.020065 | 0.018714 | 0.013595 | 0.0184 |
| 31.20833333 | 0.03518 | 0.034385 | 0.055928 | 0.041441 | 0.032689 | 0.022248 | 0.024578 | 0.074928 | 0.05767 | 0.0291 | 0.026742 | 0.045401 | 0.053646 | 0.044428 | 0.018896 | 0.019191 | 0.012643 | 0.013803 | 0.026607 | 0.021632 | 0.017156 | 0.023454 |
| 31.22222222 | 0.016433 | 0.021237 | 0.0489 | 0.035076 | 0.029513 | 0.022915 | 0.023138 | 0.054428 | 0.049337 | 0.019621 | 0.026634 | 0.035232 | 0.071104 | 0.057664 | 0.025678 | 0.022921 | 0.016536 | 0.018012 | 0.035768 | 0.027333 | 0.02297 | 0.028589 |
| 31.23611111 | 0.006655 | 0.010857 | 0.038549 | 0.024813 | 0.017603 | 0.019591 | 0.021737 | 0.039325 | 0.039731 | 0.014489 | 0.023426 | 0.025514 | 0.087991 | 0.063643 | 0.027057 | 0.023656 | 0.021367 | 0.023973 | 0.041635 | 0.03302 | 0.028068 | 0.031028 |
| 31.25 | 0.007226 | 0.011641 | 0.023989 | 0.012471 | 0.005484 | 0.010024 | 0.014425 | 0.025066 | 0.031279 | 0.013698 | 0.016255 | 0.017796 | 0.097645 | 0.060208 | 0.026081 | 0.021088 | 0.023942 | 0.027751 | 0.039668 | 0.034524 | 0.03011 | 0.029686 |
| 31.26388889 | 0.009154 | 0.010781 | 0.01016 | 0.006453 | 0.001344 | 0.003807 | 0.008834 | 0.012416 | 0.020411 | 0.009975 | 0.01178 | 0.012435 | 0.094381 | 0.051516 | 0.027366 | 0.016806 | 0.021699 | 0.025741 | 0.031539 | 0.030055 | 0.027838 | 0.024583 |
| 31.27777778 | 0.007058 | 0.004888 | 0.003743 | 0.005696 | 0.001909 | 0.003438 | 0.010143 | 0.009093 | 0.00942 | 0.004172 | 0.012746 | 0.009794 | 0.077187 | 0.04028 | 0.027003 | 0.011497 | 0.015406 | 0.018277 | 0.022553 | 0.022068 | 0.021544 | 0.016997 |
| 31.29166667 | 0.007511 | 0.003843 | 0.004996 | 0.005123 | 0.002541 | 0.004288 | 0.009681 | 0.007986 | 0.003528 | 0.001277 | 0.01378 | 0.009783 | 0.056956 | 0.028016 | 0.019986 | 0.005915 | 0.00874 | 0.009909 | 0.016786 | 0.014938 | 0.014232 | 0.009988 |
| 31.30555556 | 0.014058 | 0.004938 | 0.007677 | 0.004088 | 0.00272 | 0.005255 | 0.005544 | 0.003485 | 0.003171 | 0.002106 | 0.013236 | 0.010108 | 0.047498 | 0.020702 | 0.013096 | 0.002587 | 0.00498 | 0.005593 | 0.015758 | 0.011786 | 0.010135 | 0.006864 |
| 31.31944444 | 0.022403 | 0.003427 | 0.005993 | 0.00239 | 0.006278 | 0.004354 | 0.005259 | 0.004216 | 0.005211 | 0.005192 | 0.014877 | 0.007571 | 0.050017 | 0.021757 | 0.014913 | 0.002845 | 0.004582 | 0.006558 | 0.018242 | 0.012272 | 0.010403 | 0.007931 |
| 31.33333333 | 0.026807 | 0.003078 | 0.004229 | 0.002418 | 0.012725 | 0.003799 | 0.007884 | 0.014825 | 0.007129 | 0.006049 | 0.01506 | 0.003937 | 0.052293 | 0.02409 | 0.01936 | 0.003727 | 0.005144 | 0.008669 | 0.020277 | 0.01261 | 0.011537 | 0.009469 |
| 31.34722222 | 0.020738 | 0.006995 | 0.008359 | 0.00784 | 0.014868 | 0.009305 | 0.009947 | 0.025399 | 0.013747 | 0.005925 | 0.011194 | 0.002824 | 0.041142 | 0.017496 | 0.01594 | 0.002497 | 0.004397 | 0.007895 | 0.018396 | 0.009735 | 0.009862 | 0.007996 |
| 31.36111111 | 0.014565 | 0.013312 | 0.017086 | 0.016157 | 0.014337 | 0.016797 | 0.010995 | 0.024589 | 0.023957 | 0.012474 | 0.011356 | 0.002525 | 0.024406 | 0.009808 | 0.007845 | 0.000787 | 0.002537 | 0.005363 | 0.012898 | 0.00523 | 0.005734 | 0.004692 |
| 31.375 | 0.026767 | 0.02335 | 0.028652 | 0.022281 | 0.018581 | 0.018959 | 0.013245 | 0.018946 | 0.030234 | 0.023376 | 0.016372 | 0.004076 | 0.027171 | 0.015302 | 0.004678 | 0.000305 | 0.00139 | 0.002937 | 0.006356 | 0.003962 | 0.002356 | 0.002402 |
| 31.38888889 | 0.046605 | 0.037069 | 0.04047 | 0.026095 | 0.027303 | 0.020349 | 0.019897 | 0.024825 | 0.03479 | 0.029434 | 0.02058 | 0.01249 | 0.049318 | 0.026842 | 0.00768 | 0.001193 | 0.002598 | 0.002917 | 0.002428 | 0.007146 | 0.001735 | 0.001682 |
| 31.40277778 | 0.055572 | 0.047993 | 0.050874 | 0.032049 | 0.036025 | 0.027594 | 0.028236 | 0.04002 | 0.045083 | 0.032373 | 0.024219 | 0.02753 | 0.067746 | 0.037293 | 0.013748 | 0.004603 | 0.006589 | 0.009071 | 0.003279 | 0.011232 | 0.003572 | 0.004175 |
| 31.41666667 | 0.053965 | 0.053201 | 0.061979 | 0.042516 | 0.044492 | 0.03668 | 0.035198 | 0.04885 | 0.061486 | 0.040841 | 0.028097 | 0.046275 | 0.078961 | 0.051894 | 0.021835 | 0.011381 | 0.012933 | 0.019838 | 0.007466 | 0.01554 | 0.008588 | 0.01235 |
| 31.43055556 | 0.050758 | 0.056929 | 0.074509 | 0.054496 | 0.054001 | 0.043698 | 0.045017 | 0.050947 | 0.076297 | 0.053682 | 0.033492 | 0.06792 | 0.093705 | 0.070396 | 0.030007 | 0.020616 | 0.019773 | 0.029863 | 0.014611 | 0.022568 | 0.017255 | 0.023553 |
| 31.44444444 | 0.054232 | 0.064107 | 0.088264 | 0.066349 | 0.064773 | 0.054043 | 0.061191 | 0.057325 | 0.087907 | 0.06575 | 0.044803 | 0.089178 | 0.111399 | 0.087955 | 0.03652 | 0.030943 | 0.024967 | 0.036211 | 0.024618 | 0.032964 | 0.026396 | 0.032702 |
| 31.45833333 | 0.06656 | 0.079219 | 0.106794 | 0.083723 | 0.080704 | 0.073638 | 0.081585 | 0.070244 | 0.103409 | 0.081709 | 0.062639 | 0.105676 | 0.126408 | 0.104543 | 0.043668 | 0.042016 | 0.030184 | 0.041846 | 0.036897 | 0.045786 | 0.033773 | 0.041008 |
| 31.47222222 | 0.085851 | 0.103845 | 0.136604 | 0.11518 | 0.105077 | 0.098571 | 0.103363 | 0.083309 | 0.124784 | 0.108169 | 0.083133 | 0.117555 | 0.140522 | 0.124489 | 0.054894 | 0.054298 | 0.039997 | 0.05477 | 0.053162 | 0.060614 | 0.042065 | 0.054185 |
| 31.48611111 | 0.109975 | 0.133008 | 0.174171 | 0.160677 | 0.13275 | 0.11642 | 0.119605 | 0.093601 | 0.149245 | 0.139726 | 0.10781 | 0.125606 | 0.160566 | 0.151432 | 0.070138 | 0.070475 | 0.053569 | 0.076644 | 0.07158 | 0.076639 | 0.053305 | 0.071933 |
| 31.5 | 0.133738 | 0.160719 | 0.199037 | 0.197551 | 0.150869 | 0.120387 | 0.127909 | 0.104905 | 0.171687 | 0.166008 | 0.139842 | 0.133695 | 0.190633 | 0.187663 | 0.085535 | 0.090759 | 0.062951 | 0.096978 | 0.085759 | 0.091295 | 0.06553 | 0.089242 |
| 31.51388889 | 0.146968 | 0.183775 | 0.199603 | 0.204146 | 0.154012 | 0.121641 | 0.141138 | 0.119977 | 0.184358 | 0.185569 | 0.173496 | 0.148184 | 0.22065 | 0.233126 | 0.099032 | 0.109026 | 0.063282 | 0.105826 | 0.093484 | 0.104092 | 0.077801 | 0.104925 |
| 31.52777778 | 0.137665 | 0.192005 | 0.182461 | 0.187447 | 0.156909 | 0.129491 | 0.155178 | 0.135562 | 0.189672 | 0.196669 | 0.199178 | 0.161353 | 0.229757 | 0.274459 | 0.10973 | 0.121234 | 0.056622 | 0.104893 | 0.098561 | 0.117762 | 0.089332 | 0.120676 |
| 31.54166667 | 0.11183 | 0.173318 | 0.146008 | 0.152689 | 0.150938 | 0.123787 | 0.137705 | 0.14217 | 0.188149 | 0.183577 | 0.201405 | 0.14915 | 0.200744 | 0.268403 | 0.115426 | 0.12067 | 0.049051 | 0.100799 | 0.100888 | 0.127587 | 0.097075 | 0.131485 |
| 31.55555556 | 0.072888 | 0.119901 | 0.083001 | 0.095945 | 0.098348 | 0.078531 | 0.080982 | 0.124046 | 0.153141 | 0.121932 | 0.151372 | 0.097528 | 0.132844 | 0.179233 | 0.100985 | 0.095293 | 0.044483 | 0.094751 | 0.094669 | 0.116955 | 0.095229 | 0.120991 |
| 31.56944444 | 0.040564 | 0.055218 | 0.036797 | 0.043555 | 0.03765 | 0.030992 | 0.028373 | 0.085441 | 0.080318 | 0.050853 | 0.080432 | 0.037448 | 0.099164 | 0.109174 | 0.062753 | 0.05362 | 0.041004 | 0.085609 | 0.07704 | 0.073431 | 0.072961 | 0.083151 |
| 31.58333333 | 0.03347 | 0.029725 | 0.032182 | 0.020141 | 0.019841 | 0.023731 | 0.026766 | 0.065912 | 0.035388 | 0.037183 | 0.04678 | 0.020896 | 0.129985 | 0.125089 | 0.036541 | 0.02122 | 0.034959 | 0.070859 | 0.052904 | 0.023433 | 0.035435 | 0.039866 |
| 31.59722222 | 0.045198 | 0.066114 | 0.033092 | 0.034069 | 0.030603 | 0.050077 | 0.075568 | 0.085319 | 0.068605 | 0.072661 | 0.064179 | 0.042461 | 0.115235 | 0.101635 | 0.032298 | 0.009563 | 0.025745 | 0.048404 | 0.034533 | 0.011675 | 0.013524 | 0.013992 |
| 31.61111111 | 0.086476 | 0.148934 | 0.057344 | 0.082956 | 0.062979 | 0.094139 | 0.12381 | 0.113952 | 0.147401 | 0.139324 | 0.129987 | 0.062213 | 0.06267 | 0.055348 | 0.025839 | 0.01384 | 0.019589 | 0.030031 | 0.024368 | 0.033346 | 0.015679 | 0.00461 |
| 31.625 | 0.108801 | 0.206452 | 0.089862 | 0.113362 | 0.093785 | 0.111602 | 0.122727 | 0.114991 | 0.204186 | 0.194355 | 0.173931 | 0.066275 | 0.072034 | 0.086555 | 0.023374 | 0.022188 | 0.018326 | 0.025413 | 0.016781 | 0.051508 | 0.017243 | 0.007058 |
| 31.63888889 | 0.084717 | 0.193838 | 0.076086 | 0.092354 | 0.097953 | 0.09154 | 0.082504 | 0.081706 | 0.187957 | 0.163839 | 0.14362 | 0.050198 | 0.102584 | 0.135313 | 0.036471 | 0.027944 | 0.014492 | 0.023076 | 0.013617 | 0.054708 | 0.018944 | 0.021197 |
| 31.65277778 | 0.057569 | 0.142235 | 0.042831 | 0.051959 | 0.077372 | 0.059398 | 0.039228 | 0.046579 | 0.110924 | 0.096228 | 0.090277 | 0.028287 | 0.092826 | 0.144535 | 0.054777 | 0.029968 | 0.007821 | 0.014216 | 0.015211 | 0.044645 | 0.036858 | 0.044205 |
| 31.66666667 | 0.034693 | 0.076915 | 0.021661 | 0.03039 | 0.053267 | 0.026874 | 0.012213 | 0.036792 | 0.059811 | 0.089412 | 0.058075 | 0.01564 | 0.058416 | 0.114481 | 0.067309 | 0.024358 | 0.004741 | 0.006371 | 0.01756 | 0.029992 | 0.054513 | 0.064927 |
| 31.68055556 | 0.014544 | 0.030153 | 0.011594 | 0.028595 | 0.046927 | 0.012187 | 0.006741 | 0.037484 | 0.053274 | 0.087528 | 0.038387 | 0.022488 | 0.027532 | 0.063755 | 0.058007 | 0.013635 | 0.006343 | 0.010818 | 0.021379 | 0.022153 | 0.054333 | 0.062461 |
| 31.69444444 | 0.005296 | 0.01921 | 0.010224 | 0.02492 | 0.038533 | 0.01065 | 0.009568 | 0.031063 | 0.043652 | 0.05738 | 0.025106 | 0.049351 | 0.027676 | 0.050267 | 0.031786 | 0.00579 | 0.009704 | 0.022322 | 0.028161 | 0.021071 | 0.038356 | 0.039262 |
| 31.70833333 | 0.005984 | 0.018232 | 0.012819 | 0.029312 | 0.025463 | 0.006115 | 0.012427 | 0.021962 | 0.044683 | 0.067167 | 0.017781 | 0.05644 | 0.051613 | 0.077208 | 0.023203 | 0.003722 | 0.012147 | 0.022735 | 0.027271 | 0.027027 | 0.018401 | 0.018588 |
| 31.72222222 | 0.009338 | 0.012444 | 0.011475 | 0.034613 | 0.025234 | 0.003432 | 0.014187 | 0.015928 | 0.046147 | 0.074819 | 0.027017 | 0.033029 | 0.056212 | 0.073569 | 0.035868 | 0.005692 | 0.011358 | 0.016008 | 0.018631 | 0.02456 | 0.015784 | 0.017782 |
| 31.73611111 | 0.010945 | 0.019216 | 0.007166 | 0.025347 | 0.031751 | 0.00627 | 0.012892 | 0.021638 | 0.033576 | 0.054925 | 0.058862 | 0.010889 | 0.040686 | 0.050214 | 0.037559 | 0.012157 | 0.007355 | 0.016868 | 0.020205 | 0.015376 | 0.028759 | 0.028196 |
| 31.75 | 0.009451 | 0.044355 | 0.007472 | 0.028246 | 0.047411 | 0.008358 | 0.017189 | 0.036607 | 0.026997 | 0.046606 | 0.066941 | 0.013796 | 0.040595 | 0.071964 | 0.031029 | 0.023499 | 0.004345 | 0.019184 | 0.027961 | 0.014666 | 0.030755 | 0.027761 |
| 31.76388889 | 0.004933 | 0.047715 | 0.00962 | 0.031124 | 0.048249 | 0.006545 | 0.018259 | 0.035629 | 0.027917 | 0.03553 | 0.073671 | 0.024617 | 0.050818 | 0.107139 | 0.050194 | 0.031654 | 0.003501 | 0.015804 | 0.024004 | 0.013511 | 0.016037 | 0.033593 |
| 31.77777778 | 0.00432 | 0.035613 | 0.009547 | 0.017232 | 0.020287 | 0.003581 | 0.011953 | 0.026451 | 0.033528 | 0.031882 | 0.117242 | 0.025439 | 0.039259 | 0.084617 | 0.067257 | 0.026991 | 0.004731 | 0.017322 | 0.014268 | 0.013758 | 0.00567 | 0.054709 |
| 31.79166667 | 0.011049 | 0.046783 | 0.007886 | 0.01102 | 0.002548 | 0.002501 | 0.011919 | 0.023165 | 0.035907 | 0.037288 | 0.106179 | 0.030323 | 0.023165 | 0.049765 | 0.052674 | 0.020556 | 0.00706 | 0.023591 | 0.011677 | 0.016267 | 0.010305 | 0.06179 |
| 31.80555556 | 0.012993 | 0.045778 | 0.008213 | 0.014491 | 0.007609 | 0.003559 | 0.018706 | 0.019748 | 0.045874 | 0.023691 | 0.06378 | 0.034722 | 0.020462 | 0.059578 | 0.05352 | 0.024043 | 0.006079 | 0.018931 | 0.013581 | 0.012552 | 0.020175 | 0.046942 |
| 31.81944444 | 0.00974 | 0.023128 | 0.010113 | 0.01303 | 0.032558 | 0.006104 | 0.027103 | 0.02577 | 0.051633 | 0.022582 | 0.088653 | 0.033061 | 0.01433 | 0.059864 | 0.076533 | 0.026182 | 0.005624 | 0.010911 | 0.017819 | 0.007961 | 0.024522 | 0.02602 |
| 31.83333333 | 0.013346 | 0.01623 | 0.009977 | 0.008006 | 0.046619 | 0.008033 | 0.025 | 0.028195 | 0.047145 | 0.046576 | 0.104935 | 0.024407 | 0.007508 | 0.031681 | 0.06701 | 0.022549 | 0.012989 | 0.016819 | 0.024066 | 0.009207 | 0.01845 | 0.01816 |
| 31.84722222 | 0.020717 | 0.032299 | 0.017614 | 0.017854 | 0.022674 | 0.010496 | 0.021319 | 0.02721 | 0.060254 | 0.050293 | 0.080676 | 0.019649 | 0.008308 | 0.017394 | 0.036568 | 0.022637 | 0.022816 | 0.029219 | 0.025528 | 0.022777 | 0.010743 | 0.027112 |
| 31.86111111 | 0.019628 | 0.051192 | 0.023456 | 0.036181 | 0.016748 | 0.012729 | 0.026511 | 0.04543 | 0.066311 | 0.031072 | 0.094632 | 0.021662 | 0.014315 | 0.024012 | 0.032779 | 0.022515 | 0.020185 | 0.027094 | 0.02076 | 0.035667 | 0.012649 | 0.032739 |
| 31.875 | 0.021611 | 0.055893 | 0.02622 | 0.032446 | 0.048521 | 0.010364 | 0.0287 | 0.055777 | 0.048075 | 0.015967 | 0.122352 | 0.021673 | 0.036836 | 0.031093 | 0.032903 | 0.013896 | 0.013641 | 0.017559 | 0.015808 | 0.02723 | 0.013586 | 0.025177 |
| 31.88888889 | 0.036736 | 0.070535 | 0.038276 | 0.013475 | 0.063372 | 0.011147 | 0.030572 | 0.038973 | 0.02404 | 0.020708 | 0.095388 | 0.027765 | 0.074873 | 0.045969 | 0.017002 | 0.008997 | 0.019547 | 0.023215 | 0.01546 | 0.011514 | 0.006573 | 0.020791 |
| 31.90277778 | 0.032961 | 0.070786 | 0.03108 | 0.005674 | 0.041658 | 0.013223 | 0.026115 | 0.018612 | 0.01294 | 0.046472 | 0.068162 | 0.025251 | 0.097797 | 0.069424 | 0.006364 | 0.012998 | 0.022615 | 0.027442 | 0.025746 | 0.01425 | 0.006295 | 0.02033 |
| 31.91666667 | 0.018068 | 0.058845 | 0.014941 | 0.011907 | 0.023196 | 0.013302 | 0.028862 | 0.01172 | 0.017298 | 0.058361 | 0.074489 | 0.017341 | 0.097414 | 0.079627 | 0.016236 | 0.020069 | 0.016546 | 0.01985 | 0.042166 | 0.035772 | 0.020128 | 0.020448 |
| 31.93055556 | 0.028861 | 0.08746 | 0.015879 | 0.018779 | 0.032974 | 0.021627 | 0.044412 | 0.010943 | 0.019959 | 0.045986 | 0.061931 | 0.021445 | 0.096673 | 0.104126 | 0.057844 | 0.025006 | 0.021774 | 0.029363 | 0.047168 | 0.051905 | 0.039144 | 0.042332 |
| 31.94444444 | 0.045454 | 0.094101 | 0.021605 | 0.026447 | 0.038025 | 0.027427 | 0.041561 | 0.006394 | 0.018915 | 0.033102 | 0.072642 | 0.03216 | 0.103149 | 0.138846 | 0.090887 | 0.021786 | 0.043077 | 0.059886 | 0.033065 | 0.050044 | 0.049088 | 0.065593 |
| 31.95833333 | 0.037717 | 0.048533 | 0.037089 | 0.054211 | 0.031678 | 0.02555 | 0.038676 | 0.011171 | 0.021131 | 0.033528 | 0.142016 | 0.0324 | 0.104778 | 0.128078 | 0.064036 | 0.018019 | 0.057358 | 0.075504 | 0.017596 | 0.035849 | 0.040842 | 0.054612 |
| 31.97222222 | 0.018931 | 0.025341 | 0.051938 | 0.081987 | 0.042255 | 0.035719 | 0.064072 | 0.040165 | 0.047866 | 0.06876 | 0.195538 | 0.032602 | 0.112754 | 0.099734 | 0.022827 | 0.02472 | 0.049562 | 0.063136 | 0.018979 | 0.031684 | 0.026625 | 0.036635 |
| 31.98611111 | 0.029356 | 0.068458 | 0.037585 | 0.070089 | 0.07203 | 0.05934 | 0.100523 | 0.083065 | 0.096579 | 0.113123 | 0.195232 | 0.065338 | 0.151469 | 0.116871 | 0.029988 | 0.041506 | 0.036805 | 0.050129 | 0.03232 | 0.051379 | 0.026349 | 0.042438 |
| 32 | 0.078422 | 0.140771 | 0.02151 | 0.043959 | 0.08955 | 0.075232 | 0.118257 | 0.130056 | 0.144421 | 0.127162 | 0.17331 | 0.102768 | 0.184091 | 0.15892 | 0.076391 | 0.060252 | 0.042645 | 0.062542 | 0.055377 | 0.078311 | 0.042942 | 0.064011 |
| 32.01388889 | 0.122776 | 0.161646 | 0.045927 | 0.049603 | 0.079389 | 0.086434 | 0.110106 | 0.177163 | 0.1911 | 0.118679 | 0.158048 | 0.101749 | 0.150483 | 0.158144 | 0.121703 | 0.074285 | 0.060591 | 0.092271 | 0.073481 | 0.095184 | 0.063415 | 0.080536 |
| 32.02777778 | 0.13412 | 0.131521 | 0.103628 | 0.0865 | 0.073252 | 0.107751 | 0.101455 | 0.195841 | 0.213062 | 0.100075 | 0.148355 | 0.064026 | 0.080286 | 0.109554 | 0.130334 | 0.081052 | 0.066125 | 0.111713 | 0.067703 | 0.094658 | 0.071773 | 0.075668 |
| 32.04166667 | 0.128872 | 0.11932 | 0.163504 | 0.132288 | 0.088664 | 0.129501 | 0.108218 | 0.172697 | 0.194237 | 0.090985 | 0.14735 | 0.035073 | 0.056701 | 0.076414 | 0.10269 | 0.076935 | 0.059466 | 0.113574 | 0.055272 | 0.081803 | 0.067158 | 0.061699 |
| 32.05555556 | 0.139672 | 0.148998 | 0.187415 | 0.162541 | 0.115048 | 0.130132 | 0.106599 | 0.133979 | 0.166793 | 0.09603 | 0.154042 | 0.055315 | 0.102369 | 0.094993 | 0.076938 | 0.06608 | 0.054025 | 0.103369 | 0.051203 | 0.069308 | 0.059422 | 0.057616 |
| 32.06944444 | 0.174852 | 0.187238 | 0.158715 | 0.157061 | 0.13174 | 0.098783 | 0.077584 | 0.094879 | 0.146147 | 0.092918 | 0.137075 | 0.092956 | 0.136335 | 0.12452 | 0.069608 | 0.058235 | 0.046295 | 0.080778 | 0.04674 | 0.059775 | 0.051636 | 0.058003 |
| 32.08333333 | 0.183237 | 0.169078 | 0.104651 | 0.122399 | 0.110243 | 0.058754 | 0.038655 | 0.055719 | 0.115592 | 0.071897 | 0.092487 | 0.088758 | 0.091536 | 0.106371 | 0.061405 | 0.048524 | 0.031541 | 0.049468 | 0.036118 | 0.046757 | 0.041405 | 0.05308 |
| 32.09722222 | 0.13924 | 0.092184 | 0.075196 | 0.087752 | 0.054886 | 0.040811 | 0.015173 | 0.027363 | 0.080712 | 0.046587 | 0.054492 | 0.04527 | 0.027451 | 0.052977 | 0.04364 | 0.031764 | 0.016301 | 0.021568 | 0.024989 | 0.032927 | 0.028153 | 0.041801 |

| | | | | | | | | | | | | | | | | | | | | | |
|---|---|---|---|---|---|---|---|---|---|---|---|---|---|---|---|---|---|---|---|---|---|
| 32.11111111 | 0.08993 | 0.034597 | 0.08918 | 0.075493 | 0.01463 | 0.044654 | 0.016389 | 0.023009 | 0.064309 | 0.033807 | 0.046438 | 0.015996 | 0.004352 | 0.014705 | 0.027573 | 0.014586 | 0.007007 | 0.006245 | 0.019404 | 0.026116 | 0.014588 | 0.028382 |
| 32.125 | 0.067676 | 0.028028 | 0.111997 | 0.081276 | 0.017623 | 0.052675 | 0.034501 | 0.040712 | 0.06367 | 0.033239 | 0.061573 | 0.020142 | 0.006511 | 0.005104 | 0.018697 | 0.004218 | 0.004214 | 0.002966 | 0.018355 | 0.024496 | 0.006001 | 0.018756 |
| 32.13888889 | 0.062707 | 0.042854 | 0.11572 | 0.087659 | 0.045658 | 0.0547 | 0.055276 | 0.060906 | 0.063905 | 0.037511 | 0.073532 | 0.037501 | 0.014727 | 0.013344 | 0.015401 | 0.002224 | 0.004979 | 0.005553 | 0.018327 | 0.023072 | 0.004724 | 0.014937 |
| 32.15277778 | 0.060151 | 0.052386 | 0.104697 | 0.086116 | 0.067669 | 0.050672 | 0.065606 | 0.069744 | 0.060615 | 0.043356 | 0.070101 | 0.049718 | 0.023737 | 0.027869 | 0.017032 | 0.004801 | 0.006611 | 0.008459 | 0.017921 | 0.020874 | 0.007326 | 0.014796 |
| 32.16666667 | 0.052363 | 0.050712 | 0.085886 | 0.072166 | 0.066055 | 0.042897 | 0.061751 | 0.074947 | 0.054664 | 0.044869 | 0.061139 | 0.050931 | 0.029585 | 0.035587 | 0.019605 | 0.007447 | 0.008131 | 0.010246 | 0.018097 | 0.019187 | 0.010122 | 0.016014 |
| 32.18055556 | 0.039596 | 0.044206 | 0.065201 | 0.051275 | 0.050323 | 0.037188 | 0.056224 | 0.086213 | 0.049548 | 0.038708 | 0.055035 | 0.047774 | 0.031587 | 0.032344 | 0.018142 | 0.008255 | 0.009538 | 0.011482 | 0.019245 | 0.018588 | 0.011634 | 0.016477 |
| 32.19444444 | 0.029757 | 0.037745 | 0.053157 | 0.037833 | 0.037822 | 0.037066 | 0.055887 | 0.091549 | 0.047045 | 0.032112 | 0.047235 | 0.046956 | 0.033116 | 0.027739 | 0.014687 | 0.008967 | 0.011279 | 0.012929 | 0.01975 | 0.018901 | 0.011745 | 0.014808 |
| 32.20833333 | 0.027674 | 0.031612 | 0.052887 | 0.033998 | 0.033411 | 0.034313 | 0.046852 | 0.074541 | 0.041793 | 0.030194 | 0.03348 | 0.044333 | 0.038 | 0.030339 | 0.01548 | 0.012087 | 0.013807 | 0.015998 | 0.020026 | 0.022503 | 0.013067 | 0.013694 |
| 32.22222222 | 0.029166 | 0.024603 | 0.054671 | 0.032958 | 0.03185 | 0.025664 | 0.027956 | 0.047541 | 0.029995 | 0.024786 | 0.019774 | 0.034418 | 0.047465 | 0.038318 | 0.020778 | 0.016565 | 0.016862 | 0.020474 | 0.023552 | 0.030977 | 0.017896 | 0.016499 |
| 32.23611111 | 0.027966 | 0.01559 | 0.049623 | 0.028706 | 0.024481 | 0.020392 | 0.017363 | 0.033237 | 0.016642 | 0.013488 | 0.011095 | 0.022066 | 0.060717 | 0.045722 | 0.025135 | 0.019102 | 0.019069 | 0.022891 | 0.030308 | 0.039255 | 0.022943 | 0.0203 |
| 32.25 | 0.023769 | 0.007577 | 0.038339 | 0.018629 | 0.011619 | 0.017701 | 0.012826 | 0.028454 | 0.007545 | 0.005785 | 0.006803 | 0.013319 | 0.075322 | 0.048169 | 0.026393 | 0.018753 | 0.019037 | 0.020669 | 0.035084 | 0.040411 | 0.023953 | 0.020409 |
| 32.26388889 | 0.020964 | 0.005457 | 0.024899 | 0.008406 | 0.002913 | 0.011648 | 0.007244 | 0.01963 | 0.003069 | 0.003755 | 0.004216 | 0.007215 | 0.085398 | 0.044115 | 0.026359 | 0.016067 | 0.017044 | 0.015998 | 0.034505 | 0.033937 | 0.021797 | 0.017223 |
| 32.27777778 | 0.021307 | 0.009744 | 0.013149 | 0.005786 | 0.00107 | 0.008918 | 0.005155 | 0.012924 | 0.00145 | 0.006023 | 0.003403 | 0.003647 | 0.084672 | 0.035316 | 0.024943 | 0.011405 | 0.014791 | 0.012691 | 0.029815 | 0.025275 | 0.01924 | 0.013014 |
| 32.29166667 | 0.023364 | 0.013743 | 0.007132 | 0.009406 | 0.001335 | 0.014762 | 0.005279 | 0.011618 | 0.002605 | 0.010279 | 0.004548 | 0.003705 | 0.07605 | 0.026136 | 0.022175 | 0.006967 | 0.013154 | 0.011627 | 0.023511 | 0.01935 | 0.017353 | 0.009249 |
| 32.30555556 | 0.026187 | 0.010356 | 0.006999 | 0.011369 | 0.001211 | 0.020685 | 0.005245 | 0.010146 | 0.00731 | 0.009335 | 0.004588 | 0.006144 | 0.068605 | 0.020694 | 0.021096 | 0.005812 | 0.011427 | 0.01145 | 0.017417 | 0.017284 | 0.016512 | 0.007781 |
| 32.31944444 | 0.026721 | 0.004232 | 0.007731 | 0.008366 | 0.002616 | 0.017874 | 0.004105 | 0.01174 | 0.011691 | 0.004034 | 0.00387 | 0.007348 | 0.060156 | 0.017236 | 0.021462 | 0.006568 | 0.008655 | 0.010677 | 0.011919 | 0.015814 | 0.015351 | 0.008316 |
| 32.33333333 | 0.019657 | 0.002428 | 0.005798 | 0.004069 | 0.007151 | 0.009271 | 0.004977 | 0.013562 | 0.010392 | 0.003497 | 0.005434 | 0.00678 | 0.040185 | 0.011593 | 0.018104 | 0.005311 | 0.005044 | 0.008007 | 0.006275 | 0.010371 | 0.011879 | 0.00734 |
| 32.34722222 | 0.011568 | 0.005238 | 0.003996 | 0.004265 | 0.015912 | 0.004638 | 0.010653 | 0.018402 | 0.009134 | 0.008801 | 0.013333 | 0.009394 | 0.019108 | 0.009023 | 0.010583 | 0.003945 | 0.002056 | 0.004193 | 0.002555 | 0.004644 | 0.007207 | 0.004219 |
| 32.36111111 | 0.016175 | 0.015226 | 0.009809 | 0.012014 | 0.028979 | 0.007904 | 0.020092 | 0.026429 | 0.013699 | 0.014165 | 0.024429 | 0.013075 | 0.016994 | 0.013504 | 0.004545 | 0.004774 | 0.001017 | 0.002156 | 0.002312 | 0.004623 | 0.004369 | 0.002344 |
| 32.375 | 0.031544 | 0.030688 | 0.027349 | 0.02465 | 0.042697 | 0.014536 | 0.029537 | 0.026525 | 0.020654 | 0.023183 | 0.03036 | 0.016947 | 0.030335 | 0.019333 | 0.002678 | 0.005675 | 0.001866 | 0.002717 | 0.002954 | 0.007589 | 0.003437 | 0.002686 |
| 32.38888889 | 0.04772 | 0.042472 | 0.049677 | 0.036643 | 0.054023 | 0.022769 | 0.034983 | 0.028381 | 0.031189 | 0.037363 | 0.032103 | 0.026566 | 0.047042 | 0.024836 | 0.005164 | 0.007168 | 0.003967 | 0.005517 | 0.004562 | 0.011312 | 0.003387 | 0.004712 |
| 32.40277778 | 0.060771 | 0.049408 | 0.067019 | 0.045648 | 0.063126 | 0.035192 | 0.039705 | 0.046546 | 0.047404 | 0.046482 | 0.036796 | 0.040304 | 0.060497 | 0.03156 | 0.011441 | 0.010149 | 0.007386 | 0.011413 | 0.008591 | 0.016414 | 0.006897 | 0.009958 |
| 32.41666667 | 0.070593 | 0.057863 | 0.077846 | 0.054288 | 0.071799 | 0.050463 | 0.047068 | 0.070397 | 0.065956 | 0.049738 | 0.047255 | 0.051872 | 0.071562 | 0.040174 | 0.017916 | 0.013838 | 0.01203 | 0.019681 | 0.013736 | 0.021957 | 0.014568 | 0.018845 |
| 32.43055556 | 0.07653 | 0.069861 | 0.081131 | 0.063542 | 0.08147 | 0.061752 | 0.055031 | 0.080631 | 0.080788 | 0.057611 | 0.056631 | 0.060094 | 0.087611 | 0.052515 | 0.023607 | 0.019422 | 0.017338 | 0.028543 | 0.019059 | 0.027858 | 0.022516 | 0.028534 |
| 32.44444444 | 0.081311 | 0.082227 | 0.077364 | 0.072043 | 0.093261 | 0.067786 | 0.06395 | 0.078213 | 0.089405 | 0.071612 | 0.058898 | 0.069361 | 0.108006 | 0.070062 | 0.031777 | 0.028428 | 0.023322 | 0.038262 | 0.025422 | 0.03545 | 0.029742 | 0.036736 |
| 32.45833333 | 0.095498 | 0.095836 | 0.081434 | 0.083764 | 0.109646 | 0.077751 | 0.076345 | 0.082204 | 0.101449 | 0.085807 | 0.063299 | 0.081111 | 0.130223 | 0.093517 | 0.043424 | 0.041019 | 0.030282 | 0.050404 | 0.034645 | 0.046683 | 0.039358 | 0.045825 |
| 32.47222222 | 0.123025 | 0.115843 | 0.107055 | 0.103757 | 0.129152 | 0.097442 | 0.090679 | 0.104235 | 0.128138 | 0.103401 | 0.080026 | 0.093957 | 0.153839 | 0.118889 | 0.054905 | 0.05701 | 0.03869 | 0.064804 | 0.048051 | 0.061368 | 0.052205 | 0.058776 |
| 32.48611111 | 0.154688 | 0.148396 | 0.154557 | 0.134546 | 0.149161 | 0.123906 | 0.106824 | 0.135125 | 0.166289 | 0.130328 | 0.107868 | 0.113396 | 0.173629 | 0.140125 | 0.063623 | 0.074617 | 0.049383 | 0.07907 | 0.064995 | 0.077554 | 0.063763 | 0.073693 |
| 32.5 | 0.185692 | 0.196876 | 0.205585 | 0.170596 | 0.173607 | 0.151261 | 0.128741 | 0.155745 | 0.198449 | 0.160475 | 0.141963 | 0.139551 | 0.193773 | 0.157016 | 0.073437 | 0.0913 | 0.059096 | 0.091031 | 0.081378 | 0.096147 | 0.069518 | 0.087105 |
| 32.51388889 | 0.208345 | 0.240196 | 0.223635 | 0.193899 | 0.200088 | 0.171451 | 0.157226 | 0.157908 | 0.215918 | 0.188224 | 0.16987 | 0.156835 | 0.218806 | 0.173408 | 0.086829 | 0.105334 | 0.062508 | 0.099719 | 0.091256 | 0.115705 | 0.069151 | 0.097432 |
| 32.52777778 | 0.196942 | 0.234587 | 0.199714 | 0.19926 | 0.20658 | 0.177321 | 0.180025 | 0.153202 | 0.23024 | 0.212711 | 0.18941 | 0.151618 | 0.248018 | 0.189472 | 0.100792 | 0.109735 | 0.057882 | 0.100077 | 0.088601 | 0.127011 | 0.06934 | 0.105607 |
| 32.54166667 | 0.148143 | 0.172345 | 0.159842 | 0.186399 | 0.168449 | 0.158071 | 0.165569 | 0.159001 | 0.239266 | 0.207757 | 0.212167 | 0.123401 | 0.278334 | 0.201888 | 0.116459 | 0.102969 | 0.049253 | 0.093037 | 0.080159 | 0.132946 | 0.081216 | 0.11547 |
| 32.55555556 | 0.088857 | 0.100138 | 0.10342 | 0.135444 | 0.092069 | 0.106599 | 0.105469 | 0.169054 | 0.208891 | 0.135622 | 0.199828 | 0.087641 | 0.255767 | 0.184889 | 0.130141 | 0.092848 | 0.044394 | 0.090923 | 0.080501 | 0.130214 | 0.095055 | 0.119848 |
| 32.56944444 | 0.053552 | 0.051762 | 0.060348 | 0.068963 | 0.027967 | 0.050169 | 0.047334 | 0.145962 | 0.121569 | 0.059016 | 0.117635 | 0.066852 | 0.165593 | 0.120842 | 0.123953 | 0.076108 | 0.043883 | 0.090334 | 0.078511 | 0.101179 | 0.083472 | 0.100074 |
| 32.58333333 | 0.047632 | 0.03899 | 0.064473 | 0.032239 | 0.020624 | 0.019307 | 0.031291 | 0.083527 | 0.038083 | 0.037406 | 0.044383 | 0.077122 | 0.111657 | 0.07063 | 0.090177 | 0.052453 | 0.041339 | 0.076987 | 0.057341 | 0.062069 | 0.049265 | 0.061923 |
| 32.59722222 | 0.047586 | 0.069937 | 0.067967 | 0.032612 | 0.059624 | 0.014237 | 0.0428 | 0.03612 | 0.015953 | 0.043343 | 0.064808 | 0.109875 | 0.091449 | 0.053182 | 0.057717 | 0.035674 | 0.031672 | 0.053446 | 0.028428 | 0.037095 | 0.025207 | 0.04055 |
| 32.61111111 | 0.06208 | 0.114612 | 0.08678 | 0.071516 | 0.08383 | 0.033434 | 0.056497 | 0.042269 | 0.06297 | 0.103948 | 0.124824 | 0.109536 | 0.067035 | 0.043174 | 0.058525 | 0.029758 | 0.020384 | 0.037778 | 0.013425 | 0.02725 | 0.02659 | 0.055973 |
| 32.625 | 0.075885 | 0.115224 | 0.130163 | 0.115317 | 0.07749 | 0.064955 | 0.078058 | 0.081883 | 0.152652 | 0.188981 | 0.131251 | 0.070033 | 0.098609 | 0.07351 | 0.06818 | 0.028559 | 0.014554 | 0.039121 | 0.010937 | 0.021275 | 0.033183 | 0.078555 |
| 32.63888889 | 0.060351 | 0.079156 | 0.125606 | 0.120879 | 0.091113 | 0.085305 | 0.109034 | 0.110385 | 0.188024 | 0.191115 | 0.096392 | 0.047908 | 0.138875 | 0.121793 | 0.054333 | 0.026044 | 0.011382 | 0.04272 | 0.008178 | 0.010716 | 0.025118 | 0.062013 |
| 32.65277778 | 0.042815 | 0.071164 | 0.084705 | 0.088687 | 0.099589 | 0.083226 | 0.107311 | 0.091645 | 0.133733 | 0.113822 | 0.088196 | 0.041678 | 0.120688 | 0.129825 | 0.035398 | 0.022977 | 0.013406 | 0.042505 | 0.008887 | 0.005408 | 0.01123 | 0.039452 |
| 32.66666667 | 0.038621 | 0.087161 | 0.054335 | 0.044002 | 0.068964 | 0.05554 | 0.059961 | 0.049948 | 0.080358 | 0.058315 | 0.102401 | 0.043337 | 0.068678 | 0.090328 | 0.028095 | 0.026603 | 0.01668 | 0.043604 | 0.013772 | 0.00601 | 0.006754 | 0.042361 |
| 32.68055556 | 0.03071 | 0.076102 | 0.029494 | 0.020112 | 0.041053 | 0.020502 | 0.020489 | 0.044015 | 0.060883 | 0.047271 | 0.084568 | 0.046548 | 0.031427 | 0.049857 | 0.025907 | 0.030273 | 0.011421 | 0.034526 | 0.019969 | 0.005914 | 0.019114 | 0.037485 |
| 32.69444444 | 0.016592 | 0.047599 | 0.009817 | 0.019552 | 0.041904 | 0.008264 | 0.009394 | 0.054126 | 0.03721 | 0.029483 | 0.057485 | 0.039336 | 0.032454 | 0.057996 | 0.024669 | 0.025962 | 0.007308 | 0.020682 | 0.027589 | 0.011011 | 0.039758 | 0.044114 |
| 32.70833333 | 0.010293 | 0.040965 | 0.003846 | 0.026463 | 0.040968 | 0.01076 | 0.012685 | 0.045069 | 0.019861 | 0.015789 | 0.043642 | 0.030501 | 0.038629 | 0.073751 | 0.02489 | 0.023763 | 0.011927 | 0.024891 | 0.023524 | 0.018129 | 0.044927 | 0.06431 |
| 32.72222222 | 0.015079 | 0.038015 | 0.003076 | 0.04009 | 0.045409 | 0.006278 | 0.017885 | 0.039243 | 0.020971 | 0.035386 | 0.040314 | 0.033496 | 0.023426 | 0.047606 | 0.02849 | 0.025455 | 0.01883 | 0.03882 | 0.011707 | 0.020241 | 0.02865 | 0.055758 |
| 32.73611111 | 0.017857 | 0.025577 | 0.005177 | 0.048 | 0.066381 | 0.003111 | 0.014972 | 0.038259 | 0.025148 | 0.066818 | 0.064294 | 0.04544 | 0.018902 | 0.029608 | 0.027845 | 0.020628 | 0.022638 | 0.041441 | 0.012381 | 0.016705 | 0.01689 | 0.026158 |
| 32.75 | 0.011542 | 0.036637 | 0.009337 | 0.044158 | 0.063246 | 0.004685 | 0.016257 | 0.039929 | 0.025681 | 0.059925 | 0.06491 | 0.048763 | 0.041115 | 0.045903 | 0.024809 | 0.018036 | 0.019756 | 0.02821 | 0.019959 | 0.011953 | 0.026068 | 0.008835 |
| 32.76388889 | 0.004212 | 0.05974 | 0.023 | 0.068275 | 0.080126 | 0.008052 | 0.019214 | 0.061892 | 0.045747 | 0.04019 | 0.073573 | 0.058506 | 0.049338 | 0.054898 | 0.033473 | 0.025986 | 0.011006 | 0.014033 | 0.02077 | 0.014041 | 0.036338 | 0.011662 |
| 32.77777778 | 0.006196 | 0.055148 | 0.054988 | 0.113165 | 0.110155 | 0.015444 | 0.013869 | 0.07348 | 0.073988 | 0.060256 | 0.148718 | 0.049727 | 0.033353 | 0.042694 | 0.046081 | 0.027386 | 0.008093 | 0.019024 | 0.013574 | 0.013321 | 0.030795 | 0.015877 |
| 32.79166667 | 0.01608 | 0.028047 | 0.060141 | 0.10259 | 0.083612 | 0.017673 | 0.012889 | 0.05328 | 0.072856 | 0.069575 | 0.175522 | 0.021781 | 0.025224 | 0.035724 | 0.039656 | 0.019862 | 0.01435 | 0.0322 | 0.008432 | 0.016161 | 0.018209 | 0.013683 |
| 32.80555556 | 0.021194 | 0.007613 | 0.042165 | 0.061051 | 0.051454 | 0.015021 | 0.018132 | 0.038941 | 0.049367 | 0.044 | 0.109729 | 0.017759 | 0.03262 | 0.057628 | 0.022455 | 0.019533 | 0.014021 | 0.029546 | 0.011499 | 0.034933 | 0.015805 | 0.009645 |
| 32.81944444 | 0.014765 | 0.006911 | 0.053769 | 0.063413 | 0.053931 | 0.014274 | 0.020278 | 0.031278 | 0.022122 | 0.023136 | 0.077322 | 0.036801 | 0.043442 | 0.08288 | 0.019711 | 0.020403 | 0.008923 | 0.021248 | 0.017656 | 0.046539 | 0.024011 | 0.011305 |
| 32.83333333 | 0.007476 | 0.014525 | 0.059095 | 0.065163 | 0.05169 | 0.011057 | 0.024347 | 0.025446 | 0.010311 | 0.020463 | 0.126289 | 0.067577 | 0.060866 | 0.099659 | 0.0202 | 0.01309 | 0.015933 | 0.026527 | 0.018951 | 0.036681 | 0.027546 | 0.019717 |
| 32.84722222 | 0.009408 | 0.018924 | 0.034686 | 0.03854 | 0.045644 | 0.006786 | 0.021669 | 0.035453 | 0.01837 | 0.026289 | 0.140142 | 0.065829 | 0.061737 | 0.088296 | 0.019461 | 0.009853 | 0.02476 | 0.026479 | 0.015876 | 0.017705 | 0.019973 | 0.021976 |
| 32.86111111 | 0.012943 | 0.016968 | 0.022539 | 0.025122 | 0.048136 | 0.006429 | 0.021705 | 0.034149 | 0.028261 | 0.027109 | 0.095275 | 0.047066 | 0.038961 | 0.058117 | 0.025898 | 0.011942 | 0.017888 | 0.01314 | 0.014109 | 0.007021 | 0.010633 | 0.027982 |
| 32.875 | 0.010552 | 0.008791 | 0.030633 | 0.023799 | 0.043819 | 0.006179 | 0.03627 | 0.042906 | 0.028572 | 0.032202 | 0.111515 | 0.054053 | 0.031534 | 0.076519 | 0.024539 | 0.011782 | 0.01018 | 0.011216 | 0.011152 | 0.005594 | 0.007916 | 0.048982 |
| 32.88888889 | 0.009522 | 0.006358 | 0.031765 | 0.028907 | 0.031659 | 0.007591 | 0.034599 | 0.081285 | 0.035492 | 0.033875 | 0.165444 | 0.043263 | 0.034136 | 0.100103 | 0.019042 | 0.015856 | 0.013695 | 0.015142 | 0.017208 | 0.010232 | 0.006148 | 0.05813 |
| 32.90277778 | 0.019614 | 0.011299 | 0.022419 | 0.029375 | 0.025042 | 0.014437 | 0.024187 | 0.077542 | 0.041255 | 0.035578 | 0.151061 | 0.026434 | 0.029337 | 0.06638 | 0.026912 | 0.01673 | 0.012811 | 0.01537 | 0.029738 | 0.019316 | 0.003507 | 0.043763 |
| 32.91666667 | 0.027125 | 0.019414 | 0.021028 | 0.023693 | 0.029727 | 0.016948 | 0.021867 | 0.044275 | 0.047172 | 0.038516 | 0.08207 | 0.040978 | 0.026076 | 0.033386 | 0.037276 | 0.01186 | 0.013843 | 0.025585 | 0.027999 | 0.018349 | 0.007889 | 0.025195 |
| 32.93055556 | 0.016483 | 0.032435 | 0.022848 | 0.022879 | 0.05041 | 0.025761 | 0.028234 | 0.046471 | 0.057949 | 0.035198 | 0.050929 | 0.057525 | 0.041227 | 0.05171 | 0.032857 | 0.015903 | 0.027236 | 0.039862 | 0.022684 | 0.012769 | 0.022385 | 0.022056 |
| 32.94444444 | 0.012977 | 0.050325 | 0.03402 | 0.022859 | 0.089116 | 0.059321 | 0.058806 | 0.042564 | 0.053861 | 0.06404 | 0.063751 | 0.064402 | 0.082034 | 0.081951 | 0.022933 | 0.024277 | 0.0284 | 0.038984 | 0.024552 | 0.01881 | 0.043421 | 0.044887 |
| 32.95833333 | 0.028579 | 0.064415 | 0.086652 | 0.054631 | 0.100294 | 0.088211 | 0.098447 | 0.019546 | 0.059321 | 0.103805 | 0.055304 | 0.063761 | 0.123186 | 0.09868 | 0.032564 | 0.028724 | 0.014409 | 0.020896 | 0.02243 | 0.024949 | 0.061827 | 0.076209 |
| 32.97222222 | 0.035292 | 0.071515 | 0.149122 | 0.111477 | 0.081552 | 0.077198 | 0.103701 | 0.017095 | 0.064926 | 0.10847 | 0.071053 | 0.052028 | 0.141039 | 0.12232 | 0.066659 | 0.032206 | 0.01026 | 0.013603 | 0.016875 | 0.02771 | 0.060981 | 0.087684 |
| 32.98611111 | 0.047826 | 0.095901 | 0.163005 | 0.133375 | 0.075143 | 0.047165 | 0.071283 | 0.04001 | 0.086667 | 0.112096 | 0.14648 | 0.054327 | 0.137003 | 0.153757 | 0.105853 | 0.046038 | 0.030679 | 0.050144 | 0.019597 | 0.047817 | 0.042091 | 0.076311 |
| 33 | 0.093015 | 0.136773 | 0.121977 | 0.113197 | 0.085972 | 0.049143 | 0.059317 | 0.092593 | 0.156103 | 0.135993 | 0.213937 | 0.087678 | 0.13422 | 0.180792 | 0.130696 | 0.074029 | 0.075666 | 0.118212 | 0.037984 | 0.082243 | 0.034008 | 0.06739 |
| 33.01388889 | 0.138663 | 0.154147 | 0.061106 | 0.083987 | 0.102157 | 0.089238 | 0.093393 | 0.159524 | 0.202333 | 0.146601 | 0.230671 | 0.124353 | 0.152884 | 0.201721 | 0.12875 | 0.091607 | 0.108095 | 0.157007 | 0.065732 | 0.101297 | 0.051552 | 0.079008 |
| 33.02777778 | 0.165432 | 0.136714 | 0.030831 | 0.075102 | 0.108603 | 0.129724 | 0.137625 | 0.202149 | 0.187546 | 0.140268 | 0.214343 | 0.124424 | 0.176625 | 0.210126 | 0.10278 | 0.081829 | 0.099696 | 0.137451 | 0.087908 | 0.096173 | 0.080324 | 0.099487 |
| 33.04166667 | 0.192108 | 0.124082 | 0.054739 | 0.088851 | 0.097671 | 0.140243 | 0.15376 | 0.201546 | 0.156967 | 0.13161 | 0.197386 | 0.09889 | 0.193469 | 0.204215 | 0.080388 | 0.0618 | 0.076293 | 0.099333 | 0.096871 | 0.084995 | 0.098783 | 0.109942 |
| 33.05555556 | 0.22527 | 0.15332 | 0.114405 | 0.096116 | 0.075365 | 0.111337 | 0.116337 | 0.161965 | 0.138765 | 0.111415 | 0.18021 | 0.082297 | 0.223037 | 0.196448 | 0.077247 | 0.049035 | 0.058812 | 0.077052 | 0.090778 | 0.080221 | 0.095126 | 0.100852 |
| 33.06944444 | 0.245468 | 0.185583 | 0.171894 | 0.095832 | 0.051059 | 0.067335 | 0.048373 | 0.099337 | 0.122738 | 0.081866 | 0.14288 | 0.063698 | 0.22279 | 0.168719 | 0.076728 | 0.043208 | 0.044068 | 0.065608 | 0.070197 | 0.074709 | 0.06977 | 0.07437 |
| 33.08333333 | 0.213243 | 0.149637 | 0.198348 | 0.105065 | 0.031888 | 0.05031 | 0.01956 | 0.046557 | 0.102995 | 0.056413 | 0.109204 | 0.034279 | 0.140672 | 0.098402 | 0.061742 | 0.036489 | 0.031221 | 0.053546 | 0.044719 | 0.059913 | 0.034254 | 0.042326 |
| 33.09722222 | 0.1352 | 0.073753 | 0.183304 | 0.114539 | 0.024536 | 0.065671 | 0.039968 | 0.032352 | 0.082769 | 0.035023 | 0.105226 | 0.020568 | 0.048899 | 0.035915 | 0.036721 | 0.025535 | 0.0216 | 0.039997 | 0.026517 | 0.043043 | 0.010356 | 0.019187 |
| 33.11111111 | 0.077401 | 0.027669 | 0.144725 | 0.108901 | 0.026627 | 0.081988 | 0.053488 | 0.046846 | 0.061515 | 0.018118 | 0.097984 | 0.034379 | 0.013096 | 0.023245 | 0.019685 | 0.014959 | 0.014373 | 0.028165 | 0.018823 | 0.032936 | 0.004779 | 0.012029 |
| 33.125 | 0.065097 | 0.022058 | 0.11537 | 0.089593 | 0.036027 | 0.086352 | 0.049087 | 0.061602 | 0.046696 | 0.020242 | 0.078505 | 0.056759 | 0.007737 | 0.020856 | 0.015631 | 0.008735 | 0.009709 | 0.020434 | 0.015497 | 0.026507 | 0.005698 | 0.014552 |
| 33.13888889 | 0.073683 | 0.045581 | 0.111806 | 0.076365 | 0.05461 | 0.086171 | 0.0581 | 0.069061 | 0.047416 | 0.04322 | 0.079817 | 0.074069 | 0.010435 | 0.011749 | 0.016669 | 0.006277 | 0.008517 | 0.017555 | 0.011964 | 0.02044 | 0.008367 | 0.016275 |
| 33.15277778 | 0.083049 | 0.065338 | 0.121729 | 0.07852 | 0.074643 | 0.084102 | 0.078233 | 0.079517 | 0.055335 | 0.062613 | 0.107967 | 0.082441 | 0.02085 | 0.014848 | 0.017546 | 0.00523 | 0.00951 | 0.017081 | 0.010243 | 0.017758 | 0.009256 | 0.014491 |
| 33.16666667 | 0.083852 | 0.06707 | 0.120573 | 0.083341 | 0.079239 | 0.085317 | 0.082216 | 0.091236 | 0.057952 | 0.061163 | 0.135322 | 0.077508 | 0.033573 | 0.023868 | 0.017751 | 0.005703 | 0.010843 | 0.015857 | 0.010948 | 0.018741 | 0.006735 | 0.012532 |
| 33.18055556 | 0.079648 | 0.067526 | 0.103406 | 0.079811 | 0.065667 | 0.091603 | 0.066869 | 0.09345 | 0.054966 | 0.04792 | 0.140912 | 0.066799 | 0.043448 | 0.027187 | 0.019551 | 0.008903 | 0.01203 | 0.013672 | 0.012749 | 0.020795 | 0.005878 | 0.013695 |
| 33.19444444 | 0.076617 | 0.062566 | 0.080901 | 0.066877 | 0.04872 | 0.098182 | 0.051318 | 0.089663 | 0.053662 | 0.035429 | 0.127038 | 0.061734 | 0.052701 | 0.027122 | 0.023874 | 0.012934 | 0.013318 | 0.012121 | 0.01522 | 0.023236 | 0.008481 | 0.018193 |
| 33.20833333 | 0.070195 | 0.048399 | 0.057785 | 0.050055 | 0.03799 | 0.095641 | 0.038819 | 0.085265 | 0.055205 | 0.023934 | 0.105187 | 0.058989 | 0.065448 | 0.032107 | 0.029323 | 0.015171 | 0.014476 | 0.011834 | 0.018845 | 0.028018 | 0.012691 | 0.023438 |
| 33.22222222 | 0.058183 | 0.038463 | 0.035587 | 0.034564 | 0.030717 | 0.075318 | 0.020928 | 0.068389 | 0.047443 | 0.01175 | 0.08572 | 0.049311 | 0.083103 | 0.046174 | 0.035579 | 0.017194 | 0.015632 | 0.013145 | 0.024757 | 0.036405 | 0.017303 | 0.027675 |
| 33.23611111 | 0.046295 | 0.03312 | 0.021276 | 0.020432 | 0.019524 | 0.046276 | 0.00674 | 0.038461 | 0.026995 | 0.003807 | 0.066202 | 0.033777 | 0.101417 | 0.061488 | 0.041978 | 0.021658 | 0.016423 | 0.015455 | 0.032784 | 0.04476 | 0.022389 | 0.030555 |
| 33.25 | 0.035537 | 0.021931 | 0.015129 | 0.008846 | 0.010669 | 0.021969 | 0.005739 | 0.020537 | 0.010471 | 0.003219 | 0.040293 | 0.018925 | 0.113949 | 0.063728 | 0.044811 | 0.025372 | 0.015037 | 0.015007 | 0.038742 | 0.048311 | 0.02489 | 0.028972 |
| 33.26388889 | 0.024221 | 0.011562 | 0.010869 | 0.00465 | 0.012027 | 0.007672 | 0.01165 | 0.021116 | 0.006143 | 0.00822 | 0.017327 | 0.008537 | 0.117953 | 0.050395 | 0.041796 | 0.023218 | 0.011196 | 0.009757 | 0.038198 | 0.04613 | 0.021278 | 0.020828 |
| 33.27777778 | 0.019215 | 0.009257 | 0.008924 | 0.005 | 0.011022 | 0.004733 | 0.01645 | 0.019825 | 0.007024 | 0.014846 | 0.007506 | 0.00383 | 0.114422 | 0.033779 | 0.036145 | 0.017073 | 0.006581 | 0.003647 | 0.033451 | 0.040458 | 0.014773 | 0.012161 |
| 33.29166667 | 0.028963 | 0.012466 | 0.008852 | 0.003727 | 0.005228 | 0.008179 | 0.013638 | 0.009679 | 0.007658 | 0.015474 | 0.00607 | 0.003295 | 0.106296 | 0.023941 | 0.032312 | 0.013243 | 0.002862 | 0.000668 | 0.029756 | 0.033885 | 0.012257 | 0.009964 |
| 33.30555556 | 0.045467 | 0.015156 | 0.008731 | 0.002288 | 0.006383 | 0.00948 | 0.006508 | 0.002104 | 0.005849 | 0.010047 | 0.005063 | 0.003455 | 0.095239 | 0.019279 | 0.029799 | 0.012654 | 0.001538 | 0.000685 | 0.027075 | 0.026588 | 0.013789 | 0.011911 |
| 33.31944444 | 0.048382 | 0.012066 | 0.010583 | 0.002487 | 0.010929 | 0.010797 | 0.003926 | 0.003868 | 0.004302 | 0.007624 | 0.006438 | 0.004426 | 0.080816 | 0.013157 | 0.025011 | 0.010539 | 0.002016 | 0.0021 | 0.021858 | 0.017914 | 0.013489 | 0.011016 |
| 33.33333333 | 0.031901 | 0.008212 | 0.011052 | 0.003964 | 0.009678 | 0.016642 | 0.003789 | 0.011154 | 0.005888 | 0.008244 | 0.01265 | 0.006715 | 0.063865 | 0.00584 | 0.01911 | 0.006274 | 0.002299 | 0.002839 | 0.014416 | 0.009409 | 0.010238 | 0.006654 |
| 33.34722222 | 0.0126 | 0.007025 | 0.008642 | 0.00434 | 0.005431 | 0.017592 | 0.003752 | 0.013409 | 0.005405 | 0.007049 | 0.015829 | 0.006731 | 0.046937 | 0.001804 | 0.016689 | 0.00439 | 0.003829 | 0.004688 | 0.008439 | 0.003494 | 0.008443 | 0.004098 |
| 33.36111111 | 0.008116 | 0.010991 | 0.007023 | 0.002965 | 0.005405 | 0.014084 | 0.004235 | 0.011031 | 0.004426 | 0.00521 | 0.014338 | 0.00567 | 0.031667 | 0.001251 | 0.015965 | 0.004733 | 0.006788 | 0.009864 | 0.004416 | 0.001954 | 0.00943 | 0.004826 |
| 33.375 | 0.023732 | 0.023199 | 0.007722 | 0.003498 | 0.010338 | 0.017881 | 0.008864 | 0.02 | 0.012482 | 0.010913 | 0.022164 | 0.005703 | 0.018886 | 0.005196 | 0.011577 | 0.004324 | 0.006161 | 0.01117 | 0.00273 | 0.005447 | 0.009301 | 0.004465 |
| 33.38888889 | 0.052236 | 0.035871 | 0.014675 | 0.009563 | 0.017921 | 0.03156 | 0.020264 | 0.036365 | 0.026246 | 0.02904 | 0.046701 | 0.009471 | 0.01798 | 0.014256 | 0.00539 | 0.004784 | 0.003972 | 0.008372 | 0.005128 | 0.011654 | 0.006286 | 0.004513 |
| 33.40277778 | 0.077991 | 0.046301 | 0.02828 | 0.020163 | 0.028466 | 0.050855 | 0.031215 | 0.040693 | 0.033757 | 0.045434 | 0.076321 | 0.022179 | 0.035776 | 0.023794 | 0.003059 | 0.006188 | 0.006592 | 0.011766 | 0.009143 | 0.017236 | 0.005433 | 0.010514 |
| 33.41666667 | 0.093704 | 0.058616 | 0.04424 | 0.032096 | 0.043019 | 0.07595 | 0.041211 | 0.041035 | 0.038381 | 0.050379 | 0.100383 | 0.038865 | 0.062766 | 0.031775 | 0.005504 | 0.007462 | 0.012227 | 0.021218 | 0.013498 | 0.022197 | 0.009668 | 0.01922 |
| 33.43055556 | 0.104033 | 0.069503 | 0.061741 | 0.045753 | 0.059472 | 0.106148 | 0.055136 | 0.052747 | 0.049391 | 0.053495 | 0.121295 | 0.052742 | 0.090586 | 0.042397 | 0.011477 | 0.01225 | 0.018593 | 0.03142 | 0.019903 | 0.029093 | 0.015266 | 0.027567 |
| 33.44444444 | 0.111909 | 0.075426 | 0.080166 | 0.061972 | 0.073944 | 0.130453 | 0.070702 | 0.069157 | 0.063842 | 0.063551 | 0.142532 | 0.065717 | 0.116142 | 0.058876 | 0.020285 | 0.022662 | 0.026086 | 0.042634 | 0.028166 | 0.03889 | 0.02229 | 0.039452 |
| 33.45833333 | 0.120898 | 0.083489 | 0.101104 | 0.083553 | 0.089525 | 0.149579 | 0.086703 | 0.087362 | 0.080689 | 0.079916 | 0.171549 | 0.083464 | 0.13846 | 0.080724 | 0.031434 | 0.038023 | 0.036739 | 0.060124 | 0.039007 | 0.051717 | 0.035925 | 0.059223 |
| 33.47222222 | 0.140843 | 0.10424 | 0.129454 | 0.112197 | 0.113141 | 0.172308 | 0.105478 | 0.111612 | 0.10489 | 0.097483 | 0.221604 | 0.116458 | 0.160941 | 0.106333 | 0.045106 | 0.057943 | 0.055765 | 0.088381 | 0.052878 | 0.068519 | 0.056675 | 0.086144 |
| 33.48611111 | 0.17983 | 0.14166 | 0.164059 | 0.145062 | 0.142901 | 0.187691 | 0.129846 | 0.135811 | 0.134361 | 0.11511 | 0.281215 | 0.16966 | 0.180232 | 0.132027 | 0.059847 | 0.079322 | 0.076617 | 0.116458 | 0.067147 | 0.086501 | 0.077449 | 0.110163 |
| 33.5 | 0.232411 | 0.191835 | 0.203331 | 0.180647 | 0.170471 | 0.197723 | 0.169054 | 0.160196 | 0.167014 | 0.135943 | 0.320007 | 0.217336 | 0.192063 | 0.153014 | 0.073141 | 0.097333 | 0.079348 | 0.122173 | 0.08078 | 0.101384 | 0.090885 | 0.122802 |

| | | | | | | | | | | | | | | | | | | | | | | |
|---|---|---|---|---|---|---|---|---|---|---|---|---|---|---|---|---|---|---|---|---|---|---|
| 33.51388889 | 0.285389 | 0.243846 | 0.244685 | 0.214511 | 0.185601 | 0.213734 | 0.213344 | 0.197497 | 0.211269 | 0.162347 | 0.337883 | 0.227165 | 0.204081 | 0.168549 | 0.082608 | 0.108026 | 0.064917 | 0.104616 | 0.090747 | 0.112442 | 0.093882 | 0.128955 |
| 33.52777778 | 0.314944 | 0.281654 | 0.26751 | 0.234048 | 0.184136 | 0.198301 | 0.213063 | 0.240174 | 0.264059 | 0.188758 | 0.338916 | 0.207734 | 0.224918 | 0.172845 | 0.085703 | 0.104823 | 0.055776 | 0.094587 | 0.091762 | 0.113765 | 0.089261 | 0.137732 |
| 33.54166667 | 0.286316 | 0.281578 | 0.254903 | 0.227698 | 0.169117 | 0.137998 | 0.153864 | 0.251815 | 0.289495 | 0.196525 | 0.314849 | 0.182345 | 0.244673 | 0.159821 | 0.090921 | 0.087615 | 0.055672 | 0.103498 | 0.085742 | 0.103144 | 0.0879 | 0.151821 |
| 33.55555556 | 0.187352 | 0.219319 | 0.203253 | 0.191115 | 0.126244 | 0.073471 | 0.081254 | 0.209212 | 0.23981 | 0.164689 | 0.254961 | 0.148172 | 0.23746 | 0.137809 | 0.108917 | 0.070027 | 0.052504 | 0.110952 | 0.078371 | 0.092531 | 0.089057 | 0.154653 |
| 33.56944444 | 0.10219 | 0.112283 | 0.114014 | 0.136161 | 0.064828 | 0.033874 | 0.04451 | 0.142748 | 0.141962 | 0.09471 | 0.174004 | 0.105385 | 0.164068 | 0.100766 | 0.131447 | 0.063653 | 0.041009 | 0.098804 | 0.066186 | 0.081779 | 0.071545 | 0.122761 |
| 33.58333333 | 0.100827 | 0.034541 | 0.034213 | 0.107887 | 0.048293 | 0.038272 | 0.076981 | 0.106651 | 0.097243 | 0.043208 | 0.124345 | 0.080433 | 0.075643 | 0.054414 | 0.136025 | 0.060794 | 0.024374 | 0.065778 | 0.039752 | 0.061223 | 0.040374 | 0.073002 |
| 33.59722222 | 0.095645 | 0.041631 | 0.017248 | 0.118334 | 0.097154 | 0.069679 | 0.141735 | 0.103899 | 0.121728 | 0.061472 | 0.091878 | 0.084061 | 0.046037 | 0.04446 | 0.111668 | 0.041908 | 0.014698 | 0.04032 | 0.015218 | 0.041991 | 0.0275 | 0.047727 |
| 33.61111111 | 0.086491 | 0.103601 | 0.057691 | 0.122777 | 0.161704 | 0.082249 | 0.154496 | 0.09472 | 0.159669 | 0.119254 | 0.081389 | 0.124286 | 0.052888 | 0.08035 | 0.075685 | 0.024022 | 0.023434 | 0.047852 | 0.00941 | 0.040495 | 0.033021 | 0.050699 |
| 33.625 | 0.115462 | 0.147667 | 0.10958 | 0.115758 | 0.183678 | 0.070372 | 0.111044 | 0.081767 | 0.193212 | 0.185052 | 0.135936 | 0.158225 | 0.095401 | 0.114288 | 0.045667 | 0.026943 | 0.040944 | 0.065518 | 0.010828 | 0.045625 | 0.036587 | 0.056345 |
| 33.63888889 | 0.097936 | 0.13602 | 0.122607 | 0.108373 | 0.14711 | 0.069183 | 0.088863 | 0.087788 | 0.199362 | 0.220279 | 0.186813 | 0.113332 | 0.12216 | 0.117977 | 0.024284 | 0.027172 | 0.043741 | 0.059562 | 0.017037 | 0.041511 | 0.032151 | 0.053105 |
| 33.65277778 | 0.050328 | 0.096707 | 0.100887 | 0.096086 | 0.100281 | 0.087328 | 0.109332 | 0.092424 | 0.156235 | 0.174513 | 0.174394 | 0.041936 | 0.096453 | 0.115856 | 0.018524 | 0.024484 | 0.029237 | 0.046293 | 0.025404 | 0.038014 | 0.024158 | 0.038207 |
| 33.66666667 | 0.034507 | 0.071388 | 0.077921 | 0.080038 | 0.062543 | 0.090672 | 0.107298 | 0.080782 | 0.082401 | 0.085667 | 0.129461 | 0.024453 | 0.068228 | 0.107777 | 0.027007 | 0.027761 | 0.016759 | 0.04716 | 0.024162 | 0.032986 | 0.018002 | 0.018978 |
| 33.68055556 | 0.030224 | 0.052742 | 0.055769 | 0.059058 | 0.026357 | 0.058553 | 0.060428 | 0.055679 | 0.03466 | 0.033887 | 0.090818 | 0.066096 | 0.042447 | 0.073366 | 0.031702 | 0.026613 | 0.014356 | 0.047117 | 0.015124 | 0.021635 | 0.011384 | 0.011781 |
| 33.69444444 | 0.021684 | 0.03099 | 0.030221 | 0.037466 | 0.019532 | 0.034335 | 0.024615 | 0.032775 | 0.047014 | 0.043004 | 0.112376 | 0.154807 | 0.033274 | 0.061256 | 0.027776 | 0.024974 | 0.012868 | 0.033572 | 0.010569 | 0.032982 | 0.004013 | 0.020543 |
| 33.70833333 | 0.013666 | 0.020099 | 0.015888 | 0.027576 | 0.033209 | 0.045117 | 0.025195 | 0.04652 | 0.077912 | 0.051381 | 0.184109 | 0.241326 | 0.062628 | 0.077369 | 0.020846 | 0.020247 | 0.009662 | 0.023886 | 0.017343 | 0.064154 | 0.002761 | 0.034233 |
| 33.72222222 | 0.00813 | 0.025445 | 0.021964 | 0.044644 | 0.030281 | 0.040011 | 0.023269 | 0.066717 | 0.085423 | 0.030132 | 0.167088 | 0.204014 | 0.104989 | 0.076416 | 0.012078 | 0.014198 | 0.010333 | 0.030696 | 0.028375 | 0.071694 | 0.008438 | 0.04312 |
| 33.73611111 | 0.006403 | 0.036067 | 0.023714 | 0.050458 | 0.015867 | 0.018151 | 0.023645 | 0.052228 | 0.067438 | 0.029605 | 0.083231 | 0.085302 | 0.111878 | 0.065027 | 0.010473 | 0.014083 | 0.013778 | 0.036451 | 0.027102 | 0.048778 | 0.01114 | 0.040666 |
| 33.75 | 0.01033 | 0.032301 | 0.016969 | 0.035277 | 0.020371 | 0.012069 | 0.04222 | 0.043546 | 0.045686 | 0.05063 | 0.065719 | 0.034195 | 0.084168 | 0.051577 | 0.022002 | 0.016003 | 0.014721 | 0.031375 | 0.017354 | 0.030022 | 0.008692 | 0.034461 |
| 33.76388889 | 0.014782 | 0.022194 | 0.017869 | 0.030095 | 0.028883 | 0.016007 | 0.046834 | 0.04311 | 0.035969 | 0.080214 | 0.060622 | 0.02801 | 0.092347 | 0.074848 | 0.033314 | 0.022273 | 0.011374 | 0.022339 | 0.01369 | 0.026636 | 0.010746 | 0.046157 |
| 33.77777778 | 0.030913 | 0.040063 | 0.039834 | 0.039392 | 0.047652 | 0.031296 | 0.050515 | 0.035592 | 0.061526 | 0.138115 | 0.036834 | 0.033966 | 0.130736 | 0.143476 | 0.030516 | 0.019862 | 0.00843 | 0.022435 | 0.008377 | 0.017535 | 0.011825 | 0.054717 |
| 33.79166667 | 0.058344 | 0.084183 | 0.100805 | 0.082075 | 0.095664 | 0.036559 | 0.05455 | 0.056123 | 0.124707 | 0.184242 | 0.037791 | 0.067904 | 0.129705 | 0.180232 | 0.015789 | 0.010615 | 0.008177 | 0.027107 | 0.002315 | 0.015272 | 0.009101 | 0.037949 |
| 33.80555556 | 0.055794 | 0.099357 | 0.130378 | 0.107909 | 0.109328 | 0.025376 | 0.043029 | 0.079465 | 0.137801 | 0.144984 | 0.071051 | 0.094489 | 0.094618 | 0.145685 | 0.006904 | 0.011949 | 0.006094 | 0.020508 | 0.001761 | 0.034178 | 0.013508 | 0.021041 |
| 33.81944444 | 0.030116 | 0.059942 | 0.092242 | 0.072455 | 0.074551 | 0.030289 | 0.046779 | 0.068286 | 0.092484 | 0.062903 | 0.130496 | 0.091316 | 0.053083 | 0.074475 | 0.017253 | 0.014052 | 0.003116 | 0.012969 | 0.006137 | 0.047482 | 0.01908 | 0.018206 |
| 33.83333333 | 0.025913 | 0.017213 | 0.084021 | 0.045863 | 0.039449 | 0.0363 | 0.048293 | 0.051147 | 0.069076 | 0.043503 | 0.139727 | 0.080802 | 0.032568 | 0.03925 | 0.031057 | 0.012349 | 0.003089 | 0.012087 | 0.021187 | 0.046911 | 0.014915 | 0.019849 |
| 33.84722222 | 0.030236 | 0.004058 | 0.111236 | 0.066634 | 0.035083 | 0.030181 | 0.032744 | 0.043807 | 0.049586 | 0.056859 | 0.093562 | 0.053087 | 0.050444 | 0.0745 | 0.02714 | 0.009128 | 0.006062 | 0.011419 | 0.041583 | 0.040177 | 0.014812 | 0.015606 |
| 33.86111111 | 0.031885 | 0.012554 | 0.094765 | 0.084553 | 0.054668 | 0.03526 | 0.03077 | 0.035722 | 0.028875 | 0.057854 | 0.103463 | 0.034625 | 0.079879 | 0.140183 | 0.027815 | 0.009127 | 0.008994 | 0.010616 | 0.045222 | 0.024549 | 0.029471 | 0.00834 |
| 33.875 | 0.046991 | 0.040728 | 0.048238 | 0.074898 | 0.050271 | 0.034475 | 0.035538 | 0.034529 | 0.035354 | 0.061088 | 0.144974 | 0.05446 | 0.122685 | 0.205462 | 0.059677 | 0.020555 | 0.008646 | 0.010755 | 0.026284 | 0.009792 | 0.037305 | 0.009312 |
| 33.88888889 | 0.046616 | 0.053486 | 0.028732 | 0.049988 | 0.031057 | 0.025154 | 0.039045 | 0.040986 | 0.043217 | 0.041613 | 0.11236 | 0.051544 | 0.184759 | 0.250548 | 0.094289 | 0.02606 | 0.008324 | 0.014937 | 0.010292 | 0.006488 | 0.025229 | 0.023837 |
| 33.90277778 | 0.030784 | 0.049612 | 0.049277 | 0.048546 | 0.025096 | 0.031689 | 0.036156 | 0.041053 | 0.071233 | 0.02832 | 0.064719 | 0.020097 | 0.227203 | 0.234248 | 0.097793 | 0.018612 | 0.012187 | 0.027583 | 0.00876 | 0.006581 | 0.008999 | 0.040669 |
| 33.91666667 | 0.037775 | 0.069267 | 0.081758 | 0.096126 | 0.0194 | 0.030231 | 0.03931 | 0.034695 | 0.137123 | 0.056509 | 0.095007 | 0.005463 | 0.233307 | 0.16712 | 0.077007 | 0.013264 | 0.019023 | 0.041718 | 0.012798 | 0.011683 | 0.005381 | 0.040309 |
| 33.93055556 | 0.043062 | 0.079955 | 0.088621 | 0.117207 | 0.018526 | 0.018533 | 0.070411 | 0.029694 | 0.147134 | 0.068989 | 0.171281 | 0.030226 | 0.230777 | 0.114719 | 0.057119 | 0.011258 | 0.022435 | 0.044796 | 0.024217 | 0.027786 | 0.019939 | 0.042826 |
| 33.94444444 | 0.069949 | 0.100881 | 0.060801 | 0.077058 | 0.018384 | 0.024557 | 0.081091 | 0.049081 | 0.096745 | 0.04131 | 0.229975 | 0.076274 | 0.247503 | 0.107837 | 0.05494 | 0.018816 | 0.015323 | 0.038649 | 0.045486 | 0.039346 | 0.048808 | 0.068657 |
| 33.95833333 | 0.149721 | 0.15029 | 0.036698 | 0.042414 | 0.013721 | 0.038527 | 0.048997 | 0.096919 | 0.102773 | 0.021775 | 0.204448 | 0.080933 | 0.281643 | 0.130491 | 0.067353 | 0.037171 | 0.00548 | 0.031646 | 0.063656 | 0.041873 | 0.066155 | 0.080076 |
| 33.97222222 | 0.191457 | 0.123197 | 0.068453 | 0.044261 | 0.012423 | 0.051321 | 0.029302 | 0.119943 | 0.130797 | 0.033469 | 0.100762 | 0.050976 | 0.287608 | 0.148092 | 0.078372 | 0.052987 | 0.005787 | 0.026949 | 0.051623 | 0.040302 | 0.052435 | 0.059514 |
| 33.98611111 | 0.158556 | 0.051504 | 0.141589 | 0.081646 | 0.016451 | 0.078064 | 0.058576 | 0.097895 | 0.095809 | 0.061772 | 0.032206 | 0.048608 | 0.237069 | 0.137111 | 0.08159 | 0.058425 | 0.014545 | 0.027931 | 0.018354 | 0.040219 | 0.031018 | 0.047149 |
| 34 | 0.127207 | 0.068556 | 0.184429 | 0.130833 | 0.030926 | 0.116292 | 0.110331 | 0.066604 | 0.052858 | 0.088741 | 0.048855 | 0.085709 | 0.160726 | 0.105676 | 0.080691 | 0.053568 | 0.024048 | 0.040765 | 0.013378 | 0.054429 | 0.03102 | 0.072731 |
| 34.01388889 | 0.149699 | 0.149166 | 0.17582 | 0.15781 | 0.04688 | 0.14215 | 0.146572 | 0.06307 | 0.075099 | 0.103111 | 0.152442 | 0.128336 | 0.082815 | 0.074943 | 0.080348 | 0.050033 | 0.03528 | 0.067432 | 0.048453 | 0.078378 | 0.053841 | 0.119918 |
| 34.02777778 | 0.202471 | 0.201 | 0.138583 | 0.149464 | 0.060114 | 0.140262 | 0.151401 | 0.124746 | 0.136279 | 0.102758 | 0.283619 | 0.148396 | 0.024455 | 0.067774 | 0.085418 | 0.057761 | 0.052363 | 0.096709 | 0.093248 | 0.099425 | 0.082631 | 0.151716 |
| 34.04166667 | 0.241505 | 0.208116 | 0.094635 | 0.112233 | 0.068882 | 0.13994 | 0.132328 | 0.220169 | 0.172972 | 0.092694 | 0.315893 | 0.151336 | 0.015975 | 0.084984 | 0.091973 | 0.067889 | 0.065391 | 0.11007 | 0.121975 | 0.110892 | 0.103686 | 0.15088 |
| 34.05555556 | 0.277827 | 0.207432 | 0.106048 | 0.106377 | 0.066717 | 0.191916 | 0.112702 | 0.224161 | 0.163741 | 0.083354 | 0.261771 | 0.158487 | 0.061403 | 0.103491 | 0.090878 | 0.069789 | 0.066446 | 0.108845 | 0.125968 | 0.110368 | 0.103959 | 0.125983 |
| 34.06944444 | 0.300636 | 0.20685 | 0.17543 | 0.135197 | 0.046435 | 0.263238 | 0.080808 | 0.120881 | 0.125842 | 0.07316 | 0.186582 | 0.162033 | 0.106162 | 0.101034 | 0.07666 | 0.066405 | 0.065859 | 0.106705 | 0.107389 | 0.099036 | 0.076186 | 0.089258 |
| 34.08333333 | 0.241522 | 0.158006 | 0.193799 | 0.127052 | 0.017227 | 0.277411 | 0.04296 | 0.042119 | 0.089687 | 0.050804 | 0.11826 | 0.127306 | 0.08675 | 0.070967 | 0.053552 | 0.057554 | 0.066087 | 0.096442 | 0.07733 | 0.07791 | 0.038569 | 0.052022 |
| 34.09722222 | 0.126329 | 0.069016 | 0.131579 | 0.083318 | 0.009299 | 0.22703 | 0.032086 | 0.027648 | 0.071593 | 0.023176 | 0.087844 | 0.069744 | 0.036536 | 0.033635 | 0.034191 | 0.041136 | 0.0559 | 0.068517 | 0.051581 | 0.05382 | 0.015443 | 0.02825 |
| 34.11111111 | 0.052577 | 0.016661 | 0.072039 | 0.052732 | 0.018472 | 0.147282 | 0.02759 | 0.032213 | 0.05723 | 0.008092 | 0.07142 | 0.037432 | 0.016529 | 0.012757 | 0.023839 | 0.025247 | 0.035591 | 0.034501 | 0.037703 | 0.036824 | 0.012318 | 0.023129 |
| 34.125 | 0.037767 | 0.017364 | 0.060022 | 0.049325 | 0.026042 | 0.074108 | 0.013965 | 0.051283 | 0.038236 | 0.011393 | 0.045488 | 0.0314 | 0.02074 | 0.008224 | 0.018432 | 0.014234 | 0.015652 | 0.011167 | 0.030935 | 0.02808 | 0.018078 | 0.027413 |
| 34.13888889 | 0.051632 | 0.035431 | 0.074941 | 0.066698 | 0.043219 | 0.046419 | 0.013342 | 0.089234 | 0.038402 | 0.026227 | 0.047873 | 0.035115 | 0.019618 | 0.010201 | 0.015222 | 0.006498 | 0.003948 | 0.003841 | 0.025447 | 0.024676 | 0.021022 | 0.029533 |
| 34.15277778 | 0.086961 | 0.046845 | 0.089233 | 0.085482 | 0.073004 | 0.070078 | 0.02926 | 0.126501 | 0.056128 | 0.04287 | 0.101739 | 0.047338 | 0.018493 | 0.015575 | 0.015707 | 0.00384 | 0.002344 | 0.008094 | 0.021941 | 0.025811 | 0.019917 | 0.027392 |
| 34.16666667 | 0.129421 | 0.049017 | 0.100053 | 0.084591 | 0.088804 | 0.100621 | 0.04594 | 0.142413 | 0.065542 | 0.056123 | 0.162041 | 0.058233 | 0.030954 | 0.022282 | 0.01887 | 0.005992 | 0.005989 | 0.015934 | 0.020576 | 0.027788 | 0.017839 | 0.024834 |
| 34.18055556 | 0.146689 | 0.050006 | 0.111979 | 0.072586 | 0.079509 | 0.095058 | 0.046173 | 0.132499 | 0.056924 | 0.060173 | 0.184506 | 0.059632 | 0.042257 | 0.02722 | 0.019951 | 0.00754 | 0.00902 | 0.019992 | 0.019316 | 0.027948 | 0.014451 | 0.023518 |
| 34.19444444 | 0.129525 | 0.052468 | 0.102311 | 0.056185 | 0.062032 | 0.054213 | 0.035095 | 0.109002 | 0.038183 | 0.051493 | 0.168012 | 0.056836 | 0.045474 | 0.031771 | 0.019086 | 0.006641 | 0.010601 | 0.019771 | 0.019948 | 0.02929 | 0.010858 | 0.023699 |
| 34.20833333 | 0.098126 | 0.051984 | 0.06086 | 0.032769 | 0.045788 | 0.017931 | 0.02663 | 0.088726 | 0.029436 | 0.038806 | 0.128763 | 0.053024 | 0.047967 | 0.038522 | 0.0203 | 0.008369 | 0.012715 | 0.020236 | 0.024945 | 0.034336 | 0.011726 | 0.02691 |
| 34.22222222 | 0.070839 | 0.045517 | 0.022873 | 0.017259 | 0.026423 | 0.006751 | 0.018358 | 0.06141 | 0.031188 | 0.028824 | 0.085458 | 0.045415 | 0.050749 | 0.045657 | 0.023126 | 0.01433 | 0.01504 | 0.024082 | 0.032125 | 0.041362 | 0.017767 | 0.032158 |
| 34.23611111 | 0.048547 | 0.035344 | 0.013609 | 0.012313 | 0.008723 | 0.004561 | 0.010637 | 0.026573 | 0.027764 | 0.017043 | 0.048485 | 0.030318 | 0.052358 | 0.048598 | 0.023718 | 0.018571 | 0.015198 | 0.027536 | 0.037764 | 0.047807 | 0.023295 | 0.033981 |
| 34.25 | 0.026561 | 0.023633 | 0.019472 | 0.016572 | 0.002198 | 0.011403 | 0.010044 | 0.013995 | 0.021444 | 0.008688 | 0.020831 | 0.016709 | 0.053195 | 0.045065 | 0.021603 | 0.017173 | 0.011136 | 0.023348 | 0.039361 | 0.051493 | 0.02258 | 0.027291 |
| 34.26388889 | 0.009479 | 0.014896 | 0.033888 | 0.039578 | 0.003208 | 0.032238 | 0.014209 | 0.022526 | 0.021906 | 0.009112 | 0.010273 | 0.018362 | 0.055817 | 0.037126 | 0.019827 | 0.013084 | 0.005577 | 0.013543 | 0.036157 | 0.050104 | 0.016141 | 0.01395 |
| 34.27777778 | 0.002546 | 0.01321 | 0.045176 | 0.0555 | 0.006807 | 0.047001 | 0.015497 | 0.022914 | 0.026729 | 0.008913 | 0.019551 | 0.026659 | 0.059205 | 0.027706 | 0.019212 | 0.009412 | 0.0045 | 0.011125 | 0.030461 | 0.043103 | 0.010366 | 0.004125 |
| 34.29166667 | 0.001498 | 0.013428 | 0.031054 | 0.03697 | 0.012158 | 0.039976 | 0.015208 | 0.013363 | 0.028653 | 0.007108 | 0.035964 | 0.030511 | 0.057605 | 0.01742 | 0.017612 | 0.006047 | 0.007721 | 0.019006 | 0.024942 | 0.032336 | 0.008549 | 0.002726 |
| 34.30555556 | 0.001087 | 0.008779 | 0.011425 | 0.010035 | 0.014788 | 0.022348 | 0.015328 | 0.011473 | 0.024528 | 0.005691 | 0.04489 | 0.032571 | 0.050182 | 0.008317 | 0.014957 | 0.004064 | 0.010676 | 0.02564 | 0.020257 | 0.021233 | 0.007209 | 0.004633 |
| 34.31944444 | 0.002787 | 0.002704 | 0.00831 | 0.008239 | 0.01435 | 0.019352 | 0.013378 | 0.013868 | 0.018637 | 0.00522 | 0.044894 | 0.033775 | 0.040615 | 0.005046 | 0.013039 | 0.004045 | 0.013016 | 0.02722 | 0.016184 | 0.012613 | 0.0041 | 0.007395 |
| 34.33333333 | 0.014973 | 0.002984 | 0.011748 | 0.027494 | 0.01568 | 0.048716 | 0.010979 | 0.013185 | 0.016783 | 0.010074 | 0.046292 | 0.03318 | 0.026933 | 0.008827 | 0.010967 | 0.00493 | 0.018696 | 0.031726 | 0.01137 | 0.006716 | 0.00239 | 0.011048 |
| 34.34722222 | 0.039488 | 0.013593 | 0.011348 | 0.04256 | 0.021565 | 0.095451 | 0.012927 | 0.010399 | 0.016945 | 0.015746 | 0.053593 | 0.031767 | 0.015762 | 0.016868 | 0.007194 | 0.008561 | 0.030248 | 0.043955 | 0.006621 | 0.00572 | 0.003169 | 0.013613 |
| 34.36111111 | 0.065312 | 0.032644 | 0.011547 | 0.050761 | 0.033771 | 0.118612 | 0.019454 | 0.008551 | 0.012396 | 0.0197 | 0.060399 | 0.030209 | 0.021916 | 0.0246 | 0.003301 | 0.013005 | 0.041599 | 0.055991 | 0.007659 | 0.011262 | 0.0028 | 0.014012 |
| 34.375 | 0.089526 | 0.050128 | 0.026322 | 0.065815 | 0.047542 | 0.112129 | 0.025823 | 0.019619 | 0.011556 | 0.028366 | 0.066958 | 0.032354 | 0.041245 | 0.027678 | 0.002581 | 0.013292 | 0.046712 | 0.057851 | 0.015774 | 0.019589 | 0.00099 | 0.013872 |
| 34.38888889 | 0.113488 | 0.061691 | 0.048193 | 0.077838 | 0.055269 | 0.111468 | 0.032898 | 0.04257 | 0.027657 | 0.042949 | 0.080673 | 0.04189 | 0.063799 | 0.027704 | 0.006217 | 0.013695 | 0.050166 | 0.05567 | 0.02477 | 0.028702 | 0.000926 | 0.015727 |
| 34.40277778 | 0.129708 | 0.076261 | 0.057078 | 0.076941 | 0.062925 | 0.134582 | 0.046788 | 0.054404 | 0.04514 | 0.055308 | 0.099305 | 0.054475 | 0.085573 | 0.029836 | 0.012213 | 0.020066 | 0.057401 | 0.06085 | 0.031309 | 0.038941 | 0.003522 | 0.020022 |
| 34.41666667 | 0.147393 | 0.096791 | 0.063392 | 0.078691 | 0.079542 | 0.164257 | 0.064984 | 0.045839 | 0.050602 | 0.059755 | 0.115747 | 0.066808 | 0.102154 | 0.0358 | 0.019608 | 0.029087 | 0.066965 | 0.074176 | 0.037345 | 0.049228 | 0.008476 | 0.026812 |
| 34.43055556 | 0.180272 | 0.113215 | 0.08551 | 0.099573 | 0.103891 | 0.172343 | 0.078773 | 0.038902 | 0.056099 | 0.061953 | 0.130938 | 0.084087 | 0.115903 | 0.042886 | 0.028646 | 0.03544 | 0.073333 | 0.090506 | 0.045697 | 0.059072 | 0.016523 | 0.037882 |
| 34.44444444 | 0.220511 | 0.121525 | 0.113664 | 0.129488 | 0.12886 | 0.157062 | 0.08735 | 0.053251 | 0.07271 | 0.071404 | 0.153074 | 0.111875 | 0.129229 | 0.049253 | 0.038402 | 0.040941 | 0.073743 | 0.106477 | 0.057058 | 0.069193 | 0.028276 | 0.05406 |
| 34.45833333 | 0.261329 | 0.131818 | 0.129148 | 0.138652 | 0.140529 | 0.149616 | 0.102791 | 0.084389 | 0.100643 | 0.089131 | 0.188121 | 0.149892 | 0.142143 | 0.060502 | 0.048693 | 0.050524 | 0.074335 | 0.123942 | 0.070031 | 0.081008 | 0.043702 | 0.074952 |
| 34.47222222 | 0.306999 | 0.15319 | 0.1393 | 0.126382 | 0.135344 | 0.17031 | 0.132686 | 0.115248 | 0.133 | 0.111198 | 0.245156 | 0.185453 | 0.157635 | 0.083499 | 0.060439 | 0.062838 | 0.079945 | 0.140352 | 0.083825 | 0.097177 | 0.062625 | 0.101328 |
| 34.48611111 | 0.356514 | 0.181634 | 0.175095 | 0.134665 | 0.137074 | 0.2044 | 0.165436 | 0.138024 | 0.161903 | 0.13747 | 0.337226 | 0.212193 | 0.17621 | 0.117117 | 0.074458 | 0.072039 | 0.086202 | 0.144966 | 0.096757 | 0.118458 | 0.081621 | 0.129768 |
| 34.5 | 0.391875 | 0.208442 | 0.226187 | 0.17598 | 0.165709 | 0.216973 | 0.182663 | 0.164239 | 0.188321 | 0.162713 | 0.441278 | 0.237898 | 0.203329 | 0.153337 | 0.090041 | 0.078748 | 0.083964 | 0.135781 | 0.098841 | 0.130441 | 0.094741 | 0.150455 |
| 34.51388889 | 0.405565 | 0.231969 | 0.239463 | 0.197385 | 0.192013 | 0.19555 | 0.179777 | 0.199989 | 0.215429 | 0.174358 | 0.510257 | 0.257297 | 0.240696 | 0.182738 | 0.103675 | 0.087108 | 0.068014 | 0.126487 | 0.086634 | 0.120712 | 0.100329 | 0.16306 |
| 34.52777778 | 0.388742 | 0.25102 | 0.205667 | 0.162411 | 0.169856 | 0.168732 | 0.164212 | 0.232257 | 0.235866 | 0.170057 | 0.533482 | 0.270403 | 0.266621 | 0.201377 | 0.112491 | 0.092915 | 0.052056 | 0.123535 | 0.075501 | 0.102625 | 0.101466 | 0.169715 |
| 34.54166667 | 0.31055 | 0.245773 | 0.158425 | 0.107554 | 0.117598 | 0.156456 | 0.140796 | 0.246074 | 0.248594 | 0.154353 | 0.507944 | 0.266691 | 0.263163 | 0.196165 | 0.112487 | 0.086047 | 0.044189 | 0.10902 | 0.071808 | 0.086015 | 0.100105 | 0.160682 |
| 34.55555556 | 0.190994 | 0.173236 | 0.119034 | 0.081117 | 0.083369 | 0.133684 | 0.110509 | 0.21062 | 0.226482 | 0.113292 | 0.387739 | 0.215049 | 0.220726 | 0.152464 | 0.103082 | 0.062507 | 0.035354 | 0.074904 | 0.065743 | 0.066402 | 0.090317 | 0.122181 |
| 34.56944444 | 0.144145 | 0.082968 | 0.090801 | 0.082673 | 0.064601 | 0.079382 | 0.077204 | 0.119058 | 0.135843 | 0.055533 | 0.192675 | 0.132314 | 0.136467 | 0.08383 | 0.085939 | 0.034672 | 0.025902 | 0.040425 | 0.046362 | 0.041433 | 0.064328 | 0.060653 |
| 34.58333333 | 0.161389 | 0.051952 | 0.088173 | 0.102158 | 0.048682 | 0.026106 | 0.066489 | 0.048523 | 0.047316 | 0.024811 | 0.048874 | 0.068287 | 0.09127 | 0.031307 | 0.061769 | 0.020669 | 0.017056 | 0.025519 | 0.029284 | 0.019035 | 0.032225 | 0.026944 |
| 34.59722222 | 0.125315 | 0.064282 | 0.109458 | 0.131534 | 0.057284 | 0.012558 | 0.086888 | 0.033751 | 0.03459 | 0.0442 | 0.015203 | 0.067018 | 0.106699 | 0.027069 | 0.046576 | 0.025815 | 0.008338 | 0.027058 | 0.030872 | 0.008532 | 0.015554 | 0.028315 |
| 34.61111111 | 0.134878 | 0.147912 | 0.114884 | 0.134377 | 0.086928 | 0.029602 | 0.091708 | 0.058128 | 0.092421 | 0.111924 | 0.042657 | 0.107788 | 0.096102 | 0.063635 | 0.047145 | 0.036864 | 0.005019 | 0.028503 | 0.032925 | 0.012346 | 0.02189 | 0.037612 |
| 34.625 | 0.213881 | 0.242031 | 0.088142 | 0.095339 | 0.094377 | 0.042209 | 0.077037 | 0.128105 | 0.196085 | 0.16802 | 0.114472 | 0.130103 | 0.105257 | 0.117234 | 0.041229 | 0.033884 | 0.006982 | 0.030693 | 0.035618 | 0.028596 | 0.037384 | 0.059704 |
| 34.63888889 | 0.18685 | 0.221754 | 0.054649 | 0.053905 | 0.073098 | 0.042354 | 0.085403 | 0.171476 | 0.247051 | 0.146675 | 0.227457 | 0.142979 | 0.166377 | 0.145852 | 0.020937 | 0.020492 | 0.011031 | 0.038072 | 0.041031 | 0.041089 | 0.042373 | 0.060756 |
| 34.65277778 | 0.086735 | 0.150199 | 0.046842 | 0.057493 | 0.070313 | 0.043561 | 0.097901 | 0.145877 | 0.186742 | 0.084312 | 0.286442 | 0.133234 | 0.192767 | 0.139028 | 0.007611 | 0.012041 | 0.012909 | 0.037614 | 0.037347 | 0.037208 | 0.037302 | 0.03203 |
| 34.66666667 | 0.035791 | 0.096908 | 0.059066 | 0.087743 | 0.083661 | 0.0381 | 0.075611 | 0.106196 | 0.095242 | 0.04994 | 0.245557 | 0.0785 | 0.167238 | 0.13246 | 0.005964 | 0.007325 | 0.00946 | 0.023575 | 0.02941 | 0.026952 | 0.031305 | 0.015506 |
| 34.68055556 | 0.022732 | 0.049112 | 0.061316 | 0.090758 | 0.068805 | 0.020186 | 0.040826 | 0.061278 | 0.03803 | 0.038106 | 0.156571 | 0.029883 | 0.110929 | 0.101297 | 0.008755 | 0.003303 | 0.006168 | 0.009872 | 0.020479 | 0.019673 | 0.021642 | 0.024756 |
| 34.69444444 | 0.015573 | 0.021474 | 0.046455 | 0.059503 | 0.039802 | 0.00792 | 0.034892 | 0.036909 | 0.016818 | 0.018603 | 0.09177 | 0.024935 | 0.062156 | 0.061862 | 0.011976 | 0.003905 | 0.004541 | 0.003705 | 0.019397 | 0.013315 | 0.010202 | 0.036316 |
| 34.70833333 | 0.016448 | 0.020921 | 0.02356 | 0.033426 | 0.02524 | 0.010112 | 0.038331 | 0.059206 | 0.011166 | 0.020855 | 0.103743 | 0.0396 | 0.073309 | 0.062207 | 0.00943 | 0.007728 | 0.003929 | 0.006571 | 0.02828 | 0.016801 | 0.008219 | 0.036287 |
| 34.72222222 | 0.020838 | 0.033705 | 0.013918 | 0.047008 | 0.043188 | 0.011885 | 0.025618 | 0.062376 | 0.016192 | 0.035823 | 0.124292 | 0.036324 | 0.111185 | 0.063372 | 0.014408 | 0.009711 | 0.007613 | 0.011684 | 0.027216 | 0.025256 | 0.013486 | 0.051744 |
| 34.73611111 | 0.017939 | 0.033963 | 0.021852 | 0.065585 | 0.0768 | 0.015742 | 0.019347 | 0.04419 | 0.030973 | 0.027498 | 0.090217 | 0.02717 | 0.125873 | 0.053002 | 0.032067 | 0.018786 | 0.010596 | 0.012338 | 0.023687 | 0.025998 | 0.011678 | 0.073065 |
| 34.75 | 0.021539 | 0.031813 | 0.022604 | 0.046774 | 0.077532 | 0.032531 | 0.018958 | 0.04535 | 0.037969 | 0.031454 | 0.047209 | 0.028195 | 0.093476 | 0.035531 | 0.049439 | 0.036821 | 0.00792 | 0.01531 | 0.03149 | 0.044696 | 0.008609 | 0.059061 |
| 34.76388889 | 0.054558 | 0.076695 | 0.019754 | 0.029881 | 0.052393 | 0.03602 | 0.010179 | 0.033459 | 0.030678 | 0.039508 | 0.045859 | 0.04313 | 0.054884 | 0.019166 | 0.052096 | 0.046617 | 0.007531 | 0.013706 | 0.027631 | 0.068542 | 0.017559 | 0.026282 |
| 34.77777778 | 0.110981 | 0.161095 | 0.018942 | 0.038615 | 0.034108 | 0.017431 | 0.017535 | 0.019894 | 0.018546 | 0.025665 | 0.055477 | 0.066809 | 0.084937 | 0.019141 | 0.032339 | 0.038104 | 0.012768 | 0.005367 | 0.013601 | 0.050886 | 0.027321 | 0.017001 |
| 34.79166667 | 0.145153 | 0.216851 | 0.021124 | 0.038654 | 0.039594 | 0.010452 | 0.036825 | 0.036903 | 0.020367 | 0.039144 | 0.040242 | 0.067909 | 0.119471 | 0.025205 | 0.021881 | 0.02312 | 0.012715 | 0.000971 | 0.008993 | 0.018446 | 0.023238 | 0.018077 |
| 34.80555556 | 0.12221 | 0.199711 | 0.061155 | 0.040365 | 0.062533 | 0.019186 | 0.030076 | 0.054038 | 0.042035 | 0.09001 | 0.040543 | 0.043373 | 0.084993 | 0.031334 | 0.045904 | 0.023266 | 0.007546 | 0.005894 | 0.008948 | 0.00649 | 0.020262 | 0.009546 |
| 34.81944444 | 0.071338 | 0.122833 | 0.121231 | 0.063002 | 0.059956 | 0.024756 | 0.016244 | 0.058996 | 0.06673 | 0.098652 | 0.078556 | 0.033961 | 0.04118 | 0.026364 | 0.065885 | 0.028771 | 0.007704 | 0.015339 | 0.012538 | 0.009951 | 0.032758 | 0.012634 |
| 34.83333333 | 0.046983 | 0.057121 | 0.12231 | 0.069123 | 0.034528 | 0.028595 | 0.019894 | 0.069924 | 0.068694 | 0.060732 | 0.125572 | 0.042672 | 0.028586 | 0.014063 | 0.046751 | 0.020452 | 0.01132 | 0.015901 | 0.013104 | 0.022137 | 0.037545 | 0.020564 |
| 34.84722222 | 0.045801 | 0.045573 | 0.078191 | 0.045094 | 0.023767 | 0.026508 | 0.019316 | 0.073889 | 0.05116 | 0.043754 | 0.152367 | 0.068294 | 0.017459 | 0.013233 | 0.016934 | 0.012249 | 0.011749 | 0.017522 | 0.016921 | 0.036874 | 0.022458 | 0.016701 |
| 34.86111111 | 0.059069 | 0.061008 | 0.075078 | 0.028909 | 0.036757 | 0.024525 | 0.009427 | 0.074702 | 0.046493 | 0.042637 | 0.171675 | 0.104765 | 0.009266 | 0.021064 | 0.008375 | 0.017043 | 0.01286 | 0.032299 | 0.038058 | 0.04829 | 0.012792 | 0.019562 |
| 34.875 | 0.101637 | 0.096596 | 0.083788 | 0.02327 | 0.064915 | 0.037068 | 0.004351 | 0.087055 | 0.043237 | 0.054333 | 0.143937 | 0.093583 | 0.017179 | 0.018433 | 0.007251 | 0.025997 | 0.012568 | 0.033293 | 0.052944 | 0.04474 | 0.021302 | 0.021511 |
| 34.88888889 | 0.111143 | 0.103955 | 0.04704 | 0.014395 | 0.063894 | 0.050685 | 0.009003 | 0.082817 | 0.025878 | 0.066554 | 0.109458 | 0.07936 | 0.041729 | 0.020097 | 0.010093 | 0.023412 | 0.007686 | 0.027181 | 0.048584 | 0.029214 | 0.029067 | 0.013673 |
| 34.90277778 | 0.066782 | 0.060453 | 0.026695 | 0.016644 | 0.033176 | 0.047013 | 0.026947 | 0.05321 | 0.026584 | 0.047159 | 0.199068 | 0.085397 | 0.077634 | 0.048783 | 0.024998 | 0.016834 | 0.010064 | 0.039984 | 0.034532 | 0.016915 | 0.026648 | 0.020575 |

| | | | | | | | | | | | | | | | | | | | | | | |
|---|---|---|---|---|---|---|---|---|---|---|---|---|---|---|---|---|---|---|---|---|---|---|
| 34.91666667 | 0.033329 | 0.025021 | 0.07232 | 0.025289 | 0.034211 | 0.030753 | 0.041545 | 0.038378 | 0.066348 | 0.036493 | 0.26034 | 0.073535 | 0.108795 | 0.077727 | 0.027456 | 0.016594 | 0.013454 | 0.036392 | 0.025892 | 0.012803 | 0.027749 | 0.037396 |
| 34.93055556 | 0.054443 | 0.024234 | 0.128602 | 0.033714 | 0.046748 | 0.035468 | 0.047788 | 0.086792 | 0.158633 | 0.058468 | 0.21673 | 0.061672 | 0.128583 | 0.090371 | 0.021595 | 0.014389 | 0.01206 | 0.029301 | 0.026677 | 0.014219 | 0.035975 | 0.038809 |
| 34.94444444 | 0.101761 | 0.044204 | 0.116143 | 0.046459 | 0.03613 | 0.070596 | 0.079675 | 0.148948 | 0.235841 | 0.064741 | 0.248274 | 0.04812 | 0.14181 | 0.104235 | 0.038331 | 0.022676 | 0.017162 | 0.052399 | 0.025425 | 0.013304 | 0.039192 | 0.030937 |
| 34.95833333 | 0.096553 | 0.060039 | 0.073344 | 0.050731 | 0.049279 | 0.101854 | 0.122644 | 0.126396 | 0.191177 | 0.041711 | 0.245206 | 0.063759 | 0.152211 | 0.118182 | 0.058752 | 0.04202 | 0.018182 | 0.055164 | 0.037303 | 0.009851 | 0.034978 | 0.038125 |
| 34.97222222 | 0.064829 | 0.071904 | 0.075646 | 0.035913 | 0.073645 | 0.108228 | 0.129903 | 0.078601 | 0.096035 | 0.019044 | 0.103525 | 0.11194 | 0.164815 | 0.11992 | 0.066168 | 0.055905 | 0.014514 | 0.02871 | 0.058574 | 0.014352 | 0.035606 | 0.056474 |
| 34.98611111 | 0.102004 | 0.100031 | 0.07086 | 0.017331 | 0.061957 | 0.096521 | 0.106237 | 0.064923 | 0.05742 | 0.008999 | 0.029618 | 0.122353 | 0.175431 | 0.103579 | 0.068083 | 0.059146 | 0.014137 | 0.018677 | 0.072569 | 0.038085 | 0.043432 | 0.068987 |
| 35 | 0.220189 | 0.137656 | 0.06438 | 0.0271 | 0.037751 | 0.09322 | 0.095381 | 0.052254 | 0.049991 | 0.016766 | 0.092094 | 0.079488 | 0.158169 | 0.075245 | 0.053595 | 0.045169 | 0.010502 | 0.017846 | 0.090776 | 0.075433 | 0.049443 | 0.080897 |
| 35.01388889 | 0.316399 | 0.15648 | 0.117365 | 0.078392 | 0.038101 | 0.122112 | 0.13556 | 0.08307 | 0.076271 | 0.048047 | 0.193851 | 0.047293 | 0.108915 | 0.060837 | 0.030464 | 0.025916 | 0.009983 | 0.023916 | 0.093812 | 0.094826 | 0.052745 | 0.093586 |
| 35.02777778 | 0.304832 | 0.168002 | 0.177113 | 0.14965 | 0.068729 | 0.186915 | 0.215352 | 0.146981 | 0.147208 | 0.095869 | 0.27067 | 0.078993 | 0.064715 | 0.075818 | 0.028152 | 0.031761 | 0.028626 | 0.064525 | 0.058021 | 0.084351 | 0.054606 | 0.097088 |
| 35.04166667 | 0.237432 | 0.18958 | 0.198591 | 0.202466 | 0.113817 | 0.262612 | 0.266494 | 0.173959 | 0.187207 | 0.130781 | 0.298525 | 0.148495 | 0.068457 | 0.107629 | 0.051283 | 0.061455 | 0.074137 | 0.130508 | 0.022013 | 0.076135 | 0.057297 | 0.094894 |
| 35.05555556 | 0.20113 | 0.183649 | 0.195812 | 0.192196 | 0.118 | 0.298868 | 0.237753 | 0.146802 | 0.165913 | 0.133005 | 0.298758 | 0.192766 | 0.116402 | 0.129807 | 0.076249 | 0.084984 | 0.117155 | 0.185121 | 0.025039 | 0.086431 | 0.064268 | 0.092518 |
| 35.06944444 | 0.228029 | 0.130362 | 0.194519 | 0.134649 | 0.06754 | 0.287306 | 0.149588 | 0.085823 | 0.129211 | 0.111274 | 0.232072 | 0.177274 | 0.149008 | 0.122419 | 0.081081 | 0.086269 | 0.126291 | 0.186639 | 0.045624 | 0.090114 | 0.069116 | 0.085889 |
| 35.08333333 | 0.257625 | 0.075682 | 0.198948 | 0.121426 | 0.047504 | 0.259451 | 0.06469 | 0.04224 | 0.10007 | 0.078815 | 0.099456 | 0.125758 | 0.132621 | 0.089447 | 0.068958 | 0.068961 | 0.103753 | 0.128647 | 0.054053 | 0.074139 | 0.063938 | 0.074389 |
| 35.09722222 | 0.221083 | 0.060161 | 0.197585 | 0.174616 | 0.075181 | 0.221294 | 0.029502 | 0.039824 | 0.081261 | 0.05144 | 0.052988 | 0.089706 | 0.096975 | 0.052379 | 0.054617 | 0.046935 | 0.066595 | 0.057121 | 0.051876 | 0.055366 | 0.051269 | 0.059877 |
| 35.11111111 | 0.171399 | 0.068641 | 0.172413 | 0.177143 | 0.076531 | 0.162055 | 0.028346 | 0.068125 | 0.076553 | 0.038224 | 0.086967 | 0.080136 | 0.06715 | 0.031341 | 0.043967 | 0.030843 | 0.038895 | 0.017207 | 0.049437 | 0.046551 | 0.038644 | 0.045068 |
| 35.125 | 0.170701 | 0.068322 | 0.105441 | 0.095042 | 0.043207 | 0.084645 | 0.022464 | 0.091034 | 0.072197 | 0.036393 | 0.088549 | 0.081534 | 0.044944 | 0.029287 | 0.036104 | 0.022637 | 0.026859 | 0.007469 | 0.047389 | 0.043325 | 0.030649 | 0.034961 |
| 35.13888889 | 0.201716 | 0.066586 | 0.051384 | 0.03715 | 0.022889 | 0.024424 | 0.009245 | 0.086525 | 0.068191 | 0.044354 | 0.073736 | 0.084836 | 0.033157 | 0.032119 | 0.02895 | 0.019879 | 0.0189 | 0.004014 | 0.043801 | 0.040716 | 0.026767 | 0.030316 |
| 35.15277778 | 0.224213 | 0.080794 | 0.063268 | 0.046013 | 0.040567 | 0.012554 | 0.010138 | 0.07433 | 0.076223 | 0.057955 | 0.086339 | 0.082546 | 0.031047 | 0.030104 | 0.022763 | 0.018884 | 0.010085 | 0.001917 | 0.041465 | 0.040511 | 0.023293 | 0.02769 |
| 35.16666667 | 0.212523 | 0.094524 | 0.106431 | 0.10202 | 0.092945 | 0.037186 | 0.027292 | 0.063895 | 0.087717 | 0.071631 | 0.118158 | 0.076063 | 0.033294 | 0.027996 | 0.019513 | 0.0181 | 0.003855 | 0.001459 | 0.040303 | 0.042195 | 0.02023 | 0.024882 |
| 35.18055556 | 0.166755 | 0.087896 | 0.124905 | 0.155254 | 0.129615 | 0.083542 | 0.043916 | 0.058274 | 0.084411 | 0.079226 | 0.134948 | 0.070426 | 0.035889 | 0.032148 | 0.018261 | 0.016418 | 0.003272 | 0.002634 | 0.038302 | 0.042749 | 0.02072 | 0.023466 |
| 35.19444444 | 0.109893 | 0.064831 | 0.100762 | 0.170017 | 0.117806 | 0.132631 | 0.049285 | 0.04882 | 0.054255 | 0.071014 | 0.119731 | 0.06389 | 0.037498 | 0.039349 | 0.016296 | 0.013748 | 0.007997 | 0.009639 | 0.037932 | 0.043753 | 0.024484 | 0.025093 |
| 35.20833333 | 0.061394 | 0.035972 | 0.065168 | 0.155388 | 0.083601 | 0.156025 | 0.039341 | 0.027818 | 0.021529 | 0.047687 | 0.087108 | 0.057044 | 0.037658 | 0.041134 | 0.014877 | 0.015845 | 0.016898 | 0.02211 | 0.041268 | 0.047789 | 0.028454 | 0.029201 |
| 35.22222222 | 0.031882 | 0.01824 | 0.047558 | 0.113939 | 0.050437 | 0.143962 | 0.024119 | 0.028207 | 0.012962 | 0.022953 | 0.068316 | 0.053967 | 0.036643 | 0.034432 | 0.016002 | 0.026234 | 0.028129 | 0.035822 | 0.044418 | 0.051685 | 0.030322 | 0.033697 |
| 35.23611111 | 0.036803 | 0.022363 | 0.041796 | 0.066032 | 0.025878 | 0.112739 | 0.018038 | 0.060554 | 0.016543 | 0.00867 | 0.065975 | 0.048605 | 0.035186 | 0.022117 | 0.017042 | 0.036442 | 0.035867 | 0.044511 | 0.044998 | 0.052974 | 0.029226 | 0.034633 |
| 35.25 | 0.073949 | 0.031087 | 0.030125 | 0.036066 | 0.018482 | 0.076431 | 0.014702 | 0.068245 | 0.01859 | 0.007576 | 0.052252 | 0.030888 | 0.033948 | 0.0099 | 0.016776 | 0.036676 | 0.033481 | 0.041676 | 0.045189 | 0.052957 | 0.025711 | 0.028219 |
| 35.26388889 | 0.11178 | 0.031668 | 0.017378 | 0.035117 | 0.035855 | 0.044202 | 0.015004 | 0.045418 | 0.018608 | 0.013193 | 0.030985 | 0.016421 | 0.032439 | 0.004784 | 0.017853 | 0.029442 | 0.020508 | 0.027643 | 0.045771 | 0.050845 | 0.020988 | 0.01543 |
| 35.27777778 | 0.124505 | 0.025779 | 0.017419 | 0.051545 | 0.060006 | 0.043913 | 0.028509 | 0.04821 | 0.028473 | 0.020487 | 0.030045 | 0.024616 | 0.030381 | 0.008982 | 0.018683 | 0.022893 | 0.009064 | 0.012712 | 0.044442 | 0.04405 | 0.016495 | 0.006124 |
| 35.29166667 | 0.117732 | 0.019288 | 0.017413 | 0.050454 | 0.0641 | 0.066314 | 0.037572 | 0.059268 | 0.045422 | 0.027224 | 0.040656 | 0.039922 | 0.027859 | 0.019777 | 0.015257 | 0.018953 | 0.010055 | 0.008144 | 0.038992 | 0.033118 | 0.012403 | 0.007231 |
| 35.30555556 | 0.109106 | 0.015819 | 0.014505 | 0.034685 | 0.057059 | 0.071825 | 0.033714 | 0.052895 | 0.048888 | 0.03053 | 0.046449 | 0.044675 | 0.021988 | 0.034538 | 0.009961 | 0.015213 | 0.016845 | 0.0128 | 0.031375 | 0.022832 | 0.007753 | 0.012154 |
| 35.31944444 | 0.095167 | 0.010278 | 0.017659 | 0.033692 | 0.059799 | 0.062469 | 0.029818 | 0.041894 | 0.03738 | 0.03298 | 0.051118 | 0.044031 | 0.012195 | 0.046862 | 0.009609 | 0.011995 | 0.021814 | 0.018517 | 0.02511 | 0.016414 | 0.005257 | 0.015747 |
| 35.33333333 | 0.060051 | 0.004393 | 0.019471 | 0.044334 | 0.066541 | 0.065817 | 0.028702 | 0.029209 | 0.02784 | 0.042392 | 0.055244 | 0.045765 | 0.006955 | 0.046007 | 0.013606 | 0.010069 | 0.023932 | 0.021978 | 0.020192 | 0.011458 | 0.005824 | 0.017676 |
| 35.34722222 | 0.022632 | 0.011029 | 0.01816 | 0.050718 | 0.065255 | 0.100027 | 0.031681 | 0.03133 | 0.040849 | 0.061575 | 0.059591 | 0.049644 | 0.010544 | 0.032753 | 0.014684 | 0.006999 | 0.024298 | 0.025132 | 0.014647 | 0.006821 | 0.004512 | 0.016765 |
| 35.36111111 | 0.010662 | 0.034401 | 0.026556 | 0.061665 | 0.062697 | 0.170343 | 0.046029 | 0.053853 | 0.069494 | 0.080657 | 0.074052 | 0.052536 | 0.01167 | 0.024093 | 0.010288 | 0.00567 | 0.02961 | 0.033619 | 0.008667 | 0.007498 | 0.001708 | 0.014634 |
| 35.375 | 0.021471 | 0.05526 | 0.045162 | 0.085549 | 0.065261 | 0.246951 | 0.069421 | 0.071891 | 0.082976 | 0.091254 | 0.102071 | 0.054627 | 0.009561 | 0.029338 | 0.004993 | 0.01088 | 0.044242 | 0.051524 | 0.006796 | 0.014243 | 0.001499 | 0.012974 |
| 35.38888889 | 0.050241 | 0.056417 | 0.060593 | 0.102721 | 0.065419 | 0.268609 | 0.086558 | 0.064276 | 0.076204 | 0.099151 | 0.128662 | 0.059525 | 0.016966 | 0.037476 | 0.004593 | 0.017658 | 0.062775 | 0.075402 | 0.009984 | 0.020565 | 0.003128 | 0.011171 |
| 35.40277778 | 0.091797 | 0.050132 | 0.072142 | 0.105022 | 0.059278 | 0.232769 | 0.092571 | 0.048597 | 0.076325 | 0.110336 | 0.142762 | 0.069438 | 0.03208 | 0.036754 | 0.008632 | 0.02219 | 0.076043 | 0.093545 | 0.013523 | 0.024919 | 0.005177 | 0.01086 |
| 35.41666667 | 0.137962 | 0.058249 | 0.086282 | 0.112931 | 0.057062 | 0.202164 | 0.101019 | 0.062981 | 0.100663 | 0.12103 | 0.157857 | 0.082344 | 0.047828 | 0.034445 | 0.013529 | 0.025908 | 0.083517 | 0.102499 | 0.01804 | 0.031353 | 0.009947 | 0.016483 |
| 35.43055556 | 0.178864 | 0.085268 | 0.103174 | 0.136935 | 0.071187 | 0.200853 | 0.113903 | 0.094786 | 0.123133 | 0.123578 | 0.188039 | 0.097133 | 0.060913 | 0.044849 | 0.01802 | 0.028858 | 0.092973 | 0.114374 | 0.025819 | 0.040949 | 0.019179 | 0.029063 |
| 35.44444444 | 0.21036 | 0.122061 | 0.121772 | 0.166582 | 0.099598 | 0.214693 | 0.126153 | 0.109057 | 0.119231 | 0.121055 | 0.227211 | 0.117516 | 0.068204 | 0.067218 | 0.022403 | 0.032867 | 0.10664 | 0.132416 | 0.035825 | 0.051726 | 0.03203 | 0.044943 |
| 35.45833333 | 0.24938 | 0.155003 | 0.137108 | 0.189094 | 0.128408 | 0.228638 | 0.148619 | 0.101568 | 0.104648 | 0.129652 | 0.267963 | 0.148031 | 0.073562 | 0.090767 | 0.026892 | 0.042126 | 0.119212 | 0.151427 | 0.049477 | 0.063339 | 0.047935 | 0.061536 |
| 35.47222222 | 0.304421 | 0.174038 | 0.146458 | 0.193224 | 0.142078 | 0.226027 | 0.185454 | 0.084958 | 0.098256 | 0.143831 | 0.318766 | 0.181481 | 0.082826 | 0.102389 | 0.030748 | 0.056579 | 0.122668 | 0.161272 | 0.06993 | 0.076726 | 0.063821 | 0.080148 |
| 35.48611111 | 0.360707 | 0.186756 | 0.156691 | 0.183548 | 0.143141 | 0.223113 | 0.226327 | 0.082003 | 0.104459 | 0.147806 | 0.386413 | 0.204782 | 0.095641 | 0.093169 | 0.032664 | 0.069422 | 0.105771 | 0.145339 | 0.095634 | 0.092151 | 0.07266 | 0.10022 |
| 35.5 | 0.406933 | 0.20927 | 0.168267 | 0.185772 | 0.156029 | 0.246954 | 0.269511 | 0.100678 | 0.123016 | 0.149092 | 0.46511 | 0.21525 | 0.107663 | 0.076244 | 0.033185 | 0.072626 | 0.076468 | 0.111297 | 0.116111 | 0.109826 | 0.073123 | 0.11654 |
| 35.51388889 | 0.445488 | 0.246659 | 0.176498 | 0.205638 | 0.195986 | 0.26399 | 0.295697 | 0.131192 | 0.153465 | 0.15535 | 0.544975 | 0.224526 | 0.115025 | 0.07687 | 0.035536 | 0.067942 | 0.061016 | 0.095485 | 0.118504 | 0.125397 | 0.071742 | 0.130153 |
| 35.52777778 | 0.480376 | 0.277643 | 0.183021 | 0.21586 | 0.22679 | 0.23677 | 0.274003 | 0.162712 | 0.186385 | 0.16479 | 0.575358 | 0.242129 | 0.119526 | 0.097592 | 0.042696 | 0.068199 | 0.058997 | 0.103229 | 0.103134 | 0.132373 | 0.074981 | 0.147477 |
| 35.54166667 | 0.468026 | 0.265127 | 0.183366 | 0.192447 | 0.198735 | 0.192769 | 0.213084 | 0.174266 | 0.191005 | 0.161011 | 0.511967 | 0.241517 | 0.126551 | 0.116149 | 0.053977 | 0.073 | 0.056901 | 0.10118 | 0.081945 | 0.123214 | 0.086568 | 0.162376 |
| 35.55555556 | 0.318835 | 0.18503 | 0.164374 | 0.149285 | 0.143788 | 0.143797 | 0.131083 | 0.145981 | 0.134896 | 0.124721 | 0.37106 | 0.204548 | 0.141673 | 0.113203 | 0.061447 | 0.067608 | 0.052612 | 0.080529 | 0.068157 | 0.094818 | 0.096202 | 0.159914 |
| 35.56944444 | 0.155038 | 0.086538 | 0.134125 | 0.123924 | 0.120718 | 0.083575 | 0.0556 | 0.091278 | 0.054286 | 0.078243 | 0.188074 | 0.171871 | 0.142921 | 0.087305 | 0.059704 | 0.053502 | 0.04343 | 0.06088 | 0.063702 | 0.057995 | 0.083044 | 0.125931 |
| 35.58333333 | 0.128822 | 0.038388 | 0.115239 | 0.122266 | 0.124827 | 0.048034 | 0.034405 | 0.040969 | 0.030597 | 0.063424 | 0.051036 | 0.176403 | 0.10237 | 0.047792 | 0.056509 | 0.042947 | 0.02631 | 0.045741 | 0.057323 | 0.02821 | 0.050383 | 0.066216 |
| 35.59722222 | 0.117147 | 0.038676 | 0.109363 | 0.121408 | 0.129054 | 0.040666 | 0.066725 | 0.026393 | 0.073107 | 0.089492 | 0.086955 | 0.184989 | 0.048962 | 0.019986 | 0.05782 | 0.044221 | 0.013156 | 0.023563 | 0.043612 | 0.020346 | 0.022815 | 0.019311 |
| 35.61111111 | 0.120116 | 0.077549 | 0.098224 | 0.105391 | 0.137576 | 0.043404 | 0.091605 | 0.063902 | 0.128184 | 0.122534 | 0.266894 | 0.142612 | 0.021063 | 0.026012 | 0.055967 | 0.054149 | 0.015138 | 0.011892 | 0.024073 | 0.021323 | 0.012055 | 0.004923 |
| 35.625 | 0.192883 | 0.145294 | 0.076945 | 0.077975 | 0.14737 | 0.064511 | 0.086442 | 0.098748 | 0.13309 | 0.121963 | 0.361977 | 0.076144 | 0.026099 | 0.043268 | 0.050653 | 0.054288 | 0.02663 | 0.026871 | 0.008394 | 0.015212 | 0.013944 | 0.004091 |
| 35.63888889 | 0.240889 | 0.215034 | 0.07397 | 0.068039 | 0.126264 | 0.08237 | 0.084539 | 0.080035 | 0.078474 | 0.083006 | 0.295003 | 0.065166 | 0.052259 | 0.052025 | 0.048102 | 0.041419 | 0.033518 | 0.0505 | 0.005585 | 0.016978 | 0.021398 | 0.013022 |
| 35.65277778 | 0.209473 | 0.220063 | 0.091024 | 0.083188 | 0.081988 | 0.078429 | 0.084809 | 0.050456 | 0.038771 | 0.062903 | 0.214209 | 0.085147 | 0.083099 | 0.07524 | 0.038339 | 0.025554 | 0.023653 | 0.064005 | 0.014976 | 0.024016 | 0.029856 | 0.033257 |
| 35.66666667 | 0.10831 | 0.143864 | 0.091651 | 0.091396 | 0.051229 | 0.052276 | 0.069609 | 0.045587 | 0.048923 | 0.112181 | 0.184679 | 0.069835 | 0.118646 | 0.113583 | 0.020445 | 0.014028 | 0.012192 | 0.057551 | 0.02953 | 0.027572 | 0.0326 | 0.035498 |
| 35.68055556 | 0.025597 | 0.075735 | 0.068714 | 0.076221 | 0.039106 | 0.022515 | 0.051133 | 0.043382 | 0.059489 | 0.134646 | 0.151952 | 0.034287 | 0.124088 | 0.110802 | 0.013124 | 0.010677 | 0.011241 | 0.036082 | 0.034072 | 0.025876 | 0.028104 | 0.018101 |
| 35.69444444 | 0.006778 | 0.041205 | 0.036446 | 0.046425 | 0.034914 | 0.014929 | 0.032626 | 0.036629 | 0.053151 | 0.071405 | 0.089036 | 0.022226 | 0.073734 | 0.065145 | 0.020572 | 0.012199 | 0.01123 | 0.024994 | 0.025318 | 0.021556 | 0.029117 | 0.01883 |
| 35.70833333 | 0.013985 | 0.029009 | 0.016631 | 0.022816 | 0.028537 | 0.013653 | 0.020209 | 0.029672 | 0.092812 | 0.054118 | 0.069131 | 0.066838 | 0.042402 | 0.049154 | 0.025807 | 0.017684 | 0.007157 | 0.038076 | 0.013315 | 0.029464 | 0.039835 | 0.052161 |
| 35.72222222 | 0.018558 | 0.033903 | 0.029749 | 0.028142 | 0.015926 | 0.014323 | 0.020832 | 0.023673 | 0.146925 | 0.091036 | 0.123966 | 0.111486 | 0.05735 | 0.050405 | 0.022424 | 0.022868 | 0.003289 | 0.041791 | 0.007873 | 0.044239 | 0.038757 | 0.083363 |
| 35.73611111 | 0.023619 | 0.042511 | 0.047001 | 0.044884 | 0.014956 | 0.021058 | 0.024221 | 0.029765 | 0.121328 | 0.098769 | 0.152095 | 0.101862 | 0.05489 | 0.027141 | 0.023892 | 0.023354 | 0.005457 | 0.033115 | 0.011849 | 0.05206 | 0.022731 | 0.071128 |
| 35.75 | 0.048169 | 0.077239 | 0.041247 | 0.036786 | 0.029571 | 0.024807 | 0.026996 | 0.036926 | 0.107122 | 0.122723 | 0.115169 | 0.109061 | 0.029986 | 0.009323 | 0.023896 | 0.016774 | 0.009404 | 0.040679 | 0.02064 | 0.053256 | 0.016727 | 0.04437 |
| 35.76388889 | 0.101367 | 0.129485 | 0.041887 | 0.021841 | 0.028174 | 0.029142 | 0.027173 | 0.031998 | 0.147224 | 0.124549 | 0.087837 | 0.157528 | 0.012478 | 0.008845 | 0.016032 | 0.012956 | 0.008198 | 0.041305 | 0.025126 | 0.048172 | 0.020742 | 0.041378 |
| 35.77777778 | 0.176756 | 0.190728 | 0.043245 | 0.020872 | 0.01776 | 0.037893 | 0.033617 | 0.023849 | 0.115852 | 0.076397 | 0.122683 | 0.210952 | 0.015616 | 0.011532 | 0.013361 | 0.025549 | 0.005496 | 0.035674 | 0.021319 | 0.043146 | 0.02134 | 0.038652 |
| 35.79166667 | 0.218719 | 0.237228 | 0.033604 | 0.026273 | 0.018024 | 0.04156 | 0.043219 | 0.027205 | 0.054831 | 0.058096 | 0.167181 | 0.210048 | 0.040535 | 0.012417 | 0.012258 | 0.038151 | 0.007092 | 0.040867 | 0.027472 | 0.060634 | 0.015222 | 0.021612 |
| 35.80555556 | 0.188269 | 0.19803 | 0.058886 | 0.048709 | 0.020635 | 0.02743 | 0.052652 | 0.039773 | 0.092046 | 0.049132 | 0.128618 | 0.133158 | 0.075423 | 0.020297 | 0.00819 | 0.039196 | 0.008699 | 0.038984 | 0.046004 | 0.089417 | 0.009745 | 0.008069 |
| 35.81944444 | 0.113645 | 0.118308 | 0.114713 | 0.074134 | 0.032082 | 0.014814 | 0.057148 | 0.03331 | 0.137428 | 0.062798 | 0.056225 | 0.067325 | 0.094012 | 0.034839 | 0.0092 | 0.037108 | 0.006124 | 0.028207 | 0.049645 | 0.087156 | 0.01505 | 0.008466 |
| 35.83333333 | 0.049325 | 0.086981 | 0.140288 | 0.073344 | 0.044381 | 0.028361 | 0.054994 | 0.019477 | 0.11048 | 0.137208 | 0.039247 | 0.04175 | 0.065061 | 0.033184 | 0.021926 | 0.028594 | 0.003199 | 0.018333 | 0.034909 | 0.057358 | 0.023437 | 0.022707 |
| 35.84722222 | 0.049218 | 0.076089 | 0.109379 | 0.048482 | 0.043941 | 0.040969 | 0.070101 | 0.030458 | 0.142856 | 0.196376 | 0.04993 | 0.038051 | 0.026267 | 0.016976 | 0.038353 | 0.013128 | 0.003273 | 0.009991 | 0.023431 | 0.031252 | 0.0251 | 0.029403 |
| 35.86111111 | 0.080858 | 0.09064 | 0.083725 | 0.03723 | 0.035576 | 0.033129 | 0.072669 | 0.054039 | 0.199907 | 0.156446 | 0.121166 | 0.042841 | 0.029582 | 0.004453 | 0.043182 | 0.007958 | 0.004022 | 0.006047 | 0.023924 | 0.018007 | 0.02 | 0.023737 |
| 35.875 | 0.090501 | 0.157667 | 0.097045 | 0.049283 | 0.029529 | 0.029763 | 0.039652 | 0.057929 | 0.150014 | 0.070735 | 0.278956 | 0.04096 | 0.047228 | 0.004406 | 0.030777 | 0.025203 | 0.002767 | 0.005841 | 0.022533 | 0.011618 | 0.013658 | 0.026348 |
| 35.88888889 | 0.095633 | 0.190994 | 0.079055 | 0.044285 | 0.030385 | 0.04057 | 0.01772 | 0.042113 | 0.128813 | 0.071759 | 0.305834 | 0.0754 | 0.04368 | 0.016792 | 0.019004 | 0.046887 | 0.002936 | 0.013062 | 0.016546 | 0.011878 | 0.016582 | 0.024621 |
| 35.90277778 | 0.089928 | 0.130766 | 0.039023 | 0.036133 | 0.026424 | 0.056663 | 0.035665 | 0.054191 | 0.183655 | 0.121658 | 0.232321 | 0.135733 | 0.026539 | 0.031814 | 0.028426 | 0.052694 | 0.008691 | 0.032505 | 0.024975 | 0.016248 | 0.031546 | 0.020846 |
| 35.91666667 | 0.070163 | 0.073698 | 0.041781 | 0.05167 | 0.025549 | 0.078424 | 0.086815 | 0.081594 | 0.155093 | 0.118491 | 0.250035 | 0.135899 | 0.020899 | 0.02986 | 0.038536 | 0.04715 | 0.016702 | 0.043173 | 0.044107 | 0.012307 | 0.054662 | 0.039005 |
| 35.93055556 | 0.089543 | 0.088327 | 0.077295 | 0.052771 | 0.054472 | 0.077028 | 0.116464 | 0.064285 | 0.090986 | 0.110717 | 0.235098 | 0.114838 | 0.045419 | 0.018289 | 0.026934 | 0.034023 | 0.018822 | 0.029403 | 0.067671 | 0.020758 | 0.069792 | 0.058598 |
| 35.94444444 | 0.120411 | 0.085953 | 0.093258 | 0.026816 | 0.097239 | 0.059826 | 0.10139 | 0.040475 | 0.140302 | 0.12337 | 0.258193 | 0.143639 | 0.09245 | 0.021875 | 0.009697 | 0.024253 | 0.018802 | 0.008932 | 0.103234 | 0.056315 | 0.052737 | 0.054854 |
| 35.95833333 | 0.088575 | 0.064704 | 0.082541 | 0.015575 | 0.11489 | 0.063899 | 0.098936 | 0.068118 | 0.236527 | 0.10014 | 0.354048 | 0.152325 | 0.122822 | 0.02216 | 0.00671 | 0.022453 | 0.023658 | 0.010331 | 0.137783 | 0.09865 | 0.019449 | 0.04633 |
| 35.97222222 | 0.039172 | 0.101974 | 0.114506 | 0.033332 | 0.133465 | 0.057671 | 0.085482 | 0.101837 | 0.222512 | 0.061541 | 0.298336 | 0.130629 | 0.096928 | 0.020341 | 0.016323 | 0.030341 | 0.020392 | 0.034758 | 0.153172 | 0.139076 | 0.010019 | 0.047142 |
| 35.98611111 | 0.045483 | 0.156709 | 0.171378 | 0.058564 | 0.166475 | 0.065023 | 0.070054 | 0.095068 | 0.13431 | 0.074041 | 0.148179 | 0.117068 | 0.058093 | 0.044474 | 0.03304 | 0.055111 | 0.008078 | 0.055758 | 0.149327 | 0.154369 | 0.031223 | 0.064548 |
| 36 | 0.107476 | 0.171546 | 0.139049 | 0.061737 | 0.140632 | 0.129273 | 0.142384 | 0.072489 | 0.128124 | 0.128508 | 0.065711 | 0.101384 | 0.083285 | 0.077482 | 0.050127 | 0.07186 | 0.010624 | 0.059938 | 0.140361 | 0.13784 | 0.072328 | 0.109009 |
| 36.01388889 | 0.15845 | 0.148291 | 0.072713 | 0.040796 | 0.073085 | 0.198145 | 0.235027 | 0.085754 | 0.210768 | 0.14568 | 0.056542 | 0.130661 | 0.13905 | 0.085952 | 0.055453 | 0.072057 | 0.020215 | 0.043038 | 0.131761 | 0.114612 | 0.11151 | 0.155126 |
| 36.02777778 | 0.166771 | 0.116653 | 0.097408 | 0.037389 | 0.039604 | 0.208274 | 0.233252 | 0.173582 | 0.284814 | 0.117021 | 0.108566 | 0.22104 | 0.159547 | 0.0667 | 0.046244 | 0.064308 | 0.015564 | 0.019195 | 0.123222 | 0.094605 | 0.110247 | 0.164232 |
| 36.04166667 | 0.190379 | 0.116949 | 0.159251 | 0.079434 | 0.052307 | 0.175536 | 0.172012 | 0.270756 | 0.277429 | 0.108792 | 0.204042 | 0.291984 | 0.151244 | 0.054909 | 0.031024 | 0.047679 | 0.010602 | 0.014912 | 0.125965 | 0.081018 | 0.073707 | 0.137373 |
| 36.05555556 | 0.246379 | 0.131308 | 0.185748 | 0.155979 | 0.116008 | 0.174589 | 0.139316 | 0.292882 | 0.224512 | 0.14314 | 0.309386 | 0.279338 | 0.140246 | 0.073322 | 0.022193 | 0.035637 | 0.020309 | 0.034948 | 0.143757 | 0.075107 | 0.051989 | 0.110779 |
| 36.06944444 | 0.285204 | 0.12801 | 0.19328 | 0.234954 | 0.191452 | 0.244867 | 0.139199 | 0.266684 | 0.209847 | 0.157711 | 0.356521 | 0.202028 | 0.124524 | 0.095728 | 0.025715 | 0.03555 | 0.039924 | 0.079024 | 0.150839 | 0.077315 | 0.061204 | 0.100259 |
| 36.08333333 | 0.282145 | 0.109291 | 0.192969 | 0.24839 | 0.197023 | 0.293762 | 0.133237 | 0.242408 | 0.211383 | 0.112543 | 0.293088 | 0.132148 | 0.094722 | 0.088218 | 0.036679 | 0.039312 | 0.052239 | 0.110765 | 0.12727 | 0.079485 | 0.076304 | 0.092265 |
| 36.09722222 | 0.259221 | 0.076511 | 0.164146 | 0.15492 | 0.113964 | 0.22805 | 0.102041 | 0.233939 | 0.180091 | 0.055282 | 0.225314 | 0.123521 | 0.063081 | 0.054031 | 0.04492 | 0.037388 | 0.048777 | 0.097366 | 0.09252 | 0.06892 | 0.079958 | 0.079042 |
| 36.11111111 | 0.248155 | 0.050785 | 0.132975 | 0.04713 | 0.039128 | 0.131628 | 0.052764 | 0.198634 | 0.113888 | 0.022195 | 0.223833 | 0.149473 | 0.042909 | 0.023527 | 0.040585 | 0.03234 | 0.040541 | 0.071251 | 0.067424 | 0.050254 | 0.07258 | 0.063629 |
| 36.125 | 0.269154 | 0.049996 | 0.142612 | 0.041176 | 0.060615 | 0.089476 | 0.028604 | 0.123254 | 0.046258 | 0.01175 | 0.210354 | 0.156099 | 0.033377 | 0.01113 | 0.026032 | 0.030029 | 0.035091 | 0.059248 | 0.052325 | 0.034701 | 0.058564 | 0.049189 |
| 36.13888889 | 0.280915 | 0.055014 | 0.173286 | 0.115478 | 0.131911 | 0.097439 | 0.040719 | 0.074423 | 0.019759 | 0.02173 | 0.172516 | 0.142257 | 0.030146 | 0.01072 | 0.012793 | 0.023608 | 0.032379 | 0.056274 | 0.042421 | 0.023825 | 0.041512 | 0.03688 |
| 36.15277778 | 0.229728 | 0.041329 | 0.188001 | 0.167161 | 0.151126 | 0.125957 | 0.058265 | 0.07339 | 0.031729 | 0.050165 | 0.160277 | 0.12148 | 0.029478 | 0.015025 | 0.008611 | 0.015105 | 0.031957 | 0.056217 | 0.037285 | 0.018821 | 0.025403 | 0.026334 |
| 36.16666667 | 0.161243 | 0.033462 | 0.170192 | 0.160319 | 0.10955 | 0.147903 | 0.069548 | 0.099046 | 0.053095 | 0.078498 | 0.168336 | 0.093651 | 0.02982 | 0.020145 | 0.013964 | 0.0181 | 0.033522 | 0.057487 | 0.036185 | 0.020241 | 0.014376 | 0.01798 |
| 36.18055556 | 0.137812 | 0.060702 | 0.143159 | 0.151207 | 0.08363 | 0.171439 | 0.075968 | 0.132914 | 0.066212 | 0.086952 | 0.172485 | 0.06873 | 0.032124 | 0.025156 | 0.020309 | 0.028229 | 0.036321 | 0.059395 | 0.033787 | 0.021862 | 0.009017 | 0.012745 |
| 36.19444444 | 0.142831 | 0.096927 | 0.127639 | 0.166473 | 0.094169 | 0.196807 | 0.067744 | 0.139321 | 0.064848 | 0.080924 | 0.158153 | 0.059584 | 0.036473 | 0.030865 | 0.020422 | 0.031488 | 0.03984 | 0.062086 | 0.028892 | 0.02034 | 0.006295 | 0.010025 |
| 36.20833333 | 0.121287 | 0.097621 | 0.10557 | 0.166815 | 0.097035 | 0.176949 | 0.04662 | 0.091564 | 0.046351 | 0.069323 | 0.129132 | 0.062016 | 0.041289 | 0.037597 | 0.017037 | 0.02604 | 0.045038 | 0.066862 | 0.027357 | 0.019451 | 0.004774 | 0.009502 |
| 36.22222222 | 0.066561 | 0.0627 | 0.061324 | 0.126782 | 0.074435 | 0.095188 | 0.04603 | 0.058391 | 0.024357 | 0.049764 | 0.102331 | 0.057106 | 0.045935 | 0.045892 | 0.016806 | 0.019021 | 0.054101 | 0.077808 | 0.034099 | 0.02175 | 0.005825 | 0.012061 |
| 36.23611111 | 0.035423 | 0.033441 | 0.025035 | 0.06757 | 0.041831 | 0.031044 | 0.062887 | 0.087234 | 0.019277 | 0.027489 | 0.071666 | 0.036717 | 0.0502 | 0.055015 | 0.020047 | 0.017765 | 0.068724 | 0.099988 | 0.045994 | 0.023833 | 0.009244 | 0.01623 |
| 36.25 | 0.049156 | 0.026969 | 0.025318 | 0.024925 | 0.019539 | 0.03086 | 0.053488 | 0.094628 | 0.023285 | 0.011387 | 0.0417 | 0.024598 | 0.05209 | 0.059573 | 0.021584 | 0.020867 | 0.08124 | 0.121054 | 0.053498 | 0.022205 | 0.013384 | 0.017454 |
| 36.26388889 | 0.075783 | 0.025605 | 0.036693 | 0.014438 | 0.022312 | 0.05224 | 0.026988 | 0.063001 | 0.025374 | 0.006672 | 0.046396 | 0.032089 | 0.052568 | 0.055404 | 0.021073 | 0.021276 | 0.076895 | 0.119097 | 0.050126 | 0.018377 | 0.017823 | 0.014334 |
| 36.27777778 | 0.1021 | 0.017783 | 0.04411 | 0.029661 | 0.041653 | 0.077948 | 0.026542 | 0.046008 | 0.030496 | 0.015855 | 0.080403 | 0.036707 | 0.052933 | 0.044286 | 0.020781 | 0.016448 | 0.056573 | 0.096028 | 0.039216 | 0.015903 | 0.022957 | 0.010748 |
| 36.29166667 | 0.125805 | 0.015125 | 0.054775 | 0.053072 | 0.061374 | 0.121229 | 0.064761 | 0.062003 | 0.042775 | 0.03336 | 0.106388 | 0.037474 | 0.04917 | 0.030064 | 0.020179 | 0.009657 | 0.038983 | 0.065356 | 0.028124 | 0.015125 | 0.028959 | 0.010788 |
| 36.30555556 | 0.14136 | 0.023802 | 0.067293 | 0.071637 | 0.075193 | 0.181781 | 0.117429 | 0.106273 | 0.061125 | 0.050102 | 0.114193 | 0.049495 | 0.040564 | 0.015041 | 0.018182 | 0.005909 | 0.03301 | 0.034833 | 0.020185 | 0.013922 | 0.03401 | 0.013604 |

| | | | | | | | | | | | | | | | | | | | | | | |
|---|---|---|---|---|---|---|---|---|---|---|---|---|---|---|---|---|---|---|---|---|---|---|
| 36.31944444 | 0.153827 | 0.031768 | 0.080633 | 0.083675 | 0.081628 | 0.241118 | 0.158469 | 0.14862 | 0.077125 | 0.060402 | 0.108461 | 0.061741 | 0.031183 | 0.007221 | 0.014652 | 0.006395 | 0.030789 | 0.012968 | 0.013925 | 0.01112 | 0.034409 | 0.013759 |
| 36.33333333 | 0.160372 | 0.033748 | 0.089464 | 0.090485 | 0.082079 | 0.26531 | 0.1801 | 0.162471 | 0.087688 | 0.065762 | 0.090214 | 0.056706 | 0.02323 | 0.012264 | 0.010358 | 0.008269 | 0.024655 | 0.004401 | 0.00904 | 0.007217 | 0.029016 | 0.008725 |
| 36.34722222 | 0.151379 | 0.032128 | 0.083326 | 0.085413 | 0.074562 | 0.24602 | 0.180979 | 0.150585 | 0.101621 | 0.068697 | 0.070307 | 0.044035 | 0.016161 | 0.021009 | 0.006736 | 0.007072 | 0.017243 | 0.009378 | 0.012447 | 0.003421 | 0.020467 | 0.004908 |
| 36.36111111 | 0.133645 | 0.031462 | 0.062686 | 0.066127 | 0.058384 | 0.216755 | 0.171362 | 0.152987 | 0.119748 | 0.067766 | 0.067476 | 0.045013 | 0.009562 | 0.025968 | 0.004577 | 0.005471 | 0.023988 | 0.028062 | 0.025164 | 0.001935 | 0.011313 | 0.007249 |
| 36.375 | 0.126216 | 0.041213 | 0.04853 | 0.05582 | 0.048107 | 0.210455 | 0.168881 | 0.179493 | 0.129698 | 0.065926 | 0.085088 | 0.057929 | 0.004514 | 0.027075 | 0.003081 | 0.009864 | 0.050762 | 0.056527 | 0.036261 | 0.00417 | 0.004976 | 0.011295 |
| 36.38888889 | 0.135737 | 0.063278 | 0.063341 | 0.080814 | 0.063155 | 0.23139 | 0.183231 | 0.176094 | 0.122743 | 0.068172 | 0.107958 | 0.064904 | 0.002271 | 0.025557 | 0.001767 | 0.01515 | 0.079692 | 0.082965 | 0.041516 | 0.008988 | 0.00443 | 0.014462 |
| 36.40277778 | 0.151063 | 0.085783 | 0.103434 | 0.130467 | 0.097622 | 0.25508 | 0.209822 | 0.146436 | 0.113538 | 0.077352 | 0.125918 | 0.064277 | 0.003576 | 0.023886 | 0.001975 | 0.016683 | 0.098402 | 0.098244 | 0.045713 | 0.015342 | 0.007214 | 0.018475 |
| 36.41666667 | 0.154768 | 0.097533 | 0.125505 | 0.160135 | 0.119436 | 0.2651 | 0.246224 | 0.144627 | 0.122858 | 0.092003 | 0.143713 | 0.072258 | 0.007389 | 0.024153 | 0.004441 | 0.019041 | 0.110444 | 0.107086 | 0.053912 | 0.021502 | 0.010977 | 0.023641 |
| 36.43055556 | 0.141624 | 0.102459 | 0.104865 | 0.147019 | 0.110849 | 0.273991 | 0.296203 | 0.172893 | 0.144656 | 0.102614 | 0.171817 | 0.095165 | 0.012406 | 0.026914 | 0.007387 | 0.025338 | 0.119139 | 0.119604 | 0.067622 | 0.025504 | 0.016952 | 0.030937 |
| 36.44444444 | 0.134933 | 0.111095 | 0.080674 | 0.12174 | 0.095609 | 0.285342 | 0.325895 | 0.199721 | 0.159195 | 0.101409 | 0.211243 | 0.117817 | 0.01871 | 0.03171 | 0.009093 | 0.033276 | 0.123012 | 0.136493 | 0.08104 | 0.029334 | 0.026728 | 0.044163 |
| 36.45833333 | 0.163028 | 0.131721 | 0.085185 | 0.125063 | 0.107082 | 0.294655 | 0.314328 | 0.195263 | 0.15628 | 0.093955 | 0.252168 | 0.12396 | 0.025783 | 0.038754 | 0.009821 | 0.037897 | 0.120473 | 0.149032 | 0.08754 | 0.038017 | 0.040807 | 0.065332 |
| 36.47222222 | 0.229934 | 0.170335 | 0.116016 | 0.167622 | 0.151475 | 0.298007 | 0.303896 | 0.15071 | 0.139814 | 0.092811 | 0.287657 | 0.118514 | 0.035008 | 0.049131 | 0.011792 | 0.038069 | 0.116265 | 0.153824 | 0.093846 | 0.055639 | 0.058186 | 0.091368 |
| 36.48611111 | 0.319393 | 0.222384 | 0.158059 | 0.222387 | 0.208639 | 0.286349 | 0.310288 | 0.11162 | 0.129713 | 0.103898 | 0.33199 | 0.12502 | 0.049505 | 0.064455 | 0.018059 | 0.039982 | 0.11152 | 0.163675 | 0.117045 | 0.081661 | 0.078906 | 0.120589 |
| 36.5 | 0.399355 | 0.274011 | 0.178117 | 0.249793 | 0.249722 | 0.263588 | 0.31648 | 0.138266 | 0.147693 | 0.123316 | 0.417067 | 0.160286 | 0.065781 | 0.079678 | 0.027461 | 0.049132 | 0.104303 | 0.185336 | 0.152538 | 0.109959 | 0.101341 | 0.153542 |
| 36.51388889 | 0.436966 | 0.29962 | 0.159496 | 0.224454 | 0.24102 | 0.236774 | 0.316014 | 0.218154 | 0.199519 | 0.145782 | 0.512595 | 0.215569 | 0.0773 | 0.08161 | 0.034344 | 0.062567 | 0.105698 | 0.200258 | 0.173801 | 0.132917 | 0.117733 | 0.178743 |
| 36.52777778 | 0.437258 | 0.271006 | 0.127521 | 0.151097 | 0.171624 | 0.198909 | 0.297028 | 0.261282 | 0.24149 | 0.169642 | 0.559161 | 0.265048 | 0.083959 | 0.071011 | 0.036059 | 0.068893 | 0.109601 | 0.188703 | 0.170748 | 0.148544 | 0.123494 | 0.186702 |
| 36.54166667 | 0.432644 | 0.209366 | 0.108324 | 0.085162 | 0.100091 | 0.158056 | 0.26015 | 0.214844 | 0.208176 | 0.176878 | 0.555384 | 0.277085 | 0.096477 | 0.066502 | 0.035555 | 0.061494 | 0.091153 | 0.149524 | 0.153979 | 0.159335 | 0.122726 | 0.183714 |
| 36.55555556 | 0.370785 | 0.143505 | 0.101489 | 0.078277 | 0.094114 | 0.131629 | 0.217159 | 0.121011 | 0.116376 | 0.142612 | 0.455332 | 0.232786 | 0.130087 | 0.070021 | 0.037866 | 0.050213 | 0.056824 | 0.095948 | 0.127042 | 0.155792 | 0.115769 | 0.167409 |
| 36.56944444 | 0.227735 | 0.073702 | 0.105941 | 0.113893 | 0.145513 | 0.097227 | 0.153716 | 0.043592 | 0.046514 | 0.085442 | 0.244409 | 0.161767 | 0.169193 | 0.064902 | 0.043106 | 0.044391 | 0.03359 | 0.055006 | 0.076487 | 0.118292 | 0.09659 | 0.132081 |
| 36.58333333 | 0.144116 | 0.023703 | 0.117284 | 0.152739 | 0.209099 | 0.047056 | 0.091982 | 0.030726 | 0.041847 | 0.059191 | 0.076774 | 0.140099 | 0.173851 | 0.051371 | 0.047705 | 0.048661 | 0.029965 | 0.050254 | 0.036519 | 0.059455 | 0.065032 | 0.079852 |
| 36.59722222 | 0.123689 | 0.025174 | 0.12178 | 0.193262 | 0.268901 | 0.030381 | 0.081549 | 0.064526 | 0.080311 | 0.088674 | 0.098405 | 0.211992 | 0.15248 | 0.043203 | 0.050235 | 0.060581 | 0.030083 | 0.067621 | 0.035306 | 0.030681 | 0.033337 | 0.033247 |
| 36.61111111 | 0.12896 | 0.113809 | 0.133114 | 0.223143 | 0.309741 | 0.061368 | 0.111632 | 0.09453 | 0.147563 | 0.1292 | 0.290046 | 0.303564 | 0.147901 | 0.042749 | 0.050797 | 0.065181 | 0.028646 | 0.076594 | 0.051731 | 0.05466 | 0.013797 | 0.015545 |
| 36.625 | 0.226573 | 0.252055 | 0.149957 | 0.2042 | 0.295102 | 0.090921 | 0.128516 | 0.104251 | 0.217967 | 0.136845 | 0.455771 | 0.315388 | 0.160653 | 0.044602 | 0.050757 | 0.057497 | 0.028541 | 0.07112 | 0.083877 | 0.093209 | 0.008168 | 0.013243 |
| 36.63888889 | 0.303838 | 0.311418 | 0.140645 | 0.144297 | 0.205397 | 0.090646 | 0.104625 | 0.077378 | 0.209728 | 0.116026 | 0.420453 | 0.249923 | 0.165085 | 0.045374 | 0.044784 | 0.04022 | 0.01973 | 0.052846 | 0.090273 | 0.094235 | 0.012988 | 0.015441 |
| 36.65277778 | 0.274053 | 0.266243 | 0.111541 | 0.082661 | 0.088055 | 0.070745 | 0.068582 | 0.044478 | 0.126214 | 0.084363 | 0.253069 | 0.143882 | 0.155594 | 0.039963 | 0.026761 | 0.023147 | 0.008308 | 0.026405 | 0.052633 | 0.054031 | 0.02067 | 0.024761 |
| 36.66666667 | 0.199292 | 0.194936 | 0.094984 | 0.051347 | 0.030915 | 0.042298 | 0.034867 | 0.039108 | 0.056982 | 0.083379 | 0.106607 | 0.064215 | 0.150196 | 0.03075 | 0.008385 | 0.021008 | 0.003275 | 0.009007 | 0.023008 | 0.017671 | 0.026525 | 0.036333 |
| 36.68055556 | 0.110648 | 0.12878 | 0.103381 | 0.060106 | 0.043194 | 0.019364 | 0.012132 | 0.033585 | 0.03518 | 0.113802 | 0.040974 | 0.053523 | 0.156941 | 0.035988 | 0.006207 | 0.021023 | 0.004667 | 0.006265 | 0.019513 | 0.006867 | 0.030694 | 0.040009 |
| 36.69444444 | 0.038126 | 0.077231 | 0.109916 | 0.082018 | 0.07306 | 0.008538 | 0.016163 | 0.019536 | 0.03895 | 0.105888 | 0.053938 | 0.04397 | 0.157719 | 0.062644 | 0.017785 | 0.022775 | 0.008033 | 0.030225 | 0.021393 | 0.007224 | 0.032547 | 0.030712 |
| 36.70833333 | 0.00921 | 0.050664 | 0.086415 | 0.088373 | 0.098624 | 0.004509 | 0.031531 | 0.016019 | 0.048233 | 0.122089 | 0.075182 | 0.033065 | 0.137546 | 0.086574 | 0.032439 | 0.045236 | 0.005637 | 0.067865 | 0.017877 | 0.014277 | 0.030038 | 0.01908 |
| 36.72222222 | 0.0098 | 0.029789 | 0.044796 | 0.067879 | 0.099145 | 0.004654 | 0.0375 | 0.030073 | 0.049157 | 0.20222 | 0.056517 | 0.056423 | 0.089992 | 0.080826 | 0.042688 | 0.061372 | 0.003391 | 0.072864 | 0.026282 | 0.033186 | 0.020085 | 0.027637 |
| 36.73611111 | 0.025736 | 0.026039 | 0.031442 | 0.042674 | 0.074294 | 0.009263 | 0.03185 | 0.041267 | 0.043398 | 0.190409 | 0.054661 | 0.085407 | 0.046007 | 0.047558 | 0.037798 | 0.045274 | 0.006186 | 0.074516 | 0.061454 | 0.076434 | 0.010628 | 0.050485 |
| 36.75 | 0.06059 | 0.049231 | 0.074764 | 0.060485 | 0.080824 | 0.019625 | 0.027532 | 0.031117 | 0.032277 | 0.09023 | 0.067845 | 0.094476 | 0.048834 | 0.02563 | 0.019697 | 0.023533 | 0.015264 | 0.10482 | 0.09237 | 0.12079 | 0.012309 | 0.051171 |
| 36.76388889 | 0.10394 | 0.074503 | 0.135152 | 0.093179 | 0.106649 | 0.028776 | 0.025532 | 0.014445 | 0.021393 | 0.045432 | 0.087052 | 0.079169 | 0.071317 | 0.04402 | 0.006811 | 0.025777 | 0.025611 | 0.111897 | 0.071196 | 0.105058 | 0.017118 | 0.028394 |
| 36.77777778 | 0.133605 | 0.078318 | 0.132436 | 0.072237 | 0.07949 | 0.025615 | 0.02515 | 0.022586 | 0.036936 | 0.060718 | 0.150691 | 0.084232 | 0.063759 | 0.061157 | 0.006793 | 0.035963 | 0.021872 | 0.074663 | 0.039921 | 0.057079 | 0.014606 | 0.009324 |
| 36.79166667 | 0.132087 | 0.054294 | 0.083084 | 0.041551 | 0.042666 | 0.012942 | 0.038542 | 0.047229 | 0.051949 | 0.05894 | 0.219116 | 0.111572 | 0.038802 | 0.042532 | 0.018234 | 0.029786 | 0.016849 | 0.060669 | 0.050493 | 0.054792 | 0.009569 | 0.005619 |
| 36.80555556 | 0.102225 | 0.049129 | 0.087455 | 0.049453 | 0.051756 | 0.004175 | 0.051185 | 0.047182 | 0.064368 | 0.054029 | 0.208226 | 0.099358 | 0.035582 | 0.028607 | 0.03884 | 0.028092 | 0.029312 | 0.099032 | 0.050581 | 0.07657 | 0.01063 | 0.010064 |
| 36.81944444 | 0.065468 | 0.083722 | 0.106337 | 0.047676 | 0.050781 | 0.002573 | 0.038299 | 0.058162 | 0.115856 | 0.076222 | 0.172951 | 0.084573 | 0.037036 | 0.030401 | 0.045034 | 0.039647 | 0.037137 | 0.108728 | 0.024239 | 0.067528 | 0.016672 | 0.010705 |
| 36.83333333 | 0.033294 | 0.092073 | 0.070563 | 0.031652 | 0.023538 | 0.00784 | 0.023591 | 0.101084 | 0.138177 | 0.09318 | 0.240522 | 0.090665 | 0.021261 | 0.024268 | 0.02666 | 0.038188 | 0.024036 | 0.059818 | 0.011325 | 0.036415 | 0.022781 | 0.011093 |
| 36.84722222 | 0.009807 | 0.065507 | 0.051723 | 0.053786 | 0.033779 | 0.022377 | 0.042504 | 0.106254 | 0.097605 | 0.112481 | 0.266126 | 0.062554 | 0.021561 | 0.025818 | 0.009672 | 0.023049 | 0.008941 | 0.020095 | 0.010954 | 0.013464 | 0.026954 | 0.02681 |
| 36.86111111 | 0.03422 | 0.061747 | 0.080361 | 0.09884 | 0.074014 | 0.025959 | 0.0579 | 0.067296 | 0.092121 | 0.114952 | 0.190426 | 0.047876 | 0.052669 | 0.036601 | 0.006974 | 0.010838 | 0.006685 | 0.018025 | 0.011688 | 0.016667 | 0.02819 | 0.050742 |
| 36.875 | 0.11234 | 0.075147 | 0.107854 | 0.093563 | 0.103745 | 0.014165 | 0.048677 | 0.073888 | 0.130989 | 0.083542 | 0.207087 | 0.053667 | 0.078808 | 0.034491 | 0.020879 | 0.019152 | 0.009839 | 0.02016 | 0.020462 | 0.046502 | 0.023721 | 0.057747 |
| 36.88888889 | 0.173233 | 0.073924 | 0.108052 | 0.076638 | 0.162506 | 0.008452 | 0.061427 | 0.145596 | 0.113567 | 0.09739 | 0.291502 | 0.048143 | 0.076484 | 0.01941 | 0.041142 | 0.038996 | 0.012597 | 0.009958 | 0.026242 | 0.060668 | 0.013677 | 0.039969 |
| 36.90277778 | 0.168196 | 0.046608 | 0.069089 | 0.09104 | 0.185882 | 0.006727 | 0.066199 | 0.148768 | 0.061462 | 0.131323 | 0.298472 | 0.091177 | 0.054161 | 0.00796 | 0.036926 | 0.032426 | 0.014474 | 0.014015 | 0.022216 | 0.034178 | 0.011426 | 0.023697 |
| 36.91666667 | 0.118268 | 0.019279 | 0.045337 | 0.066636 | 0.102373 | 0.008237 | 0.063491 | 0.092194 | 0.05227 | 0.108847 | 0.291653 | 0.131903 | 0.025513 | 0.00606 | 0.02268 | 0.014541 | 0.01516 | 0.043114 | 0.025658 | 0.010724 | 0.028415 | 0.045527 |
| 36.93055556 | 0.066091 | 0.028322 | 0.081169 | 0.027023 | 0.033634 | 0.02759 | 0.084223 | 0.104091 | 0.065041 | 0.09361 | 0.324893 | 0.097535 | 0.006605 | 0.008863 | 0.030316 | 0.019165 | 0.022269 | 0.058774 | 0.029258 | 0.022777 | 0.054514 | 0.084526 |
| 36.94444444 | 0.030343 | 0.060174 | 0.125209 | 0.020081 | 0.024579 | 0.057969 | 0.083639 | 0.11421 | 0.099013 | 0.089426 | 0.288341 | 0.075295 | 0.010234 | 0.015793 | 0.032735 | 0.024404 | 0.02356 | 0.047697 | 0.038093 | 0.065149 | 0.0635 | 0.08273 |
| 36.95833333 | 0.028251 | 0.087513 | 0.118913 | 0.025931 | 0.026993 | 0.067555 | 0.063085 | 0.064604 | 0.105129 | 0.093002 | 0.175569 | 0.081885 | 0.039801 | 0.02278 | 0.02321 | 0.023537 | 0.017535 | 0.040788 | 0.06434 | 0.095767 | 0.046835 | 0.043882 |
| 36.97222222 | 0.085014 | 0.114283 | 0.071433 | 0.067013 | 0.071817 | 0.0539 | 0.065713 | 0.028195 | 0.06207 | 0.115574 | 0.097293 | 0.056 | 0.083056 | 0.024261 | 0.039845 | 0.050403 | 0.026678 | 0.037526 | 0.091218 | 0.072879 | 0.027858 | 0.013394 |
| 36.98611111 | 0.187153 | 0.145585 | 0.064503 | 0.149622 | 0.129876 | 0.047658 | 0.132573 | 0.052543 | 0.062386 | 0.14439 | 0.136601 | 0.026354 | 0.119031 | 0.034066 | 0.076402 | 0.099802 | 0.055268 | 0.032696 | 0.103603 | 0.02764 | 0.028749 | 0.019049 |
| 37 | 0.250349 | 0.138545 | 0.104046 | 0.187643 | 0.114618 | 0.057663 | 0.191344 | 0.103421 | 0.124761 | 0.212137 | 0.24544 | 0.047559 | 0.135275 | 0.067386 | 0.096086 | 0.133697 | 0.076895 | 0.044013 | 0.095458 | 0.018063 | 0.050359 | 0.068433 |
| 37.01388889 | 0.240123 | 0.083136 | 0.106487 | 0.123773 | 0.08295 | 0.06467 | 0.140384 | 0.132092 | 0.21998 | 0.201836 | 0.30259 | 0.157619 | 0.132472 | 0.096222 | 0.082842 | 0.122229 | 0.057792 | 0.072318 | 0.066295 | 0.049163 | 0.079942 | 0.11867 |
| 37.02777778 | 0.237765 | 0.05257 | 0.082387 | 0.037805 | 0.107442 | 0.07223 | 0.060307 | 0.132458 | 0.277184 | 0.106696 | 0.236972 | 0.286241 | 0.12504 | 0.095771 | 0.051876 | 0.070551 | 0.025359 | 0.0986 | 0.037759 | 0.092091 | 0.092661 | 0.111938 |
| 37.04166667 | 0.28854 | 0.101315 | 0.088875 | 0.038443 | 0.135417 | 0.134622 | 0.082644 | 0.118147 | 0.233618 | 0.08242 | 0.12489 | 0.302604 | 0.121816 | 0.089149 | 0.030241 | 0.029141 | 0.029283 | 0.103584 | 0.04087 | 0.121059 | 0.079594 | 0.069516 |
| 37.05555556 | 0.339079 | 0.17692 | 0.123721 | 0.119129 | 0.159573 | 0.276914 | 0.185049 | 0.125614 | 0.147861 | 0.105953 | 0.111886 | 0.238425 | 0.119195 | 0.094943 | 0.026757 | 0.032183 | 0.078229 | 0.108015 | 0.070608 | 0.120287 | 0.066886 | 0.056467 |
| 37.06944444 | 0.323284 | 0.15779 | 0.186064 | 0.191921 | 0.199645 | 0.406188 | 0.240682 | 0.140099 | 0.103033 | 0.100368 | 0.167161 | 0.219706 | 0.111667 | 0.098184 | 0.036377 | 0.065326 | 0.143015 | 0.133352 | 0.100796 | 0.103237 | 0.067096 | 0.076046 |
| 37.08333333 | 0.241681 | 0.076095 | 0.21992 | 0.197306 | 0.205888 | 0.425751 | 0.21366 | 0.103919 | 0.075771 | 0.061138 | 0.215048 | 0.249269 | 0.091859 | 0.079493 | 0.050628 | 0.091666 | 0.168575 | 0.137184 | 0.110094 | 0.081751 | 0.068723 | 0.0883 |
| 37.09722222 | 0.155155 | 0.064866 | 0.149744 | 0.125708 | 0.152194 | 0.330265 | 0.153455 | 0.05825 | 0.043999 | 0.025014 | 0.251303 | 0.239134 | 0.062665 | 0.042899 | 0.055951 | 0.087629 | 0.139229 | 0.095989 | 0.093756 | 0.054046 | 0.066045 | 0.08355 |
| 37.11111111 | 0.100545 | 0.096478 | 0.084266 | 0.052456 | 0.076934 | 0.16763 | 0.089419 | 0.052128 | 0.060278 | 0.024229 | 0.256578 | 0.178311 | 0.03784 | 0.015643 | 0.049226 | 0.067398 | 0.101079 | 0.050447 | 0.073252 | 0.031458 | 0.061103 | 0.077915 |
| 37.125 | 0.0883 | 0.082475 | 0.091122 | 0.041448 | 0.036471 | 0.042199 | 0.047717 | 0.042979 | 0.087726 | 0.038827 | 0.226394 | 0.120088 | 0.024086 | 0.011342 | 0.04132 | 0.047051 | 0.081375 | 0.026444 | 0.067117 | 0.025169 | 0.057756 | 0.081431 |
| 37.13888889 | 0.120661 | 0.057669 | 0.076738 | 0.038871 | 0.046198 | 0.01698 | 0.051548 | 0.044278 | 0.081358 | 0.042914 | 0.175547 | 0.078386 | 0.019168 | 0.013204 | 0.034922 | 0.027337 | 0.068029 | 0.016538 | 0.071665 | 0.031279 | 0.056599 | 0.08757 |
| 37.15277778 | 0.162891 | 0.087822 | 0.06513 | 0.040742 | 0.083784 | 0.051773 | 0.092868 | 0.094578 | 0.127762 | 0.062792 | 0.124199 | 0.054695 | 0.01885 | 0.009956 | 0.026588 | 0.016067 | 0.052318 | 0.012746 | 0.071076 | 0.03637 | 0.053424 | 0.082412 |
| 37.16666667 | 0.183532 | 0.130806 | 0.105961 | 0.091238 | 0.127075 | 0.088554 | 0.139479 | 0.177068 | 0.232708 | 0.110519 | 0.083392 | 0.047802 | 0.019471 | 0.005062 | 0.018678 | 0.016646 | 0.036272 | 0.011081 | 0.058612 | 0.032622 | 0.045192 | 0.064768 |
| 37.18055556 | 0.186161 | 0.147863 | 0.151939 | 0.1517 | 0.156225 | 0.1039 | 0.152685 | 0.253831 | 0.272104 | 0.129161 | 0.058995 | 0.044986 | 0.018773 | 0.00381 | 0.016285 | 0.023449 | 0.022852 | 0.008677 | 0.041398 | 0.024486 | 0.036292 | 0.048976 |
| 37.19444444 | 0.176776 | 0.151798 | 0.166077 | 0.181817 | 0.163528 | 0.112359 | 0.138574 | 0.280281 | 0.221774 | 0.095375 | 0.061509 | 0.045133 | 0.017586 | 0.011342 | 0.017402 | 0.026983 | 0.011593 | 0.005371 | 0.029708 | 0.018743 | 0.033979 | 0.045672 |
| 37.20833333 | 0.157857 | 0.140336 | 0.156389 | 0.180935 | 0.14983 | 0.142243 | 0.131425 | 0.279423 | 0.15681 | 0.055293 | 0.086296 | 0.052859 | 0.018238 | 0.02602 | 0.015932 | 0.018885 | 0.005992 | 0.00681 | 0.029709 | 0.017359 | 0.03947 | 0.052228 |
| 37.22222222 | 0.132346 | 0.112971 | 0.132873 | 0.149061 | 0.106206 | 0.179132 | 0.122315 | 0.268001 | 0.108047 | 0.040665 | 0.112177 | 0.060967 | 0.022075 | 0.040164 | 0.012161 | 0.006299 | 0.013247 | 0.020501 | 0.037802 | 0.018538 | 0.046912 | 0.060506 |
| 37.23611111 | 0.092666 | 0.070541 | 0.093012 | 0.096217 | 0.046762 | 0.171678 | 0.08655 | 0.19918 | 0.06082 | 0.037603 | 0.100927 | 0.047657 | 0.028315 | 0.048012 | 0.009177 | 0.005125 | 0.03627 | 0.047522 | 0.045747 | 0.018817 | 0.049157 | 0.064065 |
| 37.25 | 0.044573 | 0.028275 | 0.051095 | 0.043797 | 0.011757 | 0.120388 | 0.040853 | 0.104223 | 0.034397 | 0.031263 | 0.058963 | 0.025056 | 0.034781 | 0.049213 | 0.009442 | 0.018285 | 0.069929 | 0.079667 | 0.050383 | 0.01733 | 0.043521 | 0.060497 |
| 37.26388889 | 0.013151 | 0.008002 | 0.021487 | 0.016437 | 0.006173 | 0.065268 | 0.024214 | 0.0438 | 0.053349 | 0.041976 | 0.040433 | 0.02318 | 0.039281 | 0.045615 | 0.013017 | 0.033392 | 0.102921 | 0.10289 | 0.053555 | 0.01649 | 0.032351 | 0.049679 |
| 37.27777778 | 0.010015 | 0.005447 | 0.007753 | 0.023454 | 0.011294 | 0.037143 | 0.034567 | 0.024745 | 0.082685 | 0.075247 | 0.040139 | 0.034986 | 0.040631 | 0.038079 | 0.015351 | 0.036789 | 0.125002 | 0.107597 | 0.054386 | 0.015821 | 0.020195 | 0.03298 |
| 37.29166667 | 0.021415 | 0.006026 | 0.009812 | 0.048109 | 0.030553 | 0.054248 | 0.037102 | 0.024795 | 0.097914 | 0.104961 | 0.025724 | 0.045034 | 0.041151 | 0.028979 | 0.017888 | 0.030162 | 0.138037 | 0.095802 | 0.051966 | 0.014618 | 0.012162 | 0.016414 |
| 37.30555556 | 0.037451 | 0.005893 | 0.031105 | 0.079427 | 0.059077 | 0.089411 | 0.033475 | 0.035951 | 0.115236 | 0.118071 | 0.011574 | 0.059428 | 0.043296 | 0.022129 | 0.025098 | 0.024792 | 0.148092 | 0.076349 | 0.048483 | 0.013284 | 0.011186 | 0.006721 |
| 37.31944444 | 0.061526 | 0.013057 | 0.064614 | 0.108823 | 0.073961 | 0.11764 | 0.047311 | 0.099589 | 0.143341 | 0.119666 | 0.01794 | 0.080871 | 0.043935 | 0.015793 | 0.029586 | 0.0215 | 0.140302 | 0.054472 | 0.041938 | 0.010298 | 0.013593 | 0.003048 |
| 37.33333333 | 0.091924 | 0.034856 | 0.089464 | 0.123111 | 0.076628 | 0.157008 | 0.087704 | 0.201101 | 0.180183 | 0.122945 | 0.046339 | 0.094462 | 0.038756 | 0.010512 | 0.025226 | 0.018137 | 0.098499 | 0.029854 | 0.030303 | 0.007026 | 0.011757 | 0.004394 |
| 37.34722222 | 0.11988 | 0.062455 | 0.093858 | 0.12234 | 0.08541 | 0.210319 | 0.146945 | 0.2623 | 0.208623 | 0.137218 | 0.079381 | 0.088484 | 0.028683 | 0.01495 | 0.016176 | 0.016319 | 0.05139 | 0.018821 | 0.017093 | 0.006315 | 0.007611 | 0.014786 |
| 37.36111111 | 0.132229 | 0.077821 | 0.086041 | 0.115255 | 0.100036 | 0.251012 | 0.201743 | 0.262898 | 0.216208 | 0.159835 | 0.099688 | 0.073874 | 0.017373 | 0.027926 | 0.007369 | 0.013035 | 0.049129 | 0.036808 | 0.007682 | 0.005413 | 0.012183 | 0.032146 |
| 37.375 | 0.127213 | 0.075458 | 0.091683 | 0.117869 | 0.110523 | 0.270509 | 0.238101 | 0.238825 | 0.219158 | 0.181913 | 0.113294 | 0.069503 | 0.008293 | 0.040043 | 0.003439 | 0.011009 | 0.088293 | 0.06453 | 0.006597 | 0.005144 | 0.026559 | 0.048784 |
| 37.38888889 | 0.123739 | 0.068403 | 0.116581 | 0.135847 | 0.120764 | 0.28378 | 0.269542 | 0.21603 | 0.208424 | 0.192043 | 0.135307 | 0.079429 | 0.005042 | 0.047739 | 0.005559 | 0.020684 | 0.12008 | 0.085512 | 0.013311 | 0.010949 | 0.041953 | 0.05943 |
| 37.40277778 | 0.135345 | 0.069336 | 0.139679 | 0.155936 | 0.13288 | 0.291845 | 0.29572 | 0.200543 | 0.173295 | 0.18778 | 0.161167 | 0.095809 | 0.004917 | 0.049589 | 0.008687 | 0.035262 | 0.130562 | 0.103728 | 0.023383 | 0.018898 | 0.054344 | 0.06464 |
| 37.41666667 | 0.160614 | 0.08465 | 0.156784 | 0.175978 | 0.140092 | 0.293163 | 0.298012 | 0.188594 | 0.155944 | 0.17894 | 0.179061 | 0.105181 | 0.006558 | 0.049224 | 0.009951 | 0.038482 | 0.140011 | 0.123826 | 0.03275 | 0.024324 | 0.061 | 0.067471 |
| 37.43055556 | 0.198973 | 0.122592 | 0.179378 | 0.202065 | 0.152538 | 0.302757 | 0.301905 | 0.181387 | 0.179081 | 0.172537 | 0.199206 | 0.095905 | 0.015719 | 0.050475 | 0.01107 | 0.032168 | 0.160873 | 0.144967 | 0.038122 | 0.030718 | 0.056346 | 0.066174 |
| 37.44444444 | 0.239863 | 0.16339 | 0.201504 | 0.229788 | 0.178511 | 0.311083 | 0.318056 | 0.184272 | 0.205506 | 0.170234 | 0.222337 | 0.075562 | 0.029419 | 0.050733 | 0.01419 | 0.030879 | 0.185444 | 0.157206 | 0.040545 | 0.041698 | 0.043429 | 0.060595 |
| 37.45833333 | 0.266683 | 0.174235 | 0.214505 | 0.260542 | 0.215746 | 0.296376 | 0.320347 | 0.174018 | 0.179451 | 0.161833 | 0.237334 | 0.067542 | 0.042389 | 0.054335 | 0.019105 | 0.042063 | 0.184315 | 0.145691 | 0.048673 | 0.054678 | 0.034923 | 0.061657 |
| 37.47222222 | 0.281428 | 0.166737 | 0.214509 | 0.286913 | 0.258811 | 0.271375 | 0.320725 | 0.149967 | 0.112109 | 0.136856 | 0.259841 | 0.09227 | 0.052872 | 0.06403 | 0.025085 | 0.060764 | 0.146856 | 0.116243 | 0.068275 | 0.066147 | 0.038377 | 0.079443 |
| 37.48611111 | 0.288389 | 0.162508 | 0.183277 | 0.274301 | 0.270594 | 0.2315 | 0.311187 | 0.139025 | 0.108185 | 0.123158 | 0.298173 | 0.1357 | 0.060676 | 0.073441 | 0.032821 | 0.077339 | 0.11545 | 0.104802 | 0.087628 | 0.076993 | 0.050101 | 0.10997 |
| 37.5 | 0.29874 | 0.165474 | 0.125911 | 0.204546 | 0.219892 | 0.172383 | 0.264745 | 0.123052 | 0.156301 | 0.138513 | 0.310435 | 0.143648 | 0.069688 | 0.079006 | 0.038934 | 0.080578 | 0.110632 | 0.118842 | 0.092429 | 0.087578 | 0.063316 | 0.137028 |
| 37.51388889 | 0.322108 | 0.160216 | 0.077326 | 0.114383 | 0.130599 | 0.140592 | 0.224564 | 0.076429 | 0.150329 | 0.180804 | 0.270447 | 0.115349 | 0.076534 | 0.077271 | 0.038304 | 0.066157 | 0.094538 | 0.118181 | 0.086145 | 0.097725 | 0.073379 | 0.146905 |
| 37.52777778 | 0.341915 | 0.147915 | 0.06298 | 0.079185 | 0.0916 | 0.151238 | 0.234746 | 0.038004 | 0.134958 | 0.221147 | 0.236665 | 0.116487 | 0.076188 | 0.067063 | 0.032261 | 0.040299 | 0.063334 | 0.098978 | 0.07727 | 0.103447 | 0.076266 | 0.143566 |
| 37.54166667 | 0.358689 | 0.158893 | 0.086771 | 0.143915 | 0.172847 | 0.15827 | 0.25541 | 0.064624 | 0.167511 | 0.170727 | 0.25234 | 0.161364 | 0.078019 | 0.055718 | 0.028476 | 0.017791 | 0.039514 | 0.091699 | 0.06547 | 0.091506 | 0.075593 | 0.135598 |
| 37.55555556 | 0.335707 | 0.177708 | 0.110143 | 0.254364 | 0.313361 | 0.134464 | 0.222857 | 0.110087 | 0.188499 | 0.081581 | 0.271169 | 0.180187 | 0.096867 | 0.049775 | 0.034664 | 0.013592 | 0.026947 | 0.070442 | 0.046075 | 0.056973 | 0.074421 | 0.116519 |
| 37.56944444 | 0.223446 | 0.164512 | 0.090922 | 0.287055 | 0.389075 | 0.089115 | 0.157647 | 0.097322 | 0.143899 | 0.064124 | 0.228397 | 0.125557 | 0.114492 | 0.045381 | 0.048329 | 0.02843 | 0.028471 | 0.041845 | 0.028348 | 0.027348 | 0.061724 | 0.078768 |
| 37.58333333 | 0.155649 | 0.113007 | 0.045467 | 0.215595 | 0.341784 | 0.041137 | 0.124164 | 0.048041 | 0.067604 | 0.100709 | 0.137062 | 0.058953 | 0.096149 | 0.038782 | 0.061312 | 0.050335 | 0.038365 | 0.046552 | 0.022471 | 0.0241 | 0.037303 | 0.047438 |
| 37.59722222 | 0.161685 | 0.060693 | 0.021282 | 0.134011 | 0.216802 | 0.012033 | 0.10595 | 0.023859 | 0.033428 | 0.15576 | 0.086983 | 0.056301 | 0.063441 | 0.038623 | 0.07555 | 0.06416 | 0.040779 | 0.040863 | 0.037764 | 0.043316 | 0.019755 | 0.052433 |
| 37.61111111 | 0.123474 | 0.058097 | 0.04145 | 0.073199 | 0.100344 | 0.010806 | 0.078715 | 0.031724 | 0.06353 | 0.154222 | 0.087333 | 0.109938 | 0.061515 | 0.046469 | 0.094396 | 0.069004 | 0.038581 | 0.028249 | 0.080437 | 0.080491 | 0.021725 | 0.078089 |
| 37.625 | 0.140189 | 0.106302 | 0.085472 | 0.029011 | 0.066016 | 0.025645 | 0.057338 | 0.05515 | 0.097545 | 0.101629 | 0.072456 | 0.153231 | 0.083672 | 0.050302 | 0.107151 | 0.07084 | 0.047324 | 0.058045 | 0.104419 | 0.097664 | 0.034131 | 0.082973 |
| 37.63888889 | 0.236345 | 0.148782 | 0.117964 | 0.031875 | 0.132593 | 0.033141 | 0.042893 | 0.067704 | 0.094608 | 0.084463 | 0.094402 | 0.158745 | 0.100327 | 0.044837 | 0.096965 | 0.066119 | 0.050303 | 0.091185 | 0.079685 | 0.070777 | 0.037031 | 0.052471 |
| 37.65277778 | 0.291847 | 0.163387 | 0.125536 | 0.061281 | 0.196982 | 0.030659 | 0.03407 | 0.070388 | 0.088189 | 0.071205 | 0.220707 | 0.163398 | 0.105992 | 0.034032 | 0.064816 | 0.047267 | 0.034322 | 0.081404 | 0.041044 | 0.035921 | 0.026096 | 0.036966 |
| 37.66666667 | 0.269698 | 0.174488 | 0.109861 | 0.080352 | 0.177094 | 0.030765 | 0.024175 | 0.100024 | 0.083045 | 0.047056 | 0.254394 | 0.135696 | 0.102757 | 0.021517 | 0.029951 | 0.027717 | 0.018721 | 0.051558 | 0.018752 | 0.01499 | 0.012225 | 0.066762 |
| 37.68055556 | 0.188378 | 0.17601 | 0.077001 | 0.086628 | 0.127933 | 0.034411 | 0.01363 | 0.106135 | 0.077014 | 0.049952 | 0.141466 | 0.070237 | 0.088997 | 0.012987 | 0.010346 | 0.027425 | 0.027636 | 0.055077 | 0.018742 | 0.011954 | 0.004974 | 0.094645 |
| 37.69444444 | 0.099991 | 0.140222 | 0.050818 | 0.080297 | 0.08633 | 0.037583 | 0.008934 | 0.064005 | 0.070093 | 0.053222 | 0.149403 | 0.039195 | 0.067991 | 0.012628 | 0.006682 | 0.025225 | 0.049995 | 0.066074 | 0.020474 | 0.031547 | 0.014907 | 0.075692 |
| 37.70833333 | 0.042572 | 0.073389 | 0.055727 | 0.066821 | 0.044815 | 0.035055 | 0.010841 | 0.029264 | 0.041006 | 0.052559 | 0.274877 | 0.031341 | 0.040623 | 0.01104 | 0.006049 | 0.012874 | 0.049468 | 0.055407 | 0.019623 | 0.04908 | 0.04451 | 0.058265 |

| | | | | | | | | | | | | | | | | | | | | | | |
|---|---|---|---|---|---|---|---|---|---|---|---|---|---|---|---|---|---|---|---|---|---|---|
| 37.72222222 | 0.045692 | 0.025721 | 0.08028 | 0.053442 | 0.015818 | 0.025712 | 0.024254 | 0.019453 | 0.011088 | 0.059272 | 0.319244 | 0.038093 | 0.014613 | 0.004589 | 0.008267 | 0.018258 | 0.03769 | 0.072684 | 0.040928 | 0.053764 | 0.060731 | 0.09709 |
| 37.73611111 | 0.099111 | 0.027024 | 0.093175 | 0.04424 | 0.00699 | 0.016717 | 0.052559 | 0.026294 | 0.005122 | 0.057354 | 0.244892 | 0.056992 | 0.003981 | 0.003167 | 0.019577 | 0.034059 | 0.034573 | 0.089219 | 0.062715 | 0.058609 | 0.041361 | 0.110777 |
| 37.75 | 0.107553 | 0.041438 | 0.073399 | 0.036052 | 0.007728 | 0.01007 | 0.066796 | 0.037356 | 0.010577 | 0.055084 | 0.128245 | 0.048171 | 0.005016 | 0.006511 | 0.031806 | 0.038403 | 0.024573 | 0.070391 | 0.051196 | 0.04465 | 0.014893 | 0.063307 |
| 37.76388889 | 0.084164 | 0.056929 | 0.047295 | 0.025503 | 0.010404 | 0.006599 | 0.0488 | 0.041219 | 0.012768 | 0.058012 | 0.070555 | 0.039694 | 0.004675 | 0.007965 | 0.033924 | 0.034032 | 0.013603 | 0.044781 | 0.037254 | 0.030436 | 0.006824 | 0.03731 |
| 37.77777778 | 0.125656 | 0.077218 | 0.062788 | 0.0257 | 0.013378 | 0.008877 | 0.03306 | 0.037754 | 0.020007 | 0.049546 | 0.100958 | 0.071965 | 0.001594 | 0.008617 | 0.027301 | 0.022575 | 0.024262 | 0.031804 | 0.050467 | 0.054085 | 0.013587 | 0.049598 |
| 37.79166667 | 0.174791 | 0.054185 | 0.111536 | 0.027526 | 0.019347 | 0.018979 | 0.044109 | 0.027926 | 0.049539 | 0.025437 | 0.102536 | 0.089715 | 0.00619 | 0.00858 | 0.017883 | 0.009078 | 0.053705 | 0.050002 | 0.057691 | 0.077311 | 0.015761 | 0.053312 |
| 37.80555556 | 0.210362 | 0.014747 | 0.148158 | 0.017105 | 0.040592 | 0.034911 | 0.067076 | 0.018227 | 0.068729 | 0.025298 | 0.073857 | 0.05853 | 0.021017 | 0.007122 | 0.013678 | 0.003605 | 0.074332 | 0.078233 | 0.039448 | 0.059105 | 0.014128 | 0.060813 |
| 37.81944444 | 0.246597 | 0.013303 | 0.126381 | 0.008253 | 0.051631 | 0.039053 | 0.071653 | 0.023075 | 0.04878 | 0.06974 | 0.087422 | 0.051837 | 0.031543 | 0.007762 | 0.013914 | 0.004921 | 0.074249 | 0.077268 | 0.025484 | 0.024183 | 0.023338 | 0.081609 |
| 37.83333333 | 0.20159 | 0.047471 | 0.070267 | 0.006977 | 0.04352 | 0.031567 | 0.046422 | 0.037151 | 0.034242 | 0.136556 | 0.096429 | 0.062954 | 0.03064 | 0.010227 | 0.011088 | 0.006195 | 0.056158 | 0.057619 | 0.026977 | 0.013314 | 0.03072 | 0.069545 |
| 37.84722222 | 0.10287 | 0.095066 | 0.062247 | 0.010134 | 0.058531 | 0.027623 | 0.030129 | 0.042756 | 0.041748 | 0.160175 | 0.137017 | 0.035088 | 0.025754 | 0.009933 | 0.014286 | 0.013913 | 0.038218 | 0.06182 | 0.026492 | 0.020475 | 0.025666 | 0.038766 |
| 37.86111111 | 0.067469 | 0.092323 | 0.086667 | 0.019459 | 0.07687 | 0.02358 | 0.069725 | 0.043311 | 0.04007 | 0.113792 | 0.235555 | 0.031122 | 0.025946 | 0.009291 | 0.031764 | 0.036383 | 0.056318 | 0.083228 | 0.02532 | 0.031681 | 0.013578 | 0.040997 |
| 37.875 | 0.106759 | 0.066864 | 0.081546 | 0.028491 | 0.076483 | 0.033392 | 0.136511 | 0.055765 | 0.031385 | 0.111194 | 0.27965 | 0.063268 | 0.036597 | 0.015283 | 0.045534 | 0.053266 | 0.070494 | 0.073606 | 0.030241 | 0.037127 | 0.004849 | 0.05798 |
| 37.88888889 | 0.127216 | 0.084744 | 0.049931 | 0.022911 | 0.089437 | 0.068266 | 0.161637 | 0.071527 | 0.031596 | 0.175976 | 0.243983 | 0.055537 | 0.051853 | 0.028866 | 0.033934 | 0.048449 | 0.038996 | 0.037122 | 0.034135 | 0.031388 | 0.006362 | 0.057078 |
| 37.90277778 | 0.099183 | 0.081213 | 0.033368 | 0.015412 | 0.097647 | 0.073803 | 0.116381 | 0.057953 | 0.02907 | 0.160027 | 0.187057 | 0.025486 | 0.059132 | 0.041819 | 0.017093 | 0.033962 | 0.007956 | 0.018709 | 0.035486 | 0.038027 | 0.014821 | 0.069686 |
| 37.91666667 | 0.123996 | 0.060119 | 0.077237 | 0.032218 | 0.085441 | 0.059813 | 0.066529 | 0.03184 | 0.015685 | 0.113887 | 0.117029 | 0.016798 | 0.058868 | 0.046452 | 0.022112 | 0.037749 | 0.02099 | 0.037854 | 0.034458 | 0.0501 | 0.022469 | 0.091715 |
| 37.93055556 | 0.20156 | 0.058573 | 0.149649 | 0.06446 | 0.058436 | 0.068219 | 0.059283 | 0.035224 | 0.011678 | 0.149594 | 0.114867 | 0.01548 | 0.060904 | 0.048508 | 0.041589 | 0.069007 | 0.07183 | 0.066331 | 0.029127 | 0.04184 | 0.019914 | 0.068816 |
| 37.94444445 | 0.183736 | 0.04249 | 0.166738 | 0.079221 | 0.026876 | 0.04817 | 0.052387 | 0.043424 | 0.016931 | 0.12653 | 0.154905 | 0.015055 | 0.062271 | 0.041709 | 0.057129 | 0.09161 | 0.110994 | 0.066985 | 0.038813 | 0.047014 | 0.028689 | 0.050223 |
| 37.95833334 | 0.091374 | 0.030005 | 0.112033 | 0.056107 | 0.016672 | 0.011902 | 0.056208 | 0.032042 | 0.015964 | 0.039052 | 0.113921 | 0.013653 | 0.055123 | 0.03057 | 0.052388 | 0.079237 | 0.112453 | 0.060721 | 0.075126 | 0.088691 | 0.073185 | 0.102354 |
| 37.97222223 | 0.04726 | 0.073661 | 0.045195 | 0.020039 | 0.031275 | 0.01471 | 0.063275 | 0.01878 | 0.030368 | 0.044727 | 0.074484 | 0.027281 | 0.04616 | 0.026709 | 0.033376 | 0.050876 | 0.086502 | 0.075051 | 0.105488 | 0.100276 | 0.115782 | 0.156579 |
| 37.98611112 | 0.048148 | 0.180591 | 0.019068 | 0.017906 | 0.078674 | 0.036481 | 0.063342 | 0.01871 | 0.109009 | 0.153658 | 0.122536 | 0.067505 | 0.049527 | 0.03568 | 0.016291 | 0.030749 | 0.044414 | 0.0711 | 0.116489 | 0.062868 | 0.109814 | 0.141607 |

**Supplementary Table S6.** 1 hour integrated motion data in **(b)** 100 mM NaCl treatment on Rocket

| IntervalMid | Control1 | Control2 | Control3 | Control4 | Control5 | Control6 | Control7 | Control8 | Control9 | Control10 | Control11 | Stress1 | Stress2 | Stress3 | Stress4 | Stress5 | Stress6 | Stress7 | Stress8 | Stress9 | Stress10 | Stress11 | Stress12 |
|---|---|---|---|---|---|---|---|---|---|---|---|---|---|---|---|---|---|---|---|---|---|---|---|
| 18.1111111 | 0.01681 | 0.014586 | 0.01033 | 0.012359 | 0.000934 | 0.019917 | 0.007289 | 0.001004 | 0.003816 | 0.036615 | 0.000819 | 0.016871 | 0.015034 | 0.009743 | 0.001676 | 0.000737 | 0.013764 | 0.013182 | 0.000748 | 0.028475 | 0.012458 | 0.02468 | 0.029952 |
| 18.125 | 0.010258 | 0.008447 | 0.005423 | 0.008167 | 0.000548 | 0.009391 | 0.004999 | 0.000783 | 0.003123 | 0.021819 | 0.000912 | 0.016394 | 0.012349 | 0.003995 | 0.001885 | 0.001158 | 0.006409 | 0.029577 | 0.000674 | 0.021047 | 0.010385 | 0.028685 | 0.01137 |
| 18.1388889 | 0.003696 | 0.002858 | 0.004012 | 0.003237 | 0.000573 | 0.002132 | 0.001977 | 0.001374 | 0.001338 | 0.005738 | 0.000867 | 0.014229 | 0.009361 | 0.003017 | 0.001617 | 0.002255 | 0.006109 | 0.045899 | 0.001198 | 0.009524 | 0.007231 | 0.018379 | 0.008694 |
| 18.1527778 | 0.002527 | 0.002122 | 0.005812 | 0.001616 | 0.001117 | 0.000546 | 0.001078 | 0.002374 | 0.0008 | 0.001702 | 0.002018 | 0.014706 | 0.014896 | 0.005197 | 0.002677 | 0.004105 | 0.010938 | 0.038347 | 0.003278 | 0.004137 | 0.003398 | 0.012136 | 0.010869 |
| 18.1666667 | 0.003266 | 0.00336 | 0.006644 | 0.001829 | 0.001904 | 0.000952 | 0.001014 | 0.00301 | 0.00133 | 0.003043 | 0.003573 | 0.015782 | 0.016637 | 0.007323 | 0.005217 | 0.006379 | 0.014413 | 0.03104 | 0.005789 | 0.005626 | 0.003031 | 0.016861 | 0.010401 |
| 18.1805556 | 0.003459 | 0.0037 | 0.005952 | 0.001463 | 0.001764 | 0.003576 | 0.001522 | 0.002752 | 0.001468 | 0.003301 | 0.003739 | 0.015379 | 0.009096 | 0.008117 | 0.007126 | 0.007714 | 0.014576 | 0.049965 | 0.006971 | 0.009611 | 0.00635 | 0.026989 | 0.010766 |
| 18.1944444 | 0.004553 | 0.003022 | 0.00488 | 0.00092 | 0.001114 | 0.00815 | 0.003239 | 0.002639 | 0.001434 | 0.002108 | 0.00299 | 0.014118 | 0.004854 | 0.007979 | 0.00827 | 0.008263 | 0.013283 | 0.067322 | 0.007162 | 0.013429 | 0.00844 | 0.025773 | 0.012889 |
| 18.2083333 | 0.007574 | 0.00327 | 0.004825 | 0.001528 | 0.001312 | 0.012435 | 0.005694 | 0.004498 | 0.002581 | 0.001839 | 0.003133 | 0.011991 | 0.008351 | 0.008247 | 0.010079 | 0.009032 | 0.011663 | 0.057692 | 0.007567 | 0.017319 | 0.008766 | 0.011983 | 0.015432 |
| 18.2222222 | 0.010677 | 0.004662 | 0.004102 | 0.002281 | 0.001795 | 0.01368 | 0.00699 | 0.006134 | 0.003427 | 0.001952 | 0.003655 | 0.007865 | 0.010798 | 0.008622 | 0.012362 | 0.009801 | 0.008664 | 0.030797 | 0.008018 | 0.020238 | 0.008915 | 0.006543 | 0.017734 |
| 18.2361111 | 0.01057 | 0.004819 | 0.002812 | 0.002083 | 0.001381 | 0.011351 | 0.004996 | 0.004338 | 0.002251 | 0.00141 | 0.003094 | 0.003619 | 0.008903 | 0.008805 | 0.014422 | 0.011301 | 0.006671 | 0.010544 | 0.008541 | 0.021492 | 0.009114 | 0.012049 | 0.023193 |
| 18.25 | 0.007676 | 0.003915 | 0.003889 | 0.002204 | 0.000656 | 0.008507 | 0.002198 | 0.001336 | 0.00138 | 0.001279 | 0.002451 | 0.003326 | 0.008942 | 0.010089 | 0.016722 | 0.015037 | 0.010539 | 0.009422 | 0.010943 | 0.021981 | 0.010945 | 0.021858 | 0.032485 |
| 18.2638889 | 0.004229 | 0.00352 | 0.005866 | 0.002834 | 0.000484 | 0.006871 | 0.002003 | 0.000267 | 0.002462 | 0.001596 | 0.002616 | 0.006063 | 0.012399 | 0.012401 | 0.018625 | 0.01923 | 0.017251 | 0.015535 | 0.014969 | 0.022462 | 0.012413 | 0.031056 | 0.036334 |
| 18.2777778 | 0.002279 | 0.003209 | 0.004857 | 0.00237 | 0.000889 | 0.005099 | 0.002656 | 0.000291 | 0.003715 | 0.001529 | 0.002192 | 0.006282 | 0.014626 | 0.013467 | 0.018568 | 0.020105 | 0.019096 | 0.023756 | 0.017216 | 0.022336 | 0.011741 | 0.035975 | 0.028317 |
| 18.2916667 | 0.003048 | 0.002321 | 0.002016 | 0.001484 | 0.00105 | 0.003087 | 0.00209 | 0.00016 | 0.003552 | 0.001033 | 0.001109 | 0.004811 | 0.013041 | 0.011504 | 0.015879 | 0.016459 | 0.013914 | 0.045783 | 0.015215 | 0.019978 | 0.009433 | 0.034 | 0.016646 |
| 18.3055556 | 0.004011 | 0.001482 | 0.000831 | 0.002025 | 0.000704 | 0.002049 | 0.001928 | 0.000275 | 0.002875 | 0.000884 | 0.000614 | 0.0069 | 0.008647 | 0.006991 | 0.011008 | 0.010401 | 0.006999 | 0.072482 | 0.009919 | 0.014866 | 0.005365 | 0.022086 | 0.009549 |
| 18.3194444 | 0.003661 | 0.001324 | 0.00157 | 0.00327 | 0.000471 | 0.002003 | 0.002978 | 0.000784 | 0.002842 | 0.001035 | 0.000337 | 0.009437 | 0.004526 | 0.002772 | 0.006209 | 0.00545 | 0.00353 | 0.074151 | 0.005109 | 0.00964 | 0.002863 | 0.009073 | 0.005494 |
| 18.3333333 | 0.004015 | 0.001352 | 0.002083 | 0.002966 | 0.000408 | 0.001782 | 0.004249 | 0.000939 | 0.002919 | 0.001443 | 0.000308 | 0.007888 | 0.003001 | 0.001465 | 0.004021 | 0.003864 | 0.002979 | 0.043085 | 0.003559 | 0.007015 | 0.005453 | 0.00798 | 0.002629 |
| 18.3472222 | 0.006104 | 0.001134 | 0.001611 | 0.001551 | 0.000661 | 0.001512 | 0.005637 | 0.000855 | 0.002332 | 0.00305 | 0.001164 | 0.004899 | 0.003472 | 0.002997 | 0.004952 | 0.005408 | 0.005545 | 0.017927 | 0.00469 | 0.006979 | 0.009862 | 0.016393 | 0.001507 |
| 18.3611111 | 0.008457 | 0.00134 | 0.001605 | 0.000931 | 0.001465 | 0.002623 | 0.006673 | 0.001603 | 0.001309 | 0.006075 | 0.002304 | 0.00314 | 0.00426 | 0.005787 | 0.008132 | 0.008484 | 0.010424 | 0.01723 | 0.006821 | 0.008642 | 0.012541 | 0.017259 | 0.002911 |
| 18.375 | 0.009607 | 0.001655 | 0.002332 | 0.000674 | 0.001821 | 0.003873 | 0.006215 | 0.002777 | 0.000709 | 0.009169 | 0.002647 | 0.001977 | 0.00474 | 0.008205 | 0.011684 | 0.010812 | 0.013866 | 0.014776 | 0.008348 | 0.0106 | 0.012283 | 0.00837 | 0.006324 |
| 18.3888889 | 0.009164 | 0.001278 | 0.0023 | 0.000307 | 0.001226 | 0.003267 | 0.004683 | 0.003121 | 0.001008 | 0.010859 | 0.002462 | 0.001106 | 0.006115 | 0.010293 | 0.014248 | 0.011734 | 0.014899 | 0.011641 | 0.009151 | 0.011611 | 0.009293 | 0.005874 | 0.008506 |
| 18.4027778 | 0.007277 | 0.000839 | 0.001463 | 0.00045 | 0.000589 | 0.002854 | 0.003781 | 0.002877 | 0.001617 | 0.012071 | 0.002572 | 0.00074 | 0.00909 | 0.01365 | 0.016753 | 0.013152 | 0.014765 | 0.021253 | 0.011431 | 0.011992 | 0.00735 | 0.007467 | 0.008768 |
| 18.4166667 | 0.006246 | 0.000817 | 0.001689 | 0.000789 | 0.000632 | 0.005515 | 0.004057 | 0.003387 | 0.001577 | 0.01462 | 0.002016 | 0.002181 | 0.010439 | 0.01689 | 0.01887 | 0.014472 | 0.014438 | 0.031737 | 0.013966 | 0.011199 | 0.005948 | 0.012195 | 0.00858 |
| 18.4305556 | 0.008012 | 0.001159 | 0.003541 | 0.000652 | 0.001389 | 0.008651 | 0.004543 | 0.004939 | 0.000928 | 0.018133 | 0.000758 | 0.009222 | 0.007189 | 0.015905 | 0.017209 | 0.012034 | 0.013387 | 0.027871 | 0.01076 | 0.007261 | 0.004497 | 0.025955 | 0.006522 |
| 18.4444444 | 0.009699 | 0.001613 | 0.005178 | 0.000731 | 0.002258 | 0.008509 | 0.005093 | 0.006691 | 0.000841 | 0.020623 | 0.000414 | 0.02211 | 0.004711 | 0.010259 | 0.010411 | 0.006392 | 0.010767 | 0.014439 | 0.007605 | 0.003432 | 0.011486 | 0.02915 | 0.006164 |
| 18.4583333 | 0.007916 | 0.002014 | 0.005711 | 0.001841 | 0.002937 | 0.008767 | 0.006512 | 0.008583 | 0.001946 | 0.02202 | 0.000912 | 0.030995 | 0.008194 | 0.004389 | 0.004245 | 0.002768 | 0.00698 | 0.007115 | 0.012635 | 0.003706 | 0.024194 | 0.025578 | 0.009581 |
| 18.4722222 | 0.00449 | 0.002469 | 0.005844 | 0.002704 | 0.003408 | 0.012889 | 0.00828 | 0.009745 | 0.003037 | 0.022925 | 0.001792 | 0.025737 | 0.013272 | 0.00154 | 0.004037 | 0.002409 | 0.004154 | 0.007947 | 0.016416 | 0.004246 | 0.022241 | 0.044491 | 0.009613 |
| 18.4861111 | 0.003627 | 0.002729 | 0.005994 | 0.002566 | 0.003611 | 0.016666 | 0.009336 | 0.008176 | 0.003274 | 0.021555 | 0.003454 | 0.012048 | 0.017185 | 0.001226 | 0.007387 | 0.002539 | 0.002966 | 0.009012 | 0.016807 | 0.002987 | 0.012496 | 0.056954 | 0.006955 |
| 18.5 | 0.006847 | 0.003133 | 0.006739 | 0.00293 | 0.004384 | 0.017384 | 0.010365 | 0.005898 | 0.004321 | 0.017833 | 0.005269 | 0.004746 | 0.022287 | 0.003379 | 0.012478 | 0.003851 | 0.00227 | 0.008927 | 0.022241 | 0.00346 | 0.019992 | 0.032963 | 0.004807 |
| 18.5138889 | 0.009746 | 0.003591 | 0.006954 | 0.003688 | 0.005334 | 0.015884 | 0.011306 | 0.006861 | 0.006477 | 0.013556 | 0.005242 | 0.010685 | 0.028513 | 0.008221 | 0.019452 | 0.008854 | 0.004914 | 0.012108 | 0.031033 | 0.008678 | 0.033564 | 0.022684 | 0.002392 |
| 18.5277778 | 0.007401 | 0.002941 | 0.004906 | 0.003041 | 0.004494 | 0.012512 | 0.008859 | 0.00951 | 0.006969 | 0.008666 | 0.003478 | 0.022497 | 0.030961 | 0.011059 | 0.024155 | 0.014234 | 0.012508 | 0.021818 | 0.035649 | 0.016446 | 0.028732 | 0.056193 | 0.005133 |
| 18.5416667 | 0.004199 | 0.001594 | 0.004062 | 0.002641 | 0.003691 | 0.010274 | 0.007392 | 0.009889 | 0.008076 | 0.004573 | 0.002907 | 0.026581 | 0.023044 | 0.010223 | 0.020666 | 0.011905 | 0.015537 | 0.040818 | 0.02676 | 0.01887 | 0.013078 | 0.080516 | 0.013587 |
| 18.5555556 | 0.004091 | 0.001454 | 0.006446 | 0.004126 | 0.004624 | 0.011548 | 0.011839 | 0.009069 | 0.013191 | 0.003743 | 0.003104 | 0.019112 | 0.010225 | 0.012898 | 0.012165 | 0.006609 | 0.010048 | 0.057254 | 0.014835 | 0.012623 | 0.009242 | 0.065739 | 0.016878 |
| 18.5694444 | 0.003442 | 0.003324 | 0.006989 | 0.004663 | 0.004255 | 0.017392 | 0.01182 | 0.012519 | 0.016224 | 0.006015 | 0.002424 | 0.011187 | 0.004585 | 0.016893 | 0.012253 | 0.007534 | 0.007283 | 0.050019 | 0.013906 | 0.005979 | 0.016846 | 0.043173 | 0.011864 |
| 18.5833333 | 0.002684 | 0.007061 | 0.004898 | 0.003246 | 0.002208 | 0.028971 | 0.006706 | 0.022668 | 0.013897 | 0.011512 | 0.002271 | 0.013057 | 0.005022 | 0.014206 | 0.023149 | 0.007056 | 0.010572 | 0.026136 | 0.015453 | 0.007328 | 0.017693 | 0.0288 | 0.007808 |
| 18.5972222 | 0.004136 | 0.013565 | 0.005146 | 0.001596 | 0.001024 | 0.039381 | 0.006685 | 0.033405 | 0.01161 | 0.021229 | 0.002608 | 0.017027 | 0.006209 | 0.01023 | 0.028269 | 0.004881 | 0.015166 | 0.009453 | 0.020403 | 0.013068 | 0.010373 | 0.017322 | 0.00808 |
| 18.6111111 | 0.006048 | 0.021221 | 0.008954 | 0.001076 | 0.001494 | 0.043049 | 0.008949 | 0.037069 | 0.013227 | 0.03113 | 0.003172 | 0.012748 | 0.008171 | 0.010765 | 0.020507 | 0.010714 | 0.018048 | 0.006664 | 0.023375 | 0.01539 | 0.006636 | 0.0107 | 0.009132 |
| 18.625 | 0.006288 | 0.023915 | 0.014058 | 0.002351 | 0.002953 | 0.042987 | 0.009549 | 0.034426 | 0.018721 | 0.035286 | 0.004798 | 0.007857 | 0.008764 | 0.011627 | 0.010293 | 0.02086 | 0.016318 | 0.00888 | 0.014319 | 0.011554 | 0.005397 | 0.00968 | 0.010684 |
| 18.6388889 | 0.004572 | 0.021066 | 0.018579 | 0.004281 | 0.004211 | 0.039436 | 0.010383 | 0.031843 | 0.023802 | 0.032646 | 0.007021 | 0.01056 | 0.008347 | 0.008985 | 0.004872 | 0.02809 | 0.009693 | 0.008935 | 0.004208 | 0.00661 | 0.004797 | 0.011971 | 0.015276 |
| 18.6527778 | 0.003322 | 0.01982 | 0.022058 | 0.005941 | 0.004674 | 0.034104 | 0.011908 | 0.033776 | 0.022956 | 0.02499 | 0.00912 | 0.017111 | 0.014383 | 0.004529 | 0.004547 | 0.031467 | 0.00526 | 0.010359 | 0.003479 | 0.007651 | 0.008771 | 0.017196 | 0.023984 |
| 18.6666667 | 0.003307 | 0.022506 | 0.024573 | 0.007265 | 0.004225 | 0.03281 | 0.012975 | 0.040003 | 0.016411 | 0.014898 | 0.01034 | 0.02591 | 0.025382 | 0.00297 | 0.005976 | 0.032729 | 0.009477 | 0.015099 | 0.01098 | 0.01271 | 0.014727 | 0.024473 | 0.034489 |
| 18.6805556 | 0.005236 | 0.02493 | 0.024976 | 0.007905 | 0.002925 | 0.035494 | 0.012621 | 0.04719 | 0.009984 | 0.007191 | 0.00996 | 0.035903 | 0.031104 | 0.005897 | 0.004559 | 0.033177 | 0.015744 | 0.014294 | 0.017859 | 0.015686 | 0.017989 | 0.030633 | 0.042943 |
| 18.6944444 | 0.010321 | 0.023812 | 0.022023 | 0.00768 | 0.001404 | 0.035916 | 0.010939 | 0.053164 | 0.007469 | 0.005547 | 0.008394 | 0.041868 | 0.028941 | 0.010173 | 0.003431 | 0.031534 | 0.018364 | 0.009239 | 0.019119 | 0.015469 | 0.01672 | 0.033301 | 0.047236 |
| 18.7083333 | 0.014087 | 0.019145 | 0.017036 | 0.006768 | 0.000459 | 0.029706 | 0.008123 | 0.056346 | 0.007187 | 0.005053 | 0.006662 | 0.042866 | 0.024238 | 0.01407 | 0.008465 | 0.026767 | 0.019984 | 0.01067 | 0.018787 | 0.01461 | 0.013451 | 0.033521 | 0.048526 |
| 18.7222222 | 0.013568 | 0.014242 | 0.012623 | 0.006077 | 0.000265 | 0.018601 | 0.004699 | 0.054875 | 0.007101 | 0.003756 | 0.005559 | 0.040326 | 0.021996 | 0.01745 | 0.015509 | 0.020497 | 0.022531 | 0.014747 | 0.021055 | 0.015242 | 0.012569 | 0.033399 | 0.048886 |
| 18.7361111 | 0.011497 | 0.011899 | 0.010256 | 0.006556 | 0.00082 | 0.007855 | 0.002198 | 0.049166 | 0.008215 | 0.006071 | 0.005408 | 0.033309 | 0.022592 | 0.019949 | 0.016363 | 0.015247 | 0.025063 | 0.0167 | 0.022718 | 0.017187 | 0.01412 | 0.033475 | 0.048409 |
| 18.75 | 0.009513 | 0.011054 | 0.009797 | 0.007433 | 0.002274 | 0.002279 | 0.001171 | 0.041037 | 0.010833 | 0.008999 | 0.005881 | 0.02491 | 0.023676 | 0.022538 | 0.010509 | 0.011353 | 0.026257 | 0.020112 | 0.018808 | 0.019512 | 0.015771 | 0.033097 | 0.046482 |
| 18.7638889 | 0.006845 | 0.009582 | 0.010072 | 0.007151 | 0.003984 | 0.001469 | 0.000899 | 0.031548 | 0.012211 | 0.008299 | 0.006259 | 0.020308 | 0.023029 | 0.024882 | 0.005001 | 0.007478 | 0.026107 | 0.025843 | 0.011748 | 0.021712 | 0.016058 | 0.031144 | 0.04289 |
| 18.7777778 | 0.003483 | 0.007112 | 0.010846 | 0.005747 | 0.004666 | 0.001329 | 0.001071 | 0.020077 | 0.010308 | 0.004934 | 0.006016 | 0.017487 | 0.01964 | 0.023238 | 0.005325 | 0.005747 | 0.024196 | 0.02814 | 0.007245 | 0.022472 | 0.013919 | 0.0267 | 0.037003 |
| 18.7916667 | 0.001343 | 0.004808 | 0.012175 | 0.004401 | 0.003632 | 0.00125 | 0.001063 | 0.010677 | 0.00659 | 0.002003 | 0.005111 | 0.010991 | 0.01353 | 0.016738 | 0.008629 | 0.007929 | 0.019683 | 0.021707 | 0.006712 | 0.019818 | 0.008751 | 0.019717 | 0.028631 |
| 18.8055556 | 0.001223 | 0.003455 | 0.012382 | 0.003494 | 0.001708 | 0.001425 | 0.001043 | 0.011902 | 0.003815 | 0.001521 | 0.003651 | 0.007005 | 0.006452 | 0.008906 | 0.008743 | 0.009458 | 0.012946 | 0.011274 | 0.005689 | 0.013405 | 0.004219 | 0.011608 | 0.019022 |
| 18.8194444 | 0.001348 | 0.00288 | 0.009505 | 0.002778 | 0.000495 | 0.001454 | 0.002139 | 0.021091 | 0.00261 | 0.00223 | 0.002029 | 0.013544 | 0.00212 | 0.003574 | 0.005682 | 0.006758 | 0.006011 | 0.007129 | 0.005363 | 0.006393 | 0.00419 | 0.005186 | 0.009895 |
| 18.8333333 | 0.00129 | 0.002775 | 0.005368 | 0.001915 | 0.00053 | 0.003198 | 0.003385 | 0.026806 | 0.00193 | 0.002738 | 0.000941 | 0.02037 | 0.001224 | 0.002398 | 0.005066 | 0.004517 | 0.001872 | 0.01161 | 0.011111 | 0.003268 | 0.005194 | 0.003248 | 0.004439 |
| 18.8472222 | 0.001324 | 0.00213 | 0.004683 | 0.00146 | 0.001286 | 0.007314 | 0.003725 | 0.027533 | 0.001176 | 0.005462 | 0.001121 | 0.018174 | 0.000874 | 0.00391 | 0.007688 | 0.006683 | 0.001043 | 0.016869 | 0.016358 | 0.003805 | 0.004467 | 0.006564 | 0.005168 |
| 18.8611111 | 0.000803 | 0.001098 | 0.007732 | 0.002468 | 0.001999 | 0.011656 | 0.003436 | 0.028561 | 0.000681 | 0.010834 | 0.002425 | 0.012415 | 0.000615 | 0.005816 | 0.009624 | 0.010101 | 0.000908 | 0.018323 | 0.015441 | 0.004415 | 0.004302 | 0.012447 | 0.008778 |
| 18.875 | 0.000499 | 0.000947 | 0.009971 | 0.003537 | 0.00183 | 0.013737 | 0.003121 | 0.029012 | 0.000547 | 0.012677 | 0.002985 | 0.010959 | 0.001974 | 0.004753 | 0.010574 | 0.012665 | 0.000684 | 0.016276 | 0.012782 | 0.004344 | 0.006034 | 0.015922 | 0.01103 |
| 18.8888889 | 0.000756 | 0.00133 | 0.008375 | 0.00317 | 0.001114 | 0.013565 | 0.004461 | 0.025412 | 0.000323 | 0.010192 | 0.002237 | 0.015342 | 0.005283 | 0.002578 | 0.011793 | 0.01372 | 0.001378 | 0.011724 | 0.011904 | 0.004434 | 0.009215 | 0.013907 | 0.011671 |
| 18.9027778 | 0.002244 | 0.001402 | 0.004736 | 0.00273 | 0.00141 | 0.013501 | 0.0079 | 0.020392 | 0.000375 | 0.0106 | 0.002426 | 0.019381 | 0.0088 | 0.003454 | 0.013133 | 0.012156 | 0.002854 | 0.007455 | 0.013665 | 0.00514 | 0.01365 | 0.007846 | 0.012752 |
| 18.9166667 | 0.005741 | 0.00166 | 0.003861 | 0.003348 | 0.002527 | 0.014766 | 0.010422 | 0.019025 | 0.001381 | 0.01609 | 0.004419 | 0.018265 | 0.01132 | 0.00475 | 0.015639 | 0.010603 | 0.00522 | 0.005551 | 0.017674 | 0.007366 | 0.019145 | 0.005183 | 0.016939 |
| 18.9305556 | 0.00848 | 0.003411 | 0.006312 | 0.004452 | 0.003065 | 0.017169 | 0.009934 | 0.02089 | 0.003768 | 0.021491 | 0.00638 | 0.017077 | 0.014261 | 0.00489 | 0.020171 | 0.013362 | 0.009423 | 0.005929 | 0.022006 | 0.01146 | 0.025611 | 0.011417 | 0.024405 |
| 18.9444444 | 0.008493 | 0.006141 | 0.007803 | 0.005924 | 0.003688 | 0.019879 | 0.009013 | 0.022472 | 0.007457 | 0.023958 | 0.007121 | 0.023007 | 0.019236 | 0.00609 | 0.025941 | 0.020183 | 0.015138 | 0.007122 | 0.026917 | 0.016371 | 0.033409 | 0.021228 | 0.032304 |
| 18.9583333 | 0.007758 | 0.008424 | 0.007036 | 0.007668 | 0.004984 | 0.021117 | 0.009926 | 0.022294 | 0.010844 | 0.025377 | 0.007098 | 0.032958 | 0.026477 | 0.008875 | 0.032246 | 0.026633 | 0.020344 | 0.007977 | 0.034505 | 0.021042 | 0.04073 | 0.029432 | 0.039204 |
| 18.9722222 | 0.008067 | 0.010891 | 0.005021 | 0.008933 | 0.005703 | 0.021292 | 0.011878 | 0.02167 | 0.012214 | 0.026397 | 0.006924 | 0.039456 | 0.035768 | 0.012199 | 0.036945 | 0.030995 | 0.023987 | 0.008585 | 0.042531 | 0.025516 | 0.044146 | 0.037756 | 0.045727 |
| 18.9861111 | 0.008861 | 0.014698 | 0.003348 | 0.009805 | 0.005645 | 0.022251 | 0.01436 | 0.023025 | 0.012177 | 0.025397 | 0.007084 | 0.044086 | 0.045827 | 0.015879 | 0.037699 | 0.034649 | 0.026511 | 0.008804 | 0.046846 | 0.02928 | 0.043681 | 0.048504 | 0.050334 |
| 19 | 0.00952 | 0.018032 | 0.004928 | 0.01054 | 0.006171 | 0.023498 | 0.016617 | 0.026017 | 0.012064 | 0.022134 | 0.007937 | 0.050794 | 0.052677 | 0.019696 | 0.036271 | 0.036623 | 0.026916 | 0.006922 | 0.045619 | 0.030447 | 0.041889 | 0.060167 | 0.049627 |
| 19.0138889 | 0.010348 | 0.019004 | 0.009136 | 0.010586 | 0.007353 | 0.024012 | 0.017178 | 0.027052 | 0.011547 | 0.019829 | 0.009461 | 0.055456 | 0.053684 | 0.02215 | 0.034532 | 0.035452 | 0.024098 | 0.004476 | 0.039628 | 0.02862 | 0.039918 | 0.068663 | 0.044282 |
| 19.0277778 | 0.011666 | 0.019219 | 0.013767 | 0.009566 | 0.007659 | 0.025084 | 0.015903 | 0.024806 | 0.011271 | 0.020865 | 0.01158 | 0.055557 | 0.051012 | 0.023088 | 0.031208 | 0.03341 | 0.020451 | 0.005067 | 0.033614 | 0.025892 | 0.037957 | 0.072076 | 0.03951 |
| 19.0416667 | 0.014937 | 0.020166 | 0.019865 | 0.008455 | 0.007616 | 0.032087 | 0.014181 | 0.023398 | 0.01611 | 0.022728 | 0.013419 | 0.053347 | 0.046255 | 0.023266 | 0.025933 | 0.031894 | 0.018348 | 0.006044 | 0.031259 | 0.022998 | 0.034909 | 0.073564 | 0.037939 |
| 19.0555556 | 0.022036 | 0.019591 | 0.026646 | 0.007705 | 0.008898 | 0.046086 | 0.011578 | 0.026203 | 0.026386 | 0.022396 | 0.013613 | 0.044675 | 0.038122 | 0.019355 | 0.018793 | 0.027419 | 0.014963 | 0.005213 | 0.029062 | 0.017959 | 0.029074 | 0.075804 | 0.037853 |
| 19.0694444 | 0.027796 | 0.013537 | 0.026436 | 0.00559 | 0.00843 | 0.051663 | 0.007471 | 0.028559 | 0.028093 | 0.019058 | 0.01259 | 0.02647 | 0.028588 | 0.013627 | 0.012715 | 0.018176 | 0.008991 | 0.004457 | 0.022067 | 0.010893 | 0.020558 | 0.076293 | 0.035195 |
| 19.0833333 | 0.020879 | 0.008424 | 0.015815 | 0.002973 | 0.006696 | 0.03568 | 0.006105 | 0.023239 | 0.0154 | 0.013443 | 0.011291 | 0.00923 | 0.01792 | 0.018274 | 0.016887 | 0.009222 | 0.006322 | 0.006239 | 0.012161 | 0.005282 | 0.011023 | 0.076039 | 0.026586 |
| 19.0972222 | 0.010532 | 0.015444 | 0.005186 | 0.002959 | 0.009244 | 0.019283 | 0.008424 | 0.011737 | 0.003767 | 0.008522 | 0.008488 | 0.002076 | 0.009432 | 0.029862 | 0.029582 | 0.004484 | 0.007866 | 0.018028 | 0.008455 | 0.005146 | 0.00723 | 0.071841 | 0.014103 |
| 19.1111111 | 0.012837 | 0.025698 | 0.001532 | 0.004251 | 0.010002 | 0.021968 | 0.009937 | 0.003247 | 0.001759 | 0.006794 | 0.004554 | 0.003247 | 0.008815 | 0.029907 | 0.036583 | 0.00251 | 0.007127 | 0.044419 | 0.016042 | 0.010672 | 0.012382 | 0.053615 | 0.005205 |
| 19.125 | 0.016135 | 0.021757 | 0.00072 | 0.00451 | 0.005258 | 0.024285 | 0.010475 | 0.001272 | 0.003731 | 0.00663 | 0.003367 | 0.010687 | 0.009488 | 0.016694 | 0.035559 | 0.002981 | 0.003488 | 0.069499 | 0.031179 | 0.015853 | 0.015698 | 0.026949 | 0.004485 |
| 19.1388889 | 0.012341 | 0.009748 | 0.001278 | 0.004973 | 0.002682 | 0.016207 | 0.011649 | 0.001671 | 0.004884 | 0.004605 | 0.005328 | 0.020131 | 0.0102 | 0.005224 | 0.032265 | 0.005011 | 0.001389 | 0.066211 | 0.047746 | 0.016098 | 0.010928 | 0.008895 | 0.008231 |
| 19.1527778 | 0.00894 | 0.00557 | 0.003427 | 0.005895 | 0.003145 | 0.008313 | 0.011314 | 0.002197 | 0.003677 | 0.001653 | 0.006384 | 0.023688 | 0.014828 | 0.002509 | 0.025087 | 0.004554 | 0.001859 | 0.038036 | 0.052132 | 0.011335 | 0.005649 | 0.003483 | 0.010209 |
| 19.1666667 | 0.006895 | 0.007633 | 0.003932 | 0.005216 | 0.004032 | 0.0039 | 0.007534 | 0.00204 | 0.002287 | 0.001318 | 0.004958 | 0.019284 | 0.017549 | 0.002659 | 0.01302 | 0.003209 | 0.00309 | 0.018076 | 0.037193 | 0.006026 | 0.007397 | 0.003203 | 0.007061 |
| 19.1805556 | 0.004071 | 0.009617 | 0.003125 | 0.002847 | 0.004941 | 0.003286 | 0.004027 | 0.001401 | 0.002434 | 0.003514 | 0.00313 | 0.012526 | 0.015291 | 0.003185 | 0.003672 | 0.004508 | 0.003571 | 0.012078 | 0.017276 | 0.005143 | 0.013323 | 0.004864 | 0.00296 |
| 19.1944444 | 0.001986 | 0.010644 | 0.004822 | 0.000936 | 0.004765 | 0.004883 | 0.005704 | 0.002097 | 0.003422 | 0.00568 | 0.001866 | 0.008742 | 0.01255 | 0.007448 | 0.001716 | 0.004647 | 0.003186 | 0.009252 | 0.01002 | 0.005354 | 0.016344 | 0.007042 | 0.003339 |
| 19.2083333 | 0.002341 | 0.010009 | 0.007552 | 0.000367 | 0.003027 | 0.006478 | 0.01018 | 0.003361 | 0.005728 | 0.006379 | 0.001398 | 0.006168 | 0.011319 | 0.012243 | 0.004502 | 0.003204 | 0.004263 | 0.014538 | 0.019173 | 0.004109 | 0.017103 | 0.008789 | 0.006664 |
| 19.2222222 | 0.002888 | 0.007615 | 0.007347 | 0.000478 | 0.001378 | 0.008714 | 0.012009 | 0.003265 | 0.006358 | 0.00641 | 0.002647 | 0.004309 | 0.009516 | 0.011572 | 0.010814 | 0.004612 | 0.00848 | 0.020934 | 0.032953 | 0.004762 | 0.019731 | 0.013004 | 0.011485 |
| 19.2361111 | 0.001975 | 0.004656 | 0.005241 | 0.000778 | 0.001267 | 0.011887 | 0.010459 | 0.001853 | 0.004102 | 0.00729 | 0.003838 | 0.006161 | 0.007406 | 0.006734 | 0.016937 | 0.007665 | 0.012376 | 0.020086 | 0.039474 | 0.007768 | 0.022888 | 0.019121 | 0.015938 |
| 19.25 | 0.000915 | 0.003228 | 0.004457 | 0.00104 | 0.001316 | 0.014135 | 0.007734 | 0.000626 | 0.00439 | 0.00947 | 0.003002 | 0.008386 | 0.006068 | 0.002484 | 0.017815 | 0.008238 | 0.011659 | 0.017822 | 0.037112 | 0.011068 | 0.022895 | 0.022513 | 0.016638 |
| 19.2638889 | 0.000715 | 0.003471 | 0.004912 | 0.001712 | 0.001068 | 0.014638 | 0.004602 | 0.000483 | 0.008622 | 0.011564 | 0.001614 | 0.008264 | 0.003936 | 0.001449 | 0.014605 | 0.006657 | 0.007888 | 0.016203 | 0.030241 | 0.014281 | 0.020325 | 0.022757 | 0.014343 |
| 19.2777778 | 0.000779 | 0.002852 | 0.003837 | 0.002574 | 0.00176 | 0.014658 | 0.00279 | 0.001255 | 0.010798 | 0.011001 | 0.001598 | 0.008164 | 0.001794 | 0.004135 | 0.010224 | 0.006347 | 0.006333 | 0.018363 | 0.024622 | 0.017476 | 0.018255 | 0.024121 | 0.012023 |
| 19.2916667 | 0.000585 | 0.001885 | 0.002259 | 0.002413 | 0.003439 | 0.014025 | 0.004757 | 0.002951 | 0.00831 | 0.007111 | 0.002255 | 0.008455 | 0.00173 | 0.008153 | 0.006079 | 0.007843 | 0.009237 | 0.028763 | 0.021934 | 0.019435 | 0.017539 | 0.028125 | 0.010799 |
| 19.3055556 | 0.000511 | 0.002664 | 0.002232 | 0.001556 | 0.004912 | 0.011927 | 0.007931 | 0.005007 | 0.005673 | 0.004936 | 0.001956 | 0.007977 | 0.001787 | 0.009495 | 0.003246 | 0.008815 | 0.013174 | 0.038636 | 0.019967 | 0.018923 | 0.016624 | 0.03122 | 0.009166 |
| 19.3194444 | 0.000813 | 0.002912 | 0.001896 | 0.001346 | 0.00471 | 0.008995 | 0.008637 | 0.005918 | 0.005375 | 0.007128 | 0.001186 | 0.007139 | 0.001575 | 0.006899 | 0.001565 | 0.008056 | 0.014119 | 0.033043 | 0.0172 | 0.01604 | 0.014549 | 0.029254 | 0.005507 |
| 19.3333333 | 0.001787 | 0.001617 | 0.001844 | 0.001574 | 0.002743 | 0.005684 | 0.007691 | 0.005617 | 0.005395 | 0.008408 | 0.001042 | 0.007558 | 0.003569 | 0.003969 | 0.001095 | 0.007415 | 0.013801 | 0.020402 | 0.014535 | 0.013012 | 0.012782 | 0.023506 | 0.001734 |
| 19.3472222 | 0.003941 | 0.00117 | 0.004324 | 0.001437 | 0.001056 | 0.002801 | 0.007617 | 0.005816 | 0.00368 | 0.008546 | 0.001163 | 0.009039 | 0.007791 | 0.004512 | 0.003141 | 0.009129 | 0.016154 | 0.024908 | 0.01373 | 0.012283 | 0.012145 | 0.018991 | 0.000352 |
| 19.3611111 | 0.006764 | 0.002429 | 0.007881 | 0.001576 | 0.001008 | 0.002023 | 0.008732 | 0.007178 | 0.002141 | 0.010678 | 0.001902 | 0.007647 | 0.009062 | 0.005571 | 0.004762 | 0.010399 | 0.017773 | 0.04456 | 0.011792 | 0.01103 | 0.009294 | 0.014462 | 0.00339 |
| 19.375 | 0.009532 | 0.004422 | 0.009096 | 0.001963 | 0.001558 | 0.003323 | 0.009835 | 0.008793 | 0.004537 | 0.013768 | 0.00388 | 0.00764 | 0.008448 | 0.00465 | 0.00584 | 0.008076 | 0.013198 | 0.061036 | 0.00927 | 0.008296 | 0.009361 | 0.011408 | 0.014266 |
| 19.3888889 | 0.012819 | 0.006778 | 0.00881 | 0.002108 | 0.00215 | 0.006107 | 0.011183 | 0.010946 | 0.010696 | 0.017589 | 0.006124 | 0.015157 | 0.012894 | 0.004103 | 0.010941 | 0.005927 | 0.008635 | 0.067504 | 0.012342 | 0.0088 | 0.0199 | 0.01824 | 0.027713 |
| 19.4027778 | 0.015903 | 0.010055 | 0.011681 | 0.002883 | 0.003526 | 0.010419 | 0.013555 | 0.014784 | 0.018045 | 0.023503 | 0.007611 | 0.021286 | 0.01734 | 0.00392 | 0.016419 | 0.006202 | 0.009492 | 0.064291 | 0.017341 | 0.011037 | 0.031699 | 0.031322 | 0.033629 |
| 19.4166667 | 0.017119 | 0.013595 | 0.017799 | 0.004195 | 0.004878 | 0.015335 | 0.015914 | 0.019134 | 0.024677 | 0.031074 | 0.00754 | 0.023604 | 0.019196 | 0.007323 | 0.02137 | 0.008864 | 0.012594 | 0.050448 | 0.022359 | 0.014664 | 0.040196 | 0.0456 | 0.036787 |
| 19.4305556 | 0.016078 | 0.016018 | 0.021689 | 0.005148 | 0.004816 | 0.02044 | 0.016638 | 0.021733 | 0.029147 | 0.037342 | 0.00593 | 0.027783 | 0.023377 | 0.016686 | 0.028265 | 0.015844 | 0.01717 | 0.026922 | 0.030253 | 0.022467 | 0.048893 | 0.062099 | 0.045043 |
| 19.4444444 | 0.012847 | 0.017249 | 0.021554 | 0.005816 | 0.003815 | 0.02657 | 0.01582 | 0.022315 | 0.031146 | 0.040684 | 0.004255 | 0.029531 | 0.025353 | 0.022625 | 0.030181 | 0.022885 | 0.018821 | 0.007285 | 0.033712 | 0.026838 | 0.052395 | 0.073726 | 0.051531 |
| 19.4583333 | 0.00905 | 0.018847 | 0.021039 | 0.005771 | 0.002445 | 0.03338 | 0.01416 | 0.020391 | 0.031031 | 0.042158 | 0.002834 | 0.026213 | 0.021975 | 0.018044 | 0.025274 | 0.024959 | 0.015022 | 0.005085 | 0.029523 | 0.022355 | 0.047932 | 0.077048 | 0.050293 |
| 19.4722222 | 0.007308 | 0.024168 | 0.023741 | 0.004724 | 0.001471 | 0.038566 | 0.011883 | 0.015108 | 0.028126 | 0.041819 | 0.001557 | 0.022399 | 0.018454 | 0.008429 | 0.021256 | 0.024103 | 0.012467 | 0.020036 | 0.025578 | 0.015708 | 0.039724 | 0.077573 | 0.04465 |
| 19.4861111 | 0.009578 | 0.032468 | 0.026155 | 0.005051 | 0.001817 | 0.04222 | 0.011333 | 0.011409 | 0.023531 | 0.041143 | 0.002668 | 0.019969 | 0.016538 | 0.002049 | 0.020083 | 0.022969 | 0.012468 | 0.034783 | 0.024045 | 0.01139 | 0.03097 | 0.076837 | 0.035921 |
| 19.5 | 0.01522 | 0.03544 | 0.021288 | 0.008503 | 0.003243 | 0.045687 | 0.014199 | 0.016592 | 0.020638 | 0.042694 | 0.007009 | 0.018063 | 0.015222 | 0.001715 | 0.018845 | 0.023211 | 0.010958 | 0.027879 | 0.021612 | 0.008153 | 0.024067 | 0.072762 | 0.025085 |
| 19.5138889 | 0.018462 | 0.030051 | 0.012635 | 0.011855 | 0.003997 | 0.044638 | 0.015747 | 0.028547 | 0.019179 | 0.041434 | 0.010626 | 0.017166 | 0.01514 | 0.00581 | 0.016915 | 0.026169 | 0.008918 | 0.018145 | 0.018871 | 0.006507 | 0.0199 | 0.06383 | 0.017697 |
| 19.5277778 | 0.014392 | 0.021891 | 0.007302 | 0.010642 | 0.003533 | 0.03518 | 0.010929 | 0.035999 | 0.014986 | 0.031954 | 0.009269 | 0.015576 | 0.011833 | 0.010342 | 0.01248 | 0.026671 | 0.006648 | 0.024395 | 0.014169 | 0.005482 | 0.013406 | 0.04964 | 0.013883 |
| 19.5416667 | 0.007522 | 0.014561 | 0.008691 | 0.005771 | 0.005087 | 0.022408 | 0.005861 | 0.032428 | 0.008028 | 0.01918 | 0.005067 | 0.011143 | 0.012857 | 0.010588 | 0.00883 | 0.019061 | 0.009668 | 0.024245 | 0.014795 | 0.009016 | 0.011888 | 0.030966 | 0.011183 |
| 19.5555556 | 0.004312 | 0.010742 | 0.013377 | 0.00182 | 0.007521 | 0.01522 | 0.006189 | 0.024923 | 0.00338 | 0.010484 | 0.003545 | 0.008977 | 0.027176 | 0.011617 | 0.010721 | 0.011236 | 0.02446 | 0.015181 | 0.027199 | 0.020919 | 0.023878 | 0.014206 | 0.013107 |

| | | | | | | | | | | | | | | | | | | | | | | | |
|---|---|---|---|---|---|---|---|---|---|---|---|---|---|---|---|---|---|---|---|---|---|---|---|
| 19.5694444 | 0.004515 | 0.015208 | 0.014487 | 0.002478 | 0.005865 | 0.017177 | 0.006877 | 0.026243 | 0.003629 | 0.007842 | 0.00361 | 0.010824 | 0.028535 | 0.015663 | 0.014029 | 0.010253 | 0.034708 | 0.016352 | 0.027603 | 0.026393 | 0.02687 | 0.011736 | 0.013414 |
| 19.5833333 | 0.007252 | 0.027147 | 0.014092 | 0.008304 | 0.003245 | 0.02494 | 0.00791 | 0.039078 | 0.009879 | 0.01016 | 0.002996 | 0.009615 | 0.018792 | 0.01233 | 0.023166 | 0.015414 | 0.027327 | 0.021786 | 0.022048 | 0.016978 | 0.015809 | 0.024831 | 0.011348 |
| 19.5972222 | 0.010376 | 0.036316 | 0.014686 | 0.01429 | 0.004573 | 0.033072 | 0.010553 | 0.051058 | 0.016139 | 0.016555 | 0.006784 | 0.012594 | 0.025511 | 0.006912 | 0.038272 | 0.024341 | 0.012721 | 0.016147 | 0.034397 | 0.006594 | 0.013335 | 0.037069 | 0.017036 |
| 19.6111111 | 0.010263 | 0.034328 | 0.015229 | 0.015764 | 0.006806 | 0.035778 | 0.011506 | 0.05143 | 0.013531 | 0.024606 | 0.013965 | 0.032468 | 0.033178 | 0.009558 | 0.043347 | 0.028575 | 0.0044 | 0.010873 | 0.03886 | 0.006014 | 0.020249 | 0.035573 | 0.021772 |
| 19.625 | 0.008964 | 0.0245 | 0.015678 | 0.013513 | 0.006701 | 0.03078 | 0.011877 | 0.042922 | 0.00744 | 0.027236 | 0.016731 | 0.053096 | 0.030426 | 0.013317 | 0.033972 | 0.025922 | 0.004374 | 0.016141 | 0.024652 | 0.011497 | 0.026987 | 0.026495 | 0.018575 |
| 19.6388889 | 0.009676 | 0.015318 | 0.014573 | 0.009697 | 0.004275 | 0.022254 | 0.012073 | 0.031678 | 0.004766 | 0.01881 | 0.014978 | 0.049453 | 0.022955 | 0.010094 | 0.019828 | 0.018759 | 0.005533 | 0.019354 | 0.011676 | 0.014395 | 0.025612 | 0.020533 | 0.011938 |
| 19.6527778 | 0.011182 | 0.009727 | 0.012747 | 0.005616 | 0.00207 | 0.01411 | 0.009435 | 0.020618 | 0.004915 | 0.007526 | 0.011677 | 0.027935 | 0.014253 | 0.007889 | 0.017914 | 0.012045 | 0.009287 | 0.013193 | 0.009991 | 0.010956 | 0.018646 | 0.01703 | 0.007761 |
| 19.6666667 | 0.010707 | 0.005722 | 0.01163 | 0.002386 | 0.002229 | 0.007895 | 0.004636 | 0.011921 | 0.004946 | 0.003579 | 0.008201 | 0.017631 | 0.014876 | 0.017225 | 0.032739 | 0.015921 | 0.018759 | 0.008591 | 0.017445 | 0.009261 | 0.022962 | 0.011238 | 0.013782 |
| 19.6805556 | 0.0068 | 0.003191 | 0.009724 | 0.000885 | 0.002547 | 0.004392 | 0.001346 | 0.008332 | 0.006274 | 0.003134 | 0.006188 | 0.025229 | 0.023348 | 0.027167 | 0.044307 | 0.026268 | 0.025586 | 0.014147 | 0.021898 | 0.015524 | 0.035258 | 0.00959 | 0.025207 |
| 19.6944444 | 0.003845 | 0.002811 | 0.007942 | 0.000953 | 0.00171 | 0.003456 | 0.00073 | 0.012431 | 0.01311 | 0.001782 | 0.00638 | 0.029044 | 0.027078 | 0.025745 | 0.042845 | 0.031213 | 0.02377 | 0.018758 | 0.018579 | 0.021058 | 0.038919 | 0.018457 | 0.030501 |
| 19.7083333 | 0.00607 | 0.003142 | 0.007953 | 0.00149 | 0.001371 | 0.003797 | 0.001484 | 0.022598 | 0.01869 | 0.001221 | 0.008927 | 0.020801 | 0.023918 | 0.016868 | 0.039866 | 0.028602 | 0.016294 | 0.015281 | 0.012654 | 0.019803 | 0.034738 | 0.026392 | 0.025335 |
| 19.7222222 | 0.007891 | 0.003867 | 0.006805 | 0.001577 | 0.001623 | 0.003575 | 0.00305 | 0.032471 | 0.015427 | 0.002159 | 0.011491 | 0.011534 | 0.018527 | 0.007859 | 0.044743 | 0.020536 | 0.008138 | 0.00838 | 0.008191 | 0.013786 | 0.029086 | 0.020712 | 0.015682 |
| 19.7361111 | 0.005331 | 0.005581 | 0.003976 | 0.001283 | 0.001278 | 0.00291 | 0.005263 | 0.036664 | 0.007487 | 0.003172 | 0.010411 | 0.010588 | 0.012325 | 0.002772 | 0.04794 | 0.010129 | 0.00426 | 0.002826 | 0.006461 | 0.0064 | 0.021197 | 0.015918 | 0.014598 |
| 19.75 | 0.003286 | 0.006181 | 0.003638 | 0.001658 | 0.001472 | 0.002749 | 0.006466 | 0.03246 | 0.004985 | 0.00324 | 0.006132 | 0.01412 | 0.006137 | 0.002405 | 0.034107 | 0.003969 | 0.005006 | 0.002066 | 0.007031 | 0.002188 | 0.011418 | 0.0267 | 0.020574 |
| 19.7638889 | 0.004389 | 0.005128 | 0.004005 | 0.002759 | 0.002141 | 0.002775 | 0.004632 | 0.019627 | 0.011171 | 0.003306 | 0.002359 | 0.01255 | 0.002874 | 0.004456 | 0.013466 | 0.004788 | 0.005649 | 0.005109 | 0.00737 | 0.002645 | 0.004298 | 0.032001 | 0.017856 |
| 19.7777778 | 0.004824 | 0.006452 | 0.005272 | 0.003281 | 0.002381 | 0.002074 | 0.002411 | 0.011702 | 0.016289 | 0.005068 | 0.001383 | 0.006941 | 0.00464 | 0.005043 | 0.003861 | 0.008509 | 0.005807 | 0.007632 | 0.005569 | 0.006413 | 0.00221 | 0.020104 | 0.009633 |
| 19.7916667 | 0.003251 | 0.009466 | 0.010891 | 0.00268 | 0.003099 | 0.001095 | 0.002777 | 0.021267 | 0.011819 | 0.008057 | 0.002551 | 0.004784 | 0.008214 | 0.006409 | 0.003984 | 0.01211 | 0.006568 | 0.00795 | 0.004288 | 0.011578 | 0.002198 | 0.01278 | 0.010131 |
| 19.8055556 | 0.002856 | 0.007913 | 0.014175 | 0.001695 | 0.003992 | 0.00092 | 0.002801 | 0.035561 | 0.00614 | 0.007992 | 0.003638 | 0.010424 | 0.007798 | 0.010479 | 0.008298 | 0.013299 | 0.005284 | 0.010064 | 0.003799 | 0.01314 | 0.003255 | 0.022771 | 0.017939 |
| 19.8194444 | 0.004176 | 0.004336 | 0.011194 | 0.001298 | 0.003782 | 0.00108 | 0.00242 | 0.038787 | 0.007456 | 0.006482 | 0.002954 | 0.018777 | 0.005455 | 0.009486 | 0.014143 | 0.009631 | 0.004099 | 0.015977 | 0.003882 | 0.010165 | 0.009986 | 0.03801 | 0.026353 |
| 19.8333333 | 0.004505 | 0.004667 | 0.009916 | 0.001247 | 0.002306 | 0.001511 | 0.004157 | 0.029929 | 0.007162 | 0.01042 | 0.002398 | 0.022132 | 0.008814 | 0.007579 | 0.020439 | 0.006206 | 0.00899 | 0.022217 | 0.008581 | 0.011856 | 0.022794 | 0.046676 | 0.030891 |
| 19.8472222 | 0.003389 | 0.005278 | 0.013337 | 0.001536 | 0.001174 | 0.002493 | 0.00553 | 0.01773 | 0.003758 | 0.016494 | 0.004111 | 0.024116 | 0.019084 | 0.017991 | 0.028749 | 0.012347 | 0.019652 | 0.02808 | 0.016987 | 0.022005 | 0.036507 | 0.048668 | 0.03164 |
| 19.8611111 | 0.002647 | 0.003239 | 0.014173 | 0.00243 | 0.001498 | 0.002999 | 0.004616 | 0.010118 | 0.004929 | 0.018425 | 0.005793 | 0.032854 | 0.033986 | 0.037022 | 0.041908 | 0.029507 | 0.033655 | 0.03859 | 0.028045 | 0.035795 | 0.050067 | 0.051833 | 0.037875 |
| 19.875 | 0.002152 | 0.002307 | 0.009162 | 0.00242 | 0.001987 | 0.002812 | 0.002959 | 0.006649 | 0.009271 | 0.017573 | 0.005654 | 0.045133 | 0.051021 | 0.054561 | 0.059491 | 0.051534 | 0.049187 | 0.055759 | 0.042337 | 0.050829 | 0.064719 | 0.060185 | 0.054149 |
| 19.8888889 | 0.001653 | 0.004654 | 0.003502 | 0.00155 | 0.001675 | 0.002077 | 0.001854 | 0.005356 | 0.012096 | 0.015292 | 0.00442 | 0.051041 | 0.064926 | 0.062942 | 0.072798 | 0.069006 | 0.061118 | 0.074792 | 0.05635 | 0.06259 | 0.077387 | 0.065767 | 0.070031 |
| 19.9027778 | 0.002251 | 0.007168 | 0.004163 | 0.001677 | 0.001148 | 0.001337 | 0.001746 | 0.006997 | 0.01056 | 0.009883 | 0.002659 | 0.051047 | 0.072167 | 0.058602 | 0.074398 | 0.075234 | 0.064923 | 0.093195 | 0.065641 | 0.065196 | 0.084314 | 0.065236 | 0.074159 |
| 19.9166667 | 0.002361 | 0.00816 | 0.008976 | 0.003292 | 0.001124 | 0.001909 | 0.003885 | 0.01194 | 0.005832 | 0.003591 | 0.002243 | 0.05258 | 0.073523 | 0.048586 | 0.067824 | 0.071823 | 0.061189 | 0.100265 | 0.066708 | 0.060002 | 0.08521 | 0.065954 | 0.069504 |
| 19.9305556 | 0.001849 | 0.008818 | 0.009176 | 0.006448 | 0.001223 | 0.003386 | 0.007318 | 0.017895 | 0.002871 | 0.002618 | 0.004564 | 0.05571 | 0.069724 | 0.041865 | 0.059921 | 0.063984 | 0.056987 | 0.085789 | 0.060506 | 0.053164 | 0.081707 | 0.071937 | 0.065605 |
| 19.9444444 | 0.00313 | 0.009262 | 0.006742 | 0.009729 | 0.001233 | 0.004883 | 0.008098 | 0.02219 | 0.002907 | 0.009461 | 0.006931 | 0.054721 | 0.059709 | 0.039225 | 0.05058 | 0.053993 | 0.061531 | 0.063762 | 0.051334 | 0.047517 | 0.07499 | 0.075908 | 0.063802 |
| 19.9583333 | 0.003624 | 0.009726 | 0.006673 | 0.010161 | 0.001463 | 0.005839 | 0.006505 | 0.023611 | 0.002893 | 0.020526 | 0.007096 | 0.045036 | 0.042823 | 0.034254 | 0.03548 | 0.039988 | 0.066503 | 0.042863 | 0.038198 | 0.03868 | 0.062861 | 0.066836 | 0.057011 |
| 19.9722222 | 0.002966 | 0.00986 | 0.005569 | 0.007043 | 0.001199 | 0.005982 | 0.005761 | 0.02143 | 0.001574 | 0.028356 | 0.005522 | 0.028283 | 0.021658 | 0.022347 | 0.018296 | 0.021498 | 0.051282 | 0.021092 | 0.020687 | 0.022218 | 0.043244 | 0.042698 | 0.040973 |
| 19.9861111 | 0.004213 | 0.010443 | 0.005672 | 0.003037 | 0.000554 | 0.007187 | 0.006348 | 0.01727 | 0.00126 | 0.029803 | 0.003332 | 0.013436 | 0.006841 | 0.009922 | 0.010417 | 0.007368 | 0.022567 | 0.006238 | 0.010127 | 0.008941 | 0.021967 | 0.016751 | 0.022732 |
| 20 | 0.004581 | 0.013547 | 0.016241 | 0.000866 | 0.000967 | 0.010825 | 0.007871 | 0.015165 | 0.003455 | 0.029354 | 0.002176 | 0.007739 | 0.003091 | 0.008312 | 0.010154 | 0.003731 | 0.004572 | 0.005235 | 0.009902 | 0.006932 | 0.008503 | 0.007305 | 0.013427 |
| 20.0138889 | 0.002391 | 0.017636 | 0.030232 | 0.000513 | 0.00256 | 0.01569 | 0.011011 | 0.01567 | 0.006156 | 0.029349 | 0.003073 | 0.010023 | 0.004147 | 0.016221 | 0.007879 | 0.005887 | 0.001002 | 0.009602 | 0.007857 | 0.006478 | 0.005551 | 0.017563 | 0.014178 |
| 20.0277778 | 0.001342 | 0.021332 | 0.03198 | 0.001581 | 0.004807 | 0.023466 | 0.015498 | 0.017459 | 0.007266 | 0.028446 | 0.005989 | 0.014305 | 0.008574 | 0.020692 | 0.007282 | 0.010459 | 0.001816 | 0.008729 | 0.003115 | 0.005123 | 0.006671 | 0.034762 | 0.017325 |
| 20.0416667 | 0.003994 | 0.025723 | 0.030561 | 0.00551 | 0.009835 | 0.041755 | 0.019822 | 0.022586 | 0.008208 | 0.030356 | 0.010459 | 0.013351 | 0.013056 | 0.016297 | 0.011645 | 0.011544 | 0.001991 | 0.011051 | 0.001674 | 0.004171 | 0.005345 | 0.048945 | 0.017484 |
| 20.0555556 | 0.008786 | 0.027764 | 0.037871 | 0.012522 | 0.015258 | 0.064991 | 0.020678 | 0.032438 | 0.010566 | 0.033761 | 0.013149 | 0.012889 | 0.010953 | 0.013197 | 0.01225 | 0.009791 | 0.005447 | 0.023825 | 0.004812 | 0.003131 | 0.005265 | 0.061971 | 0.012701 |
| 20.0694444 | 0.012957 | 0.022858 | 0.042108 | 0.020843 | 0.012294 | 0.07377 | 0.014389 | 0.043167 | 0.012127 | 0.025664 | 0.010434 | 0.027594 | 0.008844 | 0.029735 | 0.014532 | 0.016501 | 0.021604 | 0.041228 | 0.014556 | 0.007064 | 0.013028 | 0.078323 | 0.009137 |
| 20.0833333 | 0.017575 | 0.012933 | 0.033585 | 0.025458 | 0.007103 | 0.064546 | 0.006341 | 0.048211 | 0.012572 | 0.011013 | 0.004905 | 0.047171 | 0.01661 | 0.067069 | 0.033623 | 0.028394 | 0.049601 | 0.056716 | 0.026584 | 0.020089 | 0.027742 | 0.090988 | 0.014862 |
| 20.0972222 | 0.021947 | 0.005048 | 0.021978 | 0.02109 | 0.010815 | 0.045174 | 0.004464 | 0.043731 | 0.01535 | 0.006131 | 0.001205 | 0.050587 | 0.026085 | 0.088222 | 0.051982 | 0.032448 | 0.067812 | 0.062791 | 0.032765 | 0.039057 | 0.041831 | 0.085934 | 0.020561 |
| 20.1111111 | 0.020424 | 0.003385 | 0.021603 | 0.011848 | 0.013571 | 0.022555 | 0.005049 | 0.033029 | 0.018843 | 0.008995 | 0.000953 | 0.038636 | 0.026663 | 0.065913 | 0.045229 | 0.026338 | 0.057203 | 0.054184 | 0.030386 | 0.049724 | 0.04469 | 0.065803 | 0.016583 |
| 20.125 | 0.012435 | 0.005045 | 0.027008 | 0.00465 | 0.008011 | 0.010814 | 0.004141 | 0.021647 | 0.017506 | 0.013228 | 0.003486 | 0.023947 | 0.020077 | 0.028195 | 0.024615 | 0.015349 | 0.028256 | 0.034131 | 0.022885 | 0.042075 | 0.033534 | 0.046279 | 0.008578 |
| 20.1388889 | 0.005694 | 0.00455 | 0.024893 | 0.001575 | 0.003383 | 0.017156 | 0.004747 | 0.01157 | 0.011025 | 0.014782 | 0.006554 | 0.01243 | 0.011477 | 0.00946 | 0.011518 | 0.007043 | 0.00938 | 0.013608 | 0.015825 | 0.022514 | 0.01665 | 0.032233 | 0.006074 |
| 20.1527778 | 0.005734 | 0.003612 | 0.018767 | 0.002017 | 0.00426 | 0.027614 | 0.00582 | 0.004463 | 0.004606 | 0.012074 | 0.007708 | 0.005765 | 0.004927 | 0.008341 | 0.008518 | 0.007616 | 0.010305 | 0.003204 | 0.010247 | 0.009996 | 0.006694 | 0.020948 | 0.011992 |
| 20.1666667 | 0.008891 | 0.007436 | 0.014979 | 0.002694 | 0.006591 | 0.027458 | 0.004353 | 0.00179 | 0.002127 | 0.008274 | 0.005785 | 0.005805 | 0.00212 | 0.011496 | 0.007188 | 0.013325 | 0.016113 | 0.003927 | 0.005674 | 0.013741 | 0.008995 | 0.011029 | 0.01959 |
| 20.1805556 | 0.011312 | 0.01277 | 0.013067 | 0.001686 | 0.008368 | 0.016605 | 0.002586 | 0.002458 | 0.001929 | 0.004676 | 0.00294 | 0.007765 | 0.002991 | 0.012259 | 0.005779 | 0.015548 | 0.016229 | 0.00972 | 0.004467 | 0.018314 | 0.013811 | 0.006028 | 0.022248 |
| 20.1944444 | 0.012324 | 0.014629 | 0.012445 | 0.0006 | 0.009376 | 0.011362 | 0.004163 | 0.00393 | 0.002819 | 0.003109 | 0.002467 | 0.006438 | 0.007498 | 0.008343 | 0.008042 | 0.012204 | 0.010193 | 0.01943 | 0.006316 | 0.013727 | 0.013528 | 0.009728 | 0.020494 |
| 20.2083333 | 0.010056 | 0.014994 | 0.012886 | 0.00046 | 0.008203 | 0.021981 | 0.008477 | 0.004589 | 0.004629 | 0.004082 | 0.003556 | 0.004283 | 0.01177 | 0.004006 | 0.010741 | 0.008211 | 0.004509 | 0.035162 | 0.009008 | 0.007299 | 0.012447 | 0.01573 | 0.019699 |
| 20.2222222 | 0.005554 | 0.016957 | 0.012345 | 0.000675 | 0.004662 | 0.0337 | 0.011243 | 0.004416 | 0.004405 | 0.003915 | 0.00359 | 0.004579 | 0.010895 | 0.004073 | 0.007743 | 0.007451 | 0.00481 | 0.04997 | 0.012433 | 0.004712 | 0.014575 | 0.018155 | 0.021742 |
| 20.2361111 | 0.00334 | 0.018472 | 0.009904 | 0.000901 | 0.001329 | 0.037023 | 0.009126 | 0.003322 | 0.004125 | 0.002347 | 0.00255 | 0.005978 | 0.008692 | 0.005632 | 0.003397 | 0.007216 | 0.005398 | 0.06122 | 0.014181 | 0.004896 | 0.016536 | 0.016205 | 0.020014 |
| 20.25 | 0.004013 | 0.01606 | 0.014769 | 0.00094 | 0.001105 | 0.035861 | 0.00493 | 0.001889 | 0.007052 | 0.001308 | 0.001875 | 0.006002 | 0.011369 | 0.004378 | 0.003439 | 0.004805 | 0.003874 | 0.069671 | 0.010496 | 0.005415 | 0.013639 | 0.009844 | 0.01139 |
| 20.2638889 | 0.00526 | 0.010914 | 0.031439 | 0.001422 | 0.002681 | 0.030867 | 0.004017 | 0.001854 | 0.0095 | 0.000746 | 0.001574 | 0.00497 | 0.01677 | 0.002055 | 0.004455 | 0.003109 | 0.003153 | 0.058957 | 0.004797 | 0.005991 | 0.007402 | 0.003882 | 0.004248 |
| 20.2777778 | 0.007256 | 0.00619 | 0.042541 | 0.00252 | 0.002964 | 0.02249 | 0.005571 | 0.002757 | 0.008808 | 0.001244 | 0.001135 | 0.004017 | 0.018913 | 0.002105 | 0.004038 | 0.003839 | 0.002388 | 0.033673 | 0.002535 | 0.007286 | 0.00361 | 0.003125 | 0.003961 |
| 20.2916667 | 0.009284 | 0.002824 | 0.031533 | 0.003041 | 0.003185 | 0.014756 | 0.007063 | 0.00261 | 0.006985 | 0.003671 | 0.001092 | 0.002658 | 0.018629 | 0.004057 | 0.002894 | 0.005588 | 0.00102 | 0.027373 | 0.001896 | 0.009102 | 0.003069 | 0.005213 | 0.007539 |
| 20.3055556 | 0.008582 | 0.001787 | 0.017661 | 0.002975 | 0.004695 | 0.010194 | 0.008866 | 0.001749 | 0.006323 | 0.007493 | 0.00109 | 0.001068 | 0.020666 | 0.00664 | 0.002662 | 0.007772 | 0.000286 | 0.039039 | 0.001649 | 0.010241 | 0.002349 | 0.006018 | 0.01248 |
| 20.3194444 | 0.005191 | 0.004183 | 0.024782 | 0.00282 | 0.005727 | 0.006248 | 0.010963 | 0.001752 | 0.006847 | 0.010729 | 0.000674 | 0.001035 | 0.022581 | 0.008669 | 0.005288 | 0.008341 | 0.000156 | 0.039169 | 0.003146 | 0.009471 | 0.001611 | 0.006925 | 0.016102 |
| 20.3333333 | 0.004165 | 0.008162 | 0.036073 | 0.002728 | 0.006476 | 0.004102 | 0.013109 | 0.00319 | 0.007421 | 0.013759 | 0.000574 | 0.005057 | 0.021053 | 0.010551 | 0.010171 | 0.005689 | 0.000942 | 0.021912 | 0.004149 | 0.006375 | 0.00453 | 0.013576 | 0.018361 |
| 20.3472222 | 0.008608 | 0.011007 | 0.036121 | 0.00305 | 0.00803 | 0.006937 | 0.015622 | 0.005461 | 0.007752 | 0.018567 | 0.000925 | 0.01358 | 0.020149 | 0.014698 | 0.017126 | 0.002679 | 0.004173 | 0.007542 | 0.002981 | 0.004965 | 0.013271 | 0.029039 | 0.023129 |
| 20.3611111 | 0.014397 | 0.012133 | 0.029493 | 0.003893 | 0.009279 | 0.010663 | 0.018217 | 0.007352 | 0.007821 | 0.023856 | 0.001194 | 0.021065 | 0.021441 | 0.01961 | 0.02528 | 0.002799 | 0.0091 | 0.006178 | 0.002116 | 0.007283 | 0.022343 | 0.048956 | 0.029809 |
| 20.375 | 0.018803 | 0.013746 | 0.020373 | 0.005384 | 0.008676 | 0.014575 | 0.020584 | 0.009159 | 0.008712 | 0.027247 | 0.001269 | 0.023122 | 0.02212 | 0.021421 | 0.031623 | 0.006857 | 0.013142 | 0.00851 | 0.003991 | 0.007499 | 0.026823 | 0.066075 | 0.034792 |
| 20.3888889 | 0.023327 | 0.01743 | 0.013393 | 0.007475 | 0.006974 | 0.022446 | 0.022205 | 0.012358 | 0.01285 | 0.02955 | 0.001444 | 0.023754 | 0.022786 | 0.020187 | 0.034074 | 0.014751 | 0.015702 | 0.013004 | 0.009193 | 0.005464 | 0.030381 | 0.07745 | 0.037571 |
| 20.4027778 | 0.026893 | 0.020812 | 0.007733 | 0.009641 | 0.005973 | 0.030823 | 0.021631 | 0.015959 | 0.021002 | 0.032669 | 0.001907 | 0.027107 | 0.024957 | 0.018366 | 0.032924 | 0.026503 | 0.017503 | 0.026758 | 0.017674 | 0.007422 | 0.035879 | 0.084127 | 0.038647 |
| 20.4166667 | 0.027464 | 0.022227 | 0.003952 | 0.012428 | 0.00543 | 0.034109 | 0.020174 | 0.018734 | 0.031993 | 0.037267 | 0.002931 | 0.030722 | 0.027014 | 0.017675 | 0.02935 | 0.039006 | 0.017809 | 0.046675 | 0.025405 | 0.013213 | 0.041637 | 0.087588 | 0.038142 |
| 20.4305556 | 0.02637 | 0.022635 | 0.008276 | 0.016715 | 0.004757 | 0.035557 | 0.02052 | 0.021444 | 0.042844 | 0.043501 | 0.004505 | 0.030093 | 0.026192 | 0.018256 | 0.024409 | 0.043735 | 0.015244 | 0.046733 | 0.026603 | 0.017836 | 0.046022 | 0.088127 | 0.037682 |
| 20.4444444 | 0.023828 | 0.021203 | 0.022094 | 0.019749 | 0.004079 | 0.041432 | 0.021176 | 0.022308 | 0.046646 | 0.050185 | 0.005812 | 0.025131 | 0.022112 | 0.017902 | 0.020636 | 0.034509 | 0.011223 | 0.022724 | 0.022146 | 0.018332 | 0.04715 | 0.085933 | 0.0394 |
| 20.4583333 | 0.01917 | 0.017514 | 0.038859 | 0.018796 | 0.003072 | 0.048273 | 0.019713 | 0.019512 | 0.037934 | 0.056689 | 0.005688 | 0.019211 | 0.018397 | 0.014921 | 0.021035 | 0.017563 | 0.008825 | 0.008125 | 0.017159 | 0.014933 | 0.04355 | 0.08482 | 0.044094 |
| 20.4722222 | 0.015933 | 0.015189 | 0.045716 | 0.017894 | 0.002414 | 0.051855 | 0.018145 | 0.015775 | 0.021959 | 0.064062 | 0.004235 | 0.015311 | 0.01892 | 0.010961 | 0.025946 | 0.005236 | 0.008604 | 0.007998 | 0.014692 | 0.01147 | 0.038365 | 0.088666 | 0.049676 |
| 20.4861111 | 0.016186 | 0.016878 | 0.03624 | 0.020656 | 0.003375 | 0.053834 | 0.019815 | 0.014886 | 0.010025 | 0.064358 | 0.005013 | 0.016798 | 0.022476 | 0.009557 | 0.032442 | 0.004018 | 0.009587 | 0.007678 | 0.016323 | 0.013057 | 0.035332 | 0.091504 | 0.052085 |
| 20.5 | 0.016543 | 0.020595 | 0.019436 | 0.023511 | 0.005834 | 0.047969 | 0.02294 | 0.018093 | 0.006878 | 0.050604 | 0.008676 | 0.023729 | 0.025329 | 0.012994 | 0.03689 | 0.013686 | 0.010994 | 0.004911 | 0.02097 | 0.019932 | 0.033777 | 0.086173 | 0.05012 |
| 20.5138889 | 0.013629 | 0.022385 | 0.010863 | 0.023395 | 0.008535 | 0.031759 | 0.022479 | 0.022573 | 0.009383 | 0.036727 | 0.010209 | 0.028993 | 0.025608 | 0.019477 | 0.038316 | 0.028976 | 0.01195 | 0.002348 | 0.02424 | 0.027342 | 0.033055 | 0.075628 | 0.045114 |
| 20.5277778 | 0.008001 | 0.01868 | 0.010594 | 0.020406 | 0.009608 | 0.016725 | 0.015975 | 0.023145 | 0.01232 | 0.032492 | 0.008221 | 0.022327 | 0.020115 | 0.020776 | 0.032534 | 0.035596 | 0.011954 | 0.006607 | 0.019748 | 0.026142 | 0.027856 | 0.056631 | 0.031414 |
| 20.5416667 | 0.005422 | 0.011988 | 0.0128 | 0.014511 | 0.00845 | 0.008821 | 0.008656 | 0.018291 | 0.012091 | 0.028729 | 0.005465 | 0.02081 | 0.027089 | 0.020439 | 0.02566 | 0.030477 | 0.02579 | 0.029578 | 0.024168 | 0.02405 | 0.025171 | 0.037701 | 0.026003 |
| 20.5555556 | 0.007677 | 0.010243 | 0.017174 | 0.007992 | 0.005994 | 0.00573 | 0.007221 | 0.012903 | 0.008188 | 0.020361 | 0.003408 | 0.043098 | 0.059316 | 0.03533 | 0.035919 | 0.034482 | 0.06111 | 0.066684 | 0.053796 | 0.041483 | 0.043142 | 0.04256 | 0.052159 |
| 20.5694444 | 0.007304 | 0.010867 | 0.019616 | 0.005976 | 0.003792 | 0.005949 | 0.006533 | 0.013202 | 0.003374 | 0.012489 | 0.002602 | 0.053686 | 0.066998 | 0.050882 | 0.044643 | 0.040656 | 0.078782 | 0.079665 | 0.067053 | 0.059508 | 0.05771 | 0.046589 | 0.068566 |
| 20.5833333 | 0.005673 | 0.013724 | 0.029432 | 0.010987 | 0.004739 | 0.011997 | 0.005163 | 0.020473 | 0.001251 | 0.010634 | 0.006582 | 0.03156 | 0.039175 | 0.041264 | 0.031823 | 0.02759 | 0.055191 | 0.054034 | 0.040517 | 0.050226 | 0.039526 | 0.034761 | 0.046412 |
| 20.5972222 | 0.008632 | 0.024846 | 0.045075 | 0.018627 | 0.009926 | 0.021817 | 0.009694 | 0.030941 | 0.002252 | 0.015146 | 0.016933 | 0.012139 | 0.028333 | 0.019246 | 0.023881 | 0.015543 | 0.022884 | 0.021713 | 0.017909 | 0.026796 | 0.015714 | 0.042072 | 0.030585 |
| 20.6111111 | 0.011377 | 0.032727 | 0.05103 | 0.024006 | 0.016063 | 0.027347 | 0.014828 | 0.041107 | 0.004872 | 0.022763 | 0.026761 | 0.018829 | 0.032794 | 0.006455 | 0.027171 | 0.018143 | 0.009544 | 0.006772 | 0.017829 | 0.01018 | 0.015649 | 0.057481 | 0.041658 |
| 20.625 | 0.011237 | 0.030753 | 0.051886 | 0.026515 | 0.018703 | 0.027755 | 0.014241 | 0.048899 | 0.008177 | 0.028883 | 0.030213 | 0.03235 | 0.02754 | 0.004865 | 0.022023 | 0.020783 | 0.009396 | 0.007041 | 0.016779 | 0.005803 | 0.027153 | 0.049815 | 0.049584 |
| 20.6388889 | 0.010413 | 0.02675 | 0.052069 | 0.024592 | 0.016219 | 0.024971 | 0.010369 | 0.050972 | 0.010376 | 0.028034 | 0.02819 | 0.030904 | 0.015635 | 0.007524 | 0.011016 | 0.019686 | 0.007946 | 0.010092 | 0.009018 | 0.009644 | 0.030383 | 0.027237 | 0.039361 |
| 20.6527778 | 0.008728 | 0.023615 | 0.040154 | 0.017884 | 0.010502 | 0.017466 | 0.006546 | 0.04543 | 0.010437 | 0.019356 | 0.021918 | 0.01974 | 0.006382 | 0.009608 | 0.004815 | 0.017691 | 0.003608 | 0.010286 | 0.003617 | 0.013717 | 0.02186 | 0.011177 | 0.020573 |
| 20.6666667 | 0.005811 | 0.02116 | 0.024974 | 0.010054 | 0.004951 | 0.009339 | 0.005092 | 0.035901 | 0.010018 | 0.00911 | 0.01358 | 0.013556 | 0.004745 | 0.011914 | 0.004482 | 0.012792 | 0.001568 | 0.010356 | 0.001761 | 0.014443 | 0.009664 | 0.009315 | 0.007157 |
| 20.6805556 | 0.003326 | 0.018772 | 0.033431 | 0.004165 | 0.002719 | 0.007119 | 0.006545 | 0.0274 | 0.010515 | 0.00286 | 0.007897 | 0.015918 | 0.006184 | 0.016262 | 0.006563 | 0.006499 | 0.001899 | 0.01129 | 0.001729 | 0.01307 | 0.003079 | 0.009893 | 0.004843 |
| 20.6944444 | 0.00414 | 0.014797 | 0.05186 | 0.002142 | 0.003785 | 0.00614 | 0.008575 | 0.021298 | 0.010597 | 0.001124 | 0.004782 | 0.01926 | 0.004768 | 0.01958 | 0.007711 | 0.004119 | 0.002035 | 0.010936 | 0.003477 | 0.011203 | 0.003431 | 0.008882 | 0.005114 |
| 20.7083333 | 0.007483 | 0.008299 | 0.049812 | 0.004274 | 0.005142 | 0.005209 | 0.007979 | 0.018621 | 0.008064 | 0.002727 | 0.002909 | 0.017307 | 0.002751 | 0.016189 | 0.006279 | 0.004604 | 0.001228 | 0.00823 | 0.005357 | 0.009164 | 0.00666 | 0.018584 | 0.005404 |
| 20.7222222 | 0.008568 | 0.003251 | 0.029572 | 0.007523 | 0.004171 | 0.012014 | 0.004603 | 0.020886 | 0.004372 | 0.005498 | 0.00632 | 0.012062 | 0.003489 | 0.007953 | 0.003689 | 0.004209 | 0.000462 | 0.004612 | 0.005257 | 0.006357 | 0.009656 | 0.040531 | 0.014661 |
| 20.7361111 | 0.005708 | 0.002466 | 0.024833 | 0.009211 | 0.002624 | 0.019289 | 0.001861 | 0.026115 | 0.003207 | 0.00704 | 0.012469 | 0.008367 | 0.003538 | 0.003993 | 0.002978 | 0.005645 | 0.000339 | 0.00264 | 0.003908 | 0.003274 | 0.009972 | 0.056457 | 0.030822 |
| 20.75 | 0.004418 | 0.00413 | 0.050657 | 0.007146 | 0.003889 | 0.014319 | 0.001176 | 0.029366 | 0.00439 | 0.008626 | 0.011775 | 0.007119 | 0.002387 | 0.008399 | 0.004176 | 0.009357 | 0.000761 | 0.004348 | 0.00293 | 0.001852 | 0.007572 | 0.049241 | 0.037363 |
| 20.7638889 | 0.008209 | 0.00721 | 0.065895 | 0.002658 | 0.004511 | 0.007591 | 0.001682 | 0.026694 | 0.005391 | 0.013559 | 0.007664 | 0.00525 | 0.003192 | 0.016198 | 0.006272 | 0.009169 | 0.001458 | 0.007555 | 0.002502 | 0.002465 | 0.006289 | 0.025695 | 0.026353 |
| 20.7777778 | 0.009907 | 0.007716 | 0.051546 | 0.000833 | 0.002615 | 0.012416 | 0.00408 | 0.017128 | 0.006133 | 0.019489 | 0.010267 | 0.003602 | 0.003941 | 0.018303 | 0.010145 | 0.00614 | 0.002054 | 0.008645 | 0.002207 | 0.003123 | 0.008778 | 0.014499 | 0.015404 |
| 20.7916667 | 0.00741 | 0.00479 | 0.032435 | 0.001373 | 0.002405 | 0.01965 | 0.006741 | 0.009862 | 0.006634 | 0.019777 | 0.015742 | 0.006498 | 0.003956 | 0.013178 | 0.014878 | 0.007746 | 0.002522 | 0.007506 | 0.003289 | 0.003144 | 0.014161 | 0.032985 | 0.024134 |
| 20.8055556 | 0.008142 | 0.003994 | 0.017022 | 0.001708 | 0.004018 | 0.018647 | 0.007688 | 0.014785 | 0.005913 | 0.013769 | 0.014136 | 0.011811 | 0.005423 | 0.012154 | 0.016809 | 0.011968 | 0.002557 | 0.006169 | 0.004206 | 0.003127 | 0.0203 | 0.064651 | 0.041334 |
| 20.8194444 | 0.011229 | 0.006073 | 0.017255 | 0.002179 | 0.004654 | 0.01197 | 0.007891 | 0.023476 | 0.005192 | 0.00782 | 0.007022 | 0.014536 | 0.007149 | 0.017667 | 0.01407 | 0.011438 | 0.00169 | 0.005642 | 0.002903 | 0.003049 | 0.0256 | 0.07722 | 0.047853 |
| 20.8333333 | 0.010967 | 0.007518 | 0.047893 | 0.003169 | 0.003762 | 0.005705 | 0.007827 | 0.02554 | 0.005956 | 0.006307 | 0.003053 | 0.013665 | 0.006601 | 0.016822 | 0.009787 | 0.006788 | 0.001543 | 0.005419 | 0.00203 | 0.003535 | 0.028858 | 0.056464 | 0.04337 |
| 20.8472222 | 0.007429 | 0.008509 | 0.079443 | 0.003668 | 0.002579 | 0.003778 | 0.006253 | 0.022605 | 0.0082 | 0.008531 | 0.002859 | 0.010786 | 0.004945 | 0.010715 | 0.009026 | 0.004175 | 0.004154 | 0.005283 | 0.003064 | 0.005733 | 0.029373 | 0.026353 | 0.035658 |
| 20.8611111 | 0.003595 | 0.010851 | 0.076445 | 0.00541 | 0.002674 | 0.004279 | 0.004099 | 0.018905 | 0.012497 | 0.012556 | 0.005694 | 0.008473 | 0.006354 | 0.012209 | 0.012619 | 0.006301 | 0.007834 | 0.005386 | 0.005373 | 0.00915 | 0.027782 | 0.017373 | 0.027707 |
| 20.875 | 0.002148 | 0.013354 | 0.044376 | 0.008693 | 0.003328 | 0.007397 | 0.005024 | 0.015472 | 0.01718 | 0.017294 | 0.013199 | 0.007972 | 0.011665 | 0.019139 | 0.016755 | 0.011082 | 0.009028 | 0.004393 | 0.007488 | 0.011892 | 0.025008 | 0.02489 | 0.018704 |
| 20.8888889 | 0.003691 | 0.01393 | 0.02156 | 0.008039 | 0.002447 | 0.018955 | 0.007958 | 0.010747 | 0.018192 | 0.023338 | 0.018912 | 0.006879 | 0.01745 | 0.021639 | 0.017839 | 0.015852 | 0.006772 | 0.003241 | 0.007065 | 0.012795 | 0.022114 | 0.025672 | 0.011045 |
| 20.9027778 | 0.005787 | 0.012287 | 0.025628 | 0.003934 | 0.001227 | 0.028674 | 0.010215 | 0.005014 | 0.015228 | 0.031154 | 0.017178 | 0.003928 | 0.021298 | 0.016149 | 0.015275 | 0.019016 | 0.00326 | 0.004723 | 0.004492 | 0.012882 | 0.021238 | 0.018471 | 0.00885 |
| 20.9166667 | 0.005576 | 0.009772 | 0.028201 | 0.002335 | 0.00243 | 0.026248 | 0.012681 | 0.002037 | 0.00979 | 0.037614 | 0.013137 | 0.002001 | 0.022473 | 0.008637 | 0.010482 | 0.01938 | 0.001115 | 0.005048 | 0.002392 | 0.014373 | 0.022344 | 0.023694 | 0.013611 |
| 20.9305556 | 0.003897 | 0.008726 | 0.020581 | 0.004324 | 0.005804 | 0.019433 | 0.014128 | 0.004109 | 0.0052 | 0.037861 | 0.012404 | 0.00414 | 0.020812 | 0.008002 | 0.006032 | 0.017273 | 0.000524 | 0.005391 | 0.001879 | 0.016865 | 0.022144 | 0.045376 | 0.023645 |
| 20.9444444 | 0.005256 | 0.010077 | 0.015948 | 0.007355 | 0.009309 | 0.016282 | 0.013437 | 0.008891 | 0.004792 | 0.031968 | 0.01387 | 0.010214 | 0.018831 | 0.011719 | 0.006081 | 0.015145 | 0.000395 | 0.012114 | 0.00153 | 0.017186 | 0.019591 | 0.065077 | 0.034974 |
| 20.9583333 | 0.010569 | 0.011682 | 0.011783 | 0.009755 | 0.012506 | 0.01773 | 0.013454 | 0.012909 | 0.004487 | 0.025701 | 0.014638 | 0.01708 | 0.020948 | 0.012591 | 0.011858 | 0.015556 | 0.000904 | 0.018266 | 0.001191 | 0.014989 | 0.018287 | 0.068776 | 0.042301 |
| 20.9722222 | 0.014826 | 0.012367 | 0.006917 | 0.010598 | 0.015035 | 0.018448 | 0.014765 | 0.014849 | 0.002975 | 0.022348 | 0.011958 | 0.023458 | 0.028006 | 0.008749 | 0.017058 | 0.017466 | 0.001917 | 0.016823 | 0.002188 | 0.013187 | 0.019372 | 0.05773 | 0.042665 |
| 20.9861111 | 0.015164 | 0.014926 | 0.004145 | 0.009497 | 0.015037 | 0.019125 | 0.017123 | 0.016032 | 0.004337 | 0.022515 | 0.0066 | 0.02944 | 0.037151 | 0.00584 | 0.016636 | 0.018006 | 0.002147 | 0.012047 | 0.004841 | 0.012701 | 0.019413 | 0.04378 | 0.03827 |
| 21 | 0.013025 | 0.020358 | 0.006497 | 0.009354 | 0.012931 | 0.026618 | 0.021446 | 0.01917 | 0.007651 | 0.027933 | 0.004018 | 0.031225 | 0.045651 | 0.009911 | 0.015799 | 0.01809 | 0.002981 | 0.008512 | 0.010034 | 0.012831 | 0.019282 | 0.039004 | 0.033981 |
| 21.0138889 | 0.01016 | 0.023916 | 0.018504 | 0.011997 | 0.011975 | 0.036487 | 0.025581 | 0.024065 | 0.010363 | 0.035581 | 0.005442 | 0.025201 | 0.047167 | 0.015548 | 0.019403 | 0.01876 | 0.006744 | 0.005861 | 0.014542 | 0.013521 | 0.021552 | 0.045625 | 0.03206 |
| 21.0277778 | 0.007858 | 0.022987 | 0.032888 | 0.013993 | 0.012804 | 0.041947 | 0.027674 | 0.027763 | 0.012186 | 0.040808 | 0.007382 | 0.015375 | 0.038201 | 0.01835 | 0.02121 | 0.018258 | 0.01179 | 0.004115 | 0.011338 | 0.012878 | 0.021902 | 0.057054 | 0.029486 |
| 21.0416667 | 0.007348 | 0.020868 | 0.04092 | 0.013524 | 0.014099 | 0.04785 | 0.027495 | 0.030456 | 0.014482 | 0.04784 | 0.009148 | 0.007789 | 0.027963 | 0.020165 | 0.015105 | 0.013887 | 0.01595 | 0.006968 | 0.006147 | 0.008311 | 0.015586 | 0.066018 | 0.023016 |

| | | | | | | | | | | | | | | | | | | | | | | | |
|---|---|---|---|---|---|---|---|---|---|---|---|---|---|---|---|---|---|---|---|---|---|---|---|
| 21.0555556 | 0.009714 | 0.020475 | 0.044039 | 0.013328 | 0.016997 | 0.060723 | 0.02583 | 0.033784 | 0.019783 | 0.058295 | 0.011598 | 0.008794 | 0.02476 | 0.01616 | 0.006971 | 0.009107 | 0.019691 | 0.014415 | 0.008766 | 0.00421 | 0.006488 | 0.069002 | 0.013947 |
| 21.0694444 | 0.015961 | 0.020423 | 0.042865 | 0.015369 | 0.019783 | 0.076454 | 0.023799 | 0.034823 | 0.028413 | 0.06036 | 0.012847 | 0.020934 | 0.024806 | 0.010137 | 0.005547 | 0.011503 | 0.022869 | 0.020662 | 0.012849 | 0.006253 | 0.002578 | 0.064217 | 0.007041 |
| 21.0833333 | 0.024992 | 0.017481 | 0.03511 | 0.017594 | 0.01734 | 0.077109 | 0.019973 | 0.030341 | 0.035223 | 0.049579 | 0.010252 | 0.031604 | 0.022382 | 0.016677 | 0.00842 | 0.015222 | 0.023063 | 0.020638 | 0.010033 | 0.011863 | 0.005545 | 0.051325 | 0.006258 |
| 21.0972222 | 0.028745 | 0.010968 | 0.02377 | 0.016173 | 0.010492 | 0.05421 | 0.013697 | 0.02426 | 0.034585 | 0.036211 | 0.005773 | 0.028633 | 0.017809 | 0.03179 | 0.008792 | 0.010978 | 0.019207 | 0.013771 | 0.004955 | 0.017098 | 0.009335 | 0.037907 | 0.009151 |
| 21.1111111 | 0.020263 | 0.004548 | 0.016387 | 0.010079 | 0.004376 | 0.024631 | 0.006865 | 0.021545 | 0.026415 | 0.026235 | 0.005171 | 0.015894 | 0.010971 | 0.040704 | 0.008228 | 0.004572 | 0.013763 | 0.006241 | 0.005092 | 0.019187 | 0.008622 | 0.032507 | 0.009012 |
| 21.125 | 0.009259 | 0.002968 | 0.014727 | 0.003578 | 0.001354 | 0.010024 | 0.00328 | 0.02114 | 0.014448 | 0.019108 | 0.008308 | 0.005084 | 0.005726 | 0.040568 | 0.009224 | 0.003069 | 0.009561 | 0.004682 | 0.007771 | 0.016586 | 0.004787 | 0.032104 | 0.005704 |
| 21.1388889 | 0.007985 | 0.006184 | 0.01441 | 0.002475 | 0.000778 | 0.012762 | 0.003975 | 0.02037 | 0.00606 | 0.014462 | 0.009683 | 0.002512 | 0.007538 | 0.035533 | 0.009468 | 0.00423 | 0.006837 | 0.006975 | 0.008532 | 0.010629 | 0.001741 | 0.031033 | 0.004339 |
| 21.1527778 | 0.01223 | 0.008547 | 0.01338 | 0.005597 | 0.001422 | 0.015744 | 0.003627 | 0.016668 | 0.005977 | 0.011817 | 0.007706 | 0.005805 | 0.010094 | 0.025872 | 0.008098 | 0.006858 | 0.004527 | 0.009441 | 0.008692 | 0.005447 | 0.00185 | 0.028512 | 0.004562 |
| 21.1666667 | 0.014266 | 0.007424 | 0.011794 | 0.006203 | 0.002646 | 0.01186 | 0.002256 | 0.009616 | 0.009103 | 0.010303 | 0.004106 | 0.010802 | 0.008955 | 0.012956 | 0.006677 | 0.01102 | 0.002511 | 0.011604 | 0.009079 | 0.005411 | 0.004427 | 0.024447 | 0.003841 |
| 21.1805556 | 0.013572 | 0.007017 | 0.008988 | 0.003234 | 0.003627 | 0.007641 | 0.003859 | 0.003675 | 0.008888 | 0.008689 | 0.001886 | 0.015562 | 0.007085 | 0.005505 | 0.007753 | 0.014597 | 0.003506 | 0.013009 | 0.010322 | 0.009219 | 0.00699 | 0.018599 | 0.002523 |
| 21.1944444 | 0.011388 | 0.007415 | 0.005158 | 0.001012 | 0.004036 | 0.00532 | 0.006931 | 0.002049 | 0.005008 | 0.006477 | 0.002563 | 0.020329 | 0.004571 | 0.008167 | 0.013311 | 0.017656 | 0.010087 | 0.015226 | 0.014448 | 0.014201 | 0.010109 | 0.012752 | 0.003408 |
| 21.2083333 | 0.007612 | 0.004775 | 0.002394 | 0.000607 | 0.004641 | 0.003353 | 0.010451 | 0.003789 | 0.002333 | 0.004617 | 0.003606 | 0.024024 | 0.001823 | 0.013939 | 0.020574 | 0.020275 | 0.017136 | 0.017856 | 0.020309 | 0.018036 | 0.013213 | 0.007864 | 0.007508 |
| 21.2222222 | 0.003593 | 0.002722 | 0.002775 | 0.001731 | 0.00538 | 0.002363 | 0.014029 | 0.006694 | 0.003844 | 0.003931 | 0.003731 | 0.022795 | 0.003465 | 0.014305 | 0.023451 | 0.019187 | 0.016035 | 0.016546 | 0.021459 | 0.016527 | 0.01135 | 0.005237 | 0.008854 |
| 21.2361111 | 0.002256 | 0.004983 | 0.005076 | 0.004376 | 0.005191 | 0.005496 | 0.015205 | 0.008743 | 0.006542 | 0.004574 | 0.003546 | 0.015637 | 0.011718 | 0.008502 | 0.020599 | 0.013142 | 0.008776 | 0.011252 | 0.015371 | 0.010195 | 0.006142 | 0.007041 | 0.005131 |
| 21.25 | 0.003618 | 0.006807 | 0.006138 | 0.004702 | 0.004291 | 0.012731 | 0.01324 | 0.009299 | 0.00817 | 0.006287 | 0.002897 | 0.00711 | 0.021074 | 0.004466 | 0.016353 | 0.006305 | 0.003955 | 0.006899 | 0.012644 | 0.005321 | 0.004141 | 0.007961 | 0.00166 |
| 21.2638889 | 0.005447 | 0.005541 | 0.004741 | 0.002267 | 0.003876 | 0.020081 | 0.0096 | 0.008964 | 0.008778 | 0.007513 | 0.001806 | 0.00231 | 0.025349 | 0.007932 | 0.013437 | 0.002419 | 0.00326 | 0.00631 | 0.013774 | 0.004547 | 0.003807 | 0.005313 | 0.000552 |
| 21.2777778 | 0.007749 | 0.005025 | 0.002294 | 0.00118 | 0.003883 | 0.023764 | 0.005342 | 0.008101 | 0.007887 | 0.006655 | 0.001616 | 0.001885 | 0.025077 | 0.013185 | 0.010562 | 0.001531 | 0.005457 | 0.008677 | 0.009814 | 0.005288 | 0.002788 | 0.004809 | 0.000524 |
| 21.2916667 | 0.012333 | 0.005944 | 0.001184 | 0.001741 | 0.003298 | 0.023066 | 0.002491 | 0.006966 | 0.005475 | 0.003933 | 0.002745 | 0.002475 | 0.024333 | 0.012721 | 0.006347 | 0.002041 | 0.008081 | 0.01186 | 0.007748 | 0.006017 | 0.003885 | 0.006905 | 0.001171 |
| 21.3055556 | 0.018636 | 0.004961 | 0.002417 | 0.003436 | 0.001864 | 0.01976 | 0.003136 | 0.005784 | 0.002881 | 0.00179 | 0.002848 | 0.002986 | 0.025388 | 0.007344 | 0.005166 | 0.002147 | 0.006663 | 0.012705 | 0.008782 | 0.005878 | 0.006087 | 0.007687 | 0.003194 |
| 21.3194444 | 0.021864 | 0.002583 | 0.004263 | 0.005505 | 0.001013 | 0.015397 | 0.005572 | 0.004199 | 0.00229 | 0.003047 | 0.001986 | 0.00534 | 0.026772 | 0.003096 | 0.010709 | 0.00244 | 0.002761 | 0.010022 | 0.007003 | 0.004288 | 0.005729 | 0.005722 | 0.006739 |
| 21.3333333 | 0.017227 | 0.001448 | 0.004535 | 0.005382 | 0.002036 | 0.010189 | 0.007576 | 0.002394 | 0.003693 | 0.0087 | 0.002714 | 0.007689 | 0.025035 | 0.001762 | 0.017734 | 0.005198 | 0.001728 | 0.007464 | 0.004329 | 0.00292 | 0.003743 | 0.002774 | 0.010267 |
| 21.3472222 | 0.009721 | 0.001837 | 0.002887 | 0.003395 | 0.004422 | 0.005702 | 0.009331 | 0.001047 | 0.00509 | 0.017854 | 0.004056 | 0.00784 | 0.020977 | 0.001773 | 0.023895 | 0.007016 | 0.004175 | 0.007548 | 0.005445 | 0.002186 | 0.002392 | 0.002537 | 0.013324 |
| 21.3611111 | 0.010275 | 0.004401 | 0.002404 | 0.003275 | 0.007533 | 0.006367 | 0.012011 | 0.001226 | 0.006652 | 0.027698 | 0.003322 | 0.008709 | 0.019266 | 0.005315 | 0.028156 | 0.005316 | 0.00548 | 0.00777 | 0.010826 | 0.001681 | 0.003578 | 0.007128 | 0.016919 |
| 21.375 | 0.01975 | 0.007947 | 0.006336 | 0.00679 | 0.010202 | 0.012476 | 0.0164 | 0.003613 | 0.00921 | 0.036605 | 0.001476 | 0.012468 | 0.020655 | 0.015775 | 0.024018 | 0.00393 | 0.005382 | 0.005444 | 0.013116 | 0.001651 | 0.008391 | 0.015479 | 0.021402 |
| 21.3888889 | 0.028853 | 0.008512 | 0.012784 | 0.011698 | 0.010888 | 0.017747 | 0.021761 | 0.006139 | 0.011696 | 0.044409 | 0.000643 | 0.016412 | 0.021436 | 0.030111 | 0.017011 | 0.005589 | 0.009581 | 0.003165 | 0.013357 | 0.001501 | 0.013799 | 0.02454 | 0.026363 |
| 21.4027778 | 0.029546 | 0.00646 | 0.018279 | 0.015235 | 0.010425 | 0.018934 | 0.02594 | 0.007085 | 0.013733 | 0.047843 | 0.000658 | 0.018445 | 0.019405 | 0.03886 | 0.019069 | 0.010627 | 0.016436 | 0.004858 | 0.016605 | 0.002433 | 0.01882 | 0.0301 | 0.030938 |
| 21.4166667 | 0.022753 | 0.00525 | 0.020906 | 0.015839 | 0.010404 | 0.017883 | 0.028113 | 0.007787 | 0.01651 | 0.046555 | 0.001874 | 0.020377 | 0.015537 | 0.035557 | 0.025503 | 0.015822 | 0.020051 | 0.0086 | 0.018129 | 0.005373 | 0.026279 | 0.030524 | 0.033898 |
| 21.4305556 | 0.016106 | 0.005638 | 0.020005 | 0.014305 | 0.010839 | 0.017867 | 0.029854 | 0.009707 | 0.02121 | 0.04713 | 0.004637 | 0.023638 | 0.011704 | 0.021183 | 0.026501 | 0.01879 | 0.020188 | 0.010791 | 0.015432 | 0.008625 | 0.036698 | 0.030904 | 0.034074 |
| 21.4444444 | 0.013418 | 0.006329 | 0.017211 | 0.01214 | 0.01102 | 0.022363 | 0.031047 | 0.011949 | 0.026735 | 0.052395 | 0.00665 | 0.026499 | 0.009562 | 0.008388 | 0.022031 | 0.019133 | 0.018975 | 0.010693 | 0.010462 | 0.011005 | 0.047236 | 0.037805 | 0.031801 |
| 21.4583333 | 0.012582 | 0.006772 | 0.015097 | 0.009155 | 0.009844 | 0.030628 | 0.02993 | 0.012685 | 0.029108 | 0.060167 | 0.005493 | 0.028296 | 0.011087 | 0.005784 | 0.017714 | 0.016457 | 0.018991 | 0.009502 | 0.004935 | 0.013261 | 0.055078 | 0.052412 | 0.029856 |
| 21.4722222 | 0.011859 | 0.00824 | 0.014783 | 0.007374 | 0.007225 | 0.036729 | 0.027194 | 0.011953 | 0.025998 | 0.068085 | 0.002444 | 0.030693 | 0.017123 | 0.007337 | 0.019942 | 0.013447 | 0.022071 | 0.008551 | 0.001309 | 0.016672 | 0.058472 | 0.070623 | 0.031344 |
| 21.4861111 | 0.012726 | 0.012637 | 0.016553 | 0.010776 | 0.005917 | 0.037429 | 0.025862 | 0.012917 | 0.020099 | 0.074474 | 0.001596 | 0.031207 | 0.02527 | 0.016555 | 0.028729 | 0.012737 | 0.02586 | 0.008397 | 0.000465 | 0.02022 | 0.056016 | 0.084348 | 0.036281 |
| 21.5 | 0.015952 | 0.019349 | 0.01995 | 0.019479 | 0.007544 | 0.037423 | 0.027143 | 0.017719 | 0.015459 | 0.08046 | 0.004374 | 0.025948 | 0.031818 | 0.036763 | 0.036782 | 0.015264 | 0.026921 | 0.009088 | 0.001432 | 0.022587 | 0.050575 | 0.087768 | 0.04176 |
| 21.5138889 | 0.017528 | 0.023323 | 0.023128 | 0.026644 | 0.008797 | 0.039557 | 0.027188 | 0.022718 | 0.01201 | 0.080461 | 0.007277 | 0.015406 | 0.033225 | 0.050678 | 0.038505 | 0.020489 | 0.025264 | 0.009336 | 0.002961 | 0.024112 | 0.04674 | 0.083549 | 0.044888 |
| 21.5277778 | 0.013579 | 0.019846 | 0.025081 | 0.024786 | 0.007126 | 0.040325 | 0.021864 | 0.023804 | 0.007325 | 0.067013 | 0.006958 | 0.009555 | 0.02331 | 0.042955 | 0.031368 | 0.022281 | 0.01919 | 0.00709 | 0.003851 | 0.021811 | 0.040966 | 0.074284 | 0.041056 |
| 21.5416667 | 0.00698 | 0.011191 | 0.026562 | 0.016507 | 0.005095 | 0.041562 | 0.012841 | 0.022085 | 0.004702 | 0.049465 | 0.004395 | 0.017858 | 0.019016 | 0.024541 | 0.02088 | 0.01795 | 0.015363 | 0.006483 | 0.010549 | 0.016197 | 0.027136 | 0.053631 | 0.027684 |
| 21.5555556 | 0.002949 | 0.005549 | 0.025252 | 0.009603 | 0.00409 | 0.045262 | 0.00745 | 0.020474 | 0.008349 | 0.037005 | 0.002648 | 0.028996 | 0.036795 | 0.017751 | 0.019102 | 0.018668 | 0.025164 | 0.014233 | 0.0271 | 0.018336 | 0.019396 | 0.027591 | 0.018683 |
| 21.5694444 | 0.003126 | 0.005259 | 0.017985 | 0.00798 | 0.002566 | 0.043049 | 0.006292 | 0.020614 | 0.009685 | 0.024557 | 0.003401 | 0.026469 | 0.038961 | 0.019787 | 0.019852 | 0.022397 | 0.033821 | 0.021554 | 0.034243 | 0.025092 | 0.025853 | 0.013464 | 0.016507 |
| 21.5833333 | 0.007214 | 0.011005 | 0.011632 | 0.011312 | 0.001294 | 0.036275 | 0.007145 | 0.025011 | 0.006398 | 0.015899 | 0.007413 | 0.016679 | 0.027452 | 0.015963 | 0.023163 | 0.018054 | 0.024502 | 0.016431 | 0.025357 | 0.019651 | 0.023953 | 0.012898 | 0.015827 |
| 21.5972222 | 0.012333 | 0.023697 | 0.014553 | 0.016354 | 0.003055 | 0.034121 | 0.01356 | 0.035491 | 0.007831 | 0.019791 | 0.013603 | 0.015266 | 0.042353 | 0.017292 | 0.03793 | 0.019862 | 0.010596 | 0.00825 | 0.026184 | 0.008806 | 0.015238 | 0.025434 | 0.02825 |
| 21.6111111 | 0.015402 | 0.033393 | 0.025455 | 0.020241 | 0.009706 | 0.033065 | 0.019811 | 0.048328 | 0.011039 | 0.026601 | 0.017895 | 0.02282 | 0.059184 | 0.025466 | 0.044936 | 0.035683 | 0.008922 | 0.010606 | 0.040959 | 0.008285 | 0.024388 | 0.04776 | 0.043776 |
| 21.625 | 0.016837 | 0.034397 | 0.041434 | 0.020745 | 0.017939 | 0.031712 | 0.02026 | 0.056037 | 0.011386 | 0.02839 | 0.019724 | 0.033316 | 0.052777 | 0.022576 | 0.035287 | 0.049358 | 0.014513 | 0.019939 | 0.046976 | 0.015392 | 0.046942 | 0.06781 | 0.050043 |
| 21.6388889 | 0.016399 | 0.030385 | 0.058287 | 0.017683 | 0.020945 | 0.034961 | 0.018649 | 0.054644 | 0.010393 | 0.031342 | 0.02222 | 0.037223 | 0.033715 | 0.02039 | 0.021748 | 0.051386 | 0.01652 | 0.026771 | 0.040393 | 0.023141 | 0.058371 | 0.080344 | 0.050114 |
| 21.6527778 | 0.013227 | 0.026761 | 0.066708 | 0.015598 | 0.018383 | 0.039884 | 0.01588 | 0.045449 | 0.010544 | 0.034926 | 0.026504 | 0.030616 | 0.014831 | 0.028746 | 0.011533 | 0.044329 | 0.015447 | 0.028011 | 0.033463 | 0.028157 | 0.050697 | 0.083755 | 0.041792 |
| 21.6666667 | 0.008896 | 0.026903 | 0.059678 | 0.016587 | 0.014542 | 0.038271 | 0.010762 | 0.033642 | 0.015058 | 0.032695 | 0.030152 | 0.024021 | 0.00752 | 0.027329 | 0.006571 | 0.03374 | 0.013485 | 0.026324 | 0.033145 | 0.028688 | 0.036635 | 0.078082 | 0.029748 |
| 21.6805556 | 0.004801 | 0.027814 | 0.039145 | 0.01673 | 0.010946 | 0.027933 | 0.005866 | 0.026027 | 0.019141 | 0.02499 | 0.028769 | 0.022476 | 0.012917 | 0.015679 | 0.006101 | 0.022528 | 0.010891 | 0.025023 | 0.039689 | 0.0261 | 0.02449 | 0.065715 | 0.022781 |
| 21.6944444 | 0.003256 | 0.025552 | 0.019704 | 0.012392 | 0.007562 | 0.014079 | 0.004884 | 0.024499 | 0.015745 | 0.016409 | 0.021575 | 0.021241 | 0.021155 | 0.009698 | 0.009082 | 0.011113 | 0.009058 | 0.023195 | 0.046407 | 0.021905 | 0.0128 | 0.045884 | 0.021163 |
| 21.7083333 | 0.006721 | 0.020273 | 0.019466 | 0.005818 | 0.004248 | 0.004884 | 0.006831 | 0.025385 | 0.008073 | 0.009472 | 0.012111 | 0.016037 | 0.023381 | 0.008689 | 0.012826 | 0.004307 | 0.007753 | 0.018345 | 0.047923 | 0.016537 | 0.005546 | 0.022992 | 0.019868 |
| 21.7222222 | 0.011921 | 0.016809 | 0.035399 | 0.002268 | 0.001814 | 0.002673 | 0.008557 | 0.026294 | 0.00399 | 0.005263 | 0.00526 | 0.008544 | 0.01587 | 0.012636 | 0.013529 | 0.004215 | 0.006211 | 0.011811 | 0.0439 | 0.011888 | 0.005142 | 0.010614 | 0.016763 |
| 21.7361111 | 0.014188 | 0.017728 | 0.046096 | 0.001849 | 0.001542 | 0.002633 | 0.009395 | 0.028274 | 0.00657 | 0.006298 | 0.003845 | 0.005489 | 0.009868 | 0.023919 | 0.009765 | 0.004517 | 0.004874 | 0.007074 | 0.035486 | 0.009007 | 0.004014 | 0.008711 | 0.014333 |
| 21.75 | 0.012205 | 0.019315 | 0.042151 | 0.002956 | 0.002572 | 0.004806 | 0.009015 | 0.03054 | 0.011184 | 0.012059 | 0.005587 | 0.007209 | 0.018018 | 0.033528 | 0.004309 | 0.005429 | 0.00464 | 0.004905 | 0.028982 | 0.00636 | 0.001229 | 0.009759 | 0.014733 |
| 21.7638889 | 0.007079 | 0.017963 | 0.030451 | 0.004397 | 0.002559 | 0.00791 | 0.007203 | 0.02878 | 0.011251 | 0.016988 | 0.007386 | 0.006815 | 0.025735 | 0.032554 | 0.001449 | 0.009298 | 0.005177 | 0.003171 | 0.025602 | 0.003794 | 0.00113 | 0.015088 | 0.015547 |
| 21.7777778 | 0.002426 | 0.013456 | 0.017486 | 0.003723 | 0.002287 | 0.008 | 0.004641 | 0.020226 | 0.007165 | 0.017978 | 0.009653 | 0.00779 | 0.018359 | 0.022357 | 0.002897 | 0.009892 | 0.004758 | 0.001916 | 0.02042 | 0.003312 | 0.003446 | 0.019754 | 0.013759 |
| 21.7916667 | 0.000997 | 0.007638 | 0.010481 | 0.001902 | 0.004043 | 0.009346 | 0.00272 | 0.010406 | 0.003481 | 0.017653 | 0.013026 | 0.01634 | 0.007771 | 0.021453 | 0.007857 | 0.006313 | 0.002847 | 0.002894 | 0.014466 | 0.005739 | 0.005712 | 0.018757 | 0.009065 |
| 21.8055556 | 0.001483 | 0.003261 | 0.015711 | 0.000858 | 0.005569 | 0.014335 | 0.002205 | 0.008615 | 0.002428 | 0.019103 | 0.015989 | 0.021275 | 0.006689 | 0.034709 | 0.014099 | 0.005792 | 0.001728 | 0.004806 | 0.010251 | 0.008899 | 0.005872 | 0.012242 | 0.004988 |
| 21.8194444 | 0.001924 | 0.002247 | 0.026478 | 0.000845 | 0.004237 | 0.01932 | 0.001911 | 0.013255 | 0.004765 | 0.021315 | 0.015958 | 0.014332 | 0.008483 | 0.035146 | 0.01784 | 0.009188 | 0.00221 | 0.006433 | 0.007728 | 0.009566 | 0.00455 | 0.007166 | 0.005929 |
| 21.8333333 | 0.002078 | 0.00344 | 0.0282 | 0.003275 | 0.001969 | 0.022255 | 0.001402 | 0.016011 | 0.009298 | 0.02153 | 0.011872 | 0.005223 | 0.007159 | 0.018329 | 0.015963 | 0.010697 | 0.002283 | 0.008156 | 0.005245 | 0.007047 | 0.004622 | 0.010967 | 0.009622 |
| 21.8472222 | 0.001929 | 0.003875 | 0.017292 | 0.007906 | 0.002216 | 0.020719 | 0.002762 | 0.015736 | 0.012893 | 0.019214 | 0.00671 | 0.003617 | 0.005345 | 0.005718 | 0.009622 | 0.008076 | 0.002806 | 0.010269 | 0.003231 | 0.003717 | 0.00836 | 0.013361 | 0.012067 |
| 21.8611111 | 0.002412 | 0.003234 | 0.009329 | 0.010663 | 0.005101 | 0.015013 | 0.007782 | 0.015883 | 0.013579 | 0.015063 | 0.003441 | 0.009052 | 0.003376 | 0.006402 | 0.003582 | 0.003801 | 0.005103 | 0.012597 | 0.005092 | 0.002581 | 0.014386 | 0.008582 | 0.014371 |
| 21.875 | 0.004143 | 0.003383 | 0.014152 | 0.009422 | 0.0072 | 0.009777 | 0.012507 | 0.017604 | 0.010997 | 0.011888 | 0.002781 | 0.016036 | 0.001609 | 0.017243 | 0.001409 | 0.001594 | 0.007475 | 0.014415 | 0.009376 | 0.003906 | 0.019223 | 0.005994 | 0.017435 |
| 21.8888889 | 0.005913 | 0.005009 | 0.024453 | 0.006727 | 0.006531 | 0.009348 | 0.012538 | 0.019849 | 0.007302 | 0.015187 | 0.006263 | 0.020231 | 0.004716 | 0.025634 | 0.003579 | 0.003738 | 0.009086 | 0.016339 | 0.01082 | 0.006348 | 0.021532 | 0.007486 | 0.020174 |
| 21.9027778 | 0.007247 | 0.007181 | 0.035044 | 0.005281 | 0.004786 | 0.015487 | 0.01046 | 0.020837 | 0.008803 | 0.024202 | 0.013819 | 0.019541 | 0.01413 | 0.018994 | 0.008127 | 0.008406 | 0.010281 | 0.018975 | 0.007808 | 0.010395 | 0.023357 | 0.007497 | 0.021943 |
| 21.9166667 | 0.007726 | 0.008334 | 0.043553 | 0.005887 | 0.005034 | 0.024979 | 0.010557 | 0.019598 | 0.017018 | 0.032267 | 0.021096 | 0.01412 | 0.023115 | 0.009649 | 0.012242 | 0.011345 | 0.010324 | 0.020392 | 0.003254 | 0.015483 | 0.026996 | 0.010851 | 0.022413 |
| 21.9305556 | 0.006931 | 0.007817 | 0.043083 | 0.008474 | 0.008837 | 0.032352 | 0.013524 | 0.017445 | 0.024 | 0.036475 | 0.024229 | 0.008944 | 0.026687 | 0.011998 | 0.01549 | 0.012337 | 0.010163 | 0.019658 | 0.002035 | 0.019206 | 0.0332 | 0.023517 | 0.023079 |
| 21.9444444 | 0.006276 | 0.007312 | 0.031373 | 0.012069 | 0.013827 | 0.035041 | 0.017384 | 0.016289 | 0.025658 | 0.037869 | 0.023775 | 0.01119 | 0.026677 | 0.015174 | 0.019054 | 0.014195 | 0.013159 | 0.018603 | 0.006596 | 0.021642 | 0.03963 | 0.040317 | 0.025478 |
| 21.9583333 | 0.006391 | 0.008652 | 0.015794 | 0.014324 | 0.016023 | 0.033059 | 0.020153 | 0.017131 | 0.022973 | 0.040201 | 0.021652 | 0.020691 | 0.026678 | 0.012323 | 0.022115 | 0.016574 | 0.017414 | 0.017392 | 0.01472 | 0.02396 | 0.041528 | 0.054973 | 0.028764 |
| 21.9722222 | 0.005574 | 0.011984 | 0.005497 | 0.012978 | 0.014797 | 0.027026 | 0.021607 | 0.018424 | 0.015887 | 0.045329 | 0.018197 | 0.028564 | 0.028621 | 0.007919 | 0.022331 | 0.018036 | 0.016005 | 0.013933 | 0.021199 | 0.024558 | 0.037745 | 0.063998 | 0.031211 |
| 21.9861111 | 0.003855 | 0.017853 | 0.002606 | 0.010053 | 0.012562 | 0.018525 | 0.022066 | 0.017598 | 0.007341 | 0.048105 | 0.013038 | 0.029051 | 0.033037 | 0.00438 | 0.01964 | 0.019505 | 0.008611 | 0.008817 | 0.024332 | 0.023181 | 0.032624 | 0.064534 | 0.032349 |
| 22 | 0.004192 | 0.025165 | 0.003104 | 0.011531 | 0.012126 | 0.012544 | 0.022775 | 0.015708 | 0.002307 | 0.04502 | 0.007457 | 0.025818 | 0.039678 | 0.005743 | 0.017134 | 0.02248 | 0.003484 | 0.006531 | 0.026978 | 0.024174 | 0.030792 | 0.057119 | 0.034413 |
| 22.0138889 | 0.00706 | 0.030321 | 0.006261 | 0.019462 | 0.015314 | 0.015589 | 0.025763 | 0.015288 | 0.001648 | 0.04219 | 0.0045 | 0.024833 | 0.047158 | 0.017834 | 0.017524 | 0.02738 | 0.005874 | 0.008667 | 0.02743 | 0.029151 | 0.031938 | 0.049692 | 0.038048 |
| 22.0277778 | 0.009774 | 0.03223 | 0.012505 | 0.027884 | 0.021016 | 0.026701 | 0.030414 | 0.017062 | 0.002875 | 0.047567 | 0.005065 | 0.027106 | 0.053038 | 0.039829 | 0.021821 | 0.03392 | 0.012747 | 0.012198 | 0.025297 | 0.034238 | 0.034922 | 0.050938 | 0.041294 |
| 22.0416667 | 0.011247 | 0.033113 | 0.020807 | 0.031644 | 0.025994 | 0.041446 | 0.034074 | 0.02034 | 0.0062 | 0.061982 | 0.007419 | 0.029358 | 0.057242 | 0.059661 | 0.029955 | 0.040417 | 0.019841 | 0.01578 | 0.027344 | 0.038027 | 0.04189 | 0.061505 | 0.047252 |
| 22.0555556 | 0.01235 | 0.035353 | 0.03066 | 0.033751 | 0.028603 | 0.061502 | 0.035461 | 0.021228 | 0.014562 | 0.075493 | 0.010086 | 0.026689 | 0.060458 | 0.06266 | 0.038586 | 0.042342 | 0.023848 | 0.017771 | 0.03506 | 0.04045 | 0.050096 | 0.07484 | 0.058562 |
| 22.0694444 | 0.014539 | 0.038837 | 0.037538 | 0.035983 | 0.029313 | 0.080309 | 0.033247 | 0.01503 | 0.025941 | 0.074726 | 0.011543 | 0.016817 | 0.059697 | 0.043649 | 0.041311 | 0.034956 | 0.021154 | 0.013694 | 0.041144 | 0.035938 | 0.050585 | 0.079163 | 0.065632 |
| 22.0833333 | 0.018132 | 0.035658 | 0.036791 | 0.032911 | 0.026843 | 0.076169 | 0.025638 | 0.006309 | 0.03423 | 0.062744 | 0.009974 | 0.008091 | 0.053275 | 0.022588 | 0.035299 | 0.02145 | 0.014868 | 0.006651 | 0.040213 | 0.022566 | 0.042753 | 0.068152 | 0.06092 |
| 22.0972222 | 0.019026 | 0.022018 | 0.030911 | 0.022418 | 0.019735 | 0.046977 | 0.014627 | 0.004553 | 0.035041 | 0.048737 | 0.005816 | 0.005387 | 0.043677 | 0.026318 | 0.025049 | 0.011098 | 0.012606 | 0.00453 | 0.034746 | 0.008995 | 0.033413 | 0.051575 | 0.050141 |
| 22.1111111 | 0.013966 | 0.012308 | 0.024763 | 0.011011 | 0.011548 | 0.018269 | 0.005565 | 0.009228 | 0.028331 | 0.034293 | 0.003545 | 0.00355 | 0.032202 | 0.042413 | 0.016745 | 0.006234 | 0.013505 | 0.006344 | 0.02864 | 0.00217 | 0.023888 | 0.039457 | 0.038449 |
| 22.125 | 0.006896 | 0.01036 | 0.020686 | 0.005031 | 0.005956 | 0.004589 | 0.001665 | 0.013241 | 0.020371 | 0.024116 | 0.005216 | 0.002439 | 0.01809 | 0.043692 | 0.012081 | 0.003122 | 0.011183 | 0.007745 | 0.022889 | 0.000404 | 0.013779 | 0.030758 | 0.0263 |
| 22.1388889 | 0.002461 | 0.010956 | 0.019161 | 0.003727 | 0.002707 | 0.001329 | 0.001122 | 0.014348 | 0.015613 | 0.020789 | 0.006369 | 0.002049 | 0.007725 | 0.031594 | 0.00993 | 0.000927 | 0.005962 | 0.007399 | 0.018727 | 0.000433 | 0.005938 | 0.0236 | 0.014886 |
| 22.1527778 | 0.001097 | 0.017941 | 0.018935 | 0.004786 | 0.001014 | 0.001224 | 0.000875 | 0.015076 | 0.012155 | 0.020401 | 0.004857 | 0.0015 | 0.005591 | 0.017654 | 0.011183 | 0.001575 | 0.00217 | 0.005203 | 0.018839 | 0.002621 | 0.002173 | 0.018179 | 0.00662 |
| 22.1666667 | 0.001627 | 0.023922 | 0.018808 | 0.007758 | 0.00064 | 0.001535 | 0.000624 | 0.017056 | 0.00763 | 0.018888 | 0.002829 | 0.003936 | 0.005562 | 0.010169 | 0.017401 | 0.006484 | 0.003715 | 0.005507 | 0.024877 | 0.008613 | 0.003667 | 0.015446 | 0.003086 |
| 22.1805556 | 0.001787 | 0.02348 | 0.018462 | 0.009242 | 0.001152 | 0.003386 | 0.001395 | 0.018865 | 0.003376 | 0.015035 | 0.001703 | 0.010337 | 0.006988 | 0.015867 | 0.024519 | 0.013203 | 0.012862 | 0.011084 | 0.033058 | 0.016351 | 0.010498 | 0.014773 | 0.002264 |
| 22.1944444 | 0.001319 | 0.021312 | 0.017033 | 0.006244 | 0.002143 | 0.006412 | 0.003172 | 0.018654 | 0.002307 | 0.010616 | 0.001167 | 0.015976 | 0.009564 | 0.027139 | 0.025319 | 0.016236 | 0.023976 | 0.015096 | 0.035915 | 0.02098 | 0.016634 | 0.012318 | 0.002307 |
| 22.2083333 | 0.001996 | 0.016475 | 0.014521 | 0.002084 | 0.003414 | 0.008306 | 0.005076 | 0.016835 | 0.003748 | 0.007598 | 0.000695 | 0.016847 | 0.007828 | 0.029438 | 0.019381 | 0.013716 | 0.029047 | 0.012534 | 0.030581 | 0.020189 | 0.017005 | 0.007346 | 0.004594 |
| 22.2222222 | 0.003155 | 0.008965 | 0.012134 | 0.000486 | 0.00454 | 0.007933 | 0.006545 | 0.014903 | 0.005612 | 0.006628 | 0.00049 | 0.015478 | 0.004029 | 0.021559 | 0.01356 | 0.009486 | 0.026578 | 0.007089 | 0.022318 | 0.015635 | 0.014861 | 0.003359 | 0.007521 |
| 22.2361111 | 0.003552 | 0.00359 | 0.009847 | 0.001162 | 0.004845 | 0.005444 | 0.007545 | 0.012657 | 0.006973 | 0.006397 | 0.000904 | 0.0165 | 0.003657 | 0.012649 | 0.012521 | 0.007954 | 0.020159 | 0.003967 | 0.017955 | 0.010225 | 0.015296 | 0.001812 | 0.006568 |
| 22.25 | 0.002786 | 0.002765 | 0.006542 | 0.003093 | 0.004235 | 0.003495 | 0.008023 | 0.008787 | 0.006929 | 0.005395 | 0.002196 | 0.019237 | 0.005733 | 0.007496 | 0.013515 | 0.008804 | 0.013986 | 0.004644 | 0.017728 | 0.005735 | 0.017303 | 0.002882 | 0.004307 |
| 22.2638889 | 0.001342 | 0.004316 | 0.003481 | 0.004947 | 0.003604 | 0.005536 | 0.007947 | 0.00429 | 0.005863 | 0.004469 | 0.00287 | 0.018751 | 0.008065 | 0.004459 | 0.011904 | 0.008915 | 0.009985 | 0.007321 | 0.017841 | 0.002946 | 0.016285 | 0.006363 | 0.00732 |
| 22.2777778 | 0.000498 | 0.006175 | 0.00346 | 0.005482 | 0.003627 | 0.009436 | 0.007398 | 0.001694 | 0.00536 | 0.004393 | 0.002231 | 0.014102 | 0.011482 | 0.003902 | 0.007772 | 0.007789 | 0.008182 | 0.010553 | 0.017487 | 0.002326 | 0.012053 | 0.008502 | 0.014532 |
| 22.2916667 | 0.000354 | 0.008249 | 0.004909 | 0.004546 | 0.003761 | 0.010577 | 0.006772 | 0.001346 | 0.006215 | 0.003745 | 0.002475 | 0.009185 | 0.0158 | 0.00631 | 0.004181 | 0.007042 | 0.007211 | 0.013519 | 0.018484 | 0.003252 | 0.008103 | 0.006991 | 0.019638 |
| 22.3055556 | 0.000422 | 0.010354 | 0.004394 | 0.002502 | 0.002812 | 0.007969 | 0.006502 | 0.002143 | 0.007657 | 0.002113 | 0.003796 | 0.00662 | 0.019634 | 0.008095 | 0.002207 | 0.006311 | 0.005051 | 0.014386 | 0.019761 | 0.003867 | 0.005878 | 0.003761 | 0.017717 |
| 22.3194444 | 0.001536 | 0.012516 | 0.002283 | 0.000804 | 0.00171 | 0.004044 | 0.005359 | 0.00374 | 0.008481 | 0.000997 | 0.00367 | 0.004437 | 0.022816 | 0.007217 | 0.00125 | 0.004508 | 0.004517 | 0.012627 | 0.016929 | 0.002735 | 0.003834 | 0.001843 | 0.010438 |
| 22.3333333 | 0.005367 | 0.014431 | 0.001031 | 0.000387 | 0.002312 | 0.001549 | 0.003851 | 0.00565 | 0.00848 | 0.000641 | 0.002312 | 0.004602 | 0.025847 | 0.004861 | 0.001669 | 0.004735 | 0.008864 | 0.013976 | 0.010043 | 0.002009 | 0.002389 | 0.001596 | 0.007779 |
| 22.3472222 | 0.010386 | 0.013663 | 0.001252 | 0.000819 | 0.003542 | 0.000848 | 0.005482 | 0.006645 | 0.008296 | 0.001485 | 0.002393 | 0.012287 | 0.028399 | 0.003204 | 0.003849 | 0.009339 | 0.012988 | 0.027361 | 0.004101 | 0.004513 | 0.003901 | 0.003087 | 0.014761 |
| 22.3611111 | 0.012265 | 0.008704 | 0.001245 | 0.002394 | 0.004499 | 0.000752 | 0.009866 | 0.005994 | 0.008699 | 0.005709 | 0.003996 | 0.023613 | 0.029184 | 0.004553 | 0.007257 | 0.014723 | 0.012591 | 0.045115 | 0.002303 | 0.007982 | 0.007224 | 0.009101 | 0.024512 |
| 22.375 | 0.00946 | 0.004749 | 0.000921 | 0.005086 | 0.005954 | 0.001263 | 0.013879 | 0.003694 | 0.010091 | 0.012878 | 0.005289 | 0.031991 | 0.027615 | 0.009927 | 0.011302 | 0.017514 | 0.009814 | 0.043047 | 0.003915 | 0.009276 | 0.010013 | 0.018437 | 0.030103 |
| 22.3888889 | 0.005413 | 0.005966 | 0.00198 | 0.007589 | 0.00785 | 0.00349 | 0.016702 | 0.002206 | 0.012389 | 0.020756 | 0.005705 | 0.03591 | 0.024833 | 0.018999 | 0.015311 | 0.017073 | 0.007881 | 0.021303 | 0.005825 | 0.008238 | 0.01163 | 0.026382 | 0.030358 |
| 22.4027778 | 0.006235 | 0.007946 | 0.004847 | 0.009279 | 0.009838 | 0.007687 | 0.019173 | 0.004055 | 0.015151 | 0.029255 | 0.006006 | 0.036035 | 0.021915 | 0.02923 | 0.018419 | 0.013934 | 0.007793 | 0.00486 | 0.006011 | 0.005777 | 0.011396 | 0.028366 | 0.026545 |
| 22.4166667 | 0.013092 | 0.009058 | 0.008929 | 0.011063 | 0.011911 | 0.0133 | 0.022248 | 0.006832 | 0.017651 | 0.038574 | 0.006962 | 0.033641 | 0.018482 | 0.035088 | 0.020426 | 0.009642 | 0.00828 | 0.001161 | 0.004705 | 0.002936 | 0.008937 | 0.022923 | 0.021233 |
| 22.4305556 | 0.020411 | 0.011298 | 0.013268 | 0.013749 | 0.013989 | 0.018964 | 0.025486 | 0.008098 | 0.019237 | 0.047453 | 0.008268 | 0.031084 | 0.014504 | 0.033794 | 0.021988 | 0.006397 | 0.00908 | 0.001109 | 0.005474 | 0.00104 | 0.007086 | 0.015757 | 0.018228 |
| 22.4444444 | 0.024566 | 0.013783 | 0.016479 | 0.016308 | 0.015187 | 0.023498 | 0.027222 | 0.008279 | 0.019594 | 0.054847 | 0.008532 | 0.030392 | 0.011317 | 0.028413 | 0.023334 | 0.005773 | 0.011557 | 0.001731 | 0.014174 | 0.000846 | 0.011243 | 0.012447 | 0.020129 |
| 22.4583333 | 0.025396 | 0.015261 | 0.017401 | 0.016812 | 0.013952 | 0.026974 | 0.025927 | 0.007035 | 0.018466 | 0.061713 | 0.006464 | 0.030882 | 0.010228 | 0.022715 | 0.024351 | 0.007124 | 0.015506 | 0.004624 | 0.024381 | 0.001151 | 0.021253 | 0.011376 | 0.025353 |
| 22.4722222 | 0.022517 | 0.016568 | 0.01739 | 0.015969 | 0.010588 | 0.030952 | 0.02328 | 0.004992 | 0.016949 | 0.070336 | 0.003499 | 0.031511 | 0.011868 | 0.018892 | 0.025932 | 0.00953 | 0.019045 | 0.008617 | 0.024209 | 0.00112 | 0.030922 | 0.010395 | 0.030781 |
| 22.4861111 | 0.0186 | 0.021226 | 0.019333 | 0.017689 | 0.00888 | 0.037133 | 0.024696 | 0.006903 | 0.017525 | 0.08218 | 0.003519 | 0.031257 | 0.01629 | 0.016812 | 0.028817 | 0.013247 | 0.021478 | 0.013075 | 0.023168 | 0.00225 | 0.038375 | 0.011959 | 0.03515 |
| 22.5 | 0.017958 | 0.029573 | 0.022169 | 0.022526 | 0.011573 | 0.043296 | 0.030087 | 0.014986 | 0.019524 | 0.09171 | 0.007388 | 0.029575 | 0.021596 | 0.013922 | 0.031285 | 0.01833 | 0.023402 | 0.017962 | 0.033566 | 0.004884 | 0.044894 | 0.016759 | 0.037765 |
| 22.5138889 | 0.019324 | 0.036019 | 0.022588 | 0.025777 | 0.01539 | 0.045231 | 0.032808 | 0.023139 | 0.019507 | 0.089329 | 0.011056 | 0.027142 | 0.023867 | 0.008445 | 0.030451 | 0.023667 | 0.024021 | 0.020393 | 0.040345 | 0.006715 | 0.048506 | 0.021492 | 0.036694 |
| 22.5277778 | 0.018741 | 0.035592 | 0.022184 | 0.025484 | 0.016108 | 0.044037 | 0.029644 | 0.026095 | 0.016672 | 0.075436 | 0.011526 | 0.019765 | 0.017891 | 0.006721 | 0.022709 | 0.023406 | 0.017567 | 0.017332 | 0.028192 | 0.006663 | 0.044371 | 0.020931 | 0.027945 |

| | | | | | | | | | | | | | | | | | | | | | | | |
|---|---|---|---|---|---|---|---|---|---|---|---|---|---|---|---|---|---|---|---|---|---|---|---|
| 22.5416667 | 0.014677 | 0.026215 | 0.024474 | 0.022893 | 0.013461 | 0.04696 | 0.020696 | 0.024827 | 0.012047 | 0.057581 | 0.010979 | 0.014308 | 0.01744 | 0.016821 | 0.015956 | 0.018513 | 0.015496 | 0.012484 | 0.016675 | 0.013061 | 0.02917 | 0.017492 | 0.019576 |
| 22.5555556 | 0.008832 | 0.013047 | 0.024429 | 0.017714 | 0.008901 | 0.052926 | 0.009605 | 0.021375 | 0.006315 | 0.041336 | 0.010592 | 0.022143 | 0.033247 | 0.030502 | 0.021119 | 0.020841 | 0.032527 | 0.014764 | 0.02392 | 0.03002 | 0.018102 | 0.021319 | 0.024146 |
| 22.5694444 | 0.00671 | 0.006299 | 0.016527 | 0.013926 | 0.004999 | 0.051711 | 0.004154 | 0.019883 | 0.002341 | 0.03161 | 0.008716 | 0.025632 | 0.033908 | 0.030872 | 0.021577 | 0.022559 | 0.042803 | 0.019602 | 0.027307 | 0.038788 | 0.019243 | 0.022369 | 0.023867 |
| 22.5833333 | 0.013526 | 0.012528 | 0.009888 | 0.01999 | 0.004041 | 0.043431 | 0.008956 | 0.025866 | 0.004567 | 0.034502 | 0.008176 | 0.017062 | 0.02345 | 0.018031 | 0.018981 | 0.016049 | 0.028387 | 0.014222 | 0.019518 | 0.027633 | 0.015643 | 0.011301 | 0.019645 |
| 22.5972222 | 0.023702 | 0.03024 | 0.013801 | 0.032783 | 0.005573 | 0.033052 | 0.019436 | 0.037132 | 0.008642 | 0.038917 | 0.011652 | 0.015656 | 0.034725 | 0.009293 | 0.033983 | 0.018415 | 0.011435 | 0.007098 | 0.023209 | 0.012497 | 0.013689 | 0.011675 | 0.036106 |
| 22.6111111 | 0.022932 | 0.043029 | 0.024201 | 0.038178 | 0.009145 | 0.023834 | 0.026665 | 0.045883 | 0.006376 | 0.029412 | 0.016153 | 0.022422 | 0.051213 | 0.010374 | 0.048271 | 0.033115 | 0.009272 | 0.011855 | 0.033854 | 0.009748 | 0.030125 | 0.041222 | 0.055173 |
| 22.625 | 0.012548 | 0.039814 | 0.035327 | 0.03238 | 0.015509 | 0.024393 | 0.027702 | 0.050201 | 0.002182 | 0.021455 | 0.02057 | 0.027692 | 0.052228 | 0.011784 | 0.047076 | 0.04584 | 0.017813 | 0.021242 | 0.039921 | 0.017815 | 0.05638 | 0.071241 | 0.060907 |
| 22.6388889 | 0.00753 | 0.029383 | 0.043932 | 0.023716 | 0.023164 | 0.037461 | 0.024751 | 0.051455 | 0.002758 | 0.029992 | 0.026055 | 0.029237 | 0.03838 | 0.010213 | 0.037076 | 0.048323 | 0.026506 | 0.026587 | 0.04289 | 0.028176 | 0.073082 | 0.061479 | 0.059804 |
| 22.6527778 | 0.008554 | 0.021541 | 0.046992 | 0.018558 | 0.028301 | 0.051105 | 0.019856 | 0.048605 | 0.005345 | 0.047264 | 0.031337 | 0.025798 | 0.0188 | 0.008856 | 0.021429 | 0.042768 | 0.027267 | 0.02874 | 0.042529 | 0.03583 | 0.071484 | 0.044558 | 0.050685 |
| 22.6666667 | 0.008637 | 0.021499 | 0.043936 | 0.017566 | 0.027695 | 0.053117 | 0.012975 | 0.042808 | 0.005496 | 0.059071 | 0.034721 | 0.019047 | 0.008433 | 0.008752 | 0.008209 | 0.033551 | 0.019321 | 0.028743 | 0.042 | 0.038795 | 0.060884 | 0.064491 | 0.036015 |
| 22.6805556 | 0.013508 | 0.030625 | 0.035653 | 0.019557 | 0.021947 | 0.042901 | 0.005813 | 0.03791 | 0.00628 | 0.053686 | 0.03344 | 0.013375 | 0.01232 | 0.009368 | 0.003316 | 0.022113 | 0.010464 | 0.025947 | 0.042924 | 0.037884 | 0.049364 | 0.090423 | 0.022843 |
| 22.6944444 | 0.022674 | 0.039173 | 0.02539 | 0.022182 | 0.014574 | 0.025165 | 0.00196 | 0.036043 | 0.012336 | 0.03502 | 0.02568 | 0.011476 | 0.018953 | 0.008994 | 0.002392 | 0.010642 | 0.005565 | 0.020244 | 0.042417 | 0.034155 | 0.034006 | 0.095751 | 0.014114 |
| 22.7083333 | 0.022426 | 0.036995 | 0.0166 | 0.020731 | 0.007509 | 0.010732 | 0.001834 | 0.037333 | 0.018405 | 0.016147 | 0.014873 | 0.01154 | 0.017788 | 0.007578 | 0.002437 | 0.003869 | 0.002743 | 0.013202 | 0.039569 | 0.027552 | 0.018293 | 0.087437 | 0.007919 |
| 22.7222222 | 0.013088 | 0.0275 | 0.009581 | 0.013728 | 0.002854 | 0.007539 | 0.004947 | 0.040944 | 0.017601 | 0.006774 | 0.006018 | 0.009502 | 0.012244 | 0.010299 | 0.004408 | 0.003742 | 0.000993 | 0.007348 | 0.032942 | 0.019164 | 0.015507 | 0.074782 | 0.003328 |
| 22.7361111 | 0.007277 | 0.015339 | 0.005209 | 0.006611 | 0.002292 | 0.007262 | 0.009422 | 0.042811 | 0.010812 | 0.00949 | 0.00156 | 0.005681 | 0.012604 | 0.016172 | 0.006287 | 0.005192 | 0.000656 | 0.004041 | 0.025385 | 0.010928 | 0.025472 | 0.065079 | 0.001952 |
| 22.75 | 0.011857 | 0.006085 | 0.006643 | 0.004467 | 0.002853 | 0.004169 | 0.010371 | 0.038743 | 0.00486 | 0.015066 | 0.000414 | 0.005513 | 0.019719 | 0.018652 | 0.005241 | 0.004994 | 0.001159 | 0.002288 | 0.025559 | 0.005306 | 0.0341 | 0.057549 | 0.003143 |
| 22.7638889 | 0.021844 | 0.003566 | 0.012936 | 0.005335 | 0.002077 | 0.00251 | 0.007119 | 0.030181 | 0.005368 | 0.015918 | 0.00092 | 0.010726 | 0.021934 | 0.016251 | 0.005417 | 0.003749 | 0.002004 | 0.001583 | 0.028499 | 0.004178 | 0.034636 | 0.046858 | 0.00481 |
| 22.7777778 | 0.026565 | 0.003643 | 0.017796 | 0.0044 | 0.001782 | 0.005116 | 0.004026 | 0.021395 | 0.010324 | 0.013053 | 0.002751 | 0.0149 | 0.014162 | 0.010506 | 0.009711 | 0.002146 | 0.002779 | 0.001748 | 0.021151 | 0.005109 | 0.029638 | 0.028841 | 0.006231 |
| 22.7916667 | 0.020888 | 0.004097 | 0.019019 | 0.002026 | 0.002892 | 0.011226 | 0.005042 | 0.014499 | 0.014645 | 0.010001 | 0.005414 | 0.012608 | 0.006388 | 0.008322 | 0.009879 | 0.001601 | 0.003103 | 0.00196 | 0.009777 | 0.004181 | 0.023949 | 0.015497 | 0.008239 |
| 22.8055556 | 0.010926 | 0.004959 | 0.018565 | 0.000778 | 0.003364 | 0.018088 | 0.00917 | 0.009707 | 0.014518 | 0.009987 | 0.008753 | 0.006764 | 0.006393 | 0.011175 | 0.008936 | 0.00247 | 0.002909 | 0.002096 | 0.007457 | 0.002617 | 0.019943 | 0.022477 | 0.011804 |
| 22.8194444 | 0.007004 | 0.00856 | 0.016828 | 0.001145 | 0.002247 | 0.024225 | 0.010488 | 0.006399 | 0.009349 | 0.013572 | 0.011713 | 0.004397 | 0.007282 | 0.009929 | 0.016221 | 0.004031 | 0.003153 | 0.001956 | 0.009967 | 0.003492 | 0.01703 | 0.038905 | 0.015583 |
| 22.8333333 | 0.009642 | 0.012316 | 0.014021 | 0.003019 | 0.001616 | 0.029293 | 0.008477 | 0.004402 | 0.004366 | 0.019106 | 0.012678 | 0.008875 | 0.004913 | 0.00559 | 0.023475 | 0.006108 | 0.00517 | 0.002261 | 0.011182 | 0.004718 | 0.01601 | 0.049086 | 0.017599 |
| 22.8472222 | 0.013458 | 0.012441 | 0.011476 | 0.003762 | 0.002494 | 0.03281 | 0.006934 | 0.006215 | 0.00329 | 0.022979 | 0.011195 | 0.016646 | 0.002874 | 0.006156 | 0.023611 | 0.007837 | 0.008451 | 0.004392 | 0.01198 | 0.003811 | 0.01758 | 0.048535 | 0.01739 |
| 22.8611111 | 0.015703 | 0.010535 | 0.010175 | 0.002468 | 0.00303 | 0.030568 | 0.005864 | 0.011638 | 0.003231 | 0.021805 | 0.00826 | 0.020139 | 0.002897 | 0.012413 | 0.019585 | 0.007922 | 0.011356 | 0.007575 | 0.012164 | 0.003544 | 0.019717 | 0.033392 | 0.015413 |
| 22.875 | 0.013821 | 0.007843 | 0.010095 | 0.001452 | 0.003624 | 0.018343 | 0.005098 | 0.016666 | 0.003876 | 0.016125 | 0.006867 | 0.015907 | 0.005306 | 0.022452 | 0.013821 | 0.007199 | 0.013818 | 0.010066 | 0.012477 | 0.00533 | 0.021436 | 0.016464 | 0.012932 |
| 22.8888889 | 0.008188 | 0.006824 | 0.010892 | 0.002108 | 0.005869 | 0.006778 | 0.006107 | 0.020047 | 0.006965 | 0.010603 | 0.008762 | 0.008821 | 0.009233 | 0.028361 | 0.008397 | 0.007966 | 0.01692 | 0.011185 | 0.014174 | 0.00741 | 0.024486 | 0.015246 | 0.011516 |
| 22.9027778 | 0.005722 | 0.009763 | 0.012141 | 0.005204 | 0.008449 | 0.004296 | 0.009057 | 0.021274 | 0.008795 | 0.009115 | 0.012725 | 0.005553 | 0.013099 | 0.021654 | 0.006149 | 0.011064 | 0.020883 | 0.011272 | 0.01514 | 0.010128 | 0.028509 | 0.021029 | 0.011708 |
| 22.9166667 | 0.009427 | 0.014238 | 0.013406 | 0.008876 | 0.009501 | 0.005789 | 0.013465 | 0.020082 | 0.006647 | 0.013328 | 0.016973 | 0.008443 | 0.016994 | 0.011598 | 0.008381 | 0.015357 | 0.025194 | 0.011125 | 0.013269 | 0.014438 | 0.030538 | 0.021436 | 0.013649 |
| 22.9305556 | 0.015229 | 0.015983 | 0.014616 | 0.011297 | 0.009673 | 0.012724 | 0.019137 | 0.01914 | 0.003342 | 0.023069 | 0.019872 | 0.015688 | 0.022189 | 0.010125 | 0.013369 | 0.018903 | 0.030863 | 0.010881 | 0.012183 | 0.019309 | 0.03046 | 0.016926 | 0.017425 |
| 22.9444444 | 0.01853 | 0.015342 | 0.015991 | 0.012503 | 0.012429 | 0.023169 | 0.024189 | 0.019941 | 0.003388 | 0.034643 | 0.019552 | 0.023328 | 0.028032 | 0.011241 | 0.01772 | 0.019612 | 0.036429 | 0.010125 | 0.020481 | 0.022487 | 0.031584 | 0.009246 | 0.022098 |
| 22.9583333 | 0.01587 | 0.015309 | 0.01697 | 0.012138 | 0.017485 | 0.026456 | 0.025939 | 0.019061 | 0.008649 | 0.042225 | 0.015222 | 0.02631 | 0.031354 | 0.008058 | 0.01837 | 0.016562 | 0.034705 | 0.008623 | 0.040319 | 0.021943 | 0.034329 | 0.003968 | 0.025102 |
| 22.9722222 | 0.008996 | 0.015808 | 0.015271 | 0.010444 | 0.018637 | 0.021202 | 0.025456 | 0.012776 | 0.014241 | 0.04202 | 0.008892 | 0.022729 | 0.029258 | 0.004129 | 0.01483 | 0.011101 | 0.024945 | 0.006248 | 0.054005 | 0.0184 | 0.033686 | 0.005624 | 0.024281 |
| 22.9861111 | 0.003396 | 0.016157 | 0.010176 | 0.009597 | 0.01497 | 0.016208 | 0.026231 | 0.005137 | 0.015068 | 0.038173 | 0.003845 | 0.017457 | 0.023581 | 0.00319 | 0.010249 | 0.00656 | 0.015526 | 0.003985 | 0.050839 | 0.015616 | 0.027041 | 0.011495 | 0.02197 |
| 23 | 0.00257 | 0.018975 | 0.005937 | 0.012955 | 0.013705 | 0.018932 | 0.028762 | 0.003375 | 0.014328 | 0.039813 | 0.002432 | 0.015757 | 0.021741 | 0.008095 | 0.009059 | 0.006029 | 0.011677 | 0.003476 | 0.049513 | 0.015802 | 0.020898 | 0.019967 | 0.023439 |
| 23.0138889 | 0.00552 | 0.023988 | 0.007099 | 0.020365 | 0.017141 | 0.028037 | 0.030862 | 0.00614 | 0.015895 | 0.04842 | 0.005054 | 0.016767 | 0.026918 | 0.018058 | 0.012734 | 0.009059 | 0.011622 | 0.00436 | 0.055469 | 0.017254 | 0.02193 | 0.027524 | 0.028837 |
| 23.0277778 | 0.009143 | 0.027449 | 0.012945 | 0.026665 | 0.02003 | 0.037206 | 0.030772 | 0.008726 | 0.018508 | 0.054635 | 0.009413 | 0.016034 | 0.030744 | 0.02526 | 0.017011 | 0.011555 | 0.01081 | 0.004773 | 0.051312 | 0.01758 | 0.025925 | 0.027943 | 0.032443 |
| 23.0416667 | 0.015348 | 0.03309 | 0.020783 | 0.030528 | 0.022823 | 0.046652 | 0.0301 | 0.01383 | 0.02147 | 0.055966 | 0.013973 | 0.011364 | 0.028555 | 0.02509 | 0.015776 | 0.010567 | 0.007169 | 0.00387 | 0.036628 | 0.015927 | 0.025222 | 0.024061 | 0.032834 |
| 23.0555556 | 0.028168 | 0.04524 | 0.029769 | 0.0363 | 0.030962 | 0.061896 | 0.031583 | 0.023618 | 0.025824 | 0.063611 | 0.020787 | 0.005941 | 0.028925 | 0.021837 | 0.010786 | 0.007226 | 0.003112 | 0.002626 | 0.024825 | 0.012197 | 0.021124 | 0.023057 | 0.034689 |
| 23.0694444 | 0.04361 | 0.053276 | 0.035587 | 0.041886 | 0.038789 | 0.078036 | 0.030752 | 0.029245 | 0.03036 | 0.076541 | 0.030103 | 0.003404 | 0.03759 | 0.020805 | 0.009349 | 0.004305 | 0.003054 | 0.002651 | 0.019583 | 0.00664 | 0.021459 | 0.024133 | 0.040386 |
| 23.0833333 | 0.051316 | 0.044121 | 0.03292 | 0.039764 | 0.035636 | 0.07615 | 0.022161 | 0.022983 | 0.032812 | 0.081326 | 0.03469 | 0.003793 | 0.047674 | 0.018176 | 0.012251 | 0.003521 | 0.00763 | 0.004807 | 0.016494 | 0.002164 | 0.026624 | 0.023707 | 0.04887 |
| 23.0972222 | 0.044457 | 0.024175 | 0.024419 | 0.028574 | 0.023679 | 0.049319 | 0.010223 | 0.0111 | 0.032367 | 0.071046 | 0.026896 | 0.008335 | 0.049407 | 0.010679 | 0.014944 | 0.004872 | 0.012194 | 0.008771 | 0.01297 | 0.000986 | 0.028264 | 0.022946 | 0.05533 |
| 23.1111111 | 0.028419 | 0.015028 | 0.016096 | 0.014678 | 0.012604 | 0.021082 | 0.003985 | 0.004666 | 0.028578 | 0.049942 | 0.014149 | 0.016403 | 0.040779 | 0.004253 | 0.015984 | 0.007303 | 0.01363 | 0.012706 | 0.00804 | 0.001644 | 0.022819 | 0.02336 | 0.052622 |
| 23.125 | 0.016686 | 0.016792 | 0.011273 | 0.005998 | 0.007858 | 0.008135 | 0.006045 | 0.006808 | 0.022105 | 0.033054 | 0.009635 | 0.022378 | 0.026191 | 0.005255 | 0.016702 | 0.009233 | 0.012781 | 0.015021 | 0.003912 | 0.00404 | 0.014763 | 0.024661 | 0.04313 |
| 23.1388889 | 0.010832 | 0.012729 | 0.00936 | 0.003627 | 0.007285 | 0.003007 | 0.009566 | 0.010361 | 0.01536 | 0.026713 | 0.011126 | 0.02333 | 0.012125 | 0.013656 | 0.017526 | 0.010166 | 0.010808 | 0.016406 | 0.006508 | 0.008227 | 0.00842 | 0.026362 | 0.035058 |
| 23.1527778 | 0.006143 | 0.010747 | 0.007364 | 0.002739 | 0.005892 | 0.000836 | 0.007754 | 0.008792 | 0.009883 | 0.023963 | 0.011033 | 0.020263 | 0.003756 | 0.018326 | 0.01773 | 0.011329 | 0.008203 | 0.017743 | 0.013998 | 0.012063 | 0.004014 | 0.028001 | 0.030004 |
| 23.1666667 | 0.004968 | 0.019443 | 0.004335 | 0.002365 | 0.003769 | 0.000703 | 0.005347 | 0.004717 | 0.005736 | 0.019787 | 0.009117 | 0.014178 | 0.002061 | 0.012113 | 0.016503 | 0.012493 | 0.005332 | 0.018247 | 0.016872 | 0.012694 | 0.002237 | 0.027828 | 0.026612 |
| 23.1805556 | 0.005478 | 0.028729 | 0.00175 | 0.00468 | 0.003328 | 0.001013 | 0.00843 | 0.002094 | 0.00287 | 0.015171 | 0.006725 | 0.006931 | 0.004437 | 0.006268 | 0.013866 | 0.011489 | 0.003255 | 0.016767 | 0.012512 | 0.00912 | 0.004592 | 0.022629 | 0.023529 |
| 23.1944444 | 0.003754 | 0.031939 | 0.001161 | 0.007775 | 0.003928 | 0.001637 | 0.011111 | 0.001716 | 0.001305 | 0.012437 | 0.004192 | 0.002097 | 0.005957 | 0.010842 | 0.011159 | 0.008482 | 0.003467 | 0.013928 | 0.007053 | 0.00449 | 0.008463 | 0.01417 | 0.018572 |
| 23.2083333 | 0.002953 | 0.027723 | 0.002382 | 0.007925 | 0.003603 | 0.001971 | 0.009736 | 0.003972 | 0.001493 | 0.011489 | 0.002805 | 0.000744 | 0.004529 | 0.019938 | 0.009741 | 0.006603 | 0.004806 | 0.012211 | 0.003494 | 0.002493 | 0.009808 | 0.009448 | 0.013221 |
| 23.2222222 | 0.008079 | 0.018238 | 0.003679 | 0.005445 | 0.002499 | 0.002891 | 0.007972 | 0.007371 | 0.003265 | 0.0109 | 0.003135 | 0.001449 | 0.003524 | 0.02577 | 0.009033 | 0.007007 | 0.005724 | 0.012392 | 0.001349 | 0.003219 | 0.007193 | 0.010627 | 0.011039 |
| 23.2361111 | 0.017102 | 0.00869 | 0.003893 | 0.003199 | 0.002282 | 0.005083 | 0.007747 | 0.010037 | 0.004742 | 0.009611 | 0.003684 | 0.002676 | 0.003355 | 0.024847 | 0.007587 | 0.007732 | 0.011845 | 0.013069 | 0.000929 | 0.004722 | 0.003705 | 0.012575 | 0.011246 |
| 23.25 | 0.022541 | 0.00441 | 0.003485 | 0.002084 | 0.003218 | 0.00552 | 0.008617 | 0.011729 | 0.004954 | 0.008166 | 0.003192 | 0.012378 | 0.003011 | 0.017169 | 0.006101 | 0.008315 | 0.021783 | 0.014009 | 0.001033 | 0.006699 | 0.004258 | 0.011084 | 0.011925 |
| 23.2638889 | 0.019527 | 0.007091 | 0.003952 | 0.00143 | 0.00424 | 0.003248 | 0.009166 | 0.011963 | 0.004772 | 0.007771 | 0.001842 | 0.026459 | 0.005028 | 0.008015 | 0.006287 | 0.009364 | 0.024129 | 0.015792 | 0.001123 | 0.009488 | 0.008561 | 0.007792 | 0.013511 |
| 23.2777778 | 0.011479 | 0.01234 | 0.005691 | 0.0015 | 0.004858 | 0.001986 | 0.008579 | 0.010467 | 0.00482 | 0.008108 | 0.001219 | 0.029593 | 0.009132 | 0.002118 | 0.00695 | 0.008726 | 0.019501 | 0.017045 | 0.001577 | 0.010685 | 0.012147 | 0.004783 | 0.015239 |
| 23.2916667 | 0.008799 | 0.017068 | 0.006527 | 0.002356 | 0.005125 | 0.003038 | 0.007096 | 0.008237 | 0.004875 | 0.00705 | 0.002371 | 0.033183 | 0.012937 | 0.000413 | 0.006154 | 0.005164 | 0.013536 | 0.016661 | 0.001502 | 0.008252 | 0.012059 | 0.002351 | 0.015783 |
| 23.3055556 | 0.014389 | 0.017841 | 0.00454 | 0.003401 | 0.00458 | 0.004023 | 0.005406 | 0.005801 | 0.004622 | 0.00425 | 0.003797 | 0.040373 | 0.015187 | 0.001138 | 0.005363 | 0.001853 | 0.008546 | 0.016294 | 0.001849 | 0.004912 | 0.008998 | 0.001078 | 0.016862 |
| 23.3194444 | 0.021285 | 0.011294 | 0.002911 | 0.004472 | 0.002783 | 0.003599 | 0.003761 | 0.003382 | 0.004187 | 0.002903 | 0.003937 | 0.04064 | 0.016893 | 0.003349 | 0.00578 | 0.001025 | 0.007354 | 0.016124 | 0.003897 | 0.003361 | 0.005233 | 0.000766 | 0.019602 |
| 23.3333333 | 0.024371 | 0.005019 | 0.004458 | 0.004976 | 0.001442 | 0.002413 | 0.002303 | 0.002876 | 0.003923 | 0.004123 | 0.002641 | 0.033345 | 0.020045 | 0.005588 | 0.005641 | 0.000902 | 0.010468 | 0.01453 | 0.005607 | 0.002558 | 0.004322 | 0.00079 | 0.020915 |
| 23.3472222 | 0.020959 | 0.005974 | 0.005521 | 0.004267 | 0.002432 | 0.001802 | 0.002398 | 0.004394 | 0.004316 | 0.005965 | 0.001701 | 0.019067 | 0.023145 | 0.004665 | 0.004594 | 0.000577 | 0.014642 | 0.012711 | 0.005056 | 0.002049 | 0.007054 | 0.000988 | 0.018789 |
| 23.3611111 | 0.011654 | 0.008189 | 0.004322 | 0.003148 | 0.00444 | 0.002984 | 0.004601 | 0.005522 | 0.005675 | 0.009223 | 0.003264 | 0.009382 | 0.024455 | 0.00339 | 0.00395 | 0.000611 | 0.01344 | 0.012358 | 0.003301 | 0.002375 | 0.008899 | 0.002703 | 0.014583 |
| 23.375 | 0.004493 | 0.006027 | 0.003565 | 0.003551 | 0.005968 | 0.005974 | 0.00715 | 0.005818 | 0.007614 | 0.01554 | 0.005928 | 0.013735 | 0.024789 | 0.007007 | 0.003405 | 0.000948 | 0.006794 | 0.012413 | 0.001821 | 0.002401 | 0.008758 | 0.0069 | 0.009207 |
| 23.3888889 | 0.004231 | 0.004396 | 0.004931 | 0.006154 | 0.007573 | 0.009718 | 0.009204 | 0.005617 | 0.009506 | 0.023022 | 0.007468 | 0.022317 | 0.023897 | 0.012998 | 0.002652 | 0.001664 | 0.002219 | 0.011709 | 0.001178 | 0.001714 | 0.008583 | 0.011235 | 0.003898 |
| 23.4027778 | 0.004768 | 0.00832 | 0.007895 | 0.009847 | 0.010025 | 0.013672 | 0.011554 | 0.004859 | 0.011487 | 0.028468 | 0.008063 | 0.025884 | 0.020775 | 0.017212 | 0.002048 | 0.002155 | 0.002958 | 0.011043 | 0.00208 | 0.001584 | 0.008246 | 0.013159 | 0.000902 |
| 23.4166667 | 0.007255 | 0.013855 | 0.010876 | 0.013051 | 0.012768 | 0.018477 | 0.014832 | 0.004439 | 0.014032 | 0.031789 | 0.00849 | 0.025222 | 0.015941 | 0.019833 | 0.002144 | 0.002776 | 0.006352 | 0.011314 | 0.003772 | 0.001927 | 0.007795 | 0.013892 | 0.00042 |
| 23.4305556 | 0.020036 | 0.01703 | 0.012924 | 0.014605 | 0.014699 | 0.02433 | 0.018867 | 0.004501 | 0.016725 | 0.036042 | 0.009431 | 0.023215 | 0.011289 | 0.022704 | 0.004934 | 0.005184 | 0.008525 | 0.011259 | 0.004076 | 0.001726 | 0.008381 | 0.016423 | 0.00219 |
| 23.4444444 | 0.035845 | 0.018051 | 0.01399 | 0.015244 | 0.015351 | 0.030445 | 0.022729 | 0.004379 | 0.018732 | 0.043659 | 0.01152 | 0.021429 | 0.008515 | 0.026038 | 0.009775 | 0.00834 | 0.008345 | 0.010265 | 0.002471 | 0.001508 | 0.010222 | 0.020408 | 0.006738 |
| 23.4583333 | 0.040436 | 0.017035 | 0.014377 | 0.015243 | 0.014136 | 0.034418 | 0.025105 | 0.003829 | 0.019504 | 0.05545 | 0.014367 | 0.018707 | 0.007395 | 0.027294 | 0.011985 | 0.008567 | 0.008542 | 0.010888 | 0.000841 | 0.001492 | 0.011308 | 0.02116 | 0.012803 |
| 23.4722222 | 0.032902 | 0.015845 | 0.015328 | 0.014278 | 0.011884 | 0.034249 | 0.026038 | 0.003629 | 0.019182 | 0.071211 | 0.017513 | 0.015163 | 0.0072 | 0.024022 | 0.010737 | 0.00493 | 0.010945 | 0.013876 | 0.000805 | 0.001934 | 0.010302 | 0.015721 | 0.018999 |
| 23.4861111 | 0.024452 | 0.018473 | 0.018473 | 0.015069 | 0.011601 | 0.033414 | 0.028671 | 0.006322 | 0.019271 | 0.086411 | 0.02043 | 0.013285 | 0.008905 | 0.017521 | 0.010545 | 0.001271 | 0.013496 | 0.015299 | 0.001346 | 0.003013 | 0.010185 | 0.008669 | 0.02587 |
| 23.5 | 0.020425 | 0.025221 | 0.02306 | 0.01938 | 0.014487 | 0.036587 | 0.03457 | 0.01288 | 0.021059 | 0.092405 | 0.021705 | 0.013801 | 0.013764 | 0.01238 | 0.014084 | 0.001327 | 0.012857 | 0.012349 | 0.00312 | 0.002748 | 0.014018 | 0.007359 | 0.03349 |
| 23.5138889 | 0.013897 | 0.031005 | 0.025718 | 0.024561 | 0.017961 | 0.041266 | 0.0382 | 0.018948 | 0.023015 | 0.086212 | 0.019713 | 0.015192 | 0.019766 | 0.010677 | 0.019285 | 0.005263 | 0.008699 | 0.007357 | 0.008993 | 0.001842 | 0.019798 | 0.012112 | 0.040155 |
| 23.5277778 | 0.008013 | 0.031362 | 0.026348 | 0.026633 | 0.018678 | 0.041702 | 0.033622 | 0.019599 | 0.02244 | 0.07199 | 0.013684 | 0.015766 | 0.020483 | 0.011144 | 0.020615 | 0.008856 | 0.00466 | 0.004277 | 0.015681 | 0.002289 | 0.022197 | 0.015497 | 0.042153 |
| 23.5416667 | 0.010604 | 0.026197 | 0.029092 | 0.024557 | 0.01682 | 0.038937 | 0.021952 | 0.015698 | 0.020602 | 0.054816 | 0.006135 | 0.012306 | 0.015077 | 0.012245 | 0.01385 | 0.008356 | 0.005703 | 0.005982 | 0.014832 | 0.006046 | 0.016258 | 0.011852 | 0.032911 |
| 23.5555556 | 0.010995 | 0.017949 | 0.030011 | 0.02021 | 0.013772 | 0.03465 | 0.009792 | 0.011266 | 0.019147 | 0.040737 | 0.002097 | 0.005415 | 0.012353 | 0.009679 | 0.006712 | 0.009058 | 0.009152 | 0.013828 | 0.008283 | 0.014789 | 0.010521 | 0.007624 | 0.01626 |
| 23.5694444 | 0.011108 | 0.012491 | 0.021629 | 0.016826 | 0.009698 | 0.028688 | 0.006917 | 0.008896 | 0.017743 | 0.038775 | 0.005427 | 0.006681 | 0.011805 | 0.006741 | 0.006493 | 0.012202 | 0.007374 | 0.022955 | 0.005976 | 0.021956 | 0.011687 | 0.008813 | 0.008951 |
| 23.5833333 | 0.029086 | 0.020103 | 0.012248 | 0.02094 | 0.007794 | 0.031563 | 0.018613 | 0.010129 | 0.020642 | 0.054531 | 0.01672 | 0.028587 | 0.011136 | 0.008976 | 0.015212 | 0.011503 | 0.002625 | 0.023829 | 0.012541 | 0.020044 | 0.011094 | 0.016509 | 0.02283 |
| 23.5972222 | 0.051936 | 0.039511 | 0.013936 | 0.03133 | 0.00998 | 0.042379 | 0.036549 | 0.014839 | 0.029489 | 0.065583 | 0.026517 | 0.057034 | 0.012865 | 0.008508 | 0.034658 | 0.014167 | 0.002198 | 0.014768 | 0.027612 | 0.01182 | 0.013431 | 0.029409 | 0.050288 |
| 23.6111111 | 0.056862 | 0.051303 | 0.02549 | 0.036525 | 0.013709 | 0.042089 | 0.043505 | 0.021449 | 0.0321 | 0.050247 | 0.02598 | 0.060932 | 0.015283 | 0.004528 | 0.047815 | 0.026597 | 0.008073 | 0.005951 | 0.045325 | 0.004524 | 0.026635 | 0.036261 | 0.06869 |
| 23.625 | 0.046594 | 0.046234 | 0.041786 | 0.034471 | 0.019793 | 0.037931 | 0.033439 | 0.028075 | 0.02261 | 0.031008 | 0.024915 | 0.037385 | 0.022085 | 0.002764 | 0.043186 | 0.038022 | 0.014541 | 0.00449 | 0.056948 | 0.002228 | 0.044422 | 0.038202 | 0.070566 |
| 23.6388889 | 0.028517 | 0.034232 | 0.057271 | 0.02981 | 0.028684 | 0.046253 | 0.018148 | 0.032296 | 0.012888 | 0.027768 | 0.028338 | 0.016252 | 0.026854 | 0.004816 | 0.032318 | 0.039478 | 0.012642 | 0.005492 | 0.058502 | 0.00478 | 0.058082 | 0.044499 | 0.063323 |
| 23.6527778 | 0.013979 | 0.023705 | 0.064208 | 0.023139 | 0.034873 | 0.054597 | 0.008062 | 0.0331 | 0.00711 | 0.034651 | 0.02649 | 0.012151 | 0.01946 | 0.014034 | 0.022293 | 0.033477 | 0.006778 | 0.004781 | 0.050246 | 0.009583 | 0.061258 | 0.050321 | 0.050799 |
| 23.6666667 | 0.013851 | 0.018167 | 0.060168 | 0.016256 | 0.035713 | 0.051388 | 0.004191 | 0.031971 | 0.004449 | 0.045774 | 0.020298 | 0.011172 | 0.00848 | 0.027588 | 0.013124 | 0.026131 | 0.005861 | 0.004325 | 0.032534 | 0.014507 | 0.053706 | 0.05014 | 0.037416 |
| 23.6805556 | 0.015845 | 0.020239 | 0.049967 | 0.010887 | 0.033308 | 0.039643 | 0.004683 | 0.031416 | 0.005982 | 0.054007 | 0.017854 | 0.013773 | 0.004867 | 0.035282 | 0.006595 | 0.019105 | 0.006812 | 0.004963 | 0.013715 | 0.017614 | 0.041762 | 0.047886 | 0.029045 |
| 23.6944444 | 0.013806 | 0.028531 | 0.038166 | 0.007453 | 0.02829 | 0.02217 | 0.007368 | 0.032269 | 0.007294 | 0.049108 | 0.019048 | 0.022465 | 0.008296 | 0.029797 | 0.003834 | 0.010444 | 0.007336 | 0.004904 | 0.003899 | 0.016952 | 0.030118 | 0.044211 | 0.02701 |
| 23.7083333 | 0.021604 | 0.036453 | 0.025969 | 0.008283 | 0.020836 | 0.009118 | 0.011657 | 0.034078 | 0.012154 | 0.02945 | 0.019689 | 0.025286 | 0.010346 | 0.015315 | 0.003708 | 0.004246 | 0.00753 | 0.004032 | 0.001332 | 0.013413 | 0.018409 | 0.03332 | 0.026867 |
| 23.7222222 | 0.034551 | 0.036853 | 0.014858 | 0.012923 | 0.012669 | 0.010143 | 0.016766 | 0.036333 | 0.023455 | 0.012105 | 0.016575 | 0.019357 | 0.006956 | 0.004749 | 0.004624 | 0.004209 | 0.014423 | 0.004671 | 0.00135 | 0.010226 | 0.008497 | 0.01753 | 0.024683 |
| 23.7361111 | 0.032324 | 0.028688 | 0.007129 | 0.016213 | 0.005921 | 0.017846 | 0.019507 | 0.03748 | 0.029162 | 0.011145 | 0.010204 | 0.01151 | 0.004575 | 0.005288 | 0.005352 | 0.004131 | 0.033467 | 0.006742 | 0.004615 | 0.008795 | 0.006348 | 0.009913 | 0.020765 |
| 23.75 | 0.018189 | 0.016497 | 0.005157 | 0.014437 | 0.003223 | 0.022546 | 0.016271 | 0.036254 | 0.0226 | 0.017111 | 0.005261 | 0.011721 | 0.008275 | 0.015786 | 0.006115 | 0.00367 | 0.041764 | 0.00732 | 0.012951 | 0.007234 | 0.013518 | 0.013603 | 0.014847 |
| 23.7638889 | 0.012441 | 0.006735 | 0.007759 | 0.008862 | 0.004303 | 0.020709 | 0.010478 | 0.032571 | 0.010995 | 0.017715 | 0.004266 | 0.019072 | 0.013325 | 0.033748 | 0.008749 | 0.007198 | 0.026686 | 0.005779 | 0.022538 | 0.004302 | 0.022857 | 0.012921 | 0.009037 |
| 23.7777778 | 0.02183 | 0.003558 | 0.010607 | 0.00399 | 0.00431 | 0.013008 | 0.010909 | 0.027037 | 0.004953 | 0.012299 | 0.005356 | 0.018264 | 0.014296 | 0.044951 | 0.013775 | 0.011008 | 0.013123 | 0.005149 | 0.02521 | 0.002258 | 0.025625 | 0.00875 | 0.009854 |
| 23.7916667 | 0.036757 | 0.00406 | 0.012915 | 0.003022 | 0.003224 | 0.005628 | 0.015364 | 0.020294 | 0.007864 | 0.007252 | 0.00656 | 0.009628 | 0.010685 | 0.032864 | 0.017447 | 0.012563 | 0.012253 | 0.007022 | 0.019071 | 0.003159 | 0.018904 | 0.013896 | 0.013957 |
| 23.8055556 | 0.042608 | 0.003152 | 0.015171 | 0.004744 | 0.004868 | 0.003755 | 0.014805 | 0.01287 | 0.011393 | 0.008029 | 0.007986 | 0.004391 | 0.006651 | 0.018965 | 0.014758 | 0.011753 | 0.010232 | 0.008982 | 0.010351 | 0.005165 | 0.008346 | 0.019128 | 0.012905 |
| 23.8194444 | 0.034371 | 0.001125 | 0.015946 | 0.005842 | 0.007335 | 0.007113 | 0.009159 | 0.005941 | 0.009949 | 0.013797 | 0.009697 | 0.004741 | 0.008059 | 0.022274 | 0.008237 | 0.007252 | 0.007332 | 0.008941 | 0.0055 | 0.005599 | 0.001961 | 0.014993 | 0.01048 |
| 23.8333333 | 0.019659 | 0.001323 | 0.014017 | 0.004887 | 0.007358 | 0.011339 | 0.003749 | 0.002035 | 0.006917 | 0.019918 | 0.010031 | 0.011386 | 0.011761 | 0.026158 | 0.006278 | 0.002473 | 0.012809 | 0.008102 | 0.0079 | 0.004494 | 0.000484 | 0.007149 | 0.012937 |
| 23.8472222 | 0.007859 | 0.003302 | 0.010533 | 0.002729 | 0.005015 | 0.012137 | 0.001425 | 0.002603 | 0.004959 | 0.021745 | 0.007861 | 0.016935 | 0.010096 | 0.029471 | 0.009664 | 0.000774 | 0.018717 | 0.008357 | 0.012703 | 0.003121 | 0.000501 | 0.002558 | 0.01655 |
| 23.8611111 | 0.004387 | 0.00324 | 0.00825 | 0.000929 | 0.002554 | 0.011119 | 0.00125 | 0.005586 | 0.004362 | 0.018255 | 0.004334 | 0.01254 | 0.006176 | 0.035356 | 0.0126 | 0.000723 | 0.015025 | 0.009919 | 0.01405 | 0.002129 | 0.001773 | 0.002898 | 0.014433 |
| 23.875 | 0.006397 | 0.00124 | 0.008963 | 0.001307 | 0.003021 | 0.016023 | 0.001857 | 0.008574 | 0.00572 | 0.013506 | 0.002135 | 0.005441 | 0.008227 | 0.032714 | 0.012589 | 0.001371 | 0.01239 | 0.011259 | 0.014806 | 0.001562 | 0.005534 | 0.004961 | 0.007373 |
| 23.8888889 | 0.006541 | 0.00175 | 0.01283 | 0.004814 | 0.006083 | 0.028448 | 0.004575 | 0.010525 | 0.009217 | 0.012183 | 0.003054 | 0.003618 | 0.015357 | 0.021388 | 0.010089 | 0.001776 | 0.016894 | 0.010445 | 0.016087 | 0.001023 | 0.009888 | 0.005207 | 0.003026 |
| 23.9027778 | 0.006129 | 0.007457 | 0.01759 | 0.009644 | 0.008284 | 0.036744 | 0.009617 | 0.011556 | 0.013264 | 0.016296 | 0.006482 | 0.00739 | 0.018198 | 0.015802 | 0.006548 | 0.00119 | 0.01527 | 0.007719 | 0.015494 | 0.000915 | 0.011112 | 0.003745 | 0.005107 |
| 23.9166667 | 0.007428 | 0.016228 | 0.020111 | 0.012729 | 0.007837 | 0.033805 | 0.015292 | 0.012277 | 0.016005 | 0.024676 | 0.012207 | 0.020608 | 0.012491 | 0.026313 | 0.004444 | 0.000962 | 0.008148 | 0.00553 | 0.015284 | 0.001457 | 0.009669 | 0.003621 | 0.009865 |
| 23.9305556 | 0.006882 | 0.021977 | 0.019545 | 0.013174 | 0.005201 | 0.026658 | 0.019994 | 0.013161 | 0.017738 | 0.034482 | 0.019046 | 0.036685 | 0.009622 | 0.040838 | 0.00514 | 0.0016 | 0.007878 | 0.005072 | 0.018012 | 0.001398 | 0.009634 | 0.00481 | 0.012636 |
| 23.9444444 | 0.007604 | 0.020481 | 0.017727 | 0.012049 | 0.003274 | 0.020603 | 0.021719 | 0.013727 | 0.019203 | 0.043658 | 0.023025 | 0.04249 | 0.01753 | 0.040349 | 0.007273 | 0.00275 | 0.01252 | 0.004832 | 0.020939 | 0.00075 | 0.011812 | 0.004972 | 0.013154 |
| 23.9583333 | 0.009364 | 0.015115 | 0.016309 | 0.010002 | 0.005676 | 0.016188 | 0.020181 | 0.01217 | 0.018644 | 0.049982 | 0.021682 | 0.039459 | 0.023464 | 0.028578 | 0.008244 | 0.004699 | 0.015651 | 0.003652 | 0.022548 | 0.000738 | 0.012956 | 0.004997 | 0.014166 |
| 23.9722222 | 0.008168 | 0.012235 | 0.014879 | 0.007848 | 0.011274 | 0.015717 | 0.018006 | 0.007624 | 0.014996 | 0.051352 | 0.016316 | 0.035262 | 0.019418 | 0.017095 | 0.006562 | 0.006201 | 0.018821 | 0.002731 | 0.028897 | 0.001095 | 0.01083 | 0.006952 | 0.016662 |
| 23.9861111 | 0.009338 | 0.014681 | 0.013612 | 0.008817 | 0.015945 | 0.021054 | 0.018314 | 0.002963 | 0.011395 | 0.048808 | 0.011851 | 0.032873 | 0.014869 | 0.008668 | 0.00456 | 0.006087 | 0.021576 | 0.002342 | 0.041965 | 0.001002 | 0.008034 | 0.009384 | 0.019644 |
| 24 | 0.015135 | 0.021198 | 0.014254 | 0.015407 | 0.018517 | 0.029518 | 0.021979 | 0.001372 | 0.011265 | 0.04793 | 0.010893 | 0.032722 | 0.018174 | 0.006611 | 0.006465 | 0.00688 | 0.018102 | 0.002397 | 0.051912 | 0.002104 | 0.009651 | 0.008318 | 0.023931 |
| 24.0138889 | 0.021776 | 0.027237 | 0.017007 | 0.023316 | 0.020434 | 0.036157 | 0.026086 | 0.002121 | 0.013673 | 0.054283 | 0.012321 | 0.034665 | 0.026718 | 0.010477 | 0.012025 | 0.010593 | 0.00991 | 0.00465 | 0.053566 | 0.00575 | 0.015911 | 0.004778 | 0.030185 |

| | | | | | | | | | | | | | | | | | | | | | | | |
|---|---|---|---|---|---|---|---|---|---|---|---|---|---|---|---|---|---|---|---|---|---|---|---|
| 24.0277778 | 0.027619 | 0.031746 | 0.02048 | 0.027378 | 0.023654 | 0.040453 | 0.027816 | 0.004532 | 0.015542 | 0.066509 | 0.016791 | 0.036106 | 0.033883 | 0.017731 | 0.018125 | 0.015675 | 0.005605 | 0.01 | 0.052617 | 0.011109 | 0.023789 | 0.006888 | 0.037983 |
| 24.0416667 | 0.03518 | 0.040937 | 0.026148 | 0.029938 | 0.030399 | 0.050035 | 0.028427 | 0.00804 | 0.01709 | 0.078755 | 0.026159 | 0.035821 | 0.042503 | 0.029011 | 0.026043 | 0.023017 | 0.009964 | 0.019895 | 0.055774 | 0.019462 | 0.035545 | 0.022728 | 0.052954 |
| 24.0555556 | 0.049936 | 0.056034 | 0.036225 | 0.035135 | 0.039682 | 0.073159 | 0.029813 | 0.011124 | 0.021107 | 0.089953 | 0.041984 | 0.033458 | 0.055566 | 0.043515 | 0.035255 | 0.031793 | 0.014235 | 0.030388 | 0.060906 | 0.027568 | 0.050232 | 0.047291 | 0.075611 |
| 24.0694444 | 0.066766 | 0.063385 | 0.044221 | 0.040597 | 0.045082 | 0.096437 | 0.029191 | 0.012184 | 0.026616 | 0.094391 | 0.056418 | 0.026111 | 0.063254 | 0.05256 | 0.035807 | 0.031741 | 0.012664 | 0.02901 | 0.057042 | 0.024064 | 0.054787 | 0.06049 | 0.087328 |
| 24.0833333 | 0.067943 | 0.049103 | 0.041921 | 0.038381 | 0.039105 | 0.089098 | 0.021545 | 0.01136 | 0.028905 | 0.080684 | 0.051429 | 0.0194 | 0.058818 | 0.050312 | 0.025494 | 0.020574 | 0.013202 | 0.016139 | 0.0432 | 0.012125 | 0.044576 | 0.054334 | 0.079013 |
| 24.0972222 | 0.050351 | 0.029246 | 0.032663 | 0.026773 | 0.023479 | 0.048893 | 0.010189 | 0.010583 | 0.026189 | 0.053299 | 0.027105 | 0.022051 | 0.048992 | 0.044062 | 0.016848 | 0.011599 | 0.015979 | 0.007774 | 0.028482 | 0.009629 | 0.03307 | 0.043353 | 0.06588 |
| 24.1111111 | 0.027528 | 0.027491 | 0.023015 | 0.013537 | 0.009264 | 0.01687 | 0.004712 | 0.009206 | 0.020369 | 0.028291 | 0.012031 | 0.029735 | 0.040771 | 0.040686 | 0.016167 | 0.011312 | 0.017624 | 0.010067 | 0.018922 | 0.016884 | 0.028975 | 0.039277 | 0.057959 |
| 24.125 | 0.011321 | 0.033887 | 0.015011 | 0.007464 | 0.002758 | 0.012942 | 0.006891 | 0.007283 | 0.013579 | 0.013732 | 0.017654 | 0.034477 | 0.033322 | 0.039293 | 0.018702 | 0.013513 | 0.016979 | 0.016133 | 0.016694 | 0.020223 | 0.028788 | 0.040119 | 0.053163 |
| 24.1388889 | 0.007542 | 0.032623 | 0.00985 | 0.008026 | 0.00132 | 0.015533 | 0.010165 | 0.006928 | 0.007469 | 0.006292 | 0.025308 | 0.033072 | 0.024828 | 0.034088 | 0.020467 | 0.013269 | 0.013214 | 0.020401 | 0.016261 | 0.017021 | 0.028578 | 0.039026 | 0.049739 |
| 24.1527778 | 0.010975 | 0.022143 | 0.007813 | 0.006235 | 0.001366 | 0.014561 | 0.009547 | 0.007056 | 0.00339 | 0.002686 | 0.026302 | 0.027606 | 0.015984 | 0.02251 | 0.020557 | 0.011818 | 0.00893 | 0.021927 | 0.012103 | 0.013404 | 0.025817 | 0.033562 | 0.045997 |
| 24.1666667 | 0.015163 | 0.012255 | 0.00731 | 0.002733 | 0.002435 | 0.011284 | 0.005595 | 0.005743 | 0.002642 | 0.002025 | 0.022314 | 0.0219 | 0.008122 | 0.011038 | 0.018724 | 0.012196 | 0.005396 | 0.021168 | 0.008397 | 0.011736 | 0.01884 | 0.028235 | 0.040563 |
| 24.1805556 | 0.020345 | 0.015399 | 0.006783 | 0.002166 | 0.003694 | 0.006508 | 0.003548 | 0.004644 | 0.004457 | 0.001892 | 0.014069 | 0.015202 | 0.003696 | 0.008238 | 0.01434 | 0.01356 | 0.003248 | 0.017626 | 0.00758 | 0.009482 | 0.01068 | 0.02634 | 0.035091 |
| 24.1944444 | 0.023114 | 0.024178 | 0.005845 | 0.002207 | 0.003753 | 0.00396 | 0.006504 | 0.006827 | 0.004981 | 0.003366 | 0.006611 | 0.015327 | 0.007515 | 0.017115 | 0.011984 | 0.011365 | 0.005878 | 0.012499 | 0.00803 | 0.008285 | 0.008594 | 0.027611 | 0.032041 |
| 24.2083333 | 0.017695 | 0.027305 | 0.004945 | 0.002315 | 0.002651 | 0.003227 | 0.011516 | 0.012227 | 0.003859 | 0.006445 | 0.004705 | 0.044361 | 0.021509 | 0.037339 | 0.022849 | 0.007054 | 0.022968 | 0.016848 | 0.009764 | 0.019249 | 0.022031 | 0.029156 | 0.02594 |
| 24.2222222 | 0.00867 | 0.025349 | 0.006912 | 0.003759 | 0.001896 | 0.00325 | 0.014888 | 0.016506 | 0.003569 | 0.007754 | 0.006068 | 0.098848 | 0.026447 | 0.054656 | 0.031161 | 0.009812 | 0.047171 | 0.030559 | 0.014277 | 0.040917 | 0.046378 | 0.028778 | 0.021603 |
| 24.2361111 | 0.008197 | 0.018356 | 0.01887 | 0.005477 | 0.002512 | 0.003297 | 0.013986 | 0.016006 | 0.003602 | 0.006591 | 0.005812 | 0.114185 | 0.019458 | 0.054424 | 0.030106 | 0.015197 | 0.059307 | 0.03017 | 0.019437 | 0.040957 | 0.051759 | 0.035571 | 0.024822 |
| 24.25 | 0.020886 | 0.009983 | 0.035801 | 0.006897 | 0.003522 | 0.005313 | 0.009662 | 0.012875 | 0.004114 | 0.005417 | 0.005163 | 0.067041 | 0.018931 | 0.040769 | 0.040118 | 0.010877 | 0.064579 | 0.020496 | 0.017601 | 0.019815 | 0.032236 | 0.041769 | 0.020099 |
| 24.2638889 | 0.032313 | 0.00869 | 0.036528 | 0.006548 | 0.004758 | 0.012506 | 0.006282 | 0.010879 | 0.004409 | 0.007395 | 0.007648 | 0.030893 | 0.013902 | 0.020513 | 0.045437 | 0.003647 | 0.062904 | 0.01986 | 0.010449 | 0.016003 | 0.020602 | 0.029084 | 0.009869 |
| 24.2777778 | 0.02849 | 0.00956 | 0.020472 | 0.004363 | 0.006299 | 0.020132 | 0.005681 | 0.009926 | 0.002792 | 0.012258 | 0.011621 | 0.030416 | 0.00444 | 0.008583 | 0.03751 | 0.003334 | 0.045743 | 0.01891 | 0.006736 | 0.027468 | 0.024464 | 0.014779 | 0.004843 |
| 24.2916667 | 0.018774 | 0.006445 | 0.007223 | 0.004088 | 0.006629 | 0.021565 | 0.006219 | 0.008798 | 0.001573 | 0.015484 | 0.012613 | 0.030248 | 0.003319 | 0.009174 | 0.030793 | 0.007153 | 0.026682 | 0.015746 | 0.008042 | 0.037686 | 0.026008 | 0.014226 | 0.003987 |
| 24.3055556 | 0.012567 | 0.005475 | 0.003158 | 0.005134 | 0.005148 | 0.017961 | 0.005775 | 0.00809 | 0.002535 | 0.014541 | 0.01032 | 0.022951 | 0.006055 | 0.009269 | 0.026669 | 0.010273 | 0.017984 | 0.015058 | 0.010781 | 0.044566 | 0.02182 | 0.017763 | 0.007894 |
| 24.3194444 | 0.007266 | 0.008374 | 0.003167 | 0.005141 | 0.002985 | 0.013871 | 0.003849 | 0.009257 | 0.004316 | 0.011373 | 0.007908 | 0.012727 | 0.008305 | 0.006171 | 0.024347 | 0.011519 | 0.016158 | 0.015133 | 0.012265 | 0.045084 | 0.017262 | 0.019642 | 0.013406 |
| 24.3333333 | 0.00463 | 0.010402 | 0.003788 | 0.004124 | 0.001863 | 0.009939 | 0.001901 | 0.011943 | 0.005455 | 0.006827 | 0.005836 | 0.005073 | 0.007482 | 0.00367 | 0.023291 | 0.012517 | 0.015427 | 0.016461 | 0.012286 | 0.039272 | 0.014721 | 0.020728 | 0.017458 |
| 24.3472222 | 0.009272 | 0.009612 | 0.005589 | 0.003054 | 0.00322 | 0.006691 | 0.00118 | 0.013619 | 0.004726 | 0.004929 | 0.00357 | 0.006543 | 0.009033 | 0.004734 | 0.02403 | 0.015631 | 0.010849 | 0.019146 | 0.011186 | 0.032644 | 0.015814 | 0.025657 | 0.023454 |
| 24.3611111 | 0.018489 | 0.006194 | 0.009001 | 0.005114 | 0.006074 | 0.008762 | 0.00111 | 0.012573 | 0.00318 | 0.01106 | 0.003611 | 0.029413 | 0.034327 | 0.013486 | 0.03054 | 0.026551 | 0.008546 | 0.022513 | 0.012219 | 0.037392 | 0.028134 | 0.052856 | 0.04439 |
| 24.375 | 0.026452 | 0.004196 | 0.012258 | 0.011855 | 0.008047 | 0.015278 | 0.001804 | 0.009156 | 0.004342 | 0.021431 | 0.006933 | 0.076974 | 0.07721 | 0.031618 | 0.046271 | 0.043135 | 0.011069 | 0.023808 | 0.019581 | 0.052965 | 0.049197 | 0.10338 | 0.07316 |
| 24.3888889 | 0.029633 | 0.00741 | 0.014702 | 0.019025 | 0.007191 | 0.020075 | 0.004246 | 0.005239 | 0.00852 | 0.030392 | 0.010781 | 0.094975 | 0.081165 | 0.039784 | 0.052699 | 0.043783 | 0.010629 | 0.020814 | 0.02473 | 0.053296 | 0.051099 | 0.115664 | 0.070217 |
| 24.4027778 | 0.03052 | 0.012027 | 0.016555 | 0.021991 | 0.004416 | 0.021991 | 0.007765 | 0.004798 | 0.013016 | 0.034913 | 0.013692 | 0.059358 | 0.050073 | 0.027171 | 0.033063 | 0.024743 | 0.012862 | 0.015581 | 0.019744 | 0.03145 | 0.026671 | 0.076923 | 0.036863 |
| 24.4166667 | 0.033323 | 0.015192 | 0.018027 | 0.021146 | 0.005095 | 0.023667 | 0.01149 | 0.009034 | 0.01599 | 0.035151 | 0.016734 | 0.029848 | 0.044076 | 0.018581 | 0.013495 | 0.011713 | 0.015563 | 0.011062 | 0.01701 | 0.015886 | 0.007515 | 0.062678 | 0.016041 |
| 24.4305556 | 0.034083 | 0.017182 | 0.020332 | 0.019417 | 0.010966 | 0.026245 | 0.014792 | 0.01355 | 0.017214 | 0.035433 | 0.020253 | 0.025393 | 0.050008 | 0.019653 | 0.011082 | 0.010292 | 0.012589 | 0.008147 | 0.020583 | 0.014127 | 0.007228 | 0.077989 | 0.013646 |
| 24.4444444 | 0.029894 | 0.017158 | 0.022679 | 0.017774 | 0.016454 | 0.027964 | 0.016386 | 0.015395 | 0.017362 | 0.037622 | 0.021551 | 0.020256 | 0.038097 | 0.015274 | 0.008805 | 0.007277 | 0.009842 | 0.0074 | 0.0177 | 0.011443 | 0.011504 | 0.077947 | 0.011566 |
| 24.4583333 | 0.023575 | 0.015998 | 0.021854 | 0.014894 | 0.01804 | 0.027276 | 0.014684 | 0.013763 | 0.015574 | 0.040926 | 0.01907 | 0.011222 | 0.02257 | 0.009636 | 0.006076 | 0.004709 | 0.011592 | 0.011447 | 0.009606 | 0.006522 | 0.010188 | 0.060237 | 0.007506 |
| 24.4722222 | 0.01648 | 0.015849 | 0.017726 | 0.011009 | 0.016718 | 0.02451 | 0.010439 | 0.008946 | 0.010514 | 0.04618 | 0.016011 | 0.004283 | 0.011299 | 0.011113 | 0.010807 | 0.006453 | 0.016825 | 0.02055 | 0.006583 | 0.003207 | 0.00753 | 0.039171 | 0.00729 |
| 24.4861111 | 0.013476 | 0.018791 | 0.014455 | 0.011173 | 0.016102 | 0.022716 | 0.01068 | 0.006553 | 0.006944 | 0.053577 | 0.015763 | 0.002395 | 0.003766 | 0.013811 | 0.013416 | 0.005999 | 0.019194 | 0.025888 | 0.006681 | 0.002517 | 0.00769 | 0.028509 | 0.009425 |
| 24.5 | 0.019422 | 0.025547 | 0.015118 | 0.018842 | 0.020729 | 0.026351 | 0.019403 | 0.011032 | 0.010371 | 0.06285 | 0.019467 | 0.005714 | 0.003242 | 0.010143 | 0.008827 | 0.004655 | 0.016224 | 0.021341 | 0.004019 | 0.004353 | 0.006771 | 0.031252 | 0.007149 |
| 24.5138889 | 0.026831 | 0.031614 | 0.017039 | 0.027414 | 0.026993 | 0.033097 | 0.029004 | 0.017415 | 0.016367 | 0.068843 | 0.022008 | 0.012269 | 0.006964 | 0.005415 | 0.004328 | 0.007884 | 0.013324 | 0.013449 | 0.000972 | 0.005768 | 0.007164 | 0.038398 | 0.002963 |
| 24.5277778 | 0.02849 | 0.033223 | 0.018731 | 0.029151 | 0.027858 | 0.040519 | 0.032612 | 0.021292 | 0.019733 | 0.062876 | 0.016255 | 0.018108 | 0.00743 | 0.008406 | 0.006535 | 0.011835 | 0.017565 | 0.012038 | 0.002472 | 0.005867 | 0.007396 | 0.04355 | 0.002798 |
| 24.5416667 | 0.027106 | 0.0294 | 0.026131 | 0.026084 | 0.02575 | 0.053747 | 0.029418 | 0.023619 | 0.020892 | 0.052074 | 0.007272 | 0.018041 | 0.010474 | 0.02215 | 0.021051 | 0.010471 | 0.034945 | 0.026031 | 0.013664 | 0.013244 | 0.010282 | 0.044405 | 0.011186 |
| 24.5555556 | 0.023091 | 0.019104 | 0.03516 | 0.021241 | 0.022834 | 0.063087 | 0.019818 | 0.023308 | 0.01943 | 0.046218 | 0.004058 | 0.013616 | 0.021888 | 0.039888 | 0.042553 | 0.007723 | 0.056561 | 0.050815 | 0.032288 | 0.031239 | 0.025335 | 0.038566 | 0.027836 |
| 24.5694444 | 0.019308 | 0.011681 | 0.031626 | 0.014903 | 0.01668 | 0.054808 | 0.012292 | 0.020813 | 0.015879 | 0.045142 | 0.007407 | 0.014661 | 0.022203 | 0.044203 | 0.048951 | 0.009473 | 0.058324 | 0.059782 | 0.040761 | 0.040951 | 0.035754 | 0.027564 | 0.03637 |
| 24.5833333 | 0.028287 | 0.021393 | 0.021931 | 0.01283 | 0.012199 | 0.047528 | 0.019542 | 0.021707 | 0.017463 | 0.04394 | 0.01921 | 0.02174 | 0.012569 | 0.030612 | 0.033568 | 0.009849 | 0.038359 | 0.04399 | 0.031086 | 0.03181 | 0.025286 | 0.017432 | 0.027426 |
| 24.5972222 | 0.045404 | 0.045226 | 0.021219 | 0.018174 | 0.013523 | 0.05105 | 0.037721 | 0.025944 | 0.027039 | 0.03711 | 0.033988 | 0.025393 | 0.013567 | 0.014946 | 0.015313 | 0.00586 | 0.018387 | 0.025631 | 0.01646 | 0.018028 | 0.01013 | 0.013042 | 0.012652 |
| 24.6111111 | 0.047243 | 0.063027 | 0.027316 | 0.022447 | 0.015218 | 0.044381 | 0.047231 | 0.026771 | 0.03399 | 0.03246 | 0.034193 | 0.025392 | 0.015992 | 0.00618 | 0.00553 | 0.002668 | 0.007768 | 0.016559 | 0.006664 | 0.010467 | 0.006686 | 0.015688 | 0.005072 |
| 24.625 | 0.034175 | 0.061826 | 0.035568 | 0.019498 | 0.015453 | 0.031916 | 0.039571 | 0.024459 | 0.031091 | 0.037019 | 0.021718 | 0.026577 | 0.012856 | 0.002599 | 0.002658 | 0.002501 | 0.00312 | 0.010945 | 0.002656 | 0.007001 | 0.01013 | 0.020702 | 0.006975 |
| 24.6388889 | 0.020602 | 0.048496 | 0.045067 | 0.016391 | 0.019942 | 0.038872 | 0.025519 | 0.02382 | 0.021621 | 0.042424 | 0.013023 | 0.026657 | 0.010478 | 0.00268 | 0.003984 | 0.003485 | 0.0022 | 0.006056 | 0.003411 | 0.005673 | 0.011803 | 0.021397 | 0.010706 |
| 24.6527778 | 0.016716 | 0.035109 | 0.052303 | 0.018392 | 0.027 | 0.0605 | 0.015596 | 0.024805 | 0.01209 | 0.038093 | 0.012129 | 0.022984 | 0.00904 | 0.003378 | 0.005322 | 0.003291 | 0.002579 | 0.003669 | 0.004385 | 0.01019 | 0.010307 | 0.017519 | 0.010975 |
| 24.6666667 | 0.025973 | 0.024743 | 0.056074 | 0.020291 | 0.029722 | 0.07256 | 0.009944 | 0.025179 | 0.008998 | 0.031051 | 0.014527 | 0.017879 | 0.007737 | 0.002745 | 0.004778 | 0.002353 | 0.002037 | 0.003394 | 0.003581 | 0.016623 | 0.007298 | 0.012862 | 0.009151 |
| 24.6805556 | 0.027388 | 0.016068 | 0.058154 | 0.018004 | 0.028745 | 0.067879 | 0.005581 | 0.024983 | 0.008225 | 0.033191 | 0.017312 | 0.013386 | 0.00626 | 0.001563 | 0.003163 | 0.001814 | 0.000991 | 0.003402 | 0.002545 | 0.014168 | 0.004422 | 0.009176 | 0.006737 |
| 24.6944444 | 0.020349 | 0.009535 | 0.057487 | 0.014043 | 0.025576 | 0.051855 | 0.003872 | 0.02437 | 0.007802 | 0.03686 | 0.022657 | 0.009825 | 0.004718 | 0.000614 | 0.001682 | 0.00153 | 0.000601 | 0.002731 | 0.001754 | 0.006336 | 0.002779 | 0.00622 | 0.004357 |
| 24.7083333 | 0.03016 | 0.005988 | 0.050723 | 0.010882 | 0.019548 | 0.030602 | 0.006586 | 0.023209 | 0.016445 | 0.031557 | 0.029103 | 0.00669 | 0.003058 | 0.000859 | 0.001023 | 0.001342 | 0.001405 | 0.001815 | 0.001425 | 0.003087 | 0.002085 | 0.003405 | 0.002403 |
| 24.7222222 | 0.045827 | 0.005997 | 0.038508 | 0.008854 | 0.013123 | 0.01495 | 0.00987 | 0.023052 | 0.028817 | 0.020211 | 0.028287 | 0.004285 | 0.003684 | 0.003641 | 0.001643 | 0.002815 | 0.004445 | 0.003286 | 0.00311 | 0.004483 | 0.002311 | 0.002589 | 0.002442 |
| 24.7361111 | 0.038367 | 0.009396 | 0.025576 | 0.00676 | 0.008267 | 0.016721 | 0.010027 | 0.021865 | 0.032539 | 0.010728 | 0.017374 | 0.00526 | 0.00892 | 0.008383 | 0.004544 | 0.007024 | 0.009349 | 0.008247 | 0.007448 | 0.008104 | 0.005337 | 0.006403 | 0.005312 |
| 24.75 | 0.016879 | 0.014316 | 0.016079 | 0.004005 | 0.004562 | 0.033089 | 0.008228 | 0.016613 | 0.024512 | 0.007986 | 0.00796 | 0.008797 | 0.014413 | 0.011995 | 0.007834 | 0.011044 | 0.012965 | 0.012967 | 0.011494 | 0.01219 | 0.010007 | 0.011696 | 0.00735 |
| 24.7638889 | 0.007942 | 0.01738 | 0.010717 | 0.003104 | 0.002562 | 0.045918 | 0.005787 | 0.011613 | 0.011572 | 0.007921 | 0.010182 | 0.009487 | 0.015624 | 0.012312 | 0.008138 | 0.011638 | 0.013234 | 0.013485 | 0.012154 | 0.013385 | 0.01148 | 0.012836 | 0.005258 |
| 24.7777778 | 0.016685 | 0.016345 | 0.00738 | 0.003603 | 0.002554 | 0.043944 | 0.003741 | 0.012193 | 0.006342 | 0.005514 | 0.017236 | 0.006232 | 0.01263 | 0.009529 | 0.005168 | 0.008652 | 0.010295 | 0.009908 | 0.009049 | 0.010181 | 0.00814 | 0.008751 | 0.003407 |
| 24.7916667 | 0.032566 | 0.011806 | 0.004829 | 0.002768 | 0.003229 | 0.030345 | 0.005075 | 0.016759 | 0.011603 | 0.005387 | 0.020339 | 0.004726 | 0.007235 | 0.004977 | 0.003004 | 0.004385 | 0.005364 | 0.004895 | 0.004699 | 0.005308 | 0.004019 | 0.003917 | 0.006662 |
| 24.8055556 | 0.041201 | 0.006406 | 0.00529 | 0.001183 | 0.003161 | 0.015217 | 0.009691 | 0.019758 | 0.017886 | 0.007374 | 0.017883 | 0.006935 | 0.002978 | 0.001309 | 0.003677 | 0.001689 | 0.001419 | 0.001351 | 0.002578 | 0.002441 | 0.002661 | 0.001832 | 0.010002 |
| 24.8194444 | 0.036885 | 0.005338 | 0.008928 | 0.000938 | 0.002502 | 0.007037 | 0.01395 | 0.016382 | 0.017205 | 0.00664 | 0.012239 | 0.007823 | 0.002309 | 0.000581 | 0.003229 | 0.001026 | 0.000488 | 0.000809 | 0.002335 | 0.001981 | 0.002004 | 0.0014 | 0.008592 |
| 24.8333333 | 0.022869 | 0.009953 | 0.011764 | 0.001818 | 0.002861 | 0.006319 | 0.015388 | 0.008366 | 0.011176 | 0.007298 | 0.007228 | 0.007037 | 0.003013 | 0.001109 | 0.001562 | 0.000877 | 0.000836 | 0.001857 | 0.002228 | 0.002646 | 0.001092 | 0.001261 | 0.006145 |
| 24.8472222 | 0.009746 | 0.01372 | 0.012195 | 0.001908 | 0.002715 | 0.008539 | 0.014027 | 0.00251 | 0.005627 | 0.01269 | 0.00424 | 0.008479 | 0.002403 | 0.001524 | 0.002472 | 0.001413 | 0.001302 | 0.002261 | 0.003325 | 0.00275 | 0.002892 | 0.002017 | 0.007345 |
| 24.8611111 | 0.008978 | 0.012887 | 0.011352 | 0.001082 | 0.001559 | 0.01142 | 0.011734 | 0.000938 | 0.005065 | 0.018557 | 0.002511 | 0.012089 | 0.002019 | 0.003362 | 0.005852 | 0.004097 | 0.003306 | 0.003311 | 0.006238 | 0.003515 | 0.007734 | 0.005233 | 0.012075 |
| 24.875 | 0.01994 | 0.010073 | 0.010659 | 0.000466 | 0.001421 | 0.015452 | 0.010063 | 0.00137 | 0.009393 | 0.023376 | 0.0014 | 0.014817 | 0.003058 | 0.005948 | 0.008705 | 0.006939 | 0.006083 | 0.005515 | 0.008774 | 0.006928 | 0.011414 | 0.007847 | 0.016359 |
| 24.8888889 | 0.031519 | 0.00791 | 0.011029 | 0.00058 | 0.002511 | 0.019288 | 0.01011 | 0.0015 | 0.012583 | 0.027845 | 0.001114 | 0.014297 | 0.003249 | 0.006992 | 0.008879 | 0.007136 | 0.007354 | 0.005777 | 0.008612 | 0.009245 | 0.010209 | 0.00619 | 0.016951 |
| 24.9027778 | 0.03009 | 0.007934 | 0.012199 | 0.002458 | 0.005697 | 0.020953 | 0.012067 | 0.001766 | 0.010669 | 0.030003 | 0.001869 | 0.012014 | 0.002082 | 0.0066 | 0.00749 | 0.005911 | 0.006934 | 0.004478 | 0.006908 | 0.008245 | 0.006907 | 0.00367 | 0.014607 |
| 24.9166667 | 0.017408 | 0.010516 | 0.013861 | 0.006504 | 0.009674 | 0.020689 | 0.014756 | 0.004788 | 0.010318 | 0.029619 | 0.003916 | 0.011326 | 0.001779 | 0.006513 | 0.007202 | 0.005954 | 0.006583 | 0.004816 | 0.006715 | 0.006934 | 0.006767 | 0.004922 | 0.012697 |
| 24.9305556 | 0.009402 | 0.01475 | 0.016147 | 0.010988 | 0.011175 | 0.02045 | 0.016882 | 0.009674 | 0.01864 | 0.02899 | 0.007663 | 0.012152 | 0.002934 | 0.007389 | 0.008218 | 0.007148 | 0.007284 | 0.006343 | 0.007984 | 0.006738 | 0.008963 | 0.009032 | 0.013247 |
| 24.9444444 | 0.01235 | 0.019908 | 0.017537 | 0.01495 | 0.010651 | 0.021997 | 0.017277 | 0.01268 | 0.027865 | 0.029716 | 0.013062 | 0.012662 | 0.003684 | 0.007486 | 0.008534 | 0.007362 | 0.007072 | 0.006424 | 0.008308 | 0.006133 | 0.00998 | 0.012351 | 0.014692 |
| 24.9583333 | 0.021165 | 0.025352 | 0.015701 | 0.018562 | 0.01138 | 0.026428 | 0.017244 | 0.011984 | 0.029806 | 0.034711 | 0.017619 | 0.012604 | 0.002681 | 0.005185 | 0.00726 | 0.005747 | 0.00439 | 0.004342 | 0.006789 | 0.004034 | 0.00943 | 0.013508 | 0.014859 |
| 24.9722222 | 0.032974 | 0.0306 | 0.012925 | 0.021262 | 0.01568 | 0.03416 | 0.021085 | 0.011033 | 0.027697 | 0.046219 | 0.017891 | 0.011128 | 0.002423 | 0.002656 | 0.004946 | 0.003484 | 0.002275 | 0.003141 | 0.00414 | 0.002866 | 0.006569 | 0.011898 | 0.013729 |
| 24.9861111 | 0.040829 | 0.03414 | 0.014182 | 0.023576 | 0.023567 | 0.041013 | 0.029163 | 0.013409 | 0.024578 | 0.056799 | 0.013721 | 0.007707 | 0.005329 | 0.002679 | 0.00259 | 0.002792 | 0.002996 | 0.005004 | 0.002617 | 0.00504 | 0.003261 | 0.00762 | 0.012043 |
| 25 | 0.037464 | 0.033858 | 0.01889 | 0.025995 | 0.030484 | 0.040728 | 0.034046 | 0.014383 | 0.02006 | 0.057963 | 0.008284 | 0.005971 | 0.007439 | 0.002783 | 0.002707 | 0.002735 | 0.003153 | 0.005518 | 0.002339 | 0.006254 | 0.003017 | 0.00521 | 0.011427 |
| 25.0138889 | 0.032127 | 0.030986 | 0.022053 | 0.025499 | 0.029867 | 0.034955 | 0.032135 | 0.009806 | 0.013466 | 0.054523 | 0.004571 | 0.009496 | 0.006798 | 0.004455 | 0.008154 | 0.002918 | 0.004518 | 0.006004 | 0.00327 | 0.005727 | 0.00597 | 0.008389 | 0.015311 |
| 25.0277778 | 0.037021 | 0.029158 | 0.022164 | 0.02252 | 0.0235 | 0.032551 | 0.029906 | 0.006426 | 0.007596 | 0.053584 | 0.002322 | 0.015931 | 0.011206 | 0.015746 | 0.020025 | 0.008168 | 0.016577 | 0.013969 | 0.009691 | 0.011528 | 0.01227 | 0.014009 | 0.026609 |
| 25.0416667 | 0.049291 | 0.03162 | 0.023318 | 0.02285 | 0.023295 | 0.041179 | 0.031436 | 0.011836 | 0.00905 | 0.058912 | 0.005152 | 0.022382 | 0.022787 | 0.033144 | 0.036888 | 0.019398 | 0.037861 | 0.027882 | 0.02235 | 0.025328 | 0.019586 | 0.01803 | 0.040926 |
| 25.0555556 | 0.061472 | 0.038653 | 0.028451 | 0.026845 | 0.034052 | 0.060015 | 0.031882 | 0.019832 | 0.019007 | 0.075375 | 0.019787 | 0.028839 | 0.030696 | 0.043071 | 0.049757 | 0.029659 | 0.051402 | 0.04076 | 0.034489 | 0.03771 | 0.027099 | 0.020359 | 0.047434 |
| 25.0694444 | 0.069854 | 0.04369 | 0.033789 | 0.031182 | 0.042021 | 0.077633 | 0.02616 | 0.021164 | 0.02983 | 0.086624 | 0.039672 | 0.032072 | 0.029057 | 0.03968 | 0.048289 | 0.031731 | 0.046062 | 0.043947 | 0.036762 | 0.038137 | 0.0332 | 0.020009 | 0.04152 |
| 25.0833333 | 0.066143 | 0.038847 | 0.034957 | 0.032047 | 0.036223 | 0.07589 | 0.015012 | 0.019301 | 0.034948 | 0.07138 | 0.044872 | 0.029341 | 0.023421 | 0.028767 | 0.036466 | 0.027062 | 0.030719 | 0.035272 | 0.029032 | 0.027903 | 0.032618 | 0.015593 | 0.030096 |
| 25.0972222 | 0.048567 | 0.022958 | 0.030511 | 0.027135 | 0.022448 | 0.047478 | 0.006335 | 0.017831 | 0.033475 | 0.043906 | 0.027827 | 0.024376 | 0.019583 | 0.019154 | 0.025715 | 0.021816 | 0.019226 | 0.024384 | 0.019668 | 0.017584 | 0.02697 | 0.010501 | 0.021804 |
| 25.1111111 | 0.02639 | 0.00918 | 0.019999 | 0.019195 | 0.009762 | 0.019928 | 0.00629 | 0.015478 | 0.0259 | 0.026692 | 0.010731 | 0.020535 | 0.017343 | 0.013577 | 0.020054 | 0.017457 | 0.013872 | 0.017405 | 0.012613 | 0.01168 | 0.020545 | 0.007674 | 0.017592 |
| 25.125 | 0.011201 | 0.008716 | 0.00842 | 0.00995 | 0.003259 | 0.017851 | 0.00909 | 0.014058 | 0.015759 | 0.023112 | 0.010722 | 0.017669 | 0.014981 | 0.010157 | 0.017723 | 0.013464 | 0.011278 | 0.013296 | 0.007668 | 0.008461 | 0.014701 | 0.007005 | 0.015346 |
| 25.1388889 | 0.010862 | 0.011128 | 0.002493 | 0.004605 | 0.002247 | 0.026616 | 0.009308 | 0.014283 | 0.008141 | 0.024588 | 0.017661 | 0.015897 | 0.013086 | 0.007634 | 0.016043 | 0.010496 | 0.009339 | 0.010918 | 0.004305 | 0.006705 | 0.010197 | 0.00814 | 0.014306 |
| 25.1527778 | 0.018448 | 0.009269 | 0.001313 | 0.00793 | 0.002215 | 0.028918 | 0.007408 | 0.012928 | 0.003582 | 0.020118 | 0.023181 | 0.015929 | 0.012532 | 0.005973 | 0.013896 | 0.009026 | 0.007388 | 0.009776 | 0.002482 | 0.006265 | 0.007778 | 0.010689 | 0.014062 |
| 25.1666667 | 0.025069 | 0.007826 | 0.001686 | 0.013253 | 0.002015 | 0.020205 | 0.00422 | 0.008098 | 0.001789 | 0.010749 | 0.024815 | 0.017646 | 0.013383 | 0.006044 | 0.013296 | 0.009066 | 0.007259 | 0.010127 | 0.00317 | 0.007791 | 0.008275 | 0.015081 | 0.015414 |
| 25.1805556 | 0.029562 | 0.007772 | 0.003431 | 0.013934 | 0.002432 | 0.010138 | 0.00316 | 0.003303 | 0.003038 | 0.006791 | 0.020223 | 0.019928 | 0.015003 | 0.008293 | 0.01583 | 0.010039 | 0.009789 | 0.012267 | 0.006368 | 0.011053 | 0.011195 | 0.02091 | 0.018101 |
| 25.1944444 | 0.030209 | 0.005778 | 0.005698 | 0.009484 | 0.003765 | 0.011934 | 0.006294 | 0.001534 | 0.004357 | 0.009294 | 0.012514 | 0.021543 | 0.015743 | 0.011123 | 0.019171 | 0.010824 | 0.011665 | 0.014678 | 0.009881 | 0.01343 | 0.014011 | 0.025269 | 0.01909 |
| 25.2083333 | 0.025281 | 0.003628 | 0.007153 | 0.005105 | 0.005495 | 0.016625 | 0.009985 | 0.001967 | 0.003775 | 0.011308 | 0.005779 | 0.020918 | 0.013099 | 0.011499 | 0.019383 | 0.009516 | 0.009863 | 0.014158 | 0.010356 | 0.011817 | 0.012847 | 0.024107 | 0.016235 |
| 25.2222222 | 0.01554 | 0.005735 | 0.007508 | 0.00563 | 0.00564 | 0.014135 | 0.011181 | 0.003844 | 0.002168 | 0.010694 | 0.003267 | 0.017536 | 0.007311 | 0.007791 | 0.015783 | 0.005666 | 0.00535 | 0.009778 | 0.006649 | 0.0069 | 0.007606 | 0.016205 | 0.011205 |
| 25.2361111 | 0.006511 | 0.007486 | 0.00706 | 0.006552 | 0.003492 | 0.009885 | 0.01064 | 0.005426 | 0.00117 | 0.00836 | 0.005422 | 0.014447 | 0.003392 | 0.002932 | 0.011499 | 0.002883 | 0.002091 | 0.005261 | 0.002731 | 0.003922 | 0.00378 | 0.007023 | 0.008644 |
| 25.25 | 0.004091 | 0.005061 | 0.006055 | 0.004992 | 0.002306 | 0.009478 | 0.009443 | 0.005979 | 0.002016 | 0.005104 | 0.009677 | 0.014527 | 0.004534 | 0.001455 | 0.009841 | 0.002209 | 0.001553 | 0.004352 | 0.002983 | 0.004386 | 0.002518 | 0.00267 | 0.012506 |
| 25.2638889 | 0.00867 | 0.00182 | 0.004436 | 0.003833 | 0.004806 | 0.012351 | 0.007735 | 0.006946 | 0.003588 | 0.004278 | 0.014264 | 0.016831 | 0.007852 | 0.003273 | 0.011473 | 0.001971 | 0.001553 | 0.00668 | 0.005686 | 0.004602 | 0.001946 | 0.002246 | 0.019504 |
| 25.2777778 | 0.01571 | 0.000625 | 0.002812 | 0.003464 | 0.008009 | 0.015945 | 0.005573 | 0.008927 | 0.004389 | 0.006653 | 0.017135 | 0.018434 | 0.00913 | 0.004433 | 0.013979 | 0.002857 | 0.00174 | 0.008835 | 0.005455 | 0.00429 | 0.002571 | 0.003272 | 0.020895 |
| 25.2916667 | 0.016705 | 0.001452 | 0.002252 | 0.002544 | 0.008101 | 0.018239 | 0.003566 | 0.011886 | 0.004644 | 0.007683 | 0.016872 | 0.017391 | 0.007491 | 0.003455 | 0.014105 | 0.00335 | 0.001872 | 0.008141 | 0.003356 | 0.005714 | 0.002665 | 0.005111 | 0.014302 |
| 25.3055556 | 0.0114 | 0.002414 | 0.002538 | 0.002717 | 0.005064 | 0.018004 | 0.002204 | 0.013747 | 0.004815 | 0.005488 | 0.01502 | 0.014067 | 0.005289 | 0.002671 | 0.01015 | 0.002753 | 0.002041 | 0.005007 | 0.004749 | 0.008814 | 0.002952 | 0.005833 | 0.006368 |
| 25.3194444 | 0.008525 | 0.003175 | 0.002555 | 0.005369 | 0.003539 | 0.014782 | 0.001301 | 0.010554 | 0.003933 | 0.003237 | 0.012814 | 0.009429 | 0.003731 | 0.002761 | 0.004342 | 0.004274 | 0.003367 | 0.002509 | 0.008714 | 0.011731 | 0.005659 | 0.004784 | 0.002706 |
| 25.3333333 | 0.008522 | 0.00608 | 0.003242 | 0.008136 | 0.005461 | 0.010309 | 0.001032 | 0.005781 | 0.002184 | 0.004301 | 0.009096 | 0.005771 | 0.002182 | 0.002426 | 0.001275 | 0.007217 | 0.004401 | 0.002113 | 0.011236 | 0.012462 | 0.007656 | 0.003329 | 0.002353 |
| 25.3472222 | 0.008107 | 0.009606 | 0.00539 | 0.008949 | 0.006173 | 0.006908 | 0.001468 | 0.005813 | 0.001474 | 0.00802 | 0.005299 | 0.00716 | 0.001 | 0.001787 | 0.001922 | 0.006526 | 0.003912 | 0.002534 | 0.009413 | 0.010094 | 0.00567 | 0.003937 | 0.004058 |
| 25.3611111 | 0.007562 | 0.010559 | 0.006998 | 0.007793 | 0.00536 | 0.005167 | 0.00217 | 0.007616 | 0.002282 | 0.012852 | 0.005442 | 0.013529 | 0.001647 | 0.001852 | 0.003778 | 0.003541 | 0.002697 | 0.004011 | 0.005033 | 0.00611 | 0.002633 | 0.006635 | 0.007669 |
| 25.375 | 0.006417 | 0.010647 | 0.007631 | 0.006479 | 0.006348 | 0.008242 | 0.003725 | 0.008266 | 0.004892 | 0.019896 | 0.008322 | 0.019375 | 0.004377 | 0.002591 | 0.003941 | 0.002665 | 0.00226 | 0.005221 | 0.003402 | 0.003742 | 0.001828 | 0.006287 | 0.010428 |
| 25.3888889 | 0.00675 | 0.011751 | 0.007663 | 0.007197 | 0.009211 | 0.017591 | 0.006332 | 0.008147 | 0.00905 | 0.028659 | 0.011878 | 0.02085 | 0.005053 | 0.0031 | 0.002499 | 0.003012 | 0.003555 | 0.003761 | 0.003916 | 0.004562 | 0.001948 | 0.004261 | 0.010372 |
| 25.4027778 | 0.011001 | 0.01327 | 0.006718 | 0.010375 | 0.012991 | 0.027799 | 0.009041 | 0.005968 | 0.013053 | 0.036783 | 0.018225 | 0.019424 | 0.003817 | 0.003877 | 0.001993 | 0.004148 | 0.0057 | 0.001401 | 0.003561 | 0.006859 | 0.002921 | 0.005678 | 0.008405 |
| 25.4166667 | 0.018957 | 0.014571 | 0.006811 | 0.015339 | 0.017463 | 0.034548 | 0.011509 | 0.002671 | 0.016099 | 0.043884 | 0.025933 | 0.018545 | 0.004432 | 0.003866 | 0.002055 | 0.00427 | 0.006429 | 0.001242 | 0.003424 | 0.007382 | 0.003236 | 0.006027 | 0.007378 |
| 25.4305556 | 0.028283 | 0.015956 | 0.008807 | 0.020539 | 0.021422 | 0.036887 | 0.013917 | 0.000781 | 0.018233 | 0.050091 | 0.029915 | 0.018937 | 0.00503 | 0.002992 | 0.002604 | 0.003279 | 0.004427 | 0.003468 | 0.005132 | 0.004919 | 0.00283 | 0.00339 | 0.009587 |
| 25.4444444 | 0.037454 | 0.017941 | 0.009583 | 0.023496 | 0.022585 | 0.036326 | 0.015903 | 0.000902 | 0.018723 | 0.053641 | 0.030535 | 0.018565 | 0.006461 | 0.003551 | 0.004587 | 0.003641 | 0.001659 | 0.005775 | 0.007378 | 0.002258 | 0.004503 | 0.002745 | 0.013825 |
| 25.4583333 | 0.047975 | 0.020084 | 0.007553 | 0.023092 | 0.020936 | 0.034724 | 0.015465 | 0.002337 | 0.0159 | 0.05341 | 0.032052 | 0.017323 | 0.008899 | 0.003151 | 0.005256 | 0.002967 | 0.001137 | 0.00613 | 0.007598 | 0.001569 | 0.005714 | 0.004005 | 0.017408 |
| 25.4722222 | 0.056485 | 0.020911 | 0.006614 | 0.022598 | 0.019448 | 0.03229 | 0.010927 | 0.006075 | 0.01065 | 0.050598 | 0.034573 | 0.016597 | 0.008025 | 0.001453 | 0.00348 | 0.001205 | 0.00198 | 0.004282 | 0.005114 | 0.001781 | 0.00459 | 0.004941 | 0.018961 |
| 25.4861111 | 0.059432 | 0.021097 | 0.011233 | 0.025505 | 0.020909 | 0.032663 | 0.007439 | 0.015018 | 0.009353 | 0.049441 | 0.037003 | 0.016905 | 0.005898 | 0.002459 | 0.002424 | 0.001214 | 0.001772 | 0.002861 | 0.002097 | 0.00344 | 0.003194 | 0.006544 | 0.01885 |
| 25.5 | 0.059482 | 0.023042 | 0.019427 | 0.03013 | 0.024783 | 0.037978 | 0.01073 | 0.026949 | 0.014555 | 0.053492 | 0.040099 | 0.019573 | 0.007582 | 0.004901 | 0.00336 | 0.002677 | 0.001294 | 0.004253 | 0.001019 | 0.00651 | 0.0031 | 0.009022 | 0.017462 |

| | | | | | | | | | | | | | | | | | | | | | | | |
|---|---|---|---|---|---|---|---|---|---|---|---|---|---|---|---|---|---|---|---|---|---|---|---|
| 25.5138889 | 0.055939 | 0.024569 | 0.024625 | 0.033101 | 0.02768 | 0.044715 | 0.017424 | 0.035882 | 0.019119 | 0.058591 | 0.040888 | 0.025103 | 0.011674 | 0.005333 | 0.003563 | 0.0039 | 0.001385 | 0.007589 | 0.002015 | 0.007839 | 0.006492 | 0.011607 | 0.01438 |
| 25.5277778 | 0.046908 | 0.023658 | 0.024617 | 0.033713 | 0.028781 | 0.054784 | 0.024311 | 0.042742 | 0.018825 | 0.059658 | 0.034606 | 0.03087 | 0.012447 | 0.007517 | 0.004172 | 0.004863 | 0.004054 | 0.010451 | 0.002952 | 0.00603 | 0.012849 | 0.015722 | 0.010136 |
| 25.5416667 | 0.034258 | 0.023745 | 0.02469 | 0.033731 | 0.029251 | 0.070938 | 0.029565 | 0.048968 | 0.016624 | 0.056572 | 0.024337 | 0.031805 | 0.008704 | 0.017408 | 0.012025 | 0.005613 | 0.013776 | 0.00989 | 0.006936 | 0.007499 | 0.013935 | 0.019322 | 0.005908 |
| 25.5555556 | 0.019932 | 0.025699 | 0.024782 | 0.030676 | 0.023643 | 0.075586 | 0.027375 | 0.047166 | 0.014604 | 0.049382 | 0.018082 | 0.026875 | 0.004397 | 0.028209 | 0.024791 | 0.009166 | 0.02658 | 0.012271 | 0.017592 | 0.017246 | 0.016059 | 0.018154 | 0.003608 |
| 25.5694444 | 0.012064 | 0.029584 | 0.020639 | 0.023992 | 0.012569 | 0.059728 | 0.021127 | 0.036153 | 0.013225 | 0.043118 | 0.020217 | 0.024266 | 0.007317 | 0.026275 | 0.02877 | 0.014473 | 0.029467 | 0.02044 | 0.022802 | 0.023703 | 0.024522 | 0.016531 | 0.004458 |
| 25.5833333 | 0.02274 | 0.039641 | 0.021222 | 0.022265 | 0.010746 | 0.046494 | 0.023292 | 0.026809 | 0.013288 | 0.045865 | 0.02939 | 0.030776 | 0.02056 | 0.013944 | 0.018831 | 0.011841 | 0.019 | 0.019225 | 0.015743 | 0.01671 | 0.020355 | 0.020891 | 0.009299 |
| 25.5972222 | 0.043308 | 0.05198 | 0.030039 | 0.027543 | 0.019216 | 0.042009 | 0.034682 | 0.022303 | 0.0137 | 0.049638 | 0.035898 | 0.039691 | 0.031488 | 0.004635 | 0.007515 | 0.006388 | 0.007264 | 0.009393 | 0.010224 | 0.007541 | 0.009277 | 0.028528 | 0.016307 |
| 25.6111111 | 0.052228 | 0.054587 | 0.034895 | 0.029797 | 0.023288 | 0.029509 | 0.043316 | 0.019339 | 0.012779 | 0.043178 | 0.029729 | 0.040546 | 0.029586 | 0.001521 | 0.005437 | 0.008226 | 0.003655 | 0.006178 | 0.011577 | 0.007788 | 0.007968 | 0.032468 | 0.019064 |
| 25.625 | 0.042559 | 0.046055 | 0.034682 | 0.023392 | 0.021935 | 0.011836 | 0.041088 | 0.020228 | 0.012895 | 0.034869 | 0.015285 | 0.034648 | 0.018405 | 0.002538 | 0.008717 | 0.012148 | 0.005515 | 0.009979 | 0.012269 | 0.011934 | 0.008731 | 0.030061 | 0.018918 |
| 25.6388889 | 0.021687 | 0.034554 | 0.037091 | 0.01469 | 0.02339 | 0.008353 | 0.030389 | 0.025591 | 0.0157 | 0.034909 | 0.00514 | 0.029046 | 0.007786 | 0.008261 | 0.011943 | 0.014573 | 0.007864 | 0.015647 | 0.01105 | 0.014904 | 0.008112 | 0.023563 | 0.022492 |
| 25.6527778 | 0.010285 | 0.024276 | 0.040535 | 0.00785 | 0.023958 | 0.025312 | 0.018066 | 0.030178 | 0.017407 | 0.041587 | 0.003017 | 0.024697 | 0.004745 | 0.013818 | 0.0132 | 0.016319 | 0.008461 | 0.019953 | 0.009054 | 0.017198 | 0.008084 | 0.015325 | 0.027721 |
| 25.6666667 | 0.01864 | 0.015612 | 0.040343 | 0.003265 | 0.018823 | 0.047966 | 0.008589 | 0.030948 | 0.016021 | 0.047737 | 0.005635 | 0.018873 | 0.010483 | 0.013301 | 0.01181 | 0.016049 | 0.007165 | 0.02019 | 0.005286 | 0.017806 | 0.006707 | 0.00815 | 0.029018 |
| 25.6805556 | 0.036361 | 0.009021 | 0.040798 | 0.001952 | 0.013652 | 0.06087 | 0.00318 | 0.029234 | 0.013213 | 0.048373 | 0.010867 | 0.012834 | 0.023052 | 0.00853 | 0.009512 | 0.014311 | 0.00561 | 0.017915 | 0.001822 | 0.016939 | 0.004551 | 0.007304 | 0.026992 |
| 25.6944444 | 0.04713 | 0.004731 | 0.045825 | 0.001228 | 0.012188 | 0.061869 | 0.002077 | 0.025389 | 0.009918 | 0.042645 | 0.019444 | 0.009849 | 0.03393 | 0.004036 | 0.007599 | 0.012683 | 0.004443 | 0.016318 | 0.000564 | 0.015761 | 0.003284 | 0.015374 | 0.025484 |
| 25.7083333 | 0.033785 | 0.00245 | 0.046861 | 0.003454 | 0.011676 | 0.049397 | 0.005341 | 0.020322 | 0.00826 | 0.032532 | 0.029061 | 0.009577 | 0.034764 | 0.002526 | 0.005595 | 0.011573 | 0.002957 | 0.015427 | 0.000524 | 0.014502 | 0.002243 | 0.027179 | 0.025233 |
| 25.7222222 | 0.017996 | 0.00197 | 0.037774 | 0.010227 | 0.009062 | 0.027259 | 0.012041 | 0.017035 | 0.010523 | 0.021745 | 0.030815 | 0.009842 | 0.027187 | 0.003189 | 0.003306 | 0.011165 | 0.00134 | 0.013982 | 0.000926 | 0.012856 | 0.001419 | 0.033091 | 0.024453 |
| 25.7361111 | 0.028335 | 0.00348 | 0.024232 | 0.012371 | 0.004788 | 0.01817 | 0.018148 | 0.016211 | 0.013733 | 0.012073 | 0.020811 | 0.009867 | 0.018345 | 0.002737 | 0.001394 | 0.011573 | 0.000513 | 0.0127 | 0.001176 | 0.011012 | 0.001802 | 0.029064 | 0.022019 |
| 25.75 | 0.043708 | 0.005616 | 0.011506 | 0.00901 | 0.002117 | 0.03042 | 0.019149 | 0.016871 | 0.013682 | 0.005149 | 0.010197 | 0.010162 | 0.012124 | 0.002059 | 0.000399 | 0.011816 | 0.000392 | 0.01233 | 0.001852 | 0.009223 | 0.002197 | 0.021095 | 0.018862 |
| 25.7638889 | 0.043798 | 0.005533 | 0.00493 | 0.0093 | 0.001019 | 0.043195 | 0.015086 | 0.019679 | 0.01027 | 0.004628 | 0.009987 | 0.010769 | 0.006969 | 0.00282 | 0.000207 | 0.010448 | 0.000324 | 0.011703 | 0.003731 | 0.007123 | 0.00222 | 0.016812 | 0.016053 |
| 25.7777778 | 0.031058 | 0.007365 | 0.00841 | 0.012405 | 0.000942 | 0.046053 | 0.008707 | 0.023849 | 0.005404 | 0.010544 | 0.015264 | 0.01112 | 0.004008 | 0.002873 | 0.000609 | 0.007859 | 0.000591 | 0.010519 | 0.004941 | 0.004074 | 0.002264 | 0.017379 | 0.013155 |
| 25.7916667 | 0.016667 | 0.017383 | 0.016744 | 0.012831 | 0.0021 | 0.042823 | 0.004176 | 0.025125 | 0.003622 | 0.0134 | 0.019813 | 0.011353 | 0.003851 | 0.002939 | 0.001335 | 0.006084 | 0.001334 | 0.009952 | 0.0043 | 0.002474 | 0.001557 | 0.018494 | 0.010254 |
| 25.8055556 | 0.017524 | 0.030368 | 0.024749 | 0.01343 | 0.002527 | 0.034621 | 0.005009 | 0.021502 | 0.007278 | 0.01015 | 0.020808 | 0.011611 | 0.006232 | 0.004499 | 0.001458 | 0.00597 | 0.001616 | 0.010254 | 0.002717 | 0.004207 | 0.001009 | 0.015621 | 0.008441 |
| 25.8194444 | 0.032896 | 0.037665 | 0.029688 | 0.014039 | 0.002645 | 0.023664 | 0.008416 | 0.016177 | 0.011134 | 0.010257 | 0.014942 | 0.012161 | 0.013303 | 0.004957 | 0.000956 | 0.006704 | 0.00109 | 0.011375 | 0.001446 | 0.004822 | 0.001848 | 0.0087 | 0.008096 |
| 25.8333333 | 0.042291 | 0.034155 | 0.029473 | 0.009668 | 0.002726 | 0.012083 | 0.00919 | 0.012449 | 0.010521 | 0.015133 | 0.009074 | 0.013748 | 0.016315 | 0.003509 | 0.000825 | 0.007188 | 0.000678 | 0.013419 | 0.001008 | 0.003051 | 0.003046 | 0.002883 | 0.008171 |
| 25.8472222 | 0.034182 | 0.021469 | 0.025207 | 0.006851 | 0.001989 | 0.005063 | 0.006952 | 0.010112 | 0.007013 | 0.02014 | 0.012042 | 0.015862 | 0.009936 | 0.003275 | 0.000818 | 0.006887 | 0.000573 | 0.015002 | 0.000989 | 0.002996 | 0.004503 | 0.001826 | 0.006976 |
| 25.8611111 | 0.017066 | 0.01027 | 0.018884 | 0.010488 | 0.002311 | 0.005307 | 0.004662 | 0.007069 | 0.003777 | 0.024997 | 0.016932 | 0.016913 | 0.005303 | 0.004914 | 0.000692 | 0.006199 | 0.000367 | 0.014771 | 0.001312 | 0.004265 | 0.007139 | 0.00525 | 0.004698 |
| 25.875 | 0.005334 | 0.006515 | 0.014448 | 0.013079 | 0.003093 | 0.006147 | 0.004204 | 0.003832 | 0.002349 | 0.029588 | 0.015097 | 0.016473 | 0.009552 | 0.005186 | 0.000834 | 0.006296 | 0.000647 | 0.014046 | 0.00145 | 0.004112 | 0.009704 | 0.01311 | 0.003488 |
| 25.8888889 | 0.002131 | 0.007786 | 0.015974 | 0.010237 | 0.003556 | 0.004721 | 0.006117 | 0.003698 | 0.002595 | 0.03387 | 0.009563 | 0.014943 | 0.01468 | 0.00354 | 0.001794 | 0.007875 | 0.002186 | 0.013769 | 0.000924 | 0.002709 | 0.008627 | 0.020444 | 0.003914 |
| 25.9027778 | 0.002001 | 0.011788 | 0.022441 | 0.007395 | 0.006639 | 0.002752 | 0.009309 | 0.00475 | 0.005002 | 0.035061 | 0.00446 | 0.012284 | 0.011334 | 0.0021 | 0.003915 | 0.009474 | 0.004593 | 0.012604 | 0.000827 | 0.00133 | 0.005268 | 0.019067 | 0.003825 |
| 25.9166667 | 0.006139 | 0.018551 | 0.03093 | 0.010515 | 0.010254 | 0.003846 | 0.012248 | 0.003569 | 0.009638 | 0.029856 | 0.003803 | 0.009532 | 0.007279 | 0.001542 | 0.005311 | 0.009201 | 0.00619 | 0.011923 | 0.001583 | 0.00066 | 0.004077 | 0.010489 | 0.00332 |
| 25.9305556 | 0.012946 | 0.027493 | 0.039095 | 0.015974 | 0.009035 | 0.015177 | 0.016453 | 0.002103 | 0.01392 | 0.025697 | 0.012384 | 0.00739 | 0.012226 | 0.001788 | 0.004737 | 0.007675 | 0.006217 | 0.013866 | 0.002316 | 0.001198 | 0.003831 | 0.003363 | 0.005711 |
| 25.9444444 | 0.012022 | 0.035663 | 0.043086 | 0.017266 | 0.004615 | 0.0318 | 0.023616 | 0.003701 | 0.016969 | 0.031076 | 0.023944 | 0.004855 | 0.016699 | 0.002914 | 0.003334 | 0.006771 | 0.005445 | 0.015875 | 0.002179 | 0.00292 | 0.002872 | 0.001021 | 0.010572 |
| 25.9583333 | 0.007483 | 0.040694 | 0.043945 | 0.017173 | 0.004792 | 0.039508 | 0.031995 | 0.00844 | 0.018612 | 0.041445 | 0.027003 | 0.002112 | 0.011623 | 0.003842 | 0.001881 | 0.006972 | 0.004458 | 0.01577 | 0.001319 | 0.005504 | 0.001824 | 0.001761 | 0.014768 |
| 25.9722222 | 0.013563 | 0.042411 | 0.045051 | 0.022979 | 0.011879 | 0.037459 | 0.037653 | 0.009867 | 0.018506 | 0.04629 | 0.023137 | 0.000964 | 0.006814 | 0.002942 | 0.001662 | 0.007333 | 0.003345 | 0.014567 | 0.00128 | 0.008886 | 0.001333 | 0.003846 | 0.018151 |
| 25.9861111 | 0.028087 | 0.04158 | 0.042087 | 0.033237 | 0.016868 | 0.034318 | 0.037827 | 0.005743 | 0.018127 | 0.043907 | 0.020274 | 0.002034 | 0.012116 | 0.001829 | 0.003562 | 0.007226 | 0.002363 | 0.013228 | 0.002548 | 0.012375 | 0.001375 | 0.005112 | 0.022423 |
| 26 | 0.038223 | 0.038728 | 0.031692 | 0.040633 | 0.014659 | 0.033619 | 0.033119 | 0.002046 | 0.017339 | 0.04154 | 0.018908 | 0.003784 | 0.017981 | 0.001666 | 0.004699 | 0.007166 | 0.002384 | 0.012747 | 0.003432 | 0.013743 | 0.001716 | 0.004008 | 0.026486 |
| 26.0138889 | 0.037401 | 0.035968 | 0.018438 | 0.039479 | 0.009372 | 0.033826 | 0.026511 | 0.004475 | 0.013896 | 0.043551 | 0.016551 | 0.00544 | 0.013758 | 0.003198 | 0.004216 | 0.007915 | 0.004513 | 0.013634 | 0.002621 | 0.010612 | 0.002065 | 0.00196 | 0.025167 |
| 26.0277778 | 0.032839 | 0.036656 | 0.010118 | 0.030131 | 0.005974 | 0.034655 | 0.023982 | 0.015626 | 0.009032 | 0.050825 | 0.015273 | 0.010219 | 0.007675 | 0.006788 | 0.004968 | 0.007065 | 0.006715 | 0.013245 | 0.002394 | 0.006473 | 0.0026 | 0.004097 | 0.018768 |
| 26.0416667 | 0.034186 | 0.040481 | 0.012388 | 0.021471 | 0.010282 | 0.042145 | 0.028329 | 0.033405 | 0.009247 | 0.062856 | 0.021663 | 0.022372 | 0.015704 | 0.009121 | 0.006608 | 0.00742 | 0.007027 | 0.008587 | 0.010418 | 0.01033 | 0.007435 | 0.017625 | 0.020237 |
| 26.0555556 | 0.041232 | 0.046177 | 0.023269 | 0.022679 | 0.022381 | 0.060905 | 0.032239 | 0.049513 | 0.015291 | 0.072173 | 0.039433 | 0.035084 | 0.034964 | 0.011562 | 0.010098 | 0.013781 | 0.008281 | 0.004543 | 0.024228 | 0.02101 | 0.014701 | 0.037305 | 0.035965 |
| 26.0694444 | 0.048851 | 0.048462 | 0.03513 | 0.031732 | 0.032362 | 0.079008 | 0.02822 | 0.054609 | 0.019043 | 0.070812 | 0.05626 | 0.034117 | 0.042042 | 0.012832 | 0.014457 | 0.014886 | 0.008001 | 0.009679 | 0.028771 | 0.024426 | 0.01303 | 0.044419 | 0.049783 |
| 26.0833333 | 0.048444 | 0.03965 | 0.0426 | 0.039319 | 0.032755 | 0.075579 | 0.016444 | 0.043913 | 0.01765 | 0.061044 | 0.052688 | 0.021491 | 0.029152 | 0.009096 | 0.01381 | 0.008547 | 0.004701 | 0.01928 | 0.024731 | 0.016927 | 0.005893 | 0.03543 | 0.048171 |
| 26.0972222 | 0.036845 | 0.023882 | 0.040981 | 0.039762 | 0.025319 | 0.048217 | 0.007557 | 0.024613 | 0.014762 | 0.046758 | 0.029407 | 0.012197 | 0.012384 | 0.007195 | 0.011076 | 0.005986 | 0.003345 | 0.020759 | 0.021924 | 0.009743 | 0.003544 | 0.021981 | 0.039609 |
| 26.1111111 | 0.020907 | 0.010373 | 0.029624 | 0.030669 | 0.014252 | 0.031435 | 0.01008 | 0.01418 | 0.012341 | 0.030947 | 0.013647 | 0.010635 | 0.004635 | 0.008457 | 0.01163 | 0.004779 | 0.002779 | 0.014127 | 0.017362 | 0.008119 | 0.003866 | 0.010054 | 0.033955 |
| 26.125 | 0.009028 | 0.003827 | 0.016967 | 0.016034 | 0.005519 | 0.040279 | 0.015613 | 0.010459 | 0.010424 | 0.022904 | 0.019691 | 0.011518 | 0.007551 | 0.00712 | 0.013523 | 0.003567 | 0.002511 | 0.007324 | 0.009401 | 0.008714 | 0.004799 | 0.003363 | 0.029536 |
| 26.1388889 | 0.004827 | 0.003133 | 0.008325 | 0.005952 | 0.002195 | 0.043609 | 0.016065 | 0.008897 | 0.008876 | 0.025771 | 0.028855 | 0.011622 | 0.016154 | 0.004961 | 0.014403 | 0.004985 | 0.003772 | 0.003262 | 0.005044 | 0.00945 | 0.005038 | 0.004727 | 0.024384 |
| 26.1527778 | 0.003183 | 0.003763 | 0.003377 | 0.005006 | 0.001121 | 0.032875 | 0.012655 | 0.017918 | 0.007926 | 0.029744 | 0.029827 | 0.009337 | 0.020709 | 0.003309 | 0.014477 | 0.00526 | 0.00447 | 0.001739 | 0.005455 | 0.009543 | 0.003135 | 0.007995 | 0.019435 |
| 26.1666667 | 0.002401 | 0.004954 | 0.002109 | 0.009975 | 0.000635 | 0.01929 | 0.008395 | 0.027926 | 0.007517 | 0.027299 | 0.027318 | 0.004967 | 0.016441 | 0.002176 | 0.014649 | 0.003158 | 0.003976 | 0.0021 | 0.007218 | 0.0084 | 0.001612 | 0.006335 | 0.014777 |
| 26.1805556 | 0.00689 | 0.007041 | 0.003275 | 0.015618 | 0.000463 | 0.009114 | 0.005162 | 0.028164 | 0.006991 | 0.021128 | 0.023971 | 0.001733 | 0.009722 | 0.002782 | 0.01455 | 0.002282 | 0.003109 | 0.002279 | 0.009724 | 0.006864 | 0.002519 | 0.003021 | 0.010242 |
| 26.1944444 | 0.015079 | 0.008375 | 0.003445 | 0.017896 | 0.000585 | 0.010272 | 0.003076 | 0.020554 | 0.005939 | 0.016585 | 0.01972 | 0.000953 | 0.006692 | 0.002787 | 0.013027 | 0.004149 | 0.002232 | 0.001798 | 0.011397 | 0.005404 | 0.004586 | 0.003385 | 0.006146 |
| 26.2083333 | 0.021329 | 0.007355 | 0.003228 | 0.016079 | 0.000815 | 0.019233 | 0.00213 | 0.010254 | 0.004482 | 0.013492 | 0.014444 | 0.001443 | 0.004933 | 0.001832 | 0.010339 | 0.004785 | 0.001284 | 0.002097 | 0.009959 | 0.003566 | 0.006435 | 0.004992 | 0.00504 |
| 26.2222222 | 0.022235 | 0.004397 | 0.0066 | 0.0121 | 0.000662 | 0.024357 | 0.001561 | 0.004225 | 0.003012 | 0.008999 | 0.00926 | 0.002449 | 0.002677 | 0.002786 | 0.007981 | 0.00328 | 0.000778 | 0.003432 | 0.006109 | 0.002103 | 0.007666 | 0.003769 | 0.008852 |
| 26.2361111 | 0.018461 | 0.002389 | 0.012531 | 0.007956 | 0.000987 | 0.021452 | 0.001617 | 0.002439 | 0.001891 | 0.004657 | 0.004599 | 0.002891 | 0.002296 | 0.003638 | 0.005955 | 0.002784 | 0.000965 | 0.00429 | 0.004231 | 0.001962 | 0.008253 | 0.002128 | 0.014295 |
| 26.25 | 0.014485 | 0.00181 | 0.017288 | 0.004886 | 0.002247 | 0.014715 | 0.003352 | 0.001382 | 0.0011 | 0.004021 | 0.002748 | 0.00178 | 0.004166 | 0.003634 | 0.003538 | 0.002132 | 0.001663 | 0.003049 | 0.005829 | 0.002386 | 0.008341 | 0.002767 | 0.016959 |
| 26.2638889 | 0.013563 | 0.001897 | 0.015372 | 0.003502 | 0.00414 | 0.009527 | 0.005634 | 0.001152 | 0.000797 | 0.007083 | 0.00574 | 0.000755 | 0.007362 | 0.005189 | 0.002666 | 0.00118 | 0.002781 | 0.001569 | 0.006038 | 0.002132 | 0.00699 | 0.003424 | 0.016213 |
| 26.2777778 | 0.013624 | 0.004368 | 0.00929 | 0.002611 | 0.006372 | 0.007101 | 0.007477 | 0.002741 | 0.001276 | 0.011111 | 0.010794 | 0.000987 | 0.010909 | 0.005448 | 0.004286 | 0.001836 | 0.003744 | 0.002076 | 0.003596 | 0.001174 | 0.003993 | 0.002224 | 0.014066 |
| 26.2916667 | 0.010308 | 0.008356 | 0.008927 | 0.003772 | 0.00821 | 0.005608 | 0.008574 | 0.006362 | 0.002228 | 0.013956 | 0.014851 | 0.001209 | 0.011999 | 0.004153 | 0.005554 | 0.002083 | 0.003753 | 0.002952 | 0.002524 | 0.000479 | 0.002187 | 0.000896 | 0.012164 |
| 26.3055556 | 0.007342 | 0.011035 | 0.010924 | 0.010299 | 0.008606 | 0.004552 | 0.008497 | 0.010721 | 0.003123 | 0.01476 | 0.016107 | 0.001004 | 0.00918 | 0.003829 | 0.004995 | 0.00143 | 0.002822 | 0.002422 | 0.004168 | 0.000455 | 0.003943 | 0.001082 | 0.010148 |
| 26.3194444 | 0.01265 | 0.011305 | 0.008972 | 0.016398 | 0.007321 | 0.004164 | 0.007277 | 0.01152 | 0.003546 | 0.013437 | 0.013115 | 0.001103 | 0.005342 | 0.003424 | 0.003828 | 0.001518 | 0.001852 | 0.001333 | 0.007072 | 0.000599 | 0.006737 | 0.002149 | 0.006645 |
| 26.3333333 | 0.022918 | 0.009783 | 0.006026 | 0.015242 | 0.005621 | 0.004163 | 0.004703 | 0.007944 | 0.0033 | 0.01039 | 0.007198 | 0.001293 | 0.003814 | 0.00301 | 0.003059 | 0.001564 | 0.001243 | 0.001126 | 0.009509 | 0.000494 | 0.007874 | 0.004101 | 0.003915 |
| 26.3472222 | 0.029881 | 0.00782 | 0.004742 | 0.009035 | 0.004757 | 0.004044 | 0.002672 | 0.006364 | 0.0025 | 0.005925 | 0.003001 | 0.001347 | 0.004177 | 0.00491 | 0.002564 | 0.002162 | 0.000868 | 0.002265 | 0.009662 | 0.000347 | 0.007995 | 0.007 | 0.005561 |
| 26.3611111 | 0.028649 | 0.006161 | 0.005277 | 0.003996 | 0.003537 | 0.003195 | 0.004191 | 0.00732 | 0.001533 | 0.003583 | 0.003848 | 0.001174 | 0.004038 | 0.008274 | 0.001981 | 0.004083 | 0.00099 | 0.003775 | 0.007513 | 0.000481 | 0.0076 | 0.008904 | 0.009586 |
| 26.375 | 0.019387 | 0.005342 | 0.007812 | 0.003271 | 0.002684 | 0.00286 | 0.008492 | 0.005535 | 0.001615 | 0.008135 | 0.00773 | 0.0007 | 0.003018 | 0.008074 | 0.00168 | 0.004399 | 0.001395 | 0.004701 | 0.004733 | 0.000523 | 0.00589 | 0.008769 | 0.013885 |
| 26.3888889 | 0.009367 | 0.005685 | 0.01231 | 0.003038 | 0.005127 | 0.005838 | 0.012539 | 0.004284 | 0.003383 | 0.018218 | 0.011874 | 0.001748 | 0.002416 | 0.003963 | 0.002059 | 0.003039 | 0.001521 | 0.005937 | 0.002732 | 0.000488 | 0.003277 | 0.007623 | 0.018844 |
| 26.4027778 | 0.008268 | 0.007194 | 0.01796 | 0.004502 | 0.010022 | 0.011557 | 0.014389 | 0.008176 | 0.005665 | 0.029116 | 0.015527 | 0.006113 | 0.002458 | 0.002878 | 0.002835 | 0.002997 | 0.001207 | 0.007121 | 0.003374 | 0.0009 | 0.00139 | 0.006132 | 0.0233 |
| 26.4166667 | 0.019012 | 0.010198 | 0.02306 | 0.014964 | 0.016388 | 0.018155 | 0.014486 | 0.014056 | 0.00806 | 0.038062 | 0.018752 | 0.013007 | 0.003101 | 0.008843 | 0.003448 | 0.00357 | 0.000638 | 0.006054 | 0.007449 | 0.001626 | 0.001293 | 0.003753 | 0.026607 |
| 26.4305556 | 0.034267 | 0.014264 | 0.025098 | 0.03013 | 0.02258 | 0.024903 | 0.014702 | 0.018319 | 0.01094 | 0.045698 | 0.02217 | 0.019671 | 0.005886 | 0.014667 | 0.003383 | 0.004401 | 0.000482 | 0.003036 | 0.012012 | 0.002355 | 0.002758 | 0.00175 | 0.028563 |
| 26.4444444 | 0.045653 | 0.016518 | 0.022023 | 0.040284 | 0.025598 | 0.028877 | 0.014843 | 0.018371 | 0.013073 | 0.0505 | 0.024684 | 0.023327 | 0.011175 | 0.011323 | 0.002334 | 0.004744 | 0.001477 | 0.001714 | 0.012748 | 0.002414 | 0.003889 | 0.001731 | 0.027499 |
| 26.4583333 | 0.048578 | 0.014506 | 0.014626 | 0.043084 | 0.024934 | 0.026288 | 0.012931 | 0.01205 | 0.012477 | 0.049061 | 0.023613 | 0.022562 | 0.015237 | 0.004521 | 0.002242 | 0.003513 | 0.00392 | 0.005663 | 0.008394 | 0.00149 | 0.003829 | 0.002926 | 0.022574 |
| 26.4722222 | 0.040229 | 0.010377 | 0.008376 | 0.039559 | 0.021884 | 0.019754 | 0.008829 | 0.005173 | 0.008633 | 0.046016 | 0.020702 | 0.018324 | 0.013547 | 0.002381 | 0.00408 | 0.003415 | 0.006191 | 0.011999 | 0.003458 | 0.000996 | 0.004197 | 0.005505 | 0.015746 |
| 26.4861111 | 0.030078 | 0.009304 | 0.010132 | 0.03312 | 0.020171 | 0.019728 | 0.008238 | 0.006581 | 0.005985 | 0.049533 | 0.022184 | 0.015114 | 0.00901 | 0.003467 | 0.00473 | 0.003464 | 0.006276 | 0.012028 | 0.002067 | 0.001958 | 0.004548 | 0.006248 | 0.010738 |
| 26.5 | 0.02976 | 0.013702 | 0.019856 | 0.029022 | 0.022406 | 0.030972 | 0.016422 | 0.017977 | 0.009151 | 0.057198 | 0.027972 | 0.016915 | 0.008589 | 0.004637 | 0.003842 | 0.002577 | 0.005364 | 0.006282 | 0.001951 | 0.00301 | 0.00568 | 0.005284 | 0.009851 |
| 26.5138889 | 0.032701 | 0.020117 | 0.029832 | 0.027846 | 0.025442 | 0.043483 | 0.027163 | 0.033818 | 0.014008 | 0.057115 | 0.03027 | 0.020931 | 0.01144 | 0.004729 | 0.003567 | 0.00367 | 0.004944 | 0.003476 | 0.001603 | 0.003625 | 0.010183 | 0.008778 | 0.012339 |
| 26.5277778 | 0.030855 | 0.025002 | 0.036512 | 0.026802 | 0.025264 | 0.049614 | 0.033085 | 0.046548 | 0.015863 | 0.045753 | 0.027051 | 0.021389 | 0.012936 | 0.008032 | 0.005896 | 0.004483 | 0.006433 | 0.003738 | 0.001446 | 0.003994 | 0.012348 | 0.012051 | 0.012686 |
| 26.5416667 | 0.027235 | 0.027238 | 0.044086 | 0.028687 | 0.025136 | 0.055553 | 0.03335 | 0.055749 | 0.016732 | 0.03062 | 0.024635 | 0.01558 | 0.009921 | 0.01611 | 0.015217 | 0.007638 | 0.014472 | 0.00851 | 0.004669 | 0.008889 | 0.011522 | 0.00895 | 0.010717 |
| 26.5555556 | 0.022899 | 0.024798 | 0.047615 | 0.030255 | 0.024787 | 0.058916 | 0.02613 | 0.061 | 0.016439 | 0.020409 | 0.024307 | 0.006721 | 0.00622 | 0.023513 | 0.029808 | 0.018705 | 0.027724 | 0.022195 | 0.013408 | 0.02209 | 0.01662 | 0.005775 | 0.014074 |
| 26.5694444 | 0.015291 | 0.023523 | 0.035149 | 0.025632 | 0.018571 | 0.049799 | 0.017146 | 0.062258 | 0.012896 | 0.02571 | 0.023568 | 0.004924 | 0.015817 | 0.02365 | 0.035703 | 0.026279 | 0.033364 | 0.032643 | 0.015042 | 0.030183 | 0.021483 | 0.005019 | 0.014843 |
| 26.5833333 | 0.011175 | 0.035624 | 0.014528 | 0.026044 | 0.014357 | 0.040595 | 0.020075 | 0.066494 | 0.011111 | 0.050929 | 0.029124 | 0.013905 | 0.037567 | 0.014417 | 0.024825 | 0.019979 | 0.025012 | 0.029002 | 0.009245 | 0.022389 | 0.014771 | 0.004993 | 0.011169 |
| 26.5972222 | 0.014777 | 0.056548 | 0.004188 | 0.032323 | 0.017813 | 0.042782 | 0.033334 | 0.068388 | 0.013678 | 0.073338 | 0.039905 | 0.023629 | 0.052164 | 0.004322 | 0.01032 | 0.008481 | 0.012064 | 0.018704 | 0.00886 | 0.01012 | 0.00588 | 0.010964 | 0.018021 |
| 26.6111111 | 0.017983 | 0.065209 | 0.003639 | 0.031533 | 0.018866 | 0.037201 | 0.040901 | 0.055789 | 0.016556 | 0.068213 | 0.036568 | 0.025457 | 0.047649 | 0.001487 | 0.005546 | 0.00257 | 0.003941 | 0.009455 | 0.007572 | 0.006846 | 0.005329 | 0.020531 | 0.028854 |
| 26.625 | 0.017428 | 0.052421 | 0.006439 | 0.024449 | 0.015358 | 0.017822 | 0.036377 | 0.034279 | 0.01911 | 0.046986 | 0.019155 | 0.019652 | 0.029444 | 0.00312 | 0.00616 | 0.001514 | 0.001288 | 0.003675 | 0.00412 | 0.010469 | 0.006657 | 0.024377 | 0.031616 |
| 26.6388889 | 0.01567 | 0.02852 | 0.018366 | 0.017186 | 0.016439 | 0.009331 | 0.025444 | 0.01671 | 0.020317 | 0.029431 | 0.007453 | 0.01085 | 0.014144 | 0.00403 | 0.005945 | 0.001681 | 0.001228 | 0.002983 | 0.005628 | 0.013505 | 0.005768 | 0.020165 | 0.028783 |
| 26.6527778 | 0.010778 | 0.011535 | 0.040417 | 0.012401 | 0.024564 | 0.018933 | 0.013503 | 0.009167 | 0.017777 | 0.018855 | 0.004692 | 0.005236 | 0.007183 | 0.005983 | 0.005523 | 0.003029 | 0.001517 | 0.005515 | 0.008248 | 0.015467 | 0.005524 | 0.013396 | 0.024076 |
| 26.6666667 | 0.004917 | 0.00878 | 0.059713 | 0.009737 | 0.030173 | 0.030841 | 0.004694 | 0.010702 | 0.012907 | 0.01464 | 0.004395 | 0.004264 | 0.003246 | 0.010791 | 0.005419 | 0.004098 | 0.001496 | 0.007069 | 0.007402 | 0.01822 | 0.004938 | 0.009733 | 0.021057 |
| 26.6805556 | 0.003797 | 0.008299 | 0.064743 | 0.006065 | 0.025901 | 0.035792 | 0.001906 | 0.023839 | 0.008709 | 0.017001 | 0.00428 | 0.003783 | 0.003536 | 0.013373 | 0.004912 | 0.003154 | 0.00137 | 0.005256 | 0.00556 | 0.019687 | 0.0034 | 0.008714 | 0.02064 |
| 26.6944444 | 0.003924 | 0.005083 | 0.05787 | 0.004834 | 0.016347 | 0.035164 | 0.004207 | 0.050686 | 0.007381 | 0.021597 | 0.005562 | 0.003517 | 0.009967 | 0.010813 | 0.004074 | 0.002129 | 0.00144 | 0.002756 | 0.00698 | 0.017303 | 0.003707 | 0.008714 | 0.020006 |
| 26.7083333 | 0.00561 | 0.005386 | 0.046484 | 0.009308 | 0.009745 | 0.030602 | 0.008726 | 0.07194 | 0.00869 | 0.023637 | 0.010761 | 0.005777 | 0.016978 | 0.006085 | 0.003483 | 0.003421 | 0.001896 | 0.002773 | 0.009038 | 0.01274 | 0.005868 | 0.009512 | 0.018096 |
| 26.7222222 | 0.014291 | 0.0076 | 0.031196 | 0.011558 | 0.008634 | 0.019147 | 0.012236 | 0.068652 | 0.010123 | 0.023719 | 0.017332 | 0.008013 | 0.019557 | 0.004291 | 0.003426 | 0.005258 | 0.002349 | 0.003322 | 0.007483 | 0.008719 | 0.007866 | 0.011215 | 0.015184 |
| 26.7361111 | 0.026189 | 0.007758 | 0.016609 | 0.00973 | 0.0082 | 0.009894 | 0.012808 | 0.046493 | 0.009835 | 0.022158 | 0.020684 | 0.006478 | 0.016478 | 0.005693 | 0.003705 | 0.005344 | 0.00229 | 0.002841 | 0.003661 | 0.0055 | 0.009131 | 0.014633 | 0.011586 |
| 26.75 | 0.030149 | 0.006763 | 0.014225 | 0.010863 | 0.009809 | 0.015515 | 0.010353 | 0.021389 | 0.008158 | 0.01831 | 0.017796 | 0.004769 | 0.009389 | 0.00475 | 0.003664 | 0.003973 | 0.001918 | 0.003146 | 0.001056 | 0.003234 | 0.00993 | 0.018509 | 0.009395 |
| 26.7638889 | 0.022397 | 0.006265 | 0.02091 | 0.011992 | 0.014115 | 0.024647 | 0.005965 | 0.00808 | 0.006687 | 0.013505 | 0.011376 | 0.008909 | 0.005472 | 0.002834 | 0.003779 | 0.00203 | 0.001469 | 0.003581 | 0.000459 | 0.002776 | 0.010104 | 0.017905 | 0.00996 |
| 26.7777778 | 0.012591 | 0.005415 | 0.021603 | 0.00975 | 0.011583 | 0.024415 | 0.003127 | 0.008096 | 0.00642 | 0.010961 | 0.011559 | 0.015992 | 0.008126 | 0.004166 | 0.004916 | 0.000916 | 0.00093 | 0.002267 | 0.000664 | 0.003584 | 0.008473 | 0.013428 | 0.011494 |
| 26.7916667 | 0.007882 | 0.004359 | 0.013891 | 0.010108 | 0.007135 | 0.016638 | 0.004237 | 0.008582 | 0.006379 | 0.012567 | 0.019995 | 0.021894 | 0.013088 | 0.006317 | 0.005717 | 0.00094 | 0.000926 | 0.001015 | 0.00157 | 0.004132 | 0.005077 | 0.028569 | 0.012182 |
| 26.8055556 | 0.012104 | 0.007013 | 0.006555 | 0.016374 | 0.009494 | 0.007718 | 0.006411 | 0.010301 | 0.005003 | 0.015173 | 0.026957 | 0.025067 | 0.018009 | 0.006135 | 0.004883 | 0.000774 | 0.001654 | 0.001469 | 0.002482 | 0.003352 | 0.00214 | 0.082885 | 0.010944 |
| 26.8194444 | 0.022775 | 0.01357 | 0.003926 | 0.027671 | 0.011663 | 0.003633 | 0.007213 | 0.018887 | 0.003816 | 0.014175 | 0.026766 | 0.022199 | 0.023834 | 0.004759 | 0.002942 | 0.000329 | 0.001962 | 0.002919 | 0.002016 | 0.002046 | 0.001465 | 0.126226 | 0.009032 |
| 26.8333333 | 0.029937 | 0.018344 | 0.002791 | 0.036021 | 0.0105 | 0.002878 | 0.007978 | 0.023298 | 0.004779 | 0.008987 | 0.020772 | 0.012891 | 0.025015 | 0.006179 | 0.001286 | 0.000392 | 0.001537 | 0.004651 | 0.000979 | 0.002046 | 0.002458 | 0.103621 | 0.009178 |
| 26.8472222 | 0.03153 | 0.018836 | 0.002298 | 0.03156 | 0.007182 | 0.001522 | 0.009415 | 0.017442 | 0.005877 | 0.006479 | 0.011862 | 0.006223 | 0.018346 | 0.01032 | 0.000747 | 0.001174 | 0.001259 | 0.004889 | 0.00071 | 0.003154 | 0.003479 | 0.051004 | 0.011438 |
| 26.8611111 | 0.030758 | 0.017257 | 0.005356 | 0.017946 | 0.003696 | 0.00076 | 0.010009 | 0.008413 | 0.004828 | 0.00998 | 0.00481 | 0.009118 | 0.013653 | 0.011387 | 0.000765 | 0.002426 | 0.001209 | 0.003555 | 0.000795 | 0.004324 | 0.003518 | 0.017126 | 0.014593 |
| 26.875 | 0.027525 | 0.01415 | 0.010397 | 0.013077 | 0.003549 | 0.001323 | 0.008959 | 0.0051 | 0.003322 | 0.010339 | 0.002941 | 0.013582 | 0.011765 | 0.007365 | 0.000781 | 0.00376 | 0.000854 | 0.003425 | 0.000598 | 0.005551 | 0.003144 | 0.004903 | 0.015899 |
| 26.8888889 | 0.01933 | 0.01011 | 0.014746 | 0.023366 | 0.003761 | 0.003529 | 0.00732 | 0.011272 | 0.00445 | 0.006304 | 0.004728 | 0.013769 | 0.00684 | 0.004278 | 0.000993 | 0.004828 | 0.000388 | 0.003525 | 0.00053 | 0.007128 | 0.003689 | 0.004456 | 0.014657 |
| 26.9027778 | 0.010069 | 0.00964 | 0.018324 | 0.033256 | 0.003667 | 0.009495 | 0.006081 | 0.019872 | 0.0073 | 0.003841 | 0.007837 | 0.013032 | 0.002974 | 0.004204 | 0.00165 | 0.0052 | 0.000228 | 0.002041 | 0.000884 | 0.008744 | 0.005284 | 0.0069 | 0.013813 |
| 26.9166667 | 0.008462 | 0.015117 | 0.022742 | 0.031734 | 0.005958 | 0.017282 | 0.006245 | 0.020912 | 0.008273 | 0.005854 | 0.008753 | 0.013571 | 0.004403 | 0.003997 | 0.001592 | 0.005052 | 0.000413 | 0.000631 | 0.001487 | 0.009099 | 0.007131 | 0.008314 | 0.015146 |
| 26.9305556 | 0.015866 | 0.023064 | 0.029139 | 0.019984 | 0.011736 | 0.024512 | 0.008612 | 0.012824 | 0.006683 | 0.017002 | 0.006534 | 0.012827 | 0.012316 | 0.006146 | 0.001233 | 0.004877 | 0.00097 | 0.000401 | 0.002665 | 0.007194 | 0.008672 | 0.00778 | 0.018056 |
| 26.9444444 | 0.0243 | 0.029098 | 0.034136 | 0.011815 | 0.021639 | 0.03076 | 0.012736 | 0.006613 | 0.005096 | 0.028952 | 0.006632 | 0.008763 | 0.021581 | 0.011531 | 0.003537 | 0.004771 | 0.001725 | 0.001209 | 0.005359 | 0.004342 | 0.010043 | 0.005111 | 0.0215 |
| 26.9583333 | 0.025919 | 0.031796 | 0.033318 | 0.020689 | 0.029693 | 0.033368 | 0.016678 | 0.010156 | 0.004674 | 0.029068 | 0.00935 | 0.006802 | 0.023112 | 0.013894 | 0.010571 | 0.004821 | 0.00229 | 0.001982 | 0.009794 | 0.003561 | 0.011154 | 0.004378 | 0.026143 |
| 26.9722222 | 0.019775 | 0.03295 | 0.028268 | 0.038618 | 0.02625 | 0.031572 | 0.01893 | 0.01535 | 0.004377 | 0.022522 | 0.008826 | 0.009116 | 0.017314 | 0.010241 | 0.018774 | 0.005093 | 0.002172 | 0.001479 | 0.01454 | 0.006258 | 0.011487 | 0.008177 | 0.031342 |
| 26.9861111 | 0.010324 | 0.035326 | 0.02435 | 0.050012 | 0.013274 | 0.029693 | 0.021449 | 0.012981 | 0.004251 | 0.023049 | 0.006714 | 0.008966 | 0.010184 | 0.004704 | 0.019465 | 0.00555 | 0.001418 | 0.000558 | 0.017615 | 0.010849 | 0.010777 | 0.012554 | 0.033665 |

| | | | | | | | | | | | | | | | | | | | | | | | |
|---|---|---|---|---|---|---|---|---|---|---|---|---|---|---|---|---|---|---|---|---|---|---|---|
| 27 | 0.006144 | 0.037352 | 0.022615 | 0.051095 | 0.005469 | 0.030651 | 0.023908 | 0.009247 | 0.004075 | 0.035371 | 0.008345 | 0.004936 | 0.005175 | 0.002166 | 0.013245 | 0.005483 | 0.00149 | 0.001083 | 0.018254 | 0.013455 | 0.007705 | 0.013768 | 0.031465 |
| 27.0138889 | 0.010558 | 0.035893 | 0.021362 | 0.049046 | 0.009108 | 0.031241 | 0.023544 | 0.017938 | 0.00323 | 0.050684 | 0.011585 | 0.002296 | 0.005422 | 0.004303 | 0.010882 | 0.003682 | 0.004266 | 0.003724 | 0.016279 | 0.009933 | 0.004782 | 0.011061 | 0.026287 |
| 27.0277778 | 0.01945 | 0.034806 | 0.022086 | 0.047814 | 0.014764 | 0.030155 | 0.027602 | 0.036918 | 0.002628 | 0.059696 | 0.011688 | 0.003001 | 0.010144 | 0.007552 | 0.016681 | 0.001243 | 0.009449 | 0.007191 | 0.011956 | 0.005747 | 0.006495 | 0.008251 | 0.020977 |
| 27.0416667 | 0.030957 | 0.040854 | 0.027702 | 0.049706 | 0.019122 | 0.034955 | 0.041586 | 0.05821 | 0.006478 | 0.063001 | 0.01199 | 0.003787 | 0.01131 | 0.007563 | 0.022608 | 0.000241 | 0.013681 | 0.009494 | 0.008381 | 0.007277 | 0.008612 | 0.014272 | 0.018051 |
| 27.0555556 | 0.041465 | 0.049872 | 0.036124 | 0.060357 | 0.029104 | 0.052741 | 0.050658 | 0.071882 | 0.015019 | 0.065734 | 0.021548 | 0.004797 | 0.012298 | 0.006612 | 0.019464 | 0.001467 | 0.014872 | 0.011198 | 0.007327 | 0.007602 | 0.006845 | 0.029014 | 0.017968 |
| 27.0694444 | 0.048429 | 0.051086 | 0.040303 | 0.071274 | 0.038622 | 0.076418 | 0.042418 | 0.068704 | 0.017783 | 0.06788 | 0.035269 | 0.009918 | 0.019673 | 0.010583 | 0.009954 | 0.004592 | 0.014059 | 0.01277 | 0.006954 | 0.004232 | 0.00358 | 0.041284 | 0.02089 |
| 27.0833333 | 0.052683 | 0.043834 | 0.038377 | 0.066335 | 0.034243 | 0.080791 | 0.024991 | 0.051555 | 0.011335 | 0.060752 | 0.036596 | 0.017183 | 0.023921 | 0.017397 | 0.003165 | 0.00648 | 0.01164 | 0.010256 | 0.008155 | 0.003091 | 0.001459 | 0.043562 | 0.02827 |
| 27.0972222 | 0.049821 | 0.031224 | 0.03331 | 0.046167 | 0.018612 | 0.05119 | 0.011076 | 0.02717 | 0.004618 | 0.039844 | 0.022774 | 0.021604 | 0.018892 | 0.019807 | 0.001902 | 0.005278 | 0.008213 | 0.005125 | 0.010619 | 0.004254 | 0.001023 | 0.034708 | 0.036948 |
| 27.1111111 | 0.036739 | 0.016231 | 0.023298 | 0.022526 | 0.005827 | 0.022719 | 0.008211 | 0.012918 | 0.00243 | 0.017378 | 0.010327 | 0.018687 | 0.010601 | 0.023068 | 0.004094 | 0.003694 | 0.005871 | 0.003454 | 0.009103 | 0.003661 | 0.001537 | 0.018588 | 0.037757 |
| 27.125 | 0.020557 | 0.00675 | 0.010595 | 0.007569 | 0.002053 | 0.025291 | 0.015509 | 0.011711 | 0.002484 | 0.006778 | 0.011959 | 0.010188 | 0.006115 | 0.02315 | 0.006931 | 0.003673 | 0.005201 | 0.003049 | 0.00559 | 0.002583 | 0.003687 | 0.006259 | 0.028132 |
| 27.1388889 | 0.010661 | 0.005065 | 0.003426 | 0.006277 | 0.002968 | 0.038144 | 0.025754 | 0.010945 | 0.002739 | 0.006522 | 0.017919 | 0.004409 | 0.004553 | 0.014967 | 0.007015 | 0.004161 | 0.004872 | 0.001443 | 0.005938 | 0.004147 | 0.006867 | 0.004375 | 0.016541 |
| 27.1527778 | 0.008398 | 0.006436 | 0.003472 | 0.012957 | 0.007006 | 0.042733 | 0.032497 | 0.015121 | 0.003519 | 0.009914 | 0.019674 | 0.003297 | 0.004047 | 0.010548 | 0.004544 | 0.004261 | 0.004386 | 0.000924 | 0.008292 | 0.006254 | 0.008501 | 0.007967 | 0.009522 |
| 27.1666667 | 0.007136 | 0.007613 | 0.007672 | 0.021272 | 0.010481 | 0.034562 | 0.030776 | 0.025278 | 0.005153 | 0.01521 | 0.020168 | 0.007032 | 0.003753 | 0.010646 | 0.00248 | 0.004587 | 0.004347 | 0.002017 | 0.010888 | 0.00705 | 0.008747 | 0.013548 | 0.006883 |
| 27.1805556 | 0.006196 | 0.006901 | 0.009799 | 0.027743 | 0.007564 | 0.017491 | 0.021873 | 0.03362 | 0.007165 | 0.02129 | 0.018861 | 0.016414 | 0.002528 | 0.011563 | 0.001634 | 0.005288 | 0.004727 | 0.003251 | 0.014062 | 0.006579 | 0.008886 | 0.017128 | 0.008558 |
| 27.1944444 | 0.011814 | 0.003899 | 0.007679 | 0.029284 | 0.003722 | 0.007092 | 0.012614 | 0.036353 | 0.008941 | 0.025738 | 0.014374 | 0.022296 | 0.003045 | 0.01228 | 0.002011 | 0.005374 | 0.005059 | 0.003136 | 0.016235 | 0.004617 | 0.00859 | 0.012887 | 0.015062 |
| 27.2083333 | 0.019413 | 0.00156 | 0.007396 | 0.024748 | 0.004231 | 0.010076 | 0.005986 | 0.031549 | 0.009974 | 0.027841 | 0.009506 | 0.017112 | 0.005427 | 0.013463 | 0.005009 | 0.004446 | 0.005179 | 0.001924 | 0.016274 | 0.00206 | 0.007682 | 0.006228 | 0.02205 |
| 27.2222222 | 0.022733 | 0.002142 | 0.009402 | 0.015329 | 0.004714 | 0.01698 | 0.003234 | 0.021597 | 0.009509 | 0.026887 | 0.007325 | 0.009122 | 0.006378 | 0.012548 | 0.010619 | 0.002709 | 0.005275 | 0.001545 | 0.014032 | 0.00087 | 0.006534 | 0.00495 | 0.022874 |
| 27.2361111 | 0.024136 | 0.00395 | 0.009611 | 0.007934 | 0.003842 | 0.01689 | 0.005384 | 0.012229 | 0.007634 | 0.02206 | 0.00692 | 0.00733 | 0.004304 | 0.009383 | 0.014216 | 0.001075 | 0.005174 | 0.002085 | 0.010998 | 0.001583 | 0.004423 | 0.005752 | 0.018683 |
| 27.25 | 0.025551 | 0.004448 | 0.007937 | 0.010477 | 0.004028 | 0.011452 | 0.009823 | 0.008078 | 0.006072 | 0.016201 | 0.006193 | 0.007444 | 0.001701 | 0.007393 | 0.012413 | 0.000719 | 0.004421 | 0.001713 | 0.009638 | 0.002324 | 0.002699 | 0.004922 | 0.016594 |
| 27.2638889 | 0.02496 | 0.00287 | 0.005431 | 0.018346 | 0.004758 | 0.008233 | 0.013527 | 0.009791 | 0.006141 | 0.013768 | 0.004442 | 0.00615 | 0.002159 | 0.007259 | 0.008485 | 0.001512 | 0.002827 | 0.001693 | 0.00865 | 0.002296 | 0.004998 | 0.006408 | 0.017811 |
| 27.2777778 | 0.020149 | 0.001508 | 0.006822 | 0.023656 | 0.004005 | 0.008195 | 0.013891 | 0.013279 | 0.007472 | 0.015303 | 0.002322 | 0.007869 | 0.005545 | 0.011917 | 0.005101 | 0.003956 | 0.002592 | 0.004979 | 0.005207 | 0.003916 | 0.01132 | 0.012706 | 0.018425 |
| 27.2916667 | 0.013812 | 0.003231 | 0.020416 | 0.022924 | 0.004232 | 0.006762 | 0.009251 | 0.01228 | 0.00834 | 0.015909 | 0.001321 | 0.012061 | 0.007856 | 0.013872 | 0.002686 | 0.007356 | 0.005198 | 0.00938 | 0.002692 | 0.006615 | 0.016501 | 0.018105 | 0.016069 |
| 27.3055556 | 0.009276 | 0.007215 | 0.037452 | 0.014556 | 0.010422 | 0.003795 | 0.005347 | 0.006741 | 0.00688 | 0.010954 | 0.002023 | 0.012869 | 0.005944 | 0.008926 | 0.001799 | 0.008216 | 0.006349 | 0.009707 | 0.005554 | 0.005535 | 0.014605 | 0.01611 | 0.010668 |
| 27.3194444 | 0.006453 | 0.009616 | 0.032659 | 0.005116 | 0.017545 | 0.002162 | 0.006634 | 0.003283 | 0.003861 | 0.004025 | 0.002627 | 0.009224 | 0.003308 | 0.005846 | 0.001917 | 0.006263 | 0.003769 | 0.006826 | 0.01087 | 0.002644 | 0.007725 | 0.00974 | 0.004866 |
| 27.3333333 | 0.00821 | 0.009017 | 0.015626 | 0.001189 | 0.016789 | 0.002679 | 0.007444 | 0.005073 | 0.001697 | 0.001278 | 0.003603 | 0.004717 | 0.005151 | 0.006178 | 0.003166 | 0.003765 | 0.001308 | 0.004733 | 0.014062 | 0.002712 | 0.004415 | 0.005553 | 0.00192 |
| 27.3472222 | 0.014766 | 0.007788 | 0.013318 | 0.000623 | 0.010229 | 0.005565 | 0.006303 | 0.010063 | 0.000919 | 0.001811 | 0.007109 | 0.002103 | 0.009176 | 0.00447 | 0.004085 | 0.00181 | 0.000876 | 0.004154 | 0.014542 | 0.003004 | 0.006545 | 0.003922 | 0.00257 |
| 27.3611111 | 0.019675 | 0.006921 | 0.016278 | 0.00151 | 0.008253 | 0.008232 | 0.004951 | 0.014654 | 0.000692 | 0.004032 | 0.00897 | 0.001491 | 0.010924 | 0.004315 | 0.003715 | 0.001683 | 0.001001 | 0.005111 | 0.012848 | 0.001588 | 0.008797 | 0.002497 | 0.005727 |
| 27.375 | 0.021345 | 0.006398 | 0.015163 | 0.008656 | 0.011105 | 0.007146 | 0.002523 | 0.016025 | 0.00074 | 0.009401 | 0.005935 | 0.002383 | 0.010883 | 0.013157 | 0.003708 | 0.002686 | 0.001351 | 0.006381 | 0.010768 | 0.000793 | 0.009251 | 0.00272 | 0.011002 |
| 27.3888889 | 0.019395 | 0.008324 | 0.015352 | 0.021017 | 0.00983 | 0.005531 | 0.000802 | 0.013294 | 0.001672 | 0.016886 | 0.003531 | 0.005812 | 0.01174 | 0.02548 | 0.00502 | 0.002253 | 0.001512 | 0.005761 | 0.010672 | 0.001849 | 0.010612 | 0.005924 | 0.018049 |
| 27.4027778 | 0.01172 | 0.013153 | 0.016547 | 0.027633 | 0.005056 | 0.010107 | 0.000733 | 0.007965 | 0.003716 | 0.024529 | 0.007036 | 0.011145 | 0.013949 | 0.027431 | 0.007332 | 0.000897 | 0.00098 | 0.003881 | 0.011373 | 0.003837 | 0.013992 | 0.009955 | 0.025023 |
| 27.4166667 | 0.006083 | 0.018333 | 0.017615 | 0.026644 | 0.003429 | 0.019713 | 0.001519 | 0.00575 | 0.006268 | 0.032783 | 0.013774 | 0.016573 | 0.01568 | 0.01726 | 0.008888 | 0.000781 | 0.000583 | 0.002152 | 0.011365 | 0.005114 | 0.016989 | 0.013325 | 0.030244 |
| 27.4305556 | 0.010983 | 0.022686 | 0.019155 | 0.022951 | 0.007764 | 0.029077 | 0.003286 | 0.012152 | 0.008706 | 0.043386 | 0.021305 | 0.021282 | 0.015602 | 0.006822 | 0.008449 | 0.00156 | 0.000885 | 0.001046 | 0.011405 | 0.005675 | 0.017739 | 0.015243 | 0.033122 |
| 27.4444444 | 0.02092 | 0.025915 | 0.022246 | 0.017912 | 0.013591 | 0.034646 | 0.005283 | 0.021348 | 0.00971 | 0.053357 | 0.029179 | 0.024272 | 0.01298 | 0.004496 | 0.007134 | 0.00209 | 0.002531 | 0.002114 | 0.010014 | 0.00624 | 0.015326 | 0.01397 | 0.034029 |
| 27.4583333 | 0.023907 | 0.024974 | 0.025387 | 0.012269 | 0.014254 | 0.033915 | 0.005231 | 0.024291 | 0.007242 | 0.056949 | 0.030189 | 0.023232 | 0.008355 | 0.009272 | 0.005336 | 0.004311 | 0.006876 | 0.008351 | 0.00722 | 0.004901 | 0.010985 | 0.00941 | 0.032323 |
| 27.4722222 | 0.019139 | 0.019613 | 0.026286 | 0.008562 | 0.010165 | 0.030359 | 0.00309 | 0.024977 | 0.003556 | 0.056952 | 0.022145 | 0.018869 | 0.005226 | 0.016723 | 0.002877 | 0.008995 | 0.01231 | 0.017619 | 0.007591 | 0.002891 | 0.01076 | 0.006767 | 0.028951 |
| 27.4861111 | 0.022741 | 0.016702 | 0.026214 | 0.012178 | 0.005777 | 0.031499 | 0.004455 | 0.032692 | 0.002571 | 0.061244 | 0.018127 | 0.015977 | 0.004426 | 0.020531 | 0.002721 | 0.010661 | 0.013348 | 0.019899 | 0.009013 | 0.002928 | 0.012745 | 0.006587 | 0.028059 |
| 27.5 | 0.039327 | 0.019907 | 0.027341 | 0.02111 | 0.002965 | 0.036795 | 0.012265 | 0.045688 | 0.003857 | 0.068543 | 0.024856 | 0.015599 | 0.005346 | 0.014378 | 0.005314 | 0.006508 | 0.009596 | 0.012828 | 0.00642 | 0.002183 | 0.009689 | 0.005112 | 0.029269 |
| 27.5138889 | 0.051323 | 0.024482 | 0.029787 | 0.021448 | 0.001642 | 0.038734 | 0.020243 | 0.053334 | 0.005801 | 0.070495 | 0.030345 | 0.015363 | 0.009261 | 0.00776 | 0.006712 | 0.002485 | 0.006566 | 0.005486 | 0.004559 | 0.000734 | 0.005618 | 0.006379 | 0.028976 |
| 27.5277778 | 0.047041 | 0.026913 | 0.034821 | 0.012517 | 0.001417 | 0.039306 | 0.025586 | 0.054604 | 0.007459 | 0.063501 | 0.027248 | 0.01223 | 0.011921 | 0.012931 | 0.005319 | 0.001919 | 0.008006 | 0.004863 | 0.004681 | 0.002303 | 0.010274 | 0.010304 | 0.024505 |
| 27.5416667 | 0.035644 | 0.02658 | 0.045843 | 0.007958 | 0.002953 | 0.045737 | 0.029063 | 0.06291 | 0.009587 | 0.048074 | 0.024235 | 0.008417 | 0.00998 | 0.020855 | 0.007241 | 0.004834 | 0.015931 | 0.014342 | 0.007979 | 0.010482 | 0.032634 | 0.010532 | 0.015384 |
| 27.5555556 | 0.026402 | 0.021827 | 0.053262 | 0.008162 | 0.009208 | 0.052547 | 0.025746 | 0.074399 | 0.010228 | 0.029709 | 0.025052 | 0.009712 | 0.00891 | 0.020569 | 0.01721 | 0.012263 | 0.027587 | 0.030834 | 0.020413 | 0.023428 | 0.064702 | 0.007249 | 0.010013 |
| 27.5694444 | 0.014932 | 0.017398 | 0.038914 | 0.010754 | 0.020392 | 0.049143 | 0.017206 | 0.068883 | 0.00668 | 0.022413 | 0.020268 | 0.009843 | 0.010374 | 0.012697 | 0.023878 | 0.016515 | 0.032586 | 0.040105 | 0.028698 | 0.030108 | 0.073861 | 0.005534 | 0.009246 |
| 27.5833333 | 0.005455 | 0.024095 | 0.015415 | 0.025031 | 0.023879 | 0.041659 | 0.015701 | 0.056538 | 0.004497 | 0.037923 | 0.010299 | 0.008473 | 0.016107 | 0.005718 | 0.016973 | 0.011035 | 0.025244 | 0.033022 | 0.019318 | 0.024335 | 0.049261 | 0.005992 | 0.011271 |
| 27.5972222 | 0.00623 | 0.040738 | 0.012757 | 0.047007 | 0.020717 | 0.039152 | 0.023373 | 0.057132 | 0.007272 | 0.06299 | 0.008434 | 0.016778 | 0.024362 | 0.008594 | 0.008319 | 0.004968 | 0.012876 | 0.018072 | 0.007319 | 0.012107 | 0.018921 | 0.010188 | 0.026326 |
| 27.6111111 | 0.012761 | 0.053958 | 0.032727 | 0.051451 | 0.029698 | 0.032762 | 0.029688 | 0.064319 | 0.010325 | 0.074124 | 0.015238 | 0.028614 | 0.021205 | 0.012895 | 0.011074 | 0.005302 | 0.004287 | 0.006804 | 0.004615 | 0.003834 | 0.004083 | 0.019466 | 0.045673 |
| 27.625 | 0.01978 | 0.055671 | 0.053183 | 0.035715 | 0.042906 | 0.019543 | 0.030615 | 0.064669 | 0.012097 | 0.068484 | 0.022055 | 0.033933 | 0.016111 | 0.010158 | 0.019626 | 0.006224 | 0.000995 | 0.00186 | 0.003419 | 0.003343 | 0.001301 | 0.023552 | 0.052263 |
| 27.6388889 | 0.023738 | 0.04453 | 0.057667 | 0.019352 | 0.044739 | 0.010364 | 0.027745 | 0.051514 | 0.013547 | 0.046504 | 0.024343 | 0.034331 | 0.025086 | 0.009323 | 0.025367 | 0.006253 | 0.001086 | 0.001352 | 0.001767 | 0.006137 | 0.002029 | 0.018004 | 0.046505 |
| 27.6527778 | 0.019655 | 0.025255 | 0.049711 | 0.013853 | 0.03488 | 0.007784 | 0.021693 | 0.02763 | 0.012925 | 0.022066 | 0.019755 | 0.03228 | 0.042697 | 0.009945 | 0.024717 | 0.009668 | 0.003003 | 0.003653 | 0.002666 | 0.009333 | 0.003524 | 0.011982 | 0.038047 |
| 27.6666667 | 0.011872 | 0.009513 | 0.038371 | 0.02088 | 0.019674 | 0.00747 | 0.016682 | 0.009245 | 0.010985 | 0.011904 | 0.010989 | 0.026894 | 0.053415 | 0.007871 | 0.018031 | 0.017044 | 0.005604 | 0.007364 | 0.005767 | 0.01244 | 0.005165 | 0.008894 | 0.036772 |
| 27.6805556 | 0.007798 | 0.004509 | 0.026598 | 0.024002 | 0.009662 | 0.008461 | 0.017098 | 0.007305 | 0.011113 | 0.013554 | 0.004652 | 0.01933 | 0.041207 | 0.010623 | 0.010361 | 0.02516 | 0.006993 | 0.009857 | 0.008606 | 0.013462 | 0.005444 | 0.005298 | 0.039313 |
| 27.6944444 | 0.005276 | 0.006073 | 0.016084 | 0.013988 | 0.011242 | 0.010469 | 0.019244 | 0.01706 | 0.013509 | 0.024593 | 0.005802 | 0.014008 | 0.019468 | 0.018629 | 0.005365 | 0.029489 | 0.006163 | 0.009275 | 0.009352 | 0.011616 | 0.00415 | 0.002003 | 0.03622 |
| 27.7083333 | 0.0048 | 0.00692 | 0.010173 | 0.008109 | 0.022603 | 0.009949 | 0.016191 | 0.034158 | 0.015178 | 0.030114 | 0.012925 | 0.010574 | 0.006949 | 0.026612 | 0.003783 | 0.028811 | 0.004383 | 0.006975 | 0.009763 | 0.008592 | 0.002223 | 0.000984 | 0.027484 |
| 27.7222222 | 0.010005 | 0.005545 | 0.012618 | 0.017653 | 0.029463 | 0.006318 | 0.008416 | 0.051531 | 0.014306 | 0.021737 | 0.016821 | 0.007436 | 0.004623 | 0.028349 | 0.003891 | 0.025237 | 0.003032 | 0.005395 | 0.010447 | 0.006373 | 0.000669 | 0.000947 | 0.016106 |
| 27.7361111 | 0.013587 | 0.007062 | 0.024797 | 0.030531 | 0.022534 | 0.005882 | 0.0023 | 0.061871 | 0.011664 | 0.012203 | 0.012718 | 0.004287 | 0.011641 | 0.018175 | 0.003506 | 0.020886 | 0.002149 | 0.004908 | 0.01036 | 0.005515 | 0.000287 | 0.001452 | 0.007069 |
| 27.75 | 0.011103 | 0.013001 | 0.032467 | 0.030259 | 0.015733 | 0.011454 | 0.001105 | 0.061111 | 0.008908 | 0.012366 | 0.006931 | 0.002881 | 0.020104 | 0.010435 | 0.002413 | 0.016514 | 0.001313 | 0.004284 | 0.009232 | 0.00539 | 0.000938 | 0.003943 | 0.004584 |
| 27.7638889 | 0.014276 | 0.016182 | 0.025364 | 0.016495 | 0.014913 | 0.017952 | 0.002886 | 0.04476 | 0.006694 | 0.019498 | 0.004264 | 0.005548 | 0.021801 | 0.016017 | 0.001437 | 0.012056 | 0.000764 | 0.003466 | 0.007878 | 0.005038 | 0.002459 | 0.007724 | 0.004382 |
| 27.7777778 | 0.024552 | 0.01267 | 0.026808 | 0.004419 | 0.01224 | 0.019964 | 0.005034 | 0.022937 | 0.004469 | 0.018596 | 0.003955 | 0.008265 | 0.018381 | 0.021382 | 0.001207 | 0.007841 | 0.001042 | 0.003671 | 0.00716 | 0.00408 | 0.004342 | 0.011607 | 0.005712 |
| 27.7916667 | 0.028026 | 0.006951 | 0.044513 | 0.002119 | 0.014367 | 0.016559 | 0.005468 | 0.019717 | 0.002091 | 0.015767 | 0.006701 | 0.0077 | 0.013574 | 0.018236 | 0.002305 | 0.00479 | 0.002247 | 0.005141 | 0.006961 | 0.003309 | 0.005716 | 0.015276 | 0.0108 |
| 27.8055556 | 0.021907 | 0.006198 | 0.04909 | 0.005109 | 0.023634 | 0.010648 | 0.00395 | 0.037255 | 0.000822 | 0.028144 | 0.012279 | 0.005839 | 0.008065 | 0.013149 | 0.004808 | 0.003565 | 0.004103 | 0.006996 | 0.006643 | 0.003645 | 0.006891 | 0.01815 | 0.016327 |
| 27.8194444 | 0.012574 | 0.011979 | 0.031368 | 0.016675 | 0.027422 | 0.005393 | 0.003577 | 0.048462 | 0.001443 | 0.035679 | 0.017157 | 0.004248 | 0.004194 | 0.017298 | 0.00719 | 0.003766 | 0.00602 | 0.00863 | 0.006359 | 0.004681 | 0.007971 | 0.020875 | 0.020234 |
| 27.8333333 | 0.011713 | 0.017604 | 0.013995 | 0.033713 | 0.019939 | 0.003268 | 0.008496 | 0.037057 | 0.001892 | 0.022143 | 0.017858 | 0.002579 | 0.004962 | 0.029385 | 0.00722 | 0.004355 | 0.007298 | 0.009993 | 0.00669 | 0.005223 | 0.007892 | 0.022521 | 0.020937 |
| 27.8472222 | 0.027446 | 0.018022 | 0.013628 | 0.034893 | 0.010777 | 0.006424 | 0.016487 | 0.02081 | 0.001563 | 0.011485 | 0.014151 | 0.002145 | 0.008638 | 0.036065 | 0.004456 | 0.004808 | 0.007855 | 0.011326 | 0.008017 | 0.004901 | 0.006031 | 0.020163 | 0.016146 |
| 27.8611111 | 0.045089 | 0.014936 | 0.024088 | 0.020899 | 0.00738 | 0.011938 | 0.019501 | 0.023621 | 0.002726 | 0.018702 | 0.00951 | 0.0045 | 0.010878 | 0.030921 | 0.001986 | 0.005066 | 0.008113 | 0.012716 | 0.009957 | 0.004293 | 0.003139 | 0.013687 | 0.008249 |
| 27.875 | 0.053141 | 0.01083 | 0.032312 | 0.016373 | 0.00753 | 0.012129 | 0.015925 | 0.033463 | 0.004574 | 0.024689 | 0.007791 | 0.006279 | 0.010185 | 0.017225 | 0.002117 | 0.00468 | 0.007959 | 0.012919 | 0.010731 | 0.003673 | 0.001329 | 0.006599 | 0.004629 |
| 27.8888889 | 0.052169 | 0.006106 | 0.042845 | 0.022092 | 0.007878 | 0.008701 | 0.010019 | 0.02854 | 0.005215 | 0.016145 | 0.009599 | 0.004435 | 0.006929 | 0.010399 | 0.002636 | 0.003076 | 0.006888 | 0.010736 | 0.009214 | 0.002931 | 0.001648 | 0.00358 | 0.00823 |
| 27.9027778 | 0.039474 | 0.002027 | 0.064656 | 0.019977 | 0.009349 | 0.012338 | 0.005393 | 0.017516 | 0.00391 | 0.006589 | 0.01113 | 0.00226 | 0.004531 | 0.019285 | 0.002522 | 0.001554 | 0.005155 | 0.007727 | 0.006588 | 0.002159 | 0.003234 | 0.004505 | 0.013432 |
| 27.9166667 | 0.024075 | 0.002408 | 0.083098 | 0.011738 | 0.012255 | 0.02237 | 0.004208 | 0.019146 | 0.00352 | 0.004763 | 0.008221 | 0.002813 | 0.006752 | 0.028273 | 0.002979 | 0.002111 | 0.003508 | 0.005701 | 0.003813 | 0.001432 | 0.005277 | 0.004215 | 0.017467 |
| 27.9305556 | 0.012655 | 0.008998 | 0.080881 | 0.007579 | 0.009987 | 0.030011 | 0.005523 | 0.033062 | 0.007845 | 0.009534 | 0.00387 | 0.004501 | 0.008826 | 0.032795 | 0.003596 | 0.004243 | 0.002267 | 0.00426 | 0.002459 | 0.000795 | 0.006938 | 0.001963 | 0.020886 |
| 27.9444444 | 0.00652 | 0.018486 | 0.067302 | 0.005024 | 0.00596 | 0.033939 | 0.007703 | 0.04409 | 0.013481 | 0.022678 | 0.005261 | 0.005942 | 0.007494 | 0.036348 | 0.003772 | 0.006569 | 0.001349 | 0.003039 | 0.004587 | 0.000838 | 0.008715 | 0.001 | 0.022252 |
| 27.9583333 | 0.008316 | 0.027126 | 0.056695 | 0.005475 | 0.005797 | 0.035813 | 0.011402 | 0.042338 | 0.015936 | 0.038012 | 0.014285 | 0.007316 | 0.010281 | 0.026211 | 0.004408 | 0.008615 | 0.000792 | 0.002352 | 0.00845 | 0.002253 | 0.011125 | 0.002626 | 0.020916 |
| 27.9722222 | 0.009553 | 0.033525 | 0.049149 | 0.017449 | 0.014318 | 0.034978 | 0.016741 | 0.02897 | 0.014628 | 0.045877 | 0.026112 | 0.009159 | 0.0202 | 0.011678 | 0.006557 | 0.010629 | 0.00116 | 0.001796 | 0.011613 | 0.004745 | 0.012247 | 0.005981 | 0.018058 |
| 27.9861111 | 0.008525 | 0.037312 | 0.043467 | 0.028154 | 0.030022 | 0.031827 | 0.022231 | 0.014654 | 0.011235 | 0.045804 | 0.03276 | 0.011697 | 0.028294 | 0.007729 | 0.009529 | 0.01286 | 0.002481 | 0.000914 | 0.013945 | 0.007128 | 0.011304 | 0.008898 | 0.01629 |
| 28 | 0.013439 | 0.037369 | 0.044422 | 0.020779 | 0.036943 | 0.026028 | 0.026094 | 0.011694 | 0.007548 | 0.041774 | 0.029895 | 0.01297 | 0.029118 | 0.00923 | 0.010849 | 0.01412 | 0.00298 | 0.0009 | 0.014912 | 0.007552 | 0.009162 | 0.009188 | 0.017876 |
| 28.0138889 | 0.021576 | 0.032935 | 0.045259 | 0.009336 | 0.032006 | 0.016256 | 0.026425 | 0.014954 | 0.004375 | 0.037419 | 0.019168 | 0.012318 | 0.023662 | 0.012995 | 0.007967 | 0.013218 | 0.002916 | 0.003283 | 0.013045 | 0.005469 | 0.006375 | 0.006464 | 0.021871 |
| 28.0277778 | 0.027083 | 0.029081 | 0.042331 | 0.008778 | 0.019812 | 0.010616 | 0.026952 | 0.016174 | 0.002978 | 0.037042 | 0.008217 | 0.013215 | 0.013707 | 0.012263 | 0.003197 | 0.012122 | 0.004226 | 0.00607 | 0.01112 | 0.003652 | 0.005126 | 0.005988 | 0.026952 |
| 28.0416667 | 0.03368 | 0.033314 | 0.049561 | 0.016269 | 0.007291 | 0.023516 | 0.031628 | 0.028509 | 0.003406 | 0.045618 | 0.010155 | 0.019381 | 0.009617 | 0.006529 | 0.005273 | 0.01519 | 0.005356 | 0.005905 | 0.015618 | 0.004786 | 0.006825 | 0.017499 | 0.03729 |
| 28.0555556 | 0.043357 | 0.042251 | 0.062385 | 0.027704 | 0.005233 | 0.056003 | 0.032319 | 0.062467 | 0.004932 | 0.063634 | 0.029204 | 0.02918 | 0.02006 | 0.005637 | 0.01765 | 0.019942 | 0.005975 | 0.003699 | 0.02421 | 0.009299 | 0.012506 | 0.037415 | 0.05388 |
| 28.0694444 | 0.0486 | 0.045446 | 0.064459 | 0.036883 | 0.008196 | 0.089337 | 0.023596 | 0.0963 | 0.006286 | 0.07987 | 0.04758 | 0.03399 | 0.037915 | 0.015217 | 0.030539 | 0.019161 | 0.005962 | 0.002355 | 0.027127 | 0.013279 | 0.02 | 0.046268 | 0.064105 |
| 28.0833333 | 0.042878 | 0.041347 | 0.050998 | 0.036255 | 0.006237 | 0.09355 | 0.011708 | 0.09687 | 0.005466 | 0.076807 | 0.043771 | 0.033518 | 0.049137 | 0.033251 | 0.033153 | 0.013173 | 0.003545 | 0.001852 | 0.024851 | 0.012191 | 0.022571 | 0.042759 | 0.057938 |
| 28.0972222 | 0.031585 | 0.035182 | 0.030178 | 0.024662 | 0.004507 | 0.059582 | 0.00513 | 0.062809 | 0.003196 | 0.054239 | 0.022843 | 0.032831 | 0.042734 | 0.044917 | 0.025957 | 0.006701 | 0.001311 | 0.001282 | 0.021879 | 0.008734 | 0.020781 | 0.038973 | 0.042076 |
| 28.1111111 | 0.020311 | 0.028057 | 0.013985 | 0.011512 | 0.00752 | 0.024923 | 0.008347 | 0.025504 | 0.002161 | 0.030998 | 0.008487 | 0.024461 | 0.024841 | 0.033724 | 0.015564 | 0.004713 | 0.001178 | 0.002179 | 0.013964 | 0.005215 | 0.01558 | 0.030188 | 0.025895 |
| 28.125 | 0.010838 | 0.01901 | 0.006641 | 0.006754 | 0.008078 | 0.017205 | 0.015592 | 0.007648 | 0.003402 | 0.019358 | 0.008762 | 0.011599 | 0.017664 | 0.020902 | 0.007275 | 0.008014 | 0.003372 | 0.007031 | 0.009936 | 0.003986 | 0.009526 | 0.016745 | 0.012197 |
| 28.1388889 | 0.008312 | 0.011195 | 0.004495 | 0.006304 | 0.004258 | 0.018717 | 0.018602 | 0.005777 | 0.004228 | 0.016883 | 0.012268 | 0.007681 | 0.023013 | 0.026193 | 0.004666 | 0.011829 | 0.008308 | 0.015123 | 0.018677 | 0.008397 | 0.010579 | 0.012663 | 0.004005 |
| 28.1527778 | 0.01062 | 0.00684 | 0.003569 | 0.007157 | 0.002502 | 0.017368 | 0.016575 | 0.011218 | 0.002691 | 0.01625 | 0.012662 | 0.008945 | 0.020536 | 0.026076 | 0.006679 | 0.012847 | 0.012354 | 0.020421 | 0.024138 | 0.012817 | 0.013591 | 0.017024 | 0.001906 |
| 28.1666667 | 0.008512 | 0.00435 | 0.006524 | 0.009697 | 0.002644 | 0.015401 | 0.013675 | 0.016111 | 0.001332 | 0.013595 | 0.012836 | 0.009505 | 0.010674 | 0.015396 | 0.008079 | 0.009628 | 0.012021 | 0.018461 | 0.01861 | 0.011587 | 0.0101 | 0.016871 | 0.002257 |
| 28.1805556 | 0.004242 | 0.004807 | 0.011599 | 0.008011 | 0.003803 | 0.011141 | 0.012097 | 0.016792 | 0.001759 | 0.009842 | 0.014928 | 0.010587 | 0.008586 | 0.009258 | 0.006135 | 0.004583 | 0.008194 | 0.012059 | 0.009514 | 0.006484 | 0.004941 | 0.010773 | 0.005116 |
| 28.1944444 | 0.005973 | 0.008601 | 0.013658 | 0.004792 | 0.007853 | 0.007293 | 0.009366 | 0.015463 | 0.003169 | 0.007566 | 0.019545 | 0.010667 | 0.014745 | 0.00698 | 0.003318 | 0.003029 | 0.00386 | 0.005983 | 0.004981 | 0.002944 | 0.005046 | 0.00618 | 0.012601 |
| 28.2083333 | 0.009741 | 0.010585 | 0.012486 | 0.005937 | 0.011444 | 0.007569 | 0.004987 | 0.011818 | 0.005205 | 0.007872 | 0.025207 | 0.007405 | 0.020701 | 0.007558 | 0.002801 | 0.005851 | 0.001806 | 0.004022 | 0.009617 | 0.004137 | 0.009492 | 0.007613 | 0.022386 |
| 28.2222222 | 0.00997 | 0.008311 | 0.010022 | 0.005904 | 0.009524 | 0.009814 | 0.002164 | 0.00697 | 0.007297 | 0.00886 | 0.028006 | 0.005301 | 0.021749 | 0.016355 | 0.003262 | 0.008694 | 0.002051 | 0.00567 | 0.016279 | 0.006427 | 0.015017 | 0.0109 | 0.027607 |
| 28.2361111 | 0.012051 | 0.005033 | 0.006725 | 0.004359 | 0.005553 | 0.013311 | 0.001435 | 0.005181 | 0.008329 | 0.008261 | 0.026533 | 0.00989 | 0.01944 | 0.027429 | 0.004335 | 0.007922 | 0.002737 | 0.005255 | 0.018306 | 0.007197 | 0.019508 | 0.009299 | 0.026507 |
| 28.25 | 0.024249 | 0.0035 | 0.003435 | 0.005722 | 0.007603 | 0.020088 | 0.001119 | 0.006265 | 0.008495 | 0.00649 | 0.022944 | 0.018147 | 0.01668 | 0.029396 | 0.008507 | 0.006076 | 0.004651 | 0.003515 | 0.017615 | 0.008825 | 0.02329 | 0.007081 | 0.026206 |
| 28.2638889 | 0.040538 | 0.003365 | 0.002059 | 0.007127 | 0.015663 | 0.026126 | 0.001356 | 0.005974 | 0.009055 | 0.004697 | 0.01903 | 0.025058 | 0.012428 | 0.025233 | 0.013078 | 0.007568 | 0.00791 | 0.005113 | 0.016113 | 0.011893 | 0.026179 | 0.010585 | 0.028667 |
| 28.2777778 | 0.046045 | 0.003895 | 0.001479 | 0.008628 | 0.024786 | 0.023154 | 0.003372 | 0.003877 | 0.010166 | 0.003847 | 0.015637 | 0.025275 | 0.00741 | 0.02338 | 0.012522 | 0.009924 | 0.008356 | 0.006717 | 0.011284 | 0.012527 | 0.023375 | 0.015141 | 0.02689 |
| 28.2916667 | 0.0376 | 0.005071 | 0.000937 | 0.013082 | 0.029442 | 0.012772 | 0.006402 | 0.00248 | 0.010797 | 0.005961 | 0.011772 | 0.015859 | 0.005933 | 0.021427 | 0.008381 | 0.00763 | 0.004811 | 0.005845 | 0.006915 | 0.008616 | 0.015422 | 0.013319 | 0.019318 |
| 28.3055556 | 0.021002 | 0.006242 | 0.001577 | 0.017895 | 0.026348 | 0.007188 | 0.006562 | 0.003935 | 0.010157 | 0.010031 | 0.006844 | 0.007215 | 0.005061 | 0.013907 | 0.007105 | 0.004543 | 0.001774 | 0.007422 | 0.010046 | 0.00377 | 0.009389 | 0.006653 | 0.012393 |
| 28.3194444 | 0.010452 | 0.006574 | 0.002713 | 0.017763 | 0.01836 | 0.009781 | 0.003448 | 0.006927 | 0.008738 | 0.011842 | 0.004697 | 0.007648 | 0.002483 | 0.008693 | 0.008835 | 0.005569 | 0.001719 | 0.010753 | 0.014429 | 0.002034 | 0.007942 | 0.00251 | 0.007848 |
| 28.3333333 | 0.01702 | 0.005736 | 0.002676 | 0.013022 | 0.009924 | 0.012521 | 0.000946 | 0.007307 | 0.007243 | 0.009819 | 0.006481 | 0.009257 | 0.001792 | 0.012192 | 0.007774 | 0.005607 | 0.002936 | 0.010172 | 0.013128 | 0.003201 | 0.007032 | 0.003702 | 0.005196 |
| 28.3472222 | 0.031492 | 0.004315 | 0.00193 | 0.008342 | 0.004728 | 0.01073 | 0.000944 | 0.00455 | 0.005059 | 0.005551 | 0.007719 | 0.008264 | 0.003218 | 0.017133 | 0.004161 | 0.003267 | 0.003906 | 0.00657 | 0.008809 | 0.004517 | 0.004721 | 0.006292 | 0.006535 |
| 28.3611111 | 0.045047 | 0.003269 | 0.001573 | 0.004912 | 0.004409 | 0.005938 | 0.00282 | 0.002566 | 0.002309 | 0.003049 | 0.006898 | 0.008126 | 0.005361 | 0.022013 | 0.00221 | 0.002924 | 0.003669 | 0.00376 | 0.005279 | 0.003878 | 0.003943 | 0.006452 | 0.008493 |
| 28.375 | 0.051432 | 0.003034 | 0.002841 | 0.002629 | 0.004667 | 0.003667 | 0.004652 | 0.003282 | 0.000865 | 0.006201 | 0.005252 | 0.008437 | 0.00625 | 0.028413 | 0.002454 | 0.004449 | 0.002706 | 0.002478 | 0.004286 | 0.002303 | 0.004144 | 0.00416 | 0.007914 |
| 28.3888889 | 0.04128 | 0.003621 | 0.007982 | 0.003203 | 0.003693 | 0.008643 | 0.004619 | 0.00414 | 0.001378 | 0.013899 | 0.006596 | 0.008102 | 0.006893 | 0.030608 | 0.00443 | 0.004632 | 0.003037 | 0.001783 | 0.005416 | 0.001834 | 0.004097 | 0.001595 | 0.008558 |
| 28.4027778 | 0.021032 | 0.004232 | 0.014405 | 0.006148 | 0.005529 | 0.01761 | 0.002888 | 0.003883 | 0.002314 | 0.022835 | 0.013446 | 0.007689 | 0.011318 | 0.028029 | 0.004829 | 0.003099 | 0.003494 | 0.00188 | 0.007435 | 0.001694 | 0.007042 | 0.001425 | 0.013253 |
| 28.4166667 | 0.008139 | 0.004318 | 0.016841 | 0.009965 | 0.010151 | 0.024601 | 0.001297 | 0.005146 | 0.002704 | 0.031368 | 0.020681 | 0.00865 | 0.017542 | 0.0284 | 0.002681 | 0.00201 | 0.002361 | 0.002467 | 0.010219 | 0.001459 | 0.012325 | 0.004725 | 0.019331 |
| 28.4305556 | 0.010056 | 0.004882 | 0.016975 | 0.014461 | 0.015062 | 0.028816 | 0.001035 | 0.010422 | 0.003131 | 0.040167 | 0.025488 | 0.010989 | 0.019088 | 0.035029 | 0.001332 | 0.001869 | 0.001959 | 0.002097 | 0.014636 | 0.001524 | 0.014378 | 0.009742 | 0.023055 |
| 28.4444444 | 0.017577 | 0.005598 | 0.01825 | 0.018467 | 0.017553 | 0.028842 | 0.00165 | 0.017125 | 0.003239 | 0.049547 | 0.027524 | 0.012784 | 0.014695 | 0.041902 | 0.001582 | 0.003001 | 0.004142 | 0.002424 | 0.020422 | 0.001464 | 0.012013 | 0.013479 | 0.024286 |
| 28.4583333 | 0.019678 | 0.004198 | 0.018209 | 0.01932 | 0.014122 | 0.021358 | 0.002512 | 0.020237 | 0.003513 | 0.056544 | 0.023997 | 0.012428 | 0.0086 | 0.040068 | 0.004017 | 0.004708 | 0.007822 | 0.006137 | 0.023974 | 0.002432 | 0.007073 | 0.013244 | 0.02328 |
| 28.4722222 | 0.018975 | 0.001964 | 0.017538 | 0.015432 | 0.007471 | 0.011986 | 0.004642 | 0.020584 | 0.005367 | 0.060176 | 0.019278 | 0.010188 | 0.004792 | 0.027442 | 0.005656 | 0.004237 | 0.010187 | 0.010942 | 0.02258 | 0.004001 | 0.002575 | 0.00951 | 0.020564 |

| | | | | | | | | | | | | | | | | | | | | | | | |
|---|---|---|---|---|---|---|---|---|---|---|---|---|---|---|---|---|---|---|---|---|---|---|---|
| 28.4861111 | 0.026623 | 0.002755 | 0.024722 | 0.011065 | 0.00514 | 0.011178 | 0.006651 | 0.025166 | 0.005628 | 0.062245 | 0.020831 | 0.01152 | 0.006223 | 0.013207 | 0.005125 | 0.002979 | 0.008305 | 0.011738 | 0.019132 | 0.00402 | 0.004228 | 0.007546 | 0.022067 |
| 28.5 | 0.041349 | 0.007242 | 0.039065 | 0.01162 | 0.008089 | 0.019317 | 0.010053 | 0.037509 | 0.004872 | 0.063711 | 0.026144 | 0.019242 | 0.011749 | 0.007226 | 0.006948 | 0.003689 | 0.00373 | 0.008152 | 0.017607 | 0.003972 | 0.010514 | 0.010062 | 0.030901 |
| 28.5138889 | 0.044912 | 0.012679 | 0.049125 | 0.014023 | 0.011369 | 0.028835 | 0.019458 | 0.050332 | 0.007397 | 0.065862 | 0.030824 | 0.025081 | 0.016565 | 0.010877 | 0.009101 | 0.004835 | 0.00111 | 0.004076 | 0.018726 | 0.004932 | 0.013767 | 0.013962 | 0.039061 |
| 28.5277778 | 0.032241 | 0.016208 | 0.054489 | 0.013902 | 0.013034 | 0.036139 | 0.029447 | 0.056712 | 0.010075 | 0.068116 | 0.034959 | 0.022832 | 0.016048 | 0.015699 | 0.008463 | 0.004268 | 0.004218 | 0.00334 | 0.017984 | 0.005649 | 0.010298 | 0.015631 | 0.035576 |
| 28.5416667 | 0.018834 | 0.017146 | 0.062965 | 0.011953 | 0.014326 | 0.041968 | 0.031692 | 0.062464 | 0.010242 | 0.068363 | 0.039594 | 0.014742 | 0.01273 | 0.015746 | 0.013139 | 0.007067 | 0.015858 | 0.012255 | 0.014351 | 0.011489 | 0.008704 | 0.011983 | 0.021036 |
| 28.5555556 | 0.012913 | 0.015702 | 0.067813 | 0.011145 | 0.015049 | 0.040505 | 0.023658 | 0.064969 | 0.007913 | 0.062589 | 0.042913 | 0.009406 | 0.013325 | 0.015681 | 0.024812 | 0.016601 | 0.029518 | 0.028215 | 0.015354 | 0.023676 | 0.017223 | 0.006326 | 0.009972 |
| 28.5694444 | 0.01127 | 0.015696 | 0.055906 | 0.014851 | 0.011932 | 0.029083 | 0.01387 | 0.054033 | 0.003859 | 0.049269 | 0.034314 | 0.008119 | 0.01518 | 0.01788 | 0.026875 | 0.019474 | 0.030458 | 0.03212 | 0.015021 | 0.027571 | 0.020742 | 0.004113 | 0.008342 |
| 28.5833333 | 0.014275 | 0.025362 | 0.037486 | 0.024549 | 0.010365 | 0.023916 | 0.016079 | 0.043668 | 0.001947 | 0.047638 | 0.018623 | 0.008039 | 0.022656 | 0.030356 | 0.016977 | 0.011831 | 0.017481 | 0.018627 | 0.012147 | 0.016784 | 0.013479 | 0.005201 | 0.014165 |
| 28.5972222 | 0.022991 | 0.042247 | 0.032069 | 0.034391 | 0.019343 | 0.033182 | 0.027106 | 0.046508 | 0.003472 | 0.063514 | 0.012644 | 0.015357 | 0.035759 | 0.04699 | 0.014216 | 0.008303 | 0.006182 | 0.006979 | 0.018435 | 0.007255 | 0.012382 | 0.009388 | 0.030729 |
| 28.6111111 | 0.031997 | 0.049168 | 0.040037 | 0.031353 | 0.030465 | 0.038464 | 0.033391 | 0.050134 | 0.00604 | 0.071357 | 0.010438 | 0.024333 | 0.038391 | 0.039793 | 0.023334 | 0.01038 | 0.004825 | 0.006741 | 0.025033 | 0.009834 | 0.020116 | 0.0138 | 0.049567 |
| 28.625 | 0.035889 | 0.040036 | 0.05316 | 0.017762 | 0.030531 | 0.028833 | 0.03289 | 0.04654 | 0.007189 | 0.066935 | 0.007167 | 0.026489 | 0.032501 | 0.020764 | 0.03289 | 0.010134 | 0.006723 | 0.008813 | 0.026602 | 0.014765 | 0.025484 | 0.016694 | 0.057322 |
| 28.6388889 | 0.035386 | 0.026127 | 0.065119 | 0.00914 | 0.020046 | 0.01994 | 0.031521 | 0.039633 | 0.006773 | 0.057822 | 0.006424 | 0.026585 | 0.024611 | 0.022167 | 0.036946 | 0.008467 | 0.006692 | 0.008574 | 0.027504 | 0.014916 | 0.02379 | 0.019218 | 0.054568 |
| 28.6527778 | 0.033845 | 0.015205 | 0.071493 | 0.007571 | 0.012116 | 0.020447 | 0.031767 | 0.030873 | 0.00631 | 0.040974 | 0.009758 | 0.028428 | 0.012943 | 0.031852 | 0.0357 | 0.008887 | 0.005522 | 0.007822 | 0.022287 | 0.01279 | 0.018074 | 0.019773 | 0.048164 |
| 28.6666667 | 0.031356 | 0.008187 | 0.065855 | 0.006575 | 0.01481 | 0.018683 | 0.031228 | 0.020103 | 0.00619 | 0.020061 | 0.020219 | 0.030007 | 0.003959 | 0.024088 | 0.029956 | 0.011532 | 0.004226 | 0.006729 | 0.011881 | 0.010023 | 0.011955 | 0.016906 | 0.041854 |
| 28.6805556 | 0.0249 | 0.004487 | 0.044757 | 0.005235 | 0.01693 | 0.010921 | 0.027701 | 0.009616 | 0.006327 | 0.008102 | 0.026296 | 0.028379 | 0.003574 | 0.016729 | 0.022292 | 0.01522 | 0.003503 | 0.005683 | 0.005416 | 0.007626 | 0.006852 | 0.011327 | 0.037253 |
| 28.6944444 | 0.013902 | 0.002353 | 0.022721 | 0.011217 | 0.0115 | 0.007388 | 0.023149 | 0.004826 | 0.007551 | 0.00904 | 0.019946 | 0.024359 | 0.008043 | 0.031751 | 0.016559 | 0.018793 | 0.003712 | 0.005716 | 0.004359 | 0.006336 | 0.003072 | 0.005726 | 0.034415 |
| 28.7083333 | 0.006578 | 0.000876 | 0.010909 | 0.019831 | 0.005826 | 0.011532 | 0.017821 | 0.005131 | 0.00917 | 0.009807 | 0.010514 | 0.020878 | 0.011763 | 0.045794 | 0.013066 | 0.021006 | 0.004223 | 0.006571 | 0.004911 | 0.005813 | 0.001851 | 0.0023 | 0.030797 |
| 28.7222222 | 0.008519 | 0.000636 | 0.012638 | 0.015286 | 0.008811 | 0.015889 | 0.009728 | 0.005086 | 0.009139 | 0.009207 | 0.005897 | 0.018852 | 0.013856 | 0.039346 | 0.009727 | 0.020721 | 0.004496 | 0.007297 | 0.005721 | 0.00534 | 0.004064 | 0.001924 | 0.025172 |
| 28.7361111 | 0.014589 | 0.002129 | 0.027165 | 0.009331 | 0.022114 | 0.014588 | 0.003749 | 0.00868 | 0.007892 | 0.015227 | 0.006626 | 0.017636 | 0.016264 | 0.022852 | 0.007432 | 0.018238 | 0.004885 | 0.007614 | 0.005902 | 0.004982 | 0.007052 | 0.003666 | 0.01898 |
| 28.75 | 0.021696 | 0.003502 | 0.039087 | 0.018128 | 0.032595 | 0.008449 | 0.004801 | 0.016281 | 0.006685 | 0.024536 | 0.012498 | 0.016971 | 0.019041 | 0.010823 | 0.007112 | 0.01447 | 0.00502 | 0.006886 | 0.005189 | 0.004321 | 0.007985 | 0.004059 | 0.015332 |
| 28.7638889 | 0.024428 | 0.003314 | 0.038106 | 0.027916 | 0.031043 | 0.003403 | 0.008542 | 0.019673 | 0.004654 | 0.027722 | 0.029583 | 0.016937 | 0.01965 | 0.011585 | 0.007363 | 0.010504 | 0.004382 | 0.005311 | 0.00504 | 0.002649 | 0.006926 | 0.003402 | 0.016593 |
| 28.7777778 | 0.018441 | 0.004013 | 0.025323 | 0.022507 | 0.021284 | 0.003422 | 0.010066 | 0.018008 | 0.002486 | 0.019041 | 0.056854 | 0.016367 | 0.018643 | 0.021405 | 0.007554 | 0.007588 | 0.003749 | 0.004497 | 0.006068 | 0.000936 | 0.004967 | 0.004705 | 0.018603 |
| 28.7916667 | 0.009984 | 0.005539 | 0.011525 | 0.009271 | 0.01222 | 0.005515 | 0.008374 | 0.01509 | 0.002596 | 0.010609 | 0.072879 | 0.013132 | 0.017979 | 0.030026 | 0.006599 | 0.005104 | 0.002952 | 0.003756 | 0.005453 | 0.000968 | 0.00427 | 0.005464 | 0.013468 |
| 28.8055556 | 0.004854 | 0.00538 | 0.005405 | 0.00216 | 0.008601 | 0.005777 | 0.005029 | 0.011829 | 0.004819 | 0.012009 | 0.062144 | 0.00746 | 0.014747 | 0.029996 | 0.00497 | 0.0033 | 0.003453 | 0.003264 | 0.003981 | 0.003801 | 0.006778 | 0.004239 | 0.006919 |
| 28.8194444 | 0.003478 | 0.005344 | 0.005417 | 0.000993 | 0.009398 | 0.004748 | 0.003047 | 0.008273 | 0.008529 | 0.011207 | 0.038269 | 0.003454 | 0.008614 | 0.02036 | 0.006111 | 0.003896 | 0.006865 | 0.005243 | 0.005821 | 0.007808 | 0.009263 | 0.004098 | 0.008898 |
| 28.8333333 | 0.004302 | 0.00762 | 0.009371 | 0.003163 | 0.015235 | 0.005374 | 0.003864 | 0.004918 | 0.011453 | 0.009074 | 0.019415 | 0.004195 | 0.007105 | 0.018242 | 0.008266 | 0.004653 | 0.008678 | 0.006925 | 0.008711 | 0.009416 | 0.007168 | 0.003775 | 0.012852 |
| 28.8472222 | 0.007092 | 0.011918 | 0.015541 | 0.014675 | 0.027363 | 0.009655 | 0.007774 | 0.002769 | 0.010111 | 0.013473 | 0.008114 | 0.006569 | 0.011802 | 0.032356 | 0.007892 | 0.003419 | 0.006529 | 0.005534 | 0.00907 | 0.007849 | 0.003077 | 0.003305 | 0.01363 |
| 28.8611111 | 0.01217 | 0.016026 | 0.021105 | 0.034386 | 0.041309 | 0.015892 | 0.014051 | 0.001837 | 0.005598 | 0.012293 | 0.00438 | 0.0072 | 0.015894 | 0.040342 | 0.00689 | 0.002012 | 0.003759 | 0.003234 | 0.007458 | 0.005378 | 0.001309 | 0.005595 | 0.013508 |
| 28.875 | 0.018156 | 0.01693 | 0.022836 | 0.040435 | 0.043699 | 0.018937 | 0.019383 | 0.002388 | 0.001793 | 0.005154 | 0.009204 | 0.006312 | 0.016009 | 0.029822 | 0.007753 | 0.002103 | 0.002312 | 0.001774 | 0.005582 | 0.003805 | 0.002152 | 0.007746 | 0.01179 |
| 28.8888889 | 0.022319 | 0.015676 | 0.022434 | 0.024233 | 0.034158 | 0.017346 | 0.021276 | 0.004651 | 0.001289 | 0.003741 | 0.018052 | 0.004972 | 0.014468 | 0.013794 | 0.009916 | 0.003309 | 0.001566 | 0.0009 | 0.004519 | 0.003266 | 0.005295 | 0.007237 | 0.008274 |
| 28.9027778 | 0.022746 | 0.015085 | 0.023829 | 0.009732 | 0.025098 | 0.016616 | 0.021493 | 0.005153 | 0.002916 | 0.006328 | 0.02785 | 0.004634 | 0.014076 | 0.007704 | 0.010843 | 0.004046 | 0.000943 | 0.000671 | 0.00399 | 0.002818 | 0.009656 | 0.005374 | 0.005127 |
| 28.9166667 | 0.020651 | 0.015147 | 0.030016 | 0.010353 | 0.02054 | 0.020671 | 0.021823 | 0.005329 | 0.00384 | 0.009613 | 0.036465 | 0.006196 | 0.013177 | 0.015018 | 0.009381 | 0.003668 | 0.000756 | 0.001569 | 0.004055 | 0.001843 | 0.013913 | 0.002923 | 0.005135 |
| 28.9305556 | 0.018011 | 0.014874 | 0.041717 | 0.016267 | 0.022828 | 0.025616 | 0.020476 | 0.010408 | 0.003579 | 0.012622 | 0.035435 | 0.009743 | 0.008779 | 0.027716 | 0.006982 | 0.003254 | 0.000759 | 0.002806 | 0.005359 | 0.000863 | 0.018072 | 0.001663 | 0.00876 |
| 28.9444444 | 0.016269 | 0.01462 | 0.053999 | 0.017951 | 0.026195 | 0.025686 | 0.017051 | 0.018526 | 0.003673 | 0.014181 | 0.021904 | 0.014769 | 0.003699 | 0.03613 | 0.005007 | 0.004116 | 0.00054 | 0.002822 | 0.007766 | 0.000624 | 0.021173 | 0.004064 | 0.013928 |
| 28.9583333 | 0.016053 | 0.016284 | 0.060466 | 0.012974 | 0.022125 | 0.021711 | 0.015066 | 0.026539 | 0.005383 | 0.016254 | 0.016329 | 0.019604 | 0.002455 | 0.041313 | 0.003952 | 0.006319 | 0.000628 | 0.001727 | 0.010821 | 0.001655 | 0.0208 | 0.009155 | 0.018646 |
| 28.9722222 | 0.016507 | 0.020735 | 0.063245 | 0.006746 | 0.013838 | 0.019271 | 0.016538 | 0.034594 | 0.008193 | 0.023006 | 0.037032 | 0.022033 | 0.007673 | 0.044979 | 0.003473 | 0.008456 | 0.001235 | 0.00091 | 0.013882 | 0.003624 | 0.017358 | 0.012906 | 0.022185 |
| 28.9861111 | 0.017002 | 0.025477 | 0.070117 | 0.005027 | 0.010531 | 0.020218 | 0.018333 | 0.040941 | 0.010211 | 0.035159 | 0.060694 | 0.021162 | 0.019948 | 0.037854 | 0.002801 | 0.009008 | 0.001312 | 0.000998 | 0.015834 | 0.004738 | 0.014042 | 0.012541 | 0.024531 |
| 29 | 0.017004 | 0.027697 | 0.075083 | 0.008143 | 0.021437 | 0.021654 | 0.016624 | 0.040874 | 0.009733 | 0.038977 | 0.059482 | 0.017985 | 0.031867 | 0.025946 | 0.001698 | 0.007811 | 0.001219 | 0.002669 | 0.015753 | 0.003768 | 0.01104 | 0.008655 | 0.02596 |
| 29.0138889 | 0.015329 | 0.027823 | 0.068063 | 0.017197 | 0.036446 | 0.021695 | 0.014067 | 0.036361 | 0.006277 | 0.025202 | 0.039938 | 0.014074 | 0.035615 | 0.02262 | 0.002241 | 0.006211 | 0.003015 | 0.005795 | 0.013342 | 0.00271 | 0.006832 | 0.004677 | 0.025564 |
| 29.0277778 | 0.012777 | 0.030064 | 0.058321 | 0.030108 | 0.042822 | 0.02373 | 0.017992 | 0.043113 | 0.002258 | 0.013153 | 0.019208 | 0.015381 | 0.032567 | 0.025358 | 0.004536 | 0.007733 | 0.004453 | 0.006668 | 0.010943 | 0.002486 | 0.00333 | 0.010013 | 0.026614 |
| 29.0416667 | 0.01346 | 0.037623 | 0.060656 | 0.041352 | 0.046315 | 0.035245 | 0.027455 | 0.068891 | 0.003595 | 0.022565 | 0.011932 | 0.028397 | 0.0342 | 0.028423 | 0.009147 | 0.019323 | 0.006377 | 0.008638 | 0.014559 | 0.006992 | 0.009542 | 0.034269 | 0.040862 |
| 29.0555556 | 0.01804 | 0.046674 | 0.073675 | 0.051788 | 0.051779 | 0.057745 | 0.031396 | 0.09361 | 0.010686 | 0.052349 | 0.023143 | 0.044009 | 0.045951 | 0.02956 | 0.021018 | 0.035667 | 0.013314 | 0.015077 | 0.031265 | 0.019341 | 0.027442 | 0.066945 | 0.06864 |
| 29.0694444 | 0.024089 | 0.048023 | 0.078279 | 0.057347 | 0.053285 | 0.079087 | 0.025949 | 0.097773 | 0.01513 | 0.082635 | 0.032752 | 0.046799 | 0.052071 | 0.022215 | 0.031328 | 0.038876 | 0.015941 | 0.0142 | 0.054657 | 0.025388 | 0.036672 | 0.085016 | 0.083761 |
| 29.0833333 | 0.028095 | 0.037844 | 0.059756 | 0.050241 | 0.04703 | 0.078179 | 0.017764 | 0.090257 | 0.01285 | 0.08386 | 0.023963 | 0.039021 | 0.040759 | 0.01476 | 0.028572 | 0.027173 | 0.00916 | 0.007954 | 0.062599 | 0.017697 | 0.026882 | 0.080678 | 0.069192 |
| 29.0972222 | 0.02375 | 0.024518 | 0.029253 | 0.032135 | 0.033007 | 0.047742 | 0.012872 | 0.081067 | 0.008745 | 0.055564 | 0.011039 | 0.030926 | 0.021158 | 0.021567 | 0.020578 | 0.015147 | 0.00323 | 0.007094 | 0.044471 | 0.008676 | 0.014713 | 0.060402 | 0.043267 |
| 29.1111111 | 0.012491 | 0.015865 | 0.010244 | 0.01643 | 0.017529 | 0.020733 | 0.012596 | 0.063887 | 0.007773 | 0.028305 | 0.005572 | 0.022328 | 0.007816 | 0.023687 | 0.015872 | 0.012283 | 0.001779 | 0.006168 | 0.025873 | 0.005155 | 0.008958 | 0.036956 | 0.024955 |
| 29.125 | 0.006103 | 0.010664 | 0.006668 | 0.013907 | 0.012476 | 0.018245 | 0.013788 | 0.039548 | 0.009279 | 0.018141 | 0.004703 | 0.012491 | 0.005601 | 0.013094 | 0.013906 | 0.011747 | 0.00227 | 0.003053 | 0.033382 | 0.003987 | 0.005712 | 0.022453 | 0.015573 |
| 29.1388889 | 0.010512 | 0.00615 | 0.004963 | 0.015578 | 0.0138 | 0.017218 | 0.011795 | 0.018489 | 0.009935 | 0.015908 | 0.006789 | 0.007978 | 0.006944 | 0.009545 | 0.012853 | 0.006684 | 0.002217 | 0.001172 | 0.044408 | 0.002589 | 0.005931 | 0.016916 | 0.011738 |
| 29.1527778 | 0.017864 | 0.002524 | 0.003177 | 0.018084 | 0.014982 | 0.011759 | 0.006928 | 0.009153 | 0.009156 | 0.010562 | 0.007662 | 0.01034 | 0.005915 | 0.014273 | 0.011388 | 0.004004 | 0.001535 | 0.000582 | 0.035961 | 0.001931 | 0.008929 | 0.01618 | 0.009208 |
| 29.1666667 | 0.019862 | 0.000901 | 0.00731 | 0.023158 | 0.017587 | 0.008524 | 0.004677 | 0.008564 | 0.008435 | 0.006101 | 0.006062 | 0.01279 | 0.006999 | 0.015017 | 0.010825 | 0.007337 | 0.001852 | 0.000794 | 0.020703 | 0.002178 | 0.00848 | 0.017568 | 0.008187 |
| 29.1805556 | 0.016675 | 0.00116 | 0.015533 | 0.02143 | 0.019078 | 0.007083 | 0.005572 | 0.007073 | 0.00782 | 0.007153 | 0.006562 | 0.011899 | 0.011279 | 0.010516 | 0.013858 | 0.010452 | 0.002283 | 0.001506 | 0.010459 | 0.002185 | 0.006213 | 0.016183 | 0.013621 |
| 29.1944444 | 0.011506 | 0.004477 | 0.021065 | 0.01286 | 0.015784 | 0.007153 | 0.005716 | 0.007463 | 0.006369 | 0.007352 | 0.007082 | 0.011611 | 0.01289 | 0.011187 | 0.018702 | 0.010403 | 0.003381 | 0.002501 | 0.006386 | 0.002966 | 0.00881 | 0.010092 | 0.023324 |
| 29.2083333 | 0.005968 | 0.011111 | 0.020362 | 0.009906 | 0.008812 | 0.00675 | 0.005461 | 0.012678 | 0.004363 | 0.009478 | 0.007537 | 0.015378 | 0.011759 | 0.018447 | 0.020428 | 0.012029 | 0.005043 | 0.002284 | 0.005932 | 0.004468 | 0.012703 | 0.003562 | 0.0285 |
| 29.2222222 | 0.002044 | 0.015599 | 0.013957 | 0.01581 | 0.00322 | 0.007861 | 0.004396 | 0.016942 | 0.003392 | 0.018135 | 0.013596 | 0.020936 | 0.012203 | 0.017036 | 0.018699 | 0.014682 | 0.005871 | 0.00127 | 0.00624 | 0.00554 | 0.01373 | 0.001749 | 0.028337 |
| 29.2361111 | 0.001163 | 0.012947 | 0.009134 | 0.022351 | 0.002523 | 0.00872 | 0.003941 | 0.017469 | 0.00393 | 0.024193 | 0.020853 | 0.02377 | 0.013006 | 0.008495 | 0.016931 | 0.011485 | 0.006056 | 0.001304 | 0.008051 | 0.006142 | 0.013374 | 0.003002 | 0.028039 |
| 29.25 | 0.001267 | 0.006265 | 0.014317 | 0.024949 | 0.006952 | 0.010078 | 0.007895 | 0.013849 | 0.004458 | 0.020549 | 0.02504 | 0.024538 | 0.011018 | 0.005343 | 0.015455 | 0.005935 | 0.005177 | 0.002478 | 0.012167 | 0.005953 | 0.011 | 0.002613 | 0.02838 |
| 29.2638889 | 0.00597 | 0.00167 | 0.020623 | 0.021099 | 0.01091 | 0.014415 | 0.011298 | 0.007289 | 0.004019 | 0.01111 | 0.025842 | 0.026498 | 0.008702 | 0.008761 | 0.012863 | 0.005404 | 0.003538 | 0.005069 | 0.014976 | 0.005584 | 0.007471 | 0.001371 | 0.028817 |
| 29.2777778 | 0.018088 | 0.001149 | 0.017498 | 0.013514 | 0.008271 | 0.018362 | 0.01037 | 0.002305 | 0.002985 | 0.00537 | 0.022843 | 0.026766 | 0.00855 | 0.019303 | 0.01049 | 0.005814 | 0.00256 | 0.007449 | 0.01214 | 0.005589 | 0.006343 | 0.001744 | 0.02899 |
| 29.2916667 | 0.026787 | 0.003259 | 0.009227 | 0.012236 | 0.004375 | 0.020146 | 0.008966 | 0.002728 | 0.004325 | 0.008245 | 0.017922 | 0.021643 | 0.008396 | 0.022117 | 0.009828 | 0.005583 | 0.002299 | 0.006506 | 0.005777 | 0.004737 | 0.007652 | 0.004977 | 0.026303 |
| 29.3055556 | 0.023728 | 0.005724 | 0.003725 | 0.017568 | 0.005644 | 0.020162 | 0.008591 | 0.007194 | 0.006798 | 0.015749 | 0.01297 | 0.014102 | 0.006945 | 0.010638 | 0.009375 | 0.006737 | 0.001885 | 0.003482 | 0.002054 | 0.002562 | 0.007875 | 0.010582 | 0.019502 |
| 29.3194444 | 0.013551 | 0.006266 | 0.003111 | 0.022654 | 0.0048 | 0.015446 | 0.00825 | 0.011934 | 0.008809 | 0.025511 | 0.011063 | 0.009107 | 0.00592 | 0.002303 | 0.007931 | 0.007004 | 0.001543 | 0.002948 | 0.004029 | 0.001026 | 0.005756 | 0.012928 | 0.011479 |
| 29.3333333 | 0.007974 | 0.004262 | 0.007295 | 0.022985 | 0.013044 | 0.01628 | 0.008565 | 0.016274 | 0.029669 | 0.038479 | 0.008947 | 0.00929 | 0.00784 | 0.001364 | 0.007423 | 0.010605 | 0.003114 | 0.00886 | 0.010751 | 0.003167 | 0.005025 | 0.009732 | 0.006903 |
| 29.3472222 | 0.011545 | 0.002456 | 0.018895 | 0.019291 | 0.010707 | 0.036649 | 0.009027 | 0.019162 | 0.065333 | 0.044288 | 0.007157 | 0.016736 | 0.014723 | 0.001948 | 0.010863 | 0.017278 | 0.008338 | 0.023624 | 0.019046 | 0.010699 | 0.01078 | 0.010425 | 0.007551 |
| 29.3611111 | 0.016055 | 0.002385 | 0.032446 | 0.017492 | 0.004338 | 0.056512 | 0.007869 | 0.017674 | 0.069081 | 0.030386 | 0.010261 | 0.026005 | 0.022651 | 0.006714 | 0.017462 | 0.020165 | 0.015884 | 0.03969 | 0.024656 | 0.02024 | 0.021537 | 0.01789 | 0.009231 |
| 29.375 | 0.016542 | 0.002585 | 0.037317 | 0.018308 | 0.002128 | 0.053696 | 0.005573 | 0.011999 | 0.038148 | 0.018682 | 0.011614 | 0.02473 | 0.021127 | 0.021694 | 0.019248 | 0.017001 | 0.0195 | 0.042307 | 0.022177 | 0.023328 | 0.026552 | 0.02015 | 0.008747 |
| 29.3888889 | 0.013953 | 0.002043 | 0.033518 | 0.017625 | 0.004111 | 0.036049 | 0.003356 | 0.007261 | 0.012064 | 0.034848 | 0.01198 | 0.013152 | 0.012036 | 0.041052 | 0.012012 | 0.010204 | 0.015229 | 0.031343 | 0.013986 | 0.016916 | 0.019039 | 0.014103 | 0.011472 |
| 29.4027778 | 0.009097 | 0.002366 | 0.026299 | 0.013126 | 0.004641 | 0.019571 | 0.001921 | 0.009623 | 0.003813 | 0.061855 | 0.018026 | 0.004535 | 0.008701 | 0.045967 | 0.004833 | 0.008177 | 0.008196 | 0.019152 | 0.011531 | 0.008045 | 0.007895 | 0.00698 | 0.019463 |
| 29.4166667 | 0.006056 | 0.005235 | 0.018463 | 0.006697 | 0.003298 | 0.009981 | 0.001775 | 0.018342 | 0.005922 | 0.078316 | 0.024532 | 0.004273 | 0.011873 | 0.030356 | 0.004075 | 0.011357 | 0.004339 | 0.012858 | 0.013765 | 0.00315 | 0.003369 | 0.003488 | 0.027161 |
| 29.4305556 | 0.007753 | 0.007879 | 0.015416 | 0.003488 | 0.003346 | 0.006099 | 0.001818 | 0.027639 | 0.01043 | 0.082187 | 0.022528 | 0.006648 | 0.013992 | 0.012467 | 0.004713 | 0.009823 | 0.004671 | 0.013014 | 0.011461 | 0.003256 | 0.003238 | 0.004767 | 0.032074 |
| 29.4444444 | 0.008652 | 0.007331 | 0.017185 | 0.005282 | 0.006804 | 0.008619 | 0.001788 | 0.031457 | 0.01263 | 0.075489 | 0.014071 | 0.00791 | 0.012511 | 0.007436 | 0.003635 | 0.005299 | 0.008559 | 0.018304 | 0.008492 | 0.007405 | 0.00498 | 0.00861 | 0.034135 |
| 29.4583333 | 0.005674 | 0.005535 | 0.020564 | 0.006546 | 0.009721 | 0.014126 | 0.003522 | 0.027358 | 0.011933 | 0.060476 | 0.008597 | 0.008588 | 0.011523 | 0.010044 | 0.003003 | 0.004927 | 0.013414 | 0.025858 | 0.010724 | 0.012531 | 0.009637 | 0.012654 | 0.033563 |
| 29.4722222 | 0.00485 | 0.006705 | 0.031496 | 0.006851 | 0.008735 | 0.018705 | 0.006508 | 0.021993 | 0.009335 | 0.042529 | 0.006098 | 0.009218 | 0.01571 | 0.010095 | 0.002893 | 0.004952 | 0.01422 | 0.030068 | 0.014942 | 0.013848 | 0.012335 | 0.015253 | 0.032405 |
| 29.4861111 | 0.013776 | 0.01286 | 0.047465 | 0.015798 | 0.01188 | 0.022501 | 0.00747 | 0.023816 | 0.009399 | 0.031632 | 0.005947 | 0.012184 | 0.023035 | 0.007787 | 0.003836 | 0.005738 | 0.008974 | 0.026466 | 0.017954 | 0.009101 | 0.008973 | 0.018414 | 0.033969 |
| 29.5 | 0.031763 | 0.02257 | 0.056401 | 0.033417 | 0.026043 | 0.023511 | 0.008695 | 0.03161 | 0.014811 | 0.032575 | 0.015473 | 0.020598 | 0.030284 | 0.008675 | 0.008579 | 0.011075 | 0.004011 | 0.02001 | 0.021788 | 0.004796 | 0.006708 | 0.02449 | 0.039199 |
| 29.5138889 | 0.048072 | 0.030572 | 0.055901 | 0.044692 | 0.039386 | 0.020484 | 0.013975 | 0.036944 | 0.020416 | 0.037781 | 0.0299 | 0.031072 | 0.035381 | 0.0148 | 0.015647 | 0.015301 | 0.004977 | 0.016288 | 0.027349 | 0.00748 | 0.012227 | 0.030234 | 0.044844 |
| 29.5277778 | 0.054441 | 0.032203 | 0.04843 | 0.042304 | 0.039339 | 0.015246 | 0.019763 | 0.040912 | 0.023175 | 0.039955 | 0.037383 | 0.036166 | 0.032124 | 0.022612 | 0.020915 | 0.015398 | 0.007651 | 0.013245 | 0.030965 | 0.012099 | 0.018934 | 0.033962 | 0.046212 |
| 29.5416667 | 0.048381 | 0.029296 | 0.04085 | 0.033624 | 0.030108 | 0.011997 | 0.022448 | 0.051539 | 0.025229 | 0.041548 | 0.043049 | 0.031678 | 0.01847 | 0.023209 | 0.021658 | 0.012475 | 0.006276 | 0.010505 | 0.027308 | 0.011599 | 0.018626 | 0.036735 | 0.040083 |
| 29.5555556 | 0.03408 | 0.024865 | 0.035242 | 0.024294 | 0.019283 | 0.013201 | 0.019442 | 0.05976 | 0.024484 | 0.040587 | 0.047041 | 0.020945 | 0.005559 | 0.014299 | 0.016846 | 0.007214 | 0.004864 | 0.012449 | 0.015136 | 0.008505 | 0.0118 | 0.034141 | 0.028571 |
| 29.5694444 | 0.020871 | 0.021423 | 0.022774 | 0.015884 | 0.009887 | 0.014431 | 0.014248 | 0.05351 | 0.017929 | 0.032101 | 0.036235 | 0.011959 | 0.008535 | 0.006904 | 0.0098 | 0.003128 | 0.007832 | 0.016617 | 0.005196 | 0.009224 | 0.006122 | 0.023432 | 0.021668 |
| 29.5833333 | 0.013604 | 0.026354 | 0.012934 | 0.01097 | 0.009434 | 0.013322 | 0.016116 | 0.043597 | 0.009834 | 0.025272 | 0.020827 | 0.013084 | 0.033623 | 0.012631 | 0.00829 | 0.002903 | 0.008368 | 0.014751 | 0.009053 | 0.009978 | 0.004276 | 0.013702 | 0.02985 |
| 29.5972222 | 0.011655 | 0.040394 | 0.018213 | 0.010126 | 0.017188 | 0.015083 | 0.023612 | 0.031967 | 0.006224 | 0.028615 | 0.018068 | 0.02254 | 0.066239 | 0.029169 | 0.014439 | 0.005003 | 0.004186 | 0.00804 | 0.019626 | 0.005762 | 0.003231 | 0.009987 | 0.046039 |
| 29.6111111 | 0.010533 | 0.048502 | 0.029017 | 0.01101 | 0.01757 | 0.020267 | 0.029532 | 0.013452 | 0.007605 | 0.036611 | 0.019611 | 0.026856 | 0.074614 | 0.041261 | 0.021182 | 0.005009 | 0.001312 | 0.004126 | 0.022222 | 0.001564 | 0.004542 | 0.006599 | 0.050673 |
| 29.625 | 0.009736 | 0.042059 | 0.037816 | 0.014693 | 0.010522 | 0.02481 | 0.032861 | 0.00617 | 0.011159 | 0.044039 | 0.01324 | 0.024661 | 0.053673 | 0.039432 | 0.024435 | 0.002528 | 0.001121 | 0.00379 | 0.01782 | 0.000639 | 0.008161 | 0.002999 | 0.040937 |
| 29.6388889 | 0.0146 | 0.029969 | 0.046734 | 0.024568 | 0.010252 | 0.028239 | 0.034079 | 0.018294 | 0.013758 | 0.049172 | 0.00813 | 0.02344 | 0.031481 | 0.02878 | 0.025557 | 0.00154 | 0.002094 | 0.004544 | 0.01423 | 0.001165 | 0.011068 | 0.003166 | 0.029635 |
| 29.6527778 | 0.023559 | 0.018722 | 0.052996 | 0.034484 | 0.013446 | 0.02831 | 0.030548 | 0.030787 | 0.013092 | 0.045869 | 0.015064 | 0.021715 | 0.017649 | 0.01796 | 0.022678 | 0.004412 | 0.005249 | 0.008565 | 0.01032 | 0.004063 | 0.009287 | 0.00509 | 0.023647 |
| 29.6666667 | 0.028912 | 0.009138 | 0.046382 | 0.032504 | 0.00911 | 0.021758 | 0.020468 | 0.026079 | 0.009066 | 0.032592 | 0.026198 | 0.014221 | 0.009052 | 0.010589 | 0.014444 | 0.010587 | 0.011625 | 0.01815 | 0.008695 | 0.011445 | 0.008832 | 0.005589 | 0.020827 |
| 29.6805556 | 0.02808 | 0.004636 | 0.029366 | 0.021191 | 0.002909 | 0.011304 | 0.01031 | 0.012714 | 0.004576 | 0.019642 | 0.037496 | 0.00681 | 0.009679 | 0.011401 | 0.006223 | 0.015378 | 0.018117 | 0.029236 | 0.014822 | 0.020109 | 0.021138 | 0.009755 | 0.015859 |
| 29.6944444 | 0.02489 | 0.003621 | 0.016218 | 0.014424 | 0.001596 | 0.004301 | 0.009099 | 0.007296 | 0.003192 | 0.012649 | 0.051313 | 0.007083 | 0.013491 | 0.018382 | 0.00302 | 0.013549 | 0.020161 | 0.034306 | 0.021158 | 0.023846 | 0.042766 | 0.017831 | 0.010319 |
| 29.7083333 | 0.021457 | 0.002895 | 0.008307 | 0.014436 | 0.001612 | 0.004529 | 0.015326 | 0.009454 | 0.00463 | 0.009472 | 0.059085 | 0.007666 | 0.014116 | 0.01856 | 0.002791 | 0.007628 | 0.016384 | 0.03015 | 0.020528 | 0.020052 | 0.064202 | 0.017107 | 0.013276 |
| 29.7222222 | 0.013962 | 0.003282 | 0.006135 | 0.012791 | 0.001176 | 0.009154 | 0.019894 | 0.010773 | 0.005268 | 0.010293 | 0.051436 | 0.004342 | 0.013927 | 0.010557 | 0.003526 | 0.005217 | 0.00958 | 0.020688 | 0.016433 | 0.012142 | 0.077507 | 0.040342 | 0.017331 |
| 29.7361111 | 0.009144 | 0.002849 | 0.012589 | 0.01299 | 0.002017 | 0.016718 | 0.018292 | 0.007276 | 0.003317 | 0.014098 | 0.033323 | 0.002231 | 0.011636 | 0.004528 | 0.005862 | 0.007516 | 0.004146 | 0.011354 | 0.012971 | 0.00514 | 0.072767 | 0.094596 | 0.012561 |
| 29.75 | 0.019017 | 0.003079 | 0.015455 | 0.017537 | 0.005723 | 0.024622 | 0.015719 | 0.003468 | 0.001343 | 0.019462 | 0.01712 | 0.004888 | 0.009113 | 0.005101 | 0.008284 | 0.010471 | 0.003234 | 0.00556 | 0.011461 | 0.002746 | 0.049937 | 0.097667 | 0.011517 |
| 29.7638889 | 0.0335 | 0.006924 | 0.010247 | 0.015175 | 0.00755 | 0.027138 | 0.016778 | 0.004353 | 0.001494 | 0.024332 | 0.007515 | 0.01139 | 0.012962 | 0.007789 | 0.01025 | 0.012237 | 0.005613 | 0.004371 | 0.013102 | 0.004819 | 0.027033 | 0.05132 | 0.018998 |
| 29.7777778 | 0.036905 | 0.01003 | 0.010509 | 0.008943 | 0.00682 | 0.024504 | 0.019151 | 0.006708 | 0.001864 | 0.023917 | 0.004183 | 0.015283 | 0.018026 | 0.00633 | 0.010795 | 0.01308 | 0.006245 | 0.004107 | 0.017927 | 0.005374 | 0.014292 | 0.029056 | 0.026541 |
| 29.7916667 | 0.031431 | 0.008532 | 0.020343 | 0.010718 | 0.008294 | 0.020408 | 0.019994 | 0.006265 | 0.001201 | 0.01775 | 0.005799 | 0.014757 | 0.019809 | 0.004761 | 0.009184 | 0.008624 | 0.004794 | 0.002958 | 0.022371 | 0.003461 | 0.010145 | 0.041637 | 0.032397 |
| 29.8055556 | 0.025927 | 0.005143 | 0.028398 | 0.015439 | 0.00922 | 0.016041 | 0.018075 | 0.004462 | 0.000488 | 0.012175 | 0.007875 | 0.015467 | 0.022966 | 0.009071 | 0.00766 | 0.007616 | 0.005287 | 0.004283 | 0.020782 | 0.004214 | 0.017999 | 0.06213 | 0.036497 |
| 29.8194444 | 0.019806 | 0.005293 | 0.024644 | 0.015773 | 0.011721 | 0.01158 | 0.012623 | 0.003205 | 0.000659 | 0.010958 | 0.008111 | 0.018621 | 0.027094 | 0.01226 | 0.00667 | 0.008477 | 0.007481 | 0.007386 | 0.012776 | 0.007482 | 0.032858 | 0.073044 | 0.034357 |
| 29.8333333 | 0.010373 | 0.009176 | 0.013332 | 0.014889 | 0.017471 | 0.006901 | 0.006076 | 0.004568 | 0.002019 | 0.010301 | 0.007591 | 0.020106 | 0.028154 | 0.009225 | 0.004758 | 0.008614 | 0.008245 | 0.008194 | 0.007093 | 0.009279 | 0.040828 | 0.070573 | 0.023771 |
| 29.8472222 | 0.00339 | 0.012504 | 0.007978 | 0.014065 | 0.022135 | 0.003481 | 0.003878 | 0.008698 | 0.003762 | 0.009467 | 0.006725 | 0.018236 | 0.024624 | 0.009667 | 0.002623 | 0.00788 | 0.007243 | 0.006858 | 0.009099 | 0.008747 | 0.037371 | 0.057368 | 0.01104 |
| 29.8611111 | 0.001962 | 0.014829 | 0.017809 | 0.01079 | 0.024191 | 0.003517 | 0.007985 | 0.011587 | 0.005181 | 0.01208 | 0.004945 | 0.014976 | 0.016325 | 0.020852 | 0.002751 | 0.007873 | 0.006302 | 0.006179 | 0.013814 | 0.008076 | 0.028844 | 0.043339 | 0.003282 |
| 29.875 | 0.00165 | 0.017301 | 0.037593 | 0.006588 | 0.024377 | 0.005299 | 0.017005 | 0.011036 | 0.006015 | 0.016418 | 0.002576 | 0.012305 | 0.009136 | 0.032232 | 0.005622 | 0.008263 | 0.006049 | 0.006723 | 0.015662 | 0.008076 | 0.021238 | 0.035586 | 0.00294 |
| 29.8888889 | 0.001631 | 0.017892 | 0.048049 | 0.003507 | 0.019539 | 0.006567 | 0.026564 | 0.008288 | 0.005758 | 0.018444 | 0.001154 | 0.009739 | 0.010164 | 0.032633 | 0.008587 | 0.006897 | 0.005517 | 0.006506 | 0.01373 | 0.006807 | 0.013799 | 0.028851 | 0.007752 |
| 29.9027778 | 0.005448 | 0.015561 | 0.039168 | 0.006192 | 0.013473 | 0.006268 | 0.030856 | 0.006546 | 0.00517 | 0.017925 | 0.001421 | 0.006663 | 0.014569 | 0.025631 | 0.008712 | 0.003671 | 0.003057 | 0.004199 | 0.009164 | 0.003578 | 0.007567 | 0.018822 | 0.010164 |
| 29.9166667 | 0.014555 | 0.013965 | 0.022883 | 0.017064 | 0.016865 | 0.016867 | 0.030385 | 0.006228 | 0.006193 | 0.014733 | 0.003923 | 0.004002 | 0.016651 | 0.017323 | 0.006389 | 0.001424 | 0.001104 | 0.001505 | 0.00395 | 0.001115 | 0.006503 | 0.014141 | 0.008214 |
| 29.9305556 | 0.028049 | 0.016509 | 0.018251 | 0.024706 | 0.023914 | 0.012082 | 0.028761 | 0.006228 | 0.008635 | 0.009064 | 0.008313 | 0.002913 | 0.017094 | 0.010242 | 0.004388 | 0.002453 | 0.002033 | 0.002071 | 0.002255 | 0.002062 | 0.005849 | 0.015285 | 0.007986 |
| 29.9444444 | 0.041461 | 0.021618 | 0.027473 | 0.022971 | 0.025799 | 0.024545 | 0.027423 | 0.010982 | 0.010489 | 0.007779 | 0.012535 | 0.002388 | 0.014381 | 0.010283 | 0.003566 | 0.00607 | 0.006373 | 0.006921 | 0.005879 | 0.006363 | 0.004337 | 0.011119 | 0.011412 |
| 29.9583333 | 0.047314 | 0.026482 | 0.035758 | 0.019326 | 0.025259 | 0.037514 | 0.02787 | 0.022737 | 0.011402 | 0.01509 | 0.015519 | 0.003407 | 0.008089 | 0.020672 | 0.002505 | 0.009032 | 0.010628 | 0.011756 | 0.010304 | 0.010934 | 0.006851 | 0.006686 | 0.013823 |

| | | | | | | | | | | | | | | | | | | | | | | | |
|---|---|---|---|---|---|---|---|---|---|---|---|---|---|---|---|---|---|---|---|---|---|---|---|
| 29.9722222 | 0.04335 | 0.030742 | 0.035964 | 0.016194 | 0.025748 | 0.046342 | 0.029252 | 0.033218 | 0.011932 | 0.025451 | 0.01676 | 0.010741 | 0.003759 | 0.036469 | 0.001362 | 0.008758 | 0.011135 | 0.012219 | 0.010483 | 0.011736 | 0.0089 | 0.014449 | 0.011125 |
| 29.9861111 | 0.036888 | 0.034814 | 0.03477 | 0.011489 | 0.025317 | 0.049247 | 0.028207 | 0.033624 | 0.011714 | 0.032825 | 0.01637 | 0.022879 | 0.003747 | 0.049877 | 0.001807 | 0.005642 | 0.008295 | 0.009014 | 0.00681 | 0.008543 | 0.006272 | 0.033736 | 0.010989 |
| 30 | 0.036905 | 0.035944 | 0.039935 | 0.006006 | 0.023979 | 0.048435 | 0.024632 | 0.029352 | 0.010387 | 0.03192 | 0.015969 | 0.03364 | 0.004044 | 0.054999 | 0.003214 | 0.002613 | 0.005957 | 0.006489 | 0.003641 | 0.005138 | 0.0023 | 0.051833 | 0.017874 |
| 30.0138889 | 0.041953 | 0.032309 | 0.044126 | 0.002862 | 0.024126 | 0.050334 | 0.019948 | 0.03202 | 0.007622 | 0.022065 | 0.015358 | 0.039714 | 0.004506 | 0.052032 | 0.00369 | 0.002995 | 0.007308 | 0.008467 | 0.003788 | 0.00607 | 0.002071 | 0.053237 | 0.019253 |
| 30.0277778 | 0.041674 | 0.029882 | 0.042574 | 0.008368 | 0.027564 | 0.057353 | 0.016058 | 0.045491 | 0.004153 | 0.012407 | 0.012414 | 0.040811 | 0.009035 | 0.04298 | 0.005252 | 0.00817 | 0.011261 | 0.014382 | 0.005483 | 0.01294 | 0.007498 | 0.038123 | 0.012314 |
| 30.0416667 | 0.035831 | 0.03512 | 0.049293 | 0.026633 | 0.03886 | 0.067944 | 0.016443 | 0.074051 | 0.003541 | 0.013096 | 0.010837 | 0.037983 | 0.022628 | 0.028943 | 0.006581 | 0.016106 | 0.014906 | 0.019968 | 0.007806 | 0.02097 | 0.014584 | 0.019344 | 0.009406 |
| 30.0555556 | 0.035929 | 0.040753 | 0.063712 | 0.047998 | 0.053453 | 0.078843 | 0.019354 | 0.101909 | 0.006047 | 0.023947 | 0.015197 | 0.030793 | 0.045471 | 0.0157 | 0.005521 | 0.019785 | 0.018747 | 0.024016 | 0.014344 | 0.024836 | 0.020368 | 0.006813 | 0.01307 |
| 30.0694444 | 0.044236 | 0.034043 | 0.065429 | 0.057167 | 0.057096 | 0.081822 | 0.017858 | 0.104172 | 0.006341 | 0.037336 | 0.02044 | 0.01972 | 0.06061 | 0.013443 | 0.005216 | 0.015577 | 0.023585 | 0.027992 | 0.021335 | 0.025726 | 0.025275 | 0.004621 | 0.013368 |
| 30.0833333 | 0.049495 | 0.018782 | 0.049446 | 0.052007 | 0.053076 | 0.067715 | 0.011055 | 0.095001 | 0.003298 | 0.050877 | 0.019265 | 0.01248 | 0.054268 | 0.02625 | 0.005547 | 0.012965 | 0.025662 | 0.028728 | 0.018487 | 0.025329 | 0.025711 | 0.006372 | 0.012219 |
| 30.0972222 | 0.040327 | 0.007633 | 0.028299 | 0.04029 | 0.051371 | 0.038678 | 0.005291 | 0.093039 | 0.001128 | 0.061212 | 0.011329 | 0.013596 | 0.036309 | 0.048848 | 0.00747 | 0.019972 | 0.022683 | 0.02406 | 0.011484 | 0.02257 | 0.019242 | 0.006651 | 0.016355 |
| 30.1111111 | 0.020396 | 0.005019 | 0.011645 | 0.030752 | 0.042848 | 0.018504 | 0.004748 | 0.089124 | 0.00178 | 0.057129 | 0.003745 | 0.014261 | 0.018843 | 0.061552 | 0.010124 | 0.030558 | 0.017993 | 0.018527 | 0.012532 | 0.018565 | 0.010726 | 0.01192 | 0.020387 |
| 30.125 | 0.005836 | 0.007649 | 0.004606 | 0.023602 | 0.023698 | 0.021063 | 0.007021 | 0.06874 | 0.004758 | 0.040164 | 0.001553 | 0.009156 | 0.007516 | 0.047496 | 0.010268 | 0.035353 | 0.014678 | 0.01566 | 0.017861 | 0.014784 | 0.004709 | 0.021312 | 0.022922 |
| 30.1388889 | 0.001521 | 0.01016 | 0.00414 | 0.013095 | 0.010373 | 0.026668 | 0.008032 | 0.037105 | 0.009254 | 0.025064 | 0.002301 | 0.004111 | 0.004728 | 0.021486 | 0.007225 | 0.031215 | 0.011874 | 0.013686 | 0.021754 | 0.010691 | 0.003037 | 0.029573 | 0.0255 |
| 30.1527778 | 0.000752 | 0.010743 | 0.003356 | 0.007386 | 0.012396 | 0.020436 | 0.006929 | 0.013281 | 0.014175 | 0.01499 | 0.003314 | 0.004274 | 0.005851 | 0.009699 | 0.00455 | 0.022052 | 0.007959 | 0.009955 | 0.026599 | 0.0062 | 0.00534 | 0.035912 | 0.026651 |
| 30.1666667 | 0.000959 | 0.010886 | 0.003552 | 0.015773 | 0.018233 | 0.010332 | 0.005164 | 0.004551 | 0.017622 | 0.007544 | 0.004392 | 0.007136 | 0.005061 | 0.01921 | 0.008182 | 0.012949 | 0.003817 | 0.005464 | 0.02991 | 0.002832 | 0.007325 | 0.035624 | 0.026935 |
| 30.1805556 | 0.002142 | 0.01249 | 0.005792 | 0.030073 | 0.020557 | 0.005169 | 0.004668 | 0.007006 | 0.016311 | 0.005181 | 0.00402 | 0.008404 | 0.003753 | 0.034101 | 0.015579 | 0.006239 | 0.001274 | 0.002832 | 0.026419 | 0.001906 | 0.005593 | 0.029071 | 0.027698 |
| 30.1944444 | 0.003267 | 0.015643 | 0.00619 | 0.038427 | 0.023637 | 0.003403 | 0.005787 | 0.012917 | 0.010806 | 0.005782 | 0.002666 | 0.009454 | 0.004806 | 0.03876 | 0.020618 | 0.005168 | 0.000414 | 0.001934 | 0.02145 | 0.002033 | 0.00305 | 0.023743 | 0.029358 |
| 30.2083333 | 0.006274 | 0.018237 | 0.003882 | 0.03467 | 0.025532 | 0.005702 | 0.006521 | 0.016108 | 0.006477 | 0.006327 | 0.004763 | 0.011202 | 0.004322 | 0.033865 | 0.021232 | 0.010739 | 0.000332 | 0.001512 | 0.020369 | 0.001459 | 0.00347 | 0.021136 | 0.030481 |
| 30.2222222 | 0.012288 | 0.018512 | 0.001957 | 0.022501 | 0.022282 | 0.017302 | 0.00531 | 0.017852 | 0.005357 | 0.008029 | 0.007116 | 0.009618 | 0.004674 | 0.023138 | 0.018488 | 0.01669 | 0.001026 | 0.002107 | 0.020158 | 0.000999 | 0.005366 | 0.016333 | 0.029393 |
| 30.2361111 | 0.013085 | 0.015465 | 0.002142 | 0.010724 | 0.015639 | 0.027153 | 0.003371 | 0.019414 | 0.005833 | 0.008409 | 0.004586 | 0.004872 | 0.010109 | 0.013425 | 0.014139 | 0.017946 | 0.002798 | 0.004248 | 0.018124 | 0.001305 | 0.006747 | 0.009064 | 0.025682 |
| 30.25 | 0.010087 | 0.01007 | 0.0037 | 0.005102 | 0.008594 | 0.019887 | 0.001989 | 0.017644 | 0.006802 | 0.005192 | 0.001862 | 0.001981 | 0.012113 | 0.014991 | 0.008831 | 0.015524 | 0.004488 | 0.006086 | 0.014641 | 0.001473 | 0.006609 | 0.006296 | 0.021821 |
| 30.2638889 | 0.01721 | 0.007025 | 0.005579 | 0.005198 | 0.004981 | 0.01046 | 0.001195 | 0.011631 | 0.007189 | 0.003772 | 0.001807 | 0.001271 | 0.006704 | 0.022579 | 0.004546 | 0.010835 | 0.004088 | 0.005248 | 0.011591 | 0.001566 | 0.004465 | 0.009138 | 0.022855 |
| 30.2777778 | 0.033016 | 0.008979 | 0.008101 | 0.004769 | 0.006409 | 0.015238 | 0.000911 | 0.009439 | 0.00659 | 0.006313 | 0.001881 | 0.001428 | 0.00258 | 0.023172 | 0.003713 | 0.005892 | 0.002729 | 0.003127 | 0.010856 | 0.003313 | 0.003344 | 0.010737 | 0.029702 |
| 30.2916667 | 0.043541 | 0.011688 | 0.010166 | 0.004649 | 0.008351 | 0.020301 | 0.000973 | 0.014226 | 0.006 | 0.006687 | 0.002063 | 0.002706 | 0.002774 | 0.016422 | 0.004226 | 0.005473 | 0.003032 | 0.002762 | 0.01233 | 0.006256 | 0.004812 | 0.009031 | 0.03532 |
| 30.3055556 | 0.041163 | 0.011341 | 0.011655 | 0.008161 | 0.007214 | 0.014779 | 0.001385 | 0.012994 | 0.005969 | 0.006516 | 0.002757 | 0.003157 | 0.003293 | 0.016645 | 0.003222 | 0.01176 | 0.002962 | 0.002598 | 0.012023 | 0.006608 | 0.005448 | 0.006813 | 0.0334 |
| 30.3194444 | 0.031475 | 0.009 | 0.012973 | 0.013193 | 0.004119 | 0.006539 | 0.00178 | 0.010359 | 0.006056 | 0.010359 | 0.00534 | 0.003223 | 0.004257 | 0.024682 | 0.002203 | 0.021176 | 0.002303 | 0.002385 | 0.008374 | 0.003746 | 0.004302 | 0.00731 | 0.026983 |
| 30.3333333 | 0.020942 | 0.006371 | 0.010894 | 0.017573 | 0.00336 | 0.003695 | 0.001672 | 0.019744 | 0.005529 | 0.010496 | 0.008224 | 0.005921 | 0.006248 | 0.021749 | 0.00188 | 0.026958 | 0.003824 | 0.004617 | 0.008666 | 0.001168 | 0.004496 | 0.009847 | 0.021067 |
| 30.3472222 | 0.013927 | 0.004732 | 0.007956 | 0.022364 | 0.008187 | 0.004223 | 0.001217 | 0.025663 | 0.003867 | 0.007703 | 0.007252 | 0.008558 | 0.009302 | 0.020143 | 0.001512 | 0.023294 | 0.004861 | 0.006266 | 0.015679 | 0.000961 | 0.005093 | 0.009305 | 0.014353 |
| 30.3611111 | 0.020172 | 0.004785 | 0.010418 | 0.030767 | 0.016631 | 0.008568 | 0.001047 | 0.018128 | 0.001923 | 0.011036 | 0.009507 | 0.007903 | 0.016125 | 0.039767 | 0.001164 | 0.012618 | 0.003247 | 0.004741 | 0.020704 | 0.002207 | 0.003445 | 0.010545 | 0.009041 |
| 30.375 | 0.032397 | 0.005531 | 0.017543 | 0.039857 | 0.0259 | 0.019897 | 0.001069 | 0.017242 | 0.001553 | 0.01569 | 0.01688 | 0.00682 | 0.025319 | 0.059244 | 0.001452 | 0.006218 | 0.001208 | 0.002363 | 0.020598 | 0.002437 | 0.001805 | 0.018243 | 0.013203 |
| 30.3888889 | 0.027091 | 0.006174 | 0.026512 | 0.041765 | 0.033609 | 0.029788 | 0.000777 | 0.034124 | 0.002729 | 0.015824 | 0.017599 | 0.008334 | 0.031003 | 0.06065 | 0.003584 | 0.008136 | 0.000501 | 0.000893 | 0.023052 | 0.001215 | 0.001816 | 0.019593 | 0.025027 |
| 30.4027778 | 0.016984 | 0.007108 | 0.033951 | 0.036318 | 0.036359 | 0.027816 | 0.000749 | 0.051629 | 0.002986 | 0.01474 | 0.010097 | 0.012087 | 0.030215 | 0.053732 | 0.006054 | 0.008688 | 0.000977 | 0.000406 | 0.034344 | 0.000678 | 0.001708 | 0.015519 | 0.036001 |
| 30.4166667 | 0.028927 | 0.008357 | 0.035235 | 0.029156 | 0.033331 | 0.016805 | 0.001789 | 0.055252 | 0.001906 | 0.015385 | 0.004343 | 0.01598 | 0.025429 | 0.047589 | 0.007751 | 0.006307 | 0.001237 | 0.000465 | 0.050398 | 0.001365 | 0.001416 | 0.024588 | 0.042961 |
| 30.4305556 | 0.047842 | 0.009806 | 0.032974 | 0.023161 | 0.027671 | 0.012551 | 0.003627 | 0.047289 | 0.001406 | 0.015785 | 0.006505 | 0.017981 | 0.02225 | 0.041641 | 0.010258 | 0.009609 | 0.00076 | 0.000451 | 0.058428 | 0.001501 | 0.002638 | 0.041158 | 0.046375 |
| 30.4444444 | 0.053333 | 0.012056 | 0.034124 | 0.019924 | 0.021707 | 0.019229 | 0.004717 | 0.037201 | 0.001329 | 0.01621 | 0.011514 | 0.016927 | 0.021327 | 0.030966 | 0.012122 | 0.016762 | 0.001211 | 0.001169 | 0.05125 | 0.000736 | 0.004833 | 0.045917 | 0.047008 |
| 30.4583333 | 0.045985 | 0.014437 | 0.039778 | 0.019443 | 0.015926 | 0.021572 | 0.004104 | 0.032408 | 0.002018 | 0.020266 | 0.009881 | 0.012813 | 0.018596 | 0.01571 | 0.010592 | 0.018584 | 0.004432 | 0.004902 | 0.038839 | 0.001332 | 0.005627 | 0.037665 | 0.046287 |
| 30.4722222 | 0.034094 | 0.014206 | 0.045506 | 0.018116 | 0.014318 | 0.014196 | 0.004767 | 0.032115 | 0.005836 | 0.022317 | 0.007602 | 0.008441 | 0.015593 | 0.004753 | 0.007939 | 0.012607 | 0.008886 | 0.01075 | 0.033136 | 0.00357 | 0.00429 | 0.025011 | 0.045665 |
| 30.4861111 | 0.026792 | 0.012071 | 0.046495 | 0.0164 | 0.020387 | 0.006729 | 0.006313 | 0.036197 | 0.009257 | 0.018373 | 0.009 | 0.008456 | 0.01654 | 0.004459 | 0.007455 | 0.005663 | 0.010975 | 0.013887 | 0.028921 | 0.004387 | 0.002273 | 0.015954 | 0.044368 |
| 30.5 | 0.024428 | 0.01346 | 0.044204 | 0.020349 | 0.033003 | 0.005321 | 0.005588 | 0.046495 | 0.007119 | 0.018865 | 0.005388 | 0.013839 | 0.019774 | 0.014917 | 0.007661 | 0.004083 | 0.010081 | 0.012619 | 0.019758 | 0.003417 | 0.002007 | 0.014504 | 0.042604 |
| 30.5138889 | 0.019603 | 0.018762 | 0.043564 | 0.025747 | 0.047697 | 0.007327 | 0.005836 | 0.059308 | 0.003889 | 0.025444 | 0.001283 | 0.019438 | 0.021421 | 0.031386 | 0.006213 | 0.005941 | 0.008498 | 0.010185 | 0.012083 | 0.003333 | 0.004531 | 0.015877 | 0.040232 |
| 30.5277778 | 0.012058 | 0.02384 | 0.044525 | 0.025834 | 0.059006 | 0.00907 | 0.007195 | 0.066421 | 0.004045 | 0.028423 | 0.000724 | 0.016848 | 0.017278 | 0.041612 | 0.005199 | 0.010414 | 0.01058 | 0.011981 | 0.008587 | 0.006995 | 0.010272 | 0.014341 | 0.030606 |
| 30.5416667 | 0.011993 | 0.027624 | 0.048531 | 0.024634 | 0.063497 | 0.010113 | 0.005712 | 0.06554 | 0.005809 | 0.026831 | 0.000643 | 0.013762 | 0.015149 | 0.036643 | 0.012133 | 0.023325 | 0.022648 | 0.025269 | 0.009815 | 0.019741 | 0.024616 | 0.023017 | 0.022197 |
| 30.5555556 | 0.017932 | 0.026977 | 0.050094 | 0.023055 | 0.061649 | 0.008179 | 0.003693 | 0.057919 | 0.006099 | 0.025241 | 0.003506 | 0.024838 | 0.02723 | 0.036172 | 0.027449 | 0.043181 | 0.043059 | 0.048557 | 0.022054 | 0.041151 | 0.046148 | 0.057831 | 0.034041 |
| 30.5694444 | 0.019532 | 0.020669 | 0.038892 | 0.017279 | 0.052256 | 0.005502 | 0.003231 | 0.043279 | 0.004375 | 0.024157 | 0.007335 | 0.032502 | 0.032765 | 0.049534 | 0.033306 | 0.051557 | 0.053824 | 0.060848 | 0.033248 | 0.05422 | 0.055912 | 0.097688 | 0.047382 |
| 30.5833333 | 0.017983 | 0.018747 | 0.024577 | 0.008936 | 0.035159 | 0.005131 | 0.005829 | 0.030181 | 0.003191 | 0.02259 | 0.006172 | 0.022093 | 0.027281 | 0.046889 | 0.021863 | 0.037811 | 0.042839 | 0.047887 | 0.02472 | 0.045082 | 0.044654 | 0.104028 | 0.034769 |
| 30.5972222 | 0.014107 | 0.026418 | 0.01629 | 0.004146 | 0.021042 | 0.004985 | 0.014268 | 0.025209 | 0.00298 | 0.023663 | 0.005794 | 0.019808 | 0.03898 | 0.025802 | 0.013256 | 0.017656 | 0.022878 | 0.024904 | 0.009186 | 0.024596 | 0.027472 | 0.075656 | 0.01862 |
| 30.6111111 | 0.006858 | 0.032332 | 0.012796 | 0.004416 | 0.016366 | 0.00844 | 0.020697 | 0.020926 | 0.004344 | 0.026961 | 0.011195 | 0.032671 | 0.053106 | 0.009368 | 0.017868 | 0.00616 | 0.008923 | 0.009349 | 0.003366 | 0.009866 | 0.018328 | 0.047597 | 0.022072 |
| 30.625 | 0.003424 | 0.028836 | 0.015508 | 0.006761 | 0.022155 | 0.015114 | 0.020163 | 0.012827 | 0.00736 | 0.030093 | 0.019801 | 0.034649 | 0.04631 | 0.005779 | 0.02485 | 0.002822 | 0.003348 | 0.003449 | 0.002504 | 0.00363 | 0.01596 | 0.035928 | 0.027267 |
| 30.6388889 | 0.006572 | 0.021793 | 0.020437 | 0.007631 | 0.029432 | 0.01915 | 0.019194 | 0.008432 | 0.010462 | 0.034627 | 0.031361 | 0.026794 | 0.03404 | 0.014373 | 0.029474 | 0.004741 | 0.004293 | 0.004681 | 0.002737 | 0.00362 | 0.01465 | 0.026971 | 0.024637 |
| 30.6527778 | 0.014354 | 0.017057 | 0.027016 | 0.006052 | 0.023563 | 0.020282 | 0.019961 | 0.01162 | 0.013067 | 0.036935 | 0.035681 | 0.024569 | 0.030383 | 0.029353 | 0.032173 | 0.009689 | 0.008206 | 0.009021 | 0.0065 | 0.009386 | 0.020967 | 0.013612 | 0.022779 |
| 30.6666667 | 0.020828 | 0.015111 | 0.035199 | 0.006488 | 0.011634 | 0.019634 | 0.019455 | 0.015674 | 0.013874 | 0.032066 | 0.026989 | 0.026077 | 0.030562 | 0.043366 | 0.032403 | 0.014503 | 0.011768 | 0.012698 | 0.014384 | 0.016018 | 0.03613 | 0.004254 | 0.026584 |
| 30.6805556 | 0.015462 | 0.013865 | 0.035247 | 0.011072 | 0.004181 | 0.016588 | 0.016883 | 0.014564 | 0.012294 | 0.020737 | 0.015176 | 0.025189 | 0.028715 | 0.050399 | 0.032586 | 0.017819 | 0.014449 | 0.015073 | 0.023689 | 0.019213 | 0.042 | 0.004318 | 0.034575 |
| 30.6944444 | 0.005925 | 0.011423 | 0.025506 | 0.019275 | 0.009019 | 0.011052 | 0.014126 | 0.009464 | 0.009745 | 0.008412 | 0.006865 | 0.022117 | 0.024809 | 0.044201 | 0.032684 | 0.020767 | 0.016158 | 0.016192 | 0.03065 | 0.019821 | 0.034831 | 0.011075 | 0.038275 |
| 30.7083333 | 0.005299 | 0.007577 | 0.018842 | 0.02871 | 0.028384 | 0.00535 | 0.010935 | 0.004389 | 0.007808 | 0.002064 | 0.003219 | 0.020448 | 0.018602 | 0.028722 | 0.031097 | 0.022988 | 0.015604 | 0.015163 | 0.030185 | 0.018493 | 0.026592 | 0.01996 | 0.035136 |
| 30.7222222 | 0.008296 | 0.003486 | 0.020294 | 0.032527 | 0.042569 | 0.003247 | 0.006801 | 0.00153 | 0.007225 | 0.002158 | 0.005386 | 0.020536 | 0.010888 | 0.019209 | 0.027916 | 0.021658 | 0.011945 | 0.01172 | 0.020609 | 0.014641 | 0.023051 | 0.029985 | 0.032737 |
| 30.7361111 | 0.008592 | 0.001089 | 0.017664 | 0.026346 | 0.039102 | 0.00437 | 0.00433 | 0.0006 | 0.008023 | 0.003625 | 0.00967 | 0.020892 | 0.004919 | 0.015016 | 0.022409 | 0.017023 | 0.006288 | 0.006656 | 0.009626 | 0.008911 | 0.022481 | 0.043842 | 0.035819 |
| 30.75 | 0.012145 | 0.001175 | 0.011537 | 0.013762 | 0.030309 | 0.005514 | 0.004382 | 0.001191 | 0.008746 | 0.002875 | 0.011068 | 0.018591 | 0.001978 | 0.012087 | 0.015381 | 0.011688 | 0.002861 | 0.003268 | 0.006419 | 0.004282 | 0.019327 | 0.055636 | 0.03785 |
| 30.7638889 | 0.023463 | 0.002421 | 0.015305 | 0.005077 | 0.023707 | 0.007146 | 0.005382 | 0.002131 | 0.007772 | 0.003308 | 0.009609 | 0.011308 | 0.00214 | 0.0175 | 0.009471 | 0.006861 | 0.00402 | 0.004037 | 0.011111 | 0.003399 | 0.01234 | 0.055814 | 0.027904 |
| 30.7777778 | 0.032485 | 0.002963 | 0.022556 | 0.003588 | 0.014858 | 0.008396 | 0.00756 | 0.003134 | 0.006455 | 0.007576 | 0.006225 | 0.003962 | 0.004762 | 0.026987 | 0.005607 | 0.003408 | 0.005543 | 0.005381 | 0.015208 | 0.004284 | 0.007095 | 0.048237 | 0.013444 |
| 30.7916667 | 0.034437 | 0.00335 | 0.023503 | 0.002985 | 0.006476 | 0.006065 | 0.010071 | 0.005503 | 0.006546 | 0.010587 | 0.002822 | 0.001509 | 0.006061 | 0.027069 | 0.003255 | 0.001589 | 0.004487 | 0.004315 | 0.014045 | 0.00352 | 0.007435 | 0.042168 | 0.010055 |
| 30.8055556 | 0.03785 | 0.007158 | 0.024495 | 0.003128 | 0.001935 | 0.00345 | 0.009324 | 0.006477 | 0.006637 | 0.010065 | 0.003487 | 0.001721 | 0.005332 | 0.017233 | 0.001835 | 0.000953 | 0.002557 | 0.002505 | 0.009516 | 0.001831 | 0.010466 | 0.038909 | 0.012103 |
| 30.8194444 | 0.046856 | 0.014274 | 0.024326 | 0.006588 | 0.001287 | 0.004412 | 0.005239 | 0.009304 | 0.005446 | 0.008244 | 0.009097 | 0.002605 | 0.008243 | 0.014234 | 0.000977 | 0.001347 | 0.001341 | 0.001479 | 0.004693 | 0.001178 | 0.011508 | 0.037393 | 0.010203 |
| 30.8333333 | 0.056959 | 0.018867 | 0.015486 | 0.009479 | 0.006538 | 0.005892 | 0.002592 | 0.017187 | 0.004257 | 0.005763 | 0.011824 | 0.005404 | 0.016358 | 0.022256 | 0.00051 | 0.002051 | 0.00106 | 0.001084 | 0.002046 | 0.001249 | 0.009574 | 0.037359 | 0.006398 |
| 30.8472222 | 0.060714 | 0.018724 | 0.007265 | 0.00988 | 0.016165 | 0.005993 | 0.003707 | 0.023097 | 0.004107 | 0.002719 | 0.007728 | 0.009129 | 0.022661 | 0.026133 | 0.000319 | 0.002097 | 0.00113 | 0.00077 | 0.001649 | 0.000965 | 0.006869 | 0.035205 | 0.00511 |
| 30.8611111 | 0.054598 | 0.01738 | 0.011699 | 0.009149 | 0.020097 | 0.00511 | 0.00671 | 0.027271 | 0.004133 | 0.003351 | 0.003881 | 0.011255 | 0.021458 | 0.018346 | 0.00091 | 0.001398 | 0.000935 | 0.000483 | 0.00307 | 0.000705 | 0.005253 | 0.02804 | 0.006098 |
| 30.875 | 0.038154 | 0.017217 | 0.024384 | 0.007344 | 0.017195 | 0.00471 | 0.009835 | 0.035291 | 0.003279 | 0.011433 | 0.002744 | 0.011945 | 0.013604 | 0.009536 | 0.002137 | 0.000673 | 0.000501 | 0.00047 | 0.005885 | 0.001125 | 0.005404 | 0.018871 | 0.00551 |
| 30.8888889 | 0.017867 | 0.017578 | 0.034222 | 0.00766 | 0.018111 | 0.008187 | 0.011507 | 0.047376 | 0.001823 | 0.020064 | 0.002213 | 0.011498 | 0.006296 | 0.007742 | 0.00206 | 0.000408 | 0.000277 | 0.000452 | 0.008661 | 0.001361 | 0.006253 | 0.011479 | 0.003115 |
| 30.9027778 | 0.007535 | 0.017584 | 0.037512 | 0.009285 | 0.019161 | 0.014939 | 0.011131 | 0.057352 | 0.000669 | 0.01746 | 0.004182 | 0.009793 | 0.005964 | 0.006333 | 0.001059 | 0.00025 | 0.000507 | 0.000357 | 0.0085 | 0.001021 | 0.005214 | 0.006733 | 0.002219 |
| 30.9166667 | 0.007137 | 0.017648 | 0.03558 | 0.006674 | 0.012525 | 0.01995 | 0.010107 | 0.057238 | 0.000304 | 0.010316 | 0.008922 | 0.007488 | 0.009145 | 0.003618 | 0.001013 | 0.000383 | 0.001059 | 0.000653 | 0.005894 | 0.001001 | 0.003807 | 0.007195 | 0.00378 |
| 30.9305556 | 0.005827 | 0.019105 | 0.025434 | 0.004213 | 0.00599 | 0.020548 | 0.009047 | 0.047559 | 0.000747 | 0.008195 | 0.013015 | 0.004795 | 0.011783 | 0.003447 | 0.001733 | 0.001189 | 0.001397 | 0.00101 | 0.006358 | 0.00135 | 0.006719 | 0.012517 | 0.003977 |
| 30.9444444 | 0.003828 | 0.022899 | 0.014045 | 0.007362 | 0.00803 | 0.019842 | 0.006953 | 0.036165 | 0.001541 | 0.006256 | 0.01464 | 0.003348 | 0.015406 | 0.009946 | 0.002073 | 0.001625 | 0.001041 | 0.000875 | 0.008235 | 0.00116 | 0.013032 | 0.017552 | 0.005158 |
| 30.9583333 | 0.005047 | 0.02887 | 0.020911 | 0.011687 | 0.015697 | 0.019892 | 0.004325 | 0.031834 | 0.001896 | 0.008519 | 0.013683 | 0.00587 | 0.020444 | 0.023538 | 0.002024 | 0.001487 | 0.000492 | 0.000524 | 0.006862 | 0.000532 | 0.018918 | 0.021259 | 0.013991 |
| 30.9722222 | 0.007875 | 0.036116 | 0.041188 | 0.01168 | 0.024084 | 0.018781 | 0.003842 | 0.037211 | 0.001577 | 0.015812 | 0.010593 | 0.011277 | 0.023851 | 0.03388 | 0.003714 | 0.0021 | 0.000404 | 0.000503 | 0.00751 | 0.000206 | 0.022442 | 0.023743 | 0.026797 |
| 30.9861111 | 0.0123 | 0.043142 | 0.057274 | 0.00796 | 0.026024 | 0.016771 | 0.006283 | 0.047802 | 0.001878 | 0.020353 | 0.00761 | 0.017356 | 0.025991 | 0.035926 | 0.00696 | 0.00275 | 0.000517 | 0.000763 | 0.013753 | 0.000415 | 0.022979 | 0.023718 | 0.034017 |
| 31 | 0.019886 | 0.047286 | 0.067534 | 0.005779 | 0.018617 | 0.01584 | 0.009184 | 0.060593 | 0.003124 | 0.021526 | 0.006738 | 0.021648 | 0.02954 | 0.036187 | 0.008354 | 0.002159 | 0.001115 | 0.001413 | 0.01954 | 0.000794 | 0.021274 | 0.021823 | 0.029871 |
| 31.0138889 | 0.024509 | 0.047947 | 0.072734 | 0.004839 | 0.016882 | 0.01585 | 0.009724 | 0.072127 | 0.003056 | 0.021633 | 0.006445 | 0.022174 | 0.029842 | 0.039498 | 0.006034 | 0.001738 | 0.003189 | 0.003628 | 0.021531 | 0.002136 | 0.018024 | 0.019939 | 0.018332 |
| 31.0277778 | 0.020497 | 0.048952 | 0.072105 | 0.004113 | 0.025251 | 0.01792 | 0.008456 | 0.081416 | 0.001822 | 0.022778 | 0.005672 | 0.02389 | 0.02433 | 0.046038 | 0.002511 | 0.002713 | 0.005867 | 0.00759 | 0.025053 | 0.005132 | 0.015506 | 0.022095 | 0.008398 |
| 31.0416667 | 0.016777 | 0.052271 | 0.072158 | 0.00487 | 0.032254 | 0.026466 | 0.009743 | 0.094997 | 0.003311 | 0.030808 | 0.007582 | 0.030264 | 0.022297 | 0.049331 | 0.003276 | 0.008164 | 0.006598 | 0.01007 | 0.036378 | 0.006205 | 0.018759 | 0.032034 | 0.003443 |
| 31.0555556 | 0.020073 | 0.053376 | 0.081287 | 0.011638 | 0.042832 | 0.040497 | 0.015027 | 0.11867 | 0.009264 | 0.04748 | 0.013819 | 0.035884 | 0.033471 | 0.042621 | 0.008788 | 0.020546 | 0.005182 | 0.008417 | 0.055365 | 0.003811 | 0.027306 | 0.04302 | 0.003053 |
| 31.0694444 | 0.023824 | 0.049257 | 0.090592 | 0.025583 | 0.058253 | 0.053148 | 0.01996 | 0.137834 | 0.015357 | 0.059557 | 0.017576 | 0.034814 | 0.052071 | 0.031545 | 0.012246 | 0.030945 | 0.003945 | 0.005269 | 0.072175 | 0.001894 | 0.032659 | 0.043426 | 0.006997 |
| 31.0833333 | 0.023633 | 0.041469 | 0.08248 | 0.029689 | 0.059879 | 0.058819 | 0.020312 | 0.125783 | 0.015733 | 0.056812 | 0.012785 | 0.028517 | 0.061057 | 0.021712 | 0.00931 | 0.028044 | 0.003554 | 0.003657 | 0.075956 | 0.001871 | 0.031135 | 0.031979 | 0.01333 |
| 31.0972222 | 0.017645 | 0.027333 | 0.050781 | 0.019958 | 0.044899 | 0.047112 | 0.015194 | 0.09515 | 0.010798 | 0.042232 | 0.005781 | 0.022513 | 0.052885 | 0.014114 | 0.00501 | 0.017539 | 0.003276 | 0.003897 | 0.063058 | 0.001746 | 0.026098 | 0.016792 | 0.01588 |
| 31.1111111 | 0.009207 | 0.014228 | 0.023728 | 0.013394 | 0.028448 | 0.025927 | 0.007343 | 0.078993 | 0.006641 | 0.023092 | 0.002933 | 0.019403 | 0.032432 | 0.014737 | 0.006087 | 0.017341 | 0.00394 | 0.005731 | 0.037938 | 0.001544 | 0.02229 | 0.007607 | 0.011667 |
| 31.125 | 0.006238 | 0.012879 | 0.028726 | 0.015191 | 0.016115 | 0.023068 | 0.002358 | 0.071219 | 0.008166 | 0.009375 | 0.002217 | 0.018604 | 0.014508 | 0.01816 | 0.009728 | 0.026532 | 0.006044 | 0.007772 | 0.015798 | 0.002376 | 0.022674 | 0.004418 | 0.006698 |
| 31.1388889 | 0.005141 | 0.013808 | 0.040549 | 0.015187 | 0.009206 | 0.038047 | 0.001502 | 0.051286 | 0.013987 | 0.007716 | 0.001859 | 0.01801 | 0.008024 | 0.012659 | 0.009977 | 0.029948 | 0.007911 | 0.008737 | 0.005214 | 0.002952 | 0.022802 | 0.002152 | 0.006727 |
| 31.1527778 | 0.002896 | 0.01079 | 0.042712 | 0.010777 | 0.013031 | 0.054128 | 0.002193 | 0.025049 | 0.018632 | 0.012389 | 0.004206 | 0.016983 | 0.005766 | 0.007775 | 0.007362 | 0.027612 | 0.008355 | 0.00869 | 0.001691 | 0.002198 | 0.016955 | 0.00269 | 0.011977 |
| 31.1666667 | 0.001877 | 0.006701 | 0.042699 | 0.012961 | 0.026775 | 0.058502 | 0.004581 | 0.011097 | 0.019947 | 0.014771 | 0.006448 | 0.016982 | 0.004966 | 0.012517 | 0.004186 | 0.024227 | 0.007884 | 0.007931 | 0.001099 | 0.00112 | 0.008542 | 0.005047 | 0.020536 |
| 31.1805556 | 0.001761 | 0.00425 | 0.038393 | 0.020966 | 0.043343 | 0.039628 | 0.007983 | 0.016051 | 0.019728 | 0.014521 | 0.004313 | 0.016741 | 0.004816 | 0.020605 | 0.002832 | 0.020047 | 0.007603 | 0.007302 | 0.001517 | 0.000711 | 0.005331 | 0.004715 | 0.030307 |
| 31.1944444 | 0.002471 | 0.005557 | 0.030571 | 0.022625 | 0.054259 | 0.020481 | 0.010289 | 0.026923 | 0.018767 | 0.013571 | 0.003863 | 0.01401 | 0.003567 | 0.02605 | 0.005259 | 0.017301 | 0.008434 | 0.008107 | 0.002168 | 0.001732 | 0.006839 | 0.002766 | 0.039706 |
| 31.2083333 | 0.006625 | 0.005644 | 0.02228 | 0.020161 | 0.0535 | 0.028021 | 0.009755 | 0.029548 | 0.015789 | 0.013316 | 0.007217 | 0.010526 | 0.003764 | 0.026316 | 0.007982 | 0.015285 | 0.010209 | 0.010511 | 0.002342 | 0.004277 | 0.007122 | 0.003719 | 0.045781 |
| 31.2222222 | 0.018257 | 0.004568 | 0.014355 | 0.021113 | 0.046838 | 0.041998 | 0.006893 | 0.022947 | 0.010488 | 0.013638 | 0.007272 | 0.007539 | 0.007324 | 0.022528 | 0.007369 | 0.010828 | 0.011355 | 0.013444 | 0.002977 | 0.006825 | 0.007798 | 0.009182 | 0.04497 |
| 31.2361111 | 0.032287 | 0.005013 | 0.008921 | 0.022047 | 0.038963 | 0.045891 | 0.006129 | 0.012542 | 0.005861 | 0.011938 | 0.004149 | 0.004194 | 0.010596 | 0.016505 | 0.005076 | 0.005345 | 0.010901 | 0.015687 | 0.00451 | 0.008195 | 0.013567 | 0.011999 | 0.037395 |
| 31.25 | 0.040467 | 0.003567 | 0.009913 | 0.019342 | 0.025236 | 0.043292 | 0.009198 | 0.007178 | 0.004477 | 0.008066 | 0.004529 | 0.002181 | 0.008229 | 0.017274 | 0.003946 | 0.002062 | 0.009543 | 0.015698 | 0.003997 | 0.008163 | 0.019877 | 0.007911 | 0.027127 |
| 31.2638889 | 0.039294 | 0.001708 | 0.011975 | 0.014093 | 0.0121 | 0.032629 | 0.013319 | 0.012725 | 0.004988 | 0.004818 | 0.006161 | 0.00326 | 0.005189 | 0.026671 | 0.004121 | 0.001086 | 0.00761 | 0.012544 | 0.001855 | 0.006332 | 0.01749 | 0.004236 | 0.019541 |
| 31.2777778 | 0.028227 | 0.004072 | 0.007851 | 0.009139 | 0.011447 | 0.017354 | 0.016305 | 0.023975 | 0.00463 | 0.004558 | 0.004279 | 0.005907 | 0.00571 | 0.029159 | 0.003851 | 0.001207 | 0.005308 | 0.007887 | 0.000688 | 0.003432 | 0.009426 | 0.006125 | 0.017154 |
| 31.2916667 | 0.014645 | 0.010302 | 0.003568 | 0.006159 | 0.016777 | 0.010304 | 0.018256 | 0.031404 | 0.002817 | 0.006953 | 0.001605 | 0.006219 | 0.005277 | 0.022308 | 0.002641 | 0.001966 | 0.00387 | 0.00463 | 0.000989 | 0.001486 | 0.005723 | 0.012688 | 0.016973 |
| 31.3055556 | 0.00876 | 0.015904 | 0.005987 | 0.00462 | 0.014131 | 0.019791 | 0.017387 | 0.032628 | 0.001744 | 0.008282 | 0.000902 | 0.004756 | 0.003687 | 0.014651 | 0.001902 | 0.00519 | 0.003839 | 0.003904 | 0.003833 | 0.001397 | 0.004494 | 0.021906 | 0.013625 |
| 31.3194444 | 0.01034 | 0.018306 | 0.015247 | 0.005619 | 0.005961 | 0.036412 | 0.010768 | 0.03183 | 0.002107 | 0.006656 | 0.001936 | 0.006602 | 0.002747 | 0.01381 | 0.002607 | 0.011036 | 0.00426 | 0.004363 | 0.00906 | 0.001921 | 0.002419 | 0.029618 | 0.007321 |
| 31.3333333 | 0.009279 | 0.018869 | 0.027893 | 0.010655 | 0.003224 | 0.050296 | 0.008388 | 0.030414 | 0.002224 | 0.005196 | 0.002749 | 0.009084 | 0.001813 | 0.021176 | 0.004705 | 0.01375 | 0.003433 | 0.003465 | 0.01343 | 0.002013 | 0.002126 | 0.030031 | 0.002956 |
| 31.3472222 | 0.00663 | 0.017977 | 0.037954 | 0.016248 | 0.008917 | 0.058545 | 0.014058 | 0.026863 | 0.002225 | 0.006762 | 0.003412 | 0.0066 | 0.00205 | 0.023145 | 0.008032 | 0.009457 | 0.002462 | 0.002441 | 0.015444 | 0.003252 | 0.003019 | 0.0205 | 0.002896 |
| 31.3611111 | 0.00972 | 0.016096 | 0.041572 | 0.017625 | 0.018835 | 0.057764 | 0.012911 | 0.021629 | 0.002662 | 0.012526 | 0.007128 | 0.002184 | 0.003792 | 0.017371 | 0.011028 | 0.003689 | 0.003821 | 0.004065 | 0.018431 | 0.006247 | 0.005236 | 0.008468 | 0.004927 |
| 31.375 | 0.012839 | 0.015936 | 0.041656 | 0.014904 | 0.025411 | 0.049578 | 0.01023 | 0.013881 | 0.00236 | 0.020797 | 0.0137 | 0.001528 | 0.006325 | 0.026255 | 0.011065 | 0.001679 | 0.005157 | 0.005315 | 0.023462 | 0.008022 | 0.01121 | 0.004094 | 0.005307 |
| 31.3888889 | 0.011161 | 0.017116 | 0.039058 | 0.010113 | 0.024571 | 0.034794 | 0.016292 | 0.007797 | 0.001635 | 0.025826 | 0.015238 | 0.004776 | 0.010176 | 0.059626 | 0.007473 | 0.001627 | 0.003635 | 0.003679 | 0.024988 | 0.00636 | 0.019353 | 0.0042 | 0.00845 |
| 31.4027778 | 0.009372 | 0.015238 | 0.027007 | 0.009594 | 0.023654 | 0.021355 | 0.022153 | 0.011291 | 0.001205 | 0.024322 | 0.01055 | 0.008536 | 0.015658 | 0.092512 | 0.005525 | 0.002647 | 0.002095 | 0.00256 | 0.020434 | 0.00357 | 0.024841 | 0.003737 | 0.01903 |
| 31.4166667 | 0.009147 | 0.011092 | 0.01091 | 0.016743 | 0.028137 | 0.026596 | 0.022968 | 0.022041 | 0.000919 | 0.018632 | 0.008842 | 0.011314 | 0.021208 | 0.098913 | 0.008824 | 0.004919 | 0.003317 | 0.004136 | 0.015158 | 0.003189 | 0.026504 | 0.003932 | 0.031643 |
| 31.4305556 | 0.006102 | 0.009888 | 0.002837 | 0.023488 | 0.027667 | 0.039135 | 0.01797 | 0.033981 | 0.001421 | 0.013882 | 0.011533 | 0.012826 | 0.024573 | 0.075969 | 0.011625 | 0.007013 | 0.004351 | 0.005244 | 0.013312 | 0.003725 | 0.024326 | 0.003643 | 0.041484 |
| 31.4444444 | 0.004799 | 0.010399 | 0.002352 | 0.024949 | 0.016952 | 0.036642 | 0.010072 | 0.041615 | 0.002614 | 0.011201 | 0.013046 | 0.011545 | 0.025308 | 0.038447 | 0.010245 | 0.007175 | 0.003366 | 0.004228 | 0.01311 | 0.003362 | 0.016172 | 0.003779 | 0.046301 |

| | | | | | | | | | | | | | | | | | | | | | | | |
|---|---|---|---|---|---|---|---|---|---|---|---|---|---|---|---|---|---|---|---|---|---|---|---|
| 31.4583333 | 0.009551 | 0.007537 | 0.004783 | 0.022426 | 0.006706 | 0.021641 | 0.007698 | 0.039655 | 0.005944 | 0.006788 | 0.00974 | 0.008522 | 0.023383 | 0.018743 | 0.006542 | 0.004541 | 0.004191 | 0.005725 | 0.012111 | 0.005476 | 0.008025 | 0.00422 | 0.042176 |
| 31.4722222 | 0.010438 | 0.002788 | 0.011158 | 0.01651 | 0.005959 | 0.012216 | 0.015308 | 0.033429 | 0.010365 | 0.002279 | 0.005831 | 0.007926 | 0.01795 | 0.029433 | 0.003137 | 0.003097 | 0.008477 | 0.011201 | 0.009802 | 0.010661 | 0.008937 | 0.003612 | 0.030874 |
| 31.4861111 | 0.005051 | 0.002773 | 0.026086 | 0.016277 | 0.022351 | 0.017889 | 0.025605 | 0.033241 | 0.00941 | 0.005874 | 0.006273 | 0.012192 | 0.013411 | 0.0372 | 0.001468 | 0.006068 | 0.012077 | 0.01463 | 0.00826 | 0.014136 | 0.015106 | 0.004534 | 0.020604 |
| 31.5 | 0.004903 | 0.008034 | 0.042574 | 0.021396 | 0.055106 | 0.025311 | 0.023653 | 0.036441 | 0.004091 | 0.019878 | 0.006694 | 0.020636 | 0.014689 | 0.025448 | 0.001209 | 0.008662 | 0.012395 | 0.013877 | 0.011664 | 0.01399 | 0.018549 | 0.010432 | 0.014029 |
| 31.5138889 | 0.013349 | 0.012296 | 0.047778 | 0.015445 | 0.08052 | 0.025739 | 0.015293 | 0.036583 | 0.001039 | 0.033603 | 0.006881 | 0.026744 | 0.017787 | 0.022065 | 0.0029 | 0.007114 | 0.010454 | 0.011343 | 0.018131 | 0.011963 | 0.01796 | 0.02103 | 0.007968 |
| 31.5277778 | 0.022358 | 0.014419 | 0.045256 | 0.005156 | 0.077287 | 0.023655 | 0.012864 | 0.039345 | 0.00219 | 0.038054 | 0.016304 | 0.021097 | 0.014314 | 0.037941 | 0.00376 | 0.007186 | 0.011113 | 0.011503 | 0.016596 | 0.012438 | 0.018685 | 0.024759 | 0.007361 |
| 31.5416667 | 0.031014 | 0.017508 | 0.042461 | 0.003483 | 0.050596 | 0.022114 | 0.01418 | 0.049449 | 0.005919 | 0.036127 | 0.029544 | 0.017498 | 0.016275 | 0.037521 | 0.007943 | 0.0189 | 0.023801 | 0.024205 | 0.020157 | 0.023933 | 0.030236 | 0.023092 | 0.023952 |
| 31.5555556 | 0.039593 | 0.01799 | 0.033588 | 0.003539 | 0.021528 | 0.019534 | 0.019187 | 0.056949 | 0.006884 | 0.030728 | 0.031534 | 0.029736 | 0.037524 | 0.021751 | 0.022104 | 0.038473 | 0.045366 | 0.046985 | 0.045293 | 0.044191 | 0.051582 | 0.039232 | 0.050391 |
| 31.5694444 | 0.037113 | 0.013715 | 0.020815 | 0.003014 | 0.007119 | 0.014496 | 0.01781 | 0.051112 | 0.005858 | 0.021865 | 0.025262 | 0.030836 | 0.047469 | 0.02079 | 0.029746 | 0.043433 | 0.052441 | 0.054679 | 0.056868 | 0.052274 | 0.064445 | 0.06766 | 0.058973 |
| 31.5833333 | 0.022364 | 0.014021 | 0.015965 | 0.005454 | 0.007573 | 0.010764 | 0.009974 | 0.037198 | 0.006376 | 0.015538 | 0.018695 | 0.020479 | 0.032442 | 0.019694 | 0.019062 | 0.025611 | 0.035604 | 0.036598 | 0.034787 | 0.038062 | 0.056071 | 0.079397 | 0.038077 |
| 31.5972222 | 0.008672 | 0.023462 | 0.018051 | 0.010706 | 0.010931 | 0.008635 | 0.013718 | 0.027236 | 0.006012 | 0.018286 | 0.012838 | 0.027358 | 0.029584 | 0.013351 | 0.009248 | 0.008508 | 0.013934 | 0.014016 | 0.015217 | 0.016966 | 0.034715 | 0.068882 | 0.016343 |
| 31.6111111 | 0.005567 | 0.030332 | 0.017179 | 0.021564 | 0.014594 | 0.007129 | 0.02499 | 0.022882 | 0.007695 | 0.025854 | 0.010218 | 0.037125 | 0.046159 | 0.008737 | 0.012854 | 0.007105 | 0.00311 | 0.003497 | 0.01852 | 0.004371 | 0.01503 | 0.050154 | 0.017629 |
| 31.625 | 0.014911 | 0.026142 | 0.010376 | 0.035839 | 0.020867 | 0.010135 | 0.029353 | 0.01741 | 0.011317 | 0.029282 | 0.013613 | 0.031631 | 0.056708 | 0.005114 | 0.018508 | 0.009866 | 0.001234 | 0.001284 | 0.029769 | 0.000995 | 0.004927 | 0.036108 | 0.023115 |
| 31.6388889 | 0.028008 | 0.01789 | 0.005057 | 0.043869 | 0.02735 | 0.015338 | 0.020435 | 0.011027 | 0.01197 | 0.026766 | 0.019592 | 0.021385 | 0.061778 | 0.005985 | 0.020578 | 0.009959 | 0.001538 | 0.000785 | 0.042102 | 0.001064 | 0.004152 | 0.029511 | 0.01891 |
| 31.6527778 | 0.033046 | 0.013438 | 0.003846 | 0.037855 | 0.03278 | 0.015443 | 0.008289 | 0.007371 | 0.011945 | 0.023196 | 0.019844 | 0.015132 | 0.069765 | 0.00578 | 0.0227 | 0.009829 | 0.002121 | 0.001437 | 0.052892 | 0.001226 | 0.007864 | 0.025817 | 0.012824 |
| 31.6666667 | 0.026431 | 0.013383 | 0.004394 | 0.019692 | 0.028943 | 0.010269 | 0.009858 | 0.004934 | 0.014431 | 0.020519 | 0.012638 | 0.011587 | 0.071199 | 0.005949 | 0.024474 | 0.01009 | 0.00371 | 0.003616 | 0.050825 | 0.001949 | 0.012058 | 0.020369 | 0.011448 |
| 31.6805556 | 0.01391 | 0.015344 | 0.006871 | 0.005875 | 0.017342 | 0.009723 | 0.016658 | 0.002979 | 0.017006 | 0.016688 | 0.009661 | 0.011106 | 0.06302 | 0.012268 | 0.025025 | 0.011002 | 0.005737 | 0.005993 | 0.038069 | 0.003782 | 0.014198 | 0.011447 | 0.016334 |
| 31.6944444 | 0.005025 | 0.016081 | 0.008775 | 0.003615 | 0.019629 | 0.016968 | 0.012876 | 0.002306 | 0.015866 | 0.010249 | 0.014195 | 0.01445 | 0.053988 | 0.020883 | 0.024831 | 0.011831 | 0.007267 | 0.007578 | 0.027707 | 0.005534 | 0.012947 | 0.005813 | 0.023375 |
| 31.7083333 | 0.002076 | 0.013372 | 0.009653 | 0.006766 | 0.040979 | 0.02199 | 0.004269 | 0.00177 | 0.011583 | 0.005406 | 0.019074 | 0.018436 | 0.047921 | 0.027564 | 0.024148 | 0.011522 | 0.008023 | 0.008248 | 0.024932 | 0.006504 | 0.010303 | 0.010411 | 0.025631 |
| 31.7222222 | 0.001332 | 0.007808 | 0.011998 | 0.012717 | 0.05236 | 0.023126 | 0.002721 | 0.003301 | 0.007147 | 0.008262 | 0.02283 | 0.020908 | 0.041551 | 0.029282 | 0.023 | 0.010759 | 0.008429 | 0.008578 | 0.027341 | 0.00718 | 0.008959 | 0.018695 | 0.020975 |
| 31.7361111 | 0.003027 | 0.003477 | 0.014928 | 0.016397 | 0.040225 | 0.022858 | 0.007534 | 0.011896 | 0.004225 | 0.013868 | 0.022746 | 0.021973 | 0.030791 | 0.022946 | 0.021053 | 0.010789 | 0.008421 | 0.008732 | 0.02952 | 0.007503 | 0.008379 | 0.022553 | 0.015006 |
| 31.75 | 0.008541 | 0.002808 | 0.016937 | 0.019375 | 0.039993 | 0.019365 | 0.014173 | 0.022673 | 0.003048 | 0.012954 | 0.017663 | 0.022208 | 0.016783 | 0.013046 | 0.018238 | 0.011197 | 0.00723 | 0.007927 | 0.025689 | 0.006648 | 0.006668 | 0.021295 | 0.011657 |
| 31.7638889 | 0.012203 | 0.002578 | 0.015276 | 0.025714 | 0.061404 | 0.013109 | 0.022189 | 0.02259 | 0.002876 | 0.007737 | 0.01086 | 0.022626 | 0.011032 | 0.005982 | 0.014925 | 0.011103 | 0.004447 | 0.005393 | 0.015532 | 0.004064 | 0.004225 | 0.017579 | 0.00947 |
| 31.7777778 | 0.011016 | 0.001958 | 0.010443 | 0.02799 | 0.064551 | 0.007033 | 0.023401 | 0.011987 | 0.002227 | 0.00673 | 0.009798 | 0.021037 | 0.019147 | 0.004131 | 0.012169 | 0.010956 | 0.001732 | 0.002374 | 0.006785 | 0.001667 | 0.0044 | 0.013721 | 0.007386 |
| 31.7916667 | 0.009087 | 0.003238 | 0.009132 | 0.02344 | 0.044766 | 0.005621 | 0.014607 | 0.005501 | 0.0012 | 0.00974 | 0.014952 | 0.017981 | 0.028292 | 0.00469 | 0.011107 | 0.011072 | 0.000845 | 0.000932 | 0.005911 | 0.000967 | 0.007519 | 0.008651 | 0.007014 |
| 31.8055556 | 0.00746 | 0.00546 | 0.00917 | 0.015755 | 0.035365 | 0.009562 | 0.006495 | 0.008954 | 0.000956 | 0.012294 | 0.016904 | 0.017283 | 0.028865 | 0.002852 | 0.011281 | 0.010847 | 0.001459 | 0.001422 | 0.009042 | 0.001218 | 0.008969 | 0.005138 | 0.010015 |
| 31.8194444 | 0.006682 | 0.006443 | 0.008416 | 0.011899 | 0.034818 | 0.016009 | 0.005172 | 0.014766 | 0.000917 | 0.014078 | 0.020385 | 0.01352 | 0.023486 | 0.003685 | 0.011784 | 0.010442 | 0.003082 | 0.003225 | 0.010474 | 0.002724 | 0.007178 | 0.00578 | 0.016073 |
| 31.8333333 | 0.008881 | 0.00486 | 0.010568 | 0.016193 | 0.029112 | 0.022357 | 0.010749 | 0.015331 | 0.000706 | 0.015257 | 0.02585 | 0.007518 | 0.015044 | 0.006897 | 0.01248 | 0.010494 | 0.004833 | 0.005142 | 0.00881 | 0.004708 | 0.00586 | 0.012855 | 0.023037 |
| 31.8472222 | 0.014584 | 0.00335 | 0.011582 | 0.023831 | 0.036556 | 0.024955 | 0.015349 | 0.009685 | 0.000592 | 0.013394 | 0.020081 | 0.008315 | 0.00832 | 0.007733 | 0.01297 | 0.010698 | 0.00588 | 0.006283 | 0.005245 | 0.005867 | 0.00663 | 0.030122 | 0.027892 |
| 31.8611111 | 0.020865 | 0.006184 | 0.0116 | 0.033503 | 0.053396 | 0.021669 | 0.013206 | 0.003971 | 0.000689 | 0.007608 | 0.008169 | 0.013415 | 0.010168 | 0.009147 | 0.012504 | 0.010597 | 0.006419 | 0.006783 | 0.002379 | 0.00649 | 0.005959 | 0.046668 | 0.029238 |
| 31.875 | 0.020065 | 0.010479 | 0.01496 | 0.045871 | 0.054799 | 0.01338 | 0.013092 | 0.003332 | 0.001408 | 0.002438 | 0.002389 | 0.016132 | 0.017451 | 0.008666 | 0.011265 | 0.010113 | 0.006759 | 0.007021 | 0.001102 | 0.006954 | 0.003107 | 0.047882 | 0.028335 |
| 31.8888889 | 0.011015 | 0.010618 | 0.018383 | 0.049798 | 0.04524 | 0.008616 | 0.013827 | 0.005532 | 0.002974 | 0.001717 | 0.002158 | 0.014857 | 0.021894 | 0.007033 | 0.009939 | 0.009027 | 0.006918 | 0.007086 | 0.000981 | 0.007291 | 0.001204 | 0.035307 | 0.026617 |
| 31.9027778 | 0.003587 | 0.006843 | 0.015102 | 0.03493 | 0.03863 | 0.015732 | 0.009742 | 0.009565 | 0.00487 | 0.005777 | 0.00413 | 0.010483 | 0.021109 | 0.007559 | 0.008453 | 0.006958 | 0.006502 | 0.006621 | 0.00216 | 0.007021 | 0.001099 | 0.02018 | 0.023795 |
| 31.9166667 | 0.006533 | 0.002939 | 0.010804 | 0.017094 | 0.02754 | 0.02695 | 0.008943 | 0.018812 | 0.005546 | 0.012724 | 0.007382 | 0.005869 | 0.01614 | 0.006959 | 0.006429 | 0.004225 | 0.00551 | 0.005653 | 0.003915 | 0.005876 | 0.00175 | 0.008991 | 0.019629 |
| 31.9305556 | 0.017802 | 0.000848 | 0.015521 | 0.015291 | 0.0124 | 0.031803 | 0.01272 | 0.032523 | 0.003945 | 0.017273 | 0.01055 | 0.005645 | 0.010031 | 0.004737 | 0.004038 | 0.002006 | 0.004653 | 0.004905 | 0.004689 | 0.004347 | 0.001747 | 0.002829 | 0.015518 |
| 31.9444444 | 0.030709 | 0.000492 | 0.018785 | 0.023859 | 0.007778 | 0.030633 | 0.013222 | 0.044555 | 0.001892 | 0.015843 | 0.010651 | 0.010712 | 0.005333 | 0.002696 | 0.002111 | 0.001117 | 0.004392 | 0.004872 | 0.003437 | 0.003262 | 0.0021 | 0.000924 | 0.01296 |
| 31.9583333 | 0.045101 | 0.001713 | 0.012775 | 0.034435 | 0.012553 | 0.029458 | 0.014923 | 0.048753 | 0.001386 | 0.013677 | 0.009733 | 0.016964 | 0.003094 | 0.004399 | 0.001859 | 0.001409 | 0.00486 | 0.005624 | 0.002364 | 0.003528 | 0.004039 | 0.000999 | 0.012251 |
| 31.9722222 | 0.05436 | 0.00507 | 0.00748 | 0.046052 | 0.018568 | 0.030231 | 0.021366 | 0.044525 | 0.001501 | 0.016245 | 0.01384 | 0.021987 | 0.005107 | 0.011196 | 0.00305 | 0.002298 | 0.005928 | 0.006813 | 0.005322 | 0.005116 | 0.00565 | 0.00221 | 0.012475 |
| 31.9861111 | 0.048277 | 0.011607 | 0.006545 | 0.055498 | 0.017007 | 0.02826 | 0.018989 | 0.039742 | 0.002346 | 0.016746 | 0.018954 | 0.022006 | 0.011249 | 0.013453 | 0.003682 | 0.003128 | 0.006502 | 0.007305 | 0.012529 | 0.006235 | 0.005698 | 0.003028 | 0.011675 |
| 32 | 0.035042 | 0.019383 | 0.011183 | 0.058565 | 0.009694 | 0.021324 | 0.014478 | 0.039728 | 0.004376 | 0.011546 | 0.016767 | 0.014712 | 0.019305 | 0.007322 | 0.003669 | 0.003782 | 0.006201 | 0.007018 | 0.022203 | 0.0062 | 0.005323 | 0.002458 | 0.008994 |
| 32.0138889 | 0.029028 | 0.026552 | 0.029412 | 0.050986 | 0.007606 | 0.012153 | 0.014629 | 0.042752 | 0.005331 | 0.009612 | 0.010407 | 0.007735 | 0.027058 | 0.003635 | 0.005114 | 0.005594 | 0.006831 | 0.008129 | 0.030878 | 0.007203 | 0.007215 | 0.00347 | 0.006419 |
| 32.0277778 | 0.033213 | 0.035899 | 0.054471 | 0.03742 | 0.012323 | 0.011815 | 0.018618 | 0.055246 | 0.005594 | 0.010173 | 0.009901 | 0.010818 | 0.033856 | 0.005475 | 0.005549 | 0.005933 | 0.007077 | 0.009399 | 0.037407 | 0.007618 | 0.007945 | 0.004756 | 0.005617 |
| 32.0416667 | 0.042628 | 0.045832 | 0.070252 | 0.03429 | 0.024705 | 0.025496 | 0.040538 | 0.080954 | 0.009609 | 0.012297 | 0.011588 | 0.025821 | 0.044138 | 0.016519 | 0.008335 | 0.007539 | 0.007798 | 0.009874 | 0.051942 | 0.008664 | 0.012212 | 0.006627 | 0.012838 |
| 32.0555556 | 0.047815 | 0.047989 | 0.079403 | 0.042551 | 0.050419 | 0.03675 | 0.059677 | 0.104119 | 0.016481 | 0.021316 | 0.007492 | 0.043554 | 0.061262 | 0.040875 | 0.018018 | 0.017152 | 0.012726 | 0.012942 | 0.077943 | 0.014858 | 0.029039 | 0.011956 | 0.030426 |
| 32.0694444 | 0.03982 | 0.03957 | 0.084905 | 0.042021 | 0.069418 | 0.040194 | 0.04906 | 0.117838 | 0.019821 | 0.028001 | 0.003514 | 0.052635 | 0.078762 | 0.059452 | 0.022284 | 0.023625 | 0.014193 | 0.013703 | 0.092895 | 0.016911 | 0.040656 | 0.015064 | 0.044584 |
| 32.0833333 | 0.025862 | 0.027044 | 0.070221 | 0.029737 | 0.060655 | 0.042592 | 0.025583 | 0.120196 | 0.01894 | 0.026247 | 0.003773 | 0.051074 | 0.083644 | 0.063006 | 0.017257 | 0.019599 | 0.008007 | 0.007831 | 0.083815 | 0.009843 | 0.035568 | 0.019337 | 0.047712 |
| 32.0972222 | 0.015677 | 0.015791 | 0.036931 | 0.018027 | 0.043913 | 0.038579 | 0.016768 | 0.102624 | 0.017783 | 0.019038 | 0.00417 | 0.041378 | 0.068117 | 0.060022 | 0.01099 | 0.011357 | 0.002423 | 0.002272 | 0.063843 | 0.003126 | 0.024562 | 0.028424 | 0.043646 |
| 32.1111111 | 0.009467 | 0.007906 | 0.016056 | 0.009182 | 0.030142 | 0.025346 | 0.019487 | 0.081435 | 0.017255 | 0.011081 | 0.00245 | 0.025997 | 0.04093 | 0.048644 | 0.006232 | 0.00738 | 0.001141 | 0.001277 | 0.040428 | 0.001309 | 0.01363 | 0.033023 | 0.035 |
| 32.125 | 0.007418 | 0.006503 | 0.020679 | 0.006181 | 0.016116 | 0.010412 | 0.018235 | 0.073264 | 0.015698 | 0.009033 | 0.001684 | 0.012311 | 0.018025 | 0.032163 | 0.005142 | 0.013379 | 0.002541 | 0.003251 | 0.01917 | 0.002778 | 0.005152 | 0.032536 | 0.025571 |
| 32.1388889 | 0.009416 | 0.008399 | 0.028242 | 0.011472 | 0.012103 | 0.003767 | 0.021276 | 0.062681 | 0.012379 | 0.009685 | 0.002527 | 0.00662 | 0.006341 | 0.022457 | 0.006648 | 0.018618 | 0.004958 | 0.005959 | 0.010356 | 0.0051 | 0.001243 | 0.033433 | 0.020435 |
| 32.1527778 | 0.01619 | 0.006499 | 0.020706 | 0.020433 | 0.017569 | 0.007868 | 0.025829 | 0.038946 | 0.008341 | 0.010798 | 0.002148 | 0.007153 | 0.005506 | 0.021752 | 0.007172 | 0.016087 | 0.00664 | 0.007674 | 0.011216 | 0.006411 | 0.001001 | 0.036652 | 0.021885 |
| 32.1666667 | 0.020077 | 0.002805 | 0.012169 | 0.025106 | 0.020271 | 0.018051 | 0.01686 | 0.016907 | 0.005001 | 0.011137 | 0.001341 | 0.009455 | 0.01234 | 0.023432 | 0.005992 | 0.011557 | 0.006343 | 0.007355 | 0.012538 | 0.005955 | 0.002232 | 0.039829 | 0.027166 |
| 32.1805556 | 0.014175 | 0.002079 | 0.020298 | 0.020803 | 0.017933 | 0.028874 | 0.006422 | 0.007808 | 0.002757 | 0.01153 | 0.001469 | 0.011183 | 0.01942 | 0.022299 | 0.003637 | 0.006595 | 0.004189 | 0.005092 | 0.01176 | 0.003832 | 0.003225 | 0.040171 | 0.031228 |
| 32.1944444 | 0.005829 | 0.003492 | 0.033106 | 0.014016 | 0.01922 | 0.038617 | 0.005349 | 0.01394 | 0.001376 | 0.020118 | 0.001887 | 0.012357 | 0.018003 | 0.018749 | 0.002235 | 0.003032 | 0.001918 | 0.002699 | 0.010459 | 0.001563 | 0.002771 | 0.035769 | 0.032443 |
| 32.2083333 | 0.001938 | 0.005198 | 0.034597 | 0.01047 | 0.024588 | 0.050227 | 0.00519 | 0.032262 | 0.000653 | 0.029129 | 0.002349 | 0.013882 | 0.010285 | 0.01208 | 0.003728 | 0.003796 | 0.001102 | 0.001704 | 0.009406 | 0.000768 | 0.001625 | 0.028025 | 0.032565 |
| 32.2222222 | 0.003106 | 0.005329 | 0.031037 | 0.008855 | 0.026236 | 0.061009 | 0.006929 | 0.044295 | 0.000718 | 0.030324 | 0.003054 | 0.015734 | 0.005993 | 0.011229 | 0.00697 | 0.005622 | 0.000998 | 0.001353 | 0.006324 | 0.000791 | 0.000941 | 0.021659 | 0.03304 |
| 32.2361111 | 0.008288 | 0.008955 | 0.029002 | 0.008796 | 0.026394 | 0.061821 | 0.01148 | 0.040457 | 0.001917 | 0.029801 | 0.006593 | 0.017018 | 0.006069 | 0.026054 | 0.011315 | 0.00552 | 0.001198 | 0.001187 | 0.003801 | 0.00124 | 0.001166 | 0.019649 | 0.033727 |
| 32.25 | 0.010245 | 0.019111 | 0.026224 | 0.013125 | 0.031142 | 0.047975 | 0.013432 | 0.034495 | 0.003231 | 0.028689 | 0.011629 | 0.018254 | 0.010332 | 0.036591 | 0.016642 | 0.004999 | 0.003177 | 0.002574 | 0.007061 | 0.003605 | 0.003184 | 0.022721 | 0.033176 |
| 32.2638889 | 0.0102 | 0.022421 | 0.021677 | 0.019524 | 0.037452 | 0.026653 | 0.018293 | 0.034377 | 0.003008 | 0.021874 | 0.011564 | 0.020165 | 0.018821 | 0.026975 | 0.021072 | 0.005506 | 0.006404 | 0.005318 | 0.01277 | 0.007248 | 0.006337 | 0.031743 | 0.03116 |
| 32.2777778 | 0.0204 | 0.014951 | 0.018844 | 0.020767 | 0.039396 | 0.020463 | 0.027172 | 0.030327 | 0.001557 | 0.01093 | 0.011085 | 0.020071 | 0.022646 | 0.012127 | 0.021978 | 0.005734 | 0.008102 | 0.00692 | 0.016256 | 0.009259 | 0.008339 | 0.04543 | 0.028148 |
| 32.2916667 | 0.037468 | 0.008878 | 0.016816 | 0.016034 | 0.036771 | 0.036967 | 0.034469 | 0.017198 | 0.000904 | 0.003541 | 0.013536 | 0.015424 | 0.018467 | 0.008067 | 0.017961 | 0.004625 | 0.006037 | 0.005191 | 0.017624 | 0.007225 | 0.007936 | 0.068716 | 0.023003 |
| 32.3055556 | 0.051213 | 0.006176 | 0.013151 | 0.008525 | 0.02974 | 0.046735 | 0.032275 | 0.009265 | 0.001016 | 0.001873 | 0.011448 | 0.00796 | 0.01281 | 0.016127 | 0.0104 | 0.005216 | 0.003354 | 0.003613 | 0.018438 | 0.004123 | 0.005107 | 0.092903 | 0.015827 |
| 32.3194444 | 0.056675 | 0.00523 | 0.018833 | 0.005648 | 0.019937 | 0.039719 | 0.02663 | 0.01809 | 0.000781 | 0.002609 | 0.006515 | 0.003522 | 0.010364 | 0.029701 | 0.003945 | 0.008107 | 0.004447 | 0.00611 | 0.018241 | 0.004869 | 0.003484 | 0.082847 | 0.009257 |
| 32.3333333 | 0.051857 | 0.008118 | 0.037971 | 0.012244 | 0.02431 | 0.030349 | 0.023452 | 0.035897 | 0.001163 | 0.005246 | 0.003729 | 0.004996 | 0.010785 | 0.034951 | 0.00212 | 0.00954 | 0.006622 | 0.008863 | 0.015221 | 0.007194 | 0.006845 | 0.040861 | 0.004463 |
| 32.3472222 | 0.036155 | 0.00799 | 0.057768 | 0.016148 | 0.039527 | 0.017008 | 0.01678 | 0.053294 | 0.002805 | 0.010442 | 0.002161 | 0.007729 | 0.010837 | 0.02466 | 0.00321 | 0.009209 | 0.007162 | 0.009311 | 0.010287 | 0.007832 | 0.011695 | 0.013838 | 0.002231 |
| 32.3611111 | 0.01847 | 0.004088 | 0.070517 | 0.00925 | 0.037992 | 0.007159 | 0.012925 | 0.064232 | 0.004321 | 0.015824 | 0.001063 | 0.009141 | 0.008182 | 0.019163 | 0.003963 | 0.008833 | 0.006655 | 0.008721 | 0.005264 | 0.007253 | 0.01468 | 0.015608 | 0.003756 |
| 32.375 | 0.012119 | 0.002849 | 0.072263 | 0.008157 | 0.022969 | 0.009222 | 0.008881 | 0.064266 | 0.003789 | 0.021751 | 0.000457 | 0.009901 | 0.004198 | 0.031698 | 0.00313 | 0.008022 | 0.00504 | 0.00664 | 0.003262 | 0.005732 | 0.015801 | 0.021673 | 0.007011 |
| 32.3888889 | 0.011972 | 0.003123 | 0.058912 | 0.013021 | 0.010351 | 0.013332 | 0.005294 | 0.05189 | 0.001978 | 0.029505 | 0.000321 | 0.009685 | 0.002129 | 0.047227 | 0.002663 | 0.006022 | 0.00286 | 0.003541 | 0.007701 | 0.003498 | 0.014491 | 0.021536 | 0.009705 |
| 32.4027778 | 0.009245 | 0.005191 | 0.033491 | 0.008422 | 0.005603 | 0.016786 | 0.006444 | 0.029375 | 0.000829 | 0.036562 | 0.000293 | 0.009685 | 0.001913 | 0.060981 | 0.004125 | 0.003421 | 0.001946 | 0.001645 | 0.015113 | 0.002341 | 0.012078 | 0.023369 | 0.011478 |
| 32.4166667 | 0.004832 | 0.009106 | 0.014373 | 0.005246 | 0.009196 | 0.024953 | 0.011693 | 0.014085 | 0.00052 | 0.039275 | 0.000863 | 0.011098 | 0.001878 | 0.074563 | 0.004914 | 0.002334 | 0.002087 | 0.001035 | 0.019915 | 0.002458 | 0.011157 | 0.032426 | 0.014114 |
| 32.4305556 | 0.004923 | 0.010481 | 0.012753 | 0.011658 | 0.018658 | 0.034734 | 0.024435 | 0.020009 | 0.000788 | 0.03771 | 0.004917 | 0.012833 | 0.00186 | 0.074371 | 0.003178 | 0.003479 | 0.001553 | 0.000772 | 0.021659 | 0.001782 | 0.011678 | 0.043212 | 0.020615 |
| 32.4444444 | 0.013525 | 0.009431 | 0.012402 | 0.017815 | 0.032447 | 0.043129 | 0.034724 | 0.032585 | 0.001655 | 0.034221 | 0.012735 | 0.013434 | 0.002368 | 0.055152 | 0.001138 | 0.004525 | 0.001034 | 0.000948 | 0.022455 | 0.000985 | 0.011144 | 0.044563 | 0.028074 |
| 32.4583333 | 0.022823 | 0.007593 | 0.006679 | 0.01717 | 0.042018 | 0.04876 | 0.035558 | 0.034095 | 0.002244 | 0.028911 | 0.019848 | 0.012669 | 0.004852 | 0.029517 | 0.000875 | 0.004028 | 0.002155 | 0.002939 | 0.02449 | 0.002289 | 0.008271 | 0.034582 | 0.030332 |
| 32.4722222 | 0.021123 | 0.004104 | 0.004177 | 0.014289 | 0.03962 | 0.037873 | 0.038769 | 0.02807 | 0.002132 | 0.023427 | 0.019879 | 0.012184 | 0.007507 | 0.013134 | 0.001171 | 0.003197 | 0.003872 | 0.006298 | 0.027401 | 0.004819 | 0.004726 | 0.027394 | 0.026595 |
| 32.4861111 | 0.013576 | 0.002252 | 0.009012 | 0.015241 | 0.031494 | 0.016914 | 0.044309 | 0.030746 | 0.002804 | 0.023587 | 0.012649 | 0.014453 | 0.008847 | 0.013931 | 0.001601 | 0.002398 | 0.004252 | 0.007069 | 0.027109 | 0.006199 | 0.003153 | 0.03417 | 0.019763 |
| 32.5 | 0.014504 | 0.004502 | 0.026188 | 0.025339 | 0.021193 | 0.012601 | 0.036097 | 0.045236 | 0.004566 | 0.031763 | 0.010523 | 0.019417 | 0.011867 | 0.024706 | 0.001955 | 0.002172 | 0.005197 | 0.006669 | 0.022442 | 0.007504 | 0.00296 | 0.044075 | 0.012692 |
| 32.5138889 | 0.017954 | 0.006087 | 0.047915 | 0.033319 | 0.013727 | 0.024512 | 0.028664 | 0.060752 | 0.005323 | 0.040786 | 0.018987 | 0.022926 | 0.014886 | 0.030828 | 0.001677 | 0.003668 | 0.008041 | 0.008675 | 0.01828 | 0.010075 | 0.002053 | 0.044641 | 0.006667 |
| 32.5277778 | 0.018473 | 0.005657 | 0.056039 | 0.023391 | 0.0195 | 0.035381 | 0.033149 | 0.075061 | 0.004771 | 0.04322 | 0.027949 | 0.017355 | 0.011876 | 0.028334 | 0.005321 | 0.007772 | 0.013515 | 0.013988 | 0.013364 | 0.014987 | 0.003094 | 0.036762 | 0.0062 |
| 32.5416667 | 0.019976 | 0.007024 | 0.050512 | 0.017618 | 0.030961 | 0.037873 | 0.027854 | 0.08467 | 0.005537 | 0.038924 | 0.029603 | 0.012507 | 0.011287 | 0.026074 | 0.018812 | 0.0183 | 0.027022 | 0.027849 | 0.012403 | 0.027319 | 0.012126 | 0.023295 | 0.020991 |
| 32.5555556 | 0.017767 | 0.007818 | 0.034866 | 0.024738 | 0.034344 | 0.03258 | 0.028131 | 0.081216 | 0.005809 | 0.034156 | 0.022957 | 0.019922 | 0.021895 | 0.02514 | 0.035833 | 0.029888 | 0.042818 | 0.044728 | 0.023856 | 0.042023 | 0.024988 | 0.013279 | 0.049439 |
| 32.5694444 | 0.011296 | 0.005807 | 0.020167 | 0.023702 | 0.023206 | 0.019785 | 0.048331 | 0.066007 | 0.003905 | 0.035084 | 0.011676 | 0.020398 | 0.025359 | 0.02212 | 0.036251 | 0.026728 | 0.040765 | 0.042878 | 0.027241 | 0.040828 | 0.02419 | 0.012818 | 0.070022 |
| 32.5833333 | 0.005864 | 0.005 | 0.020829 | 0.017824 | 0.010914 | 0.011968 | 0.050552 | 0.052654 | 0.002908 | 0.043736 | 0.004909 | 0.015124 | 0.025203 | 0.030132 | 0.021451 | 0.014506 | 0.021466 | 0.022642 | 0.019401 | 0.022301 | 0.01558 | 0.021352 | 0.063983 |
| 32.5972222 | 0.006142 | 0.004357 | 0.025549 | 0.014218 | 0.007136 | 0.017757 | 0.028414 | 0.046813 | 0.002616 | 0.056002 | 0.007458 | 0.022818 | 0.04062 | 0.045177 | 0.015638 | 0.011134 | 0.008137 | 0.009014 | 0.025413 | 0.00766 | 0.019918 | 0.036092 | 0.038982 |
| 32.6111111 | 0.010209 | 0.007007 | 0.029841 | 0.008703 | 0.006327 | 0.022261 | 0.013674 | 0.037096 | 0.002585 | 0.05738 | 0.016041 | 0.034716 | 0.053021 | 0.044653 | 0.019945 | 0.014249 | 0.00675 | 0.007934 | 0.036556 | 0.005535 | 0.027925 | 0.042707 | 0.015789 |
| 32.625 | 0.010397 | 0.018652 | 0.02681 | 0.009379 | 0.010068 | 0.01964 | 0.010459 | 0.021429 | 0.003776 | 0.044492 | 0.026967 | 0.039724 | 0.050442 | 0.033425 | 0.021554 | 0.016295 | 0.006383 | 0.007446 | 0.036684 | 0.005051 | 0.026341 | 0.036742 | 0.00407 |
| 32.6388889 | 0.007598 | 0.027532 | 0.01101 | 0.015452 | 0.018893 | 0.014796 | 0.018976 | 0.009597 | 0.005847 | 0.030498 | 0.038953 | 0.036531 | 0.042803 | 0.024415 | 0.019545 | 0.015651 | 0.003791 | 0.004351 | 0.034718 | 0.002561 | 0.018634 | 0.027456 | 0.003602 |
| 32.6527778 | 0.010062 | 0.022451 | 0.006098 | 0.016983 | 0.033274 | 0.01018 | 0.04536 | 0.006239 | 0.008485 | 0.021232 | 0.04641 | 0.02835 | 0.035215 | 0.017068 | 0.016622 | 0.01188 | 0.001589 | 0.001654 | 0.036552 | 0.001797 | 0.010475 | 0.019802 | 0.010487 |
| 32.6666667 | 0.016161 | 0.012493 | 0.015462 | 0.012792 | 0.042561 | 0.007249 | 0.055145 | 0.006867 | 0.009499 | 0.015277 | 0.036572 | 0.019743 | 0.032583 | 0.012696 | 0.013774 | 0.007025 | 0.000972 | 0.000731 | 0.037011 | 0.002452 | 0.004851 | 0.013504 | 0.018651 |
| 32.6805556 | 0.016554 | 0.009408 | 0.019973 | 0.006595 | 0.034217 | 0.004691 | 0.03522 | 0.005179 | 0.007735 | 0.01571 | 0.01985 | 0.013686 | 0.034586 | 0.011612 | 0.01117 | 0.003028 | 0.001515 | 0.001166 | 0.033757 | 0.003324 | 0.001957 | 0.007819 | 0.025118 |
| 32.6944444 | 0.011016 | 0.014186 | 0.016266 | 0.00383 | 0.019021 | 0.004272 | 0.024274 | 0.002534 | 0.005933 | 0.026099 | 0.021649 | 0.01016 | 0.032887 | 0.007567 | 0.00851 | 0.000829 | 0.00228 | 0.001749 | 0.030169 | 0.004241 | 0.00141 | 0.004945 | 0.030252 |
| 32.7083333 | 0.005978 | 0.023569 | 0.017886 | 0.003661 | 0.012963 | 0.007535 | 0.025278 | 0.001985 | 0.005376 | 0.039399 | 0.035672 | 0.006714 | 0.025615 | 0.003375 | 0.00567 | 0.000369 | 0.003431 | 0.002784 | 0.030262 | 0.005395 | 0.001714 | 0.003572 | 0.032898 |
| 32.7222222 | 0.003768 | 0.026671 | 0.028566 | 0.007342 | 0.016289 | 0.012952 | 0.019677 | 0.002416 | 0.004608 | 0.042678 | 0.040991 | 0.003546 | 0.019303 | 0.004953 | 0.003567 | 0.000726 | 0.004842 | 0.00472 | 0.035105 | 0.006805 | 0.00259 | 0.002924 | 0.031616 |
| 32.7361111 | 0.005763 | 0.016966 | 0.033932 | 0.017636 | 0.021191 | 0.019252 | 0.01335 | 0.002956 | 0.003002 | 0.032911 | 0.032262 | 0.002498 | 0.018084 | 0.007182 | 0.003712 | 0.001437 | 0.005147 | 0.005705 | 0.039313 | 0.007243 | 0.003767 | 0.005968 | 0.02786 |
| 32.75 | 0.011076 | 0.009111 | 0.035237 | 0.023401 | 0.024121 | 0.024848 | 0.009535 | 0.002967 | 0.001314 | 0.017869 | 0.020851 | 0.004591 | 0.017915 | 0.007221 | 0.005527 | 0.002544 | 0.003956 | 0.004398 | 0.036982 | 0.005776 | 0.003292 | 0.01061 | 0.023585 |
| 32.7638889 | 0.014483 | 0.012728 | 0.032773 | 0.019914 | 0.021405 | 0.030306 | 0.007409 | 0.002092 | 0.000332 | 0.007142 | 0.014309 | 0.008811 | 0.012597 | 0.010752 | 0.006659 | 0.003235 | 0.002842 | 0.002615 | 0.029989 | 0.003913 | 0.002739 | 0.014383 | 0.019813 |
| 32.7777778 | 0.014671 | 0.016854 | 0.027893 | 0.013328 | 0.014399 | 0.032003 | 0.00863 | 0.001909 | 0.000122 | 0.004934 | 0.007729 | 0.011313 | 0.005255 | 0.018326 | 0.006166 | 0.002617 | 0.003038 | 0.002489 | 0.022274 | 0.003639 | 0.003535 | 0.016459 | 0.01778 |
| 32.7916667 | 0.010051 | 0.01466 | 0.032009 | 0.009841 | 0.00931 | 0.022372 | 0.010185 | 0.004198 | 0.000288 | 0.007154 | 0.002326 | 0.01069 | 0.001936 | 0.020288 | 0.005448 | 0.001815 | 0.003652 | 0.003472 | 0.015283 | 0.004601 | 0.00353 | 0.013606 | 0.017832 |
| 32.8055556 | 0.004516 | 0.008977 | 0.034435 | 0.017223 | 0.012552 | 0.009765 | 0.01732 | 0.007777 | 0.000553 | 0.007618 | 0.000635 | 0.009054 | 0.002582 | 0.016251 | 0.006417 | 0.002405 | 0.003115 | 0.003447 | 0.010407 | 0.004916 | 0.002956 | 0.007693 | 0.017477 |
| 32.8194444 | 0.008526 | 0.007738 | 0.026494 | 0.029188 | 0.028529 | 0.008044 | 0.025528 | 0.00892 | 0.000894 | 0.005137 | 0.000334 | 0.005873 | 0.0048 | 0.017822 | 0.008857 | 0.004436 | 0.001692 | 0.002047 | 0.007505 | 0.003826 | 0.00327 | 0.005582 | 0.013588 |
| 32.8333333 | 0.021152 | 0.01324 | 0.019834 | 0.036496 | 0.038526 | 0.014443 | 0.019262 | 0.007289 | 0.001237 | 0.003305 | 0.000378 | 0.003402 | 0.005234 | 0.021288 | 0.010719 | 0.006522 | 0.000861 | 0.001115 | 0.004321 | 0.002426 | 0.00338 | 0.007132 | 0.007022 |
| 32.8472222 | 0.031533 | 0.01367 | 0.02601 | 0.036315 | 0.02636 | 0.021562 | 0.01089 | 0.006392 | 0.00149 | 0.00327 | 0.000551 | 0.005281 | 0.003535 | 0.017941 | 0.010713 | 0.00662 | 0.00079 | 0.001036 | 0.002267 | 0.002098 | 0.002991 | 0.00712 | 0.002891 |
| 32.8611111 | 0.030781 | 0.007068 | 0.030551 | 0.028947 | 0.012021 | 0.027385 | 0.013504 | 0.008518 | 0.001331 | 0.004401 | 0.001341 | 0.010574 | 0.001951 | 0.014055 | 0.008884 | 0.004503 | 0.001272 | 0.001216 | 0.003435 | 0.003092 | 0.005145 | 0.004765 | 0.003739 |
| 32.875 | 0.020633 | 0.006516 | 0.030788 | 0.019786 | 0.011951 | 0.02916 | 0.015149 | 0.01126 | 0.00101 | 0.007825 | 0.002313 | 0.016825 | 0.003702 | 0.011636 | 0.006385 | 0.002033 | 0.002165 | 0.002197 | 0.004728 | 0.00436 | 0.01055 | 0.004167 | 0.006992 |
| 32.8888889 | 0.009934 | 0.016644 | 0.049033 | 0.01266 | 0.013389 | 0.028297 | 0.012361 | 0.009719 | 0.001284 | 0.012149 | 0.006898 | 0.020068 | 0.010741 | 0.008189 | 0.004973 | 0.000741 | 0.002165 | 0.002837 | 0.00467 | 0.004519 | 0.014307 | 0.007598 | 0.010882 |
| 32.9027778 | 0.003073 | 0.026607 | 0.067375 | 0.011637 | 0.011281 | 0.030836 | 0.016814 | 0.004732 | 0.001752 | 0.014775 | 0.01657 | 0.020268 | 0.015803 | 0.005204 | 0.004945 | 0.000564 | 0.001176 | 0.001838 | 0.005001 | 0.003435 | 0.013082 | 0.011851 | 0.015911 |
| 32.9166667 | 0.001074 | 0.029808 | 0.054517 | 0.019771 | 0.016028 | 0.034079 | 0.036035 | 0.002336 | 0.002268 | 0.013727 | 0.02316 | 0.019507 | 0.011668 | 0.005427 | 0.004659 | 0.001268 | 0.000441 | 0.000578 | 0.006423 | 0.00236 | 0.009291 | 0.01493 | 0.021726 |
| 32.9305556 | 0.001997 | 0.028208 | 0.025155 | 0.034187 | 0.021954 | 0.029777 | 0.061744 | 0.004231 | 0.003733 | 0.008444 | 0.024348 | 0.016023 | 0.006304 | 0.010726 | 0.003024 | 0.002304 | 0.000463 | 0.000484 | 0.008376 | 0.00234 | 0.007041 | 0.018369 | 0.02695 |

| | | | | | | | | | | | | | | | | | | | | | | | |
|---|---|---|---|---|---|---|---|---|---|---|---|---|---|---|---|---|---|---|---|---|---|---|---|
| 32.9444444 | 0.002606 | 0.022559 | 0.015343 | 0.043521 | 0.016468 | 0.016906 | 0.07116 | 0.005484 | 0.004512 | 0.00343 | 0.019286 | 0.01016 | 0.009838 | 0.016946 | 0.001371 | 0.002434 | 0.001302 | 0.001512 | 0.009979 | 0.003283 | 0.008157 | 0.024451 | 0.031027 |
| 32.9583333 | 0.004188 | 0.016654 | 0.033214 | 0.041032 | 0.010368 | 0.007421 | 0.050785 | 0.003532 | 0.002884 | 0.002133 | 0.015063 | 0.006415 | 0.017495 | 0.017911 | 0.000945 | 0.002673 | 0.002166 | 0.002537 | 0.010763 | 0.004096 | 0.010768 | 0.032918 | 0.033057 |
| 32.9722222 | 0.007127 | 0.015196 | 0.042705 | 0.03441 | 0.01616 | 0.015047 | 0.030288 | 0.004683 | 0.001596 | 0.003443 | 0.024109 | 0.007771 | 0.021027 | 0.014345 | 0.000972 | 0.003698 | 0.001815 | 0.002128 | 0.009902 | 0.003505 | 0.011336 | 0.038169 | 0.032398 |
| 32.9861111 | 0.007711 | 0.015408 | 0.028391 | 0.028438 | 0.020576 | 0.031923 | 0.045168 | 0.016453 | 0.002652 | 0.007326 | 0.033121 | 0.012654 | 0.018667 | 0.012313 | 0.001318 | 0.003637 | 0.001266 | 0.001425 | 0.007405 | 0.002225 | 0.00906 | 0.034173 | 0.030511 |
| 33 | 0.010833 | 0.013608 | 0.020902 | 0.021167 | 0.016056 | 0.035965 | 0.061793 | 0.038542 | 0.003142 | 0.009487 | 0.028506 | 0.016924 | 0.013237 | 0.011207 | 0.002547 | 0.00236 | 0.002847 | 0.003102 | 0.004662 | 0.003202 | 0.005347 | 0.02133 | 0.029804 |
| 33.0138889 | 0.021179 | 0.011606 | 0.020073 | 0.0135 | 0.014596 | 0.026423 | 0.040421 | 0.069015 | 0.003564 | 0.00559 | 0.017054 | 0.017402 | 0.011265 | 0.00938 | 0.005224 | 0.002073 | 0.006917 | 0.007695 | 0.002368 | 0.007876 | 0.003865 | 0.010595 | 0.033491 |
| 33.0277778 | 0.022224 | 0.013093 | 0.014419 | 0.009252 | 0.019157 | 0.019272 | 0.018829 | 0.108875 | 0.006453 | 0.004556 | 0.008117 | 0.016994 | 0.022802 | 0.010939 | 0.007048 | 0.002663 | 0.011245 | 0.013209 | 0.002082 | 0.012853 | 0.005856 | 0.008819 | 0.039397 |
| 33.0416667 | 0.018476 | 0.021778 | 0.020265 | 0.011849 | 0.017824 | 0.021715 | 0.018395 | 0.143935 | 0.006993 | 0.016839 | 0.00428 | 0.021385 | 0.046649 | 0.01759 | 0.007072 | 0.005231 | 0.01101 | 0.013338 | 0.008381 | 0.011432 | 0.008088 | 0.011747 | 0.037835 |
| 33.0555556 | 0.039665 | 0.033415 | 0.032797 | 0.019375 | 0.013115 | 0.030715 | 0.025955 | 0.151823 | 0.006035 | 0.038158 | 0.003884 | 0.028294 | 0.068967 | 0.029515 | 0.010928 | 0.011763 | 0.006232 | 0.006988 | 0.022727 | 0.006176 | 0.013005 | 0.022147 | 0.028032 |
| 33.0694444 | 0.073427 | 0.035559 | 0.036026 | 0.024379 | 0.01543 | 0.037436 | 0.049099 | 0.140724 | 0.008705 | 0.05417 | 0.006019 | 0.030792 | 0.083513 | 0.042661 | 0.016088 | 0.015 | 0.002915 | 0.001953 | 0.038068 | 0.004783 | 0.018064 | 0.033263 | 0.021746 |
| 33.0833333 | 0.079187 | 0.024312 | 0.046461 | 0.024462 | 0.015312 | 0.037089 | 0.060588 | 0.133687 | 0.010898 | 0.054486 | 0.007319 | 0.027223 | 0.084149 | 0.050375 | 0.015042 | 0.010764 | 0.001971 | 0.000435 | 0.045896 | 0.004141 | 0.017277 | 0.030415 | 0.025005 |
| 33.0972222 | 0.049852 | 0.011285 | 0.07028 | 0.0271 | 0.013946 | 0.029775 | 0.042846 | 0.136198 | 0.008179 | 0.038941 | 0.00657 | 0.019939 | 0.06901 | 0.050858 | 0.010991 | 0.004894 | 0.001383 | 0.000403 | 0.044766 | 0.001785 | 0.01749 | 0.022519 | 0.03206 |
| 33.1111111 | 0.018399 | 0.006729 | 0.079307 | 0.029721 | 0.018463 | 0.018093 | 0.022428 | 0.144192 | 0.003273 | 0.019018 | 0.005413 | 0.010848 | 0.051954 | 0.042422 | 0.008414 | 0.001927 | 0.001276 | 0.001216 | 0.037266 | 0.000802 | 0.021288 | 0.022203 | 0.03413 |
| 33.125 | 0.006558 | 0.006419 | 0.067686 | 0.022357 | 0.018752 | 0.007513 | 0.010519 | 0.138029 | 0.001736 | 0.010969 | 0.004324 | 0.003267 | 0.040946 | 0.028127 | 0.006776 | 0.001877 | 0.001486 | 0.00212 | 0.027502 | 0.00145 | 0.022847 | 0.031763 | 0.028585 |
| 33.1388889 | 0.003811 | 0.004721 | 0.051008 | 0.012682 | 0.019174 | 0.004664 | 0.004319 | 0.105536 | 0.00413 | 0.014005 | 0.003303 | 0.001757 | 0.035633 | 0.019165 | 0.005577 | 0.00162 | 0.001606 | 0.002875 | 0.019897 | 0.002369 | 0.022391 | 0.052764 | 0.022001 |
| 33.1527778 | 0.002067 | 0.003181 | 0.032613 | 0.008806 | 0.026227 | 0.012707 | 0.003728 | 0.066675 | 0.005567 | 0.012327 | 0.002108 | 0.004327 | 0.036687 | 0.018082 | 0.004664 | 0.000666 | 0.001453 | 0.00262 | 0.012812 | 0.002511 | 0.021448 | 0.075379 | 0.018596 |
| 33.1666667 | 0.00374 | 0.003896 | 0.030409 | 0.008066 | 0.026913 | 0.026424 | 0.005571 | 0.034579 | 0.003633 | 0.007763 | 0.001963 | 0.005155 | 0.037605 | 0.015827 | 0.003639 | 0.001024 | 0.000982 | 0.001531 | 0.005351 | 0.001784 | 0.018767 | 0.076808 | 0.014611 |
| 33.1805556 | 0.00752 | 0.007299 | 0.05876 | 0.009889 | 0.016511 | 0.038964 | 0.011081 | 0.01882 | 0.001734 | 0.00885 | 0.003789 | 0.004712 | 0.031774 | 0.009099 | 0.002155 | 0.002542 | 0.000571 | 0.001162 | 0.003439 | 0.000847 | 0.014769 | 0.053405 | 0.009664 |
| 33.1944444 | 0.011415 | 0.010751 | 0.086994 | 0.010971 | 0.005796 | 0.047978 | 0.024986 | 0.035714 | 0.001769 | 0.008779 | 0.00674 | 0.005128 | 0.023319 | 0.006146 | 0.001539 | 0.003602 | 0.001017 | 0.001793 | 0.009239 | 0.000652 | 0.010285 | 0.025892 | 0.007747 |
| 33.2083333 | 0.016765 | 0.009572 | 0.104307 | 0.008202 | 0.003514 | 0.052438 | 0.028935 | 0.075851 | 0.001943 | 0.010879 | 0.009238 | 0.004855 | 0.018293 | 0.015347 | 0.003116 | 0.004149 | 0.002587 | 0.003289 | 0.016233 | 0.001696 | 0.005982 | 0.012301 | 0.010685 |
| 33.2222222 | 0.020554 | 0.005228 | 0.12356 | 0.009113 | 0.007148 | 0.050124 | 0.02842 | 0.110941 | 0.002135 | 0.022861 | 0.009631 | 0.004493 | 0.016501 | 0.031569 | 0.00447 | 0.004672 | 0.004236 | 0.005011 | 0.017879 | 0.003197 | 0.004734 | 0.018419 | 0.015177 |
| 33.2361111 | 0.01838 | 0.002324 | 0.129784 | 0.015767 | 0.008686 | 0.042846 | 0.042631 | 0.119176 | 0.002627 | 0.037037 | 0.007572 | 0.006488 | 0.013134 | 0.041113 | 0.003422 | 0.004435 | 0.005254 | 0.005815 | 0.015378 | 0.004465 | 0.006436 | 0.033158 | 0.016678 |
| 33.25 | 0.012613 | 0.002072 | 0.115599 | 0.016528 | 0.006052 | 0.034444 | 0.04211 | 0.097268 | 0.002673 | 0.045307 | 0.004544 | 0.009458 | 0.007617 | 0.04001 | 0.001491 | 0.003077 | 0.006244 | 0.006048 | 0.012699 | 0.006012 | 0.007615 | 0.045628 | 0.016528 |
| 33.2638889 | 0.008417 | 0.003282 | 0.086132 | 0.011309 | 0.006082 | 0.026017 | 0.024533 | 0.059389 | 0.002337 | 0.044895 | 0.00292 | 0.009741 | 0.004778 | 0.034345 | 0.001718 | 0.002115 | 0.006174 | 0.005872 | 0.009902 | 0.006472 | 0.007666 | 0.055663 | 0.016417 |
| 33.2777778 | 0.005991 | 0.002912 | 0.045526 | 0.014716 | 0.014425 | 0.018616 | 0.030356 | 0.029598 | 0.00183 | 0.036247 | 0.003693 | 0.006784 | 0.003786 | 0.026291 | 0.005033 | 0.003222 | 0.004067 | 0.004258 | 0.006388 | 0.004448 | 0.006835 | 0.058954 | 0.014401 |
| 33.2916667 | 0.006424 | 0.001936 | 0.021235 | 0.027377 | 0.024658 | 0.013597 | 0.05727 | 0.030298 | 0.001558 | 0.023428 | 0.00515 | 0.003556 | 0.0019 | 0.014965 | 0.007601 | 0.003748 | 0.001907 | 0.002585 | 0.00317 | 0.002001 | 0.004437 | 0.050848 | 0.011949 |
| 33.3055556 | 0.011817 | 0.003928 | 0.035331 | 0.033207 | 0.024011 | 0.012301 | 0.056671 | 0.053707 | 0.002237 | 0.011304 | 0.005381 | 0.002483 | 0.002283 | 0.007827 | 0.00611 | 0.003865 | 0.002606 | 0.004382 | 0.002724 | 0.002575 | 0.003658 | 0.037611 | 0.013609 |
| 33.3194444 | 0.016302 | 0.006977 | 0.067 | 0.027345 | 0.012339 | 0.013939 | 0.025121 | 0.074607 | 0.003479 | 0.005909 | 0.004072 | 0.005092 | 0.00647 | 0.012791 | 0.005095 | 0.008591 | 0.007465 | 0.009948 | 0.00741 | 0.007704 | 0.006281 | 0.027611 | 0.019598 |
| 33.3333333 | 0.018403 | 0.008947 | 0.082759 | 0.020823 | 0.003145 | 0.015668 | 0.011367 | 0.092842 | 0.004399 | 0.007868 | 0.002739 | 0.011653 | 0.011881 | 0.020988 | 0.009655 | 0.016864 | 0.014738 | 0.017245 | 0.015323 | 0.015046 | 0.009467 | 0.021858 | 0.027223 |
| 33.3472222 | 0.019391 | 0.017135 | 0.068535 | 0.016899 | 0.002251 | 0.016103 | 0.020677 | 0.119048 | 0.004078 | 0.008105 | 0.003507 | 0.01875 | 0.016753 | 0.020163 | 0.015649 | 0.023714 | 0.020978 | 0.023759 | 0.021796 | 0.02107 | 0.013524 | 0.016953 | 0.033722 |
| 33.3611111 | 0.013894 | 0.033795 | 0.041276 | 0.01172 | 0.005454 | 0.012763 | 0.021636 | 0.13489 | 0.003311 | 0.005441 | 0.005743 | 0.019091 | 0.015855 | 0.014176 | 0.013687 | 0.02271 | 0.019172 | 0.02199 | 0.020857 | 0.019336 | 0.012914 | 0.013669 | 0.032385 |
| 33.375 | 0.009449 | 0.035875 | 0.021117 | 0.012581 | 0.006853 | 0.006562 | 0.0121 | 0.111863 | 0.004411 | 0.005318 | 0.006332 | 0.01287 | 0.010746 | 0.008655 | 0.010152 | 0.014596 | 0.012914 | 0.0145 | 0.012631 | 0.013351 | 0.010643 | 0.0205 | 0.019557 |
| 33.3888889 | 0.021125 | 0.029309 | 0.019825 | 0.02026 | 0.007777 | 0.00341 | 0.007462 | 0.06166 | 0.004939 | 0.006405 | 0.004455 | 0.010639 | 0.011781 | 0.005381 | 0.017623 | 0.011426 | 0.015369 | 0.015111 | 0.006487 | 0.016264 | 0.017338 | 0.033585 | 0.009457 |
| 33.4027778 | 0.036787 | 0.047776 | 0.040152 | 0.02442 | 0.010884 | 0.0031 | 0.011243 | 0.026802 | 0.002787 | 0.009624 | 0.002366 | 0.01282 | 0.017076 | 0.00936 | 0.025475 | 0.014124 | 0.021864 | 0.020799 | 0.004208 | 0.023398 | 0.019974 | 0.037501 | 0.013863 |
| 33.4166667 | 0.037218 | 0.066207 | 0.053625 | 0.023673 | 0.00906 | 0.002335 | 0.019463 | 0.030129 | 0.001363 | 0.01493 | 0.001623 | 0.012263 | 0.019923 | 0.016171 | 0.024757 | 0.01406 | 0.023448 | 0.022603 | 0.002494 | 0.025279 | 0.011904 | 0.029238 | 0.021859 |
| 33.4305556 | 0.028984 | 0.052377 | 0.05675 | 0.019251 | 0.004417 | 0.004838 | 0.018175 | 0.059052 | 0.001942 | 0.018063 | 0.002878 | 0.008717 | 0.018991 | 0.016749 | 0.021439 | 0.011537 | 0.022121 | 0.022071 | 0.003851 | 0.023963 | 0.007399 | 0.016824 | 0.024637 |
| 33.4444444 | 0.020853 | 0.027367 | 0.074303 | 0.012014 | 0.00491 | 0.01463 | 0.011291 | 0.086956 | 0.003171 | 0.019205 | 0.005464 | 0.0045 | 0.013901 | 0.015247 | 0.018394 | 0.009846 | 0.01983 | 0.021112 | 0.006498 | 0.021826 | 0.009679 | 0.013026 | 0.024245 |
| 33.4583333 | 0.015271 | 0.012348 | 0.08693 | 0.005664 | 0.005266 | 0.025914 | 0.011094 | 0.096911 | 0.003242 | 0.019308 | 0.007186 | 0.002461 | 0.00717 | 0.020416 | 0.015549 | 0.008437 | 0.017952 | 0.020873 | 0.005465 | 0.020096 | 0.009865 | 0.019681 | 0.022452 |
| 33.4722222 | 0.022594 | 0.009033 | 0.067841 | 0.003333 | 0.003354 | 0.032884 | 0.012521 | 0.092348 | 0.002611 | 0.01817 | 0.007974 | 0.005125 | 0.003067 | 0.029667 | 0.012677 | 0.005868 | 0.017036 | 0.020899 | 0.005077 | 0.01903 | 0.005795 | 0.021638 | 0.019093 |
| 33.4861111 | 0.042481 | 0.017852 | 0.036191 | 0.005264 | 0.003862 | 0.034425 | 0.015179 | 0.087921 | 0.005705 | 0.020346 | 0.008288 | 0.01184 | 0.004941 | 0.034471 | 0.008187 | 0.003943 | 0.014314 | 0.017244 | 0.010907 | 0.016006 | 0.002577 | 0.014002 | 0.012059 |
| 33.5 | 0.055344 | 0.028627 | 0.019285 | 0.010147 | 0.007335 | 0.028604 | 0.013648 | 0.094687 | 0.012317 | 0.030363 | 0.008103 | 0.019627 | 0.012407 | 0.03278 | 0.003882 | 0.006032 | 0.009083 | 0.010484 | 0.017353 | 0.010591 | 0.003207 | 0.005397 | 0.005851 |
| 33.5138889 | 0.045793 | 0.026373 | 0.014028 | 0.012042 | 0.014763 | 0.016518 | 0.008547 | 0.110403 | 0.016094 | 0.042104 | 0.007386 | 0.024804 | 0.019826 | 0.025777 | 0.004443 | 0.010158 | 0.004194 | 0.004739 | 0.018976 | 0.005149 | 0.006186 | 0.003529 | 0.008247 |
| 33.5277778 | 0.02482 | 0.016099 | 0.011852 | 0.007936 | 0.020196 | 0.006211 | 0.009418 | 0.120781 | 0.01651 | 0.047577 | 0.006172 | 0.024802 | 0.022003 | 0.01699 | 0.007238 | 0.011525 | 0.001865 | 0.001828 | 0.015809 | 0.001687 | 0.008455 | 0.007791 | 0.014496 |
| 33.5416667 | 0.014193 | 0.010867 | 0.021043 | 0.00449 | 0.016071 | 0.005311 | 0.01253 | 0.116716 | 0.017341 | 0.048812 | 0.007751 | 0.016725 | 0.016263 | 0.013069 | 0.007817 | 0.010101 | 0.006378 | 0.006178 | 0.013406 | 0.004928 | 0.012998 | 0.016296 | 0.013923 |
| 33.5555556 | 0.011343 | 0.010595 | 0.03741 | 0.00441 | 0.012336 | 0.012081 | 0.010572 | 0.102021 | 0.014464 | 0.051656 | 0.011251 | 0.01095 | 0.015044 | 0.027457 | 0.012697 | 0.014623 | 0.020607 | 0.020485 | 0.027894 | 0.018293 | 0.026733 | 0.043743 | 0.018086 |
| 33.5694444 | 0.01239 | 0.010404 | 0.043969 | 0.006995 | 0.016279 | 0.016872 | 0.008173 | 0.077575 | 0.007477 | 0.057765 | 0.0098 | 0.018321 | 0.026107 | 0.054547 | 0.023032 | 0.025413 | 0.033738 | 0.033719 | 0.066158 | 0.032329 | 0.043553 | 0.079156 | 0.043557 |
| 33.5833333 | 0.019671 | 0.012622 | 0.040352 | 0.009206 | 0.014681 | 0.016868 | 0.011892 | 0.046686 | 0.003607 | 0.065406 | 0.006143 | 0.020708 | 0.027276 | 0.058043 | 0.02166 | 0.025434 | 0.031322 | 0.031093 | 0.103089 | 0.032078 | 0.041229 | 0.078739 | 0.071975 |
| 33.5972222 | 0.018847 | 0.019154 | 0.031535 | 0.013183 | 0.008394 | 0.024455 | 0.011196 | 0.021353 | 0.003543 | 0.06882 | 0.009584 | 0.017812 | 0.022545 | 0.039824 | 0.012461 | 0.013468 | 0.017015 | 0.016362 | 0.10384 | 0.019 | 0.020263 | 0.060631 | 0.081625 |
| 33.6111111 | 0.010321 | 0.024358 | 0.018181 | 0.018023 | 0.012304 | 0.038871 | 0.004724 | 0.00807 | 0.006798 | 0.062803 | 0.018646 | 0.026451 | 0.028416 | 0.029253 | 0.013607 | 0.005442 | 0.005458 | 0.005051 | 0.076645 | 0.00765 | 0.008363 | 0.062316 | 0.072107 |
| 33.625 | 0.006765 | 0.020939 | 0.008312 | 0.016421 | 0.024608 | 0.045818 | 0.004041 | 0.006426 | 0.012876 | 0.04456 | 0.026423 | 0.036054 | 0.032194 | 0.022682 | 0.016908 | 0.005862 | 0.001342 | 0.00146 | 0.055472 | 0.003351 | 0.009494 | 0.059505 | 0.048369 |
| 33.6388889 | 0.005906 | 0.014128 | 0.008407 | 0.015014 | 0.033559 | 0.04464 | 0.005087 | 0.007546 | 0.018925 | 0.022923 | 0.028971 | 0.037271 | 0.034682 | 0.016992 | 0.015763 | 0.00939 | 0.000889 | 0.001021 | 0.040344 | 0.002831 | 0.010045 | 0.041479 | 0.022608 |
| 33.6527778 | 0.005692 | 0.014436 | 0.010335 | 0.015463 | 0.029861 | 0.039358 | 0.002314 | 0.00581 | 0.023383 | 0.016913 | 0.026289 | 0.033981 | 0.041841 | 0.03123 | 0.016244 | 0.011804 | 0.002188 | 0.001964 | 0.02233 | 0.003975 | 0.007511 | 0.026247 | 0.011236 |
| 33.6666667 | 0.00628 | 0.018628 | 0.006819 | 0.012801 | 0.017499 | 0.031011 | 0.002257 | 0.00402 | 0.025666 | 0.017872 | 0.022424 | 0.032927 | 0.048745 | 0.060092 | 0.018662 | 0.013387 | 0.004144 | 0.003775 | 0.016777 | 0.00541 | 0.004303 | 0.016284 | 0.014994 |
| 33.6805556 | 0.006607 | 0.025601 | 0.002872 | 0.007796 | 0.007675 | 0.020487 | 0.007554 | 0.005174 | 0.025958 | 0.015408 | 0.01713 | 0.036437 | 0.055777 | 0.072754 | 0.020953 | 0.014354 | 0.005631 | 0.00548 | 0.027659 | 0.008137 | 0.002971 | 0.009154 | 0.02015 |
| 33.6944444 | 0.005486 | 0.036401 | 0.00359 | 0.006322 | 0.003681 | 0.00931 | 0.009311 | 0.008718 | 0.023744 | 0.026371 | 0.010224 | 0.040908 | 0.063669 | 0.063693 | 0.022939 | 0.014686 | 0.007389 | 0.007589 | 0.039121 | 0.011231 | 0.003473 | 0.005231 | 0.020114 |
| 33.7083333 | 0.003608 | 0.041565 | 0.007239 | 0.015145 | 0.003046 | 0.002464 | 0.009901 | 0.010466 | 0.019132 | 0.050909 | 0.009083 | 0.047311 | 0.069149 | 0.04667 | 0.024735 | 0.016275 | 0.010232 | 0.010869 | 0.049135 | 0.010089 | 0.003211 | 0.004728 | 0.016852 |
| 33.7222222 | 0.007535 | 0.039897 | 0.010876 | 0.033472 | 0.006002 | 0.001439 | 0.016292 | 0.009317 | 0.013676 | 0.068464 | 0.015416 | 0.054767 | 0.069457 | 0.034778 | 0.024953 | 0.01882 | 0.013087 | 0.014106 | 0.061318 | 0.006234 | 0.003123 | 0.007166 | 0.013364 |
| 33.7361111 | 0.017146 | 0.038753 | 0.013559 | 0.042373 | 0.01074 | 0.001645 | 0.016524 | 0.008055 | 0.008415 | 0.068511 | 0.02264 | 0.053402 | 0.063968 | 0.033635 | 0.023741 | 0.020623 | 0.014747 | 0.01576 | 0.068657 | 0.006583 | 0.005897 | 0.010356 | 0.011132 |
| 33.75 | 0.019741 | 0.043472 | 0.011611 | 0.030949 | 0.009409 | 0.00339 | 0.016221 | 0.008574 | 0.003852 | 0.05551 | 0.027493 | 0.04127 | 0.051735 | 0.030788 | 0.02319 | 0.021365 | 0.015484 | 0.016152 | 0.070559 | 0.012601 | 0.009696 | 0.01222 | 0.009223 |
| 33.7638889 | 0.012873 | 0.054 | 0.007335 | 0.024949 | 0.005305 | 0.008824 | 0.021808 | 0.009002 | 0.002044 | 0.03833 | 0.026753 | 0.029521 | 0.033917 | 0.020695 | 0.023622 | 0.02174 | 0.015585 | 0.01588 | 0.070354 | 0.020774 | 0.011514 | 0.012964 | 0.006686 |
| 33.7777778 | 0.013025 | 0.055942 | 0.010671 | 0.041391 | 0.007667 | 0.012809 | 0.019449 | 0.007671 | 0.004132 | 0.022376 | 0.018295 | 0.023072 | 0.01711 | 0.010245 | 0.023627 | 0.021562 | 0.01468 | 0.014799 | 0.063727 | 0.028225 | 0.013376 | 0.013478 | 0.004042 |
| 33.7916667 | 0.022847 | 0.044868 | 0.018698 | 0.061115 | 0.011539 | 0.013707 | 0.017126 | 0.005601 | 0.006624 | 0.010807 | 0.007994 | 0.019423 | 0.0091 | 0.004295 | 0.021146 | 0.019709 | 0.012437 | 0.012745 | 0.054035 | 0.031862 | 0.017101 | 0.015128 | 0.002099 |
| 33.8055556 | 0.030041 | 0.039692 | 0.02574 | 0.061634 | 0.00904 | 0.014833 | 0.021234 | 0.004161 | 0.006318 | 0.006549 | 0.00228 | 0.016031 | 0.008706 | 0.005718 | 0.014126 | 0.015508 | 0.008745 | 0.009325 | 0.04564 | 0.025337 | 0.017491 | 0.018895 | 0.001708 |
| 33.8194444 | 0.036094 | 0.038414 | 0.033885 | 0.047112 | 0.007567 | 0.013945 | 0.024326 | 0.00311 | 0.005191 | 0.008496 | 0.000771 | 0.011642 | 0.007341 | 0.012529 | 0.006681 | 0.009489 | 0.004409 | 0.004784 | 0.034122 | 0.011008 | 0.012233 | 0.023742 | 0.004098 |
| 33.8333333 | 0.043486 | 0.025505 | 0.040832 | 0.030243 | 0.021445 | 0.008806 | 0.02548 | 0.003222 | 0.006164 | 0.011355 | 0.000629 | 0.006414 | 0.00565 | 0.024108 | 0.00488 | 0.004022 | 0.001412 | 0.001498 | 0.025283 | 0.002369 | 0.006776 | 0.027387 | 0.009225 |
| 33.8472222 | 0.053624 | 0.010796 | 0.039749 | 0.01762 | 0.042002 | 0.006975 | 0.019284 | 0.004945 | 0.007167 | 0.013269 | 0.001158 | 0.003316 | 0.009743 | 0.043125 | 0.005472 | 0.001169 | 0.000644 | 0.000849 | 0.022629 | 0.00139 | 0.003577 | 0.028457 | 0.016333 |
| 33.8611111 | 0.068175 | 0.006406 | 0.028113 | 0.012218 | 0.043313 | 0.013767 | 0.011346 | 0.005903 | 0.005294 | 0.017218 | 0.002145 | 0.005854 | 0.01483 | 0.059704 | 0.003786 | 0.000487 | 0.000938 | 0.001171 | 0.018412 | 0.001815 | 0.001964 | 0.027402 | 0.024095 |
| 33.875 | 0.078648 | 0.008208 | 0.01591 | 0.00976 | 0.026636 | 0.021012 | 0.014113 | 0.006363 | 0.002356 | 0.023306 | 0.002422 | 0.011476 | 0.012073 | 0.057729 | 0.001849 | 0.000818 | 0.001157 | 0.00122 | 0.011974 | 0.002282 | 0.002742 | 0.025199 | 0.029929 |
| 33.8888889 | 0.07832 | 0.008342 | 0.014903 | 0.008645 | 0.011818 | 0.023077 | 0.016751 | 0.008966 | 0.000895 | 0.027174 | 0.002648 | 0.01617 | 0.008155 | 0.038069 | 0.001918 | 0.001606 | 0.000868 | 0.000879 | 0.007382 | 0.00308 | 0.004926 | 0.022321 | 0.031402 |
| 33.9027778 | 0.061615 | 0.005337 | 0.026534 | 0.011187 | 0.006121 | 0.022134 | 0.01368 | 0.014945 | 0.001368 | 0.023501 | 0.004409 | 0.018869 | 0.014465 | 0.01774 | 0.002564 | 0.002287 | 0.000528 | 0.000549 | 0.004395 | 0.003536 | 0.006588 | 0.018421 | 0.027659 |
| 33.9166667 | 0.033924 | 0.00348 | 0.037102 | 0.014027 | 0.007291 | 0.020985 | 0.011537 | 0.019546 | 0.003337 | 0.014686 | 0.00697 | 0.020781 | 0.020783 | 0.011432 | 0.002715 | 0.002637 | 0.000824 | 0.000902 | 0.002386 | 0.002693 | 0.007329 | 0.013821 | 0.020419 |
| 33.9305556 | 0.012967 | 0.003852 | 0.036163 | 0.012617 | 0.008231 | 0.020923 | 0.009613 | 0.016326 | 0.005575 | 0.008951 | 0.008967 | 0.024225 | 0.015901 | 0.013246 | 0.002769 | 0.00302 | 0.001265 | 0.001579 | 0.002587 | 0.001245 | 0.00776 | 0.009901 | 0.01273 |
| 33.9444444 | 0.005702 | 0.004447 | 0.027644 | 0.007522 | 0.00849 | 0.019819 | 0.011465 | 0.008113 | 0.006977 | 0.005272 | 0.008507 | 0.029379 | 0.007976 | 0.011757 | 0.0021 | 0.003224 | 0.001291 | 0.001849 | 0.004155 | 0.000477 | 0.008616 | 0.007613 | 0.006711 |
| 33.9583333 | 0.006337 | 0.006466 | 0.015394 | 0.006676 | 0.011854 | 0.014712 | 0.018612 | 0.002289 | 0.008667 | 0.0031 | 0.005304 | 0.034441 | 0.006306 | 0.008549 | 0.001709 | 0.002374 | 0.000949 | 0.001321 | 0.005532 | 0.001002 | 0.010392 | 0.007275 | 0.002949 |
| 33.9722222 | 0.012107 | 0.010852 | 0.009984 | 0.0137 | 0.016714 | 0.009365 | 0.019371 | 0.001906 | 0.01071 | 0.006579 | 0.004736 | 0.037787 | 0.009251 | 0.006426 | 0.003975 | 0.002429 | 0.000718 | 0.000579 | 0.006473 | 0.002419 | 0.012745 | 0.008117 | 0.001482 |
| 33.9861111 | 0.023573 | 0.014072 | 0.014987 | 0.019167 | 0.020711 | 0.011466 | 0.013055 | 0.003559 | 0.009481 | 0.015704 | 0.010558 | 0.037522 | 0.014021 | 0.008599 | 0.008066 | 0.005668 | 0.000673 | 0.000506 | 0.006939 | 0.003413 | 0.014001 | 0.008228 | 0.002593 |
| 34 | 0.030773 | 0.014561 | 0.025944 | 0.018355 | 0.020544 | 0.017386 | 0.014838 | 0.004793 | 0.00469 | 0.021407 | 0.015885 | 0.031878 | 0.021067 | 0.013762 | 0.011133 | 0.008529 | 0.001238 | 0.001554 | 0.00664 | 0.002695 | 0.012557 | 0.006061 | 0.005986 |
| 34.0138889 | 0.023558 | 0.015119 | 0.045643 | 0.014875 | 0.019315 | 0.018629 | 0.01666 | 0.012331 | 0.00186 | 0.016565 | 0.012788 | 0.024235 | 0.026813 | 0.013055 | 0.009379 | 0.006801 | 0.003953 | 0.004525 | 0.005274 | 0.00212 | 0.007846 | 0.003688 | 0.010987 |
| 34.0277778 | 0.011836 | 0.017951 | 0.064211 | 0.011449 | 0.021643 | 0.017275 | 0.009985 | 0.036271 | 0.002273 | 0.012329 | 0.006742 | 0.02079 | 0.030554 | 0.007789 | 0.003769 | 0.003267 | 0.007462 | 0.007951 | 0.004691 | 0.002952 | 0.002604 | 0.003151 | 0.015258 |
| 34.0416667 | 0.007152 | 0.024513 | 0.06238 | 0.011338 | 0.025588 | 0.02311 | 0.005687 | 0.07779 | 0.005294 | 0.014324 | 0.004734 | 0.021042 | 0.040209 | 0.009277 | 0.003884 | 0.003553 | 0.006755 | 0.007489 | 0.007406 | 0.004712 | 0.002435 | 0.004019 | 0.014067 |
| 34.0555556 | 0.005916 | 0.031638 | 0.038322 | 0.013834 | 0.026019 | 0.0403 | 0.003517 | 0.119167 | 0.012397 | 0.023188 | 0.003969 | 0.020654 | 0.0561 | 0.021064 | 0.011359 | 0.007901 | 0.002893 | 0.003688 | 0.010964 | 0.008716 | 0.006836 | 0.006828 | 0.010512 |
| 34.0694444 | 0.005825 | 0.029877 | 0.013953 | 0.01124 | 0.017994 | 0.056214 | 0.002824 | 0.141457 | 0.016762 | 0.045784 | 0.004598 | 0.020499 | 0.064578 | 0.030349 | 0.016856 | 0.010627 | 0.00217 | 0.002859 | 0.010372 | 0.010269 | 0.007633 | 0.00739 | 0.020575 |
| 34.0833333 | 0.005172 | 0.017928 | 0.010199 | 0.005795 | 0.008029 | 0.056547 | 0.004156 | 0.144769 | 0.013508 | 0.066715 | 0.007448 | 0.021841 | 0.058271 | 0.027708 | 0.017162 | 0.007701 | 0.004394 | 0.005734 | 0.006307 | 0.006492 | 0.004842 | 0.004227 | 0.052694 |
| 34.0972222 | 0.005556 | 0.008484 | 0.026189 | 0.007548 | 0.00575 | 0.045753 | 0.00382 | 0.140921 | 0.007253 | 0.067443 | 0.008611 | 0.020138 | 0.044791 | 0.019592 | 0.014859 | 0.005055 | 0.005899 | 0.007299 | 0.003509 | 0.004507 | 0.006606 | 0.002532 | 0.08484 |
| 34.1111111 | 0.012379 | 0.010211 | 0.031643 | 0.013007 | 0.007748 | 0.035134 | 0.003249 | 0.145121 | 0.003296 | 0.05802 | 0.006834 | 0.012677 | 0.032322 | 0.012796 | 0.010558 | 0.007817 | 0.005373 | 0.006088 | 0.003389 | 0.007236 | 0.012963 | 0.001834 | 0.08624 |
| 34.125 | 0.019206 | 0.01593 | 0.018561 | 0.012275 | 0.007181 | 0.02725 | 0.006428 | 0.150561 | 0.003373 | 0.049207 | 0.004299 | 0.005574 | 0.023786 | 0.010982 | 0.005071 | 0.010582 | 0.004181 | 0.004399 | 0.004492 | 0.009959 | 0.017563 | 0.00088 | 0.062322 |
| 34.1388889 | 0.016919 | 0.017357 | 0.016286 | 0.007154 | 0.005589 | 0.018249 | 0.012566 | 0.130718 | 0.003599 | 0.037893 | 0.002874 | 0.00498 | 0.020209 | 0.016268 | 0.002328 | 0.010422 | 0.004369 | 0.004314 | 0.005282 | 0.009357 | 0.017239 | 0.000659 | 0.041464 |
| 34.1527778 | 0.013041 | 0.014651 | 0.018306 | 0.003977 | 0.00694 | 0.010523 | 0.015708 | 0.089343 | 0.002643 | 0.0224 | 0.001874 | 0.005883 | 0.020059 | 0.022267 | 0.004175 | 0.011294 | 0.006373 | 0.006677 | 0.004018 | 0.006076 | 0.015217 | 0.001183 | 0.033209 |
| 34.1666667 | 0.022198 | 0.008999 | 0.0141 | 0.004957 | 0.00958 | 0.012128 | 0.014066 | 0.049811 | 0.001824 | 0.008939 | 0.001068 | 0.006866 | 0.020868 | 0.021685 | 0.007069 | 0.014678 | 0.009251 | 0.010389 | 0.002506 | 0.003034 | 0.014657 | 0.003167 | 0.031812 |
| 34.1805556 | 0.035261 | 0.003094 | 0.017396 | 0.012237 | 0.013873 | 0.017093 | 0.008822 | 0.02619 | 0.00096 | 0.00514 | 0.001176 | 0.011782 | 0.020386 | 0.017744 | 0.008357 | 0.017701 | 0.011459 | 0.01293 | 0.003472 | 0.003009 | 0.013949 | 0.006098 | 0.033102 |
| 34.1944444 | 0.036003 | 0.001617 | 0.028254 | 0.020521 | 0.023824 | 0.014654 | 0.005903 | 0.035669 | 0.000612 | 0.010084 | 0.001081 | 0.013063 | 0.015319 | 0.017777 | 0.008804 | 0.018106 | 0.012529 | 0.013153 | 0.004817 | 0.00473 | 0.012079 | 0.008298 | 0.034915 |
| 34.2083333 | 0.032184 | 0.005833 | 0.033421 | 0.019853 | 0.033422 | 0.010287 | 0.010415 | 0.073776 | 0.002391 | 0.014914 | 0.000606 | 0.007401 | 0.009008 | 0.021623 | 0.009189 | 0.015261 | 0.012473 | 0.011579 | 0.005028 | 0.004844 | 0.009553 | 0.008406 | 0.034106 |
| 34.2222222 | 0.031175 | 0.01319 | 0.02844 | 0.013471 | 0.034646 | 0.012927 | 0.013803 | 0.102777 | 0.007717 | 0.013645 | 0.000806 | 0.00307 | 0.009056 | 0.020679 | 0.007853 | 0.009435 | 0.010158 | 0.008921 | 0.005245 | 0.002941 | 0.006325 | 0.005743 | 0.028842 |
| 34.2361111 | 0.024819 | 0.017734 | 0.024403 | 0.009476 | 0.029387 | 0.019232 | 0.010607 | 0.097058 | 0.013144 | 0.009531 | 0.001505 | 0.004263 | 0.010637 | 0.012019 | 0.005334 | 0.005046 | 0.006114 | 0.005183 | 0.004661 | 0.002395 | 0.003742 | 0.003703 | 0.021577 |
| 34.25 | 0.012797 | 0.015798 | 0.024327 | 0.006569 | 0.022382 | 0.022029 | 0.006144 | 0.062227 | 0.014084 | 0.011831 | 0.002522 | 0.009066 | 0.006798 | 0.00602 | 0.006635 | 0.006543 | 0.004705 | 0.003039 | 0.003154 | 0.005785 | 0.004405 | 0.00647 | 0.014525 |
| 34.2638889 | 0.00469 | 0.011511 | 0.023585 | 0.008435 | 0.01372 | 0.020611 | 0.005951 | 0.032448 | 0.01186 | 0.021612 | 0.003372 | 0.013664 | 0.002134 | 0.007994 | 0.010425 | 0.009811 | 0.006177 | 0.004386 | 0.002414 | 0.009665 | 0.00603 | 0.010244 | 0.008971 |
| 34.2777778 | 0.003736 | 0.009186 | 0.022826 | 0.020826 | 0.008356 | 0.017921 | 0.010199 | 0.042717 | 0.00959 | 0.030417 | 0.003303 | 0.012362 | 0.001279 | 0.01429 | 0.009404 | 0.008591 | 0.005175 | 0.004591 | 0.003475 | 0.008945 | 0.005158 | 0.008898 | 0.006972 |
| 34.2916667 | 0.005163 | 0.008498 | 0.021402 | 0.025691 | 0.013284 | 0.016005 | 0.013088 | 0.079755 | 0.007956 | 0.036872 | 0.0022 | 0.00758 | 0.003573 | 0.022106 | 0.005177 | 0.004957 | 0.003448 | 0.004188 | 0.008706 | 0.004883 | 0.005363 | 0.004799 | 0.007791 |
| 34.3055556 | 0.008427 | 0.007955 | 0.017356 | 0.014424 | 0.021298 | 0.015172 | 0.011082 | 0.107358 | 0.00667 | 0.041873 | 0.001108 | 0.007731 | 0.007537 | 0.02296 | 0.00372 | 0.004159 | 0.005151 | 0.006564 | 0.016947 | 0.002094 | 0.008159 | 0.0035 | 0.00805 |
| 34.3194444 | 0.02039 | 0.0067 | 0.013417 | 0.011145 | 0.024255 | 0.014391 | 0.007119 | 0.1176 | 0.004594 | 0.040132 | 0.001049 | 0.009823 | 0.009916 | 0.014914 | 0.003427 | 0.005204 | 0.006397 | 0.007873 | 0.023062 | 0.001719 | 0.009127 | 0.00382 | 0.005645 |
| 34.3333333 | 0.034744 | 0.004357 | 0.020915 | 0.032448 | 0.020618 | 0.010126 | 0.00695 | 0.119996 | 0.002031 | 0.030412 | 0.001403 | 0.008614 | 0.008376 | 0.012743 | 0.002802 | 0.005841 | 0.005641 | 0.007122 | 0.02494 | 0.002468 | 0.0082 | 0.003256 | 0.002251 |
| 34.3472222 | 0.036659 | 0.003986 | 0.042805 | 0.054373 | 0.01375 | 0.004084 | 0.010987 | 0.113025 | 0.001101 | 0.017802 | 0.002618 | 0.006344 | 0.005645 | 0.0206 | 0.00238 | 0.007721 | 0.00635 | 0.007541 | 0.023915 | 0.002266 | 0.00869 | 0.003225 | 0.000826 |
| 34.3611111 | 0.027797 | 0.005259 | 0.050677 | 0.043054 | 0.01099 | 0.002658 | 0.013951 | 0.092134 | 0.002118 | 0.008821 | 0.005482 | 0.004984 | 0.003692 | 0.024692 | 0.001356 | 0.010691 | 0.008194 | 0.008392 | 0.022494 | 0.000951 | 0.009728 | 0.003228 | 0.002073 |
| 34.375 | 0.031418 | 0.004894 | 0.031625 | 0.023432 | 0.013166 | 0.004081 | 0.01209 | 0.063537 | 0.004675 | 0.008694 | 0.007083 | 0.005974 | 0.003445 | 0.020603 | 0.000863 | 0.011942 | 0.00726 | 0.007049 | 0.022639 | 0.000963 | 0.009012 | 0.002024 | 0.005475 |
| 34.3888889 | 0.051542 | 0.007168 | 0.018907 | 0.028636 | 0.012216 | 0.003759 | 0.007736 | 0.035444 | 0.007263 | 0.013729 | 0.00485 | 0.011037 | 0.005962 | 0.014514 | 0.000886 | 0.011471 | 0.003429 | 0.004676 | 0.025055 | 0.001952 | 0.007891 | 0.001034 | 0.008522 |
| 34.4027778 | 0.060874 | 0.012525 | 0.021587 | 0.037258 | 0.006608 | 0.002294 | 0.00758 | 0.017253 | 0.007599 | 0.017467 | 0.001836 | 0.016455 | 0.006728 | 0.010526 | 0.000903 | 0.011029 | 0.000872 | 0.003768 | 0.027122 | 0.002026 | 0.007775 | 0.000613 | 0.009861 |
| 34.4166667 | 0.051091 | 0.016086 | 0.021535 | 0.026671 | 0.00389 | 0.001743 | 0.010621 | 0.018412 | 0.008018 | 0.01808 | 0.000915 | 0.017225 | 0.004 | 0.009621 | 0.001009 | 0.009834 | 0.000517 | 0.003058 | 0.025278 | 0.00237 | 0.007173 | 0.001023 | 0.011563 |

| | | | | | | | | | | | | | | | | | | | | | | | |
|---|---|---|---|---|---|---|---|---|---|---|---|---|---|---|---|---|---|---|---|---|---|---|---|
| 34.4305556 | 0.029772 | 0.016736 | 0.028351 | 0.014682 | 0.011824 | 0.002409 | 0.010161 | 0.032764 | 0.01095 | 0.01414 | 0.000843 | 0.016361 | 0.003015 | 0.0094 | 0.000963 | 0.008152 | 0.000409 | 0.001434 | 0.020557 | 0.004098 | 0.005642 | 0.002622 | 0.013293 |
| 34.4444444 | 0.010875 | 0.013255 | 0.039499 | 0.016771 | 0.024311 | 0.003835 | 0.00675 | 0.040209 | 0.012393 | 0.007213 | 0.001086 | 0.018657 | 0.005825 | 0.010803 | 0.003056 | 0.008277 | 0.000514 | 0.000747 | 0.01667 | 0.005446 | 0.004573 | 0.003433 | 0.01231 |
| 34.4583333 | 0.011412 | 0.012132 | 0.038502 | 0.014423 | 0.022268 | 0.00446 | 0.008177 | 0.034027 | 0.008725 | 0.00216 | 0.003274 | 0.02097 | 0.009812 | 0.021135 | 0.007376 | 0.009277 | 0.000853 | 0.001402 | 0.014882 | 0.00584 | 0.003242 | 0.003635 | 0.008483 |
| 34.4722222 | 0.026731 | 0.020023 | 0.034593 | 0.008231 | 0.012731 | 0.004839 | 0.015948 | 0.025506 | 0.005182 | 0.003508 | 0.006135 | 0.01974 | 0.010318 | 0.046111 | 0.010958 | 0.009494 | 0.000729 | 0.001803 | 0.014143 | 0.00562 | 0.001272 | 0.00551 | 0.00435 |
| 34.4861111 | 0.037902 | 0.024306 | 0.031814 | 0.014742 | 0.012306 | 0.011974 | 0.019532 | 0.030212 | 0.004894 | 0.017572 | 0.006002 | 0.019085 | 0.008182 | 0.061337 | 0.01287 | 0.009766 | 0.00121 | 0.001904 | 0.014199 | 0.00372 | 0.000747 | 0.006996 | 0.002132 |
| 34.5 | 0.03206 | 0.017813 | 0.027944 | 0.02486 | 0.016496 | 0.025012 | 0.015101 | 0.049327 | 0.003109 | 0.042207 | 0.005491 | 0.022575 | 0.009439 | 0.040652 | 0.012219 | 0.010882 | 0.002323 | 0.002658 | 0.015153 | 0.001346 | 0.00185 | 0.005405 | 0.00196 |
| 34.5138889 | 0.018966 | 0.011264 | 0.022982 | 0.02582 | 0.021163 | 0.031851 | 0.008634 | 0.062838 | 0.000871 | 0.059387 | 0.008157 | 0.026779 | 0.012837 | 0.022497 | 0.00879 | 0.0125 | 0.002201 | 0.002915 | 0.015616 | 0.000789 | 0.00273 | 0.003805 | 0.001804 |
| 34.5277778 | 0.016999 | 0.008748 | 0.013686 | 0.02434 | 0.025237 | 0.031265 | 0.004805 | 0.068515 | 0.003217 | 0.060051 | 0.011861 | 0.023545 | 0.011211 | 0.037118 | 0.007421 | 0.01058 | 0.004701 | 0.005231 | 0.012139 | 0.005319 | 0.005313 | 0.008082 | 0.004154 |
| 34.5416667 | 0.021754 | 0.007712 | 0.010009 | 0.035223 | 0.024784 | 0.027465 | 0.004394 | 0.076339 | 0.012179 | 0.05431 | 0.014118 | 0.017551 | 0.011335 | 0.052215 | 0.017677 | 0.012963 | 0.019214 | 0.019123 | 0.014335 | 0.020969 | 0.017649 | 0.023145 | 0.016055 |
| 34.5555556 | 0.02044 | 0.007157 | 0.016779 | 0.050741 | 0.018239 | 0.018974 | 0.003908 | 0.083806 | 0.019936 | 0.051571 | 0.011307 | 0.022018 | 0.022479 | 0.047948 | 0.03604 | 0.02853 | 0.04074 | 0.041164 | 0.032322 | 0.041985 | 0.037822 | 0.043098 | 0.033802 |
| 34.5694444 | 0.013536 | 0.005826 | 0.020658 | 0.049353 | 0.009499 | 0.011022 | 0.002164 | 0.090027 | 0.016639 | 0.05743 | 0.006193 | 0.02775 | 0.026089 | 0.03731 | 0.041679 | 0.037171 | 0.048746 | 0.050312 | 0.045621 | 0.050382 | 0.048661 | 0.051957 | 0.043391 |
| 34.5833333 | 0.007552 | 0.005822 | 0.020433 | 0.036427 | 0.006278 | 0.012882 | 0.000787 | 0.093435 | 0.011791 | 0.076474 | 0.006256 | 0.023783 | 0.017408 | 0.034919 | 0.031415 | 0.029101 | 0.040849 | 0.042743 | 0.040329 | 0.04405 | 0.045196 | 0.047202 | 0.041544 |
| 34.5972222 | 0.006904 | 0.007082 | 0.027318 | 0.022615 | 0.00649 | 0.019892 | 0.000682 | 0.096499 | 0.014586 | 0.101295 | 0.006968 | 0.014936 | 0.015688 | 0.036182 | 0.021869 | 0.019226 | 0.032485 | 0.034076 | 0.029689 | 0.036032 | 0.037958 | 0.039663 | 0.035761 |
| 34.6111111 | 0.010142 | 0.006369 | 0.035802 | 0.008709 | 0.004408 | 0.02166 | 0.00333 | 0.098554 | 0.017078 | 0.116806 | 0.005388 | 0.008879 | 0.01731 | 0.026091 | 0.016169 | 0.012655 | 0.026423 | 0.027713 | 0.02112 | 0.029328 | 0.030314 | 0.032219 | 0.027614 |
| 34.625 | 0.014512 | 0.004169 | 0.033624 | 0.007944 | 0.004008 | 0.016005 | 0.008357 | 0.090648 | 0.015154 | 0.11441 | 0.004639 | 0.013333 | 0.012426 | 0.009838 | 0.008584 | 0.006746 | 0.01845 | 0.019601 | 0.012605 | 0.020874 | 0.020977 | 0.022842 | 0.016019 |
| 34.6388889 | 0.017883 | 0.003812 | 0.02672 | 0.019612 | 0.010853 | 0.012307 | 0.012437 | 0.071417 | 0.010291 | 0.090743 | 0.002942 | 0.022822 | 0.007035 | 0.002248 | 0.002805 | 0.003082 | 0.011007 | 0.012003 | 0.006753 | 0.012951 | 0.013026 | 0.014118 | 0.006155 |
| 34.6527778 | 0.015743 | 0.006259 | 0.026097 | 0.028934 | 0.023799 | 0.021892 | 0.014338 | 0.047341 | 0.008445 | 0.049204 | 0.001549 | 0.027867 | 0.005445 | 0.004788 | 0.002542 | 0.004048 | 0.005893 | 0.006586 | 0.009314 | 0.007329 | 0.008241 | 0.007967 | 0.002202 |
| 34.6666667 | 0.010512 | 0.008414 | 0.025127 | 0.035794 | 0.027015 | 0.03792 | 0.014348 | 0.026815 | 0.015157 | 0.025577 | 0.002608 | 0.029651 | 0.005183 | 0.013236 | 0.004958 | 0.006839 | 0.002298 | 0.002762 | 0.017731 | 0.003444 | 0.005746 | 0.003992 | 0.001202 |
| 34.6805556 | 0.012457 | 0.007604 | 0.014896 | 0.040879 | 0.015459 | 0.049872 | 0.012316 | 0.012279 | 0.019972 | 0.041133 | 0.004521 | 0.029557 | 0.005253 | 0.020948 | 0.007436 | 0.00652 | 0.001344 | 0.001968 | 0.021976 | 0.002409 | 0.005777 | 0.003171 | 0.001249 |
| 34.6944444 | 0.018889 | 0.004508 | 0.006587 | 0.037085 | 0.008086 | 0.048896 | 0.008093 | 0.004749 | 0.016 | 0.046085 | 0.005307 | 0.026942 | 0.003509 | 0.021869 | 0.007059 | 0.003945 | 0.003894 | 0.005266 | 0.0179 | 0.004965 | 0.00862 | 0.005994 | 0.004083 |
| 34.7083333 | 0.016199 | 0.001841 | 0.012775 | 0.022114 | 0.009745 | 0.031101 | 0.005605 | 0.007418 | 0.010752 | 0.035203 | 0.004538 | 0.0222 | 0.00183 | 0.018197 | 0.003894 | 0.003441 | 0.007332 | 0.009785 | 0.011814 | 0.008352 | 0.011799 | 0.009594 | 0.00855 |
| 34.7222222 | 0.010716 | 0.00079 | 0.026101 | 0.008047 | 0.011897 | 0.014531 | 0.010245 | 0.014443 | 0.008151 | 0.063796 | 0.003428 | 0.016392 | 0.002328 | 0.016539 | 0.002455 | 0.003727 | 0.008616 | 0.011767 | 0.011032 | 0.009408 | 0.012148 | 0.01038 | 0.011578 |
| 34.7361111 | 0.015816 | 0.001268 | 0.033257 | 0.004483 | 0.01062 | 0.01338 | 0.015804 | 0.016737 | 0.006205 | 0.114754 | 0.002712 | 0.011093 | 0.002969 | 0.021129 | 0.00375 | 0.002159 | 0.00823 | 0.011307 | 0.014666 | 0.008968 | 0.011074 | 0.008904 | 0.013497 |
| 34.75 | 0.023776 | 0.001784 | 0.032553 | 0.005867 | 0.007676 | 0.019113 | 0.014366 | 0.012286 | 0.003733 | 0.131663 | 0.002021 | 0.008405 | 0.005574 | 0.028267 | 0.005596 | 0.00064 | 0.007898 | 0.010394 | 0.01665 | 0.009145 | 0.011878 | 0.007869 | 0.016848 |
| 34.7638889 | 0.02727 | 0.001826 | 0.025127 | 0.0064 | 0.006231 | 0.025074 | 0.008574 | 0.008517 | 0.002432 | 0.109068 | 0.001176 | 0.011179 | 0.011291 | 0.034717 | 0.004956 | 0.001547 | 0.006527 | 0.00823 | 0.015034 | 0.008229 | 0.012451 | 0.006298 | 0.020818 |
| 34.7777778 | 0.028647 | 0.003929 | 0.017312 | 0.007704 | 0.003895 | 0.032151 | 0.004319 | 0.012789 | 0.00283 | 0.074903 | 0.001327 | 0.018412 | 0.017721 | 0.039898 | 0.003461 | 0.005368 | 0.00395 | 0.00484 | 0.013219 | 0.005033 | 0.009122 | 0.003637 | 0.023184 |
| 34.7916667 | 0.026056 | 0.010138 | 0.018314 | 0.011909 | 0.001948 | 0.037006 | 0.004283 | 0.019301 | 0.002491 | 0.040299 | 0.005167 | 0.024396 | 0.020741 | 0.039539 | 0.005852 | 0.010131 | 0.003691 | 0.004236 | 0.014448 | 0.003551 | 0.004523 | 0.00303 | 0.024043 |
| 34.8055556 | 0.017401 | 0.017102 | 0.022931 | 0.016616 | 0.005039 | 0.040219 | 0.006471 | 0.022466 | 0.002842 | 0.022047 | 0.013172 | 0.026392 | 0.019898 | 0.033535 | 0.009292 | 0.014127 | 0.006944 | 0.007689 | 0.019924 | 0.006062 | 0.003646 | 0.005338 | 0.023436 |
| 34.8194444 | 0.008034 | 0.020577 | 0.028081 | 0.018325 | 0.015548 | 0.040132 | 0.007528 | 0.027199 | 0.003696 | 0.042908 | 0.018733 | 0.027591 | 0.019343 | 0.027423 | 0.010131 | 0.0176 | 0.011153 | 0.012187 | 0.028267 | 0.009872 | 0.00611 | 0.008839 | 0.020264 |
| 34.8333333 | 0.004032 | 0.022704 | 0.035459 | 0.014947 | 0.027842 | 0.027986 | 0.006811 | 0.036815 | 0.002878 | 0.089488 | 0.014159 | 0.029824 | 0.019757 | 0.023891 | 0.009628 | 0.019176 | 0.01386 | 0.015168 | 0.035025 | 0.012381 | 0.008028 | 0.011786 | 0.015068 |
| 34.8472222 | 0.003053 | 0.02445 | 0.035319 | 0.009131 | 0.027084 | 0.013461 | 0.005316 | 0.044115 | 0.002787 | 0.123431 | 0.007855 | 0.029006 | 0.01809 | 0.020162 | 0.0082 | 0.016798 | 0.013523 | 0.015105 | 0.035134 | 0.012108 | 0.007346 | 0.012223 | 0.011136 |
| 34.8611111 | 0.002422 | 0.024836 | 0.021899 | 0.007747 | 0.016021 | 0.014591 | 0.003194 | 0.040997 | 0.002863 | 0.11998 | 0.011795 | 0.021365 | 0.013485 | 0.012822 | 0.004924 | 0.011675 | 0.010263 | 0.011798 | 0.026961 | 0.008855 | 0.00441 | 0.009716 | 0.010556 |
| 34.875 | 0.002552 | 0.025805 | 0.011684 | 0.012492 | 0.01105 | 0.026178 | 0.001317 | 0.02909 | 0.002255 | 0.085172 | 0.017006 | 0.010981 | 0.008094 | 0.005654 | 0.001745 | 0.007307 | 0.006399 | 0.007443 | 0.014154 | 0.004985 | 0.001592 | 0.006133 | 0.011594 |
| 34.8888889 | 0.006004 | 0.027626 | 0.021732 | 0.016269 | 0.010354 | 0.03463 | 0.000963 | 0.016689 | 0.003482 | 0.040005 | 0.01383 | 0.006526 | 0.004041 | 0.005204 | 0.000847 | 0.005511 | 0.004252 | 0.00483 | 0.005441 | 0.002958 | 0.001519 | 0.003595 | 0.011565 |
| 34.9027778 | 0.011059 | 0.021338 | 0.040872 | 0.014684 | 0.00629 | 0.039003 | 0.00182 | 0.014152 | 0.0042 | 0.013836 | 0.006955 | 0.009044 | 0.002366 | 0.01093 | 0.001146 | 0.004547 | 0.003489 | 0.003932 | 0.005841 | 0.002224 | 0.003115 | 0.00199 | 0.010277 |
| 34.9166667 | 0.011436 | 0.010516 | 0.049515 | 0.010389 | 0.001896 | 0.042458 | 0.003694 | 0.029864 | 0.002403 | 0.025109 | 0.002591 | 0.0123 | 0.005097 | 0.017871 | 0.001317 | 0.003221 | 0.002317 | 0.002563 | 0.010582 | 0.001679 | 0.003171 | 0.001395 | 0.009444 |
| 34.9305556 | 0.008314 | 0.011355 | 0.037694 | 0.00688 | 0.00633 | 0.043888 | 0.007305 | 0.047312 | 0.001715 | 0.056156 | 0.001452 | 0.01437 | 0.010595 | 0.021659 | 0.001627 | 0.003595 | 0.001587 | 0.00138 | 0.015887 | 0.002313 | 0.002052 | 0.002872 | 0.00989 |
| 34.9444444 | 0.012177 | 0.015613 | 0.015704 | 0.007454 | 0.020199 | 0.040442 | 0.012207 | 0.043055 | 0.003588 | 0.062739 | 0.001628 | 0.017229 | 0.01232 | 0.020249 | 0.004101 | 0.005535 | 0.00282 | 0.002403 | 0.021551 | 0.004188 | 0.002669 | 0.005072 | 0.011024 |
| 34.9583333 | 0.0292 | 0.013128 | 0.011951 | 0.013568 | 0.027452 | 0.029424 | 0.01587 | 0.030928 | 0.006449 | 0.036434 | 0.002108 | 0.021097 | 0.008738 | 0.017843 | 0.008733 | 0.005853 | 0.004262 | 0.00413 | 0.026417 | 0.005689 | 0.00469 | 0.006513 | 0.011881 |
| 34.9722222 | 0.03737 | 0.010341 | 0.024482 | 0.0184 | 0.019506 | 0.015825 | 0.01717 | 0.029436 | 0.010095 | 0.019133 | 0.002198 | 0.023367 | 0.00441 | 0.020587 | 0.012726 | 0.003951 | 0.004184 | 0.004331 | 0.029403 | 0.00569 | 0.005295 | 0.006833 | 0.011402 |
| 34.9861111 | 0.024183 | 0.01045 | 0.030919 | 0.017066 | 0.0083 | 0.012349 | 0.01765 | 0.033314 | 0.014495 | 0.030681 | 0.003528 | 0.021942 | 0.003465 | 0.024926 | 0.013937 | 0.002395 | 0.002992 | 0.002989 | 0.030113 | 0.004105 | 0.004561 | 0.005715 | 0.009058 |
| 35 | 0.010757 | 0.011944 | 0.02799 | 0.012659 | 0.006908 | 0.019133 | 0.017975 | 0.031594 | 0.015826 | 0.050982 | 0.007848 | 0.018722 | 0.004608 | 0.025747 | 0.011454 | 0.003035 | 0.0016 | 0.00166 | 0.028274 | 0.00231 | 0.004666 | 0.003339 | 0.005123 |
| 35.0138889 | 0.017413 | 0.012331 | 0.024763 | 0.010514 | 0.008953 | 0.025844 | 0.015732 | 0.022154 | 0.010892 | 0.073281 | 0.009424 | 0.015262 | 0.005555 | 0.022844 | 0.006229 | 0.004545 | 0.001607 | 0.00272 | 0.023383 | 0.002737 | 0.004693 | 0.002368 | 0.003364 |
| 35.0277778 | 0.032009 | 0.012829 | 0.022791 | 0.012443 | 0.005693 | 0.028759 | 0.010074 | 0.019183 | 0.004602 | 0.104217 | 0.005361 | 0.012547 | 0.006122 | 0.018416 | 0.002028 | 0.004133 | 0.004326 | 0.007048 | 0.017463 | 0.005609 | 0.003316 | 0.004776 | 0.007922 |
| 35.0416667 | 0.028268 | 0.017962 | 0.027886 | 0.011466 | 0.003398 | 0.022488 | 0.008615 | 0.029565 | 0.003777 | 0.123255 | 0.004414 | 0.012198 | 0.010592 | 0.01921 | 0.000914 | 0.002153 | 0.009035 | 0.012404 | 0.014354 | 0.009269 | 0.003602 | 0.007872 | 0.017301 |
| 35.0555556 | 0.022393 | 0.025436 | 0.057198 | 0.006251 | 0.00427 | 0.018727 | 0.018202 | 0.0365 | 0.006621 | 0.097367 | 0.012493 | 0.01081 | 0.023157 | 0.028247 | 0.001294 | 0.001109 | 0.014289 | 0.01706 | 0.013409 | 0.013822 | 0.00813 | 0.011491 | 0.030877 |
| 35.0694444 | 0.021216 | 0.029023 | 0.095377 | 0.002996 | 0.005352 | 0.037411 | 0.029403 | 0.029683 | 0.006632 | 0.0438 | 0.02161 | 0.005964 | 0.032574 | 0.039668 | 0.001635 | 0.002355 | 0.017999 | 0.020379 | 0.011622 | 0.018035 | 0.013796 | 0.016081 | 0.048433 |
| 35.0833333 | 0.012873 | 0.027113 | 0.112705 | 0.002776 | 0.006407 | 0.062511 | 0.02731 | 0.026144 | 0.003302 | 0.014437 | 0.020365 | 0.002618 | 0.028379 | 0.041189 | 0.001459 | 0.004693 | 0.017811 | 0.019889 | 0.010601 | 0.018307 | 0.015286 | 0.017493 | 0.062578 |
| 35.0972222 | 0.007607 | 0.019473 | 0.111411 | 0.003792 | 0.008168 | 0.068819 | 0.015973 | 0.039407 | 0.001256 | 0.015117 | 0.011253 | 0.003559 | 0.017456 | 0.027422 | 0.002679 | 0.004897 | 0.013964 | 0.01513 | 0.011879 | 0.014291 | 0.012204 | 0.01397 | 0.06349 |
| 35.1111111 | 0.005163 | 0.010943 | 0.098094 | 0.004938 | 0.009542 | 0.053012 | 0.013586 | 0.054581 | 0.001511 | 0.018653 | 0.005996 | 0.004513 | 0.008783 | 0.011072 | 0.006324 | 0.005247 | 0.009283 | 0.009636 | 0.011628 | 0.009434 | 0.009072 | 0.008821 | 0.049869 |
| 35.125 | 0.005168 | 0.00974 | 0.070919 | 0.004786 | 0.008689 | 0.029323 | 0.022311 | 0.0521 | 0.00194 | 0.014511 | 0.009128 | 0.004648 | 0.004415 | 0.00407 | 0.009599 | 0.008784 | 0.006826 | 0.007018 | 0.007469 | 0.006993 | 0.009024 | 0.006015 | 0.030939 |
| 35.1388889 | 0.009506 | 0.011458 | 0.044565 | 0.004749 | 0.00781 | 0.015837 | 0.024444 | 0.031914 | 0.002145 | 0.010284 | 0.013124 | 0.00687 | 0.003559 | 0.006032 | 0.00909 | 0.011864 | 0.007566 | 0.008135 | 0.004814 | 0.007734 | 0.011596 | 0.006716 | 0.016806 |
| 35.1527778 | 0.01607 | 0.009124 | 0.039532 | 0.009405 | 0.009319 | 0.024308 | 0.015475 | 0.012679 | 0.001959 | 0.012834 | 0.013059 | 0.008116 | 0.004137 | 0.012377 | 0.005815 | 0.013807 | 0.009761 | 0.010728 | 0.006964 | 0.009926 | 0.014188 | 0.009101 | 0.010541 |
| 35.1666667 | 0.025185 | 0.009614 | 0.045969 | 0.018444 | 0.015224 | 0.046353 | 0.012805 | 0.008465 | 0.002192 | 0.019928 | 0.010957 | 0.007482 | 0.003292 | 0.017308 | 0.004876 | 0.017425 | 0.01105 | 0.011821 | 0.008065 | 0.011181 | 0.014556 | 0.010584 | 0.008875 |
| 35.1805556 | 0.031039 | 0.01519 | 0.034623 | 0.024191 | 0.020484 | 0.057625 | 0.020576 | 0.019334 | 0.003334 | 0.024115 | 0.006894 | 0.007067 | 0.002165 | 0.020564 | 0.006266 | 0.021467 | 0.010643 | 0.010643 | 0.004975 | 0.010502 | 0.012372 | 0.009933 | 0.007715 |
| 35.1944444 | 0.030274 | 0.017348 | 0.025135 | 0.020579 | 0.015809 | 0.050468 | 0.025475 | 0.040392 | 0.003171 | 0.021771 | 0.003021 | 0.00538 | 0.005053 | 0.026959 | 0.005642 | 0.021535 | 0.008917 | 0.008246 | 0.002431 | 0.008254 | 0.008561 | 0.007297 | 0.005443 |
| 35.2083333 | 0.027855 | 0.015981 | 0.05473 | 0.011062 | 0.01058 | 0.030702 | 0.028325 | 0.060373 | 0.003003 | 0.017533 | 0.001544 | 0.002699 | 0.012337 | 0.035517 | 0.003057 | 0.014842 | 0.005936 | 0.005389 | 0.003202 | 0.005244 | 0.004954 | 0.003799 | 0.002889 |
| 35.2222222 | 0.031297 | 0.01419 | 0.090691 | 0.007026 | 0.016376 | 0.016641 | 0.035995 | 0.070785 | 0.005805 | 0.018358 | 0.001611 | 0.002335 | 0.02059 | 0.036583 | 0.001667 | 0.006848 | 0.002576 | 0.00269 | 0.004959 | 0.002375 | 0.002659 | 0.001388 | 0.002352 |
| 35.2361111 | 0.044099 | 0.010941 | 0.093689 | 0.011383 | 0.021487 | 0.027234 | 0.038603 | 0.07435 | 0.010662 | 0.025982 | 0.001905 | 0.005683 | 0.02481 | 0.024283 | 0.00152 | 0.00411 | 0.000849 | 0.001057 | 0.006746 | 0.0007 | 0.001346 | 0.001074 | 0.004217 |
| 35.25 | 0.05307 | 0.008146 | 0.07896 | 0.014973 | 0.016423 | 0.050041 | 0.026654 | 0.076137 | 0.014734 | 0.035054 | 0.00349 | 0.008857 | 0.022356 | 0.009135 | 0.001205 | 0.003123 | 0.001185 | 0.00105 | 0.007908 | 0.00079 | 0.001332 | 0.001347 | 0.005118 |
| 35.2638889 | 0.042904 | 0.007735 | 0.067896 | 0.013242 | 0.014041 | 0.065284 | 0.020041 | 0.076067 | 0.014826 | 0.039864 | 0.012039 | 0.007096 | 0.017033 | 0.003703 | 0.000864 | 0.001519 | 0.002169 | 0.002596 | 0.006059 | 0.002349 | 0.0041 | 0.002002 | 0.004238 |
| 35.2777778 | 0.025113 | 0.007047 | 0.060371 | 0.009507 | 0.023059 | 0.063838 | 0.031086 | 0.072436 | 0.011108 | 0.038485 | 0.021904 | 0.003532 | 0.01164 | 0.007058 | 0.00041 | 0.002247 | 0.002784 | 0.004463 | 0.002673 | 0.004148 | 0.008718 | 0.003972 | 0.004974 |
| 35.2916667 | 0.018691 | 0.004271 | 0.05348 | 0.011908 | 0.02974 | 0.044098 | 0.037009 | 0.065409 | 0.005957 | 0.029783 | 0.022264 | 0.002711 | 0.006932 | 0.016656 | 0.000808 | 0.005397 | 0.003406 | 0.005155 | 0.001636 | 0.004542 | 0.011329 | 0.004688 | 0.006443 |
| 35.3055556 | 0.019864 | 0.002191 | 0.045353 | 0.021978 | 0.025276 | 0.020326 | 0.028604 | 0.052824 | 0.002081 | 0.015942 | 0.017754 | 0.002691 | 0.005136 | 0.027694 | 0.002315 | 0.008475 | 0.003561 | 0.004405 | 0.002458 | 0.003469 | 0.010889 | 0.003138 | 0.005421 |
| 35.3194444 | 0.015213 | 0.002803 | 0.033066 | 0.030638 | 0.015128 | 0.012745 | 0.017467 | 0.030765 | 0.002148 | 0.00797 | 0.012365 | 0.001998 | 0.004083 | 0.026532 | 0.003102 | 0.009033 | 0.002797 | 0.002658 | 0.002404 | 0.002198 | 0.00858 | 0.001603 | 0.0033 |
| 35.3333333 | 0.0124 | 0.004954 | 0.021859 | 0.034442 | 0.006532 | 0.026817 | 0.017465 | 0.014811 | 0.005783 | 0.012072 | 0.006964 | 0.001486 | 0.002071 | 0.014797 | 0.004944 | 0.007854 | 0.002287 | 0.000906 | 0.001383 | 0.001621 | 0.005684 | 0.001129 | 0.003245 |
| 35.3472222 | 0.020812 | 0.006999 | 0.015448 | 0.034072 | 0.005697 | 0.050447 | 0.032066 | 0.022309 | 0.010245 | 0.01757 | 0.00789 | 0.001467 | 0.002044 | 0.008758 | 0.010385 | 0.008112 | 0.002673 | 0.001522 | 0.002891 | 0.002115 | 0.005393 | 0.001824 | 0.004323 |
| 35.3611111 | 0.027638 | 0.008103 | 0.011577 | 0.026927 | 0.006991 | 0.065475 | 0.040854 | 0.036604 | 0.014289 | 0.015511 | 0.014328 | 0.002864 | 0.005081 | 0.010414 | 0.015828 | 0.010469 | 0.004811 | 0.005385 | 0.009147 | 0.004348 | 0.008753 | 0.004258 | 0.003294 |
| 35.375 | 0.02853 | 0.007307 | 0.008904 | 0.015117 | 0.003181 | 0.059022 | 0.028549 | 0.037703 | 0.018363 | 0.009968 | 0.014442 | 0.006475 | 0.00851 | 0.010441 | 0.018182 | 0.012544 | 0.00745 | 0.008728 | 0.01638 | 0.006981 | 0.011664 | 0.007077 | 0.001759 |
| 35.3888889 | 0.03265 | 0.005566 | 0.015939 | 0.008936 | 0.003579 | 0.037896 | 0.018454 | 0.025086 | 0.021064 | 0.011702 | 0.007304 | 0.00959 | 0.008967 | 0.009867 | 0.019955 | 0.012413 | 0.007365 | 0.008311 | 0.02021 | 0.006993 | 0.010842 | 0.007036 | 0.003416 |
| 35.4027778 | 0.039586 | 0.006206 | 0.039553 | 0.011584 | 0.00712 | 0.017459 | 0.029651 | 0.017952 | 0.019845 | 0.025041 | 0.003425 | 0.010301 | 0.006751 | 0.010841 | 0.021275 | 0.01093 | 0.00464 | 0.005433 | 0.020243 | 0.004443 | 0.007459 | 0.004014 | 0.007959 |
| 35.4166667 | 0.041371 | 0.009052 | 0.072308 | 0.013771 | 0.008337 | 0.007277 | 0.039572 | 0.031469 | 0.012624 | 0.040561 | 0.002489 | 0.009516 | 0.00396 | 0.01228 | 0.019834 | 0.009801 | 0.002494 | 0.003095 | 0.018159 | 0.002439 | 0.00376 | 0.002093 | 0.0126 |
| 35.4305556 | 0.036605 | 0.011011 | 0.098025 | 0.014198 | 0.018279 | 0.0098 | 0.034653 | 0.052073 | 0.005339 | 0.048812 | 0.002216 | 0.007993 | 0.0017 | 0.020915 | 0.019365 | 0.01005 | 0.003013 | 0.003288 | 0.015695 | 0.003011 | 0.001844 | 0.003535 | 0.015116 |
| 35.4444444 | 0.0284 | 0.009571 | 0.087652 | 0.014122 | 0.030713 | 0.017052 | 0.022404 | 0.06601 | 0.005231 | 0.047008 | 0.003087 | 0.006395 | 0.000864 | 0.031767 | 0.022852 | 0.011722 | 0.004449 | 0.005042 | 0.013314 | 0.004631 | 0.002632 | 0.005138 | 0.015491 |
| 35.4583333 | 0.021057 | 0.006197 | 0.047768 | 0.010207 | 0.026845 | 0.025852 | 0.013524 | 0.062222 | 0.007031 | 0.035925 | 0.003441 | 0.00425 | 0.000897 | 0.035615 | 0.025302 | 0.01255 | 0.005747 | 0.007176 | 0.010999 | 0.005881 | 0.004801 | 0.005707 | 0.015465 |
| 35.4722222 | 0.018994 | 0.006802 | 0.023282 | 0.011091 | 0.011411 | 0.036896 | 0.018628 | 0.038914 | 0.005486 | 0.020296 | 0.004671 | 0.00332 | 0.000656 | 0.039242 | 0.025085 | 0.012217 | 0.006942 | 0.008699 | 0.009869 | 0.006725 | 0.006077 | 0.007716 | 0.015026 |
| 35.4861111 | 0.022272 | 0.010636 | 0.027666 | 0.019445 | 0.0064 | 0.039226 | 0.030537 | 0.018137 | 0.003119 | 0.013491 | 0.005998 | 0.007976 | 0.001497 | 0.049597 | 0.025771 | 0.013248 | 0.005686 | 0.006517 | 0.010286 | 0.005663 | 0.004598 | 0.008709 | 0.011674 |
| 35.5 | 0.029927 | 0.011638 | 0.050465 | 0.018729 | 0.013385 | 0.025047 | 0.033322 | 0.010985 | 0.001701 | 0.028273 | 0.004643 | 0.016829 | 0.004731 | 0.055868 | 0.02573 | 0.014688 | 0.002606 | 0.002543 | 0.010272 | 0.002903 | 0.002817 | 0.005278 | 0.006308 |
| 35.5138889 | 0.040696 | 0.00816 | 0.075282 | 0.008956 | 0.014697 | 0.014567 | 0.022089 | 0.010676 | 0.001681 | 0.056683 | 0.002273 | 0.022072 | 0.007243 | 0.045146 | 0.021012 | 0.013252 | 0.001155 | 0.001032 | 0.007256 | 0.001319 | 0.002834 | 0.001938 | 0.003944 |
| 35.5277778 | 0.048571 | 0.003213 | 0.083197 | 0.01393 | 0.011366 | 0.023705 | 0.009066 | 0.009849 | 0.002696 | 0.077264 | 0.002365 | 0.017435 | 0.008403 | 0.028121 | 0.014302 | 0.011414 | 0.005978 | 0.00542 | 0.007629 | 0.006016 | 0.007137 | 0.007014 | 0.010144 |
| 35.5416667 | 0.044533 | 0.001644 | 0.067254 | 0.032506 | 0.015375 | 0.032197 | 0.004071 | 0.018407 | 0.003741 | 0.078468 | 0.005727 | 0.019164 | 0.019435 | 0.032641 | 0.021873 | 0.023516 | 0.025777 | 0.025408 | 0.02487 | 0.025407 | 0.027995 | 0.026769 | 0.030888 |
| 35.5555556 | 0.029353 | 0.00235 | 0.041776 | 0.038682 | 0.018023 | 0.025606 | 0.004171 | 0.045353 | 0.006072 | 0.065793 | 0.007571 | 0.041517 | 0.042264 | 0.068533 | 0.046888 | 0.04961 | 0.055199 | 0.057601 | 0.053801 | 0.05437 | 0.064811 | 0.053768 | 0.059289 |
| 35.5694444 | 0.018115 | 0.005108 | 0.02326 | 0.026515 | 0.013061 | 0.014336 | 0.007519 | 0.049356 | 0.008136 | 0.052745 | 0.005772 | 0.054844 | 0.049925 | 0.092548 | 0.056104 | 0.057452 | 0.065684 | 0.071086 | 0.068437 | 0.066216 | 0.086332 | 0.06396 | 0.06981 |
| 35.5833333 | 0.022203 | 0.014314 | 0.017359 | 0.015414 | 0.009978 | 0.013776 | 0.012021 | 0.034269 | 0.006297 | 0.052381 | 0.007128 | 0.038426 | 0.039477 | 0.072273 | 0.036322 | 0.034572 | 0.046237 | 0.05126 | 0.057058 | 0.048775 | 0.071577 | 0.047261 | 0.050972 |
| 35.5972222 | 0.023692 | 0.024118 | 0.021929 | 0.016558 | 0.00994 | 0.032367 | 0.012338 | 0.049408 | 0.004101 | 0.063105 | 0.010914 | 0.016988 | 0.048661 | 0.041324 | 0.014031 | 0.015012 | 0.020111 | 0.022486 | 0.034736 | 0.022176 | 0.037863 | 0.021582 | 0.022216 |
| 35.6111111 | 0.017498 | 0.026216 | 0.018112 | 0.016177 | 0.014981 | 0.057429 | 0.006332 | 0.061448 | 0.006646 | 0.067773 | 0.018772 | 0.014583 | 0.068382 | 0.023208 | 0.005521 | 0.016234 | 0.005983 | 0.006765 | 0.015899 | 0.00657 | 0.014853 | 0.006061 | 0.006346 |
| 35.625 | 0.018909 | 0.022594 | 0.006012 | 0.013615 | 0.024655 | 0.062742 | 0.001877 | 0.043674 | 0.008609 | 0.060827 | 0.031984 | 0.025705 | 0.057149 | 0.010936 | 0.007861 | 0.017031 | 0.003506 | 0.004413 | 0.007986 | 0.003802 | 0.012972 | 0.003153 | 0.006119 |
| 35.6388889 | 0.021968 | 0.018662 | 0.001429 | 0.013607 | 0.032364 | 0.044069 | 0.001655 | 0.024918 | 0.005472 | 0.049803 | 0.0332 | 0.036092 | 0.026431 | 0.004939 | 0.015391 | 0.010644 | 0.006933 | 0.007934 | 0.015576 | 0.007447 | 0.018196 | 0.006681 | 0.011713 |
| 35.6527778 | 0.029456 | 0.016392 | 0.002947 | 0.016946 | 0.035063 | 0.020434 | 0.001735 | 0.01548 | 0.002524 | 0.038367 | 0.020484 | 0.041835 | 0.007945 | 0.00944 | 0.024157 | 0.00694 | 0.011534 | 0.012606 | 0.029238 | 0.012375 | 0.018815 | 0.011792 | 0.017705 |
| 35.6666667 | 0.04639 | 0.012793 | 0.009213 | 0.033188 | 0.029849 | 0.007552 | 0.002962 | 0.00932 | 0.001794 | 0.025818 | 0.008086 | 0.042742 | 0.003617 | 0.028431 | 0.029016 | 0.007065 | 0.014201 | 0.015784 | 0.037361 | 0.015497 | 0.015249 | 0.015308 | 0.021603 |
| 35.6805556 | 0.061134 | 0.009458 | 0.015296 | 0.04443 | 0.022172 | 0.007341 | 0.003884 | 0.004833 | 0.001795 | 0.012578 | 0.002255 | 0.03915 | 0.005384 | 0.056585 | 0.028077 | 0.008052 | 0.014131 | 0.015528 | 0.037963 | 0.015604 | 0.009815 | 0.015917 | 0.021591 |
| 35.6944444 | 0.070178 | 0.021134 | 0.017206 | 0.034867 | 0.018258 | 0.009799 | 0.004848 | 0.004351 | 0.005419 | 0.004464 | 0.001423 | 0.033514 | 0.013763 | 0.076936 | 0.02698 | 0.009292 | 0.013028 | 0.013803 | 0.033675 | 0.014231 | 0.005636 | 0.015058 | 0.019412 |
| 35.7083333 | 0.071562 | 0.050138 | 0.024864 | 0.01956 | 0.024149 | 0.009026 | 0.00939 | 0.006669 | 0.012841 | 0.004245 | 0.001372 | 0.026872 | 0.026866 | 0.080231 | 0.027668 | 0.011252 | 0.011163 | 0.011876 | 0.028223 | 0.012022 | 0.003277 | 0.013222 | 0.016163 |
| 35.7222222 | 0.065587 | 0.070428 | 0.038554 | 0.0124 | 0.039636 | 0.007891 | 0.013259 | 0.00999 | 0.018864 | 0.008188 | 0.001414 | 0.019001 | 0.035774 | 0.072521 | 0.025101 | 0.013135 | 0.007228 | 0.007862 | 0.024461 | 0.007922 | 0.002401 | 0.009143 | 0.011654 |
| 35.7361111 | 0.060647 | 0.069119 | 0.049642 | 0.011725 | 0.044608 | 0.007108 | 0.010614 | 0.013469 | 0.020869 | 0.010604 | 0.001656 | 0.01234 | 0.035688 | 0.064725 | 0.019114 | 0.014871 | 0.003012 | 0.003471 | 0.024479 | 0.003502 | 0.003726 | 0.004211 | 0.007459 |
| 35.75 | 0.051099 | 0.058938 | 0.055789 | 0.011352 | 0.030806 | 0.004599 | 0.005972 | 0.014669 | 0.018719 | 0.01856 | 0.001652 | 0.010607 | 0.03296 | 0.059024 | 0.015157 | 0.016659 | 0.001041 | 0.001809 | 0.027199 | 0.00133 | 0.004035 | 0.001352 | 0.006084 |
| 35.7638889 | 0.036177 | 0.056849 | 0.052569 | 0.011749 | 0.014729 | 0.002018 | 0.004675 | 0.012877 | 0.014546 | 0.042116 | 0.002144 | 0.013644 | 0.033649 | 0.0456 | 0.014351 | 0.017585 | 0.000596 | 0.001315 | 0.027979 | 0.000798 | 0.003026 | 0.000864 | 0.007707 |
| 35.7777778 | 0.034239 | 0.069899 | 0.036823 | 0.010921 | 0.009812 | 0.001999 | 0.005519 | 0.01554 | 0.011278 | 0.060558 | 0.003526 | 0.017773 | 0.035424 | 0.026912 | 0.013031 | 0.016669 | 0.000594 | 0.000712 | 0.024333 | 0.0006 | 0.004385 | 0.001333 | 0.009485 |
| 35.7916667 | 0.047894 | 0.09145 | 0.021836 | 0.010502 | 0.008592 | 0.005121 | 0.006354 | 0.024279 | 0.007821 | 0.055297 | 0.003408 | 0.020006 | 0.032531 | 0.019493 | 0.00931 | 0.014208 | 0.00092 | 0.000727 | 0.017951 | 0.000788 | 0.006109 | 0.001256 | 0.008986 |
| 35.8055556 | 0.053384 | 0.120693 | 0.015753 | 0.01748 | 0.003848 | 0.012044 | 0.006962 | 0.027179 | 0.006782 | 0.03352 | 0.002296 | 0.019972 | 0.021733 | 0.016819 | 0.004981 | 0.011637 | 0.001161 | 0.001174 | 0.012083 | 0.001218 | 0.006323 | 0.000598 | 0.007406 |
| 35.8194444 | 0.035499 | 0.146287 | 0.010473 | 0.022943 | 0.003739 | 0.018944 | 0.007459 | 0.022846 | 0.009026 | 0.021629 | 0.00279 | 0.019518 | 0.012601 | 0.009984 | 0.001976 | 0.009756 | 0.001075 | 0.001195 | 0.008543 | 0.001176 | 0.006275 | 0.000719 | 0.007114 |
| 35.8333333 | 0.01375 | 0.14984 | 0.004705 | 0.018342 | 0.00971 | 0.020972 | 0.006056 | 0.018373 | 0.008051 | 0.036629 | 0.003021 | 0.018979 | 0.017912 | 0.008379 | 0.001616 | 0.007343 | 0.001252 | 0.001093 | 0.006746 | 0.001053 | 0.005383 | 0.001855 | 0.007713 |
| 35.8472222 | 0.005716 | 0.14192 | 0.004769 | 0.01194 | 0.014475 | 0.019827 | 0.005291 | 0.015116 | 0.004332 | 0.04846 | 0.002015 | 0.016741 | 0.029698 | 0.007334 | 0.004268 | 0.003836 | 0.001361 | 0.001213 | 0.005304 | 0.001068 | 0.003061 | 0.002575 | 0.006896 |
| 35.8611111 | 0.007864 | 0.134354 | 0.011432 | 0.011108 | 0.017051 | 0.018176 | 0.01123 | 0.011961 | 0.002415 | 0.03498 | 0.001102 | 0.01282 | 0.031235 | 0.003991 | 0.007027 | 0.001654 | 0.001128 | 0.001271 | 0.003706 | 0.00084 | 0.001099 | 0.002447 | 0.003961 |
| 35.875 | 0.015106 | 0.115838 | 0.016695 | 0.018511 | 0.016787 | 0.014558 | 0.019912 | 0.008271 | 0.001838 | 0.018284 | 0.000487 | 0.00891 | 0.02059 | 0.003545 | 0.007266 | 0.001538 | 0.001276 | 0.001489 | 0.002281 | 0.000773 | 0.000637 | 0.002249 | 0.001307 |
| 35.8888889 | 0.01783 | 0.085376 | 0.013985 | 0.029712 | 0.01223 | 0.009088 | 0.022306 | 0.004975 | 0.002179 | 0.019761 | 0.000334 | 0.005741 | 0.013931 | 0.00451 | 0.005714 | 0.001889 | 0.001526 | 0.001645 | 0.001289 | 0.000826 | 0.000615 | 0.002002 | 0.000856 |
| 35.9027778 | 0.011818 | 0.058903 | 0.007874 | 0.035943 | 0.011379 | 0.004331 | 0.016673 | 0.005828 | 0.00321 | 0.031123 | 0.00051 | 0.003041 | 0.019035 | 0.010817 | 0.004263 | 0.003077 | 0.001161 | 0.001269 | 0.001648 | 0.000644 | 0.001116 | 0.001438 | 0.001639 |

| | | | | | | | | | | | | | | | | | | | | | | | |
|---|---|---|---|---|---|---|---|---|---|---|---|---|---|---|---|---|---|---|---|---|---|---|---|
| 35.9166667 | 0.008272 | 0.042134 | 0.008949 | 0.032101 | 0.016483 | 0.00331 | 0.008879 | 0.012727 | 0.003523 | 0.03173 | 0.000608 | 0.002045 | 0.019692 | 0.022778 | 0.004224 | 0.006029 | 0.000691 | 0.000786 | 0.004015 | 0.000715 | 0.002566 | 0.000827 | 0.002701 |
| 35.9305556 | 0.012482 | 0.030129 | 0.020246 | 0.019742 | 0.01687 | 0.007187 | 0.006955 | 0.024963 | 0.00286 | 0.019093 | 0.001007 | 0.003989 | 0.011165 | 0.03203 | 0.005862 | 0.01001 | 0.001089 | 0.001071 | 0.007245 | 0.001436 | 0.003899 | 0.000782 | 0.00369 |
| 35.9444444 | 0.021395 | 0.020731 | 0.029857 | 0.00895 | 0.009779 | 0.010432 | 0.010388 | 0.043703 | 0.00218 | 0.01173 | 0.002095 | 0.007177 | 0.007114 | 0.029296 | 0.008499 | 0.013297 | 0.002089 | 0.002073 | 0.010111 | 0.002412 | 0.004651 | 0.001351 | 0.004387 |
| 35.9583333 | 0.033711 | 0.013475 | 0.026909 | 0.008331 | 0.003062 | 0.008383 | 0.011628 | 0.055272 | 0.002718 | 0.015948 | 0.00339 | 0.011081 | 0.014251 | 0.015199 | 0.011766 | 0.014082 | 0.002891 | 0.002965 | 0.012378 | 0.003238 | 0.0057 | 0.002059 | 0.005428 |
| 35.9722222 | 0.041928 | 0.007656 | 0.016885 | 0.011318 | 0.002766 | 0.003892 | 0.009369 | 0.041094 | 0.004289 | 0.014431 | 0.004123 | 0.017546 | 0.023706 | 0.00698 | 0.014506 | 0.011738 | 0.00295 | 0.003042 | 0.013953 | 0.003535 | 0.007489 | 0.002436 | 0.007145 |
| 35.9861111 | 0.040362 | 0.003591 | 0.008803 | 0.01111 | 0.005145 | 0.002379 | 0.006021 | 0.019668 | 0.006643 | 0.010271 | 0.004297 | 0.02218 | 0.029868 | 0.013952 | 0.014147 | 0.008313 | 0.002119 | 0.002108 | 0.014458 | 0.002901 | 0.008491 | 0.001879 | 0.008629 |
| 36 | 0.038263 | 0.002365 | 0.004566 | 0.018354 | 0.00887 | 0.007393 | 0.004006 | 0.019607 | 0.008721 | 0.019368 | 0.005438 | 0.018531 | 0.036424 | 0.025556 | 0.010933 | 0.006783 | 0.00137 | 0.001396 | 0.01393 | 0.001951 | 0.006835 | 0.001733 | 0.008739 |
| 36.0138889 | 0.048109 | 0.00362 | 0.003924 | 0.037171 | 0.021305 | 0.017827 | 0.005885 | 0.034173 | 0.008968 | 0.031621 | 0.008247 | 0.009735 | 0.033927 | 0.030864 | 0.007773 | 0.008178 | 0.002324 | 0.002834 | 0.012671 | 0.002724 | 0.004614 | 0.004574 | 0.007141 |
| 36.0277778 | 0.063949 | 0.003797 | 0.006427 | 0.054092 | 0.03857 | 0.029375 | 0.008896 | 0.039486 | 0.007637 | 0.036972 | 0.010842 | 0.003155 | 0.017931 | 0.029022 | 0.007836 | 0.011563 | 0.004192 | 0.005686 | 0.012279 | 0.004678 | 0.00497 | 0.007709 | 0.005461 |
| 36.0416667 | 0.054924 | 0.005987 | 0.008418 | 0.057066 | 0.047058 | 0.042999 | 0.013353 | 0.02744 | 0.006019 | 0.038793 | 0.013806 | 0.002561 | 0.010529 | 0.027159 | 0.015876 | 0.014838 | 0.003836 | 0.006094 | 0.015822 | 0.004134 | 0.005349 | 0.006525 | 0.007245 |
| 36.0555556 | 0.029066 | 0.015587 | 0.007989 | 0.044223 | 0.039837 | 0.063101 | 0.020127 | 0.020562 | 0.008948 | 0.036529 | 0.018642 | 0.006097 | 0.022853 | 0.031621 | 0.030463 | 0.016815 | 0.002055 | 0.003312 | 0.021963 | 0.002347 | 0.004961 | 0.004007 | 0.013211 |
| 36.0694444 | 0.028442 | 0.025134 | 0.005929 | 0.025026 | 0.023059 | 0.083708 | 0.024289 | 0.032929 | 0.015615 | 0.028231 | 0.020087 | 0.010836 | 0.034693 | 0.041539 | 0.041363 | 0.018138 | 0.001302 | 0.001065 | 0.025882 | 0.002473 | 0.004872 | 0.00381 | 0.020112 |
| 36.0833333 | 0.033168 | 0.024759 | 0.002715 | 0.012689 | 0.011215 | 0.096633 | 0.026889 | 0.046262 | 0.013935 | 0.017551 | 0.012952 | 0.016755 | 0.027726 | 0.047632 | 0.044598 | 0.01972 | 0.000948 | 0.000483 | 0.026849 | 0.002543 | 0.002804 | 0.003829 | 0.025186 |
| 36.0972222 | 0.020961 | 0.015999 | 0.000988 | 0.007687 | 0.007311 | 0.090012 | 0.026875 | 0.044653 | 0.008543 | 0.014676 | 0.005765 | 0.018022 | 0.015502 | 0.040056 | 0.042103 | 0.02165 | 0.000832 | 0.000367 | 0.025782 | 0.001539 | 0.002194 | 0.002643 | 0.025612 |
| 36.1111111 | 0.019016 | 0.007395 | 0.000867 | 0.006196 | 0.005417 | 0.054379 | 0.021224 | 0.029358 | 0.012325 | 0.035051 | 0.00712 | 0.011185 | 0.02208 | 0.023891 | 0.033648 | 0.021002 | 0.000775 | 0.000579 | 0.022368 | 0.001074 | 0.006098 | 0.00193 | 0.020869 |
| 36.125 | 0.031178 | 0.004478 | 0.00098 | 0.012866 | 0.005188 | 0.021514 | 0.02308 | 0.018939 | 0.015485 | 0.073984 | 0.010001 | 0.004962 | 0.035876 | 0.014111 | 0.021873 | 0.015308 | 0.000504 | 0.000838 | 0.01841 | 0.000955 | 0.008828 | 0.002554 | 0.017194 |
| 36.1388889 | 0.029736 | 0.009396 | 0.00148 | 0.024657 | 0.004728 | 0.024649 | 0.040409 | 0.026991 | 0.009405 | 0.08835 | 0.009181 | 0.002747 | 0.032358 | 0.015134 | 0.011192 | 0.008036 | 0.0005 | 0.001028 | 0.015453 | 0.00056 | 0.007612 | 0.002601 | 0.017976 |
| 36.1527778 | 0.015358 | 0.021276 | 0.002572 | 0.030571 | 0.009632 | 0.054291 | 0.056365 | 0.047484 | 0.00281 | 0.056763 | 0.007387 | 0.001798 | 0.017247 | 0.021377 | 0.007601 | 0.003091 | 0.000915 | 0.001614 | 0.012273 | 0.000736 | 0.006837 | 0.002128 | 0.018966 |
| 36.1666667 | 0.013154 | 0.030251 | 0.005917 | 0.027516 | 0.023377 | 0.084191 | 0.061188 | 0.066391 | 0.001052 | 0.024082 | 0.006151 | 0.003141 | 0.011222 | 0.029283 | 0.012809 | 0.001933 | 0.001862 | 0.002643 | 0.008904 | 0.001698 | 0.00831 | 0.002669 | 0.018025 |
| 36.1805556 | 0.02459 | 0.030198 | 0.010012 | 0.017707 | 0.029864 | 0.096534 | 0.062148 | 0.065721 | 0.001957 | 0.026726 | 0.004965 | 0.006663 | 0.017315 | 0.032971 | 0.016795 | 0.005837 | 0.003292 | 0.00407 | 0.007087 | 0.002946 | 0.009716 | 0.004051 | 0.015581 |
| 36.1944444 | 0.041067 | 0.023851 | 0.009412 | 0.013083 | 0.020431 | 0.094996 | 0.057681 | 0.042542 | 0.005639 | 0.046427 | 0.003446 | 0.010109 | 0.020992 | 0.027673 | 0.0132 | 0.013997 | 0.005327 | 0.006245 | 0.006959 | 0.004809 | 0.010431 | 0.006281 | 0.011395 |
| 36.2083333 | 0.058062 | 0.013197 | 0.00584 | 0.027969 | 0.007541 | 0.088826 | 0.039267 | 0.023357 | 0.011078 | 0.06609 | 0.002274 | 0.011795 | 0.018223 | 0.018702 | 0.00703 | 0.022795 | 0.007662 | 0.008828 | 0.007116 | 0.007242 | 0.01132 | 0.009077 | 0.00673 |
| 36.2222222 | 0.066156 | 0.006263 | 0.005489 | 0.047794 | 0.005208 | 0.079558 | 0.021117 | 0.038477 | 0.015435 | 0.088297 | 0.009126 | 0.011318 | 0.019171 | 0.016246 | 0.00611 | 0.02759 | 0.00907 | 0.010394 | 0.006455 | 0.008816 | 0.011808 | 0.01083 | 0.003689 |
| 36.2361111 | 0.066712 | 0.006944 | 0.006342 | 0.04801 | 0.011195 | 0.06052 | 0.028887 | 0.078131 | 0.015113 | 0.098541 | 0.021203 | 0.009374 | 0.024749 | 0.024106 | 0.011299 | 0.026815 | 0.008826 | 0.010064 | 0.00495 | 0.008466 | 0.011092 | 0.010439 | 0.003384 |
| 36.25 | 0.074429 | 0.010659 | 0.006021 | 0.031642 | 0.012803 | 0.034019 | 0.050018 | 0.094564 | 0.009035 | 0.08041 | 0.019242 | 0.007094 | 0.023249 | 0.035342 | 0.016282 | 0.02329 | 0.007667 | 0.008641 | 0.003397 | 0.007049 | 0.010228 | 0.00893 | 0.004922 |
| 36.2638889 | 0.082481 | 0.02611 | 0.00657 | 0.0159 | 0.00749 | 0.026527 | 0.053235 | 0.06615 | 0.0055 | 0.048286 | 0.012709 | 0.005207 | 0.013597 | 0.043892 | 0.016875 | 0.019918 | 0.006536 | 0.00726 | 0.002561 | 0.005794 | 0.009917 | 0.007644 | 0.006607 |
| 36.2777778 | 0.067035 | 0.044135 | 0.005589 | 0.008429 | 0.004286 | 0.039499 | 0.03615 | 0.027897 | 0.007865 | 0.021699 | 0.015159 | 0.003889 | 0.008666 | 0.048365 | 0.014222 | 0.016962 | 0.005489 | 0.006078 | 0.002097 | 0.004831 | 0.009023 | 0.006586 | 0.007799 |
| 36.2916667 | 0.034008 | 0.041985 | 0.002724 | 0.010819 | 0.006864 | 0.044695 | 0.030396 | 0.011167 | 0.011862 | 0.010014 | 0.013201 | 0.003141 | 0.010906 | 0.04587 | 0.010934 | 0.013212 | 0.004331 | 0.004862 | 0.001419 | 0.003884 | 0.007334 | 0.005419 | 0.008554 |
| 36.3055556 | 0.010961 | 0.033059 | 0.001941 | 0.020302 | 0.010956 | 0.039158 | 0.050992 | 0.010029 | 0.015082 | 0.013889 | 0.006135 | 0.002209 | 0.011957 | 0.032149 | 0.007021 | 0.008379 | 0.003079 | 0.003566 | 0.000774 | 0.002873 | 0.006262 | 0.004104 | 0.008995 |
| 36.3194444 | 0.008672 | 0.032855 | 0.002569 | 0.025279 | 0.012533 | 0.034027 | 0.064838 | 0.017896 | 0.013359 | 0.021066 | 0.002487 | 0.001343 | 0.010139 | 0.018371 | 0.004992 | 0.003984 | 0.001931 | 0.00234 | 0.000532 | 0.001967 | 0.006211 | 0.002729 | 0.00917 |
| 36.3333333 | 0.011814 | 0.035178 | 0.003126 | 0.023672 | 0.010014 | 0.029264 | 0.055301 | 0.031192 | 0.009306 | 0.02378 | 0.002273 | 0.001521 | 0.008548 | 0.020969 | 0.007474 | 0.0018 | 0.001333 | 0.001686 | 0.000491 | 0.001672 | 0.006504 | 0.001833 | 0.008621 |
| 36.3472222 | 0.011796 | 0.031516 | 0.005198 | 0.023362 | 0.005398 | 0.02201 | 0.036096 | 0.044652 | 0.008352 | 0.02003 | 0.003401 | 0.00185 | 0.00778 | 0.02929 | 0.009494 | 0.00244 | 0.001655 | 0.00207 | 0.000522 | 0.00235 | 0.007475 | 0.00213 | 0.006858 |
| 36.3611111 | 0.023699 | 0.018489 | 0.009172 | 0.026718 | 0.002741 | 0.012067 | 0.019831 | 0.055509 | 0.006891 | 0.01207 | 0.003399 | 0.002982 | 0.005984 | 0.032777 | 0.007523 | 0.005337 | 0.002876 | 0.003403 | 0.001179 | 0.003751 | 0.010588 | 0.003742 | 0.004203 |
| 36.375 | 0.040501 | 0.007219 | 0.010521 | 0.03086 | 0.002281 | 0.004507 | 0.021303 | 0.053555 | 0.003843 | 0.012026 | 0.001887 | 0.007194 | 0.003716 | 0.039427 | 0.004403 | 0.009737 | 0.004579 | 0.005175 | 0.002772 | 0.005265 | 0.015666 | 0.0063 | 0.002389 |
| 36.3888889 | 0.035176 | 0.0058 | 0.006884 | 0.026732 | 0.003345 | 0.002556 | 0.038466 | 0.034856 | 0.002205 | 0.024206 | 0.000754 | 0.013994 | 0.005336 | 0.053209 | 0.00485 | 0.015405 | 0.006507 | 0.007147 | 0.005404 | 0.006729 | 0.019884 | 0.009279 | 0.003478 |
| 36.4027778 | 0.024534 | 0.006444 | 0.003512 | 0.018509 | 0.005557 | 0.00257 | 0.05236 | 0.025456 | 0.002669 | 0.037247 | 0.00104 | 0.020234 | 0.012231 | 0.068726 | 0.01181 | 0.021274 | 0.008356 | 0.009073 | 0.0087 | 0.008125 | 0.020936 | 0.011494 | 0.00626 |
| 36.4166667 | 0.032425 | 0.007527 | 0.003356 | 0.019827 | 0.00482 | 0.003198 | 0.048166 | 0.042516 | 0.004809 | 0.040247 | 0.00156 | 0.021499 | 0.020434 | 0.071933 | 0.021675 | 0.024446 | 0.008209 | 0.009116 | 0.010372 | 0.007831 | 0.018337 | 0.010768 | 0.00686 |
| 36.4305556 | 0.031902 | 0.008409 | 0.005328 | 0.025476 | 0.002535 | 0.003183 | 0.031592 | 0.054828 | 0.007614 | 0.033419 | 0.001207 | 0.015076 | 0.026418 | 0.052159 | 0.027048 | 0.022547 | 0.00609 | 0.006641 | 0.007619 | 0.005969 | 0.012151 | 0.007483 | 0.00535 |
| 36.4444444 | 0.017079 | 0.01058 | 0.008539 | 0.027499 | 0.005152 | 0.002323 | 0.029301 | 0.045533 | 0.011941 | 0.023328 | 0.000904 | 0.007532 | 0.027735 | 0.028078 | 0.025783 | 0.016889 | 0.006796 | 0.006701 | 0.003587 | 0.007209 | 0.00584 | 0.007287 | 0.006725 |
| 36.4583333 | 0.012719 | 0.020859 | 0.00817 | 0.022953 | 0.008665 | 0.00411 | 0.03391 | 0.03069 | 0.01358 | 0.014192 | 0.002622 | 0.006992 | 0.01927 | 0.02377 | 0.021921 | 0.012743 | 0.010615 | 0.011371 | 0.002194 | 0.011277 | 0.003407 | 0.011376 | 0.010385 |
| 36.4722222 | 0.011811 | 0.031838 | 0.005745 | 0.013001 | 0.01146 | 0.008633 | 0.021255 | 0.016371 | 0.00866 | 0.010219 | 0.005715 | 0.007532 | 0.011052 | 0.029444 | 0.020593 | 0.01355 | 0.01142 | 0.013357 | 0.00275 | 0.012142 | 0.002762 | 0.012781 | 0.00966 |
| 36.4861111 | 0.012066 | 0.028381 | 0.007301 | 0.009869 | 0.022117 | 0.01506 | 0.007939 | 0.007234 | 0.004452 | 0.020534 | 0.005418 | 0.005481 | 0.013469 | 0.035057 | 0.022565 | 0.014926 | 0.008182 | 0.009373 | 0.005243 | 0.009283 | 0.001765 | 0.00932 | 0.004954 |
| 36.5 | 0.027493 | 0.015332 | 0.007113 | 0.013069 | 0.033788 | 0.020452 | 0.005445 | 0.005544 | 0.004039 | 0.046085 | 0.002645 | 0.006974 | 0.014084 | 0.046422 | 0.023625 | 0.012075 | 0.005366 | 0.005318 | 0.006776 | 0.006471 | 0.003419 | 0.005623 | 0.001964 |
| 36.5138889 | 0.047246 | 0.008661 | 0.006092 | 0.011263 | 0.032065 | 0.021357 | 0.00527 | 0.008887 | 0.004354 | 0.069994 | 0.00198 | 0.009495 | 0.014057 | 0.050519 | 0.022097 | 0.007409 | 0.004519 | 0.004306 | 0.00568 | 0.005032 | 0.007958 | 0.003914 | 0.001628 |
| 36.5277778 | 0.056863 | 0.006444 | 0.011562 | 0.009309 | 0.0187 | 0.02048 | 0.0091 | 0.020724 | 0.003961 | 0.078886 | 0.002127 | 0.008034 | 0.015107 | 0.036276 | 0.01686 | 0.005231 | 0.006225 | 0.006118 | 0.0047 | 0.006104 | 0.010969 | 0.004834 | 0.00263 |
| 36.5416667 | 0.056089 | 0.005395 | 0.016193 | 0.011087 | 0.007229 | 0.019201 | 0.018067 | 0.038818 | 0.0037 | 0.08016 | 0.002693 | 0.012741 | 0.024238 | 0.027103 | 0.012839 | 0.01186 | 0.016201 | 0.016506 | 0.010371 | 0.015401 | 0.016032 | 0.013735 | 0.009622 |
| 36.5555556 | 0.046369 | 0.010383 | 0.012837 | 0.013723 | 0.005692 | 0.013057 | 0.031158 | 0.054532 | 0.006417 | 0.075581 | 0.003163 | 0.030529 | 0.051277 | 0.039007 | 0.017661 | 0.026109 | 0.033441 | 0.035331 | 0.024744 | 0.032275 | 0.028157 | 0.030384 | 0.024438 |
| 36.5694444 | 0.027131 | 0.024687 | 0.006952 | 0.020595 | 0.006255 | 0.004528 | 0.03253 | 0.047584 | 0.009641 | 0.052139 | 0.002468 | 0.039753 | 0.05531 | 0.04577 | 0.01779 | 0.027544 | 0.039584 | 0.042789 | 0.030412 | 0.0393 | 0.035711 | 0.03824 | 0.031858 |
| 36.5833333 | 0.012347 | 0.035189 | 0.004226 | 0.022291 | 0.003823 | 0.006509 | 0.026772 | 0.031255 | 0.013643 | 0.025633 | 0.002081 | 0.026643 | 0.033781 | 0.033361 | 0.011164 | 0.021444 | 0.02484 | 0.027135 | 0.020657 | 0.025349 | 0.025033 | 0.025452 | 0.022292 |
| 36.5972222 | 0.016821 | 0.030486 | 0.004014 | 0.015523 | 0.006511 | 0.025334 | 0.039672 | 0.038754 | 0.02201 | 0.025588 | 0.005728 | 0.015244 | 0.029618 | 0.029316 | 0.015587 | 0.031965 | 0.009656 | 0.010529 | 0.014777 | 0.009778 | 0.012304 | 0.010409 | 0.014144 |
| 36.6111111 | 0.022334 | 0.026058 | 0.003879 | 0.01488 | 0.014616 | 0.049527 | 0.050044 | 0.045222 | 0.031388 | 0.042118 | 0.018623 | 0.020514 | 0.043266 | 0.043425 | 0.025032 | 0.041913 | 0.008687 | 0.009487 | 0.017725 | 0.008836 | 0.01771 | 0.010235 | 0.019722 |
| 36.625 | 0.01744 | 0.032019 | 0.002171 | 0.022225 | 0.018035 | 0.058713 | 0.042829 | 0.034274 | 0.035407 | 0.048813 | 0.030936 | 0.028795 | 0.063669 | 0.0542 | 0.02665 | 0.034136 | 0.010469 | 0.011225 | 0.019021 | 0.010622 | 0.027768 | 0.012591 | 0.026552 |
| 36.6388889 | 0.017547 | 0.039247 | 0.00366 | 0.028278 | 0.014685 | 0.04764 | 0.031526 | 0.022298 | 0.029931 | 0.043091 | 0.02857 | 0.03207 | 0.070258 | 0.052331 | 0.021204 | 0.020686 | 0.008936 | 0.009253 | 0.01763 | 0.008526 | 0.027234 | 0.010208 | 0.027025 |
| 36.6527778 | 0.021506 | 0.037454 | 0.014971 | 0.026428 | 0.014009 | 0.029358 | 0.021147 | 0.013329 | 0.019367 | 0.037381 | 0.016661 | 0.030234 | 0.045297 | 0.051085 | 0.014265 | 0.009814 | 0.006876 | 0.007085 | 0.014886 | 0.006475 | 0.016621 | 0.007786 | 0.022888 |
| 36.6666667 | 0.016937 | 0.025767 | 0.033047 | 0.017617 | 0.020319 | 0.013857 | 0.013145 | 0.006826 | 0.010366 | 0.03118 | 0.008284 | 0.023371 | 0.021902 | 0.051696 | 0.009095 | 0.003212 | 0.004115 | 0.004265 | 0.009755 | 0.00415 | 0.006788 | 0.005308 | 0.016084 |
| 36.6805556 | 0.008205 | 0.013219 | 0.051186 | 0.008836 | 0.023583 | 0.006801 | 0.006664 | 0.00313 | 0.004157 | 0.018013 | 0.009182 | 0.016069 | 0.030985 | 0.043482 | 0.005346 | 0.00127 | 0.001606 | 0.001472 | 0.004517 | 0.001666 | 0.005966 | 0.002658 | 0.010437 |
| 36.6944444 | 0.002954 | 0.009437 | 0.067124 | 0.006822 | 0.017377 | 0.010185 | 0.002691 | 0.004877 | 0.001282 | 0.007614 | 0.013188 | 0.012804 | 0.048872 | 0.038532 | 0.006743 | 0.001945 | 0.001396 | 0.00097 | 0.003 | 0.001013 | 0.005652 | 0.001808 | 0.009003 |
| 36.7083333 | 0.016239 | 0.010809 | 0.078418 | 0.010865 | 0.008266 | 0.018352 | 0.003872 | 0.011873 | 0.002497 | 0.010976 | 0.013192 | 0.013417 | 0.055691 | 0.047434 | 0.015681 | 0.00403 | 0.00352 | 0.002995 | 0.005409 | 0.002784 | 0.004327 | 0.003553 | 0.011265 |
| 36.7222222 | 0.03923 | 0.008414 | 0.079593 | 0.015734 | 0.005213 | 0.022806 | 0.009177 | 0.022488 | 0.005113 | 0.023593 | 0.009203 | 0.014301 | 0.052898 | 0.056263 | 0.023247 | 0.007409 | 0.005685 | 0.005243 | 0.007992 | 0.004962 | 0.00679 | 0.005798 | 0.013739 |
| 36.7361111 | 0.043254 | 0.004209 | 0.067891 | 0.01482 | 0.011633 | 0.017202 | 0.014134 | 0.037006 | 0.006419 | 0.036104 | 0.006588 | 0.013262 | 0.03589 | 0.054787 | 0.020935 | 0.01061 | 0.006114 | 0.005792 | 0.00822 | 0.005689 | 0.009992 | 0.006489 | 0.01402 |
| 36.75 | 0.040301 | 0.002181 | 0.044553 | 0.010767 | 0.018611 | 0.011119 | 0.013068 | 0.05419 | 0.00565 | 0.03644 | 0.012033 | 0.011474 | 0.024647 | 0.047287 | 0.012601 | 0.011901 | 0.004871 | 0.00461 | 0.006717 | 0.0048 | 0.012498 | 0.005492 | 0.012097 |
| 36.7638889 | 0.035949 | 0.001216 | 0.02169 | 0.011744 | 0.016465 | 0.014082 | 0.008877 | 0.068287 | 0.002545 | 0.02571 | 0.021973 | 0.01027 | 0.040304 | 0.038643 | 0.006832 | 0.010781 | 0.002835 | 0.002592 | 0.004251 | 0.002989 | 0.014975 | 0.003607 | 0.008844 |
| 36.7777778 | 0.024795 | 0.000508 | 0.020557 | 0.012715 | 0.010028 | 0.015258 | 0.011556 | 0.058139 | 0.002048 | 0.025352 | 0.026777 | 0.009251 | 0.05771 | 0.027812 | 0.007552 | 0.00822 | 0.001373 | 0.00121 | 0.002219 | 0.001411 | 0.01471 | 0.001743 | 0.005467 |
| 36.7916667 | 0.019341 | 0.000418 | 0.040669 | 0.010572 | 0.005844 | 0.01112 | 0.01612 | 0.05722 | 0.005992 | 0.037405 | 0.023022 | 0.008069 | 0.057813 | 0.017429 | 0.010395 | 0.005934 | 0.001576 | 0.001606 | 0.002967 | 0.001261 | 0.010042 | 0.001163 | 0.002816 |
| 36.8055556 | 0.018527 | 0.002014 | 0.059504 | 0.018425 | 0.003227 | 0.008443 | 0.013727 | 0.117791 | 0.008651 | 0.047468 | 0.01335 | 0.007082 | 0.051279 | 0.017878 | 0.00922 | 0.004988 | 0.002415 | 0.002709 | 0.005675 | 0.002124 | 0.008449 | 0.002078 | 0.001093 |
| 36.8194444 | 0.013711 | 0.005384 | 0.063905 | 0.030104 | 0.001091 | 0.008225 | 0.007677 | 0.173869 | 0.007315 | 0.050394 | 0.005116 | 0.007282 | 0.045339 | 0.022086 | 0.005611 | 0.005641 | 0.002028 | 0.00255 | 0.007594 | 0.00219 | 0.014064 | 0.002384 | 0.000577 |
| 36.8333333 | 0.007142 | 0.007225 | 0.052324 | 0.023488 | 0.000743 | 0.016035 | 0.003605 | 0.162531 | 0.004513 | 0.044913 | 0.002621 | 0.008914 | 0.043477 | 0.015293 | 0.003827 | 0.006781 | 0.001176 | 0.001538 | 0.007852 | 0.001361 | 0.017473 | 0.001624 | 0.001247 |
| 36.8472222 | 0.005149 | 0.006496 | 0.029345 | 0.008856 | 0.001659 | 0.032723 | 0.002886 | 0.120864 | 0.003262 | 0.033389 | 0.003963 | 0.010831 | 0.042489 | 0.010678 | 0.003131 | 0.006373 | 0.00152 | 0.001498 | 0.00741 | 0.001098 | 0.012666 | 0.001603 | 0.002197 |
| 36.8611111 | 0.006767 | 0.006812 | 0.009439 | 0.002071 | 0.002246 | 0.048639 | 0.003933 | 0.092226 | 0.00491 | 0.023112 | 0.008631 | 0.012571 | 0.031284 | 0.021568 | 0.002261 | 0.004312 | 0.002156 | 0.002208 | 0.007398 | 0.001355 | 0.007048 | 0.002033 | 0.002524 |
| 36.875 | 0.008551 | 0.009342 | 0.006363 | 0.001679 | 0.005173 | 0.06033 | 0.004372 | 0.075063 | 0.006774 | 0.017898 | 0.014649 | 0.013925 | 0.019529 | 0.031577 | 0.00256 | 0.002263 | 0.002118 | 0.002536 | 0.00768 | 0.001322 | 0.009089 | 0.001766 | 0.002203 |
| 36.8888889 | 0.011355 | 0.011516 | 0.021043 | 0.006228 | 0.009028 | 0.072601 | 0.003295 | 0.048143 | 0.005024 | 0.014507 | 0.02023 | 0.012804 | 0.030201 | 0.03621 | 0.003679 | 0.001872 | 0.001819 | 0.002435 | 0.007428 | 0.001058 | 0.014883 | 0.001199 | 0.001785 |
| 36.9027778 | 0.01362 | 0.009592 | 0.034284 | 0.010834 | 0.010868 | 0.084113 | 0.005508 | 0.02144 | 0.003971 | 0.016078 | 0.025266 | 0.007832 | 0.050074 | 0.042229 | 0.003852 | 0.003078 | 0.001721 | 0.002326 | 0.006426 | 0.000918 | 0.017954 | 0.001047 | 0.001819 |
| 36.9166667 | 0.01301 | 0.004417 | 0.040233 | 0.011094 | 0.020549 | 0.085992 | 0.017251 | 0.012852 | 0.007545 | 0.018846 | 0.026935 | 0.003889 | 0.061207 | 0.036135 | 0.003708 | 0.003234 | 0.001992 | 0.002396 | 0.005269 | 0.001089 | 0.01873 | 0.001394 | 0.002363 |
| 36.9305556 | 0.011318 | 0.002373 | 0.041368 | 0.007791 | 0.034652 | 0.084205 | 0.030359 | 0.010661 | 0.010141 | 0.022285 | 0.025107 | 0.004423 | 0.068448 | 0.019669 | 0.005439 | 0.002535 | 0.002336 | 0.002485 | 0.004412 | 0.001423 | 0.017787 | 0.001828 | 0.002888 |
| 36.9444444 | 0.009476 | 0.004734 | 0.024703 | 0.004549 | 0.037667 | 0.090699 | 0.029438 | 0.006094 | 0.008516 | 0.033243 | 0.025052 | 0.004886 | 0.059313 | 0.008018 | 0.009363 | 0.004328 | 0.002464 | 0.002552 | 0.003553 | 0.00158 | 0.014786 | 0.001773 | 0.002851 |
| 36.9583333 | 0.006424 | 0.005732 | 0.006575 | 0.006435 | 0.035948 | 0.091812 | 0.016615 | 0.004385 | 0.007017 | 0.03219 | 0.027217 | 0.003839 | 0.036645 | 0.006707 | 0.013427 | 0.006915 | 0.002303 | 0.002584 | 0.002499 | 0.001394 | 0.01088 | 0.001066 | 0.00214 |
| 36.9722222 | 0.002785 | 0.003182 | 0.003616 | 0.008772 | 0.036092 | 0.075352 | 0.00841 | 0.0114 | 0.007469 | 0.019373 | 0.025307 | 0.003403 | 0.029295 | 0.015839 | 0.016294 | 0.008059 | 0.001665 | 0.002116 | 0.001972 | 0.000885 | 0.00686 | 0.000518 | 0.001087 |
| 36.9861111 | 0.001587 | 0.003848 | 0.006461 | 0.006233 | 0.030775 | 0.053914 | 0.0139 | 0.026512 | 0.006572 | 0.019745 | 0.016597 | 0.003052 | 0.029491 | 0.026879 | 0.019209 | 0.010867 | 0.000874 | 0.001188 | 0.002452 | 0.000591 | 0.003643 | 0.000913 | 0.000436 |
| 37 | 0.003318 | 0.011393 | 0.007935 | 0.005179 | 0.022938 | 0.032875 | 0.023754 | 0.03542 | 0.004298 | 0.032512 | 0.008153 | 0.003061 | 0.019633 | 0.03029 | 0.023468 | 0.015456 | 0.000565 | 0.000687 | 0.002933 | 0.000815 | 0.002427 | 0.001276 | 0.000555 |
| 37.0138889 | 0.00466 | 0.01866 | 0.006509 | 0.011948 | 0.026469 | 0.016303 | 0.028509 | 0.030433 | 0.004114 | 0.03892 | 0.008171 | 0.00458 | 0.012014 | 0.023105 | 0.028356 | 0.017232 | 0.001281 | 0.001746 | 0.002588 | 0.001982 | 0.002165 | 0.001512 | 0.001493 |
| 37.0277778 | 0.00419 | 0.022579 | 0.004243 | 0.02036 | 0.034885 | 0.018065 | 0.029244 | 0.032421 | 0.004977 | 0.027089 | 0.012905 | 0.007131 | 0.009426 | 0.016146 | 0.035999 | 0.016761 | 0.002161 | 0.003622 | 0.004285 | 0.002986 | 0.002543 | 0.001722 | 0.002868 |
| 37.0416667 | 0.003421 | 0.030757 | 0.006137 | 0.018672 | 0.026493 | 0.033934 | 0.021664 | 0.051868 | 0.004727 | 0.009111 | 0.018204 | 0.011774 | 0.017297 | 0.024991 | 0.048746 | 0.02034 | 0.00322 | 0.004435 | 0.012701 | 0.003697 | 0.007009 | 0.003141 | 0.007356 |
| 37.0555556 | 0.003014 | 0.042688 | 0.017072 | 0.009899 | 0.01276 | 0.04655 | 0.013637 | 0.066795 | 0.005354 | 0.003444 | 0.025038 | 0.018762 | 0.049809 | 0.044015 | 0.061879 | 0.028665 | 0.005739 | 0.005086 | 0.024415 | 0.006268 | 0.014599 | 0.00757 | 0.016616 |
| 37.0694444 | 0.004066 | 0.048248 | 0.025653 | 0.007894 | 0.012031 | 0.048408 | 0.022478 | 0.073812 | 0.00754 | 0.004506 | 0.027274 | 0.023154 | 0.071752 | 0.051477 | 0.064276 | 0.034455 | 0.006103 | 0.004867 | 0.028737 | 0.006908 | 0.016408 | 0.00973 | 0.022976 |
| 37.0833333 | 0.006382 | 0.044078 | 0.018949 | 0.013326 | 0.011923 | 0.042651 | 0.038002 | 0.093046 | 0.007246 | 0.002989 | 0.01865 | 0.021351 | 0.050543 | 0.039431 | 0.052153 | 0.032755 | 0.0034 | 0.002916 | 0.022615 | 0.00385 | 0.009942 | 0.006907 | 0.022263 |
| 37.0972222 | 0.006063 | 0.032715 | 0.007881 | 0.014819 | 0.009636 | 0.032343 | 0.041573 | 0.109621 | 0.004357 | 0.004345 | 0.008408 | 0.01397 | 0.036483 | 0.019638 | 0.036173 | 0.024 | 0.002391 | 0.002406 | 0.012042 | 0.002086 | 0.005634 | 0.003581 | 0.018925 |
| 37.1111111 | 0.005958 | 0.020576 | 0.008071 | 0.010813 | 0.011626 | 0.017539 | 0.029015 | 0.088303 | 0.002721 | 0.01255 | 0.007541 | 0.007005 | 0.063149 | 0.011342 | 0.021191 | 0.012483 | 0.00351 | 0.00374 | 0.005011 | 0.00287 | 0.010087 | 0.001739 | 0.015651 |
| 37.125 | 0.011881 | 0.018348 | 0.017375 | 0.005994 | 0.01416 | 0.006442 | 0.016518 | 0.044497 | 0.002088 | 0.020793 | 0.009601 | 0.005494 | 0.073665 | 0.021855 | 0.009102 | 0.006726 | 0.004561 | 0.00491 | 0.005218 | 0.003996 | 0.016274 | 0.001465 | 0.012972 |
| 37.1388889 | 0.018861 | 0.028995 | 0.025662 | 0.002958 | 0.0138 | 0.009106 | 0.017472 | 0.021809 | 0.003616 | 0.024153 | 0.00713 | 0.005326 | 0.04447 | 0.03366 | 0.007059 | 0.008493 | 0.005245 | 0.005747 | 0.007853 | 0.004774 | 0.018275 | 0.002688 | 0.011385 |
| 37.1527778 | 0.020526 | 0.041928 | 0.030752 | 0.002378 | 0.013957 | 0.020591 | 0.020655 | 0.031564 | 0.0078 | 0.027592 | 0.002986 | 0.006176 | 0.025664 | 0.031448 | 0.014782 | 0.0117 | 0.006496 | 0.00714 | 0.009244 | 0.006153 | 0.01751 | 0.004704 | 0.01001 |
| 37.1666667 | 0.019248 | 0.045824 | 0.034113 | 0.00353 | 0.020416 | 0.024604 | 0.022721 | 0.050934 | 0.009117 | 0.030118 | 0.001696 | 0.011767 | 0.030686 | 0.020802 | 0.029336 | 0.014549 | 0.008658 | 0.009464 | 0.009537 | 0.008502 | 0.015966 | 0.00714 | 0.007647 |
| 37.1805556 | 0.018545 | 0.035865 | 0.028406 | 0.005136 | 0.023568 | 0.017696 | 0.029636 | 0.062291 | 0.00611 | 0.0266 | 0.004943 | 0.016447 | 0.029299 | 0.013591 | 0.045894 | 0.017542 | 0.010109 | 0.011112 | 0.007203 | 0.010208 | 0.013845 | 0.00878 | 0.005444 |
| 37.1944444 | 0.016322 | 0.01853 | 0.014276 | 0.006698 | 0.016239 | 0.013439 | 0.036352 | 0.066106 | 0.00339 | 0.01819 | 0.009911 | 0.013736 | 0.01915 | 0.012211 | 0.052045 | 0.020652 | 0.009321 | 0.010314 | 0.004639 | 0.00953 | 0.00994 | 0.007985 | 0.005296 |
| 37.2083333 | 0.010891 | 0.011706 | 0.007133 | 0.010513 | 0.012728 | 0.011193 | 0.032788 | 0.062806 | 0.002196 | 0.011438 | 0.011012 | 0.007794 | 0.013805 | 0.009382 | 0.040709 | 0.024163 | 0.007377 | 0.008084 | 0.006047 | 0.007418 | 0.004957 | 0.005524 | 0.006199 |
| 37.2222222 | 0.005431 | 0.018924 | 0.007408 | 0.013143 | 0.017993 | 0.007577 | 0.019776 | 0.051973 | 0.001683 | 0.012335 | 0.006537 | 0.004682 | 0.01168 | 0.004521 | 0.021103 | 0.02546 | 0.006114 | 0.006539 | 0.008114 | 0.005964 | 0.001567 | 0.003588 | 0.005955 |
| 37.2361111 | 0.003994 | 0.026801 | 0.006414 | 0.012711 | 0.020703 | 0.010012 | 0.016153 | 0.03849 | 0.002034 | 0.017339 | 0.002045 | 0.003411 | 0.009321 | 0.00371 | 0.012248 | 0.023103 | 0.005599 | 0.005944 | 0.00749 | 0.005423 | 0.0009 | 0.002555 | 0.004876 |
| 37.25 | 0.006504 | 0.029232 | 0.005877 | 0.012854 | 0.017905 | 0.011984 | 0.031278 | 0.03358 | 0.004298 | 0.015617 | 0.001685 | 0.001818 | 0.011028 | 0.006895 | 0.019528 | 0.020122 | 0.005308 | 0.005728 | 0.005286 | 0.005098 | 0.001234 | 0.001936 | 0.003868 |
| 37.2638889 | 0.008561 | 0.02689 | 0.006952 | 0.012337 | 0.015085 | 0.008253 | 0.049971 | 0.043299 | 0.008365 | 0.007804 | 0.003395 | 0.002166 | 0.016875 | 0.009838 | 0.029322 | 0.018726 | 0.005277 | 0.00578 | 0.00306 | 0.004914 | 0.000983 | 0.002018 | 0.002708 |
| 37.2777778 | 0.008008 | 0.019385 | 0.006458 | 0.008032 | 0.014791 | 0.005463 | 0.061525 | 0.060042 | 0.013005 | 0.003554 | 0.005387 | 0.00509 | 0.019871 | 0.008633 | 0.033252 | 0.018289 | 0.005215 | 0.00572 | 0.003173 | 0.004698 | 0.000986 | 0.00227 | 0.001852 |
| 37.2916667 | 0.006508 | 0.009279 | 0.00445 | 0.003424 | 0.014239 | 0.004215 | 0.059143 | 0.074077 | 0.017134 | 0.003624 | 0.006836 | 0.005321 | 0.019458 | 0.005708 | 0.03143 | 0.01674 | 0.00445 | 0.004948 | 0.005307 | 0.003883 | 0.001259 | 0.001682 | 0.002099 |
| 37.3055556 | 0.005139 | 0.003191 | 0.00402 | 0.002135 | 0.010776 | 0.002527 | 0.037794 | 0.084686 | 0.019741 | 0.008641 | 0.005461 | 0.002689 | 0.017514 | 0.007809 | 0.025882 | 0.012925 | 0.00299 | 0.003542 | 0.007192 | 0.002502 | 0.001018 | 0.00078 | 0.00295 |
| 37.3194444 | 0.006379 | 0.004561 | 0.00481 | 0.002726 | 0.006611 | 0.002055 | 0.027595 | 0.093155 | 0.020506 | 0.021486 | 0.002849 | 0.002121 | 0.012439 | 0.012145 | 0.01894 | 0.008125 | 0.001942 | 0.00254 | 0.008654 | 0.001513 | 0.00146 | 0.000529 | 0.003174 |
| 37.3333333 | 0.009947 | 0.008984 | 0.008743 | 0.005189 | 0.004153 | 0.002168 | 0.050075 | 0.082396 | 0.019871 | 0.031258 | 0.002637 | 0.003483 | 0.005904 | 0.01089 | 0.010596 | 0.004339 | 0.002589 | 0.00326 | 0.009871 | 0.001994 | 0.003889 | 0.001434 | 0.002188 |
| 37.3472222 | 0.010414 | 0.011841 | 0.018193 | 0.007315 | 0.002457 | 0.003335 | 0.071288 | 0.048664 | 0.019621 | 0.023311 | 0.003612 | 0.004039 | 0.002854 | 0.011327 | 0.006453 | 0.002368 | 0.004415 | 0.005324 | 0.010369 | 0.003595 | 0.007282 | 0.003307 | 0.001361 |
| 37.3611111 | 0.008036 | 0.013665 | 0.02744 | 0.007739 | 0.001615 | 0.007965 | 0.065537 | 0.036863 | 0.017749 | 0.012296 | 0.004274 | 0.002866 | 0.003052 | 0.021597 | 0.010342 | 0.001783 | 0.005061 | 0.006161 | 0.009454 | 0.0042 | 0.009289 | 0.003928 | 0.001612 |
| 37.375 | 0.006331 | 0.015414 | 0.033504 | 0.008352 | 0.002556 | 0.013601 | 0.042381 | 0.065111 | 0.012233 | 0.01864 | 0.004999 | 0.002017 | 0.002996 | 0.034673 | 0.015111 | 0.003128 | 0.003945 | 0.004871 | 0.007811 | 0.003081 | 0.009658 | 0.00271 | 0.001856 |
| 37.3888889 | 0.004364 | 0.014938 | 0.03602 | 0.007503 | 0.005061 | 0.017377 | 0.020205 | 0.094454 | 0.007466 | 0.030113 | 0.005676 | 0.003702 | 0.005457 | 0.045158 | 0.0176 | 0.007486 | 0.003111 | 0.003718 | 0.007835 | 0.002254 | 0.010493 | 0.002226 | 0.002556 |

| | | | | | | | | | | | | | | | | | | | | | | | |
|---|---|---|---|---|---|---|---|---|---|---|---|---|---|---|---|---|---|---|---|---|---|---|---|
| 37.4027778 | 0.003077 | 0.009441 | 0.031302 | 0.005074 | 0.005651 | 0.019496 | 0.01678 | 0.096811 | 0.005587 | 0.031408 | 0.006198 | 0.007246 | 0.013556 | 0.049448 | 0.02003 | 0.013286 | 0.003123 | 0.003572 | 0.009798 | 0.002476 | 0.011912 | 0.002885 | 0.004108 |
| 37.4166667 | 0.005669 | 0.004043 | 0.019352 | 0.008096 | 0.007631 | 0.021078 | 0.028225 | 0.068546 | 0.003619 | 0.02328 | 0.006833 | 0.010282 | 0.023837 | 0.047052 | 0.021894 | 0.017976 | 0.002775 | 0.003092 | 0.011864 | 0.002415 | 0.011847 | 0.002939 | 0.005313 |
| 37.4305556 | 0.009341 | 0.004161 | 0.007759 | 0.016787 | 0.016577 | 0.022573 | 0.033604 | 0.036254 | 0.002917 | 0.013083 | 0.007561 | 0.012751 | 0.035253 | 0.040555 | 0.022313 | 0.021245 | 0.001773 | 0.001927 | 0.013143 | 0.001559 | 0.00986 | 0.001944 | 0.005763 |
| 37.4444444 | 0.011634 | 0.00648 | 0.003154 | 0.020448 | 0.020679 | 0.022086 | 0.024001 | 0.041243 | 0.008609 | 0.009676 | 0.006492 | 0.0134 | 0.043819 | 0.029497 | 0.021556 | 0.022772 | 0.001621 | 0.001909 | 0.01329 | 0.001497 | 0.006026 | 0.002118 | 0.004952 |
| 37.4583333 | 0.013693 | 0.009892 | 0.00328 | 0.0139 | 0.014908 | 0.017311 | 0.013008 | 0.084541 | 0.016946 | 0.013093 | 0.004438 | 0.010225 | 0.040074 | 0.017468 | 0.022598 | 0.021878 | 0.003706 | 0.004868 | 0.012877 | 0.003405 | 0.002938 | 0.005261 | 0.00276 |
| 37.4722222 | 0.013191 | 0.013588 | 0.003259 | 0.009486 | 0.013955 | 0.011722 | 0.019318 | 0.122852 | 0.017798 | 0.012936 | 0.004174 | 0.008626 | 0.026843 | 0.013583 | 0.029494 | 0.020842 | 0.005267 | 0.007055 | 0.013862 | 0.00485 | 0.002575 | 0.007239 | 0.001039 |
| 37.4861111 | 0.008715 | 0.012432 | 0.007021 | 0.014476 | 0.01798 | 0.013997 | 0.037855 | 0.118483 | 0.014507 | 0.013849 | 0.004769 | 0.012384 | 0.014594 | 0.017862 | 0.040359 | 0.02141 | 0.003565 | 0.004771 | 0.016218 | 0.003761 | 0.002283 | 0.004753 | 0.001475 |
| 37.5 | 0.003925 | 0.007898 | 0.014722 | 0.014869 | 0.013533 | 0.02596 | 0.048206 | 0.086608 | 0.015484 | 0.025765 | 0.007409 | 0.015756 | 0.008733 | 0.019691 | 0.047412 | 0.021096 | 0.001338 | 0.001977 | 0.017676 | 0.001692 | 0.00274 | 0.001754 | 0.003529 |
| 37.5138889 | 0.001685 | 0.00728 | 0.015529 | 0.008253 | 0.008163 | 0.039543 | 0.040928 | 0.052095 | 0.016214 | 0.02999 | 0.012113 | 0.013015 | 0.008109 | 0.019751 | 0.044697 | 0.017159 | 0.000751 | 0.001283 | 0.016941 | 0.000959 | 0.008682 | 0.000925 | 0.005038 |
| 37.5277778 | 0.001684 | 0.00903 | 0.01159 | 0.006465 | 0.010772 | 0.048558 | 0.023604 | 0.02237 | 0.012756 | 0.018145 | 0.01442 | 0.009835 | 0.011395 | 0.023989 | 0.029059 | 0.013359 | 0.004984 | 0.00484 | 0.013416 | 0.006409 | 0.024032 | 0.005237 | 0.006636 |
| 37.5416667 | 0.003343 | 0.007319 | 0.011843 | 0.009699 | 0.012186 | 0.055022 | 0.009554 | 0.010491 | 0.007531 | 0.006936 | 0.015162 | 0.019417 | 0.024339 | 0.021532 | 0.020786 | 0.024827 | 0.023107 | 0.022169 | 0.016947 | 0.024902 | 0.046995 | 0.022689 | 0.017063 |
| 37.5555556 | 0.005885 | 0.003656 | 0.011124 | 0.009709 | 0.020219 | 0.054832 | 0.00633 | 0.01259 | 0.003275 | 0.006633 | 0.015219 | 0.037141 | 0.041419 | 0.012219 | 0.034133 | 0.048508 | 0.049102 | 0.048869 | 0.032293 | 0.049249 | 0.066506 | 0.04708 | 0.036283 |
| 37.5694444 | 0.00821 | 0.002765 | 0.005234 | 0.00892 | 0.038512 | 0.039501 | 0.00928 | 0.012631 | 0.002981 | 0.01973 | 0.010958 | 0.039492 | 0.036425 | 0.008795 | 0.033199 | 0.049576 | 0.056077 | 0.05718 | 0.036134 | 0.056226 | 0.066536 | 0.055176 | 0.043648 |
| 37.5833333 | 0.011641 | 0.00627 | 0.001636 | 0.009757 | 0.043446 | 0.020593 | 0.009879 | 0.013499 | 0.009353 | 0.038184 | 0.004611 | 0.022679 | 0.024066 | 0.011917 | 0.017007 | 0.027528 | 0.037713 | 0.039035 | 0.020601 | 0.039479 | 0.046181 | 0.040338 | 0.030198 |
| 37.5972222 | 0.017088 | 0.010915 | 0.001874 | 0.007175 | 0.034585 | 0.015578 | 0.012117 | 0.01494 | 0.019413 | 0.049364 | 0.002574 | 0.009205 | 0.041449 | 0.023947 | 0.015147 | 0.019362 | 0.016799 | 0.01776 | 0.00902 | 0.018595 | 0.023803 | 0.019989 | 0.01292 |
| 37.6111111 | 0.02105 | 0.012776 | 0.003636 | 0.004184 | 0.030644 | 0.025619 | 0.015118 | 0.010876 | 0.022419 | 0.055437 | 0.00725 | 0.0093 | 0.074074 | 0.034599 | 0.017872 | 0.026425 | 0.006356 | 0.007125 | 0.009466 | 0.007064 | 0.011158 | 0.007265 | 0.006632 |
| 37.625 | 0.020945 | 0.013471 | 0.009769 | 0.004715 | 0.04003 | 0.039647 | 0.010725 | 0.009641 | 0.019251 | 0.05932 | 0.016576 | 0.012979 | 0.104787 | 0.029945 | 0.017636 | 0.030025 | 0.002958 | 0.003813 | 0.010025 | 0.00318 | 0.006492 | 0.002012 | 0.008327 |
| 37.6388889 | 0.019195 | 0.016849 | 0.015458 | 0.009322 | 0.057581 | 0.042106 | 0.003936 | 0.013226 | 0.020093 | 0.050572 | 0.025322 | 0.013806 | 0.119989 | 0.018952 | 0.014207 | 0.028786 | 0.002249 | 0.003524 | 0.007374 | 0.002802 | 0.005877 | 0.00086 | 0.009544 |
| 37.6527778 | 0.018867 | 0.020103 | 0.01477 | 0.012873 | 0.055999 | 0.034194 | 0.002815 | 0.015116 | 0.021625 | 0.030367 | 0.029437 | 0.012734 | 0.10255 | 0.0124 | 0.007515 | 0.021551 | 0.002091 | 0.003755 | 0.005556 | 0.003131 | 0.006732 | 0.001149 | 0.00931 |
| 37.6666667 | 0.019148 | 0.018743 | 0.012457 | 0.010855 | 0.029358 | 0.027927 | 0.005105 | 0.013356 | 0.018154 | 0.013273 | 0.027722 | 0.01169 | 0.064493 | 0.008442 | 0.006988 | 0.010398 | 0.001953 | 0.003592 | 0.004561 | 0.003094 | 0.007476 | 0.001472 | 0.009722 |
| 37.6805556 | 0.016411 | 0.013656 | 0.011368 | 0.01081 | 0.009738 | 0.021975 | 0.005247 | 0.009216 | 0.013354 | 0.005726 | 0.022499 | 0.011566 | 0.030586 | 0.005543 | 0.015031 | 0.006478 | 0.001885 | 0.003263 | 0.003473 | 0.002963 | 0.007831 | 0.001523 | 0.010472 |
| 37.6944444 | 0.009684 | 0.008753 | 0.008285 | 0.015845 | 0.009897 | 0.018216 | 0.003668 | 0.004929 | 0.010057 | 0.005306 | 0.018472 | 0.012263 | 0.026435 | 0.010362 | 0.023316 | 0.009953 | 0.001931 | 0.003031 | 0.002113 | 0.003034 | 0.008145 | 0.001368 | 0.010939 |
| 37.7083333 | 0.00488 | 0.008152 | 0.00345 | 0.022402 | 0.016568 | 0.018786 | 0.003232 | 0.00182 | 0.006871 | 0.007289 | 0.014889 | 0.013109 | 0.057919 | 0.018179 | 0.021138 | 0.010904 | 0.00195 | 0.002701 | 0.001176 | 0.003038 | 0.008193 | 0.001073 | 0.010983 |
| 37.7222222 | 0.007173 | 0.011529 | 0.001394 | 0.026147 | 0.016502 | 0.016903 | 0.004011 | 0.001186 | 0.003309 | 0.010834 | 0.009209 | 0.013231 | 0.101972 | 0.014526 | 0.011429 | 0.007994 | 0.00154 | 0.001909 | 0.001504 | 0.00251 | 0.007195 | 0.000751 | 0.010391 |
| 37.7361111 | 0.012759 | 0.015164 | 0.003817 | 0.022132 | 0.012305 | 0.012669 | 0.0051 | 0.003035 | 0.000922 | 0.016797 | 0.005859 | 0.012839 | 0.125039 | 0.008754 | 0.005355 | 0.007324 | 0.000899 | 0.001067 | 0.00179 | 0.00143 | 0.004937 | 0.000919 | 0.009444 |
| 37.75 | 0.016363 | 0.016967 | 0.007556 | 0.012156 | 0.017398 | 0.009365 | 0.005378 | 0.005071 | 0.001122 | 0.022282 | 0.01056 | 0.011935 | 0.112471 | 0.018692 | 0.003757 | 0.009866 | 0.000778 | 0.000989 | 0.001131 | 0.000638 | 0.002392 | 0.001584 | 0.008556 |
| 37.7638889 | 0.015538 | 0.016422 | 0.008222 | 0.007334 | 0.028308 | 0.011524 | 0.003682 | 0.007727 | 0.004328 | 0.024215 | 0.021136 | 0.009926 | 0.079942 | 0.035895 | 0.004107 | 0.010369 | 0.00106 | 0.001323 | 0.001087 | 0.000644 | 0.000761 | 0.001915 | 0.007658 |
| 37.7777778 | 0.009315 | 0.013443 | 0.005155 | 0.014645 | 0.032916 | 0.021389 | 0.002635 | 0.012916 | 0.007659 | 0.021644 | 0.029553 | 0.007279 | 0.041928 | 0.043661 | 0.007755 | 0.006538 | 0.001271 | 0.001517 | 0.002416 | 0.000952 | 0.000333 | 0.001879 | 0.007136 |
| 37.7916667 | 0.005514 | 0.009698 | 0.002223 | 0.023799 | 0.029236 | 0.031253 | 0.006578 | 0.019533 | 0.006753 | 0.01674 | 0.032443 | 0.00491 | 0.021965 | 0.047067 | 0.016312 | 0.004498 | 0.001481 | 0.001767 | 0.003859 | 0.001287 | 0.000844 | 0.00202 | 0.007724 |
| 37.8055556 | 0.010731 | 0.007668 | 0.003066 | 0.024684 | 0.021599 | 0.035674 | 0.013644 | 0.023866 | 0.003377 | 0.018925 | 0.033056 | 0.003157 | 0.042037 | 0.056024 | 0.023626 | 0.010451 | 0.001619 | 0.00199 | 0.004121 | 0.001491 | 0.002004 | 0.002065 | 0.008962 |
| 37.8194444 | 0.015908 | 0.006819 | 0.006572 | 0.01999 | 0.013999 | 0.037152 | 0.018602 | 0.019155 | 0.002303 | 0.026987 | 0.033579 | 0.002051 | 0.076226 | 0.062447 | 0.025342 | 0.019888 | 0.001399 | 0.00191 | 0.003379 | 0.001267 | 0.003716 | 0.001451 | 0.009709 |
| 37.8333333 | 0.014728 | 0.005583 | 0.007202 | 0.014942 | 0.008051 | 0.037139 | 0.019847 | 0.012176 | 0.003449 | 0.025261 | 0.033645 | 0.001911 | 0.095809 | 0.065025 | 0.024561 | 0.025452 | 0.000826 | 0.001523 | 0.002787 | 0.000806 | 0.004935 | 0.000916 | 0.009514 |
| 37.8472222 | 0.012206 | 0.004196 | 0.005371 | 0.009378 | 0.003862 | 0.035528 | 0.018086 | 0.01854 | 0.00511 | 0.016281 | 0.033131 | 0.003585 | 0.10111 | 0.068143 | 0.021272 | 0.02531 | 0.000501 | 0.001005 | 0.00267 | 0.000913 | 0.008077 | 0.001491 | 0.008438 |
| 37.8611111 | 0.011205 | 0.003305 | 0.00406 | 0.011364 | 0.002694 | 0.031973 | 0.013848 | 0.027832 | 0.005902 | 0.018374 | 0.032209 | 0.006541 | 0.097315 | 0.062411 | 0.01509 | 0.020233 | 0.001035 | 0.001087 | 0.002015 | 0.002019 | 0.018342 | 0.002147 | 0.006754 |
| 37.875 | 0.011123 | 0.003321 | 0.002792 | 0.029387 | 0.005878 | 0.027081 | 0.008435 | 0.023096 | 0.005433 | 0.035468 | 0.03074 | 0.008223 | 0.087403 | 0.048337 | 0.008391 | 0.012882 | 0.00207 | 0.002149 | 0.001551 | 0.003398 | 0.030571 | 0.001933 | 0.004928 |
| 37.8888889 | 0.011689 | 0.003953 | 0.003636 | 0.049376 | 0.014754 | 0.024045 | 0.003732 | 0.012442 | 0.005463 | 0.052475 | 0.02827 | 0.007372 | 0.074238 | 0.042027 | 0.003492 | 0.007006 | 0.002895 | 0.003198 | 0.003322 | 0.004241 | 0.036698 | 0.001845 | 0.00347 |
| 37.9027778 | 0.011783 | 0.004242 | 0.004456 | 0.057875 | 0.029097 | 0.024701 | 0.0017 | 0.007318 | 0.004329 | 0.058724 | 0.025913 | 0.00489 | 0.059959 | 0.039286 | 0.001417 | 0.003903 | 0.002953 | 0.003154 | 0.005456 | 0.004019 | 0.03752 | 0.002085 | 0.00285 |
| 37.9166667 | 0.009089 | 0.004324 | 0.003359 | 0.056165 | 0.041859 | 0.029887 | 0.003187 | 0.005489 | 0.001714 | 0.056862 | 0.025386 | 0.002127 | 0.039618 | 0.029184 | 0.00238 | 0.002041 | 0.002159 | 0.002317 | 0.004859 | 0.002732 | 0.034163 | 0.001932 | 0.00362 |
| 37.9305556 | -0.00617 | -0.01002 | -0.02109 | -0.08516 | -0.08404 | -0.09378 | -0.01153 | -0.00274 | -0.00302 | -0.07526 | -0.05224 | -0.00021 | -0.04939 | -0.02271 | -0.01959 | -0.00098 | -0.00779 | -0.00957 | -0.00592 | -0.00822 | -0.03349 | -0.00973 | -0.01718 |
| 37.8611111 | -0.00594 | -0.0097 | -0.01561 | -0.09128 | -0.09406 | -0.08618 | -0.01231 | -0.00425 | -0.00175 | -0.09341 | -0.05175 | 4.81E-05 | -0.03152 | -0.02889 | -0.01297 | -0.00107 | -0.00469 | -0.00616 | -0.00422 | -0.00504 | -0.04526 | -0.00609 | -0.01377 |
| 37.875 | 0.009739 | 0.004799 | 0.011523 | 0.042023 | 0.017906 | 0.04974 | 0.004689 | 0.003136 | 0.003813 | 0.013784 | 0.027274 | 0.001564 | 0.08045 | 0.011498 | 0.020034 | 0.002675 | 0.009308 | 0.010122 | 0.009291 | 0.009763 | 0.004692 | 0.010607 | 0.012153 |
| 37.8888889 | 0.013658 | 0.003434 | 0.008082 | 0.044969 | 0.010107 | 0.052468 | 0.011057 | 0.007623 | 0.004987 | 0.012628 | 0.027844 | 0.001938 | 0.11025 | 0.010353 | 0.026091 | 0.004379 | 0.011101 | 0.011804 | 0.012797 | 0.011569 | 0.001075 | 0.012516 | 0.012747 |
| 37.9027778 | 0.01584 | 0.003488 | 0.010823 | 0.050156 | 0.016882 | 0.057291 | 0.02224 | 0.013394 | 0.008476 | 0.016578 | 0.025775 | 0.00195 | 0.114379 | 0.005949 | 0.035278 | 0.003368 | 0.010773 | 0.011354 | 0.014743 | 0.011255 | 0.000424 | 0.012115 | 0.010953 |
| 37.9166667 | 0.017917 | 0.007049 | 0.015904 | 0.054051 | 0.02679 | 0.061258 | 0.040084 | 0.016179 | 0.015098 | 0.016142 | 0.021769 | 0.003028 | 0.087829 | 0.002672 | 0.045454 | 0.002795 | 0.009701 | 0.010178 | 0.015679 | 0.0103 | 0.00153 | 0.010957 | 0.009282 |
